# Discovery of diverse chimeric peptides in a eukaryotic proteome sets the stage for the experimental proof of the mosaic translation hypothesis

**DOI:** 10.1101/2024.12.01.626167

**Authors:** Umut Çakır, Noujoud Gabed, Yunus Emre Köroğlu, Selen Kaya, Senjuti Sinharoy, Vagner A. Benedito, Marie Brunet, Xavier Roucou, Igor S. Kryvoruchko

## Abstract

The high complexity of eukaryotic organisms enabled their evolutionary success, which became possible due to the diversification of eukaryotic proteomes. Various mechanisms contributed to this process. Alternative splicing had the largest known impact among these mechanisms: tens or hundreds of protein isoforms produced from a single genetic locus. Earlier, we hypothesized that along with alternative splicing, a different but conceptually similar mechanism creates novel versions of existing proteins in all eukaryotes. However, this mechanism acts at the level of translation, where the novelty of an amino acid sequence is achieved via multiple programmed ribosomal frameshifting. This mechanism, which is termed mosaic translation, is very difficult to demonstrate even with the most up-to-date molecular tools. Thus, it remained unnoticed so far. Using only a portion of all mass spectrometry proteomic data generated from various organs of the model plant *Medicago truncatula*, we attempted the first step toward the experimental proof of this hypothesis. Our original *in silico* approach resulted in the discovery of two candidates for mosaic proteins (homologs of EF1α and RuBisCo) and 154 candidates for chimeric peptides. Chimeric peptides and polypeptides are produced in the course of one ribosomal frameshifting event and may correspond to parts of mosaic proteins. In addition, our analysis reveals the possibility of translation of chimeric peptides from five ribosomal RNA transcripts, ten long non-coding RNA transcripts, and one transfer RNA transcript. These findings are very novel and will be the basis for experimental validation in future studies. In this work, we present multiple lines of indirect evidence that support the validity of our *in silico* data.

## 1. Introduction

Recently, it has been recognized that eukaryotic transcripts have a polycistronic nature, which revolutionized our understanding of the proteome complexity (Mouilleron et al., 2016). The central point in this paradigm-shifting development was the discovery of translated alternative open reading frames (altORFs) and their inventory in the variety of organisms (Brunet et al., 2019, 2021; Leblanc et al., 2024). These regions of transcripts can be defined as relatively long stop-free sequences in any reading frame. Products of their translation, alternative proteins (altProts), may resemble some annotated proteins or be entirely unique. In both cases, they are thought to be an important source of protein novelty in the evolution (Orr et al., 2020; Riegger and Caliskan, 2022; Ardern, 2023). AltORFs often overlap with annotated coding sequences, also called reference ORFs (refORFs), or other altORFs located on the same transcript (Mouilleron et al., 2016; Orr et al., 2020). Earlier, high abundance of conserved altORFs in plant transcriptomes prompted us to hypothesize that a mechanism by which the information from overlapping ORFs is combined in continuous polypeptide molecules in the course of multiple programmed ribosomal frameshifting (PRF) may have importance in the adaptability of organisms to internal and external conditions (Çakır et al., 2023). In contrast to a chimeric peptide or polypeptide, which originates from a single PRF event per transcript, with only a few known examples in prokaryotes (Blinkowa and Walker, 1990; Flower and McHenry, 1990; Tsuchihashi and Kornberg, 1990; Chaijarasphong et al, 2016; Meydan et al., 2017) and eukaryotes (Matsufuji et al., 1995; Clark et al., 2007; Ivanov and Atkins, 2007; Ren et al., 2024), a peptide or polypeptide produced via two and more PRF events has a mosaic nature because it combines translational products of multiple reading frames (Ketteler, 2012). Thus, we refer to this mode of translation as mosaic translation. Mosaic Gag-Pro-Pol polypeptides produced by this mechanism have been described in some viruses (Jacks, 1990; Hatfield et al., 1992; Rex et al., 2019), which have to overlap their genes with high density for the more complete usage of their limited genomic space. In contrast, this way of using the coding potential of altORFs has not been anticipated to play an important role in eukaryotes, probably because of their large genomes. To the best of our knowledge, only one study has suggested the existence of mosaic proteins in eukaryotes (Ketteler, 2012). We took a long step forward and proposed why mosaic translation may be a ubiquitous phenomenon with fundamental importance in all domains of life. Our chief argument was the enormous biological advantage that should result from the manifold expansion of protein-coding capacity. The direct proof of this hypothesis is nearly impossible without a revolution in the read length of protein sequencers. The plan to develop a long-read method by which mosaic proteins can be discovered at a large scale has been mentioned by Timp and Timp (2020). It is based on the nanopore sequencing principle, which proved to be very useful in long-read sequencing of nucleic acids (Pugh, 2023). Despite very recent revolutionary developments in nanopore sequencing of proteins, the long-read version of the method is still not available due to major challenges. The first challenge is slowing down and stretching the long polypeptide molecules during their transit through the nanopore. The second challenge is discriminating among amino acids with different posttranslational modifications (Martin-Baniandres et al., 2023; Wang et al., 2024). Due to the absence of any adequate alternative to the protein nanopore sequencing, earlier, we proposed a strategy that can detect candidates for mosaic translation based on available mass spectrometry (MS) proteomic data. If conserved and/or translated altORFs overlap with refORFs, other altORFs, or both, possible PRF events that change the translation from one frame to another can be modeled. Once the frameshifted sequences are modeled, they can be used as a query database in the searches for corresponding MS peptides in biological samples. This way, we proposed to detect individual chimeric peptides in the first place. If two or more chimeric peptides validated by MS originate from the same transcript, depending on the frameshift type and the distance between PRF sites, they can be parts of a mosaic protein. Despite very large computational requirements, the principle advantage of our approach is the use of MS proteomic data. For many organisms, such data are already available in a very large amount (Perez-Riverol et al., 2022; Deutsch et al., 2023). Up to 60% of high-quality peptides derived from eukaryotes that have been identified by MS cannot be assigned to any genomic location (Ning et al., 2010; Pathan et al., 2017). A similar problem has been found in assigning the minimal proteome of the smallest culturable bacterium with only 729 ORFs (Lluch-Senar et al., 2016). Earlier, it was suggested that this unmapped “dark” proteome could be composed of translational products of altORFs (Mouilleron et al., 2016; Orr et al., 2020). A recent study in humans identified many thousands of previously unknown peptides, so-called non-canonical ORFs (ncORFs), which further emphasizes the true complexity of the proteome (Cao et al., 2024). We extend this idea to mosaic peptides and proteins, which are made of altORF/ncORF translation blocks. Novel PRF sites can be detected with ribosome profiling, or Ribo-Seq (Ingolia et al., 2011; Kiniry et al., 2019, 2021; Richardson and Eddy, 2023). While Ribo-Seq-based information about positions of the frameshifts is expensive and technically challenging to generate, it is also indirect compared to the information deduced from MS proteomic reads. Many organisms have no Ribo-Seq datasets deposited publicly, including the model plant *Medicago truncatula*. Here we demonstrate the efficient application of our MS-based approach on three selected proteomic datasets from *M. truncatula*. To our knowledge, this pilot study is the first attempt to demonstrate the existence of mosaic peptides and proteins in a non-viral biological system. Although we detected only two good candidates for mosaic translation in the selected three datasets, this approach can be extended to other datasets of *M. truncatula* and MS data from other organisms, which can ultimately lead to the discovery of the large number of mosaic proteins. Importantly, in the course of this work, we also detected many chimeric peptides previously unthought for a eukaryotic genome. We hope that the novelty of our observations will ultimately bring chimeric and mosaic proteins into the spotlight and will motivate the scientific community to apply our approach to other organisms, including humans.

## 2. Material and methods

### 2.1. *In-silico* extraction and translation of altProts

The *M. truncatula* genome assembly and annotated features (v. 5.1.7) were downloaded from the *M. truncatula* genome portal MtrunA17r5.0-ANR (Pecrix et al., 2018; https://medicago.toulouse.inra.fr/MtrunA17r5.0-ANR/). In each transcript, we identified regions that are at least 60 base pairs long and do not contain any in-frame stop codons in three reading frames. These regions were then translated into sequences of amino acids using the standard genetic code. This way, we generated a query database of peptide and polypeptide sequences (altProts), from which reference proteins (refProts) were eliminated based on their identity to sequences from the annotated proteome. Input and output sequences of the corresponding script are FASTA-formatted. The script supplies output sequences with unique identifiers that contain the following information: locus identifier, reading frame (three forward frames denoted as 1F to 3F), nucleotide coordinates of the extracted region on the transcript (first base and last base), and the length of the extracted region on the transcript in bases. The code for generating this query database will be published elsewhere (Çakır et al., preprint in preparation).

### 2.2. Sequence similarity searches using a protein reference database

Each altProt was compared with entries from the reference protein database UniProt v. 2020_02 (UniProt Consortium, 2019) using DIAMOND v. 0.9.14 (Buchfink et al., 2021). To build a DIAMOND search database using a “makedb” command, a single fasta-file was generated from the downloaded and concatenated UniProt sequences. The following parameters were used for the similarity searches: “max-target-seqs”, which is the maximum number of hits, was set to zero; this setting enabled the report of all hits per query; the maximal expect value was set to 10^−3^; to enable higher sensitivity of the searches, an option “more-sensitive” was used. For efficient downstream analysis of significant hits, the following information was recorded for each sequence with at least one hit: “qseqid”, “stitle”, “length”, “pident”, “ppos”, “qcovhsp”, “evalue”, and the number of hits. The information on taxonomy of all hits was recorded for in-depth phylogenetic analysis.

### 2.3. Validation of altProts by MS proteomics

We searched for mass spectra-derived peptides that match each *in-silico* generated altProt with the aid of SearchGUI v. 4.0.41 (Barsnes & Vaudel, 2018) and its partner tool PeptideShaker v. 2.0.33 (Vaudel et al., 2015). Three proteomic datasets publicly deposited at the ProteomeXchange database (Deutsch et al., 2023) were used for this purpose: PXD002692 (Marx et al., 2016), PXD013606 (Shin et al., 2021), and PXD022278 (Castañeda et al., 2021), which contain nine, one, and six samples respectively (Supplementary Dataset S1). ThermoRawFileParser v. 1.1.2 (Hulstaert et al., 2020) was employed for the conversion of raw data from these three datasets to Mascot Generic Format (MGF). X!Tandem, MS-GF+, OMSSA, and Comet search algorithms were used in all searches. The search database consisted of the following three components: (1) *in-silico* translated altProts; (2) all annotated refProts; and (3) the contaminant database known as cRAP (common Repository of Adventitious Proteins), which was downloaded from the GPM resource (GPM: Generalized Proteomics data Meta-analysis; https://www.thegpm.org/crap/). Validated altProts were excluded from the analysis if they grouped with a refProt or a contaminant sequence (Related Proteins). Carbamidomethylation of C was set to fixed modification, and acetylation of protein N-termini and oxidation of M were set to variable modifications. Precursor and fragment tolerance were set to 4.5 ppm and 20.0 ppm, respectively. At most, two missed cleavages were allowed. False discovery rate (FDR) of 1% was used to validate peptide-spectrum matches (PSMs), peptides, and proteins with target/decoy hit distribution. The decoy spectral library was generated by *in-silico* reversing the collection of refProt, altProt, and contaminant sequences. Software version-specific default settings were used for parameters that are not mentioned here. The results of PeptideShaker confidence classification of the chimeric peptides are available in Supplementary Dataset S1.

The inclusion of all mRNA-derived altProts in the analysis causes the inflation of the search database (more than 840,000 after the inclusion of refProts, Supplementary Table S1), which strongly affects the number of confident protein identifications (Li et al., 2016; Kumar et al., 2017). To avoid the database inflation, a two-step MS search approach was used (Jagtap et al., 2013). In the first step, the list of altProt was shuffled and split into ten chunks. Each chunk was used as a search database, and proteins validated in this first step were subsequently included in the second search. Because there were 16 organs/conditions in datasets PXD002692, PXD013606, and PXD022278 combined and ten groups of the mRNA-derived altProt sequences, 160 MS searches were conducted in the first step of the two-step approach and 16 in the second step for these three datasets. The parameters of individual searches were specified above. The two-step approach was not used for non-mRNA-derived altProts because their relatively small number does not inflate the search database. Thus, a single-step MS search protocol was used for those sequences. MS searches for ncRNA, rRNA, and tRNA-derived altProts were conducted separately. Because there were 16 organs/conditions in total in the three MS datasets combined, 16 MS searches were conducted for each group of non-mRNA-derived altProts. All validated altProts were recorded for further analysis. The MS datasets were also searched independently, for instance, validated proteins from dataset PXD002692 were not combined with those from PXD013606 or PXD022278 for the search database of the second step.

### 2.4. Modeling of chimeric proteins for MS searches

Throughout the text, we discriminate between terms “chimeric peptide” and “chimeric protein”. Namely, we reserve the former term for short amino acid sequences deduced from MS proteomics. In contrast, *in-silico* generated models for matching corresponding MS peptides are referred to as chimeric protein models even though their actual length generally does not exceed 40 amino acids in our study. This is meant to emphasize that these models may represent longer chimeric sequences or fragments of mosaic proteins. We also use the term “chimeric protein” in contexts where the length is not relevant.

Although our in-house script can use the list of any overlapping or non-overlapping ORFs on the same transcript as a starting point, regardless of their conservation and translation status, the number of chimeric protein models thus generated would be astronomic. For this reason, we modeled chimeric proteins based on two lists of altProts. The first list contained altProts validated by MS searches, and the second list contained so-called conserved altProts. The definition of “conserved” in this case refers to the presence of at least one hit with at least 70% identity (e-value below 0.001) in the global sequence similarity search using UniProt as a reference protein database.

Each MS-validated and/or conserved altProt has a corresponding altORF with coordinates on its transcript and locus information. Our chimeric protein modeling algorithm determines and *in-silico* translates products of many possible PRF events that may occur if an altORF overlaps with its refORF or other altORFs on the same transcript. We use the word combination “many possible PRF events” instead of “all possible PRF events” to emphasize that some situations were deliberately left beyond the scope of our analysis. For example, we considered only PRF values −2, −1, +1, and +2, which correspond to the backward and forward slippage of ribosomes, respectively, by one or two nucleotides, although “longer” events that cause the frameshifting by up to six bases are known (Weiss et al., 1987; Yan et al. 2015). The reason for focusing on the “shortest” PRF events and the simplest scenarios was the phenomenon of the search database inflation (Li et al., 2016; Kumar et al., 2017). In short, generating models for all theoretically possible chimeric proteins is technically feasible with our algorithm. However, it is counterproductive to include all the models into the analysis. Thus, we included only situations that would serve as the most convincing illustration of chimeric translation without inflating the MS search database. This explains why the settings for the generation of chimeric models were not uniform for all cases but were tailored to specific scenarios described in the corresponding software article (Çakır et al., preprint in preparation). Furthermore, those settings depended on the location of the frameshift relative to the involved ORFs (the 5’-end vs the 3’-end).

### 2.5. Validation of chimeric protein models by MS proteomics

The number of chimeric protein models derived from MS-validated altProts is moderate (36,536) and does not cause inflation of the search database. In contrast, the models derived from conserved altProts are numerous (533,569), which causes inflation of the search database (see Section 4.3). Thus, similarly to the MS-validation of mRNA-derived altProts, the models from the latter group were searched by the two-step MS approach, while the regular MS searches were conducted for the validation of the former group. The validation of chimeric models involved the same three MS datasets and the same pipeline as the validation of altProts. The list of chimeric protein models derived from the conserved altProts was split into ten chunks. Each chunk was searched separately in 16 organs/conditions in the first step. Then, validated chimeric protein models were searched one more time (phase two of the two-step approach). Validated models were recorded for further analysis. Because there were 16 organs/conditions in those three datasets, 160 and 16 MS searches were conducted in the first and second steps of the two-step approach, respectively, for the models derived from conserved altProts. Unlike in the altProt validation, altProts used for the modeling of chimeric proteins were also included in the search database in addition to the refProt and cRAP databases, which permitted the elimination of chimeric peptides identical to sequences of altProts. This was necessary because some chimeric peptides were different from their altProts or refProts by only one or two amino acids. If these different amino acids are indistinguishable by MS, true altProts or true refProts may be categorized as chimeric proteins. Validated chimeric protein models were excluded from the analysis if they grouped with an altProt, a refProt, or a contaminant sequence. Thus, our final dataset (Supplementary Dataset S1) is free from this ambiguity. Like with the validation of altProts, the three MS datasets were searched independently.

### 2.6. Visualization of chimeric MS peptides, chimeric protein models, mosaic proteins, and their respective transcripts

Sequences of MS-supported chimeric peptides were idenfied and manually mapped to their transcripts in Geneious^®^ v. 7.1 (Dotmatics Ltd., MA, USA, https://www.geneious.com). The location of each PRF event, its value (−2, −1, +1, or +2), type, and subtype (Supplementary Dataset S1) were deduced manually. To create images of transcripts associated with each PRF event, we calculated sizes and coordinates of each feature in Geneious^®^ and then converted them to distances in centimeters to be plotted. Transcript lengths were scaled to either 10 cm or 20 cm in the images, depending on the purpose of each figure, so that transcripts of two different lengths in nucleotides appear equal in lengths in the images. This way, we unified visualization of transcripts that ranged from 84 bp to 10,214 bp.

### 2.7. Protein folding predictions

To estimate the proportion of potentially unstructured protein sequences in our collection of putative chimeric peptides, and also to discriminate between sequences that fold into alpha-helices and beta-sheets, we predicted their folding with ColabFold v1.5.2 (Mirdita et al., 2022) and visualized it with ChimeraX v. 1.6.1 (Pettersen et al., 2004). The predictions were run in the unrelaxed mode. Because MS peptides that support chimeric models are too short for folding predictions (8-30 aa, Supplementary Dataset S1), we based the predictions on chimeric models, which are longer than MS peptides. Five predictions per sequence were made, out of which the top-score predictions, and in some cases strongly differing additional predictions with a lower score, were depicted.

### 2.8. Homology searches

To identify homology groups within the dataset, transcript and protein sequences that correspond to chimeric peptides were aligned with a Geneious^®^ alignment tool, which is a built-in option in the Geneious^®^ software, and also with Clustal Omega (Sievers et al., 2011). To identify small-scale local similarities among sequences, we employed the MEME (Multiple EM for Motif Elicitation) software v. 5.4.1 (Bailey and Elkan, 1994; Bailey et al., 2015). In addition, BLASTN and BLASTP tools (Altschul et al., 1990; Boratyn et al., 2013) and the *M. truncatula* genome portal (Pecrix et al., 2018) were used for the identification of homology groups based on shared subjects. The same tools were used for the annotation of transcripts and translational products of refORFs and altORFs involved in the production of chimeric peptides (Supplementary Dataset S1).

### 2.9. Identification of alternative sources

In searches for non-chimeric alternative sources of chimeric MS peptides, we used genomic, repeat element, and transcript data from the *M. truncatula* genome portal (Pecrix et al., 2018), 50 selected RNA-Seq datasets of *M. truncatula* deposited at the Sequence Read Archive (SRA) database (Leinonen et al., 2011; Katz et al., 2022), and the RNA-Seq-based gene expression atlas of this organism (MtExpress v. 3, Carrere et al., 2021; https://medicago.toulouse.inrae.fr/GEA). In-house scripts based on TBLASTN were used for the similarity searches in this part of the study. MtExpress was also used for transcription profiles of primary sources and the selection of the most likely alternative chimeric sources listed in Supplementary Dataset S1. In searches for chimeric alternative sources, we used a script that will be published separately as the area of its application is broad (Çakır et al., preprint in preparation). Transcript and repeat-element data from the newer version 5.1.9 of the *M. truncatula* genome browser (Pecrix et al., 2018) were used in this analysis. Each primary-source transcript was manually analyzed for differences between v. 5.1.7 and v. 5.1.9. There were no sequence or length differences found between these two genome releases for any of the primary-source transcripts.

### 2.10. Statistical analysis

To estimate the deviation from the randomness assumption in the relationships between various features of chimeric peptides and their transcripts, we mostly employed the chi-square tests for the association and the goodness-of-fit as conservative non-parametric tools. In some cases, we used Pearson and Spearman correlation analyses along with corresponding t-tests for the significance of their p-values. For the assessment of differences between BLASTP and BLASTN-related characteristics of four RNA types (mRNA, ncRNA, rRNA, and tRNA), Kholmogorov-Smirnov test was utilized in R (R Core Team, 2021). This test is an adequate non-parametric procedure suitable for the comparisons of samples that have very different variances and sizes. In cases where the application of the chi-square test was inappropriate (at least one cell with an expected value below five), three alternative tests were used in R: (1) the Fisher’s exact test for more than one level and more than two proportions; (2) the Exact Multinomial Test for one level and more than two proportions; (3) the Exact Binomial Test for one level and two proportions. In some cases, indicated in supplementary figures, the exact tests could not be run because of a large number of categories and/or very low values in specific groups. An alpha-level of 5% was used as a significance threshold in all tests. The default setting of all tests mentioned above is to calculate a two-sided p-value. Graphs for supplementary figures were generated in Microsoft Excel and in some cases in R or SPSS^®^ Statistics v. 27.0 (IBM^®^ Corporation, NY, USA, https://www.ibm.com/products/spss-statistics).

## 4. Results

### 4.1. The *M. truncatula* transcriptome contains thousands of altORFs with a conservation signature

To enable a comprehensive study with potential for novel observations, we have not limited our analysis to mRNA transcripts. We included three other RNA types that are classically defined as non-coding: ncRNA, rRNA, and tRNA, which are among the longest of all known RNA types (Palazzo and Lee, 2015; Boivin et al., 2019). With the same intention, we have not limited our definition of an open reading frame (ORF) to transcript regions that start with or contain the standard translation initiation codon AUG (Sieber et al., 2018). Here we define an ORF as a stop-free region above certain length in any forward reading frame, which conceptually has the potential to be translated to a short peptide or a longer chain of amino acids. Eventually, this ORF definition proved to be useful as it led to the discovery of many translated ORFs that do not start with an ATG or even contain no ATG at any position (results not shown). On the other hand, the inclusion of non-coding RNA types allowed us to detect chimeric peptides potentially translated from all the three groups of transcripts, including tRNA, and non-chimeric altProts translated from ncRNA.

Because we planned to use the DIAMOND BLASTP analysis as a part of our detection pipeline, we chose 60 nt as a minimal ORF length. This decision was conditioned by the intrinsic limitations of the BLASTP algorithm in handling peptide sequences shorter than 20 aa (Altschul, 1991). The annotated transcriptome v. 5.1.7 of *M. truncatula* (Pecrix et al., 2018) contains 875,356 altORFs longer than 59 nt, which correspond to putative altProts of 20 aa or longer (Supplementary Table S1). We extracted these sequences by *in-silico* translation and subjected them to the BLASTP analysis. AltProts that had at least one hit were retained for further analysis if they passed the following threshold: e-value at most 0.001, percent amino acid identity at least 70. There were 13,078 such sequences (Table 1), which we conditionally call conserved altProts, although in reality their conservation is often limited to the *M. truncatula* proteome. Table 1, Supplementary Figures S1-S5, and Supplementary Tables S1-S4 illustrate details and statistics of the BLASTP results.

**Table 1.** Statistics on altProts with at least one hit in the global BLASTP analysis after the application of the filter^1^.

| Statistic | mRNA | ncRNA | rRNA | tRNA |
| --- | --- | --- | --- | --- |
| Number of transcripts | 44,624 | 5,657 | 62 | 974 |
| Median length of transcripts, nt | 1,280 | 413 | 120 | 75 |
| Total number of altORFs<sup>2</sup> | 8,718 | 3,549 | 385 | 426 |
| altORFs per transcript | 0.20 | 0.63 | 6.21 | 0.44 |
| Transcripts per altORF | 5.12 | 1.59 | 0.16 | 2.29 |
| Median length of altProts, aa | 53.0 | 47.0 | 45.0 | 25.0 |
| Median length of alignment, aa | 38.0 | 36.0 | 40.0 | 24.0 |
| Median percent identity | 87.5 | 91.4 | 93.3 | 100.0 |
| Median percent query coverage | 82.5 | 84.5 | 94.8 | 100.0 |
| altORFs longer than 300 nt | 795 | 215 | 15 | 0 |
| altORFs longer than 600 nt | 33 | 27 | 3 | 0 |

<sup>1</sup>BLASTP filter settings: e-value ≤ 0.001; percent identity ≥ 70.
<sup>2</sup>There are 13,078 of such altORFs combined from the RNA types mentioned here.

### 4.2. Three public MS proteomic datasets of *M. truncatula* provide evidence for translation of 805 putative altProts

In parallel with the BLASTP analysis, all 875,356 putative altProts were analyzed for the presence of corresponding MS spectra in three selected proteomic datasets of *M. truncatula* (Marx et al., 2016; Shin et al., 2021; Castañeda et al., 2021). This analysis revealed 805 unique putative altProts with evidence for translation, 122 of which are also present in the list of conserved altProts. Supplementary Figures S4 and S5 show distributions of the top-hit percent identity and the number of hits, respectively, for the 122 conserved MS-supported altProts. Most of the altProts detected by MS (720 unique sequences) correspond to mRNA transcripts. However, we have also found 85 unique ncRNA-derived altProts supported by MS. No validated non-chimeric MS peptides were found for rRNA-altProts and tRNA-altProts (Supplementary Figure S6). In total, 16 biological samples contain MS peptides corresponding to altProts. More than half of these peptides (573) were detected in just four samples: seeds (155), 10-dpi nodules (147), flowers (136), and buds (135), which may point to the importance of altProts in the reproduction and symbiotic nitrogen fixation (Supplementary Figure S6).

In the BLASTP analysis, the taxonomic status of the top-hit subject sequence may reflect the degree of conservation of the query sequence among different organisms, especially if the number of significant hits is small. In our analysis, 70, 65, and 39% of altProts have between 1 and 5 hits before the application of the 70% filter, after the filter, and in the set of conserved MS-supported altProts, respectively. The corresponding median numbers of hits in these three samples are 2, 2, and 16.5, which is rather low (Supplementary Figures S2, S3, and S5). This indicates that, as a whole, altProts detected in our study do not share similarity with proteins from many species but just a few. A very large portion of these altProts have the top-similarity with proteins from *M. truncatula* itself or from other legume species (Supplementary Figure S7). At the same time, the majority of MS-supported altProts (683 out of 805, which is ca. 85%) have no similarity with any annotated protein at all. On the other hand, percent amino acid identity with the top hit is another parameter that may reflect the degree of conservation. In our dataset (Supplementary Figure S1), median values of percent identity are very high, which may point to the potential origin of many altProts from duplication events and frameshifting mutations in the *M. truncatula* genome and its common ancestors with other legume species. Percent identity reaches the maximum in tRNA-altProts (100%) followed by rRNA-altProts (93.3%), which has also the highest median number of hits (Supplementary Figure S1). Together with other parameters described in Table 1, these observations highlight a very special role of tRNA and rRNA in genome evolution, which was proposed earlier (Root-Bernstein and Root-Bernstein, 2015, 2016, 2019; Caetano-Anollés and Caetano-Anollés, 2016; de Farias et al., 2016).

### 4.3. Modeling of chimeric proteins based on the combined evidence for conservation and translation helped to validate MS peptides that match 156 chimeric models

Our next goal was to generate chimeric models for matching them to peptides present in MS proteomic samples. We searched for altORFs that overlap with annotated coding sequences (refORFs) and/or each other among altORFs of 13,780 conserved and/or translated altProts (13,078 conserved + 805 translated - 103 both conserved and translated, see Table 1 and also Supplementary Figures S4 and S6). Upon detection of such overlapping ORFs, they were used for the modeling of many possible PRF positions at which the translational switch can occur. Four PRF categories are considered in this study: −2, −1, +1, and +2. These symbols indicate the ribosomal movement back (−) or forth (+) by the corresponding number of nucleotides. To explore a possibility of unconventional PRF events that bridge closely spaced non-overlapping (adjacent) ORFs, we also included ORFs that were separated by 1-10 nt. For this category of ORF pairs, we have modeled chimeric proteins using only forward frameshifts, from +1 to +10. In some cases, the software generated models that incorporated stop codons. Such models were eliminated from the analysis or shortened by a few amino acids. In this process, 570,105 models of chimeric proteins were generated, 36,536 of which were modeled with MS-supported altProts and the remaining 533,569 ones used conserved putative altProts as the basis (redundant numbers of models shown in Supplementary Table S5). These models were subjected to the search for matching MS peptides using the same three datasets as for the identification of translated altORFs. This search delivered translation evidence for 156 putative chimeric proteins, 20 of which were modeled with MS-supported altProts and 135 with conserved putative altProts. One of these sequences, chimeric protein 78 (CP78) was modeled with an altProt that was conserved and MS-supported at the same time (Supplementary Dataset S1). None of chimeric protein models generated with non-overlapping adjacent ORFs (Scenario 3 in Çakır et al., preprint in preparation) matched validated MS peptides in our study. It should be noted that refORFs were assumed to be translated in this analysis. However, if a chimeric protein was modeled with a conserved altORF that overlaps with a refORF, it was scored as modeled with a conserved altORF. This definition was appropriate because our primary goal was to study translation of altORFs, whereas refORF translation is an expected condition. Supplementary Dataset S2 illustrates the details of each chimeric model, its matching MS peptide, position, type, and value of PRF, and positions of involved ORFs relative to their transcripts. Supplementary Dataset S3 contains sequences of 156 chimeric models, their matching MS peptides, and corresponding 145 transcripts with detailed annotation of features. Supplementary Dataset S1 provides a very comprehensive summary of many aspects of the identified chimeric proteins and permits easy cross-comparison of various categories.

After the analysis was completed, we realized the existence of two sources of redundancy among 570,105 models of chimeric proteins. Firstly, chimeric models that corresponded to 103 conserved MS-supported altORFs were counted twice because we conducted the modeling separately for conserved and translated altORFs. Secondly, chimeric models in which PRF events involved only altORFs were counted twice, so our pipeline generated two identical sets of models for each altORF. In the first set, altORF1 (the upstream one) was considered as a refORF, and in the other set, altORF2 (the downstream one) was treated as a refORF. These two sources of redundancy were subsequently eliminated from the pipeline (Çakır et al., preprint in preparation). Accordingly, corresponding non-redundant counts of chimeric models were recorded for different altORF types, PRF values, RNA types, MS proteomic studies (Supplementary Tables S6-S9), and also for different chromosomal locations (see Section 4.5.9 in Supplementary Results). In supplementary figures, expected counts based on the distribution of chimeric models were calculated using non-redundant numbers of chimeric models per category without models that come from non-overlapping ORFs (Supplementary Table S9). These models that correspond to the group “other” were excluded from the calculations of expected counts because none of them received MS-support. It should be noted that the inclusion of 99,355 duplicated chimeric models in our MS search process (ca. 17% of 570,105) is unlikely to have affected the efficiency of detection because models with identical sequences were combined by the analysis software into related protein groups.

### 4.4. The detection of eight transcripts associated with multiple MS-supported chimeric peptides enabled the discovery of first candidates for mosaic translation

Our primary analysis revealed the presence of eight transcripts each associated with more than one frameshifting event (Supplementary Dataset S4). We also conducted a nearly exhaustive search for genomic and transcript regions that can potentially produce any of the 156 detected chimeric MS peptides, with or without PRF involved. We call them alternative sources throughout the manuscript. That additional analysis indicated conservation of some chimeric peptides within the *M. truncatula* genome and pointed to other transcripts with multiple frameshifting potential. Transcripts mentioned in this section are referred to as primary sources (145 transcripts). They will be the focus of our main discussion.

Among eight transcripts with multiple frameshifting potential, six were annotated as mRNA. They have two or three MS-deduced putative PRF sites per transcript, most of which are located within the boundaries of their refORFs (except for MtrunA17_Chr5g0430341, CP90 and CP91). Two non-mRNA transcripts are MtrunA17_MTg0490971 (ncRNA, CP150 and CP151) and MtrunA17_Chr5g0422291 (rRNA, CP88 and CP89). Each of them has two putative PRF sites per transcript (Supplementary Dataset S4).

Next, we analyzed the sequence context of each chimeric peptide in this group, with the focus on the PRF value, frames involved, the distance between the PRF sites, and the presence of in-frame stop codons between the PRF sites. This analysis suggested that six transcripts out of the eight are unlikely candidates for mosaic translation unless additional MS-supported PRF sites are found for those transcripts. For example, in MtrunA17_Chr1g0185811 (mRNA, CP16, CP17, and CP18), all the three PRF sites have the same PRF value (+1). In addition, they belong to the same PRF type (1→2) and PRF subtype (a→r). Supplementary Dataset S1 explains the definitions of these categories. Such arrangement of PRF sites does not support the possibility of mosaic translation from MtrunA17_Chr1g0185811. In MtrunA17_Chr5g0430341 (mRNA, CP90 and CP91), two PRF sites are separated by 1,373 nt. Although the first PRF site has value −2 and the second one +2, both PRF events in this transcript start from frame 3, which is incompatible with the production of one continuous mosaic protein of the category that we called short round trip (Çakır et al., 2023). Furthermore, there are 25 stop codons in frame 1 between the first and the second PRF site, which would require the discovery of many additional chimeric peptides to form a continuous mosaic “bridge” between the two PRF sites. Unlike other transcripts in this dataset, MtrunA17_Chr1g0200071 (mRNA, CP23 and CP24) and MtrunA17_Chr6g0457461 (mRNA, CP98, CP99, and CP100) are good candidates for mosaic translation (Figure 1 and Supplementary Dataset S4). The annotated product of MtrunA17_Chr1g0200071 is a protein-synthesizing GTPase. The transcript of MtrunA17_Chr1g0200071 has 73.5% nucleotide identity with an alternative-source transcript of CP24 and CP101, MtrunA17_Chr6g0458111 (elongation factor 1-alpha named *MtEF1A1* in Xu et al., 2023) and the primary-source transcript of CP101, MtrunA17_Chr6g0458091. *MtEF1A1* is the key regulator of translation involved in abiotic stress responses and adaptation to the environment. MtrunA17_Chr6g0458091 is occasionally used as a housekeeping gene for RT-PCR analysis (Gomez et al., 2009; Jiang et al., 2018) despite suboptimal stability of its expression in various tissues (Kakar et al., 2008). The second candidate for mosaic translation, MtrunA17_Chr6g0457461, encodes a ribulose-bisphosphate carboxylase (RuBisCo), which is the major enzyme in photosynthesis (Chen et al., 2023). Despite completely different putative cellular functions, our study indicates that these two transcripts, MtrunA17_Chr1g0200071 and MtrunA17_Chr6g0457461, have much in common. The first PRF site in both transcripts has the same value −2, the same PRF type (2→3), and the same PRF subtype (r→a). Likewise, the second PRF site shares the same characteristics in those two transcripts: the PRF value +2, the PRF type (3→2), and the PRF subtype (a→r). The distance between the two PRF sites is only 81 nt in the transcript of MtrunA17_Chr1g0200071 and 60 nt in MtrunA17_Chr6g0457461. Neither have in-frame stop codons between the two PRF sites, which makes them perfect candidates for producing mosaic proteins of the short round trip category (Çakır et al., 2023).

**Figure 1.**
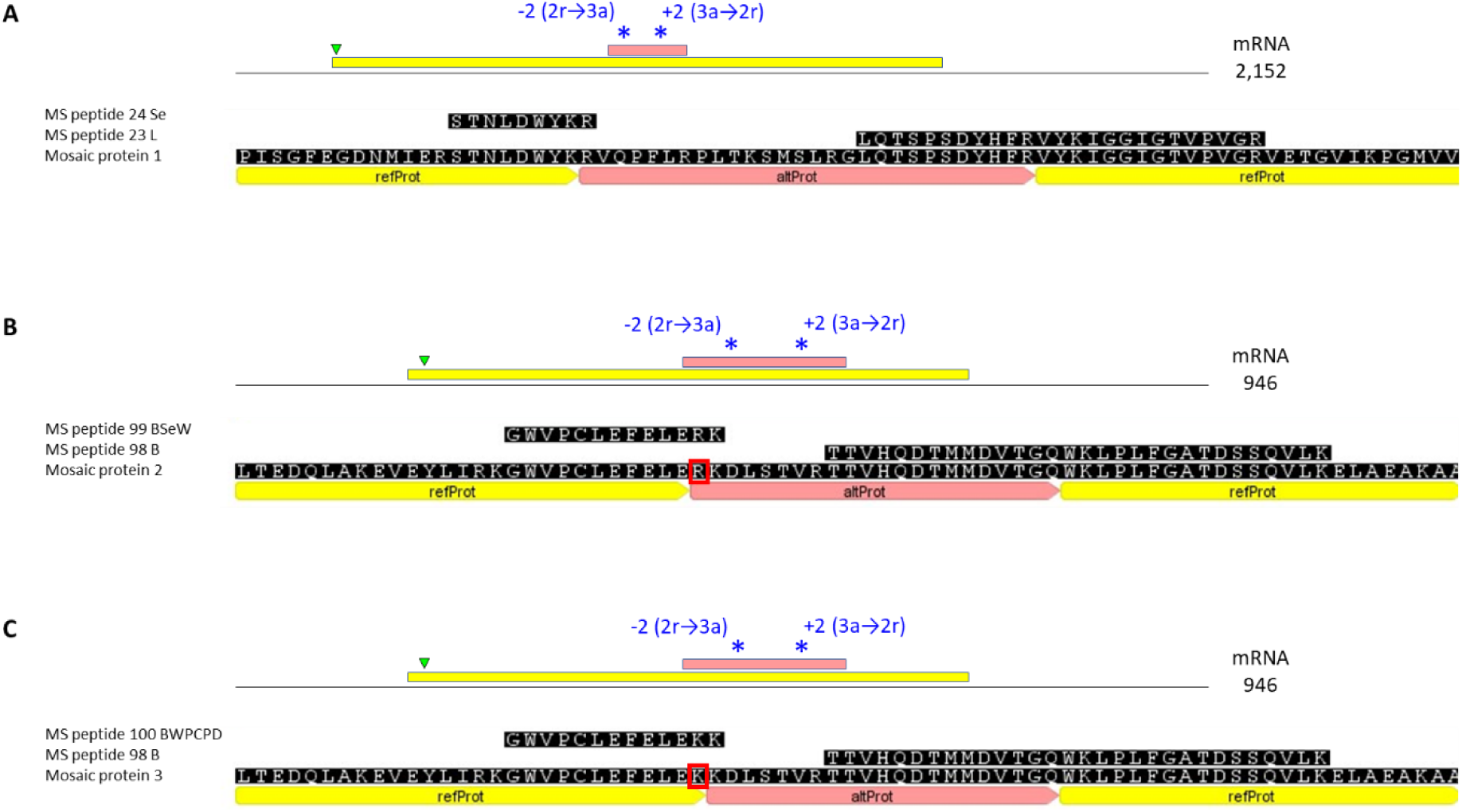
Three candidate mosaic proteins deduced from chimeric MS peptides. A, Mosaic protein 1 translated from transcript MtrunA17_Chr1g0200071 (the locus annotated as a putative protein-synthesizing GTPase). B, Mosaic protein 2 translated from transcript MtrunA17_Chr6g0457461 (the locus annotated as a putative ribulose-bisphosphate carboxylase, RuBisCo). C, Mosaic protein 3 translated from the same transcript MtrunA17_Chr6g0457461. Mosaic proteins 2 and 3 differ by one amino acid around the −2 PRF site (red-boxed R and K, respectively). The upper portion of each figure depicts a corresponding transcript with ORFs mapped and scaled relative to the whole transcript length. Asterisks represent positions of PRF events. The text in blue describes the PRF value, type, and subtype of each frameshifting event. Interpretation example: −2 (2r→3a) refers to a frameshift with value minus 2 from a refORF in frame 2 (yellow) to an altORF in frame 3 (pink). The green triangle indicates the position of the first in-frame translational start codon (AUG).

According to our study, MtrunA17_Chr1g0200071 is expected to produce one 448 aa long mosaic protein, which is of the same length as the annotated product. However, this mosaic protein contains 28 aa entirely different from the annotated protein in the middle of the sequence (Figure 1A, Supplementary Dataset S5). This segment introduced by putative mosaic translation does not disrupt the overall three-dimensional structure of the protein, as predicted by ColabFold, but rather modifies it (Supplementary Figure S8). This is not very surprising because the NCBI BLASTP analysis of the conserved altProt that donates its sequence to the mosaic protein has the top similarity to elongation factor 1-alpha from *M. truncatula*.

In contrast to MtrunA17_Chr1g0200071, the second likely candidate for mosaic translation, MtrunA17_Chr6g0457461, is expected to produce two 177 aa long mosaic proteins, which are of the same length as the annotated RuBisCo. Because of the presence of the second −2 PRF site only one nucleotide away from the first site, the two mosaic proteins of MtrunA17_Chr6g0457461 differ by just one amino acid. One mosaic isoform contains 22 aa in the middle of the sequence that are entirely different from the annotated protein (Figure 1B, Supplementary Dataset S5). The other mosaic isoform contains the same segment but one amino acid shorter at the left side (21 aa different from the annotated product, Figure 1C, Supplementary Dataset S5). Like in the case with MtrunA17_Chr1g0200071, these segments introduced by putative mosaic translation somewhat modify the three-dimensional structure of the protein, as predicted by ColabFold (Supplementary Figures S9 and S10). The translational product of the conserved altORF involved in the putative PRF events of both mosaic isoforms is similar to RuBisCo from another legume species, *Arachis duranensis* (peanut). Thus, it is possible that this altORF emerged via a frameshifting mutation, which is older than in the case of MtrunA17_Chr1g0200071. The complete ColabFold prediction files corresponding to the annotated and mosaic proteins of MtrunA17_Chr1g0200071 and MtrunA17_Chr6g0457461 can be found in Supplementary Dataset S6.

Interestingly, the putative PRF sites in transcripts mentioned in this section repeat in distinct groups. Namely, three transcripts, MtrunA17_Chr1g0200071, MtrunA17_Chr6g0457461, and MtrunA17_Chr5g0430341, have the first PRF site with the value −2 followed by a +2 PRF site. The other group, MtrunA17_Chr1g0185811, MtrunA17_MTg0490471, and MtrunA17_MTg0490971, have repeated +1 sites. The only rRNA transcript in this group, MtrunA17_Chr5g0422291, has repeated −1 sites. MtrunA17_Chr3g0144151 has no particular pattern of PRF sites, which is an exception in this subset of transcripts (Supplementary Dataset S4). These non-random observations may reflect the biological relevance of this grouping.

### 4.5. Multiple significant associations between various parameters of the dataset reject the null hypothesis of its artifactual origin

Our hypothesis about the mosaic nature of proteins produced by MtrunA17_Chr1g0200071 and MtrunA17_Chr6g0457461 relies on the validity of the entire dataset. So far, MS proteomics is the most powerful and accurate large-scale method for the discovery of new proteins (Timp and Timp, 2020; Bennett et al., 2023; Messner et al., 2023). Despite its unique status among currently available methods, MS-based detection of peptides and proteins is inherently prone to false discoveries even when the standard procedures for their control are implemented, especially in proteogenomics (Aggarwal et al., 2022; Ebadi et al., 2023). In our study, we used a standard FDR of 1% as a threshold. However, because of the very large search space used for the detection of chimeric peptides (more than half-million of chimeric protein models, Supplementary Table S5), we had to apply a two-step search procedure. Since the primary purpose of this approach is to lower the rate of false negatives (Jagtap et al., 2013), it is likely to be associated with more difficult control over FDR. To address this known weakness of the large-scale search for matching MS peptides, we conducted a very comprehensive analysis of chimeric peptide features summarized in Supplementary Dataset S1. Our intention was to detect significant relationships between features that are not compatible with the null hypothesis about the randomness of the dataset. If the various features were occurring randomly, it would suggest that the 156 validated chimeric peptides might be false discoveries. Much of the data described in Figures 2-8, Supplementary Figures S11-S62 (see Supplementary Results), and discussed below passed statistical tests. Collectively, our statistical analysis revealed a clear deviation from the randomness assumption. Thus, the detected sequences must have biological relevance.

**Figure 2.**
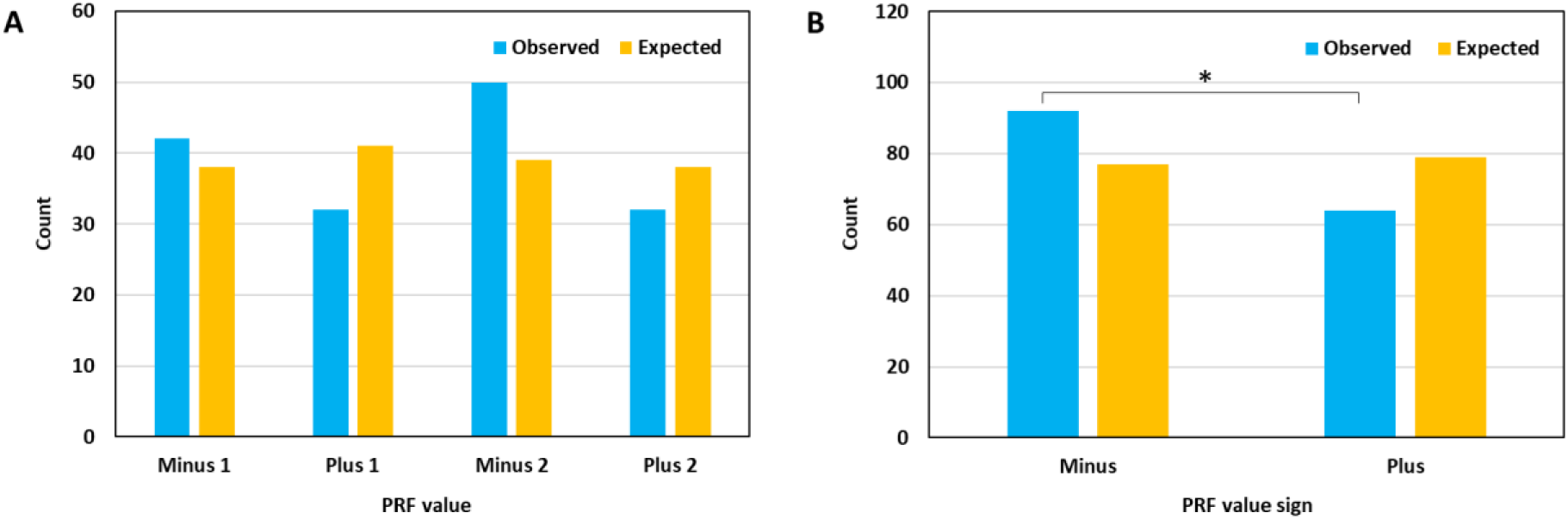
Distribution of 156 chimeric MS peptides in different groups according to PRF values. A, Four separate groups of PRF. B, Two combined groups of PRF. Expected values were calculated based on the numbers of corresponding chimeric protein models (non-redundant counts in Supplementary Table S9, 469,600 models in total). There is no significant difference between the observed and expected proportions for the separate groups (A) and a significant difference (p = 0.016; the chi-square test for goodness-of-fit) for the combined groups (B). The distribution of PRF values within individual groups in A and B is not significantly different from the uniform distribution. However, the distribution of observed values in B is significantly different from the uniform distribution (p = 0.025; the chi-square test for homogeneity): PRF values with the negative sign are significantly overrepresented in the dataset.

**Figure 3.**
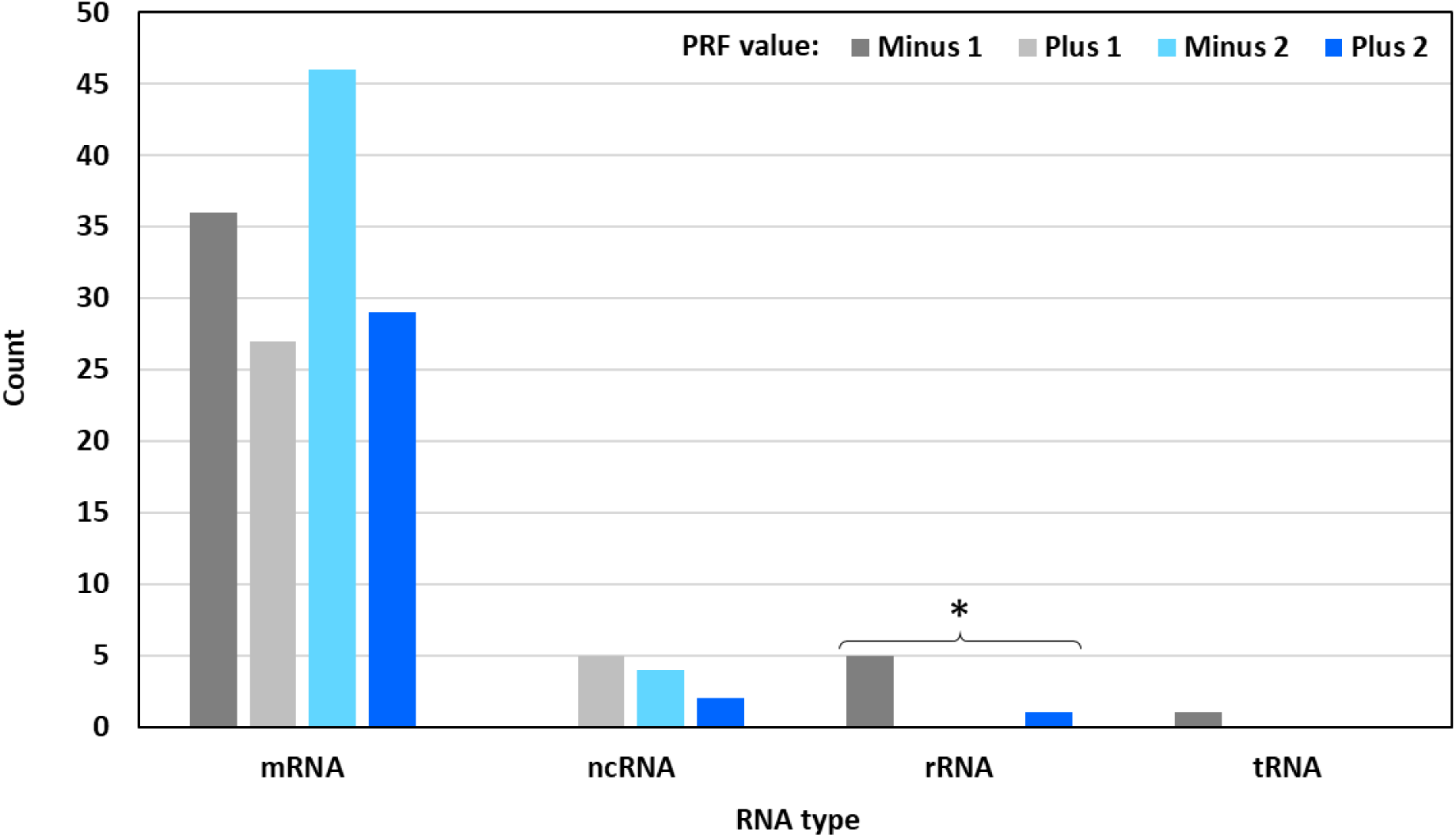
Distribution of 156 chimeric MS peptides in different groups based on the RNA type and the PRF value. The Fisher’s exact test shows significant association between the RNA type and the PRF value (p = 0.007 with tRNA included, p = 7.024E-07 with tRNA excluded). Within individual RNA groups, the distribution of PRF values is significantly different from the uniform distribution in one case indicated with an asterisk (the Exact Multinomial Test, p = 0.019). With PRF values −1 and +2 only, the difference is not significant. Note the absence of certain PRF values from ncRNA and rRNA. Remarkably, the only PRF value in tRNA is the same as the most frequent PRF value in rRNA.

**Figure 4.**
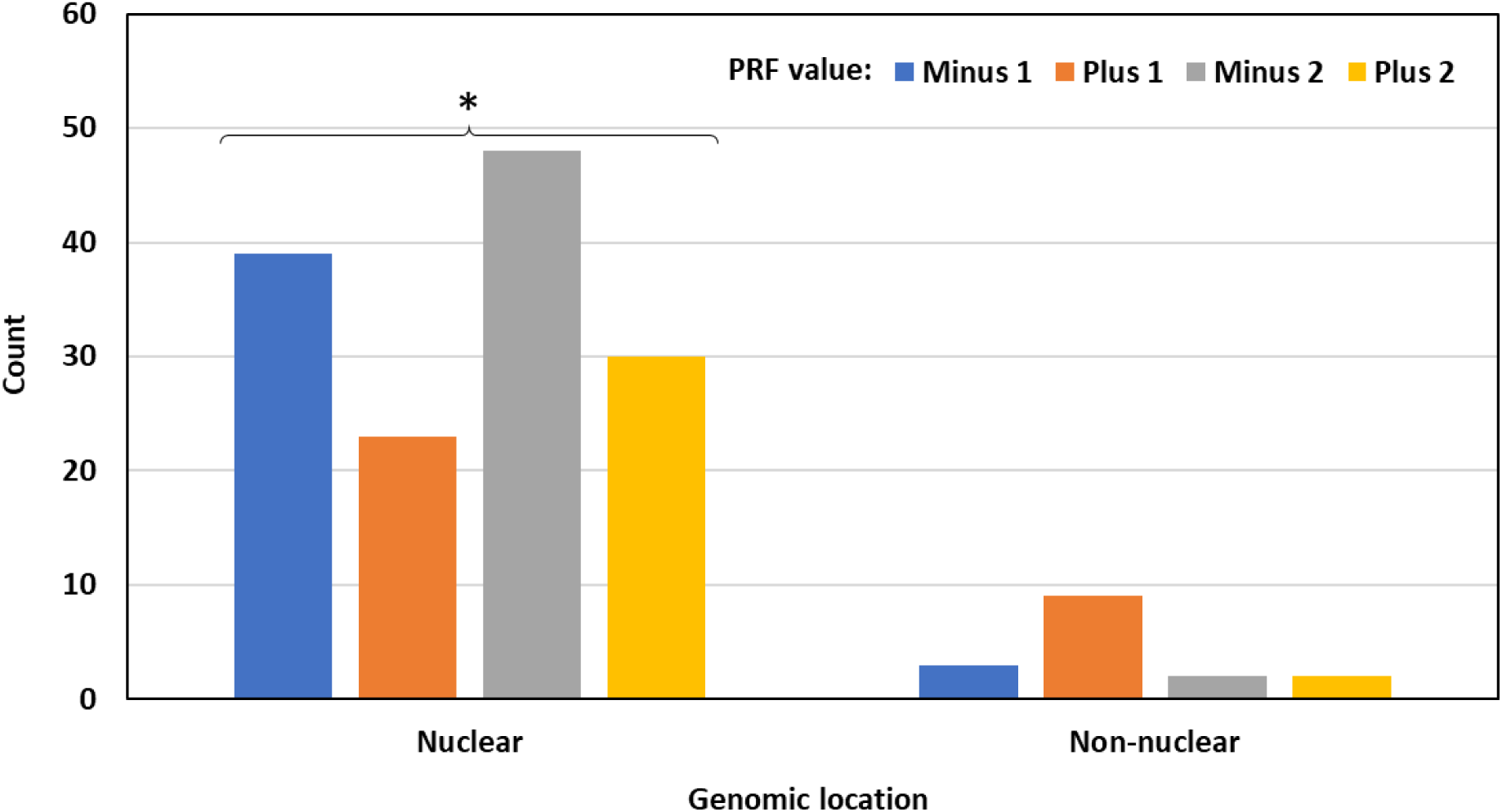
Distribution of 156 chimeric MS peptides based on the genomic location of their primary-source genes and the PRF value. “Nuclear”: Chromosome 0 (unmapped loci) to Chromosome 8. “Non-nuclear”: chloroplasts and mitochondria. There is no significant association between the genomic location and the PRF value. Within individual genomic locations, the distribution of PRF values is significantly different from the uniform distribution in one case indicated with an asterisk (the Exact Multinomial Test, p = 0.018). PRF events with value +1 are significantly underrepresented in nuclear transcripts and non-significantly overrepresented in chloroplasts and mitochondria.

**Figure 5.**
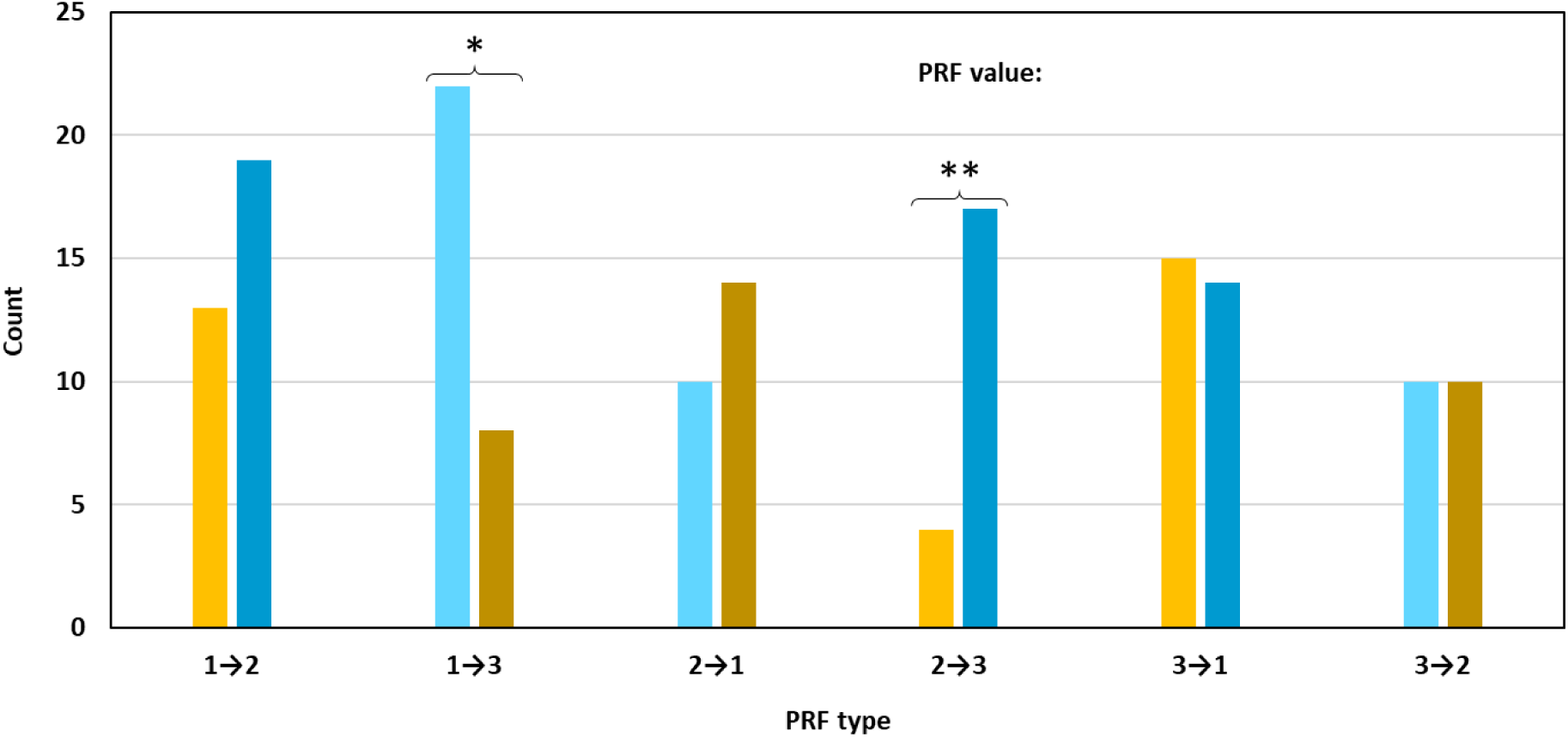
Distribution of 156 chimeric MS peptides based on the PRF type and value. Each PRF type has only two possible values: one positive and one negative. There is significant association between the PRF type and the PRF value based on the chi-square test (p = 0.007, a two-category test with categories of the same absolute value combined). Within individual groups, the distribution is significantly different from the uniform distribution in two cases indicated with asterisks: 1→3 and 2→3 (the chi-square test for homogeneity, p-values are 0.011 and 0.005, respectively). Backward PRF is more frequent for the shift from any frame to frame 3.

**Figure 6.**
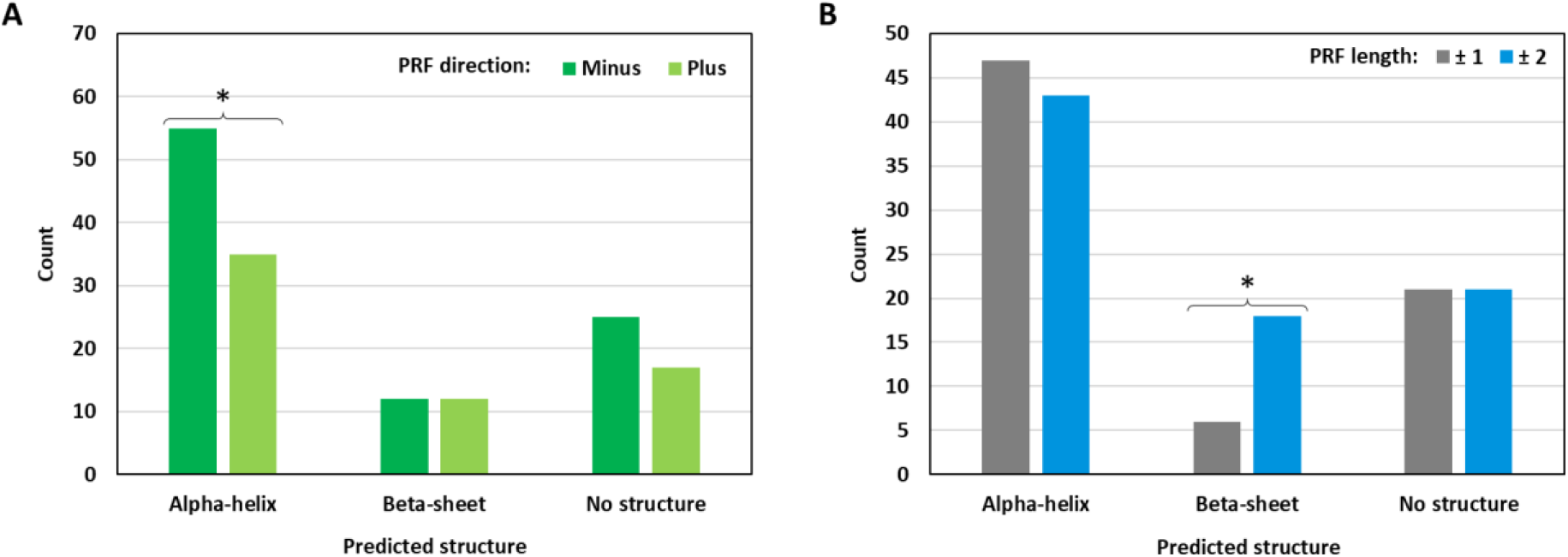
Distribution of 156 MS-supported models in three folding categories based on two aspects of the PRF value. A, grouping by the direction of PRF (minus, backward frameshifts; plus, forward frameshifts). B, grouping by the length of PRF (±1, “short” frameshifts; ±2, “long” frameshifts). There is no significant association in A and B. Within individual folding categories, the distribution is significantly different from the uniform distribution in two cases indicated with asterisks (the chi-square test for homogeneity, p-values are 0.035 and 0.014 for A and B, respectively). Backward PRF values are significantly overrepresented in chimeric peptides with predicted alpha-helices. Beta-sheet structures are three times more abundant in the models with “long” PRF values.

**Figure 7.**
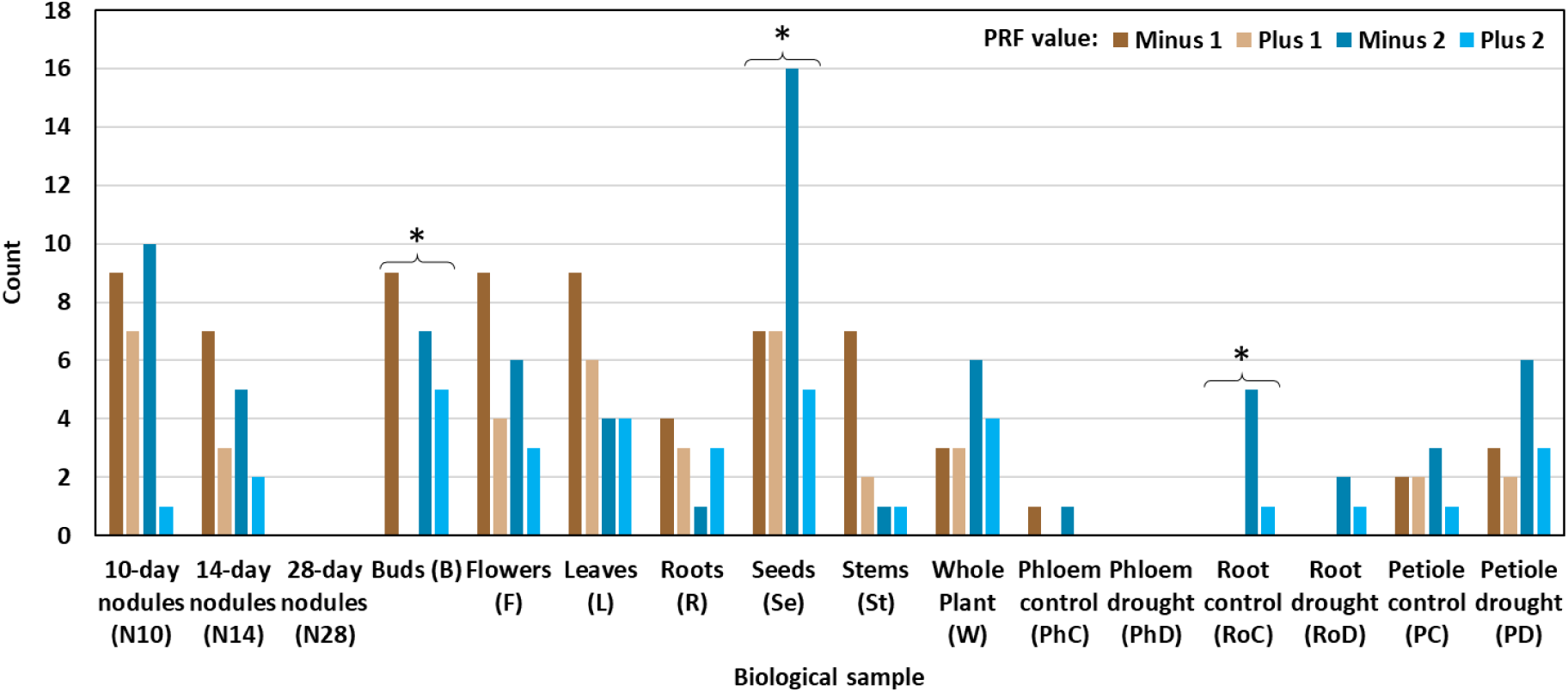
Distribution of 156 chimeric MS peptides in 16 biological samples depending on the PRF value. The counts are not additive because some chimeric peptides were identified in multiple samples. No chimeric peptides were detected in samples N28 and PhD. The PRF value is significantly associated with the sample (the Fisher’s exact test with a simulated p-value, p = 0.010). Within individual samples, the distribution of PRF values is significantly different from the uniform distribution in three cases indicated with asterisks. P-values of the chi-square test for homogeneity are as follows: 0.036 (B) and 0.040 (Se). The significance in sample RoC was assessed with the Exact Multinomial Test (p = 0.019). No chimeric peptides with PRF values −1 and +1 were found in samples RoC and RoD. Chimeric peptides with PRF values +1 and +2 are absent from sample PhC. Sample B lacks chimeric peptides with PRF value +1.

**Figure 8.**
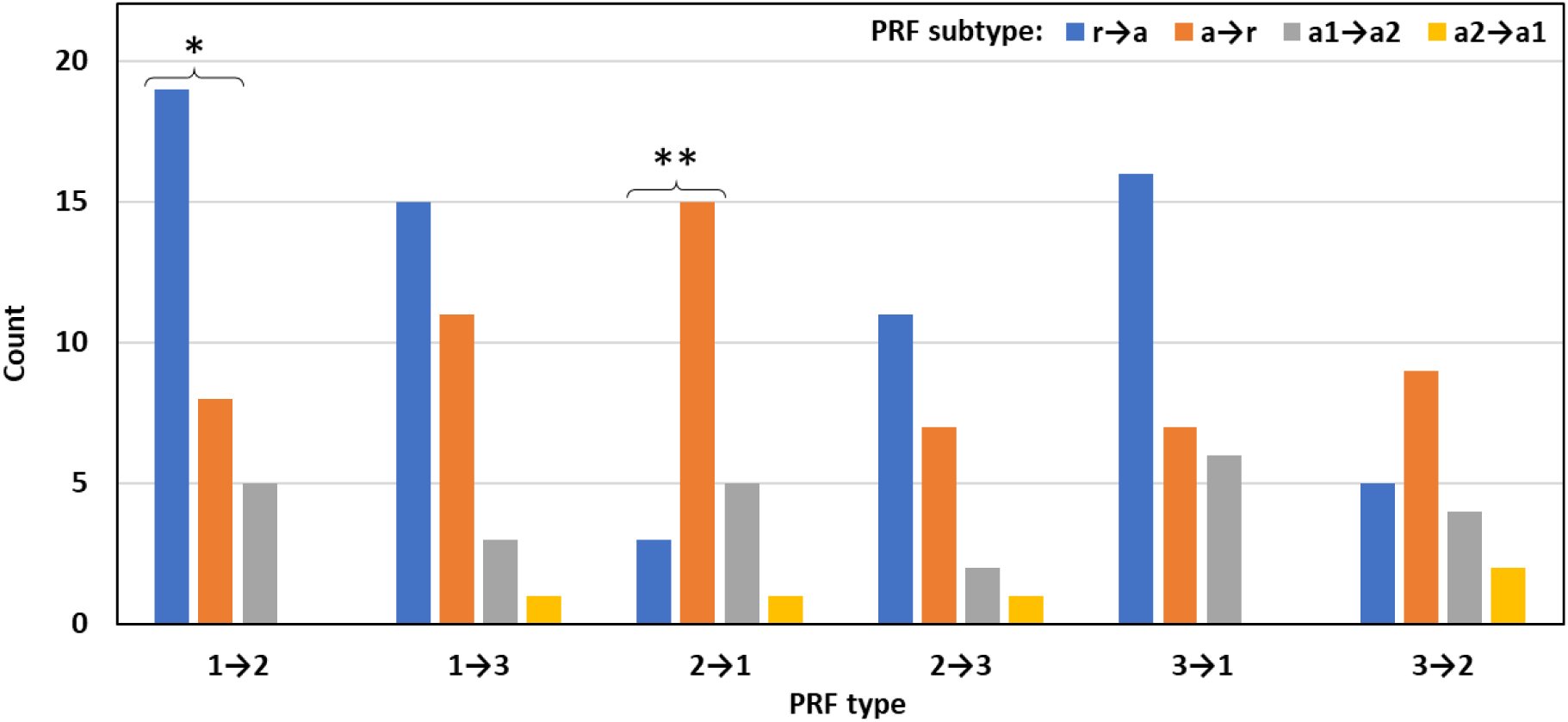
Distribution of 156 chimeric MS peptides in different PRF categories depending on the PRF type and subtype. r→a, PRF from refORF to altORF; a→r, PRF from altORF to refORF; a1→a2, PRF from altORF1 to altORF2, where altORF1 starts upstream of altORF2; a2→a1, PRF from altORF2 to altORF1, where altORF1 starts upstream of altORF2. There is no significant association between the PRF type and the PRF subtype. Proportion pairs in which the distribution is significantly different from the uniform distribution are marked with asterisks. P-values of the chi-square test for homogeneity of two proportions: *0.034; **0.005. In PRF type 1→2, PRF subtype r→a is significantly more frequent than PRF subtype a→r. In PRF type 2→1, it is the other way around, which means refORFs involved in PRF are associated with frame 1 and altORFs with frame 2 in those two categories. There is a non-significant trend of the same kind in PRF types 2→3 and 3→2, where refORFs are associated with frame 2.

## 5. Discussion

### 5.1. Old is gold: MS proteomics can help reveal the existence of mosaic proteins in the absence of methodology for long-read protein sequencing

Earlier, we proposed that multiple PRF may play an important role in the adaptability of organisms. The versatility of their proteomes may have been greatly expanded via the production of polypeptides that incorporate translation products of different reading frames. We refer to this hypothetical mechanism as mosaic translation (Çakır et al., 2023). Experimental demonstration of this mechanism is a major challenge for a number of reasons. They are associated with technical limitations of the protein detection methods. The chief group of methods is based on MS proteomics. These methods infer the presence of long continuous polypeptide sequences via the detection of their short fragments (7-35 aa, Swaney et al., 2010). Unfortunately, this methodology permits the identification of only a small fraction of non-canonical peptides actually present in a sample. Many biologically relevant peptides that are expressed at a low level and/or have small size remain undetected by MS proteomics, which does not reflect their instability or the lack of biological importance (Wacholder and Carvunis, 2023). Unequivocal demonstration of the mosaic nature of a protein requires long sequencing reads. Recently, a breakthrough in the development of nanopore-based sequencing of proteins has been reported (Martin-Baniandres et al., 2023; Wang et al., 2024). Still, the current state of this methodology is too far from offering significant help with the detection of long continuous mosaic polypeptides in a complex mixture of amino acid sequences. In the absence of any alternative to MS-based proteomics, we decided to mine the existing MS proteomic datasets for fragments of putative mosaic proteins that we modeled using an original script (Çakır et al., preprint in preparation). Two types of data were used as inputs for the script: (1) altProts that have a conservation signature based on the global sequence similarity searches and (2) altProts that have matching MS-validated peptides regardless of their conservation signature. Due to the nature of MS proteomics as a detection strategy (Wacholder and Carvunis, 2023), 156 chimeric peptides reported here likely represent only “the tip of an iceberg”, in which the hidden part is awaiting discovery.

### 5.2 Main findings and arguments for their validity

Upon the detection of 156 MS peptides that match chimeric models, we found that some of them could be mapped to the same transcripts. This way, we discovered eight transcripts associated with multiple PRF events, two or three per transcript. Based on the PRF value and type, we showed that two of these transcripts are good candidates for the production of mosaic proteins (Figure 1). One of them corresponds to a putative protein-synthesizing GTPase, which is a close homolog of elongation factor alpha (MtEF1α). The other one encodes a putative ribulose-bisphosphate carboxylase (RuBisCo), which is an enzyme central to photosynthesis. Intriguingly, both putative mosaic proteins involve the same PRF events: the first −2 frameshift changes translation from frame 2 (refORF) to frame 3 (altORF). The second +2 frameshift brings translation back to the refORF located in frame 2. We think the exact correspondence between these two PRF events in completely unrelated transcripts argues for a biological reason for this conservation. The discovery of two strikingly convergent transcripts that are likely to produce mosaic proteins in our non-viral system is highly novel and unexpected. So far, mosaic proteins produced via PRF are known exclusively in viruses (Jacks, 1990; Hatfield et al., 1992; Ketteler, 2012; Rex et al., 2019). This is the main finding of our study. The remaining six transcripts with multiple PRF sites can also be candidates for mosaic translation because additional PRF events associated with these transcripts can be found in the future.

Ideally, MS peptides matching a mosaic protein must be long enough to include both PRF sites. Due to the short read length of the current MS technology (7-35 aa, Swaney et al., 2010), putative mosaic proteins produced by MtrunA17_Chr1g0200071 and MtrunA17_Chr6g0457461 have no matching MS peptides that would join the products of individual PRF sites. However, the gap in the MS peptide coverage is fairly small: 16 aa for MtrunA17_Chr1g0200071 and only 6 aa for MtrunA17_Chr6g0457461, which may indicate that the peptides are indeed part of a continuous mosaic sequence (Figure 1, Supplementary Dataset S5). These short distances between MS peptides permit testing the effects of corresponding synthetic mosaic proteins *in vivo*. In addition, due to these short distances, generation of MS proteomic data from the same samples using different digestive enzymes could potentially yield peptide “bridges” that span the gaps in the peptide coverage. This would serve as indirect evidence for continuity. Nevertheless, it is possible that MS peptides corresponding to the PRF sites of these transcripts are translated individually as chimeric non-mosaic sequences. On one hand, this possibility is supported by the existence of alternative sources of these peptides. On the other hand, while three MS peptides that correspond to putative PRF sites in MtrunA17_Chr6g0457461 were all detected in the same organ (buds), two MS peptides of MtrunA17_Chr1g0200071 were found in two different organs (one in seeds and the other one in leaves). Given the ambiguity associated with these multi-PRF sequences, the following three scenarios are not mutually exclusive: (1) chimeric peptides are produced individually from the same transcript; (2) chimeric peptides are produced individually from two or more different transcripts; (3) chimeric peptides are produced in the course of mosaic translation as parts of a continuous amino acid sequence from one transcript or multiple transcripts. Dedicated wet lab studies are required for discrimination between these scenarios. Despite the lack of solid proof, MS peptides corresponding to MtrunA17_Chr1g0200071 and MtrunA17_Chr6g0457461 provide the first experimental hint for a possibility of mosaic translation in a non-viral system. Thus, they deserve close investigation in *M. truncatula*, which will pave the avenue toward the discovery of mosaic proteins in other organisms. Our methodology, in combination with parallel multi-enzyme digestion of protein samples for MS, is likely to reveal many more candidates for mosaic proteins.

Our study provides unique information of fundamental importance in several other respects. Firstly, the discovery of the large number of chimeric peptides in a non-viral system is another novel outcome of our work because only a few chimeric peptides have so far been known outside the viral kingdom: at least three in bacteria (Blinkowa and Walker, 1990; Flower and McHenry, 1990; Tsuchihashi and Kornberg, 1990; Chaijarasphong et al, 2016; Meydan et al., 2017) and at least three in eukaryotes (Matsufuji et al., 1995; Clark et al., 2007; Ivanov and Atkins, 2007; Ren et al., 2024). In this respect, our results agree with the prediction according to which as much as 10% of genes in a eukaryotic genome can be associated with PRF (Ketteler, 2012). A parallel study in the human published very recently reached similar conclusions (Ren et al., 2024; see Section 5.7). Secondly, the inclusion of non-mRNA transcript types into our search pipeline revealed an entirely novel possibility. It showed that MS-supported PRF events can be detected in ncRNA, rRNA, and even tRNA transcripts, all of which are traditionally considered non-coding. Although still very exotic, the ability to serve as a template for translation has already been demonstrated for ncRNA (Ingolia et al., 2011; Ruiz-Orera and Albà, 2019; Zaheed et al., 2021), rRNA (Hashimoto et al., 2001; Maximov et al., 2002), pre-miRNA (Wang et al., 2014; Couzigou et al., 2016), and pre-siRNA (Yoshikawa et al., 2016). However, no information of this type has ever been reported for tRNA. At the same time, to the best of our knowledge, PRF events have never been detected in any non-mRNA transcript so far. Thus, our work sends several important messages that require close attention.

Naturally, the presence of several non-ordinary findings in one computational MS proteomics-based study raises doubts about the validity of the entire dataset. Thus, let us set our zero-hypothesis as follows: the dataset is the collection of false-positives obtained due to the difficulty of controlling false discovery rate when the procedure involves a very large search database. The detailed manual analysis of various parameters (Supplementary Dataset S1) in conjunction with stringent statistical procedures clearly indicates that most observations are non-random and most associations are significant. This concerns not one or two parameters but several dozens of various aspects. We summarized 37 of the most obvious significant observations in Supplementary Dataset S29. Supplementary Dataset S30 lists 14 of the most relevant significant associations. Taken together, these findings speak strongly against the null-hypothesis, which indicates our data have biological relevance despite the high false discovery rate expected for an MS proteomics-based study.

### 5.3. RNA-Seq data as a support for chimeric nature of MS-validated peptides

Another major concern associated with the discovery of highly novel peptide sequences is their true origin. Can our chimeric peptides have alternative sources that do not involve PRF? Using genomic DNA as a basis, we found that one chimeric peptide in our dataset, CP80, can potentially be produced from two genomic loci with unknown translation status. However, these loci are not transcribed, which makes them unlikely non-chimeric sources of the peptide identical to CP80 (see Sections 4.7.1 and 4.7.3 in Supplementary Results). Alternative splicing events such as exon skipping or intron retention, along with the products of RNA editing and natural polymorphisms, can also potentially mimic chimeric peptides. Using the collection of individual reads from 50 relevant RNA-Seq runs, we demonstrate that only one to four MS-validated chimeric peptides (CP54, CP93, CP140, and CP148, or CP54 alone) could originate from unusual versions of native transcripts without PRF. Contrary to our predictions summarized in Supplementary Dataset S1, none of these peptides is likely to be produced via alternative splicing (Supplementary Datasets S18 and S19).

### 5.4 The existence of potential alternative sources is a challenge for the functional analysis of chimeric proteins

Because of the chimeric nature of MS-validated peptides in our dataset and their short length typical for an MS study, many of them can have multiple origins (Supplementary Datasets S1 and S22 to S24). This genetic redundancy will make it very difficult to study their functions by conventional loss-of-function methods such as insertional mutagenesis. Conceptually, RNA interference (RNAi) can be used to downregulate a group of related transcripts (Mello and Conte, 2004). Alternative sources of chimeric peptides in our study share a limited similarity, which may permit their simultaneous silencing (as can be judged from their annotation, Supplementary Dataset S24). Nevertheless, construction of multiple mutants may be required to learn the loss-of-function phenotypes of these loci. Moreover, demonstration of individual phenotypes for a refProt, altProts, and a chimeric protein(s) derived from a given transcript will require complementation of null-mutants with constructs in which these proteins are disabled differentially. Using an in-house script (Çakır et al., preprint in preparation), we conducted a comprehensive inventory of genetic loci that can potentially produce chimeric peptides identified in our study. This inventory is crucial for understanding the biological roles of chimeric proteins. At the same time, ca. 58% of chimeric peptides reported here have unique sources, which means they can be functionally characterized without generation of multiple mutants. Based on the analysis of expression profiles, only 31 chimeric peptides out of 156 (ca. 20%) can have alternative sources at least as likely as their primary sources (Supplementary Dataset S1). Thus, it is possible that most of the 145 transcripts that are denoted as primary sources in our study are indeed the main or even the only contributors to the translation of these peptides. This possibility is supported by numerous significant associations observed at the level of transcripts and genomic DNA of the primary-source loci (e.g., Figure 3, 4, 5, and 8, Supplementary Figures S11-S15 and S50-S62). These observations cannot be expected under an assumption that translation of chimeric peptides detected in this study occurs mainly or exclusively from other transcripts (alternative sources). A recent discovery of the promotive effect of synthetic complementary peptides on the expression of matching transcripts offers a unique opportunity for studying mosaic and chimeric proteins (Ormancey et al., 2023). Using synthetic versions of these proteins, one can study the gain-of-function phenotypes associated not with a specific locus but with all loci that actually contribute to the production of a mosaic or chimeric protein.

### 5.5. Four genetic loci associated with the production of chimeric peptides in *M. truncatula* have been in the focus of functional studies

In the course of our work, we realized the importance of learning if any genes listed as primary or alternative sources of chimeric peptides have already been studied by loss-of-function approaches. Like in the case of altProts, chimeric proteins are expected to introduce ambiguity to the interpretation of any genetic study. In the absence of knowledge about an altProt or a chimeric protein translated along with a refProt, naturally, any mutant phenotype linked to the locus is automatically attributed to the loss-of-function of a refProt. Incorrectly inferred phenotype-genotype relationships can have dramatic consequences for efforts to cure genetic diseases or develop a more resistant/more productive crop. To address this important task, we have conducted a nearly comprehensive analysis of more than a thousand of loss-of-function studies in *M. truncatula* that were published since the time of its advent as a model organism (Barker et al., 1990). By January 2024, these studies involved at least 627 genetic loci. To our surprise, only two genes from our primary-source list have been in the focus of such studies so far. One of them is a sucrose synthase gene *MtSUCS1* (CP80, MtrunA17_Chr4g0070011). Antisense-mediated transcript knockdown of *MtSUCS1* leads to the defects in symbiotic nitrogen fixation and arbuscular mycorrhizal symbiosis (Baier et al., 2007, 2010). The other gene is a cytokinin oxidase/dehydrogenase *MtCKX6*, which was targeted by insertional mutagenesis using tobacco retrotransposon *Tnt1*. The phenotype of the *Mtckx6* mutants was analyzed in the context of root development. It turned out to be not different from the phenotype of wild-type plants (Wang et al., 2021). One gene listed among two alternative sources of CP104 was targeted by RNAi. It encodes a pathogenesis-related protein MtPR10-5 (MtrunA17_Chr4g0067951). Downregulation of this gene together with four likely off-target loci resulted in reduced colonization and suppressed infection by an oomycete pathogen *Aphanomyces euteiches* (Colditz et al., 2007; Samac et al., 2011). Our meta-analysis of publications on 627 loci targeted by at least one loss-of-function study does not include genes analyzed exclusively via gain-of-function approaches. However, one such study is worth mentioning here because it was conducted on an alternative source of two chimeric peptides, CP24 and CP101 (MtrunA17_Chr6g0458111). This locus is among six possible sources of these CPs, all of which are annotated as protein-synthesizing GTPases, including a qRT-PCR housekeeping gene *MtEF1* (Gomez et al., 2009; Jiang et al., 2018). Overexpression of MtrunA17_Chr6g0458111 named *MtEF1A1* in the original study renders transgenic plants of *M. truncatula* and *Arabidopsis thaliana* more salt-tolerant (Xu et al., 2023). It may be interesting to clarify whether the reported phenotypes of these genes can be attributed to their refProts, chimeric peptides, or both.

The scarcity of previously studied genes in our dataset can probably be explained by the fact that most of the 145 transcripts have transcription profiles not specific to a particular biological process (Supplementary Datasets S1 and S27). In contrast, functional genomics studies naturally focus on genes with process-specific or organ-specific transcription profiles.

### 5.6. Chimeric sequences may be important for symbiotic nitrogen fixation and seed development

Nearly one quarter of chimeric peptides in our study were identified in nodule samples. Together with chimeric peptides found in seeds, they make up 41% of the dataset (64 sequences, Supplementary Dataset S1, Supplementary Figure S36). This overrepresentation of two biological processes may have technical reasons: we cannot meaningfully compare abundances of validated proteins in different samples even if they are prepared by the same group (Supplementary Figures S6 and S36). However, it may also point to important biological roles of altORFs and chimeric peptides in symbiotic nitrogen fixation and seed development along with other biological processes. Although we do not know if the detected MS peptides represent short sequences or are fragments of long proteins, it should be noted that symbiotic nitrogen fixation is known to involve short peptides for its regulation (Van de Velde et al., 2010; Djordjevic et al., 2015; Kereszt et al., 2018; Roy and Müller, 2022; Ito et al., 2024; Roy et al., 2024). Among chimeric peptides detected in multiple samples, the largest number of sequences was shared by nodule and seed samples (eight chimeric peptides, Supplementary Figure S38). This moderate association is surprising in the context of our study because seed development and biological nitrogen fixation are processes expected to have only a limited overlap in gene expression (Benedito et al., 2008; Mergaert et al., 2020). This is further supported by the fact that only two genes out of 627 targeted by at least one loss-of-function approach in *M. truncatula* (as of January 2024) are likely to have the roles in both processes: MtCYP15a (Sheokand et al., 2005) and MtNOOT2 (Magne et al., 2018; Zhang et al., 2022; Liu et al., 2023). On the other hand, root and nodule samples could be expected to have much overlap in the abundance of peptides based on similarities in gene expression (Benedito et al., 2008; Mergaert et al., 2020). Contrary to this expectation, roots share only three chimeric peptides with nodules (Supplementary Dataset S1). Observations summarized in this section warrant dedicated functional studies on our candidate loci. Differential mutagenesis-based analyses of refProts and corresponding chimeric peptides from our dataset may provide useful insights into molecular mechanisms shared by nodules and seeds.

### 5.7. Our data complement a very recent MS-based study conducted on human samples

In this section, we would like to mention a very remarkable study that was published during the preparation of our manuscript. Ren and associates (2024) discovered at least 405 unique chimeric peptides in 32 diverse human samples not associated with any pathological condition. These peptides correspond to 454 loci, all of which have naturally occurring repeat codon sequences at the putative PRF sites. This group not only provided evidence for the presence of such peptides in multiple samples but also functionally characterized one of the loci, which encodes a histone deacetylase HsHDAC1. This study is very important because it shows how widespread PRF can be in eukaryotic organisms. Although our work has a similar goal and methodology, it does not duplicate the data of Ren et al. (2024). The studies are different in a few principle points and thus are complementary. Firstly, none of 156 chimeric peptides reported here originate from loci with repeated codons at the site of putative PRF. Secondly, our dataset contains a homolog of *HsHDAC1*, which is a gene encoding a histone deacetylase MtHDA9 (MtrunA17_Chr5g0393401, CP81). It also contains a few other loci associated with histone deacetylase activity and histones in general (Supplementary Dataset S1). Transcripts of *HsHDAC1* and *MtHDA9* share only ca. 57% nucleotide identity, and their PRF sites do not coincide in the alignment. Thirdly, while the availability of many MS proteomic datasets for the human permitted the identification of chimeric peptides across very many samples, the *M. truncatula* datasets are still very limited. Thus, only 27 chimeric peptides out of 156 (ca. 17%) were identified in more than one sample. This presents a major technical limitation of our study along with the absence of wet lab validation of our candidate peptides. However, in a few other aspects, our work has a broader scope and offers additional points of novelty. For example, our methodology is conceptually open to the detection of PRF events *de novo*, with no prior knowledge about similarity of PRF sites to already characterized cases. Along with this principle difference, our dataset is not limited to chimeric peptides produced by “short” PRF events (−1 and +1). It includes many sequences that require “long” frameshifts (−2 and +2) for their generation, namely ca. 53%. Next, our dataset includes three non-mRNA transcript types, which constitute ca. 12% of the sequences. Importantly, our study had the goal of detecting multiple occurrence of PRF sites per transcript, which resulted in the discovery of 15 non-RE and eight RE loci potentially associated with multiple frameshifting. Lastly, there is a major difference in the way we considered alternative sources of chimeric sequences. Namely, along with the exclusion of sequences that correspond to refProts and non-chimeric products of short ORFs, we conducted a very comprehensive search for additional alternative sources. This search permitted the discrimination between truly chimeric sequences and those that can be produced from unusual forms of transcripts as follows from the analysis of RNA-Seq data. With an in-house script (Çakır et al., preprint in preparation), we discovered alternative chimeric sources for many peptides and highlighted the role of repeat elements as a potential origin of some chimeric peptides. In summary, the messages sent by these two studies will likely have synergistic effect, which will motivate other research groups for functional characterization of the most interesting candidate peptides.

### 5.8. Our data support the modern view on the pre-biotic evolution and suggest that the ability to undergo “short” frameshifting may be an ancestral feature of all genomes

Besides the identification of candidate loci for the production of mosaic and chimeric proteins, our results also indirectly support the modern view on the origin of genomes, and the central role of rRNA in genome evolution. According to the consensus, first genomes and enzymes were composed of single-stranded RNA. These proto-genomes were similar to the modern rRNA in several aspects. They are thought to have been formed by ligation of tRNA-like molecules that acted as monomers, or proto-genes (Root-Bernstein and Root-Bernstein, 2015, 2016, 2019; Caetano-Anollés and Caetano-Anollés, 2016; de Farias et al., 2016). This scenario is not purely hypothetical. It is based on experimental observations. Modern rRNA and tRNA molecules have striking features that point to their common evolution. rRNA molecules contain sequences very similar to tRNAs of all 20 usual proteinogenic amino acids (Root-Bernstein and Root-Bernstein, 2015). Furthermore, rRNA-like sequences were found in samples of the poly-adenylated mRNAs, which makes them likely templates for translation. So far, at least 29 such transcripts have been detected in various organisms (e.g., Chooi and Leiby 1981; Divecha and Charleston, 1995; Kermekchiev and Ivanova, 2001; Coelho et al., 2002; Scharf et al., 2005; extensively reviewed in Kong et al., 2008 and Root-Bernstein and Root-Bernstein, 2019). Many of them share similarity with rRNA transcripts detected in this study, far from the putative PRF sites. However, four such sequences make significant same-strand nucleotide alignments with regions very close to the PRF sites: *Mus musculus* testin-2 (Divecha and Charleston, 1995); *Homo sapiens* mitochondria-encoded Humanin (Hashimoto et al., 2001); *Candida albicans* mitochondrial protein Tar1p (closely related to *Saccharomyces cerevisiae* Tar1p, Coelho et al., 2002); and *Reticulitermes flavipes* rRNA-like mRNA (Scharf et al., 2005). Alignments of these transcripts are shown in Supplementary Dataset S31. Interestingly, Humanin, which is a 24 aa long peptide, is thought to be translated directly from rRNA (Maximov et al., 2002). At the amino acid level, PRF-altORFs located on one tRNA and five rRNA transcripts detected in our study have similarity with annotated proteins of 31 different types (Supplementary Datasets S1 and S31), out of which only two have previously been associated with rRNA-like mRNA transcripts: testin-2 (Divecha and Charleston, 1995) and receptor-like serine/threonine-protein kinase (Kong et al., 2008). Thus, our study expands the knowledge about rRNA-like mRNA transcripts by detecting 29 additional protein types encoded by rRNA-like sequences. So far, there are only three known rRNA-located ORFs related to rRNA-specific functions. This suggests that the majority of rRNA-ORFs with the protein coding capacity may be involved in other biological processes. According to the ribosome-first hypothesis, in rRNA-like proto-genomes, relatively short ORFs overlapped with each other in all reading frames and directions. Thus, the density of protein-coding regions was very high in contrast to modern genomes where only a portion of genes overlap (Root-Bernstein and Root-Bernstein, 2016). Our data support this hypothesis. For example, the analysis of all major transcript types in *M. truncatula* revealed that rRNA has the highest number of ORFs per transcript, when we consider ORFs that have high similarity to known proteins after the conversion of ORFs to amino acid sequences, i.e. conserved altProts. With median length of 120 nt, rRNA transcripts are on average ten times shorter than mRNA transcripts and nearly four times shorter than ncRNA transcripts. Despite the short length, rRNA transcripts contain ca. 31 and ten times more conserved altORFs compared to mRNA and ncRNA, respectively (Table 1). Another remarkable observation concerns the percent identity of ORF translations aligned with annotated proteins from the global collection. The median value of percent identity is the highest for tRNA (100%) followed by rRNA (93.3%), as can be seen in Table 1 and Supplementary Figure S1. The same concerns the median value of query coverage (100% and 94.8%, respectively; the global BLASTP analysis, Table 1). This means that ORF translations from tRNA and rRNA have the highest degree of similarity to annotated proteins, even though they are traditionally considered as non-coding RNA types. Likewise, ORF translations from rRNA have the highest median number of hits per query in the global BLASTP analysis (Supplementary Figures S2 and S3). It is ten times higher compared to mRNA. This supports observations of other researchers according to which rRNA-like sequences are overrepresented in genomes (Root-Bernstein and Root-Bernstein, 2019). However, our data confirm this overrepresentation at the protein level, which once again brings rRNA into the spotlight as a potential template for translation. This overrepresentation is also evident at the transcript level, as follows from our analysis of shared subjects in BLASTN (Supplementary Figures S63-S65). In view of these special characteristics of rRNA, it may seem to be not surprising that rRNA transcripts are significantly (25-fold) overrepresented in our dataset (Supplementary Figure S50A). Many of our chimeric models were built using conserved altProts, which were most abundant in rRNA (Table 1). The surprising point is the absence of MS-validated altProts from either rRNA or tRNA in our dataset and the presence of only MS-validated chimeric peptides associated with these transcript types (Supplementary Figure S6). This paradox suggests that some characteristics of rRNA and tRNA distinguish them from other RNA types with regard to PRF. The nearly complete prevalence of PRF value −1 in rRNA and tRNA (Figure 3) further points to the tight evolutionary link between these transcript types. This prevalence is remarkable; the only rRNA transcript associated with multiple frameshifting in our study (MtrunA17_Chr5g0422291) has both PRF sites with value −1 (Supplementary Dataset S4). PRF in tRNA and rRNA has been unknown so far. Likewise, the ability of ncRNA to produce chimeric peptides has never been described before. In our study, 11 such peptides originate from ncRNA. One of eight multi-PRF transcripts illustrated in Supplementary Dataset S4 is ncRNA from mitochondria (MtrunA17_MTg0490971).

Since many peptides in our dataset have more than one potential source in the transcriptome, there is a valid question about the true origin of tRNA- and rRNA-derived chimeric MS peptides. To address this concern, we examined the annotation of loci that are alternative sources of non-mRNA-derived chimeric peptides (Supplementary Dataset S1). This analysis revealed that both alternative sources of CP72 (tRNA) are annotated as tRNA. However, they can also be produced by the reverse complement of repeat elements that overlap those tRNA loci. All five non-repeat alternative sources of CP87 (rRNA) are annotated as rRNA. However, this peptide can also be produced by repeat elements. CP1, CP88, CP89, and CP141 have only repeat-element alternative sources. In contrast, CP114 (rRNA) has no detected alternative sources. This indicates that our study is the first one to demonstrate the presence of a unique chimeric rRNA-encoded MS peptide in a proteomic dataset. This is in line with the suggestion that the non-chimeric peptide Humanin (HN1) may be translated directly from rRNA (Hashimoto et al., 2001; Maximov et al., 2002) because the transcript of CP114 shares 67.4% nucleotide identity with the transcript of Humanin (Supplementary Dataset S31). Interestingly, CP114 is the only rRNA-chimeric peptide with PRF value +2. All other rRNA- and tRNA-peptides have PRF value −1 (Figure 3). It should be noted that no evidence for the association of repeat elements with PRF value −1 or any other PRF value can be found in our data. Furthermore, among repeat elements mentioned above, only those that correspond to CP88 and CP89 have sufficiently high expression in corresponding organs to be considered as true sources of those rRNA-derived peptides. At the same time, regions around PRF sites in the transcript of CP87, CP88 and CP89 share nucleotide identity with two different rRNA-like mRNA transcripts described earlier (Supplementary Datasets S28 and S31). Together, these observations suggest that CP114 is not the only chimeric peptide translated directly from rRNA (Supplementary Dataset S1). To the best of our knowledge, the detection of rRNA and also ncRNA loci with more than one PRF site per transcript is also unique to our study (Supplementary Datasets S4 and S28).

Is there anything special about non-nuclear transcripts when we consider them in the context of PRF? We noticed a trend toward overrepresentation of PRF value +1 in mitochondrial transcripts regardless of the RNA type (Supplementary Figure S11) and also in non-nuclear transcripts combined (Figure 4). Remarkably, the trend is opposite for nuclear transcripts, where PRF value +1 is underrepresented. Two out of eight multi-PRF transcripts shown in Supplementary Dataset S4 have mitochondrial origin, one mRNA and one ncRNA. Both transcripts contain exclusively +1 PRF sites. Chloroplast and mitochondrial transcripts are significantly overrepresented in the dataset (19-fold and 12-fold, respectively; Supplementary Figure S50). These observations reflect the origin of these organelles and highlight the PRF-related difference between nuclear (eukaryotic) and non-nuclear (prokaryotic) transcripts. The biological significance of this difference deserves close attention. It is possible that “short” frameshifts that appear to be associated with transcripts of ancient origin in our study (−1 frameshifts in rRNA and +1 frameshifts in non-nuclear transcripts) represent the ancestral feature whereas “long” frameshifts evolved later. Earlier, we emphasized that prokaryotes are likely to have tremendously benefited from mosaic translation and from PRF in general given the absence of alternative splicing and relatively small genome sizes. In this respect, they are close to viruses where PRF is essential for survival. We hypothesize that the ability to undergo PRF has been an intrinsic feature of all genomes from the very beginning. Conceivably, at some early point in the evolution, it has been harnessed for mosaic translation. This process must have diversified proteomes of primitive organisms because it utilizes the genomic space more efficiently and inventively. This “new dimension” of the proteomes probably facilitated the gradual increase in complexity and adaptability of organisms, which ensured their evolutionary success (Çakır et al., 2023).

## 6. Conclusions

We hope that our study will constitute a rich resource for many discoveries associated with PRF and translation from non-mRNA transcript types. In a broader sense, we anticipate that this work will seed a new field of genetic studies that will consider the nearly infinite protein-coding potential of transcripts due to multiple PRF events. We also hope that many genetic diseases so far not explained with traditional views on the proteome complexity will ultimately find their cures when chimeric and mosaic proteins in eukaryotes will be discovered at a large scale. Likewise, the progress in the development of more resilient and more productive crops will benefit from the discovery of chimeric and mosaic proteins involved in efficient stress responses and other agriculturally important traits. We believe, major advances in nanopore sequencing of proteins will enable the direct detection of chimeric and mosaic proteins, which will leave no doubt about their composite origin. As more evidence supporting the conclusions of our study becomes available, it is possible that our view on the protein-coding potential in higher eukaryotes will undergo a transition from two-dimensional (refProt plus altProt) to multi-dimensional (refProt plus numerous multi-frame chimeric products). We are looking forward to further development of this concept, which is currently in its infancy.

**Umut Çakır**: Conceptualization, Methodology, Software, Validation, Formal analysis, Resources, Data Curation, Writing - Original Draft, Writing - Review & Editing, Visualization, Funding acquisition. **Noujoud Gabed**: Conceptualization, Validation, Formal analysis, Resources, Writing - Review & Editing. **Yunus Emre Köroğlu**, Methodology, Software. **Selen Kaya**: Investigation, Data Curation. **Senjuti Sinharoy**: Conceptualization, Writing - Review & Editing. **Vagner A. Benedito**: Conceptualization, Writing - Review & Editing. **Marie Brunet**: Methodology. **Xavier Roucou**: Methodology, Writing - Review & Editing. **Igor S. Kryvoruchko**: Conceptualization, Methodology, Validation, Formal analysis, Resources, Data Curation, Writing - Original Draft, Writing - Review & Editing, Visualization, Supervision, Project administration, Funding acquisition.

## Declaration of Competing Interest

The authors declare no competing interest relevant to this study.

## Data availability

Sequences of MS-supported chimeric models, their corresponding MS peptides and transcripts, as well as genomic DNA, alignments, and other relevant features with detailed annotation are available in Geneious^®^ format in the online version of this article. Chimeric protein model sequences used as MS search databases, certificates of analysis, and peptide/protein report files from MS searches are available as separate supplementary datasets (Supplementary Datasets S32-S36) in the online version of this article. Corresponding files for the conservation evidence and MS-validation of altProts that were the basis for modeling of chimeric proteins will be published later in a paper with the focus on altProts. Another article will contain our meta-analysis on 627 *M. truncatula* genes studied with loss-of-function approaches and will address MS-supported and conserved altORFs located on already studied genes. Two dedicated software articles will describe the details and provide the codes of in-house scripts generated for this study: one for the modeling of chimeric peptides and the other one for the detection of alternative chimeric sources (Çakır et al., preprints in preparation). Data on more than 18,600 conserved altORFs, 805 MS-supported altORFs, 156 chimeric peptides, and their non-RE alternative sources will be integrated into the *M. truncatula* genome portal MtrunA17r5.0-ANR as a separate track. The complete set of data related to this preprint is available in the Zenodo repository.

## Acknowledgements

This work was supported by the Scientific and Technological Research Council of Turkey (TÜBİTAK) grants and Boğaziçi University standard research grant (BAP-P) to UÇ and IK (TÜBİTAK 1001 120Z514, TÜBİTAK 1002 120Z247, and BAP-P 18841), a Canada Research Chair in functional proteomics and discovery of novel proteins to XR, and a Canadian Institutes of Health Research Project Grant PJT-175322 to XR and MB. MB was supported by a Fonds de Recherche du Québec en Santé (FRQS) Junior 1 award (307936). Computational analysis was conducted using the server of the Turkish National e-Science e-Infrastructure (TRUBA) center. The completion of this study was possible due to the support of IK by United Arab Emirates University and the support of UÇ by the IMPRS-Genome Science PhD program.

## Abbreviations

altORF: alternative open reading frame
refORF: reference open reading frame
altProt: alternative protein
refProt: reference protein
CP: chimeric peptide
PRF: programmed ribosomal frameshifting
mRNA: messenger RNA
ncRNA: non-coding RNA
rRNA: ribosomal RNA
tRNA: transfer RNA
CDS: coding sequence
UTR: untranslated region
aa: amino acids
bp: base pairs
nt: nucleotides
MS: mass spectrometry

## Supporting information

This preprint contains supplementary materials: Supplementary Results (1), Supplementary Figures (67), Supplementary Tables (9), and Supplementary Datasets (36).

## Supplementary Results

Supplementary Results are available as a separate file.

## Supplementary Figures

Supplementary Figures are available as a separate file.

## Supplementary Tables

Supplementary Tables are available as a separate file.

**<u>Supplementary Table S1</u>**. Statistics on all ORFs 60 nt or longer grouped by RNA type.

**<u>Supplementary Table S2</u>**. Statistics on altProts with at least one hit in the global BLASTP analysis before the application of the filter.

**<u>Supplementary Table S3</u>**. Numbers of altProts before and after the elimination of queries with less than 70% identity to annotated proteins.

**<u>Supplementary Table S4</u>**. Distribution of transcripts in different categories based on RNA type, cellular localization, mass spectrometry (MS) support of altORFs, and conservation of altORF translation products.

**<u>Supplementary Table S5</u>**. Redundant counts of chimeric protein models, shown per altProt type, transcript type, and programmed ribosomal frameshifting (PRF) value.

**<u>Supplementary Table S6</u>**. Non-redundant counts of chimeric proteins modeled with altORFs that are MS-supported and conserved at the same time, shown per transcript type and PRF value.

**<u>Supplementary Table S7</u>**. Non-redundant counts of chimeric proteins modeled with MS-supported altORFs, shown per proteomic study and PRF value.

**<u>Supplementary Table S8</u>**. Non-redundant counts of chimeric proteins modeled with conserved altORFs, shown per transcript type and PRF value.

**<u>Supplementary Table S9</u>**. Non-redundant counts of chimeric protein models, shown per altProt type, transcript type, and PRF value.

## Supplementary Datasets

Supplementary Datasets are available as individual files

**<u>Supplementary Dataset S1</u>**. Master table. A master table that summarizes various parameters of 156 chimeric MS peptides validated in this study and charecteristics of their corresponding transcripts.

**<u>Supplementary Dataset S2</u>**. Graphical summary. A graphical summary on alignments between 156 chimeric peptide models (CPs) and corresponding MS peptides (left) together with transcript models (right) that feature the transcript type, length in nucleotides, and relative positions of ORFs (to scale) involved in the production of CPs.

**<u>Supplementary Dataset S3</u>**. Relevant sequences with detailed annotation. This Geneious^®^-formatted dataset contains sequences of 156 MS-supported chimeric models (CPs), their corresponding MS peptides and transcripts, as well as genomic DNA, alignments, and other informative features with detailed annotation.

**<u>Supplementary Dataset S4</u>**. Primary-source transcripts associated with multiple PRF events. Eight primary-source transcripts associated with multiple PRF events.

**<u>Supplementary Dataset S5</u>**. Mosaic proteins. Three candidate mosaic proteins deduced from chimeric MS peptides.

**<u>Supplementary Dataset S6</u>**. Folding of mosaic proteins.

**<u>Supplementary Dataset S7</u>**. Folding of chimeric protein models Part 1. A graphical summary on folding predictions for MS-supported chimeric peptide models (CPs) 1-50.

**<u>Supplementary Dataset S7</u>**. Folding of chimeric protein models Part 2. A graphical summary on folding predictions for MS-supported chimeric peptide models (CPs) 51-100.

**<u>Supplementary Dataset S7</u>**. Folding of chimeric protein models Part 3. A graphical summary on folding predictions for MS-supported chimeric peptide models (CPs) 101-156.

**<u>Supplementary Dataset S7</u>**. Folding of chimeric protein models Part 4 Alpha-helices only. A graphical summary on folding predictions for 90 MS-supported chimeric peptide models (CPs) that contain alpha-helices and no beta-sheets.

**<u>Supplementary Dataset S7</u>**. Folding of chimeric protein models Part 5 Beta-sheets only. A graphical summary on folding predictions for 24 MS-supported chimeric peptide models (CPs) that contain beta-sheets regardless of the presence of alpha-helices.

**<u>Supplementary Dataset S8</u>**. Alpha helices vs PRF value. Distribution of 90 alpha-helix-forming models of chimeric MS pepsample tides in four categories based on the PRF position relative to the helices.

**<u>Supplementary Dataset S9</u>**. Alpha helices inside only vs PRF value. Distribution of 72 MS-supported models based on the position of PRF sites inside alpha-helices.

**<u>Supplementary Dataset S10</u>**. Beta sheets vs PRF value. Distribution of 24 MS-supported chimeric peptide models with predicted beta-sheets in four categories based on the position of PRF sites relative to the beta-sheets.

**<u>Supplementary Dataset S11</u>**. Overlapping genes. The genomic landscape at and around the primary-source loci of 156 MS-supported chimeric peptides.

**<u>Supplementary Dataset S12</u>**. Nucleotide alignments of random RNA samples. Distribution of percent nucleotide identity values of 45 pairwise alignments among ten randomly chosen transcripts per RNA category.

**<u>Supplementary Dataset S13</u>**. Homology groups 1 to 10. Homology groups based on pairwise nucleotide alignments with percent identity values 56 and above.

**<u>Supplementary Dataset S14</u>**. Alignments among 34 primary-source transcripts with detectable similarity. A graphical summary on nucleotide alignments among all 34 primary-source transcripts that have the global or the local similarity to each other.

**<u>Supplementary Dataset S15</u>**. Summary on sequences with shared subjects in the BLASTN analysis. Summary on the primary-source transcript sequences with shared subjects in the BLASTN analysis and sequences that have direct similarity over the entire length.

**<u>Supplementary Dataset S16</u>**. Sequences with shared subjects in the BLASTN analysis. Primary-source transcript sequences with shared subjects in the BLASTN analysis.

**<u>Supplementary Dataset S17</u>**. Selected RNA-Seq runs. Fifty RNA-Seq runs selected for the identification of non-chimeric alternative sources of 156 MS-supported chimeric peptides.

**<u>Supplementary Dataset S18</u>**. Details of the RNA-Seq analysis.

**<u>Supplementary Dataset S19</u>**. RNA-Seq read alignments. A graphical summary on alignments between transcripts of 15 MS-supported chimeric peptides and RNA-Seq reads from 50 selected RNA-Seq runs.

**<u>Supplementary Dataset S20</u>**. RNA-Seq reads with exact matches in more than one run.

**<u>Supplementary Dataset S21</u>**. Sequencing adapters and primers.

**<u>Supplementary Dataset S22</u>**. Non-repeat-element alternative chimeric sources. Non-repeat-element alternative chimeric sources of 156 MS-supported chimeric peptides.

**<u>Supplementary Dataset S23</u>**. Repeat-element alternative chimeric sources. Repeat-element alternative chimeric and non-chimeric sources of 156 MS-supported chimeric peptides.

**<u>Supplementary Dataset S24</u>**. Summary on alternative sources of chimeric peptides. This dataset summarizes the information from Supplementary Datasets S22 and S23.

**<u>Supplementary Dataset S25</u>**. Non-overlapping repeat elements. A list of 70 alternative-source repeat elements that do not osample verlap with any locus at the putative PRF sites.

**<u>Supplementary Dataset S26</u>**. Expression profiles of non-overlapping repeat elements. Expression profiles of ten non-overlapping repeat elements that have log2 TMM values above zero.

**<u>Supplementary Dataset S27</u>**. Expression profiles of 156 primary-source transcripts.

**<u>Supplementary Dataset S28</u>**. Alternative-source transcripts associated with multiple PRF events.

**<u>Supplementary Dataset S29</u>**. Summary on the most relevant statistically significant observations.

**<u>Supplementary Dataset S30</u>**. Summary on the most relevant statistically significant associations.

**<u>Supplementary Dataset S31</u>**. Alignments of rRNA transcripts with known rRNA-like mRNA transcripts.

**<u>Supplementary Dataset S32</u>**. All chimeric protein models (redundant).

**<u>Supplementary Dataset S33</u>**. All chimeric protein models (unique).

**<u>Supplementary Dataset S34</u>**. Chimeric proteins modeled with conserved altProts (redundant).

**<u>Supplementary Dataset S35</u>**. Chimeric proteins modeled with MS-supported altProts (redundant).

**<u>Supplementary Dataset S36</u>**. MS reports and certificates of analysis.

---

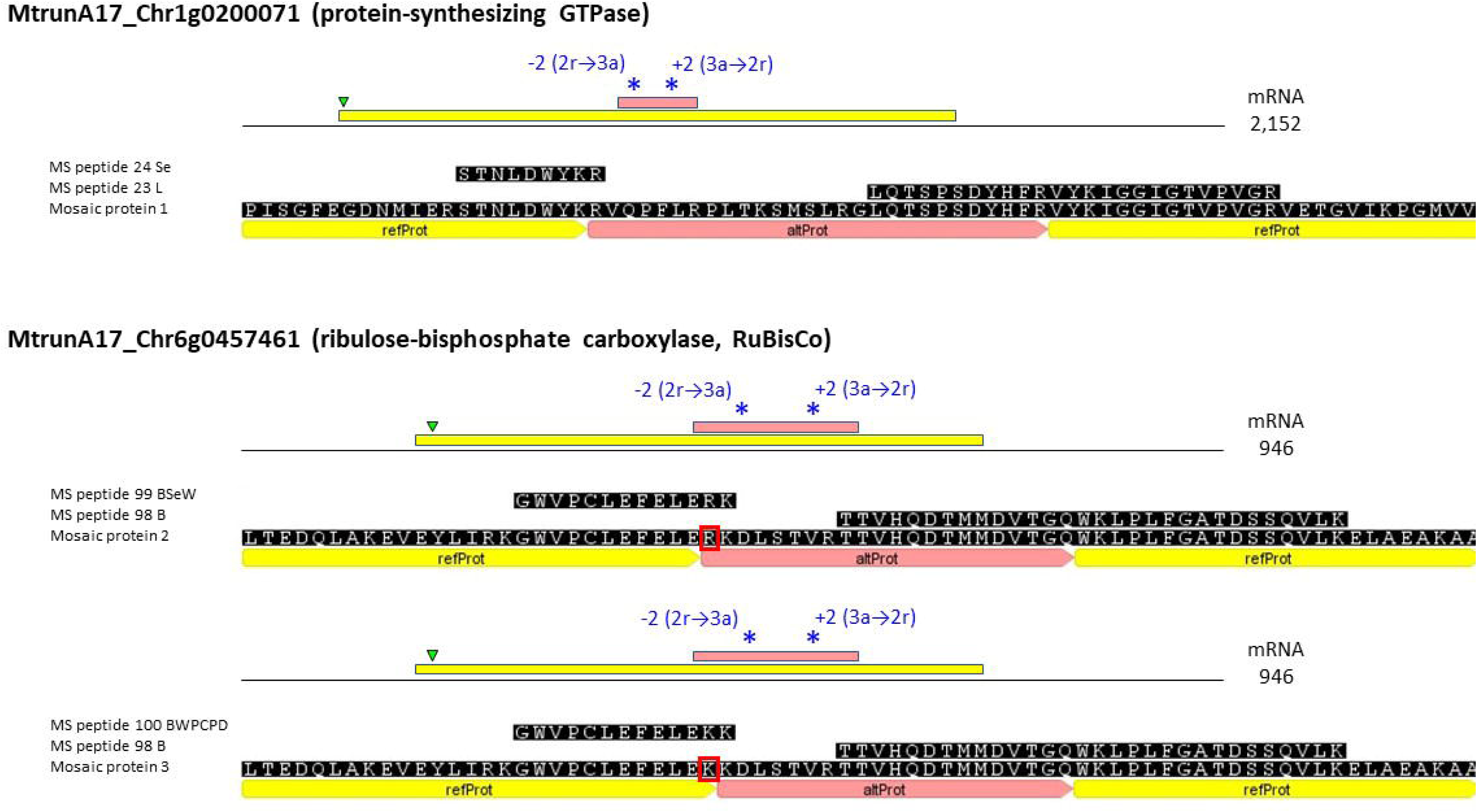

## Notes

### Competing Interest Statement

The authors have declared no competing interest.

