## Supplementary material for "Discovery of diverse chimeric peptides in a eukaryotic proteome sets the stage for the experimental proof of the mosaic translation hypothesis": Supplementary Dataset S2 Graphical summary.pdf

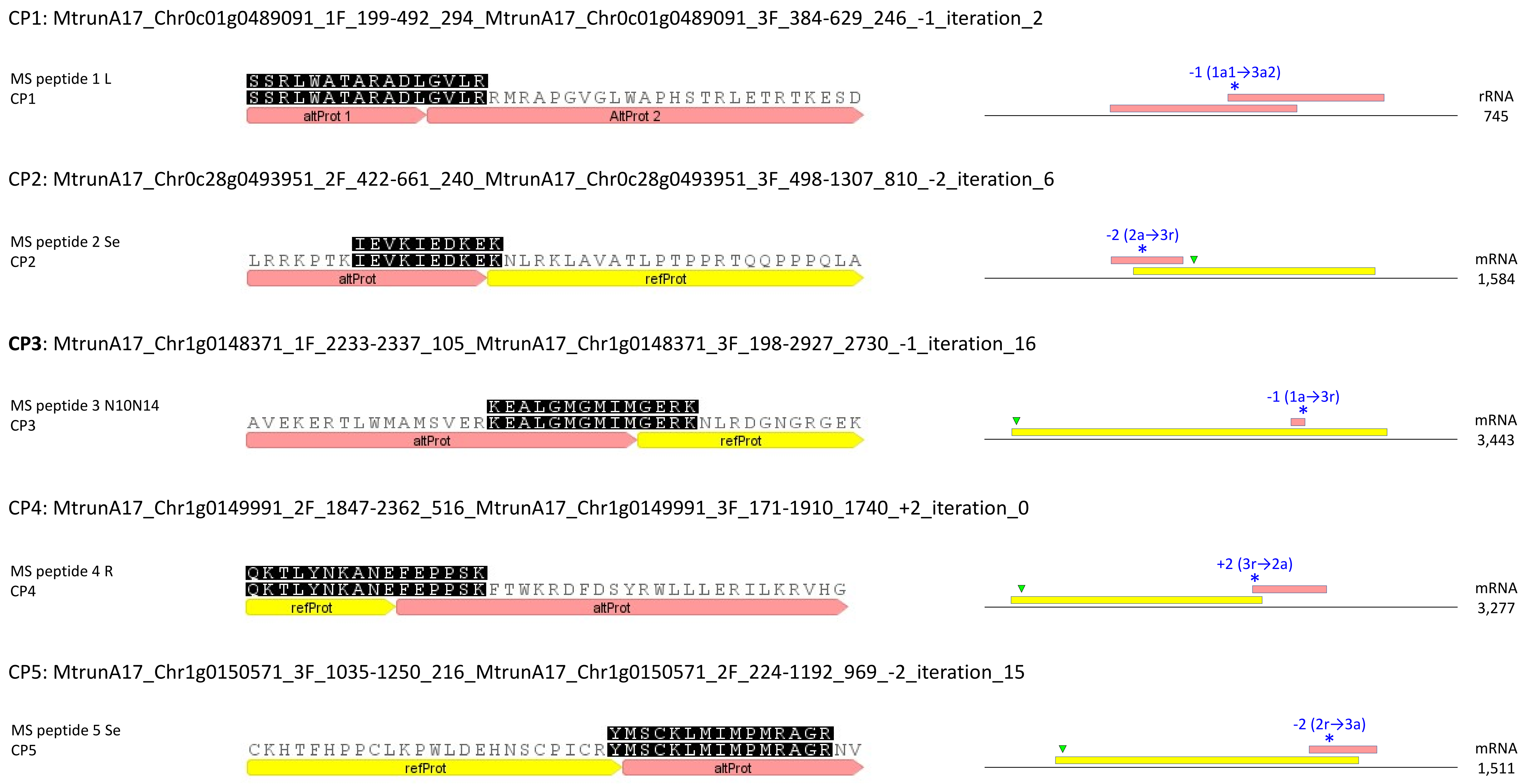
CP1: MtrunA17_Chr0c01g0489091_1F_199-492_294_MtrunA17_Chr0c01g0489091_3F_384-629_246_-1_iteration_2
MS peptide 1 L
CP1
SSRLWATARADLGVLR
SSRLWATARADLGVLRRMRAPGVGLWAPHSTRLETRTKESD
altProt 1
AltProt 2
-1 (1a1→3a2)
*
rRNA
745
CP2: MtrunA17_Chr0c28g0493951_2F_422-661_240_MtrunA17_Chr0c28g0493951_3F_498-1307_810_-2_iteration_6
MS peptide 2 Se
CP2
IEVKIEDKEK
LRRKPTKIEVKIEDKEKNLRKLAVATLPTPPRTQQPPPQLA
altProt
refProt
-2 (2a→3r)
*
mRNA
1,584
CP3: MtrunA17_Chr1g0148371_1F_2233-2337_105_MtrunA17_Chr1g0148371_3F_198-2927_2730_-1_iteration_16
MS peptide 3 N10N14
CP3
KEALGMGMIMGERK
AVEKERTLWMAMSVERKEALGMGMIMGERKNLRDGNGRGEK
altProt
refProt
-1 (1a→3r)
*
mRNA
3,443
CP4: MtrunA17_Chr1g0149991_2F_1847-2362_516_MtrunA17_Chr1g0149991_3F_171-1910_1740_+2_iteration_0
MS peptide 4 R
CP4
QKTLYNKANEFEPPSK
QKTLYNKANEFEPPSKFTWKRDFDSYRWLLLERILKRVHG
refProt
altProt
+2 (3r→2a)
*
mRNA
3,277
CP5: MtrunA17_Chr1g0150571_3F_1035-1250_216_MtrunA17_Chr1g0150571_2F_224-1192_969_-2_iteration_15
MS peptide 5 Se
CP5
YMSCKLMIMPMRAGR
CKHTFHPPCLKPWLDEHNSCPICRYMSCKLMIMPMRAGRNV
refProt
altProt
-2 (2r→3a)
*
mRNA
1,511

CP6: MtrunA17_Chr1g0152521_2F_290-436_147_MtrunA17_Chr1g0152521_1F_1-969_969_-2_iteration_20

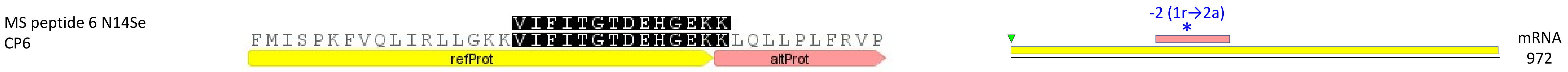


CP7: MtrunA17_Chr1g0153001_1F_178-519_342_MtrunA17_Chr1g0153001_3F_396-806_411_-1_iteration_9

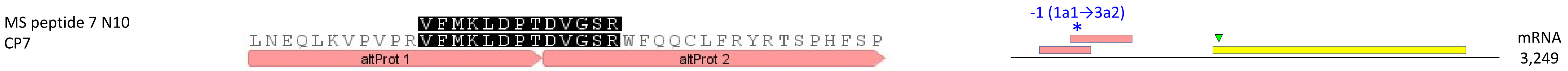


**CP8**: MtrunA17_Chr1g0155251_1F_1228-1467_240_MtrunA17_Chr1g0155251_3F_138-2090_1953_-1_iteration_16

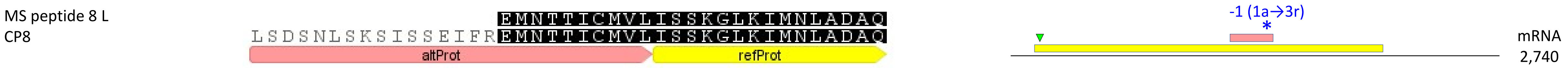


<u>CP9</u>: MtrunA17_Chr1g0158341_2F_758-859_102_MtrunA17_Chr1g0158341_1F_1-1251_1251_+2_iteration_17

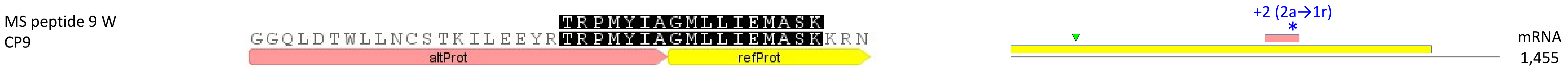


**CP10**: MtrunA17_Chr1g0162101_3F_3-221_219_MtrunA17_Chr1g0162101_1F_1-255_255_+2_iteration_1

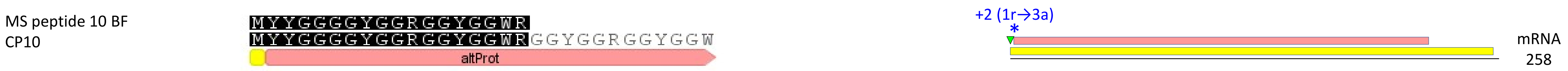

CP11: MtrunA17_Chr1g0164591_3F_3-338_336_MtrunA17_Chr1g0164591_2F_314-1369_1056_-1_iteration_2

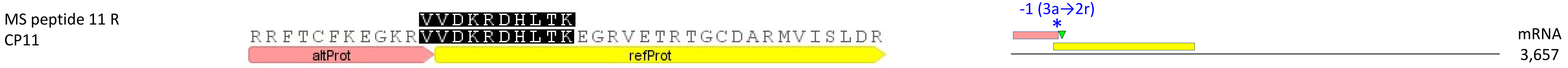


**CP12**: MtrunA17_Chr1g0178361_1F_529-618_90_MtrunA17_Chr1g0178361_3F_201-1124_924_-1_iteration_9

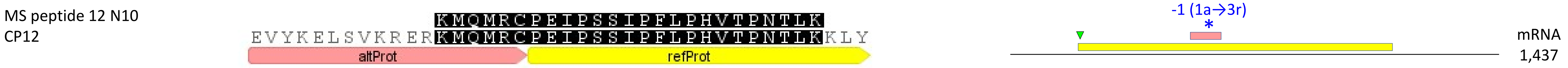


CP13: MtrunA17_Chr1g0181761_1F_445-570_126_MtrunA17_Chr1g0181761_3F_18-1262_1245_+2_iteration_0

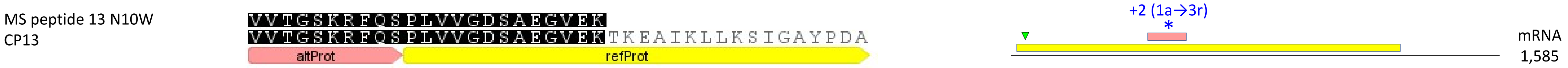


**CP14**: MtrunA17_Chr1g0182591_1F_4162-4257_96_MtrunA17_Chr1g0182591_2F_11-4228_4218_-1_iteration_11

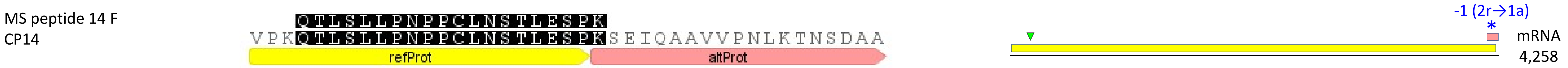


CP15: MtrunA17_Chr1g0183001_1F_2749-2943_195_MtrunA17_Chr1g0183001_3F_375-4916_4542_-2_iteration_16

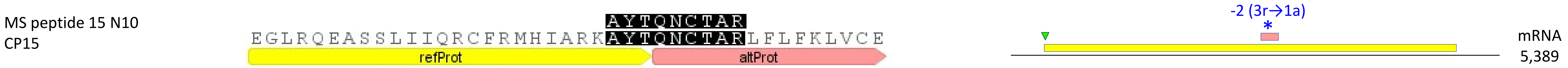

**CP16**: MtrunA17_Chr1g0185811_1F_535-621_87_MtrunA17_Chr1g0185811_2F_2-1774_1773_+1_iteration_10

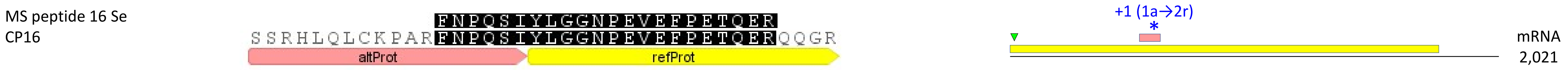


**CP17**: MtrunA17_Chr1g0185811_1F_535-621_87_MtrunA17_Chr1g0185811_2F_2-1774_1773_+1_iteration_2

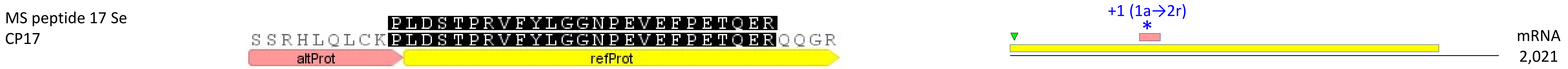


**CP18**: MtrunA17_Chr1g0185811_1F_535-621_87_MtrunA17_Chr1g0185811_2F_2-1774_1773_+1_iteration_9

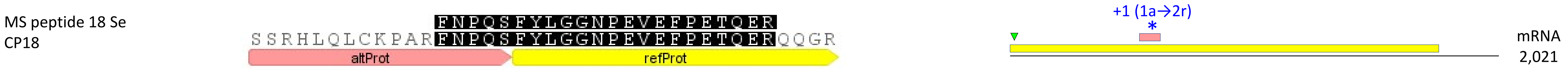


CP19: MtrunA17_Chr1g0185871_1F_1447-1602_156_MtrunA17_Chr1g0185871_3F_309-3371_3063_-2_iteration_3

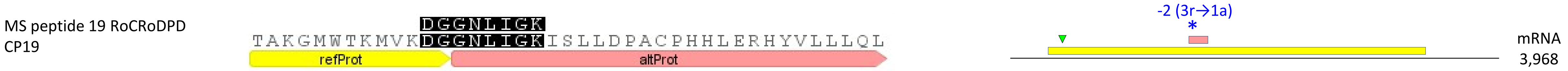


CP20: MtrunA17_Chr1g0190571_2F_62-262_201_MtrunA17_Chr1g0190571_1F_1-2274_2274_+2_iteration_10

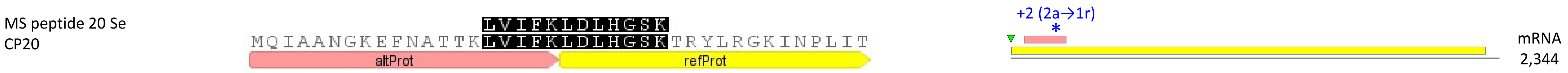

CP21: MtrunA17_Chr1g0191411_1F_1660-1869_210_MtrunA17_Chr1g0191411_2F_1601-1732_132_+2_iteration_22

MS peptide 21 St — RHVIRDLNIIIC
CP21 — LIGSDCHINFDPVQIQSMTLASSTLQIRRHVIRDLNIIIC
altProt 1 | altProt 2

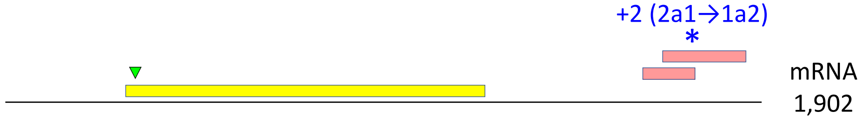


CP22: MtrunA17_Chr1g0198091_3F_255-443_189_MtrunA17_Chr1g0198091_1F_379-507_129_+1_iteration_2

MS peptide 22 W — LGPQWSNRELENSMK
CP22 — LGPQWSNRELENSMKYTESMVEIGKKVATTIRKRSIEMVE
altProt | refProt

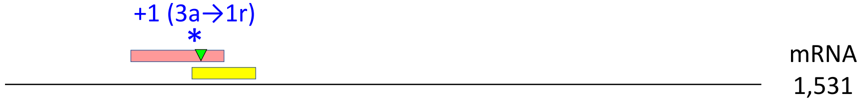


CP23: MtrunA17_Chr1g0200071_3F_828-1001_174_MtrunA17_Chr1g0200071_2F_218-1567_1350_+2_iteration_1

MS peptide 23 L — LQTSPSDYHFRVYKIGGIGTVPVGR
CP23 — LQTSPSDYHFRVYKIGGIGTVPVGRVETGVIKPGMVVTFA
altProt | refProt

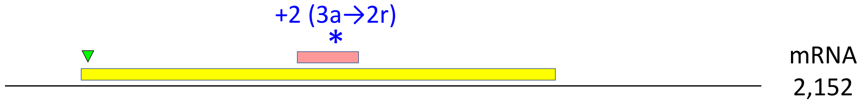


CP24: MtrunA17_Chr1g0200071_3F_828-1001_174_MtrunA17_Chr1g0200071_2F_218-1567_1350_-2_iteration_11

MS peptide 24 Se — STNLDWYKR
CP24 — PISGFEGDNMIERSTNLDWYKRVQPFLRPLTKSMSLRGLQT
refProt | altProt

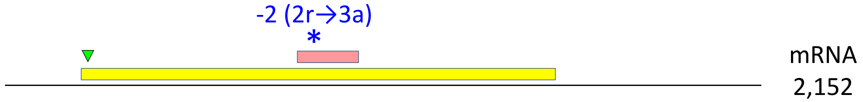


CP25: MtrunA17_Chr1g0202001_2F_149-421_273_MtrunA17_Chr1g0202001_1F_1-2331_2331_-1_iteration_0

MS peptide 25 N10N14 — LVMVQSLDLHGSYRRR
CP25 — EFNATFKLVMVQSLDLHGSYRRRLFGEINPSITELQHLTYL
altProt | refProt

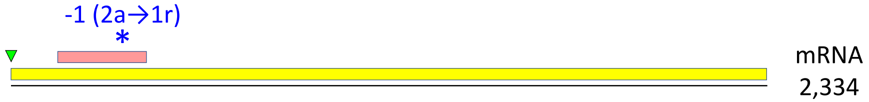

CP26: MtrunA17_Chr1g0205601_3F_321-395_75_MtrunA17_Chr1g0205601_1F_13-369_357_-1_iteration_17

MS peptide 26 FR NFMVITWDSPKE
CP26 PNTPSPIFSGNTGLEAFYARPKLKNFMVITWDSPKE
refProt altProt

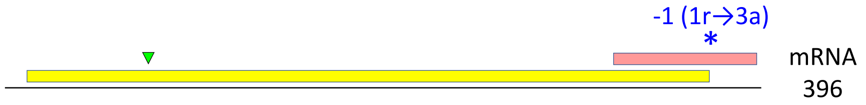


CP27: MtrunA17_Chr1g0207811_1F_1462-1824_363_MtrunA17_Chr1g0207811_3F_207-1913_1707_+1_iteration_19

MS peptide 27 L TREAGIQDIHHQINQK
CP27 ALARMEDLGLKCRDVRTREAGIQDIHHQINQKRWNLFVVI
refProt altProt

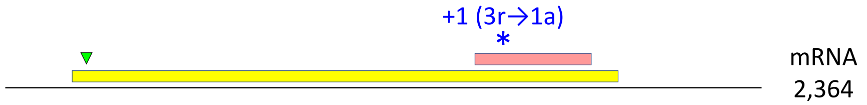


CP28: MtrunA17_Chr1g0207921_2F_260-391_132_MtrunA17_Chr1g0207921_1F_1-597_597_+1_iteration_6

MS peptide 28 N10N14 VNGAFFLQR
CP28 NKFEVLVDRVNGAFFLQRDGKRFVTFTESILVPGLPWFLF
refProt altProt

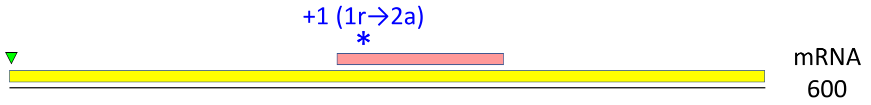


CP29: MtrunA17_Chr1g0209791_2F_2-130_129_MtrunA17_Chr1g0209791_1F_1-288_288_+2_iteration_5

MS peptide 29 Se SMNVFCSLKFCK
CP29 AGRMEKMSVGRSMNVFCSLKFCKSQALGGHMNQRETETLN
altProt refProt

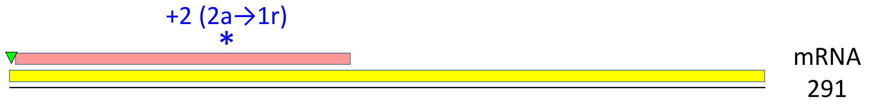


CP30: MtrunA17_Chr1g0210521_2F_782-928_147_MtrunA17_Chr1g0210521_3F_192-917_726_-1_iteration_23

MS peptide 30 N10 MWLQNLQMKLMGGRR
CP30 KTENLQFQADSFQRQGRQLRRKMWLQNLQMKLMGGRRNPYL
refProt altProt

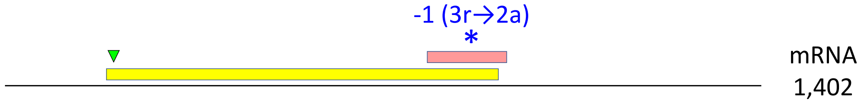

CP31: MtrunA17_Chr1g0212961_2F_617-703_87_MtrunA17_Chr1g0212961_3F_63-1136_1074_-2_iteration_2

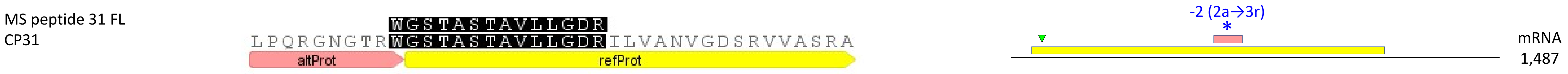


CP32: MtrunA17_Chr1g1004575_2F_299-379_81_MtrunA17_Chr1g1004575_3F_333-416_84_-2_iteration_11

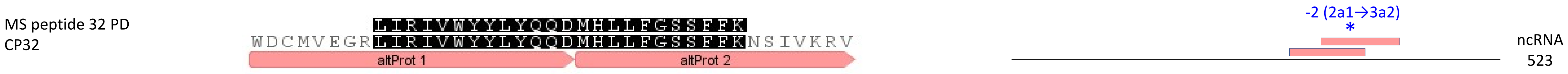


CP33: MtrunA17_Chr2g0283311_1F_766-849_84_MtrunA17_Chr2g0283311_2F_263-868_606_-2_iteration_14

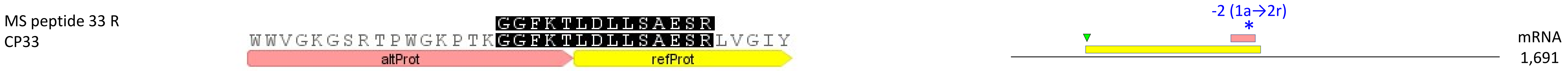


CP34: MtrunA17_Chr2g0285461_2F_1553-1795_243_MtrunA17_Chr2g0285461_1F_838-3573_2736_+2_iteration_0

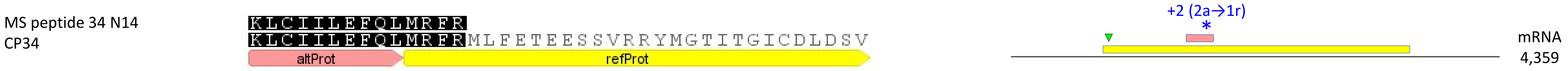


CP35: MtrunA17_Chr2g0292921_2F_1202-1453_252_MtrunA17_Chr2g0292921_1F_187-1545_1359_-2_iteration_6

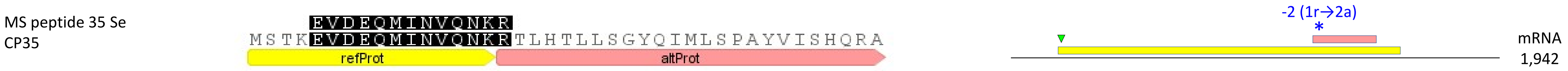

**CP36**: MtrunA17_Chr2g0298731_1F_3127-3255_129_MtrunA17_Chr2g0298731_3F_105-3674_3570_-1_iteration_7

MS peptide 36 St — YLSMNQMKVDSLDK
CP36 — RLRKSVPKYLSMNQMKVDSLDKRPYEQEADNVEGSLDFHYV
altProt refProt

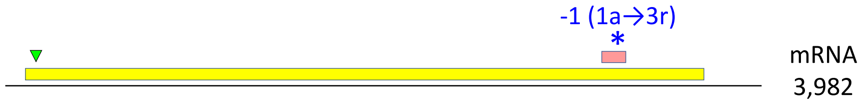


CP37: MtrunA17_Chr2g0299561_2F_3347-3535_189_MtrunA17_Chr2g0299561_1F_97-3969_3873_-1_iteration_17

MS peptide 37 N10 — FLLNLHQK
CP37 — PYQPQKQKYLWRYQIHLLQMKFLLNLHQKKILNLHIIGRFI
altProt refProt

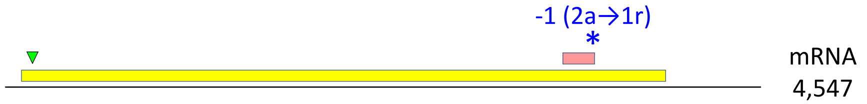


CP38: MtrunA17_Chr2g0304891_3F_627-725_99_MtrunA17_Chr2g0304891_1F_643-1512_870_-2_iteration_1

MS peptide 38 W — IPLGMNIK
CP38 — IPLGMNIKEYNQTNHSLEILVLVGEEAGNPVFGINV
altProt refProt

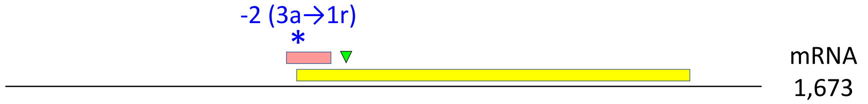


CP39: MtrunA17_Chr2g0305951_3F_3-113_111_MtrunA17_Chr2g0305951_1F_1-153_153_-1_iteration_14

MS peptide 39 N14 — SLNLQRSWLPLLILYVNDAFGTAHR
CP39 — MSLNLQRSWLPLLILYVNDAFGTAHRAHTSI
refProt altProt

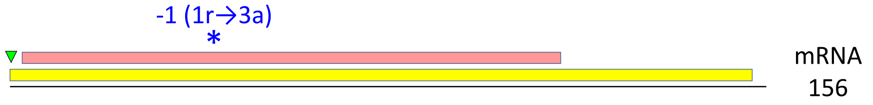


**CP40**: MtrunA17_Chr2g0309251_1F_1300-1383_84_MtrunA17_Chr2g0309251_3F_795-2144_1350_-1_iteration_5

MS peptide 40 B — RSQKSTPQPK
CP40 — GLALFKILERRSQKSTPQPKPAKPLPDILTQSGHQHAI
altProt refProt

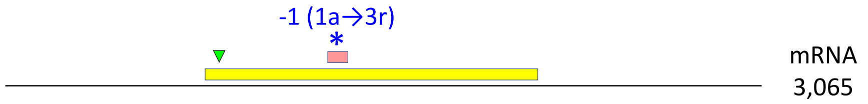

CP41: MtrunA17_Chr2g0310041_3F_3-95_93_MtrunA17_Chr2g0310041_1F_1-540_540_-2_iteration_20

MS peptide 41 RoC
CP41

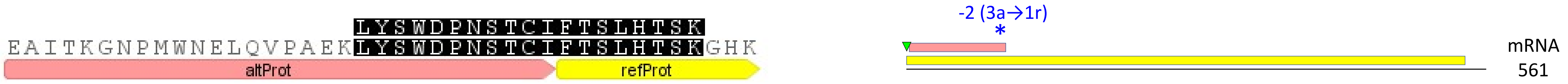


**CP42**: MtrunA17_Chr2g0312631_1F_2020-2094_75_MtrunA17_Chr2g0312631_3F_213-3398_3186_+2_iteration_11

MS peptide 42 L
CP42

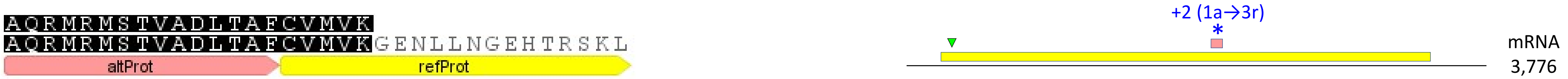


CP43: MtrunA17_Chr2g0316291_3F_1647-1745_99_MtrunA17_Chr2g0316291_2F_101-3394_3294_-2_iteration_1

MS peptide 43 W
CP43

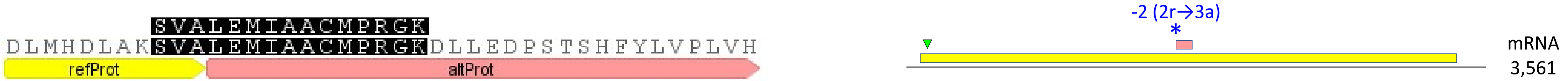


CP44: MtrunA17_Chr2g0326801_1F_2404-2493_90_MtrunA17_Chr2g0326801_3F_2394-2507_114_-2_iteration_19

MS peptide 44 N10
CP44

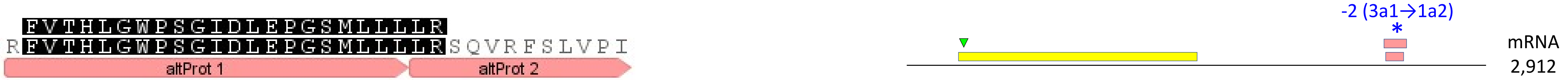


CP45: MtrunA17_Chr2g0328091_3F_87-245_159_MtrunA17_Chr2g0328091_2F_107-1243_1137_-1_iteration_5

MS peptide 45 F
CP45

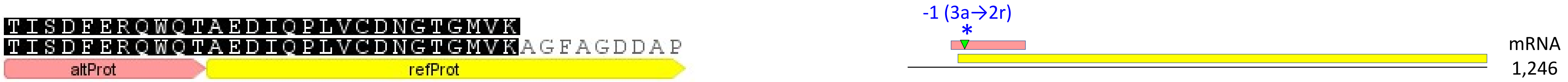

CP46: MtrunA17_Chr2g0329031_3F_1209-1379_171_MtrunA17_Chr2g0329031_2F_197-2440_2244_+2_iteration_20

MS peptide 46 B
CP46

PCMMIVHVGCQRMK
DGGSWLIGLNLEMLSGFKPCMMIVHVGCQRMKDISLAGLS
altProt refProt

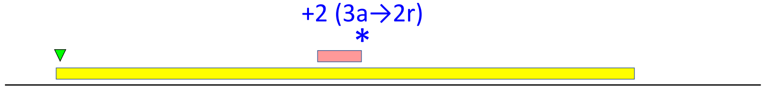


mRNA
2,935

CP47: MtrunA17_Chr3g0079521_2F_1580-1747_168_MtrunA17_Chr3g0079521_1F_25-2292_2268_-1_iteration_10

MS peptide 47 St
CP47

RTLFEYYAWDYR
TTNIFQAQERFWILQKYRRTLFEYYAWDYRVLKTFARHYST
altProt refProt

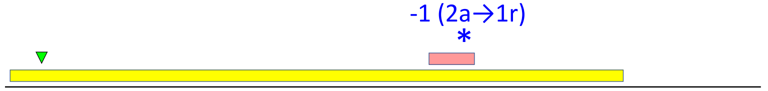


mRNA
2,795

**CP48**: MtrunA17_Chr3g0083801_1F_76-174_99_MtrunA17_Chr3g0083801_3F_12-737_726_+1_iteration_3

MS peptide 48 N10
CP48

MAMIENETKQEANVWFVYAMK
FFFVLVKMAMIENETKQEANVWFVYAMKKATLKPFVTSNA
refProt altProt

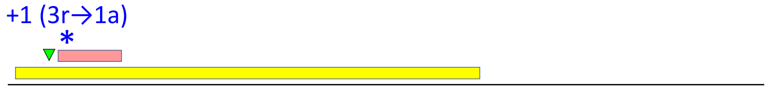


mRNA
1,182

**CP49**: MtrunA17_Chr3g0091141_1F_976-1071_96_MtrunA17_Chr3g0091141_2F_170-1891_1722_-2_iteration_11

MS peptide 49 B
CP49

SKNLALEPAPTMVK
LLQCTWRIRTVWHYNKSKNLALEPAPTMVKWIRVLYSDFTA
altProt refProt

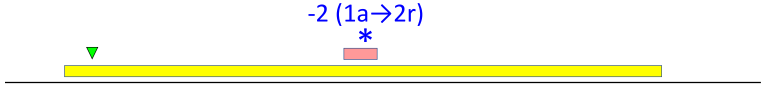


mRNA
2,180

CP50: MtrunA17_Chr3g0091671_2F_14-130_117_MtrunA17_Chr3g0091671_1F_1-153_153_-2_iteration_14

MS peptide 50 PD
CP50

NPVDHPHGGGEGRAPIGVEK
MNPVDHPHGGGEGRAPIGVEKNPQLLGVILHLEEE
refProt altProt

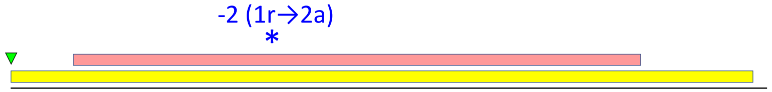


mRNA
156

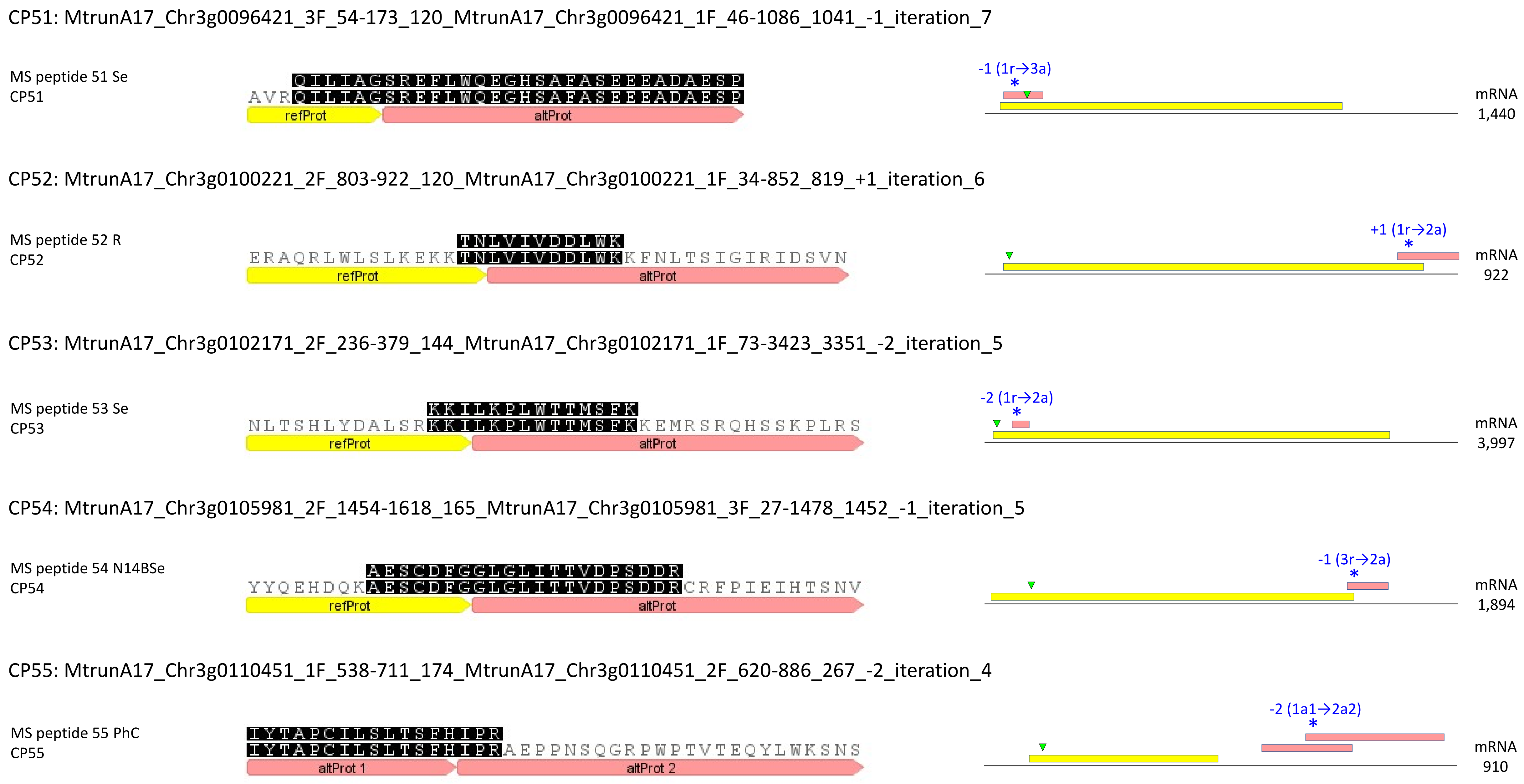
CP51: MtrunA17_Chr3g0096421_3F_54-173_120_MtrunA17_Chr3g0096421_1F_46-1086_1041_-1_iteration_7
MS peptide 51 Se
CP51
QILIAGSREEFLWQEGHSAFASEEEADAESP
AVRQILIAGSREEFLWQEGHSAFASEEEADAESP
refProt
altProt
-1 (1r→3a)
*
mRNA
1,440
CP52: MtrunA17_Chr3g0100221_2F_803-922_120_MtrunA17_Chr3g0100221_1F_34-852_819_+1_iteration_6
MS peptide 52 R
CP52
TNLVIVDDLWK
ERAQRLWLSLKEKKTNLVIVDDLWKKFNLTSIGIRIDSVN
refProt
altProt
+1 (1r→2a)
*
mRNA
922
CP53: MtrunA17_Chr3g0102171_2F_236-379_144_MtrunA17_Chr3g0102171_1F_73-3423_3351_-2_iteration_5
MS peptide 53 Se
CP53
KKILKPLWTTMSFK
NLTSHLYDALSRKKILKPLWTTMSFKKEMRSRQHSSKPLRS
refProt
altProt
-2 (1r→2a)
*
mRNA
3,997
CP54: MtrunA17_Chr3g0105981_2F_1454-1618_165_MtrunA17_Chr3g0105981_3F_27-1478_1452_-1_iteration_5
MS peptide 54 N14BSe
CP54
AESCDFGGLGLITTVDPSDDR
YYQEHDQKAESCDFGGLGLITTVDPSDDRCRFPIEIHTSNV
refProt
altProt
-1 (3r→2a)
*
mRNA
1,894
CP55: MtrunA17_Chr3g0110451_1F_538-711_174_MtrunA17_Chr3g0110451_2F_620-886_267_-2_iteration_4
MS peptide 55 PhC
CP55
IYTAPCILSLTSFHIPR
IYTAPCILSLTSFHIPRAEPPNSQGRPWPTVTEQYLWKSNS
altProt 1
altProt 2
-2 (1a1→2a2)
*
mRNA
910

CP56: MtrunA17_Chr3g0113591_2F_1295-1465_171_MtrunA17_Chr3g0113591_1F_124-2748_2625_-2_iteration_19

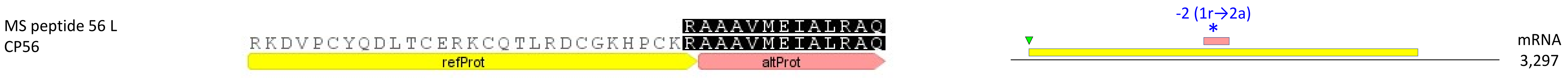


CP57: MtrunA17_Chr3g0124631_2F_1127-1249_123_MtrunA17_Chr3g0124631_1F_130-2304_2175_+2_iteration_19

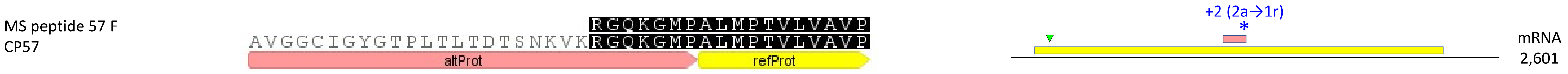


CP58: MtrunA17_Chr3g0127321_2F_1055-1303_249_MtrunA17_Chr3g0127321_1F_1-1329_1329_+2_iteration_18

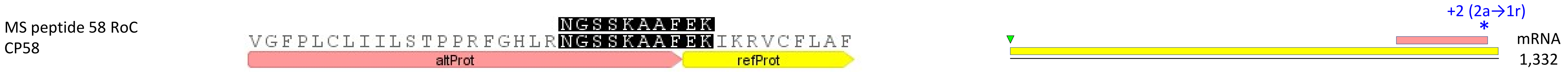


CP59: MtrunA17_Chr3g0130671_3F_1026-1190_165_MtrunA17_Chr3g0130671_1F_334-1179_846_-1_iteration_5

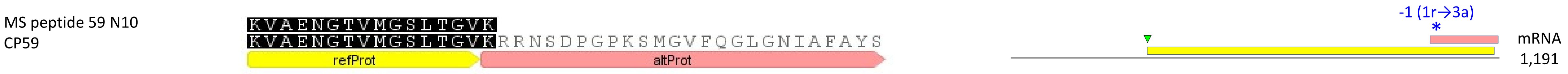


CP60: MtrunA17_Chr3g0135761_2F_734-958_225_MtrunA17_Chr3g0135761_1F_1-2115_2115_+2_iteration_0

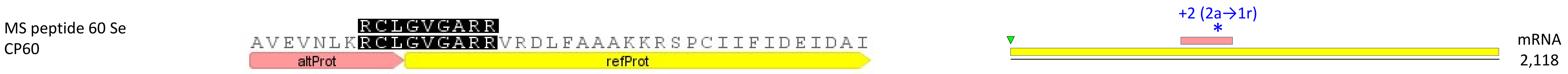

CP61: MtrunA17_Chr3g0137391_3F_915-1070_156_MtrunA17_Chr3g0137391_2F_23-1768_1746_-1_iteration_8

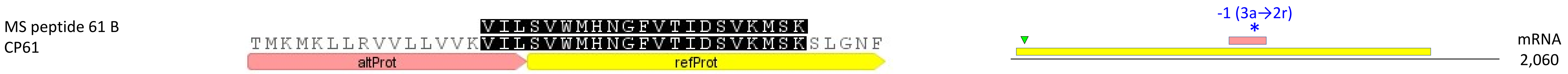


CP62: MtrunA17_Chr3g0141901_2F_1316-1585_270_MtrunA17_Chr3g0141901_1F_70-1371_1302_-2_iteration_17

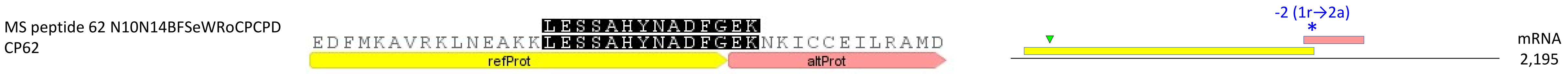


CP63: MtrunA17_Chr3g0144151_1F_1222-1311_90_MtrunA17_Chr3g0144151_3F_33-1415_1383_+2_iteration_3

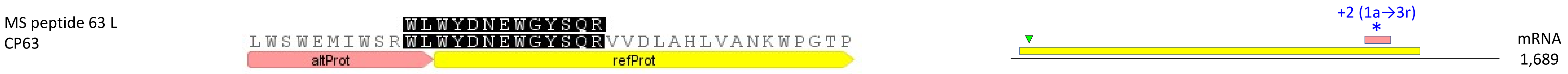


CP64: MtrunA17_Chr3g0144151_1F_1222-1311_90_MtrunA17_Chr3g0144151_3F_33-1415_1383_-1_iteration_5

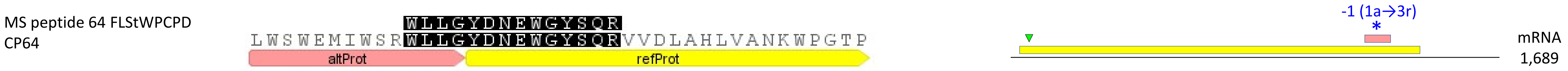


CP65: MtrunA17_Chr3g0144151_1F_1222-1311_90_MtrunA17_Chr3g0144151_3F_33-1415_1383_-1_iteration_6

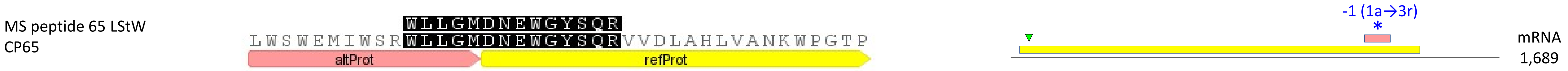

CP66: MtrunA17_Chr3g1011650_2F_26-340_315_MtrunA17_Chr3g1011650_3F_108-224_117_-2_iteration_2

MS peptide 66 RoC LSRKTVETFIDNTMSFK
CP66 LSRKTVETFIDNTMSFKKEMRSRQHSSKPLRSHMHLLSSSQ
altProt 1 altProt 2

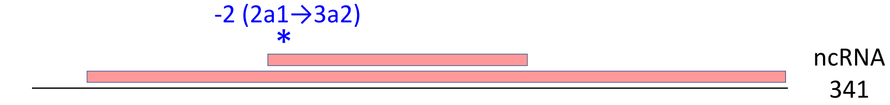


CP67: MtrunA17_Chr4g0000131_1F_1471-1644_174_MtrunA17_Chr4g0000131_3F_159-2459_2301_+1_iteration_7

MS peptide 67 N10 AVQALLAANLNDEIEPALPK
CP67 AVQALLAANLNDEIEPALPKDMISLRNLRLQRTLLEILRV
refProt altProt

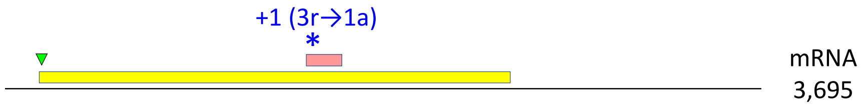


CP68: MtrunA17_Chr4g0000891_3F_3-473_471_MtrunA17_Chr4g0000891_1F_1-471_471_+2_iteration_6

MS peptide 68 N14 RLSQIFGTRIAFFHK
CP68 MKRLKRLSQIFGTRIAFFHKINNGCCTKLL
refProt altProt

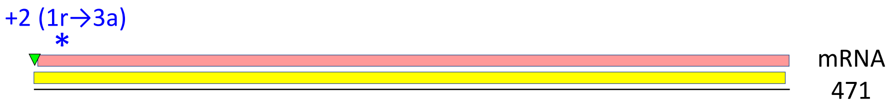


**CP69**: MtrunA17_Chr4g0004721_1F_634-813_180_MtrunA17_Chr4g0004721_3F_171-1157_987_+1_iteration_9

MS peptide 69 N10 TVALLGWEGLAQPLPR
CP69 GNWKHGDYKLTTKFSGKTVALLGWEGLAQPLPRELKALTV
refProt altProt

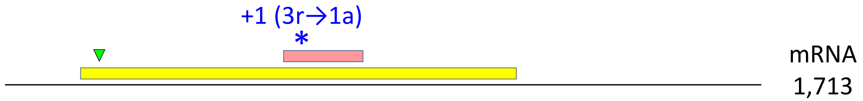


CP70: MtrunA17_Chr4g0014251_3F_45-242_198_MtrunA17_Chr4g0014251_1F_1-327_327_+1_iteration_4

MS peptide 70 N10 QPFNDHPAFFTSSPRCHRHR
CP70 QPFNDHPAFFTSSPRCHRHRLRLLEKLLILIETYLRNMNA
altProt refProt

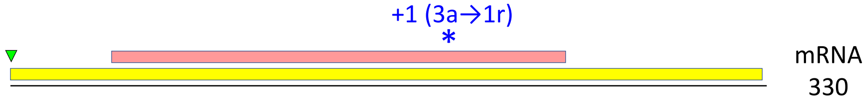

**CP71**: MtrunA17_Chr4g0022491_3F_762-923_162_MtrunA17_Chr4g0022491_2F_413-1177_765_+2_iteration_9

MS peptide 71 W LMALRDPKMAGLVTAMSLFCTR

CP71 LMALRDPKMAGLVTAMSLFCTRCNKERRFRLRSKEECRIH

altProt refProt

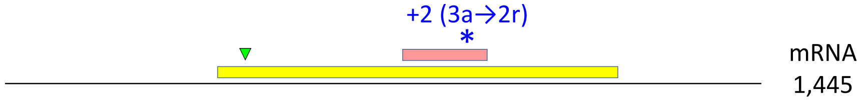


CP72: MtrunA17_Chr4g0023791_1F_1-84_84_MtrunA17_Chr4g0023791_2F_2-82_81_-1_iteration_5

MS peptide 72 F QTSKSVGLCLRR

CP72 VDMSEWLRRQTSKSVGLCLRRFESCCRR

altProt 2 altProt 1

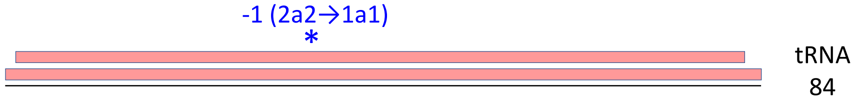


CP73: MtrunA17_Chr4g0034271_3F_15-344_330_MtrunA17_Chr4g0034271_1F_1-738_738_-1_iteration_7

MS peptide 73 PD MKYSEISHFSHPKTQTK

CP73 MKYSEISHFSHPKTQTKIRAQRISIQMRRLQRNRH

refProt altProt

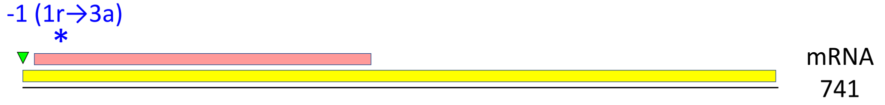


CP74: MtrunA17_Chr4g0034491_2F_1151-1708_558_MtrunA17_Chr4g0034491_1F_88-2415_2328_-2_iteration_1

MS peptide 74 F LEELPGLDTLKR

CP74 LEELPGLDTLKRSQELLMARQKLSEREITELRMHGTWQNRN

refProt altProt

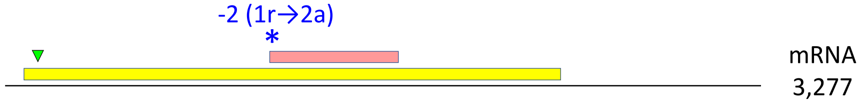


**CP75**: MtrunA17_Chr4g0037381_1F_880-1014_135_MtrunA17_Chr4g0037381_2F_422-1024_603_-1_iteration_16

MS peptide 75 B GGGGGGGYERGGGGYGGGGGGYGGWR

CP75 RRGGGGGGGYERGGGGYGGGGGGYGGWRWWLWWKRSCYSCG

refProt altProt

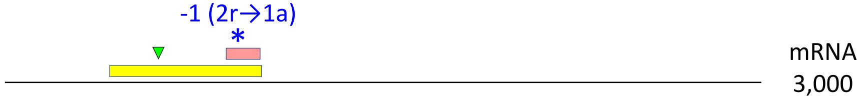

CP76: MtrunA17_Chr4g0040471_3F_2295-2474_180_MtrunA17_Chr4g0040471_1F_85-2868_2784_-1_iteration_14

MS peptide 76 N10N14FLRSeSt: LSHDTDSEVAMAAVISLGLNRRWNK
CP76: DTLSRLSHDTDSEVAMAAVISLGLNRRWNKQCSNCRHASQS
refProt altProt

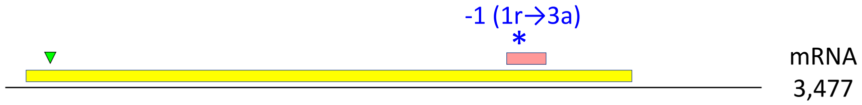


**CP77**: MtrunA17_Chr4g0055331_2F_62-127_66_MtrunA17_Chr4g0055331_1F_1-738_738_+2_iteration_9

MS peptide 77 B: IMQGTFLQPDEMQRANYFTFVMLIKMSPPD
CP77: KIMQGTFLQPDEMQRANYFTFVMLIKMSPPD
altProt refProt

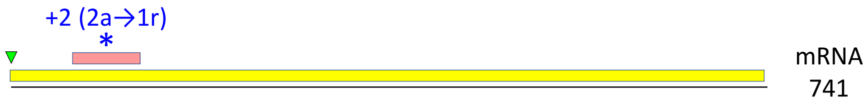


**CP78**: MtrunA17_Chr4g0059001_2F_83-277_195_MtrunA17_Chr4g0059001_1F_1-840_840_-2_iteration_11

MS peptide 78 B: KNIVLFGDSLTEQSFR
CP78: REVLVRCICLSRKSSSSDERKNIVLFGDSLTEQSFRVNGWG
refProt altProt

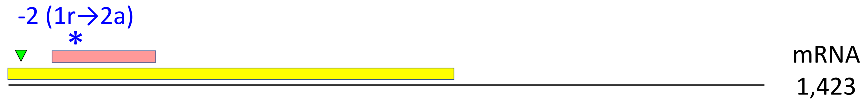


CP79: MtrunA17_Chr4g0063201_2F_446-535_90_MtrunA17_Chr4g0063201_1F_262-522_261_-2_iteration_18

MS peptide 79 Se: LLYTNCGKIR
CP79: NHSLIHLINLILQYNNYHWCGGSSTRLLYTNCGKIRFNCWD
refProt altProt

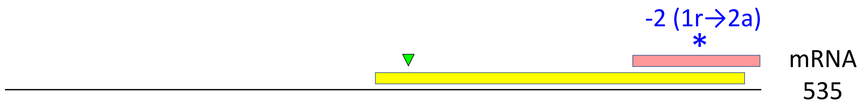


CP80: MtrunA17_Chr4g0070011_3F_1743-1832_90_MtrunA17_Chr4g0070011_1F_58-2481_2424_-1_iteration_16

MS peptide 80 N14R: PIIFHHGK
CP80: EELLYSSVENEEHICVLKDRNKPIIFHHGKVGPCEEHYRTC
refProt altProt

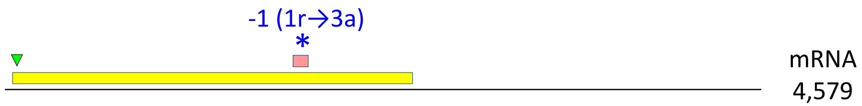

CP81: MtrunA17_Chr5g0393401_2F_1649-1765_117_MtrunA17_Chr5g0393401_3F_282-1781_1500_-2_iteration_10

MS peptide 81 N10F VRGGFNGCGWTPFIKEEQENLIDDHK

CP81 GDCMLQGDRVRGGFNGCGWTPFIKEEQENLIDDHKSR

altProt refProt

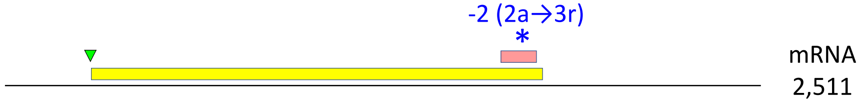


CP82: MtrunA17_Chr5g0400181_1F_859-1017_159_MtrunA17_Chr5g0400181_3F_312-2192_1881_+1_iteration_14

MS peptide 82 PD IPFDGVGR

CP82 PSPQQLLQLINEFVEKERIPFDGVGRLKYFWGETRGQVEK

refProt altProt

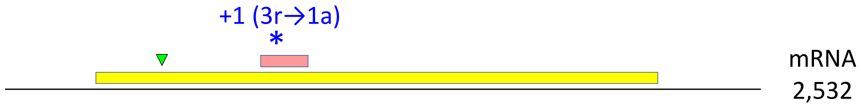


CP83: MtrunA17_Chr5g0405061_2F_182-412_231_MtrunA17_Chr5g0405061_3F_228-2174_1947_-2_iteration_20

MS peptide 83 N10 LHEVPDVHSDPPIWIHCMSLMK

CP83 KECVEKLQLLNSPVERQRRLHEVPDVHSDPPIWIHCMSLMK

altProt refProt

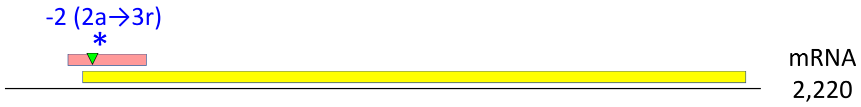


CP84: MtrunA17_Chr5g0408581_2F_2615-2860_246_MtrunA17_Chr5g0408581_1F_1-3168_3168_-2_iteration_12

MS peptide 84 RoC LVSAIGGSSHIR

CP84 MVAHVSDFGIARLVSAIGGSSHIRILRQLELKGLLAMLLQS

refProt altProt

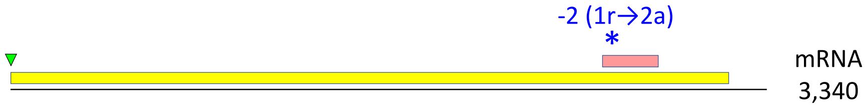


CP85: MtrunA17_Chr5g0415031_1F_931-1152_222_MtrunA17_Chr5g0415031_3F_114-4400_4287_+1_iteration_19

MS peptide 85 L KEDEAGENGNEIIE

CP85 PDQDHDGGSLRAEKERKRRMEKEKDRKEDEAGENGNEIIE

refProt altProt

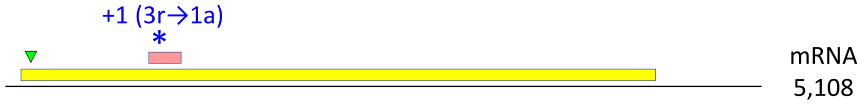

CP86: MtrunA17_Chr5g0415911_1F_3553-3948_396_MtrunA17_Chr5g0415911_3F_2832-3632_801_-2_iteration_6

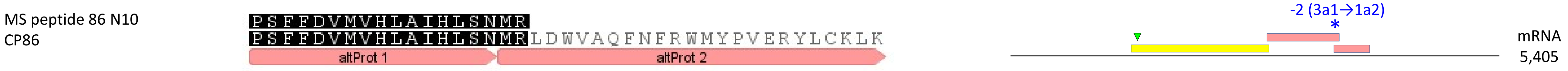


CP87: MtrunA17_Chr5g0421761_2F_1835-1918_84_MtrunA17_Chr5g0421761_3F_1818-1925_108_-1_iteration_4

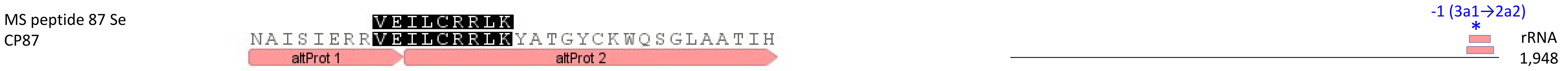


CP88: MtrunA17_Chr5g0422291_2F_3056-3259_204_MtrunA17_Chr5g0422291_3F_3120-3197_78_-1_iteration_5

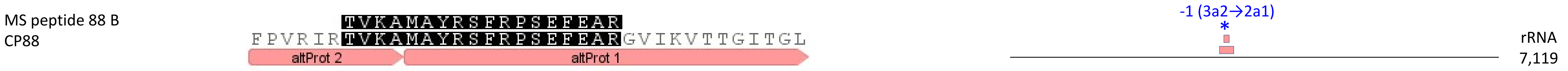


CP89: MtrunA17_Chr5g0422291_2F_4808-5617_810_MtrunA17_Chr5g0422291_1F_4921-5037_117_-1_iteration_29

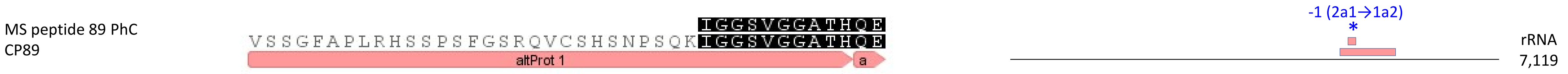


CP90: MtrunA17_Chr5g0430341_2F_2501-2623_123_MtrunA17_Chr5g0430341_3F_2298-2531_234_+2_iteration_3

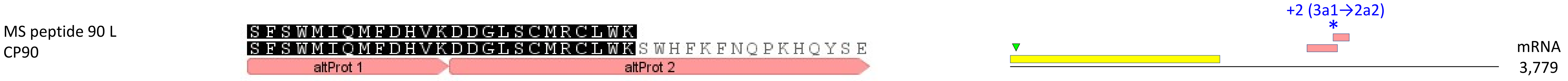

CP91: MtrunA17_Chr5g0430341_3F_1059-1172_114_MtrunA17_Chr5g0430341_1F_1-1620_1620_-2_iteration_9

MS peptide 91 W GTPIILSTKHRGVILK
CP91 FPISRGTPIILSTKHRGVILKVERGEMFPCLSSLDIWKCPK
altProt refProt

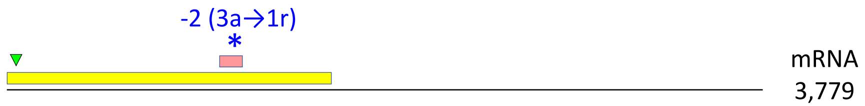


<u>CP92</u>: MtrunA17_Chr5g0431401_3F_1149-1388_240_MtrunA17_Chr5g0431401_2F_2-1354_1353_+1_iteration_18

MS peptide 92 RSt SSKLESDVVFAKEDMESLLR
CP92 VDEWKNGWKVEKSSKLESDVVFAKEDMESLLRGLWILKAK
refProt altProt

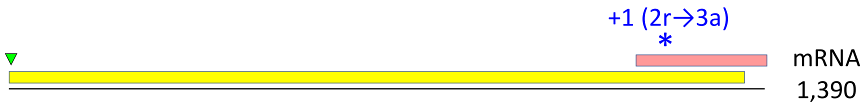


CP93: MtrunA17_Chr5g0435191_1F_883-1032_150_MtrunA17_Chr5g0435191_2F_677-979_303_-1_iteration_25

MS peptide 93 Se IEMNCLSGRVIKQIK
CP93 PLQAHEVDTARFFFQTMLSRKIEMNCLSGRVIKQIKAGFTI
refProt altProt

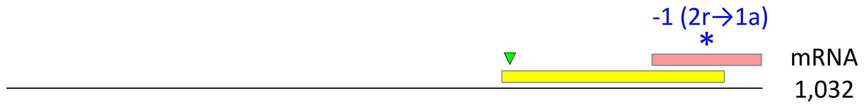


<u>CP94</u>: MtrunA17_Chr5g0444231_2F_758-967_210_MtrunA17_Chr5g0444231_3F_3-902_900_+2_iteration_11

MS peptide 94 F ILDELGNEMDSTSNRLDFVKK
CP94 EKILDELGNEMDSTSNRLDFVKKSGNGDEEGQCKGADDDD
refProt altProt

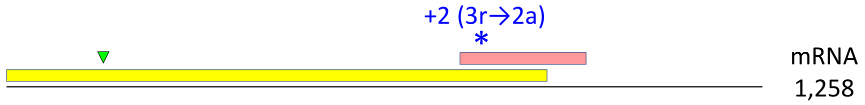


CP95: MtrunA17_Chr6g0451601_2F_434-577_144_MtrunA17_Chr6g0451601_1F_91-543_453_-2_iteration_29

MS peptide 95 F KECHCMLFLTPDNDFAGDEQTITLDEKL
CP95 GFWNCPCVPMRERKECHCMLFLTPDNDFAGDEQTITLDEKL
refProt a

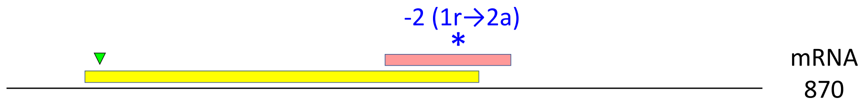

CP96: MtrunA17_Chr6g0452781_3F_4245-4397_153_MtrunA17_Chr6g0452781_1F_1-4458_4458_-1_iteration_15

MS peptide 96 F
CP96

YCDDAGNPTETYPFNVNGSPLRSSSYLFSR
APVRYCDDAGNPTETYPFNVNGSPLRSSSYLFSRRKTSCND
refProt altProt

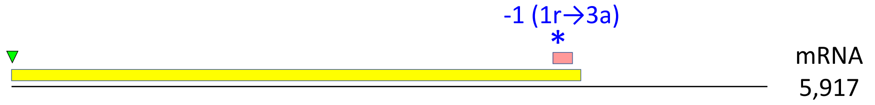


CP97: MtrunA17_Chr6g0457351_1F_406-636_231_MtrunA17_Chr6g0457351_3F_279-575_297_-2_iteration_22

MS peptide 97 RoD
CP97

DALSISQKRIP
LARLKKVPTLKCTKWHKVILFGFGQEGQKRDALSISQKRIP
altProt 1 altProt 2

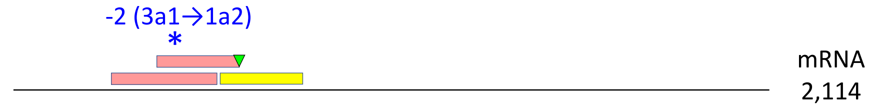


CP98: MtrunA17_Chr6g0457461_3F_438-596_159_MtrunA17_Chr6g0457461_2F_170-715_546_+2_iteration_6

MS peptide 98 B
CP98

TTVHQDTMMDVTGQWKLPLFGATDSSQVLK
VRTTVHQDTMMDVTGQWKLPLFGATDSSQVLKELAEAKAA
altProt refProt

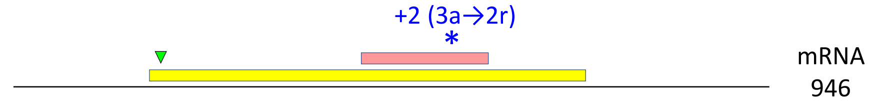


CP99: MtrunA17_Chr6g0457461_3F_438-596_159_MtrunA17_Chr6g0457461_2F_170-715_546_-2_iteration_17

MS peptide 99 BSeW
CP99

GWVPCLEFELERK
LTEDQLAKEVEYLIRKGWVPCLEFELERKDLSTVRTTVHQD
refProt altProt

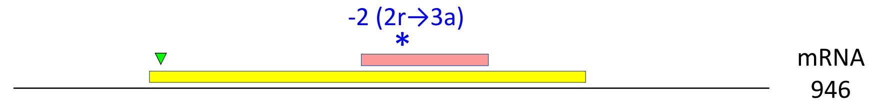


<u>CP100</u>: MtrunA17_Chr6g0457461_3F_438-596_159_MtrunA17_Chr6g0457461_2F_170-715_546_-2_iteration_18

MS peptide 100 BWPCPD
CP100

GWVPCLEFELEKK
LTEDQLAKEVEYLIRKGWVPCLEFELEKKDLSTVRTTVHQD
refProt altProt

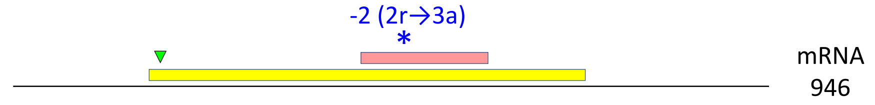

CP101: MtrunA17_Chr6g0458091_1F_1240-1362_123_MtrunA17_Chr6g0458091_2F_275-1621_1347_-2_iteration_4

MS peptide 101 N10N14Se WGYAPVLDCHTSHIAVK
CP101 PSHHHEPPWTDWKWGYAPVLDCHTSHIAVKFAELVTKIDRR

altProt refProt

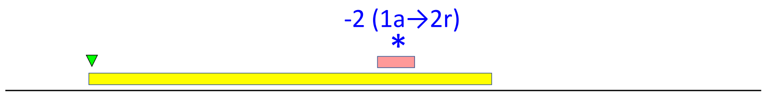


mRNA 2,527

CP102: MtrunA17_Chr6g0459481_3F_183-263_81_MtrunA17_Chr6g0459481_1F_1-363_363_-1_iteration_12

MS peptide 102 L VWITFGEHK
CP102 RIMIPLLSIDEVSLFAIRRVWITFGEHKVHCKELPCFK

refProt altProt

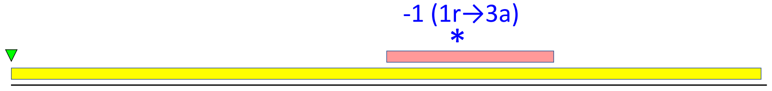


mRNA 366

CP103: MtrunA17_Chr6g0461931_3F_234-497_264_MtrunA17_Chr6g0461931_2F_86-298_213_+1_iteration_7

MS peptide 103 PC SGPSVLSR
CP103 TATNNKILTLTRTARSGPSVLSRILKLEDIKVEVKDDENV

refProt altProt

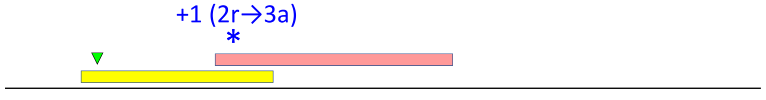


mRNA 840

CP104: MtrunA17_Chr6g0462271_3F_3-149_147_MtrunA17_Chr6g0462271_1F_1-180_180_+1_iteration_5

MS peptide 104 W AKGDAFSRPLR
CP104 PIEKEVEEGKAKGDAFSRPLRVMFWPILITTNPSSNLPSA

altProt refProt

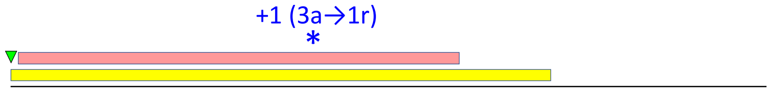


mRNA 252

CP105: MtrunA17_Chr6g0468411_3F_408-509_102_MtrunA17_Chr6g0468411_1F_1-507_507_+2_iteration_28

MS peptide 105 Se IMSWLSQGFLSMETSTMLLTILHLGI
CP105 SDKKTFNGPPEMLRIMSWLSQGFLSMETSTMLLTILHLGI

refProt a

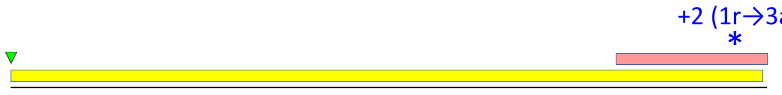


mRNA 510

CP106: MtrunA17_Chr6g0476751_3F_2274-2426_153_MtrunA17_Chr6g0476751_2F_359-2653_2295_+2_iteration_5

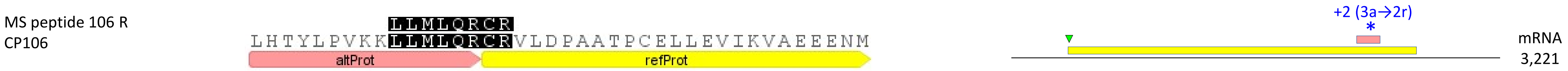


CP107: MtrunA17_Chr6g0479001_1F_3799-3981_183_MtrunA17_Chr6g0479001_2F_3701-3817_117_+2_iteration_5

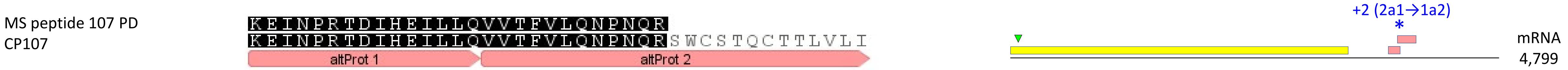


CP108: MtrunA17_Chr6g0485321_1F_1768-1881_114_MtrunA17_Chr6g0485321_3F_1674-1892_219_+2_iteration_13

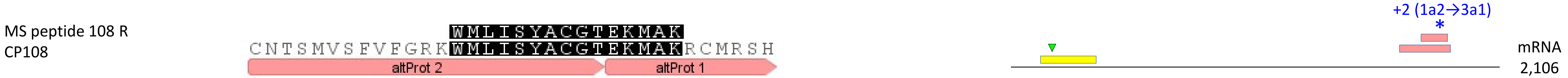


CP109: MtrunA17_Chr6g0486961_3F_1875-2120_246_MtrunA17_Chr6g0486961_1F_1-2943_2943_-2_iteration_12

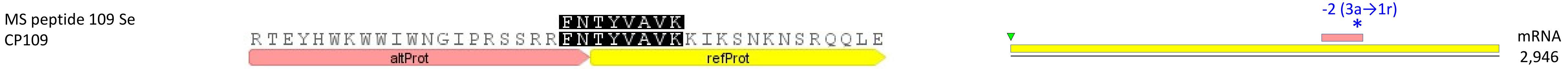


CP110: MtrunA17_Chr7g0214741_1F_340-462_123_MtrunA17_Chr7g0214741_3F_132-1478_1347_+1_iteration_8

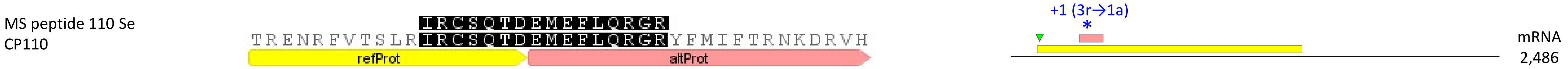

CP111: MtrunA17_Chr7g0214911_3F_183-368_186_MtrunA17_Chr7g0214911_2F_140-1999_1860_-2_iteration_6

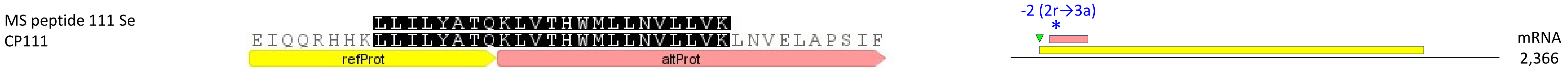


CP112: MtrunA17_Chr7g0219691_1F_1507-1602_96_MtrunA17_Chr7g0219691_3F_12-2699_2688_-2_iteration_2

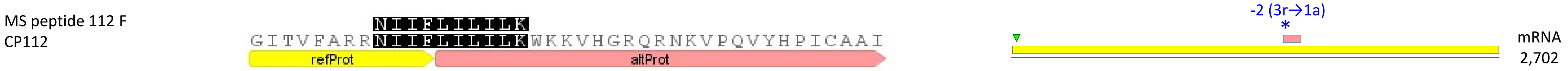


CP113: MtrunA17_Chr7g0221631_3F_3-125_123_MtrunA17_Chr7g0221631_1F_1-351_351_-1_iteration_13

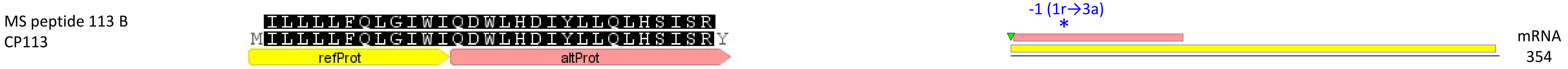


CP114: MtrunA17_Chr7g0229401_2F_2402-2755_354_MtrunA17_Chr7g0229401_3F_2268-2465_198_+2_iteration_1

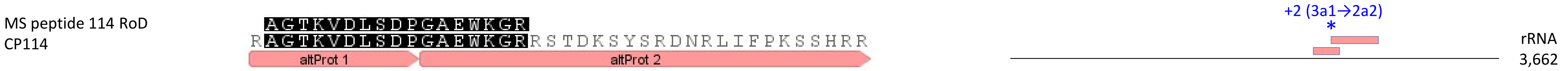


CP115: MtrunA17_Chr7g0230341_2F_2-307_306_MtrunA17_Chr7g0230341_1F_16-186_171_-1_iteration_3

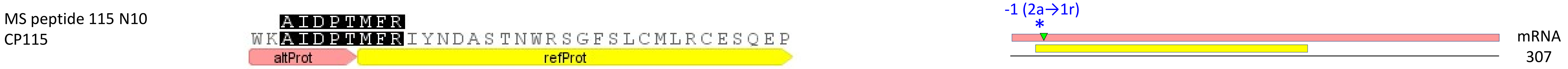

CP116: MtrunA17_Chr7g0232851_1F_502-684_183_MtrunA17_Chr7g0232851_2F_587-775_189_-2_iteration_22

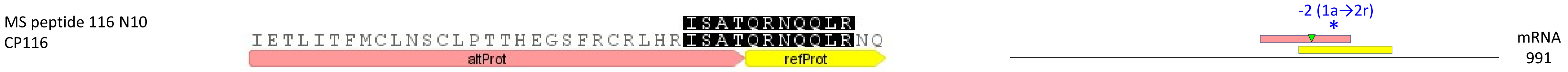


CP117: MtrunA17_Chr7g0237331_2F_1172-1372_201_MtrunA17_Chr7g0237331_1F_88-2508_2421_+2_iteration_18

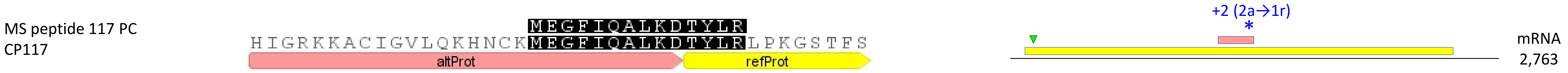


CP118: MtrunA17_Chr7g0251971_3F_3-161_159_MtrunA17_Chr7g0251971_1F_1-189_189_-1_iteration_4

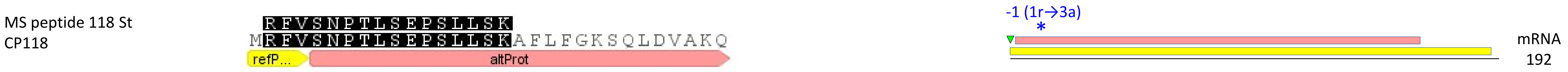


**CP119**: MtrunA17_Chr7g0259611_2F_185-292_108_MtrunA17_Chr7g0259611_3F_201-779_579_-2_iteration_6

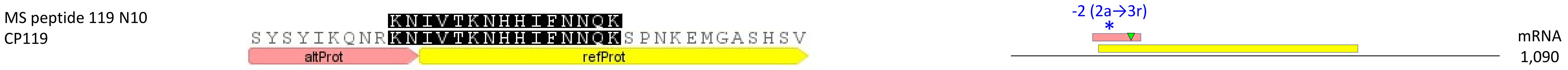


CP120: MtrunA17_Chr7g0262821_3F_1662-1847_186_MtrunA17_Chr7g0262821_2F_86-2131_2046_-1_iteration_13

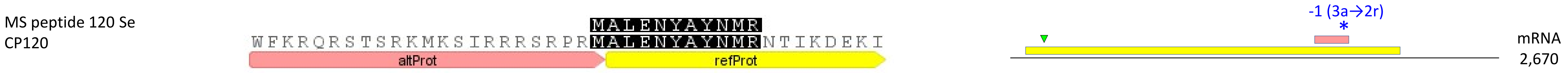

CP121: MtrunA17_Chr7g0270811_3F_1536-1718_183_MtrunA17_Chr7g0270811_2F_8-1711_1704_-2_iteration_10

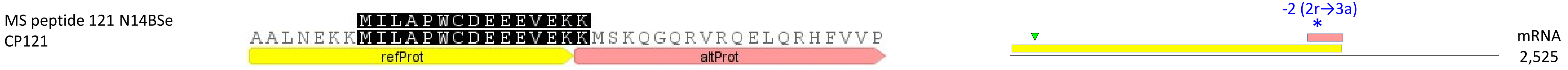


CP122: MtrunA17_Chr7g1034306_1F_502-681_180_MtrunA17_Chr7g1034306_2F_533-703_171_-2_iteration_17

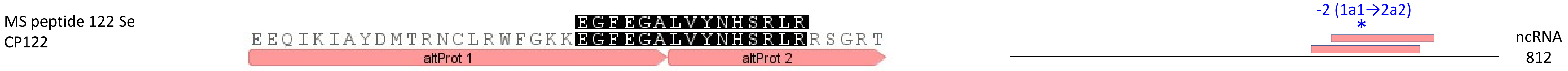


CP123: MtrunA17_Chr8g0338111_1F_436-534_99_MtrunA17_Chr8g0338111_2F_476-733_258_-2_iteration_11

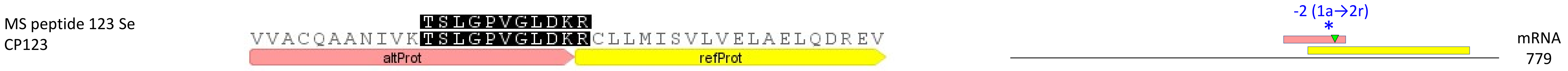


CP124: MtrunA17_Chr8g0338301_2F_1574-1681_108_MtrunA17_Chr8g0338301_1F_1-1680_1680_+1_iteration_17

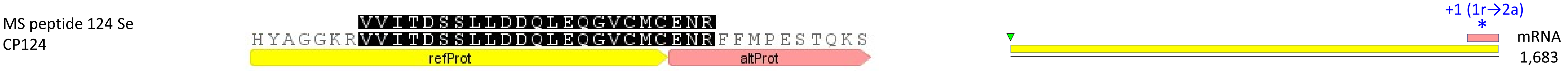


CP125: MtrunA17_Chr8g0339891_2F_998-1174_177_MtrunA17_Chr8g0339891_3F_426-1067_642_+2_iteration_11

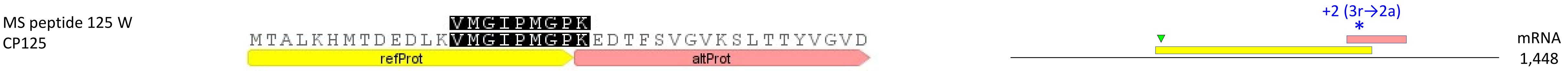

CP126: MtrunA17_Chr8g0342881_2F_656-892_237_MtrunA17_Chr8g0342881_1F_94-2667_2574_-2_iteration_1

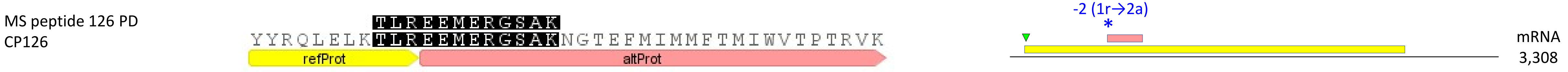


CP127: MtrunA17_Chr8g0345421_3F_2838-2963_126_MtrunA17_Chr8g0345421_2F_83-3610_3528_-2_iteration_6

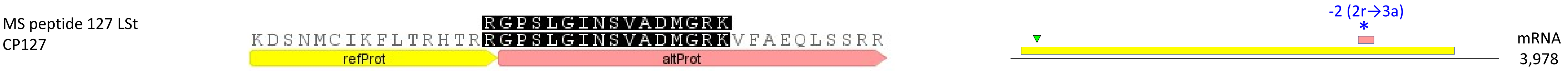


CP128: MtrunA17_Chr8g0353711_2F_110-283_174_MtrunA17_Chr8g0353711_1F_1-282_282_+1_iteration_21

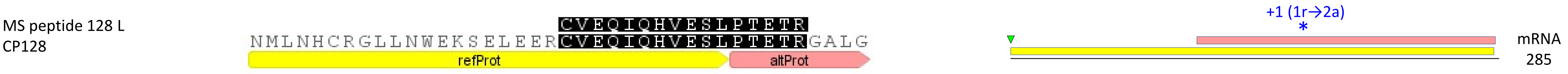


CP129: MtrunA17_Chr8g0355501_1F_1552-1761_210_MtrunA17_Chr8g0355501_3F_63-1898_1836_-2_iteration_6

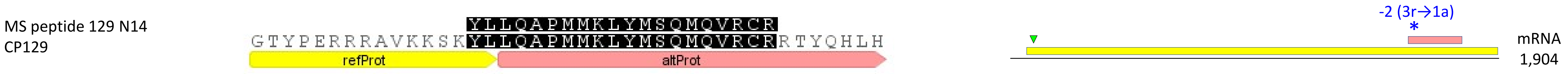


CP130: MtrunA17_Chr8g0356581_1F_1387-1464_78_MtrunA17_Chr8g0356581_2F_1394-1516_123_+1_iteration_5

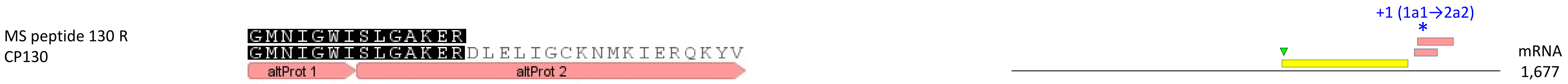

CP131: MtrunA17_Chr8g0365341_2F_773-1216_444_MtrunA17_Chr8g0365341_1F_169-2331_2163_+2_iteration_7

MS peptide 131 B
CP131

MTAKIHILDK
TTTMKTTLKTTKMTAKIHILDKTENKVPEYLKEAKVYFPS
altProt refProt

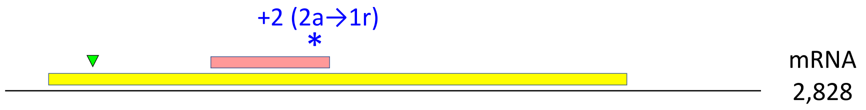


CP132: MtrunA17_Chr8g0368731_2F_1343-1540_198_MtrunA17_Chr8g0368731_1F_1-1875_1875_+1_iteration_0

MS peptide 132 PC
CP132

PLVNMSAQCFMGK
RFLLTVKPLVNMSAQCFMGKKRMFKFFTEMIPFMCCLDFI
refProt altProt

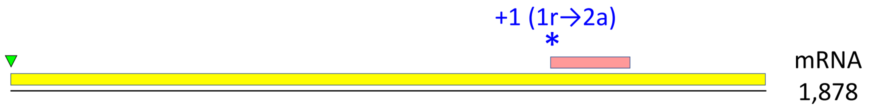


CP133: MtrunA17_Chr8g0368931_1F_889-1059_171_MtrunA17_Chr8g0368931_3F_3-1043_1041_-2_iteration_12

MS peptide 133 Se
CP133

STNEARINENFDVFDWAIPLK
LPKSTNEARINENFDVFDWAIPLKTCLLNFLRFNRQDYSGV
refProt altProt

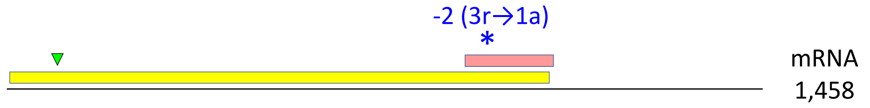


CP134: MtrunA17_Chr8g0371281_3F_696-851_156_MtrunA17_Chr8g0371281_2F_122-955_834_-2_iteration_4

MS peptide 134 N10
CP134

IATMLMYLSRMLK
HAGQRIATMLMYLSRMLKKGVRRCSQMPREILALCLGGMSF
refProt altProt

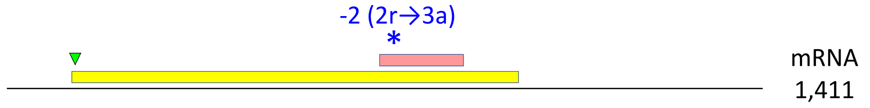


CP135: MtrunA17_Chr8g0371741_1F_691-888_198_MtrunA17_Chr8g0371741_3F_189-794_606_+1_iteration_16

MS peptide 135 L
CP135

QERNLETSLRSIPNLK
NCFFEEFVLQLMHAIQVAEKAQKKQERNLETSLRSIPNLKW
refProt altProt

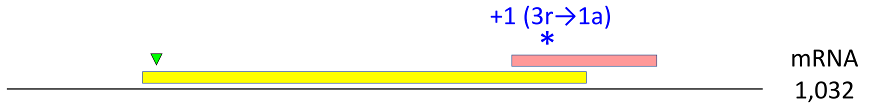

CP136: MtrunA17_Chr8g0373091_1F_457-702_246_MtrunA17_Chr8g0373091_3F_234-1106_873_+1_iteration_3

MS peptide 136 F
CP136

PSLMQPTK
VKAELGKPSLMQPTKGRSRYILTVVRLQNQLLRVEGPIGC
refProt altProt

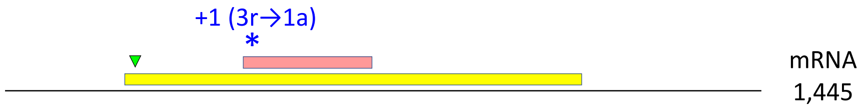


**CP137**: MtrunA17_Chr8g0376411_1F_433-582_150_MtrunA17_Chr8g0376411_3F_156-1463_1308_-1_iteration_6

MS peptide 137 BF
CP137

DWKEEMERDPCVCVMGEDVGHYGGSYK
LGMNFSFLKPCVKDWKEEMERDPCVCVMGEDVGHYGGSYKV
altProt refProt

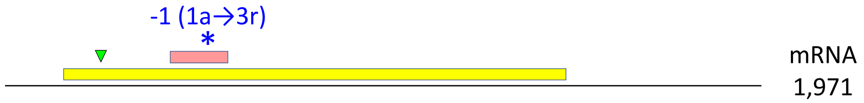


**CP138**: MtrunA17_Chr8g0377071_2F_1406-1693_288_MtrunA17_Chr8g0377071_1F_511-2715_2205_-1_iteration_17

MS peptide 138 L
CP138

VVEVLMSLRR
KRLLNLQLNLKERSTCFRVVEVLMSLRRLNKIDEGTYGVVY
altProt refProt

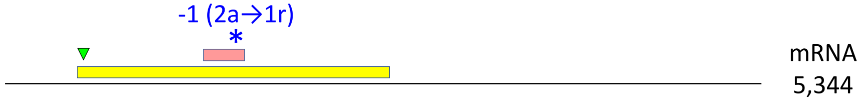


CP139: MtrunA17_Chr8g0385331_1F_1456-1719_264_MtrunA17_Chr8g0385331_3F_255-1670_1416_-2_iteration_8

MS peptide 139 B
CP139

RLMLNHAEMMTAGTLVCSVIVIR
RLMLNHAEMMTAGTLVCSVIVIRSWCHFSCTGFAFLHPNTF
refProt altProt

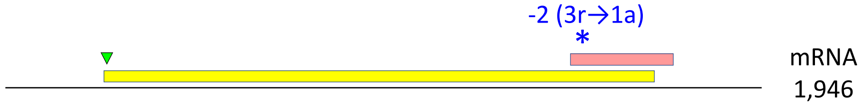


CP140: MtrunA17_Chr8g0392351_3F_96-272_177_MtrunA17_Chr8g0392351_1F_1-105_105_+2_iteration_4

MS peptide 140 PD
CP140

DMWLNGSHDIKLLFR
DMWLNGSHDIKLLFRGRSHHNSLLLNFLSVAYALHYCIGW
altProt 1 altProt 2

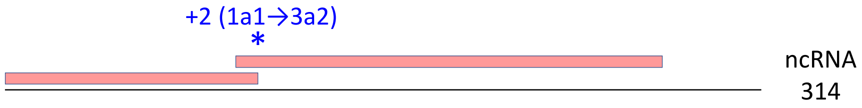

CP141: MtrunA17_CPg0492331_2F_584-874_291_MtrunA17_CPg0492331_3F_603-704_102_-1_iteration_19

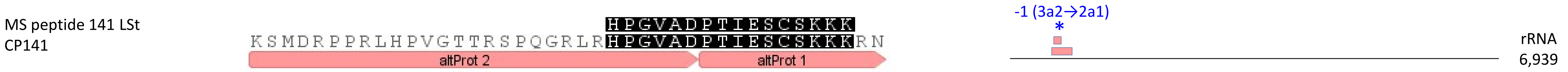


CP142: MtrunA17_CPg0492381_1F_631-750_120_MtrunA17_CPg0492381_2F_671-820_150_+1_iteration_1

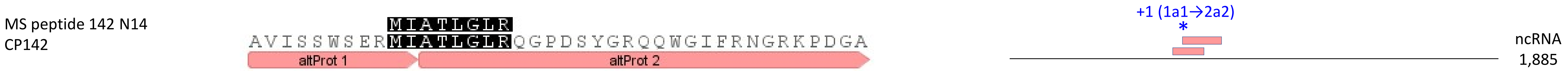


CP143: MtrunA17_CPg0492461_1F_3232-3426_195_MtrunA17_CPg0492461_3F_1413-7163_5751_+1_iteration_12

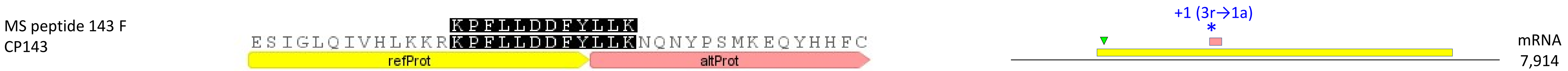


CP144: MtrunA17_CPg0492851_2F_209-328_120_MtrunA17_CPg0492851_3F_129-1157_1029_+1_iteration_6

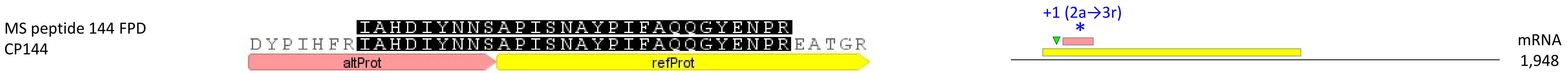


CP145: MtrunA17_CPg0492941_3F_825-947_123_MtrunA17_CPg0492941_1F_31-1572_1542_-1_iteration_2

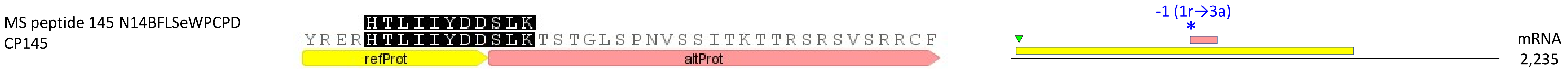

CP146: MtrunA17_CPg0493291_1F_733-804_72_MtrunA17_CPg0493291_2F_698-799_102_+2_iteration_18

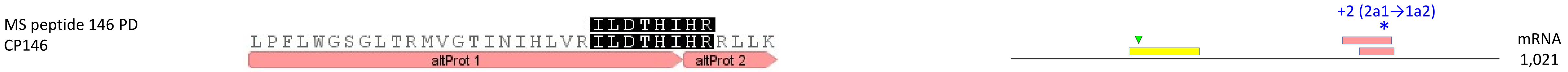


CP147: MtrunA17_CPg0493401_2F_1223-1450_228_MtrunA17_CPg0493401_3F_261-1778_1518_-2_iteration_7

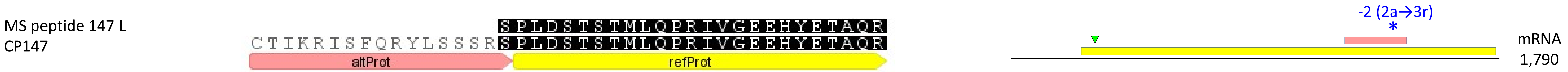


CP148: MtrunA17_MTg0490471_2F_1031-1147_117_MtrunA17_MTg0490471_1F_220-1740_1521_+1_iteration_7

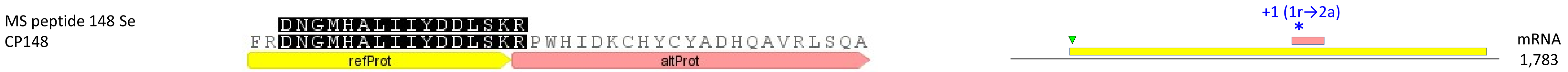


CP149: MtrunA17_MTg0490471_2F_422-538_117_MtrunA17_MTg0490471_1F_220-1740_1521_+1_iteration_16

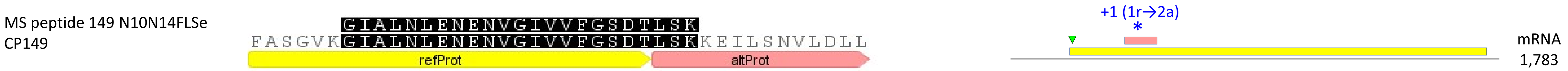


CP150: MtrunA17_MTg0490971_2F_1562-1699_138_MtrunA17_MTg0490971_3F_1542-1727_186_+1_iteration_12

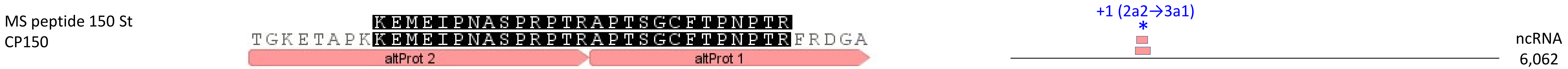

CP151: MtrunA17_MTg0490971_2F_476-628_153_MtrunA17_MTg0490971_1F_277-534_258_+1_iteration_0

MS peptide 151 W
CP151

QVMGPVRGERNGSSSHFSPQPPK
CTYSLYDKQVMGPVRGERNGSSSHFSPQPPKLKRNRGTLQ
altProt 1 altProt 2

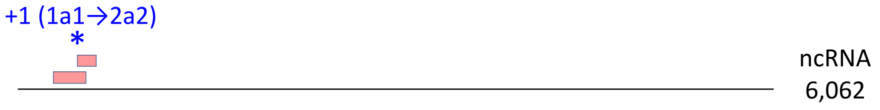


CP152: MtrunA17_MTg0491151_1F_4534-4659_126_MtrunA17_MTg0491151_2F_4511-4612_102_+2_iteration_15

MS peptide 152 Se
CP152

RSAFYLFR
AAMVRAGKHGHCIGGRRSAFYLFRSYVRPTSHRIRGT
altProt 1 altProt 2

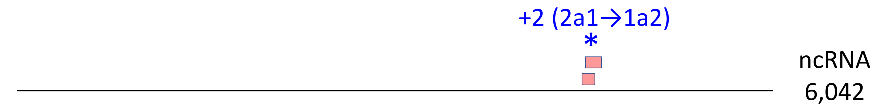


CP153: MtrunA17_MTg0491291_2F_1358-1447_90_MtrunA17_MTg0491291_3F_1329-1421_93_-1_iteration_18

MS peptide 153 B
CP153

WMGMPLAVLALIYP
SDPLSHAFLRAFGYATSHSRLSRCLRWMGMPLAVLALIYP
altProt 1 altProt 2

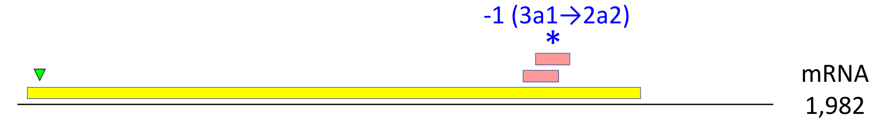


CP154: MtrunA17_MTg0491501_1F_661-825_165_MtrunA17_MTg0491501_3F_624-734_111_-2_iteration_6

MS peptide 154 PC
CP154

WKKGIAMDSYEK
MNLSRWKKGIAMDSYEKHSGTRRFNRTTNSYVVQGNKRSAS
altProt 1 altProt 2

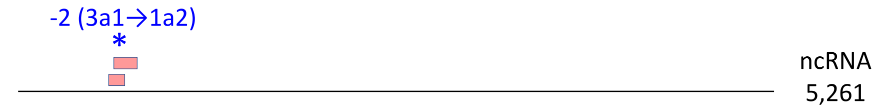


CP155: MtrunA17_MTg0491621_1F_1660-1836_177_MtrunA17_MTg0491621_3F_1377-1772_396_+1_iteration_16

MS peptide 155 N10
CP155

VLRPRFIAAAIWFTGTTK
SAPIRIKGSLAFARSREILRVLRPRFIAAAIWFTGTTKKS
altProt 1 altProt 2

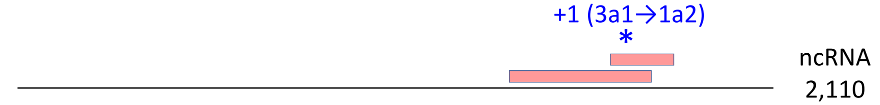

CP156: MtrunA17_MTg0491711_1F_2815-2910_96_MtrunA17_MTg0491711_3F_2664-2906_243_+1_iteration_24

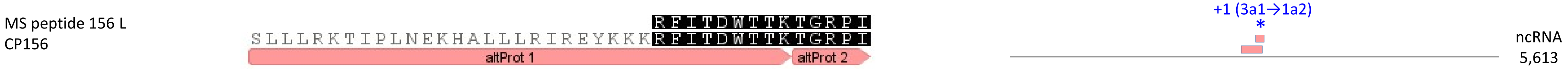

**Supplementary Dataset S2**. A graphical summary on alignments between 156 chimeric peptide models (CPs) and corresponding MS peptides (left) together with transcript models (right) that feature the transcript type, length in nucleotides, and relative positions of ORFs (to scale) involved in the production of CPs. Reference ORFs (refORFs) and reference proteins (refProts) are shown in yellow. Alternative ORFs (altORFs) and alternative proteins (altProts) are shown in pink. The first in-frame start codon (AUG) in each refORF is marked with a green triangle. If an altORF and a refORF are in the same frame, they are shown in the same level (e.g. CP7). If they are in different reading frames, they are shown in different levels (e.g. CP21). Codes of CPs modeled with MS-supported altProts are shown in a bold font. Codes of CPs with "confident" MS peptide detection in at least one sample are underlined. "Confident" is not the same as validated. For a short chimeric model sequence (mostly 40 amino acids or shorter in our dataset), it is very difficult to receive the status "confident" due to the nature of the MS method. Transcript models also show the positions and characteristics of programmed ribosomal frameshifting (PRF) events, which are mapped with an asterisk. The description of a PRF event should be interpreted as follows. For example, CP1, -1 (1a1→3a2): a minus 1 frameshift changes the translation from frame 1 to frame 3, which corresponds to the change from altORF1 to altORF2. An altORF1 in this study is defined as the one that starts earlier than an altORF2. Switches from altORF2 to altORF1 are also found in this dataset but are five times less frequent (e.g. CP108 and four other CPs). Throughout this study, unique numbers are assigned to chimeric models and their matching chimeric MS peptides. MS peptide identifiers also contain codes of biological samples in which they were detected. For example, the chimeric MS peptide of CP3 was identified in nodules of two developmental stages (10 and 14 days post inoculation). Thus, the identifier of MS peptide 3 ends with "N10N14".
