## Supplementary material for "Discovery of diverse chimeric peptides in a eukaryotic proteome sets the stage for the experimental proof of the mosaic translation hypothesis": Supplementary Dataset S4 Primary-source transcripts associated with multiple PRF events.pdf

### MtrunA17_Chr1g0185811

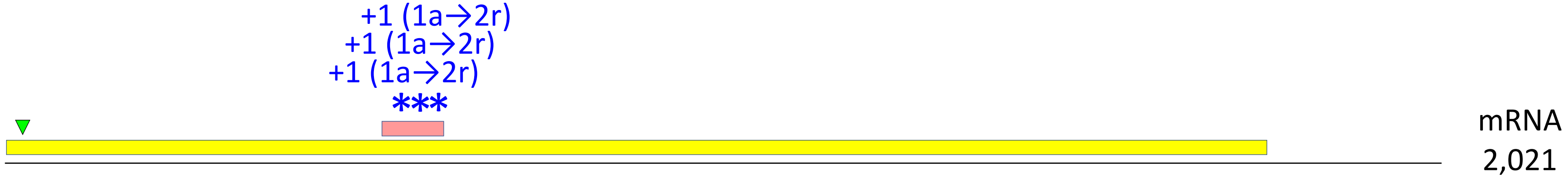


**CP17**: MtrunA17_Chr1g0185811_1F_535-621_87_MtrunA17_Chr1g0185811_2F_2-1774_1773_+1_iteration_2

MS peptide 17 Se
CP17

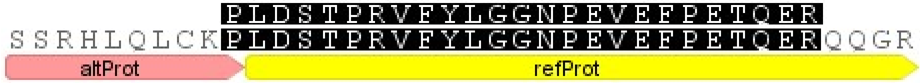


**CP18**: MtrunA17_Chr1g0185811_1F_535-621_87_MtrunA17_Chr1g0185811_2F_2-1774_1773_+1_iteration_9

MS peptide 18 Se
CP18

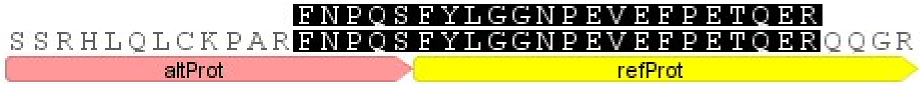


**CP16**: MtrunA17_Chr1g0185811_1F_535-621_87_MtrunA17_Chr1g0185811_2F_2-1774_1773_+1_iteration_10

MS peptide 16 Se
CP16

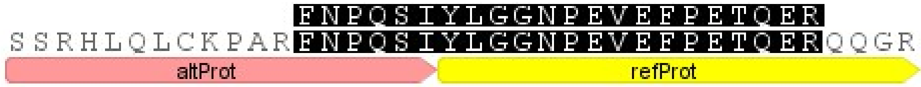

### MtrunA17_Chr1g0200071

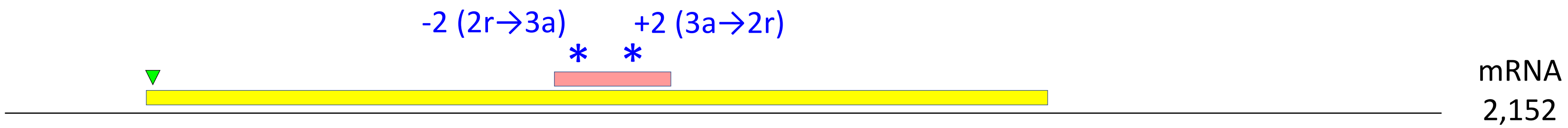


CP24: MtrunA17_Chr1g0200071_3F_828-1001_174_MtrunA17_Chr1g0200071_2F_218-1567_1350_-2_iteration_11

MS peptide 24 Se STNLDWYKR
CP24 PISGFEGDNMIERSTNLDWYKRVQPFLRPLTKSMSLRGLQT
refProt altProt

CP23: MtrunA17_Chr1g0200071_3F_828-1001_174_MtrunA17_Chr1g0200071_2F_218-1567_1350_+2_iteration_1

MS peptide 23 L LQTSPSDYHFRVYKIGGIGTVPVGR
CP23 LQTSPSDYHFRVYKIGGIGTVPVGRVETGVIKPGMVVTFA
altProt refProt

### MtrunA17_Chr3g0144151

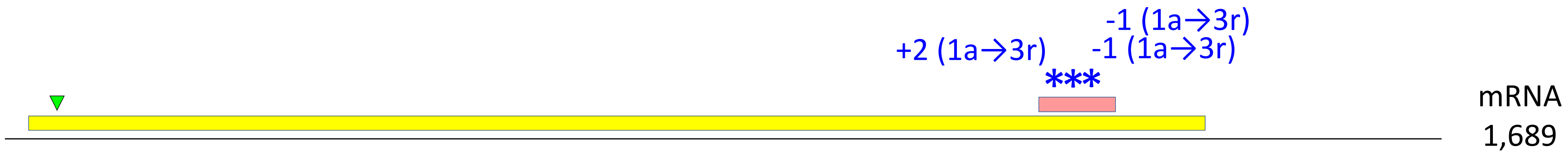


CP63: MtrunA17_Chr3g0144151_1F_1222-1311_90_MtrunA17_Chr3g0144151_3F_33-1415_1383_+2_iteration_3

MS peptide 63 L
CP63

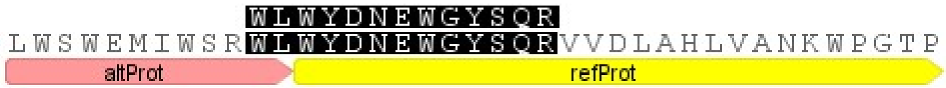


<u>CP64</u>: MtrunA17_Chr3g0144151_1F_1222-1311_90_MtrunA17_Chr3g0144151_3F_33-1415_1383_-1_iteration_5

MS peptide 64 FLStWPCPD
CP64

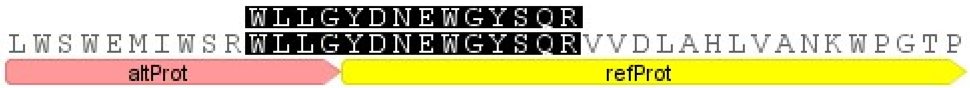


<u>CP65</u>: MtrunA17_Chr3g0144151_1F_1222-1311_90_MtrunA17_Chr3g0144151_3F_33-1415_1383_-1_iteration_6

MS peptide 65 LStW
CP65

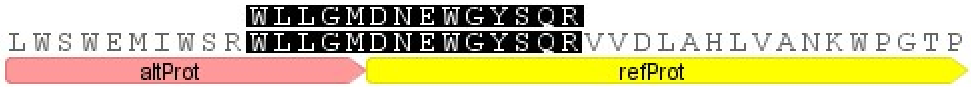

### MtrunA17_Chr5g0422291

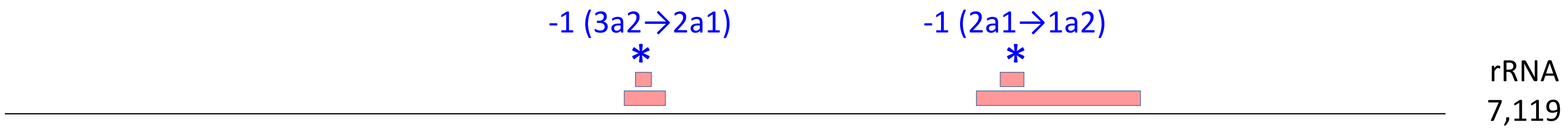


CP88: MtrunA17_Chr5g0422291_2F_3056-3259_204_MtrunA17_Chr5g0422291_3F_3120-3197_78_-1_iteration_5

MS peptide 88 B
CP88

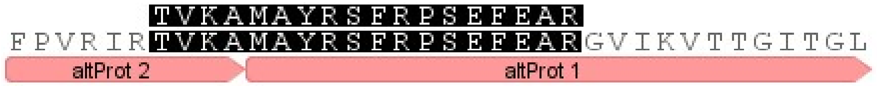


CP89: MtrunA17_Chr5g0422291_2F_4808-5617_810_MtrunA17_Chr5g0422291_1F_4921-5037_117_-1_iteration_29

MS peptide 89 PhC
CP89

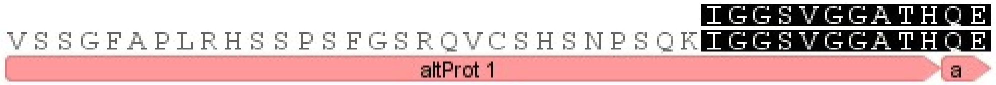

### MtrunA17_Chr5g0430341

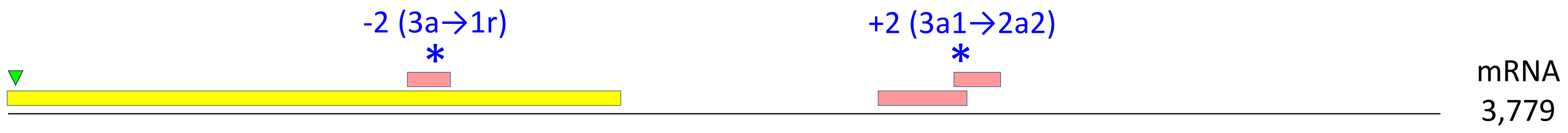


CP91: MtrunA17_Chr5g0430341_3F_1059-1172_114_MtrunA17_Chr5g0430341_1F_1-1620_1620_-2_iteration_9

MS peptide 91 W
CP91

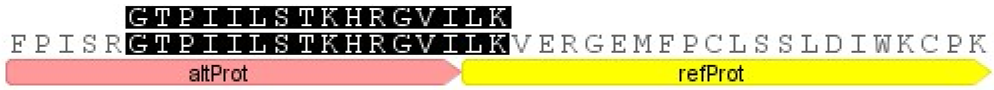


CP90: MtrunA17_Chr5g0430341_2F_2501-2623_123_MtrunA17_Chr5g0430341_3F_2298-2531_234_+2_iteration_3

MS peptide 90 L
CP90

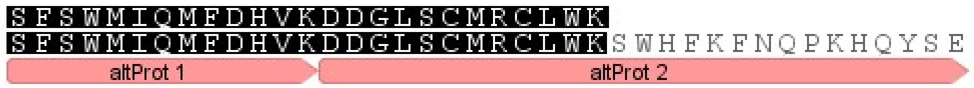

### MtrunA17_Chr6g0457461

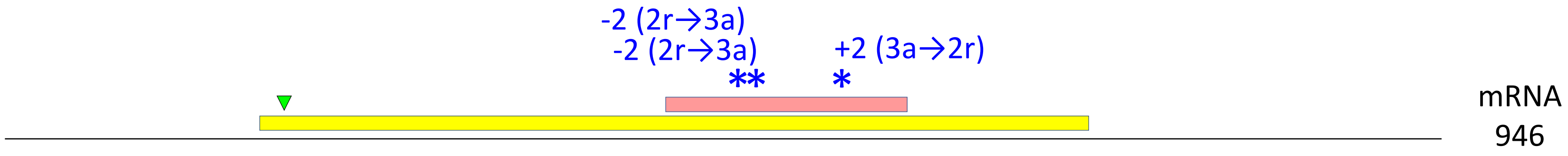


CP99: MtrunA17_Chr6g0457461_3F_438-596_159_MtrunA17_Chr6g0457461_2F_170-715_546_-2_iteration_17

MS peptide 99 BSeW
CP99

GWVPCLEFELERK
LTEDQLAKEVEYLIRKGWVPCLEFELERKDLSTVRTTVHQD
refProt altProt

CP100: MtrunA17_Chr6g0457461_3F_438-596_159_MtrunA17_Chr6g0457461_2F_170-715_546_-2_iteration_18

MS peptide 100 BWPCPD
CP100

GWVPCLEFELEKK
LTEDQLAKEVEYLIRKGWVPCLEFELEKKDLSTVRTTVHQD
refProt altProt

CP98: MtrunA17_Chr6g0457461_3F_438-596_159_MtrunA17_Chr6g0457461_2F_170-715_546_+2_iteration_6

MS peptide 98 B
CP98

TTVHQDTMMDVTGQWKLPLFGATDSSQVLK
VRTTVHQDTMMDVTGQWKLPLFGATDSSQVLKELAEAKAA
altProt refProt

### MtrunA17_MTg0490471

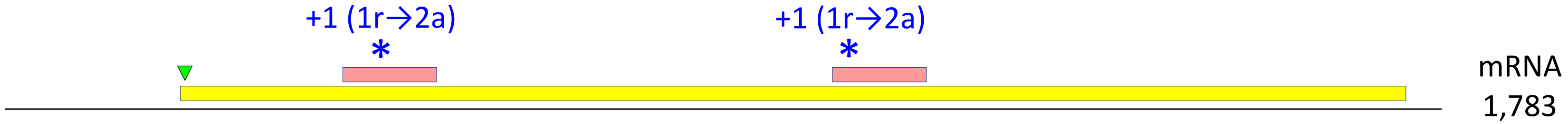


CP149: MtrunA17_MTg0490471_2F_422-538_117_MtrunA17_MTg0490471_1F_220-1740_1521_+1_iteration_16

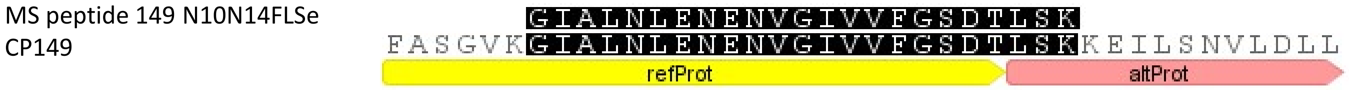


<u>CP148</u>: MtrunA17_MTg0490471_2F_1031-1147_117_MtrunA17_MTg0490471_1F_220-1740_1521_+1_iteration_7

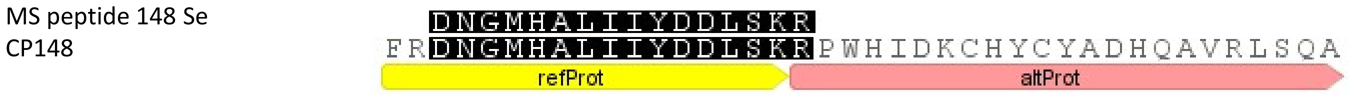

### MtrunA17_MTg0490971

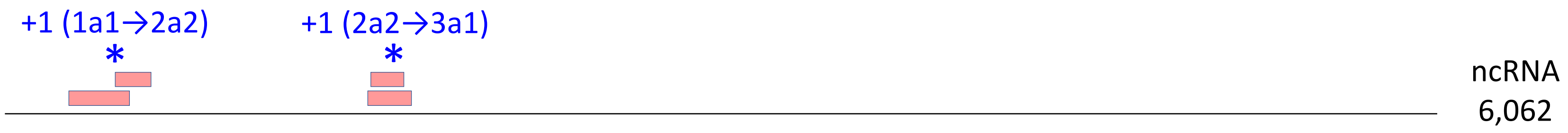


CP151: MtrunA17_MTg0490971_2F_476-628_153_MtrunA17_MTg0490971_1F_277-534_258_+1_iteration_0

MS peptide 151 W QVMGPVRGERNGSSSHFSPQPPK
CP151 CTYSLYDKQVMGPVRGERNGSSSHFSPQPPKLKRNRGTLQ
altProt 1 altProt 2

CP150: MtrunA17_MTg0490971_2F_1562-1699_138_MtrunA17_MTg0490971_3F_1542-1727_186_+1_iteration_12

MS peptide 150 St KEMEIPNASPRPTRAPTSGCFTPNPTR
CP150 TGKETAPKKEMEIPNASPRPTRAPTSGCFTPNPTRFRDGA
altProt 2 altProt 1

**Supplementary Dataset S4**. Eight primary-source transcripts associated with multiple PRF events. Chimeric peptide models (CPs) and corresponding MS peptides (bottom) are shown together with transcript models (top) that feature the transcript type, length in nucleotides, and relative positions of ORFs (to scale) involved in the production of CPs. Reference ORFs (refORFs) and reference proteins (refProts) are shown in yellow. Alternative ORFs (altORFs) and alternative proteins (altProts) are shown in pink. The first in-frame start codon (AUG) in each refORF is marked with a green triangle. Codes of CPs modeled with MS-supported altProts are shown in bold. Codes of CPs with "confident" MS peptide detection in at least one sample are underlined. Transcript models also show the positions and characteristics of PRF events, which are mapped with an asterisk. The description of a PRF event should be interpreted as follows. For example, CP88, -1 (3a2→2a1): a minus 1 frameshift changes the translation from frame 3 to frame 2, which corresponds to the change from altORF2 to altORF1. An altORF1 in this study is defined as the one that starts earlier than an altORF2. Throughout this study, unique numbers are assigned to chimeric models and their matching chimeric MS peptides. MS peptide identifiers also contain codes of biological samples in which they were detected. For example, the chimeric MS peptide of CP65 was identified in leaves, stems, and the whole plant. Thus, the identifier of MS peptide 65 ends with "LStW".
