## Supplementary material for "Discovery of diverse chimeric peptides in a eukaryotic proteome sets the stage for the experimental proof of the mosaic translation hypothesis": Supplementary Dataset S5 Mosaic proteins.pdf

### Hypothetical mosaic protein 1 translated from transcript MtrunA17_Chr1g0200071
#### Putative protein-synthesizing GTPase

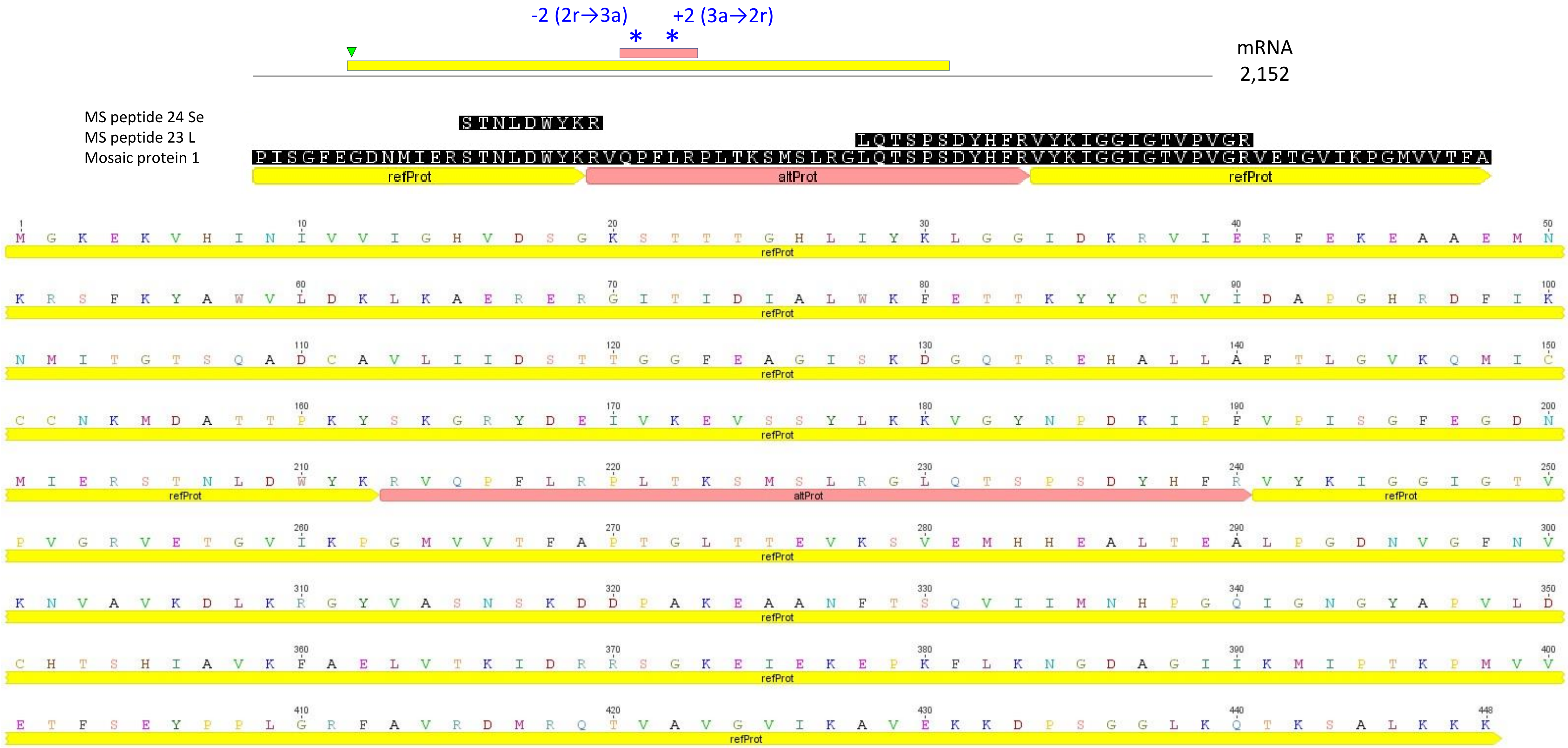

#### Hypothetical mosaic protein 2 translated from transcript MtrunA17_Chr6g0457461
#### Putative ribulose-bisphosphate carboxylase (RuBisCo)

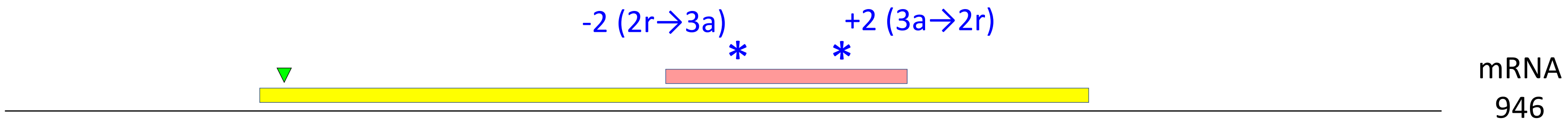


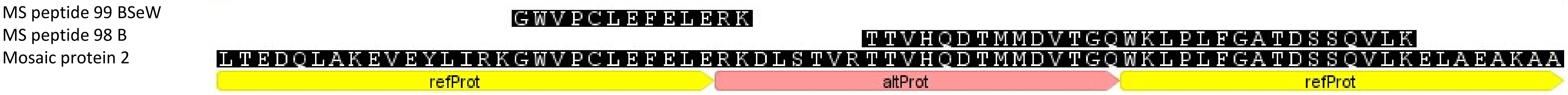


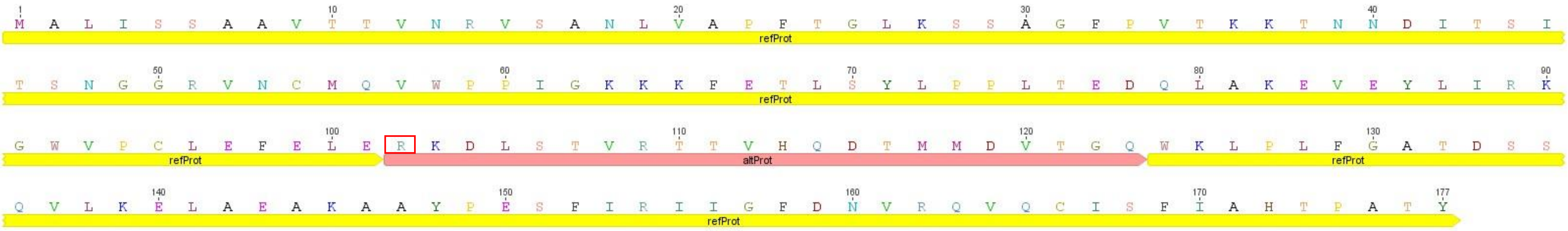

### Hypothetical mosaic protein 3 translated from transcript MtrunA17_Chr6g0457461
### Putative ribulose-bisphosphate carboxylase (RuBisCo)

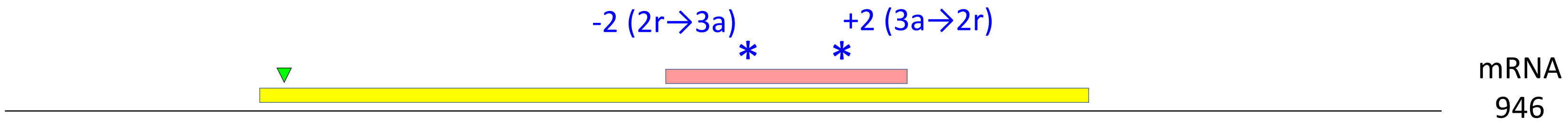


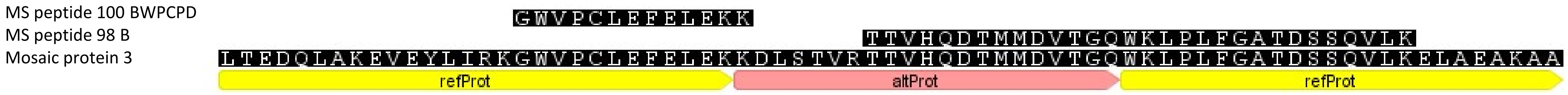


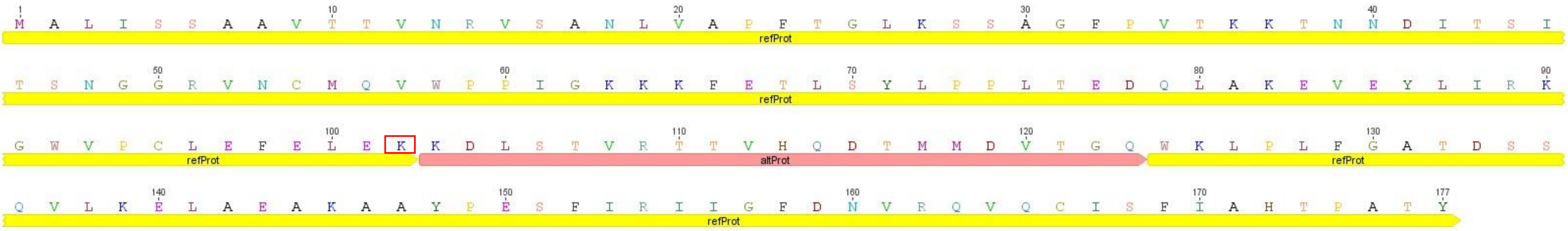

**Supplementary Dataset S5.** Three candidate mosaic proteins deduced from chimeric MS peptides. This dataset is an extended version of Figure 1. Mosaic proteins 2 and 3 differ by one amino acid around the -2 PRF site (red-boxed R and K, respectively). The upper portion of each figure depicts a corresponding transcript with ORFs mapped and scaled relative to the whole transcript length. Asterisks represent positions of PRF events. The text in blue describes the PRF value, type, and subtype of each frameshifting event. Interpretation example: -2 (2r→3a) refers to a frameshift with value minus 2 from a refORF in frame 2 (yellow) to an altORF in frame 3 (pink). The green triangle indicates the position of the first in-frame translational start codon (AUG). The complete sequence of each hypothetical mosaic protein is shown at the bottom of corresponding images.
