## Supplementary material for "Discovery of diverse chimeric peptides in a eukaryotic proteome sets the stage for the experimental proof of the mosaic translation hypothesis": Supplementary Dataset S7 Folding of chimeric protein models Part 1.pdf

### CP1: MtrunA17_Chr0c01g0489091

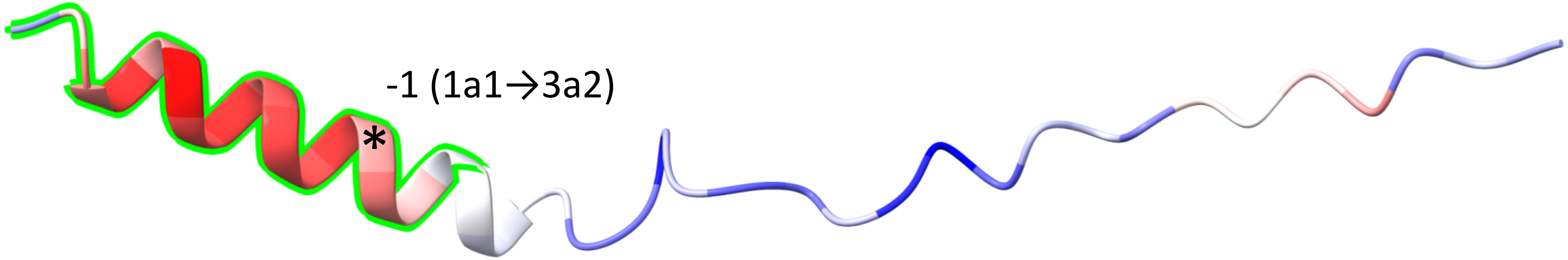


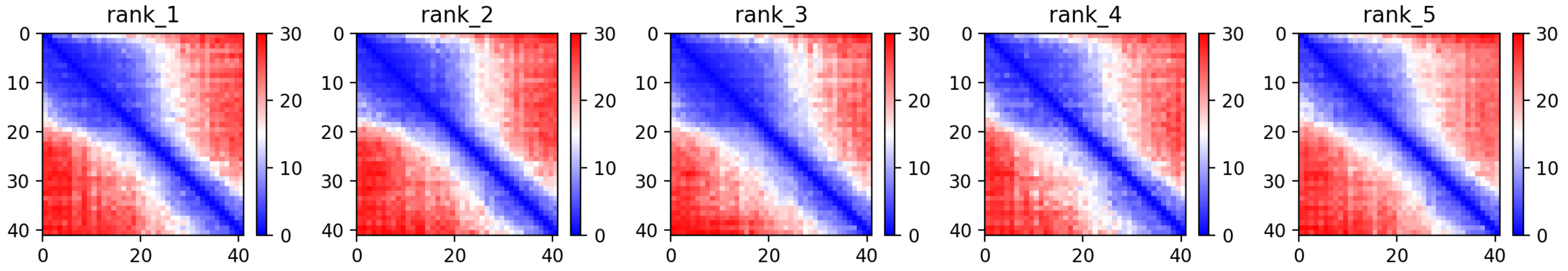

CP1: MtrunA17_Chr0c01g0489091

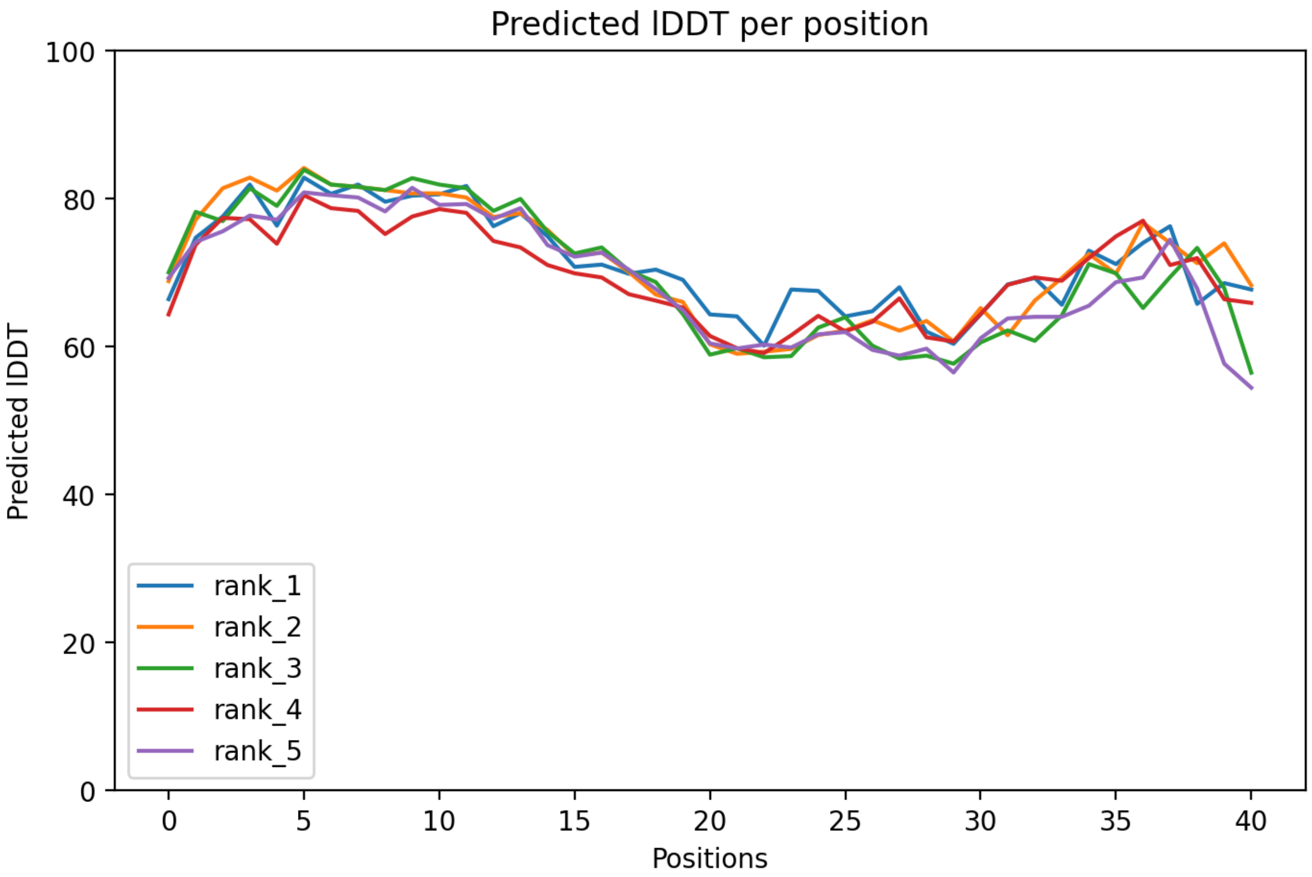

### CP1: MtrunA17_Chr0c01g0489091

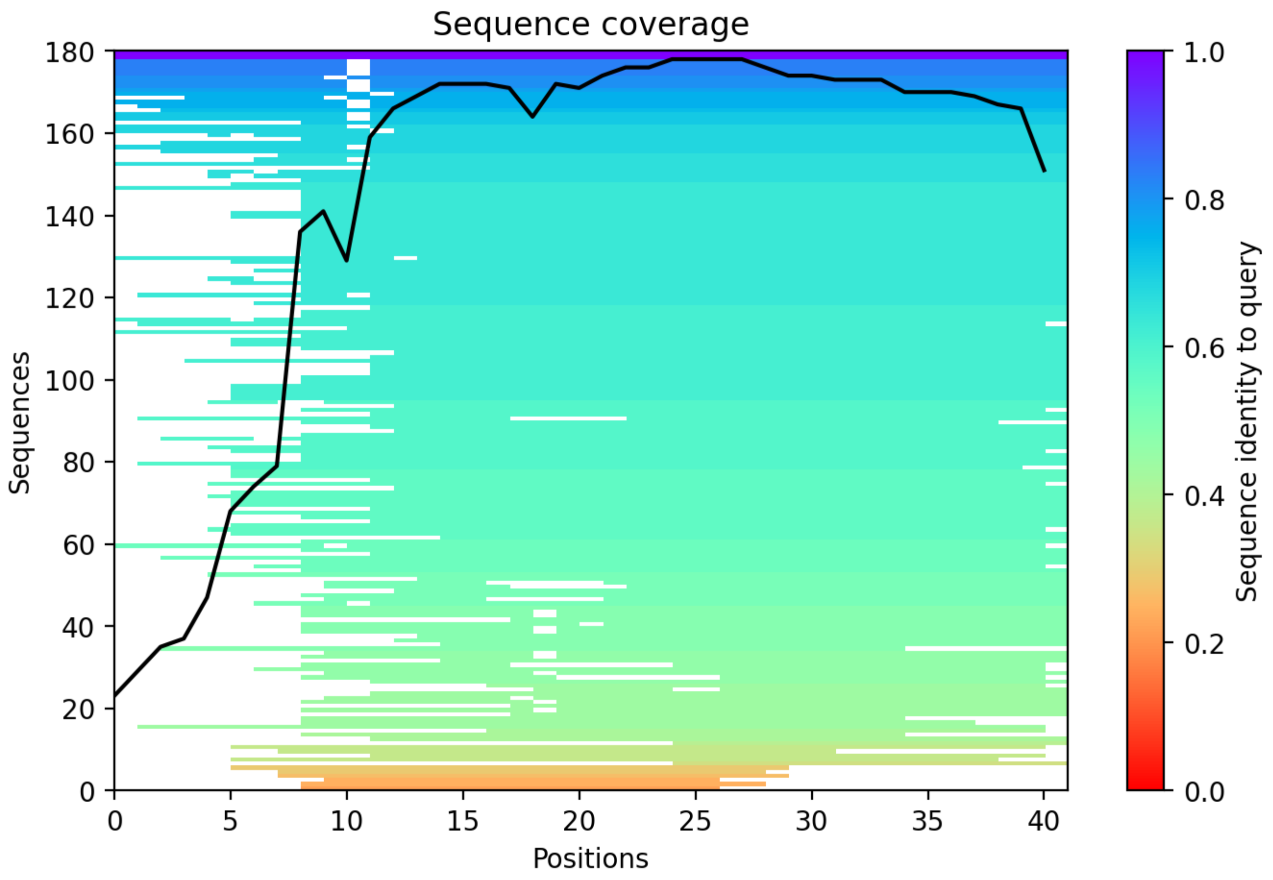

CP2: MtrunA17_Chr0c28g0493951

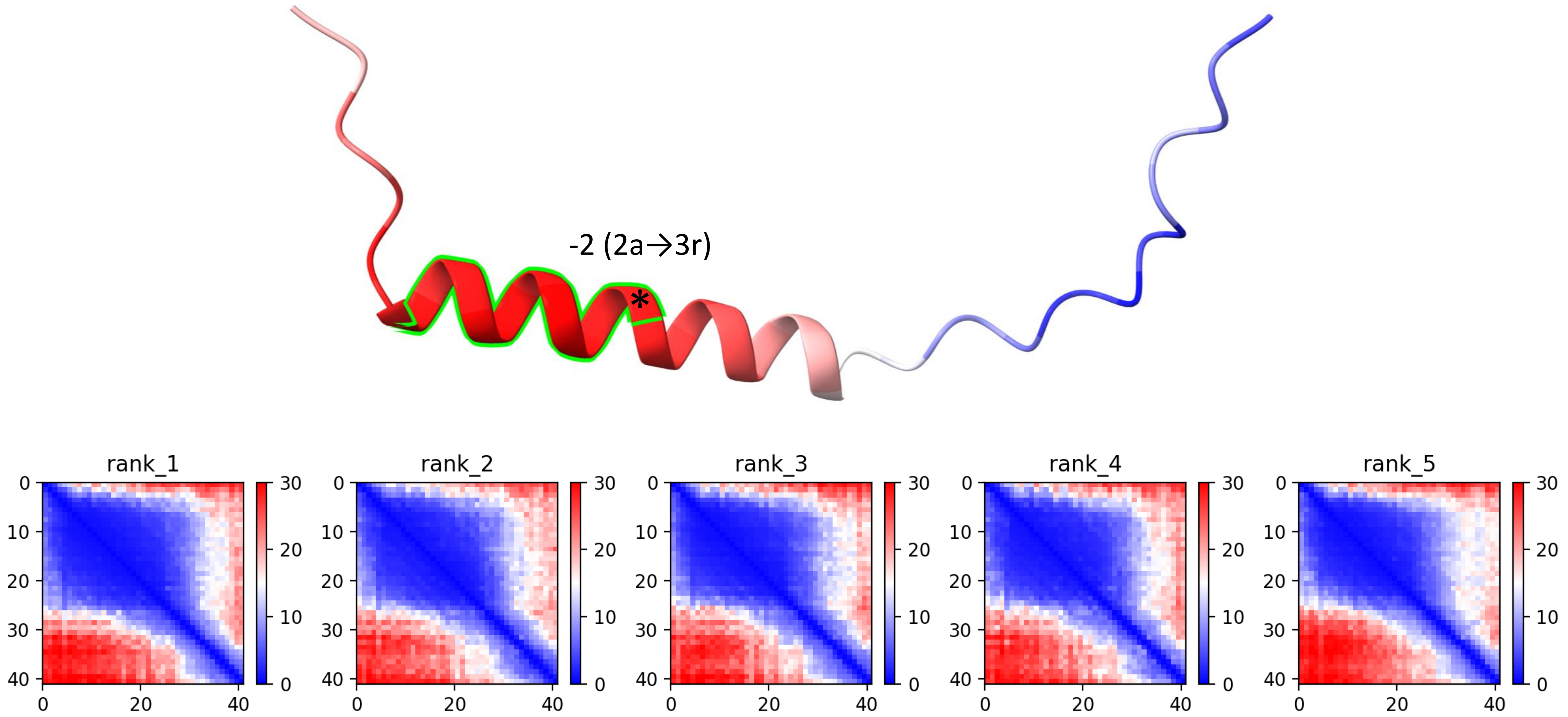

CP2: MtrunA17_Chr0c28g0493951

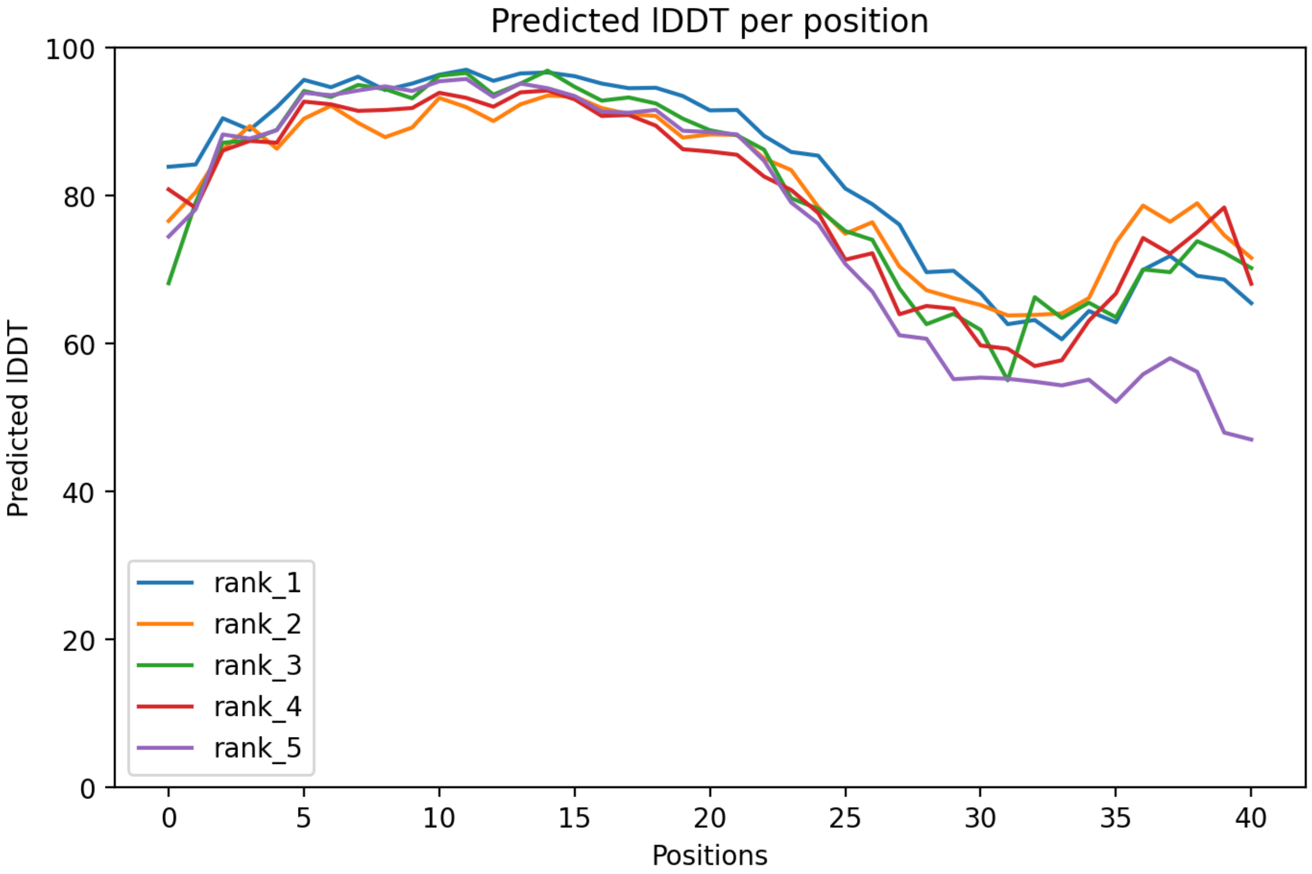

CP2: MtrunA17_Chr0c28g0493951

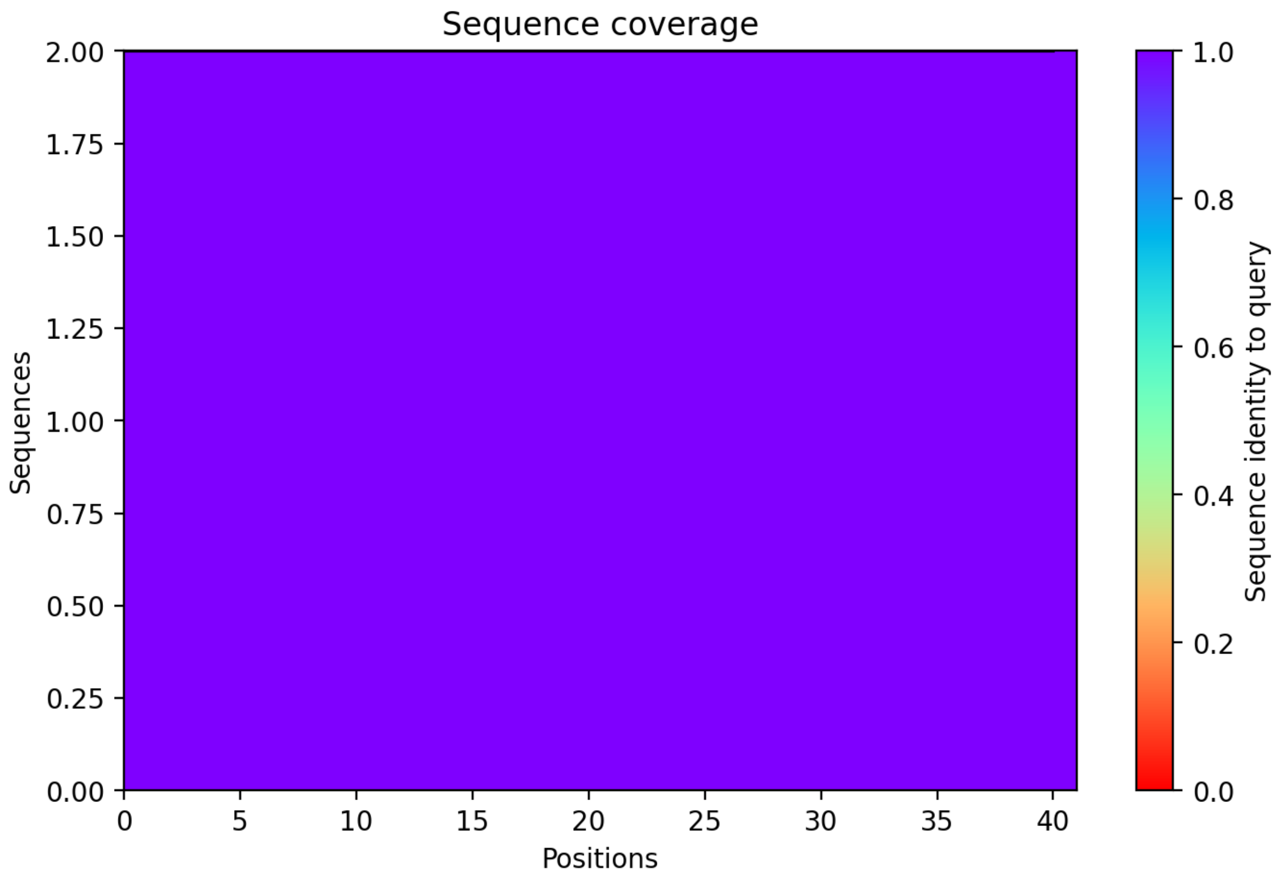

CP3: MtrunA17_Chr1g0148371

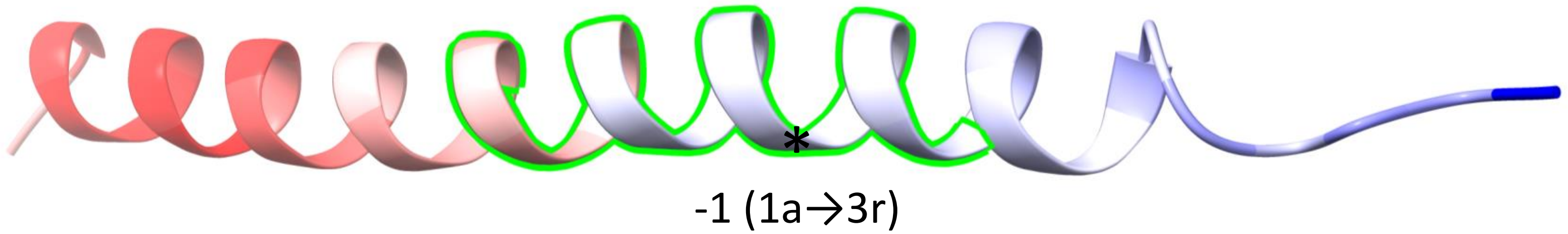


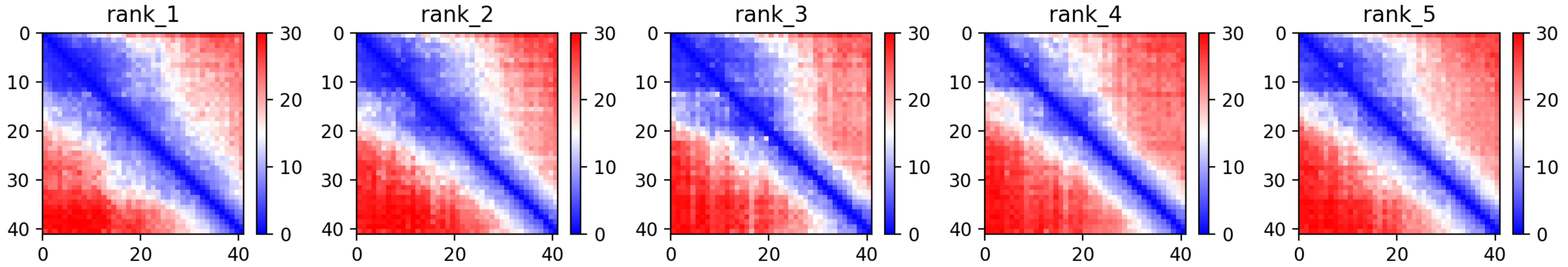

CP3: MtrunA17_Chr1g0148371

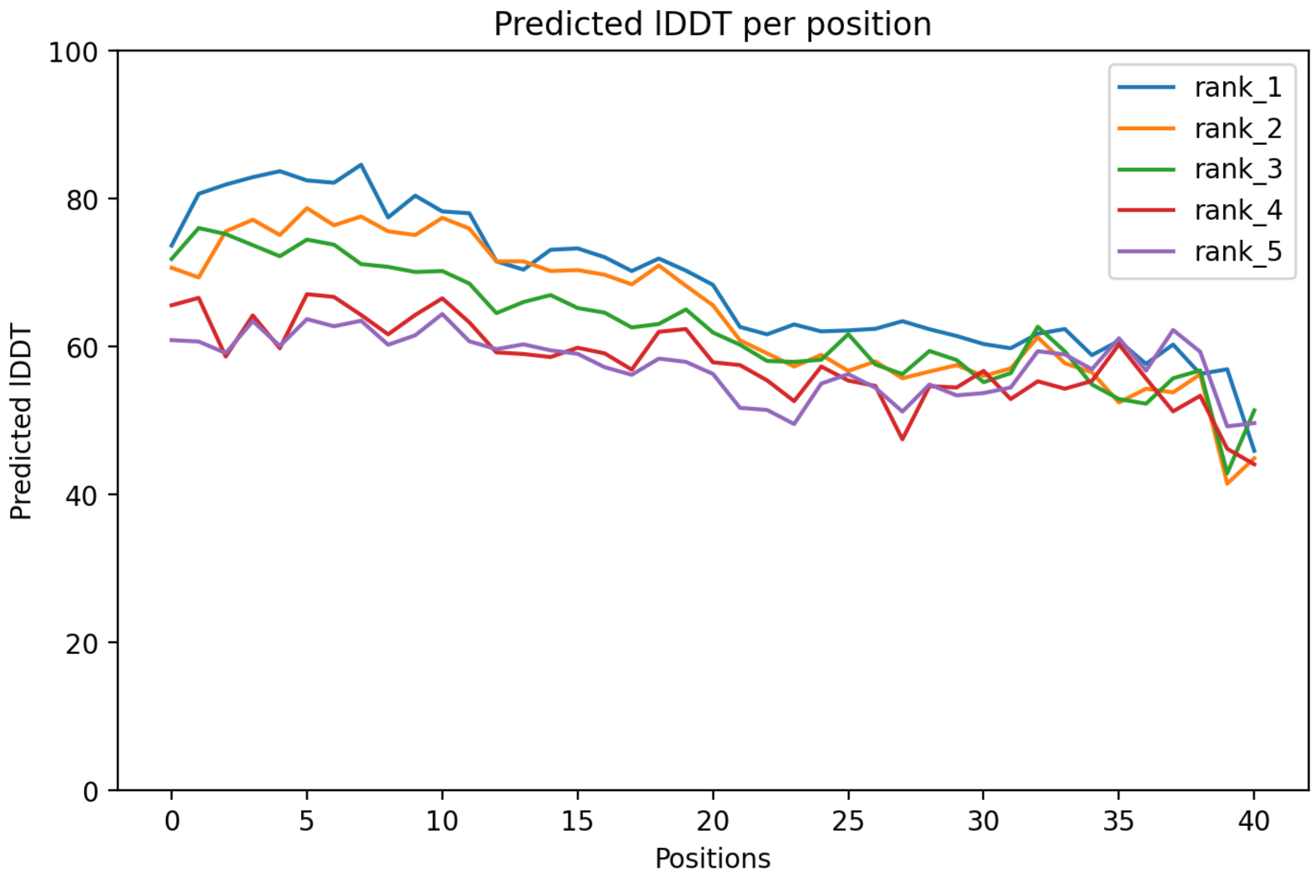

CP3: MtrunA17_Chr1g0148371

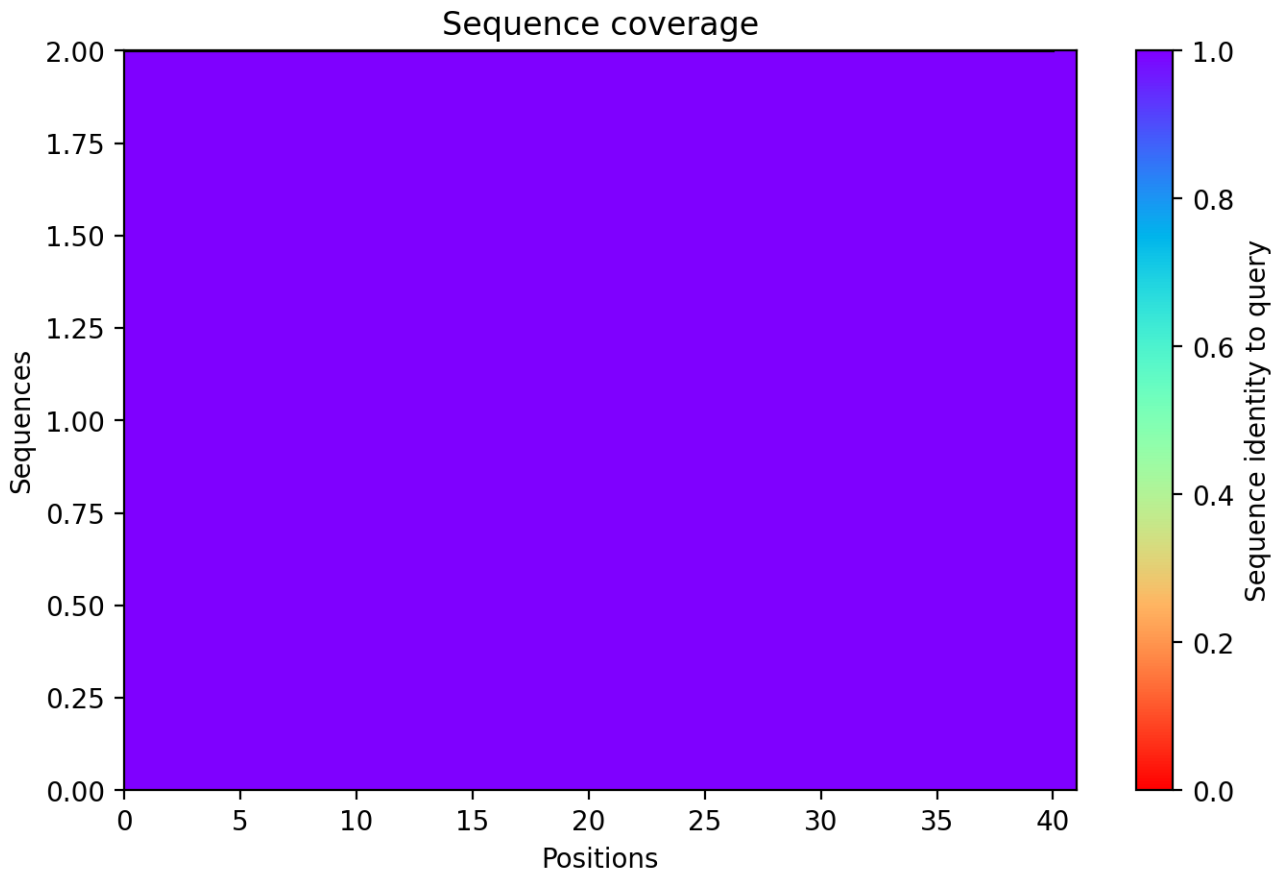

CP4: MtrunA17_Chr1g0149991

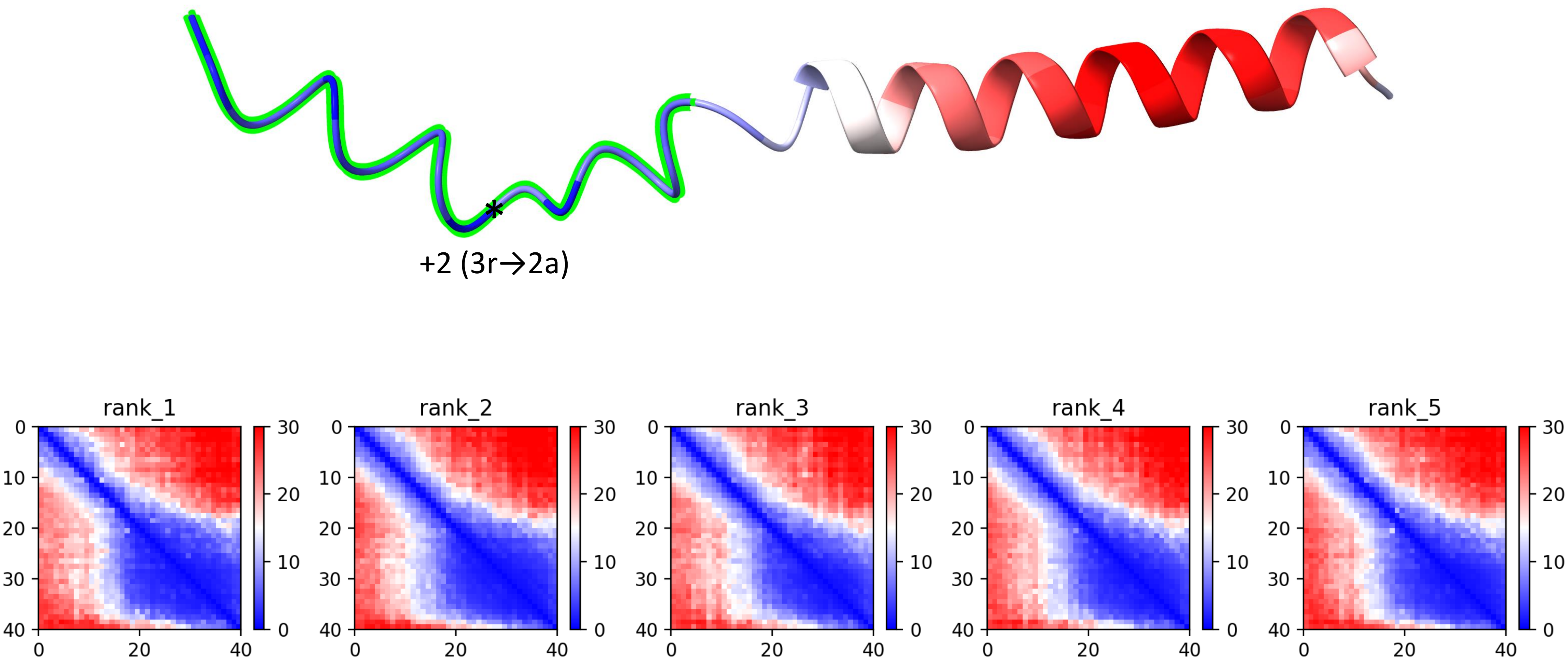

CP4: MtrunA17_Chr1g0149991

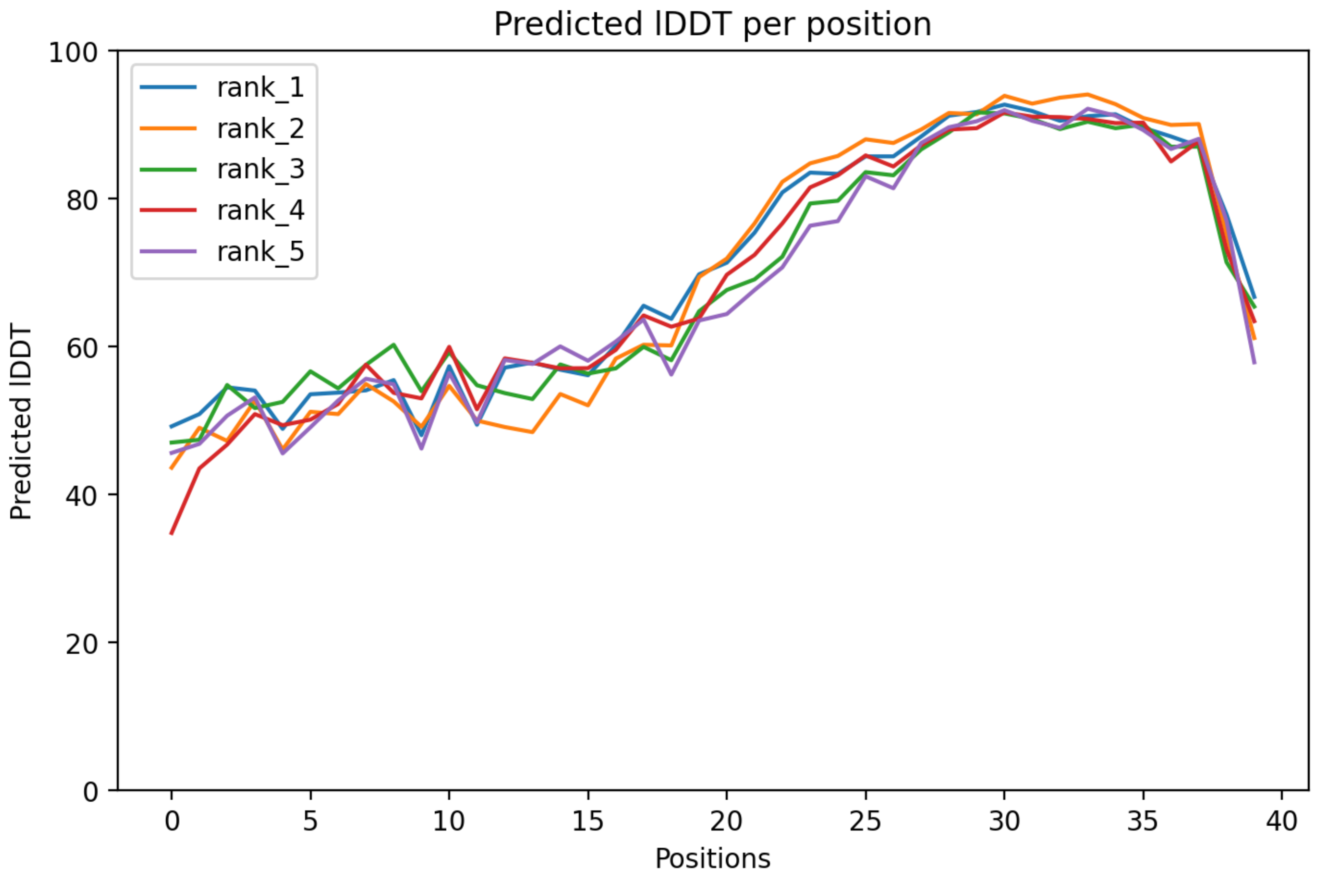

CP4: MtrunA17_Chr1g0149991

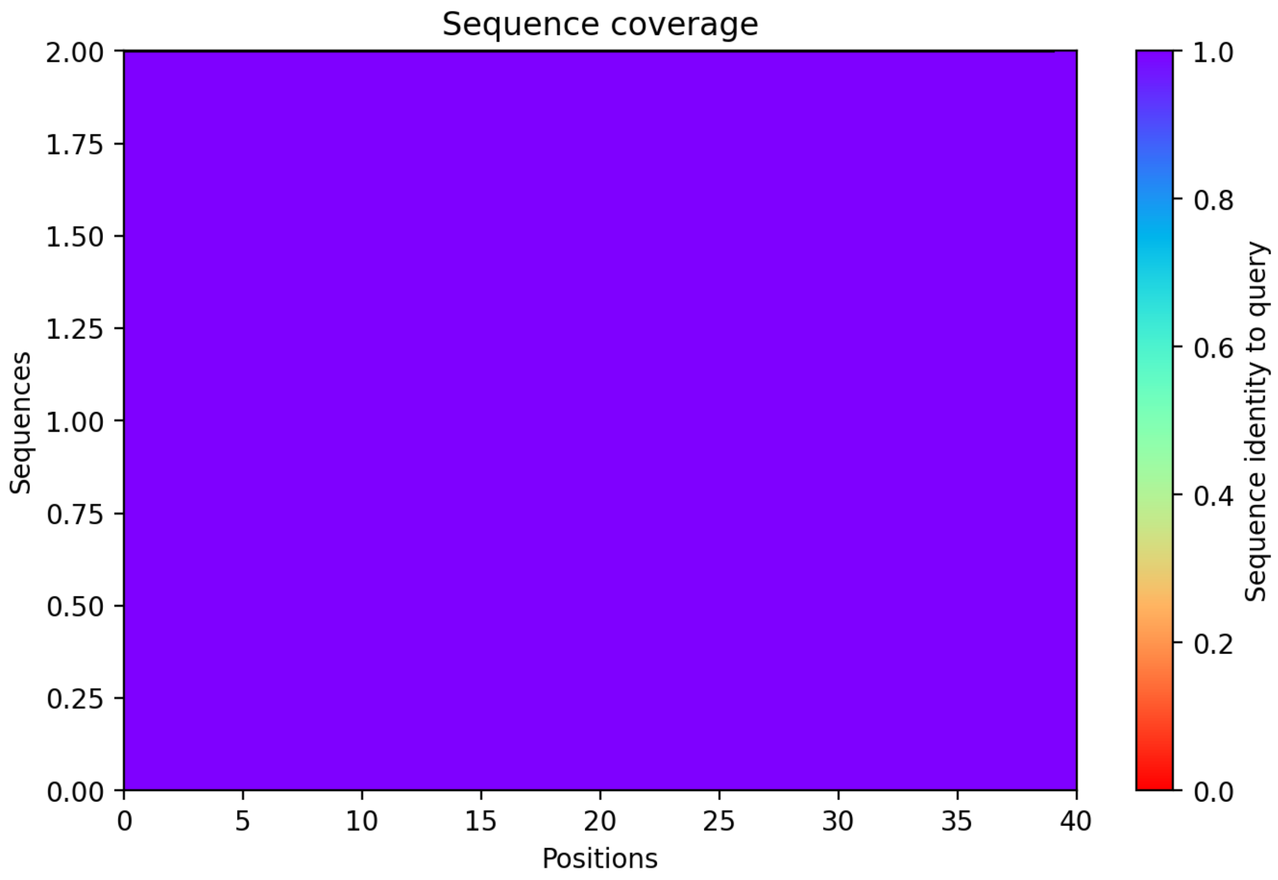

CP5: MtrunA17_Chr1g0150571

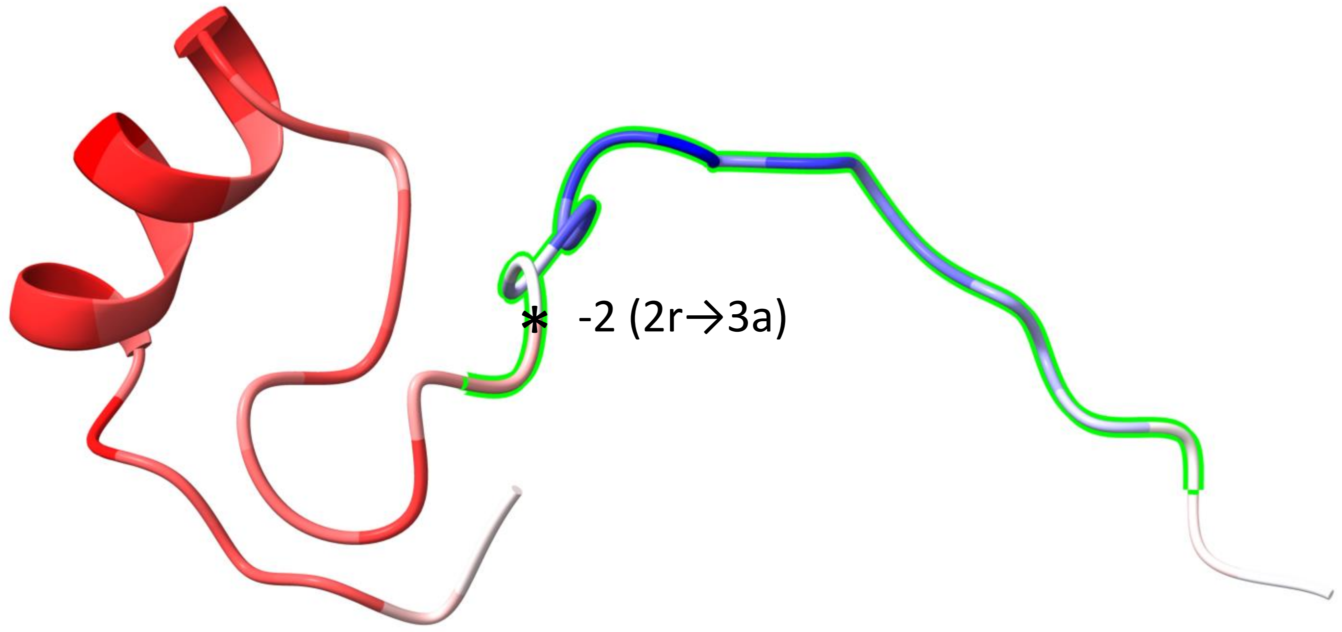


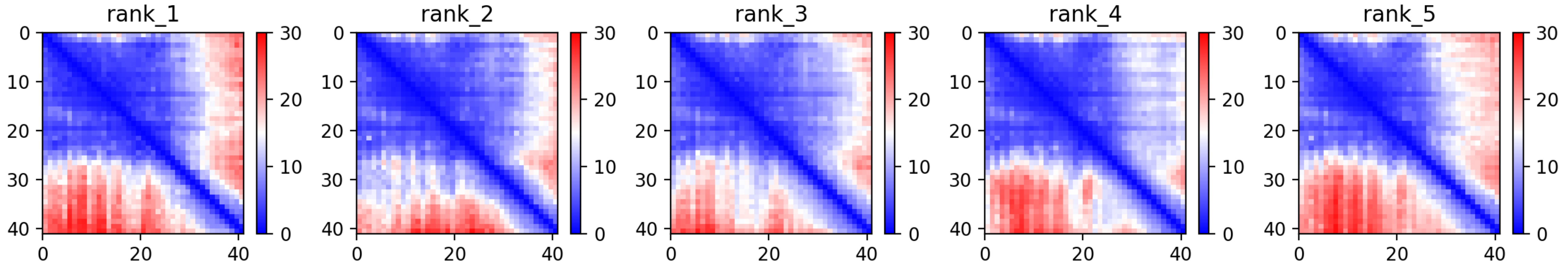

### CP5: MtrunA17_Chr1g0150571

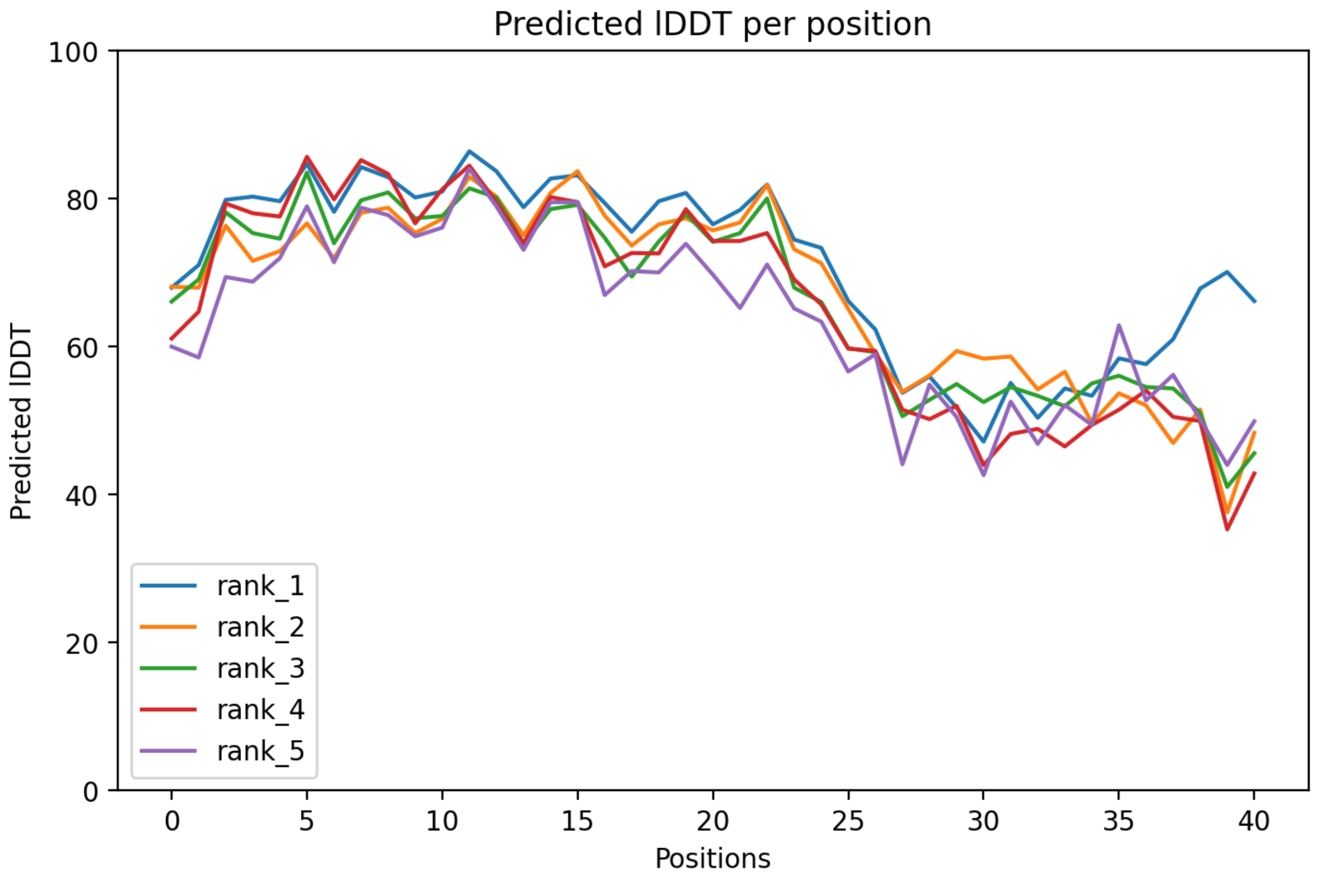

CP5: MtrunA17_Chr1g0150571

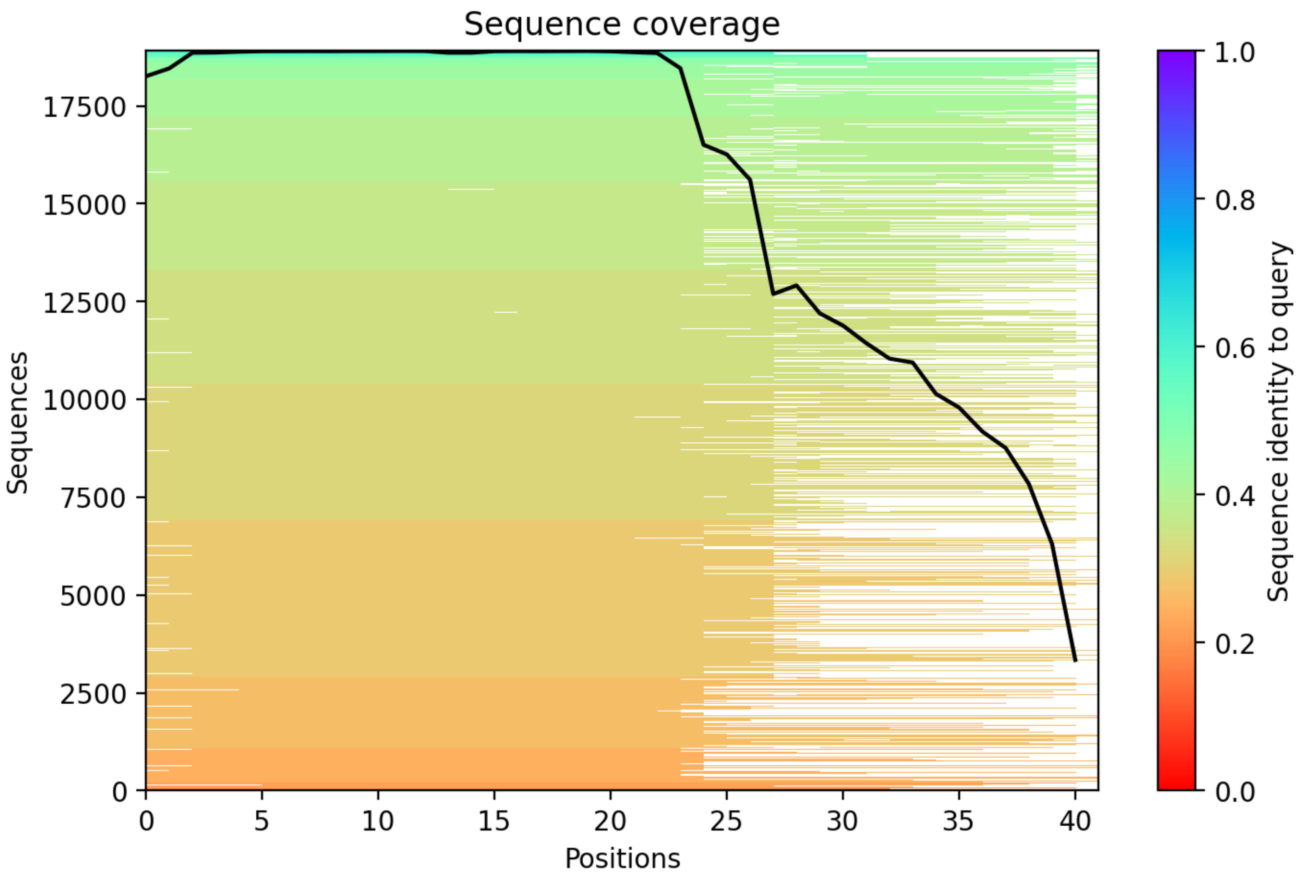

CP6: MtrunA17_Chr1g0152521

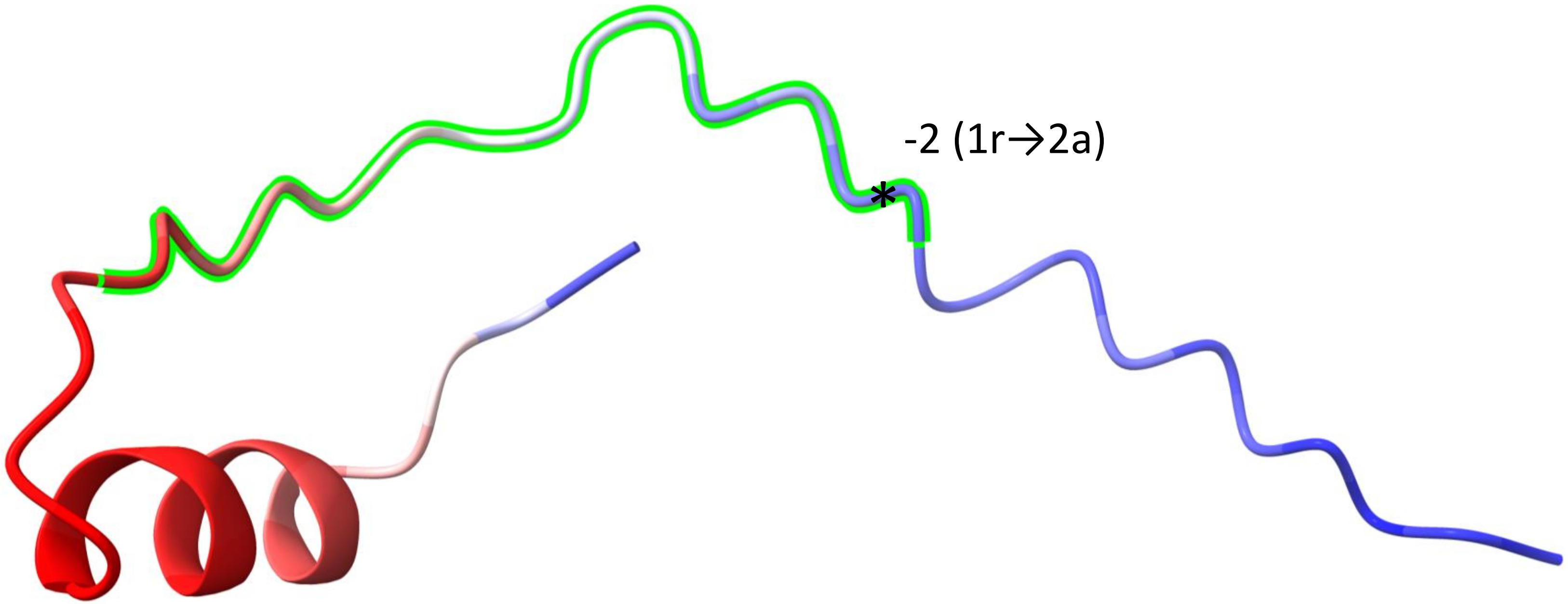


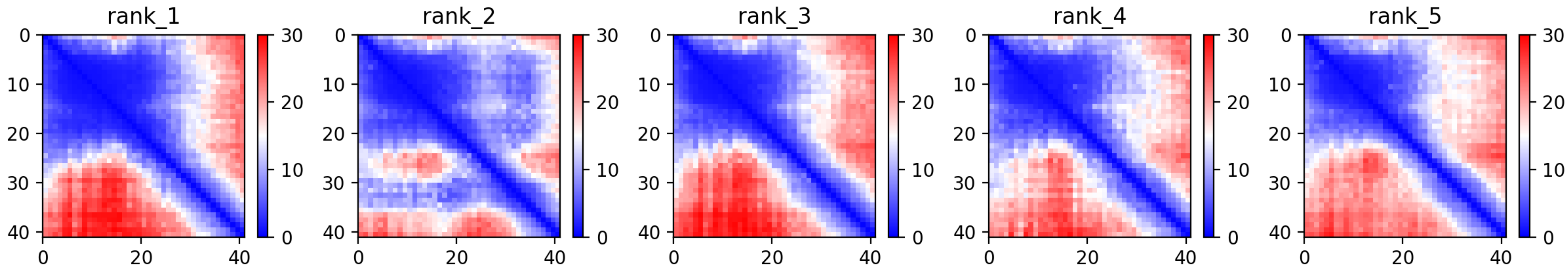

### CP6: MtrunA17_Chr1g0152521

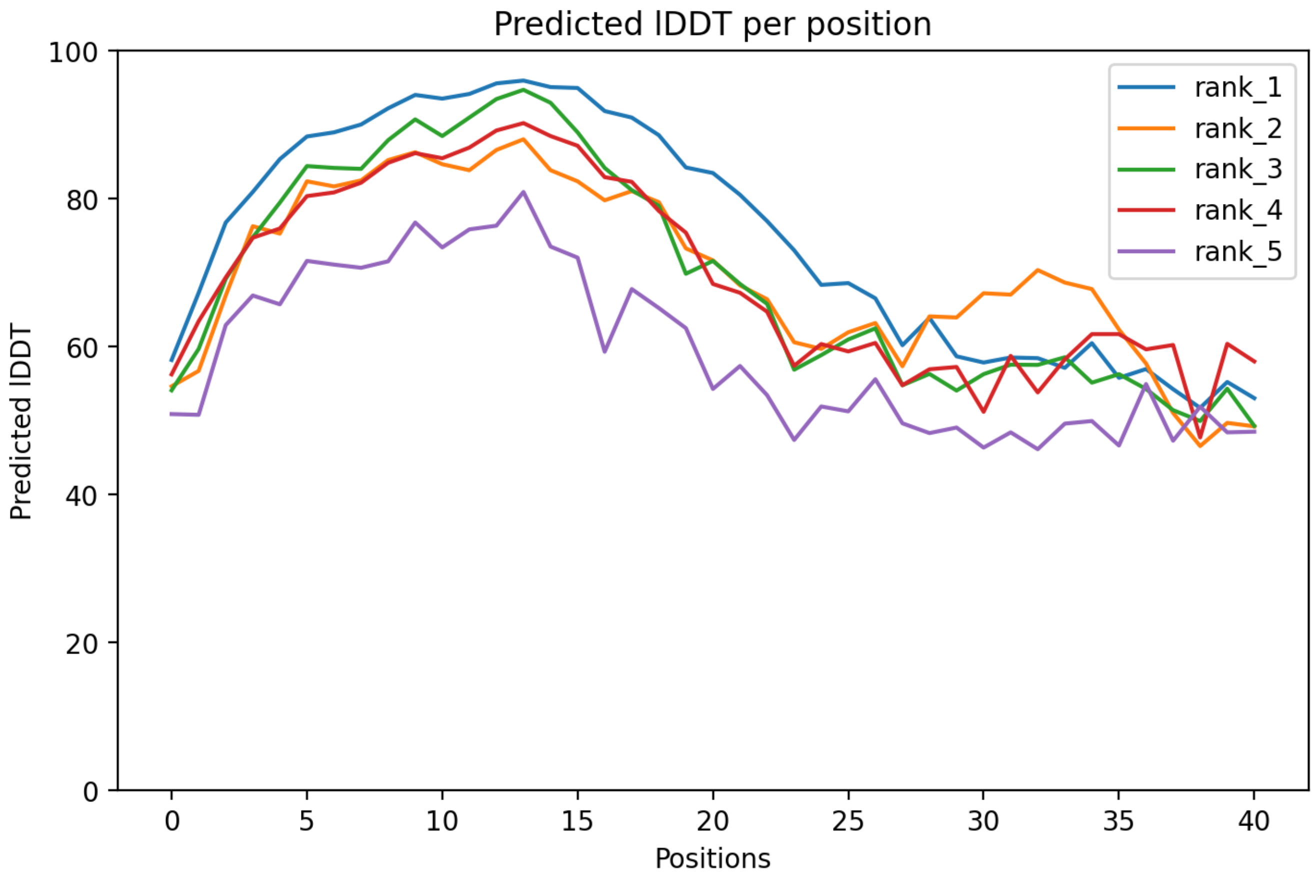

### CP6: MtrunA17_Chr1g0152521

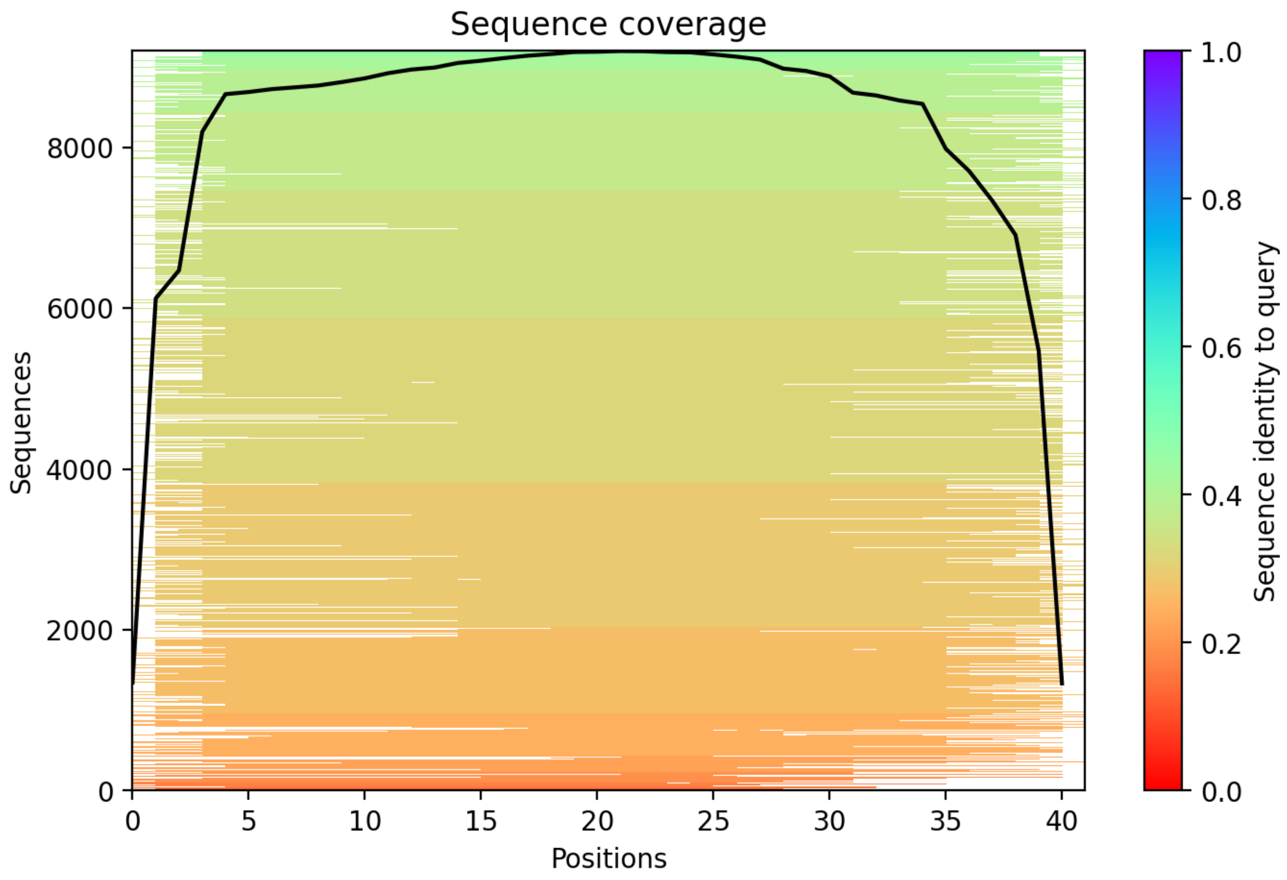

CP7: MtrunA17_Chr1g0153001

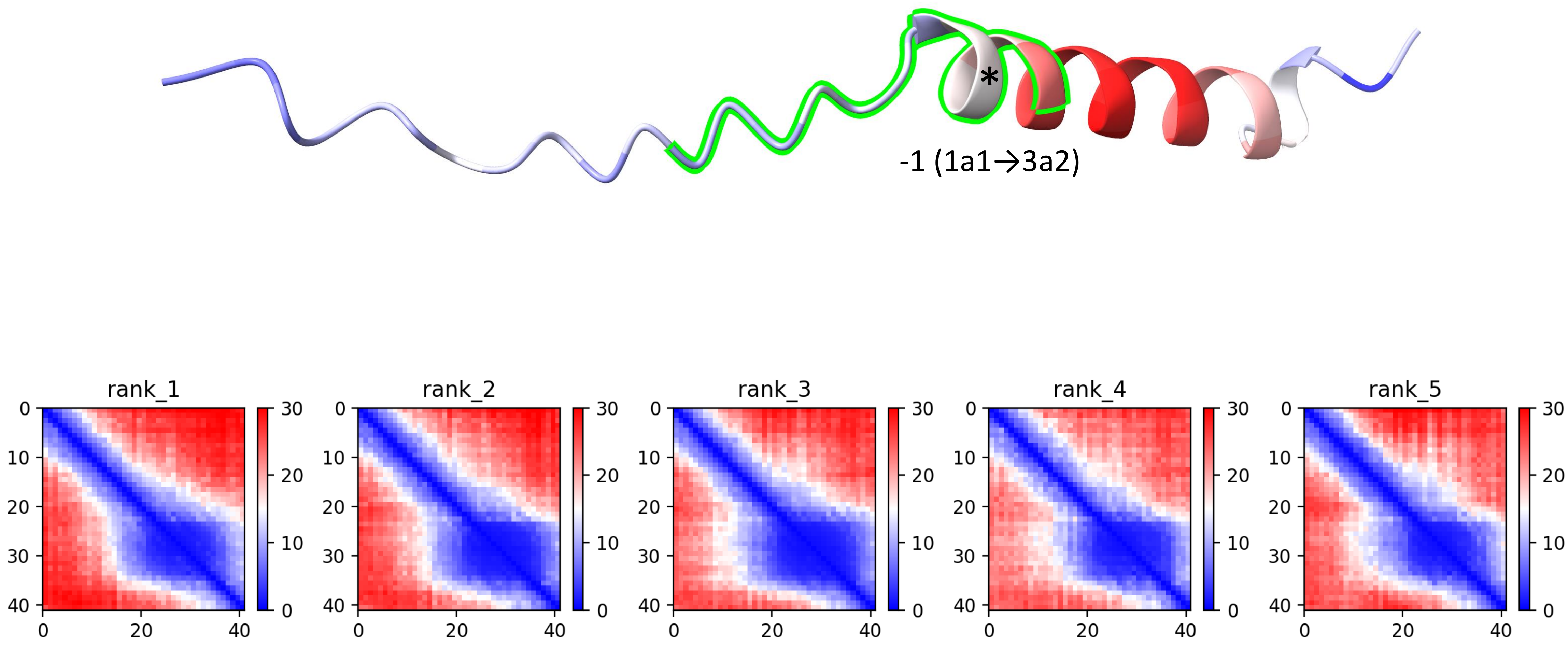

### CP7: MtrunA17_Chr1g0153001

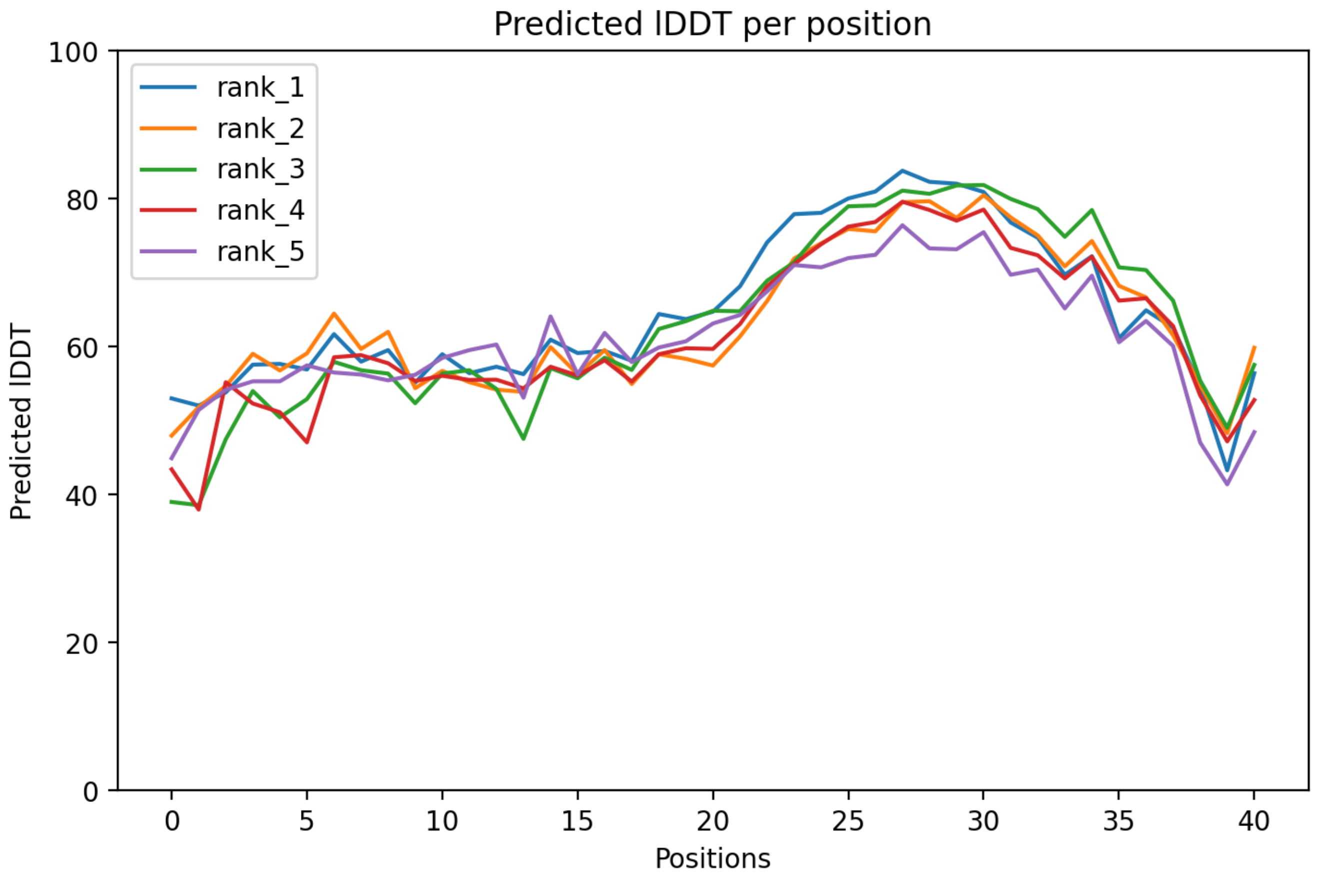

CP7: MtrunA17_Chr1g0153001

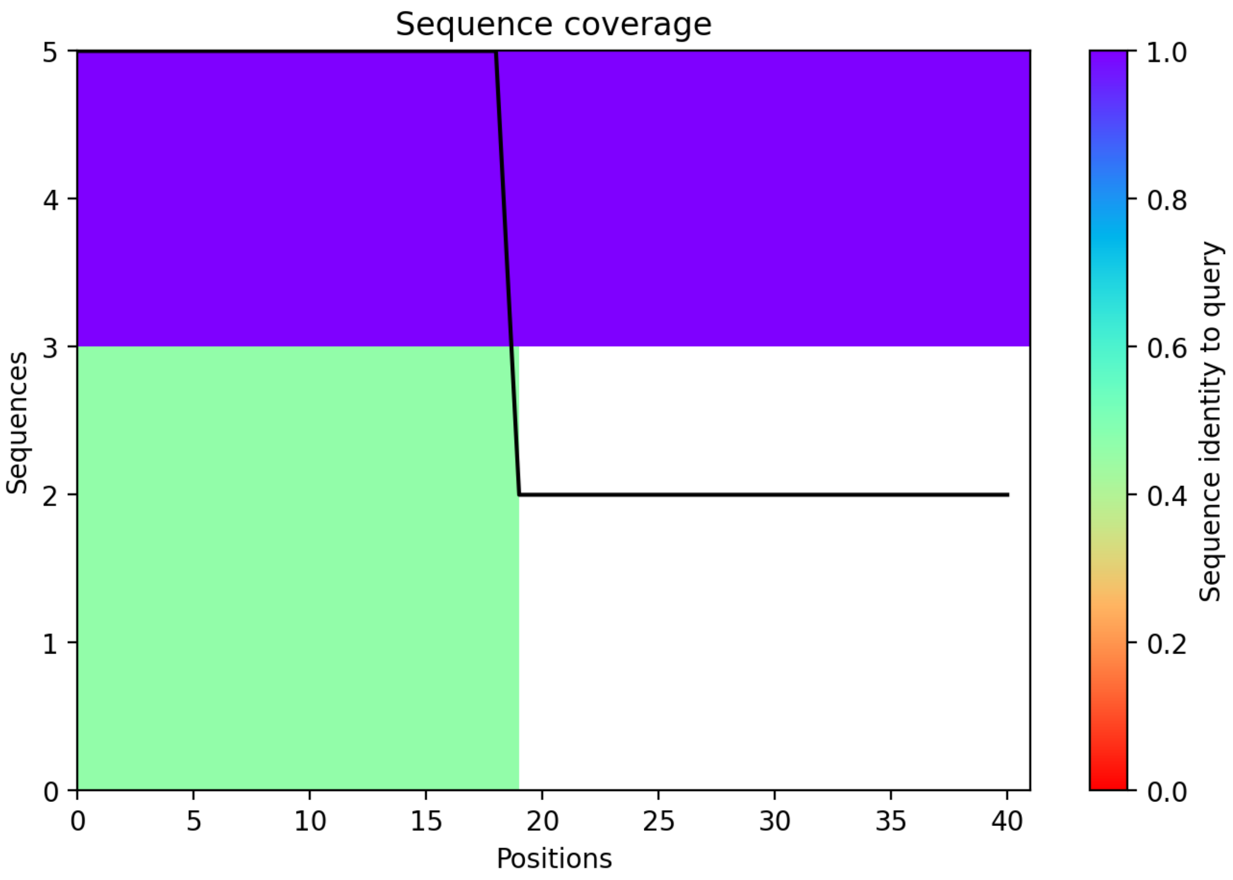

CP8: MtrunA17_Chr1g0155251

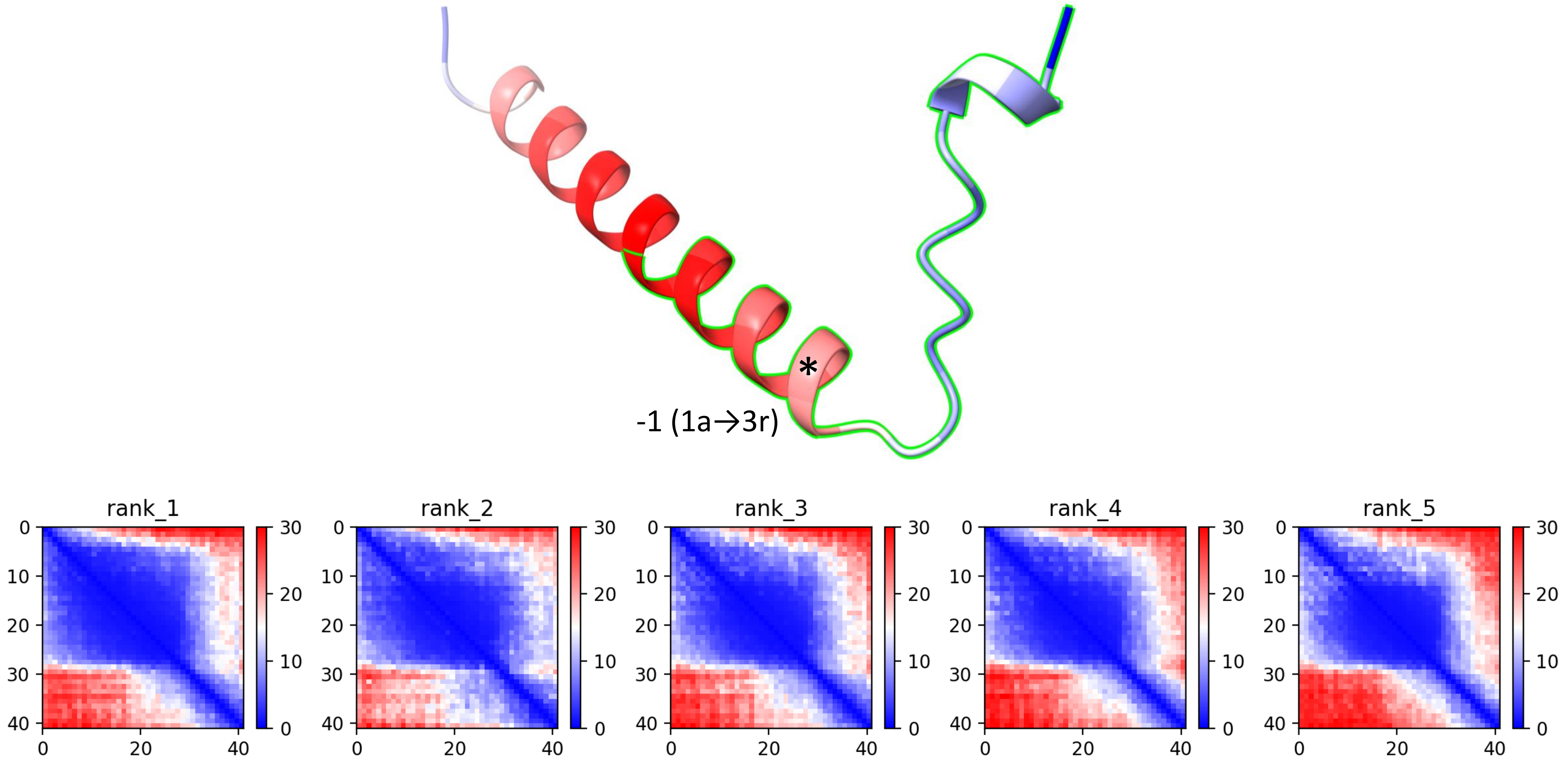

CP8: MtrunA17_Chr1g0155251

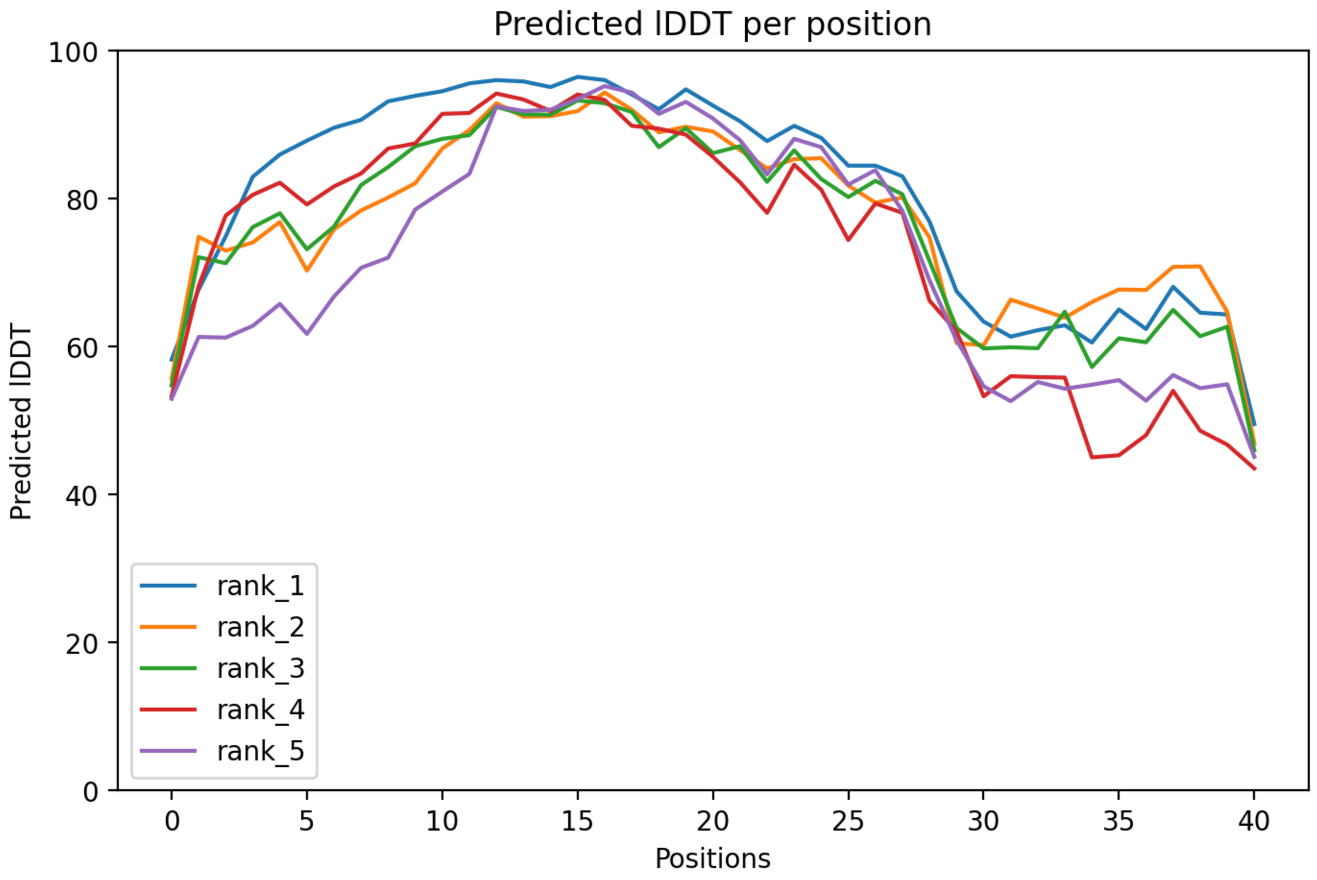

### CP8: MtrunA17_Chr1g0155251

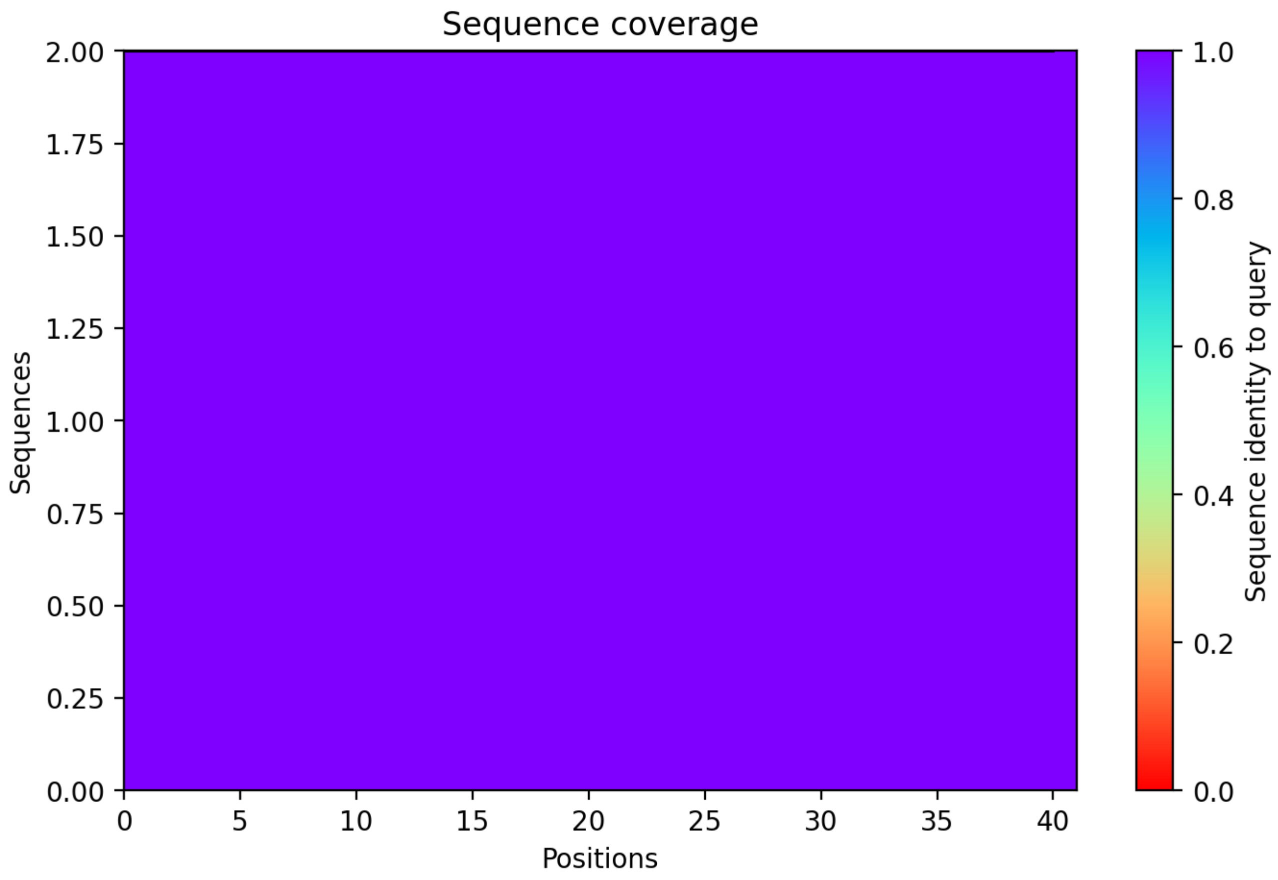

CP9: MtrunA17_Chr1g0158341

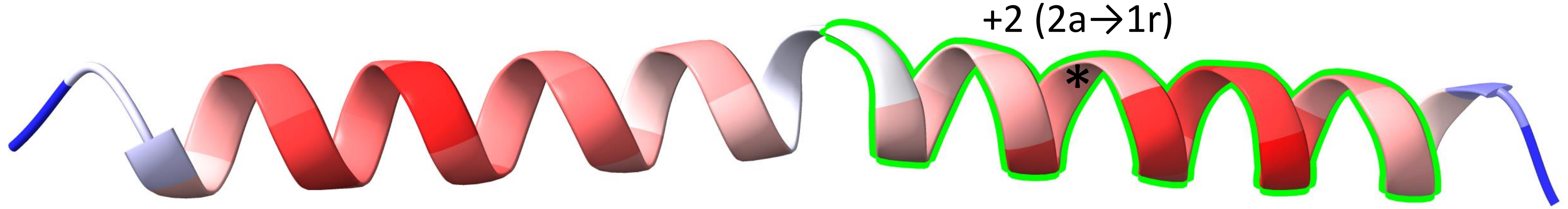


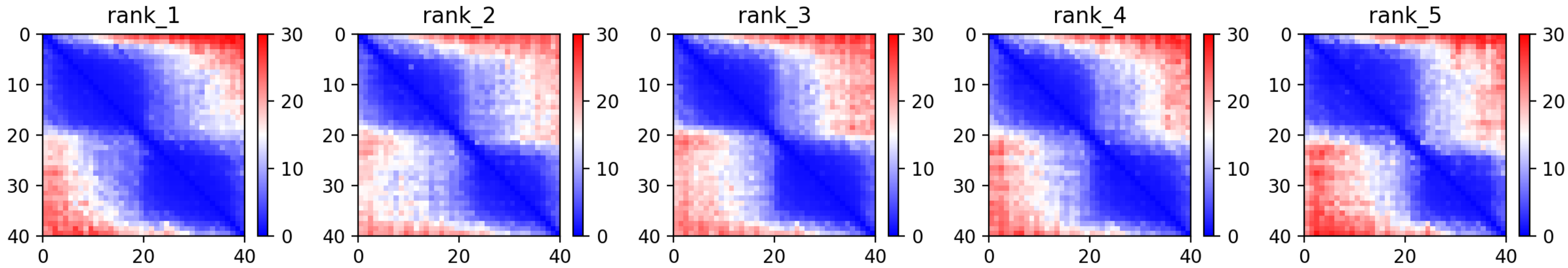

CP9: MtrunA17_Chr1g0158341

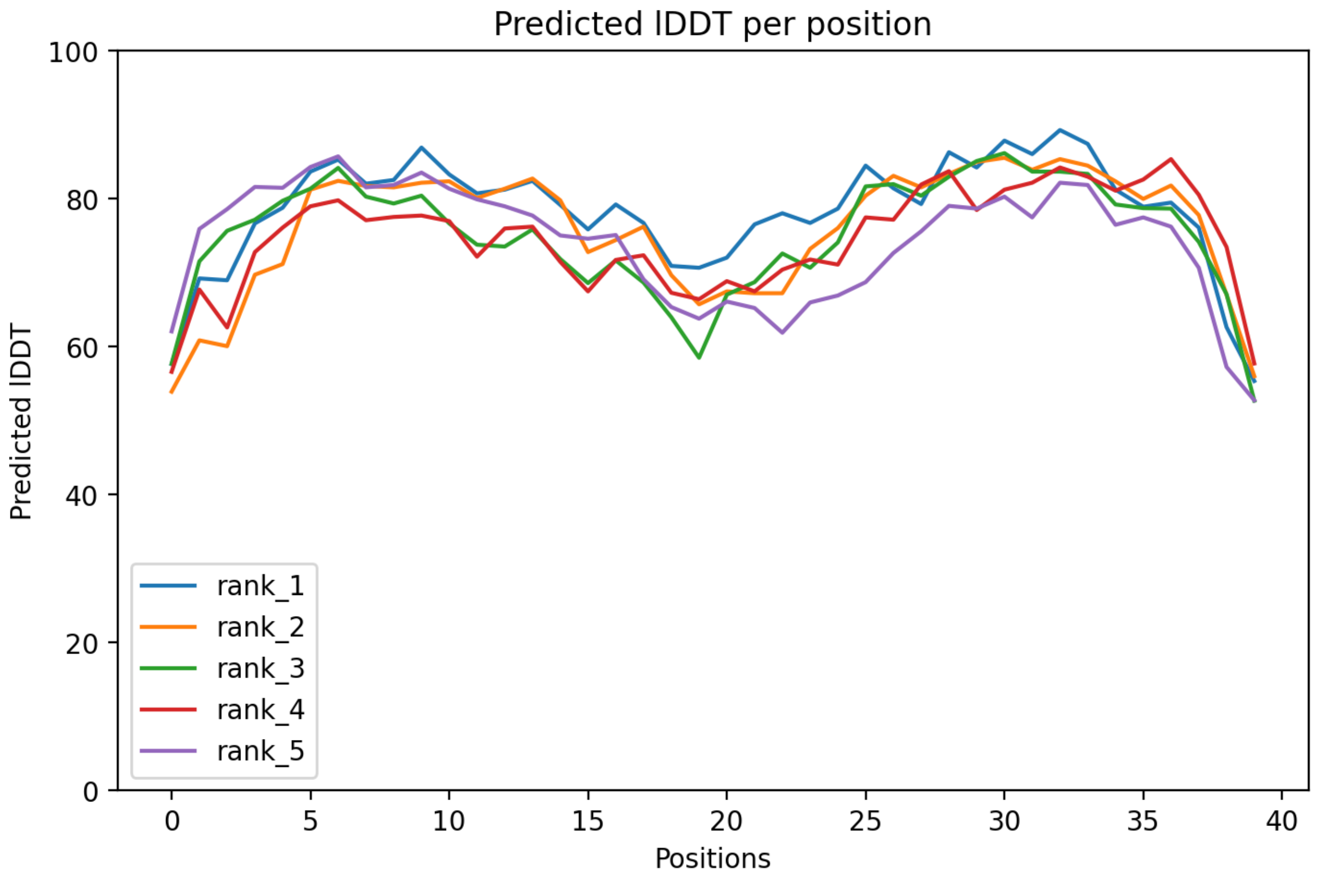

CP9: MtrunA17_Chr1g0158341

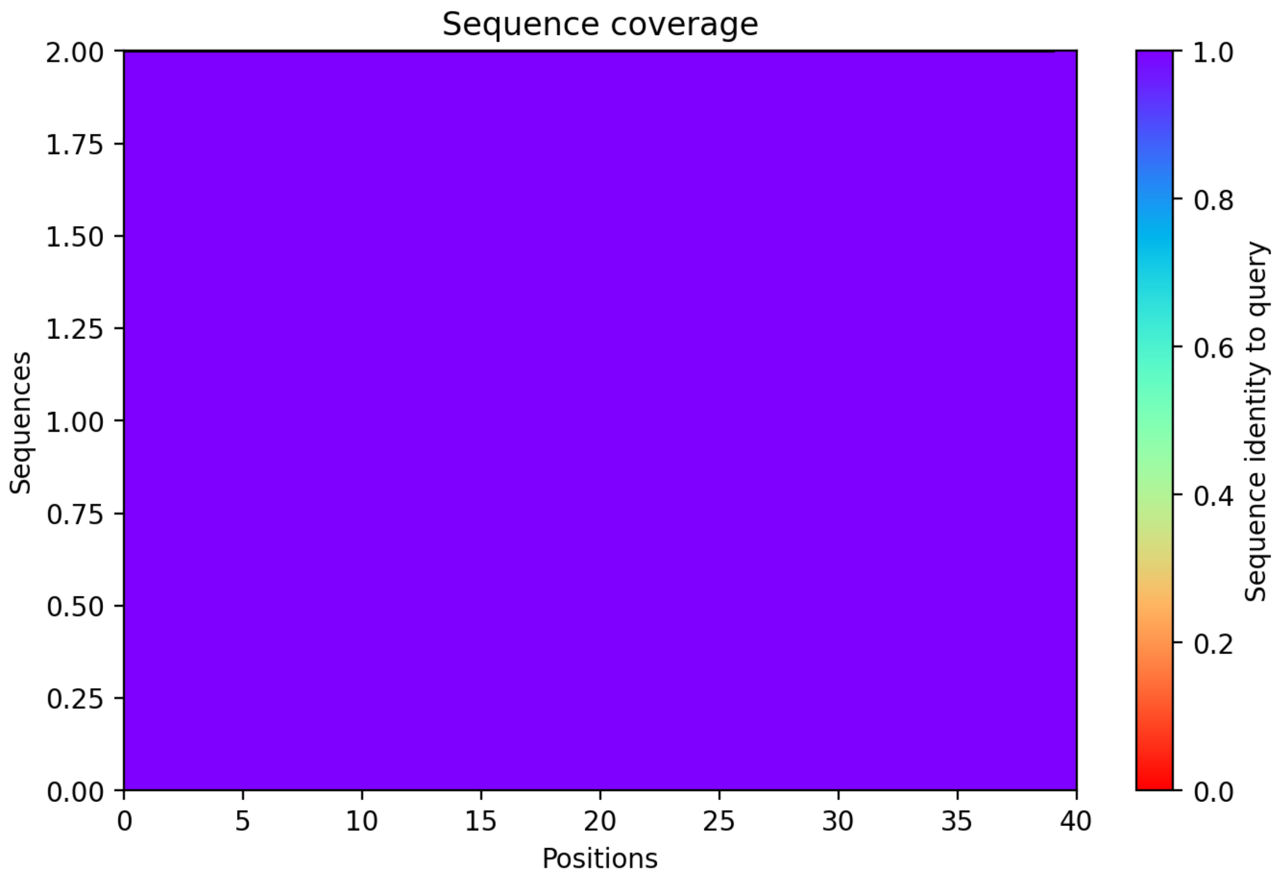

CP10: MtrunA17_Chr1g0162101

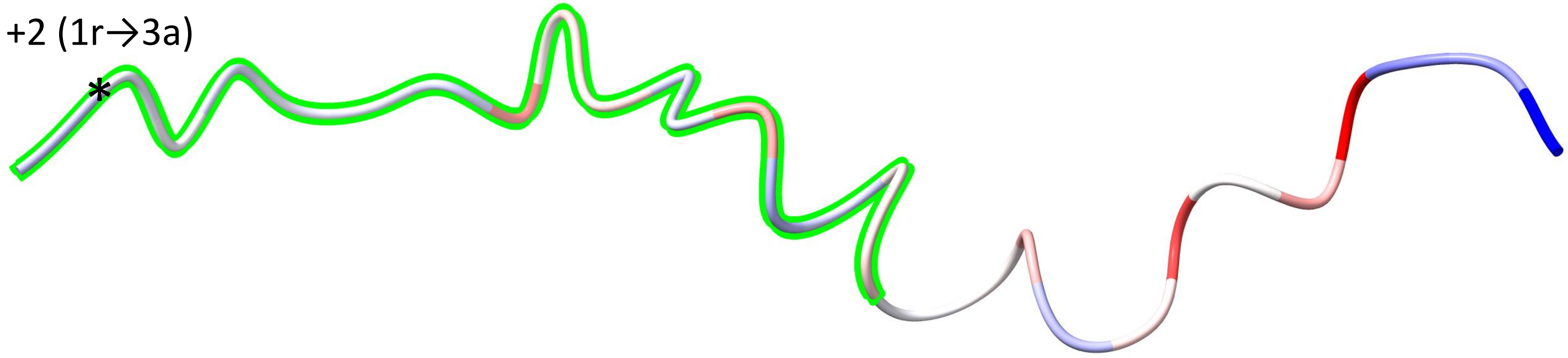


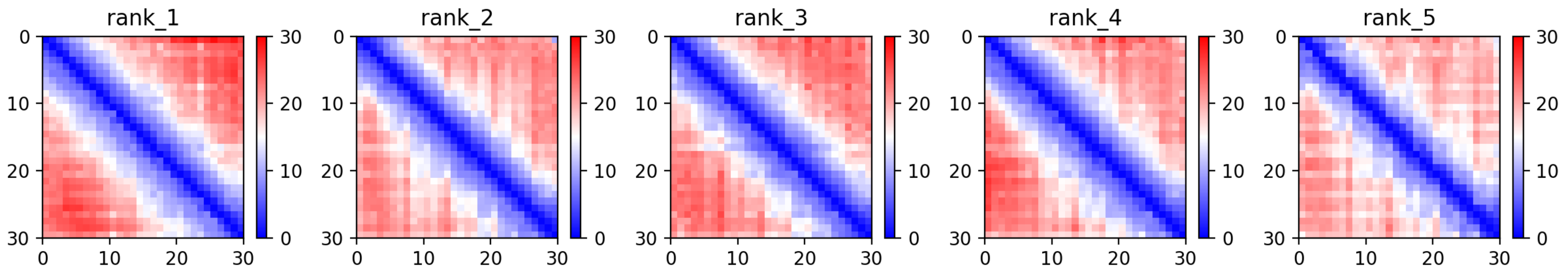

CP10: MtrunA17_Chr1g0162101

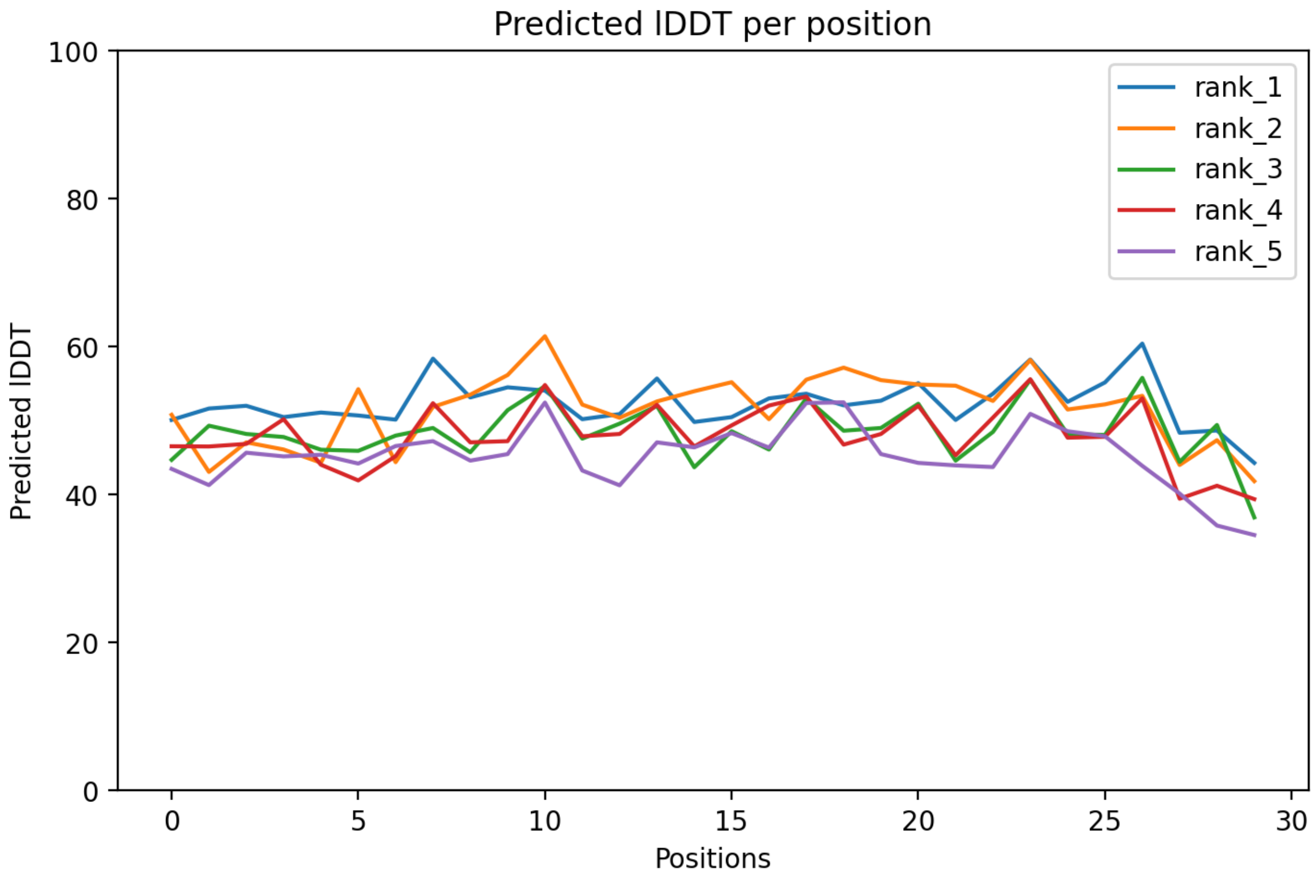

CP10: MtrunA17_Chr1g0162101

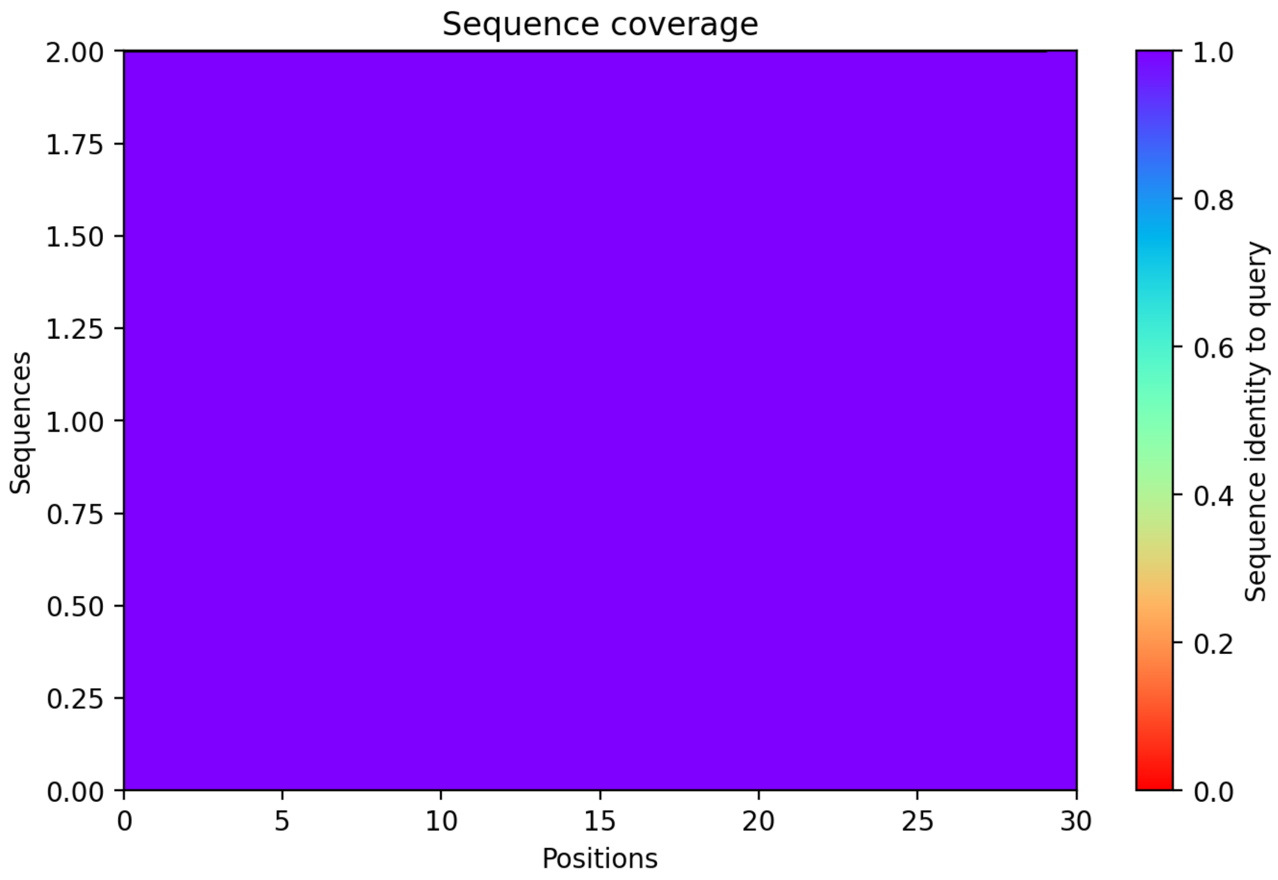

### CP11: MtrunA17_Chr1g0164591

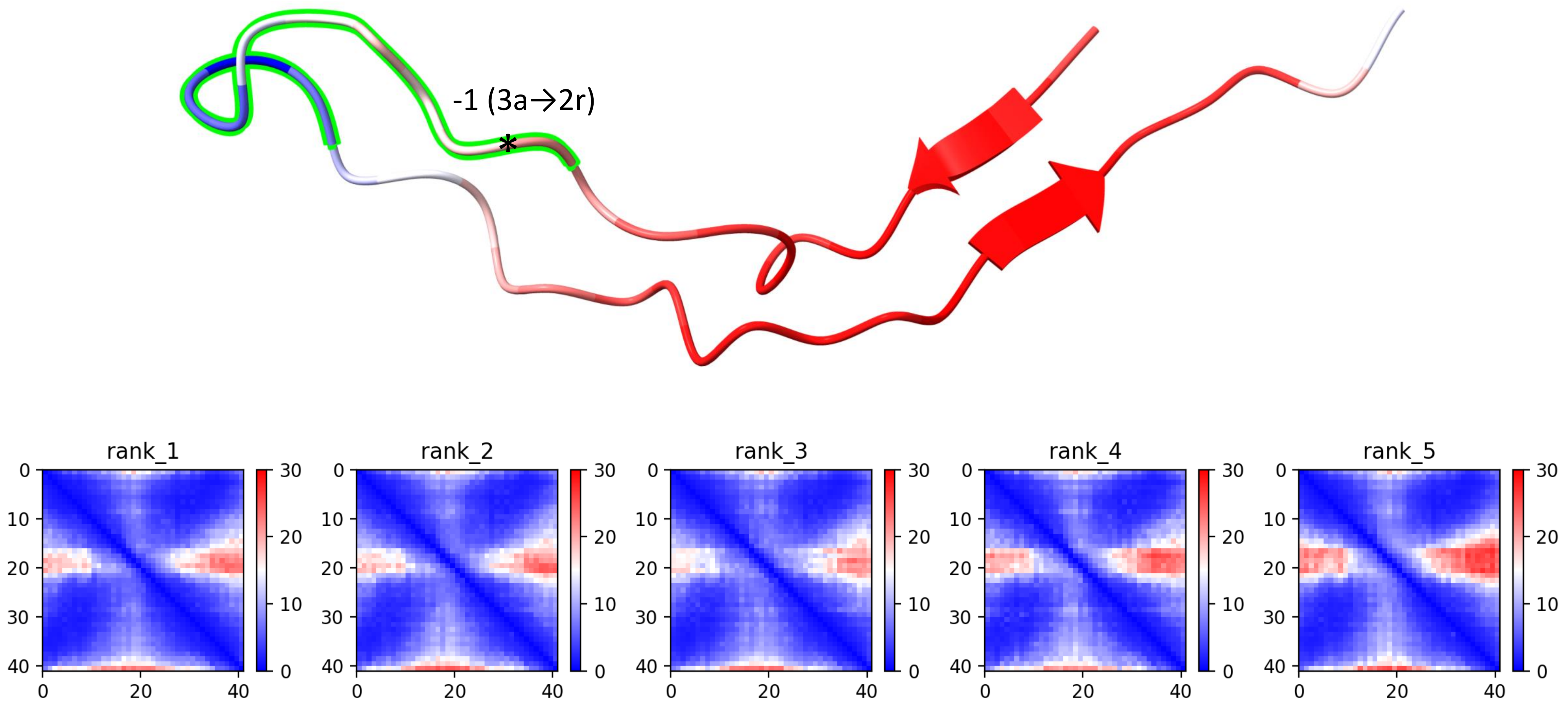

CP11: MtrunA17_Chr1g0164591

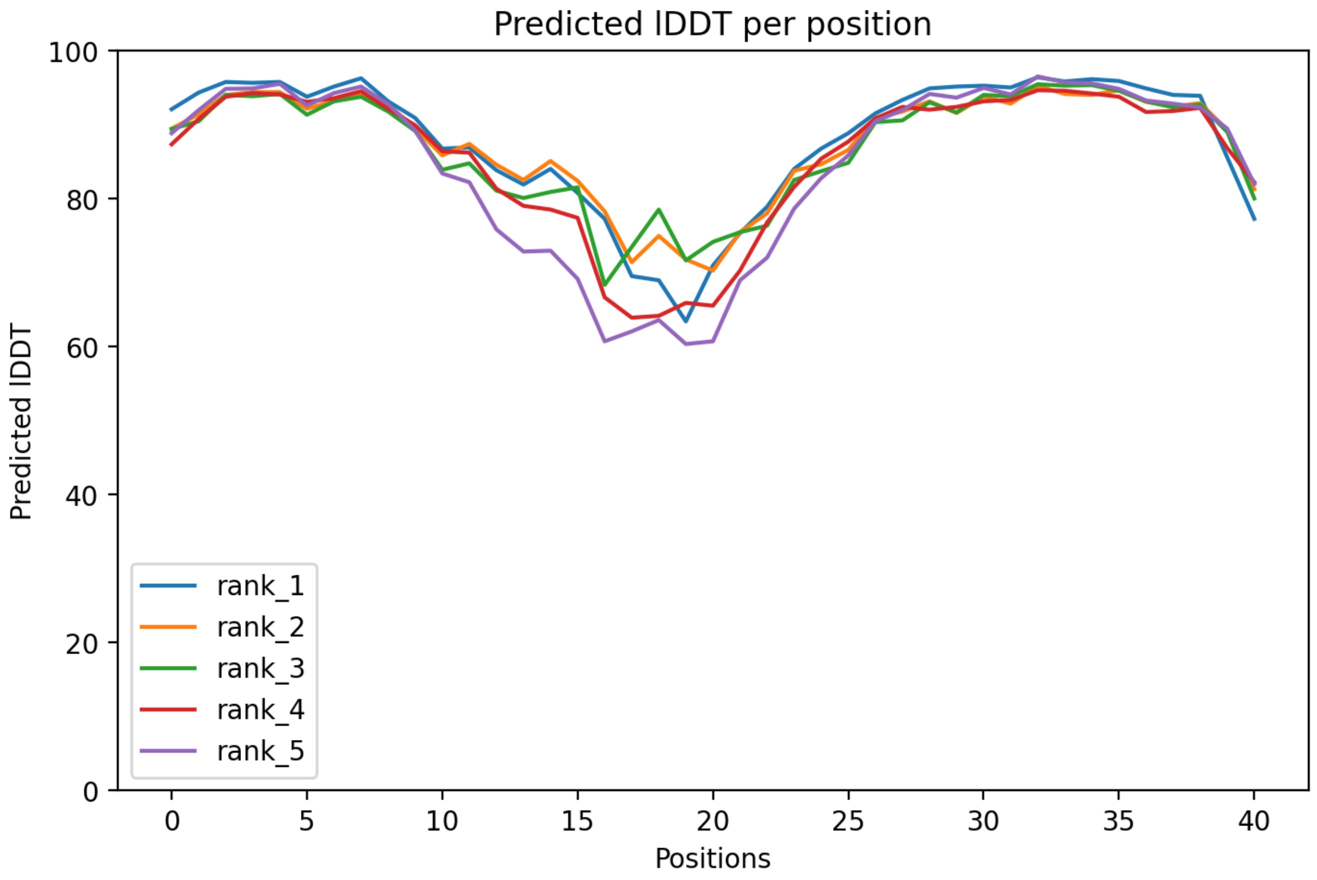

CP11: MtrunA17_Chr1g0164591

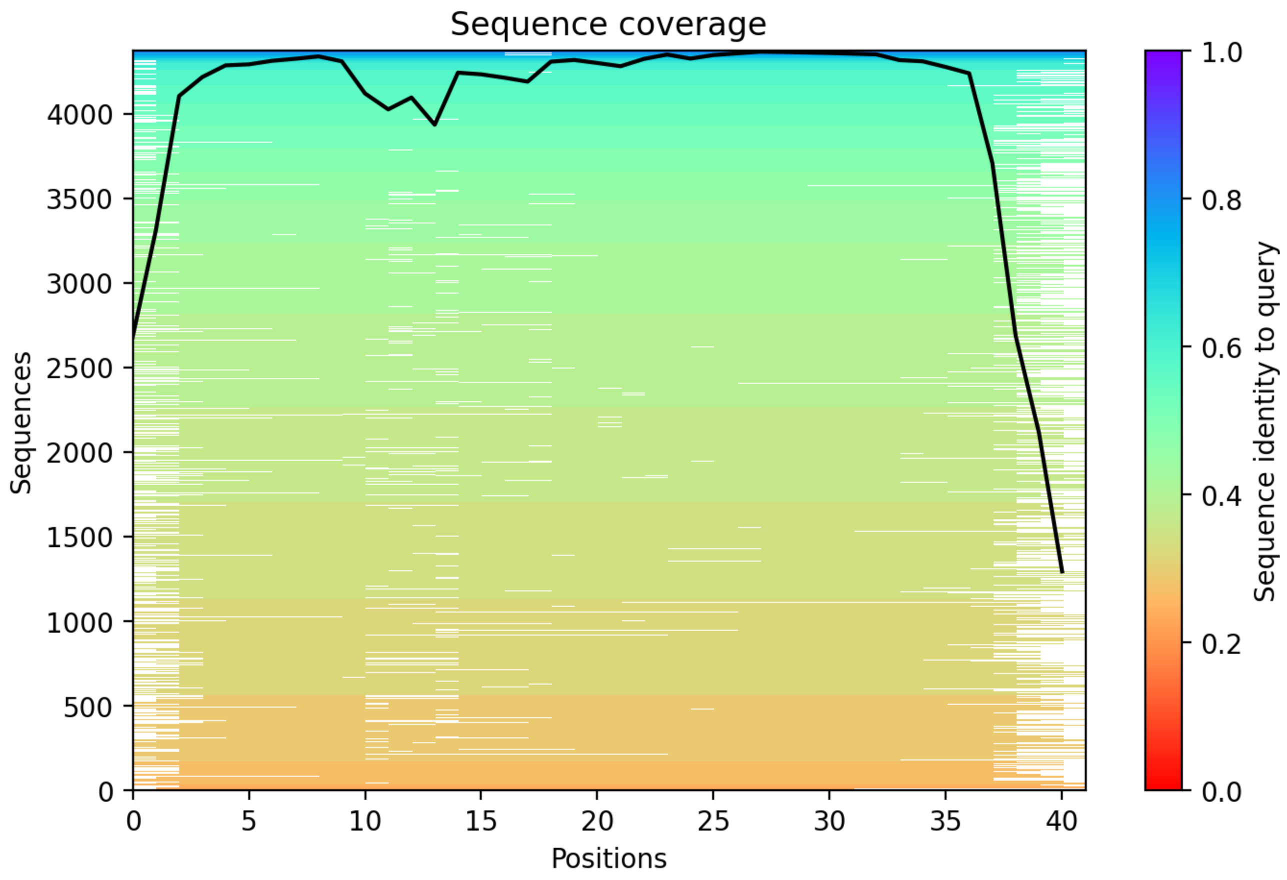

CP12: MtrunA17_Chr1g0178361

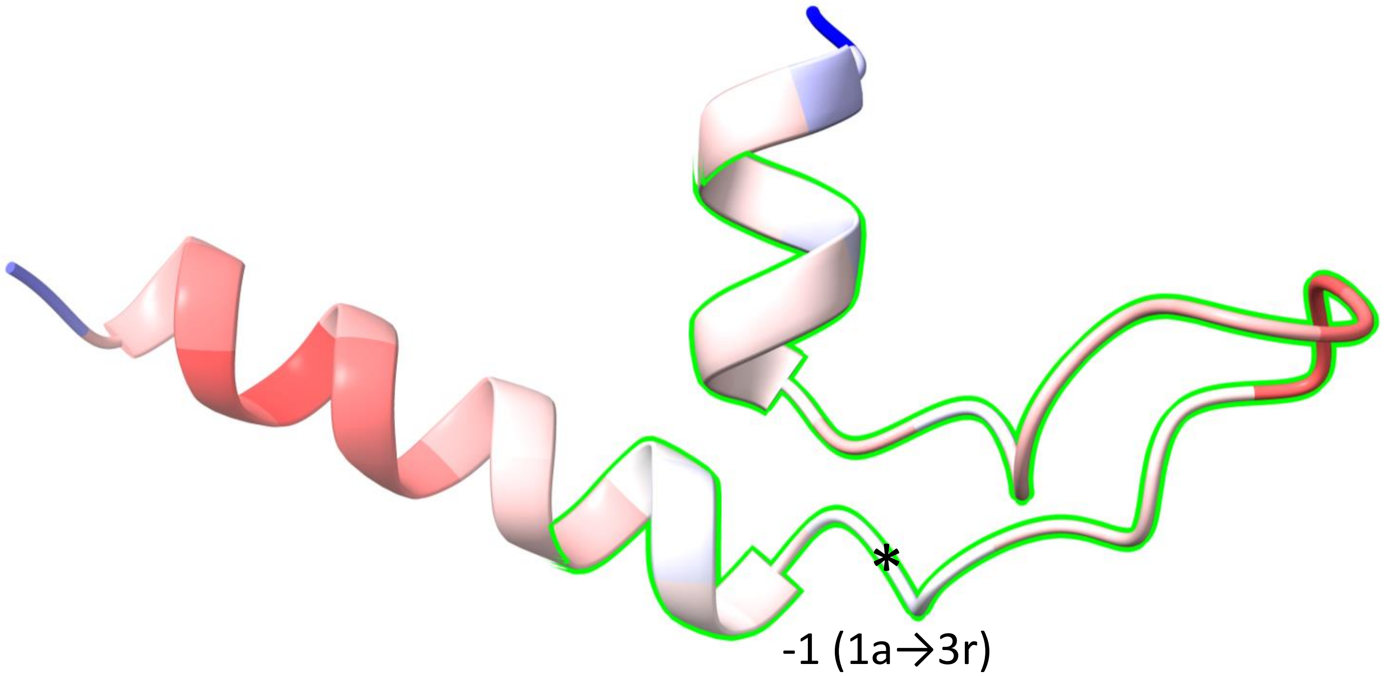


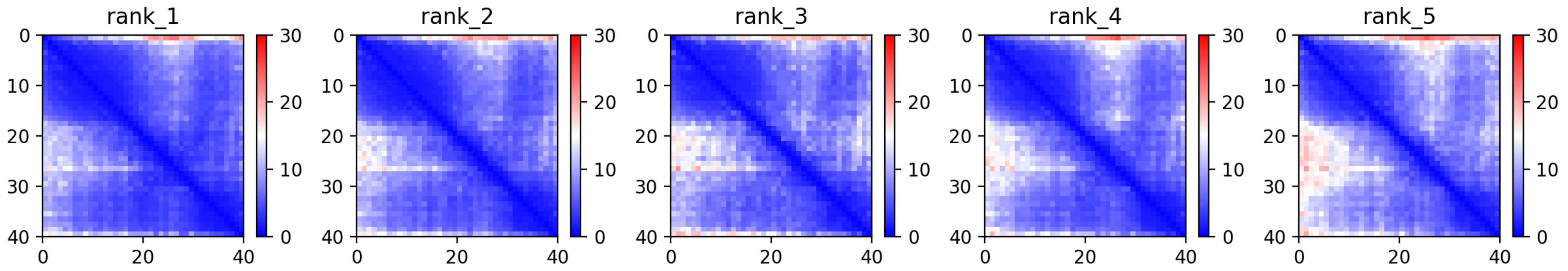

CP12: MtrunA17_Chr1g0178361

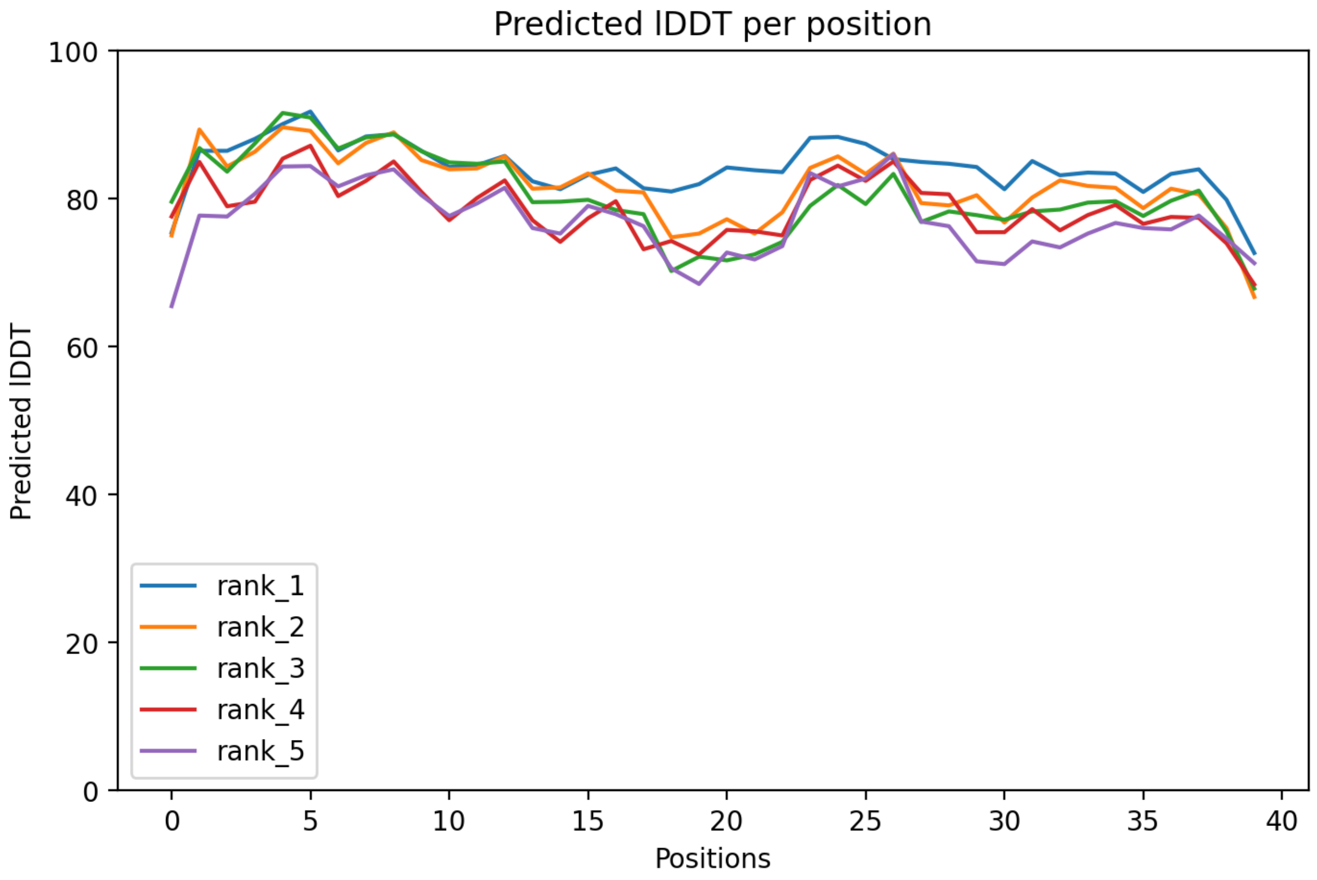

### CP12: MtrunA17_Chr1g0178361

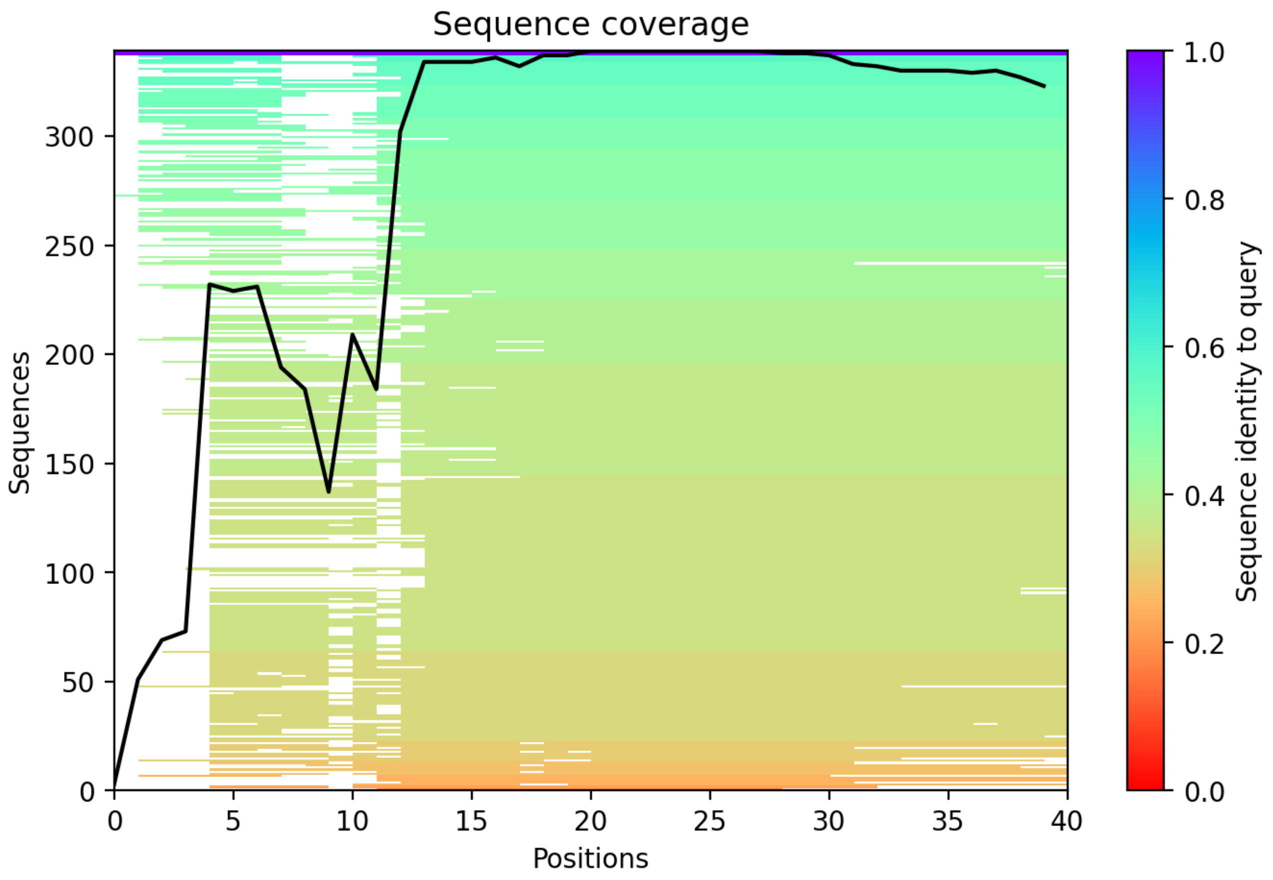

CP13: MtrunA17_Chr1g0181761

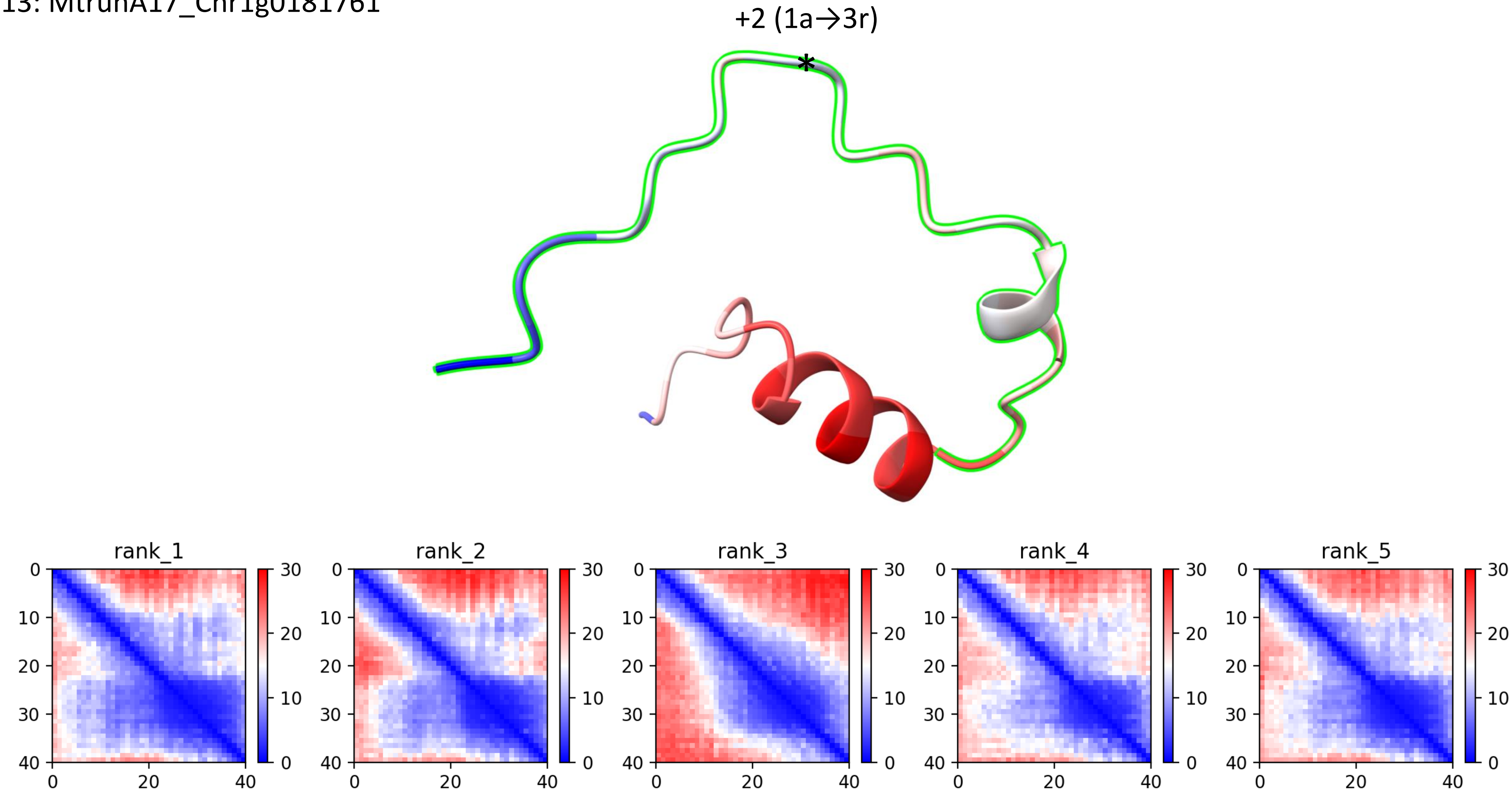

CP13: MtrunA17_Chr1g0181761

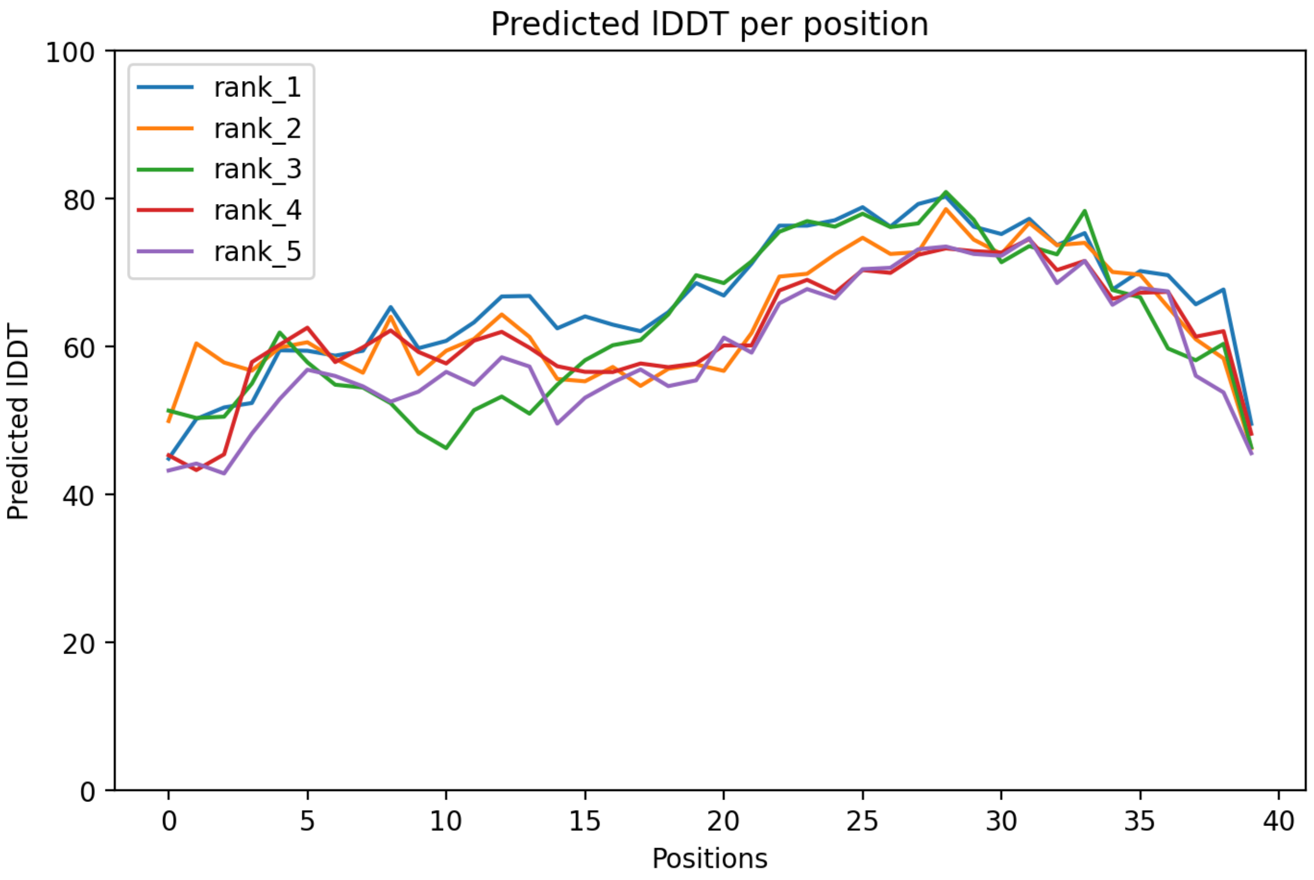

### CP13: MtrunA17_Chr1g0181761

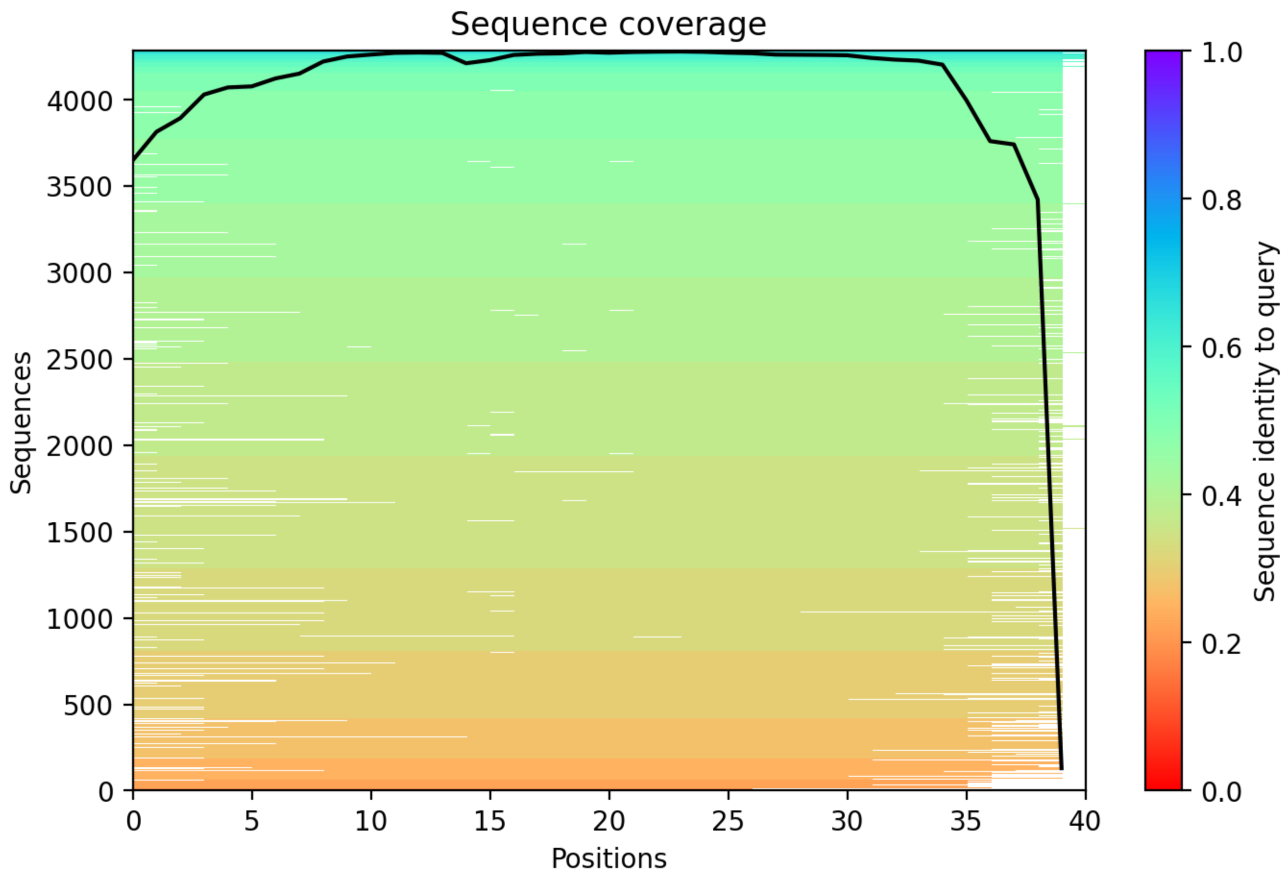

### CP14: MtrunA17_Chr1g0182591

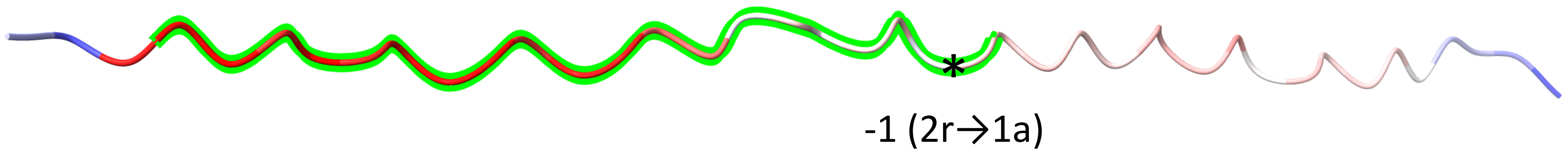


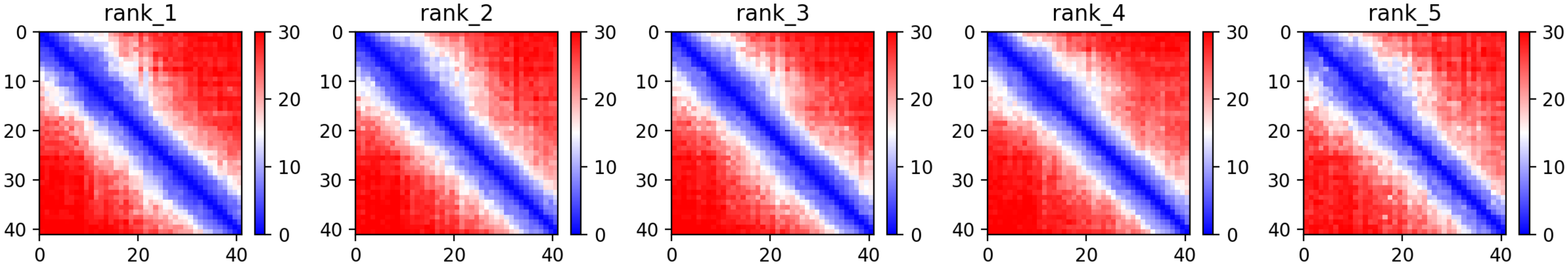

CP14: MtrunA17_Chr1g0182591

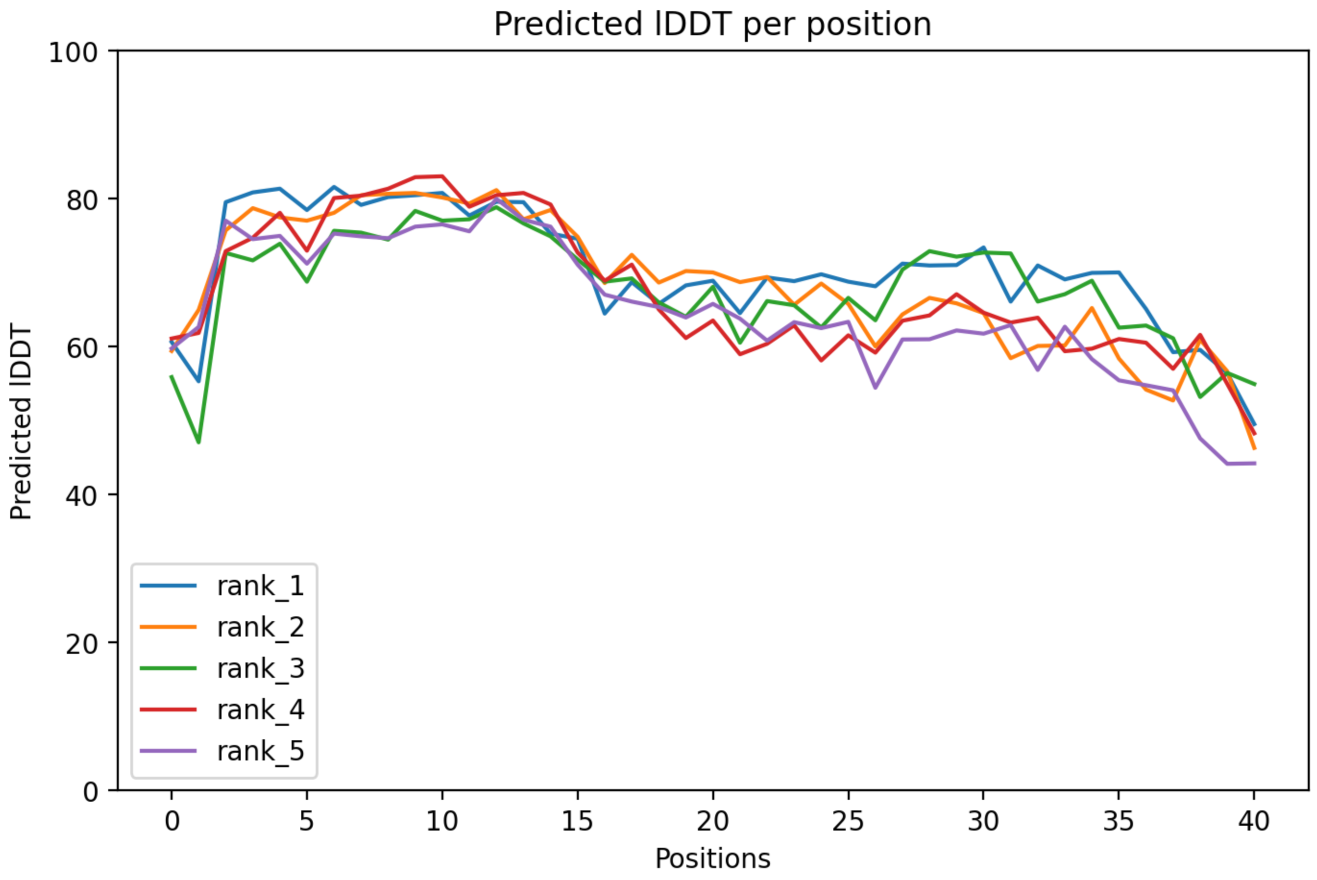

CP14: MtrunA17_Chr1g0182591

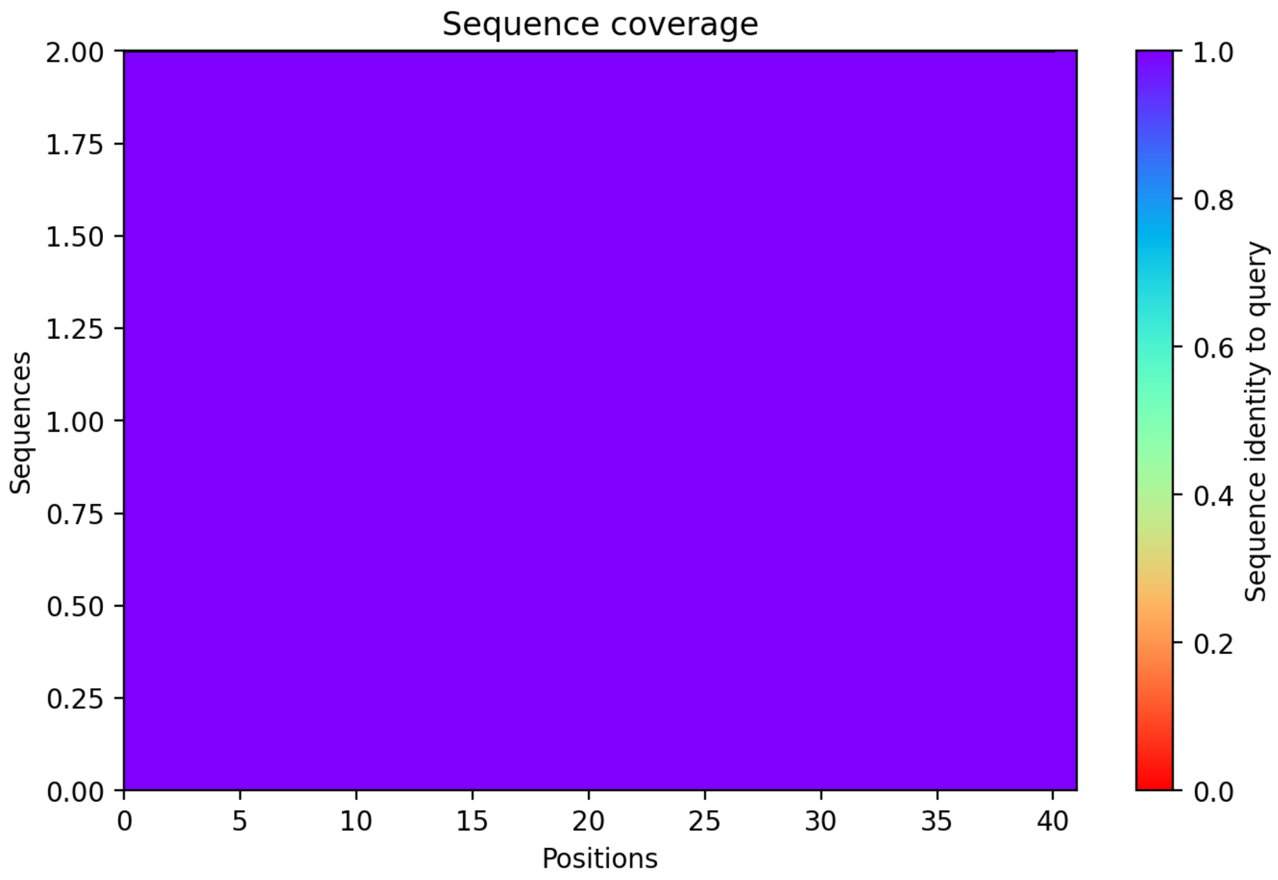

CP15: MtrunA17_Chr1g0183001

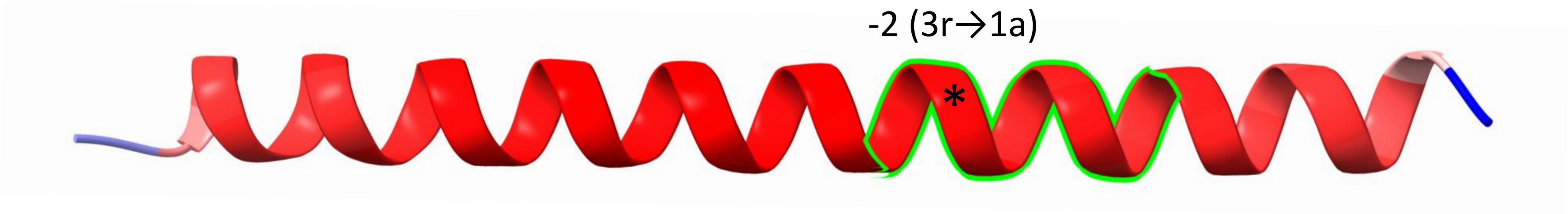


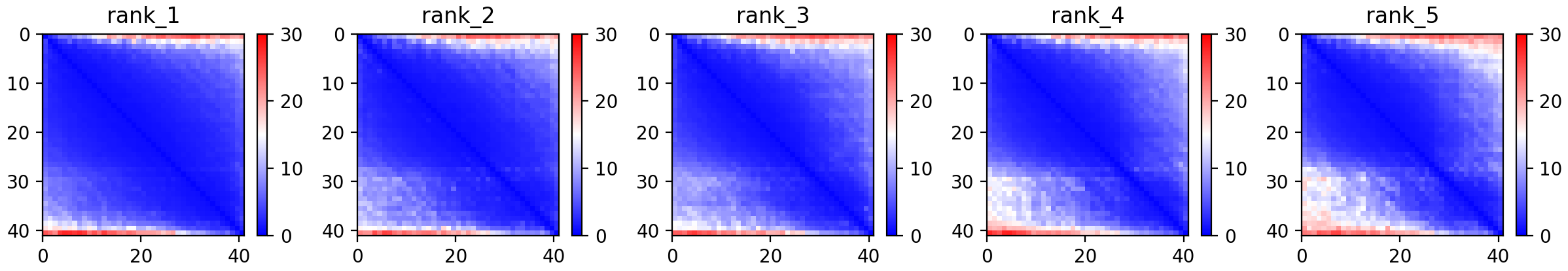

CP15: MtrunA17_Chr1g0183001

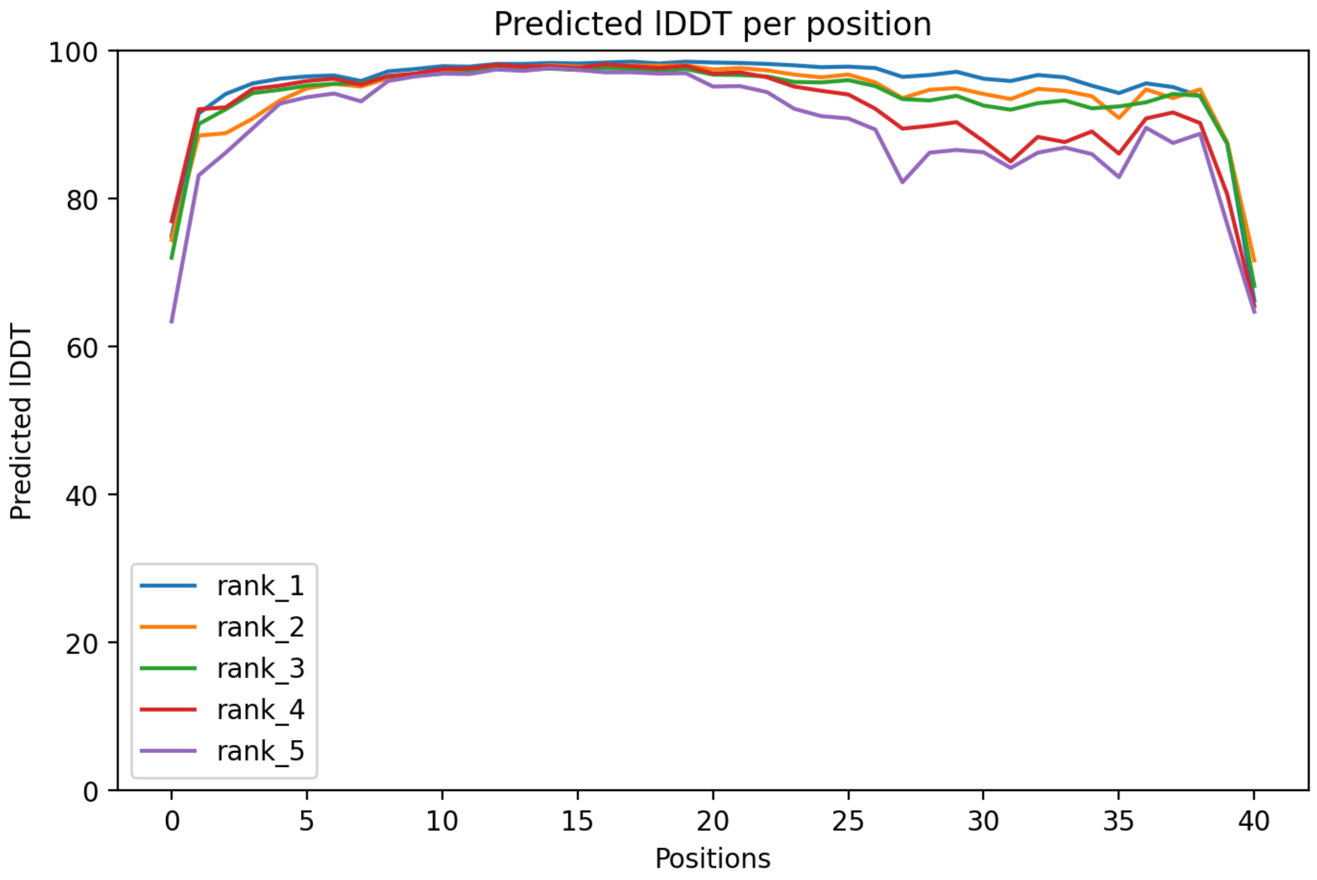

CP15: MtrunA17_Chr1g0183001

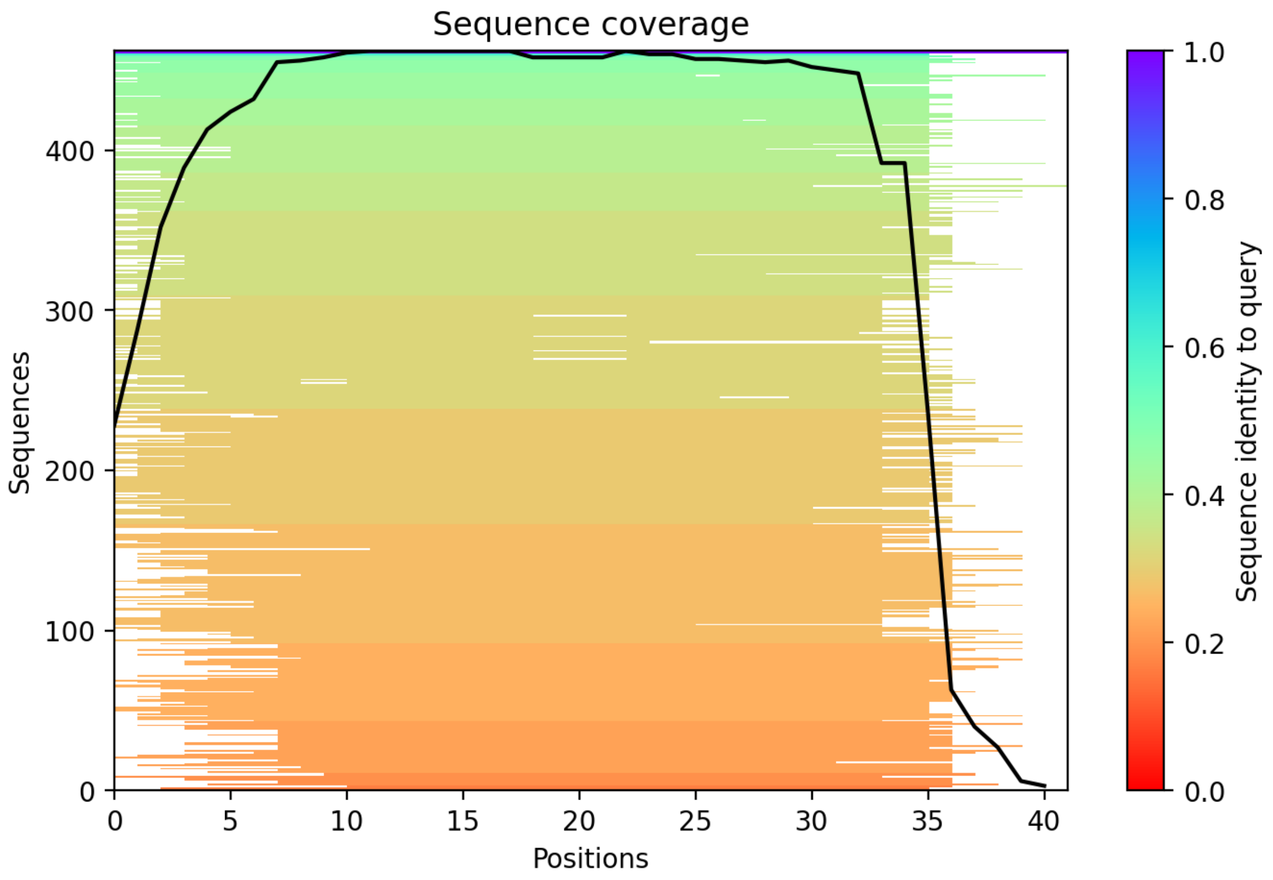

CP16: MtrunA17_Chr1g0185811

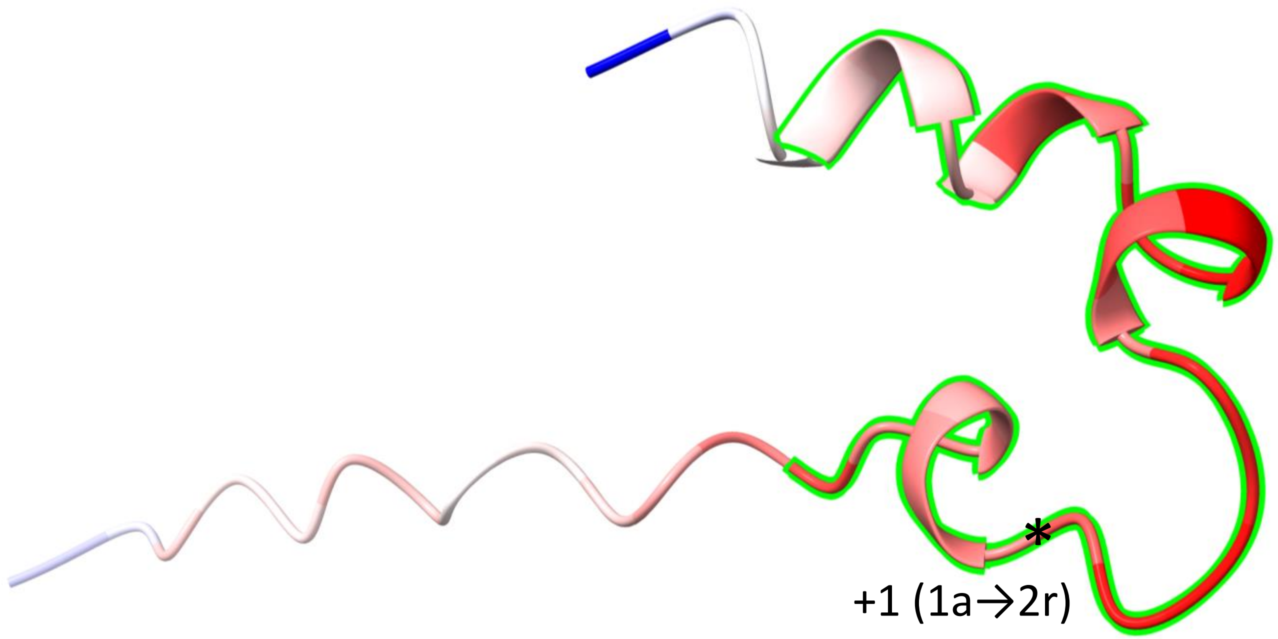


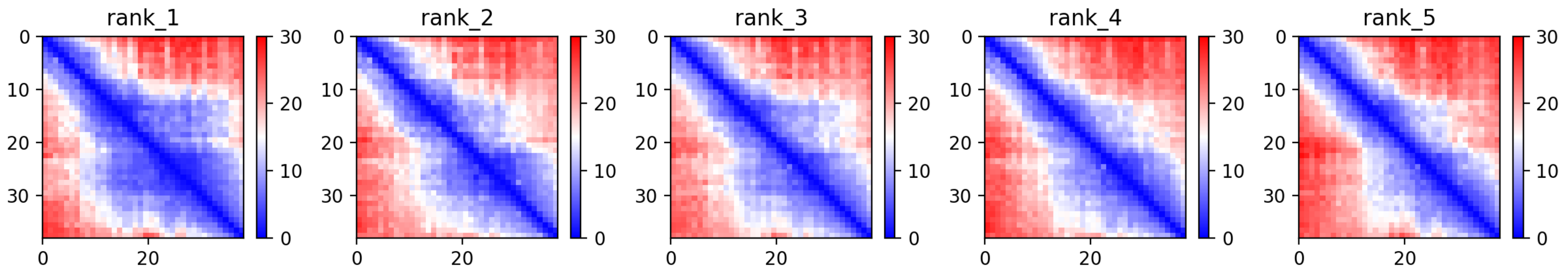

### CP16: MtrunA17_Chr1g0185811

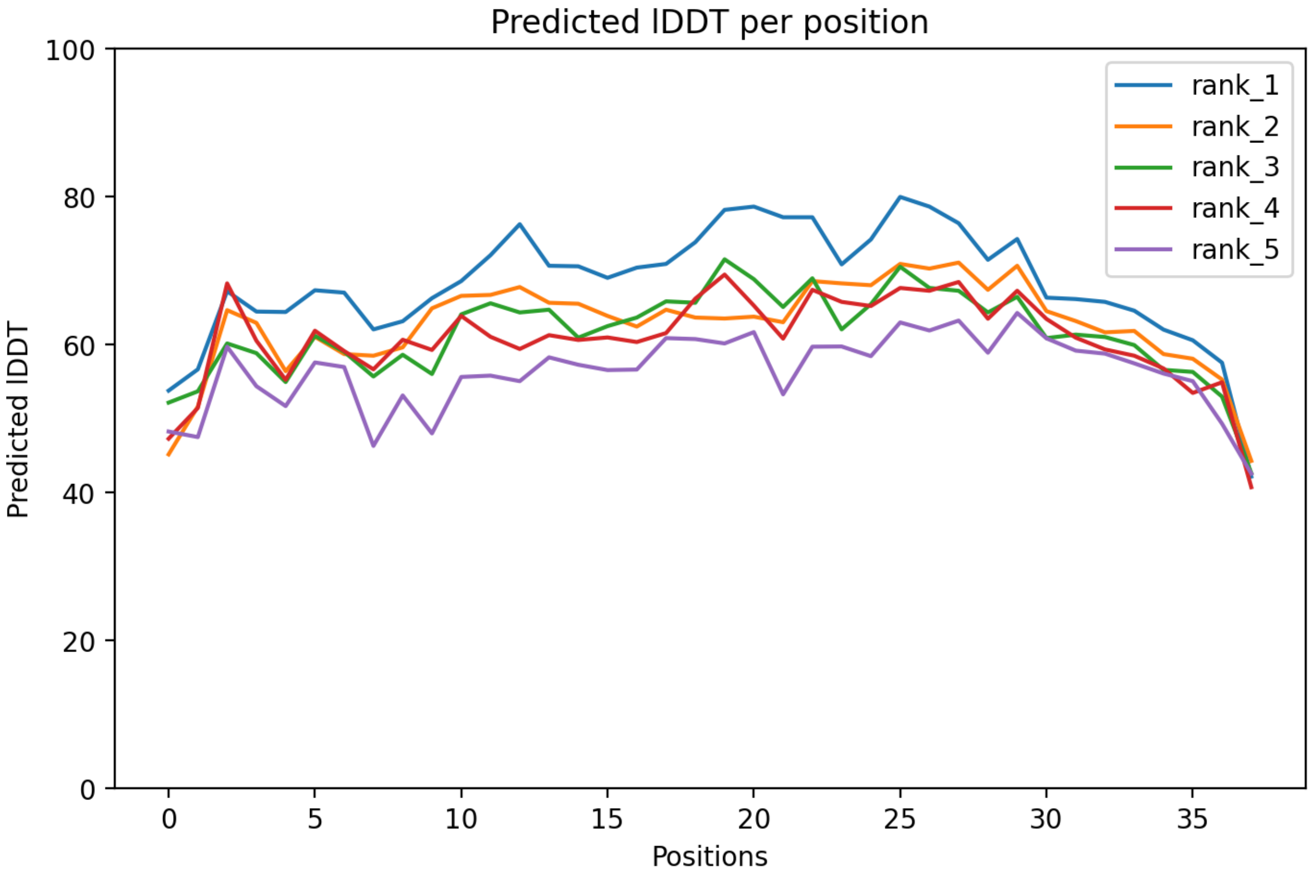

### CP16: MtrunA17_Chr1g0185811

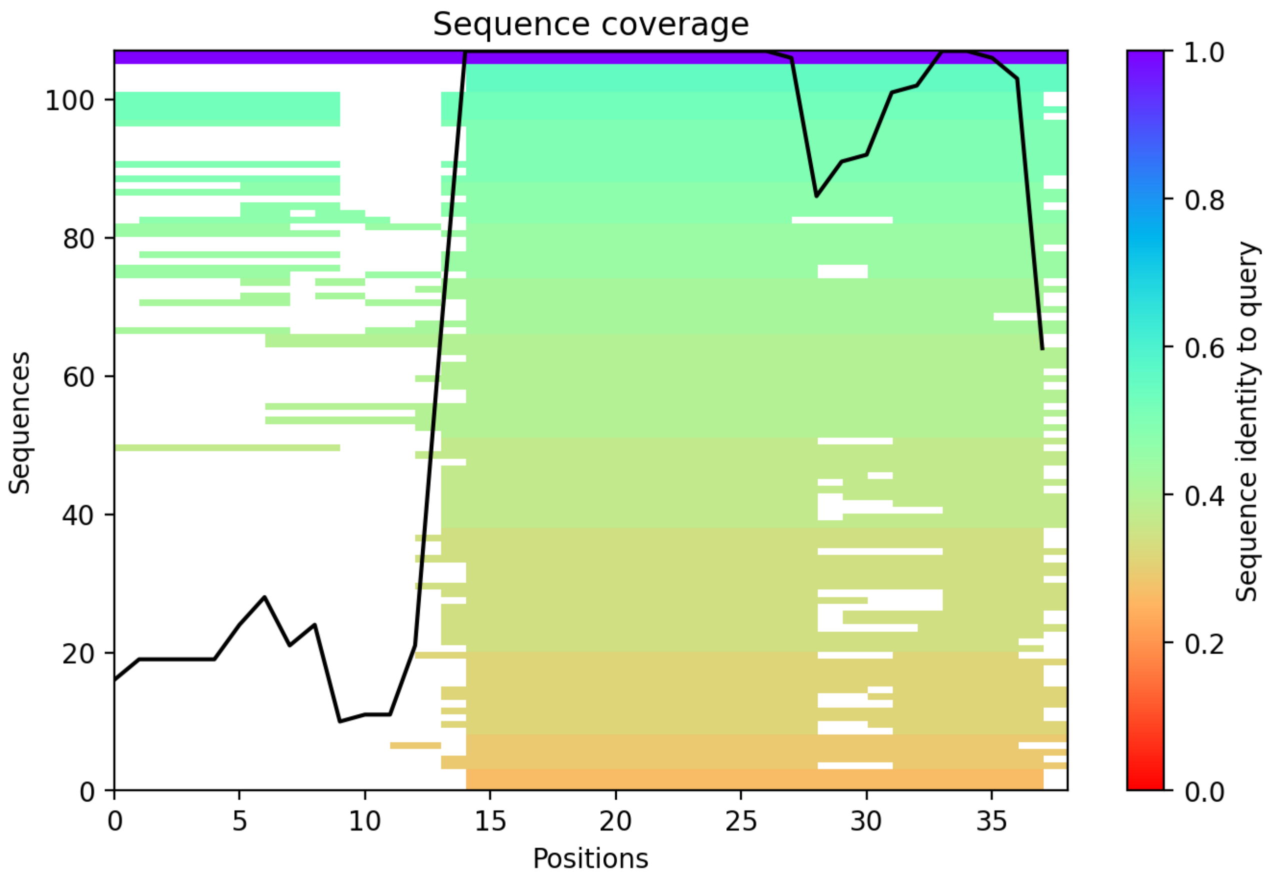

### CP17: MtrunA17_Chr1g0185811

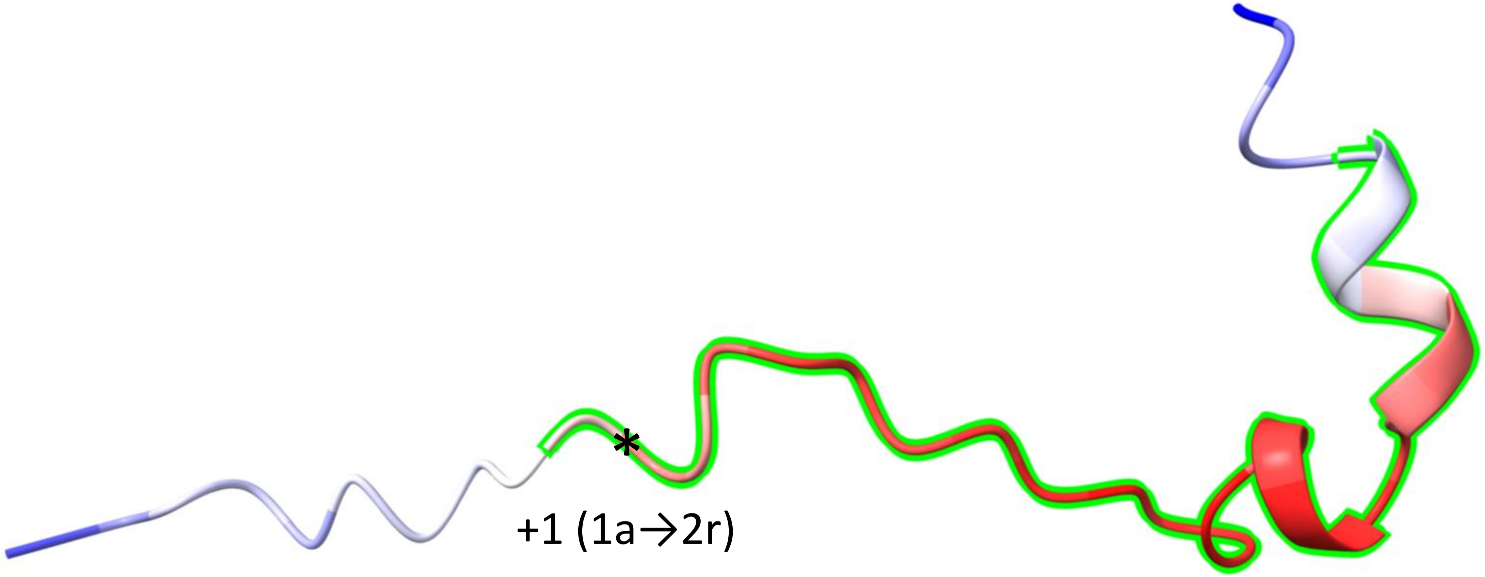


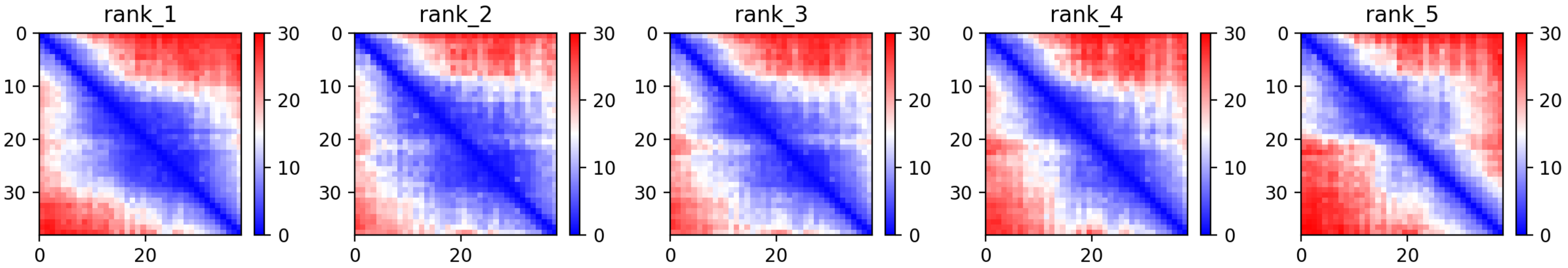

CP17: MtrunA17_Chr1g0185811

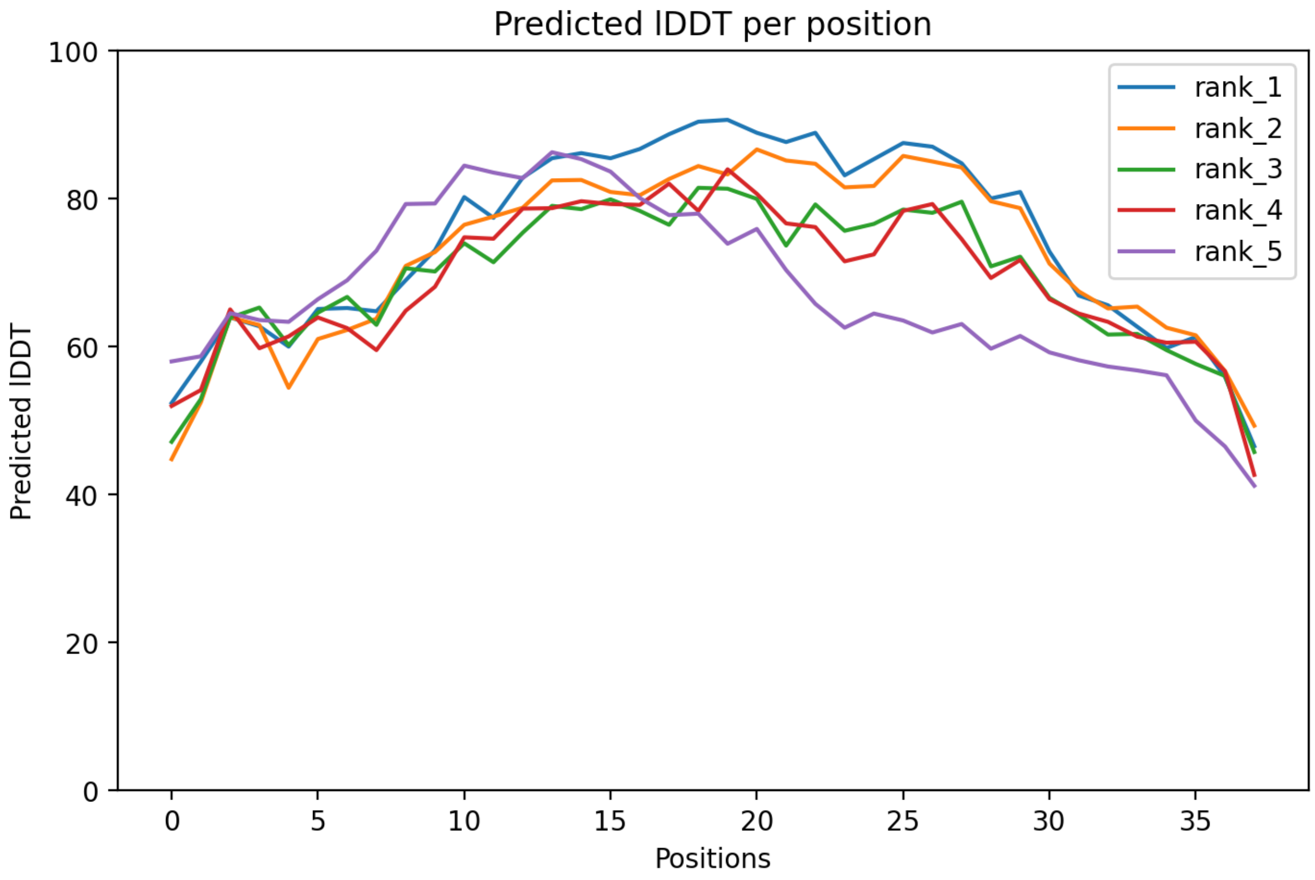

CP17: MtrunA17_Chr1g0185811

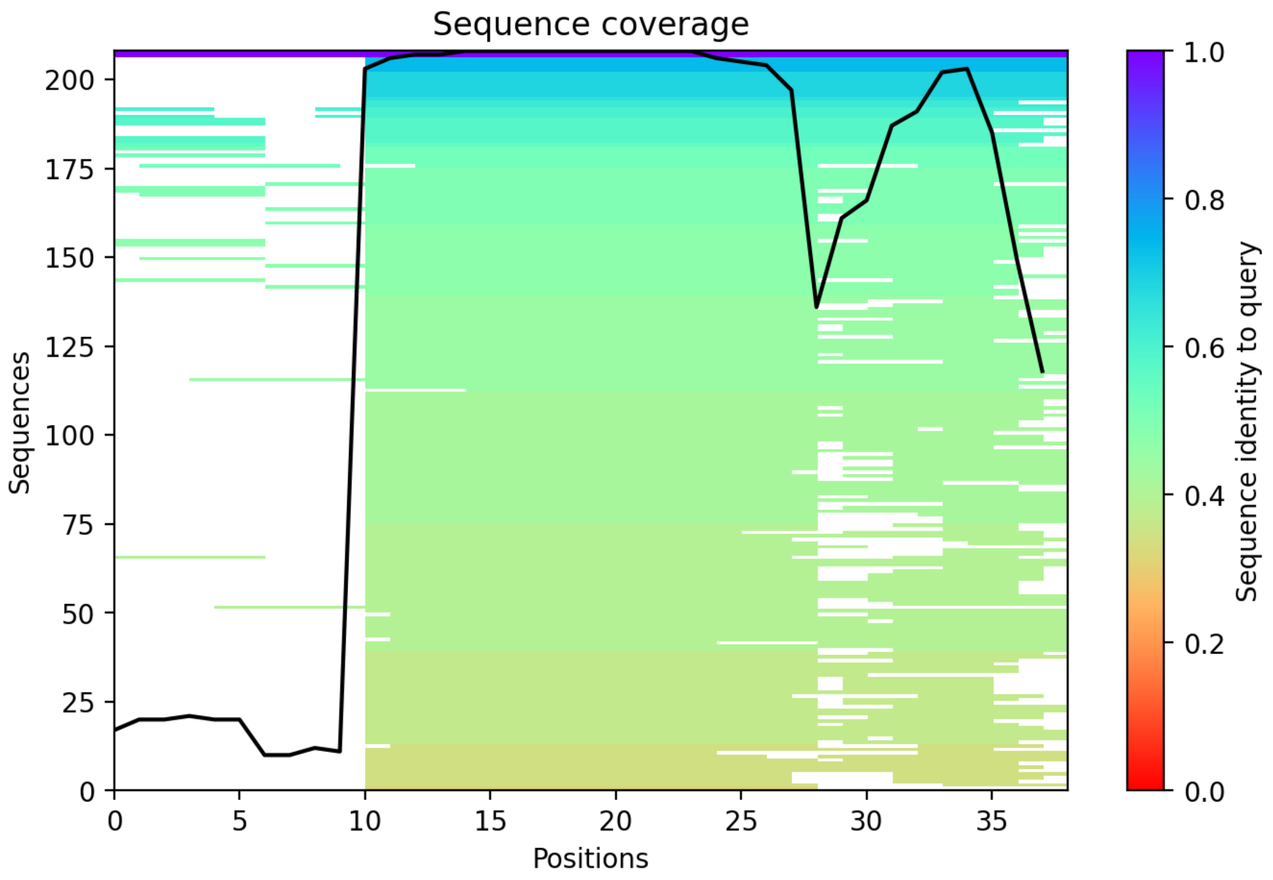

CP18: MtrunA17_Chr1g0185811

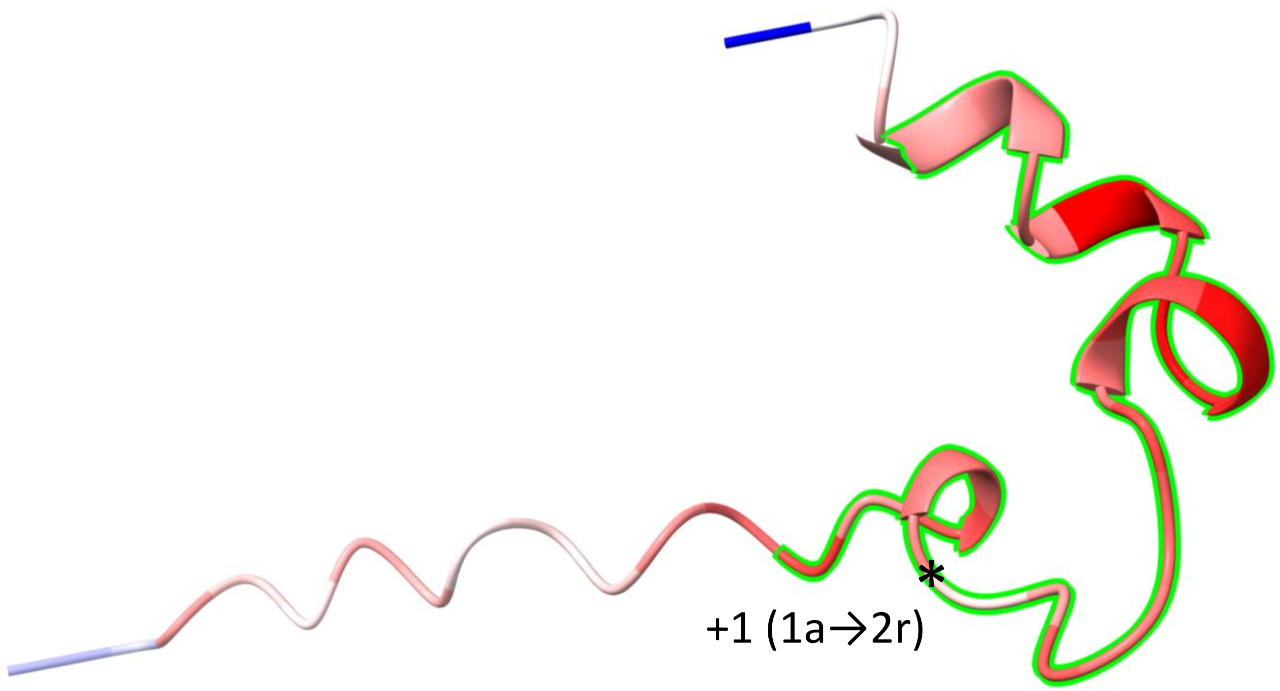


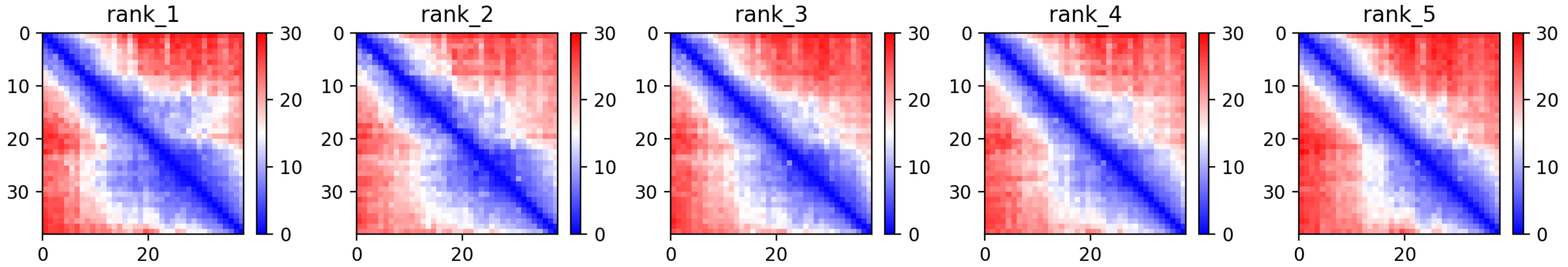

CP18: MtrunA17_Chr1g0185811

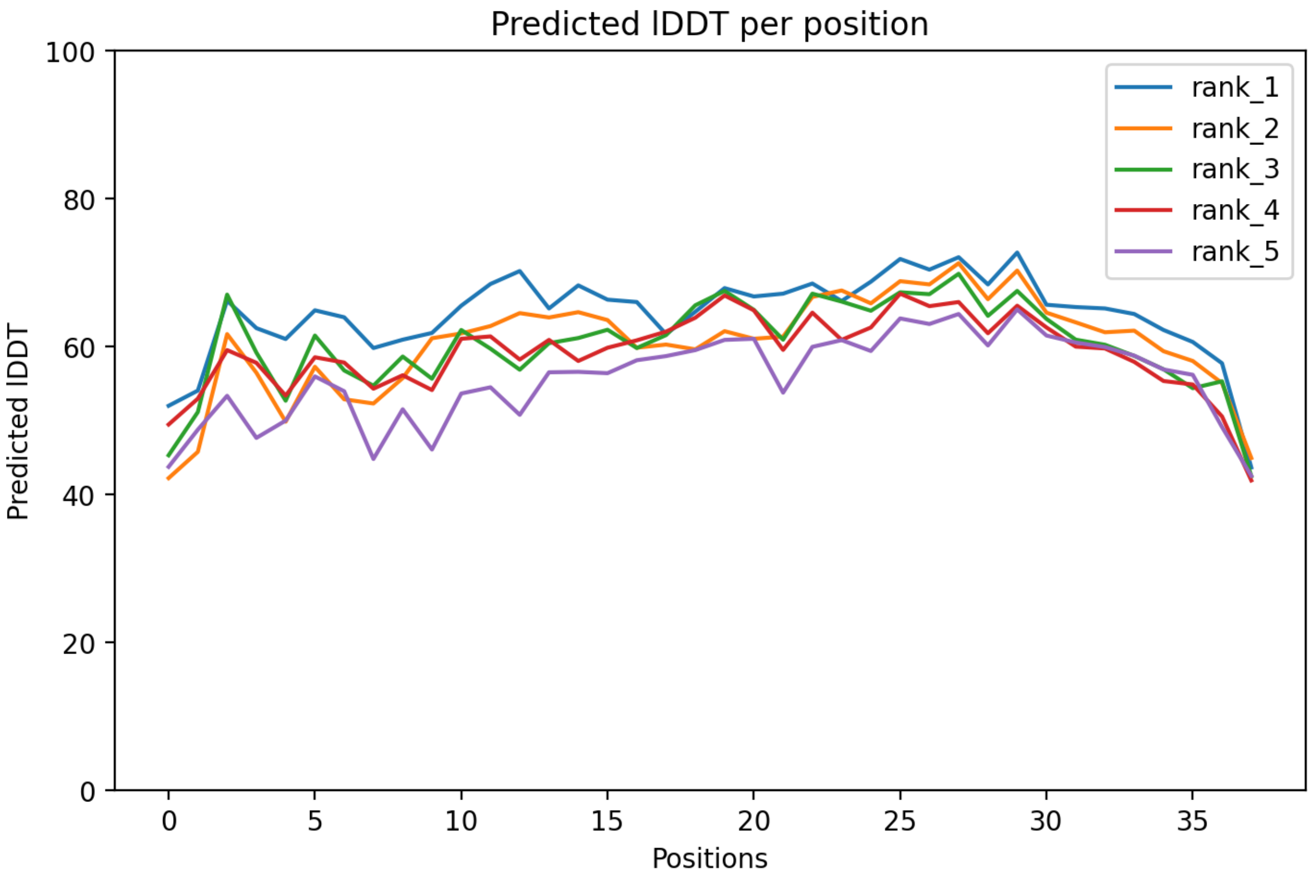

CP18: MtrunA17_Chr1g0185811

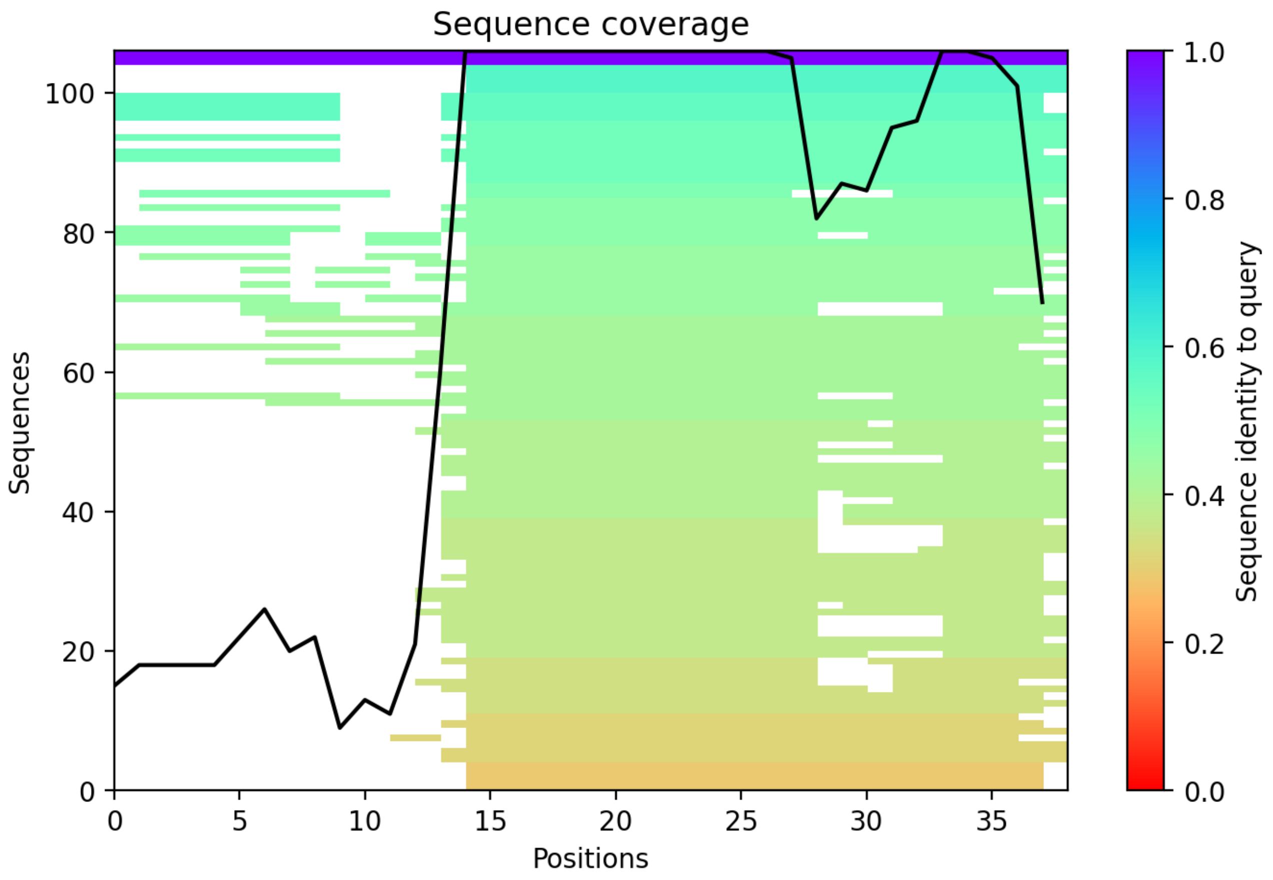

CP19: MtrunA17_Chr1g0185871

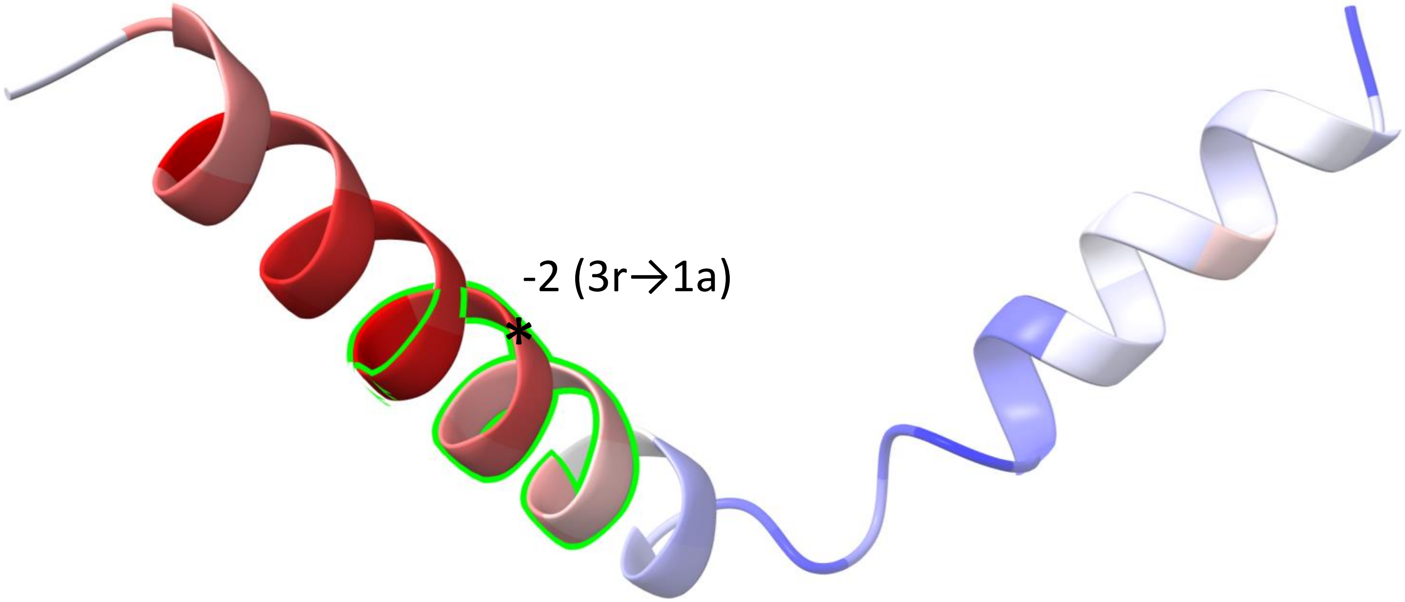


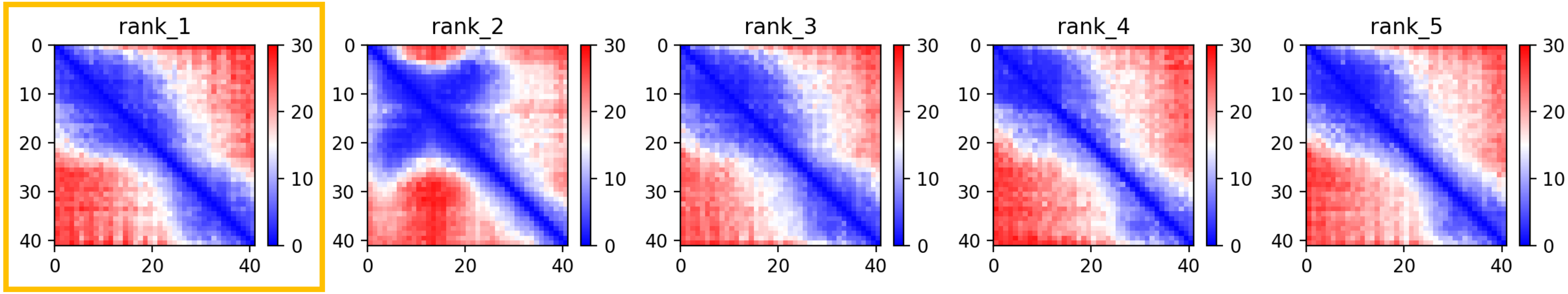

CP19: MtrunA17_Chr1g0185871

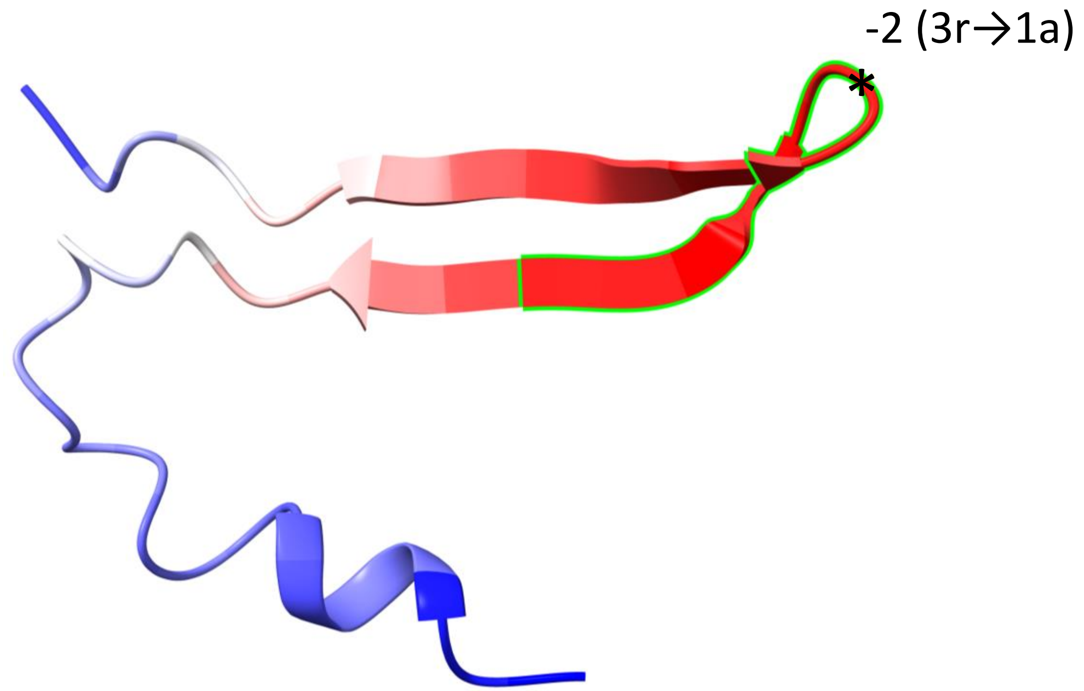


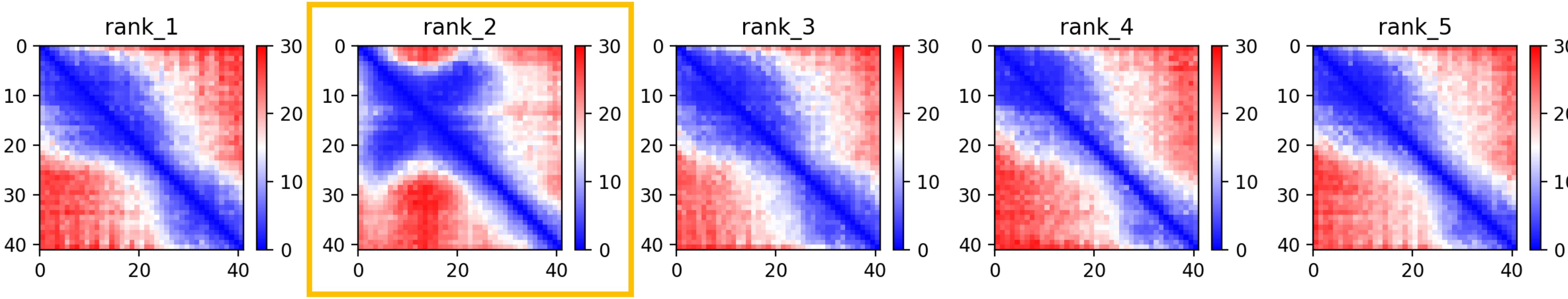

### CP19: MtrunA17_Chr1g0185871

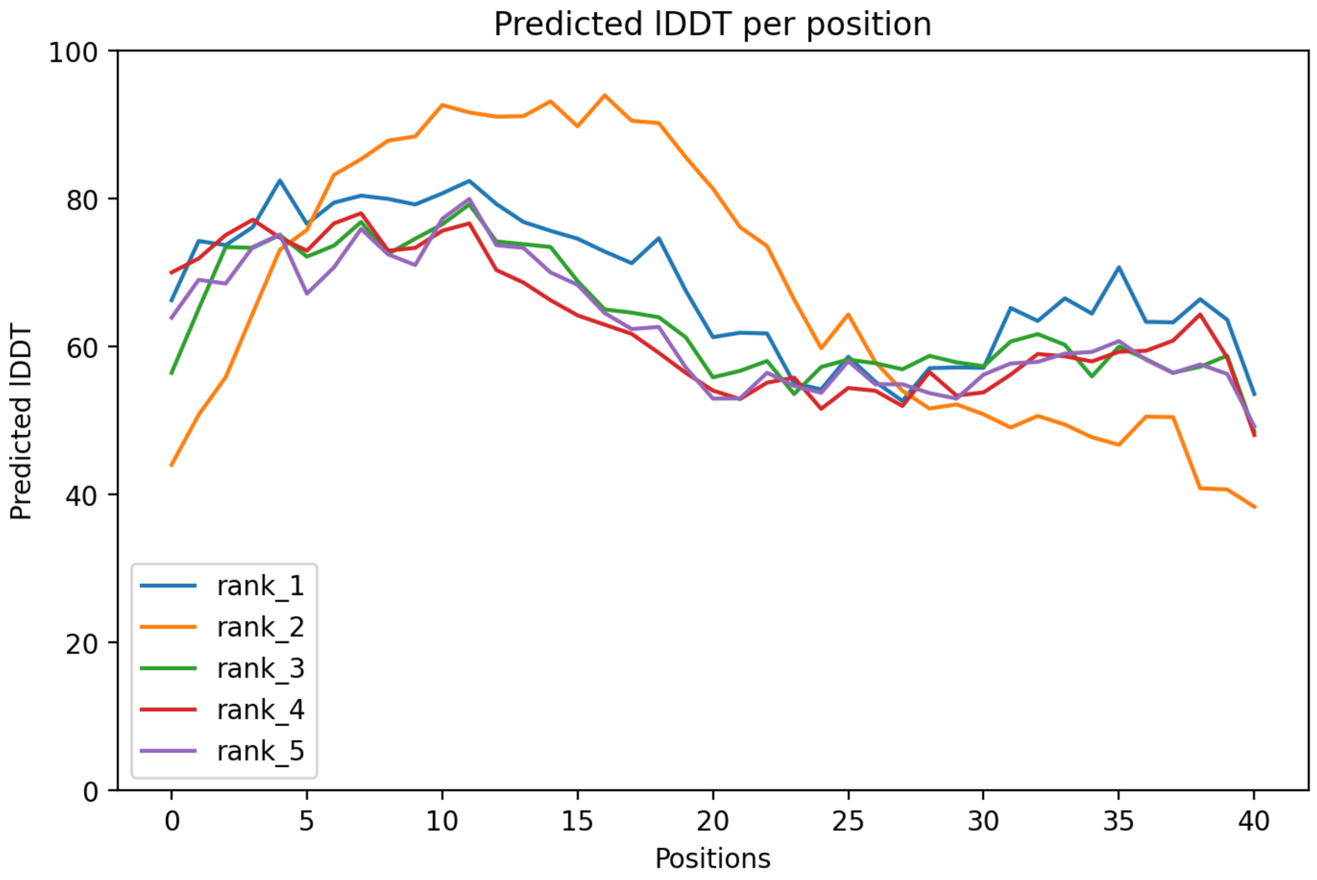

### CP19: MtrunA17_Chr1g0185871

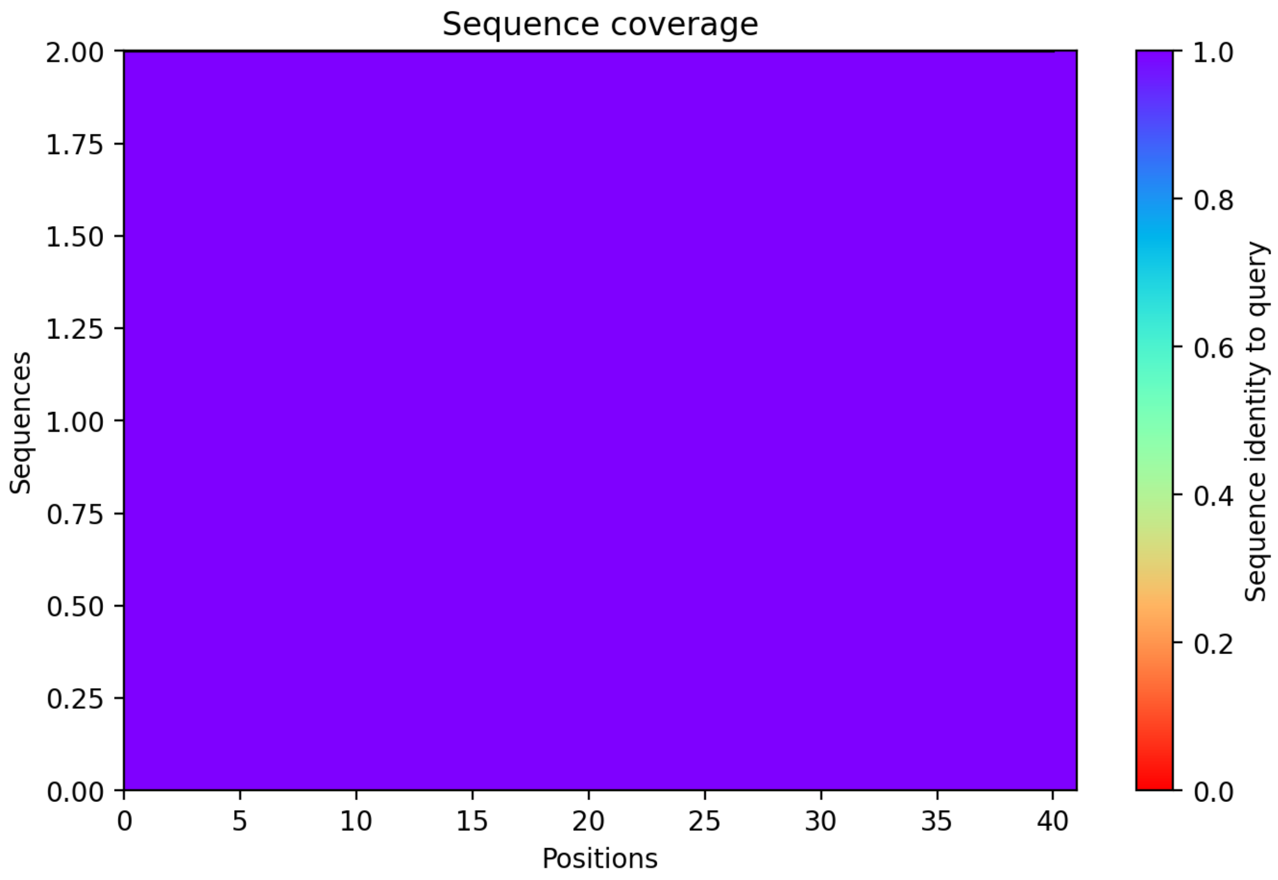

CP20: MtrunA17_Chr1g0190571

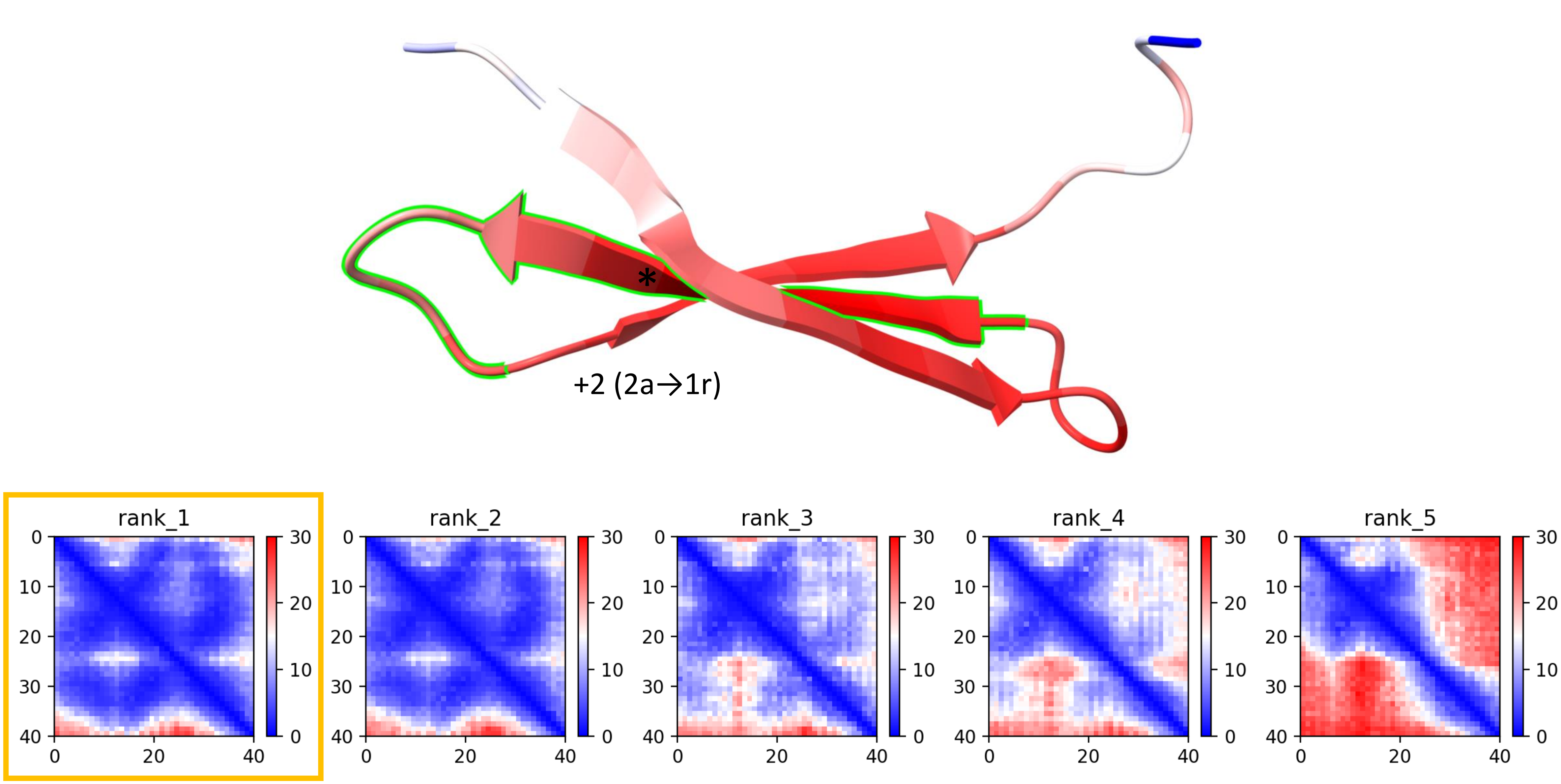

CP20: MtrunA17_Chr1g0190571

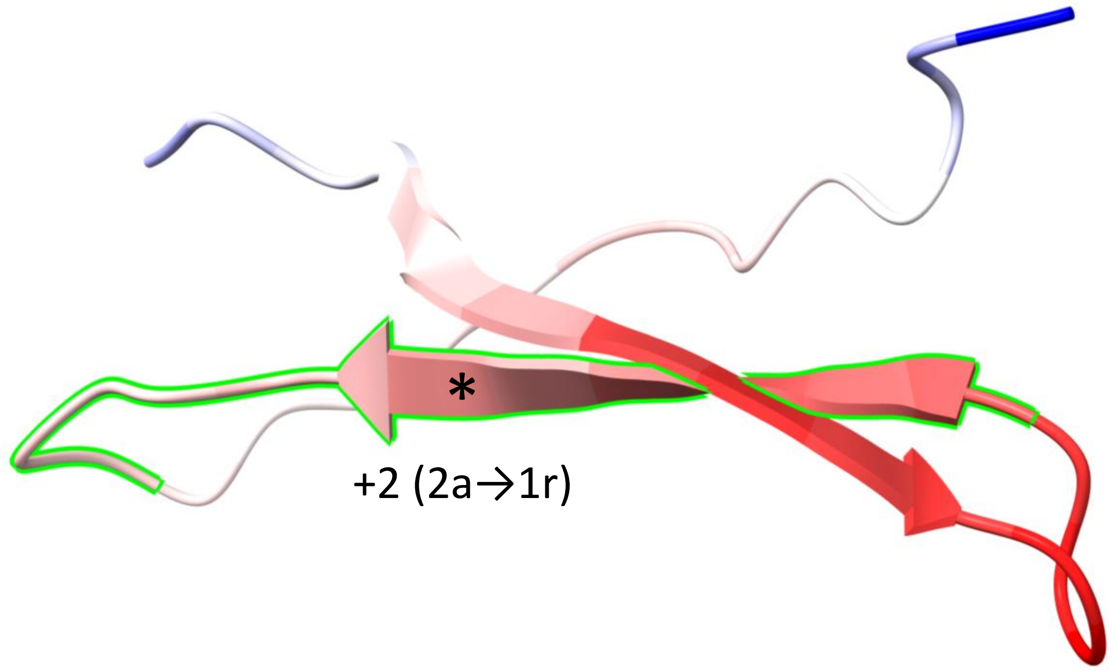


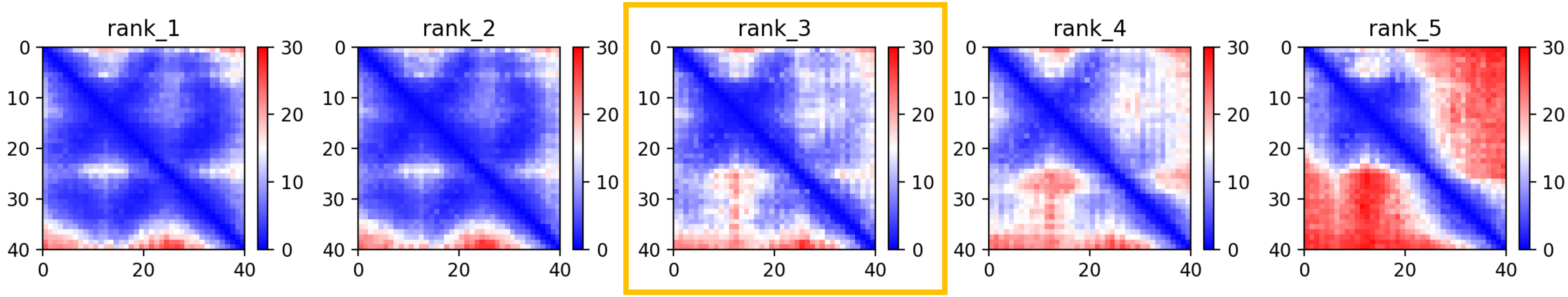

CP20: MtrunA17_Chr1g0190571

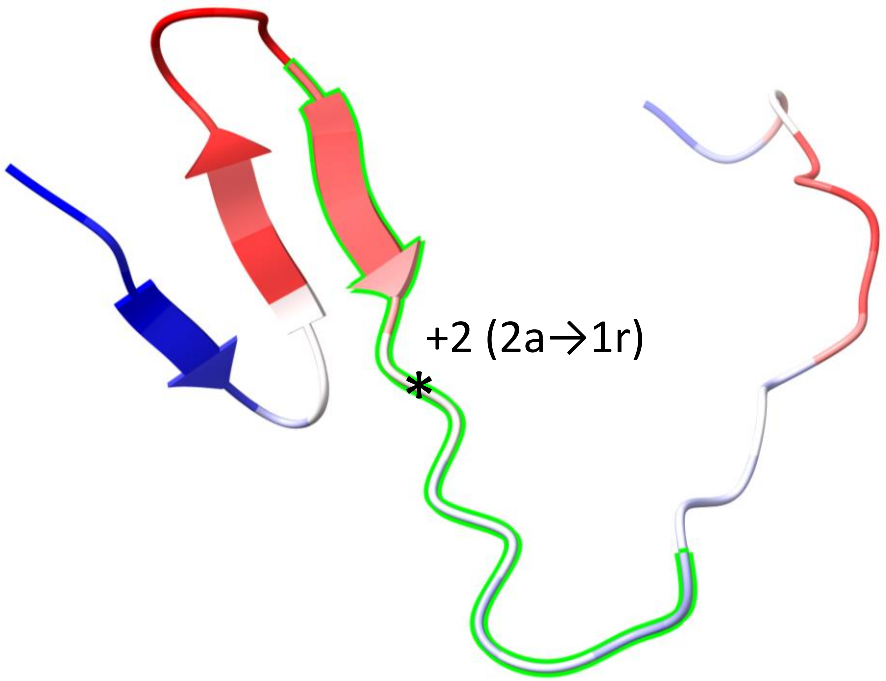


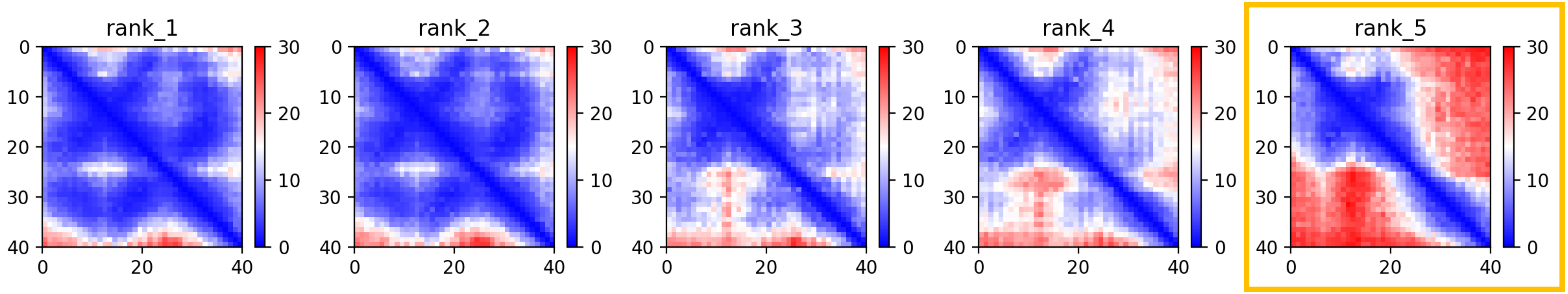

CP20: MtrunA17_Chr1g0190571

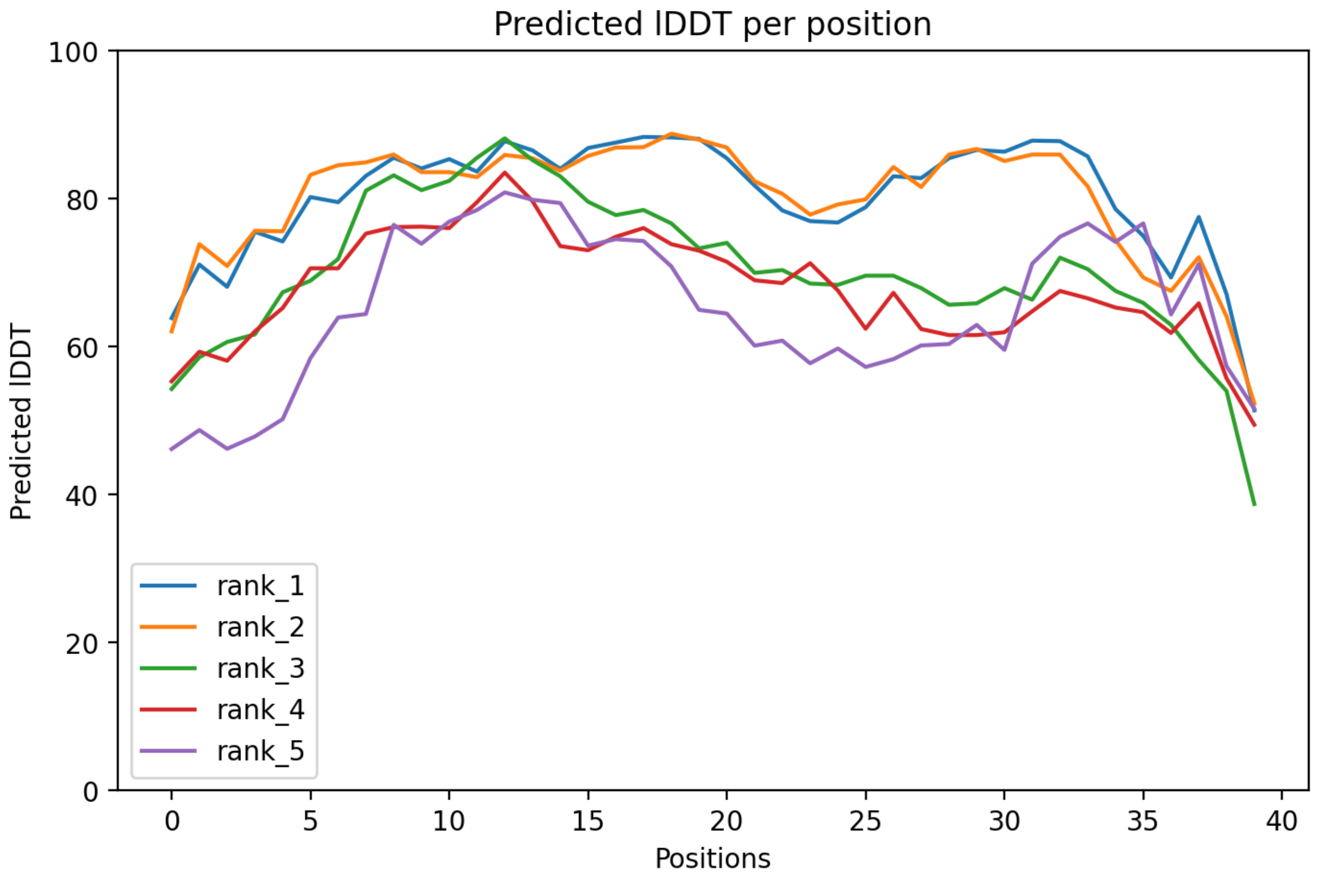

CP20: MtrunA17_Chr1g0190571

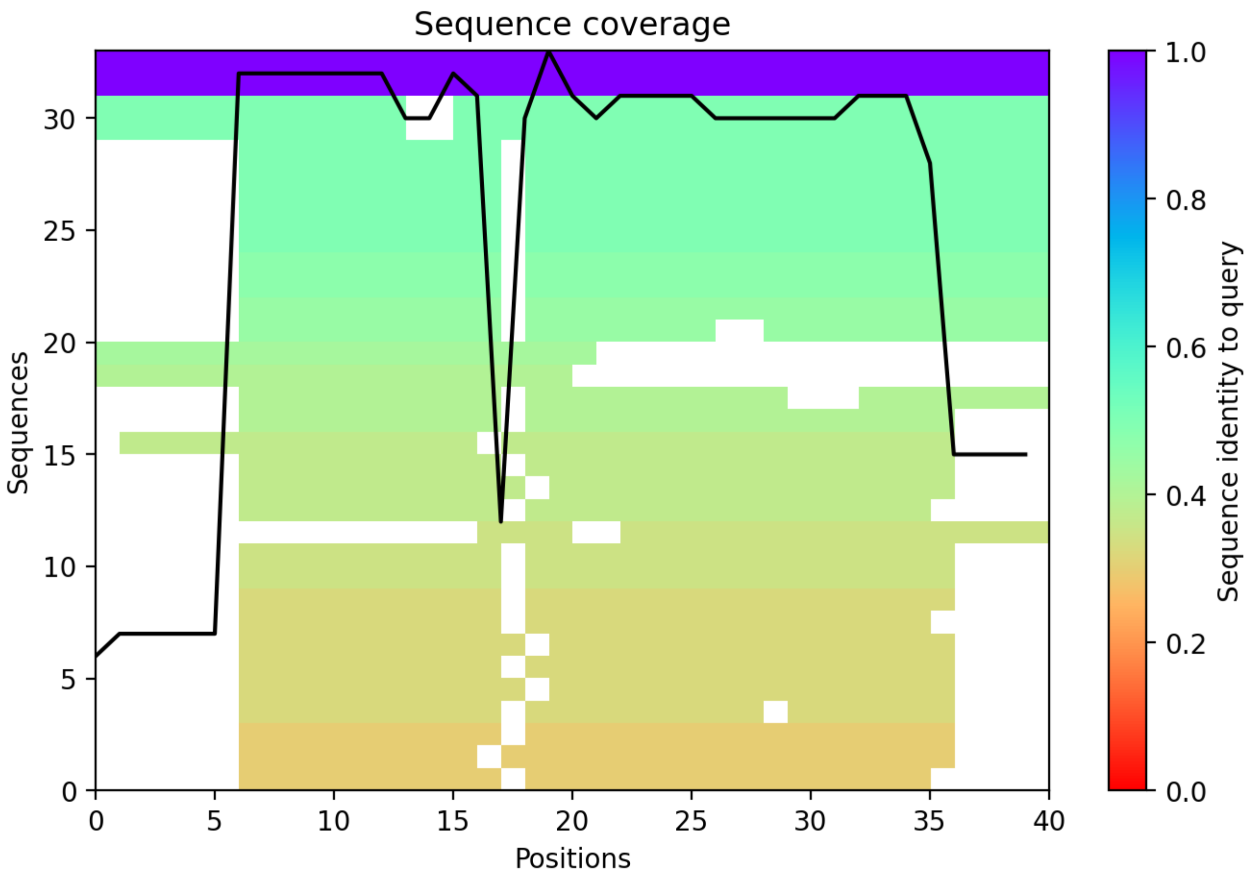

CP21: MtrunA17_Chr1g0191411

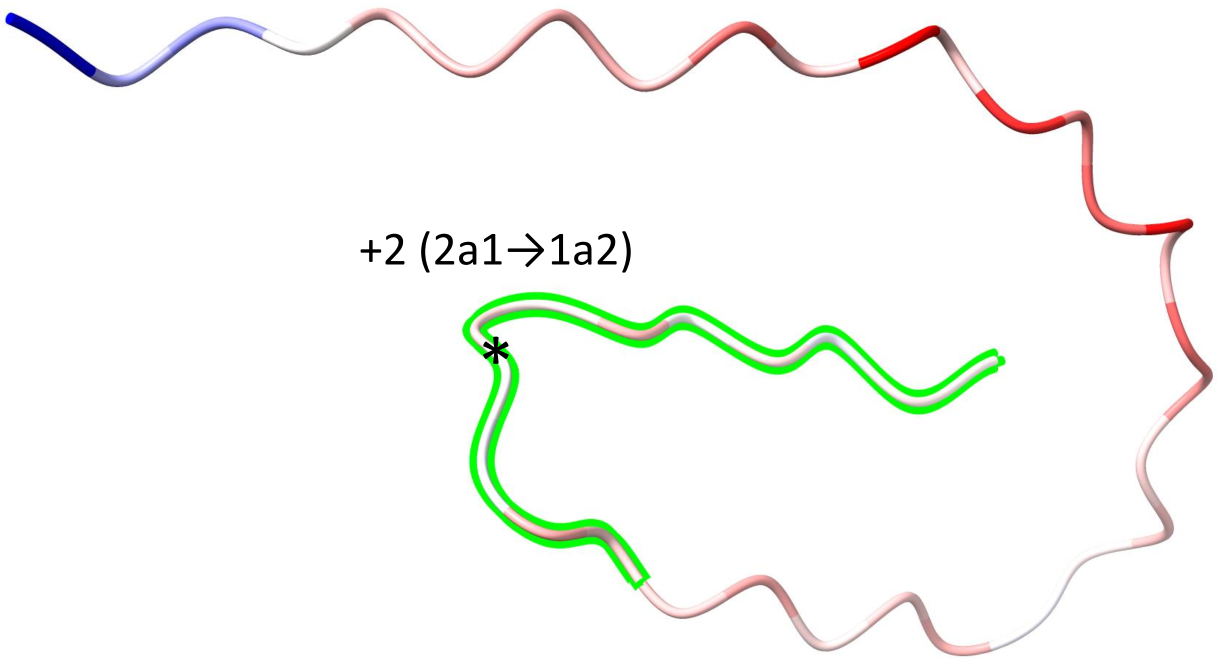


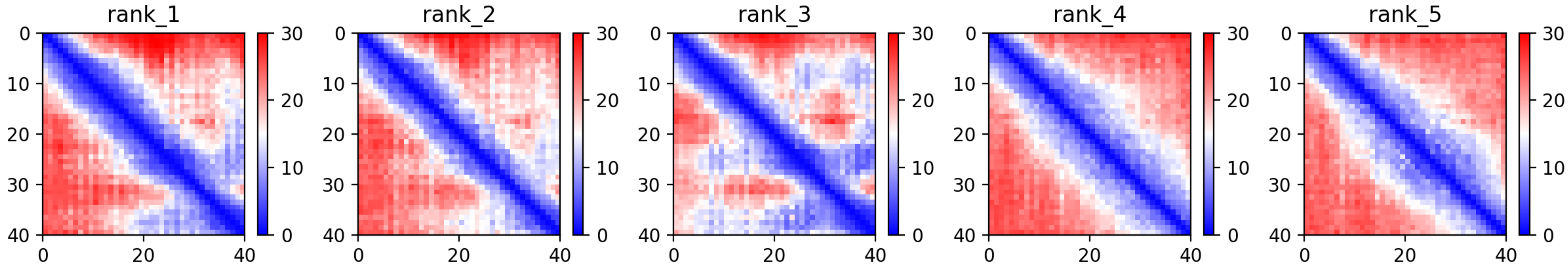

CP21: MtrunA17_Chr1g0191411

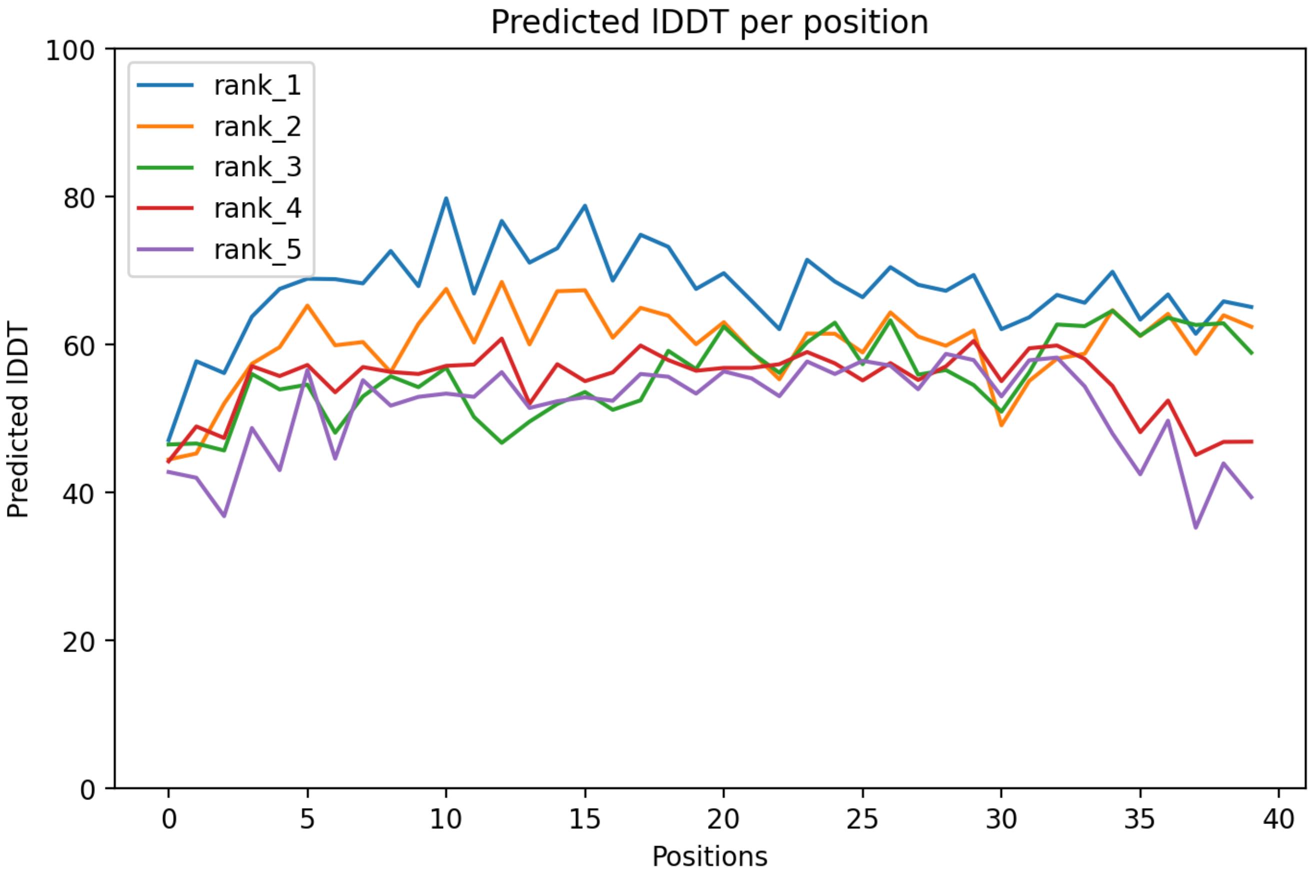

CP21: MtrunA17_Chr1g0191411

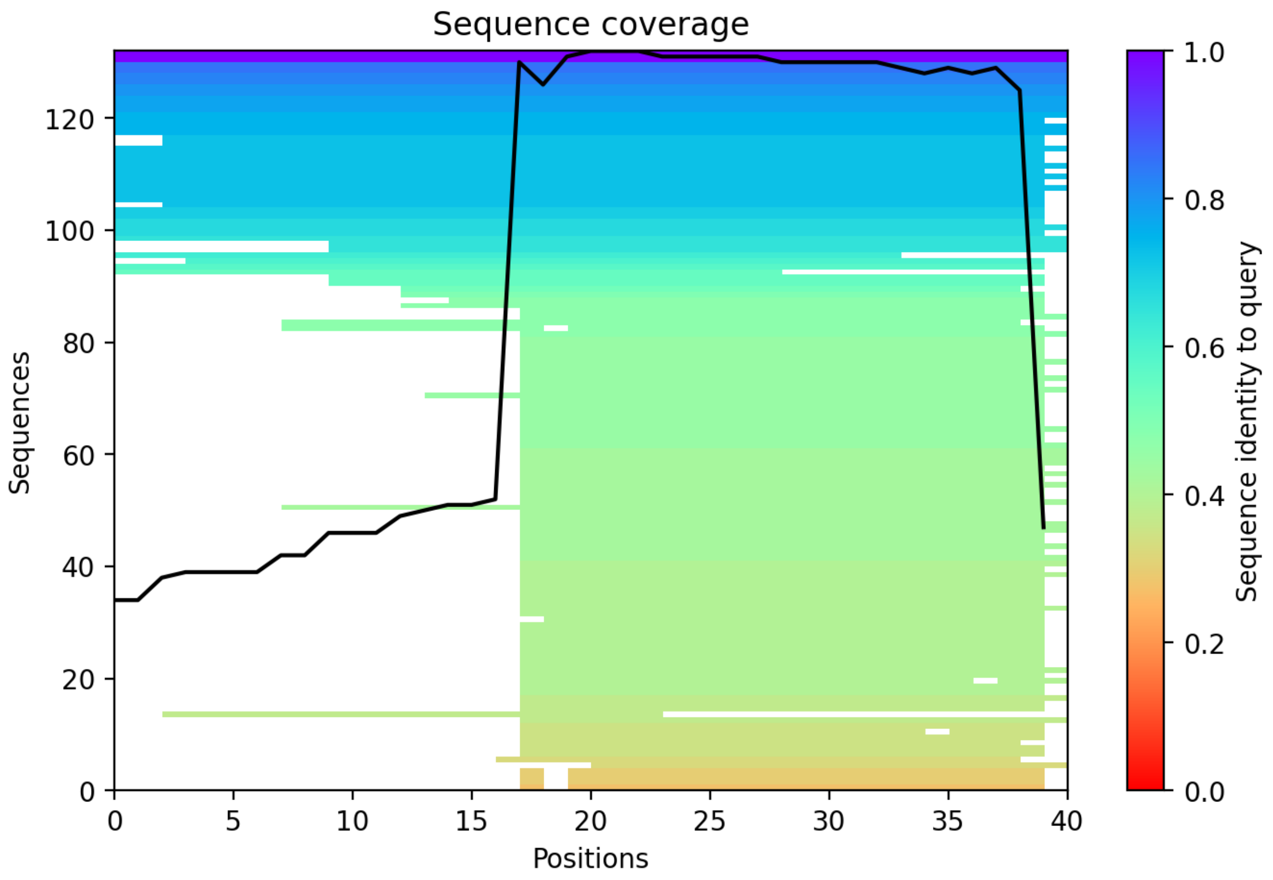

CP22: MtrunA17_Chr1g0198091

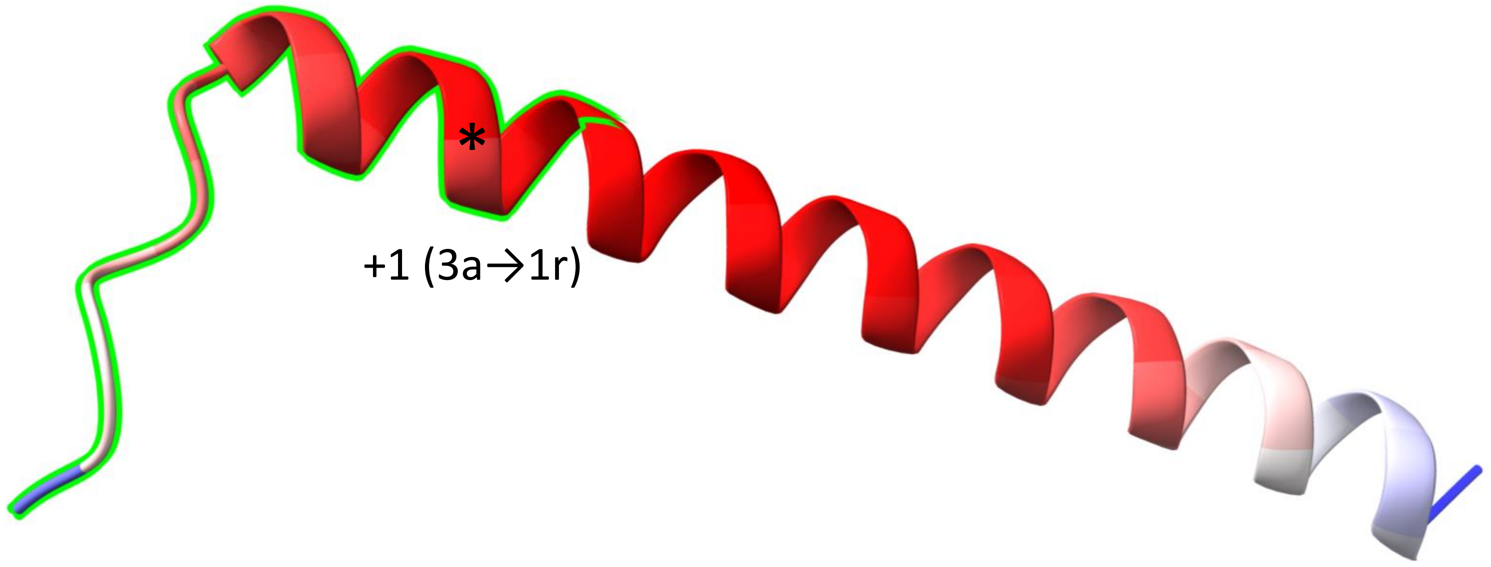


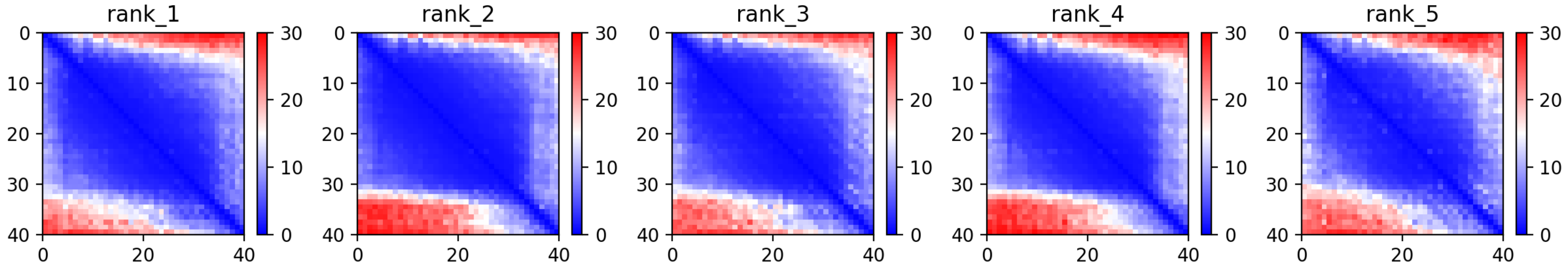

### CP22: MtrunA17_Chr1g0198091

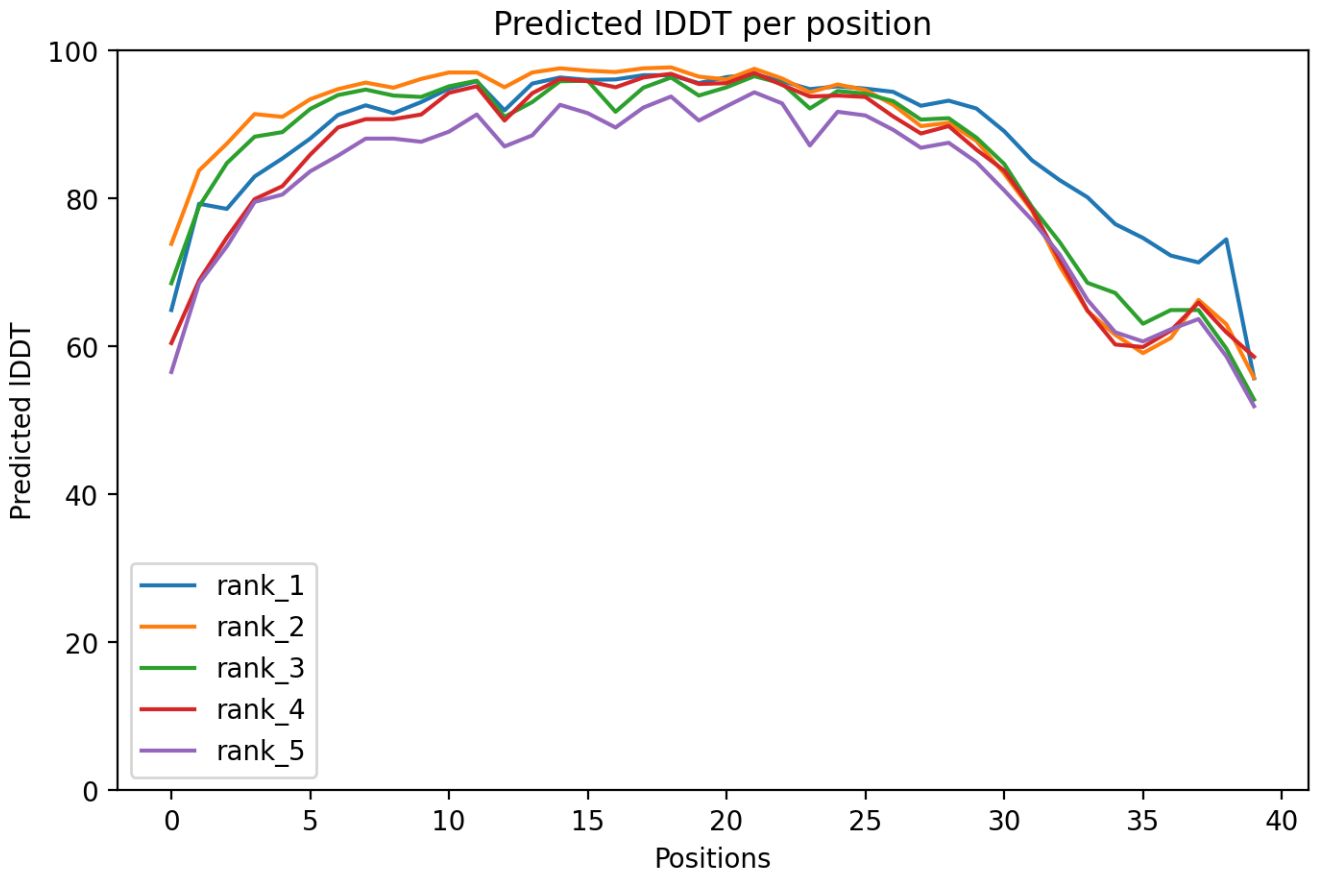

CP22: MtrunA17_Chr1g0198091

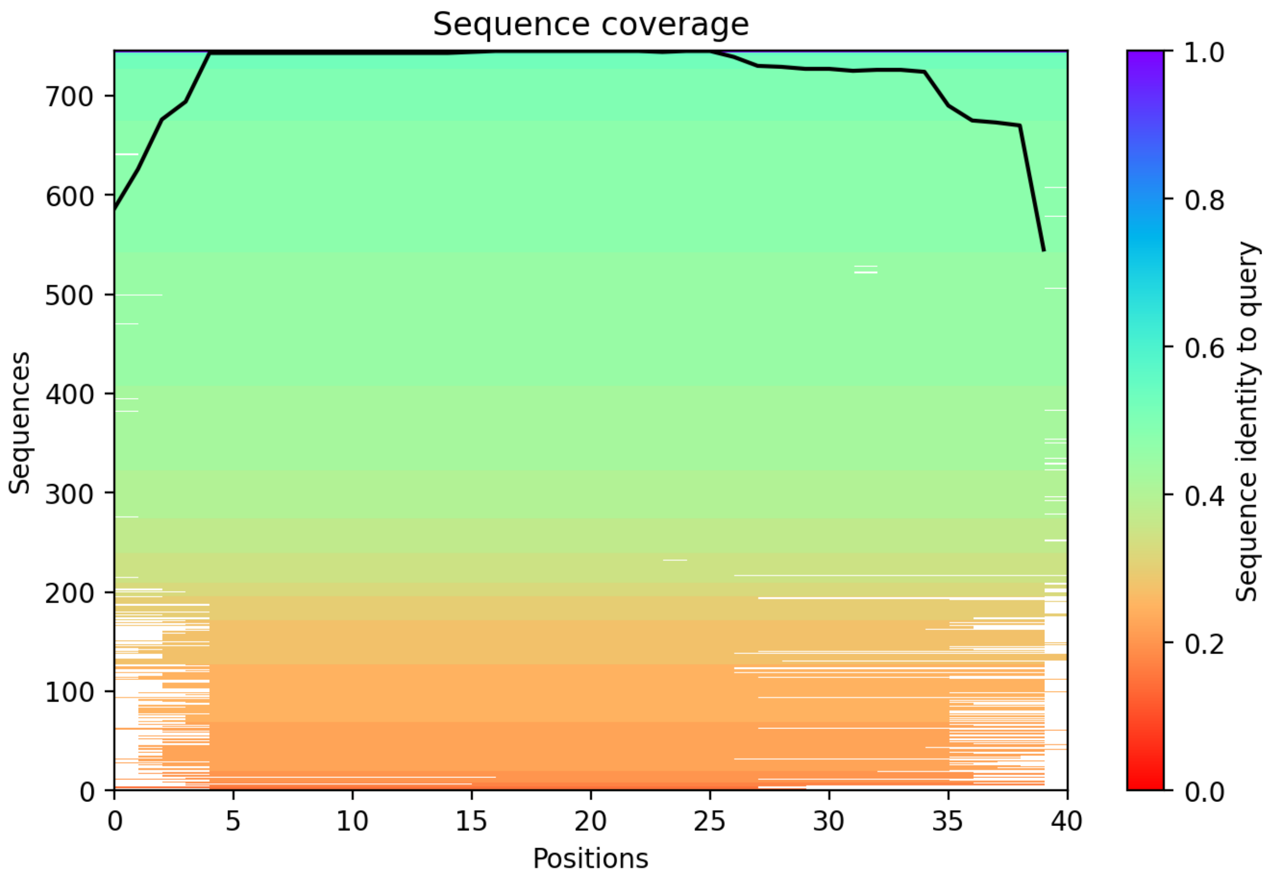

CP23: MtrunA17_Chr1g0200071

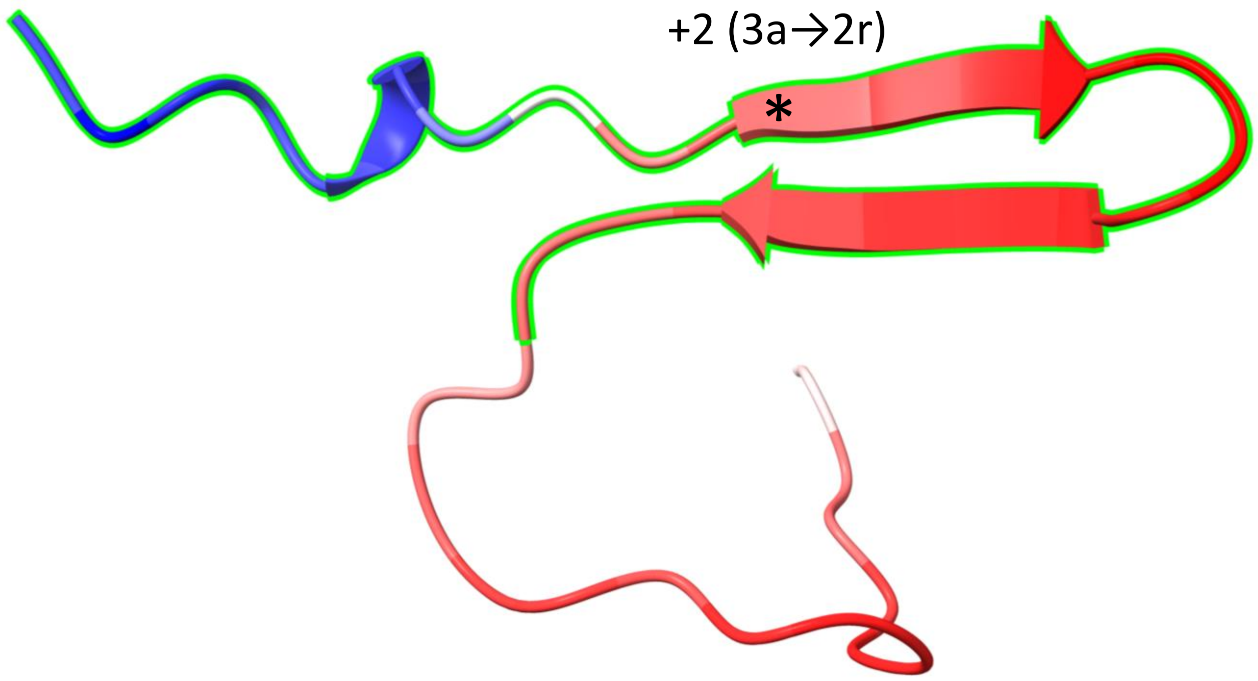


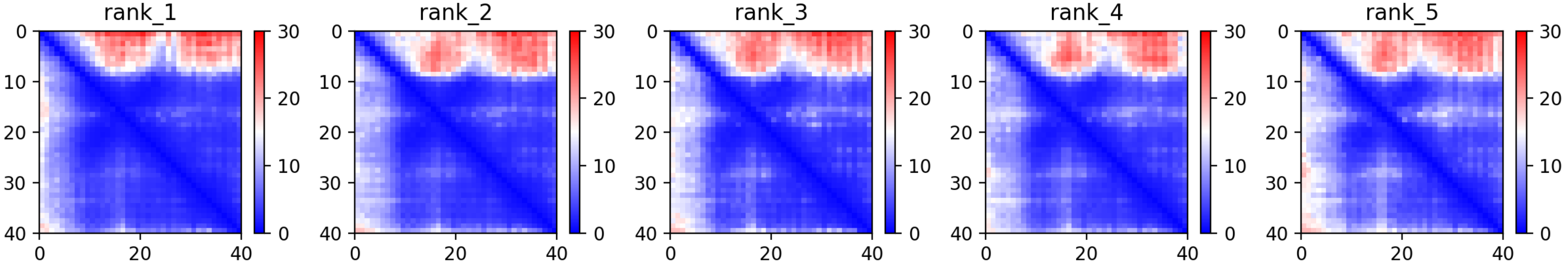

CP23: MtrunA17_Chr1g0200071

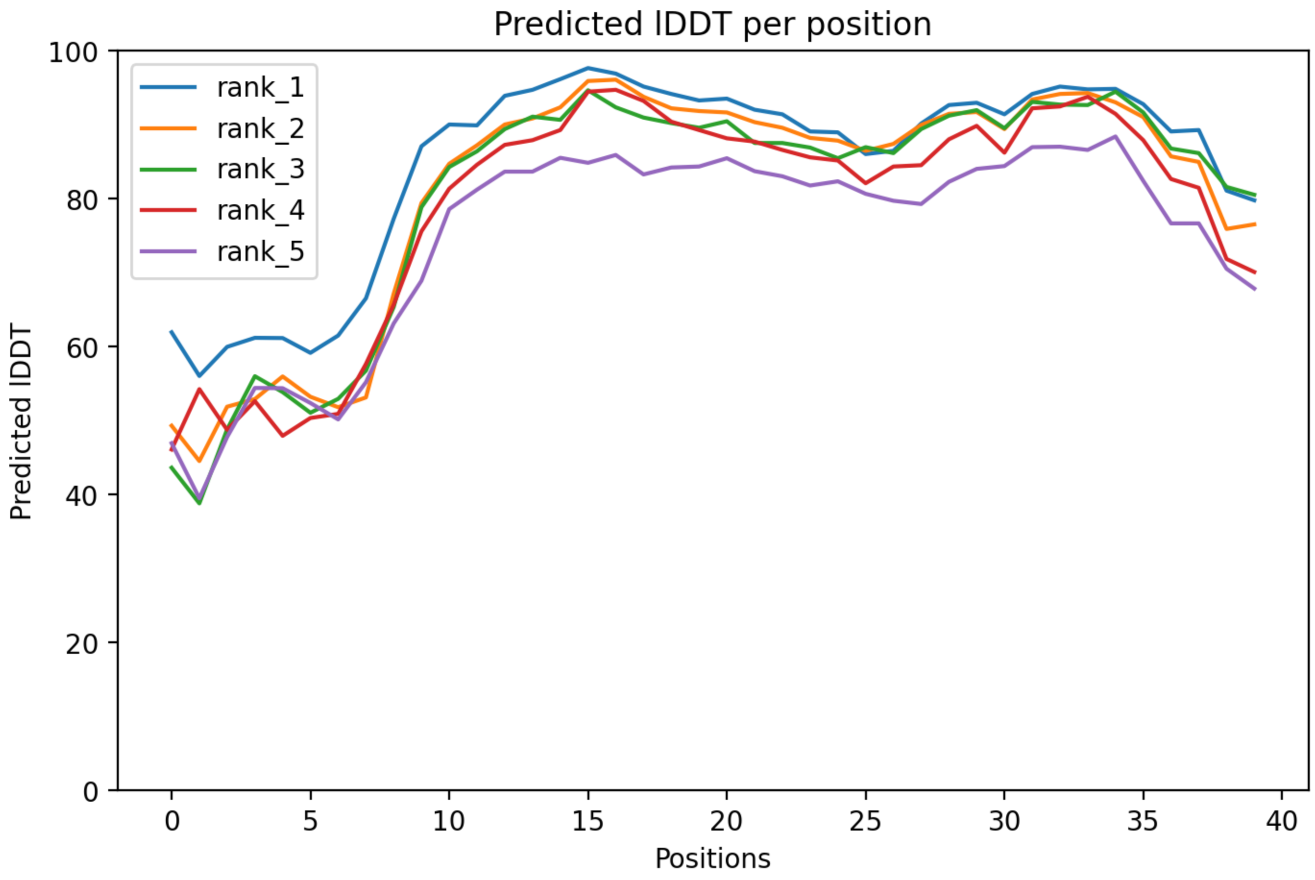

CP23: MtrunA17_Chr1g0200071

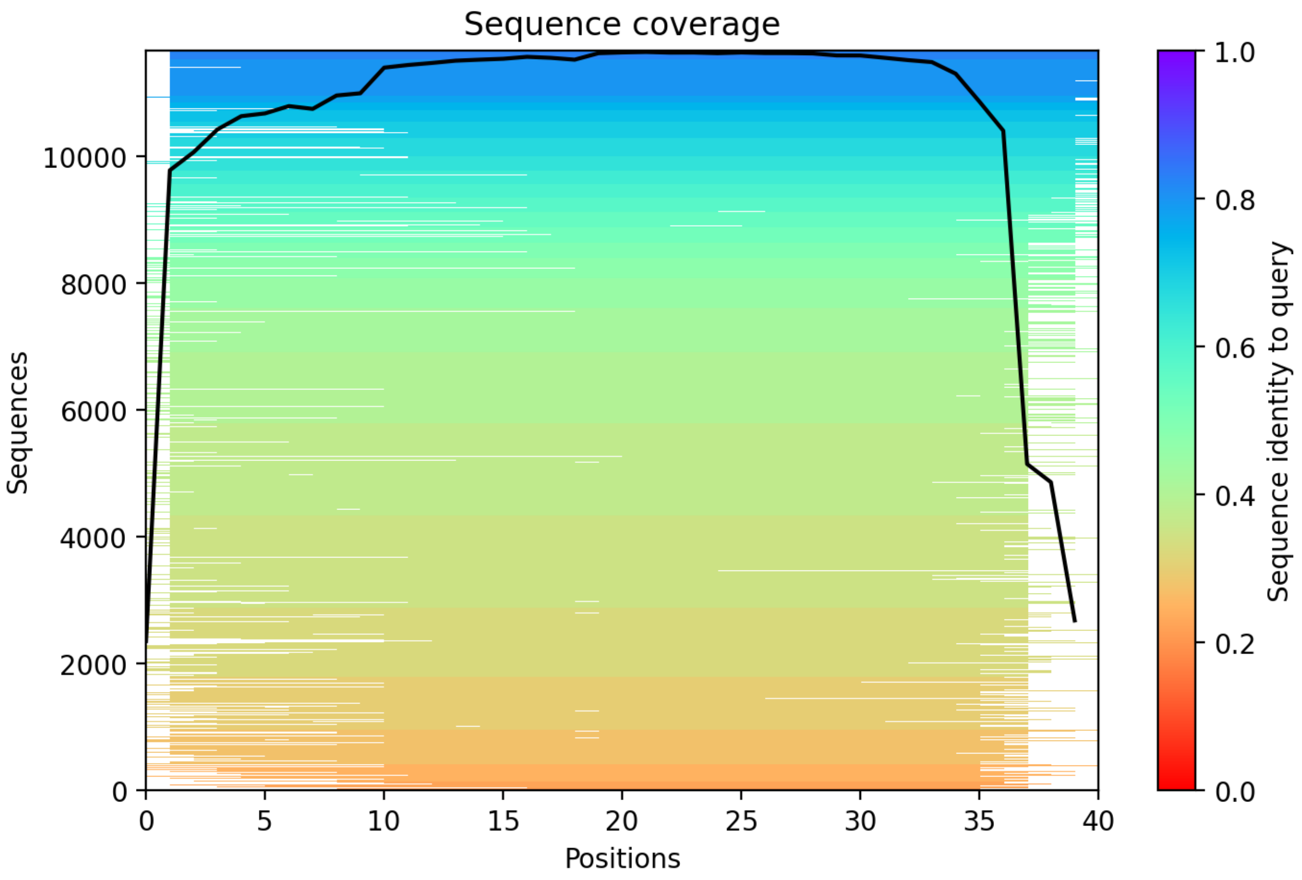

CP24: MtrunA17_Chr1g0200071

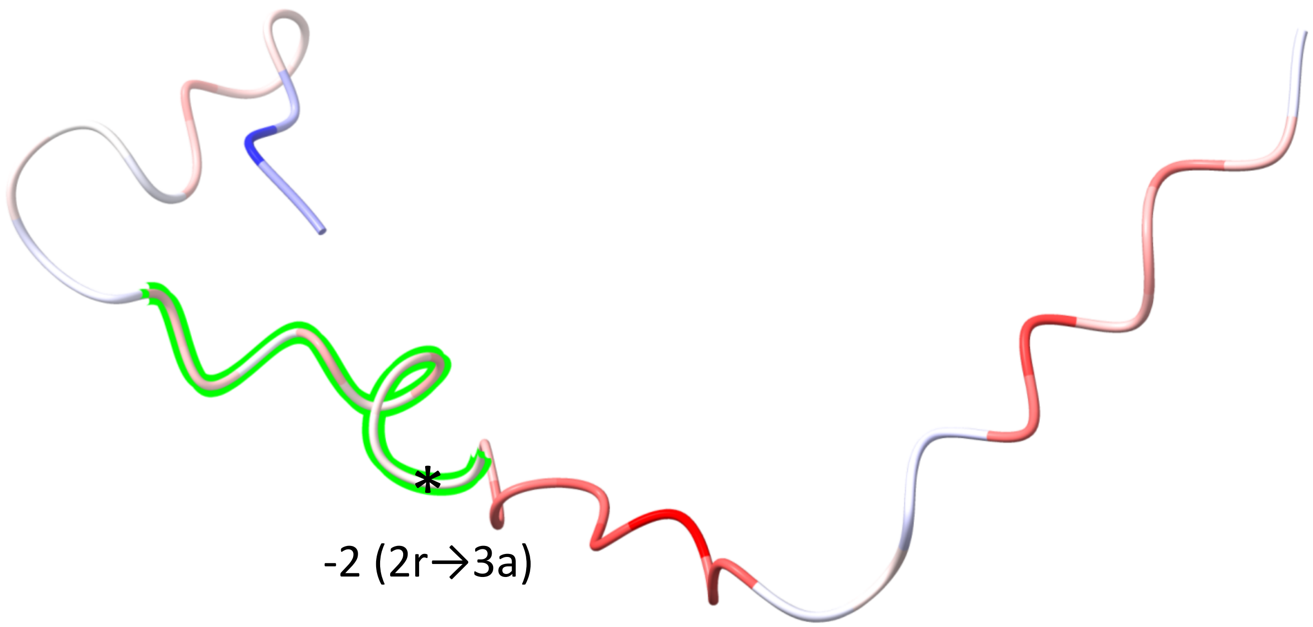


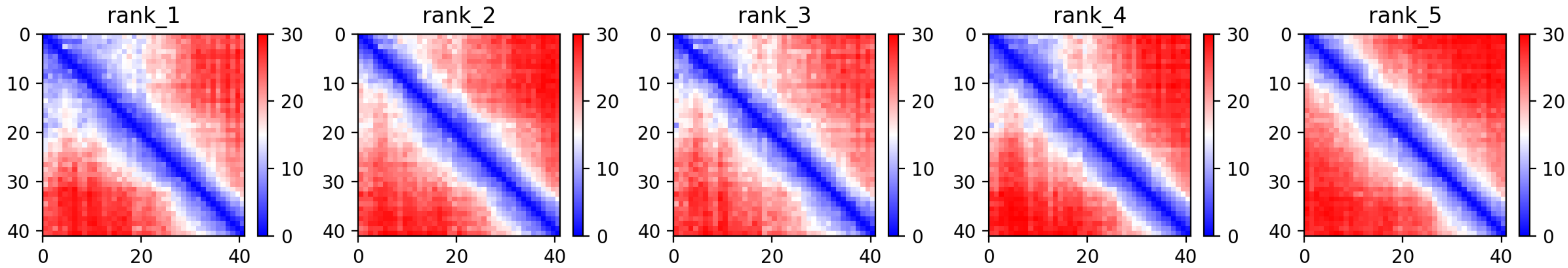

CP24: MtrunA17_Chr1g0200071

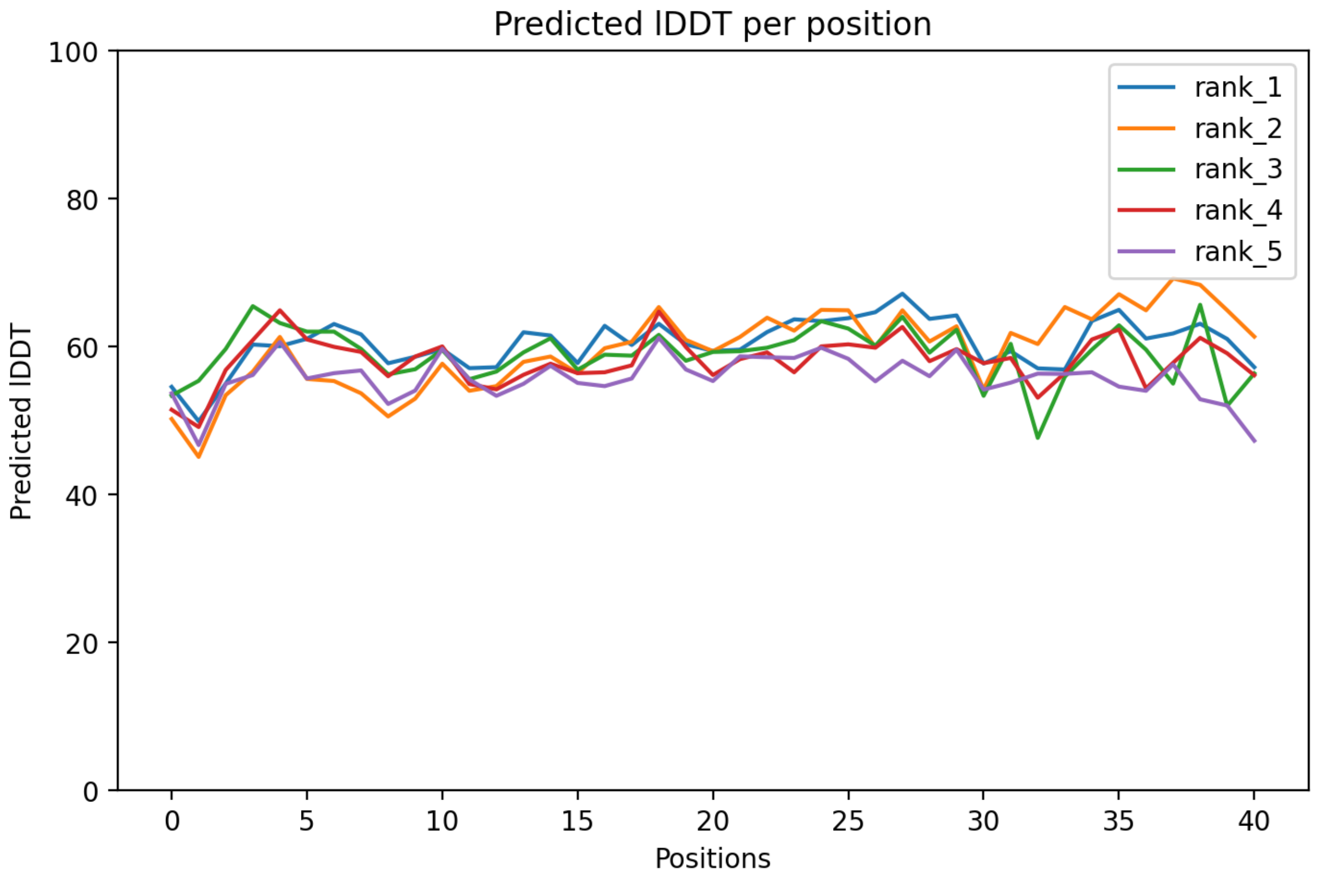

### CP24: MtrunA17_Chr1g0200071

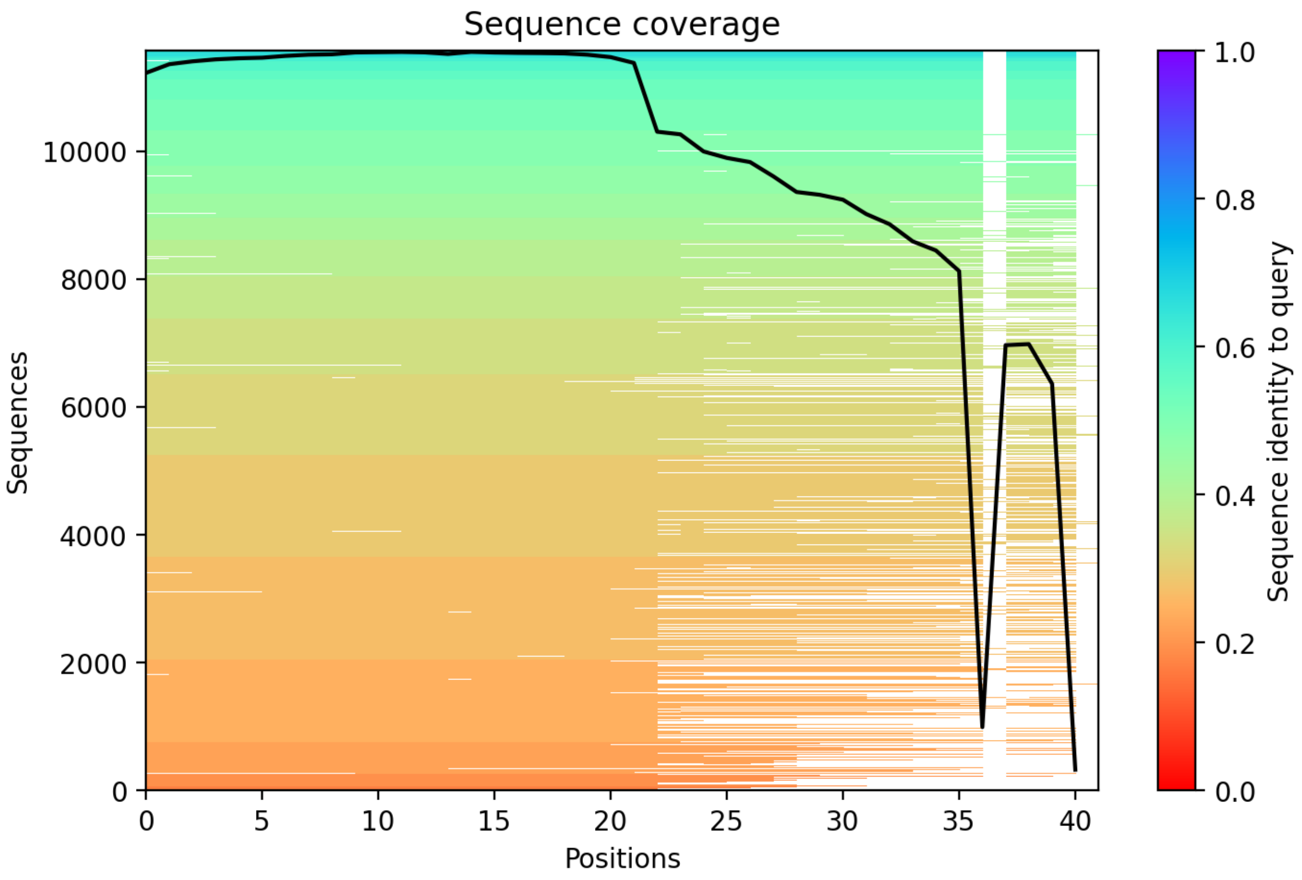

CP25: MtrunA17_Chr1g0202001

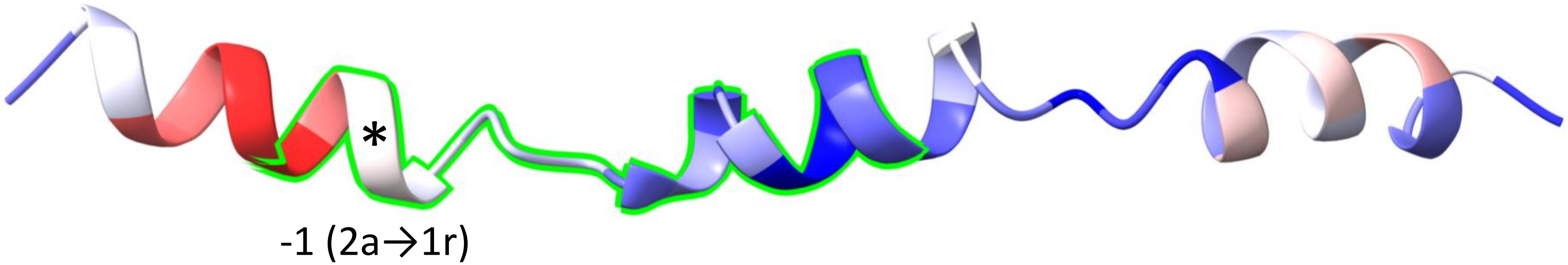


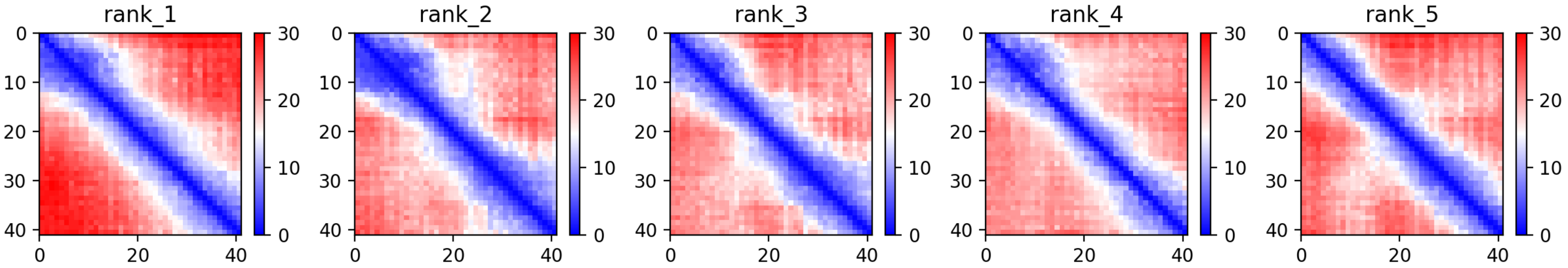

CP25: MtrunA17_Chr1g0202001

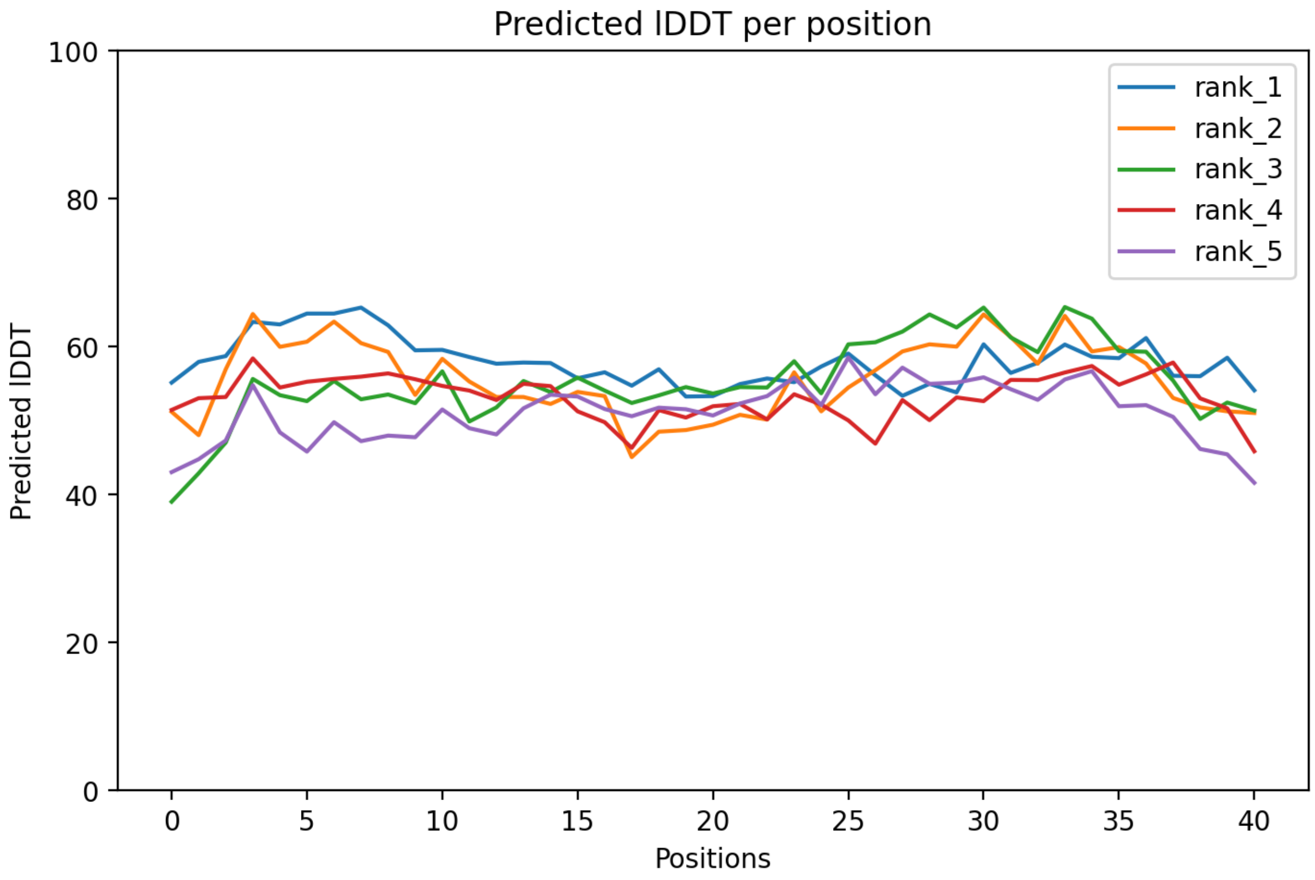

CP25: MtrunA17_Chr1g0202001

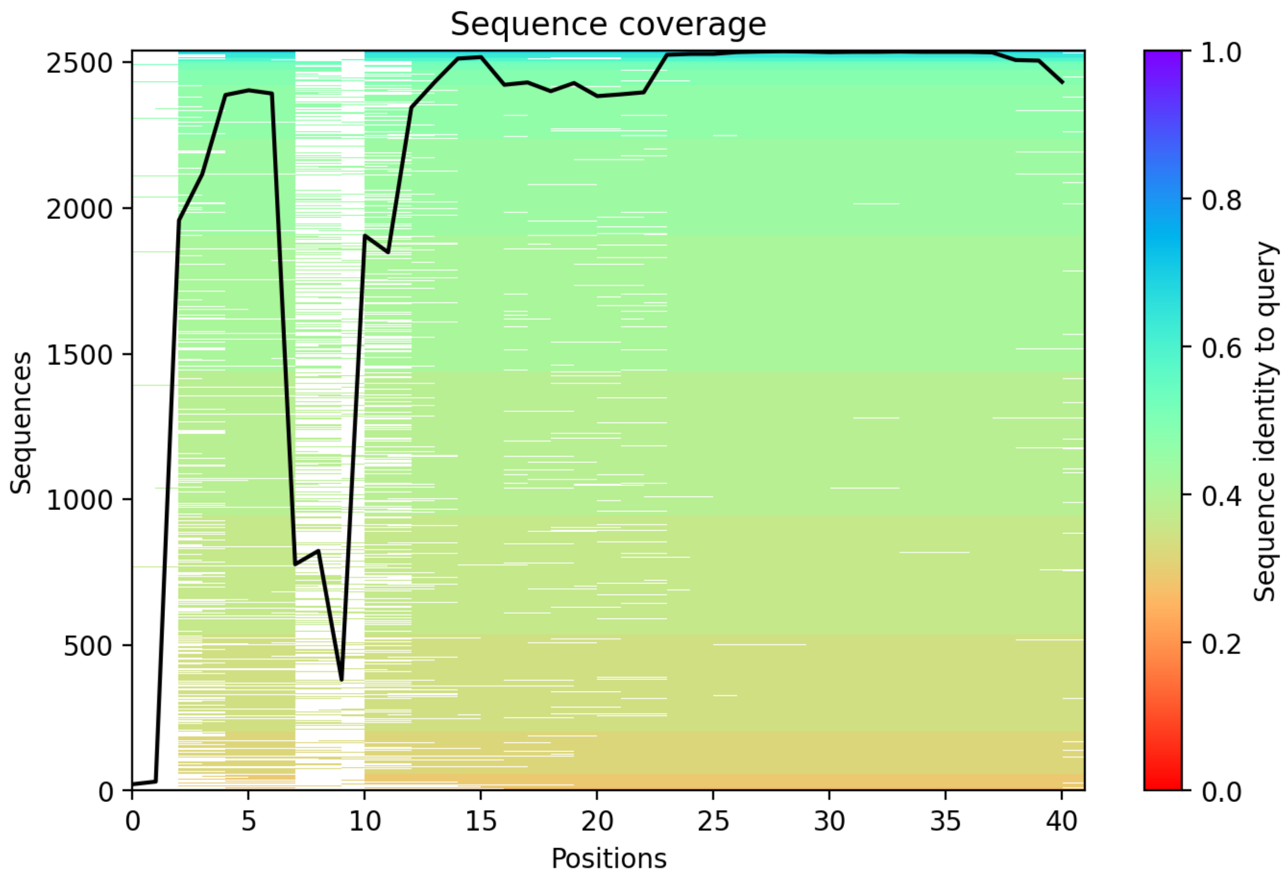

CP26: MtrunA17_Chr1g0205601

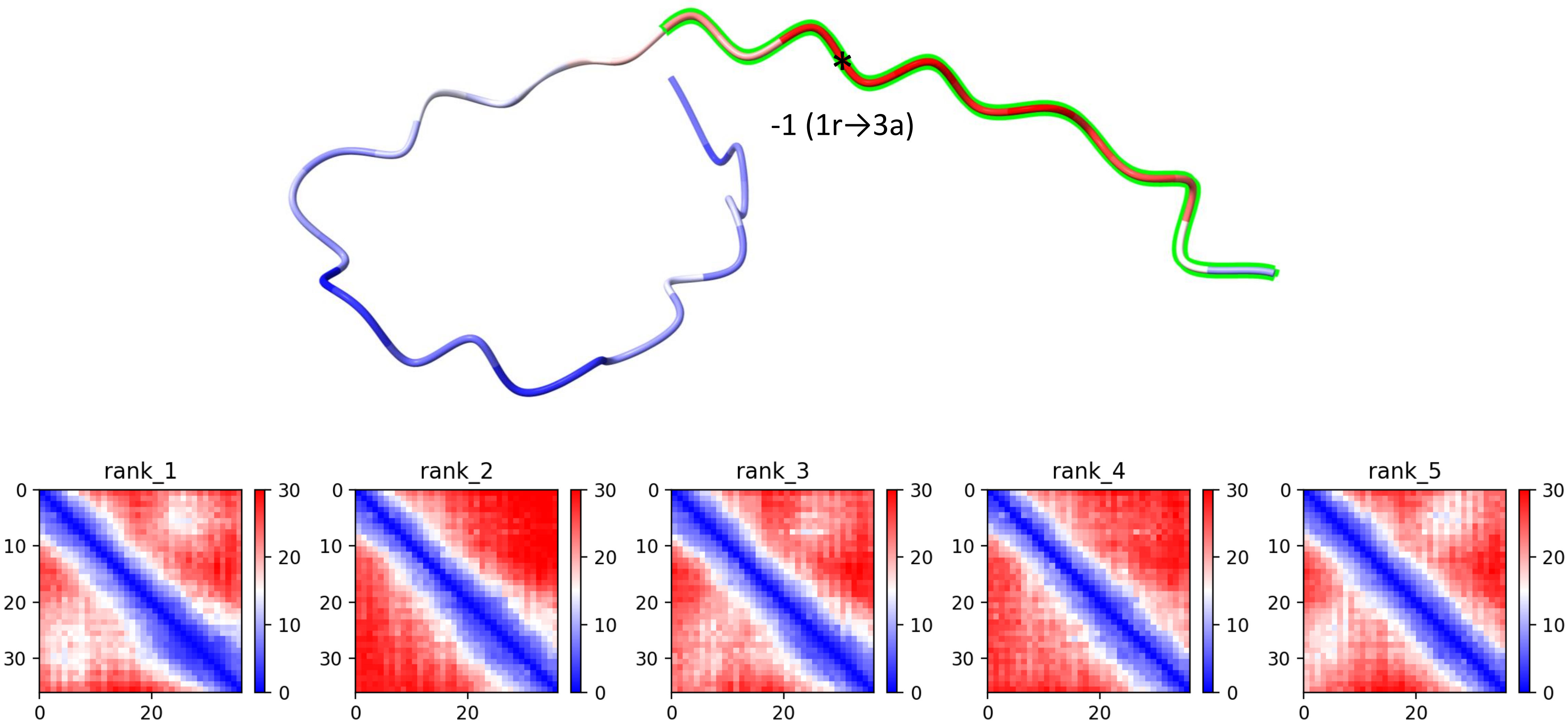

CP26: MtrunA17_Chr1g0205601

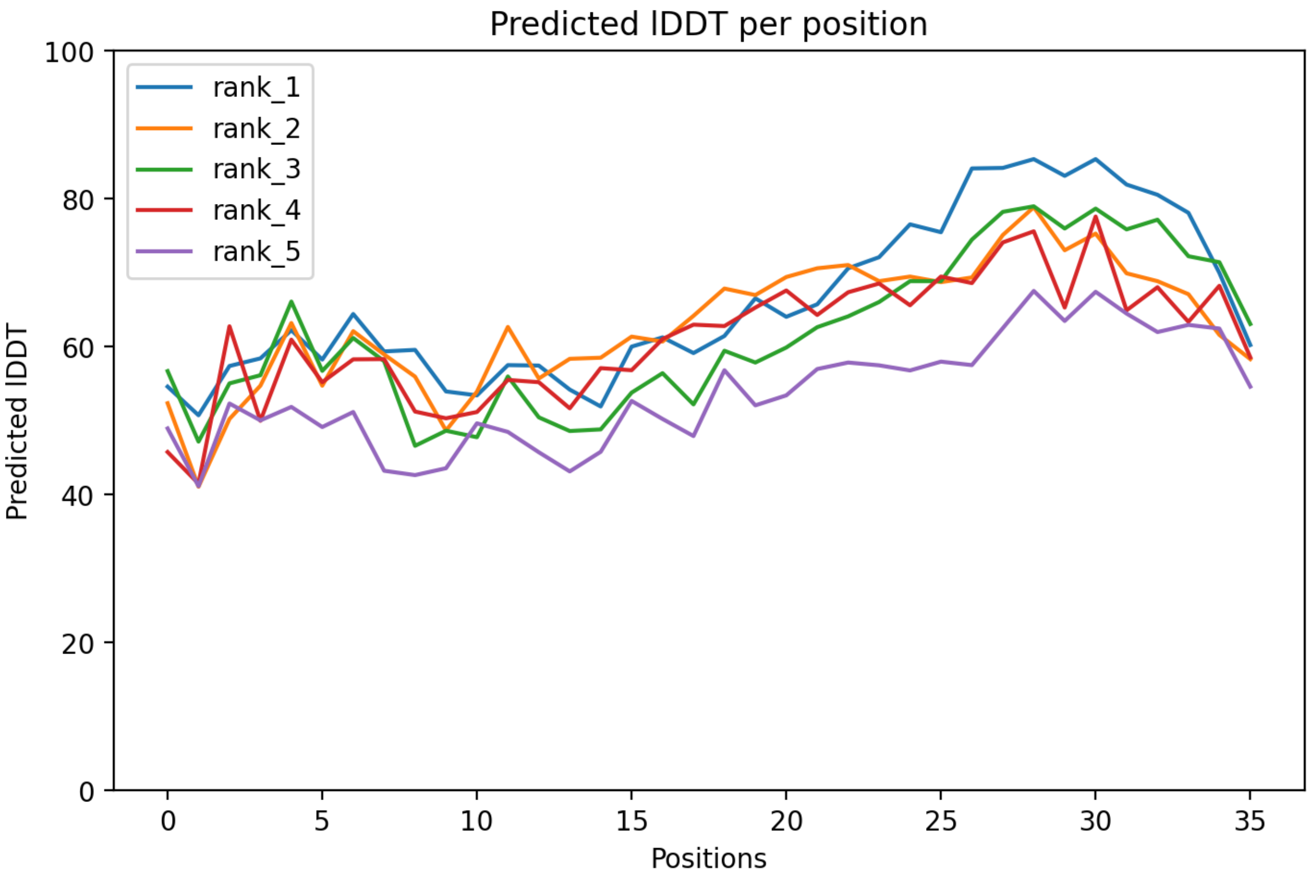

CP26: MtrunA17_Chr1g0205601

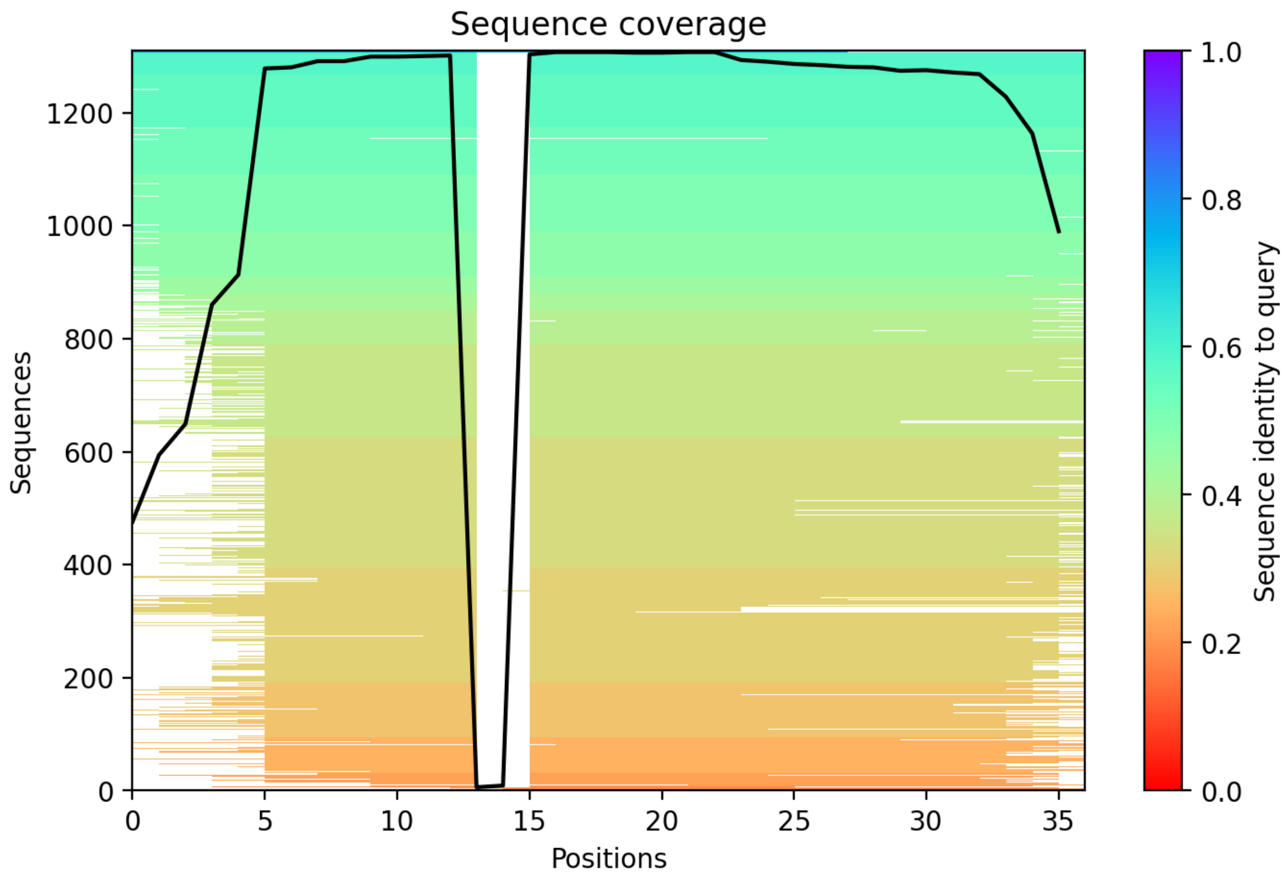

CP27: MtrunA17_Chr1g0207811

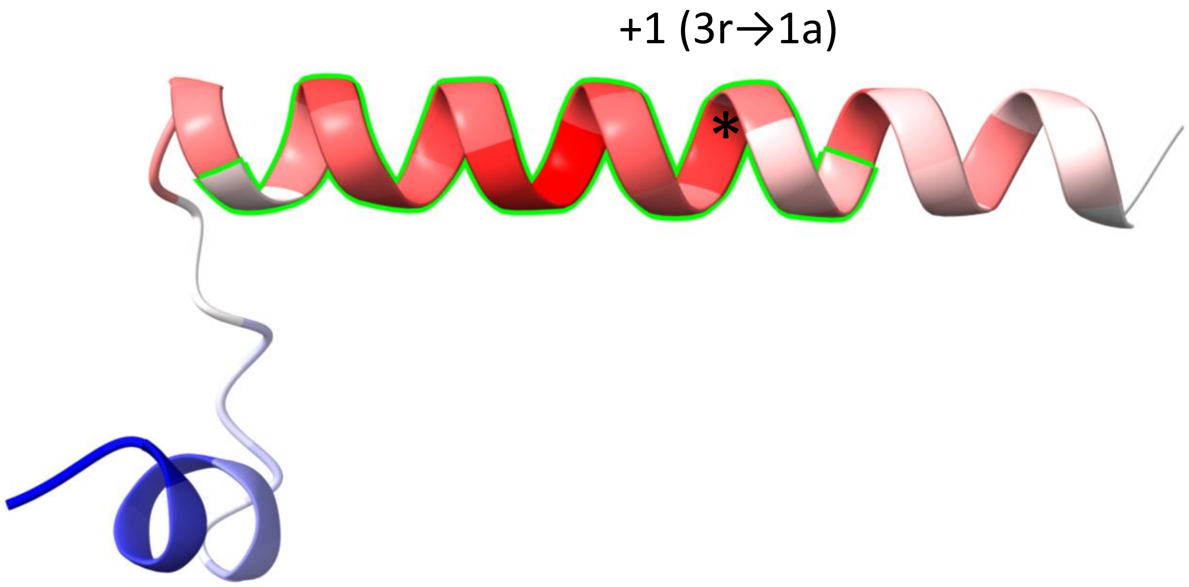


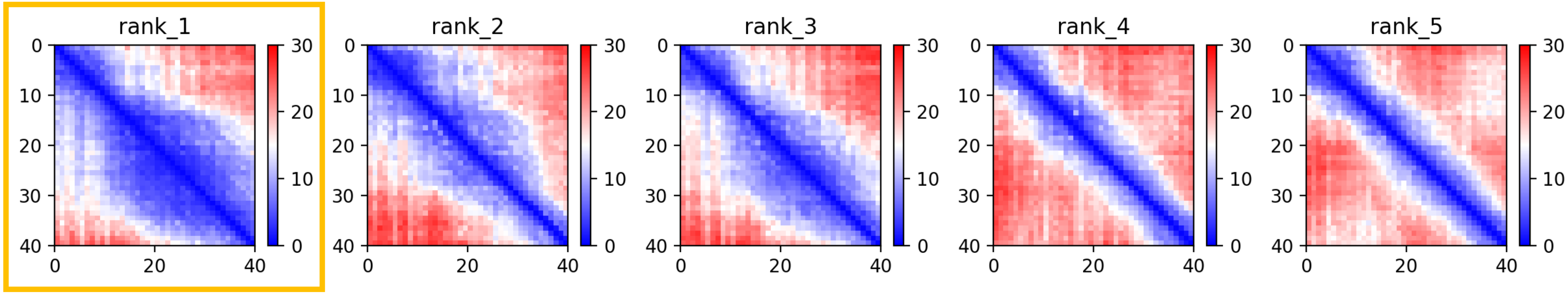

CP27: MtrunA17_Chr1g0207811

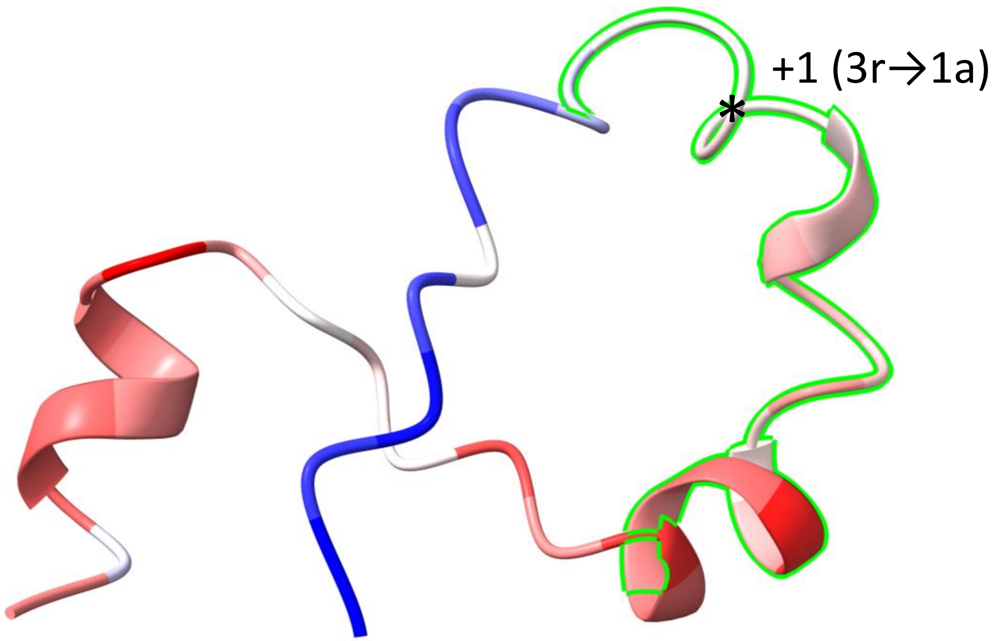


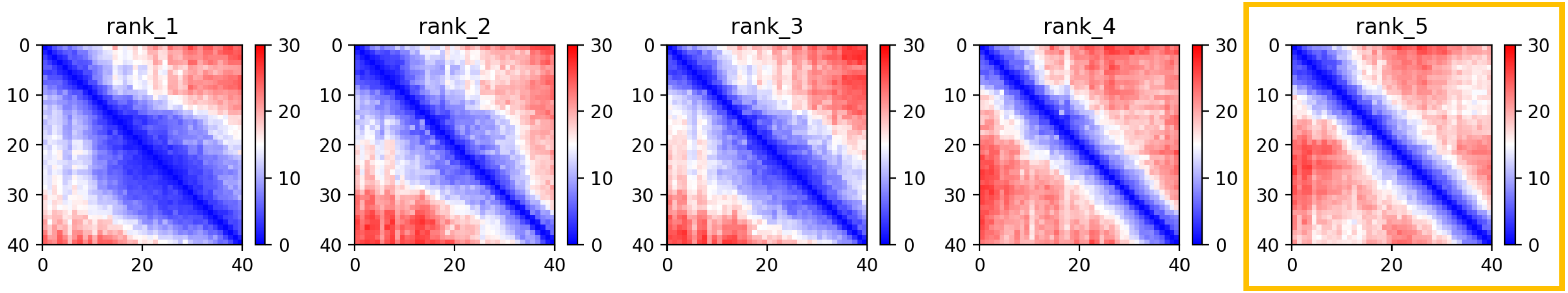

CP27: MtrunA17_Chr1g0207811

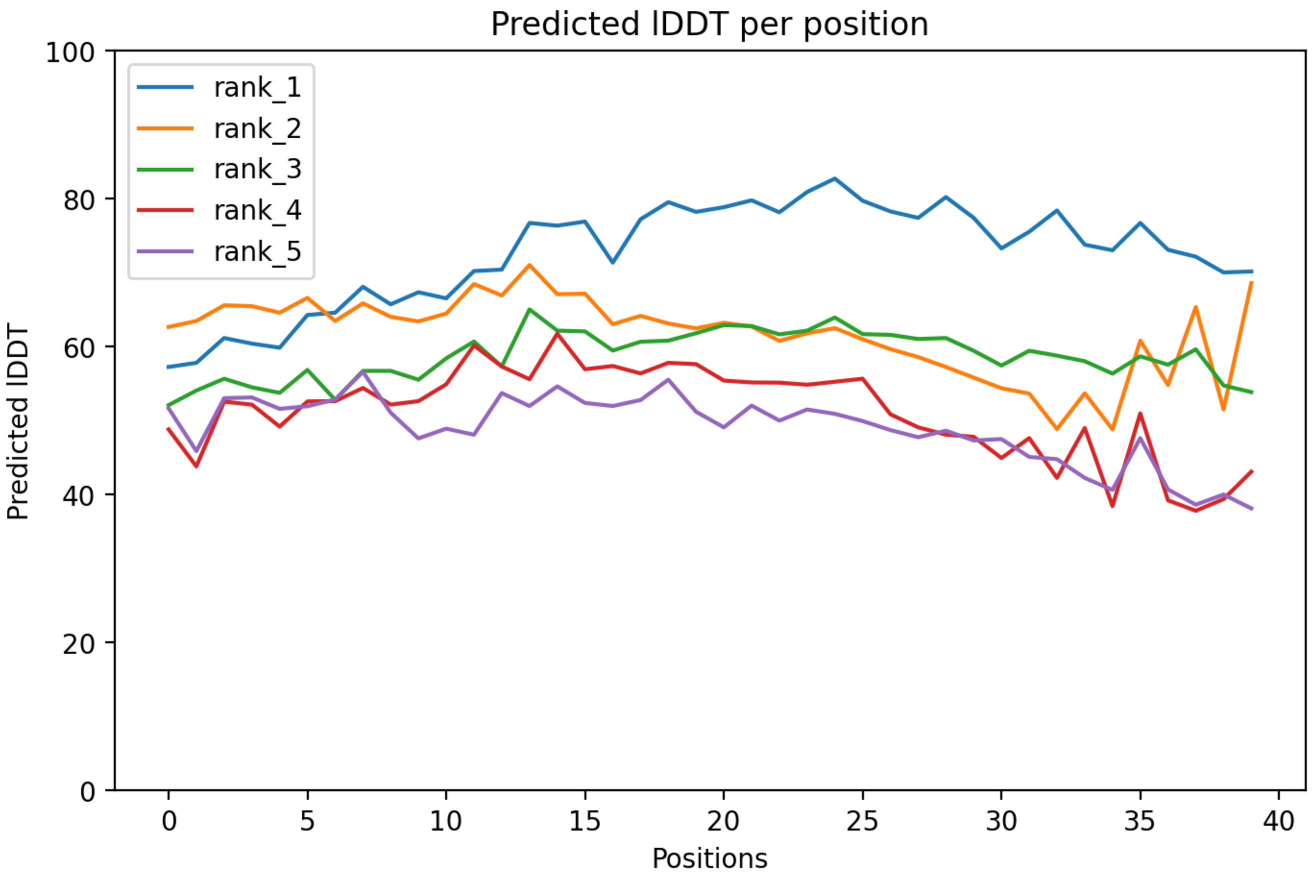

### CP27: MtrunA17_Chr1g0207811

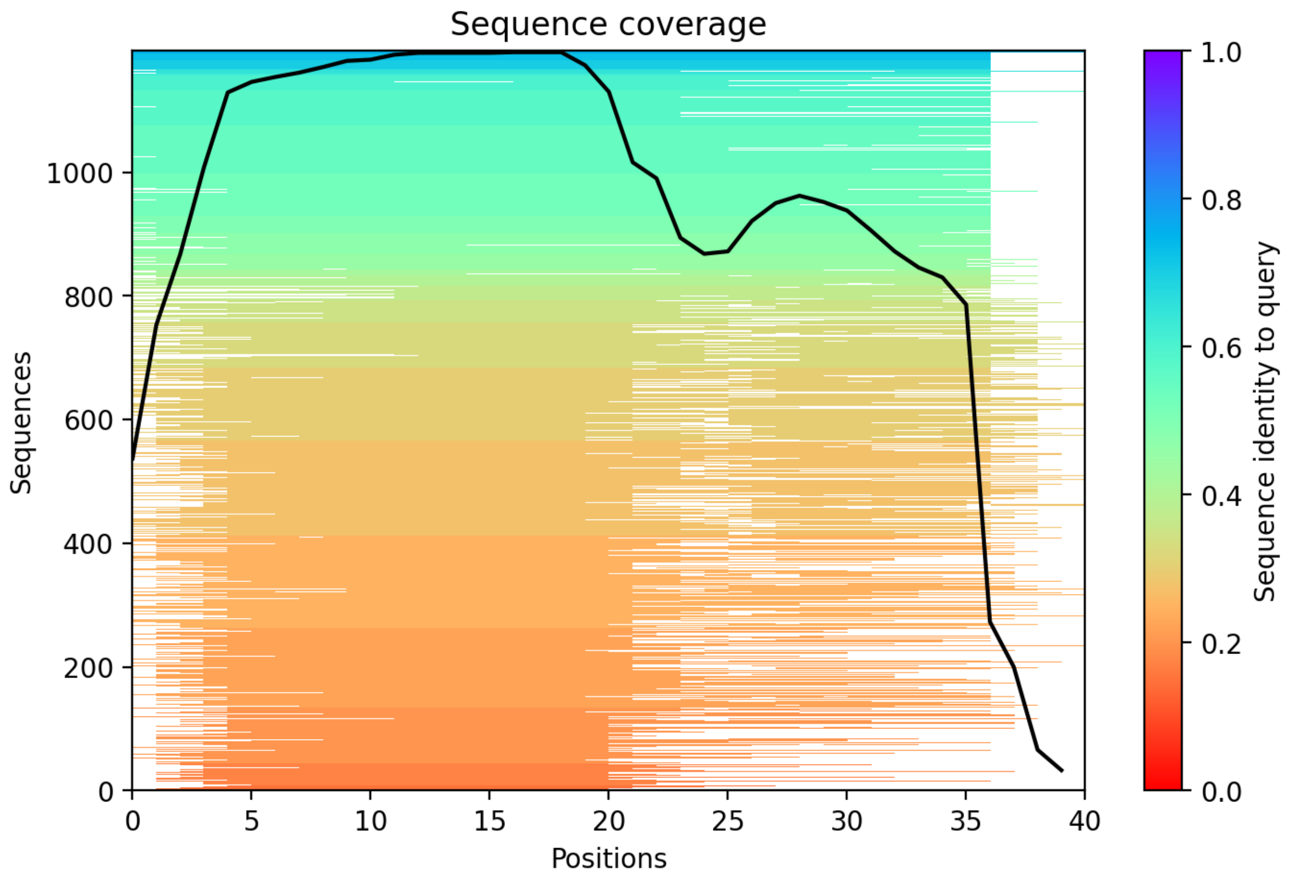

CP28: MtrunA17_Chr1g0207921

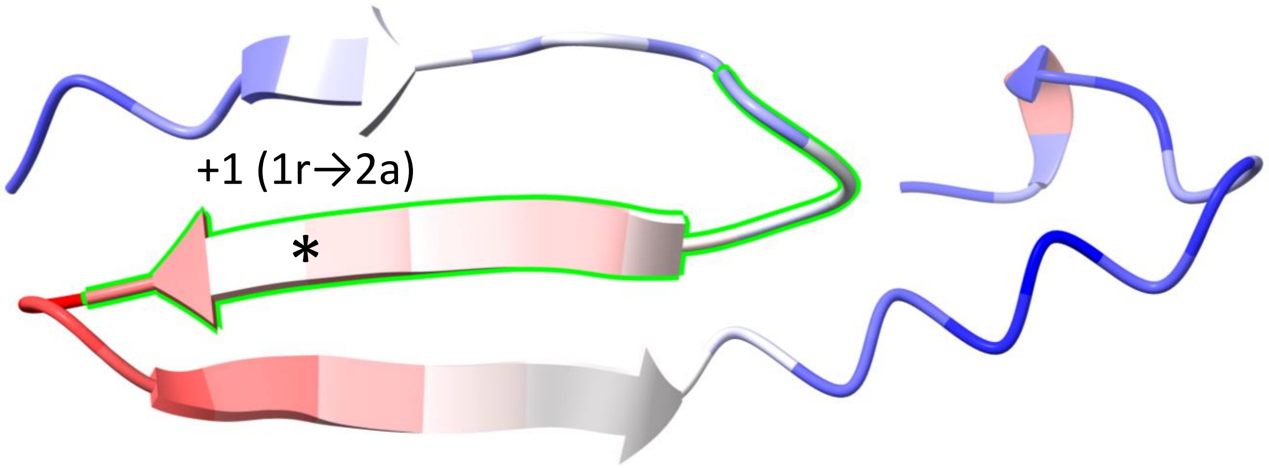


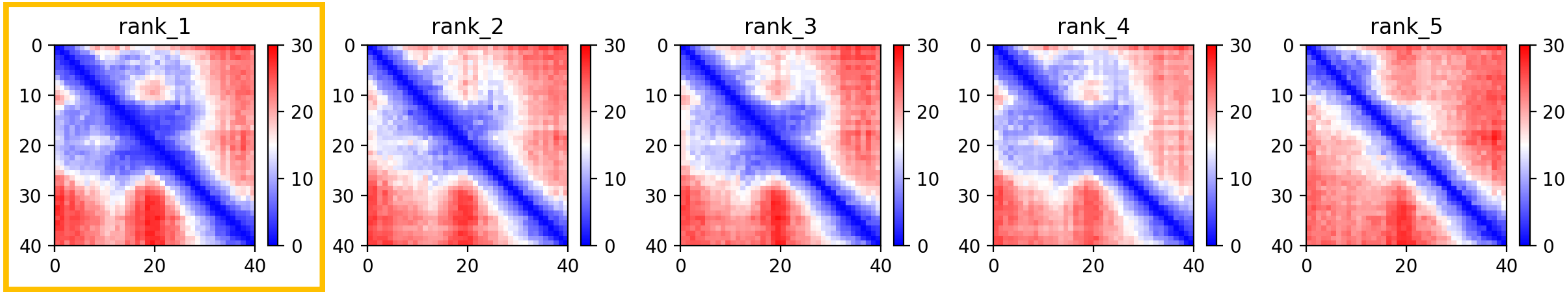

CP28: MtrunA17_Chr1g0207921

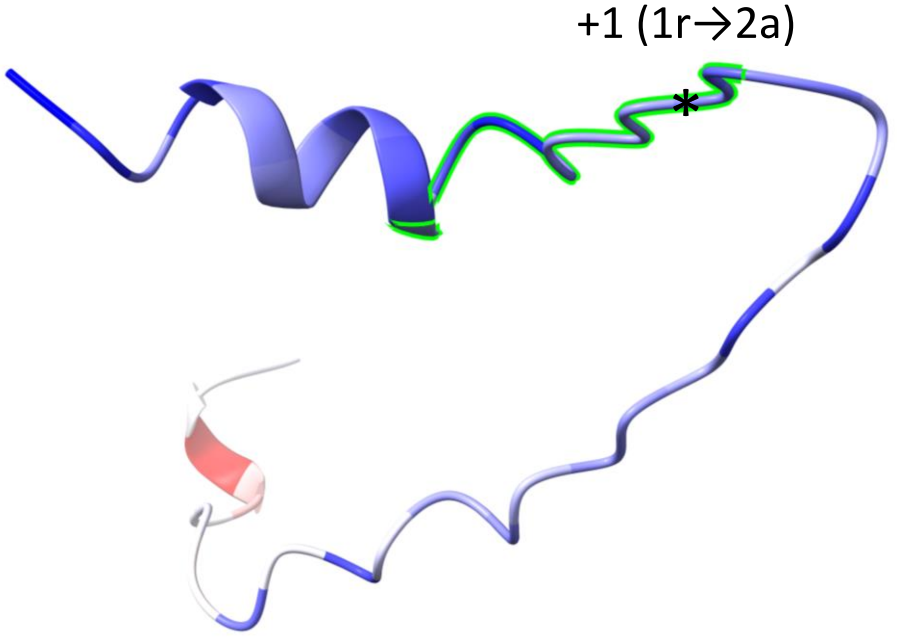


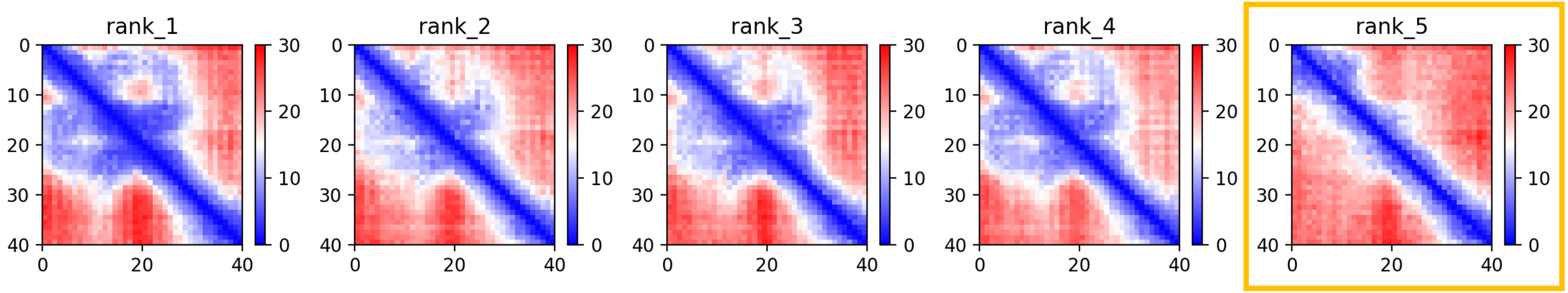

CP28: MtrunA17_Chr1g0207921

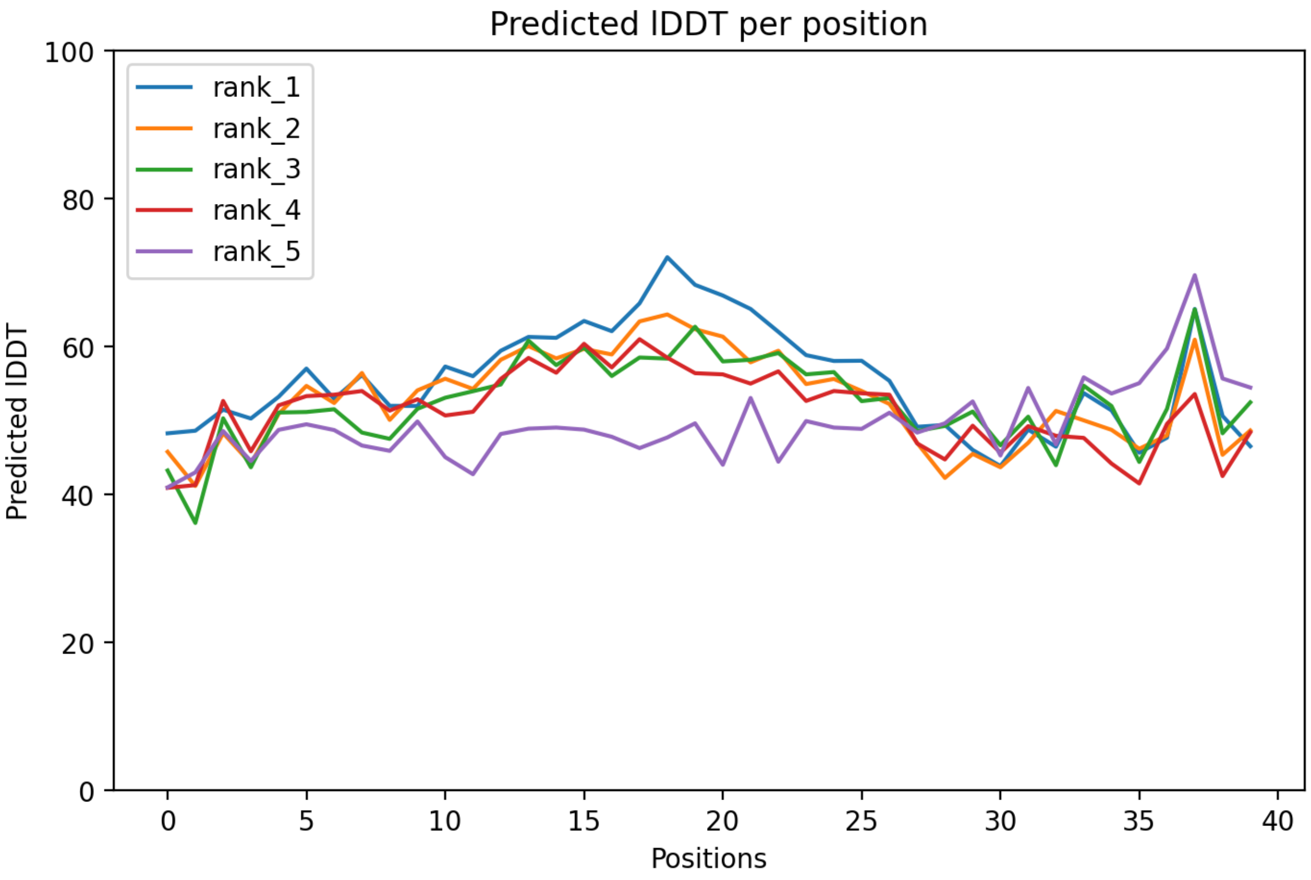

CP28: MtrunA17_Chr1g0207921

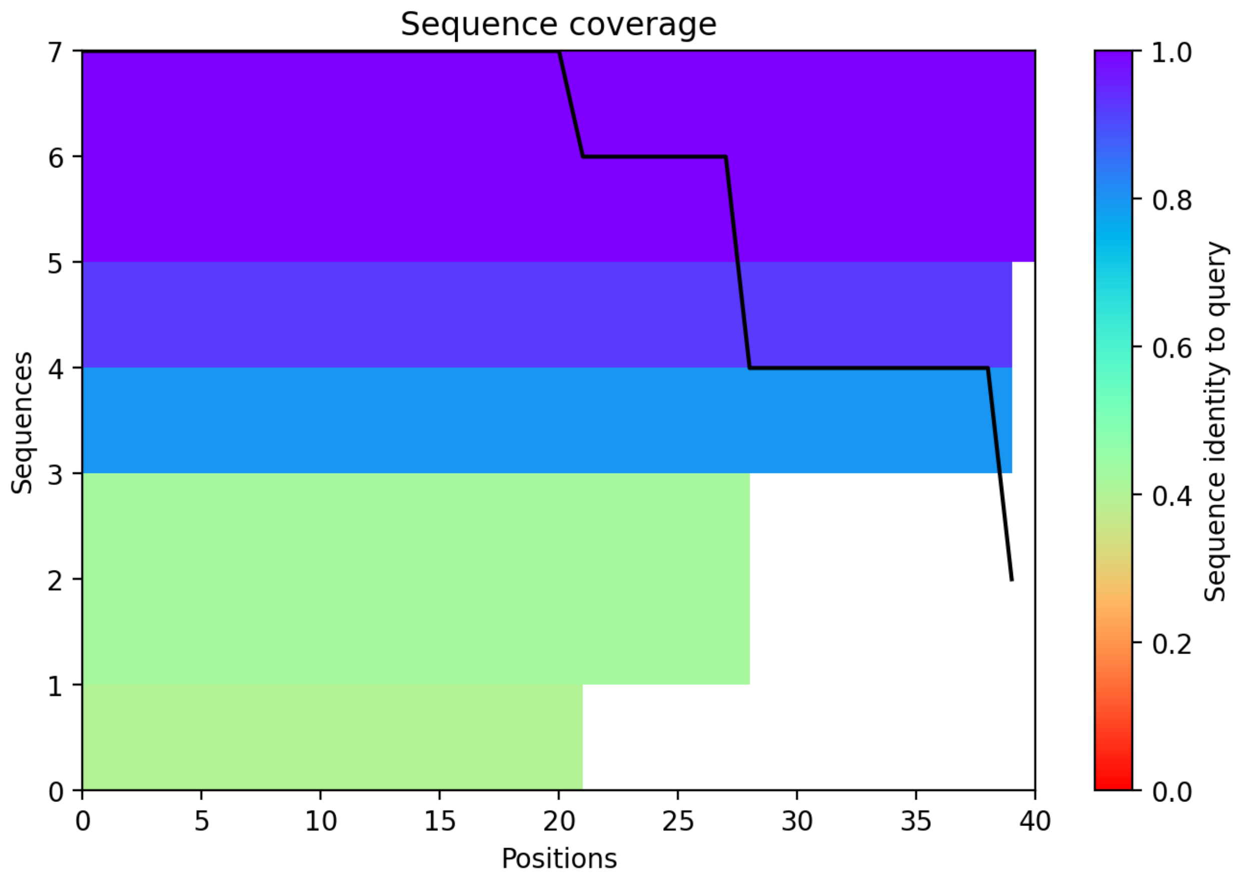

CP29: MtrunA17_Chr1g0209791

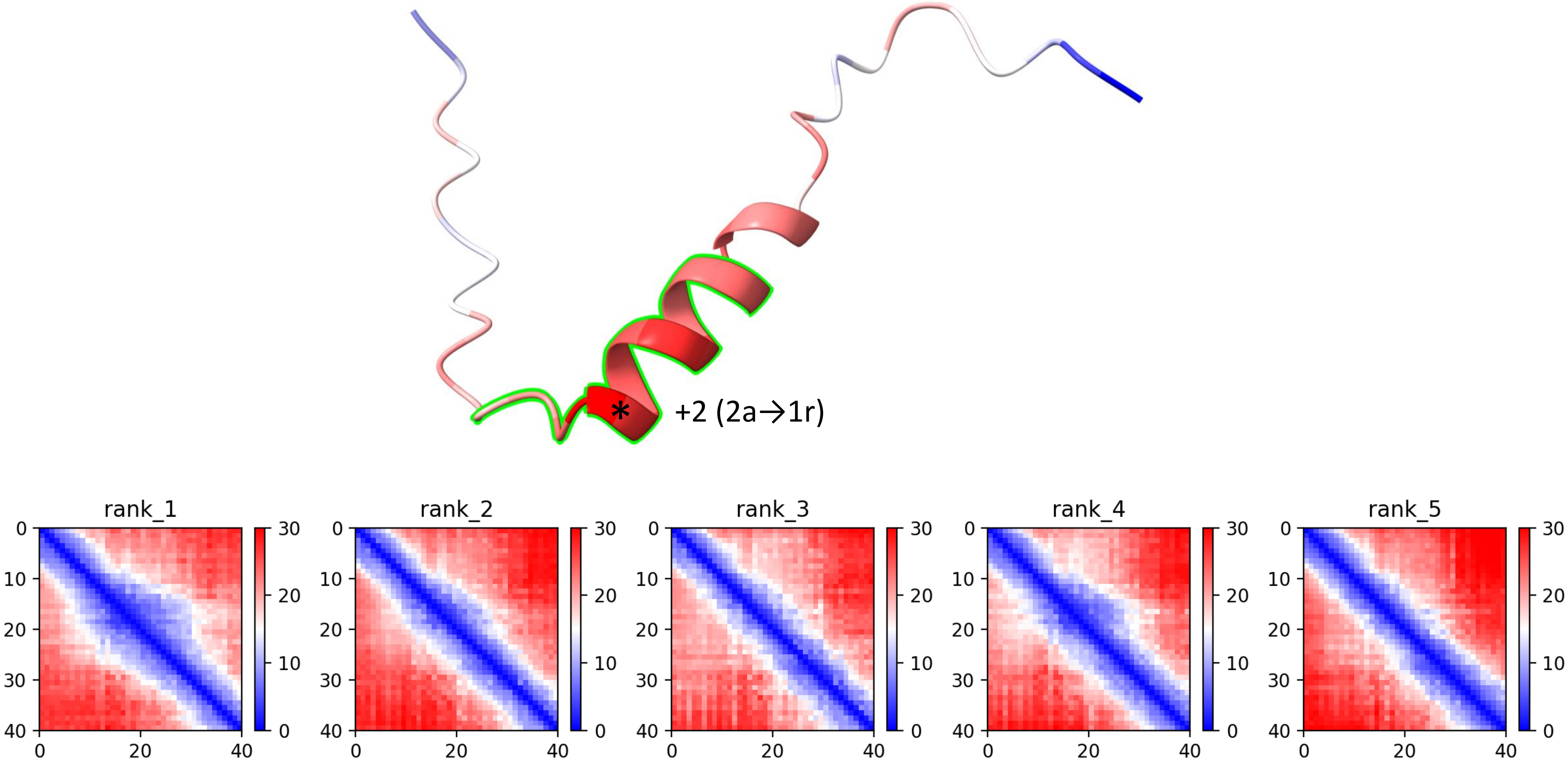

CP29: MtrunA17_Chr1g0209791

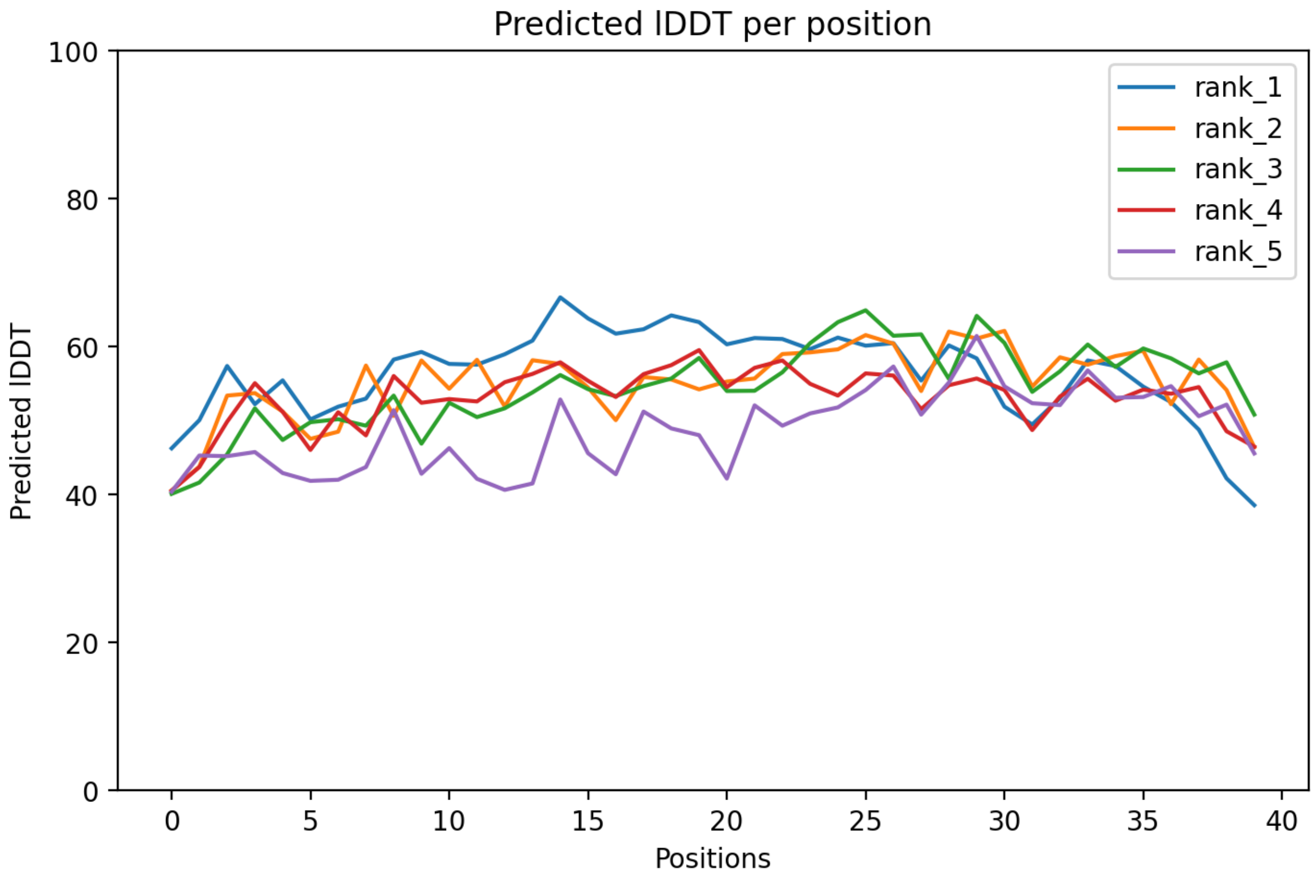

CP29: MtrunA17_Chr1g0209791

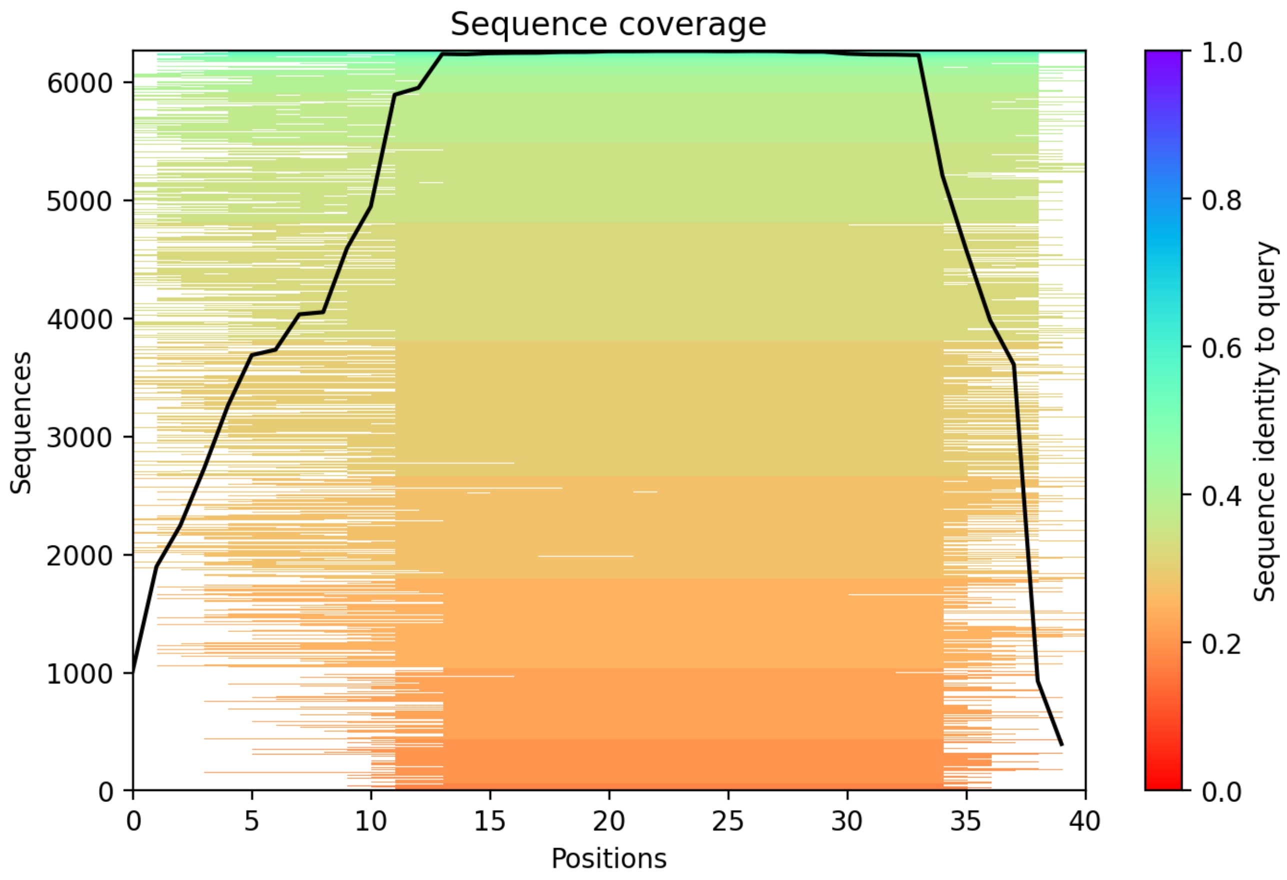

CP30: MtrunA17_Chr1g0210521

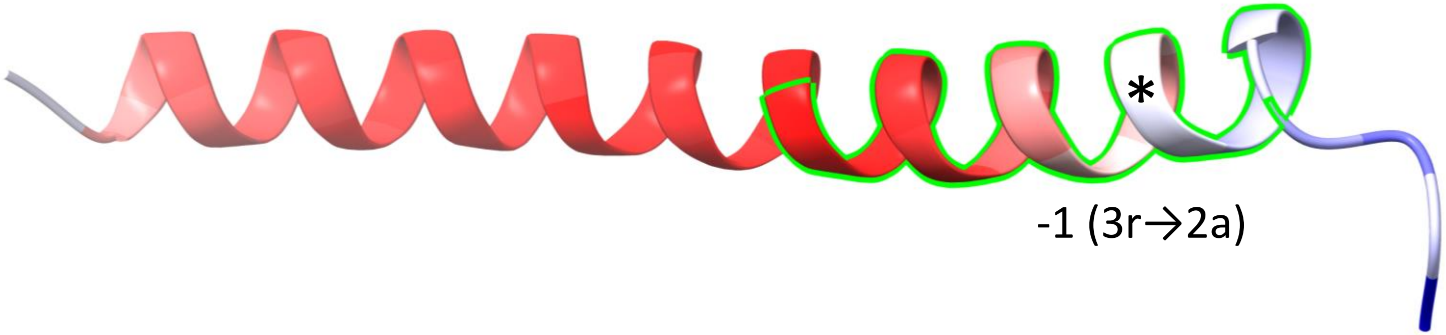


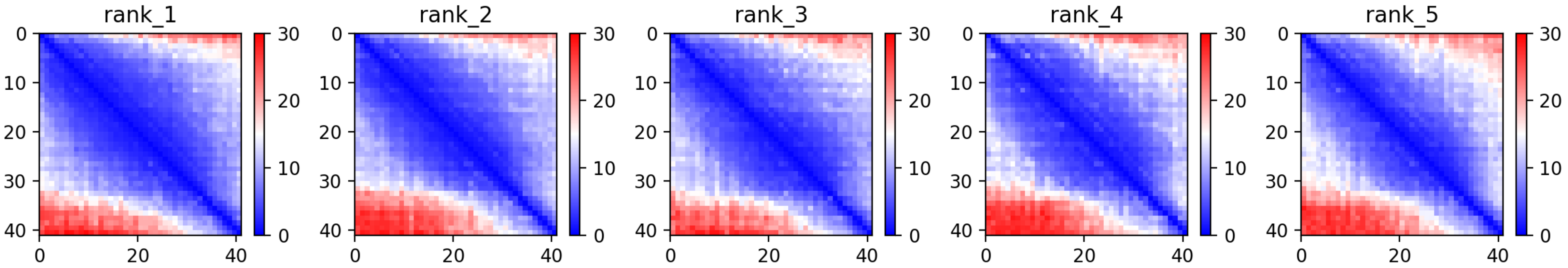

### CP30: MtrunA17_Chr1g0210521

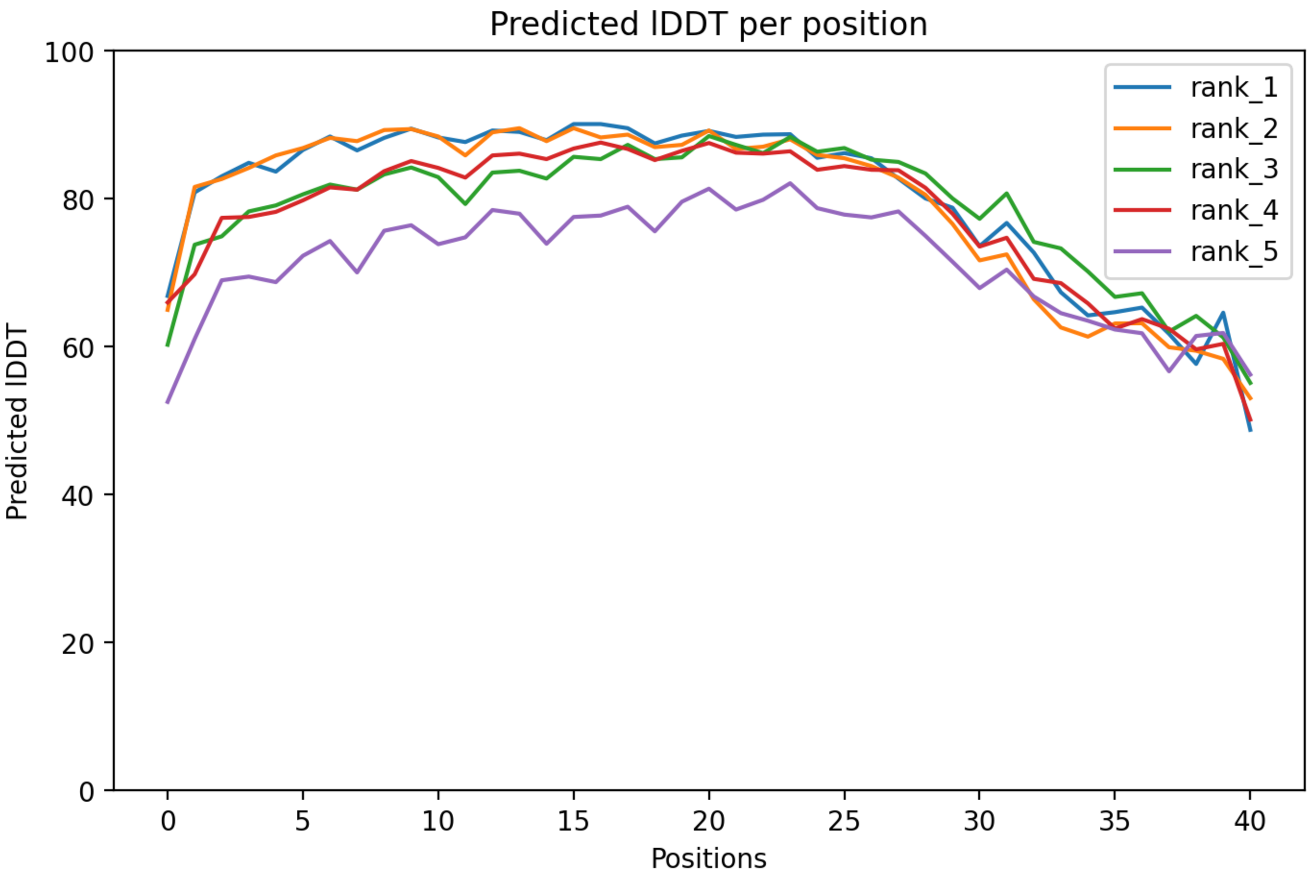

### CP30: MtrunA17_Chr1g0210521

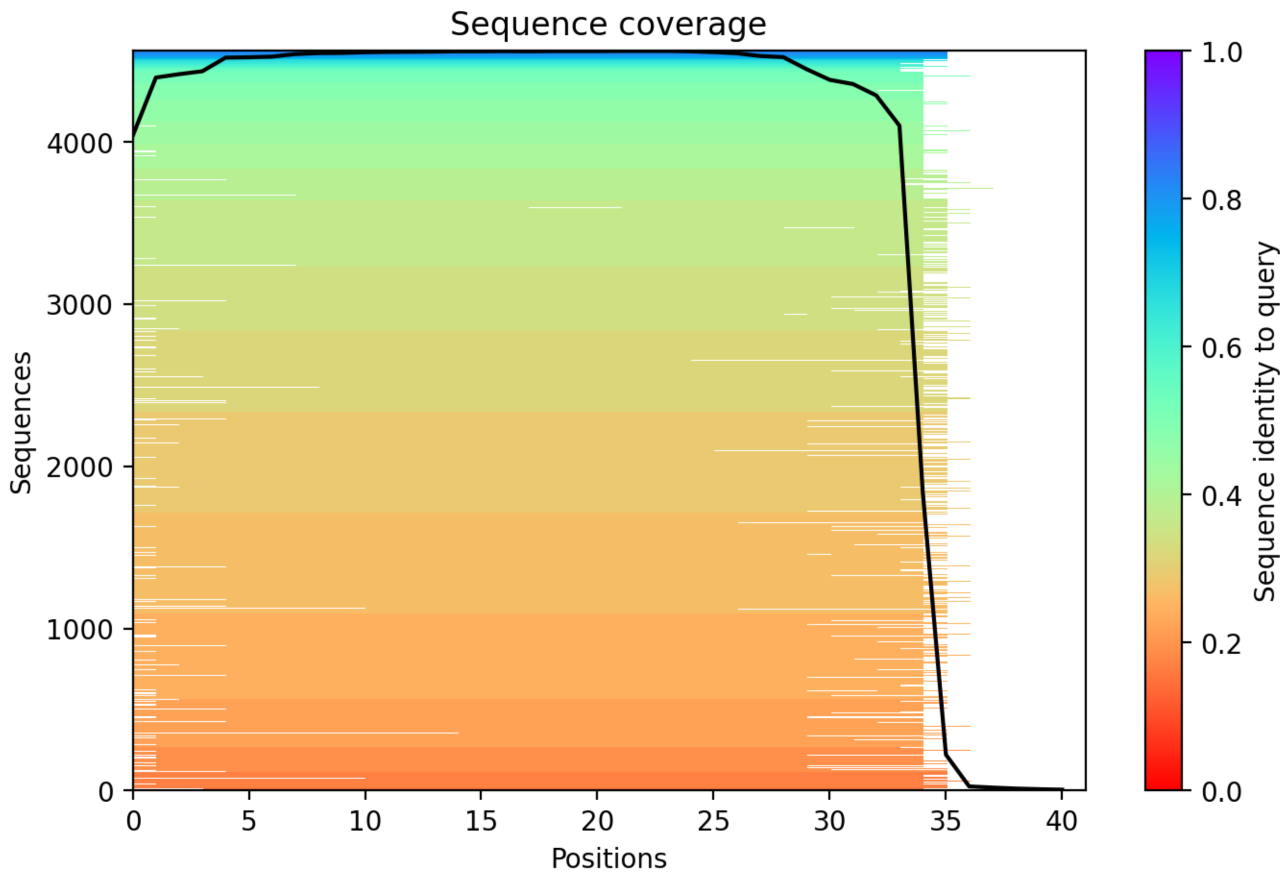

CP31: MtrunA17_Chr1g0212961

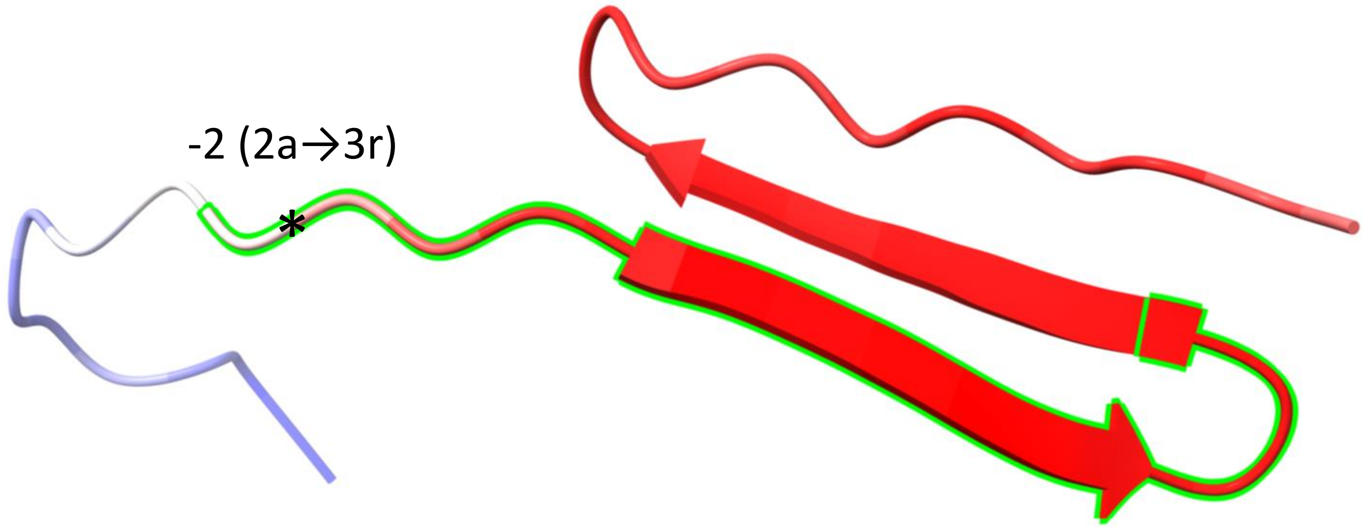


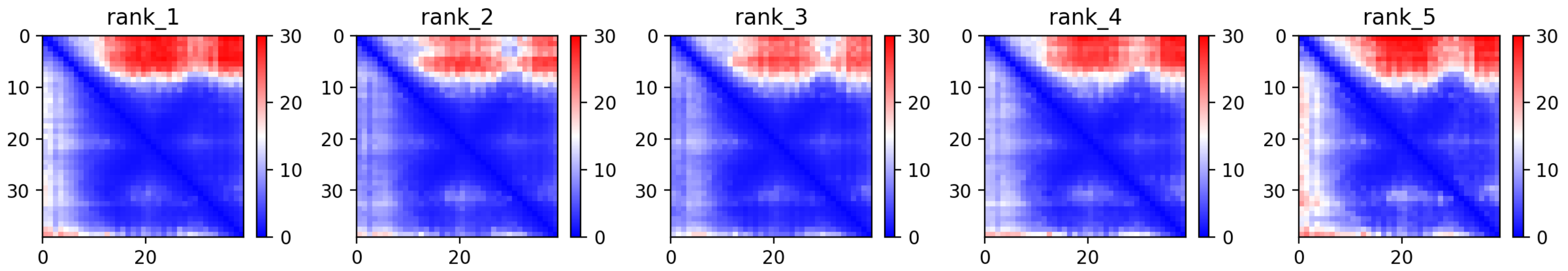

CP31: MtrunA17_Chr1g0212961

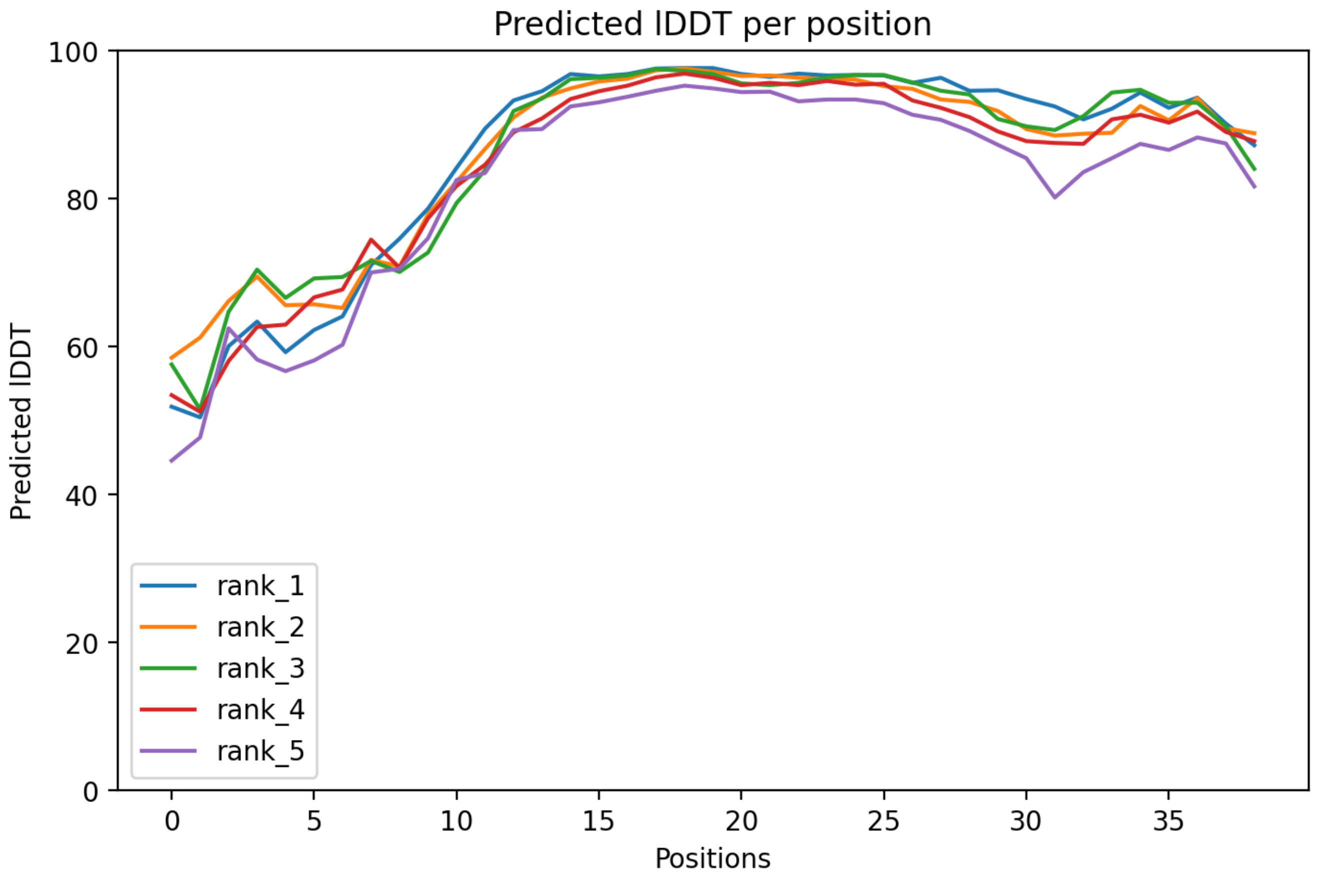

CP31: MtrunA17_Chr1g0212961

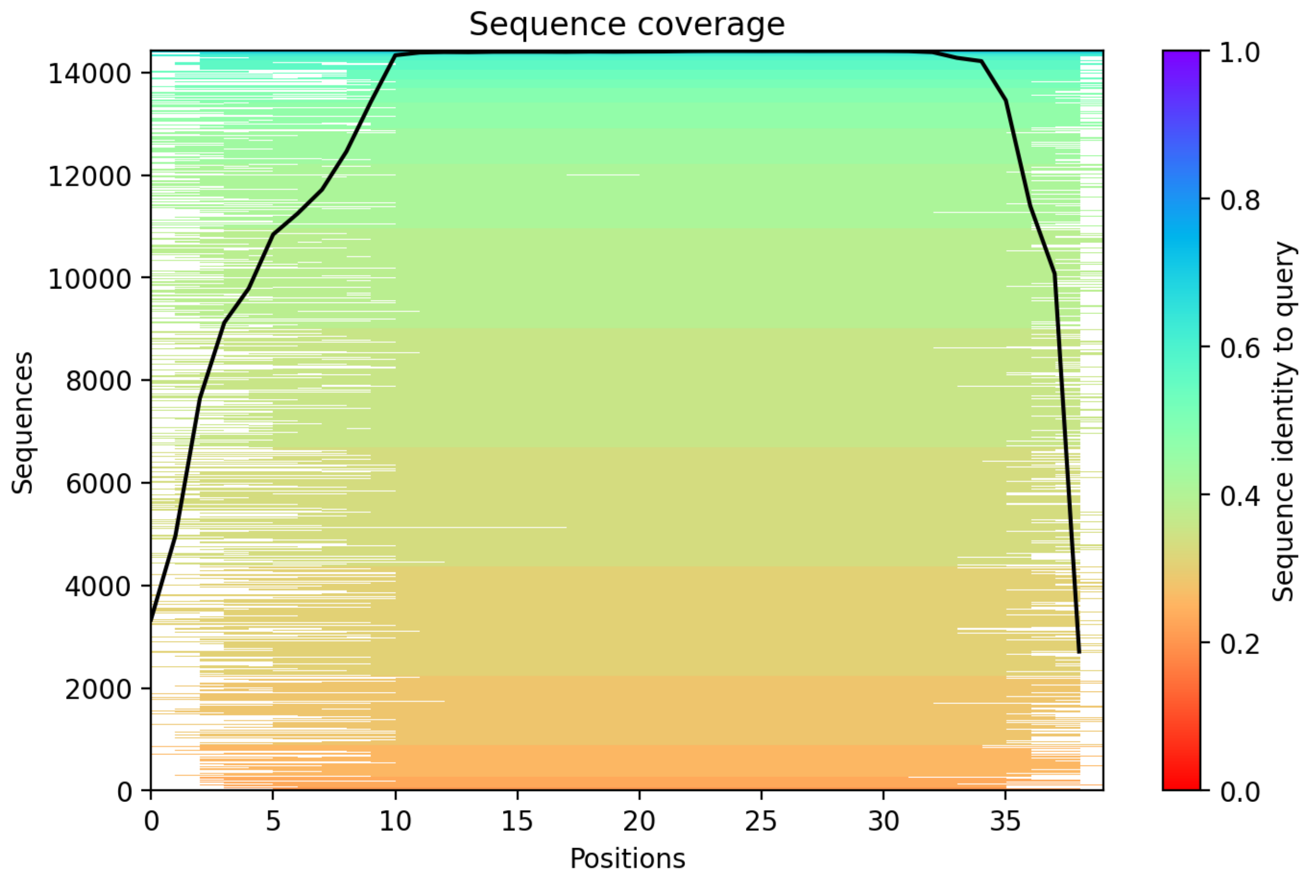

### CP32: MtrunA17_Chr1g1004575

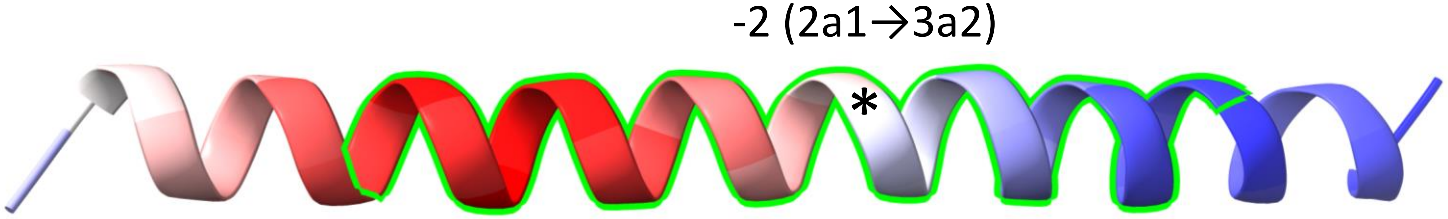


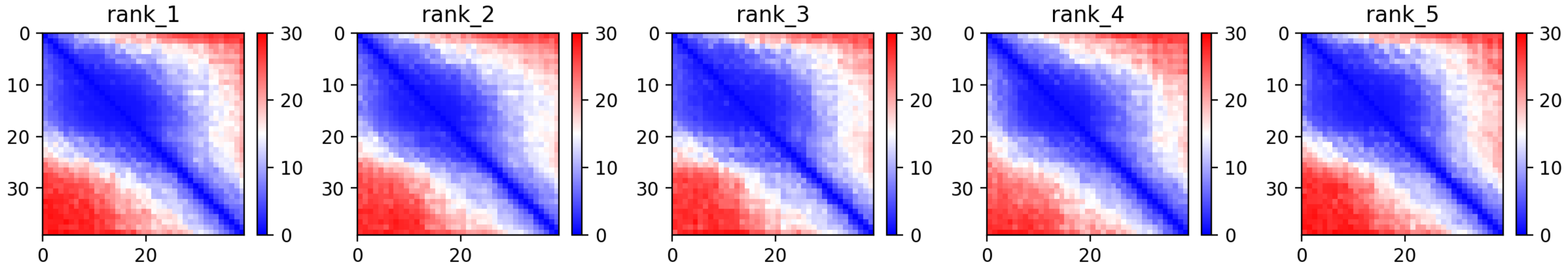

CP32: MtrunA17_Chr1g1004575

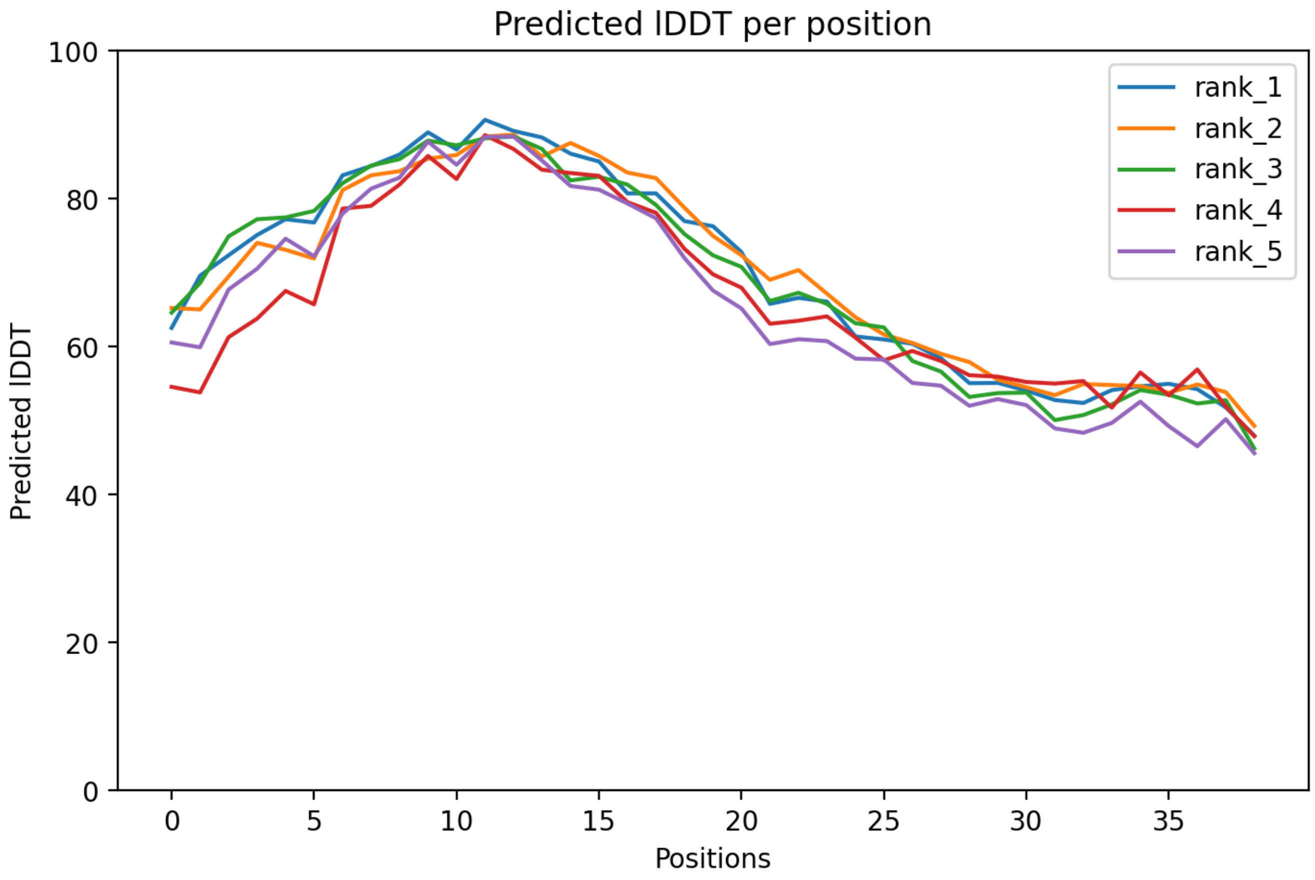

### CP32: MtrunA17_Chr1g1004575

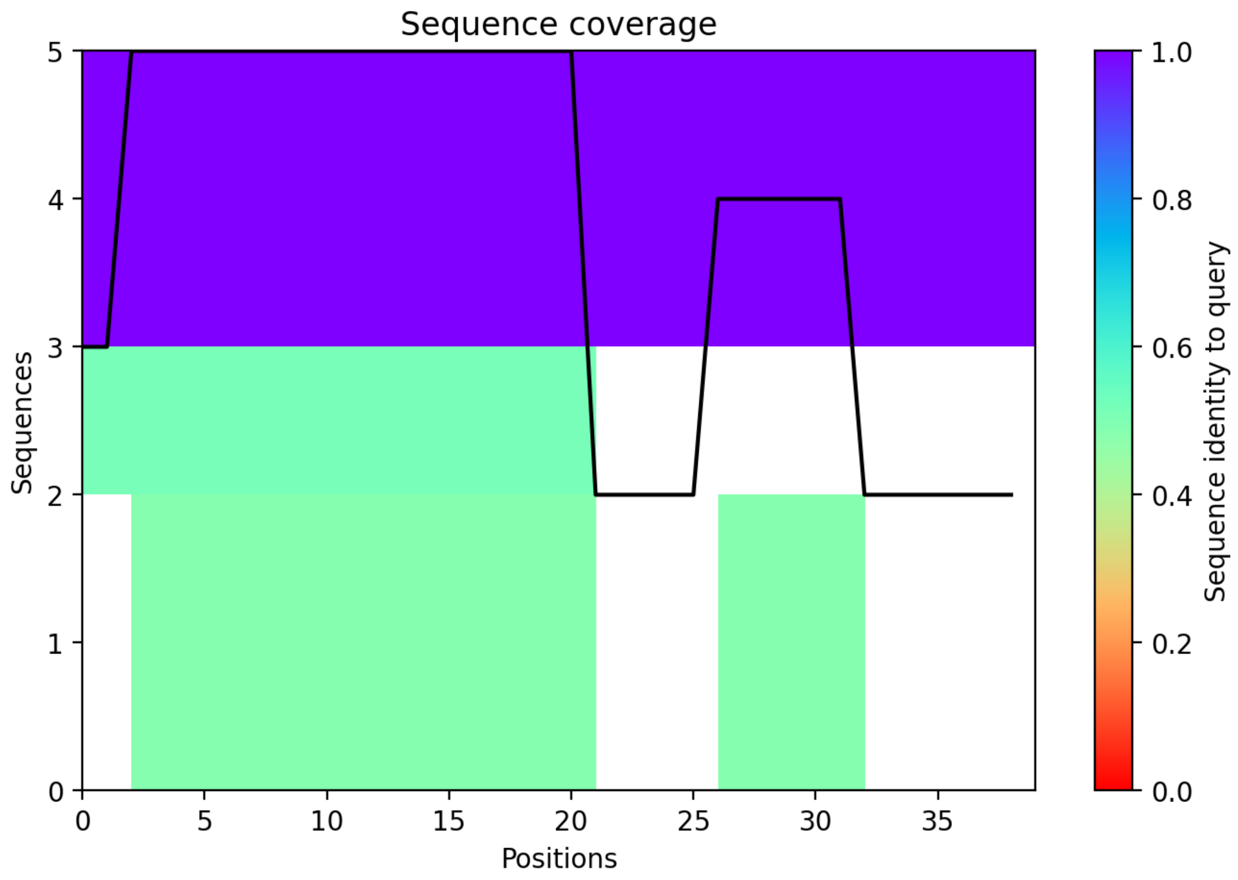

CP33: MtrunA17_Chr2g0283311

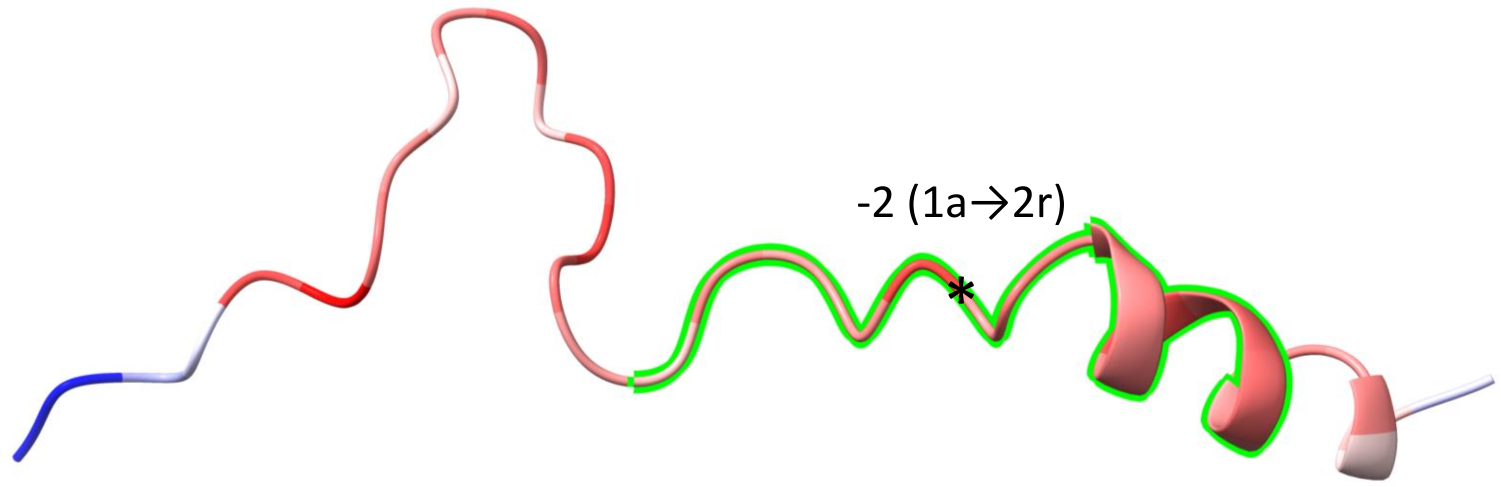


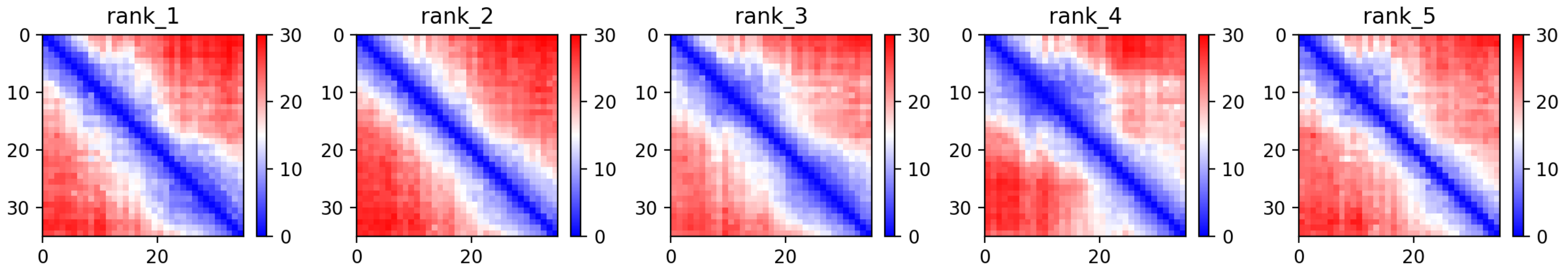

CP33: MtrunA17_Chr2g0283311

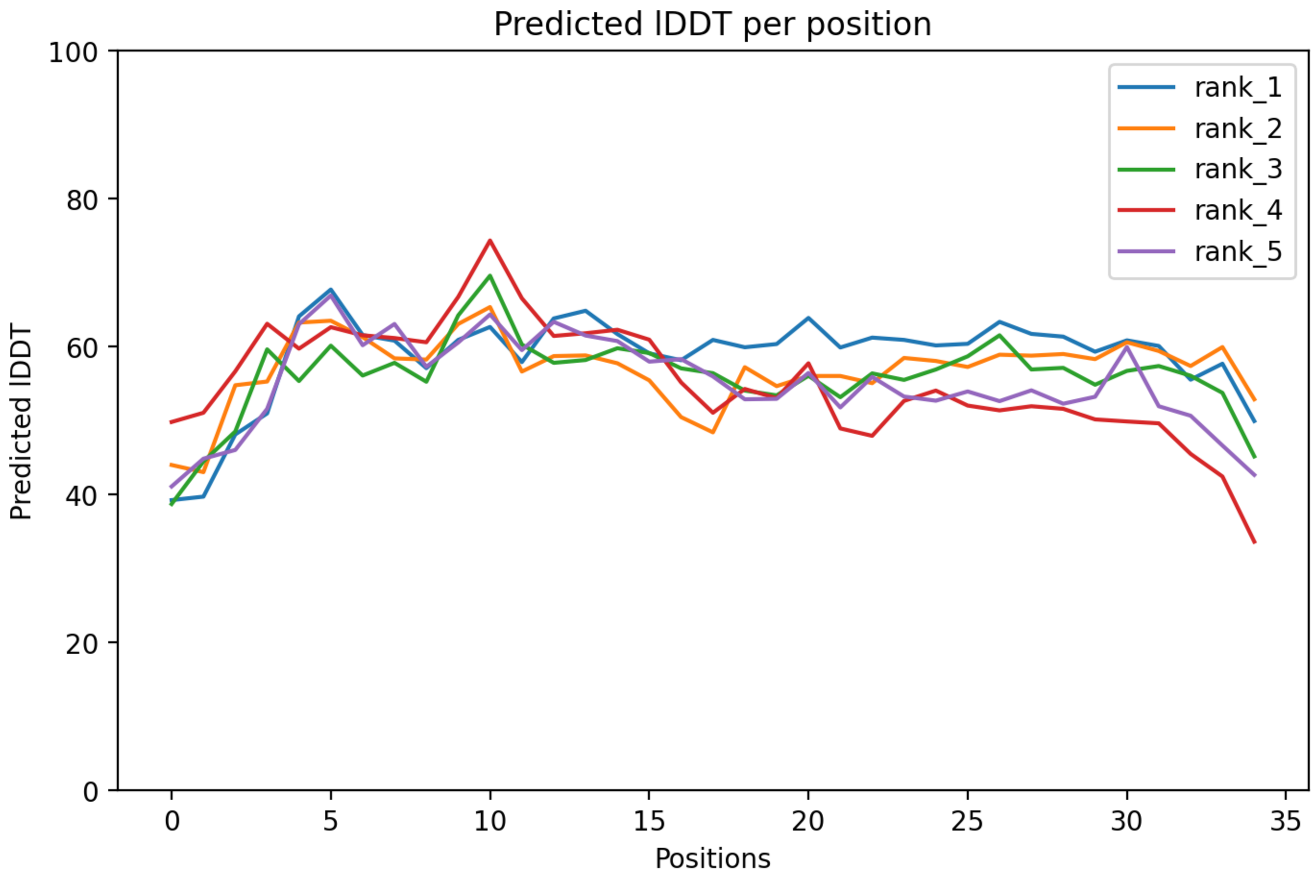

CP33: MtrunA17_Chr2g0283311

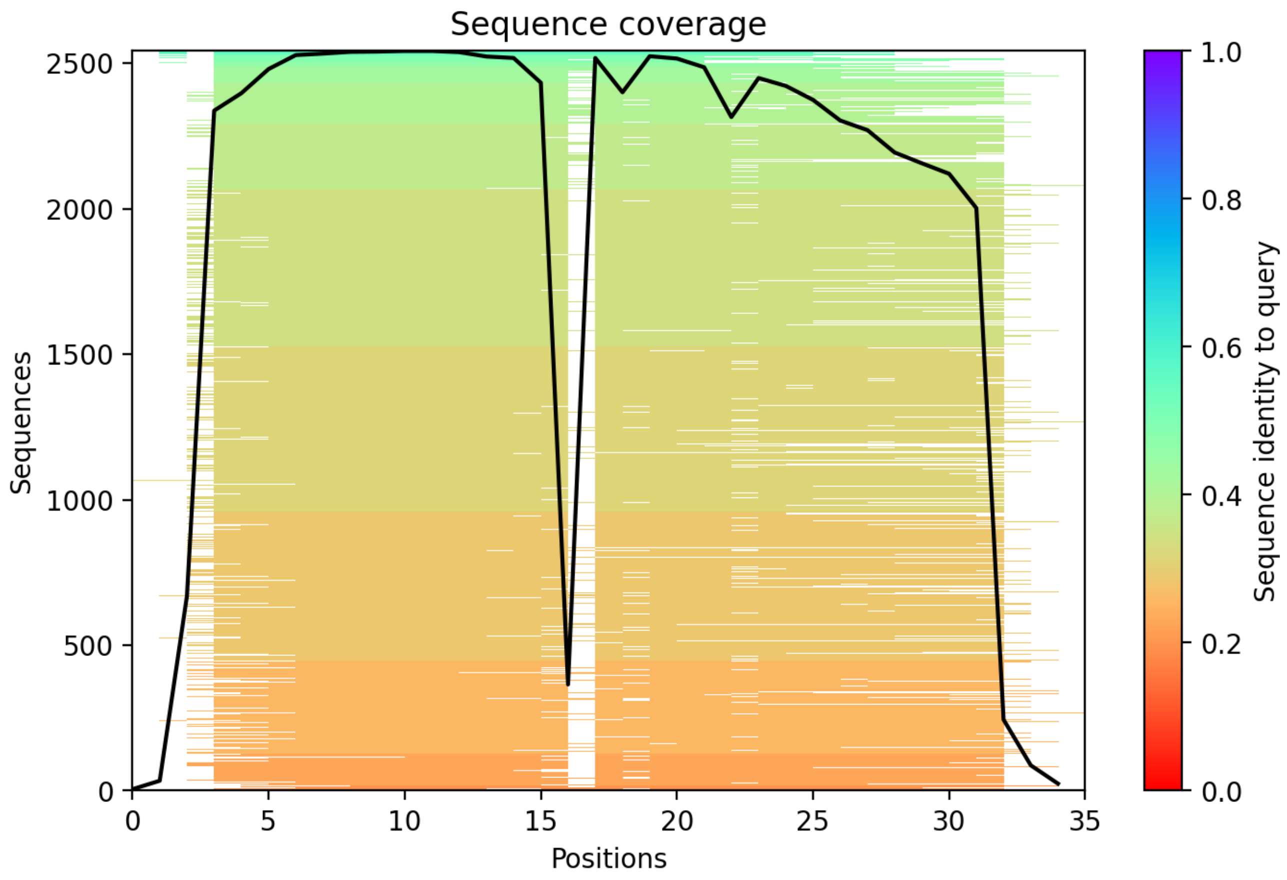

CP34: MtrunA17_Chr2g0285461

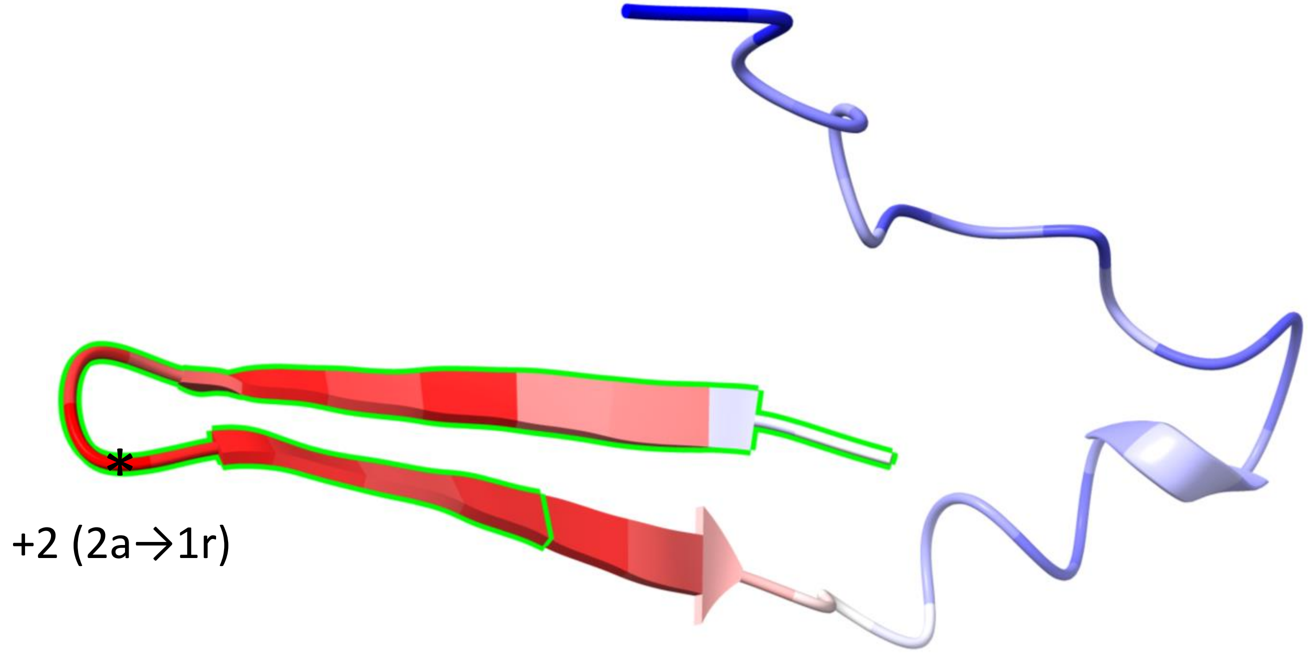


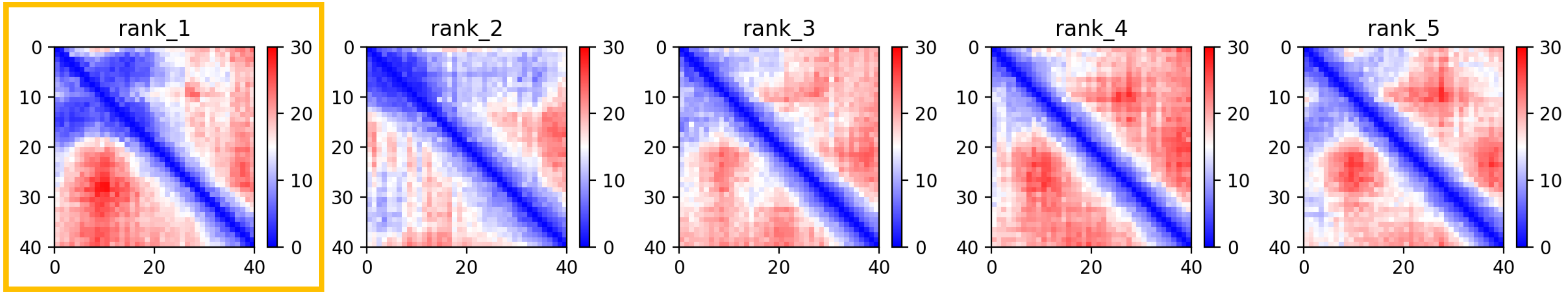

CP34: MtrunA17_Chr2g0285461

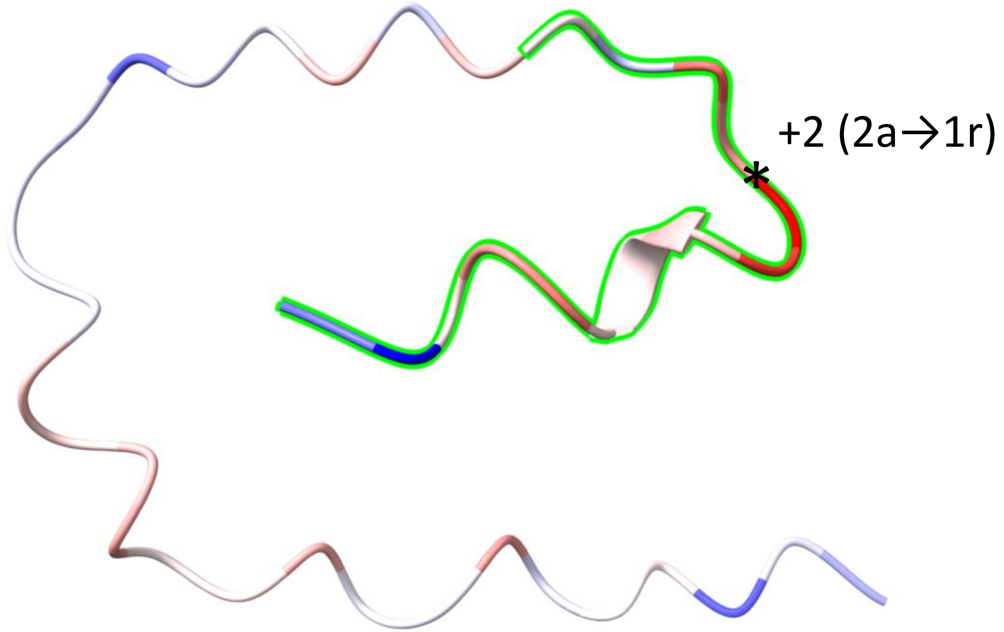


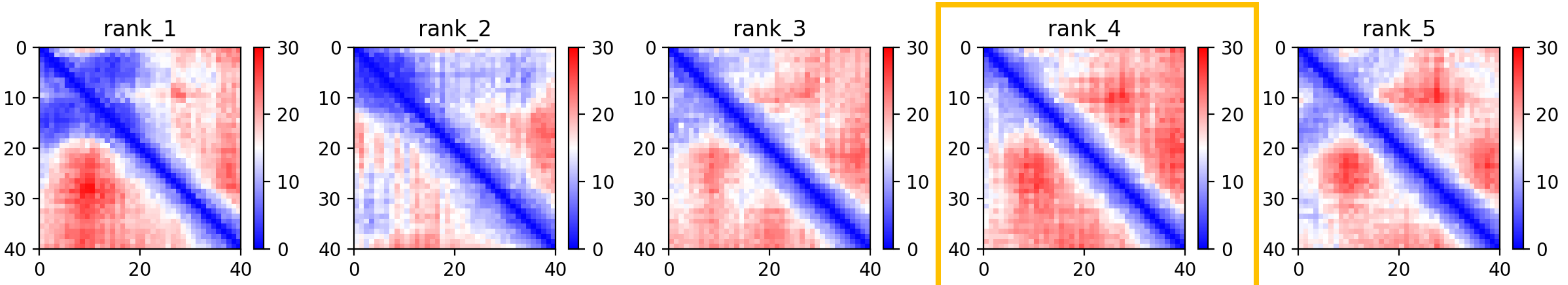

CP34: MtrunA17_Chr2g0285461

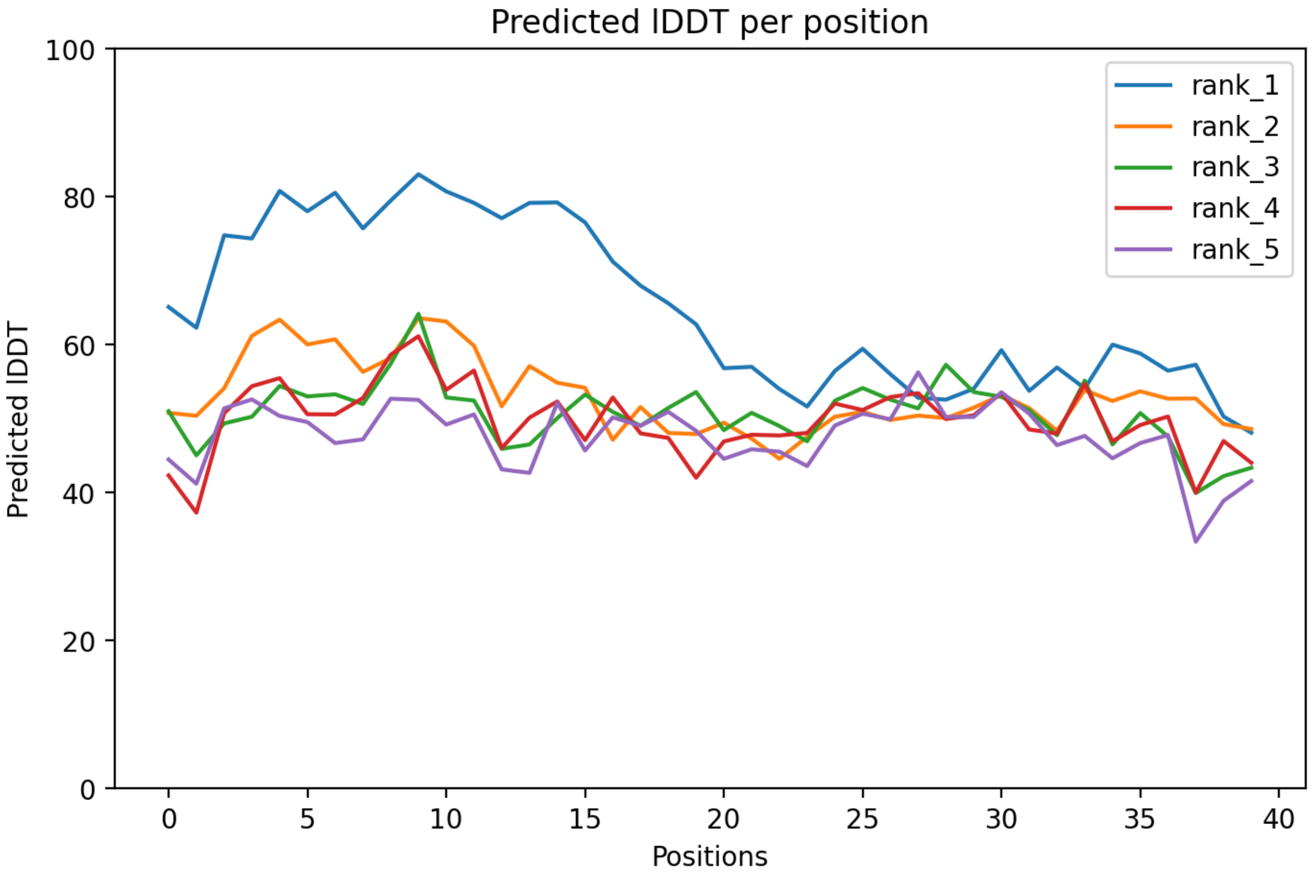

CP34: MtrunA17_Chr2g0285461

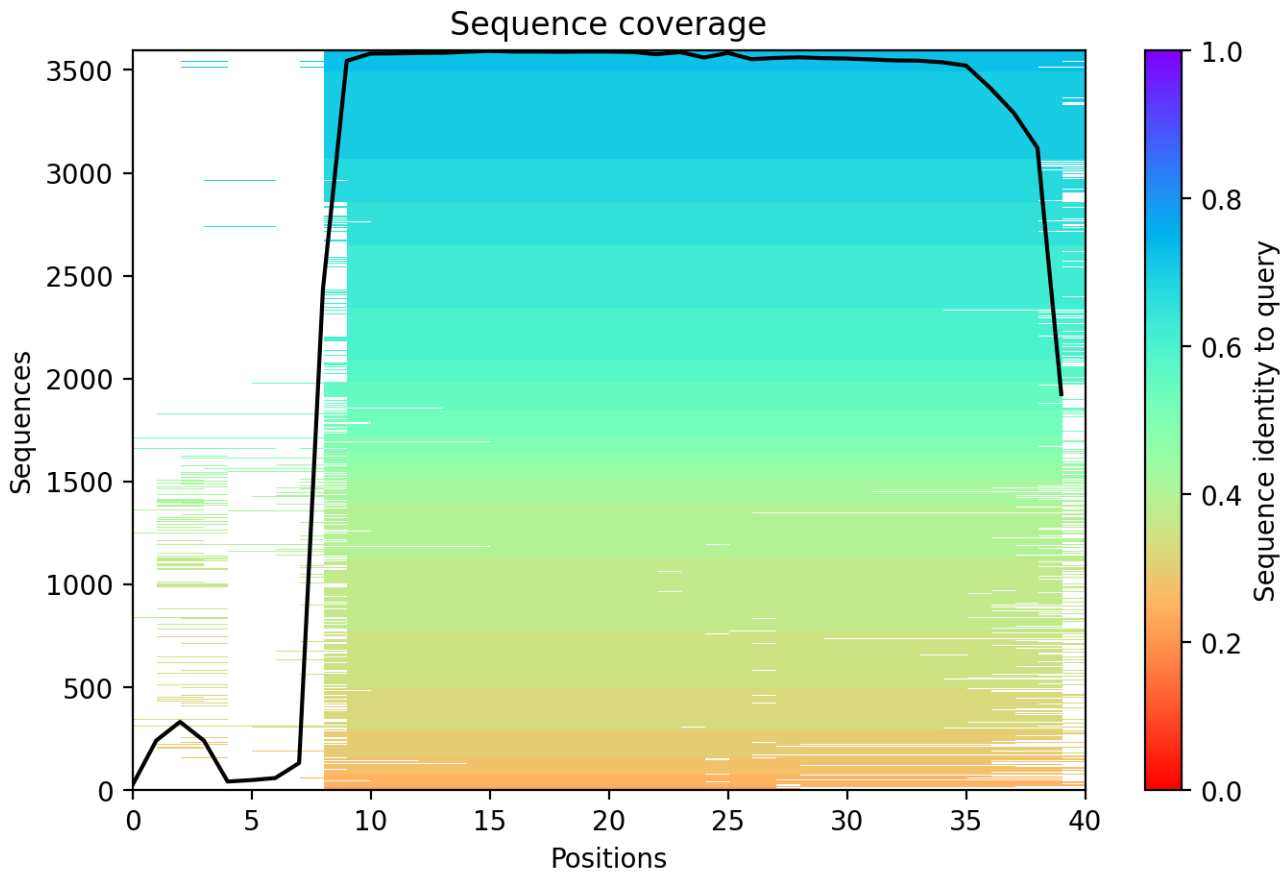

CP35: MtrunA17_Chr2g0292921

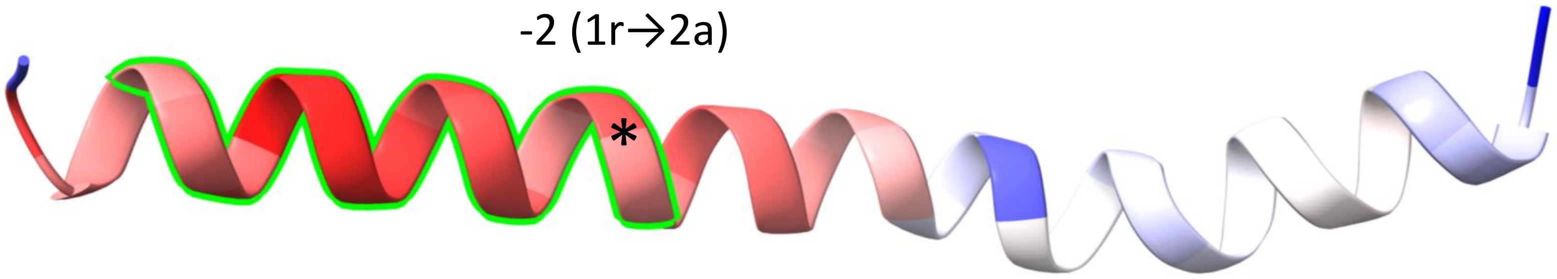


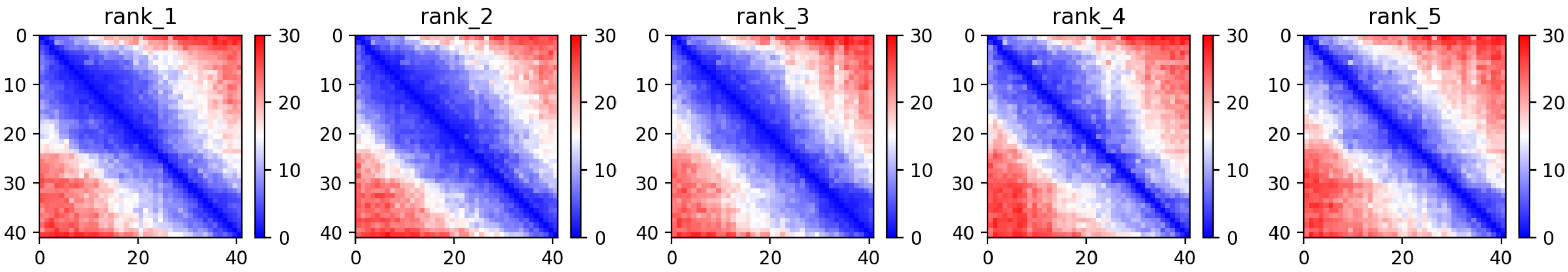

CP35: MtrunA17_Chr2g0292921

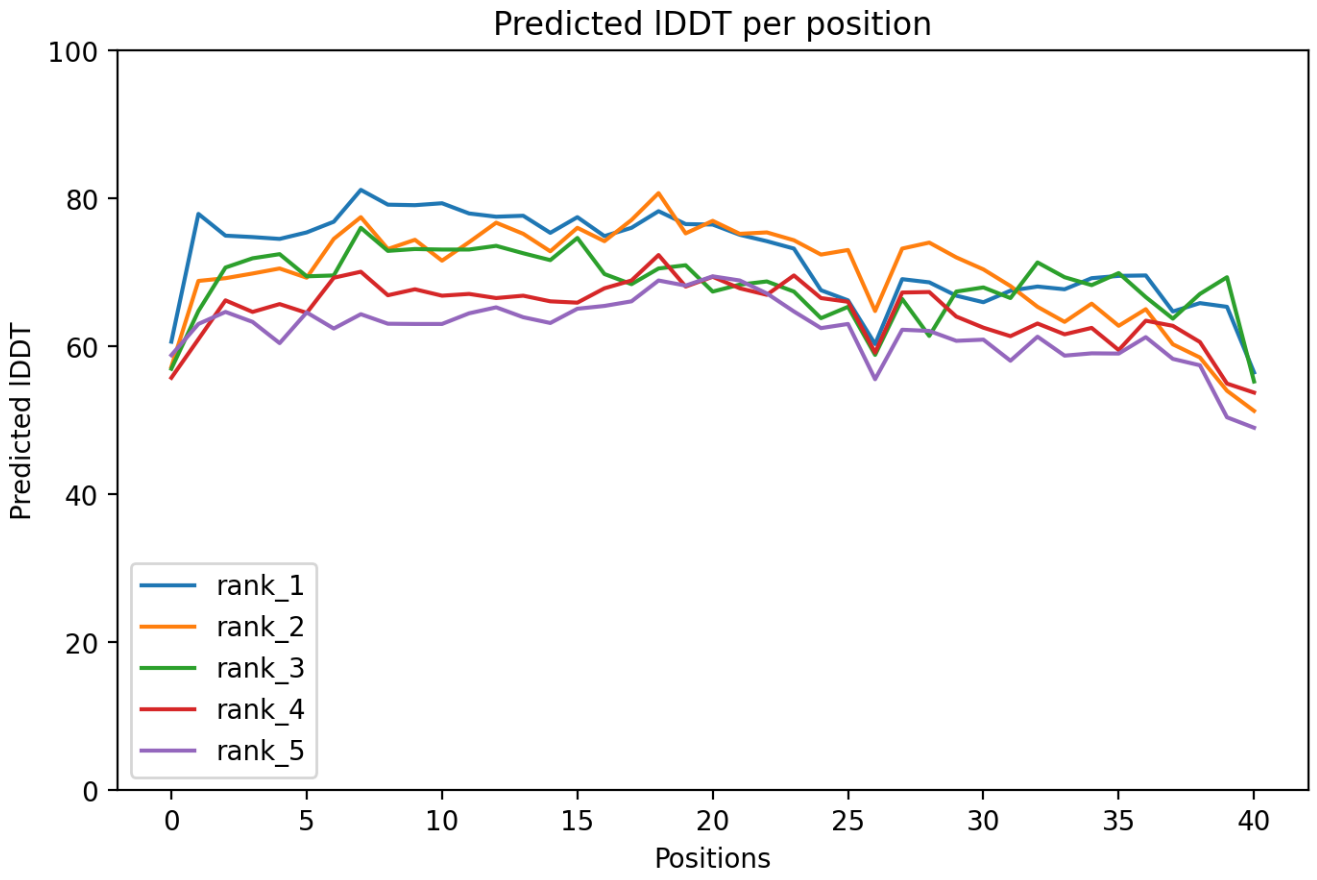

CP35: MtrunA17_Chr2g0292921

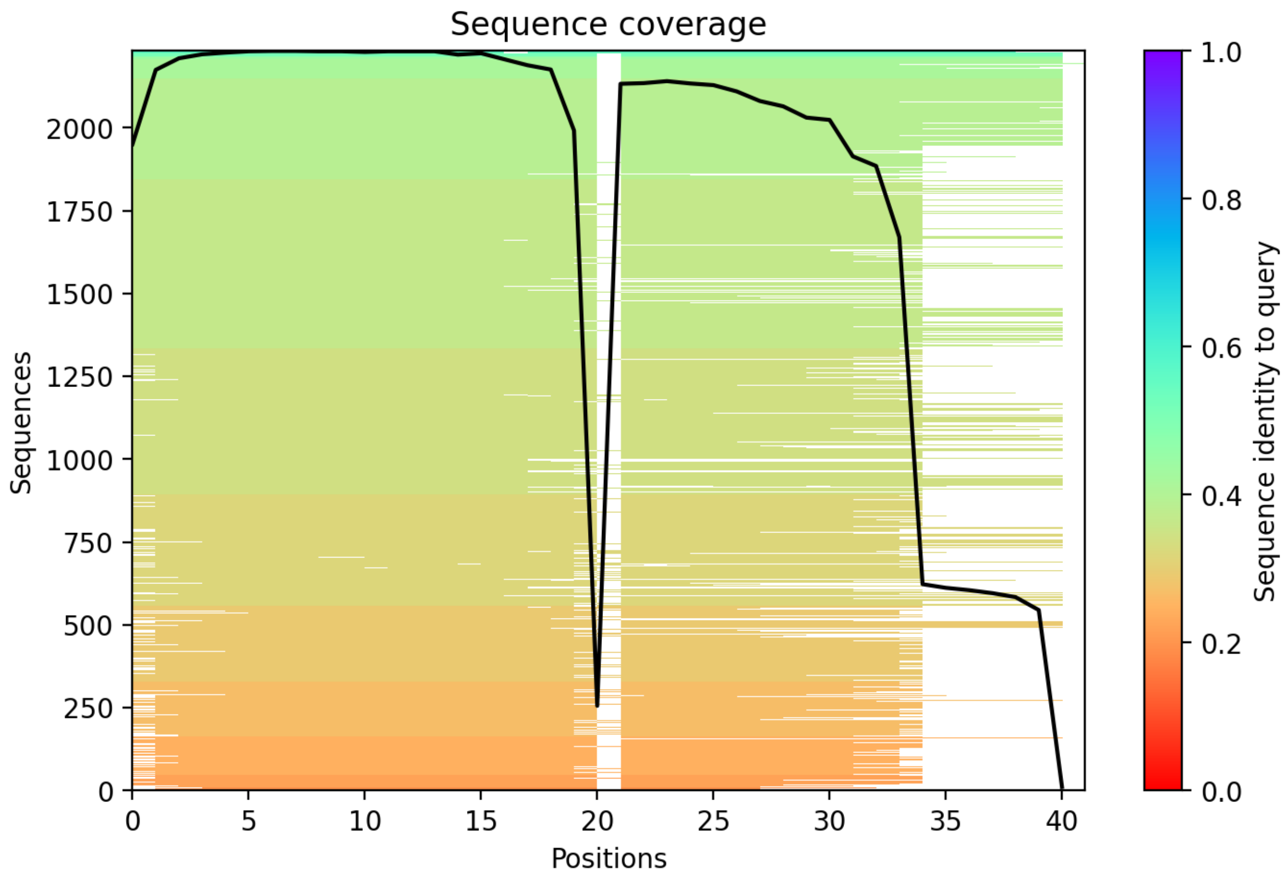

CP36: MtrunA17_Chr2g0298731

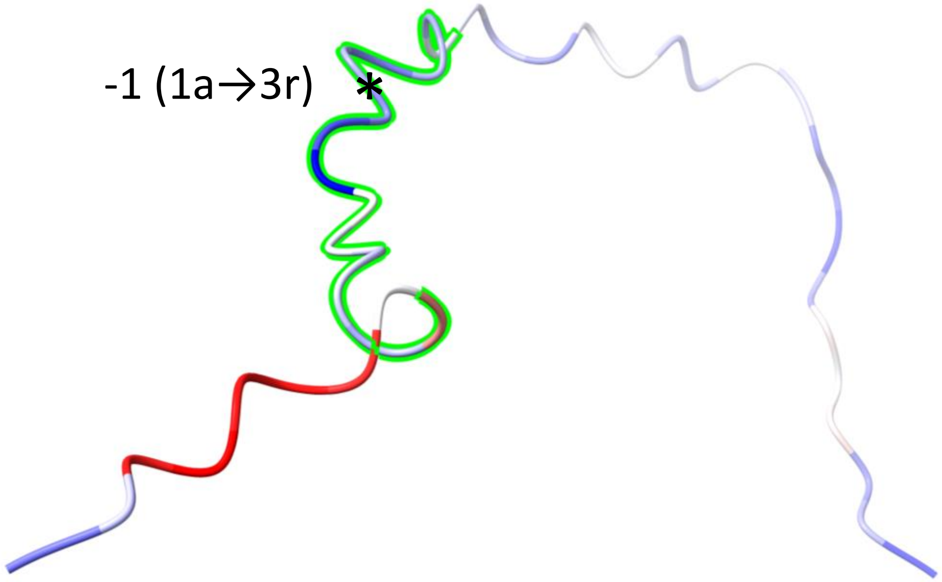


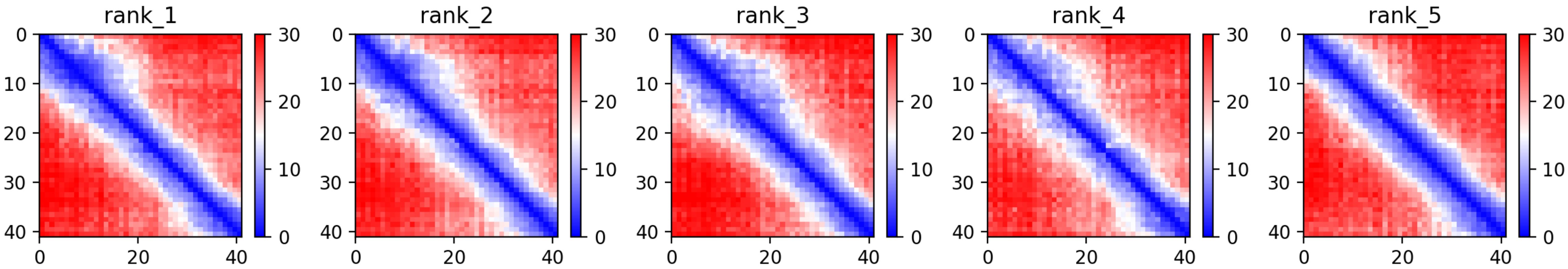

CP36: MtrunA17_Chr2g0298731

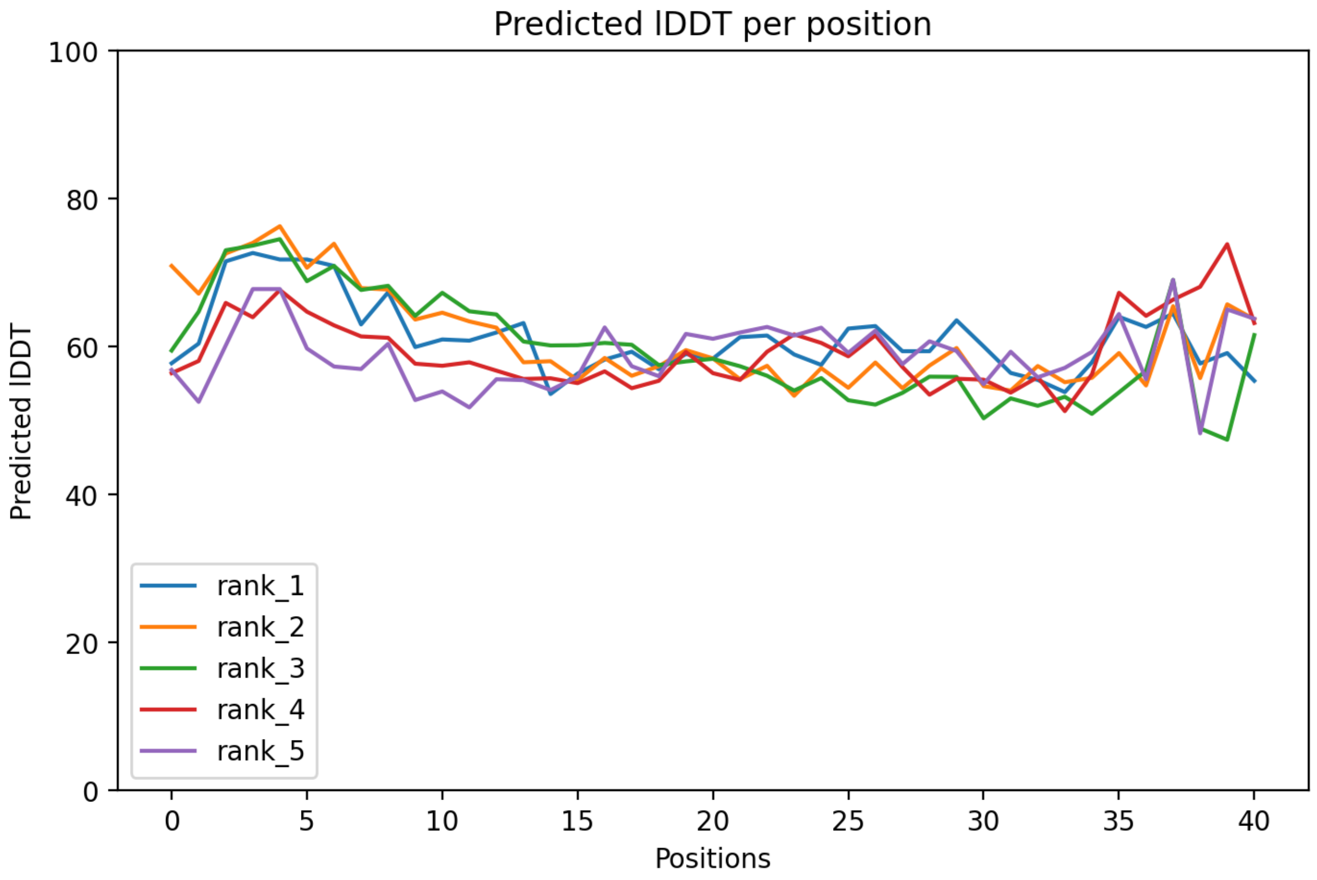

### CP36: MtrunA17_Chr2g0298731

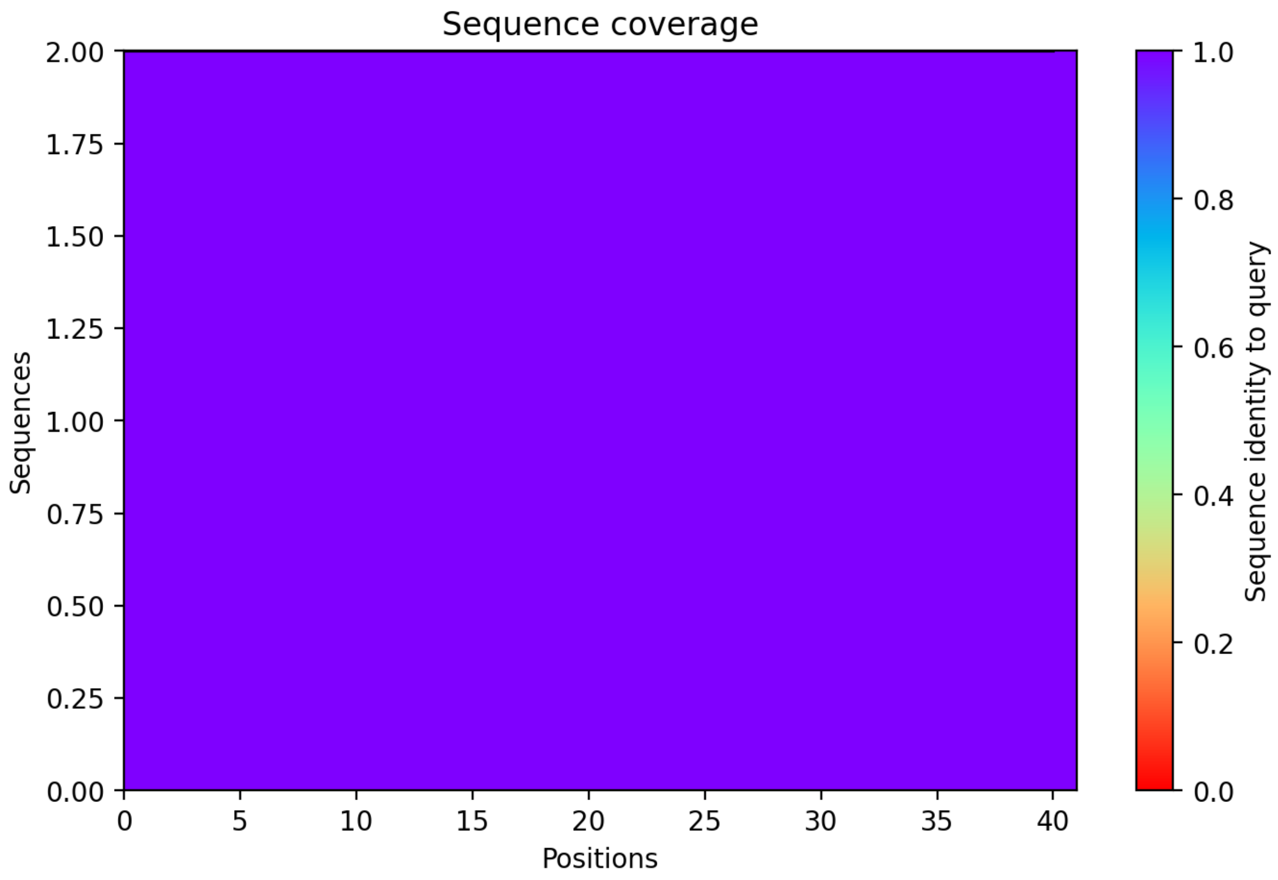

CP37: MtrunA17_Chr2g0299561

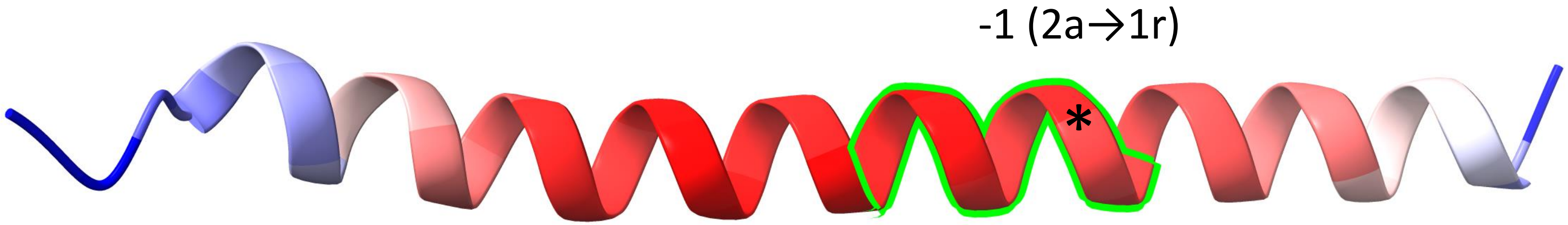


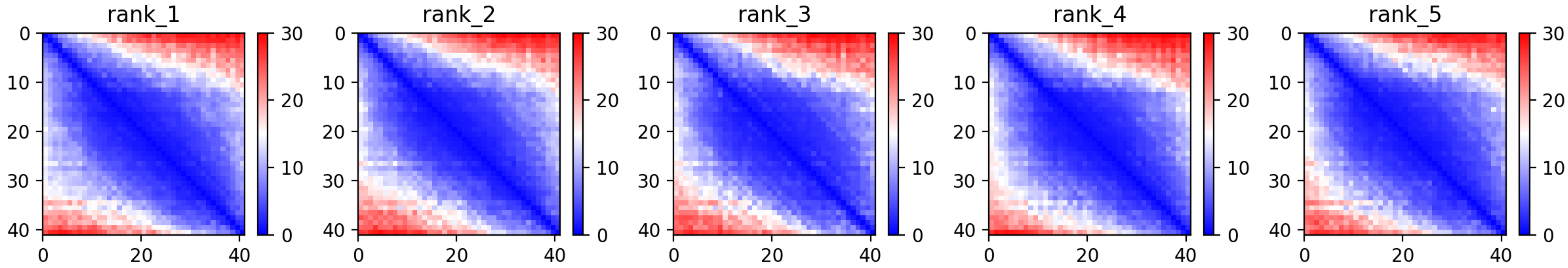

CP37: MtrunA17_Chr2g0299561

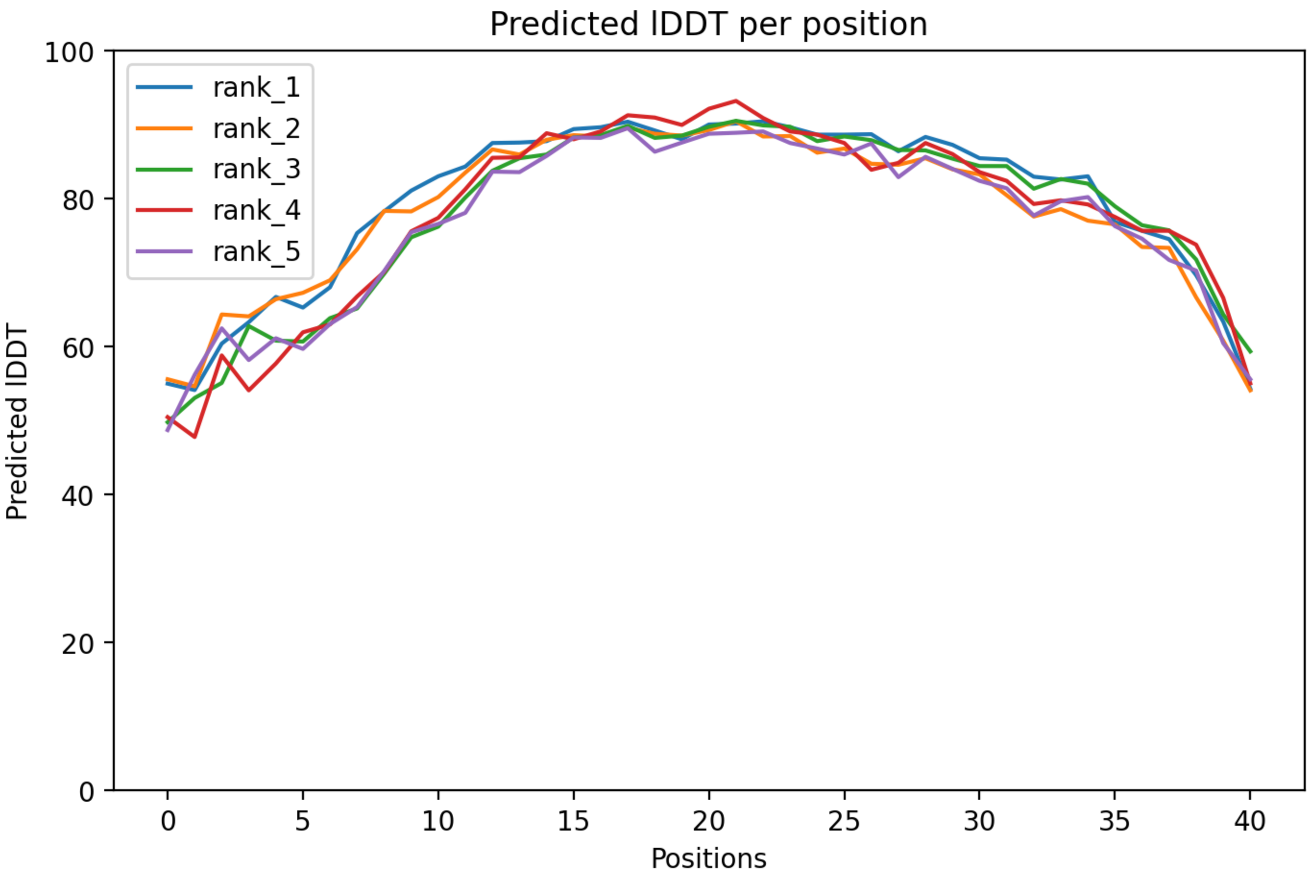

### CP37: MtrunA17_Chr2g0299561

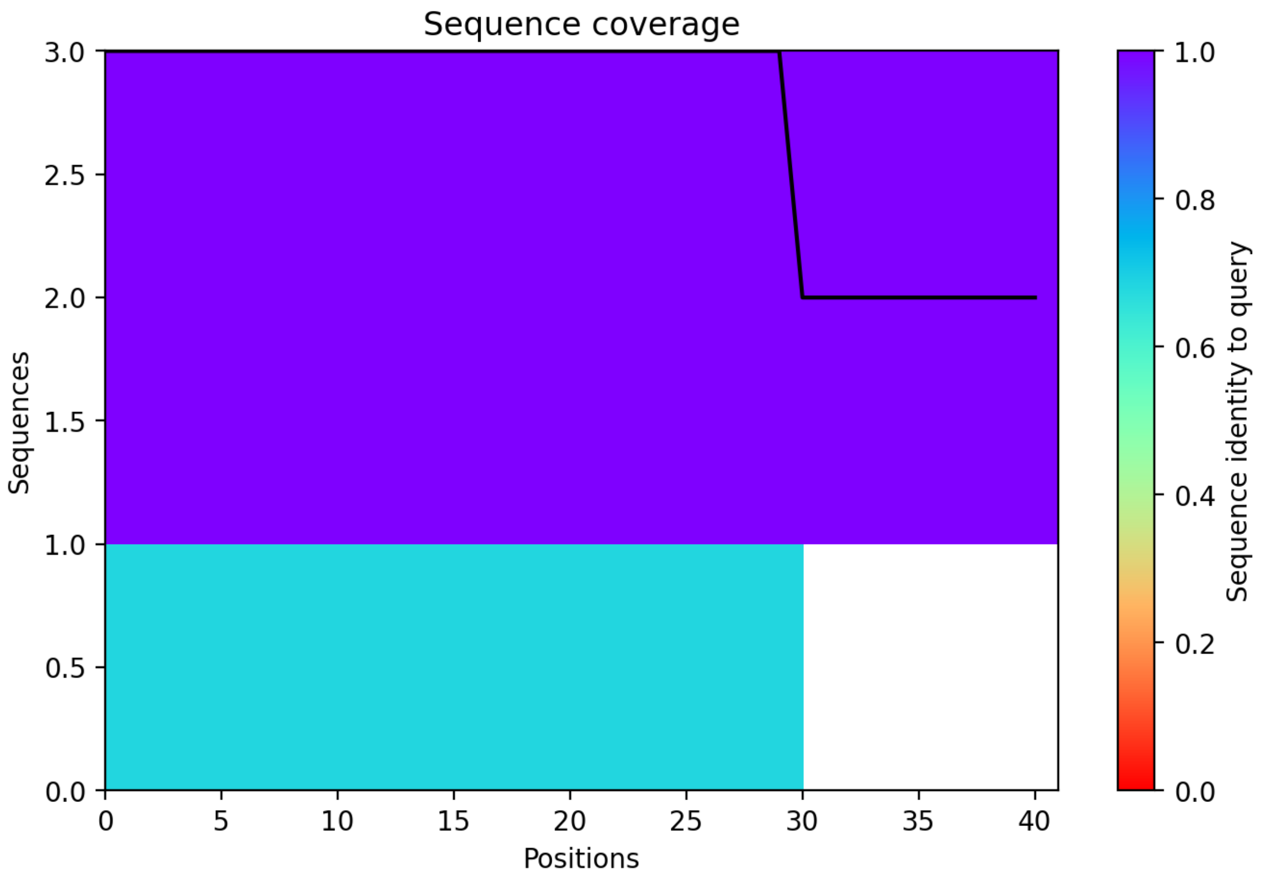

CP38: MtrunA17_Chr2g0304891

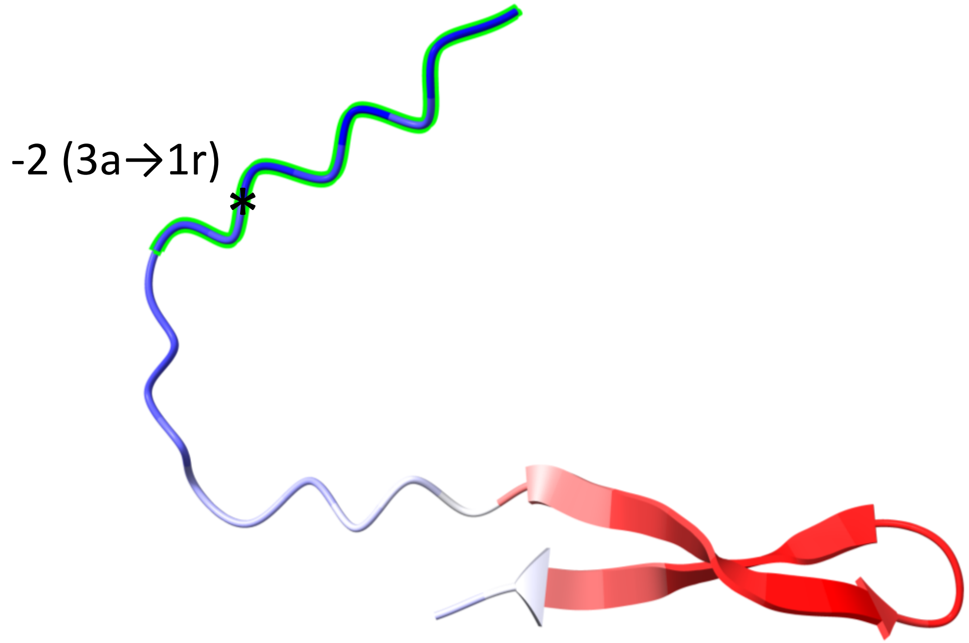


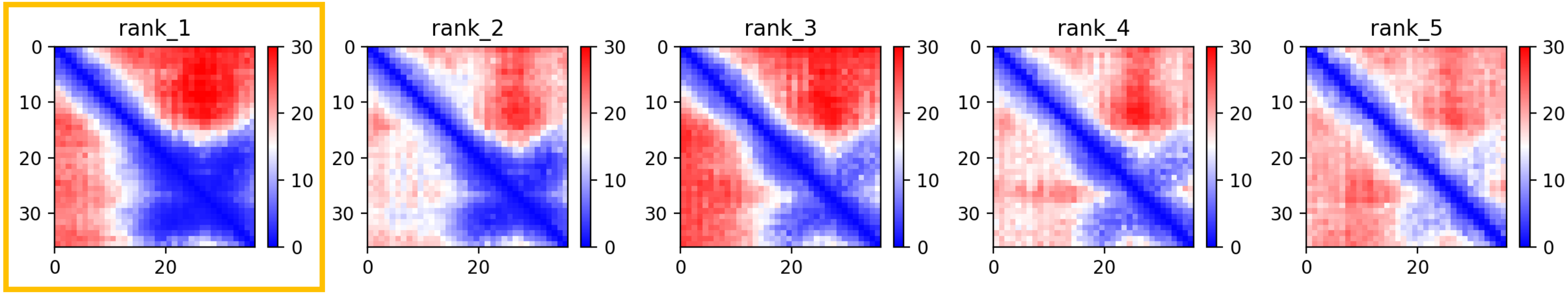

### CP38: MtrunA17_Chr2g0304891

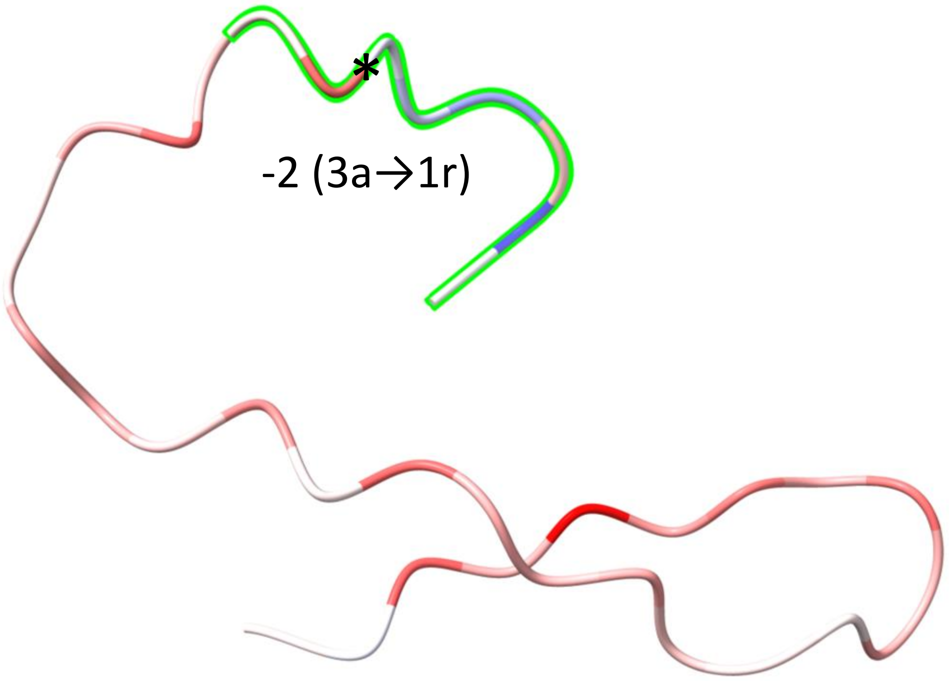


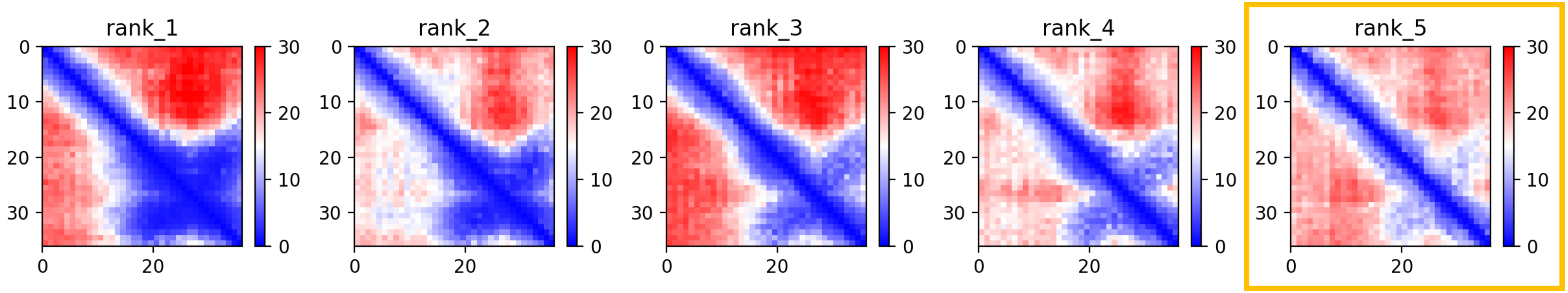

CP38: MtrunA17_Chr2g0304891

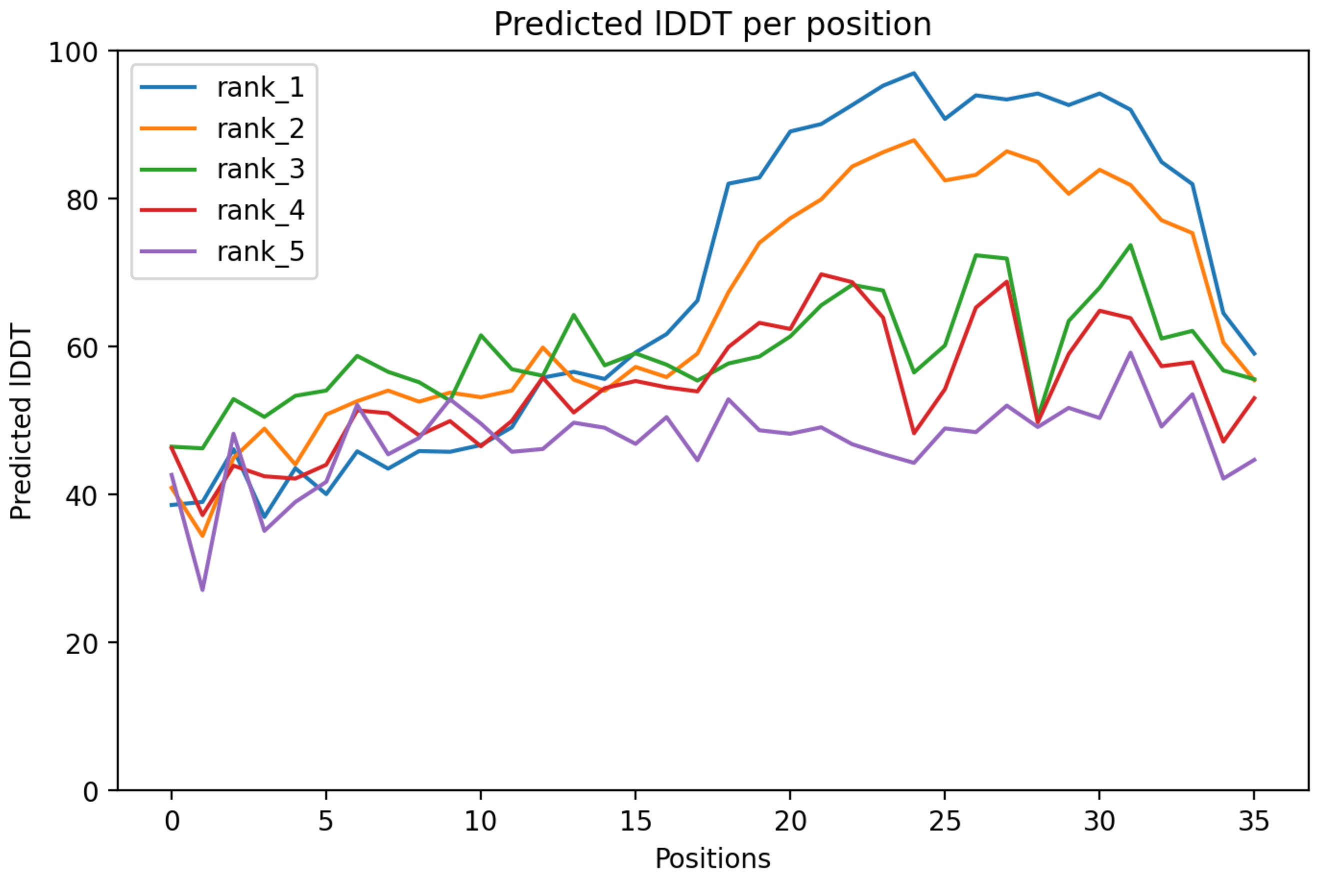

### CP38: MtrunA17_Chr2g0304891

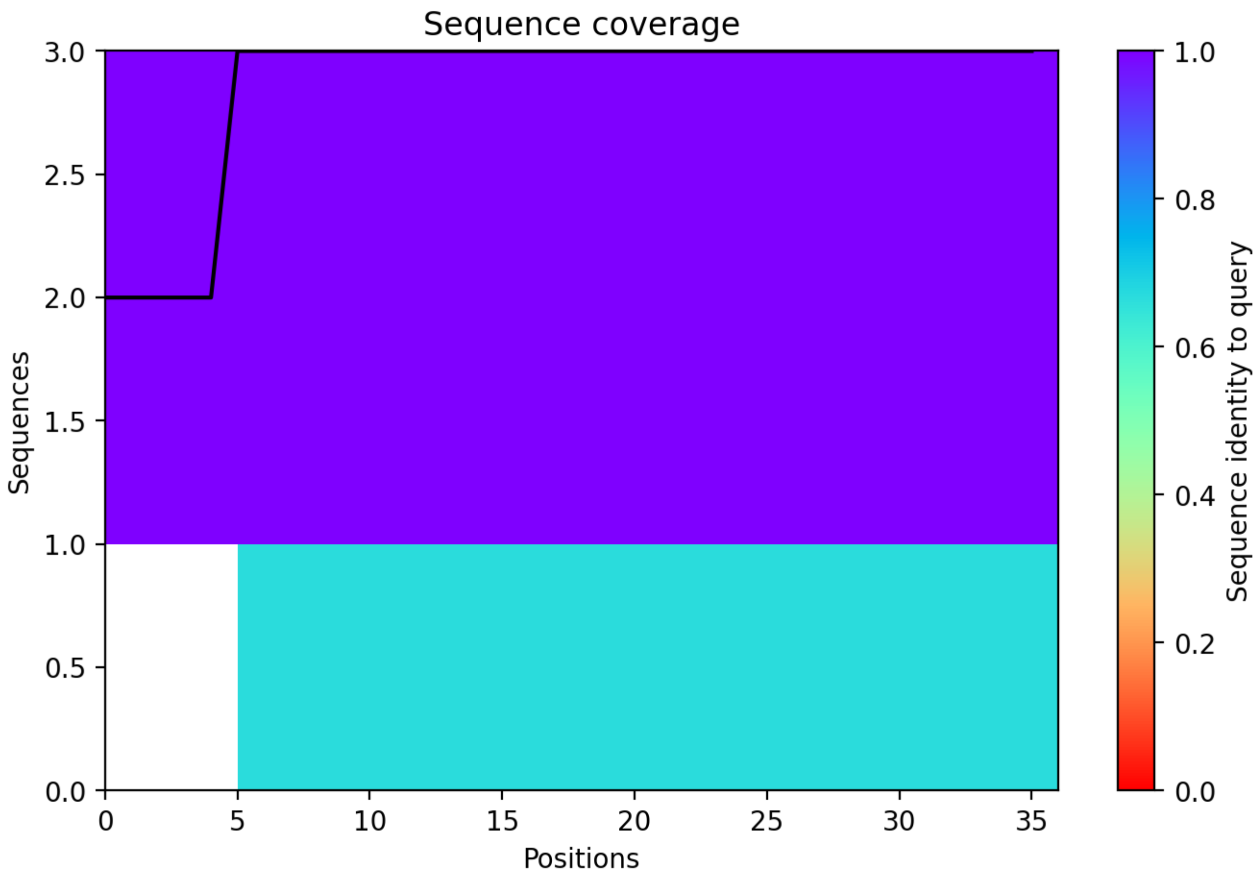

CP39: MtrunA17_Chr2g0305951

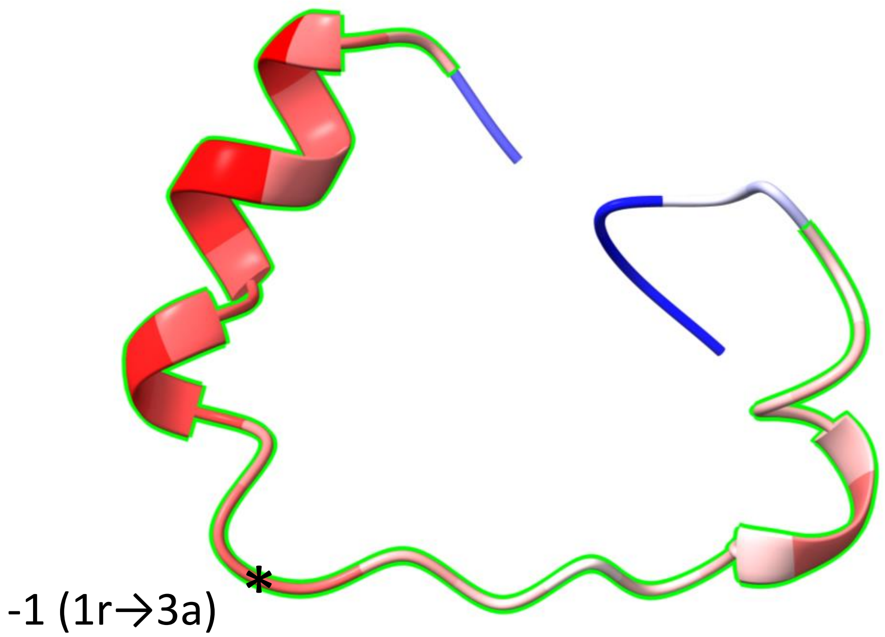


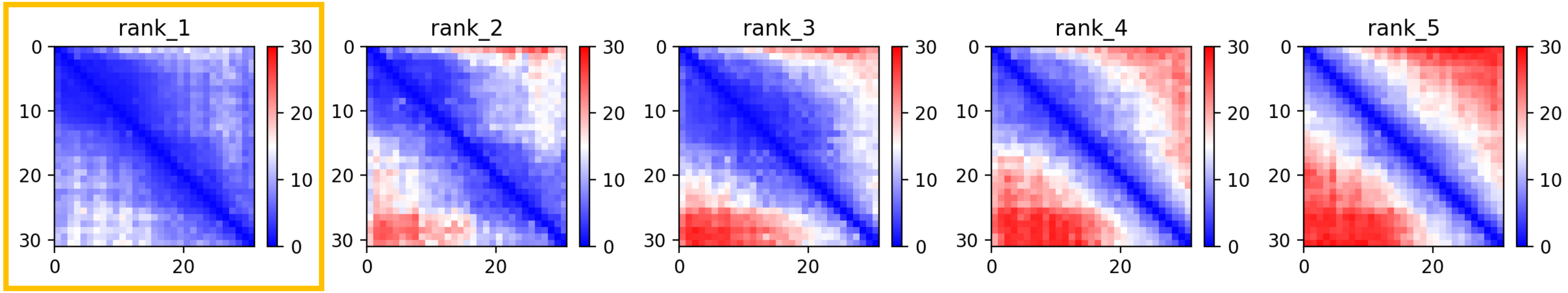

CP39: MtrunA17_Chr2g0305951

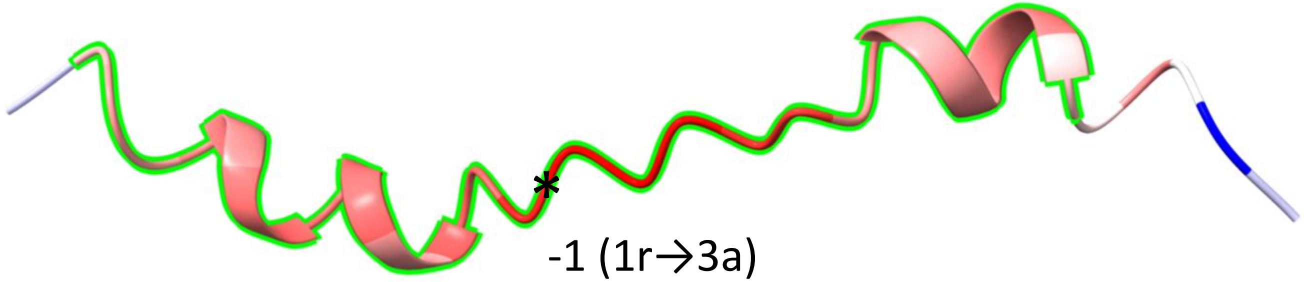


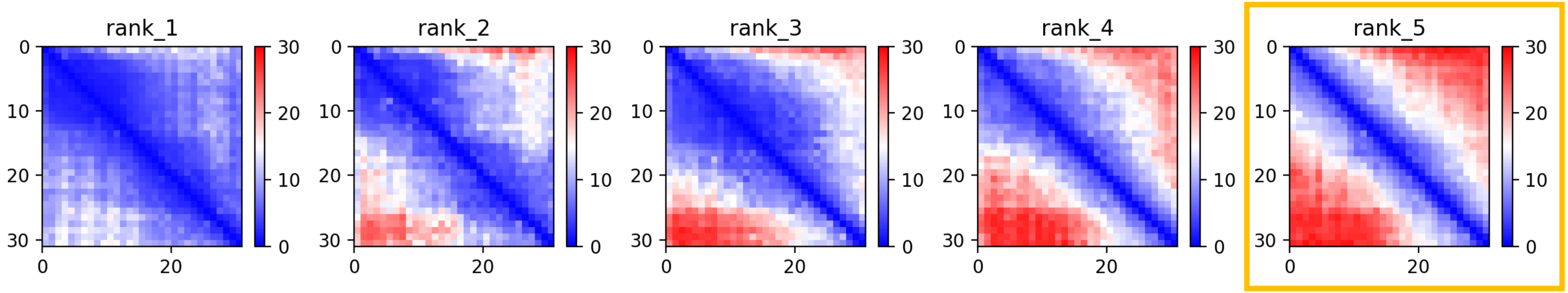

CP39: MtrunA17_Chr2g0305951

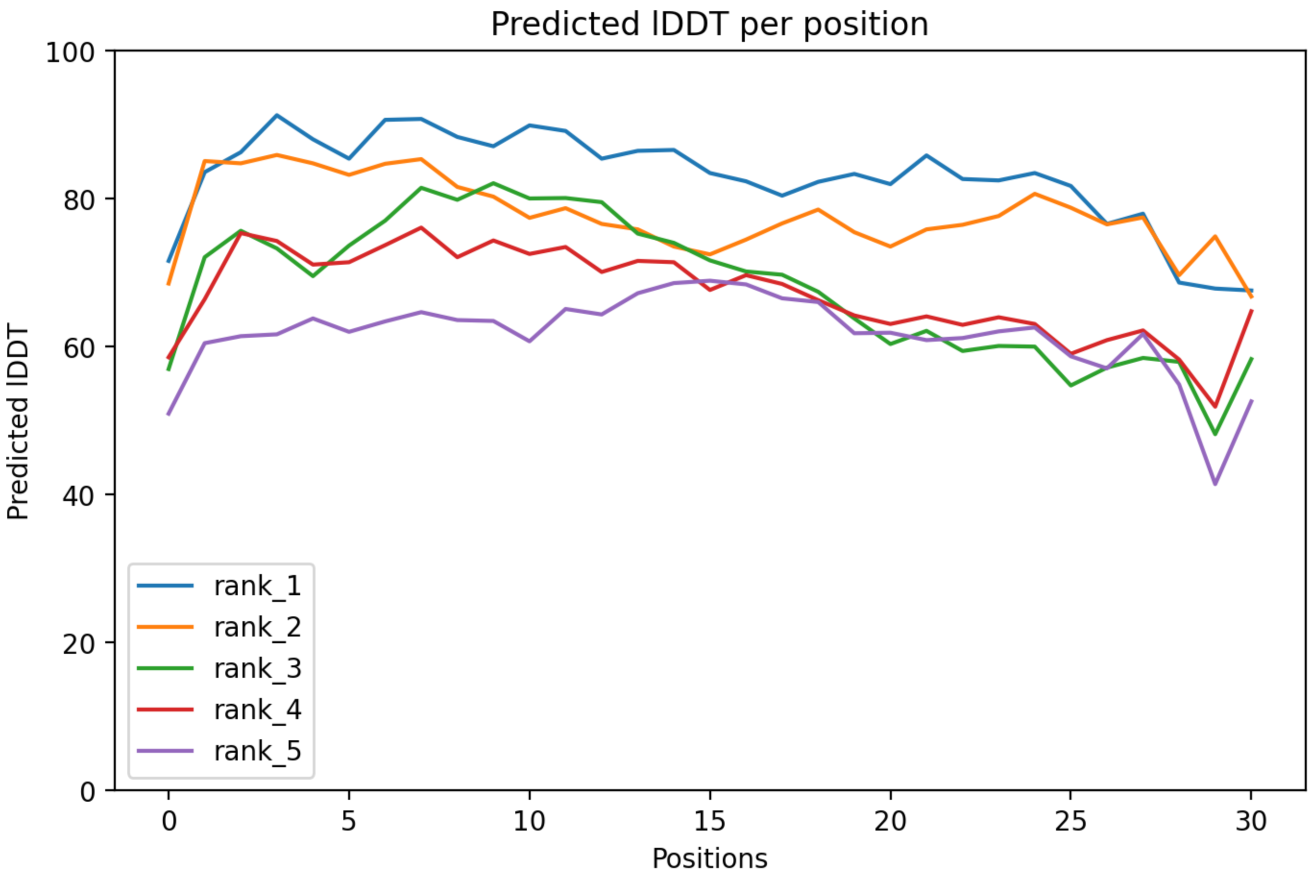

CP39: MtrunA17_Chr2g0305951

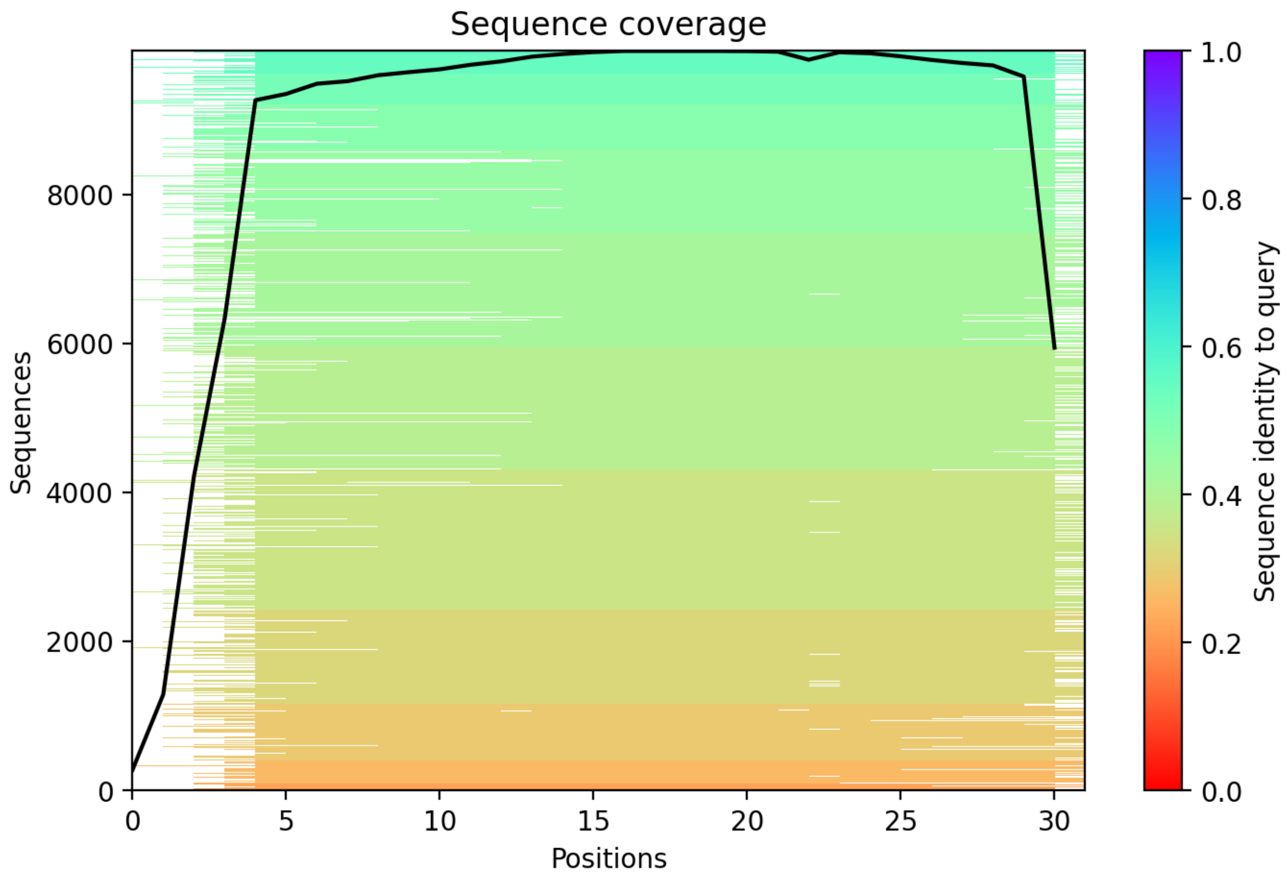

CP40: MtrunA17_Chr2g0309251

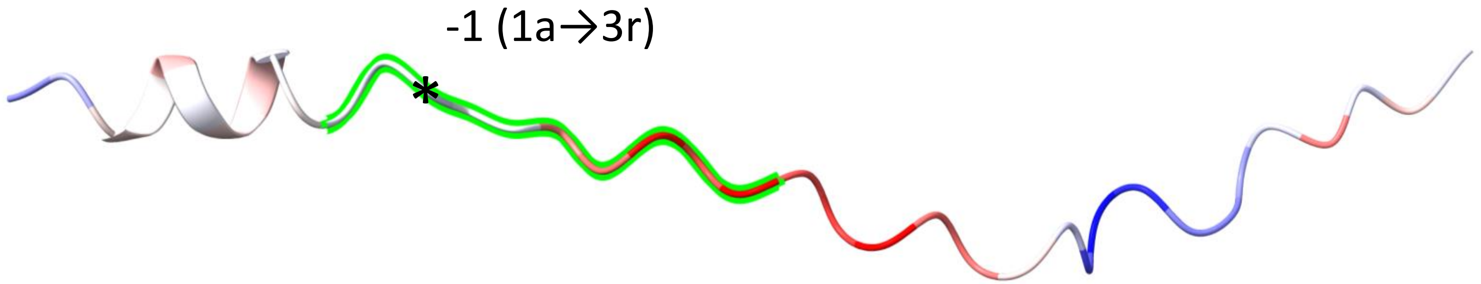


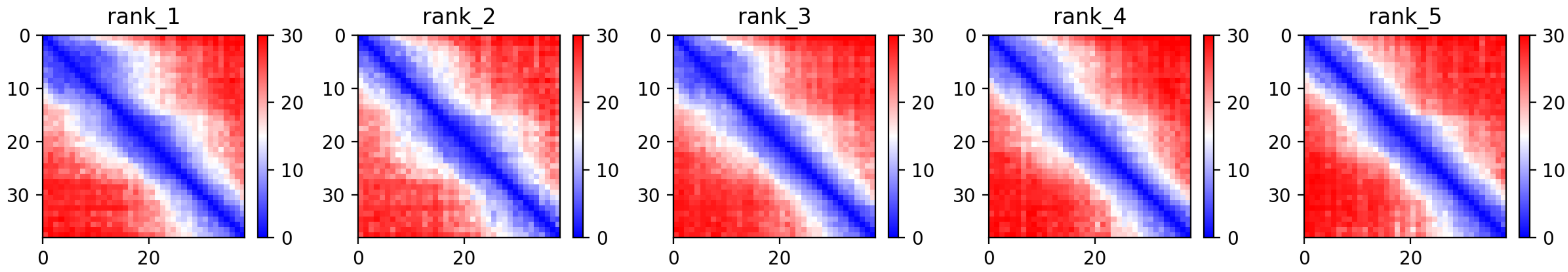

### CP40: MtrunA17_Chr2g0309251

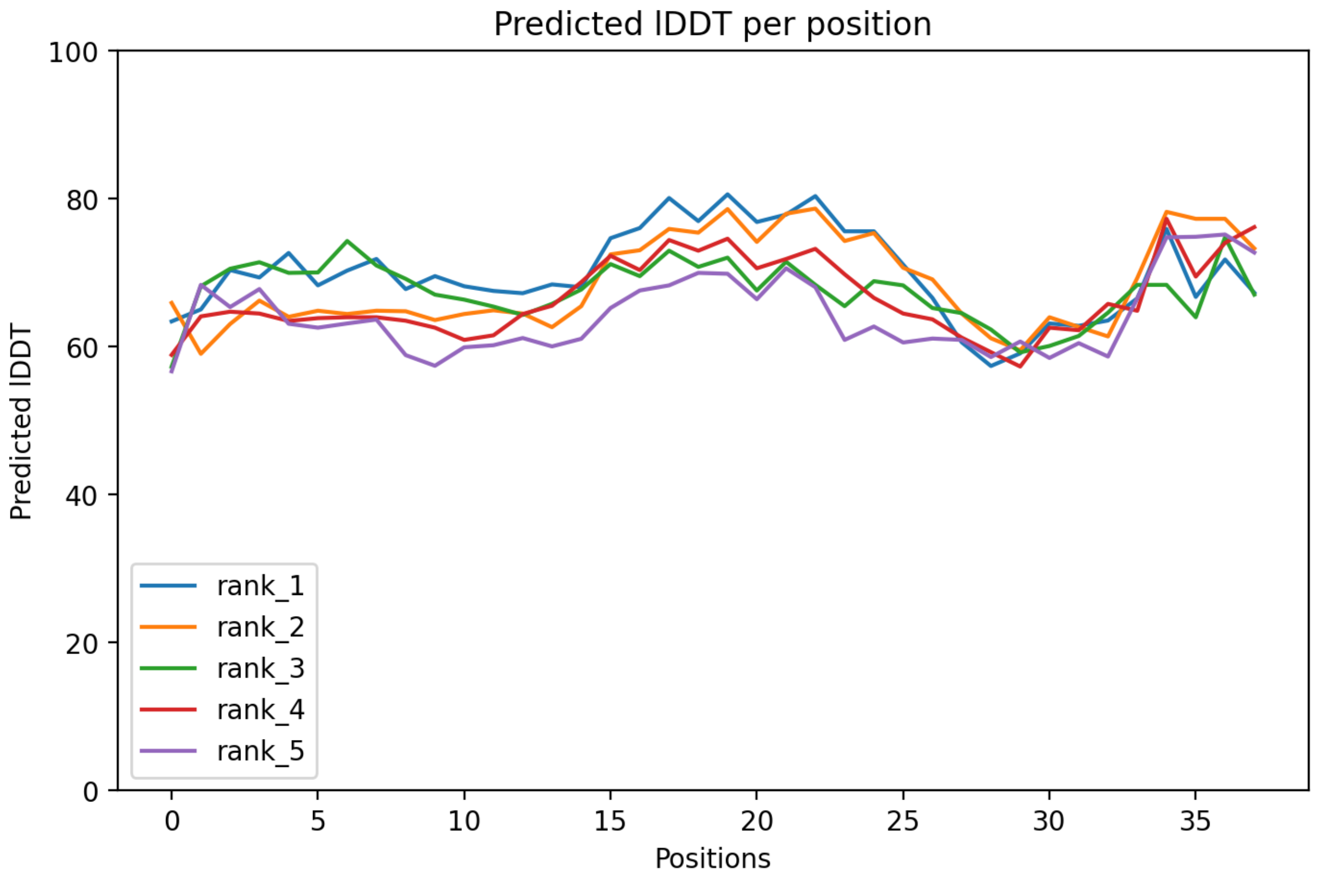

CP40: MtrunA17_Chr2g0309251

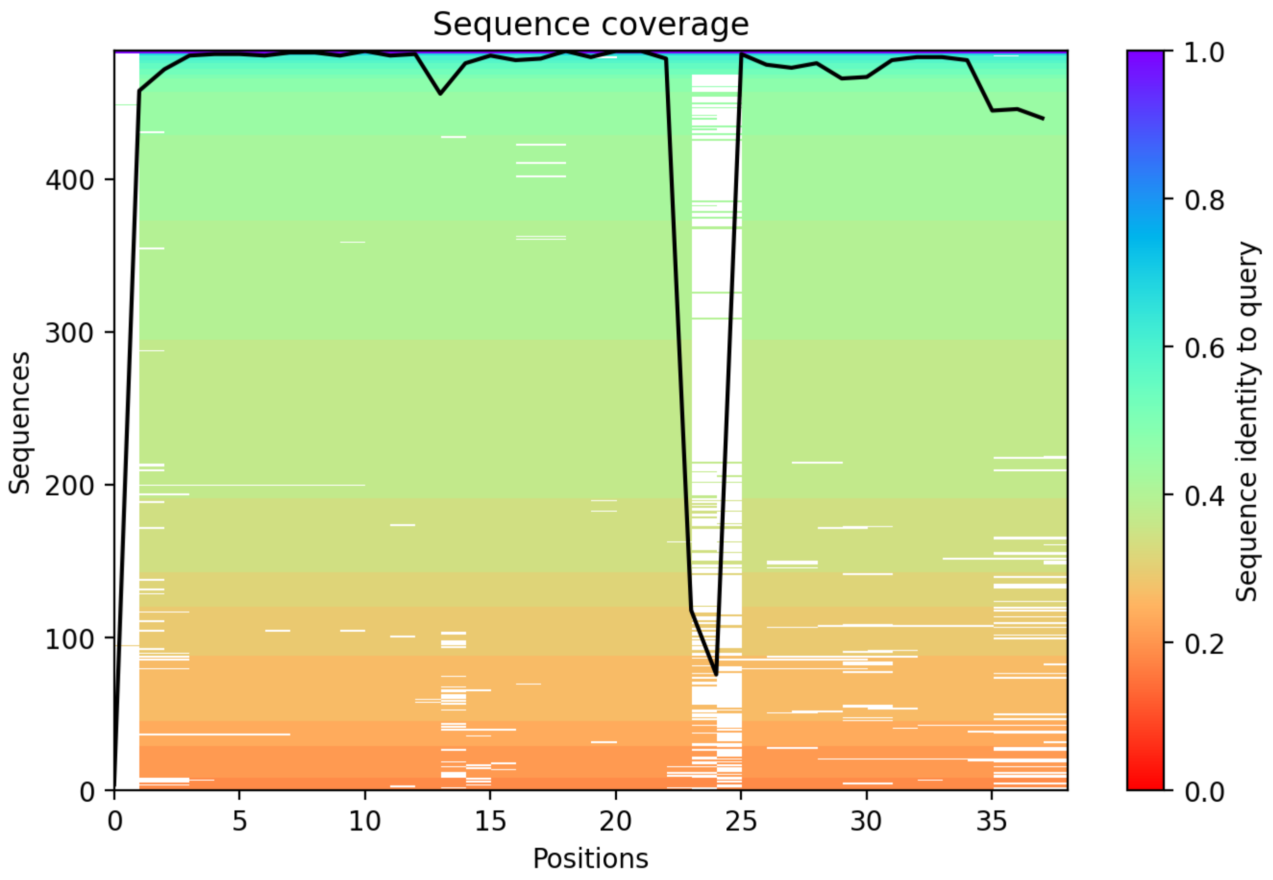

CP41: MtrunA17_Chr2g0310041

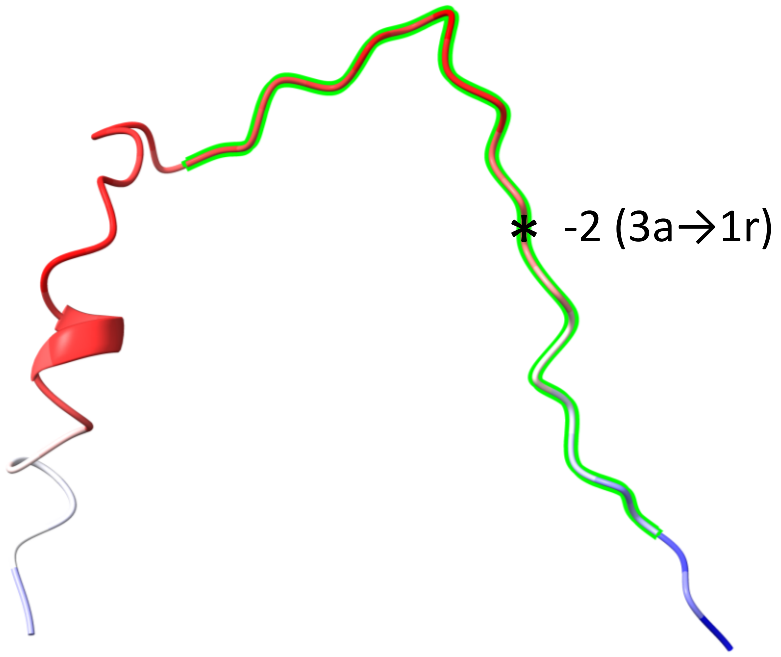


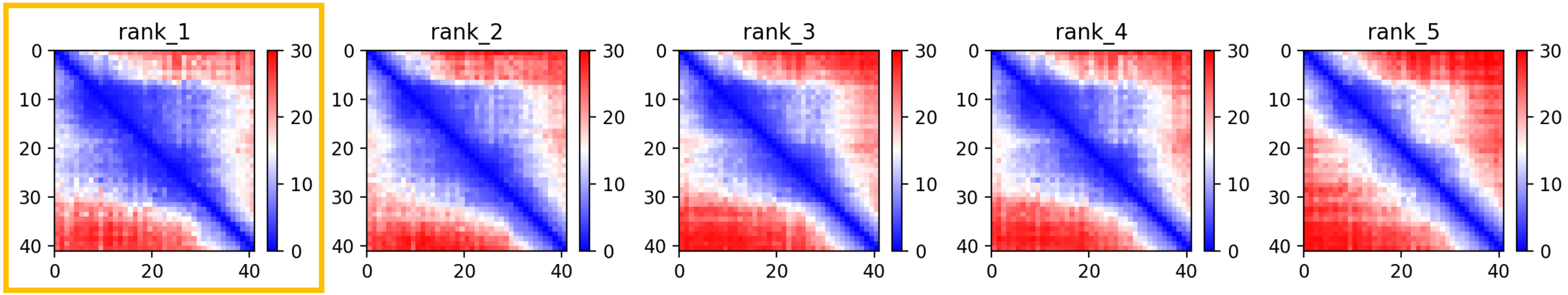

CP41: MtrunA17_Chr2g0310041

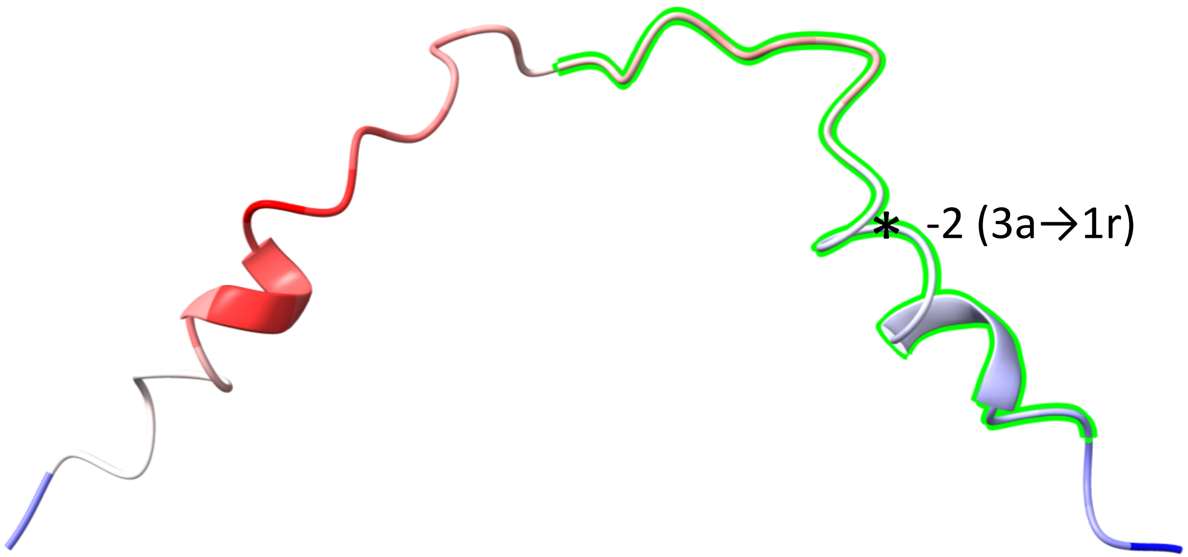


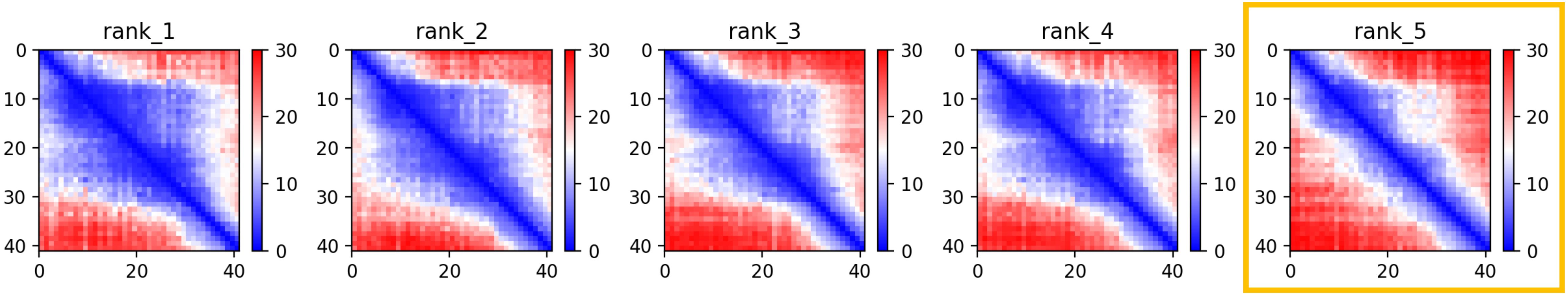

CP41: MtrunA17_Chr2g0310041

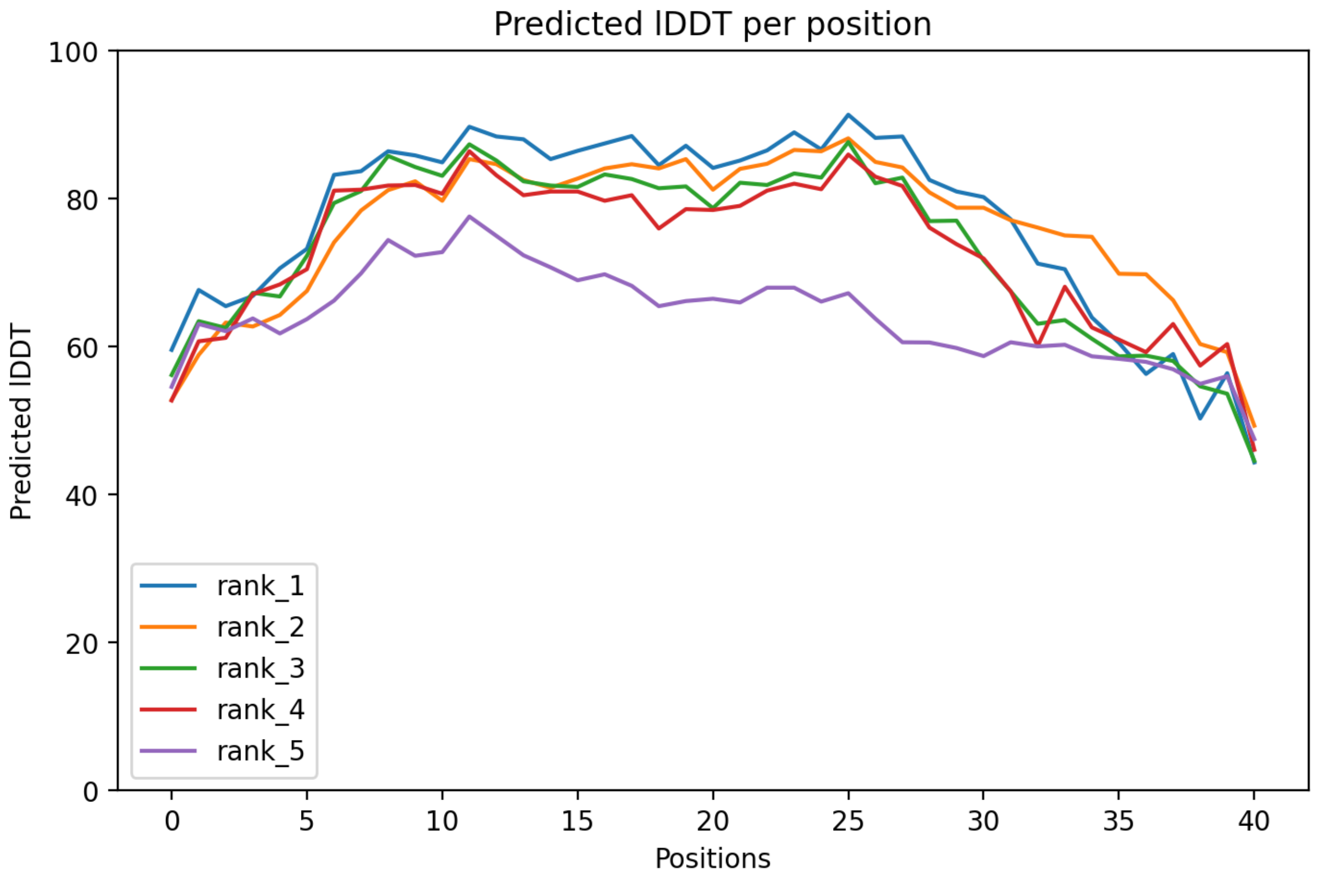

CP41: MtrunA17_Chr2g0310041

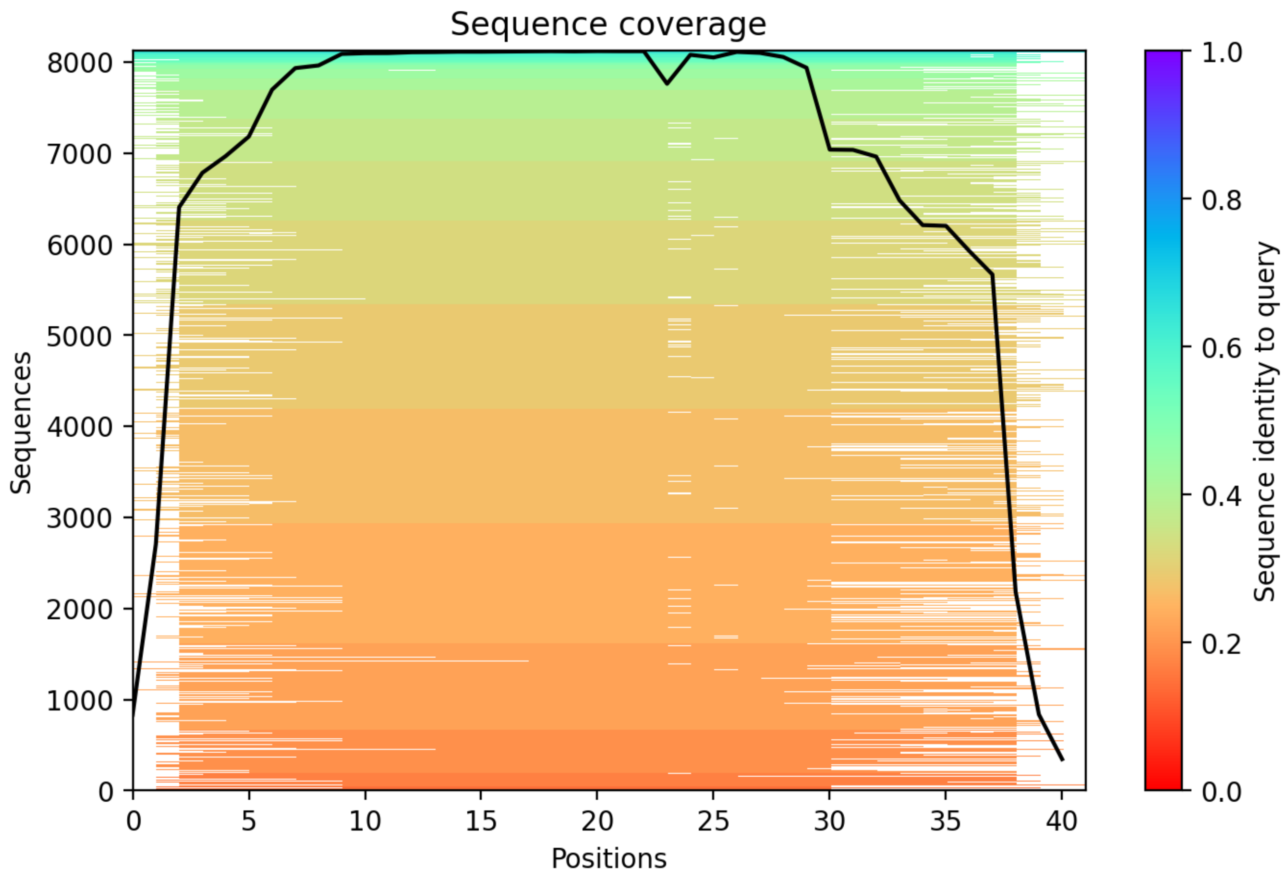

CP42: MtrunA17_Chr2g0312631

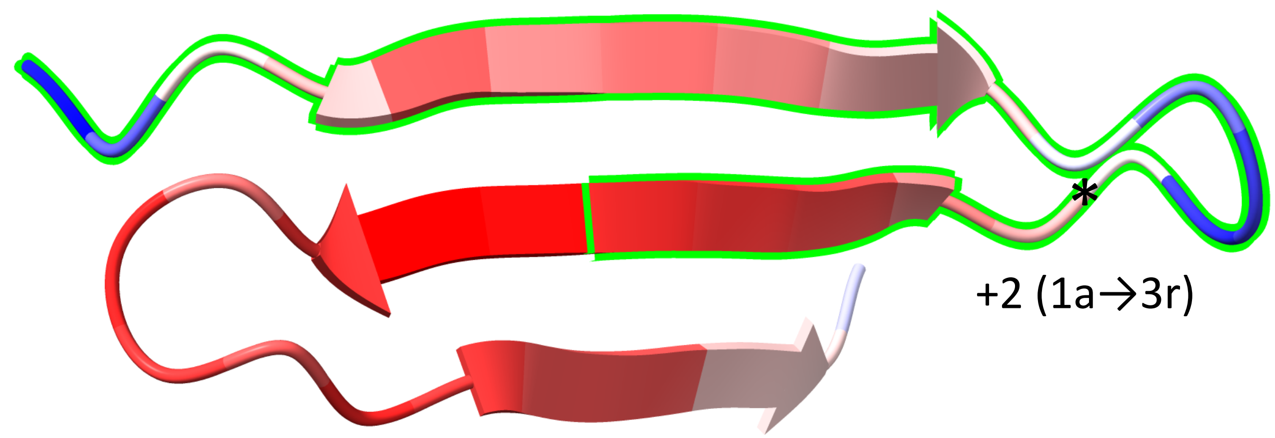


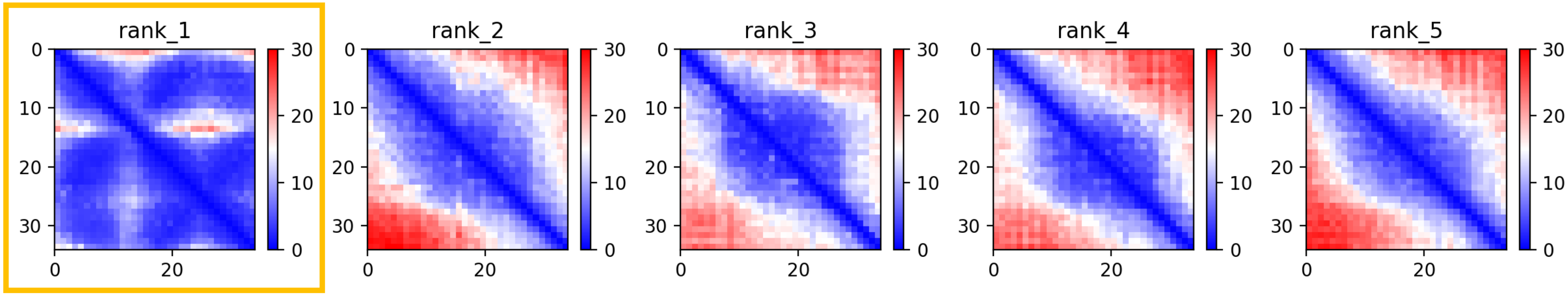

CP42: MtrunA17_Chr2g0312631

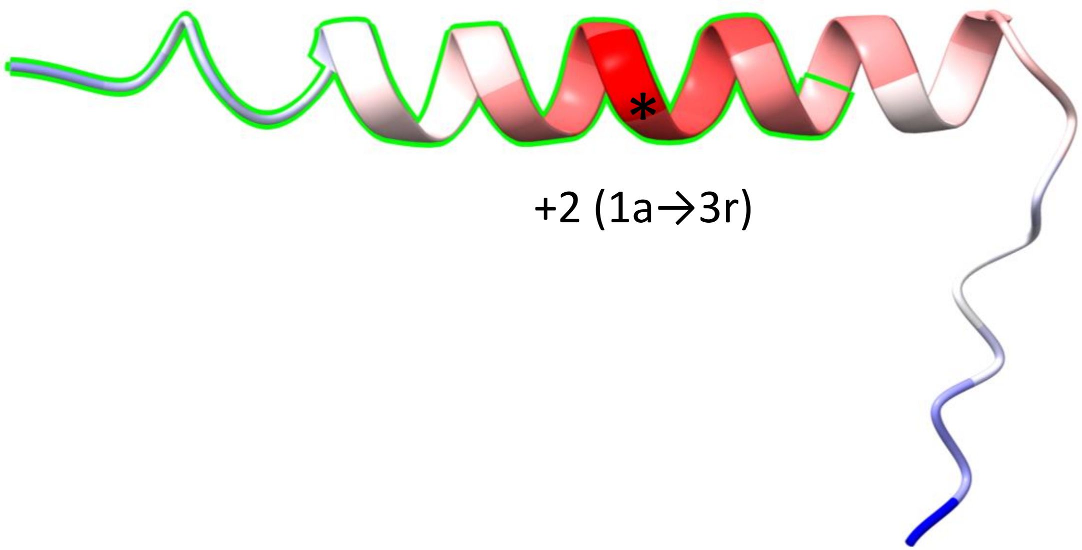


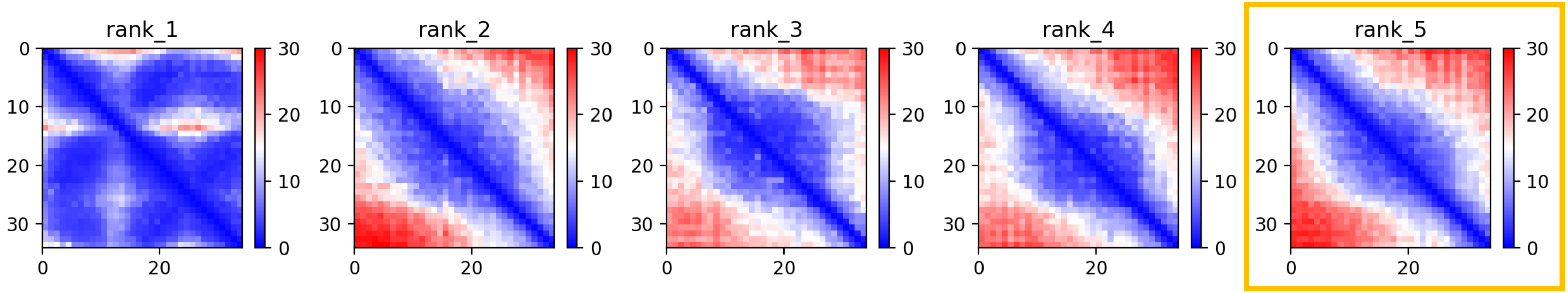

CP42: MtrunA17_Chr2g0312631

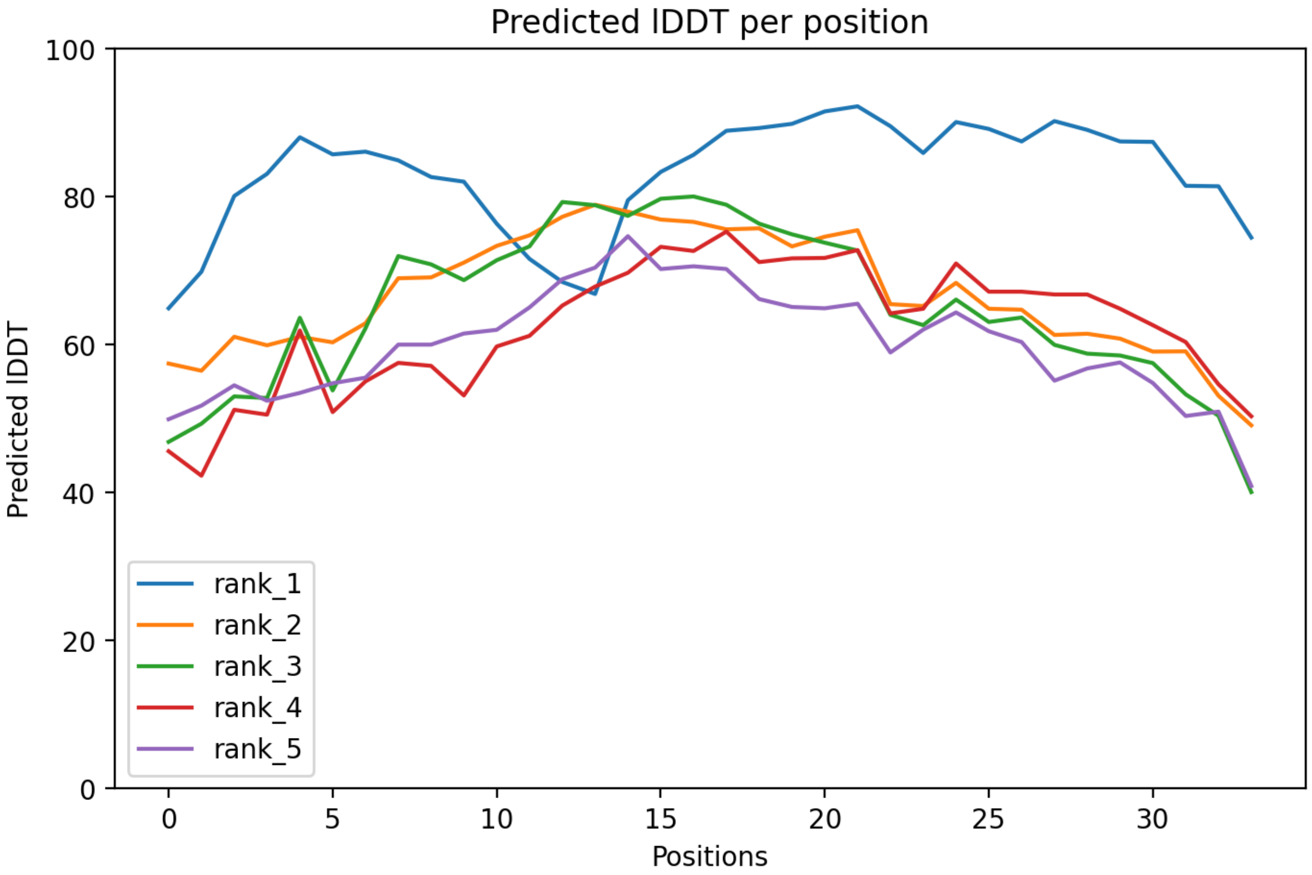

CP42: MtrunA17_Chr2g0312631

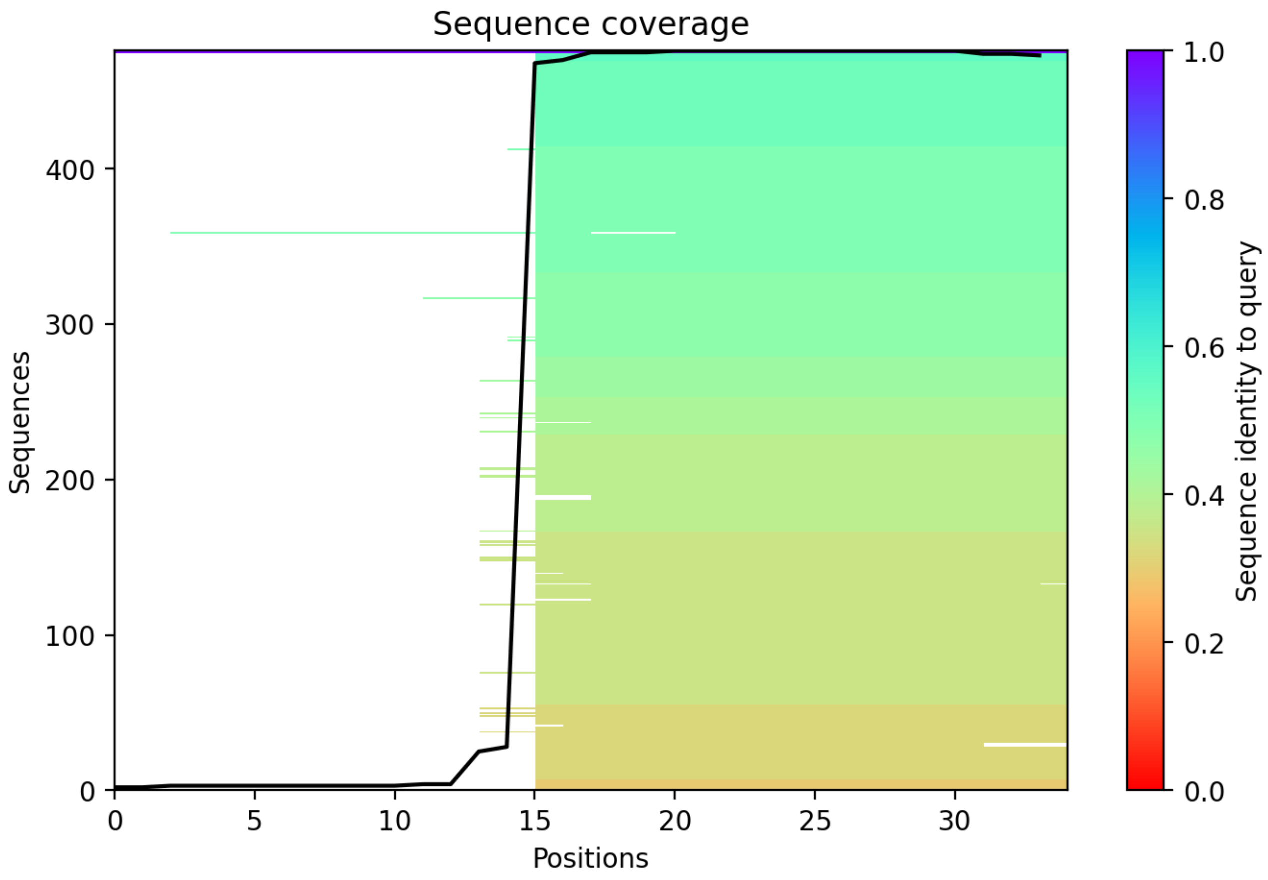

### CP43: MtrunA17_Chr2g0316291

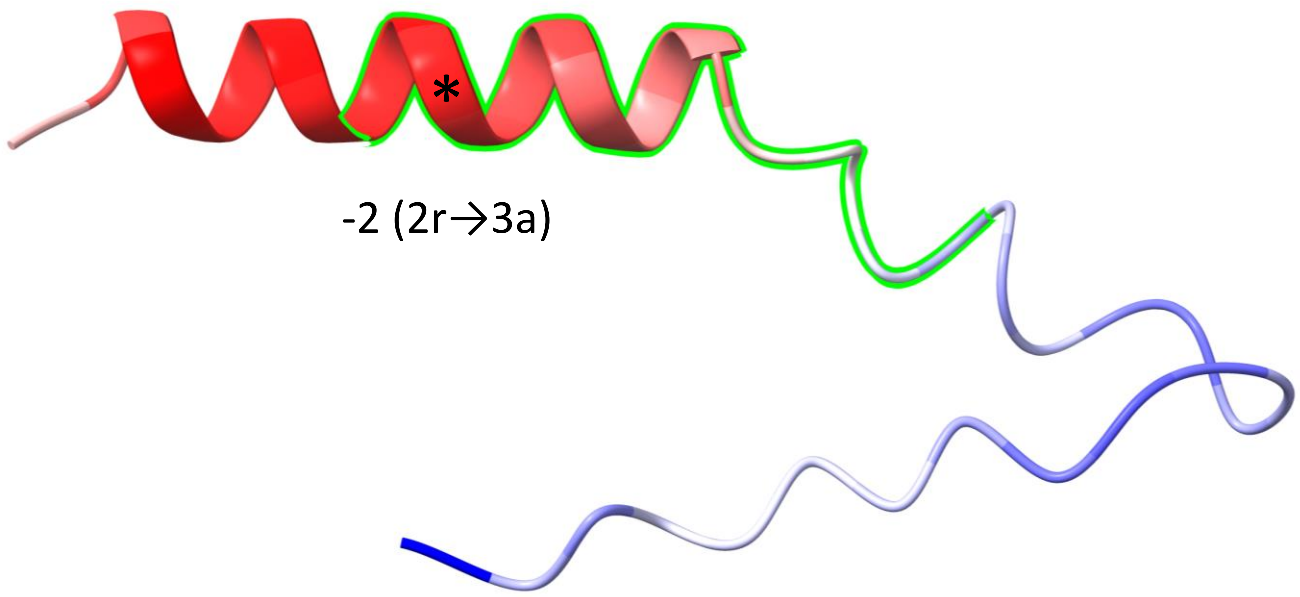


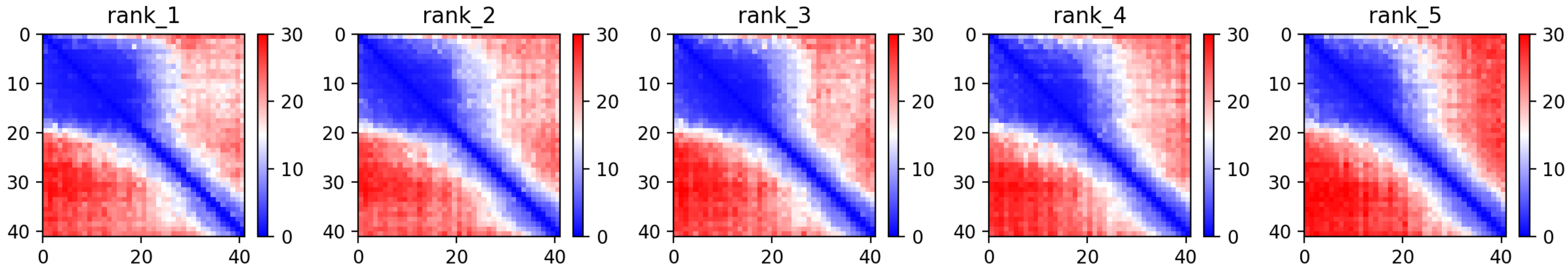

CP43: MtrunA17_Chr2g0316291

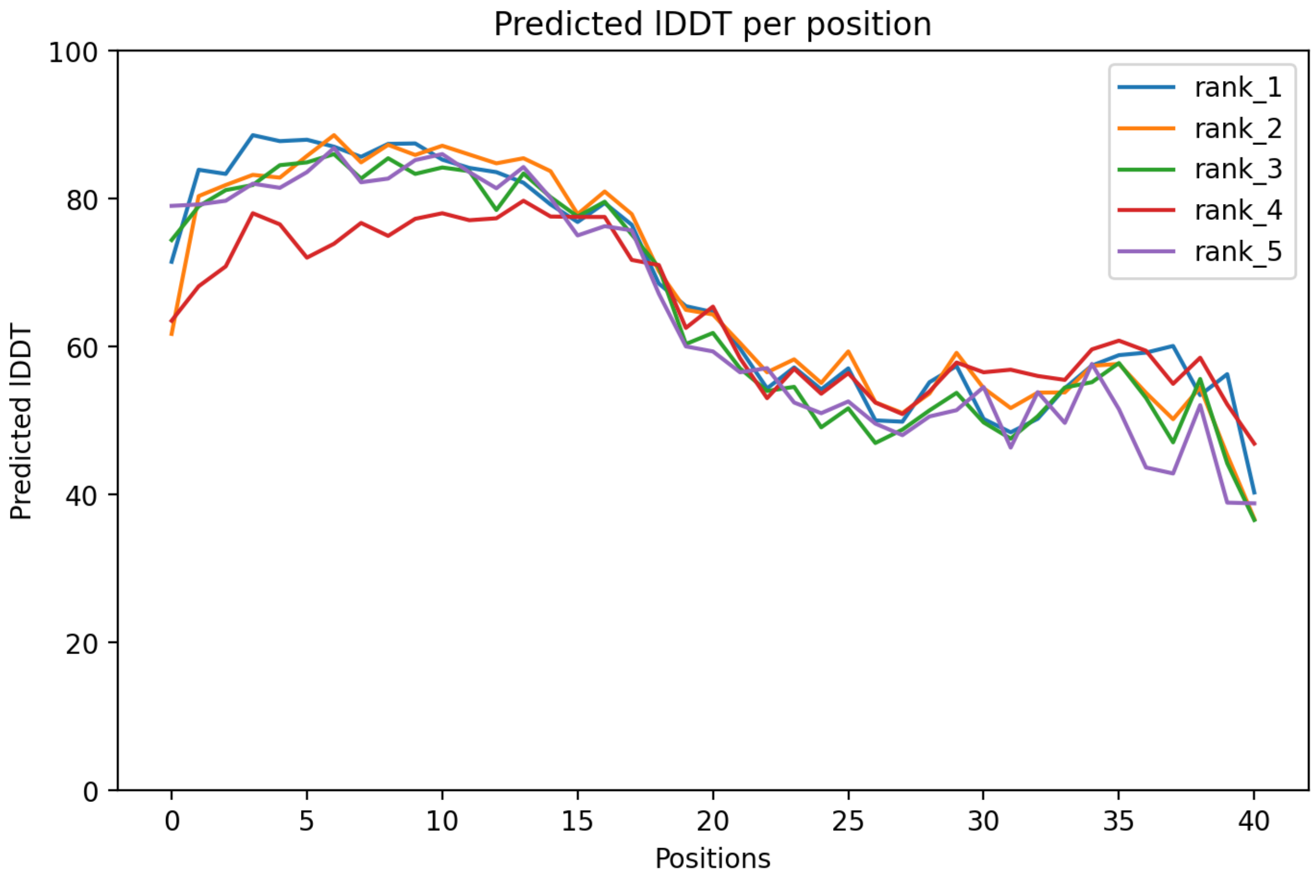

### CP43: MtrunA17_Chr2g0316291

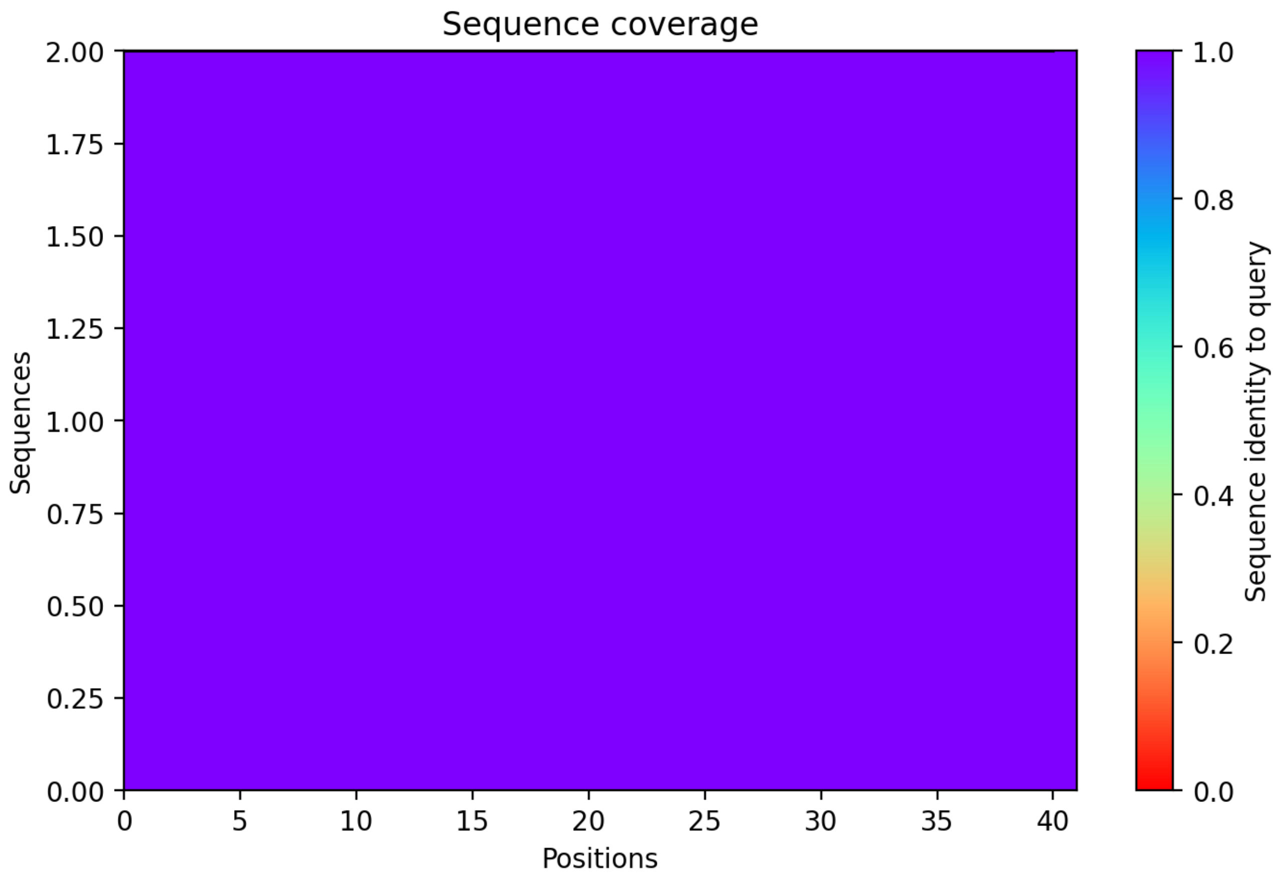

CP44: MtrunA17_Chr2g0326801

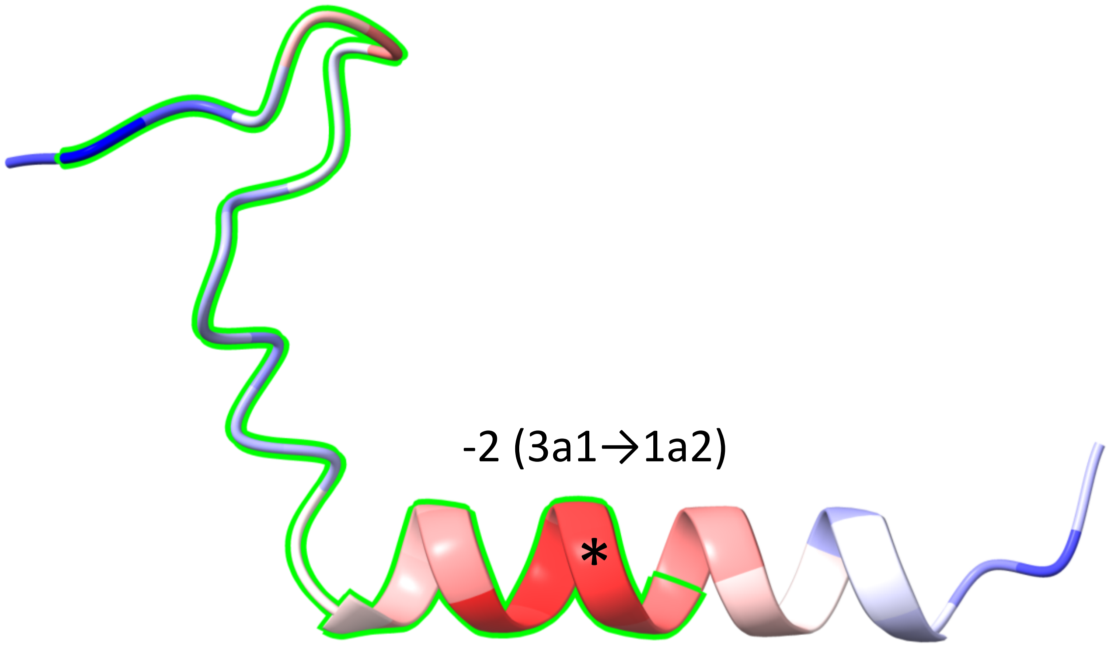


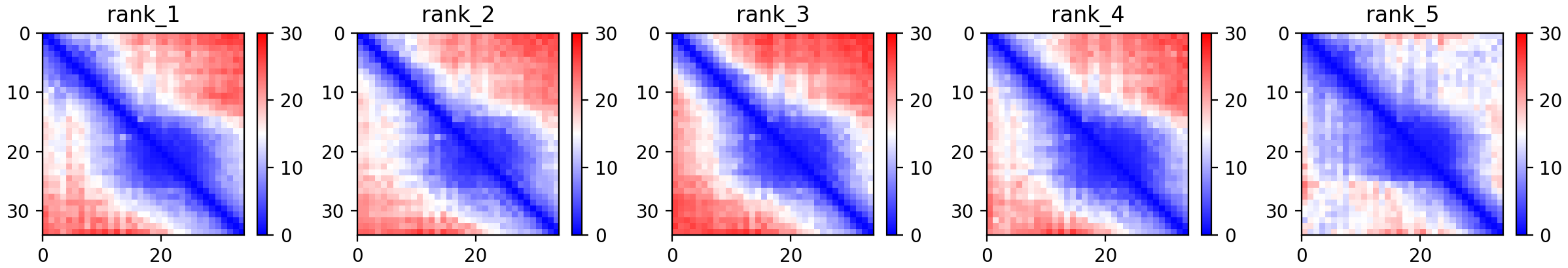

CP44: MtrunA17_Chr2g0326801

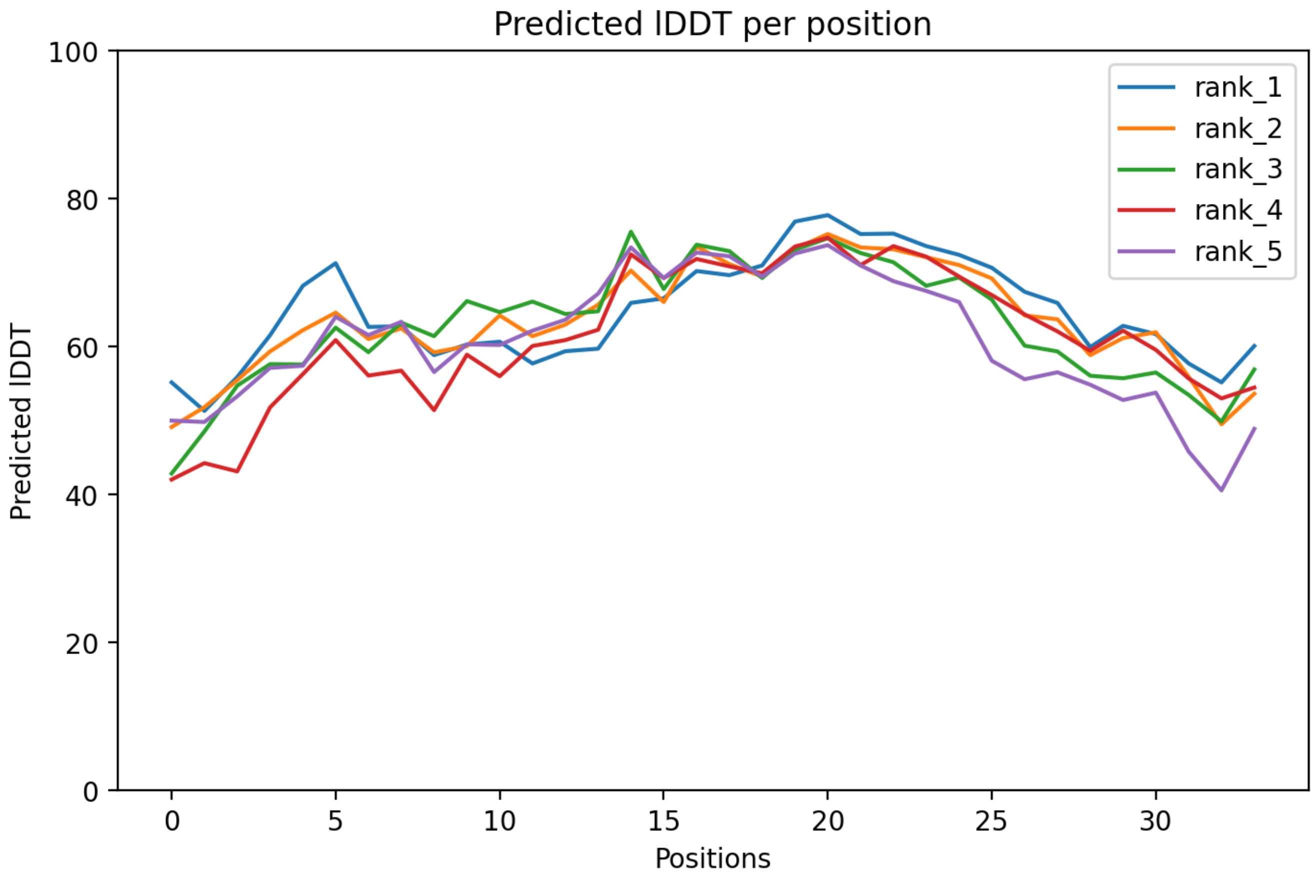

### CP44: MtrunA17_Chr2g0326801

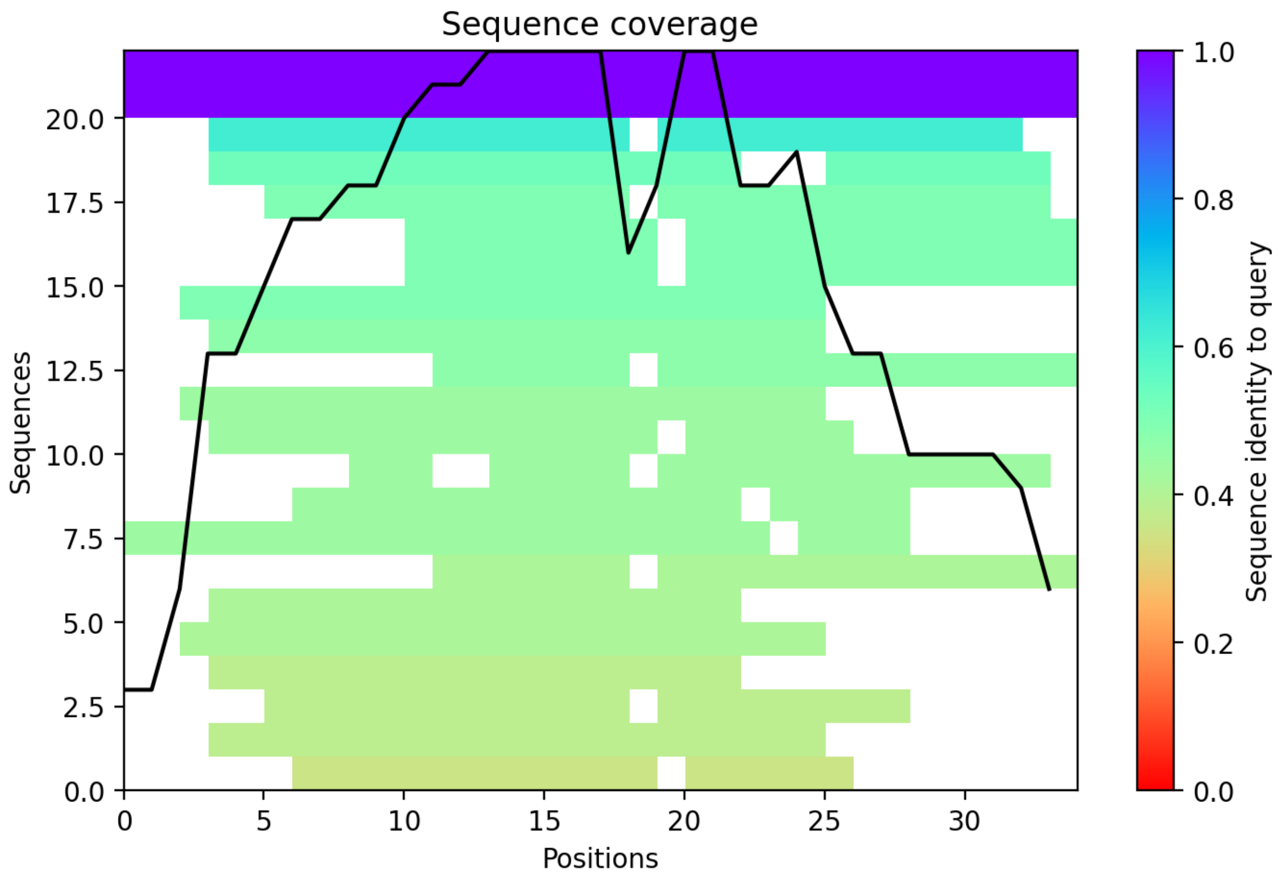

CP45: MtrunA17_Chr2g0328091

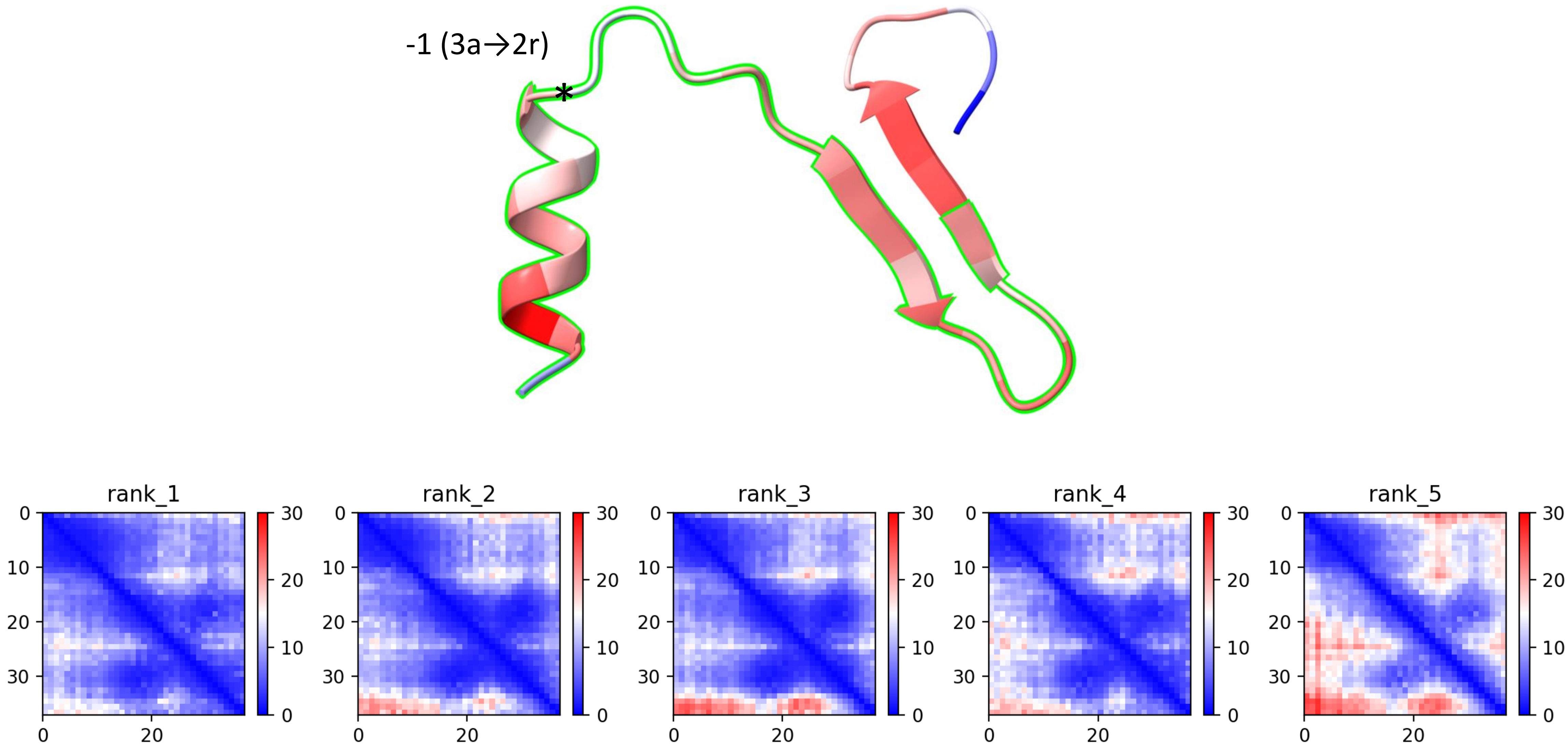

CP45: MtrunA17_Chr2g0328091

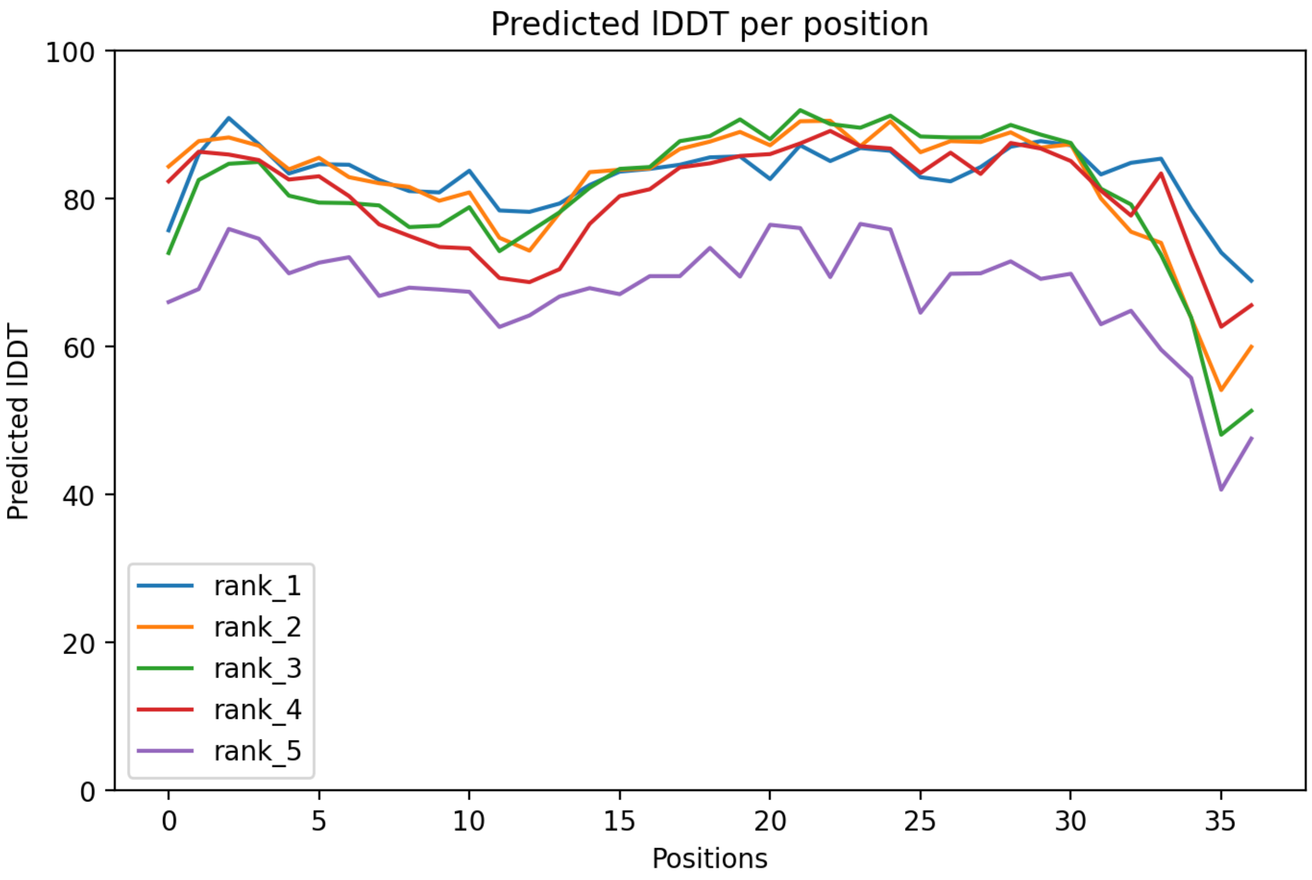

### CP45: MtrunA17_Chr2g0328091

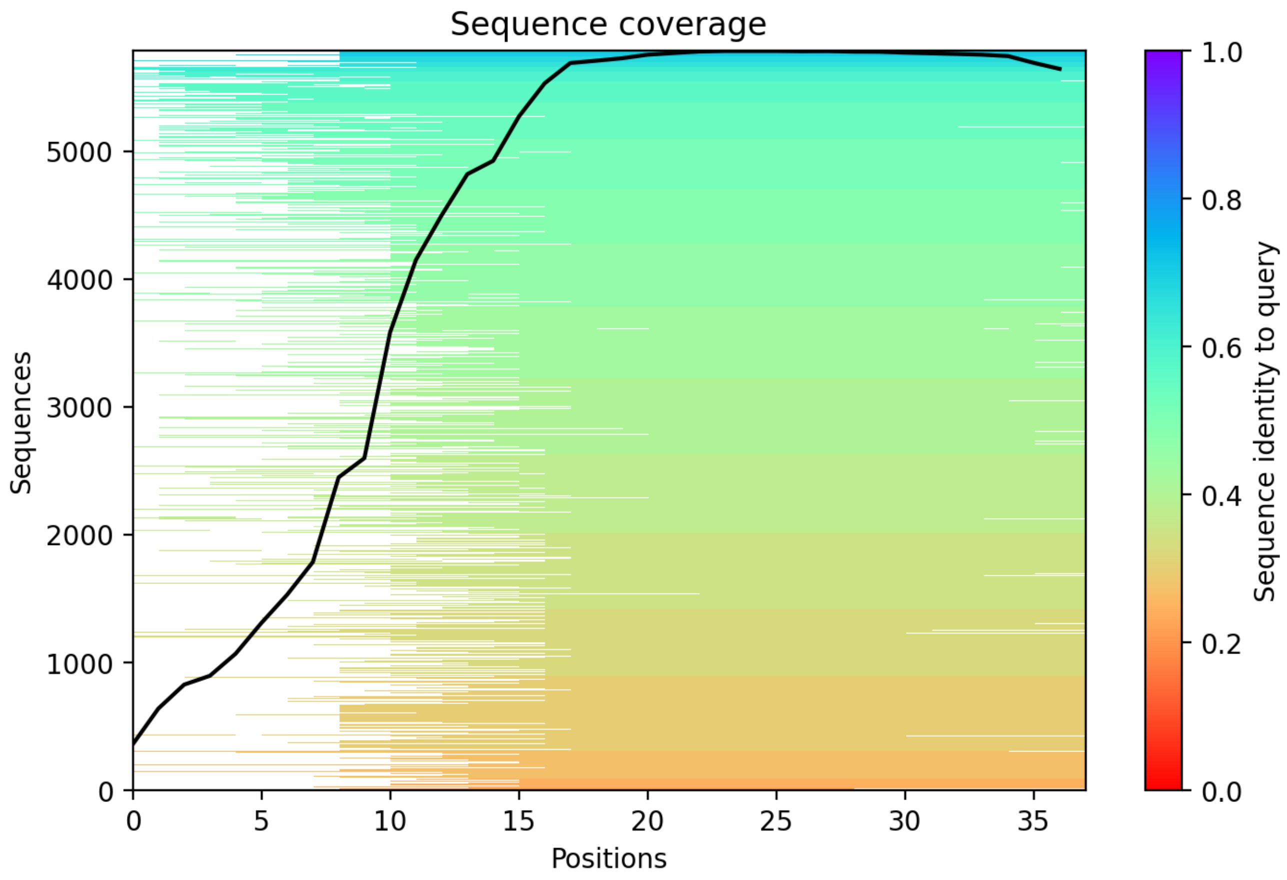

CP46: MtrunA17_Chr2g0329031

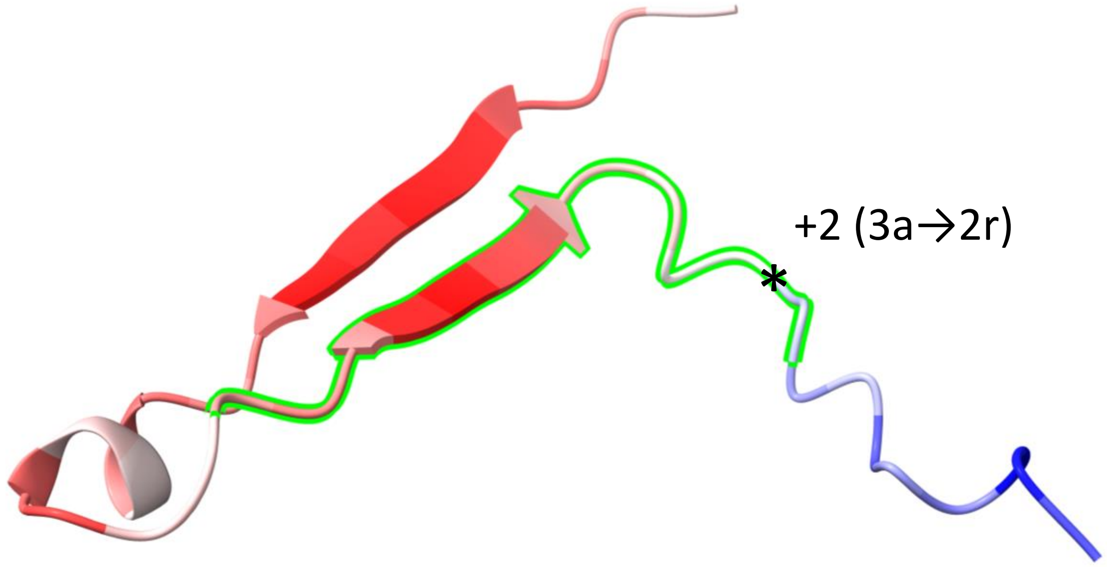


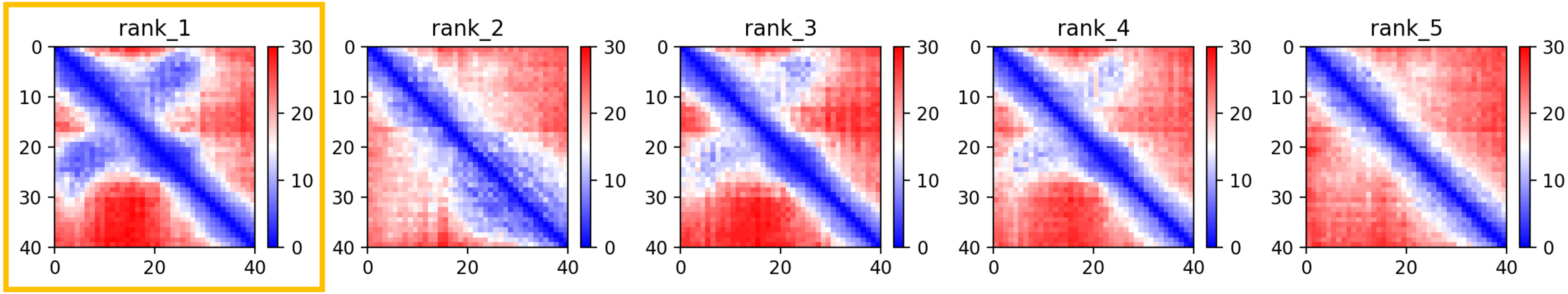

CP46: MtrunA17_Chr2g0329031

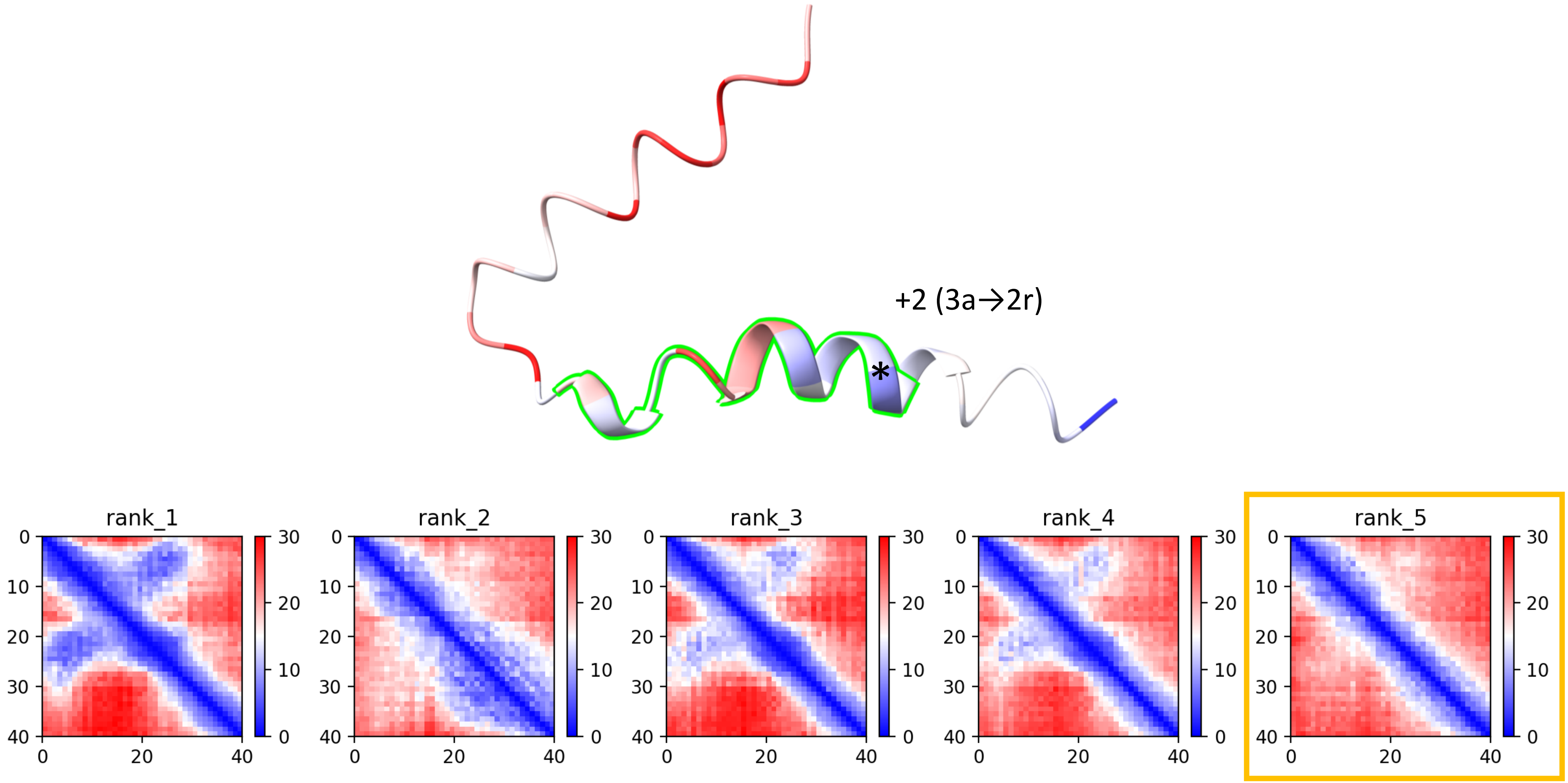

CP46: MtrunA17_Chr2g0329031

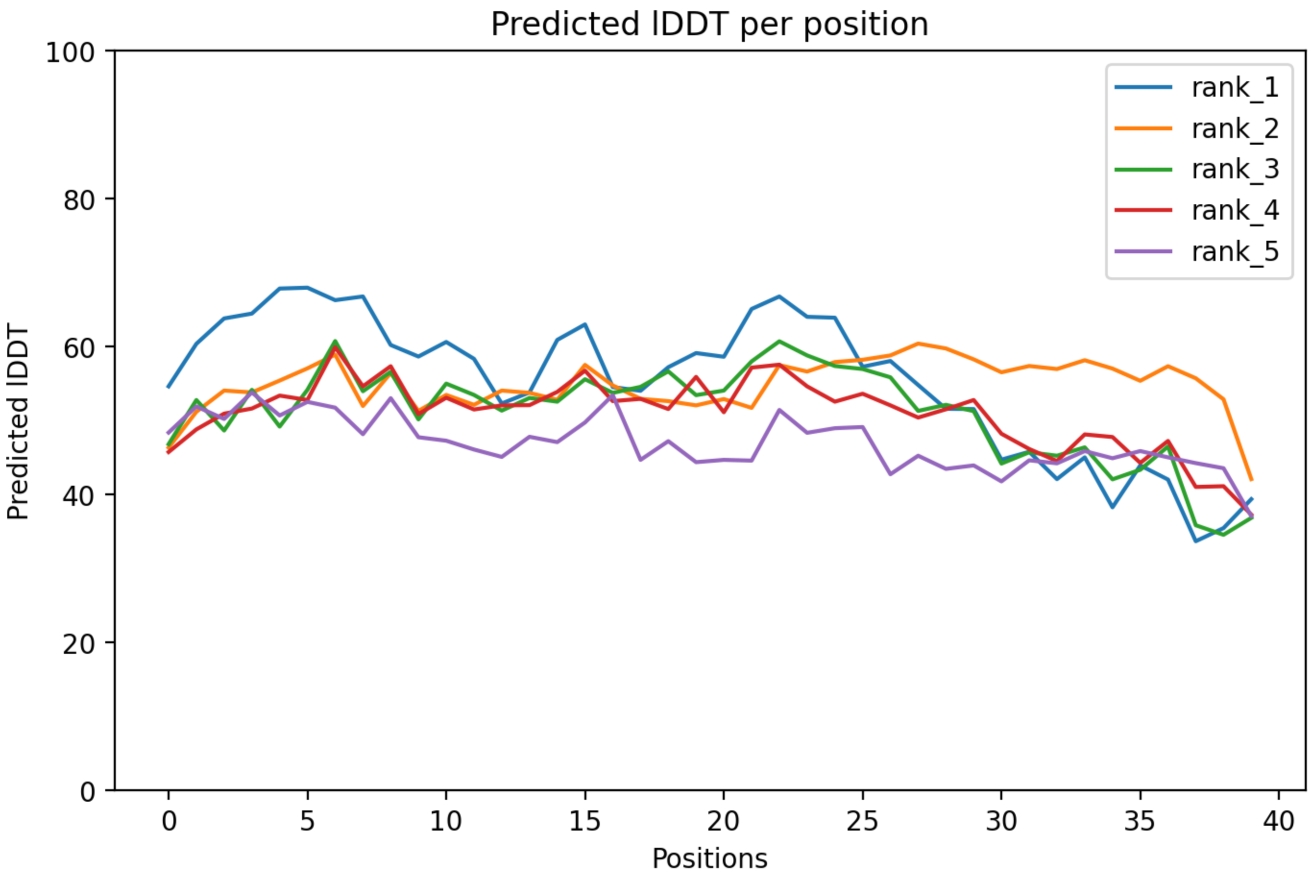

CP46: MtrunA17_Chr2g0329031

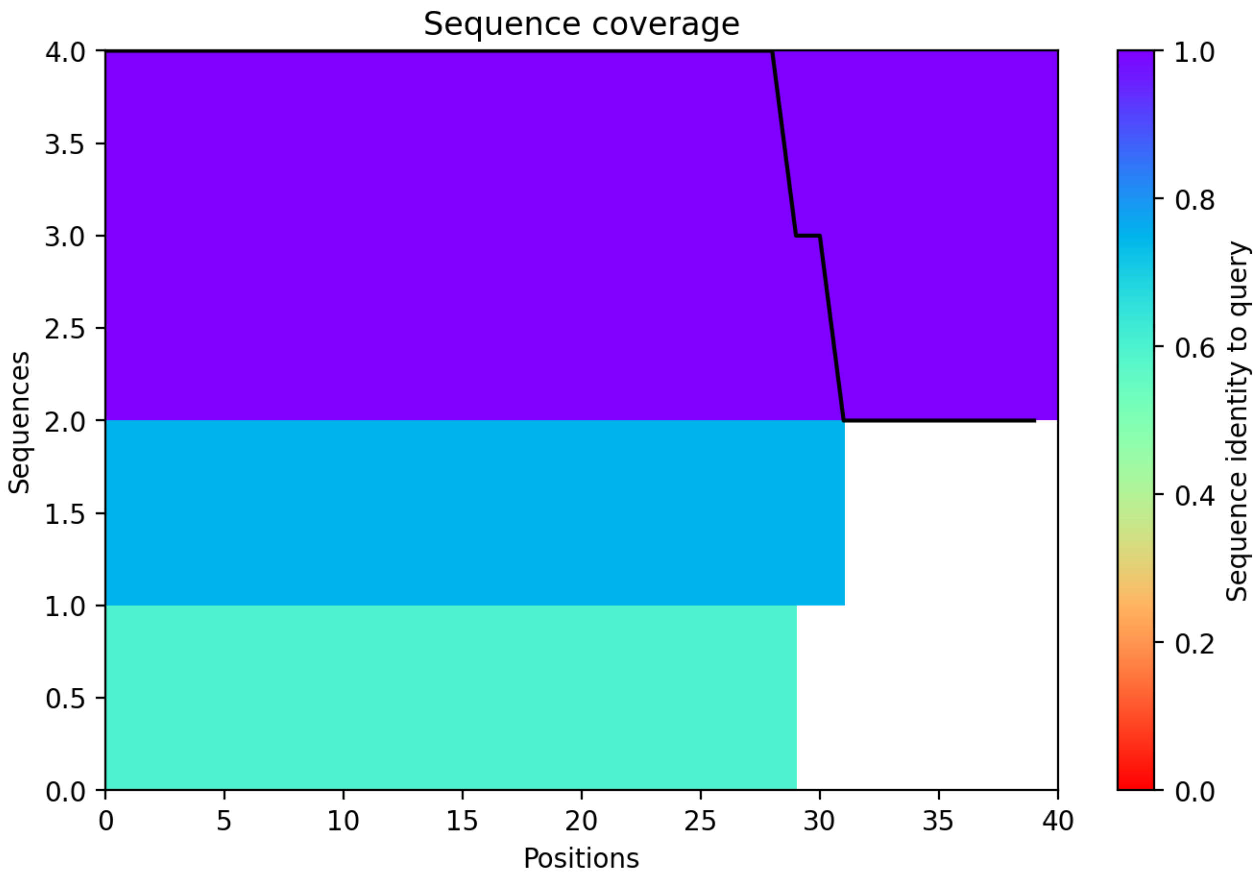

CP47: MtrunA17_Chr3g0079521

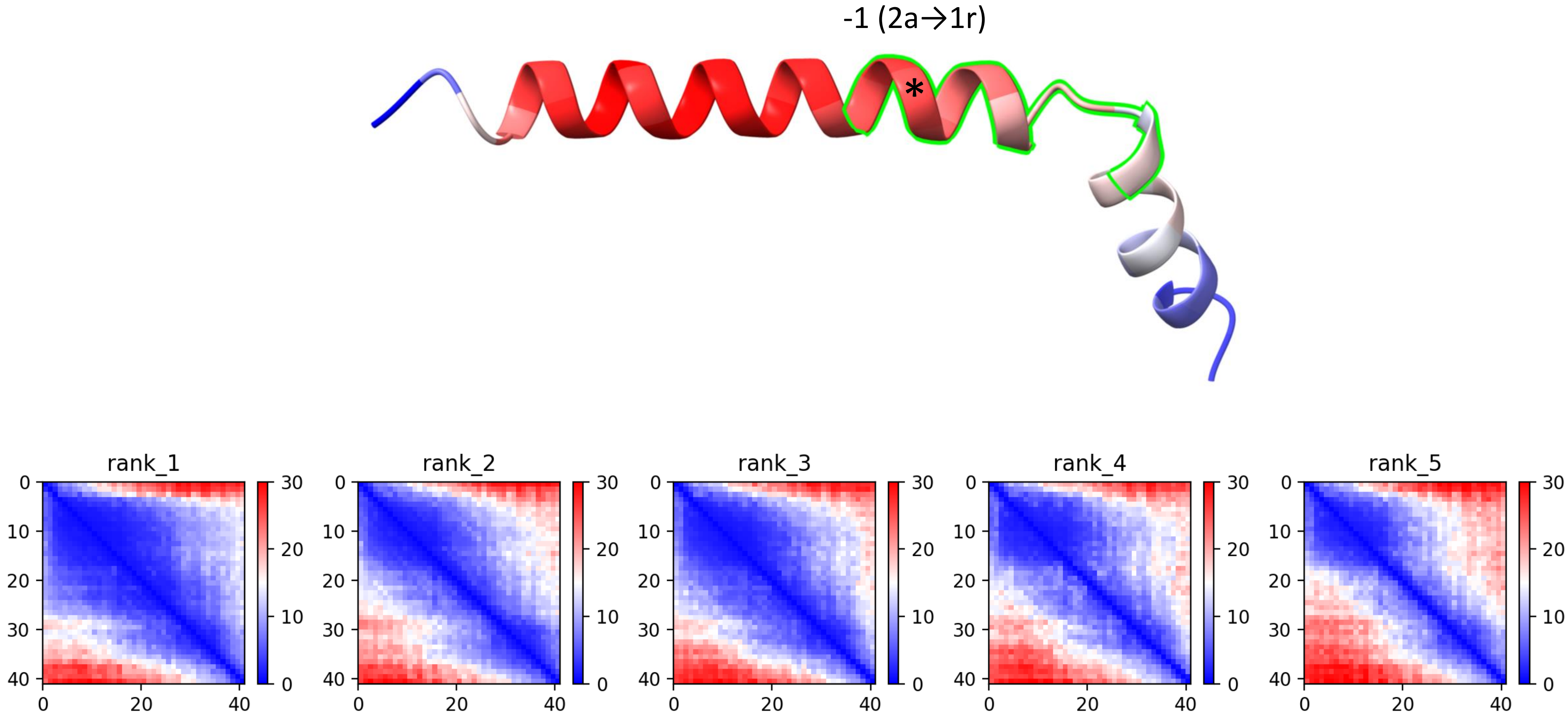

CP47: MtrunA17_Chr3g0079521

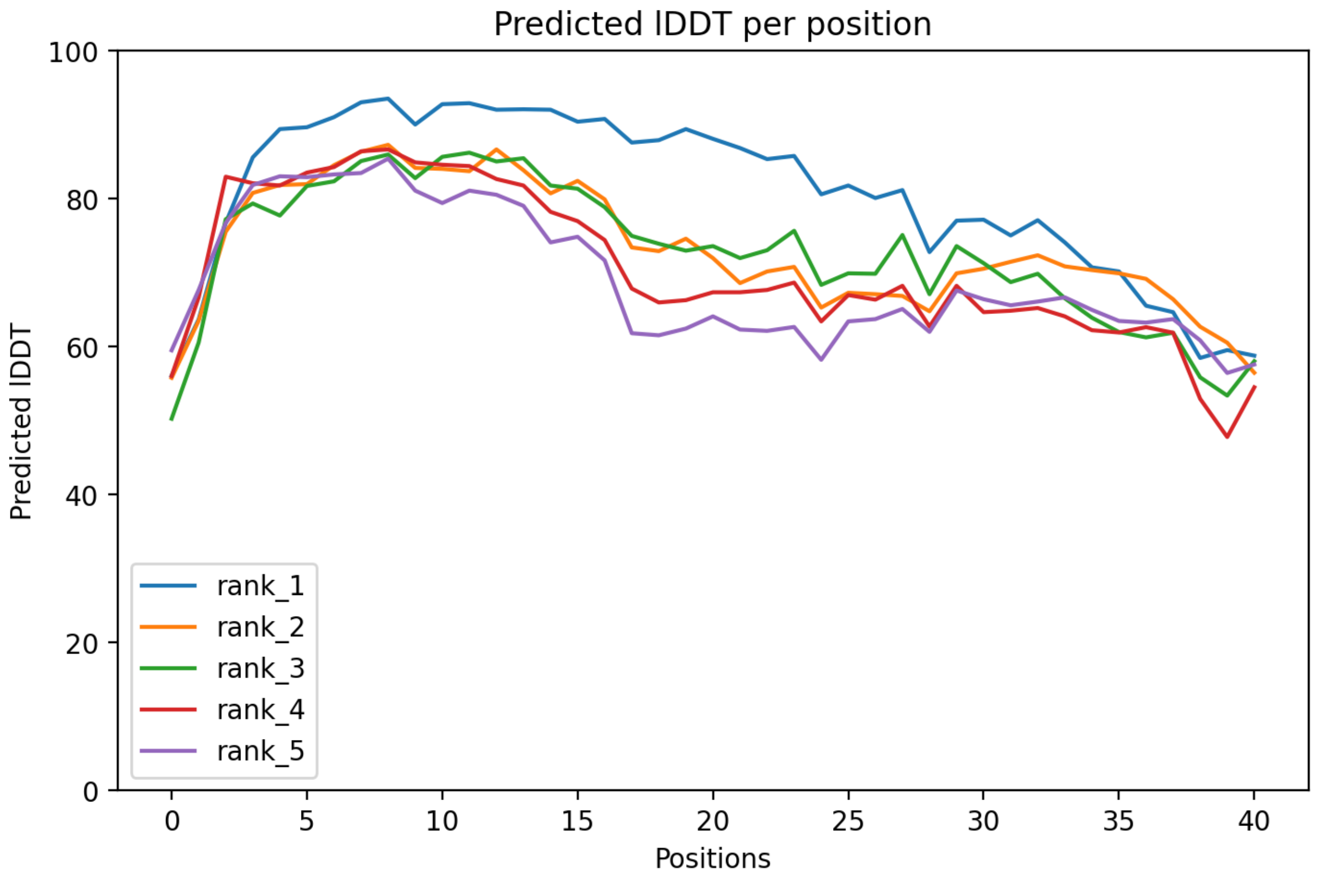

CP47: MtrunA17_Chr3g0079521

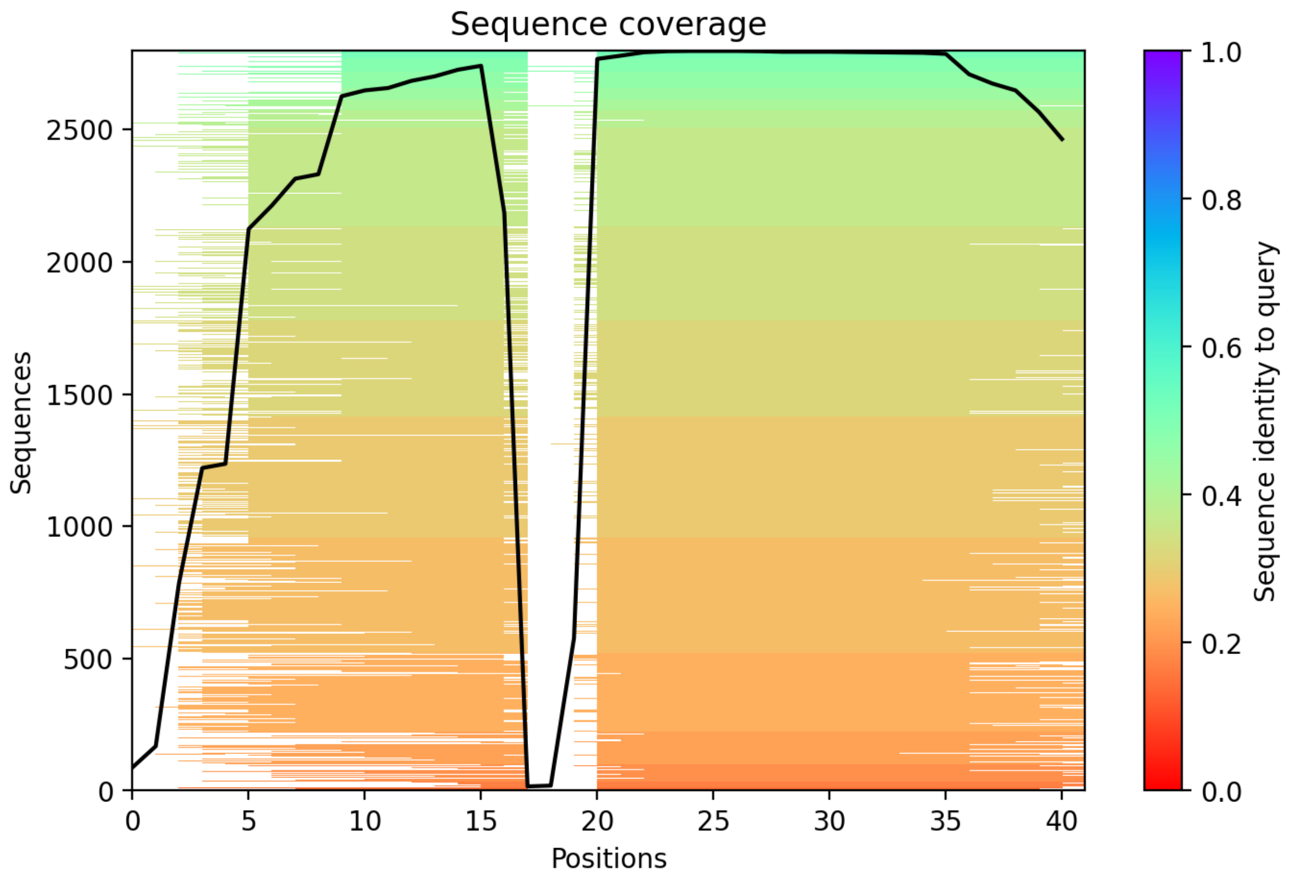

CP48: MtrunA17_Chr3g0083801

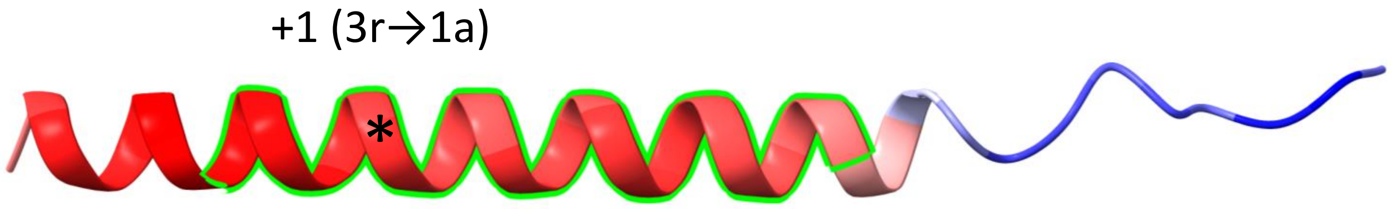


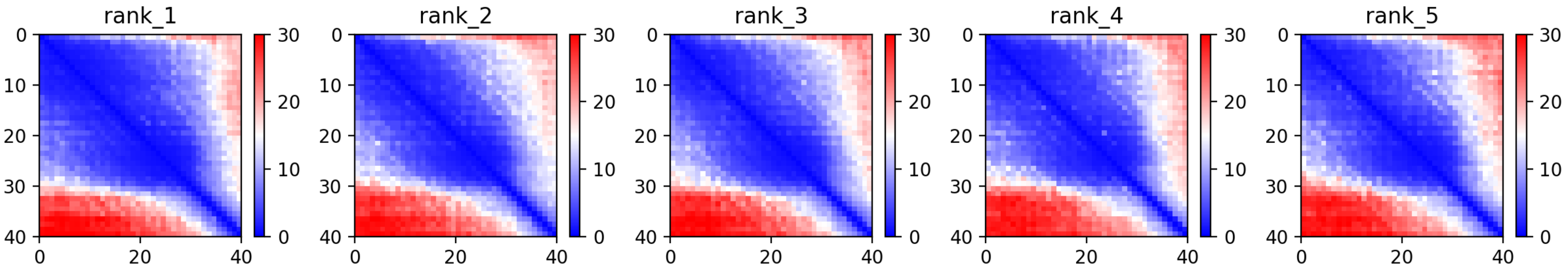

### CP48: MtrunA17_Chr3g0083801

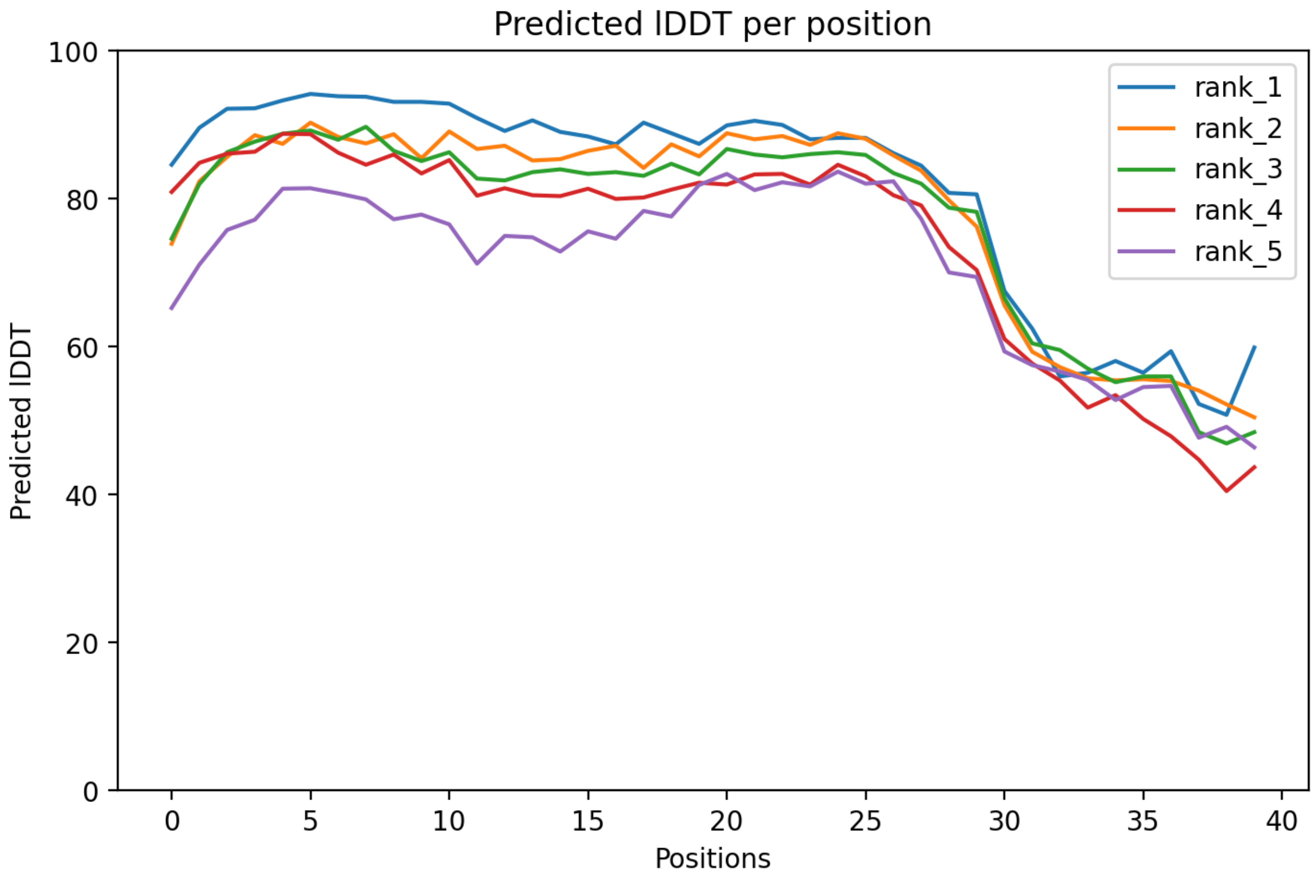

CP48: MtrunA17_Chr3g0083801

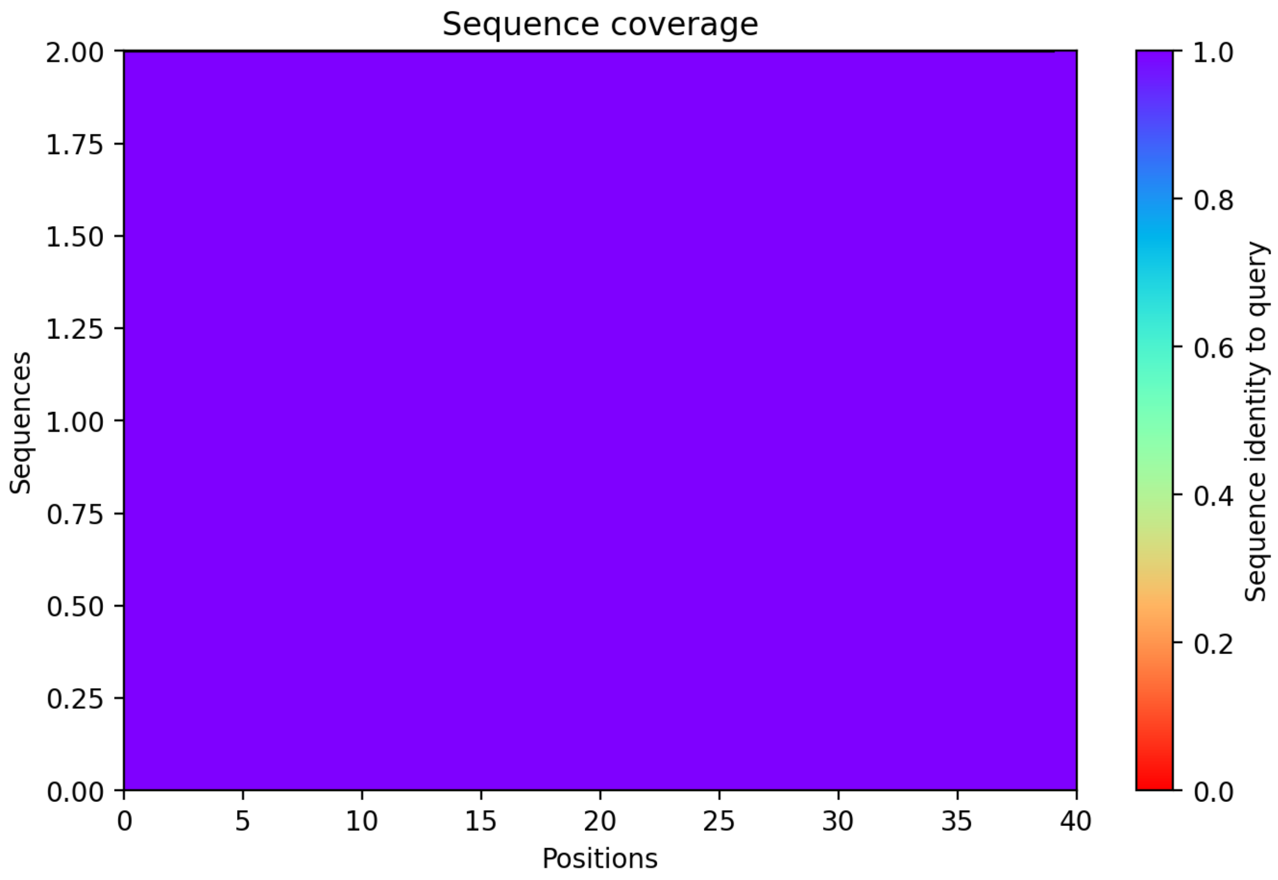

### CP49: MtrunA17_Chr3g0091141

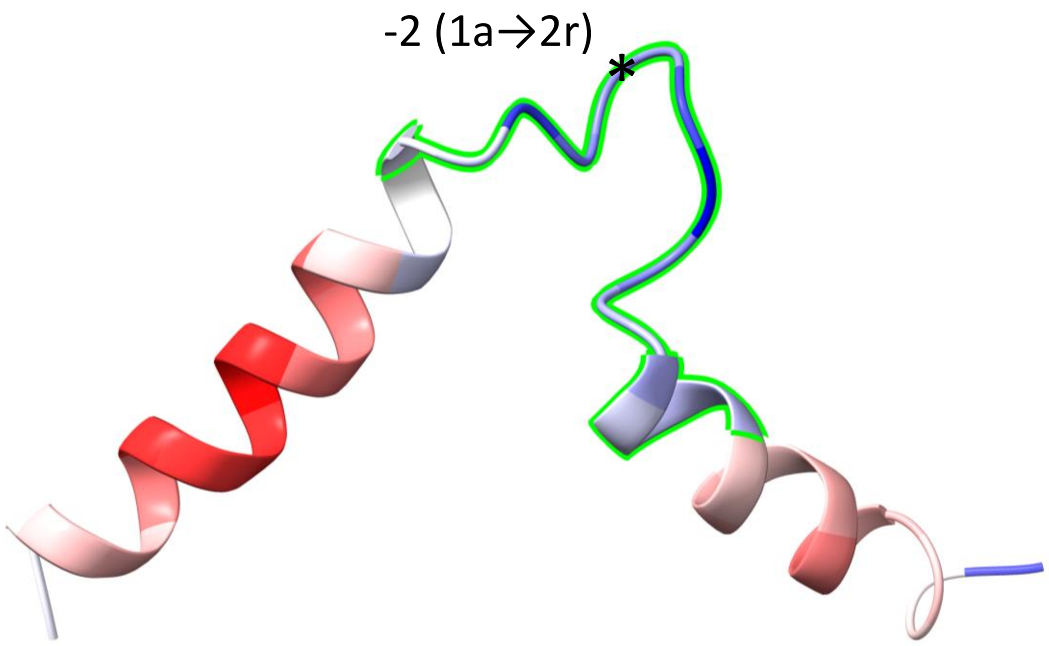


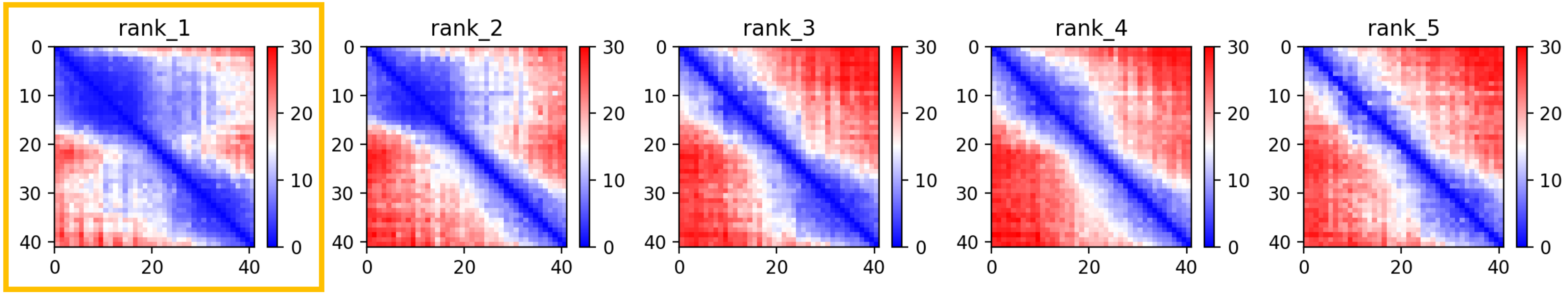

CP49: MtrunA17_Chr3g0091141

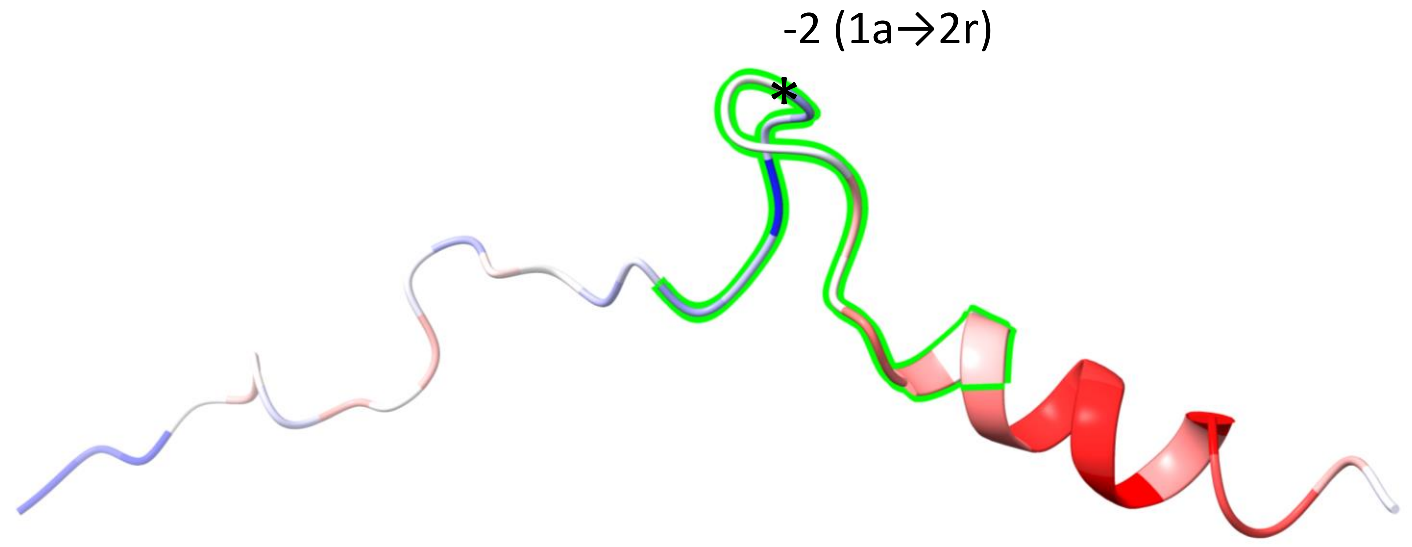


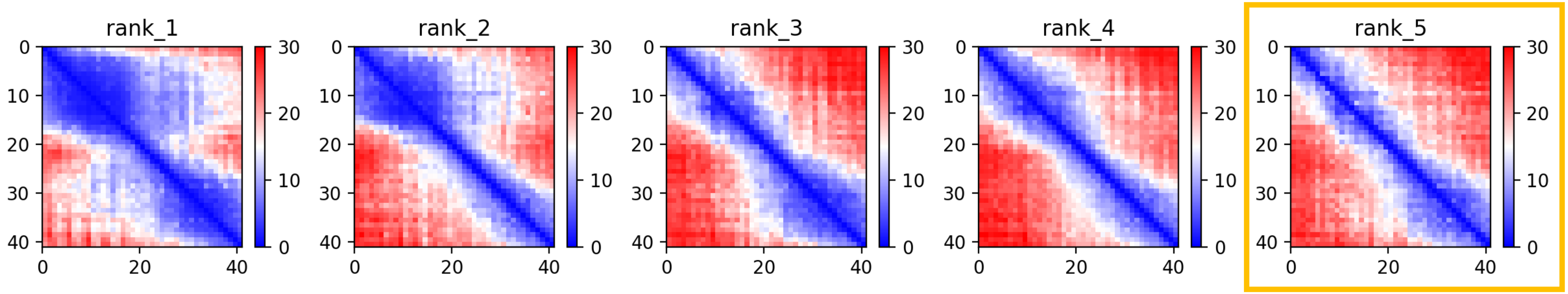

CP49: MtrunA17_Chr3g0091141

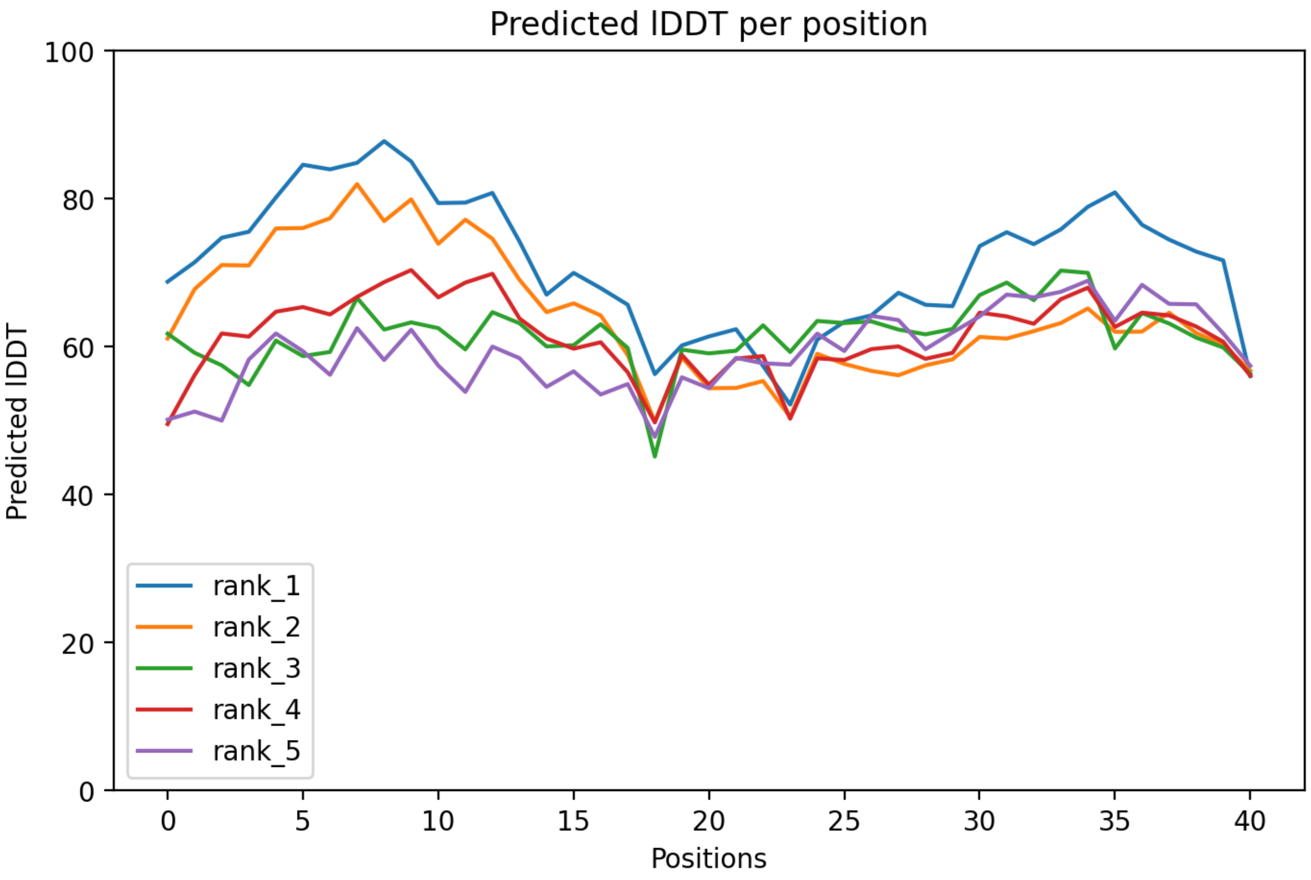

### CP49: MtrunA17_Chr3g0091141

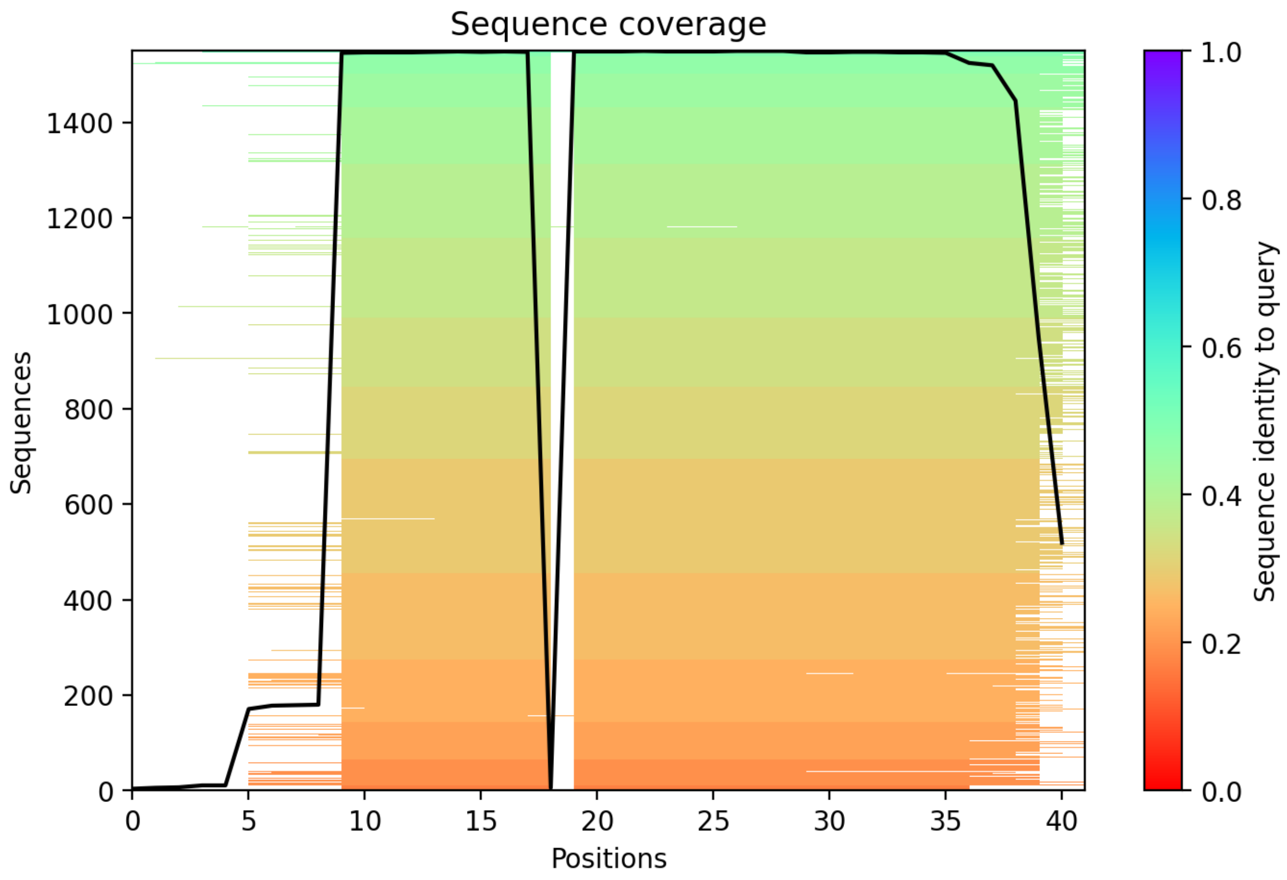

CP50: MtrunA17_Chr3g0091671

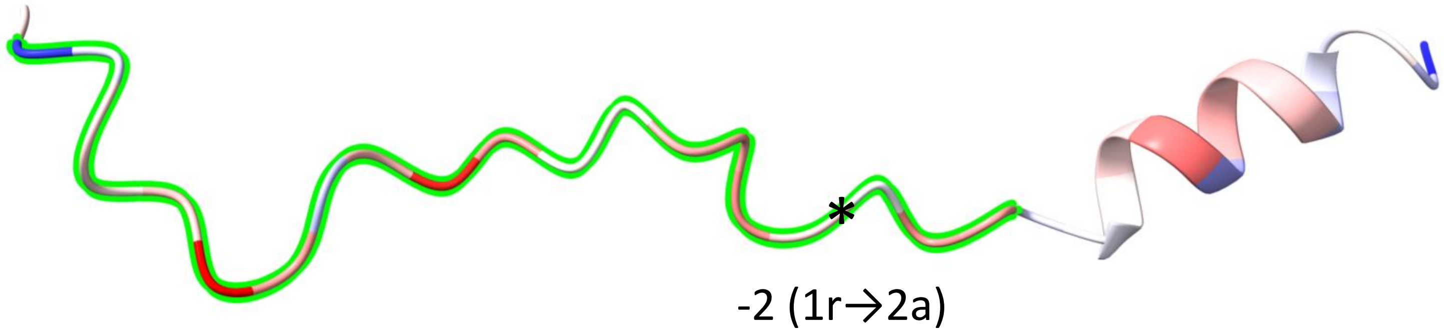


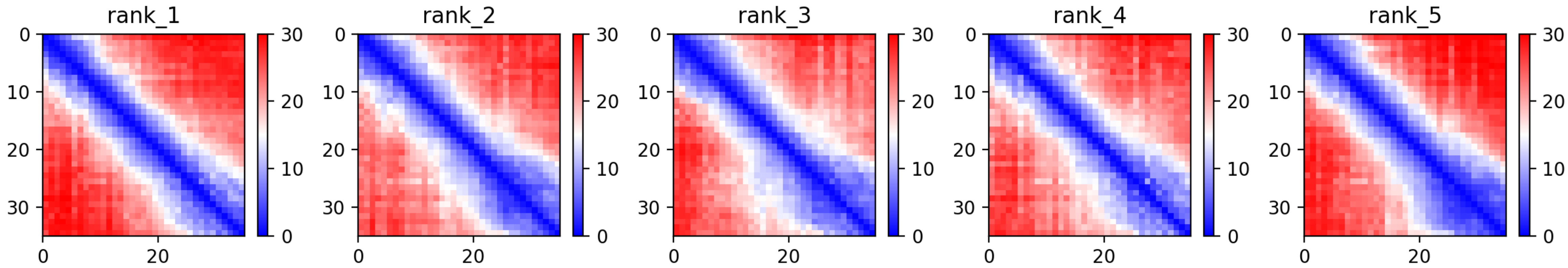

CP50: MtrunA17_Chr3g0091671

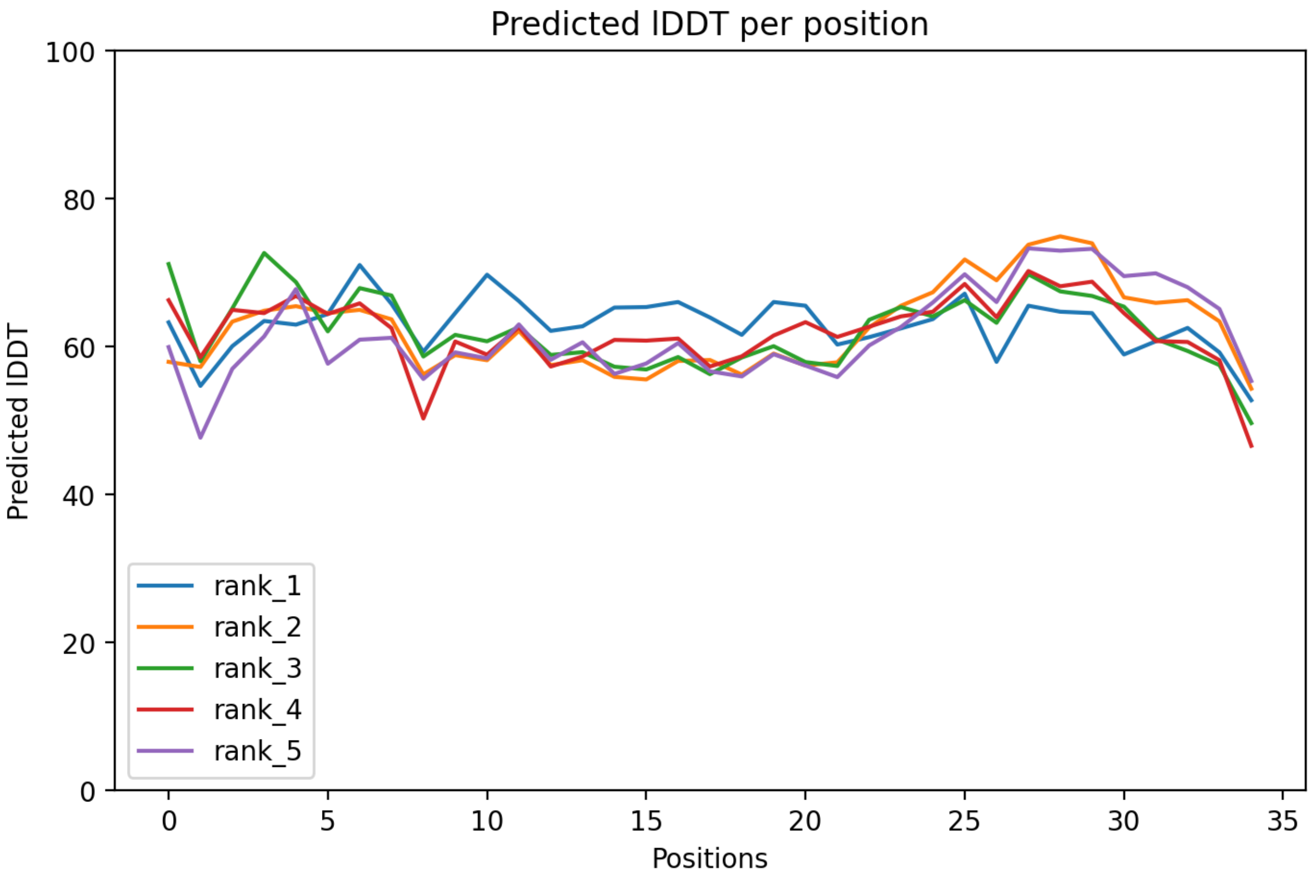

### CP50: MtrunA17_Chr3g0091671

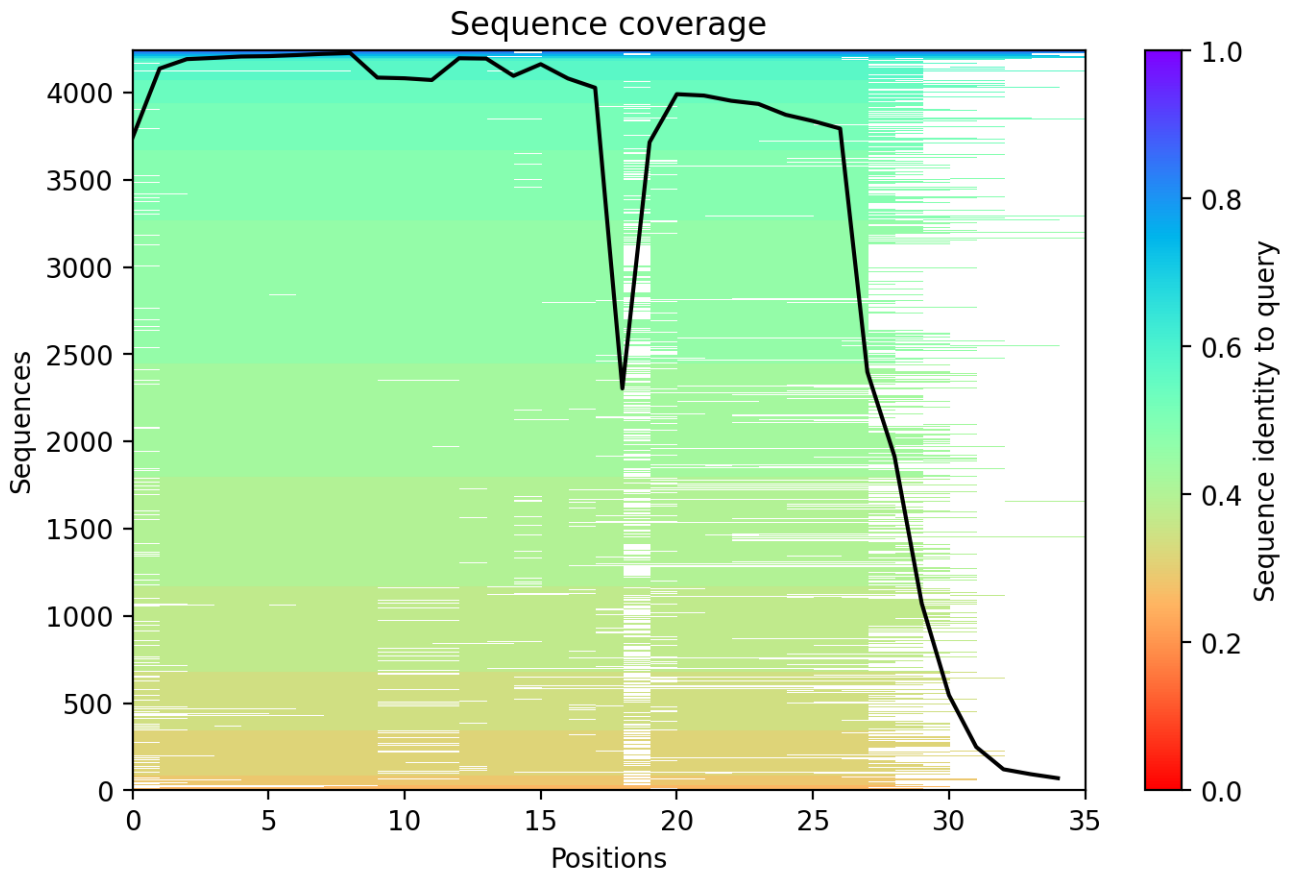

**Supplementary Dataset S7 Part 1**. A graphical summary on folding predictions for MS-supported chimeric peptide models (CPs) 1-50. Folding categories were predicted by ColabFold v. 1.5.2 and visualized with ChimeraX v. 1.6.1. For each CP, three images are displayed: (1) the predicted folding structure (top) along with heat maps of five predictions (bottom); (2) per-residue confidence score (pLDDT) for five predictions; and (3) a sequence coverage plot. By default, the structure shown corresponds to the prediction with rank 1. In cases where any prediction with a rank 2 to 5 is visibly different from the rank-1-prediction, a corresponding structure is shown in addition to the rank-1-prediction (its heat map is boxed with orange). Each structure was manually oriented in space to maximize the two-dimensional projection and to position the N-terminus on the left-hand side. The blue-white-red gradient indicates the prediction confidence based on the range of b-factor values. Blue stands for the lowest and red stands for the highest values (the highest confidence). A sequence segment highlighted with lime delimits the MS peptide-matching portion of the chimeric model. Positions of PRF events are labeled with an asterisk accompanied with PRF characteristics. The description of a PRF event should be interpreted as follows. For example, CP1, -1 (1a1→3a2): a minus 1 frameshift changes the translation from frame 1 to frame 3, which corresponds to the change from altORF1 to altORF2. An altORF1 in this study is defined as the one that starts earlier than an altORF2.
