## Supplementary material for "Discovery of diverse chimeric peptides in a eukaryotic proteome sets the stage for the experimental proof of the mosaic translation hypothesis": Supplementary Dataset S7 Folding of chimeric protein models Part 2.pdf

### CP51: MtrunA17_Chr3g0096421

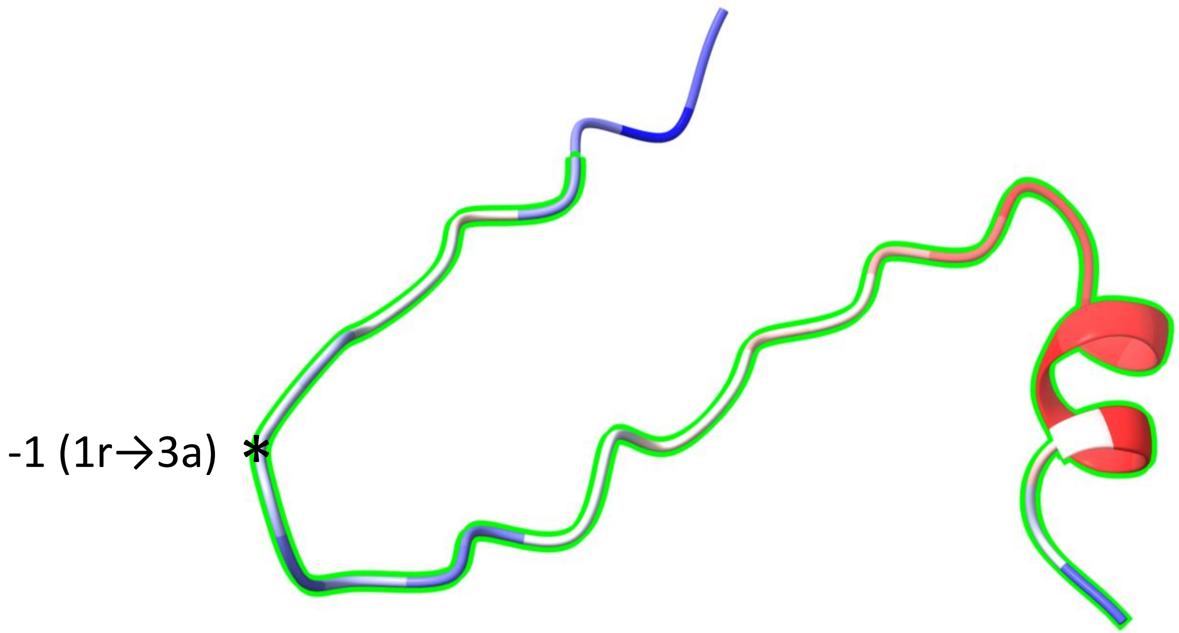


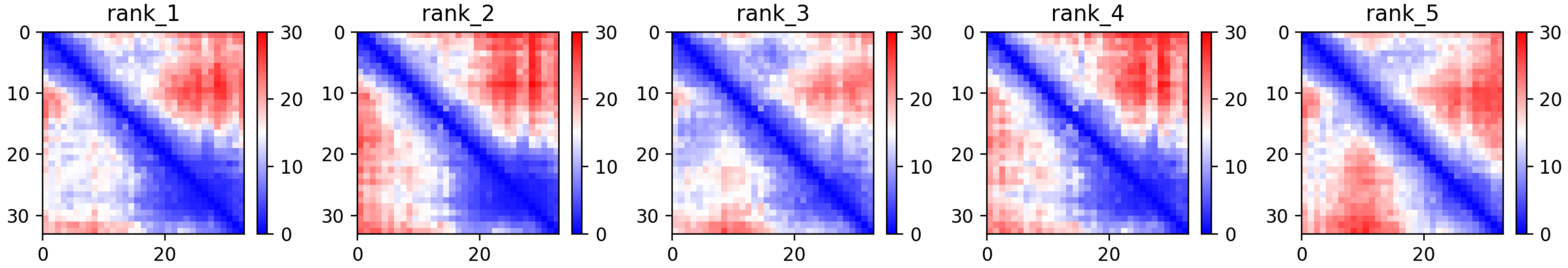

CP51: MtrunA17_Chr3g0096421

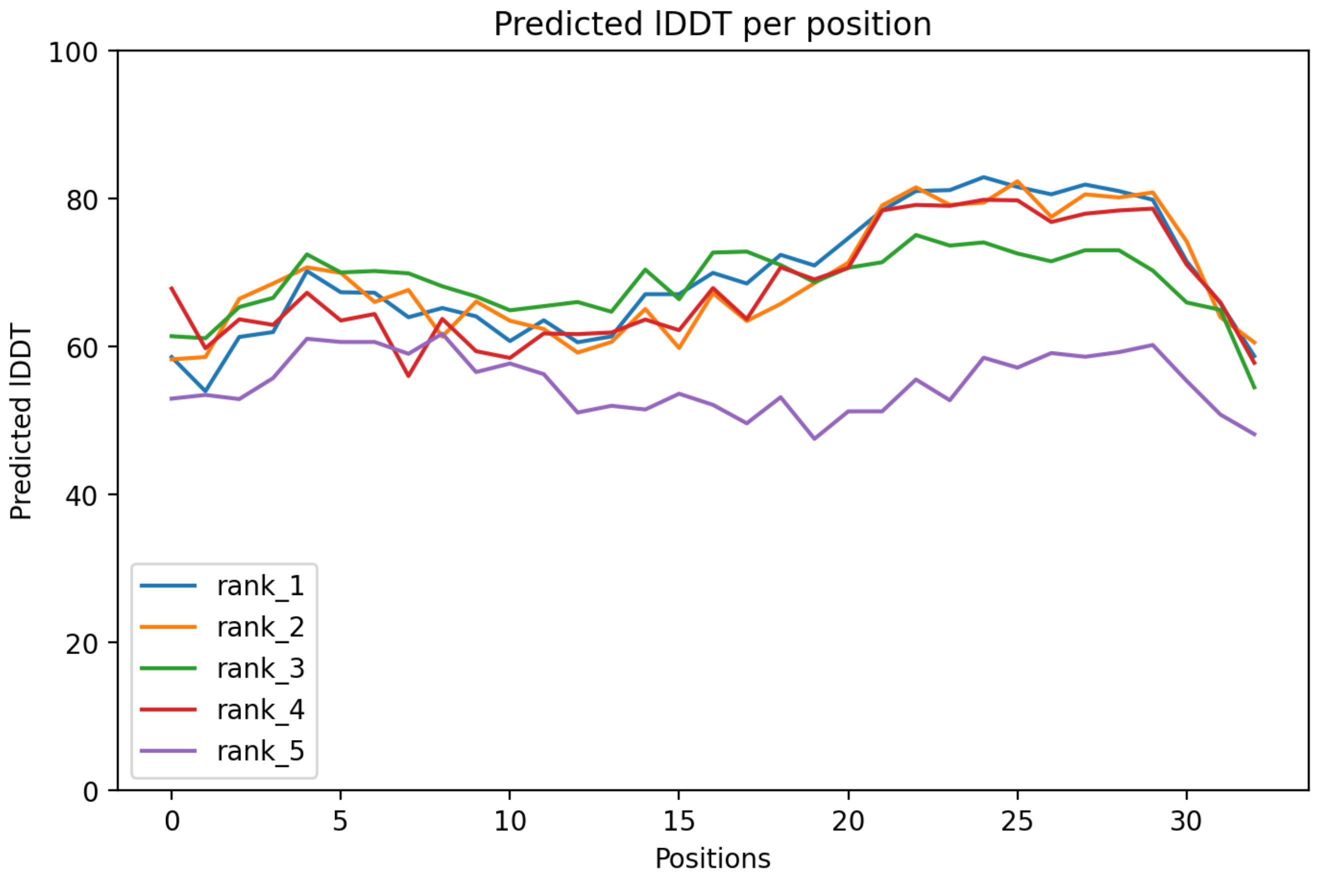

### CP51: MtrunA17_Chr3g0096421

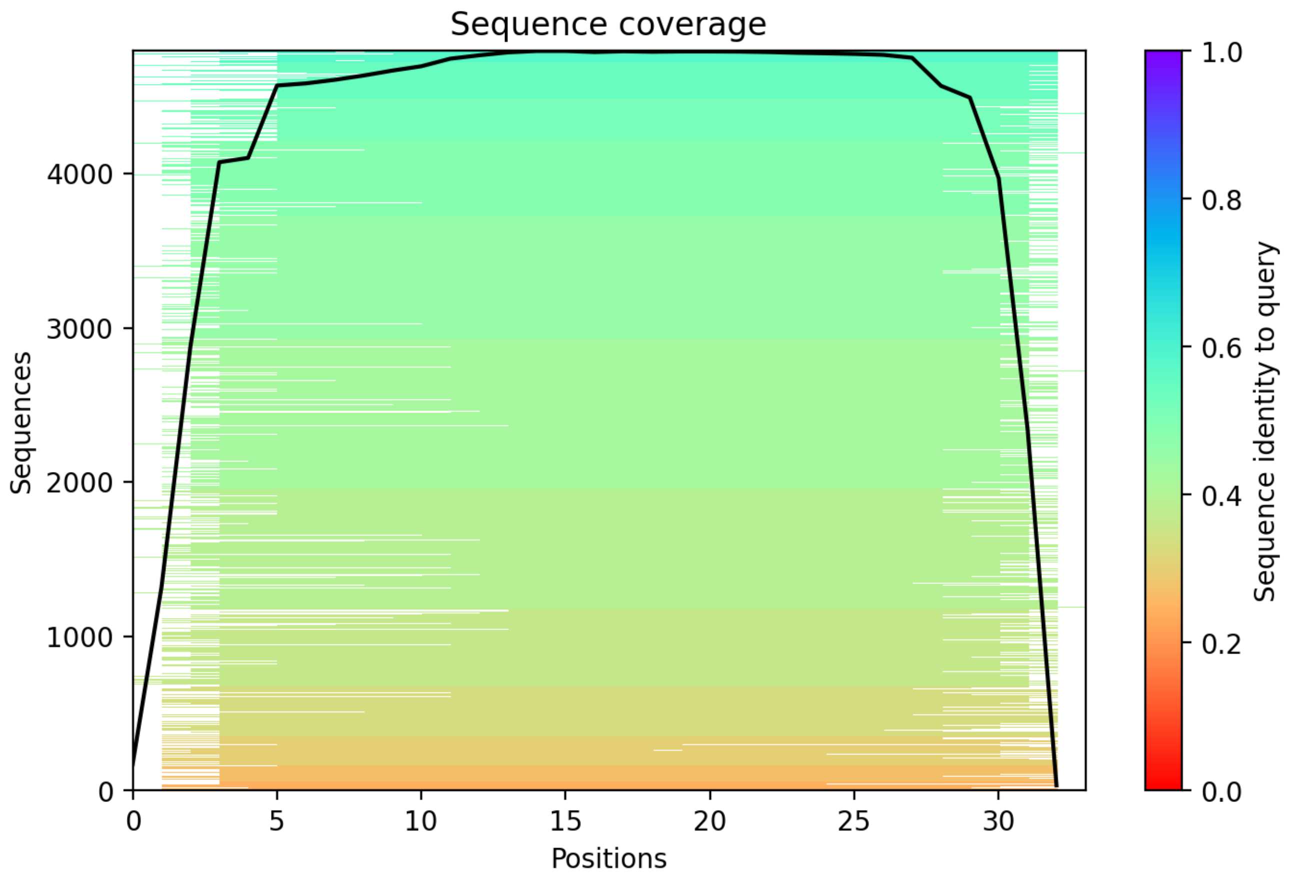

CP52: MtrunA17_Chr3g0100221

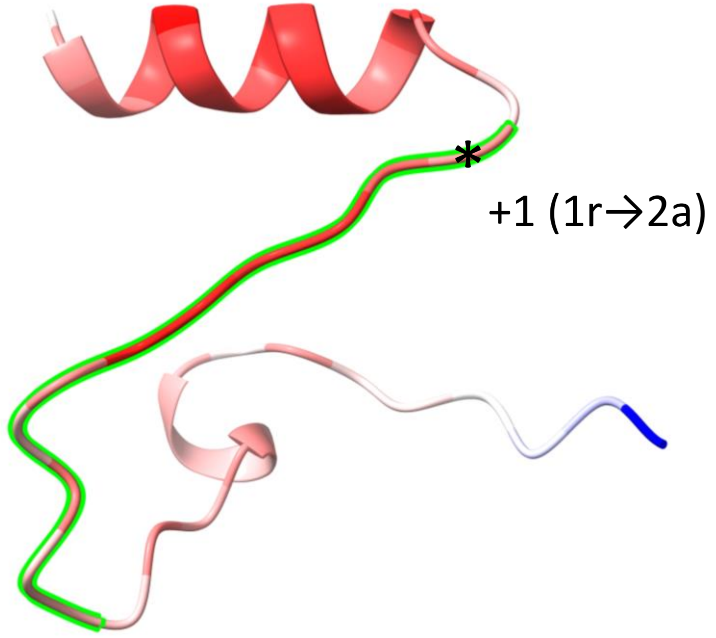


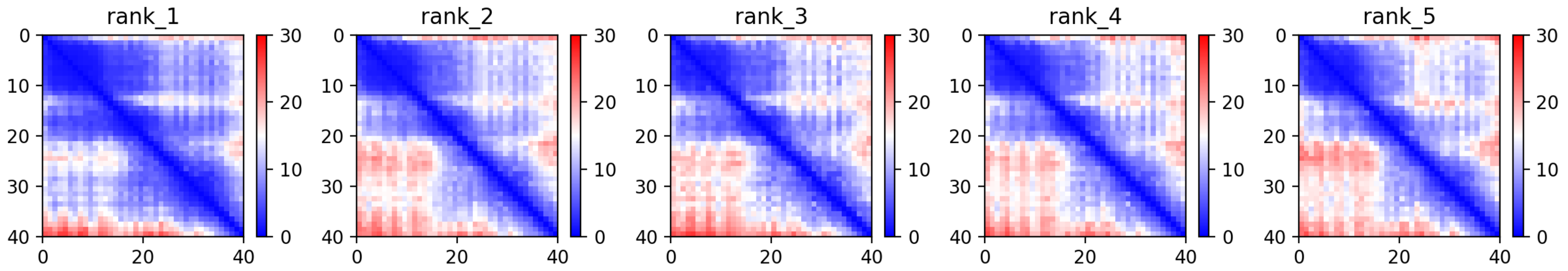

CP52: MtrunA17_Chr3g0100221

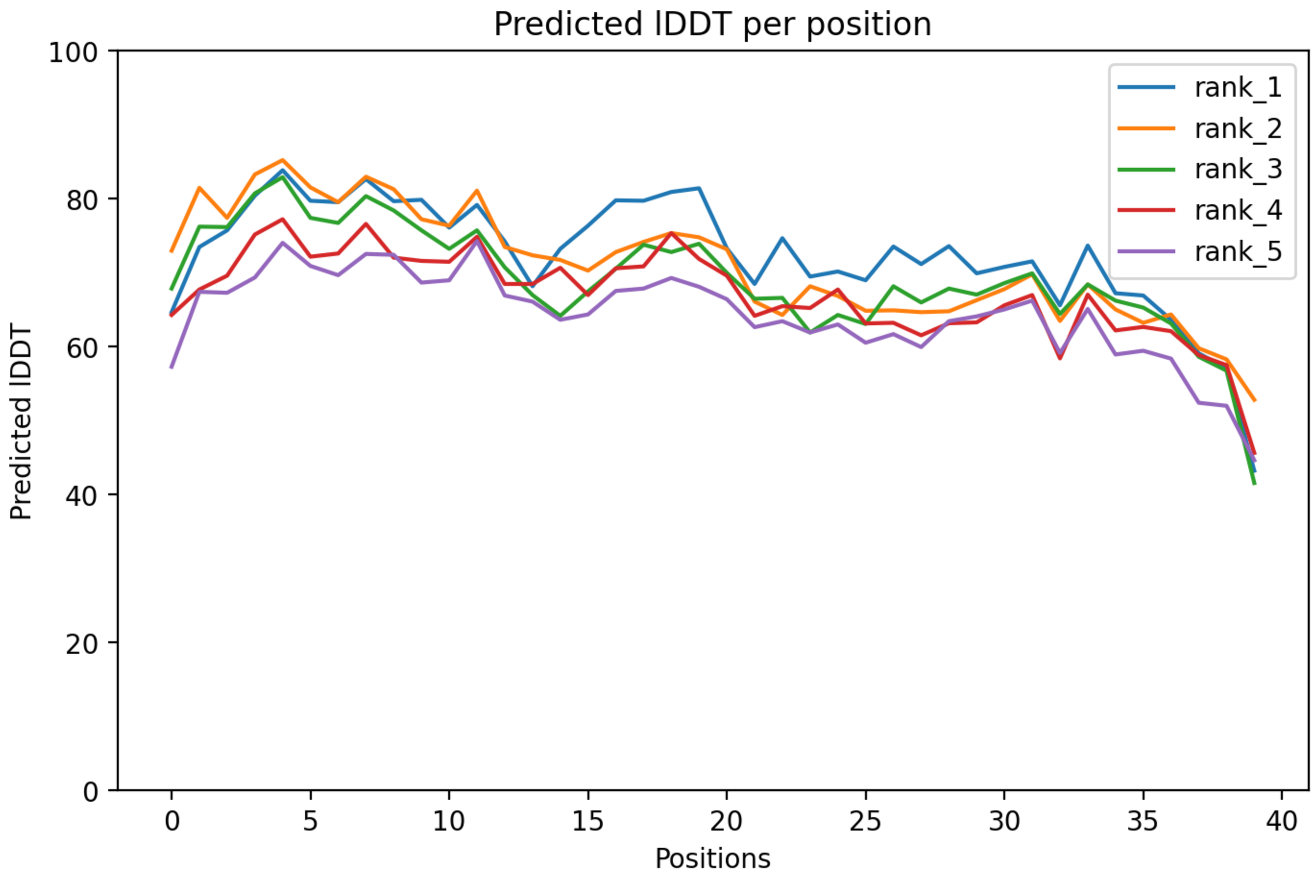

CP52: MtrunA17_Chr3g0100221

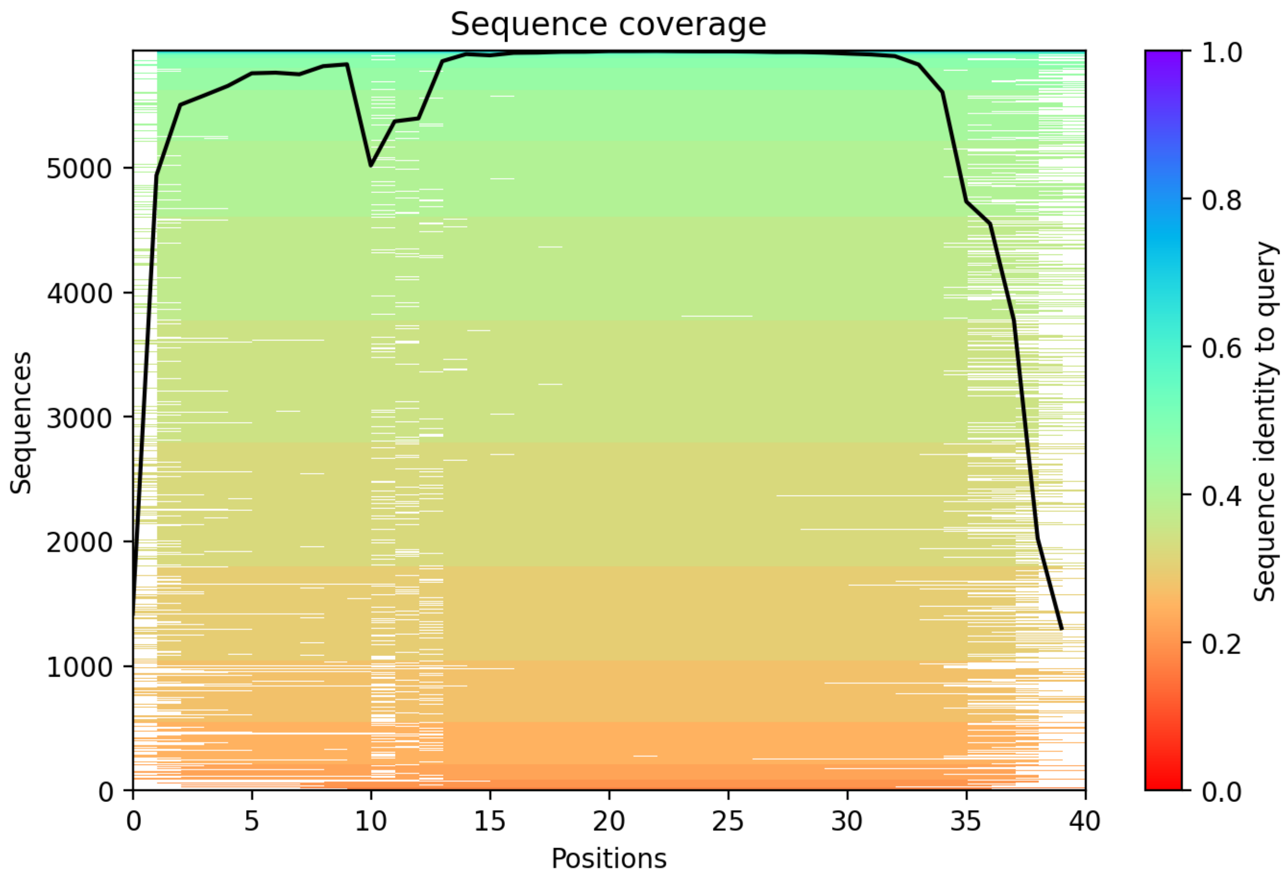

### CP53: MtrunA17_Chr3g0102171

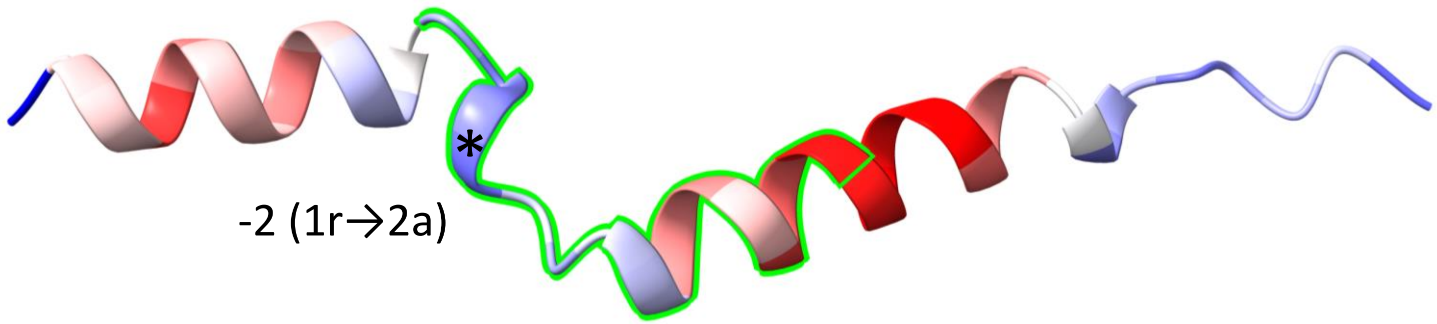


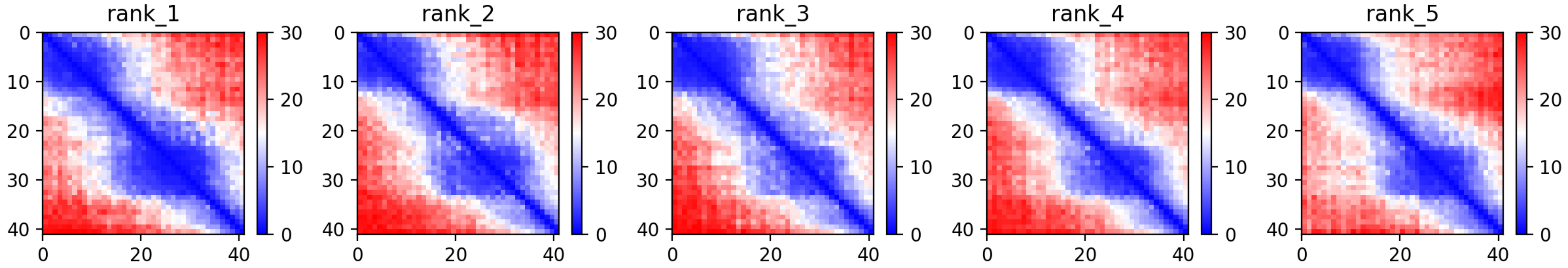

CP53: MtrunA17_Chr3g0102171

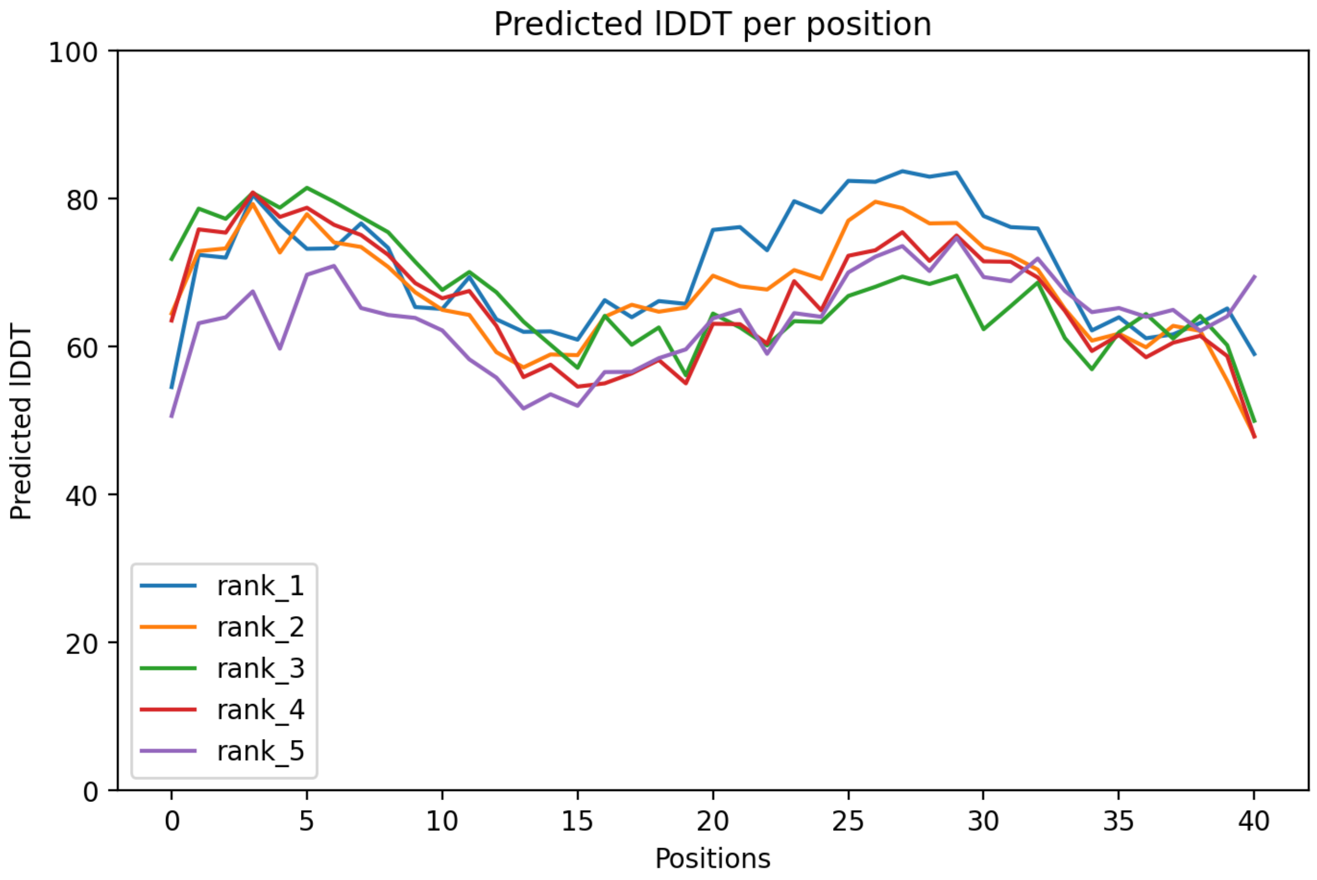

### CP53: MtrunA17_Chr3g0102171

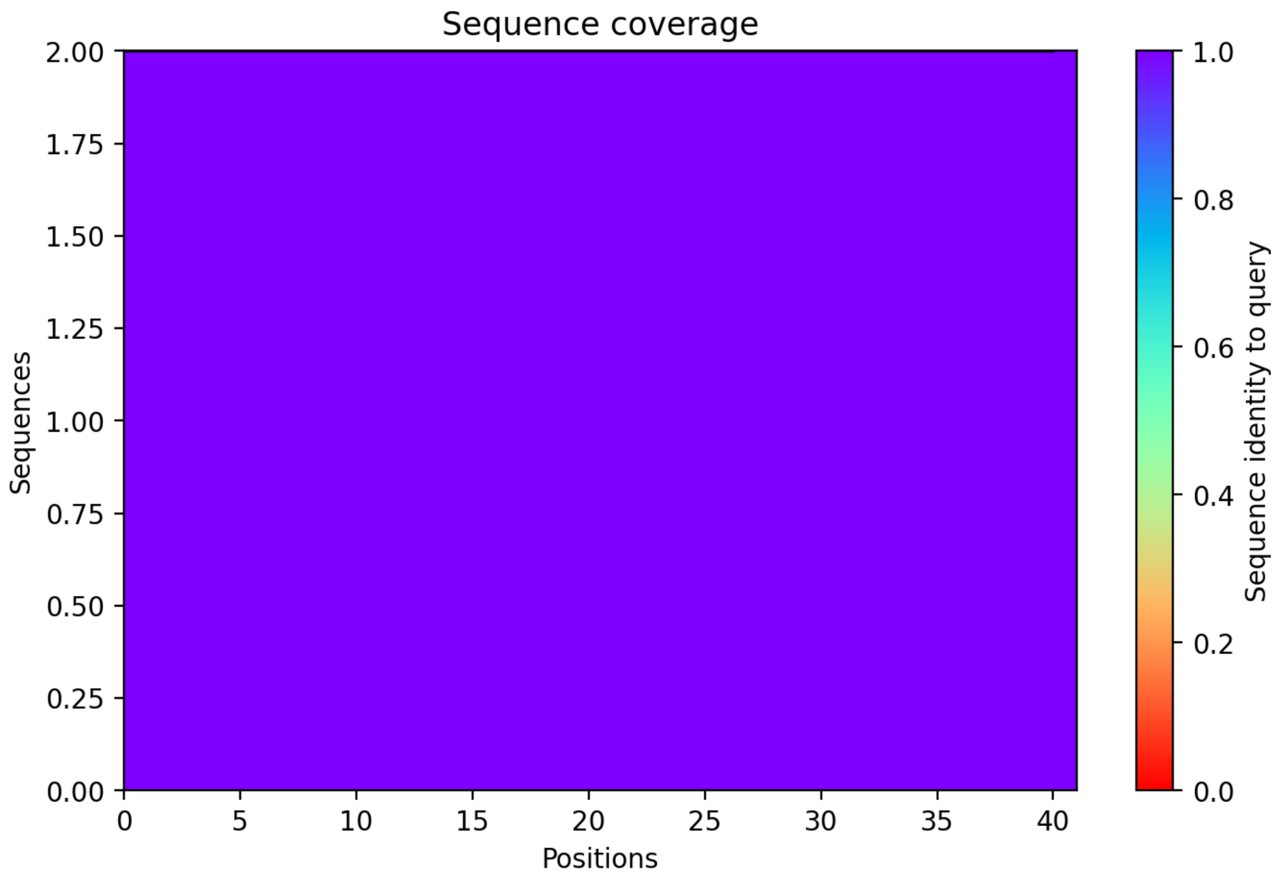

CP54: MtrunA17_Chr3g0105981

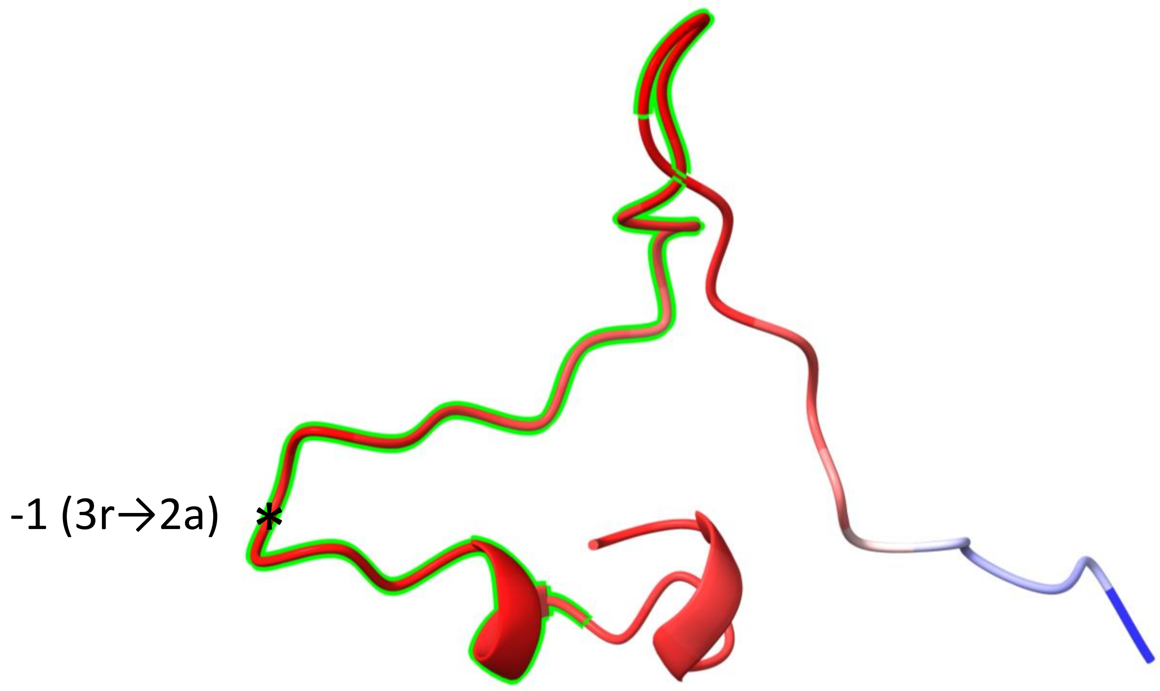


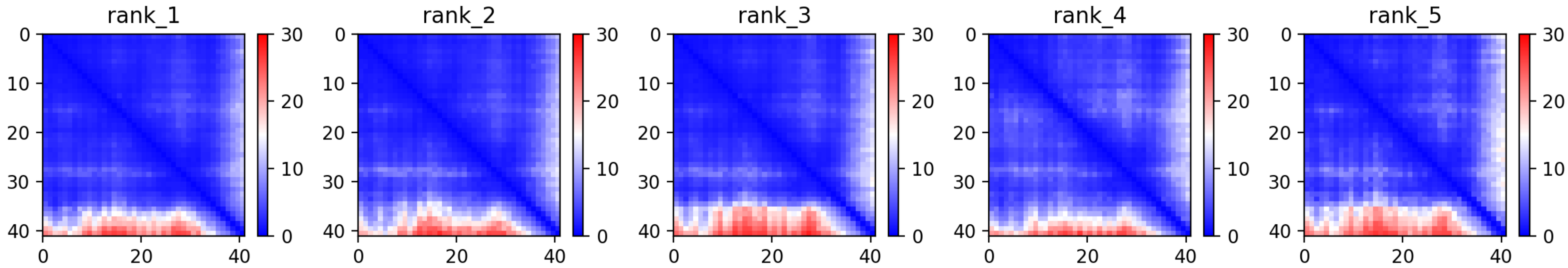

CP54: MtrunA17_Chr3g0105981

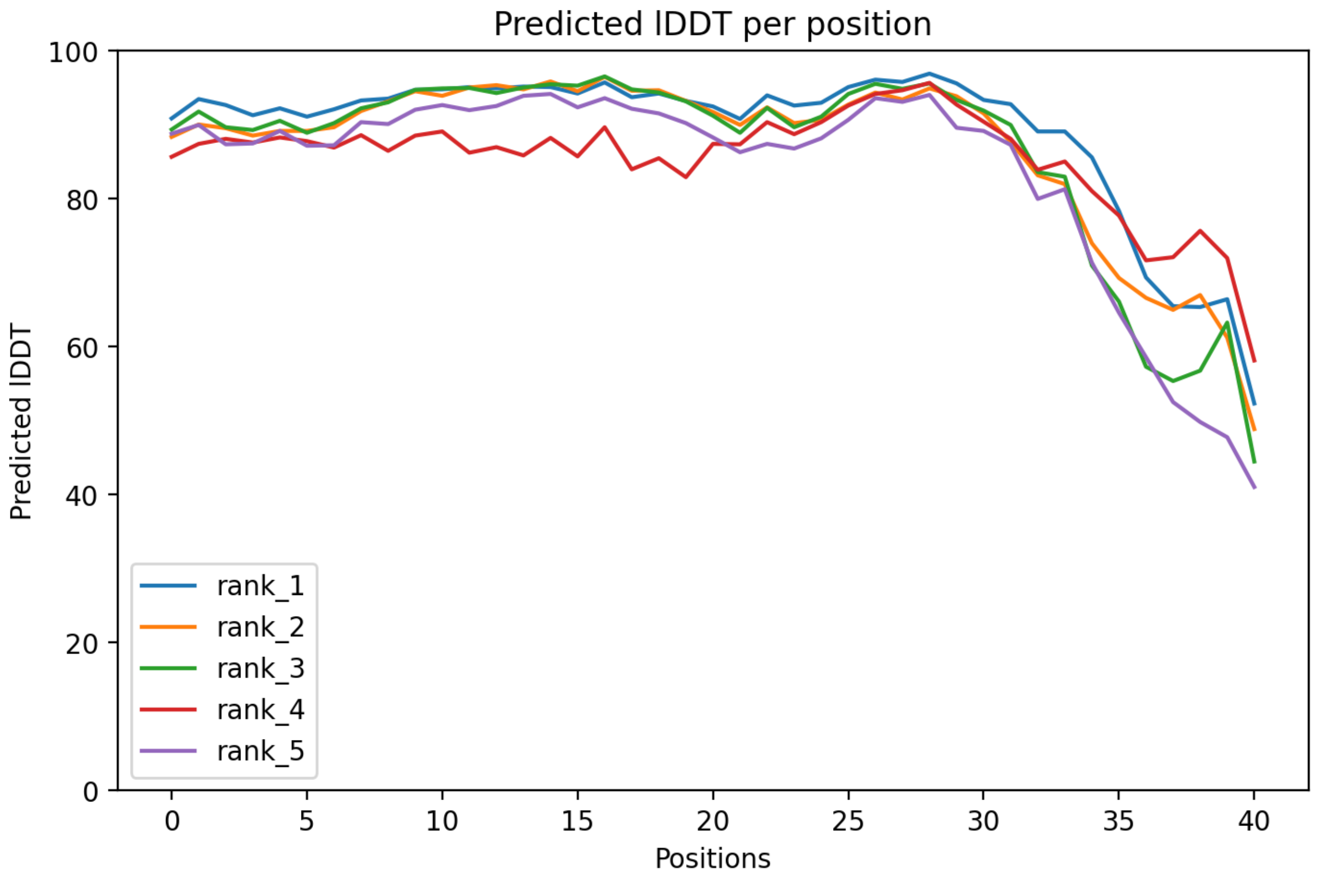

### CP54: MtrunA17_Chr3g0105981

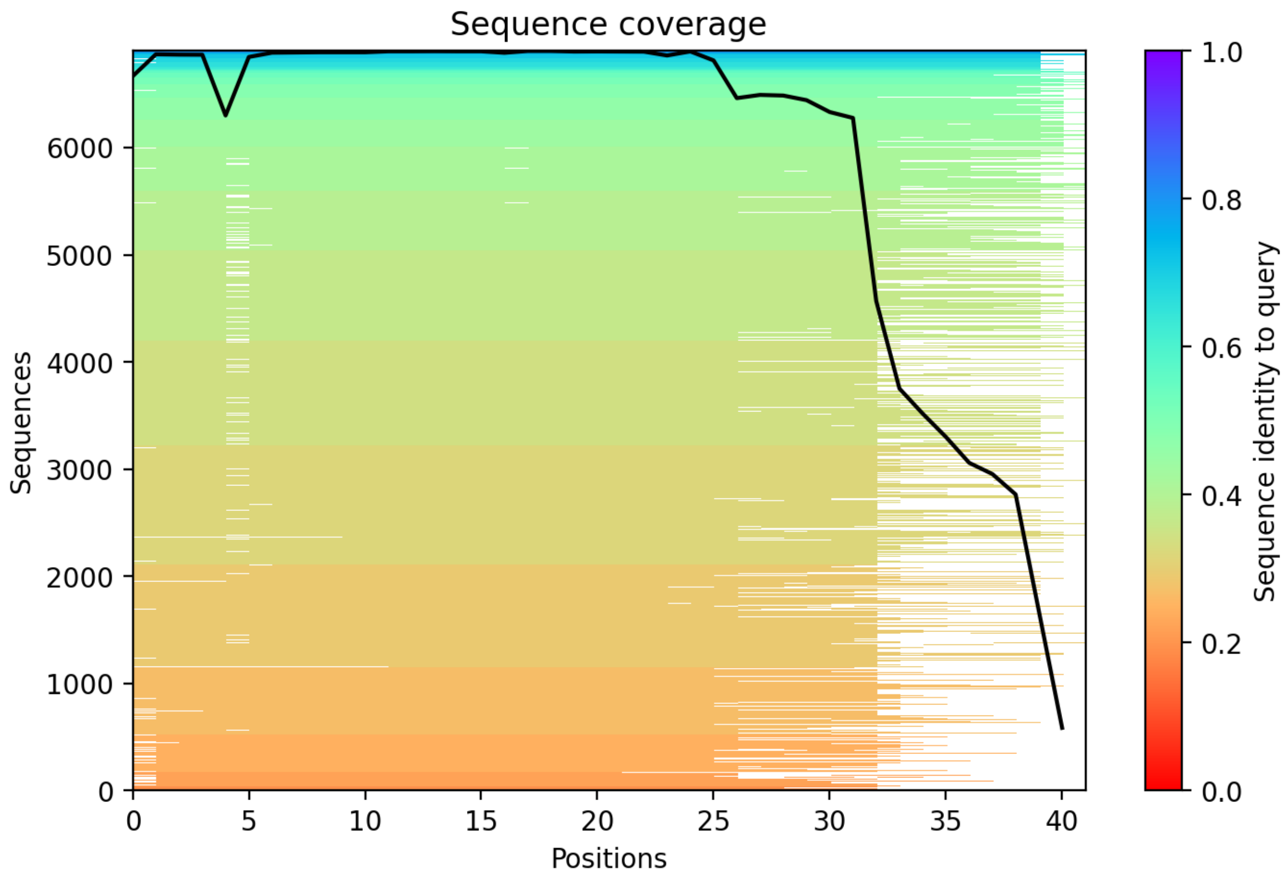

CP55: MtrunA17_Chr3g0110451

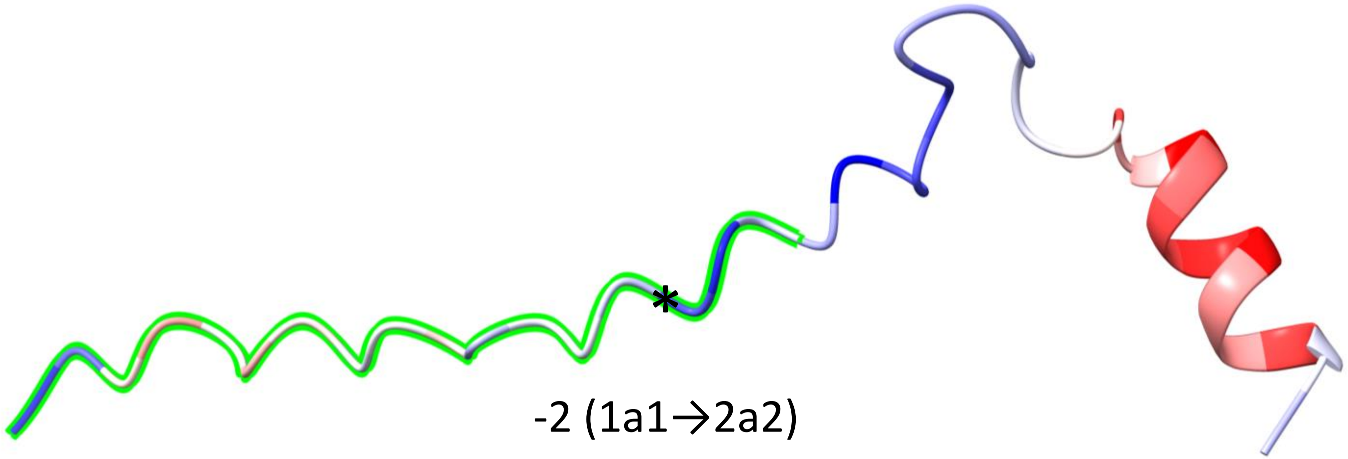


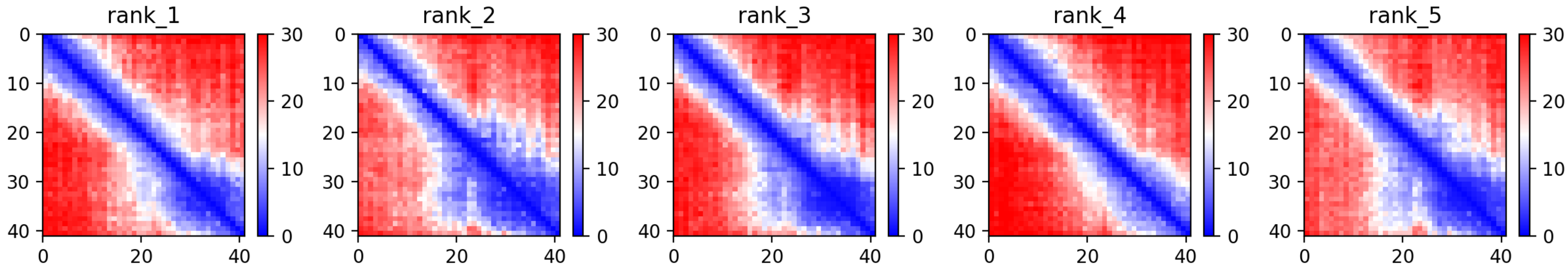

CP55: MtrunA17_Chr3g0110451

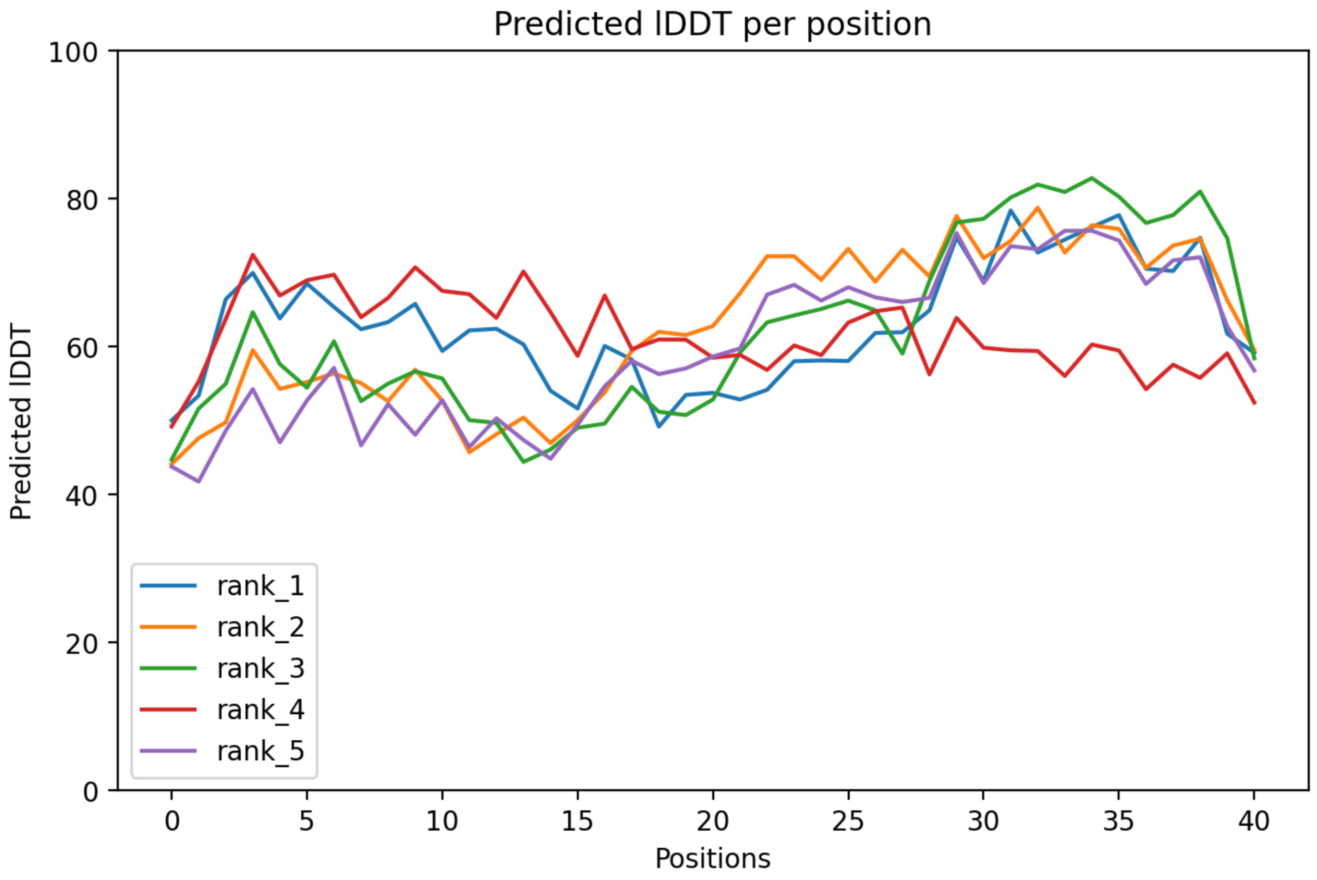

CP55: MtrunA17_Chr3g0110451

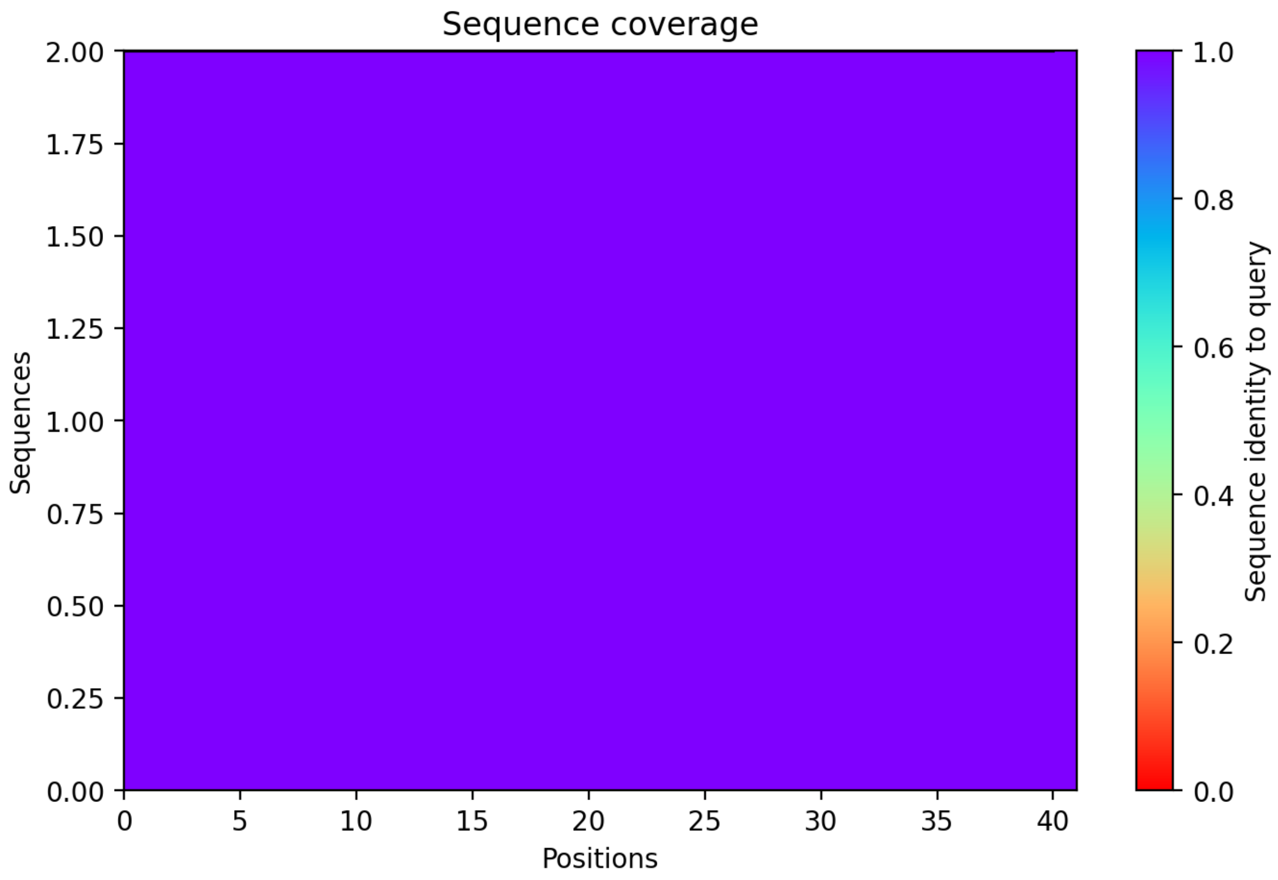

CP56: MtrunA17_Chr3g0113591

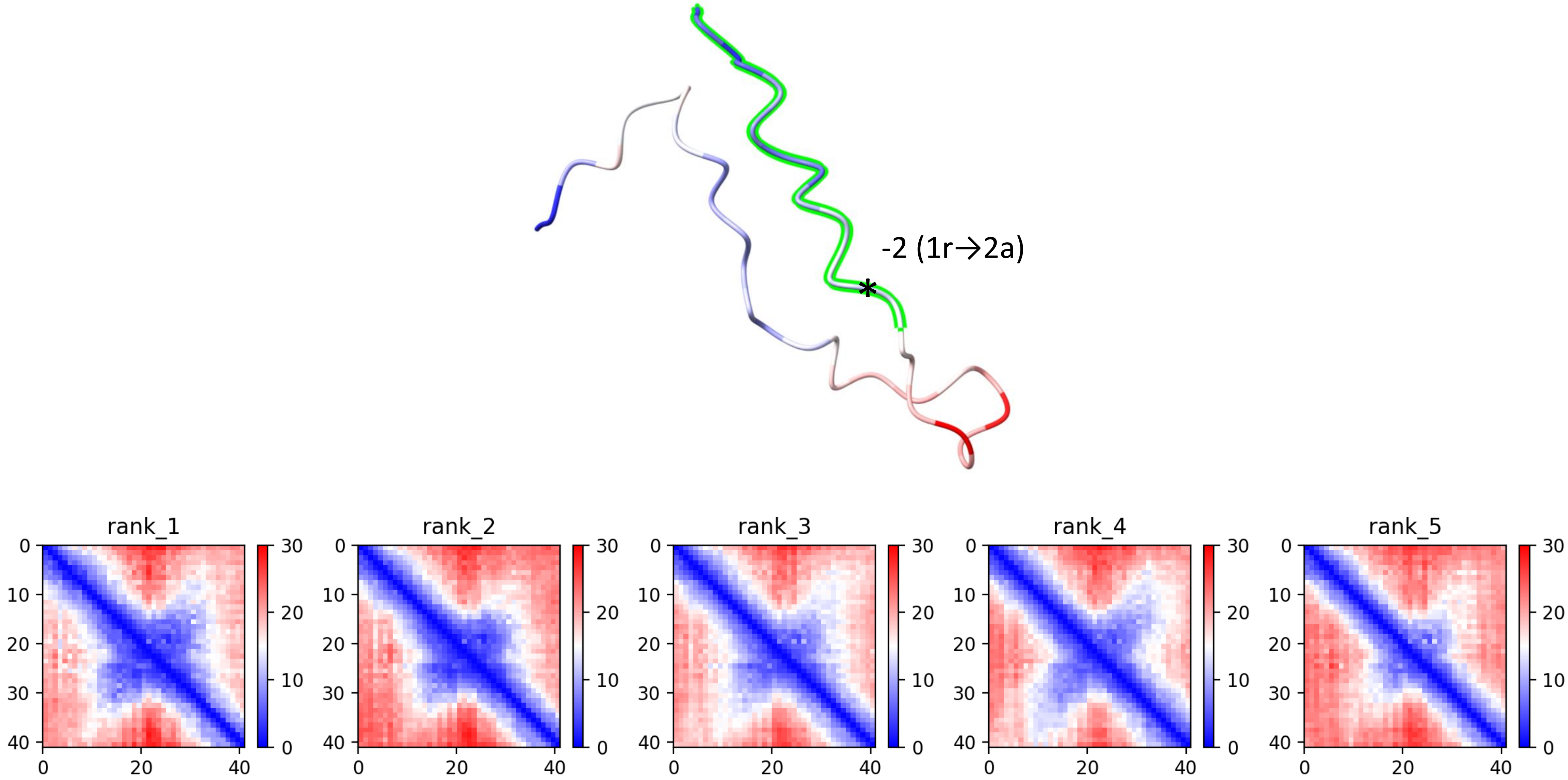

CP56: MtrunA17_Chr3g0113591

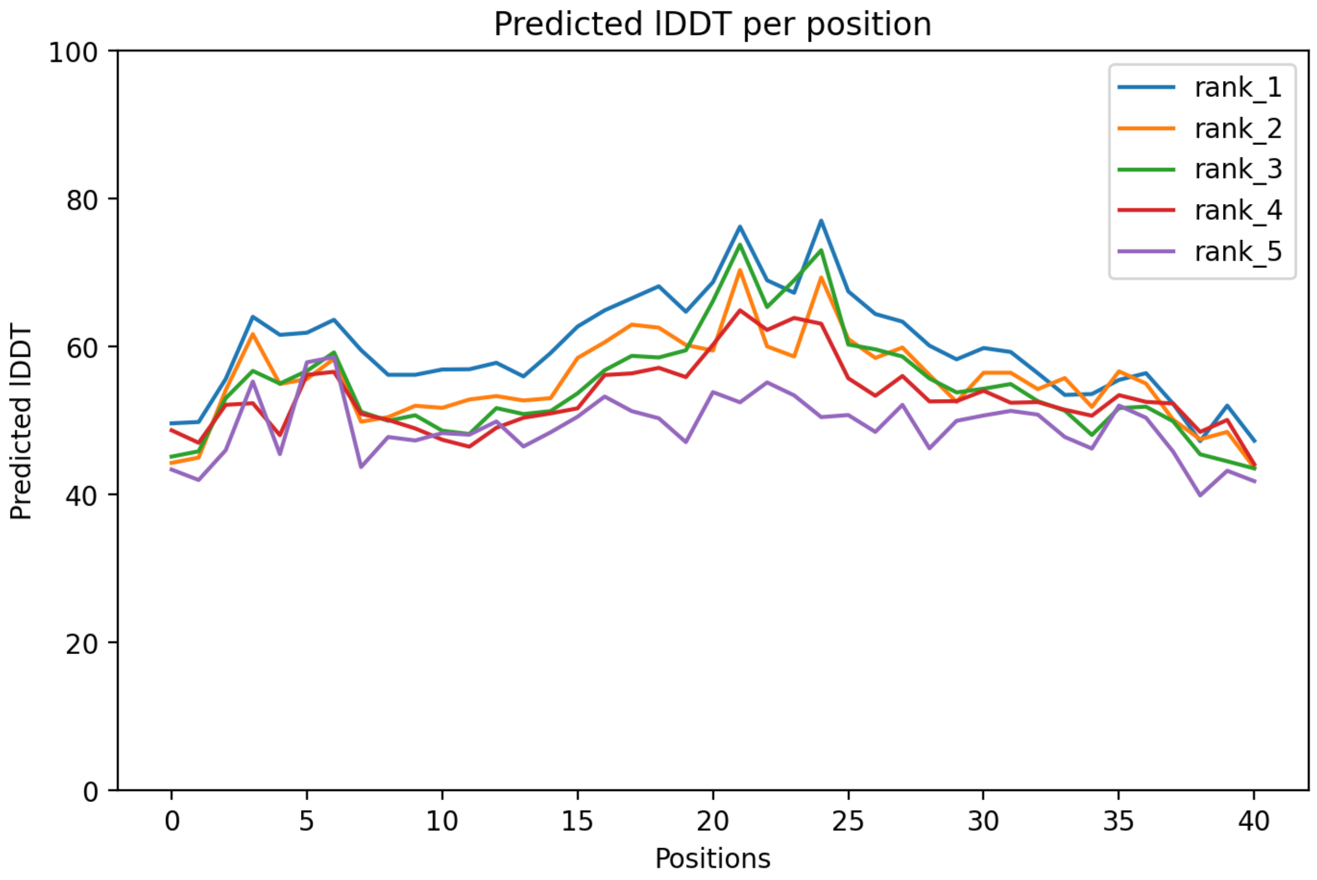

### CP56: MtrunA17_Chr3g0113591

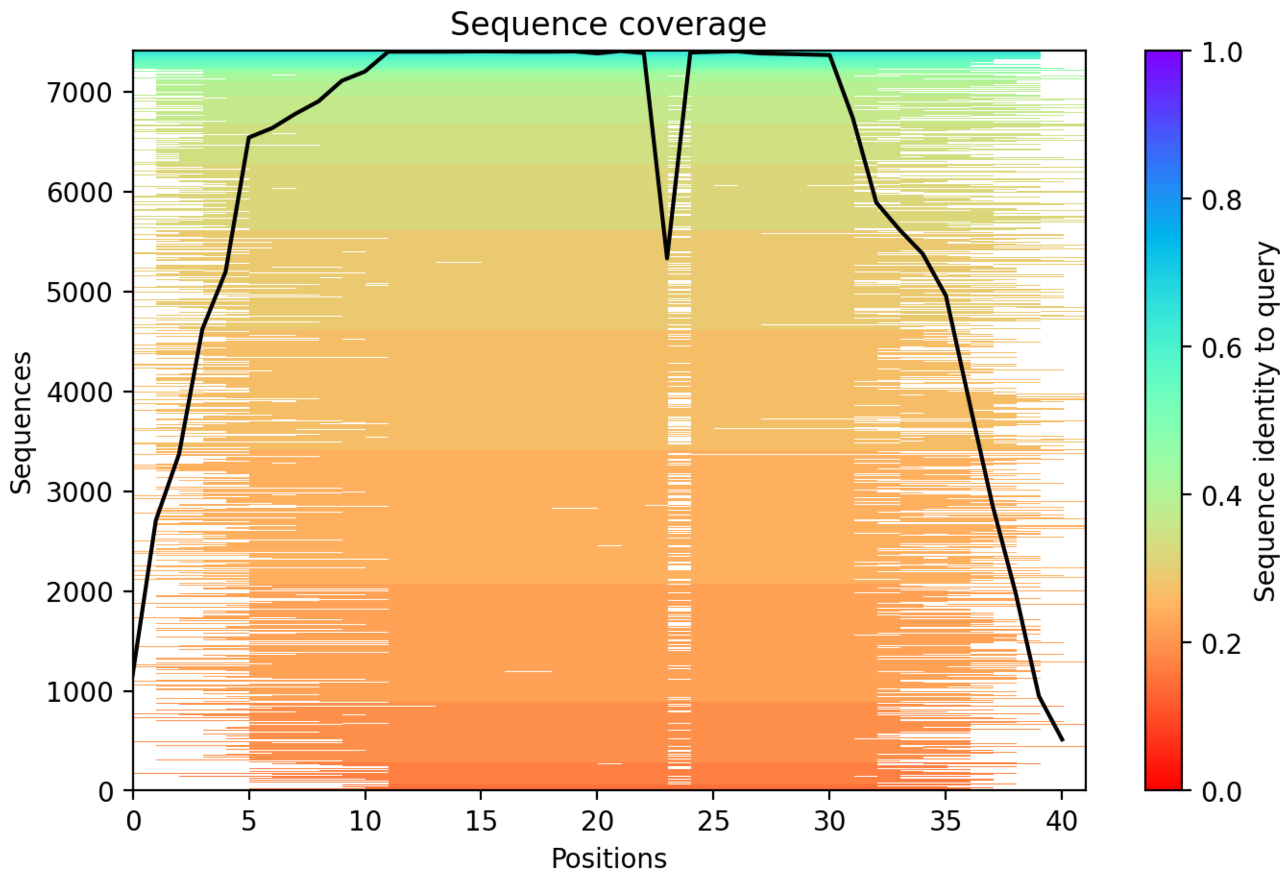

CP57: MtrunA17_Chr3g0124631

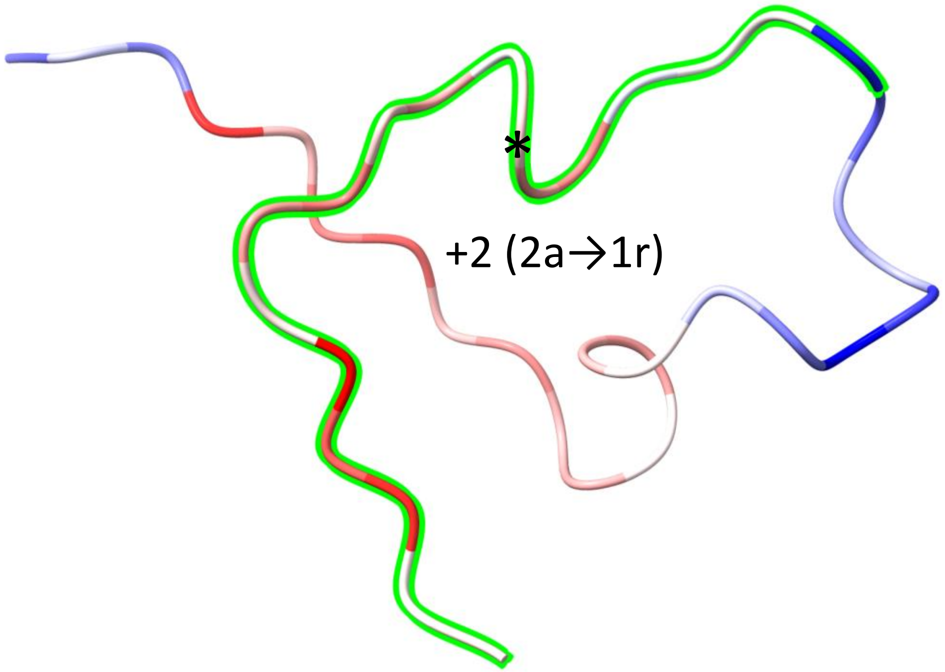


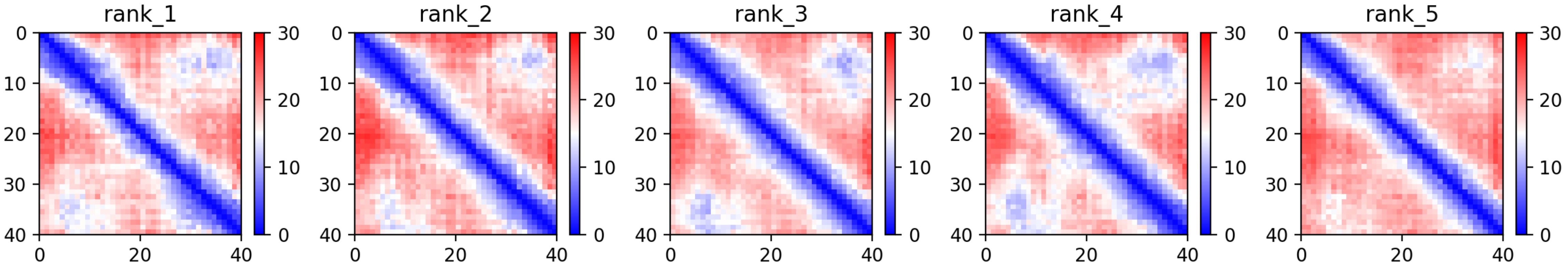

CP57: MtrunA17_Chr3g0124631

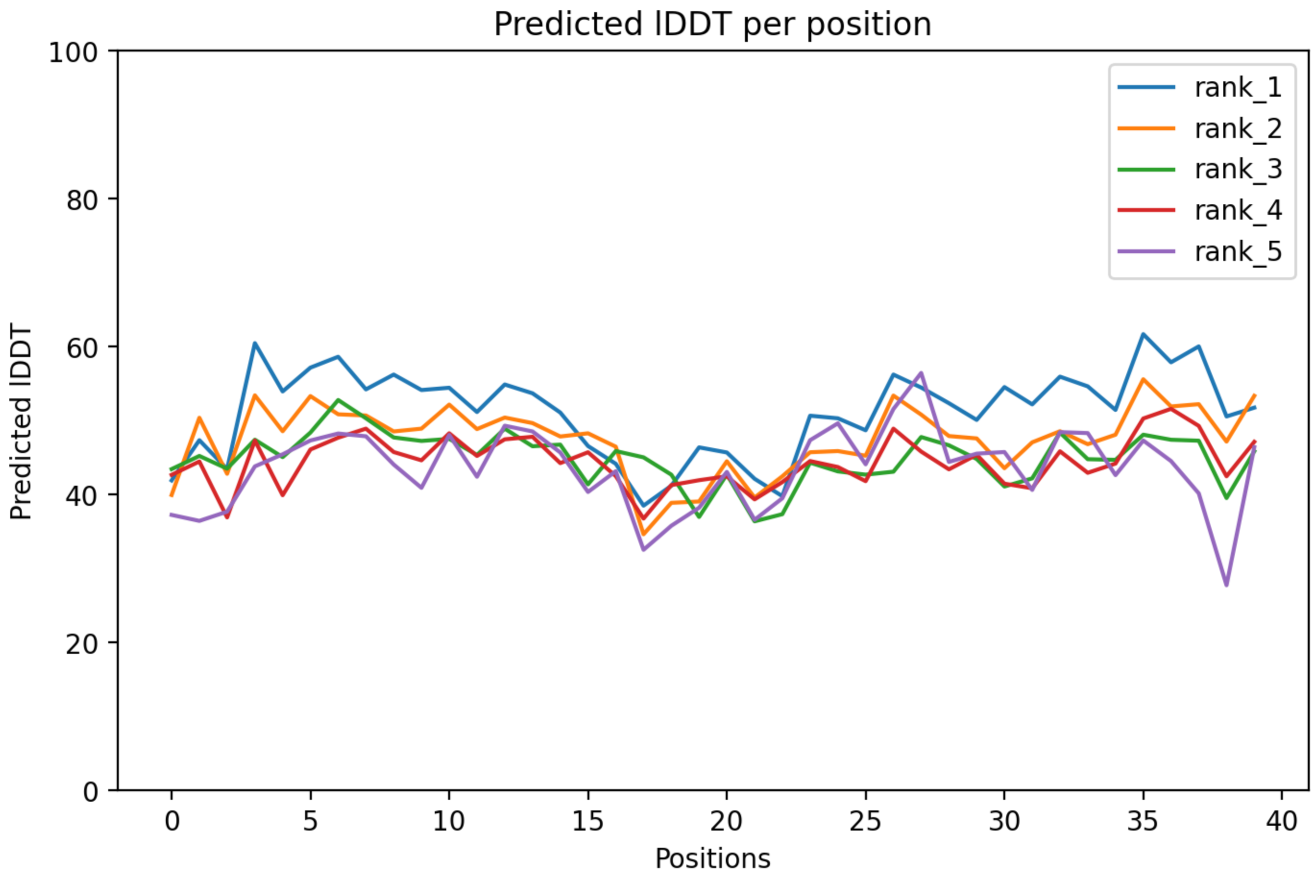

### CP57: MtrunA17_Chr3g0124631

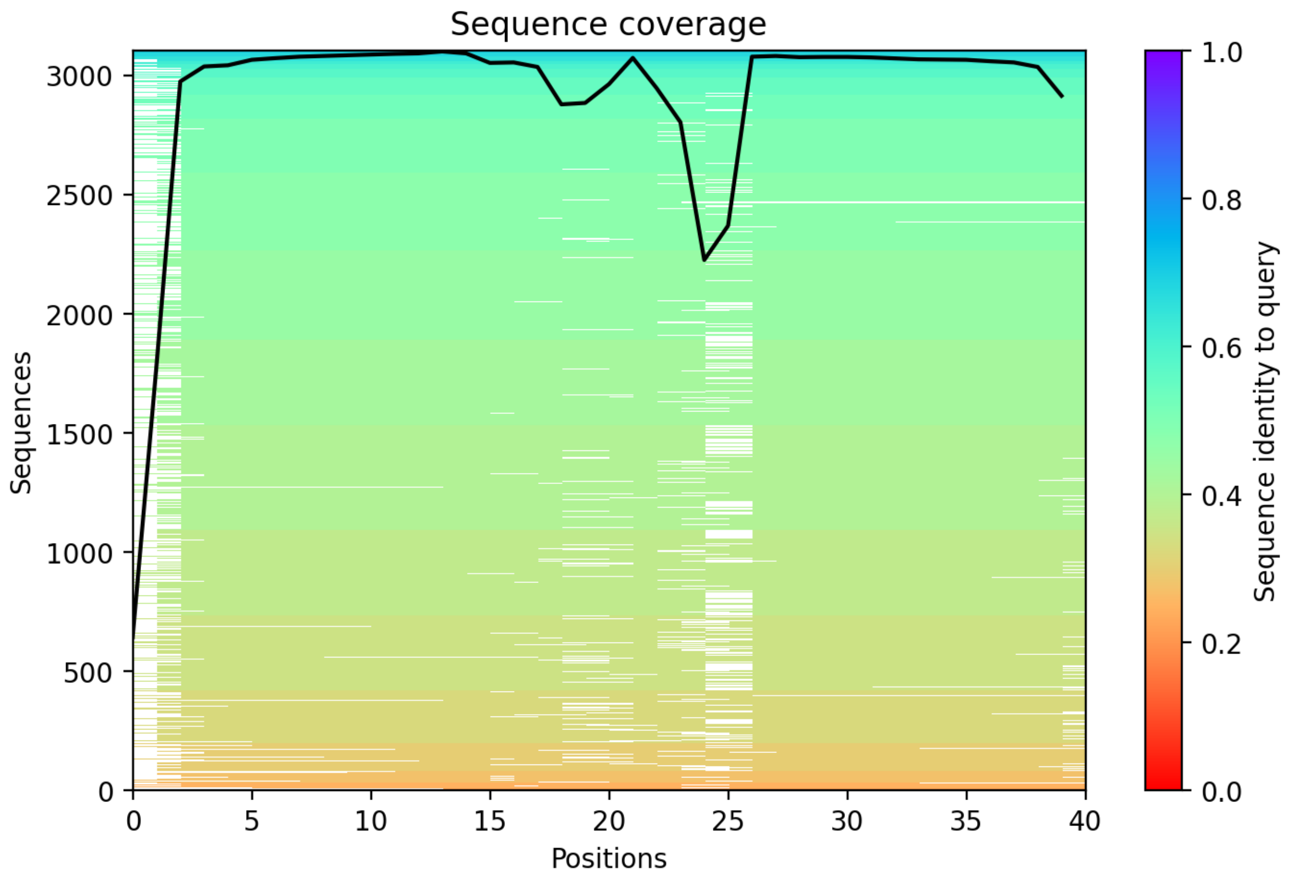

CP58: MtrunA17_Chr3g0127321

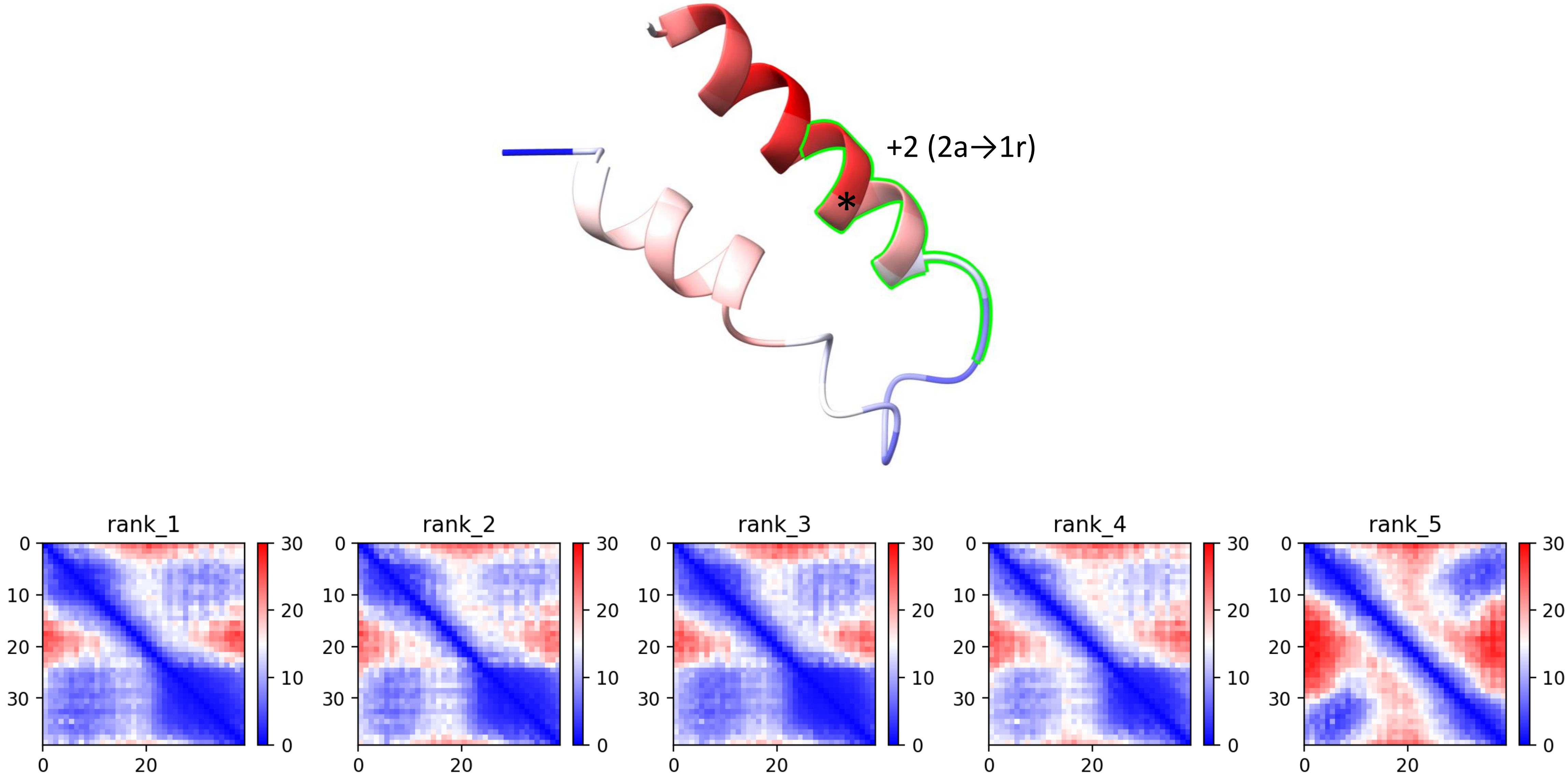

CP58: MtrunA17_Chr3g0127321

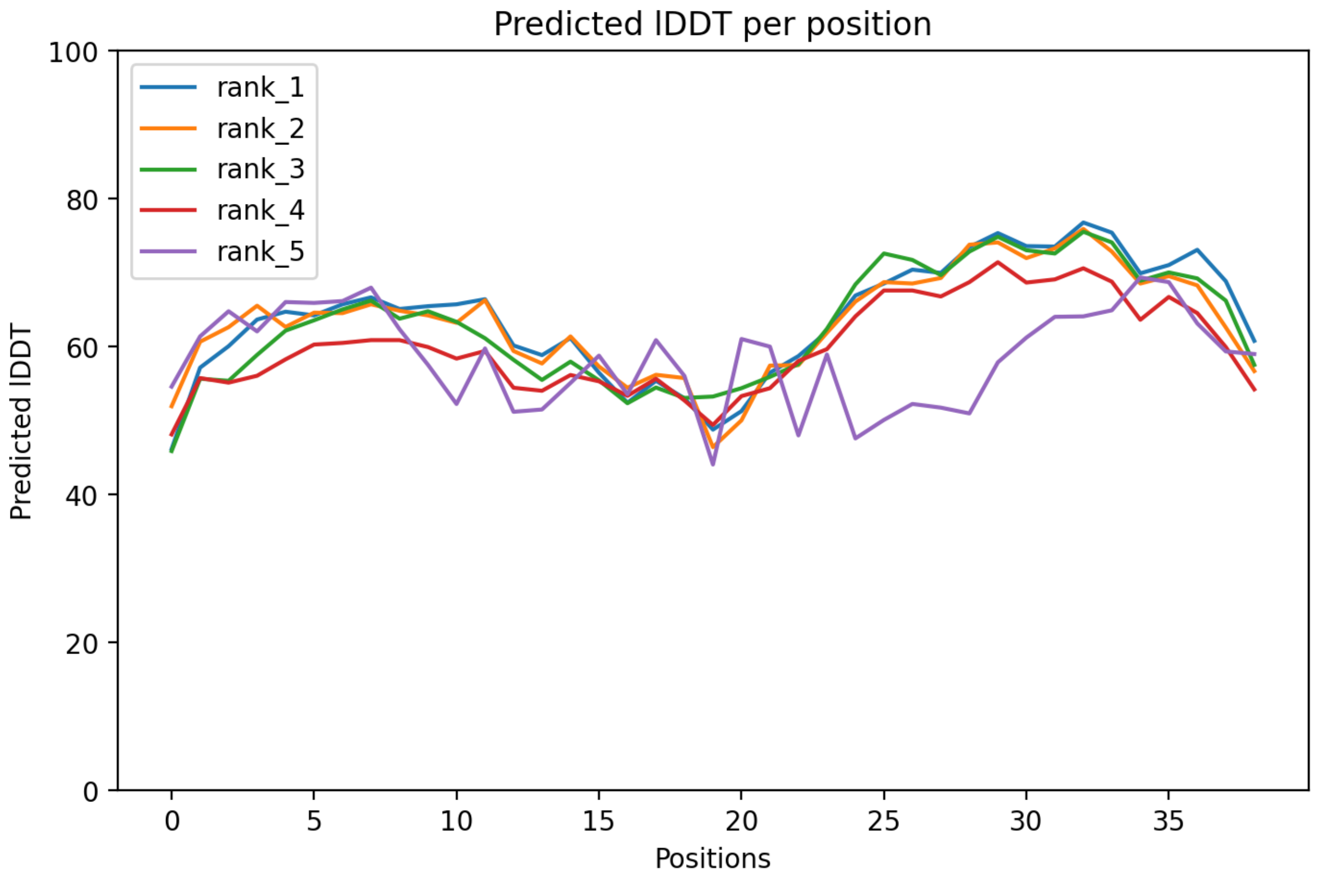

### CP58: MtrunA17_Chr3g0127321

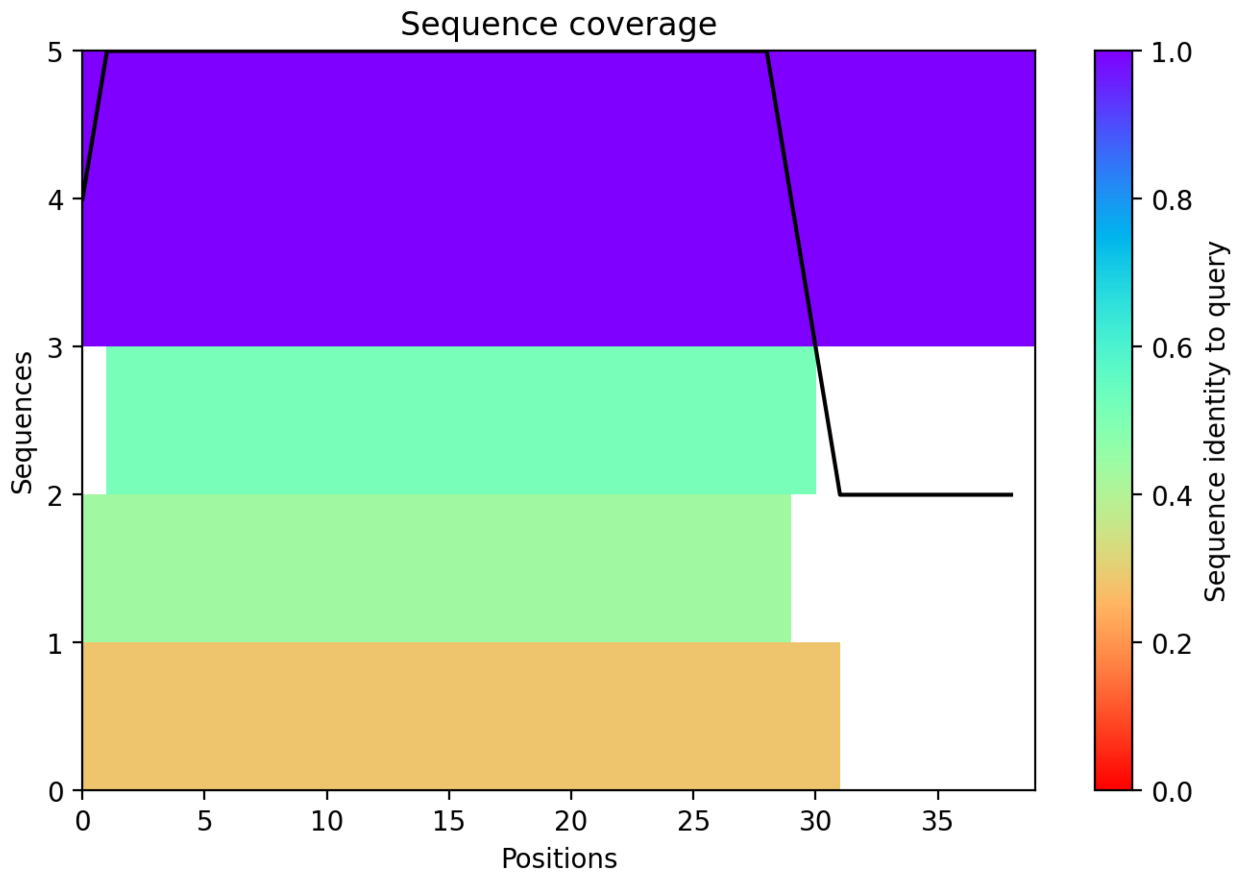

CP59: MtrunA17_Chr3g0130671

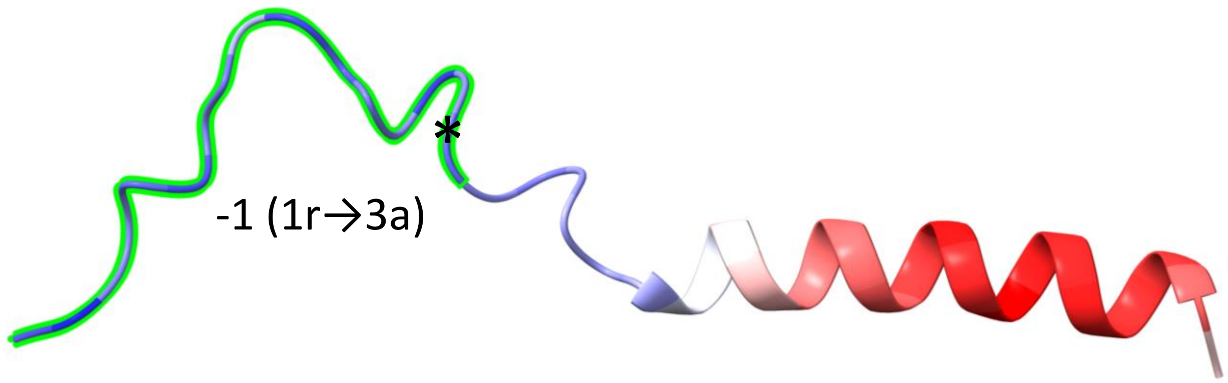


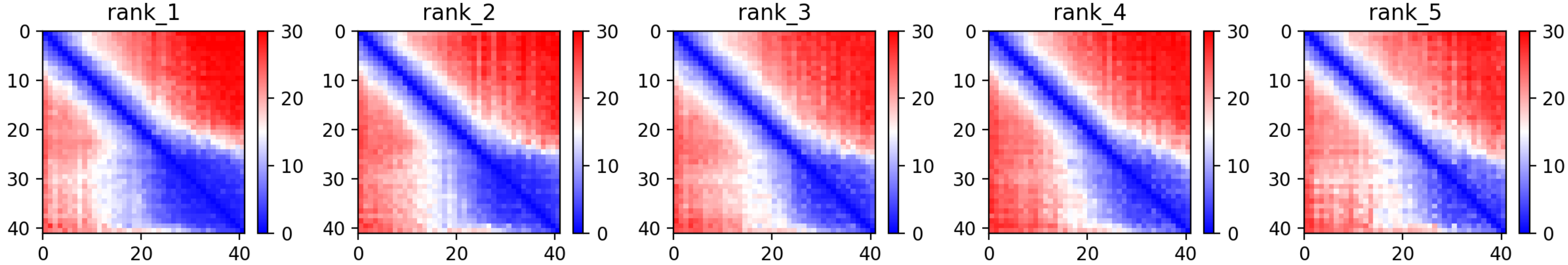

CP59: MtrunA17_Chr3g0130671

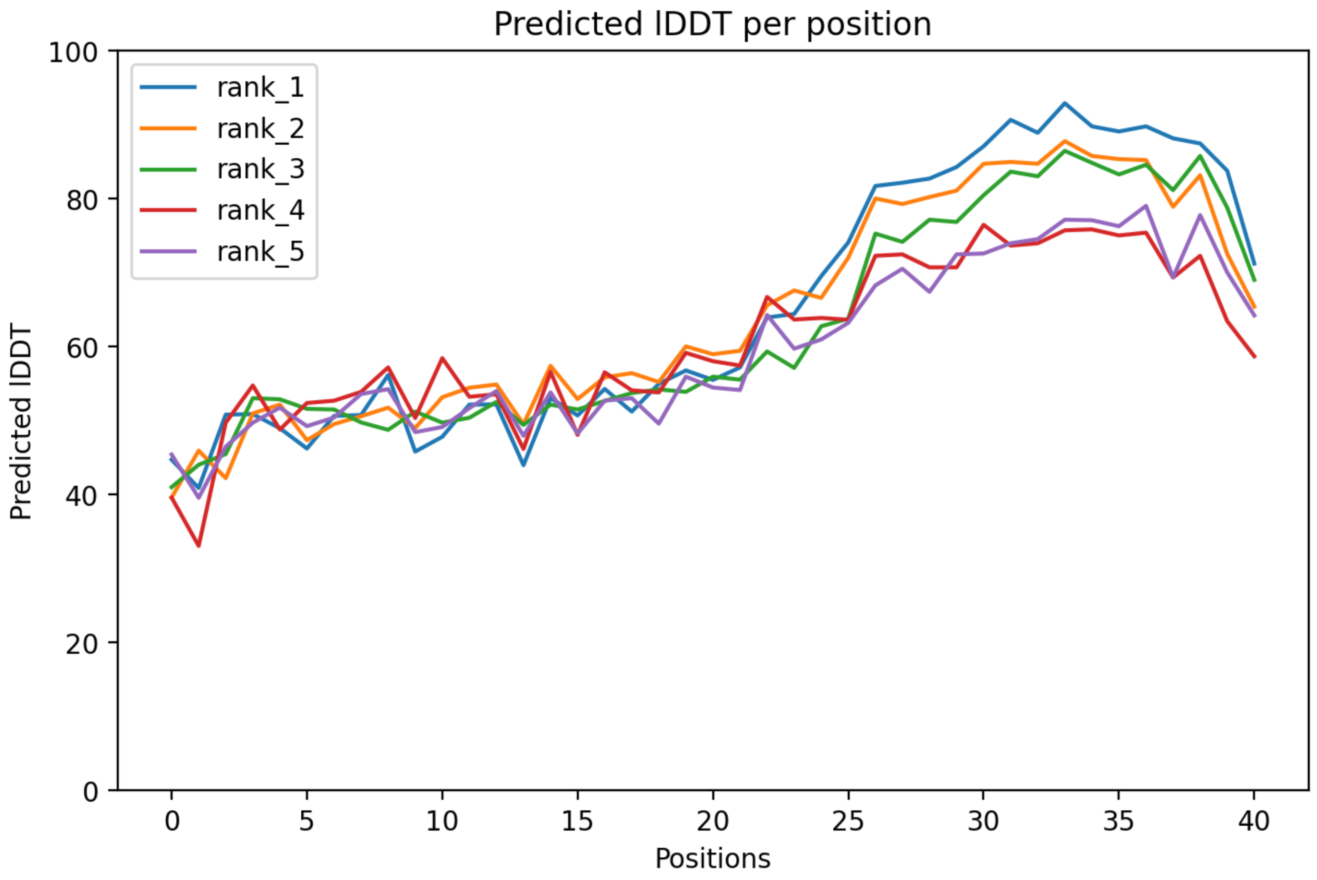

### CP59: MtrunA17_Chr3g0130671

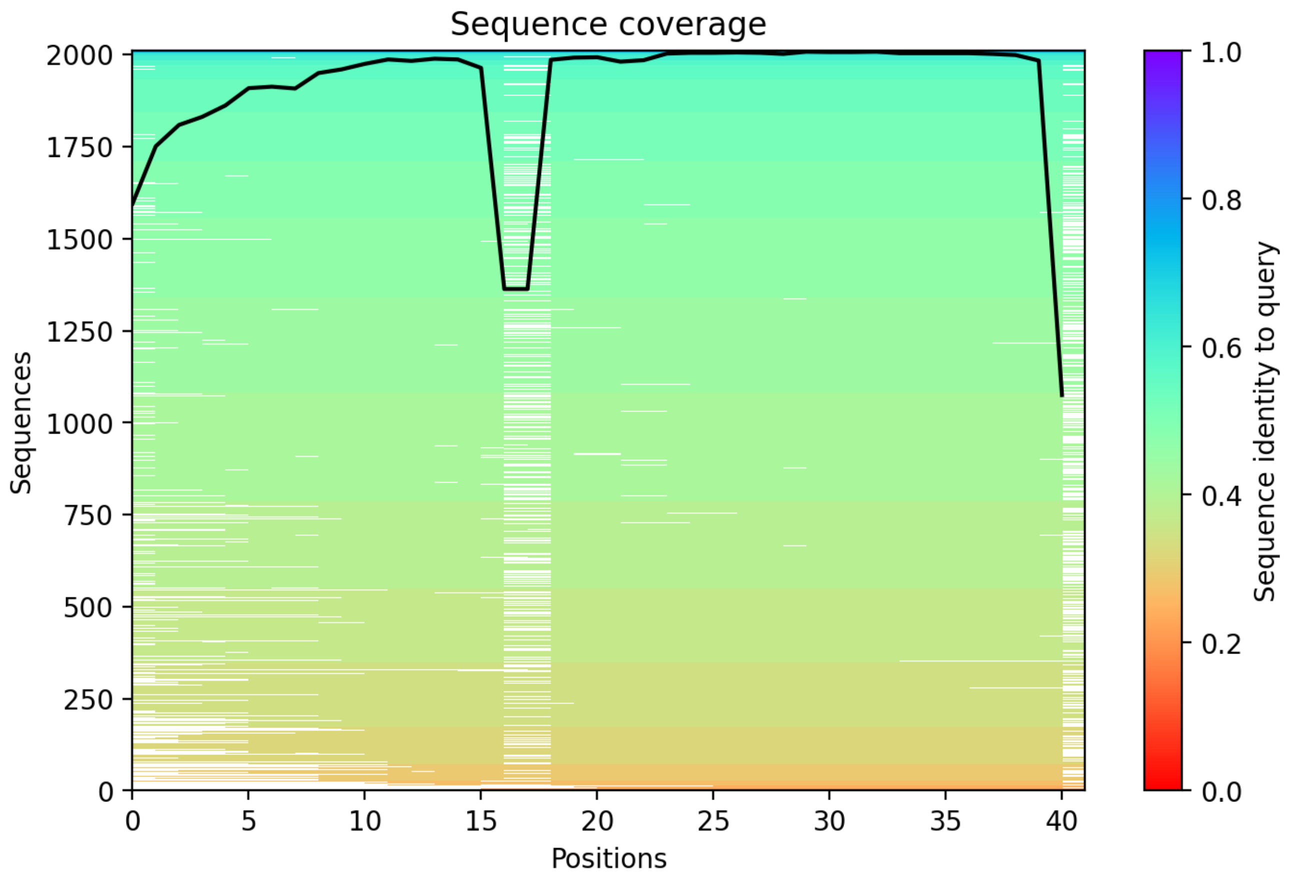

CP60: MtrunA17_Chr3g0135761

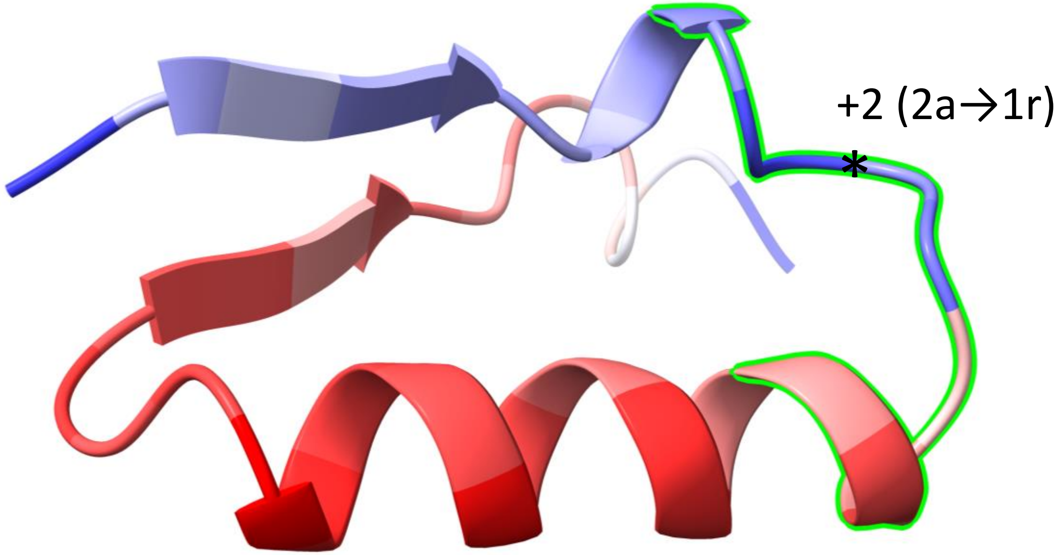


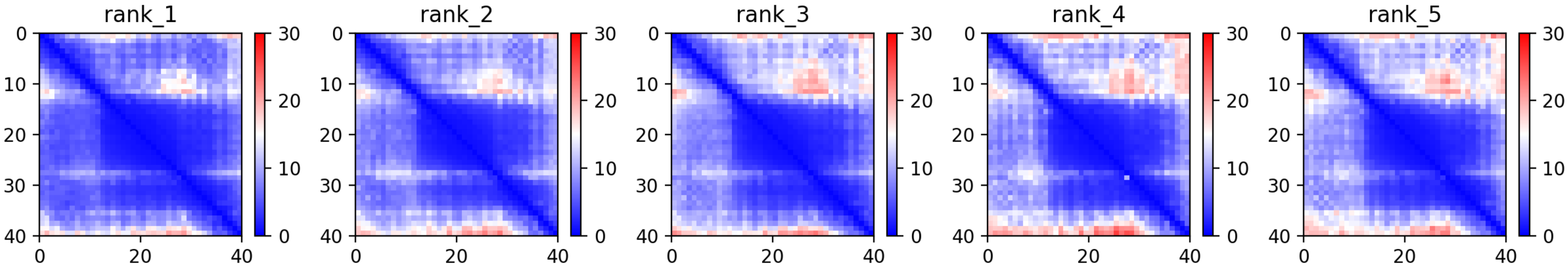

CP60: MtrunA17_Chr3g0135761

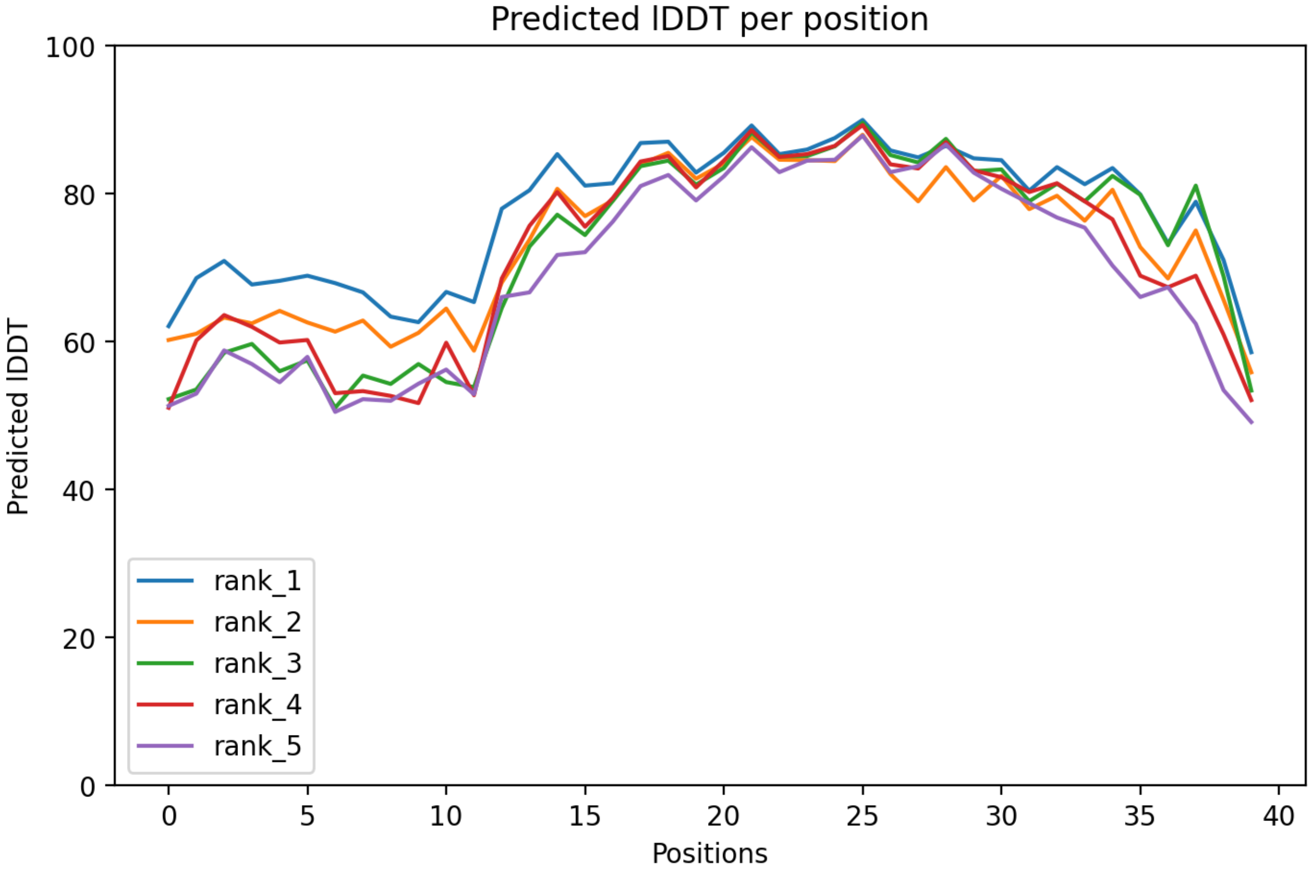

### CP60: MtrunA17_Chr3g0135761

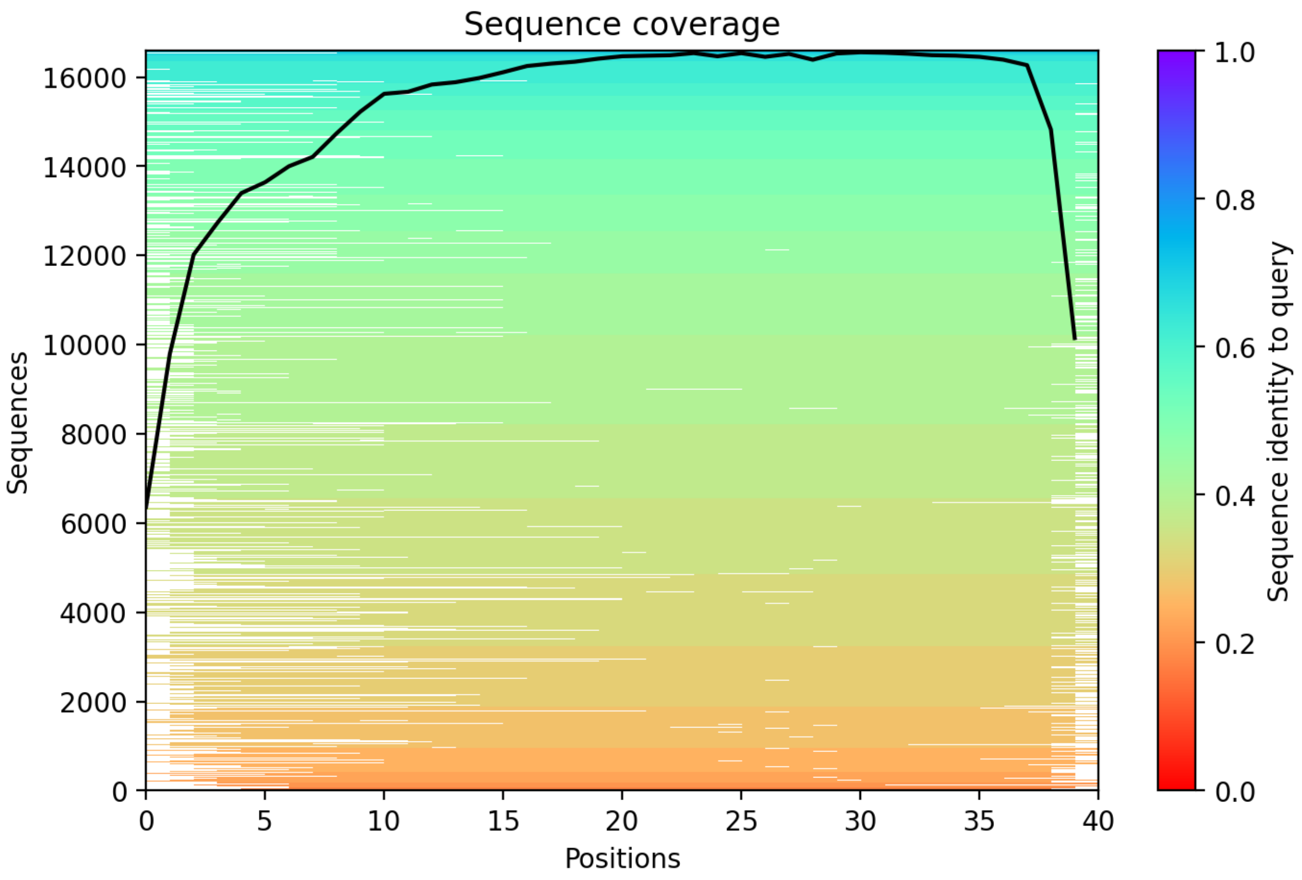

### CP61: MtrunA17_Chr3g0137391

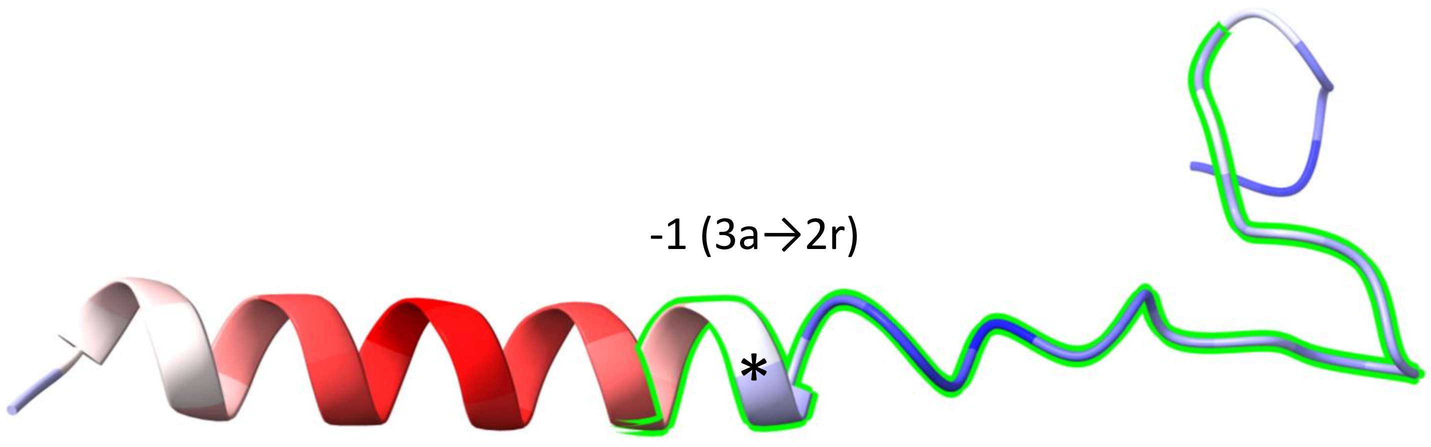


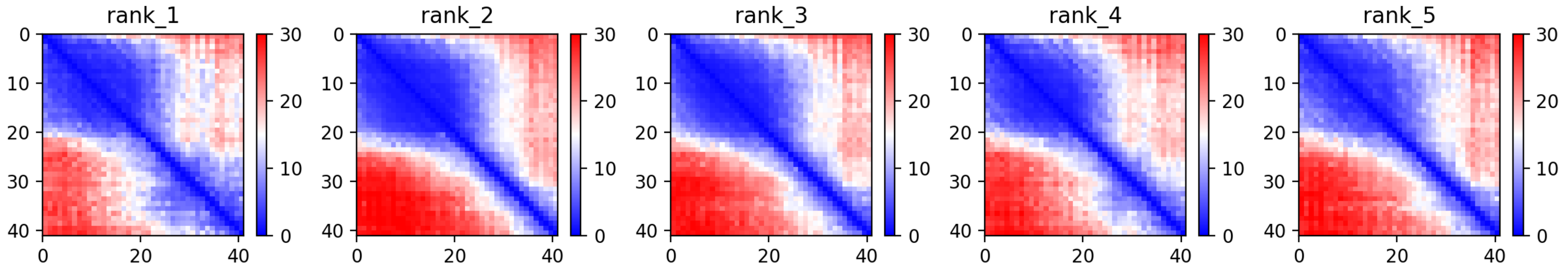

CP61: MtrunA17_Chr3g0137391

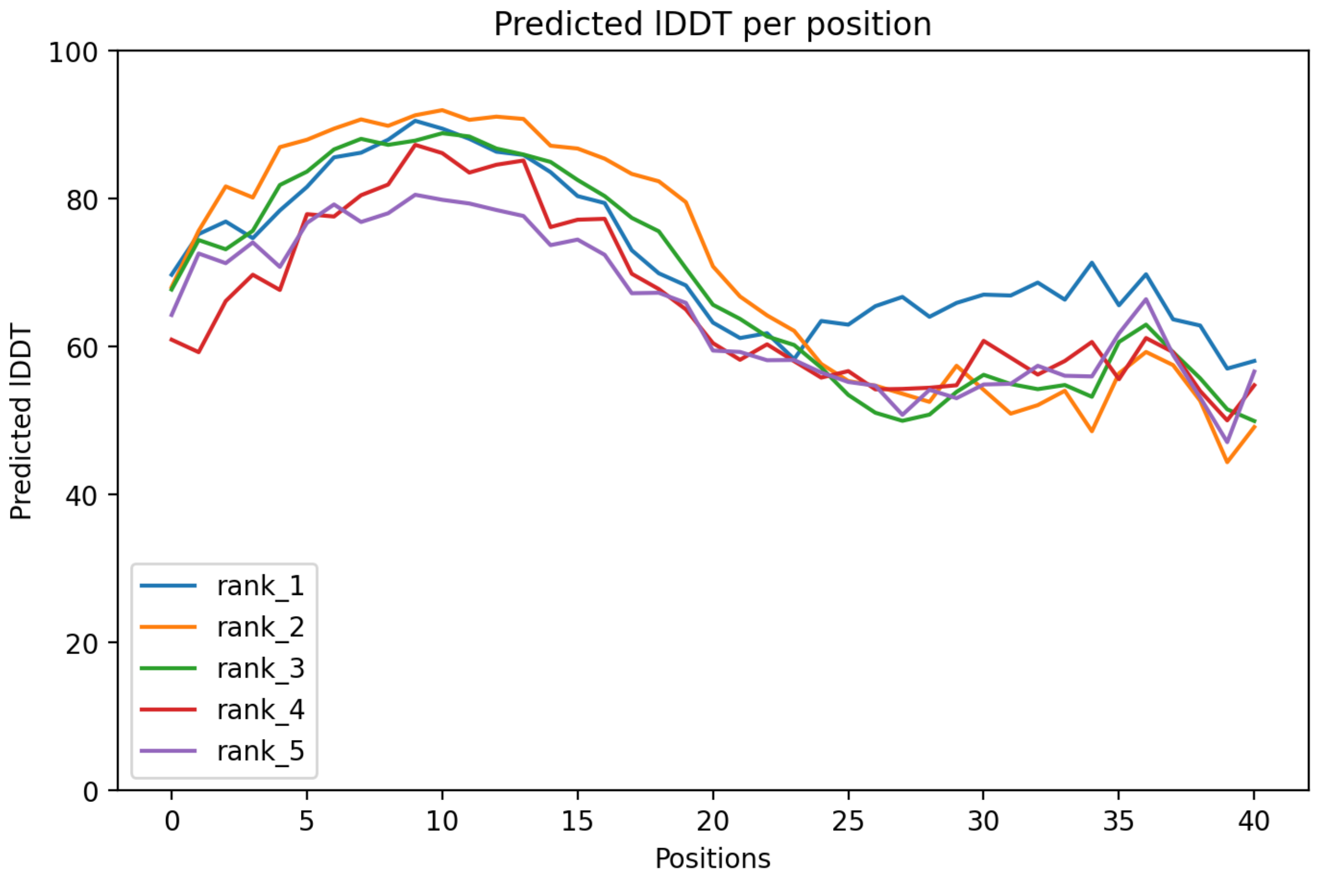

CP61: MtrunA17_Chr3g0137391

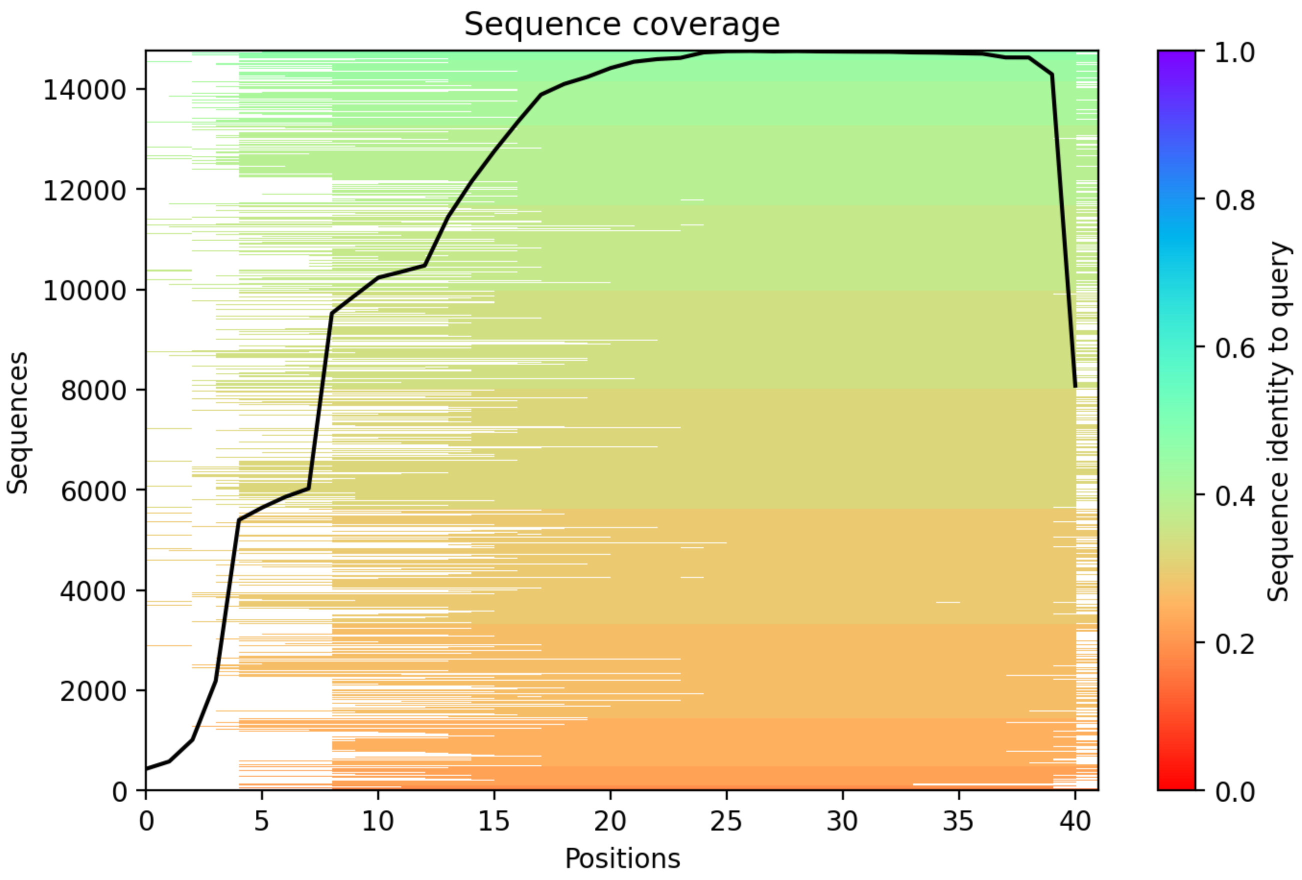

### CP62: MtrunA17_Chr3g0141901

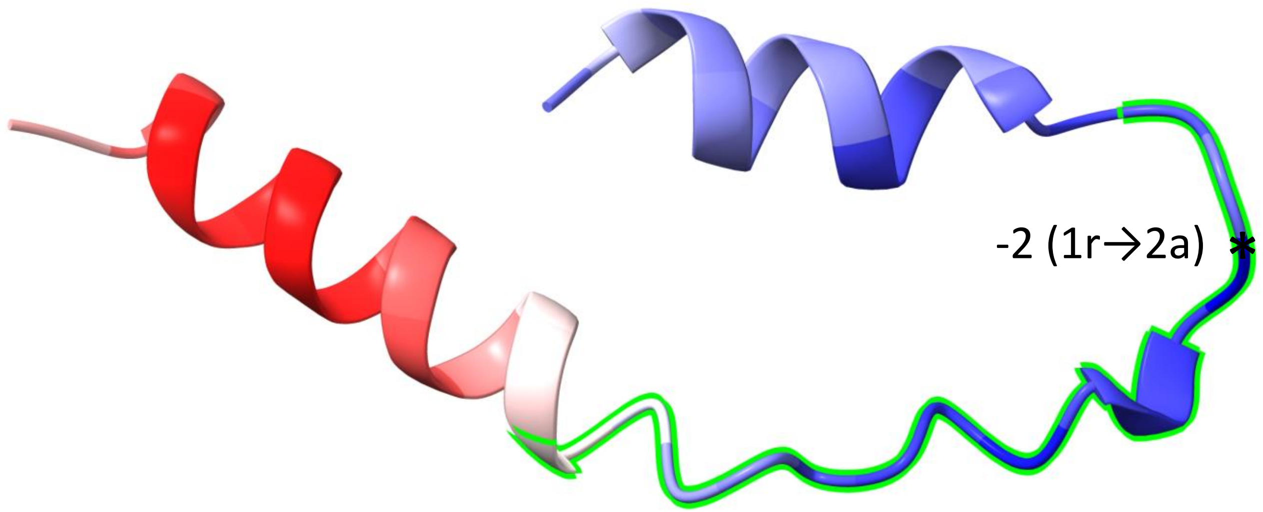


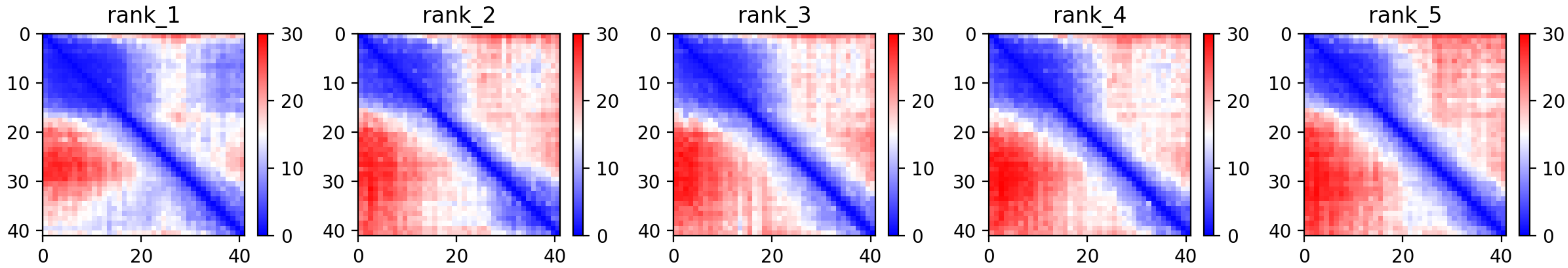

CP62: MtrunA17_Chr3g0141901

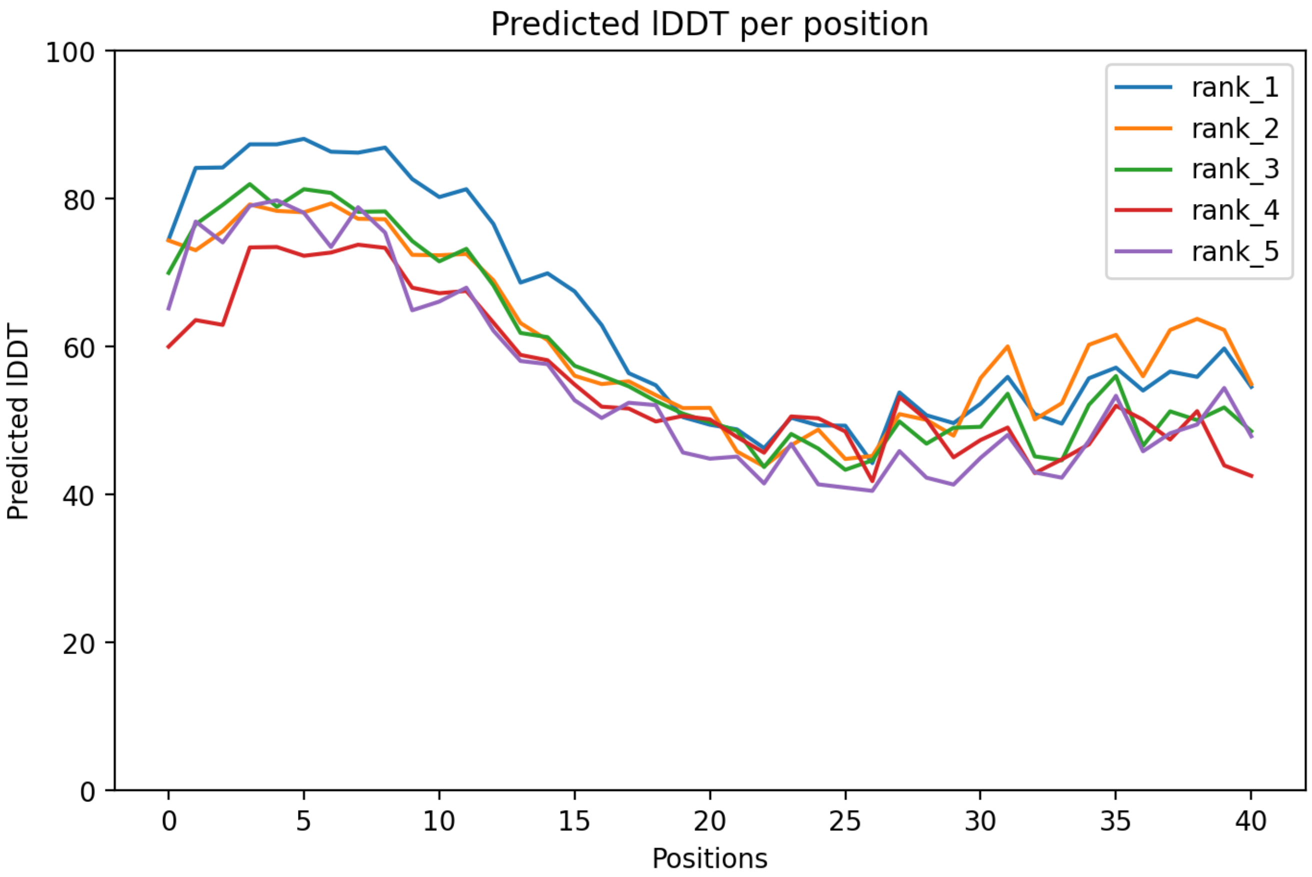

### CP62: MtrunA17_Chr3g0141901

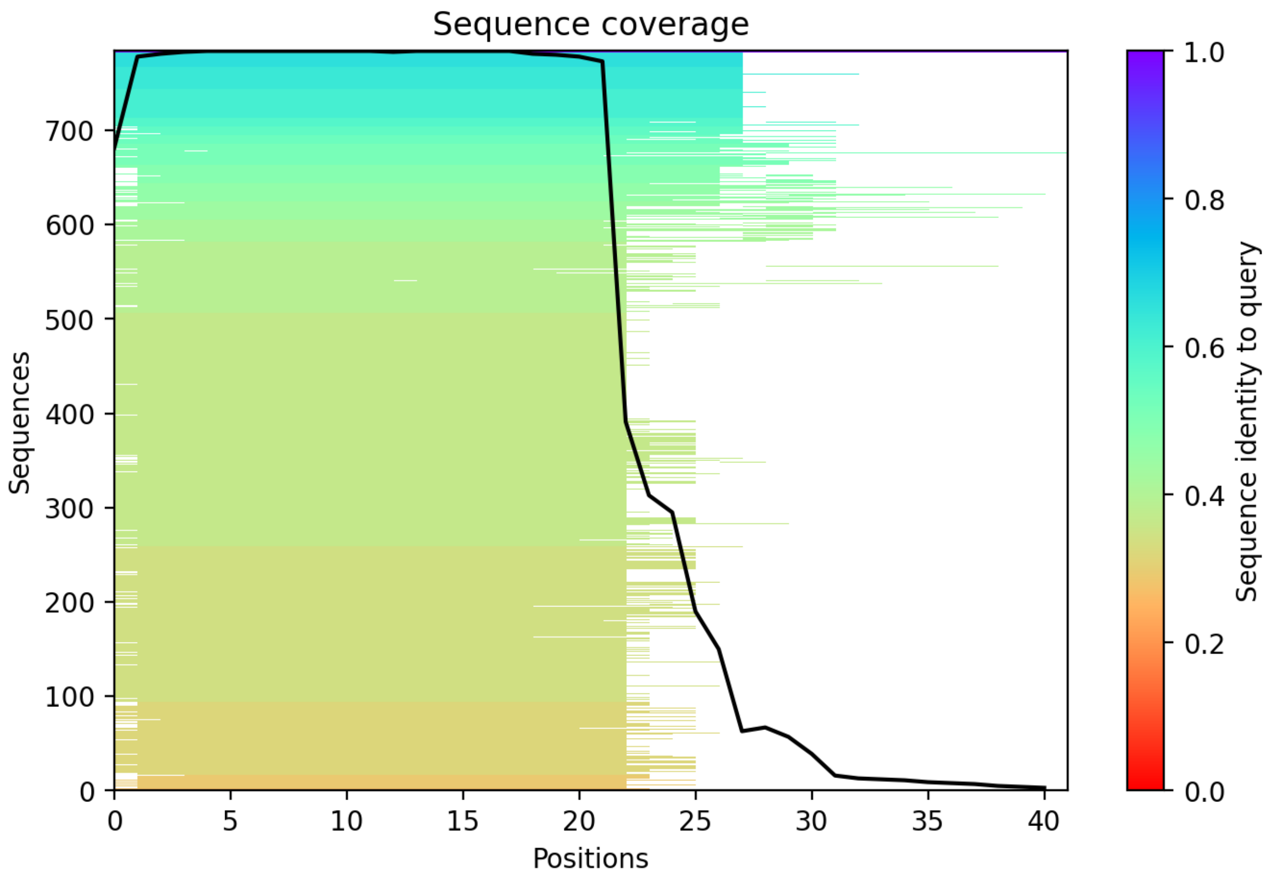

CP63: MtrunA17_Chr3g0144151

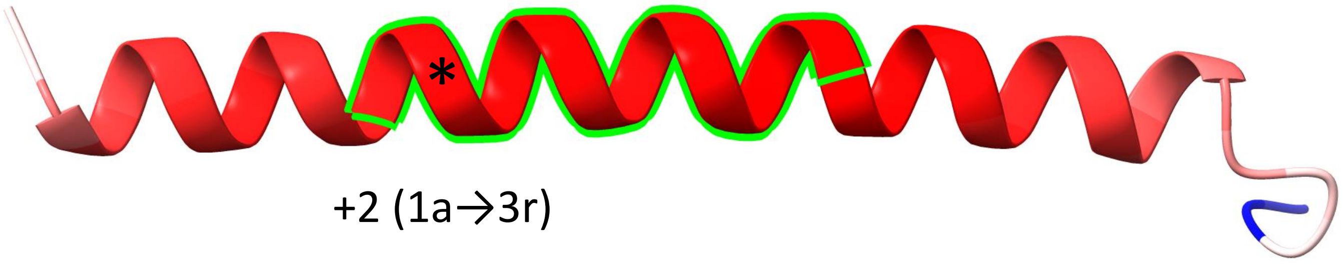


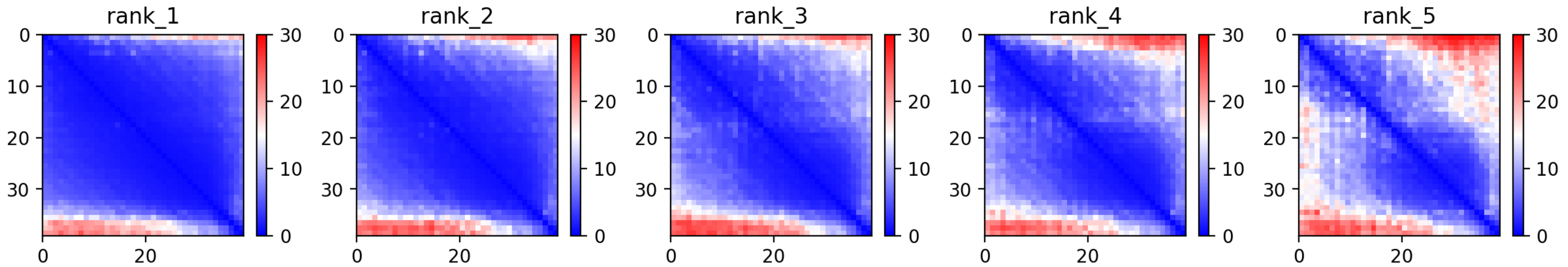

CP63: MtrunA17_Chr3g0144151

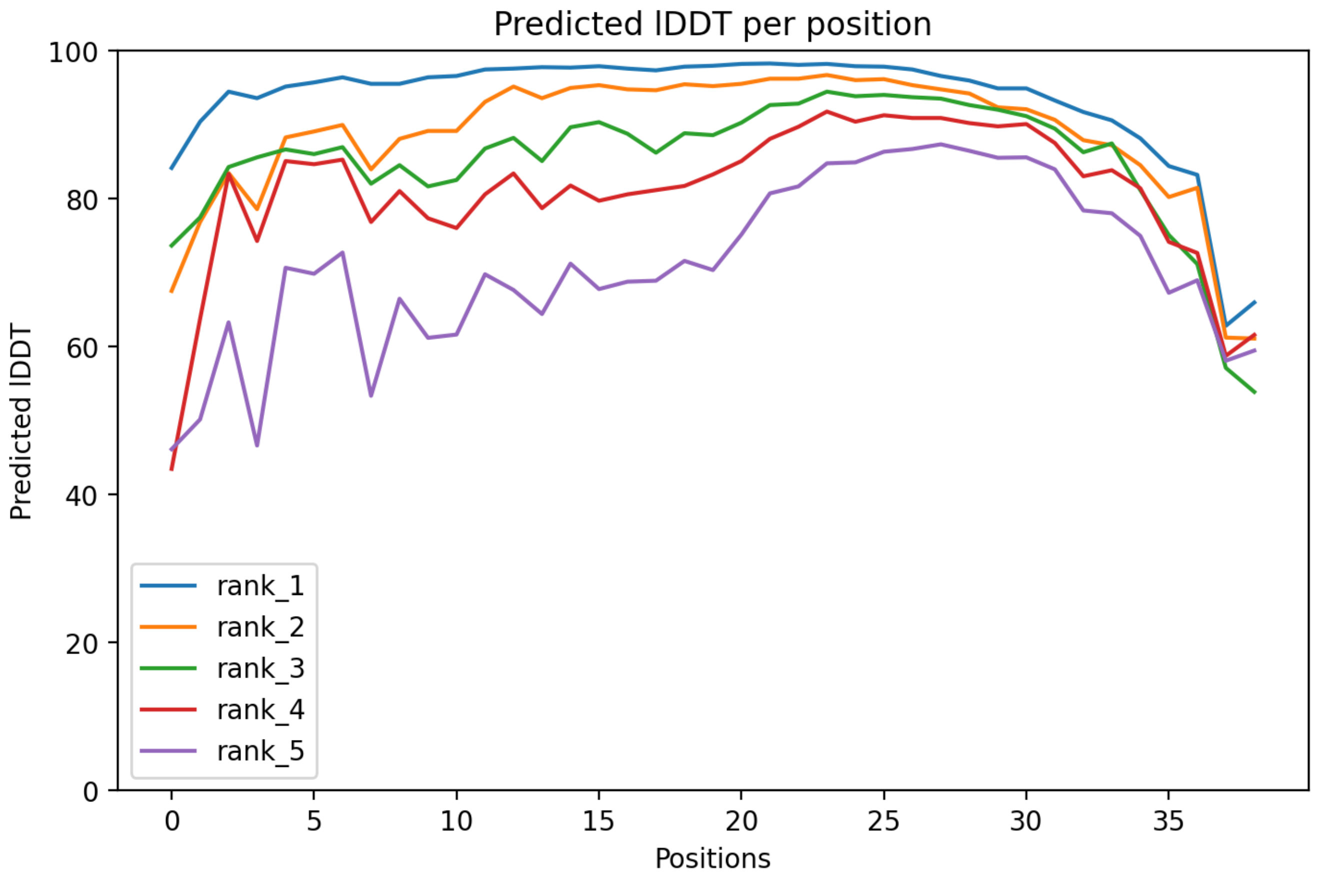

### CP63: MtrunA17_Chr3g0144151

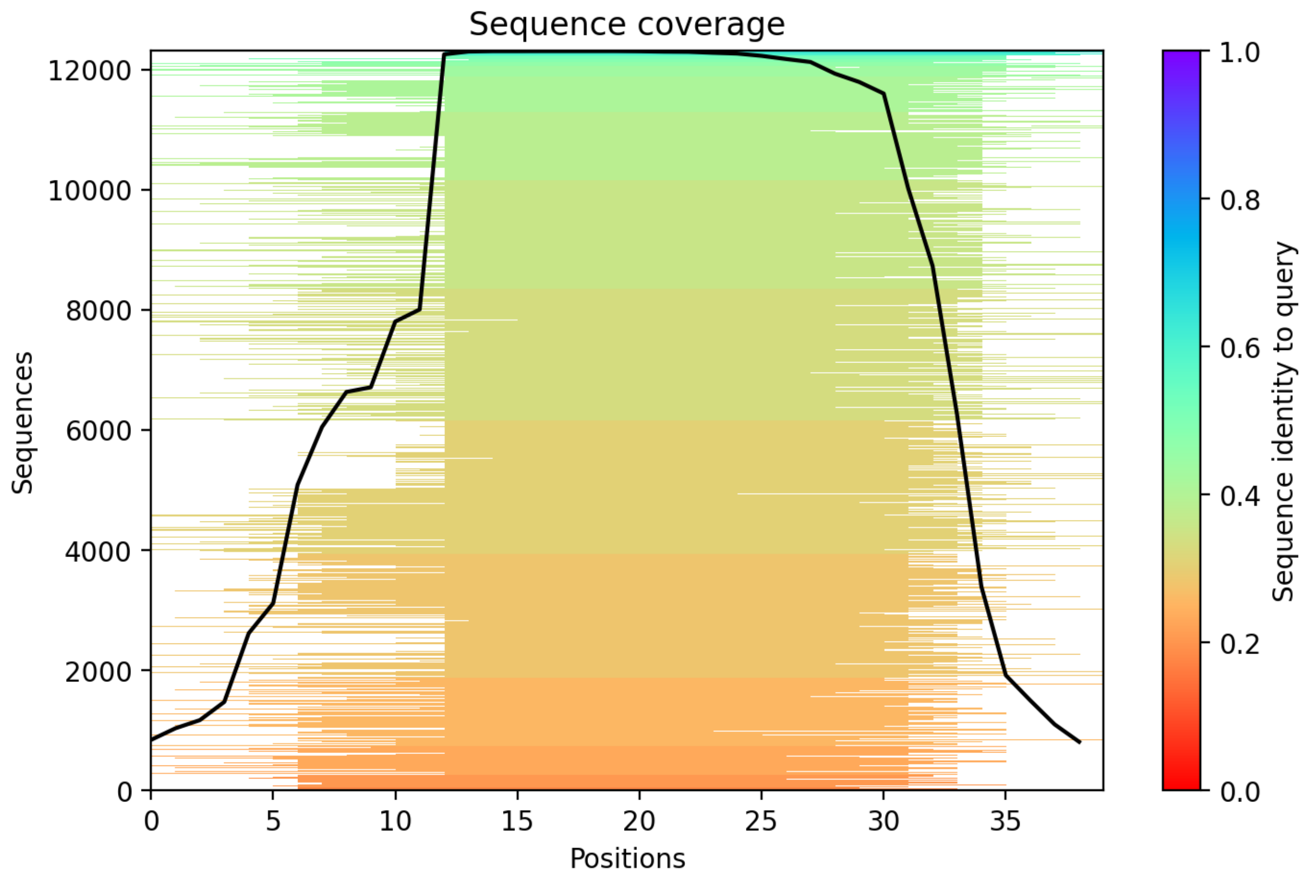

CP64: MtrunA17_Chr3g0144151

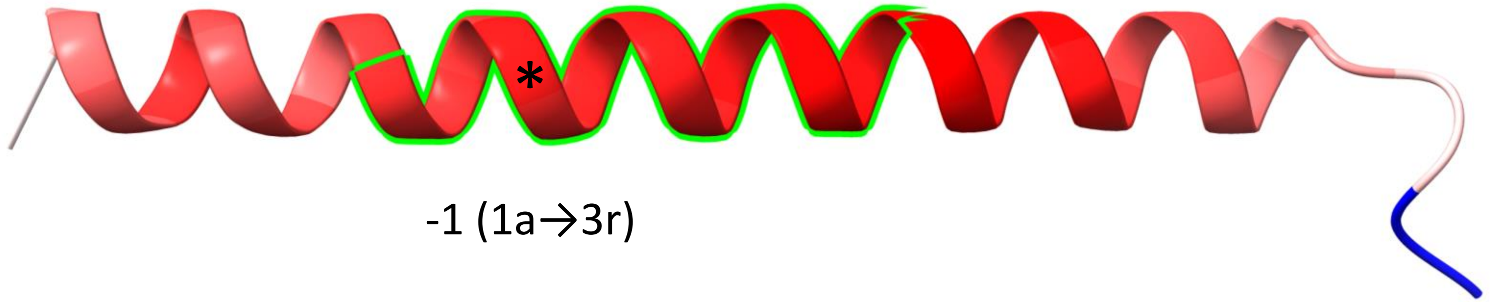


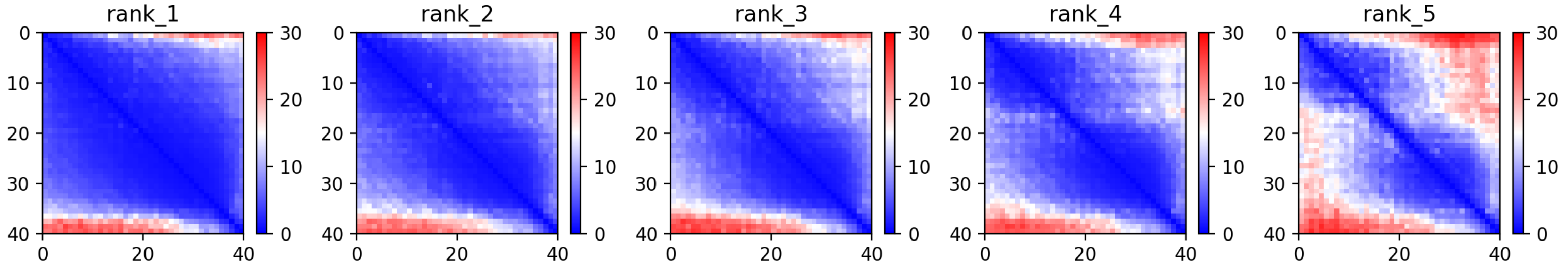

CP64: MtrunA17_Chr3g0144151

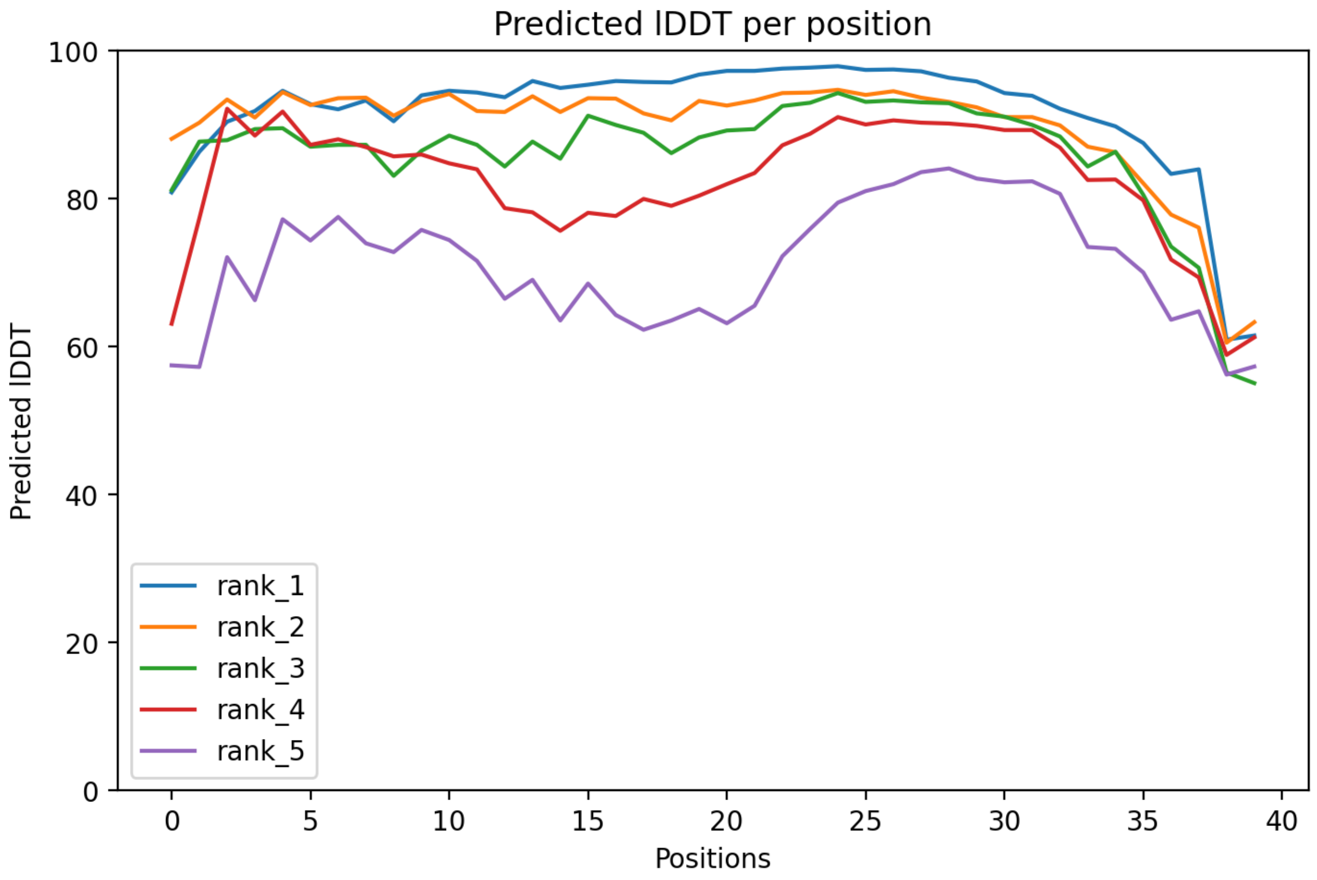

CP64: MtrunA17_Chr3g0144151

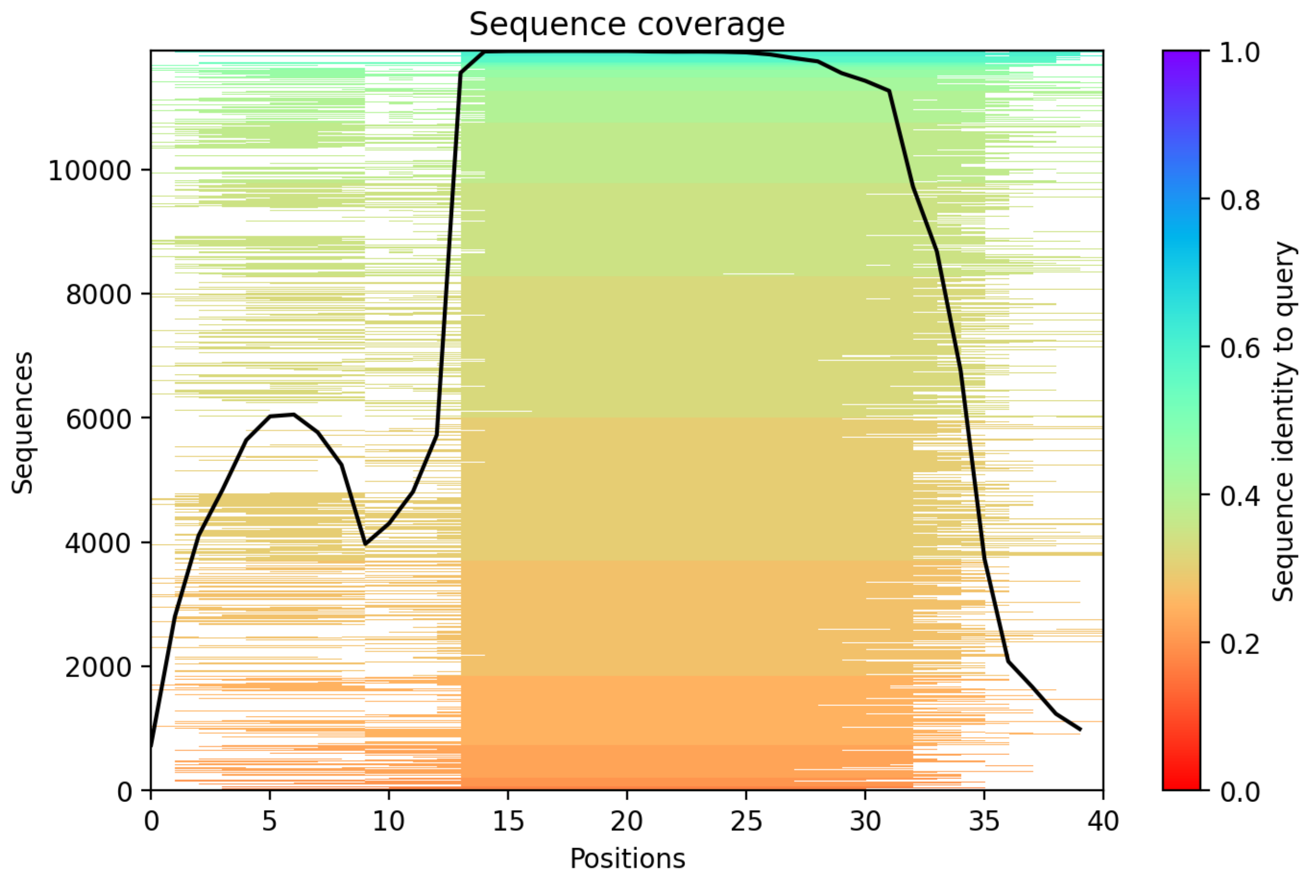

CP65: MtrunA17_Chr3g0144151

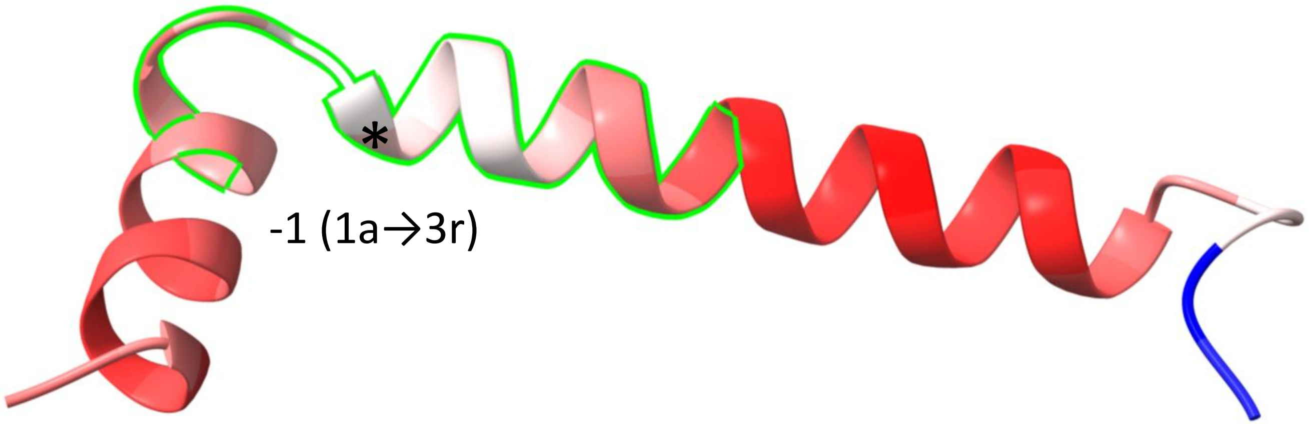


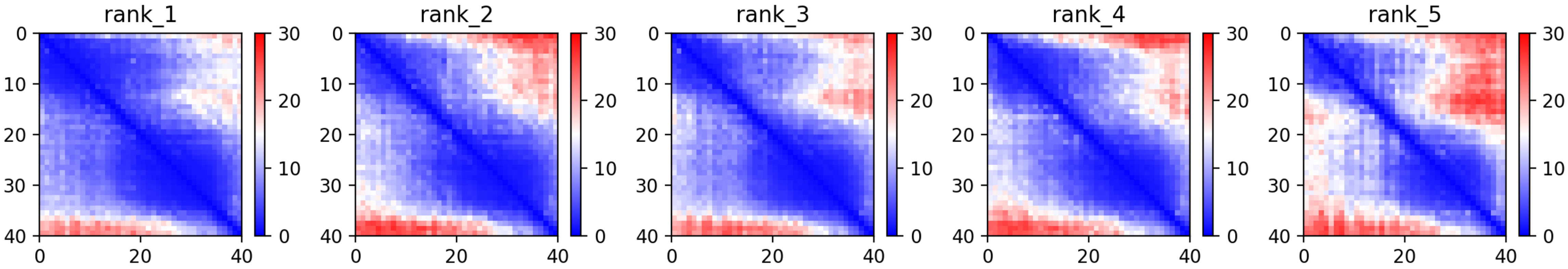

CP65: MtrunA17_Chr3g0144151

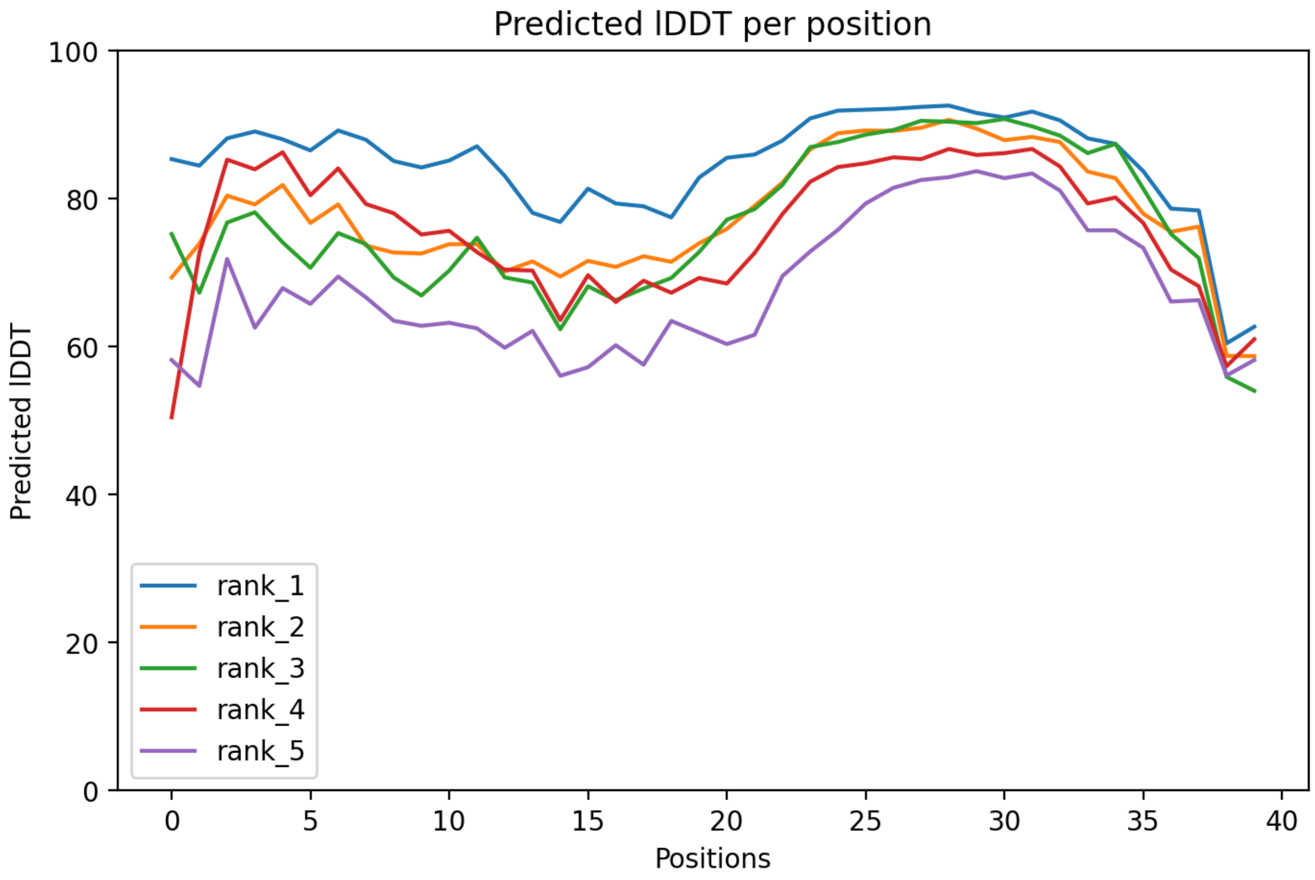

### CP65: MtrunA17_Chr3g0144151

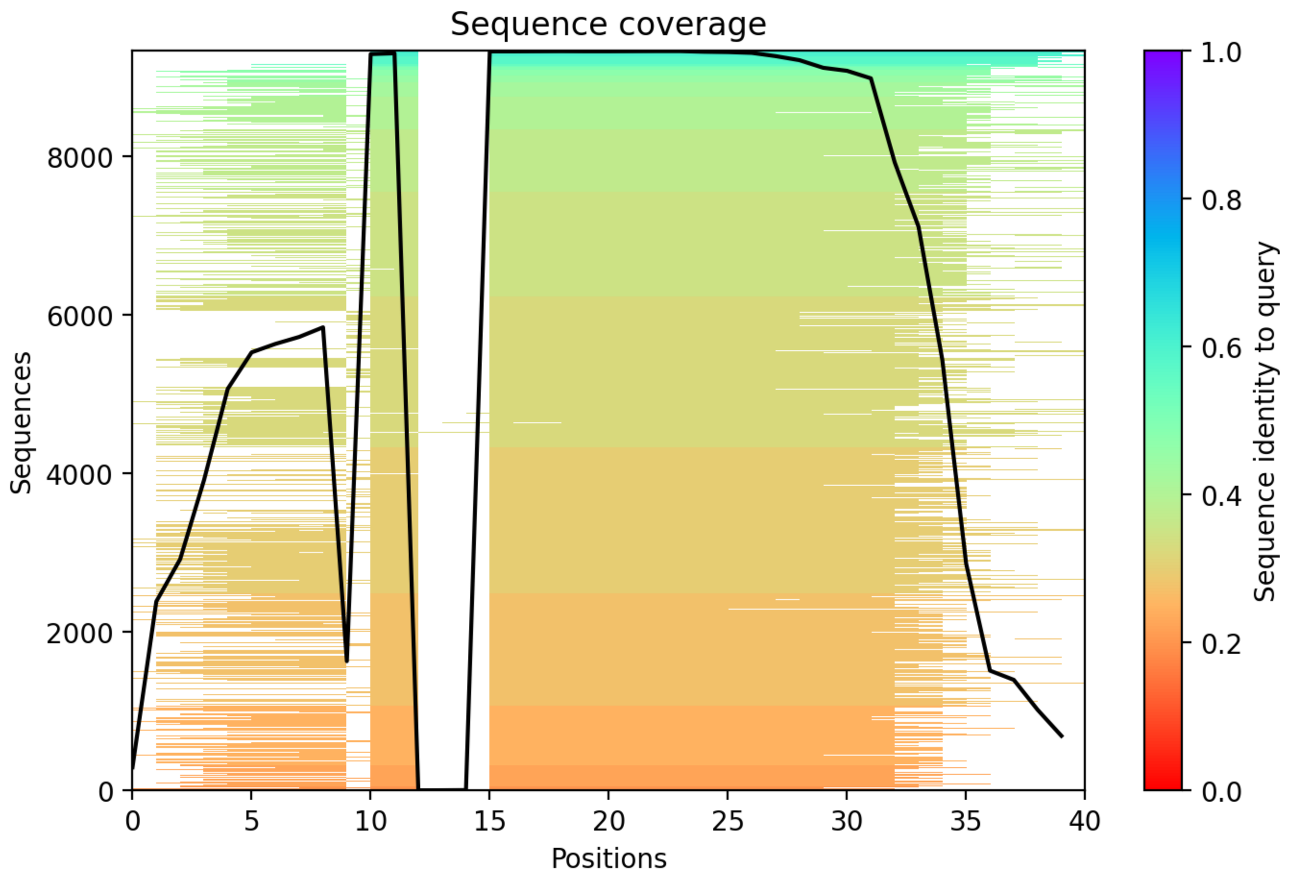

### CP66: MtrunA17_Chr3g1011650

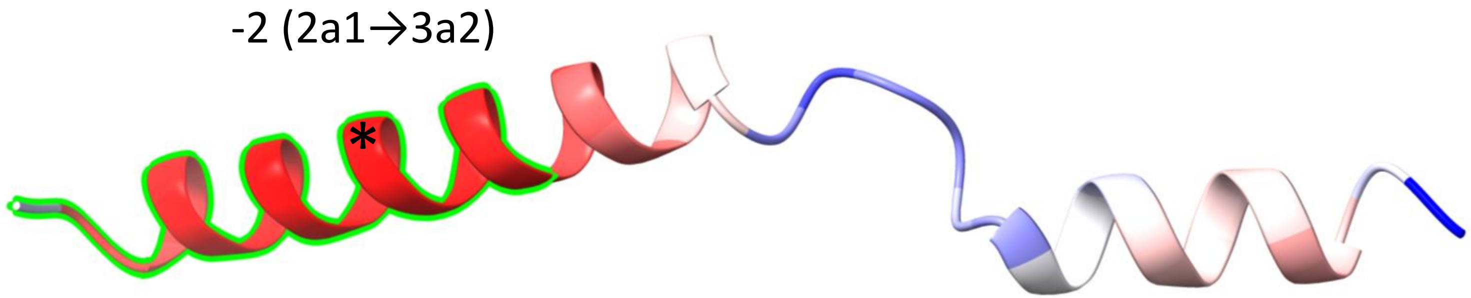


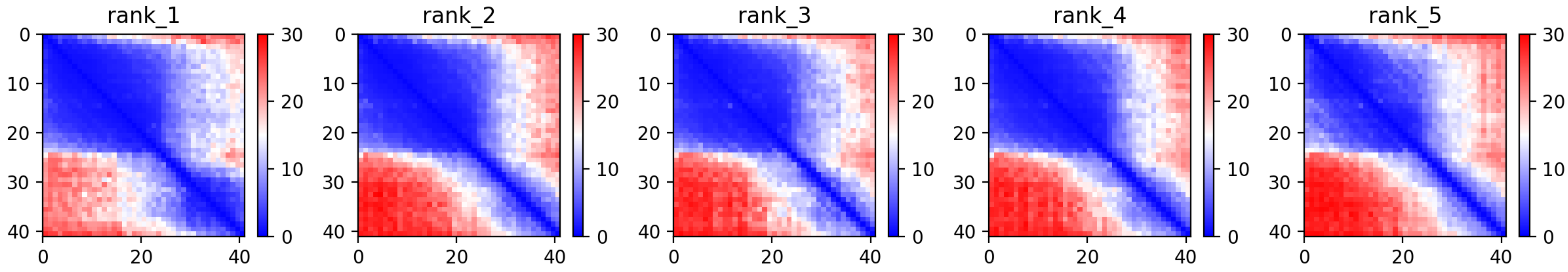

### CP66: MtrunA17_Chr3g1011650

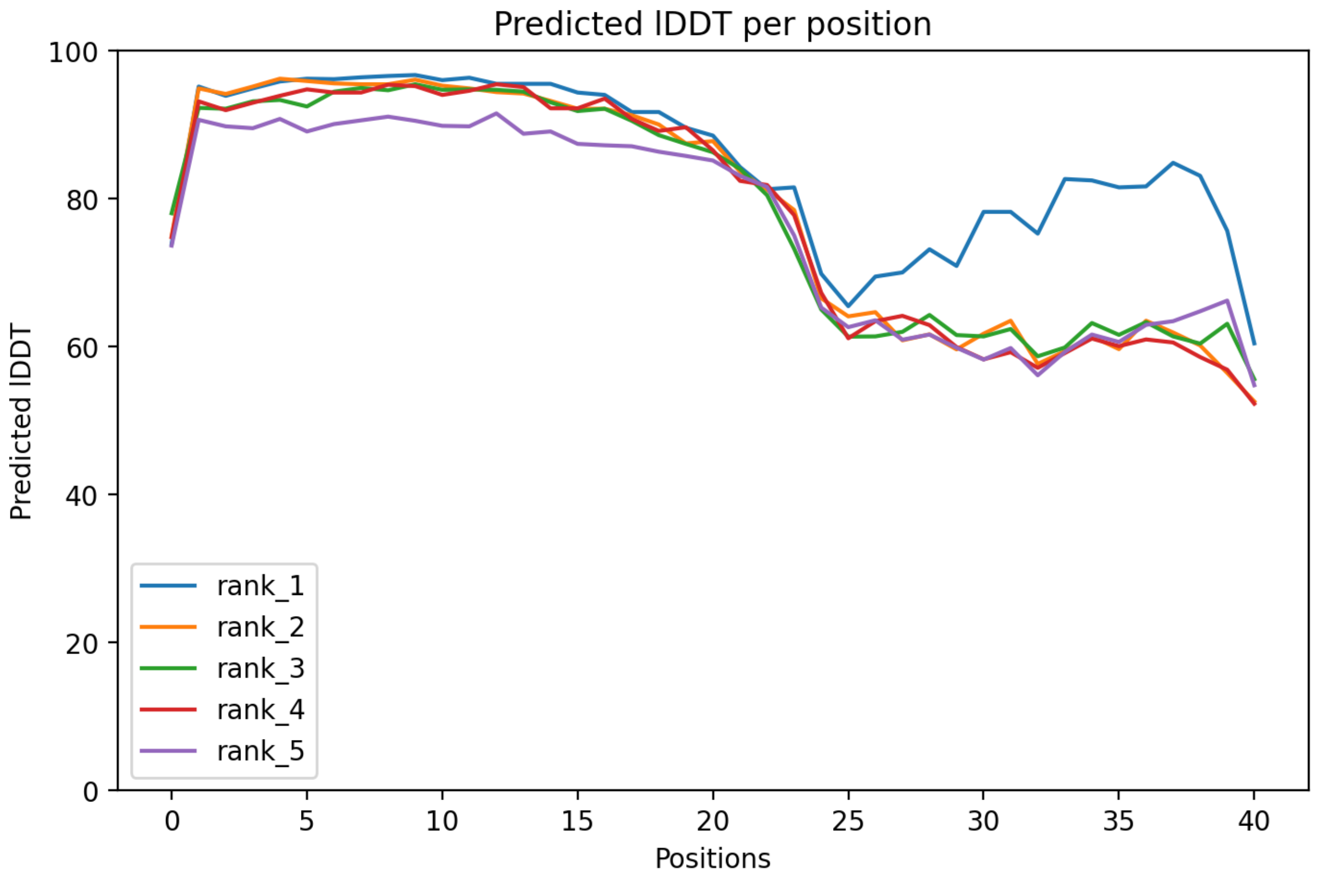

### CP66: MtrunA17_Chr3g1011650

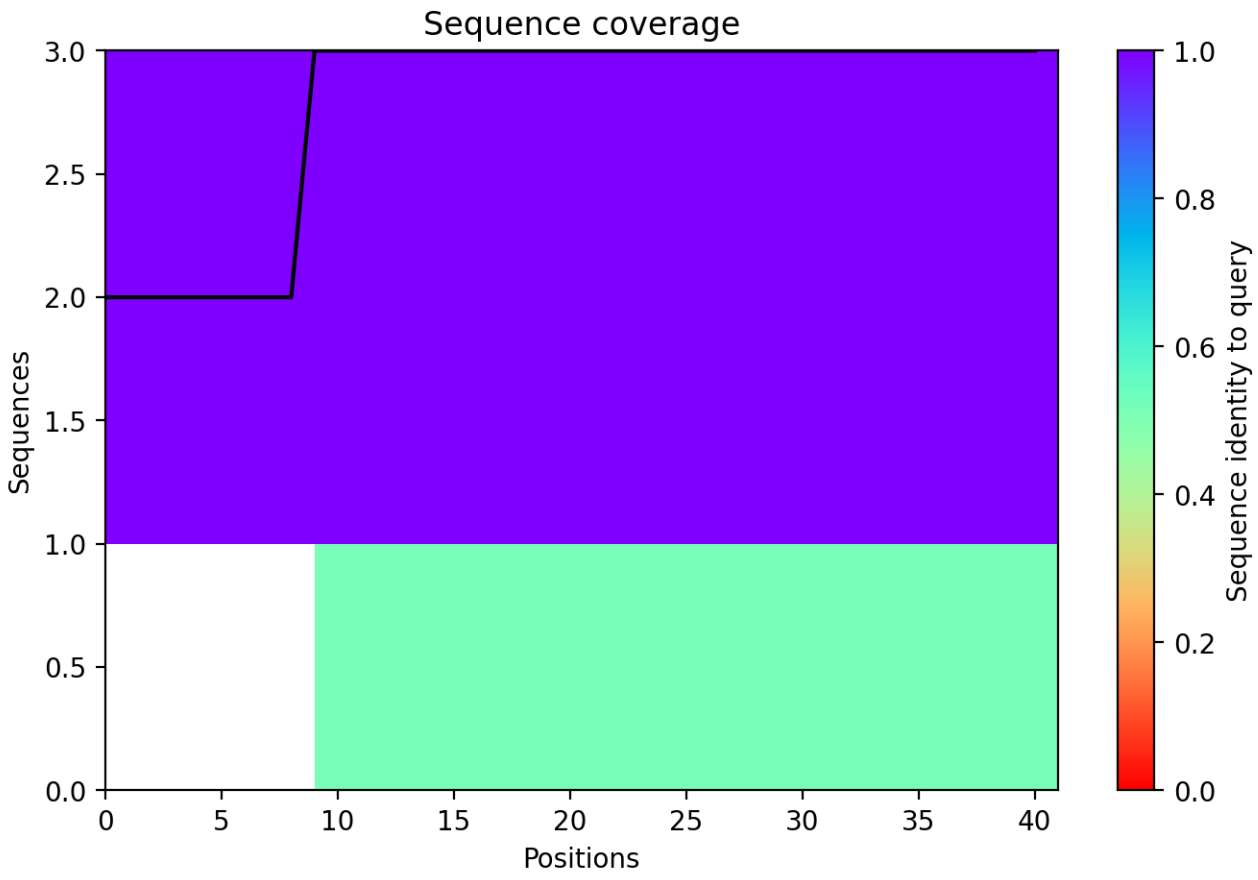

CP67: MtrunA17_Chr4g0000131

+1 (3r→1a)

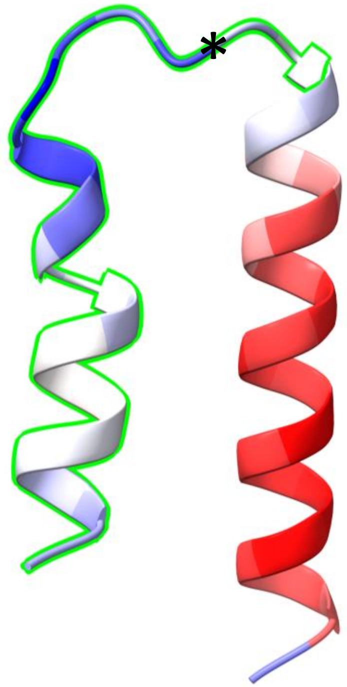


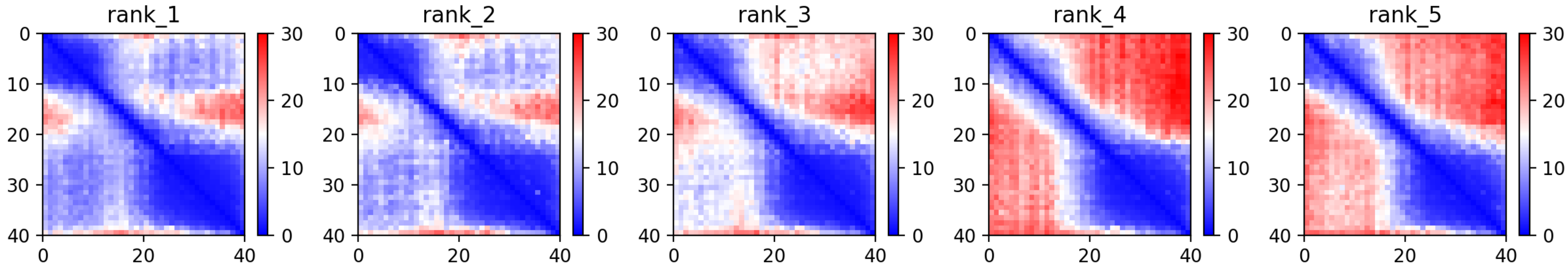

CP67: MtrunA17_Chr4g0000131

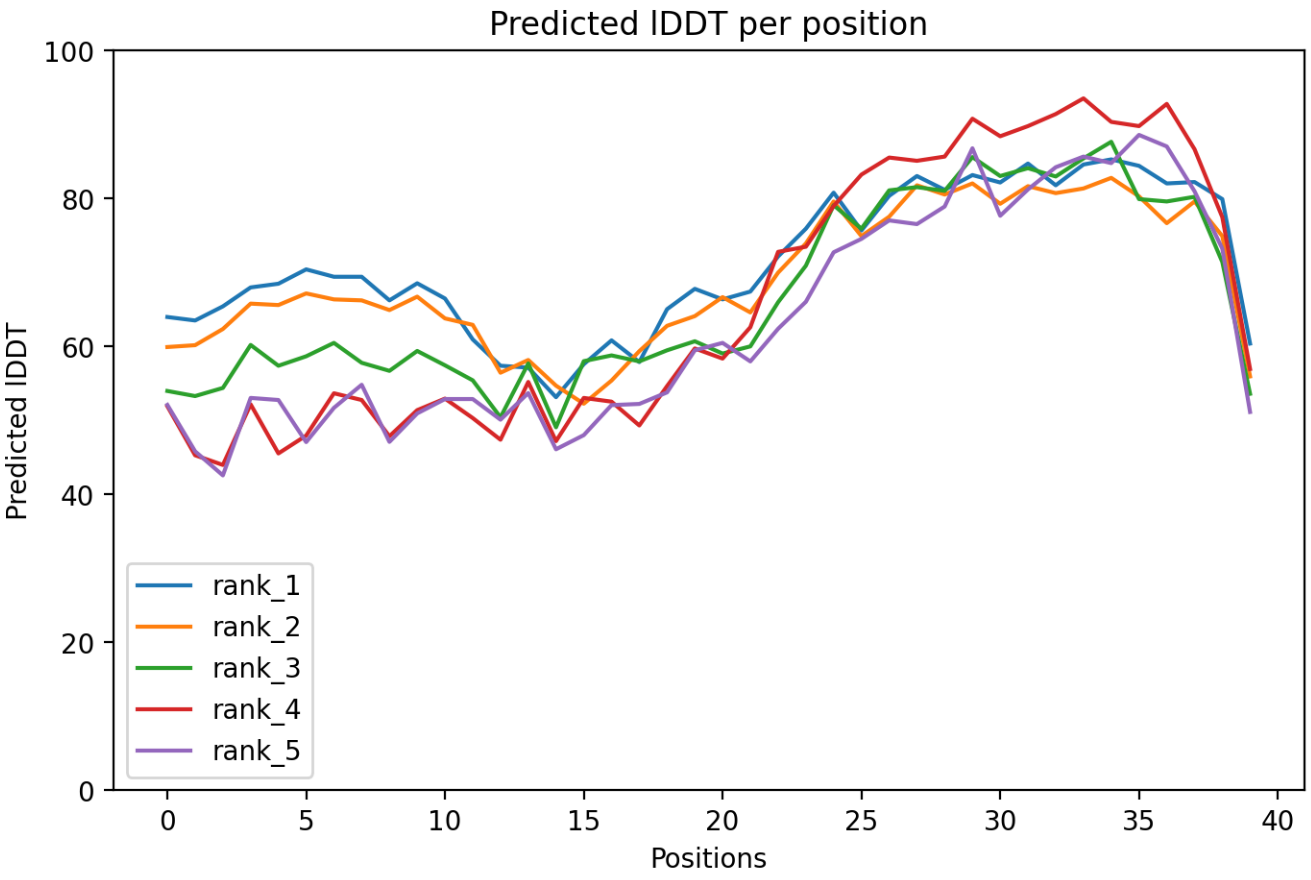

### CP67: MtrunA17_Chr4g0000131

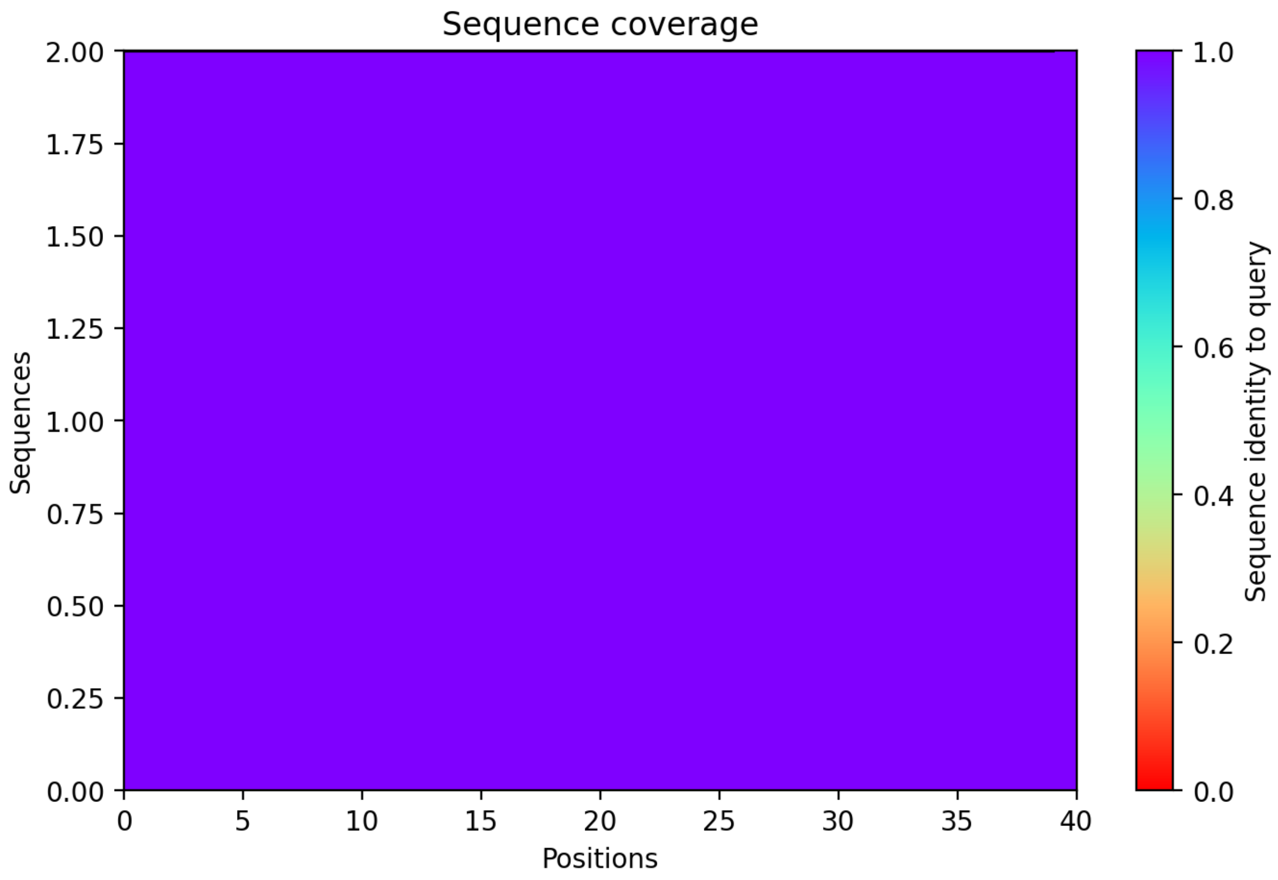

### CP68: MtrunA17_Chr4g0000891

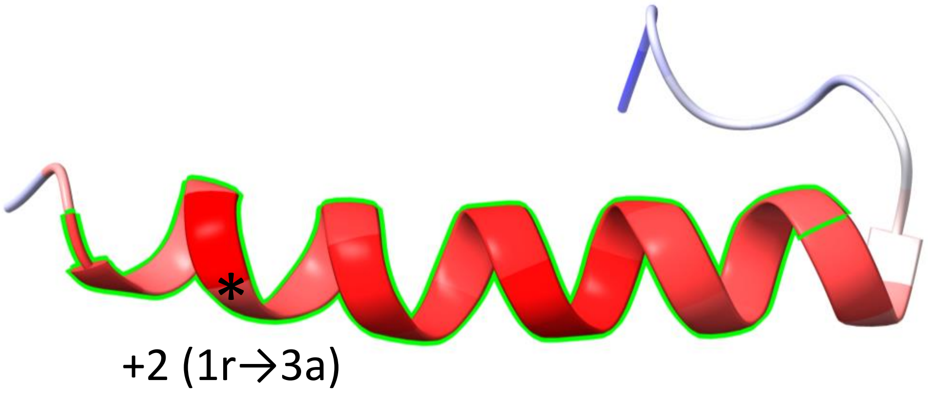


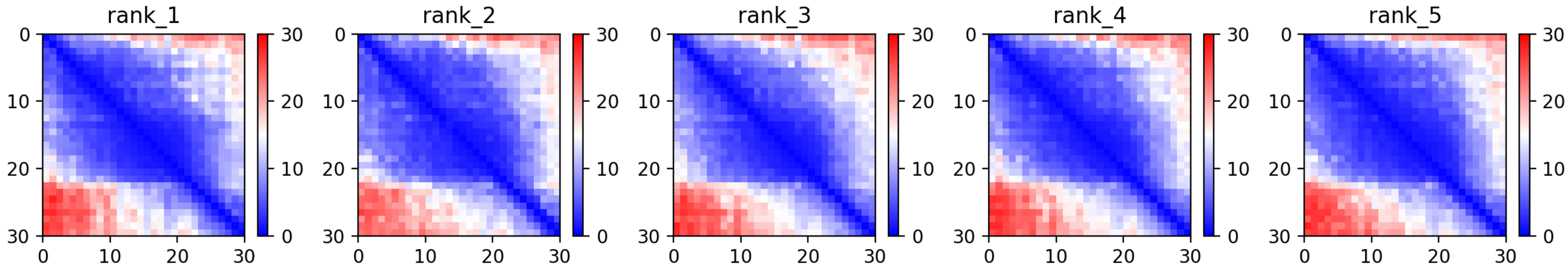

CP68: MtrunA17_Chr4g0000891

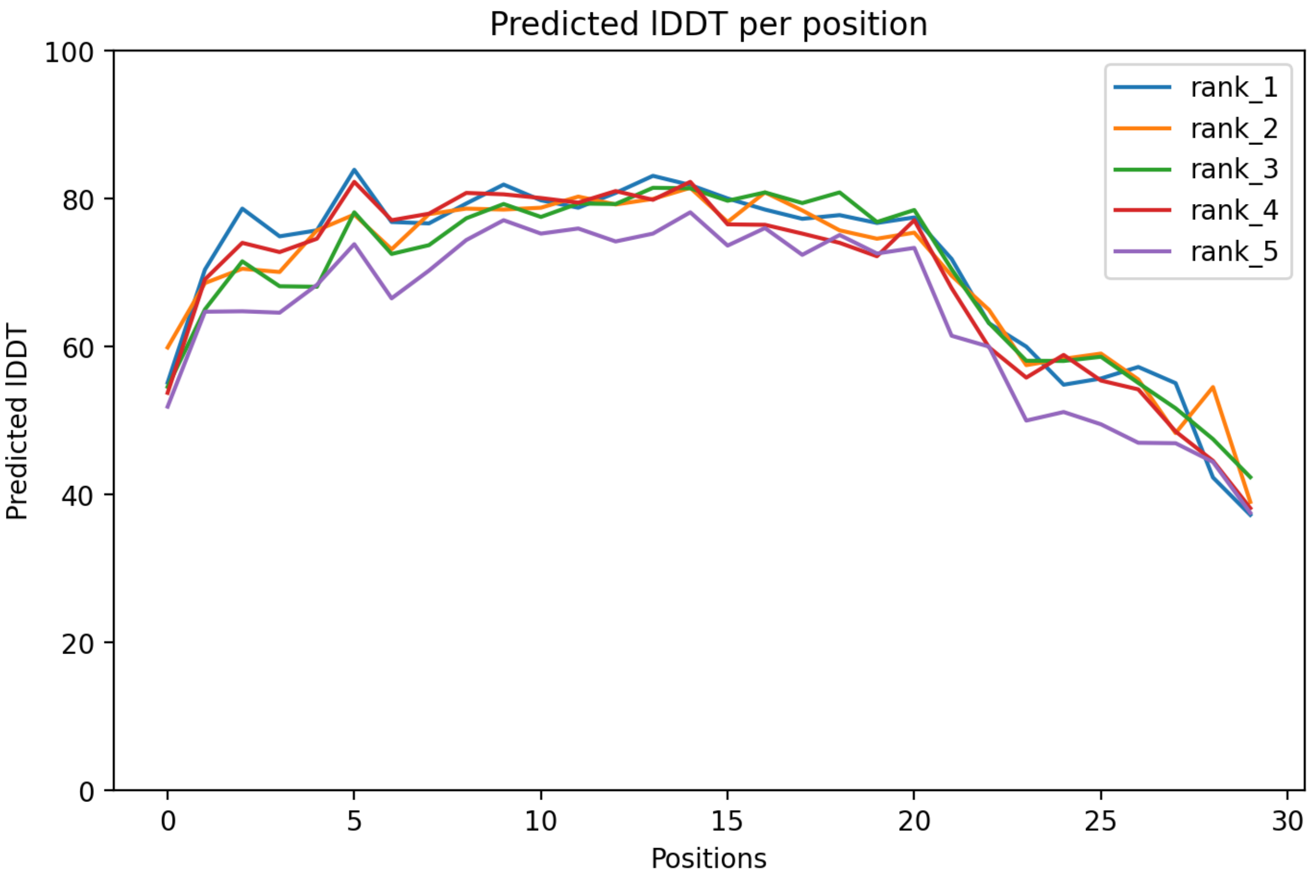

### CP68: MtrunA17_Chr4g0000891

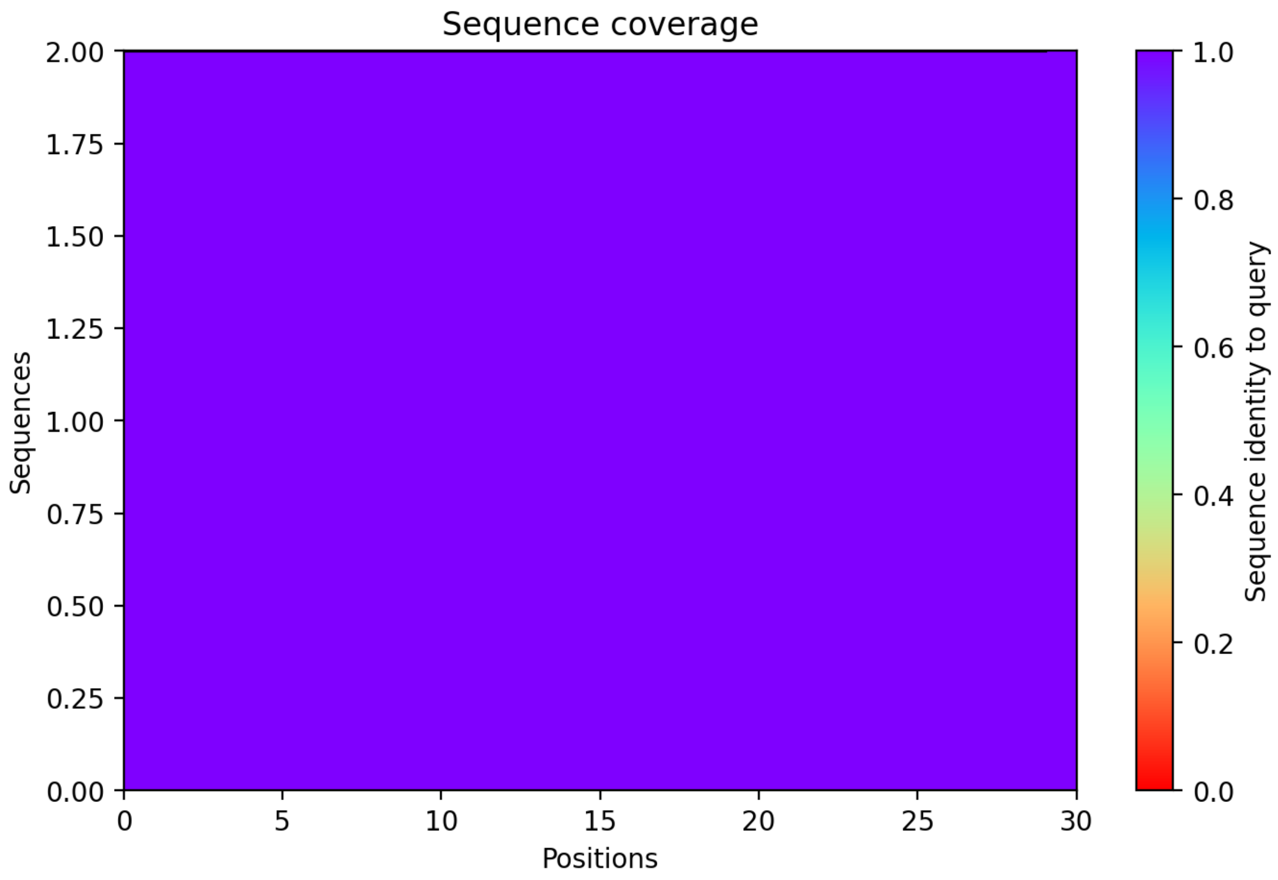

CP69: MtrunA17_Chr4g0004721

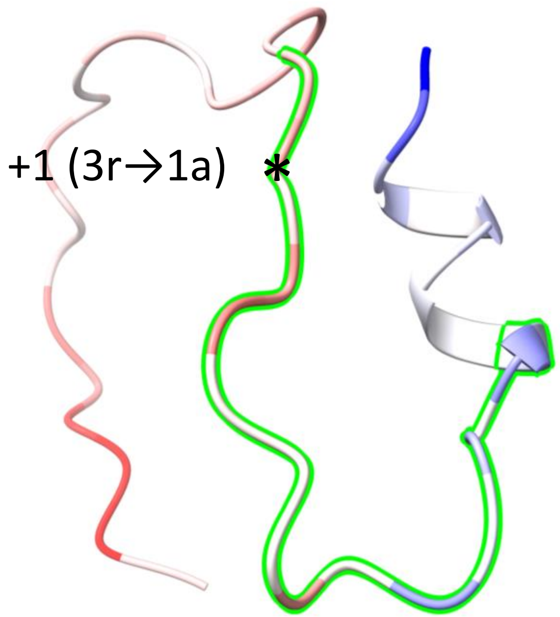


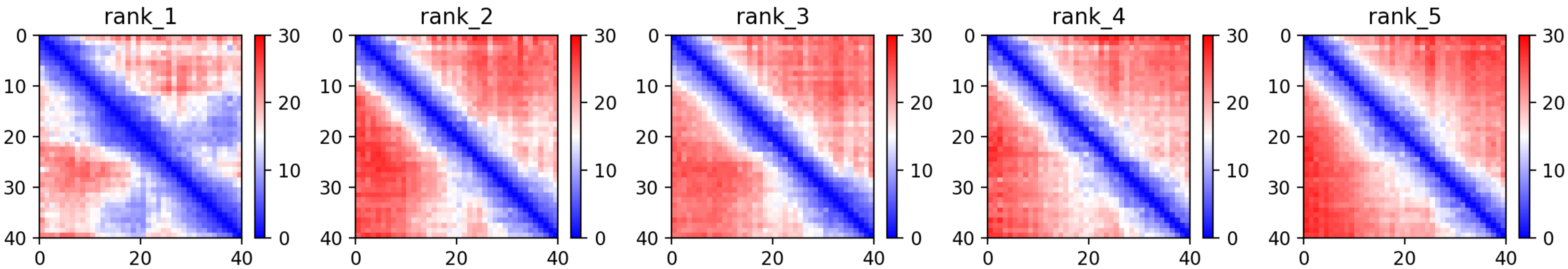

CP69: MtrunA17_Chr4g0004721

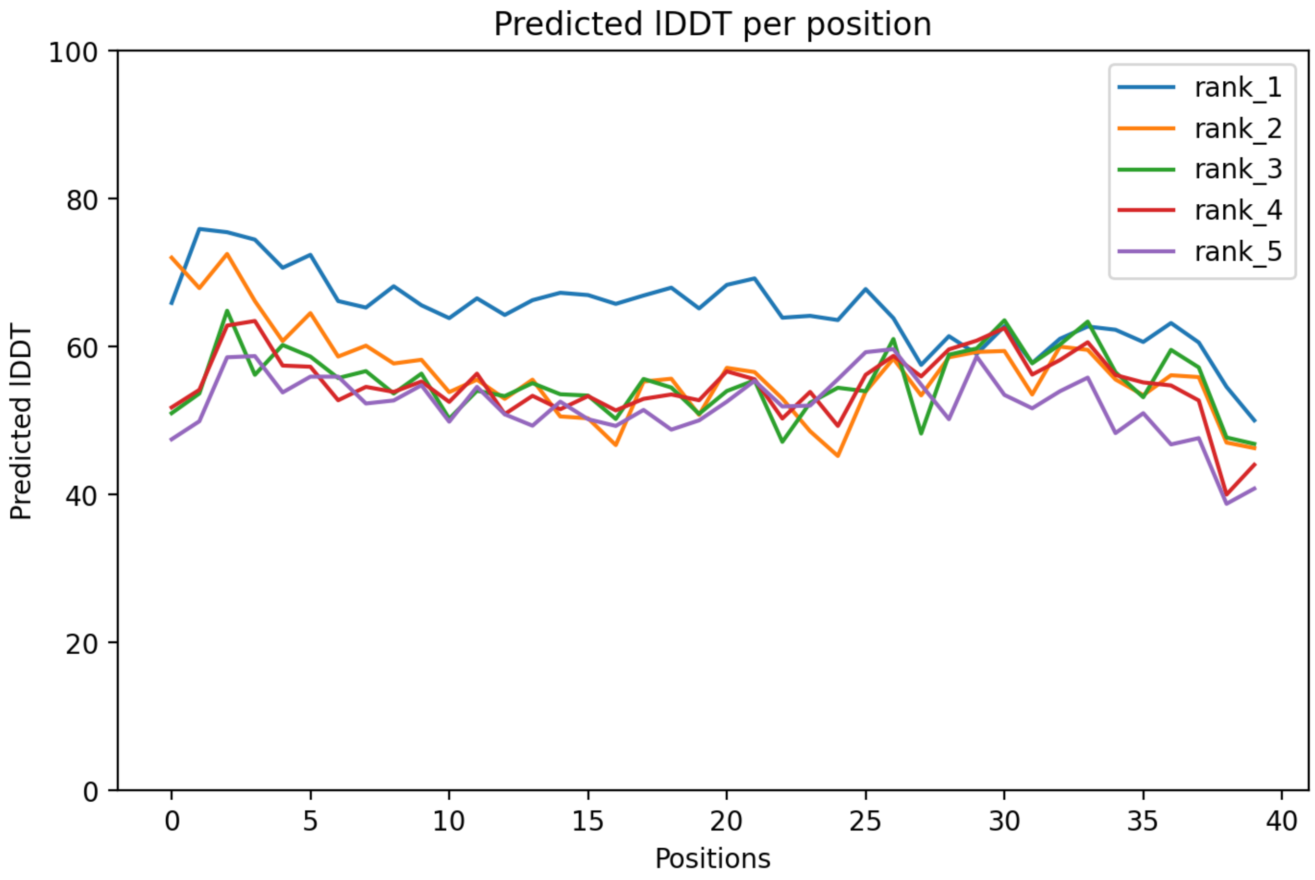

### CP69: MtrunA17_Chr4g0004721

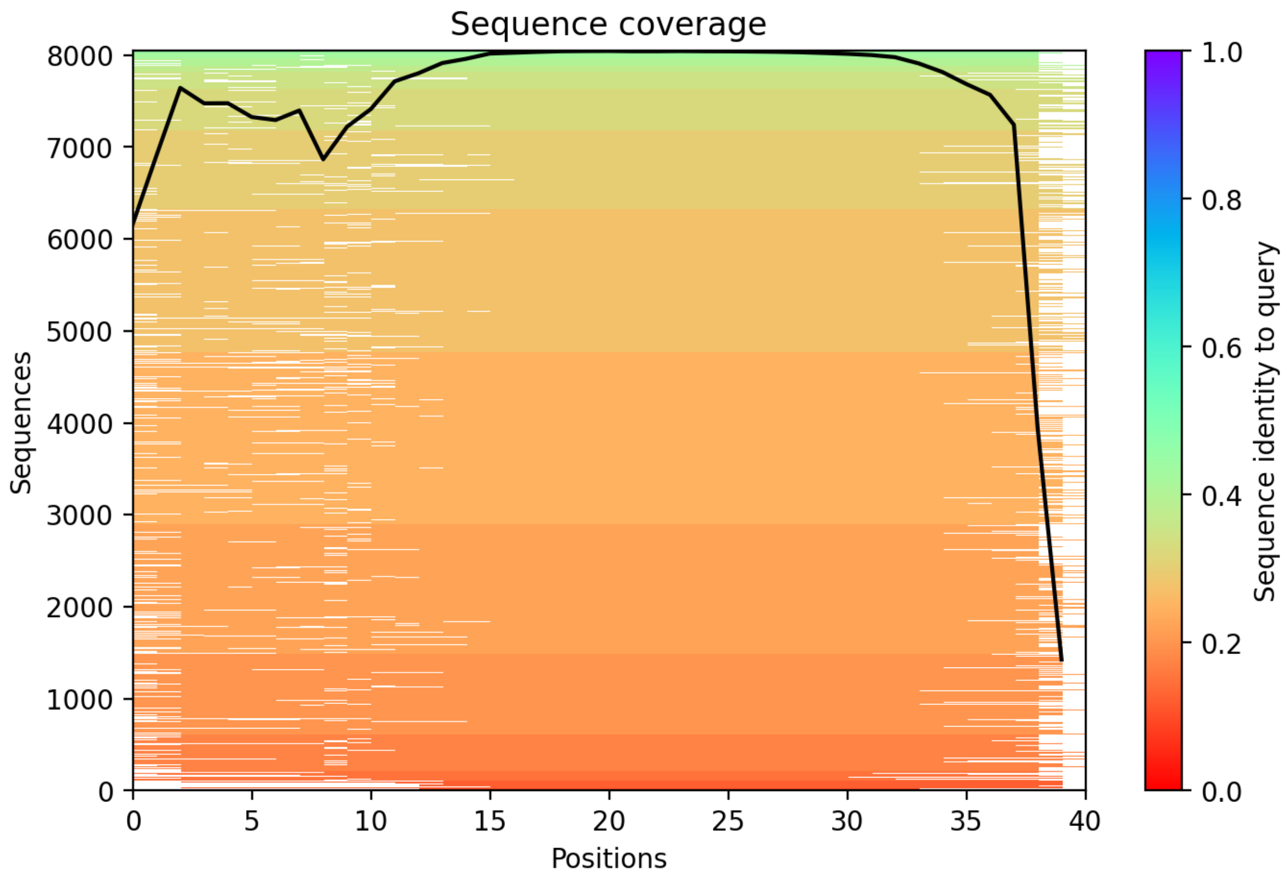

CP70: MtrunA17_Chr4g0014251

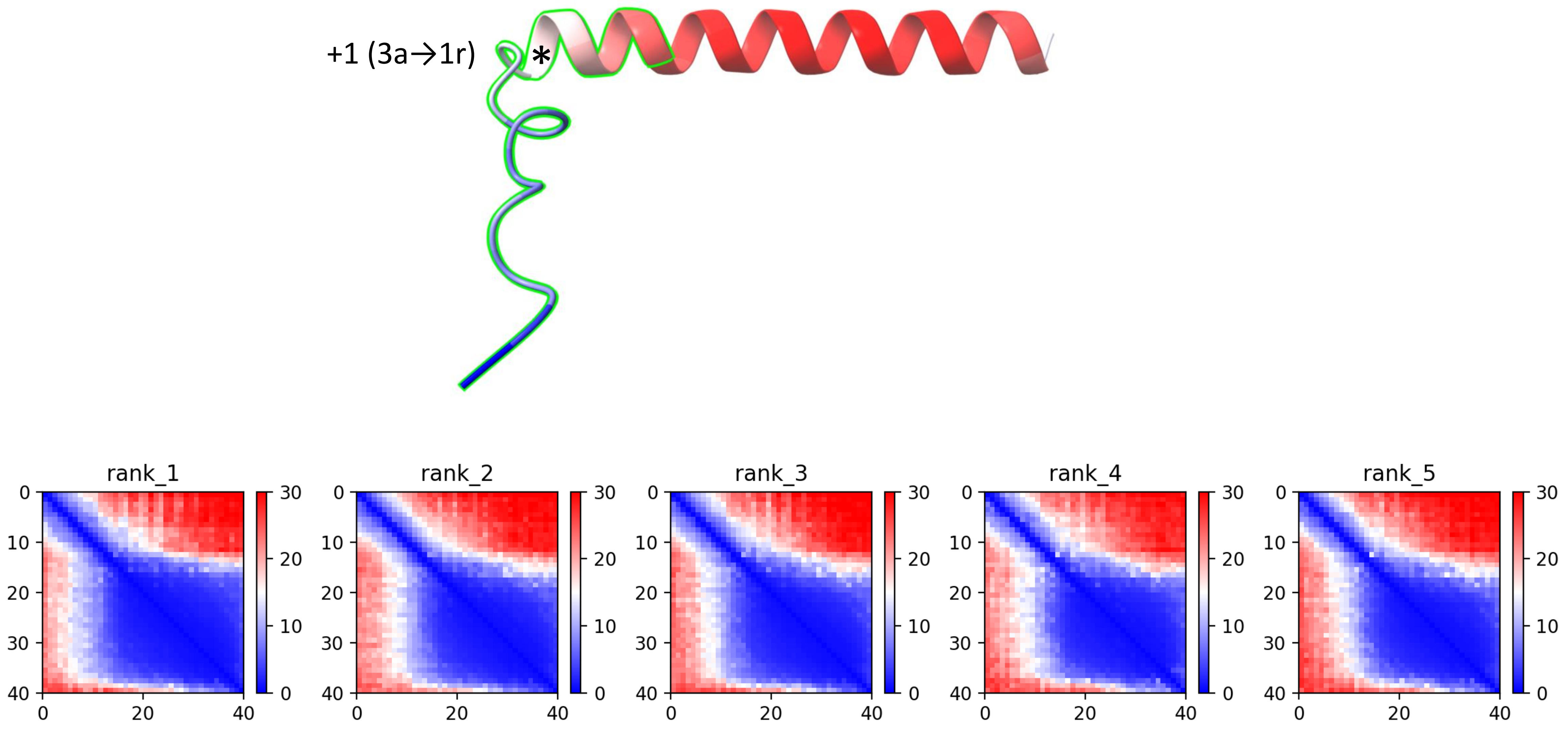

### CP70: MtrunA17_Chr4g0014251

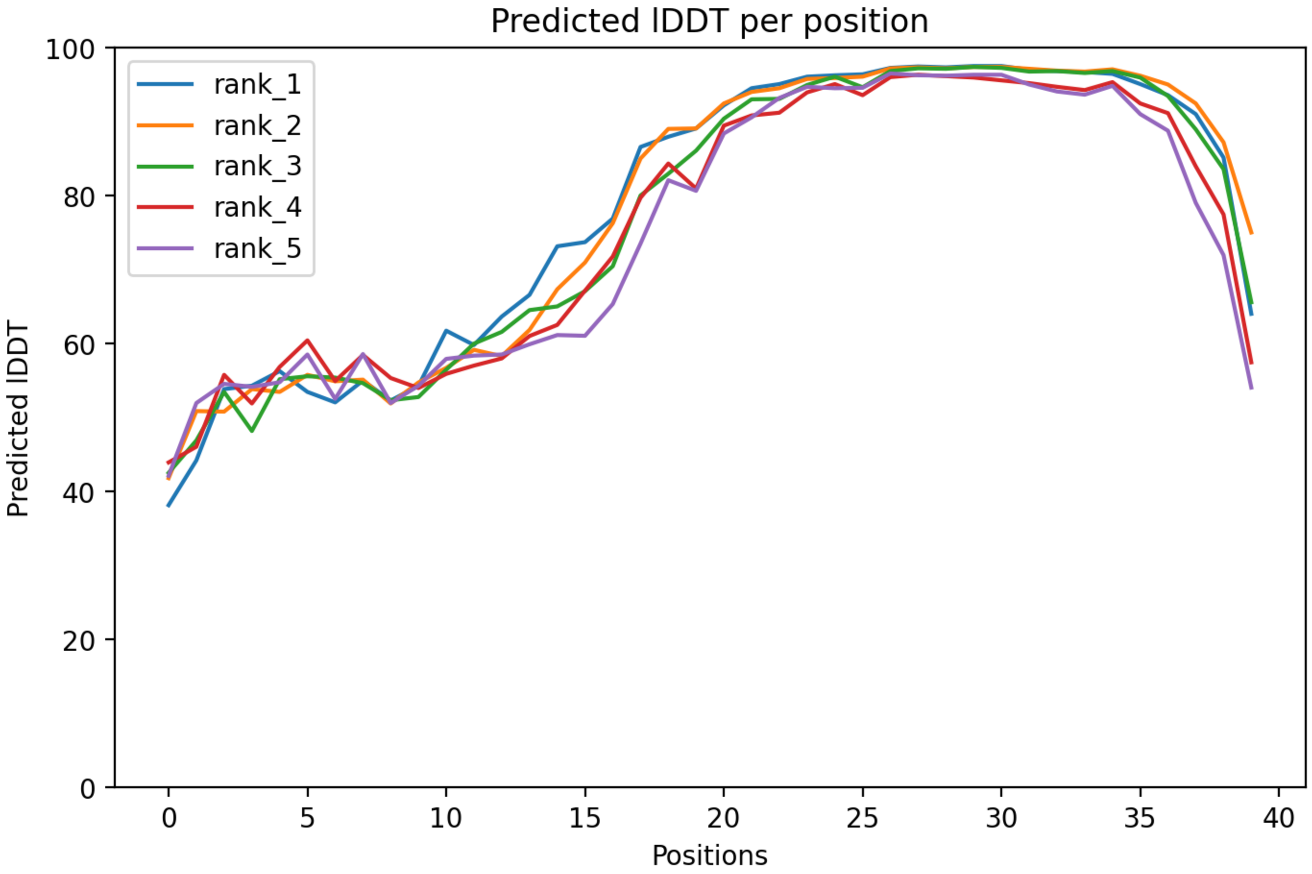

### CP70: MtrunA17_Chr4g0014251

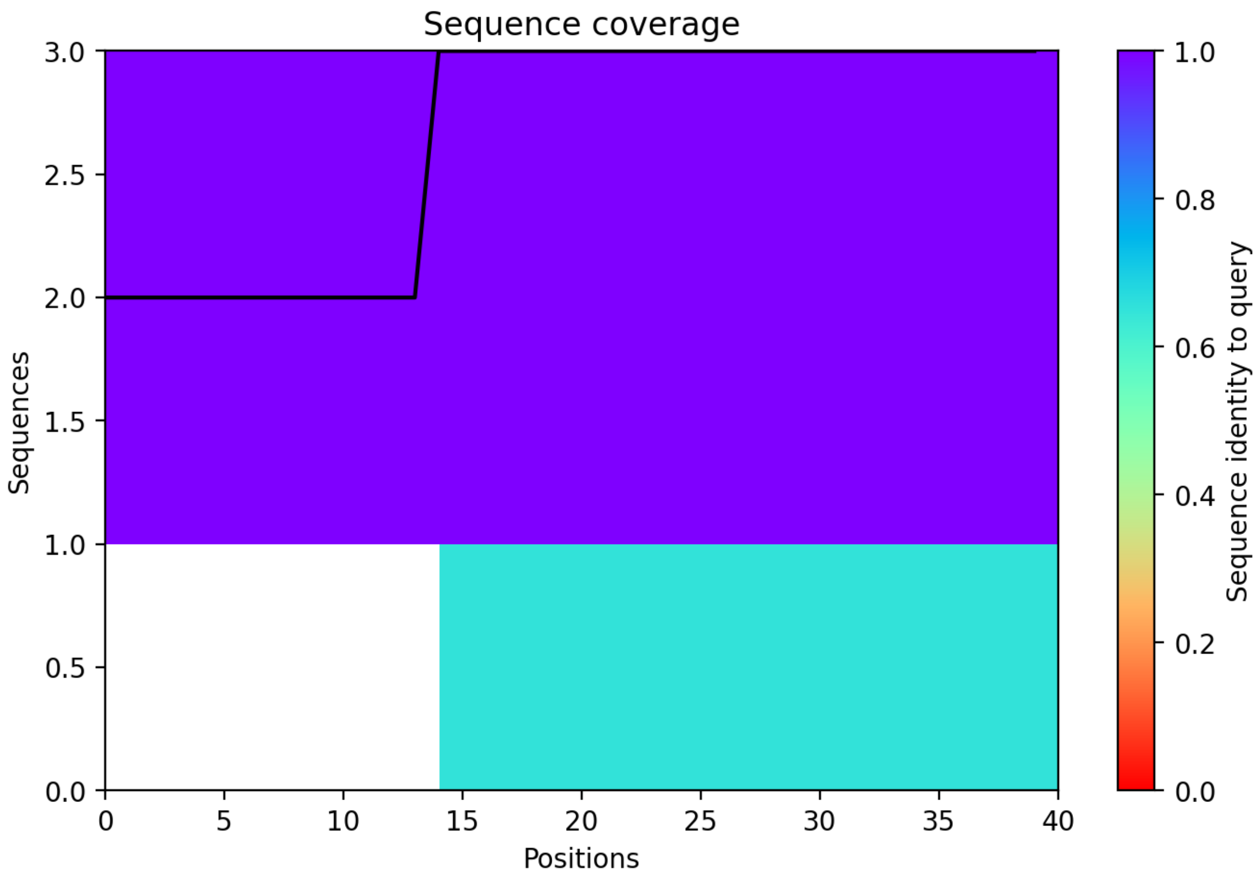

### CP71: MtrunA17_Chr4g0022491

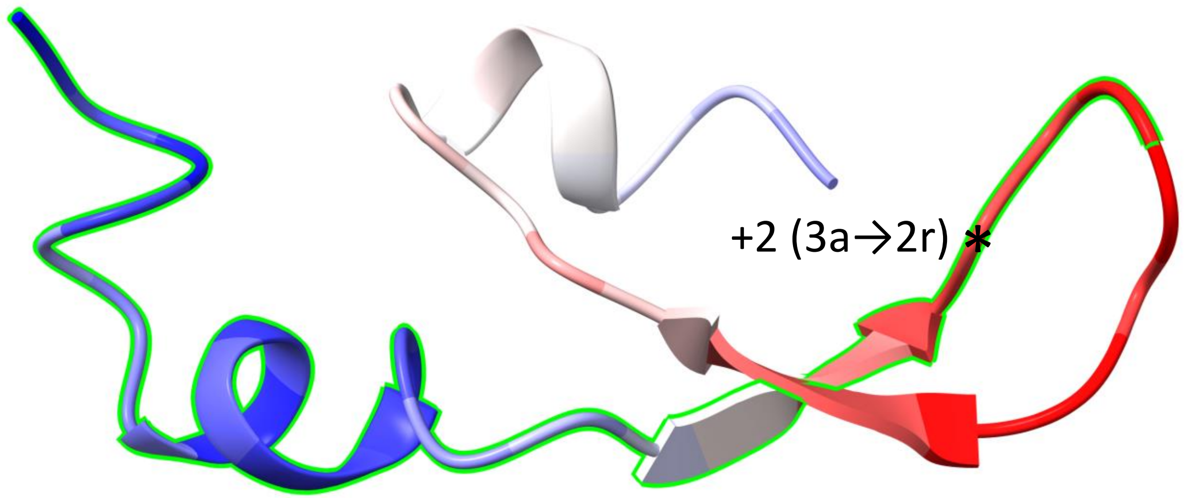


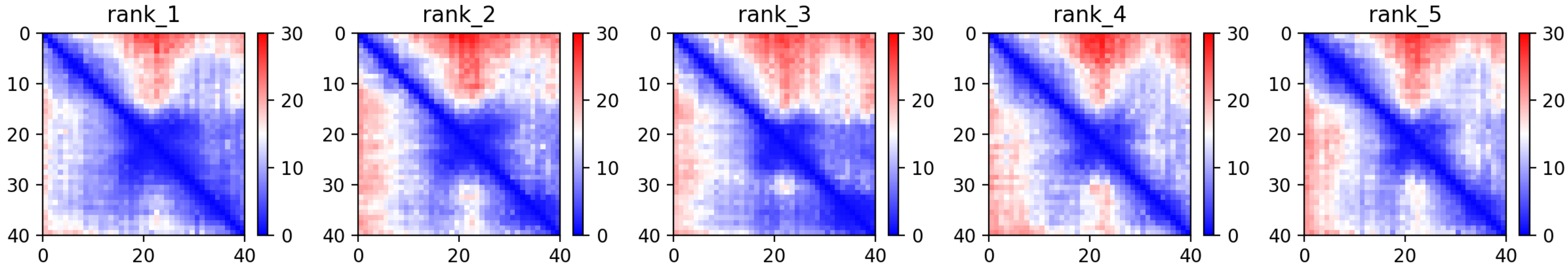

CP71: MtrunA17_Chr4g0022491

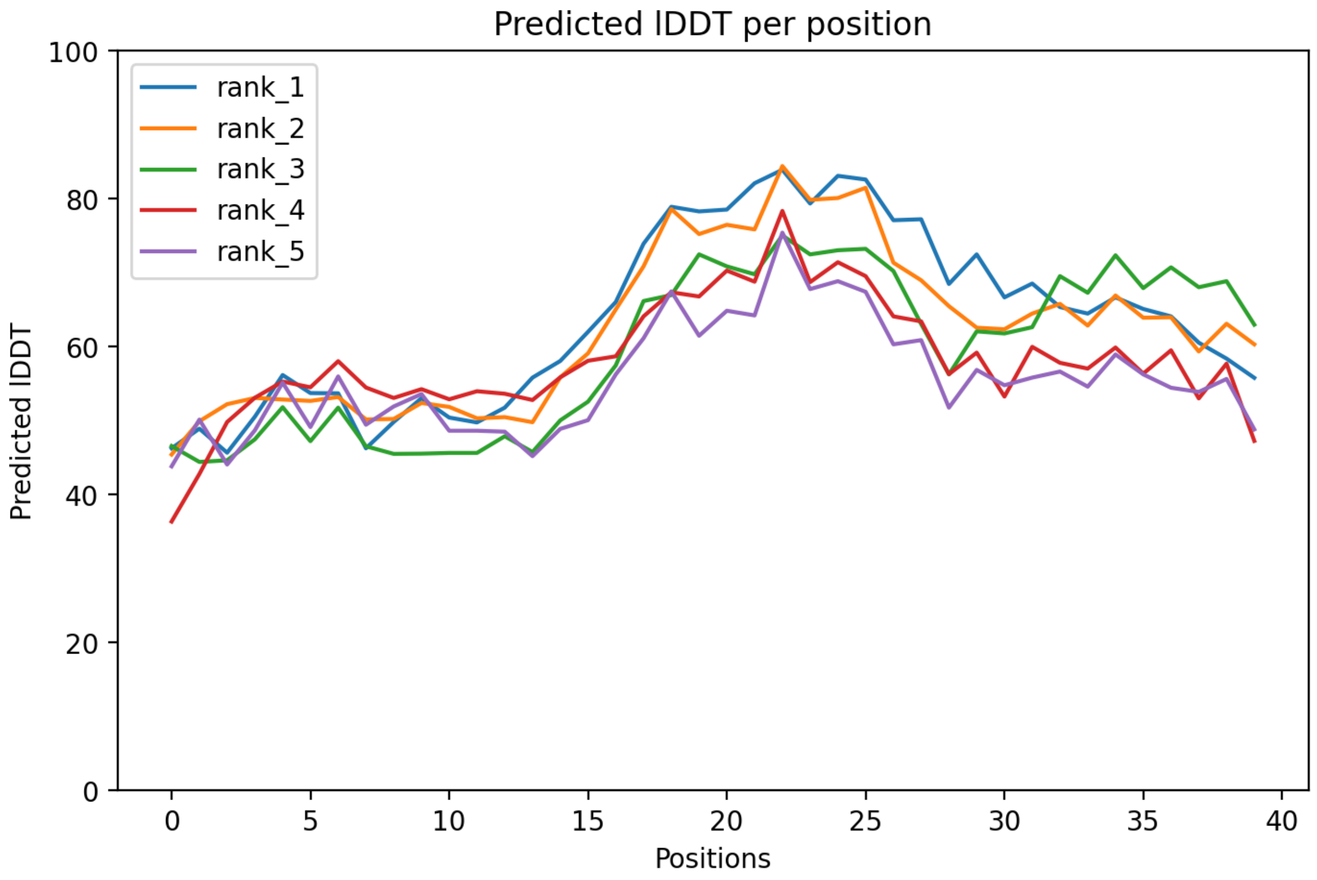

### CP71: MtrunA17_Chr4g0022491

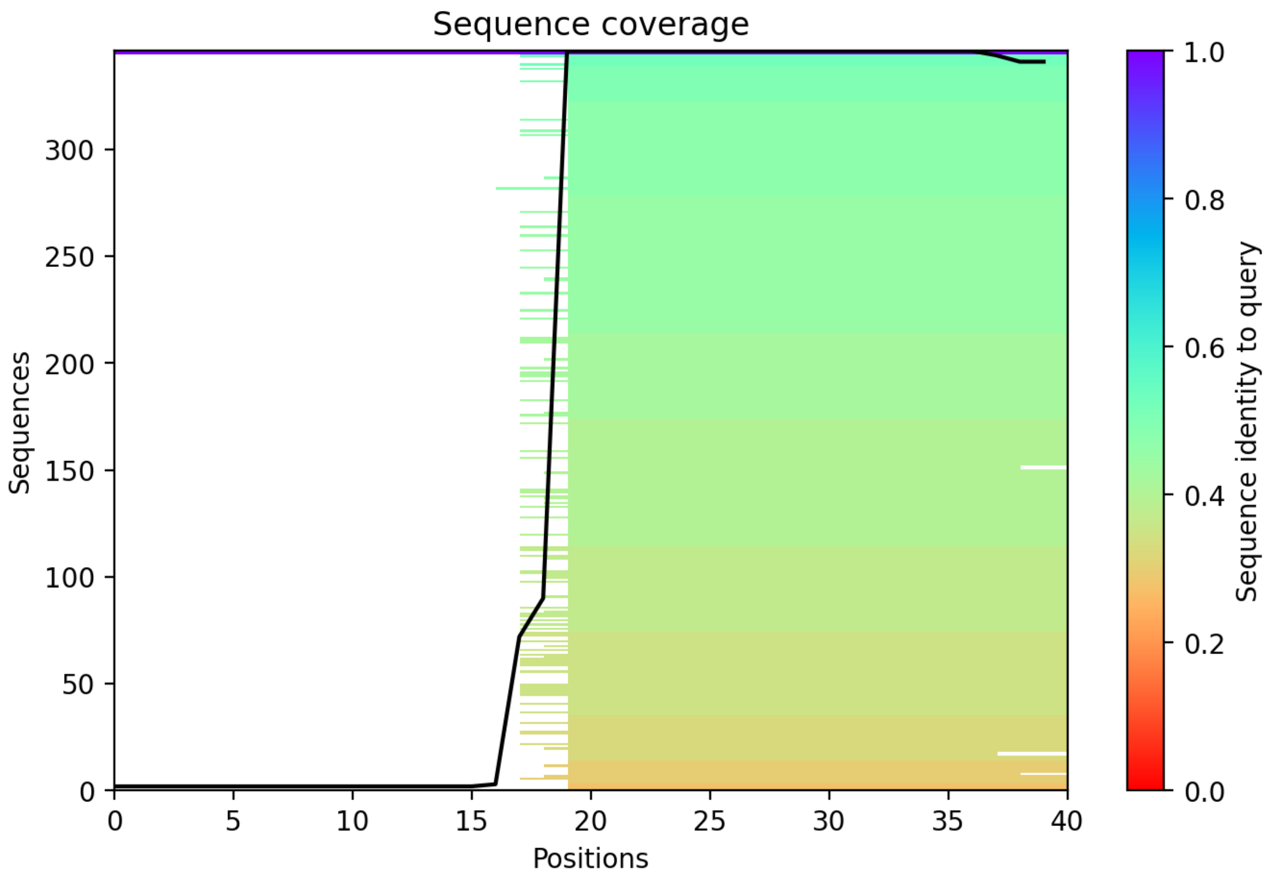

### CP72: MtrunA17_Chr4g0023791

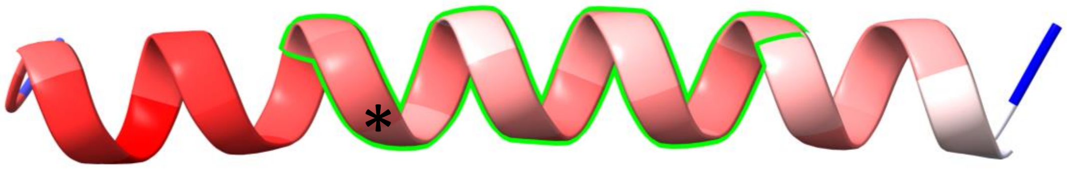


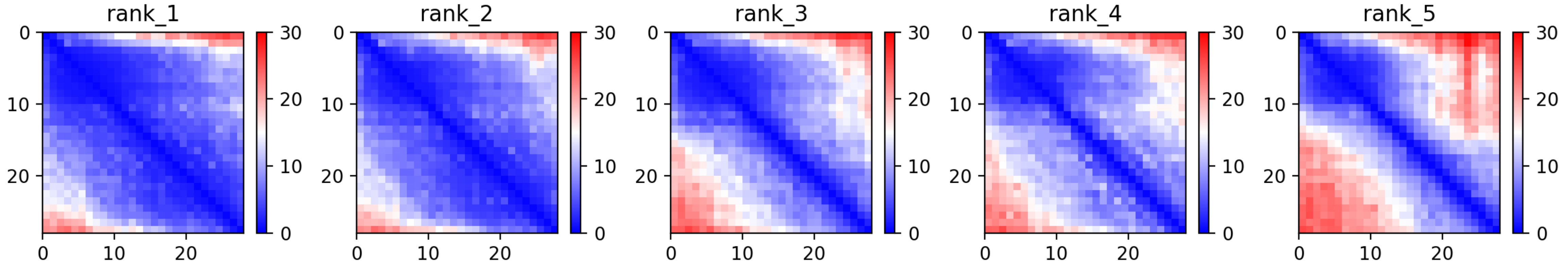

### CP72: MtrunA17_Chr4g0023791

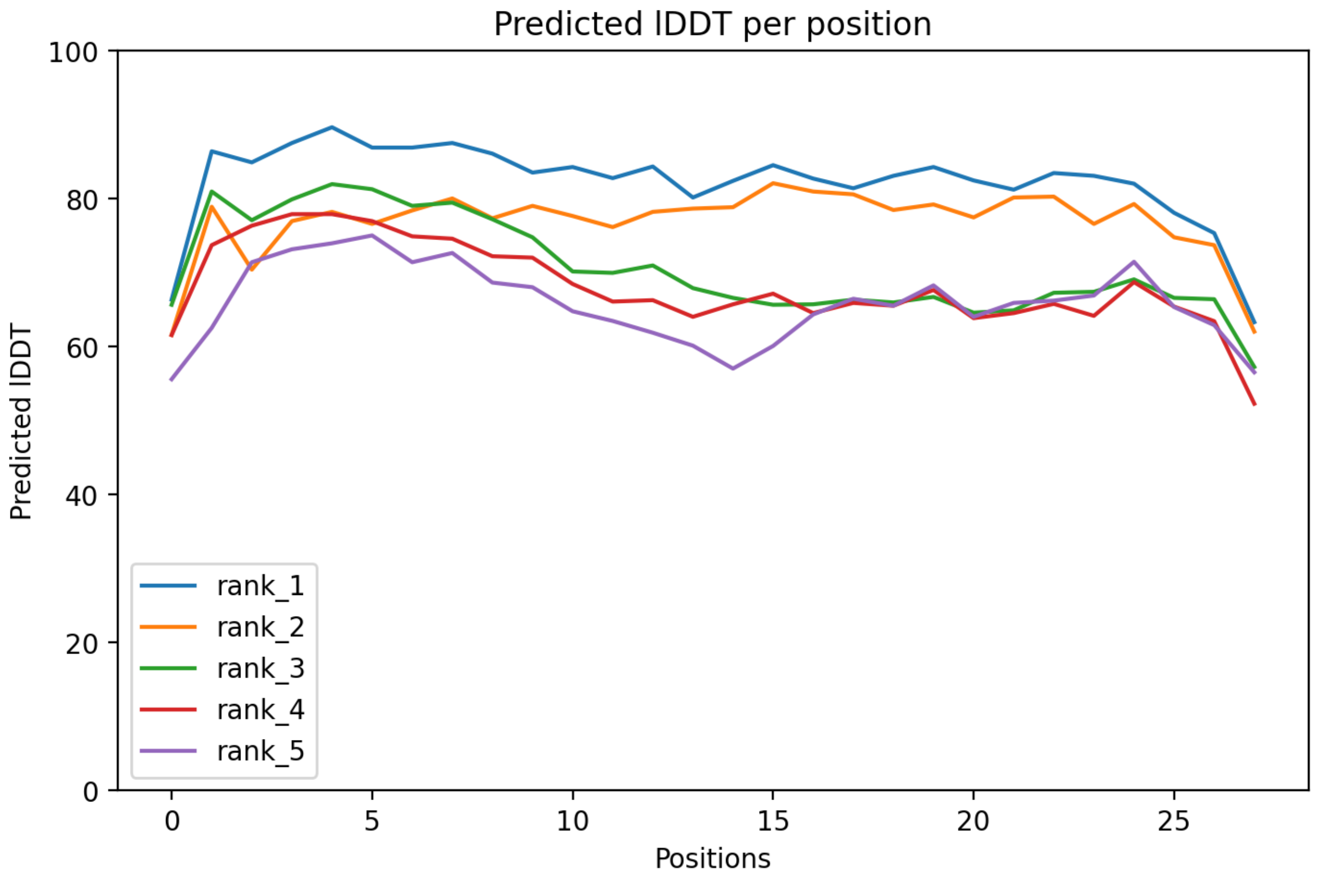

### CP72: MtrunA17_Chr4g0023791

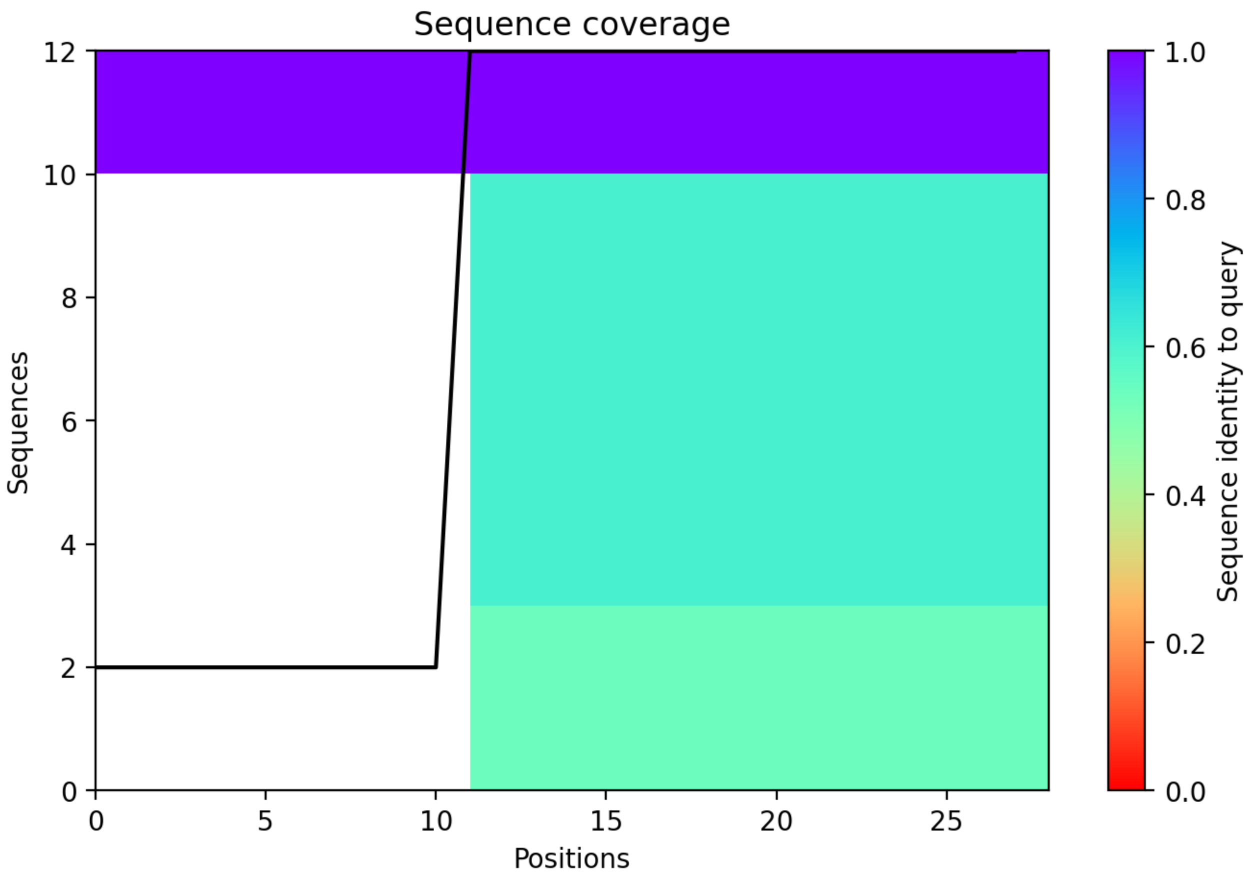

CP73: MtrunA17_Chr4g0034271

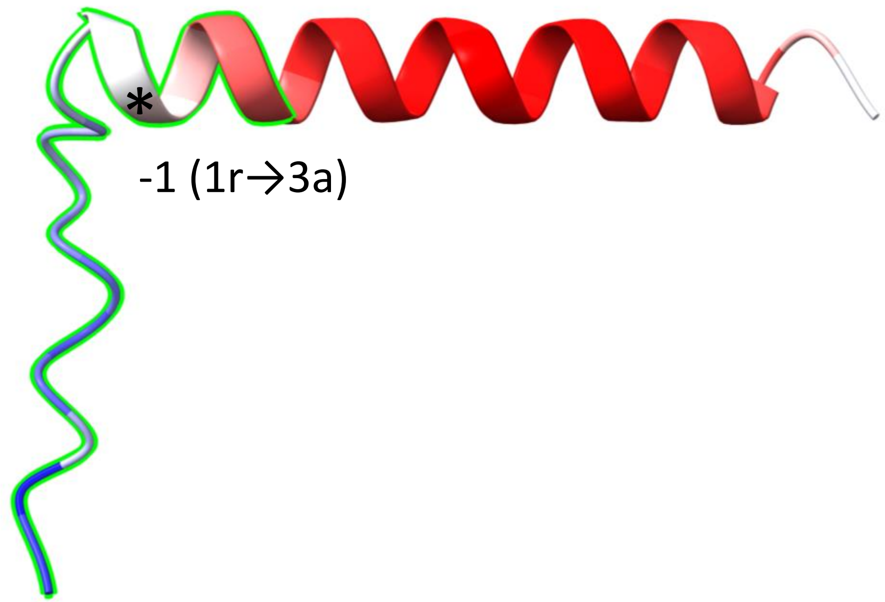


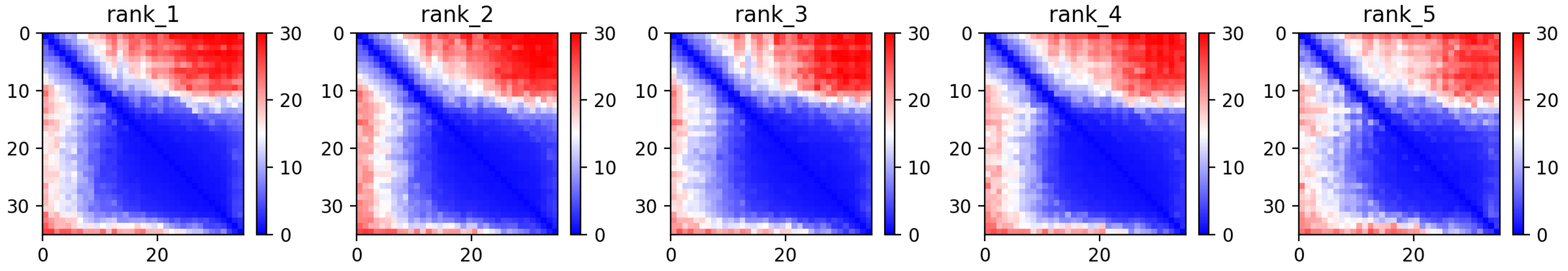

CP73: MtrunA17_Chr4g0034271

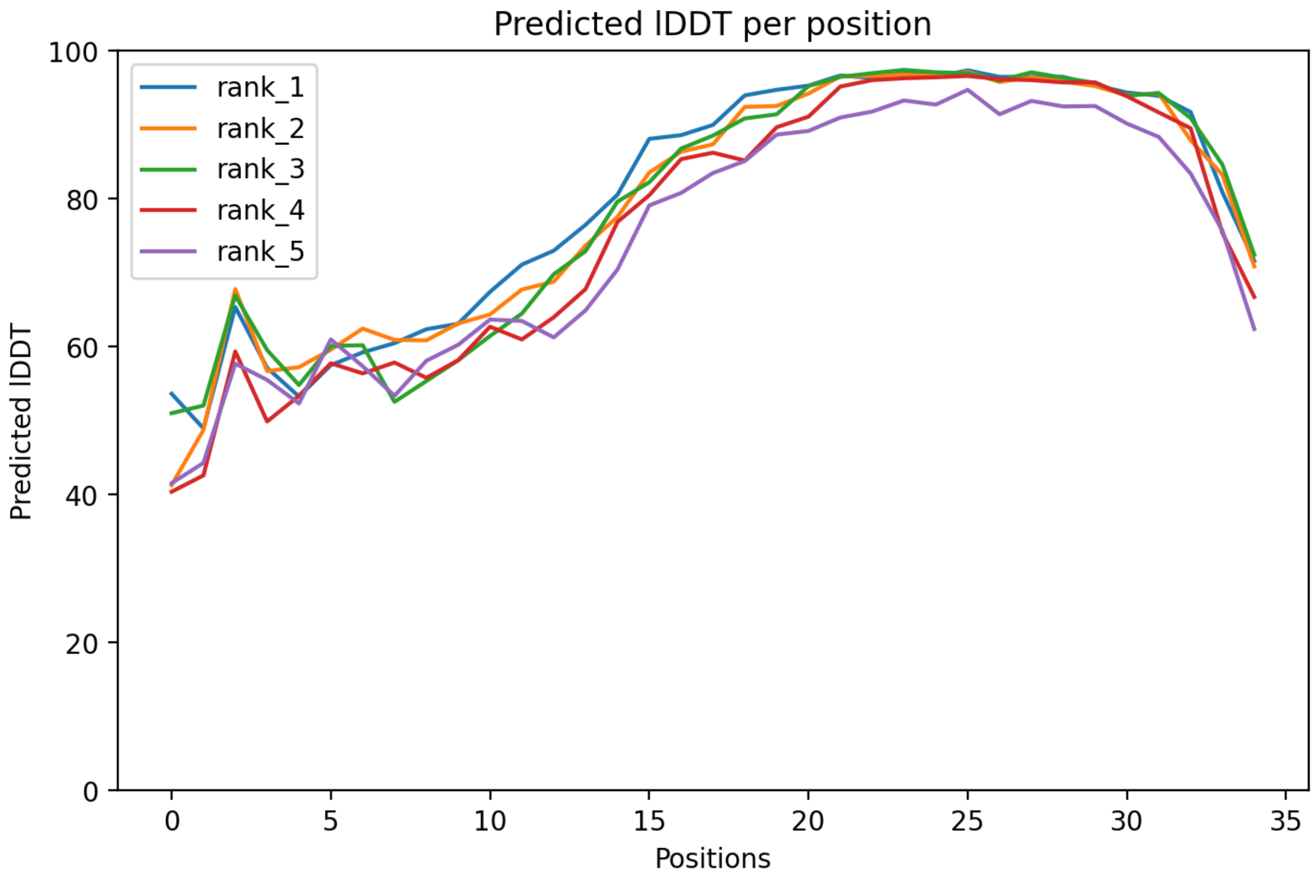

### CP73: MtrunA17_Chr4g0034271

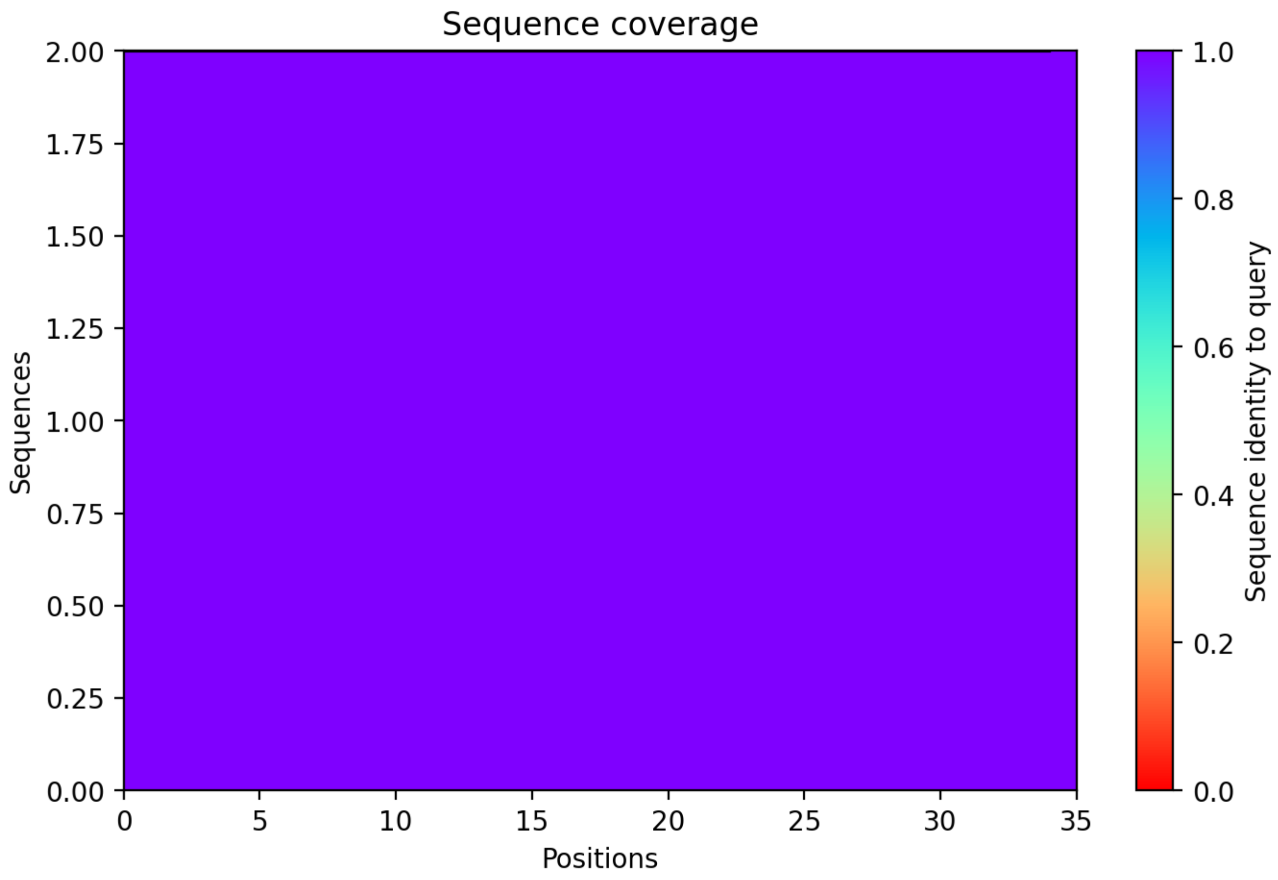

### CP74: MtrunA17_Chr4g0034491

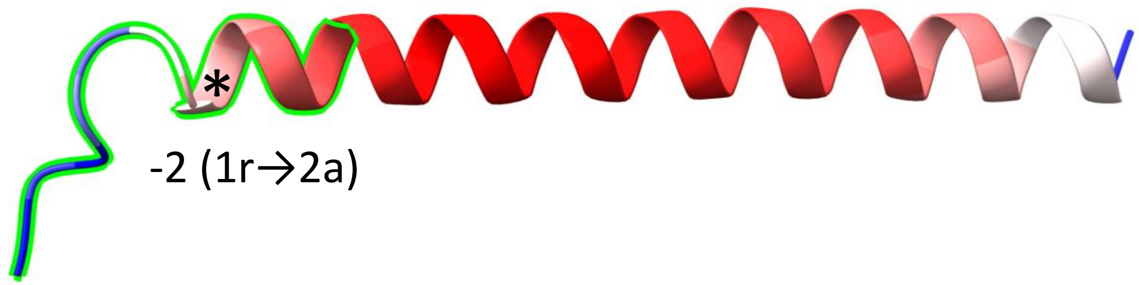


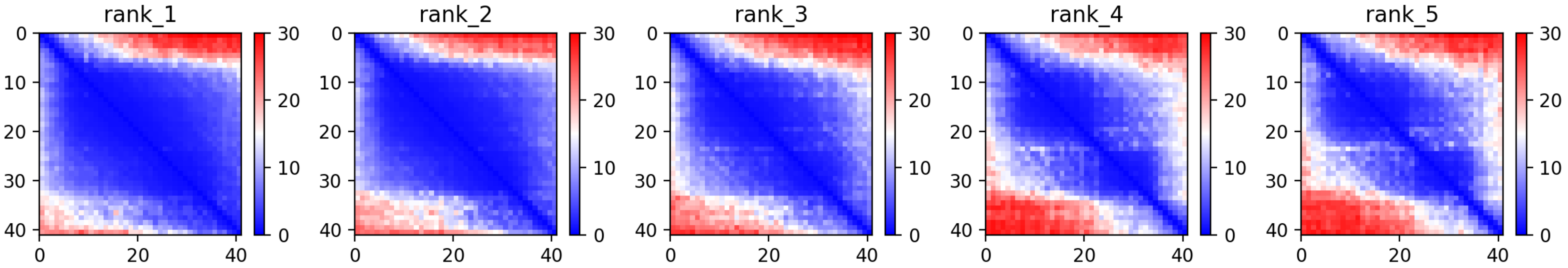

### CP74: MtrunA17_Chr4g0034491

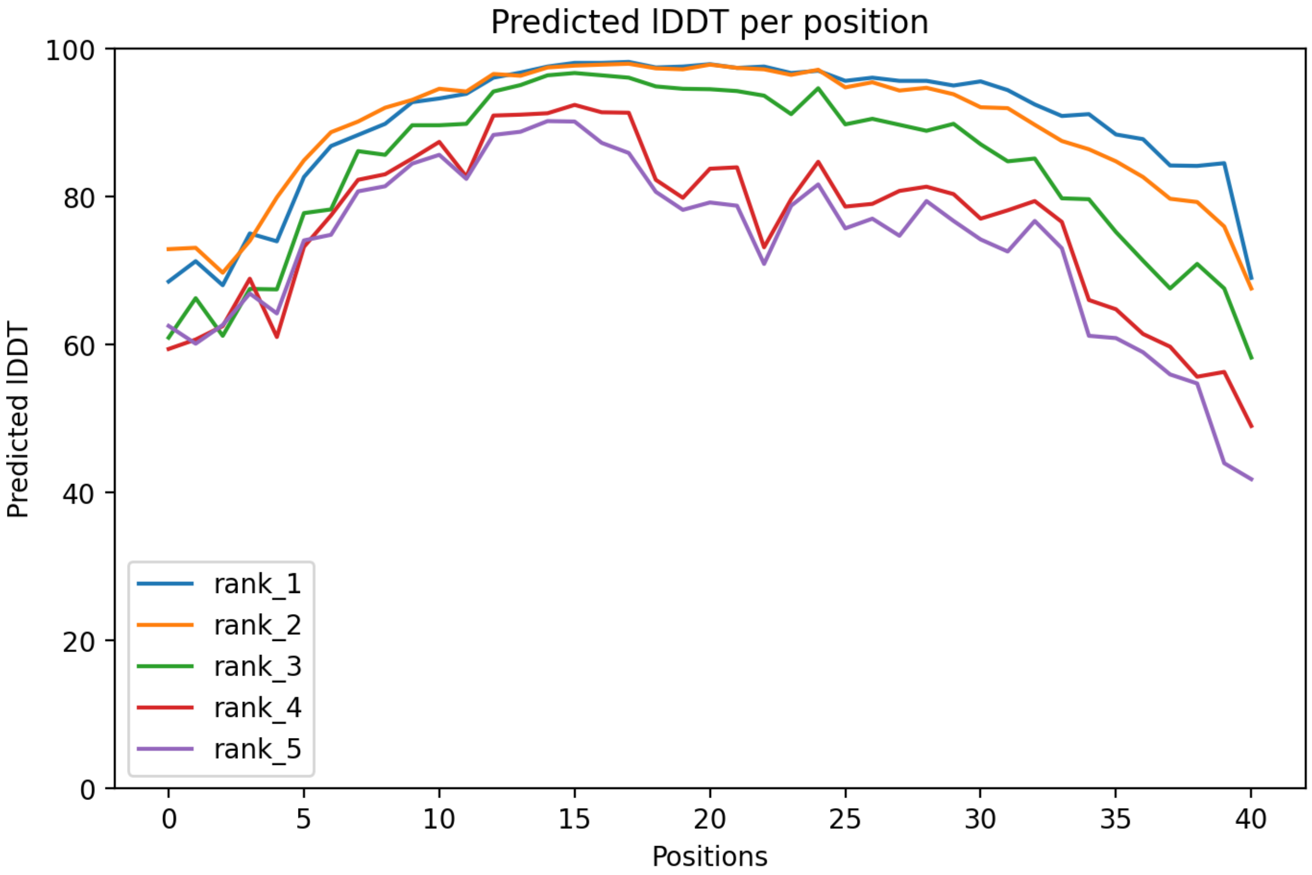

### CP74: MtrunA17_Chr4g0034491

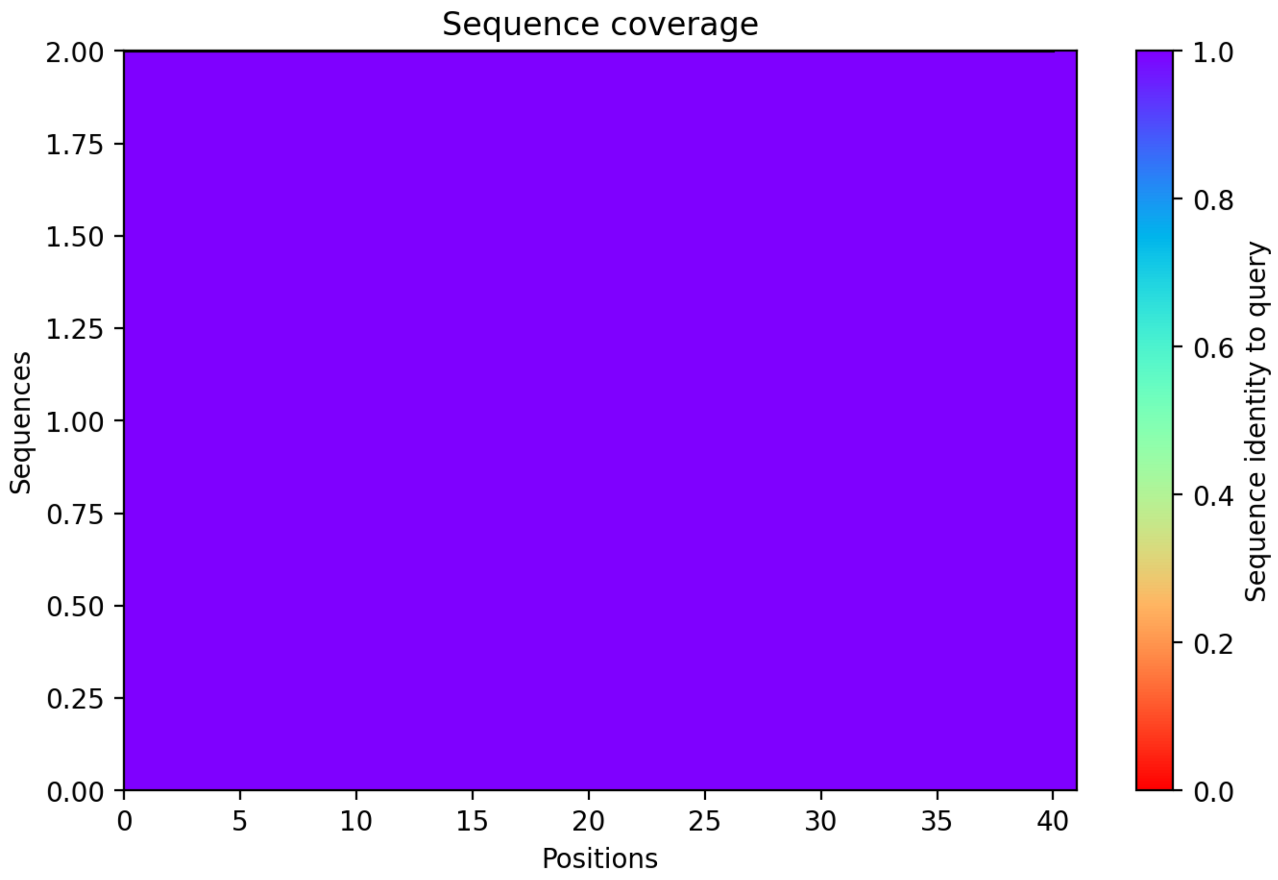

### CP75: MtrunA17_Chr4g0037381

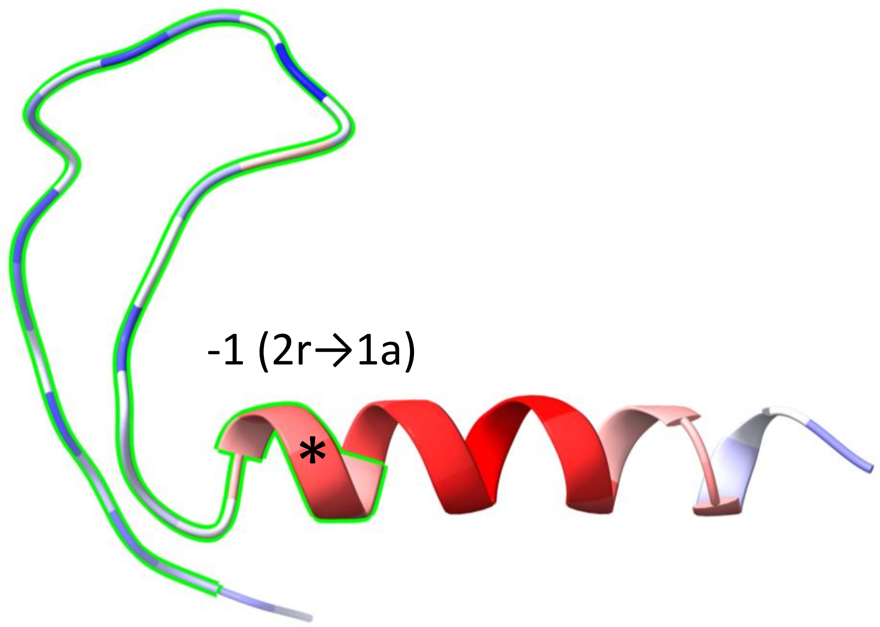


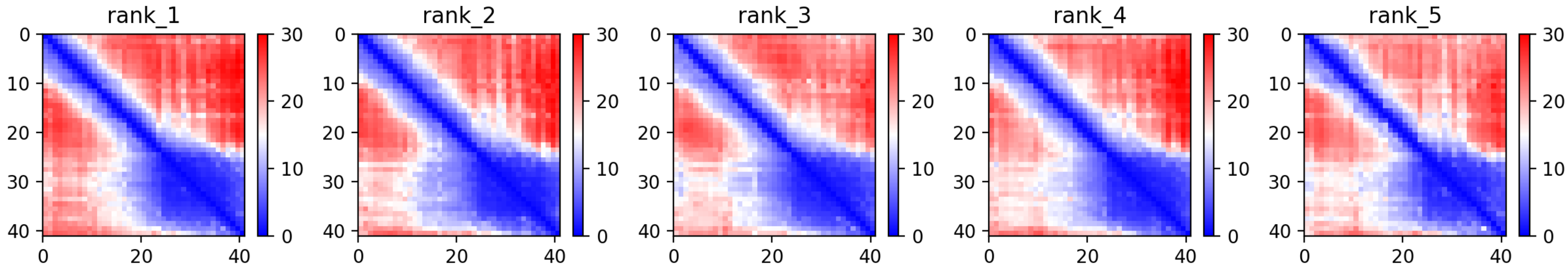

CP75: MtrunA17_Chr4g0037381

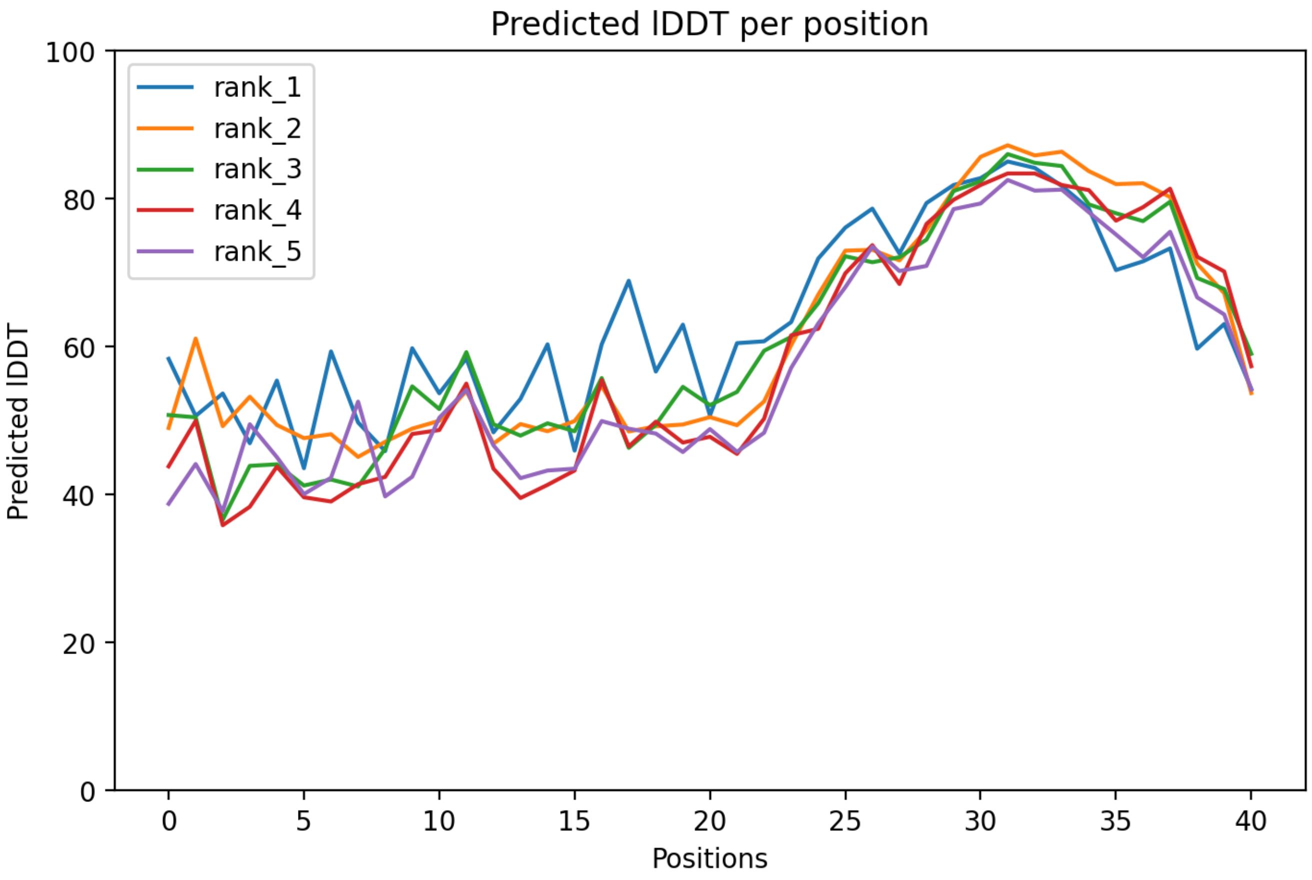

### CP75: MtrunA17_Chr4g0037381

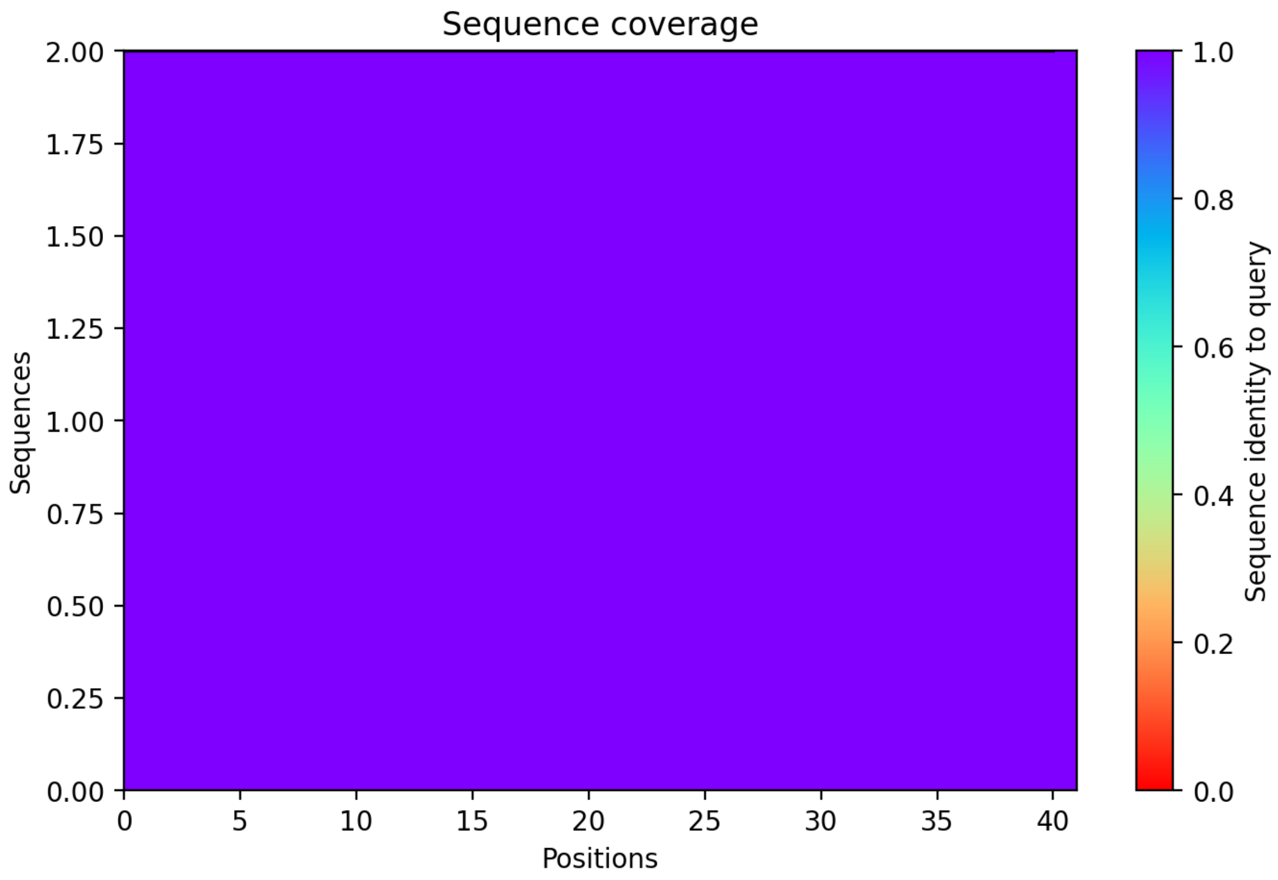

CP76: MtrunA17_Chr4g0040471

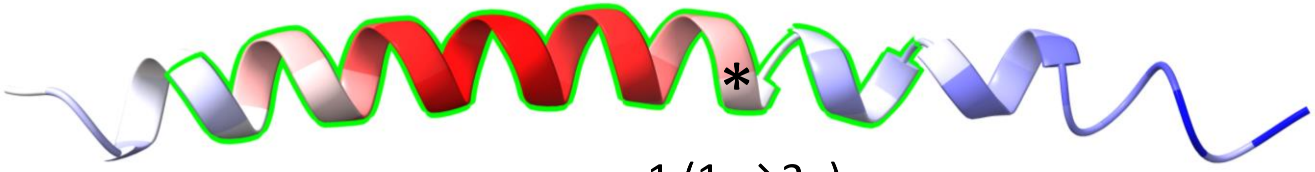


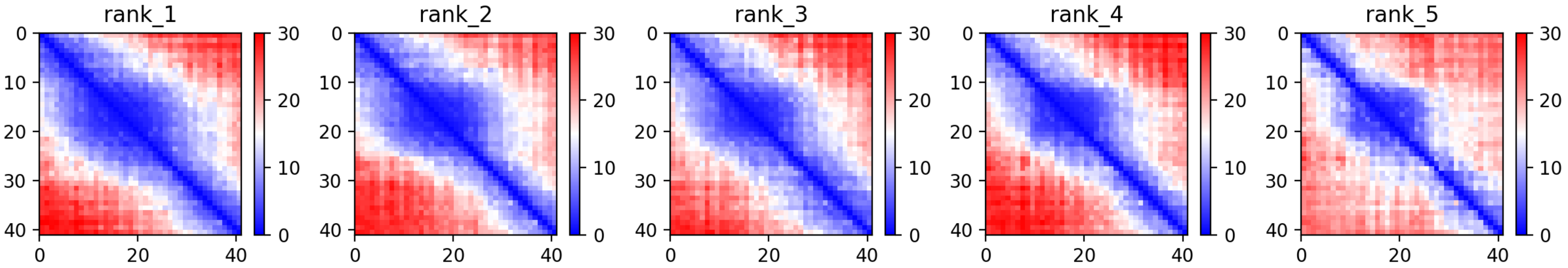

CP76: MtrunA17_Chr4g0040471

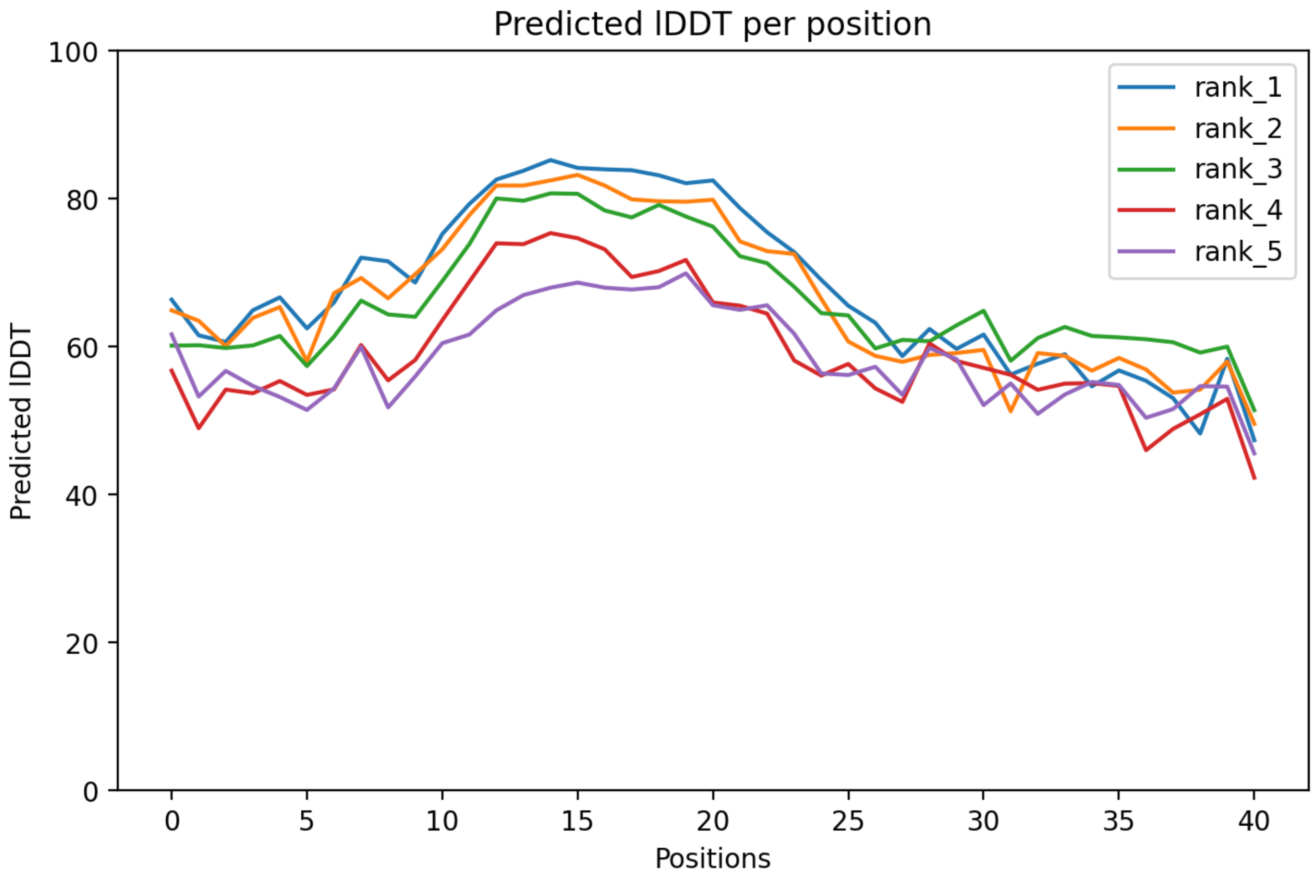

CP76: MtrunA17_Chr4g0040471

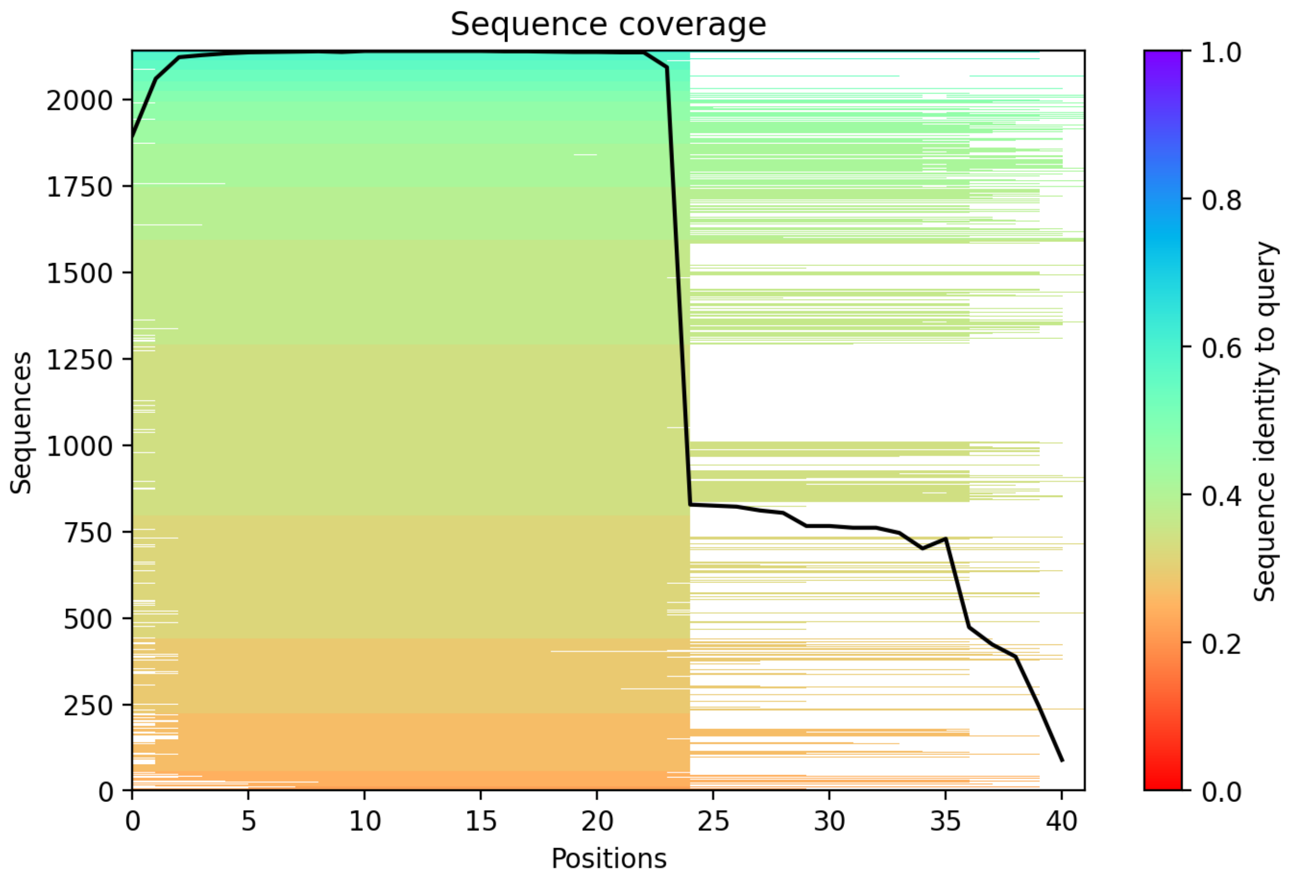

### CP77: MtrunA17_Chr4g0055331

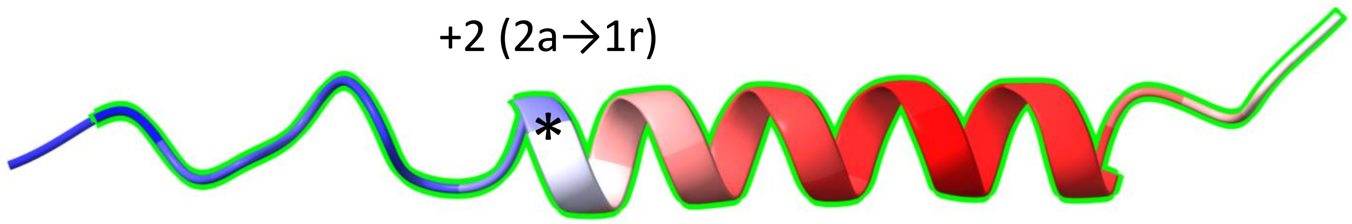


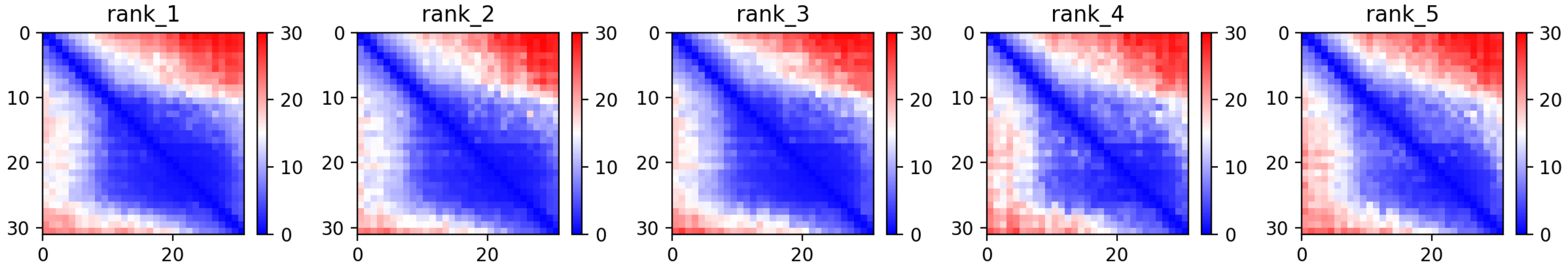

CP77: MtrunA17_Chr4g0055331

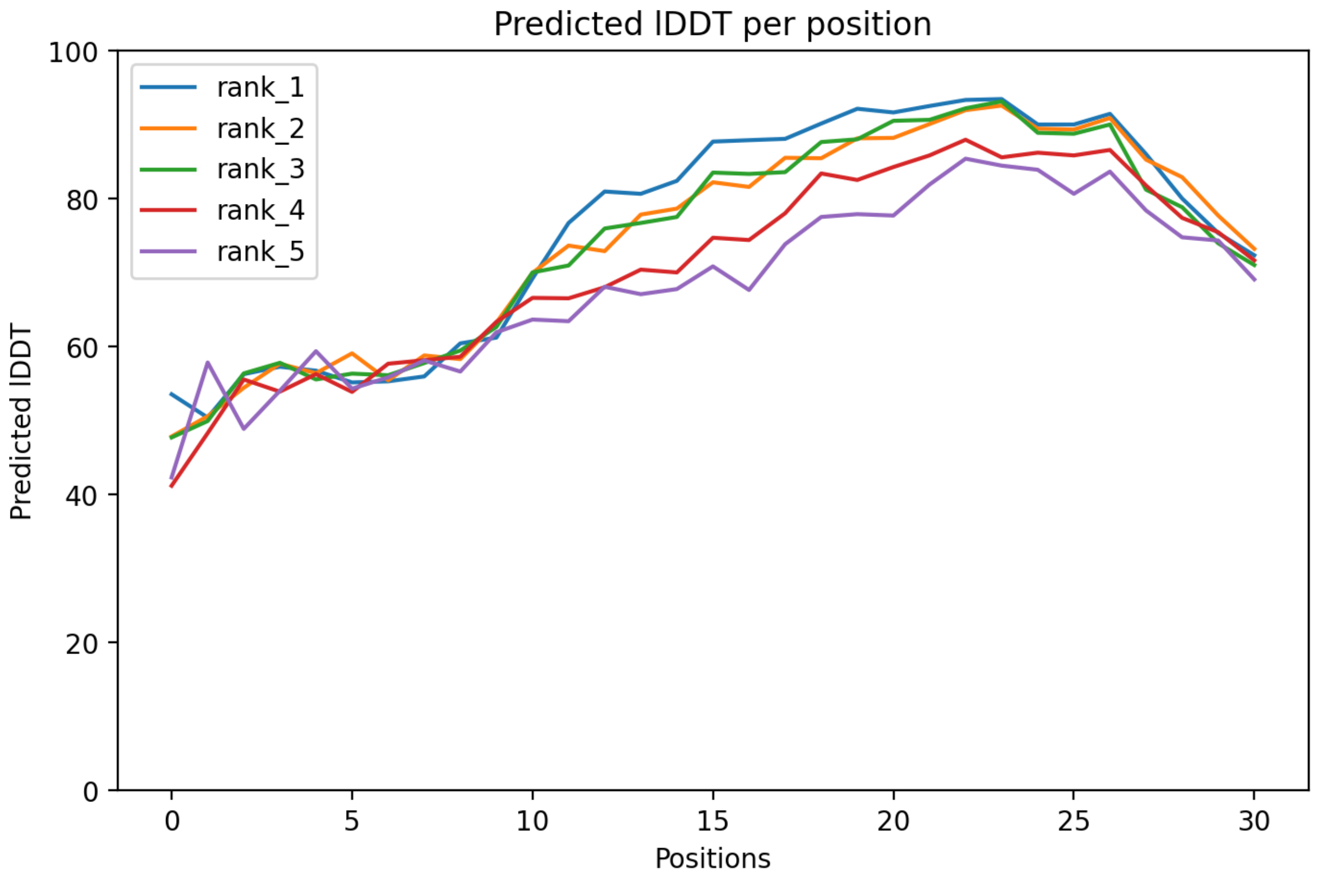

CP77: MtrunA17_Chr4g0055331

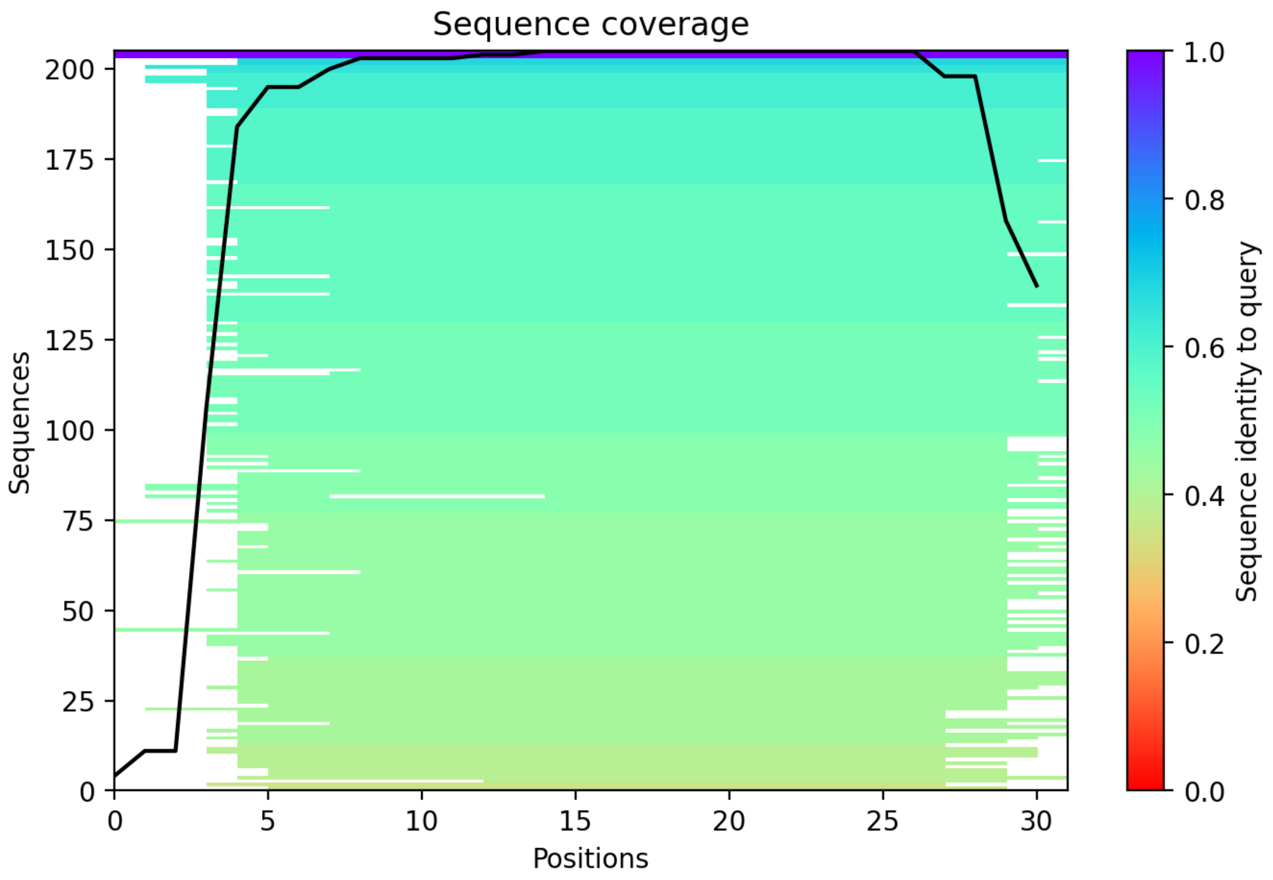

### CP78: MtrunA17_Chr4g0059001

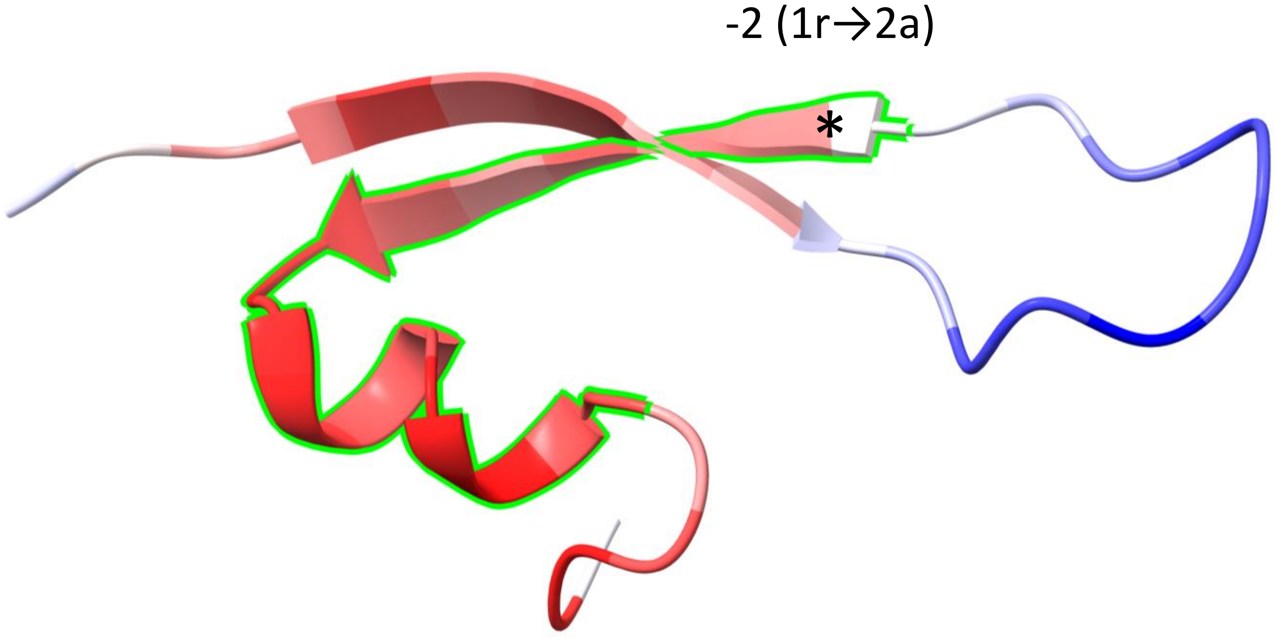


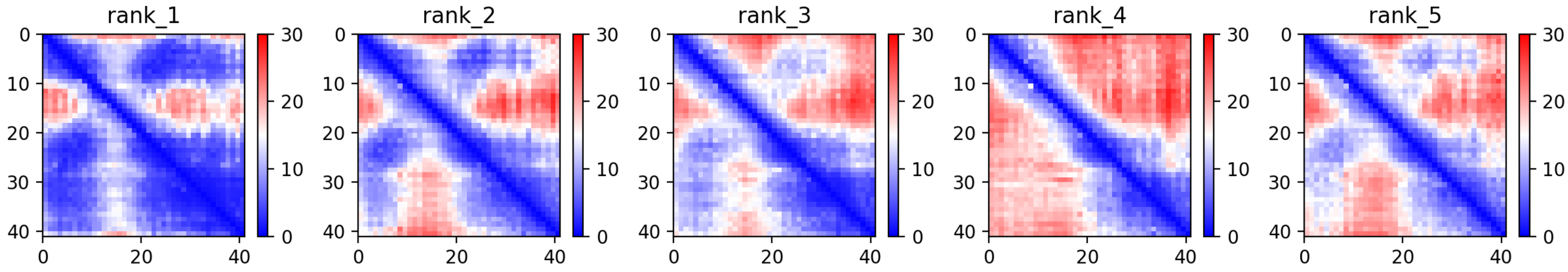

CP78: MtrunA17_Chr4g0059001

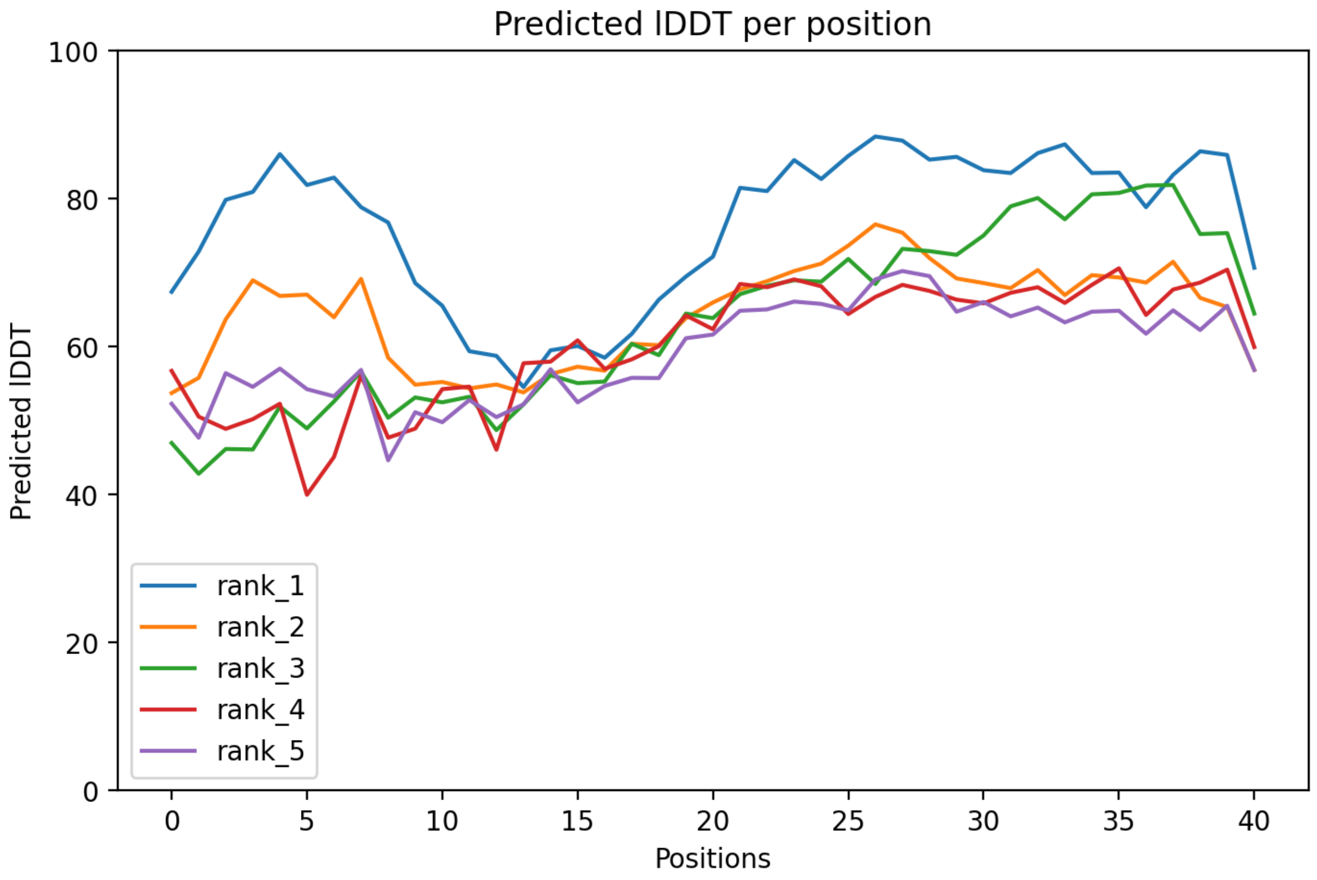

### CP78: MtrunA17_Chr4g0059001

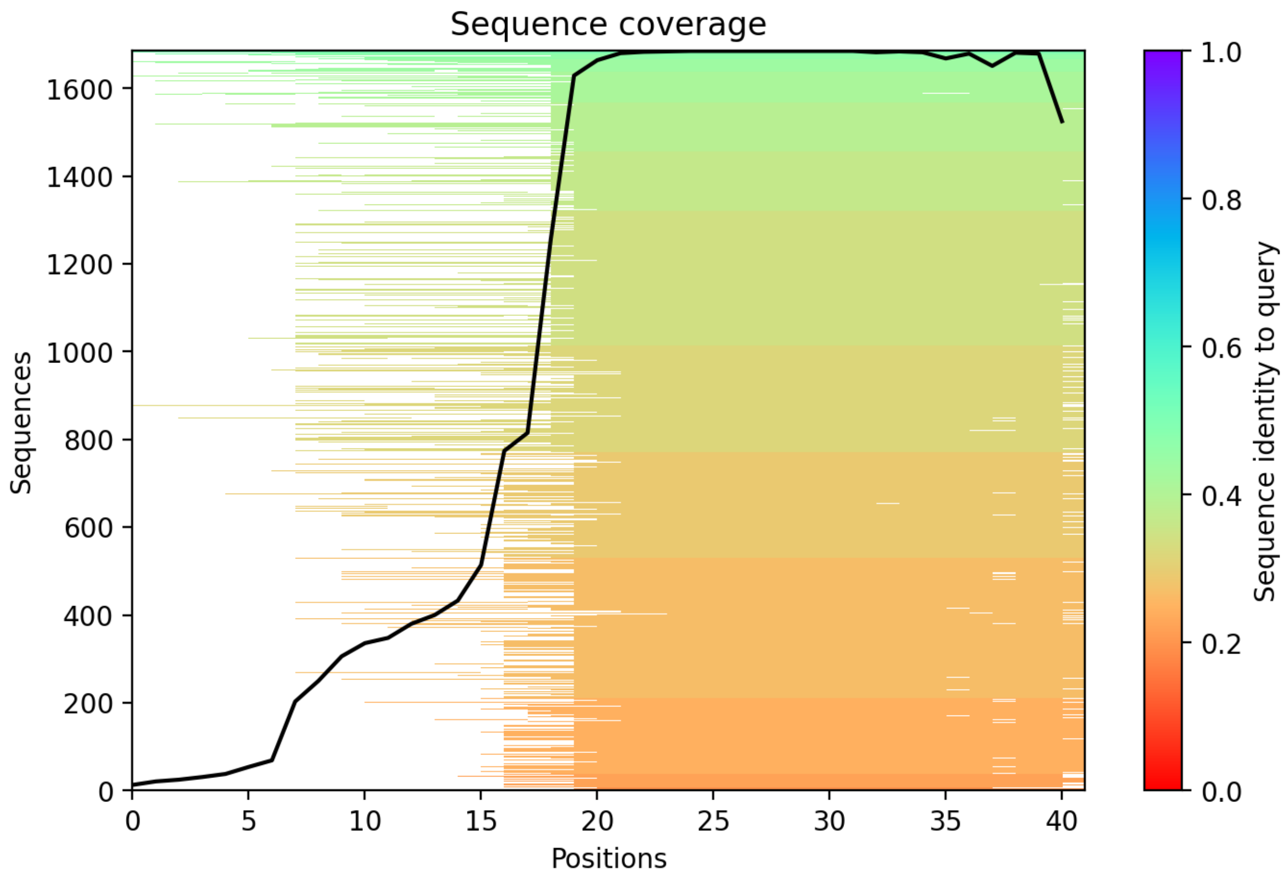

CP79: MtrunA17_Chr4g0063201

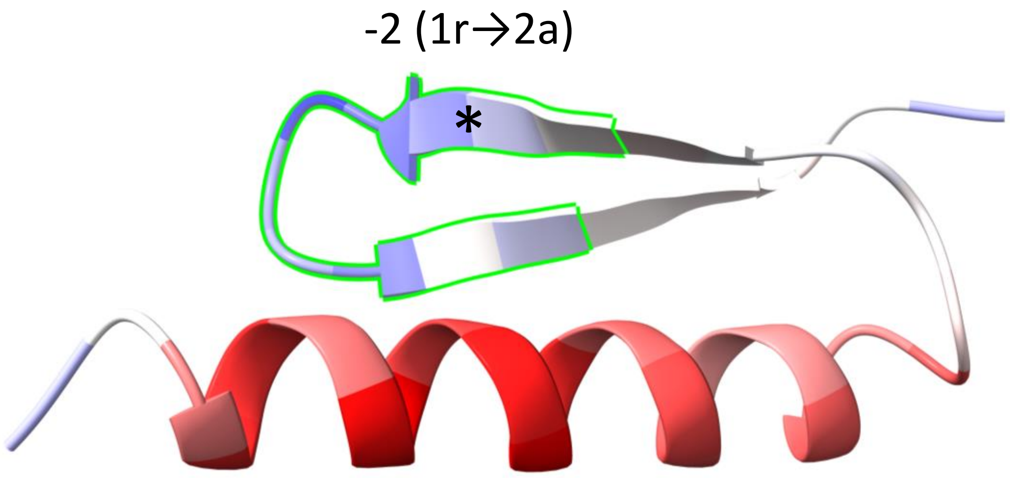


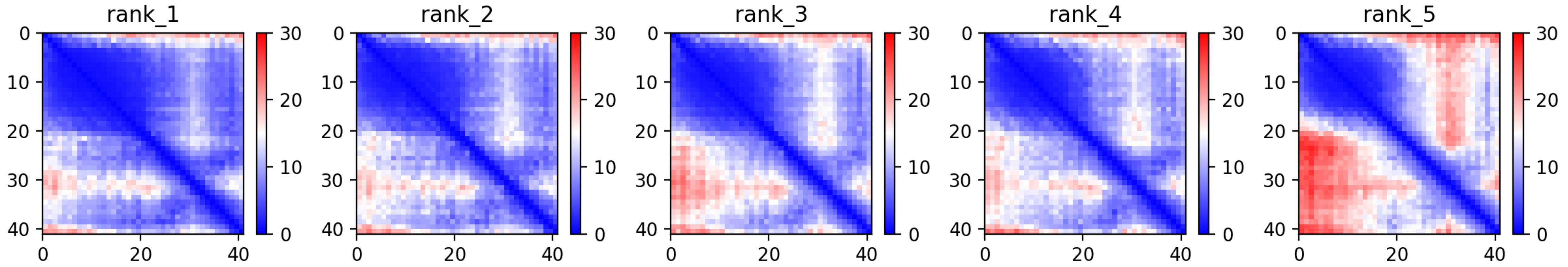

CP79: MtrunA17_Chr4g0063201

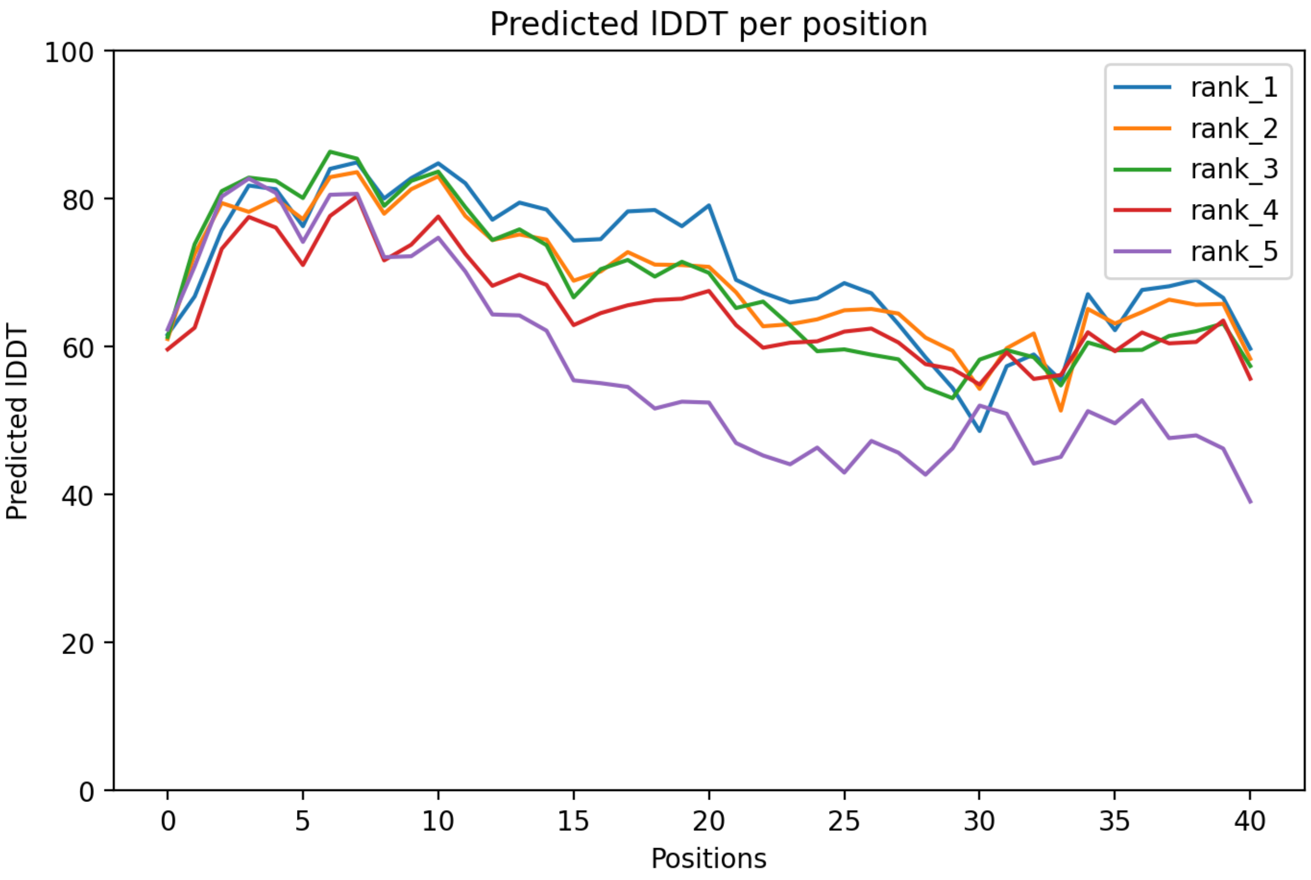

### CP79: MtrunA17_Chr4g0063201

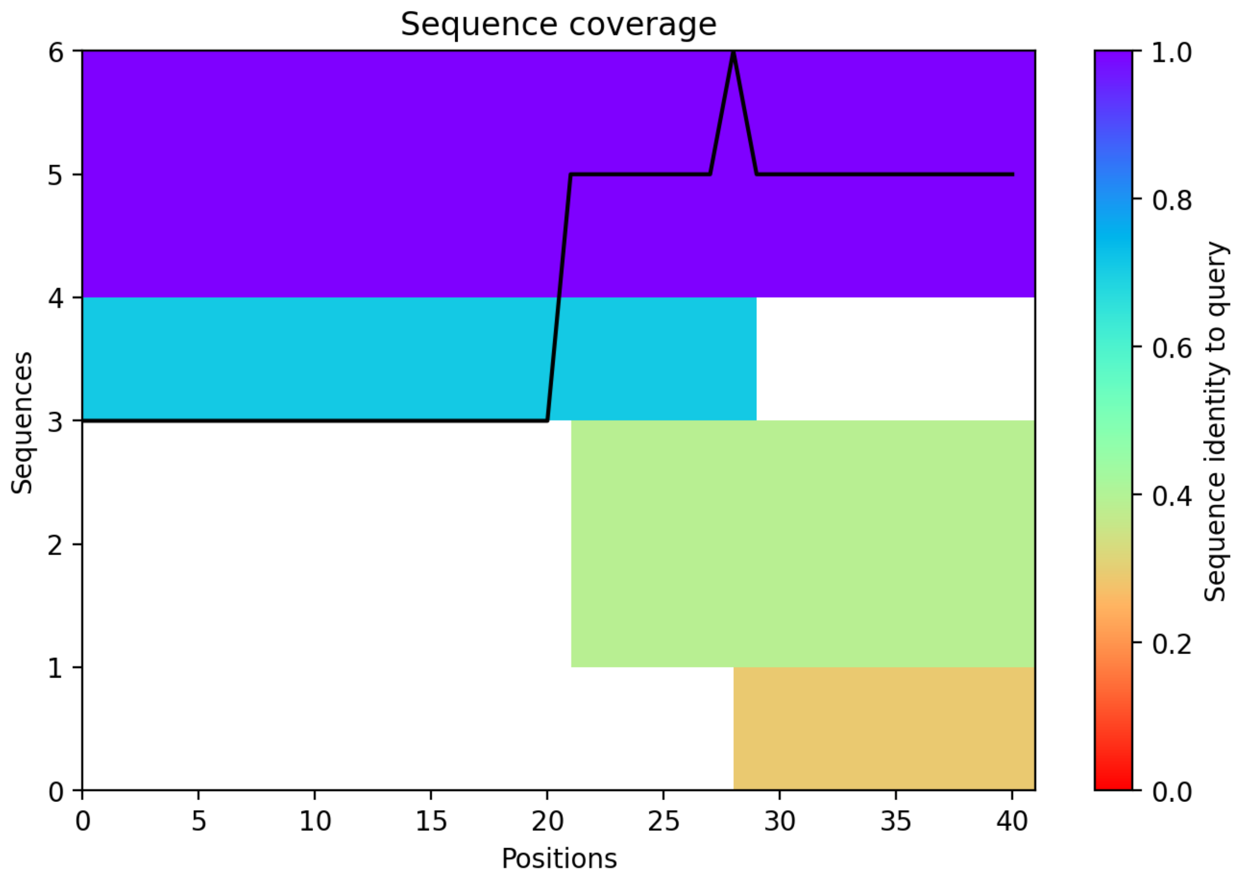

CP80: MtrunA17_Chr4g0070011

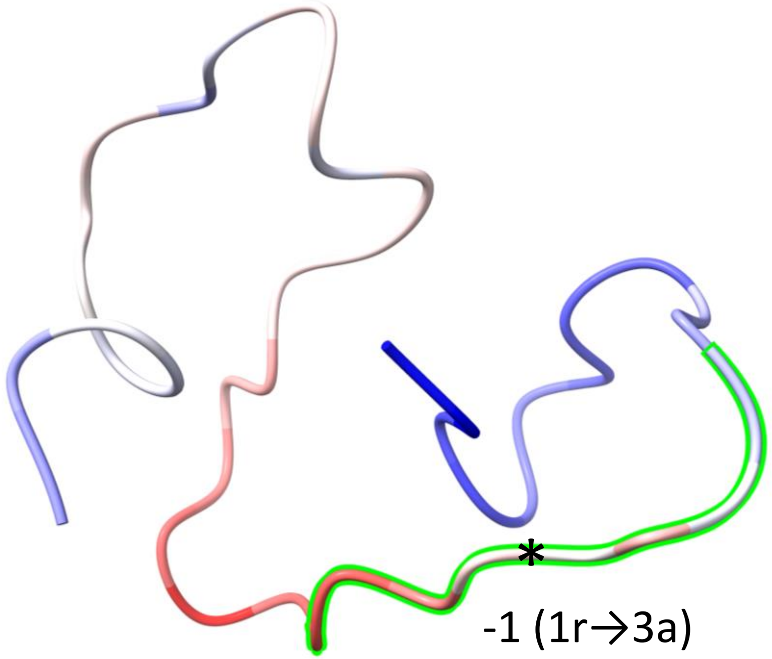


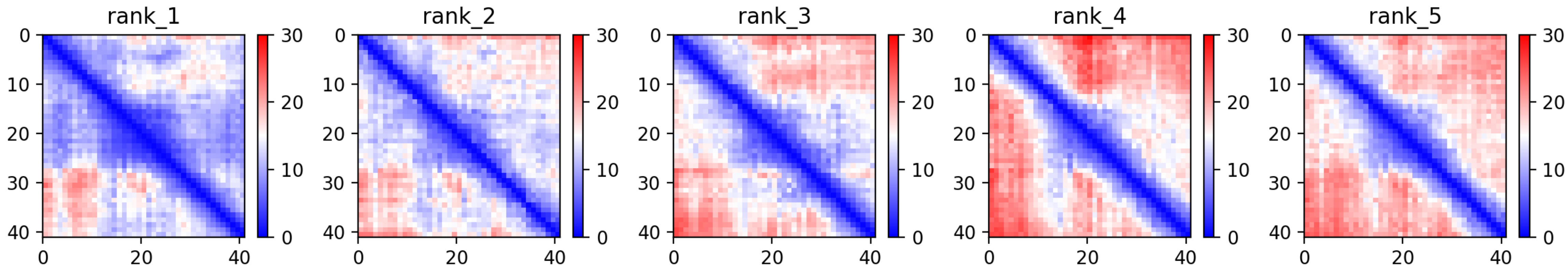

### CP80: MtrunA17_Chr4g0070011

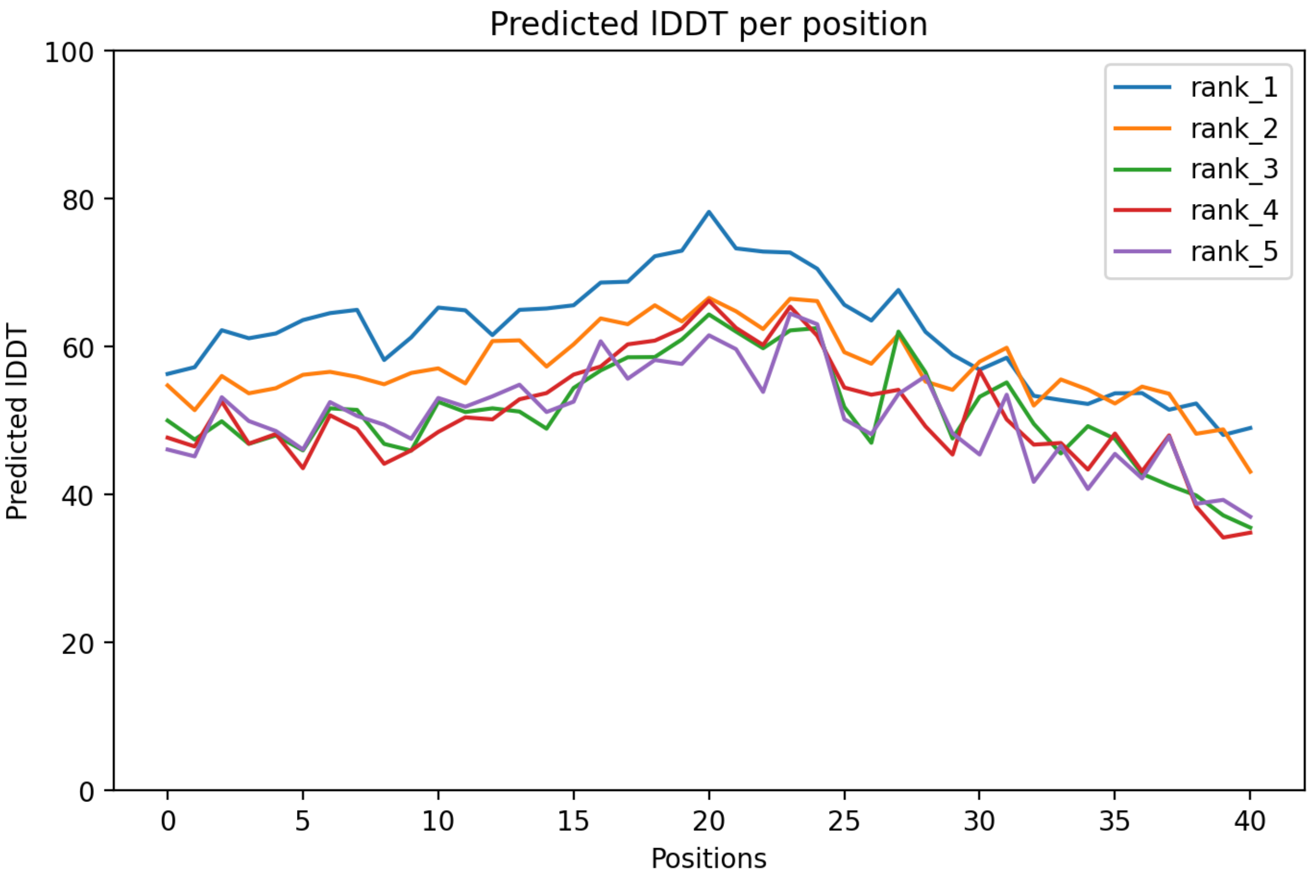

CP80: MtrunA17_Chr4g0070011

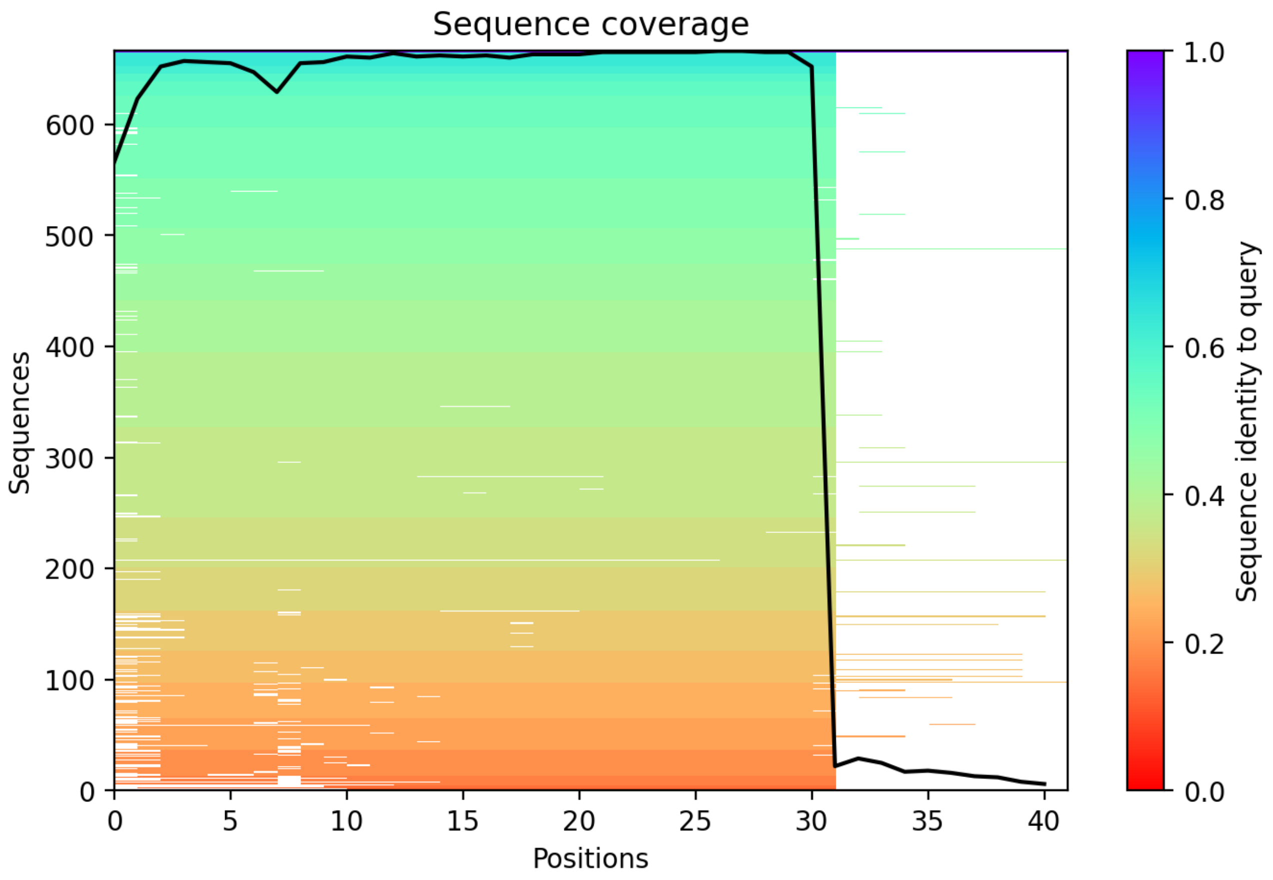

CP81: MtrunA17_Chr5g0393401

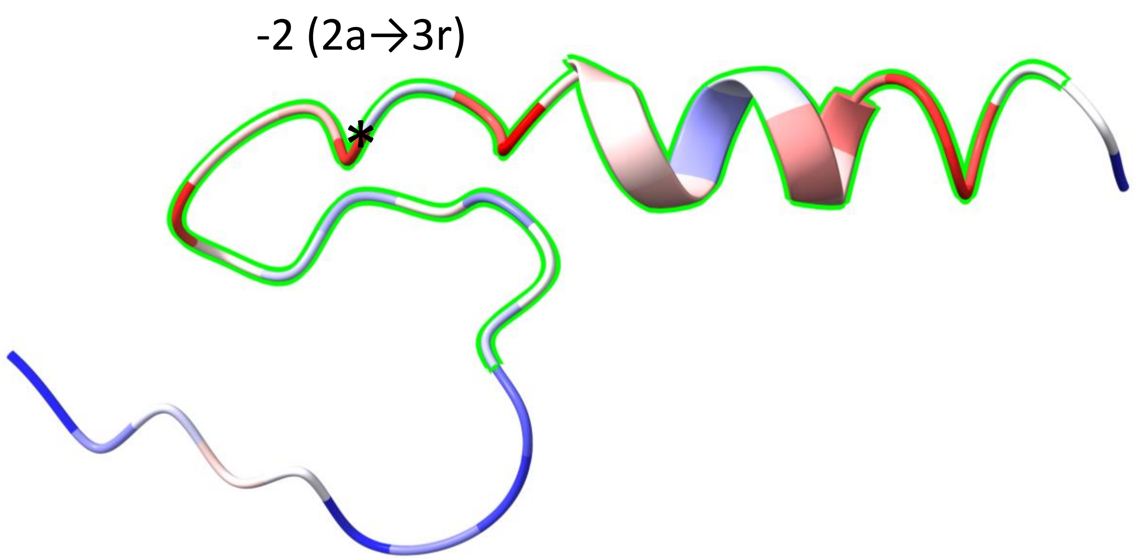


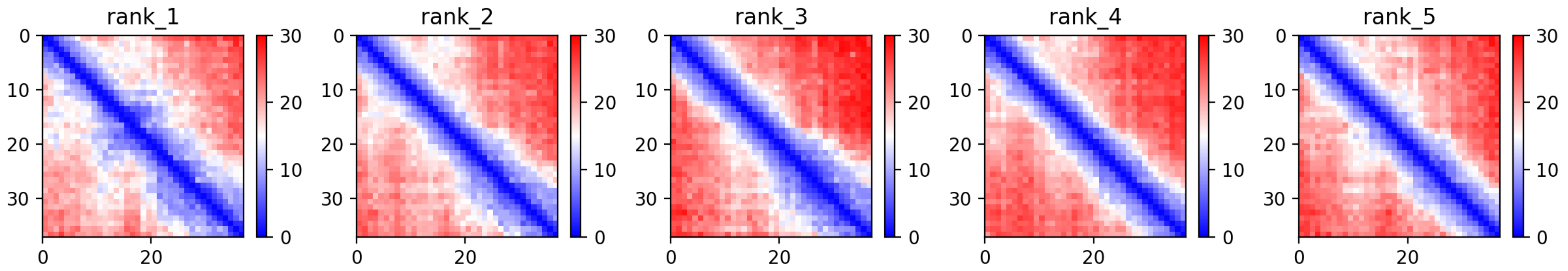

CP81: MtrunA17_Chr5g0393401

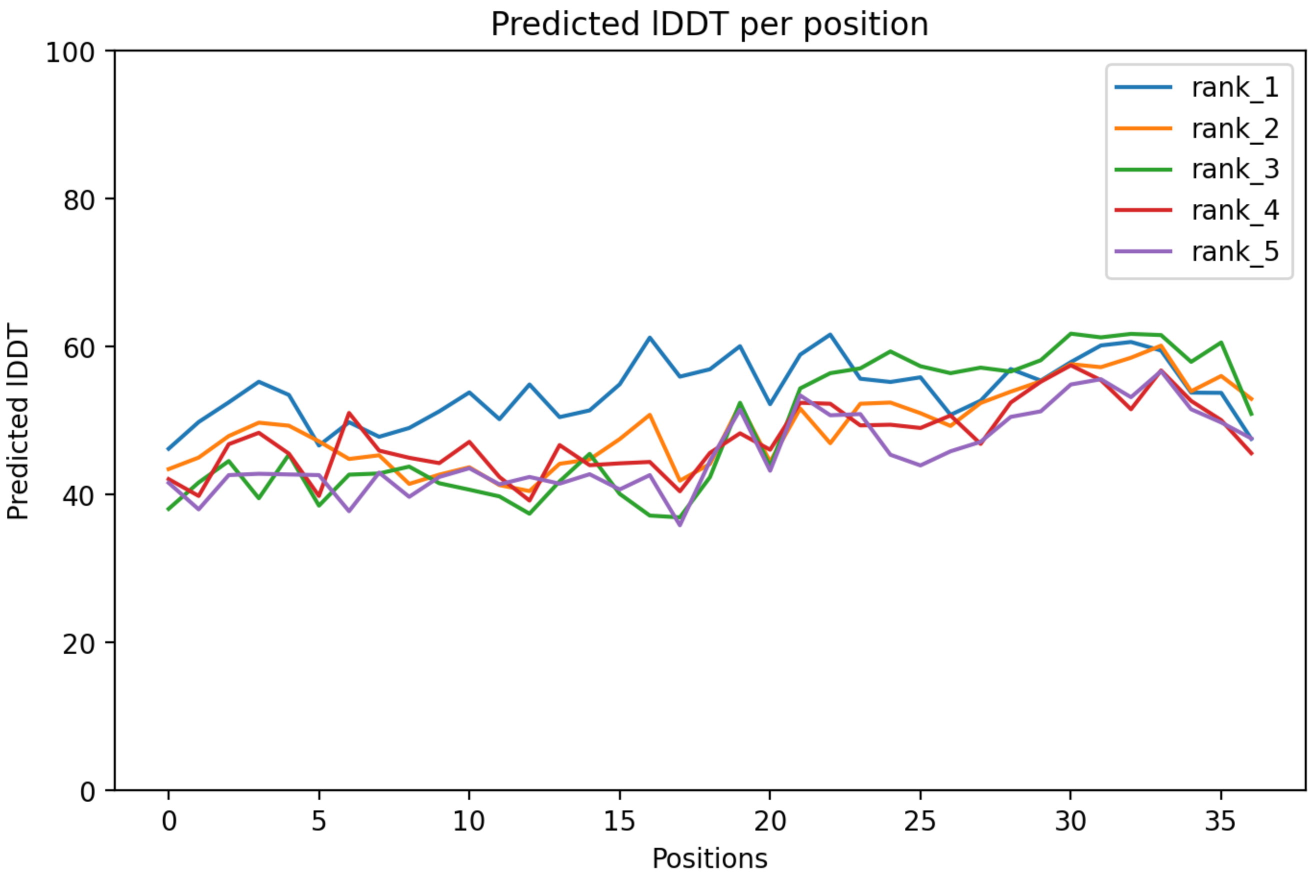

CP81: MtrunA17_Chr5g0393401

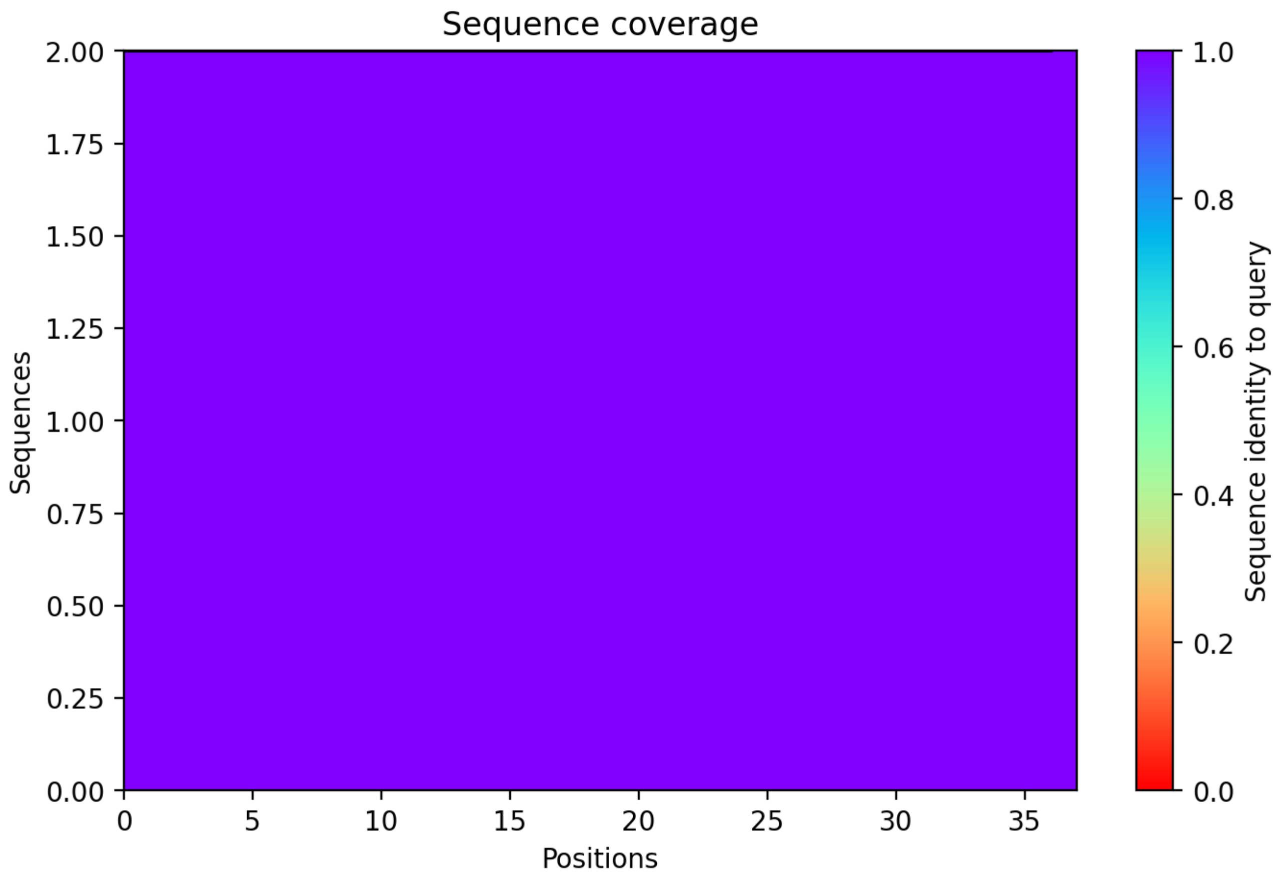

### CP82: MtrunA17_Chr5g0400181

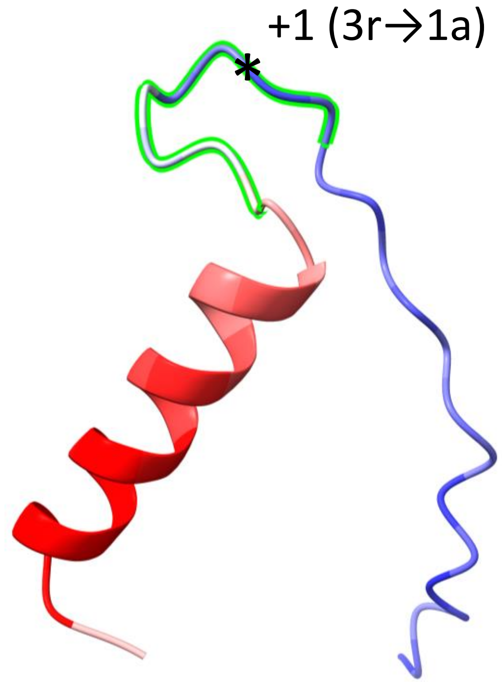


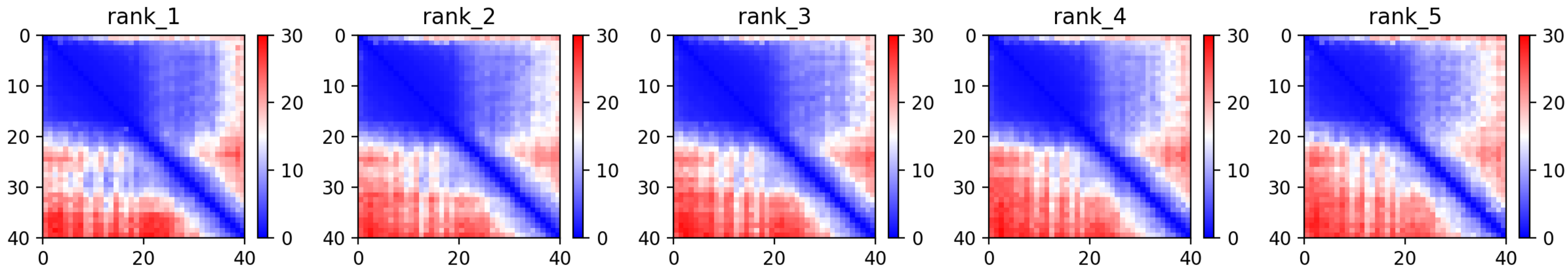

CP82: MtrunA17_Chr5g0400181

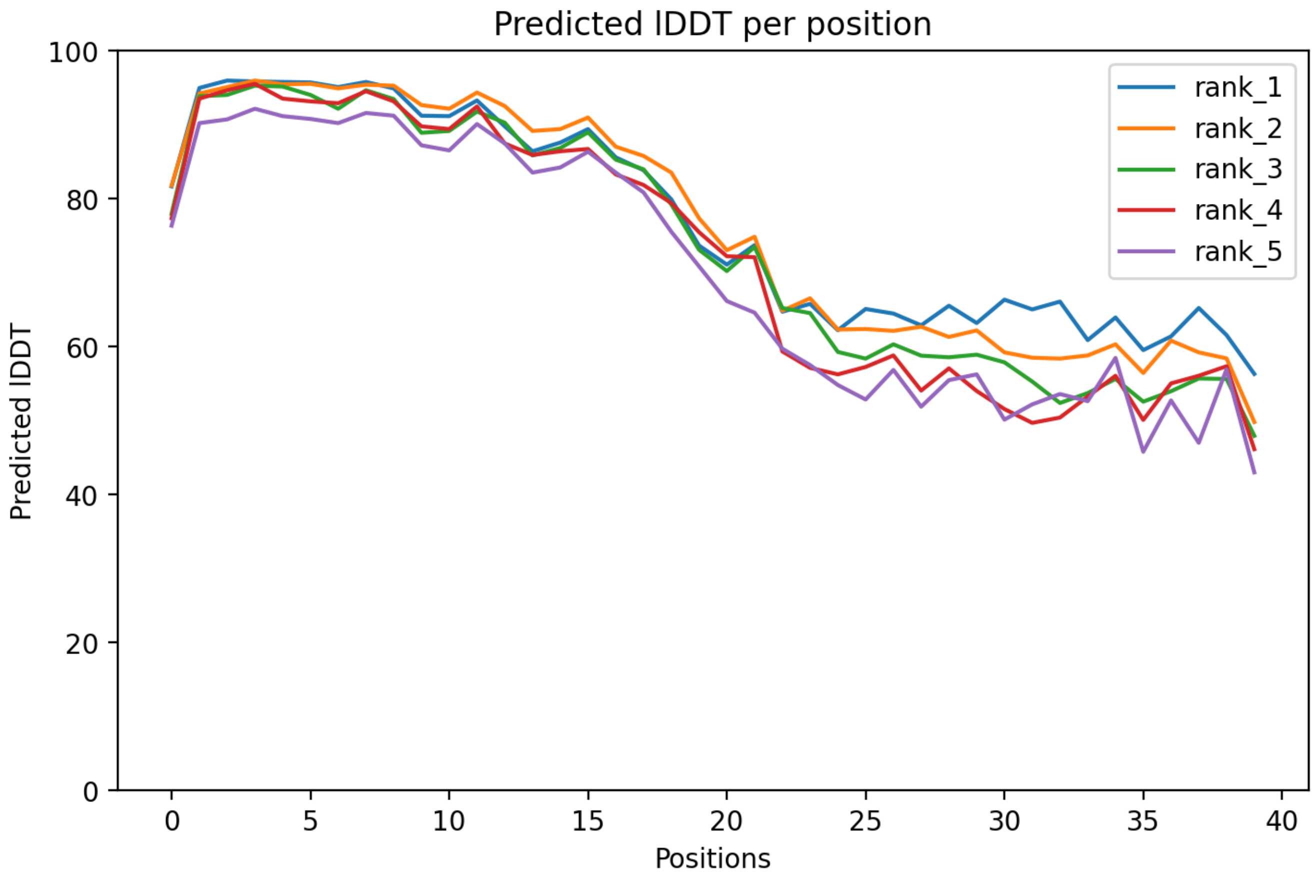

### CP82: MtrunA17_Chr5g0400181

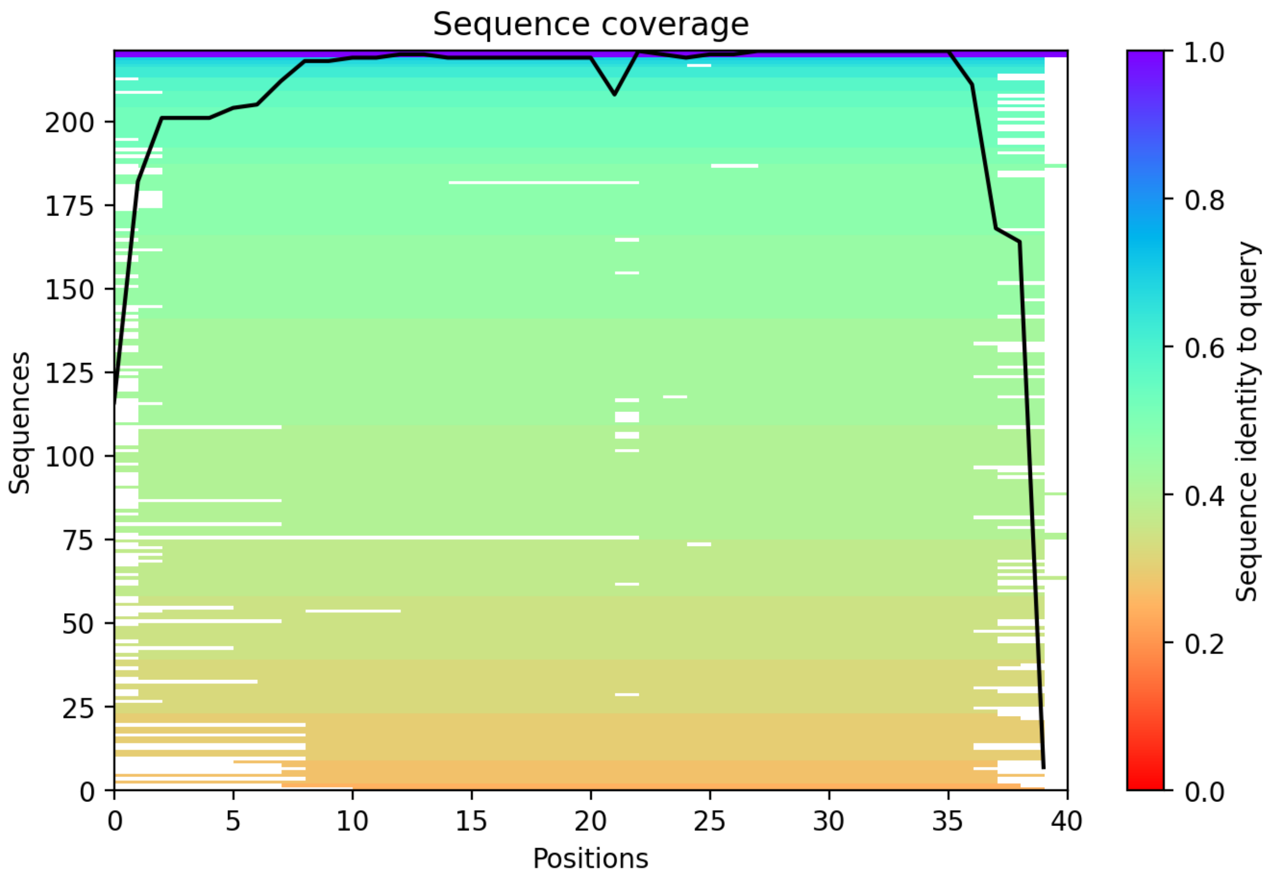

CP83: MtrunA17_Chr5g0405061

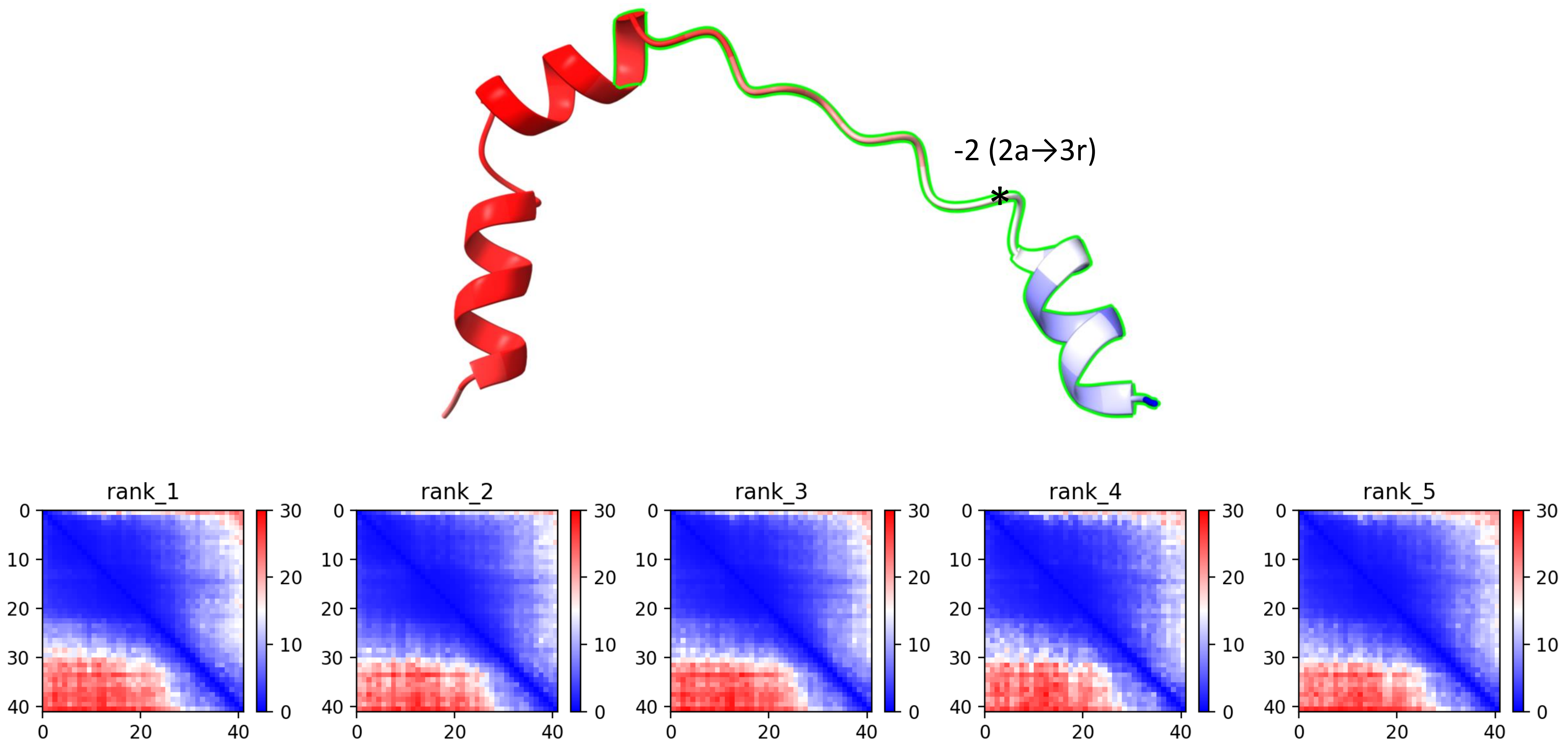

### CP83: MtrunA17_Chr5g0405061

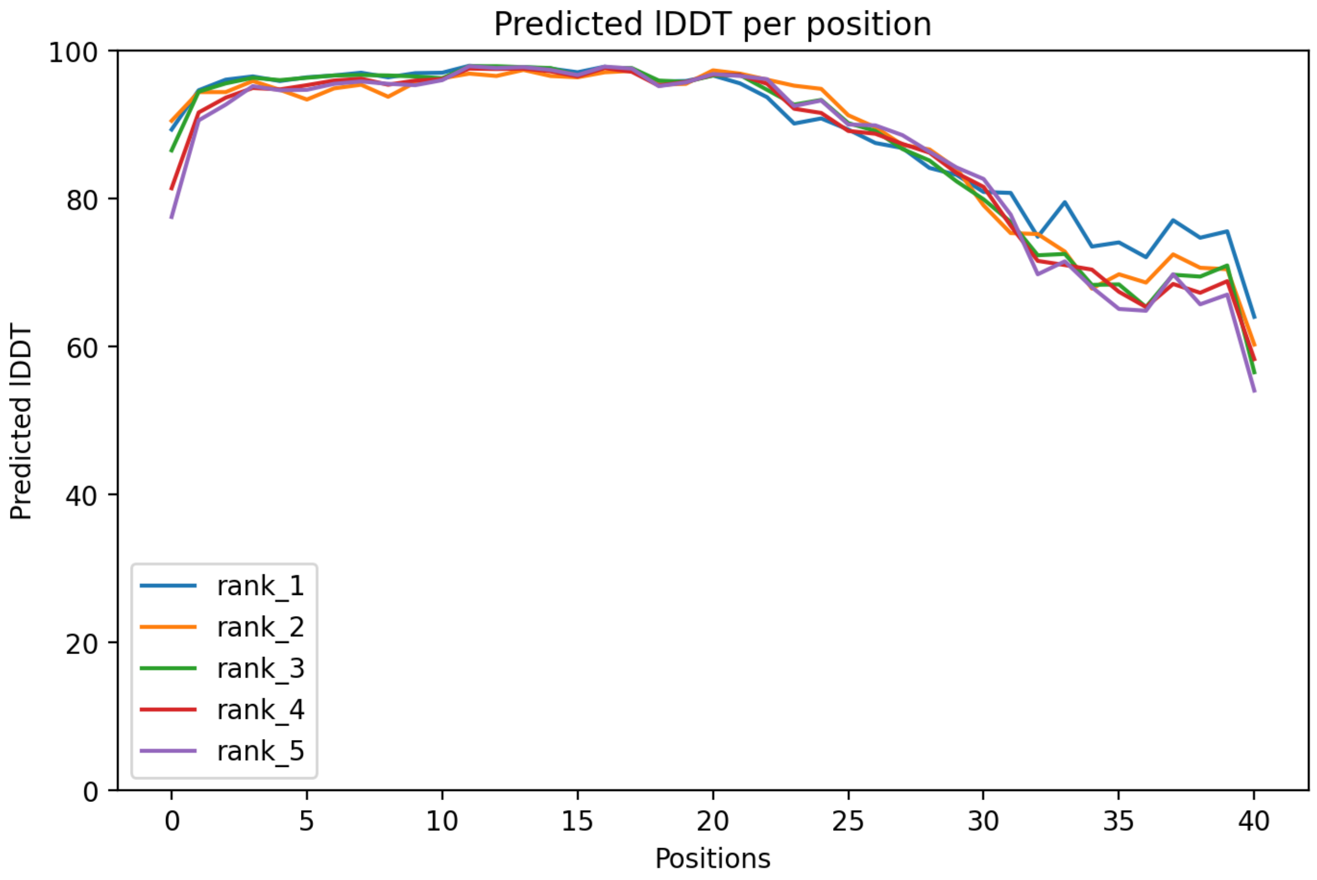

### CP83: MtrunA17_Chr5g0405061

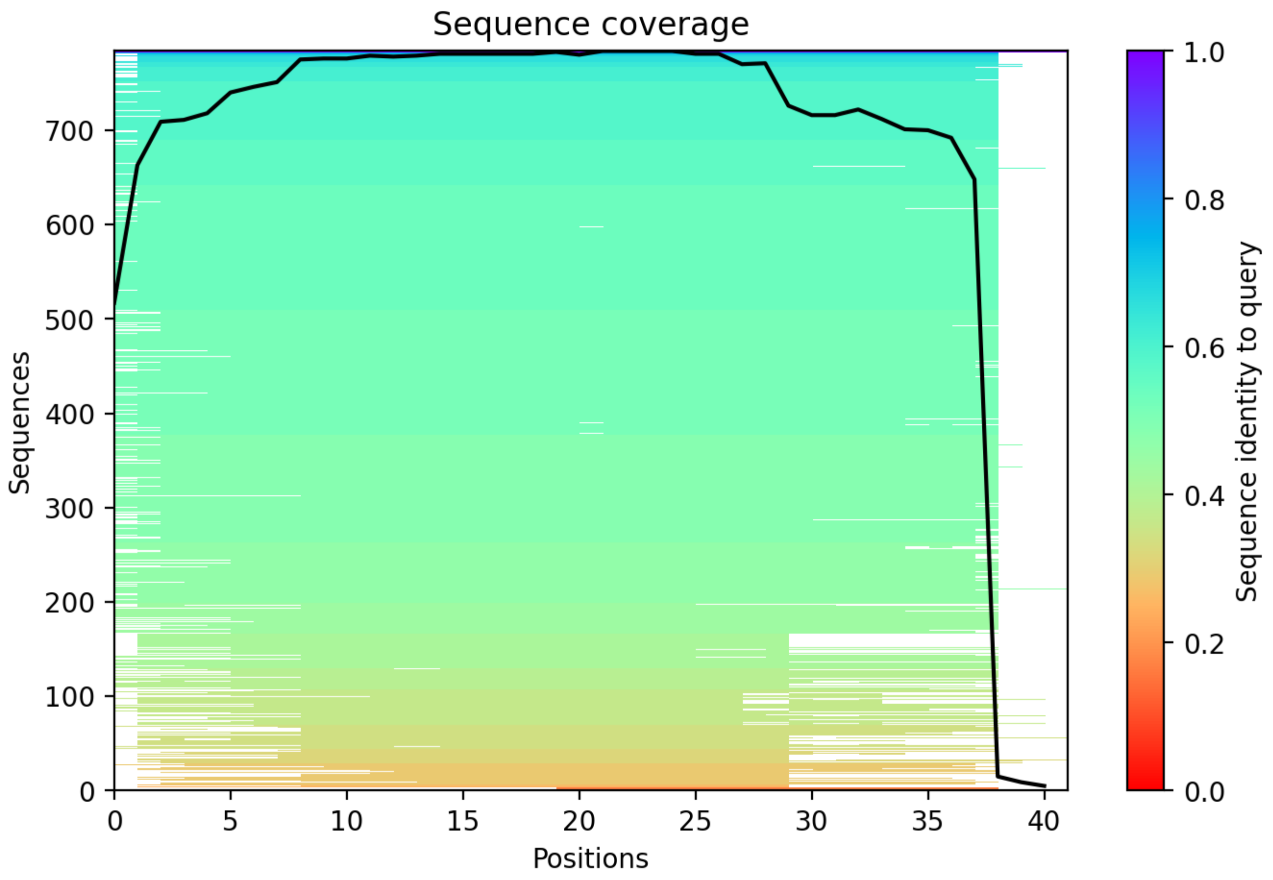

CP84: MtrunA17_Chr5g0408581

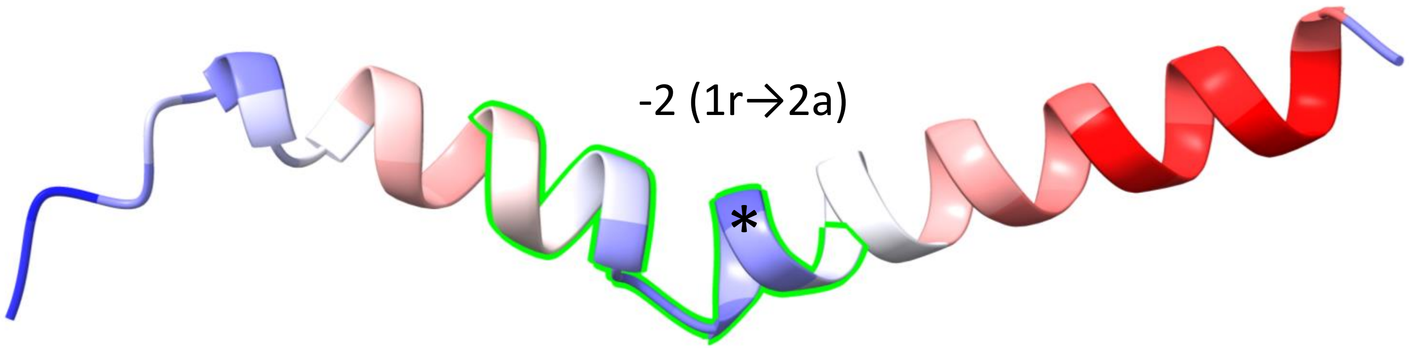


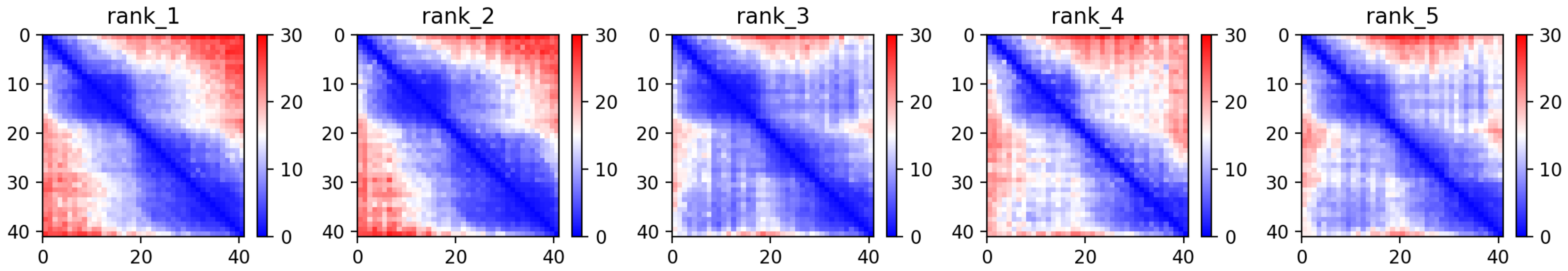

### CP84: MtrunA17_Chr5g0408581

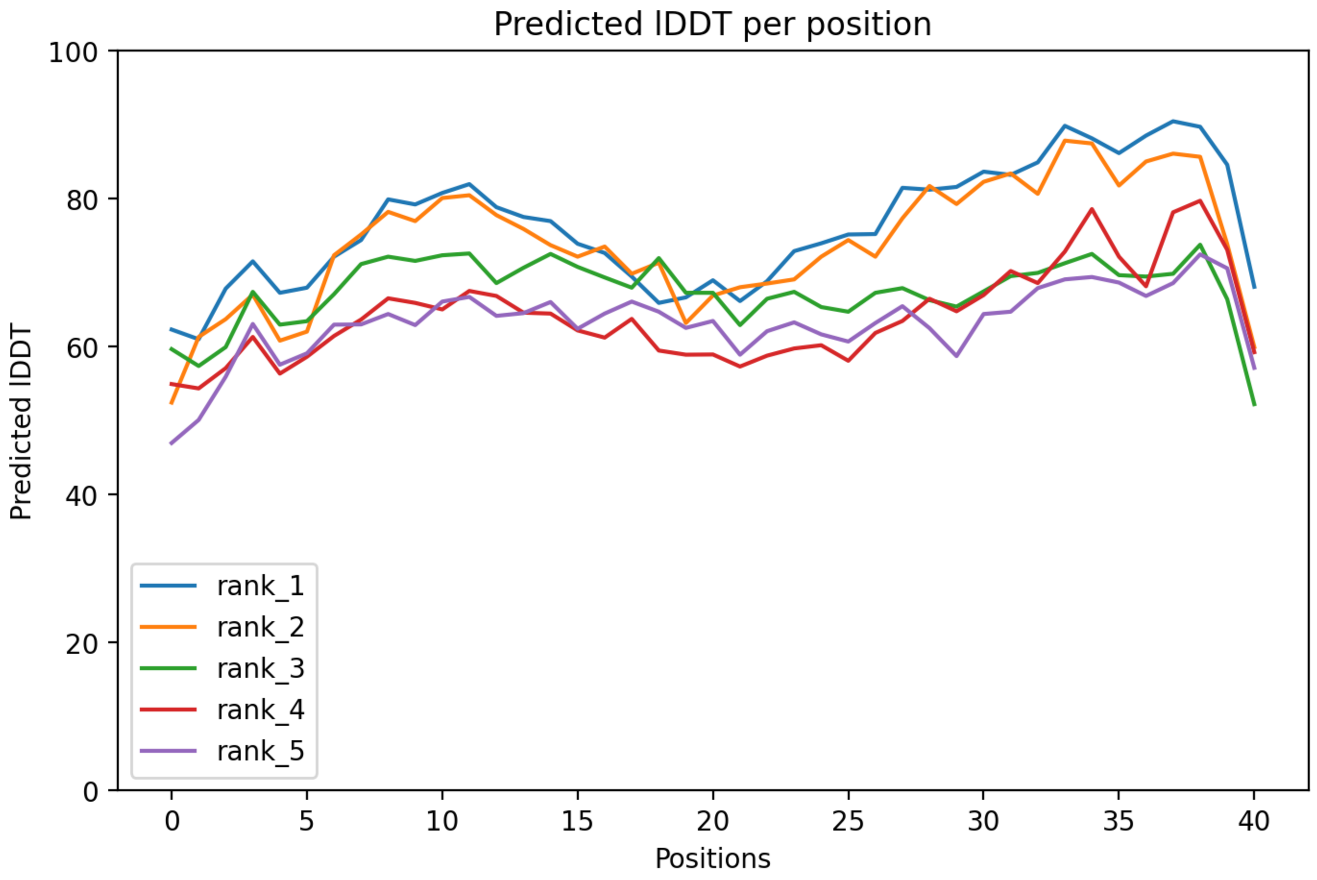

CP84: MtrunA17_Chr5g0408581

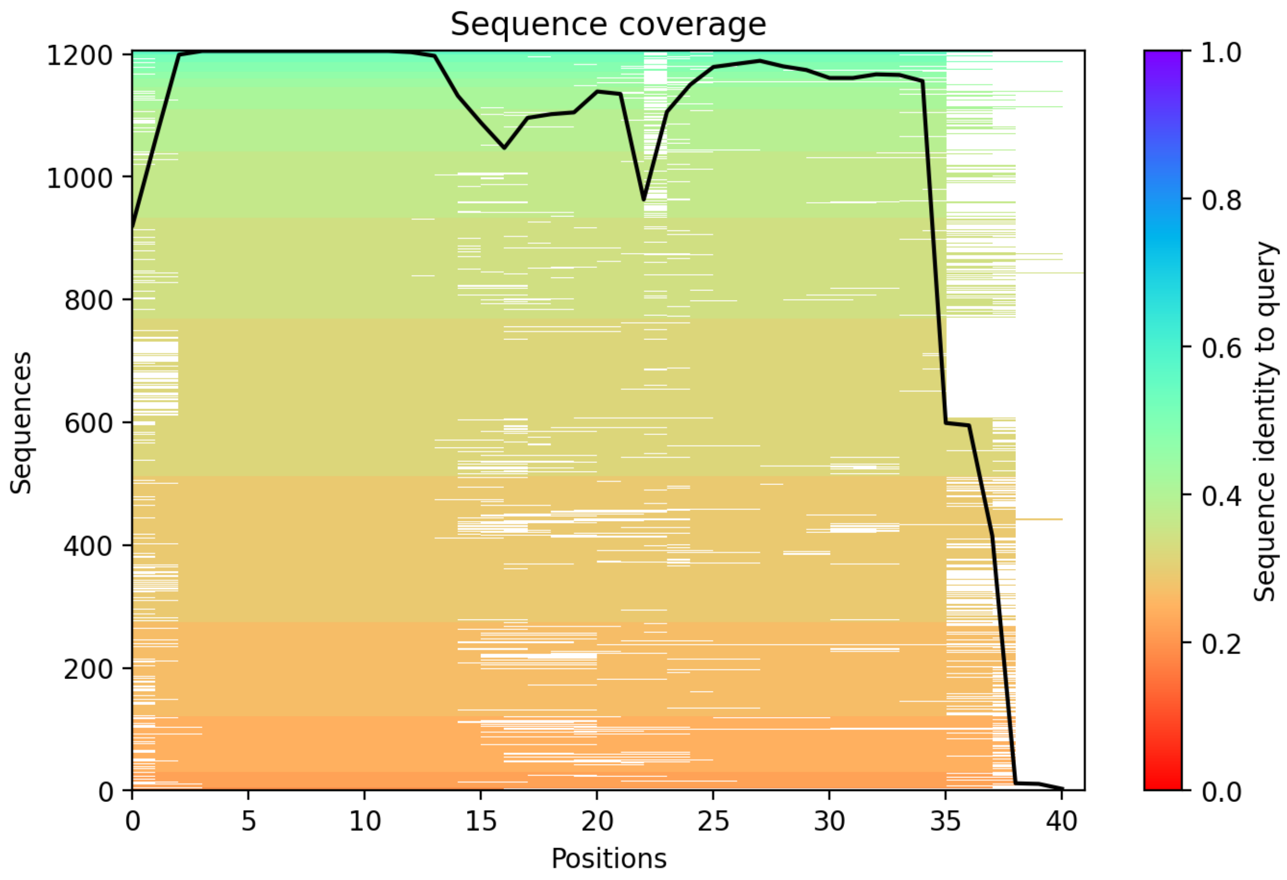

### CP85: MtrunA17_Chr5g0415031

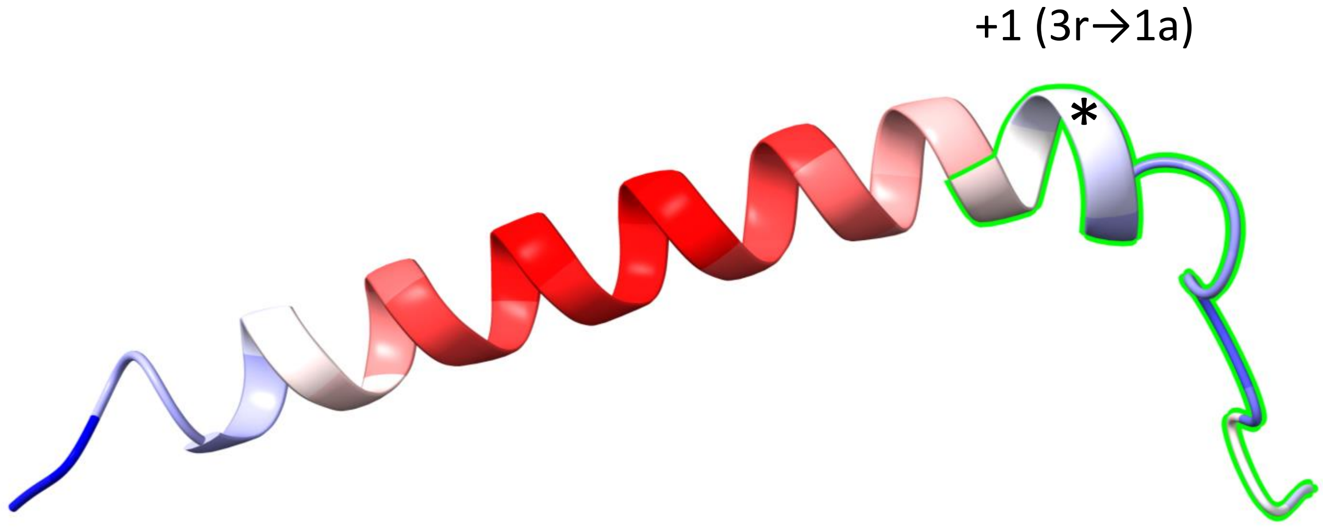


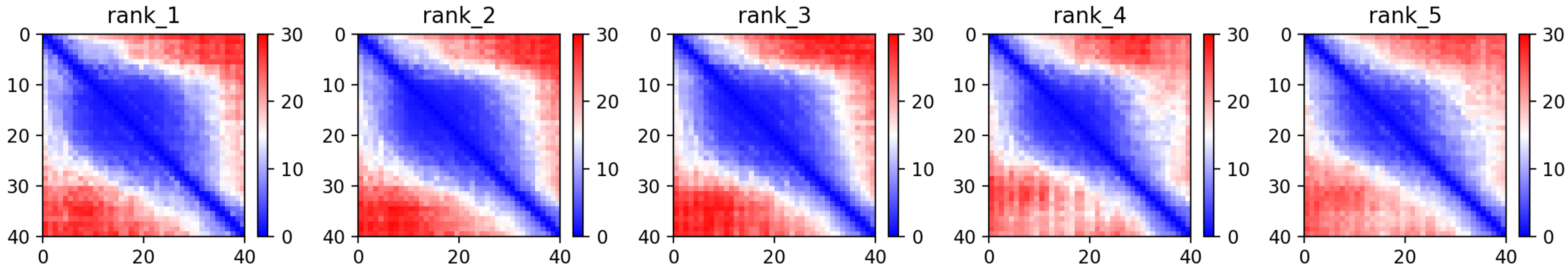

CP85: MtrunA17_Chr5g0415031

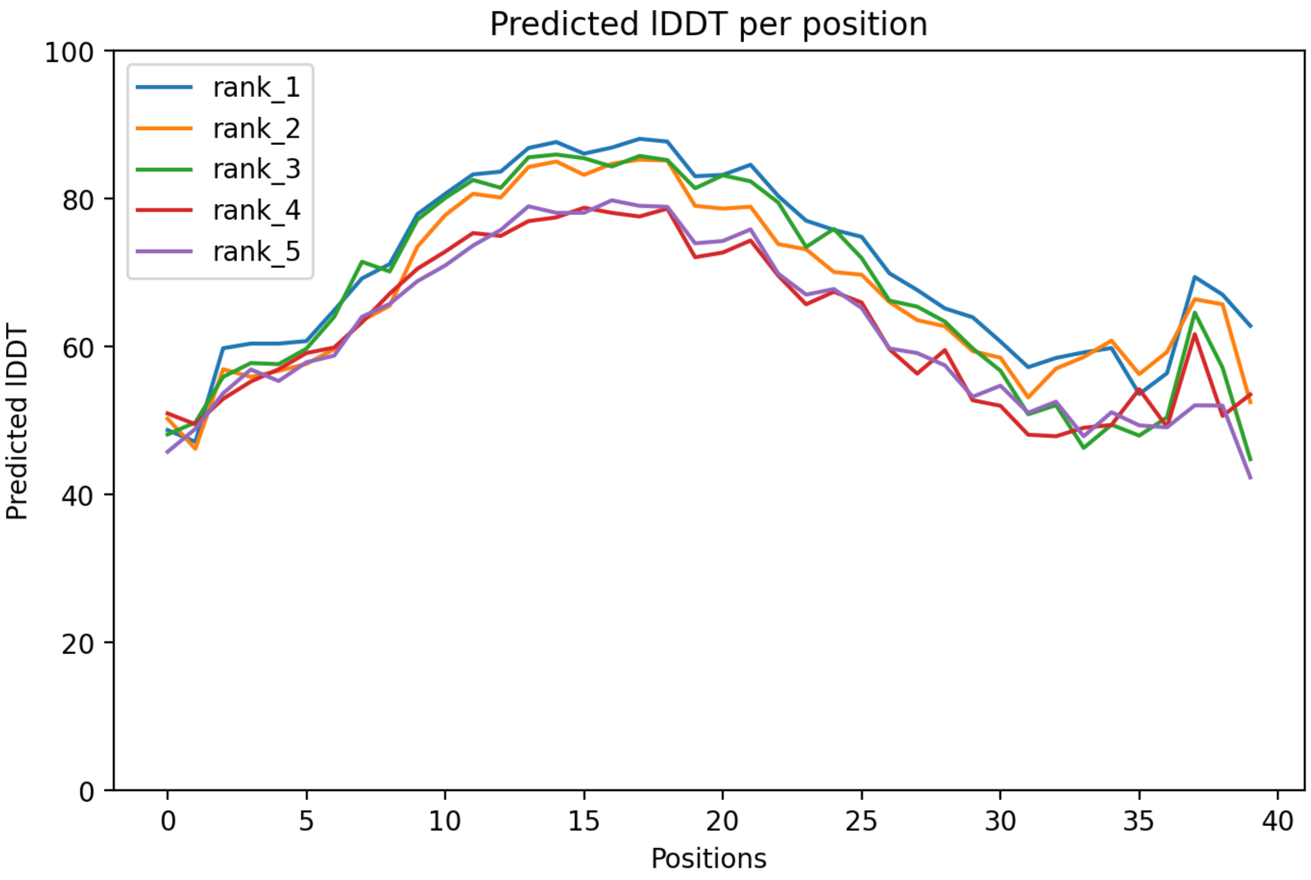

### CP85: MtrunA17_Chr5g0415031

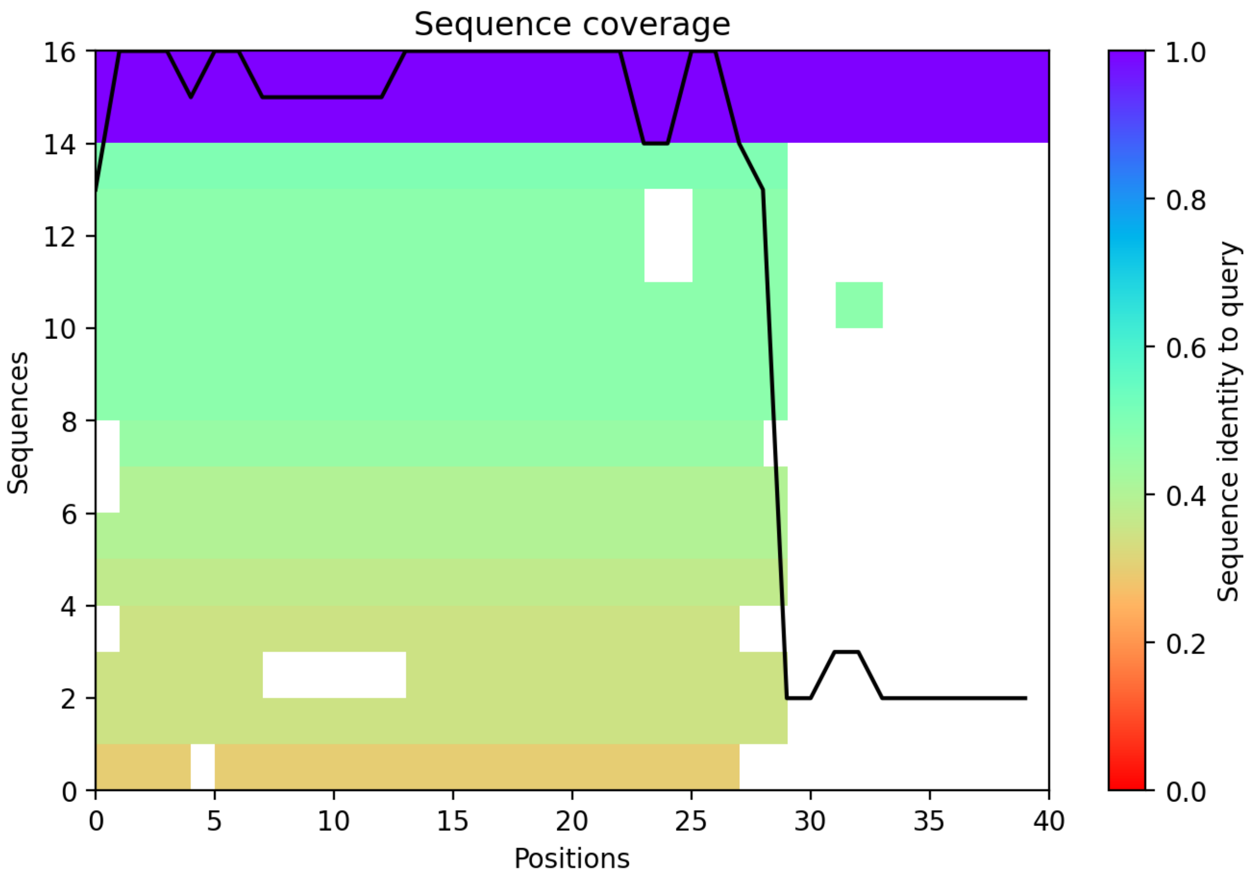

CP86: MtrunA17_Chr5g0415911

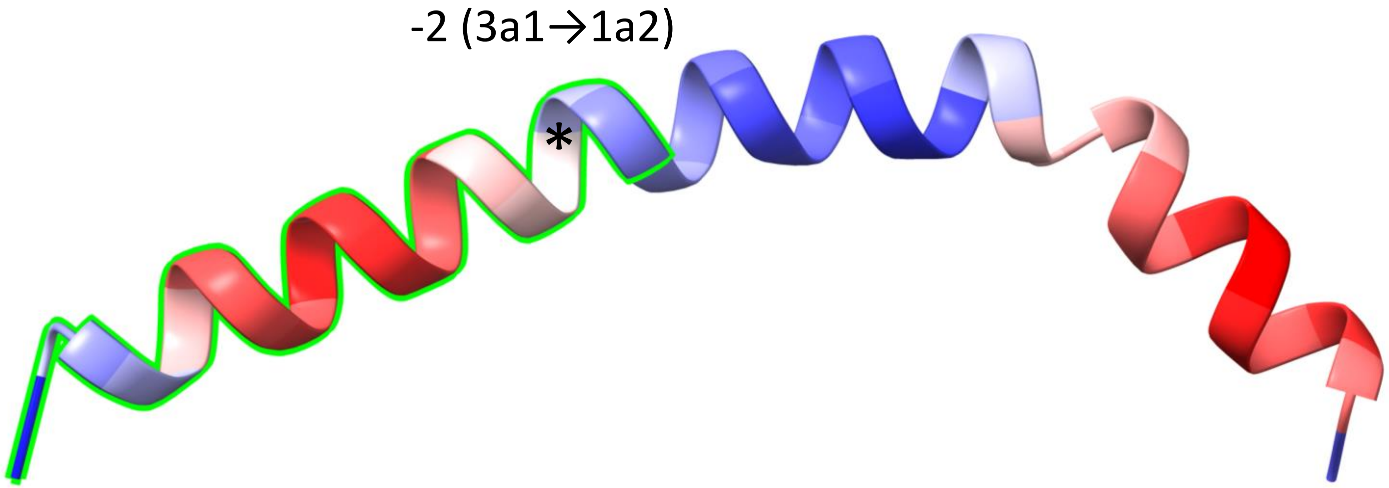


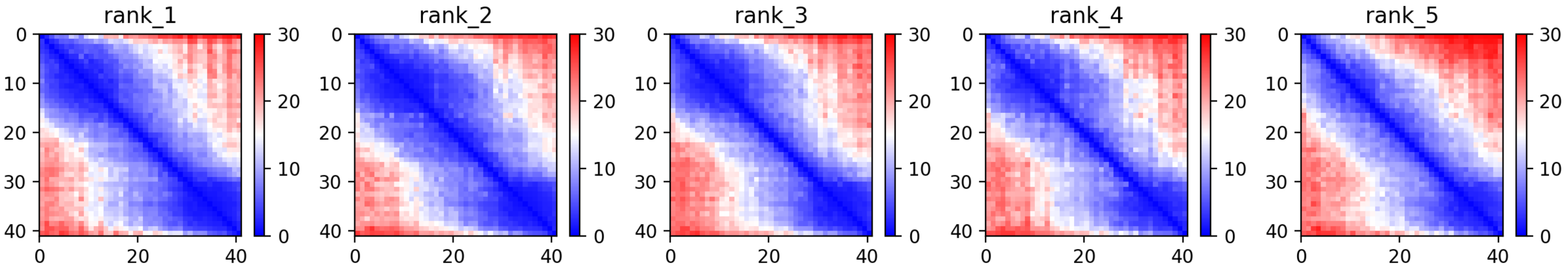

### CP86: MtrunA17_Chr5g0415911

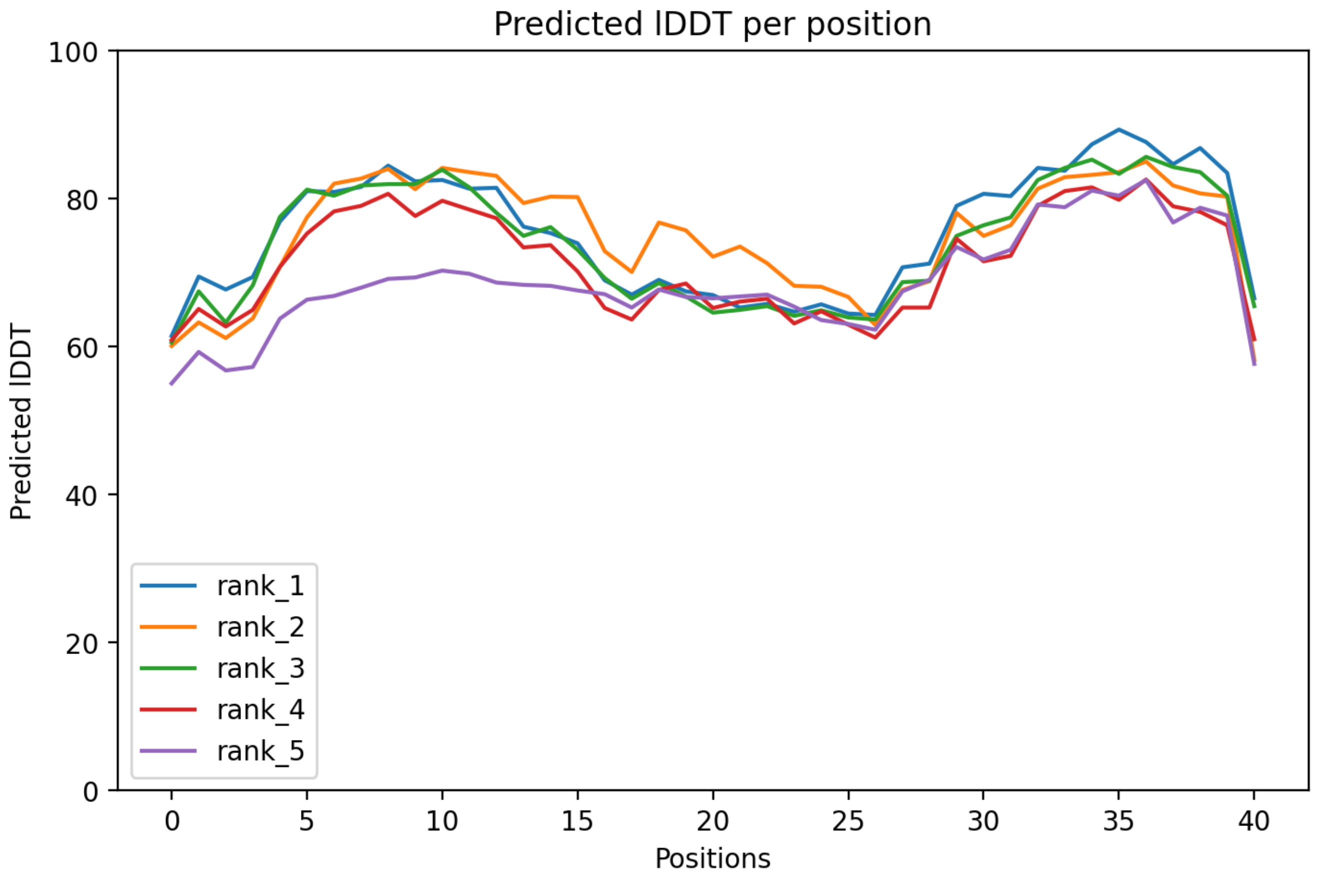

### CP86: MtrunA17_Chr5g0415911

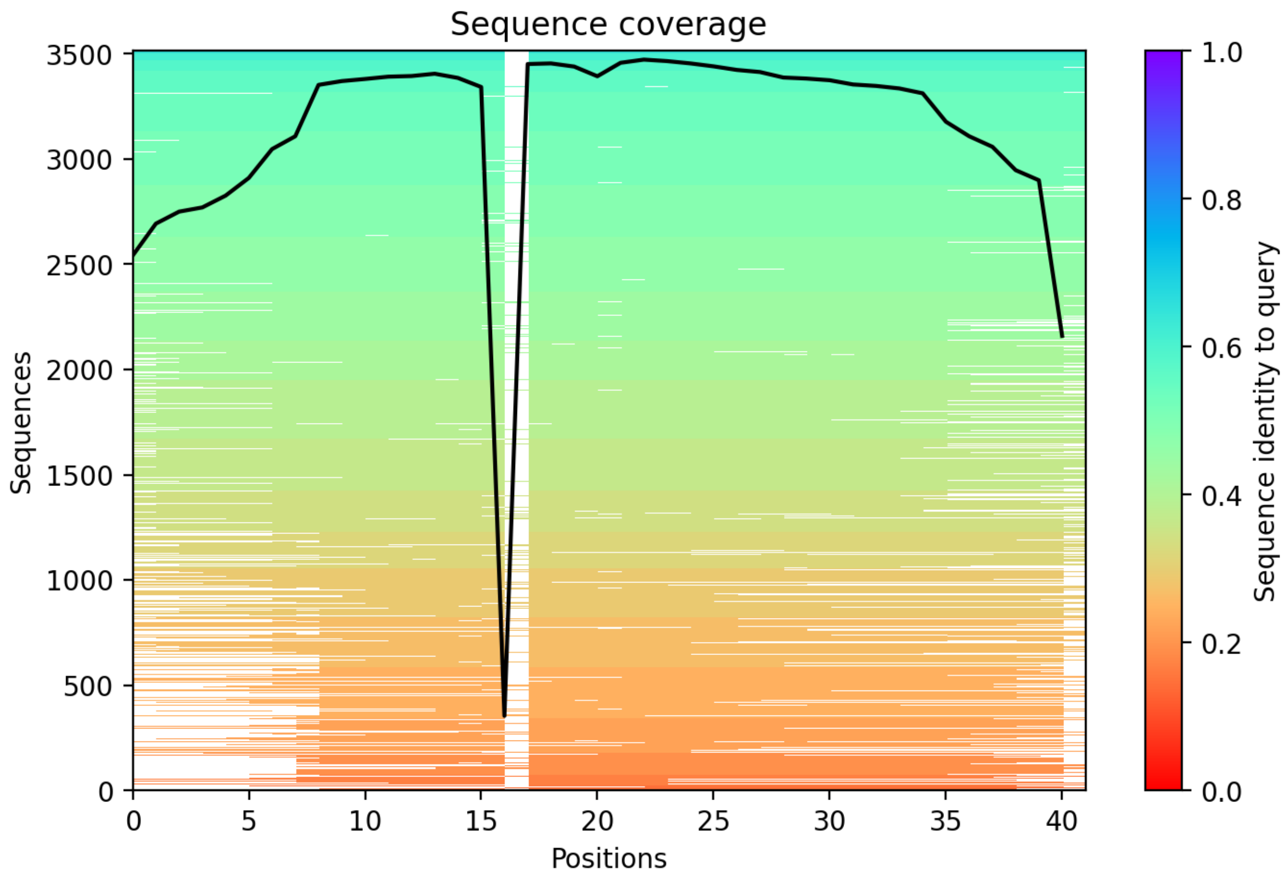

CP87: MtrunA17_Chr5g0421761

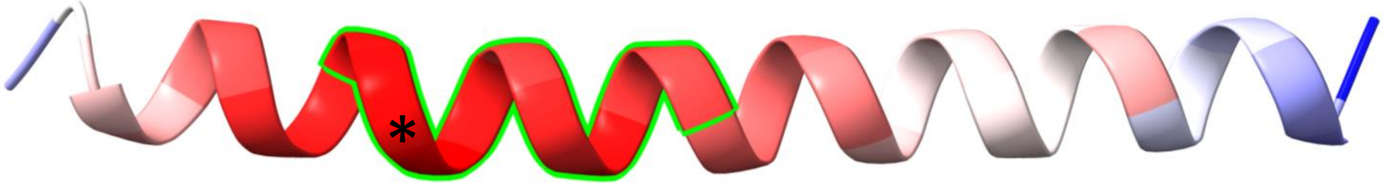


-1 (3a1→2a2)

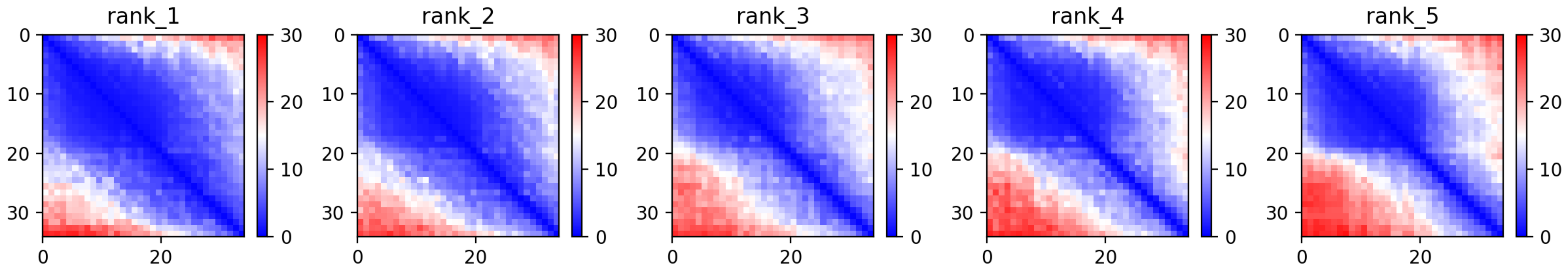

CP87: MtrunA17_Chr5g0421761

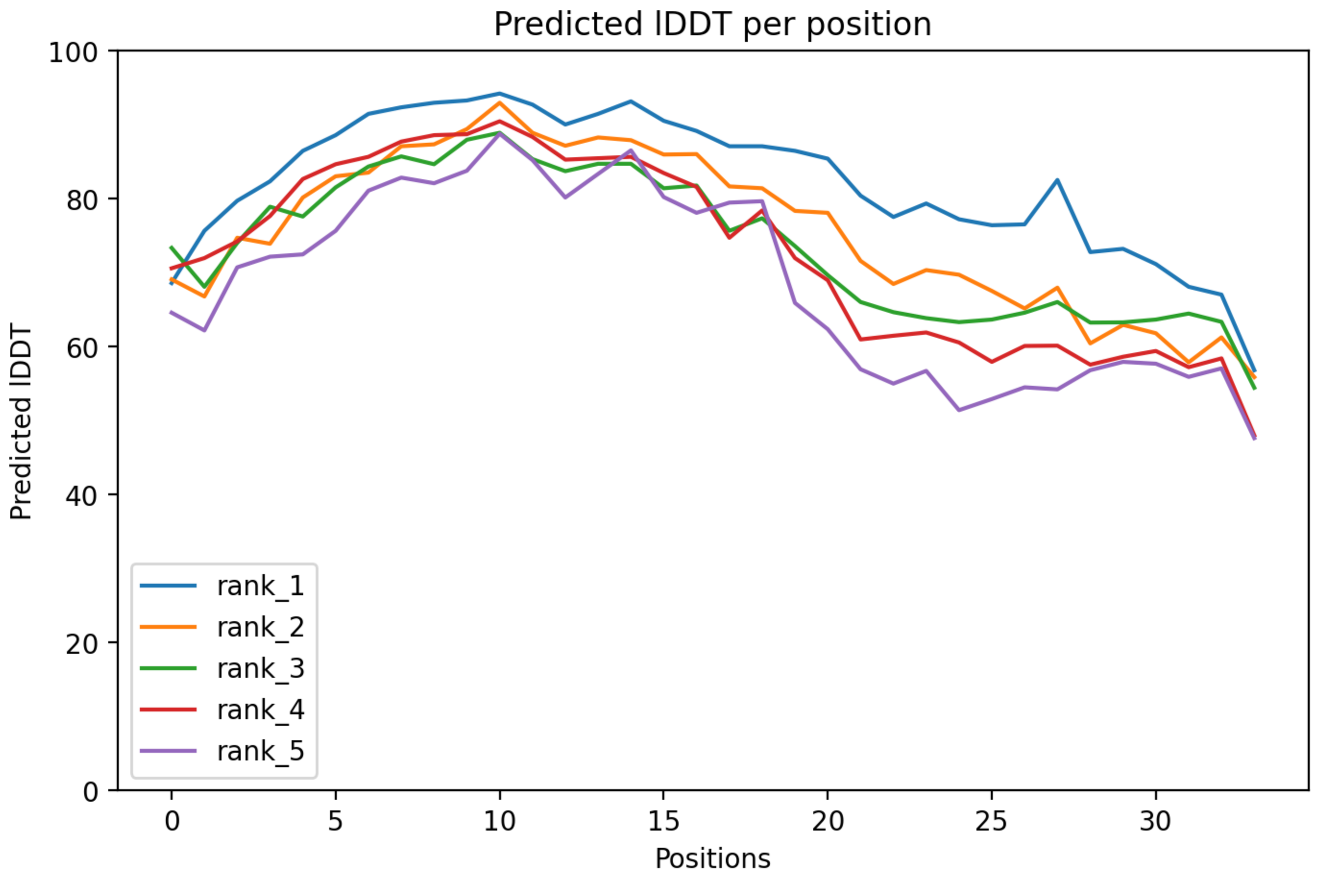

CP87: MtrunA17_Chr5g0421761

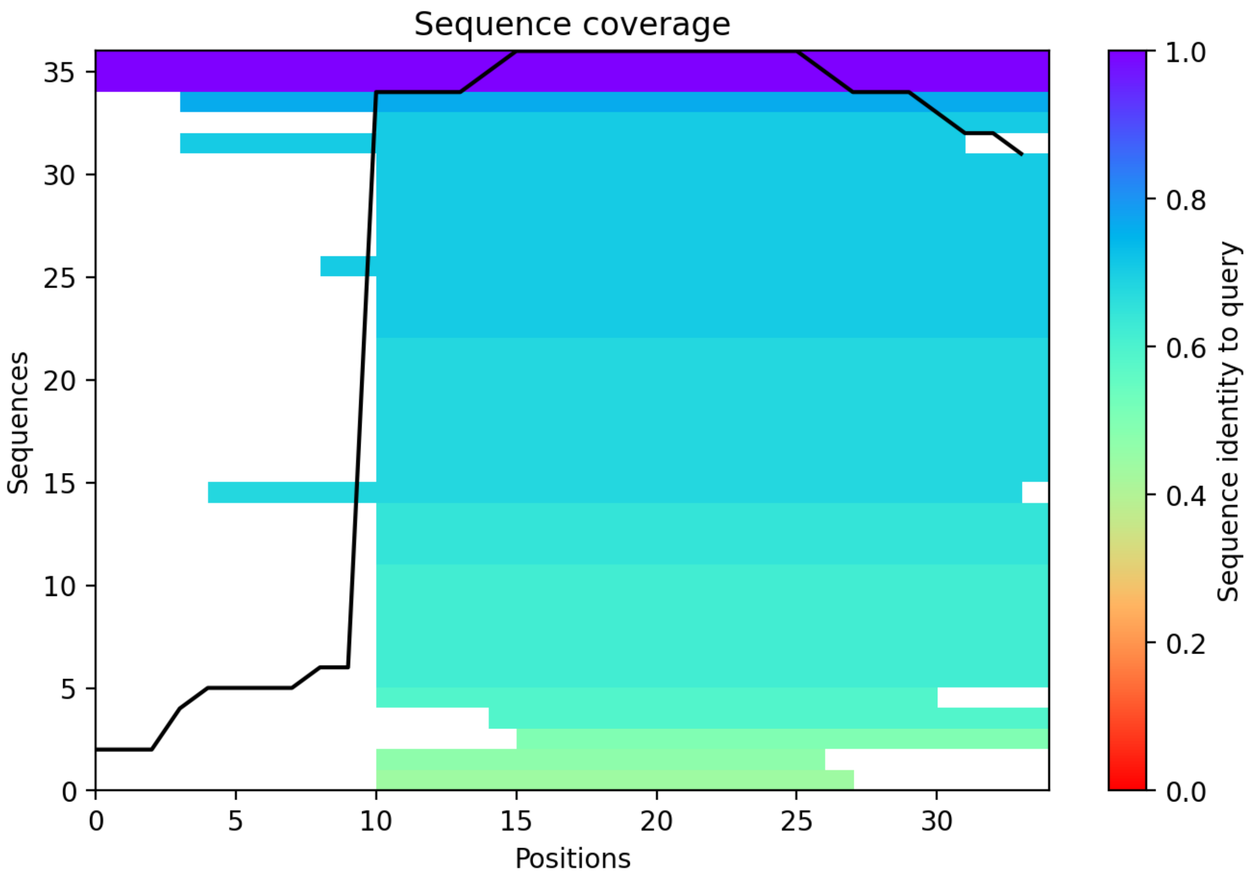

### CP88: MtrunA17_Chr5g0422291

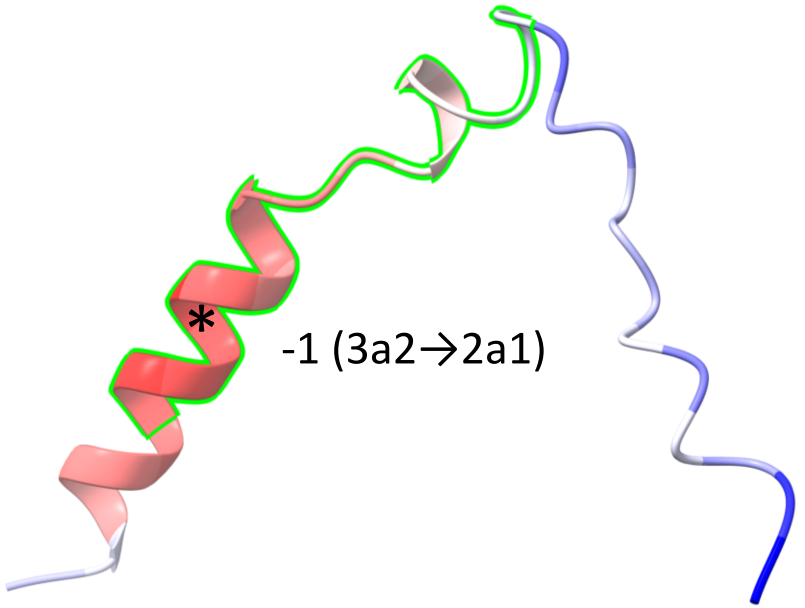


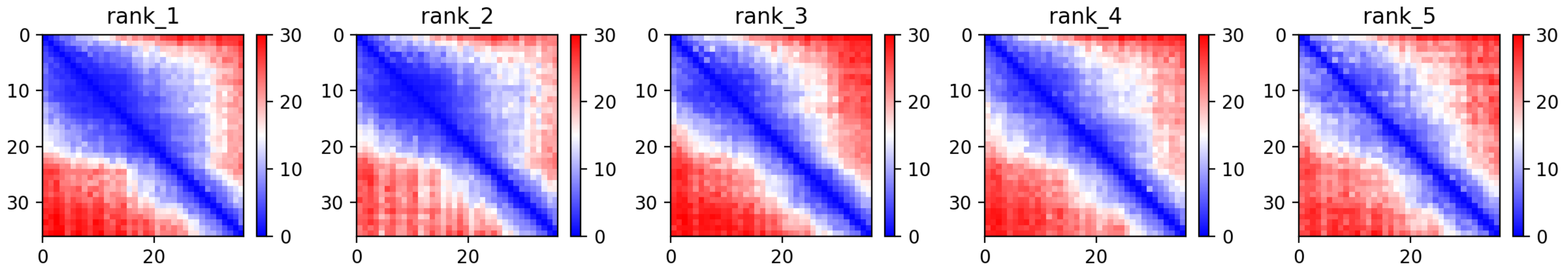

CP88: MtrunA17_Chr5g0422291

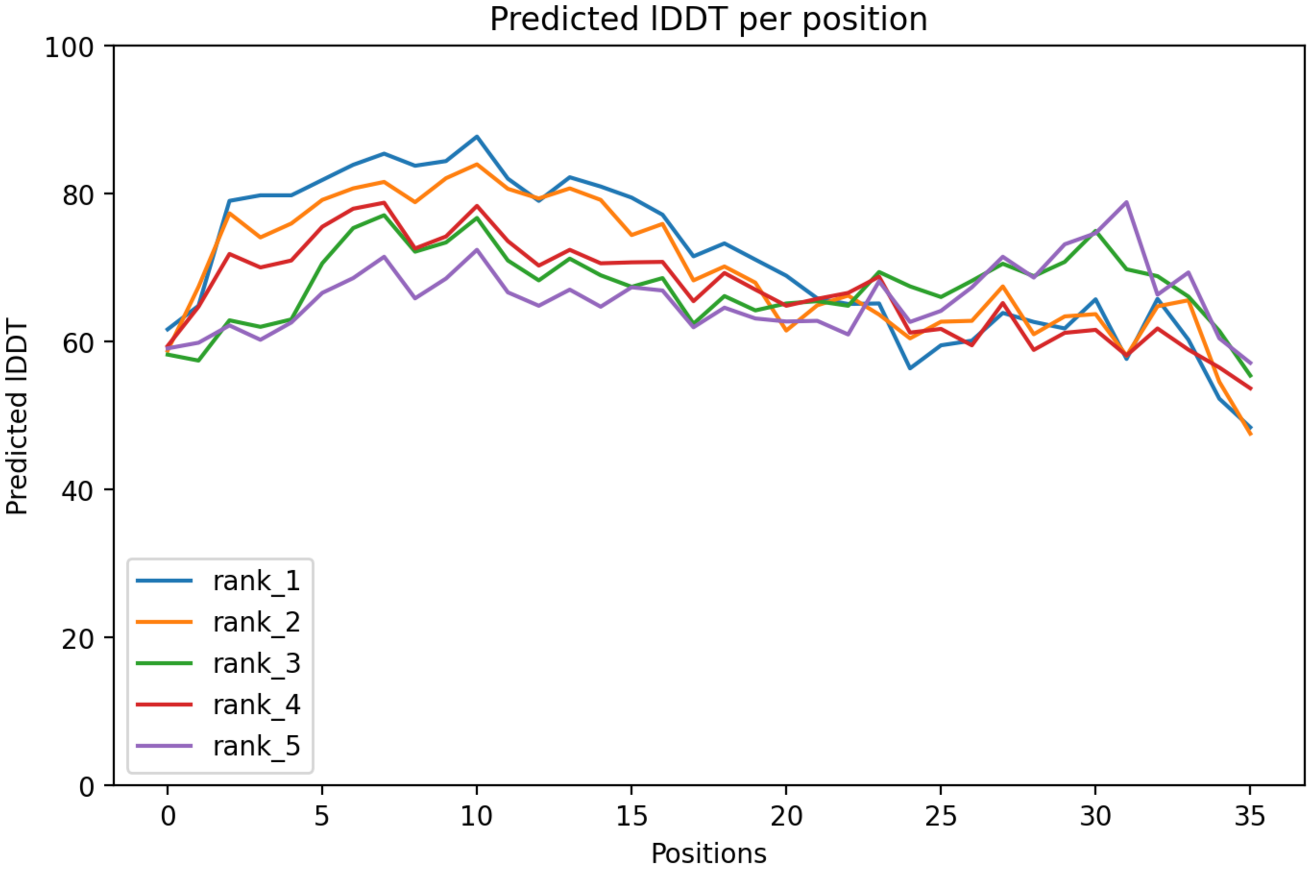

### CP88: MtrunA17_Chr5g0422291

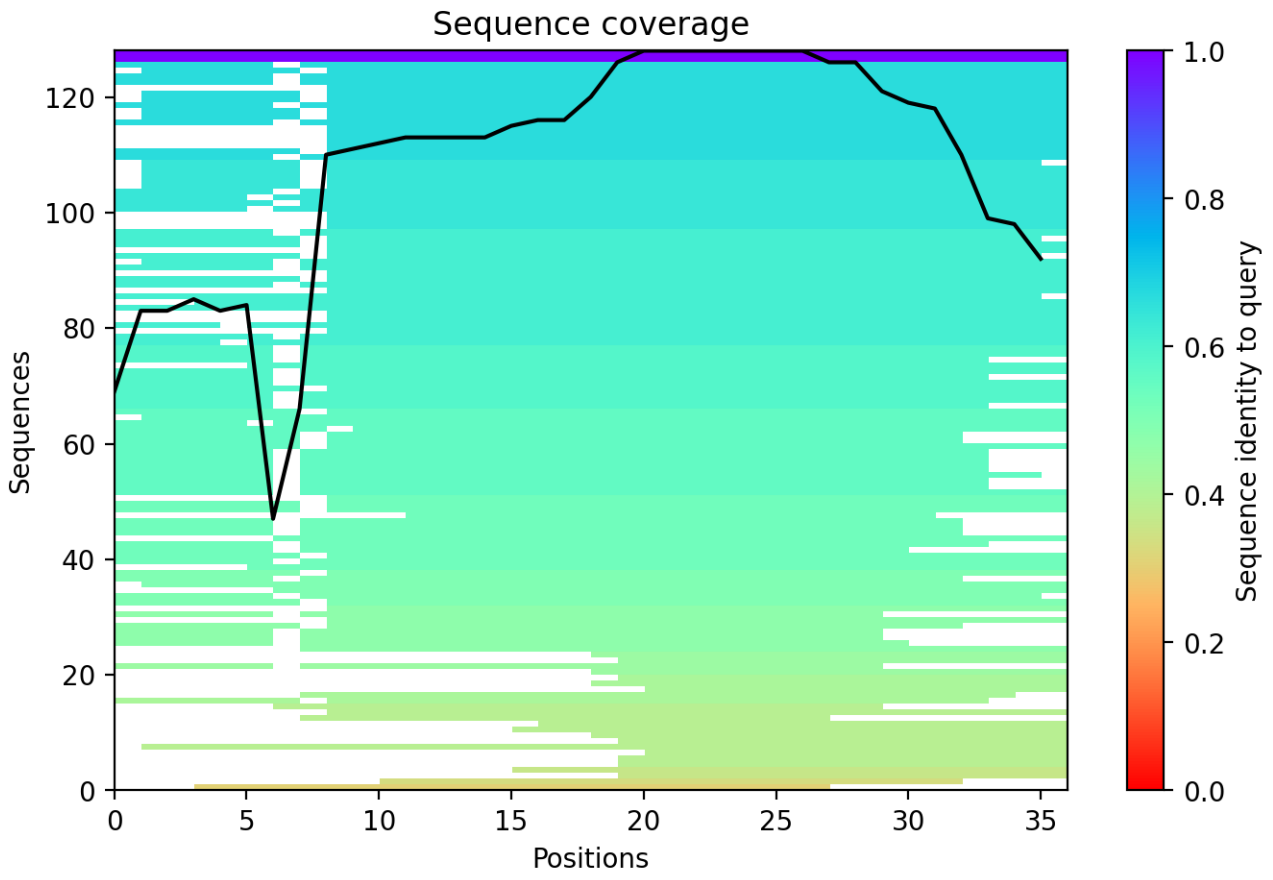

### CP89: MtrunA17_Chr5g0422291

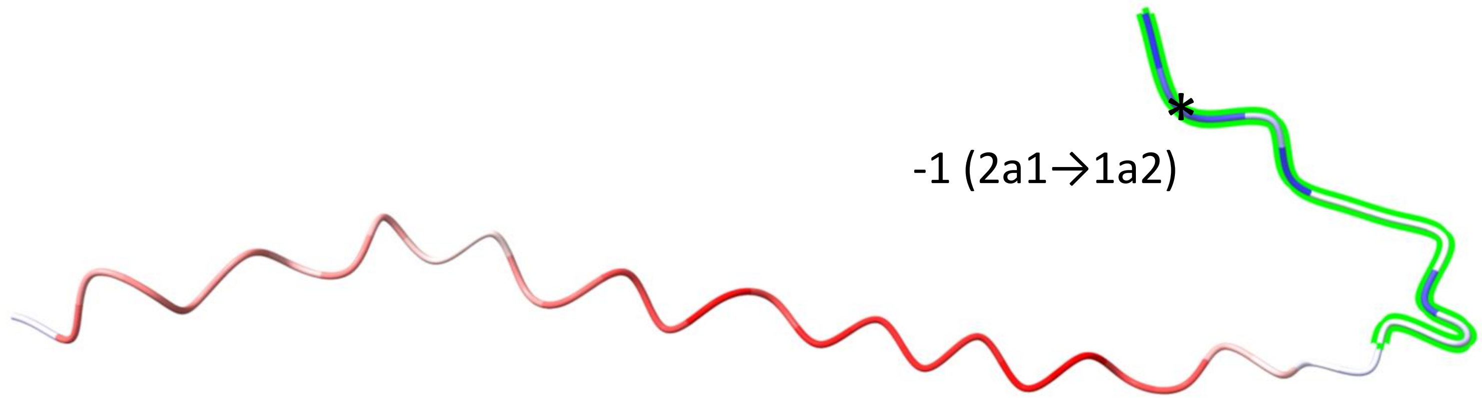


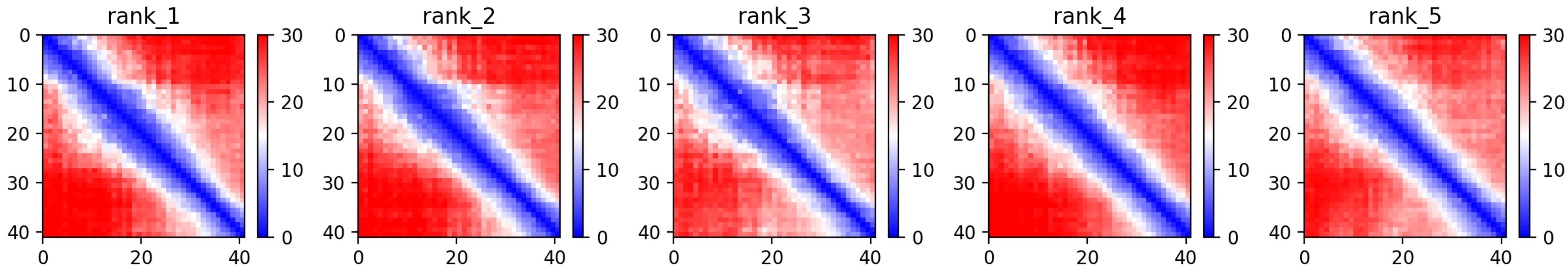

### CP89: MtrunA17_Chr5g0422291

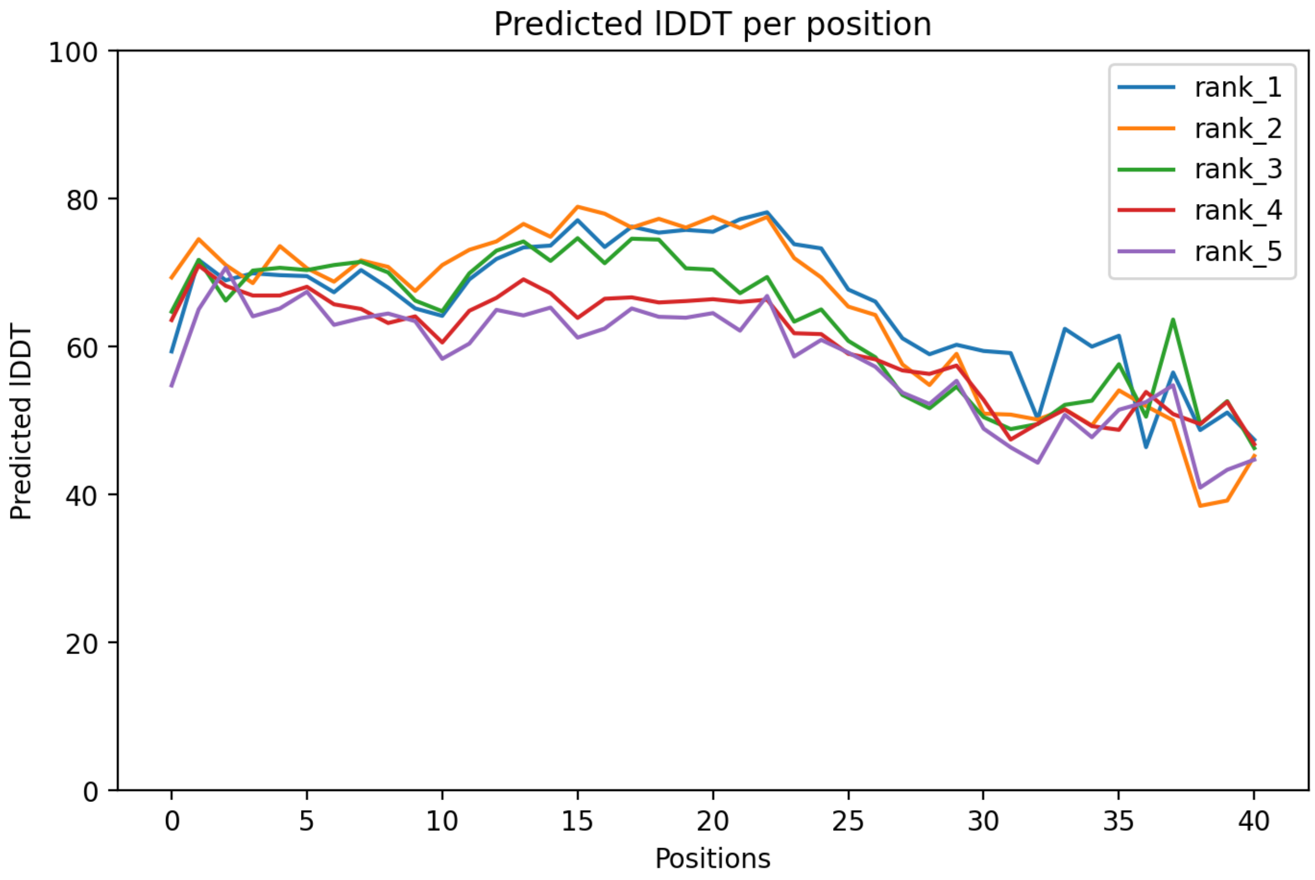

### CP89: MtrunA17_Chr5g0422291

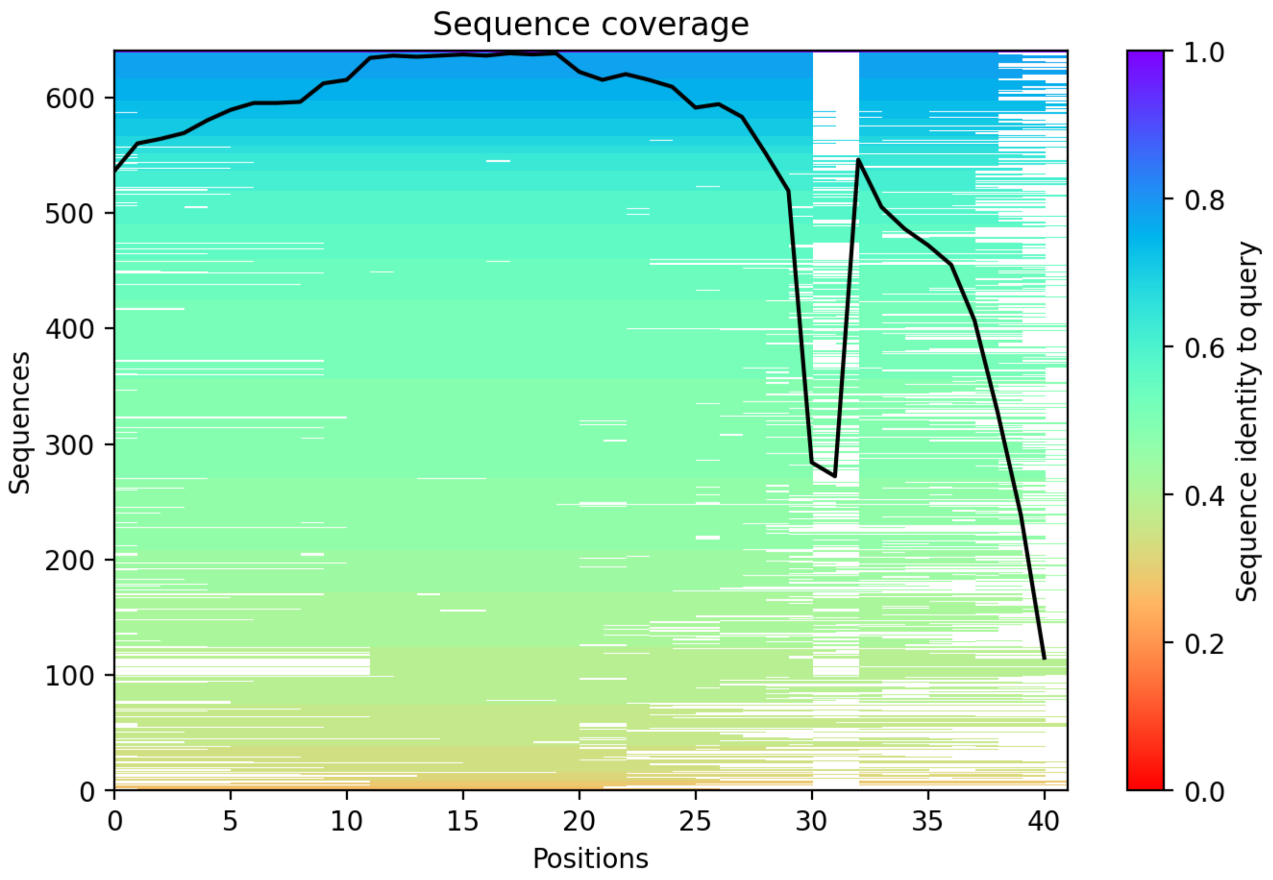

CP90: MtrunA17_Chr5g0430341

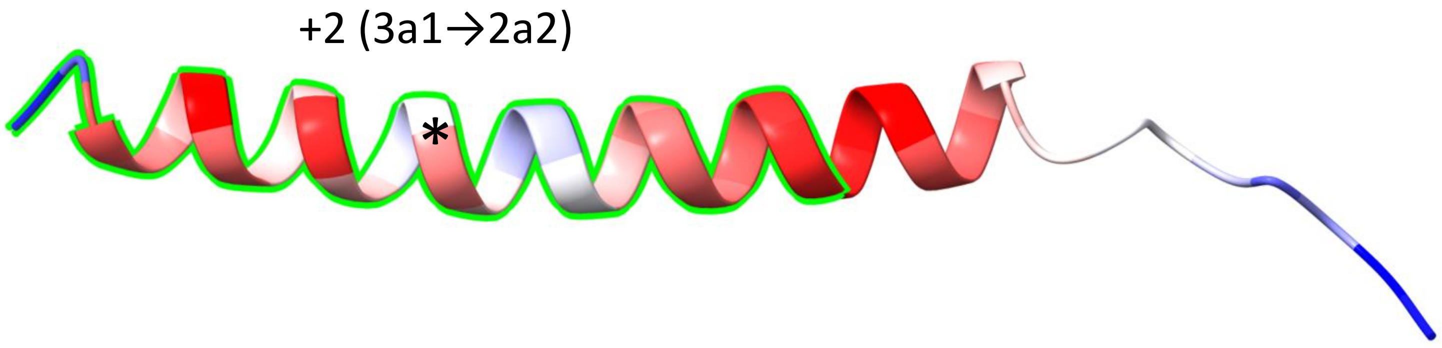


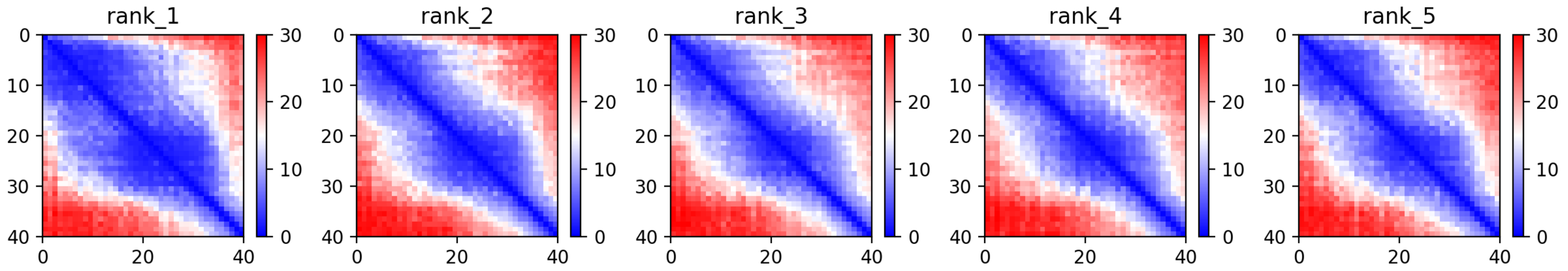

CP90: MtrunA17_Chr5g0430341

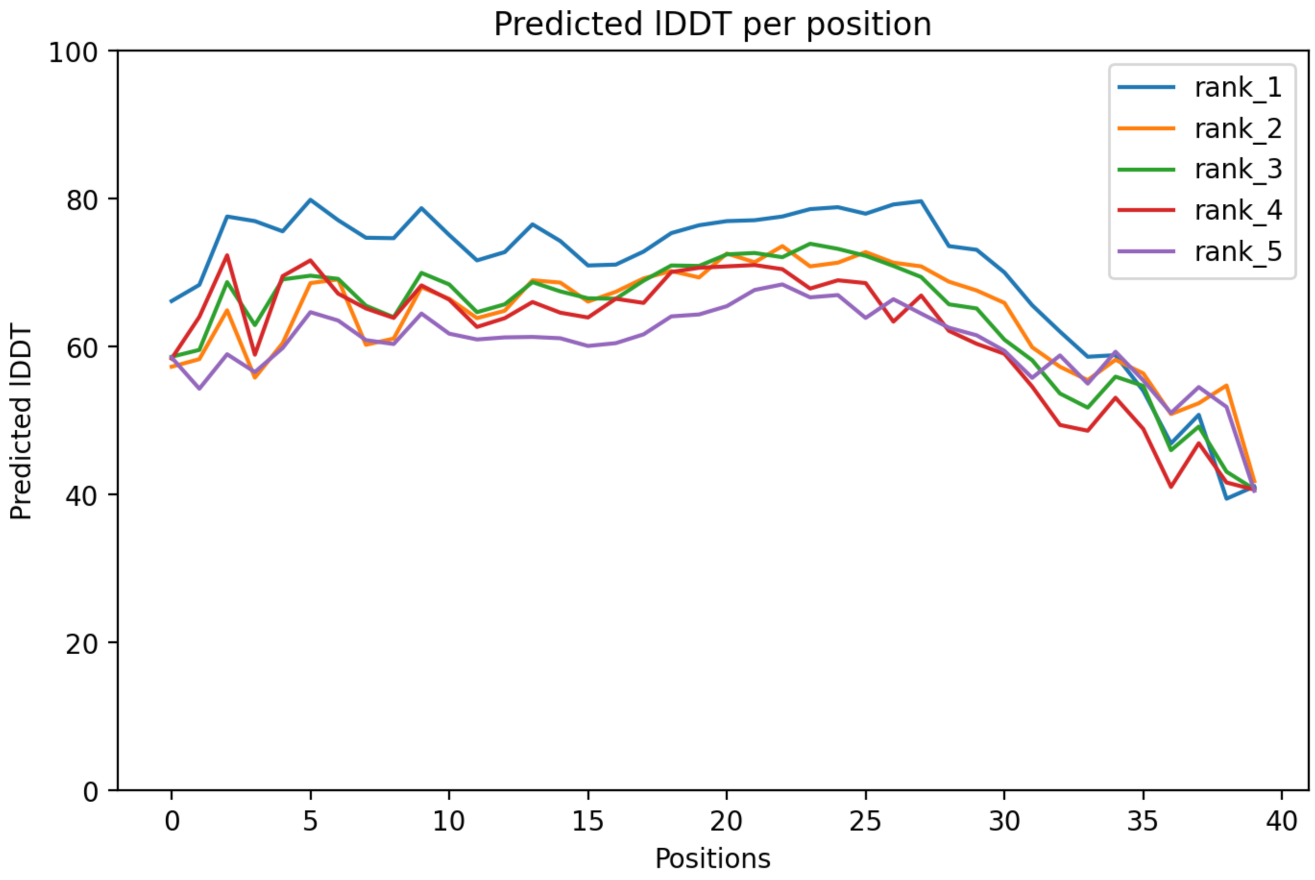

CP90: MtrunA17_Chr5g0430341

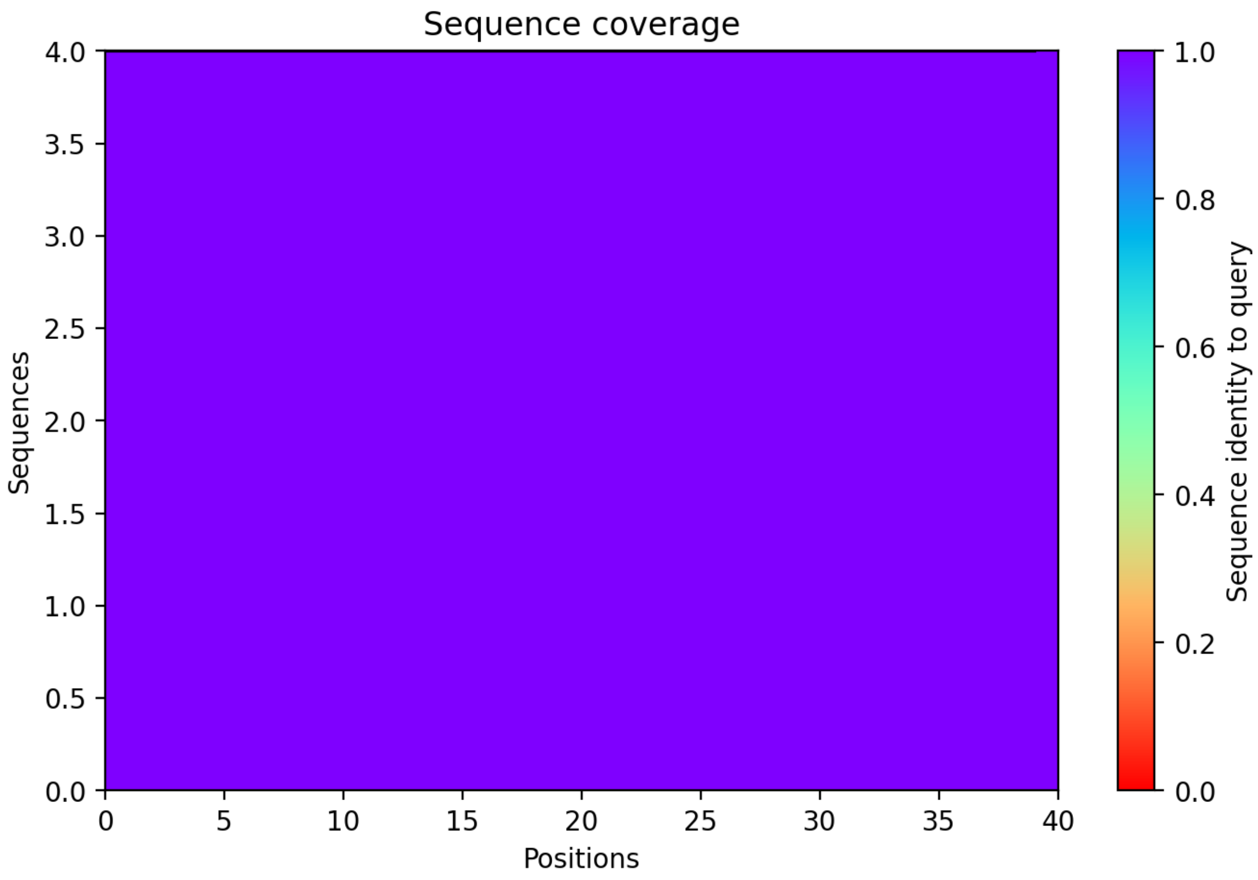

CP91: MtrunA17_Chr5g0430341

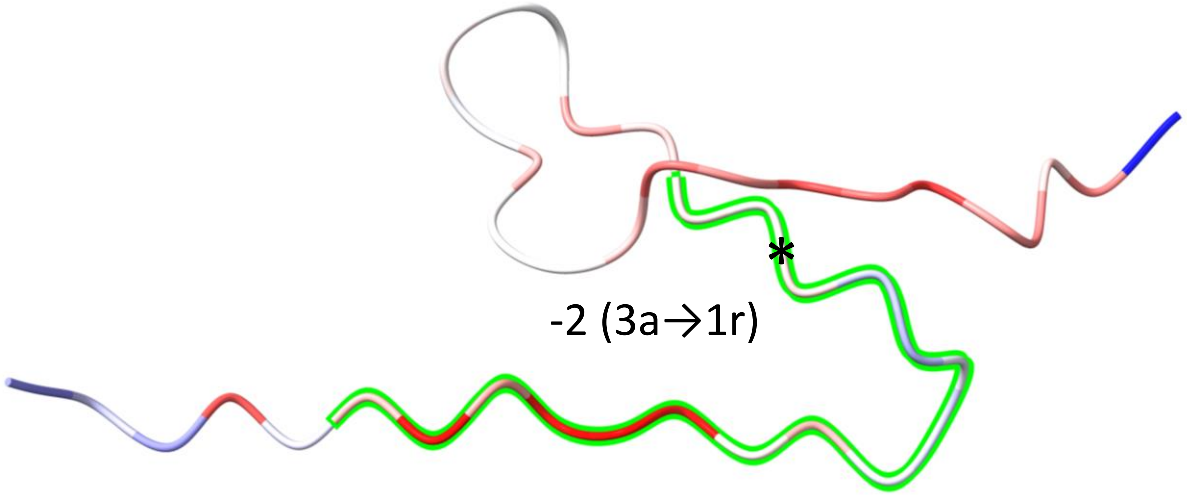


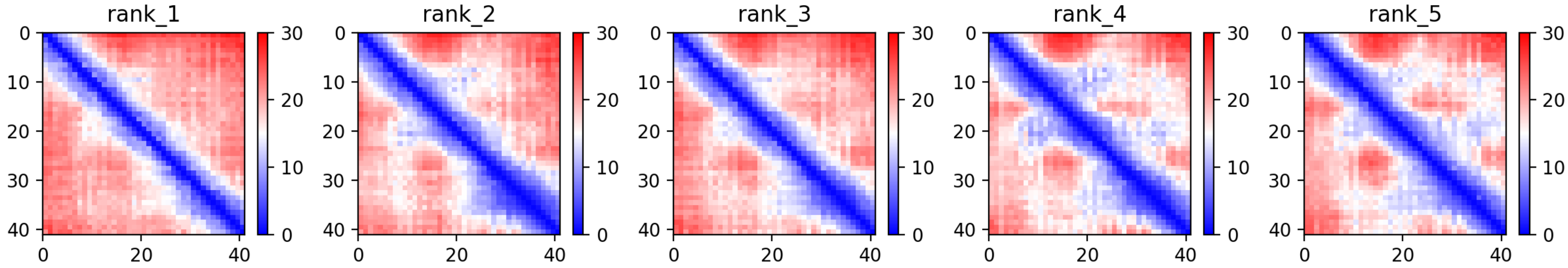

CP91: MtrunA17_Chr5g0430341

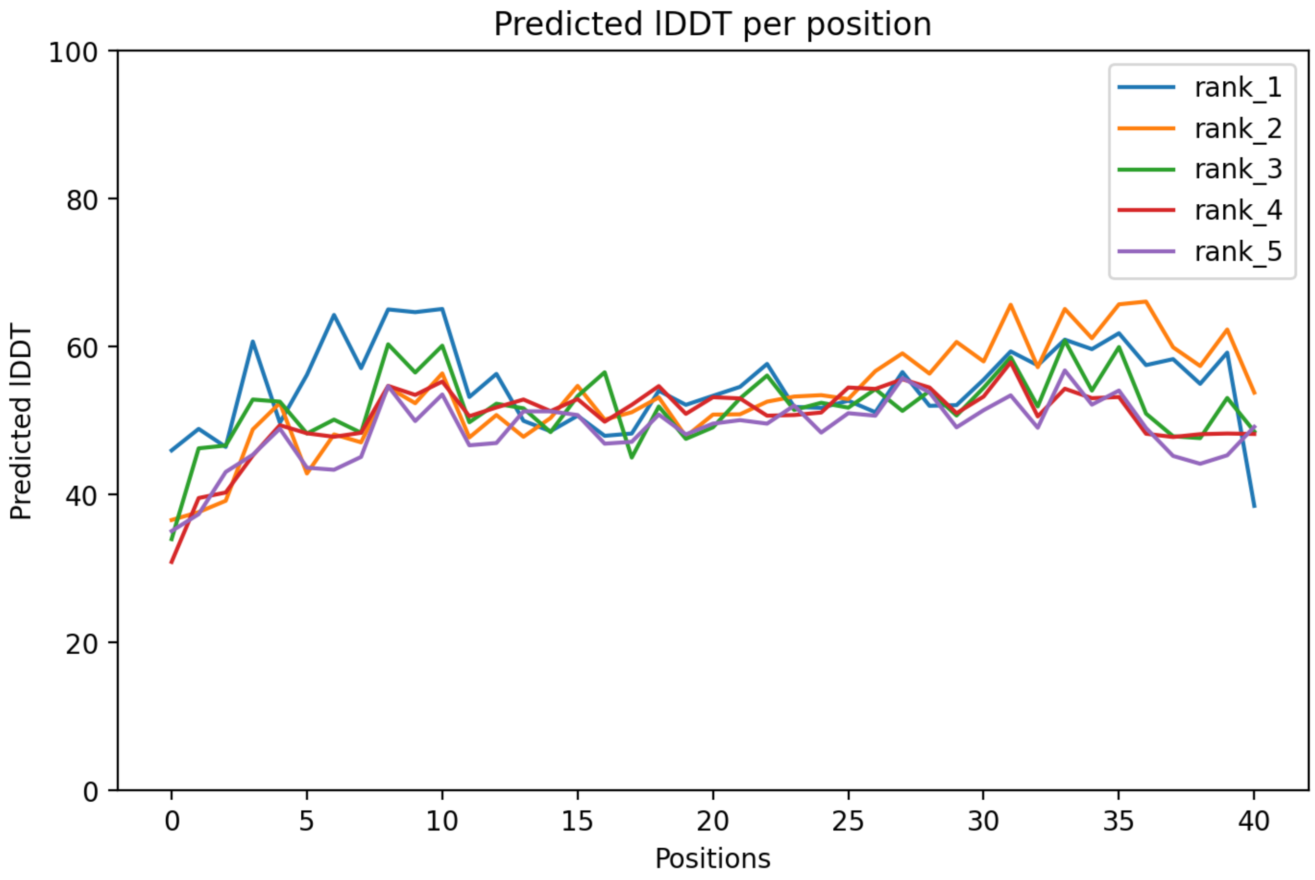

### CP91: MtrunA17_Chr5g0430341

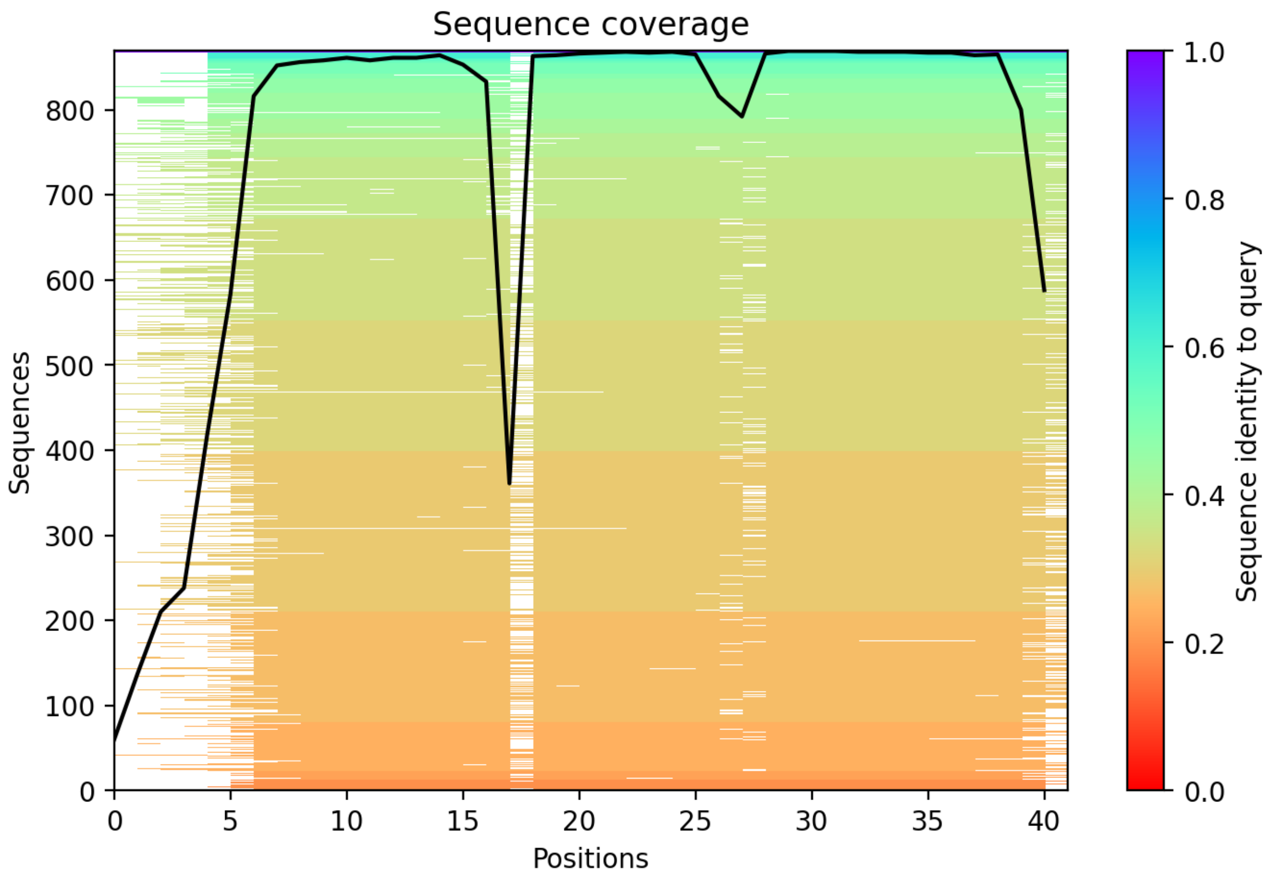

### CP92: MtrunA17_Chr5g0431401

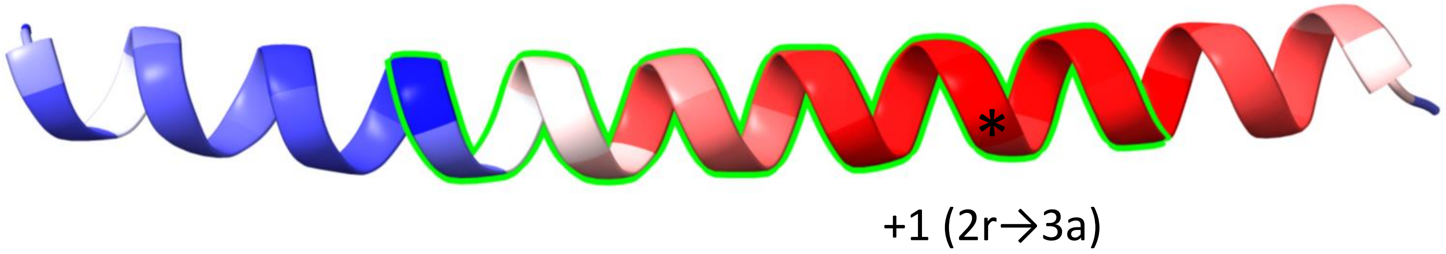


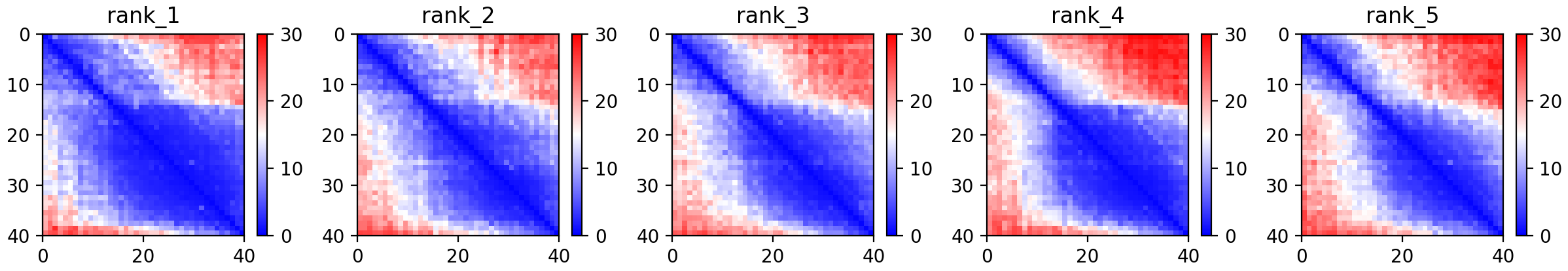

CP92: MtrunA17_Chr5g0431401

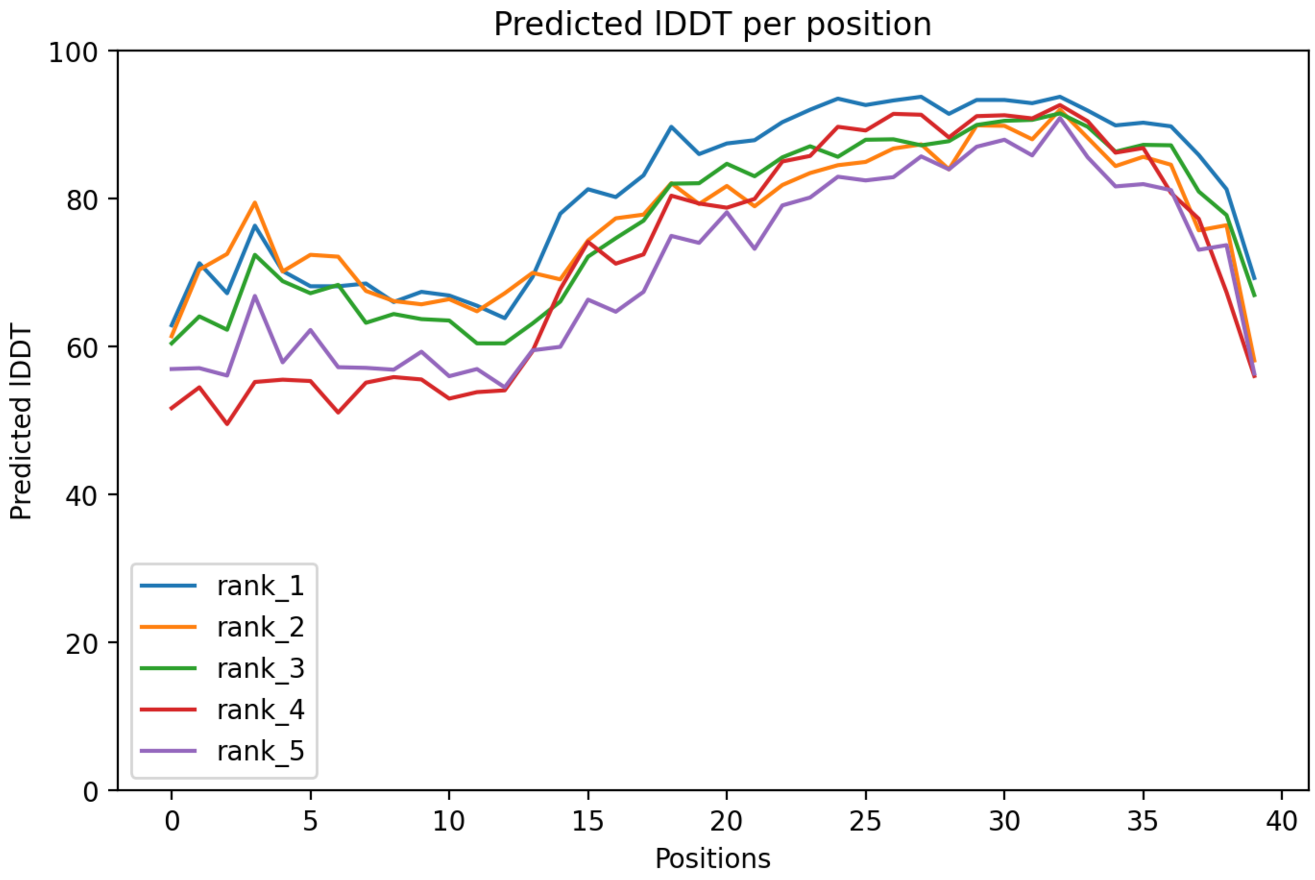

### CP92: MtrunA17_Chr5g0431401

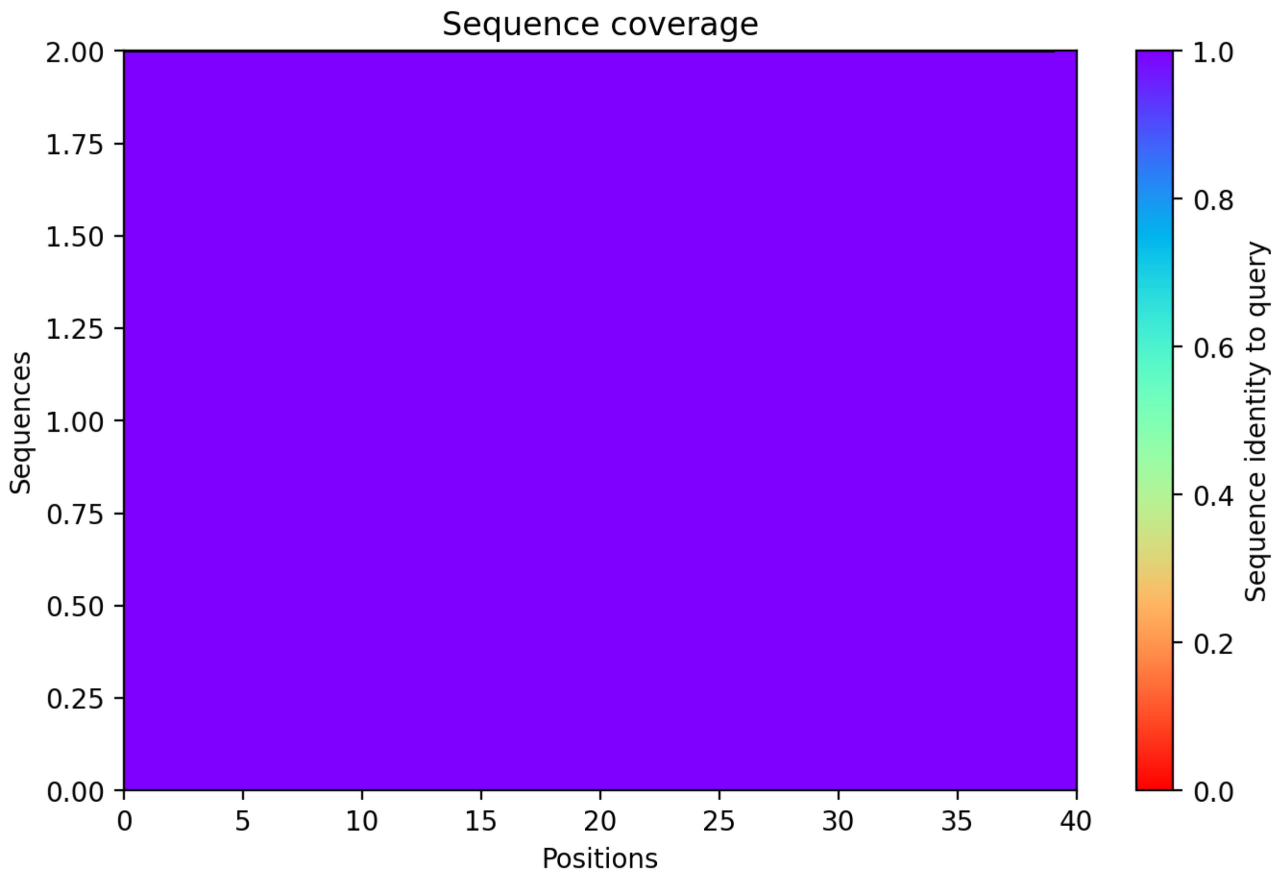

CP93: MtrunA17_Chr5g0435191

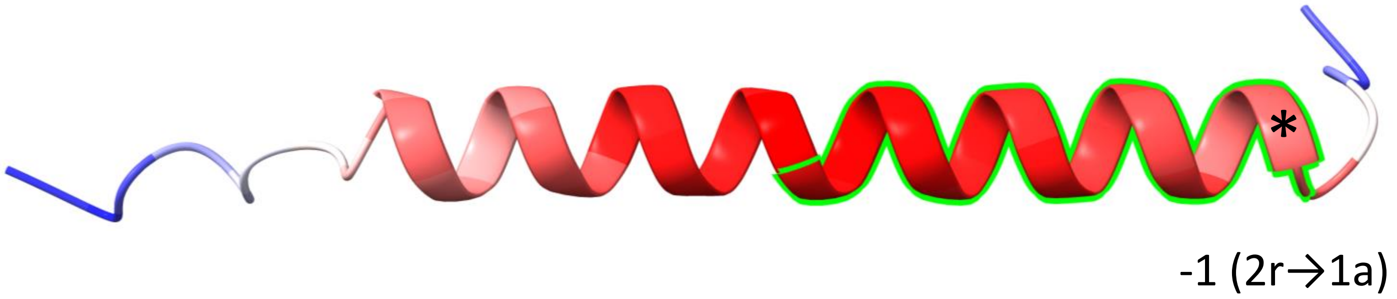


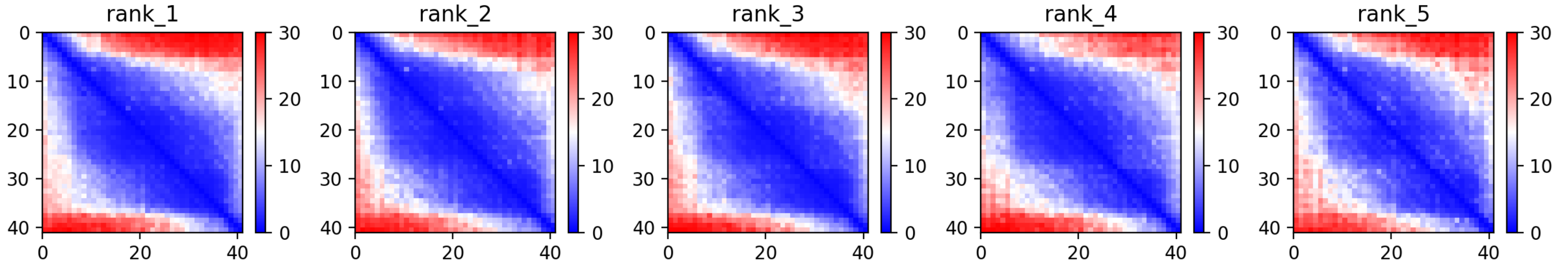

### CP93: MtrunA17_Chr5g0435191

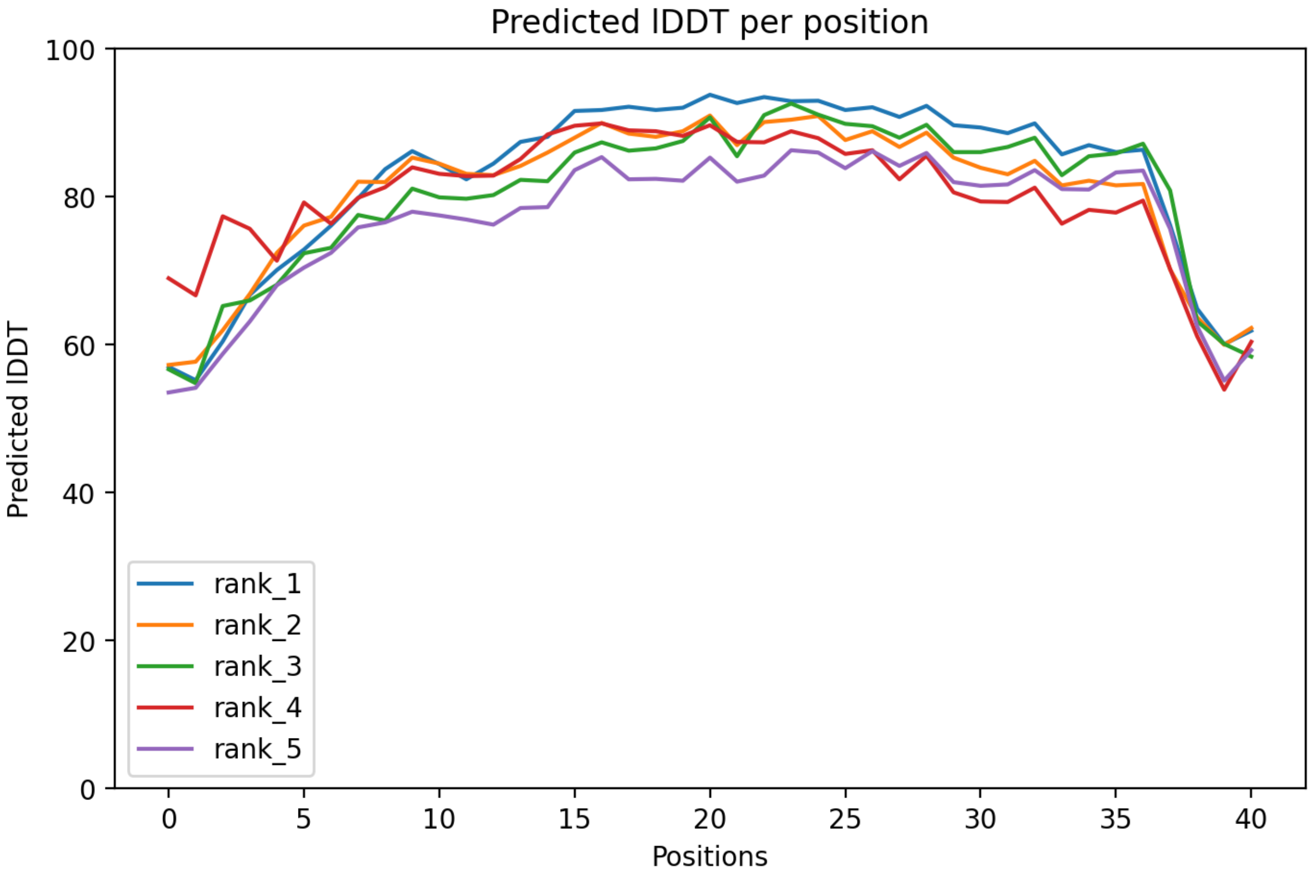

### CP93: MtrunA17_Chr5g0435191

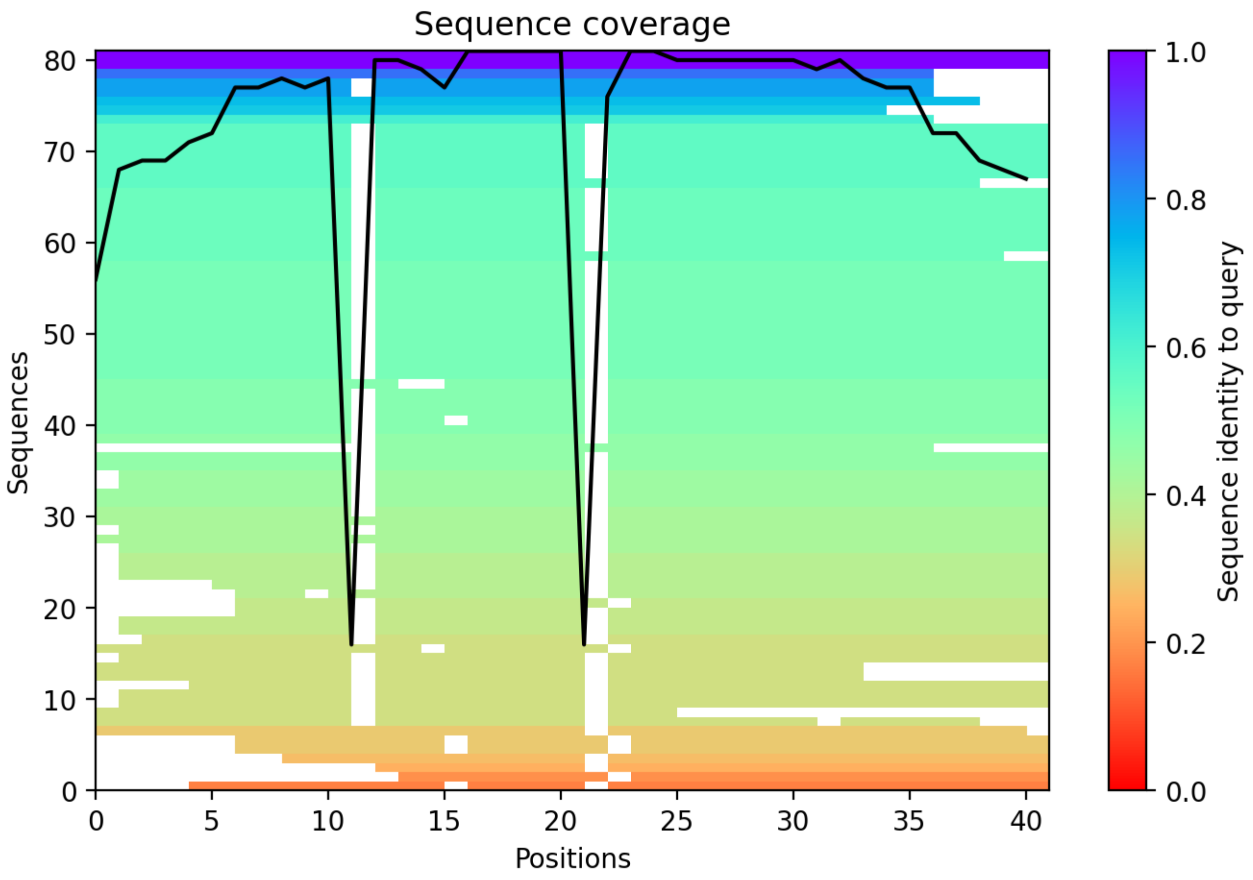

CP94: MtrunA17_Chr5g0444231

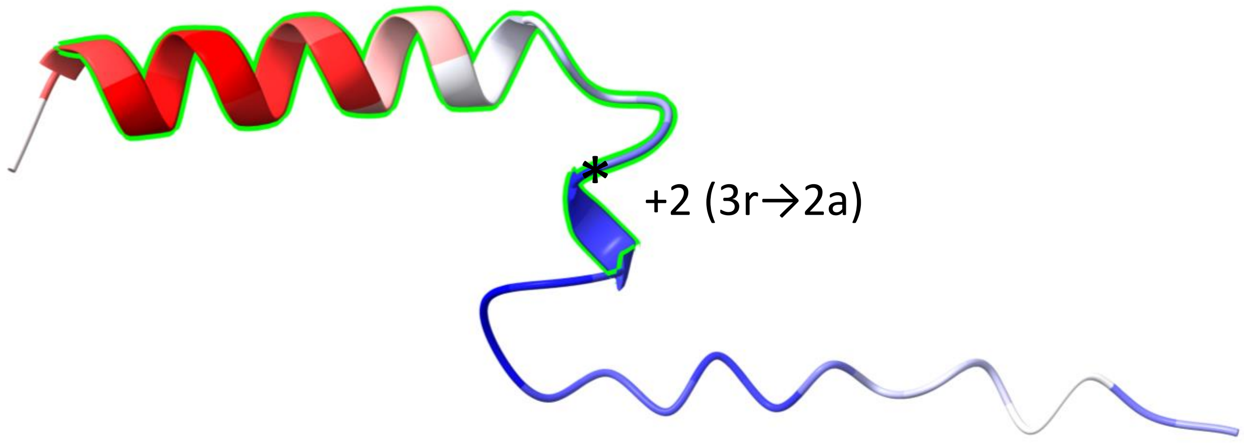


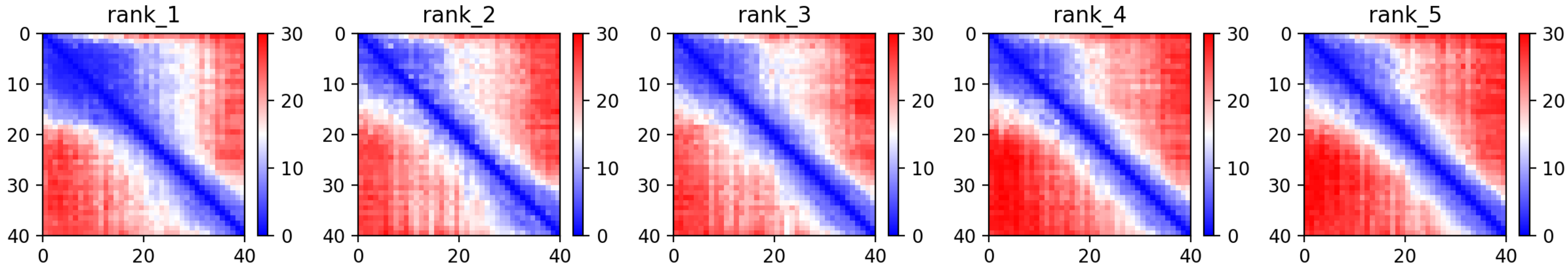

CP94: MtrunA17_Chr5g0444231

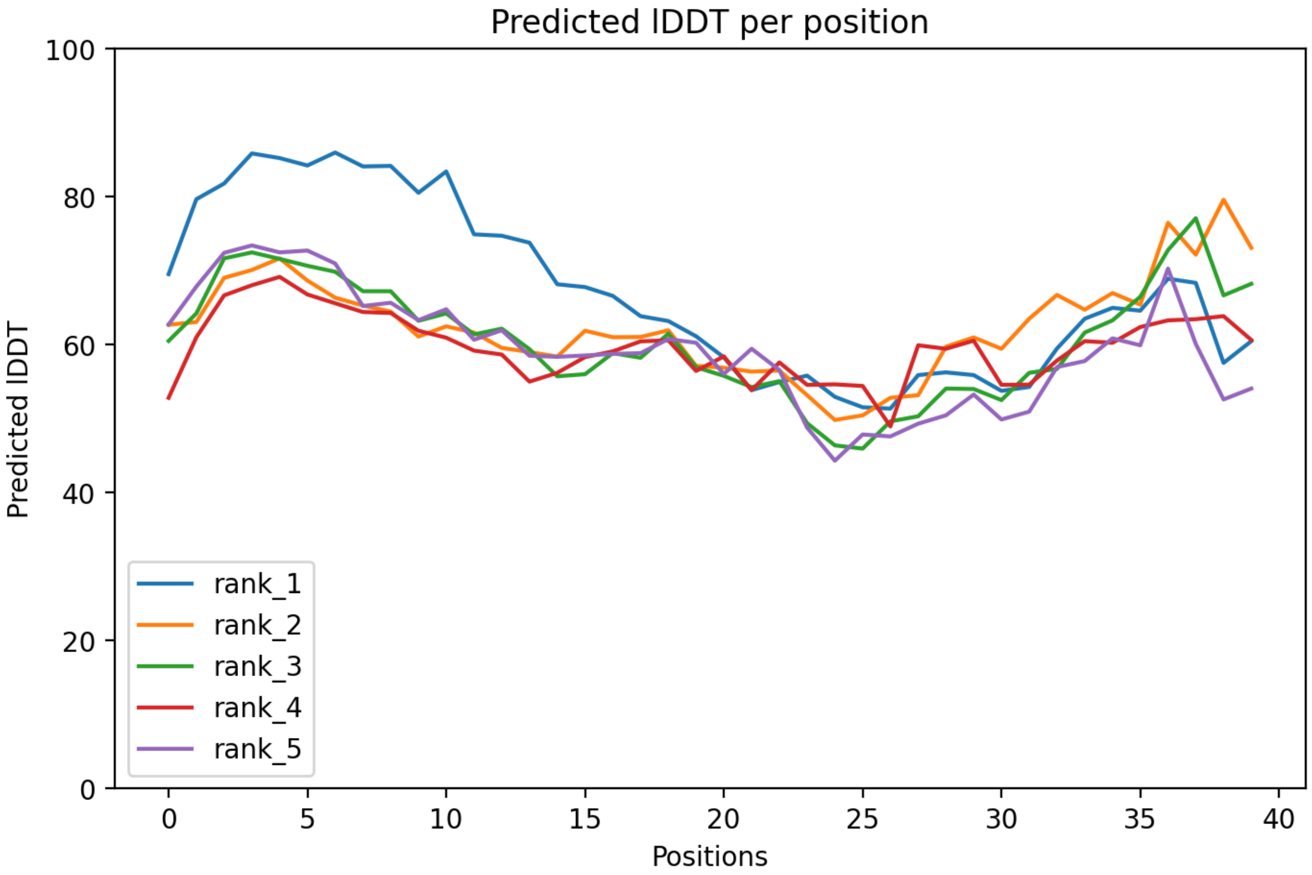

CP94: MtrunA17_Chr5g0444231

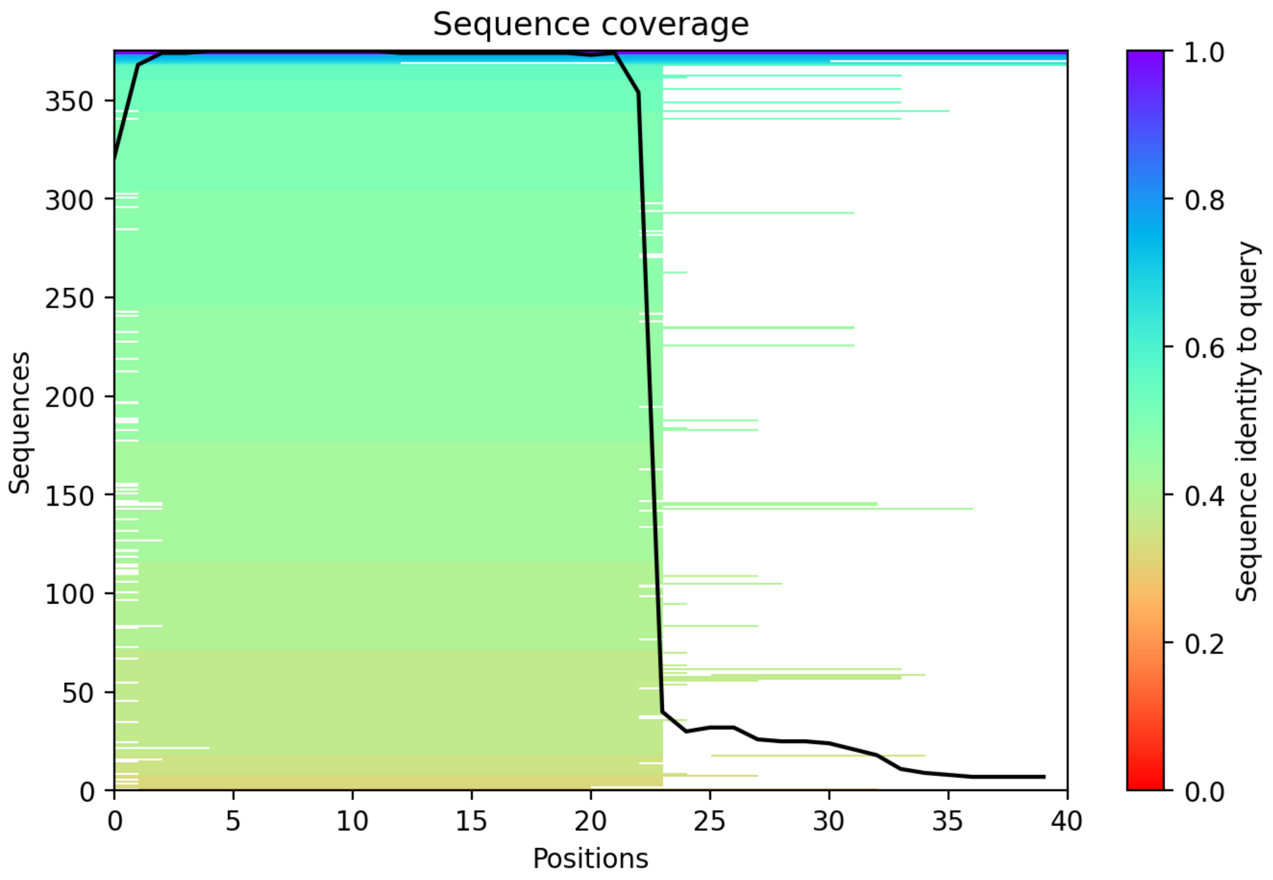

CP95: MtrunA17_Chr6g0451601

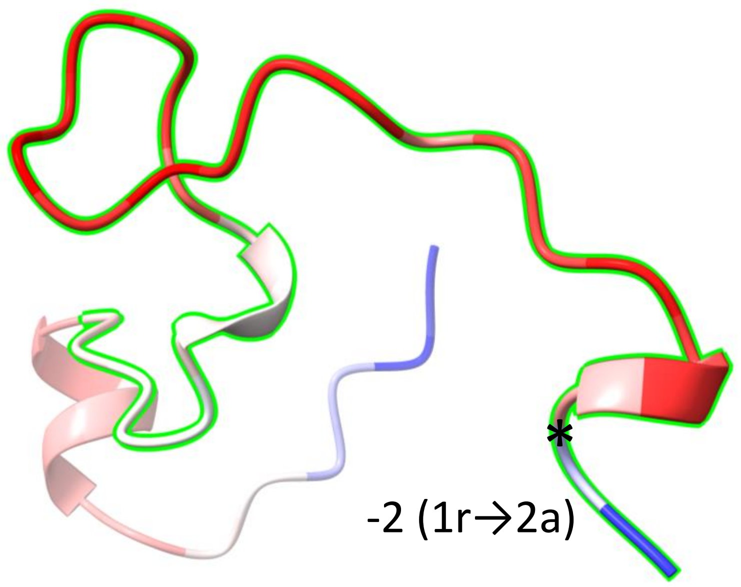


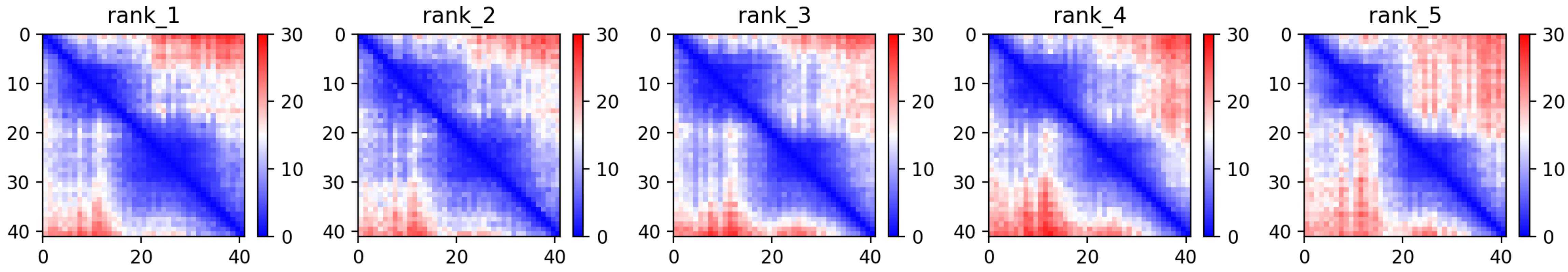

### CP95: MtrunA17_Chr6g0451601

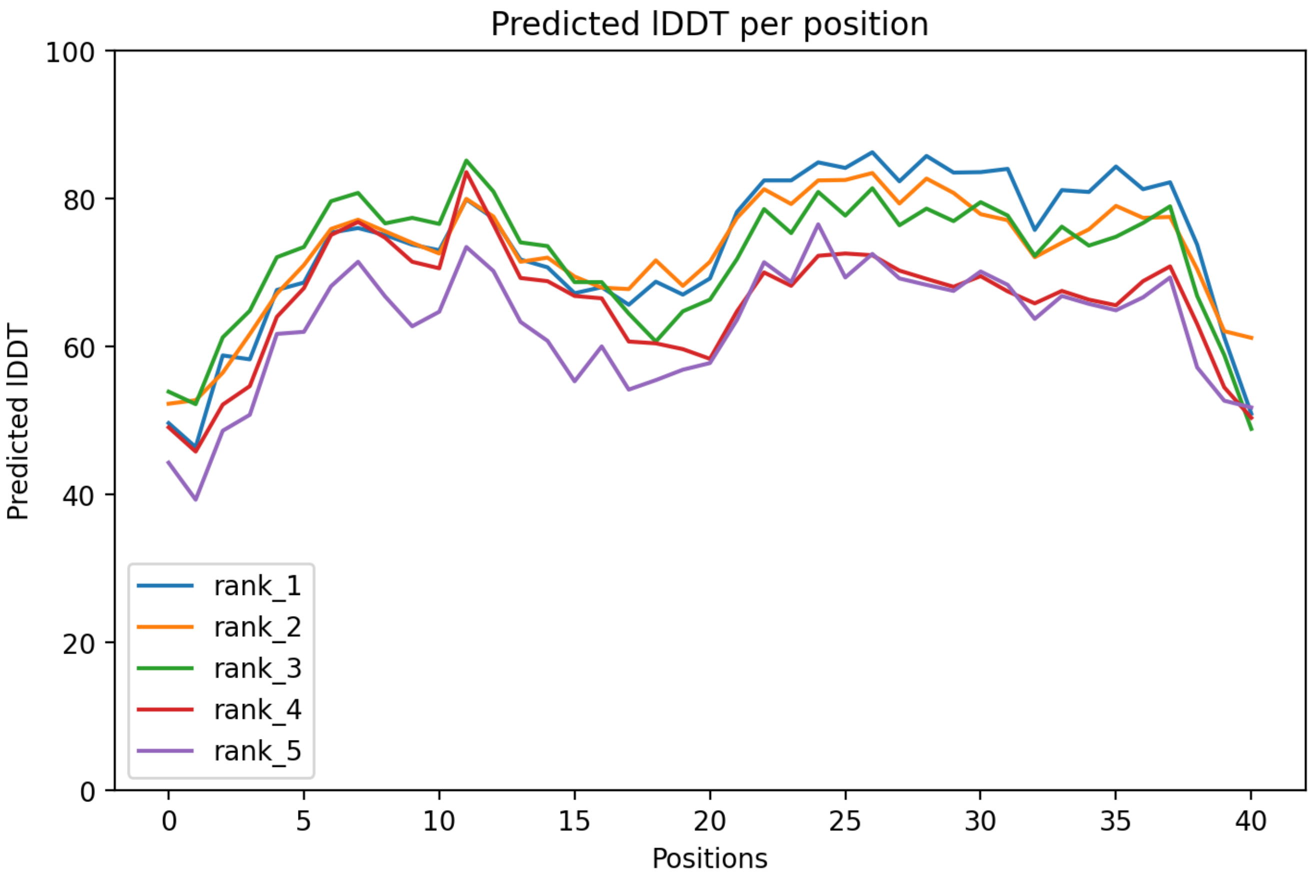

CP95: MtrunA17_Chr6g0451601

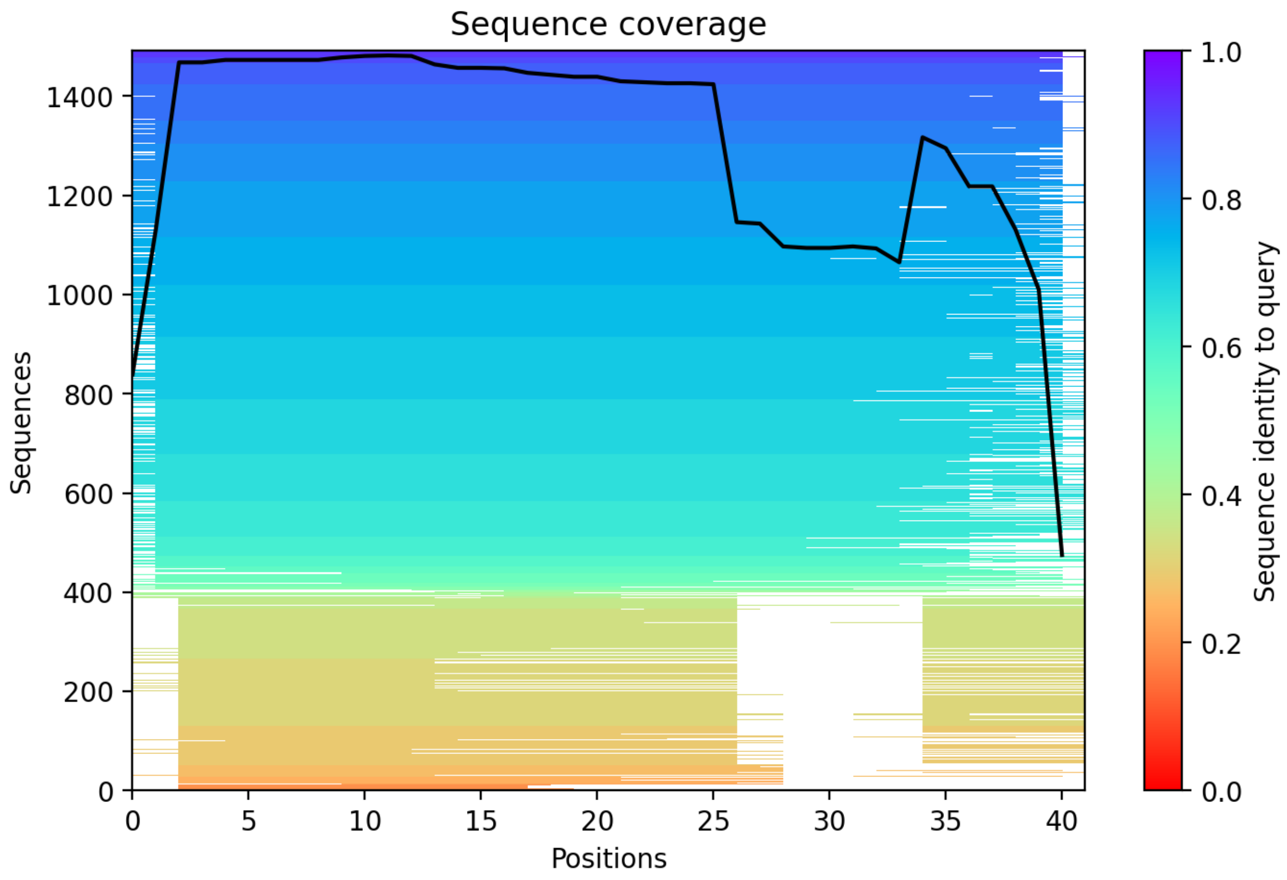

CP96: MtrunA17_Chr6g0452781

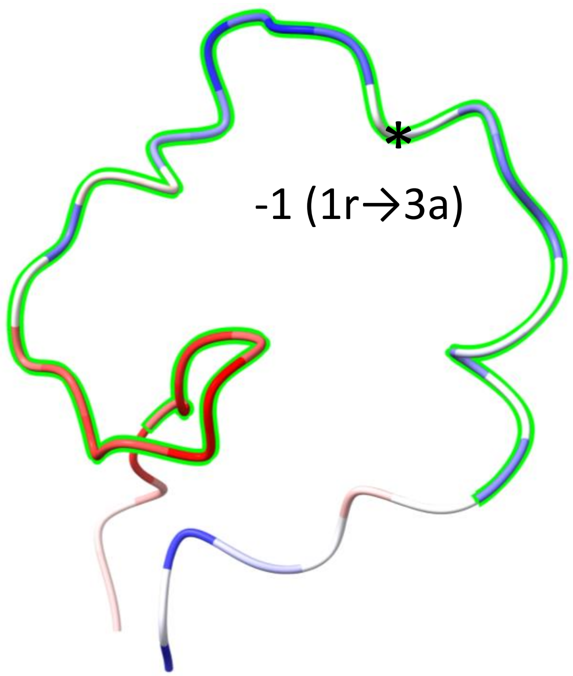


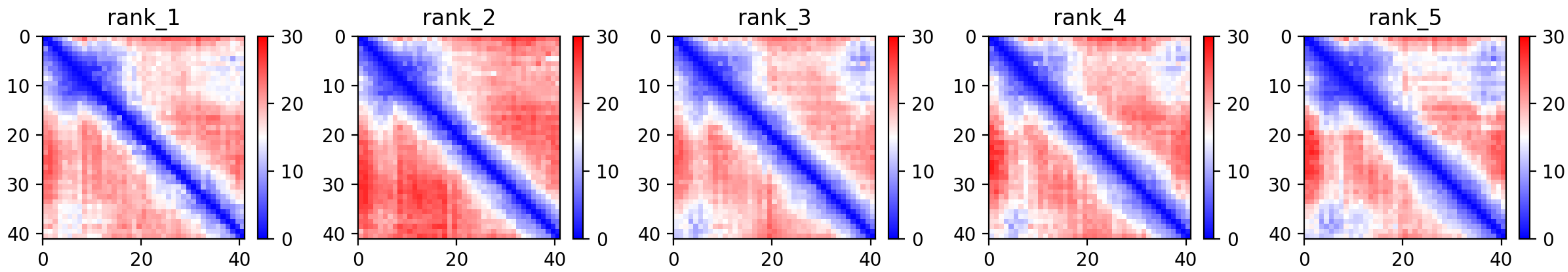

CP96: MtrunA17_Chr6g0452781

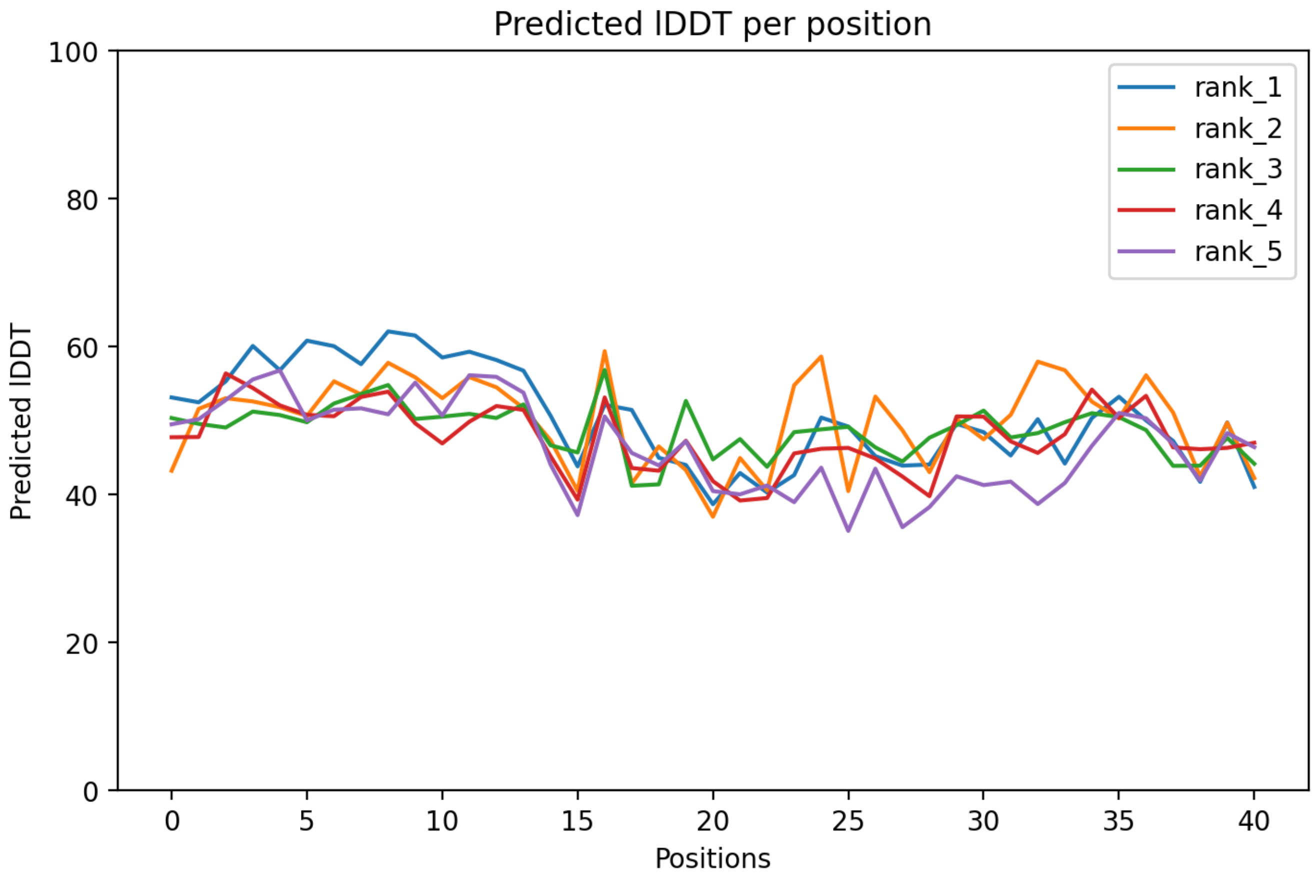

### CP96: MtrunA17_Chr6g0452781

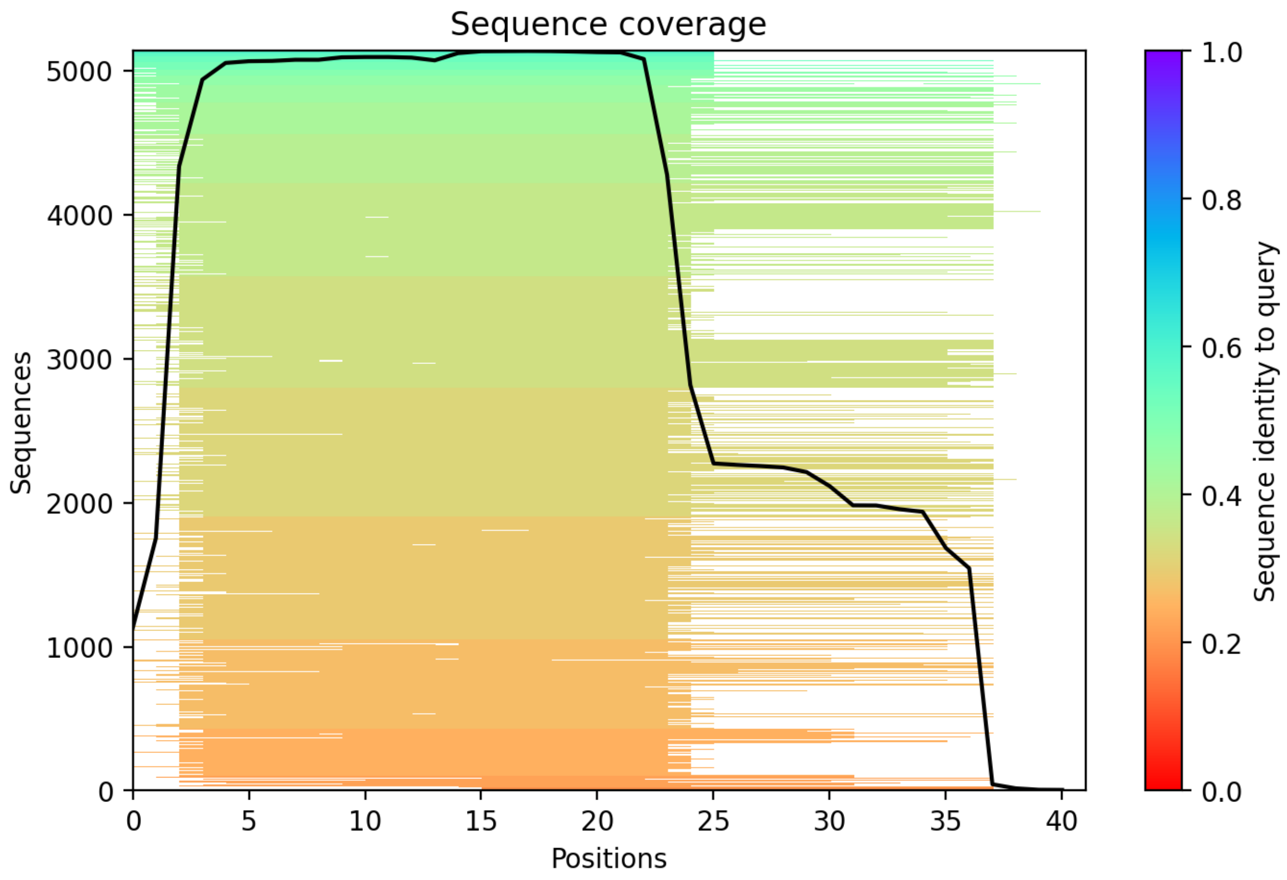

CP97: MtrunA17_Chr6g0457351

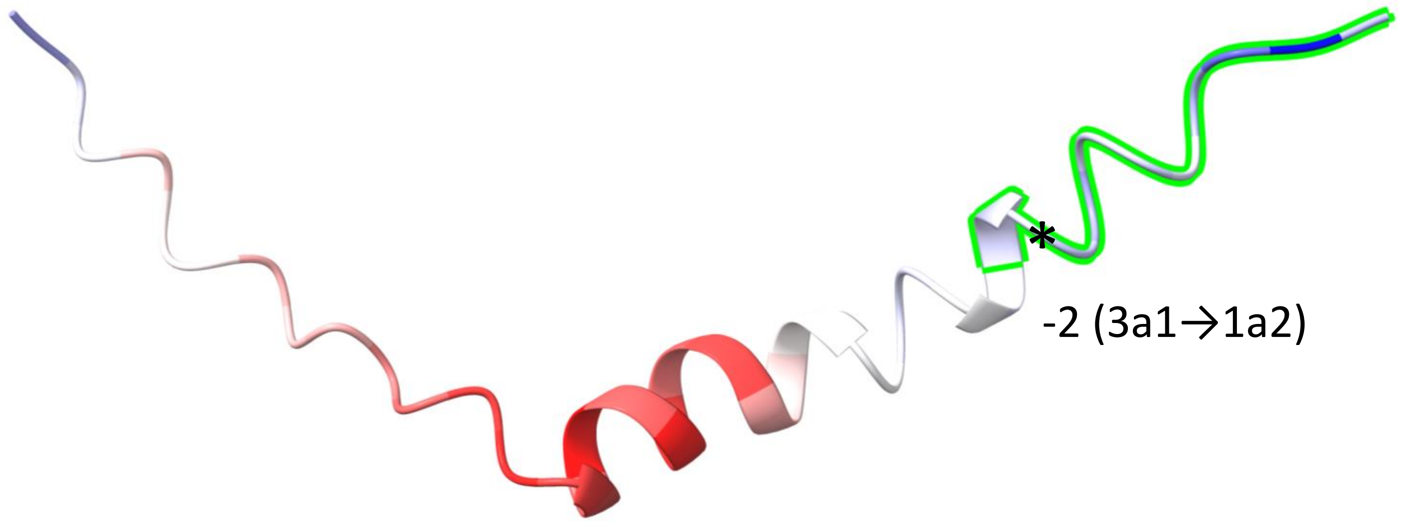


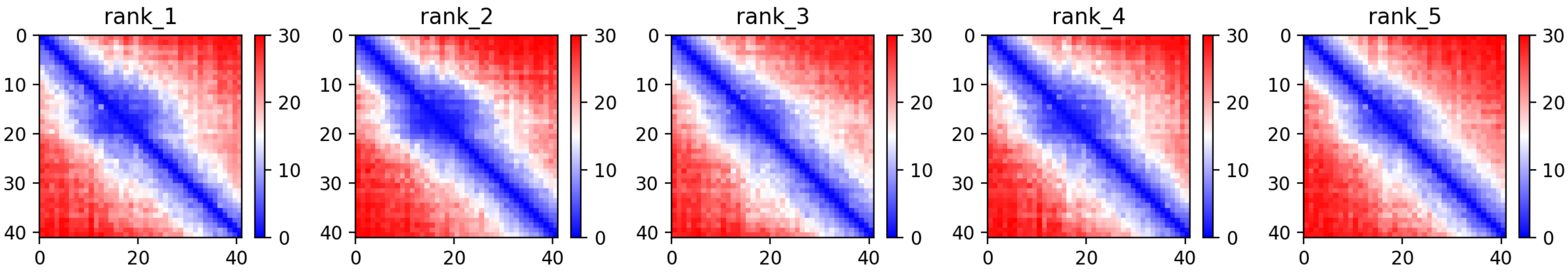

### CP97: MtrunA17_Chr6g0457351

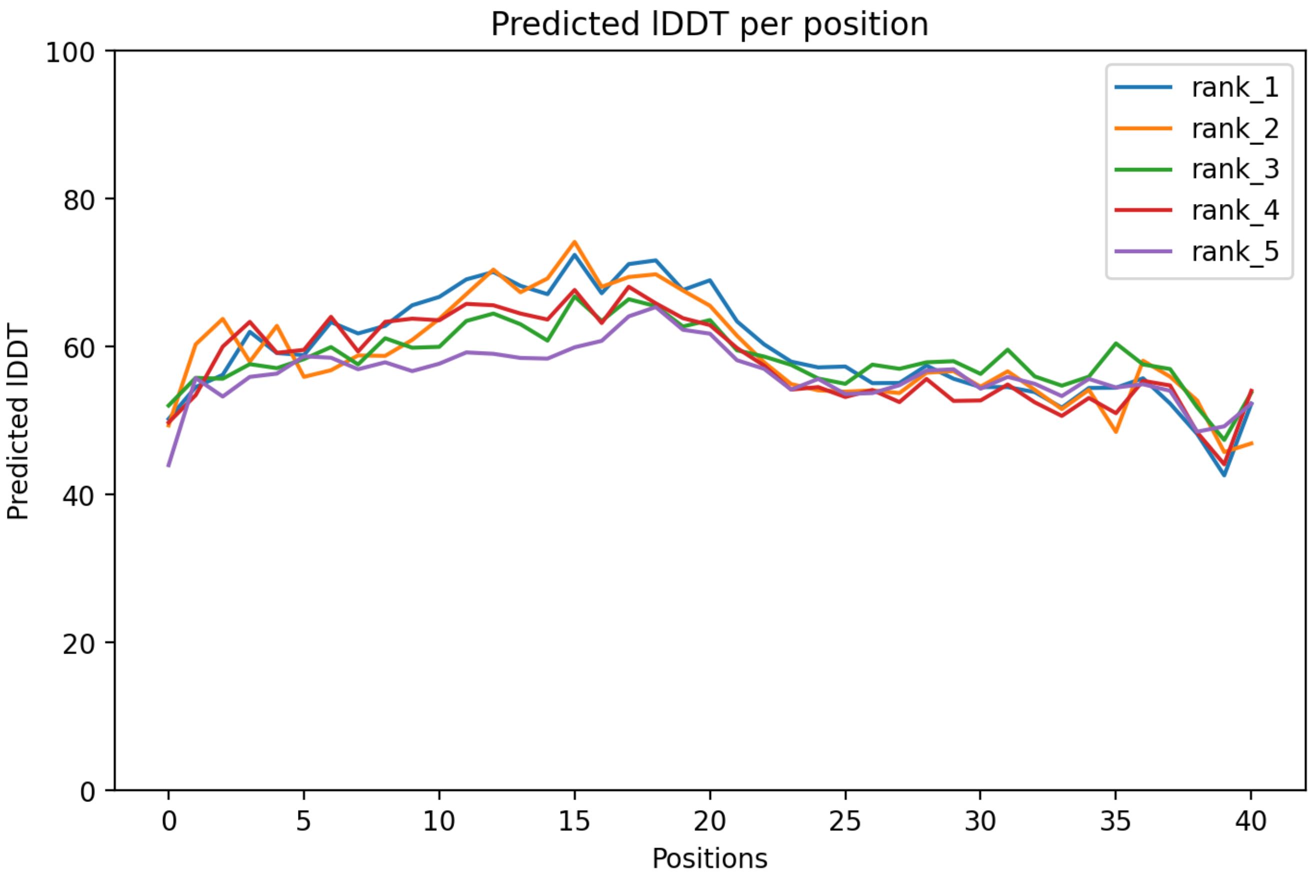

### CP97: MtrunA17_Chr6g0457351

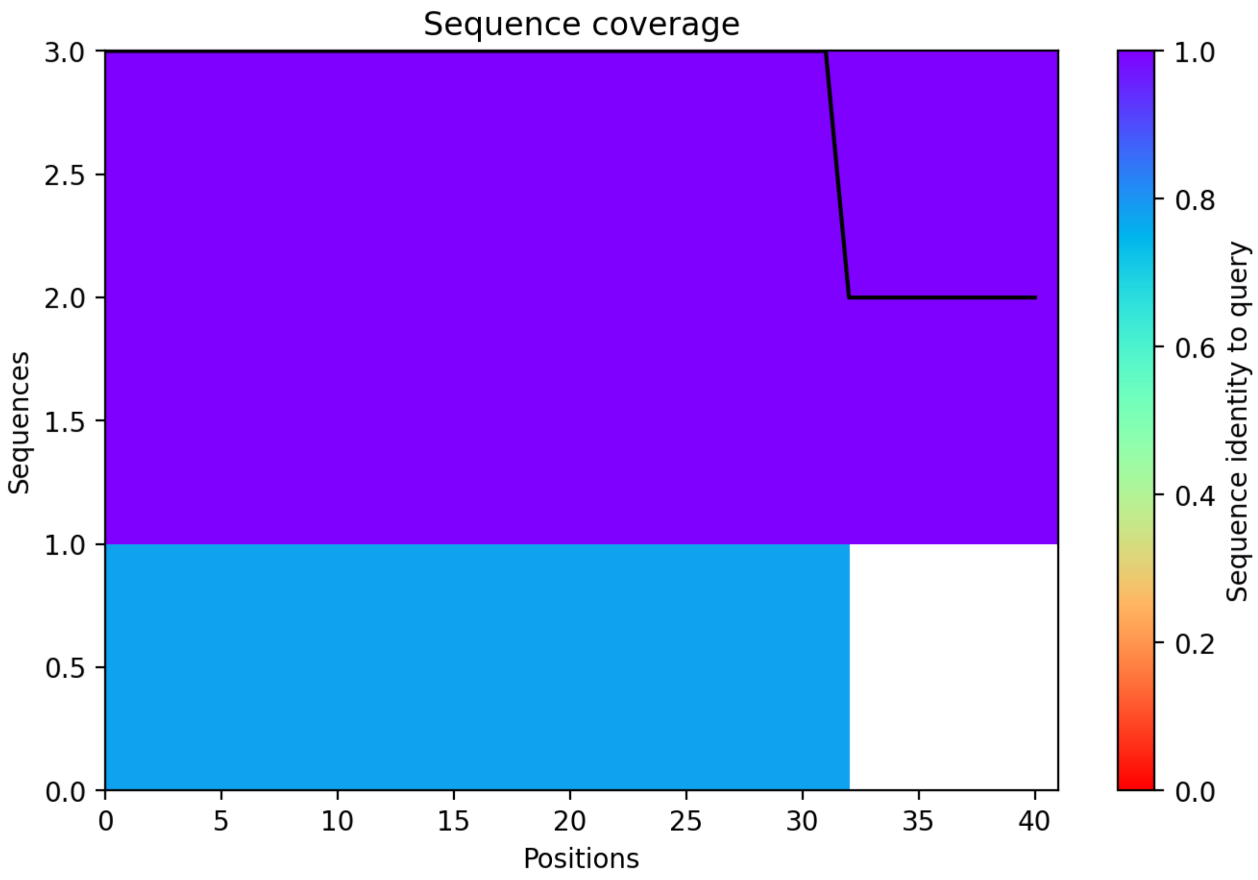

CP98: MtrunA17_Chr6g0457461

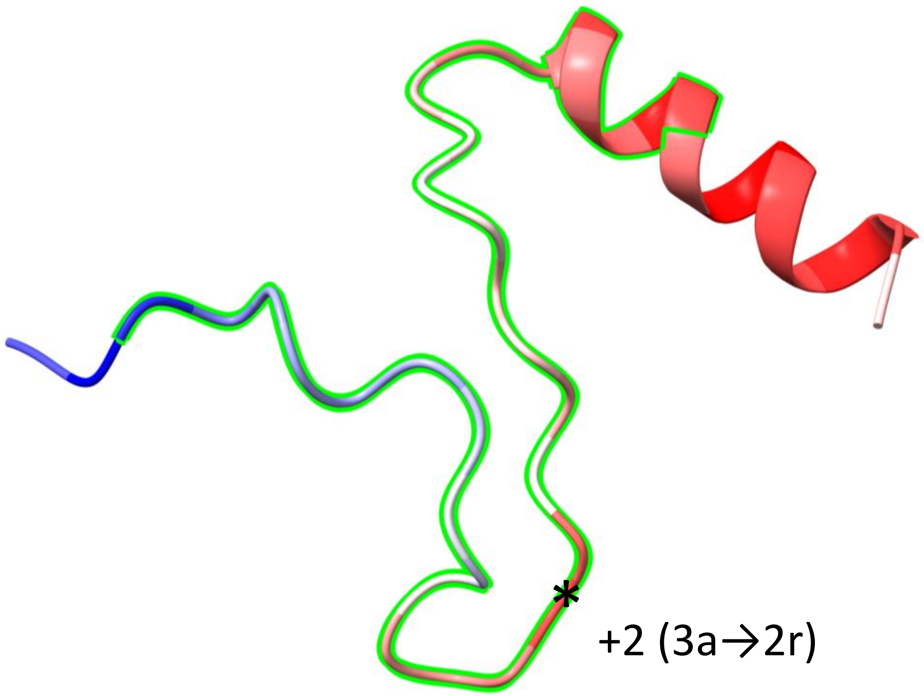


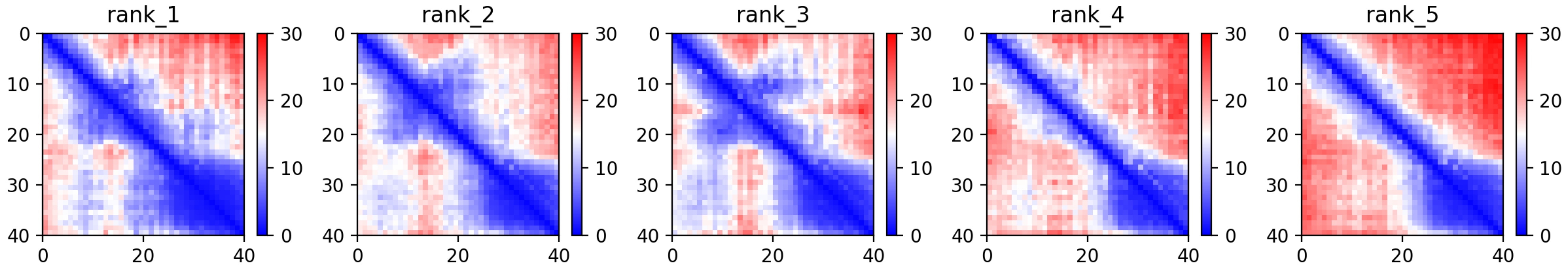

CP98: MtrunA17_Chr6g0457461

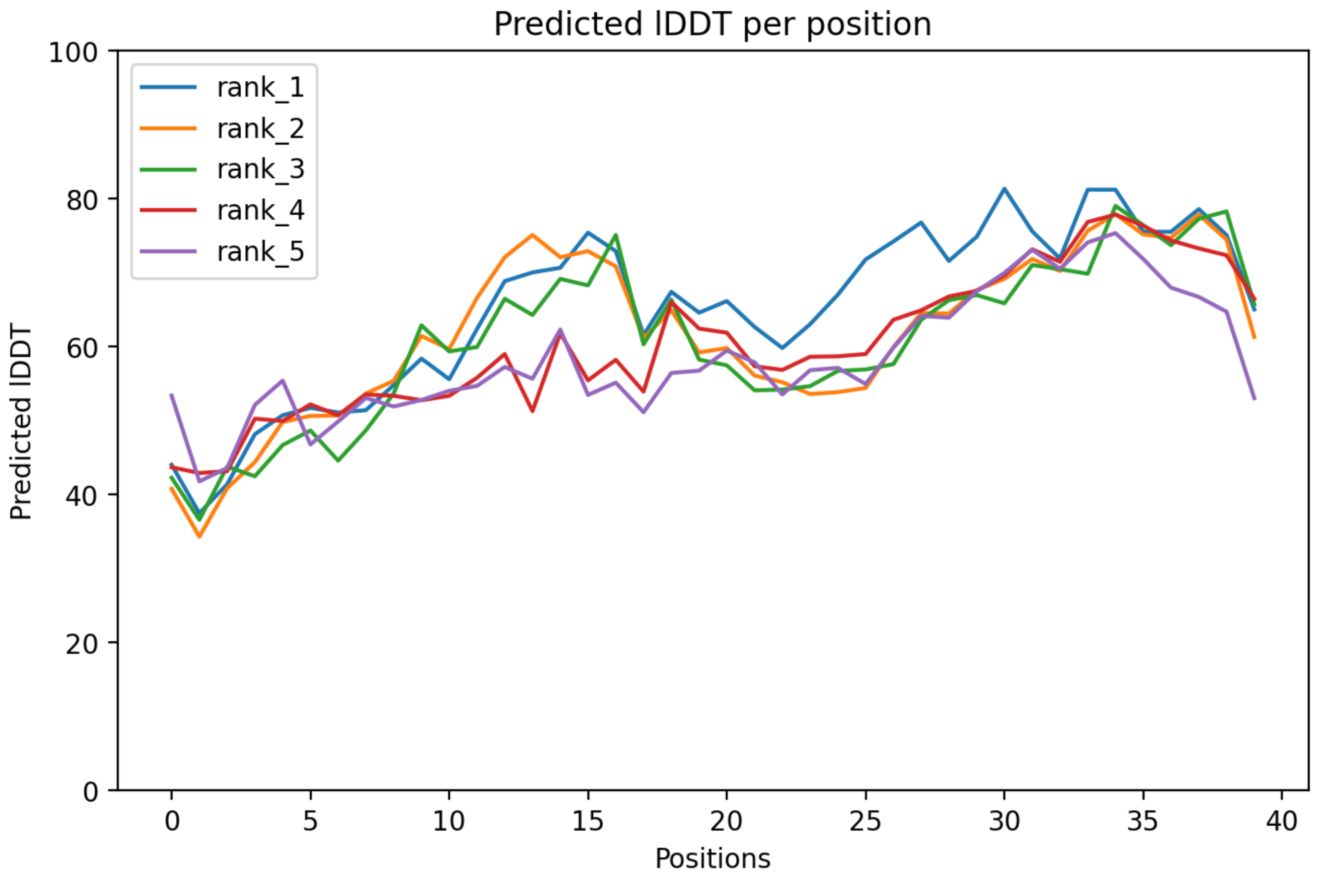

CP98: MtrunA17_Chr6g0457461

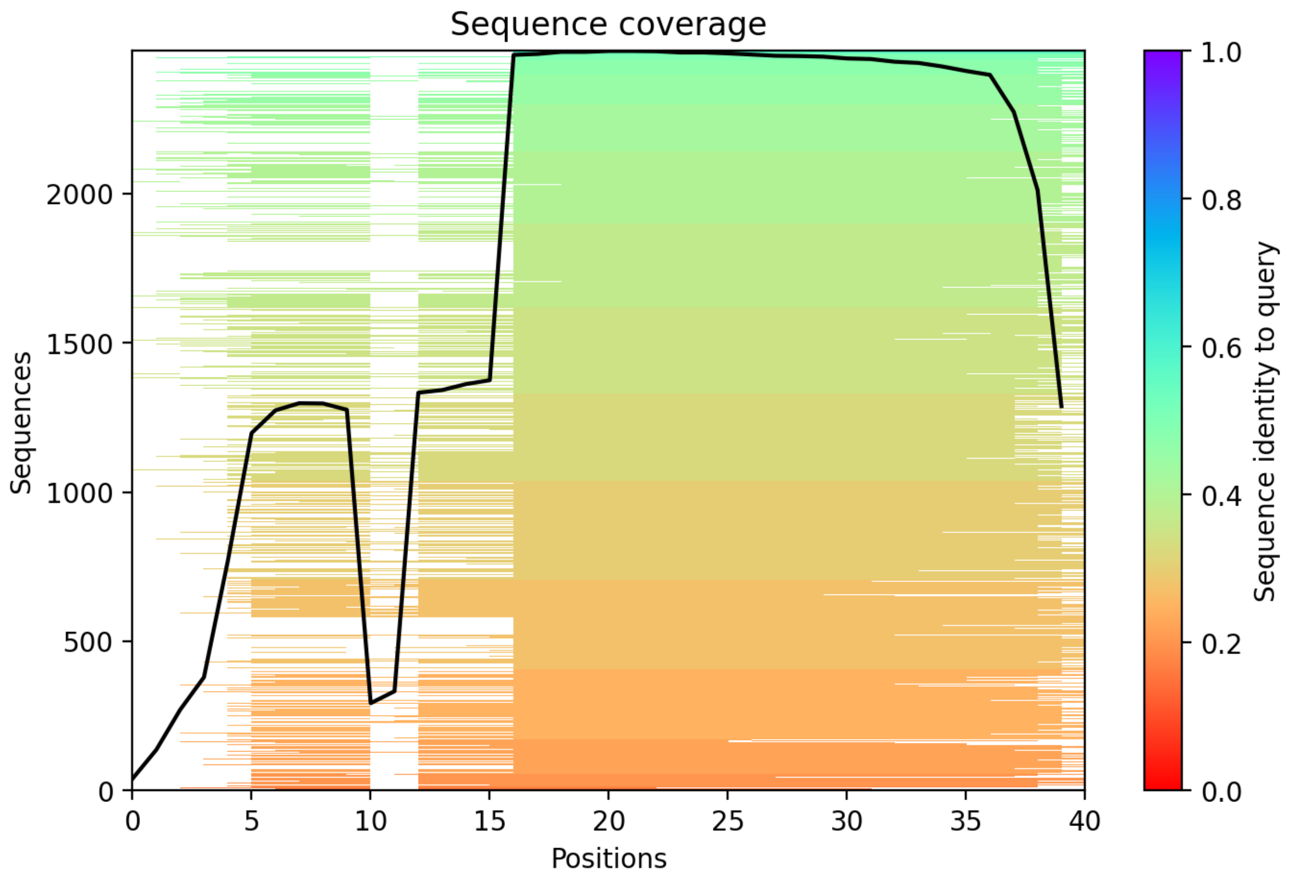

### CP99: MtrunA17_Chr6g0457461

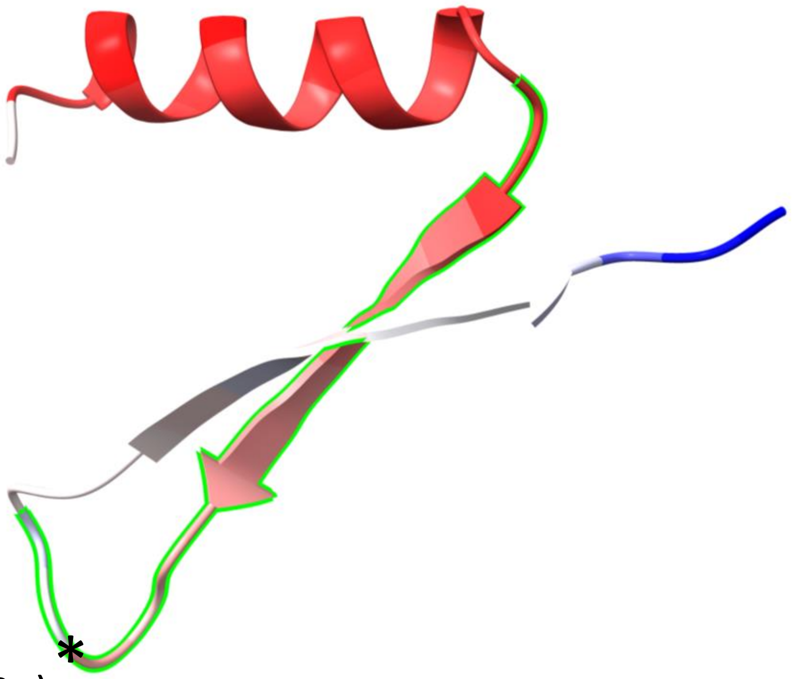


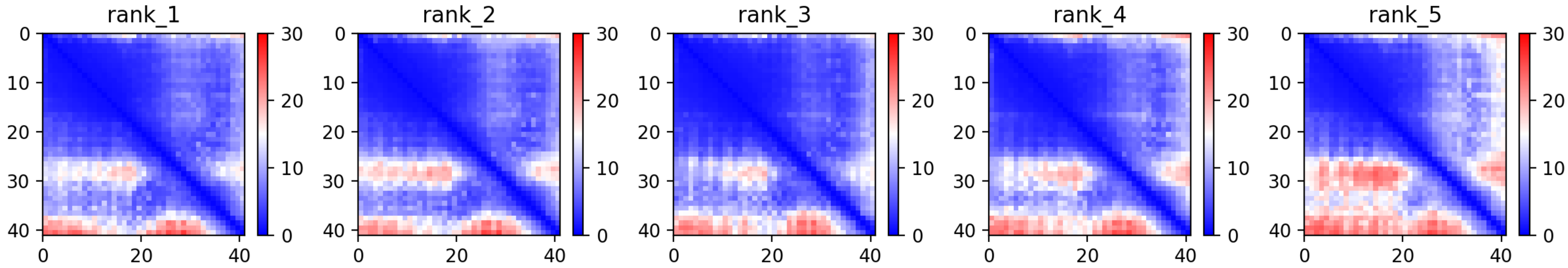

CP99: MtrunA17_Chr6g0457461

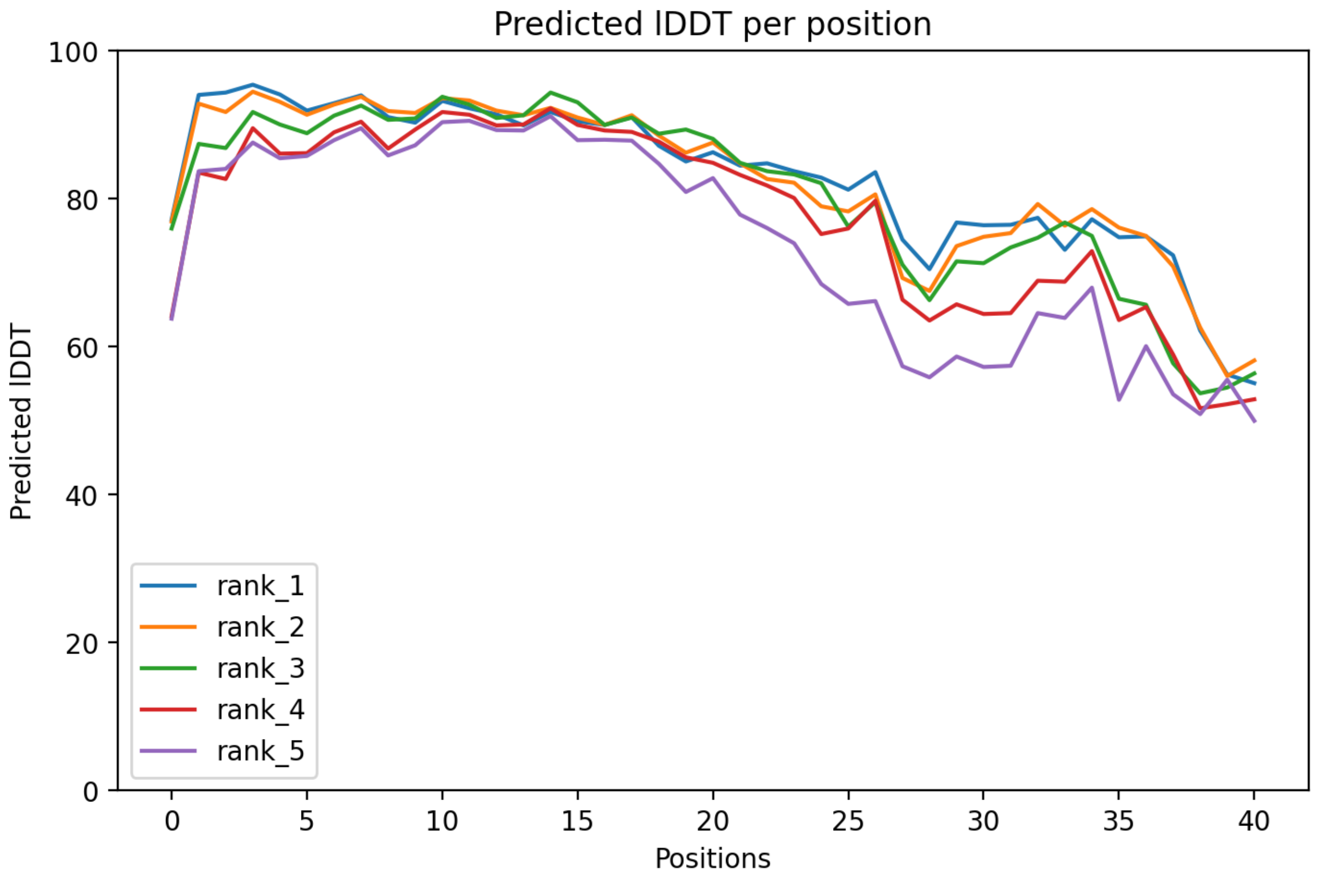

CP99: MtrunA17_Chr6g0457461

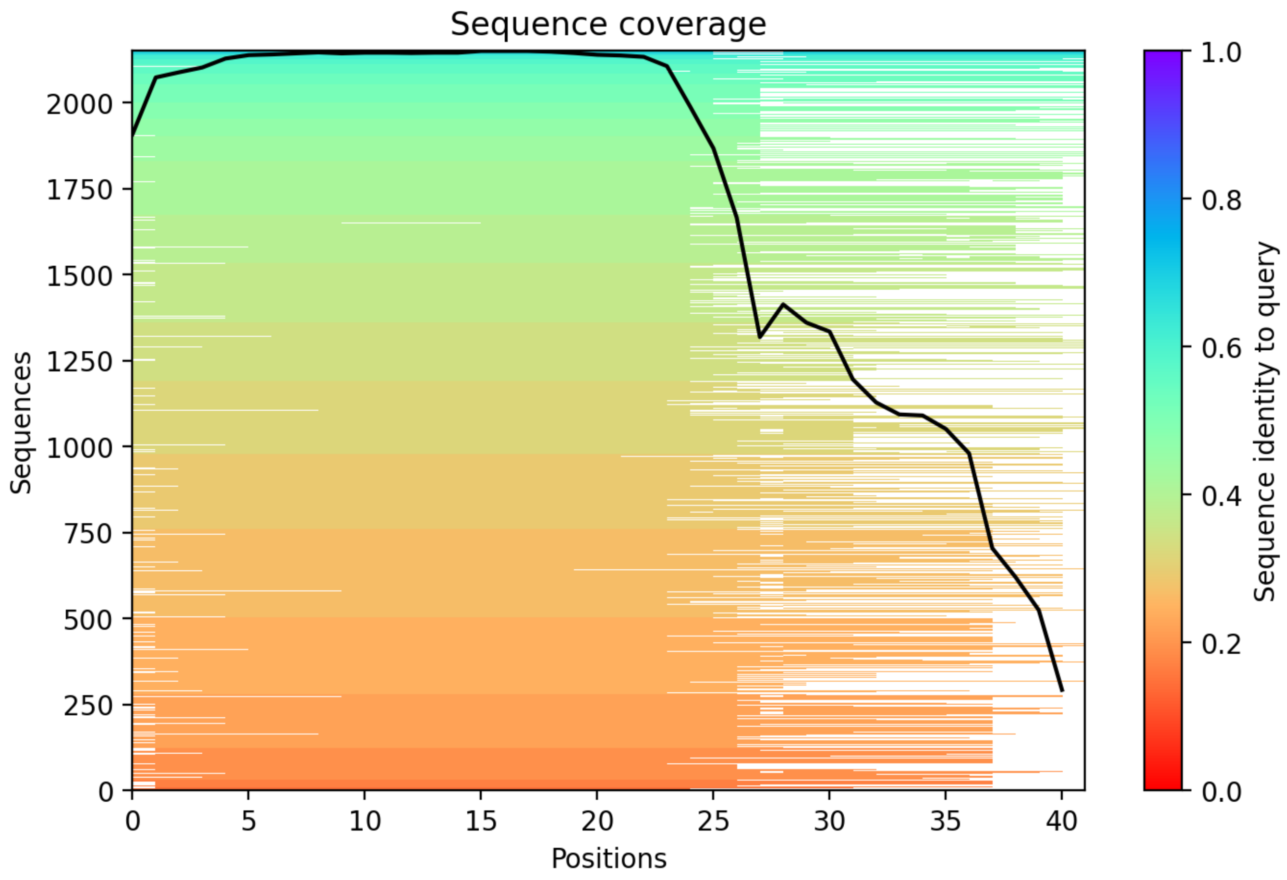

CP100: MtrunA17_Chr6g0457461

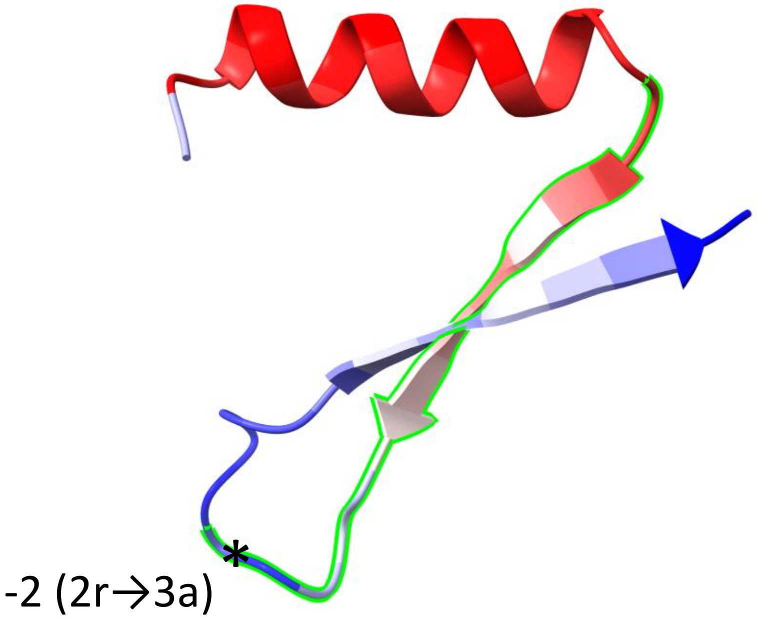


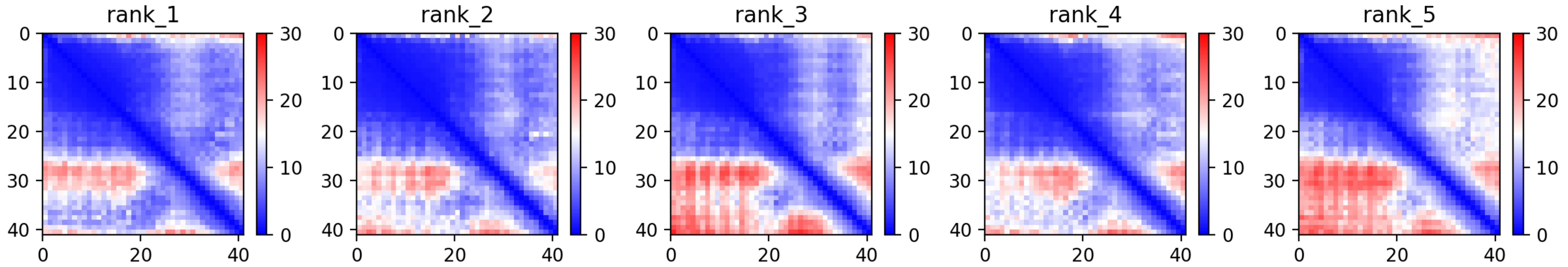

CP100: MtrunA17_Chr6g0457461

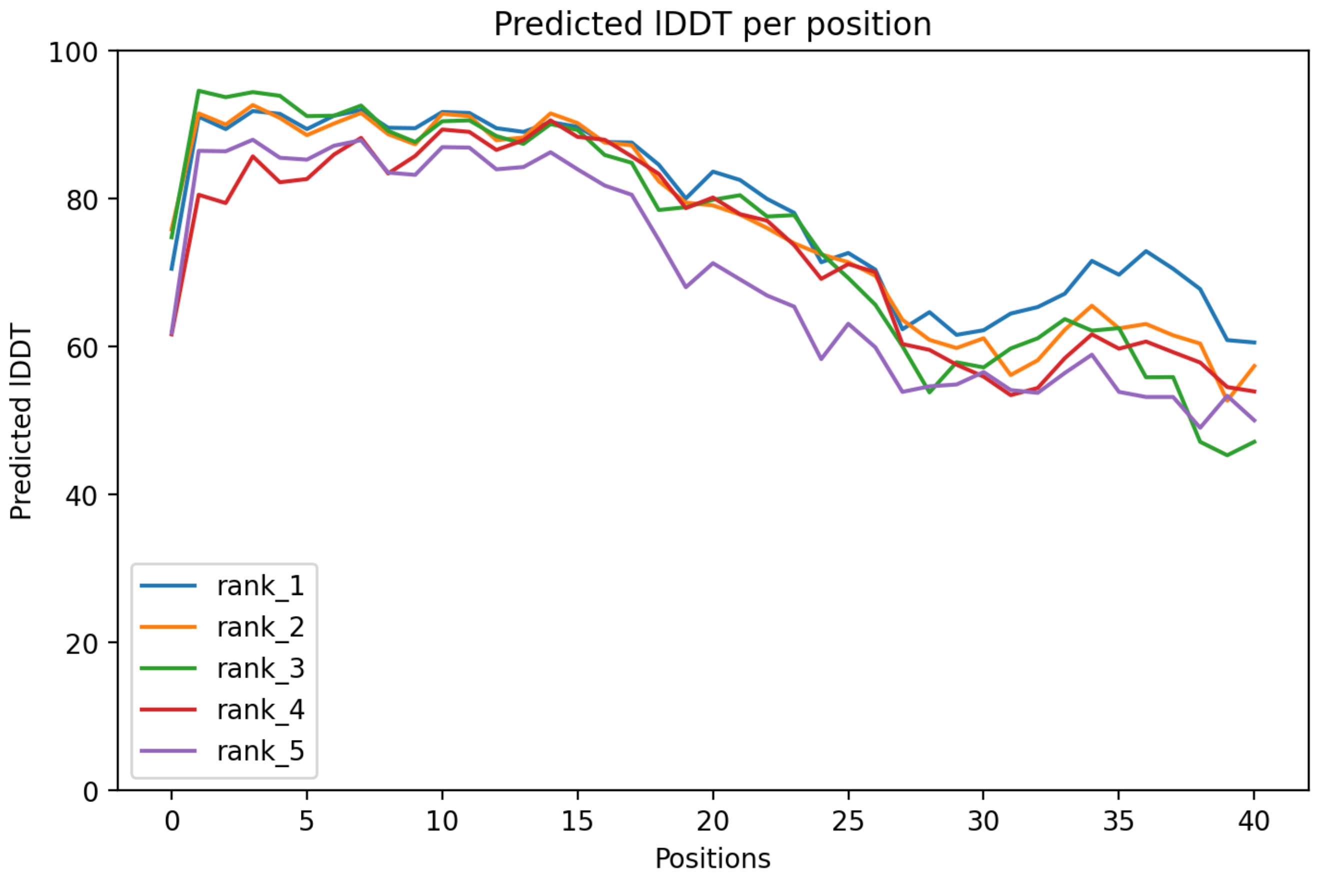

CP100: MtrunA17_Chr6g0457461

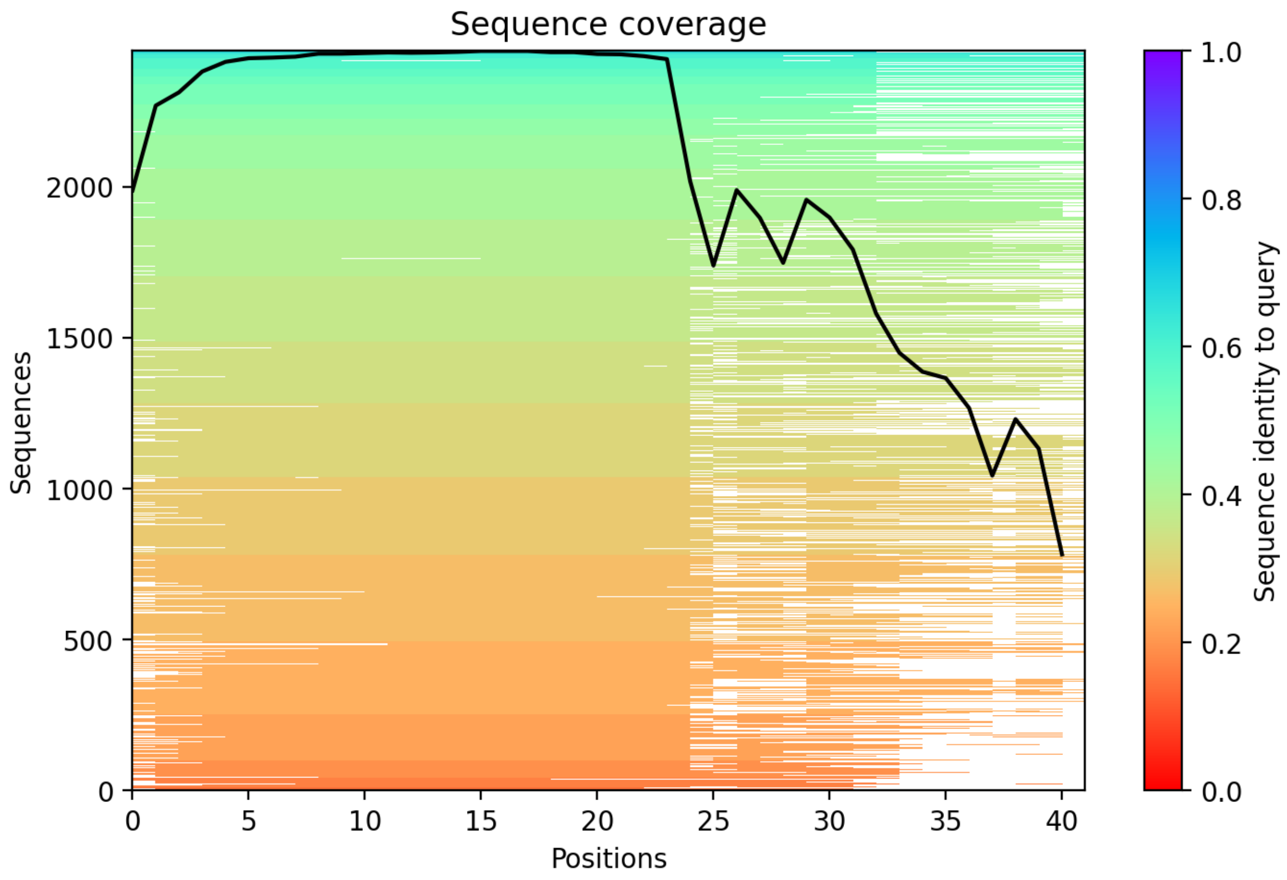

**Supplementary Dataset S7 Part 2**. A graphical summary on folding predictions for MS-supported chimeric peptide models (CPs) 51-100. The rest of the legend is the same as for Supplementary Dataset S7 Part 1.
