## Supplementary material for "Discovery of diverse chimeric peptides in a eukaryotic proteome sets the stage for the experimental proof of the mosaic translation hypothesis": Supplementary Dataset S7 Folding of chimeric protein models Part 3.pdf

CP101: MtrunA17_Chr6g0458091

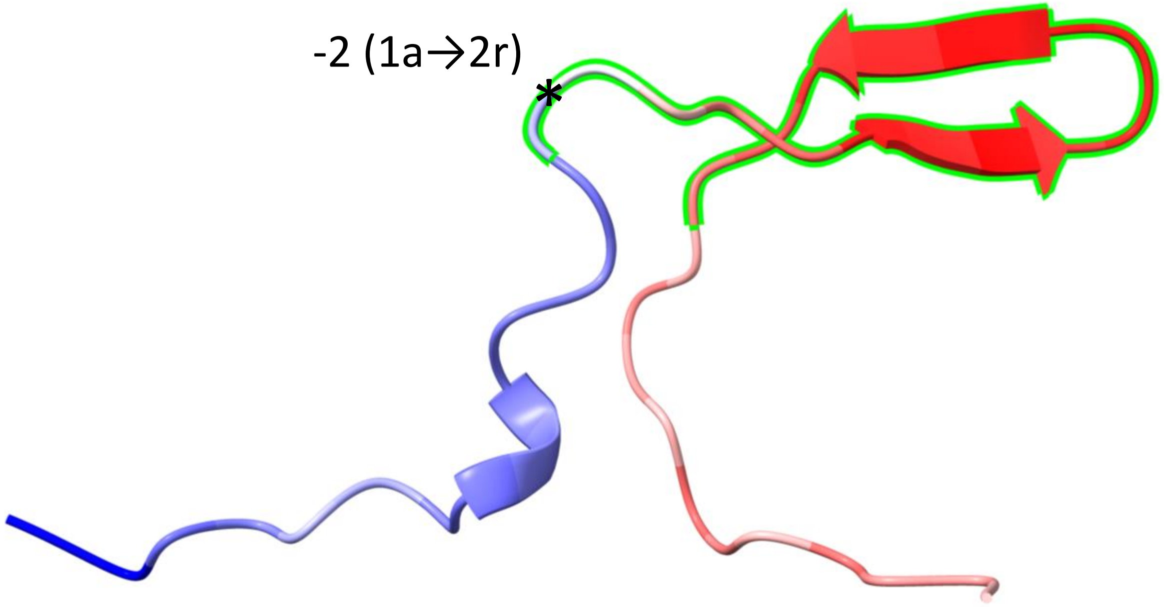


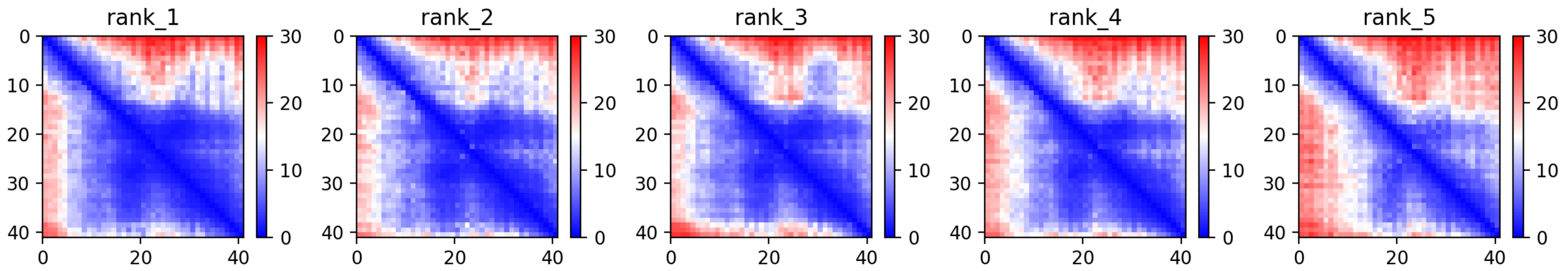

CP101: MtrunA17_Chr6g0458091

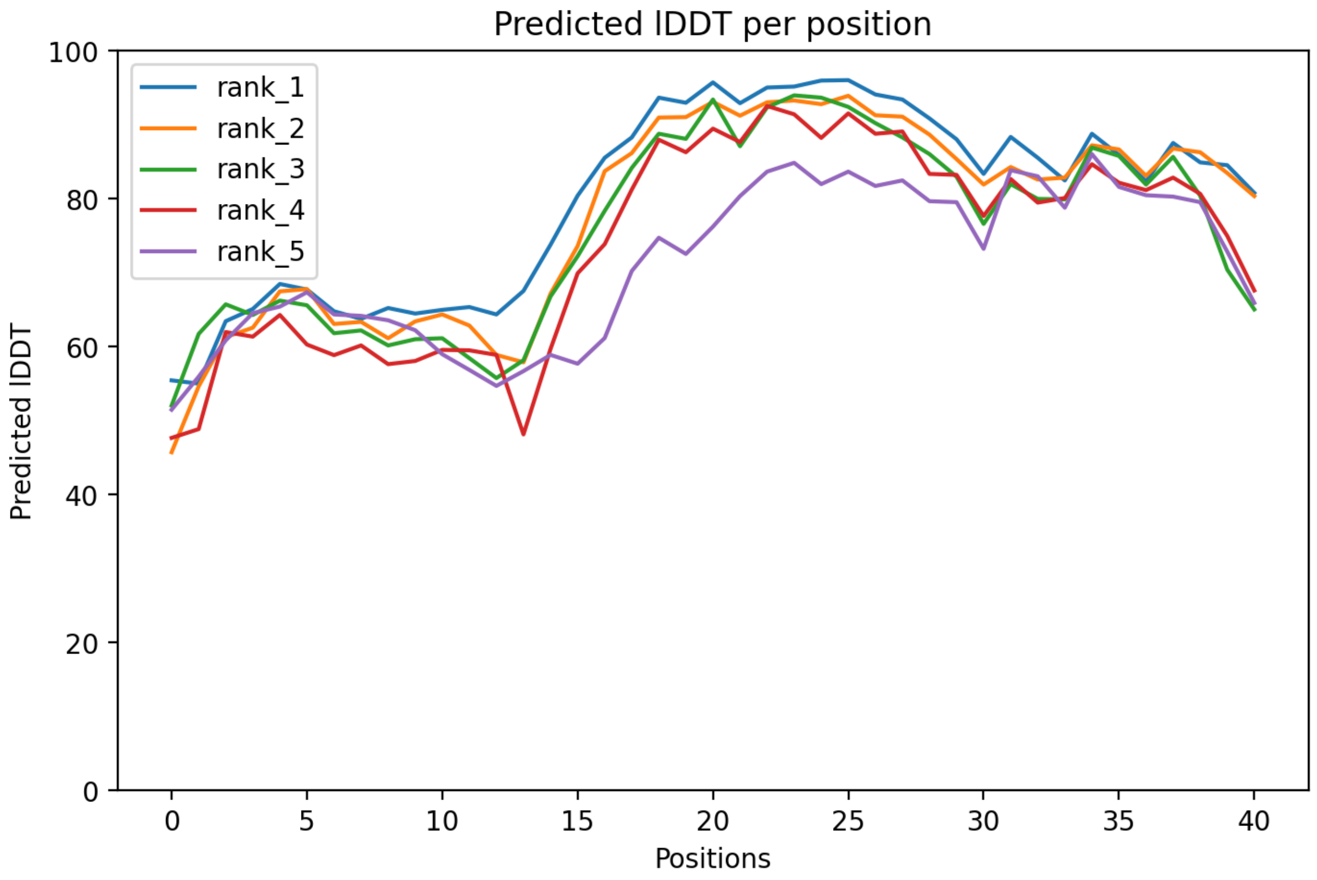

CP101: MtrunA17_Chr6g0458091

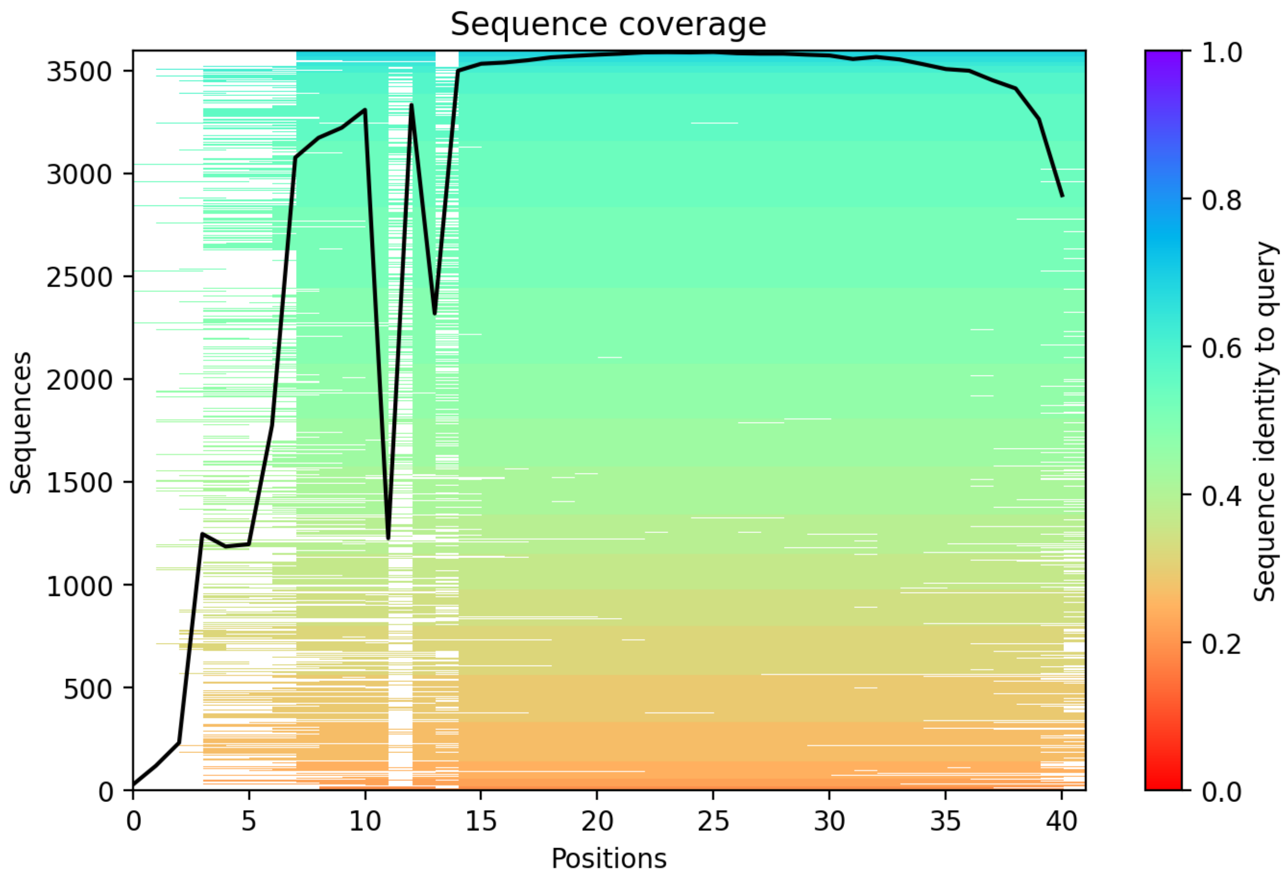

CP102: MtrunA17_Chr6g0459481

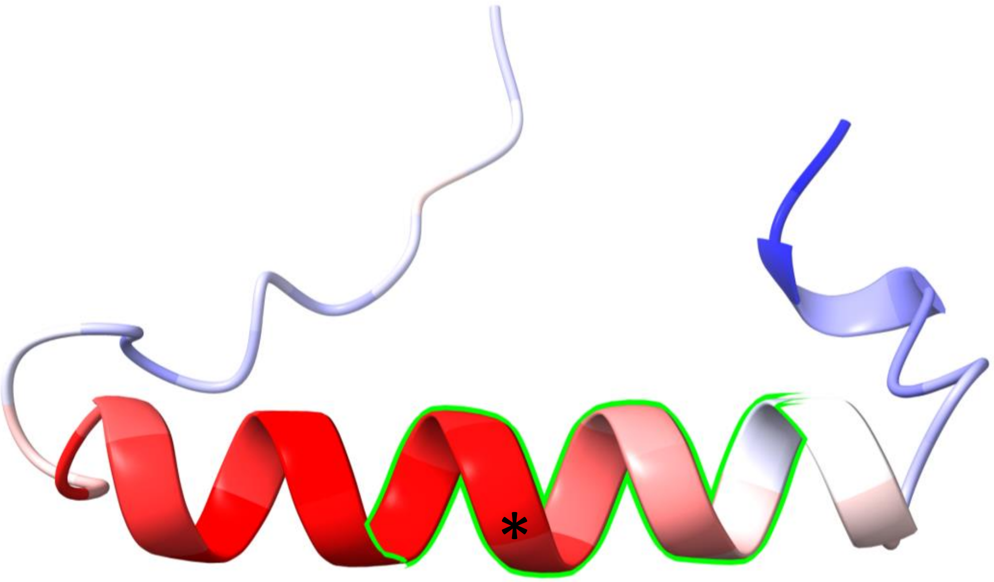


-1 (1r→3a)

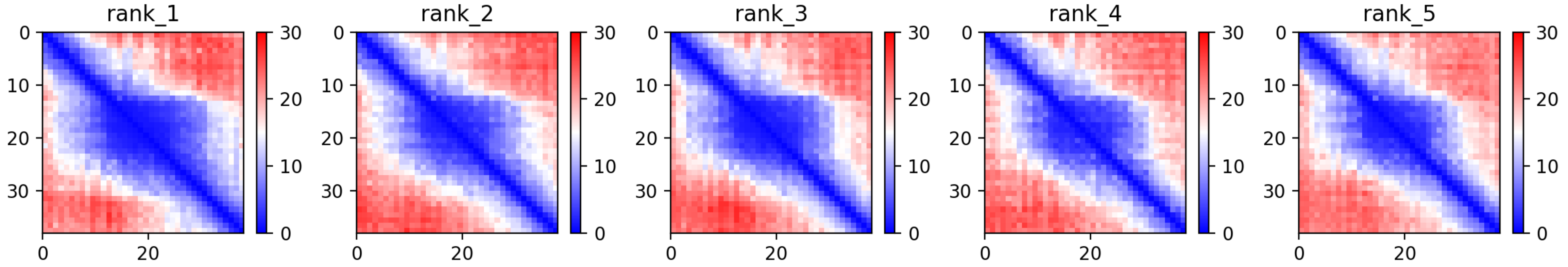

CP102: MtrunA17_Chr6g0459481

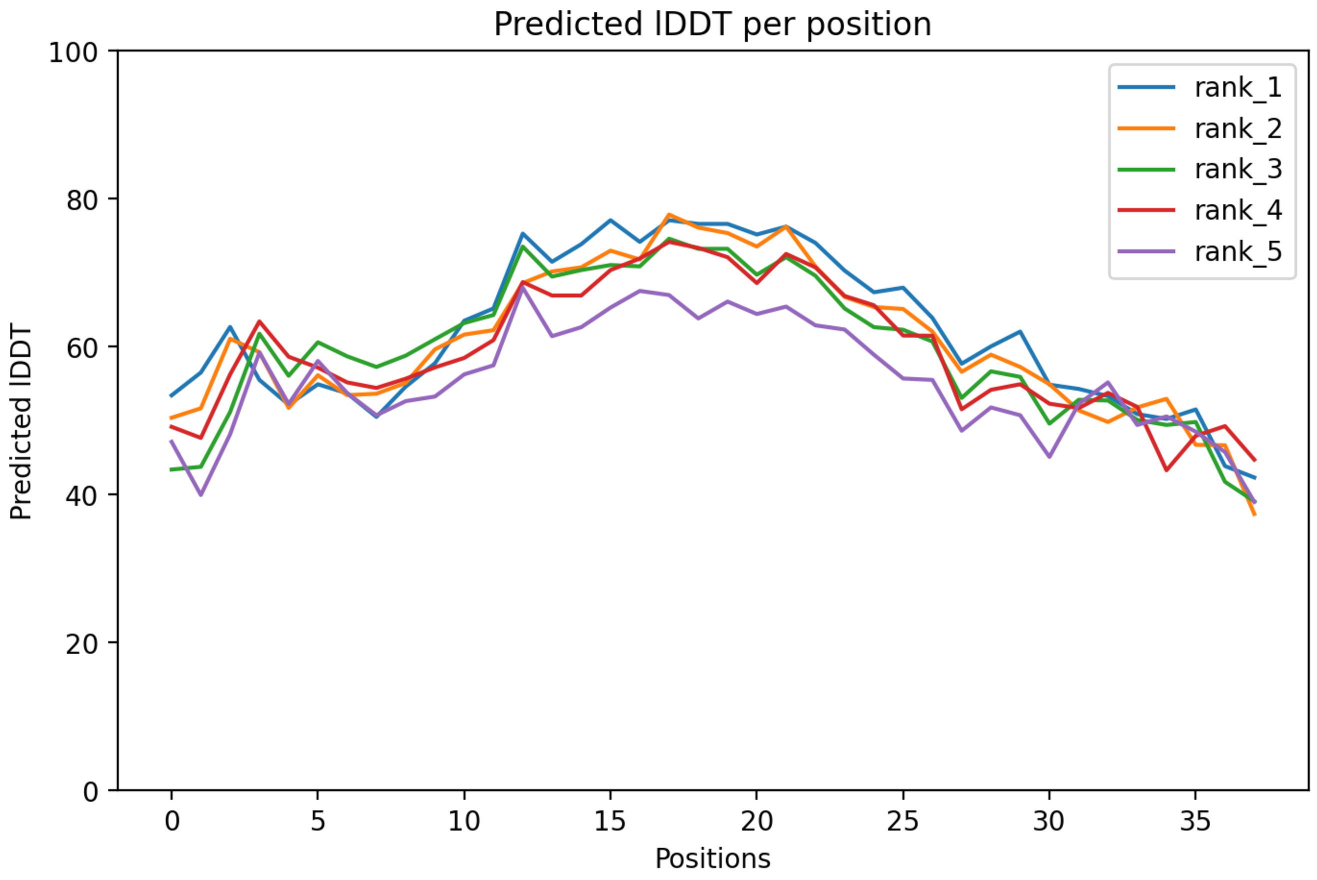

### CP102: MtrunA17_Chr6g0459481

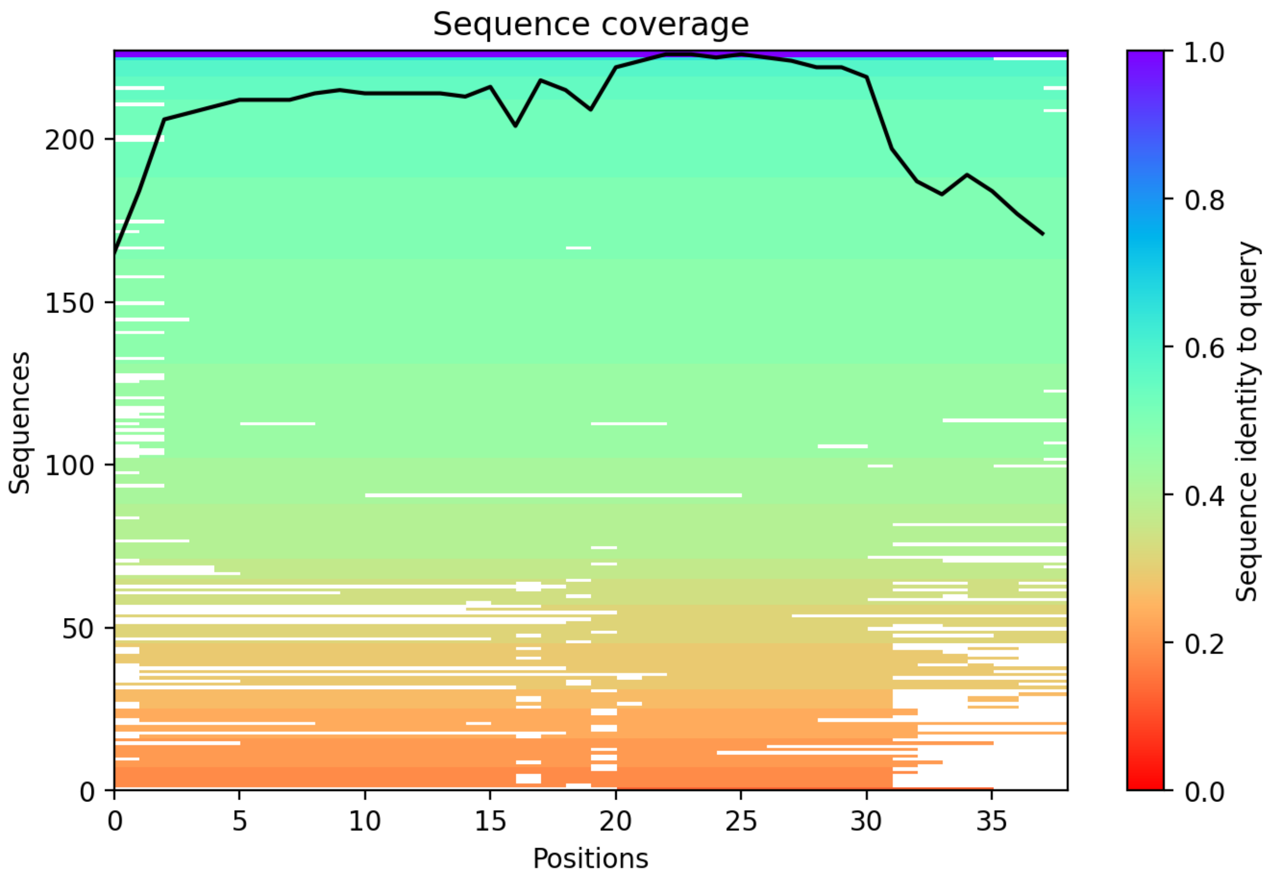

CP103: MtrunA17_Chr6g0461931

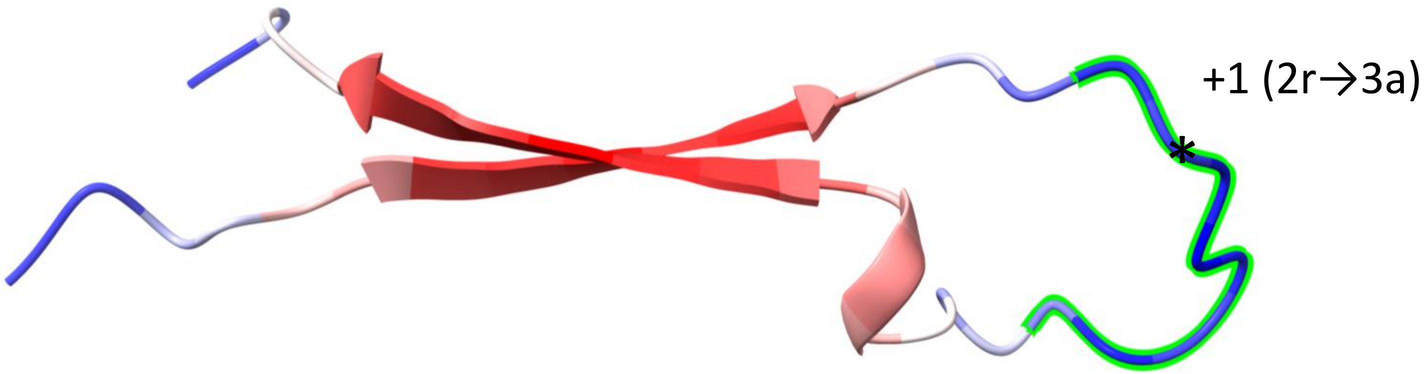


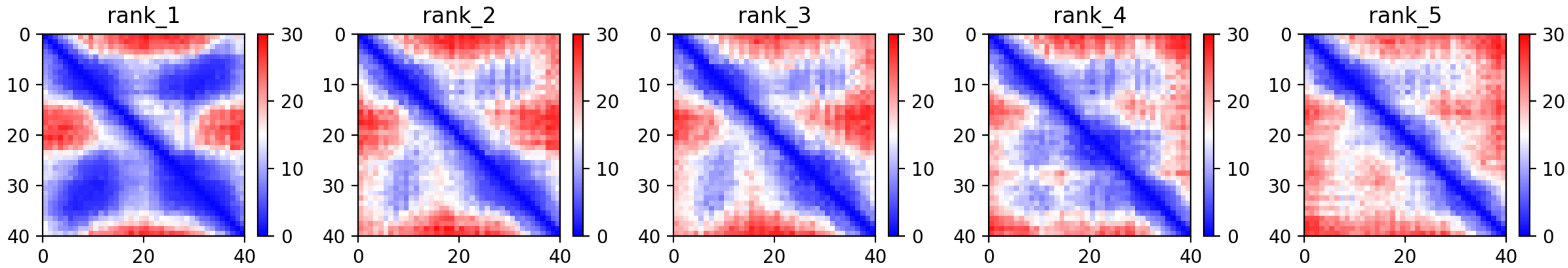

CP103: MtrunA17_Chr6g0461931

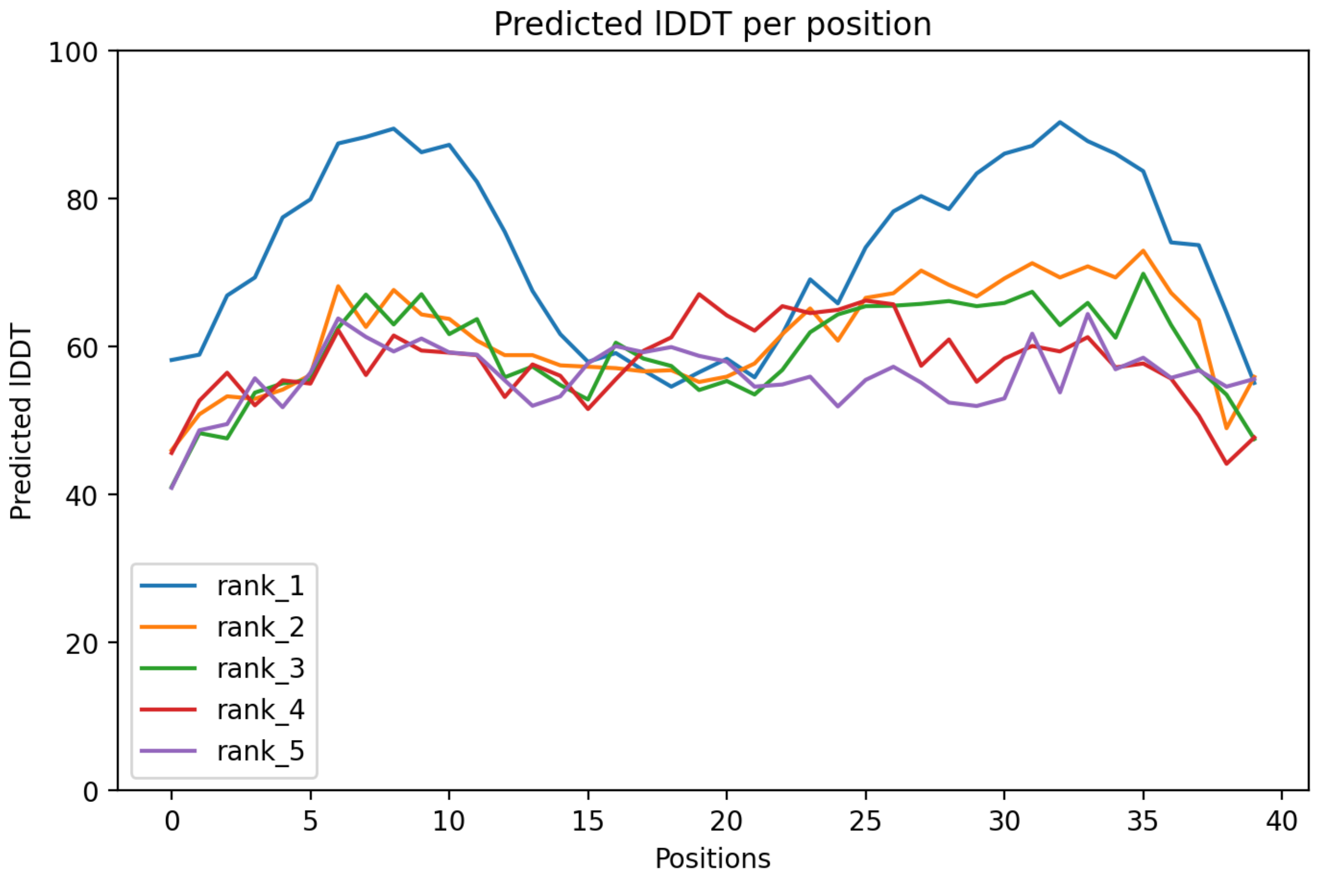

### CP103: MtrunA17_Chr6g0461931

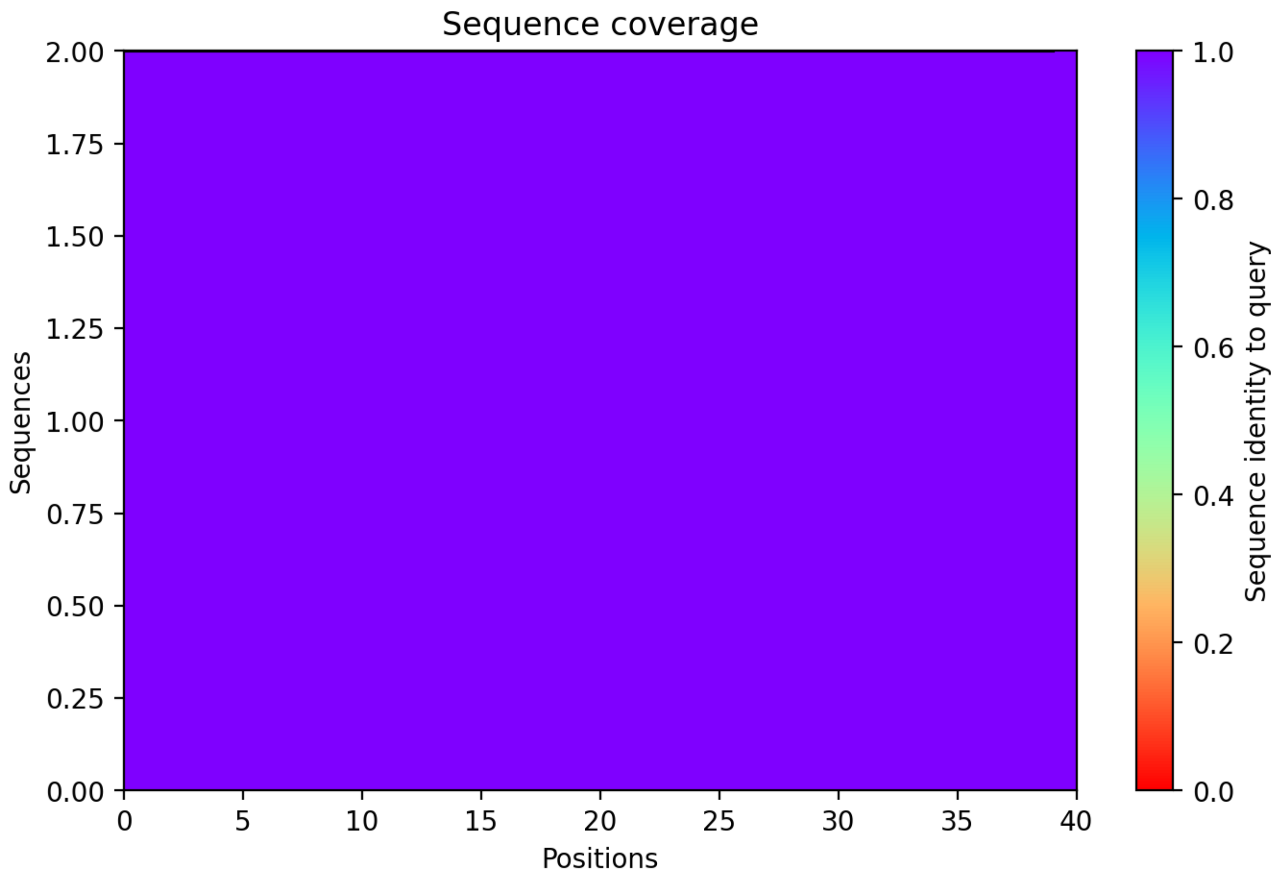

### CP104: MtrunA17_Chr6g0462271

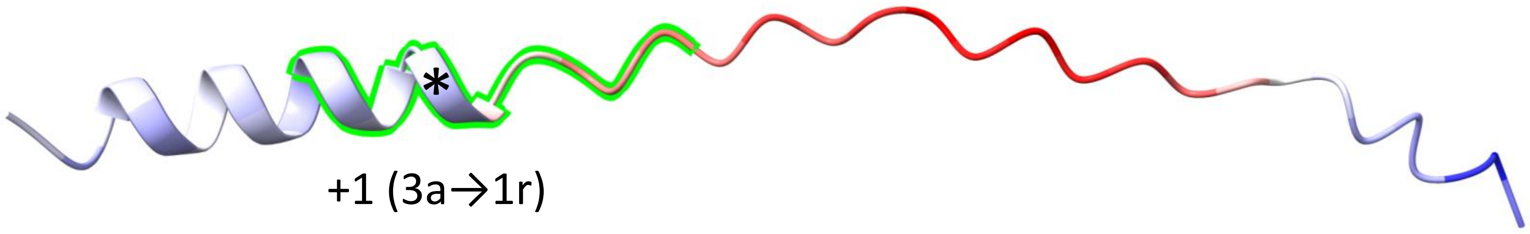


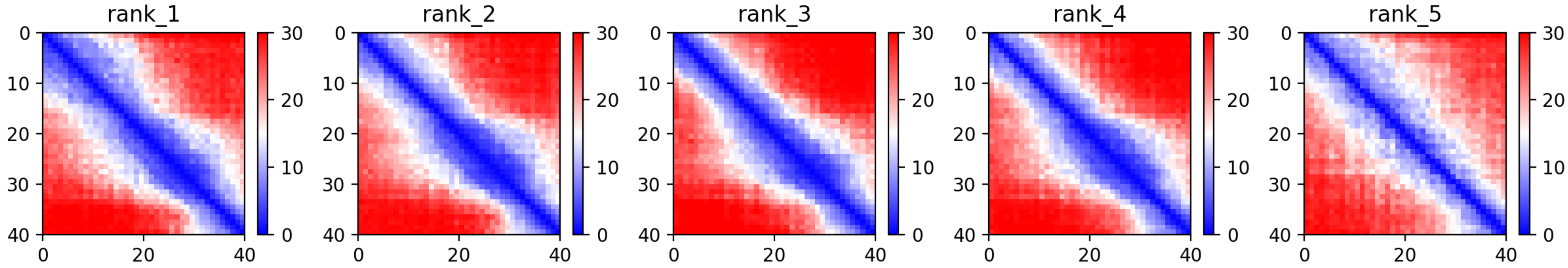

CP104: MtrunA17_Chr6g0462271

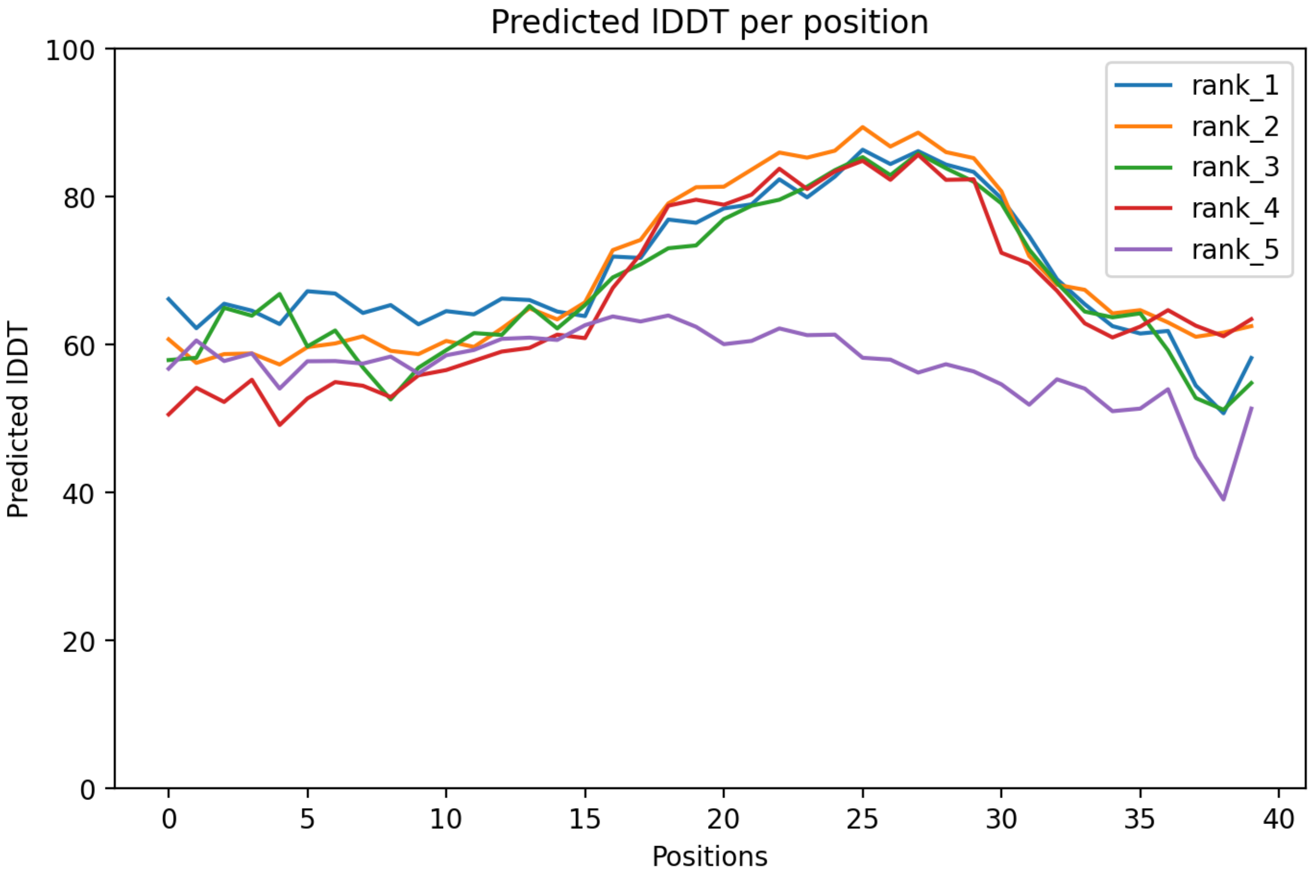

CP104: MtrunA17_Chr6g0462271

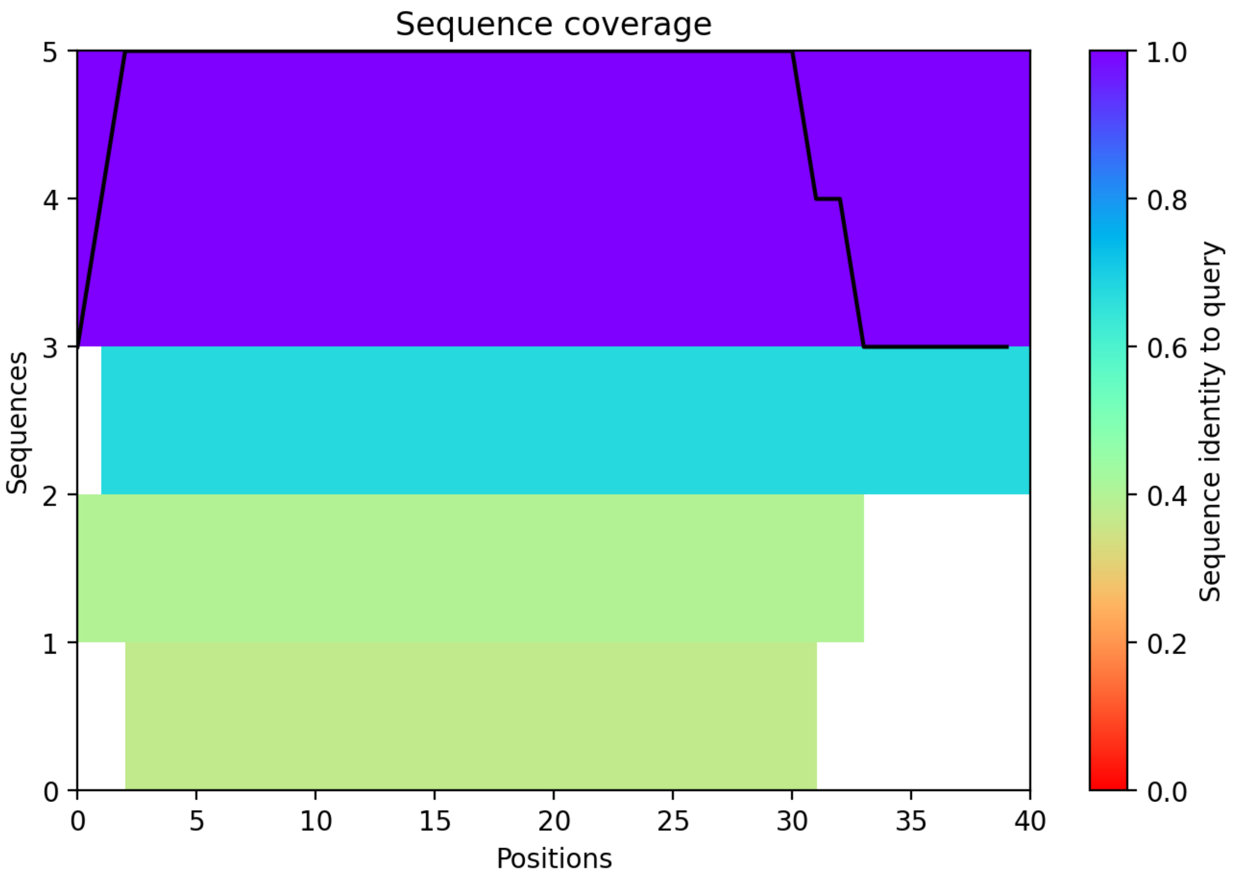

CP105: MtrunA17_Chr6g0468411

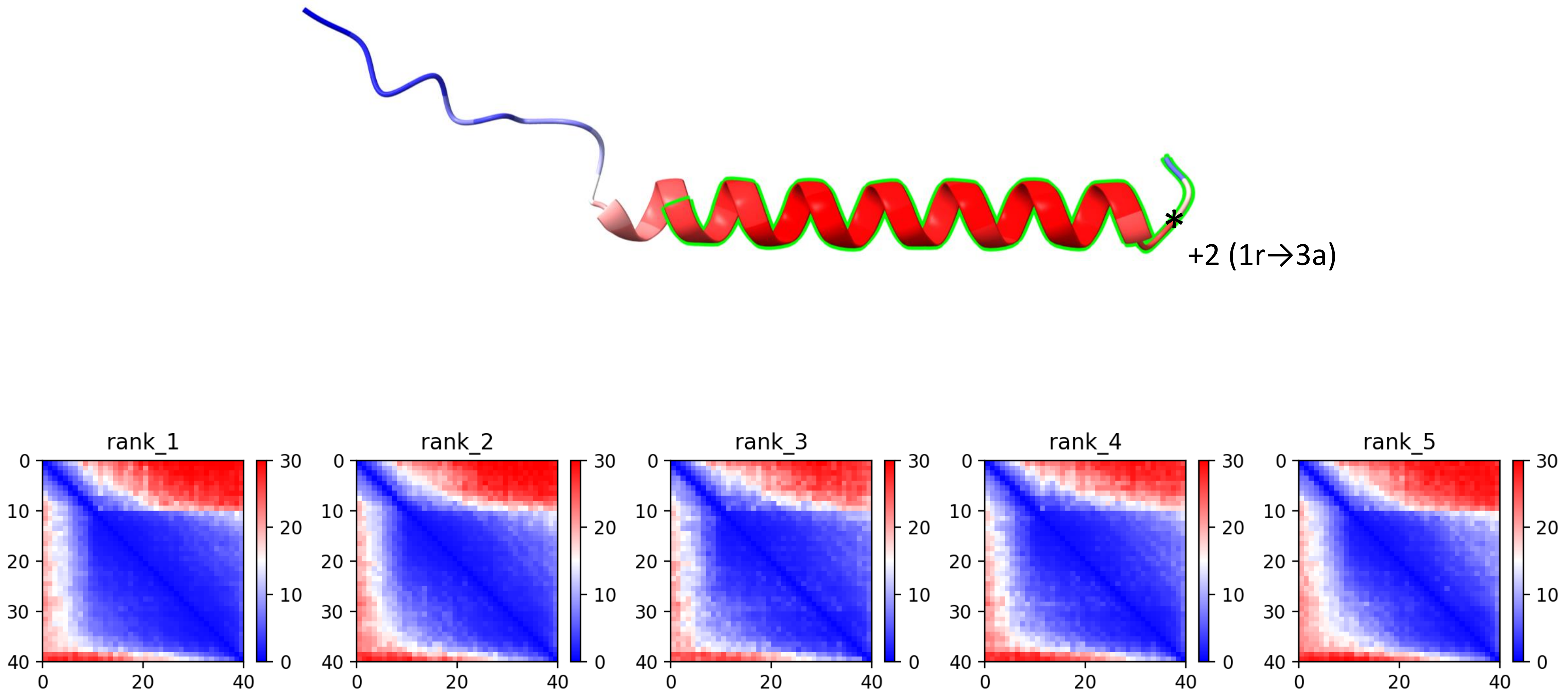

CP105: MtrunA17_Chr6g0468411

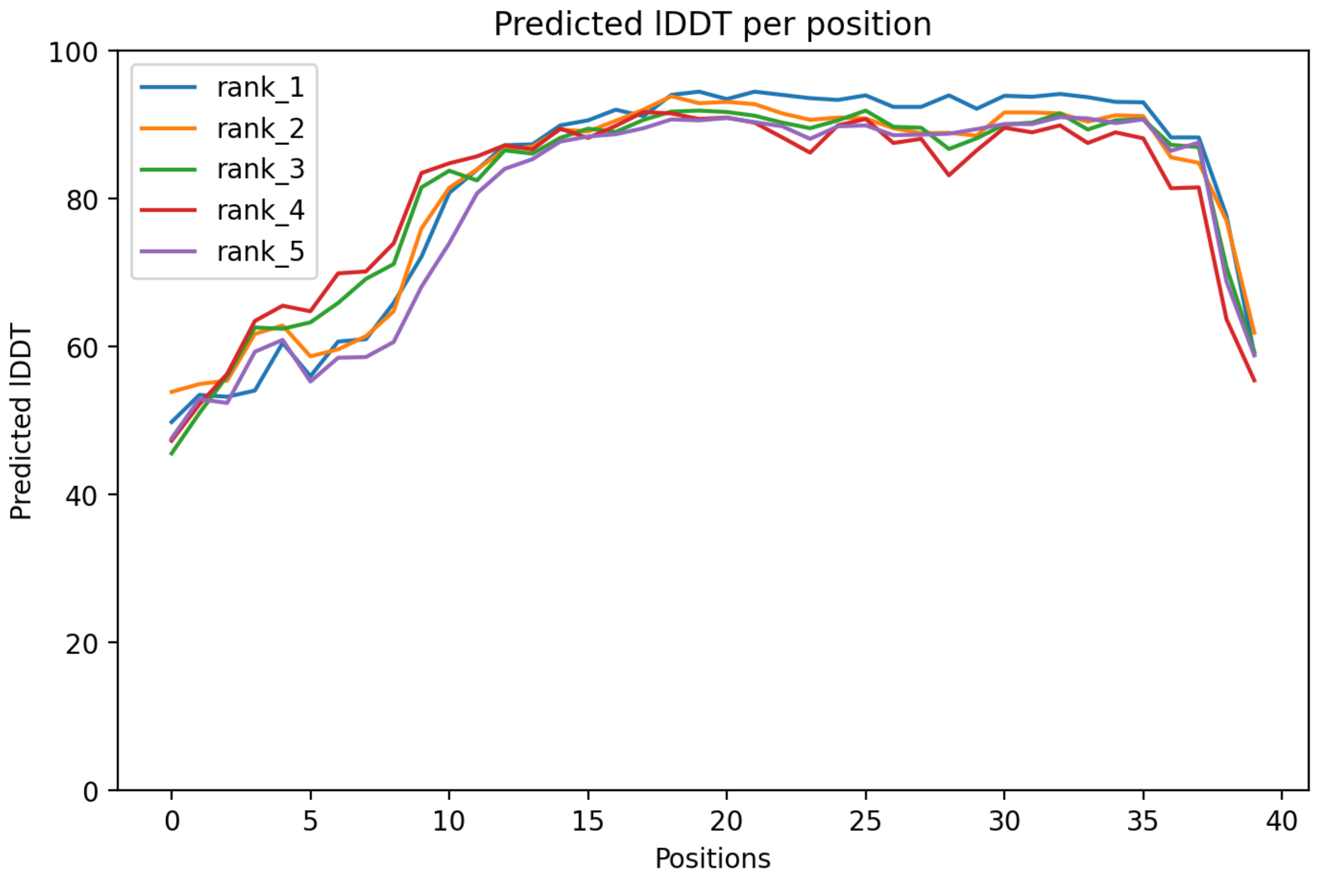

CP105: MtrunA17_Chr6g0468411

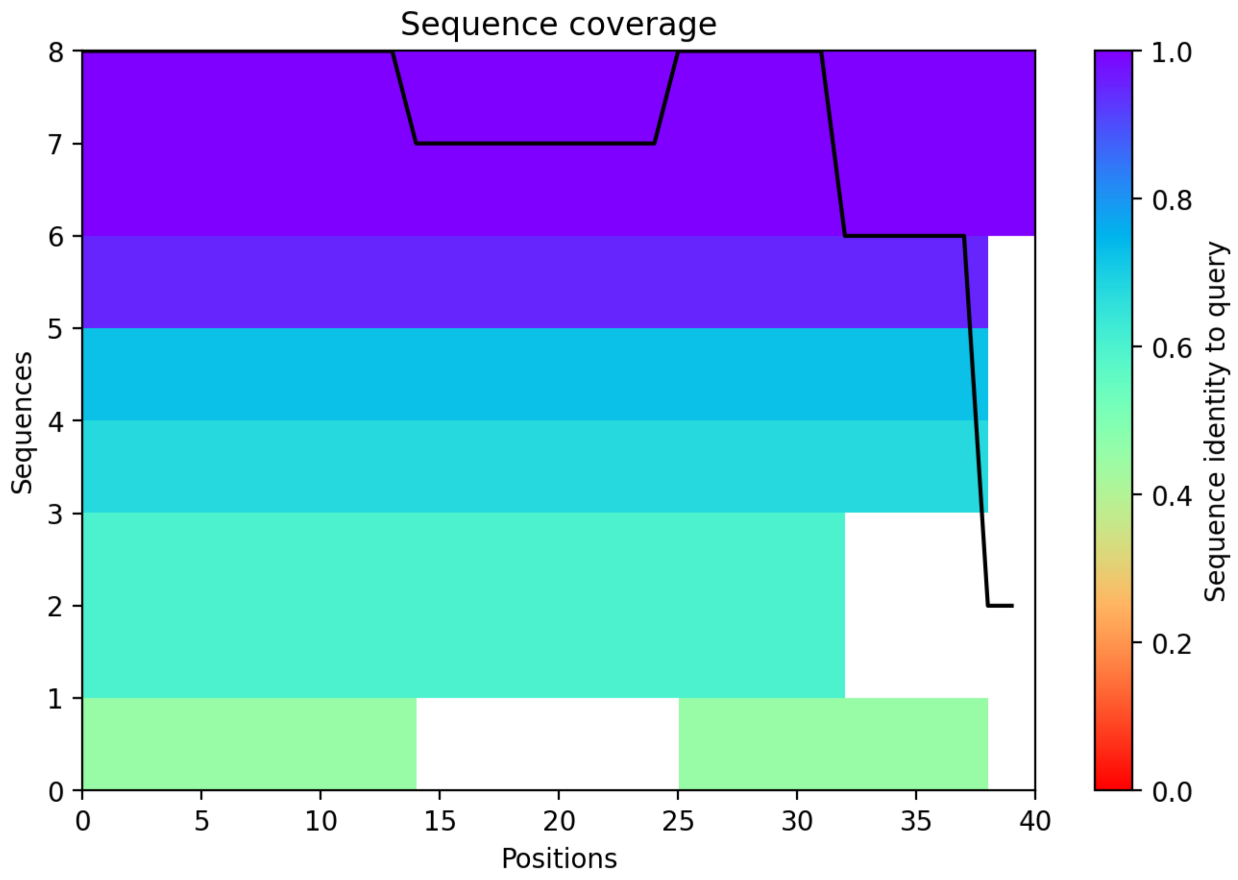

### CP106: MtrunA17_Chr6g0476751

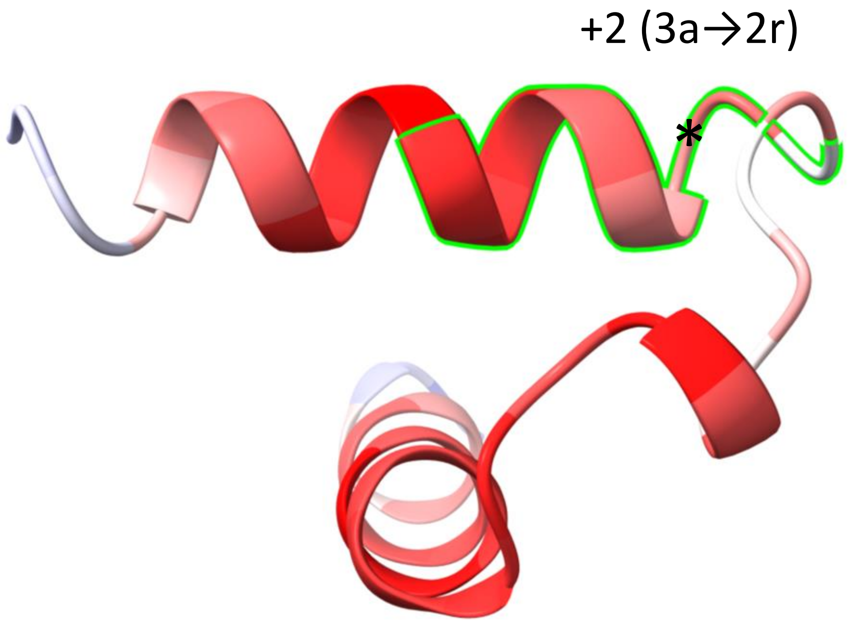


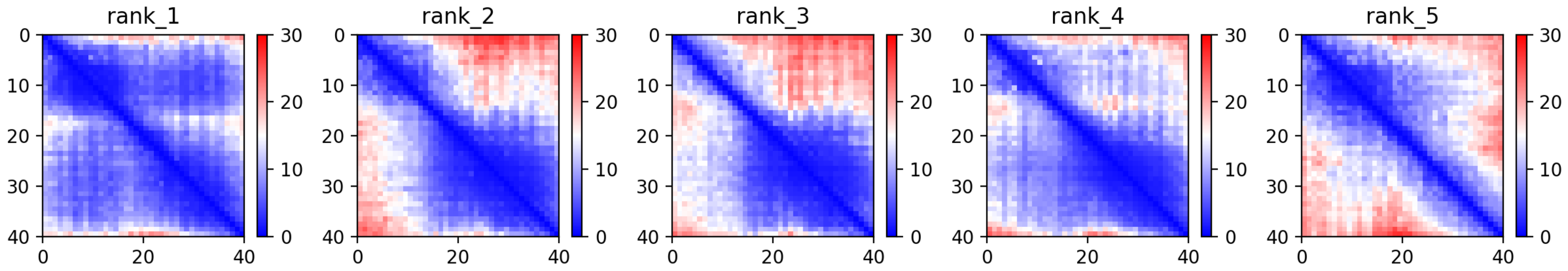

CP106: MtrunA17_Chr6g0476751

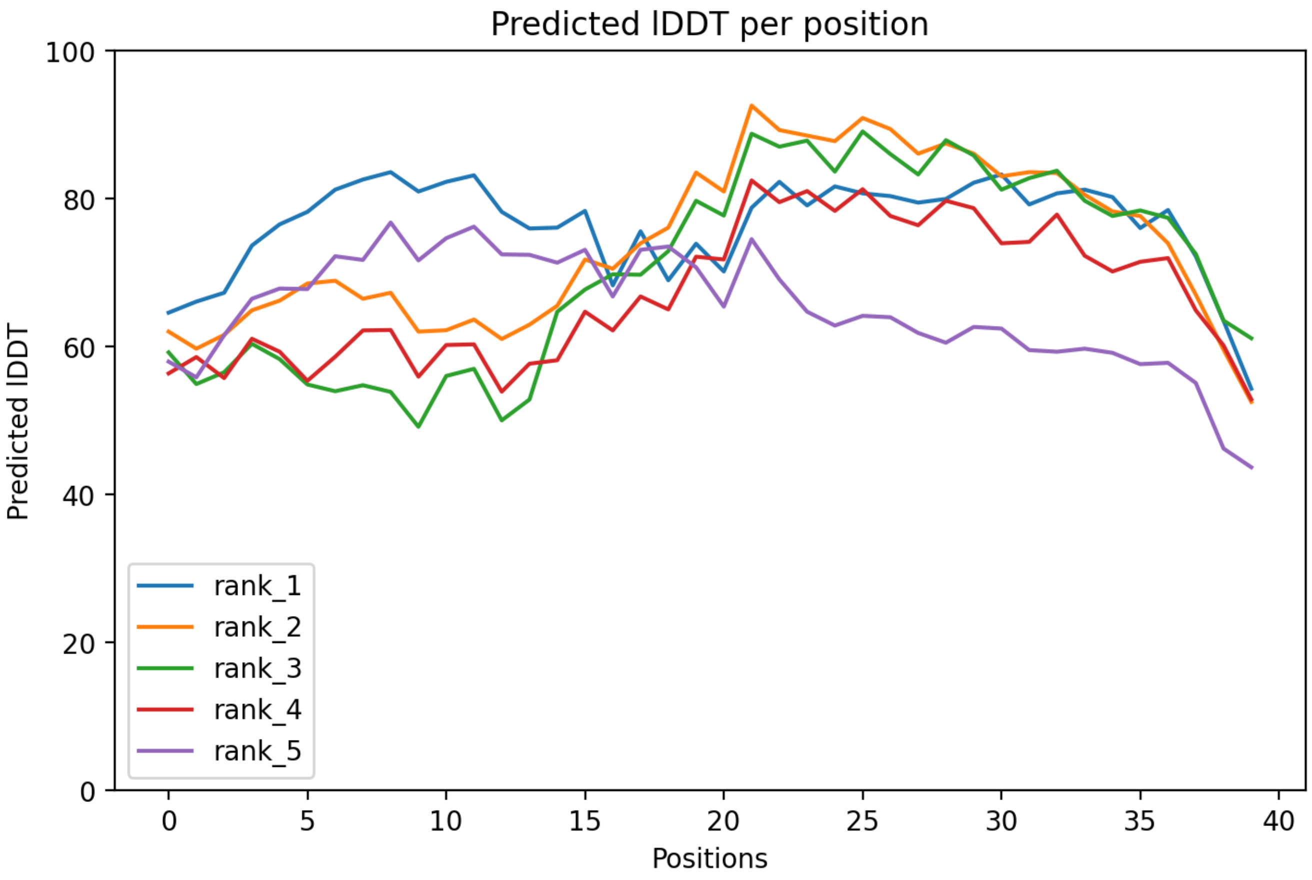

### CP106: MtrunA17_Chr6g0476751

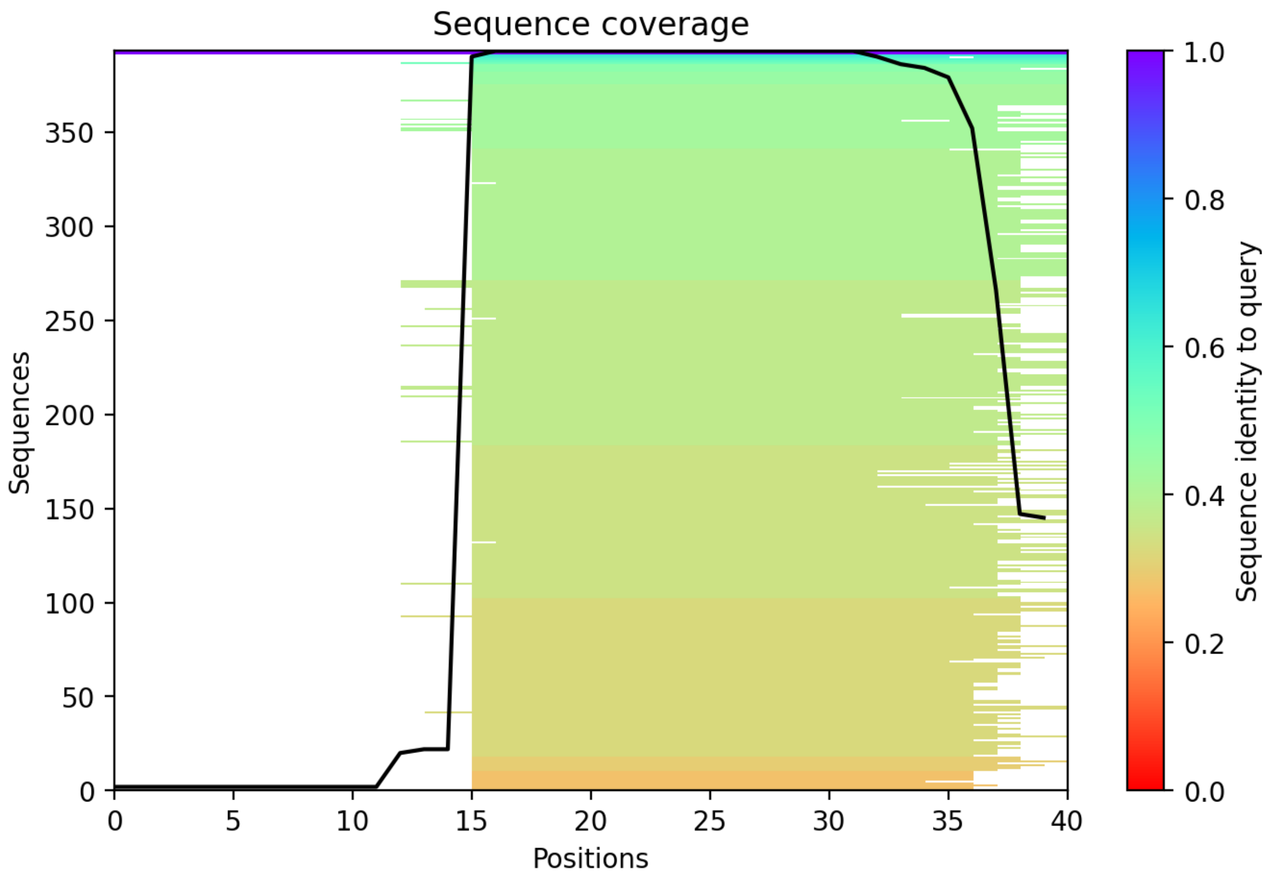

CP107: MtrunA17_Chr6g0479001

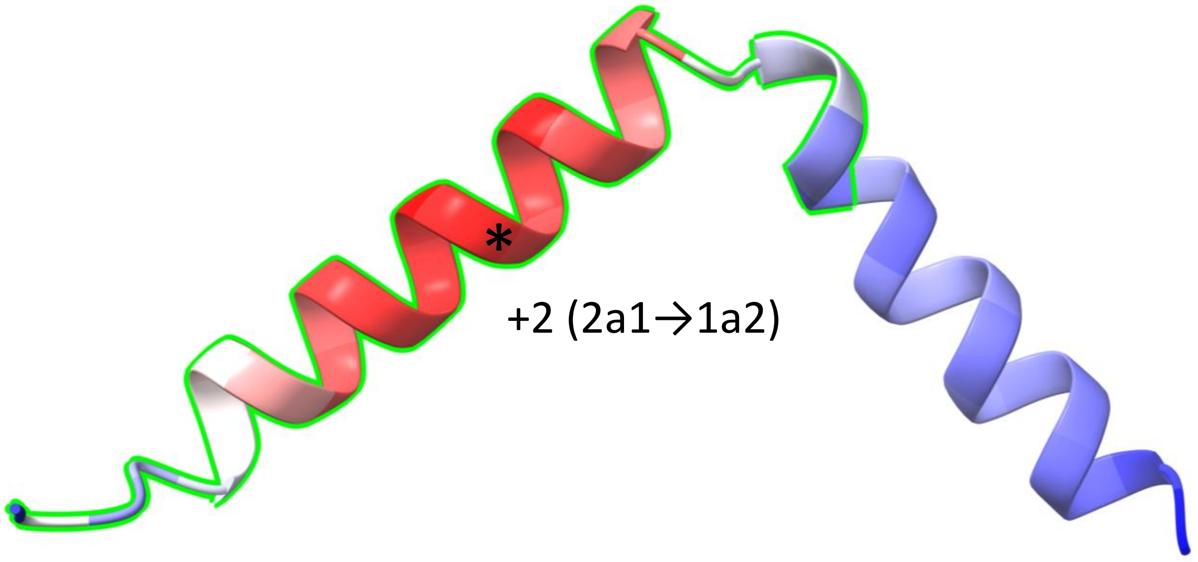


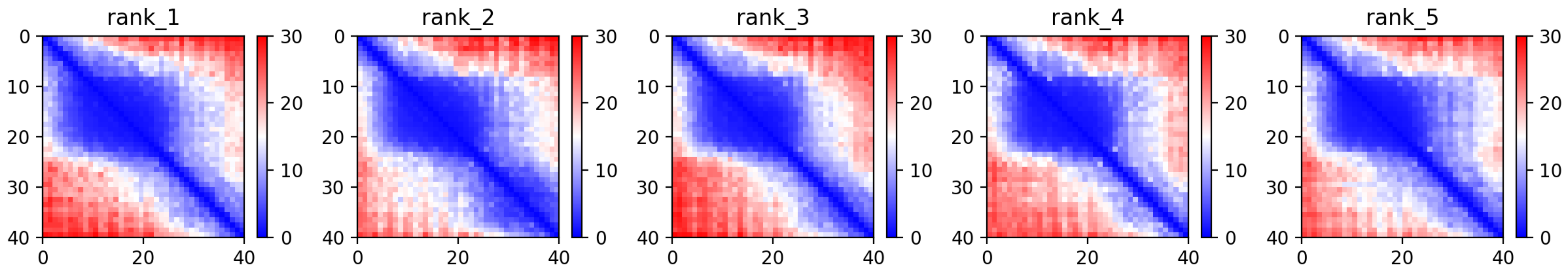

CP107: MtrunA17_Chr6g0479001

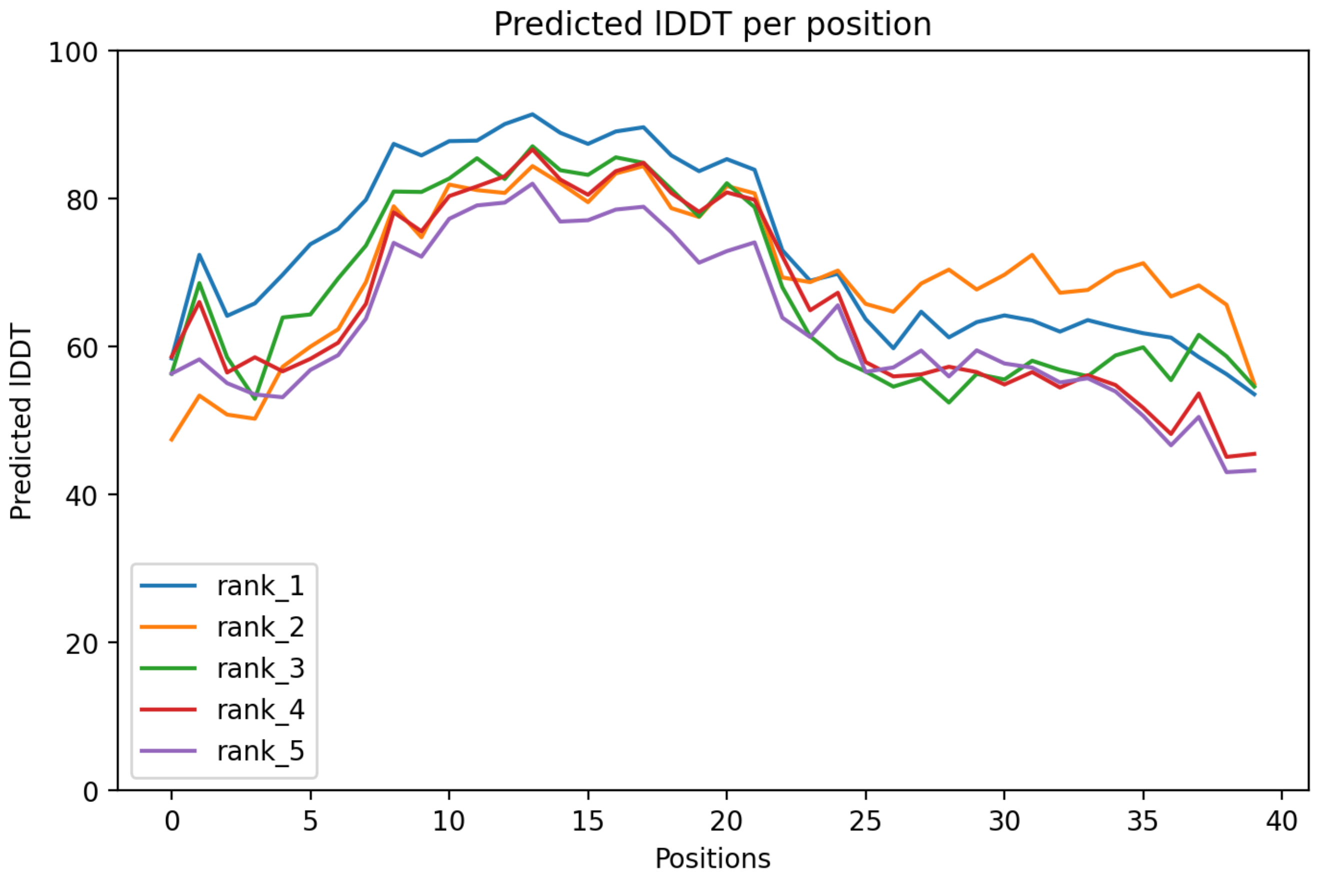

### CP107: MtrunA17_Chr6g0479001

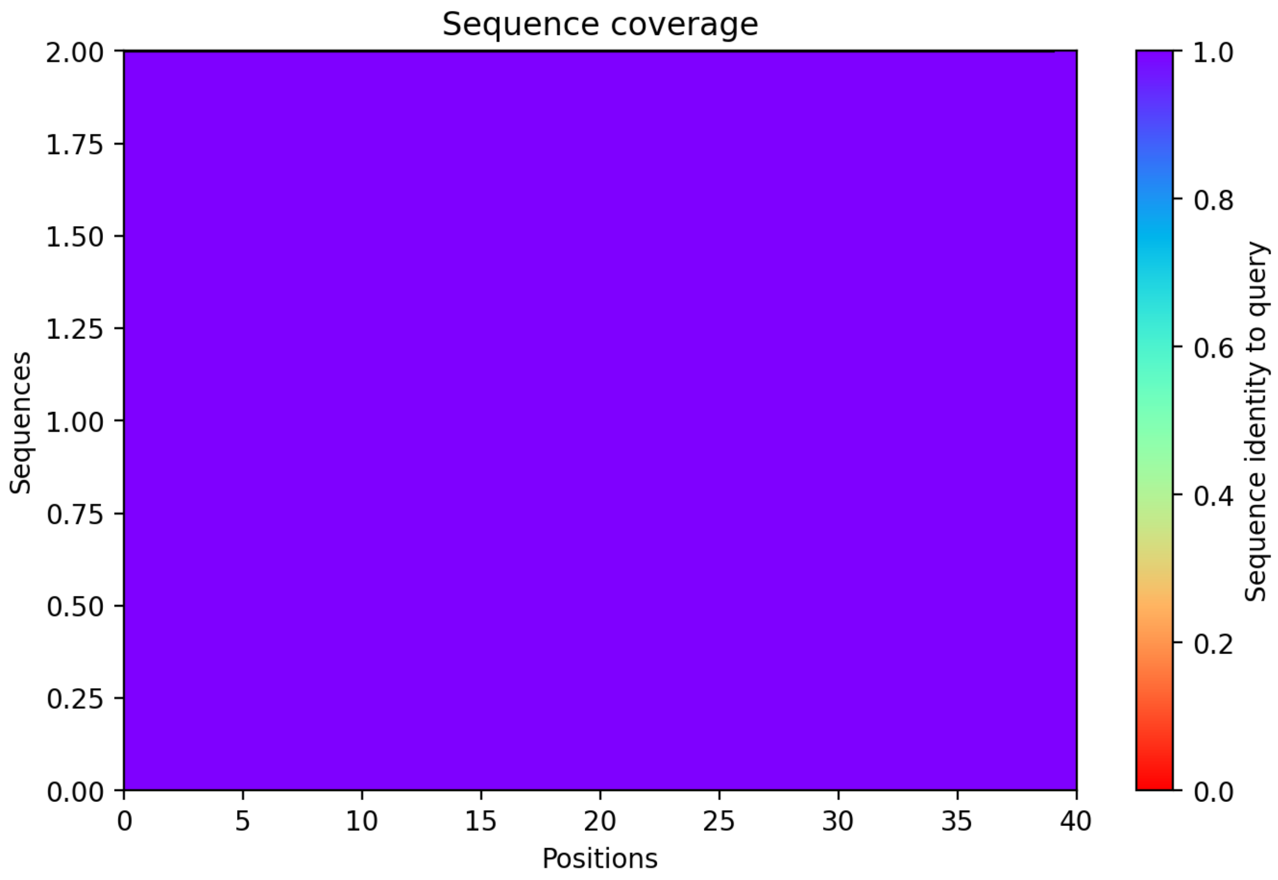

### CP108: MtrunA17_Chr6g0485321

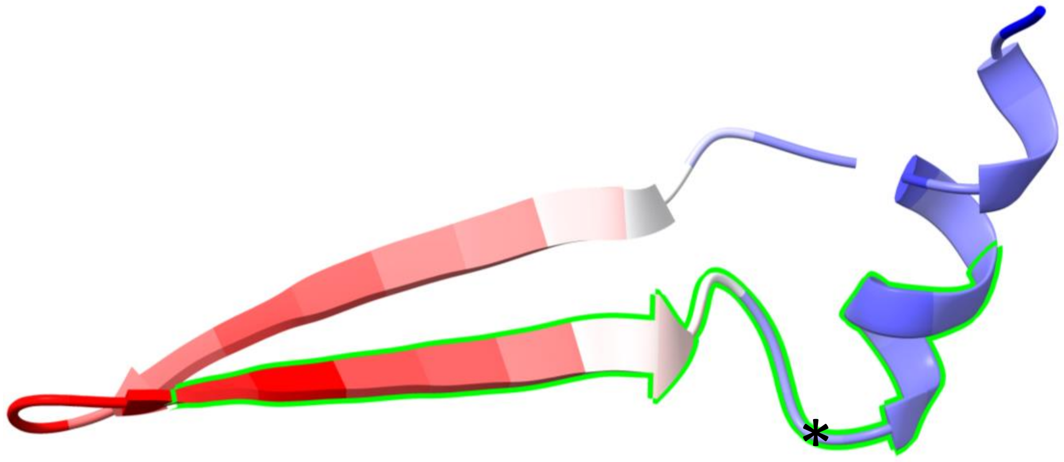


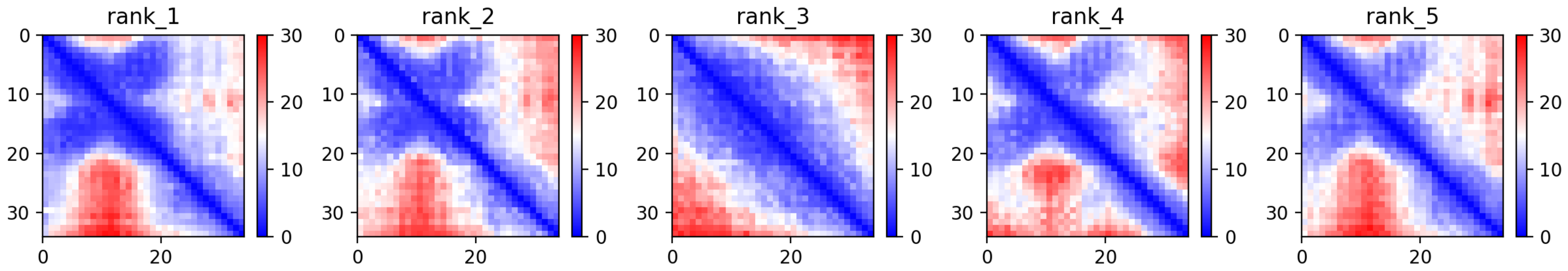

CP108: MtrunA17_Chr6g0485321

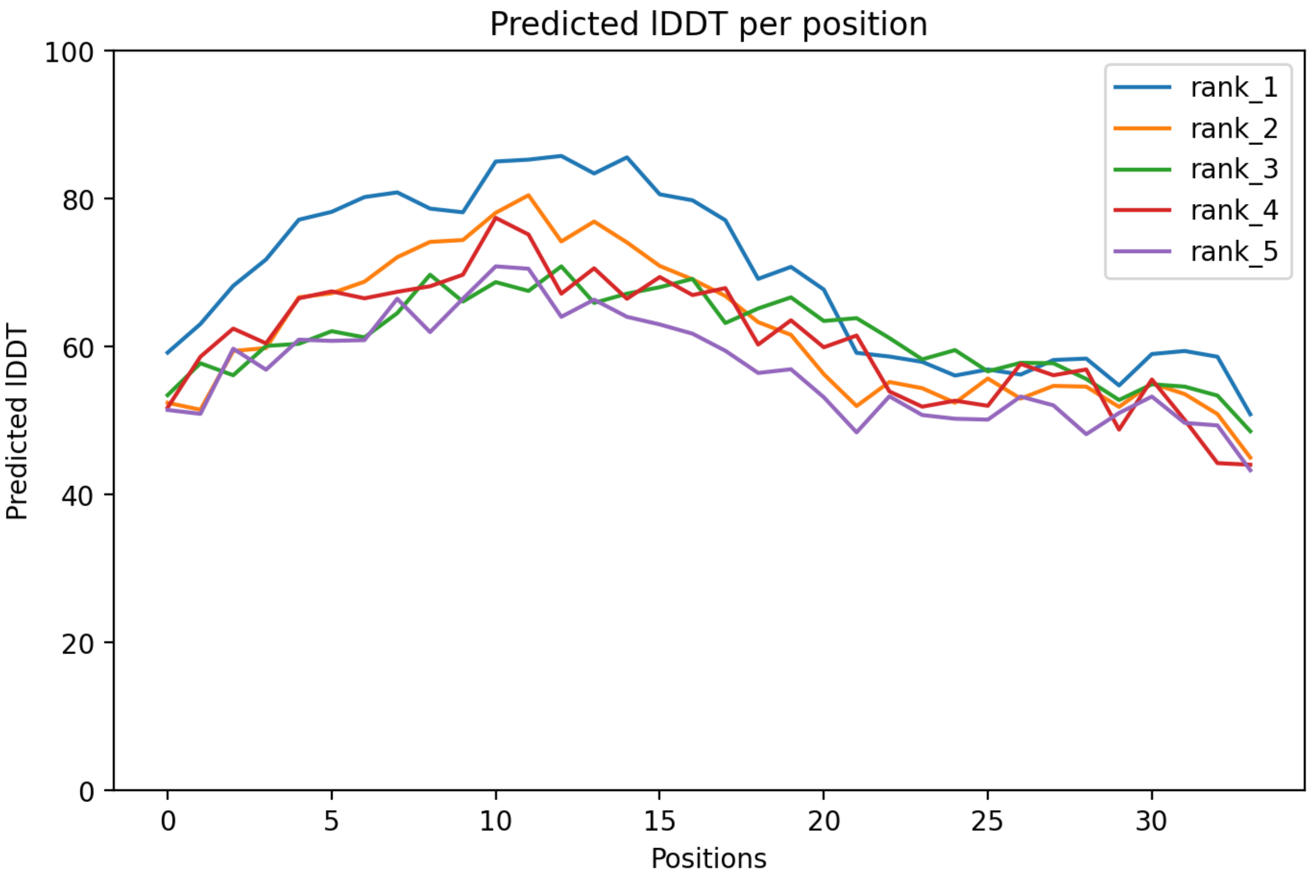

### CP108: MtrunA17_Chr6g0485321

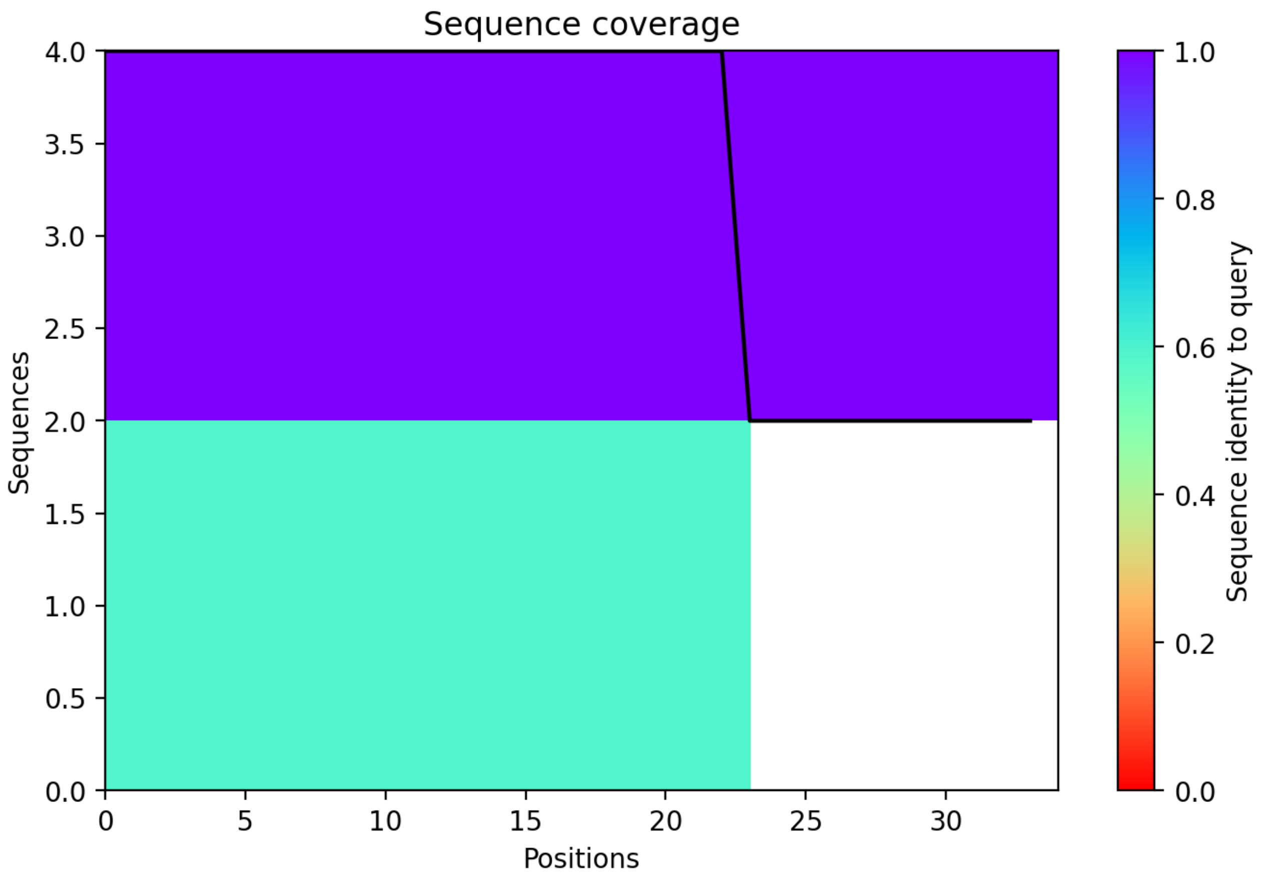

CP109: MtrunA17_Chr6g0486961

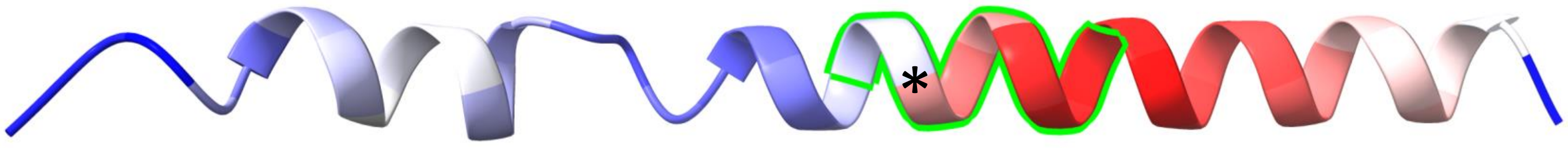


-2 (3a→1r)

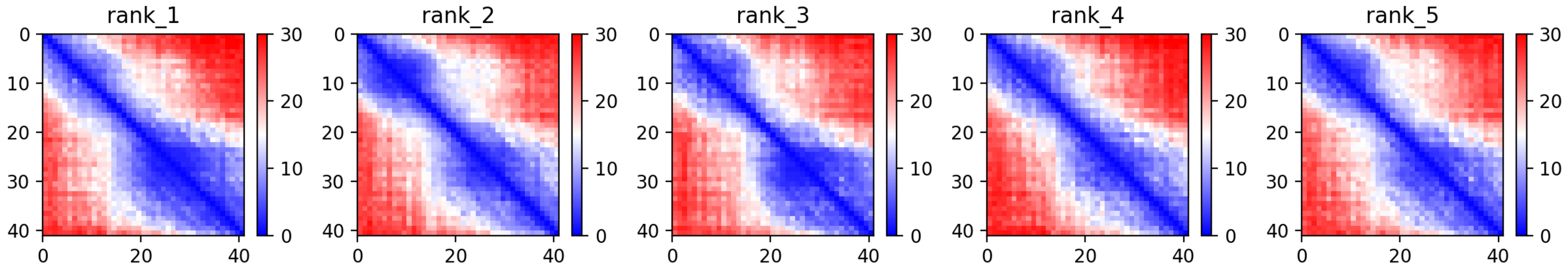

### CP109: MtrunA17_Chr6g0486961

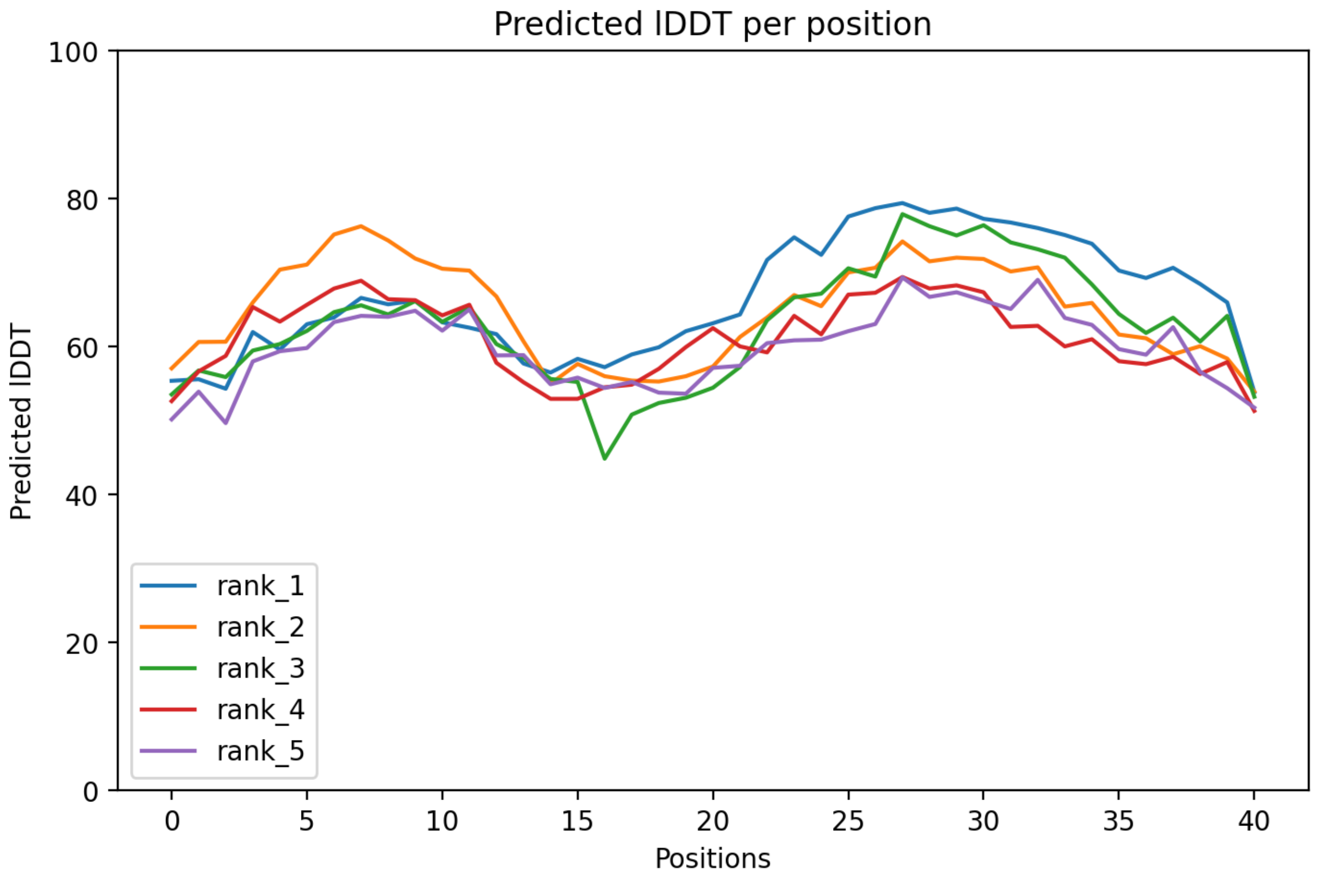

### CP109: MtrunA17_Chr6g0486961

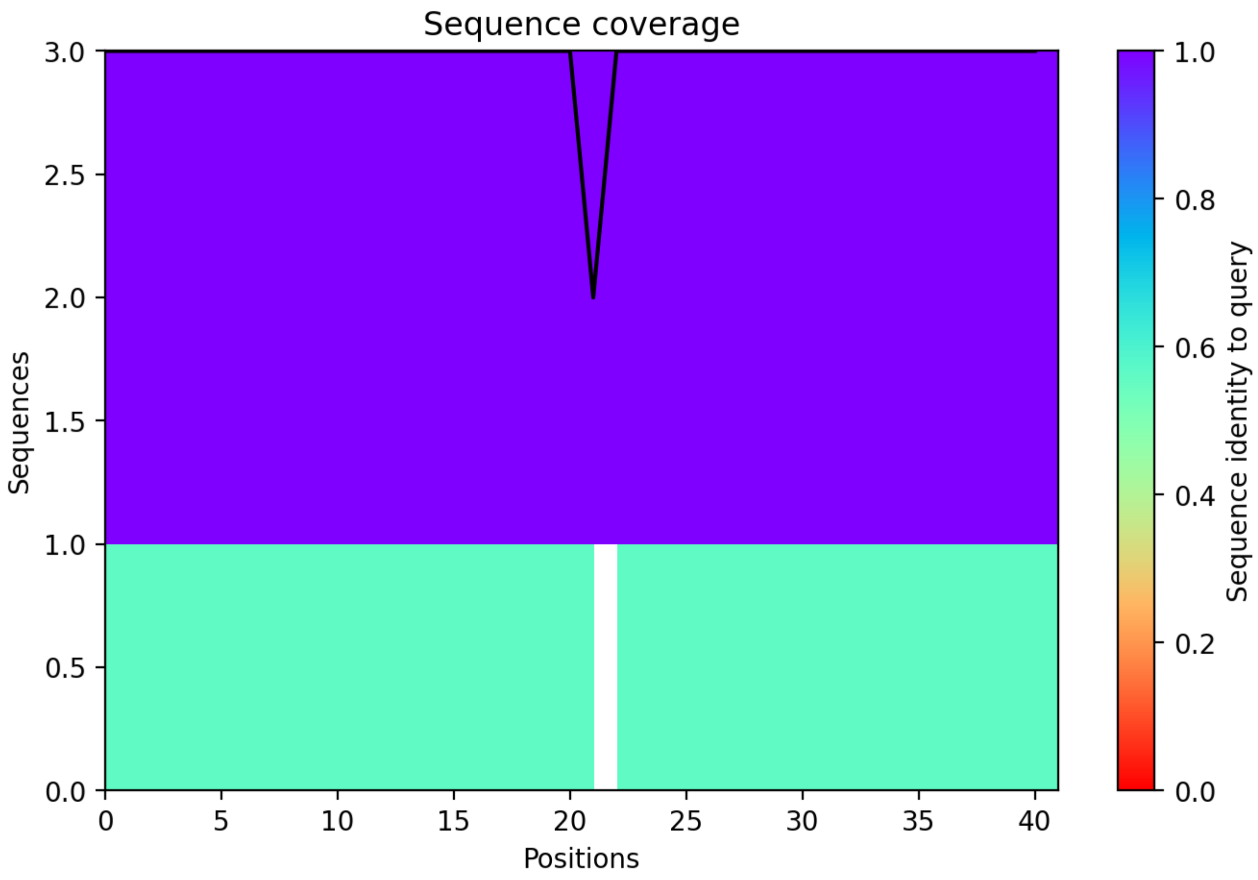

### CP110: MtrunA17_Chr7g0214741

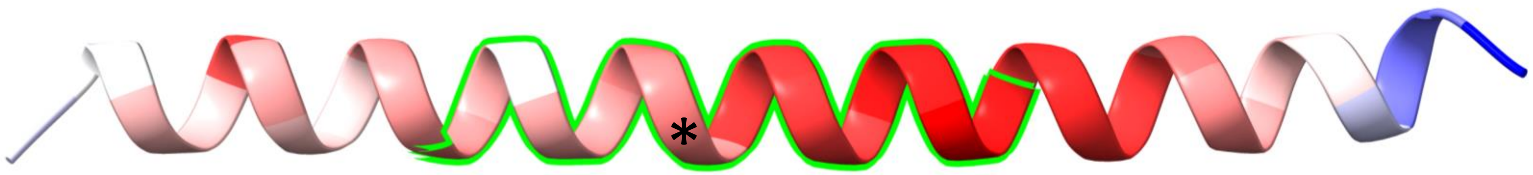


+1 (3r→1a)

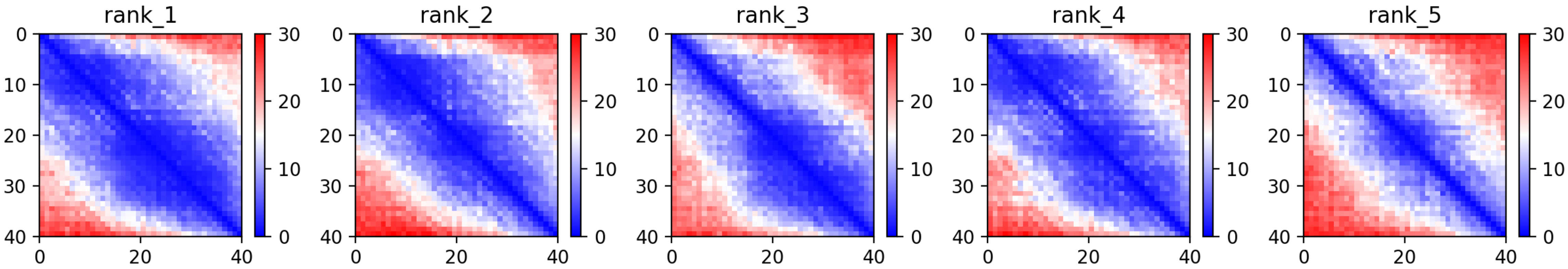

CP110: MtrunA17_Chr7g0214741

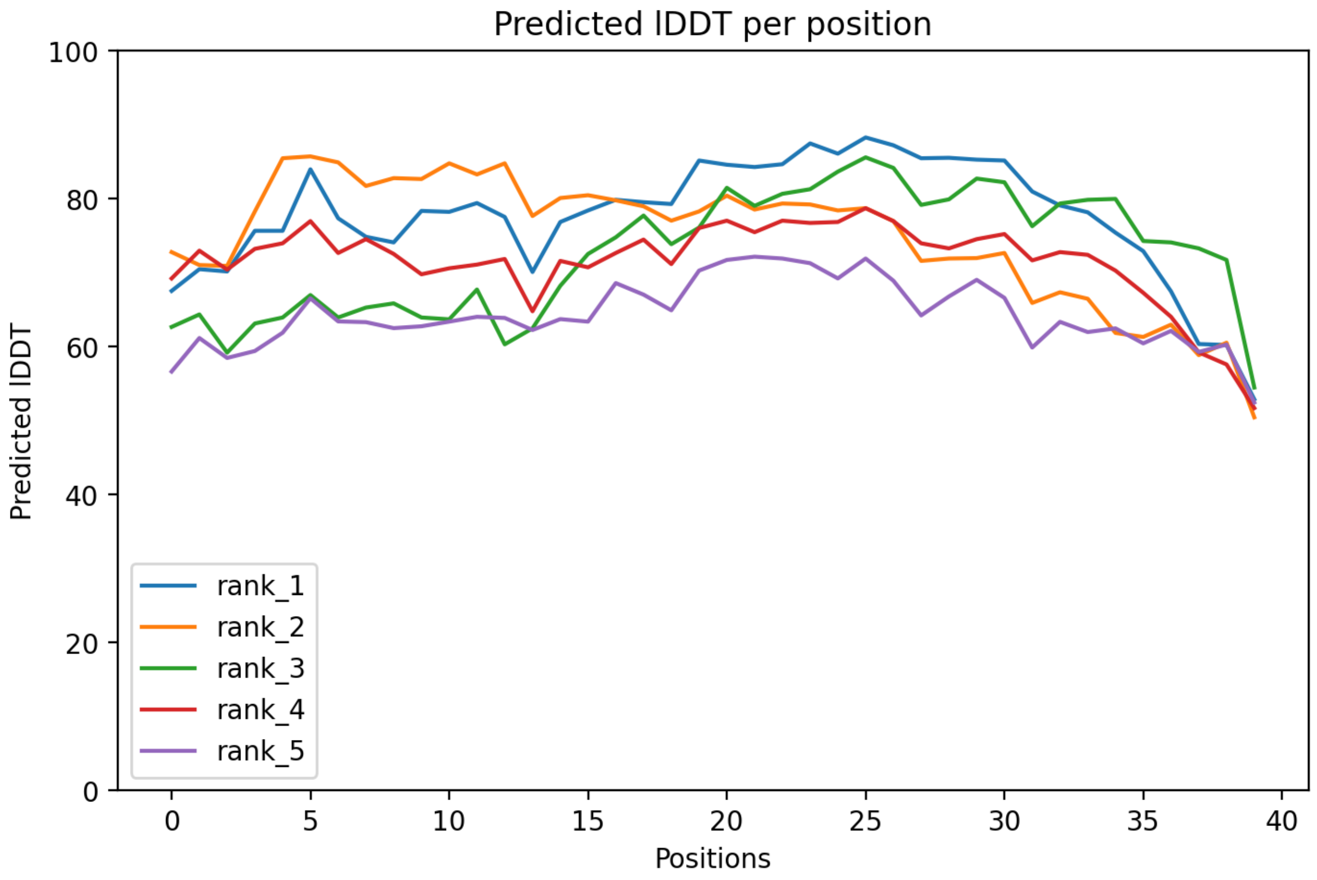

### CP110: MtrunA17_Chr7g0214741

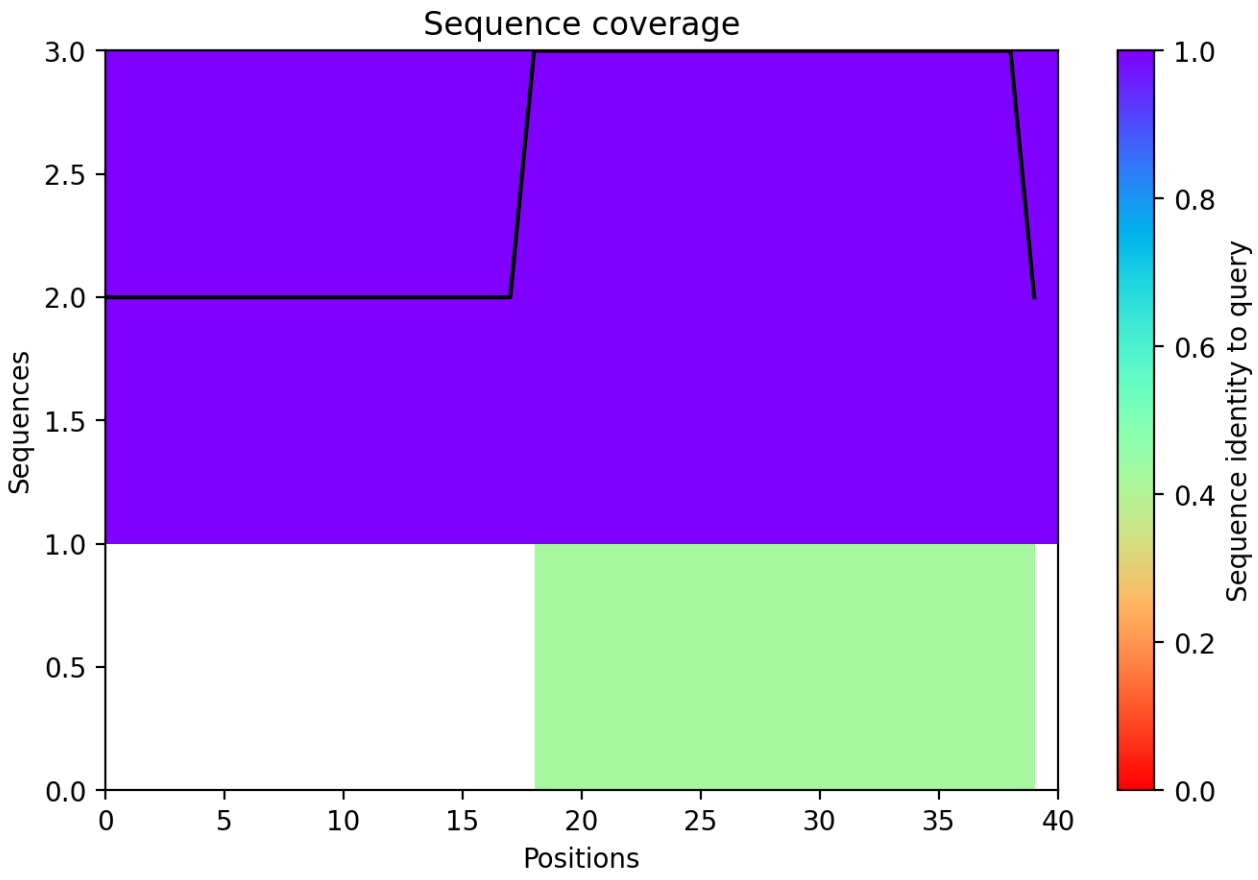

CP111: MtrunA17_Chr7g0214911

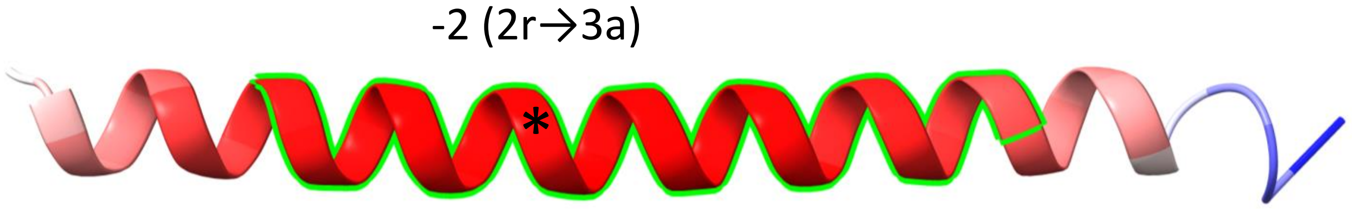


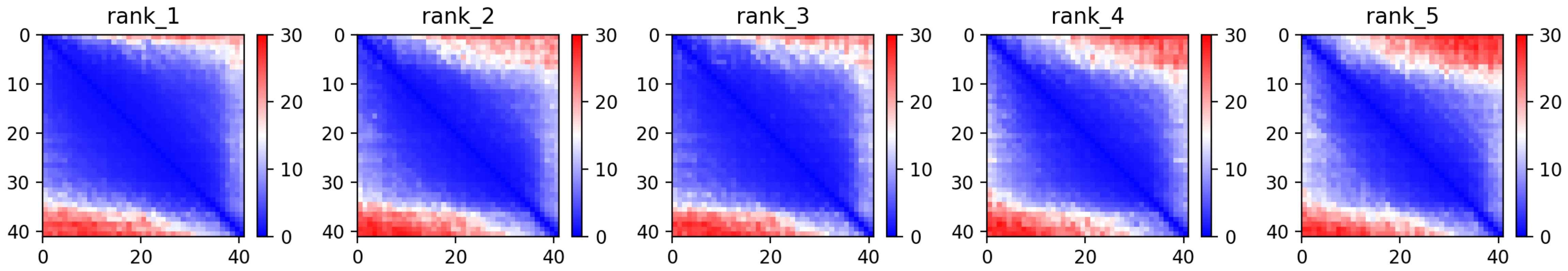

CP111: MtrunA17_Chr7g0214911

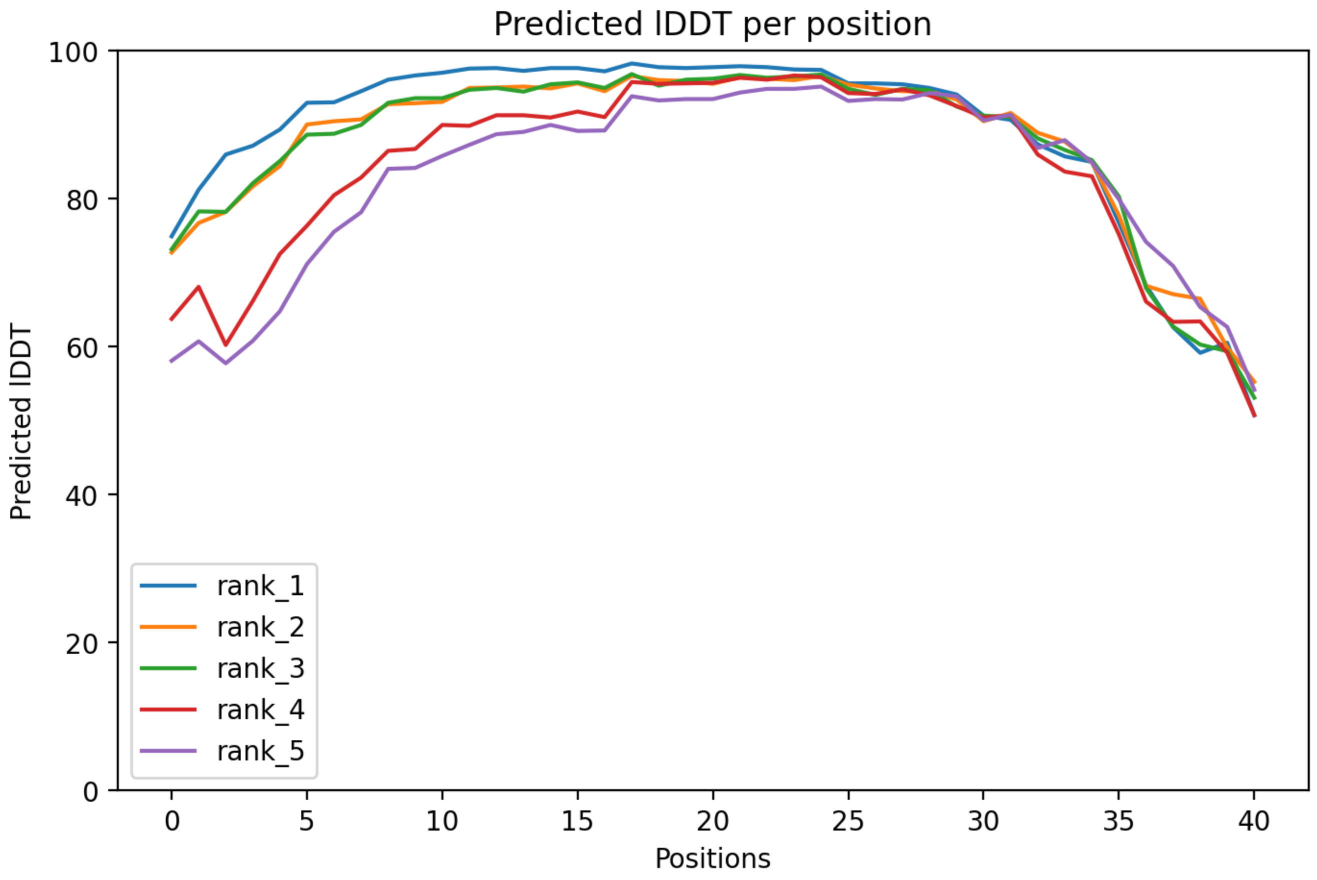

### CP111: MtrunA17_Chr7g0214911

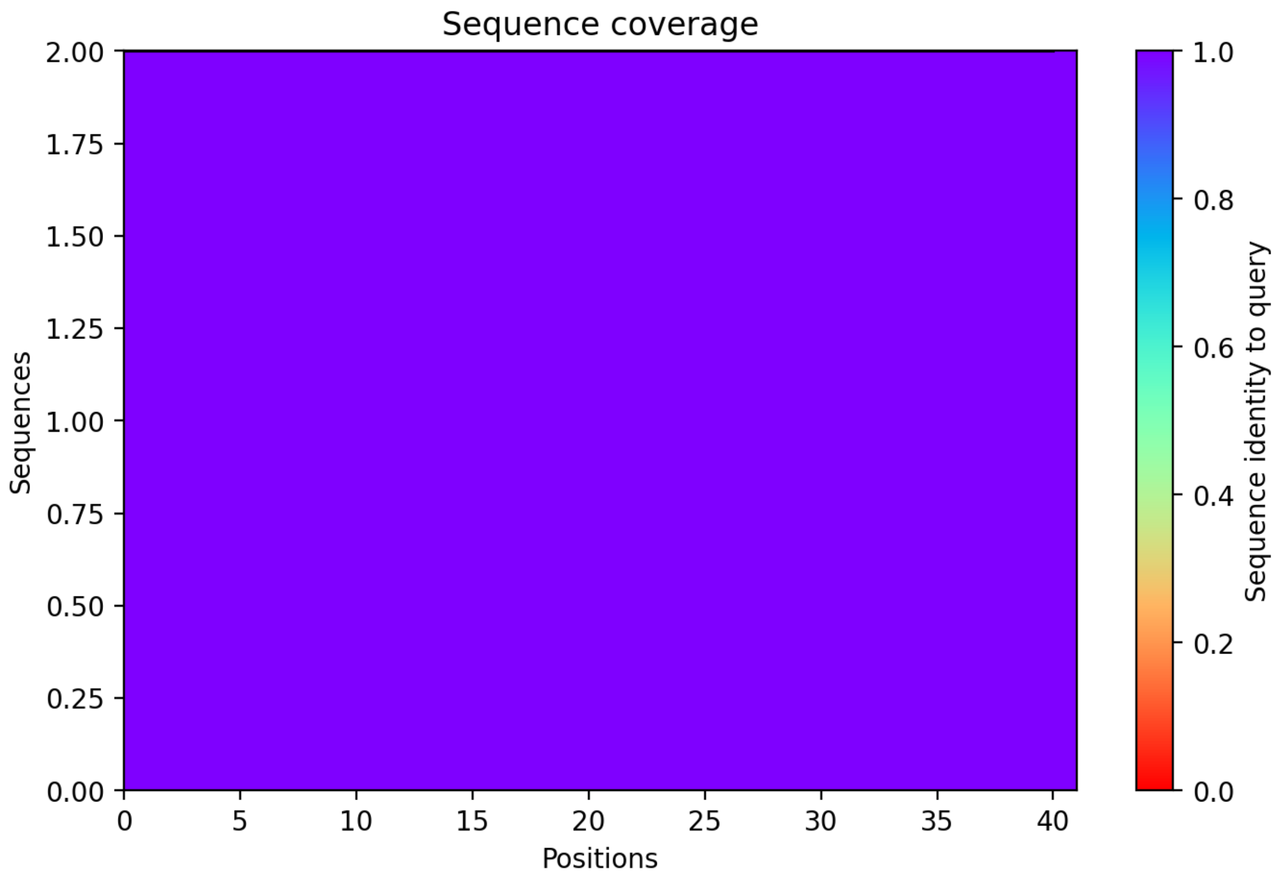

CP112: MtrunA17_Chr7g0219691

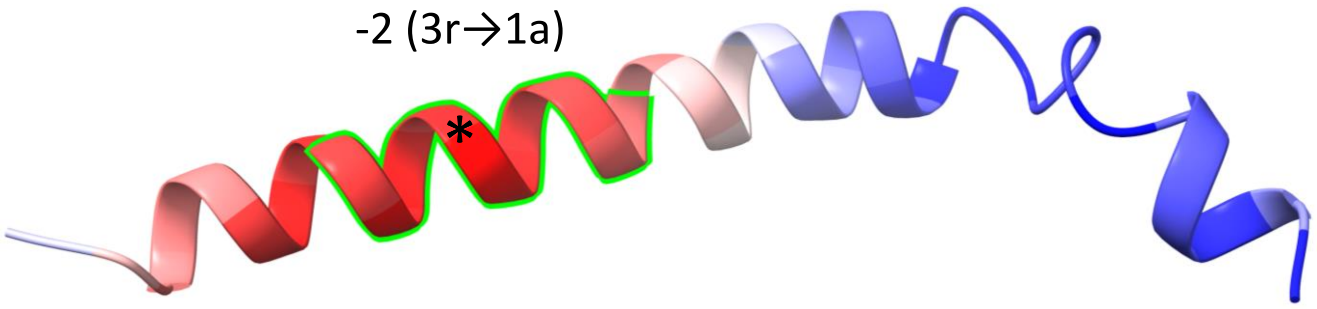


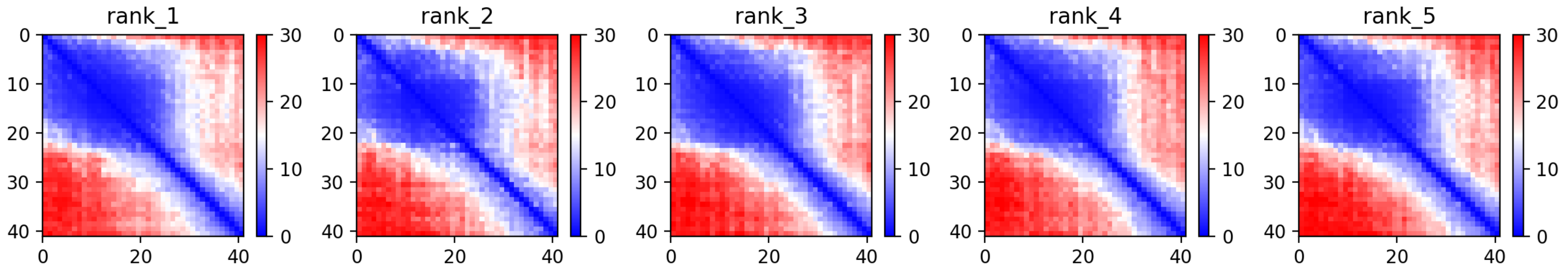

CP112: MtrunA17_Chr7g0219691

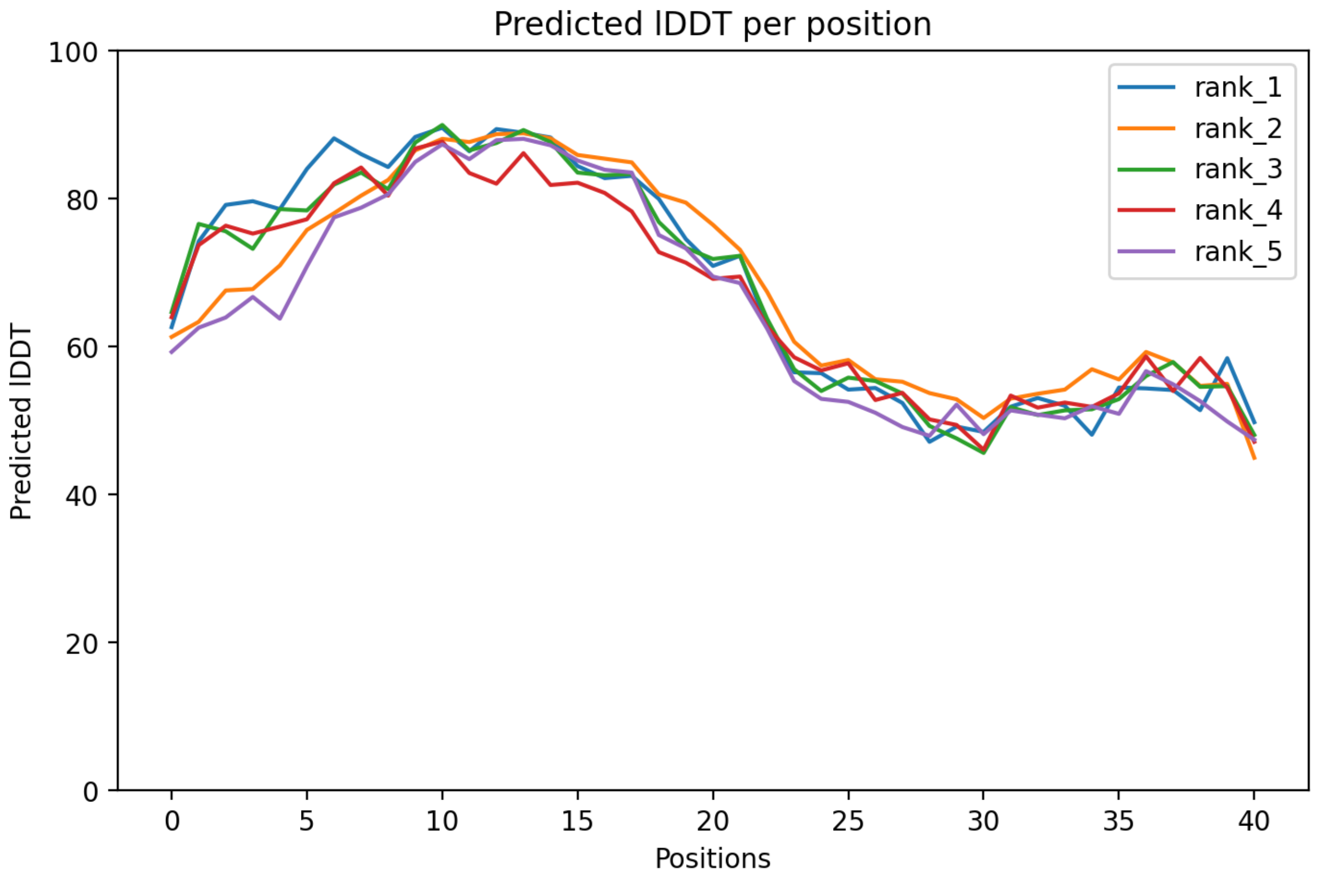

CP112: MtrunA17_Chr7g0219691

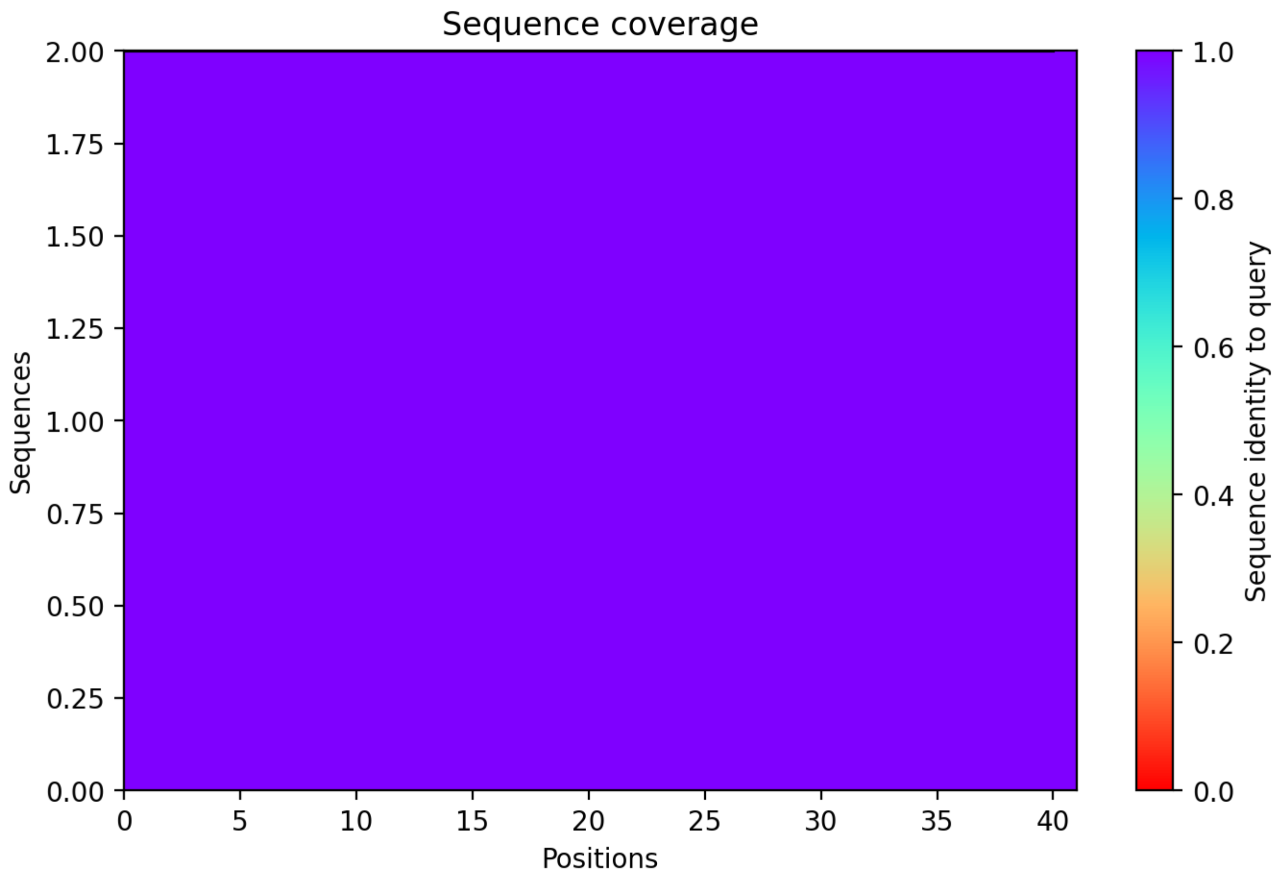

CP113: MtrunA17_Chr7g0221631

-1 (1r→3a)

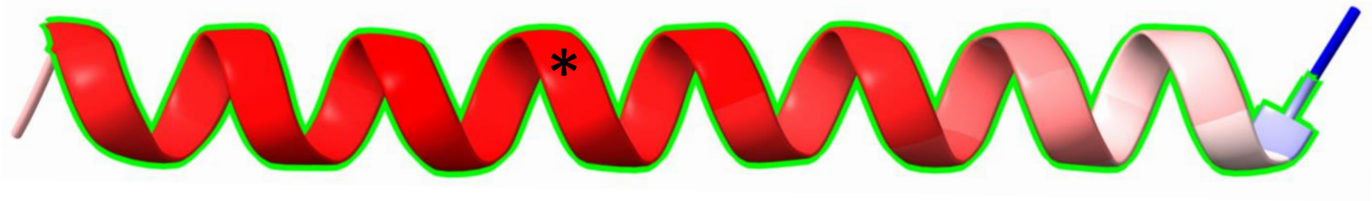


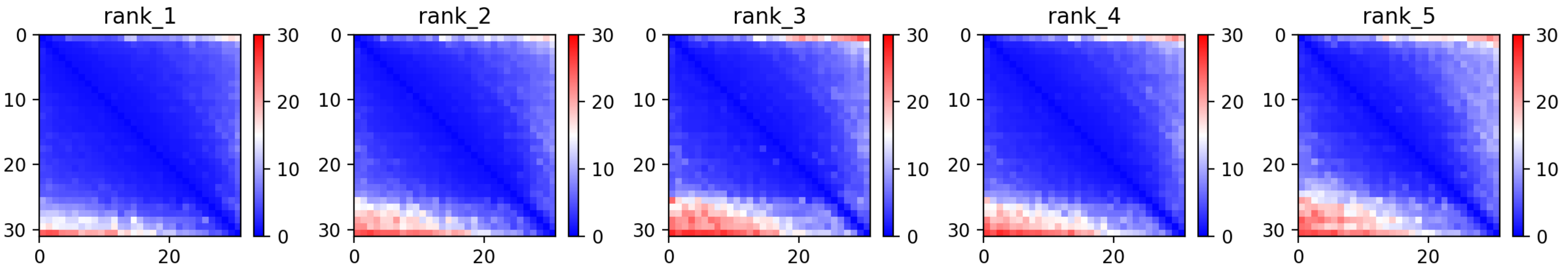

### CP113: MtrunA17_Chr7g0221631

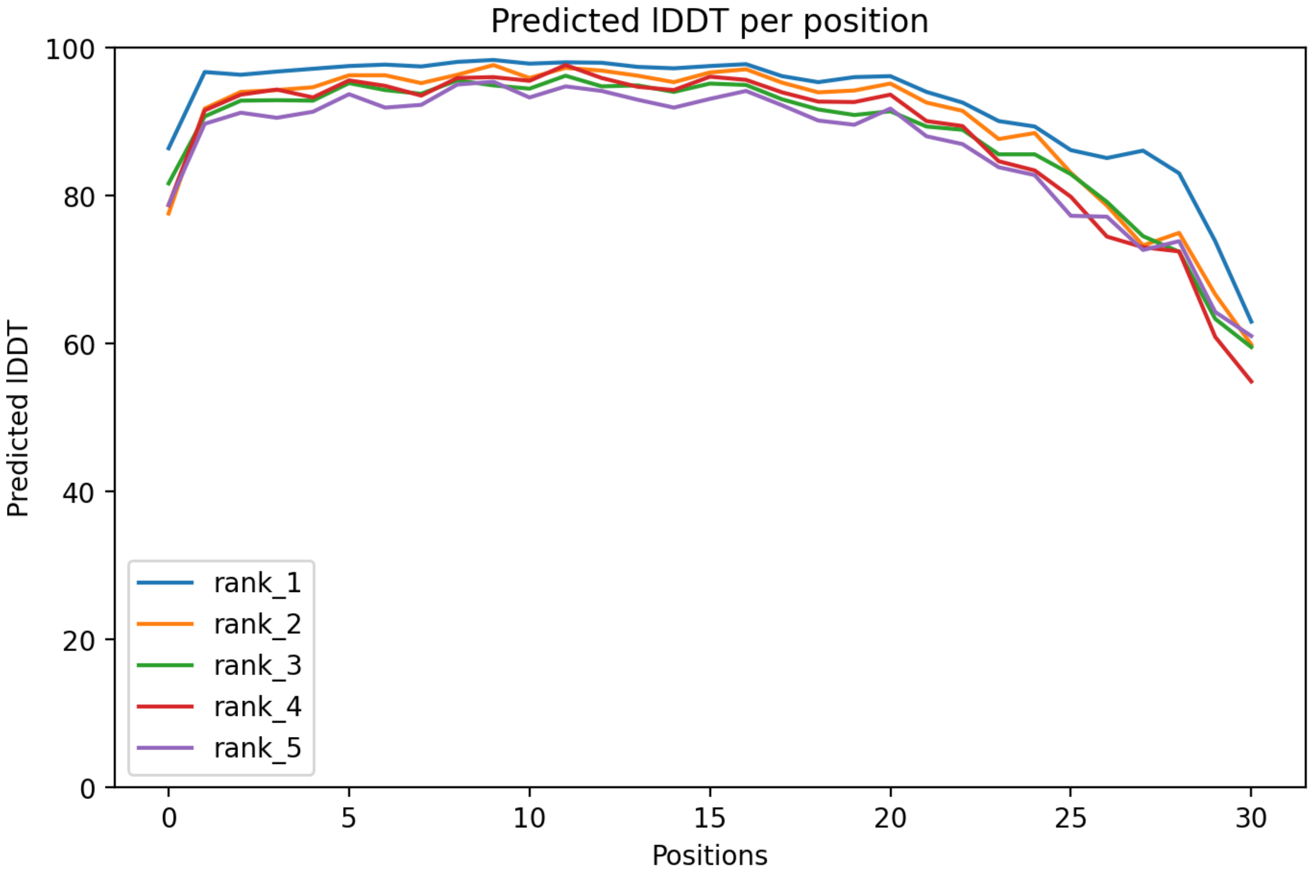

CP113: MtrunA17_Chr7g0221631

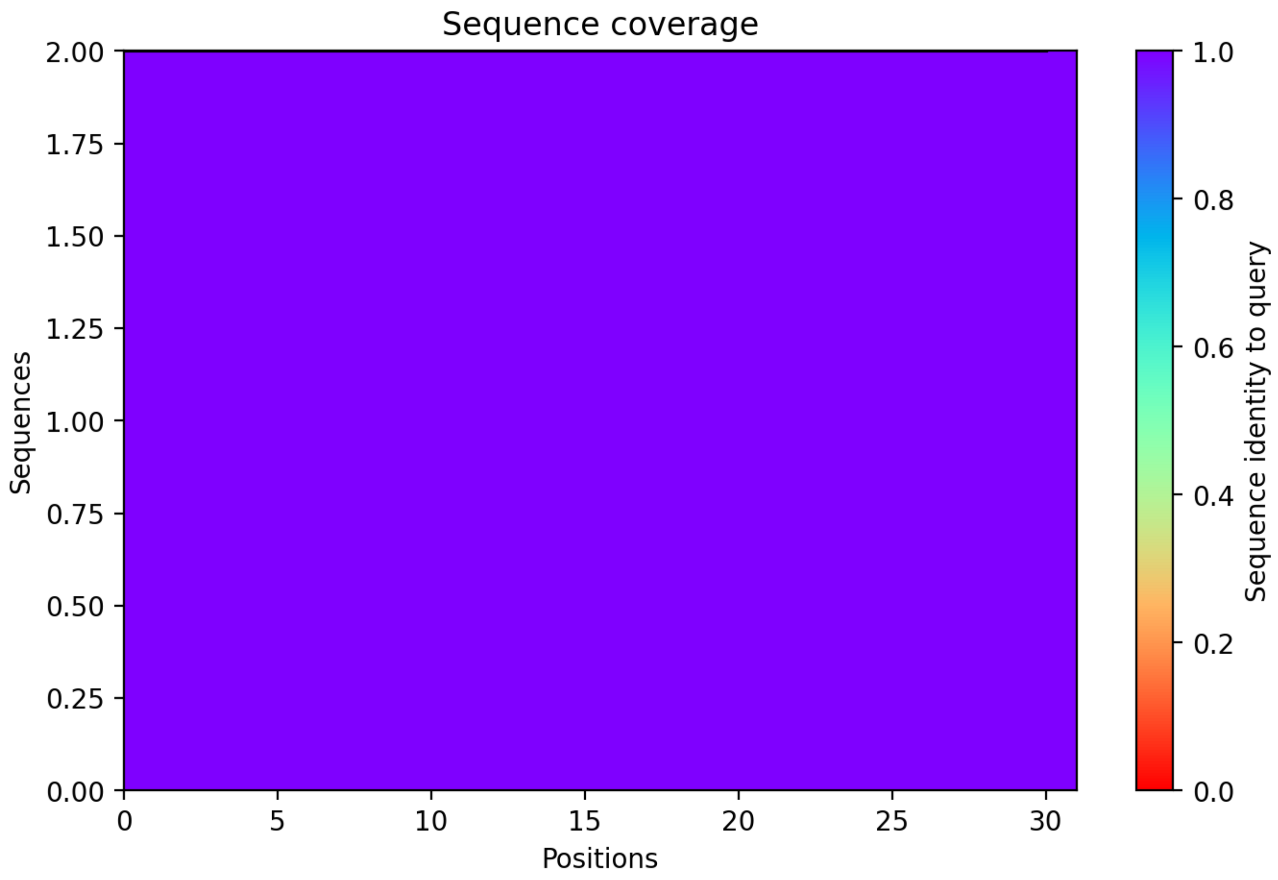

CP114: MtrunA17_Chr7g0229401

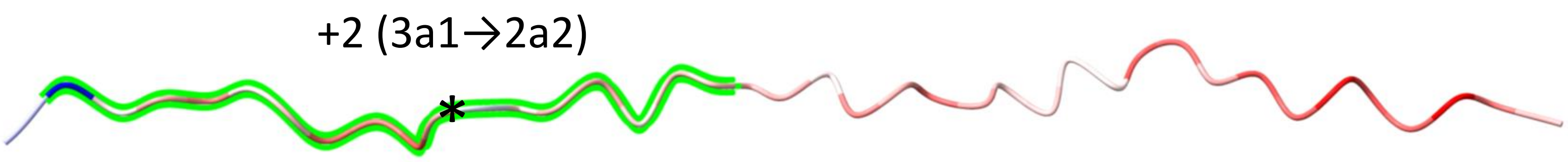


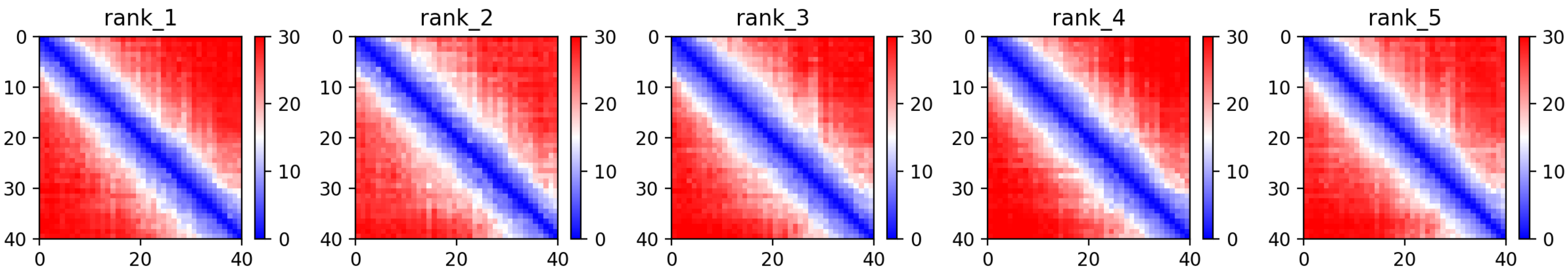

### CP114: MtrunA17_Chr7g0229401

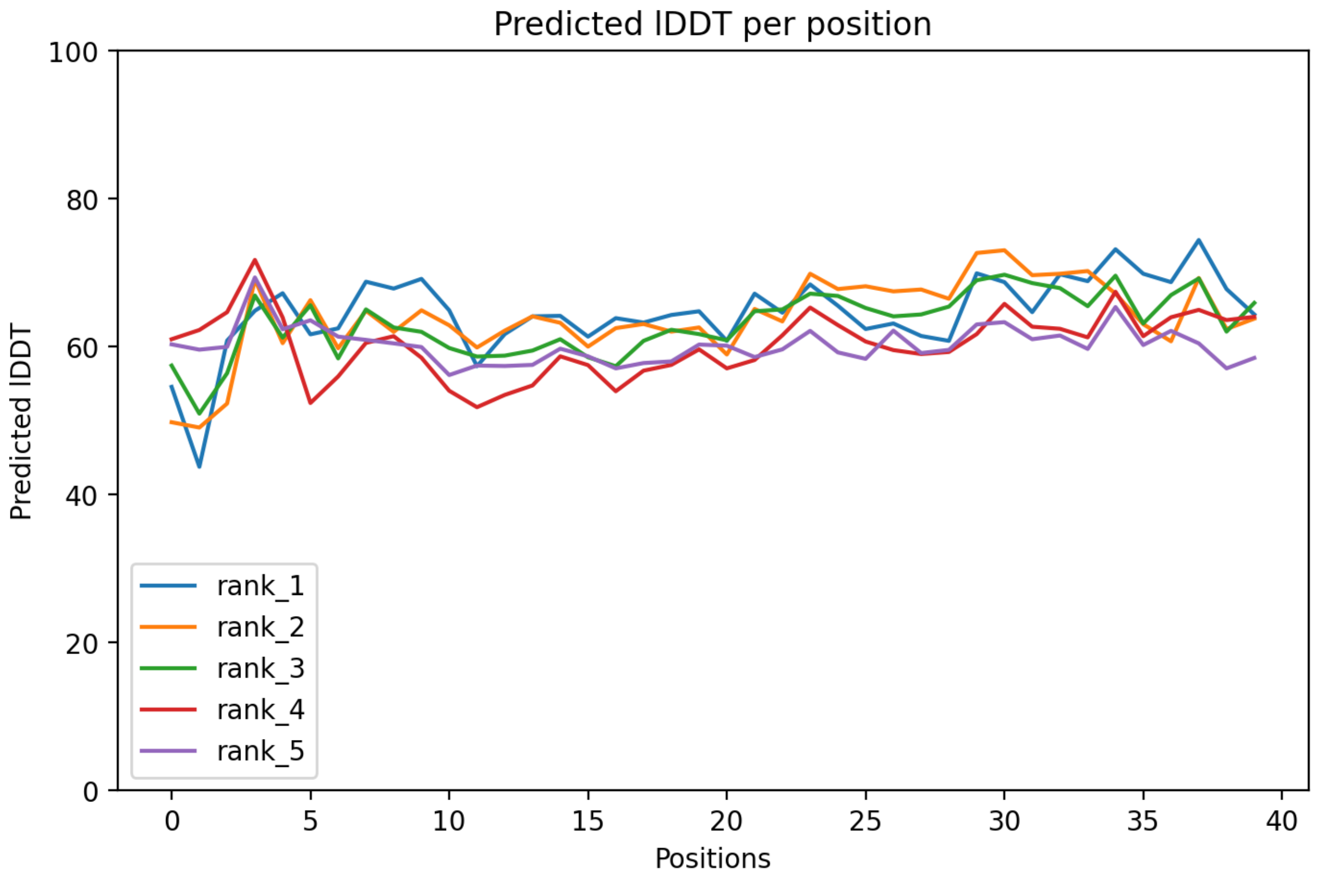

### CP114: MtrunA17_Chr7g0229401

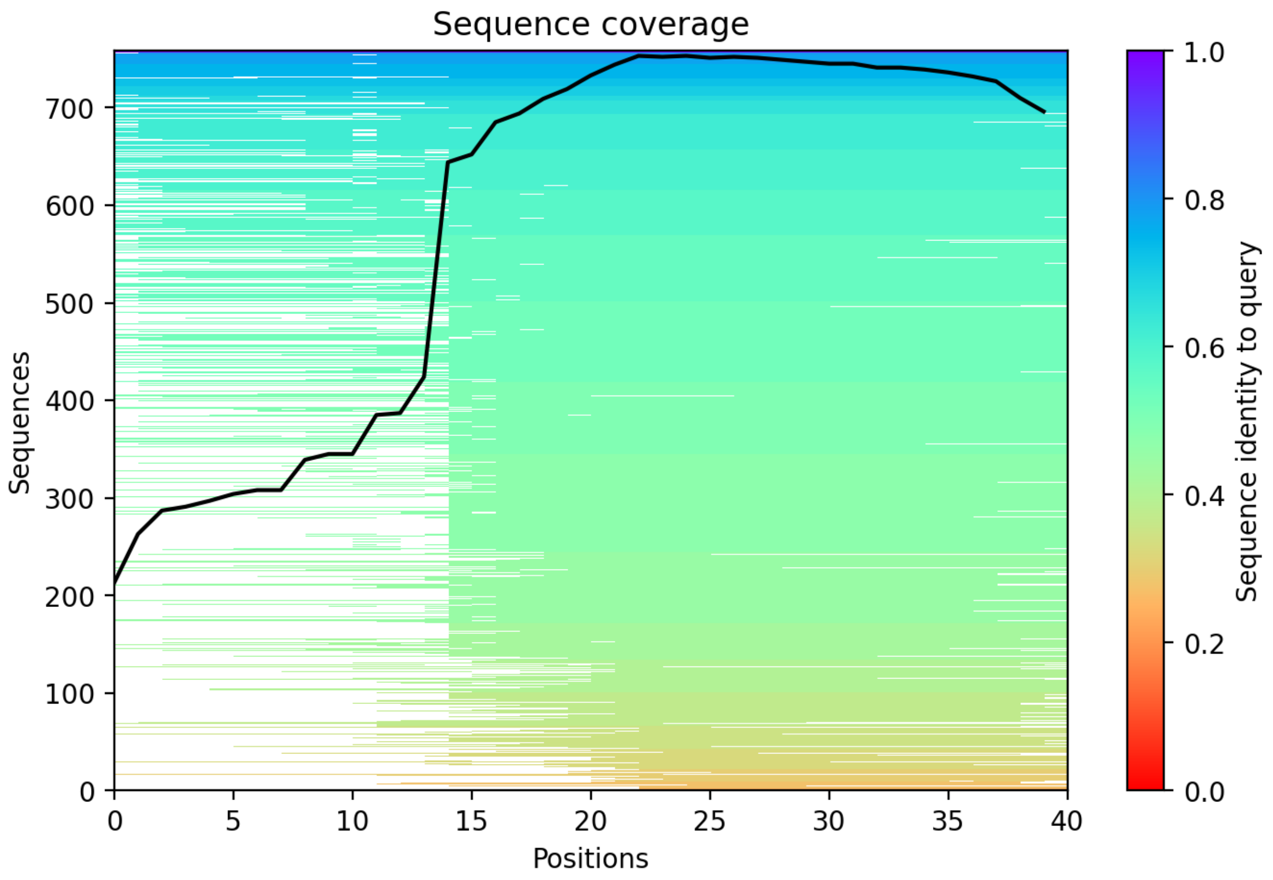

CP115: MtrunA17_Chr7g0230341

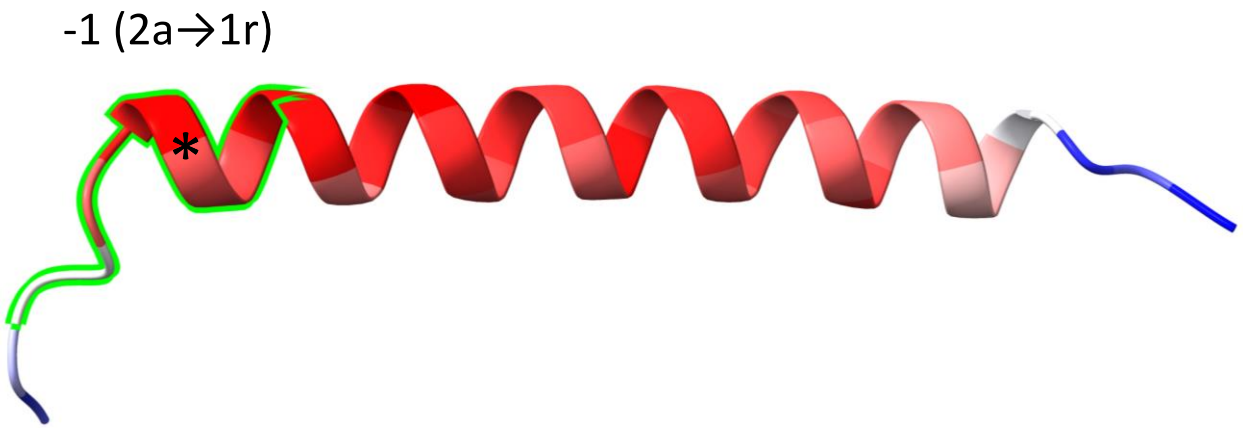


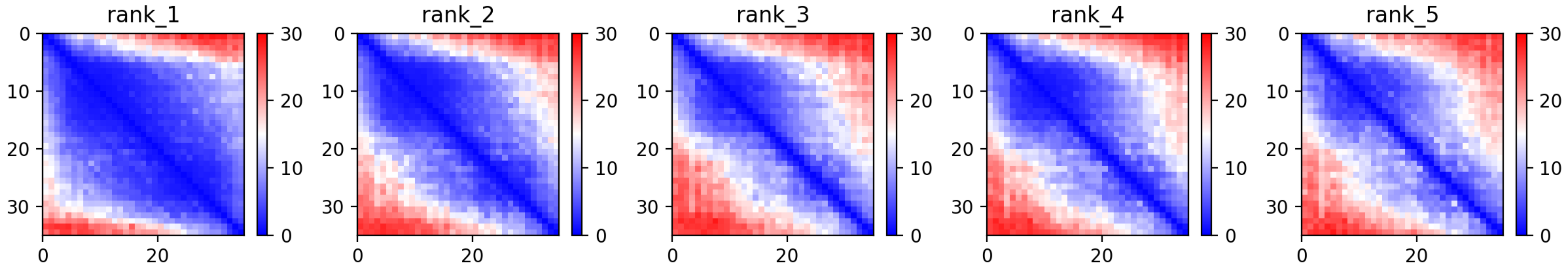

CP115: MtrunA17_Chr7g0230341

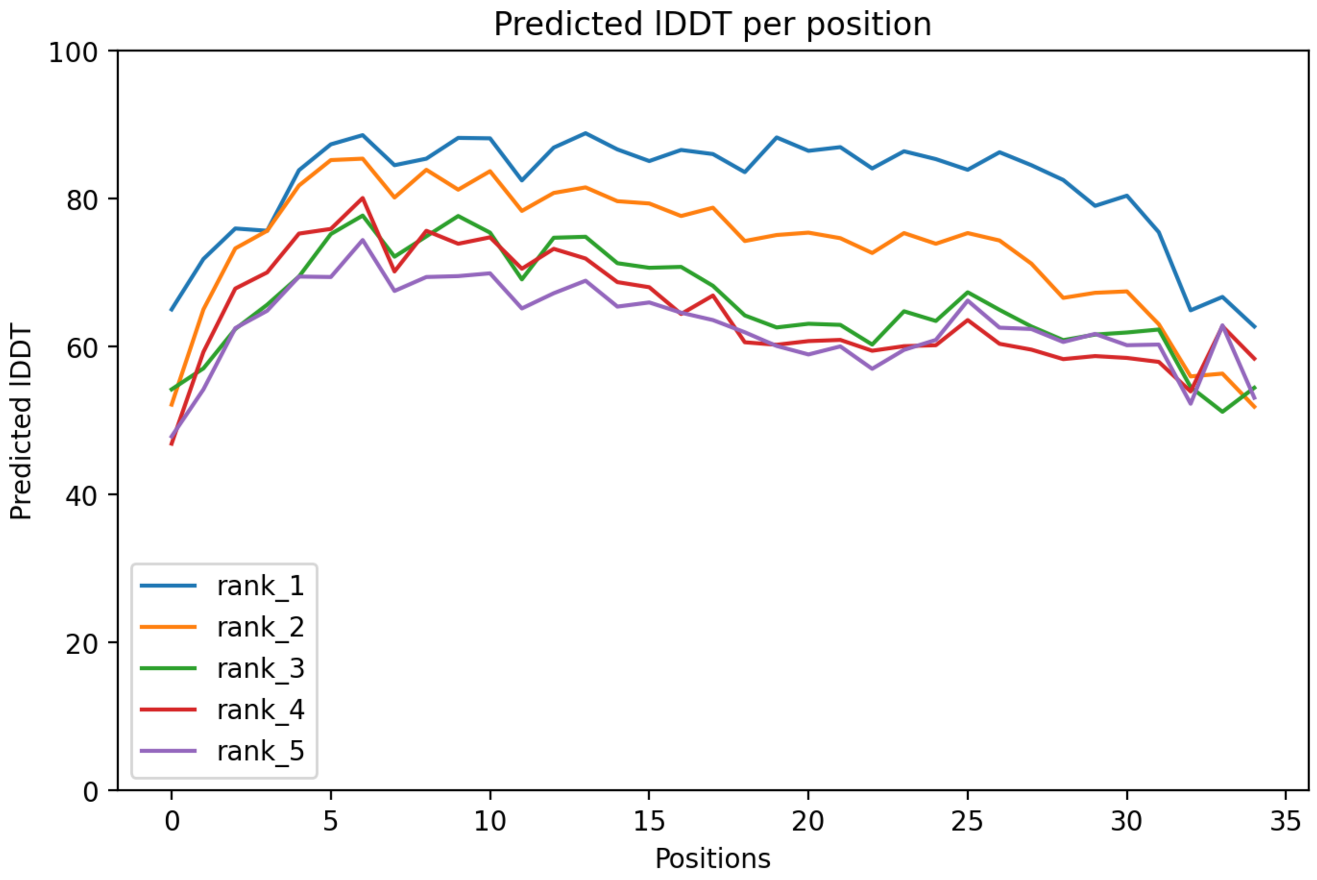

CP115: MtrunA17_Chr7g0230341

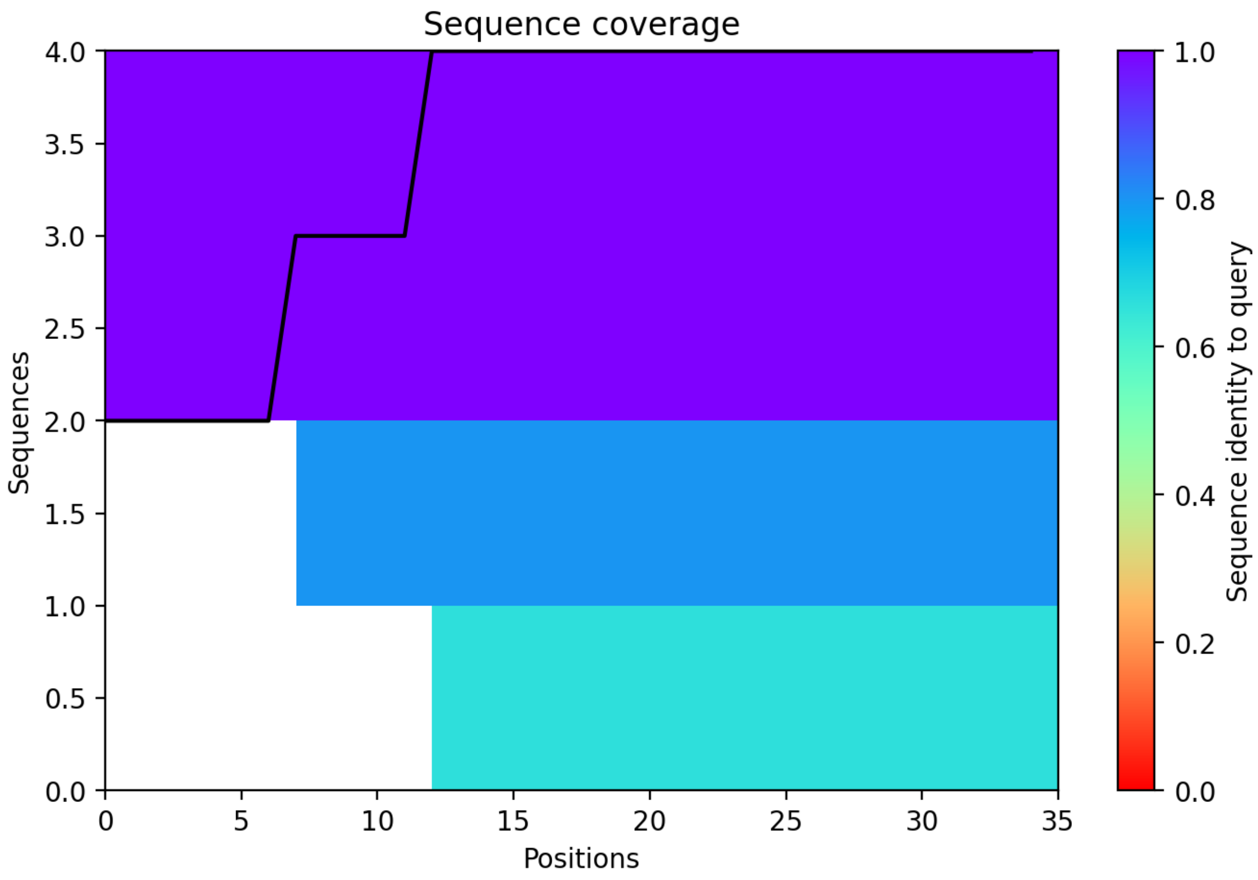

CP116: MtrunA17_Chr7g0232851

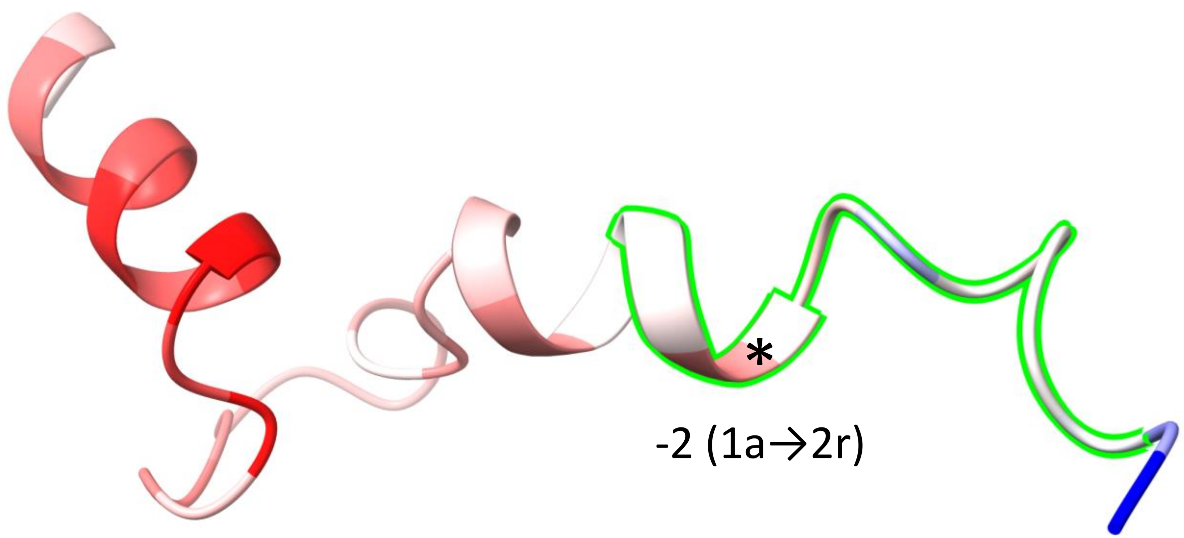


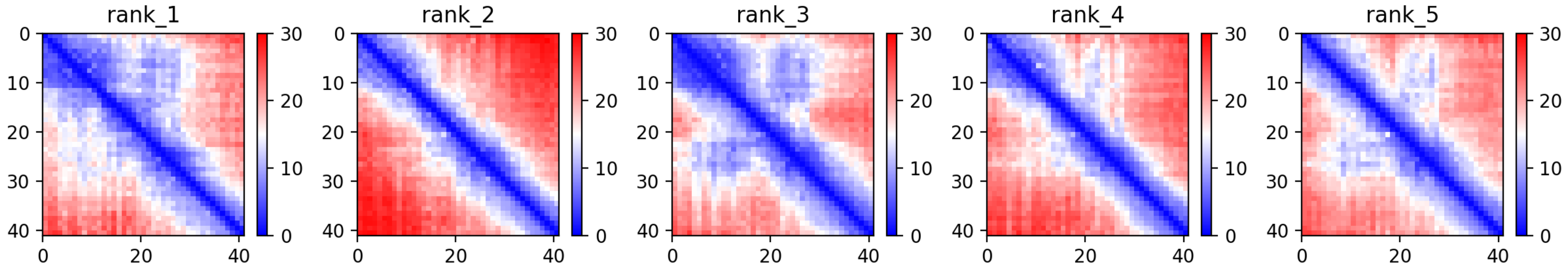

CP116: MtrunA17_Chr7g0232851

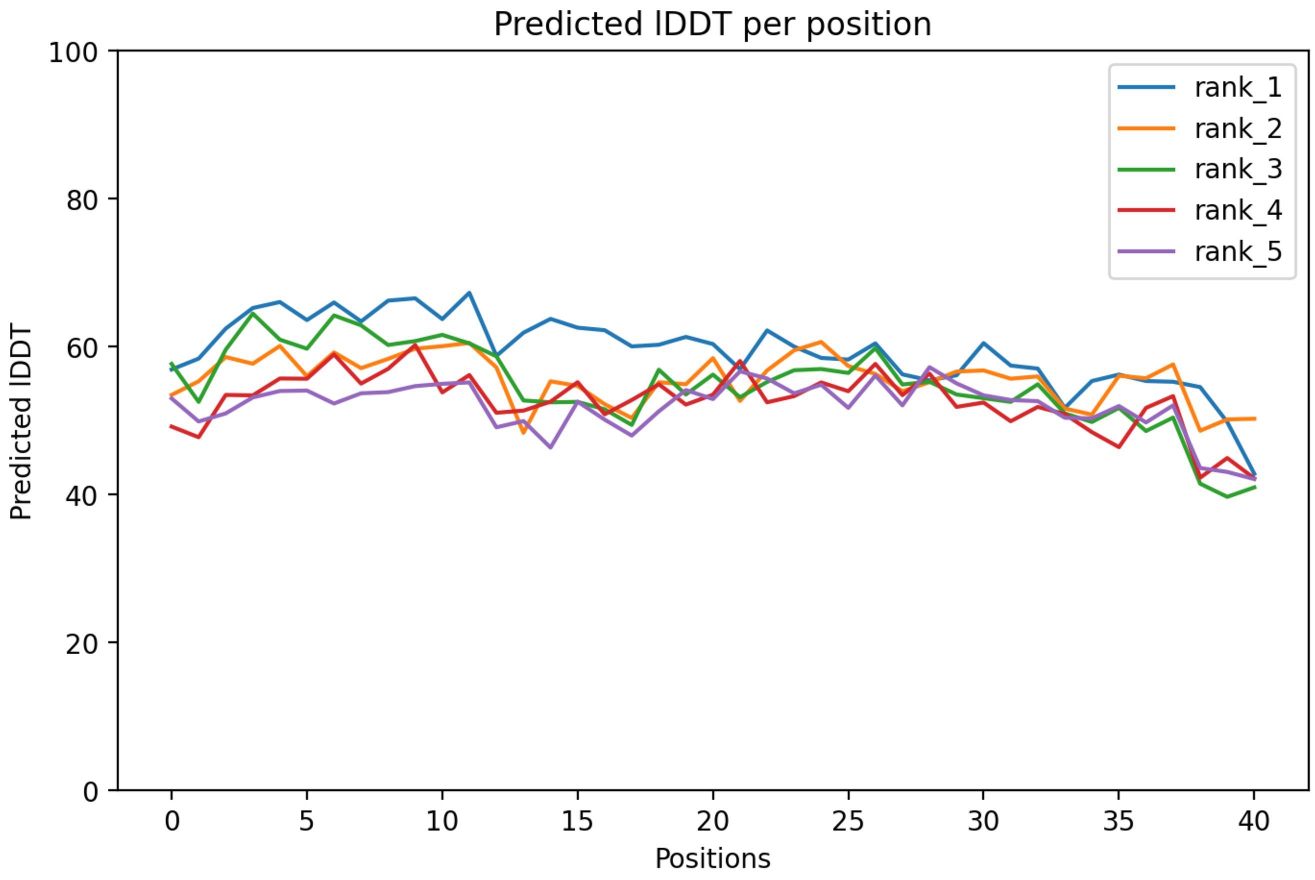

### CP116: MtrunA17_Chr7g0232851

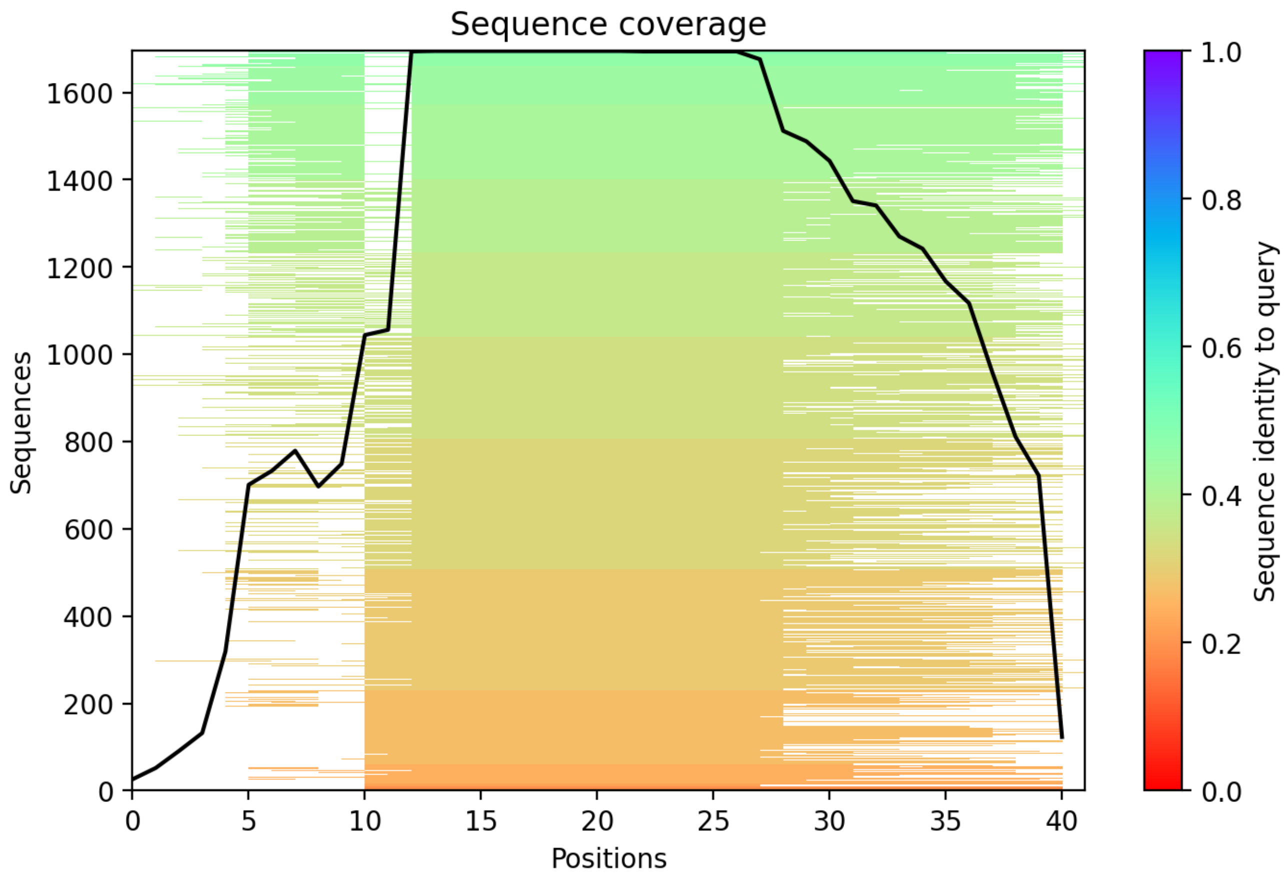

CP117: MtrunA17_Chr7g0237331

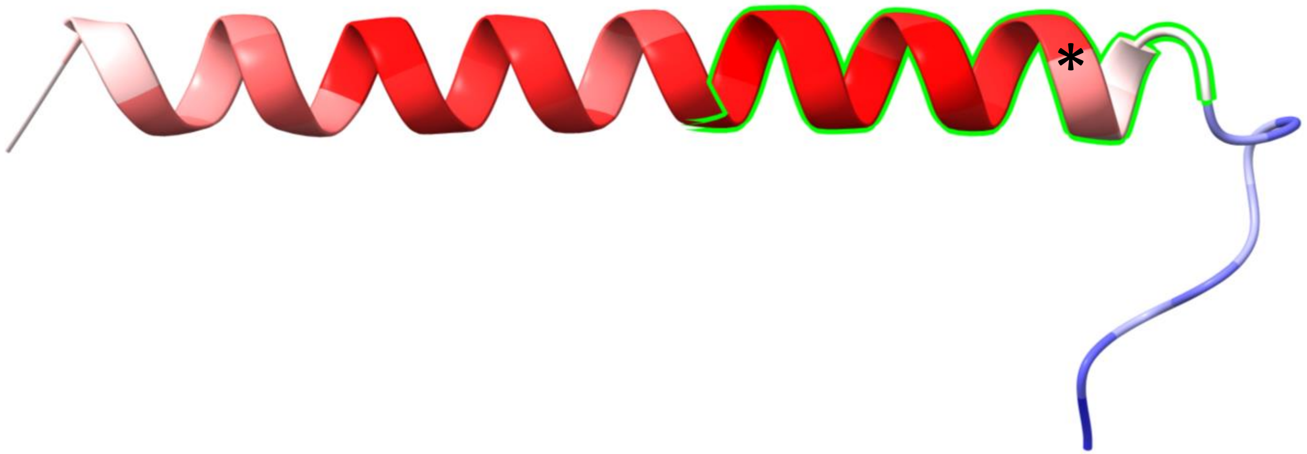


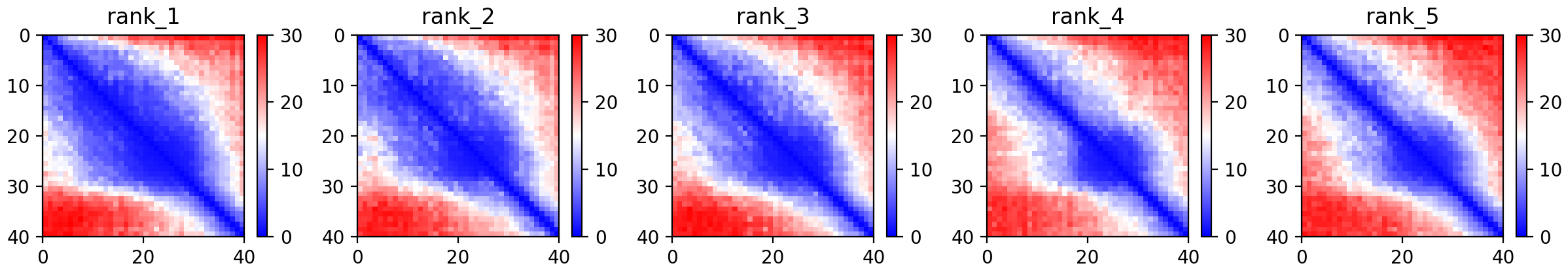

CP117: MtrunA17_Chr7g0237331

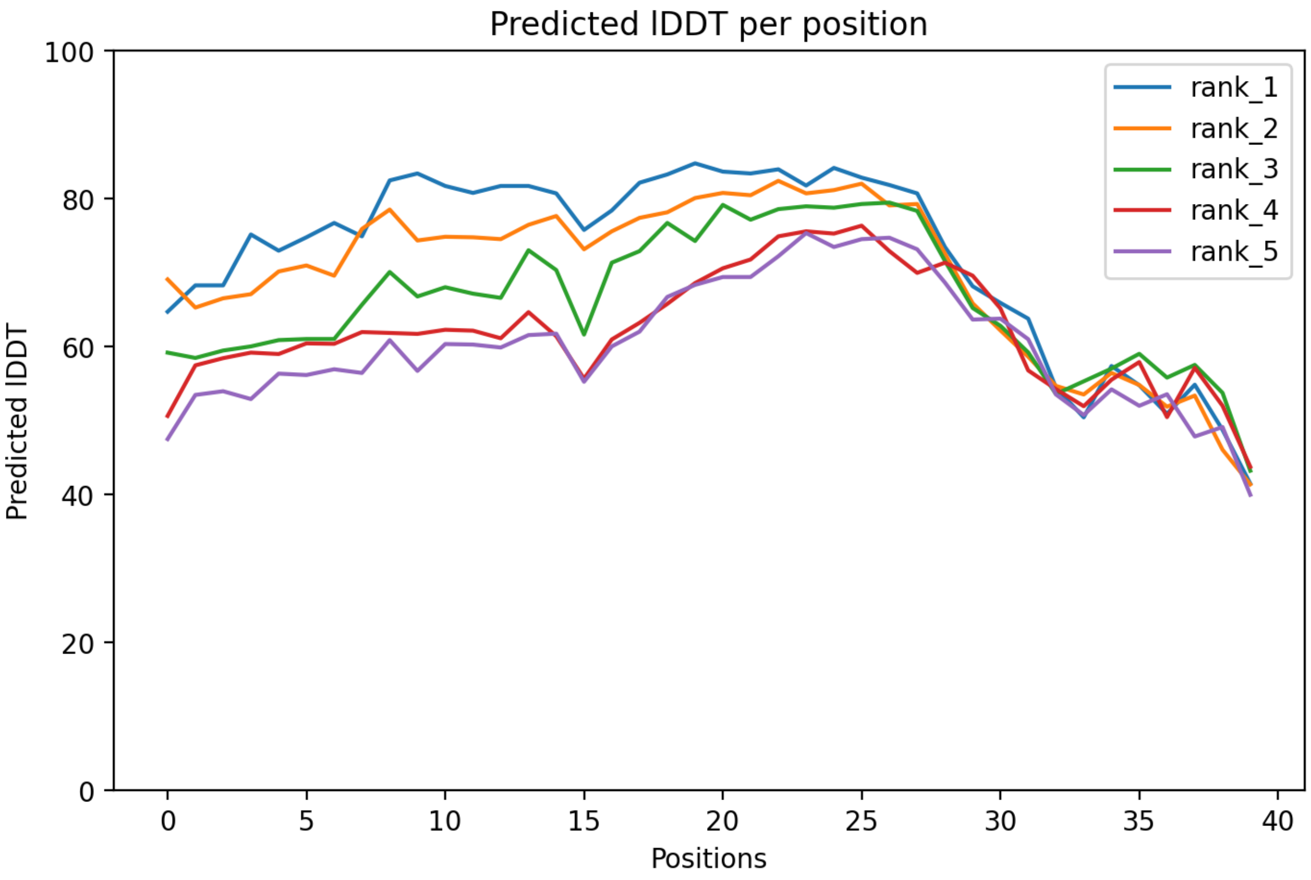

### CP117: MtrunA17_Chr7g0237331

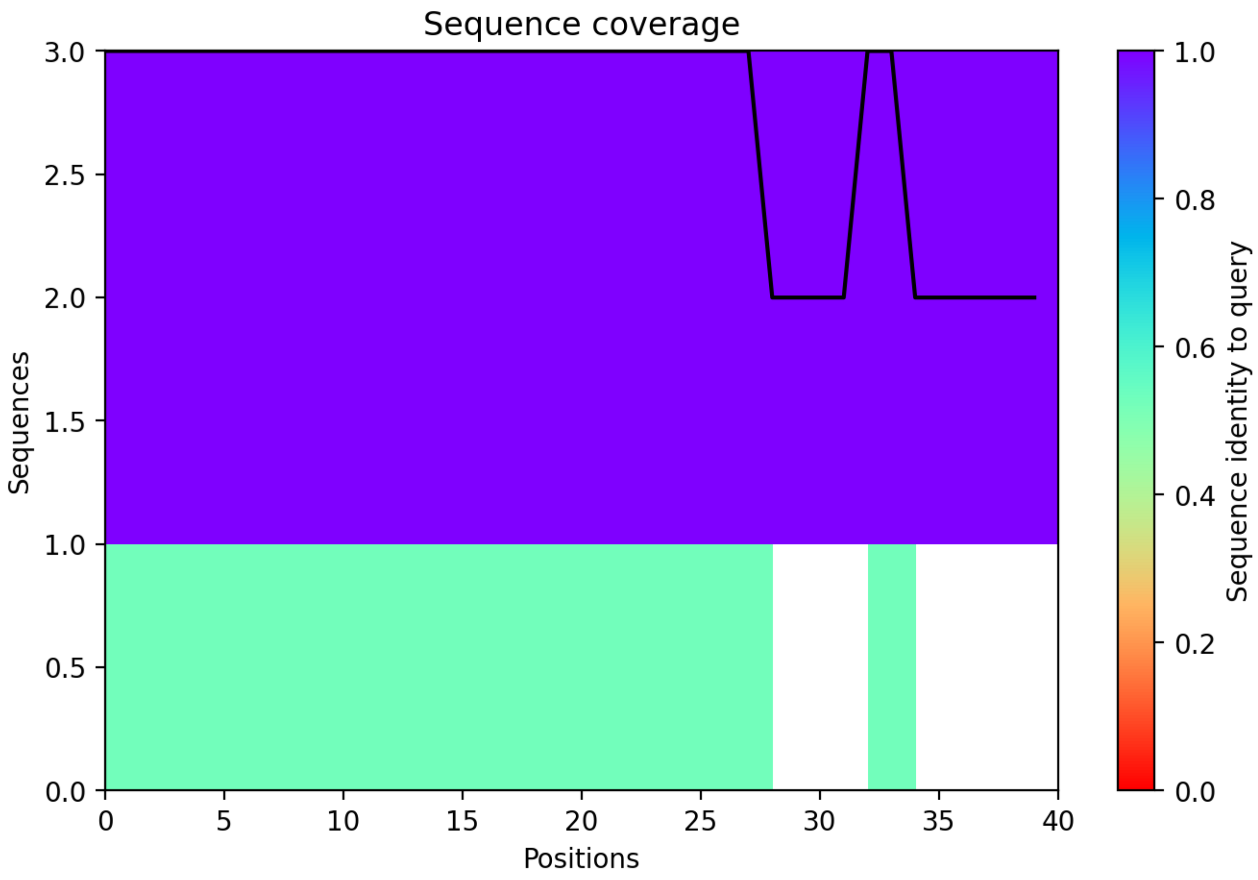

### CP118: MtrunA17_Chr7g0251971

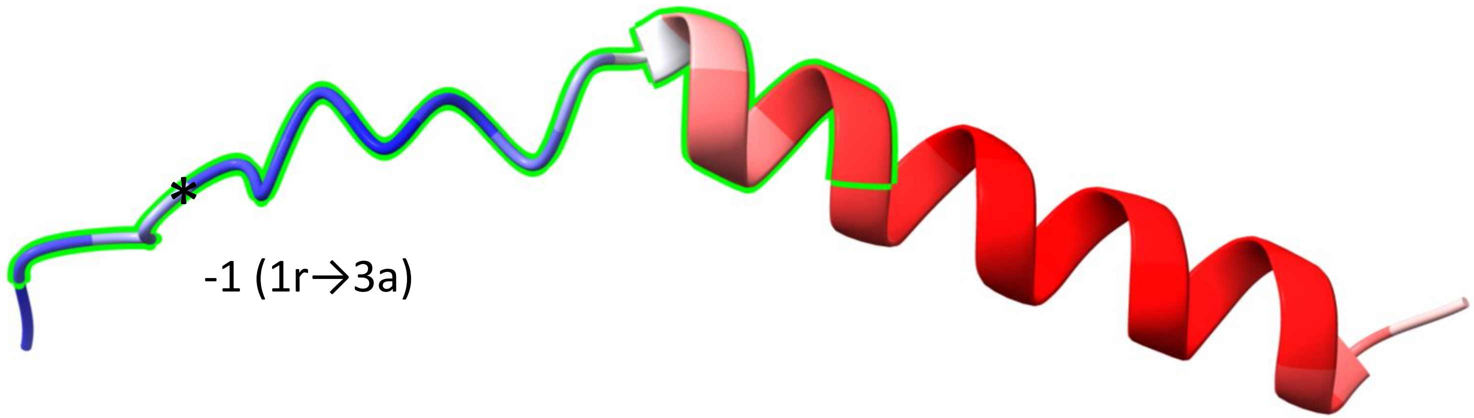


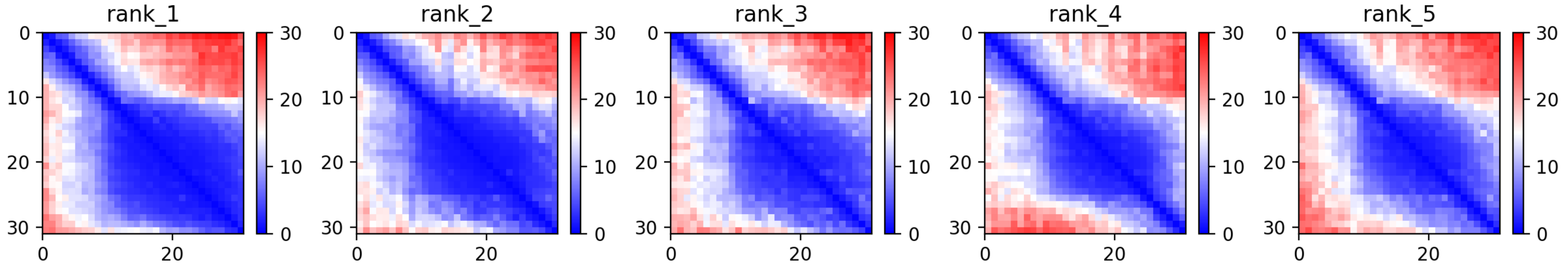

CP118: MtrunA17_Chr7g0251971

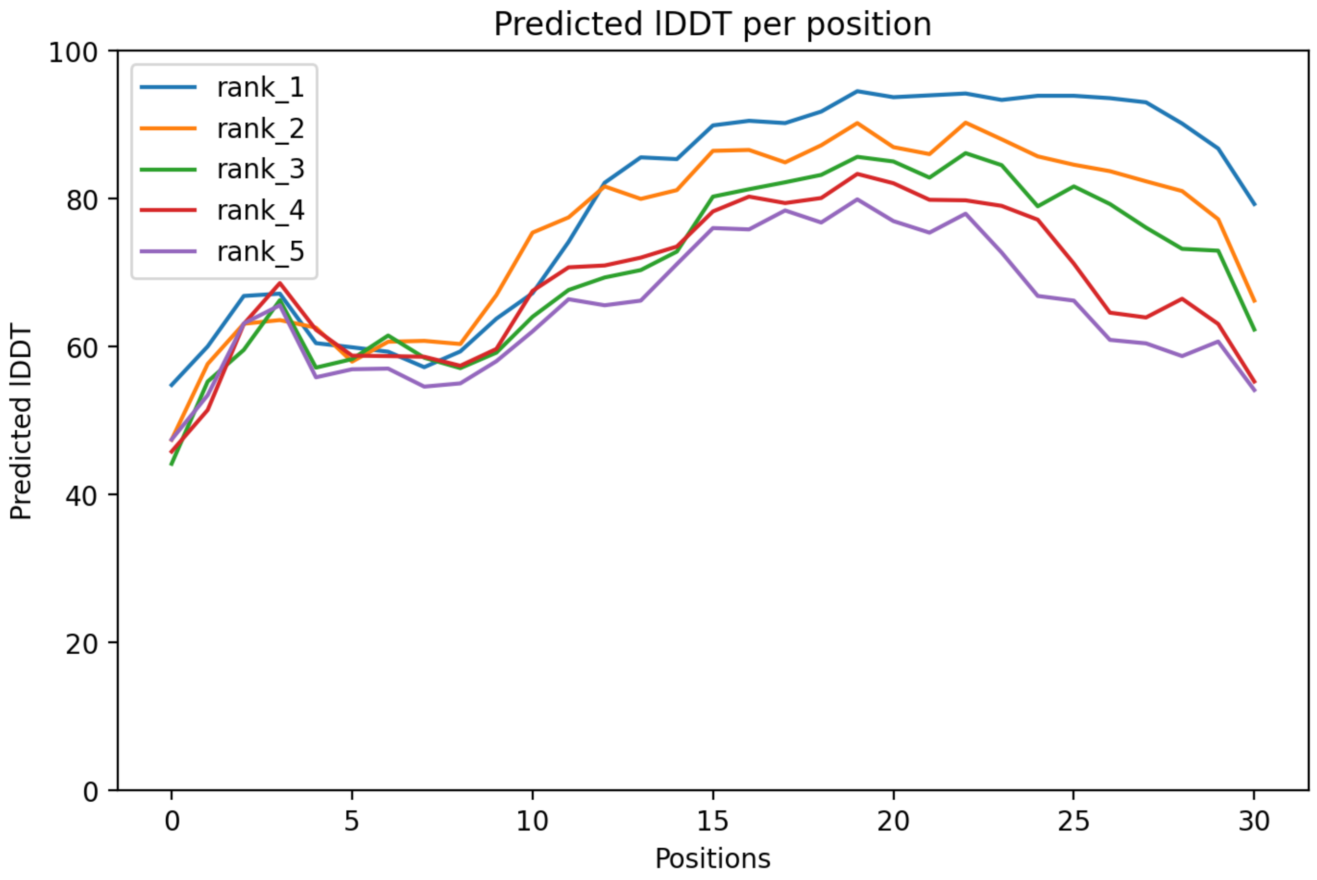

### CP118: MtrunA17_Chr7g0251971

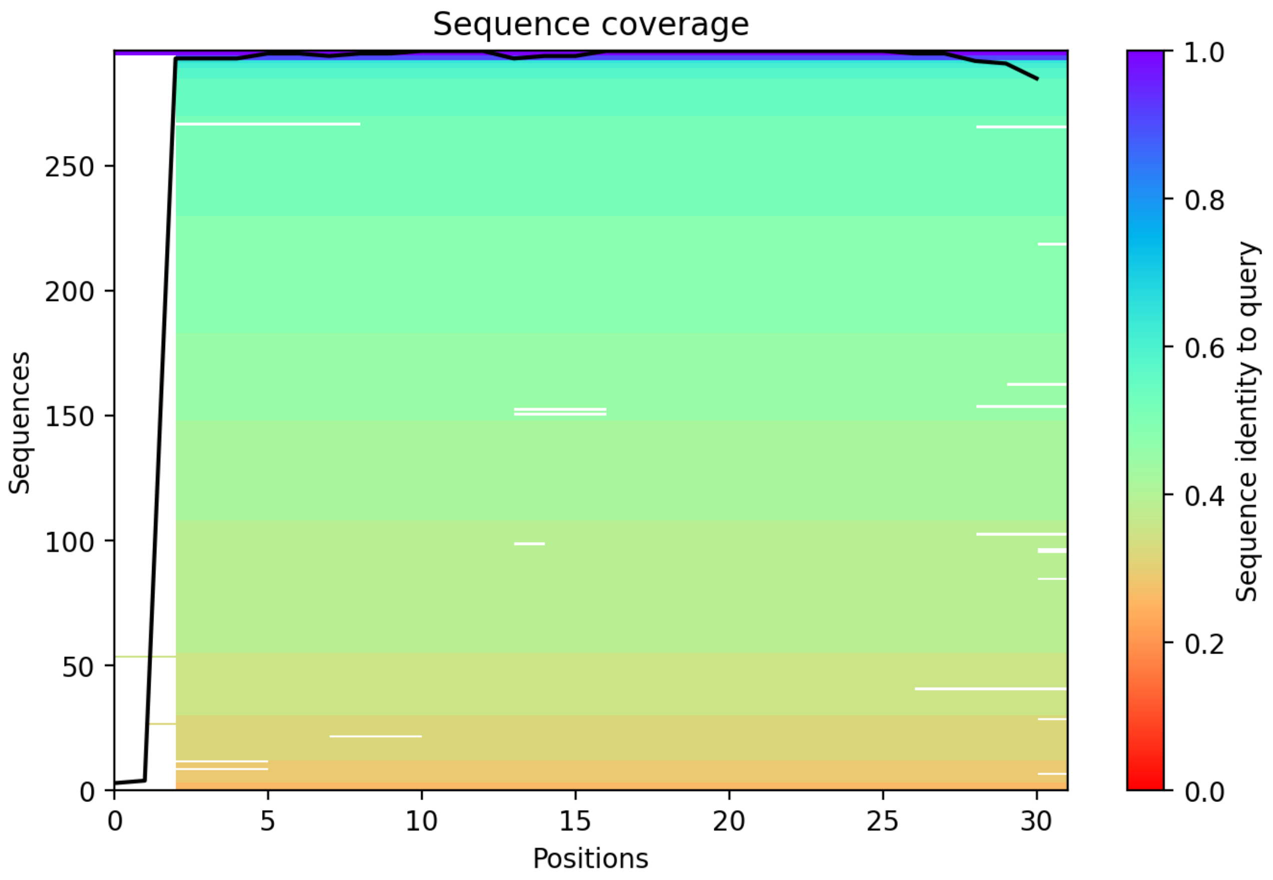

CP119: MtrunA17_Chr7g0259611

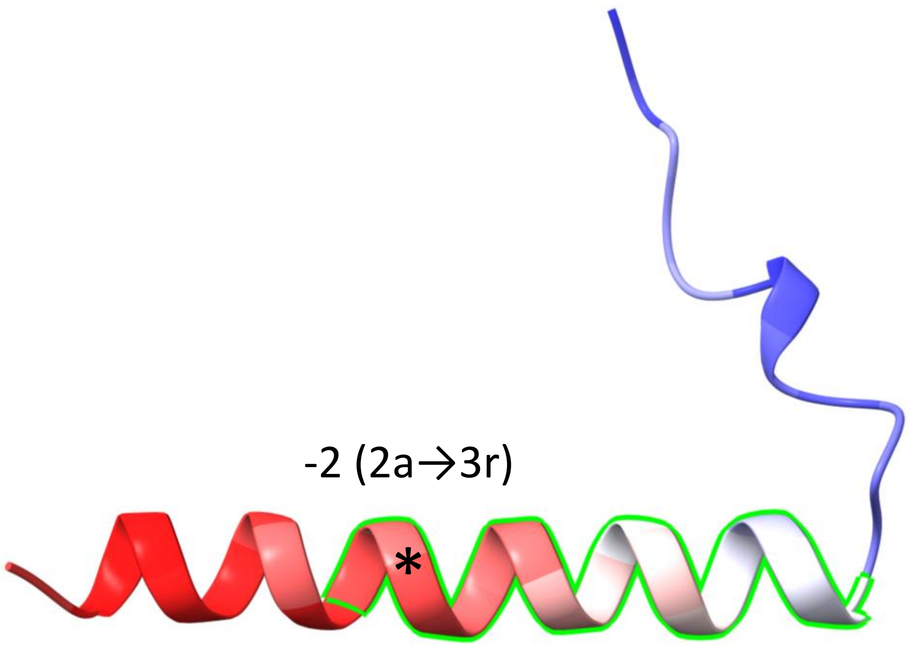


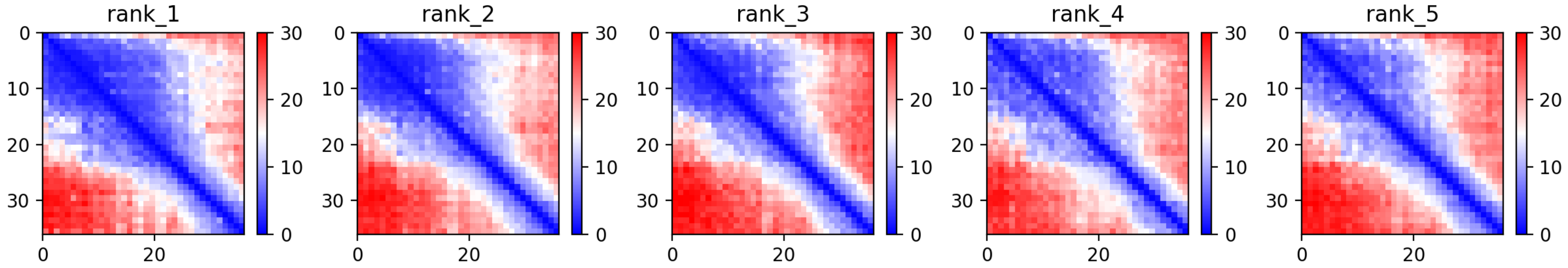

### CP119: MtrunA17_Chr7g0259611

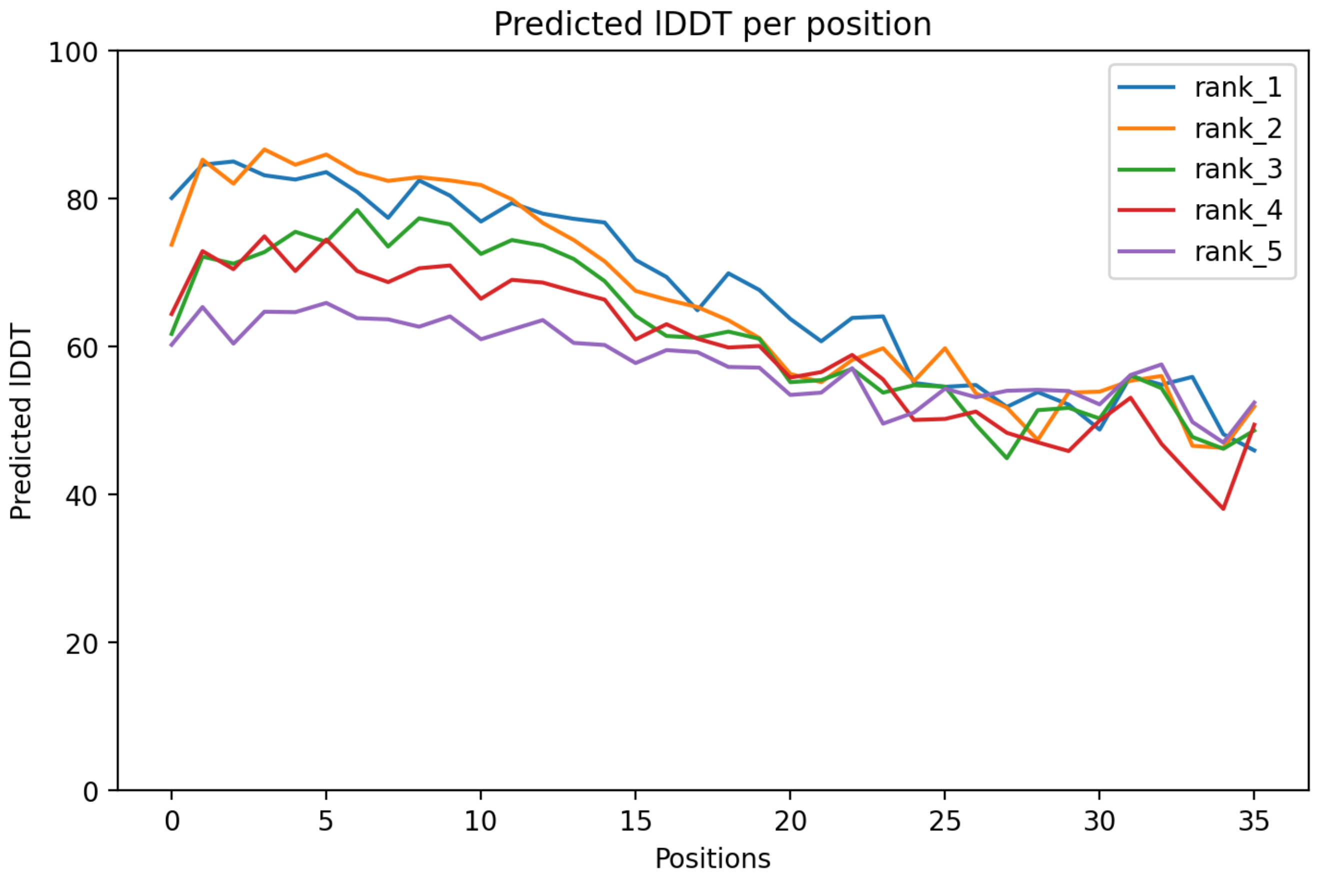

### CP119: MtrunA17_Chr7g0259611

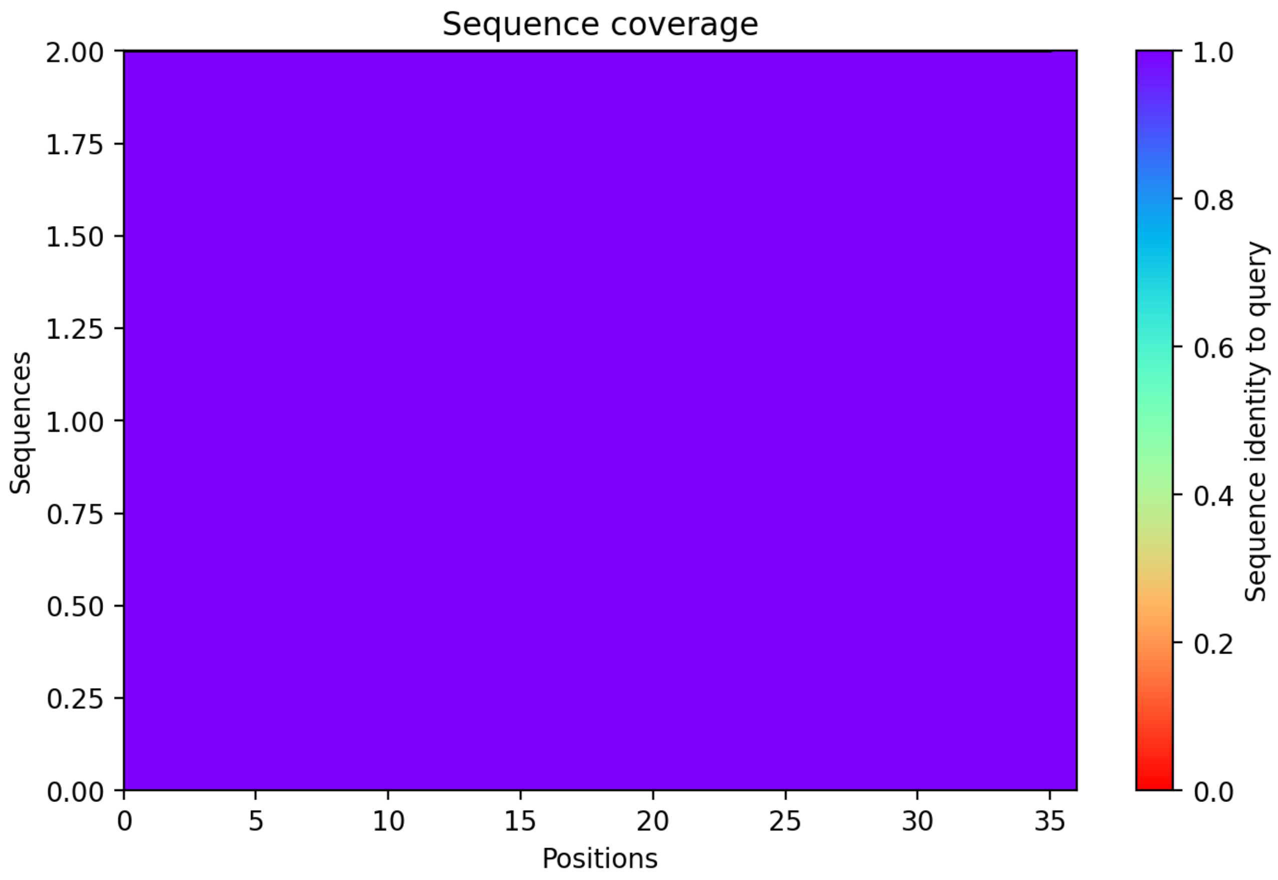

### CP120: MtrunA17_Chr7g0262821

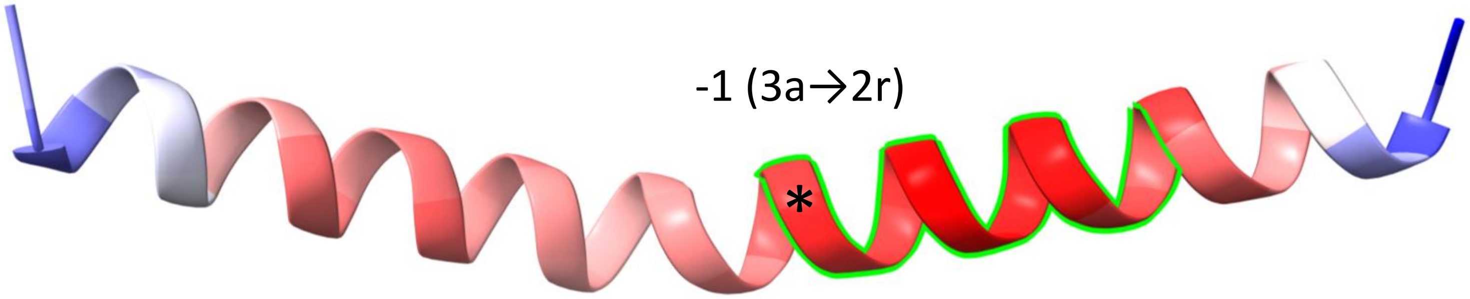


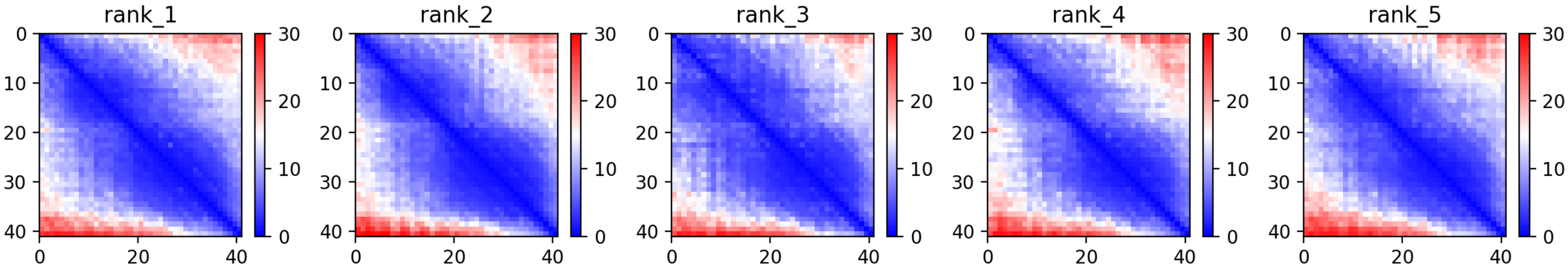

CP120: MtrunA17_Chr7g0262821

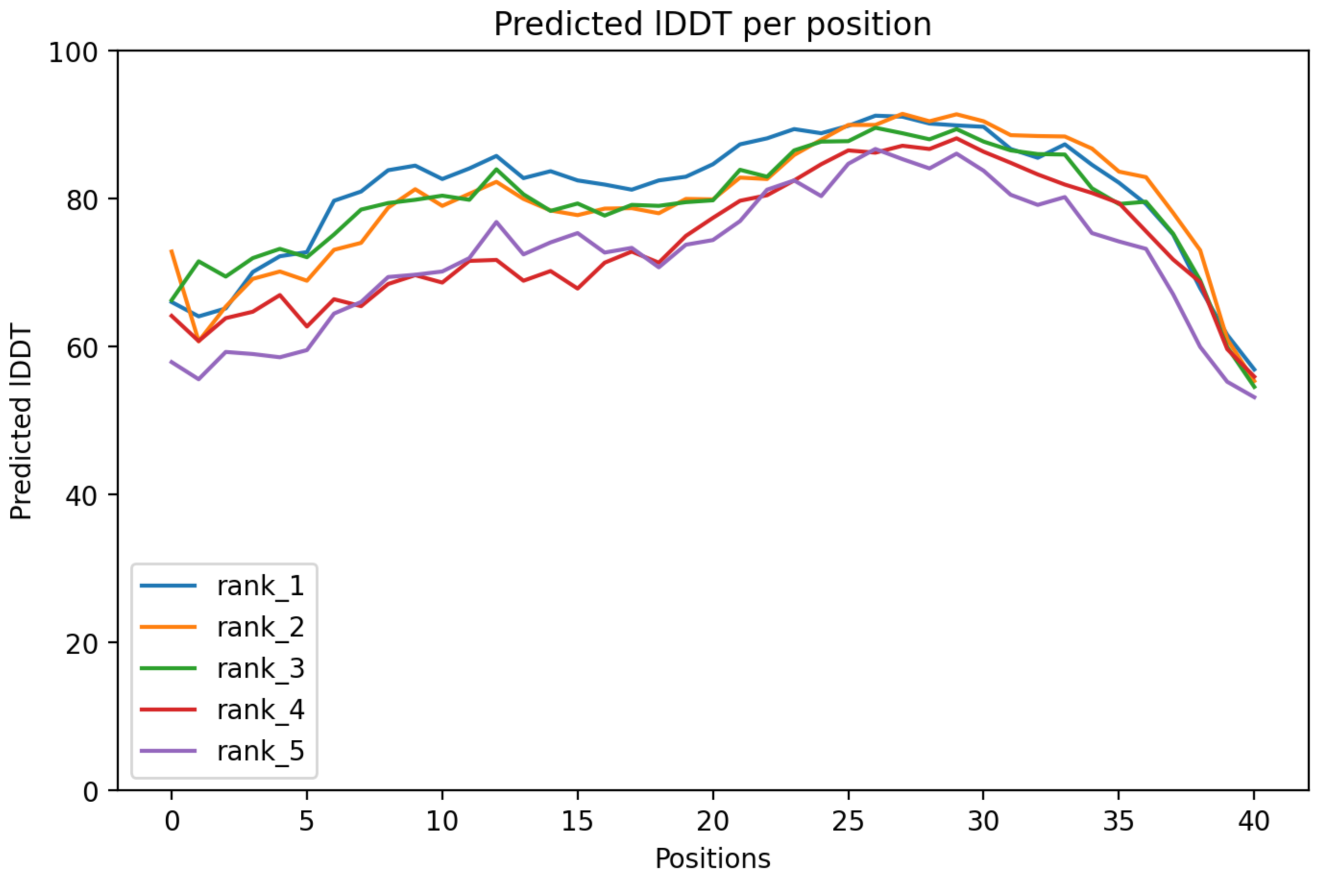

CP120: MtrunA17_Chr7g0262821

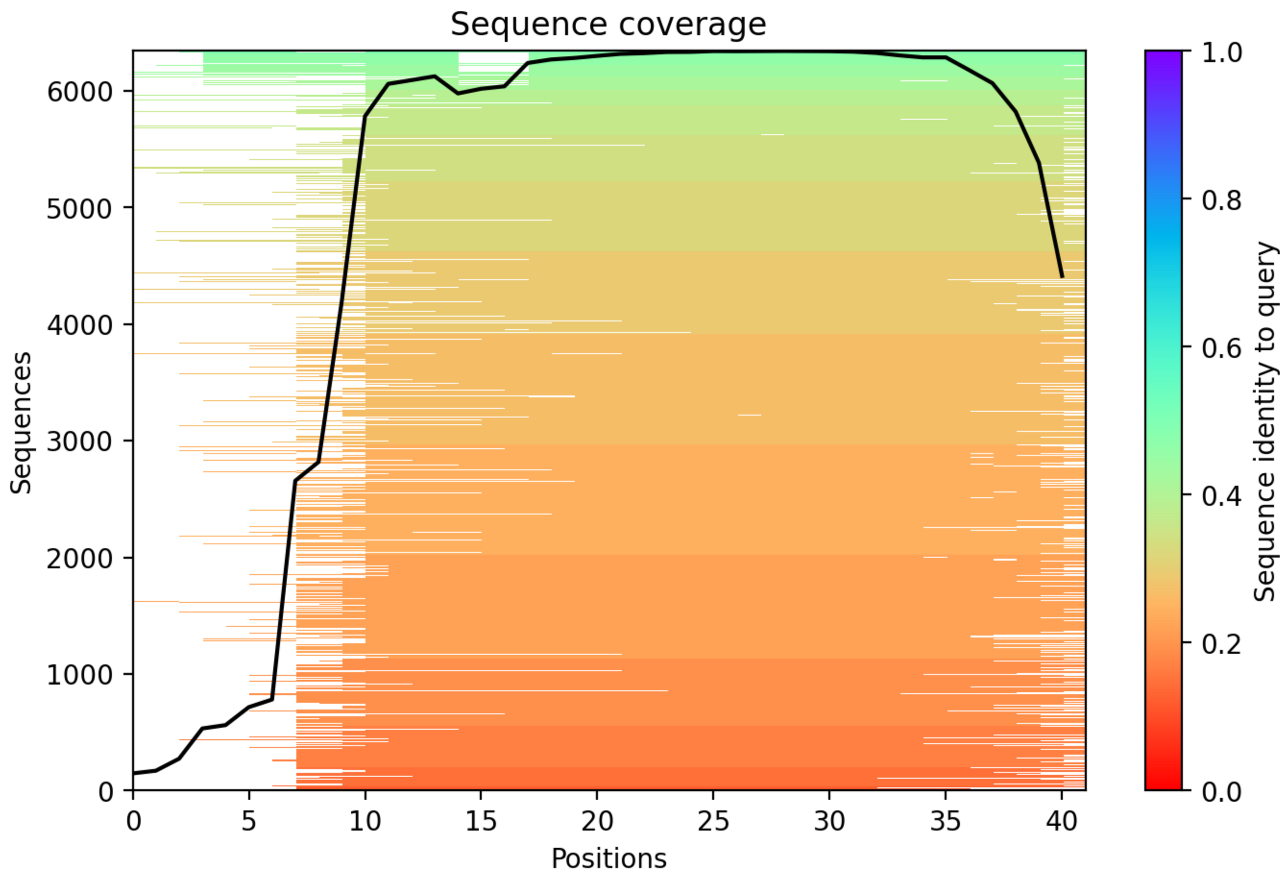

CP121: MtrunA17_Chr7g0270811

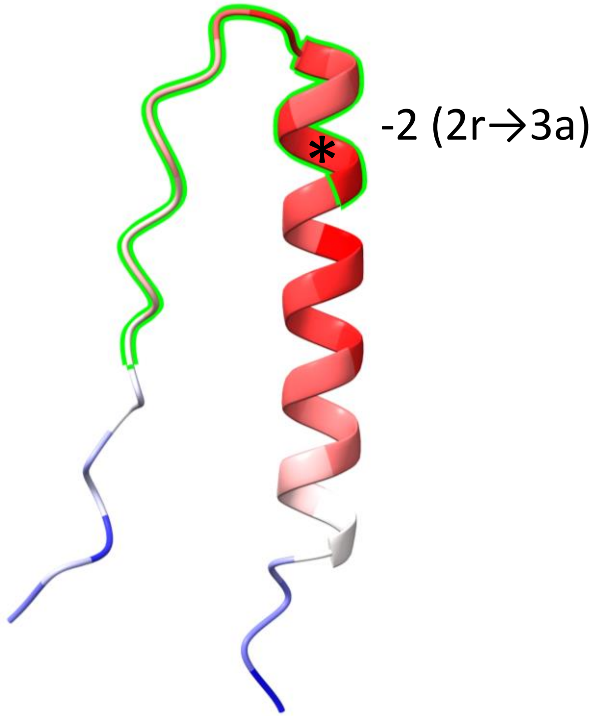


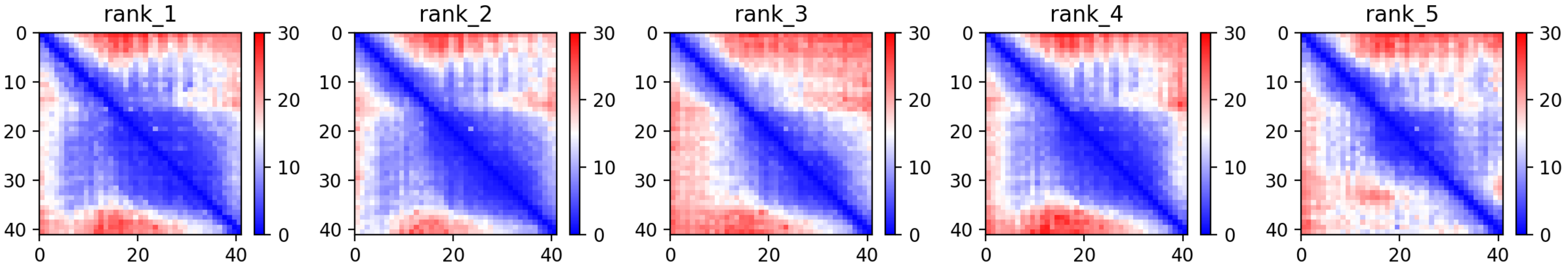

CP121: MtrunA17_Chr7g0270811

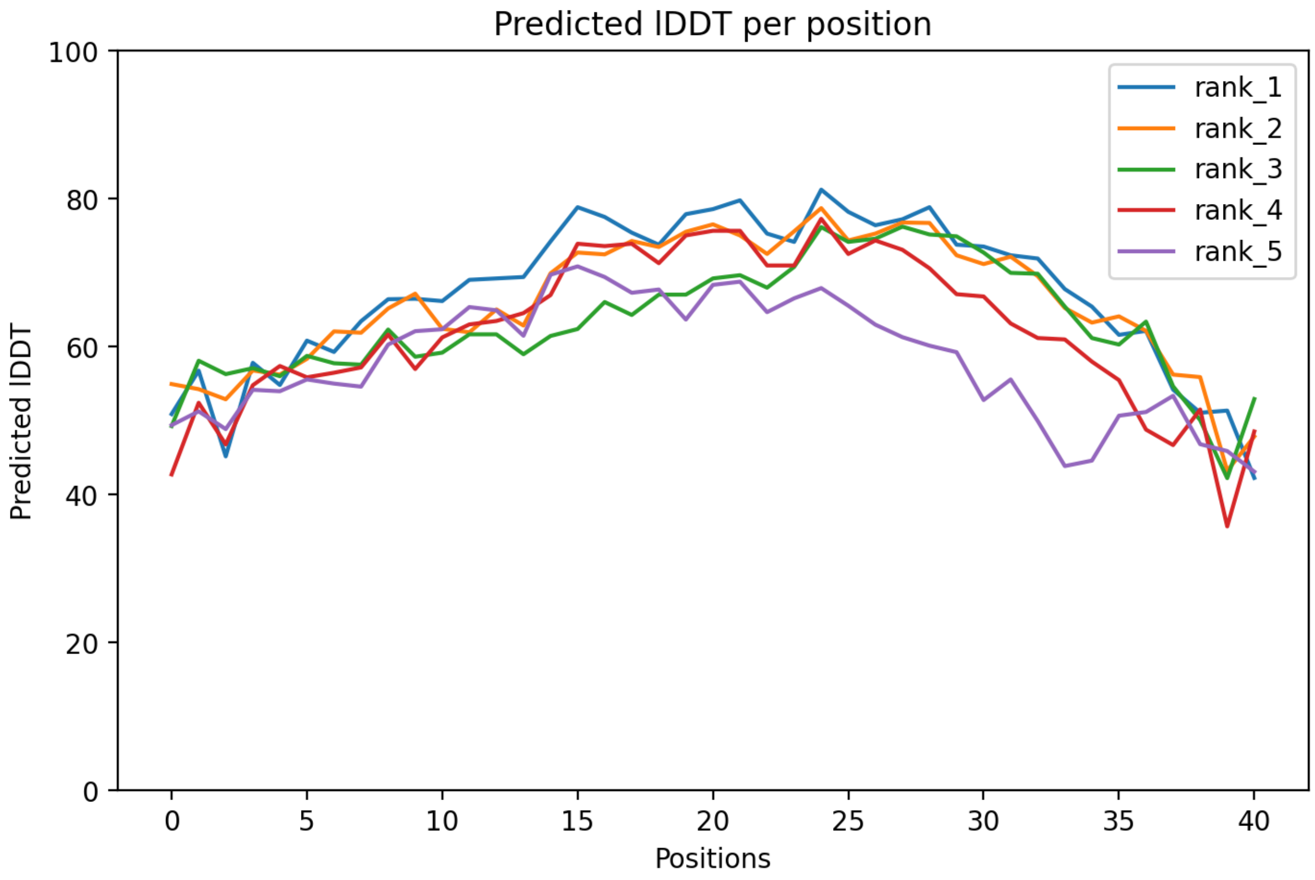

CP121: MtrunA17_Chr7g0270811

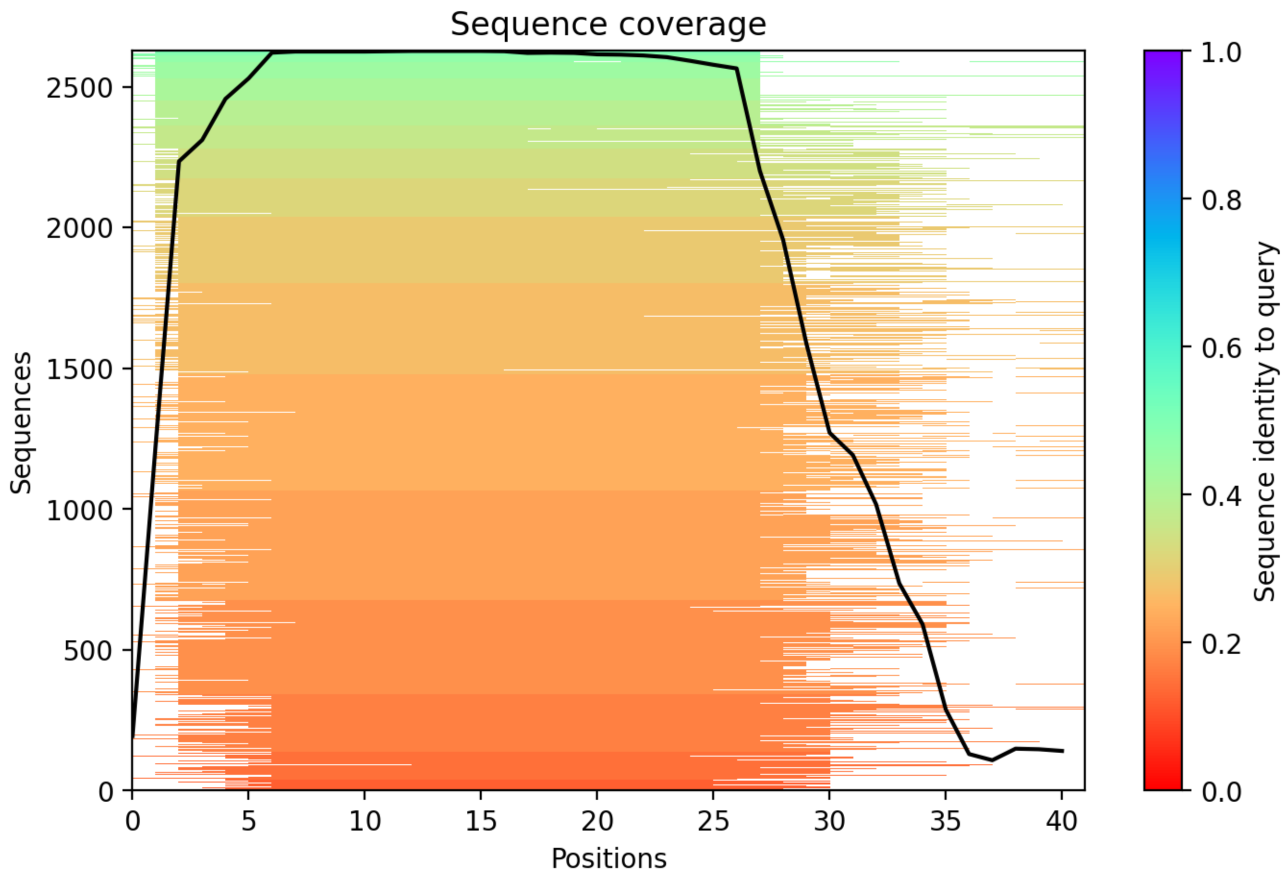

CP122: MtrunA17_Chr7g1034306

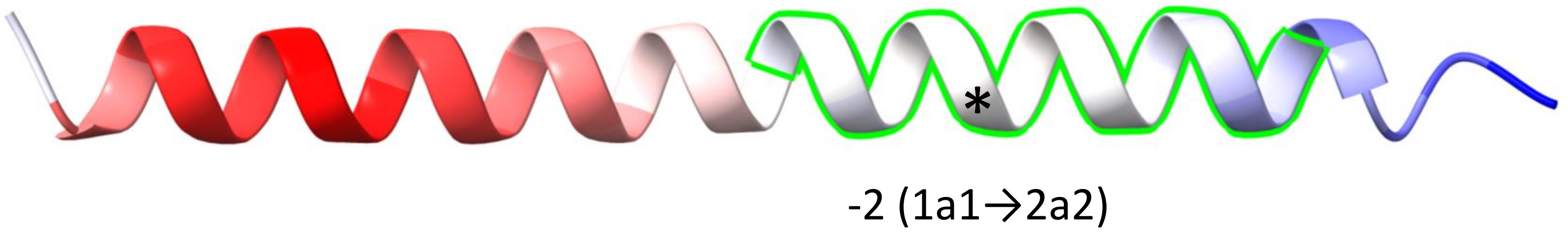


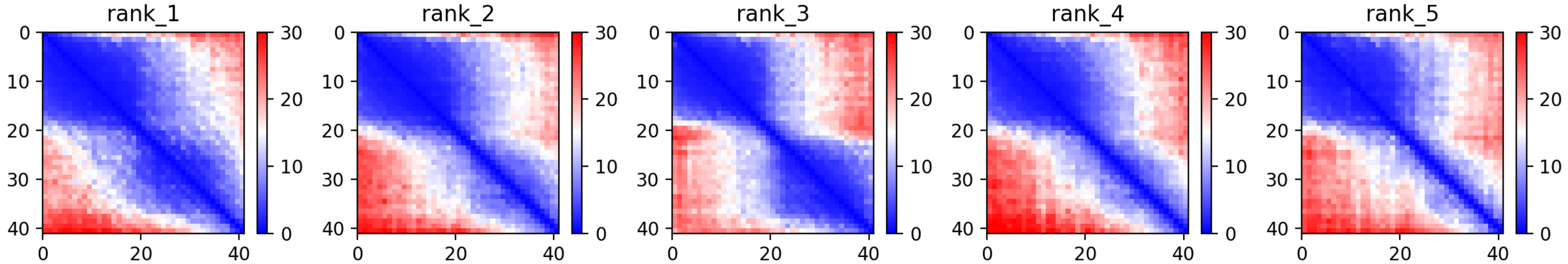

### CP122: MtrunA17_Chr7g1034306

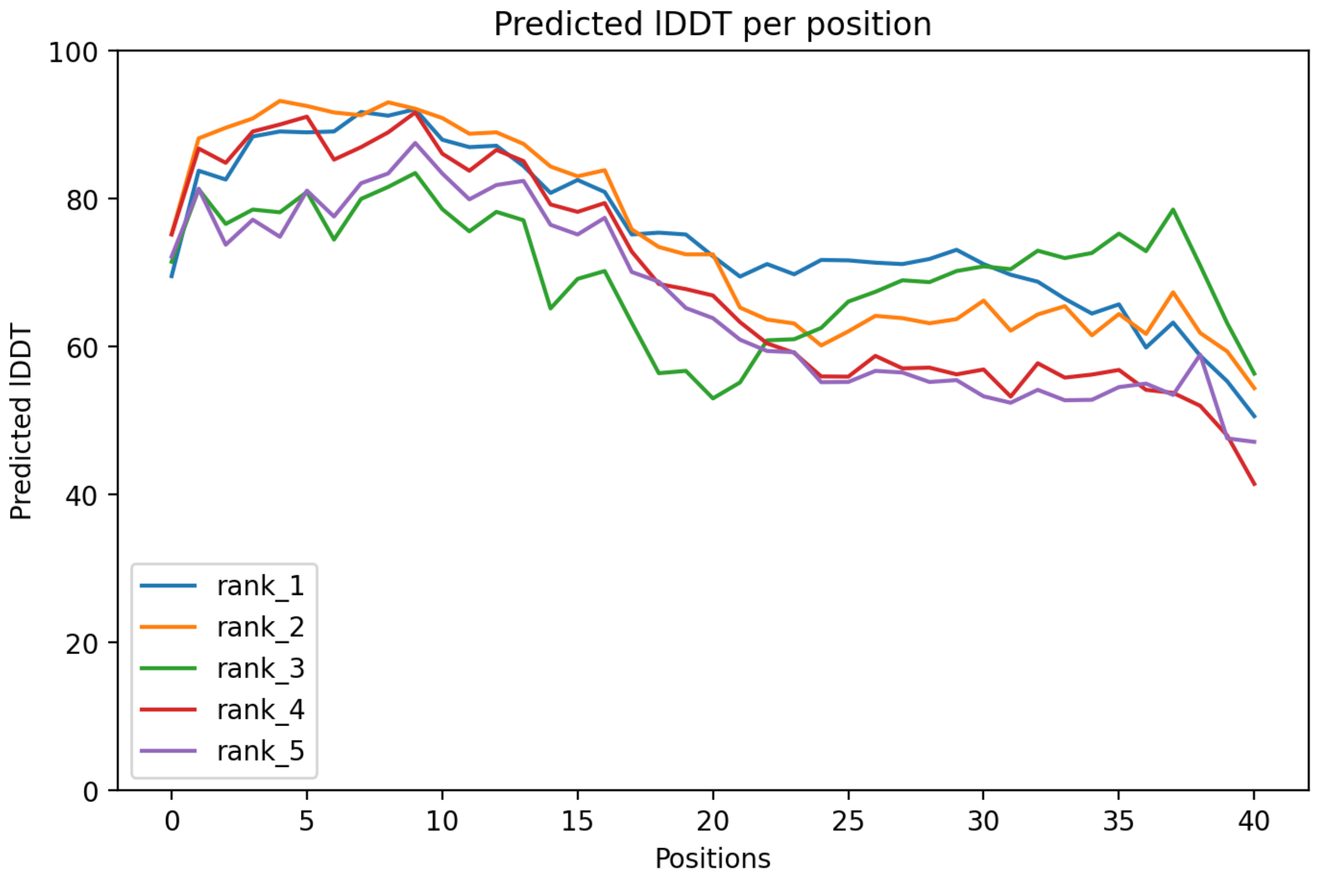

### CP122: MtrunA17_Chr7g1034306

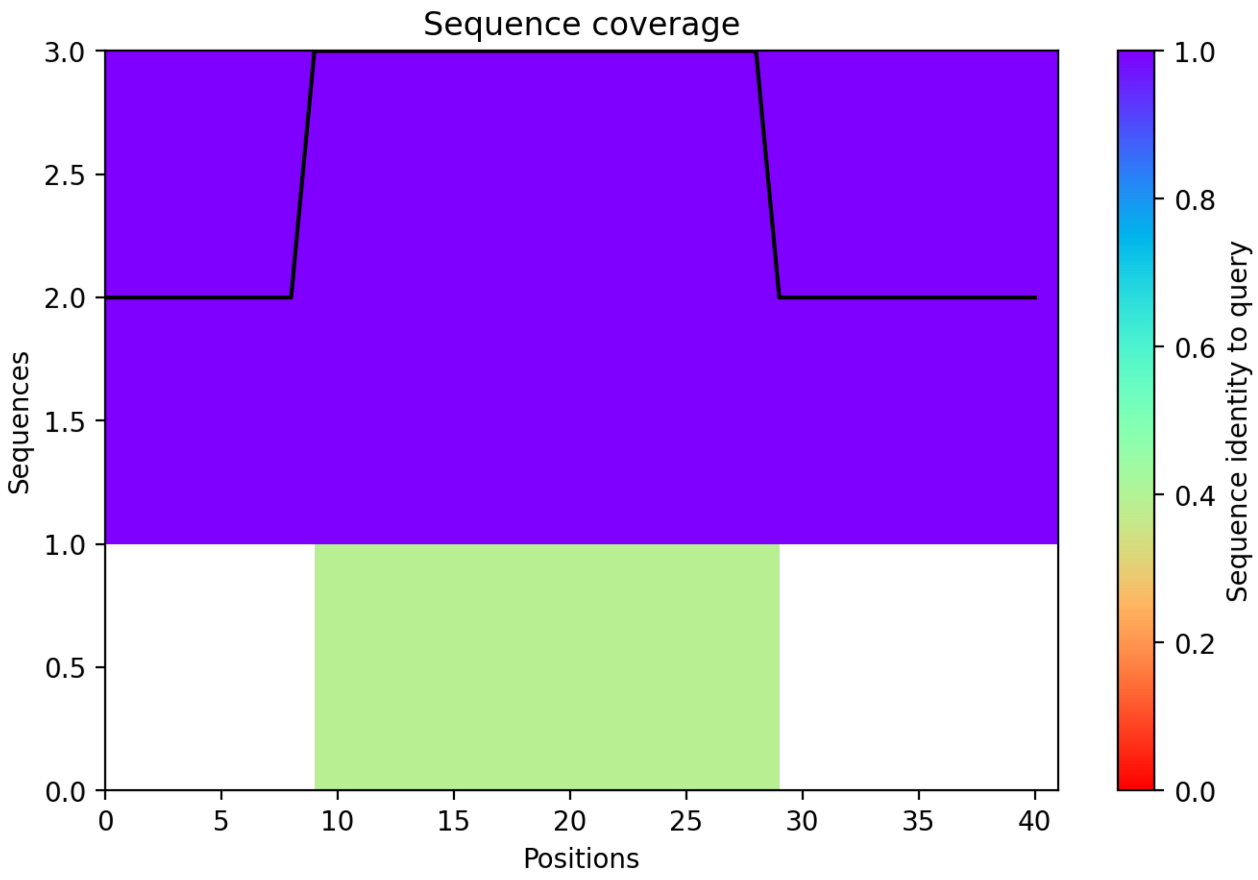

CP123: MtrunA17_Chr8g0338111

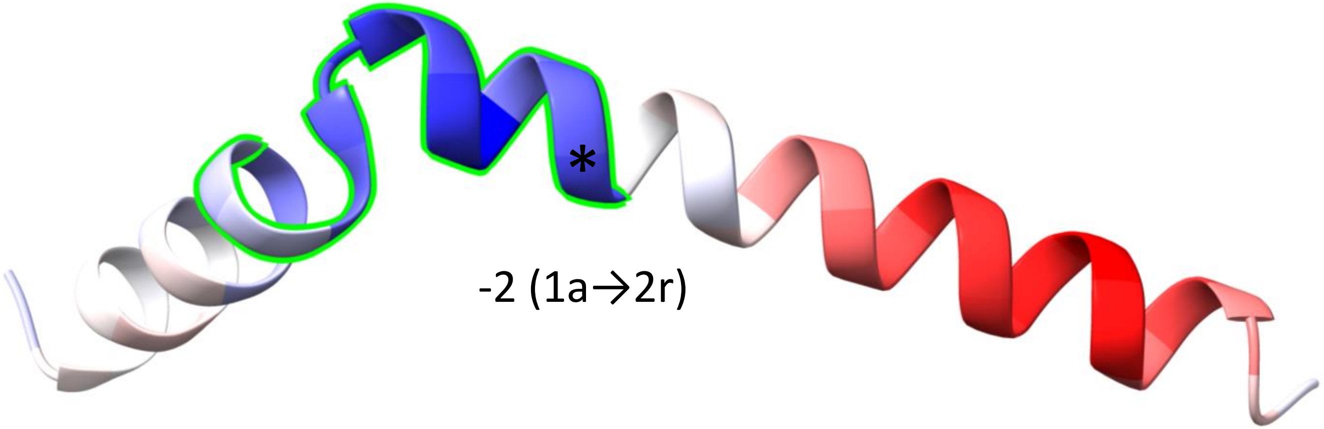


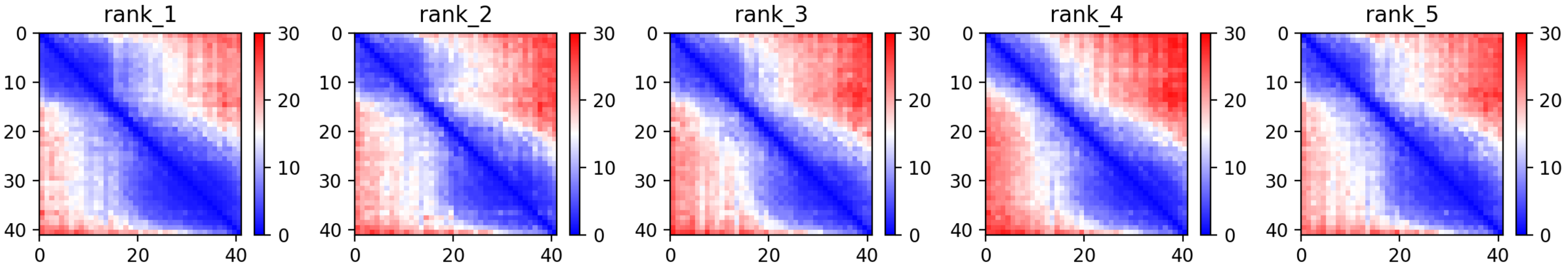

CP123: MtrunA17_Chr8g0338111

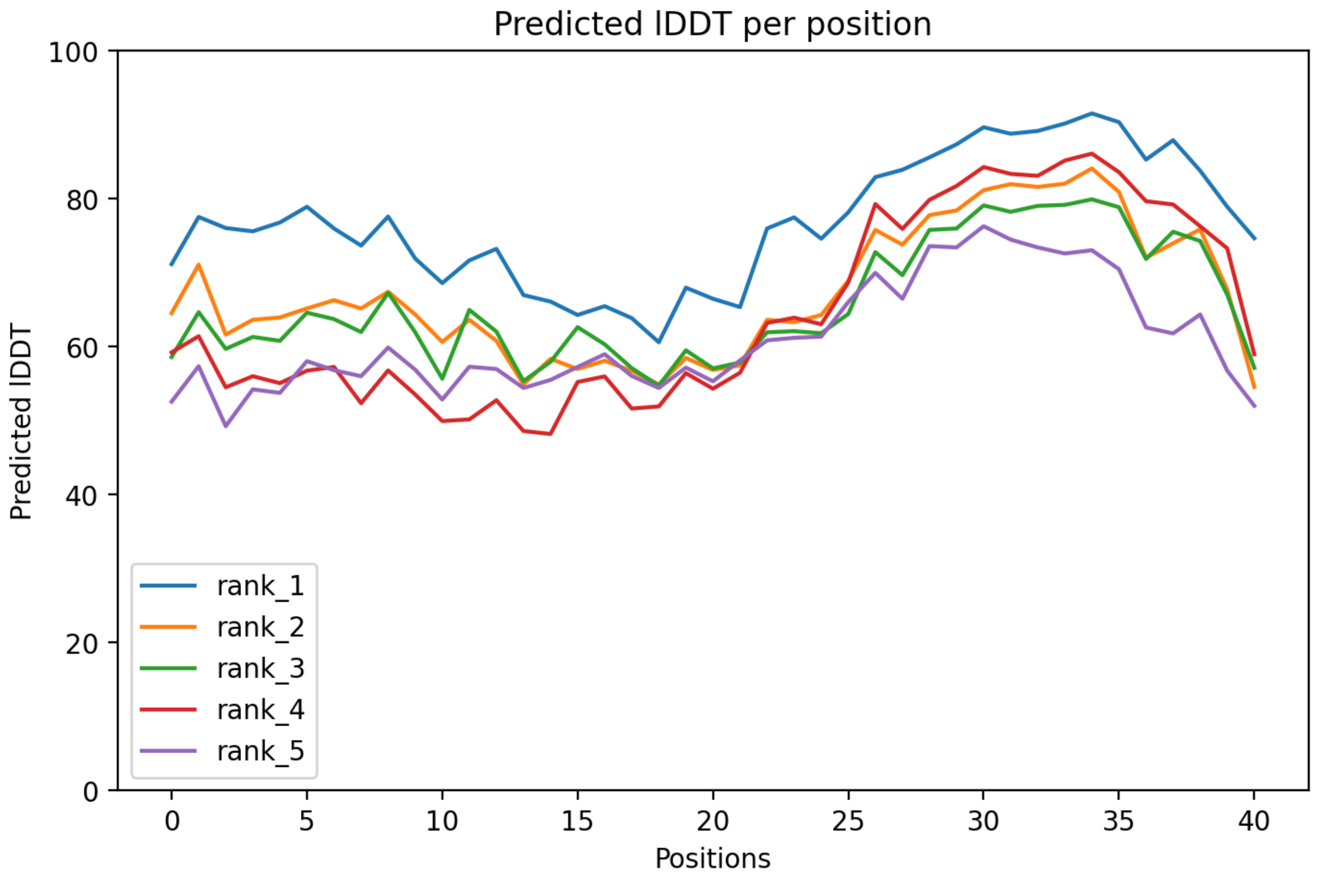

CP123: MtrunA17_Chr8g0338111

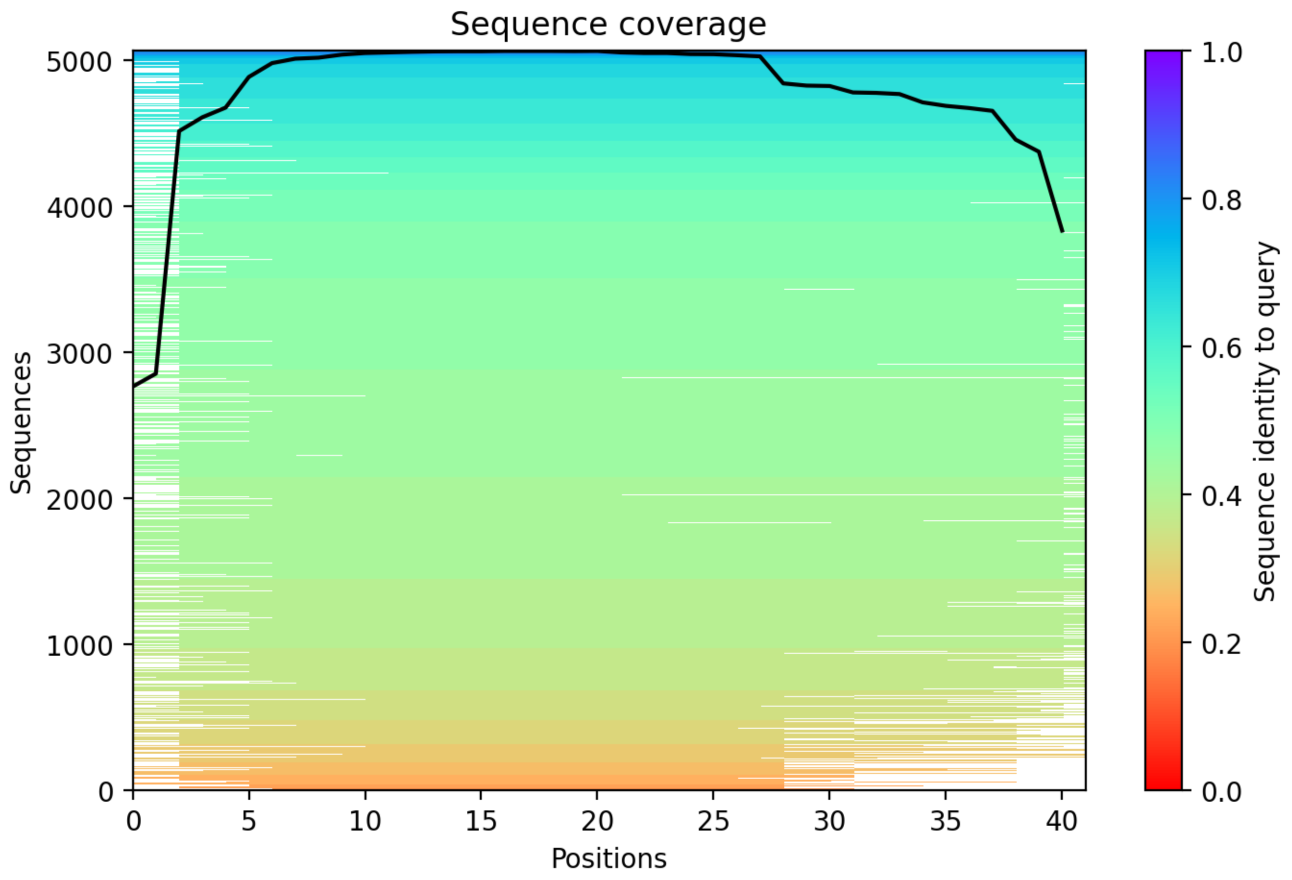

CP124: MtrunA17_Chr8g0338301

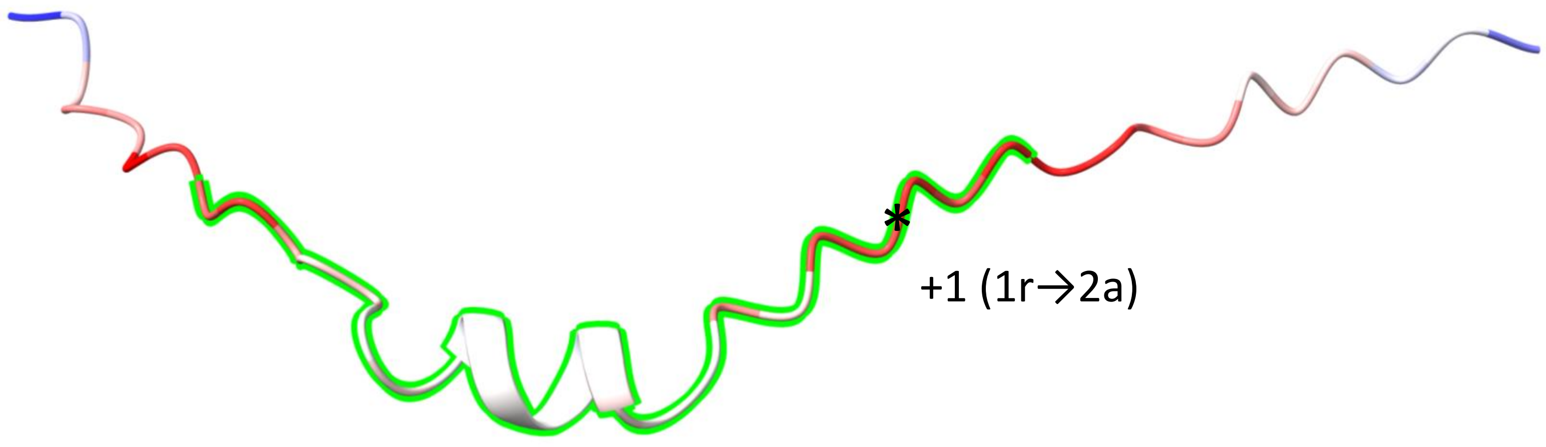


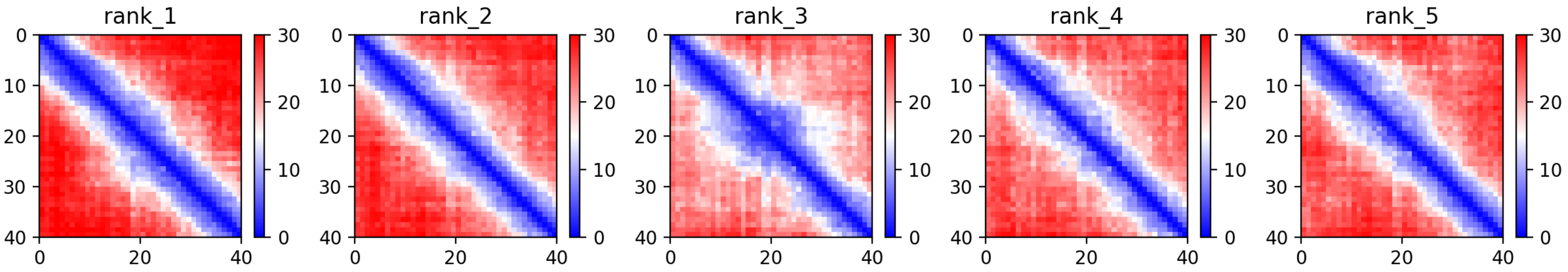

CP124: MtrunA17_Chr8g0338301

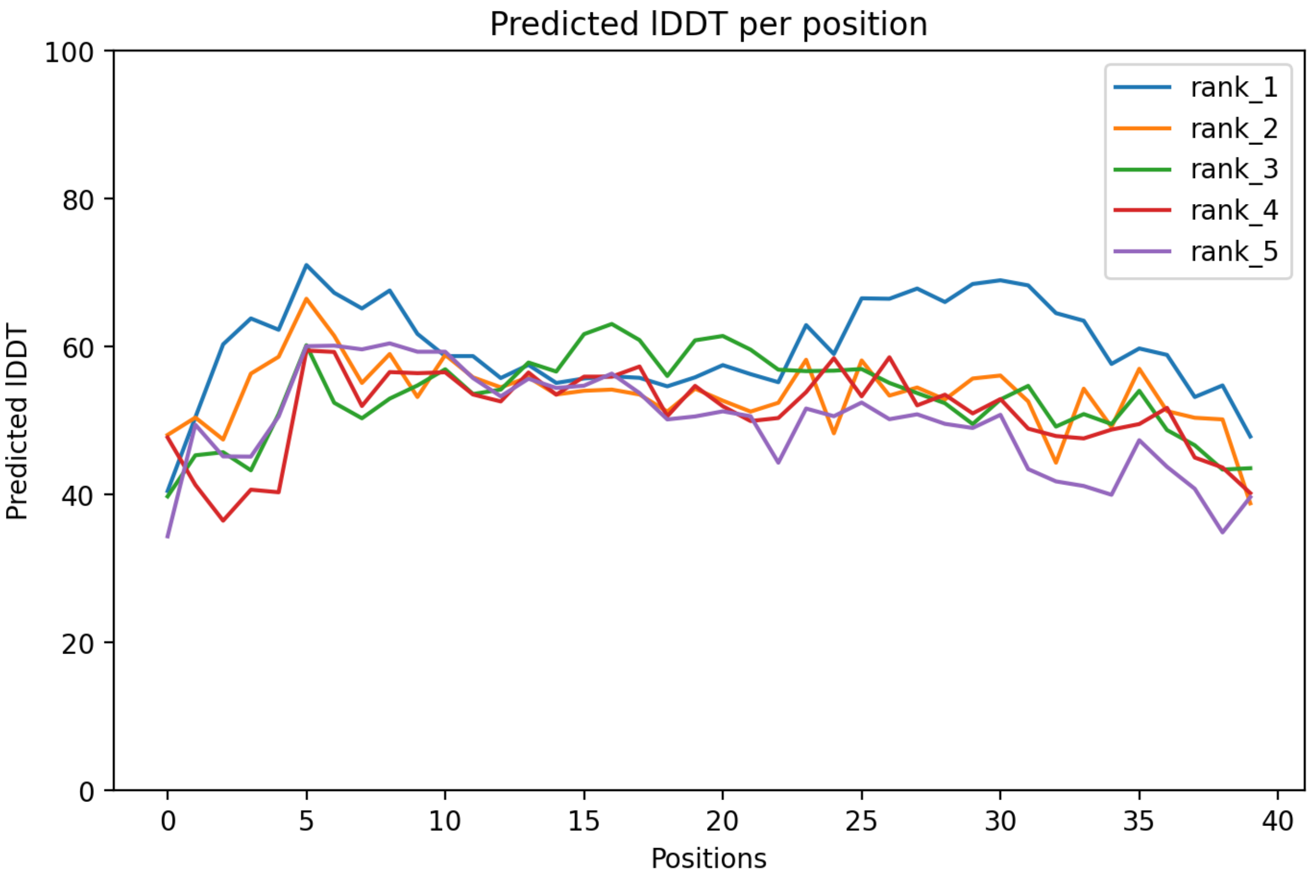

CP124: MtrunA17_Chr8g0338301

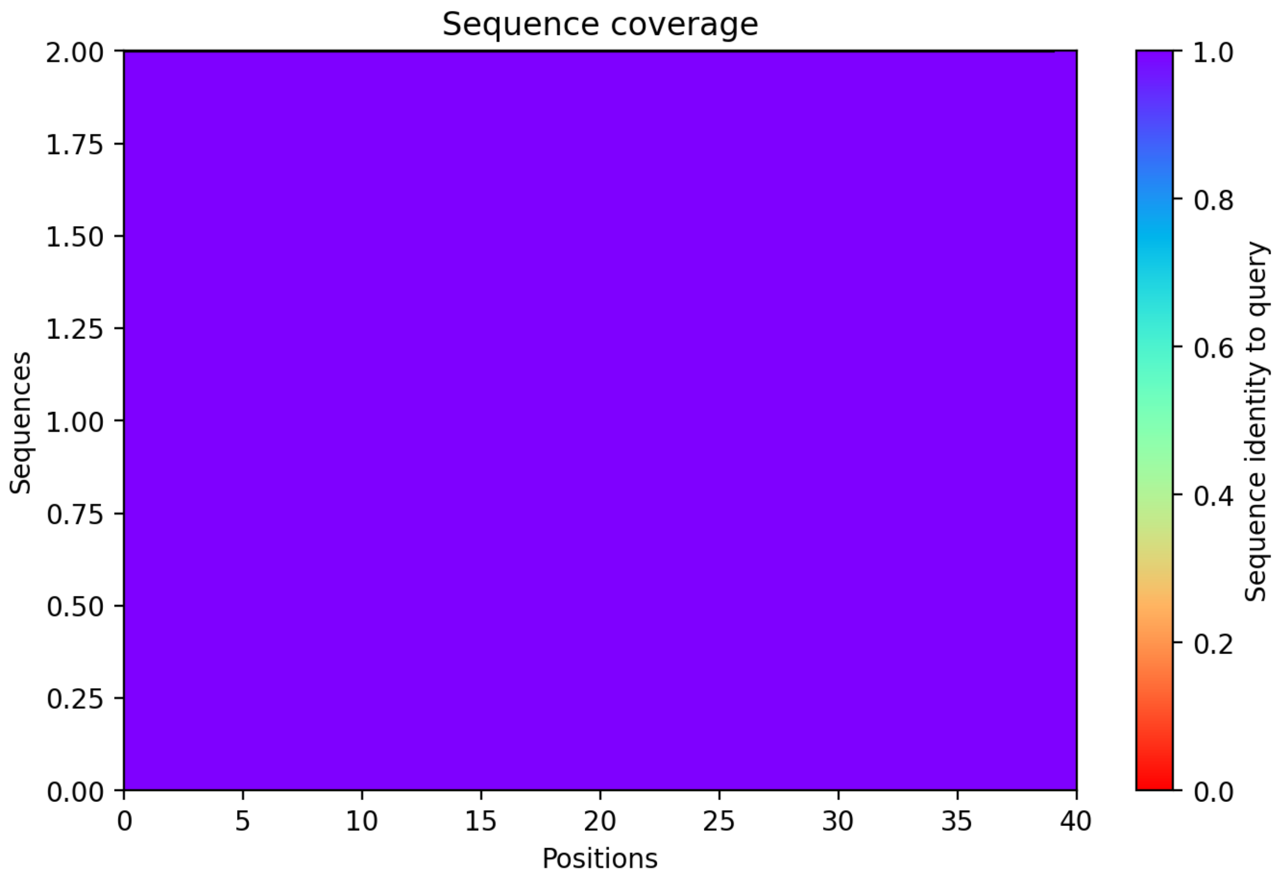

CP125: MtrunA17_Chr8g0339891

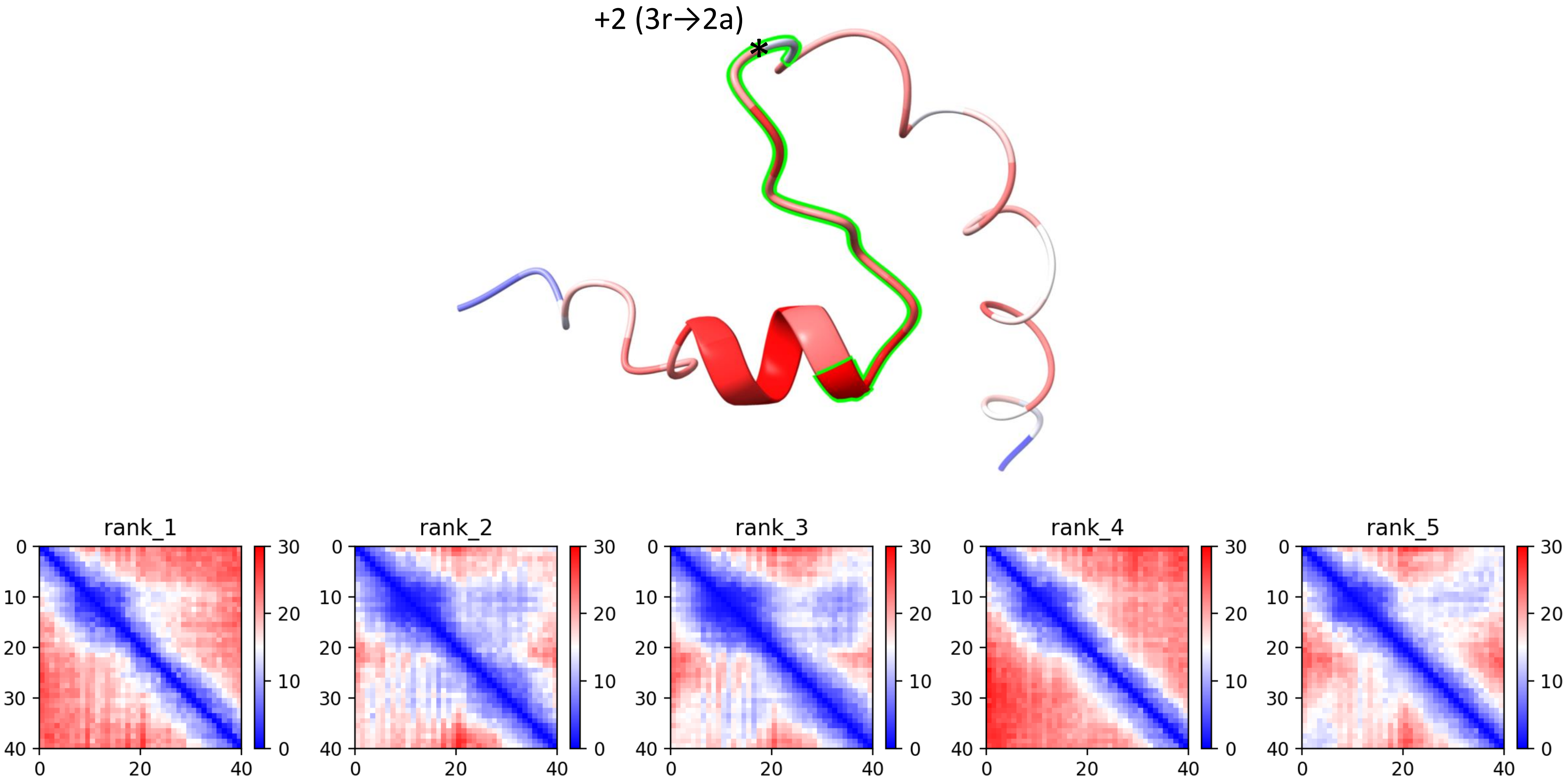

CP125: MtrunA17_Chr8g0339891

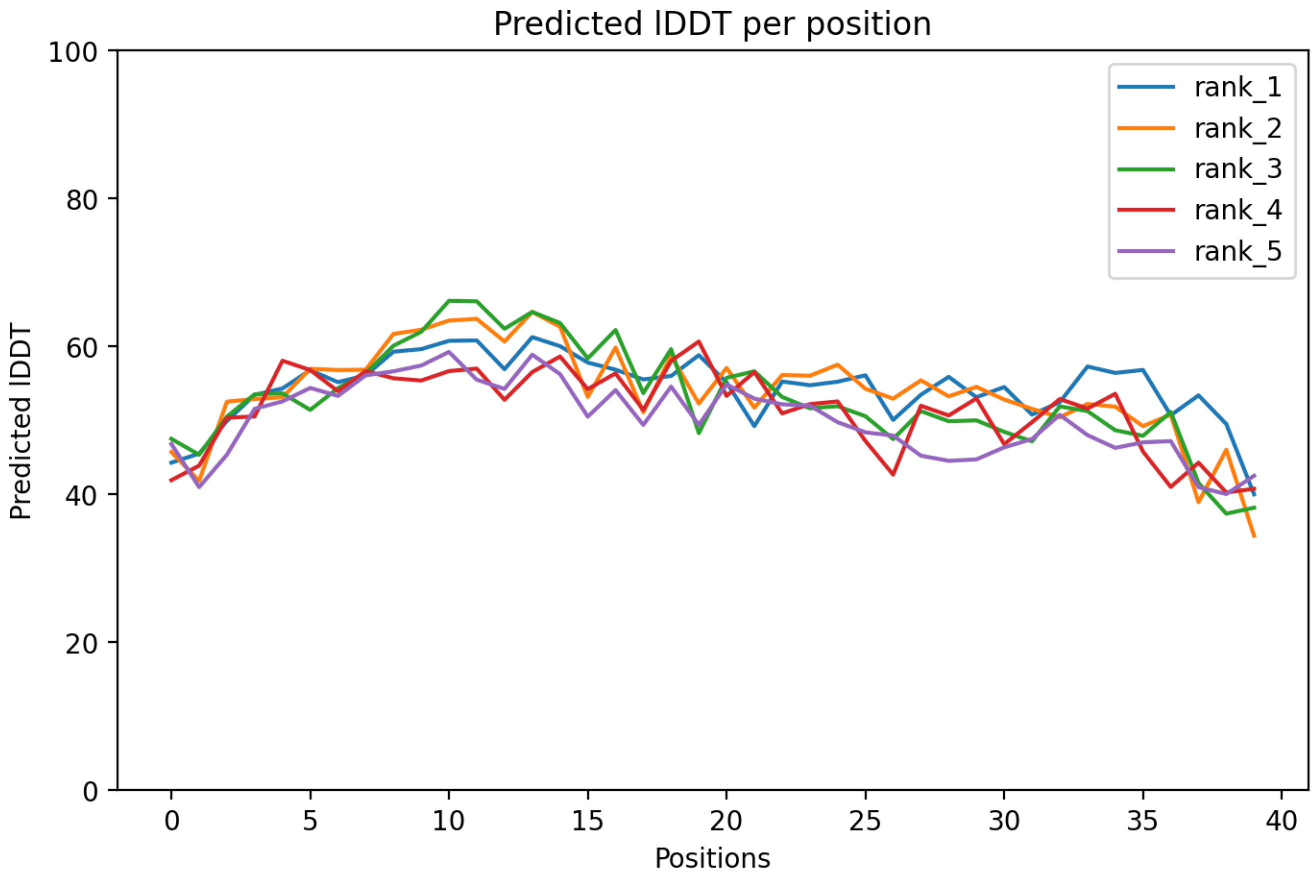

CP125: MtrunA17_Chr8g0339891

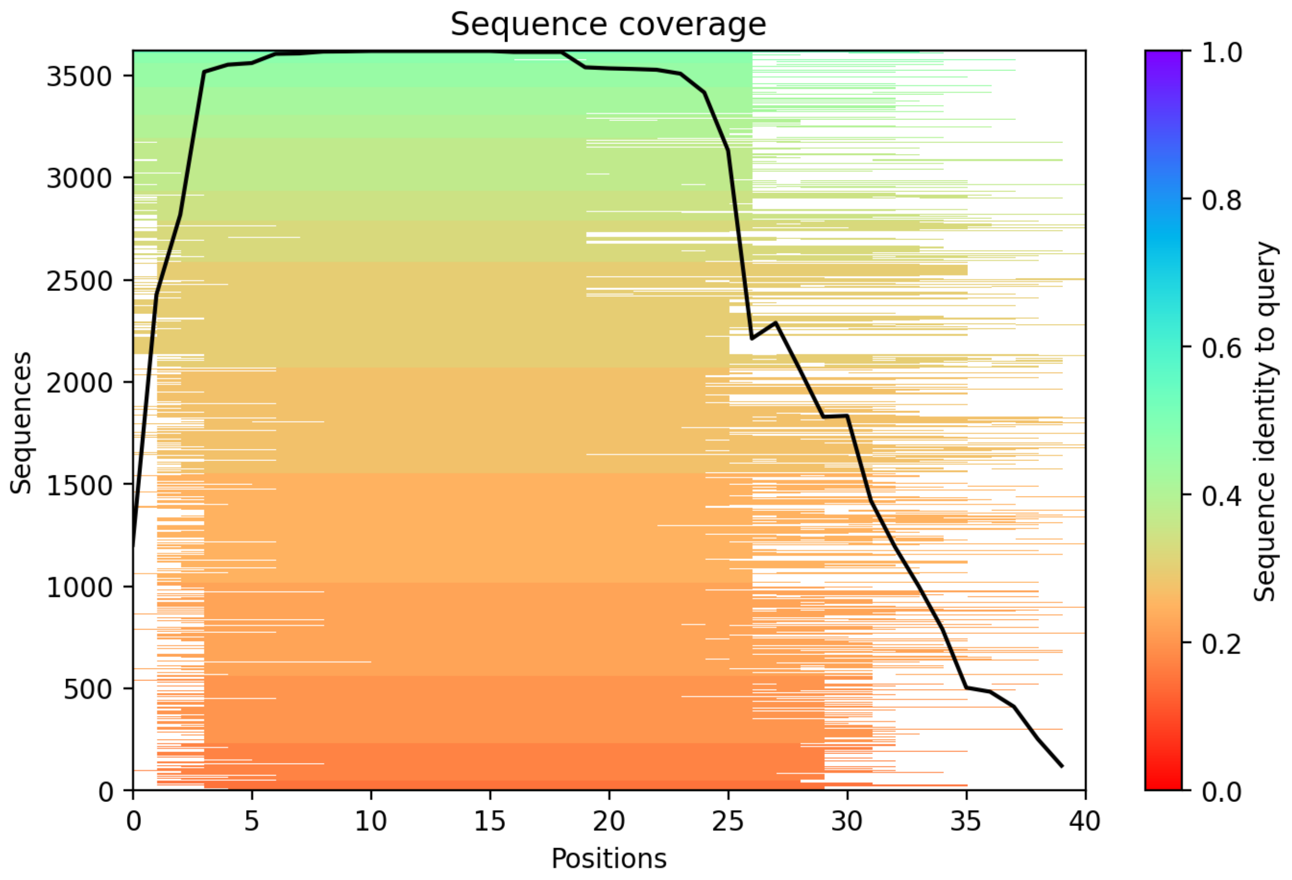

CP126: MtrunA17_Chr8g0342881

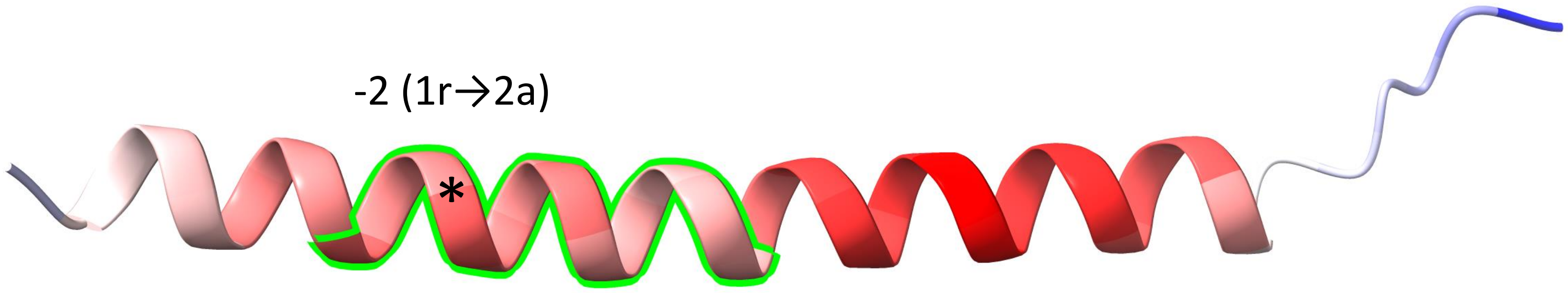


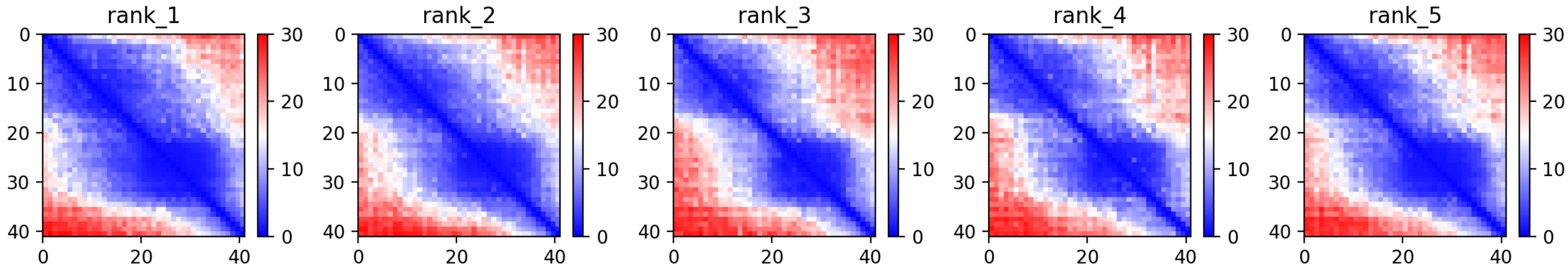

CP126: MtrunA17_Chr8g0342881

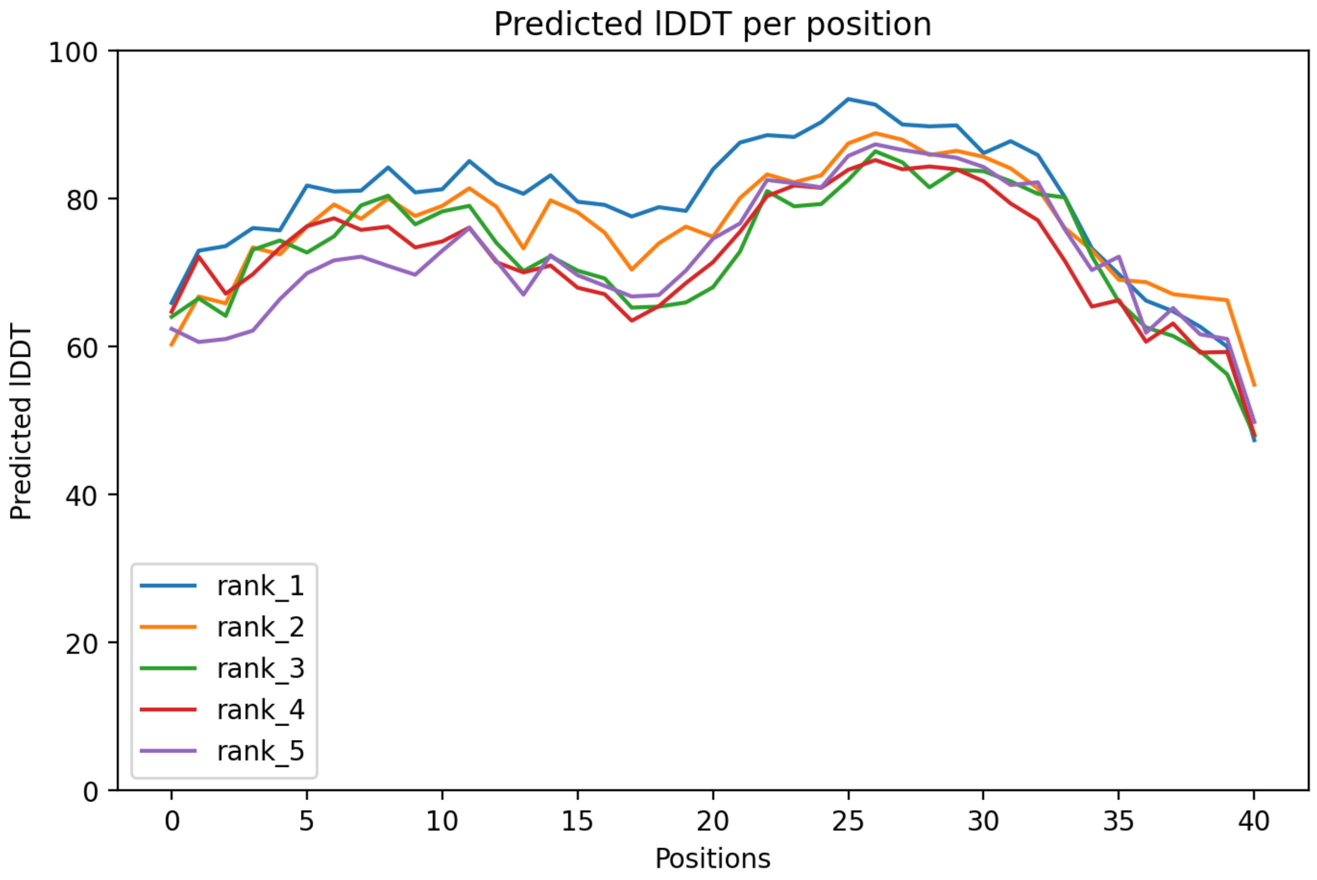

CP126: MtrunA17_Chr8g0342881

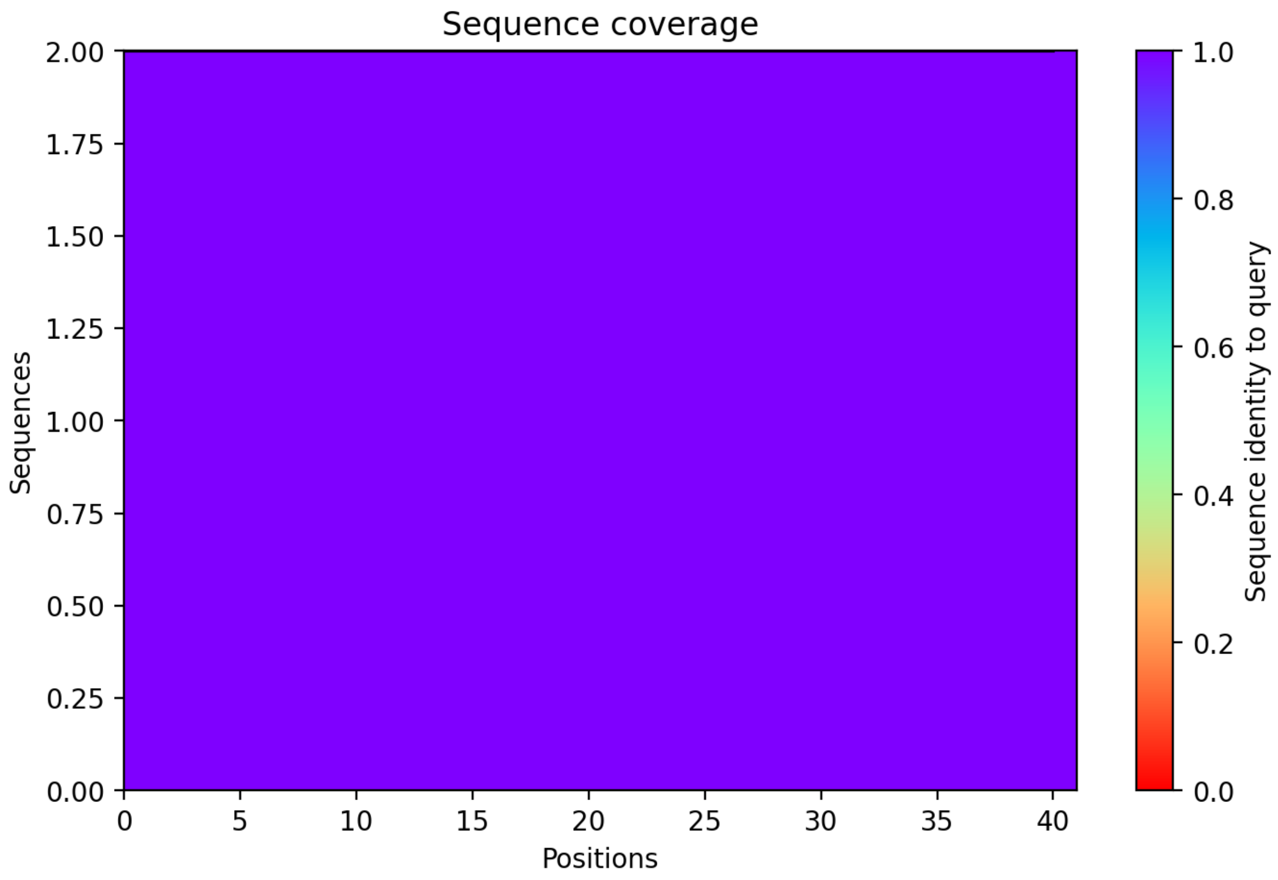

CP127: MtrunA17_Chr8g0345421

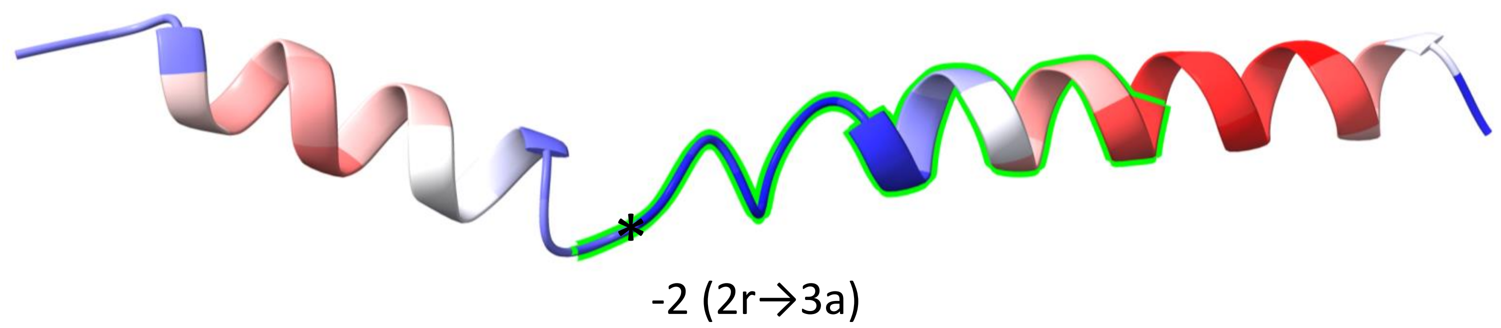


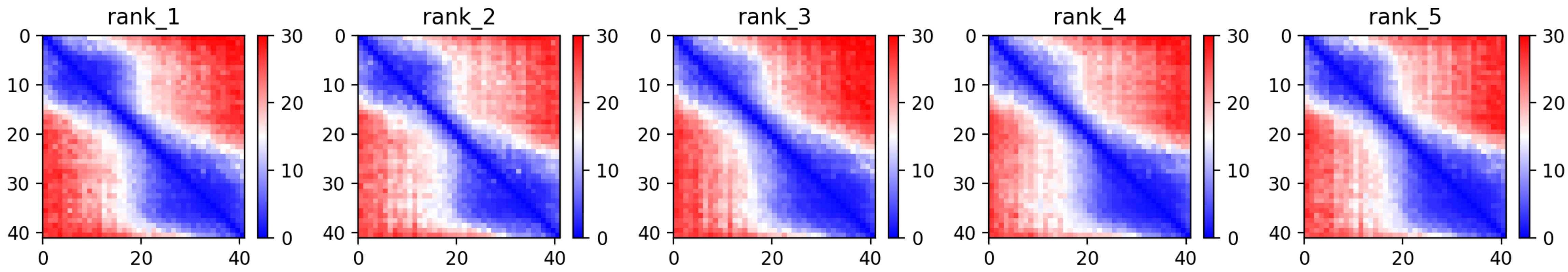

CP127: MtrunA17_Chr8g0345421

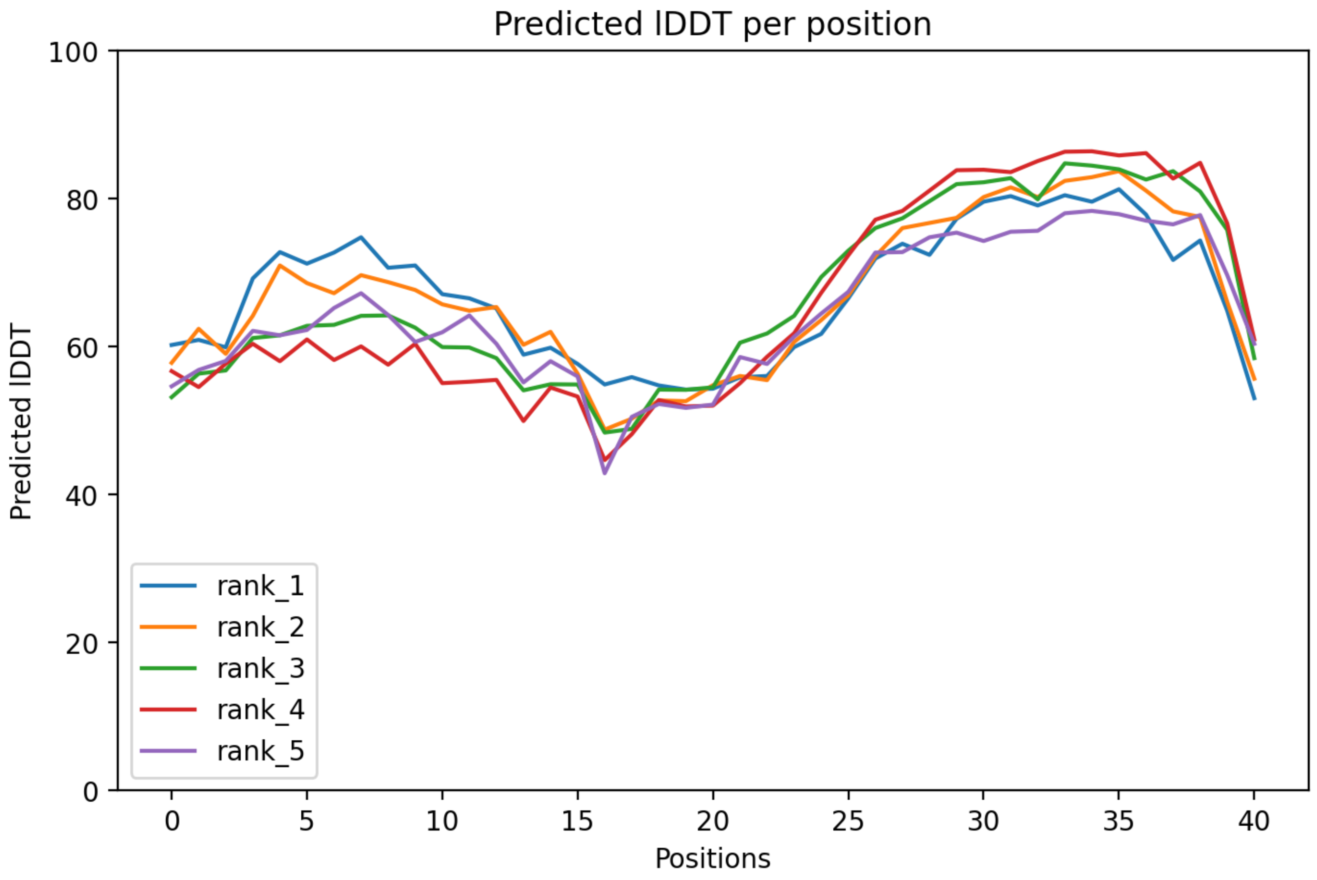

### CP127: MtrunA17_Chr8g0345421

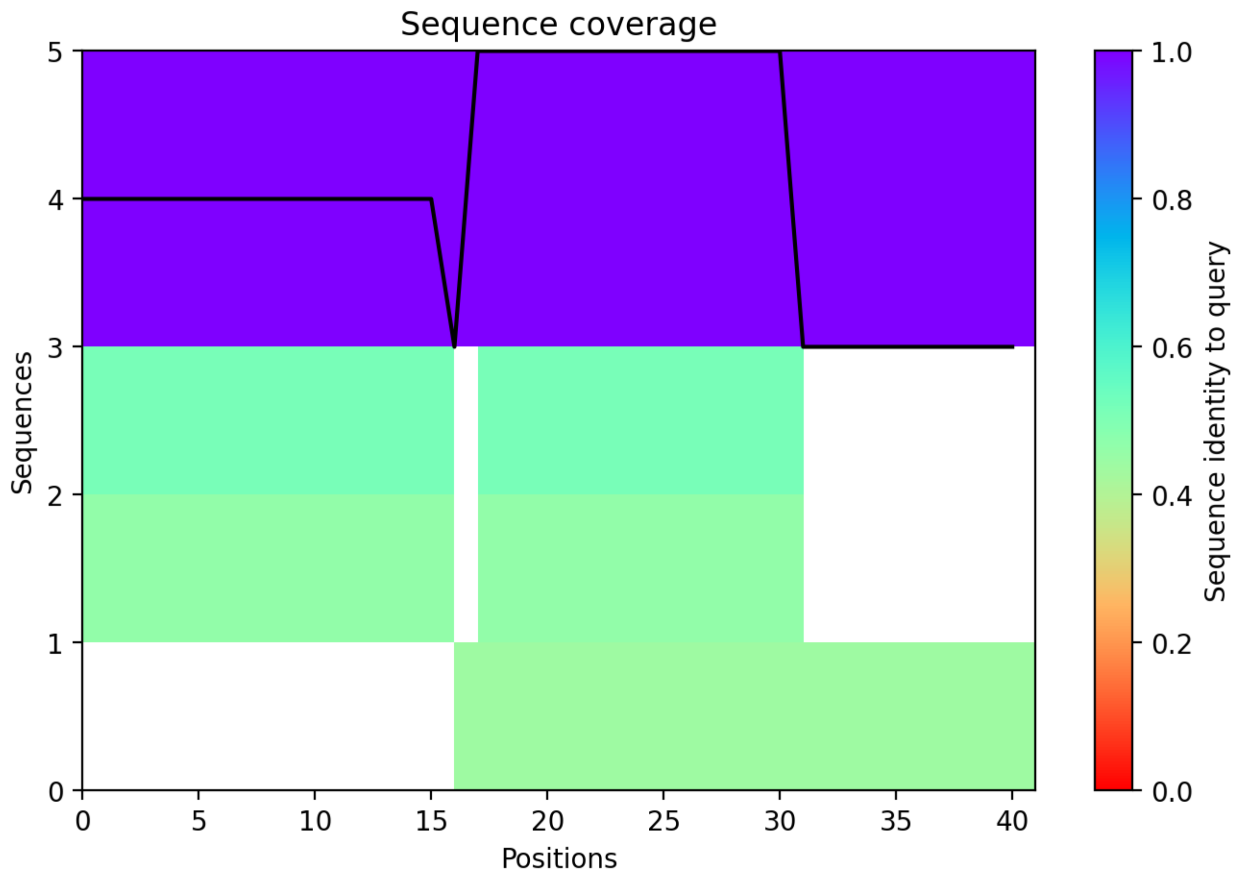

### CP128: MtrunA17_Chr8g0353711

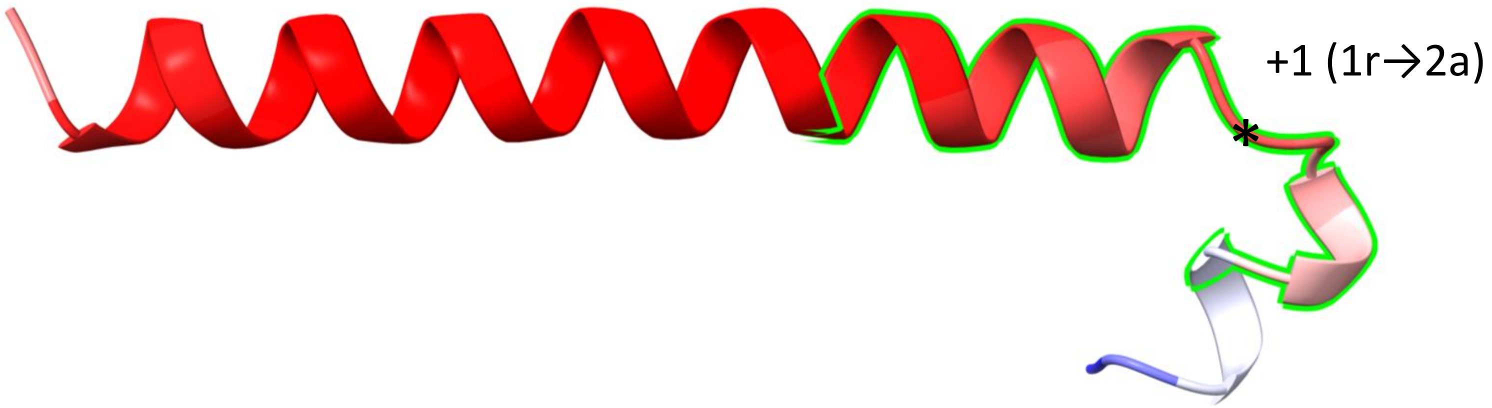


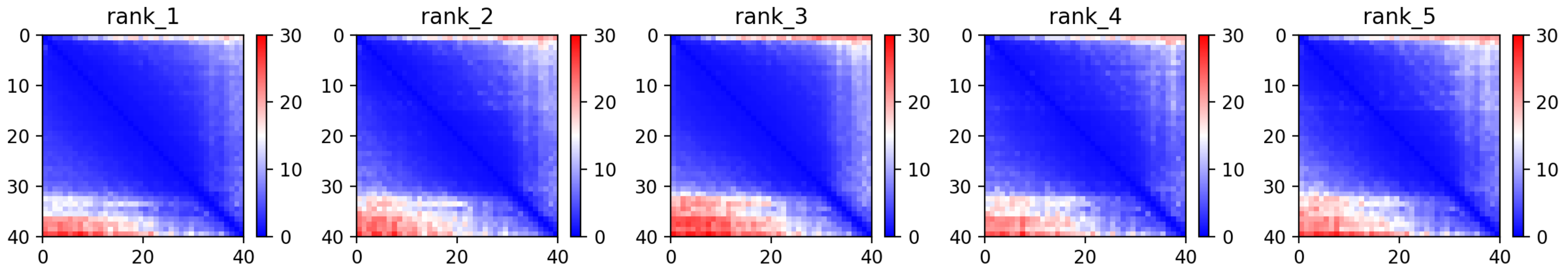

CP128: MtrunA17_Chr8g0353711

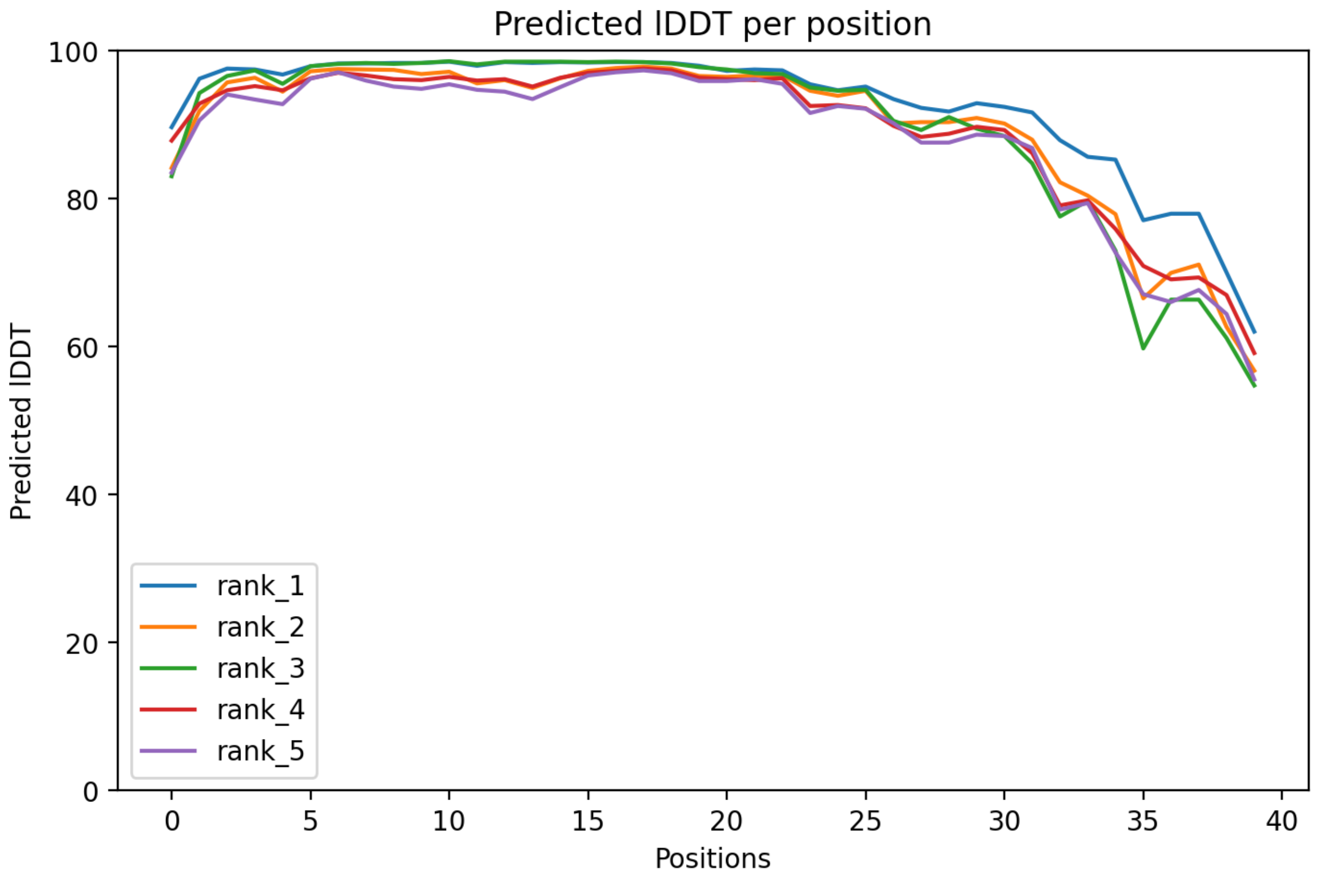

### CP128: MtrunA17_Chr8g0353711

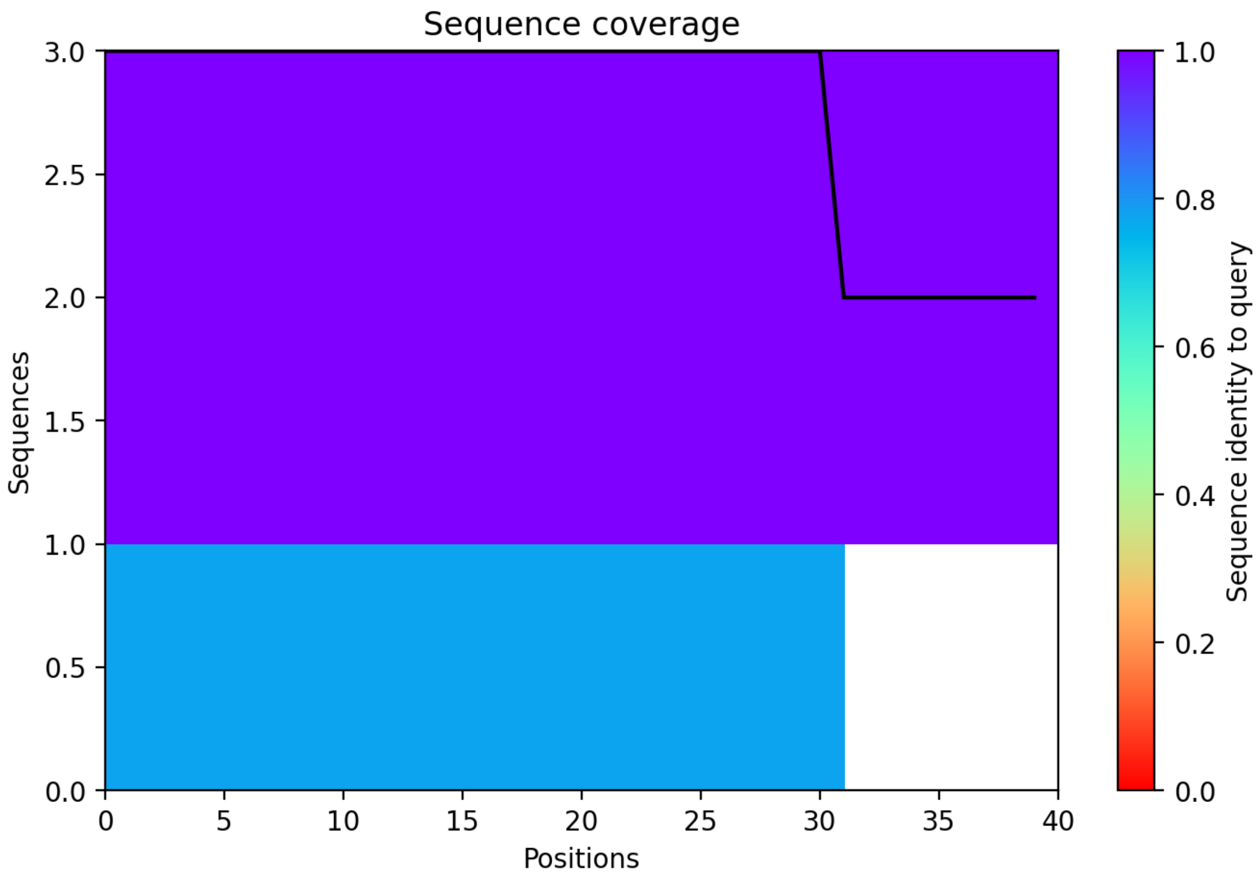

CP129: MtrunA17_Chr8g0355501

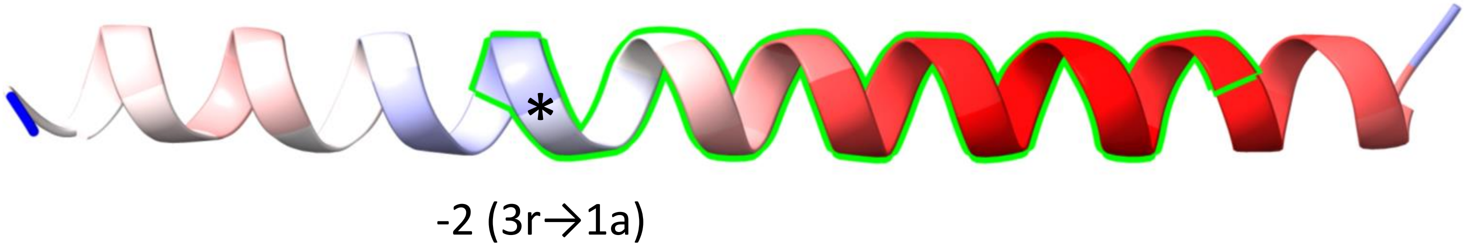


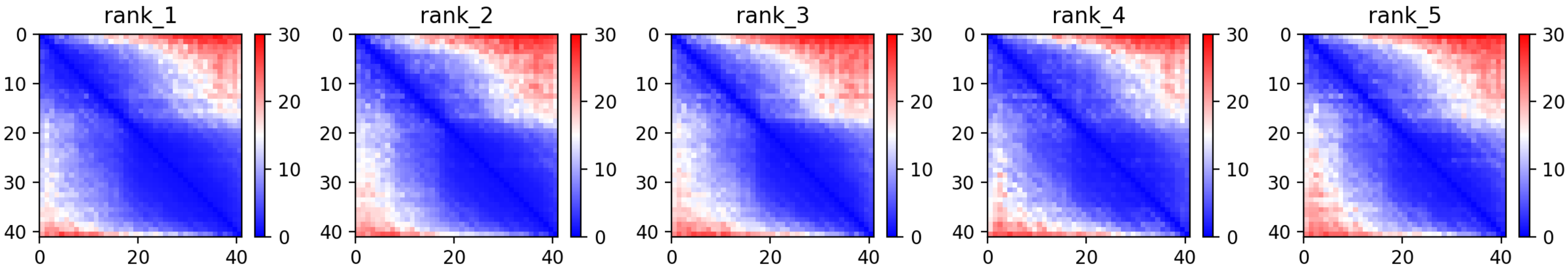

### CP129: MtrunA17_Chr8g0355501

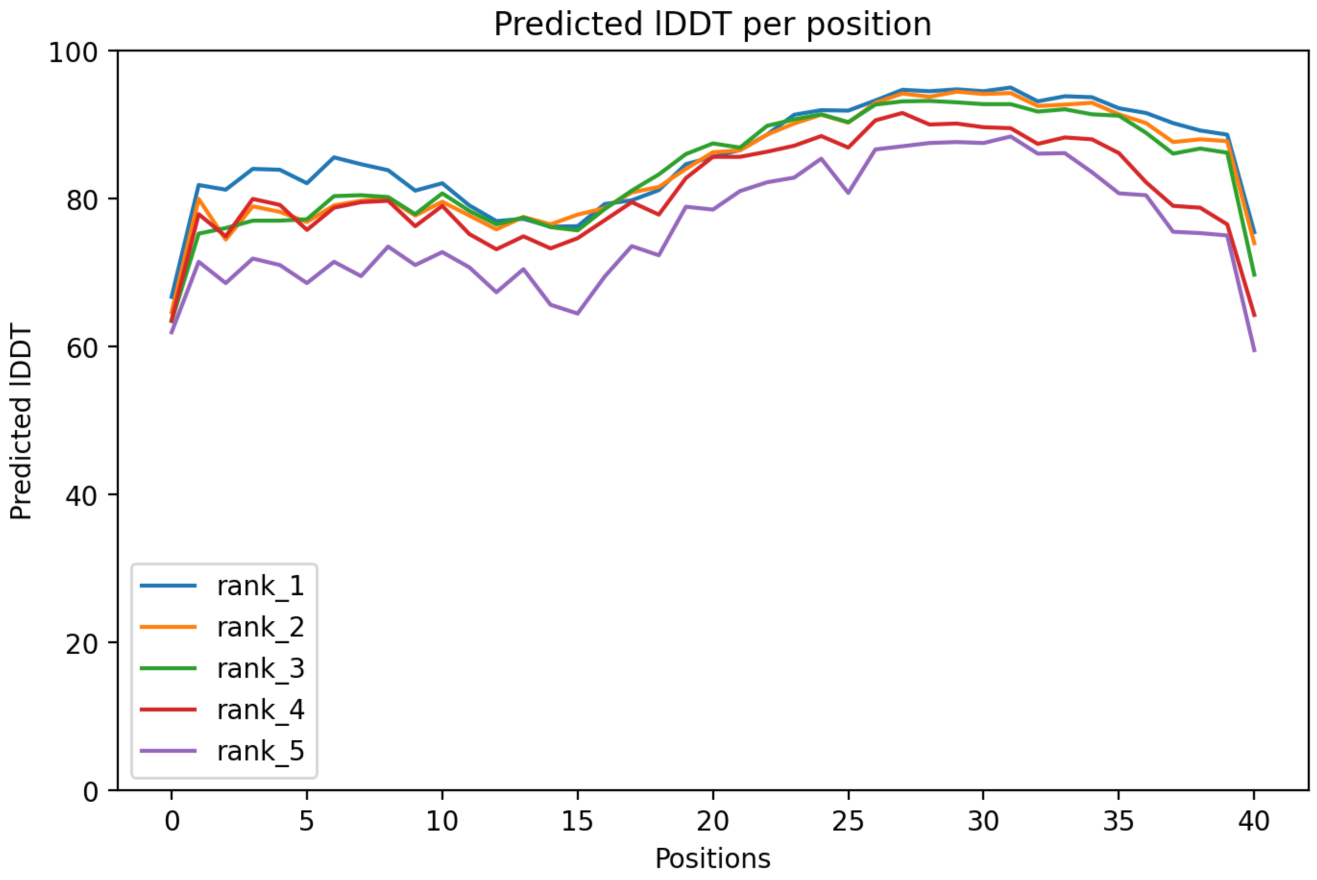

CP129: MtrunA17_Chr8g0355501

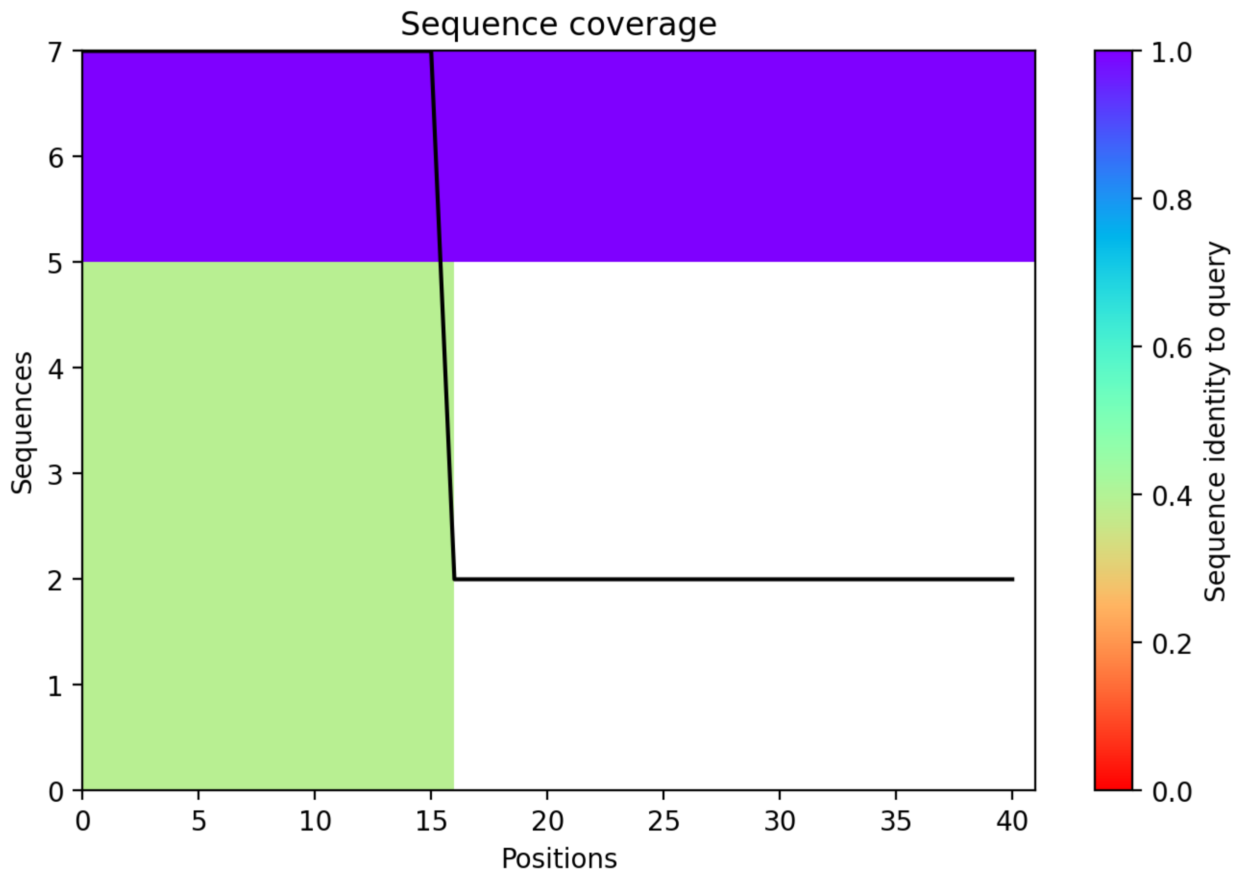

CP130: MtrunA17_Chr8g0356581

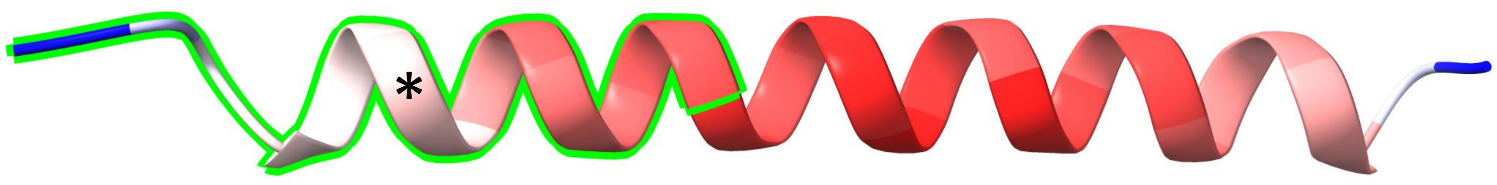


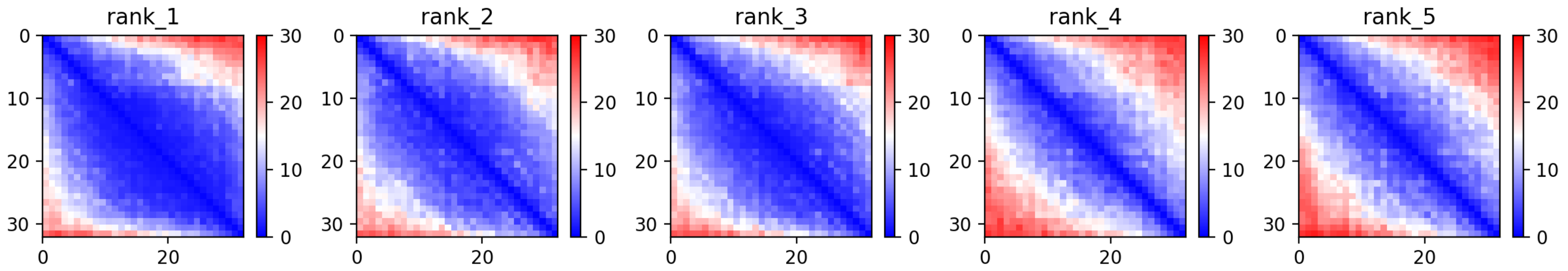

CP130: MtrunA17_Chr8g0356581

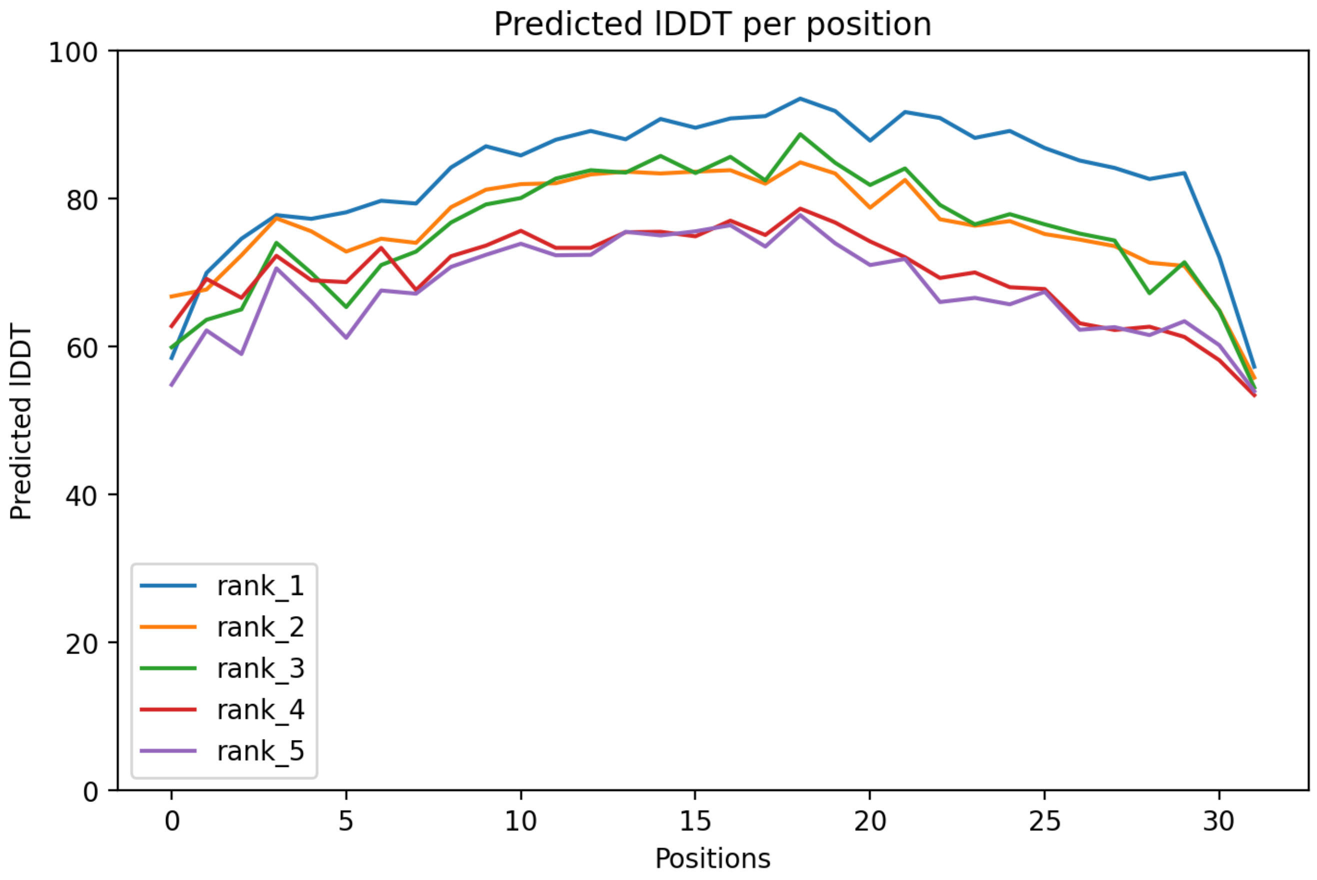

### CP130: MtrunA17_Chr8g0356581

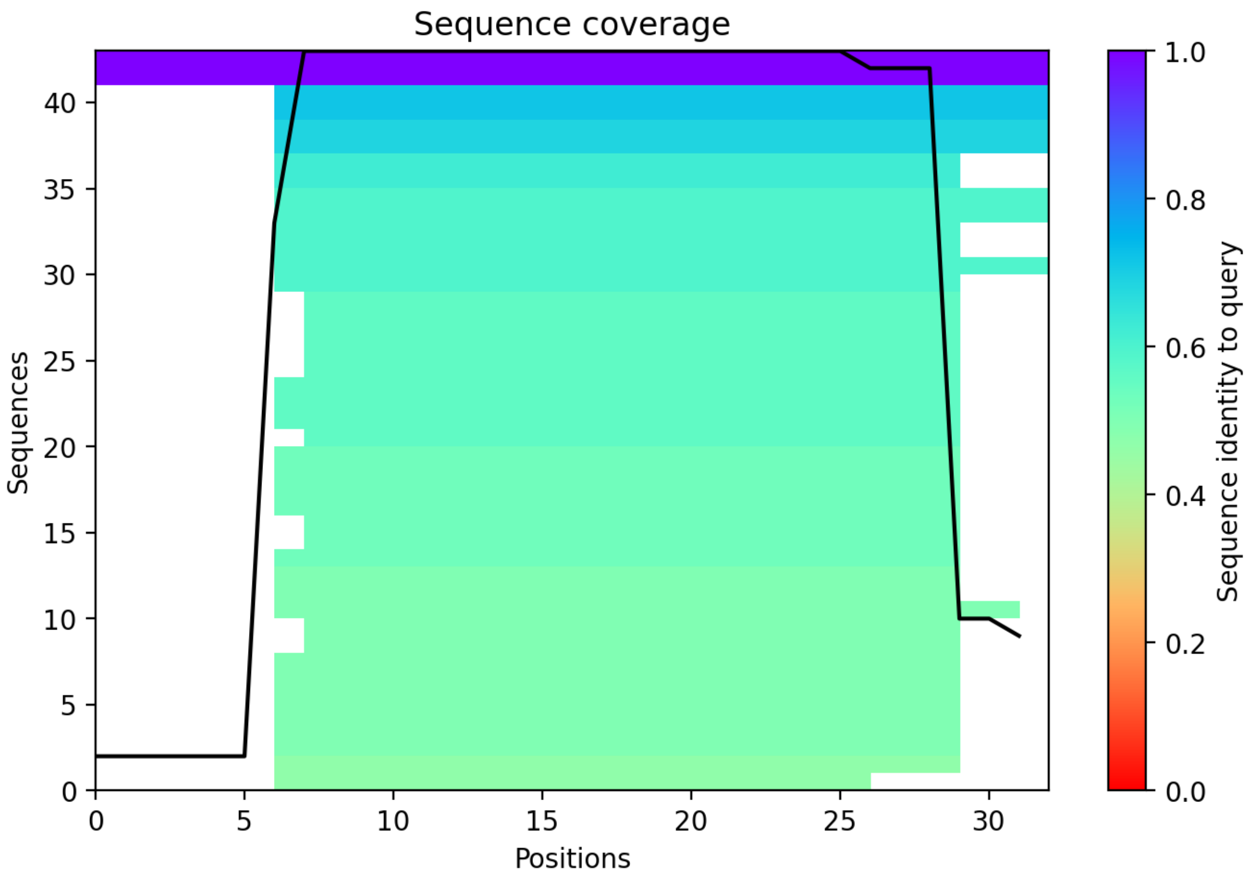

CP131: MtrunA17_Chr8g0365341

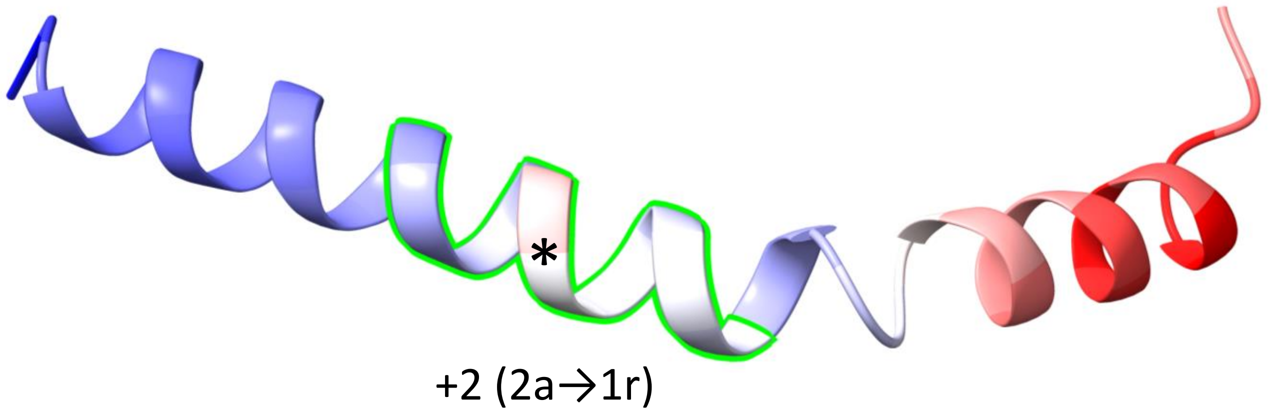


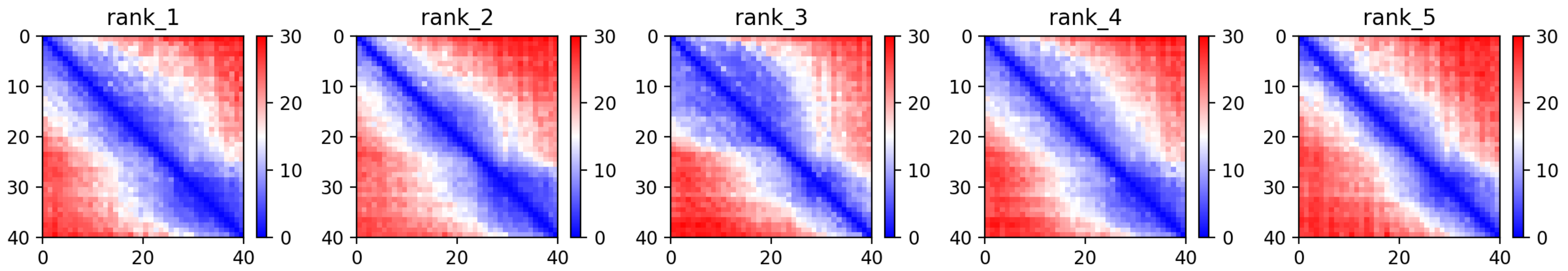

CP131: MtrunA17_Chr8g0365341

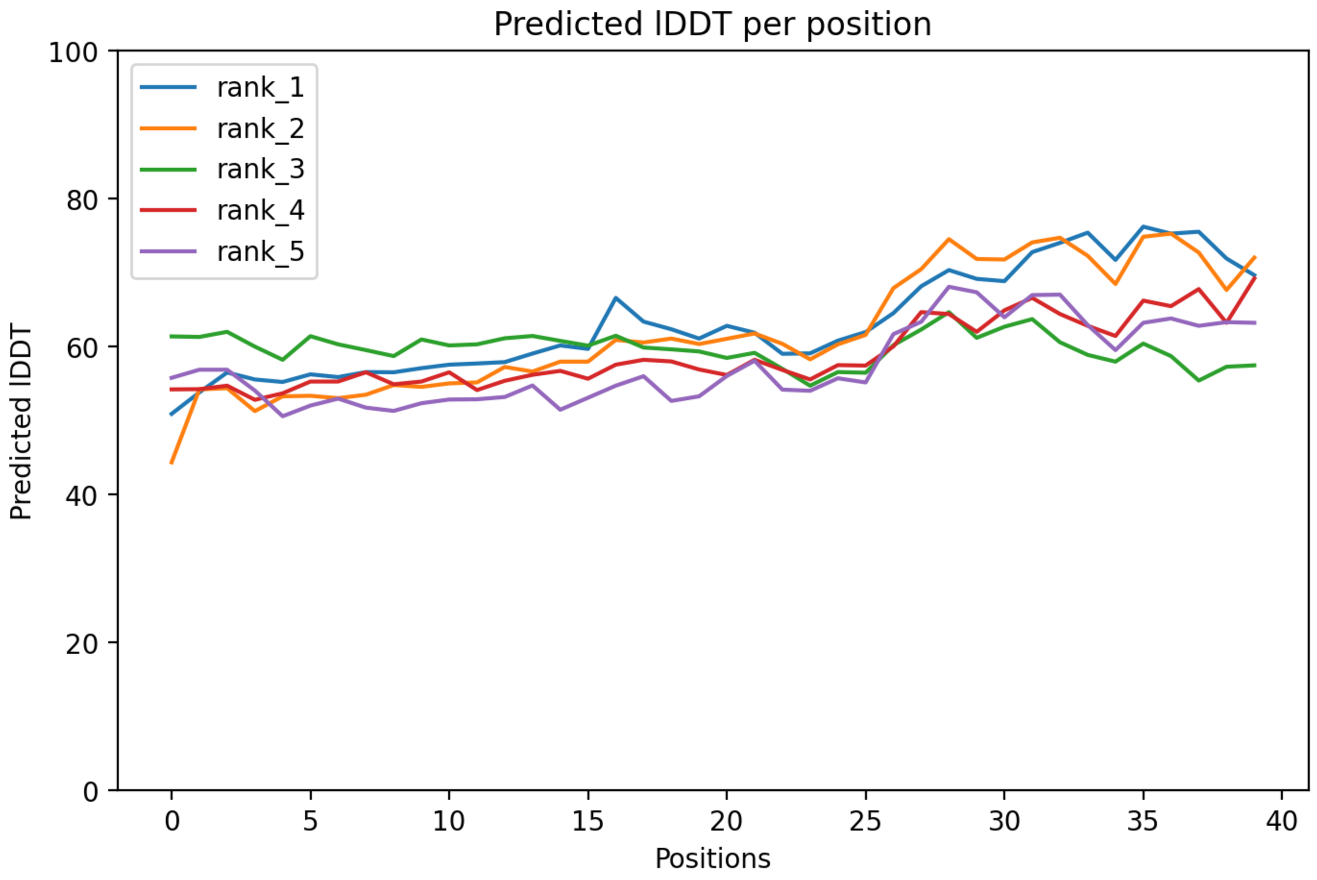

CP131: MtrunA17_Chr8g0365341

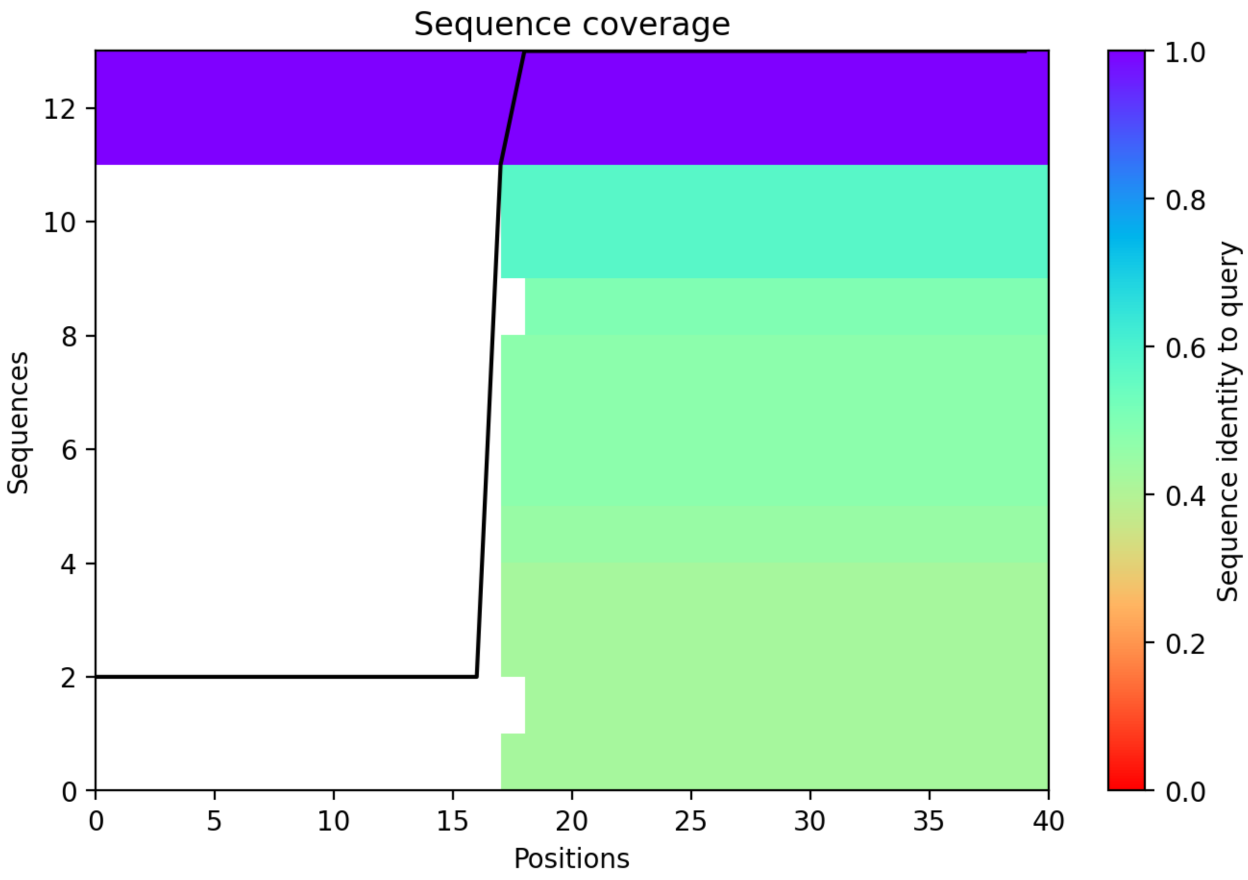

CP132: MtrunA17_Chr8g0368731

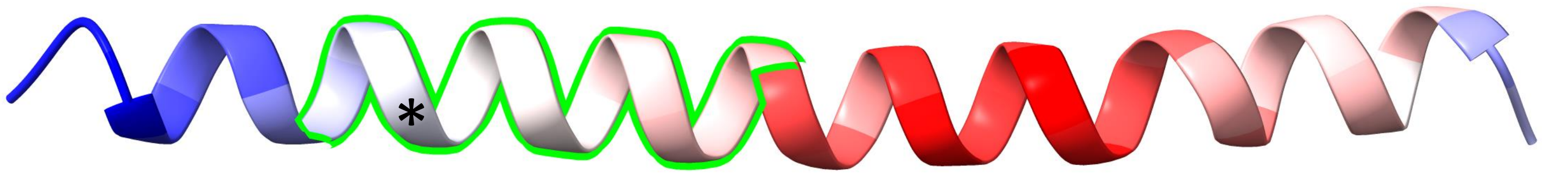


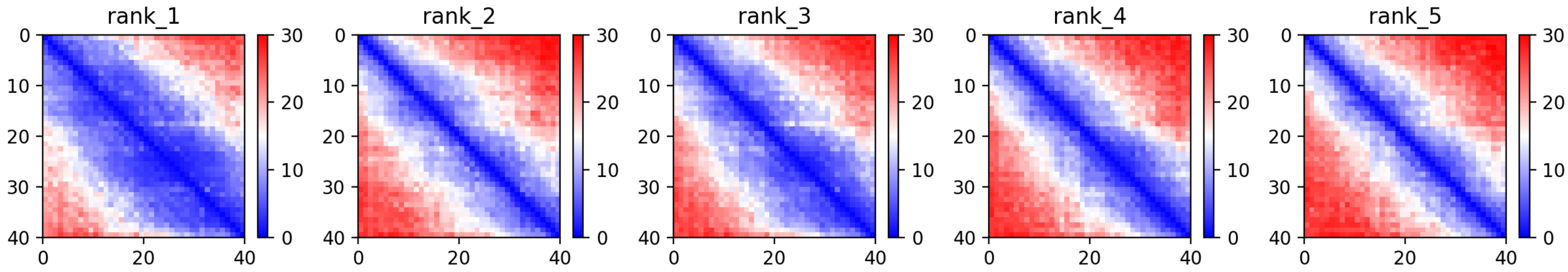

CP132: MtrunA17_Chr8g0368731

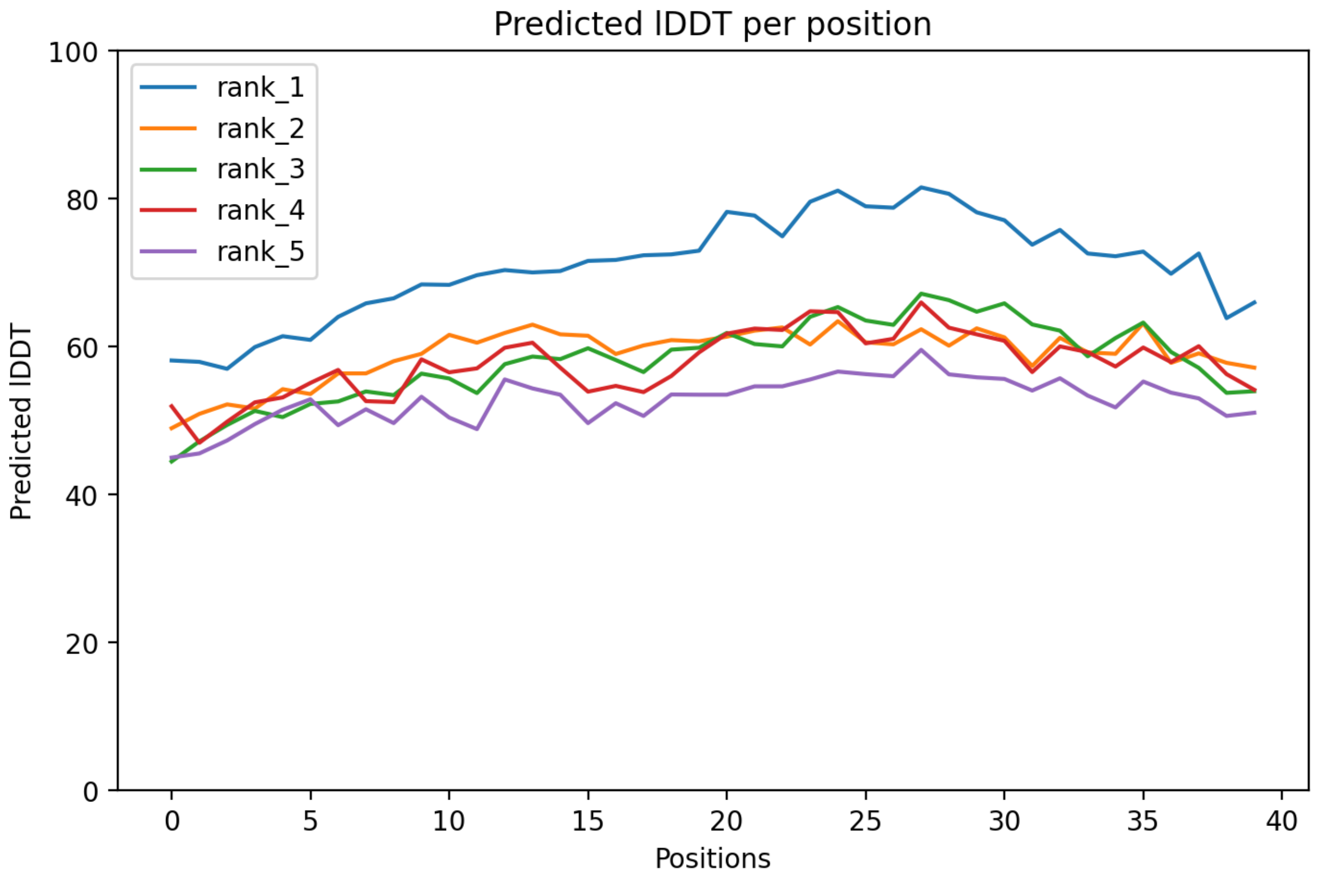

### CP132: MtrunA17_Chr8g0368731

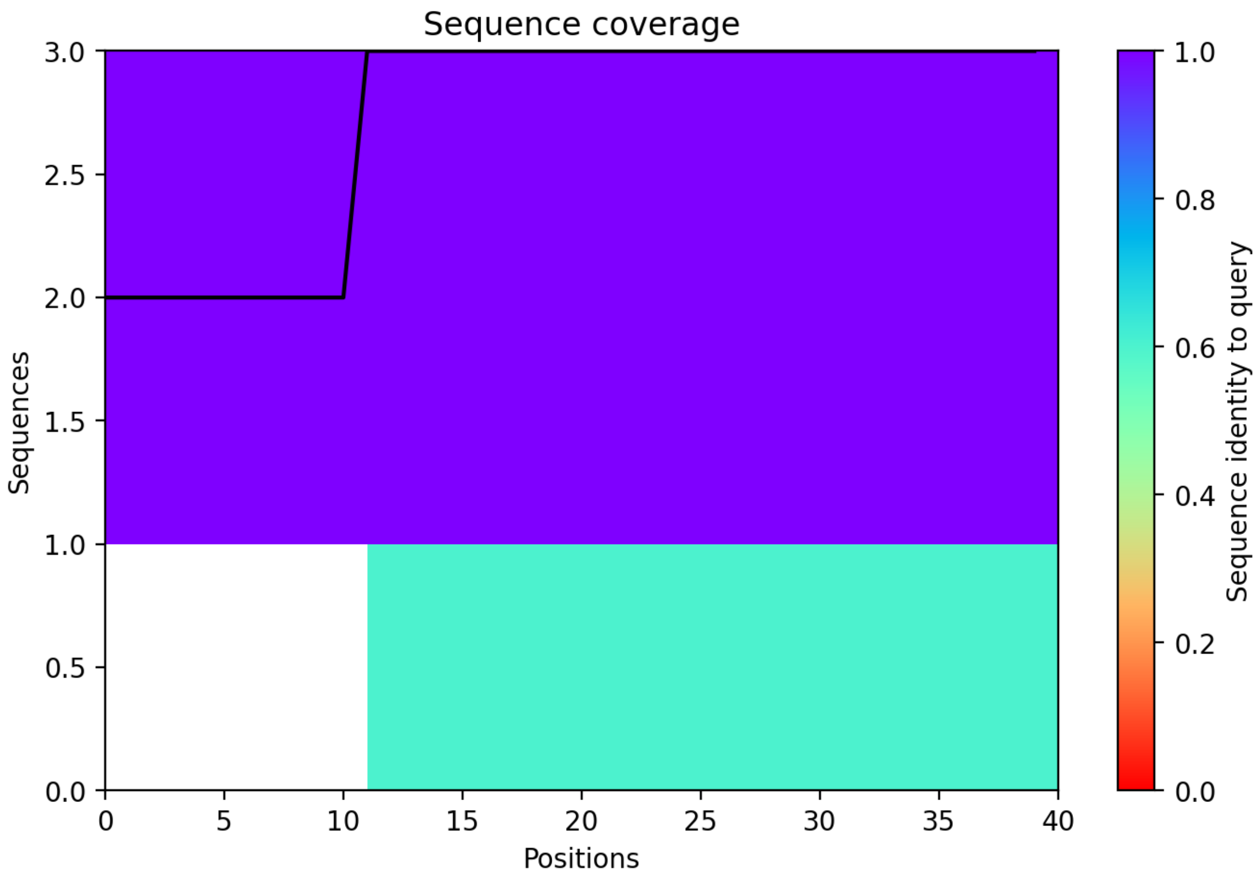

### CP133: MtrunA17_Chr8g0368931

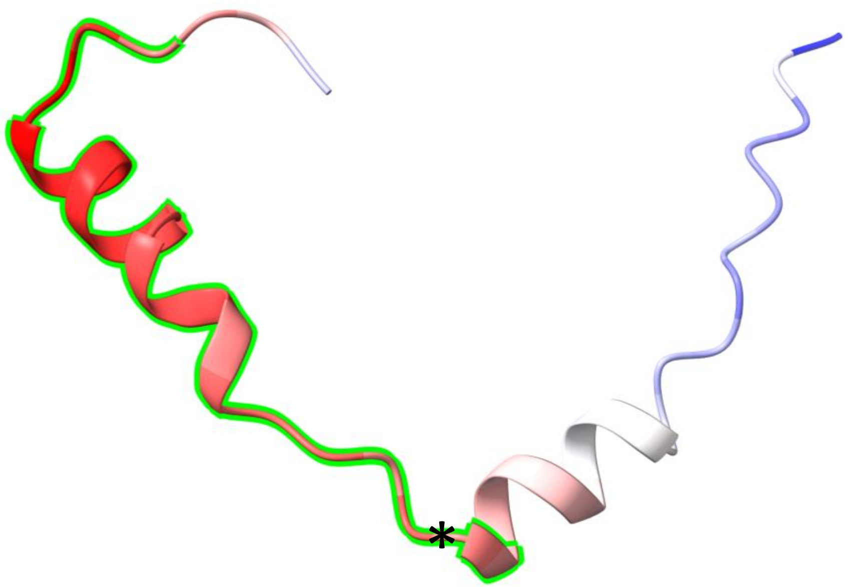


-2 (3r→1a)

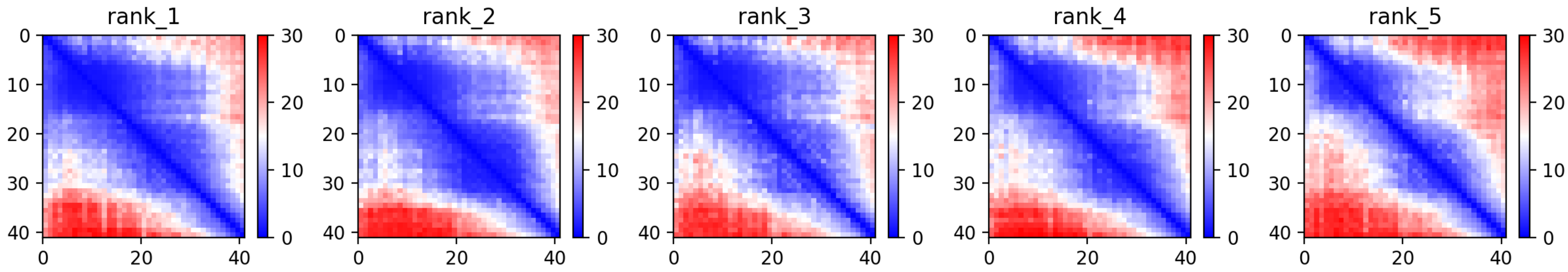

### CP133: MtrunA17_Chr8g0368931

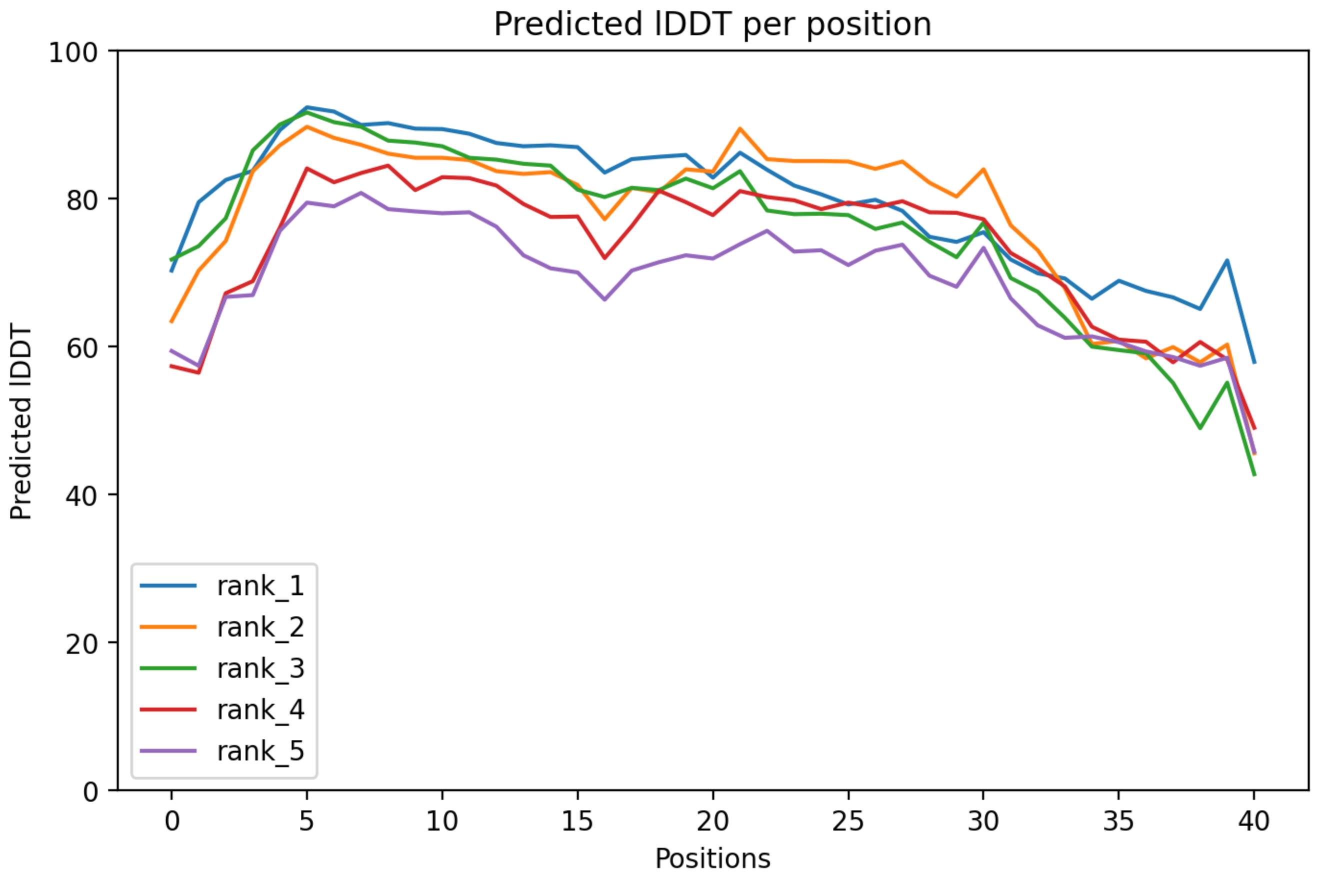

### CP133: MtrunA17_Chr8g0368931

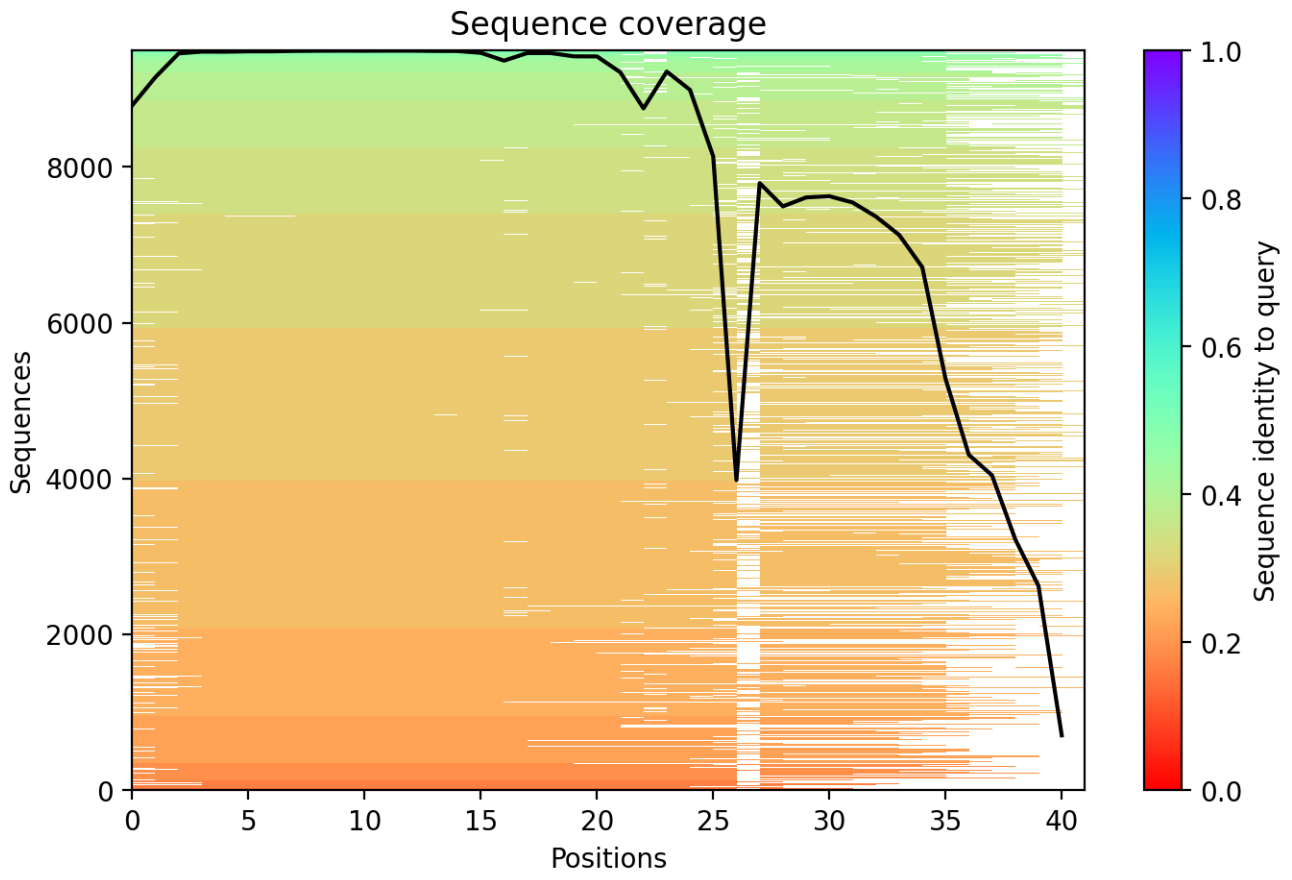

### CP134: MtrunA17_Chr8g0371281

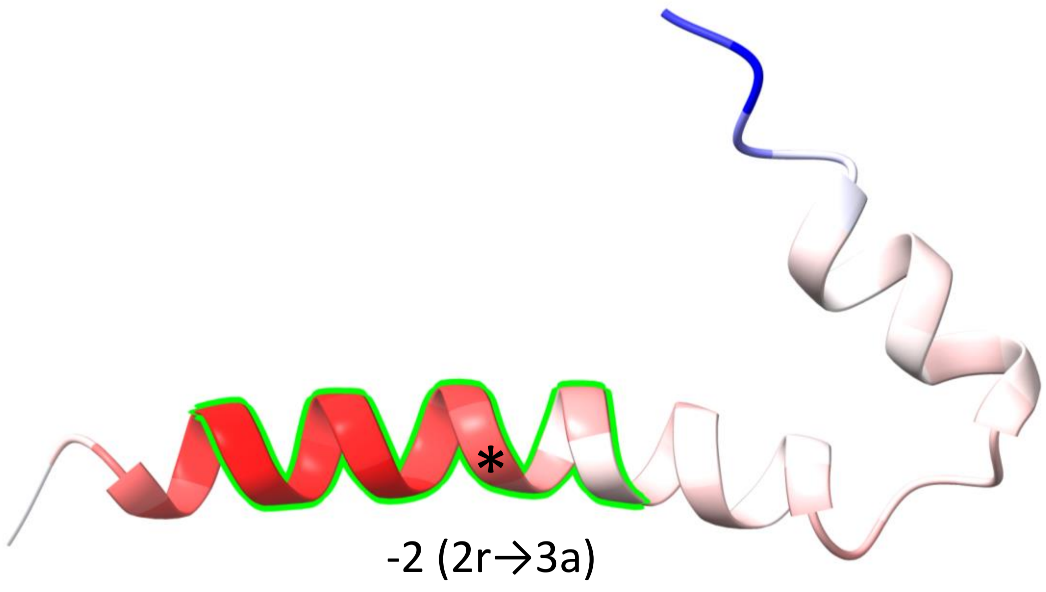


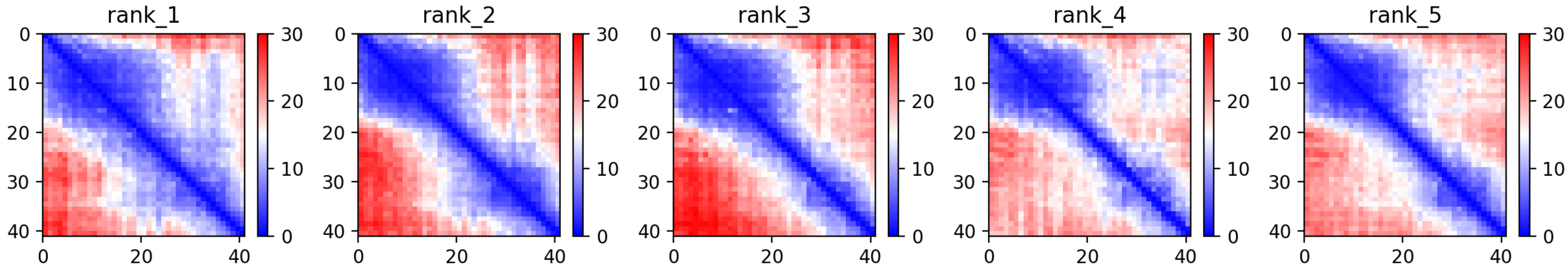

### CP134: MtrunA17_Chr8g0371281

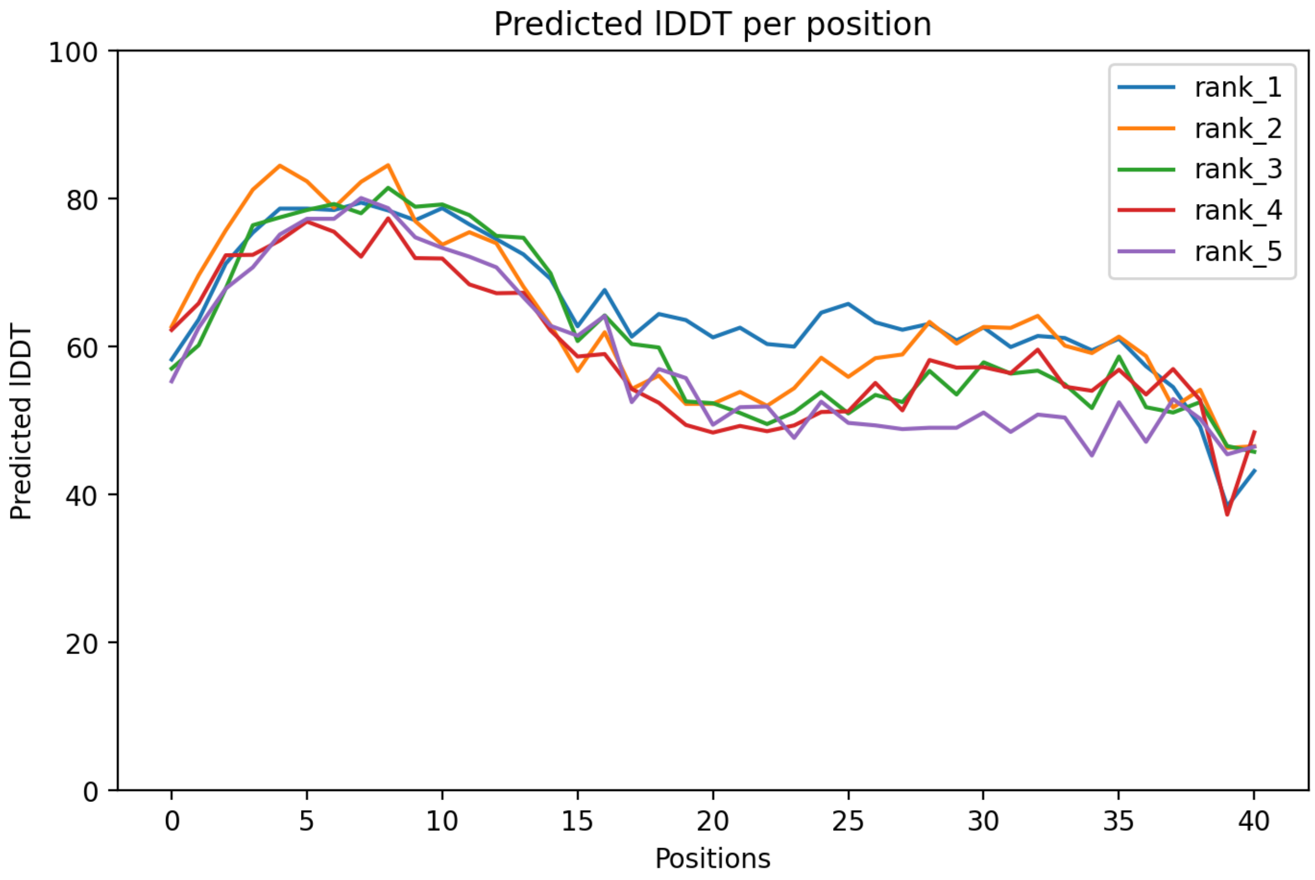

CP134: MtrunA17_Chr8g0371281

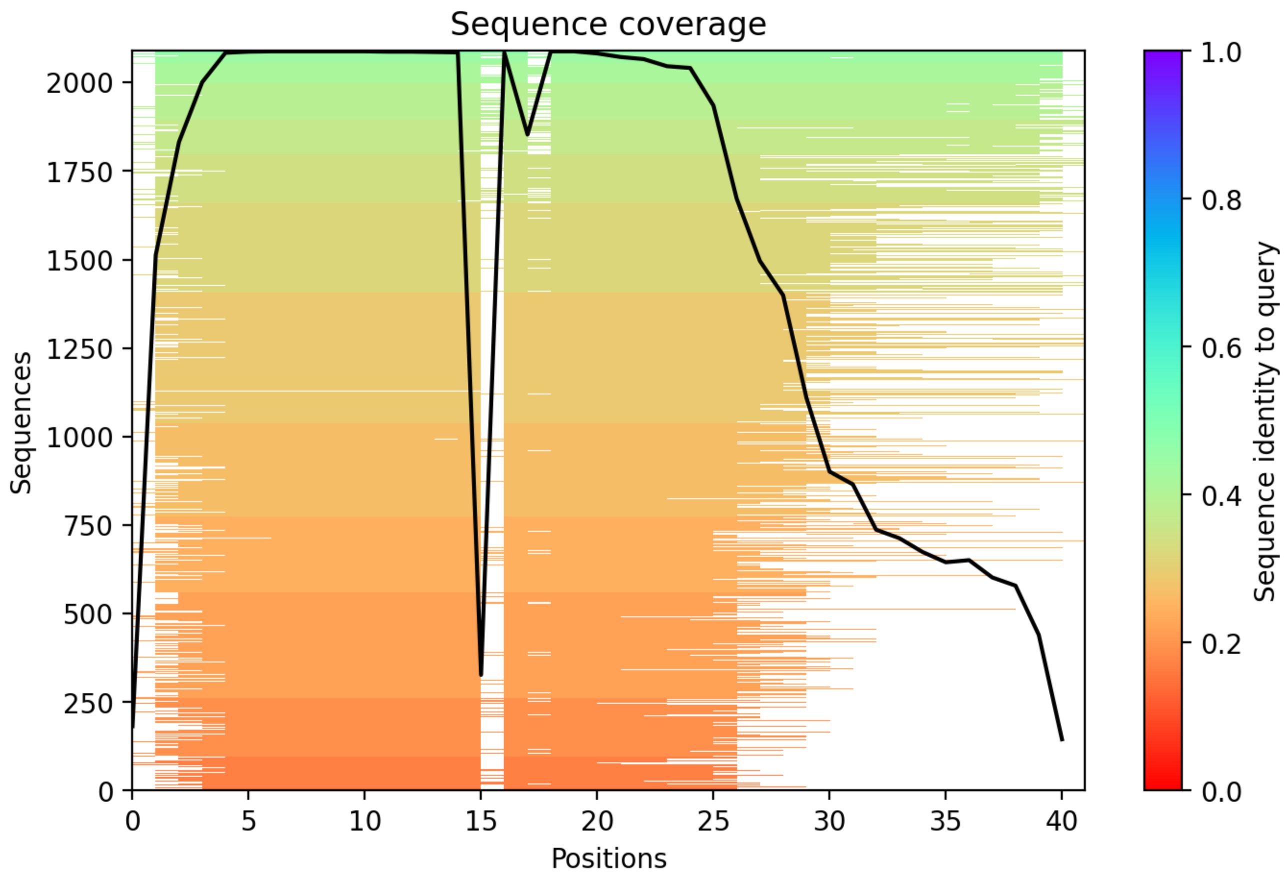

CP135: MtrunA17_Chr8g0371741

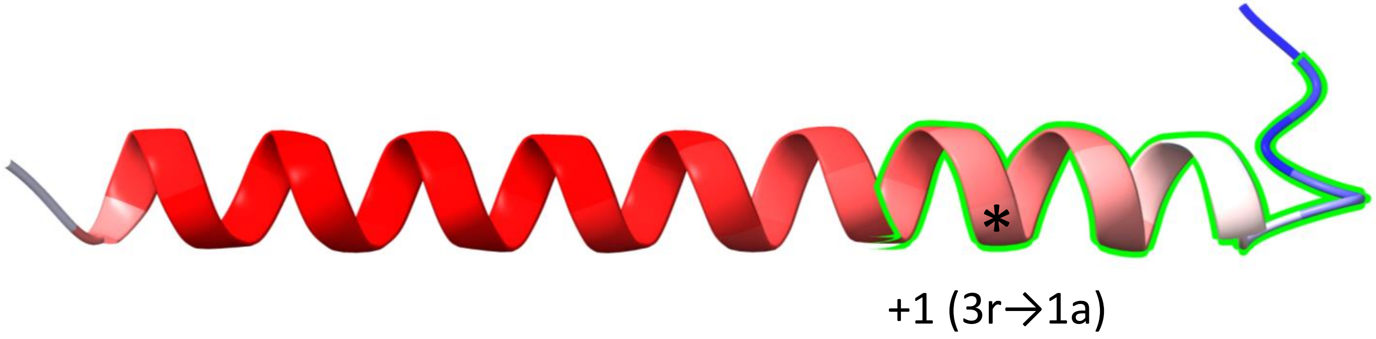


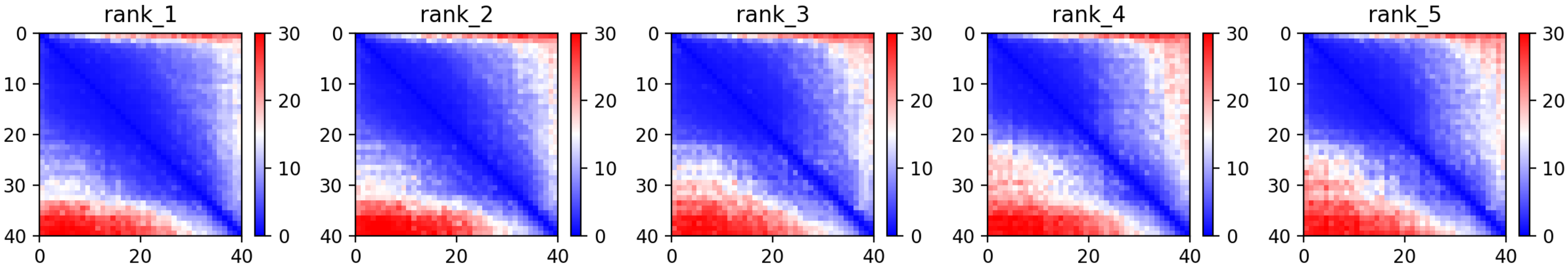

CP135: MtrunA17_Chr8g0371741

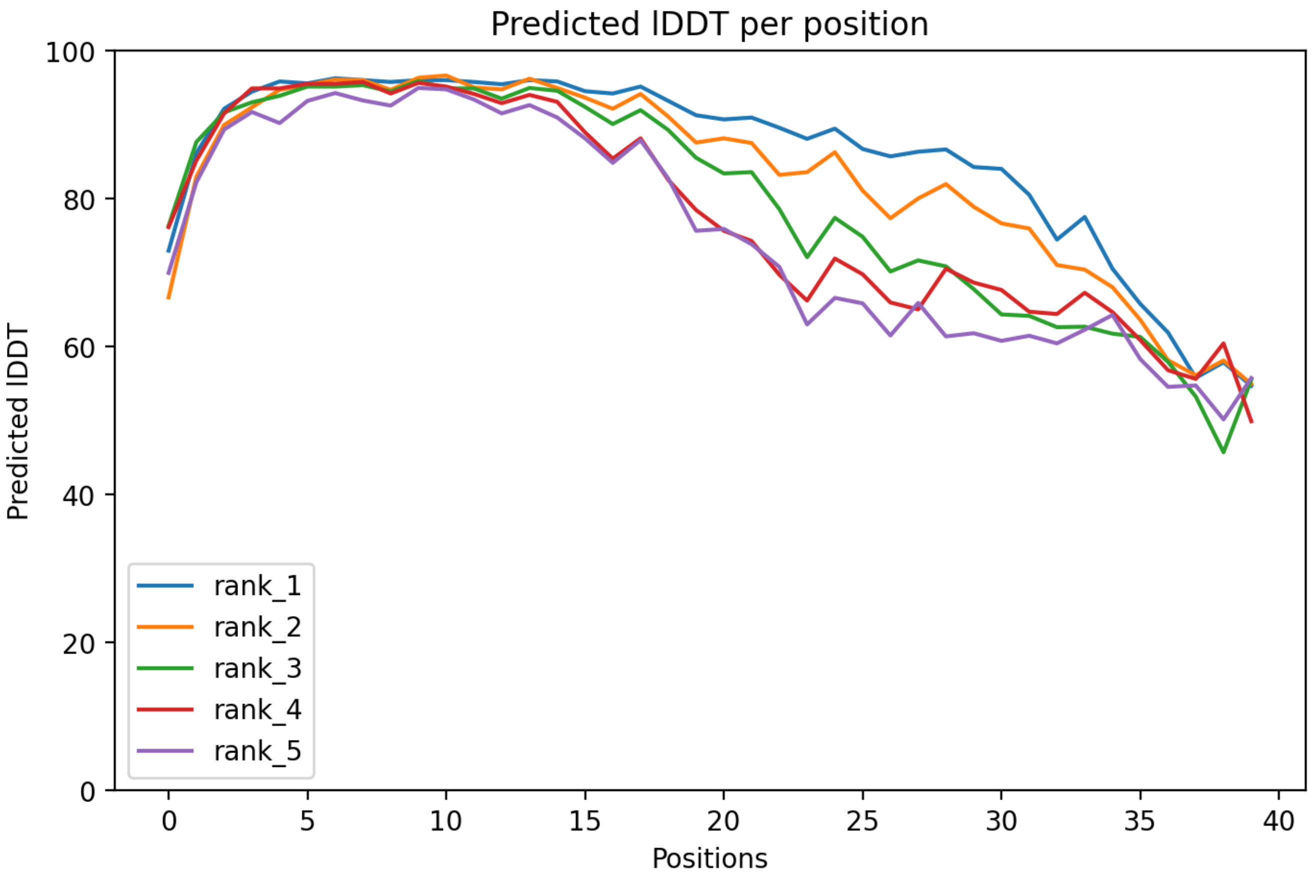

CP135: MtrunA17_Chr8g0371741

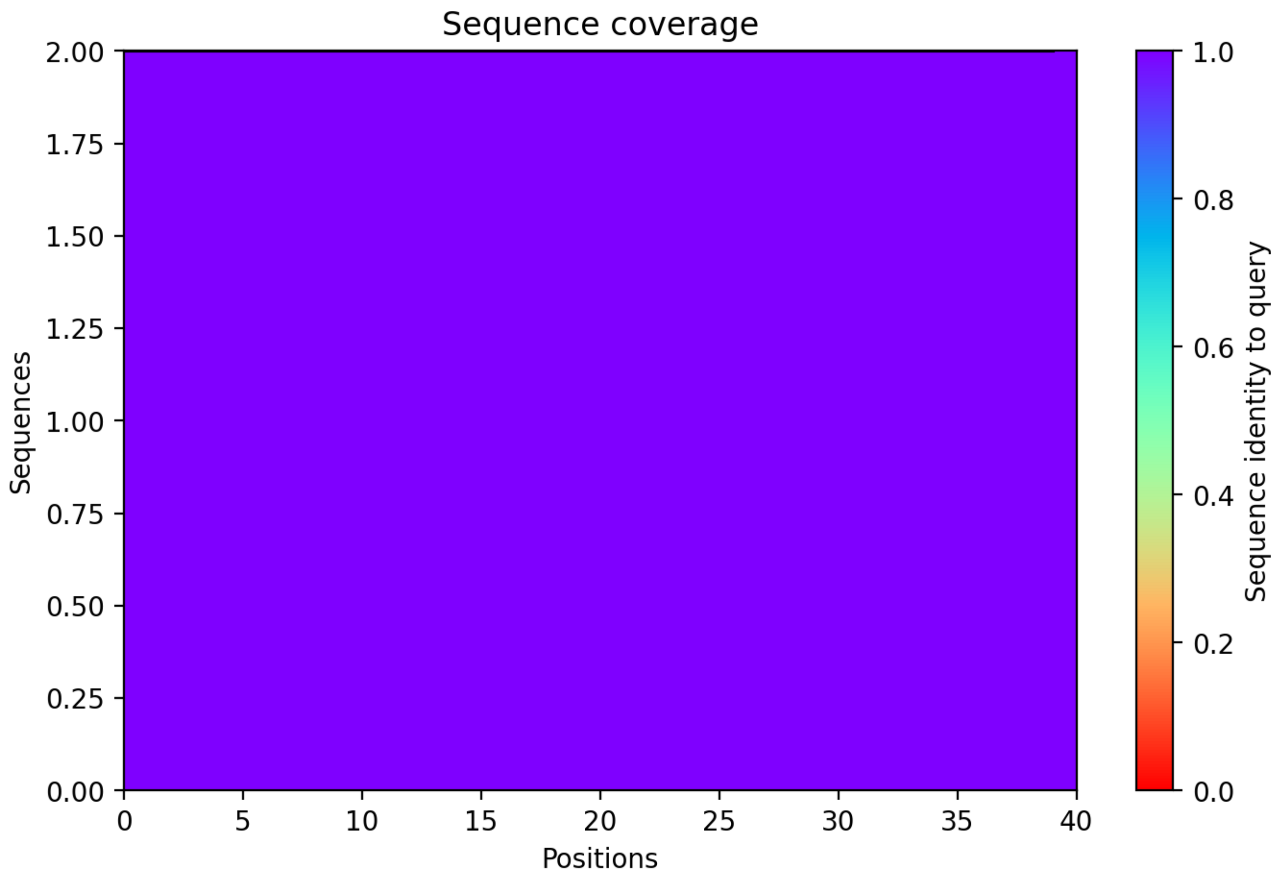

CP136: MtrunA17_Chr8g0373091

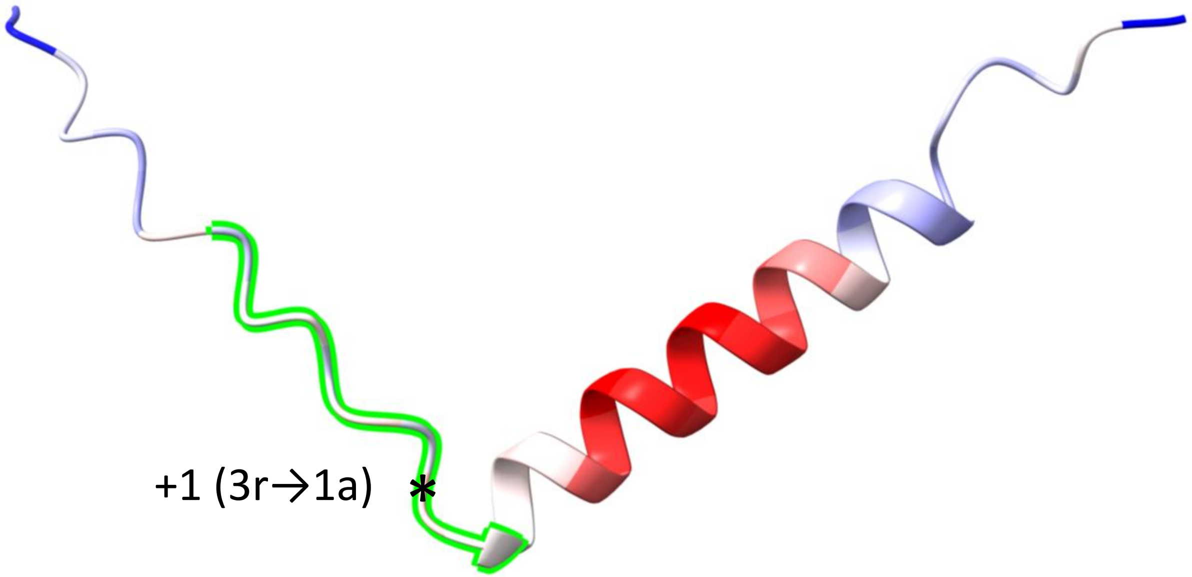


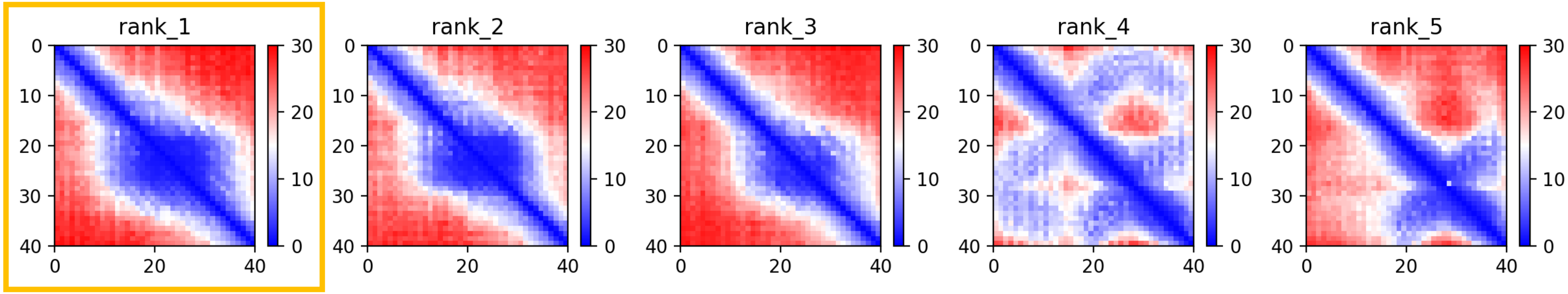

CP136: MtrunA17_Chr8g0373091

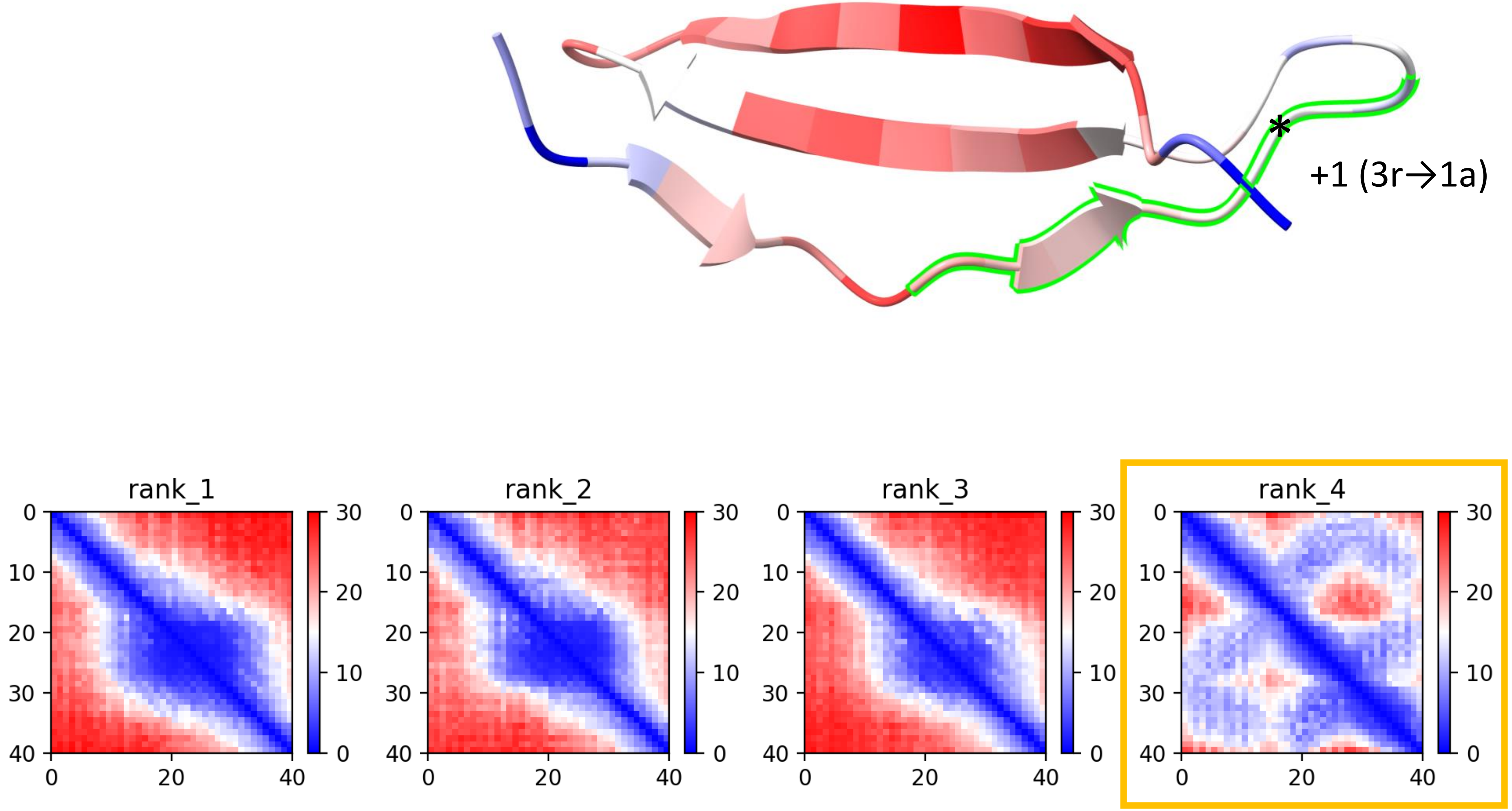

CP136: MtrunA17_Chr8g0373091

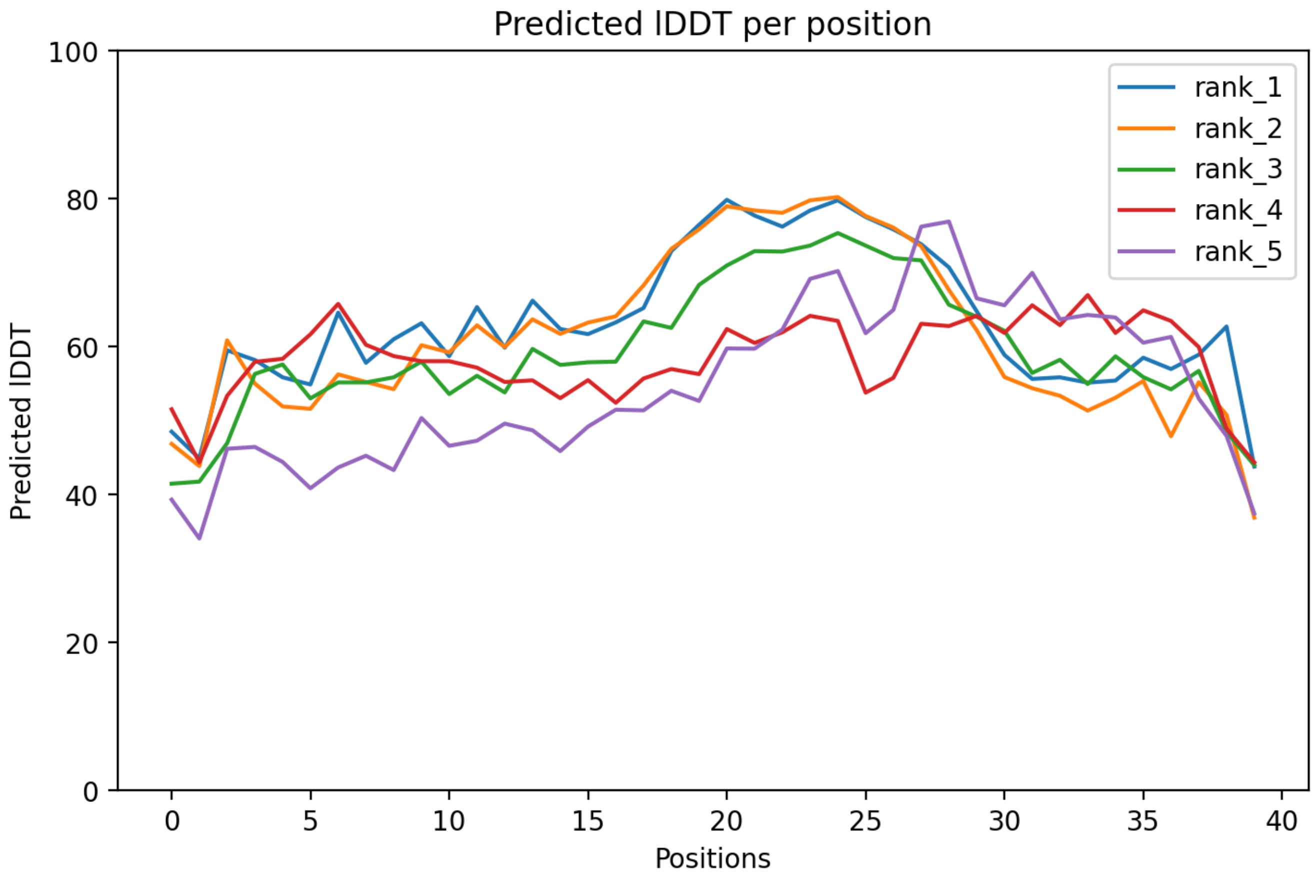

### CP136: MtrunA17_Chr8g0373091

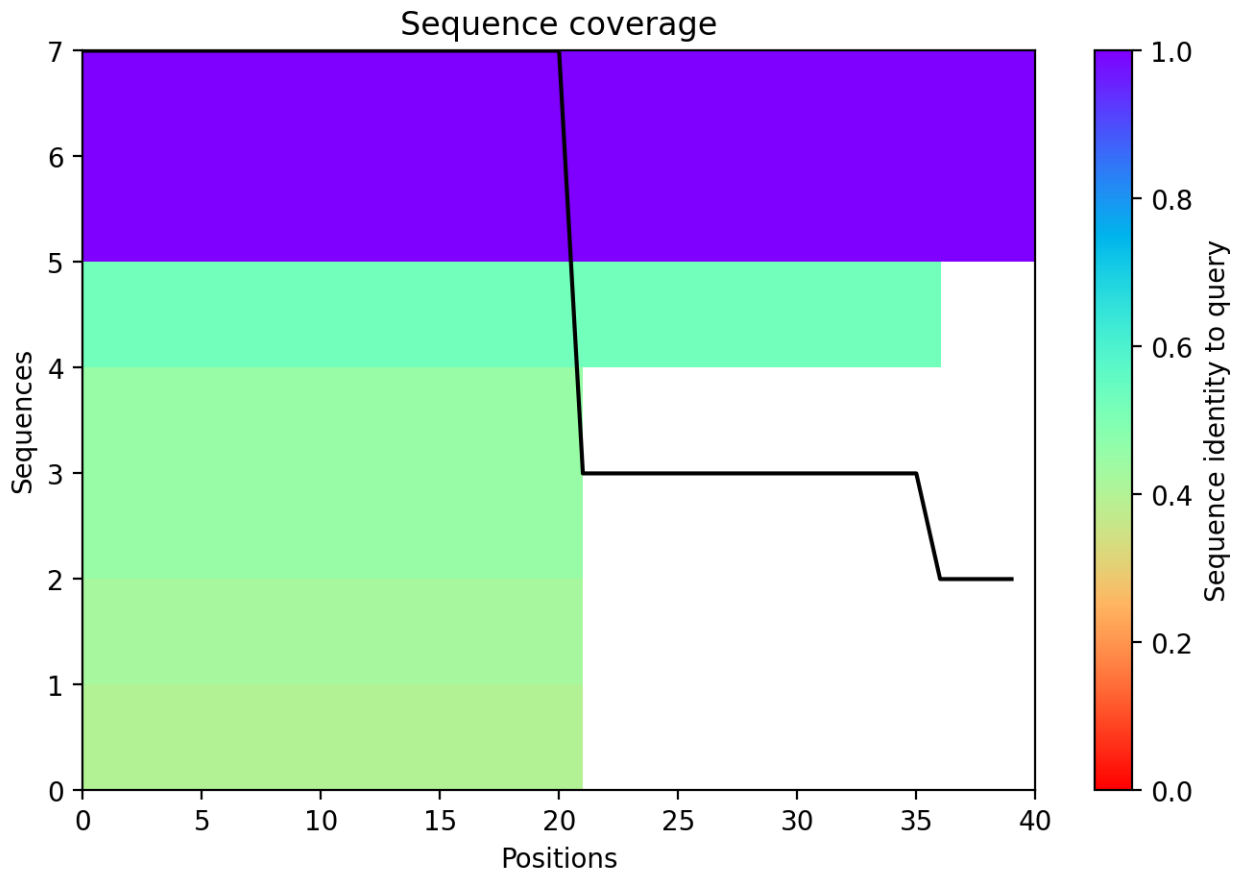

CP137: MtrunA17_Chr8g0376411

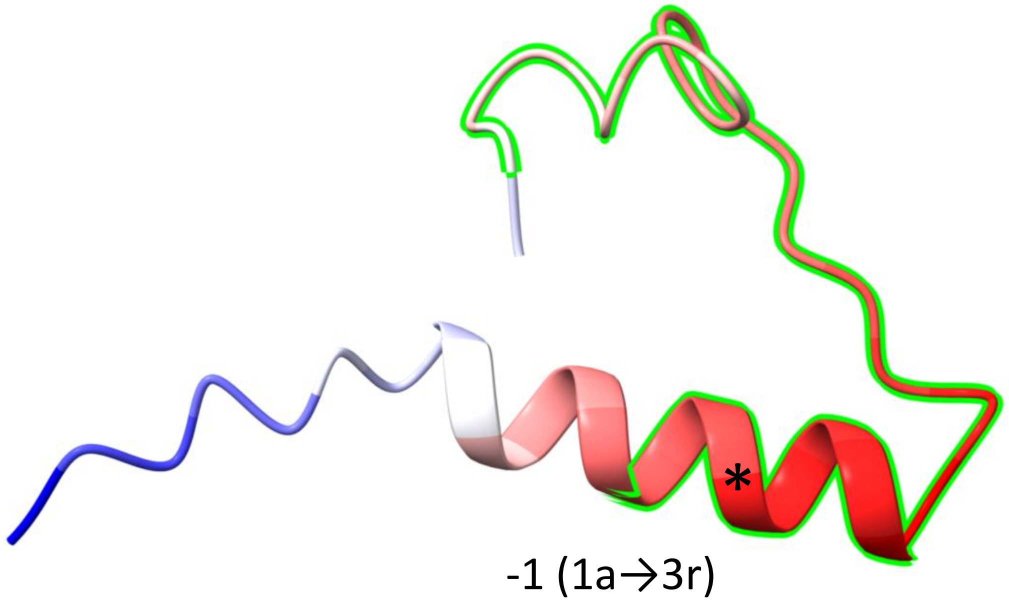


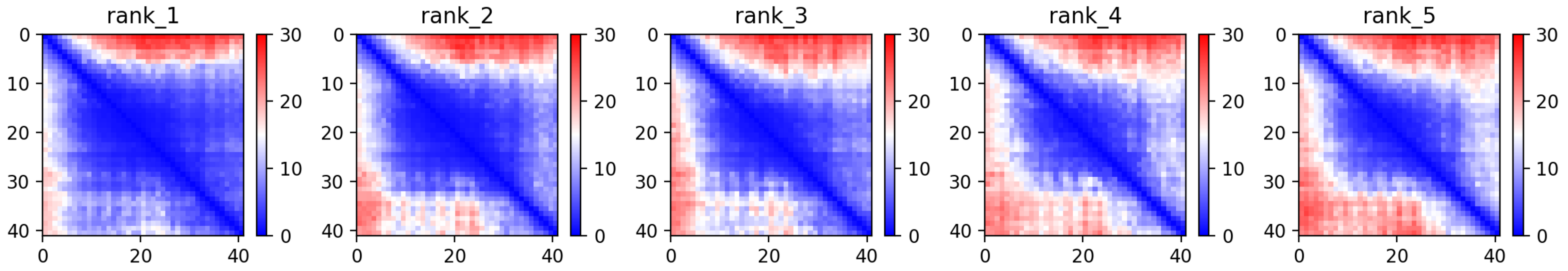

CP137: MtrunA17_Chr8g0376411

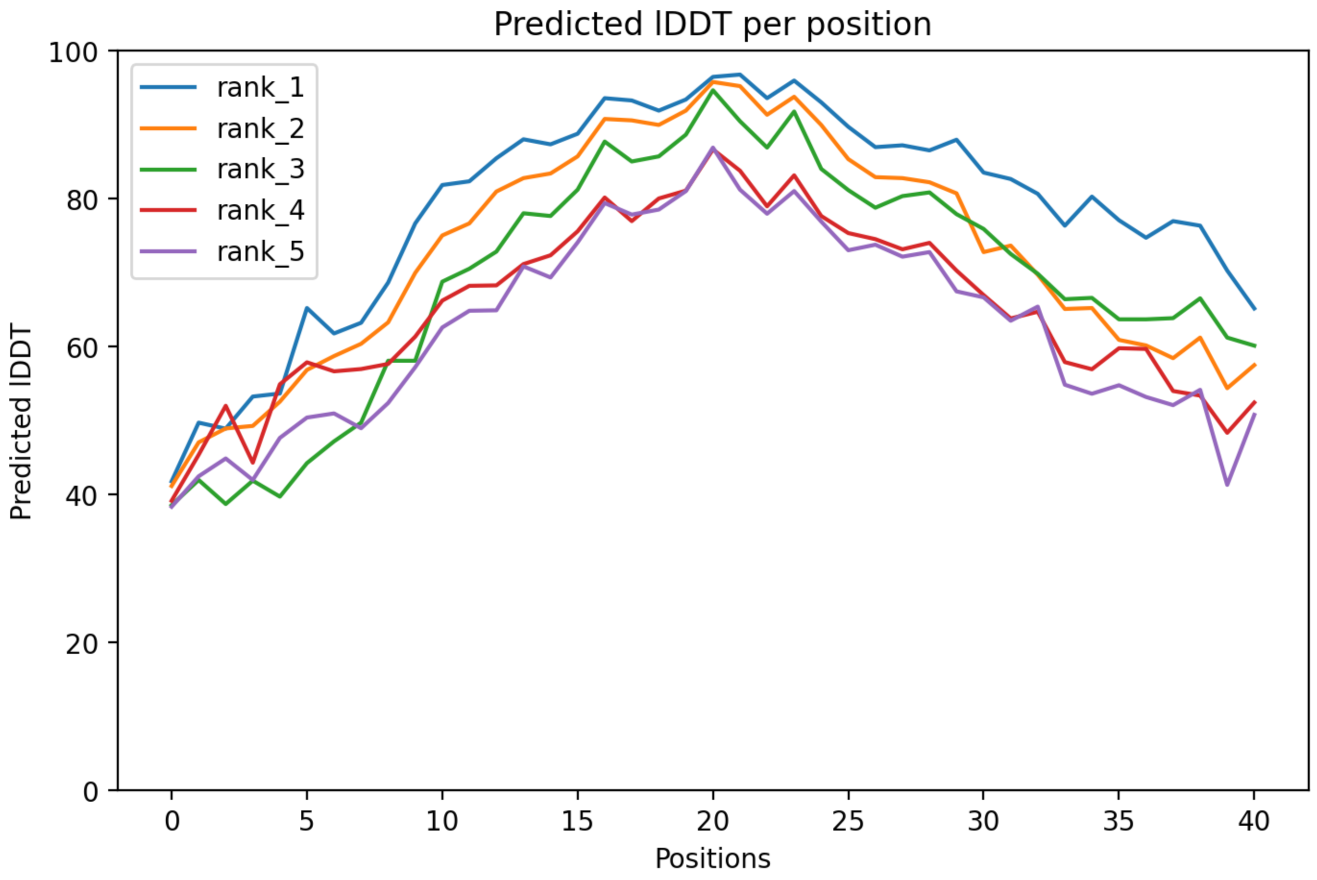

CP137: MtrunA17_Chr8g0376411

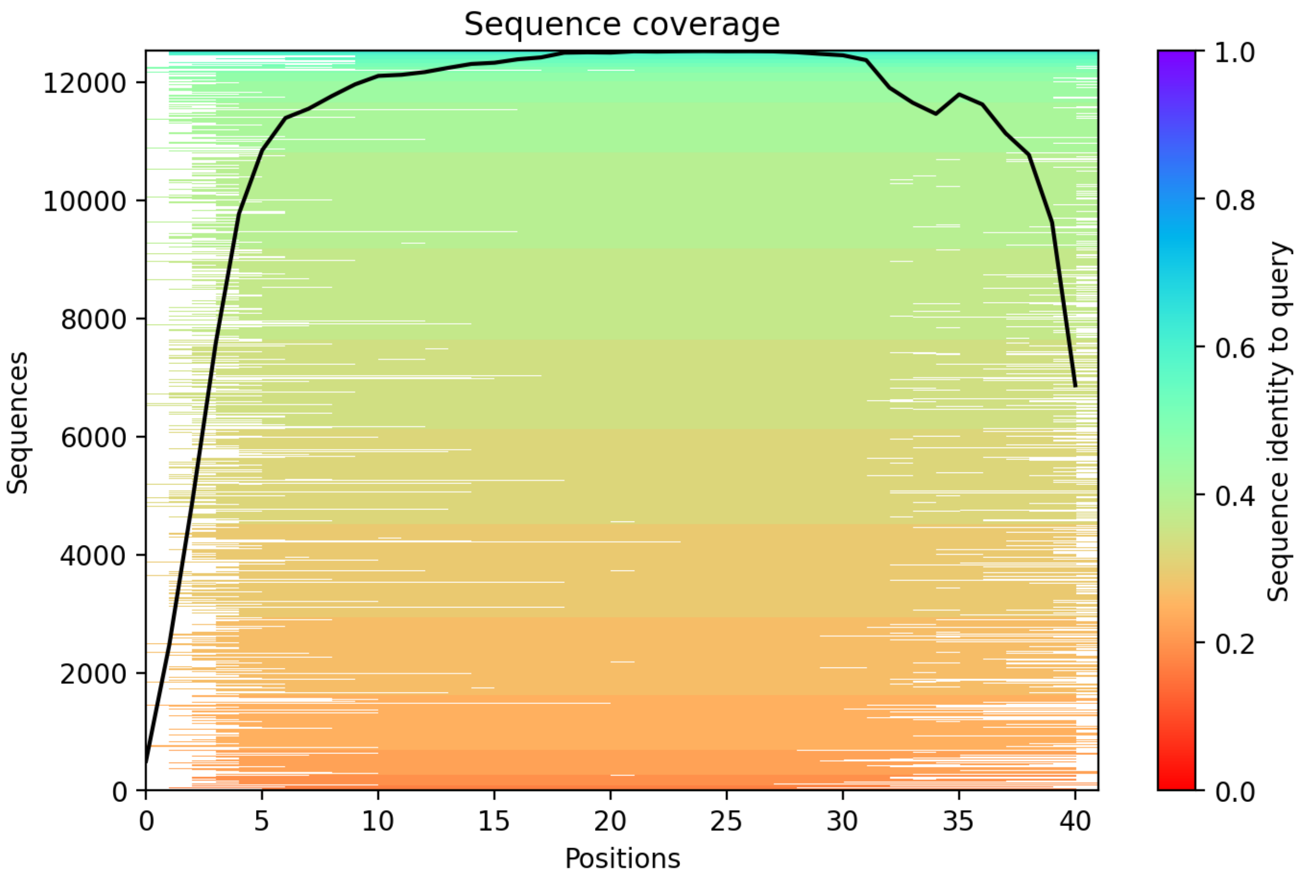

CP138: MtrunA17_Chr8g0377071

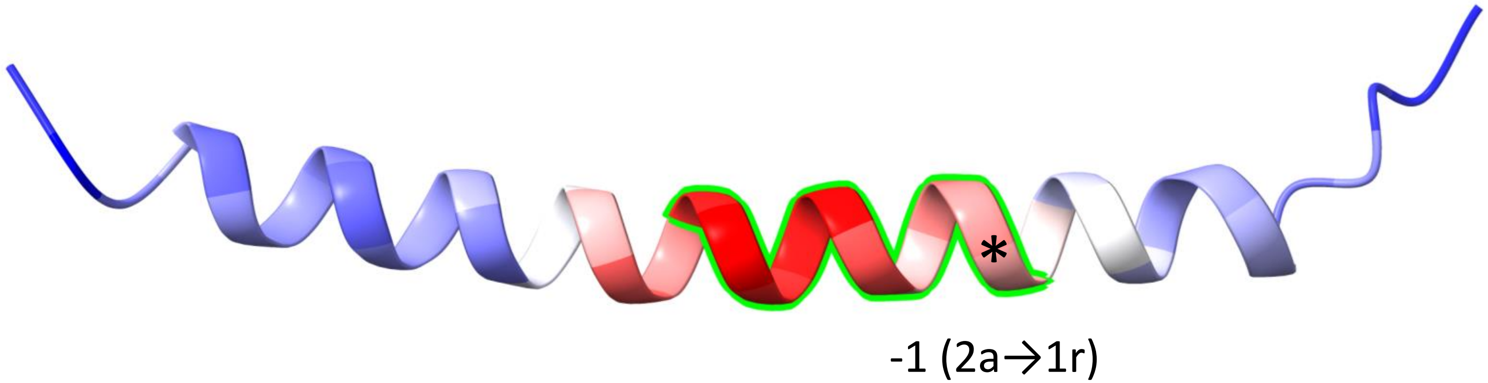


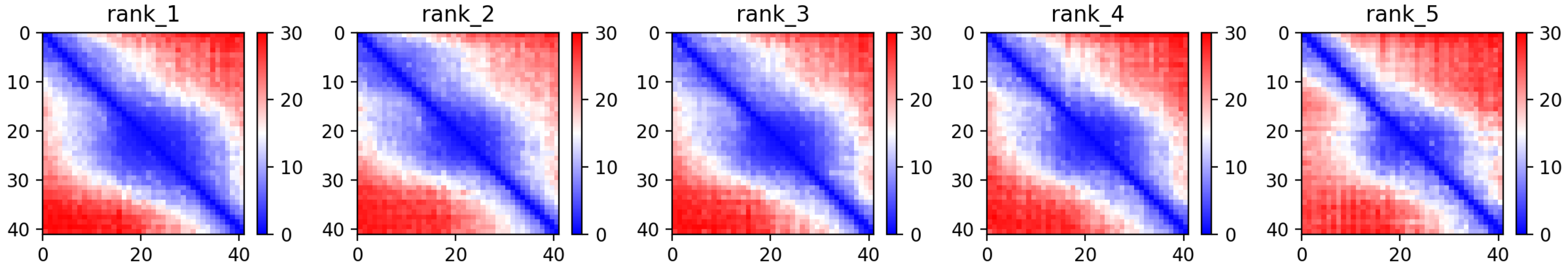

CP138: MtrunA17_Chr8g0377071

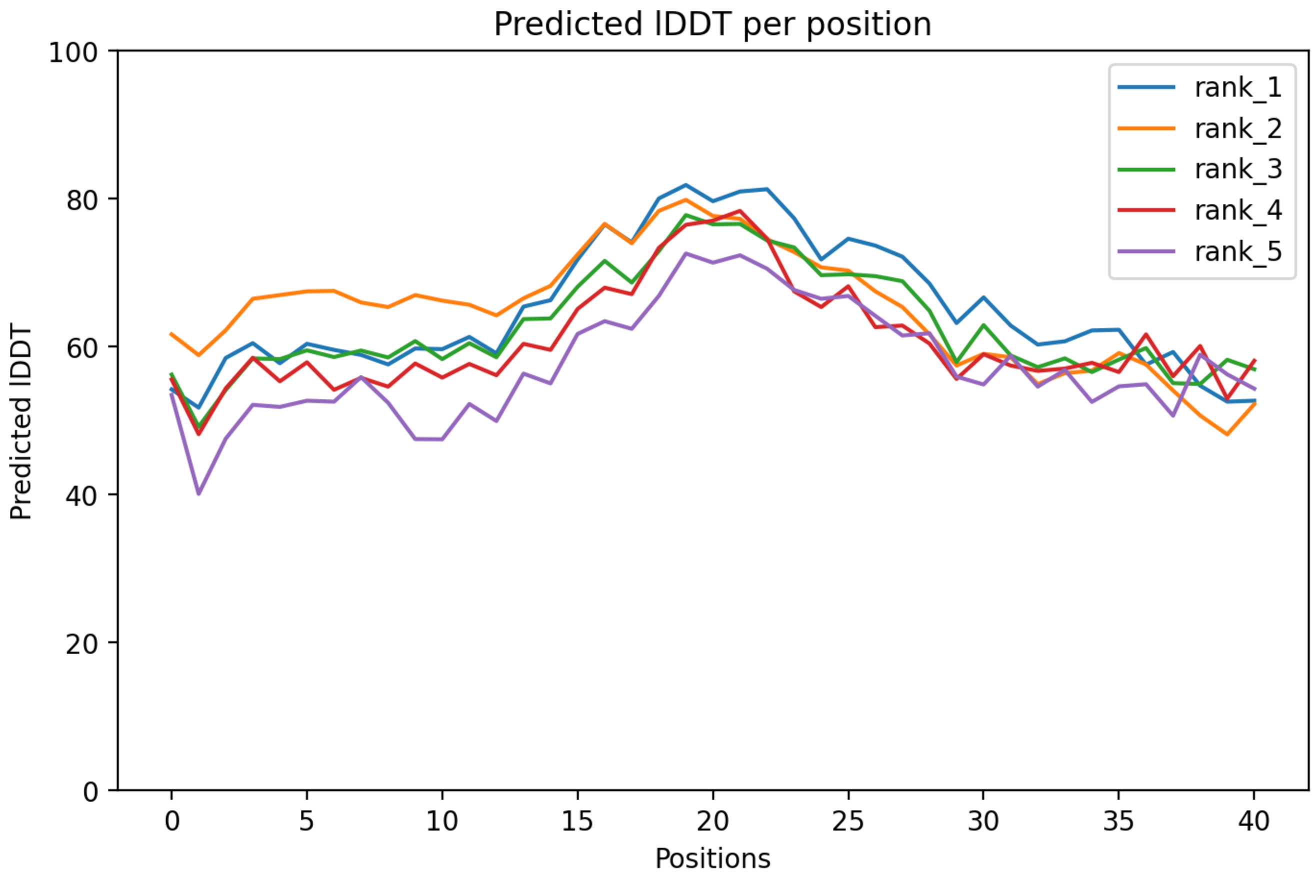

CP138: MtrunA17_Chr8g0377071

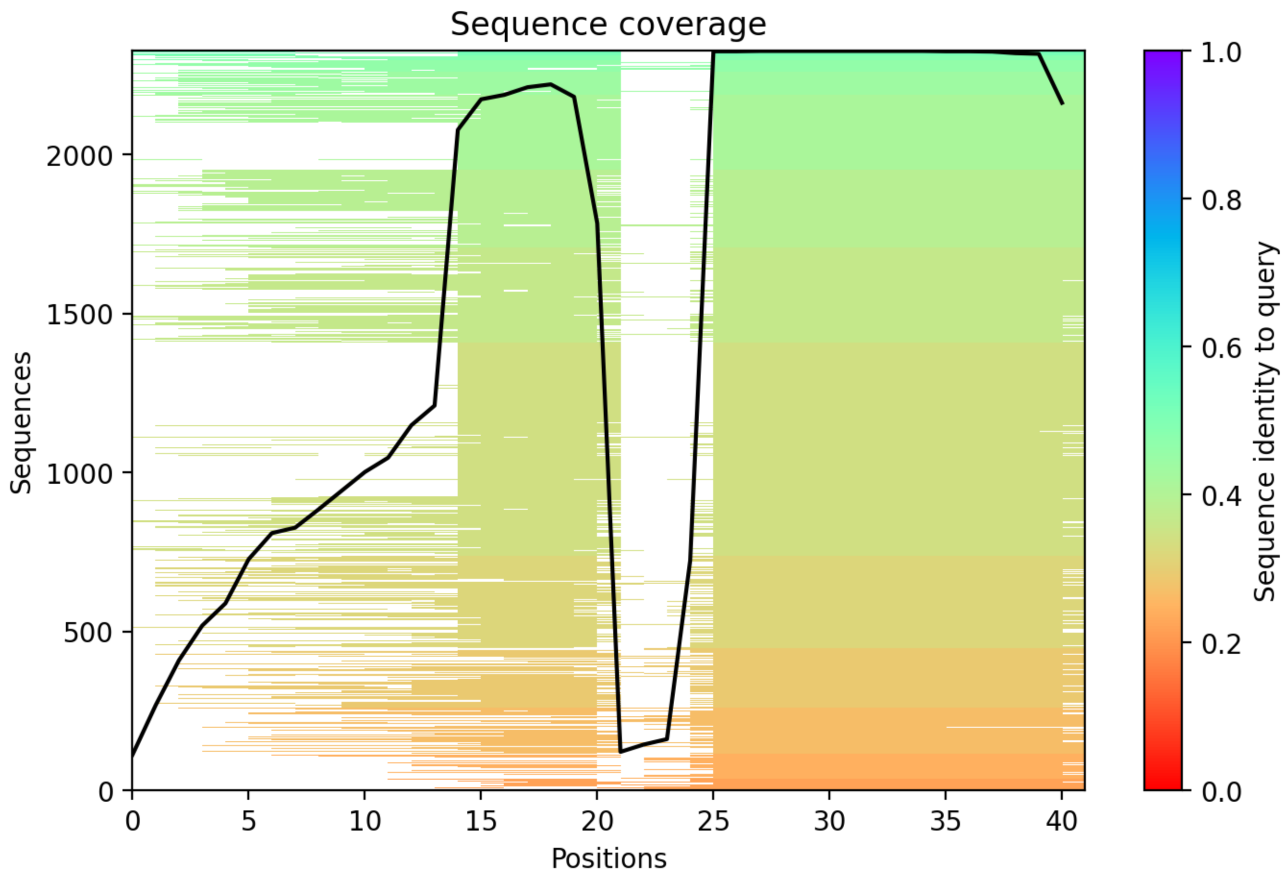

### CP139: MtrunA17_Chr8g0385331

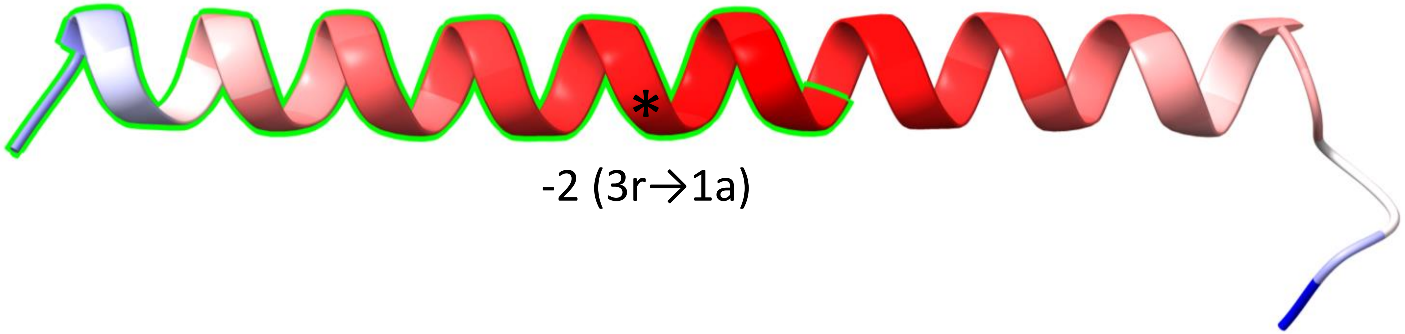


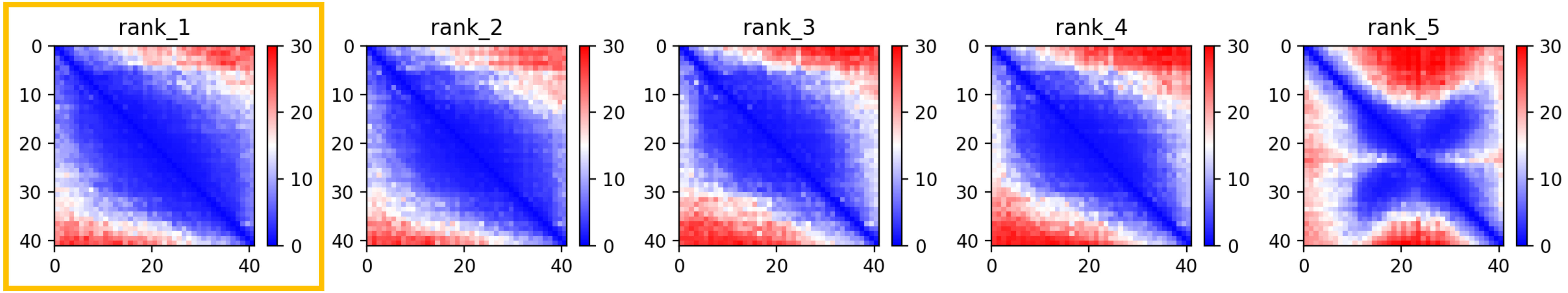

CP139: MtrunA17_Chr8g0385331

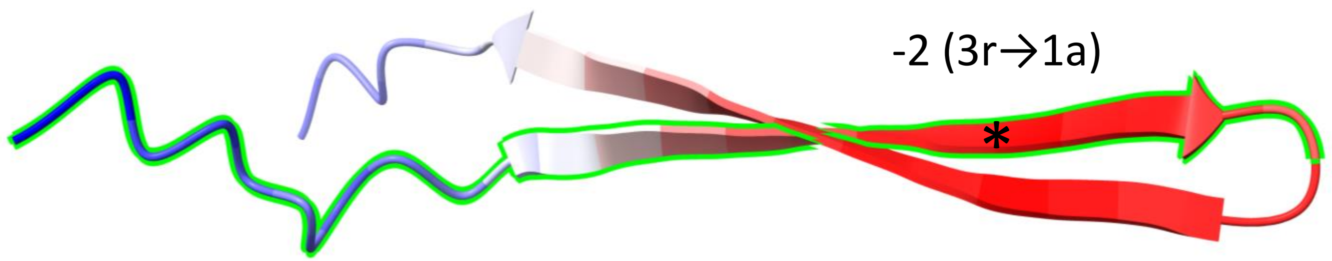


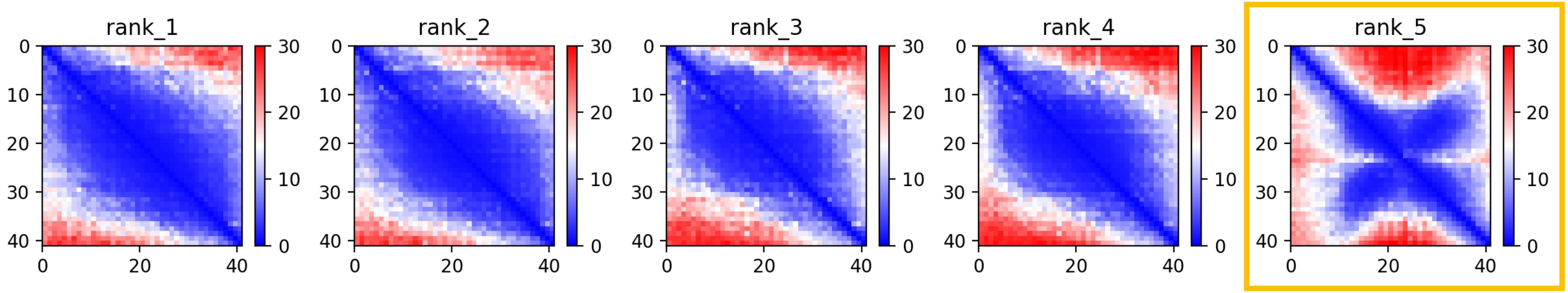

CP139: MtrunA17_Chr8g0385331

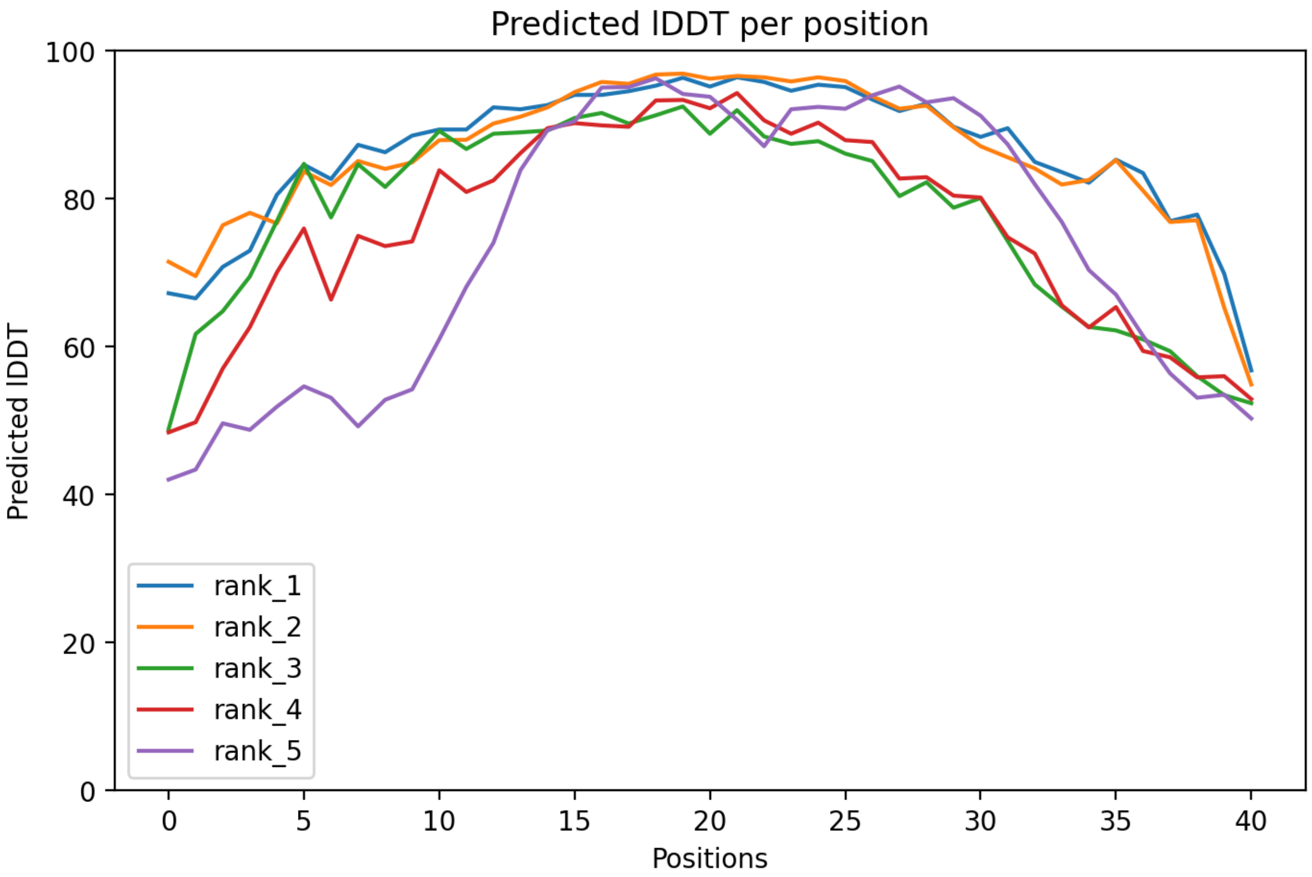

CP139: MtrunA17_Chr8g0385331

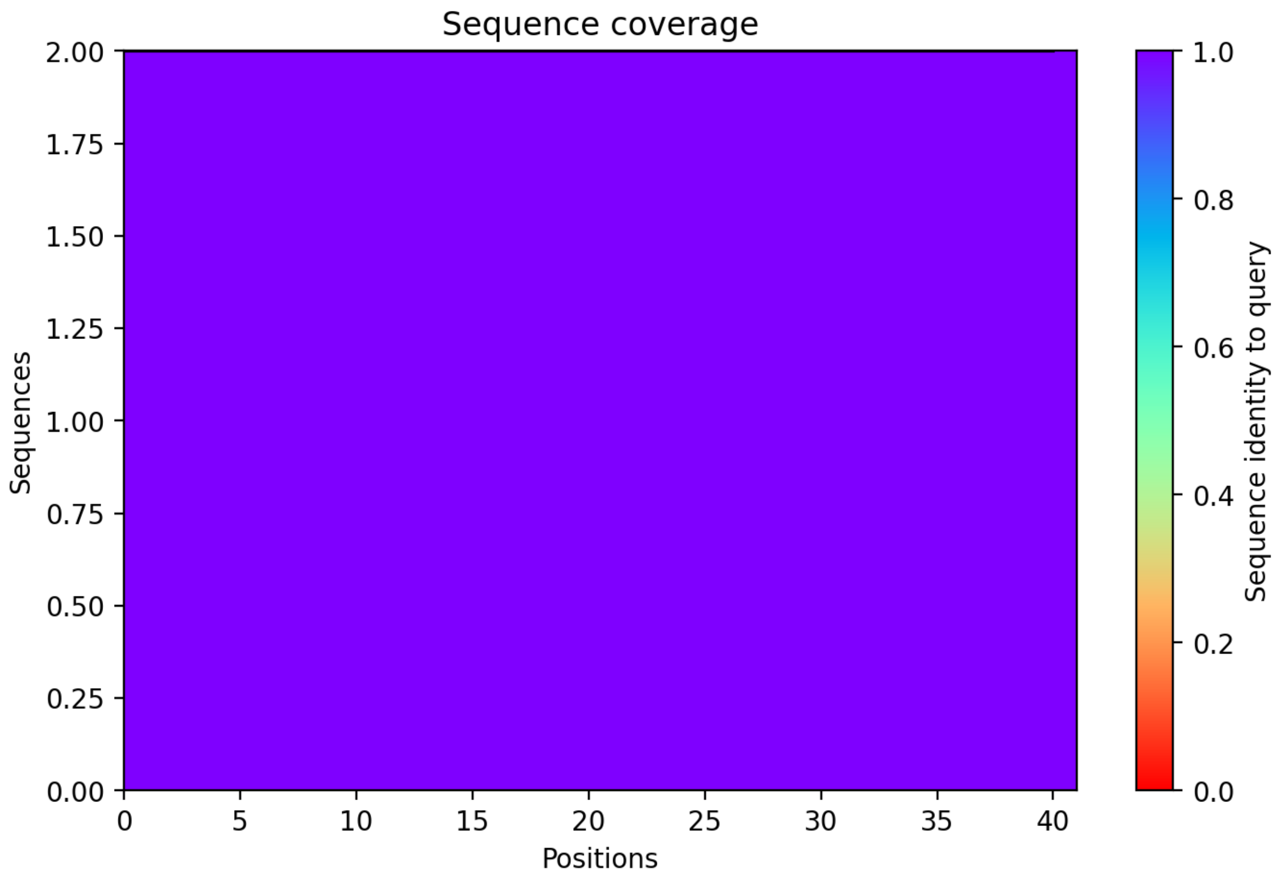

### CP140: MtrunA17_Chr8g0392351

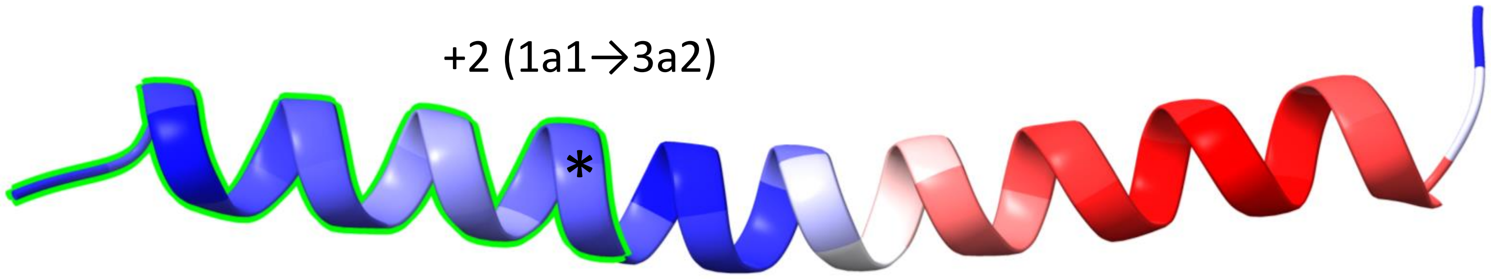


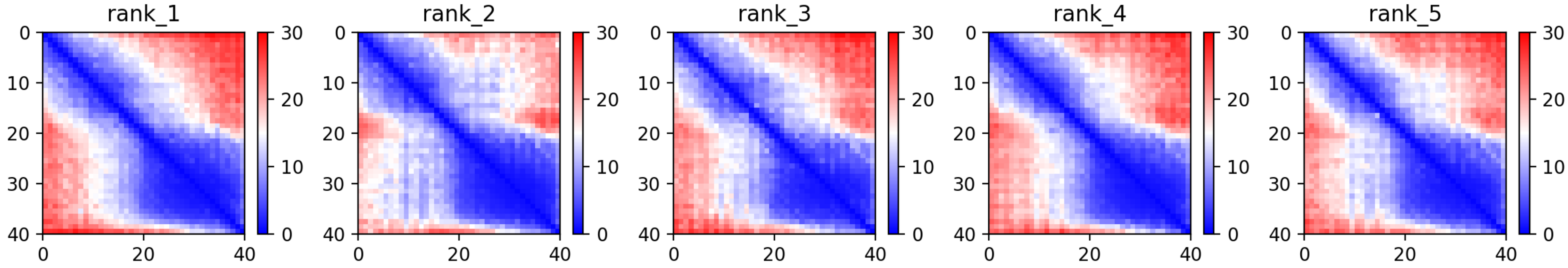

CP140: MtrunA17_Chr8g0392351

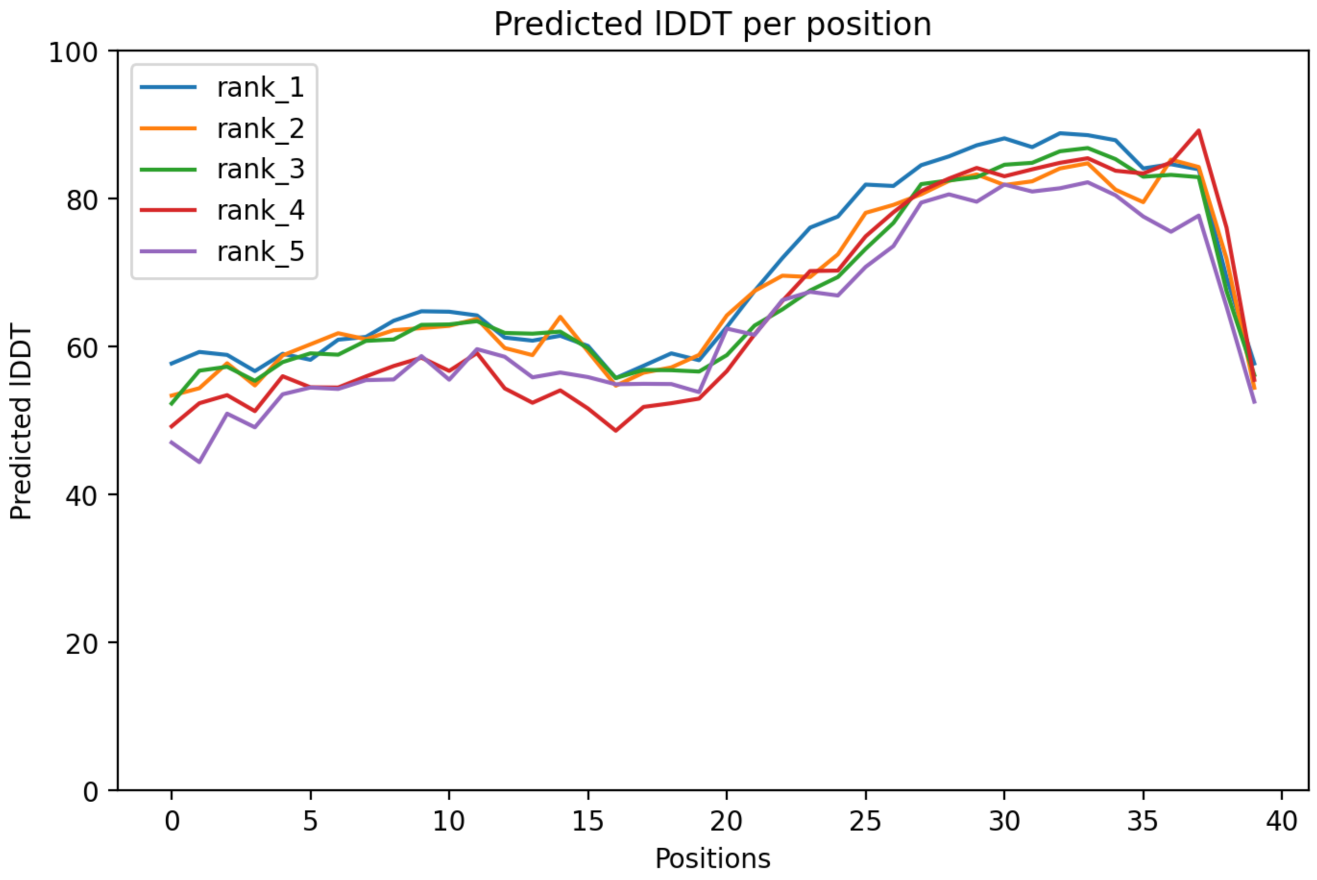

CP140: MtrunA17_Chr8g0392351

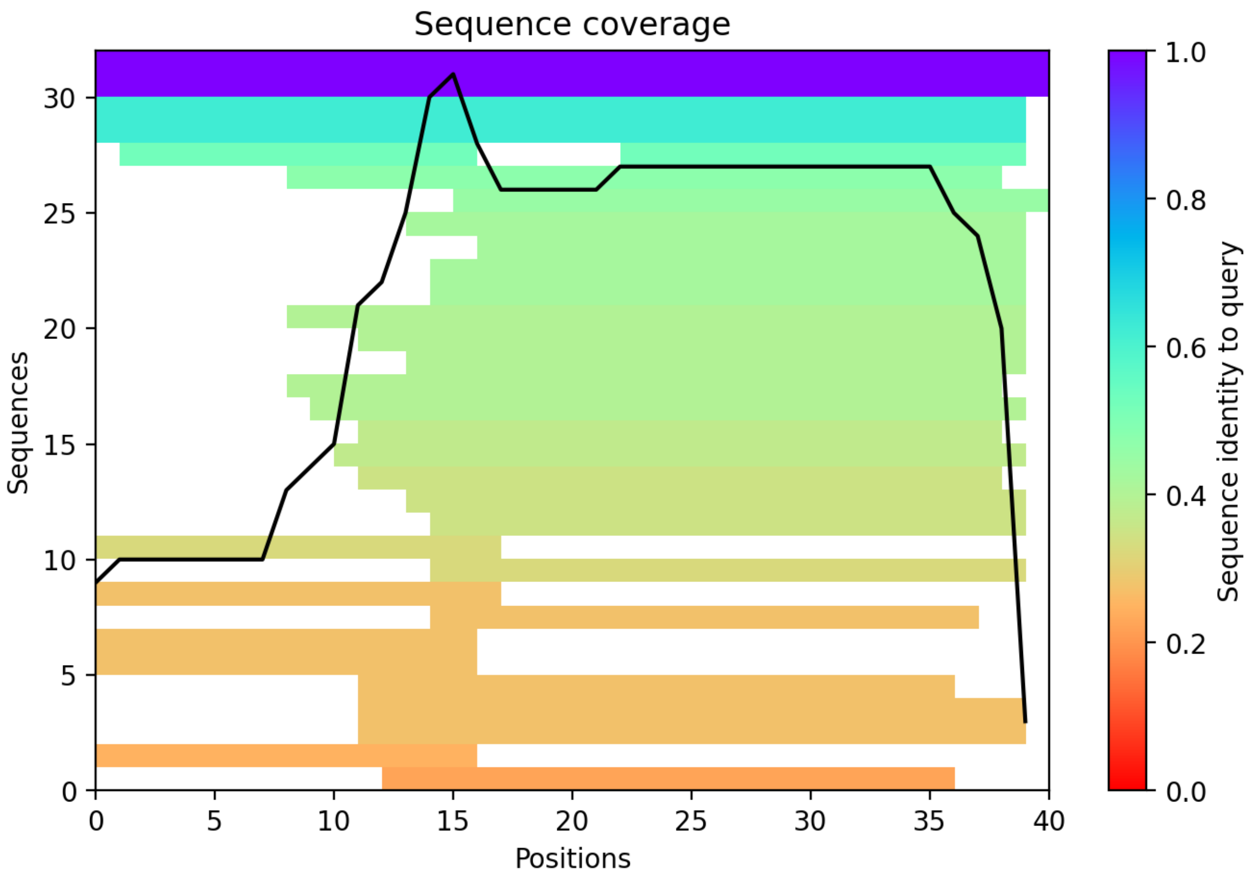

CP141: MtrunA17_CPg0492331

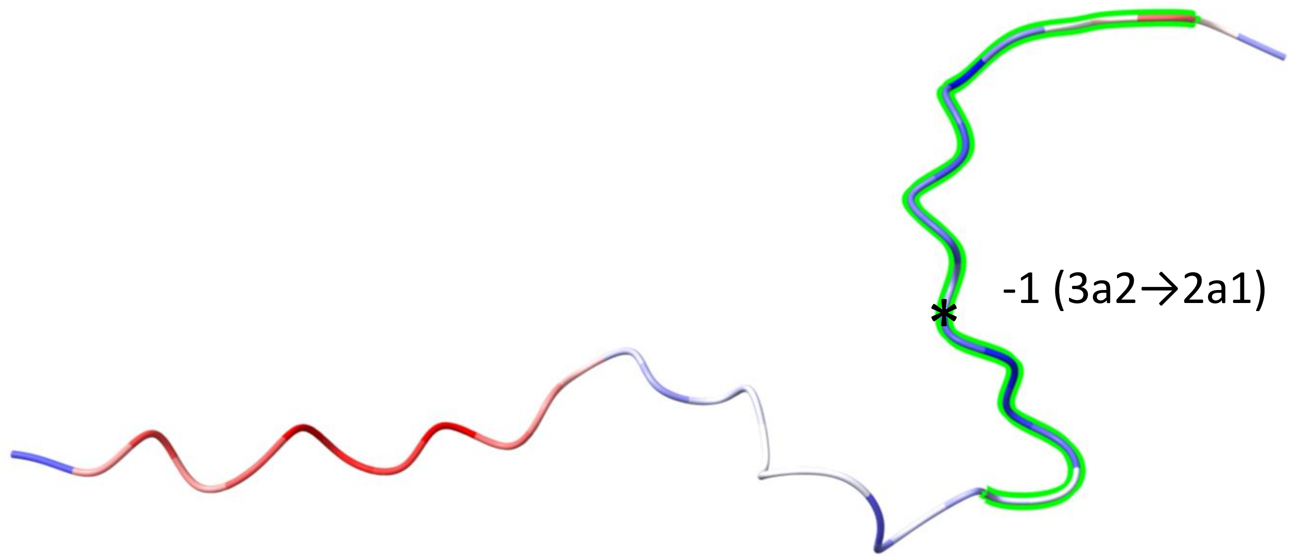


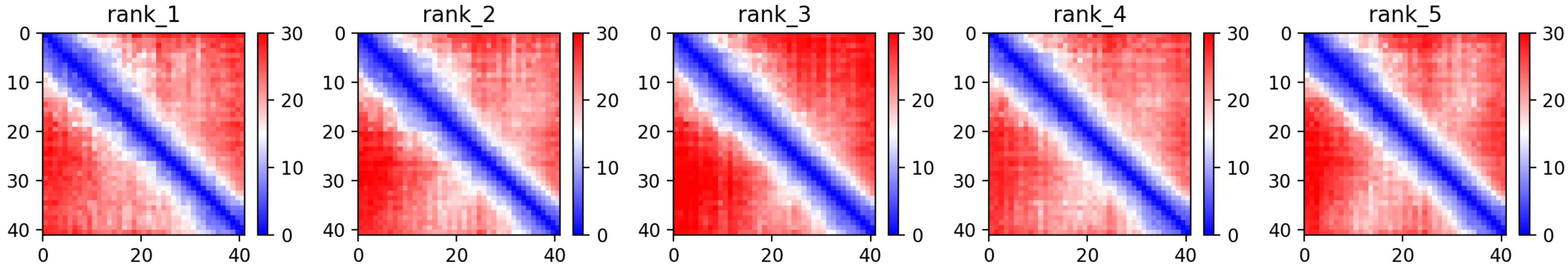

CP141: MtrunA17_CPg0492331

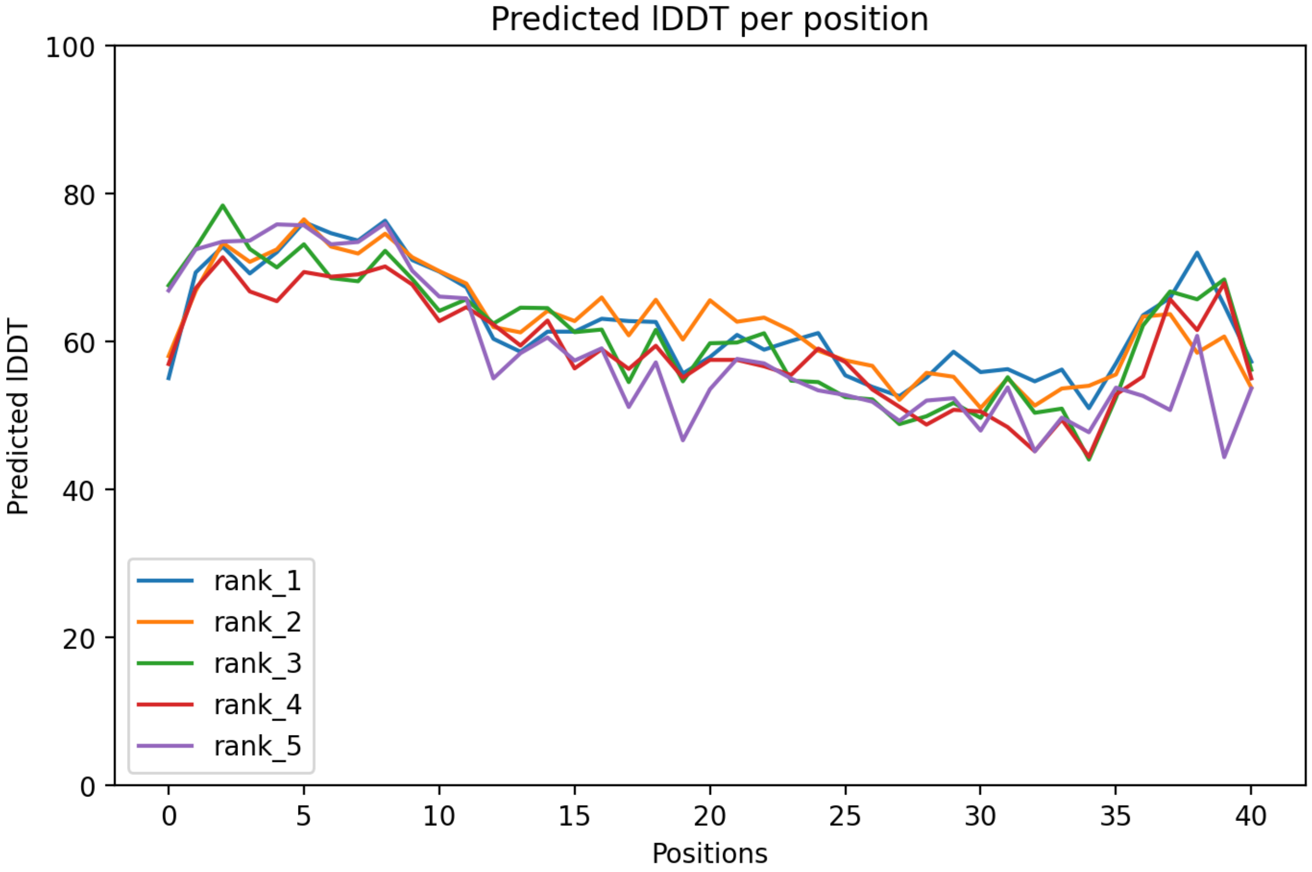

### CP141: MtrunA17_CPg0492331

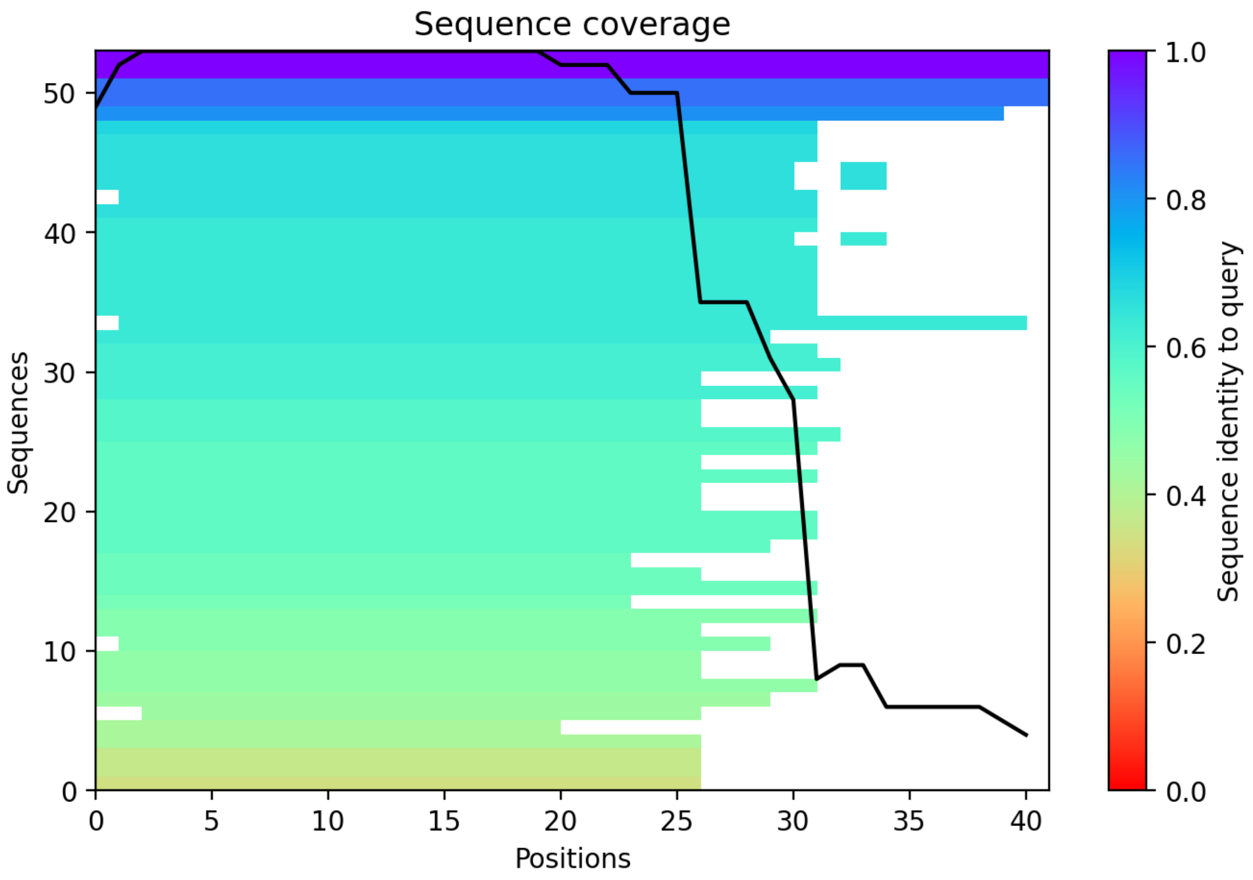

CP142: MtrunA17_CPg0492381

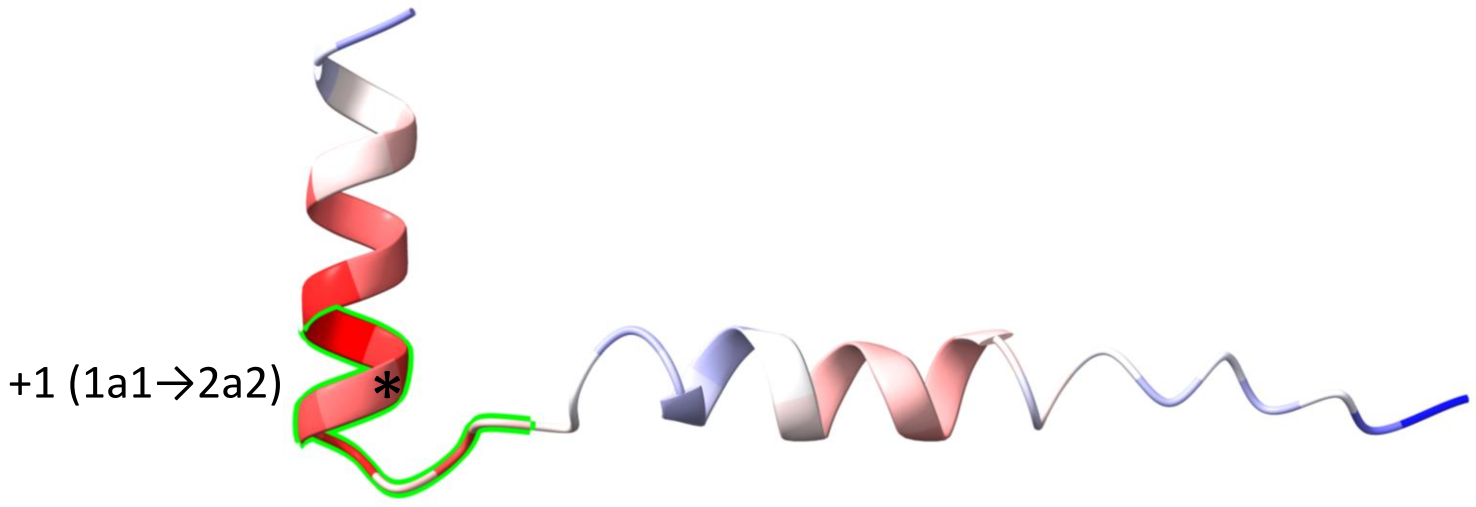


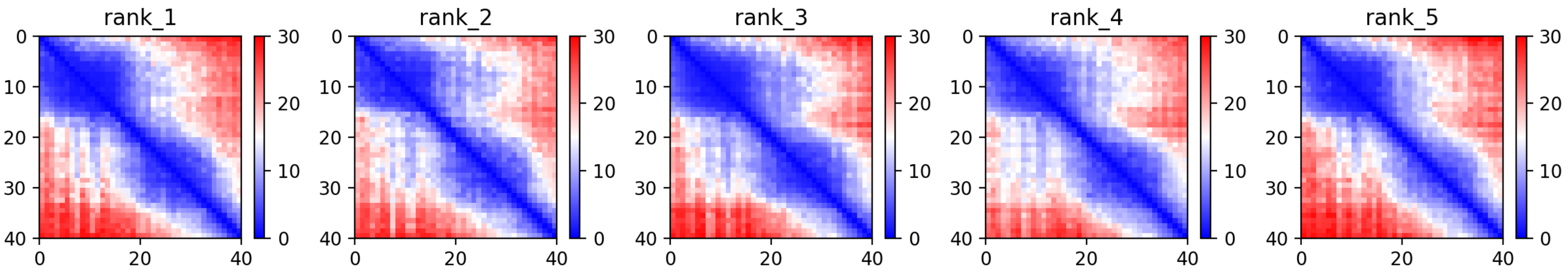

CP142: MtrunA17_CPg0492381

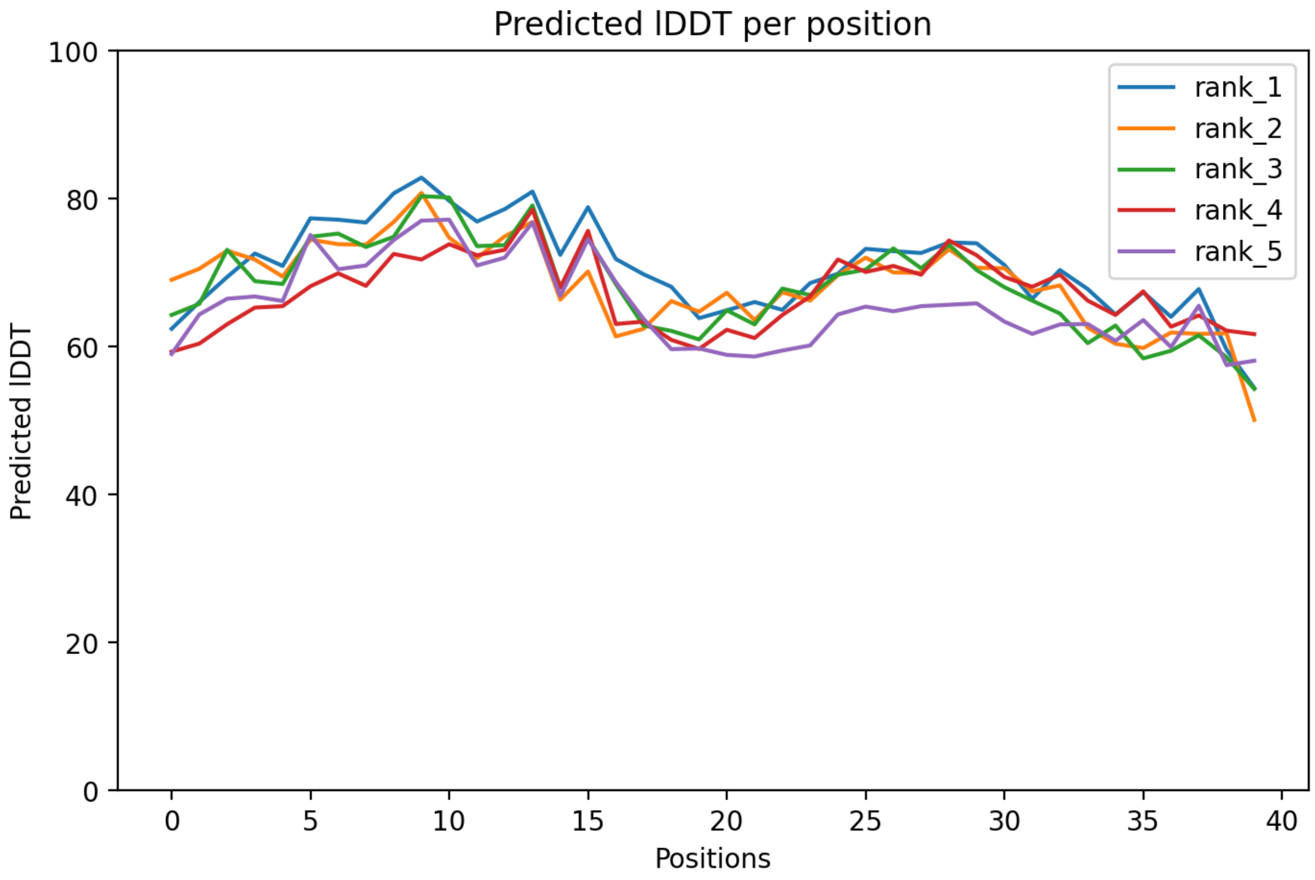

### CP142: MtrunA17_CPg0492381

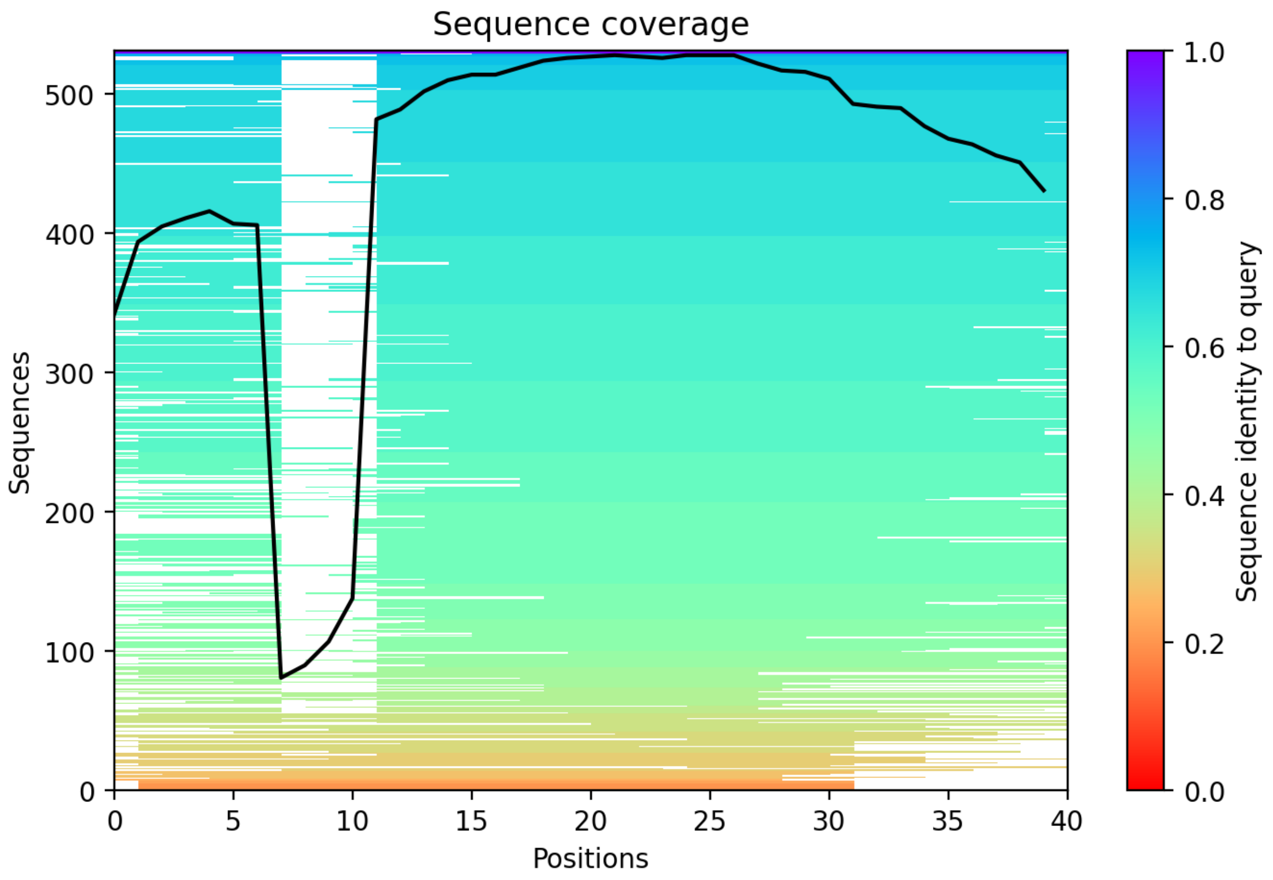

### CP143: MtrunA17_CPg0492461

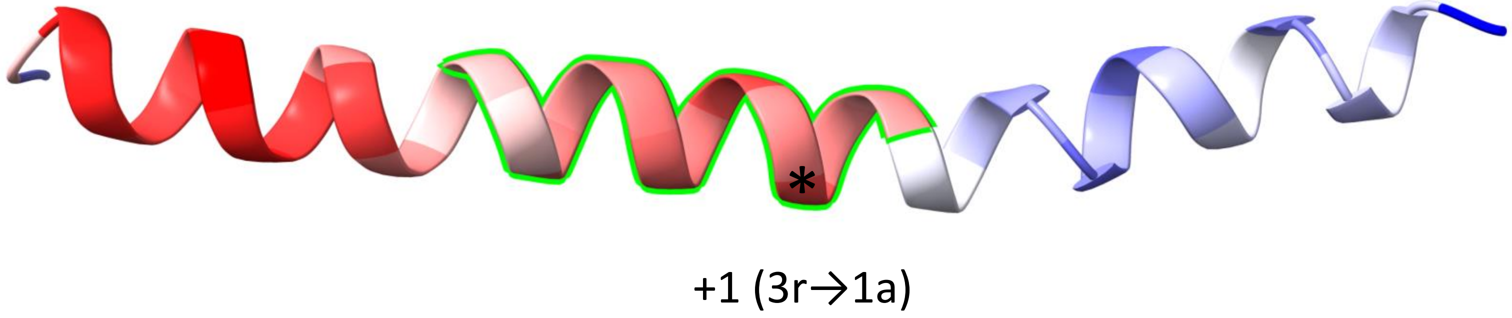


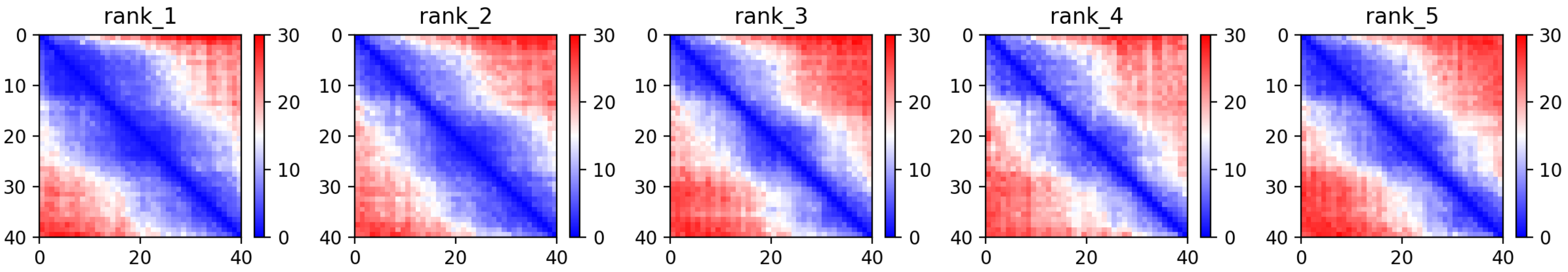

CP143: MtrunA17_CPg0492461

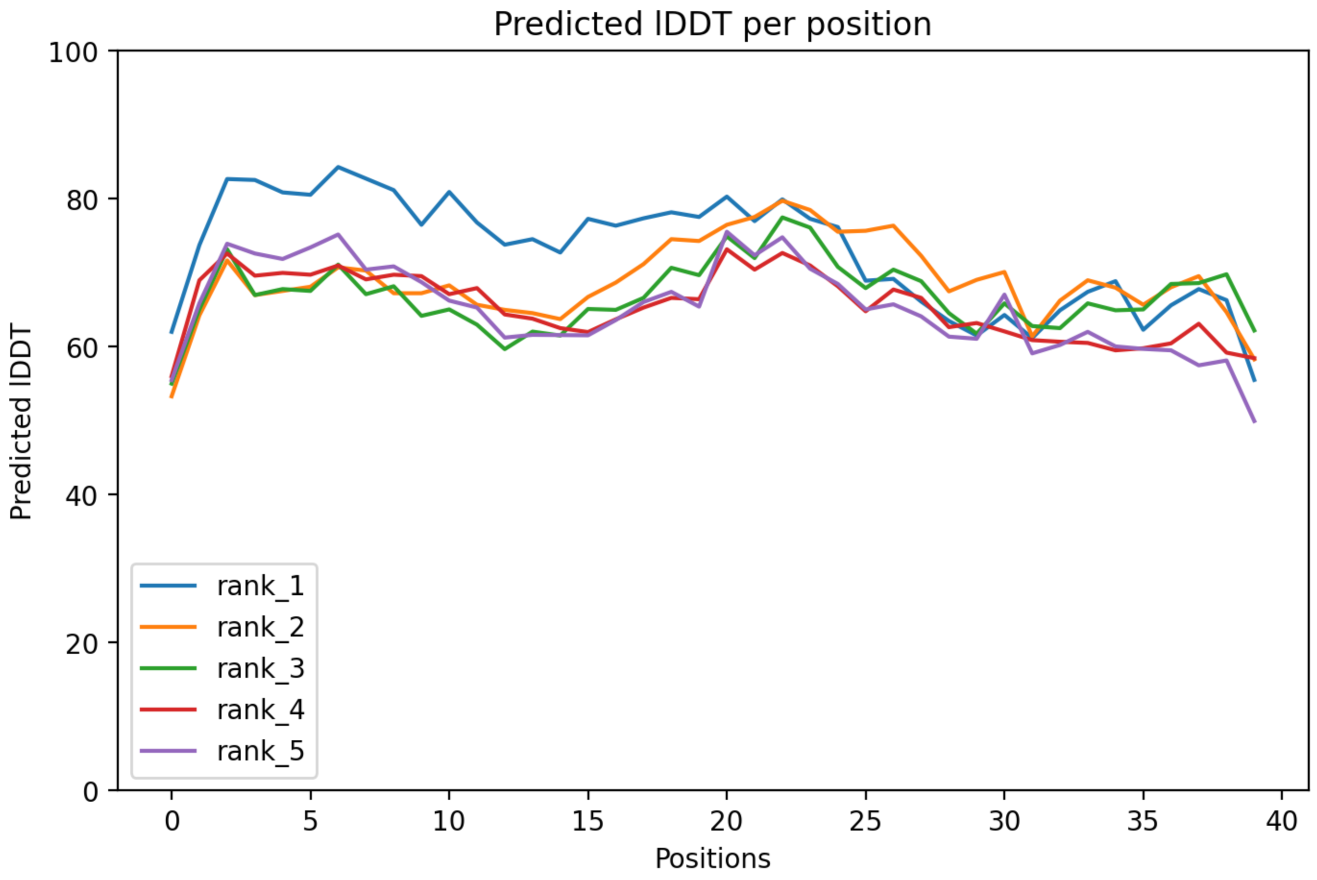

### CP143: MtrunA17_CPg0492461

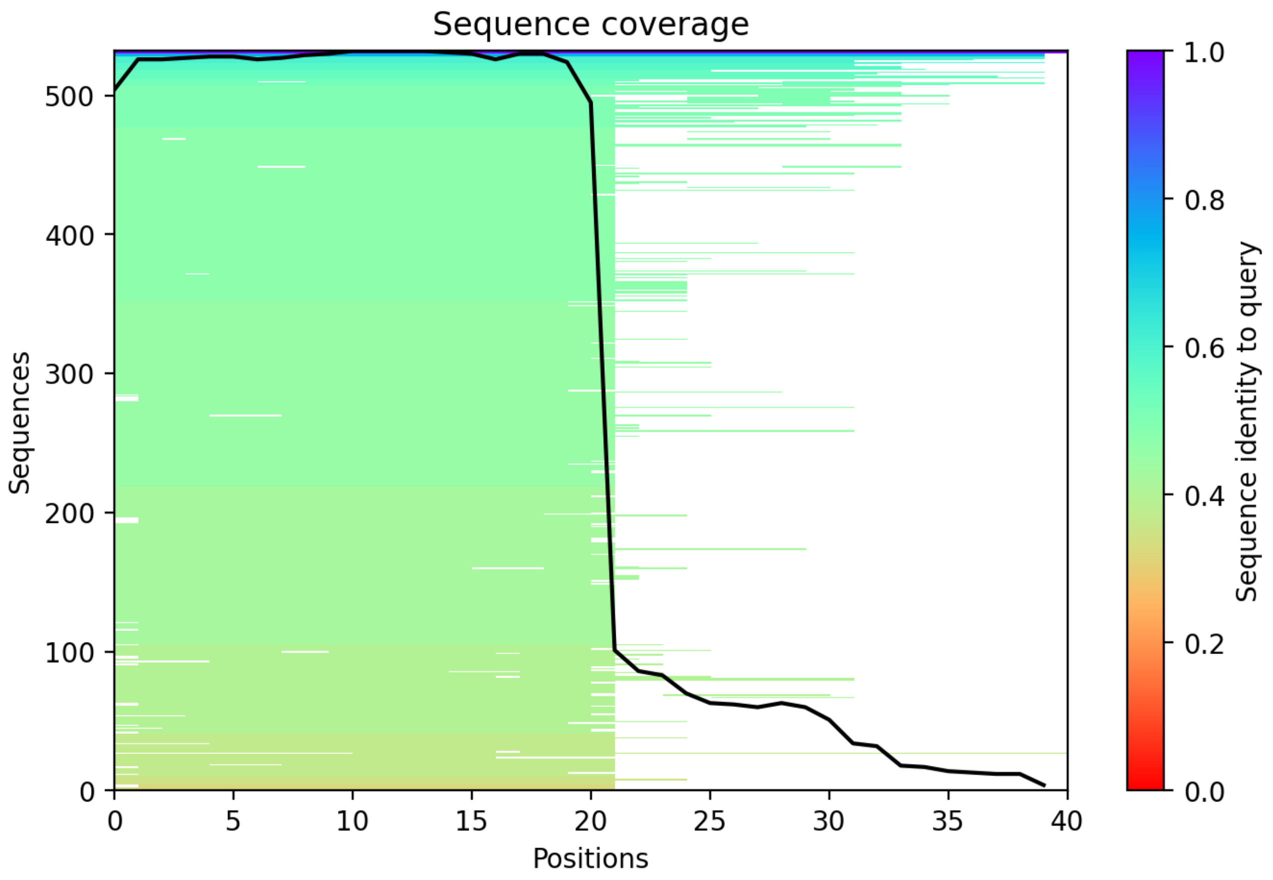

### CP144: MtrunA17_CPg0492851

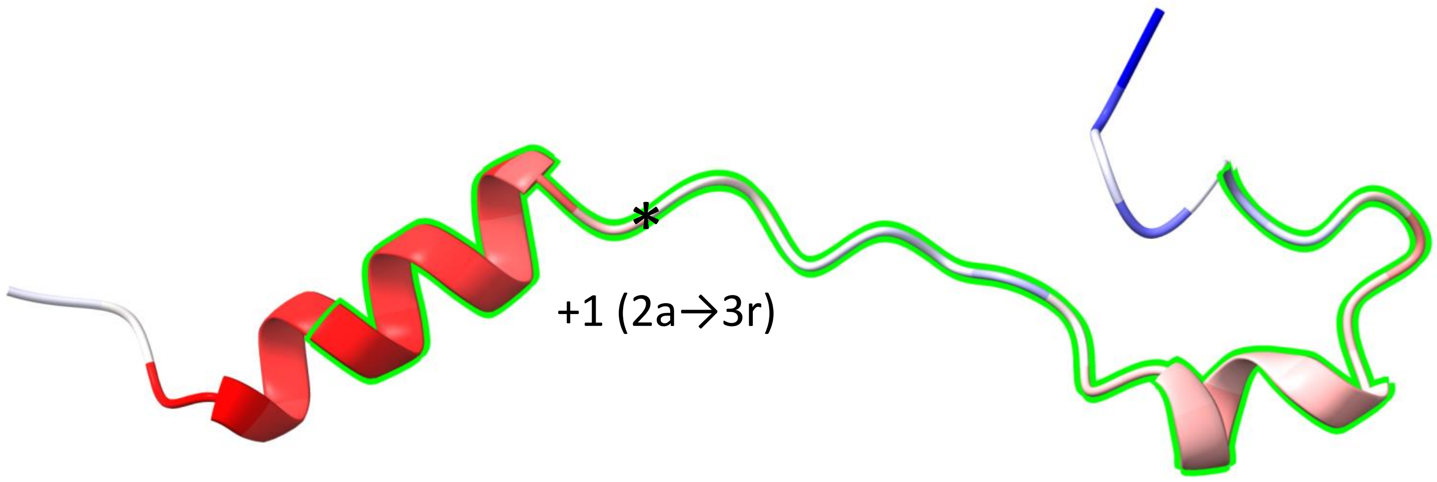


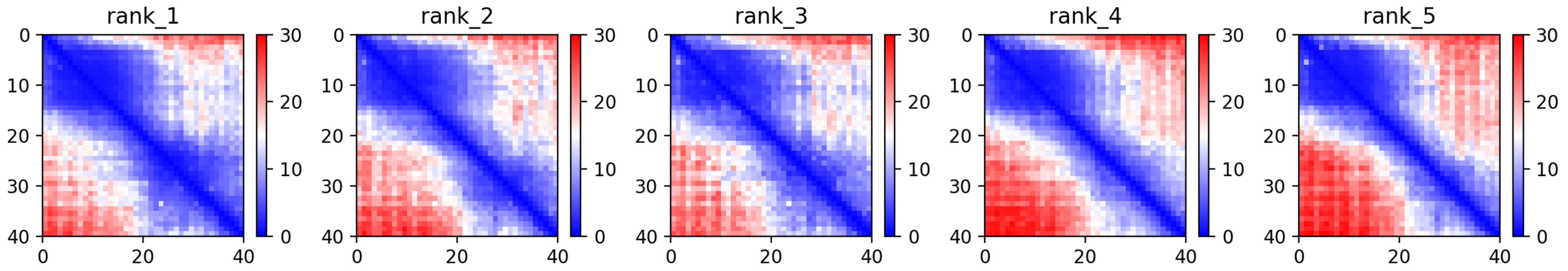

CP144: MtrunA17_CPg0492851

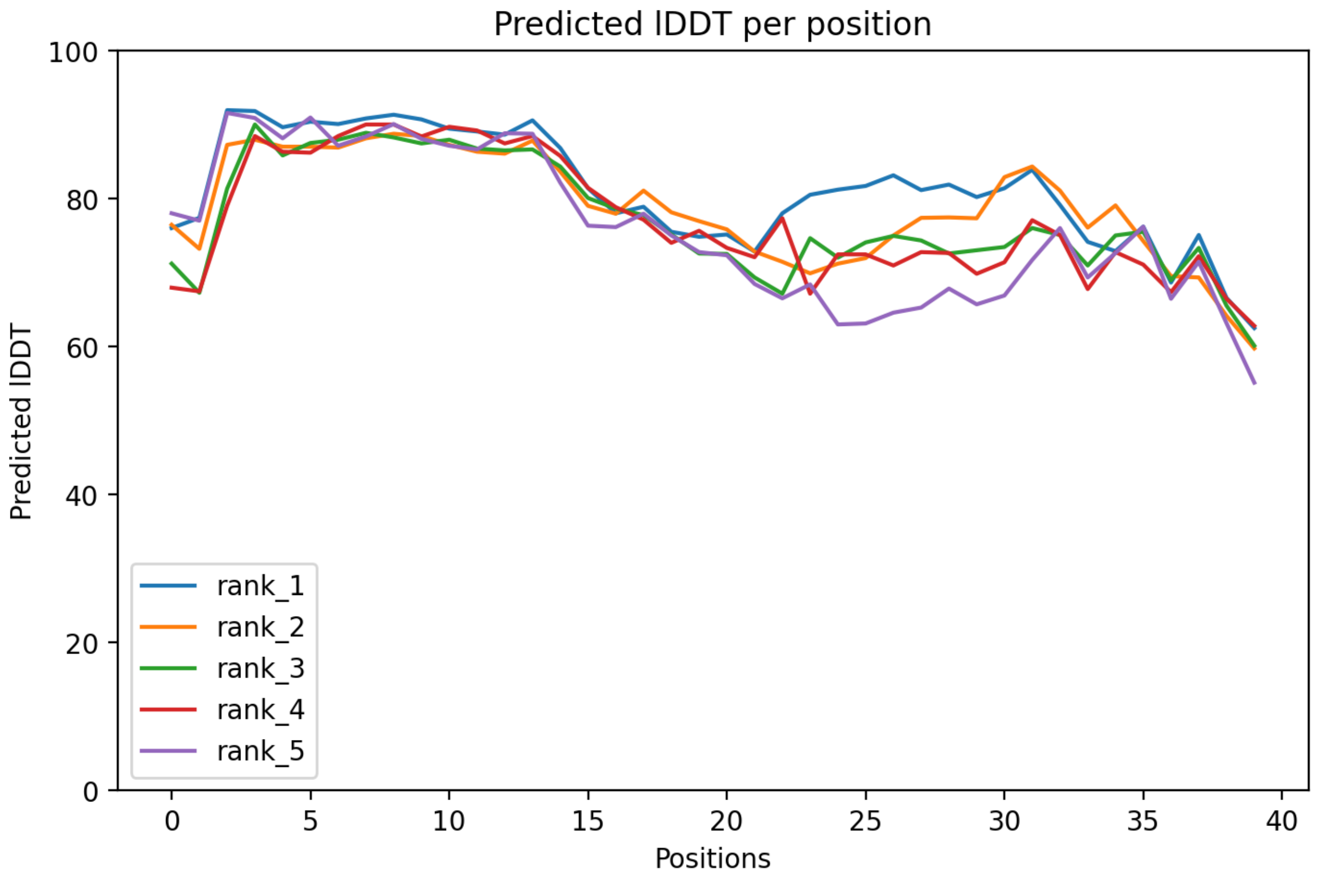

CP144: MtrunA17_CPg0492851

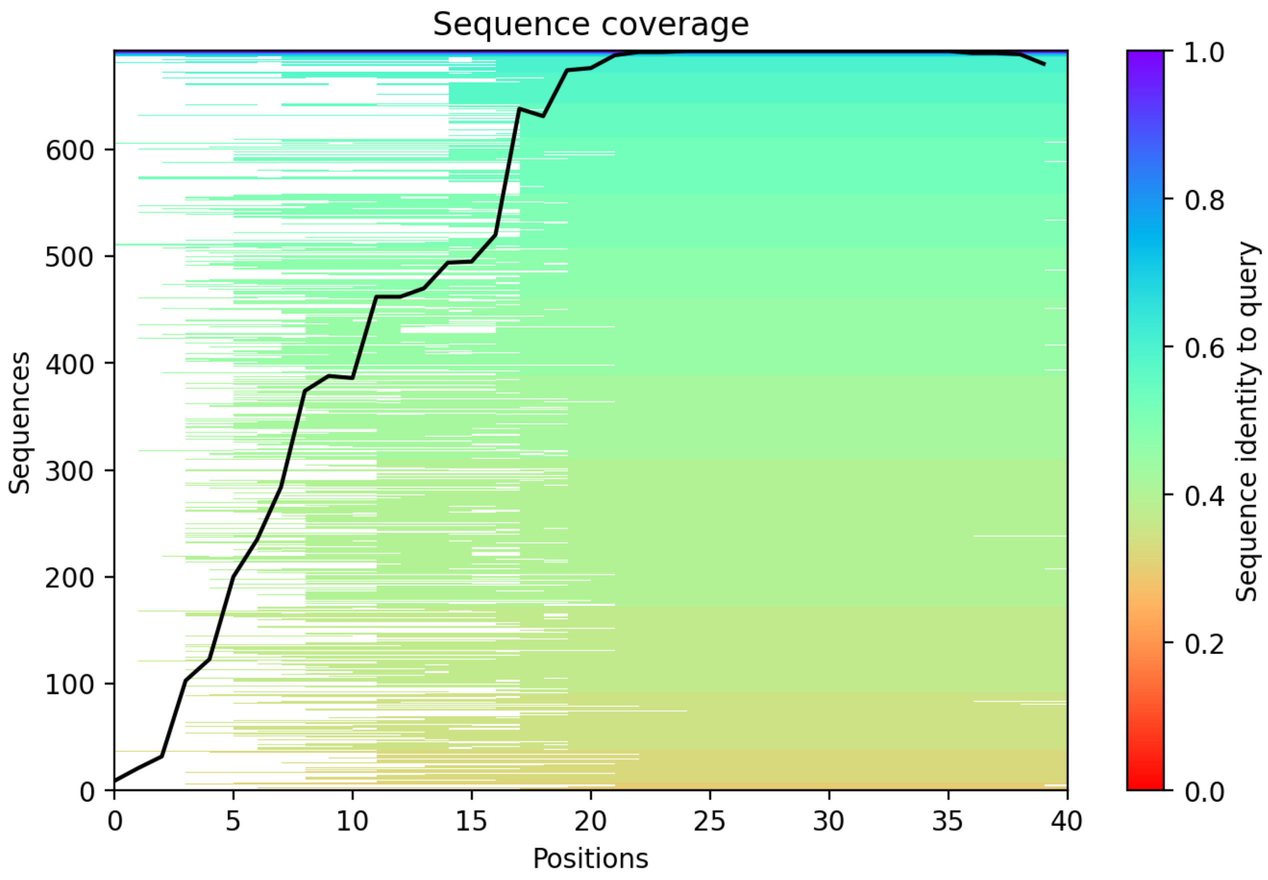

CP145: MtrunA17_CPg0492941

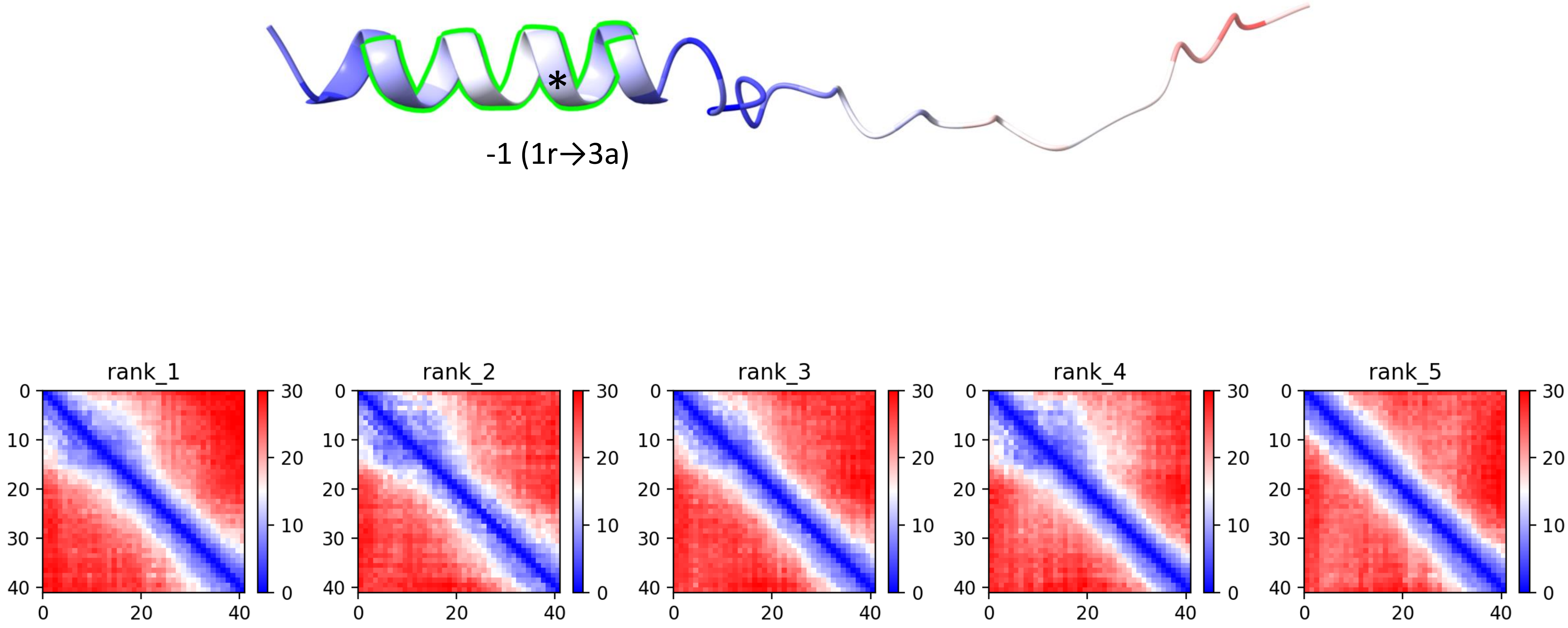

### CP145: MtrunA17_CPg0492941

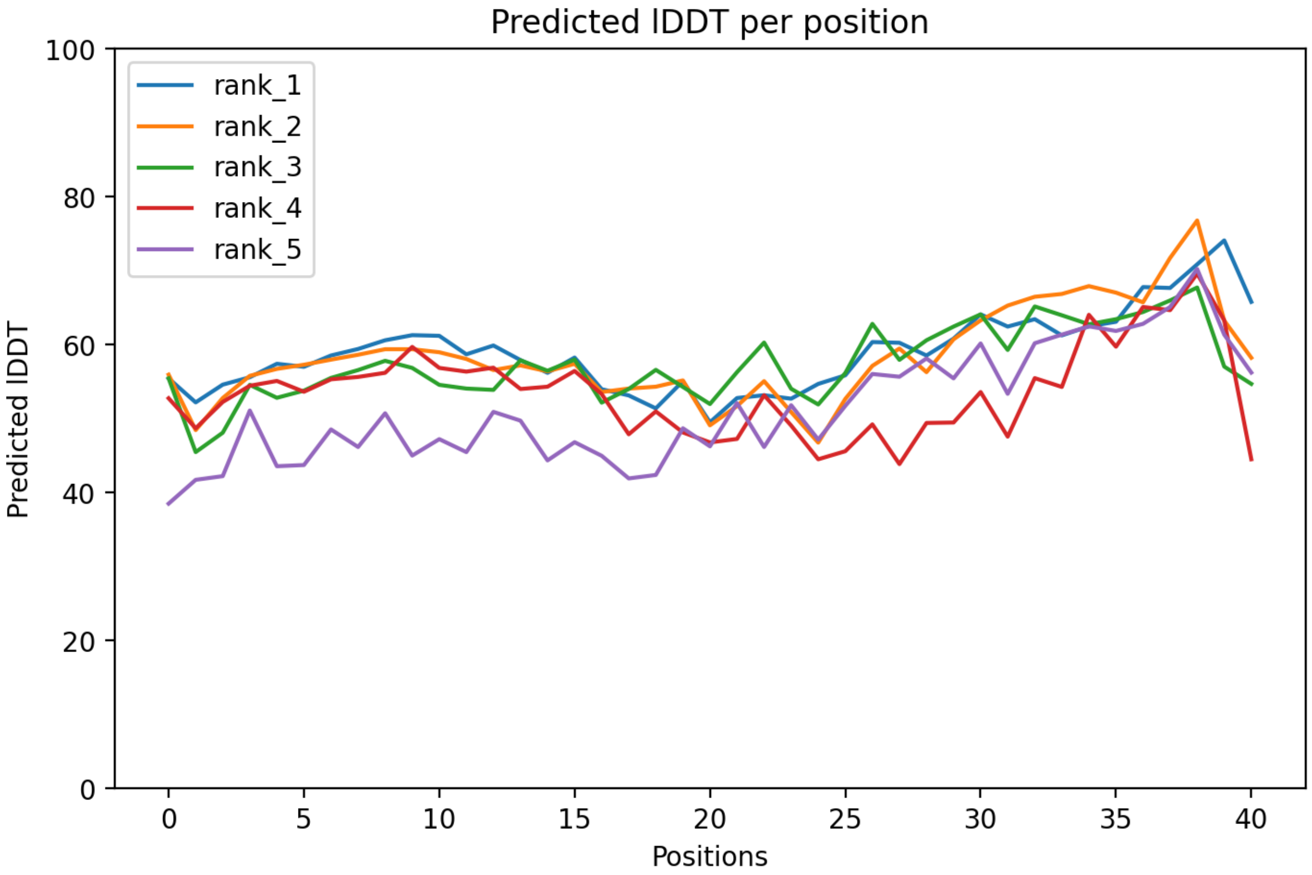

### CP145: MtrunA17_CPg0492941

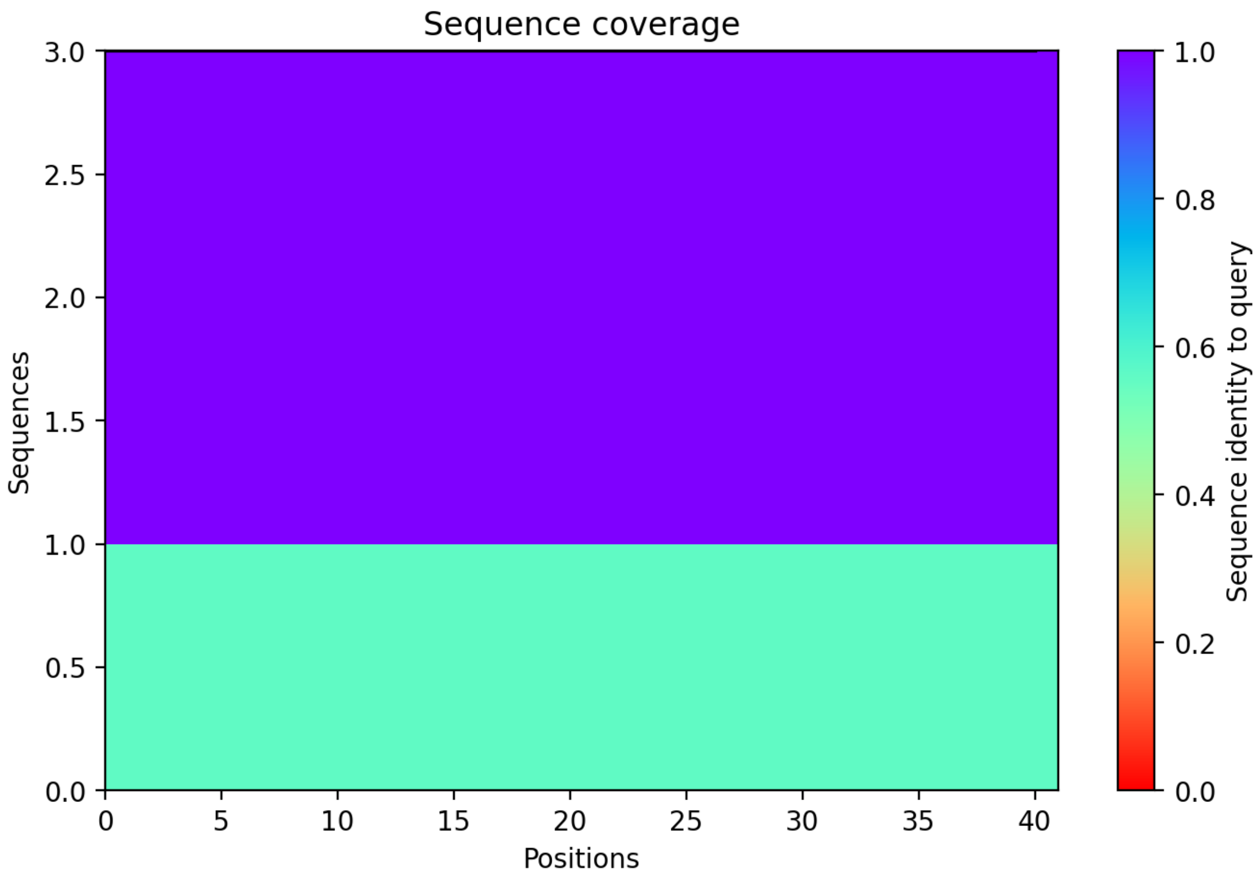

CP146: MtrunA17_CPg0493291

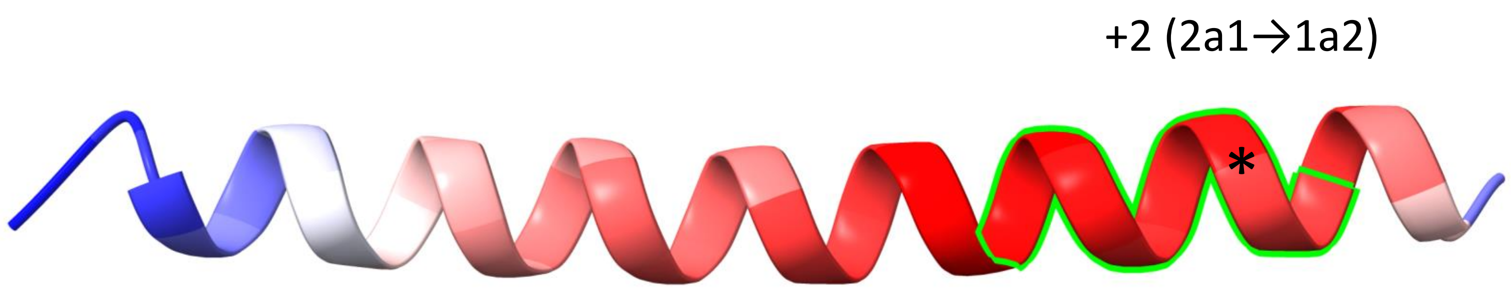


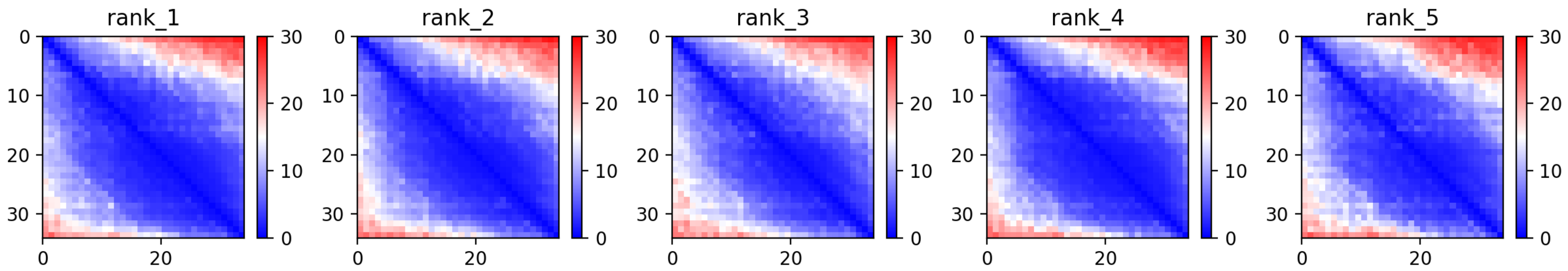

### CP146: MtrunA17_CPg0493291

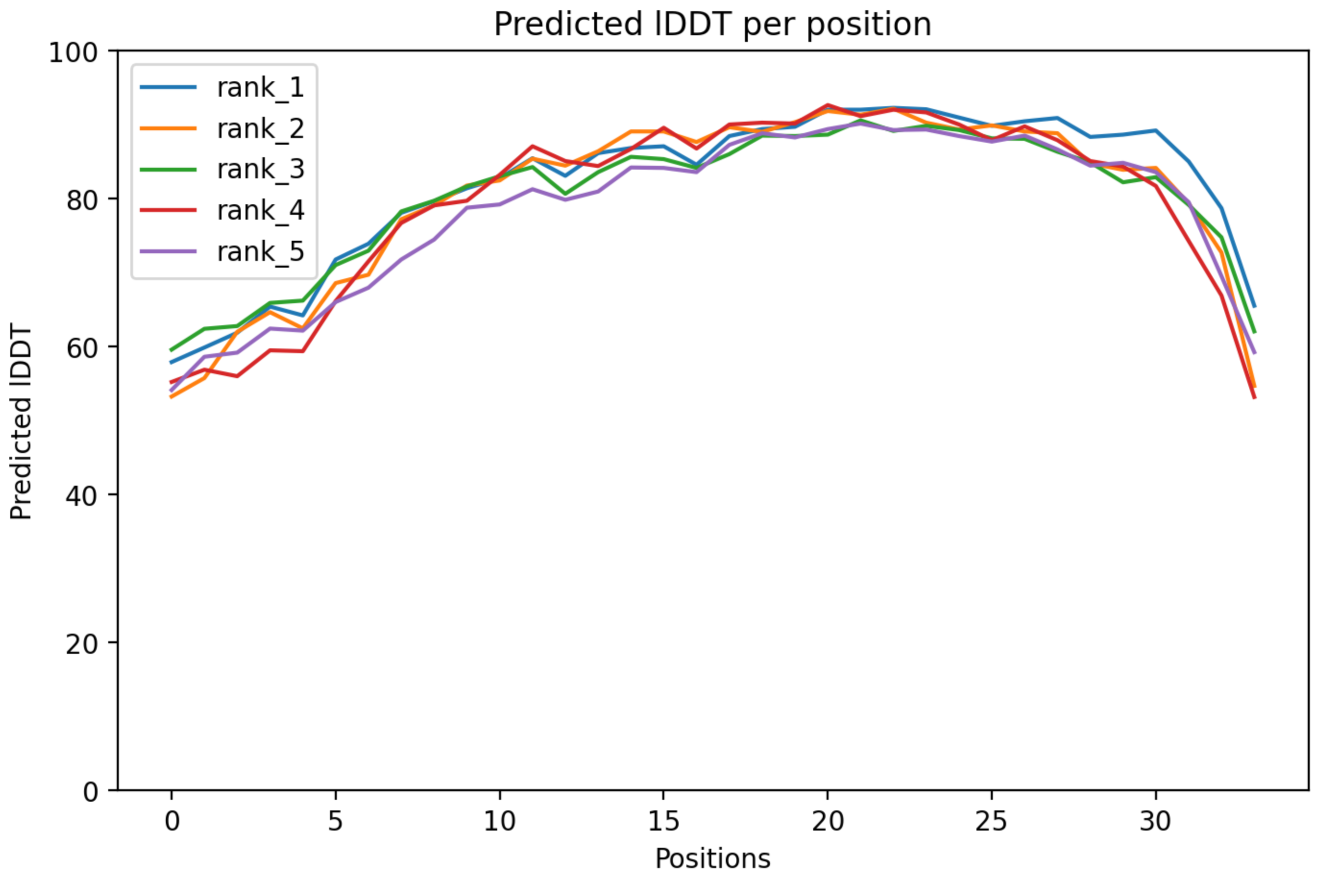

CP146: MtrunA17_CPg0493291

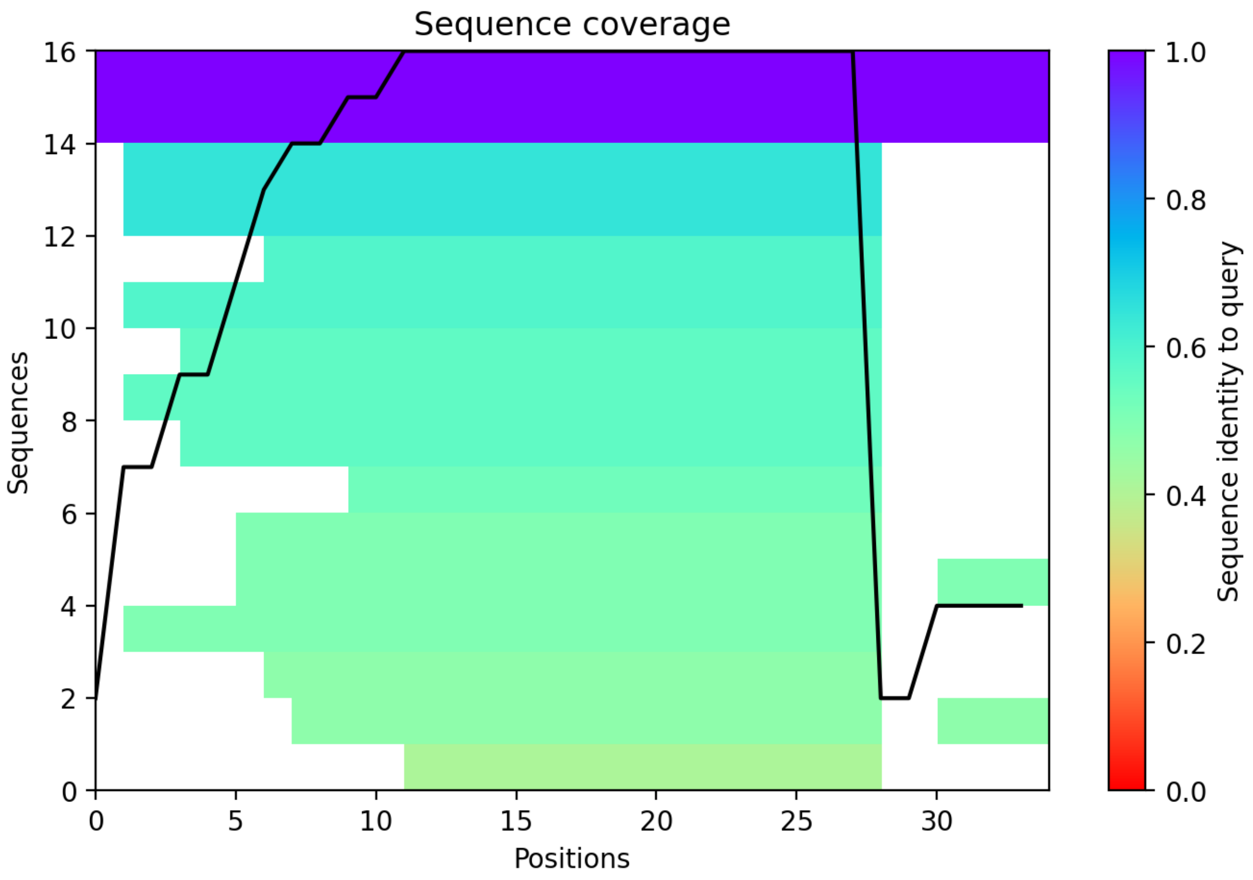

CP147: MtrunA17_CPg0493401

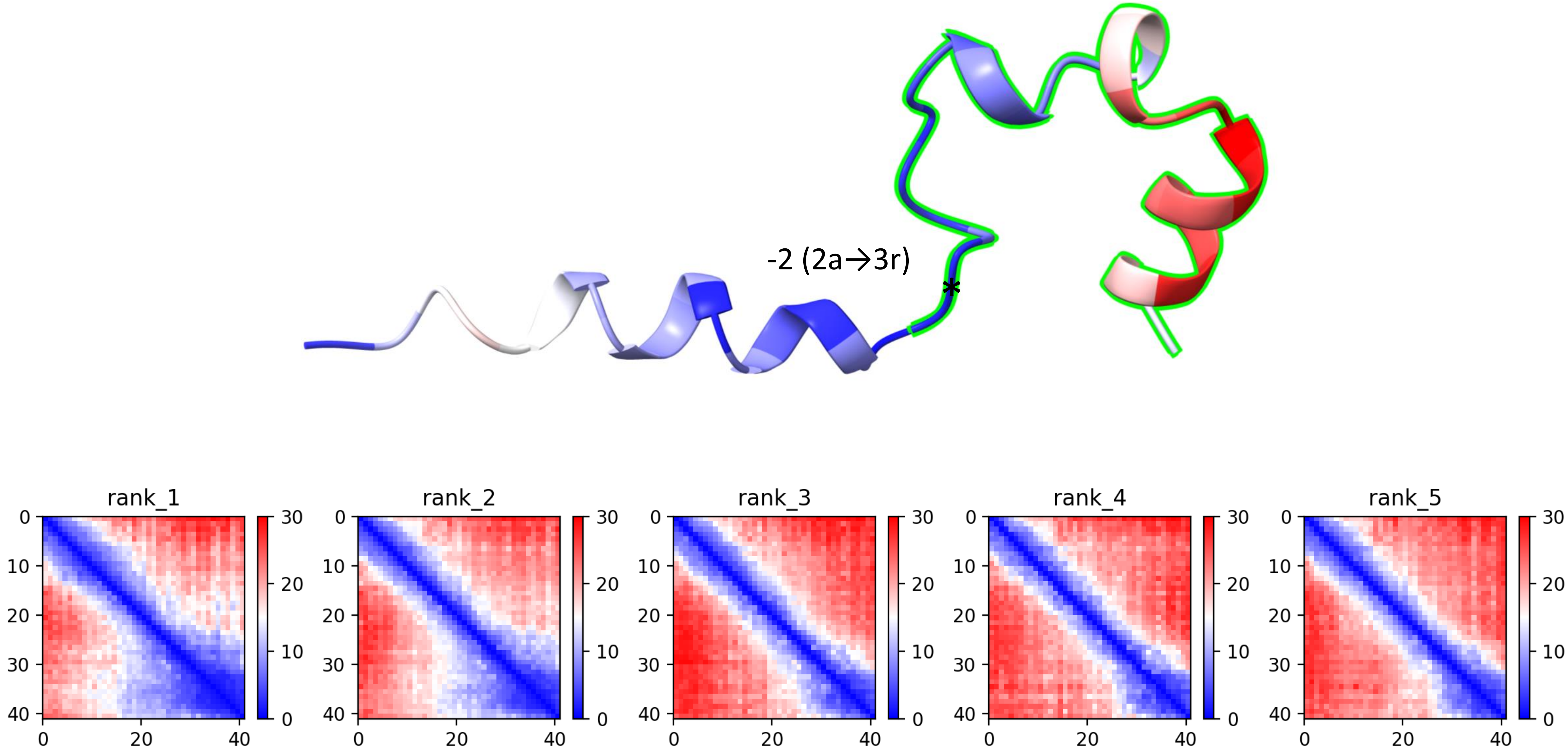

CP147: MtrunA17_CPg0493401

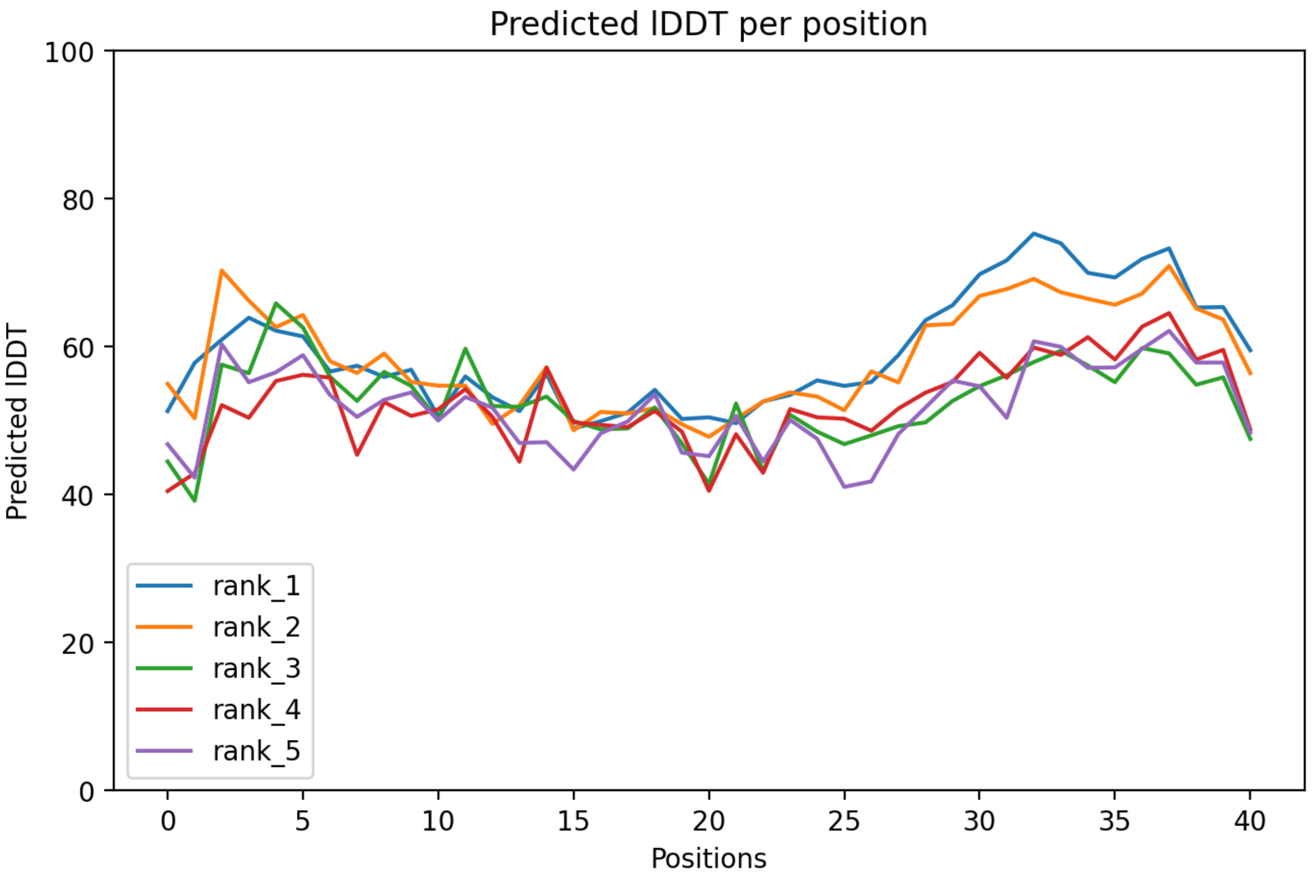

CP147: MtrunA17_CPg0493401

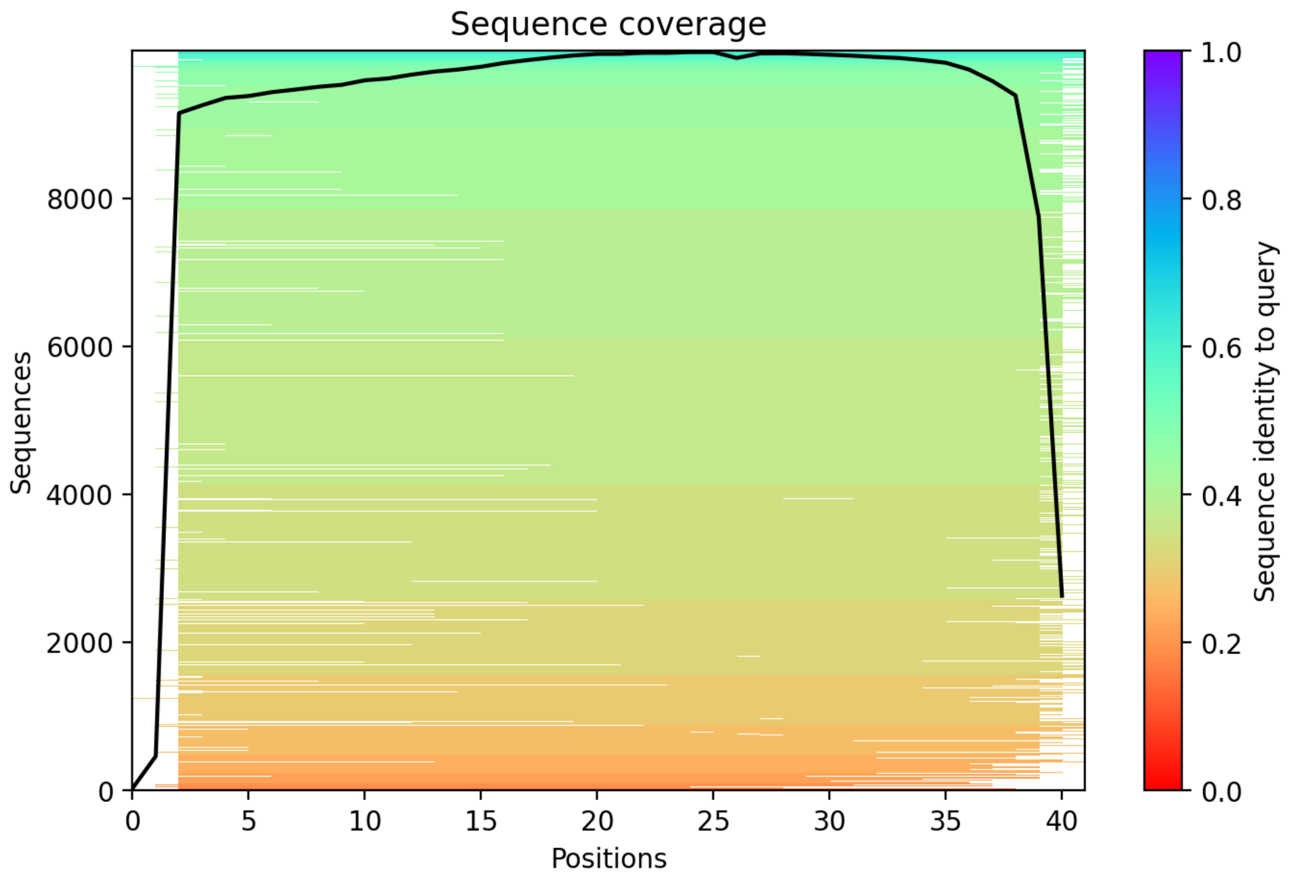

CP148: MtrunA17_MTg0490471

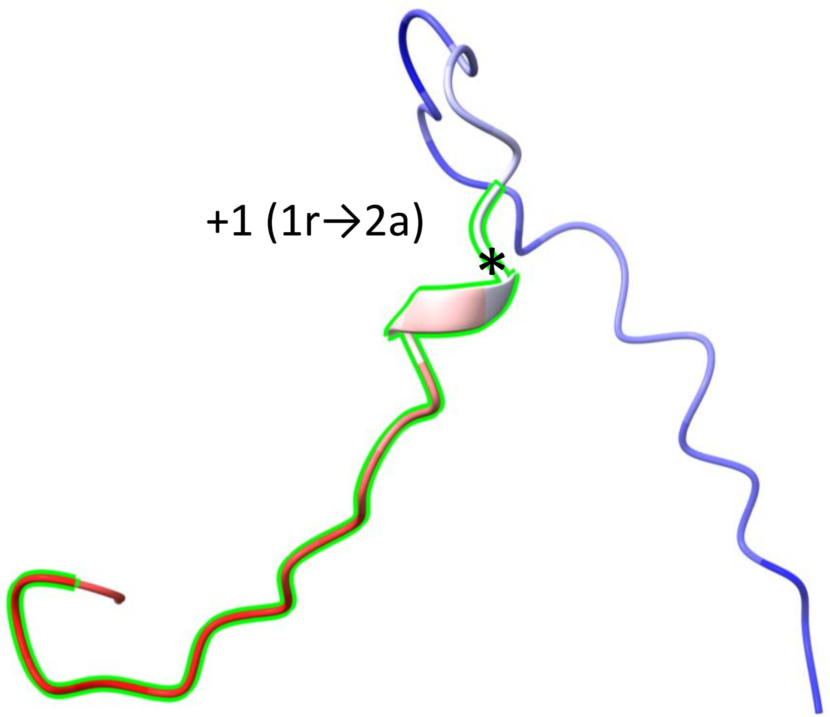


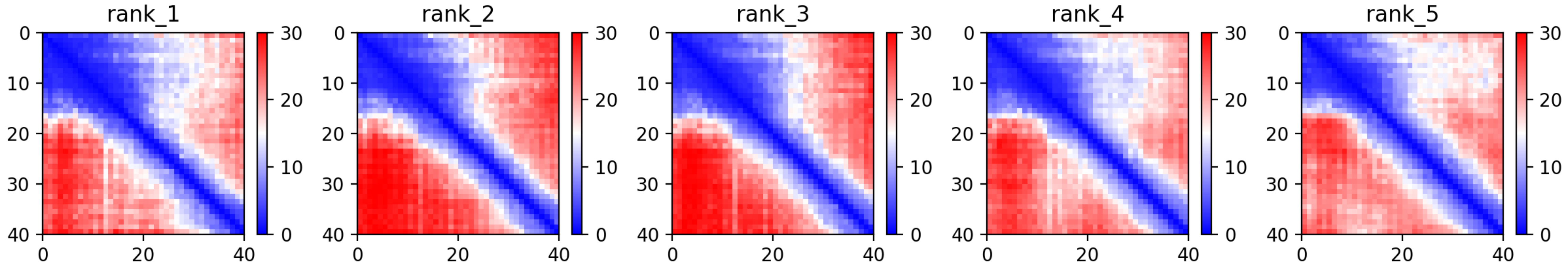

### CP148: MtrunA17_MTg0490471

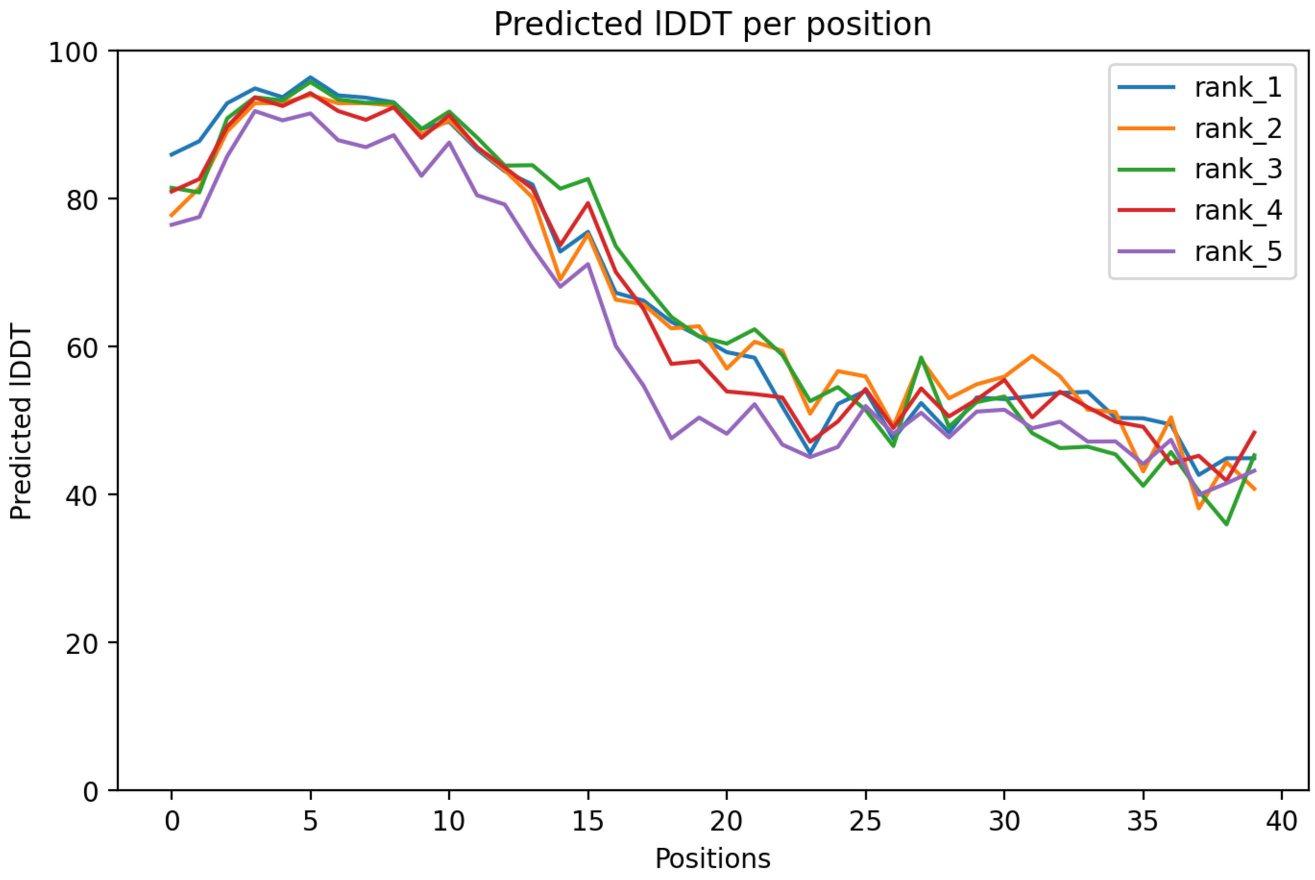

### CP148: MtrunA17_MTg0490471

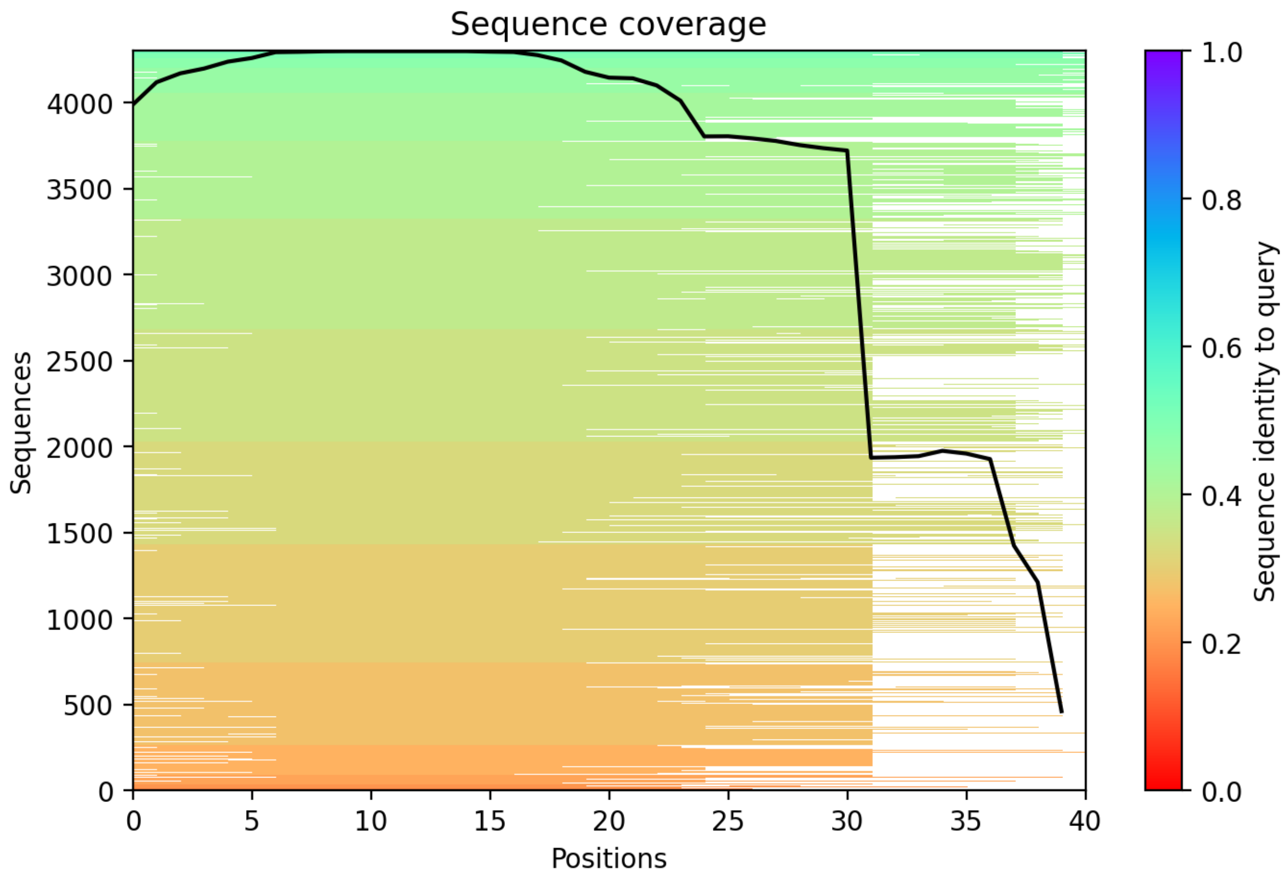

### CP149: MtrunA17_MTg0490471

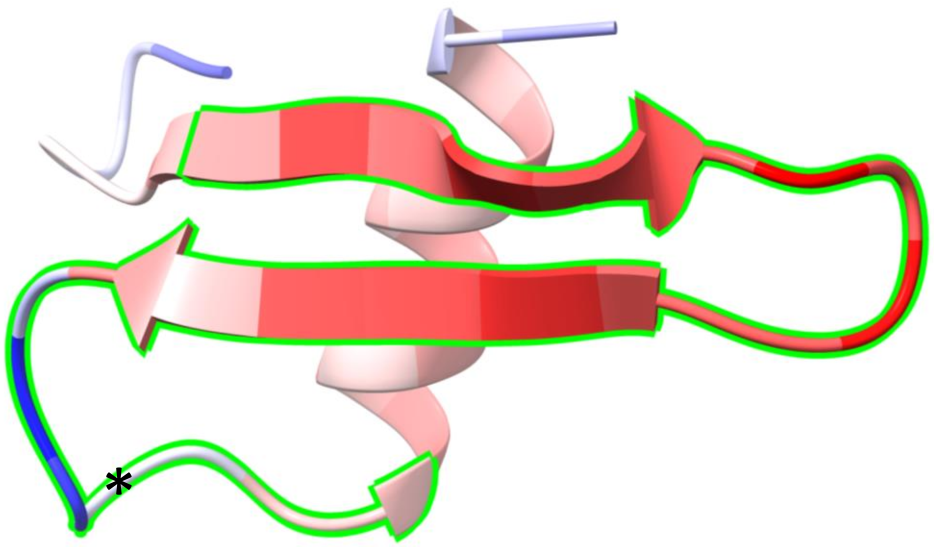


+1 (1r→2a)

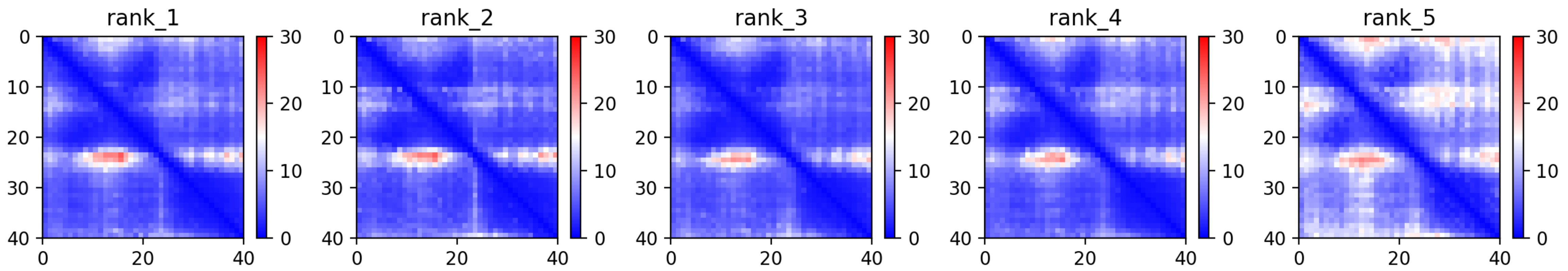

CP149: MtrunA17_MTg0490471

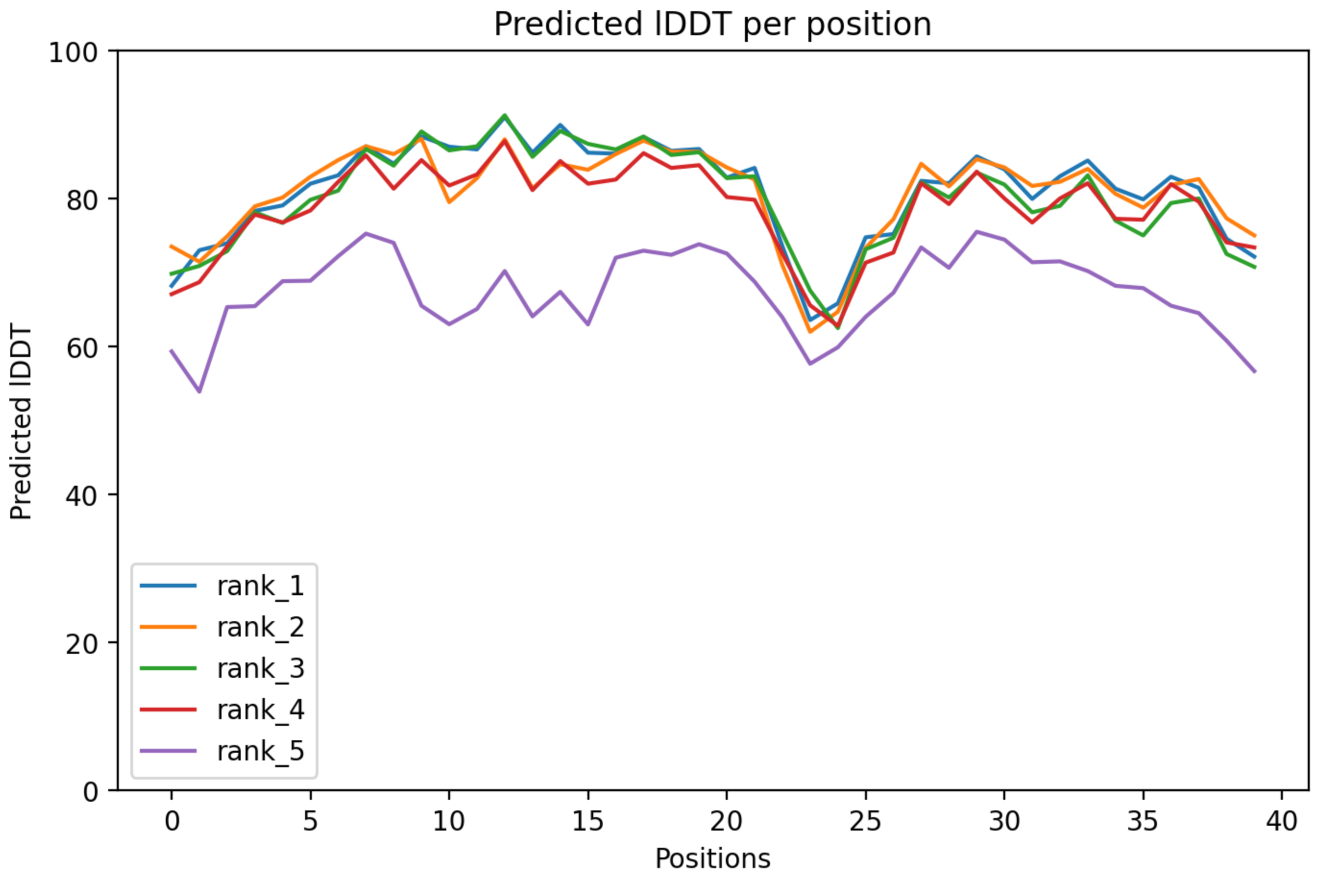

### CP149: MtrunA17_MTg0490471

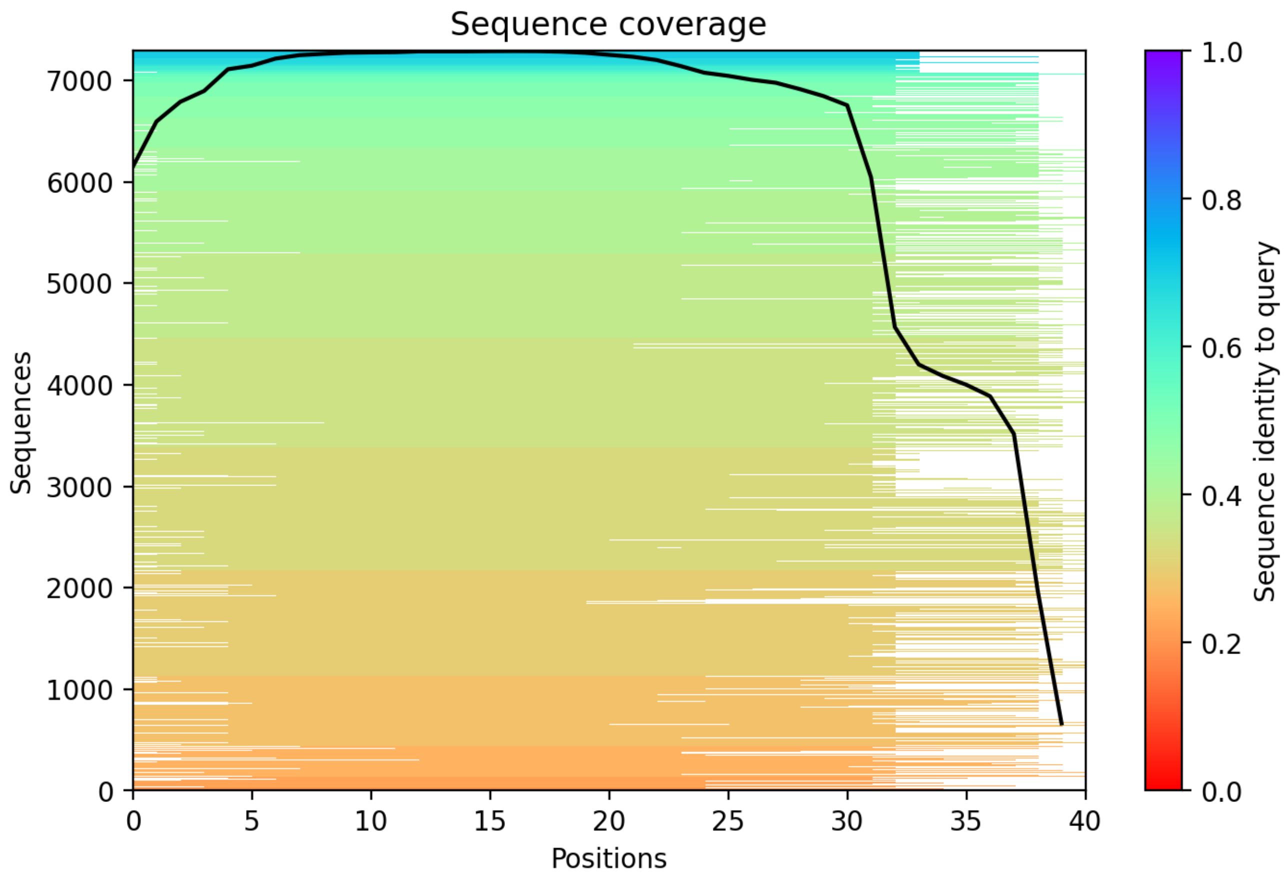

### CP150: MtrunA17_MTg0490971

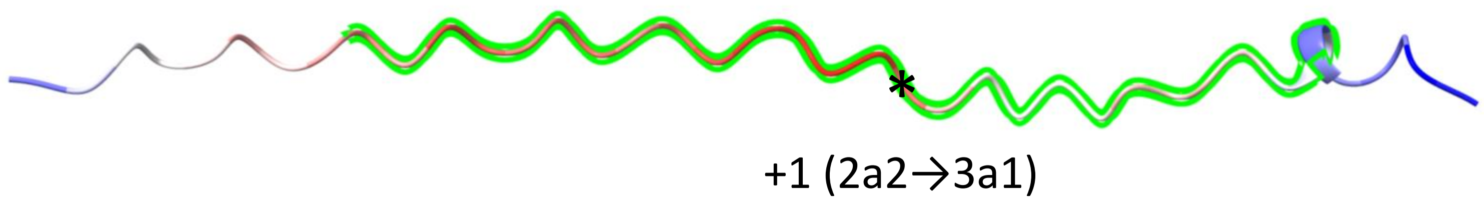


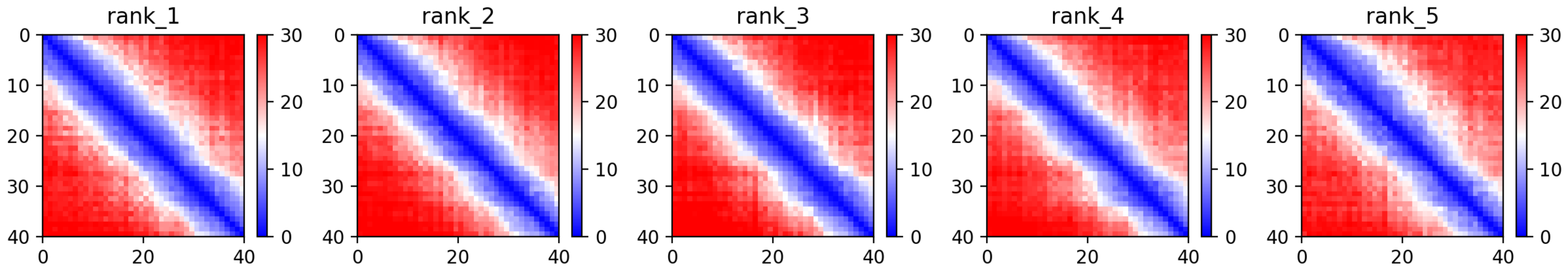

### CP150: MtrunA17_MTg0490971

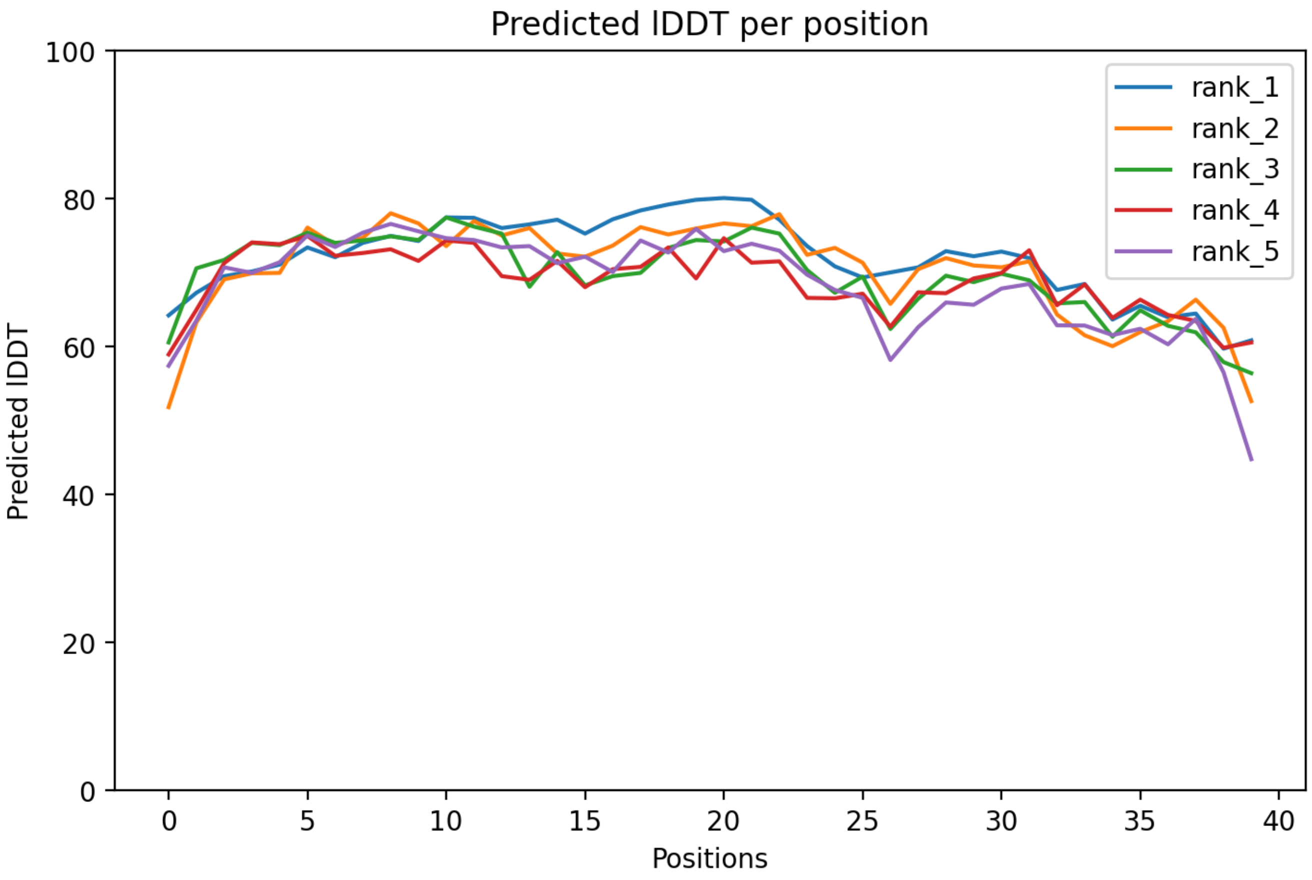

CP150: MtrunA17_MTg0490971

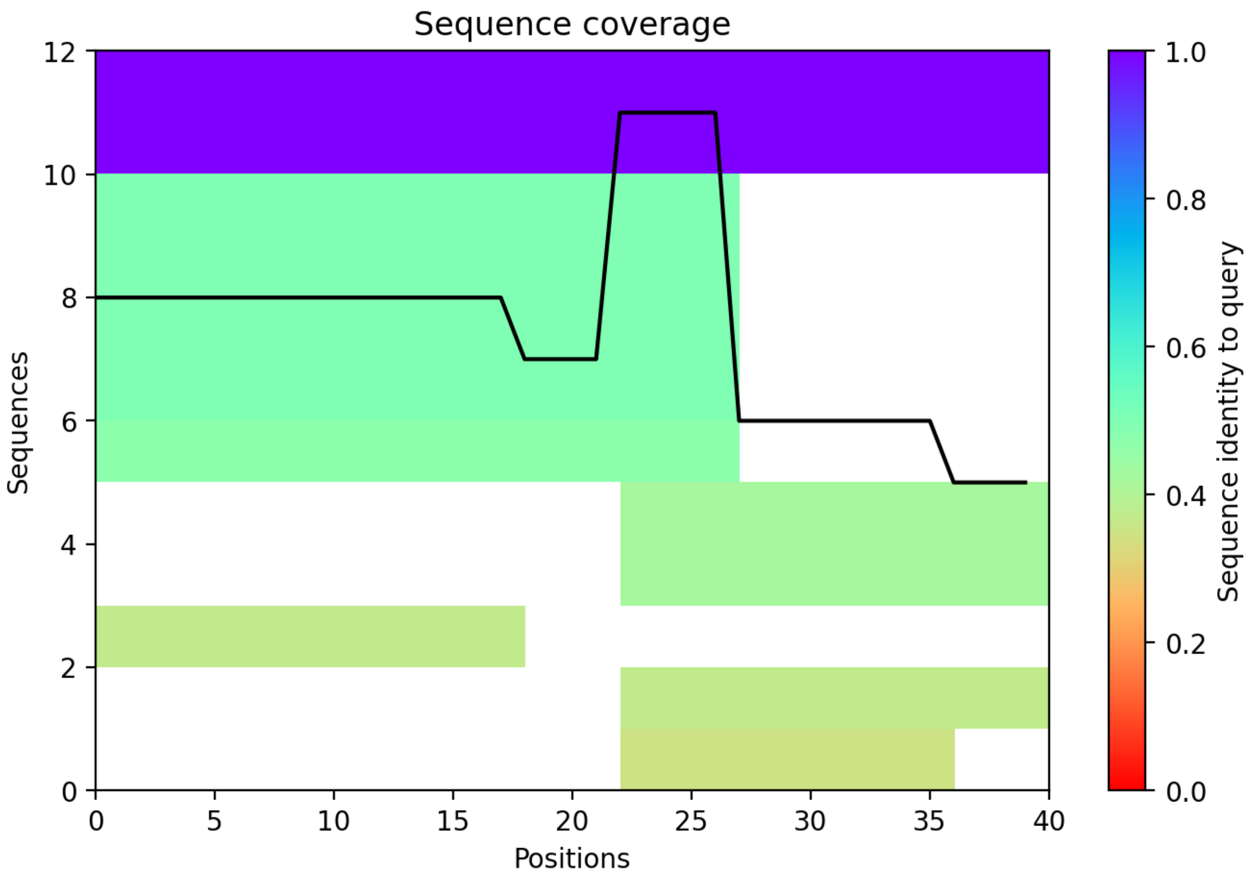

CP151: MtrunA17_MTg0490971

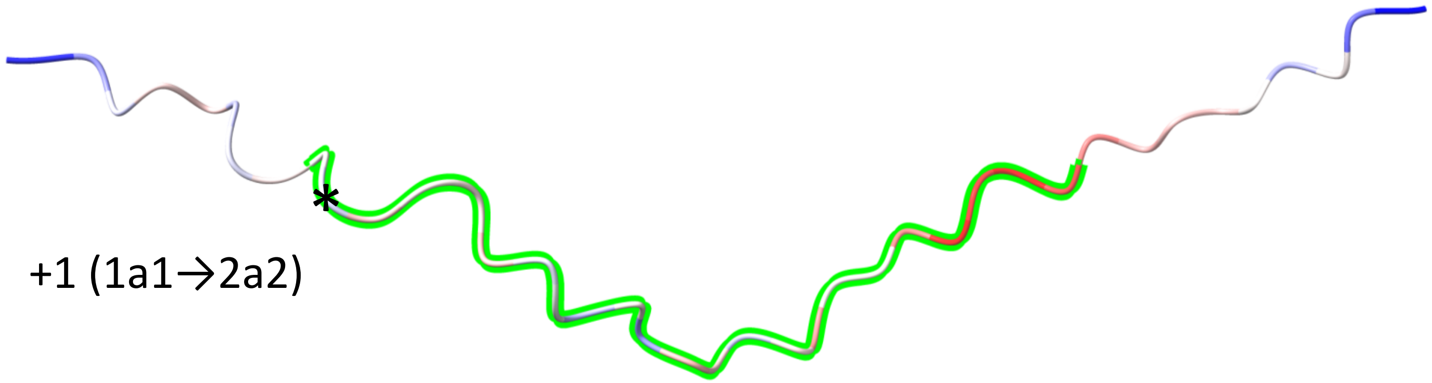


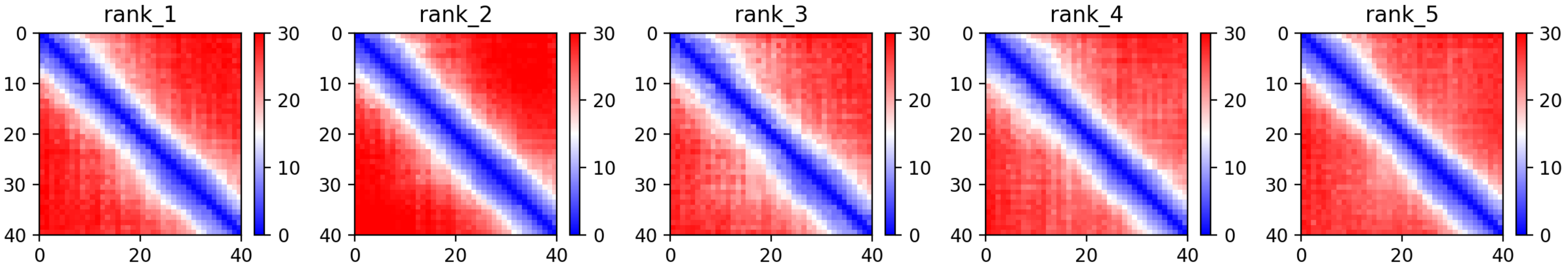

CP151: MtrunA17_MTg0490971

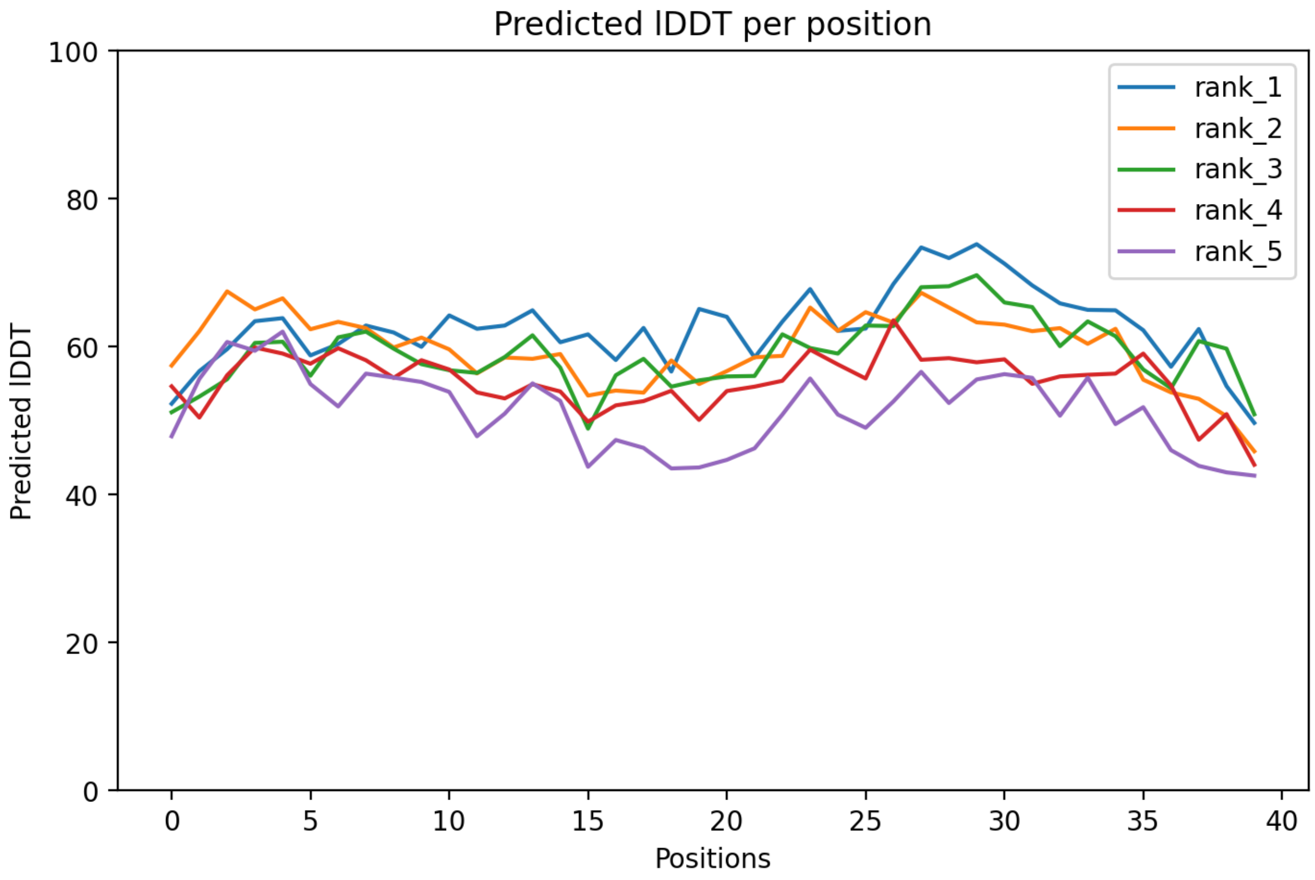

CP151: MtrunA17_MTg0490971

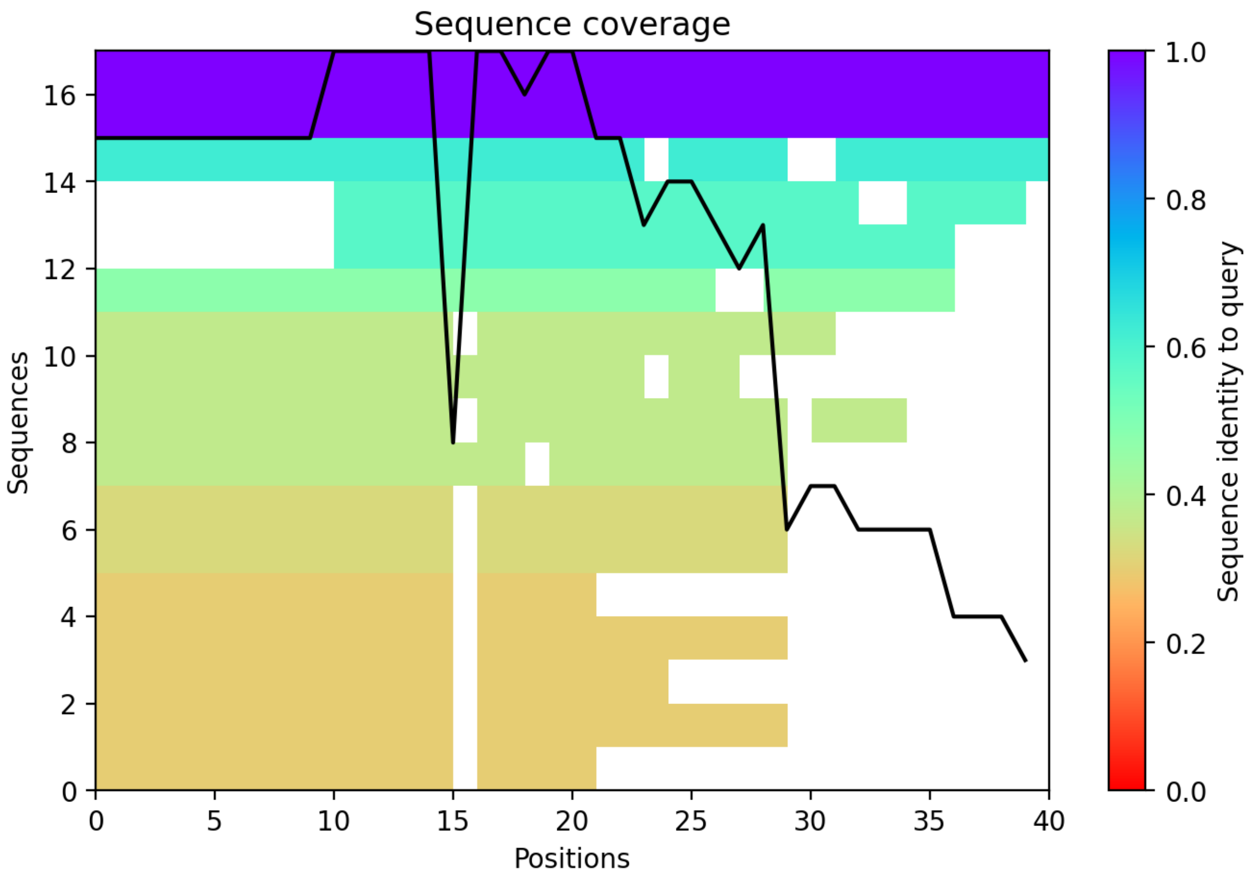

CP152: MtrunA17_MTg0491151

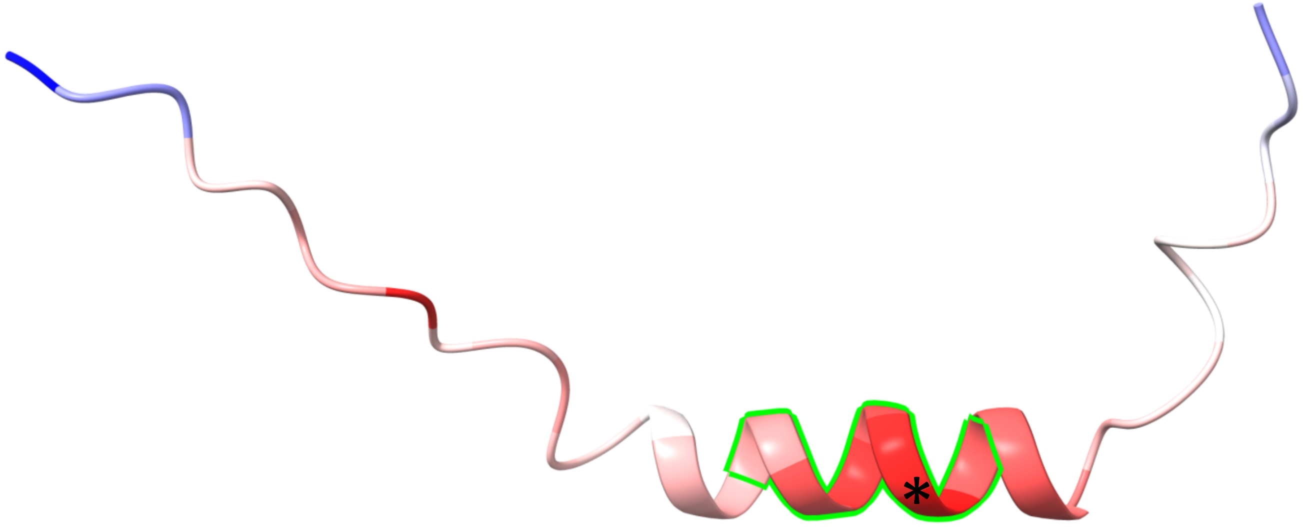


+2 (2a1→1a2)

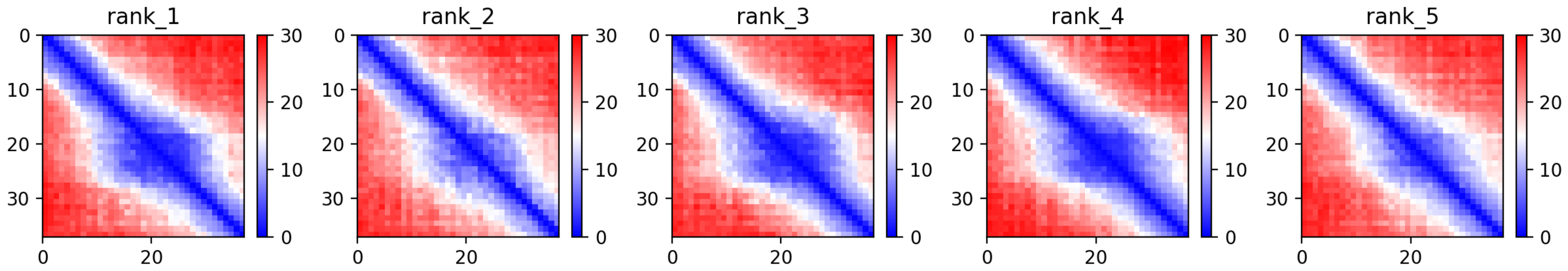

CP152: MtrunA17_MTg0491151

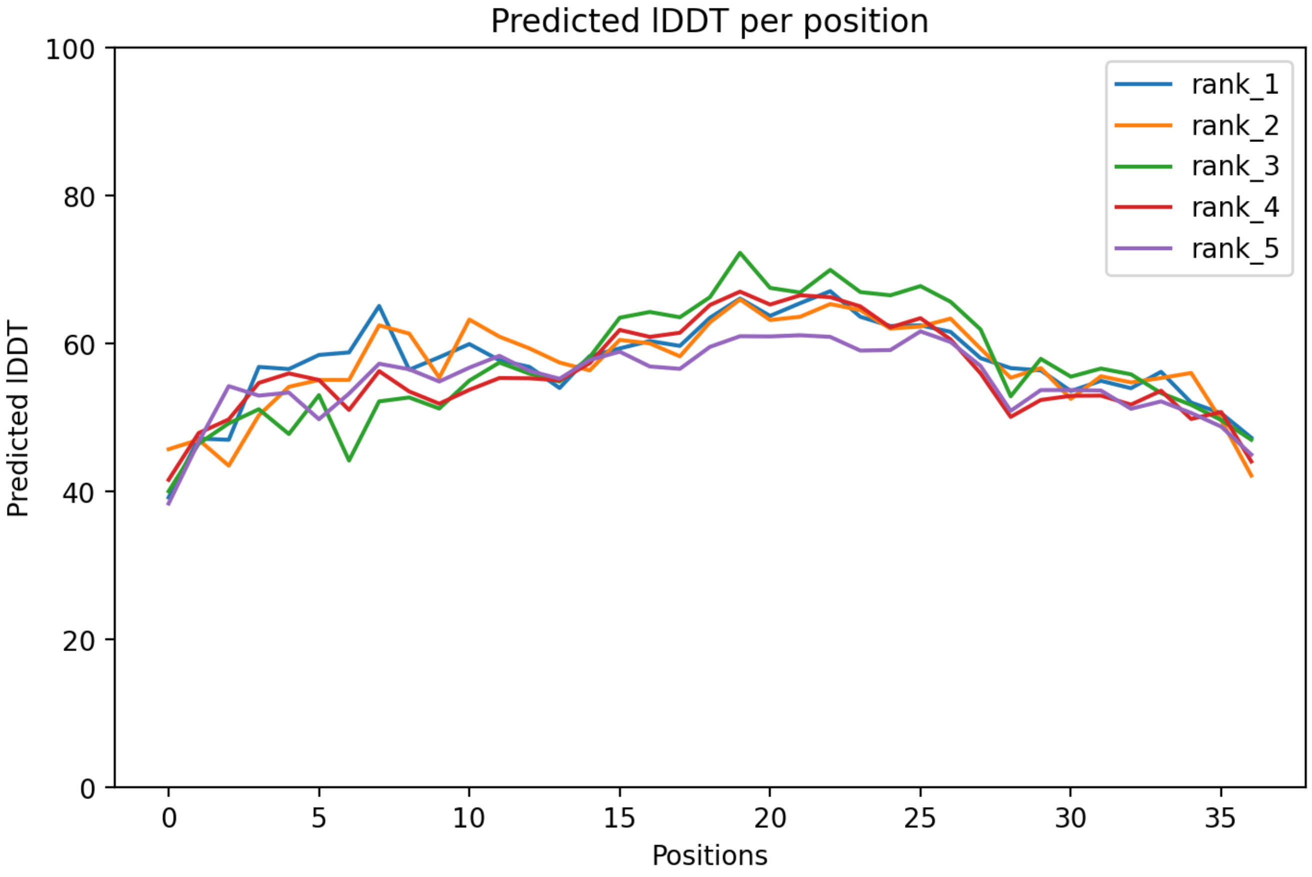

CP152: MtrunA17_MTg0491151

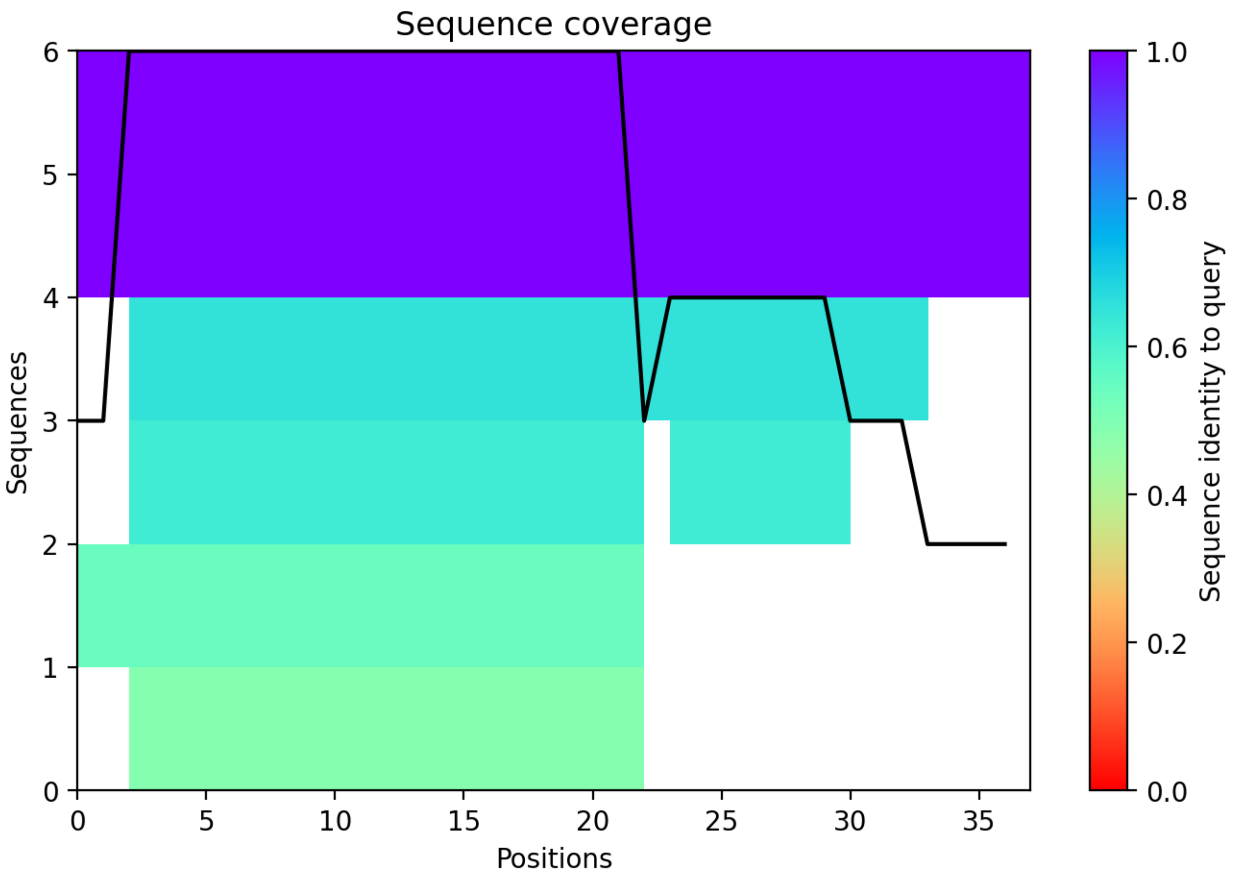

CP153: MtrunA17_MTg0491291

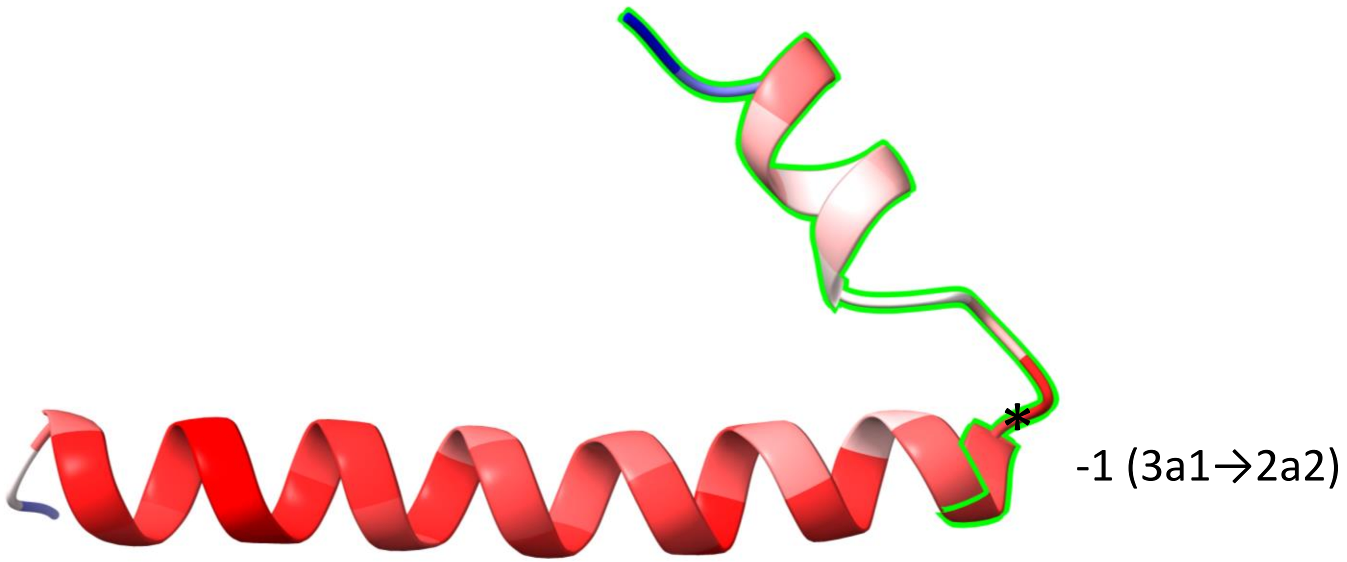


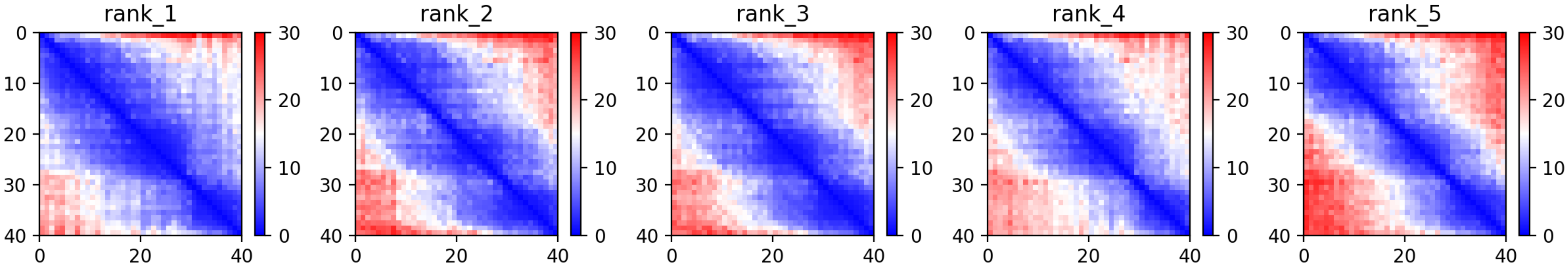

### CP153: MtrunA17_MTg0491291

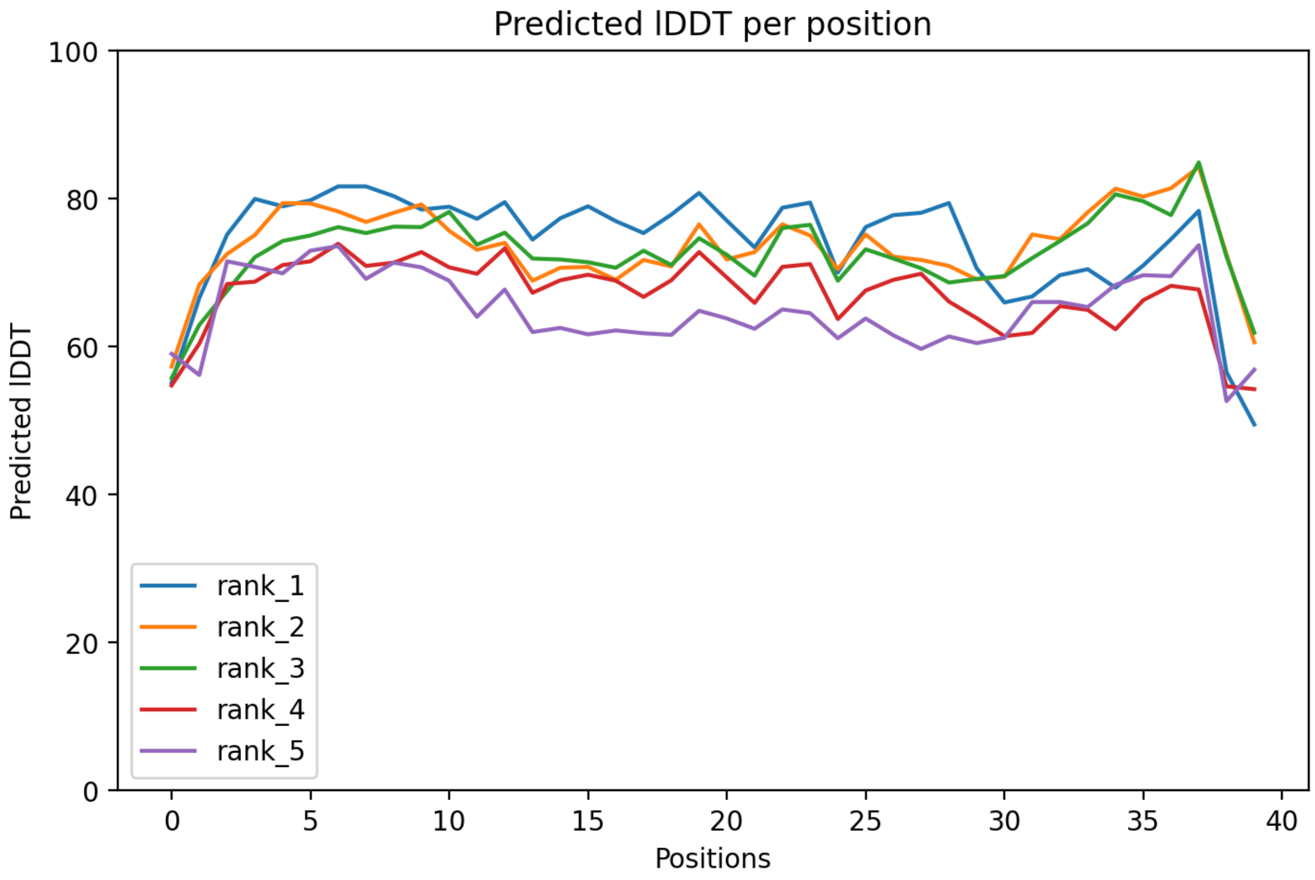

### CP153: MtrunA17_MTg0491291

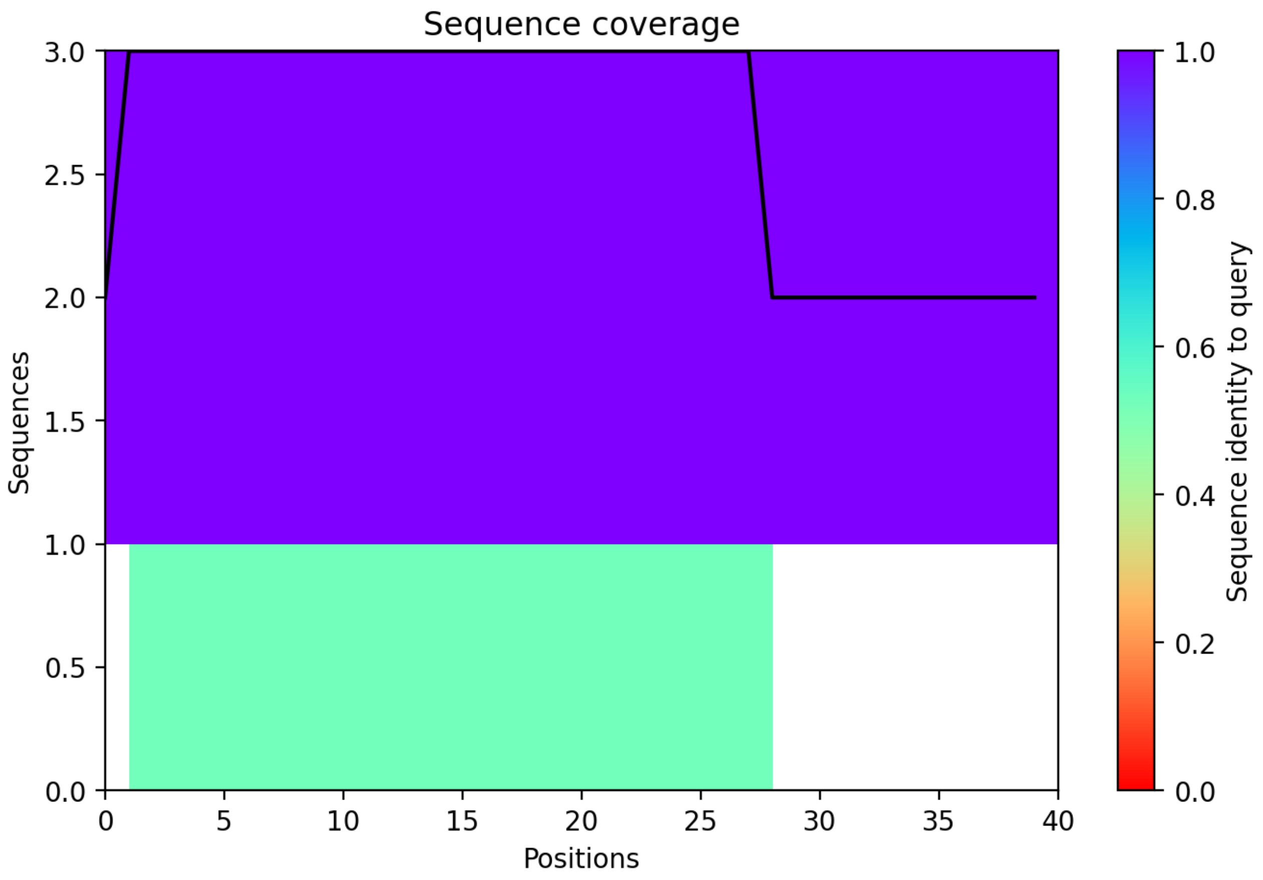

CP154: MtrunA17_MTg0491501

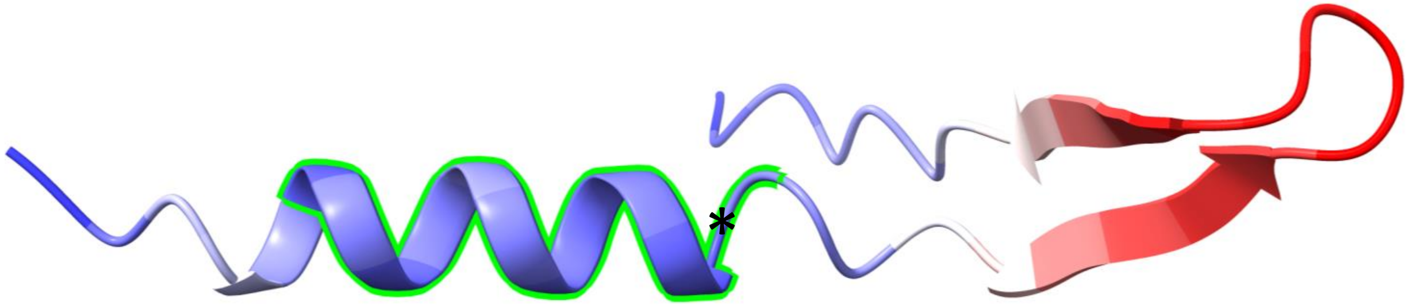


-2 (3a1→1a2)

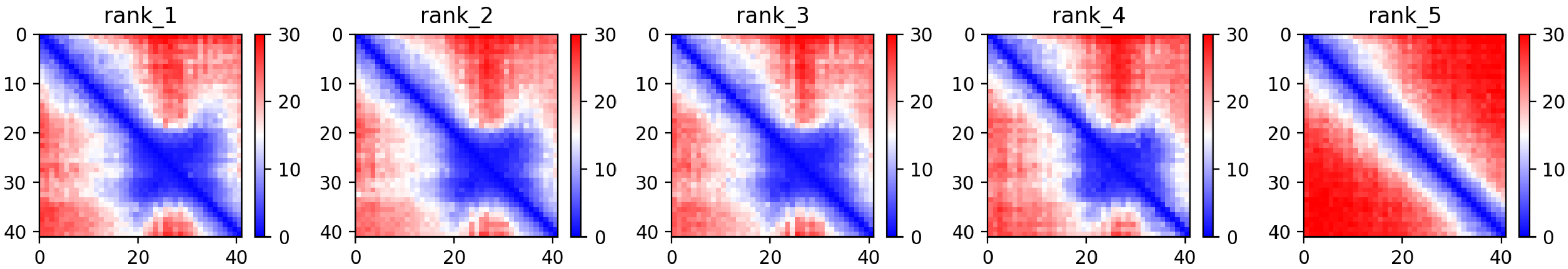

CP154: MtrunA17_MTg0491501

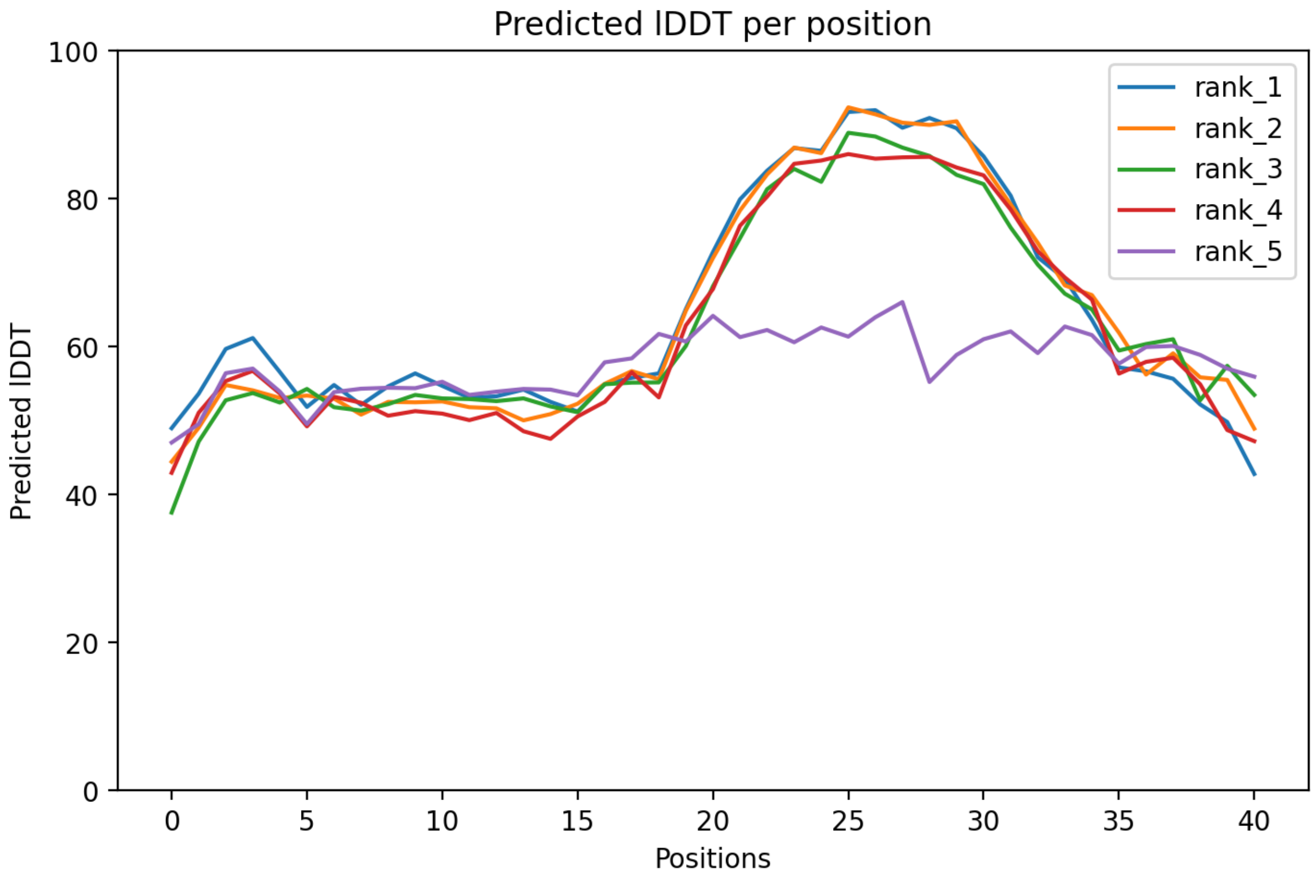

### CP154: MtrunA17_MTg0491501

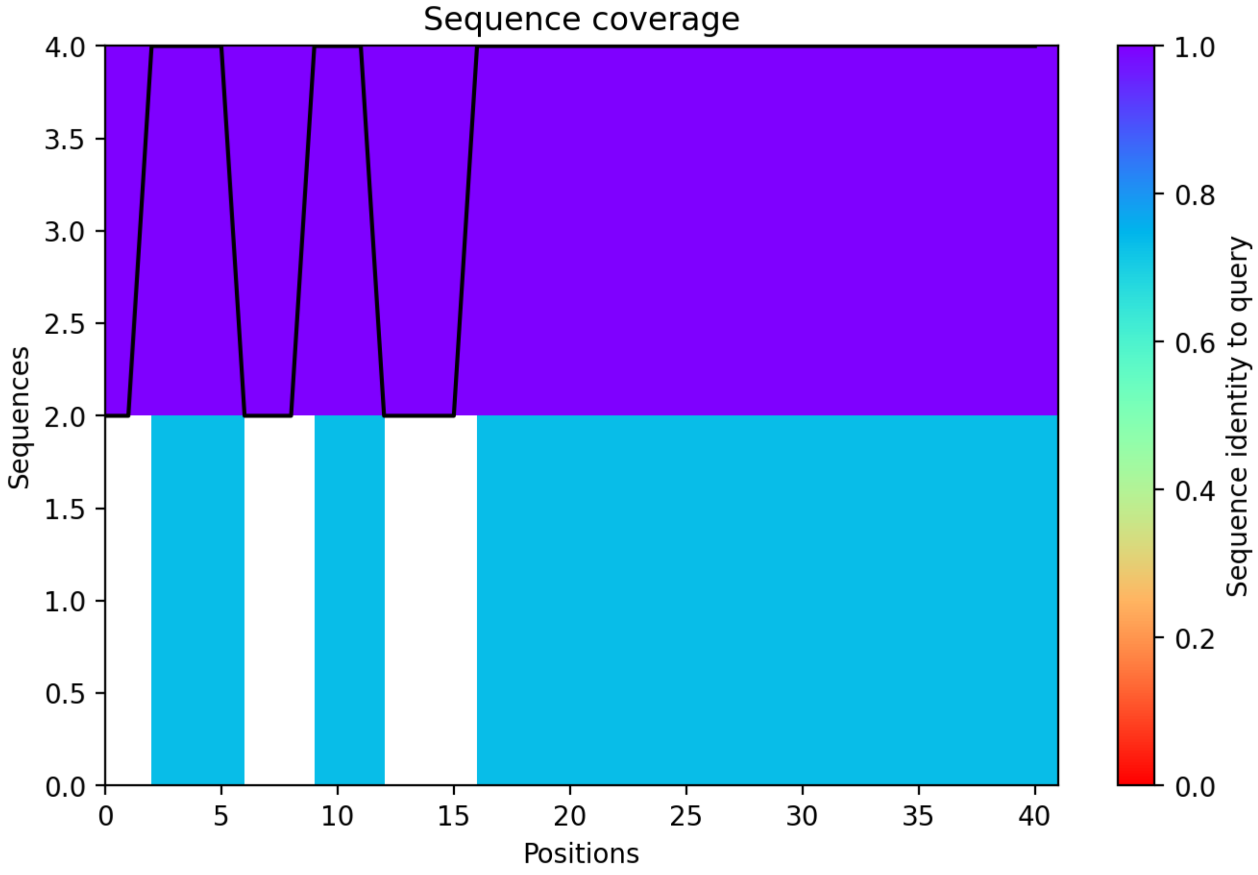

CP155: MtrunA17_MTg0491621

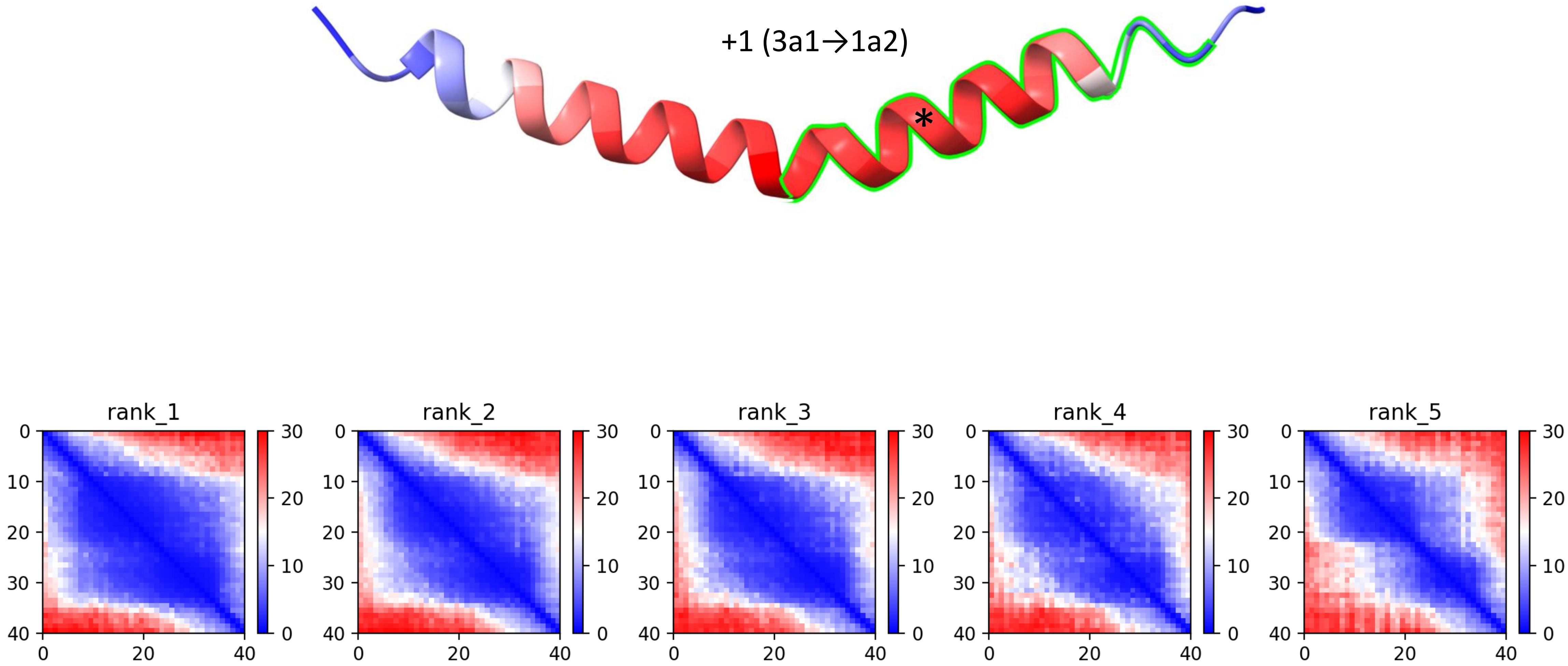

### CP155: MtrunA17_MTg0491621

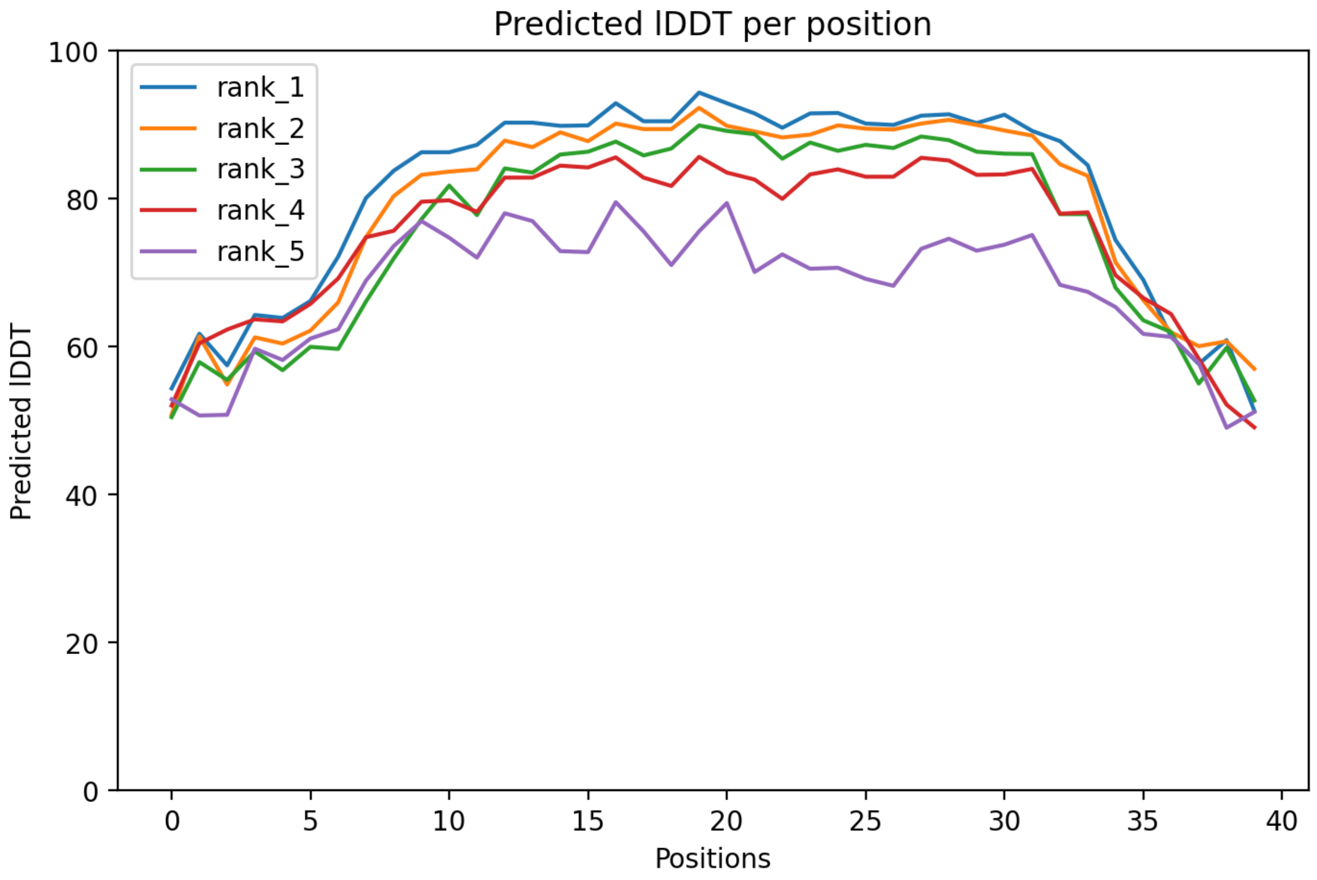

CP155: MtrunA17_MTg0491621

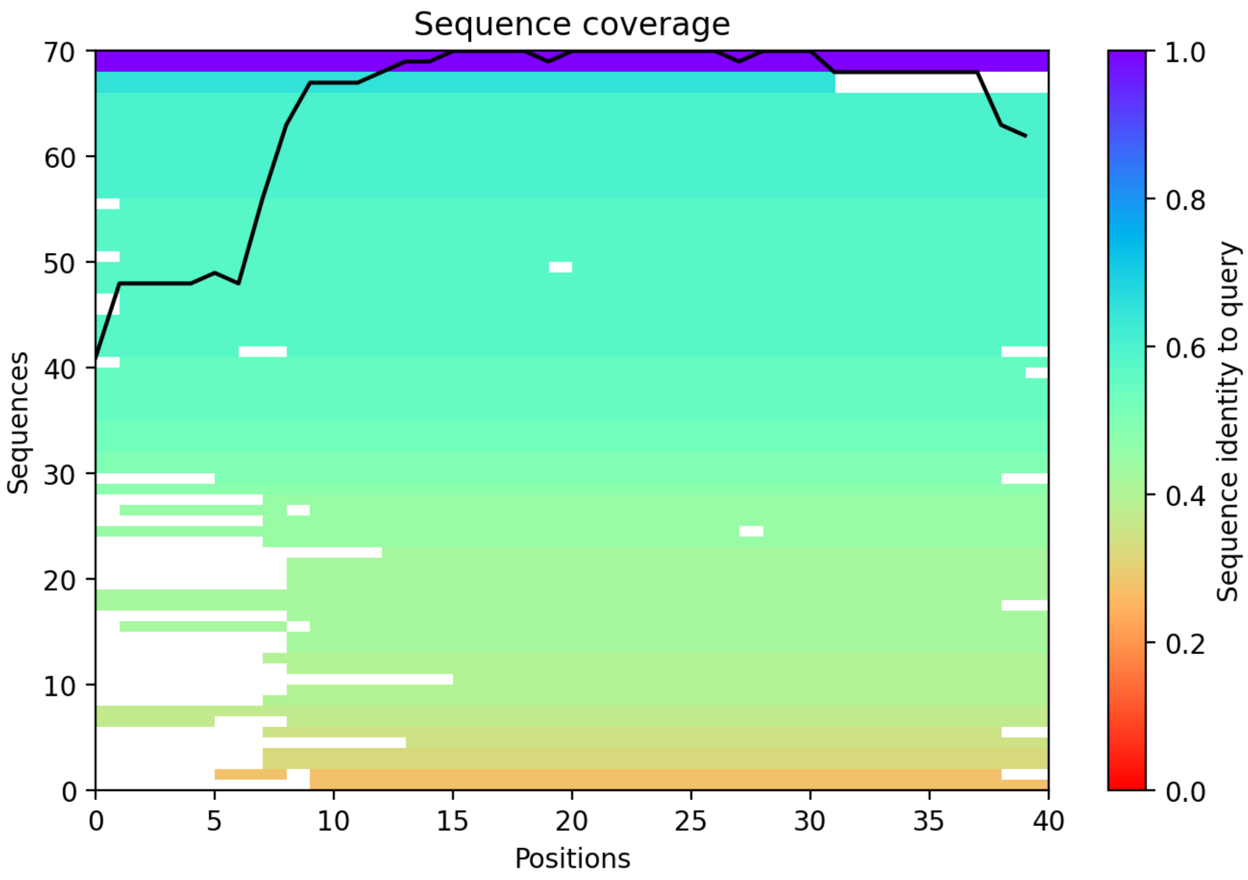

### CP156: MtrunA17_MTg0491711

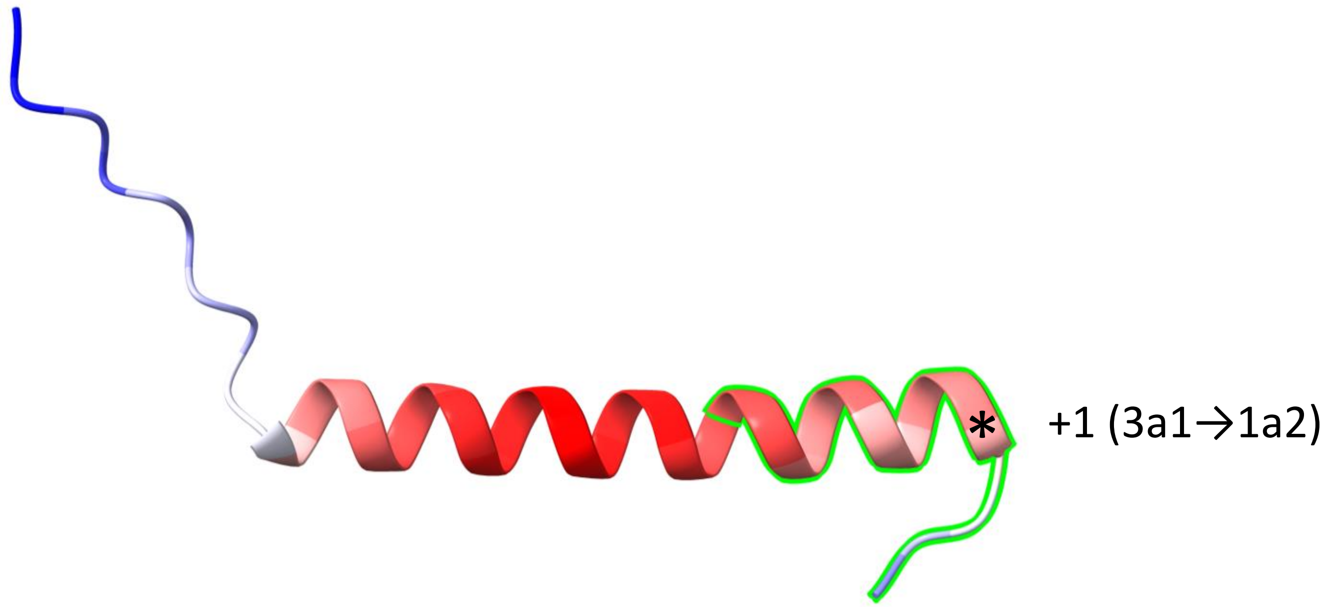


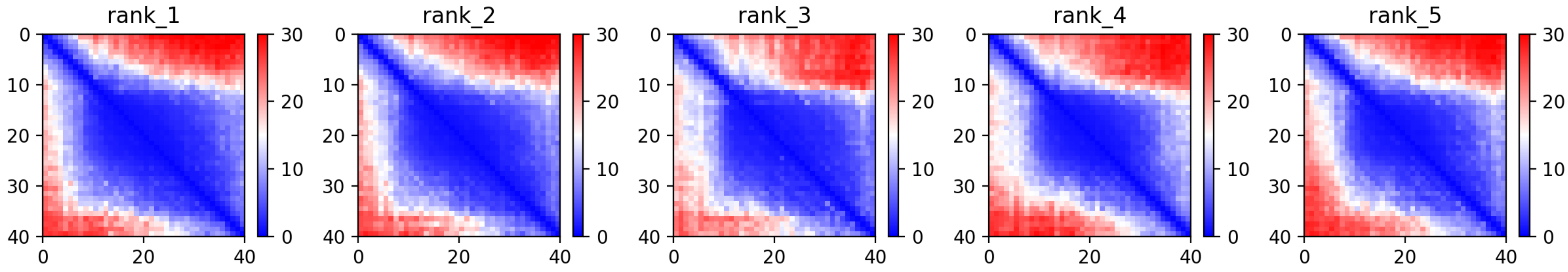

CP156: MtrunA17_MTg0491711

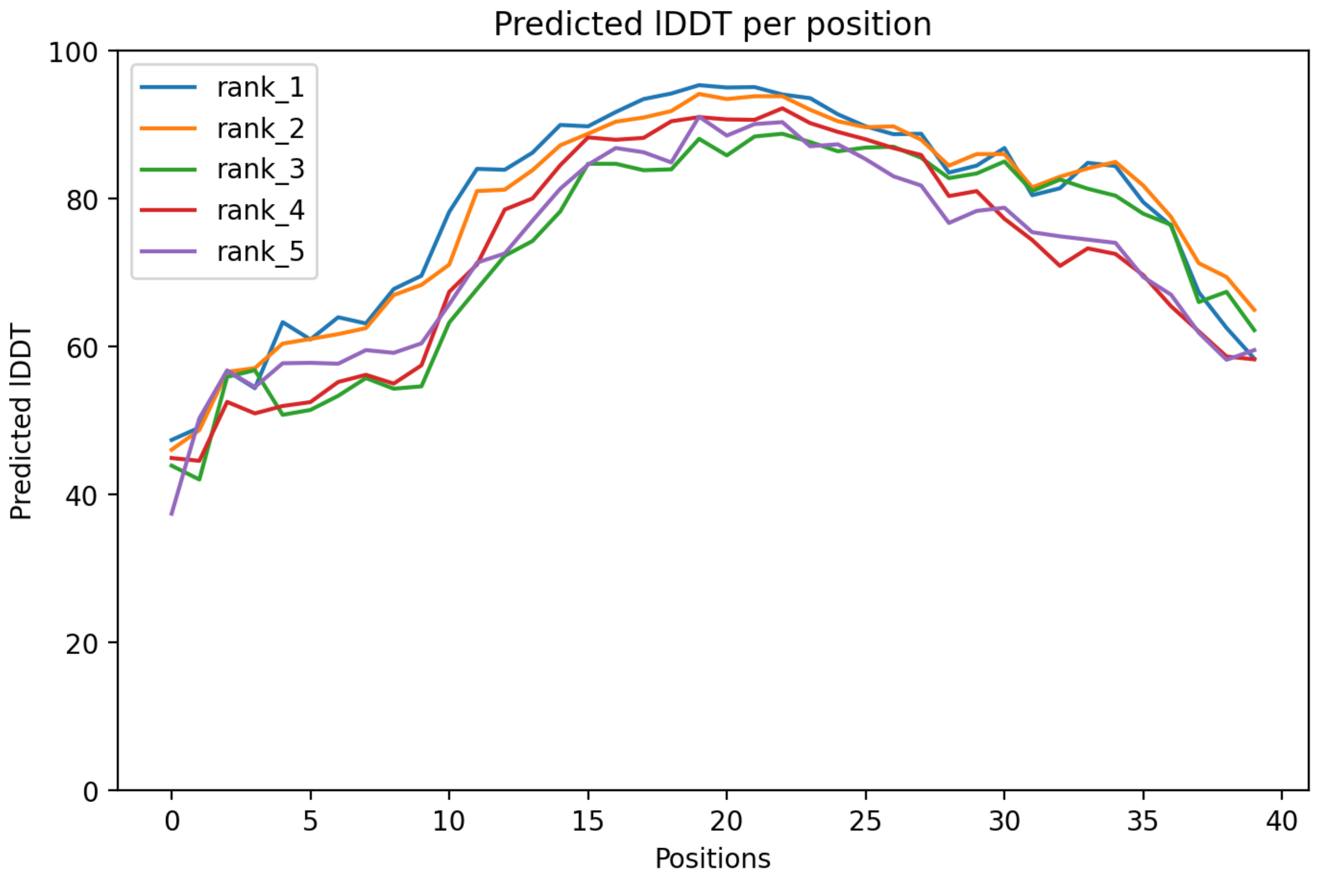

CP156: MtrunA17_MTg0491711

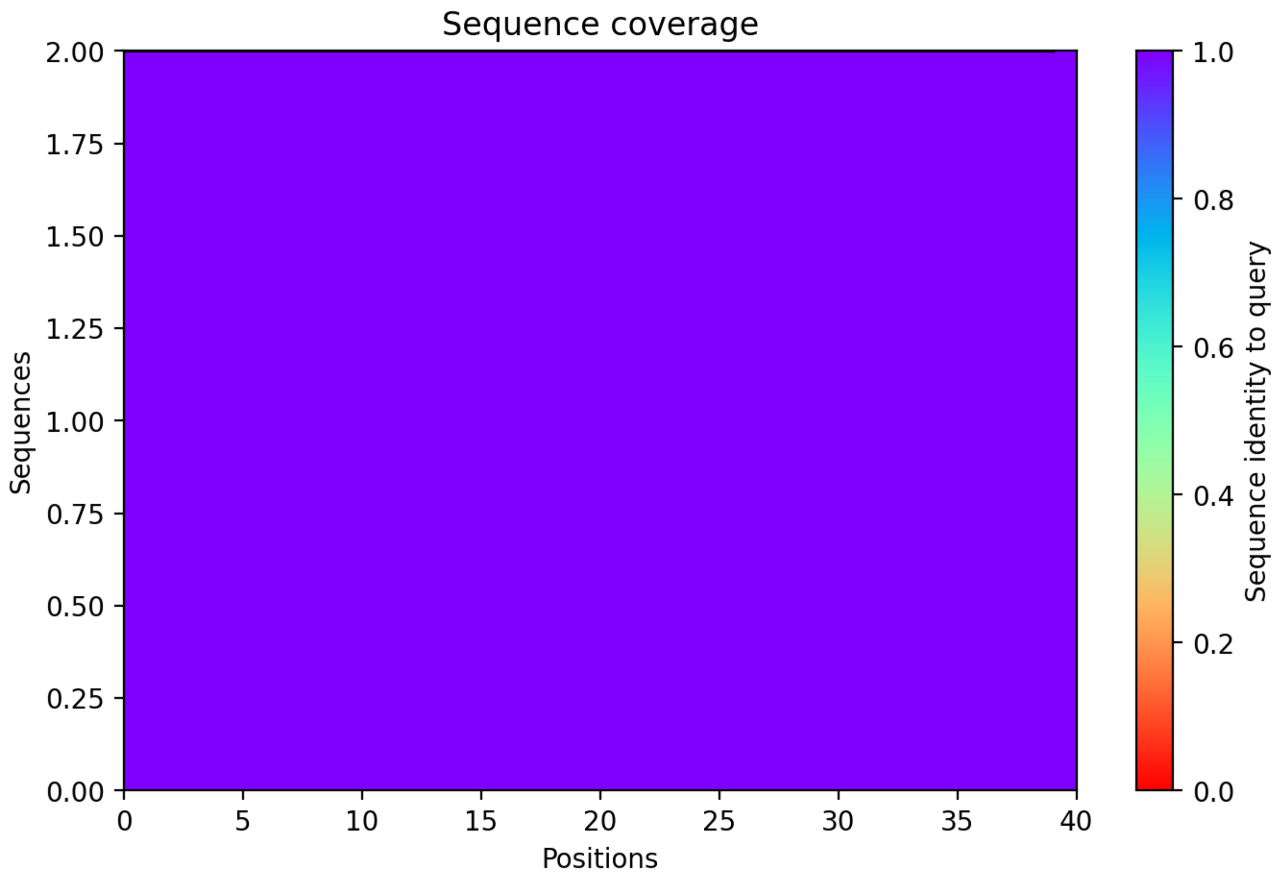

**Supplementary Dataset S7 Part 3**. A graphical summary on folding predictions for MS-supported chimeric peptide models (CPs) 101-156. The rest of the legend is the same as for Supplementary Dataset S7 Part 1.
