## Supplementary material for "Discovery of diverse chimeric peptides in a eukaryotic proteome sets the stage for the experimental proof of the mosaic translation hypothesis": Supplementary Dataset S7 Folding of chimeric protein models Part 4 Alpha-helices only.pdf

CP1: MtrunA17_Chr0c01g0489091

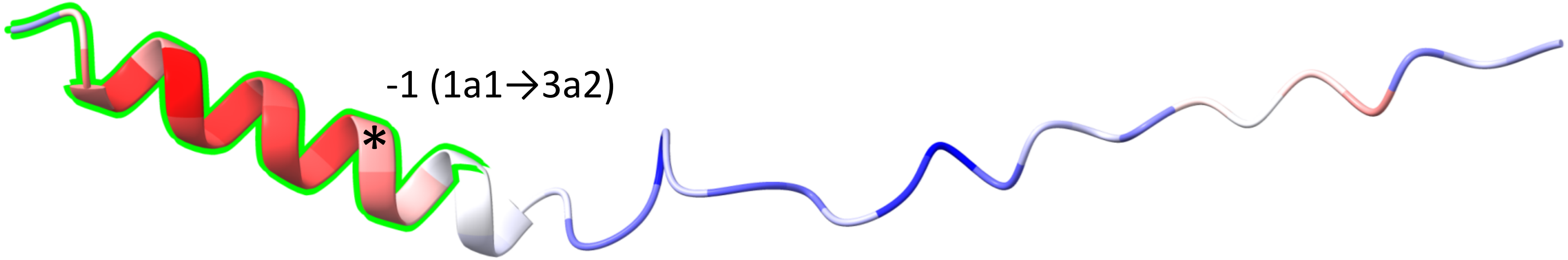


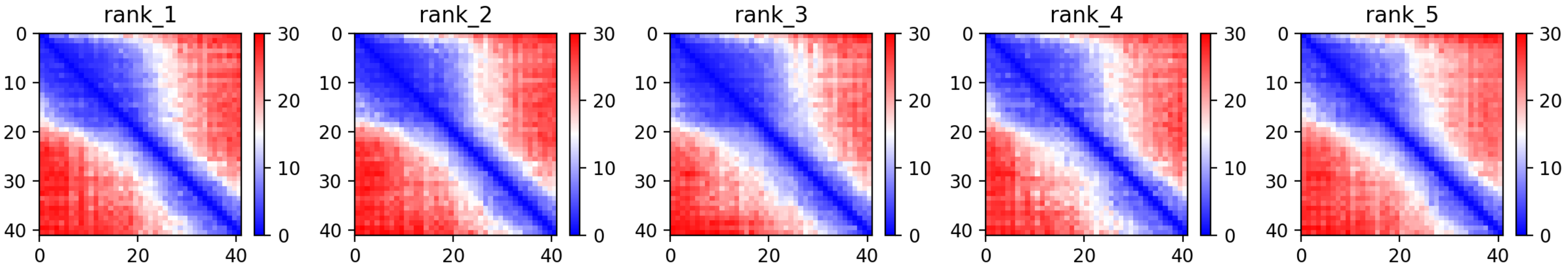

CP2: MtrunA17_Chr0c28g0493951

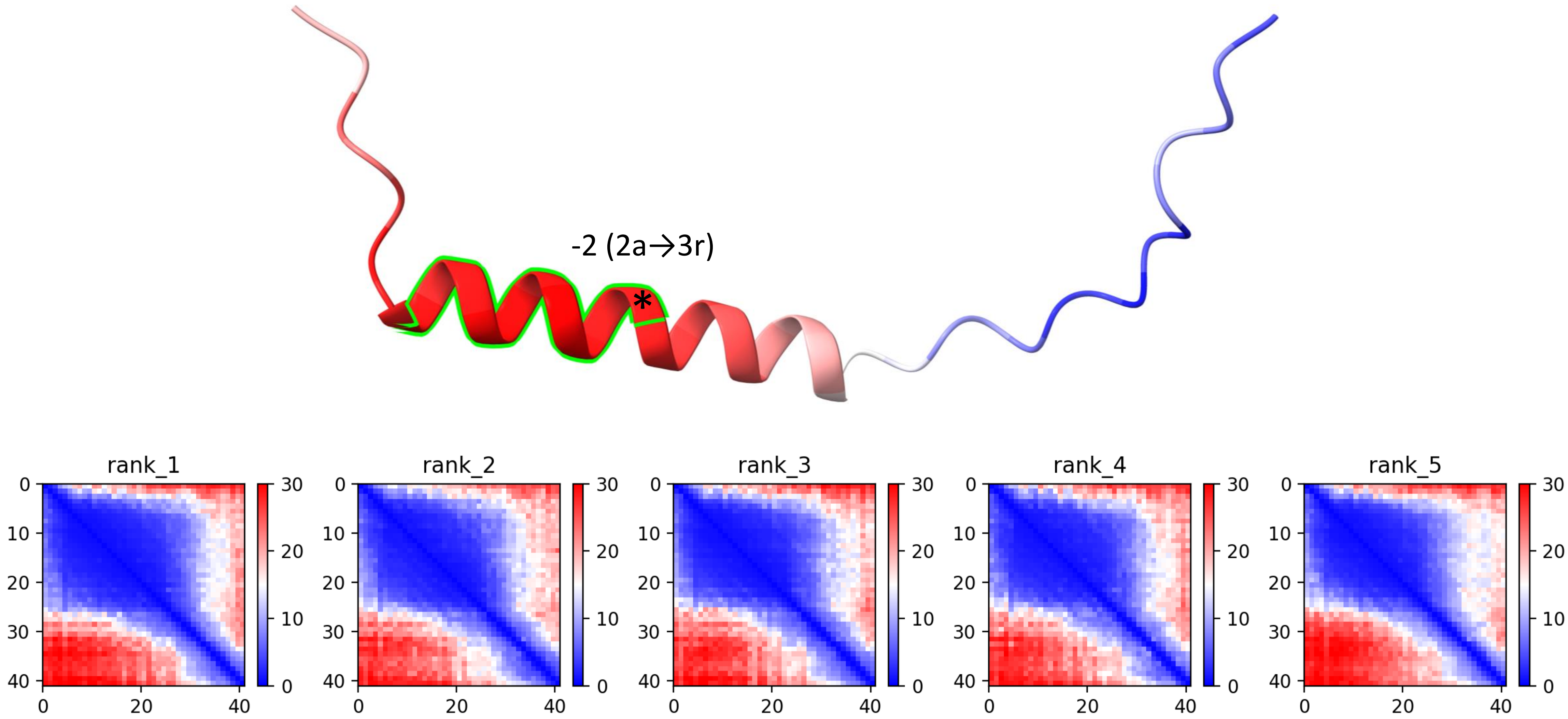

CP3: MtrunA17_Chr1g0148371

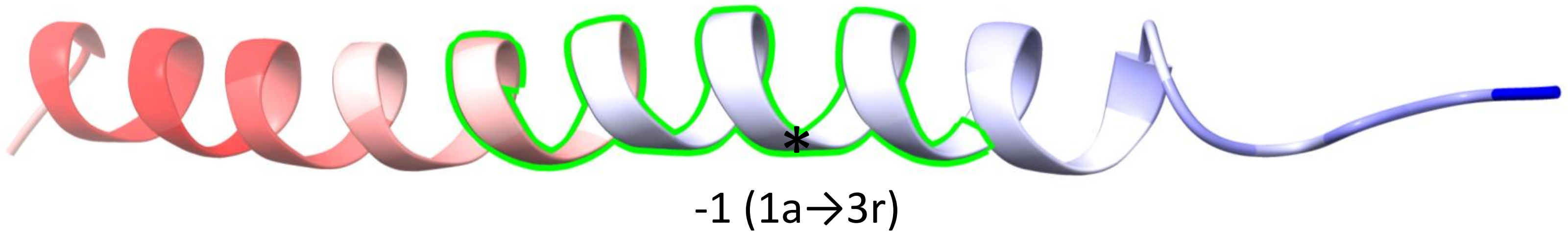


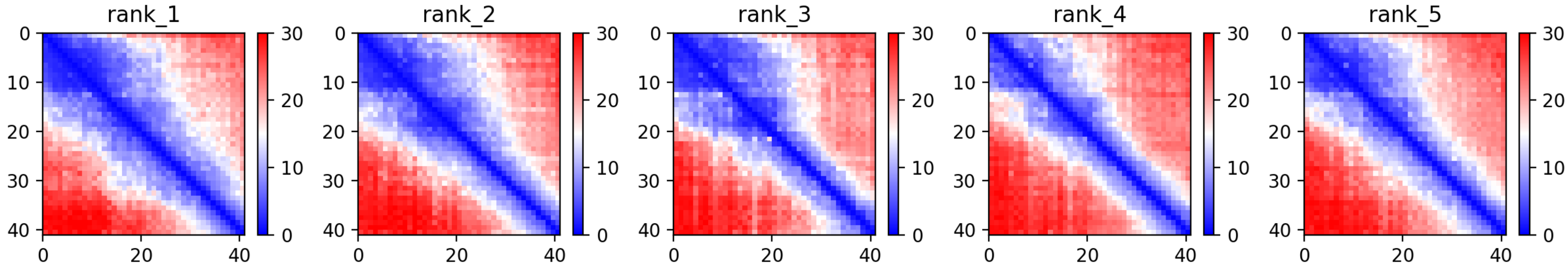

CP4: MtrunA17_Chr1g0149991

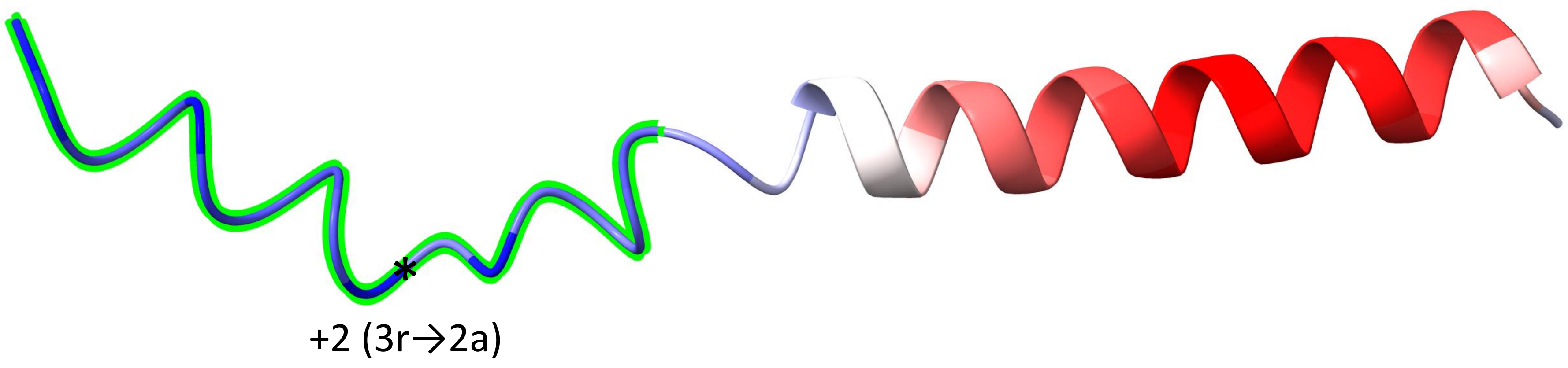


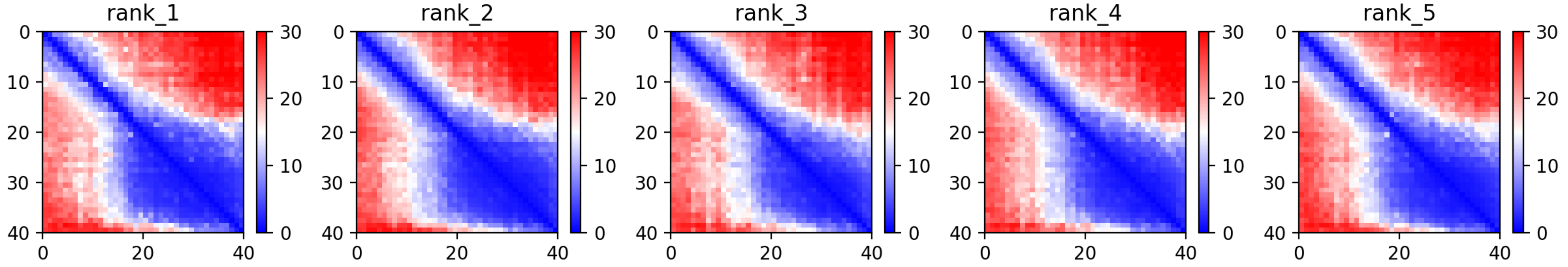

CP7: MtrunA17_Chr1g0153001

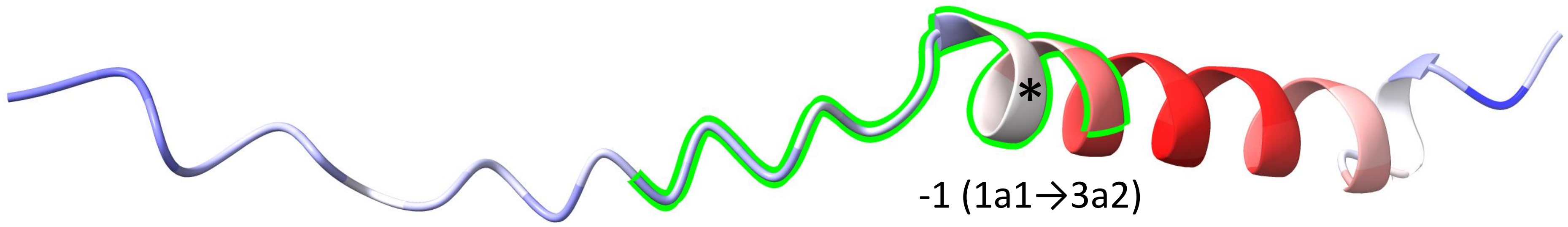


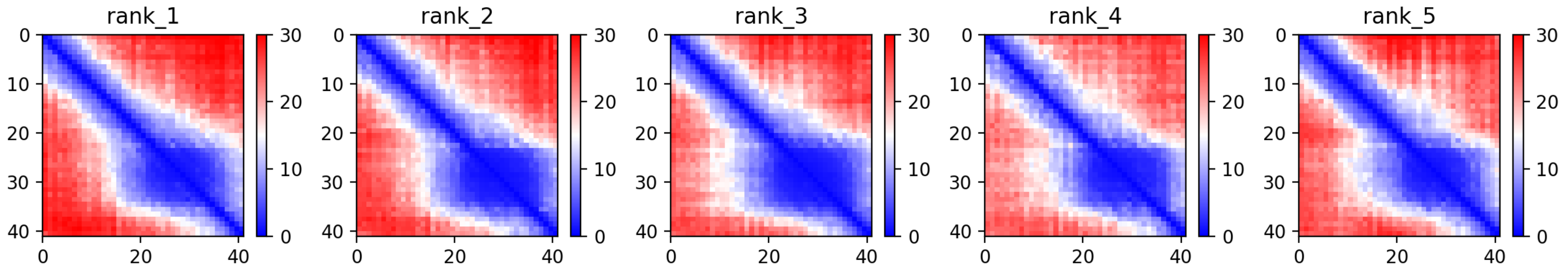

CP8: MtrunA17_Chr1g0155251

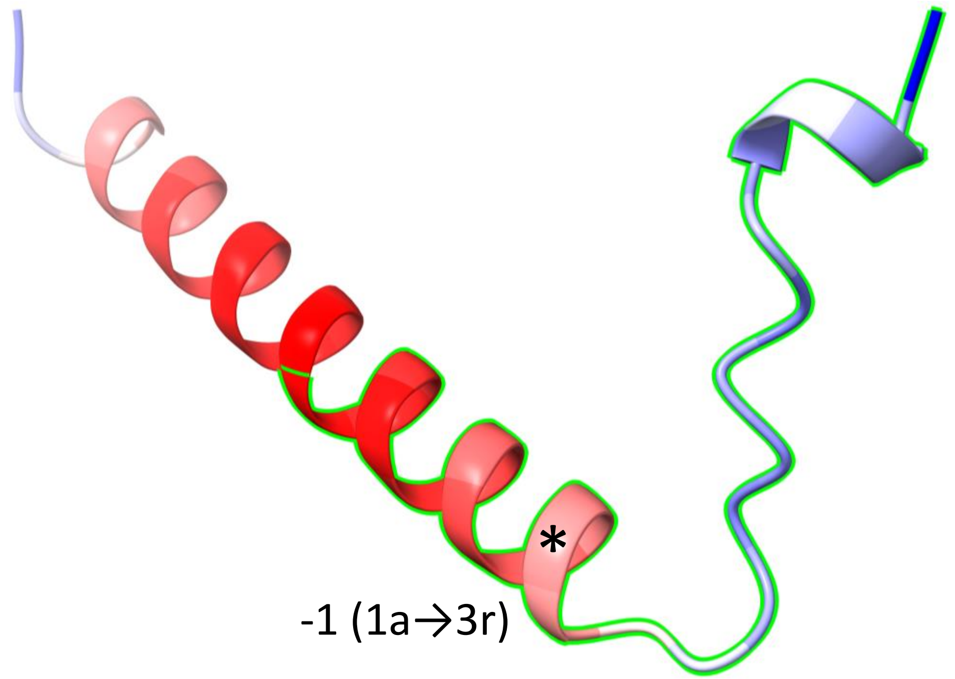


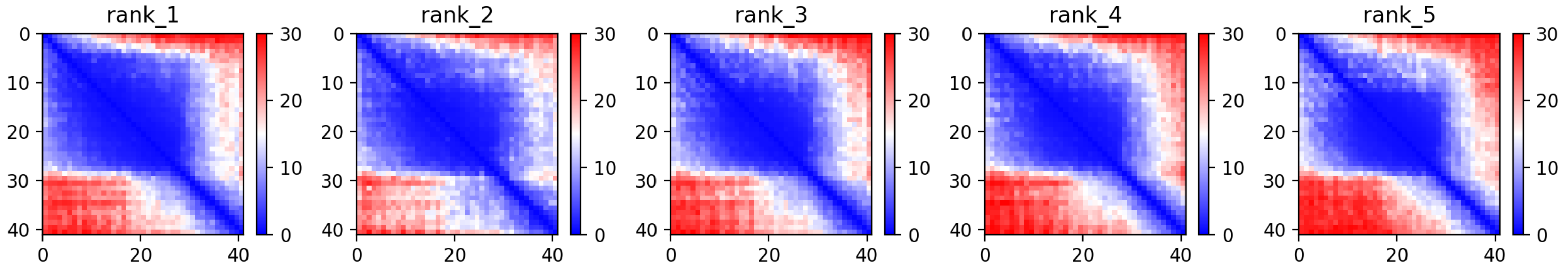

CP9: MtrunA17_Chr1g0158341

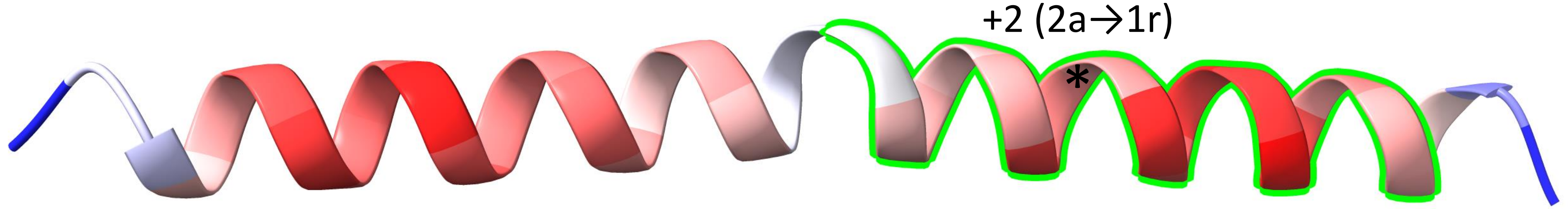


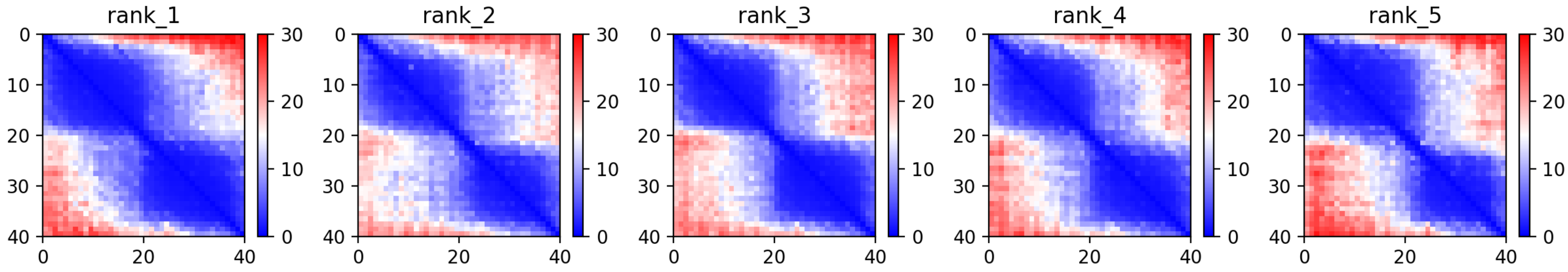

CP12: MtrunA17_Chr1g0178361

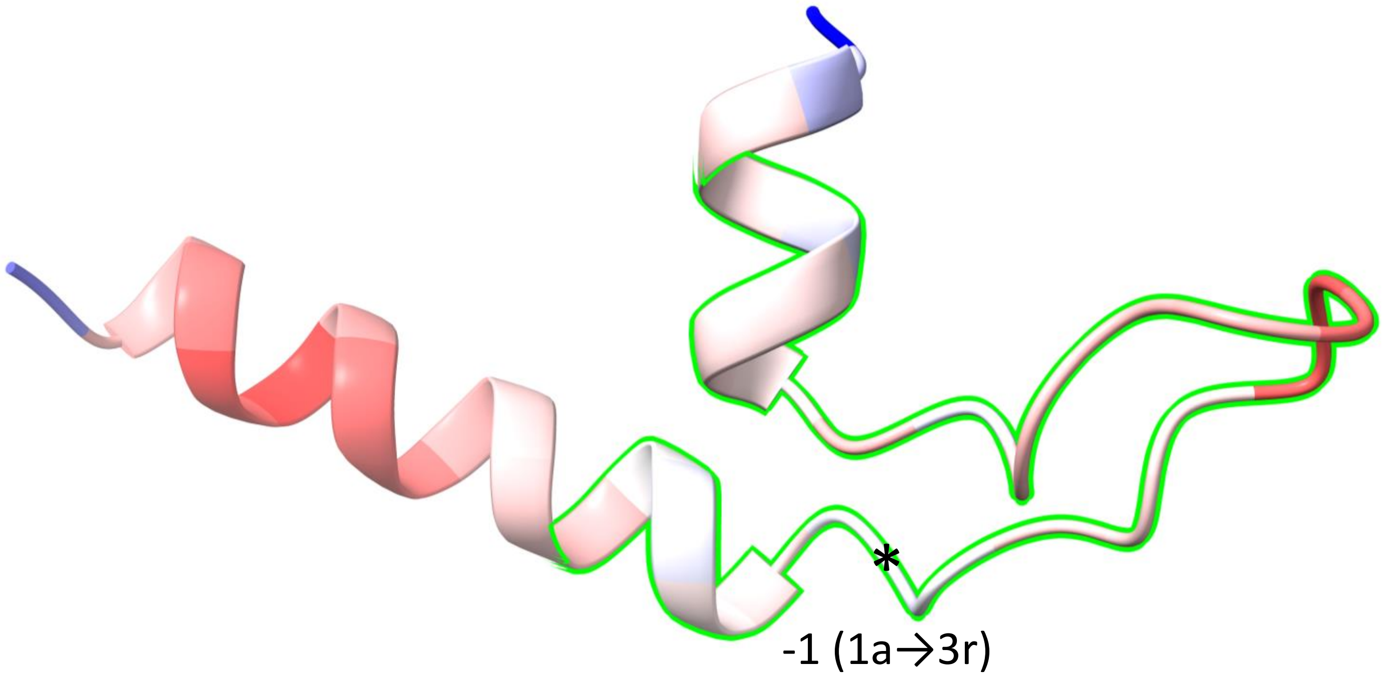


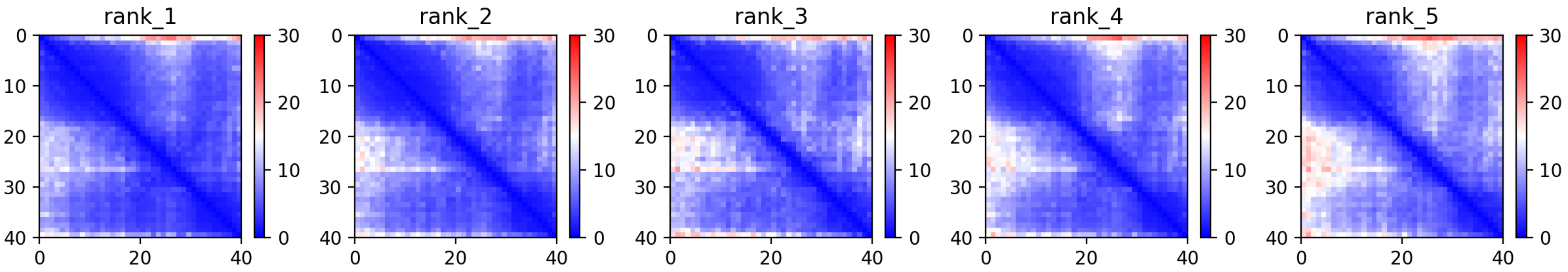

CP15: MtrunA17_Chr1g0183001

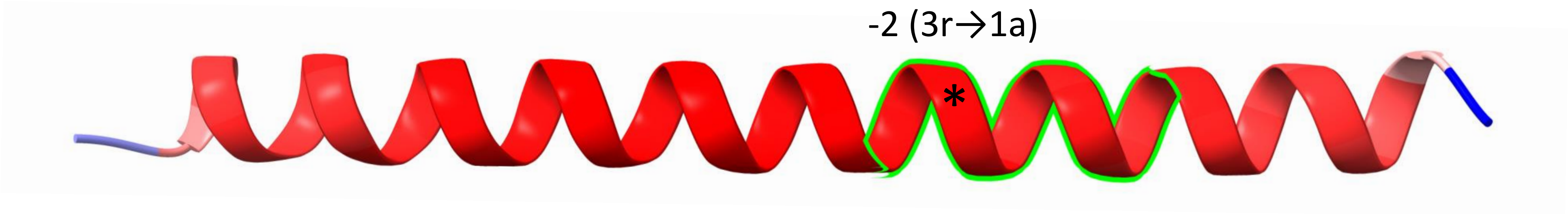


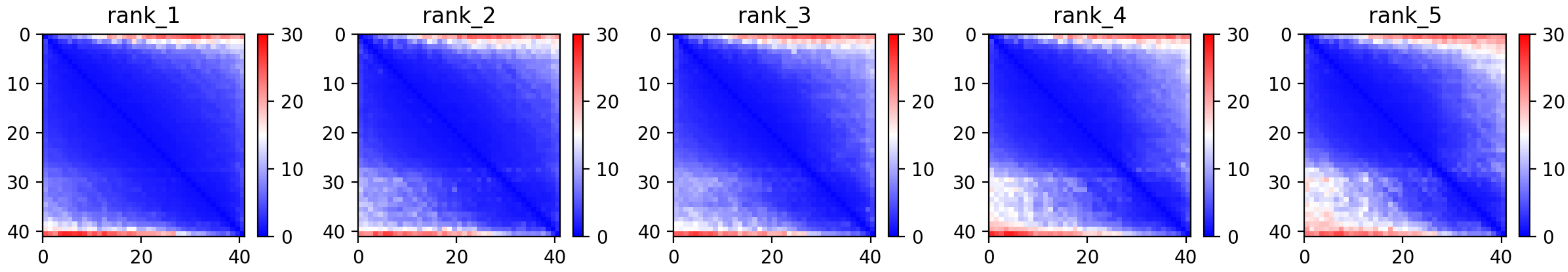

CP22: MtrunA17_Chr1g0198091

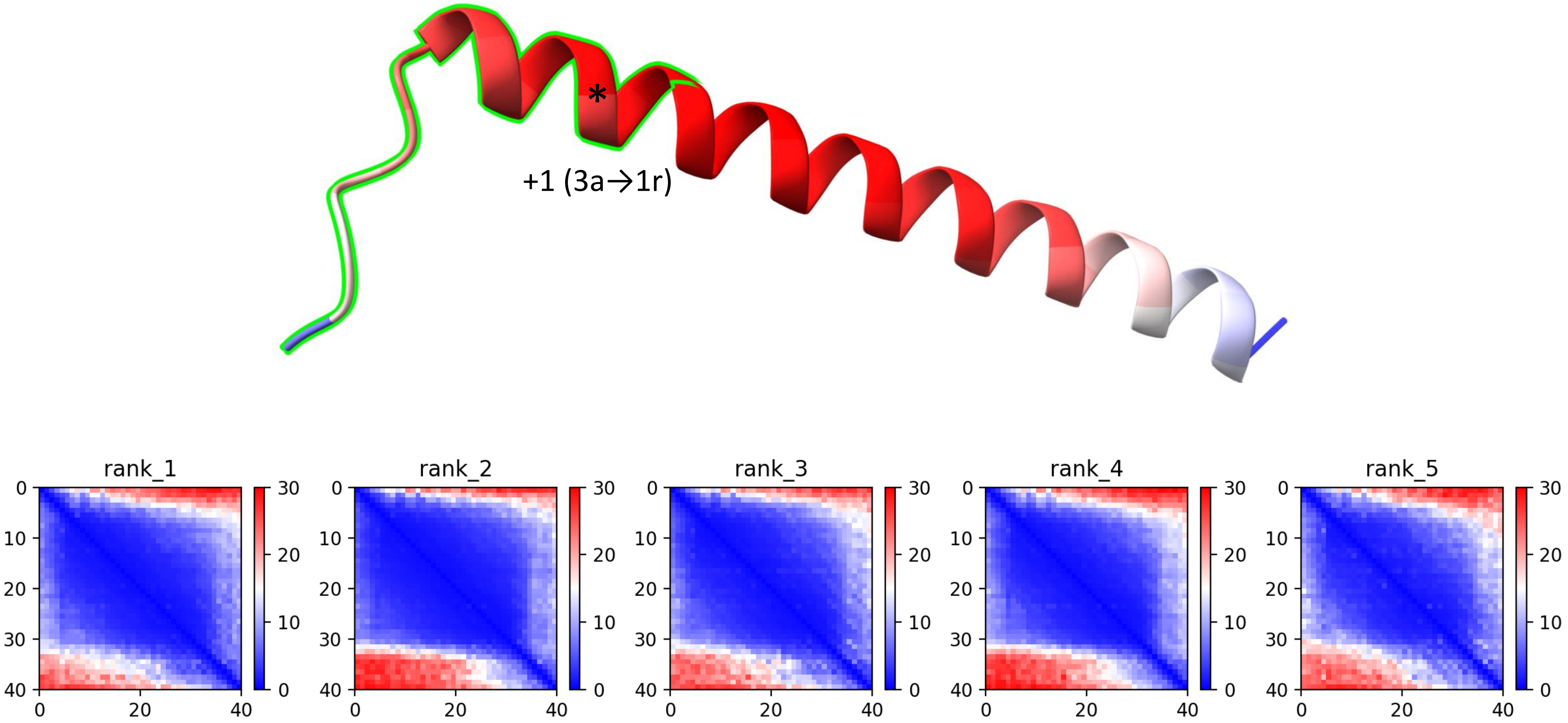

CP25: MtrunA17_Chr1g0202001

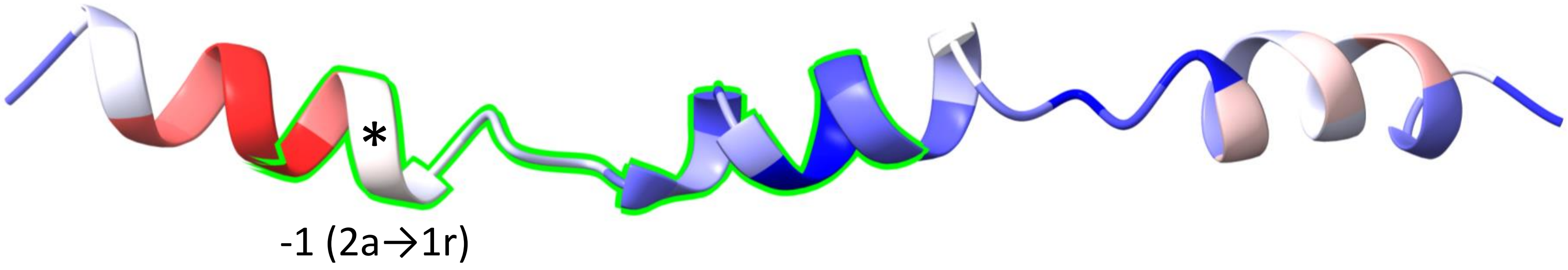


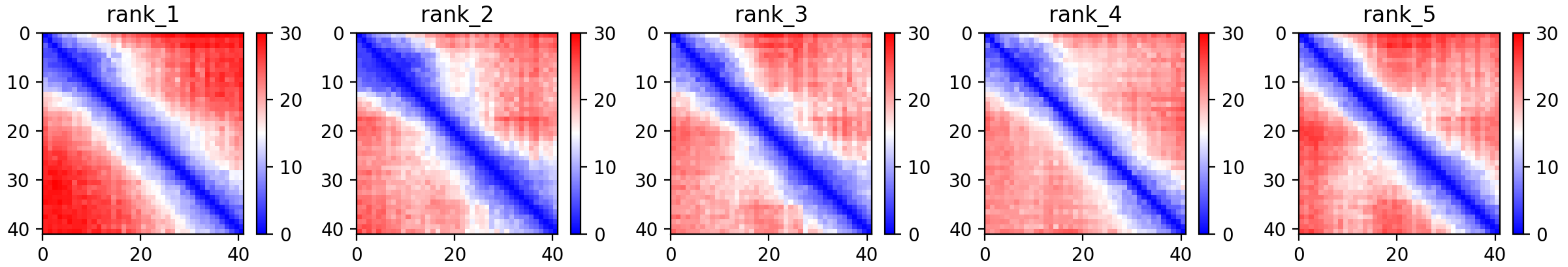

CP27: MtrunA17_Chr1g0207811

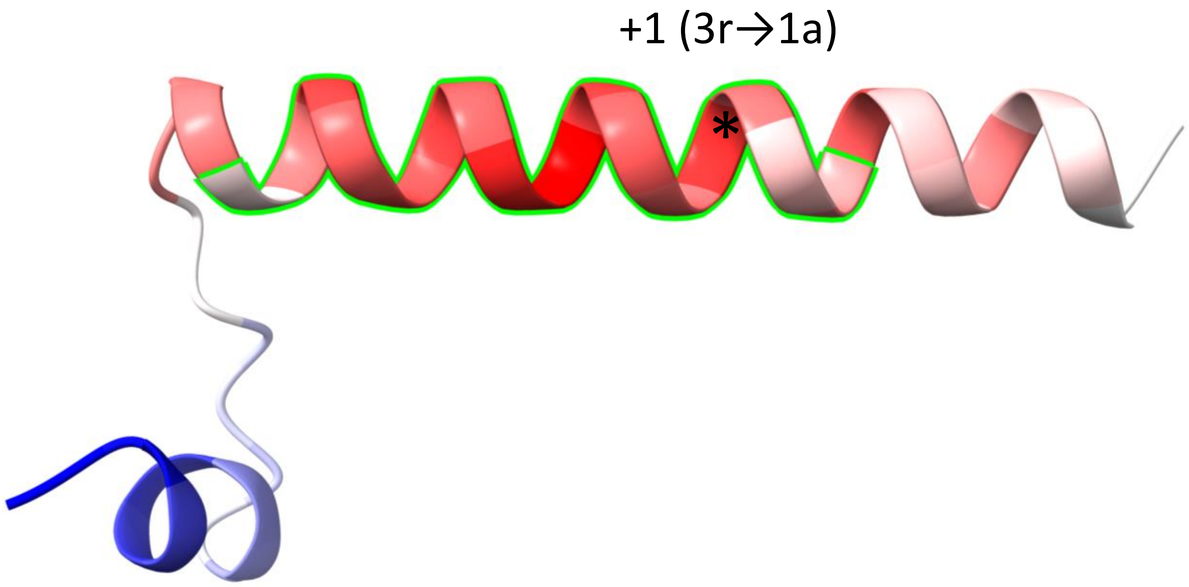


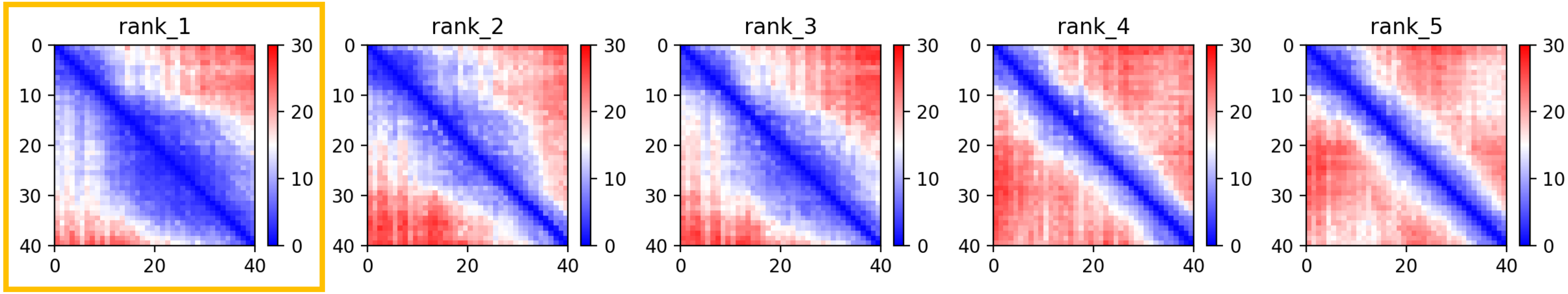

CP30: MtrunA17_Chr1g0210521

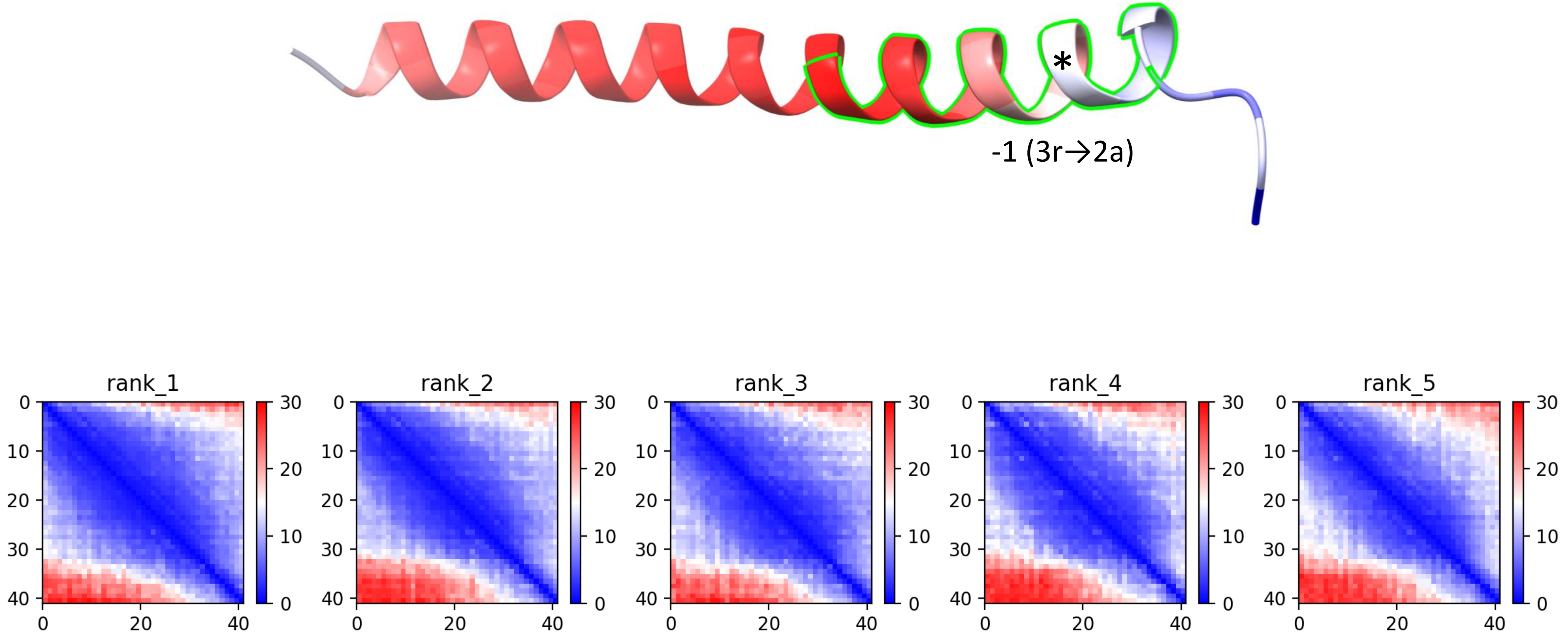

CP32: MtrunA17_Chr1g1004575

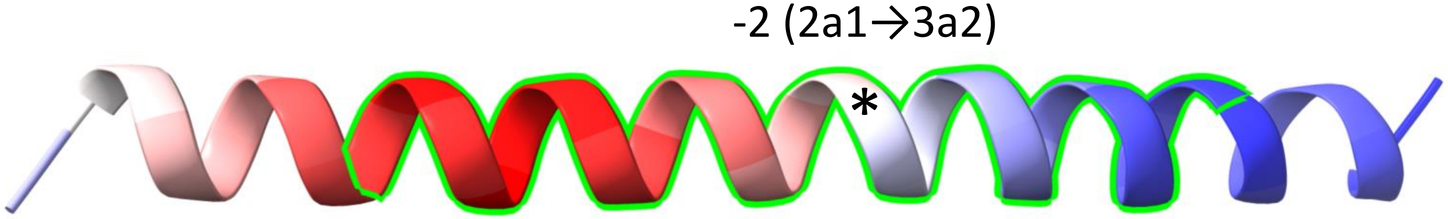


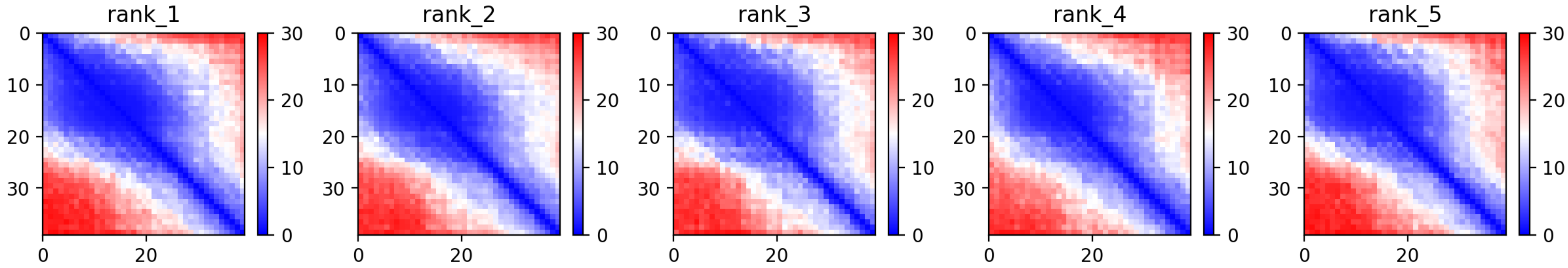

CP35: MtrunA17_Chr2g0292921

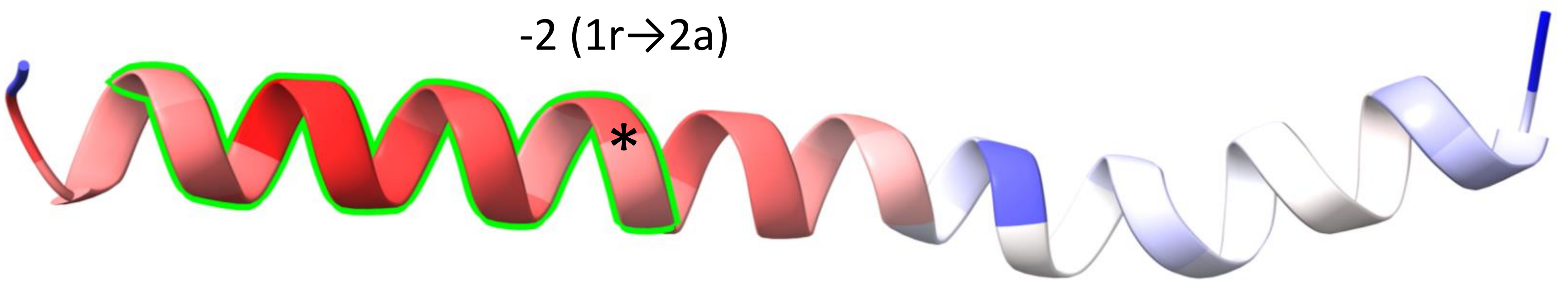


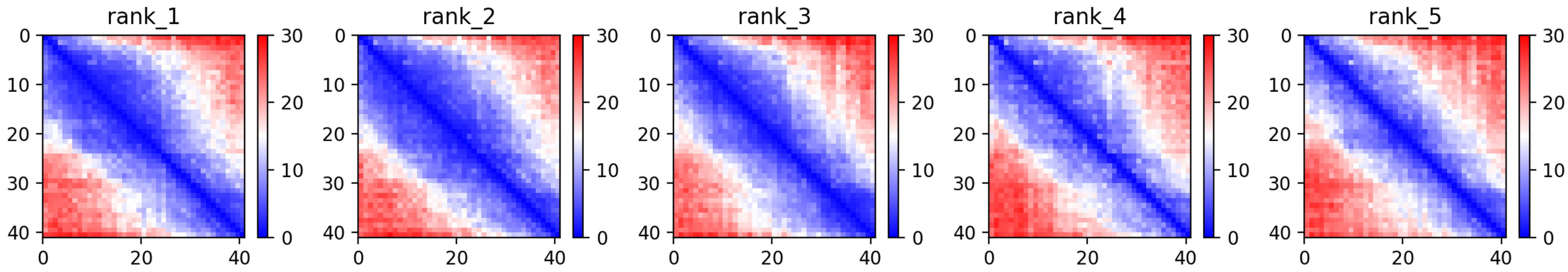

CP37: MtrunA17_Chr2g0299561

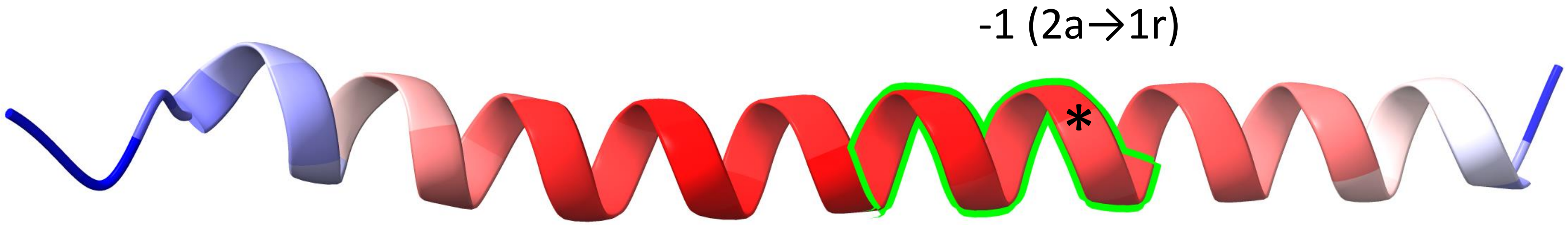


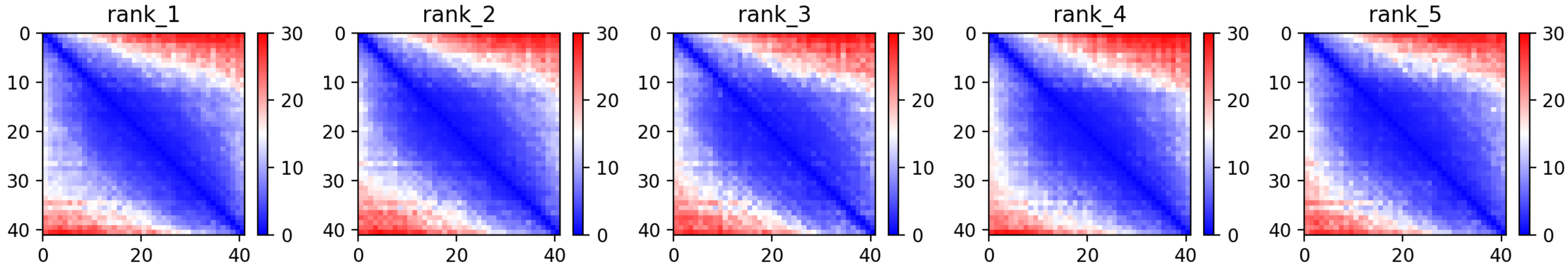

CP43: MtrunA17_Chr2g0316291

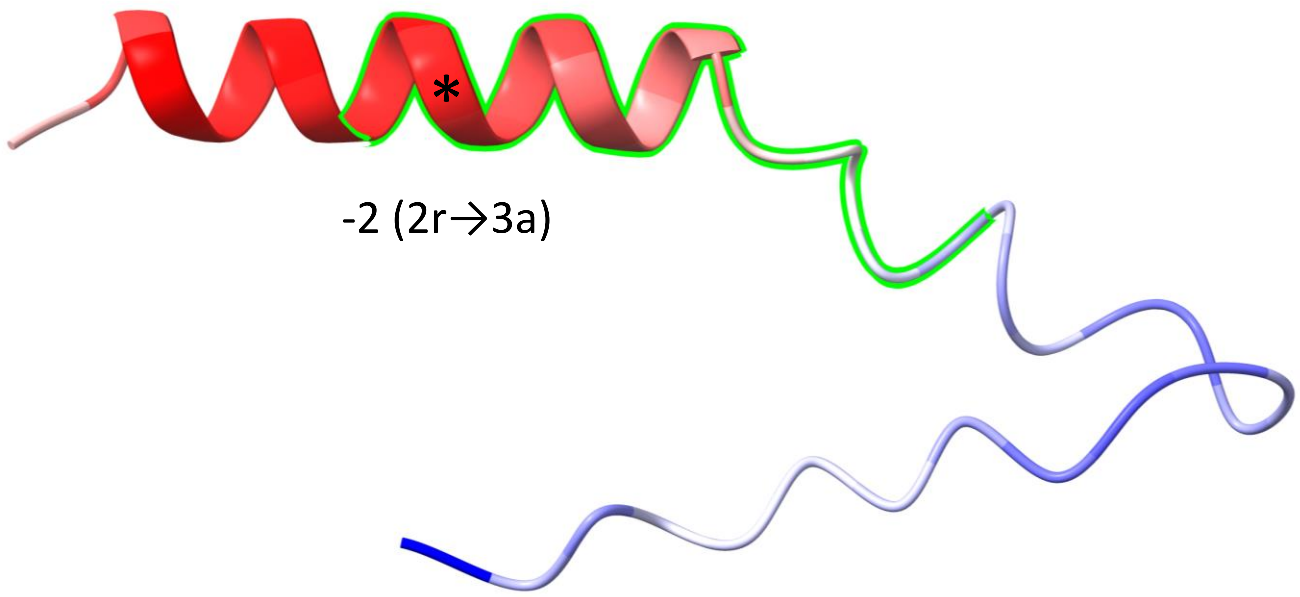


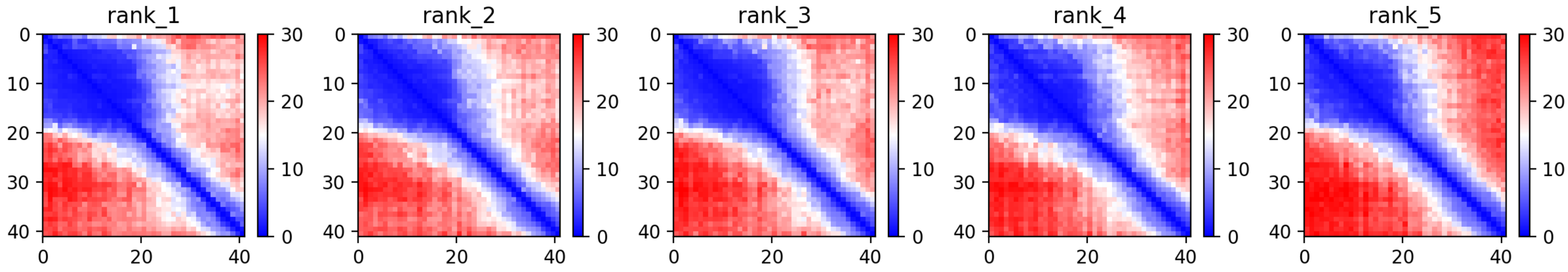

CP44: MtrunA17_Chr2g0326801

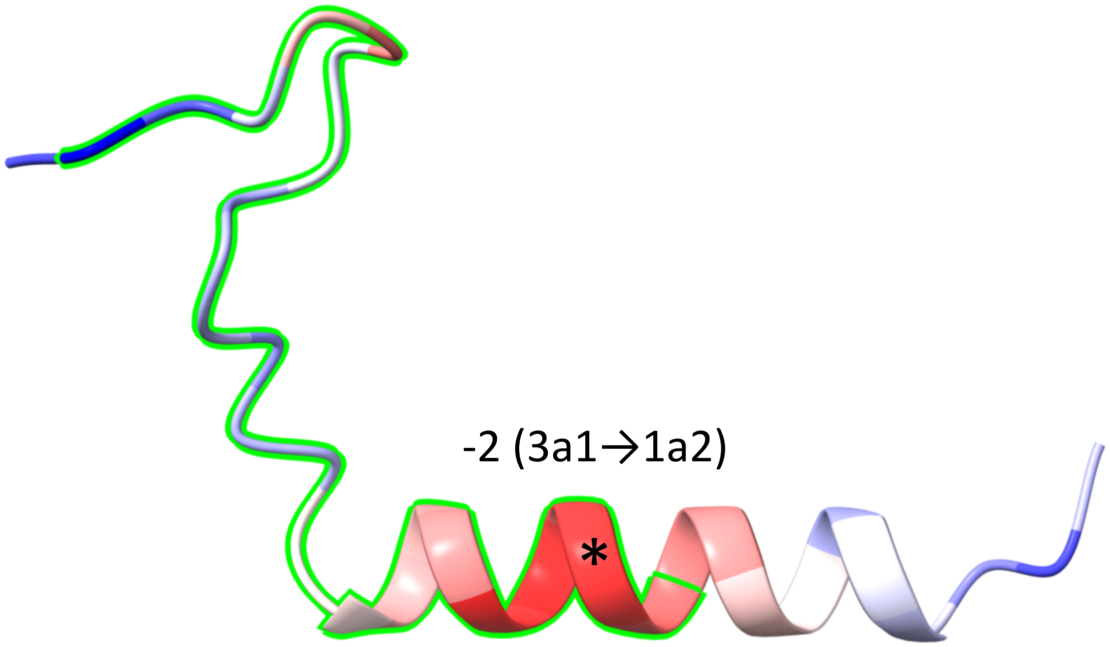


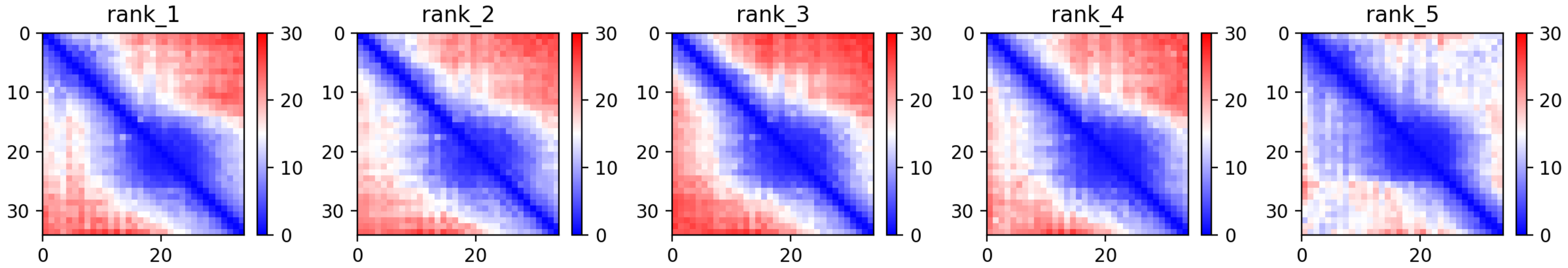

CP47: MtrunA17_Chr3g0079521

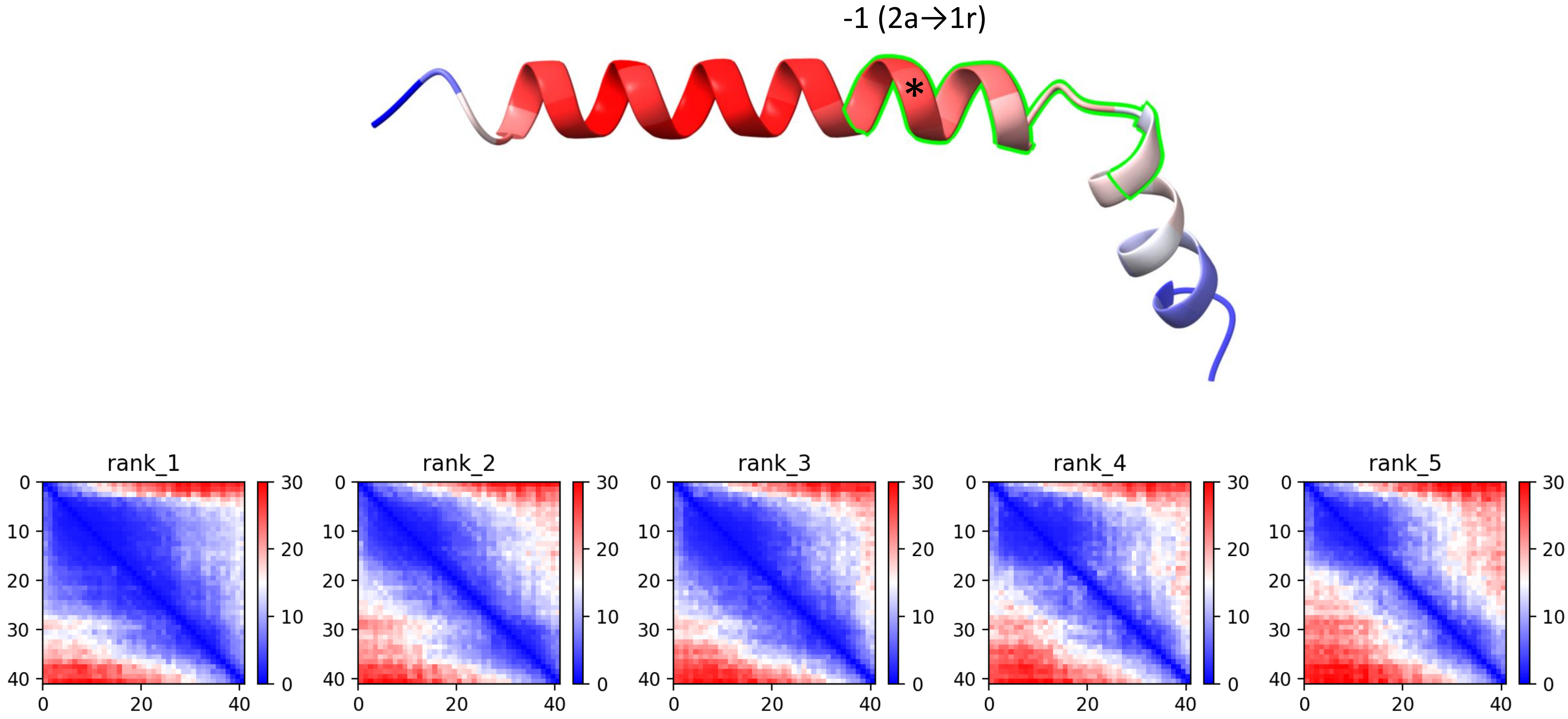

CP48: MtrunA17_Chr3g0083801

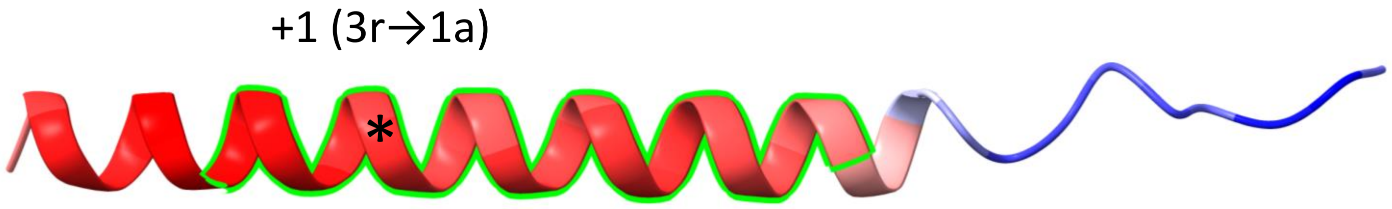


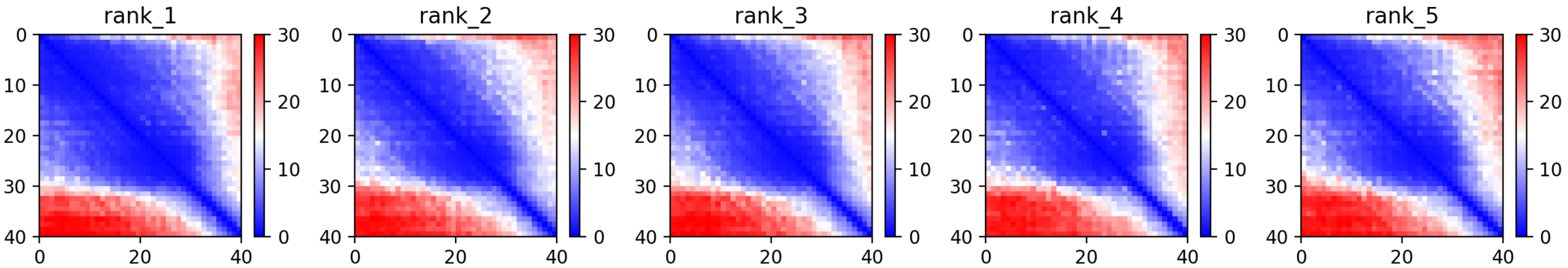

CP49: MtrunA17_Chr3g0091141

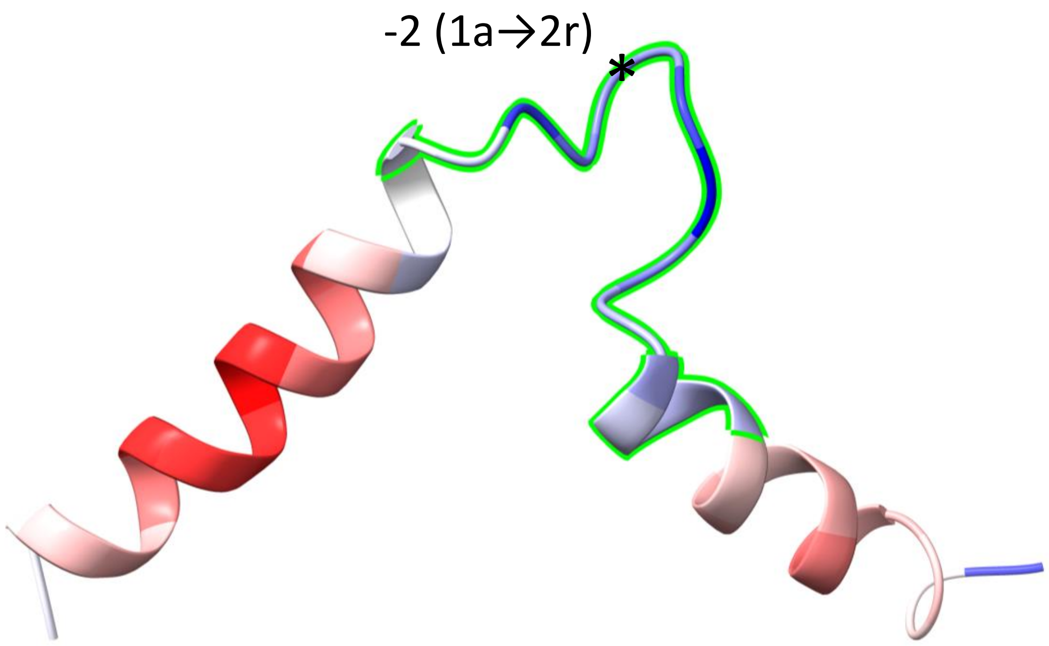


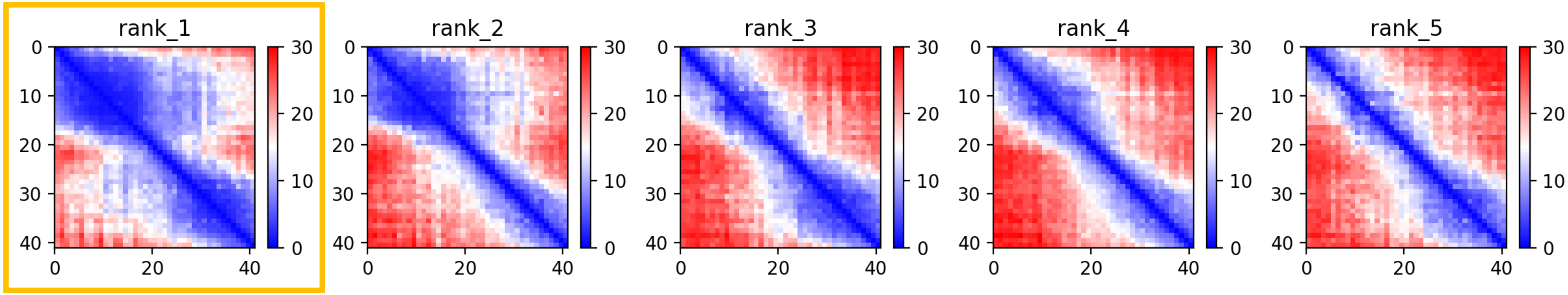

CP53: MtrunA17_Chr3g0102171

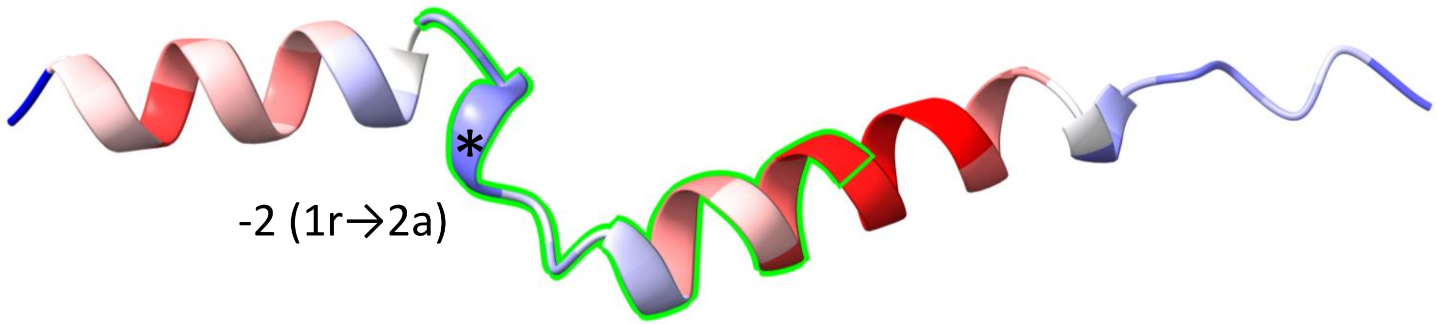


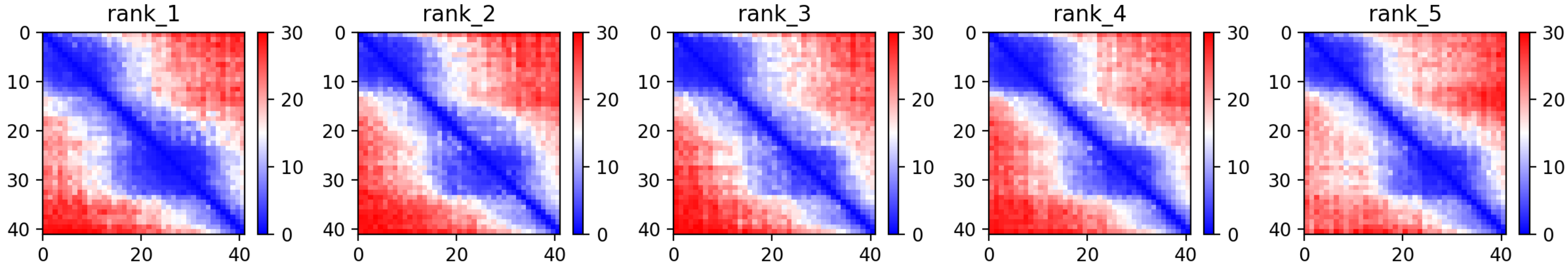

CP58: MtrunA17_Chr3g0127321

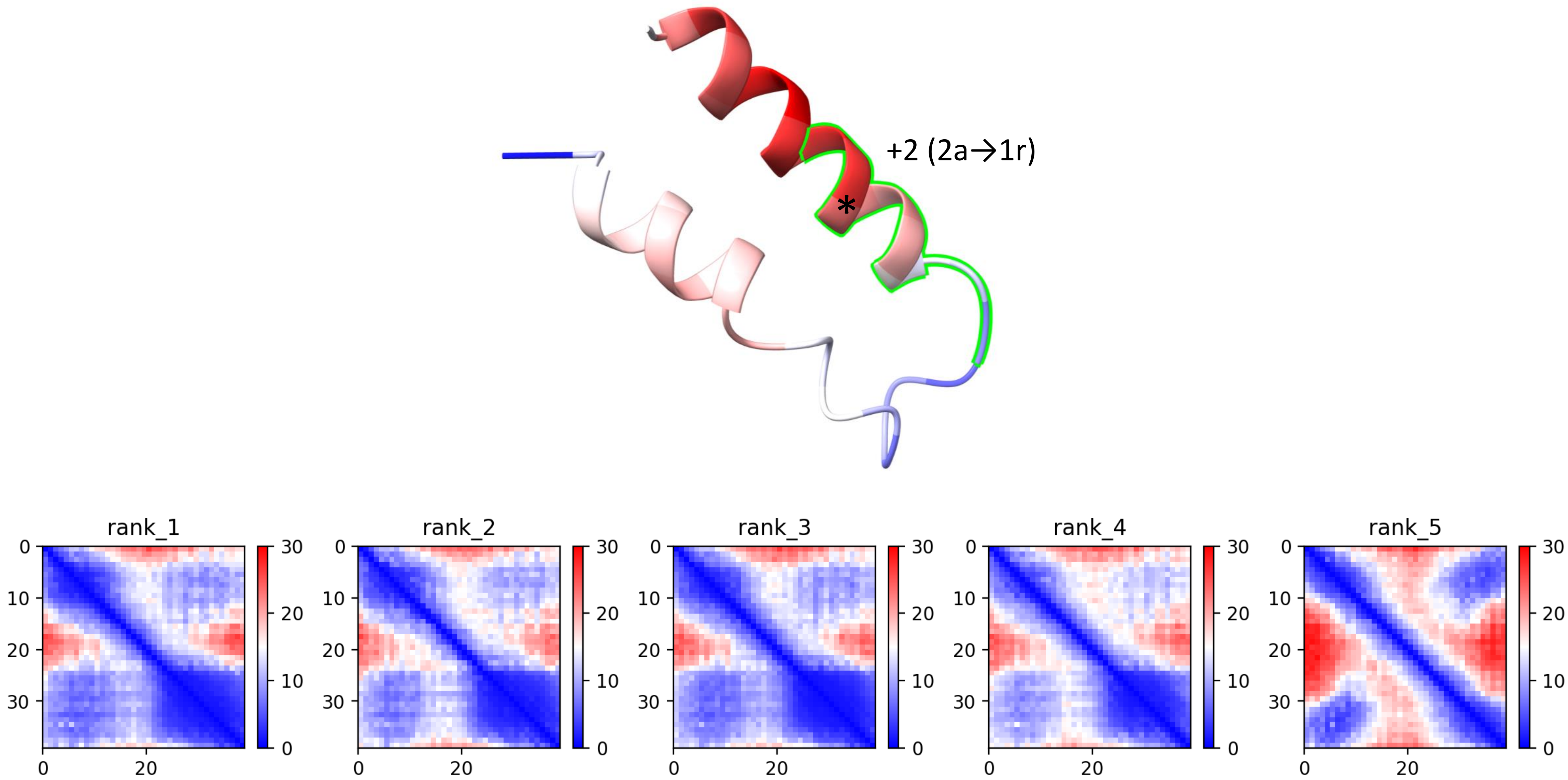

CP59: MtrunA17_Chr3g0130671

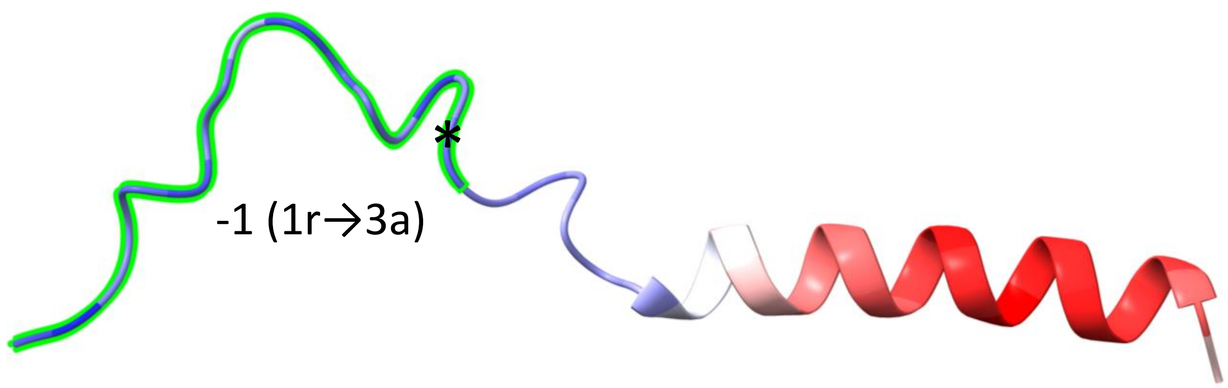


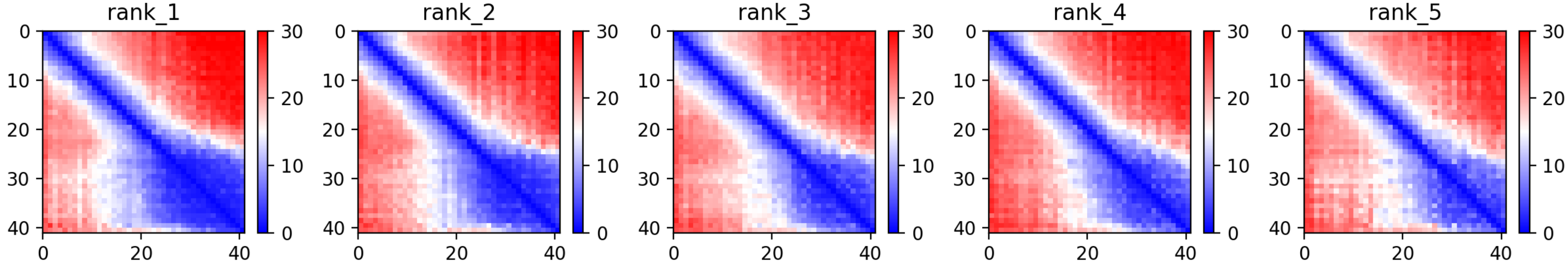

CP61: MtrunA17_Chr3g0137391

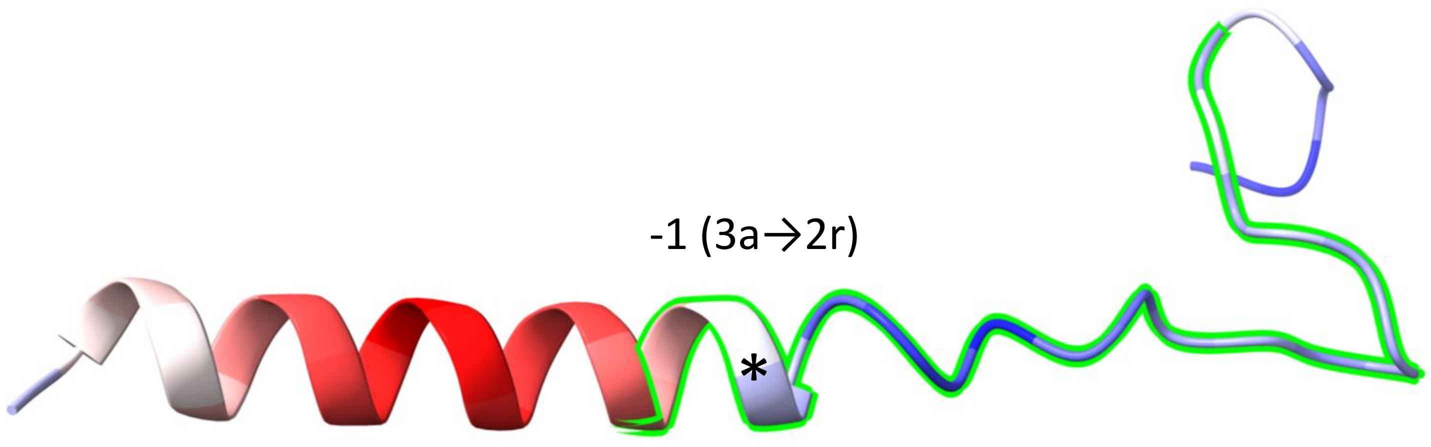


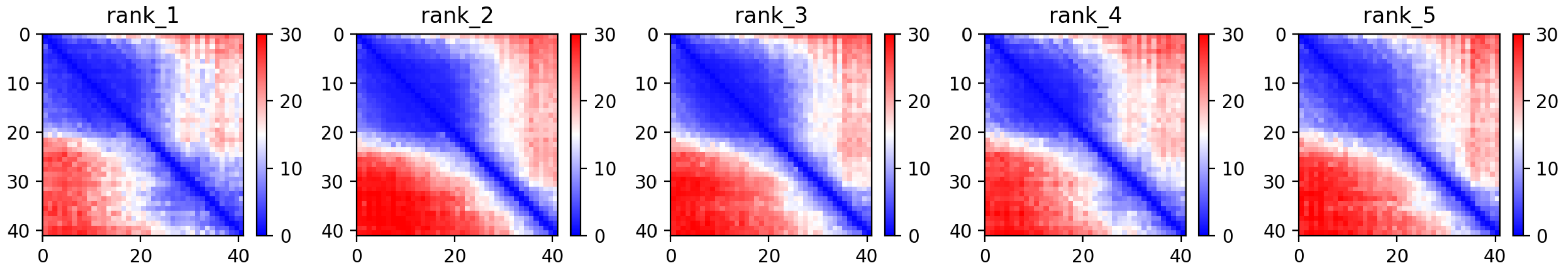

CP62: MtrunA17_Chr3g0141901

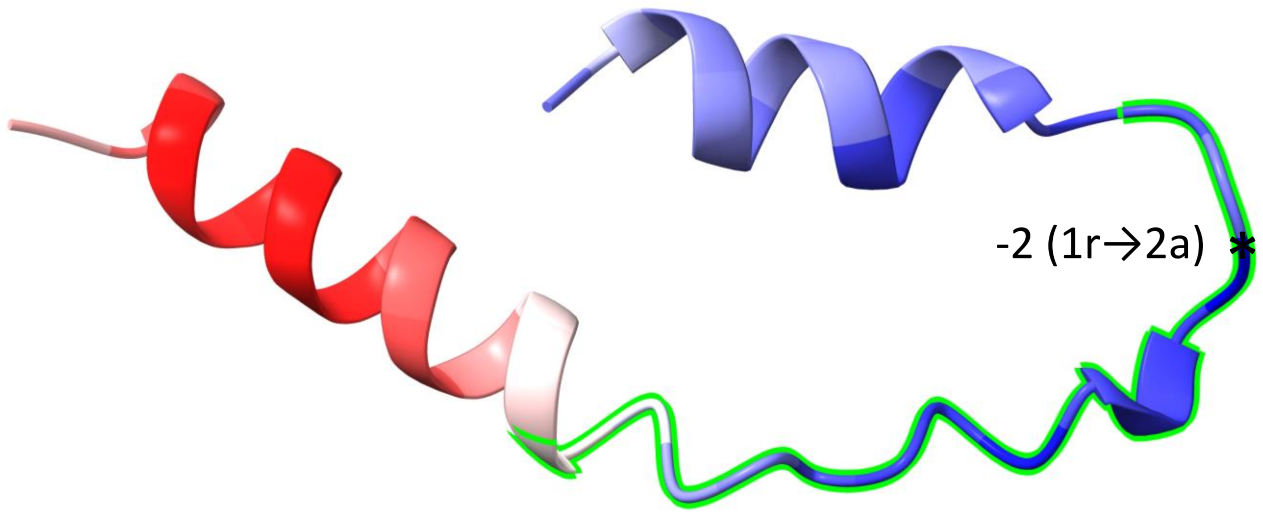


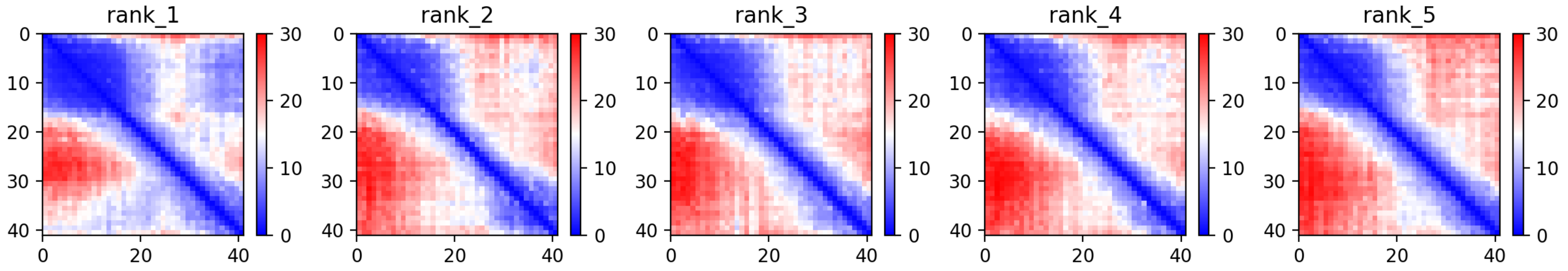

CP63: MtrunA17_Chr3g0144151

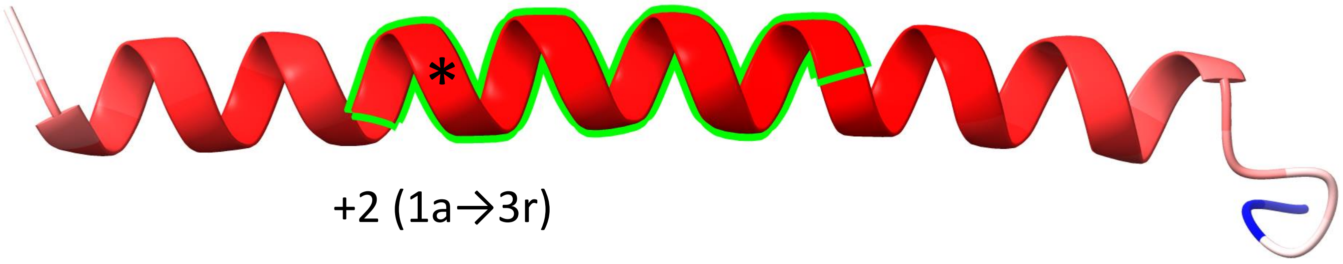


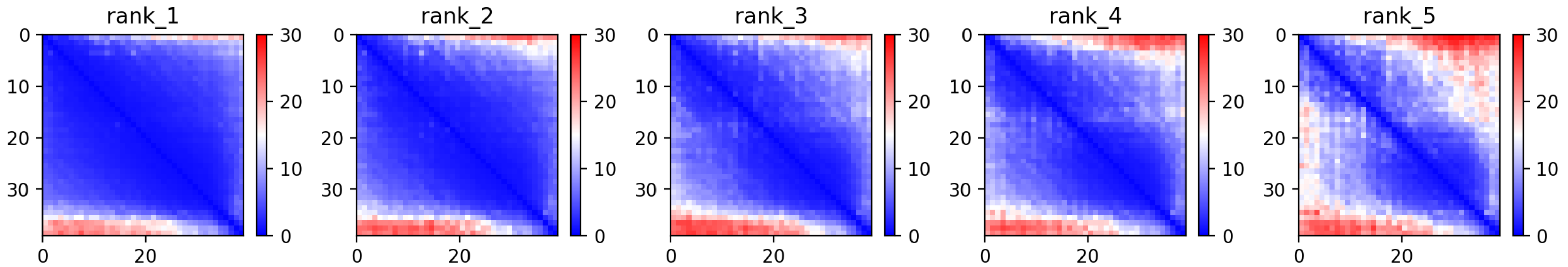

### CP64: MtrunA17_Chr3g0144151

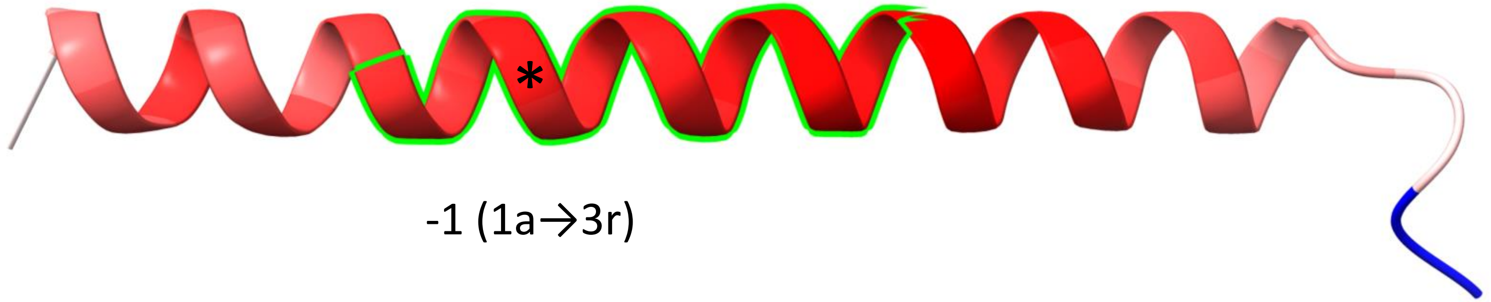


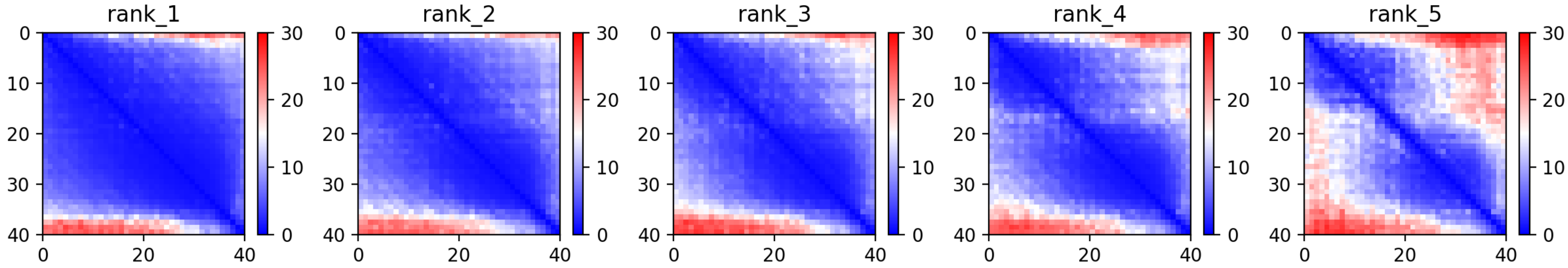

CP65: MtrunA17_Chr3g0144151

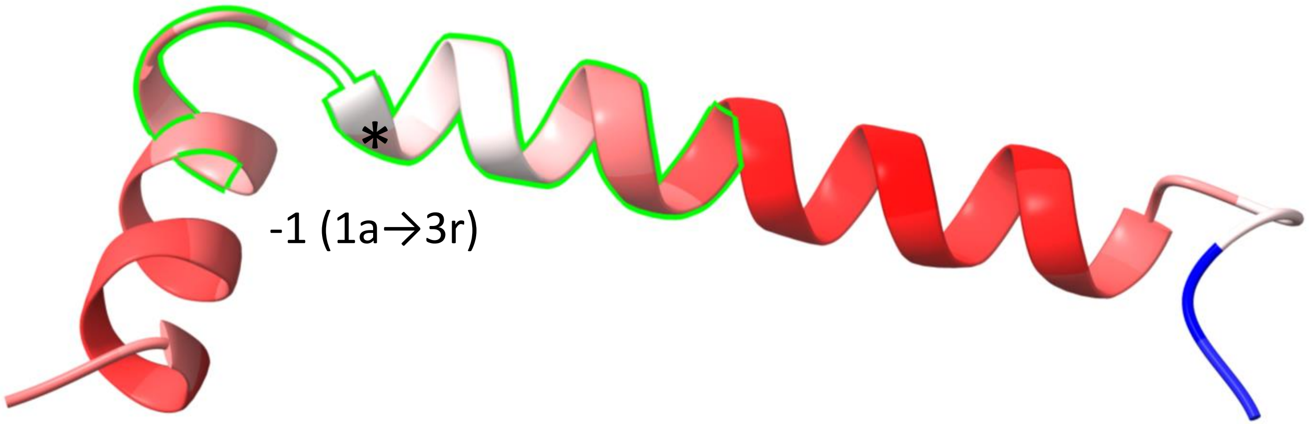


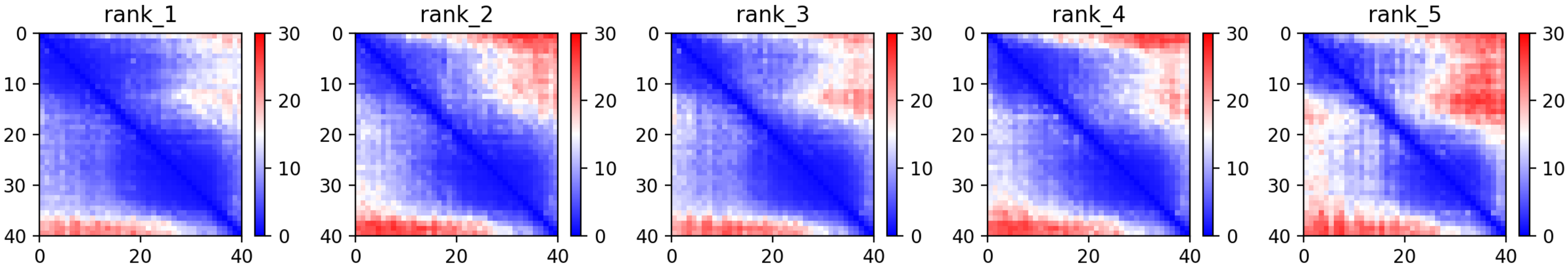

CP66: MtrunA17_Chr3g1011650

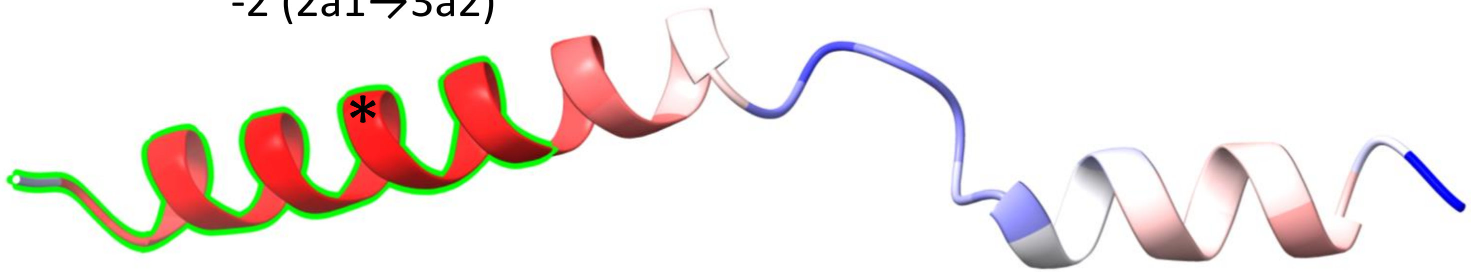


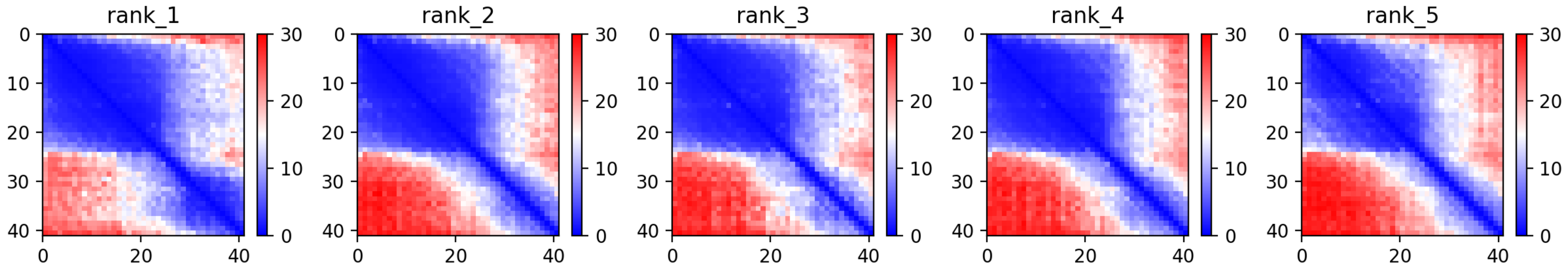

### CP67: MtrunA17_Chr4g0000131

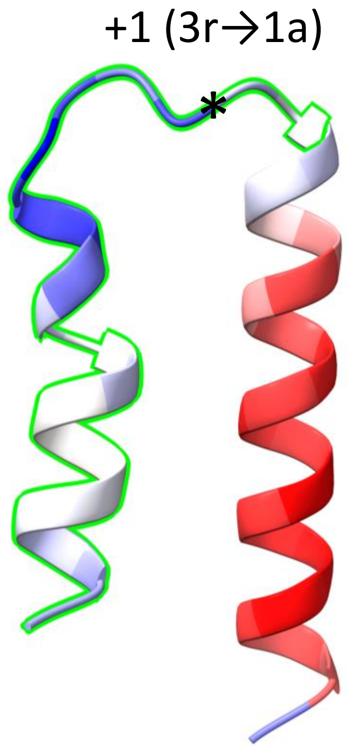


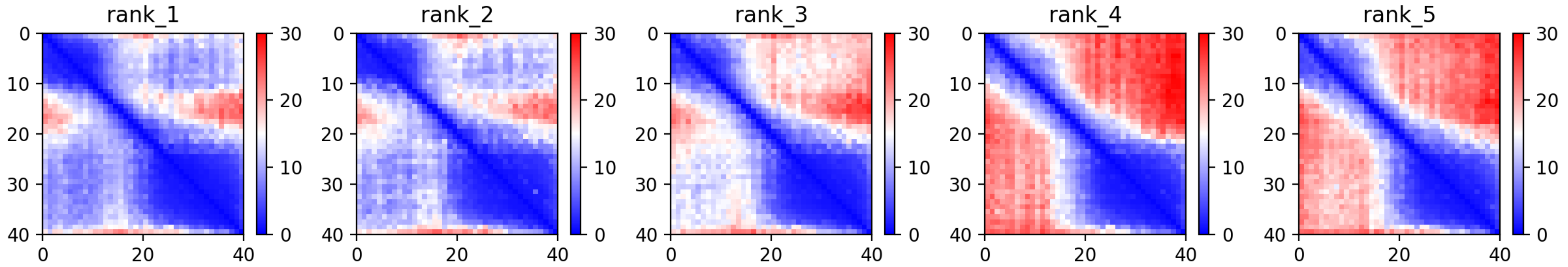

CP68: MtrunA17_Chr4g0000891

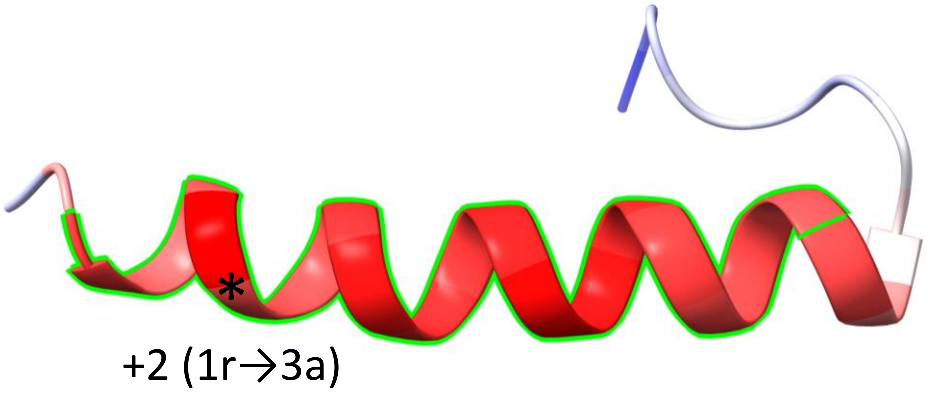


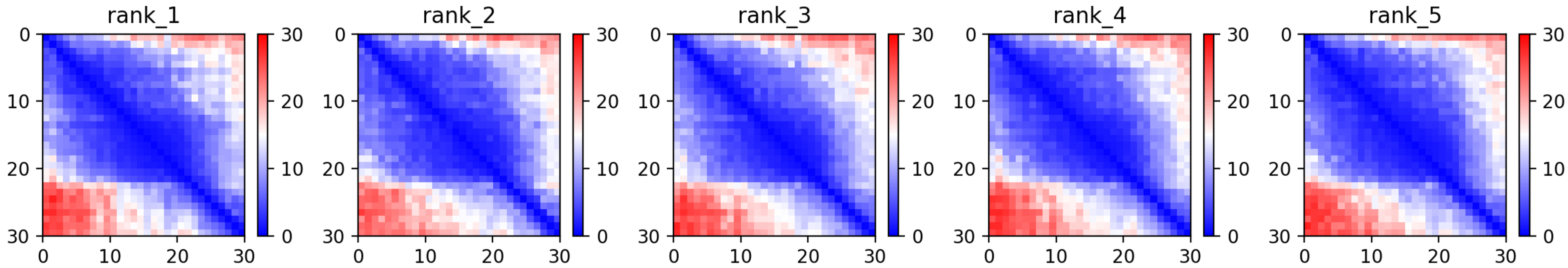

CP70: MtrunA17_Chr4g0014251

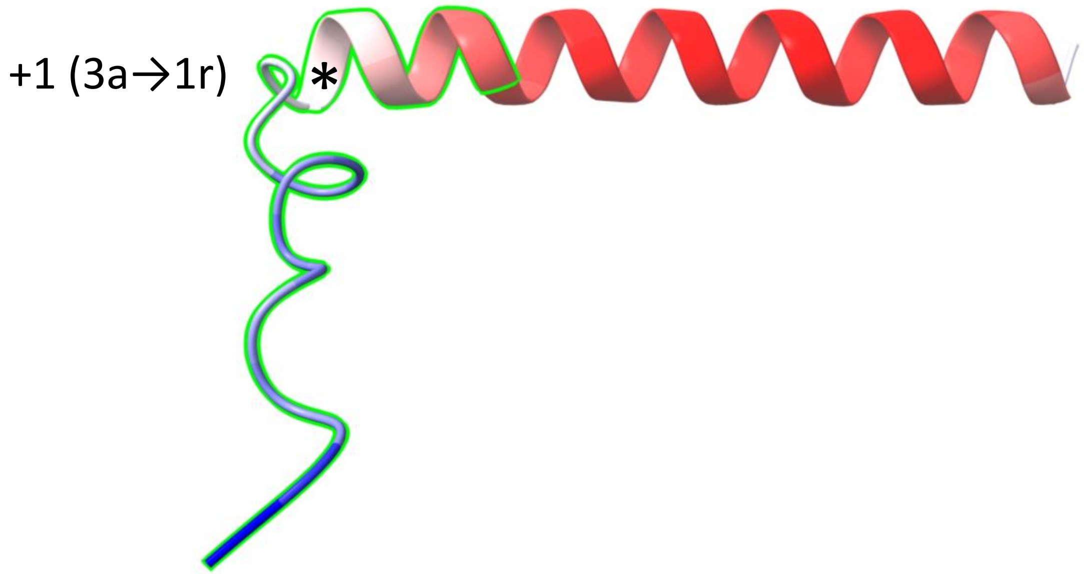


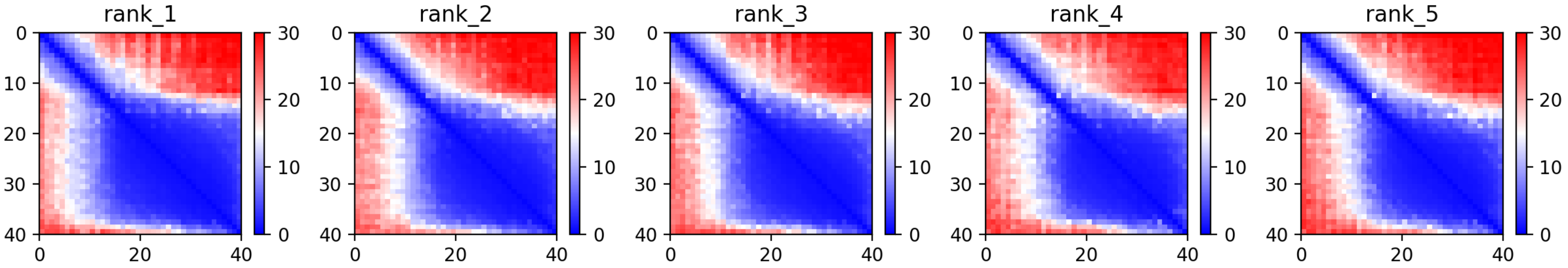

CP72: MtrunA17_Chr4g0023791

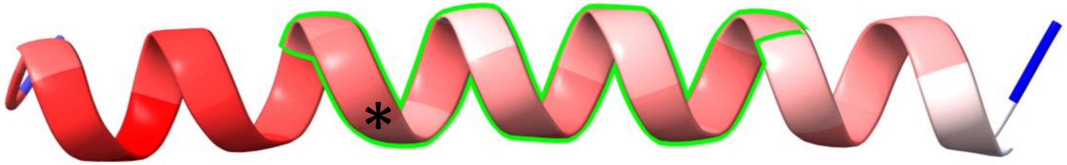


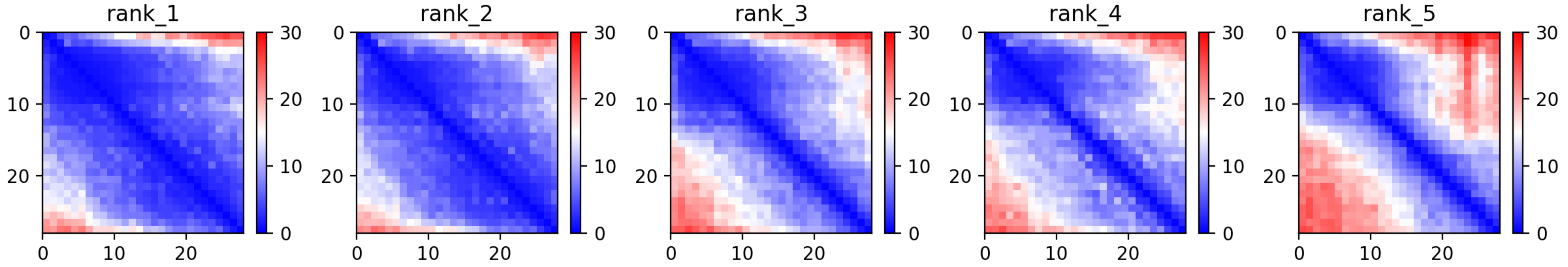

CP73: MtrunA17_Chr4g0034271

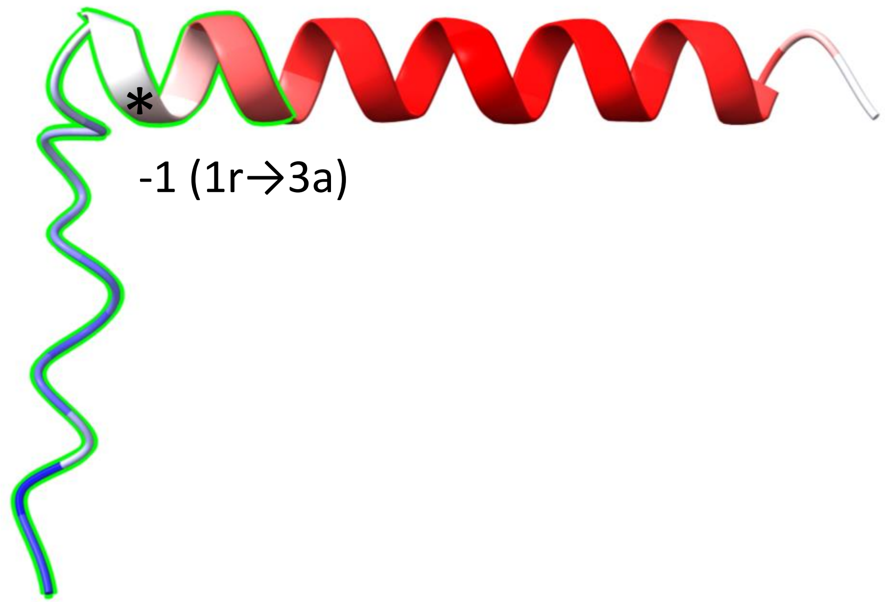


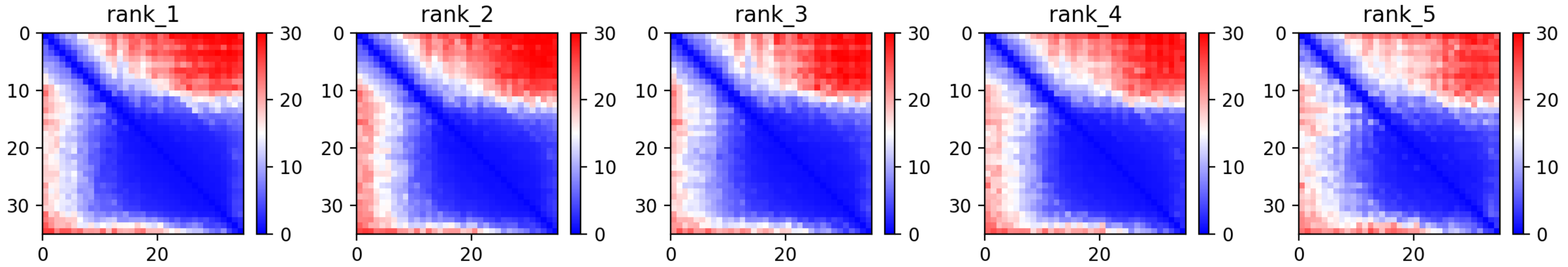

CP74: MtrunA17_Chr4g0034491

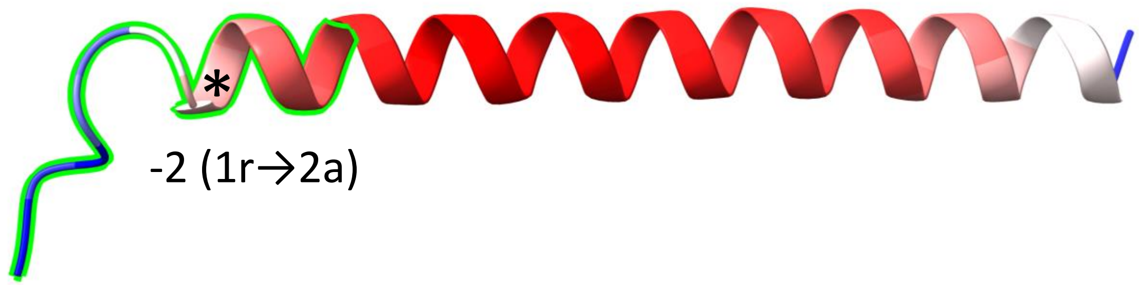


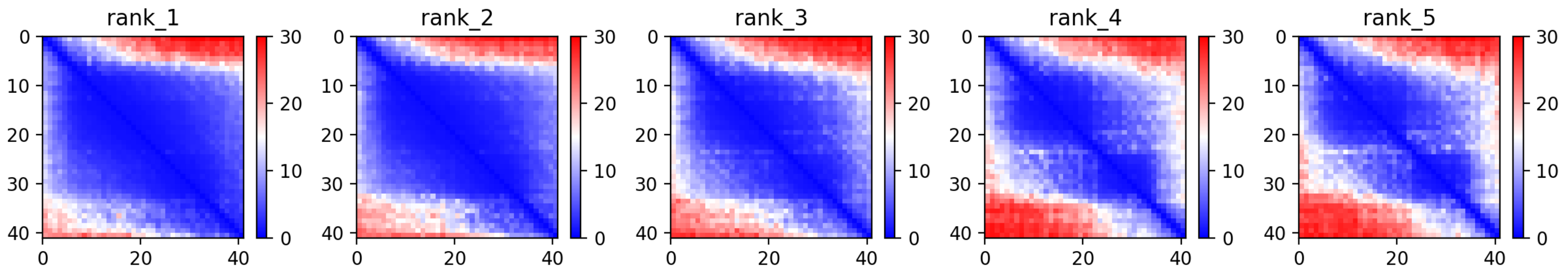

CP75: MtrunA17_Chr4g0037381

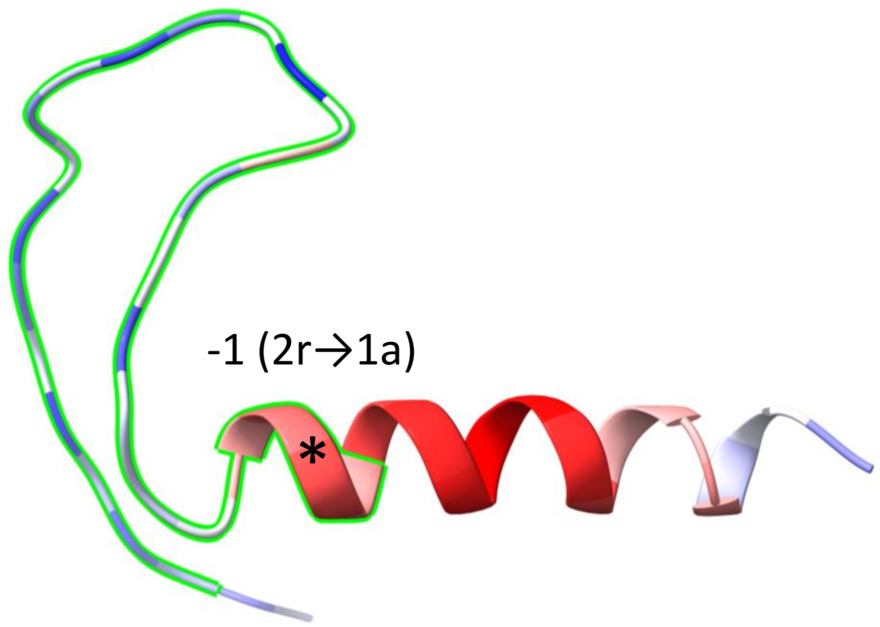


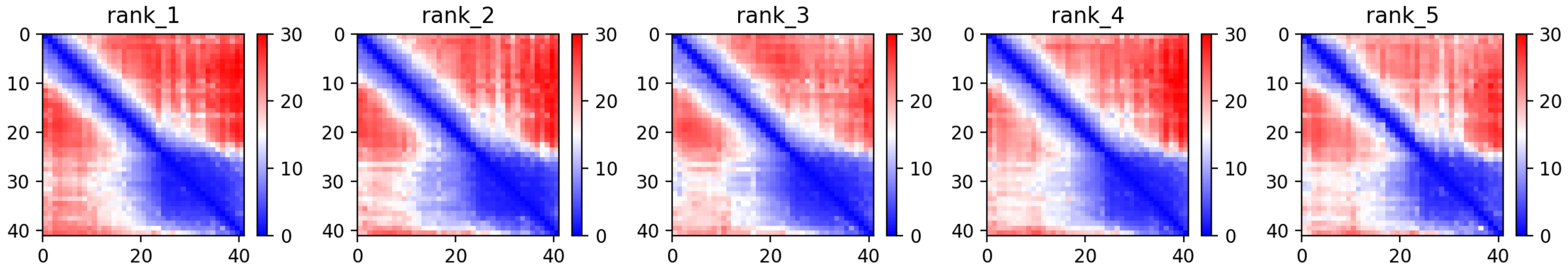

CP76: MtrunA17_Chr4g0040471

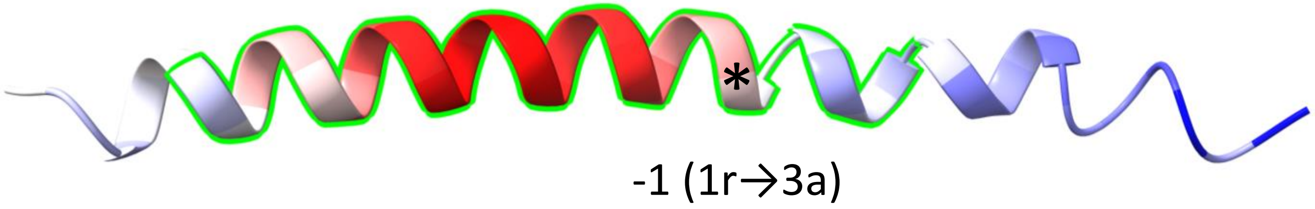


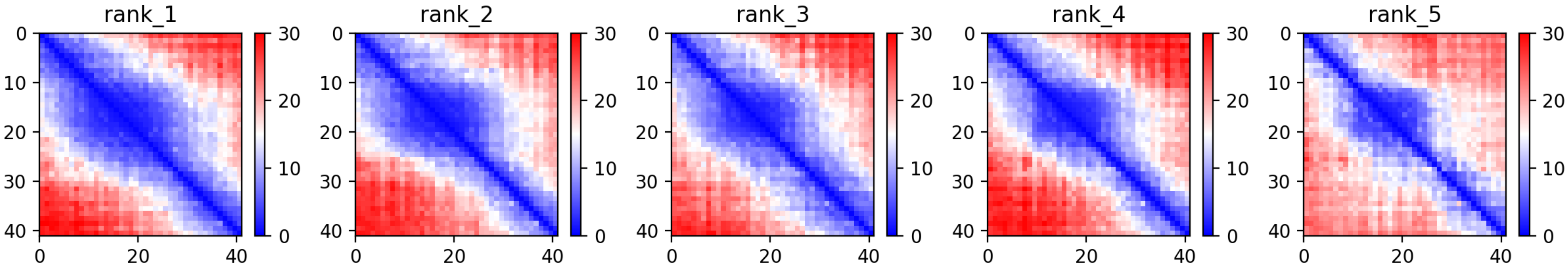

### CP77: MtrunA17_Chr4g0055331

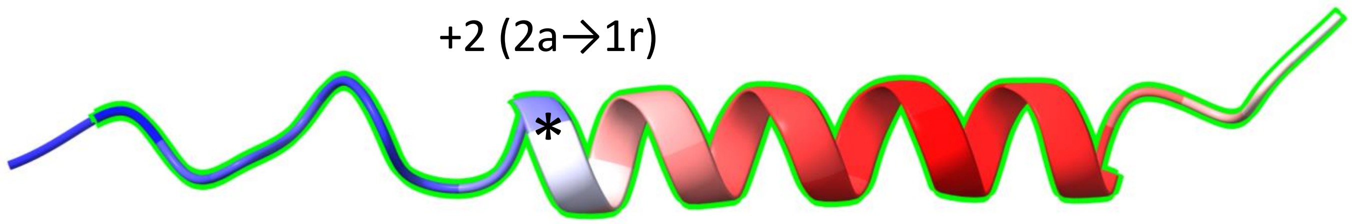


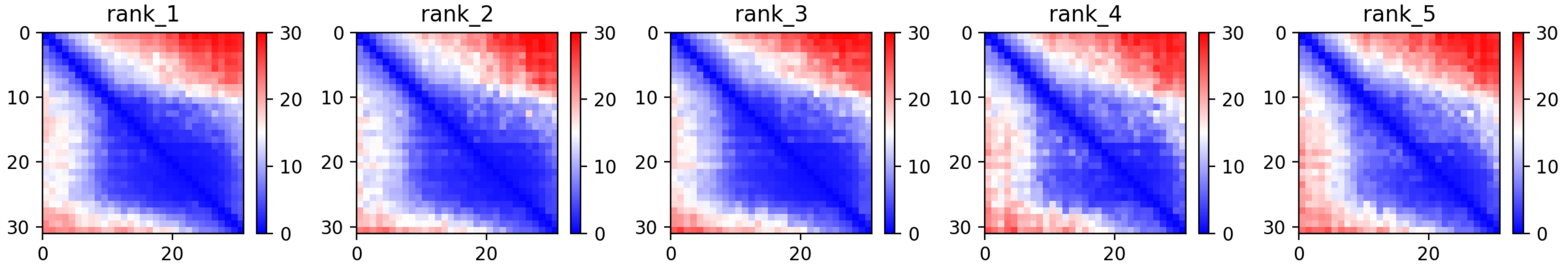

### CP82: MtrunA17_Chr5g0400181

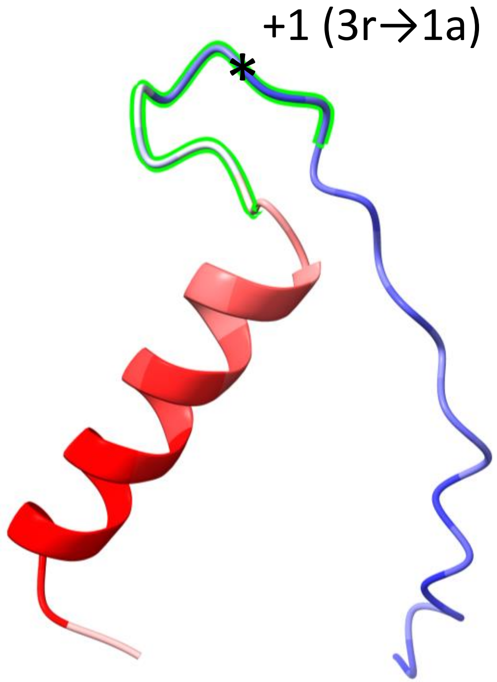


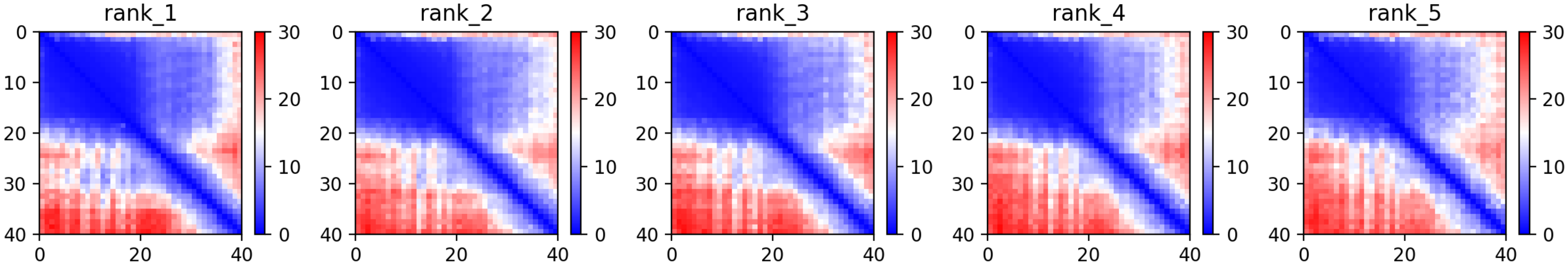

CP83: MtrunA17_Chr5g0405061

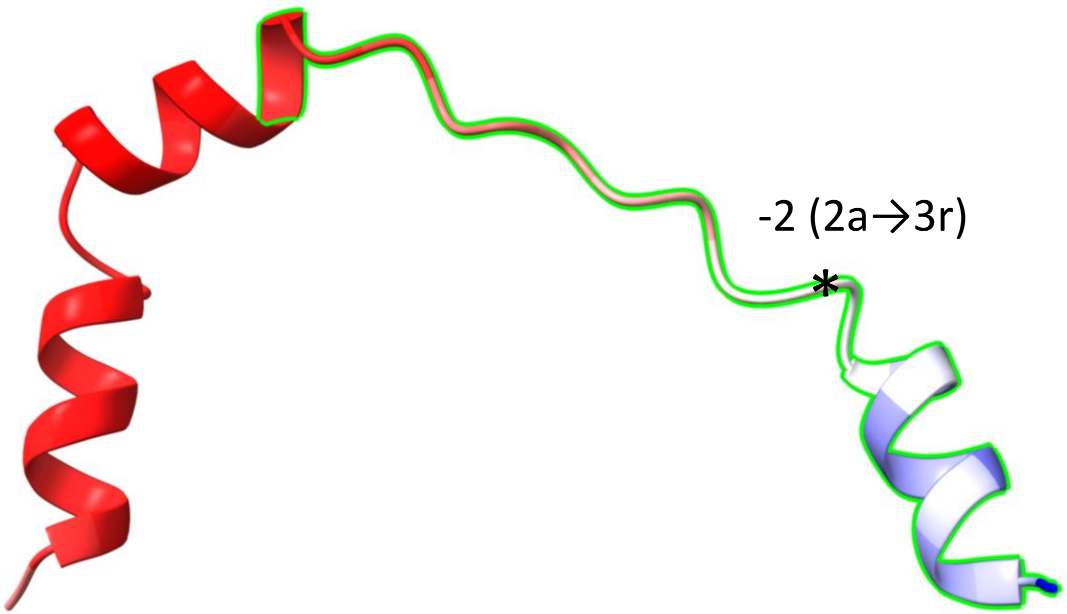


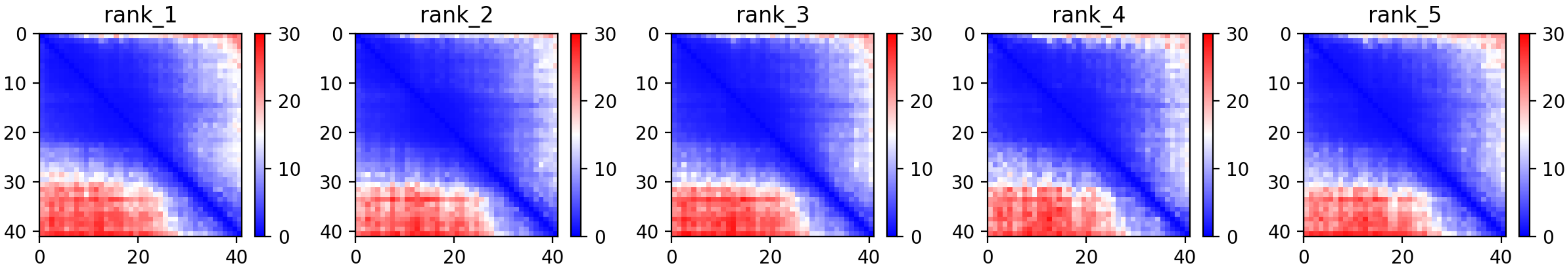

### CP84: MtrunA17_Chr5g0408581

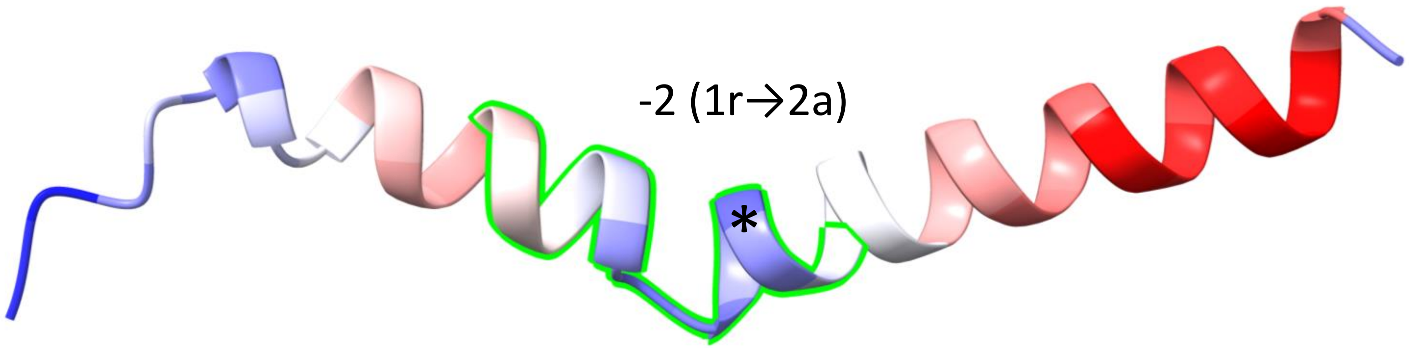


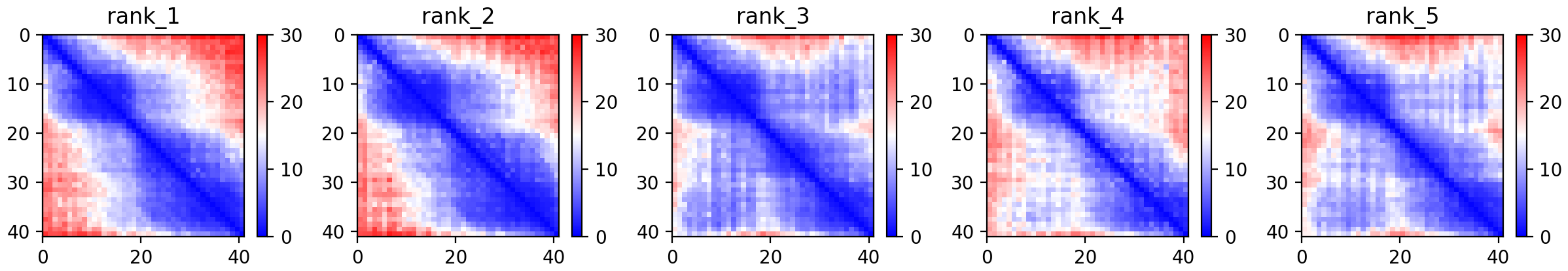

CP85: MtrunA17_Chr5g0415031

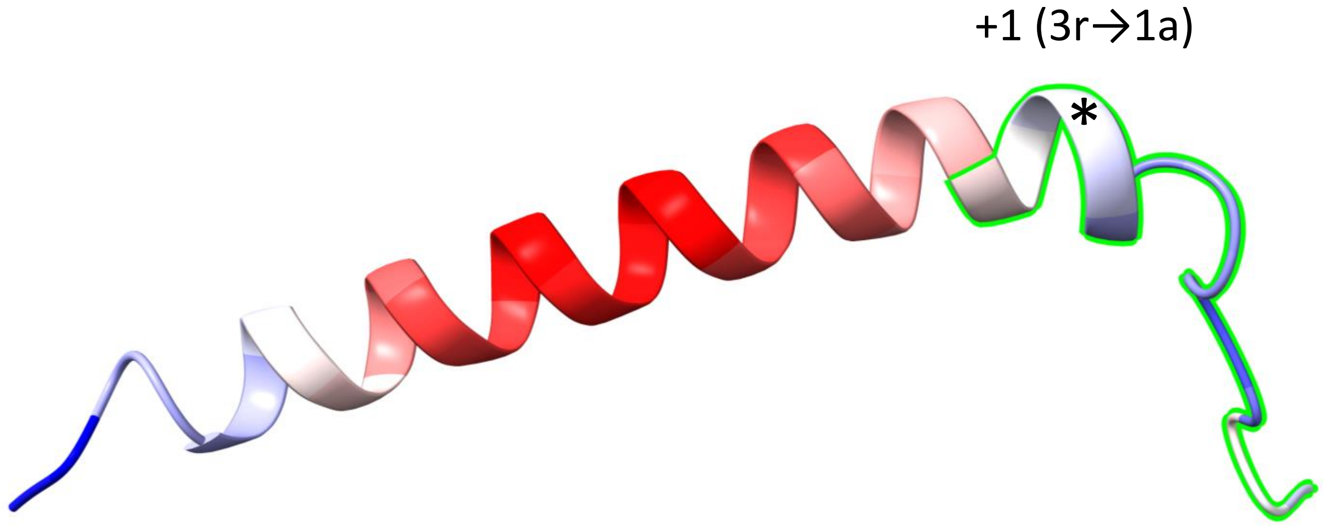


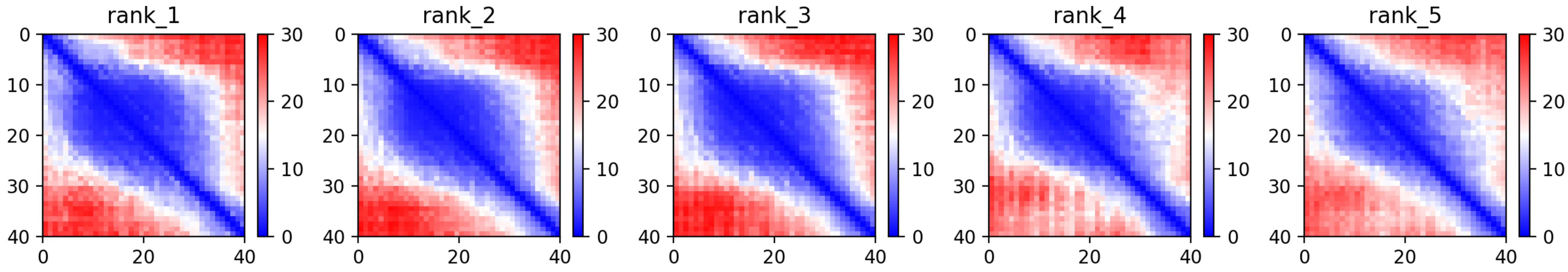

CP86: MtrunA17_Chr5g0415911

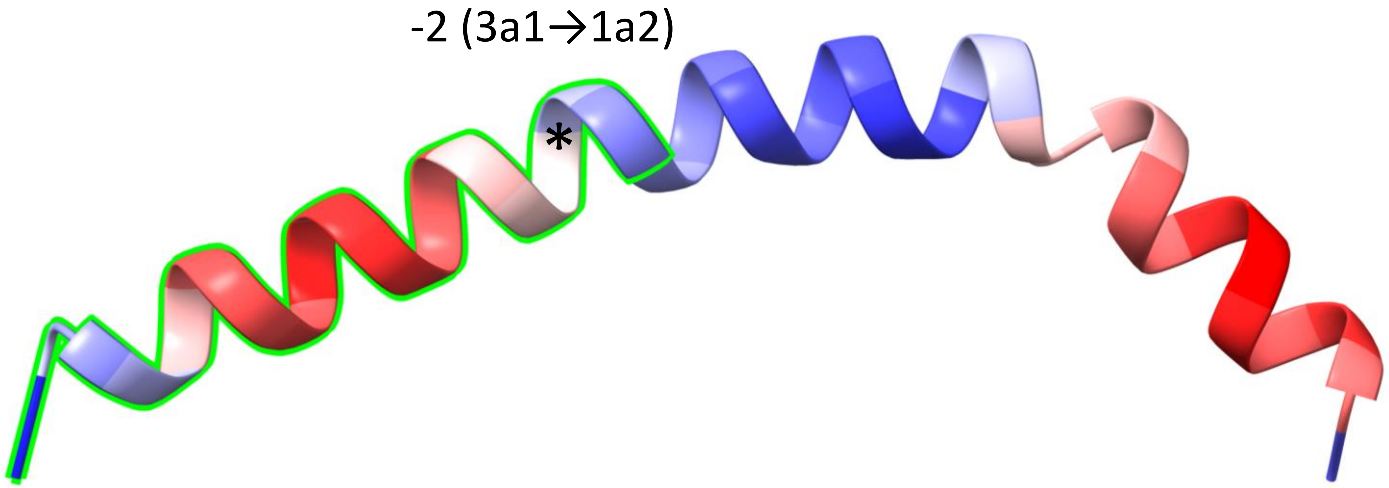


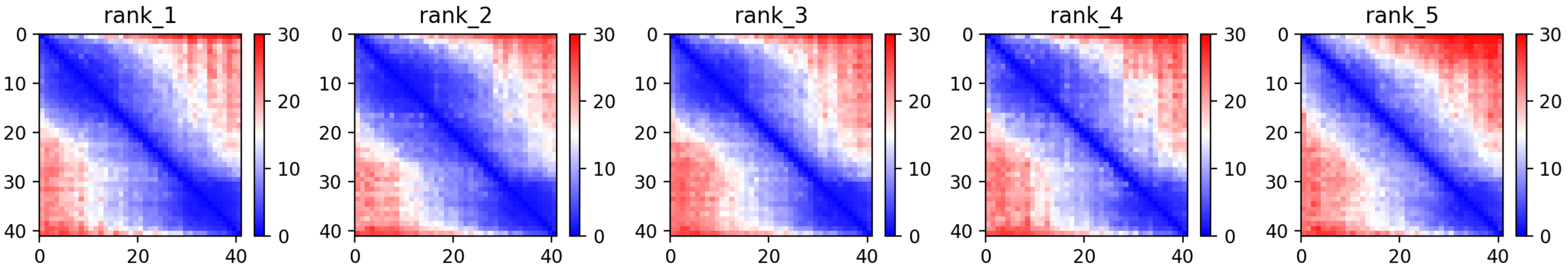

CP87: MtrunA17_Chr5g0421761

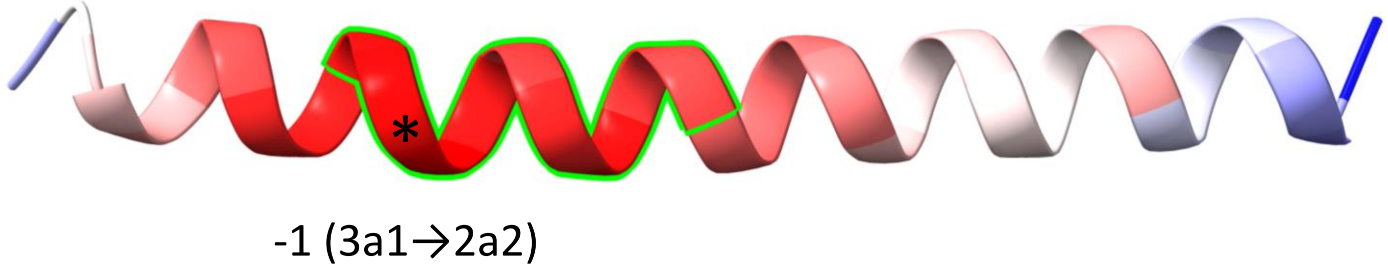


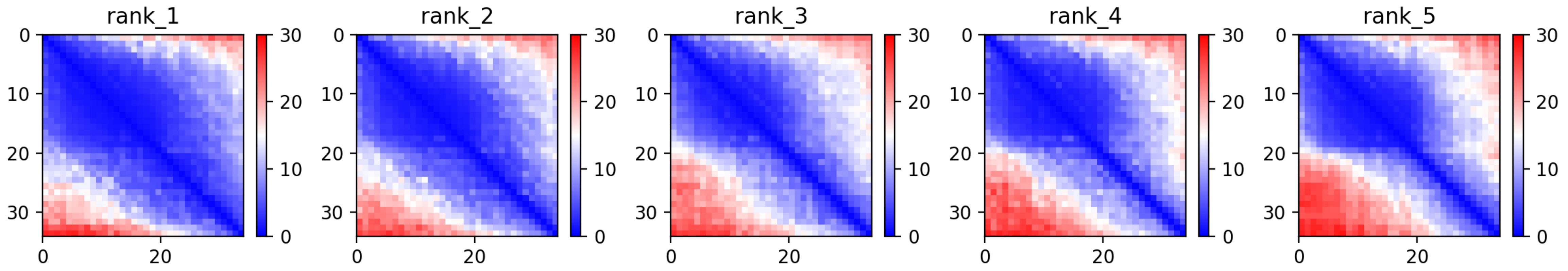

CP88: MtrunA17_Chr5g0422291

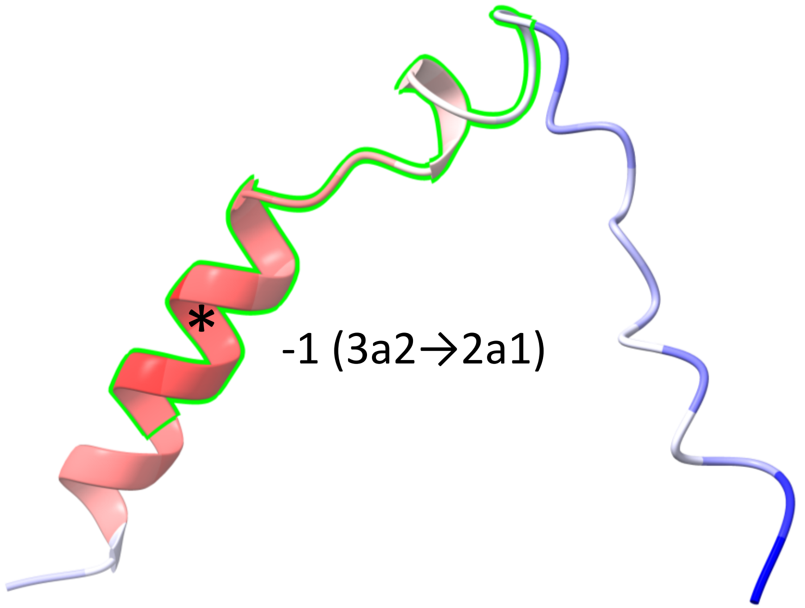


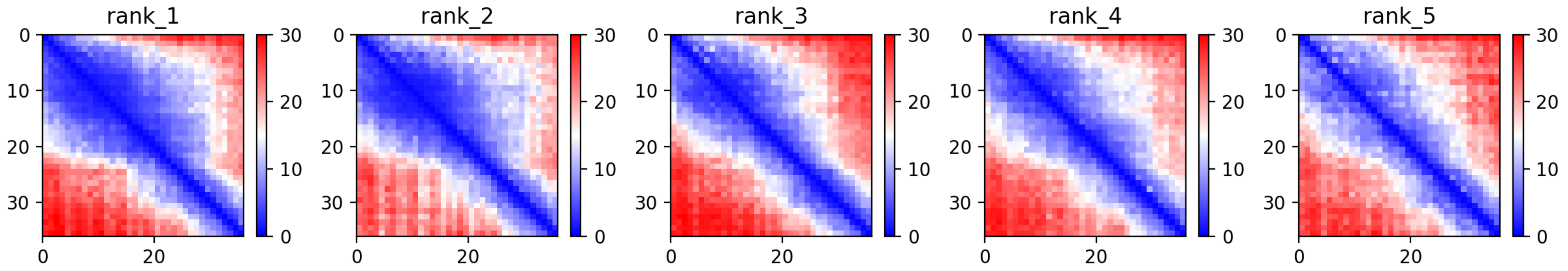

CP90: MtrunA17_Chr5g0430341

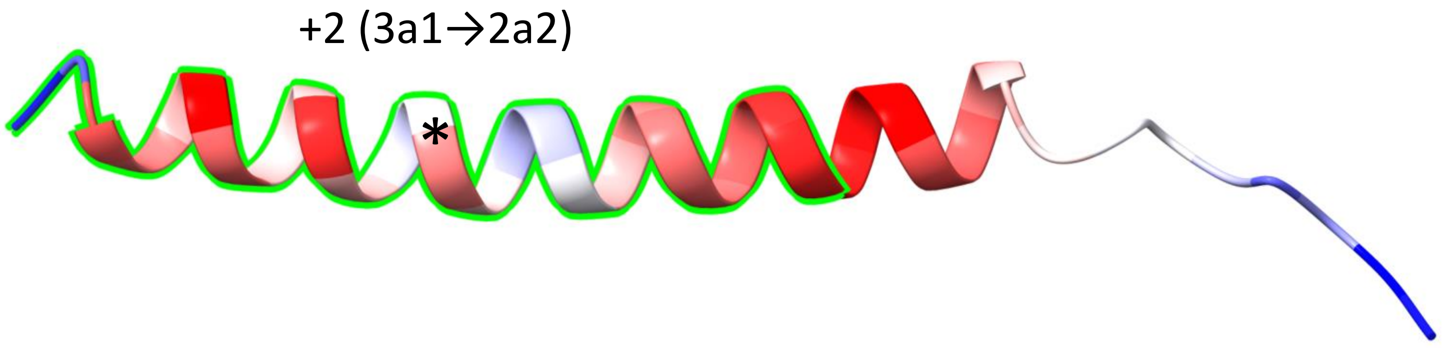


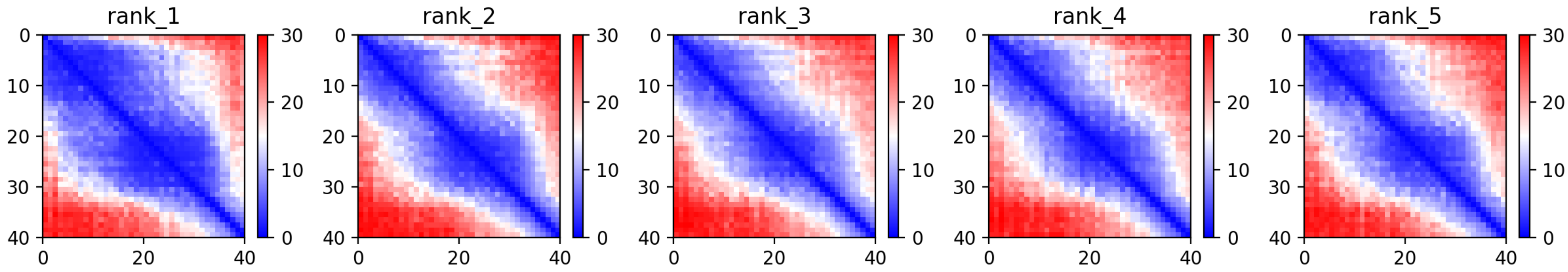

CP92: MtrunA17_Chr5g0431401

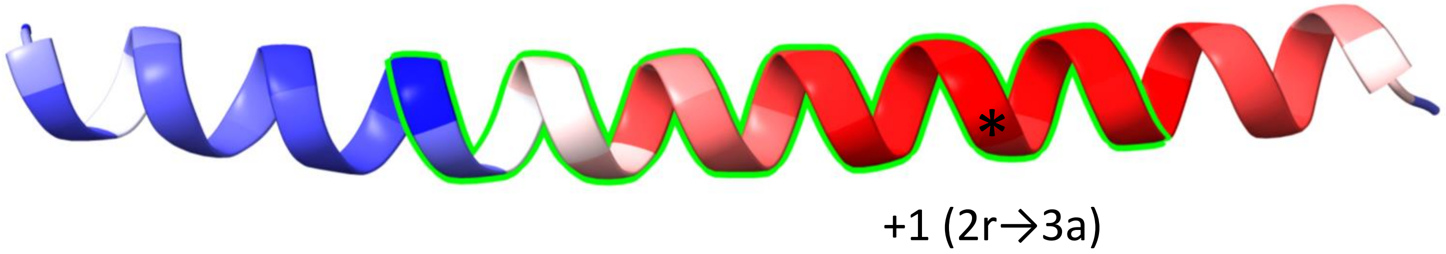


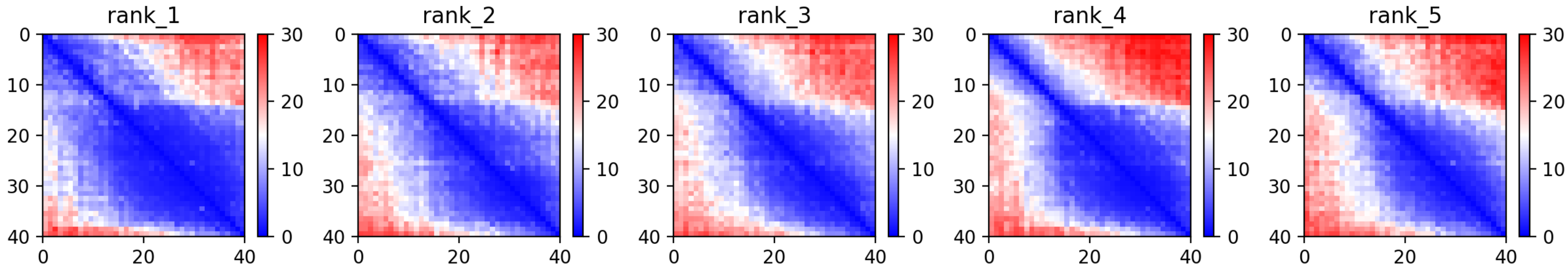

CP93: MtrunA17_Chr5g0435191

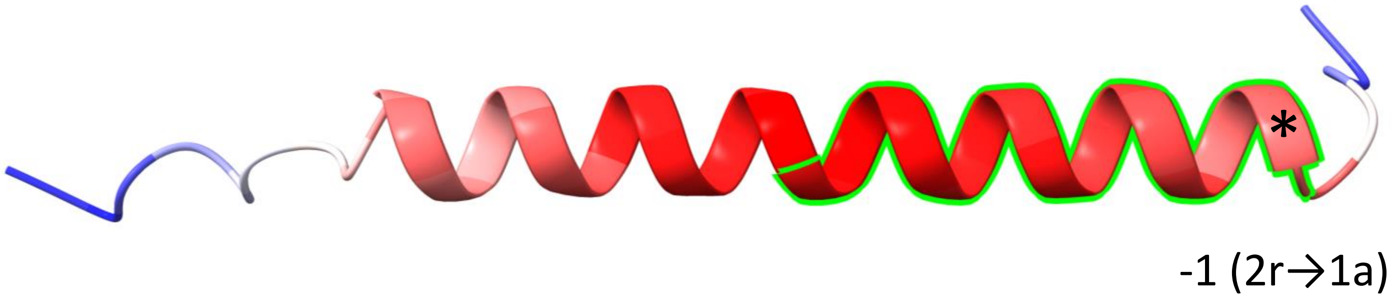


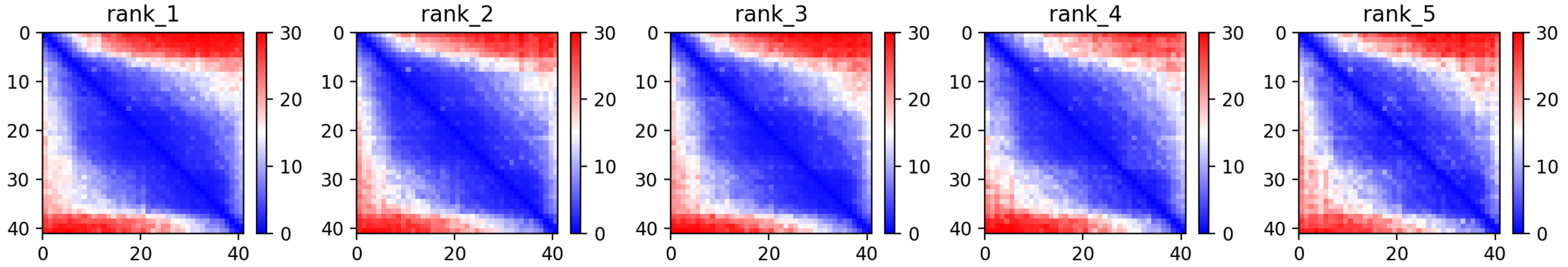

CP94: MtrunA17_Chr5g0444231

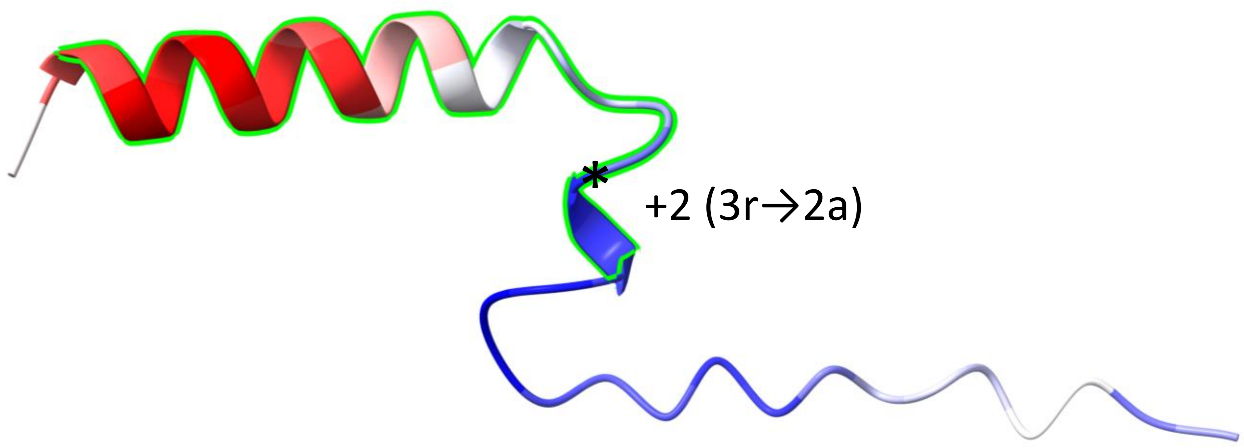


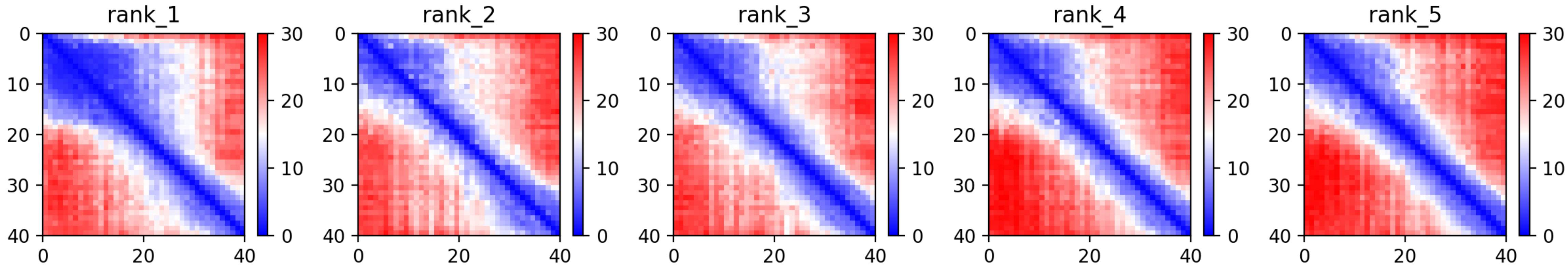

CP98: MtrunA17_Chr6g0457461

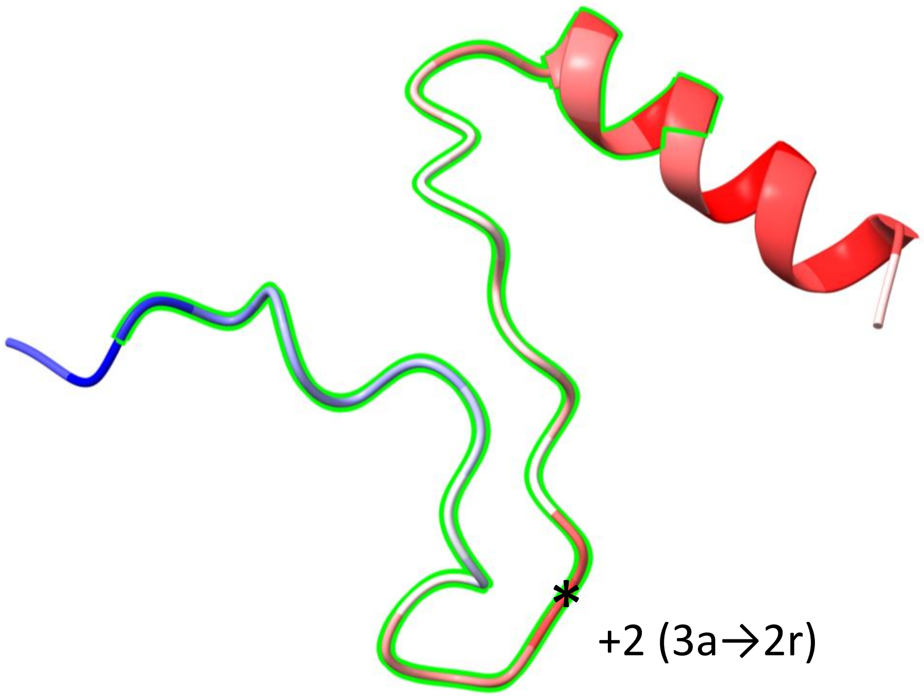


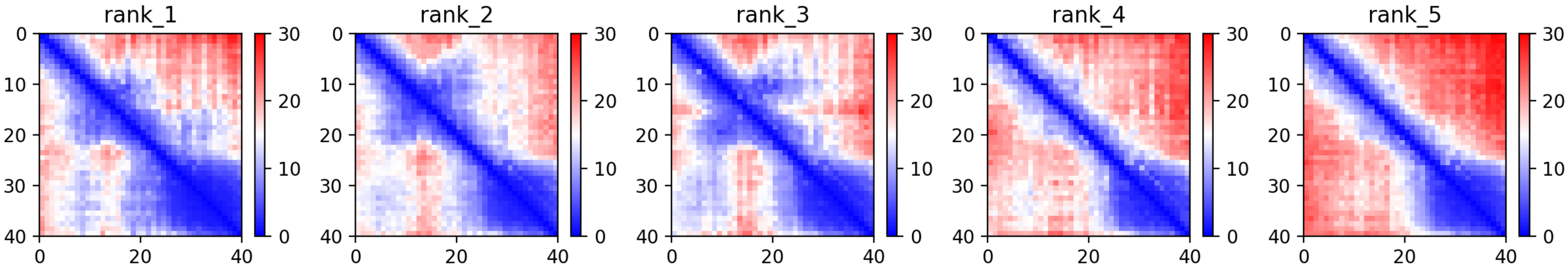

CP102: MtrunA17_Chr6g0459481

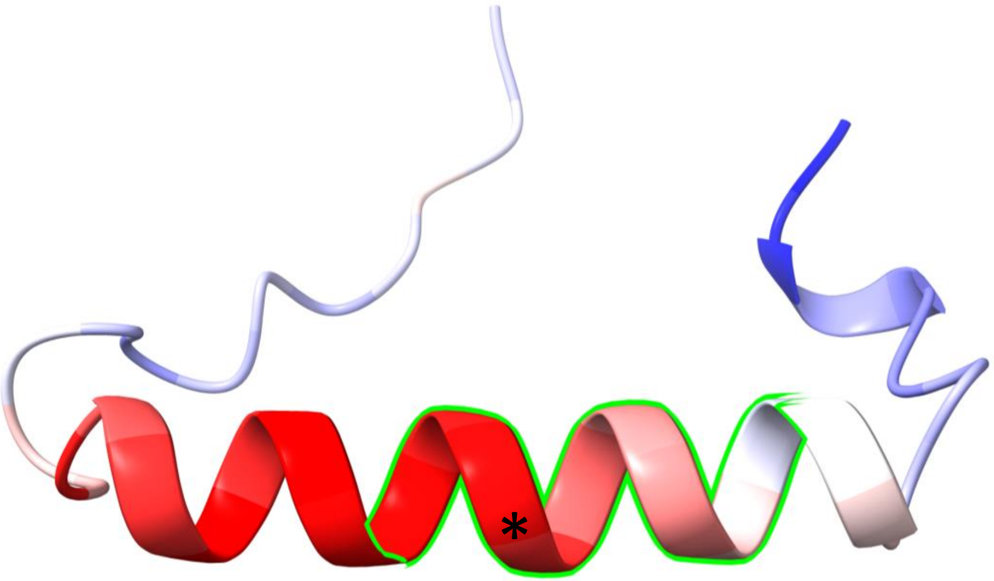


-1 (1r→3a)

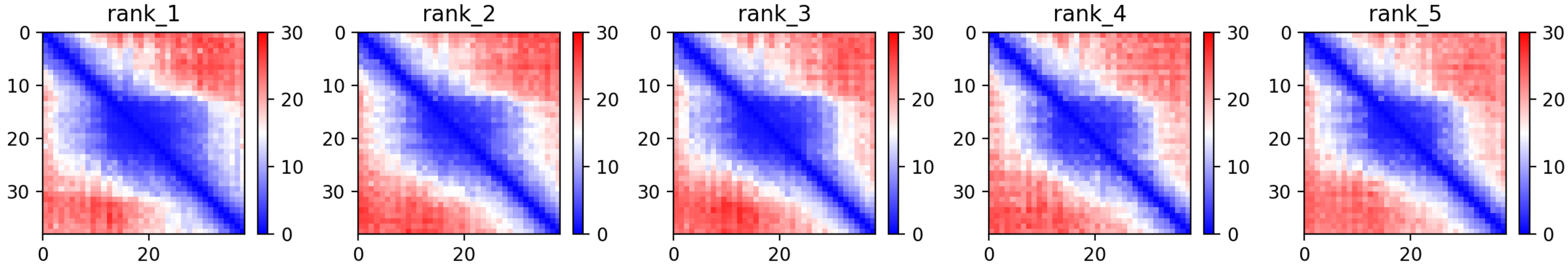

### CP104: MtrunA17_Chr6g0462271

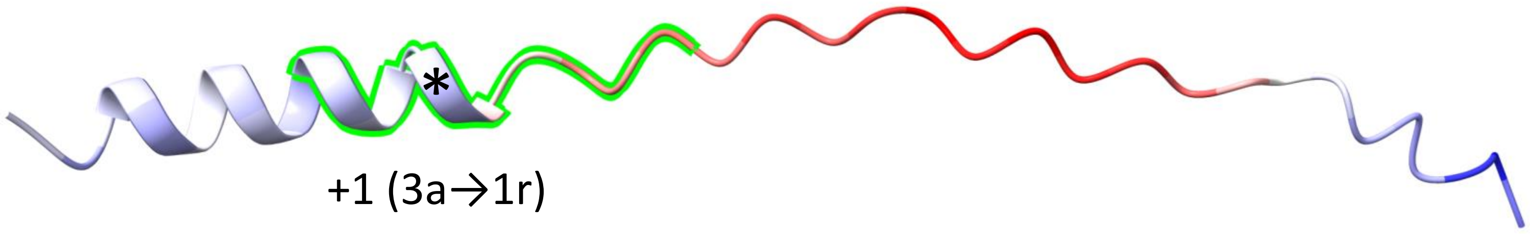


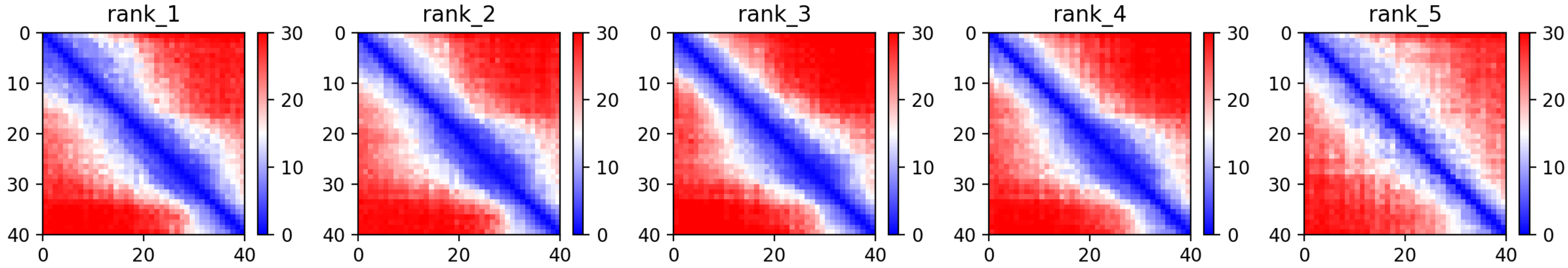

### CP105: MtrunA17_Chr6g0468411

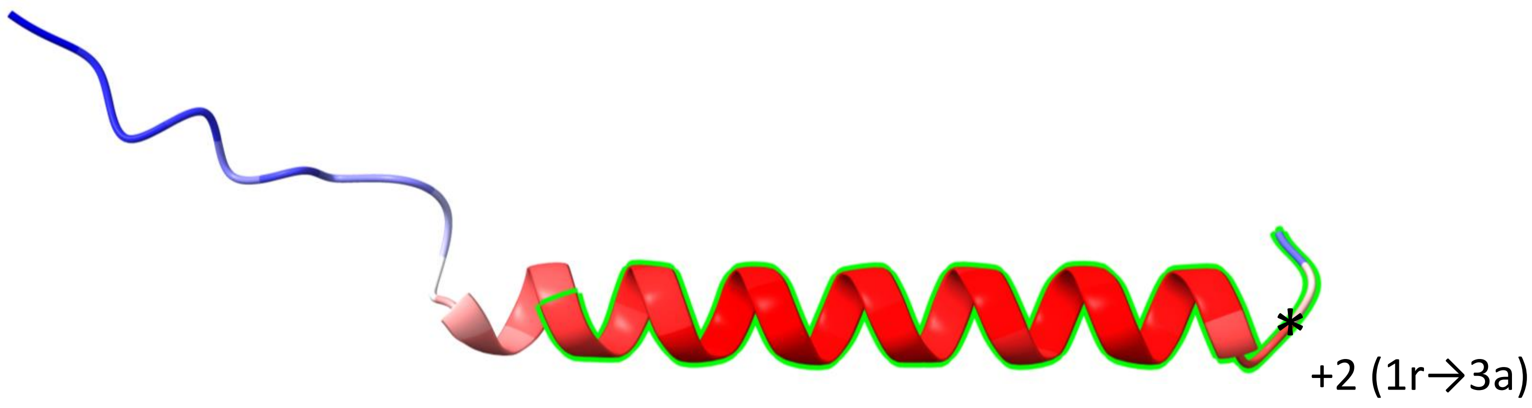


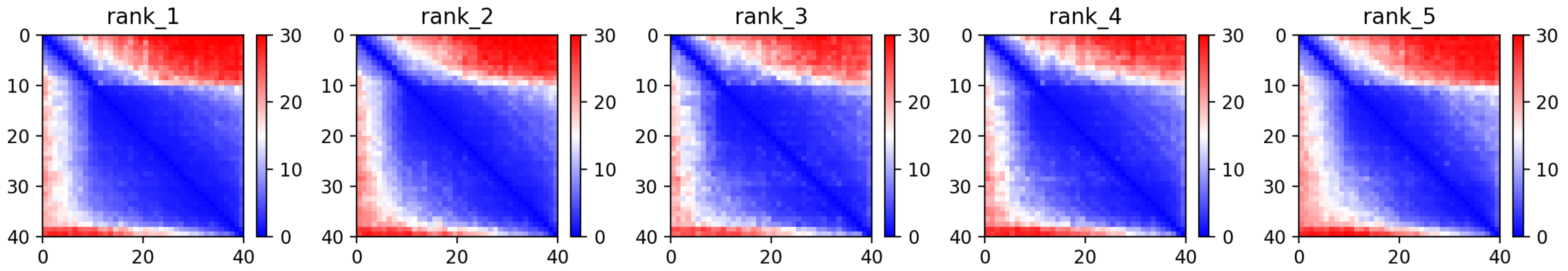

CP106: MtrunA17_Chr6g0476751

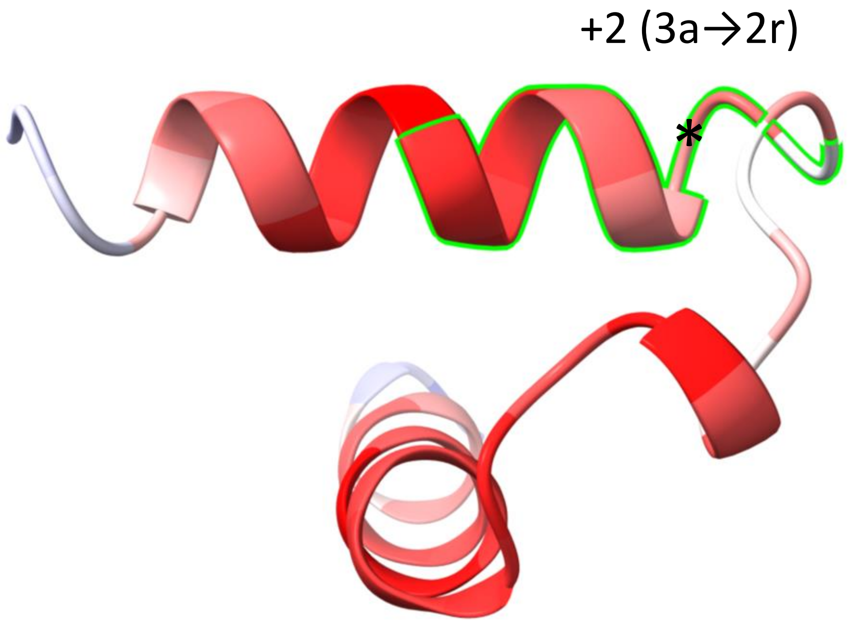


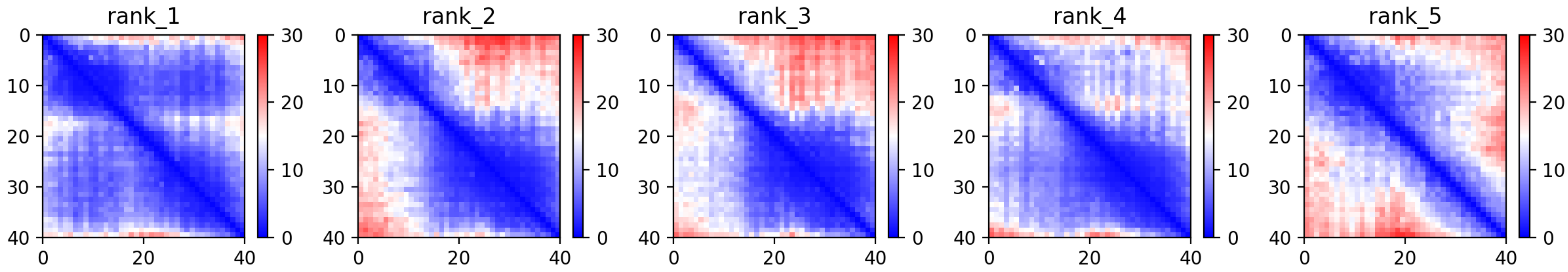

CP107: MtrunA17_Chr6g0479001

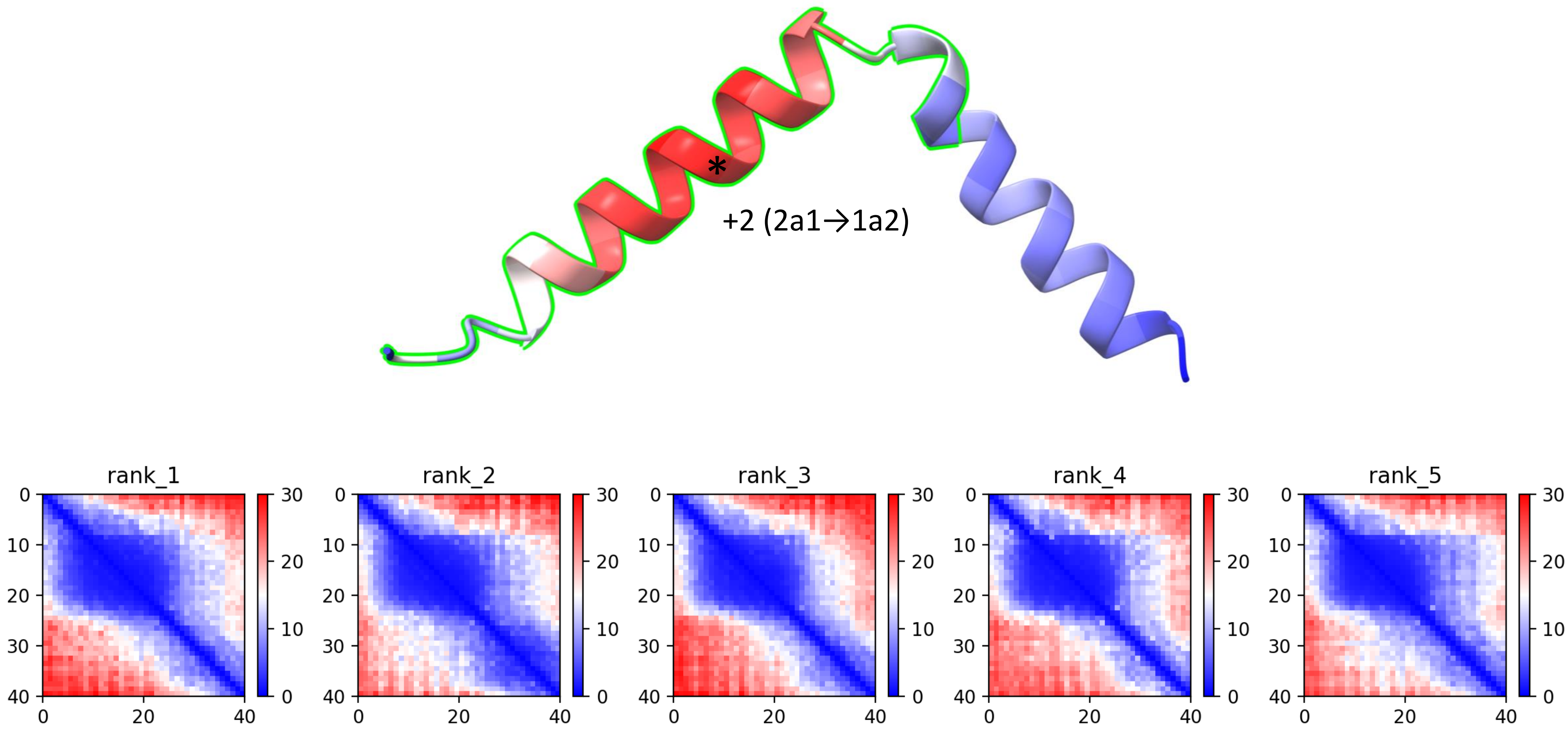

CP109: MtrunA17_Chr6g0486961

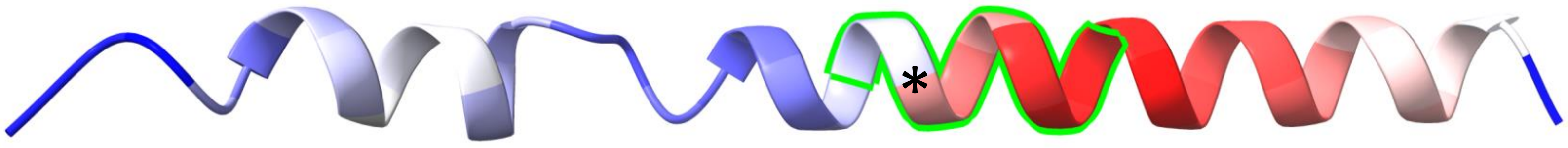


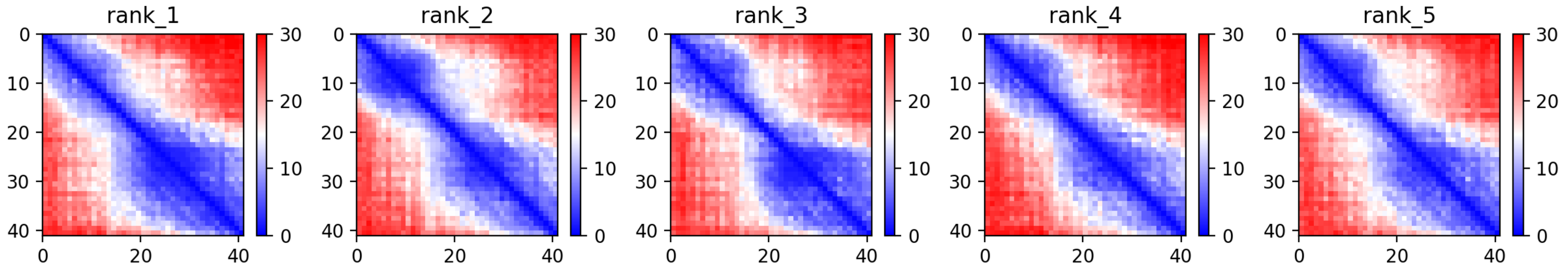

CP110: MtrunA17_Chr7g0214741

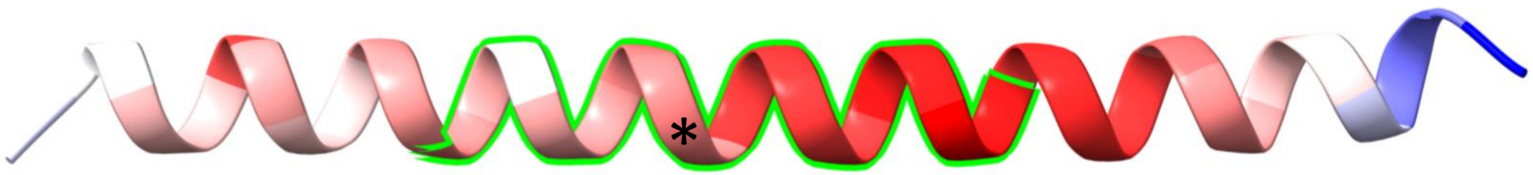


+1 (3r→1a)

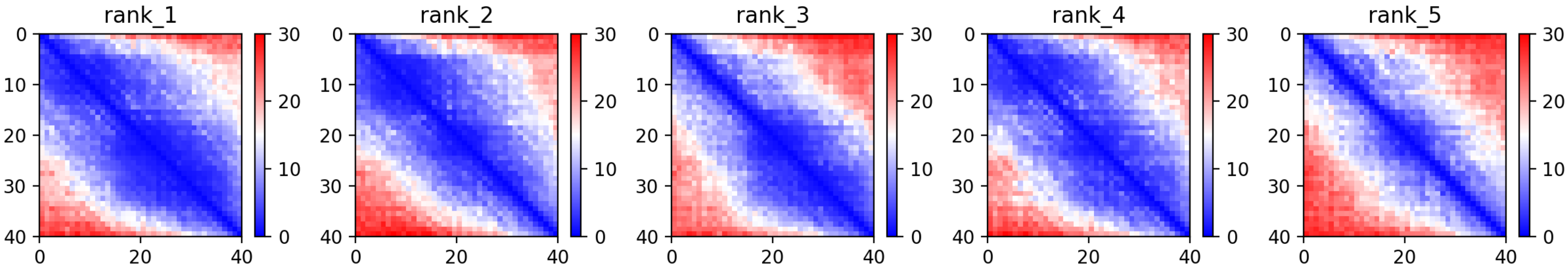

### CP111: MtrunA17_Chr7g0214911

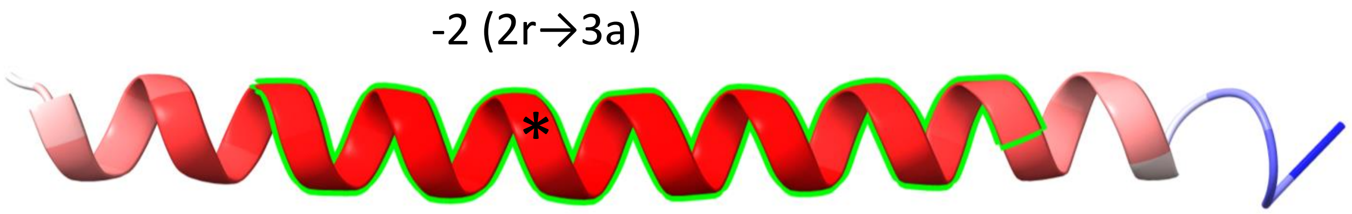


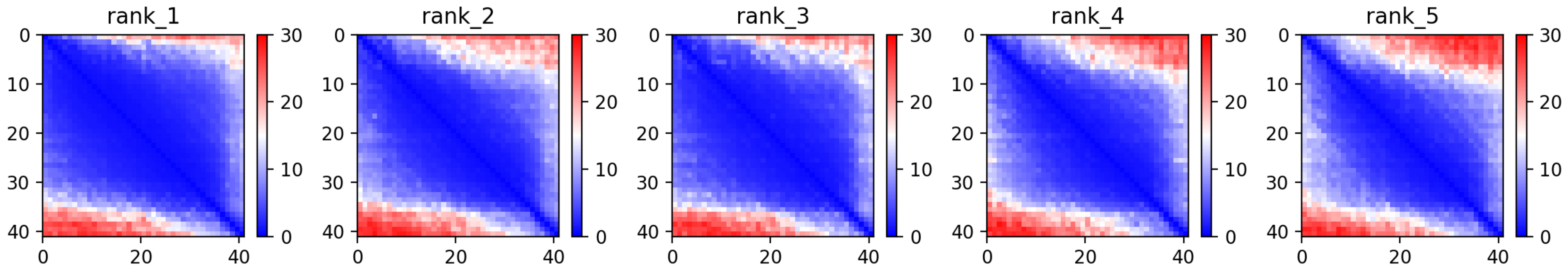

### CP112: MtrunA17_Chr7g0219691

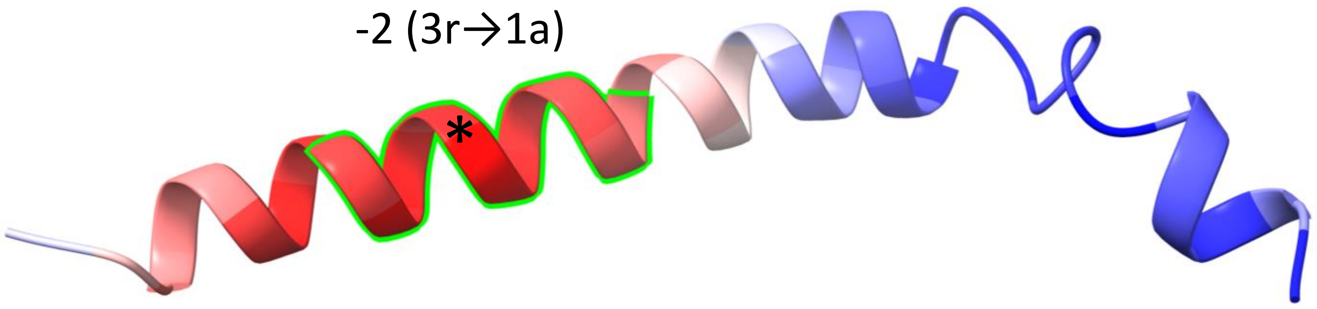


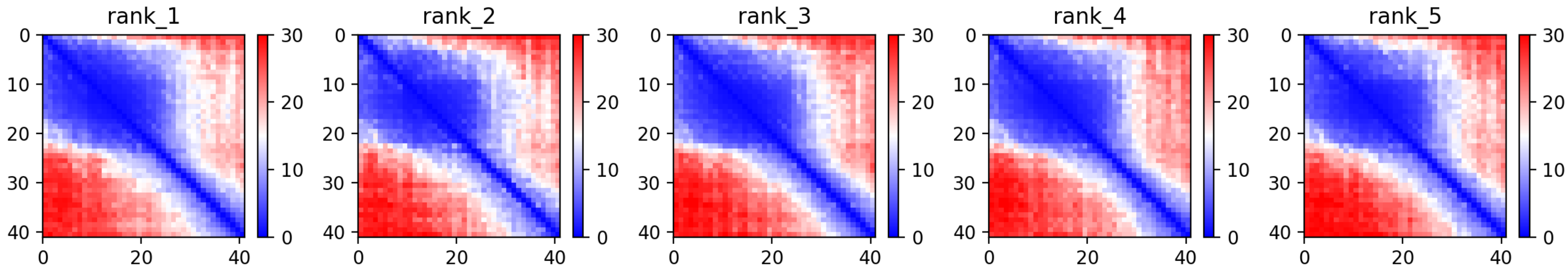

CP113: MtrunA17_Chr7g0221631

-1 (1r→3a)

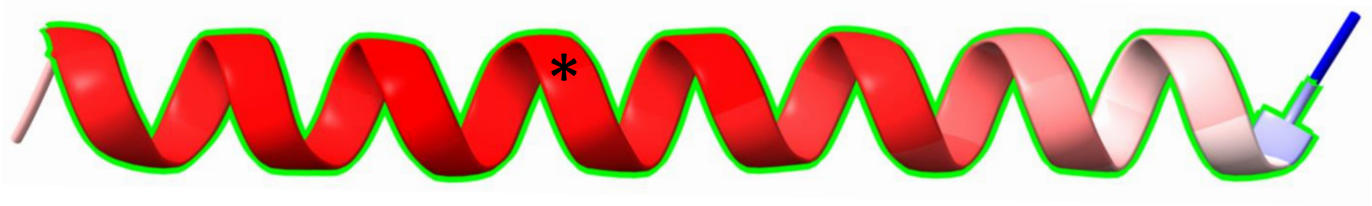


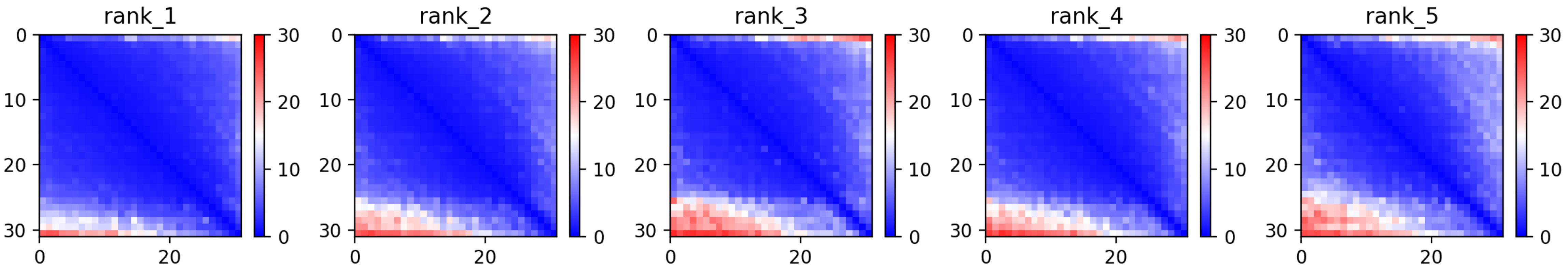

### CP115: MtrunA17_Chr7g0230341

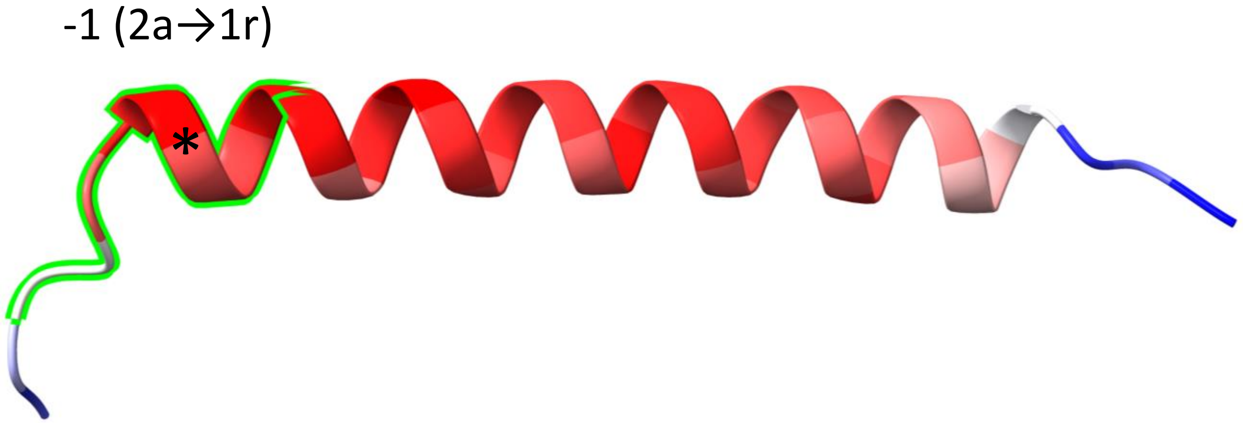


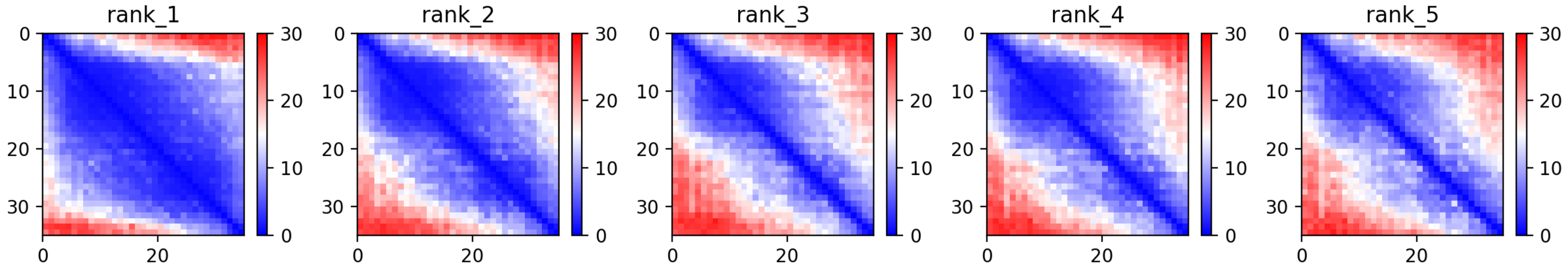

CP117: MtrunA17_Chr7g0237331

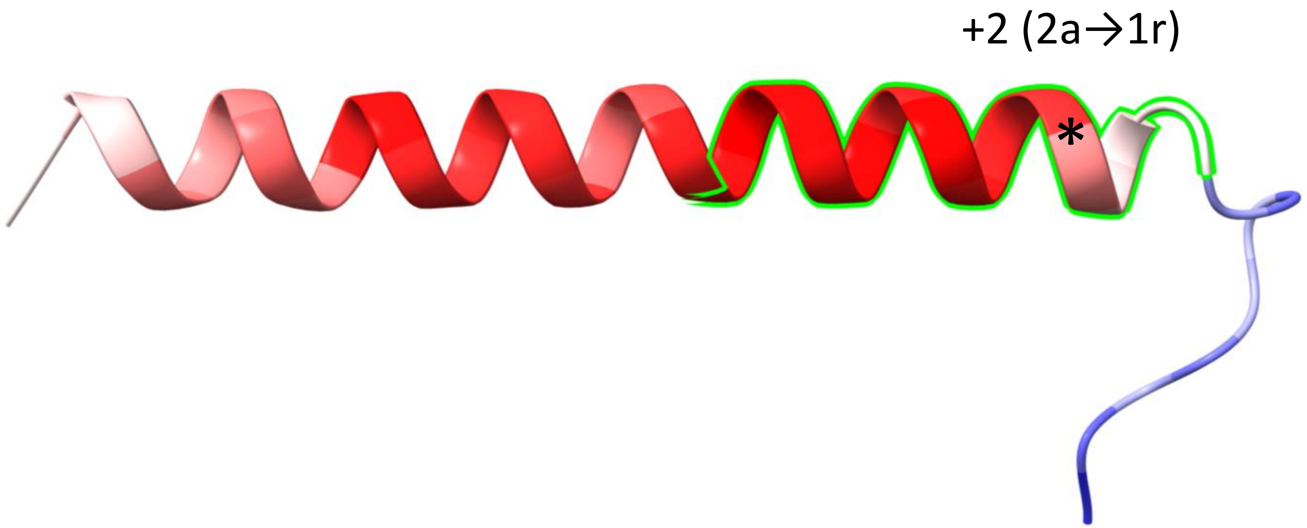


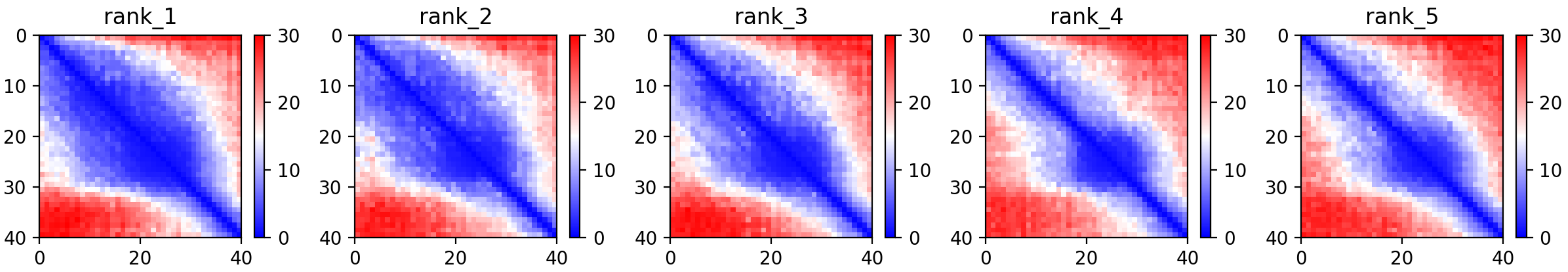

CP118: MtrunA17_Chr7g0251971

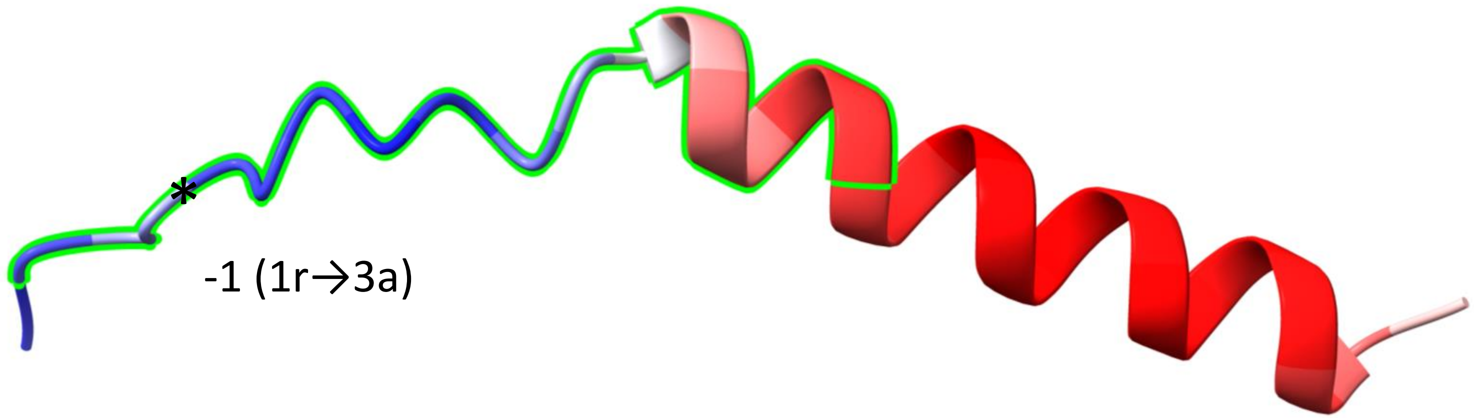


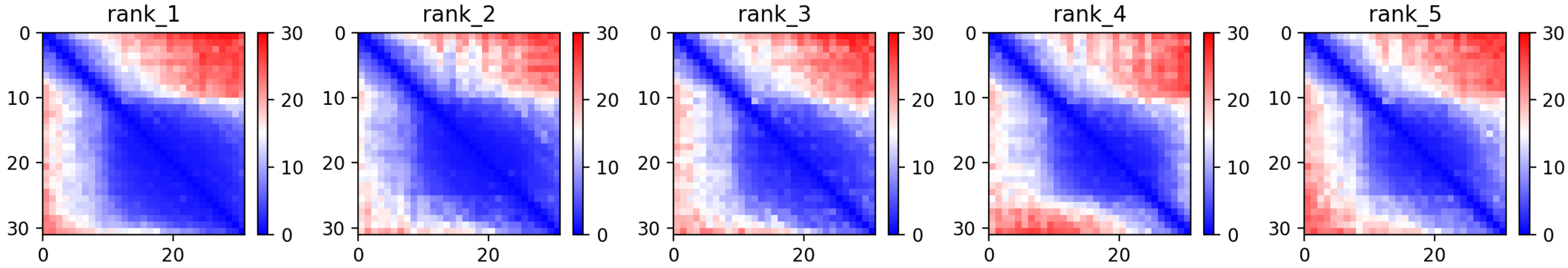

CP119: MtrunA17_Chr7g0259611

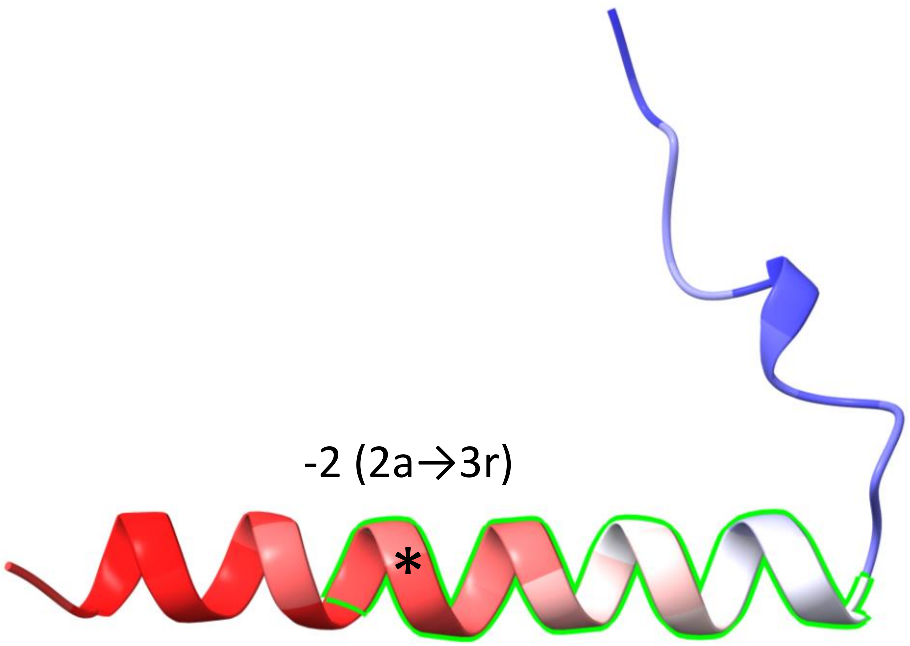


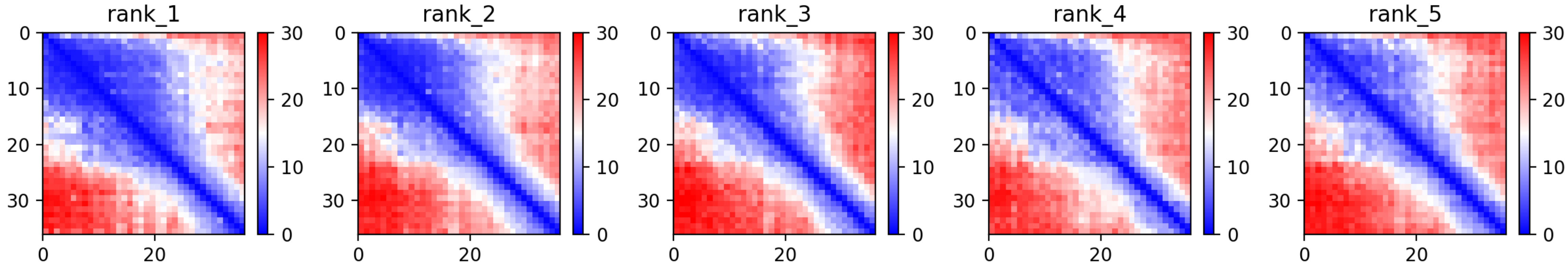

CP120: MtrunA17_Chr7g0262821

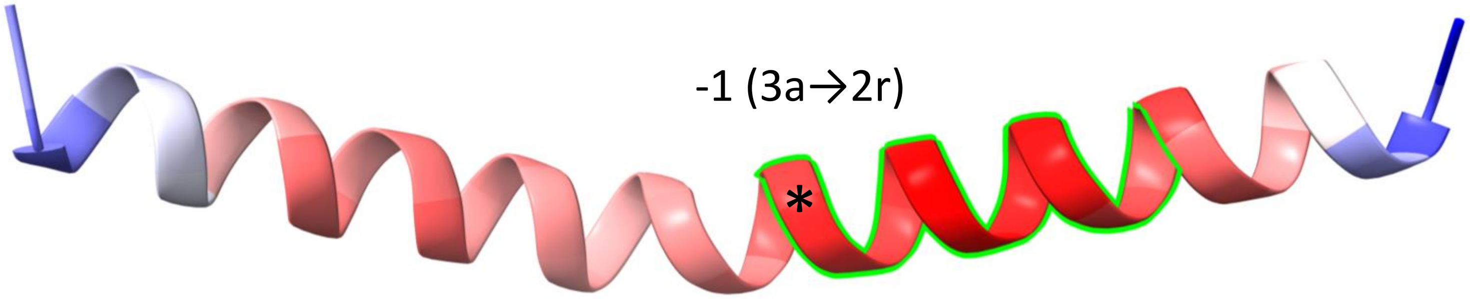


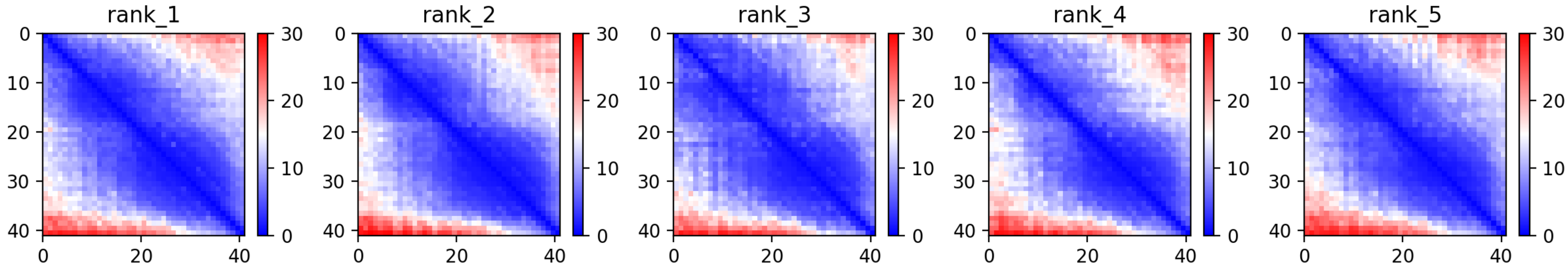

CP121: MtrunA17_Chr7g0270811

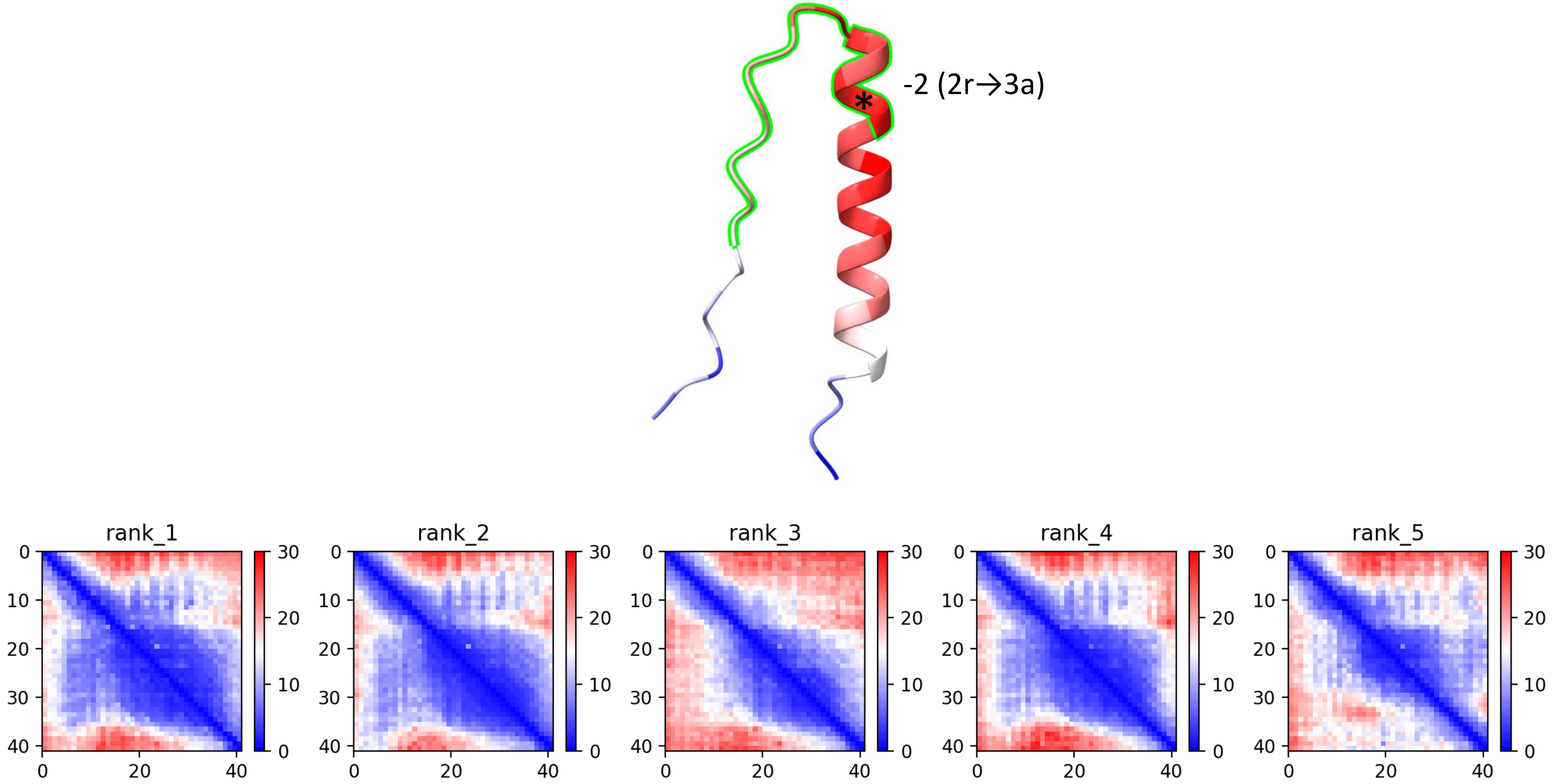

### CP122: MtrunA17_Chr7g1034306

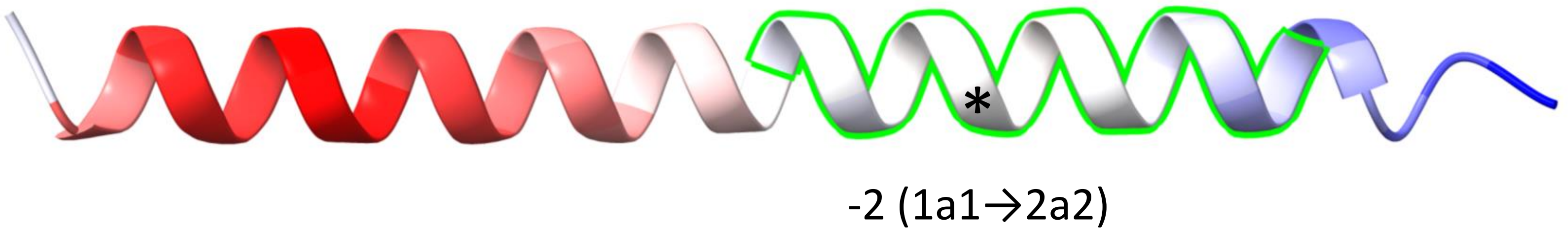


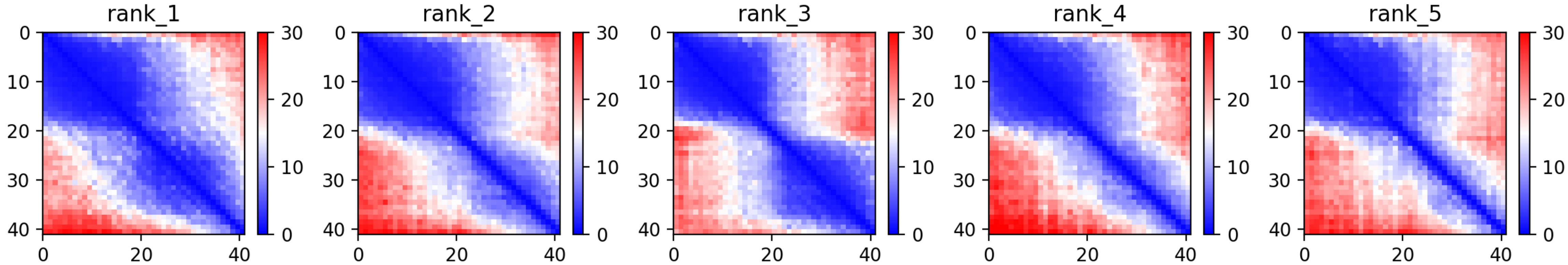

### CP123: MtrunA17_Chr8g0338111

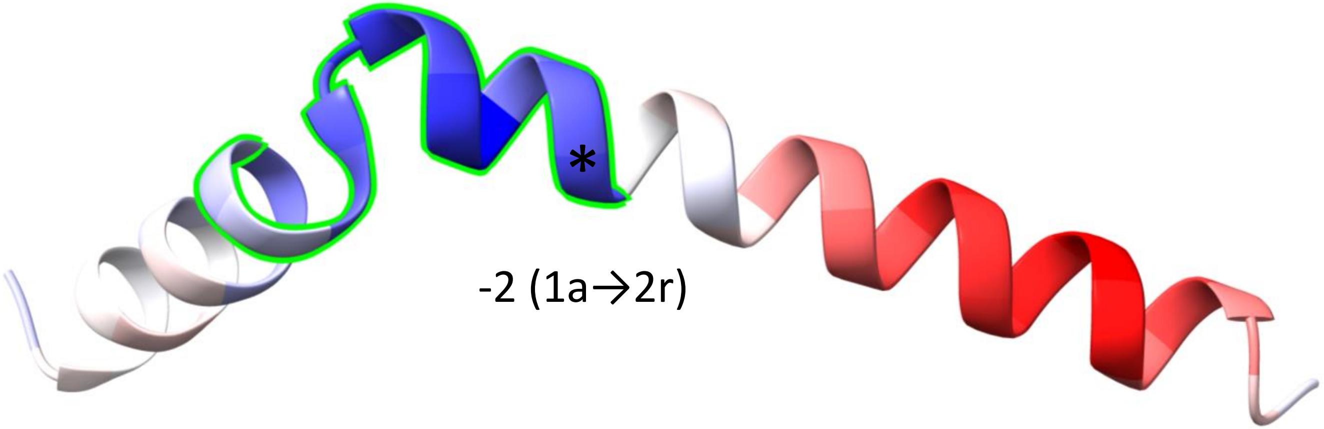


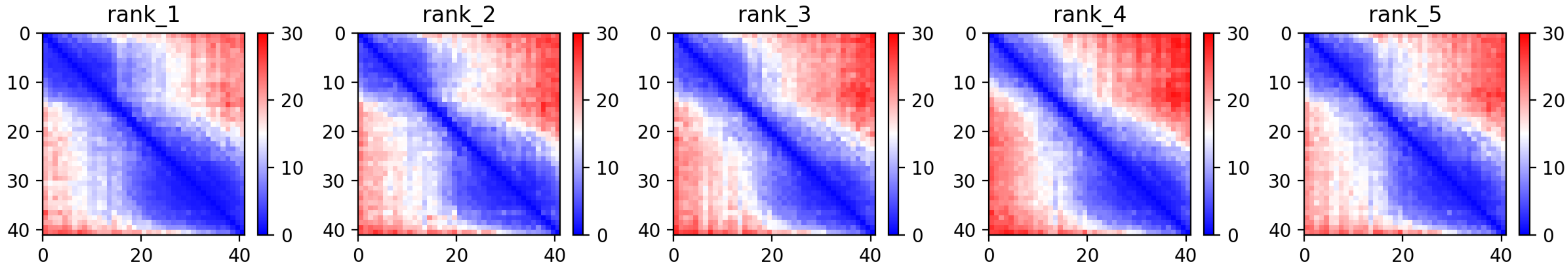

CP126: MtrunA17_Chr8g0342881

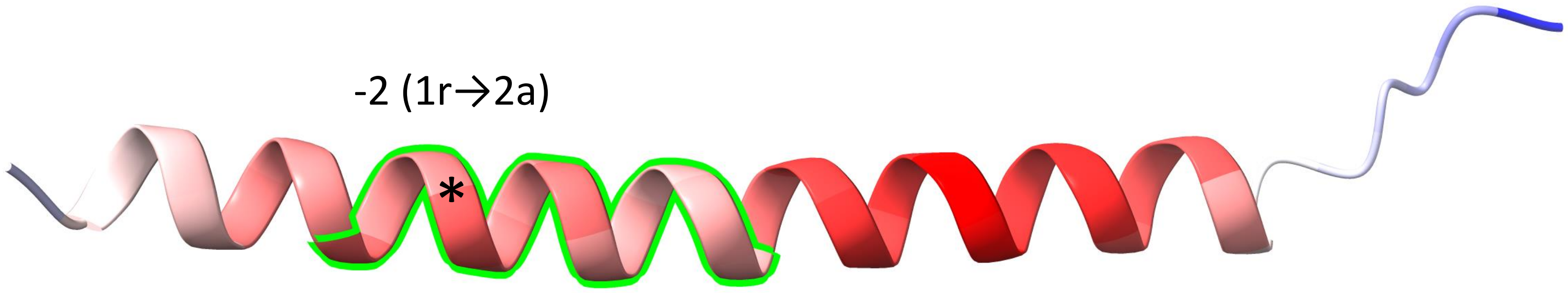


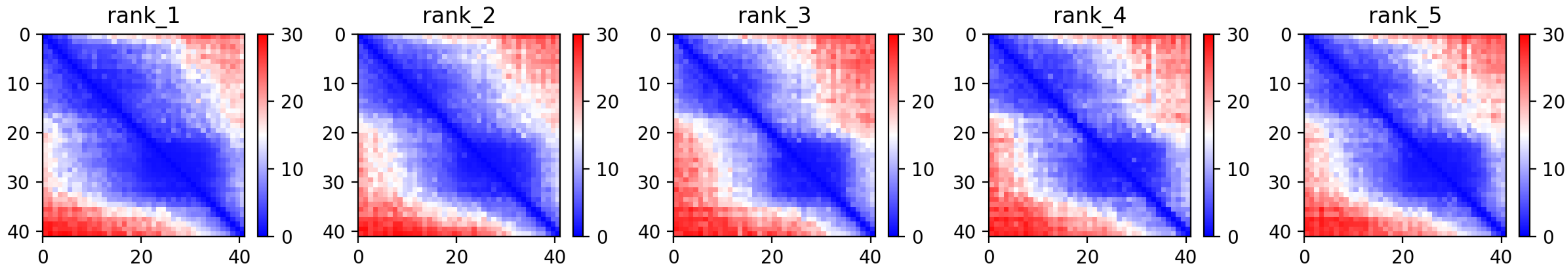

CP127: MtrunA17_Chr8g0345421

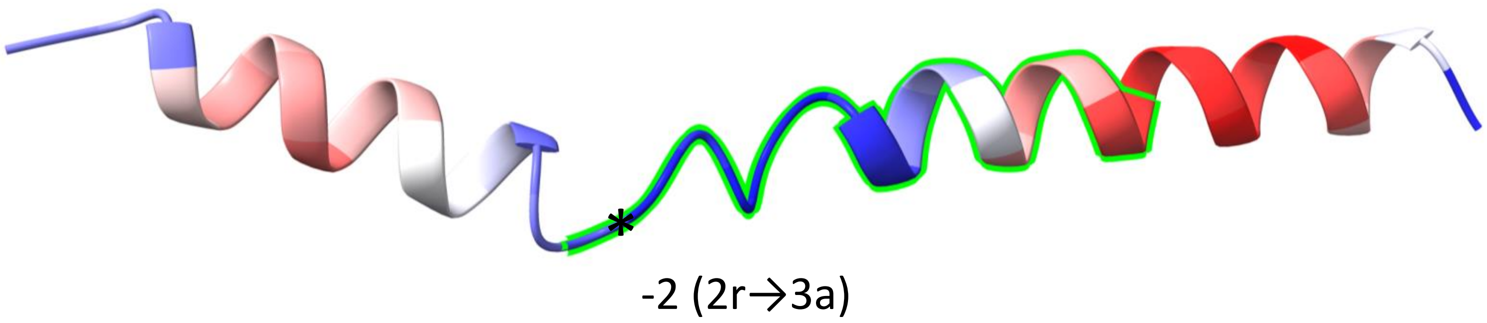


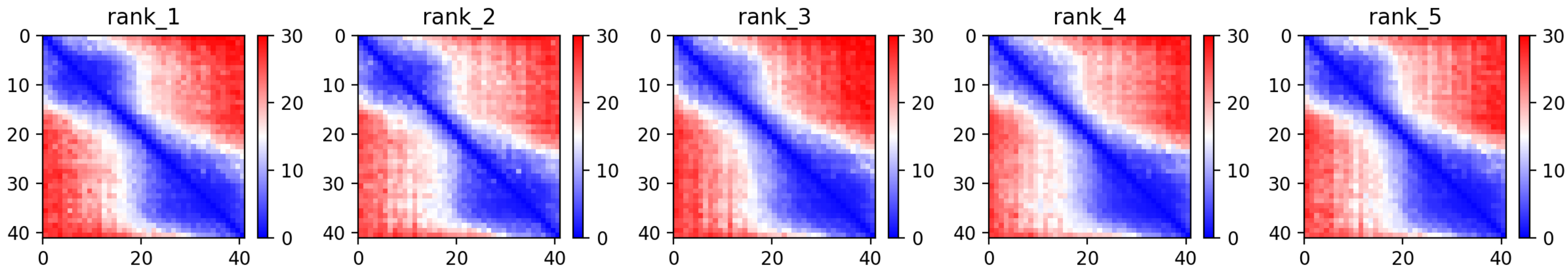

CP128: MtrunA17_Chr8g0353711

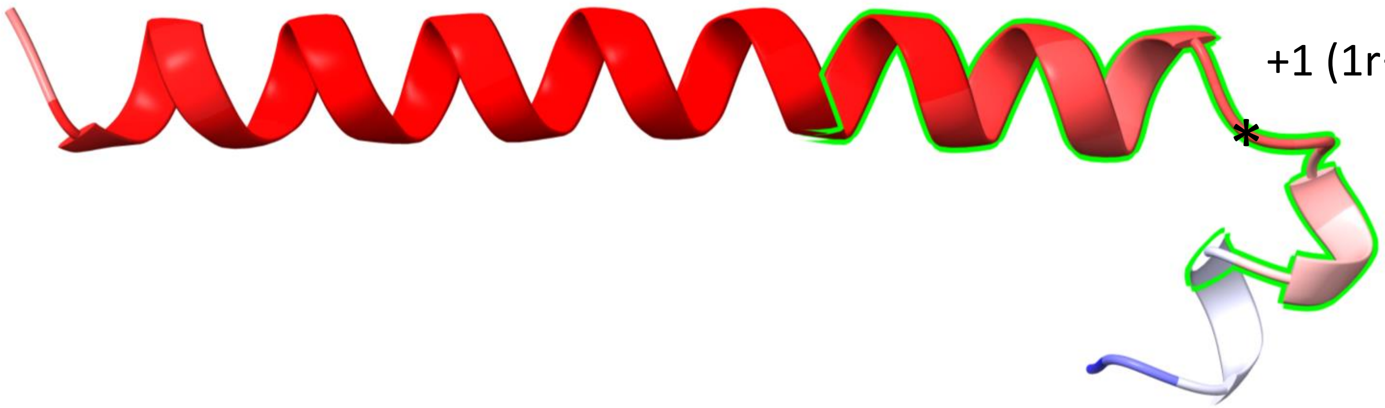


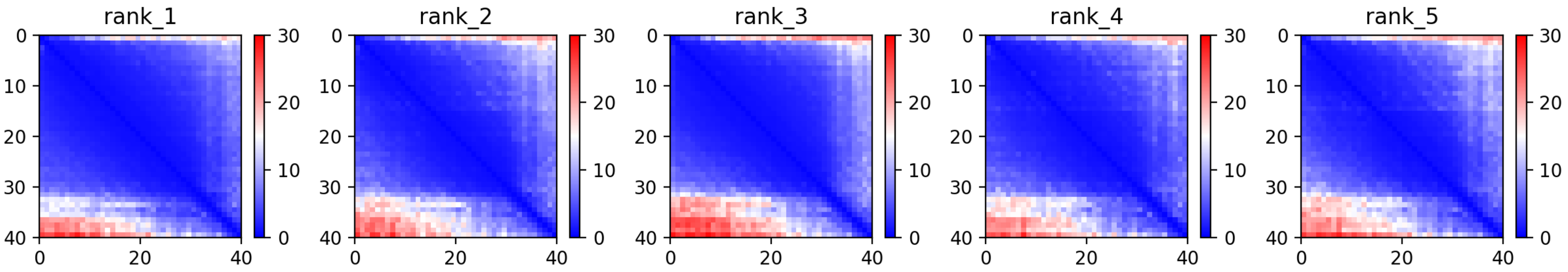

CP129: MtrunA17_Chr8g0355501

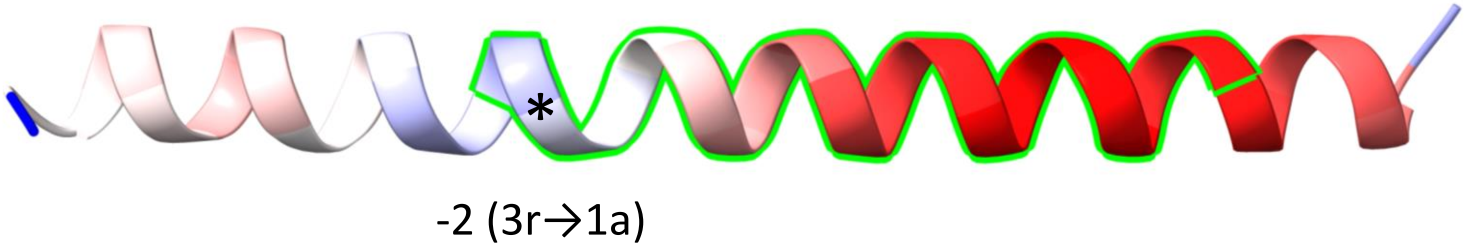


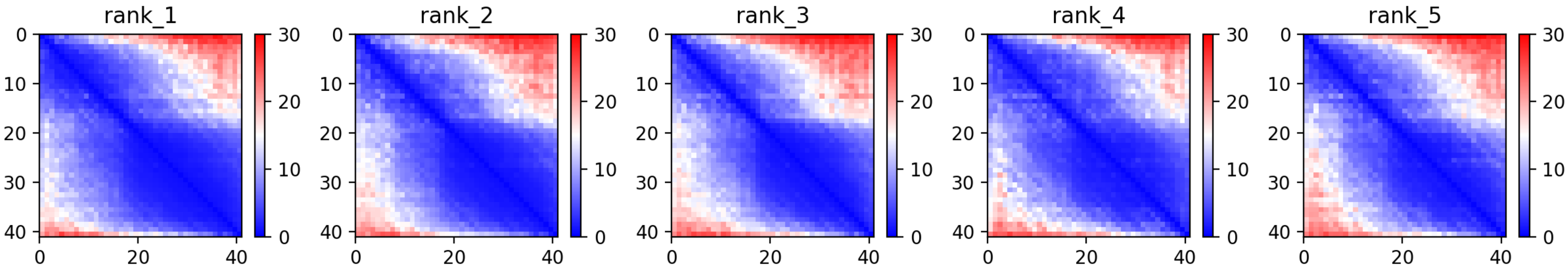

CP130: MtrunA17_Chr8g0356581

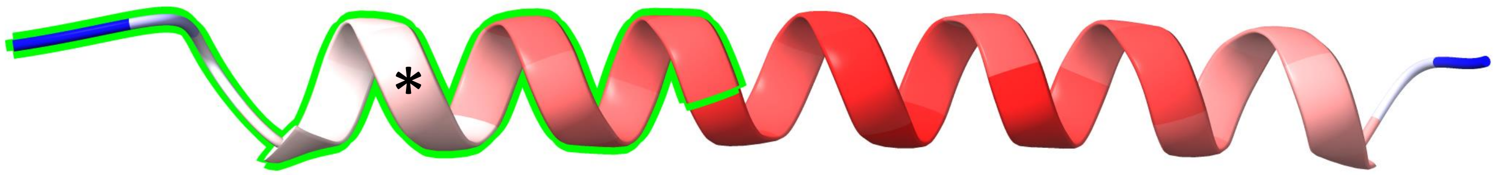

+1 (1a1→2a2)

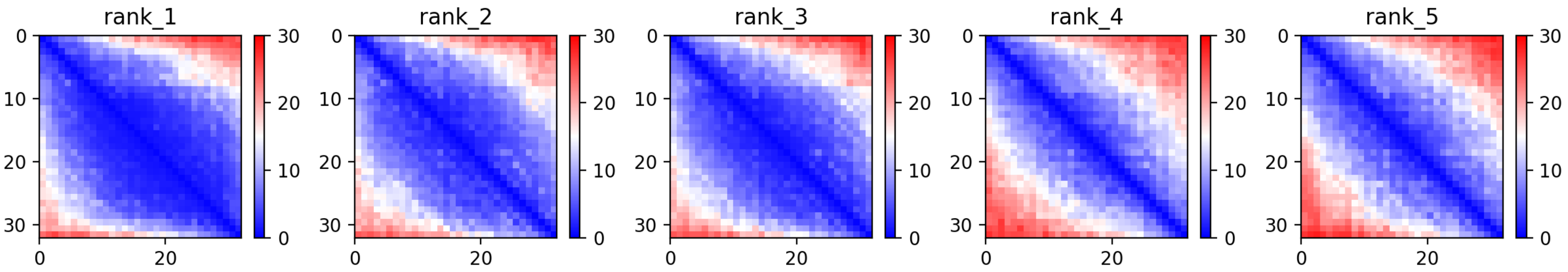

### CP131: MtrunA17_Chr8g0365341

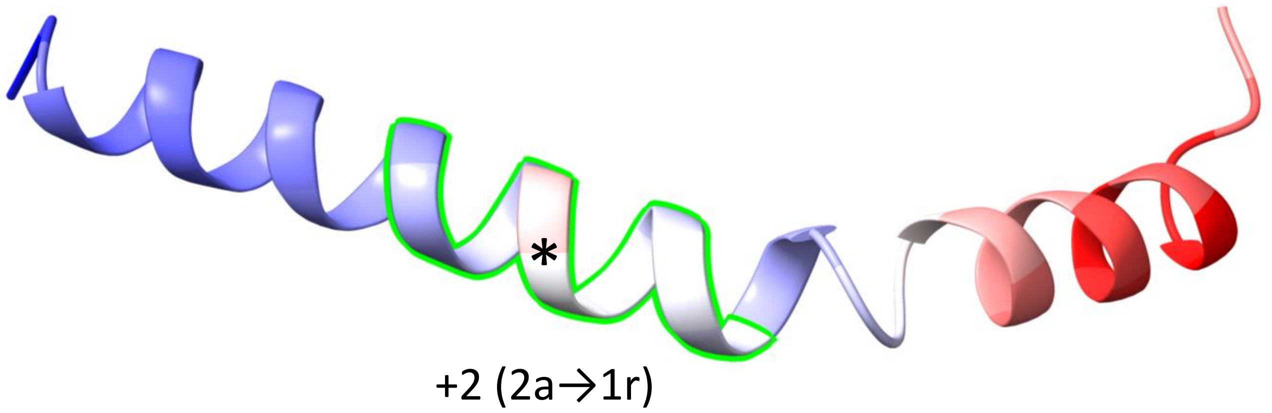


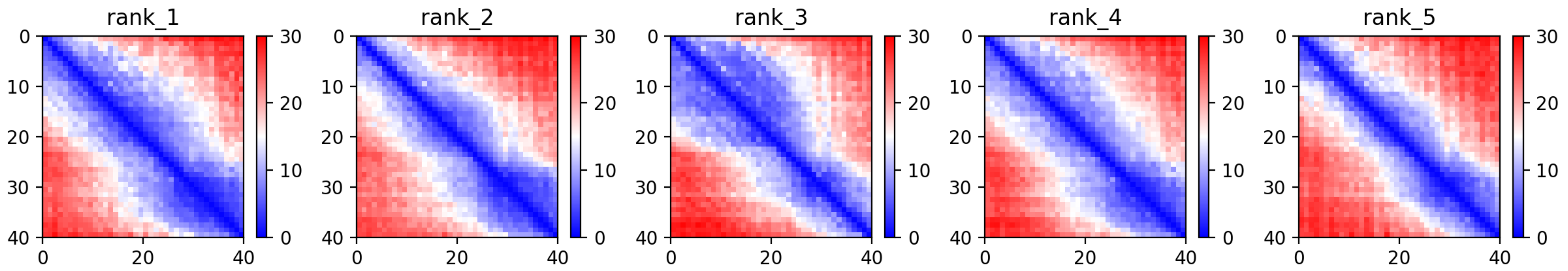

CP132: MtrunA17_Chr8g0368731

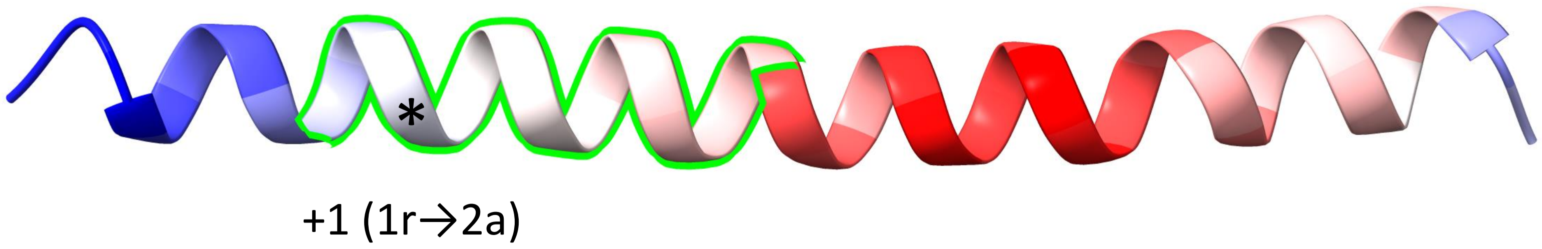


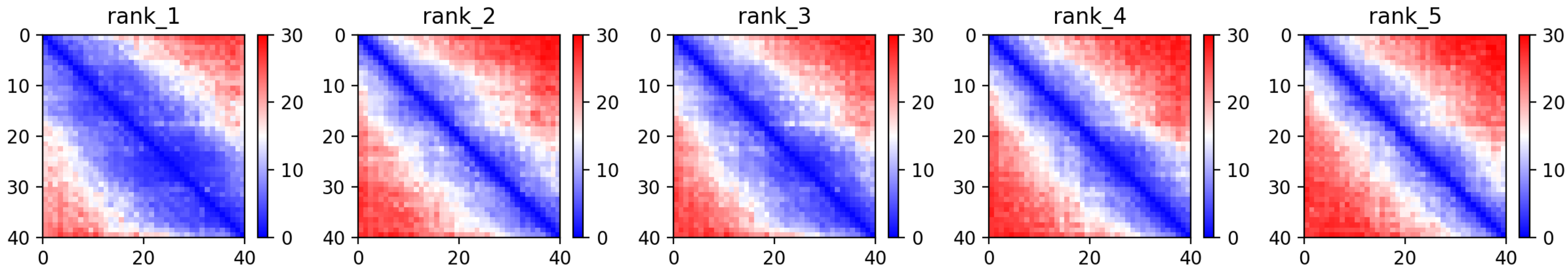

CP133: MtrunA17_Chr8g0368931

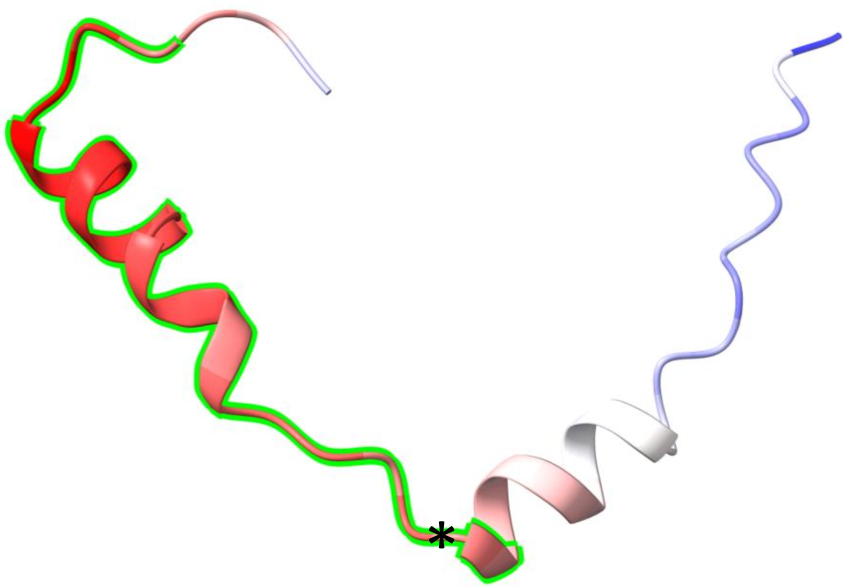


-2 (3r→1a)

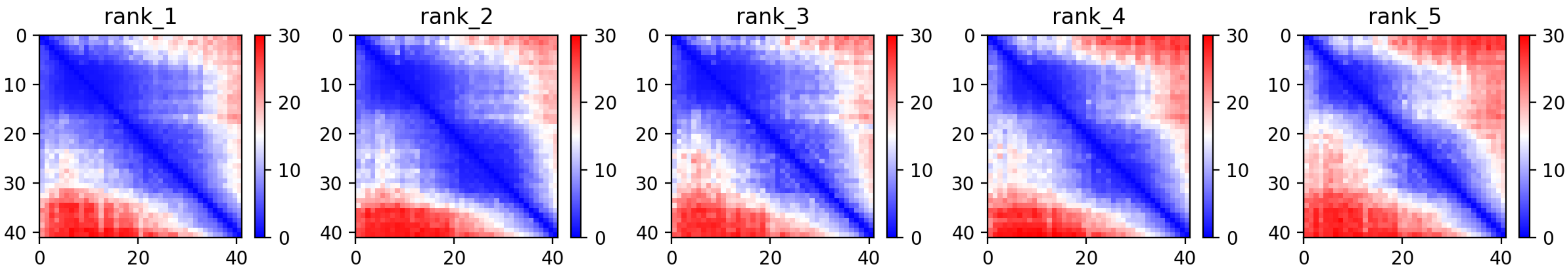

CP134: MtrunA17_Chr8g0371281

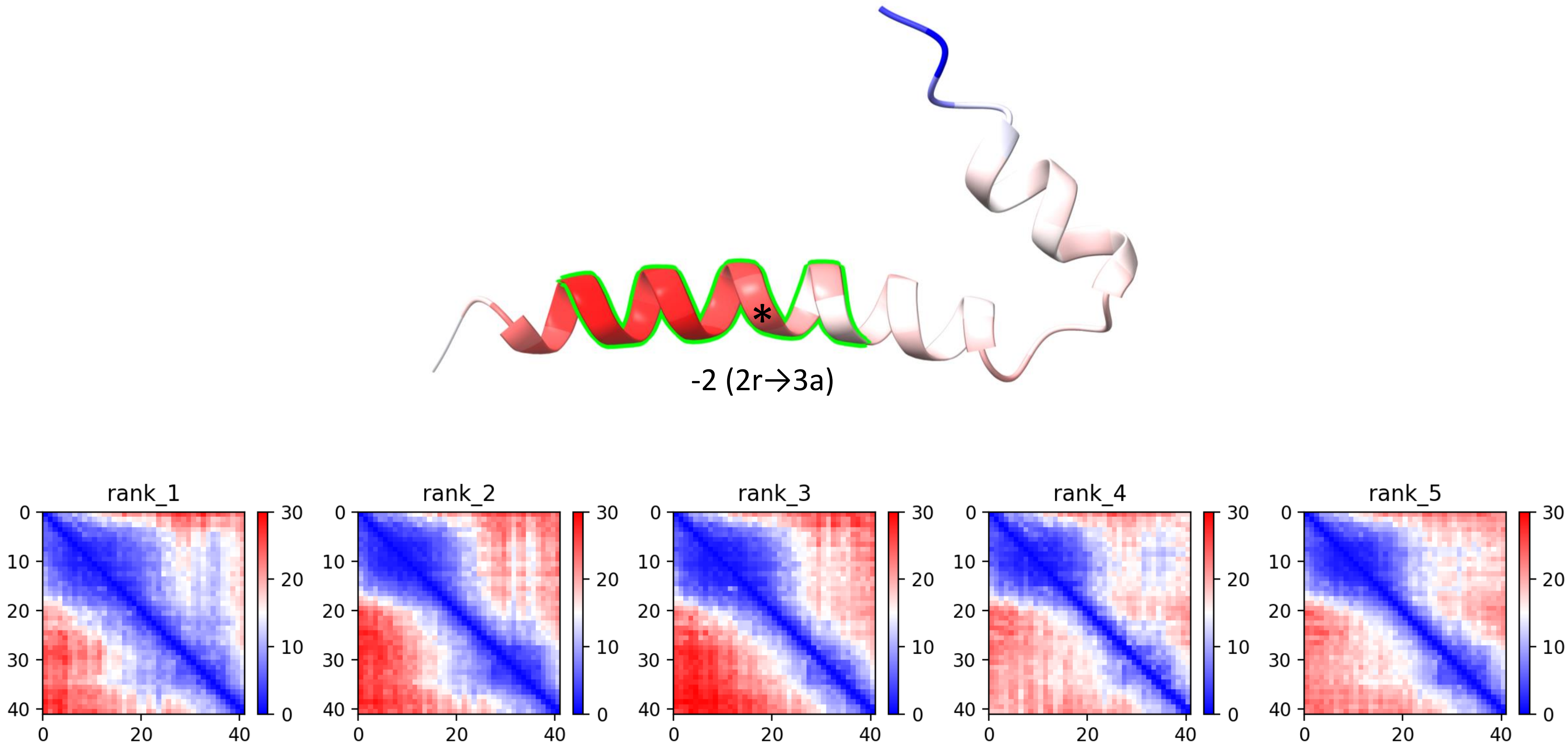

CP135: MtrunA17_Chr8g0371741

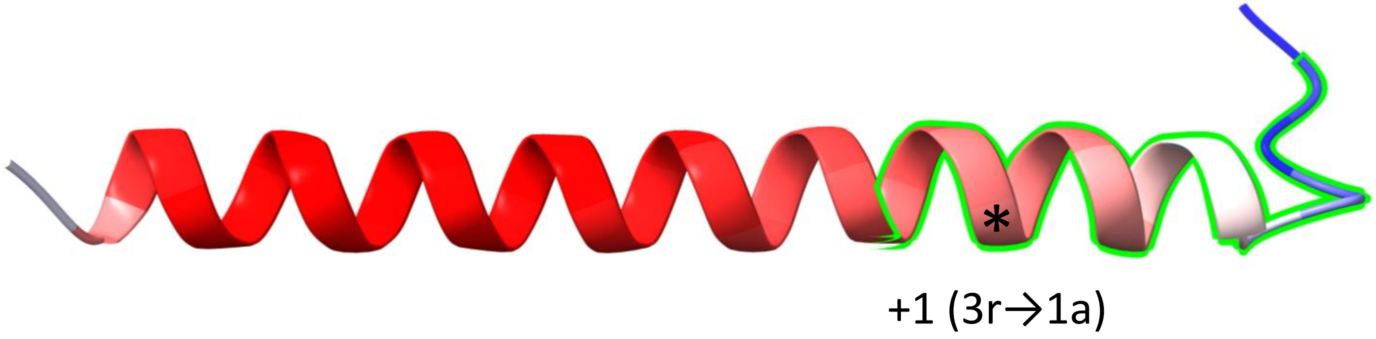


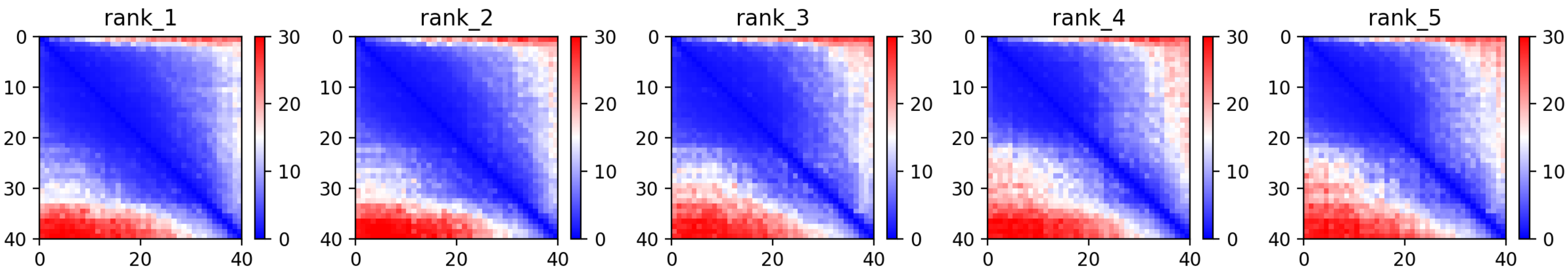

CP137: MtrunA17_Chr8g0376411

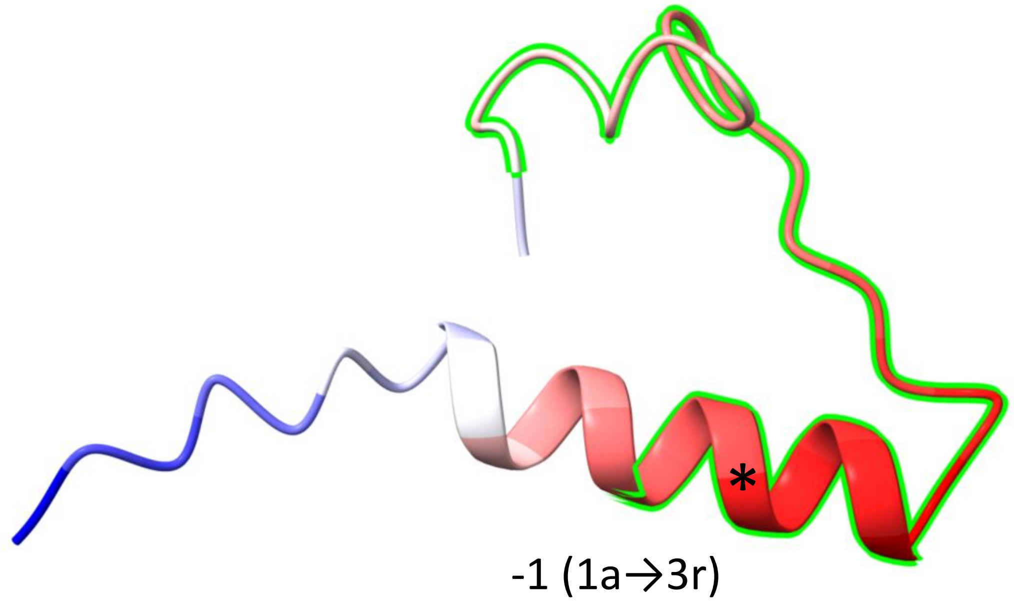


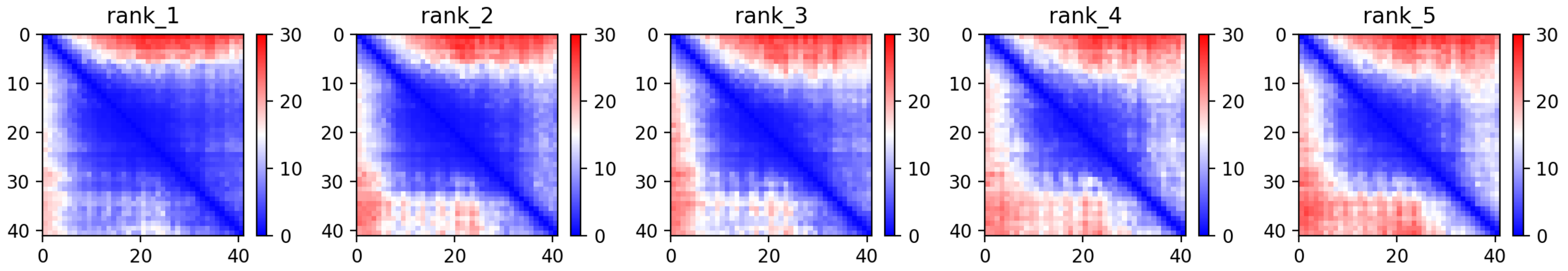

CP138: MtrunA17_Chr8g0377071

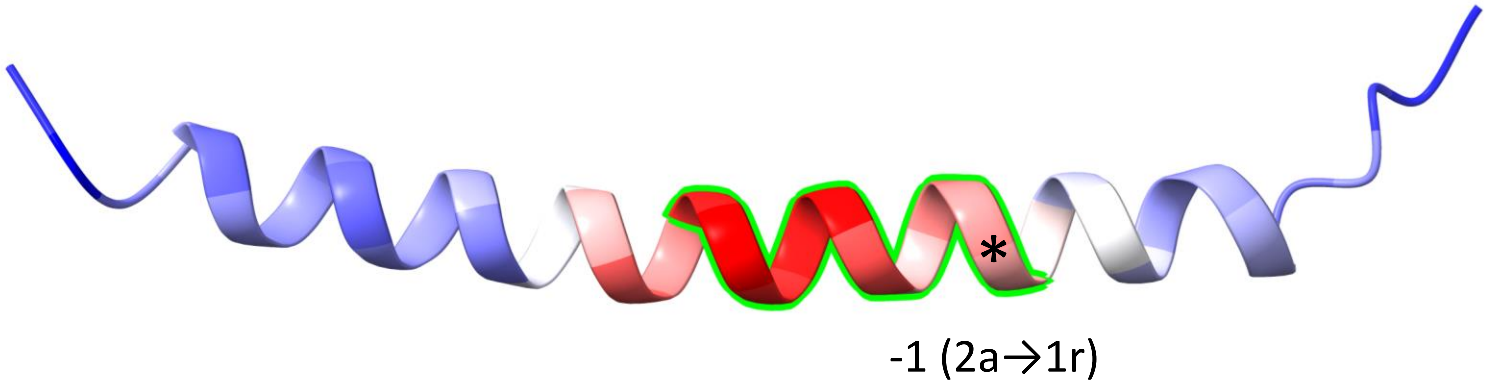


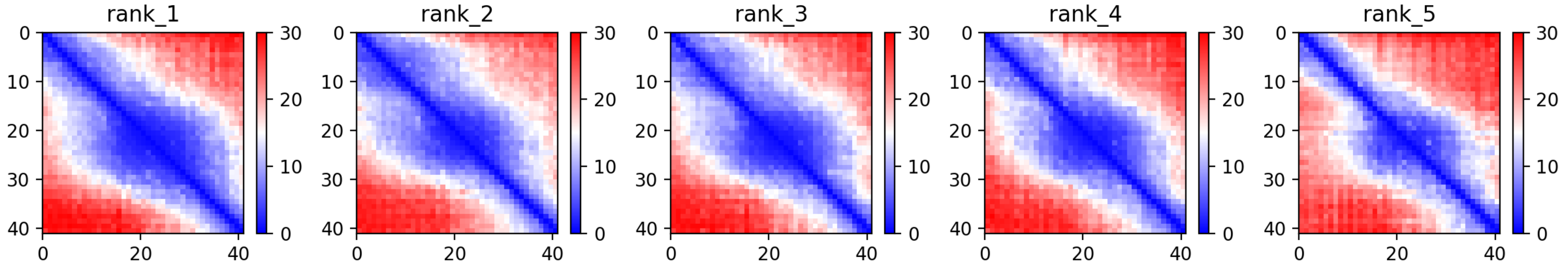

CP140: MtrunA17_Chr8g0392351

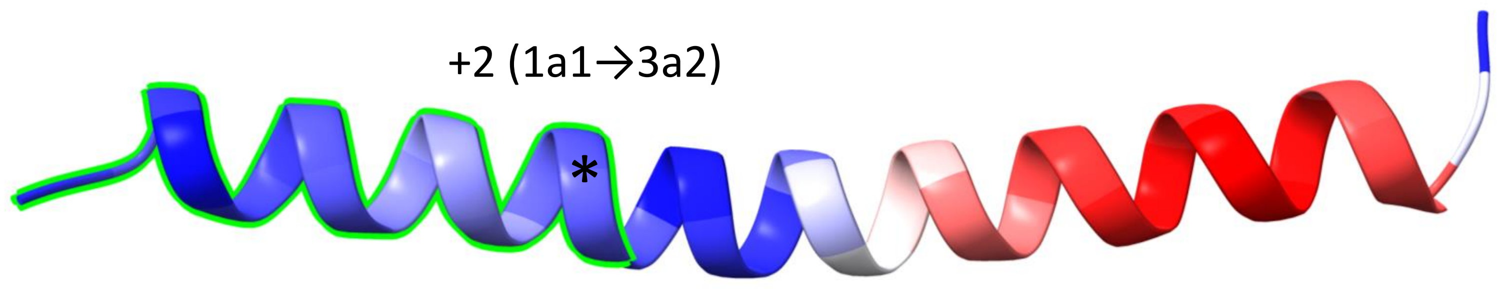


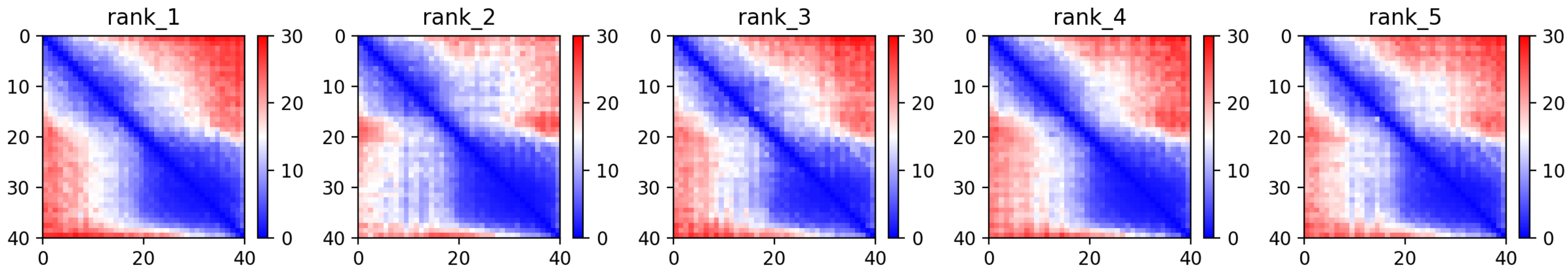

CP142: MtrunA17_CPg0492381

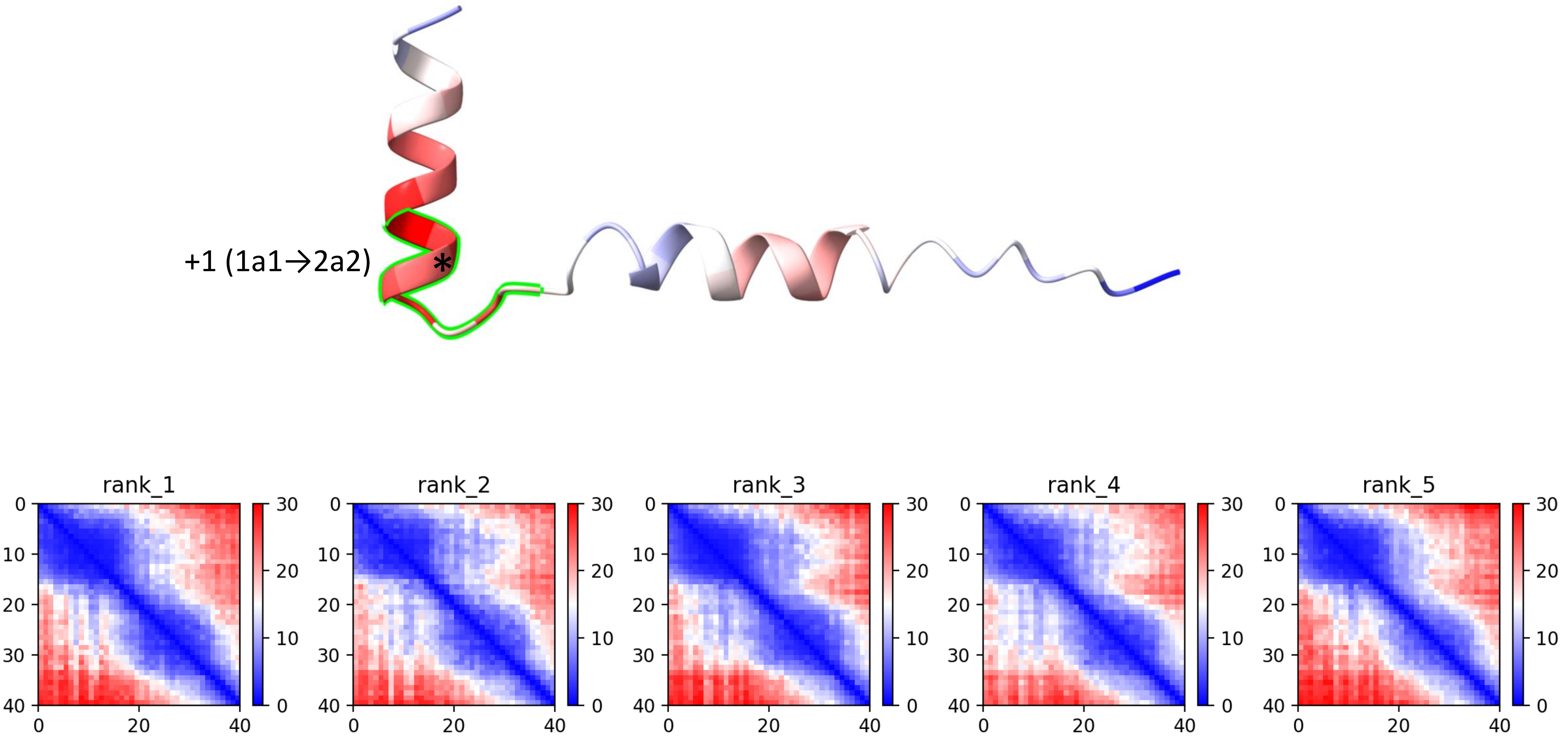

CP143: MtrunA17_CPg0492461

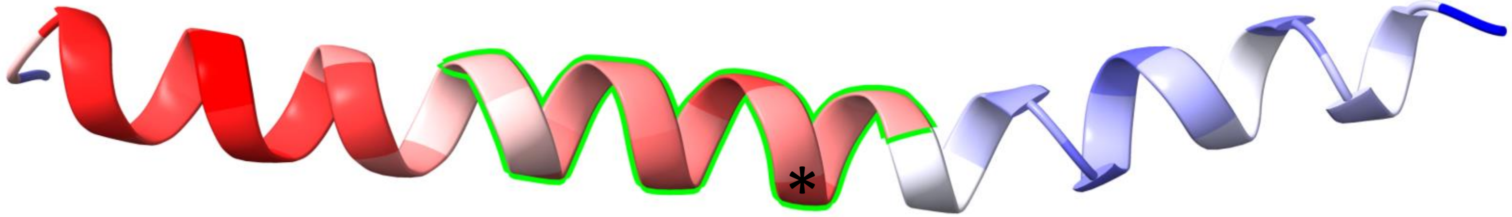

+1 (3r→1a)

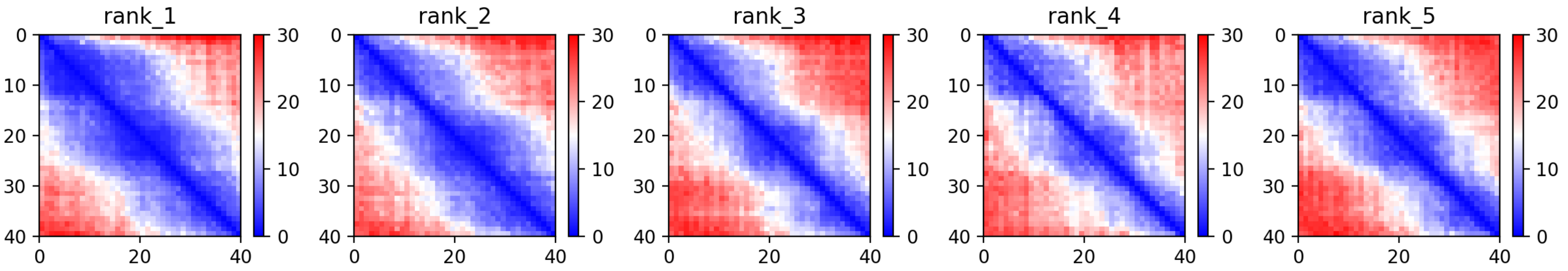

CP145: MtrunA17_CPg0492941

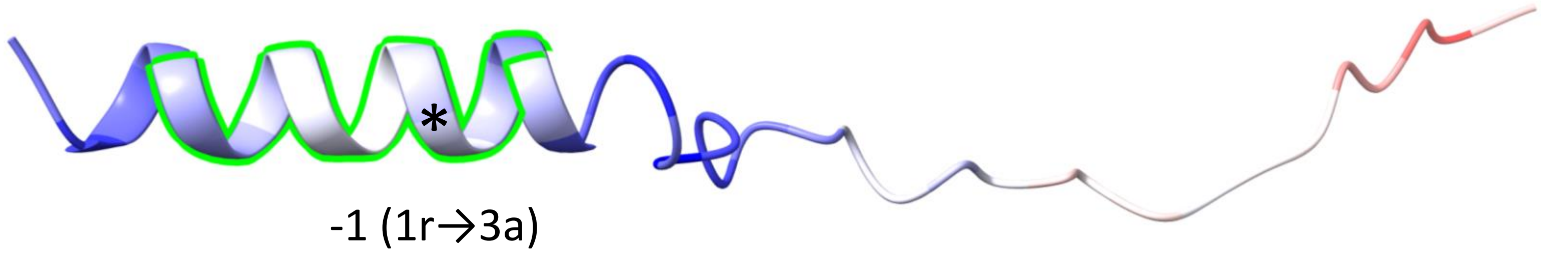


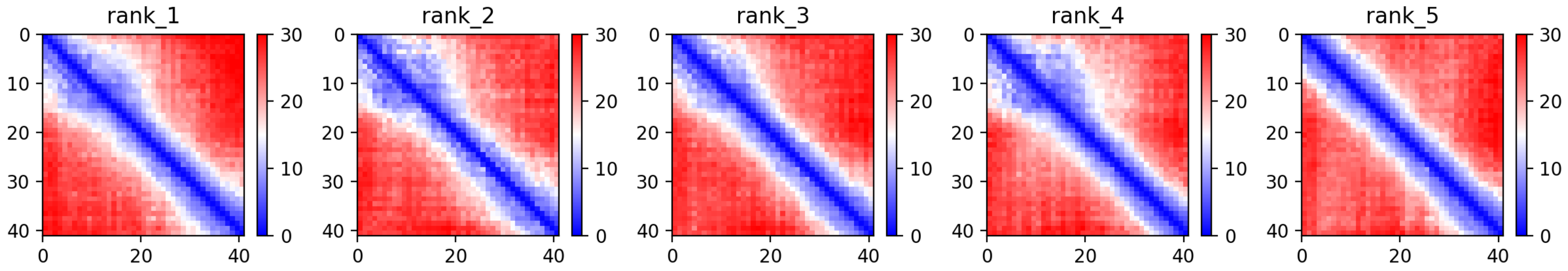

CP146: MtrunA17_CPg0493291

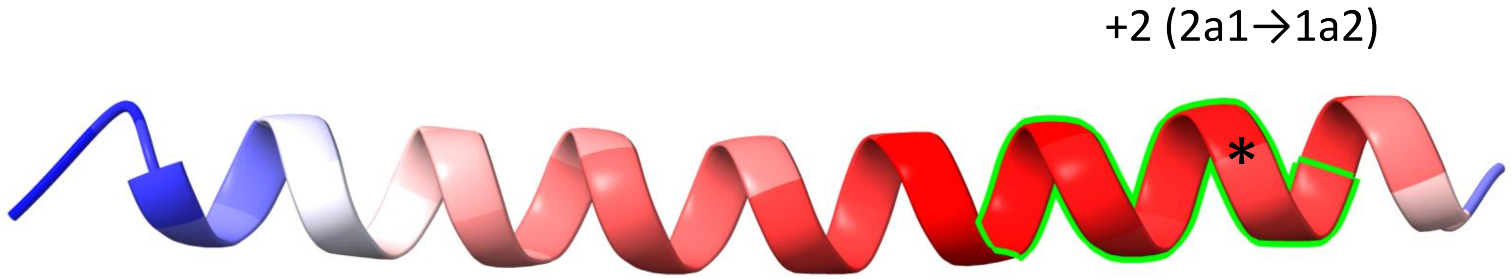


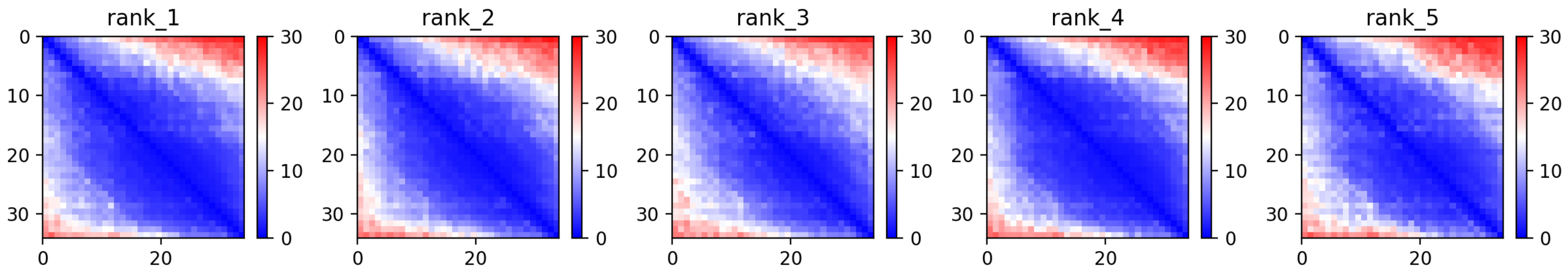

CP152: MtrunA17_MTg0491151

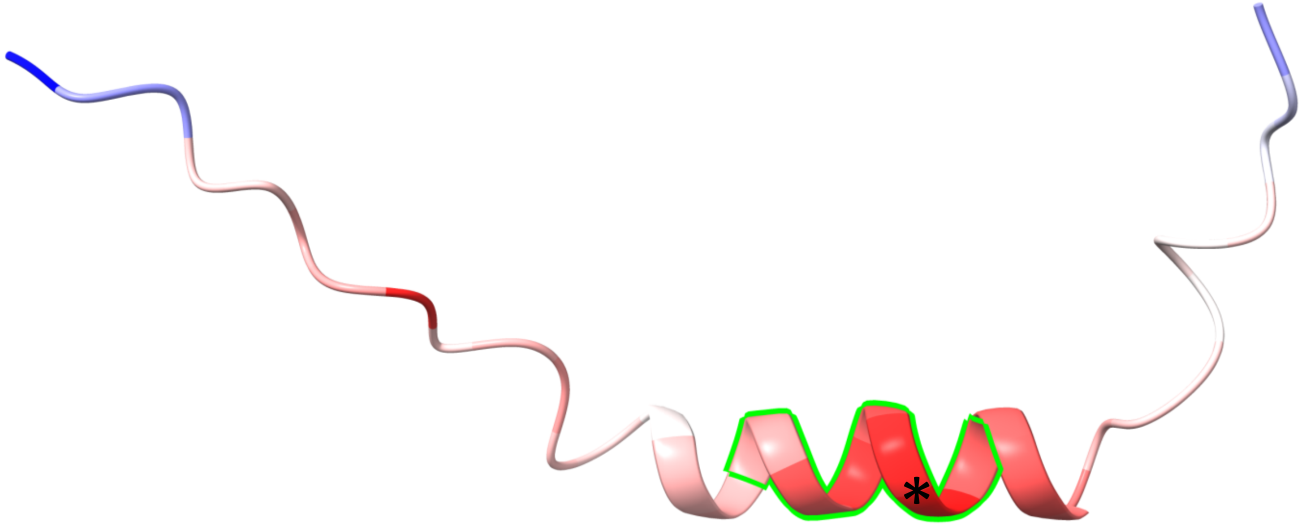


+2 (2a1→1a2)

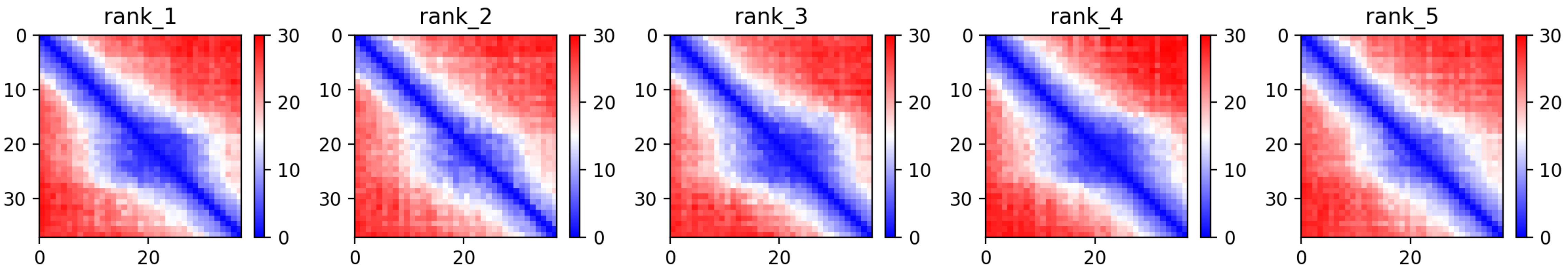

### CP153: MtrunA17_MTg0491291

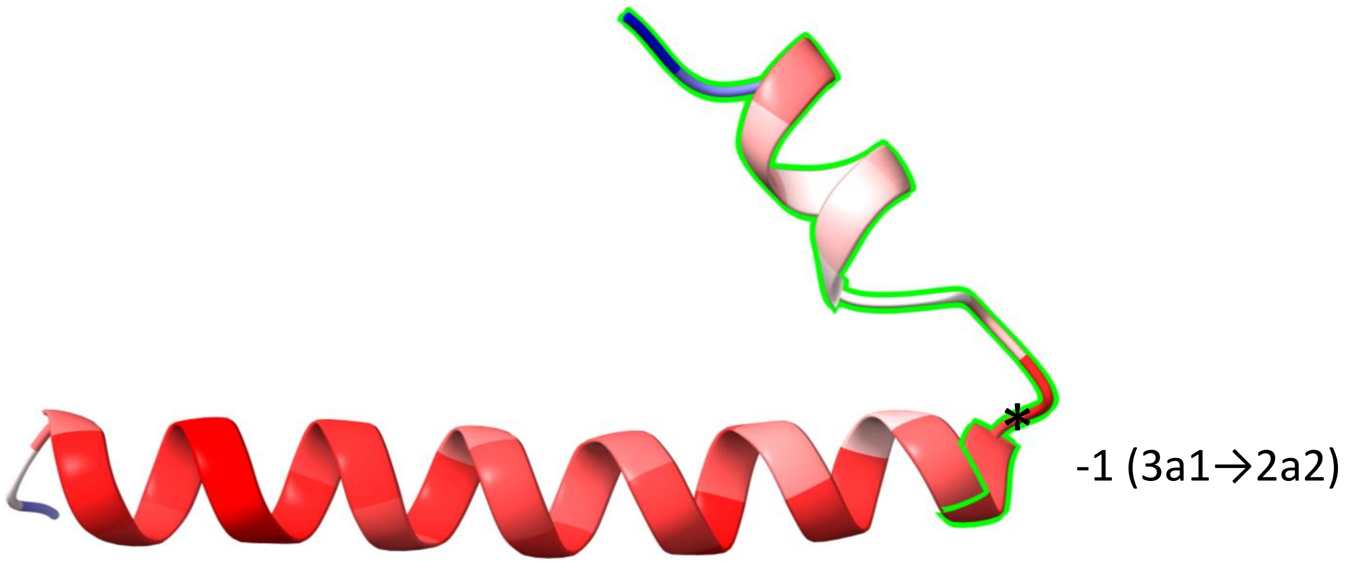


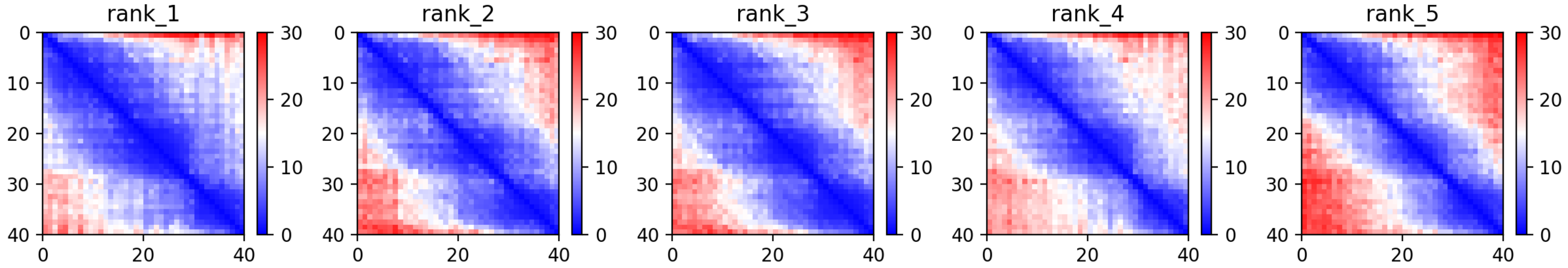

CP155: MtrunA17_MTg0491621

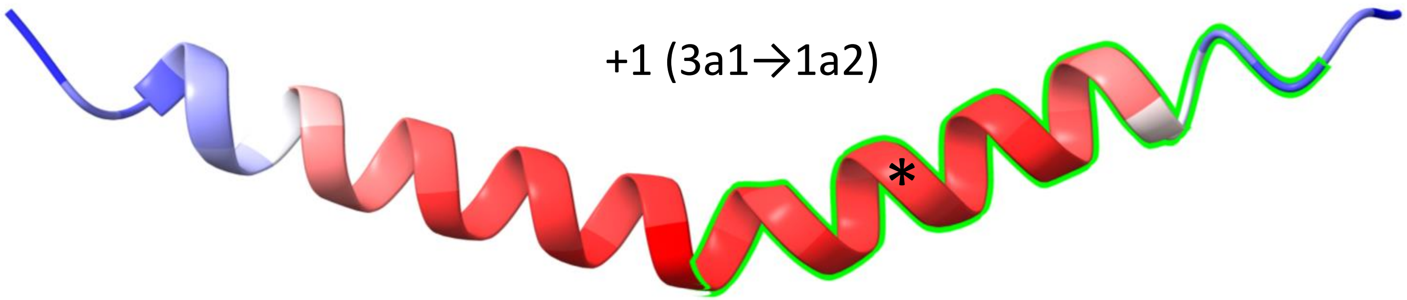


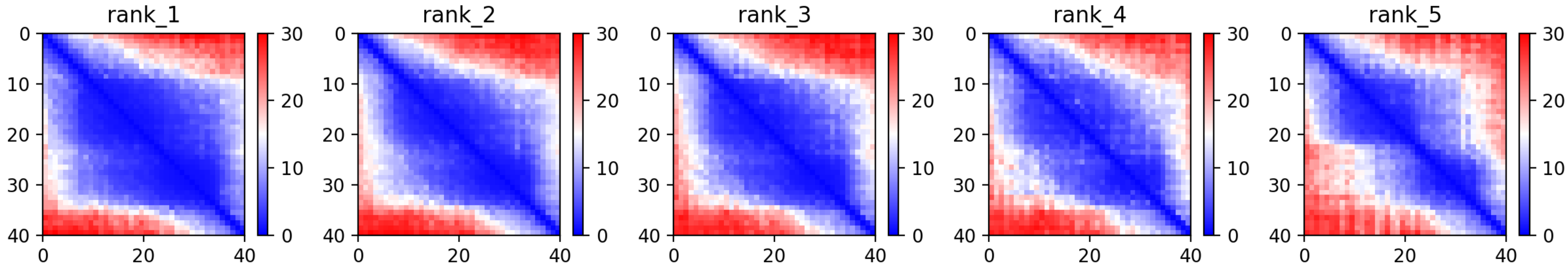

CP156: MtrunA17_MTg0491711

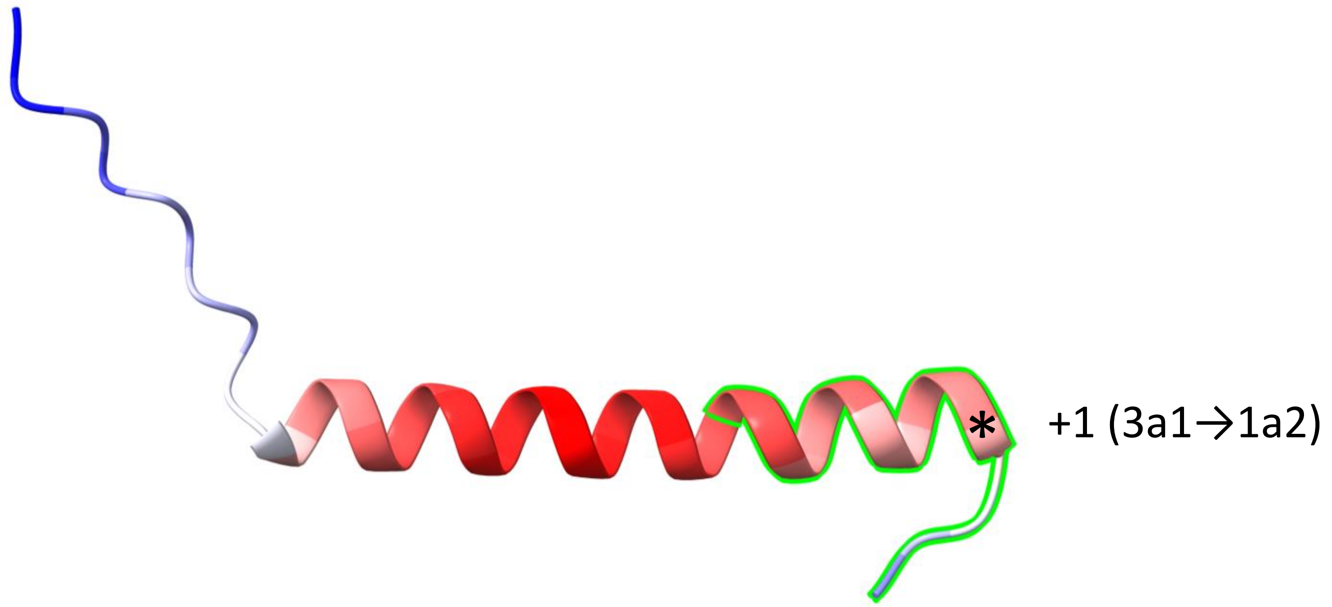


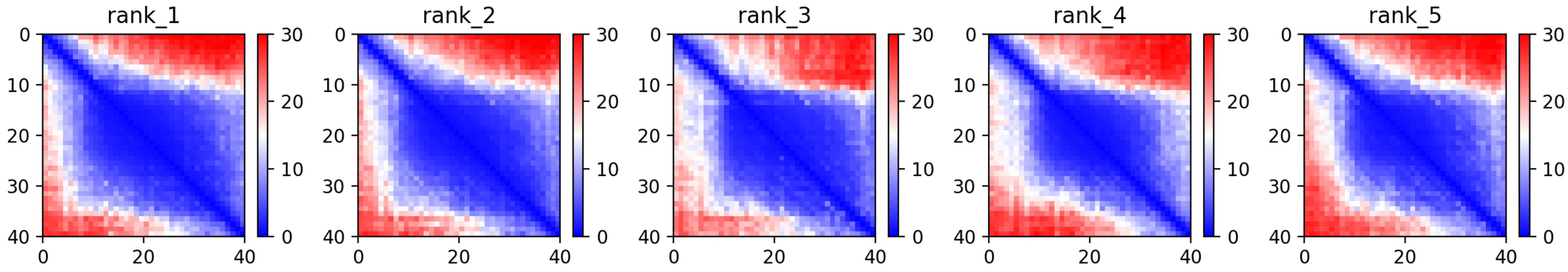

**Supplementary Dataset S7 Part 4**. A graphical summary on folding predictions for 90 MS-supported chimeric peptide models (CPs) that contain alpha-helices and no beta-sheets. For each CP, one image is displayed: the predicted folding structure (top) along with heat maps of five predictions (bottom). The rest of the legend is the same as for Supplementary Dataset S7 Part 1.
