## Supplementary material for "Discovery of diverse chimeric peptides in a eukaryotic proteome sets the stage for the experimental proof of the mosaic translation hypothesis": Supplementary Dataset S7 Folding of chimeric protein models Part 5 Beta-sheets only.pdf

CP11: MtrunA17_Chr1g0164591

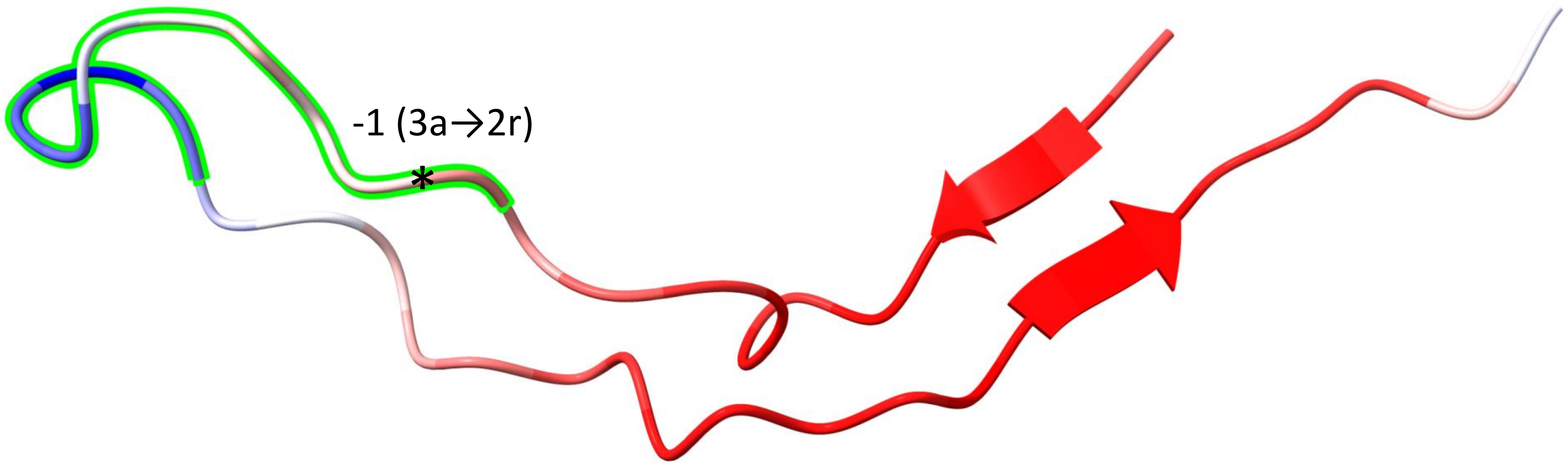


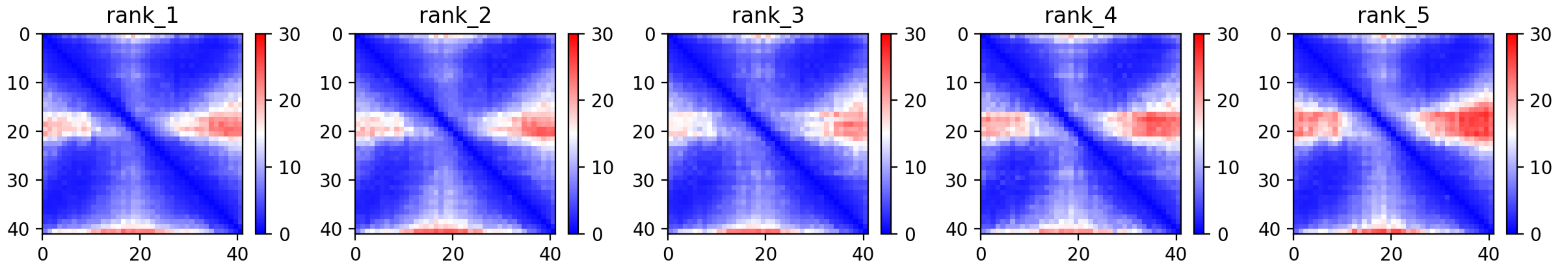

### CP19: MtrunA17_Chr1g0185871

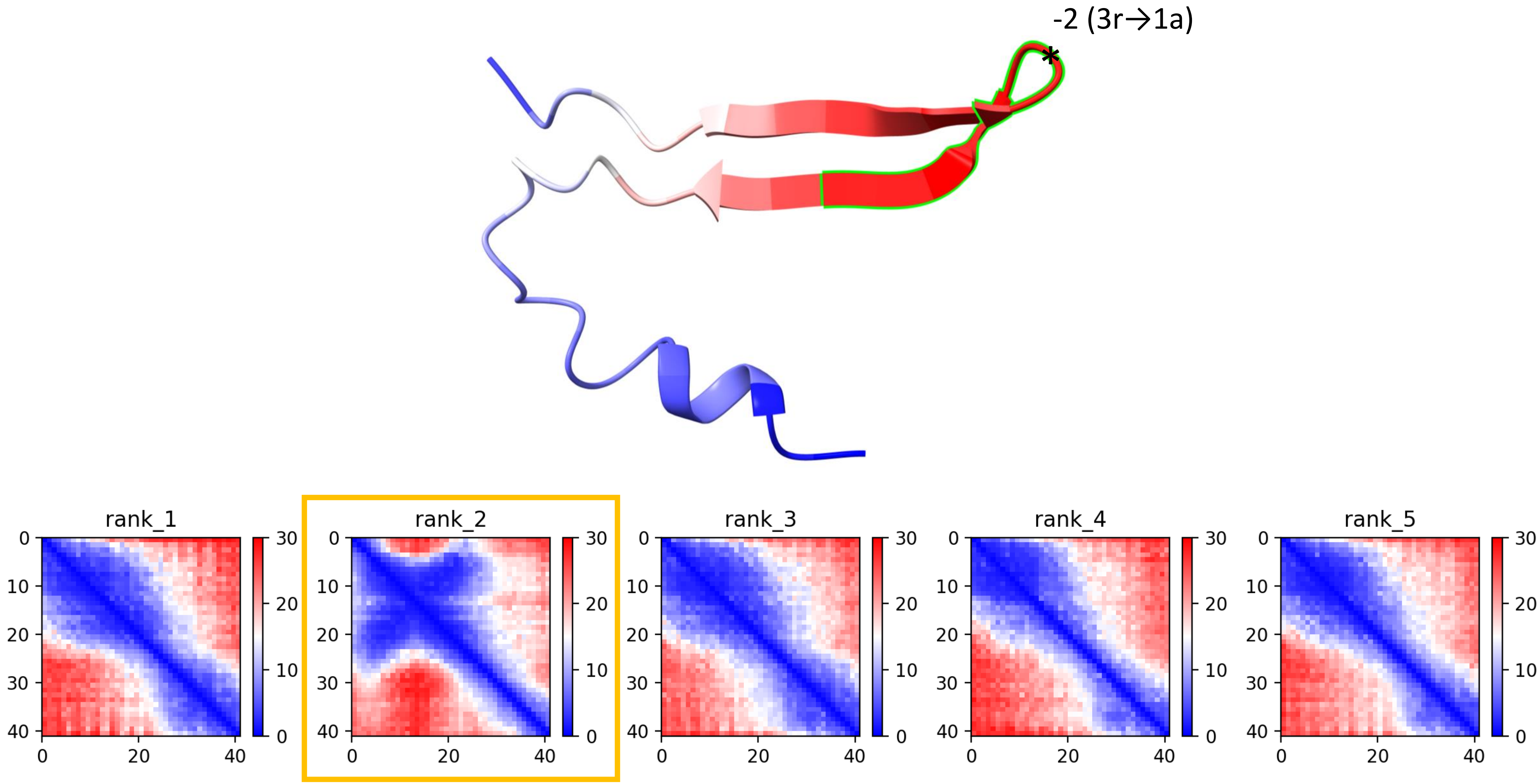

CP20: MtrunA17_Chr1g0190571

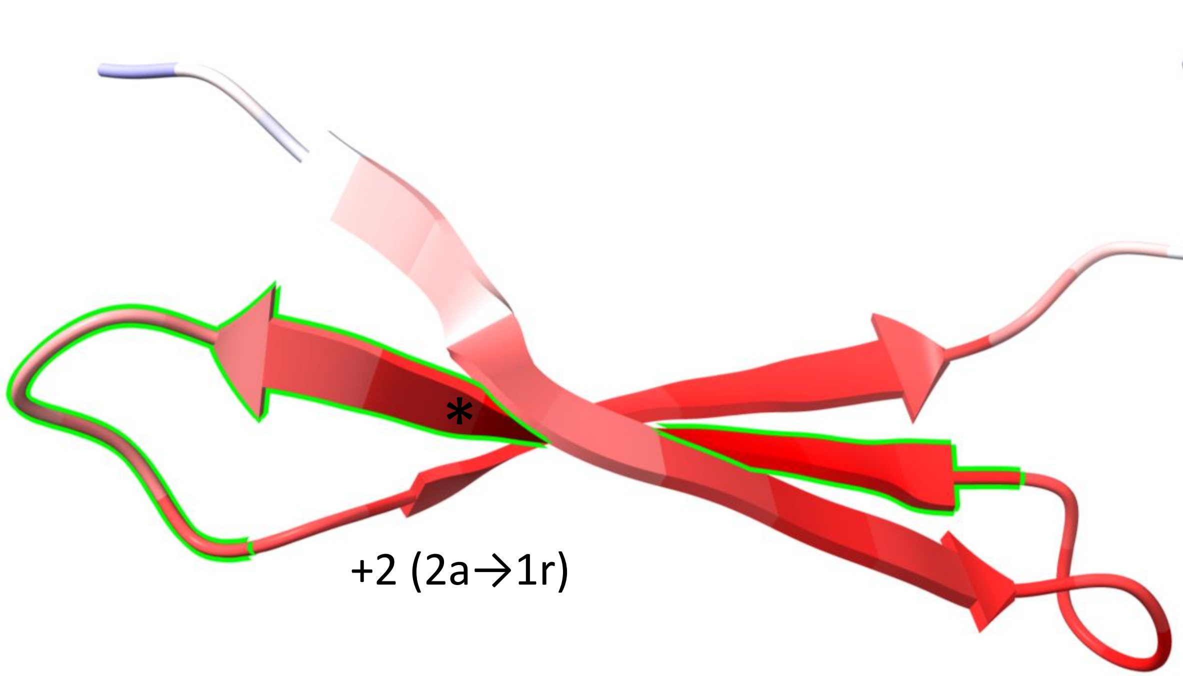


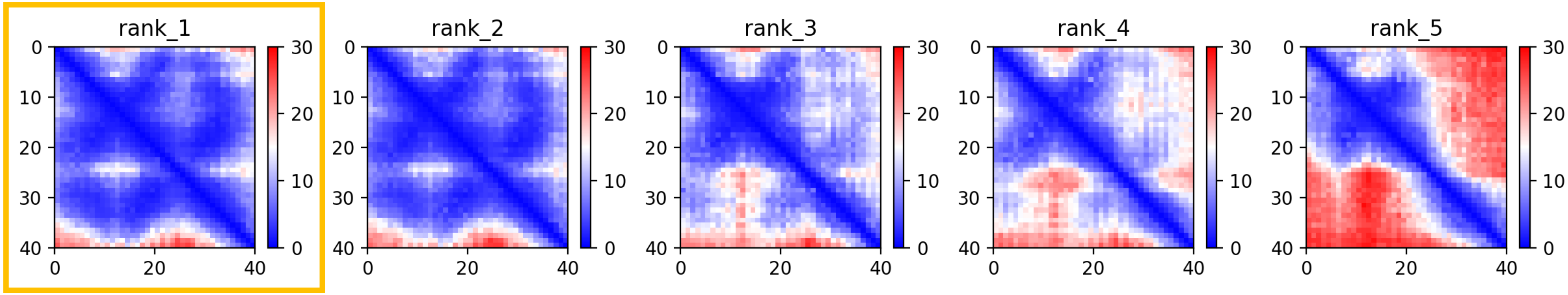

CP20: MtrunA17_Chr1g0190571

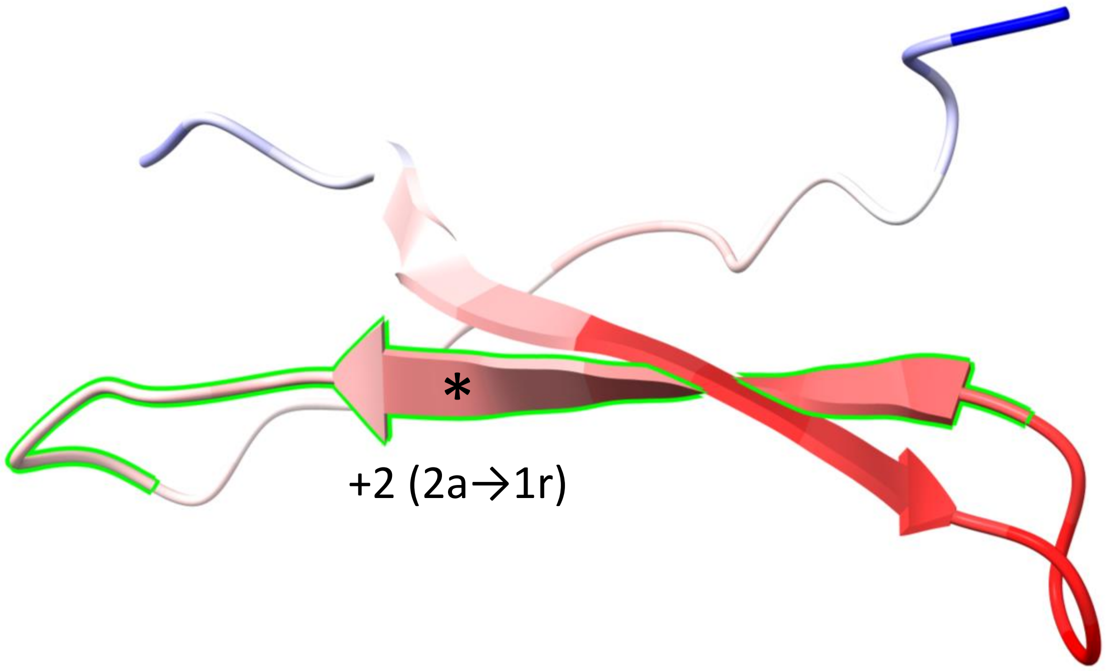


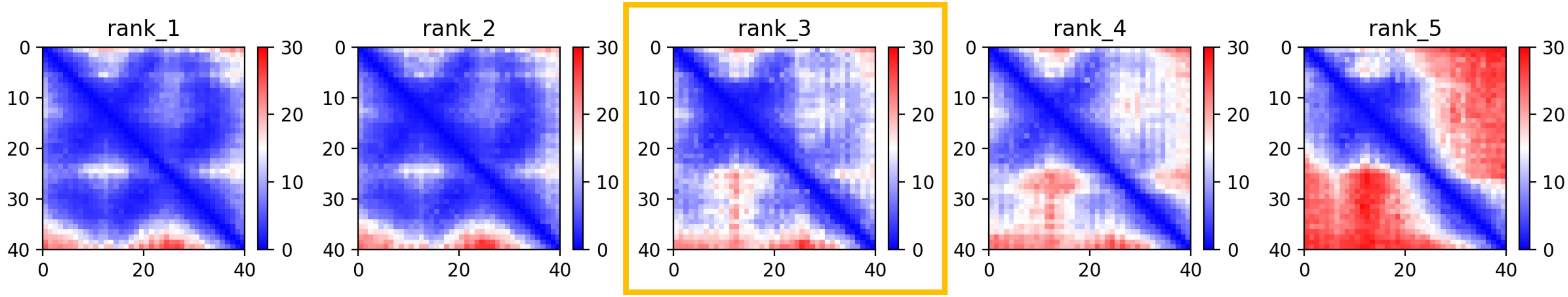

CP20: MtrunA17_Chr1g0190571

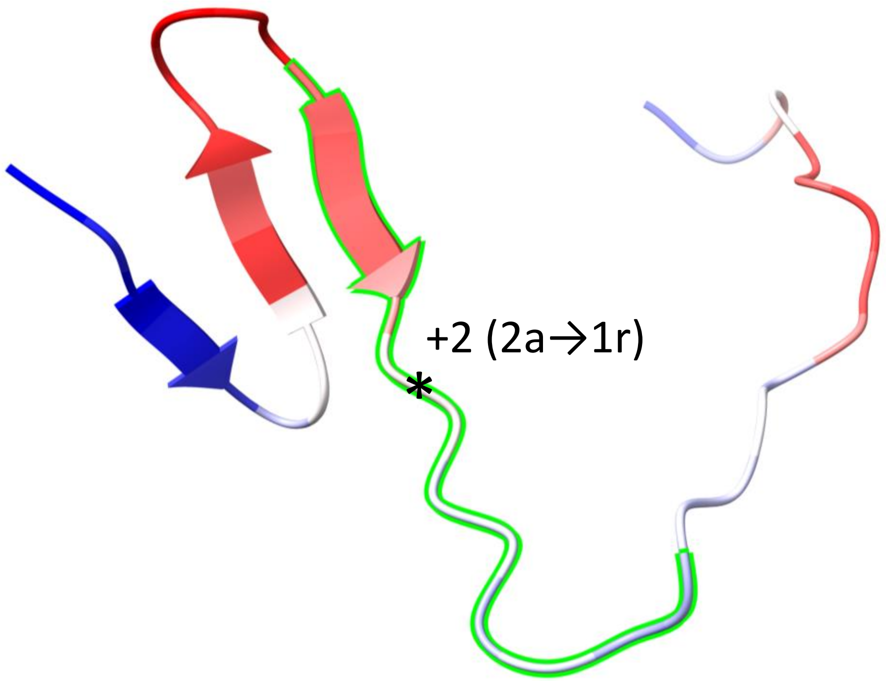


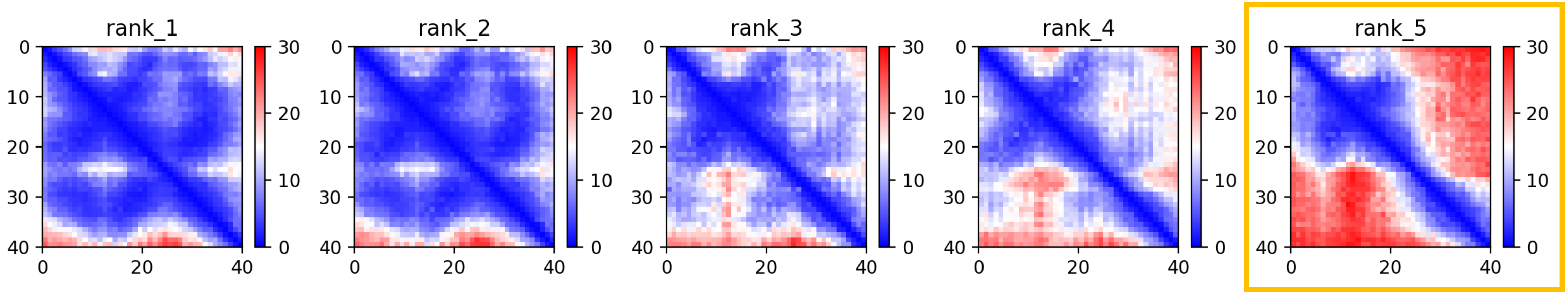

### CP23: MtrunA17_Chr1g0200071

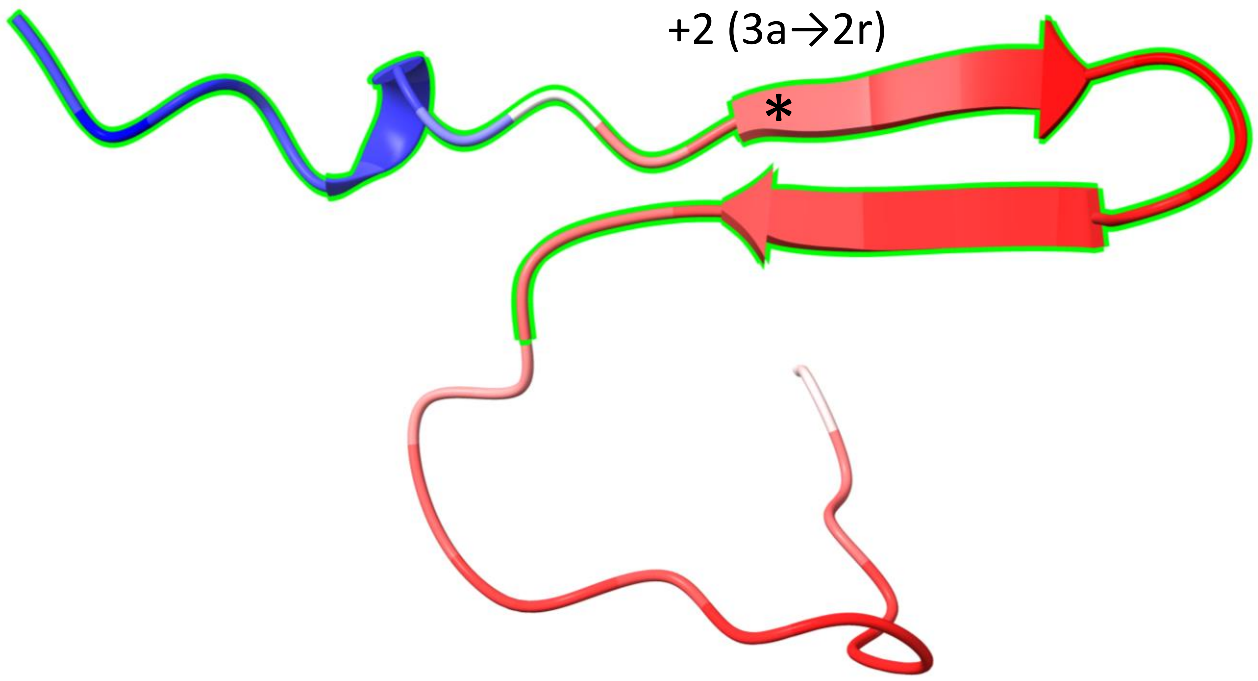


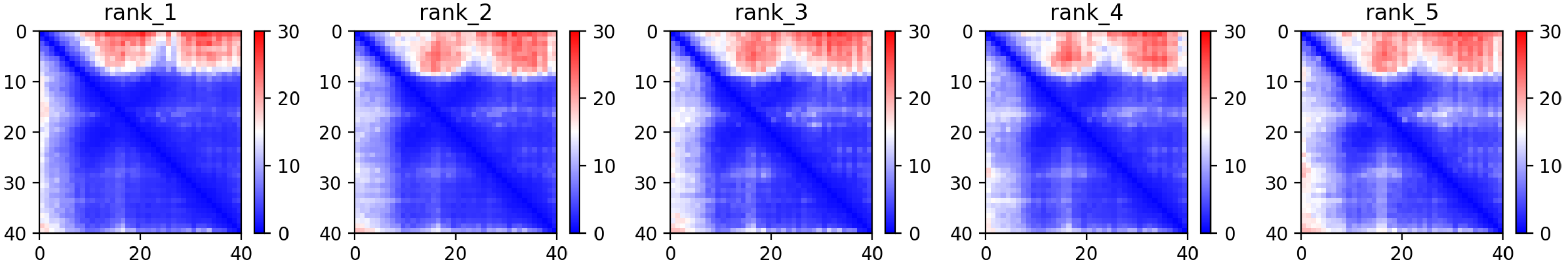

CP28: MtrunA17_Chr1g0207921

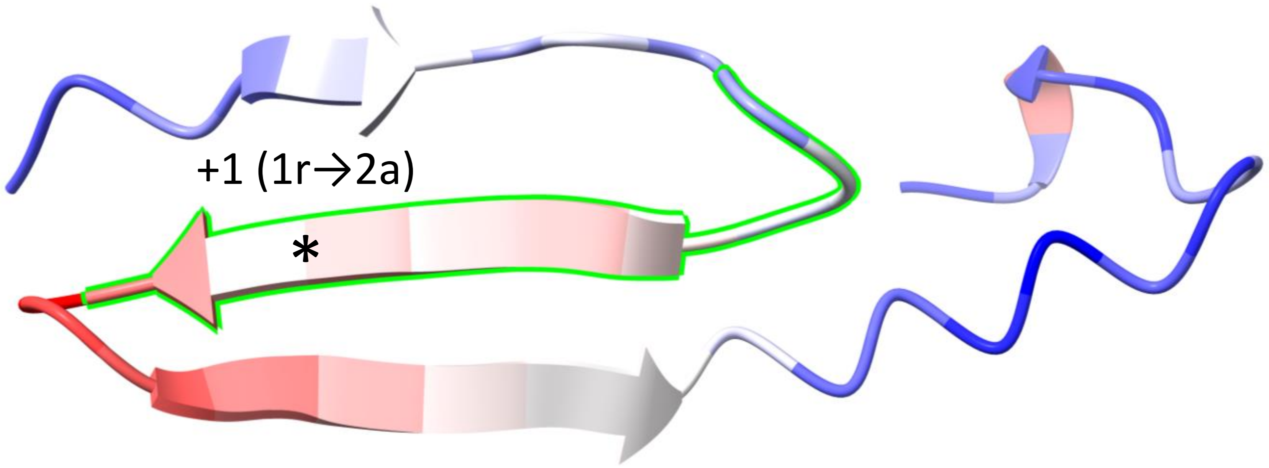


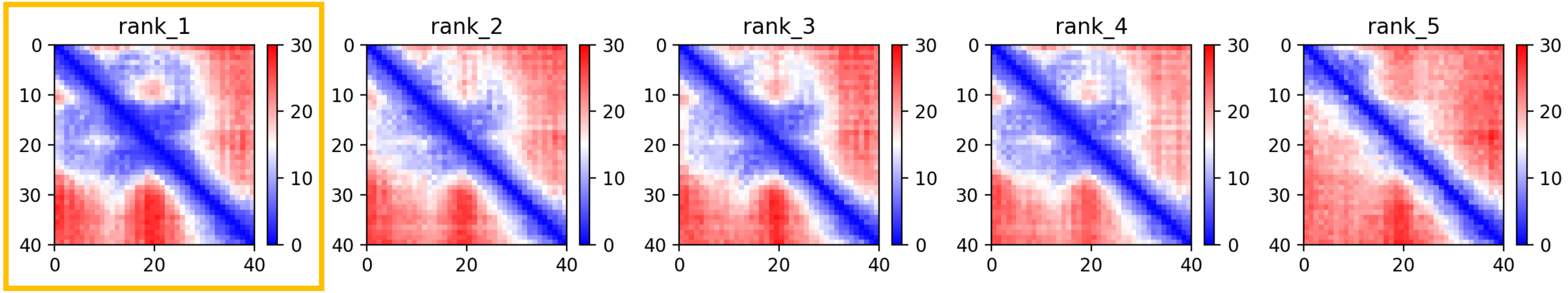

CP31: MtrunA17_Chr1g0212961

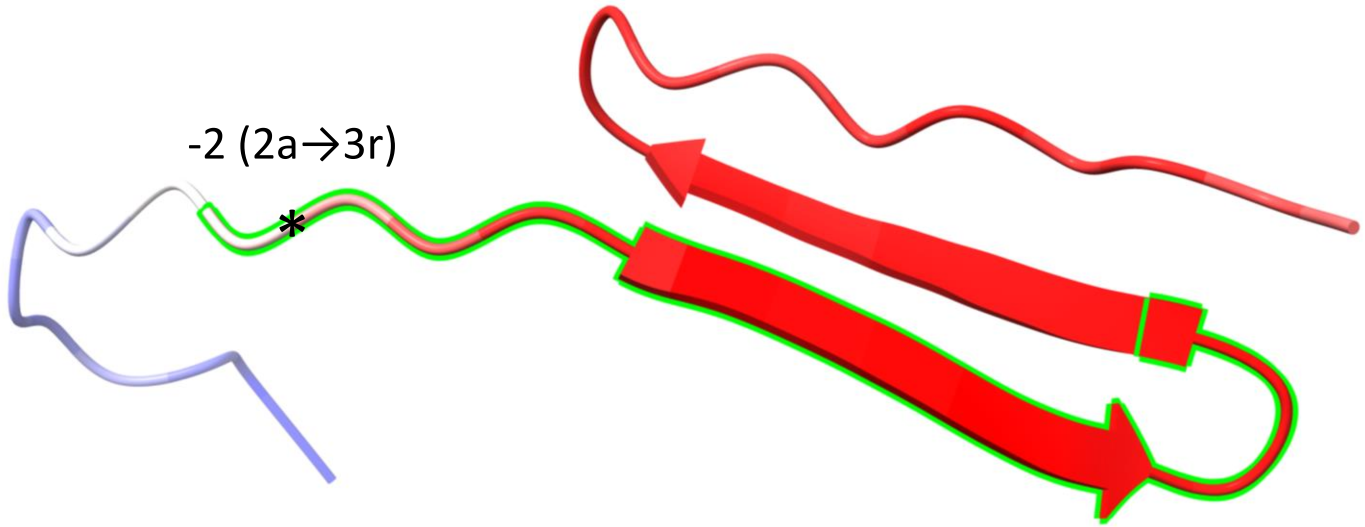


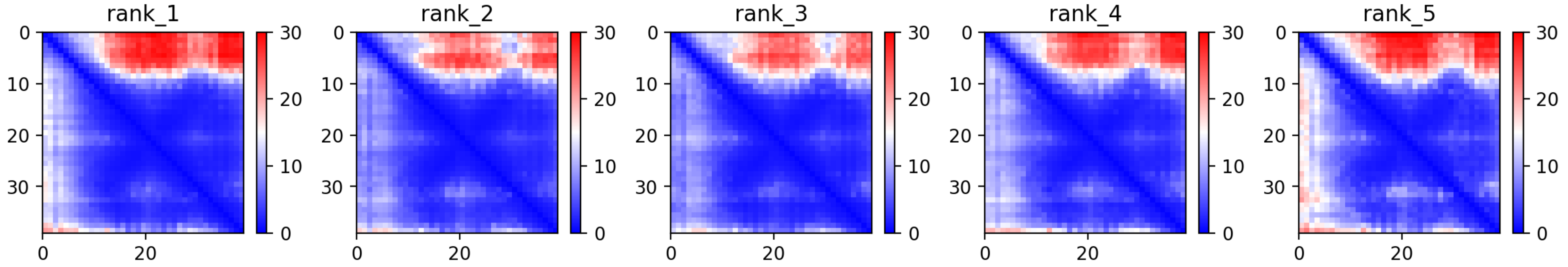

CP34: MtrunA17_Chr2g0285461

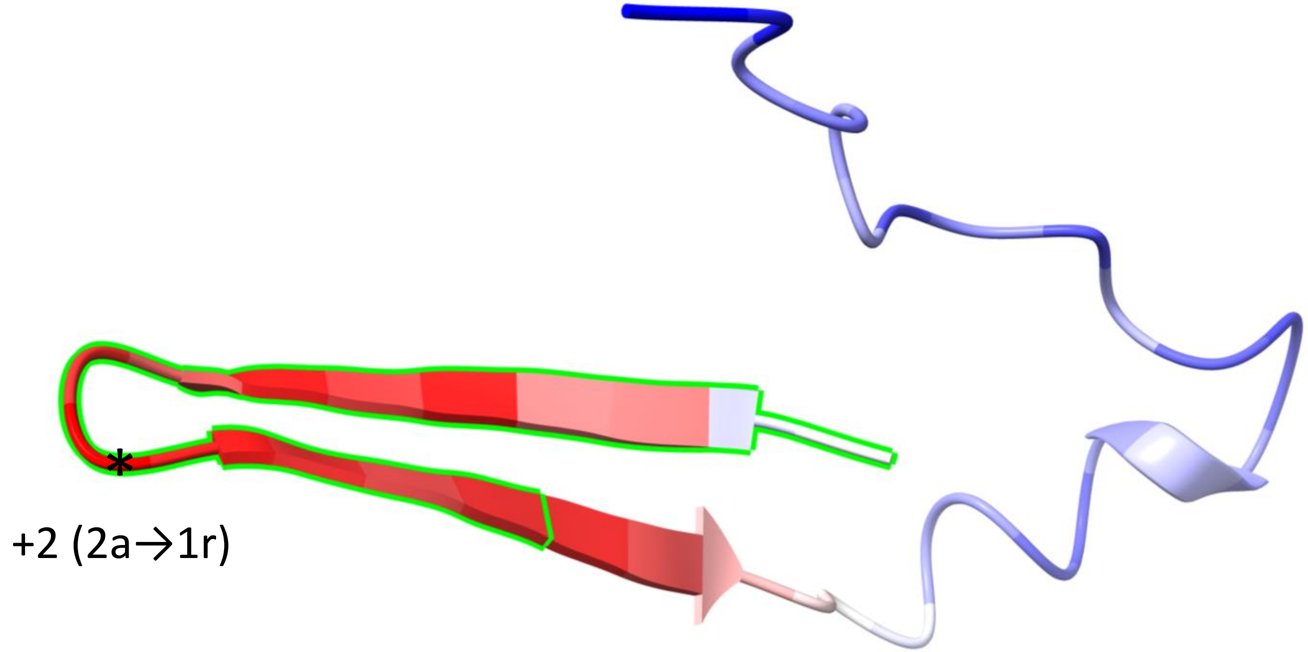


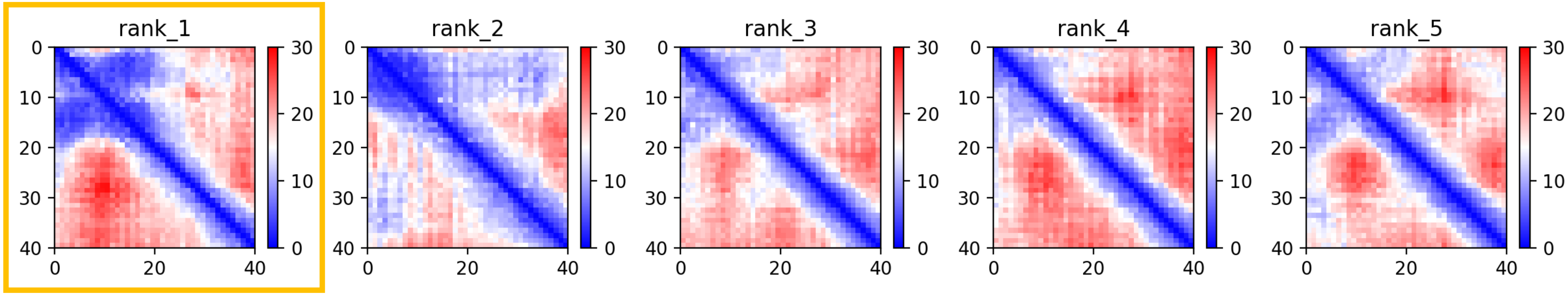

CP38: MtrunA17_Chr2g0304891

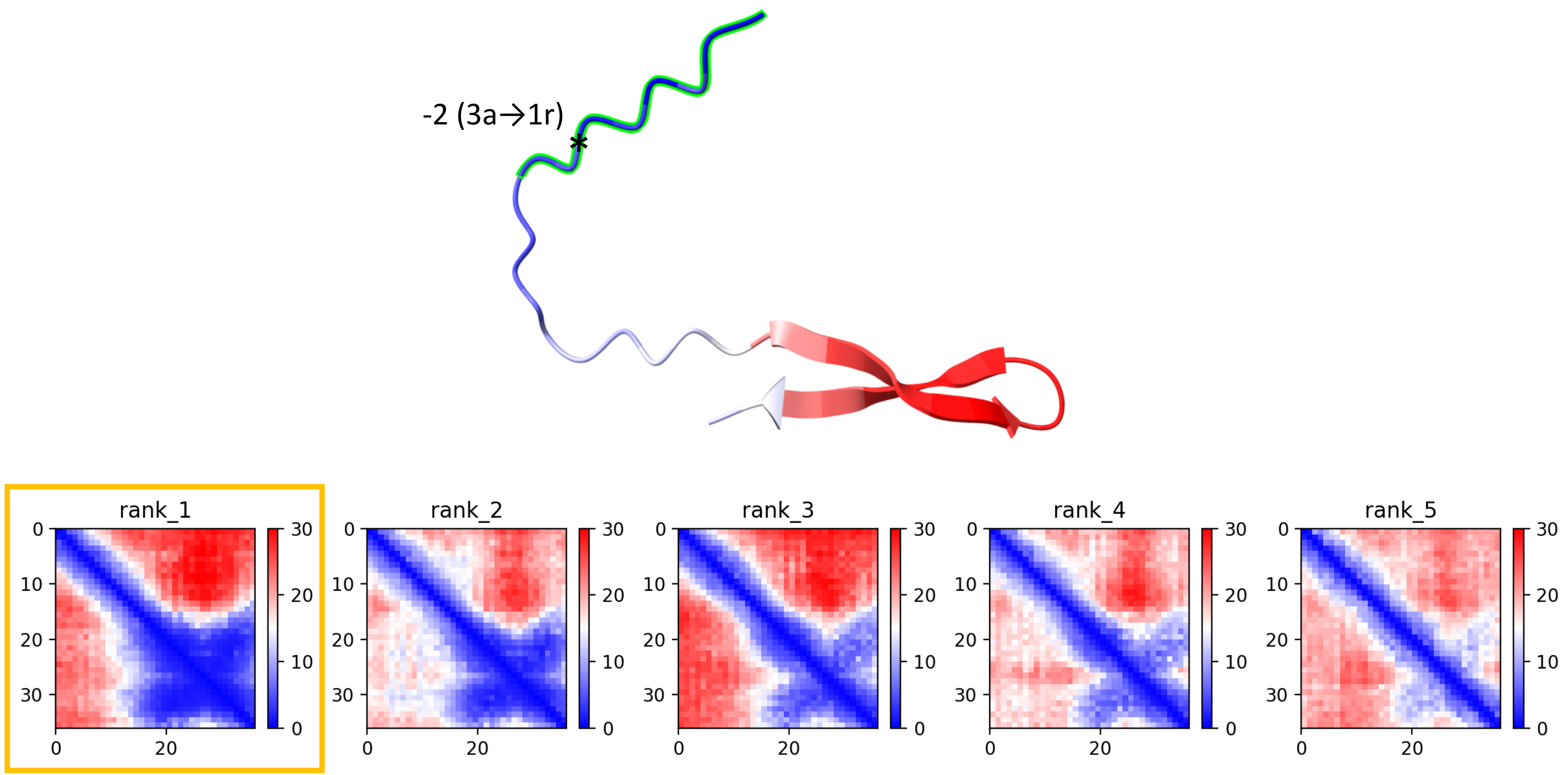

CP42: MtrunA17_Chr2g0312631

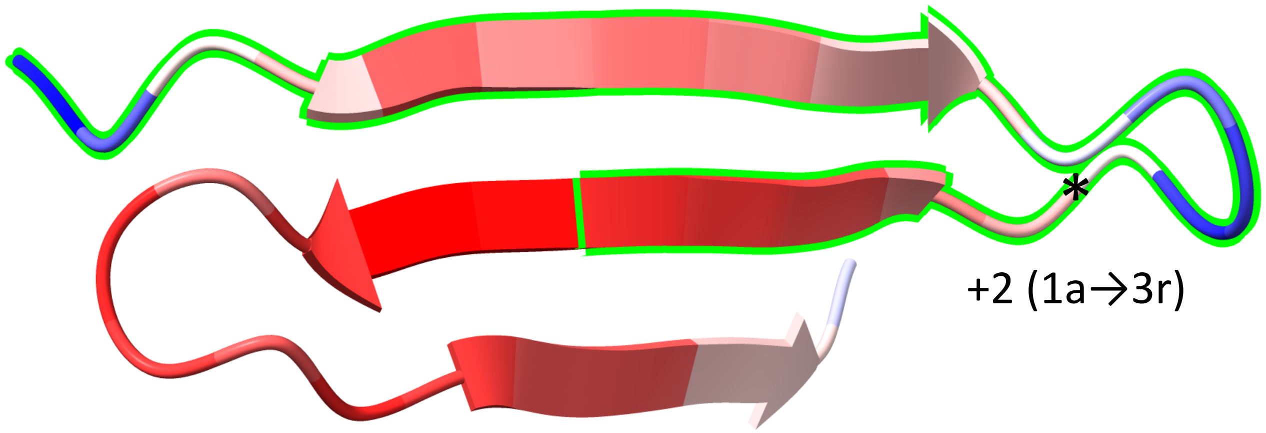


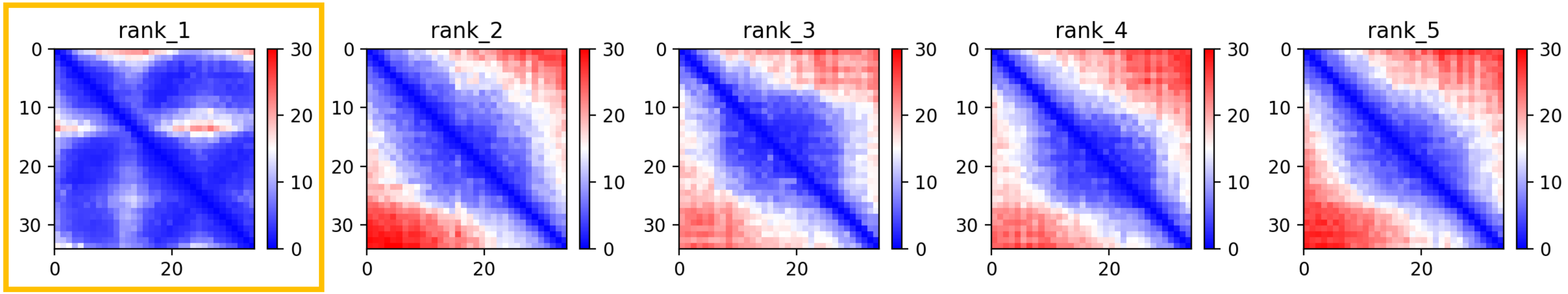

CP45: MtrunA17_Chr2g0328091

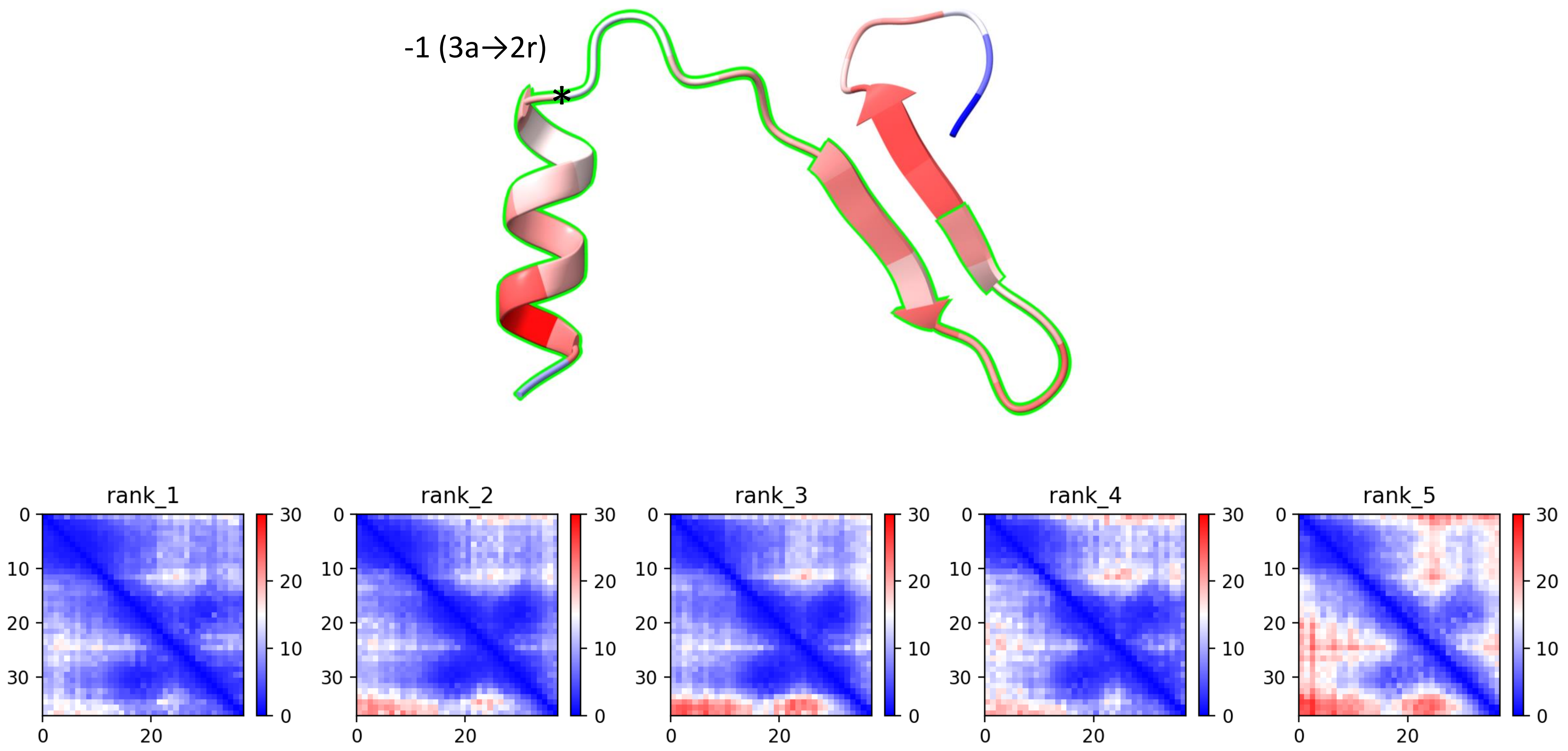

CP46: MtrunA17_Chr2g0329031

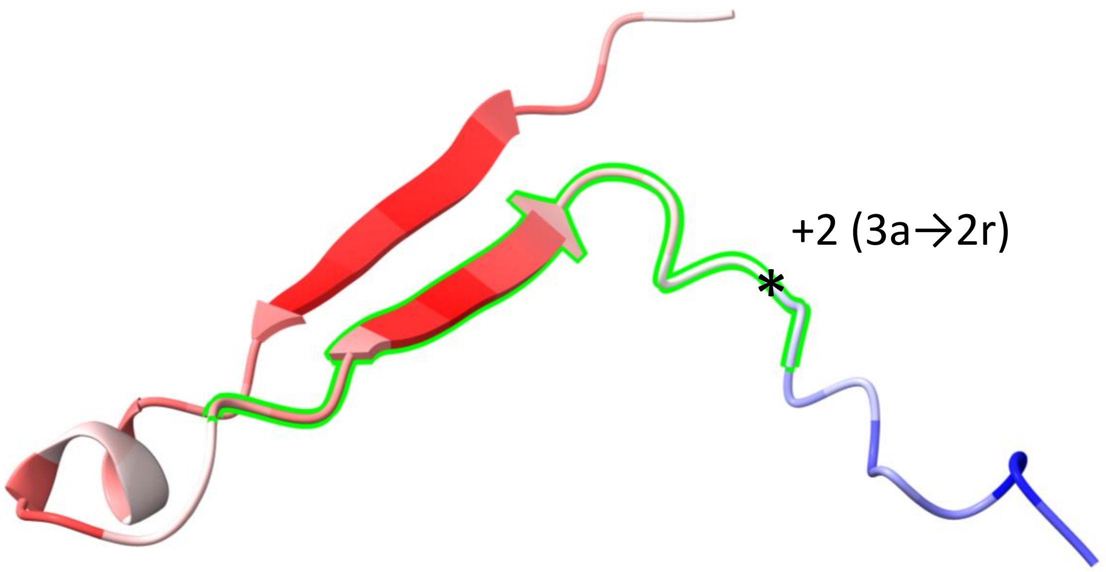


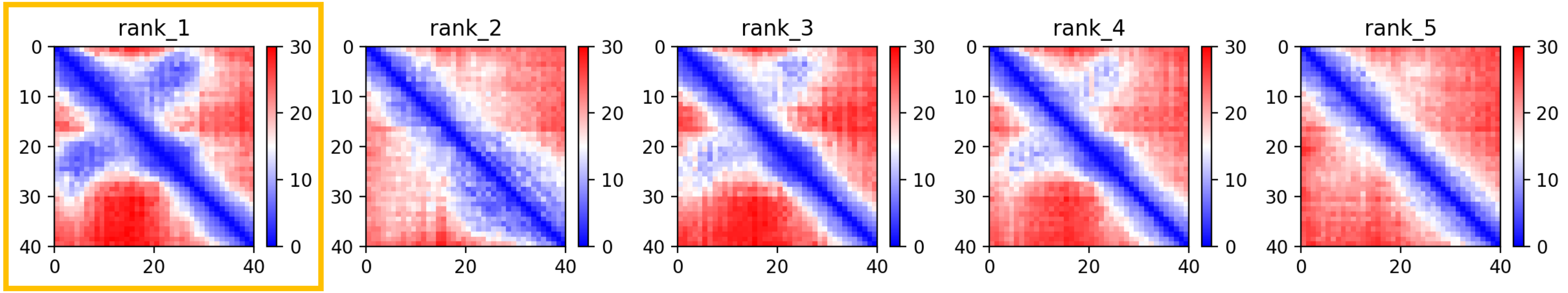

CP60: MtrunA17_Chr3g0135761

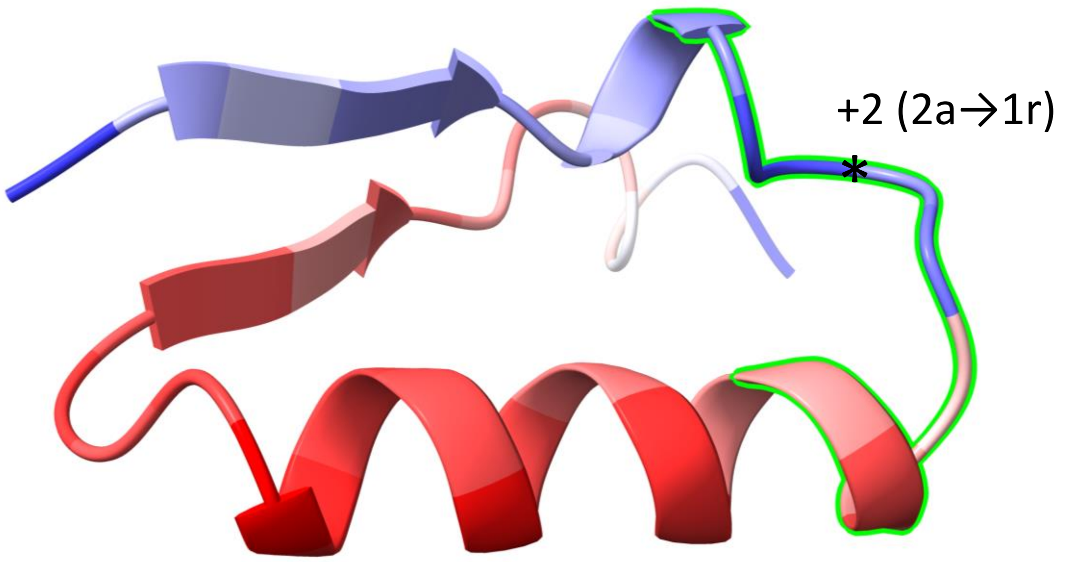


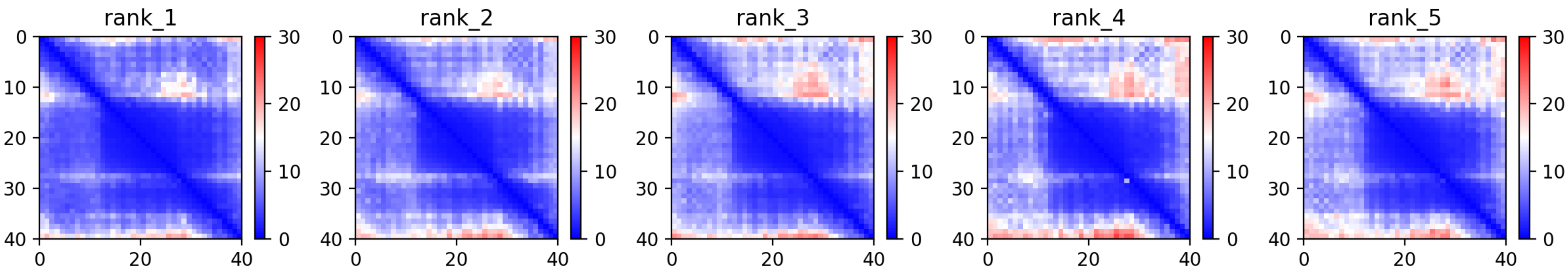

### CP71: MtrunA17_Chr4g0022491

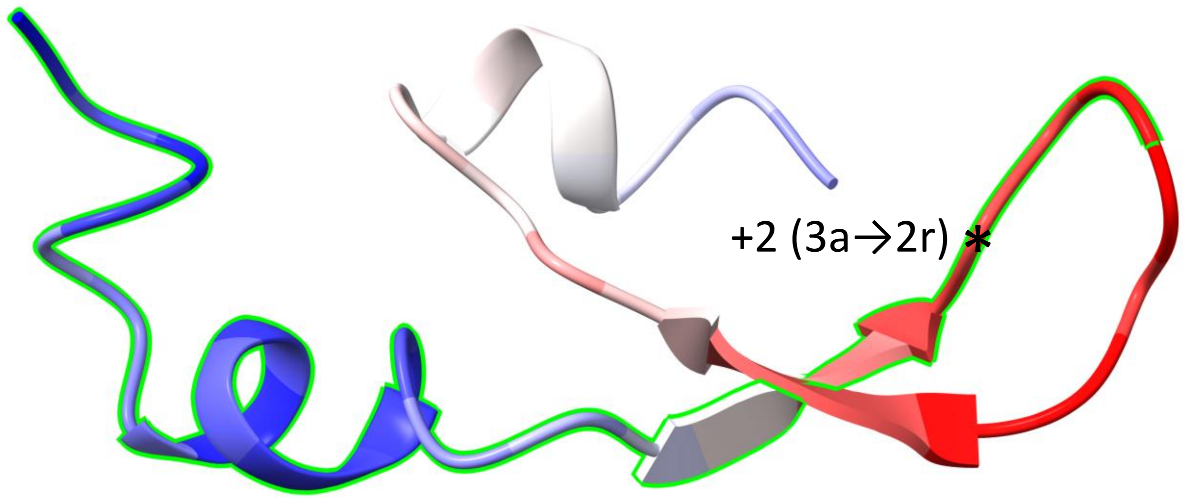


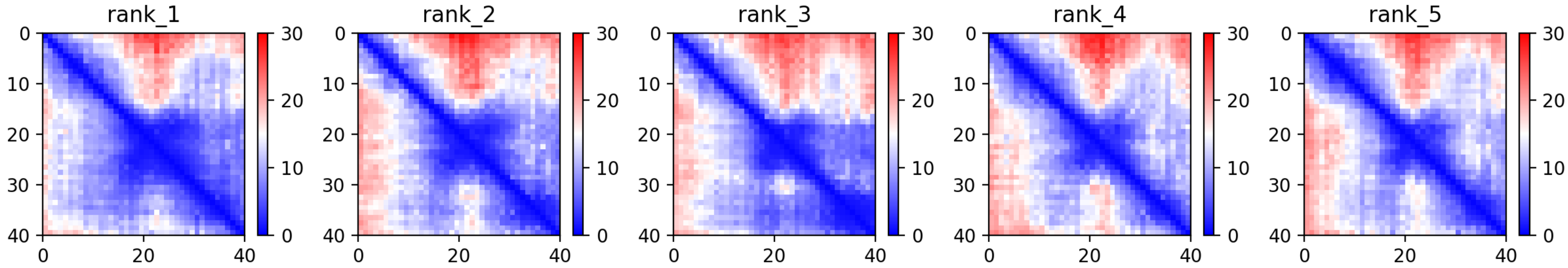

### CP78: MtrunA17_Chr4g0059001

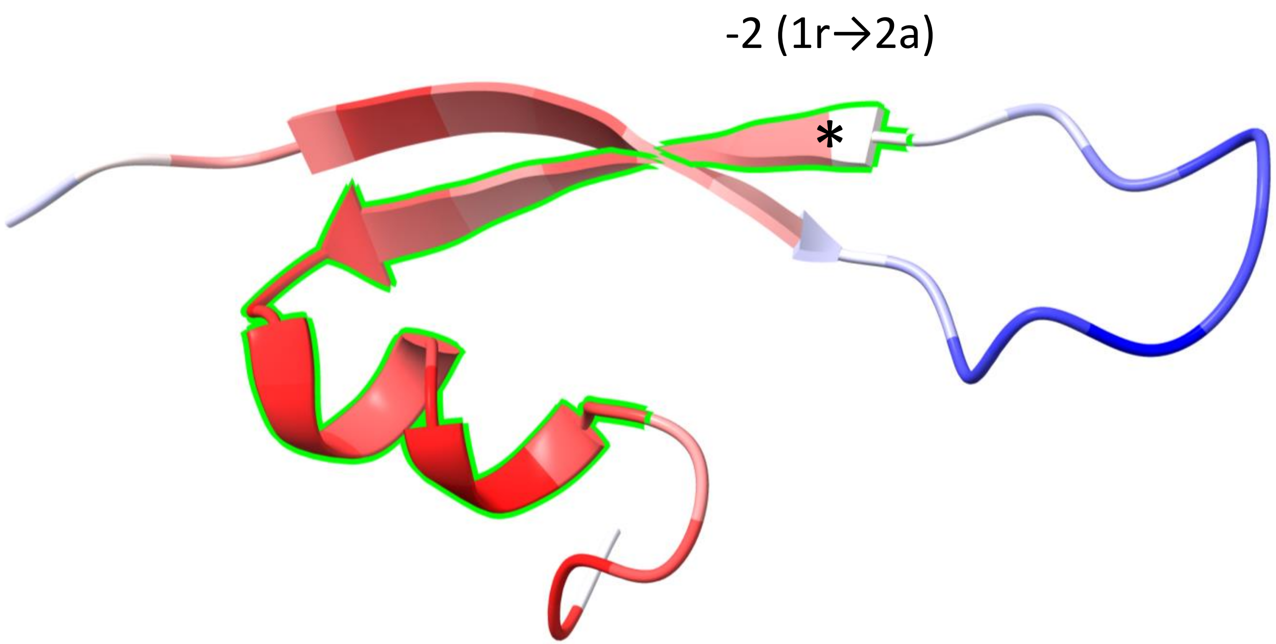


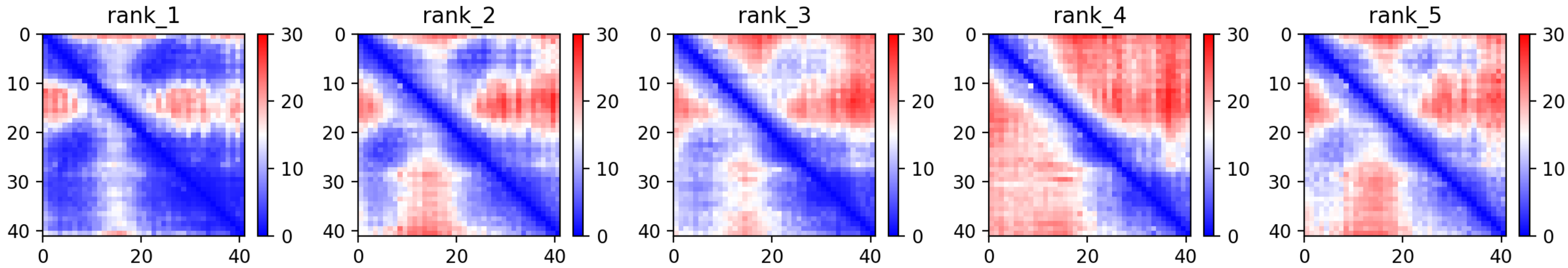

CP79: MtrunA17_Chr4g0063201

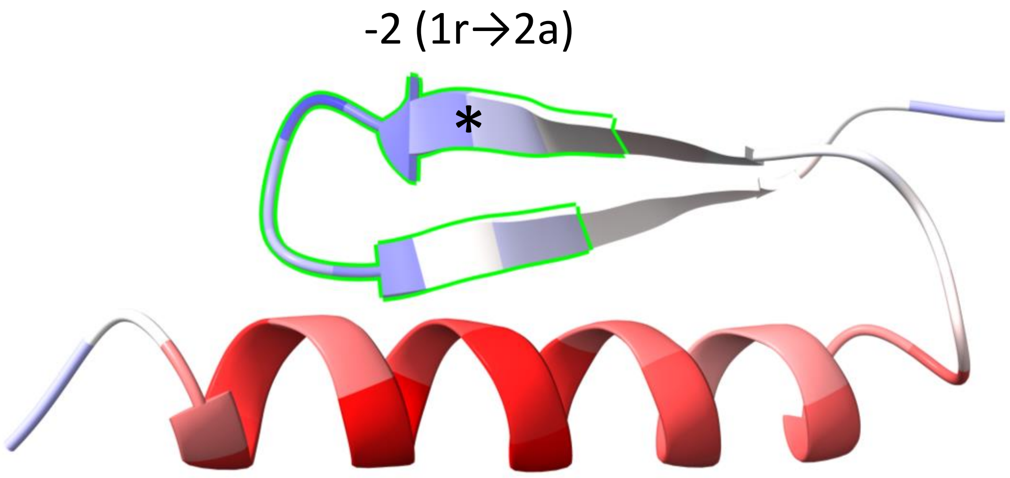


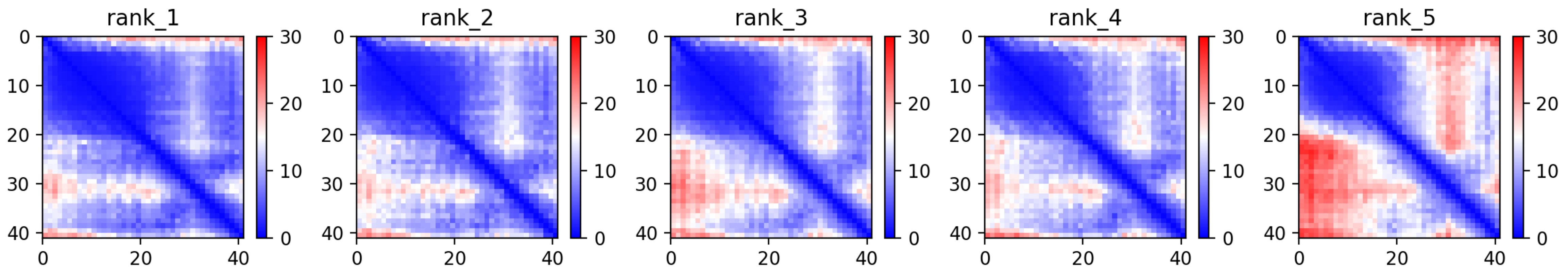

CP99: MtrunA17_Chr6g0457461

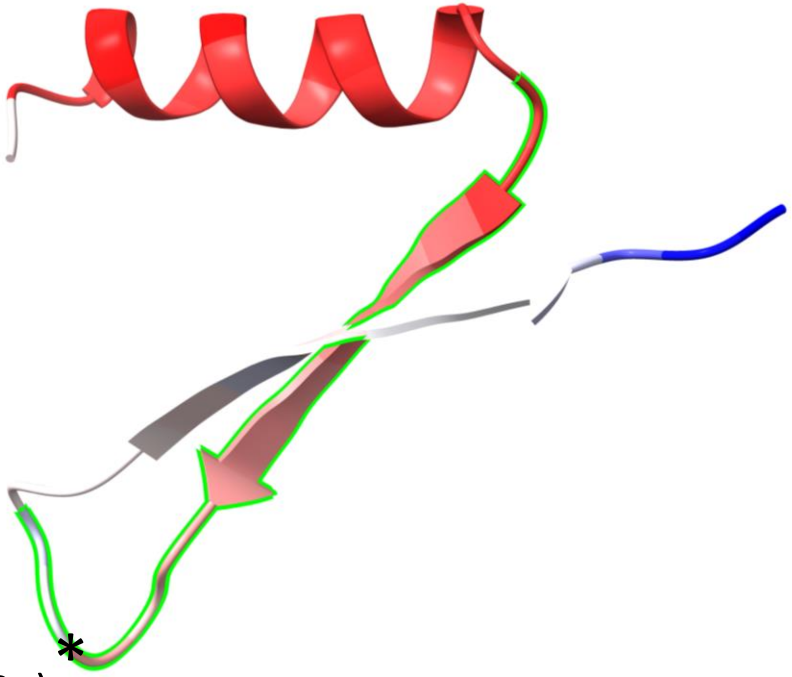


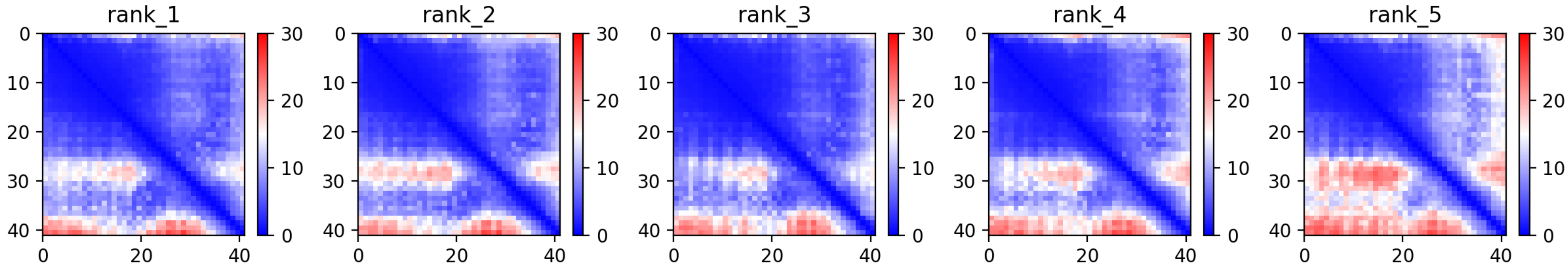

CP100: MtrunA17_Chr6g0457461

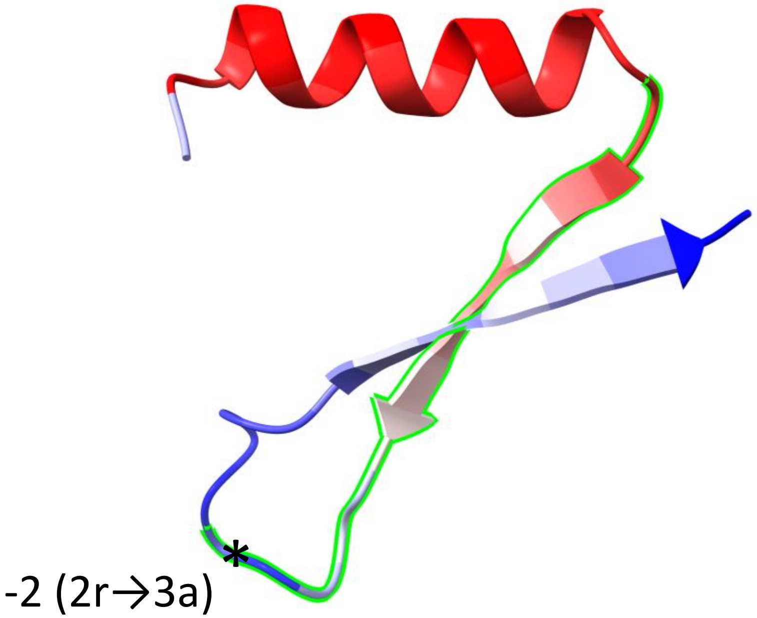


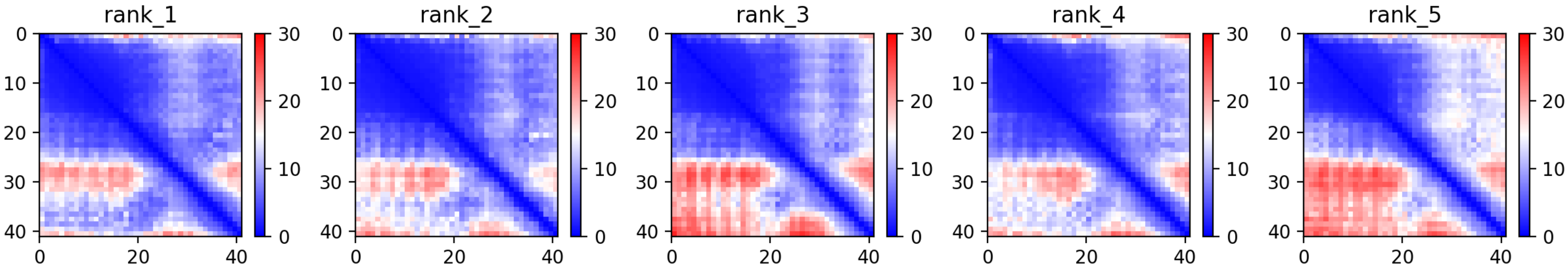

CP101: MtrunA17_Chr6g0458091

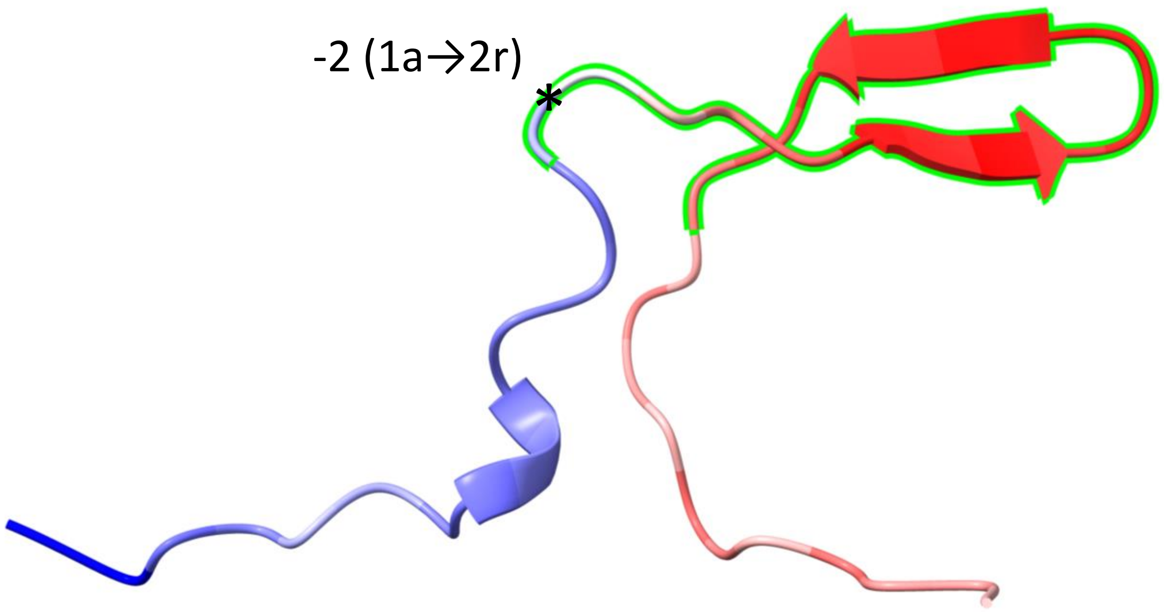


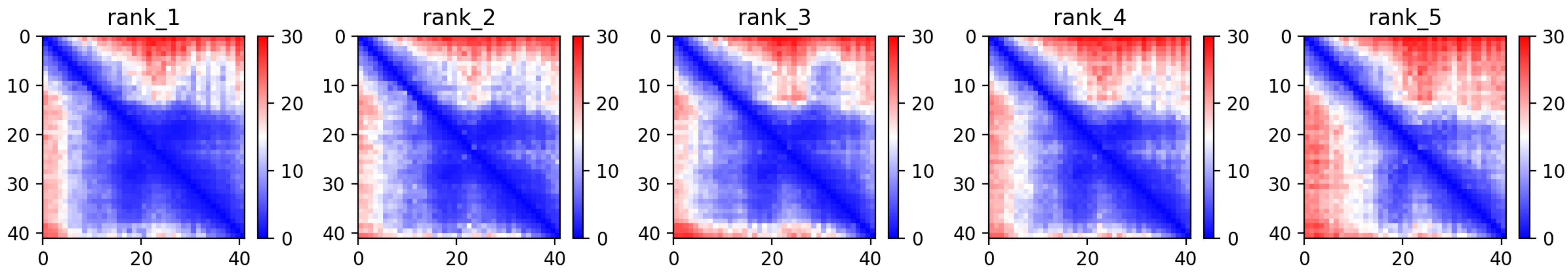

CP103: MtrunA17_Chr6g0461931

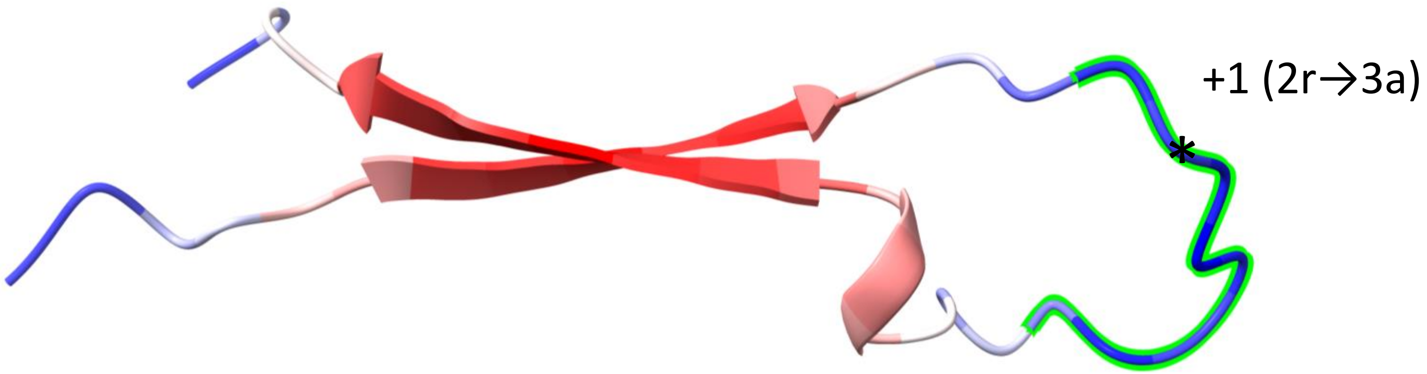


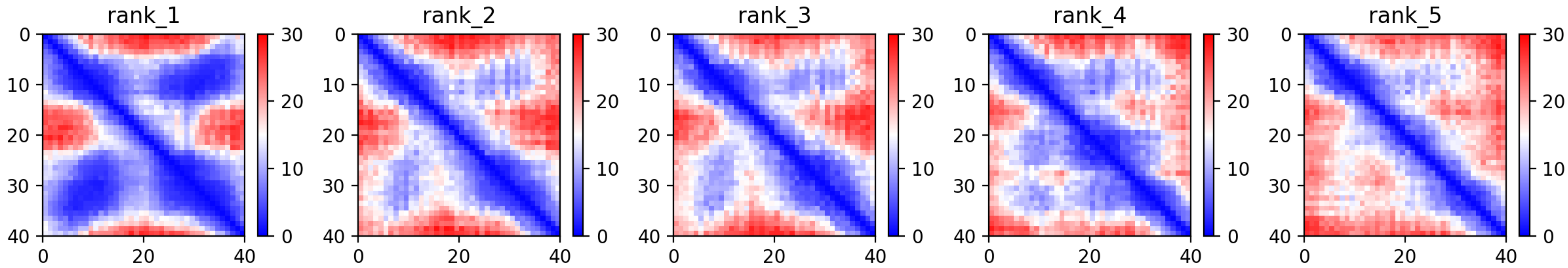

CP108: MtrunA17_Chr6g0485321

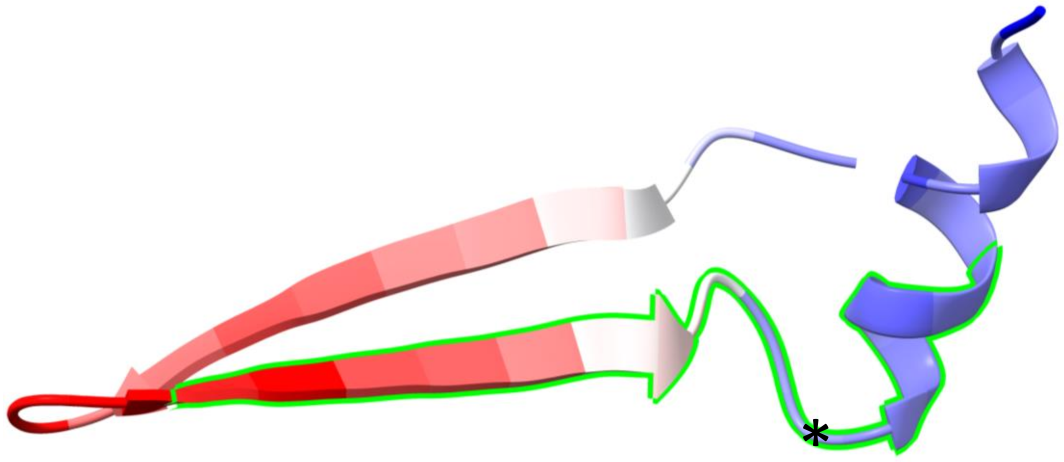


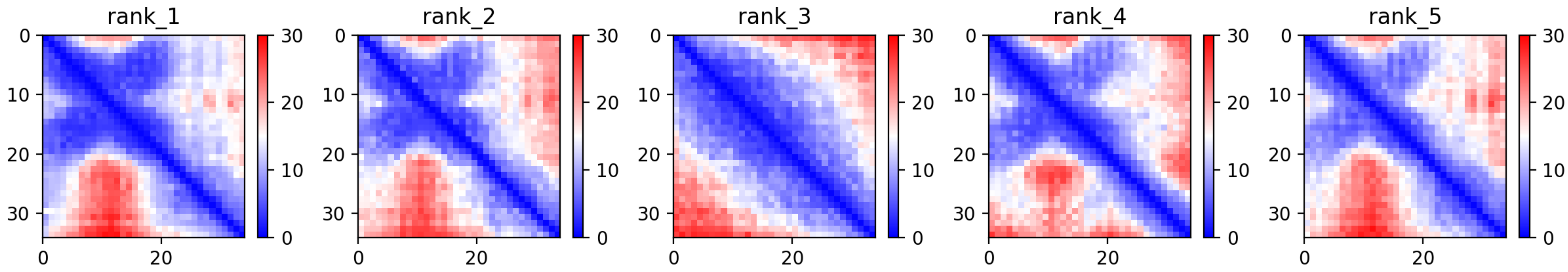

### CP136: MtrunA17_Chr8g0373091

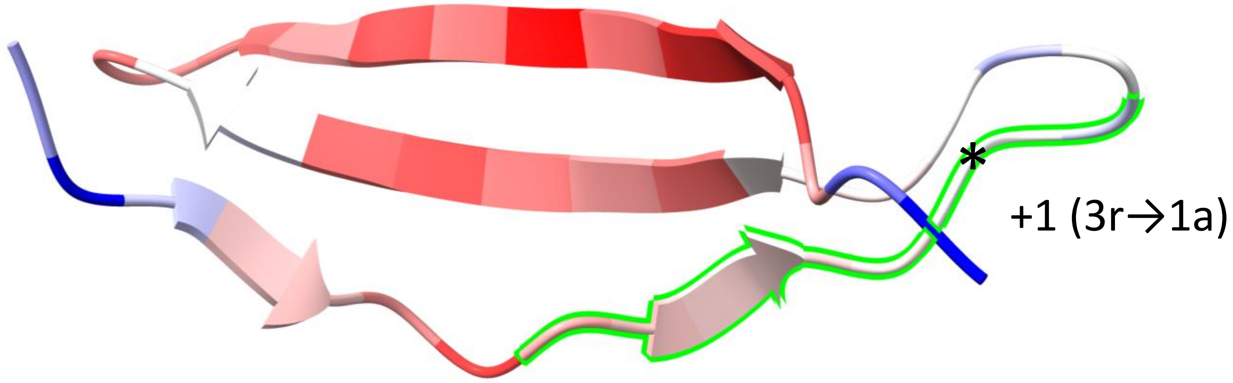


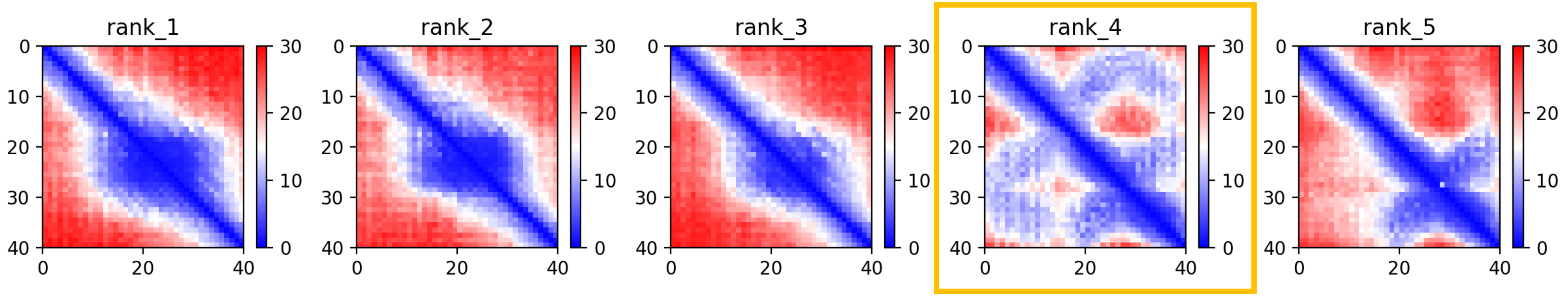

### CP139: MtrunA17_Chr8g0385331

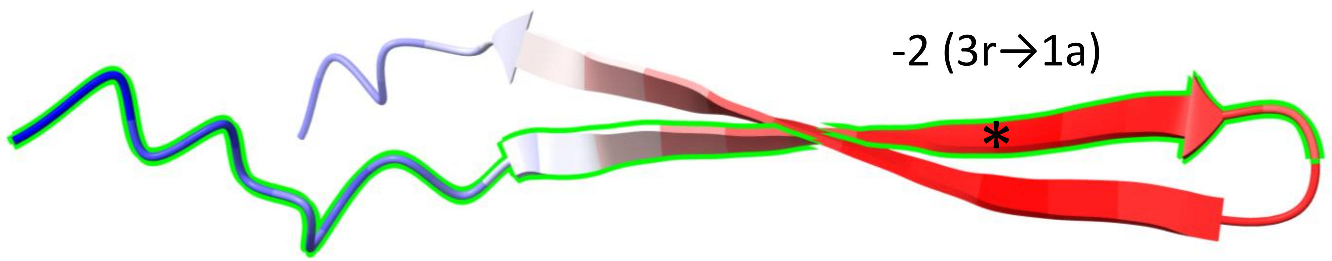


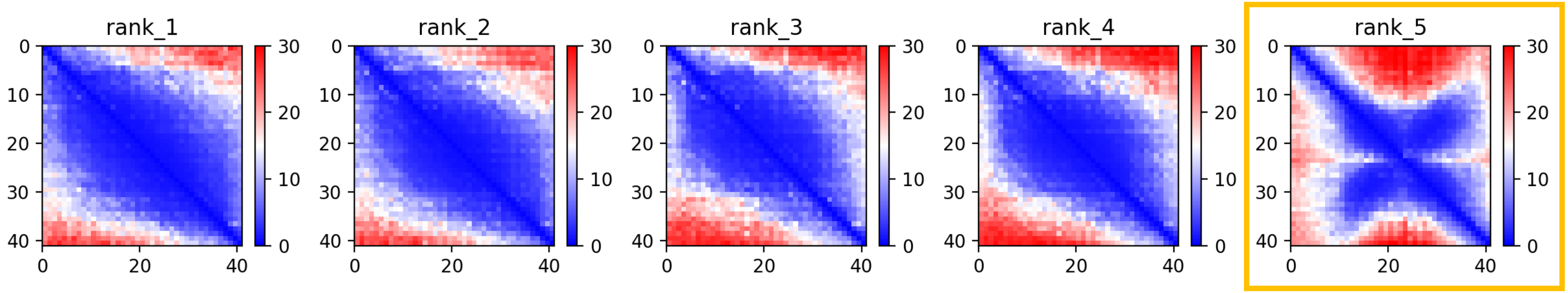

CP149: MtrunA17_MTg0490471

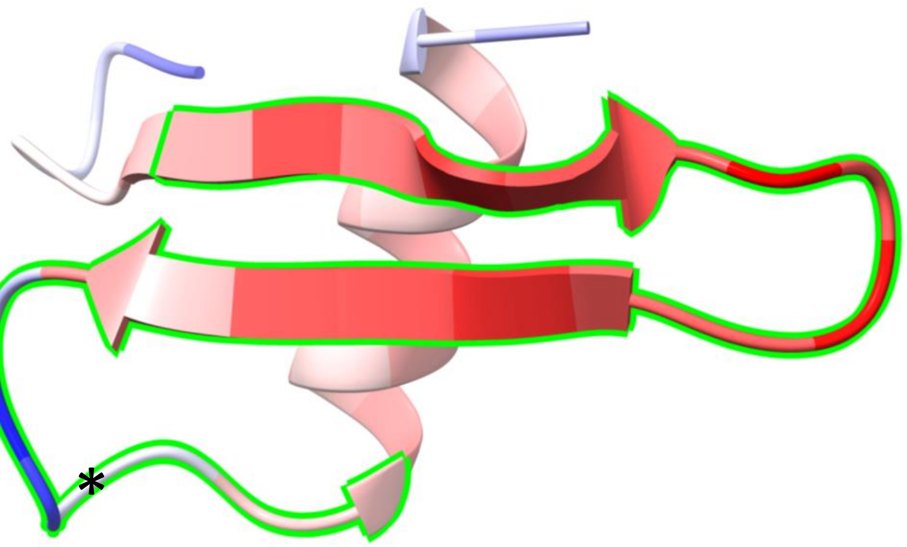


+1 (1r→2a)

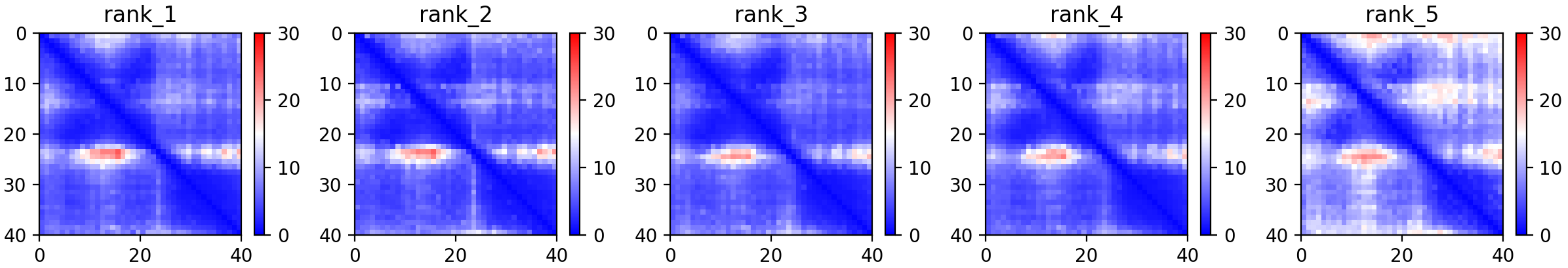

CP154: MtrunA17_MTg0491501

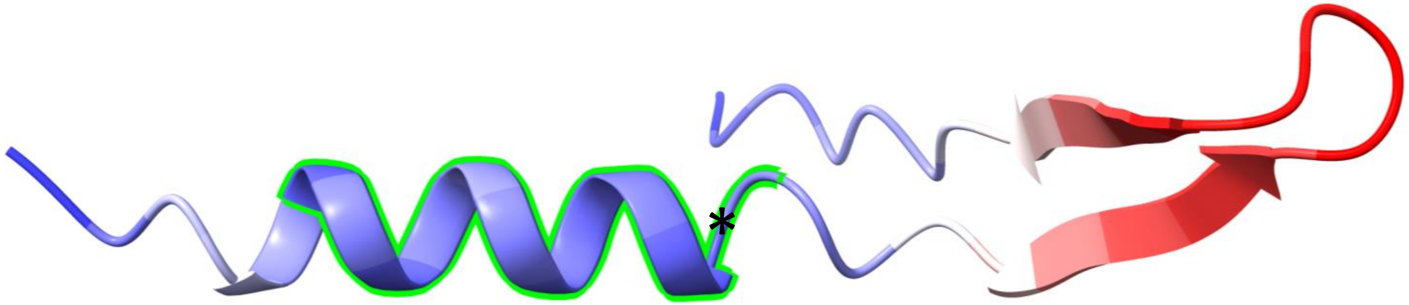


-2 (3a1→1a2)

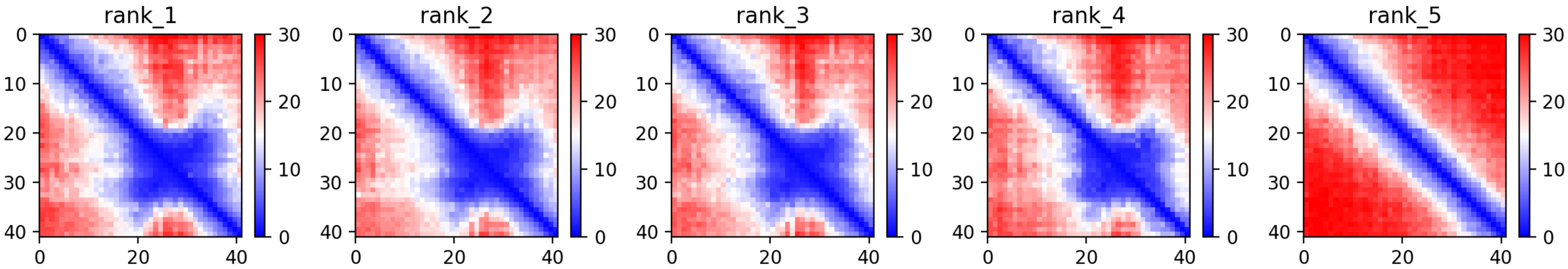

**Supplementary Dataset S7 Part 5**. A graphical summary on folding predictions for 24 MS-supported chimeric peptide models (CPs) that contain beta-sheets regardless of the presence of alpha-helices. For each CP, one image is displayed: the predicted folding structure (top) along with heat maps of five predictions (bottom). The rest of the legend is the same as for Supplementary Dataset S7 Part 1.
