## Supplementary material for "Discovery of diverse chimeric peptides in a eukaryotic proteome sets the stage for the experimental proof of the mosaic translation hypothesis": Supplementary Dataset S11 Overlapping genes.pdf

CP1: MtrunA17_Chr0c01g0489091

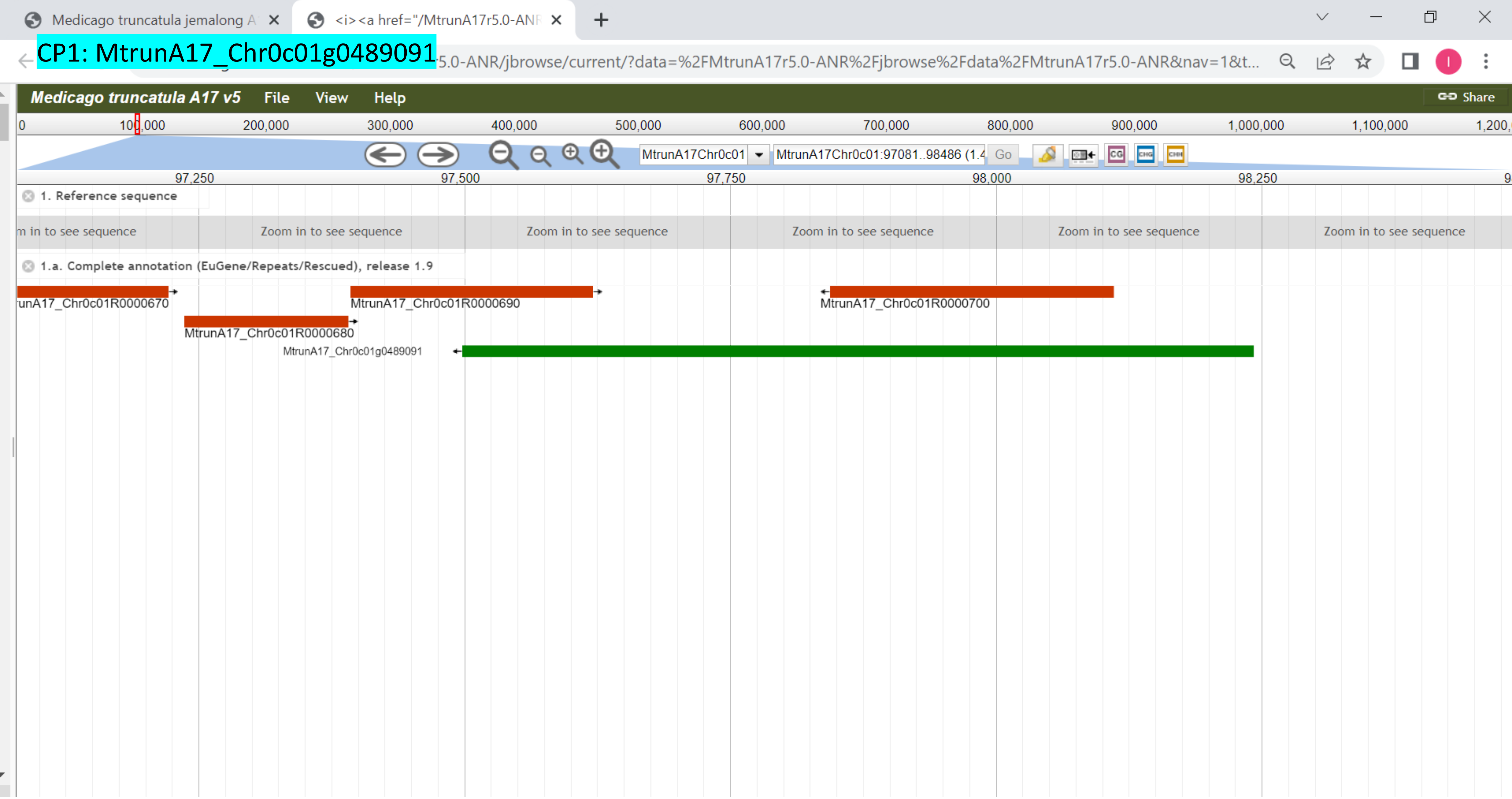

CP2: MtrunA17_Chr0c28g0493951

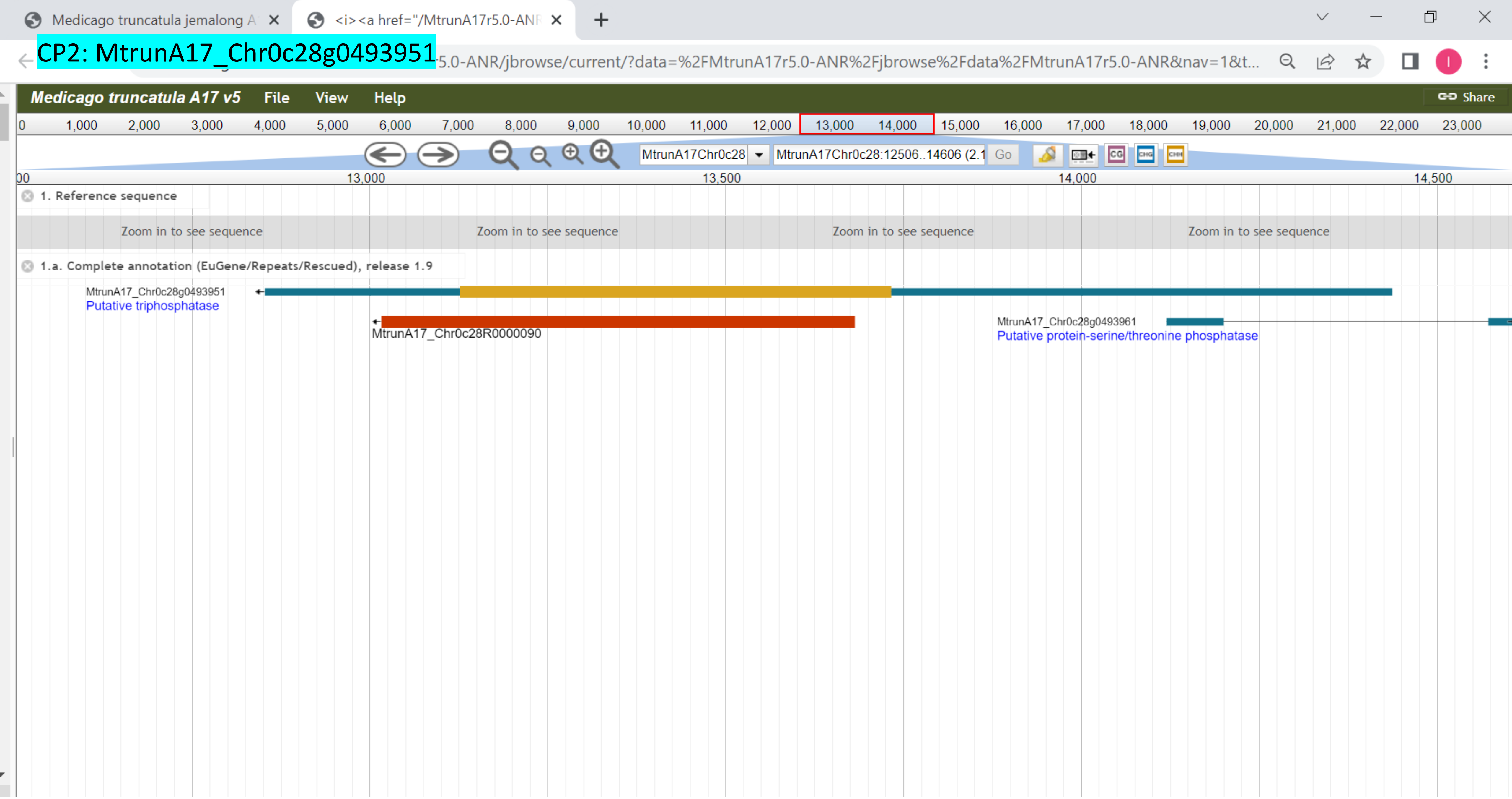

CP3: MtrunA17_Chr1g0148371

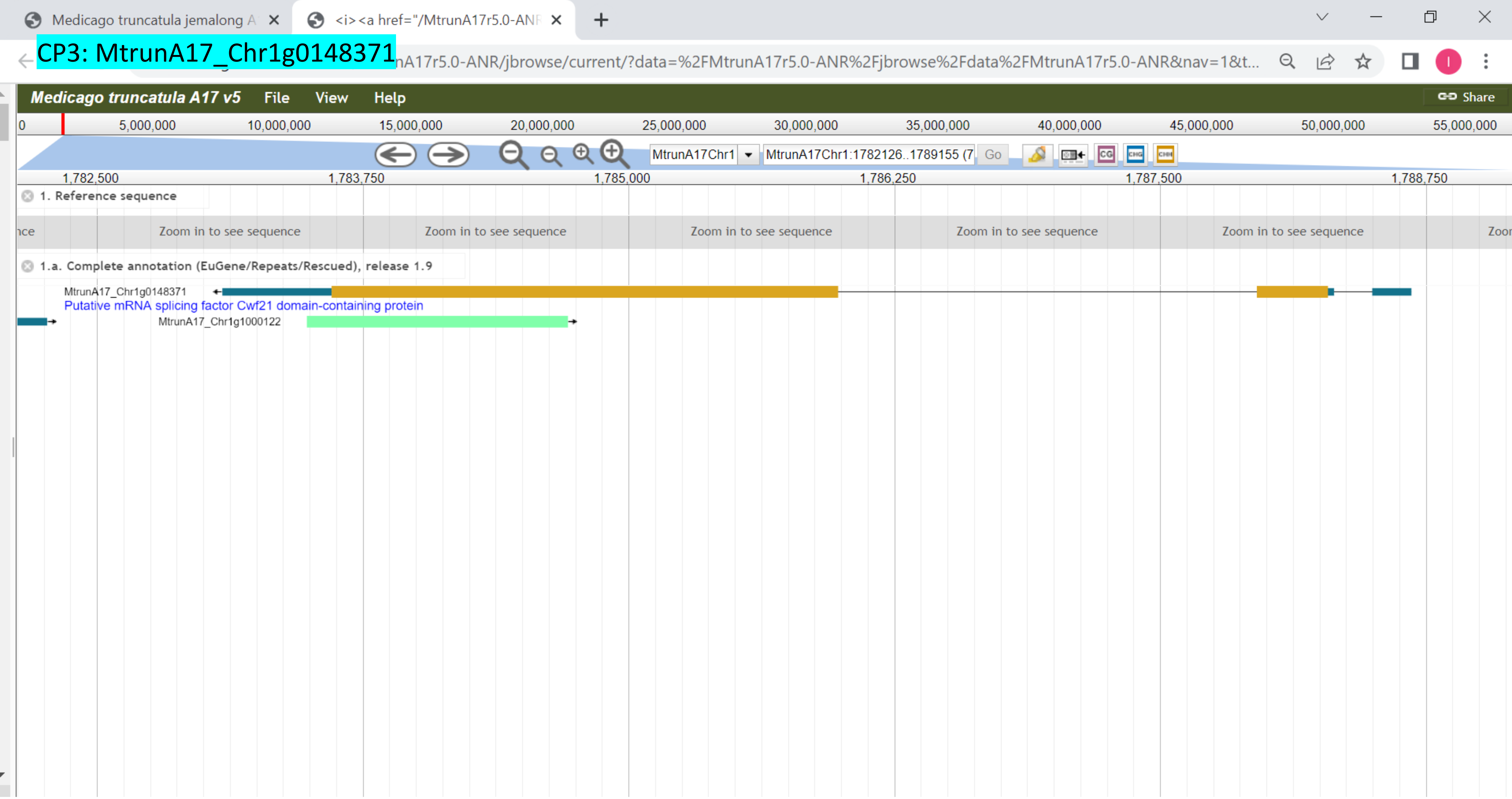

CP4: MtrunA17_Chr1g0149991

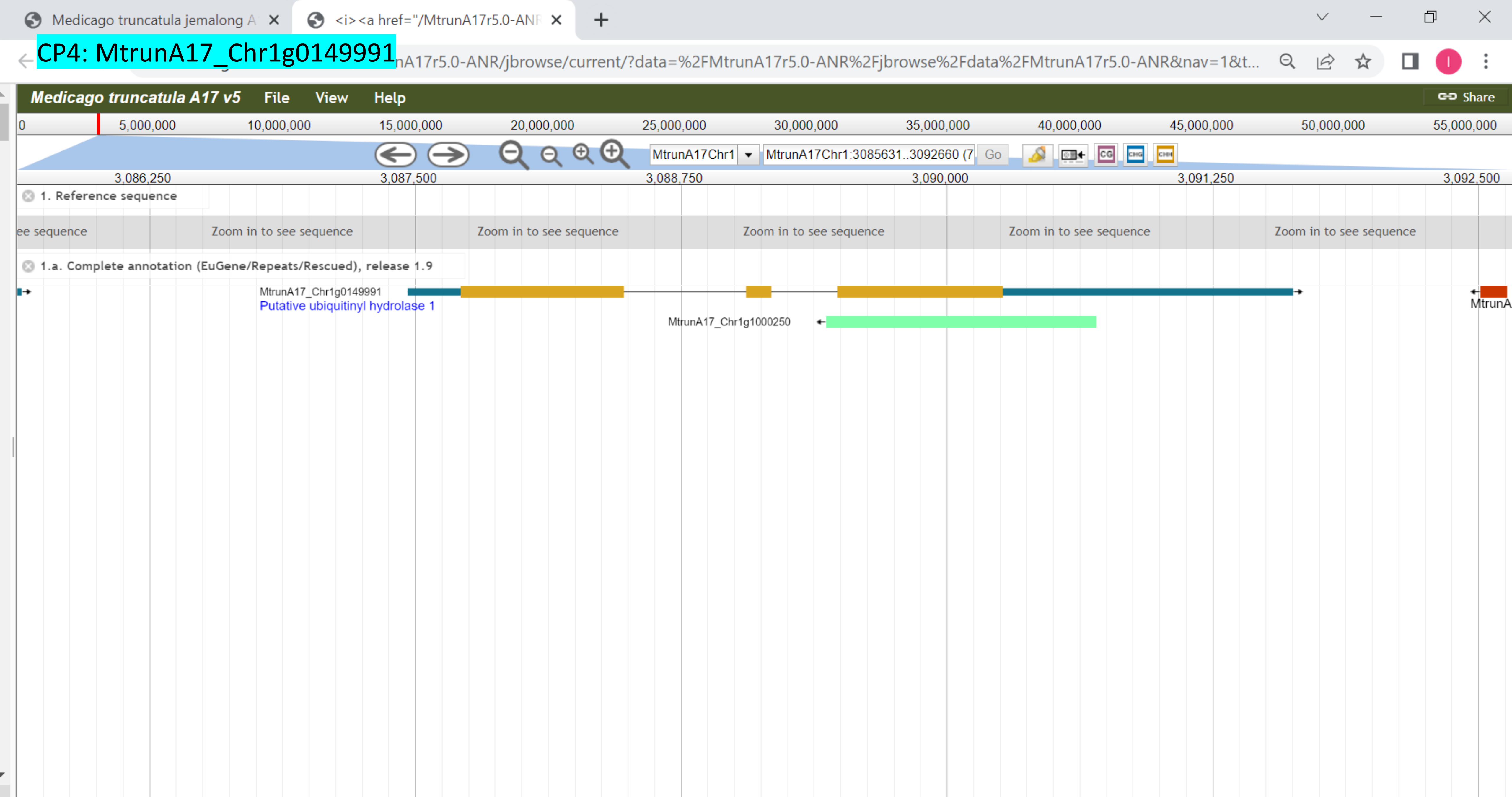

### CP5: MtrunA17_Chr1g0150571

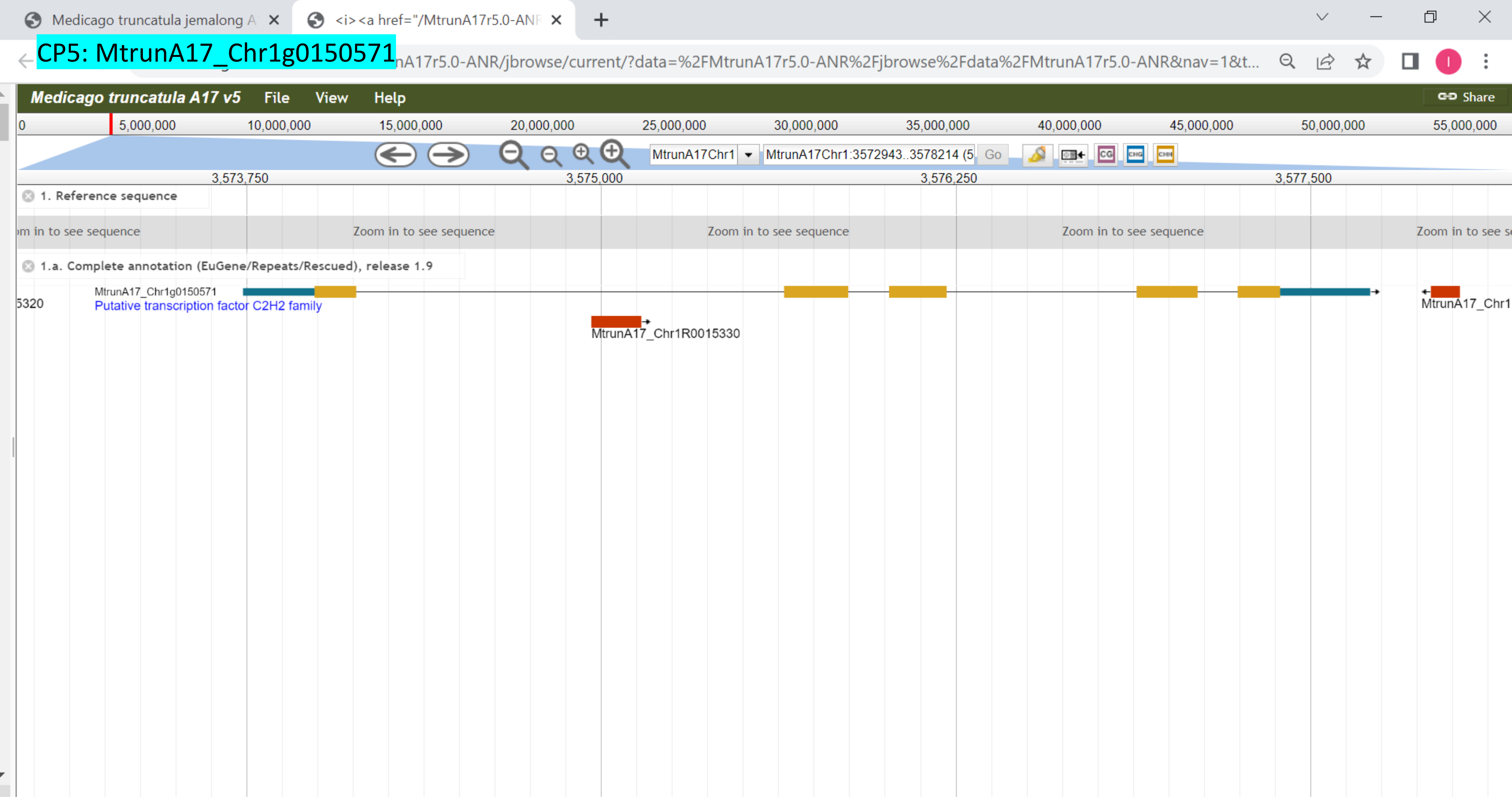

CP6: MtrunA17_Chr1g0152521

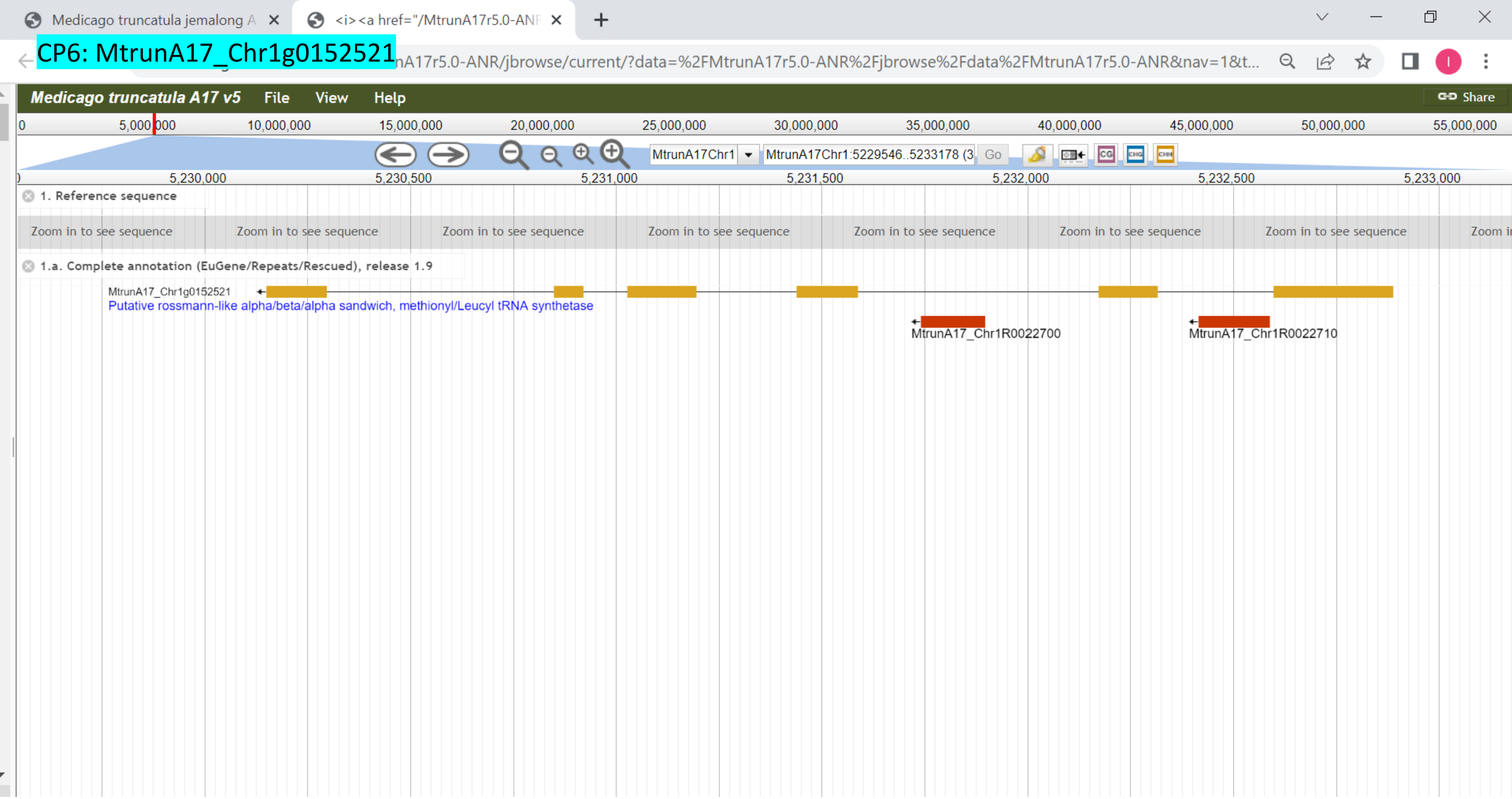

CP7: MtrunA17_Chr1g0153001

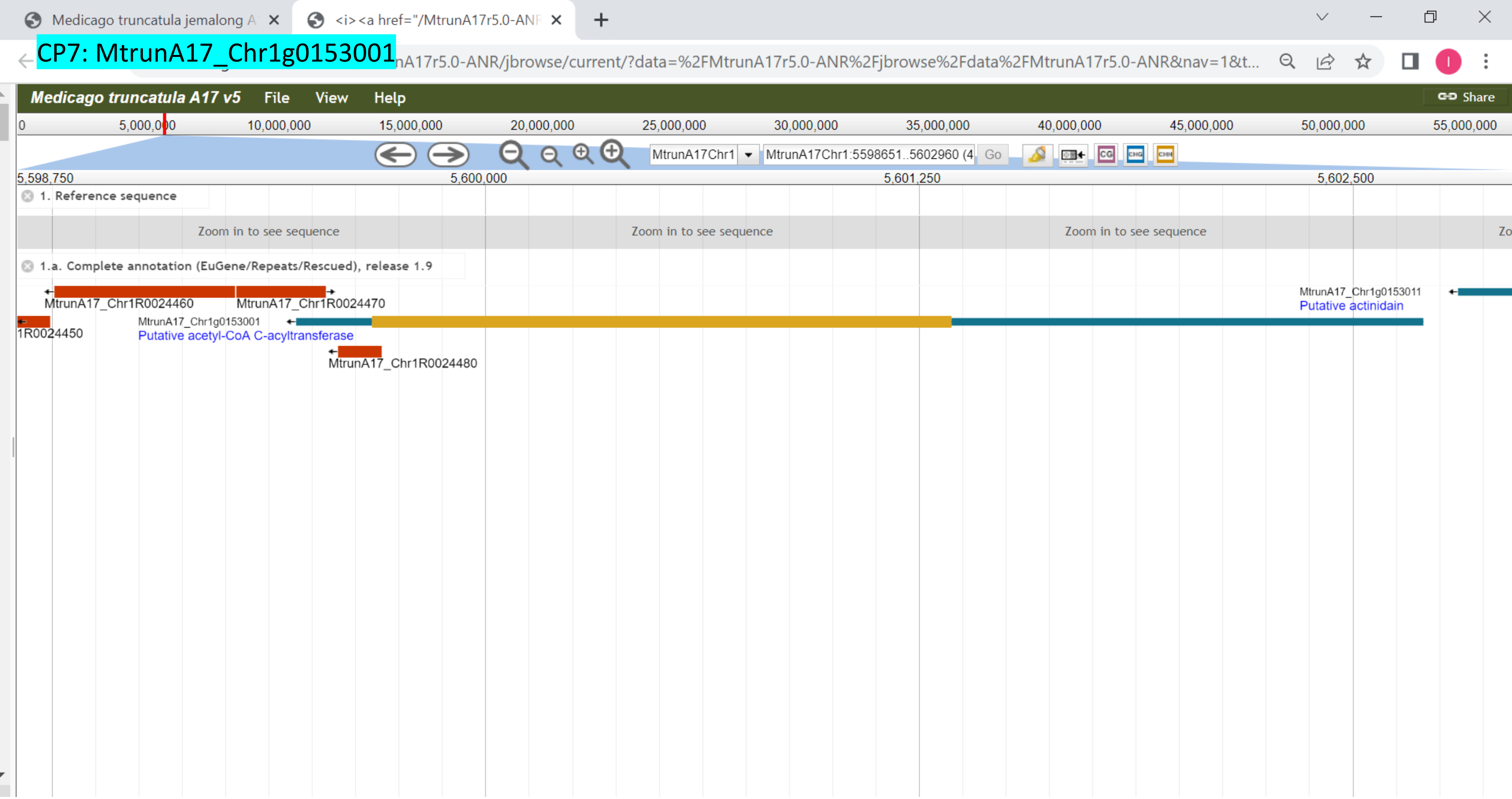

CP8: MtrunA17_Chr1g0155251

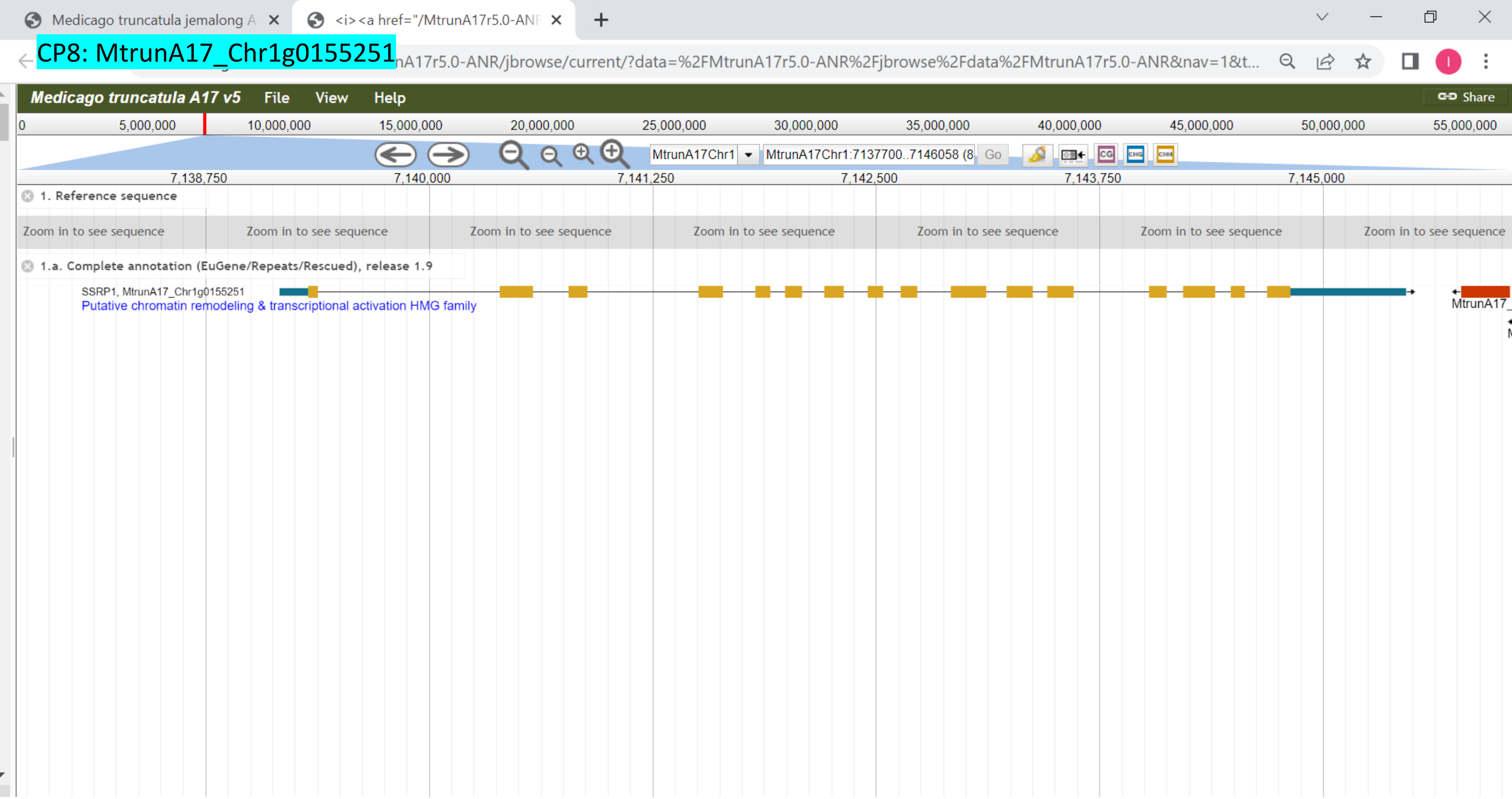

### CP9: MtrunA17_Chr1g0158341

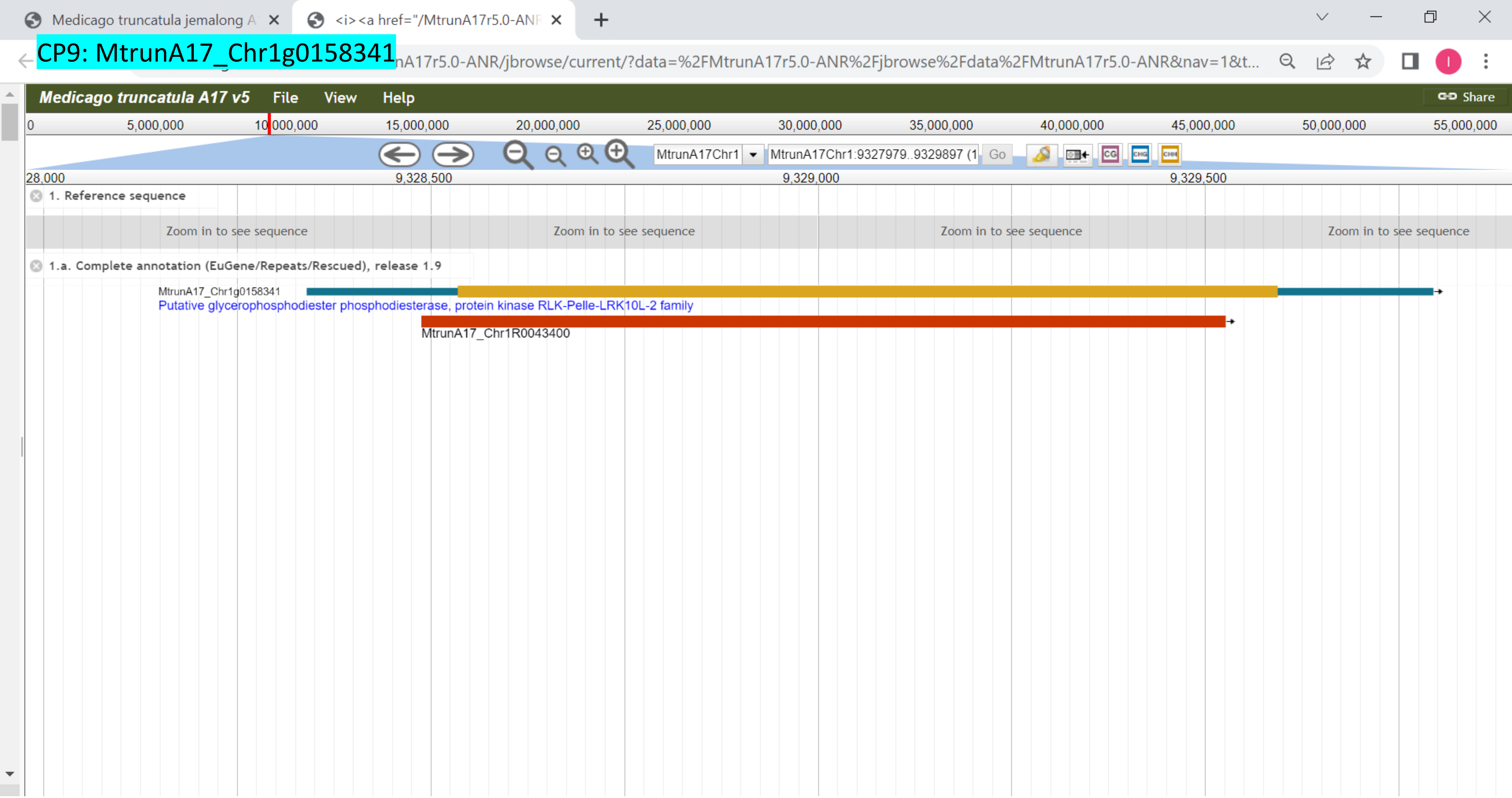

### CP10: MtrunA17_Chr1g0162101

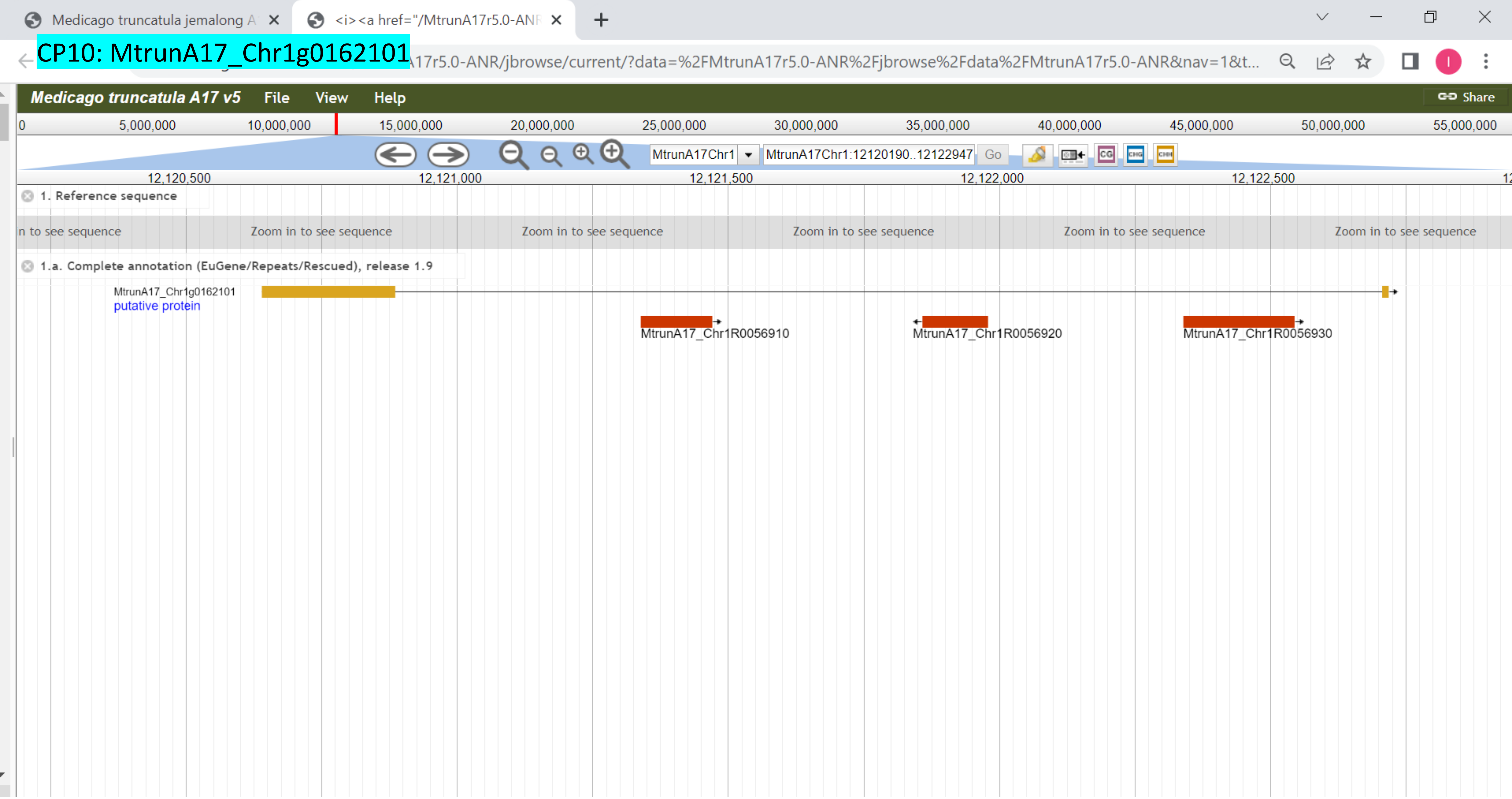

CP11: MtrunA17_Chr1g0164591

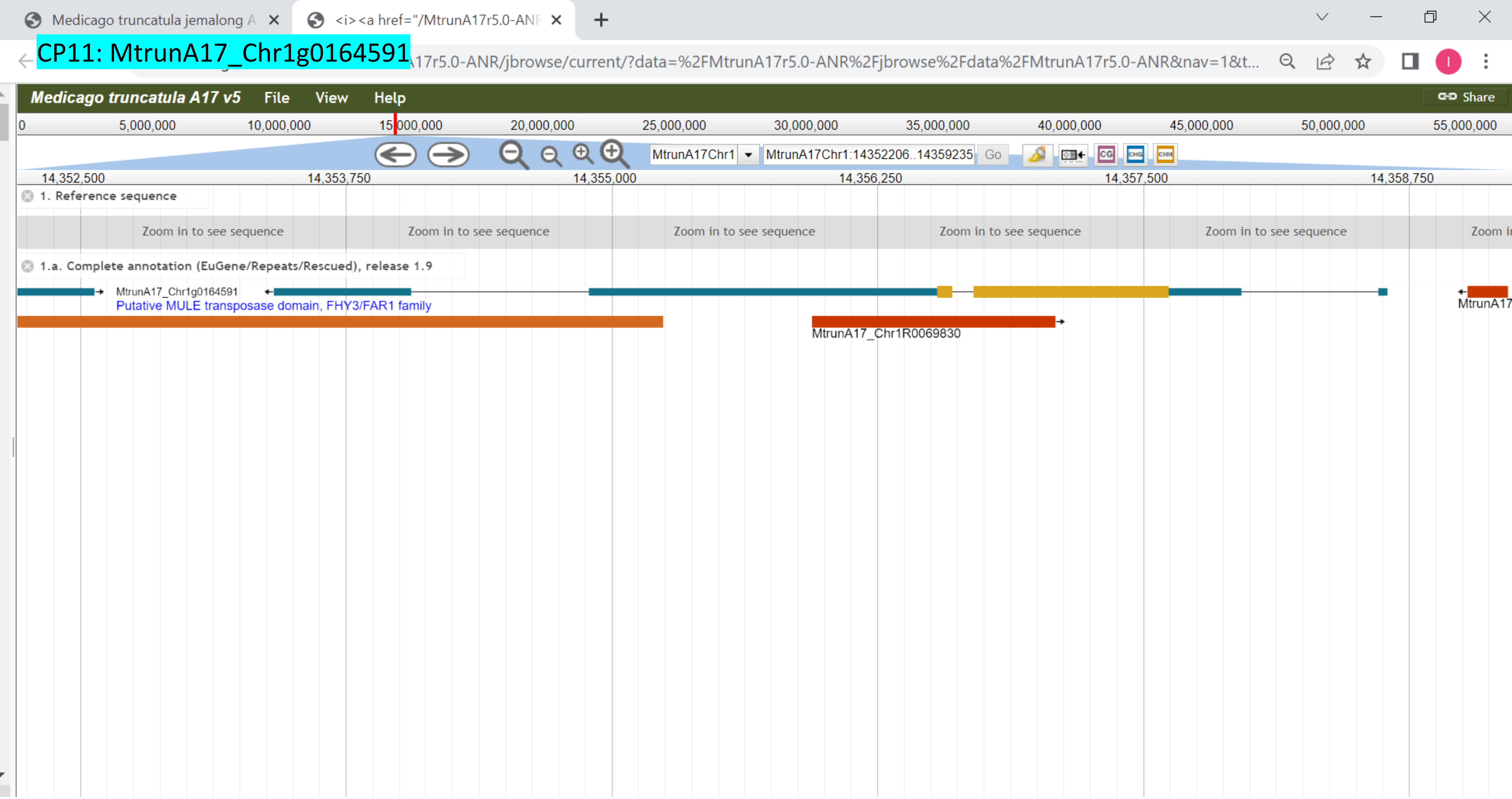

CP12: MtrunA17_Chr1g0178361

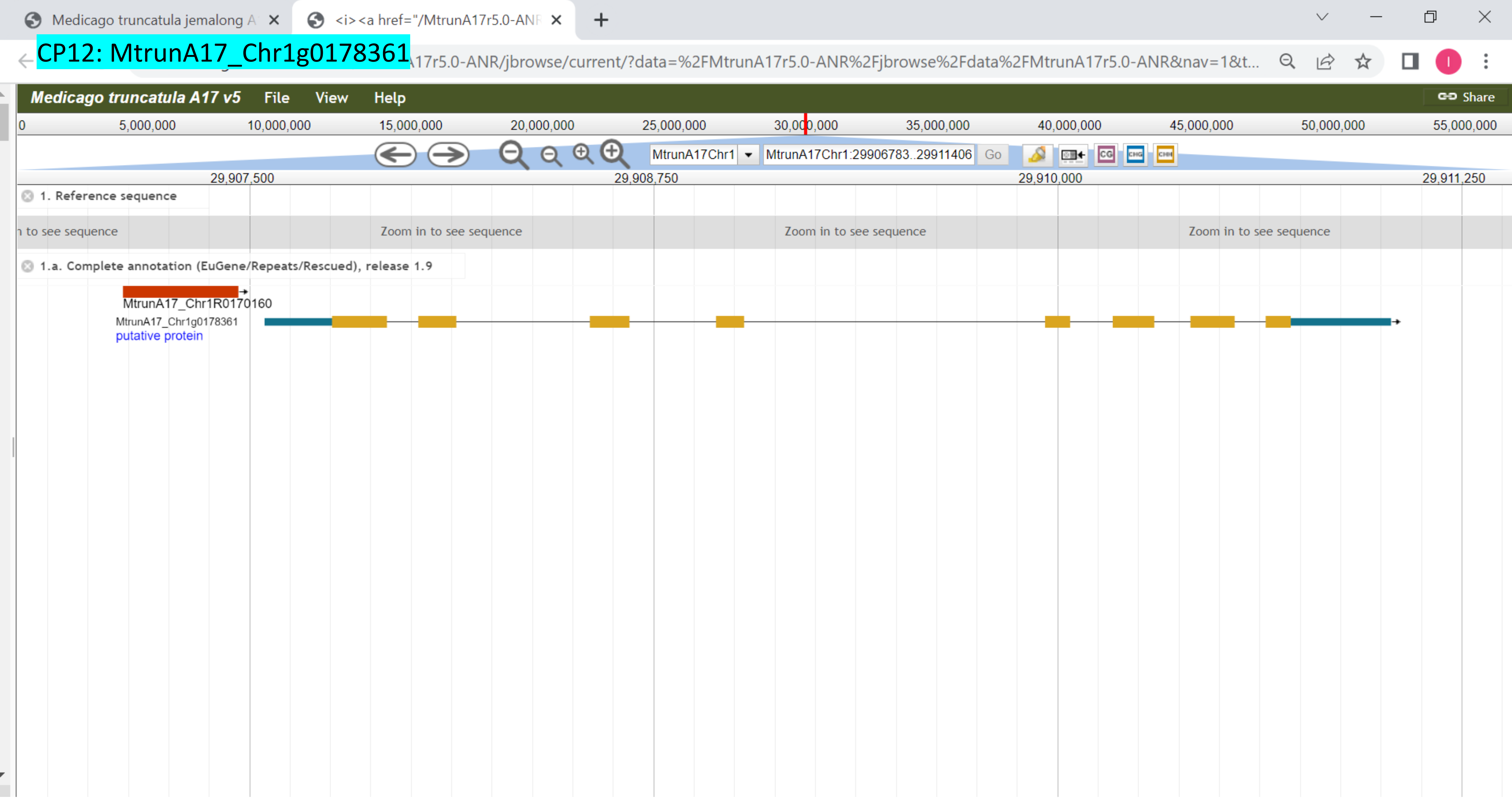

### CP13: MtrunA17_Chr1g0181761

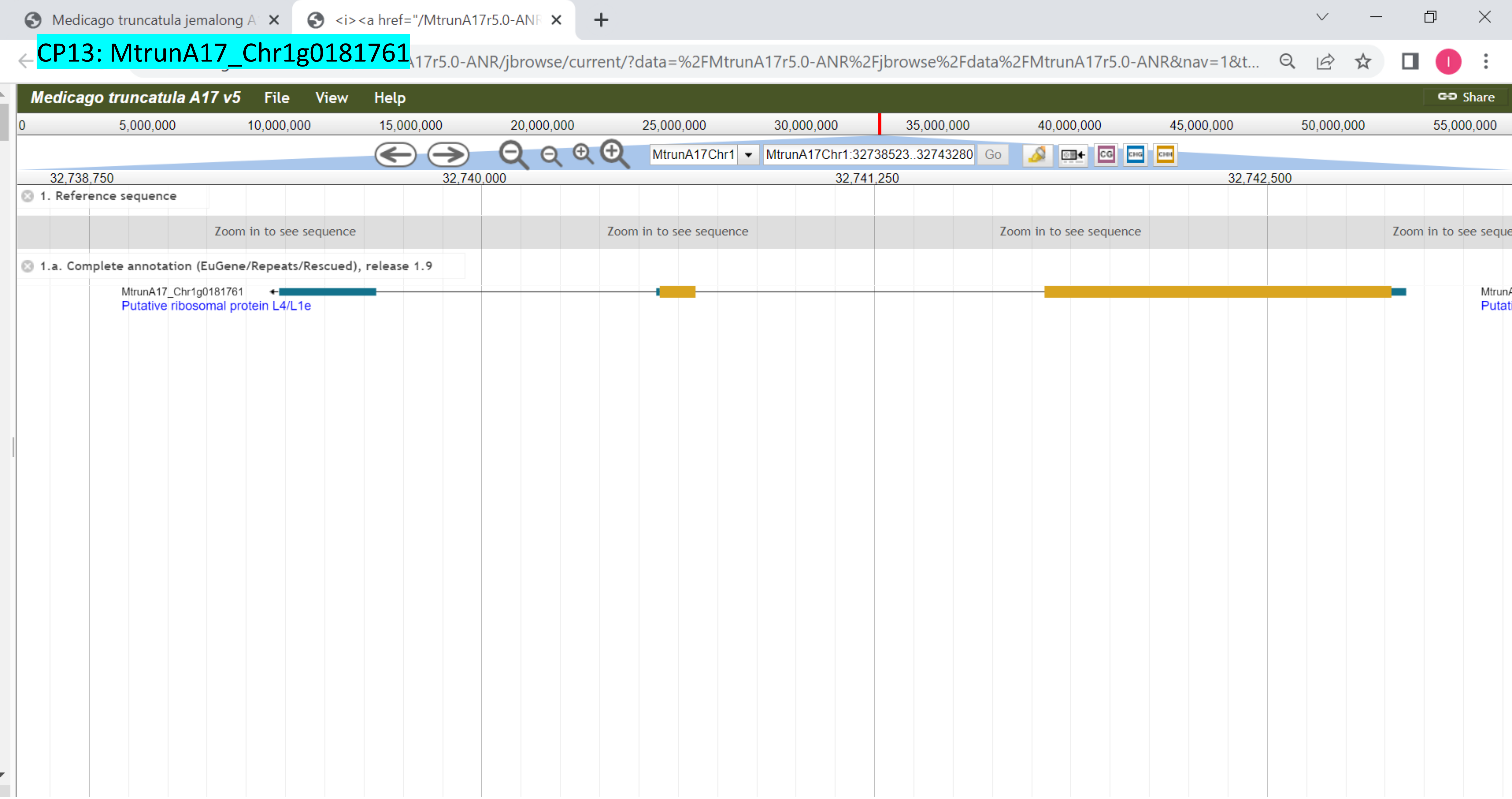

CP14: MtrunA17_Chr1g0182591

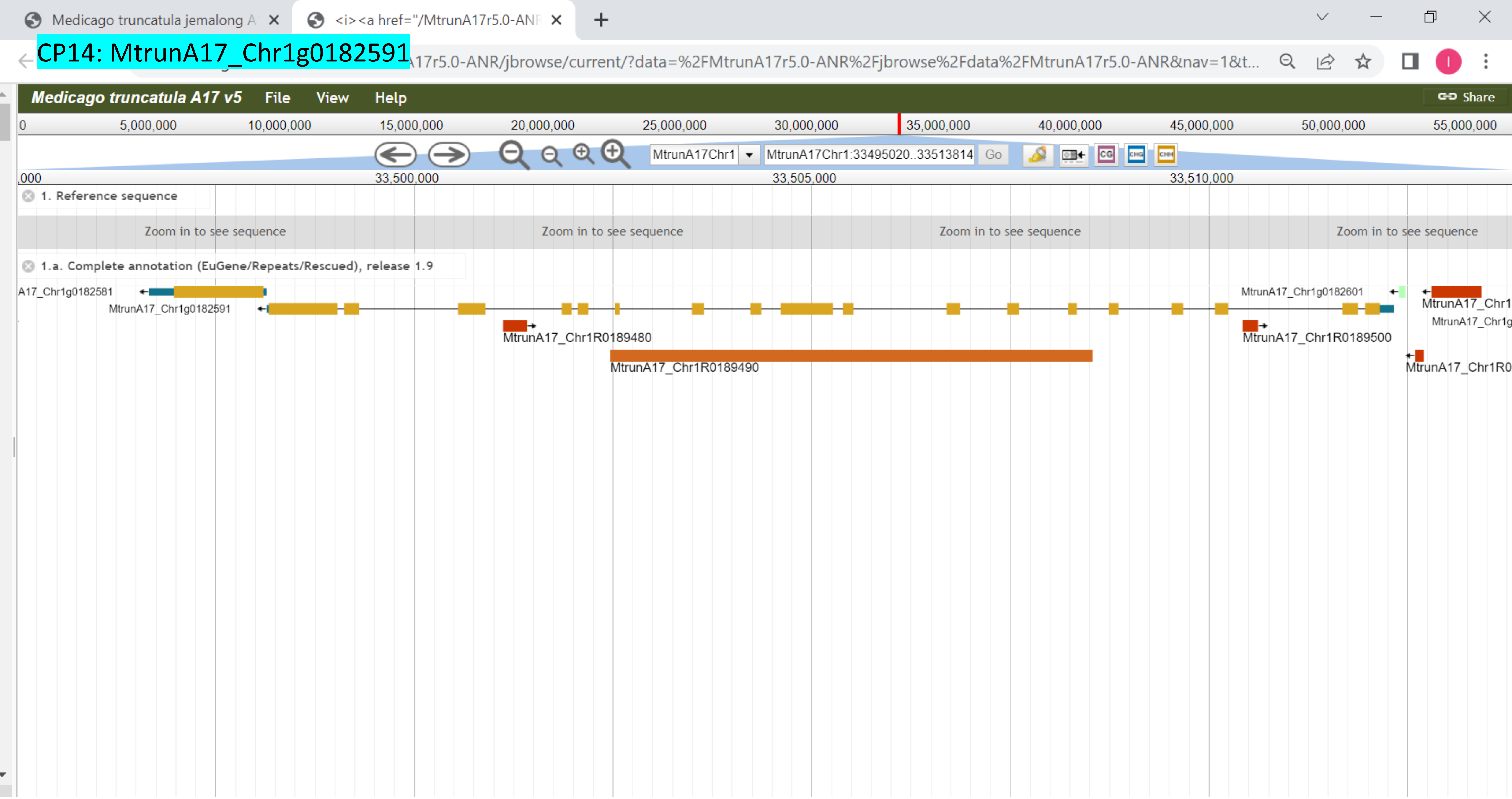

CP15: MtrunA17_Chr1g0183001

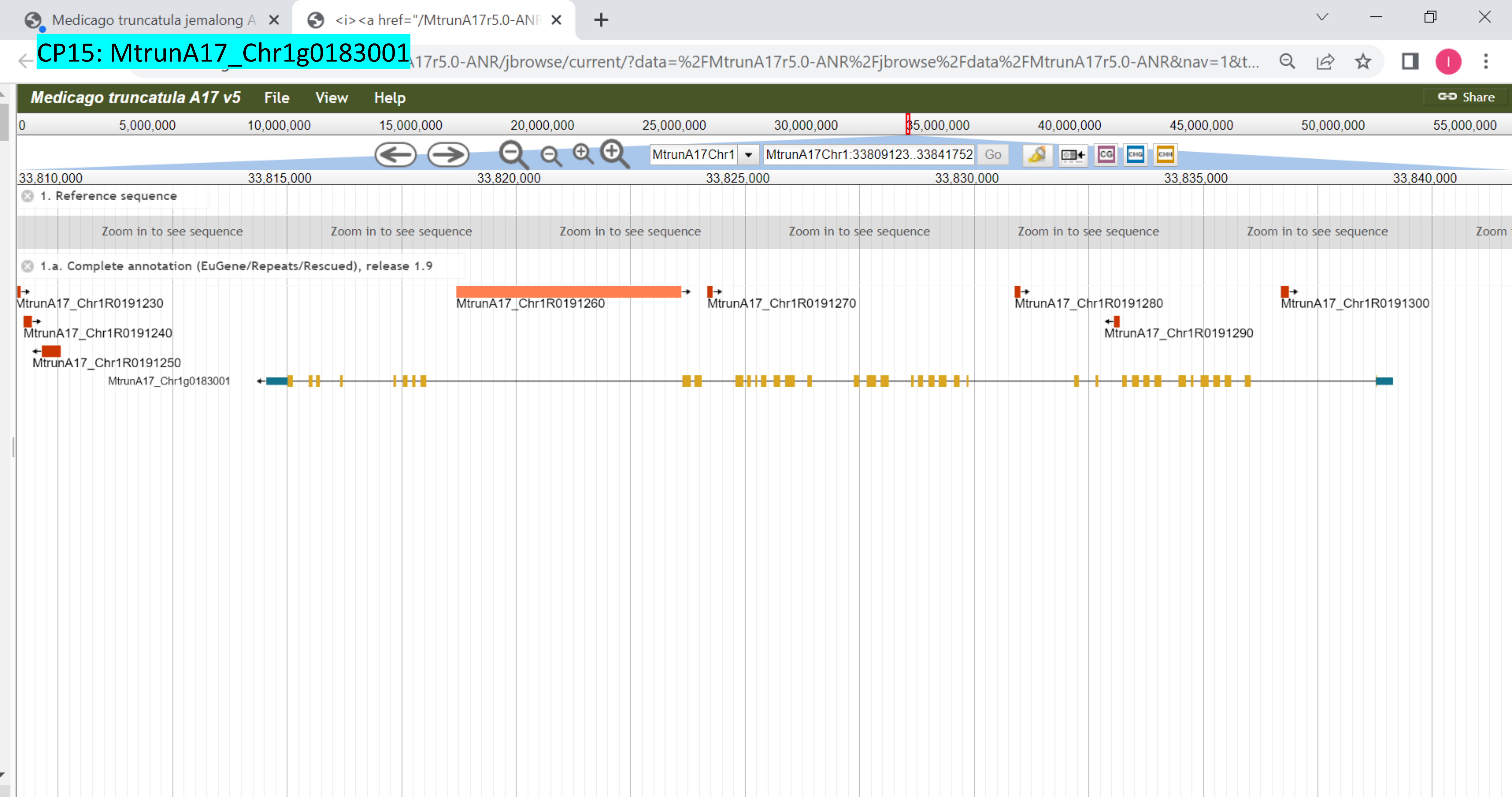

CP16: MtrunA17_Chr1g0185811

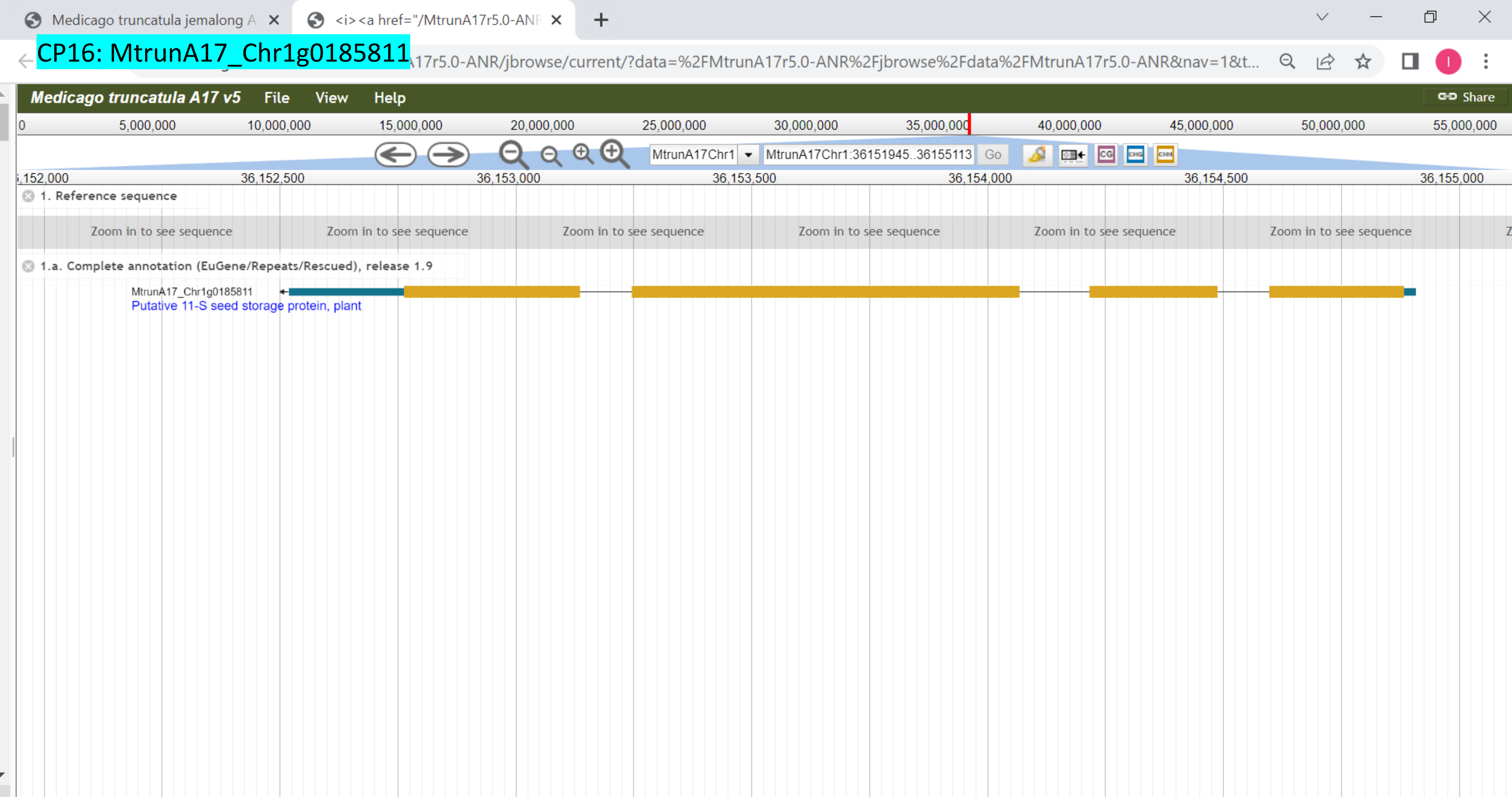

CP17: MtrunA17_Chr1g0185811

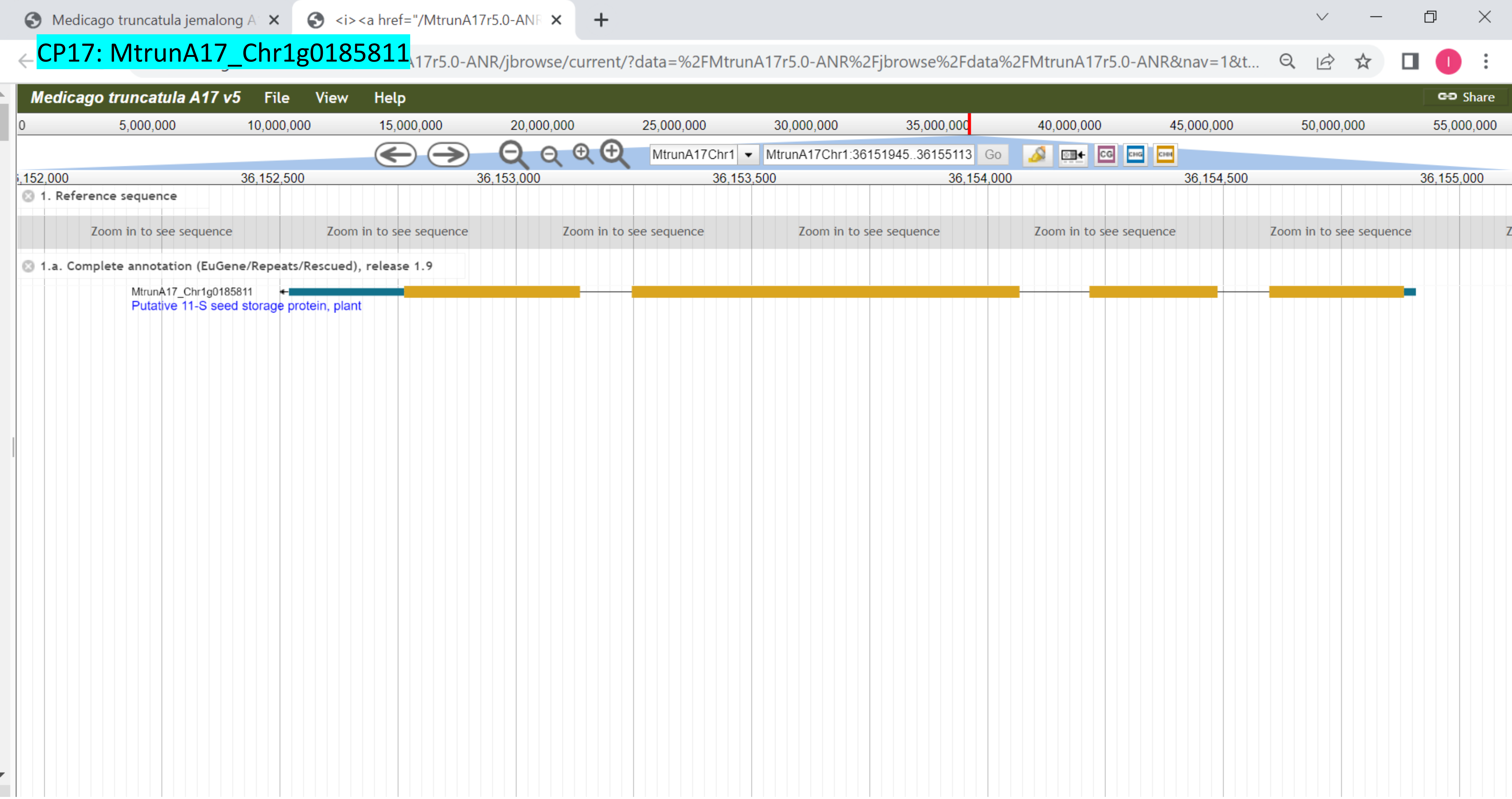

### CP18: MtrunA17_Chr1g0185811

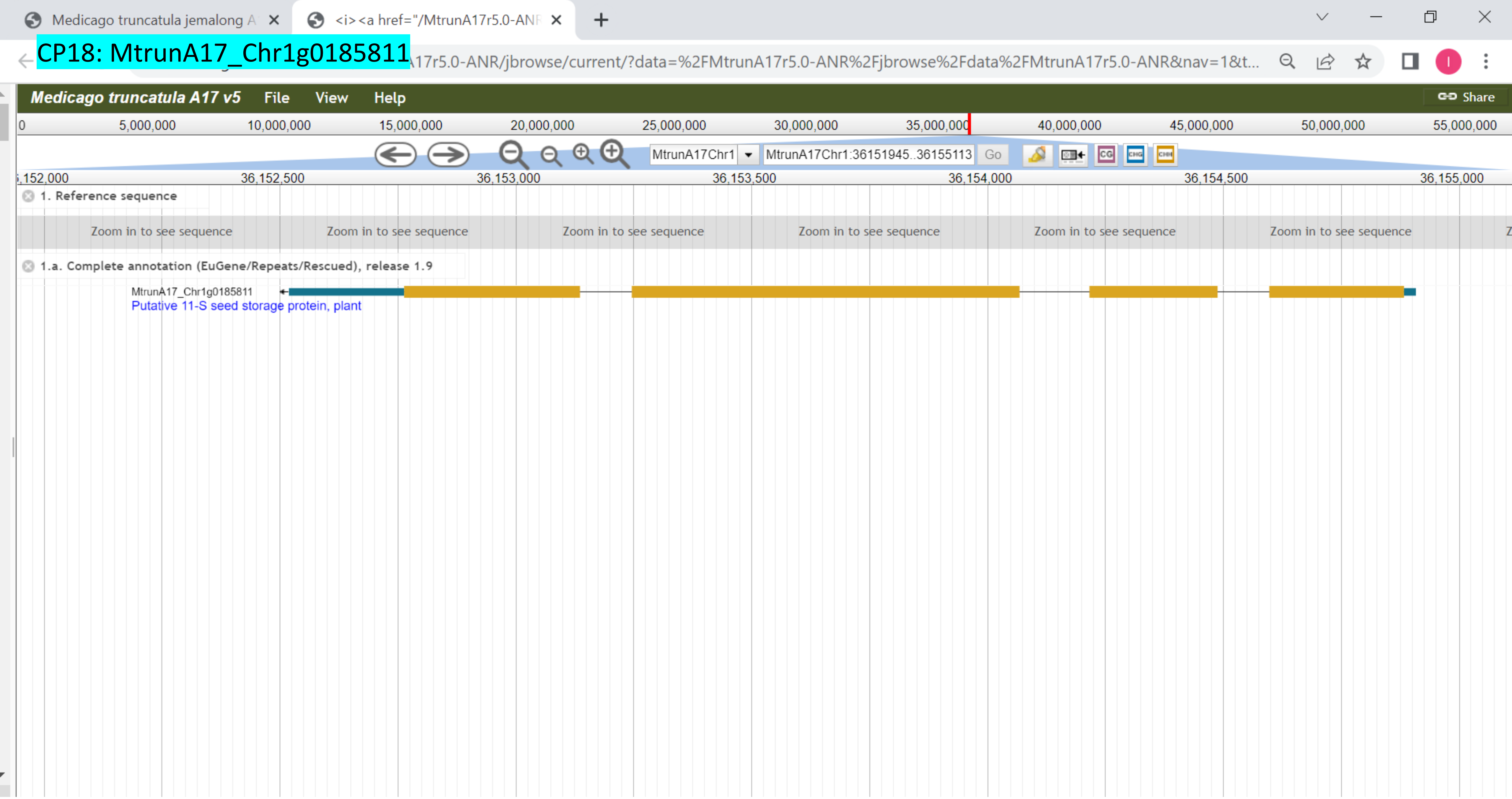

### CP19: MtrunA17_Chr1g0185871

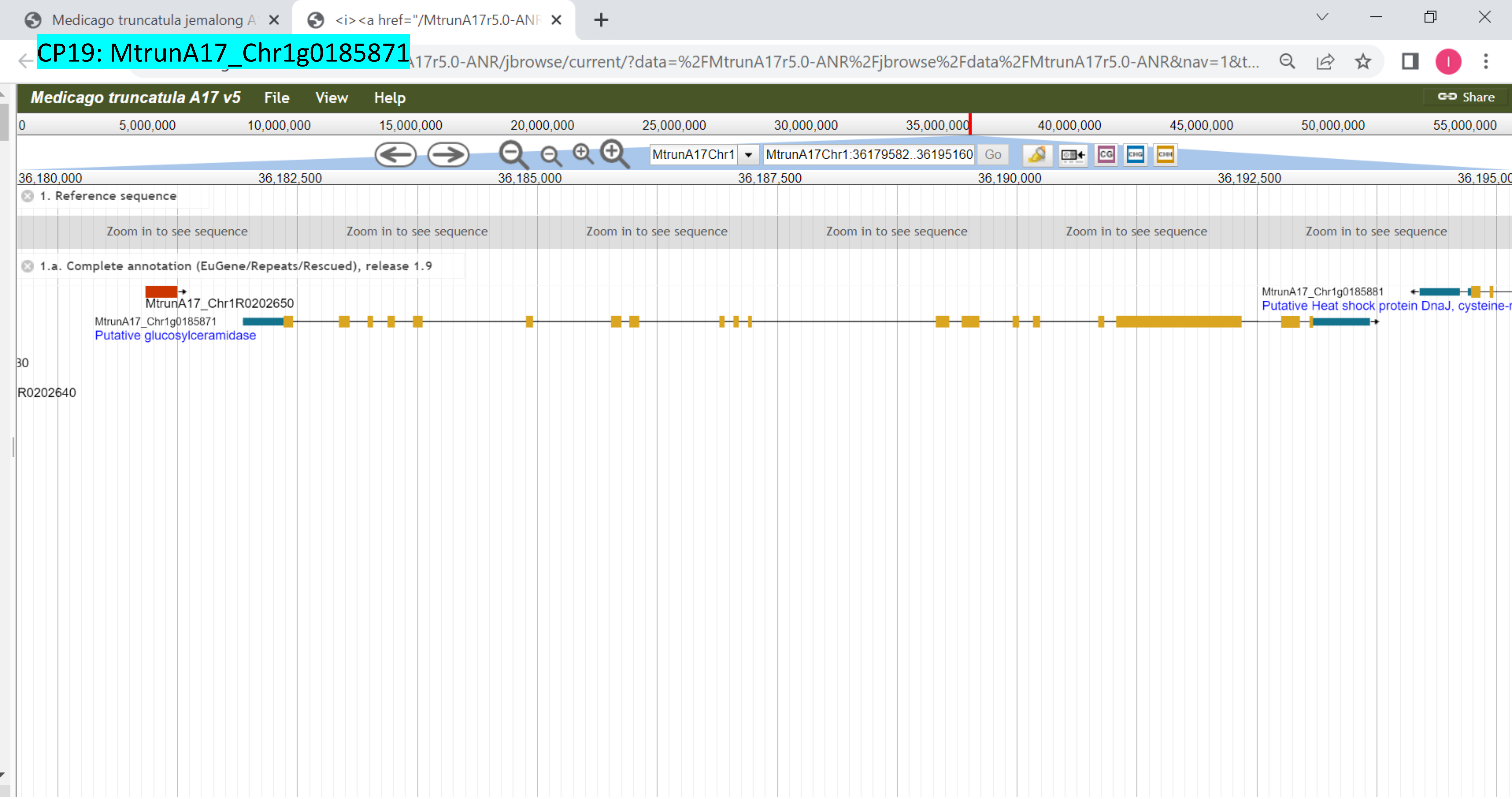

CP20: MtrunA17_Chr1g0190571

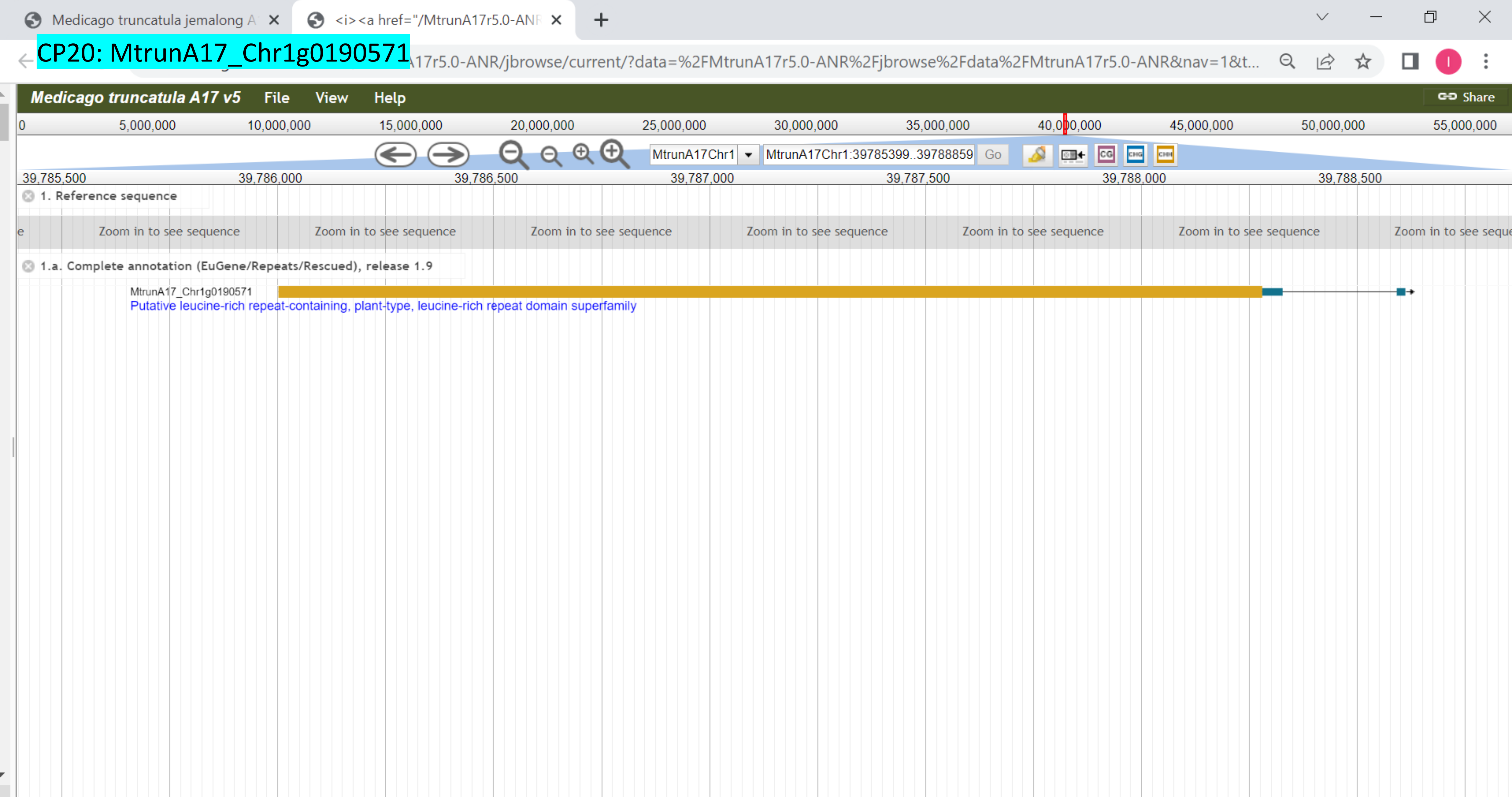

### CP21: MtrunA17_Chr1g0191411

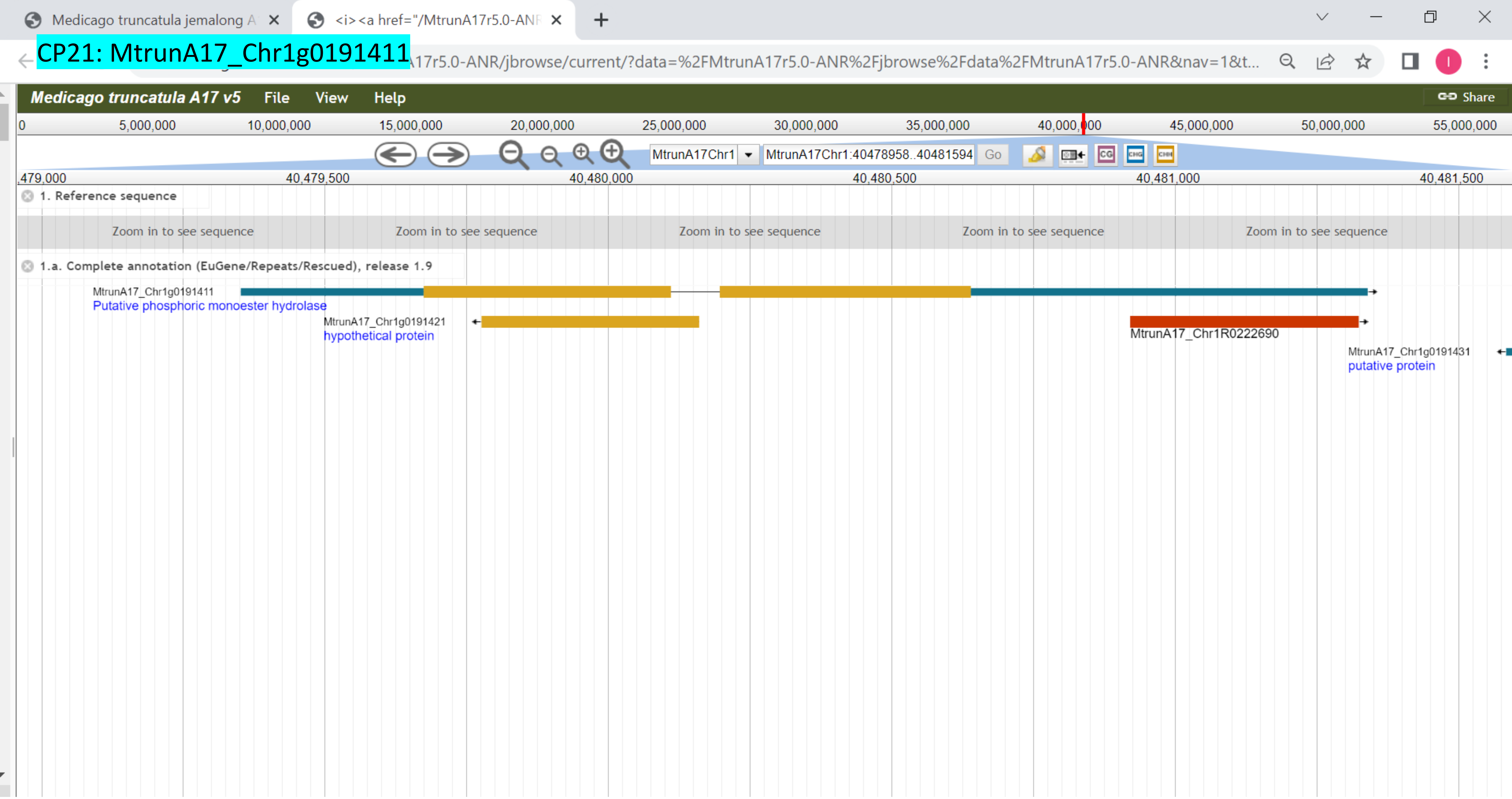

CP22: MtrunA17_Chr1g0198091

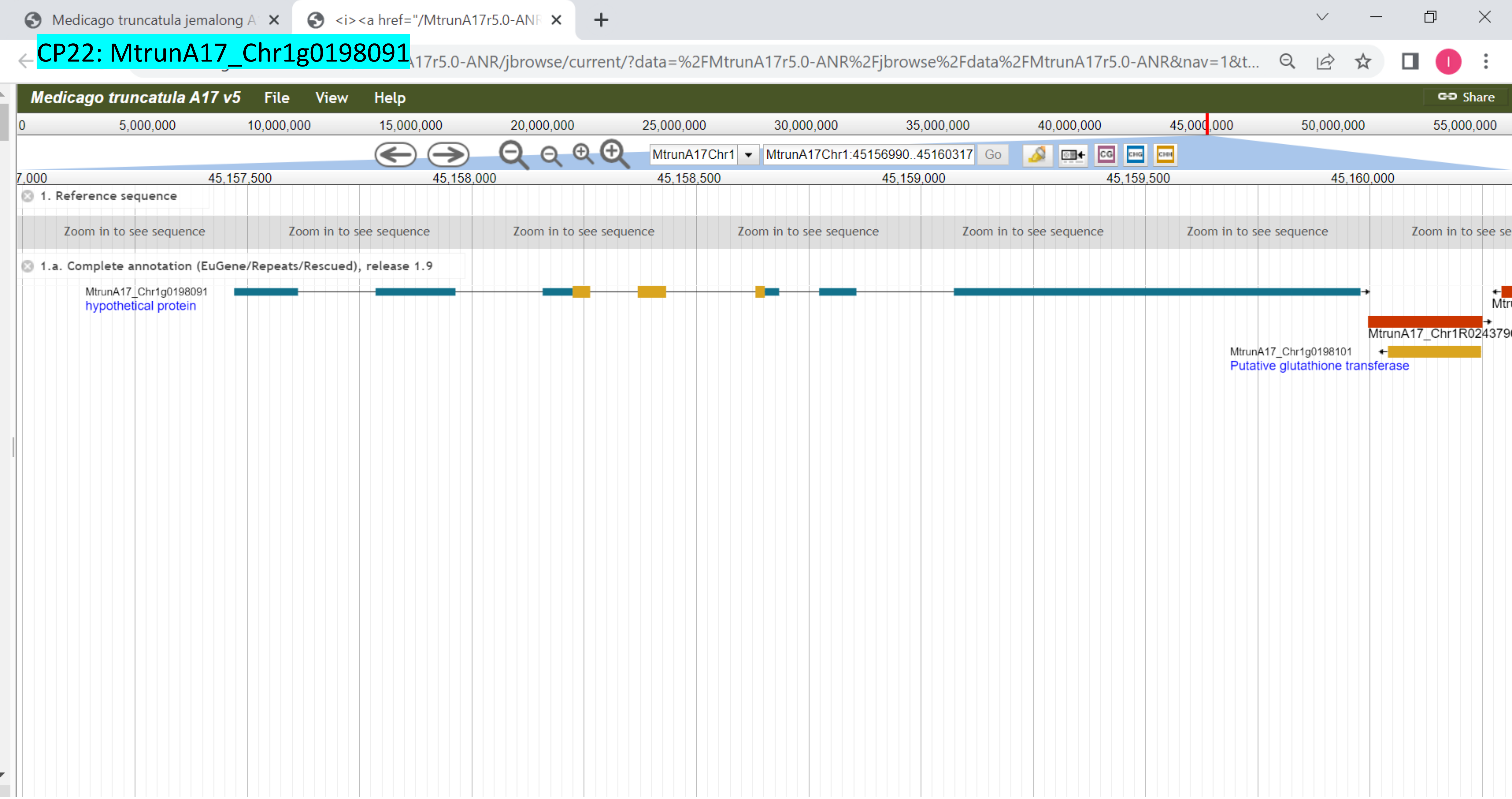

### CP23: MtrunA17_Chr1g0200071

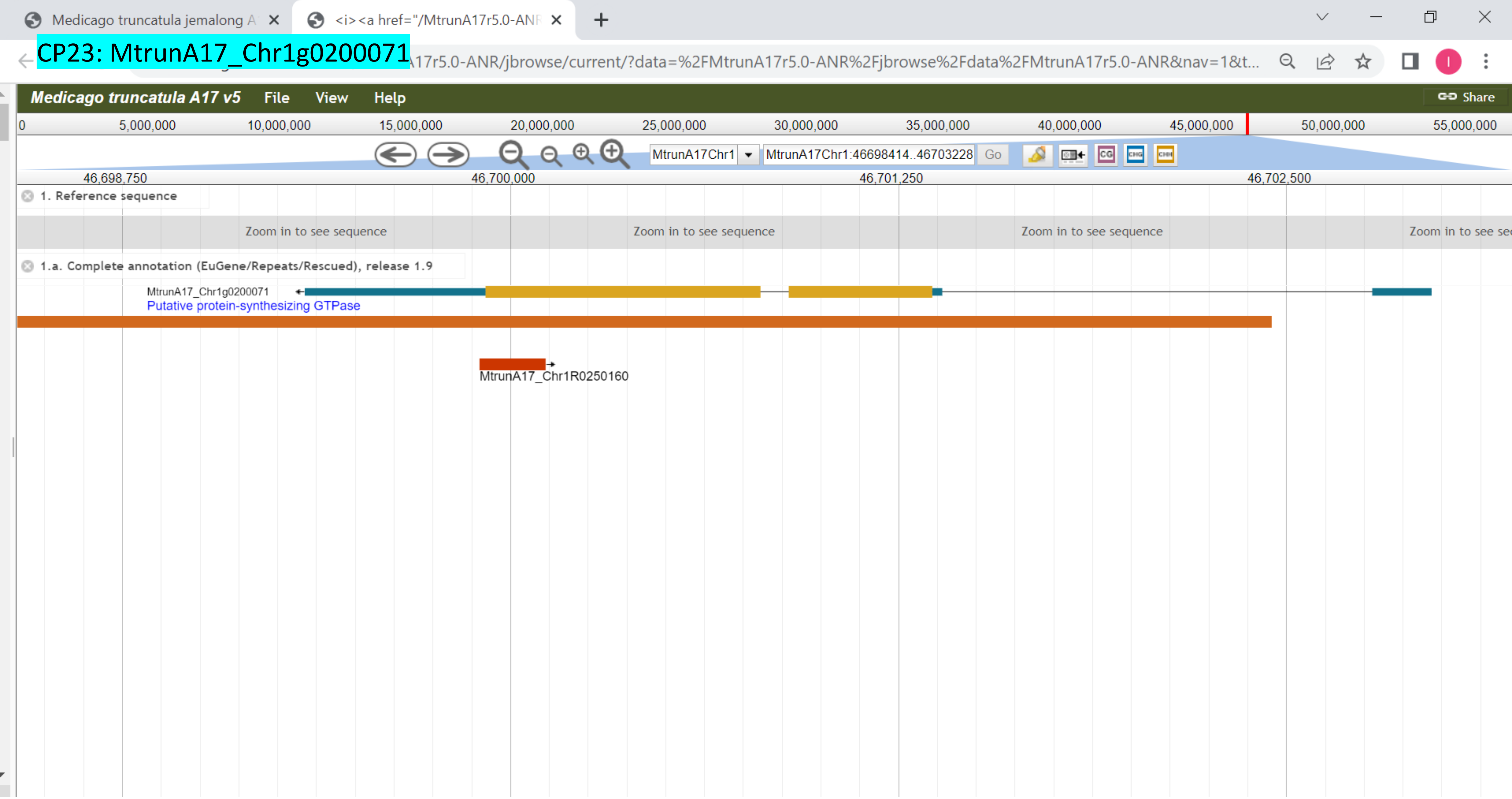

CP24: MtrunA17_Chr1g0200071

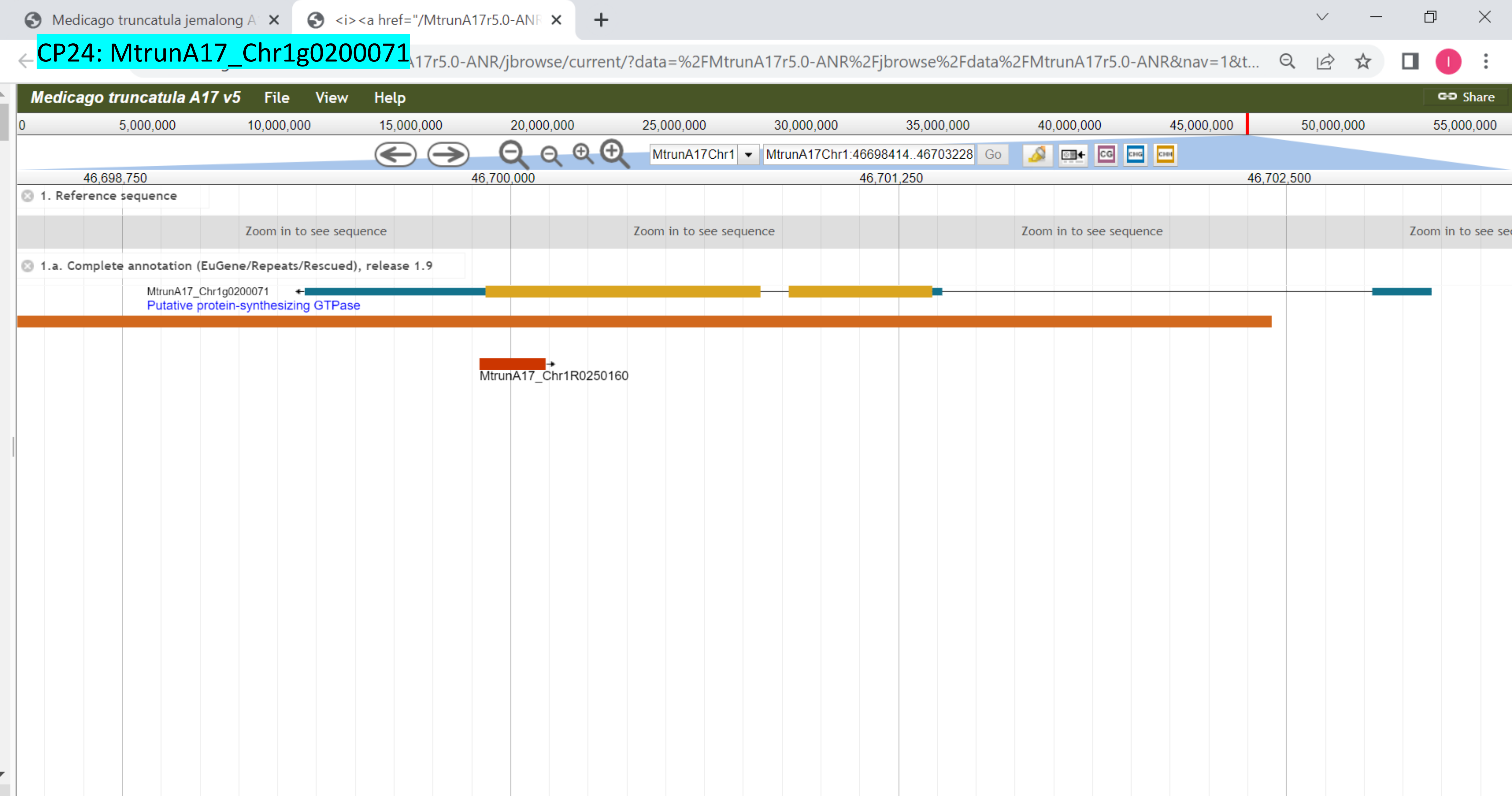

### CP25: MtrunA17_Chr1g0202001

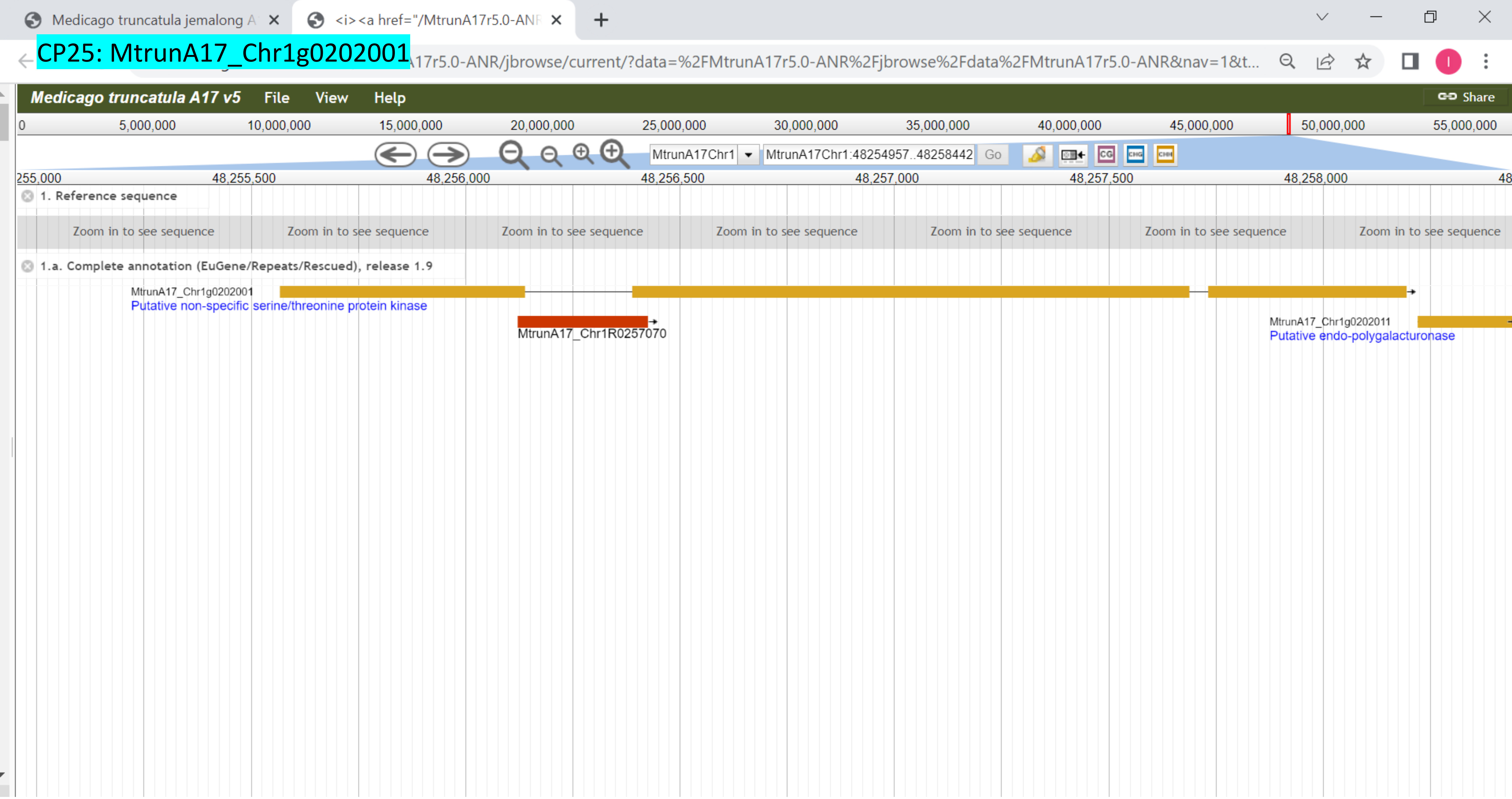

CP26: MtrunA17_Chr1g0205601

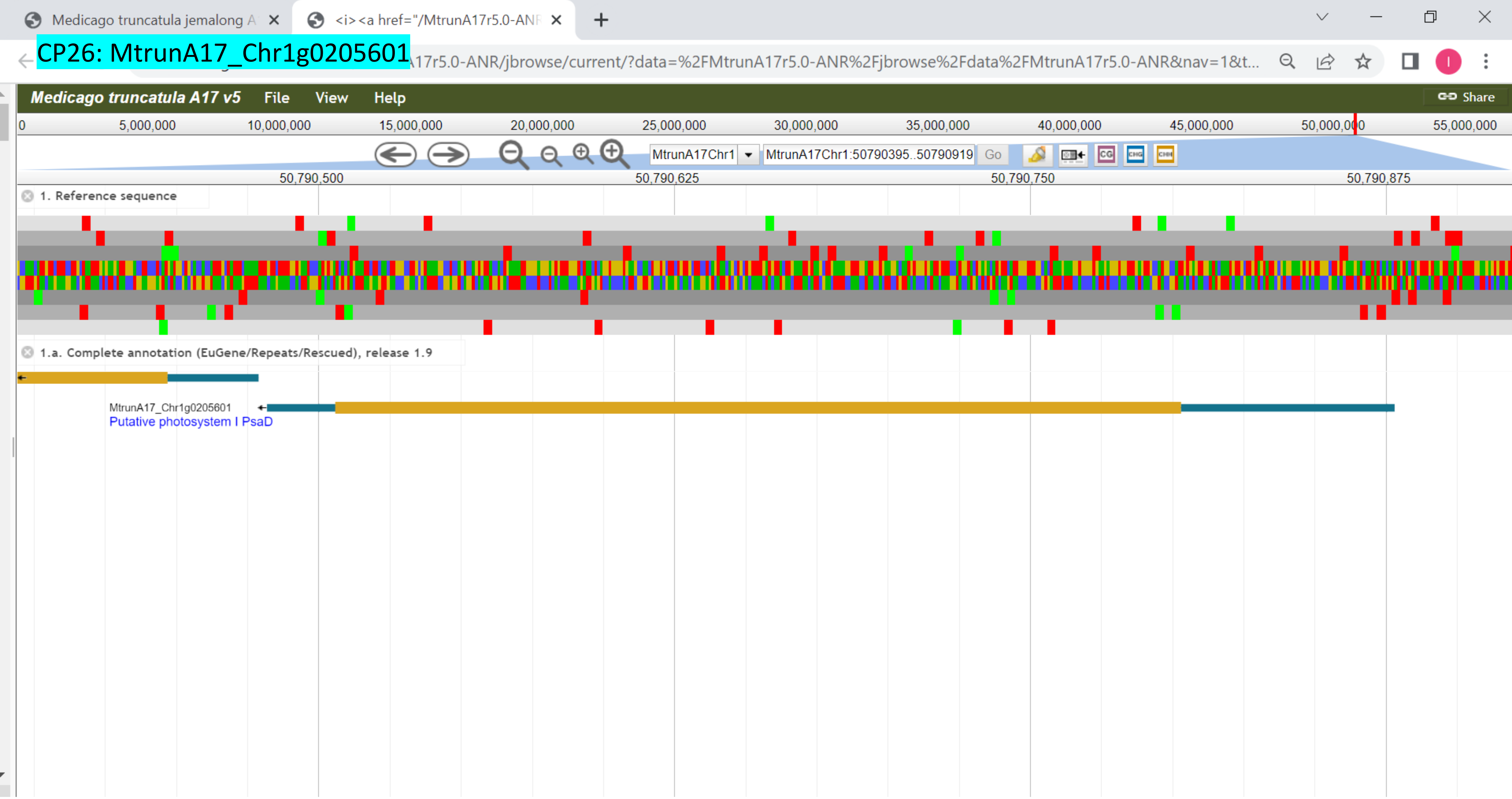

### CP27: MtrunA17_Chr1g0207811

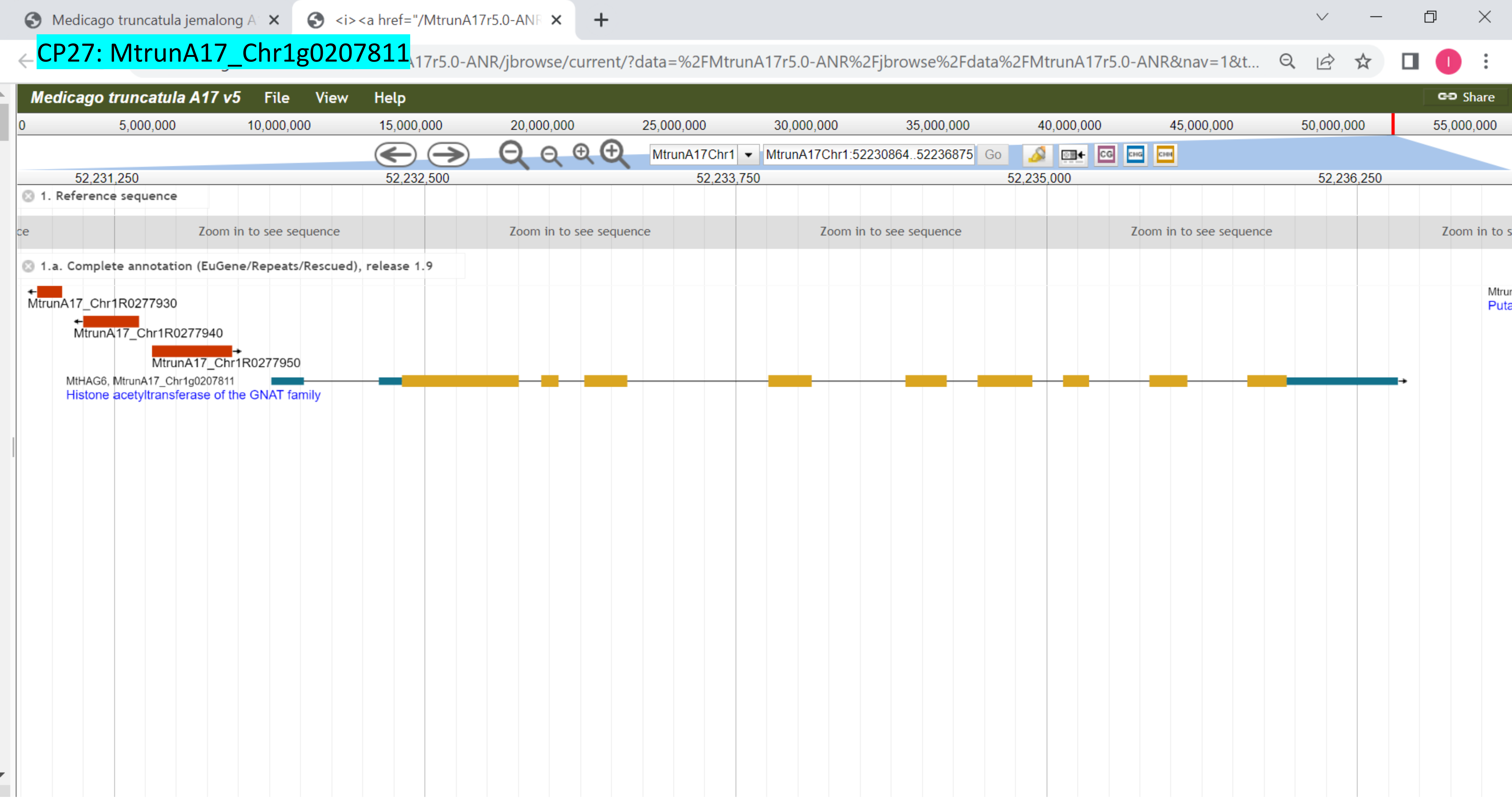

CP28: MtrunA17_Chr1g0207921

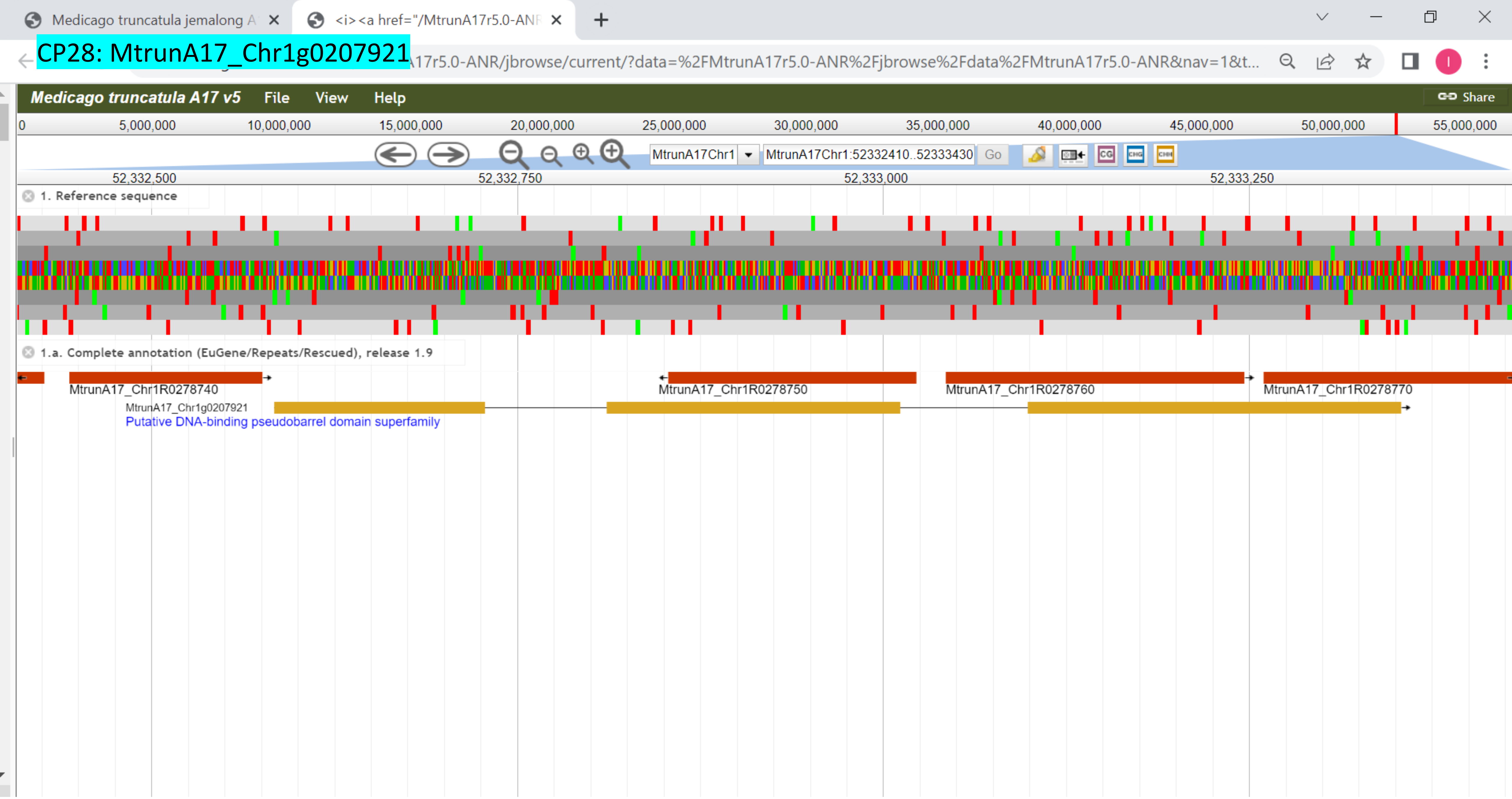

### CP29: MtrunA17_Chr1g0209791

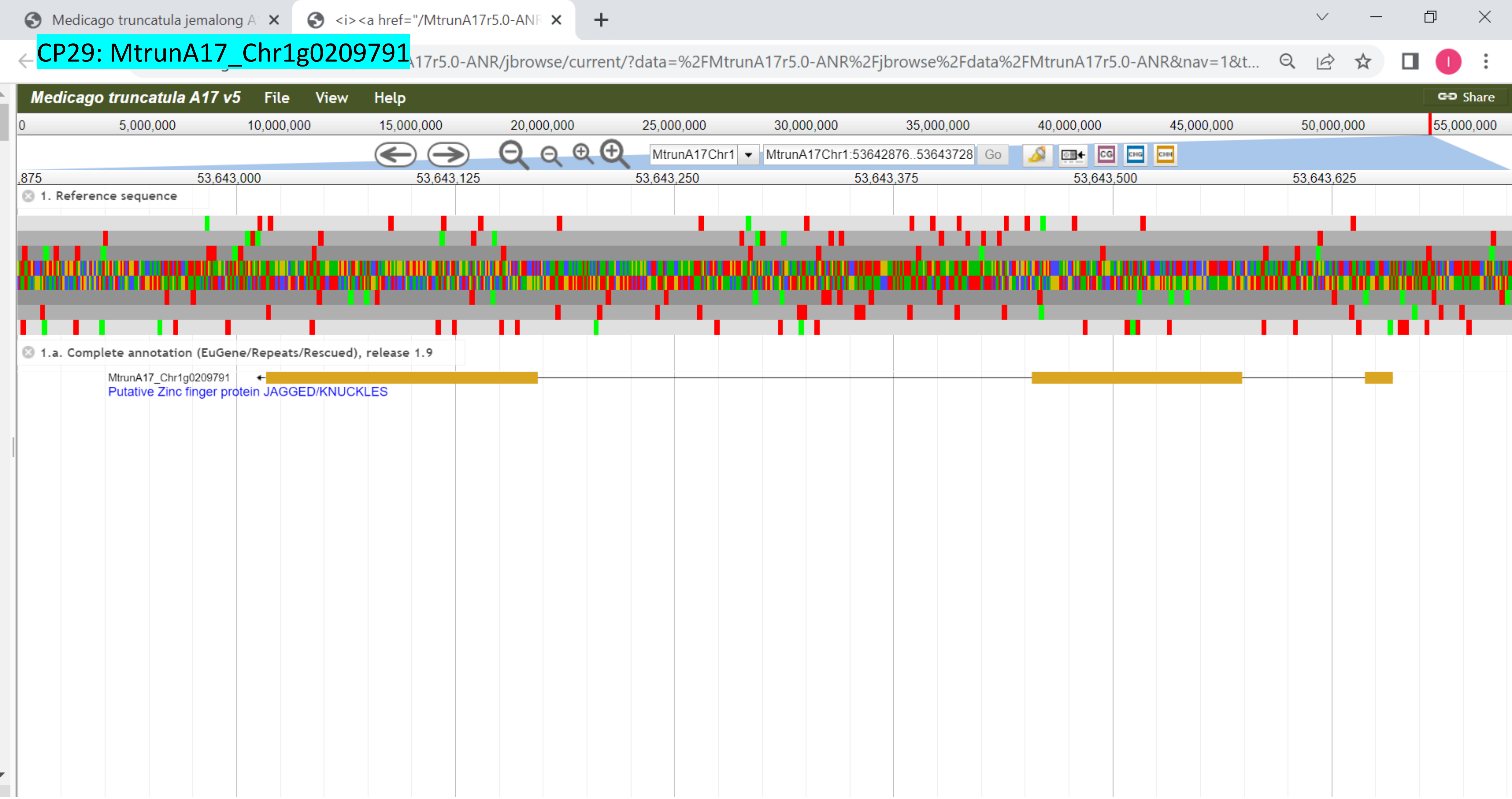

CP30: MtrunA17_Chr1g0210521

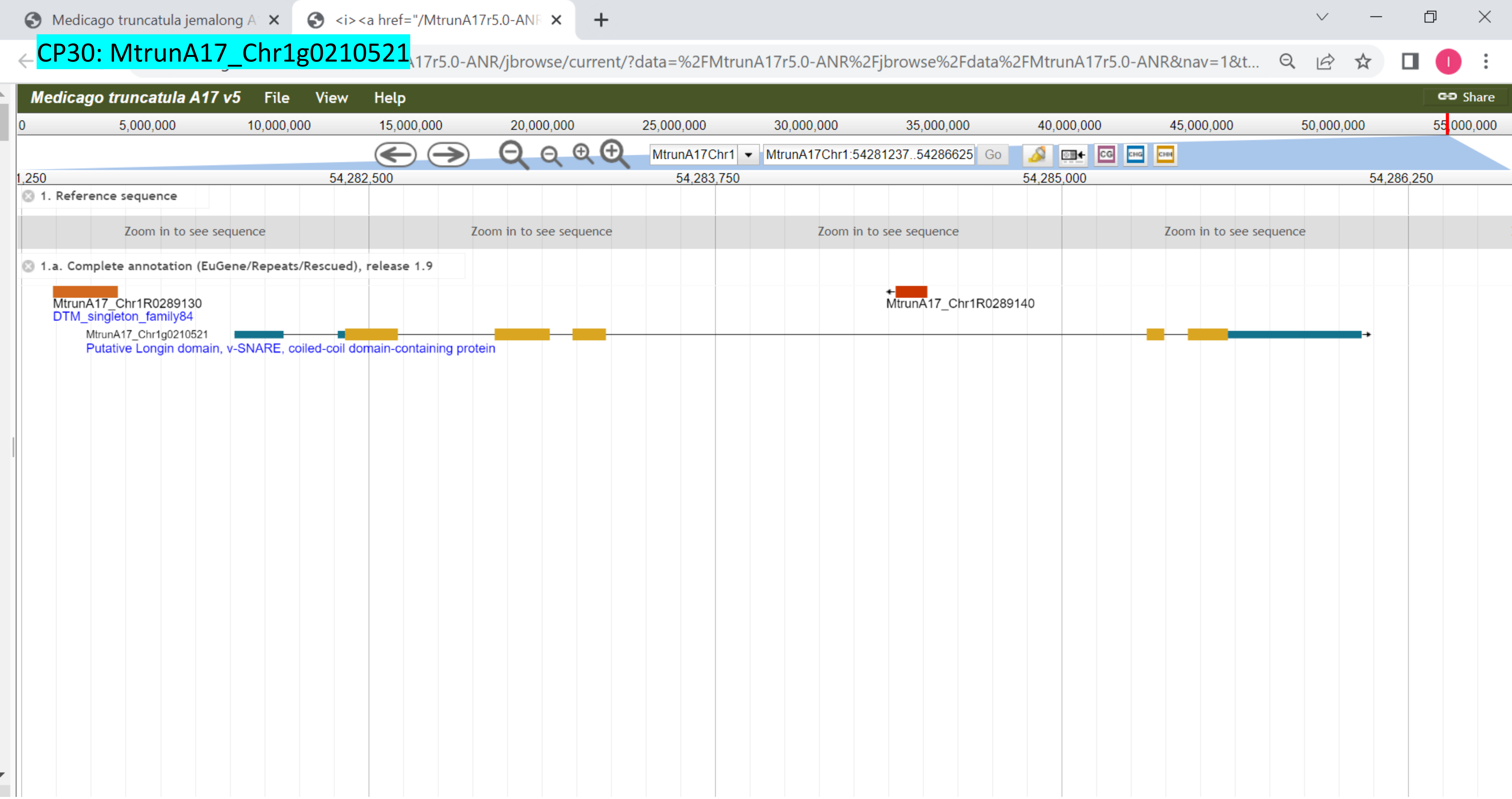

### CP31: MtrunA17_Chr1g0212961

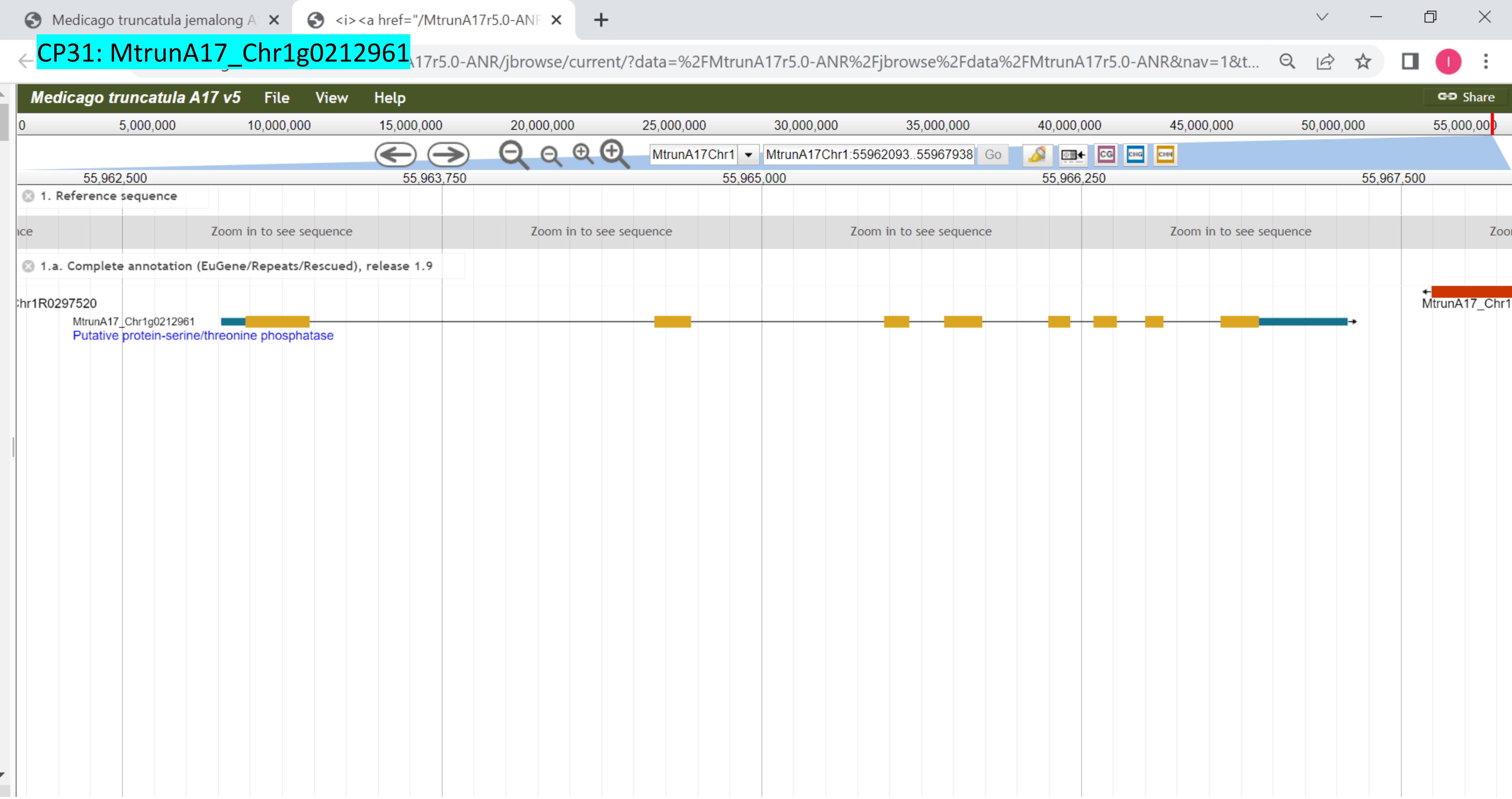

CP32: MtrunA17_Chr1g1004575

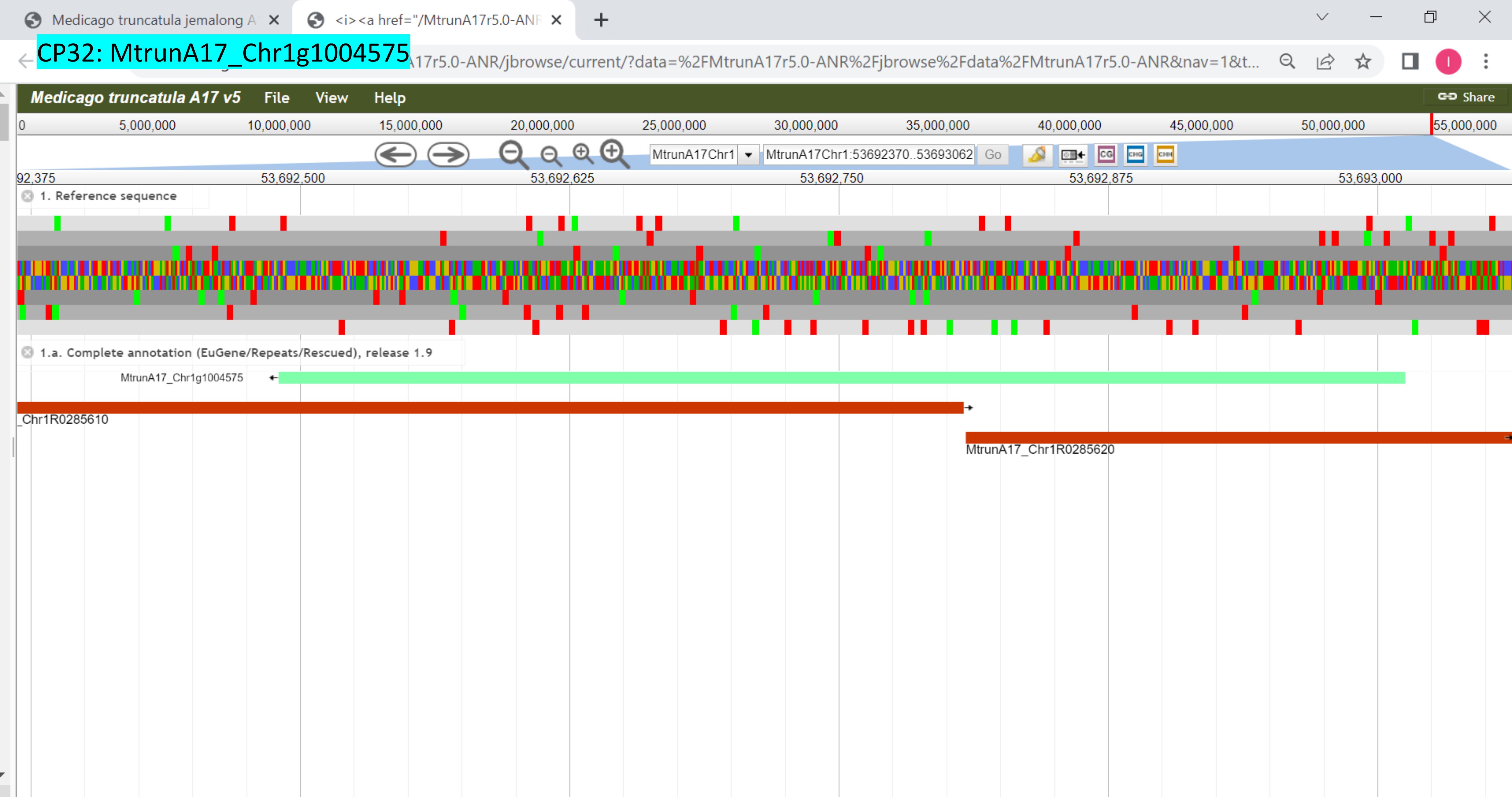

CP33: MtrunA17_Chr2g0283311

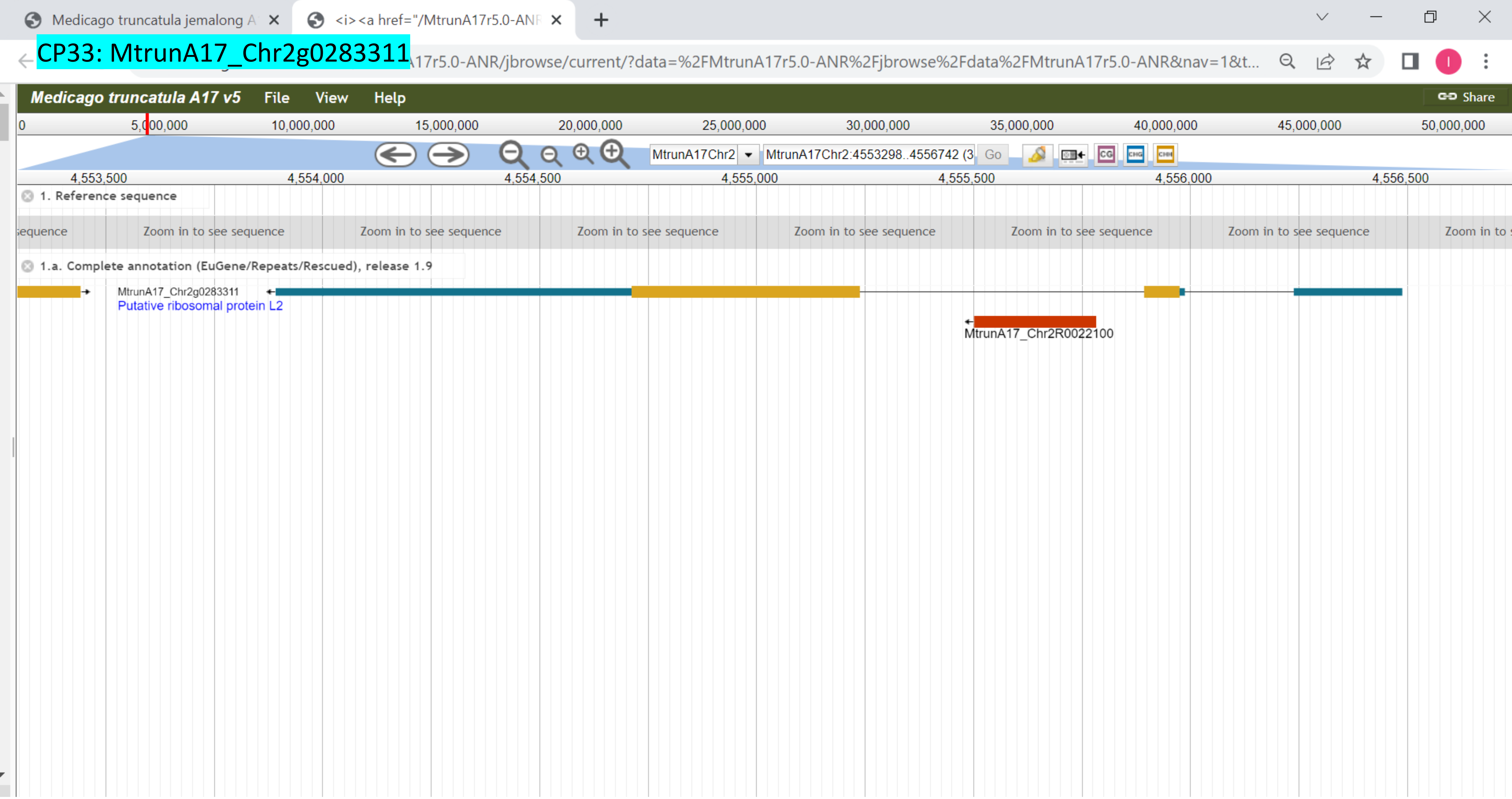

CP34: MtrunA17_Chr2g0285461

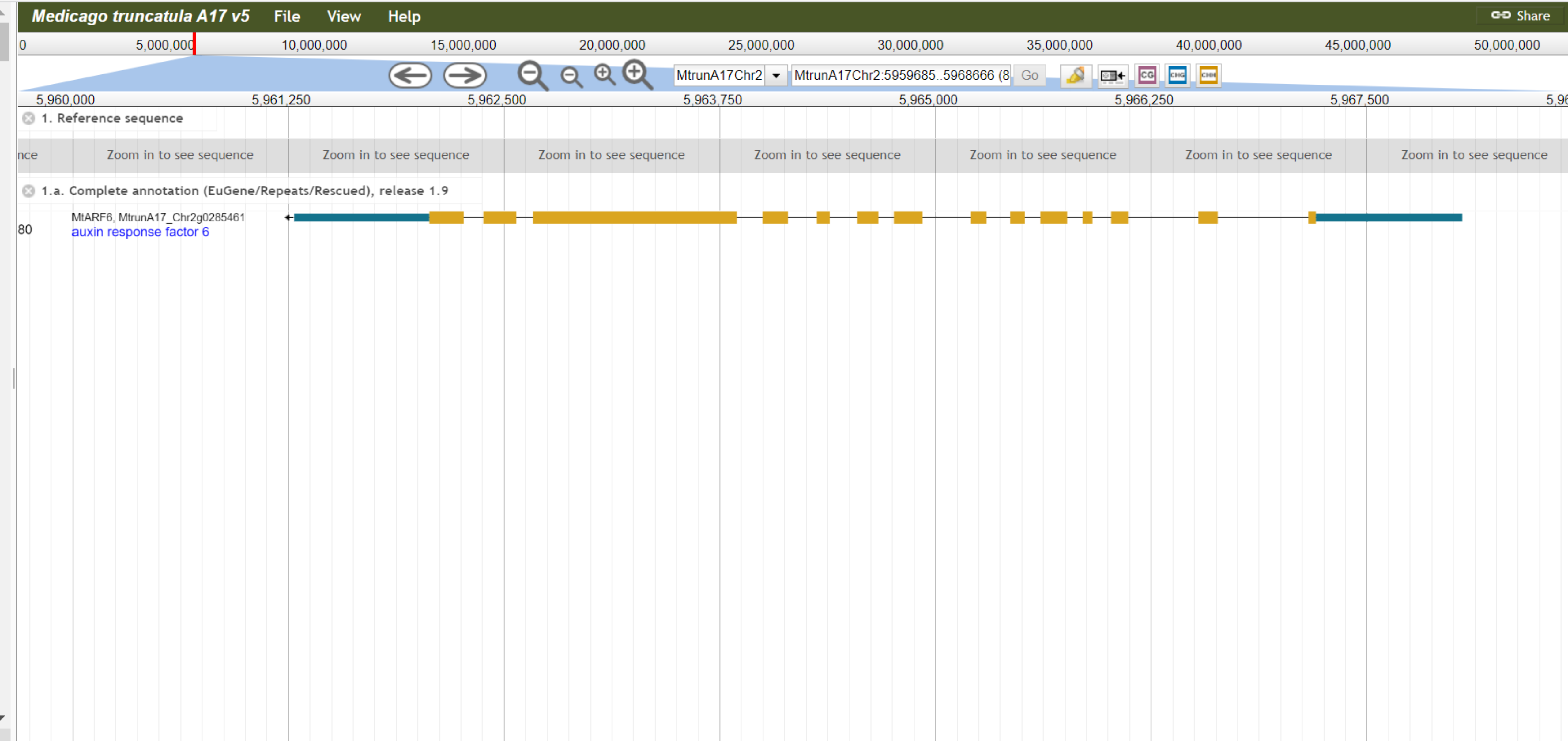

CP35: MtrunA17_Chr2g0292921

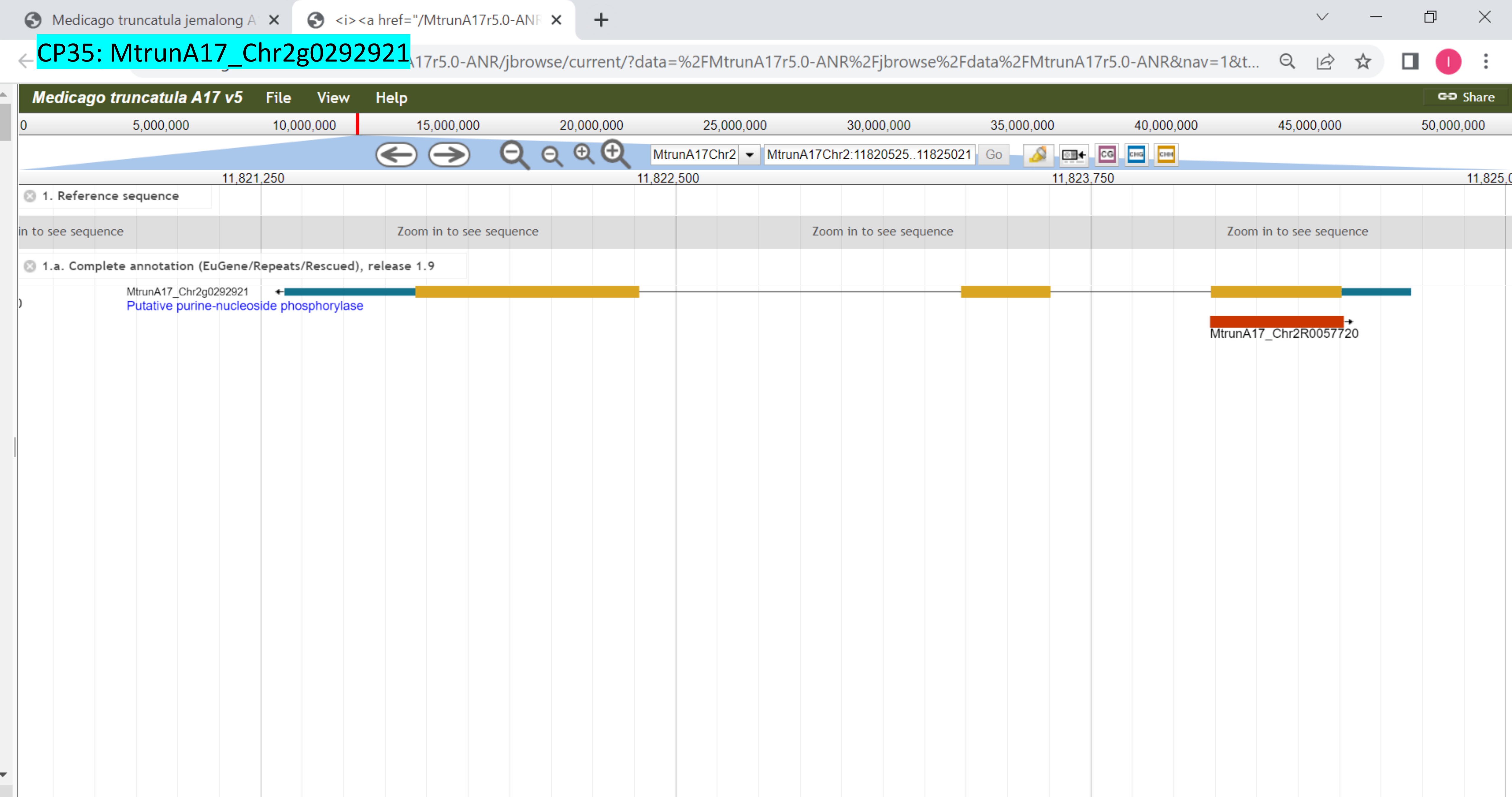

CP36: MtrunA17_Chr2g0298731

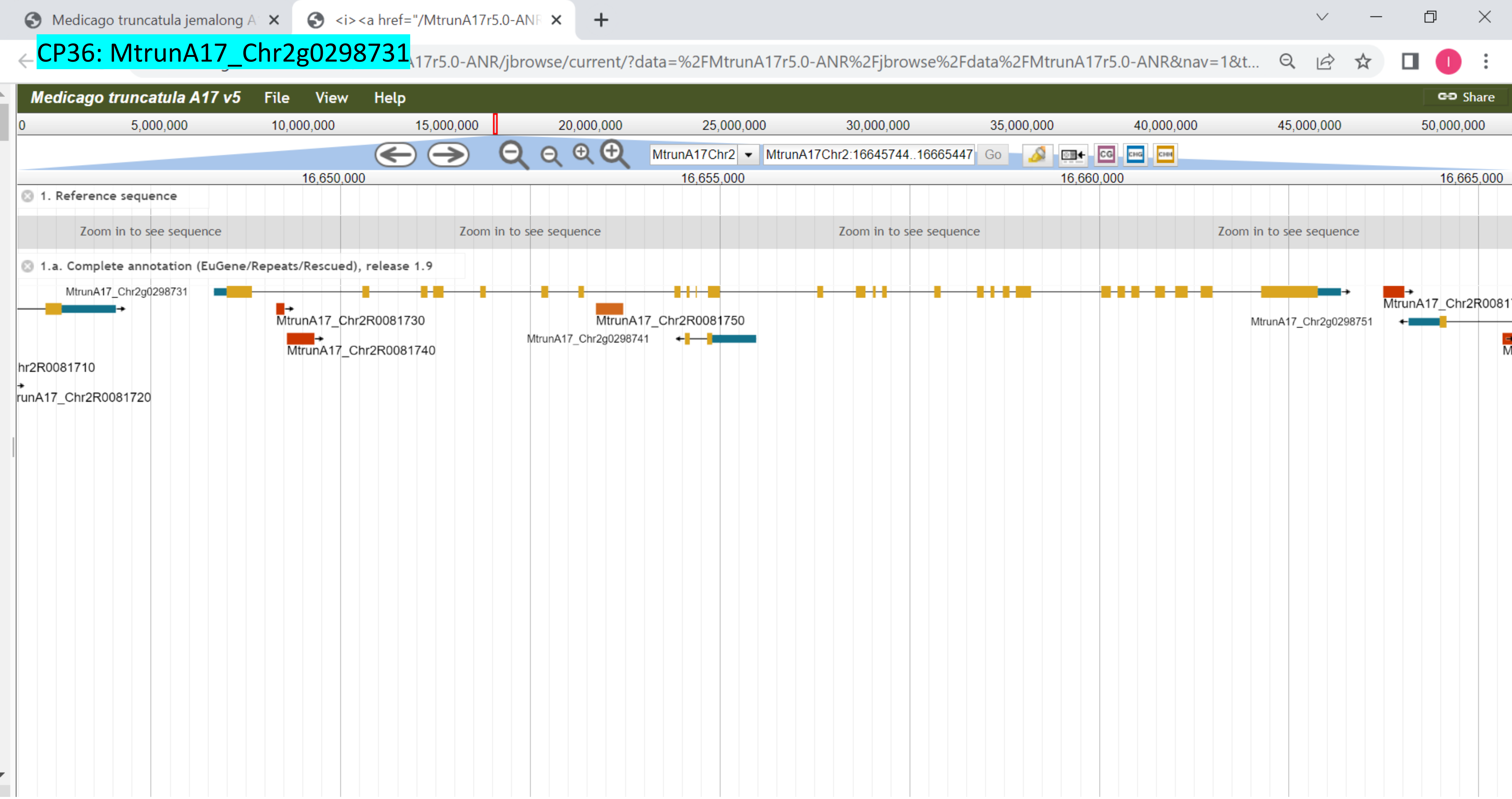

### CP37: MtrunA17_Chr2g0299561

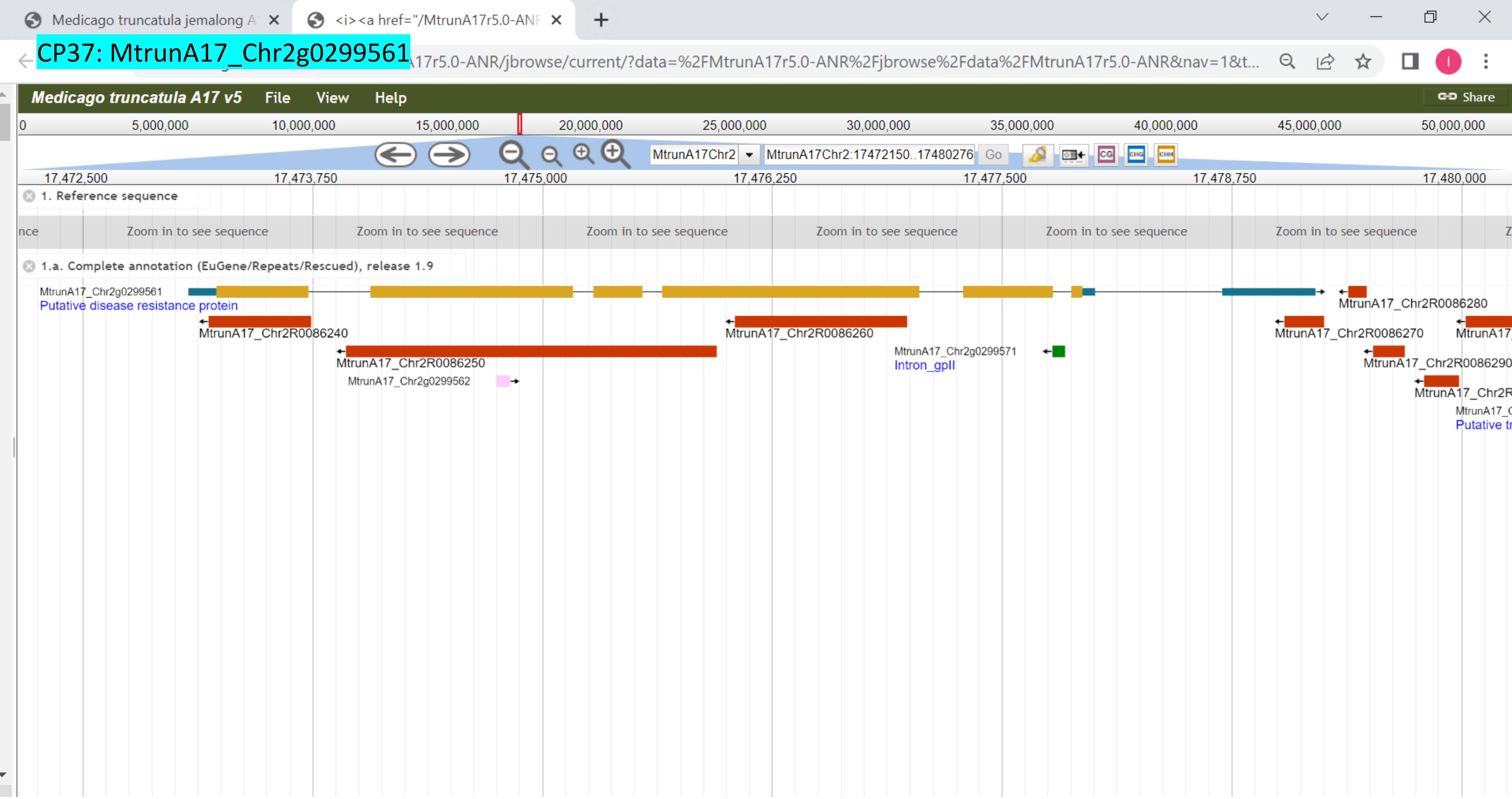

### CP38: MtrunA17_Chr2g0304891

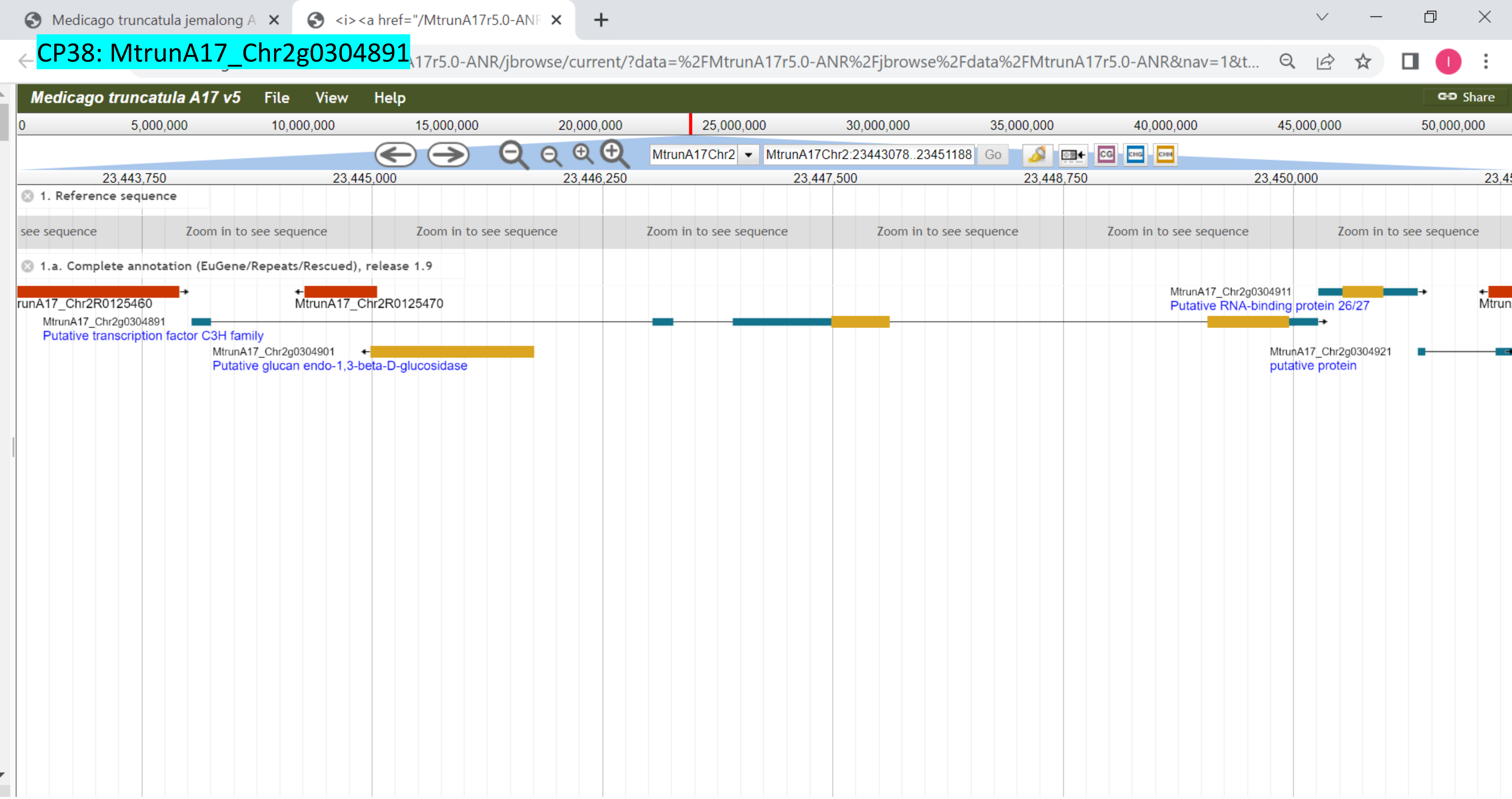

CP39: MtrunA17_Chr2g0305951

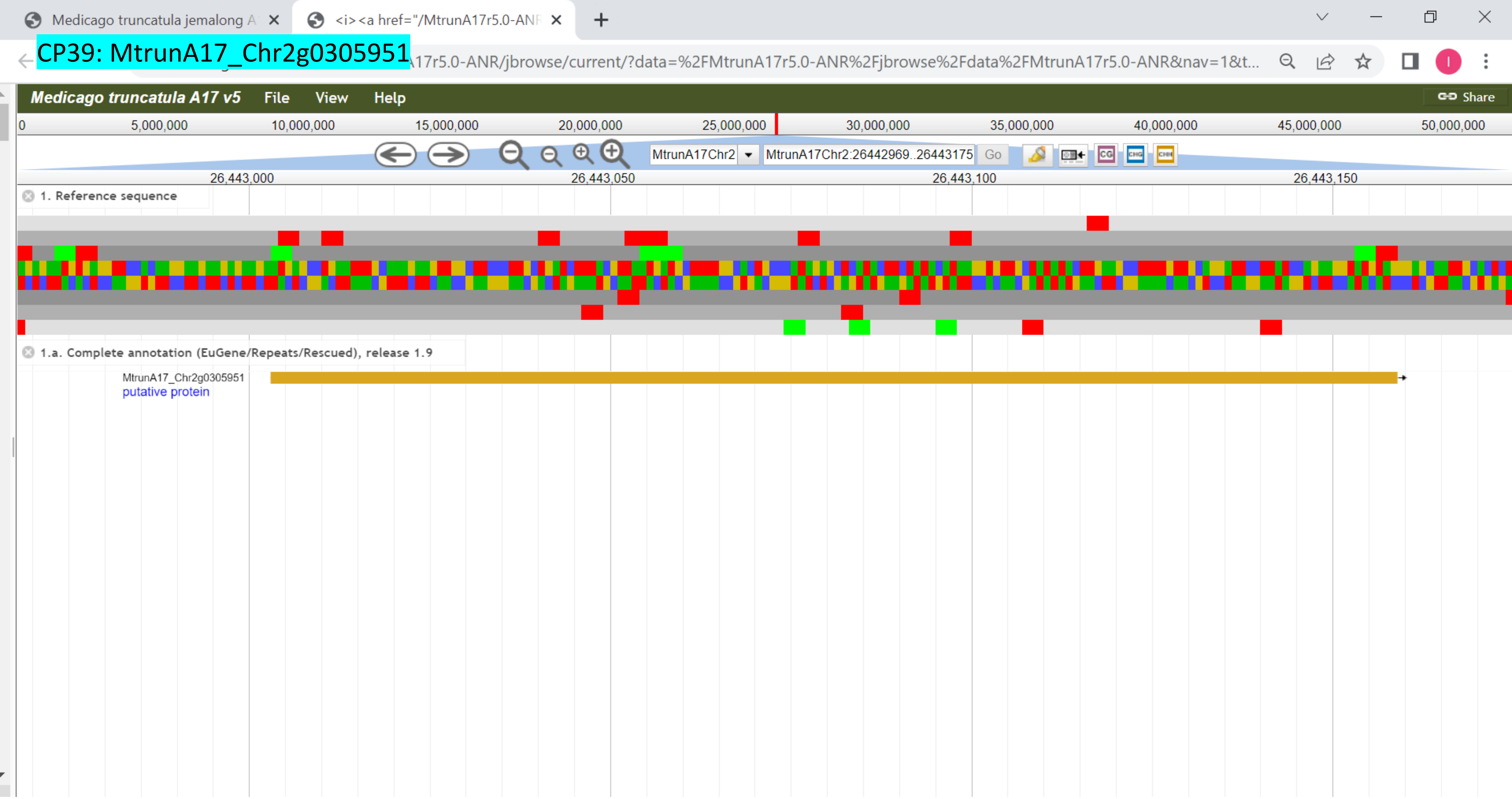

### CP40: MtrunA17_Chr2g0309251

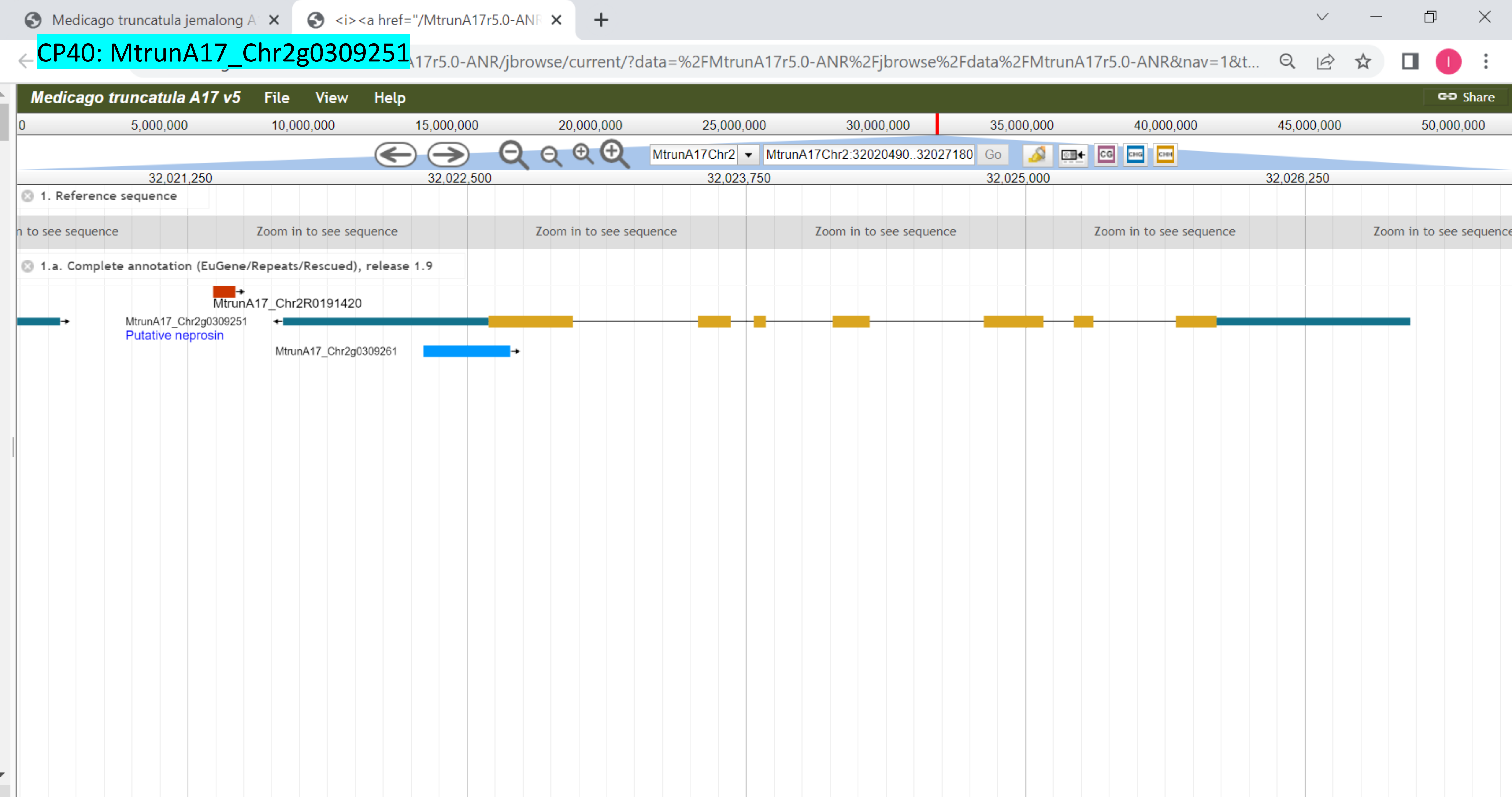

CP41: MtrunA17_Chr2g0310041

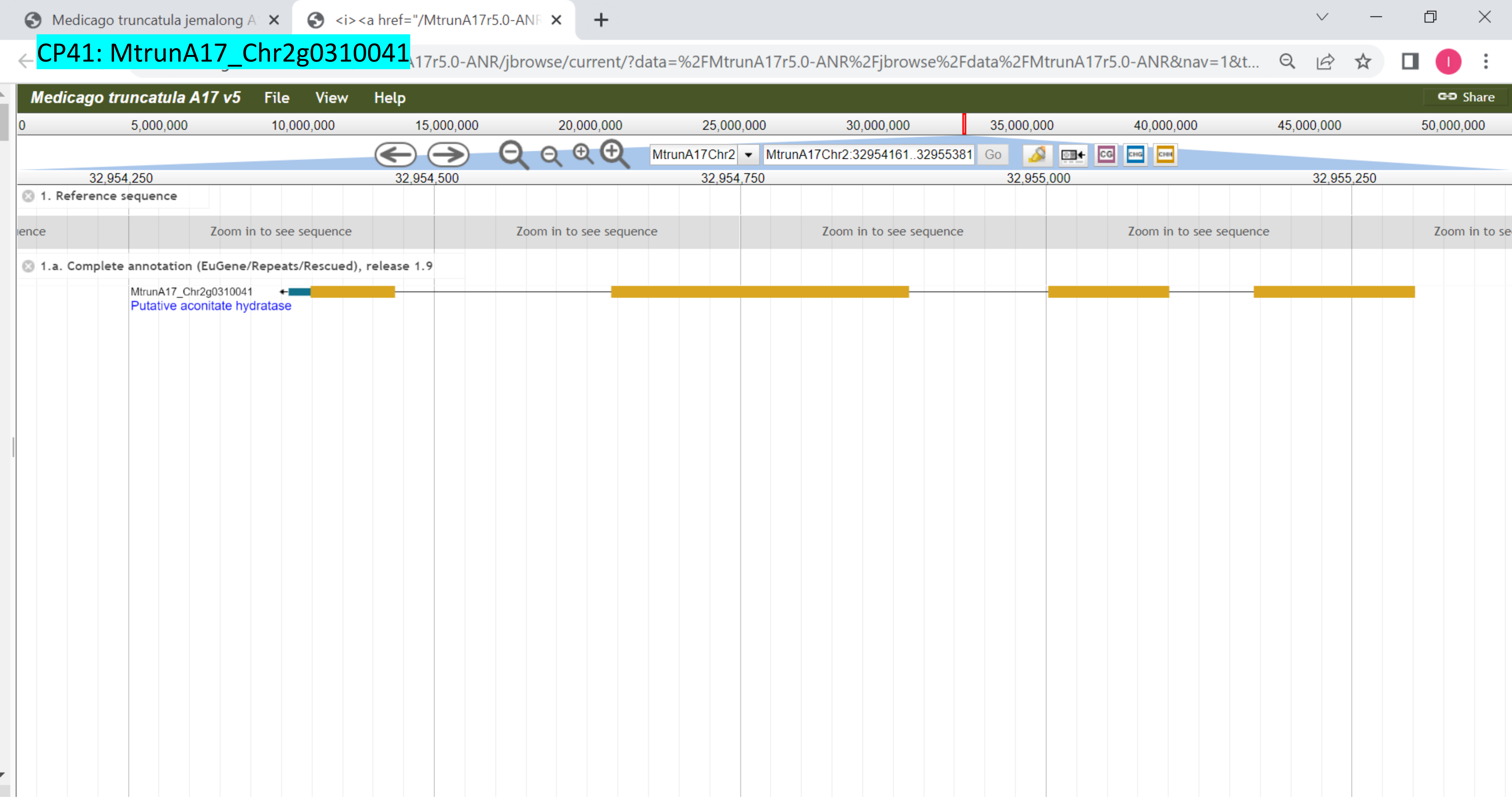

CP42: MtrunA17_Chr2g0312631

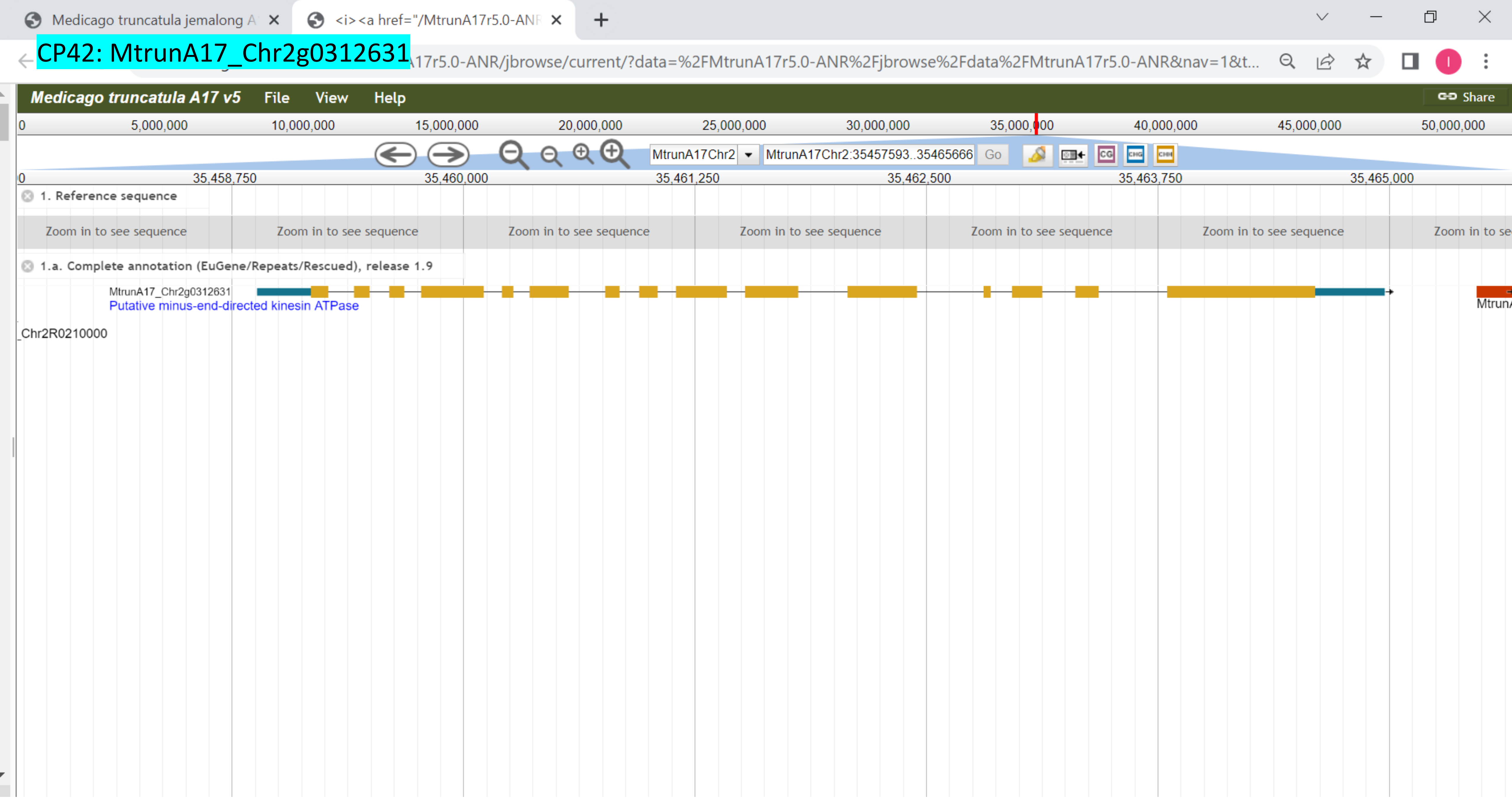

CP43: MtrunA17_Chr2g0316291

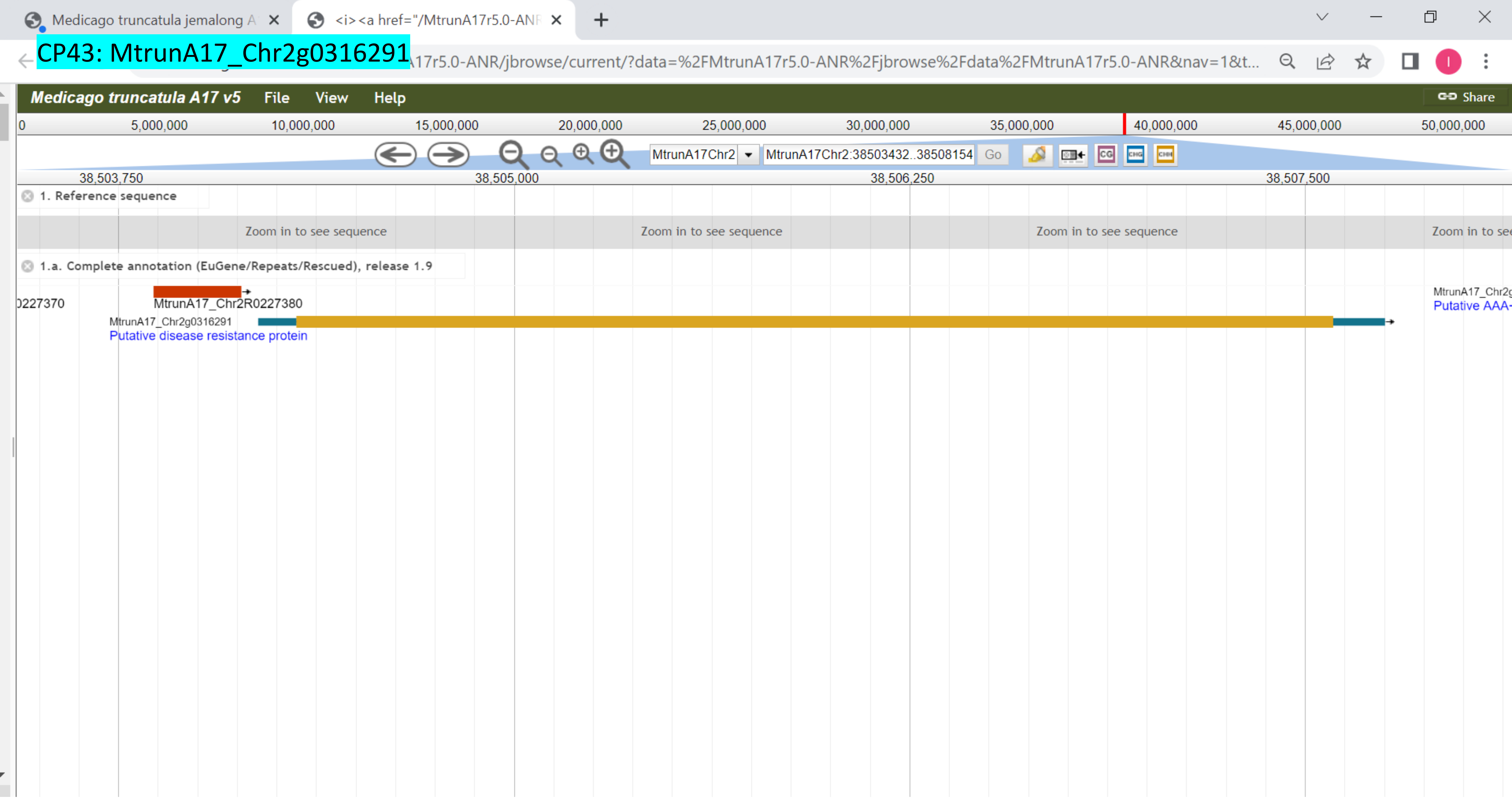

### CP44: MtrunA17_Chr2g0326801

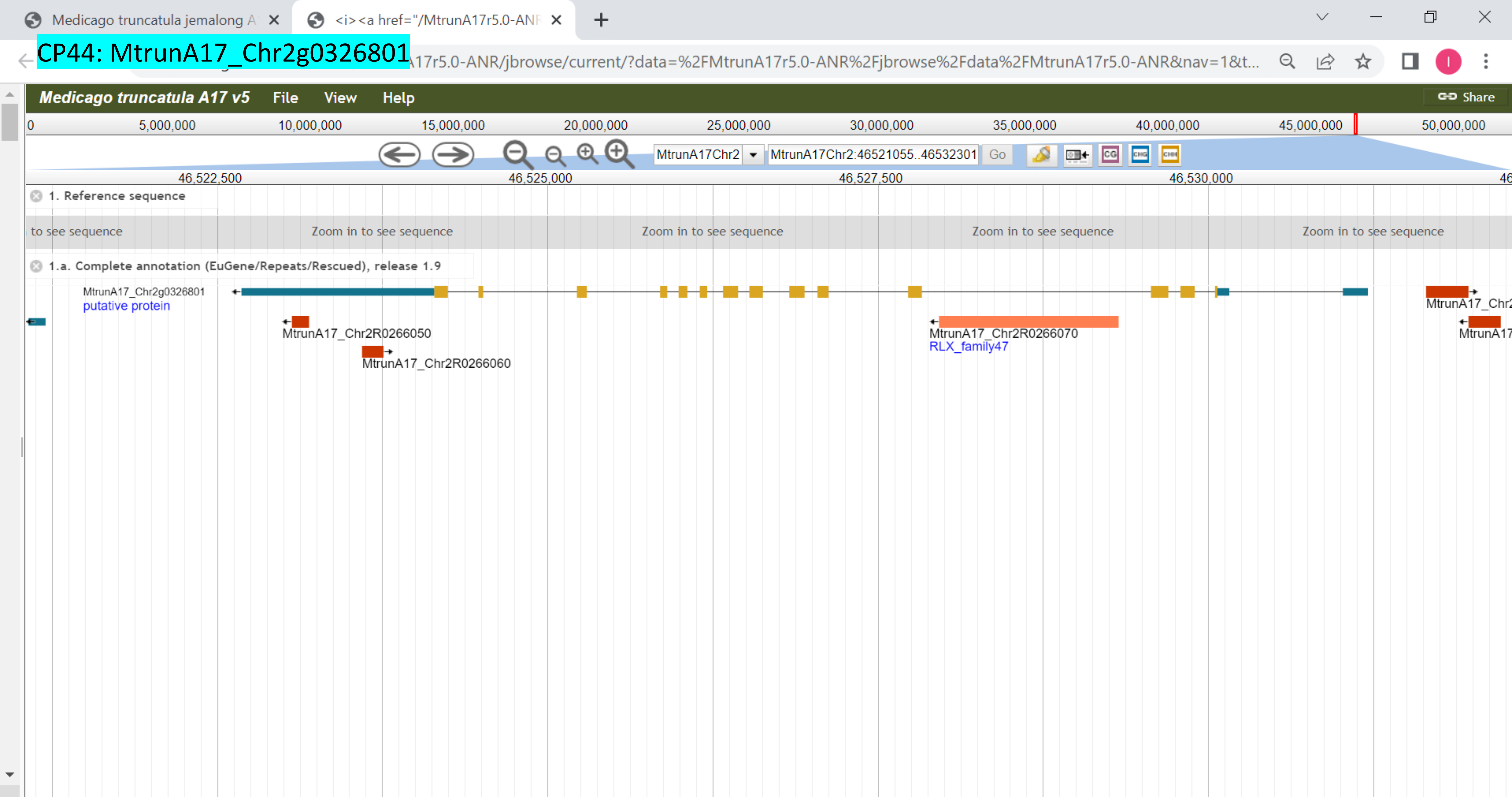

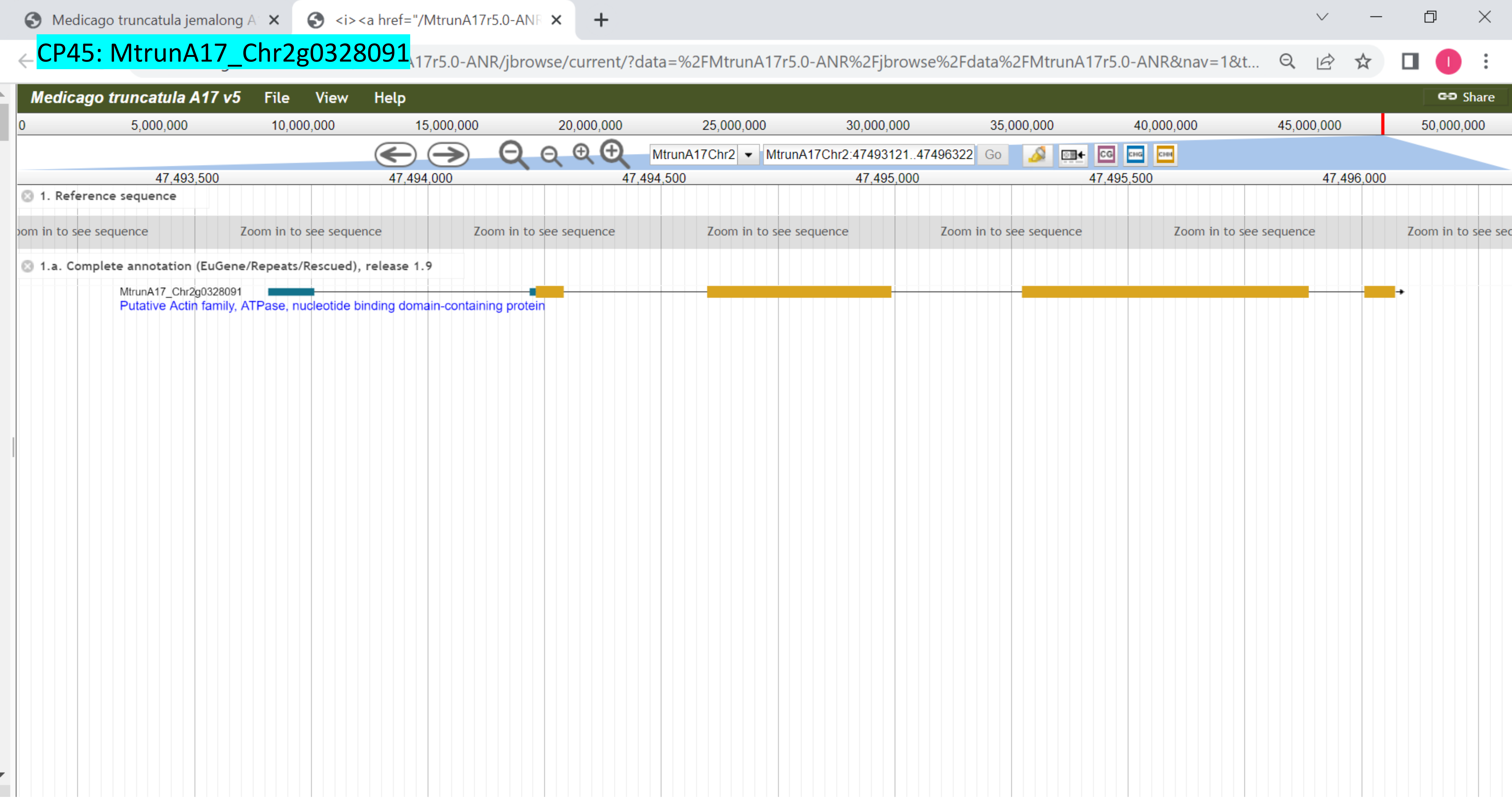

CP45: MtrunA17_Chr2g0328091
Medicago truncatula jemalong A
<i><a href="/MtrunA17r5.0-ANR
17r5.0-ANR/jbrowse/current/?data=%2FMtrunA17r5.0-ANR%2Fjbrowse%2Fdata%2FMtrunA17r5.0-ANR&nav=1&t...
Medicago truncatula A17 v5
File
View
Help
Share
0
5,000,000
10,000,000
15,000,000
20,000,000
25,000,000
30,000,000
35,000,000
40,000,000
45,000,000
50,000,000
MtrunA17Chr2
MtrunA17Chr2:47493121..47496322
Go
47,493,500
47,494,000
47,494,500
47,495,000
47,495,500
47,496,000
1. Reference sequence
Zoom in to see sequence
1.a. Complete annotation (EuGene/Repeats/Rescued), release 1.9
MtrunA17_Chr2g0328091
Putative Actin family, ATPase, nucleotide binding domain-containing protein

### CP46: MtrunA17_Chr2g0329031

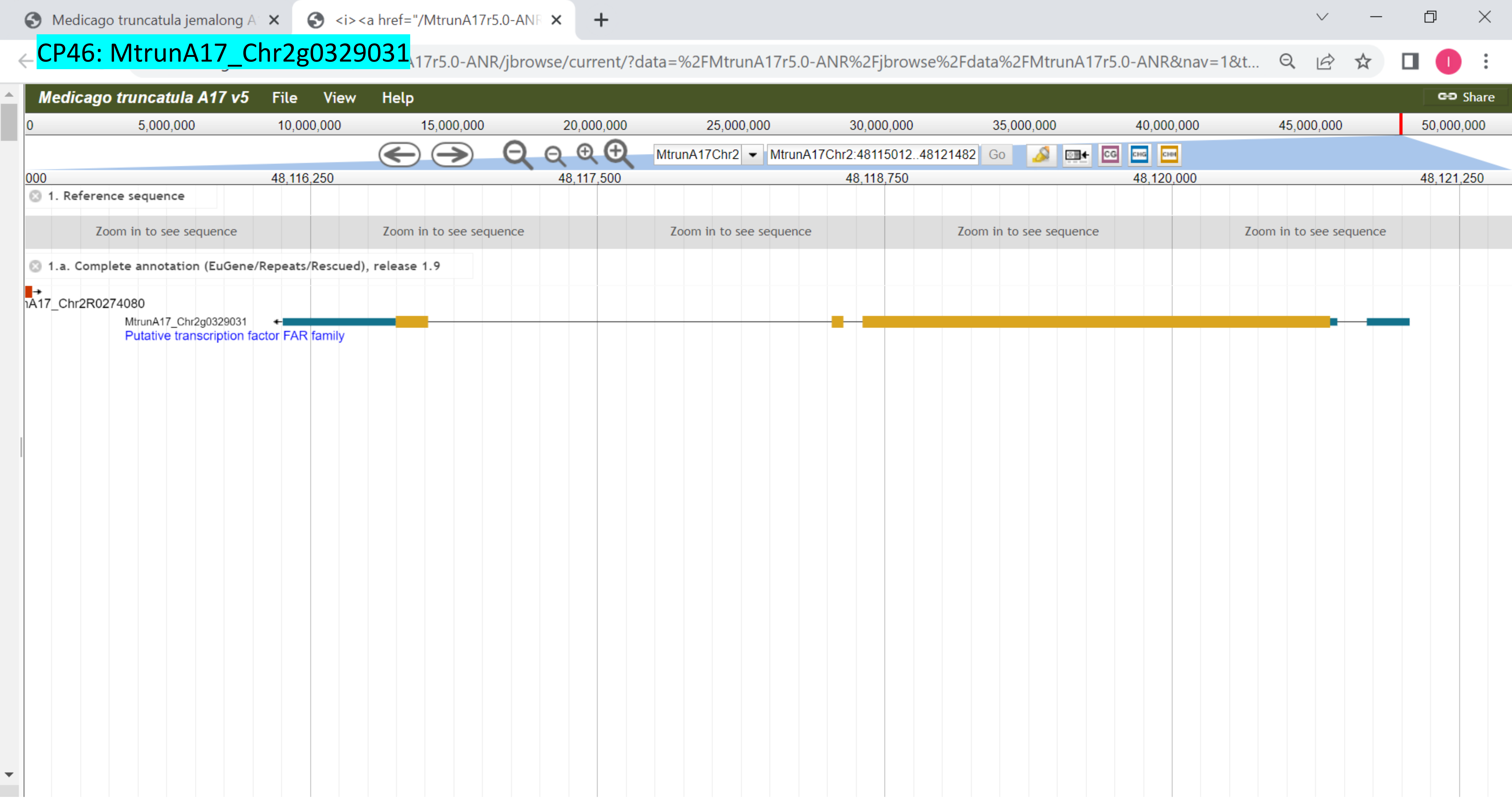

CP47: MtrunA17_Chr3g0079521

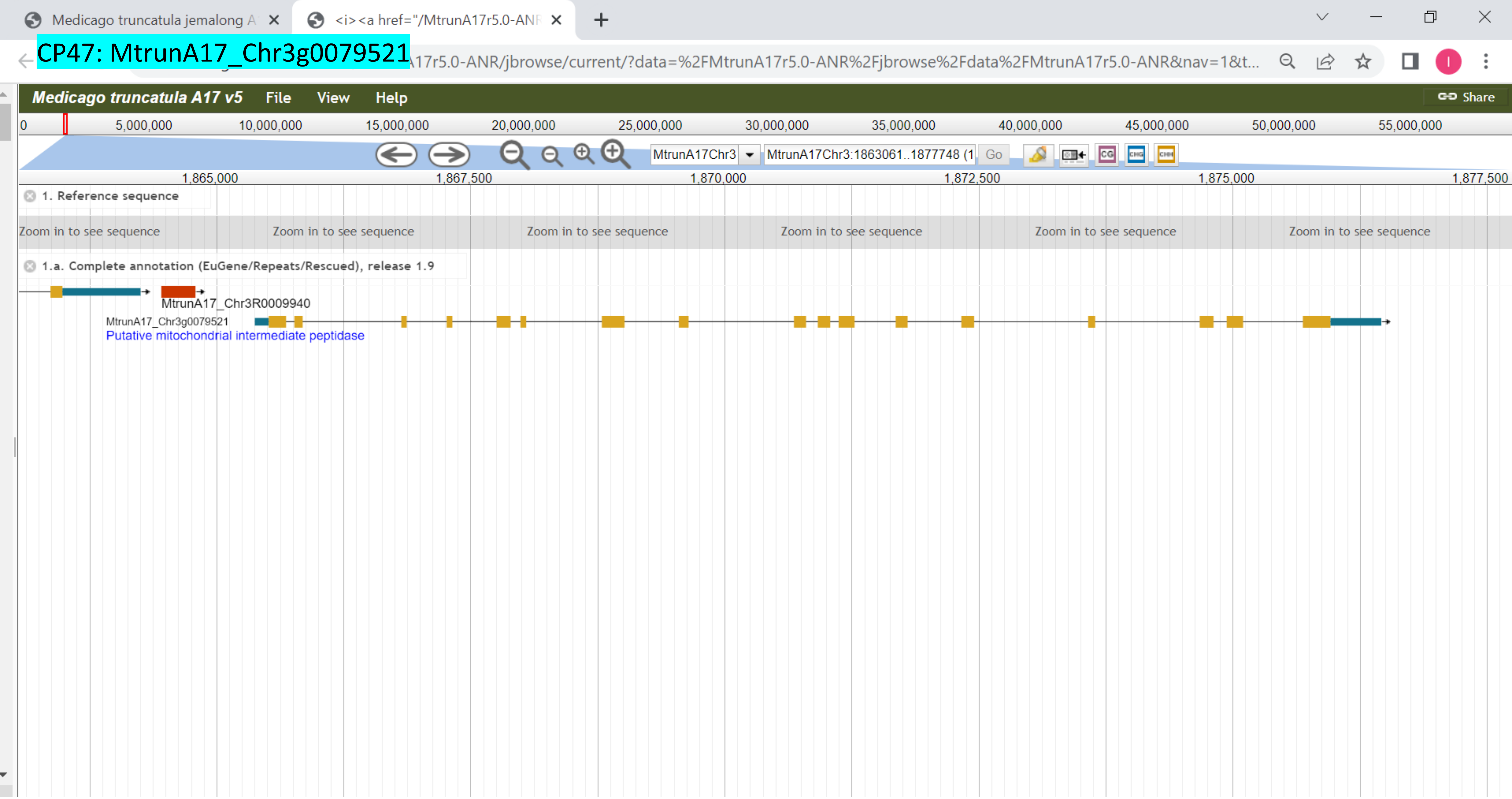

CP48: MtrunA17_Chr3g0083801

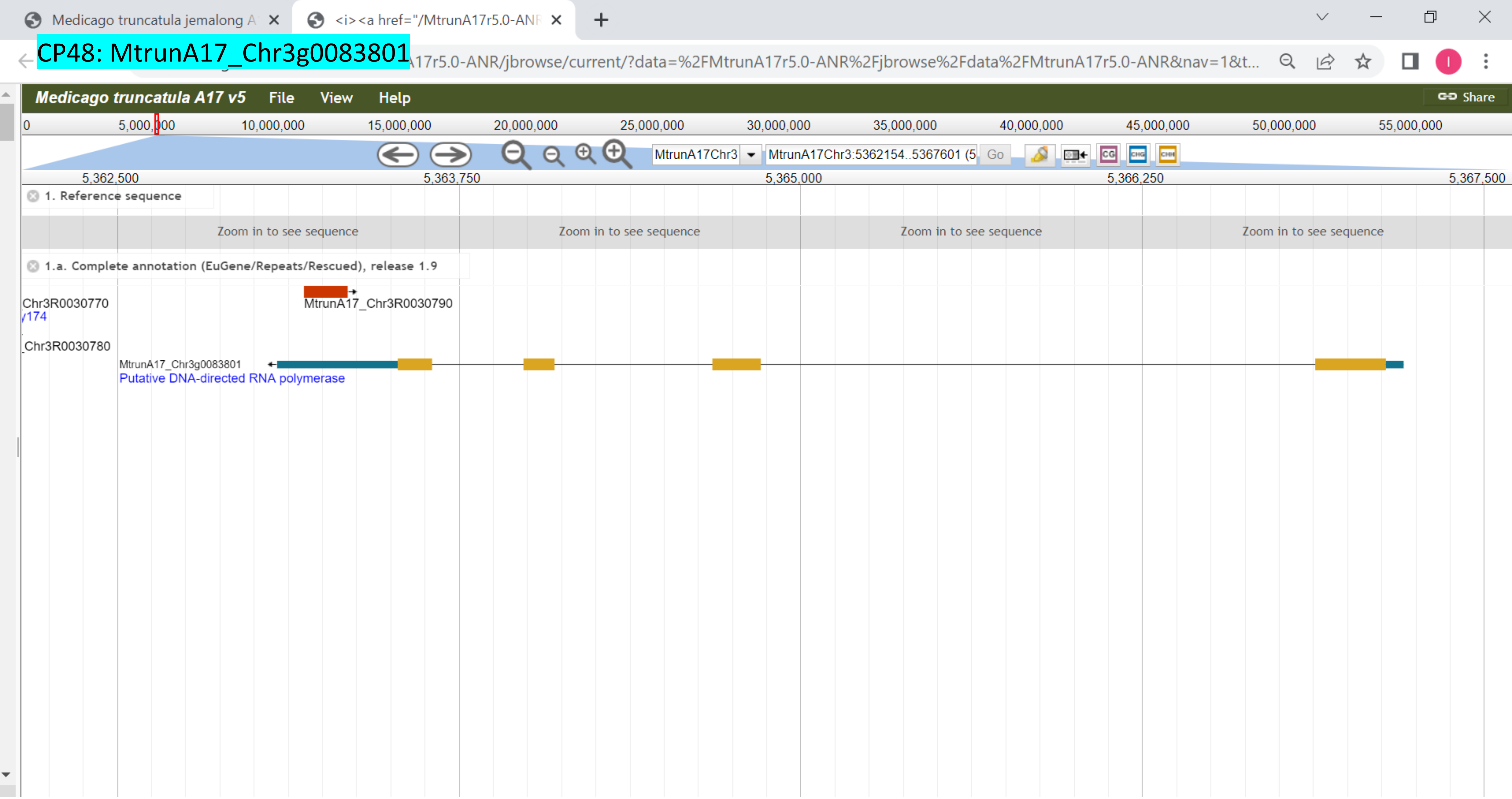

### CP49: MtrunA17_Chr3g0091141

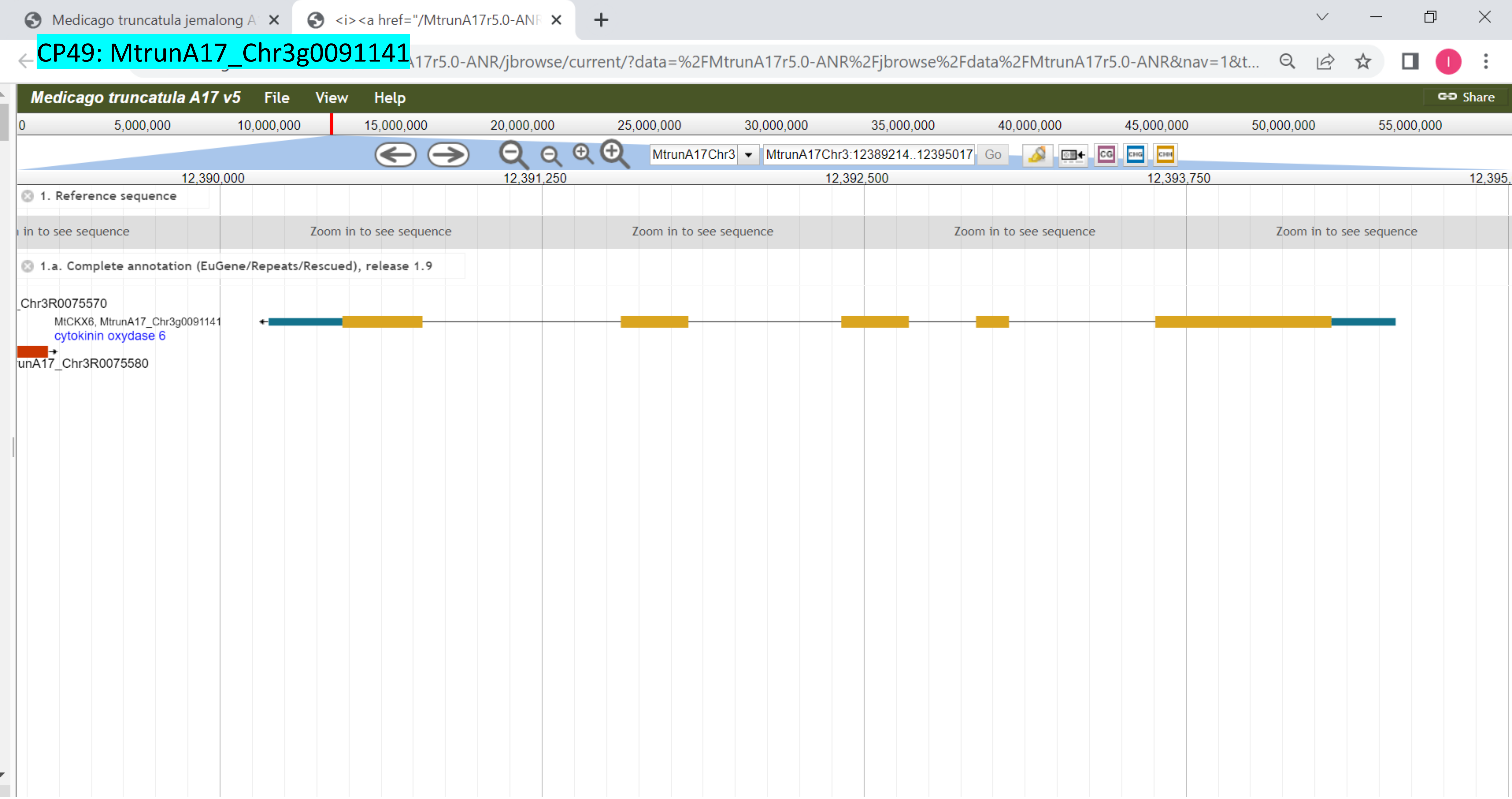

CP50: MtrunA17_Chr3g0091671

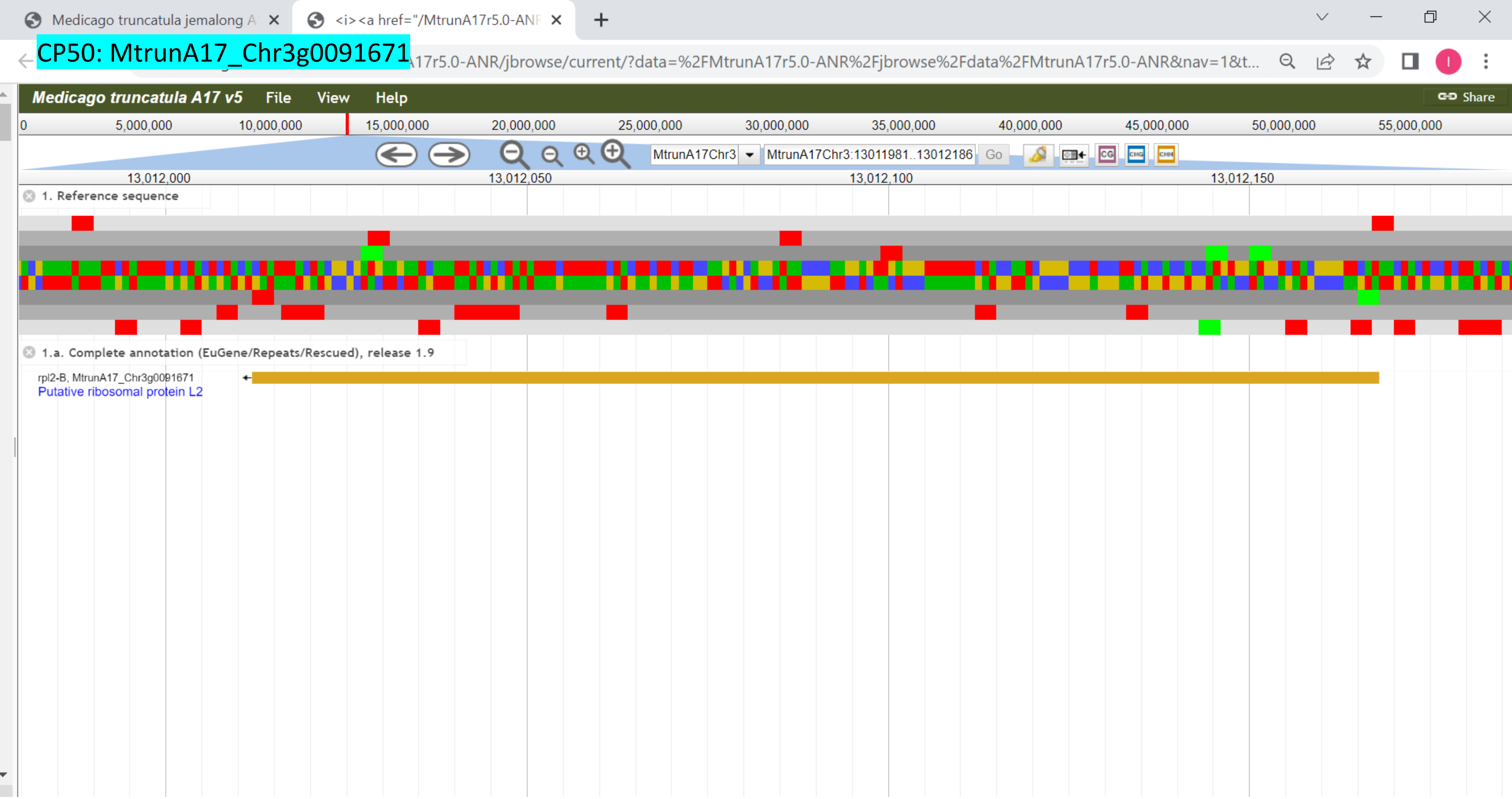

CP51: MtrunA17_Chr3g0096421

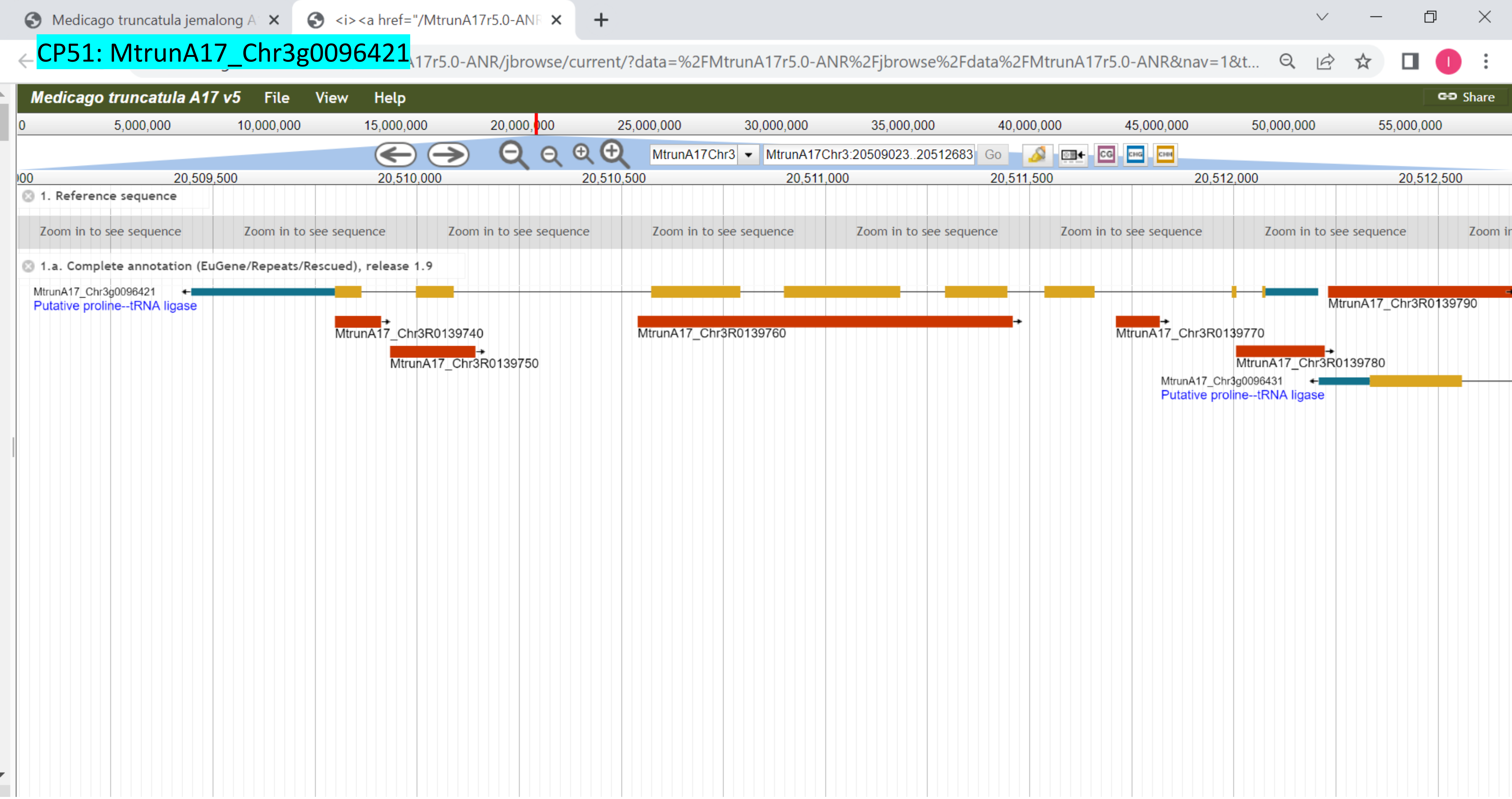

CP52: MtrunA17_Chr3g0100221

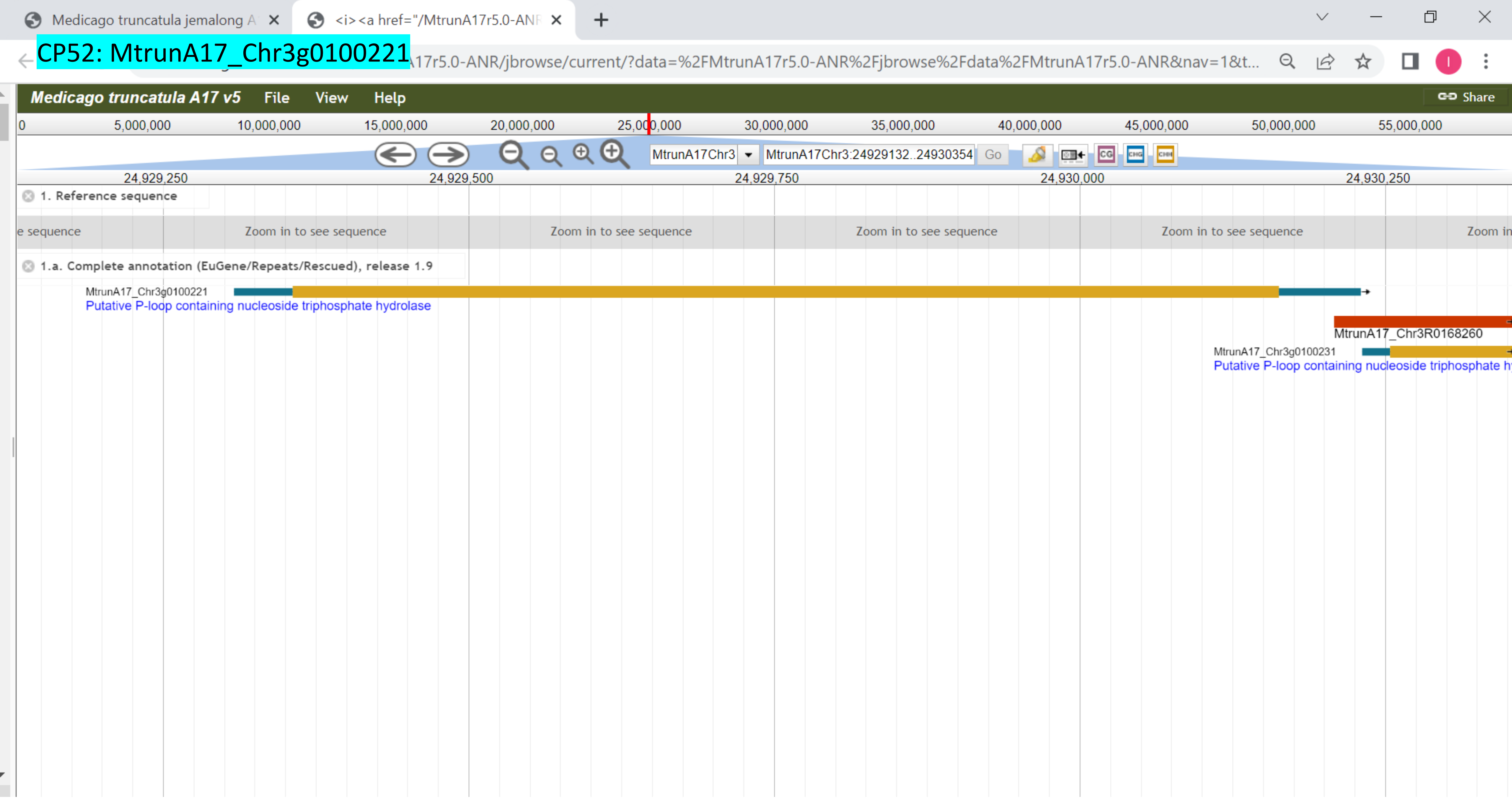

CP53: MtrunA17_Chr3g0102171

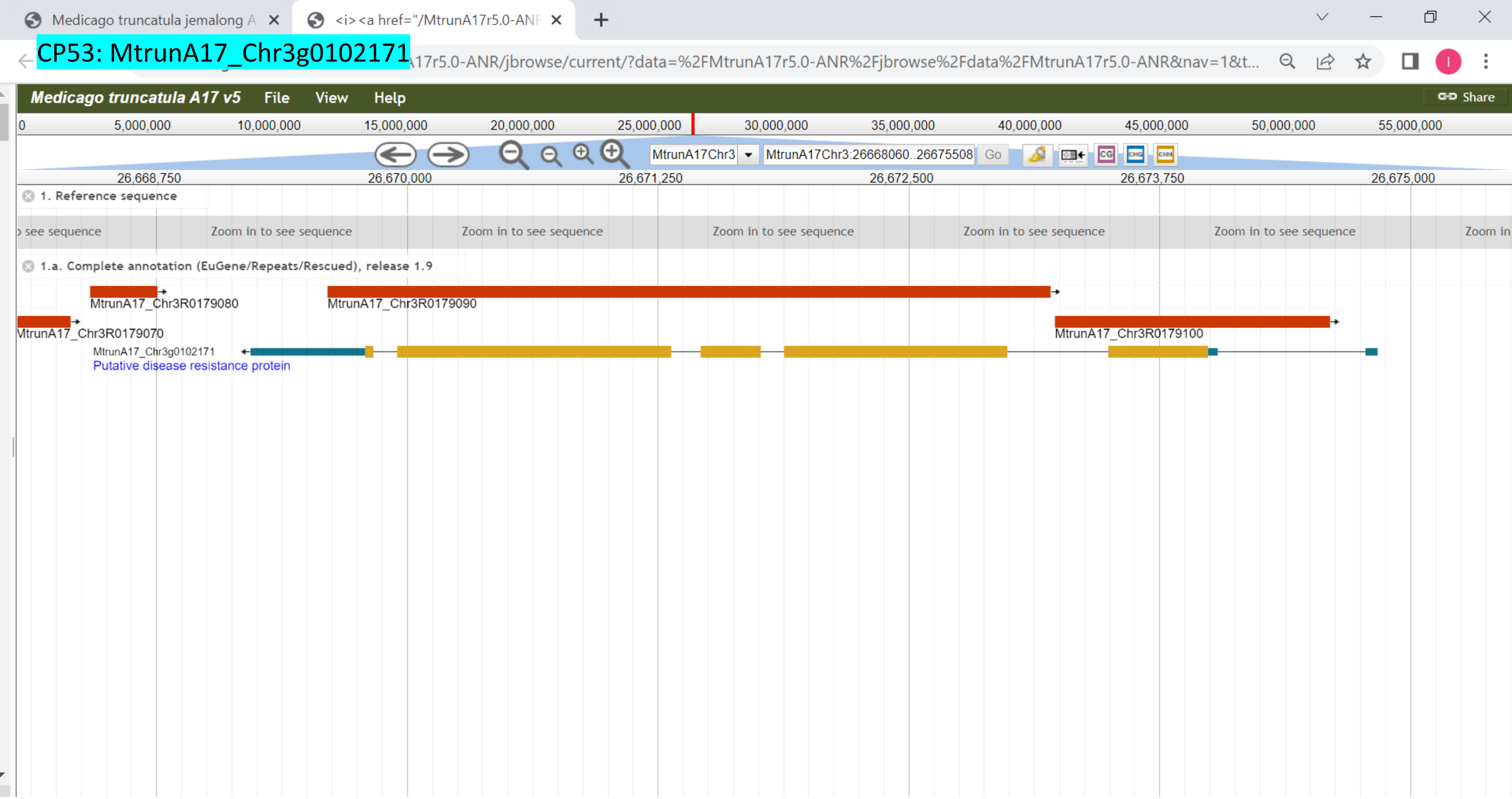

### CP54: MtrunA17_Chr3g0105981

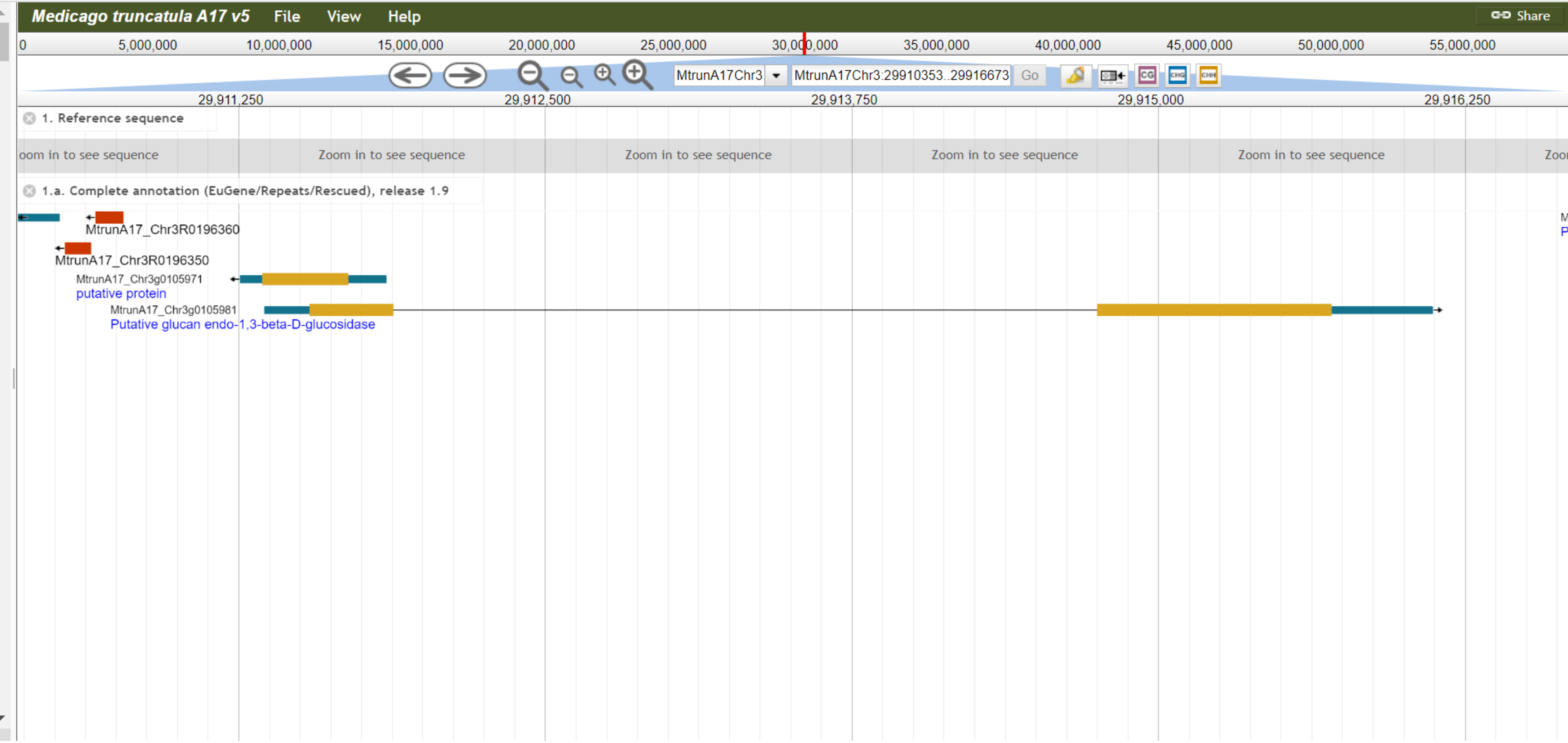

CP55: MtrunA17_Chr3g0110451

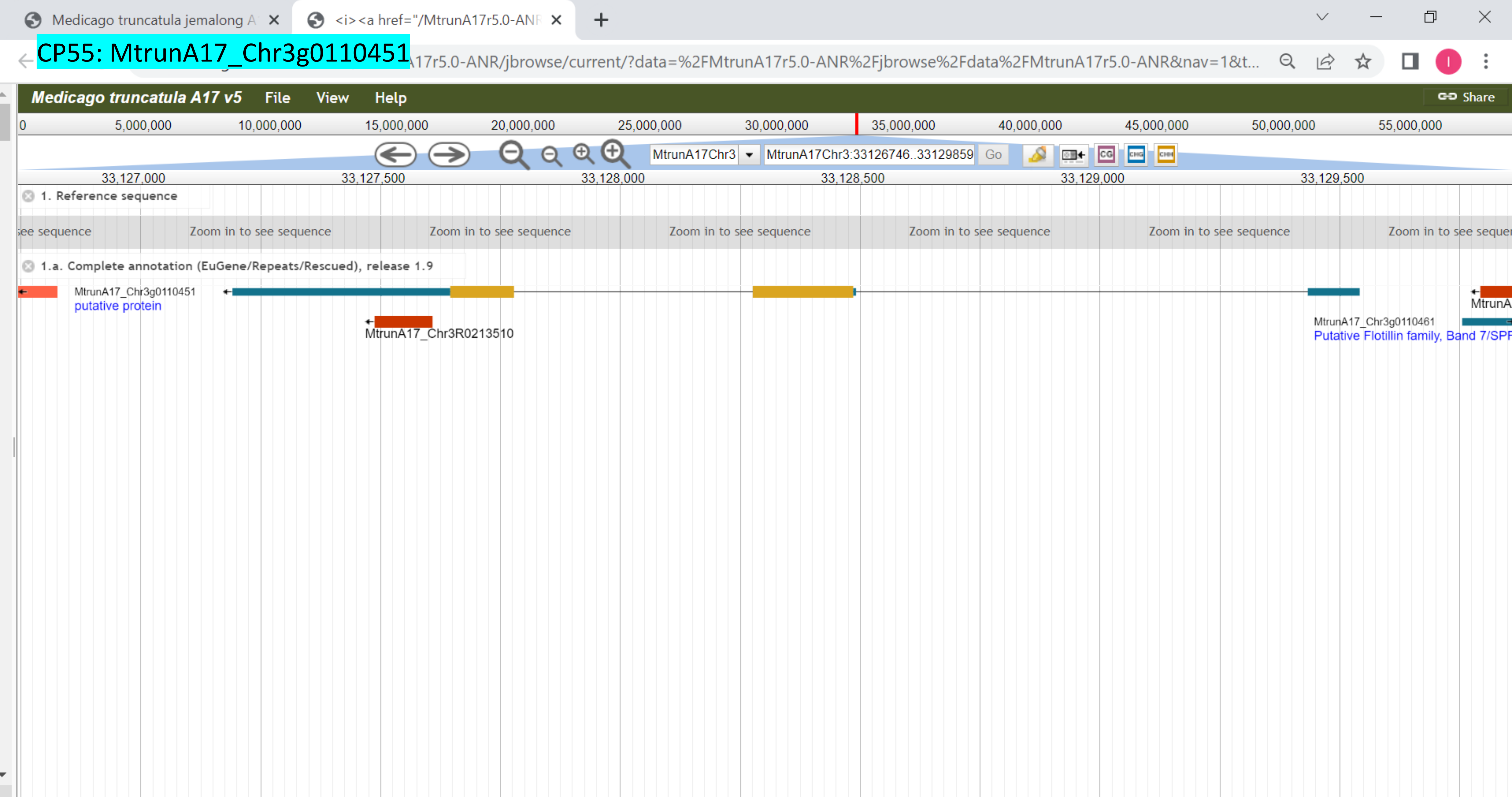

CP56: MtrunA17_Chr3g0113591

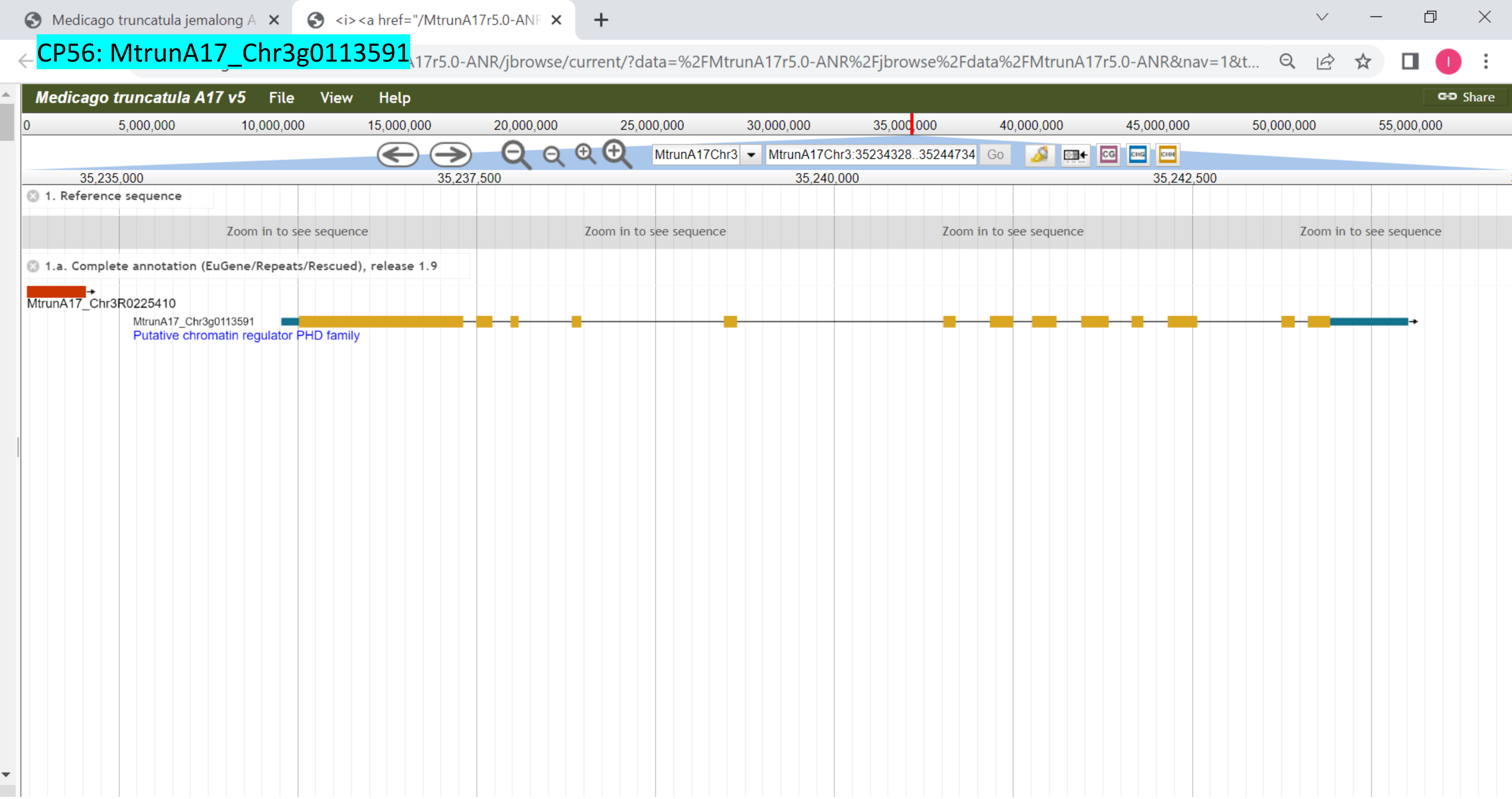

CP57: MtrunA17_Chr3g0124631

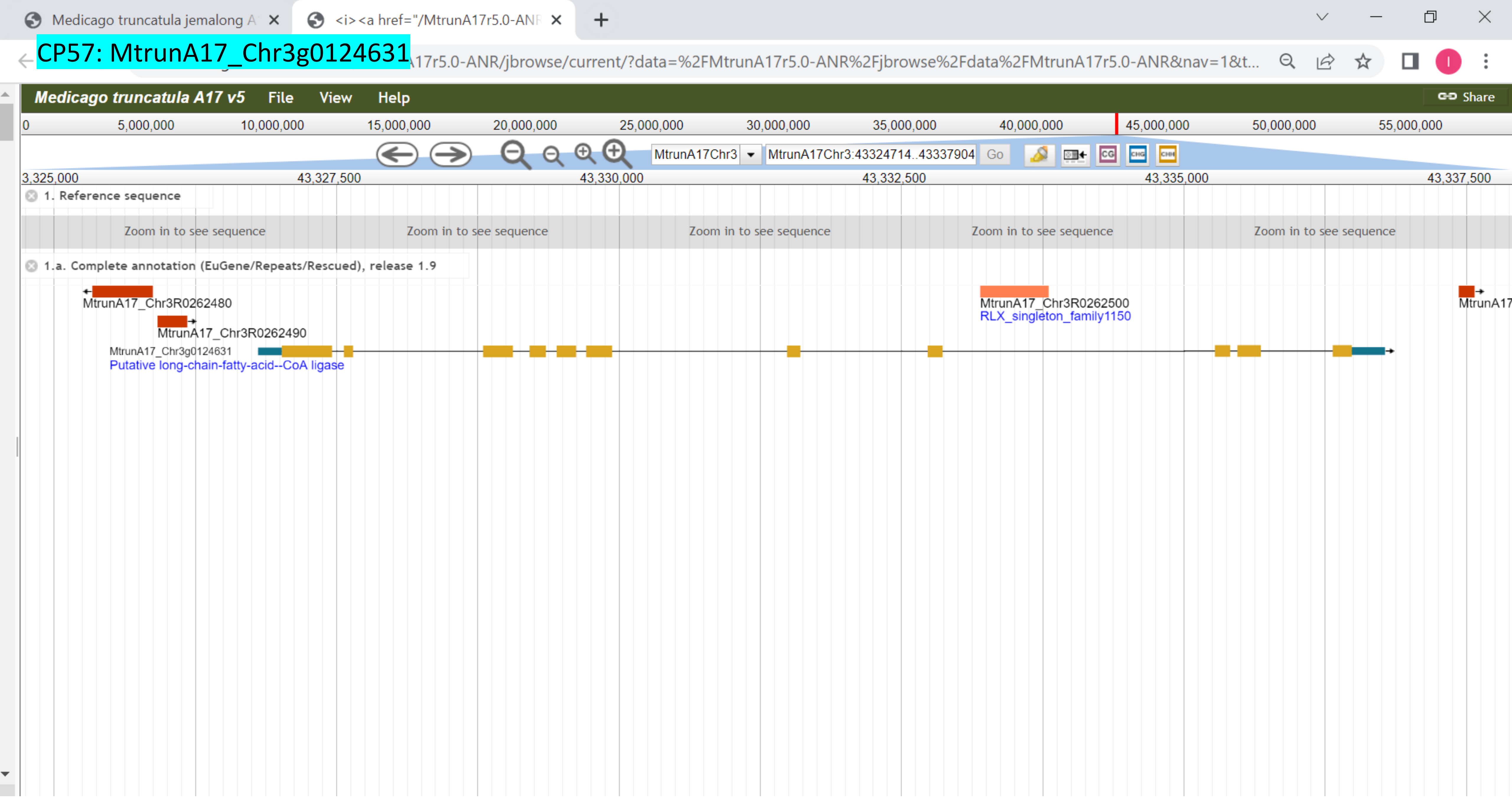

### CP58: MtrunA17_Chr3g0127321

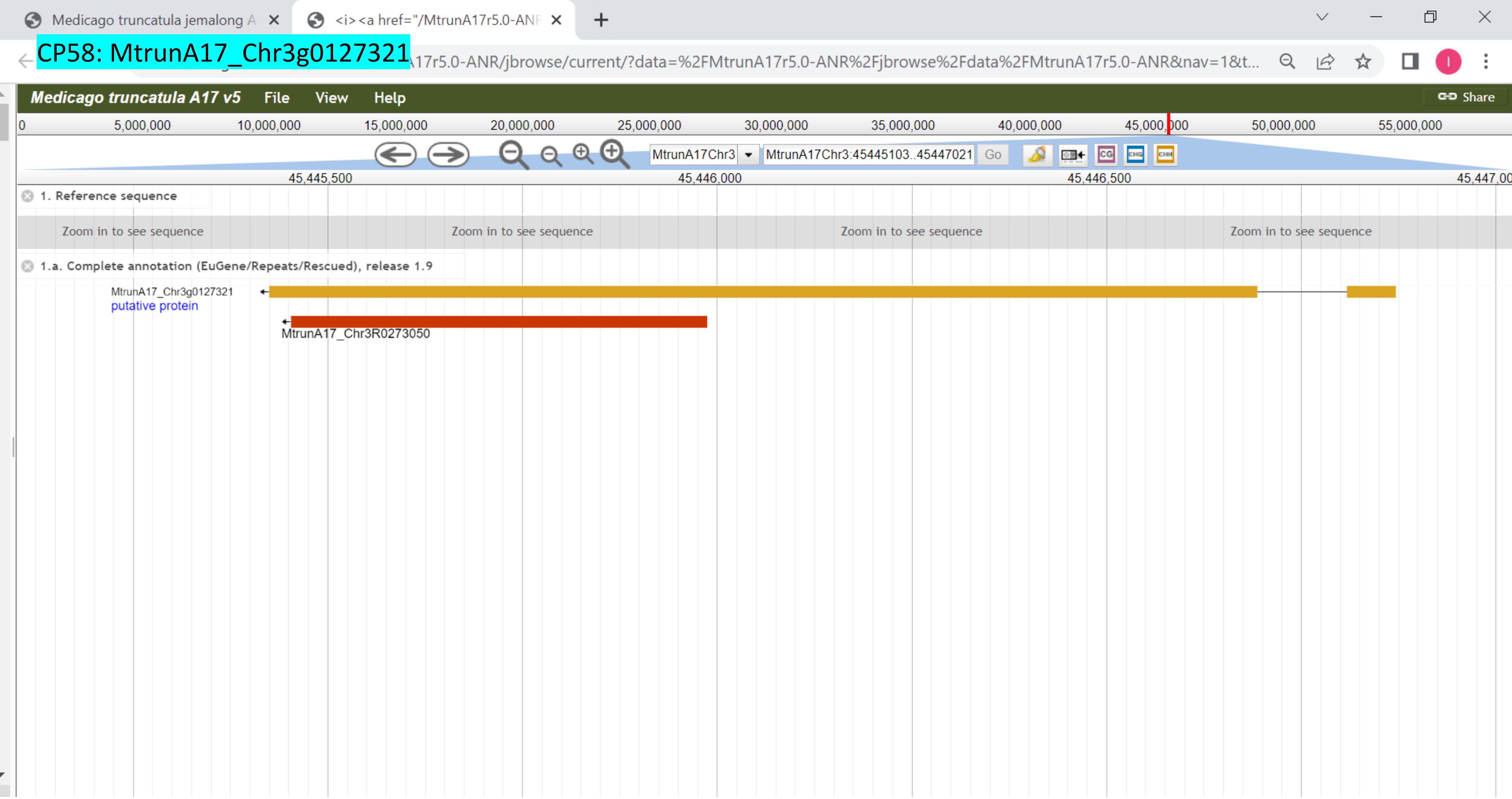

### CP59: MtrunA17_Chr3g0130671

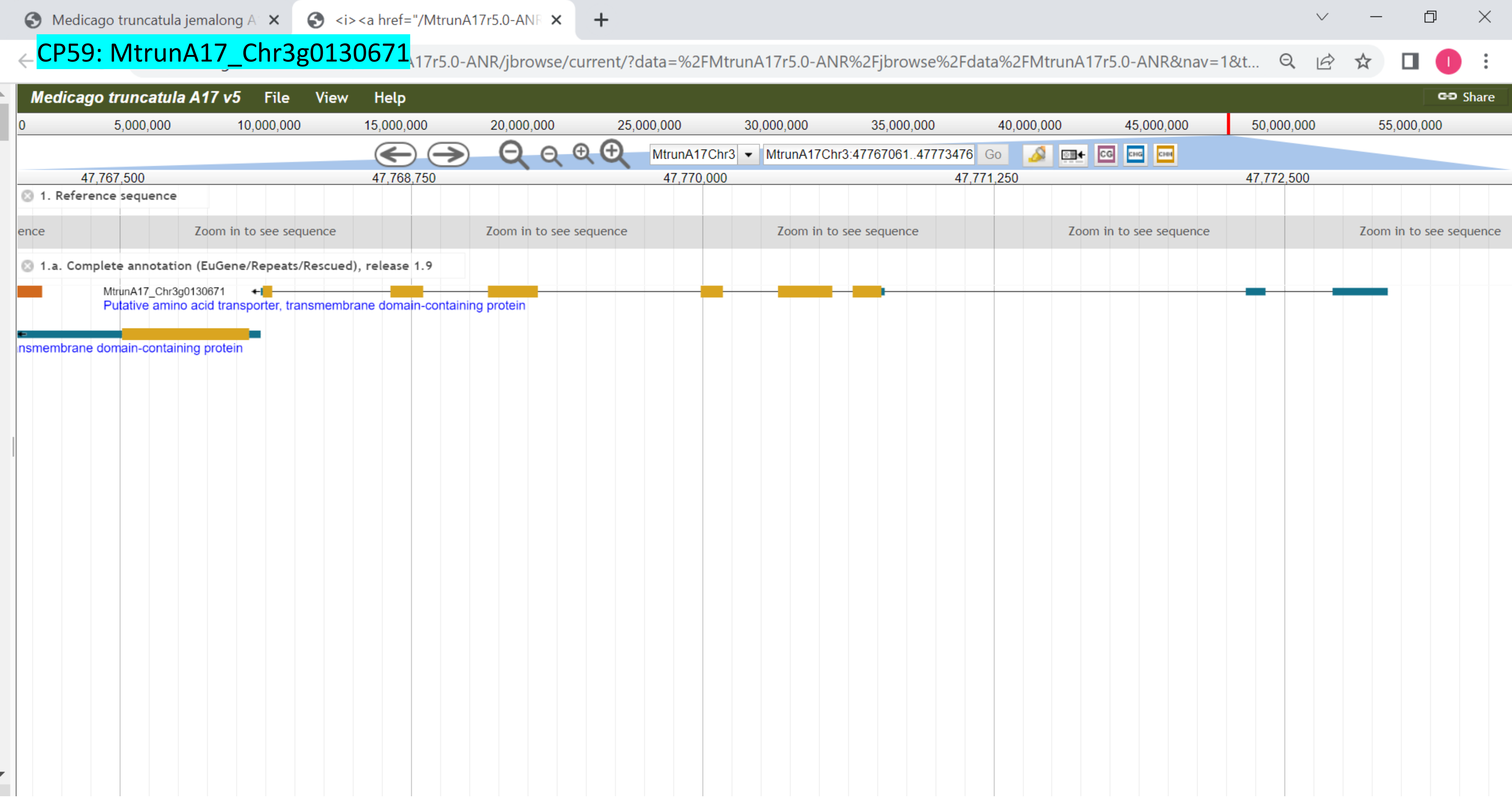

CP60: MtrunA17_Chr3g0135761

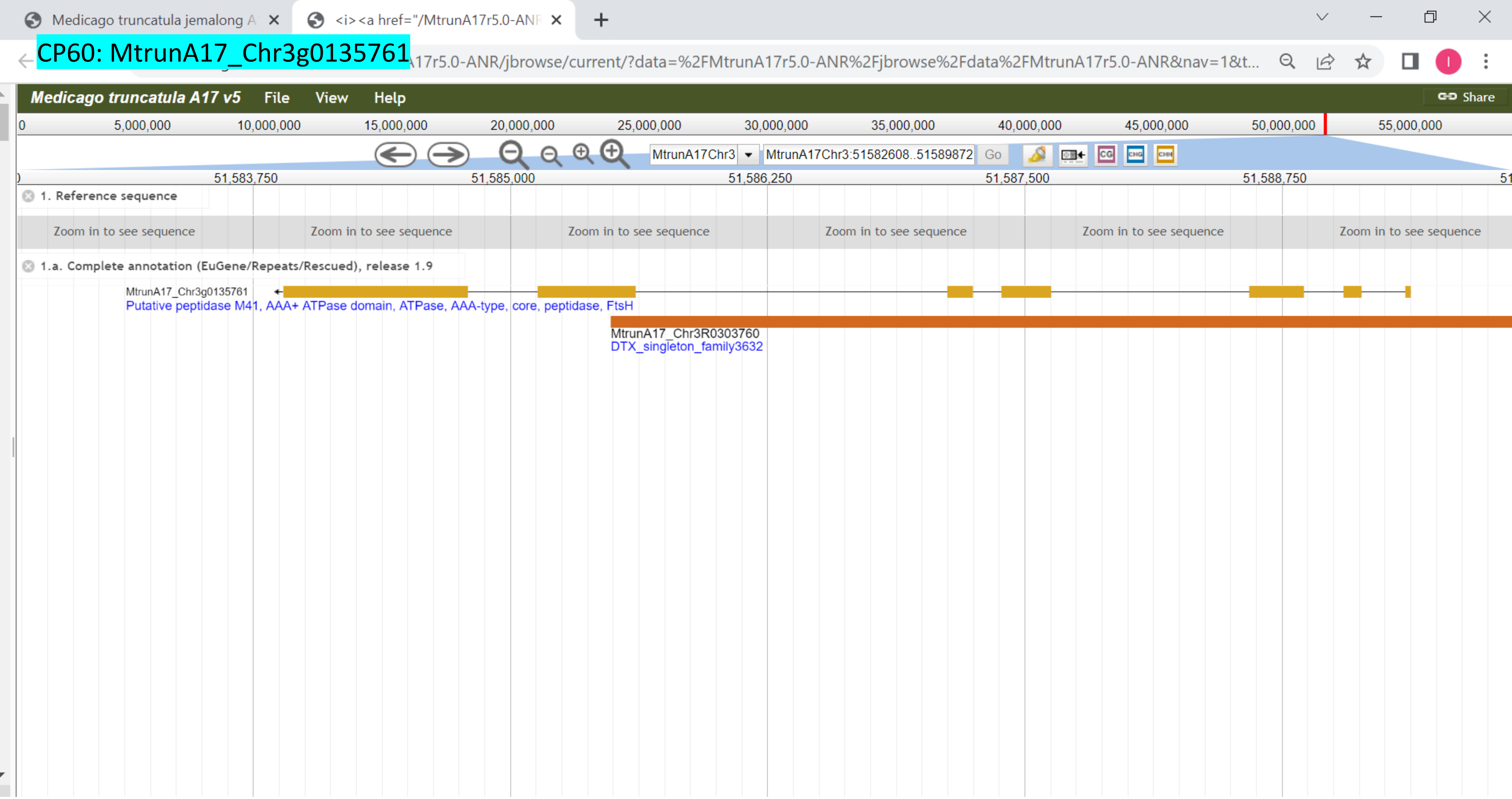

CP61: MtrunA17_Chr3g0137391

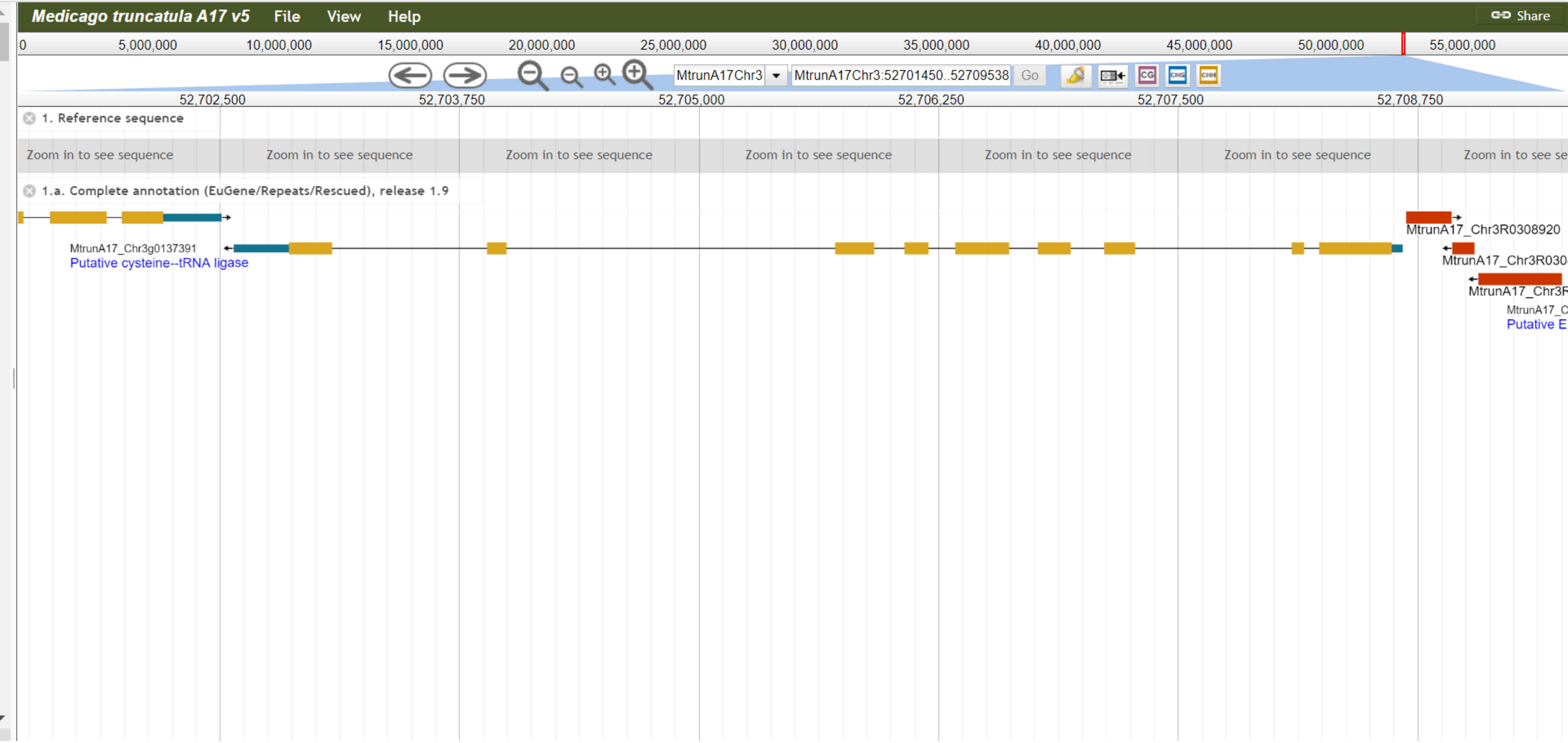

### CP62: MtrunA17_Chr3g0141901

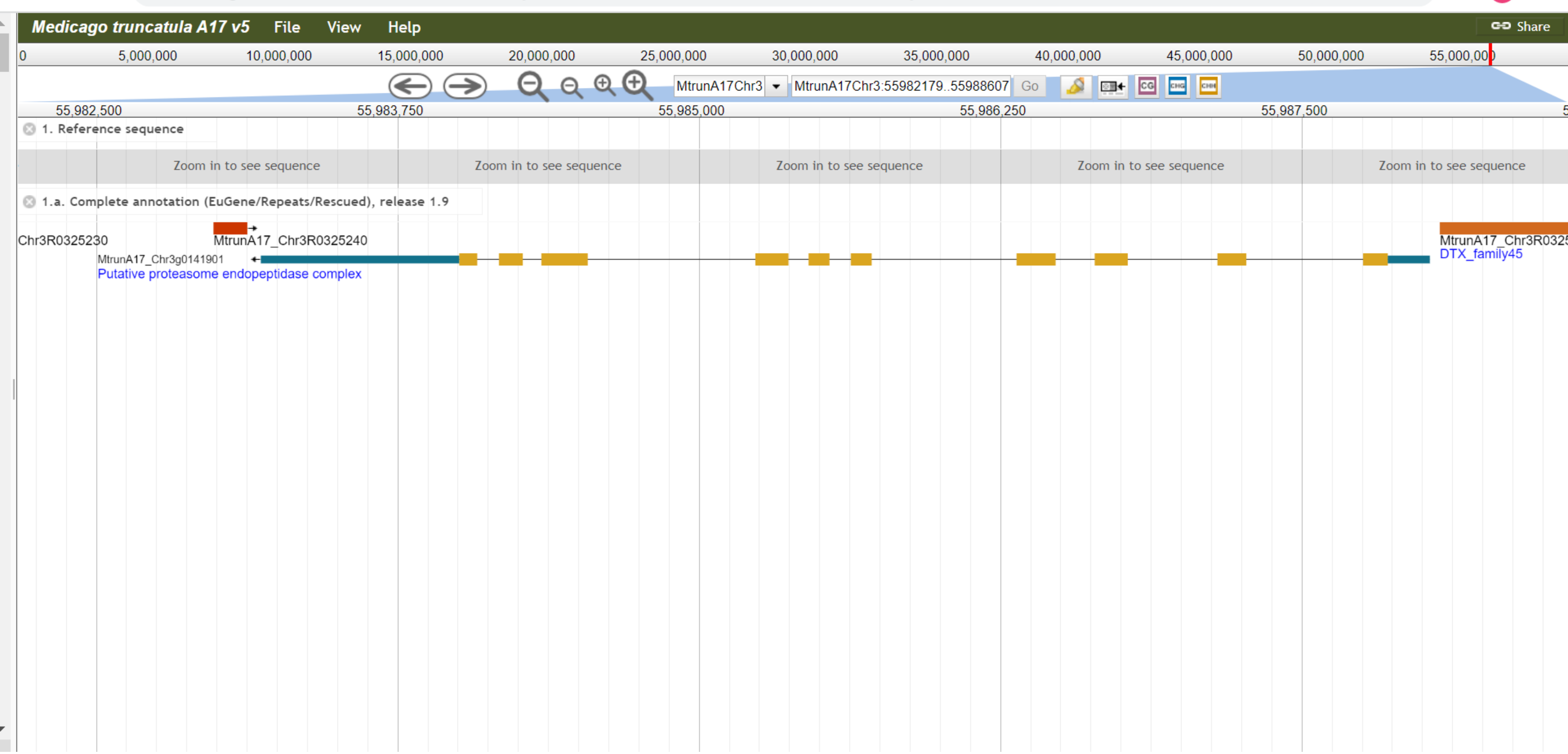

CP63: MtrunA17_Chr3g0144151

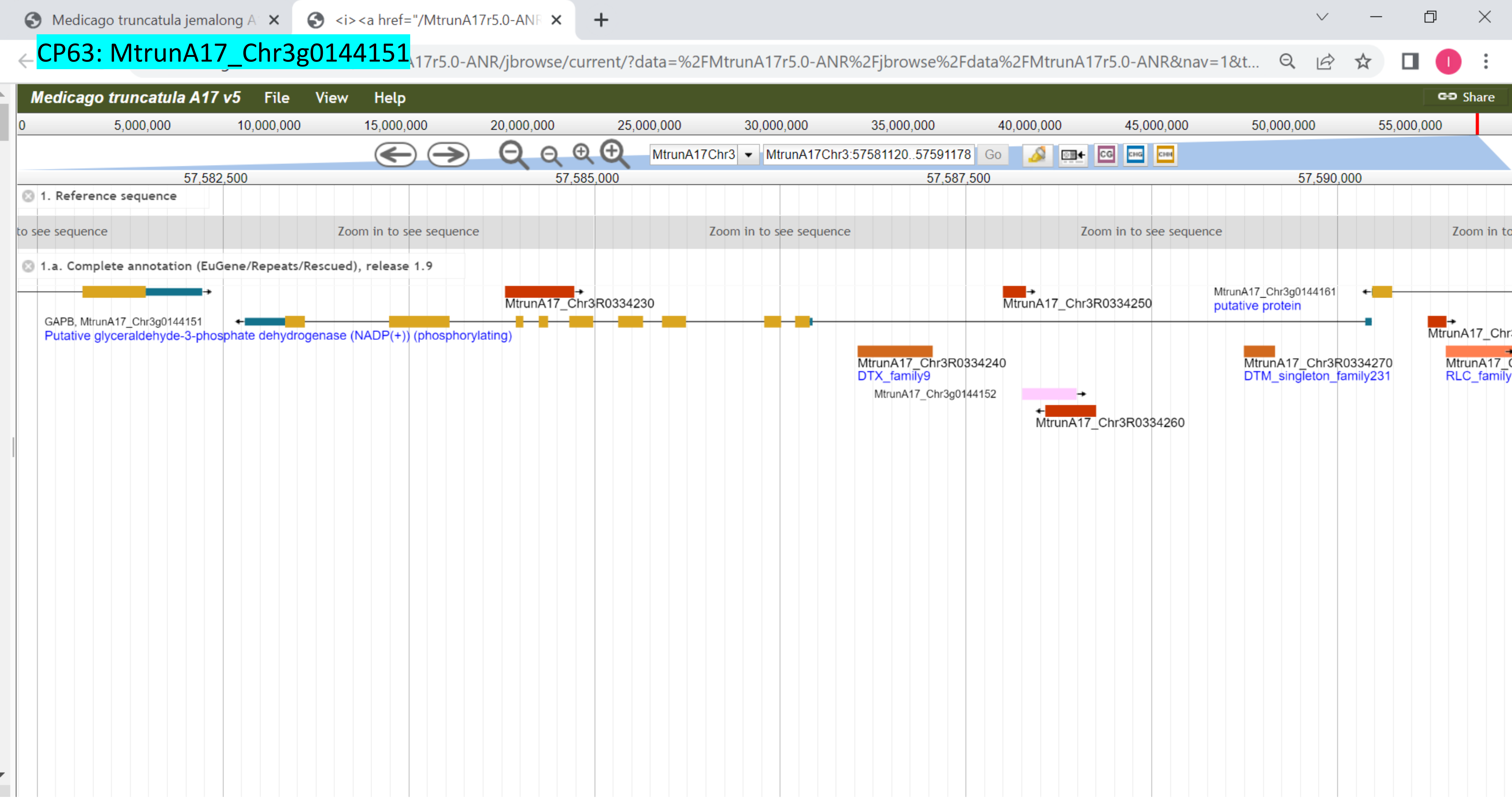

### CP64: MtrunA17_Chr3g0144151

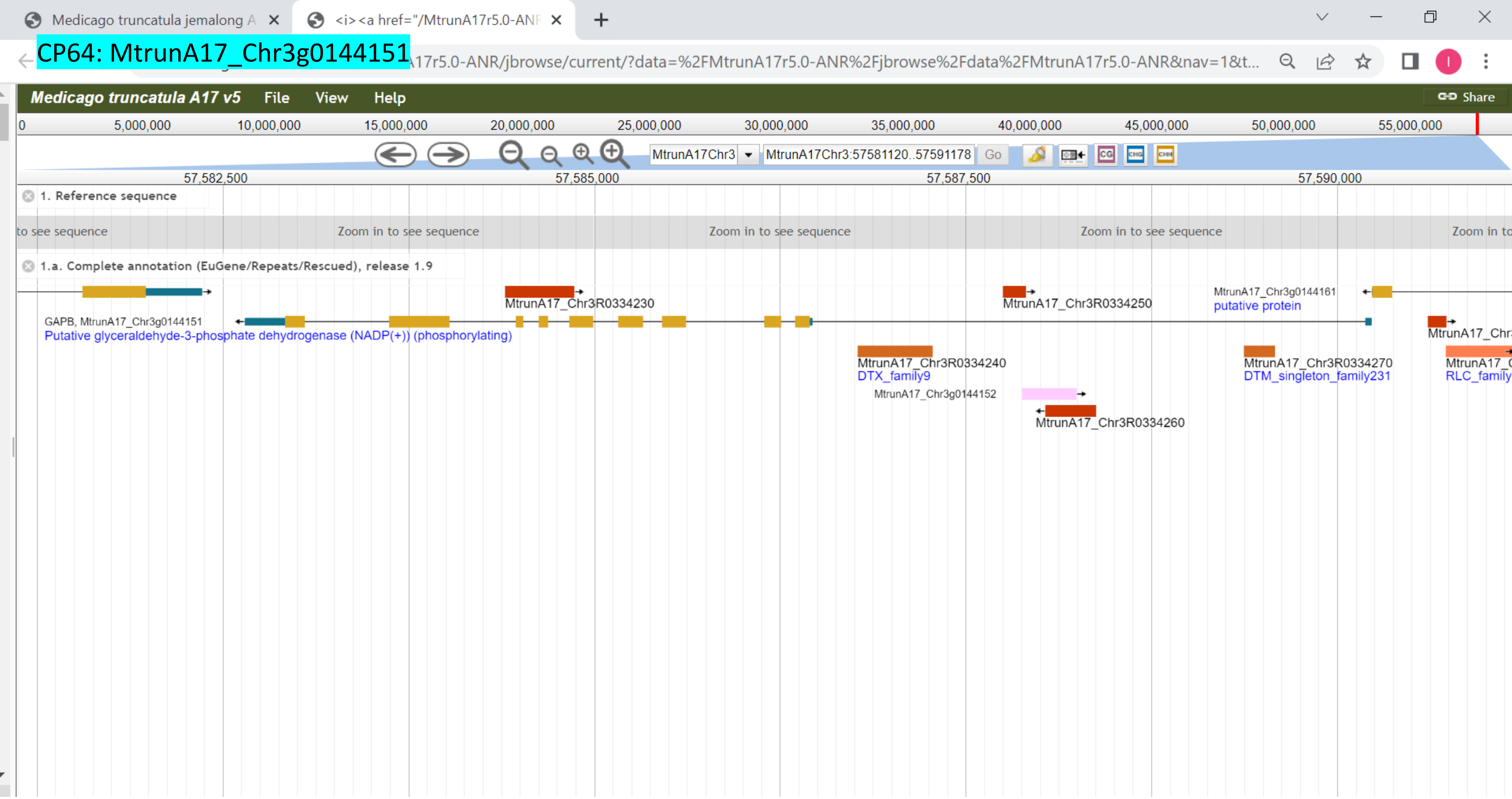

CP65: MtrunA17_Chr3g0144151

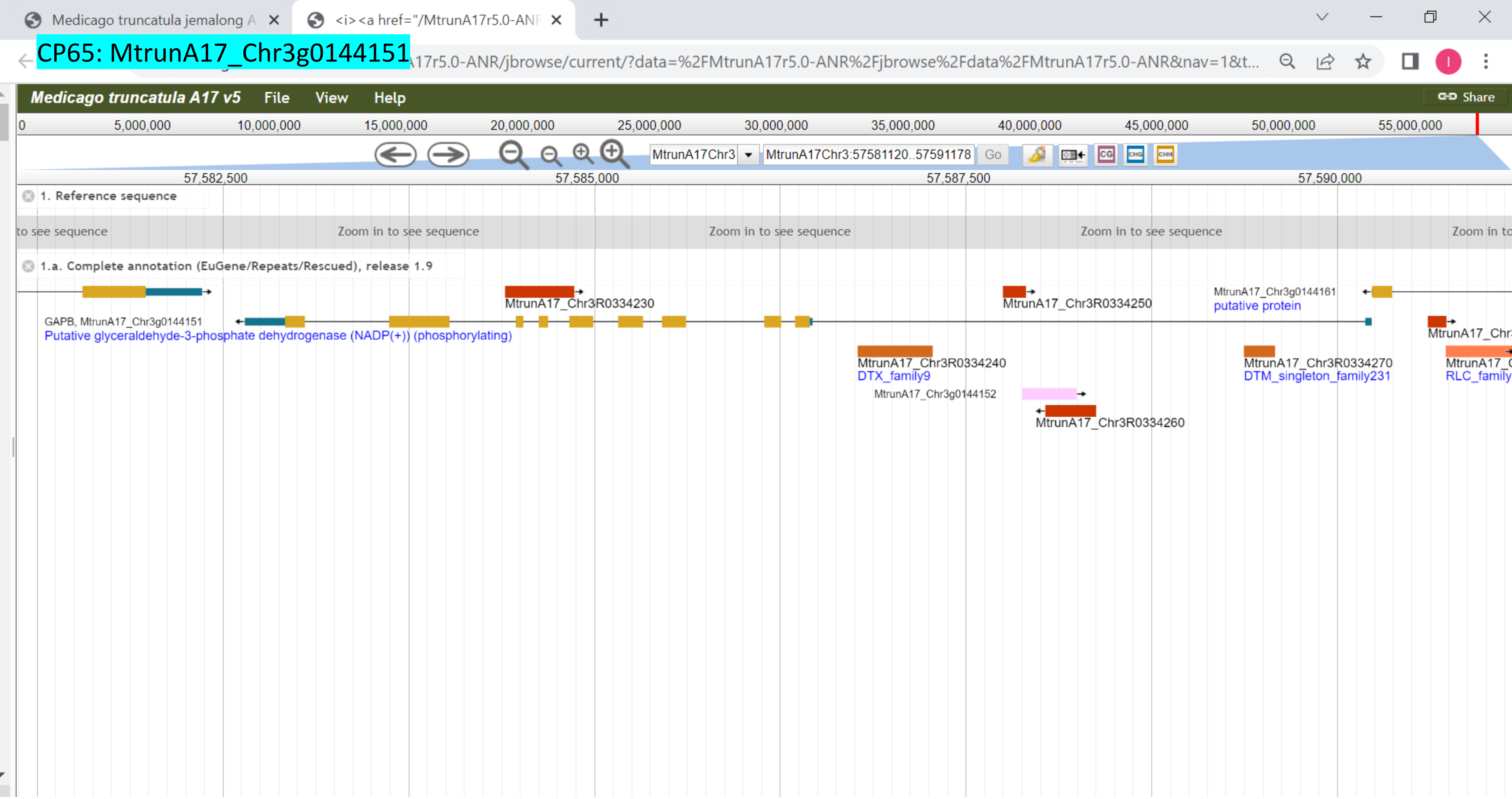

CP66: MtrunA17_Chr3g1011650

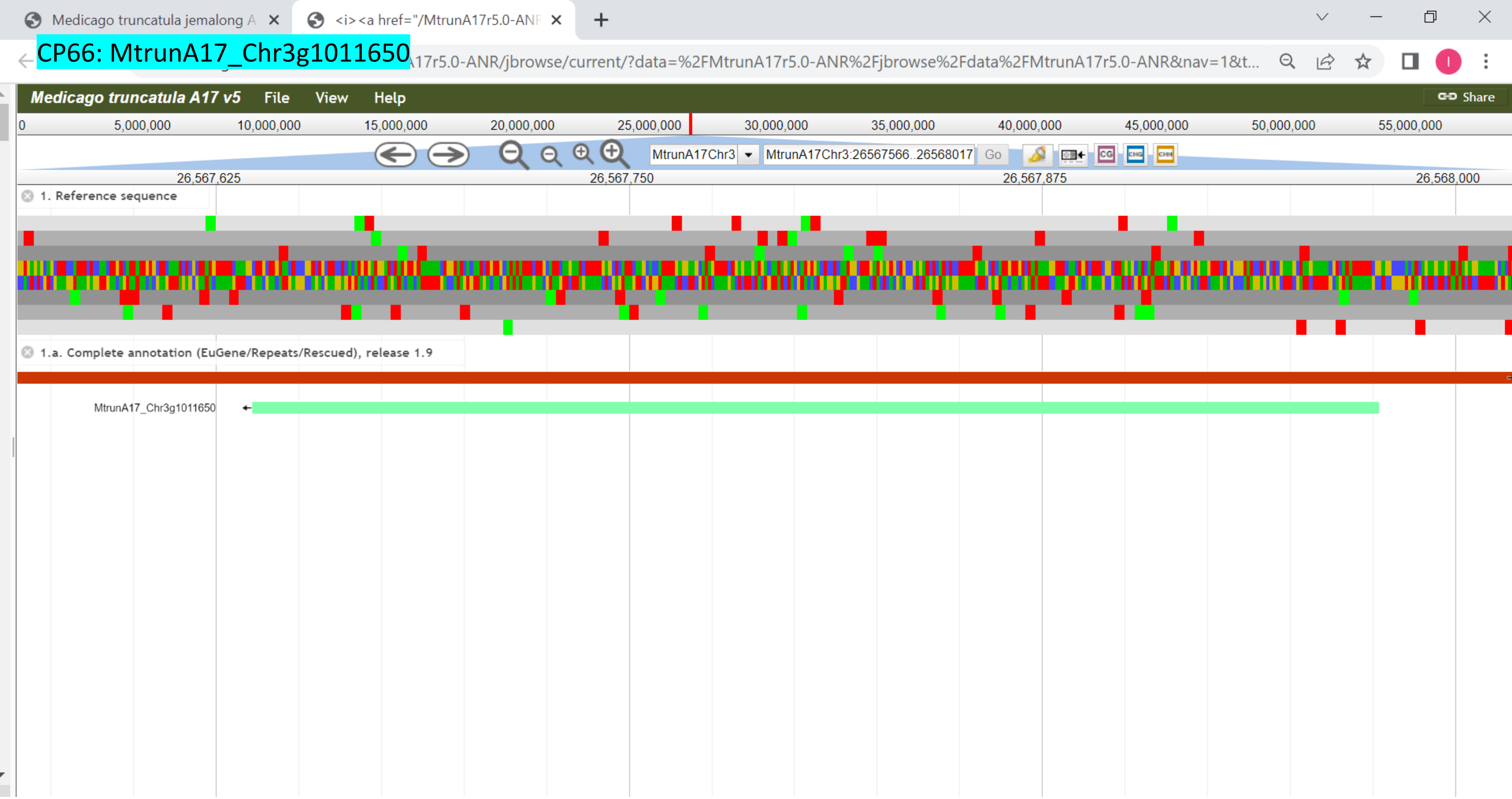

CP67: MtrunA17_Chr4g0000131

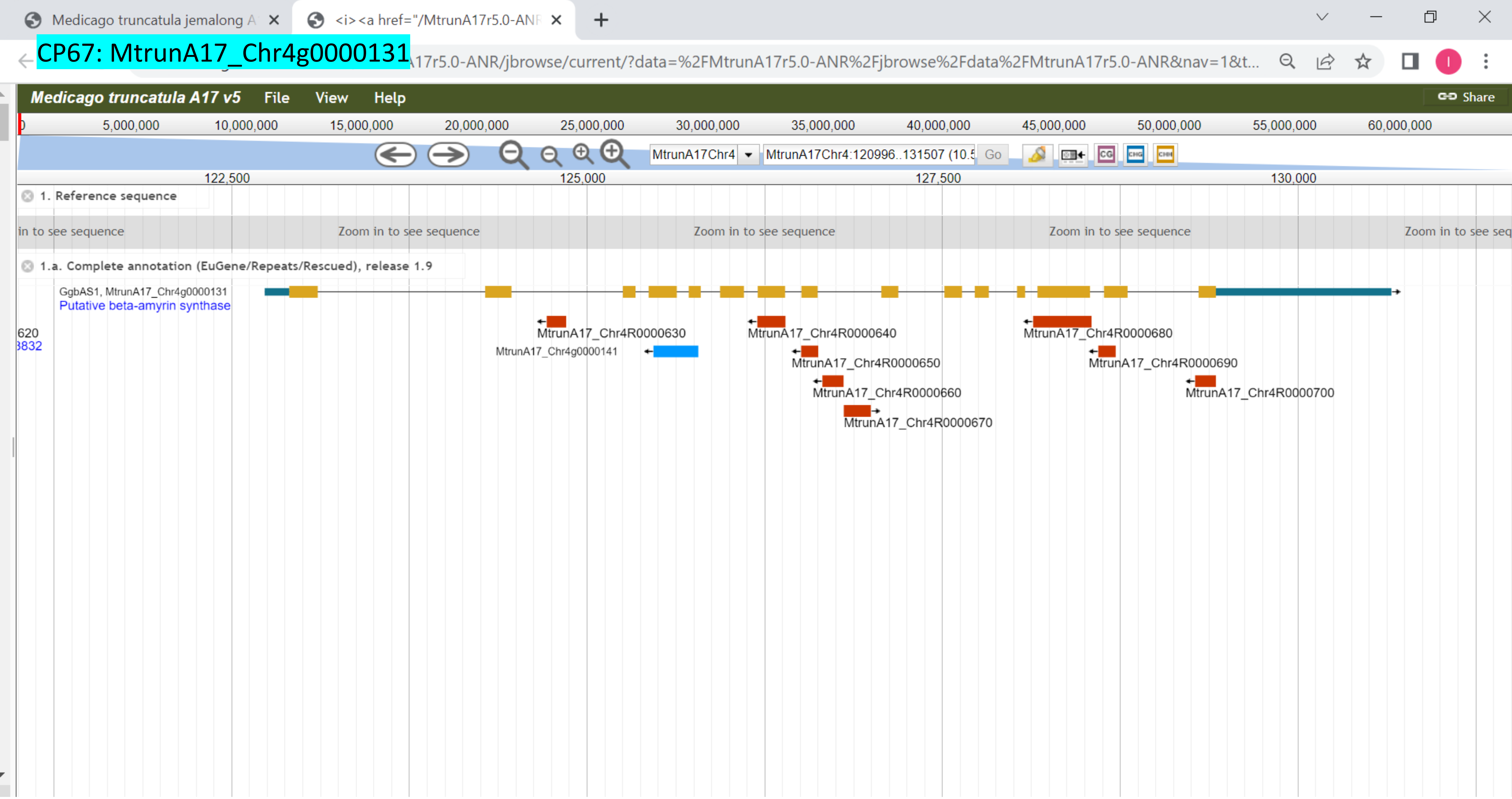

CP68: MtrunA17_Chr4g0000891

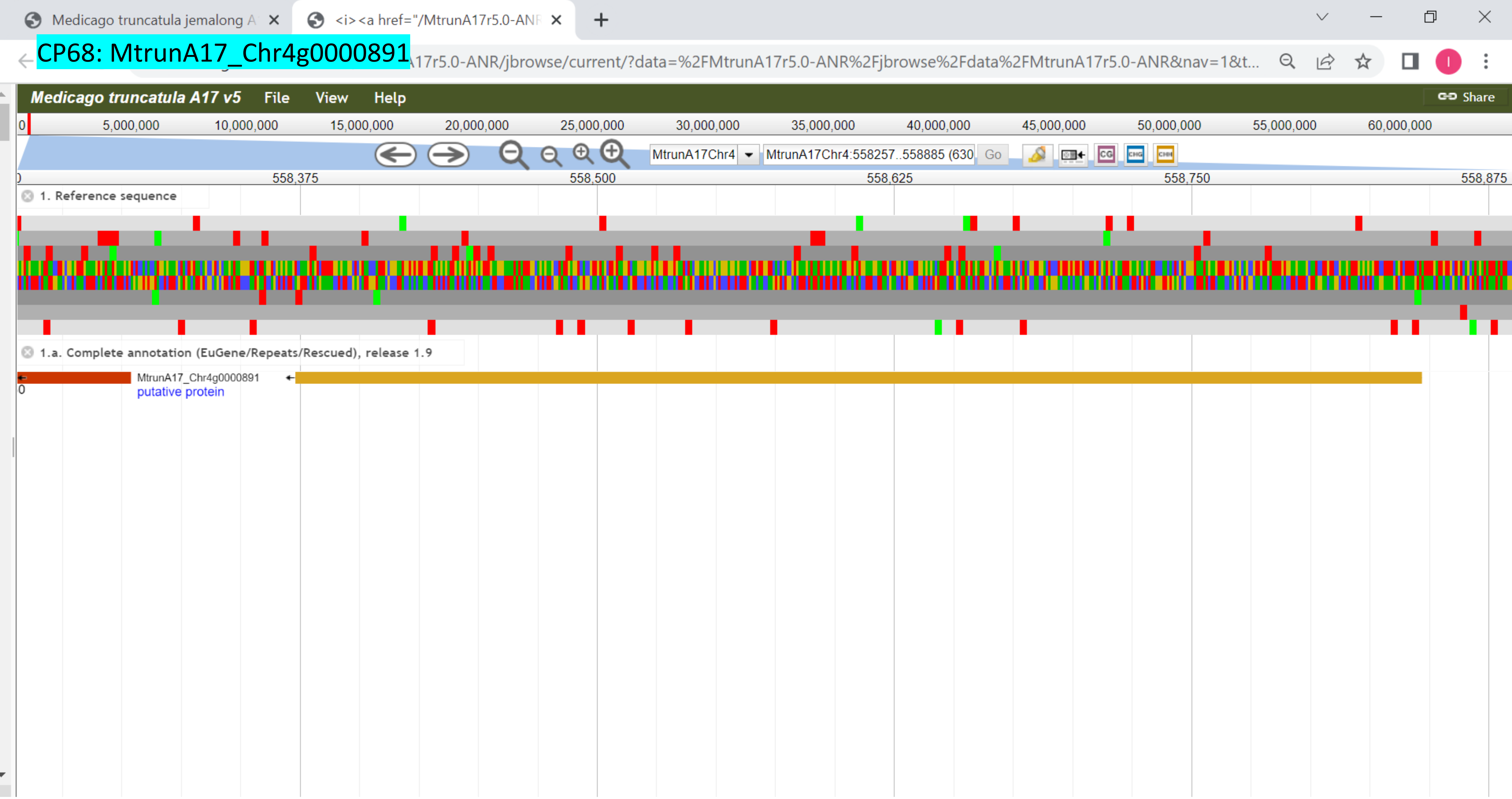

CP69: MtrunA17_Chr4g0004721

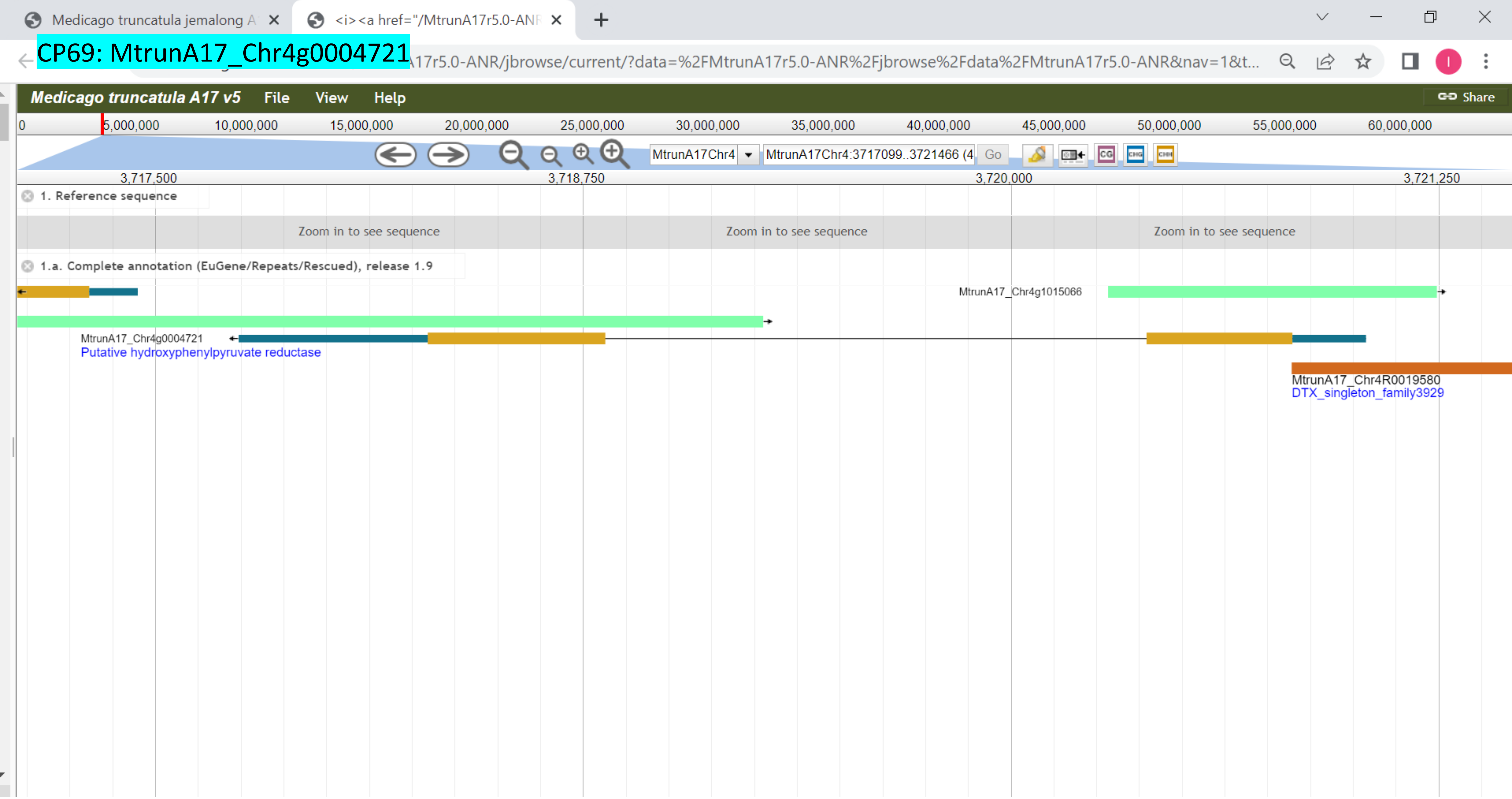

CP70: MtrunA17_Chr4g0014251

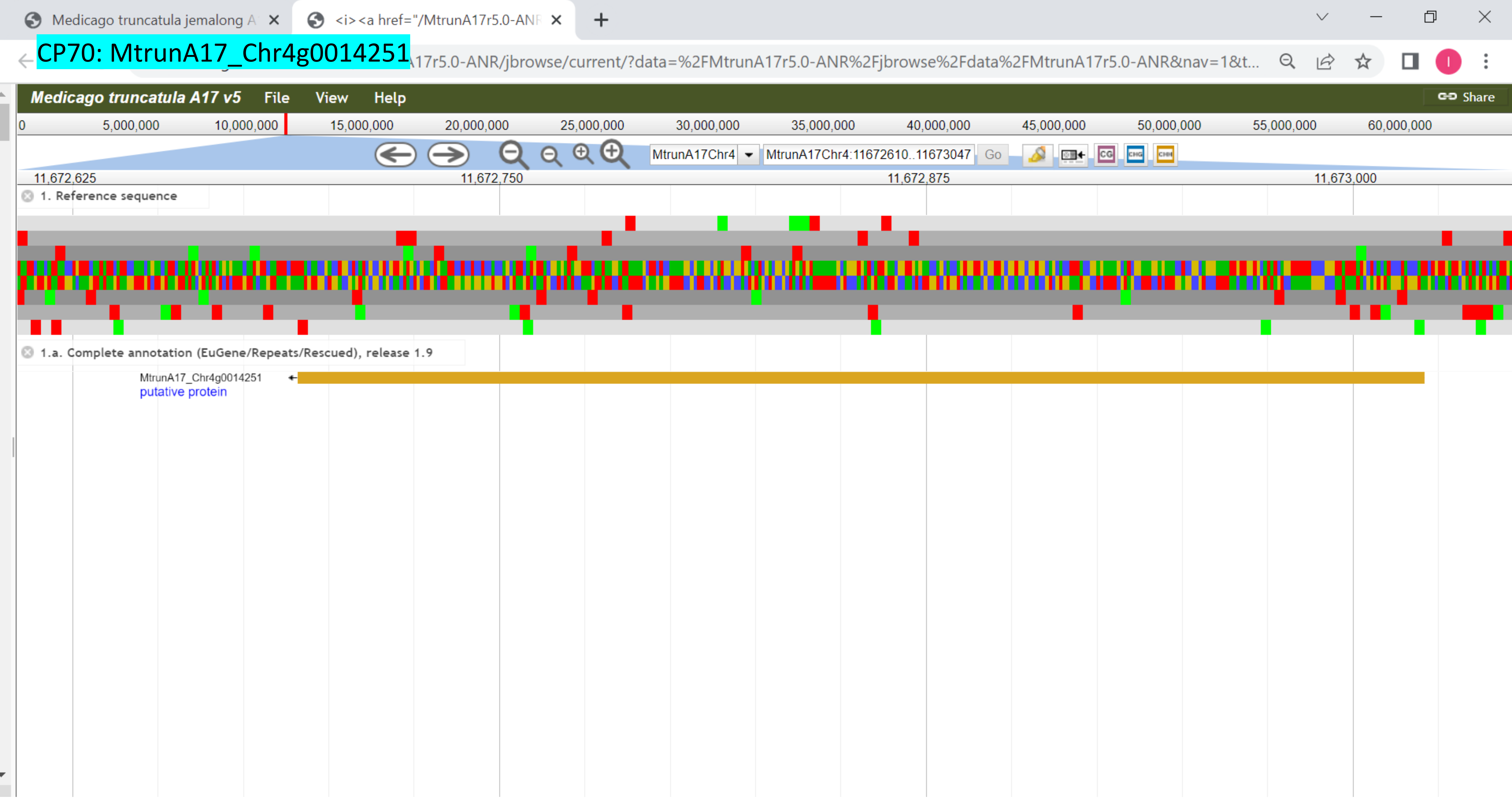

### CP71: MtrunA17_Chr4g0022491

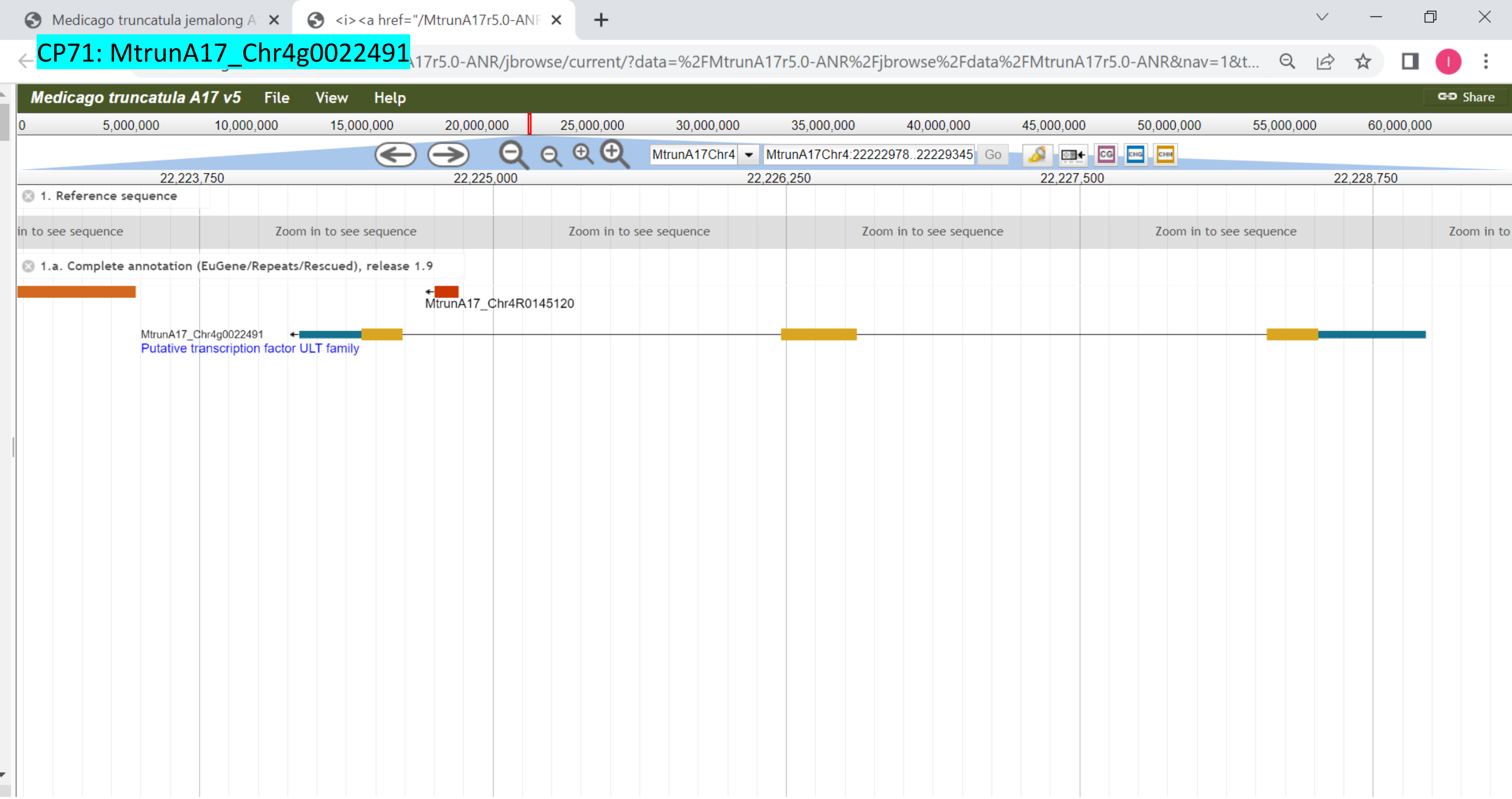

### CP72: MtrunA17_Chr4g0023791

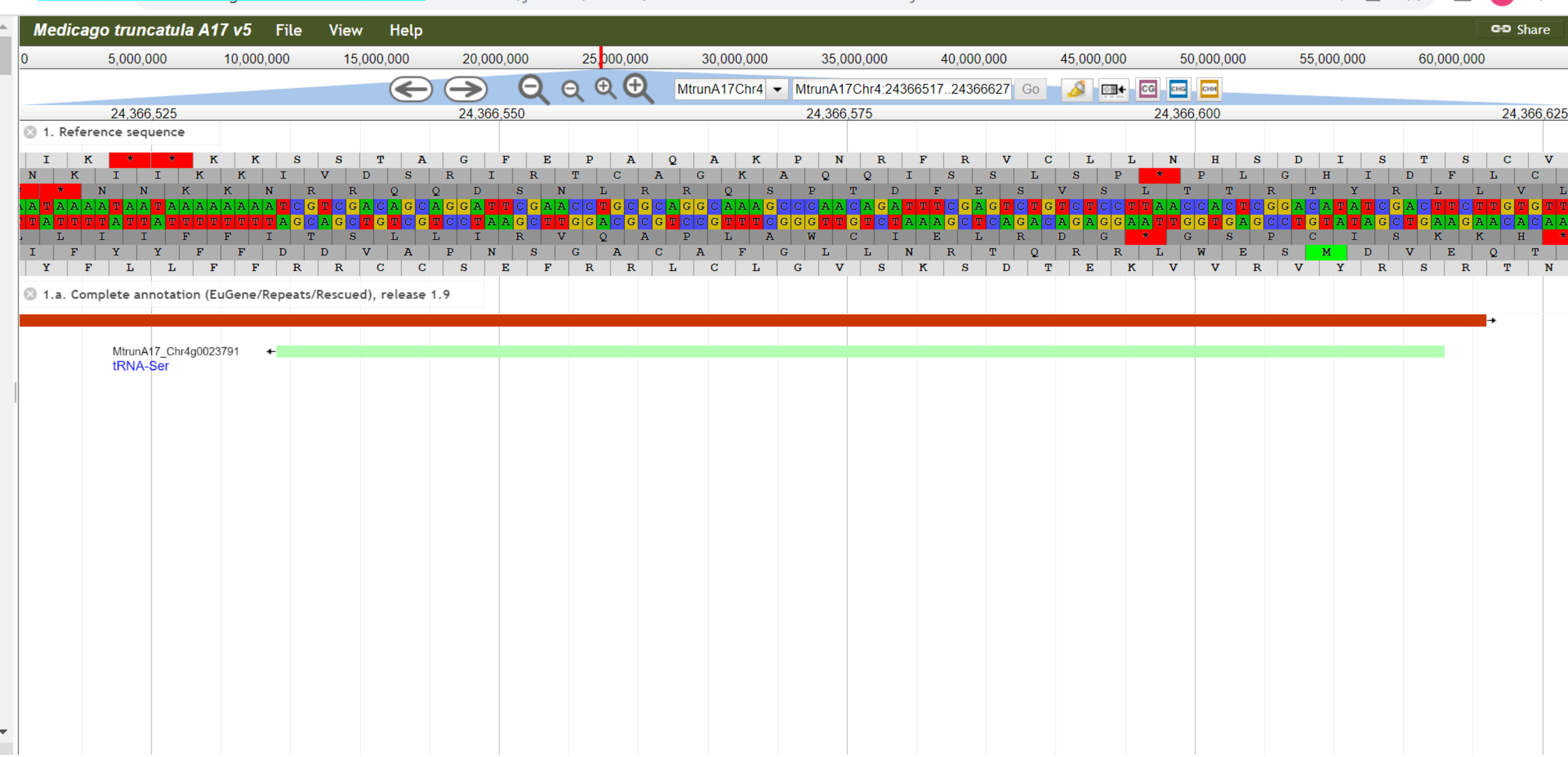

CP73: MtrunA17_Chr4g0034271

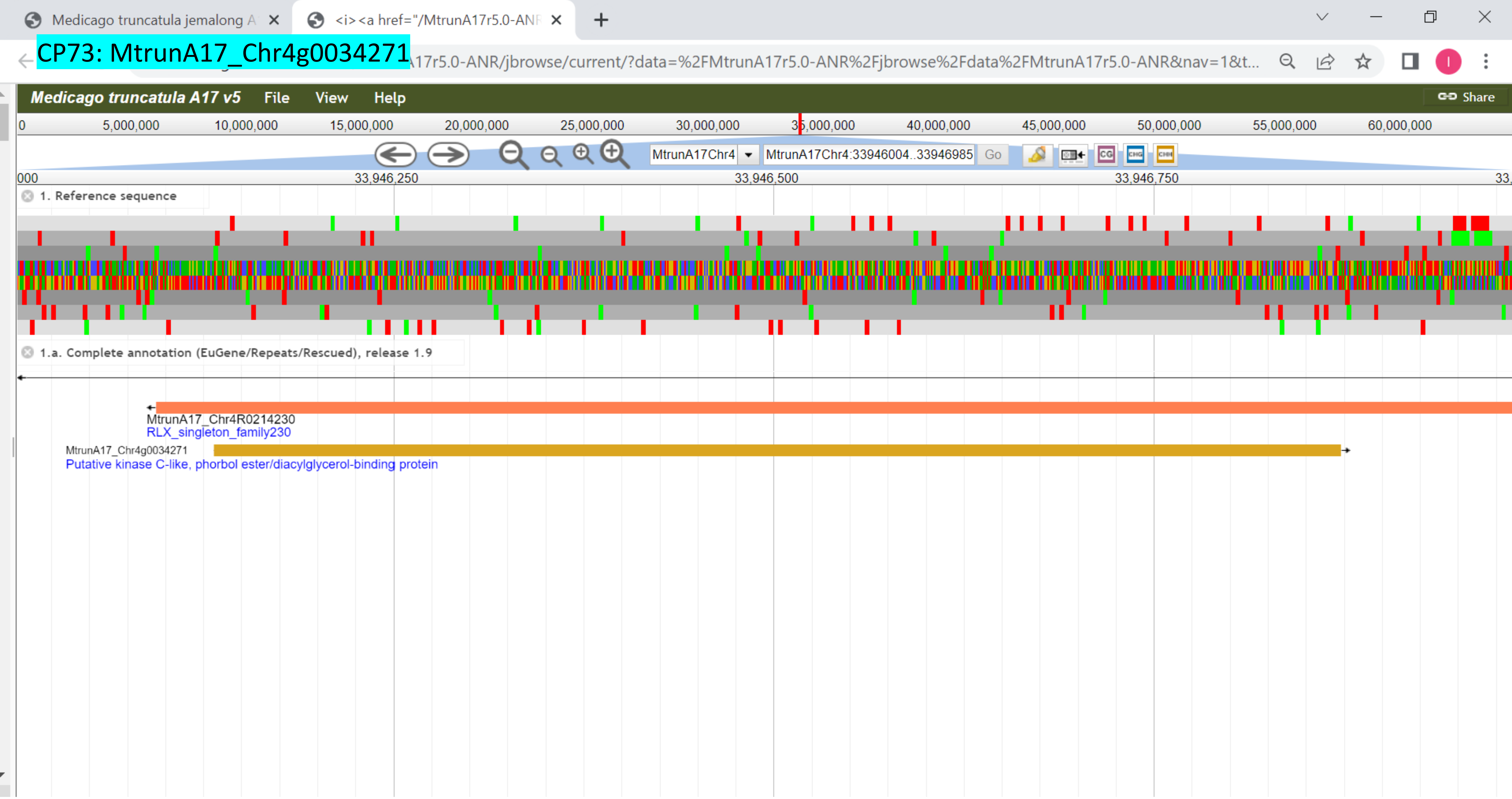

### CP74: MtrunA17_Chr4g0034491

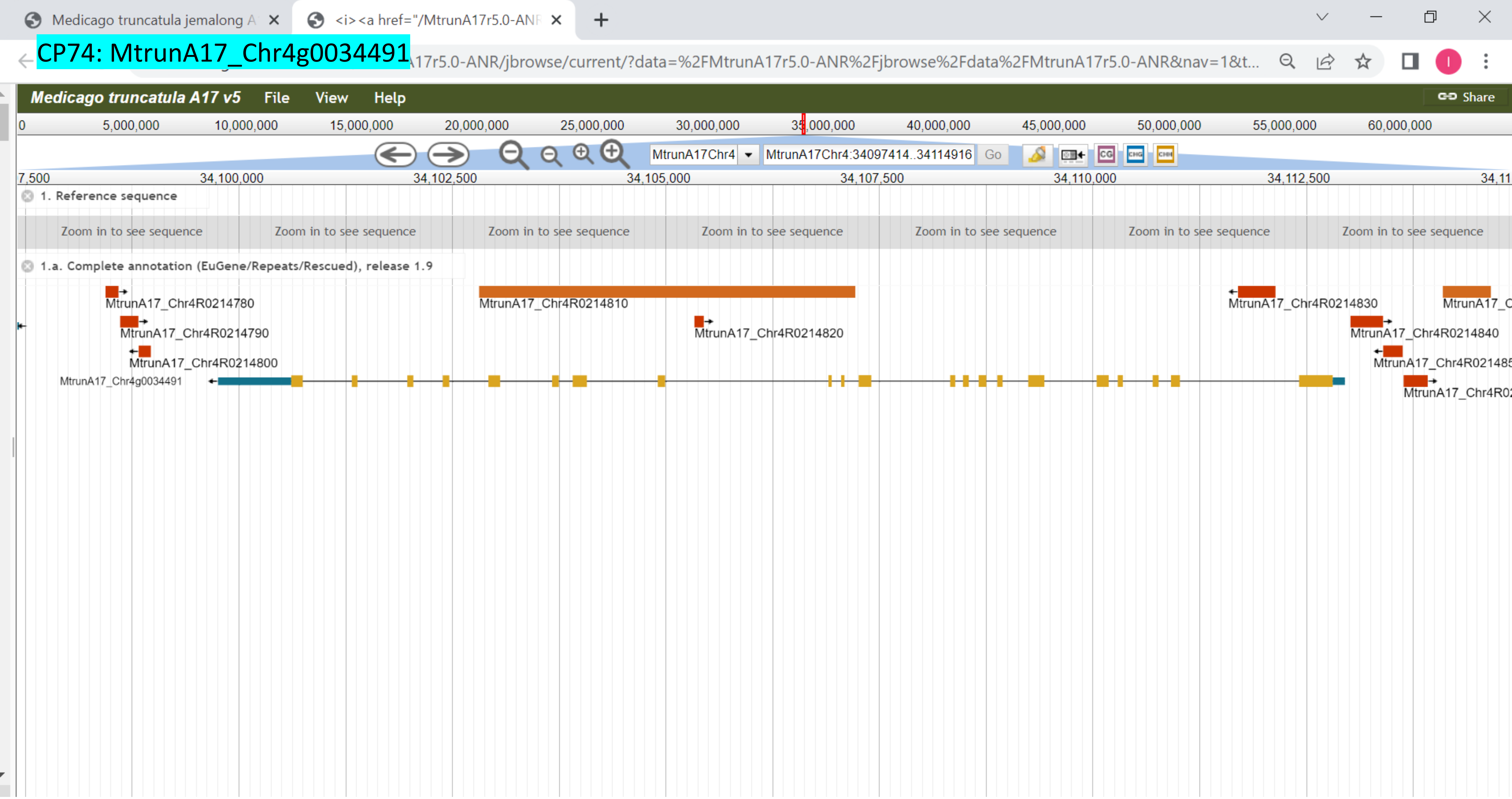

CP75: MtrunA17_Chr4g0037381

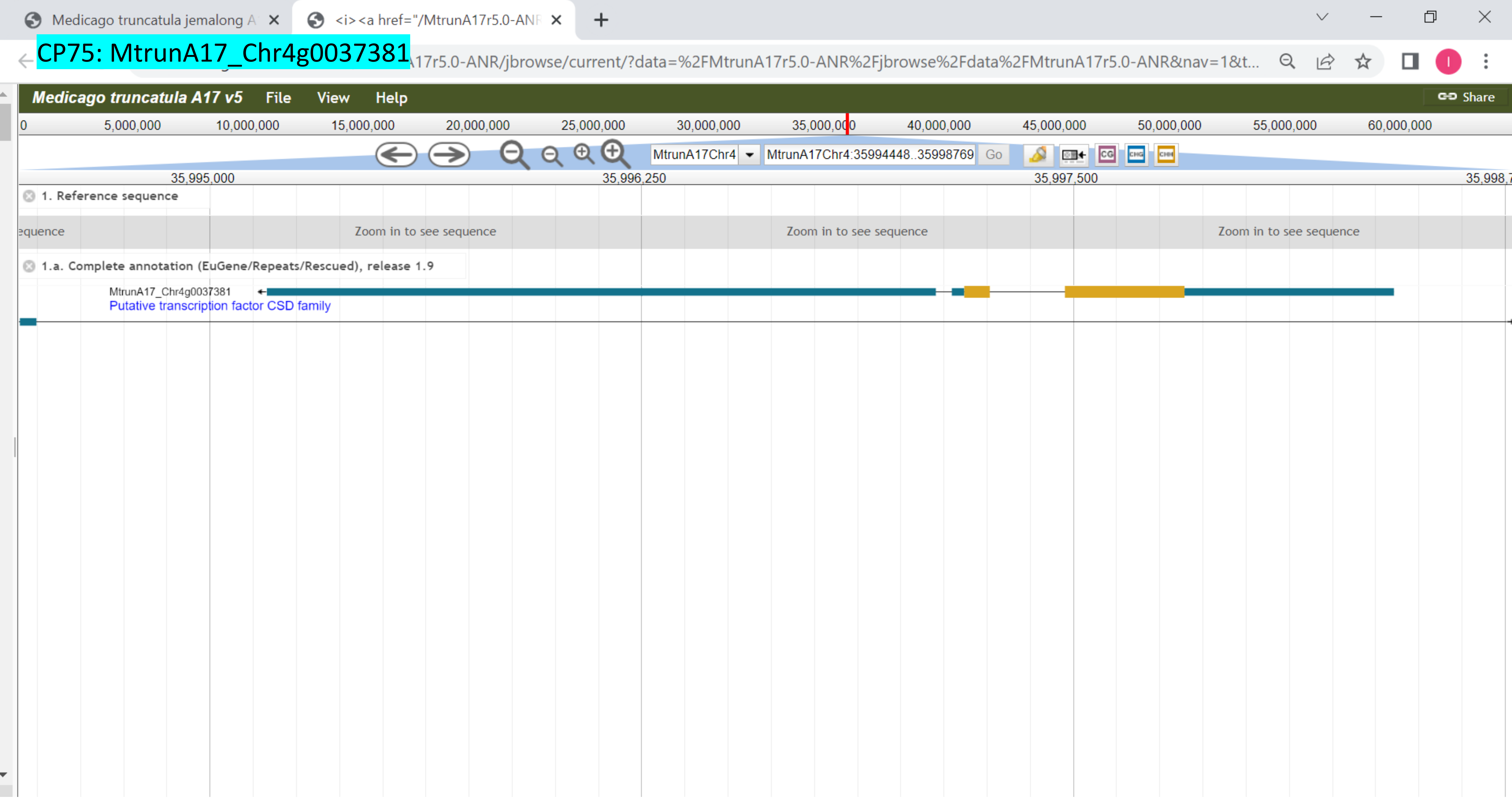

### CP76: MtrunA17_Chr4g0040471

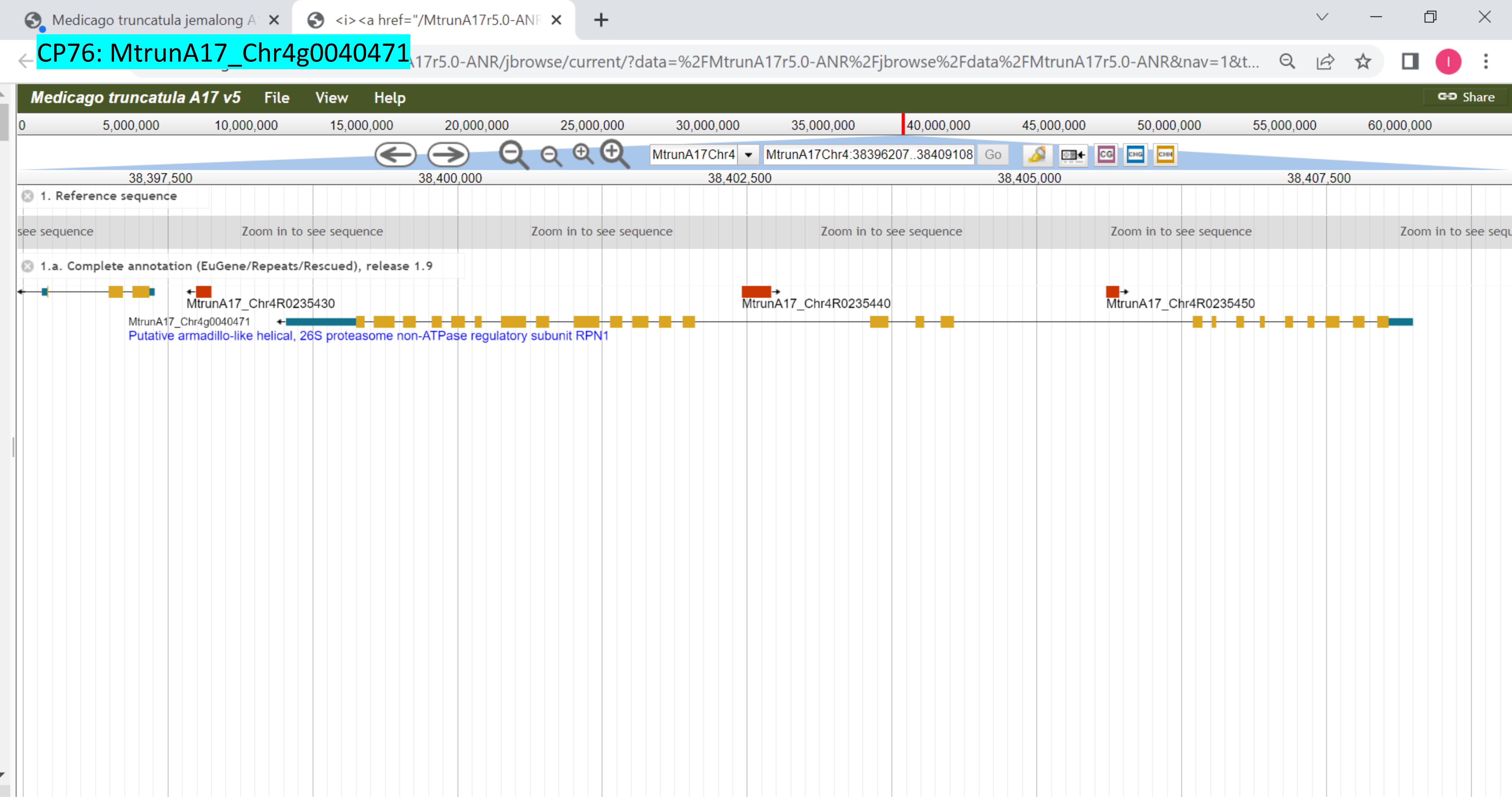

CP77: MtrunA17_Chr4g0055331

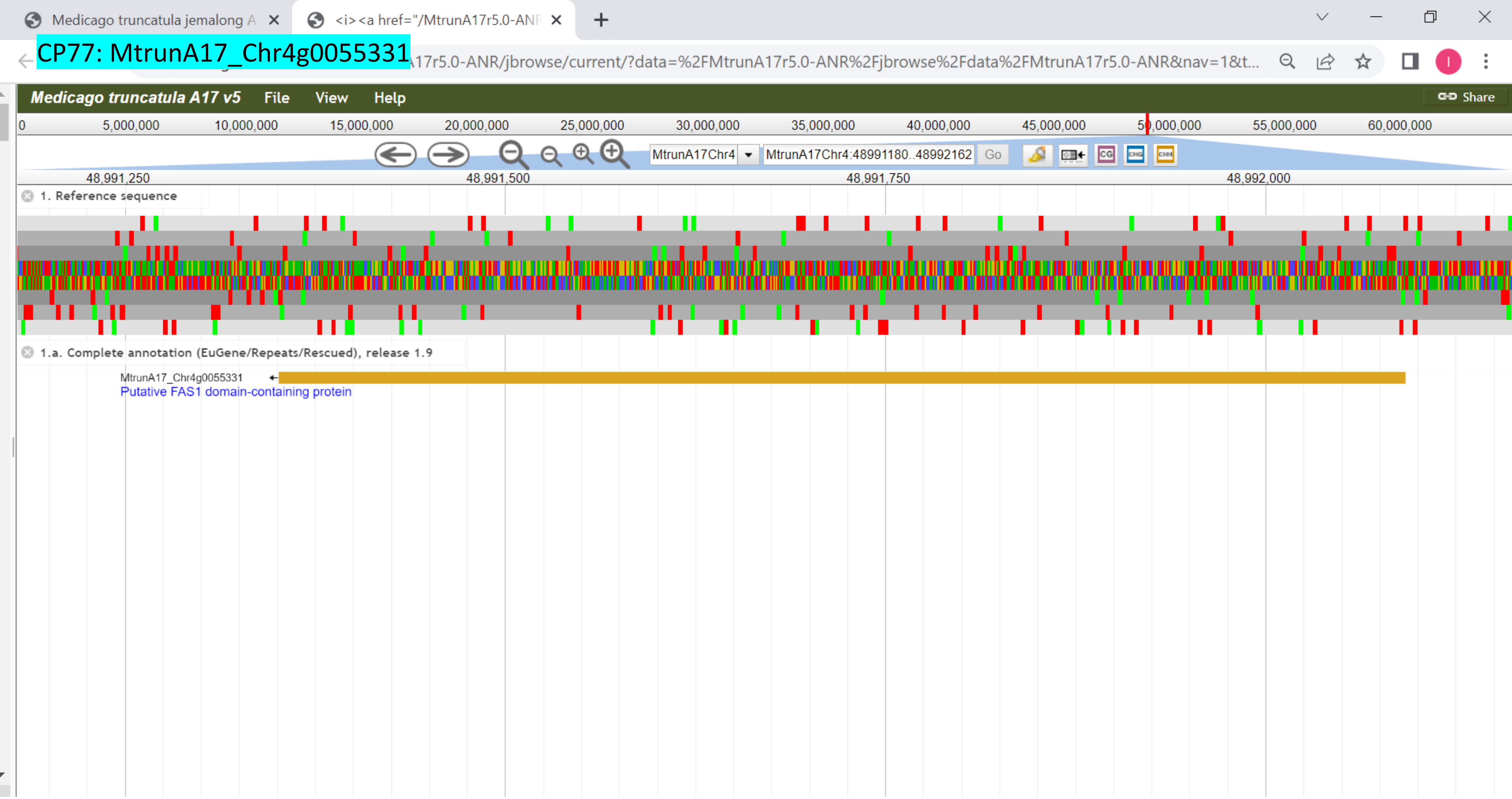

CP78: MtrunA17_Chr4g0059001

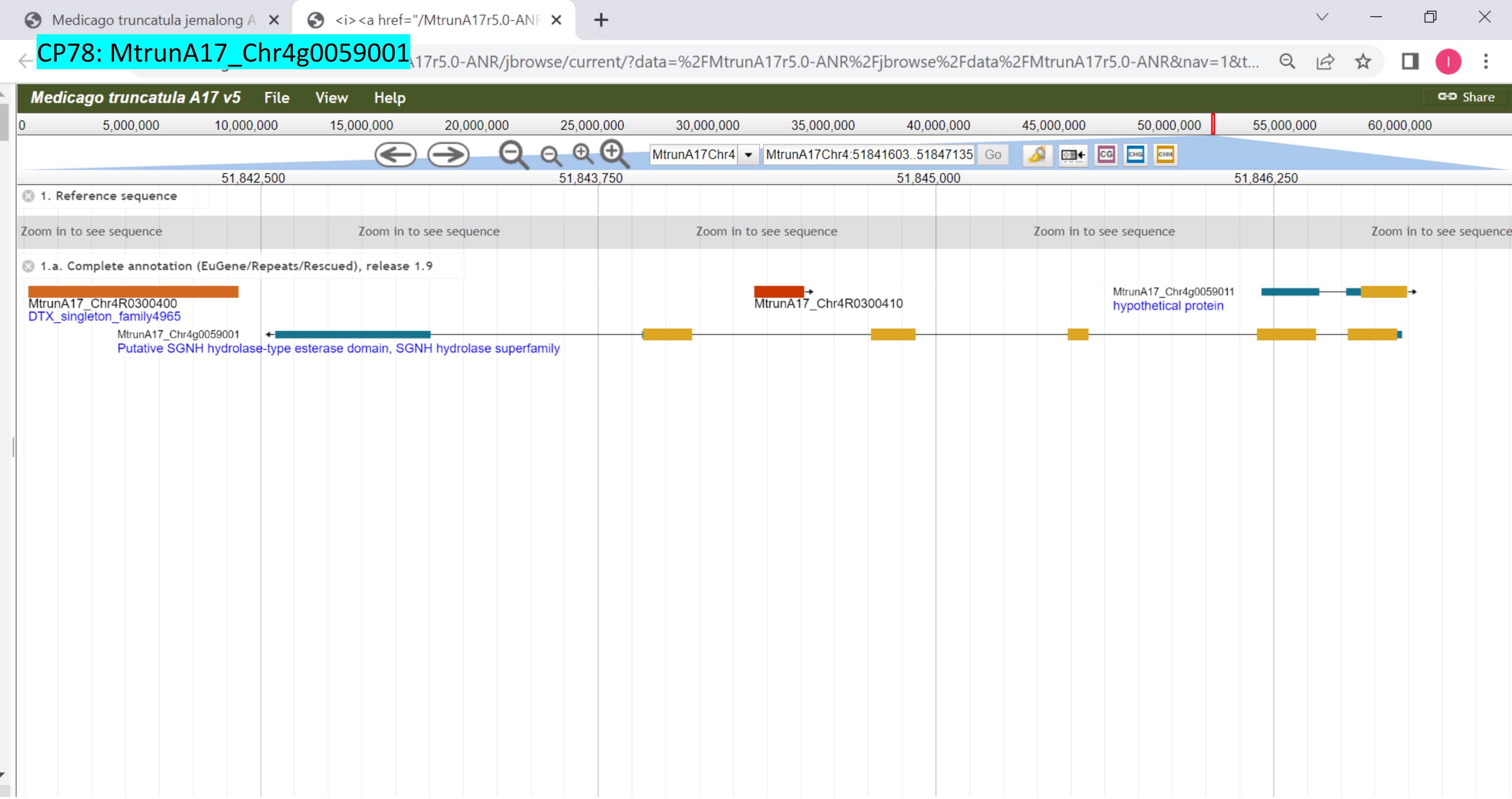

CP79: MtrunA17_Chr4g0063201

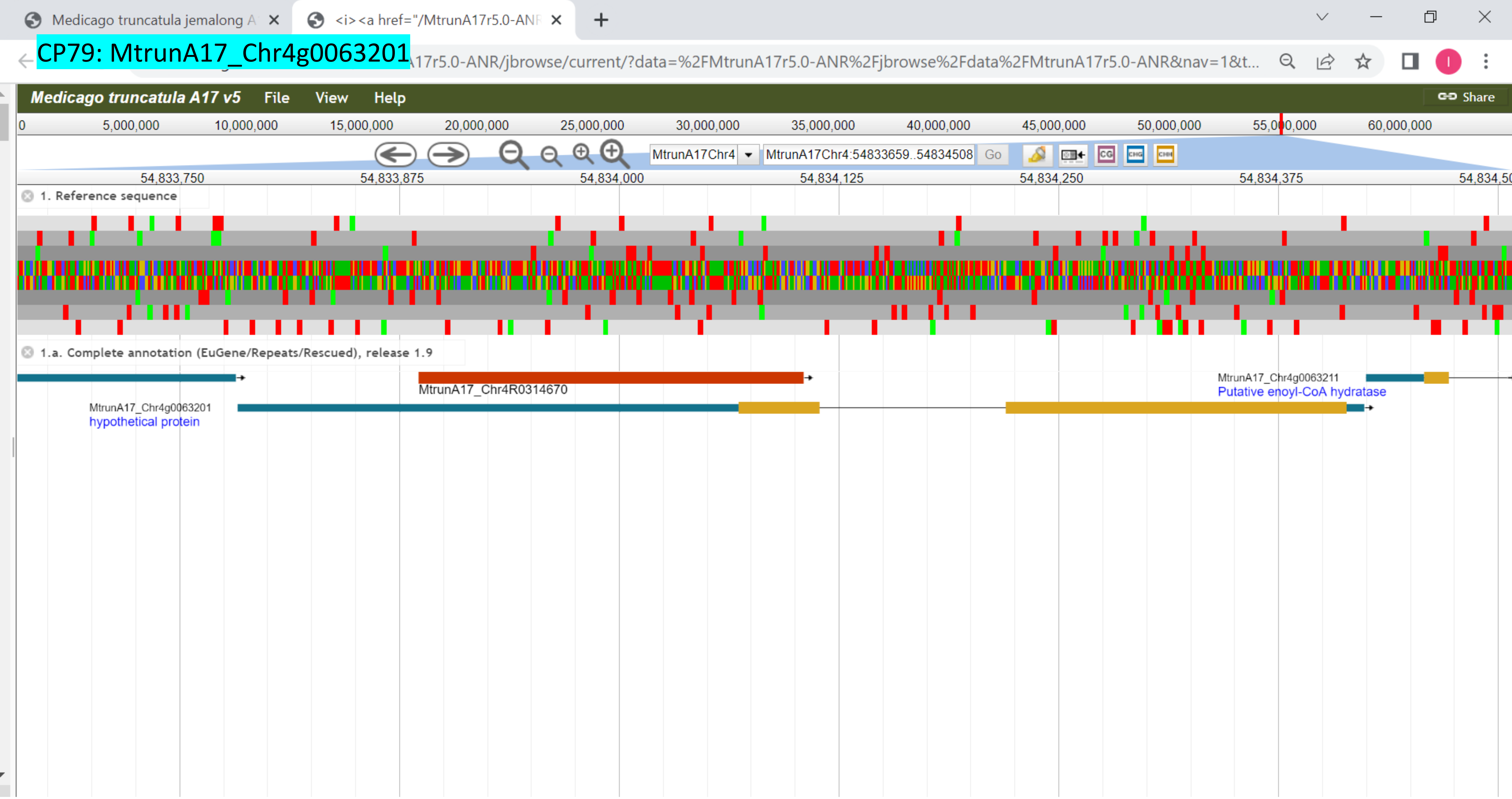

CP80: MtrunA17_Chr4g0070011

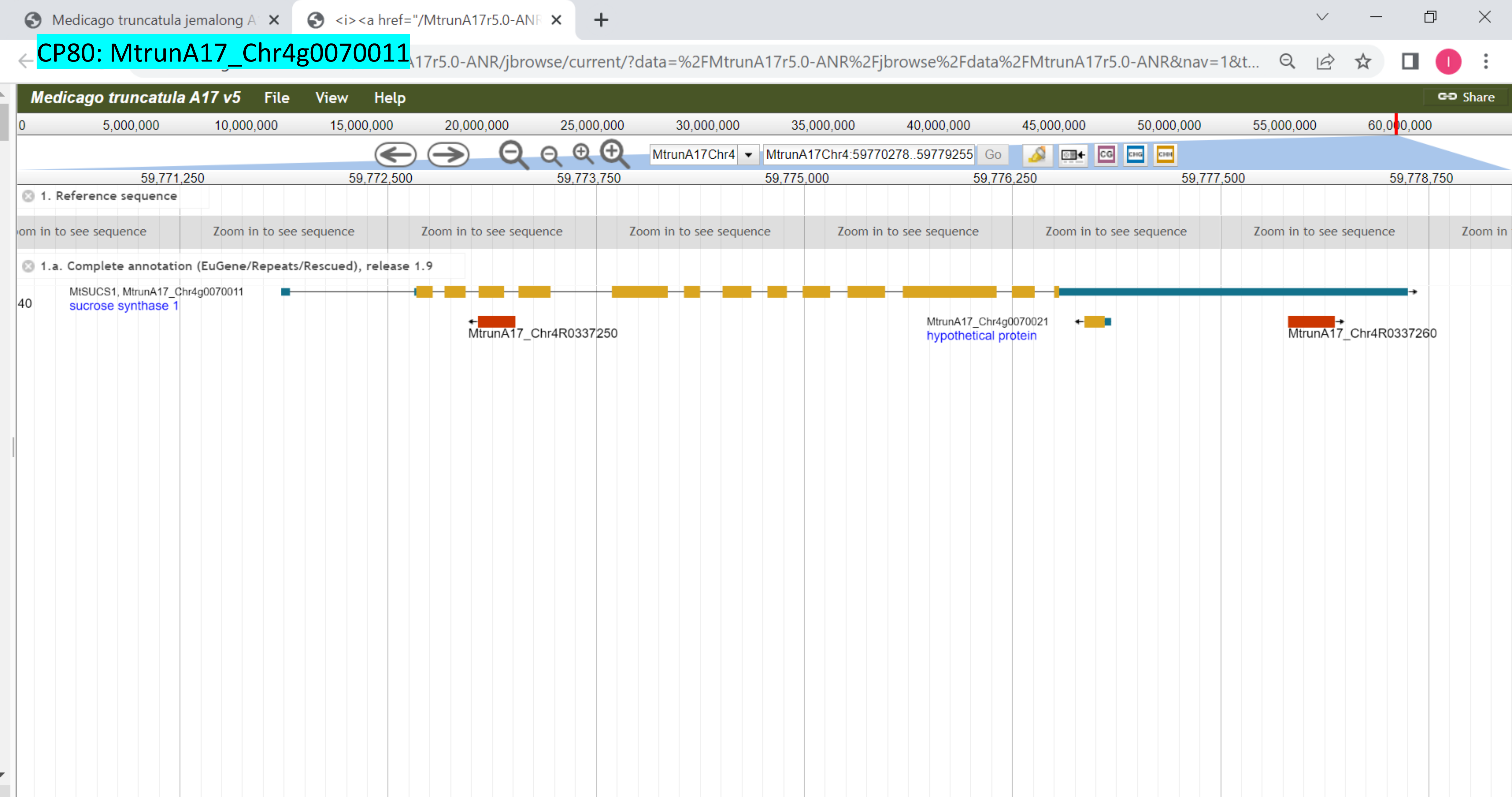

CP81: MtrunA17_Chr5g0393401

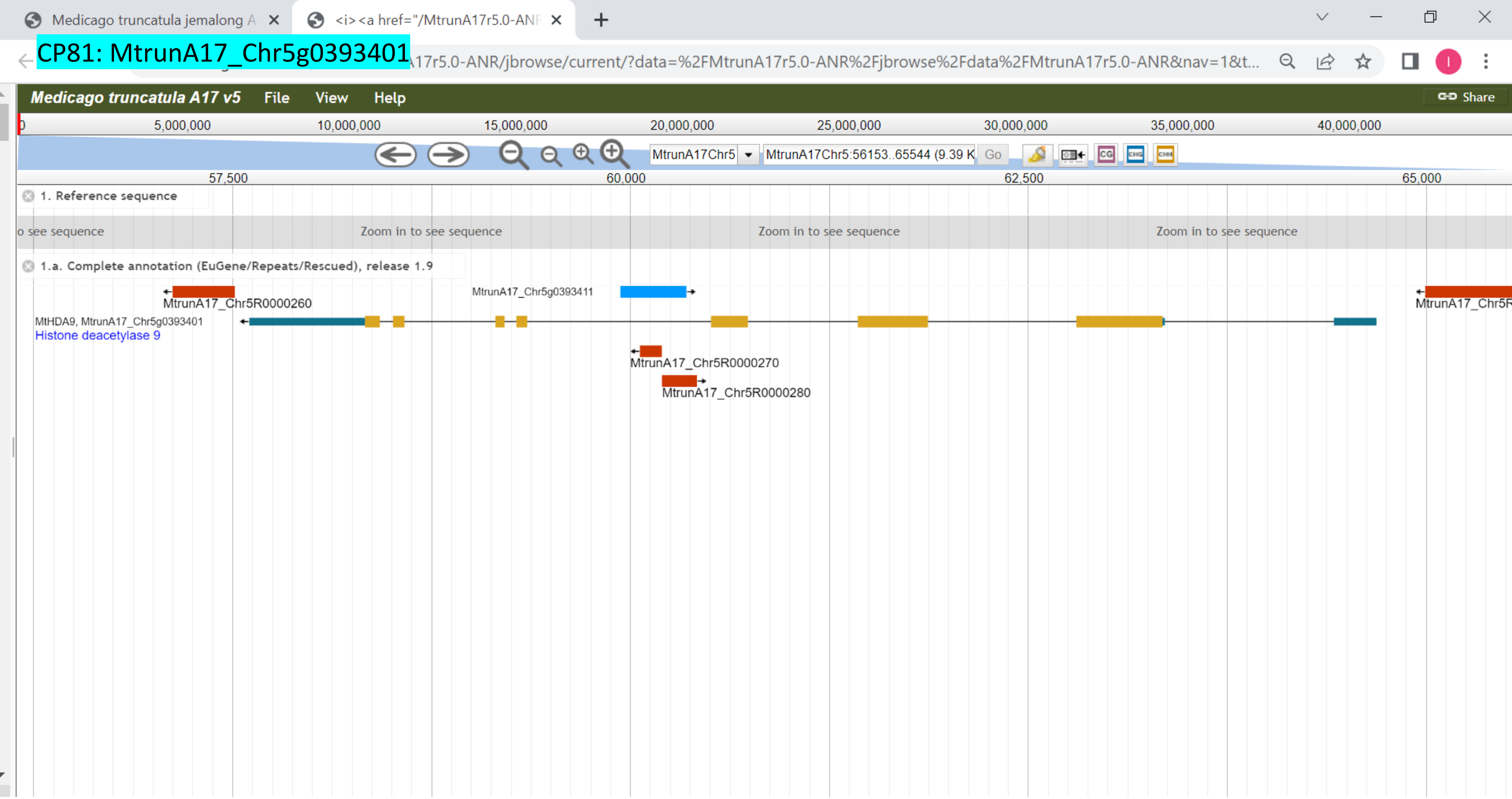

CP82: MtrunA17_Chr5g0400181

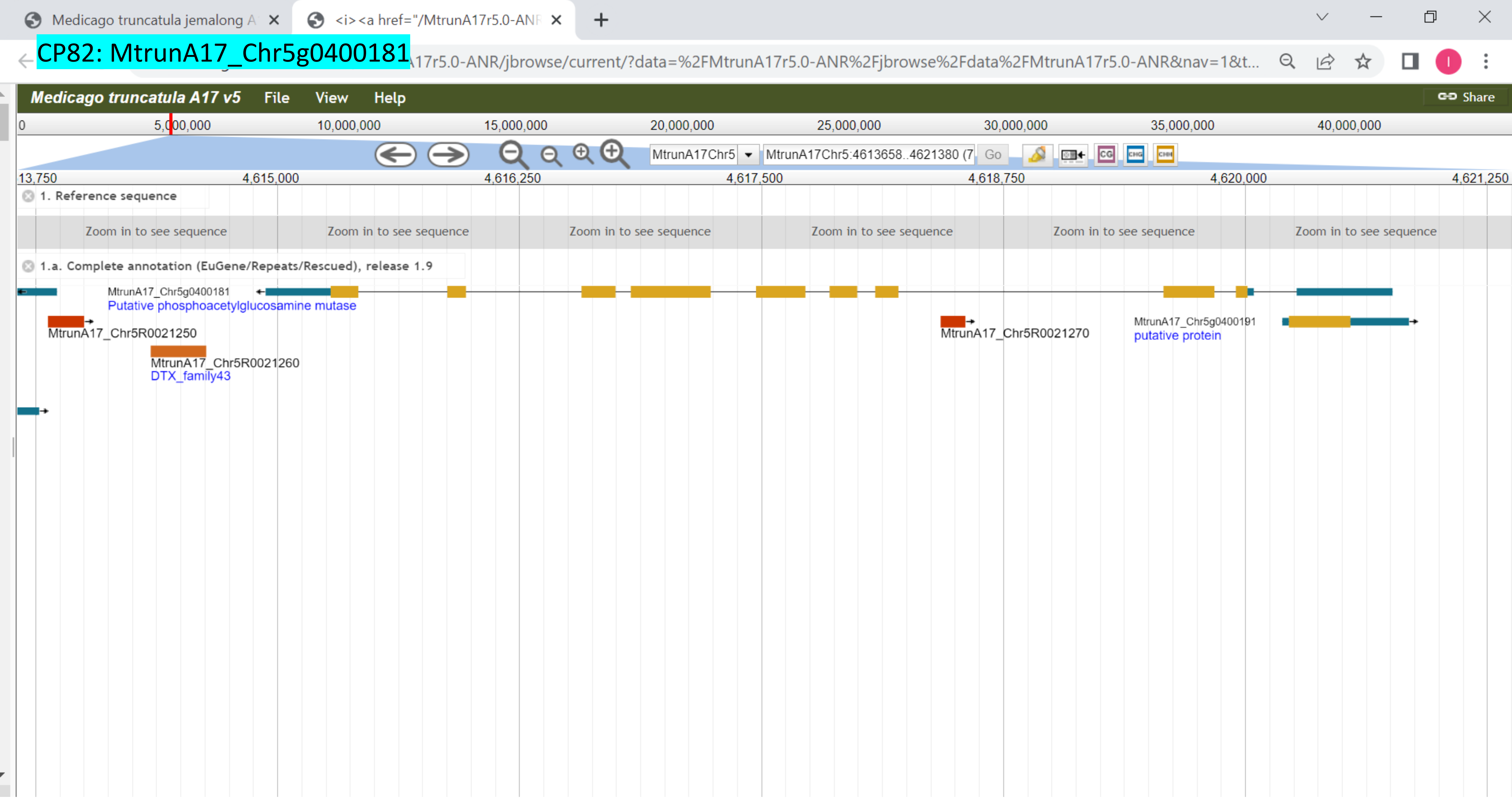

CP83: MtrunA17_Chr5g0405061

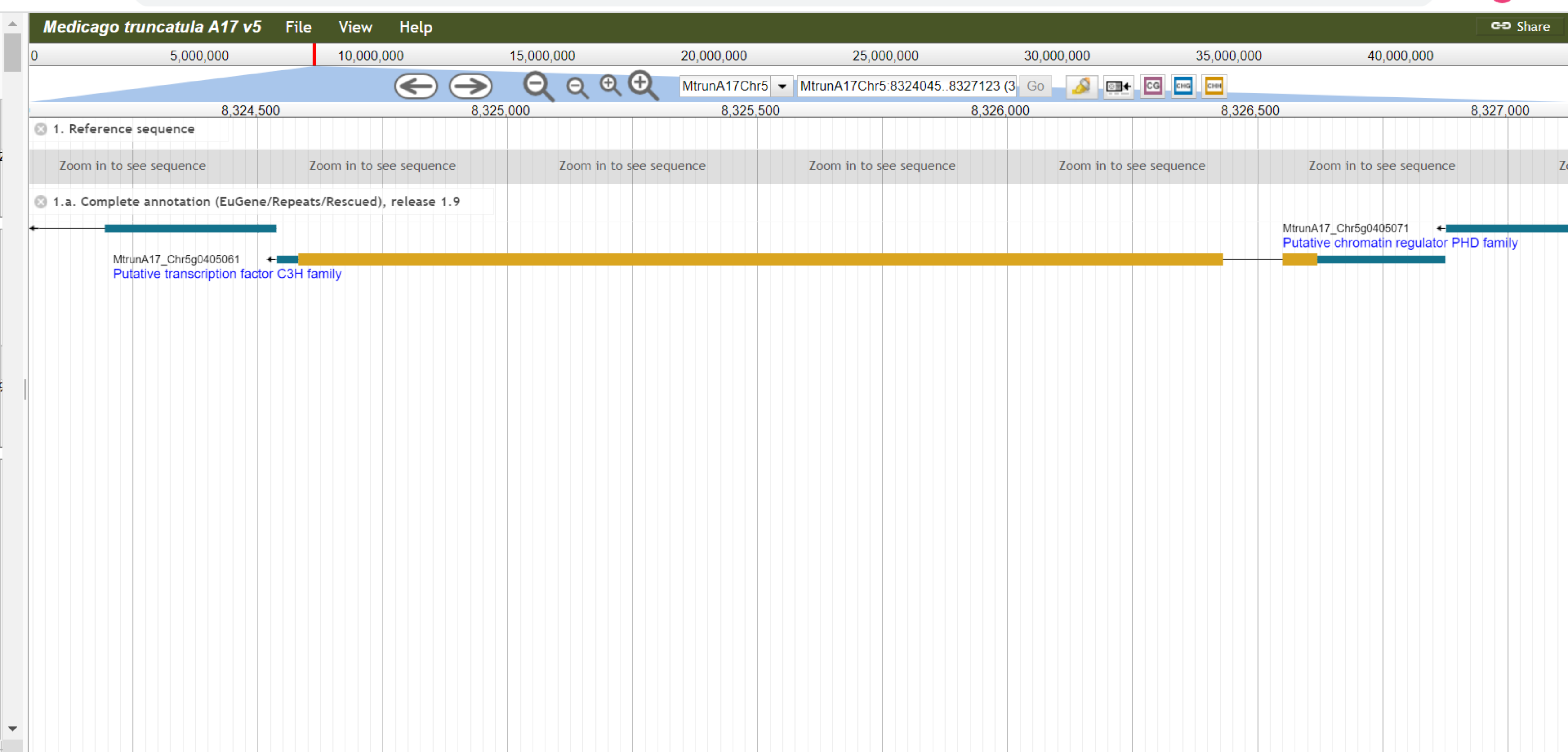

CP84: MtrunA17_Chr5g0408581

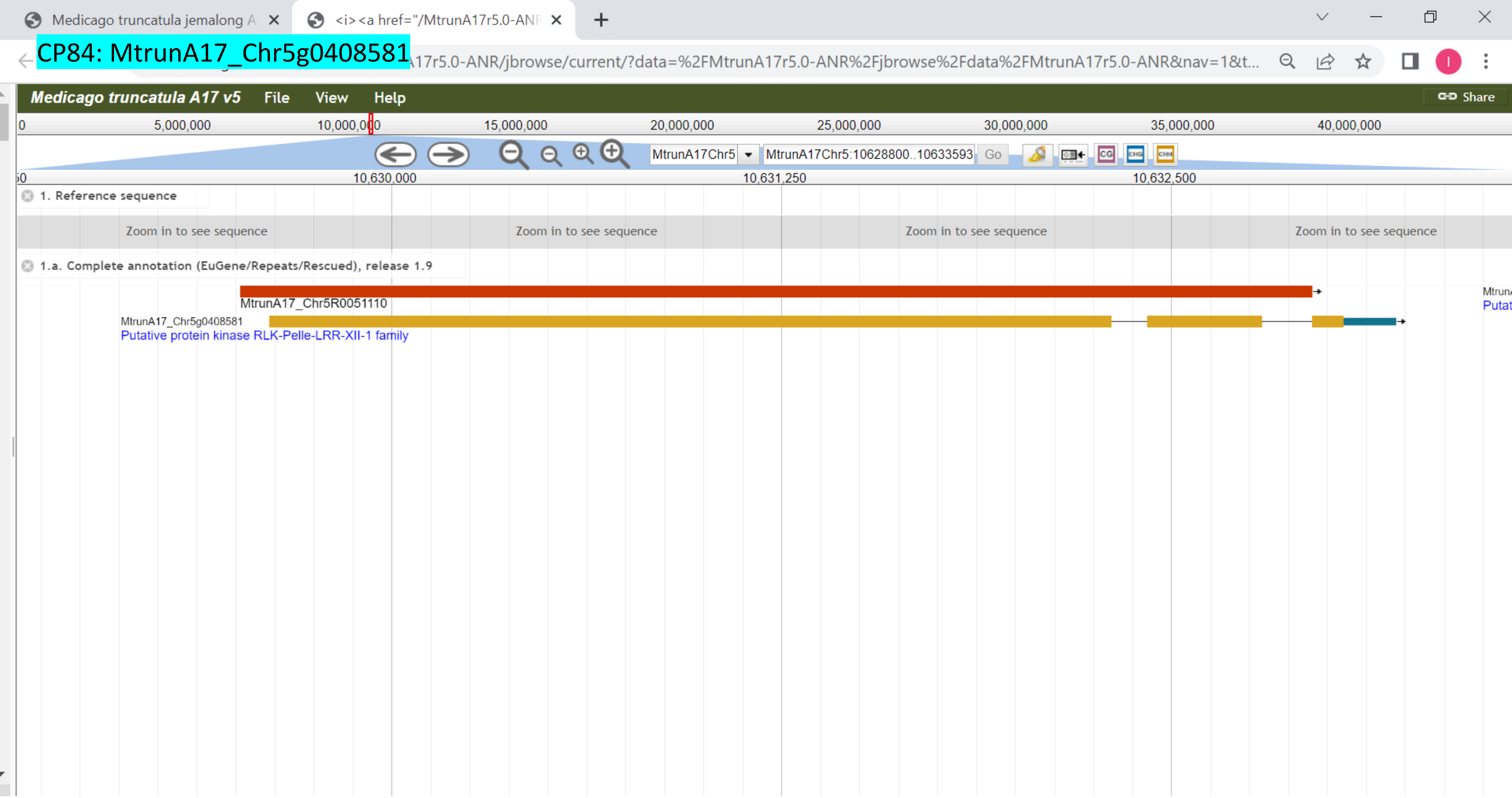

### CP85: MtrunA17_Chr5g0415031

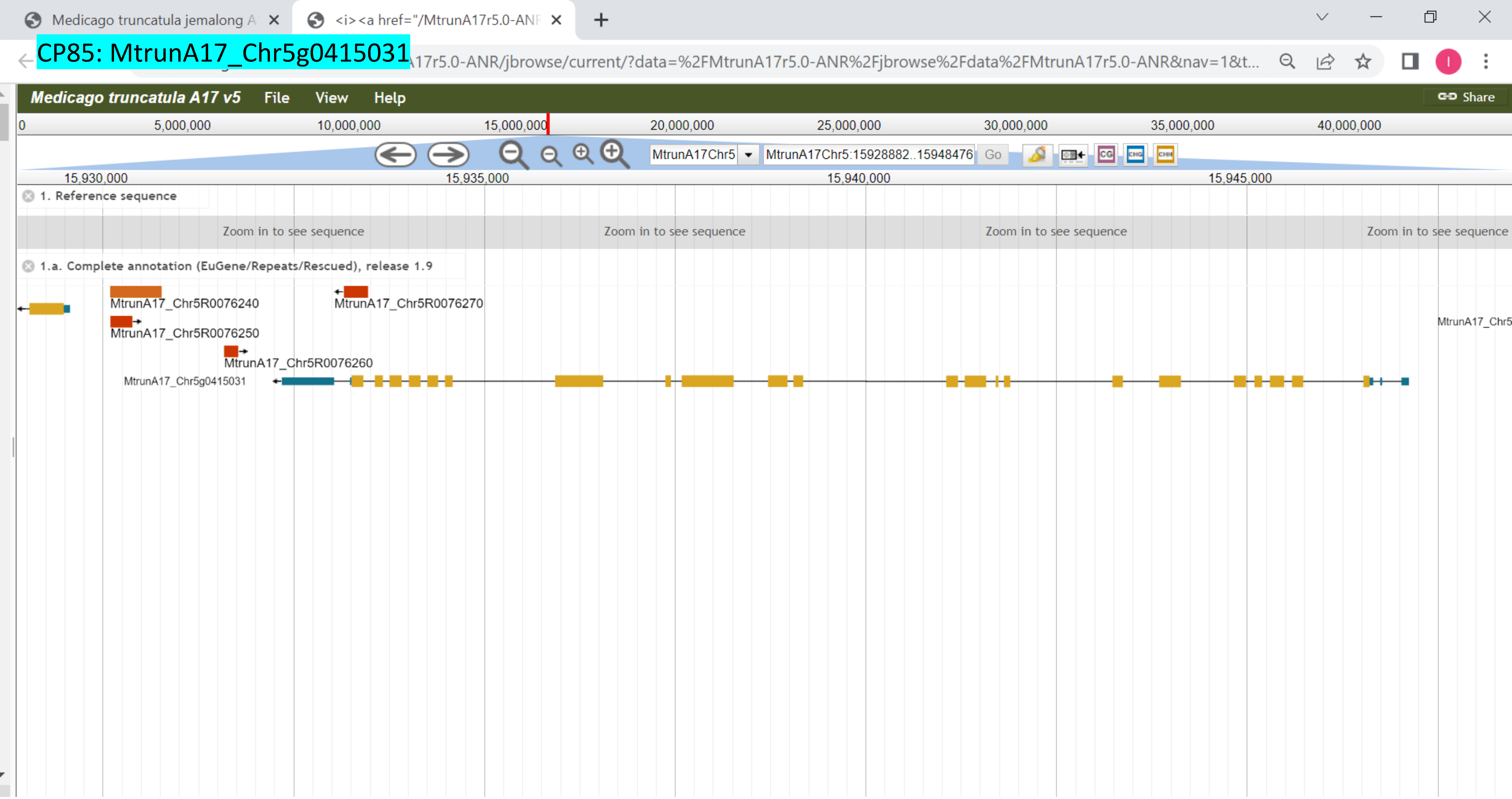

CP86: MtrunA17_Chr5g0415911

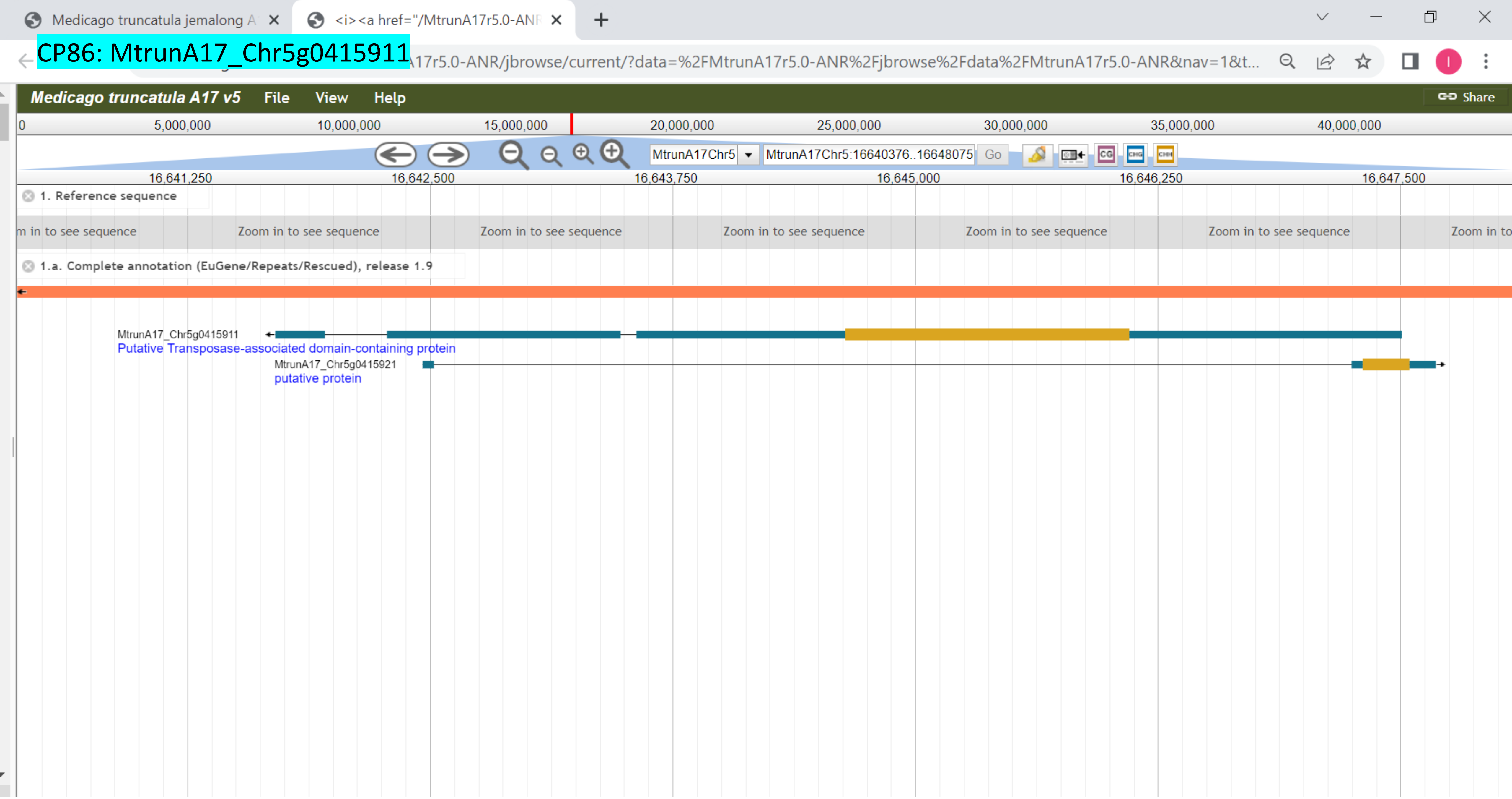

CP87: MtrunA17_Chr5g0421761

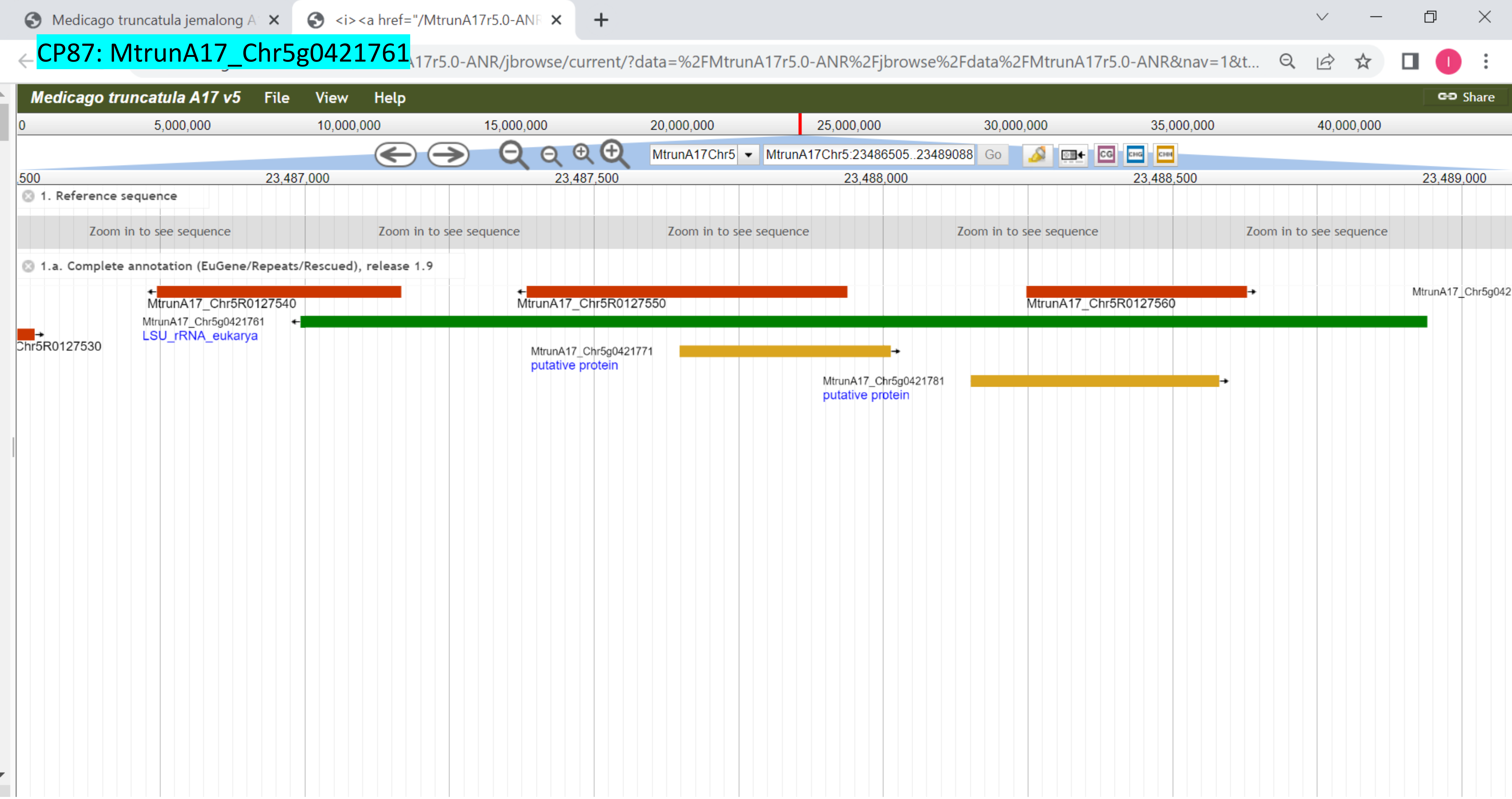

### CP88: MtrunA17_Chr5g0422291

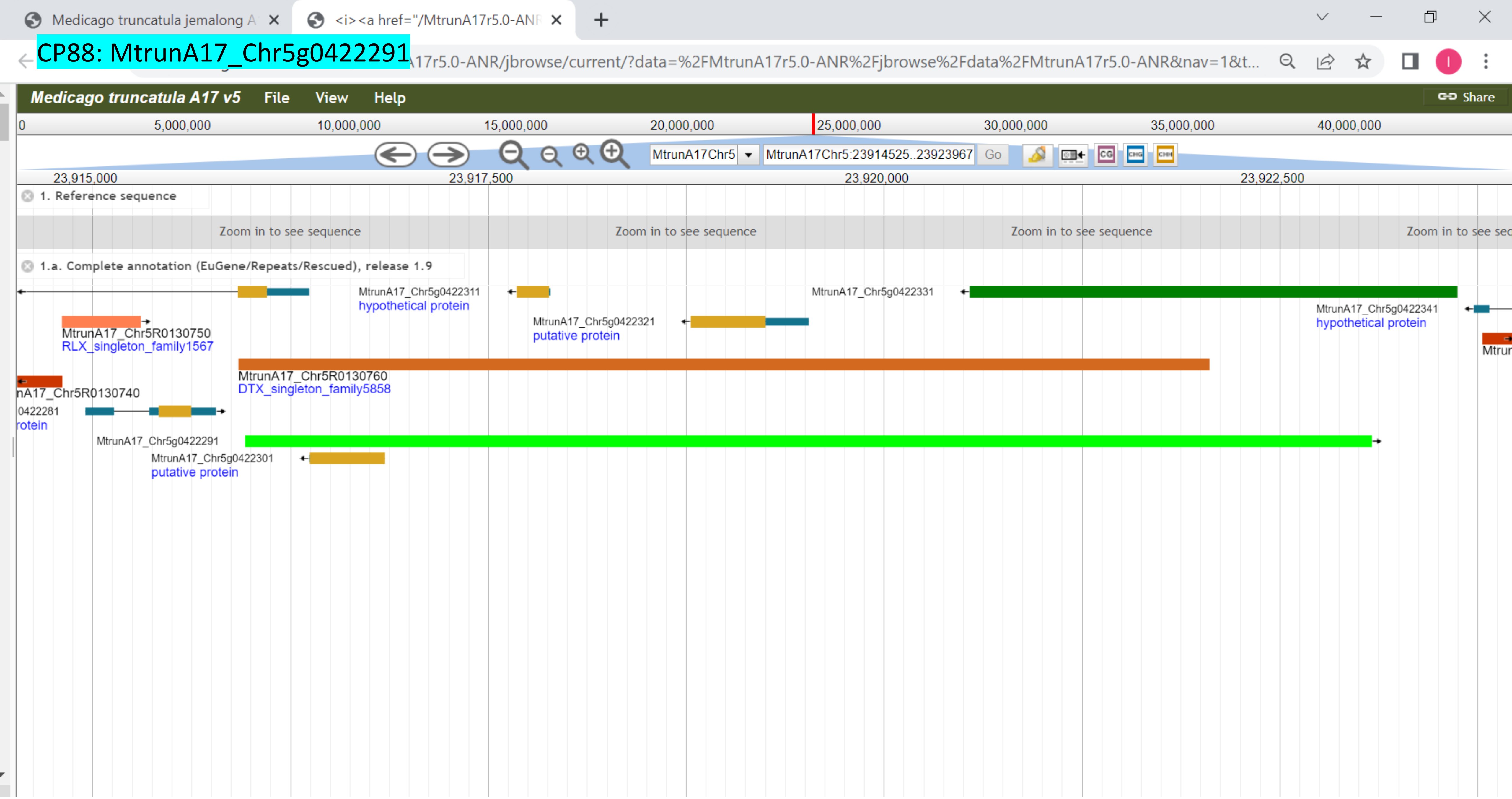

### CP89: MtrunA17_Chr5g0422291

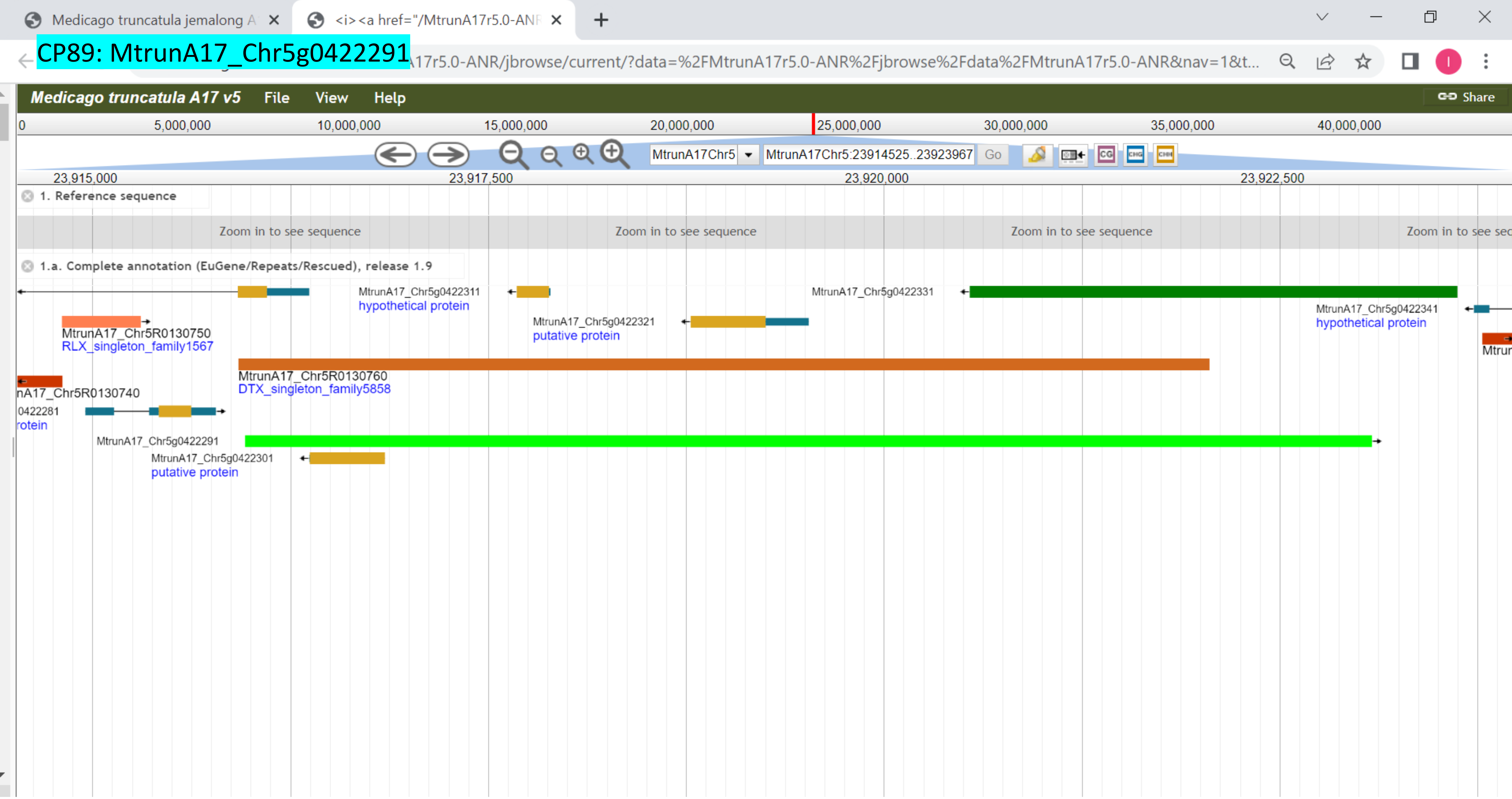

### CP90: MtrunA17_Chr5g0430341

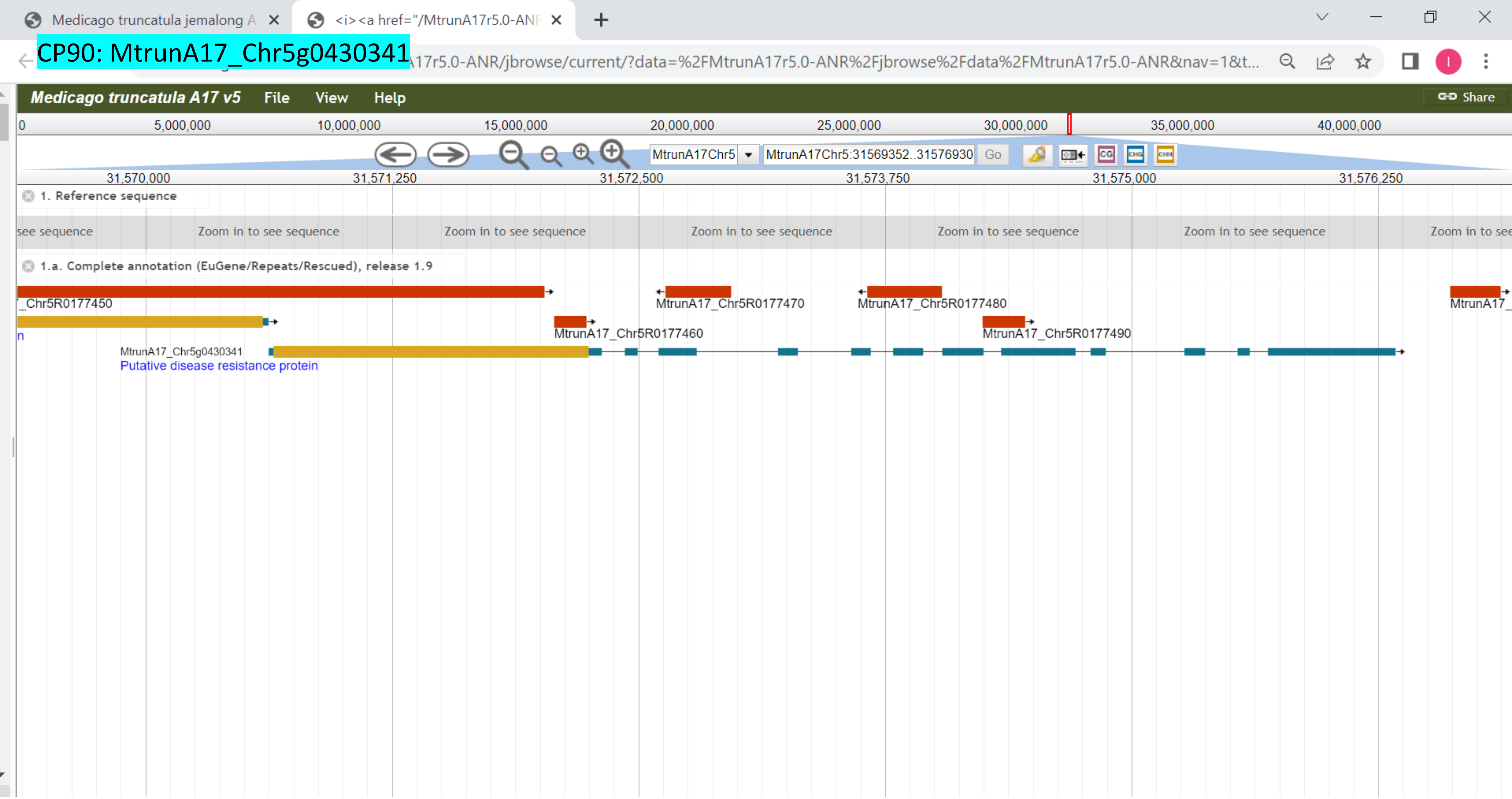

### CP91: MtrunA17_Chr5g0430341

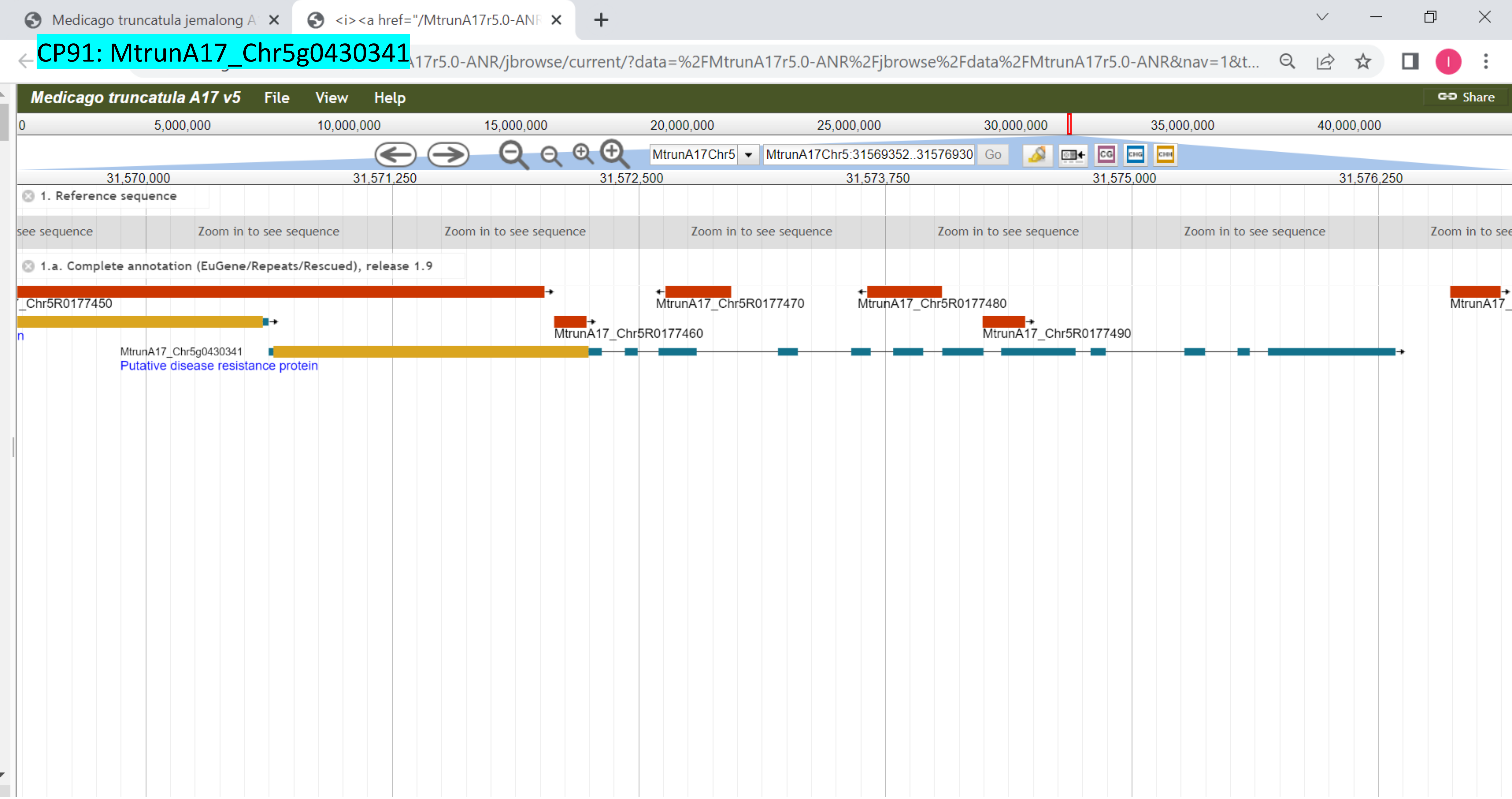

CP92: MtrunA17_Chr5g0431401

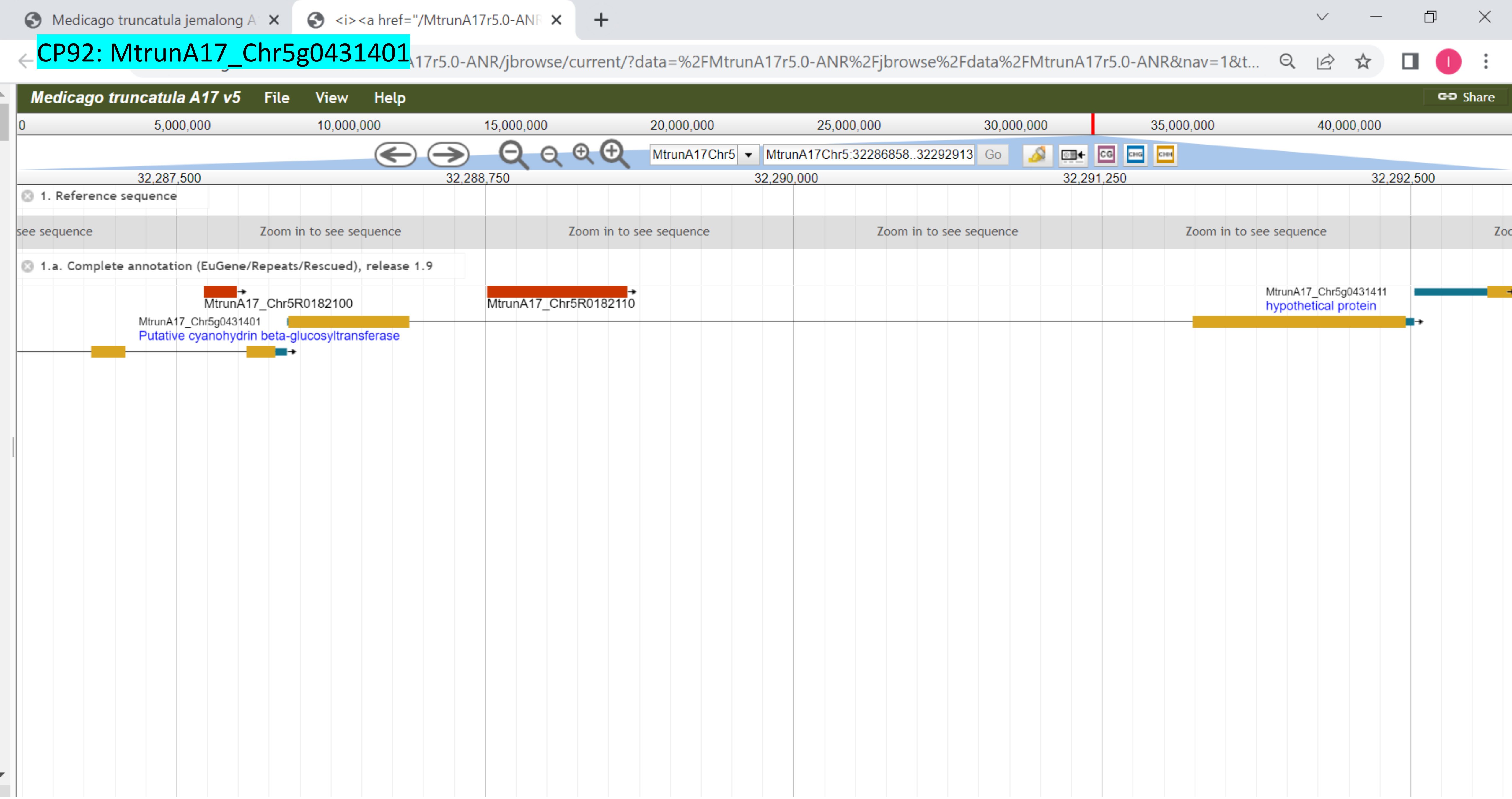

CP93: MtrunA17_Chr5g0435191

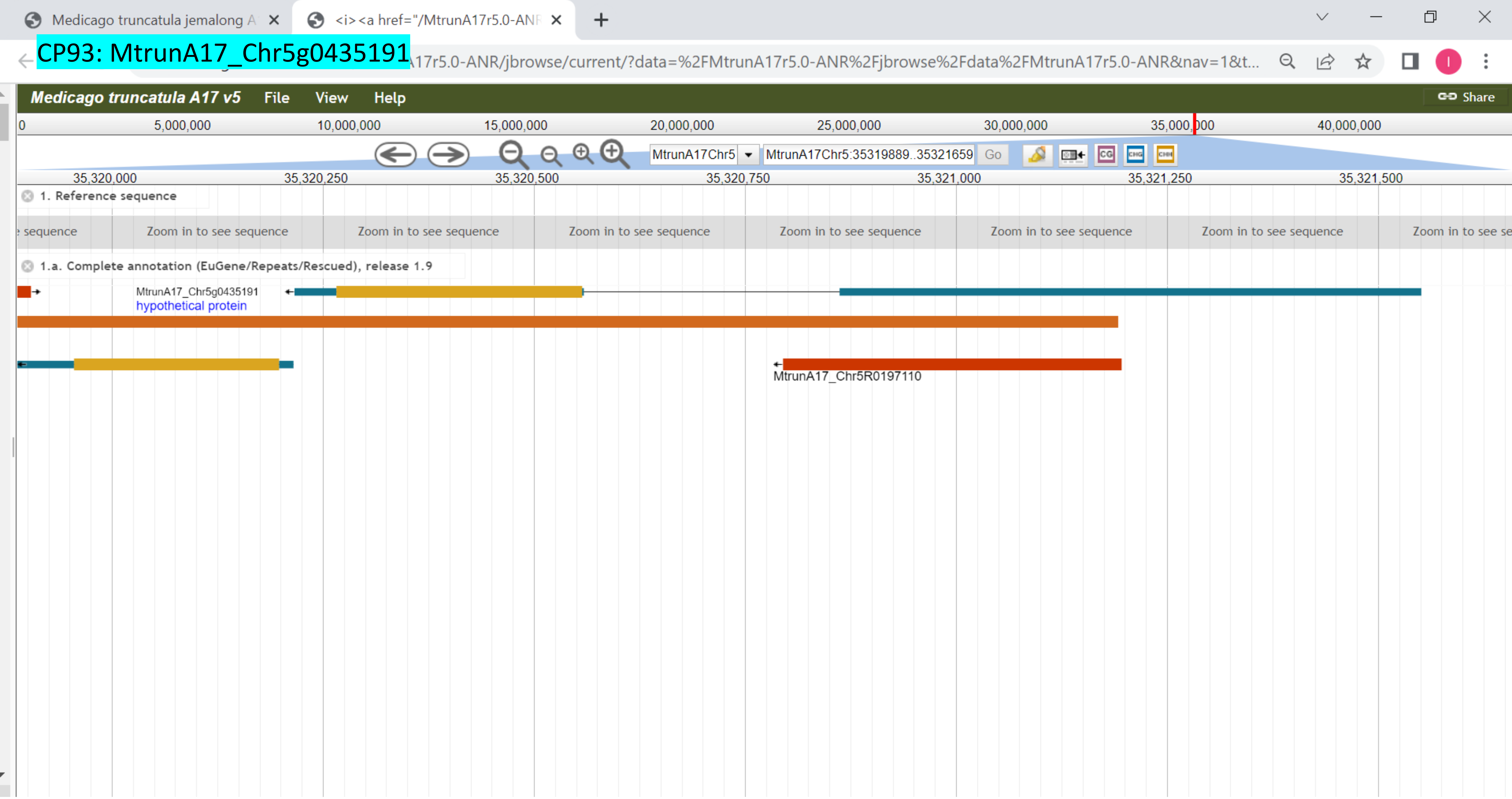

CP94: MtrunA17_Chr5g0444231

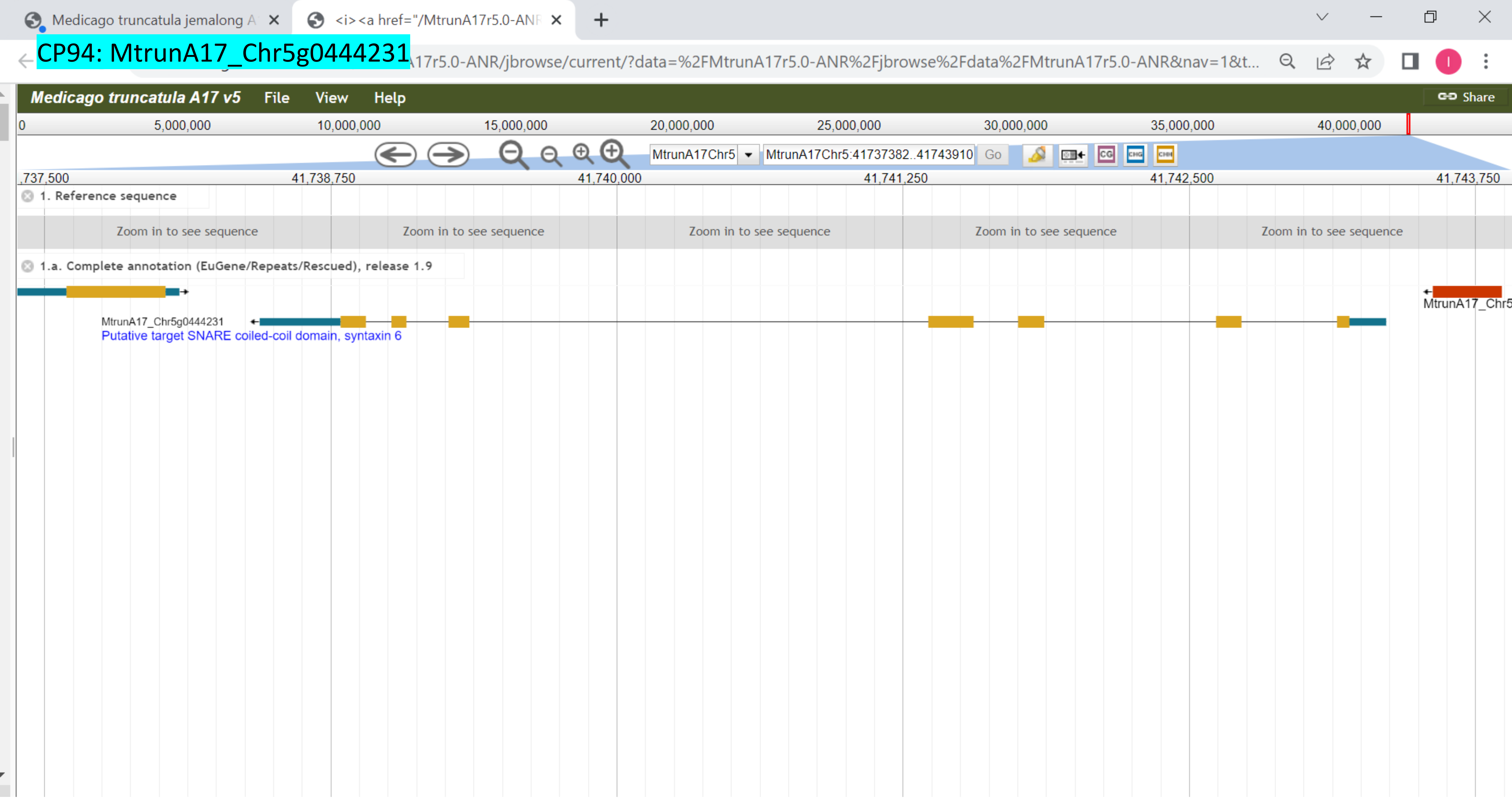

### CP95: MtrunA17_Chr6g0451601

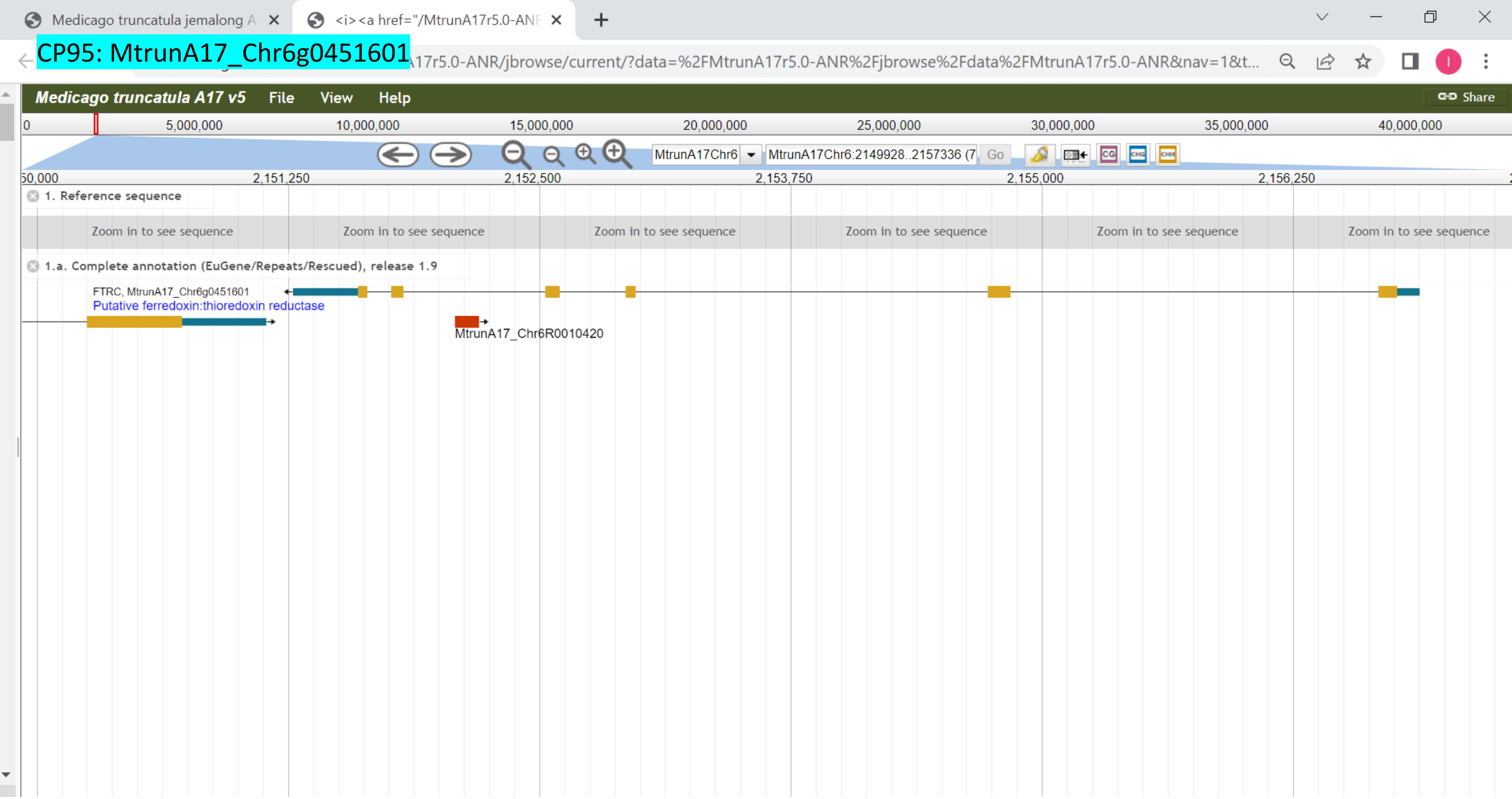

CP96: MtrunA17_Chr6g0452781

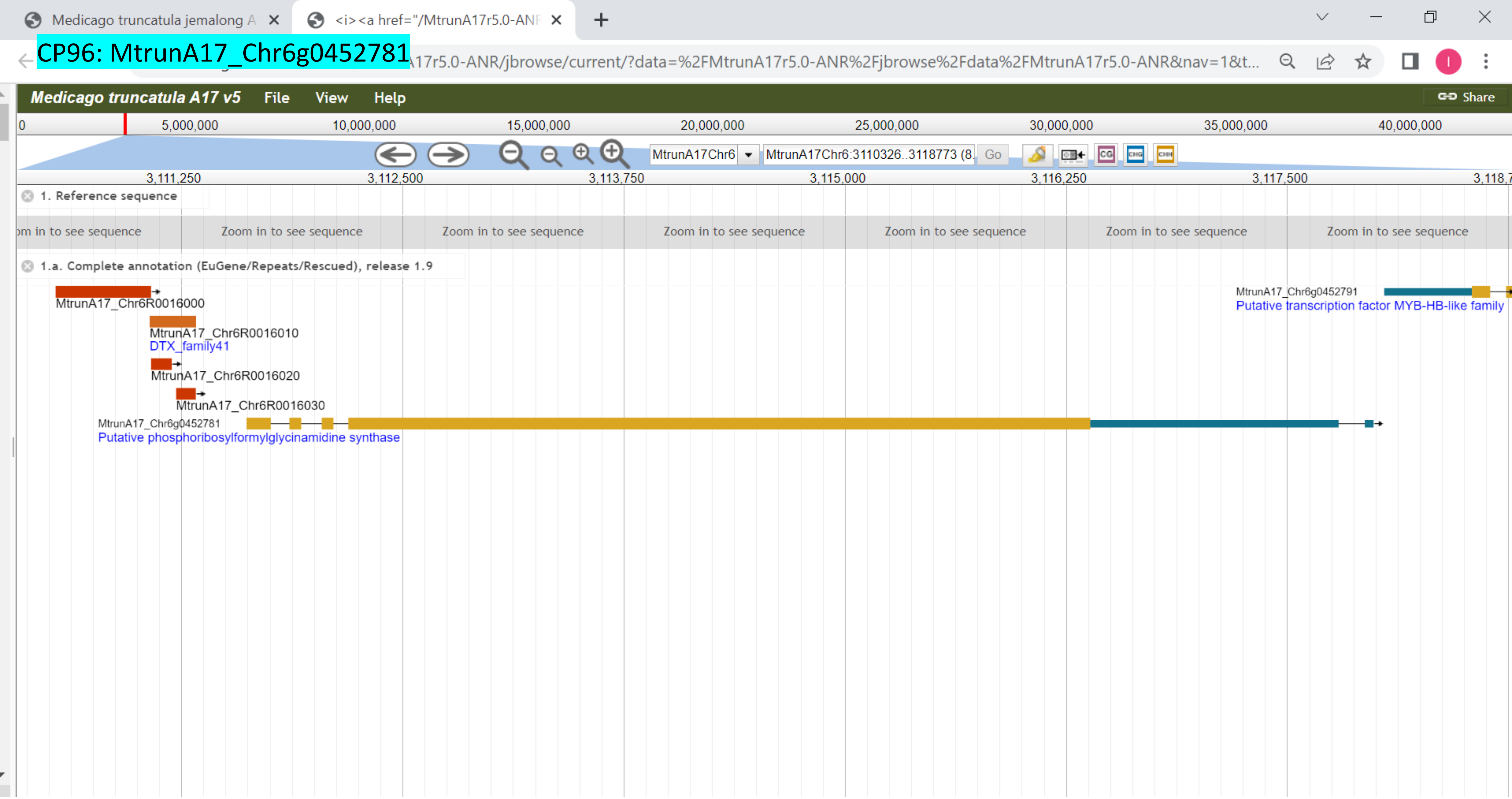

CP97: MtrunA17_Chr6g0457351

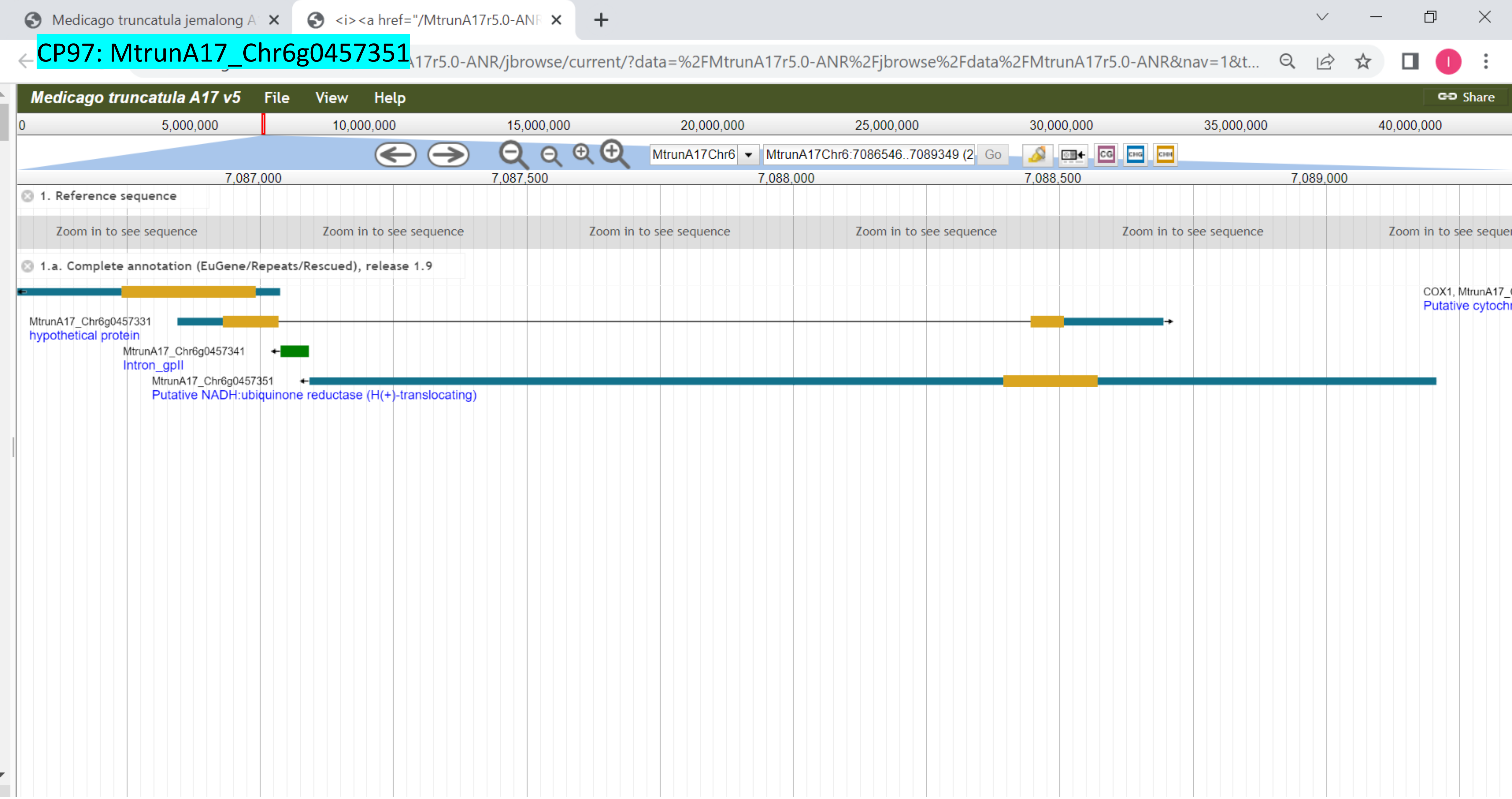

CP98: MtrunA17_Chr6g0457461

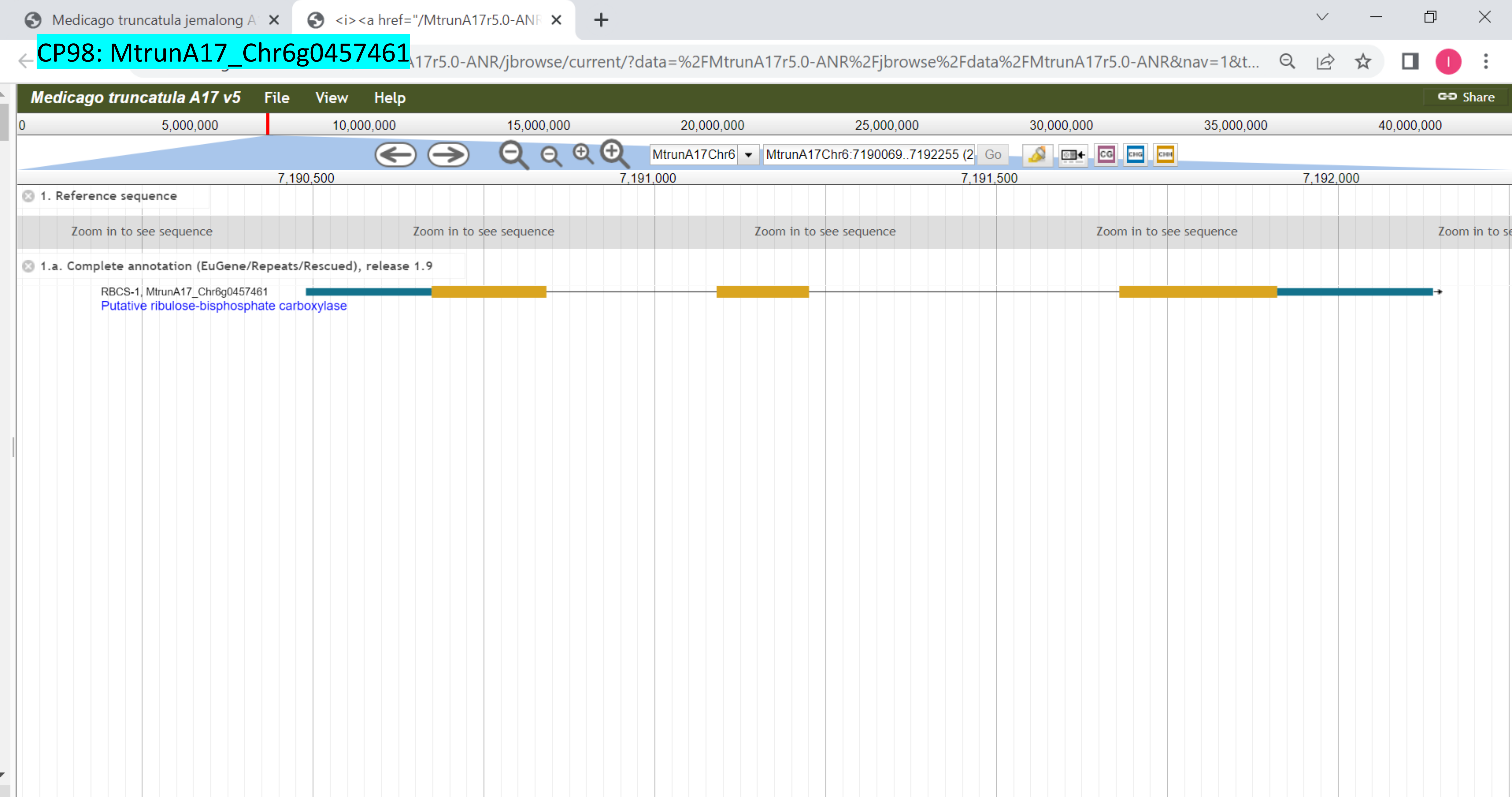

CP99: MtrunA17_Chr6g0457461

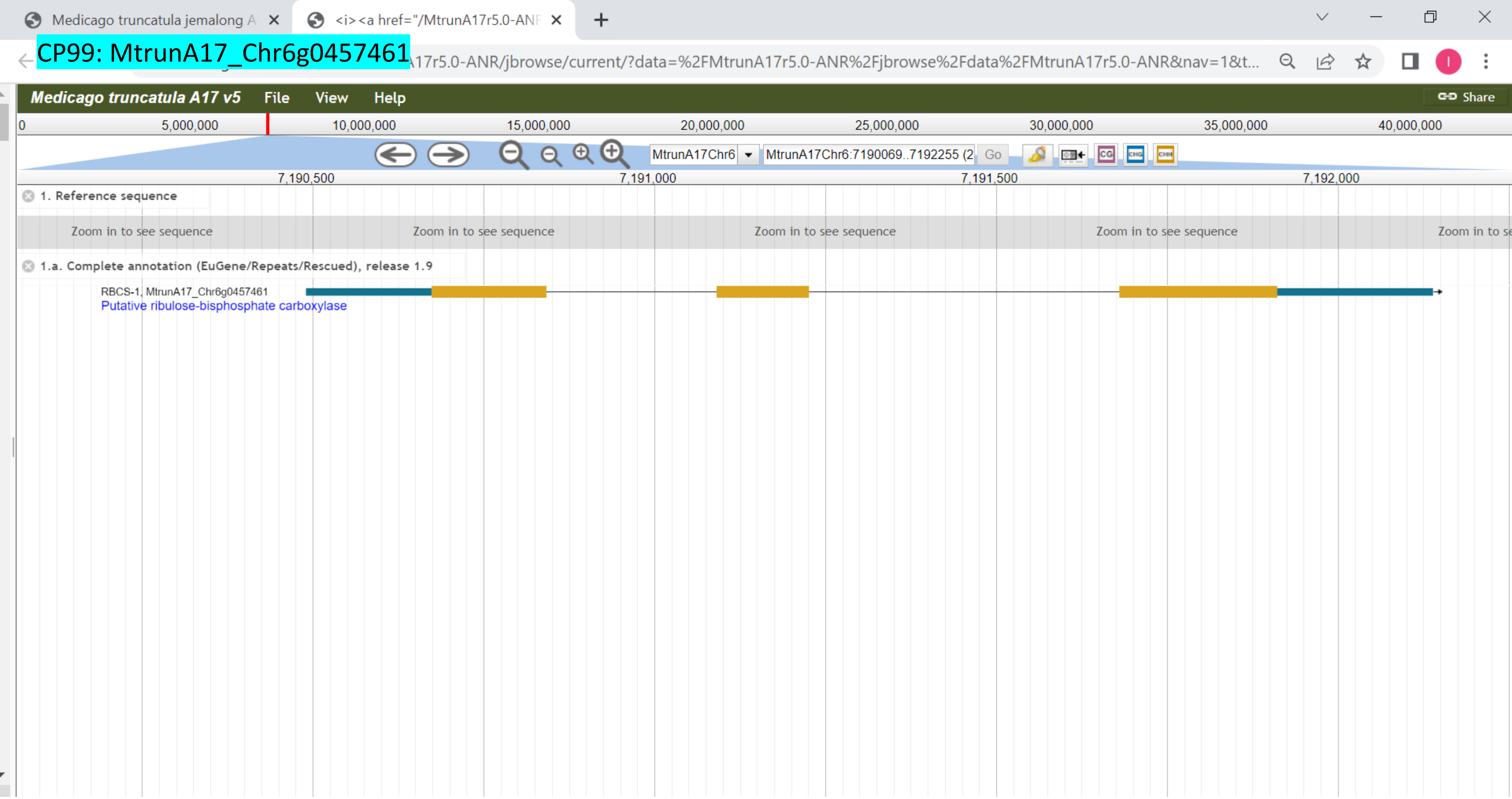

CP100: MtrunA17_Chr6g0457461

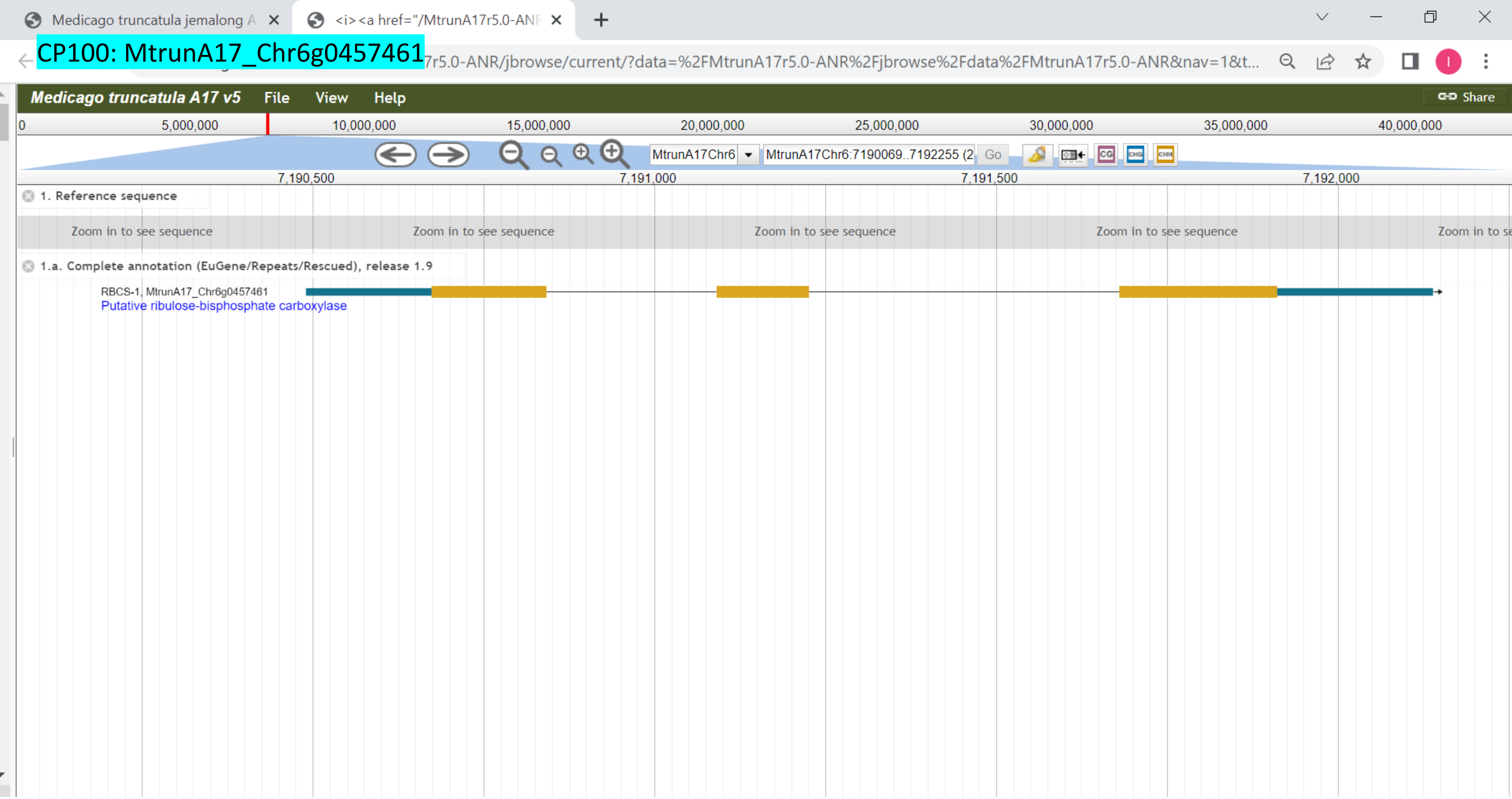

CP101: MtrunA17_Chr6g0458091

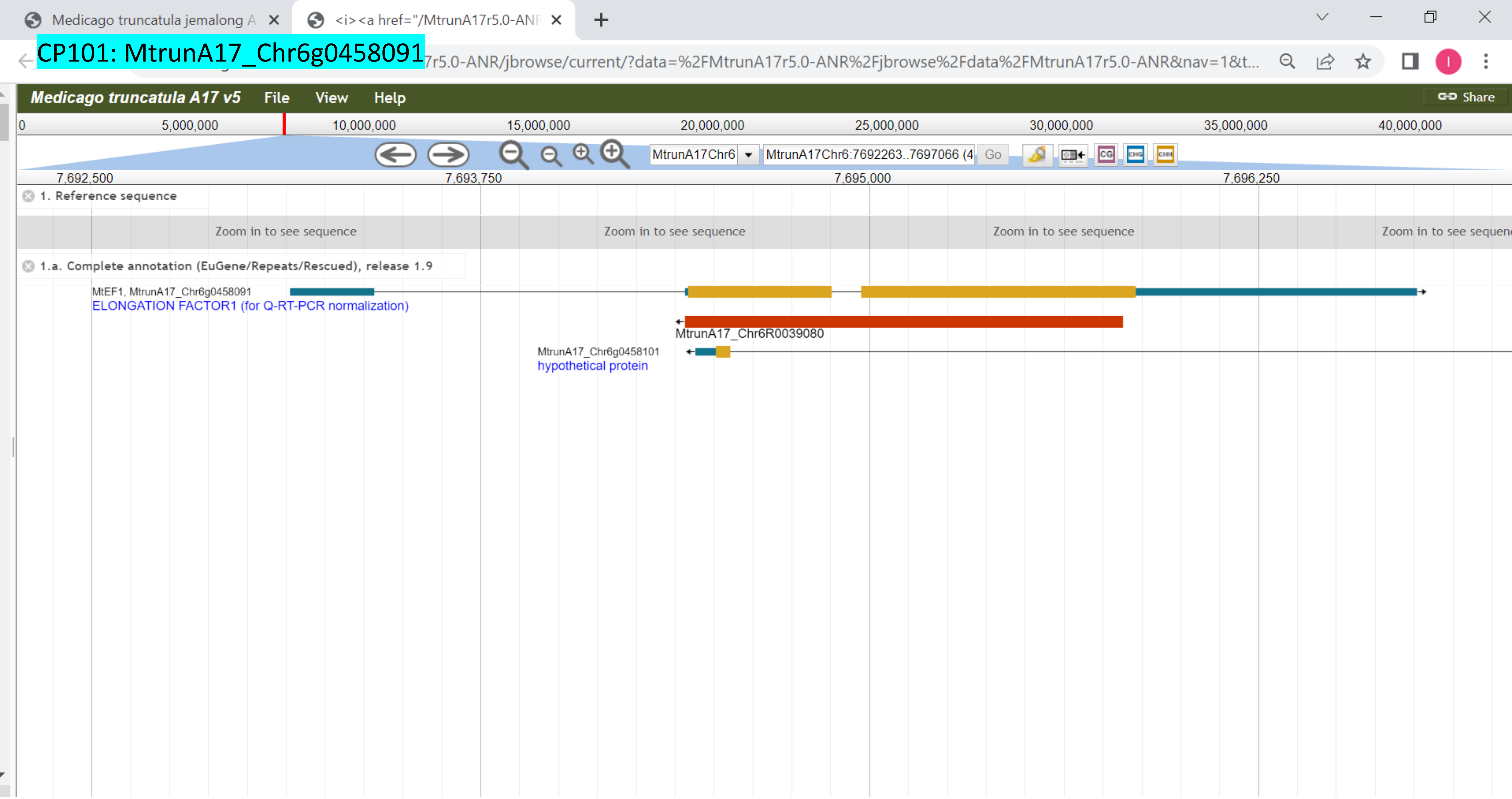

### CP102: MtrunA17_Chr6g0459481

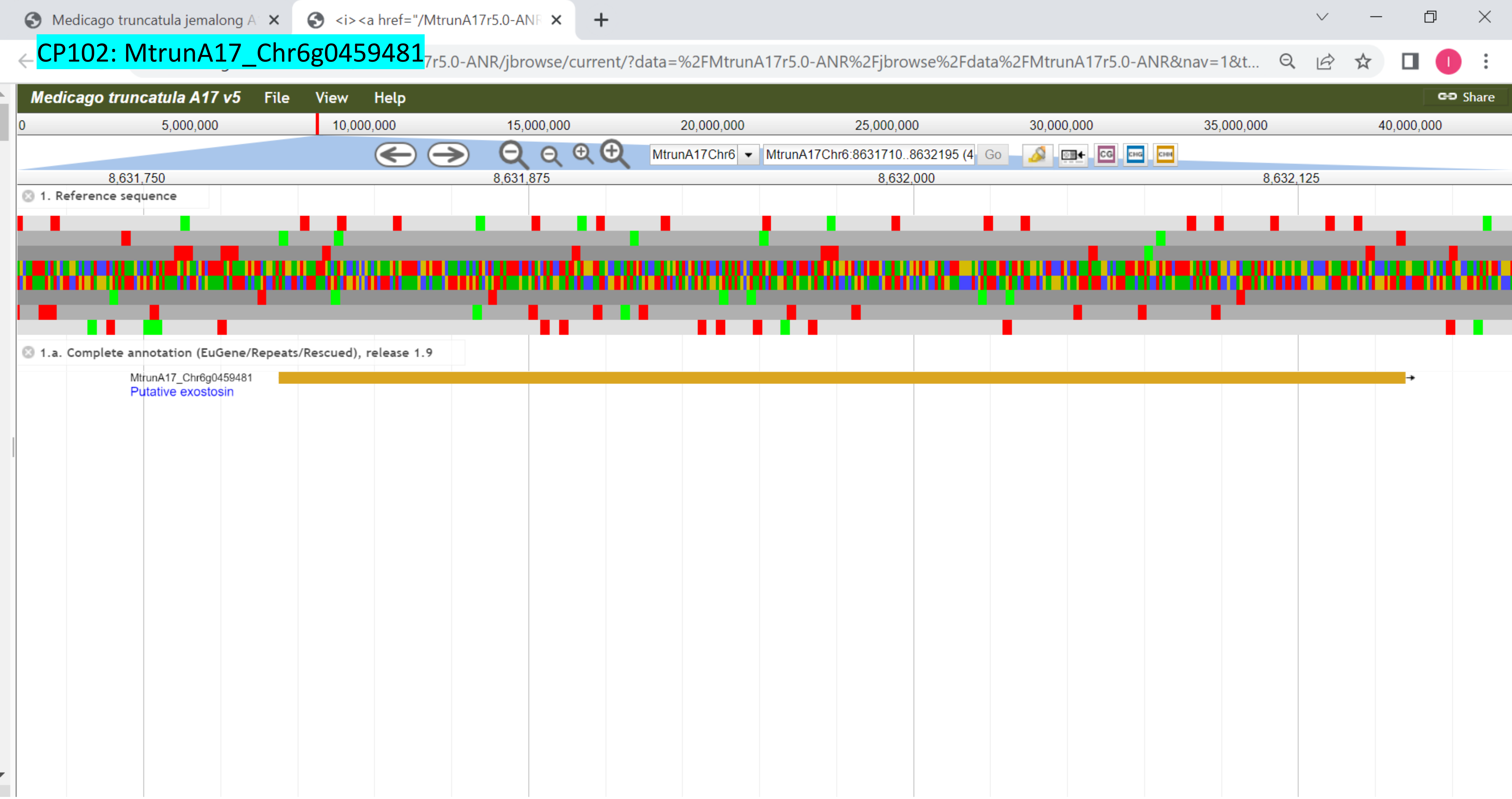

### CP103: MtrunA17_Chr6g0461931

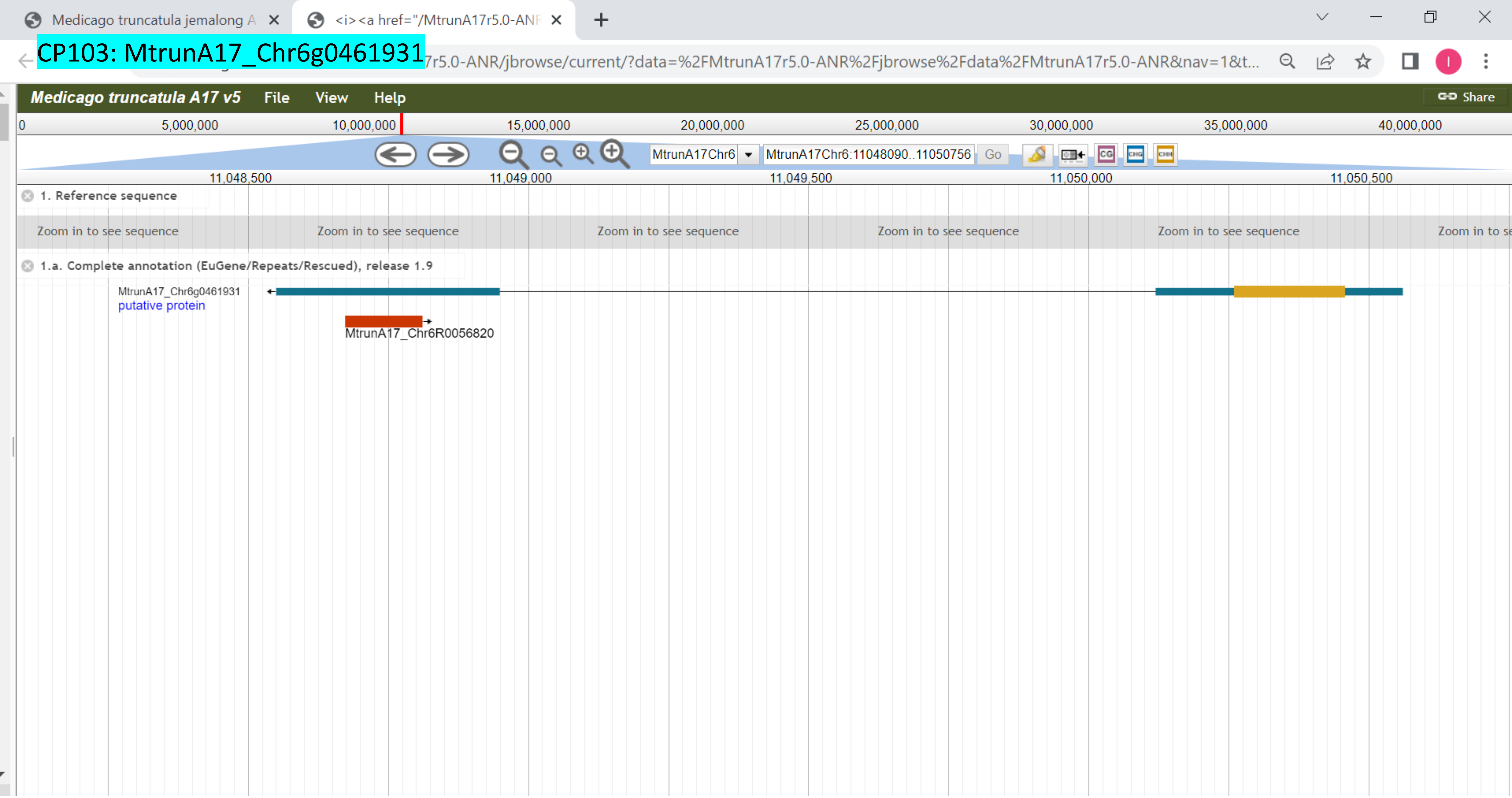

### CP104: MtrunA17_Chr6g0462271

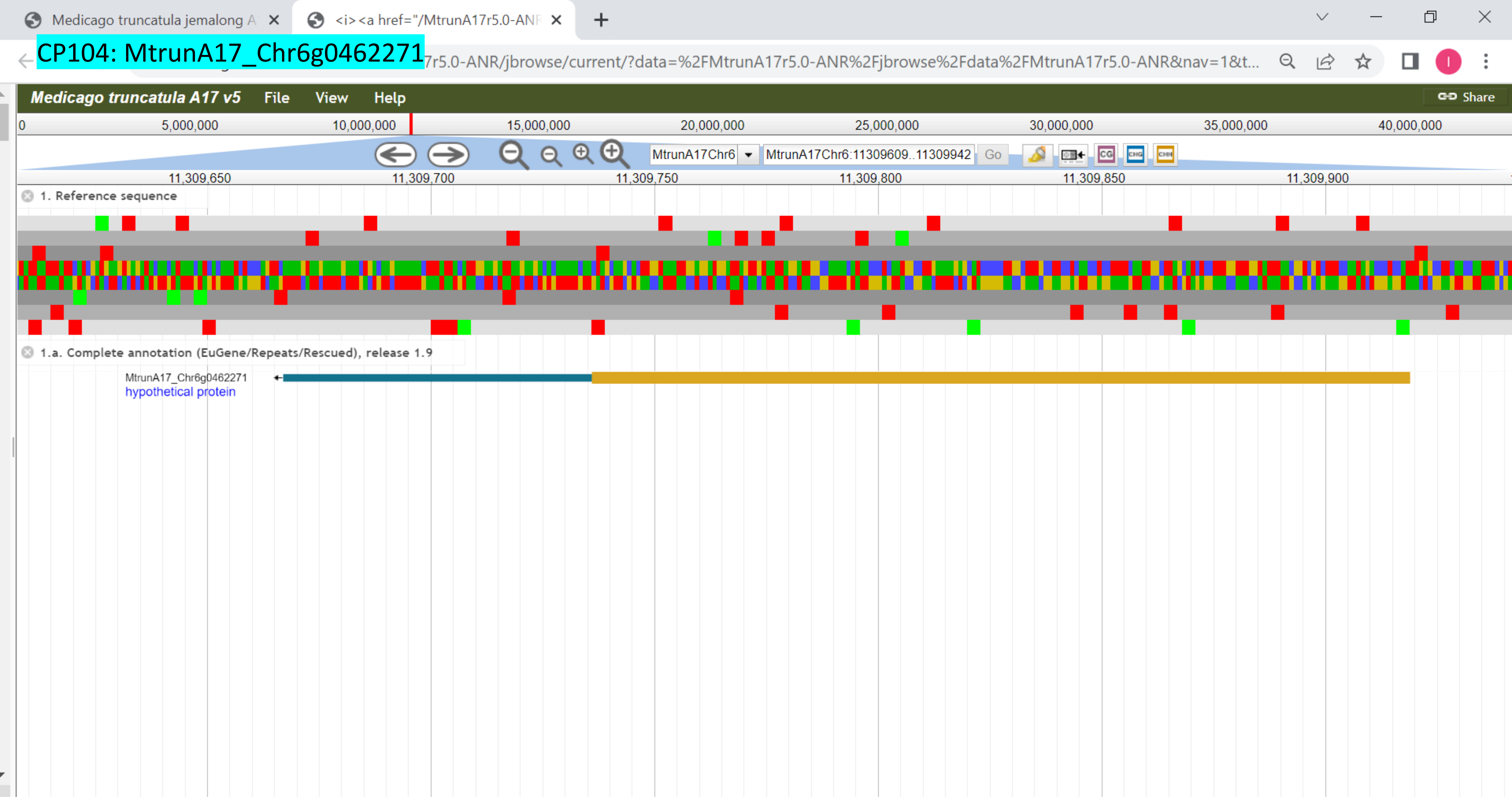

### CP105: MtrunA17_Chr6g0468411

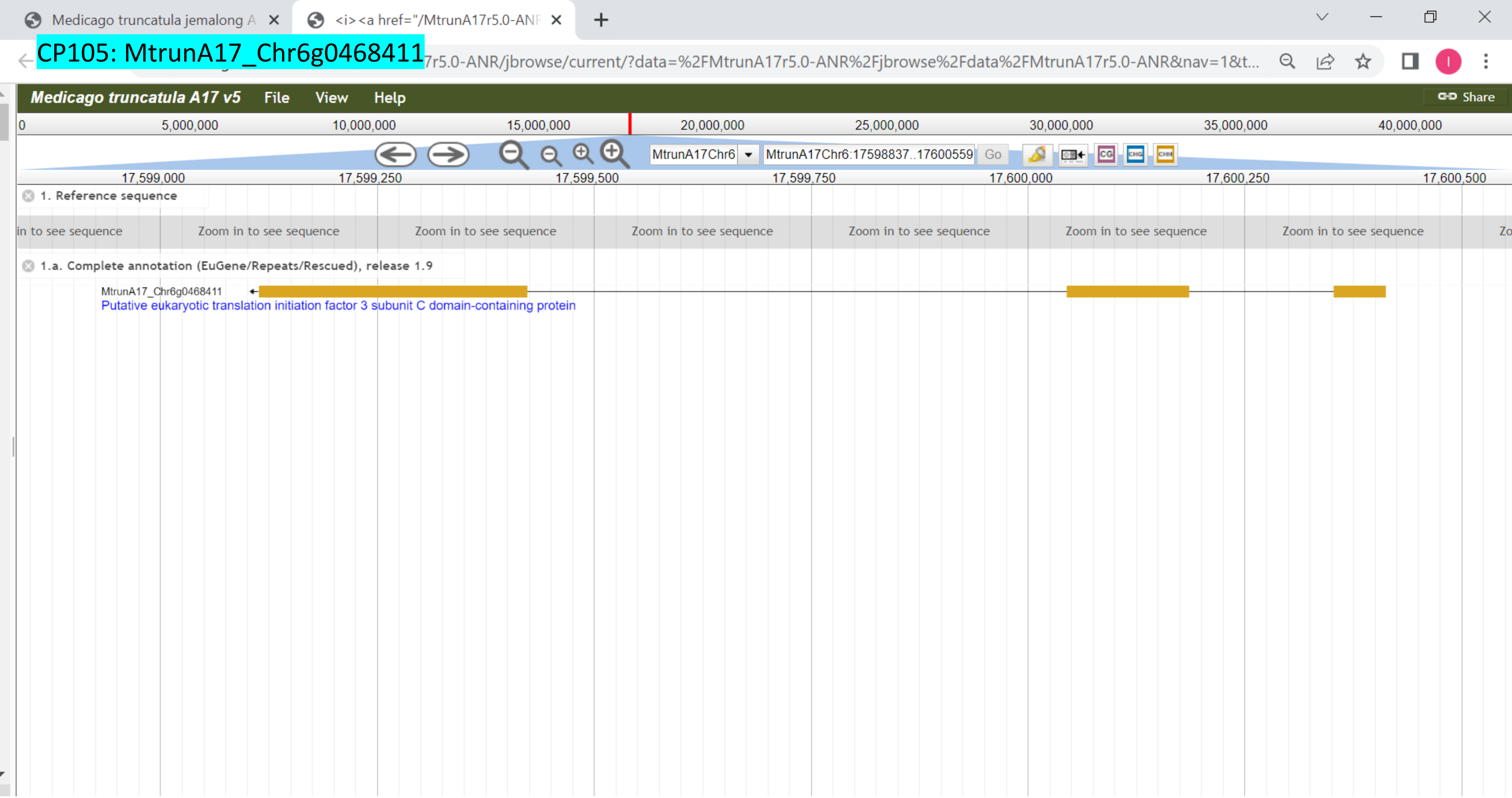

### CP106: MtrunA17_Chr6g0476751

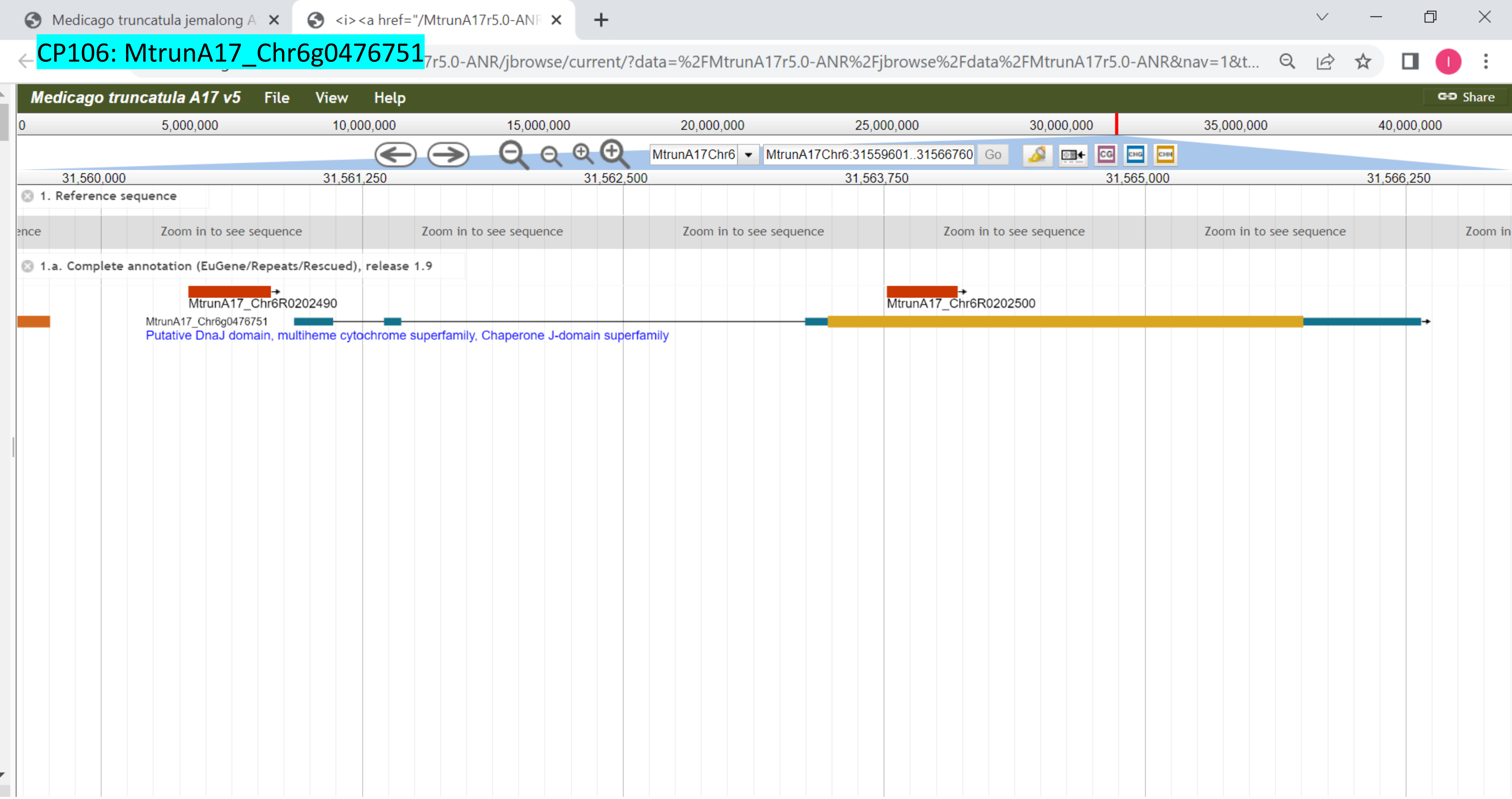

### CP107: MtrunA17_Chr6g0479001

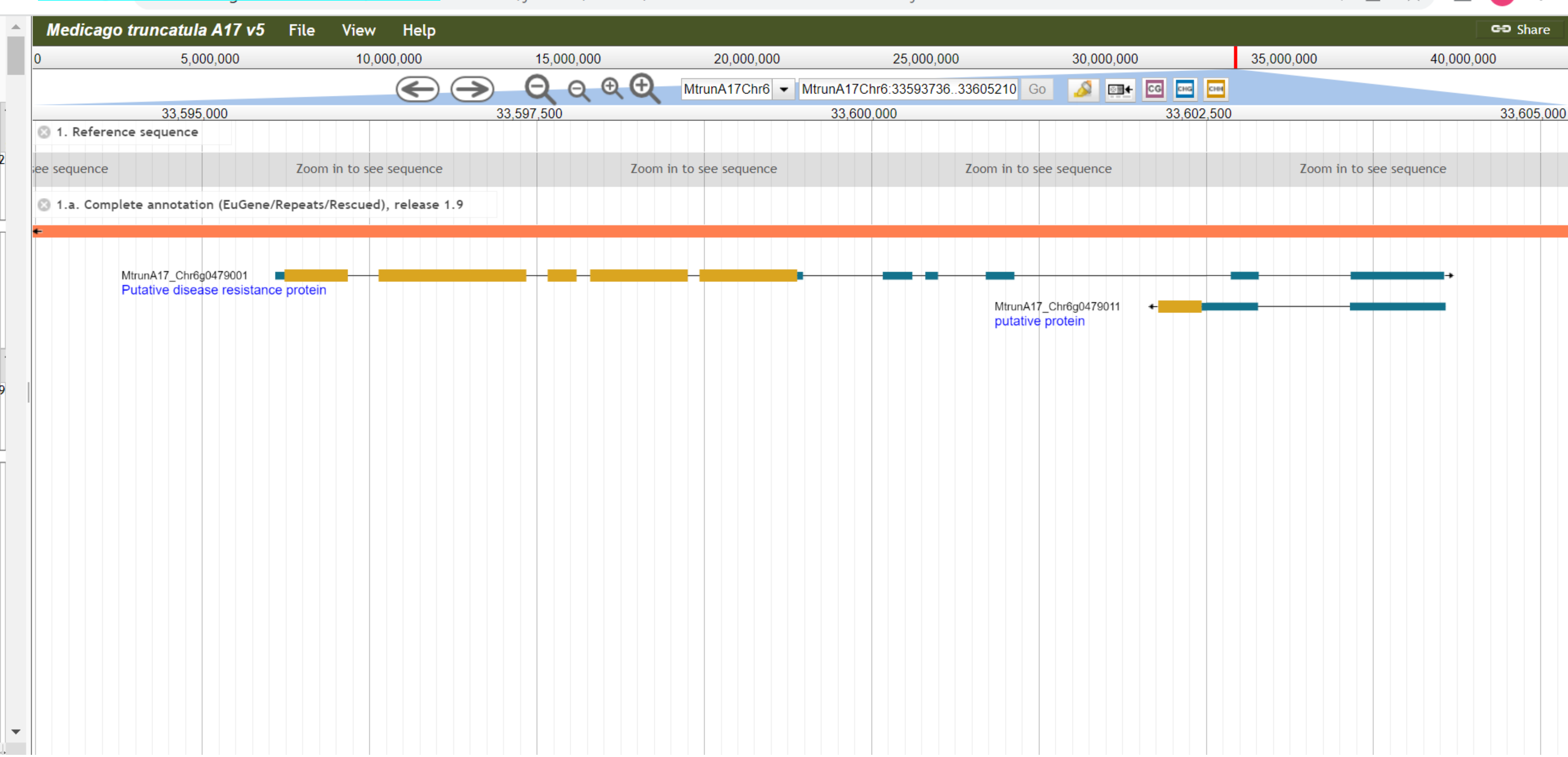

CP108: MtrunA17_Chr6g0485321

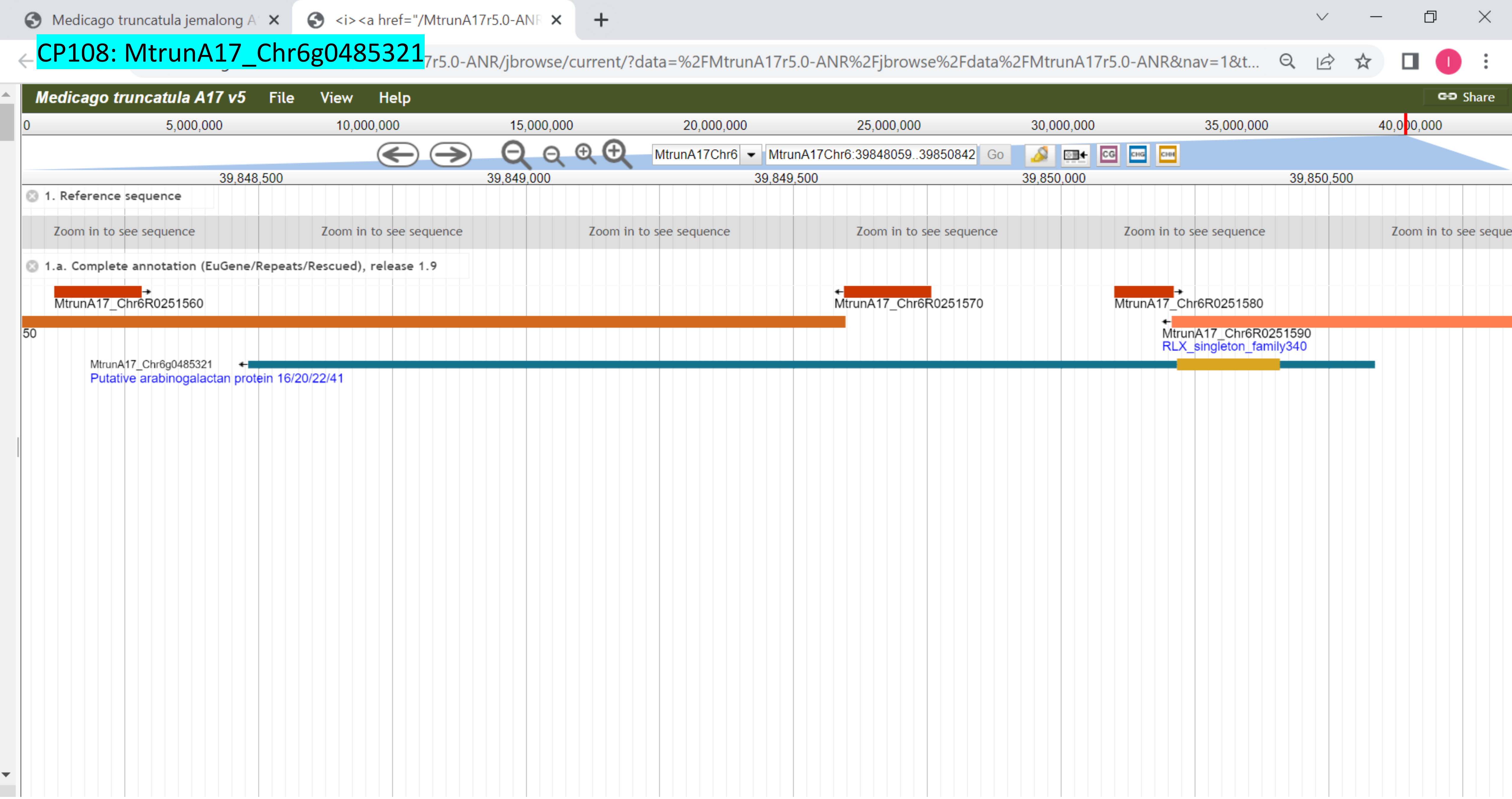

CP109: MtrunA17_Chr6g0486961

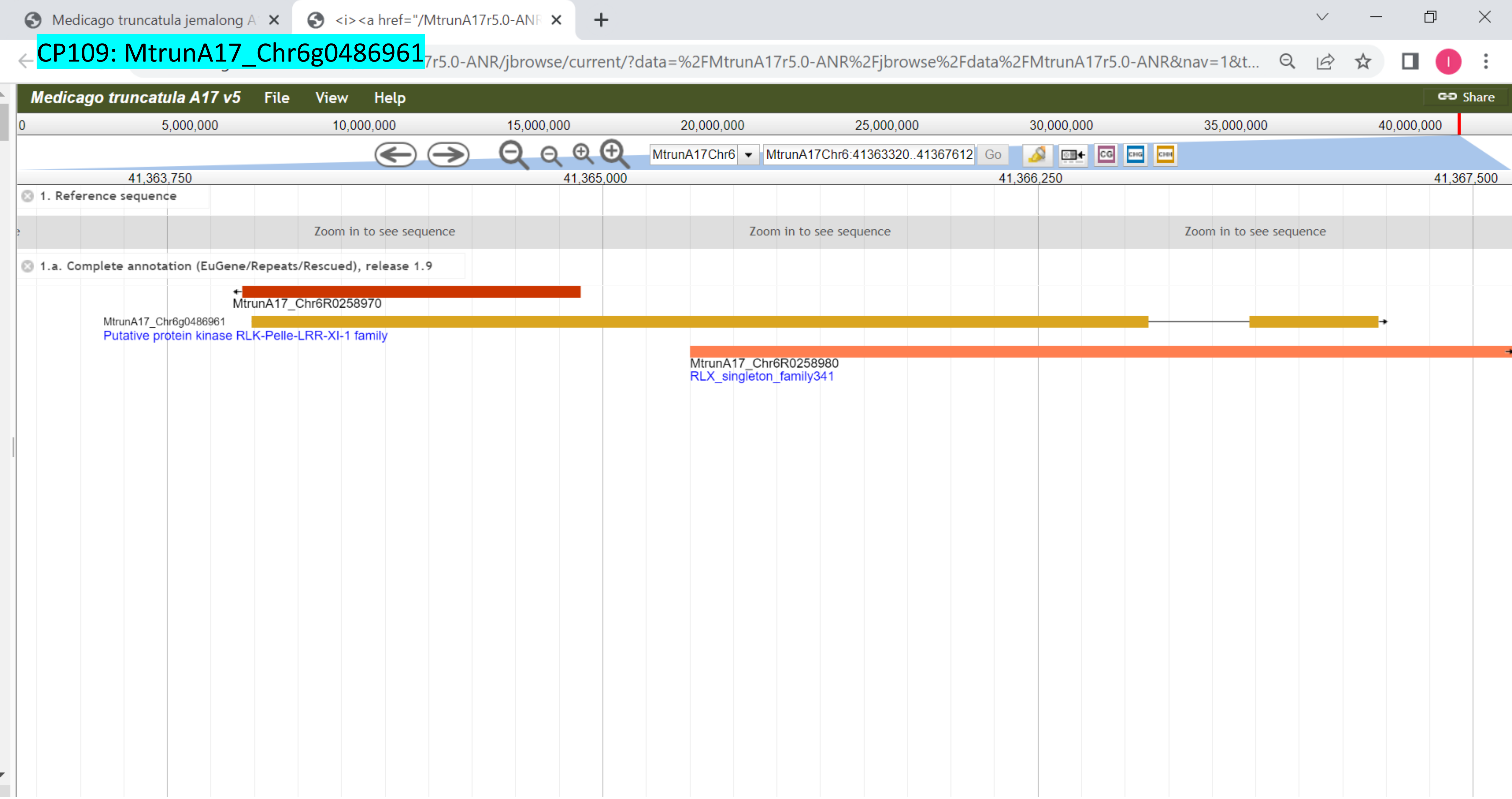

CP110: MtrunA17_Chr7g0214741

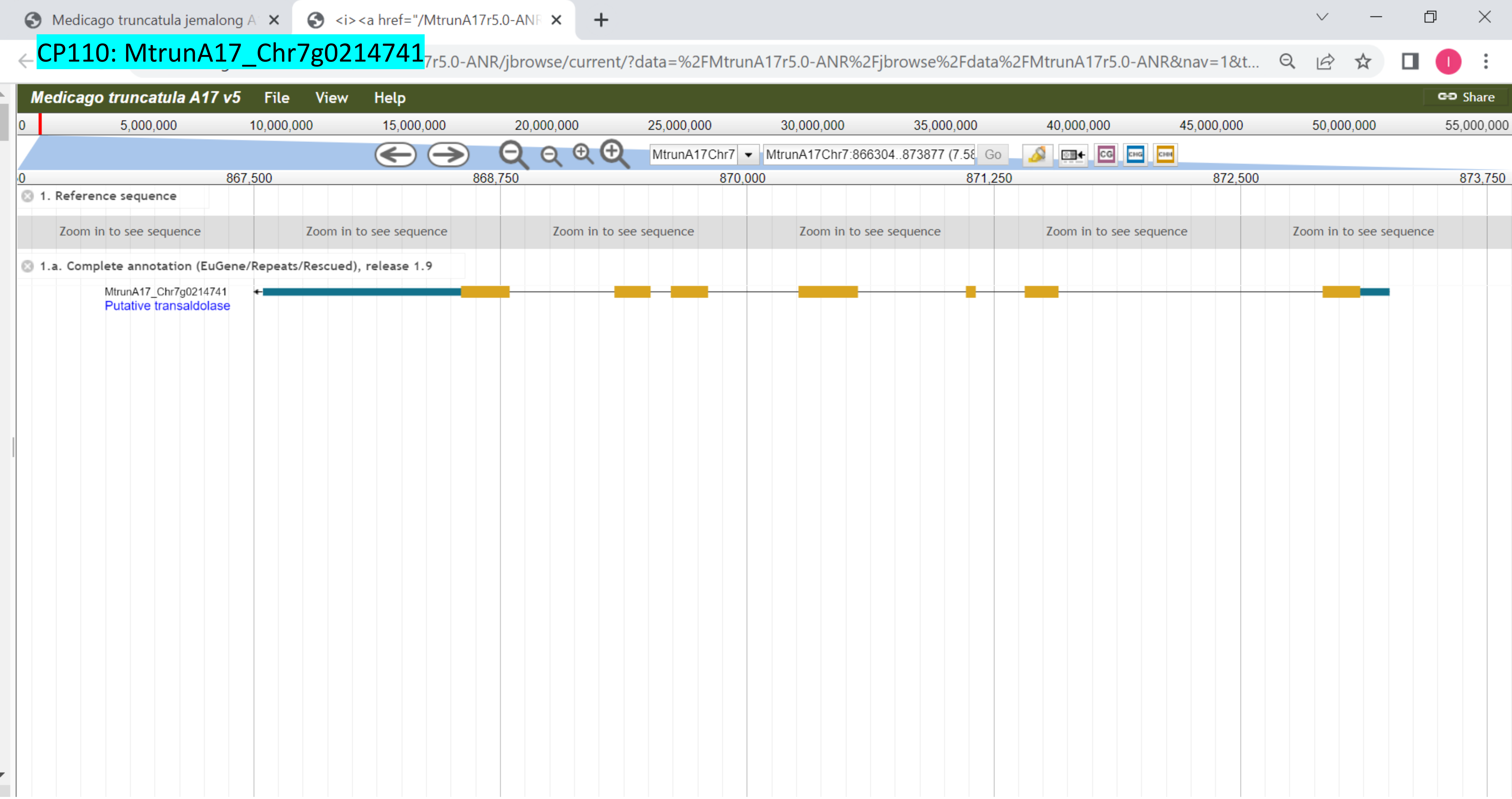

CP111: MtrunA17_Chr7g0214911

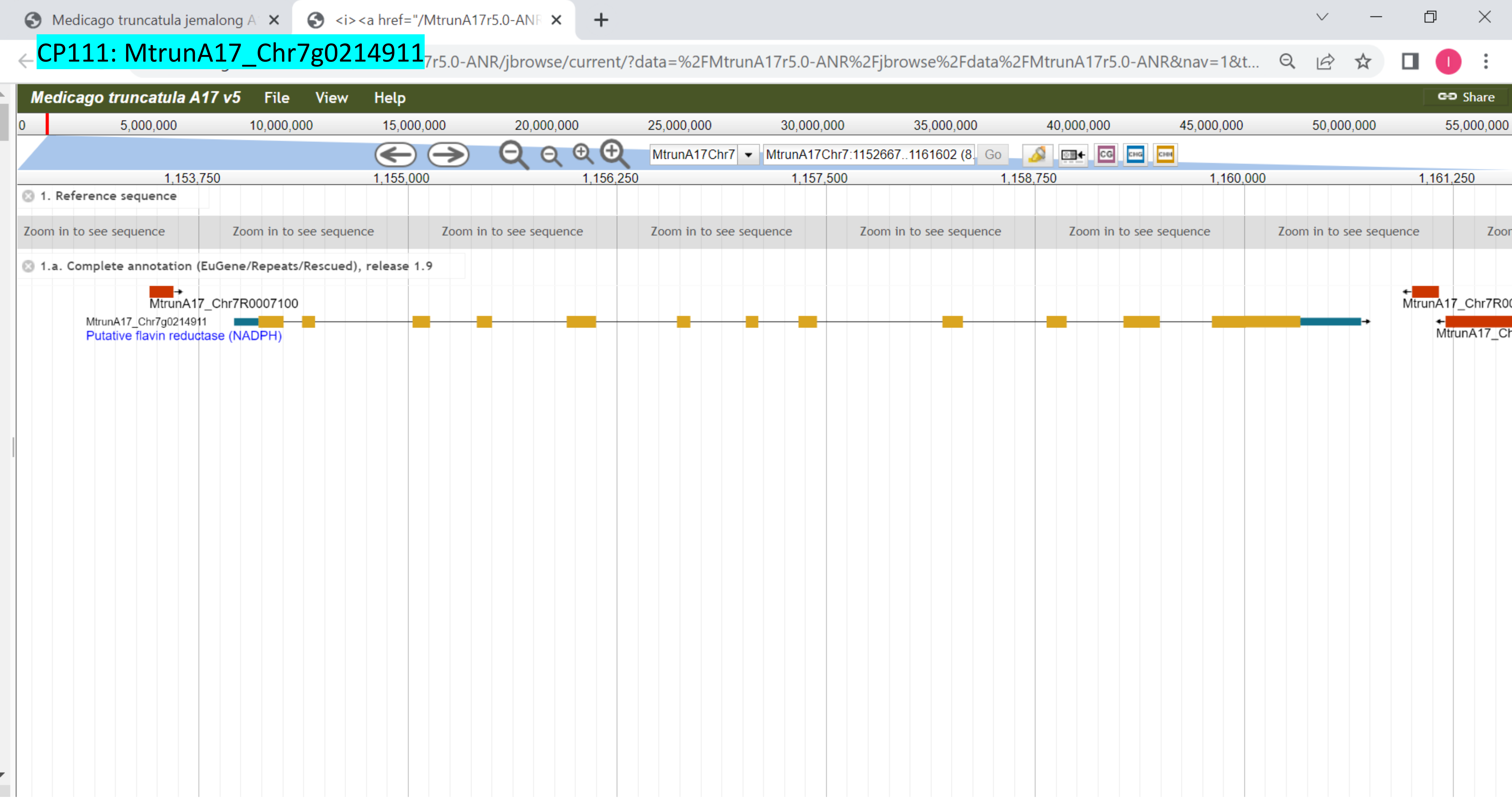

### CP112: MtrunA17_Chr7g0219691

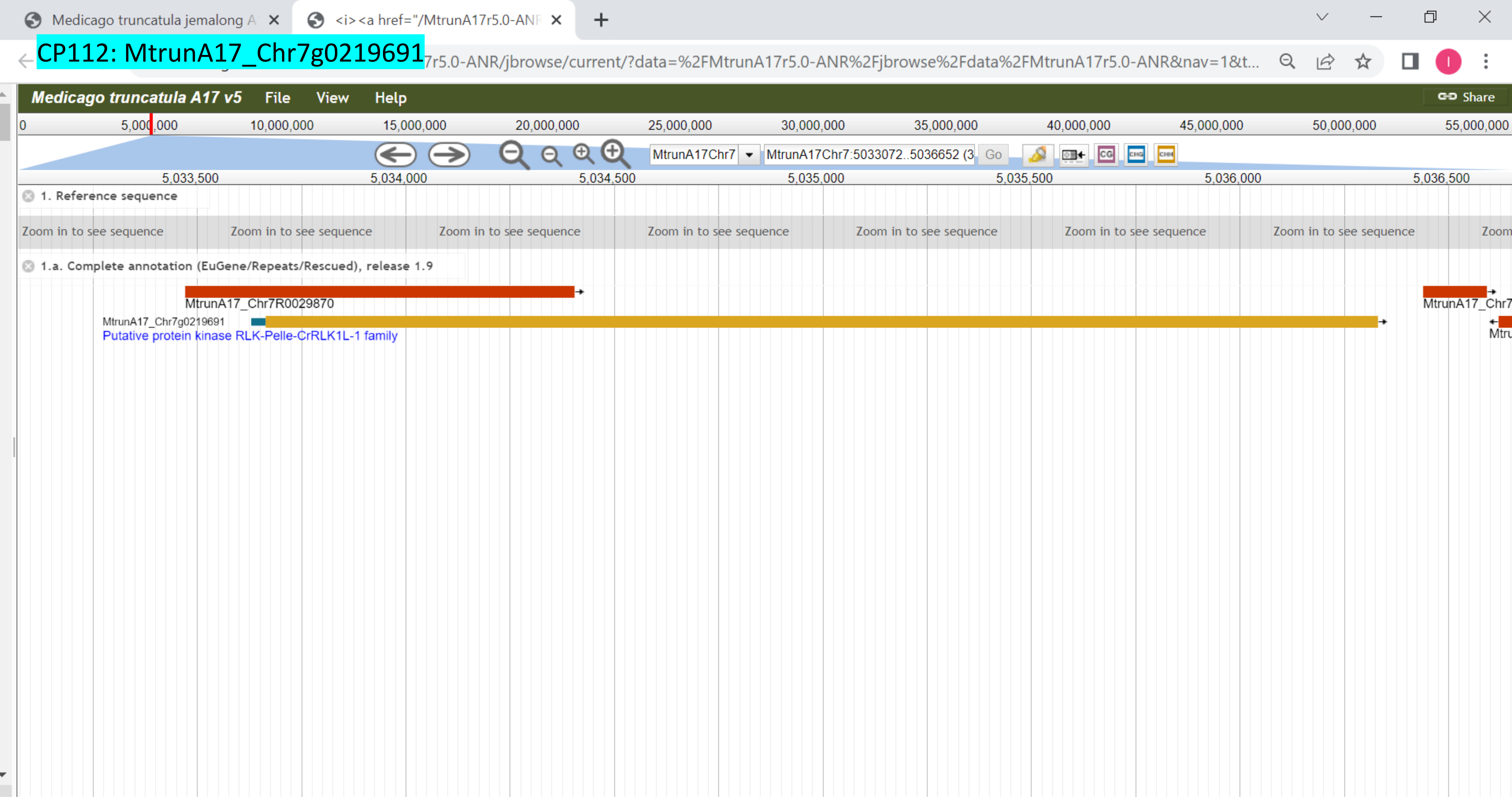

### CP113: MtrunA17_Chr7g0221631

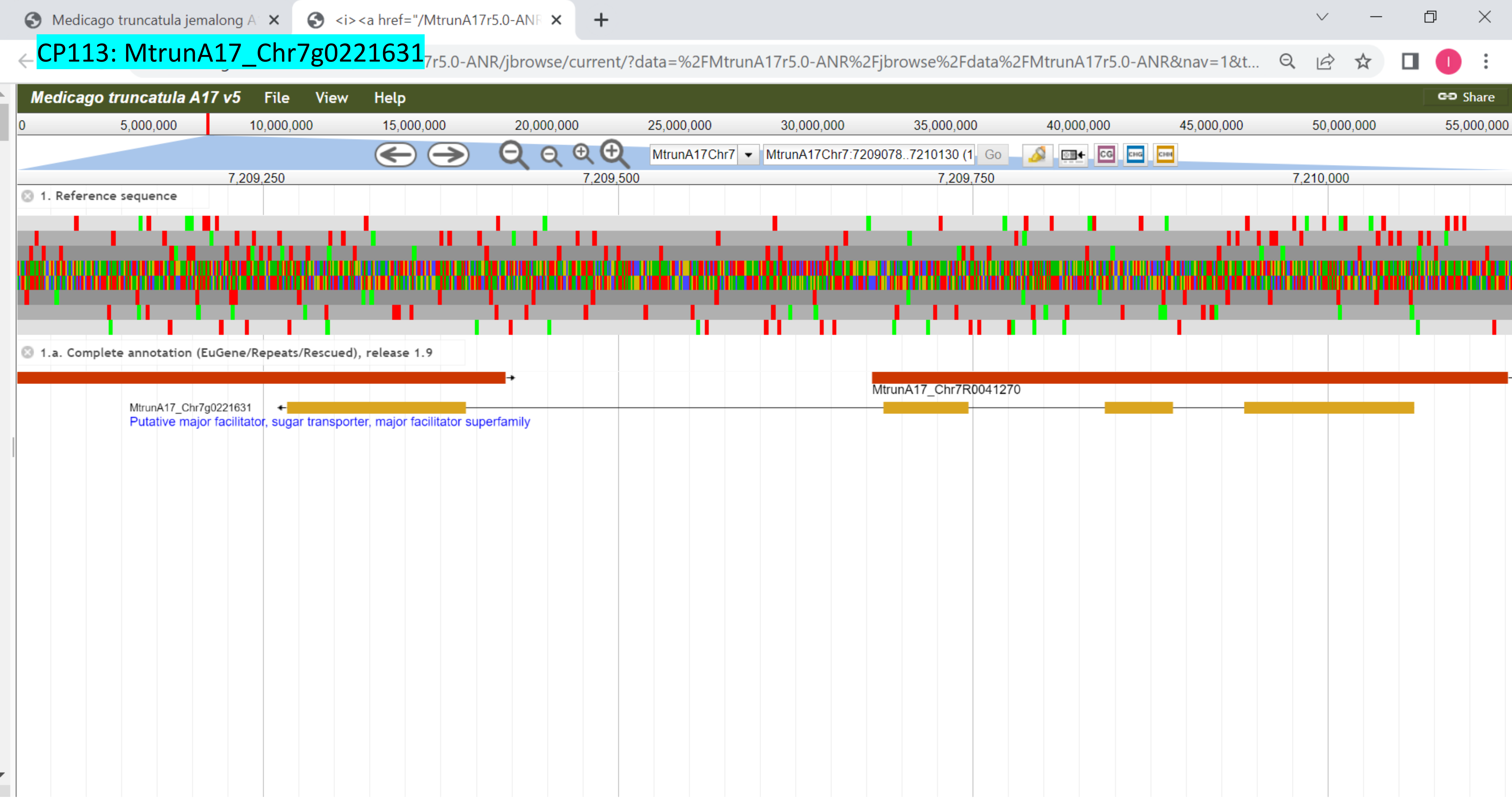

CP114: MtrunA17_Chr7g0229401

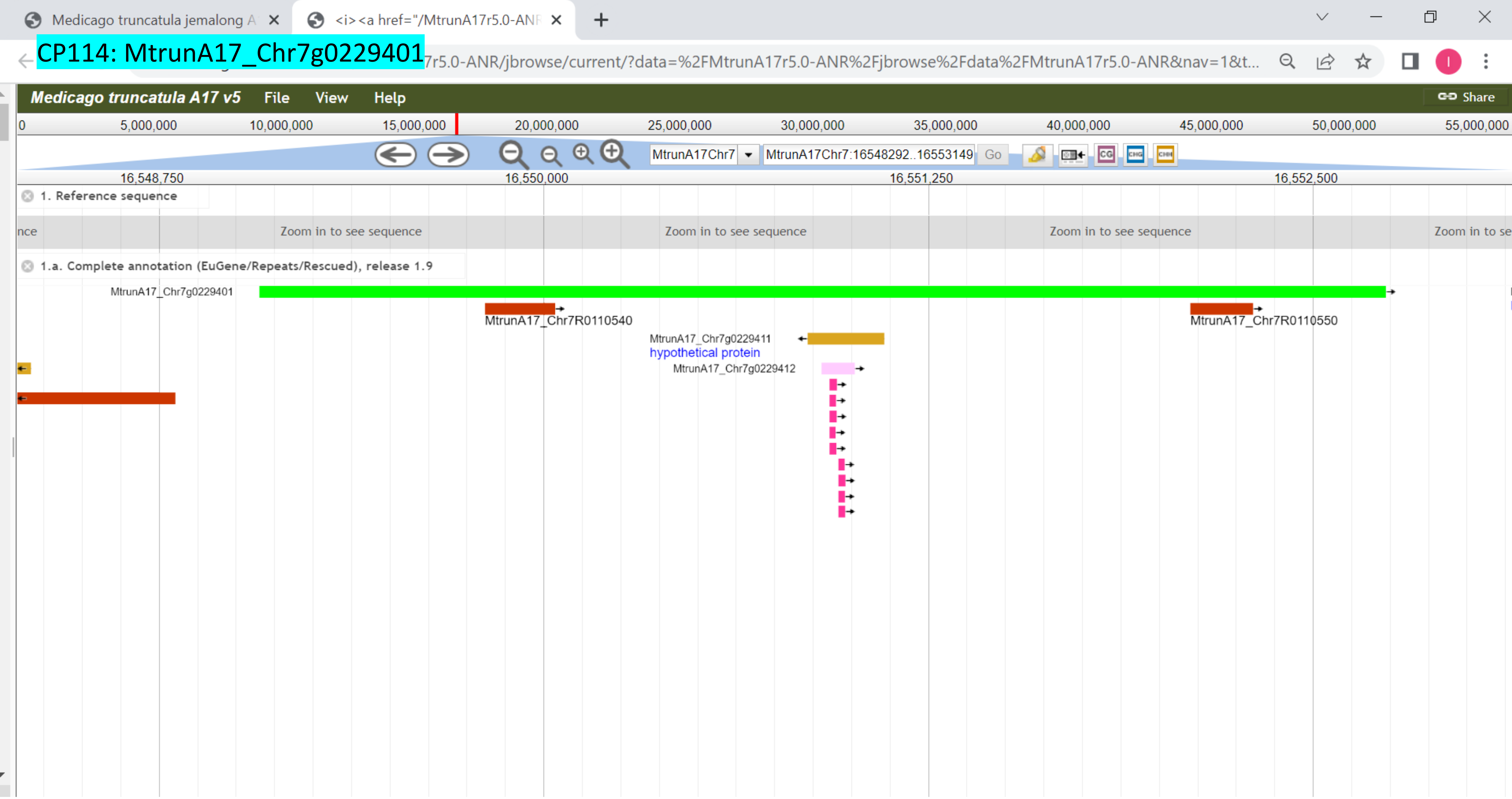

CP115: MtrunA17_Chr7g0230341

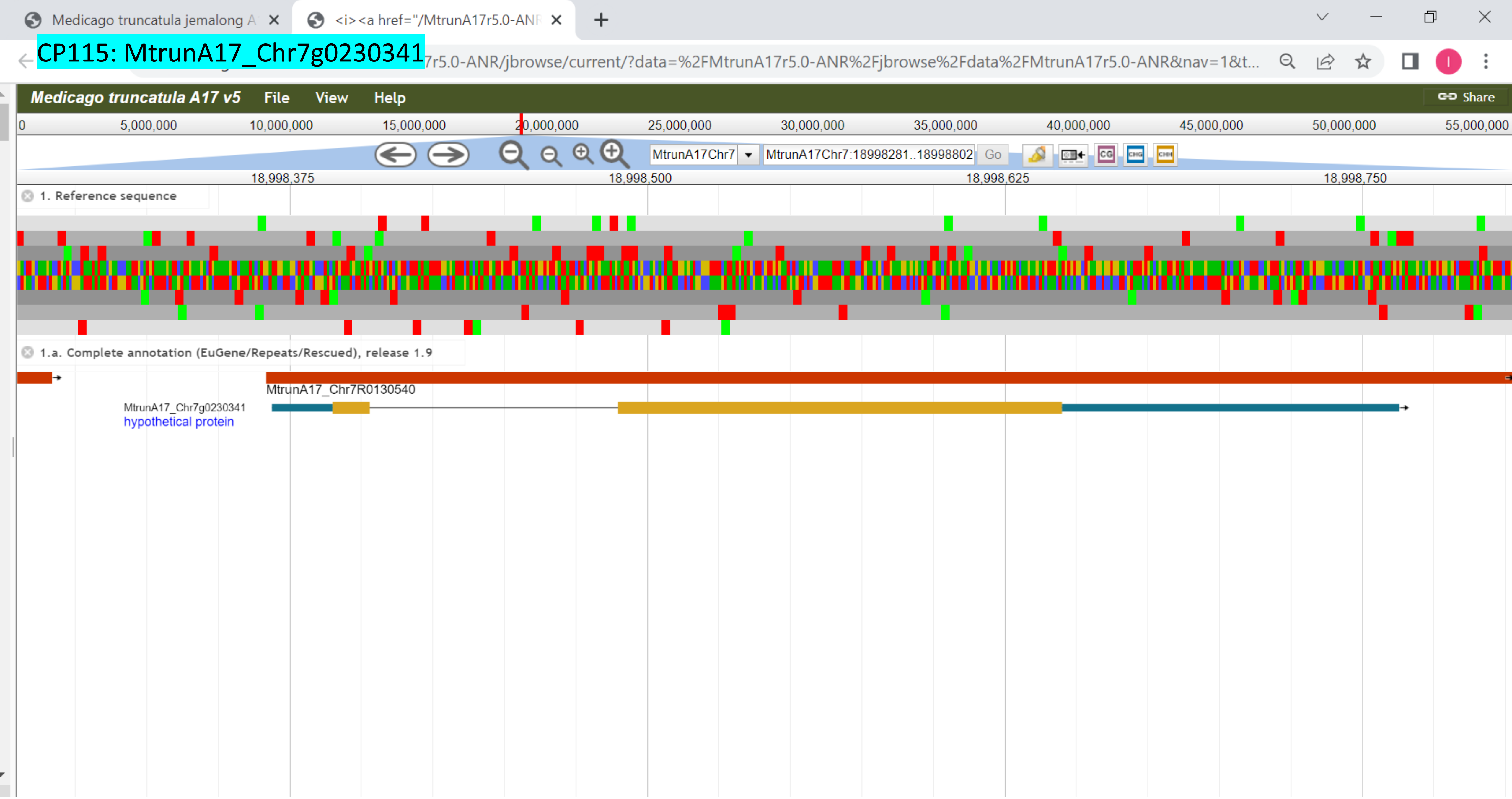

CP116: MtrunA17_Chr7g0232851

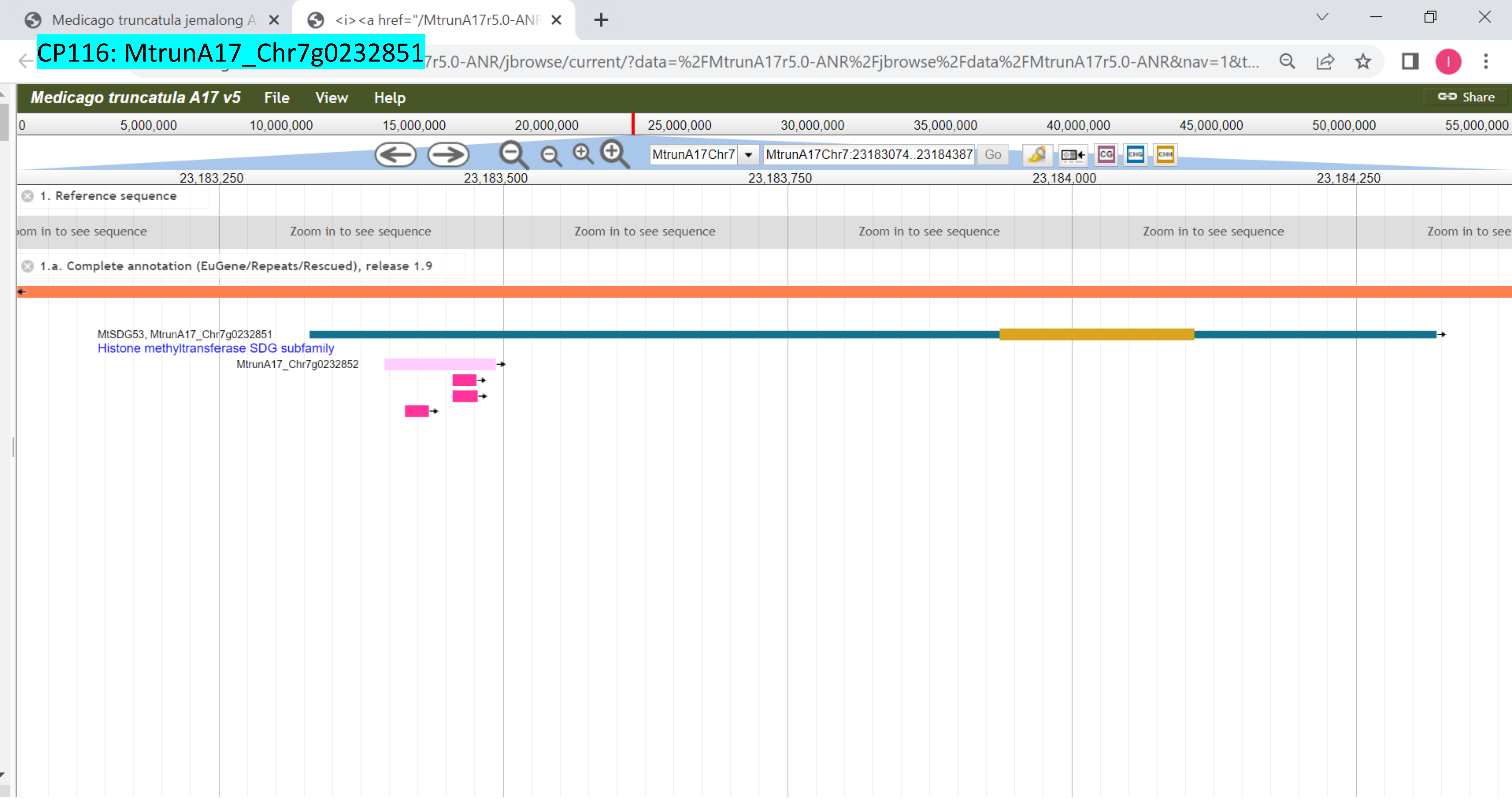

### CP117: MtrunA17_Chr7g0237331

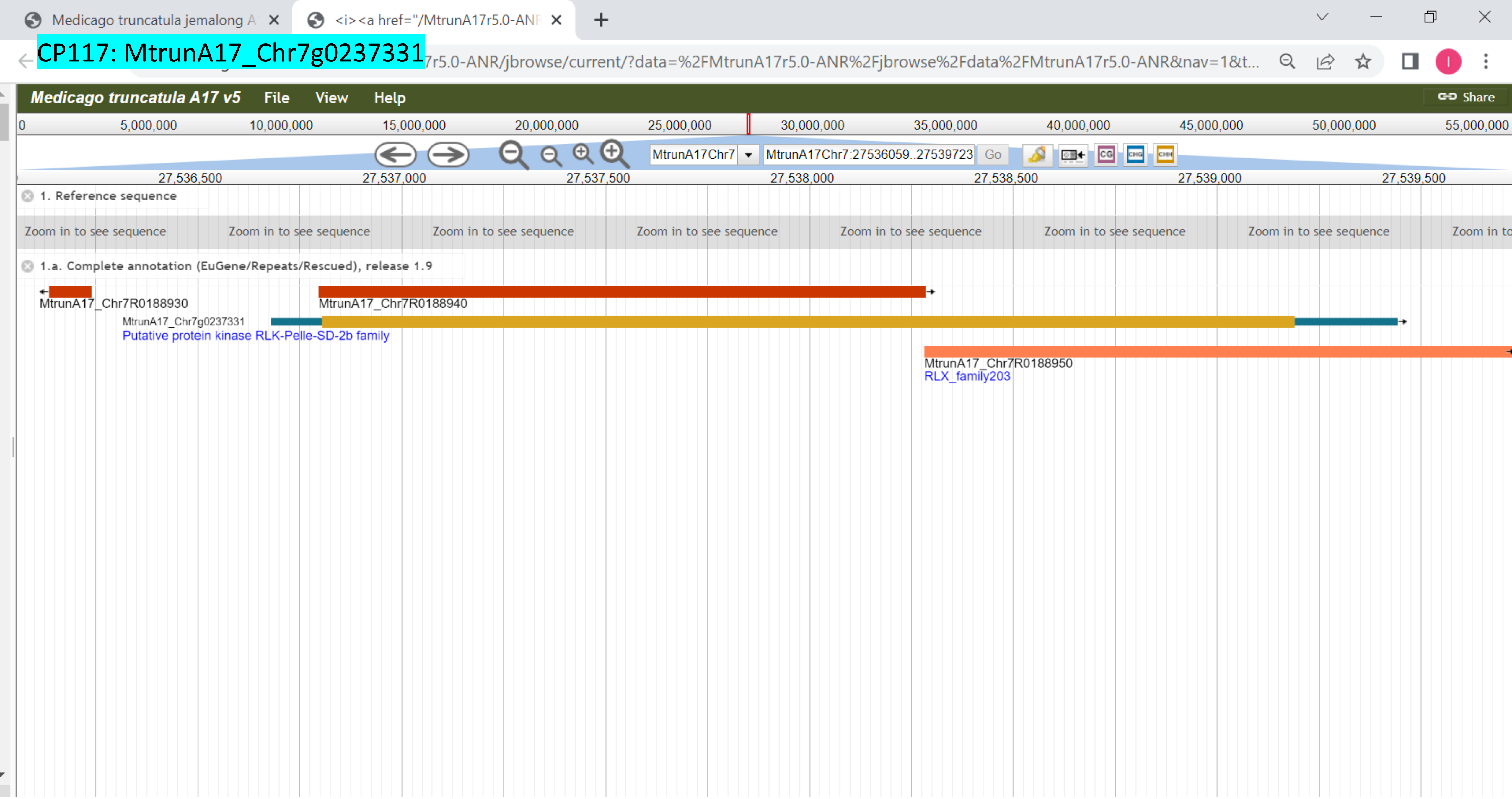

CP118: MtrunA17_Chr7g0251971

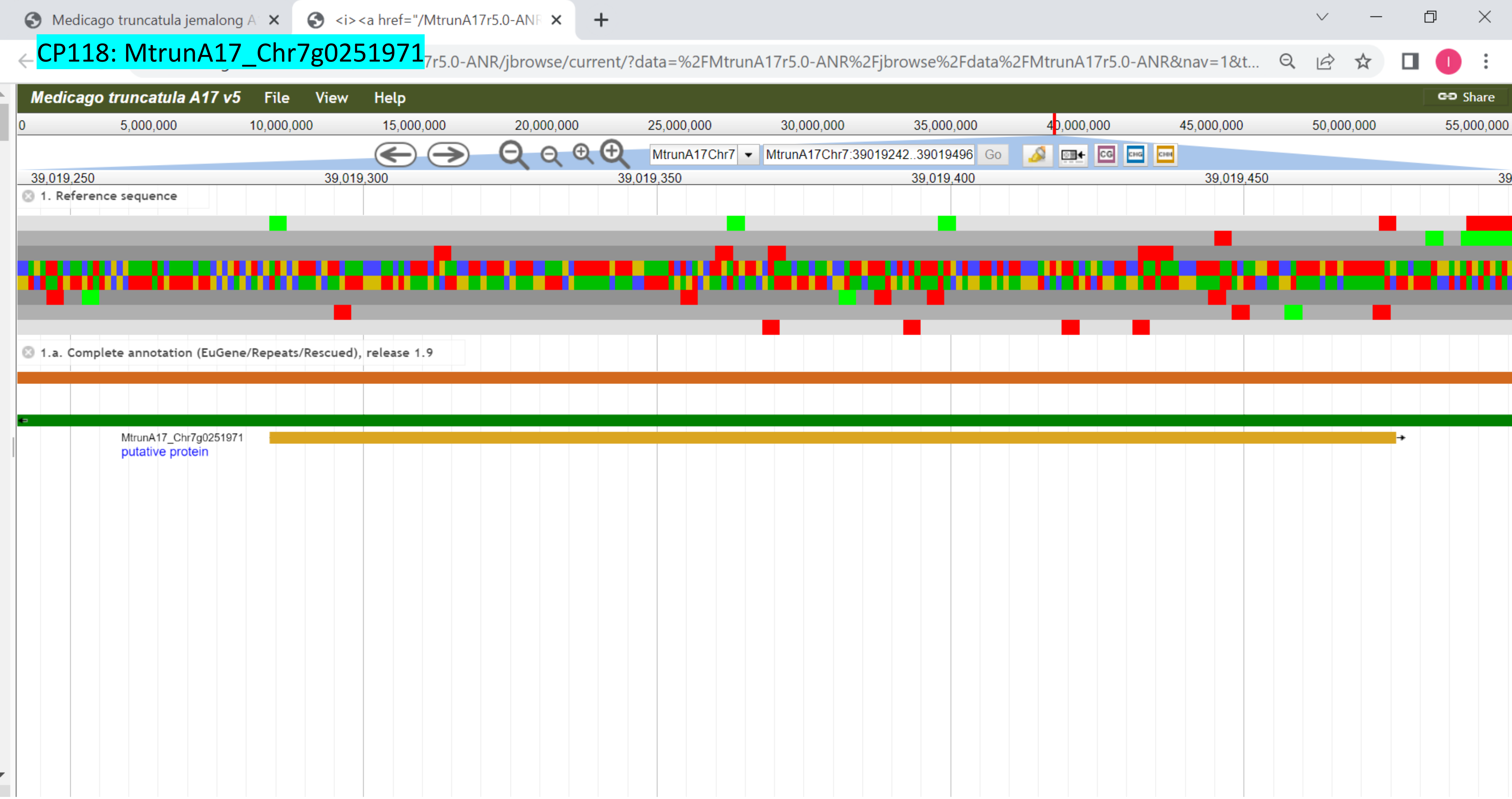

### CP119: MtrunA17_Chr7g0259611

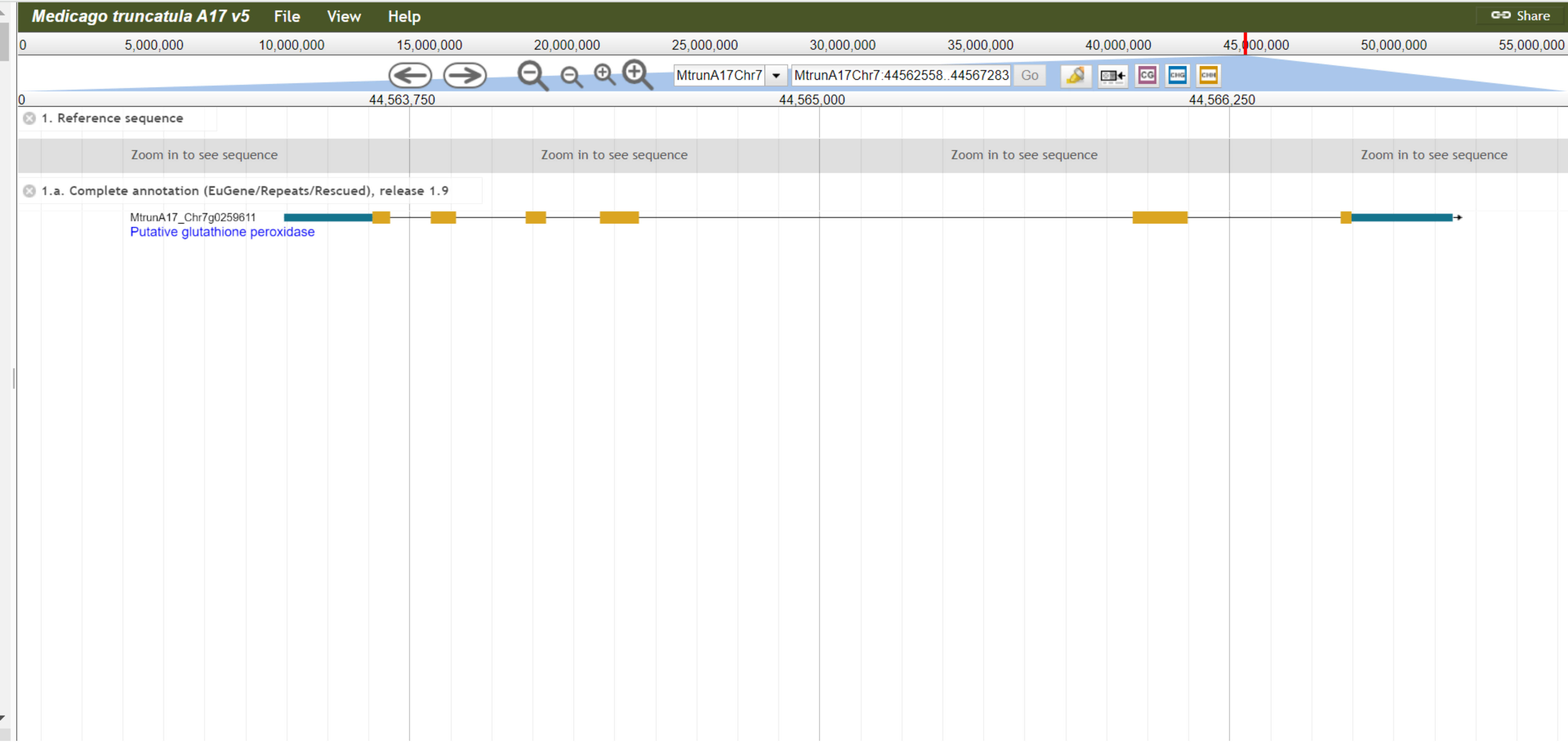

CP120: MtrunA17_Chr7g0262821

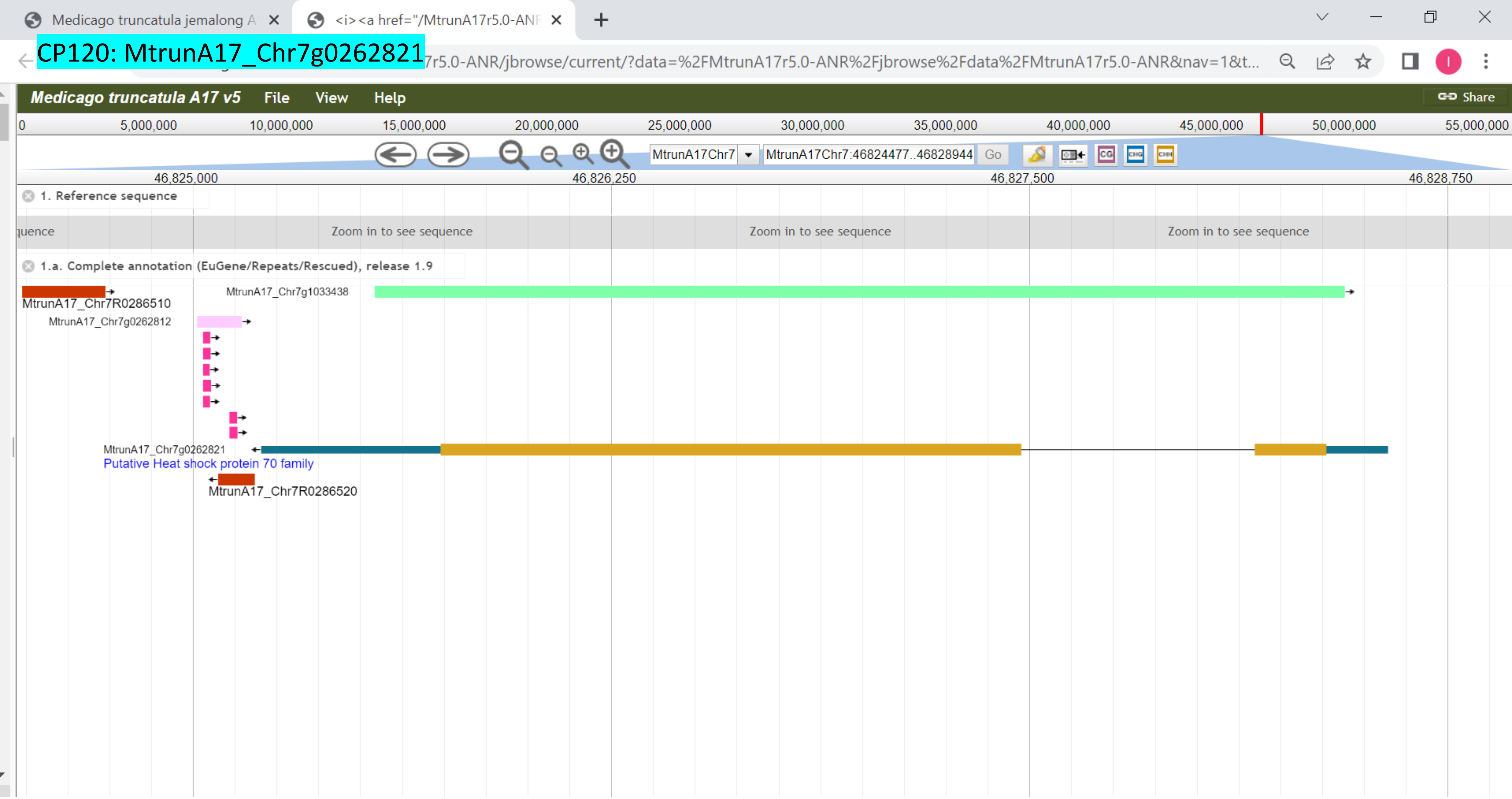

### CP121: MtrunA17_Chr7g0270811

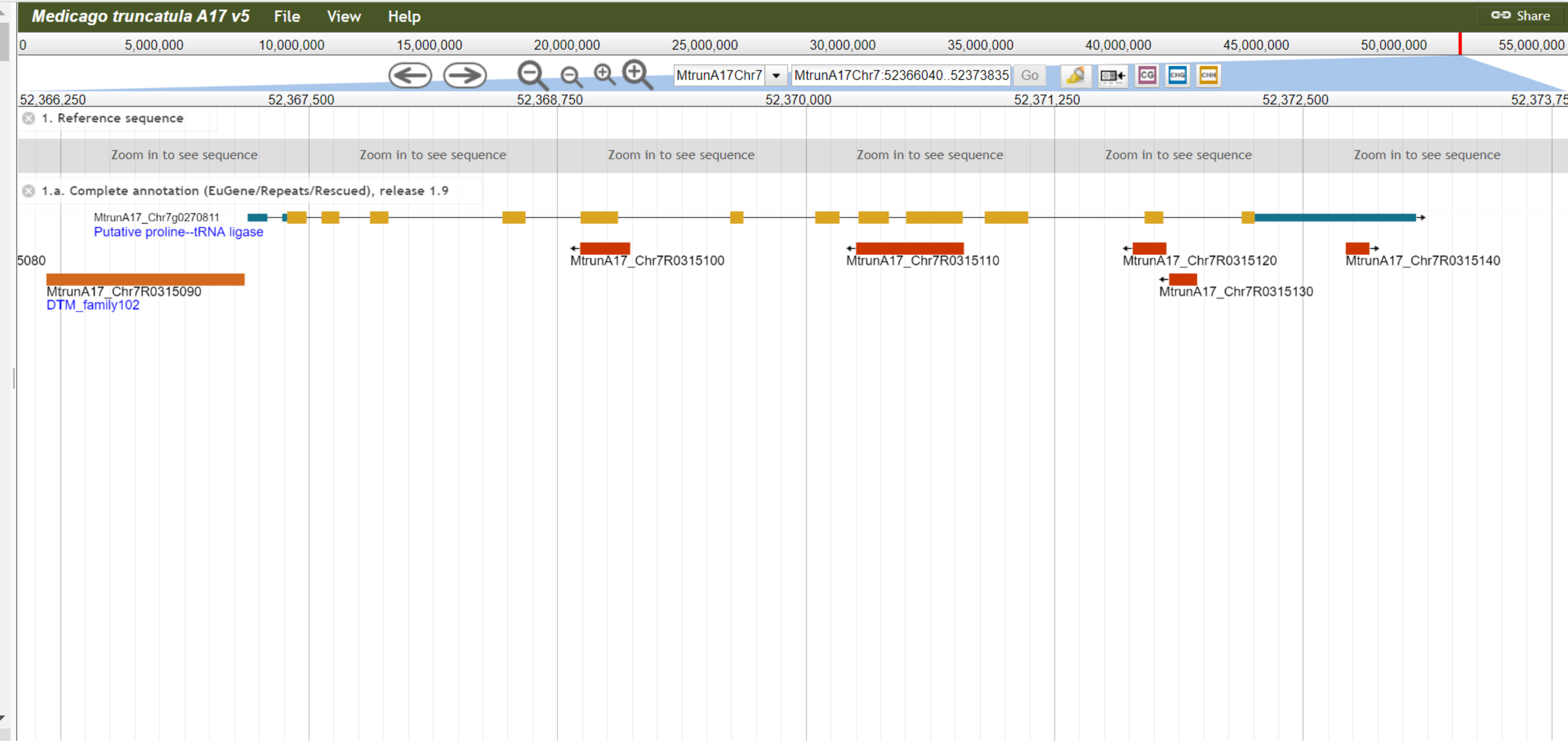

CP122: MtrunA17_Chr7g1034306

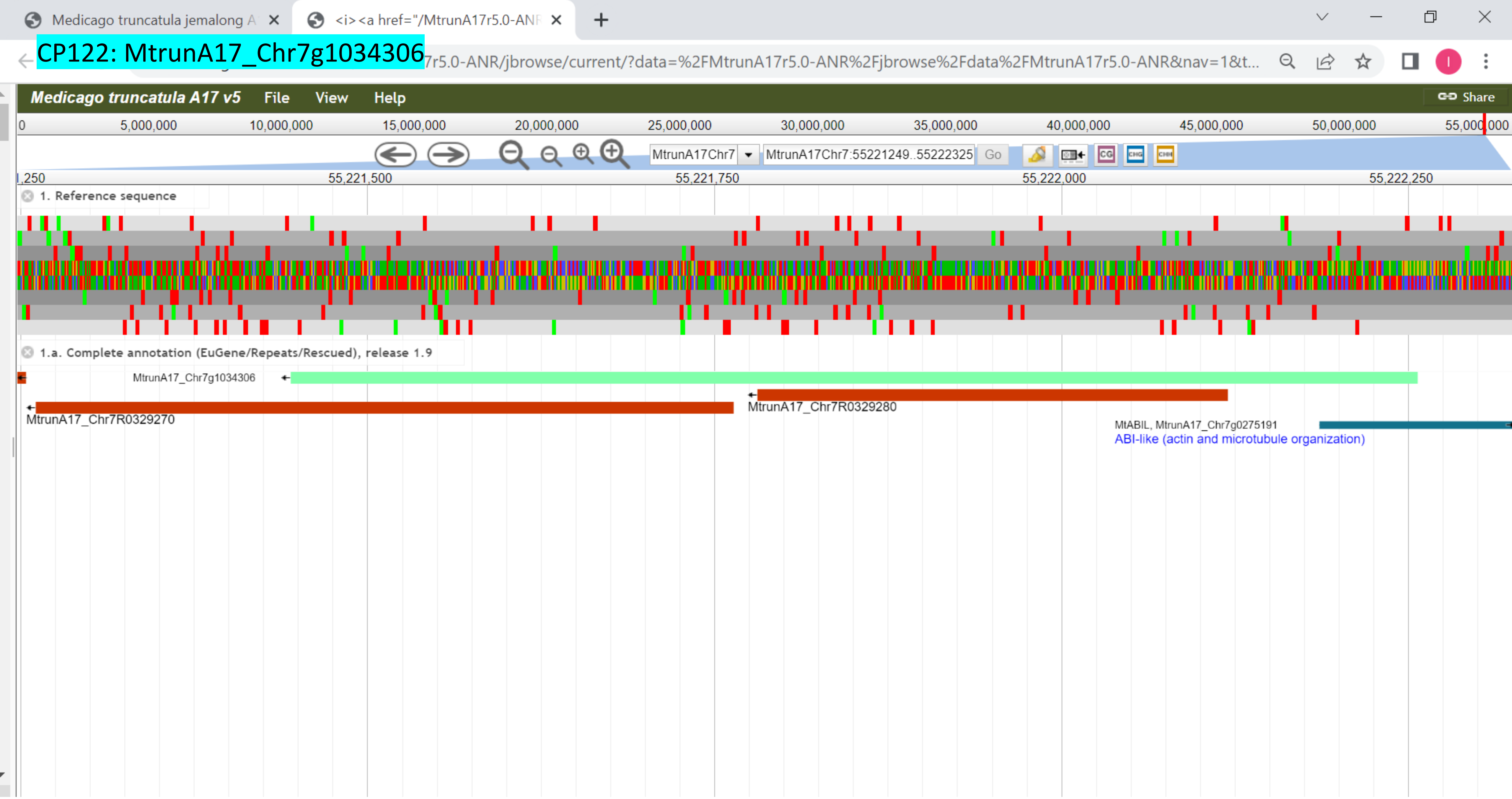

CP123: MtrunA17_Chr8g0338111

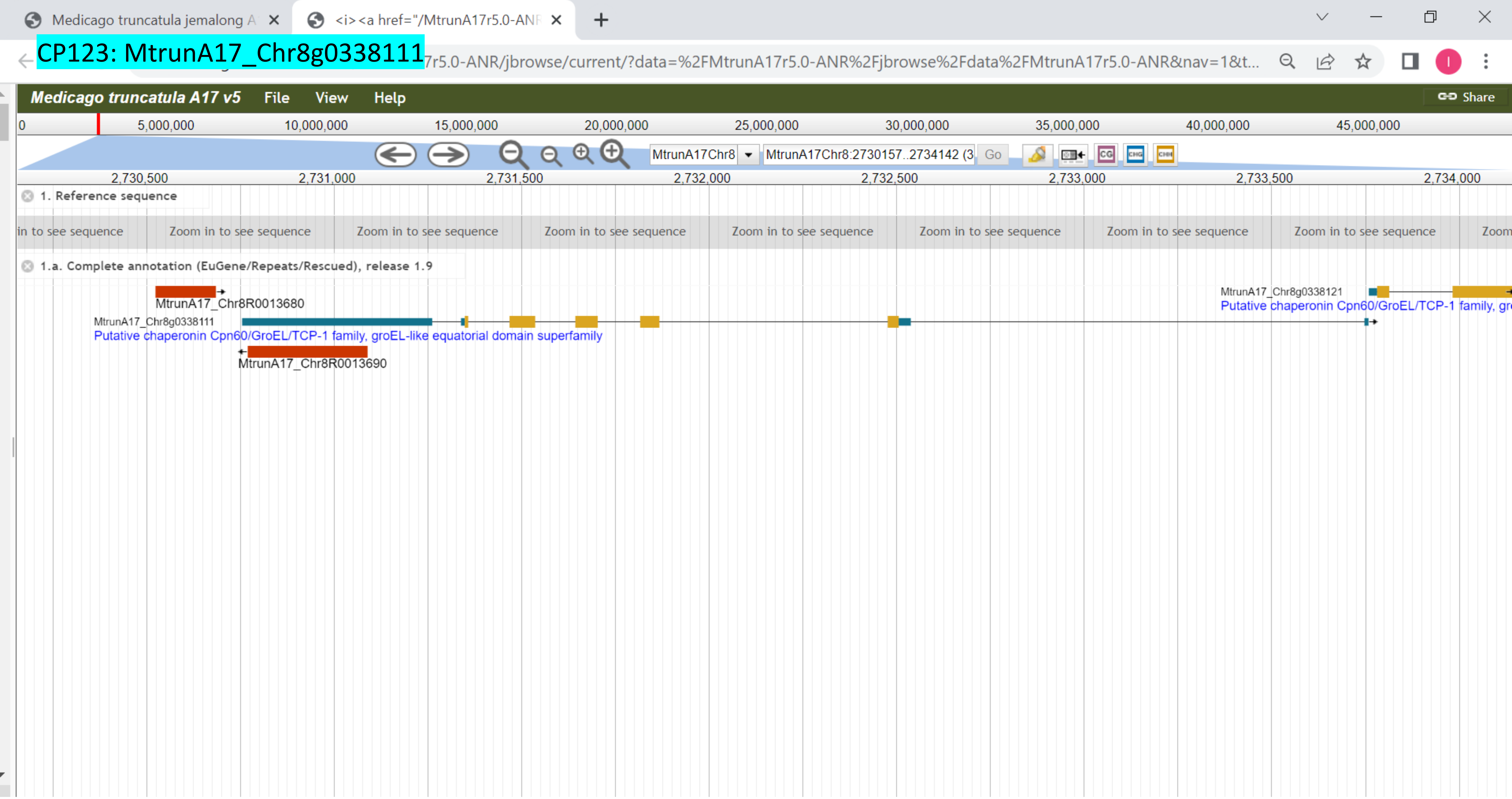

CP124: MtrunA17_Chr8g0338301

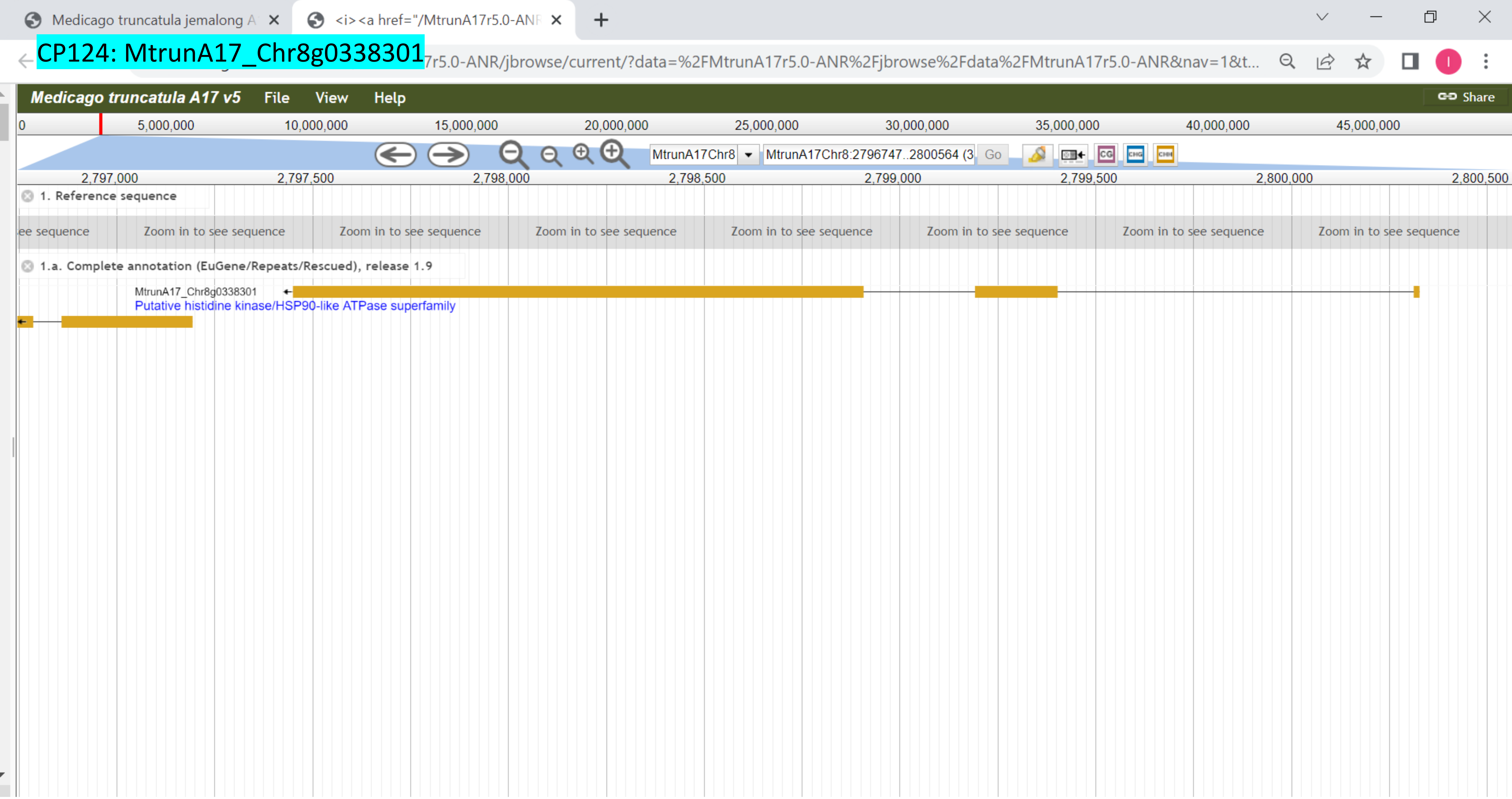

### CP125: MtrunA17_Chr8g0339891

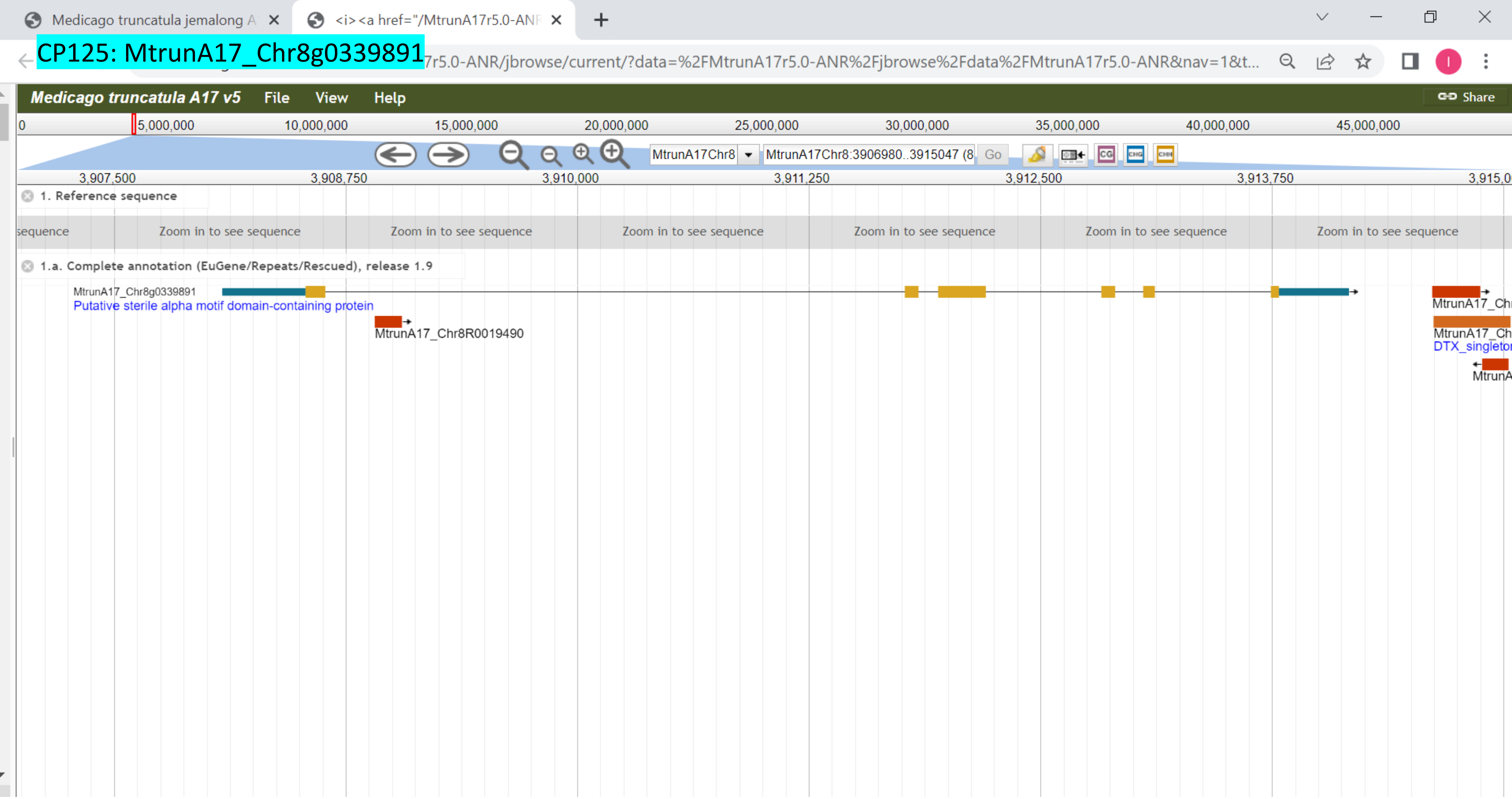

### CP126: MtrunA17_Chr8g0342881

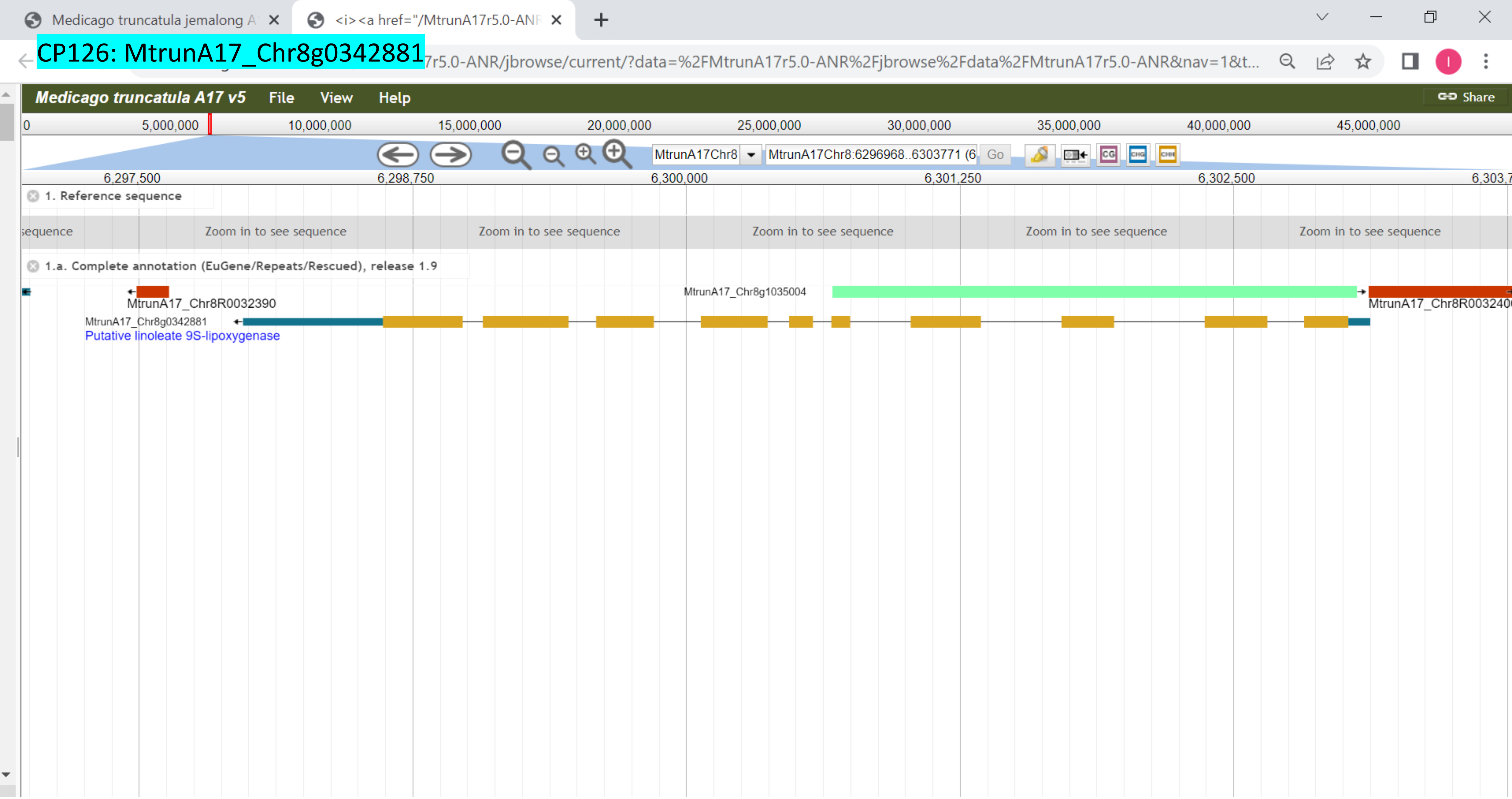

CP127: MtrunA17_Chr8g0345421

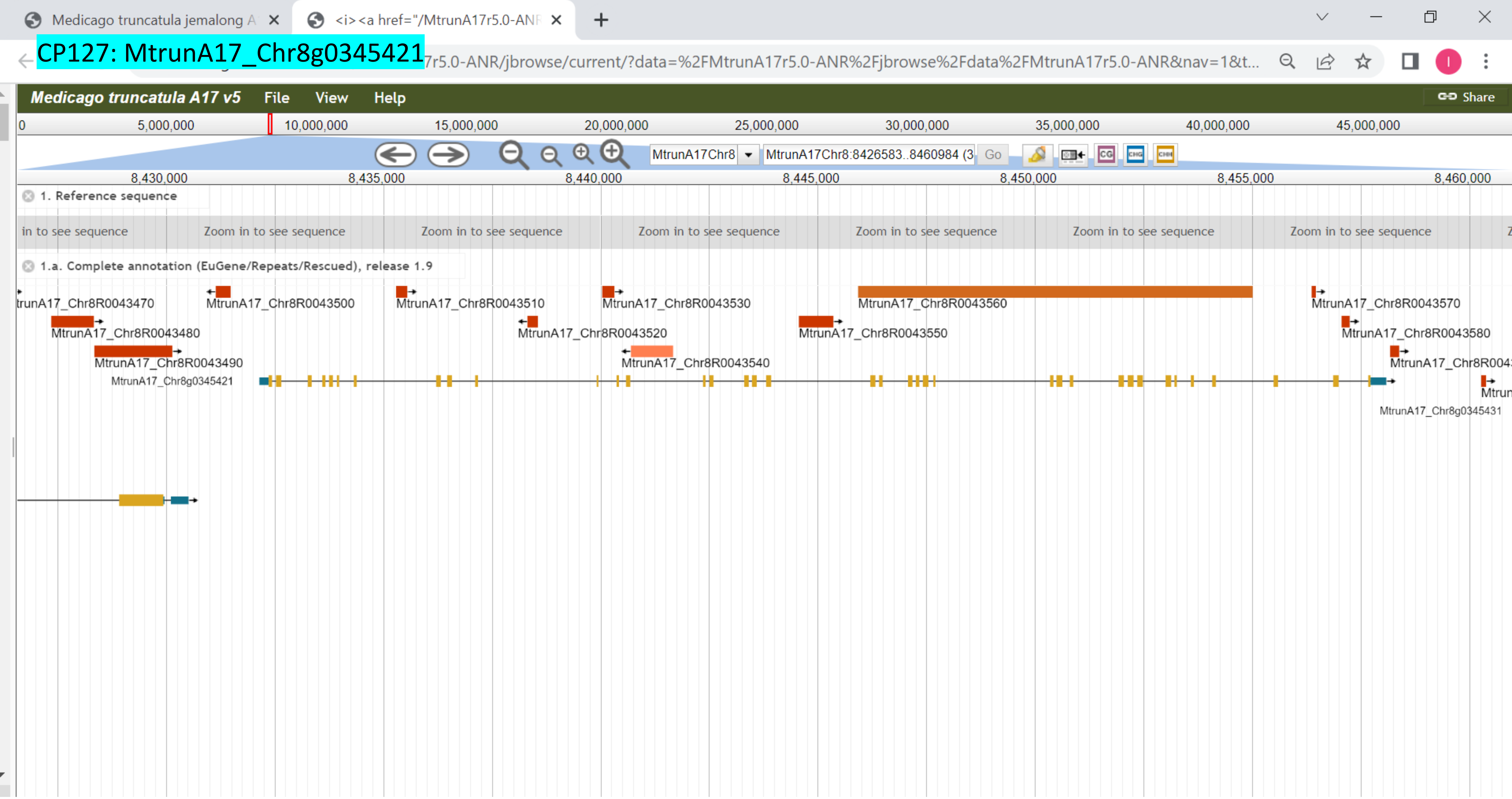

### CP128: MtrunA17_Chr8g0353711

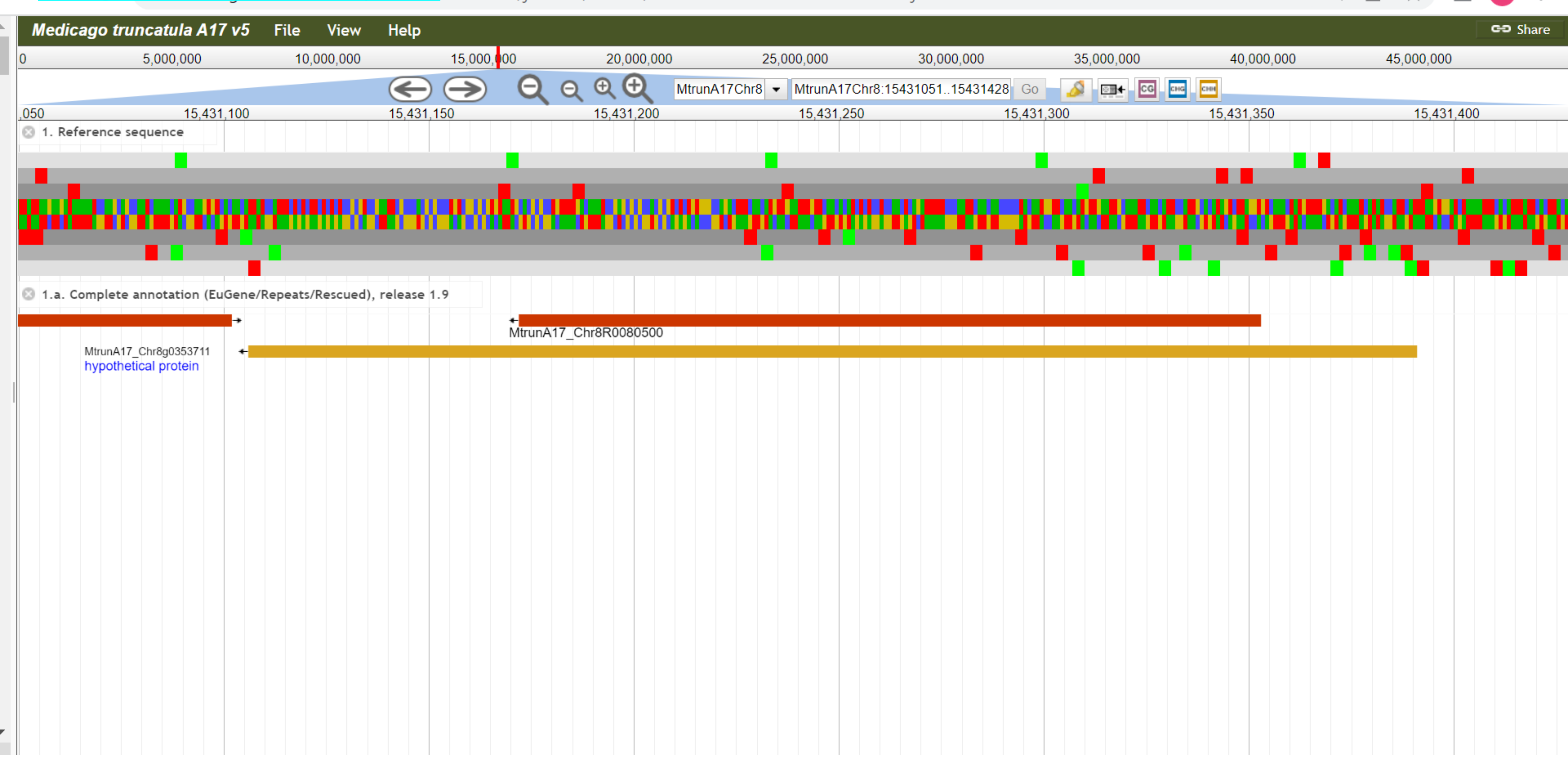

### CP129: MtrunA17_Chr8g0355501

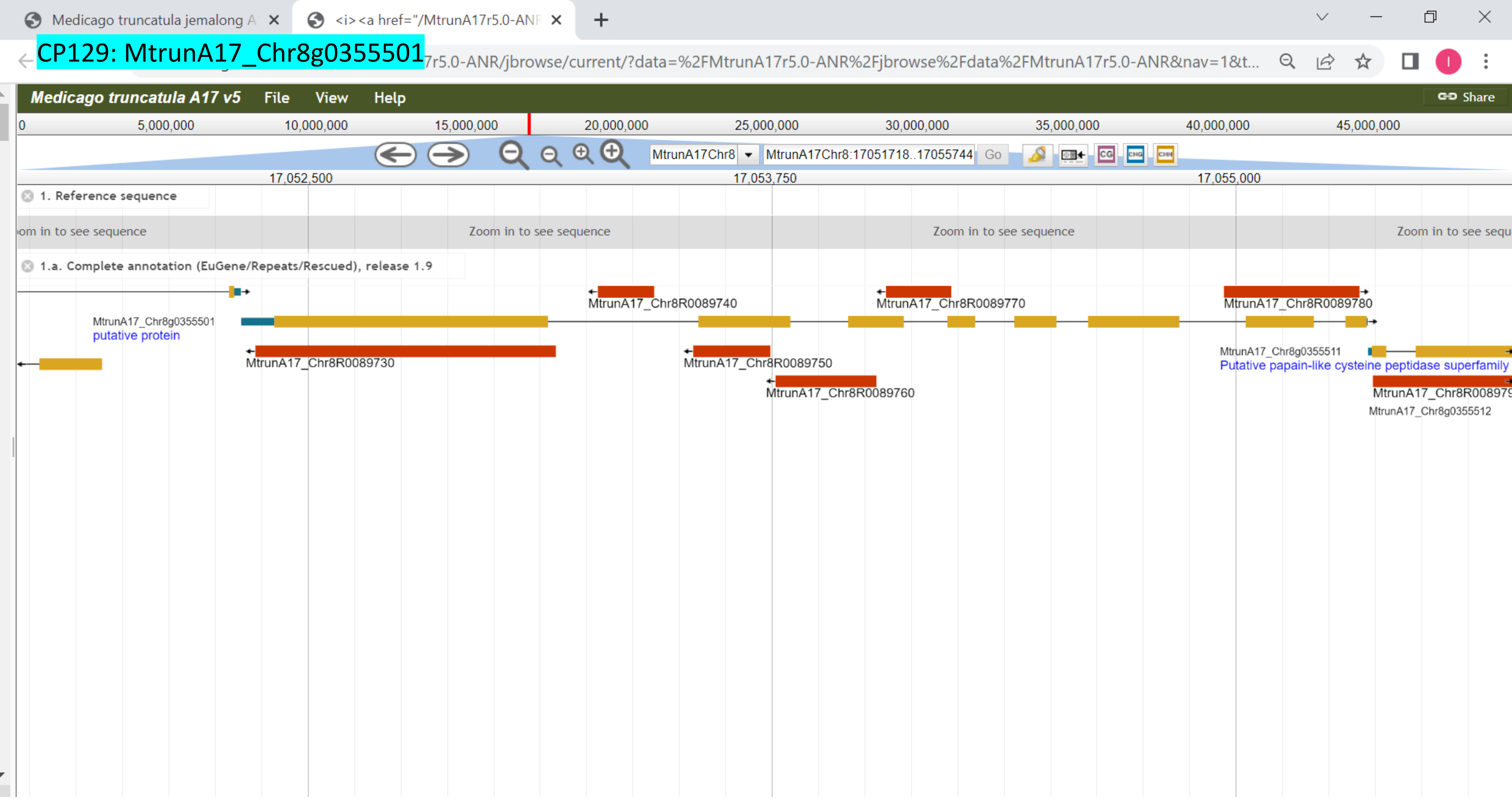

### CP130: MtrunA17_Chr8g0356581

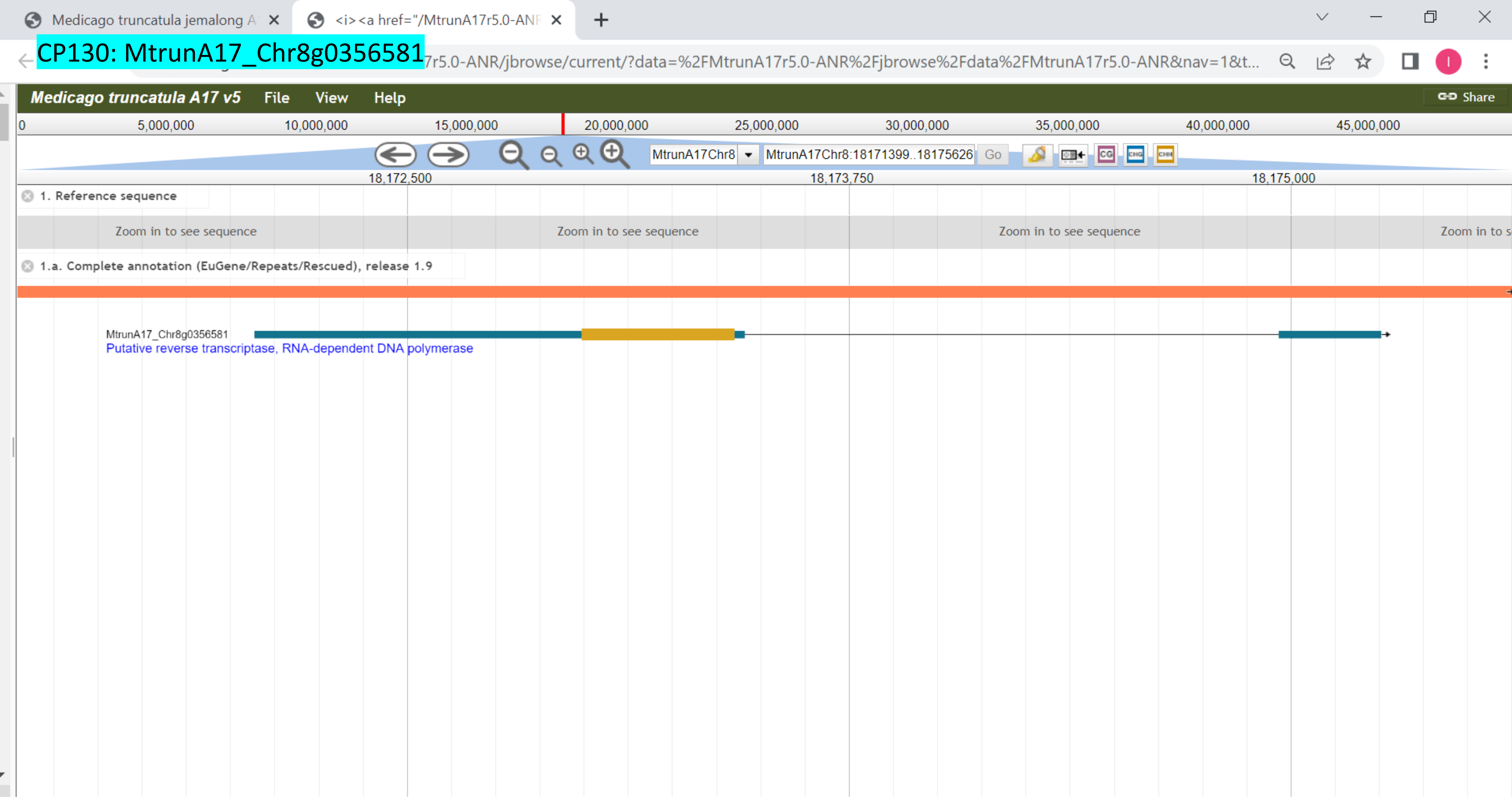

### CP131: MtrunA17_Chr8g0365341

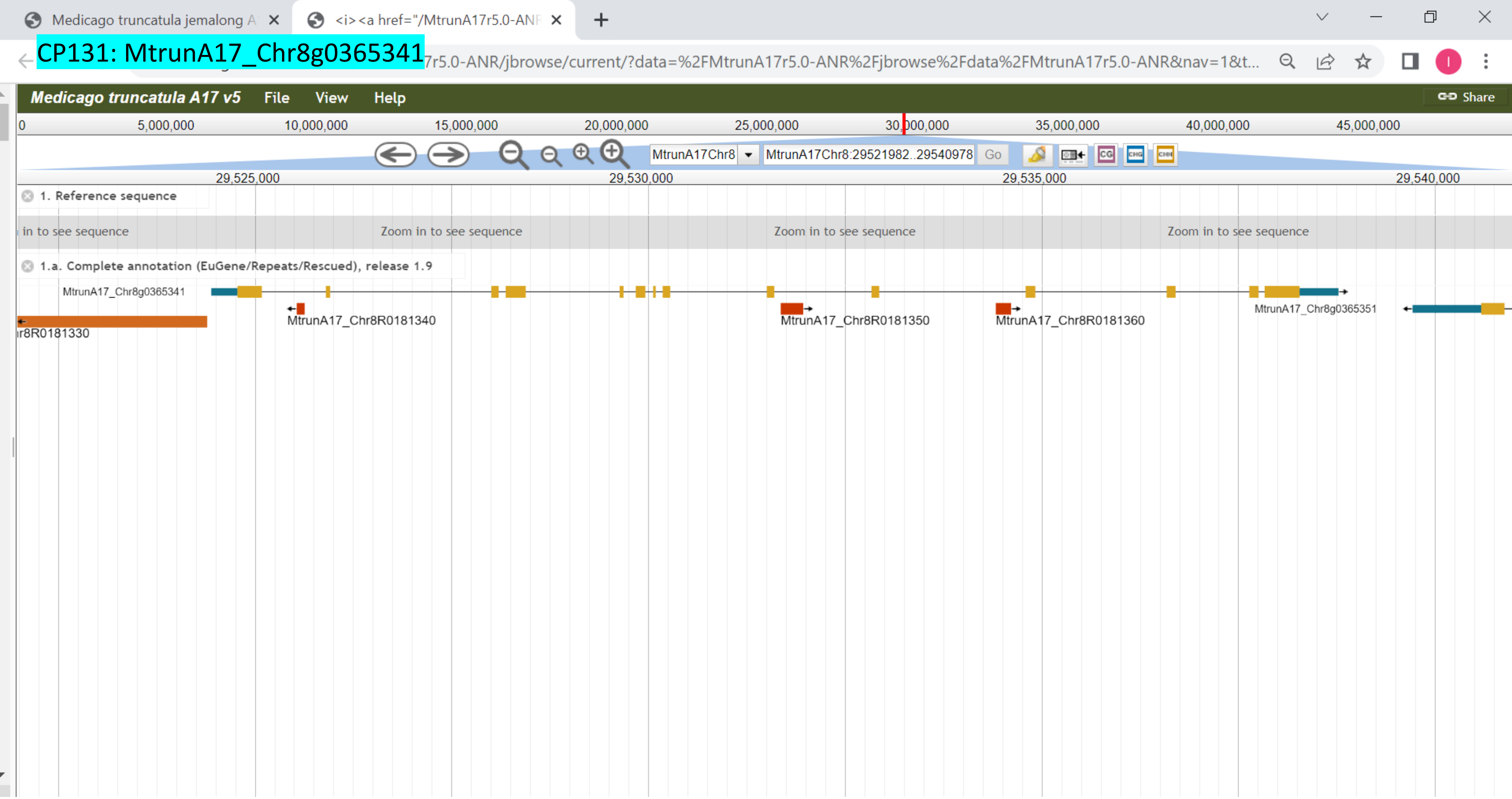

CP132: MtrunA17_Chr8g0368731

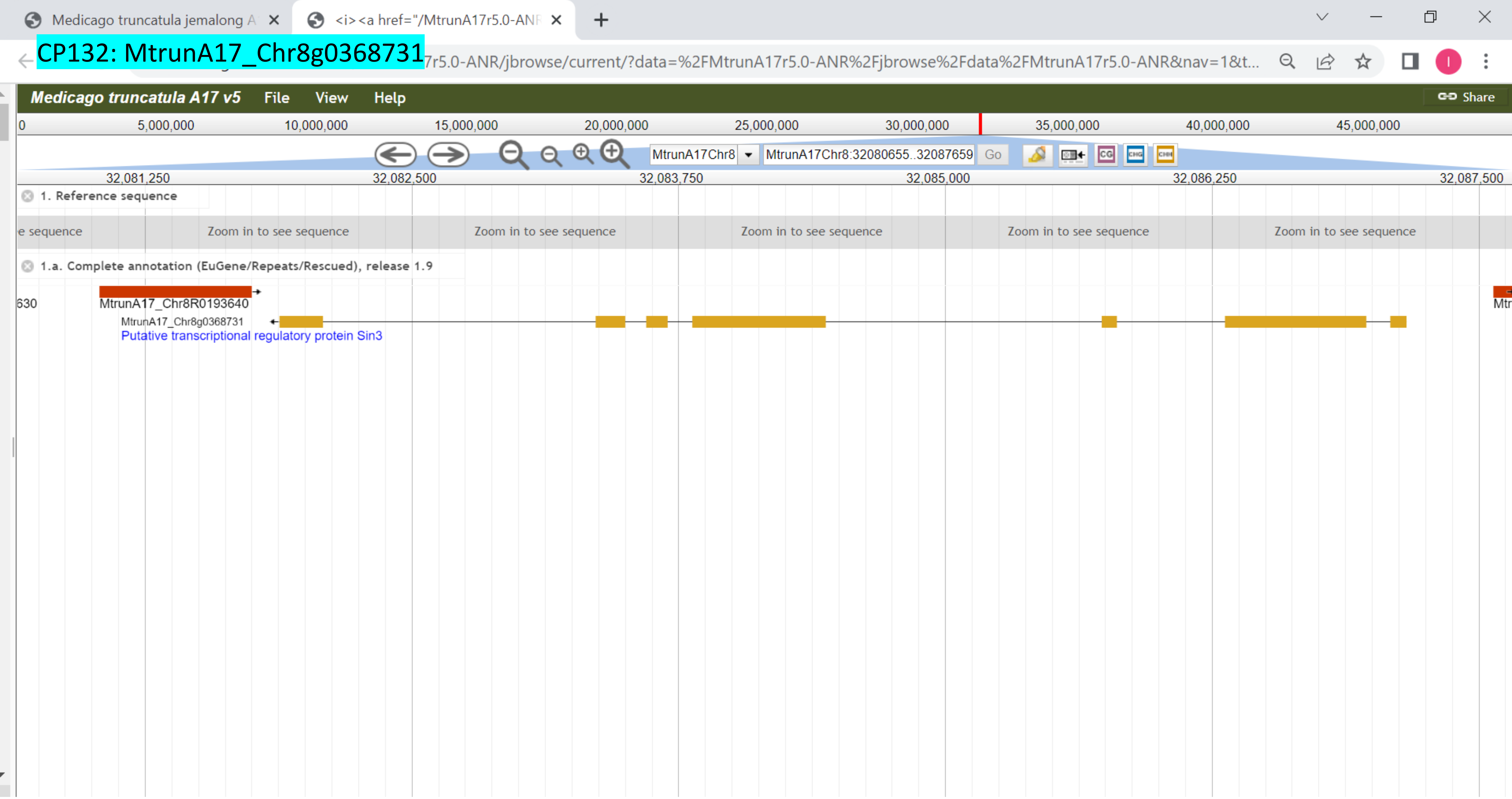

CP133: MtrunA17_Chr8g0368931

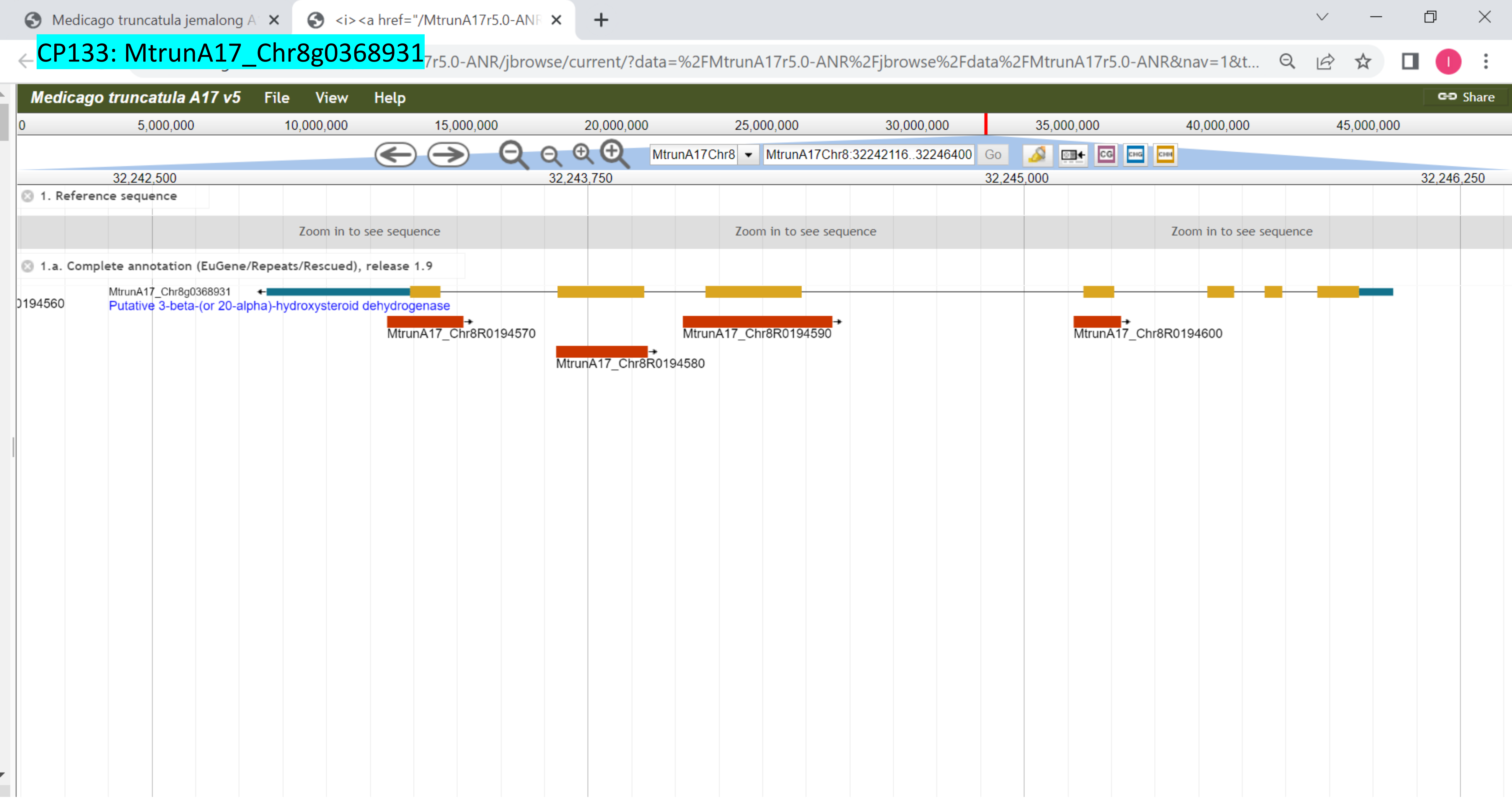

CP134: MtrunA17_Chr8g0371281

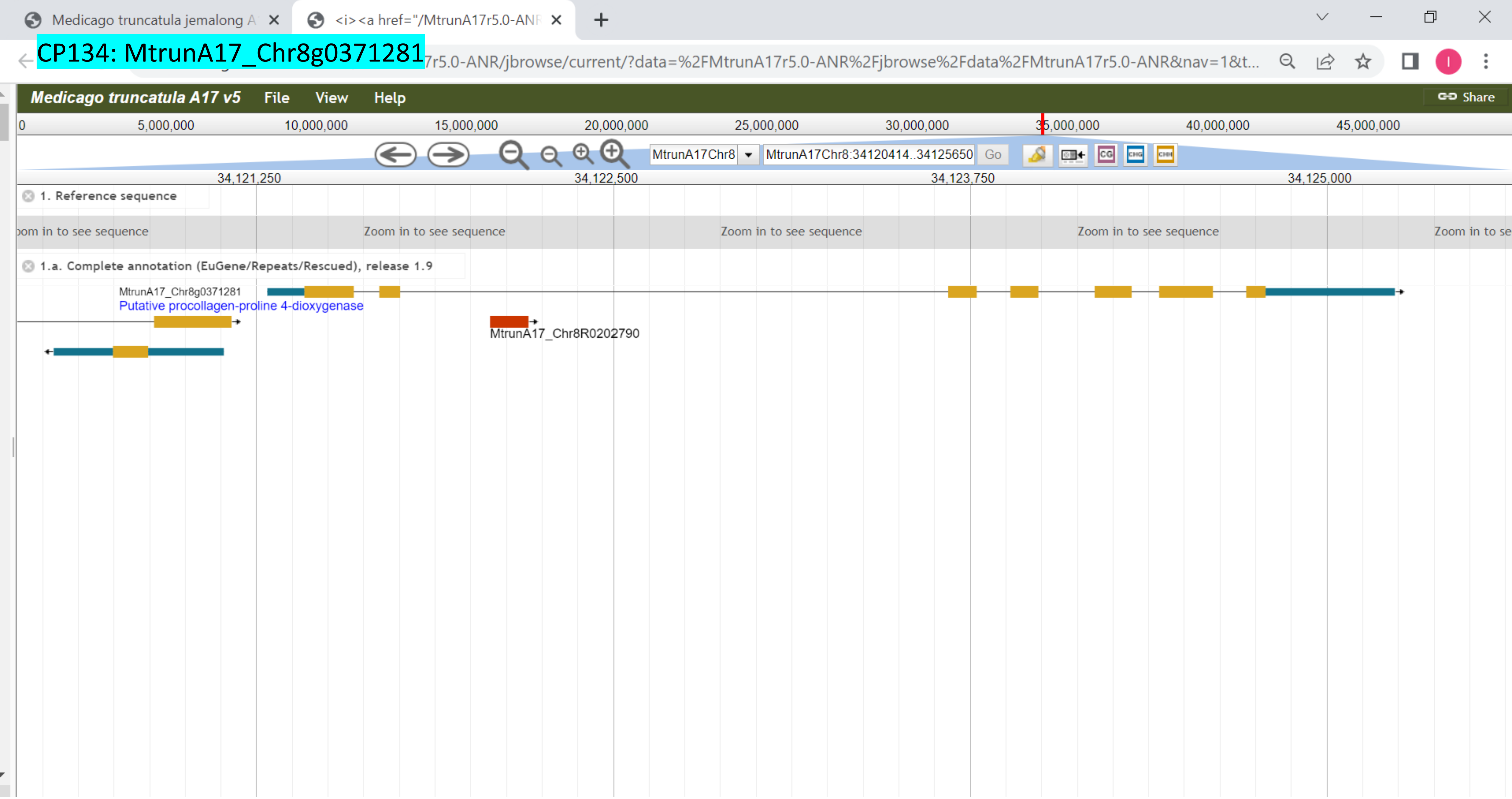

CP135: MtrunA17_Chr8g0371741

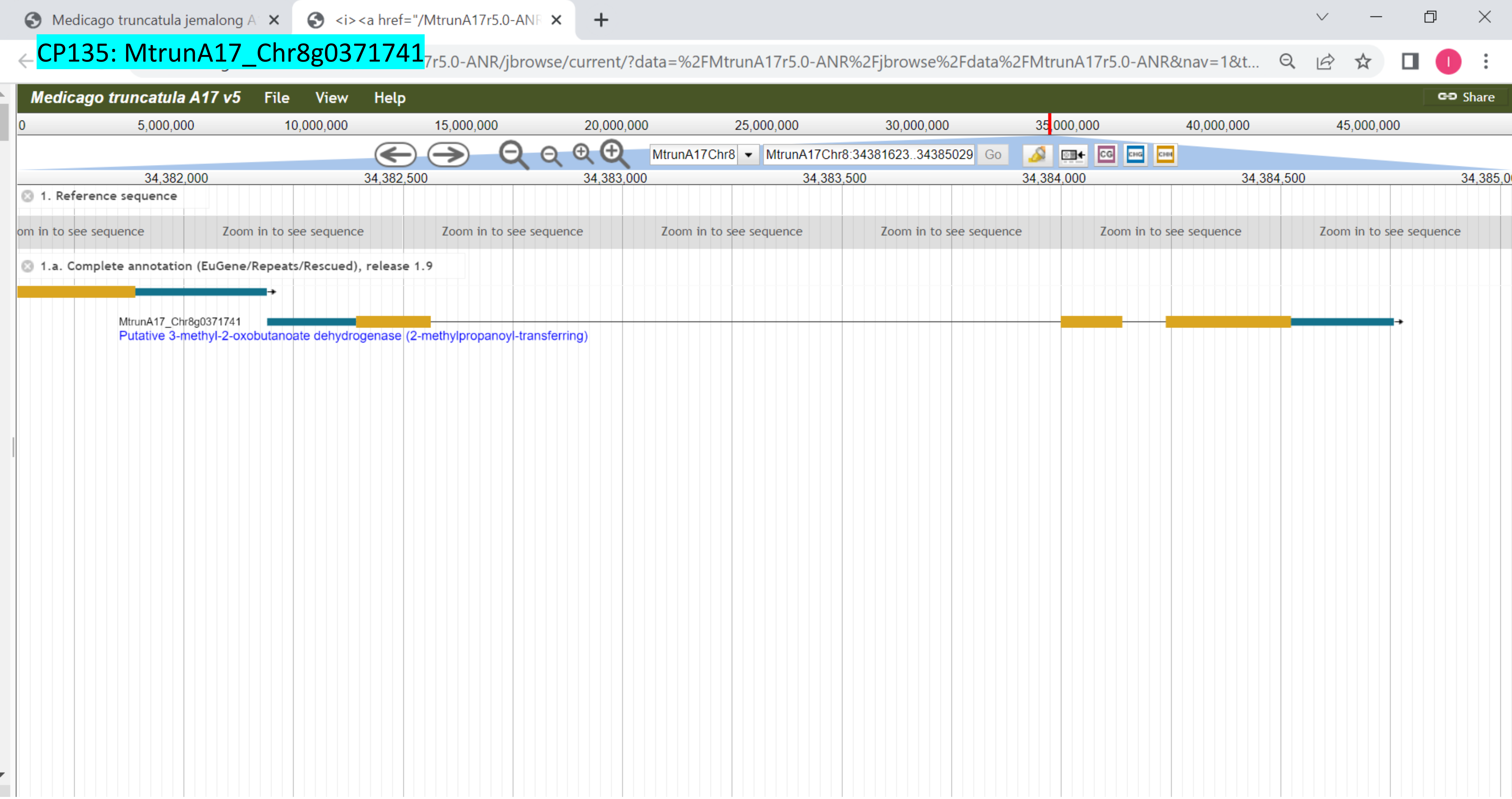

### CP136: MtrunA17_Chr8g0373091

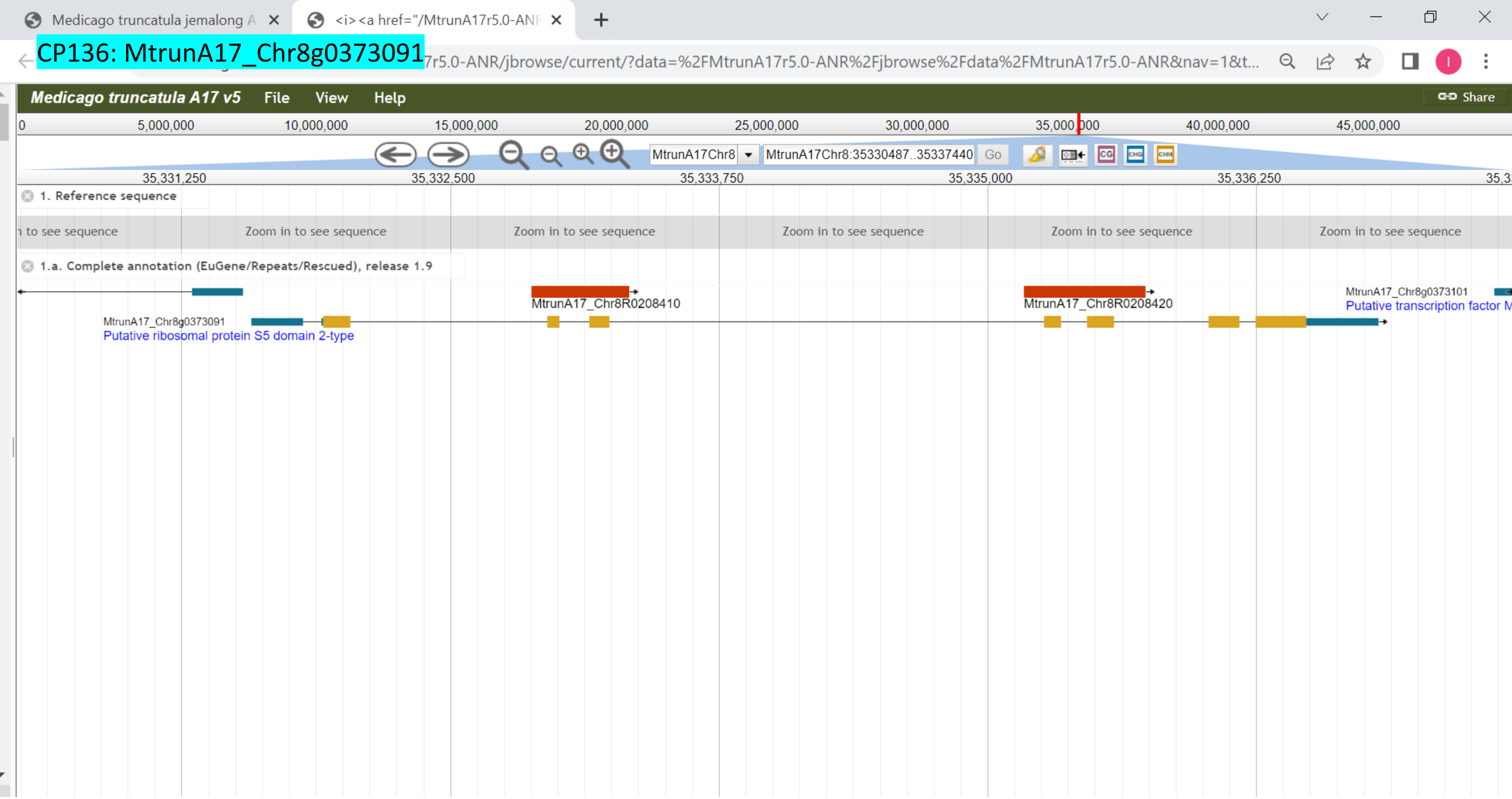

CP137: MtrunA17_Chr8g0376411

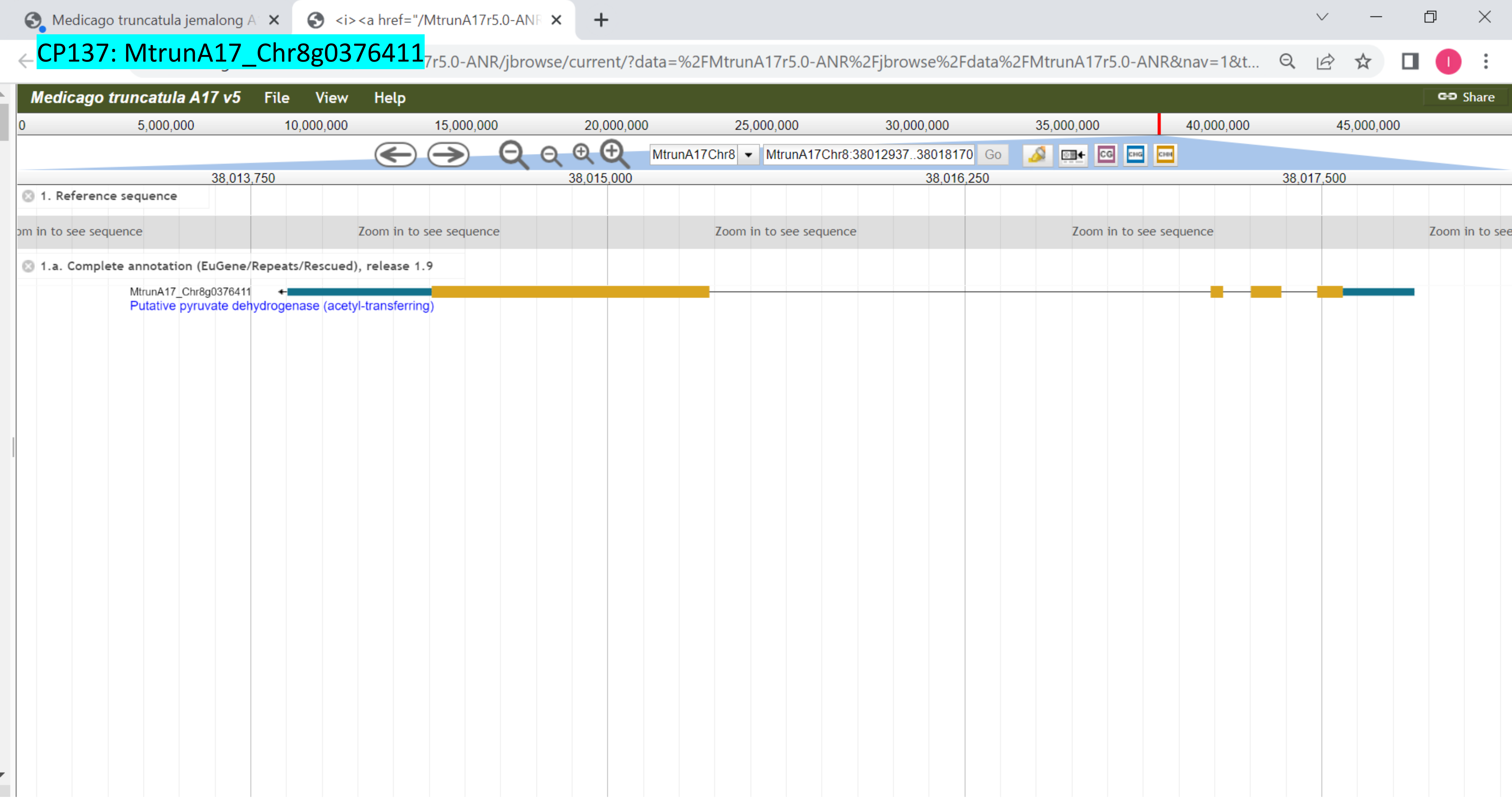

CP138: MtrunA17_Chr8g0377071

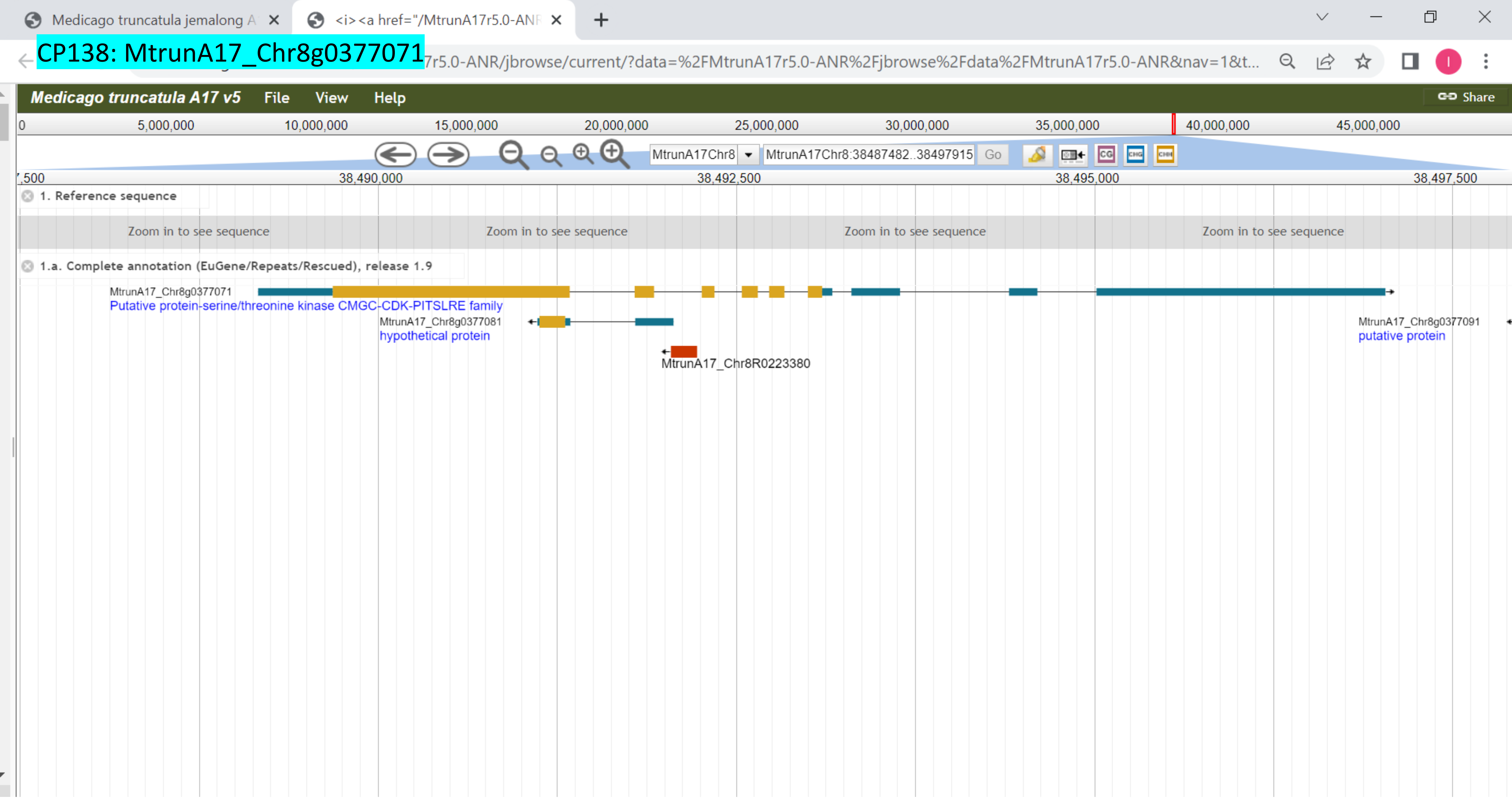

CP139: MtrunA17_Chr8g0385331

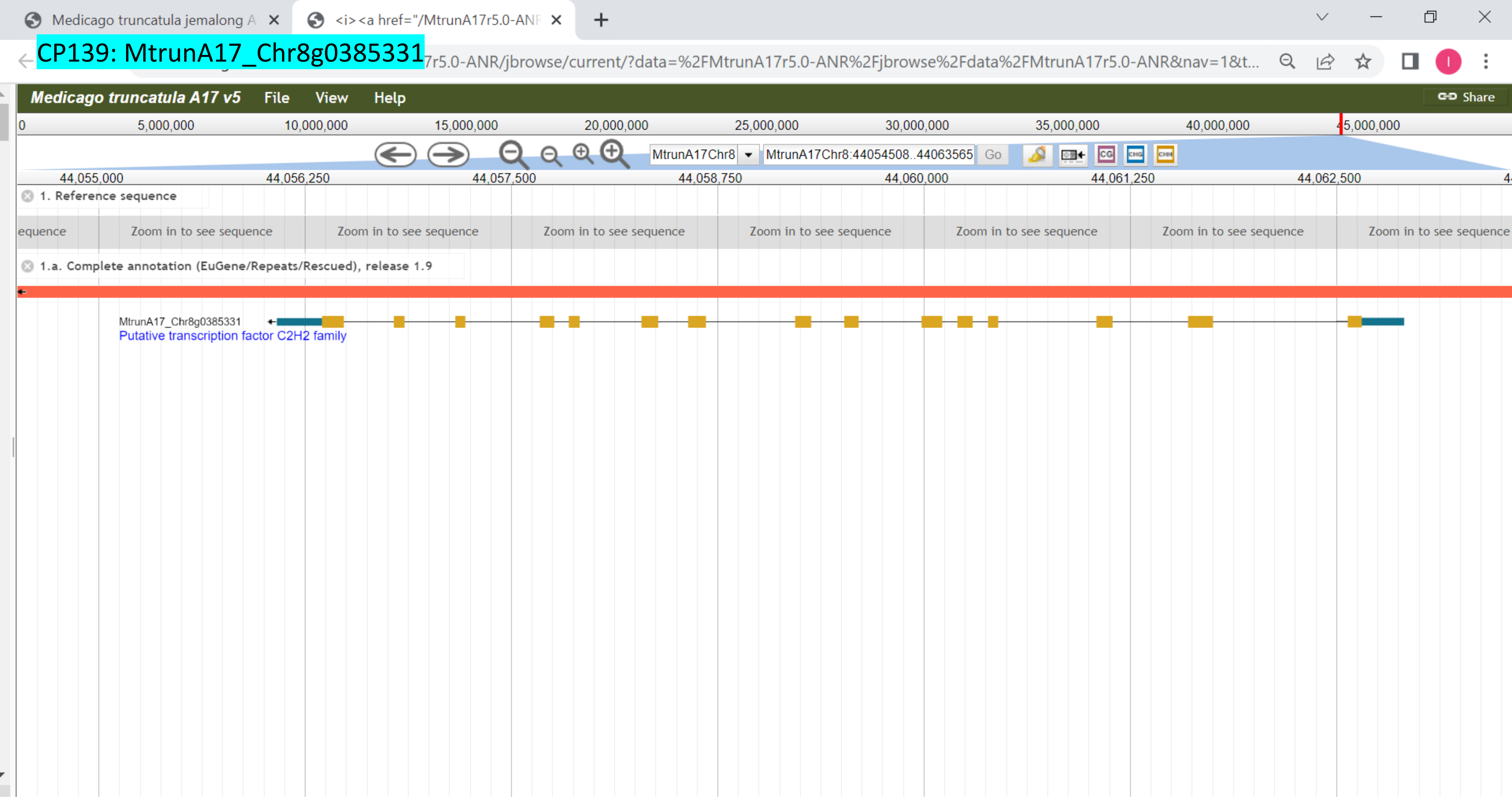

CP140: MtrunA17_Chr8g0392351

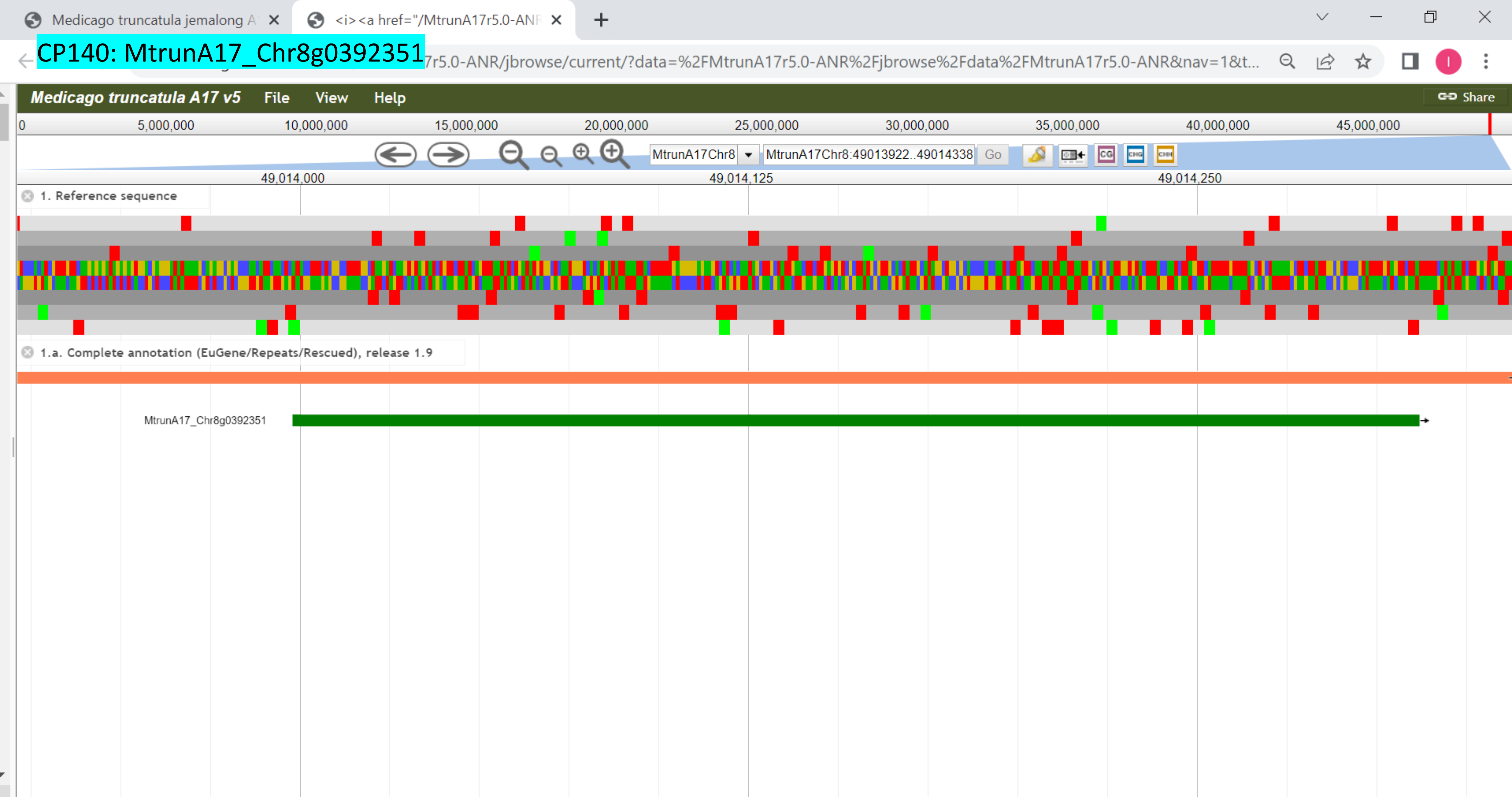

### CP141: MtrunA17_CPg0492331

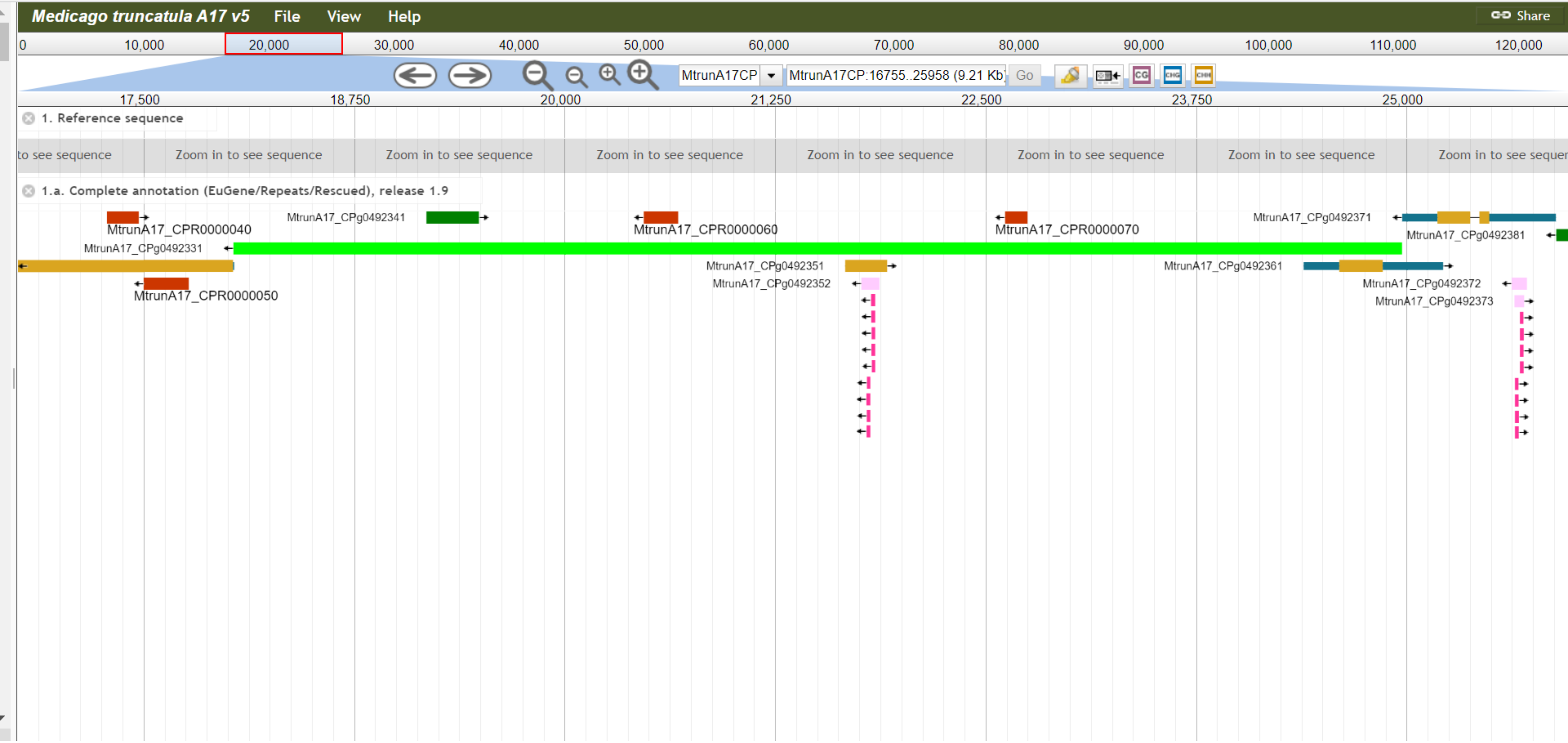

CP142: MtrunA17_CPg0492381

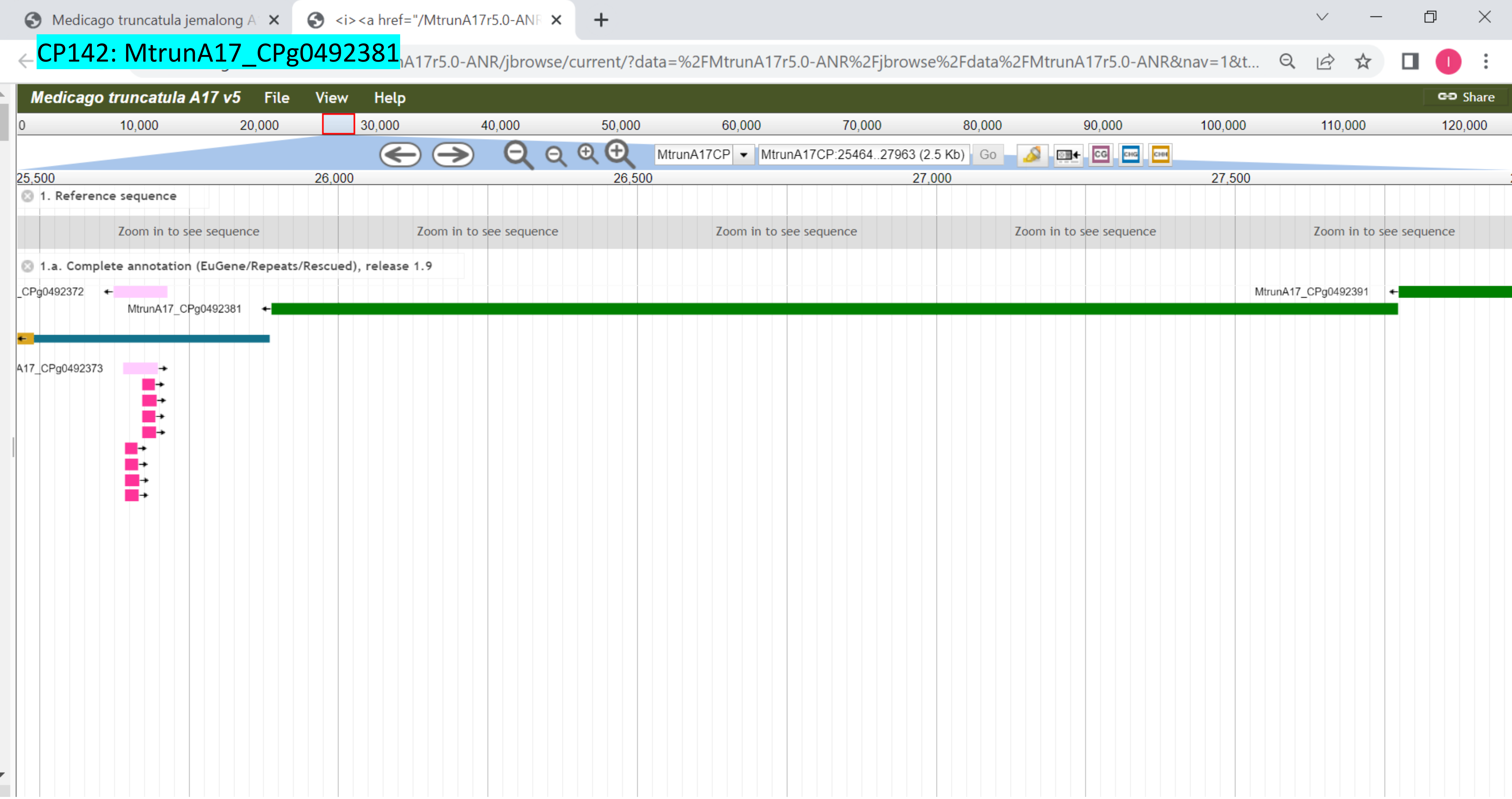

### CP143: MtrunA17_CPg0492461

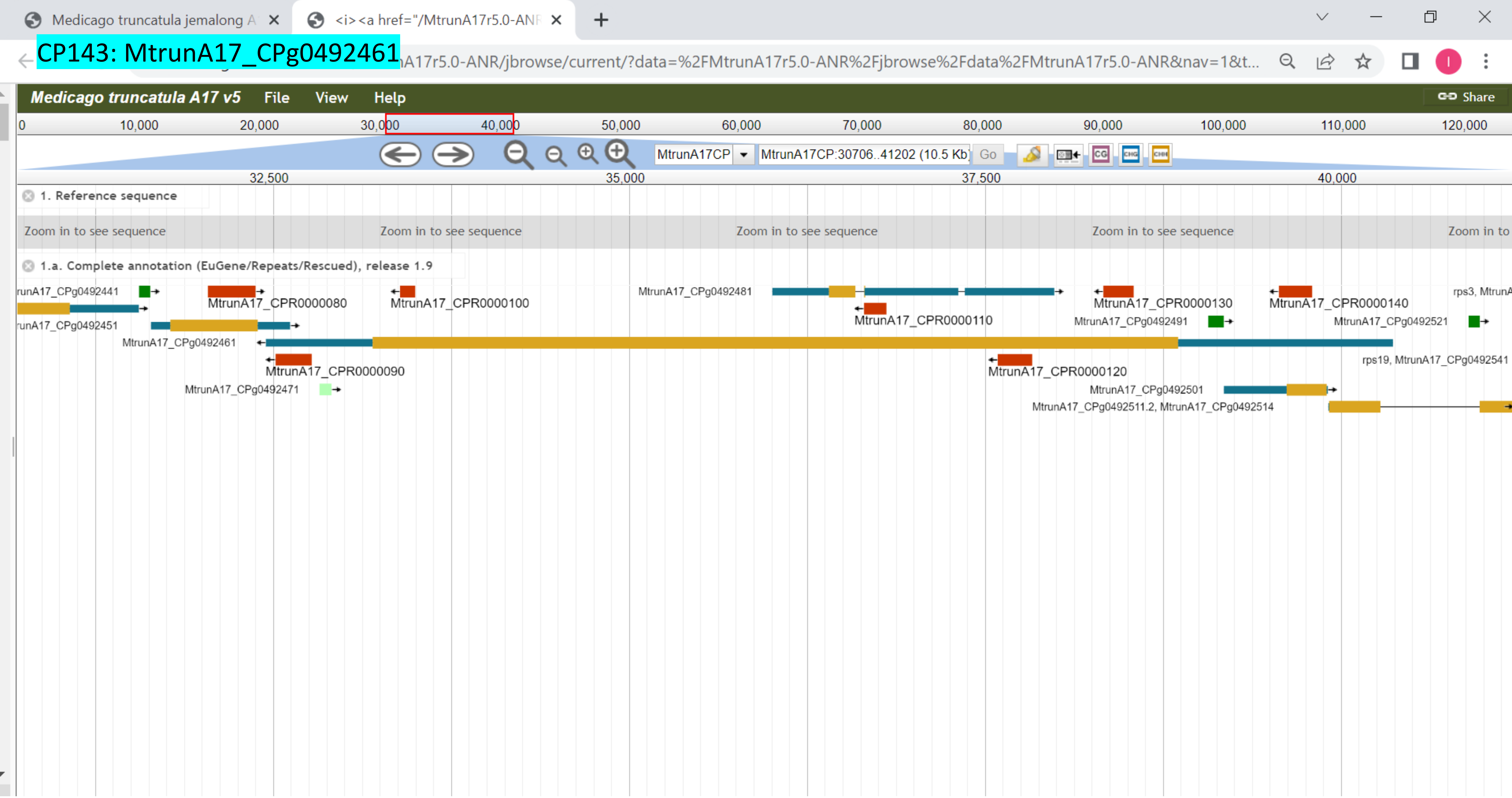

CP144: MtrunA17_CPg0492851

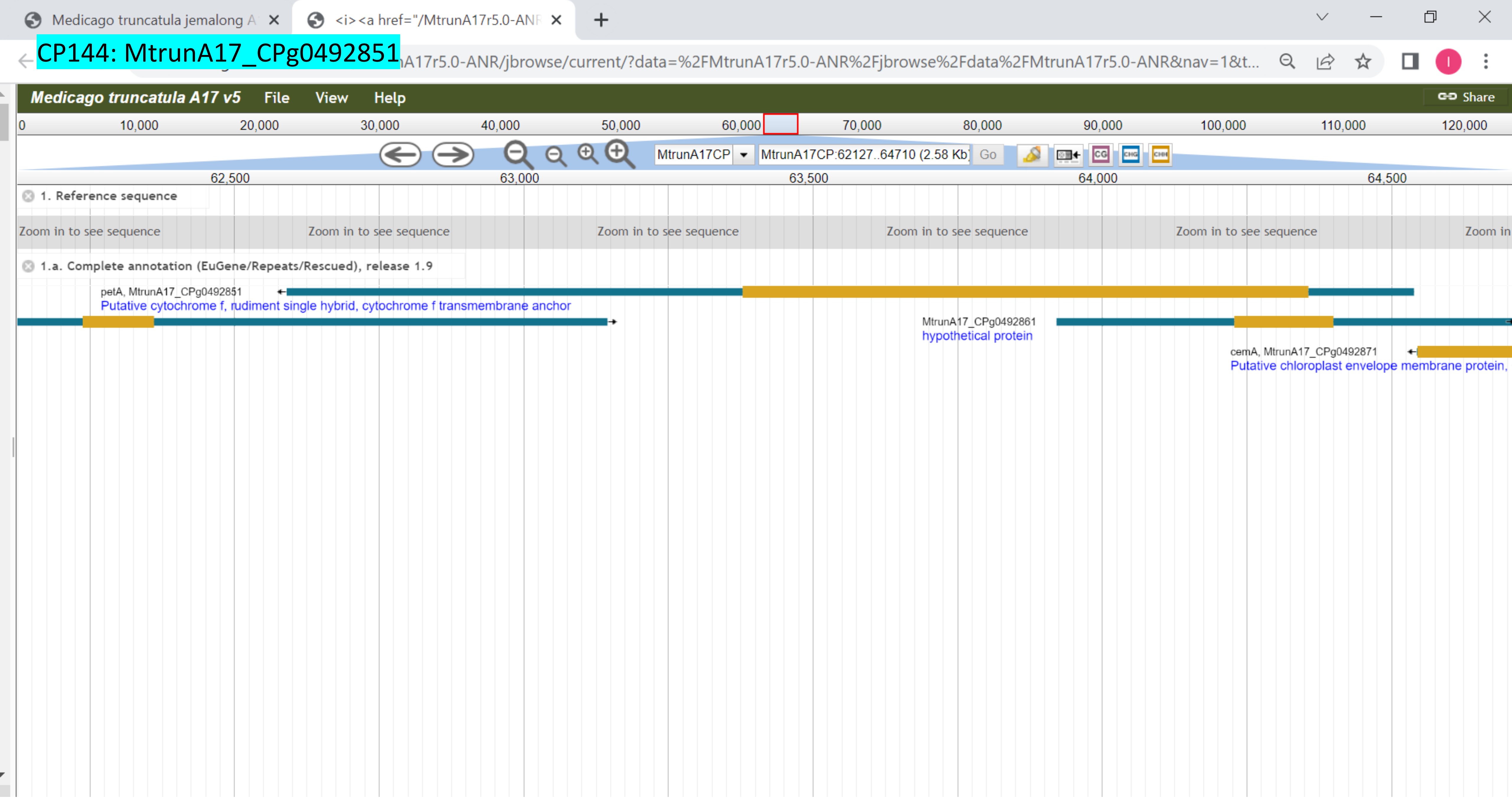

CP145: MtrunA17_CPg0492941

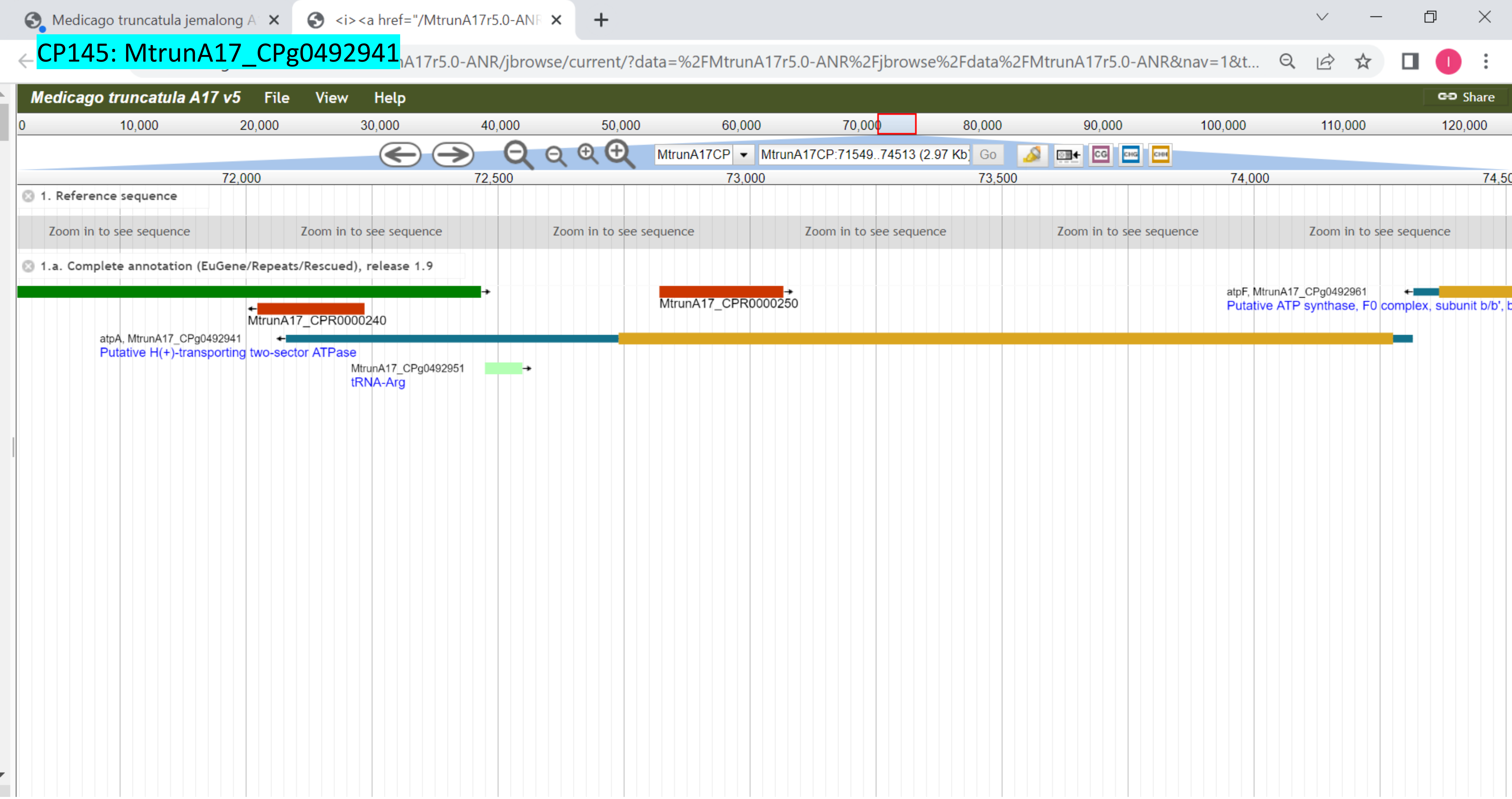

### CP146: MtrunA17_CPg0493291

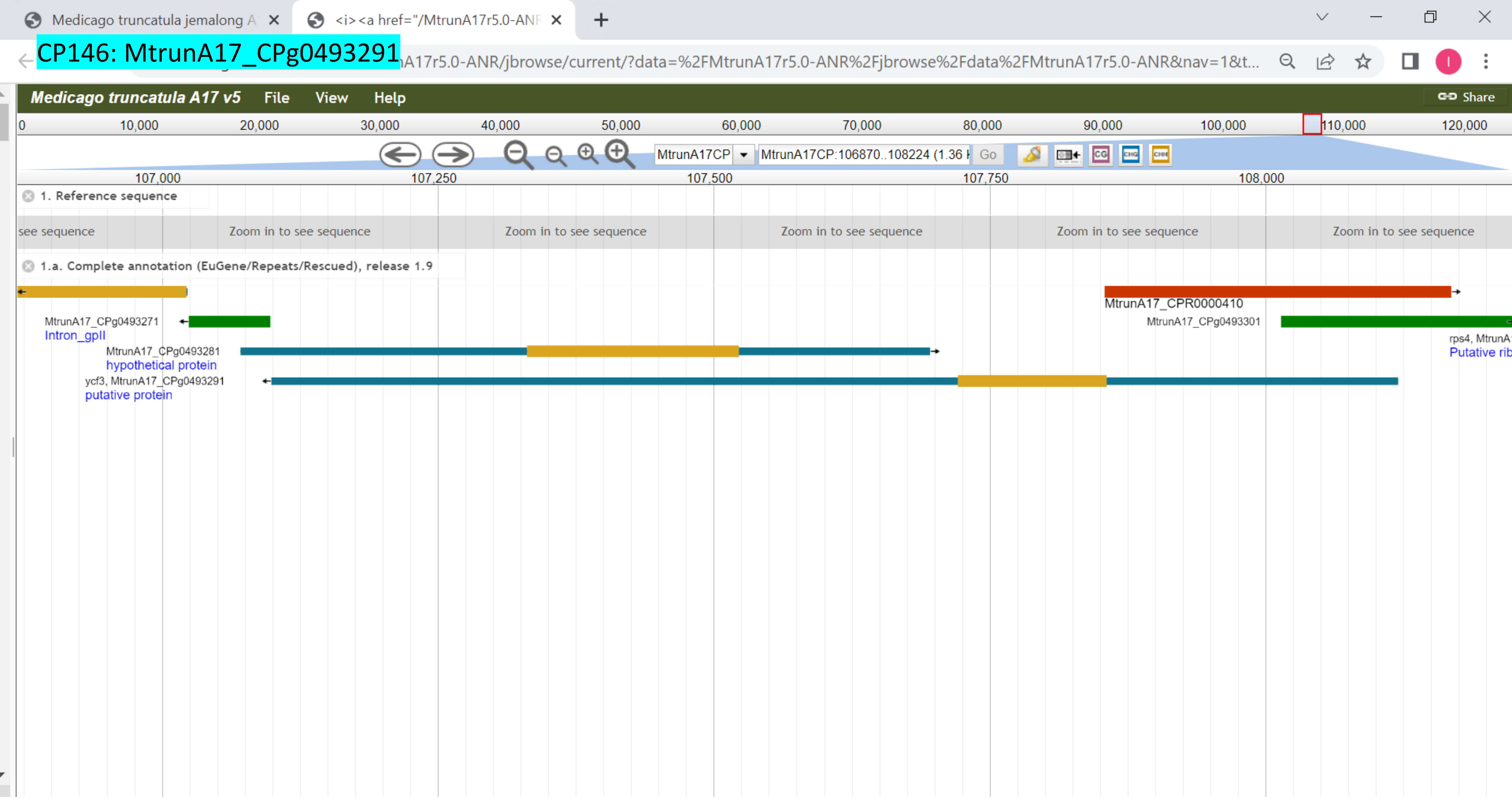

### CP147: MtrunA17_CPg0493401

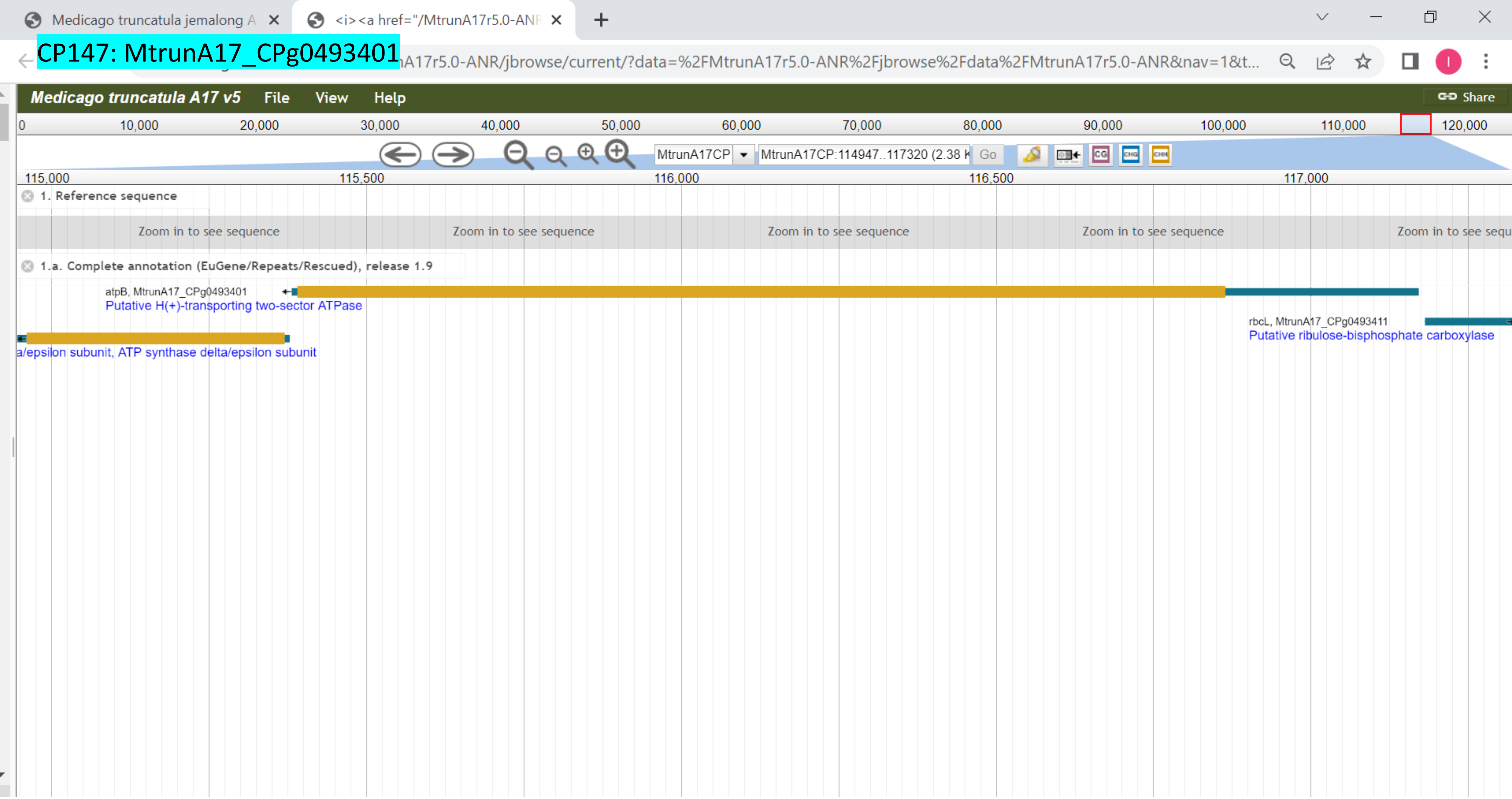

CP148: MtrunA17_MTg0490471

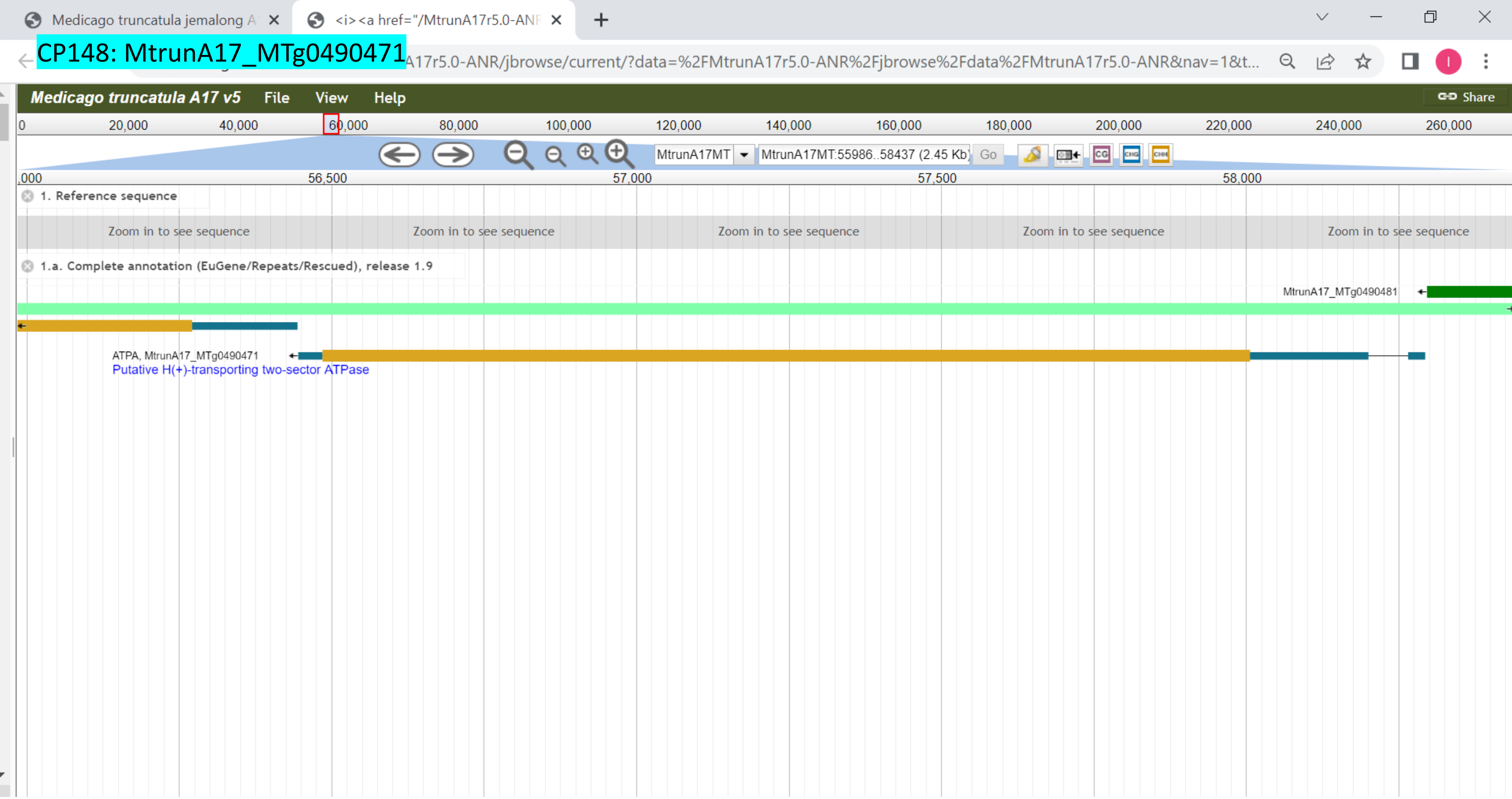

CP149: MtrunA17_MTg0490471

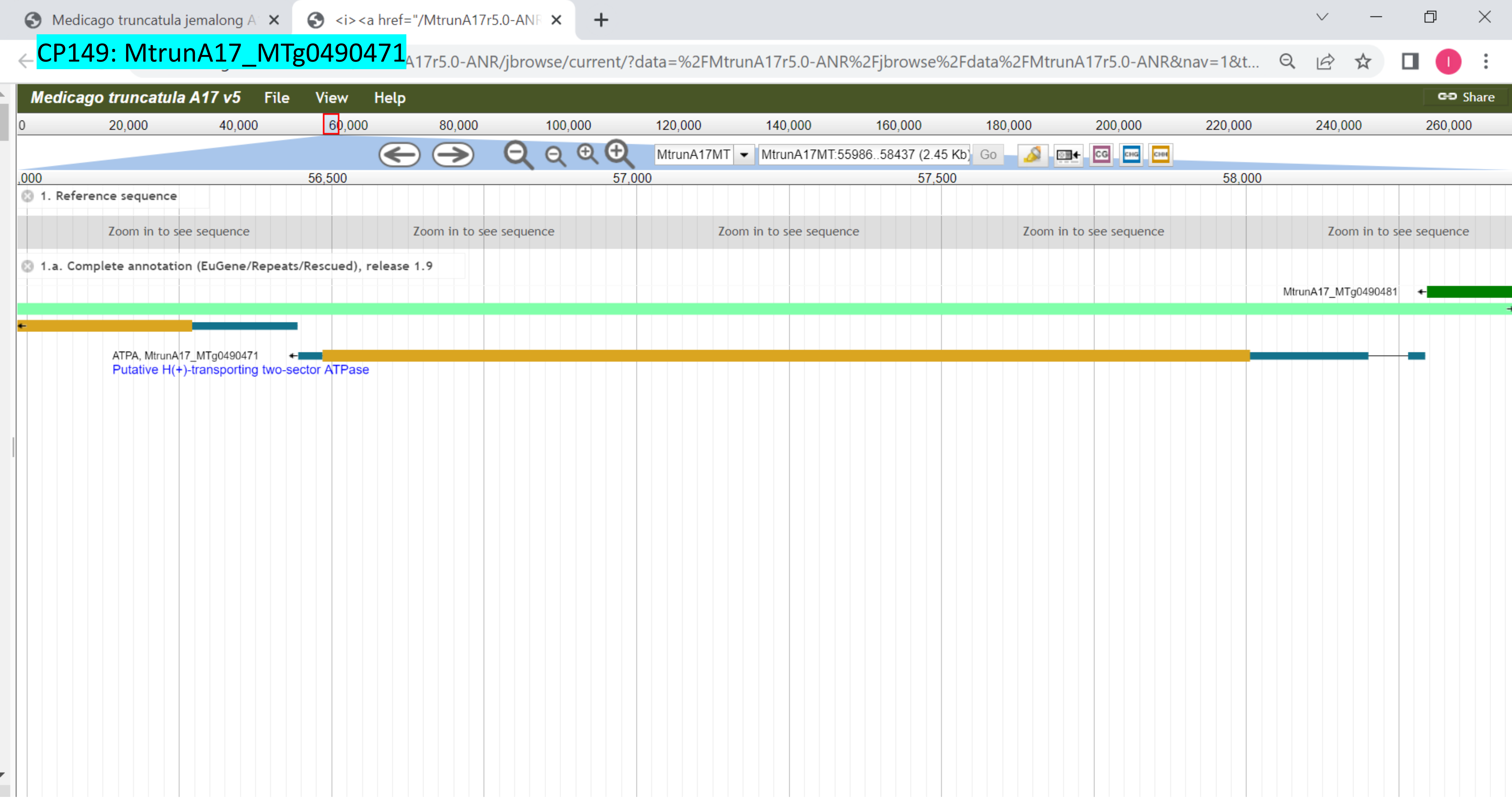

CP150: MtrunA17_MTg0490971

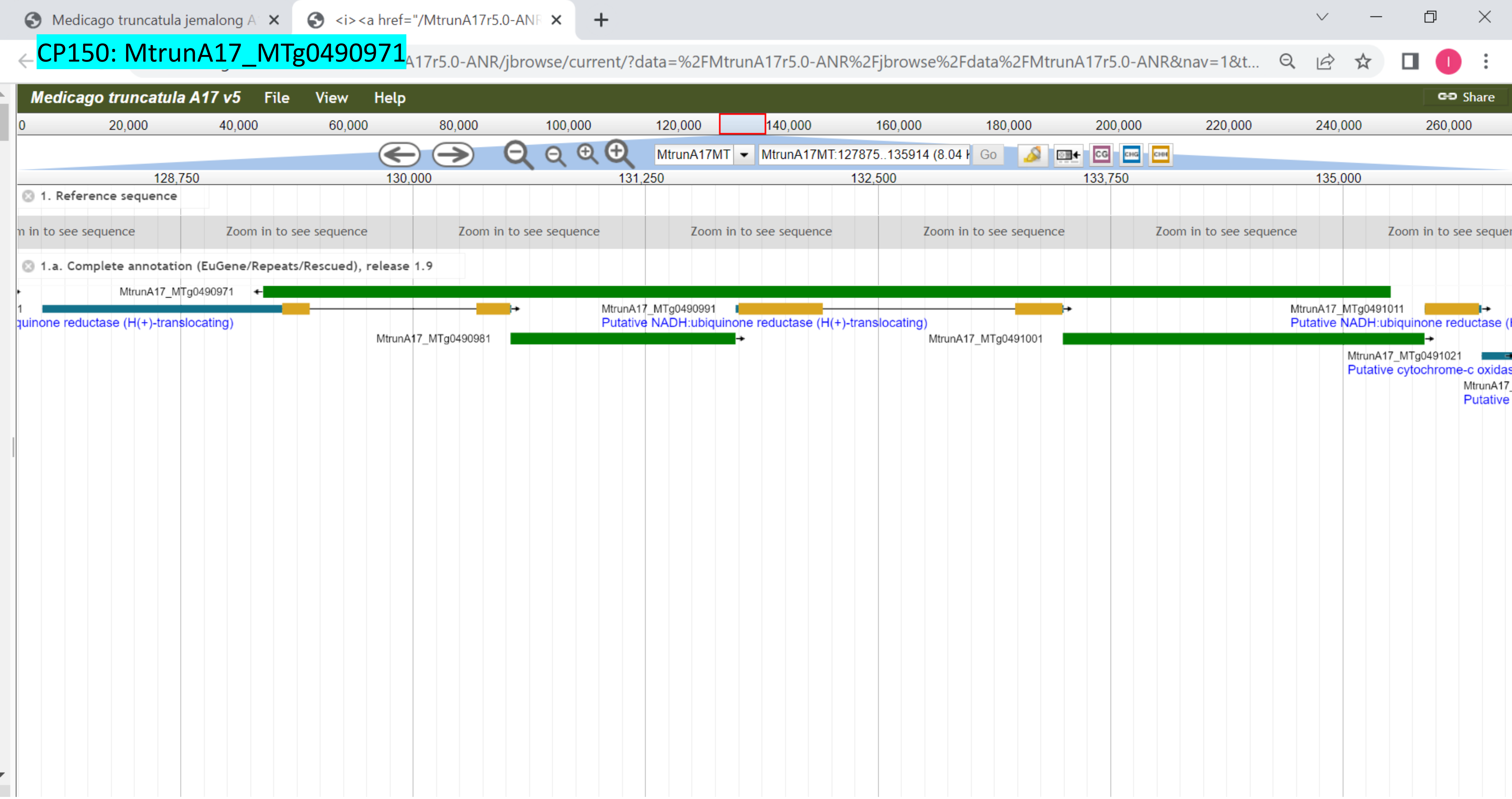

### CP151: MtrunA17_MTg0490971

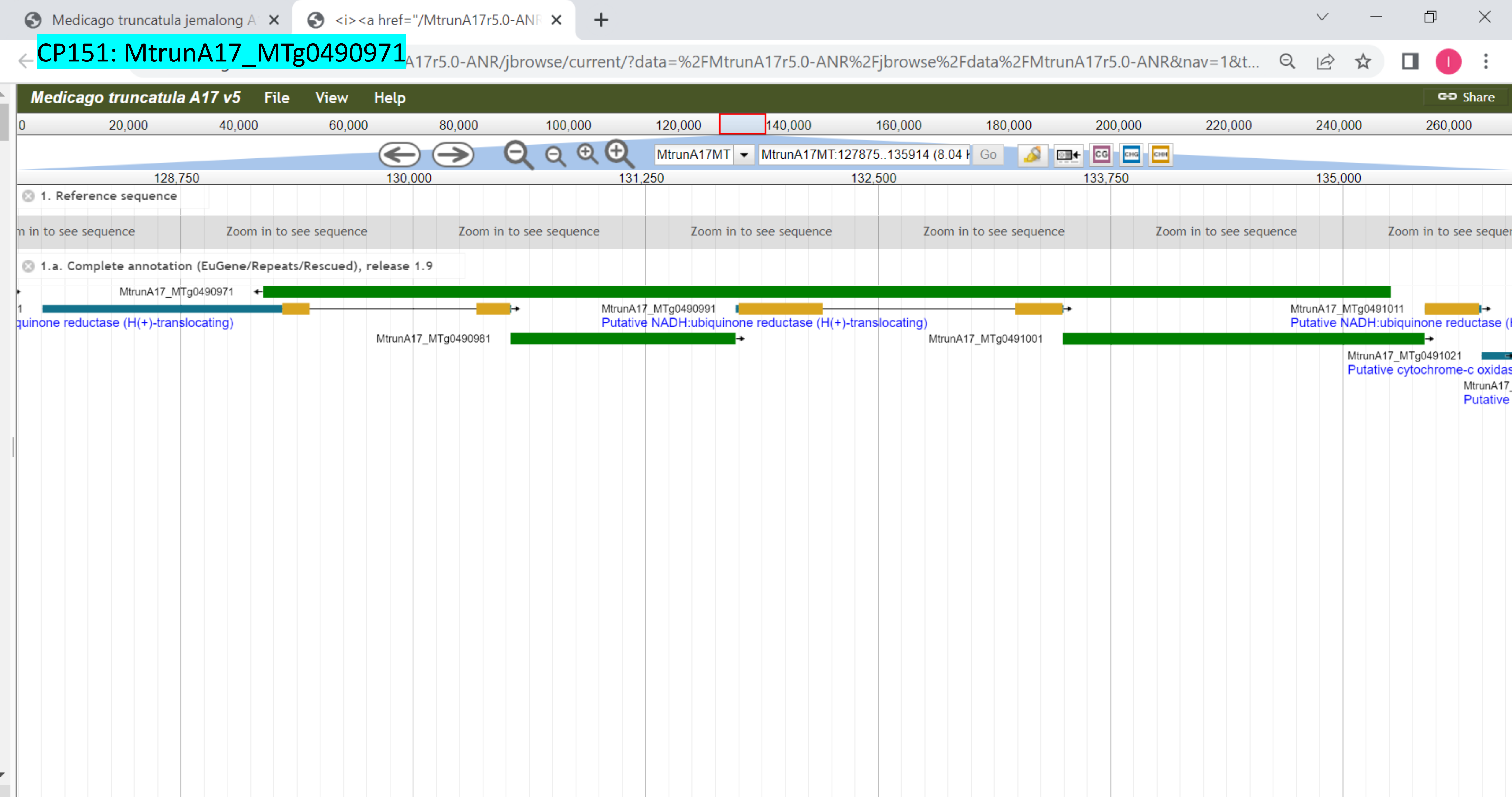

CP152: MtrunA17_MTg0491151

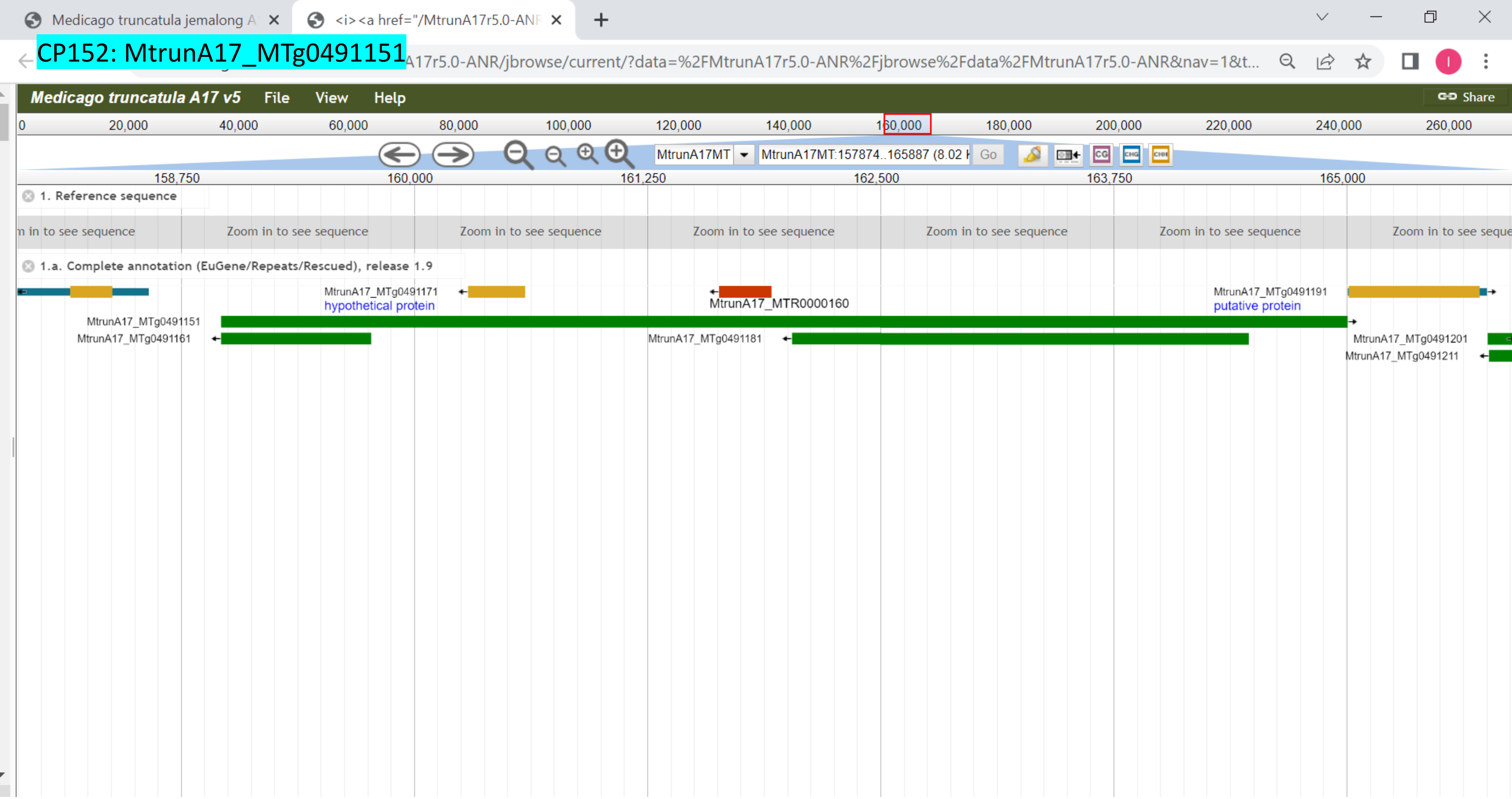

### CP153: MtrunA17_MTg0491291

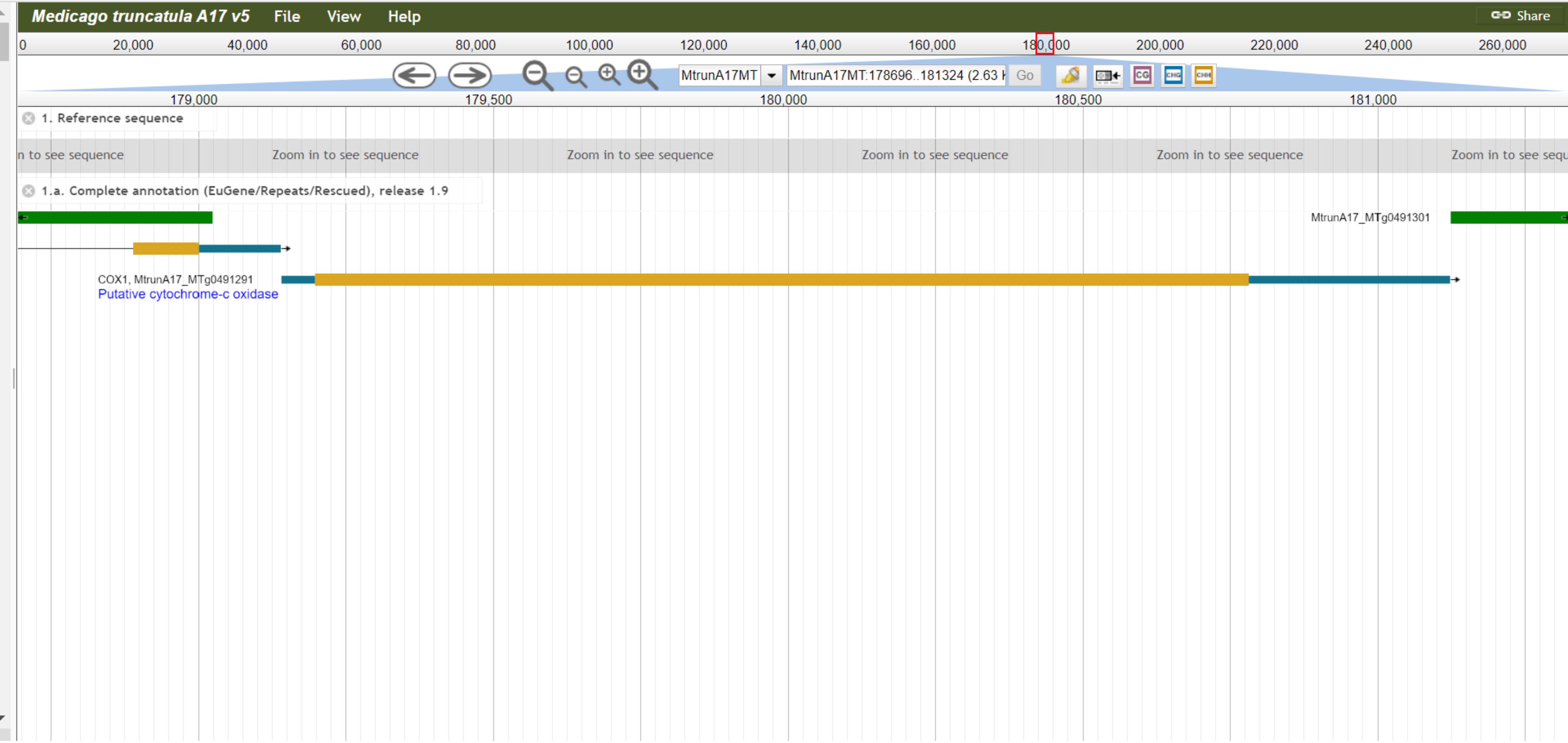

### CP154: MtrunA17_MTg0491501

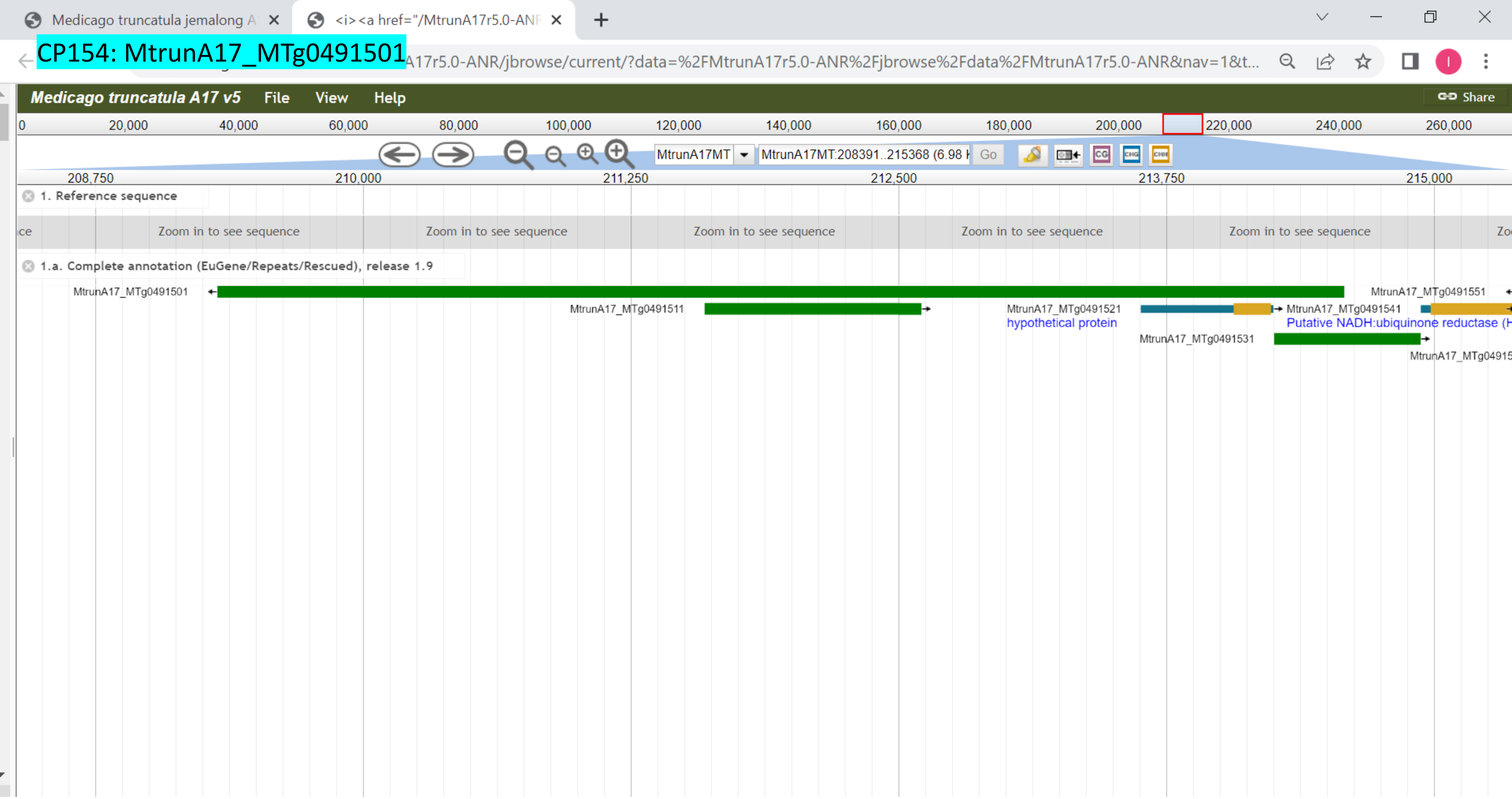

### CP155: MtrunA17_MTg0491621

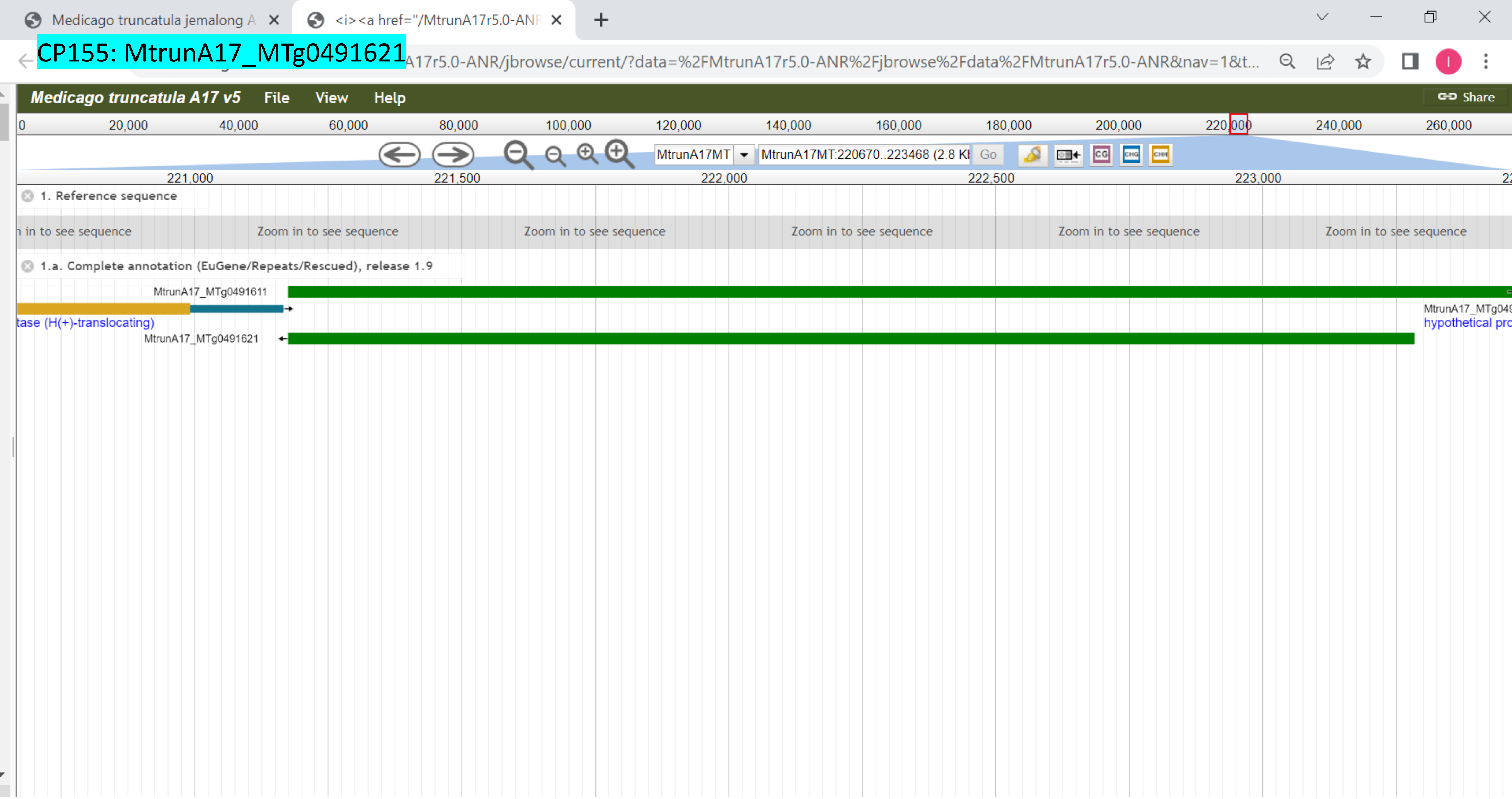

CP156: MtrunA17_MTg0491711

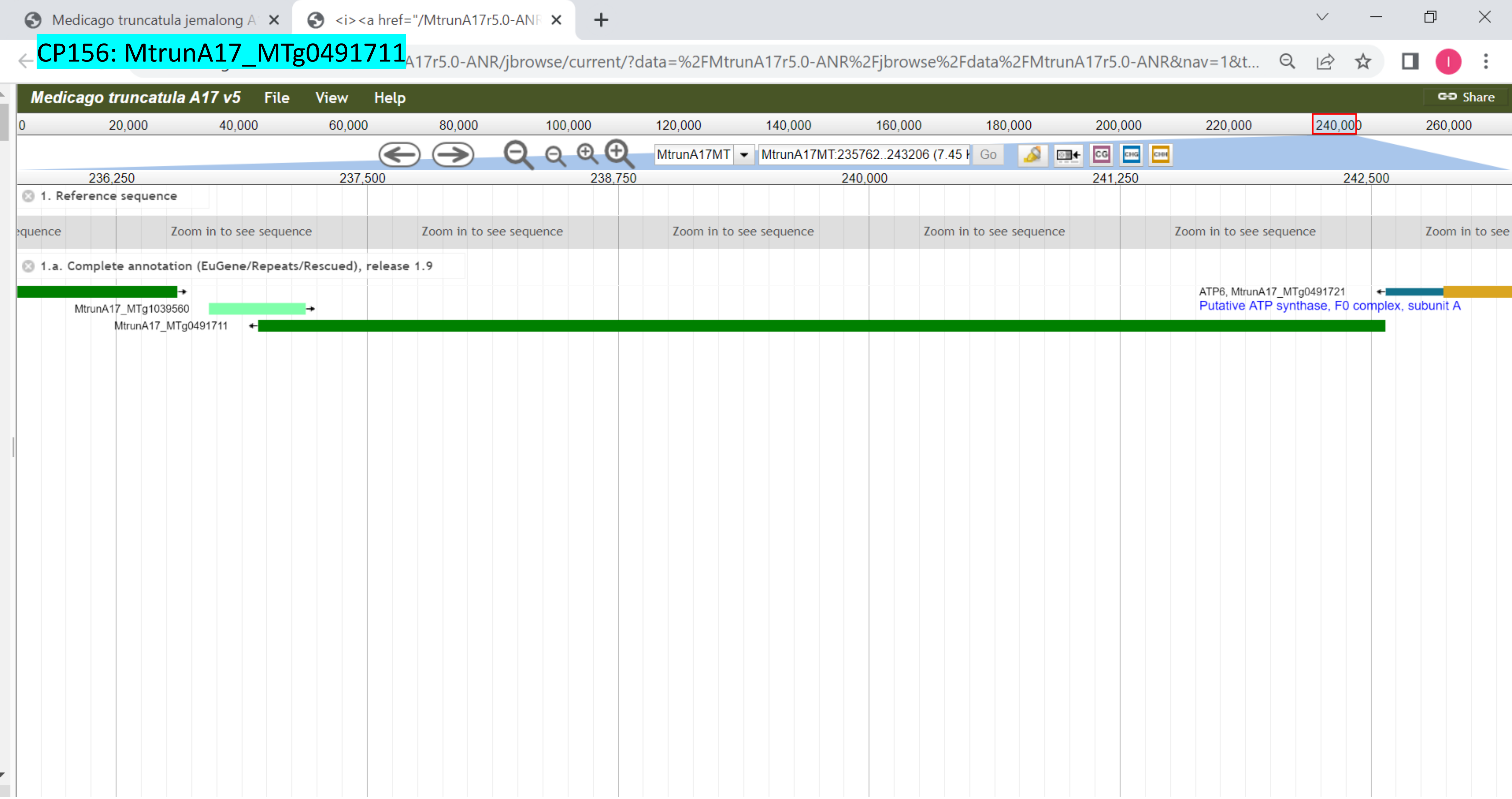

**Supplementary Dataset S11**. The genomic landscape at and around the primary-source loci of 156 MS-supported chimeric peptides. Each image represents a screenshot from the *Medicago truncatula* genome browser v. 5.1.7. Color codes for genes of different RNA types are shown below. Arrows pointing from left to right and in the opposite direction indicate loci that are located on the plus and minus strands, respectively.

- exons of mRNA genes
- introns of mRNA genes
- UTRs of mRNA genes
- ncRNA and tRNA genes
- rRNA genes
- pre-miRNA genes
- repeat elements
