## Supplementary material for "Discovery of diverse chimeric peptides in a eukaryotic proteome sets the stage for the experimental proof of the mosaic translation hypothesis": Supplementary Dataset S14 Alignments among 34 primary-source transcripts with detectable similarity.pdf

CP1: MtrunA17_Chr0c01g0489091
CP87: MtrunA17_Chr5g0421761
% identity **64.3**

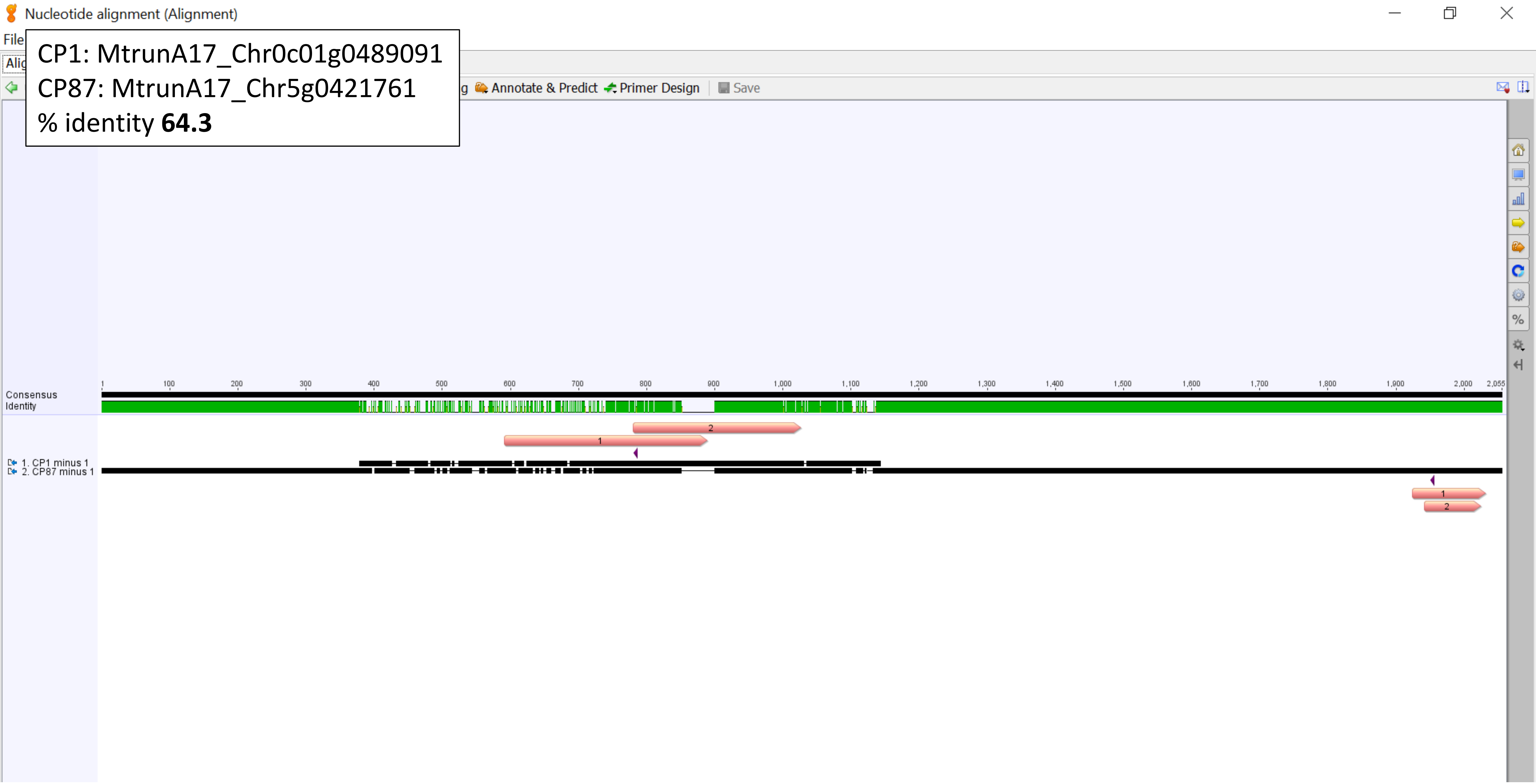

CP1: MtrunA17_Chr0c01g0489091
CP88_89: MtrunA17_Chr5g0422291
% identity **87.5**

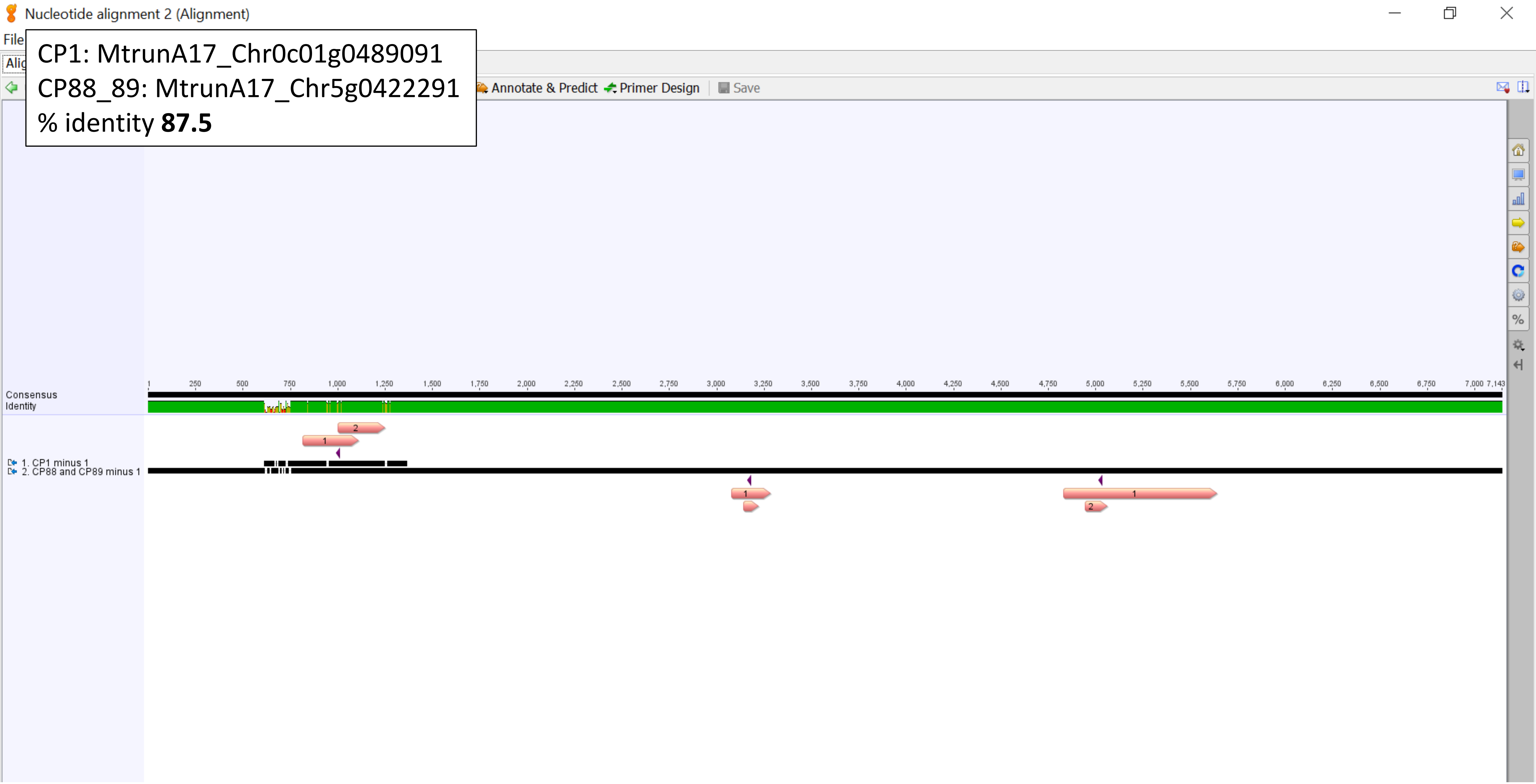

CP8: MtrunA17_Chr1g0155251
CP44: MtrunA17_Chr2g0326801
% identity **51.1**

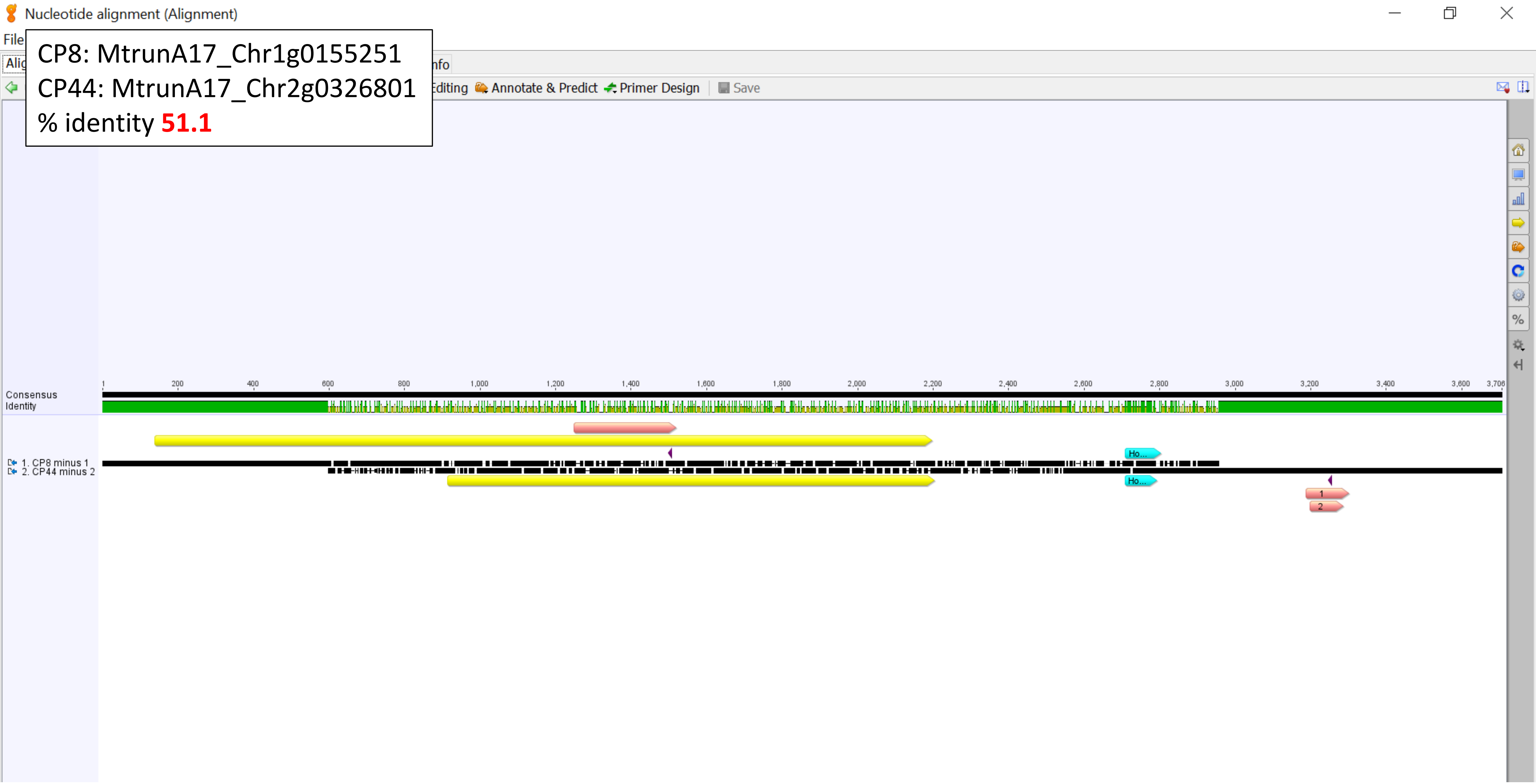

CP8: MtrunA17_Chr1g0155251
CP44: MtrunA17_Chr2g0326801
Homology region, % identity **83.9**

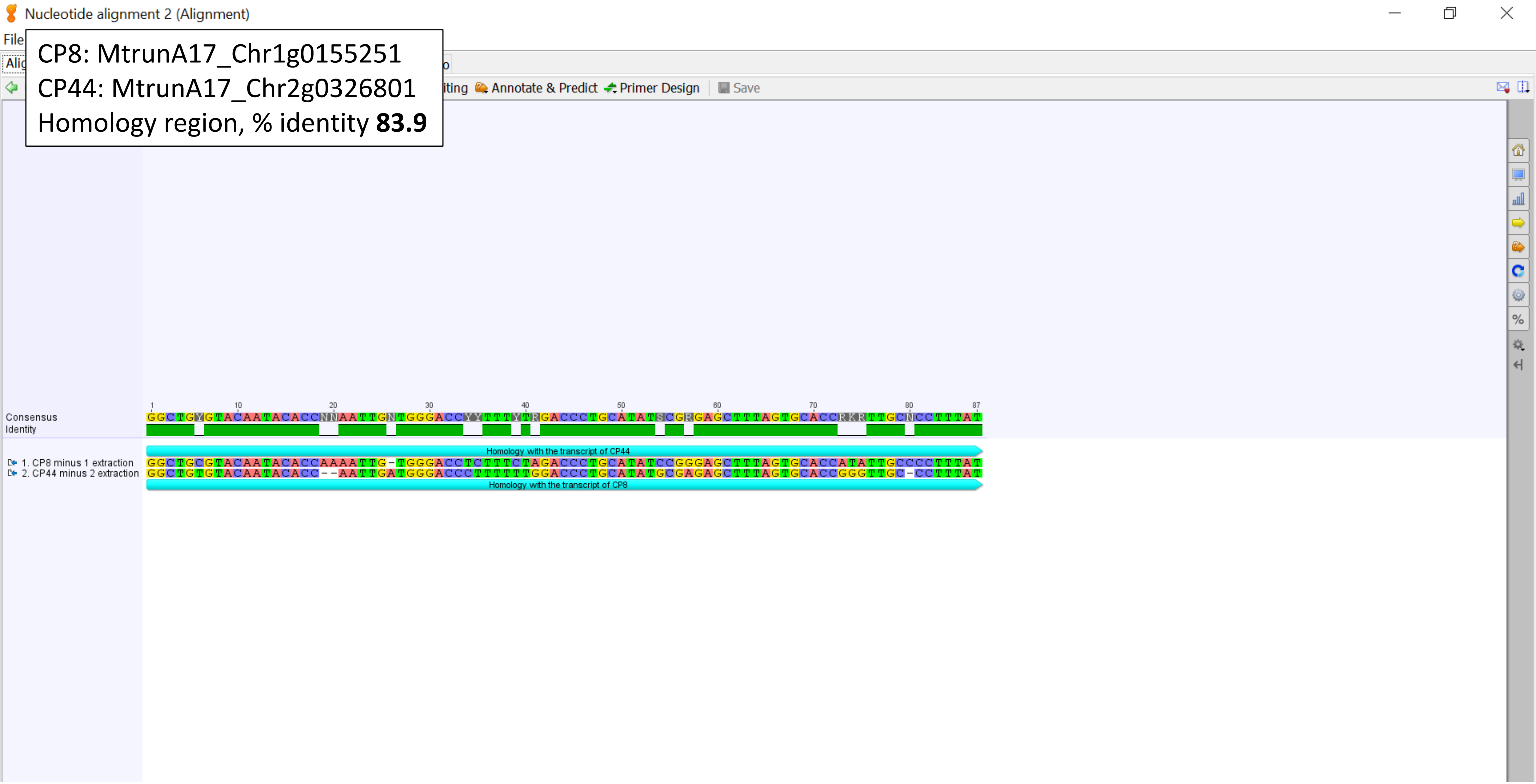

CP10: MtrunA17_Chr1g0162101
CP75: MtrunA17_Chr4g0037381
% identity **58.9**

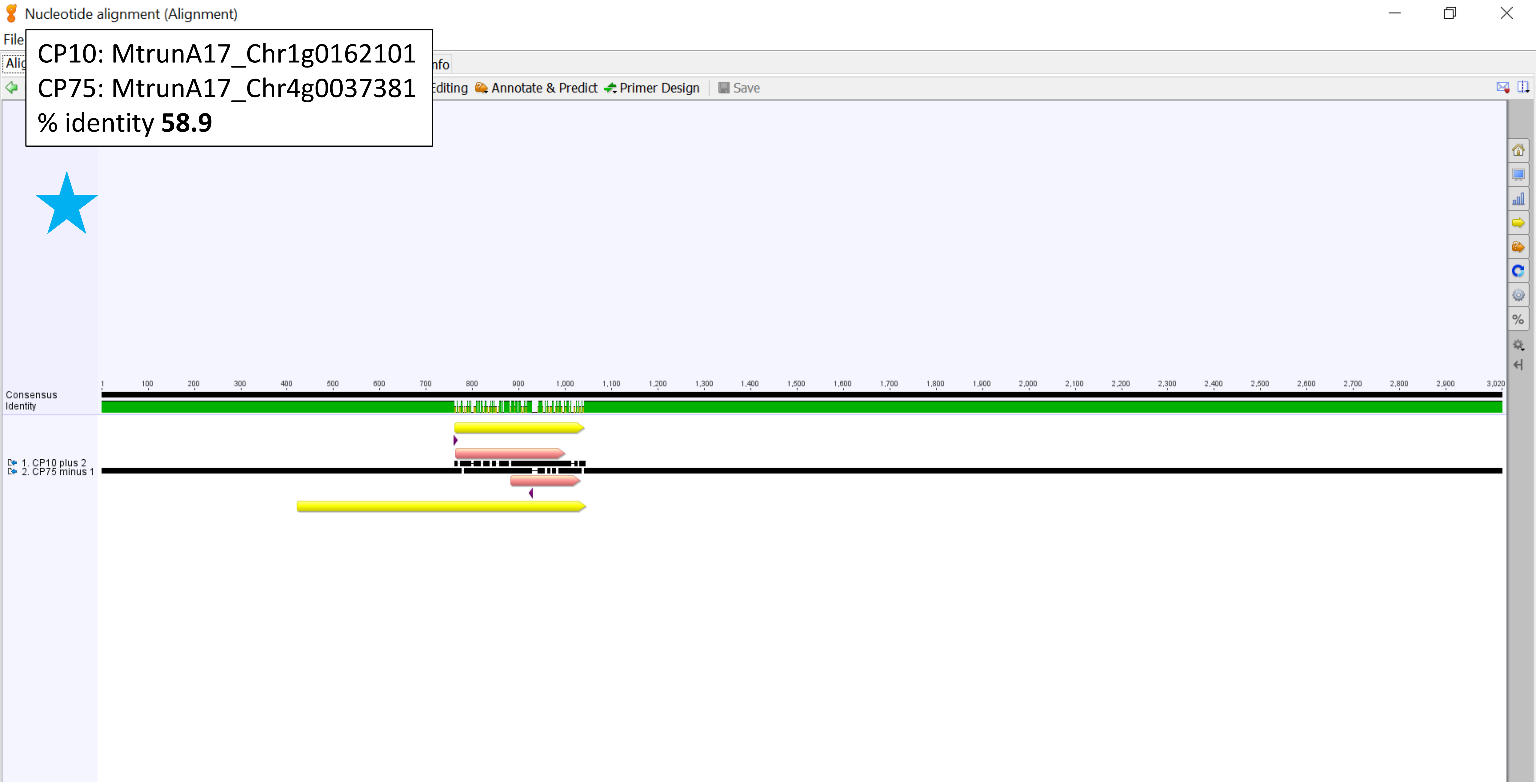

CP20: MtrunA17_Chr1g0190571
CP25: MtrunA17_Chr1g0202001
% identity **58.9**

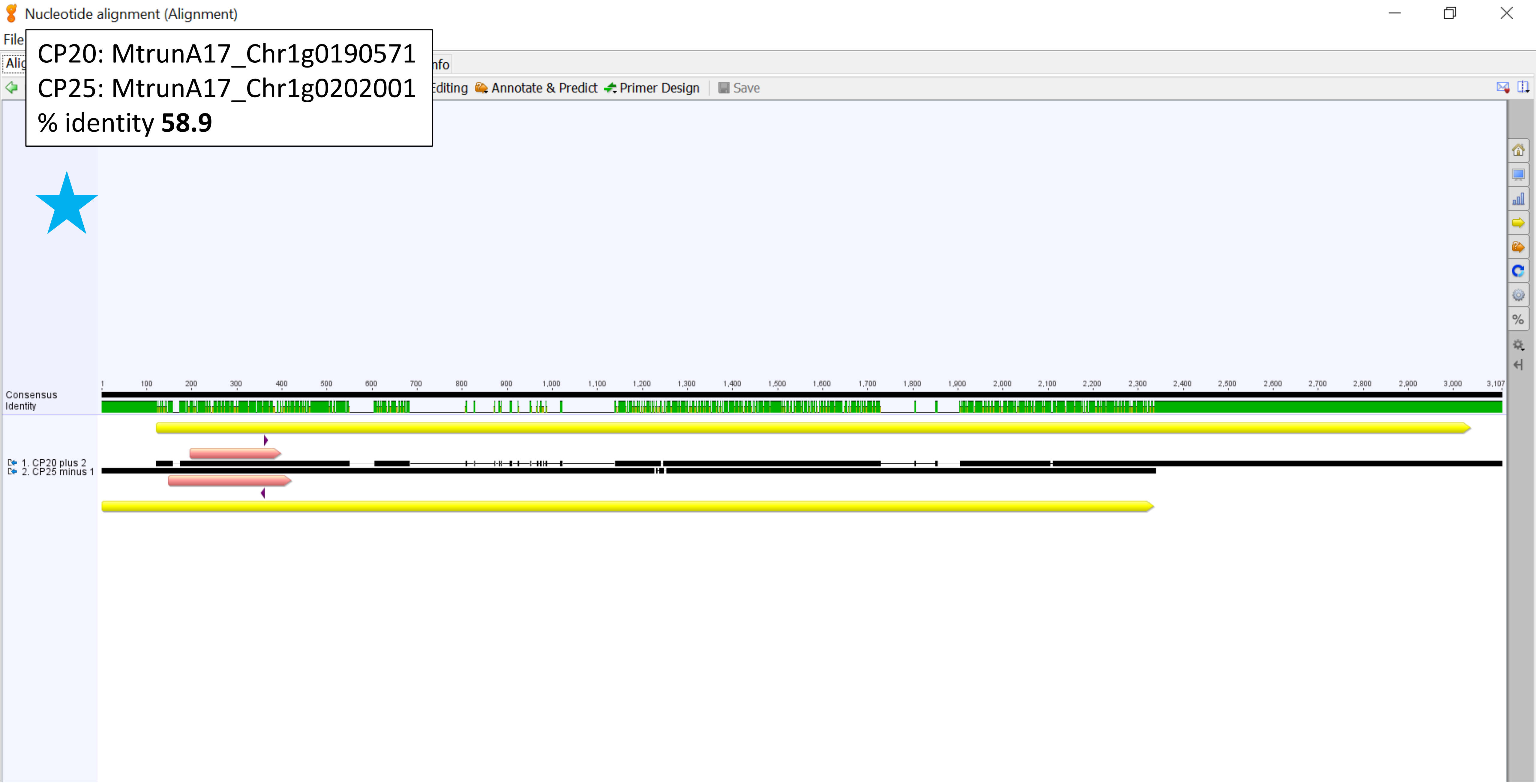

CP23_24: MtrunA17_Chr1g0200071
CP101: MtrunA17_Chr6g0458091
% identity **73.5**

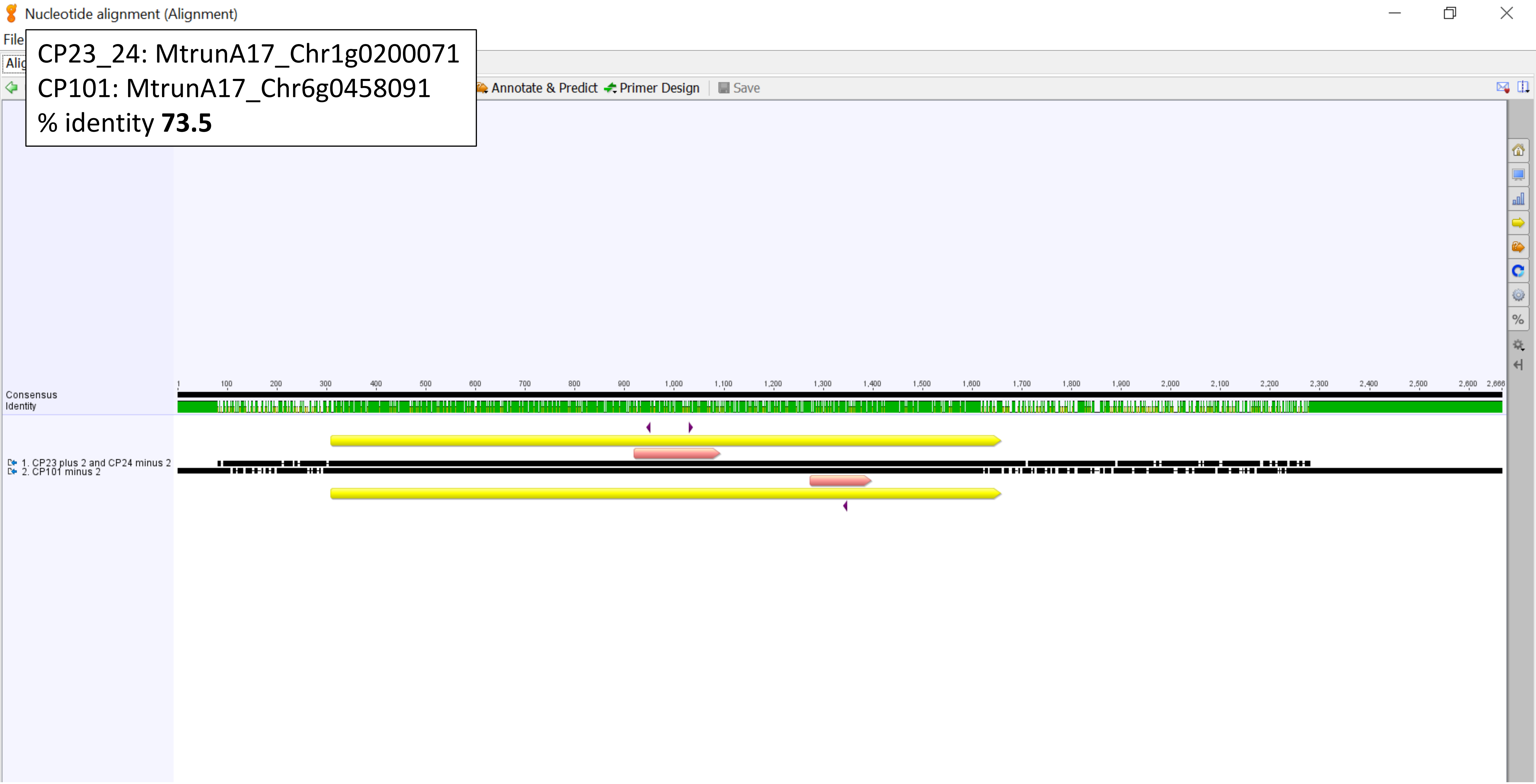

CP33: MtrunA17_Chr2g0283311
CP50: MtrunA17_Chr3g0091671
% identity **56.5**

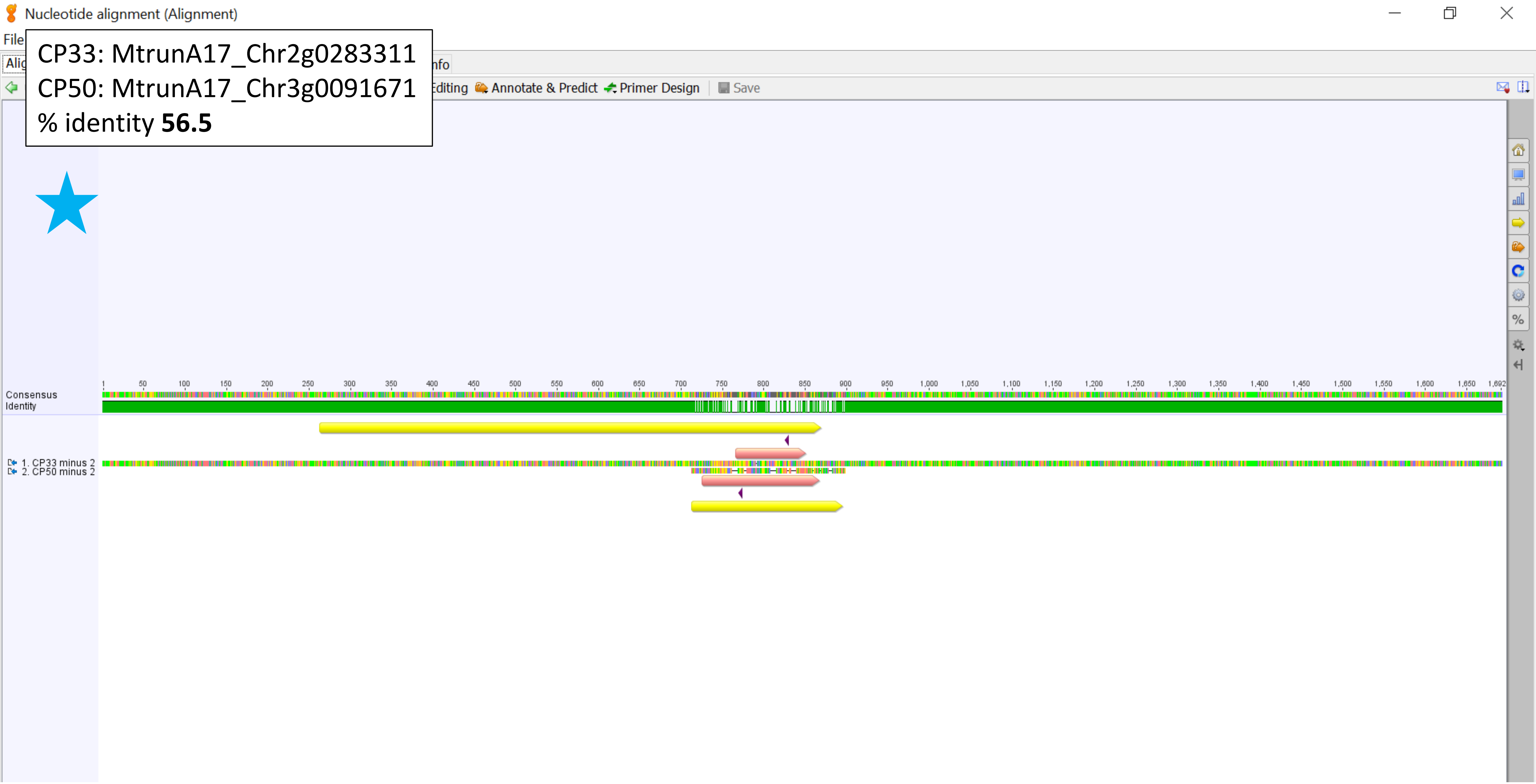

CP37: MtrunA17_Chr2g0299561
CP53: MtrunA17_Chr3g0102171
% identity **53.8**

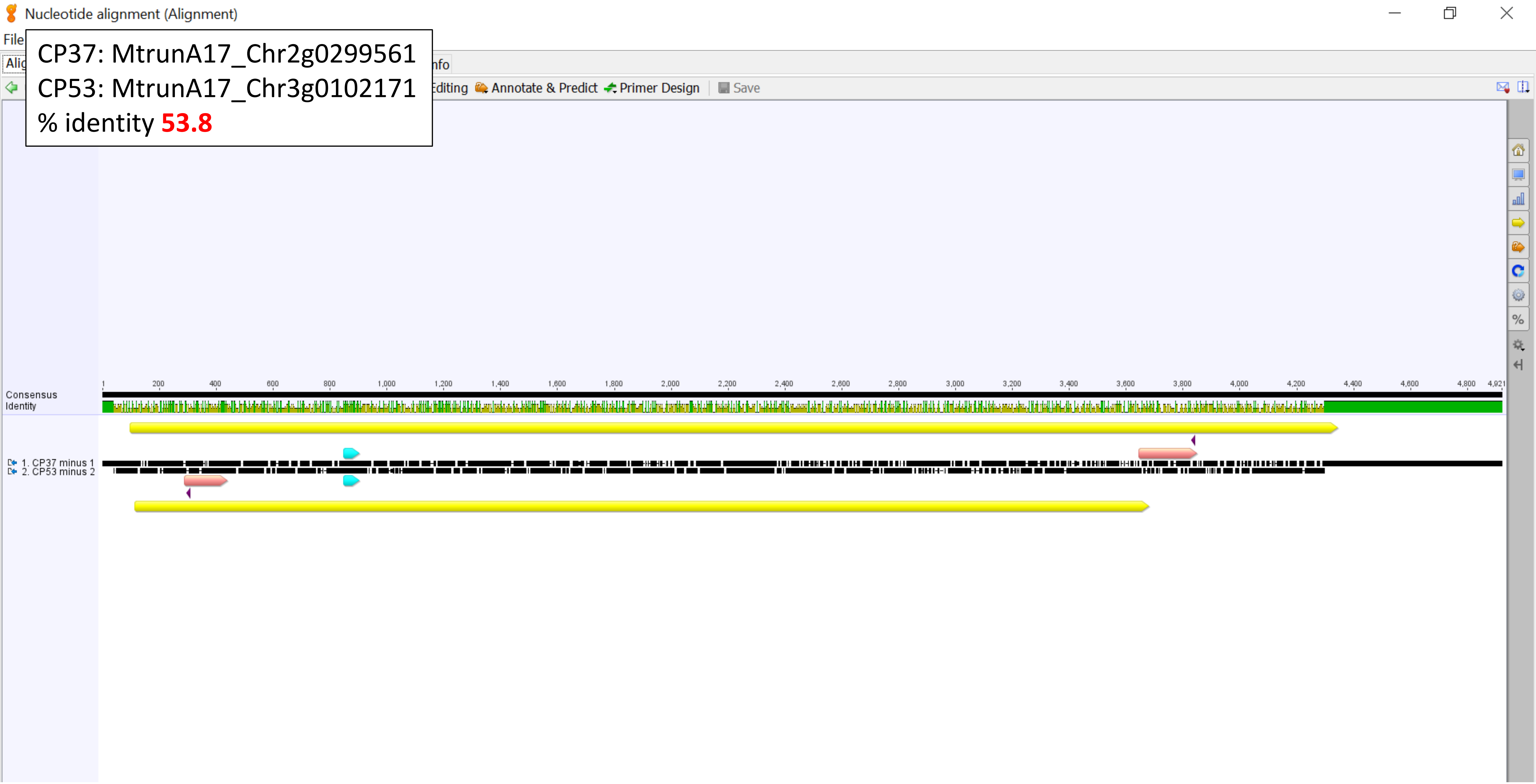

CP37: MtrunA17_Chr2g0299561
CP53: MtrunA17_Chr3g0102171
Homology region, % identity **83.6**

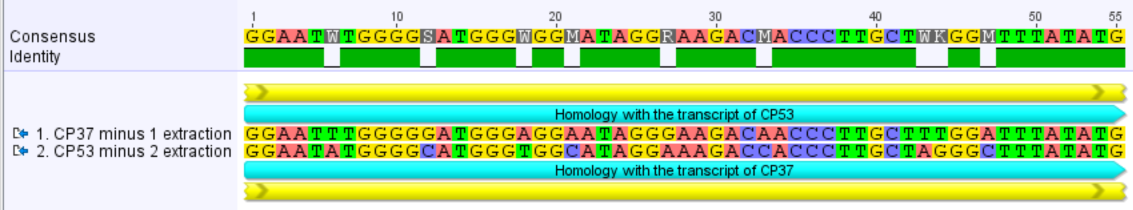

CP37: MtrunA17_Chr2g0299561
CP66: MtrunA17_Chr3g1011650
% identity **63.3**

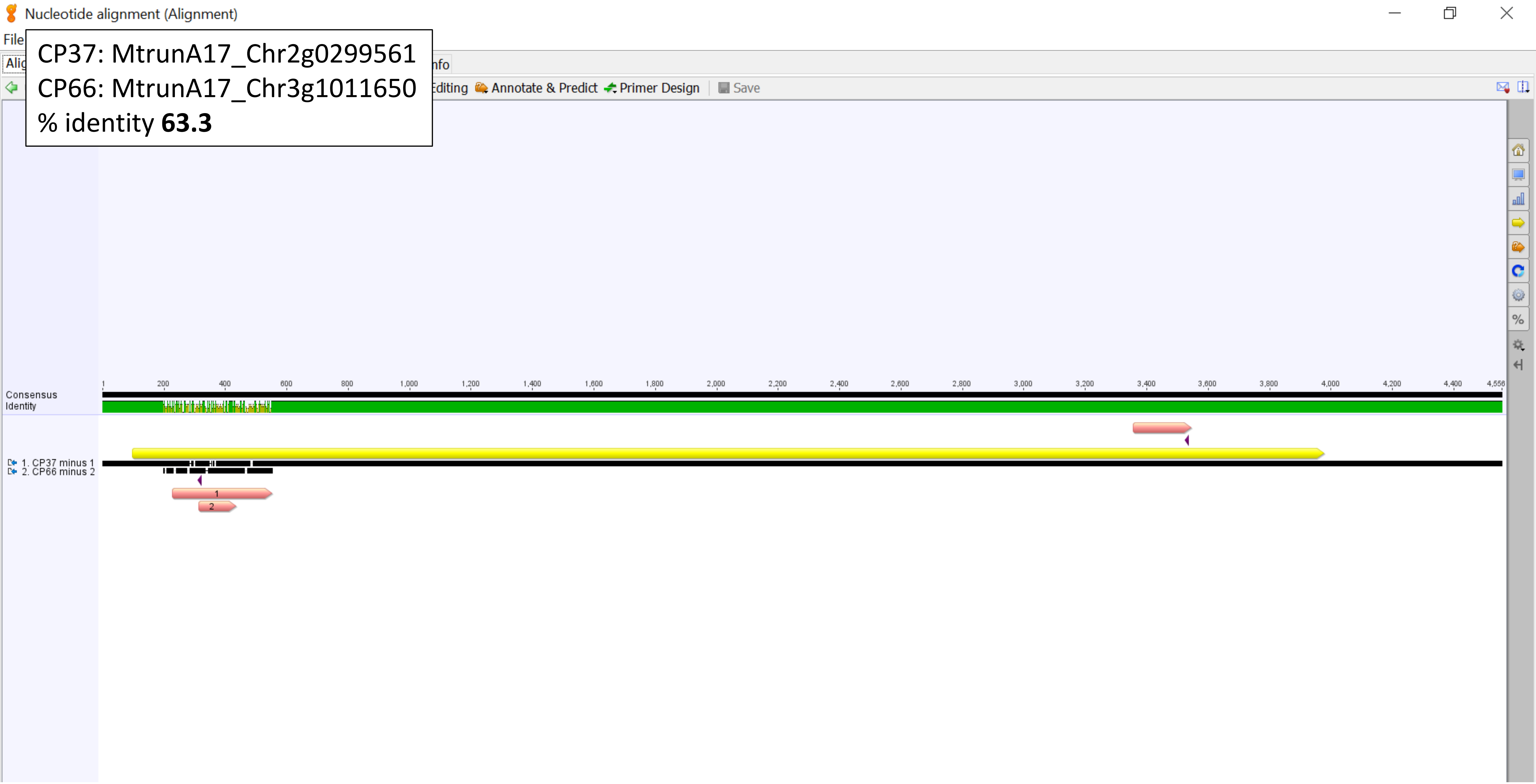

CP51: MtrunA17_Chr3g0096421
CP121: MtrunA17_Chr7g0270811
% identity **72.8**

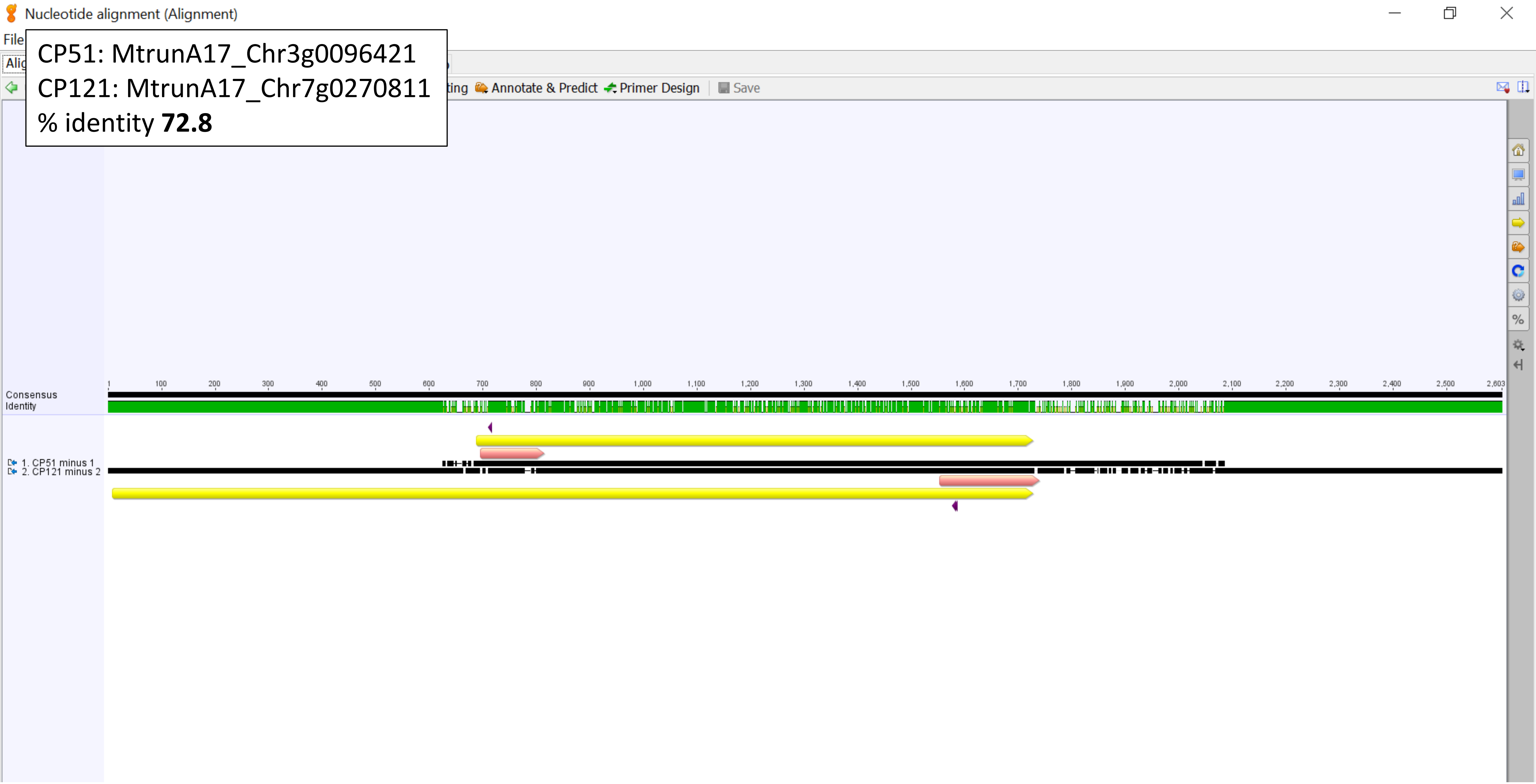

Nucleotide alignment 2 (Alignment)

CP53: MtrunA17_Chr3g0102171
CP66: MtrunA17_Chr3g1011650
% identity **93.0**

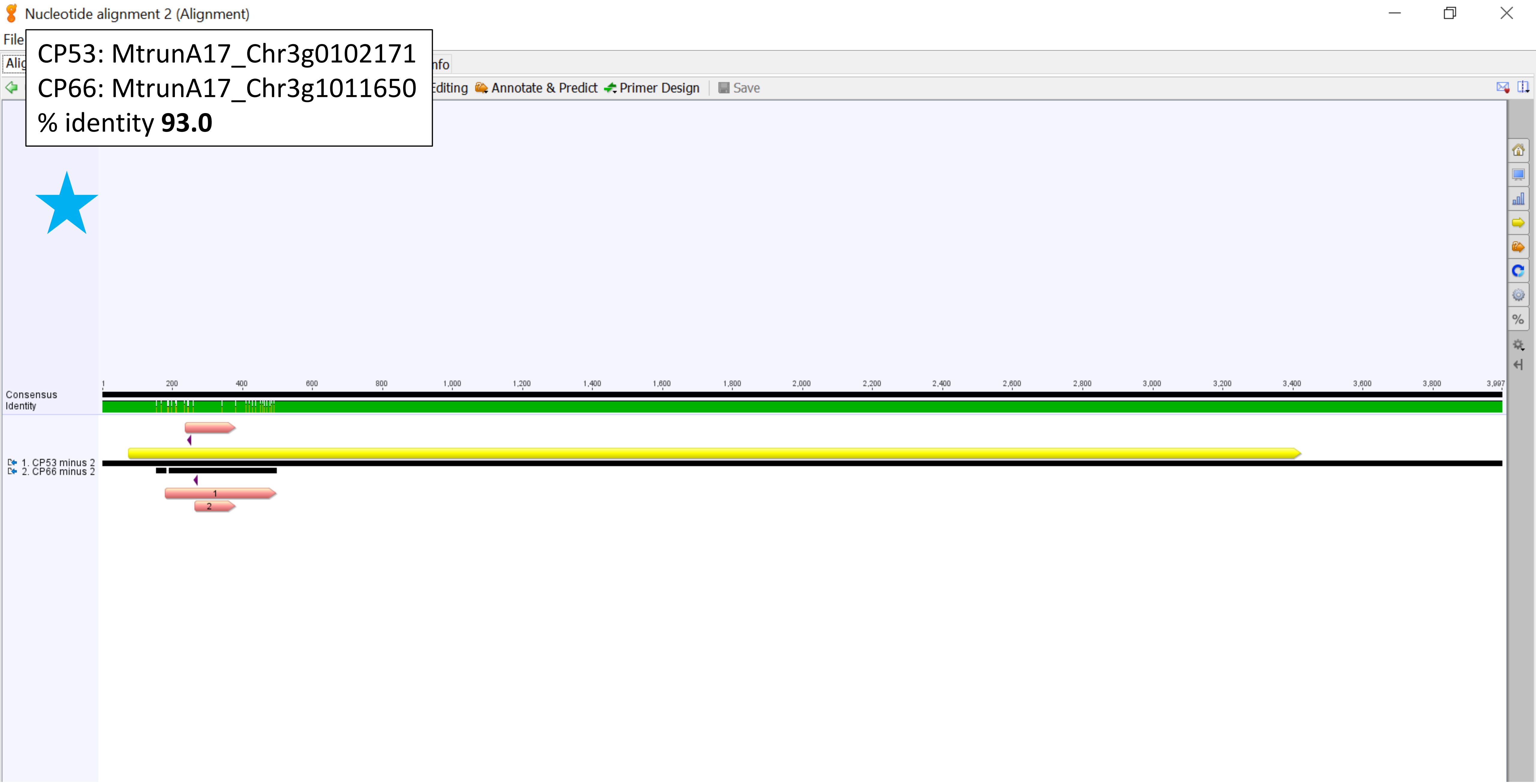

CP60: MtrunA17_Chr3g0135761
CP62: MtrunA17_Chr3g0141901
% identity **52.1**

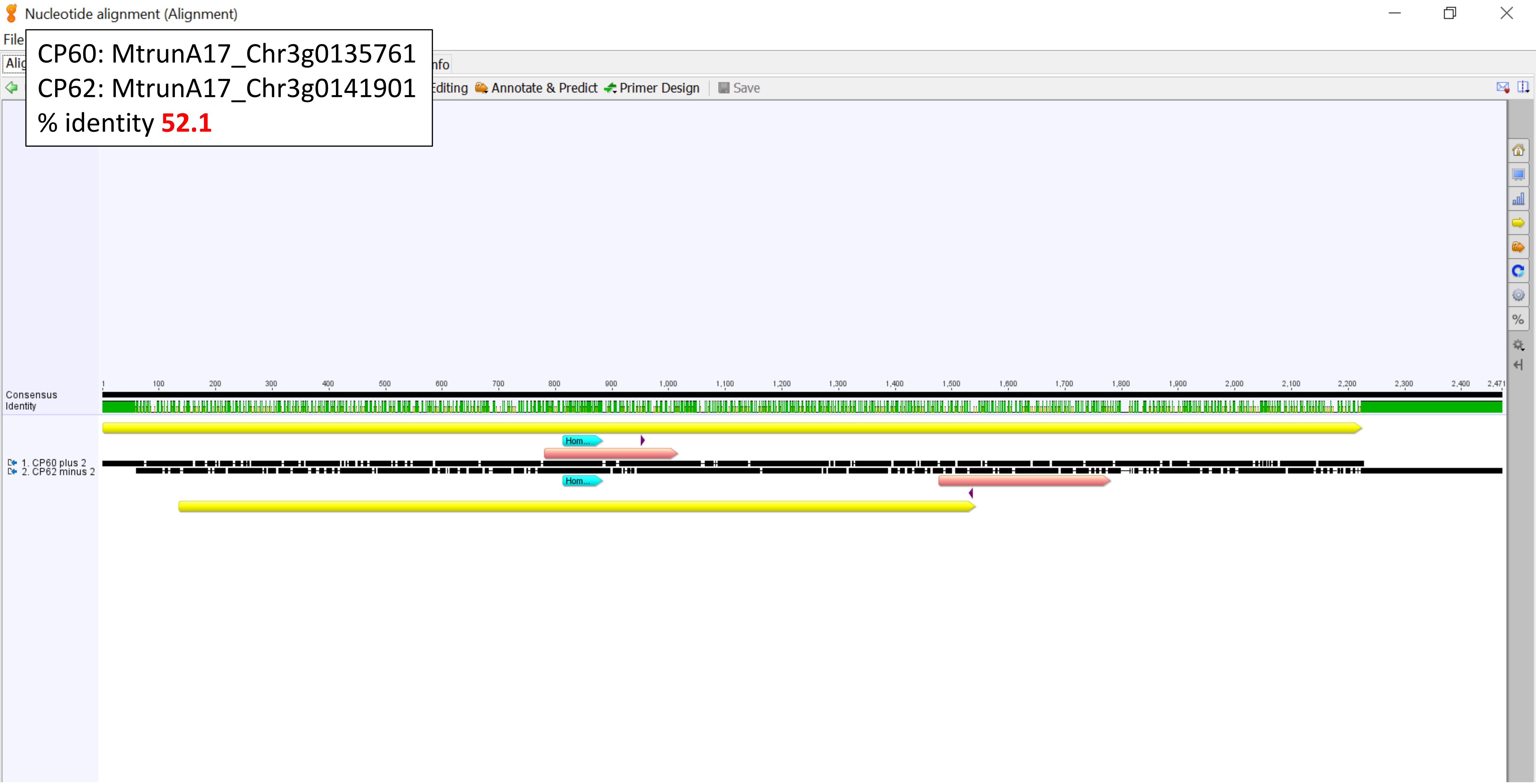

CP60: MtrunA17_Chr3g0135761
CP62: MtrunA17_Chr3g0141901
Homology region, % identity **79.2**

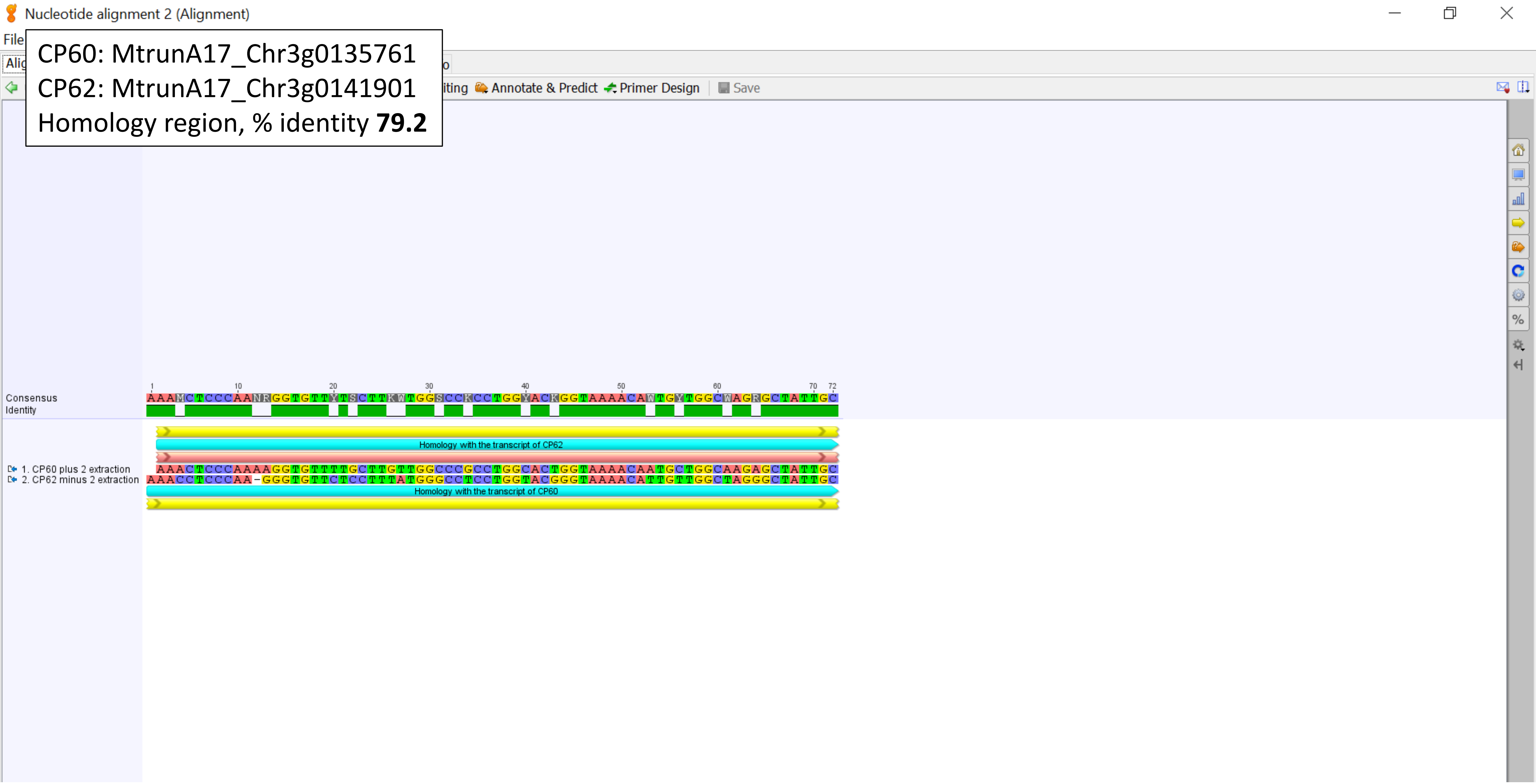

CP66: MtrunA17_Chr3g1011650
CP107: MtrunA17_Chr6g0479001
% identity **59.9**

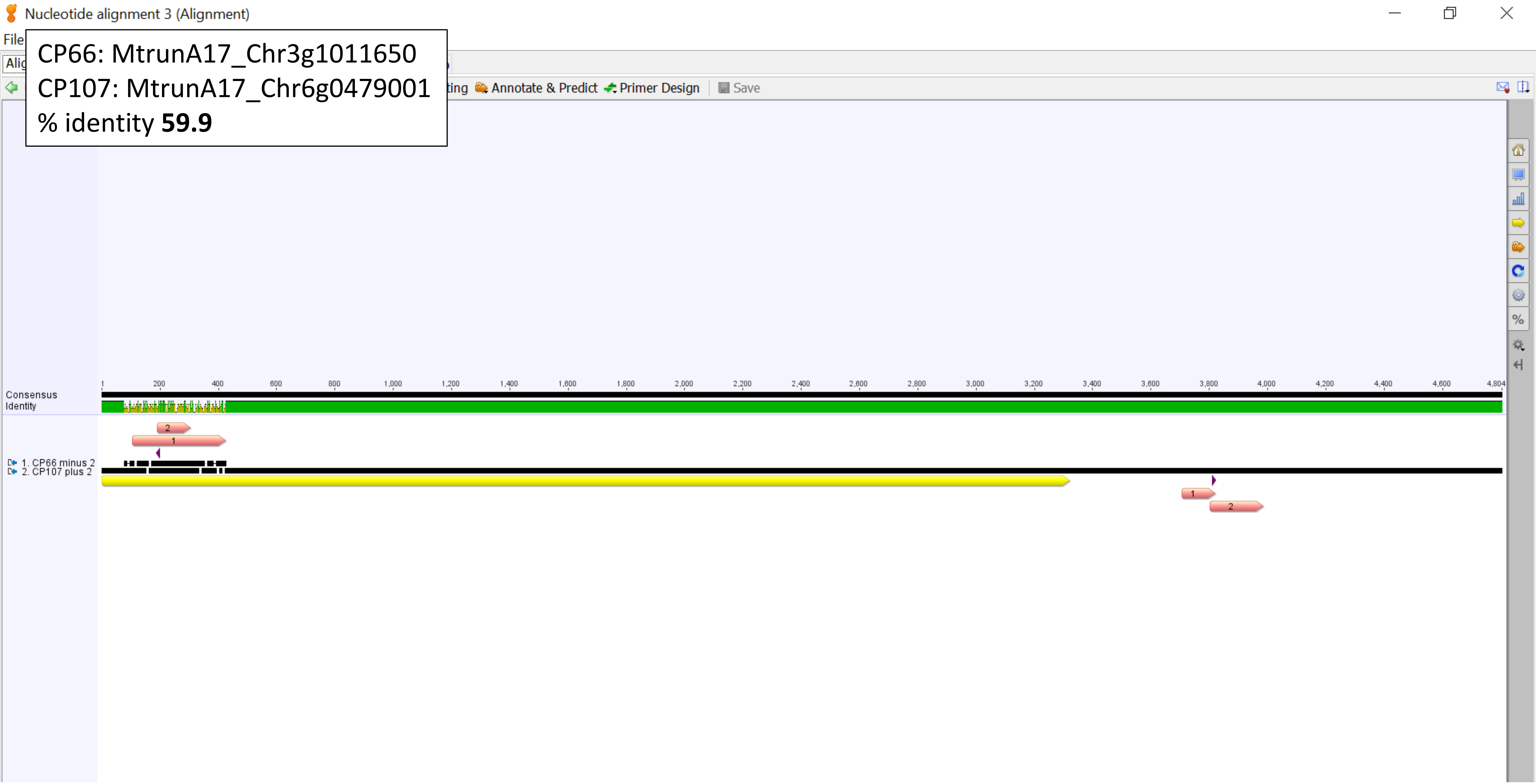

CP72: MtrunA17_Chr4g0023791
CP87: MtrunA17_Chr5g0421761
% identity **58.8**

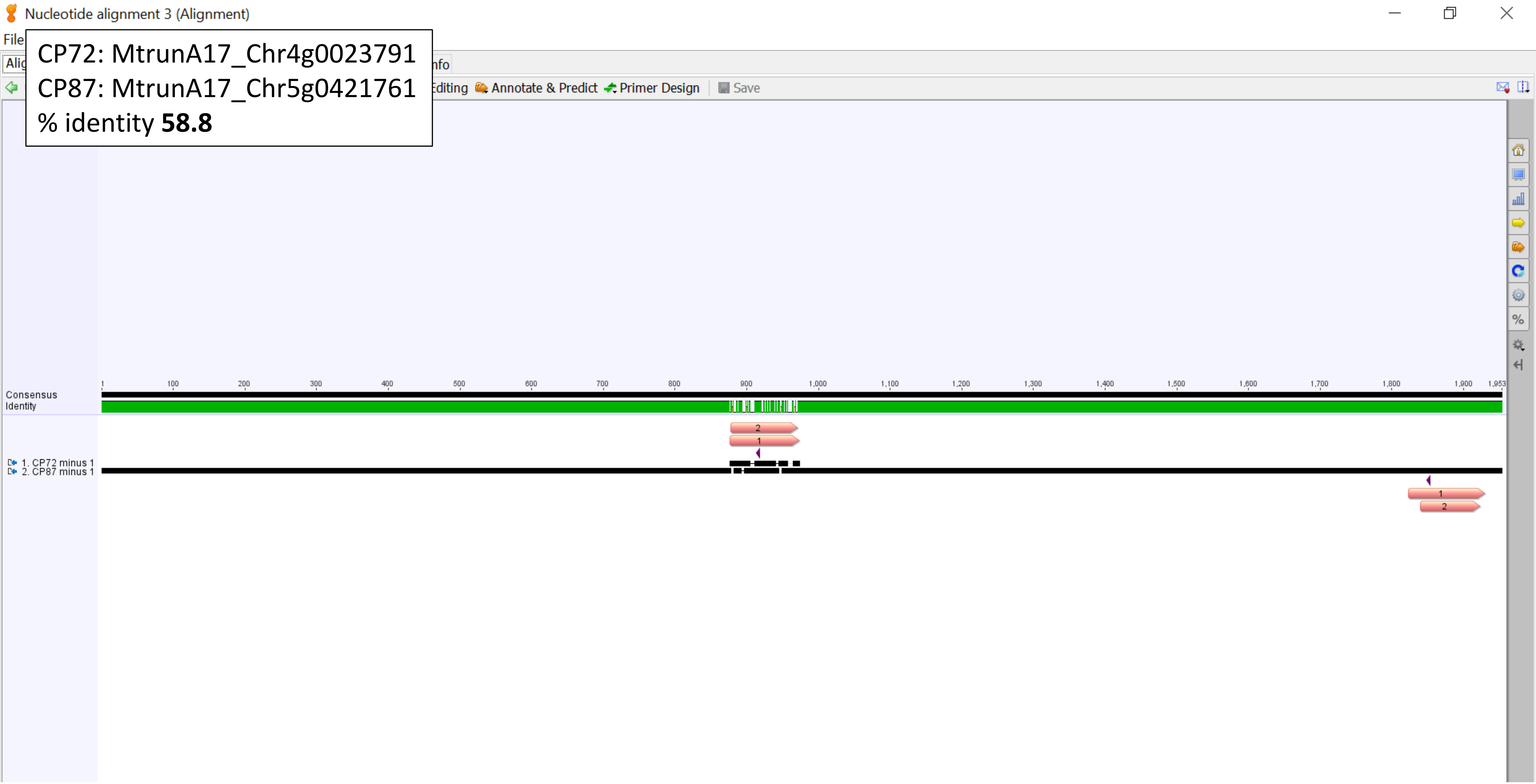

CP72: MtrunA17_Chr4g0023791
CP87: MtrunA17_Chr5g0421761
Zoom-in view

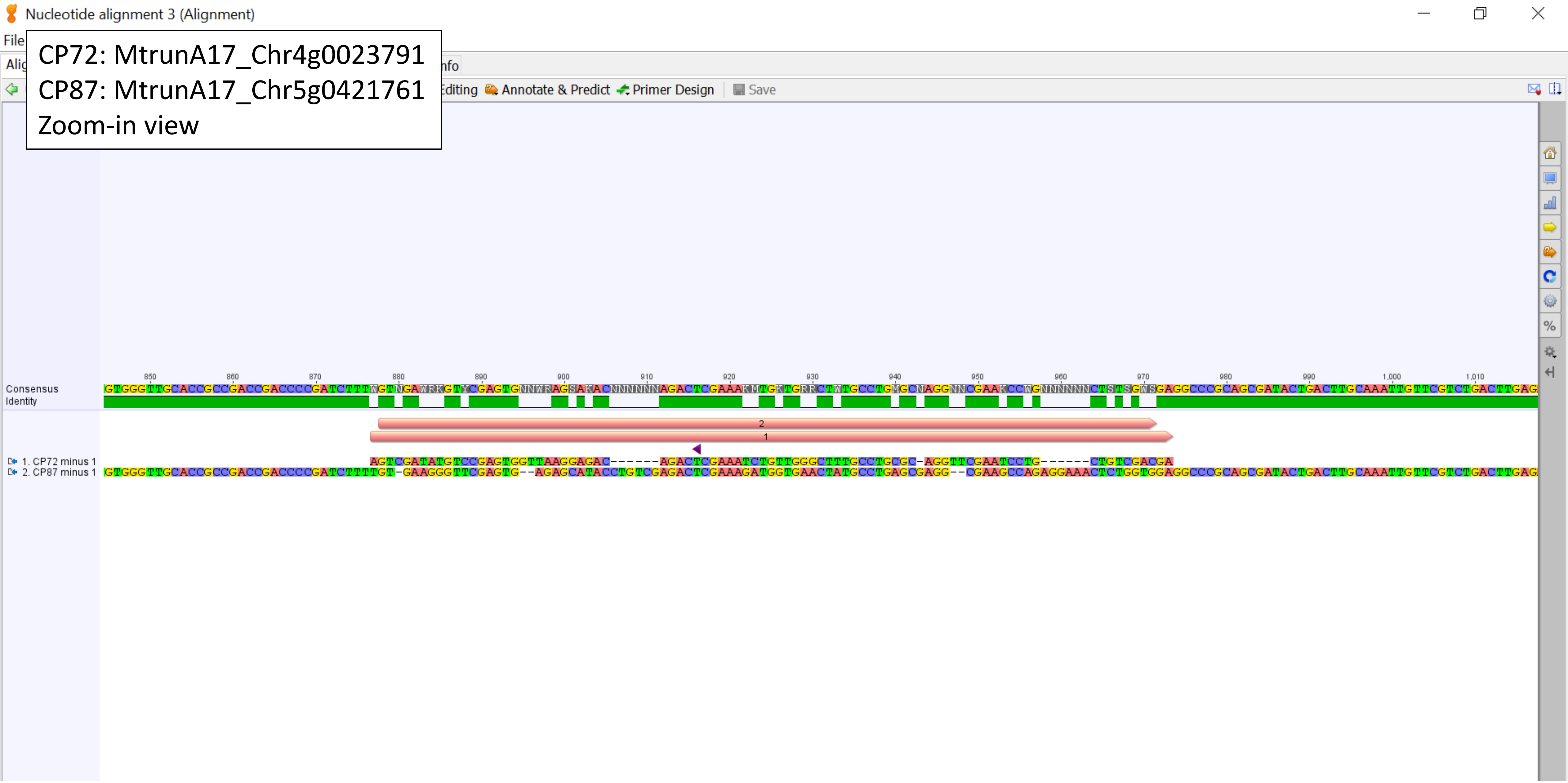

CP72: MtrunA17_Chr4g0023791
CP88_89: MtrunA17_Chr5g0422291
% identity **56.3**

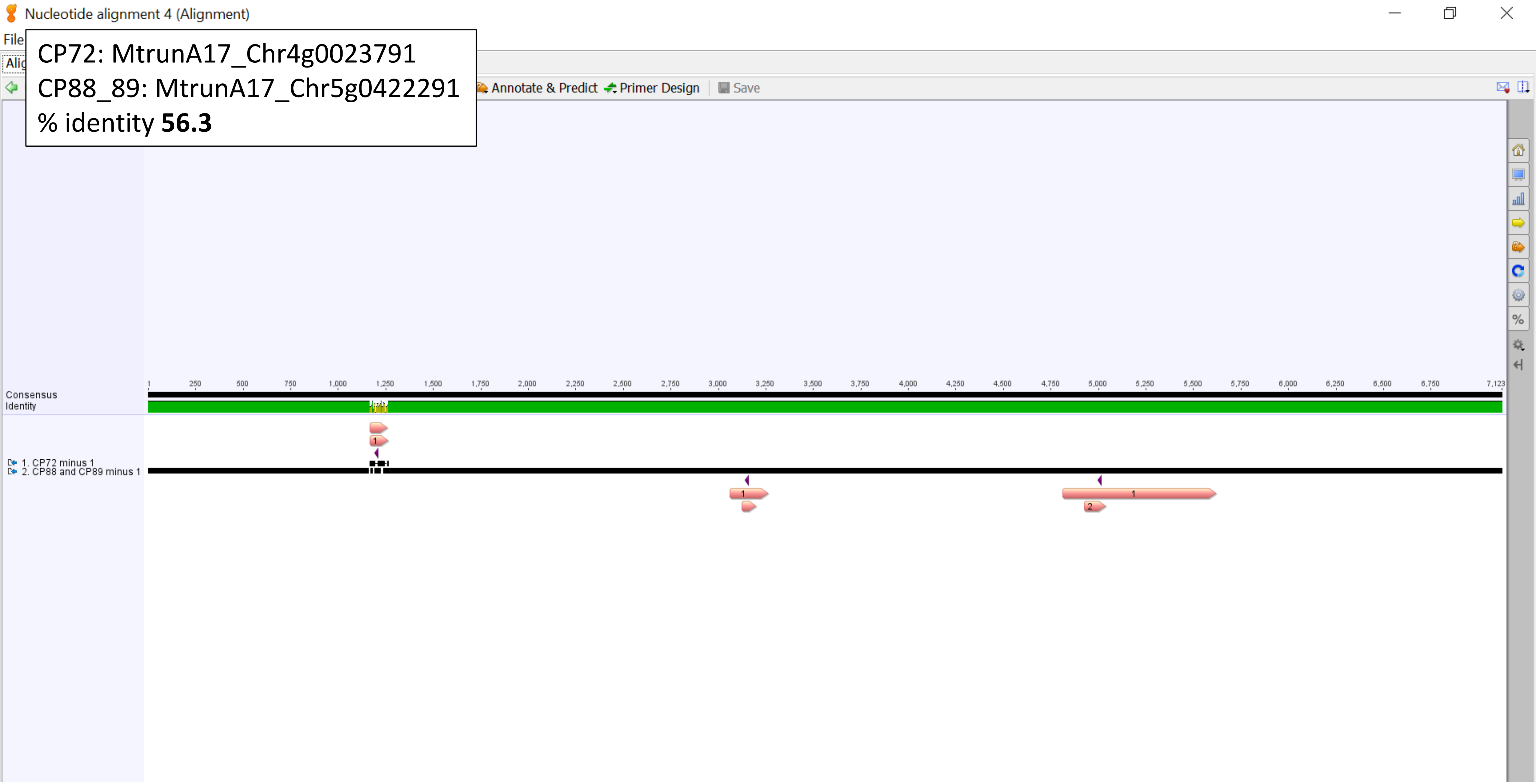

CP72: MtrunA17_Chr4g0023791
CP88_89: MtrunA17_Chr5g0422291
Zoom-in view

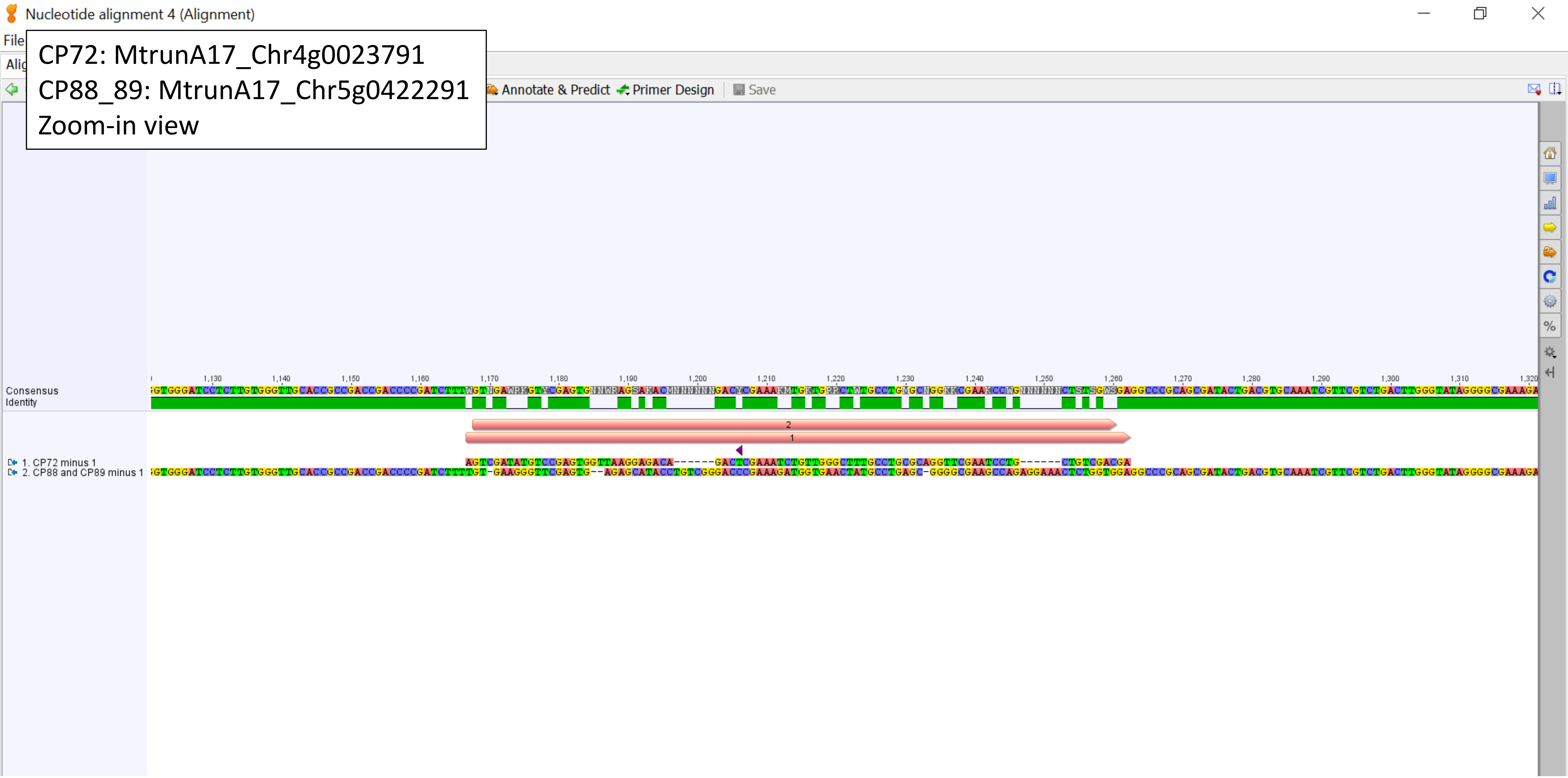

CP85: MtrunA17_Chr5g0415031
CP132: MtrunA17_Chr8g0368731
% identity **74.8**

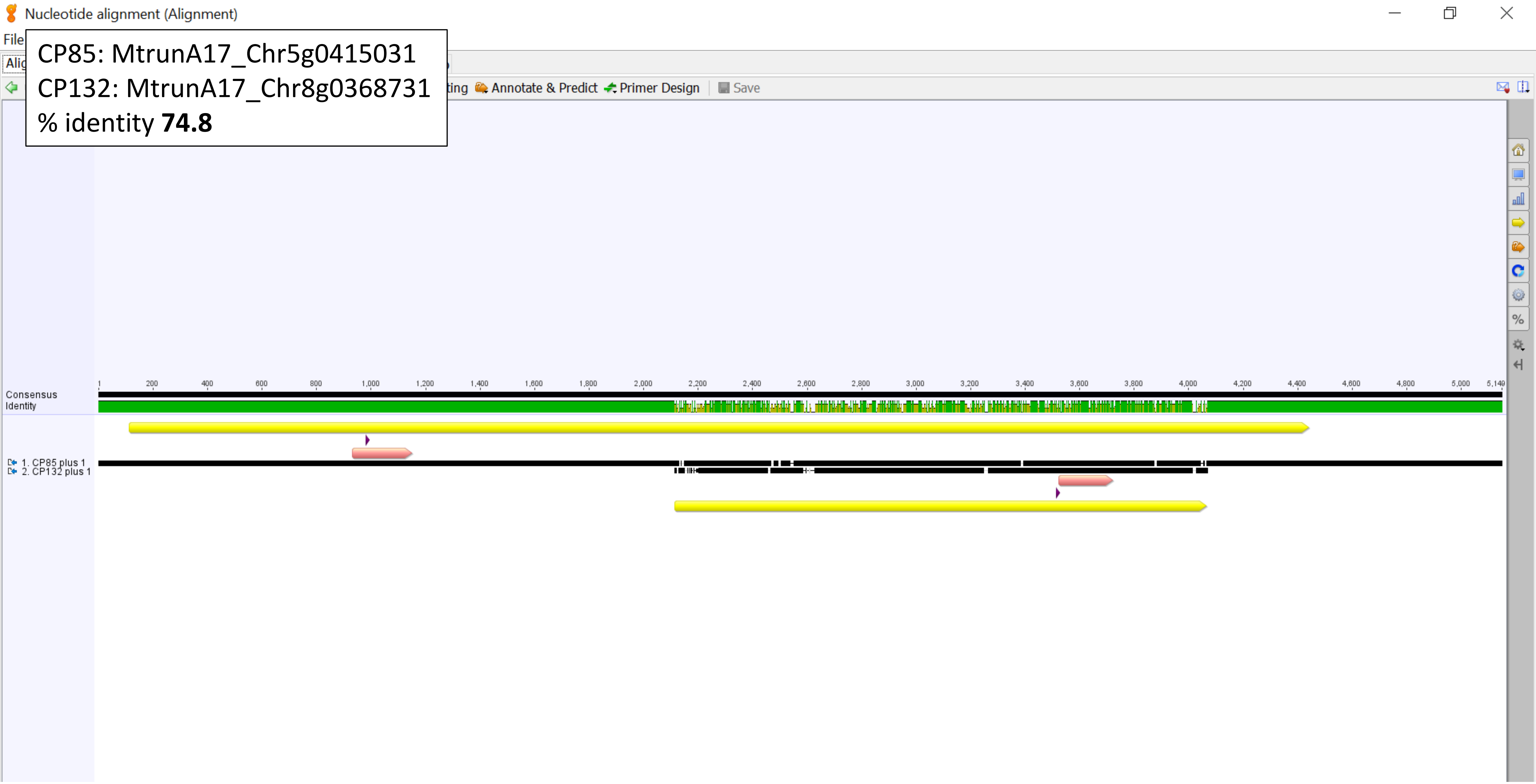

CP87: MtrunA17_Chr5g0421761

CP88_89: MtrunA17_Chr5g0422291

Geneious alignment, % identity **63.1**

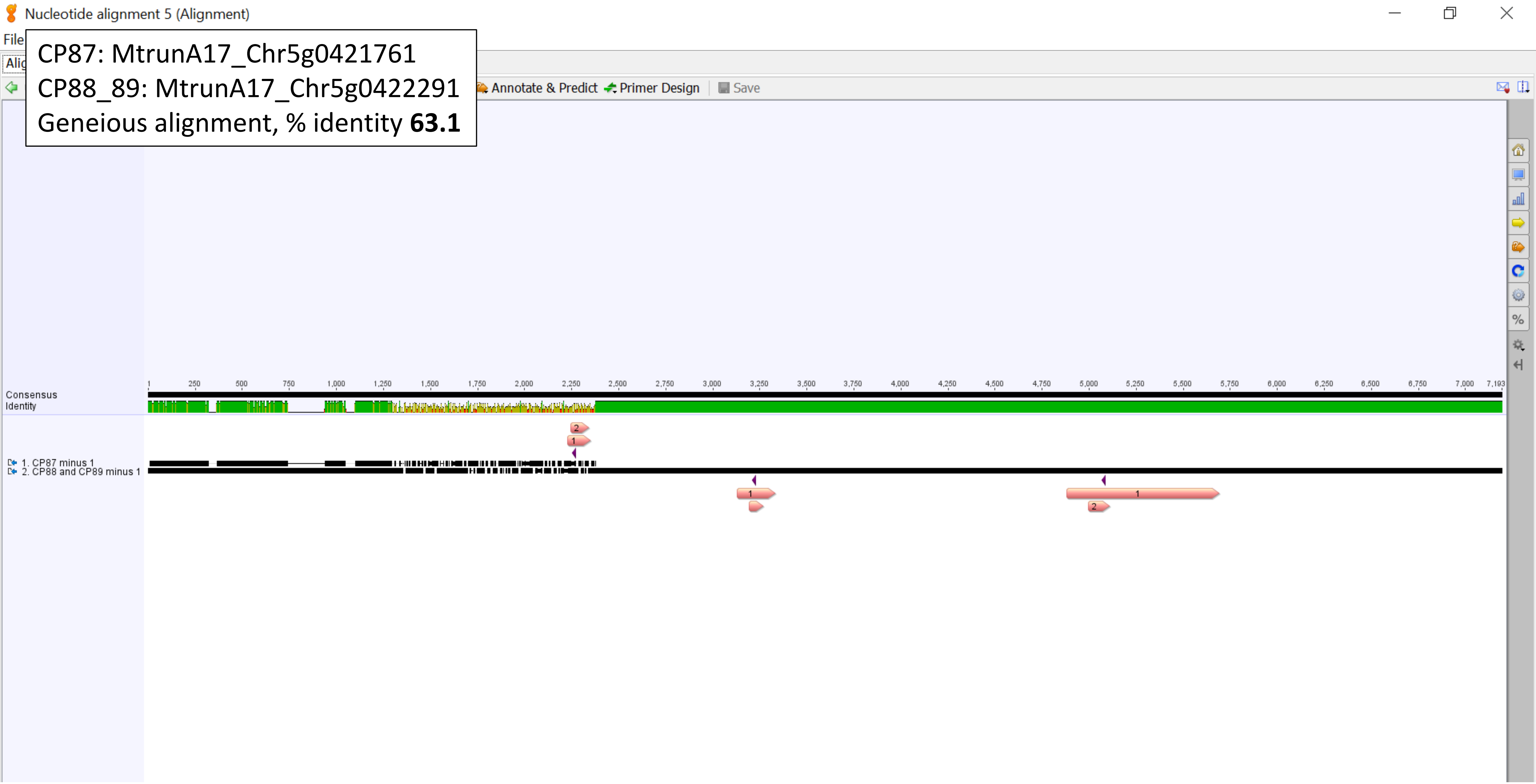

CP87: MtrunA17_Chr5g0421761
CP88_89: MtrunA17_Chr5g0422291
MUSCLE alignment, , % identity **25.9**

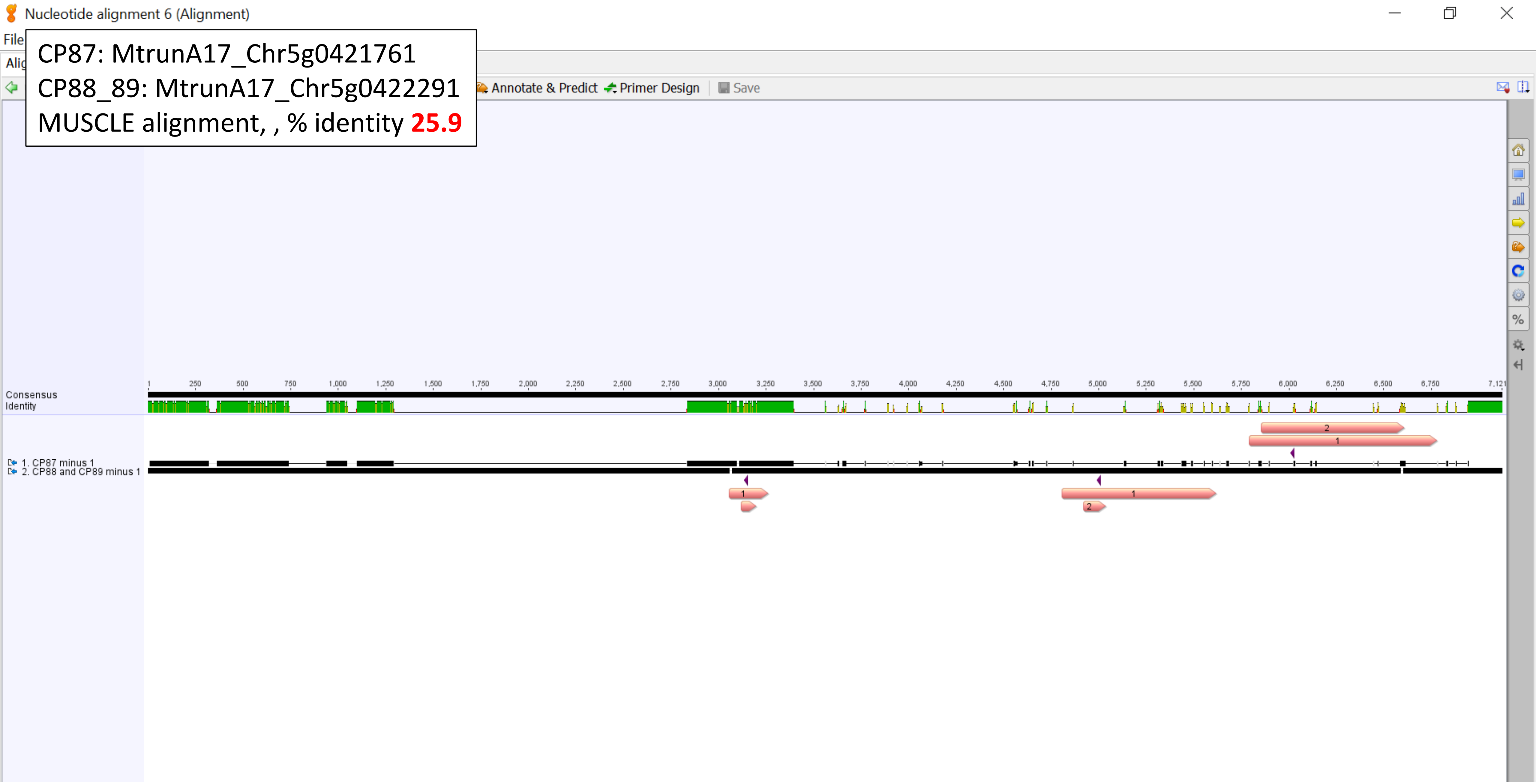

CP87: MtrunA17_Chr5g0421761
CP114: MtrunA17_Chr7g0229401
% identity **52.4**

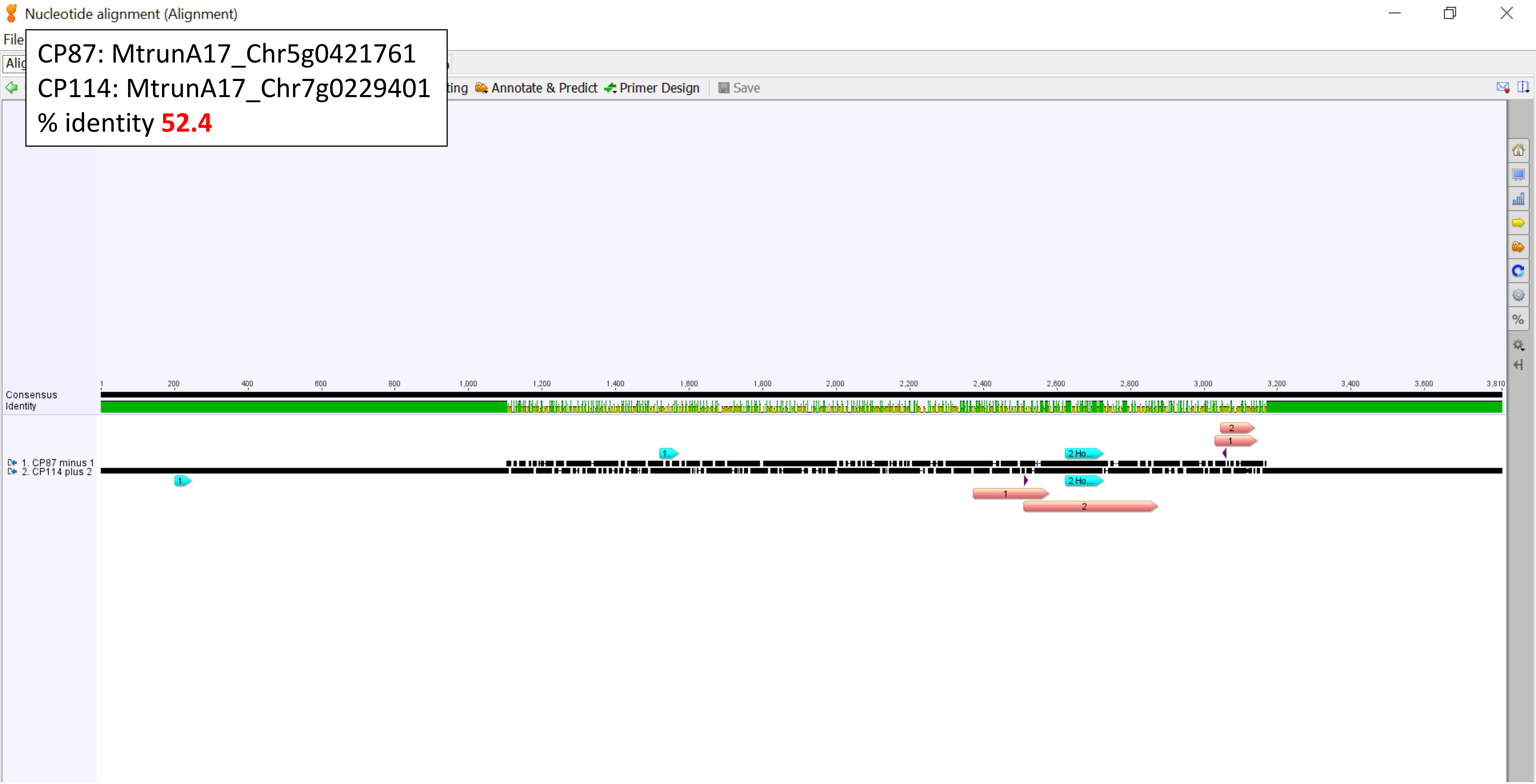

CP87: MtrunA17_Chr5g0421761
CP114: MtrunA17_Chr7g0229401
Homology region 1, % identity **87.0**

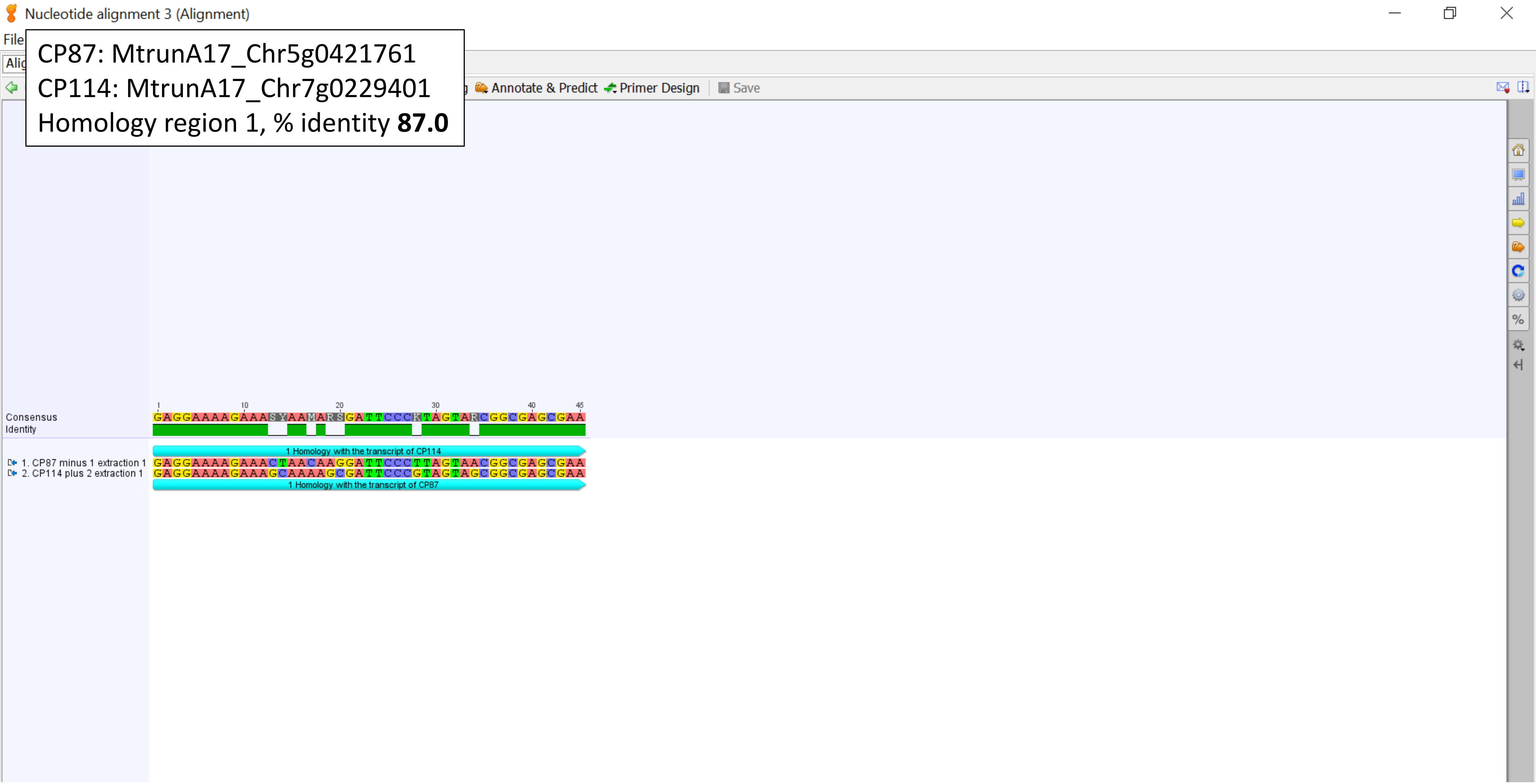

CP87: MtrunA17_Chr5g0421761
CP114: MtrunA17_Chr7g0229401
Homology region 2, % identity **77.8**

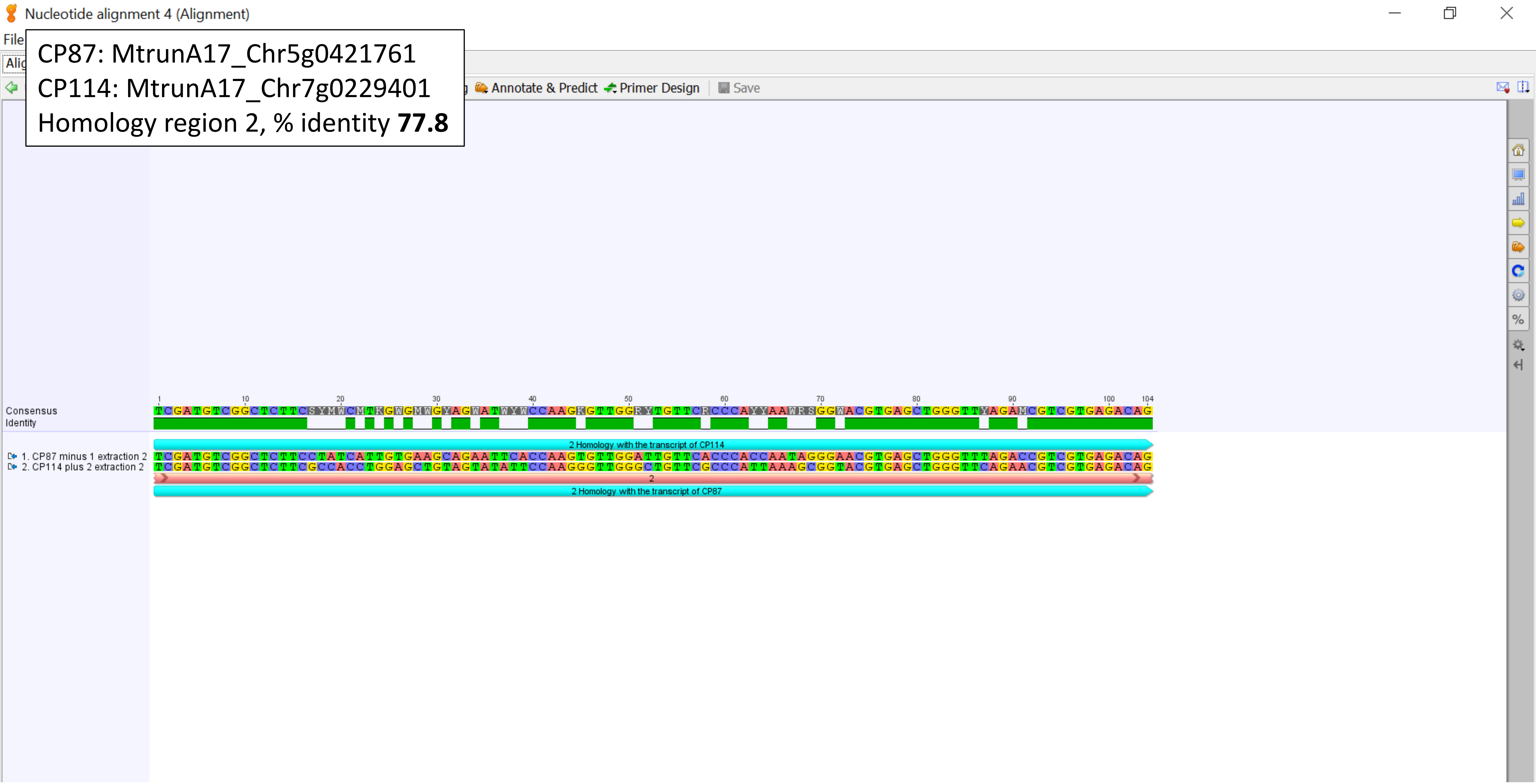

CP87: MtrunA17_Chr5g0421761
CP141: MtrunA17_CPg0492331
% identity **52.3**

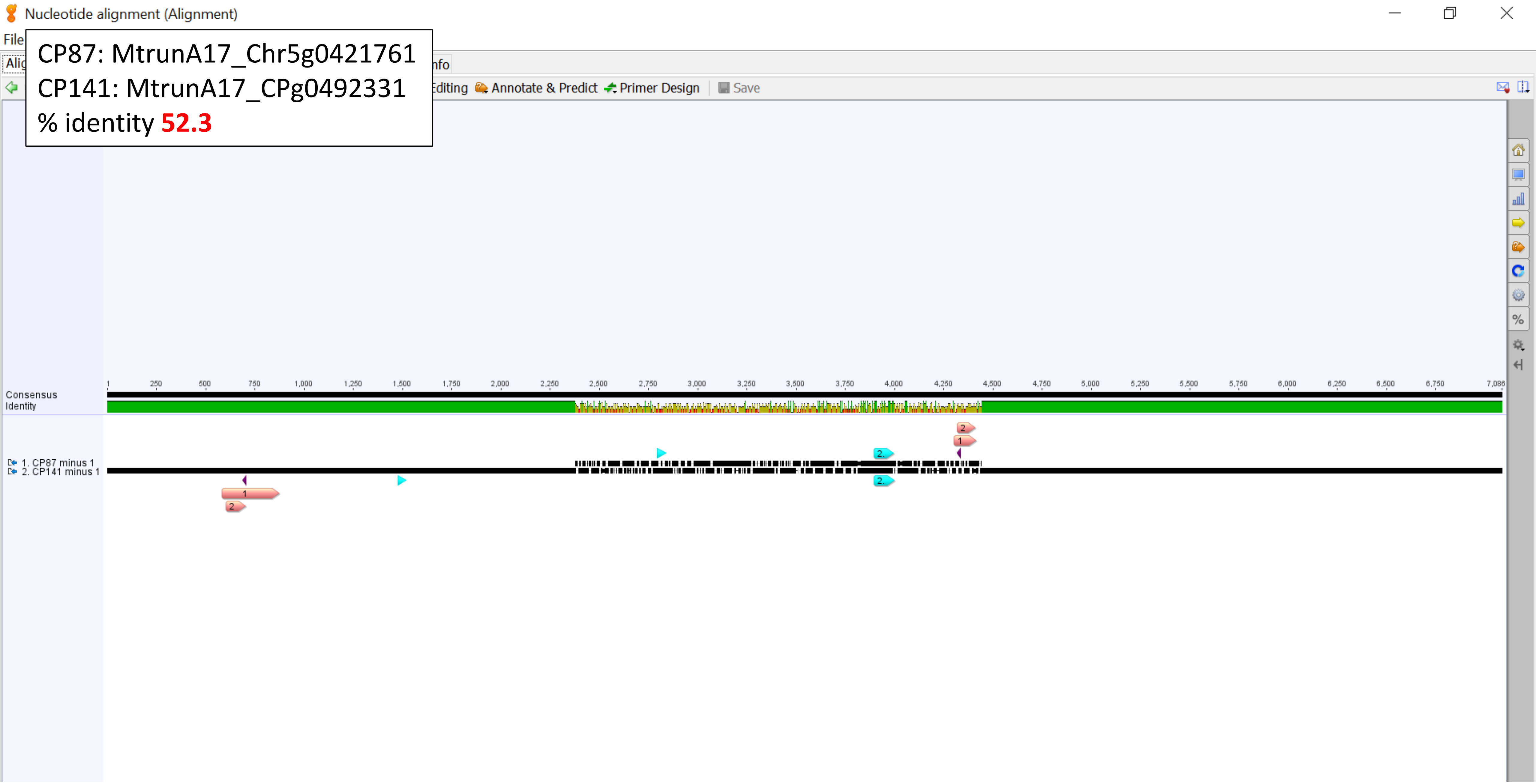

CP87: MtrunA17_Chr5g0421761
CP141: MtrunA17_CPg0492331
Homology region 1, % identity **87.0**

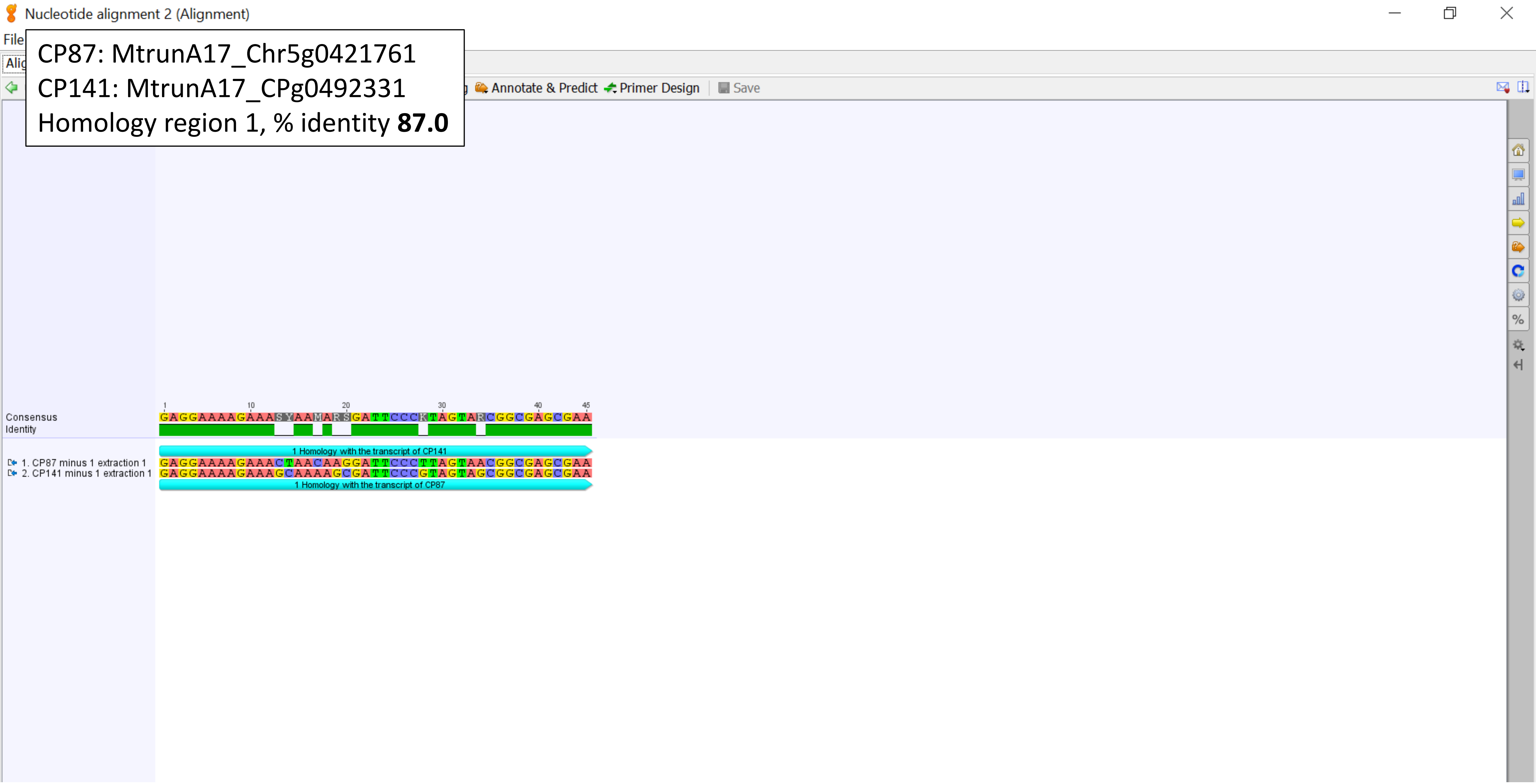

CP87: MtrunA17_Chr5g0421761
CP141: MtrunA17_CPg0492331
Homology region 2, % identity **77.4**

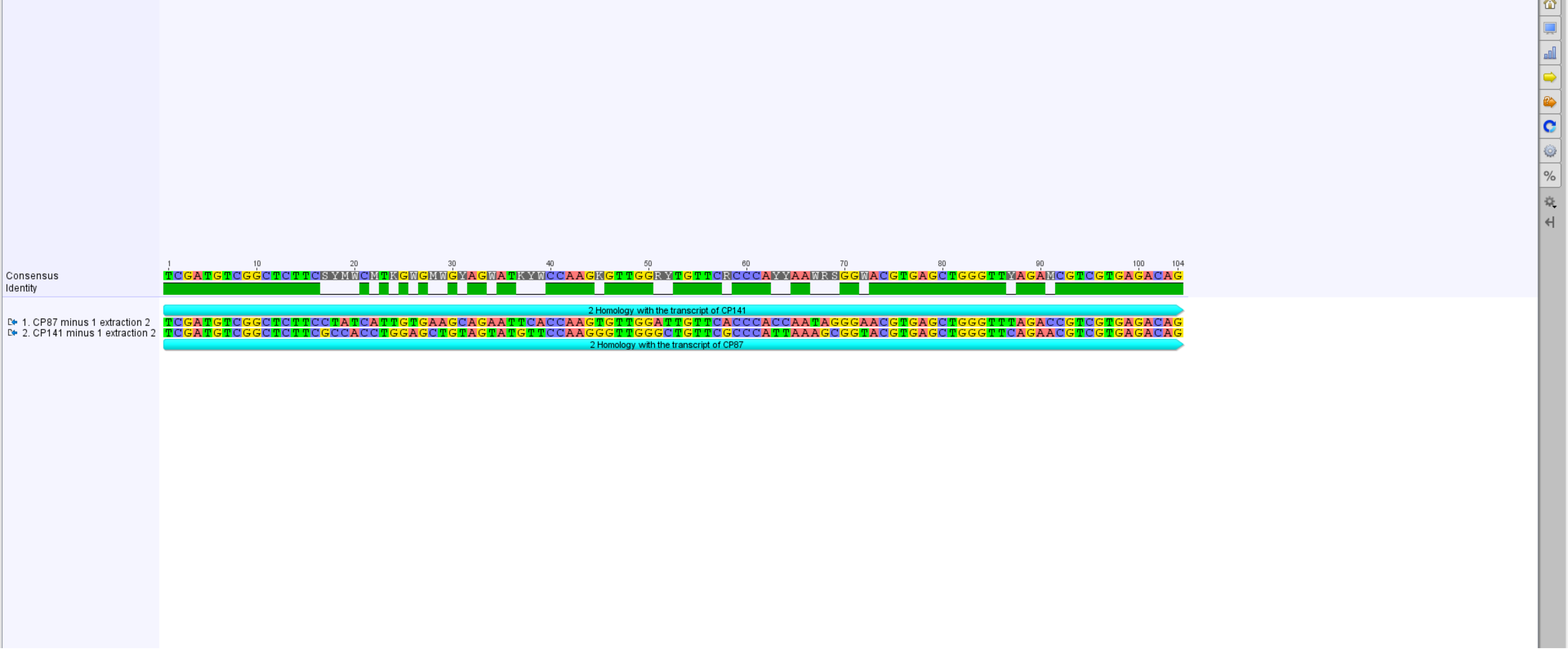

CP88_89: MtrunA17_Chr5g0422291
CP114: MtrunA17_Chr7g0229401
% identity **51.4**

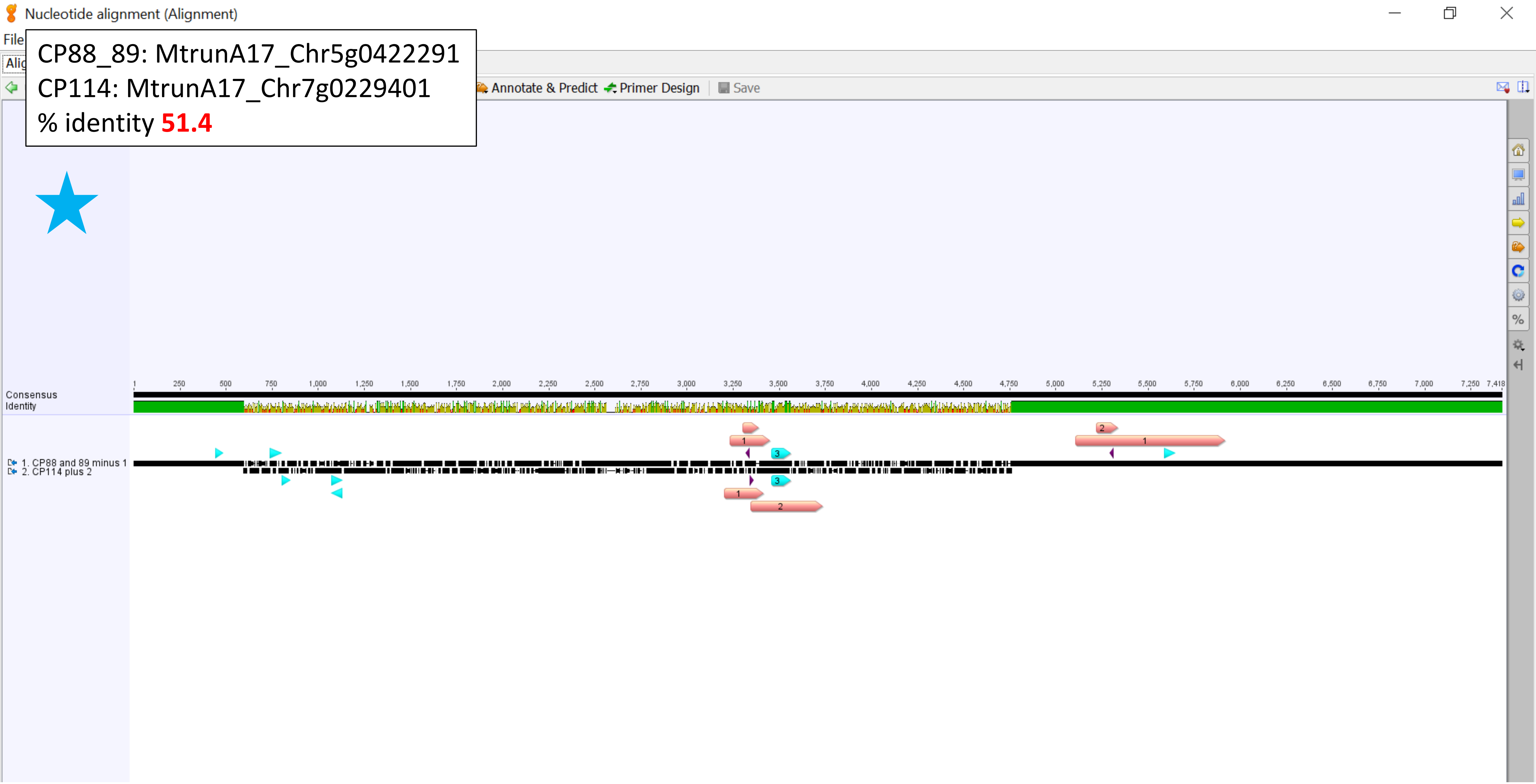

CP88_89: MtrunA17_Chr5g0422291
CP114: MtrunA17_Chr7g0229401
Zoom-in view, left side

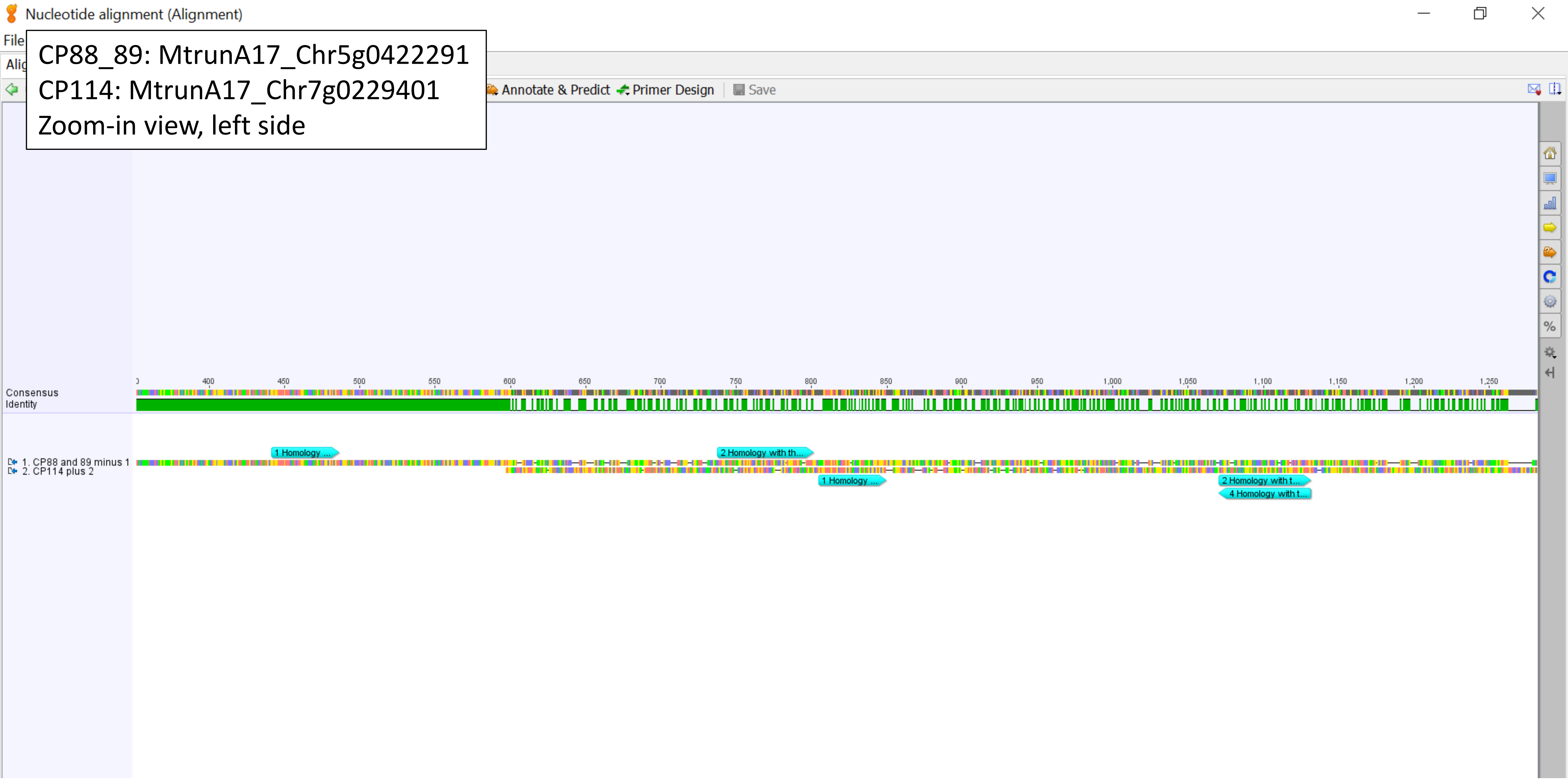

CP88_89: MtrunA17_Chr5g0422291
CP114: MtrunA17_Chr7g0229401
Zoom-in view, right side

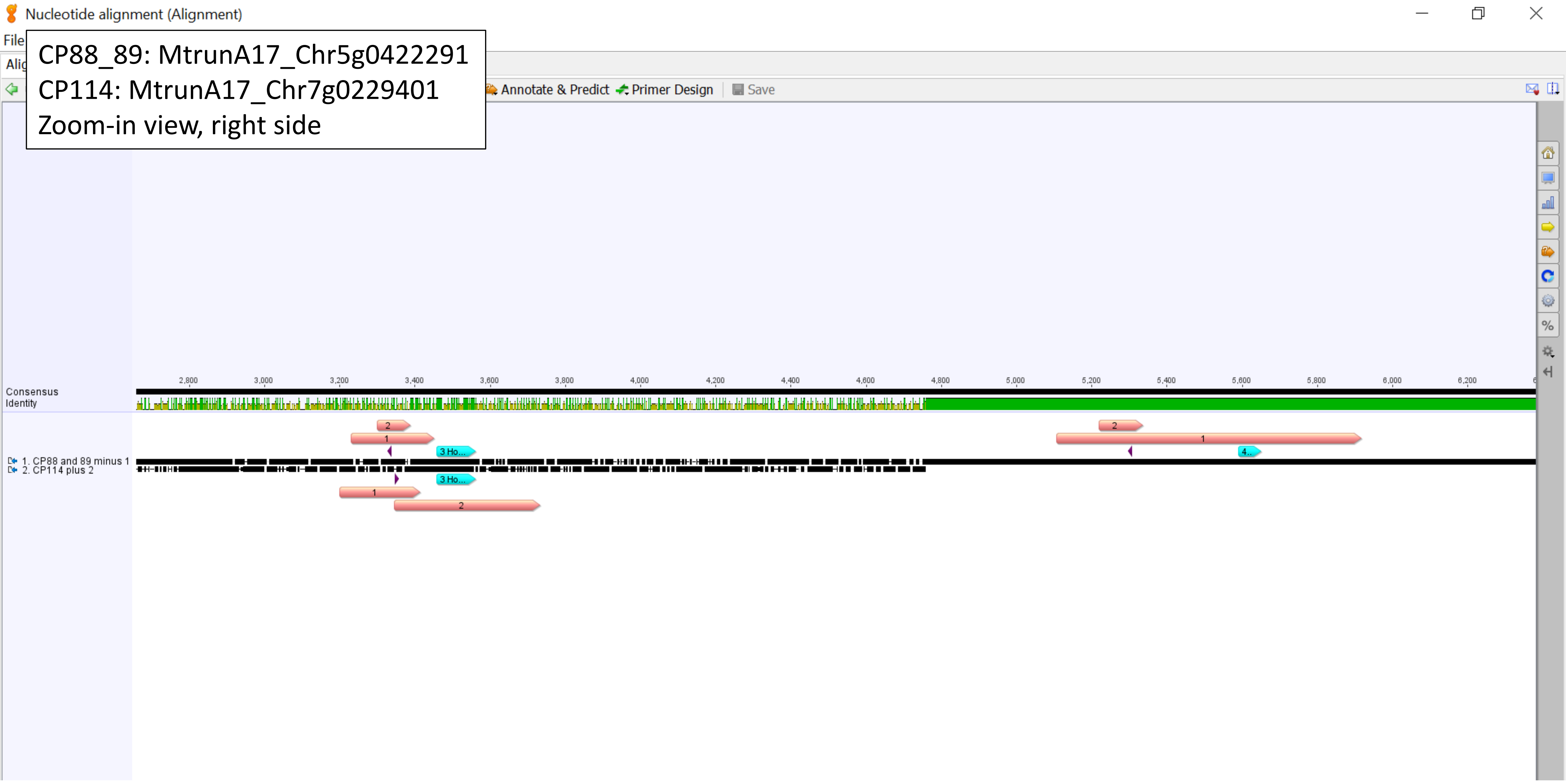

CP88_89: MtrunA17_Chr5g0422291
CP114: MtrunA17_Chr7g0229401
Homology region 1, % identity **87.0**

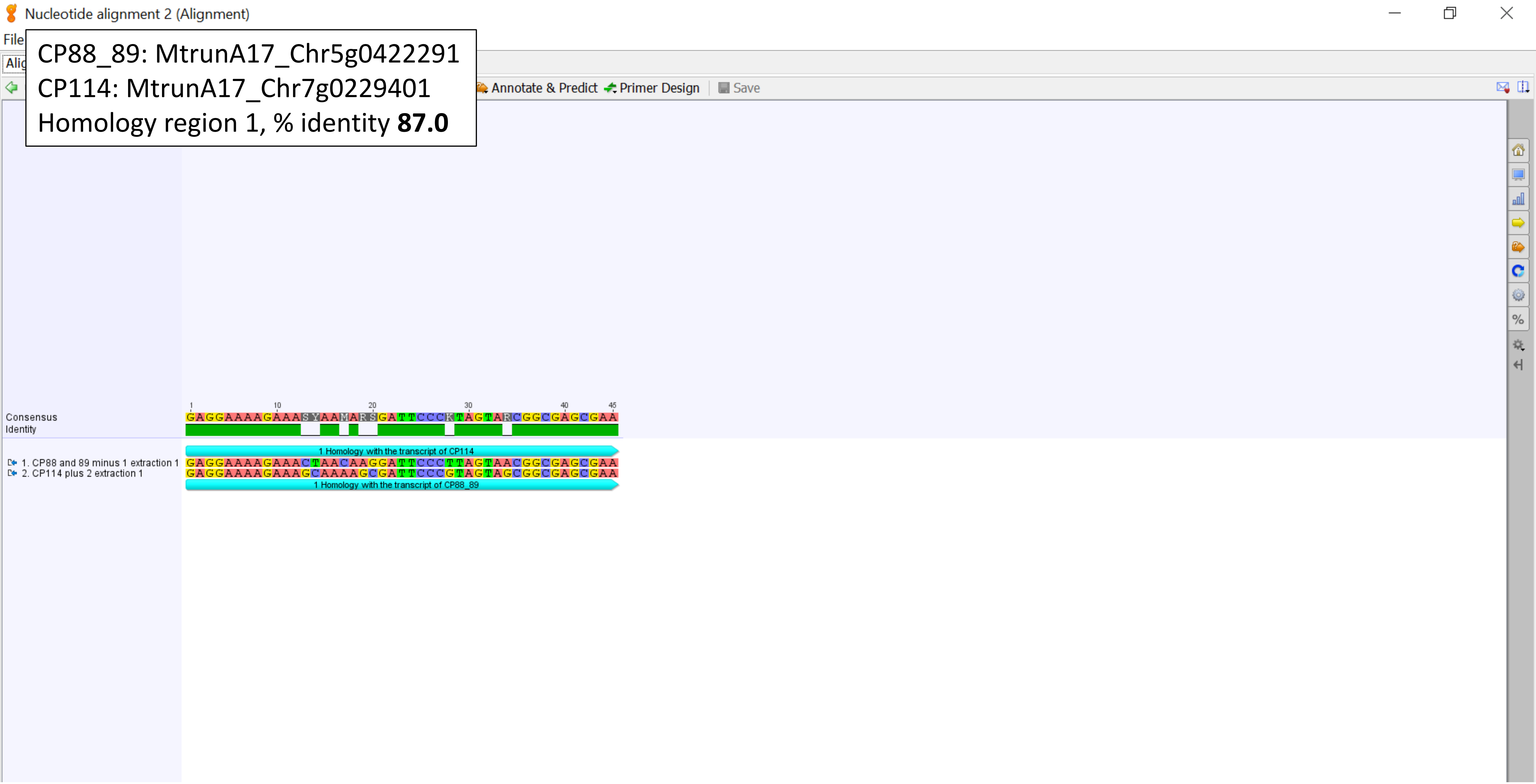

CP88_89: MtrunA17_Chr5g0422291
CP114: MtrunA17_Chr7g0229401
Homology region 2, % identity **80.3**

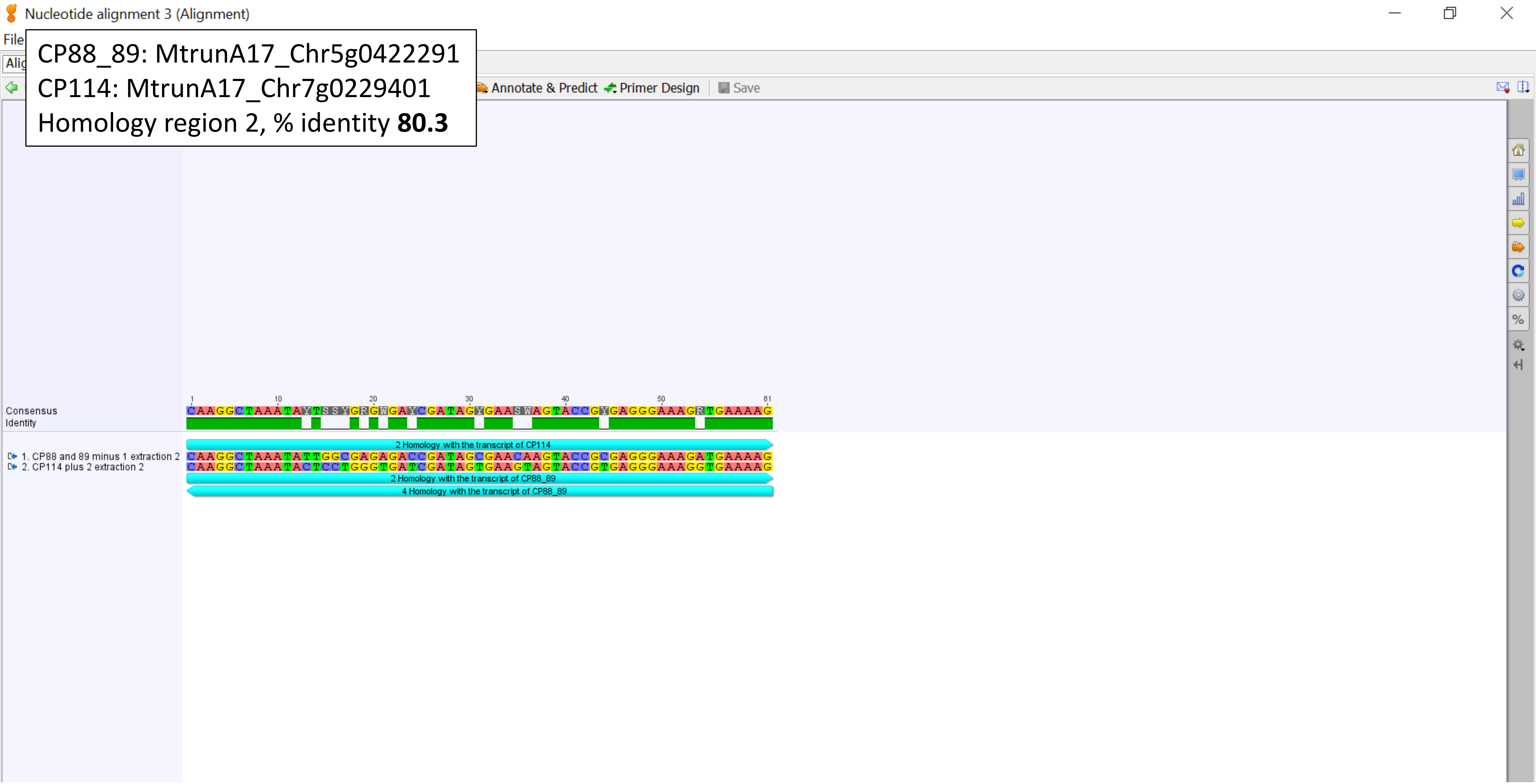

Nucleotide alignment 4 (Alignment)

CP88_89: MtrunA17_Chr5g0422291
CP114: MtrunA17_Chr7g0229401
Homology region 3, % identity **77.8**

Annotate & Predict | Primer Design | Save

Consensus
Identity

```
1        10        20        30        40        50        60        70        80        90        100 104
TCGATGTCGGCTCTTCSYMWCMTKGWGMWGYAGWATWYWCCAAGKGTTGGRYTGTTCRCCCAYYAAWRSGGWACGTGAGCTGGGTTYAGAMCGTCGTGAGACAG
```

3 Homology with the transcript of CP114

1. CP88 and 89 minus 1 extraction 3
```
TCGATGTCGGCTCTTCCTATCATTGTGAAGCAGAATTCACCAAGTGTTGGATTGTTCACCCACCAATAGGGAACGTGAGCTGGGTTTAGACCGTCGTGAGACAG
```
2. CP114 plus 2 extraction 3
```
TCGATGTCGGCTCTTCGCCACCTGGAGCTGTAGTATATTCCAAGGGTTGGGCTGTTCGCCCATTAAAGCGGTACGTGAGCTGGGTTCAGAACGTCGTGAGACAG
```

2

3 Homology with the transcript of CP88_89

CP88_89: MtrunA17_Chr5g0422291
CP114: MtrunA17_Chr7g0229401
Homology region 4, % identity **80.3**

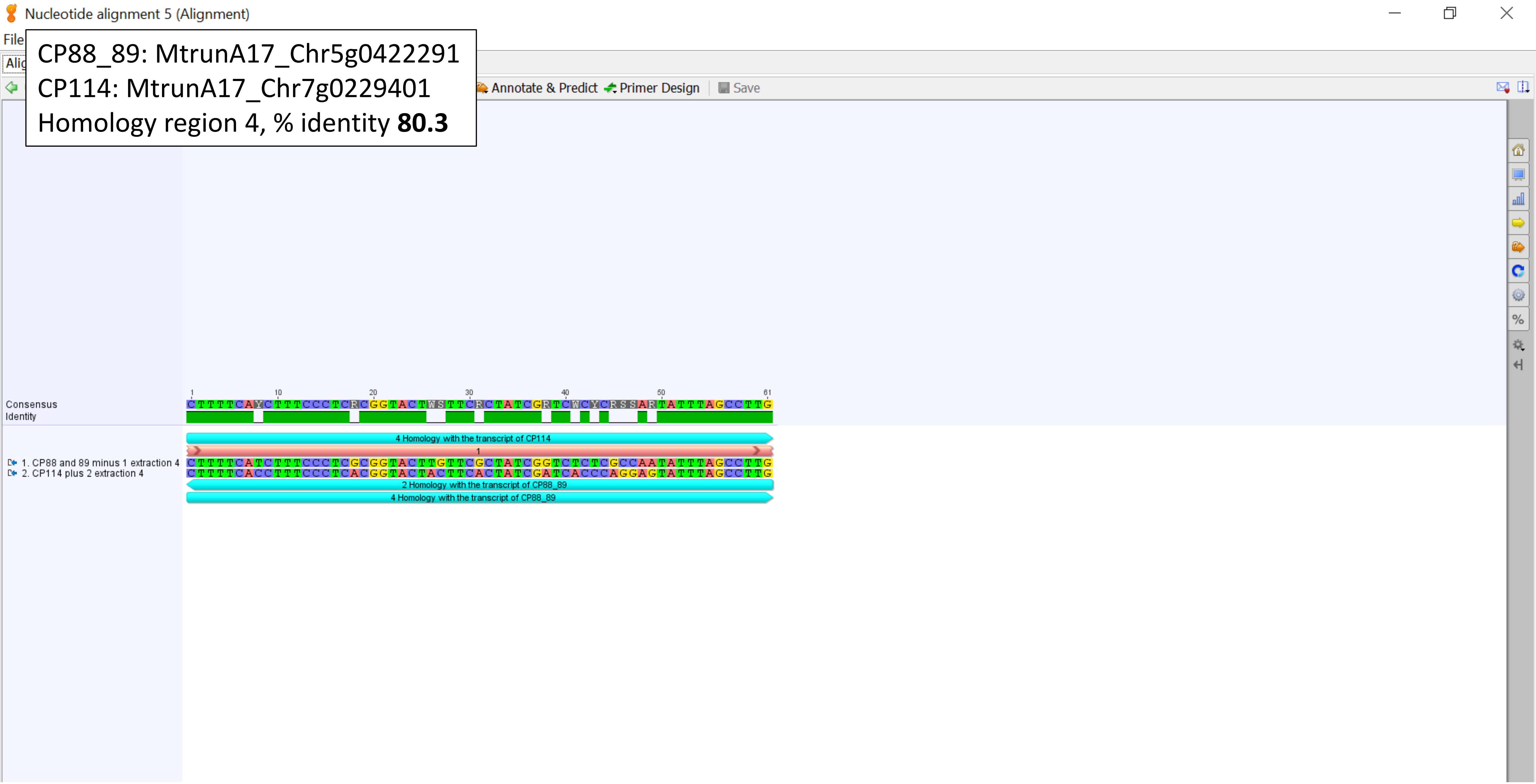

CP88_89: MtrunA17_Chr5g0422291
CP141: MtrunA17_CPg0492331
% identity **49.8**

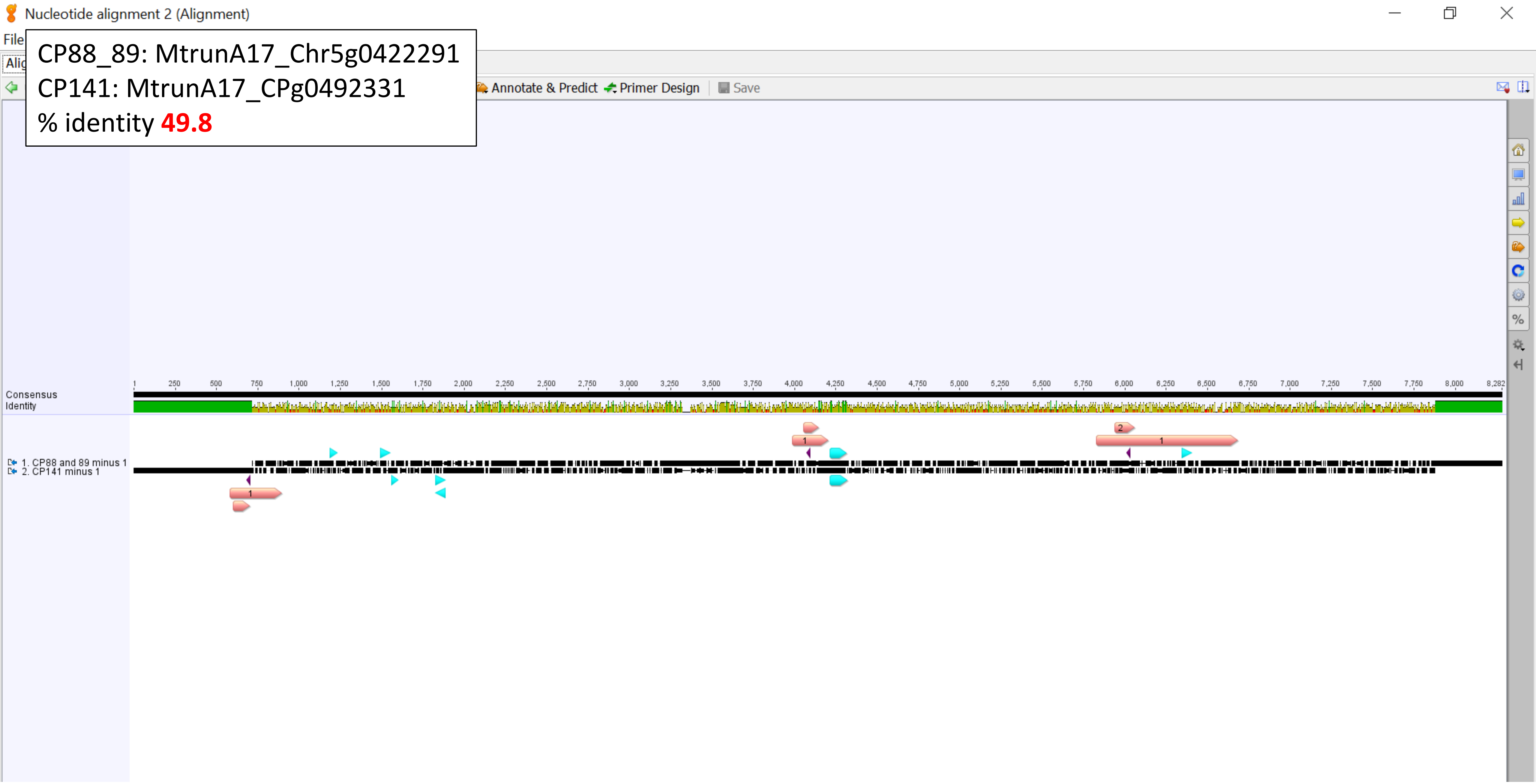

CP88_89: MtrunA17_Chr5g0422291
CP141: MtrunA17_CPg0492331
Zoom-in view, left side

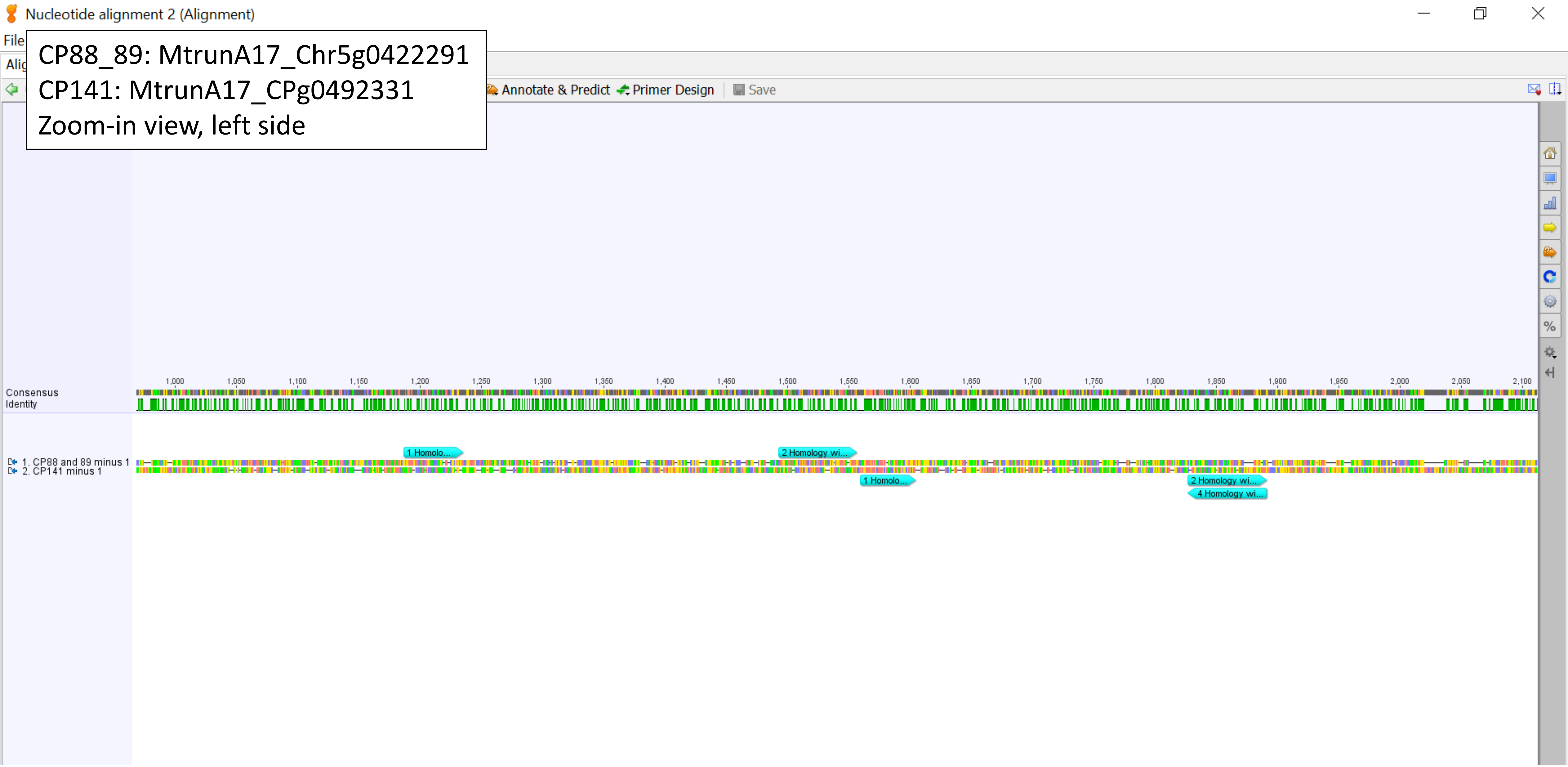

CP88_89: MtrunA17_Chr5g0422291
CP141: MtrunA17_CPg0492331
Zoom-in view, right side

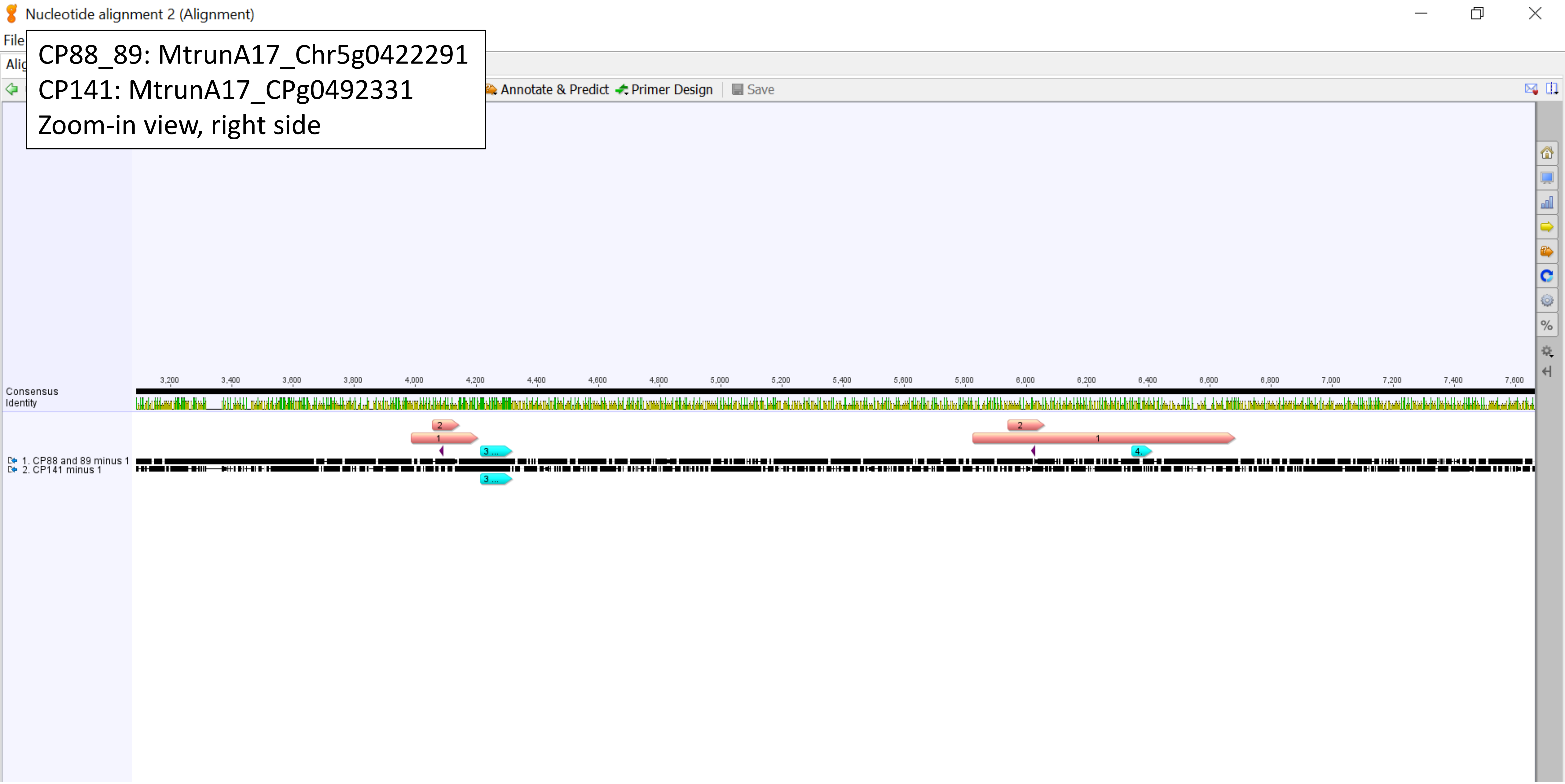

CP88_89: MtrunA17_Chr5g0422291
CP141: MtrunA17_CPg0492331
Homology region 1, % identity **87.0**

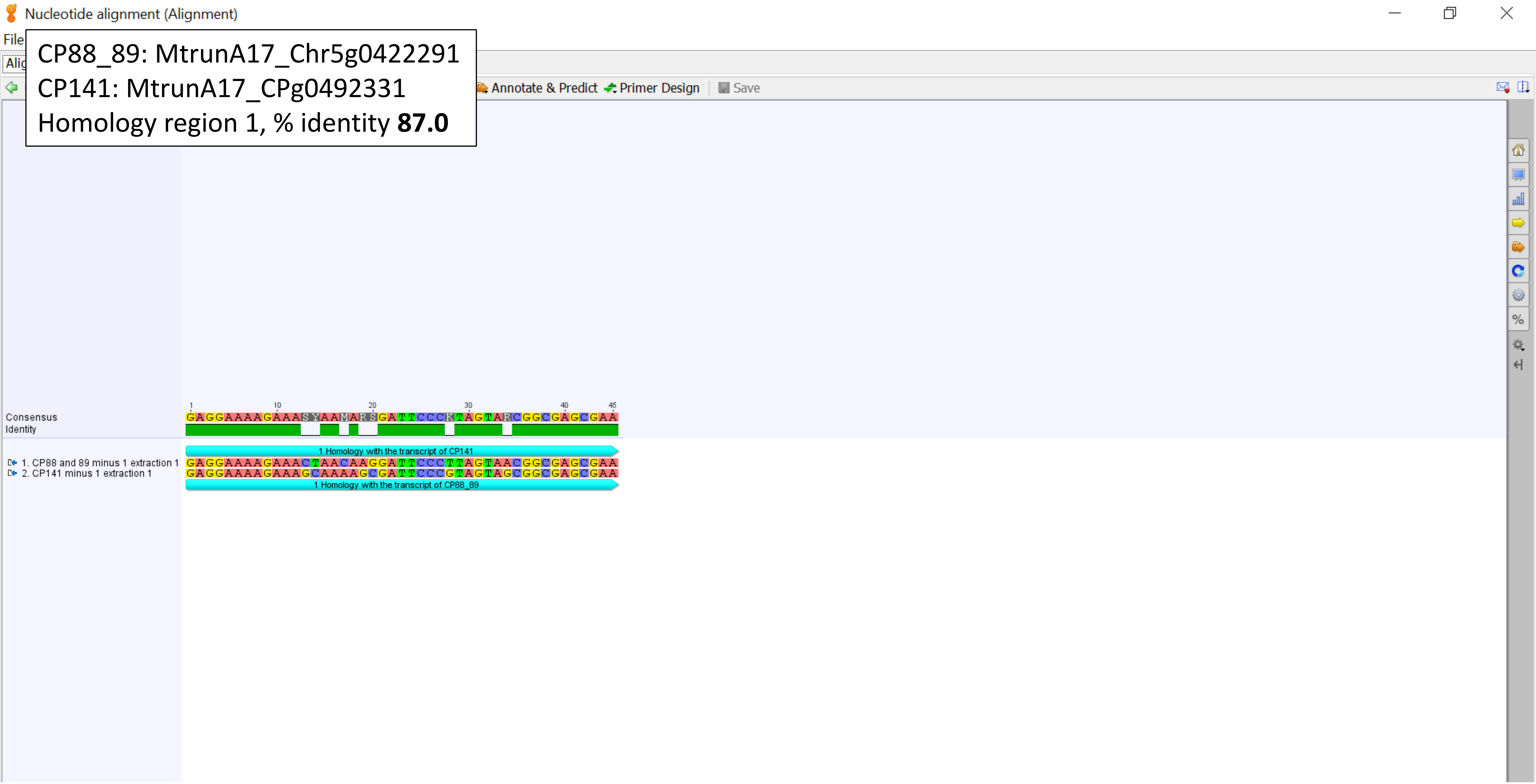

CP88_89: MtrunA17_Chr5g0422291
CP141: MtrunA17_CPg0492331
Homology region 2, % identity **82.0**

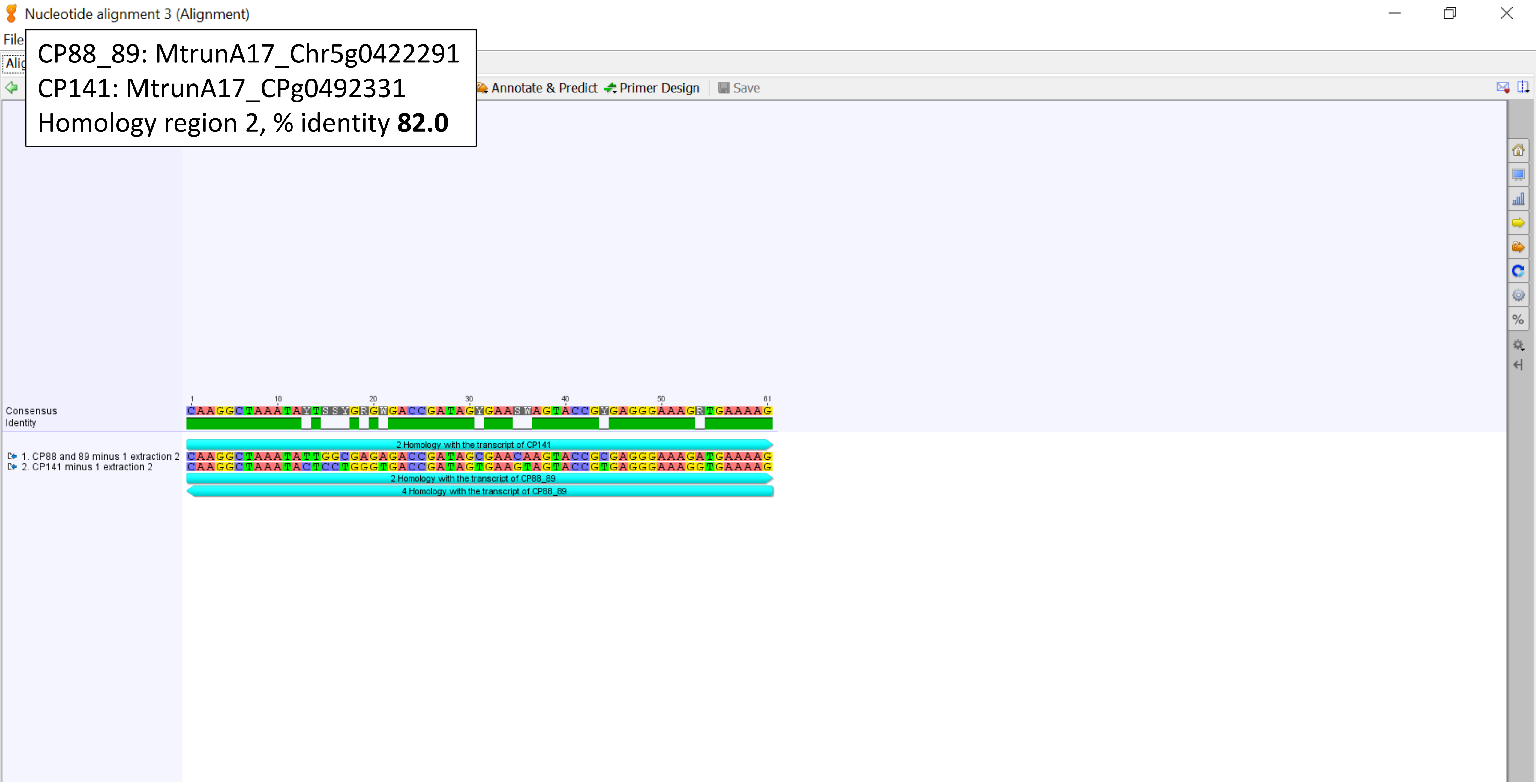

CP88_89: MtrunA17_Chr5g0422291
CP141: MtrunA17_CPg0492331
Homology region 3, % identity **77.4**

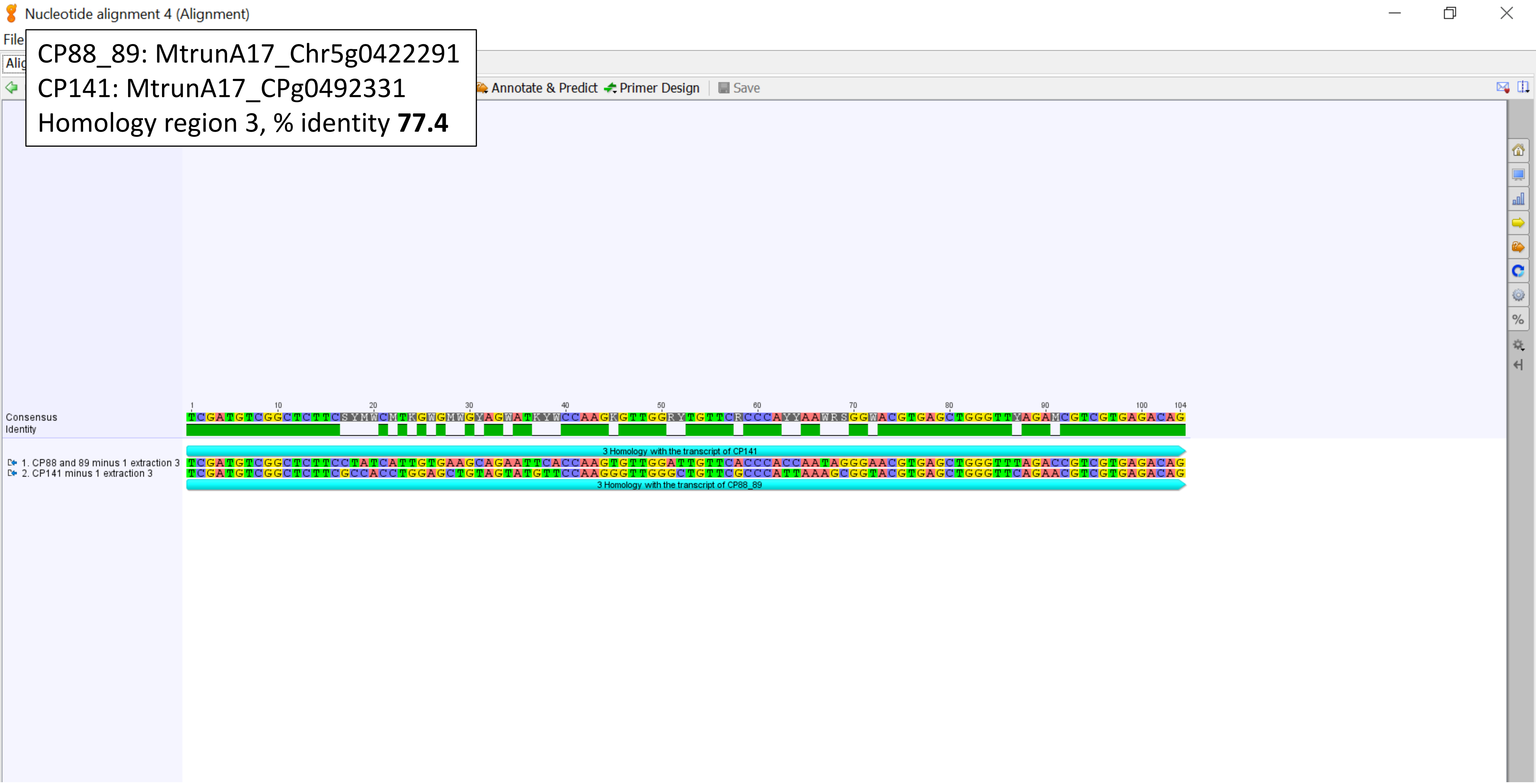

CP88_89: MtrunA17_Chr5g0422291
CP141: MtrunA17_CPg0492331
Homology region 4, % identity **82.0**

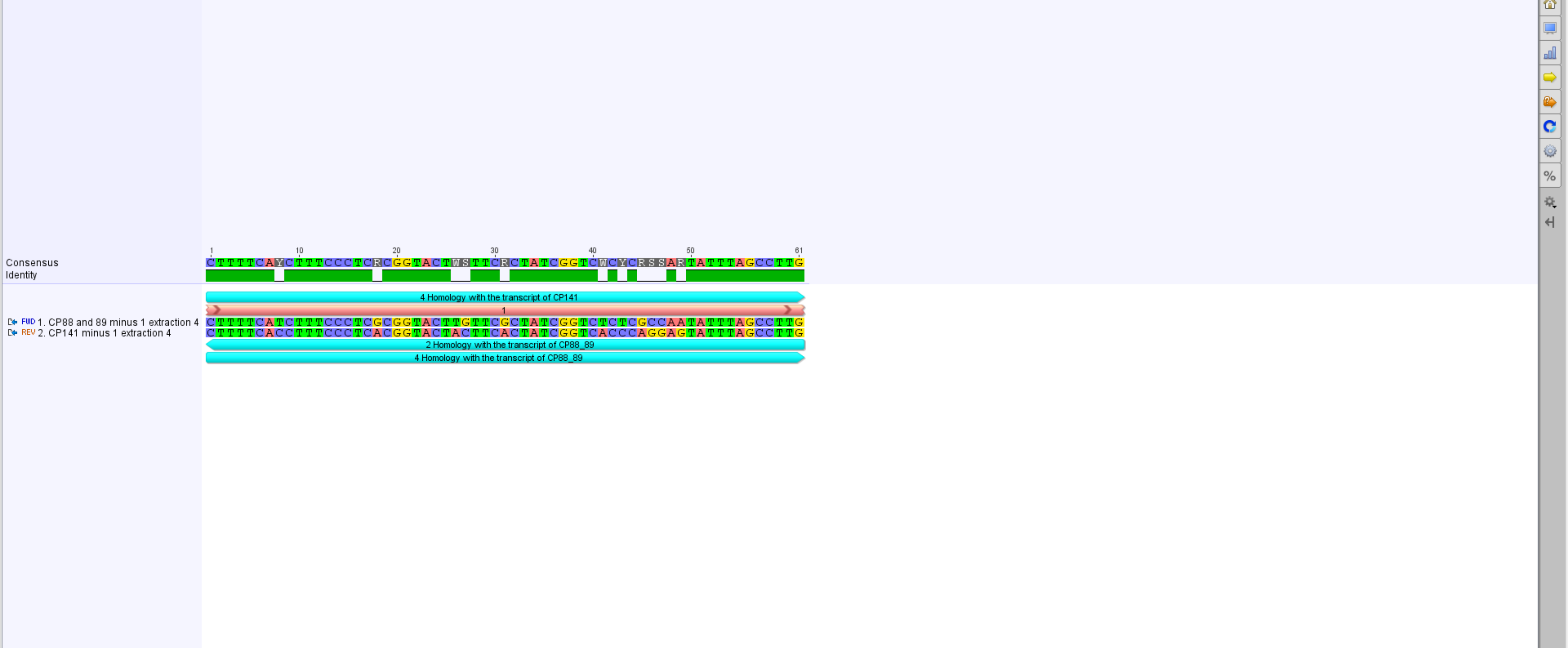

CP88_89: MtrunA17_Chr5g0422291
CP142: MtrunA17_CPg0492381
% identity **50.5**

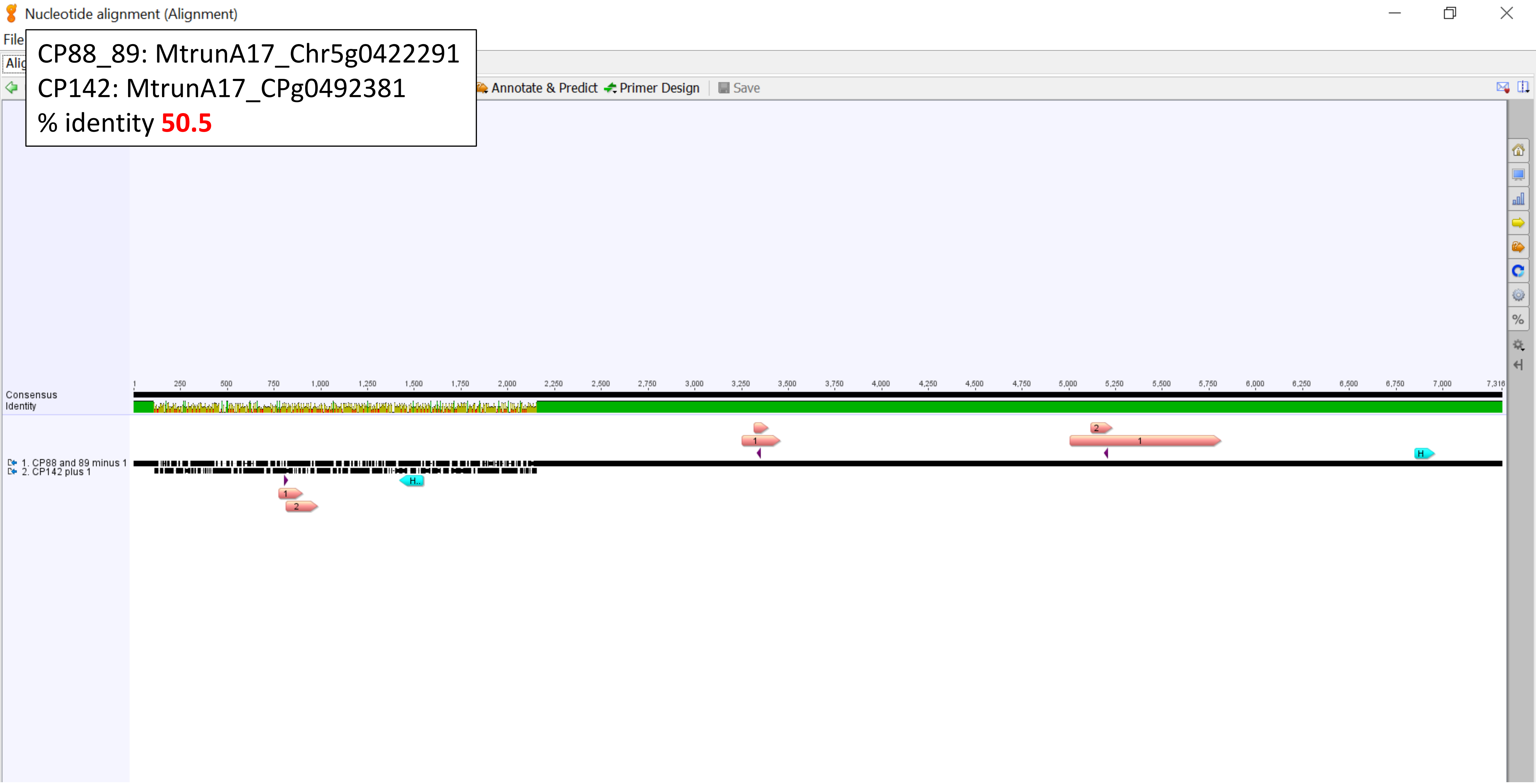

Nucleotide alignment 3 (Alignment)

CP88_89: MtrunA17_Chr5g0422291
CP142: MtrunA17_CPg0492381
Homology region, % identity **78.3**

Annotate & Predict   Primer Design   Save

```
           1        10        20        30        40        50        60        70        80        90        100  106
Consensus  CCTGGTAAGKTTCYYCGYKTTGMRTCRAATTAARCCRCAKGCTCCACYSCTKGTGNGKGCCCNNCCGTCAATTCCTTTRAGTTTCAKYCTTGCGAMCRTACTCCCC
Identity

           Homology with the transcript of CP142
FWD 1. CP88 and 89 minus 1 extraction
           CCTGGTAAGTTTCCCCGTGTTGAGTCAAATTAAGCCGCAGGCTCCACTCCTGGTG-GTGCCCTTCCGTCAATTCCTTTAAGTTTCAGCCTTGCGACCATACTCCCC
REV 2. CP142 plus 1 extraction
           CCTGGTAAGGTTCTTCGCTTTGCATCGAATTAAACCACATGCTCCACCGCTTGTGCGGGCCC--CCGTCAATTCCTTTGAGTTTCATTCTTGCGAACGTACTCCCC
           Homology with the transcript of CP88_89
```

CP93: MtrunA17_Chr5g0435191
CP122: MtrunA17_Chr7g1034306
% identity **52.1**

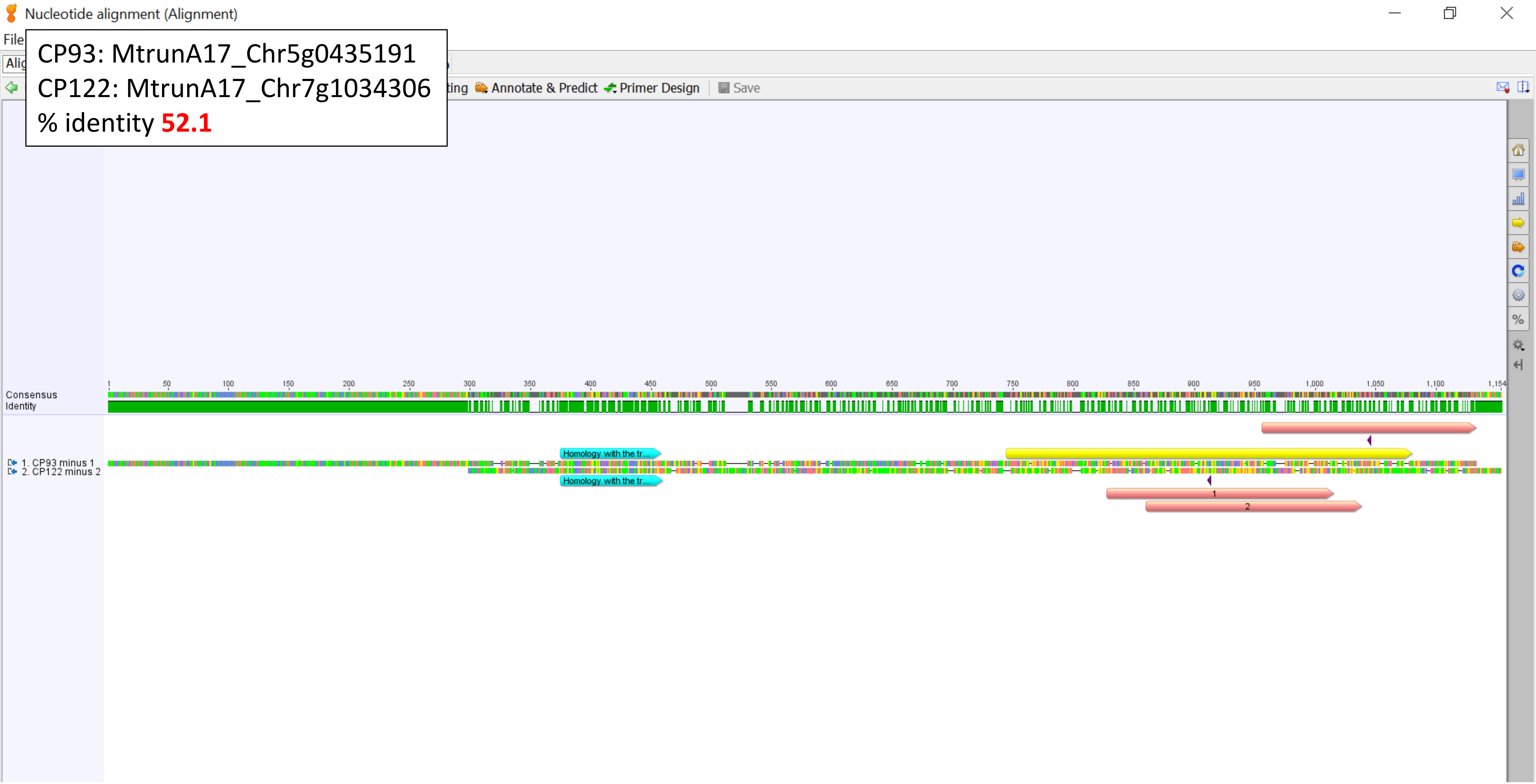

CP93: MtrunA17_Chr5g0435191
CP122: MtrunA17_Chr7g1034306
Homology region, % identity **83.5**

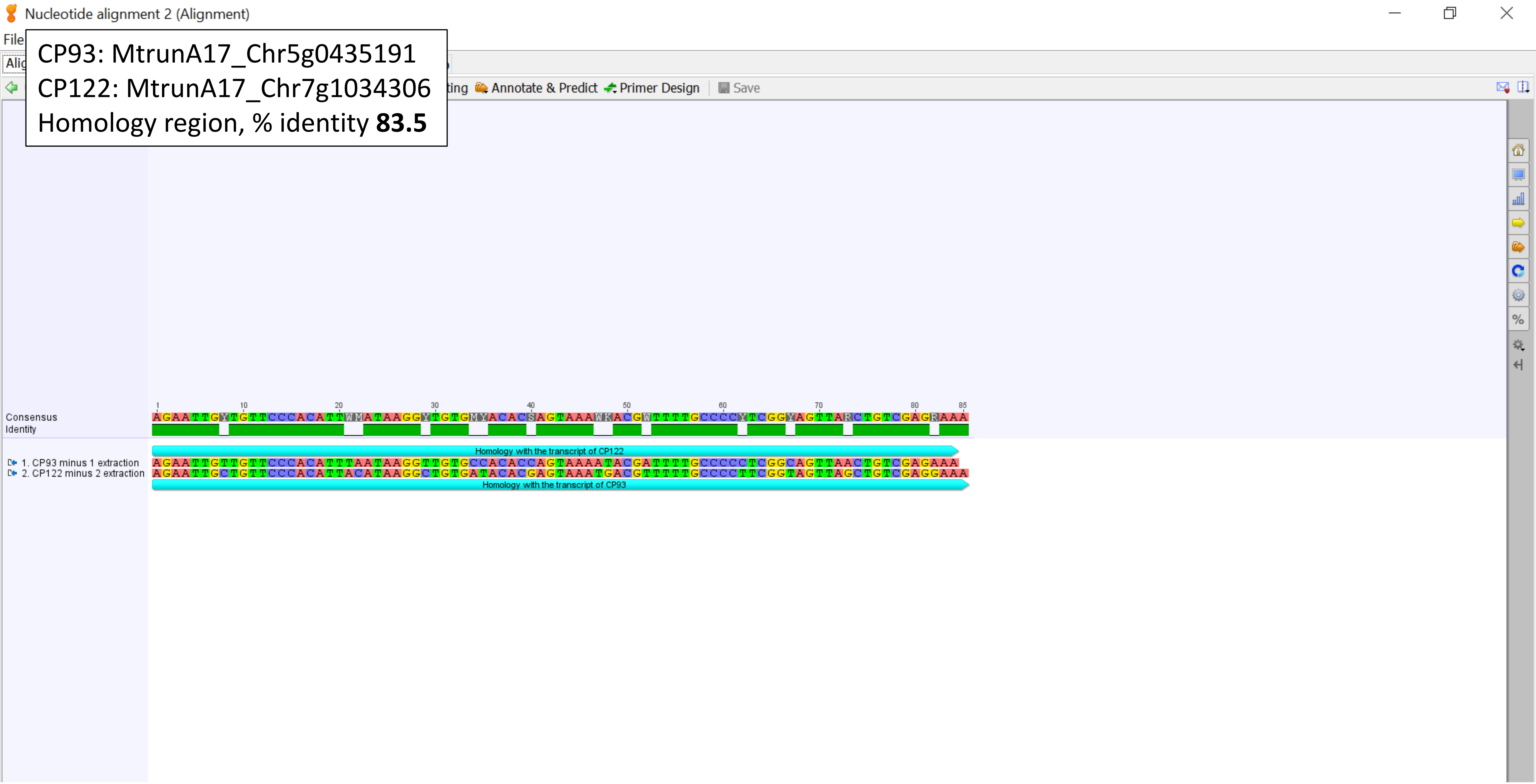

CP114: MtrunA17_Chr7g0229401
CP141: MtrunA17_CPg0492331
% identity **99.8**

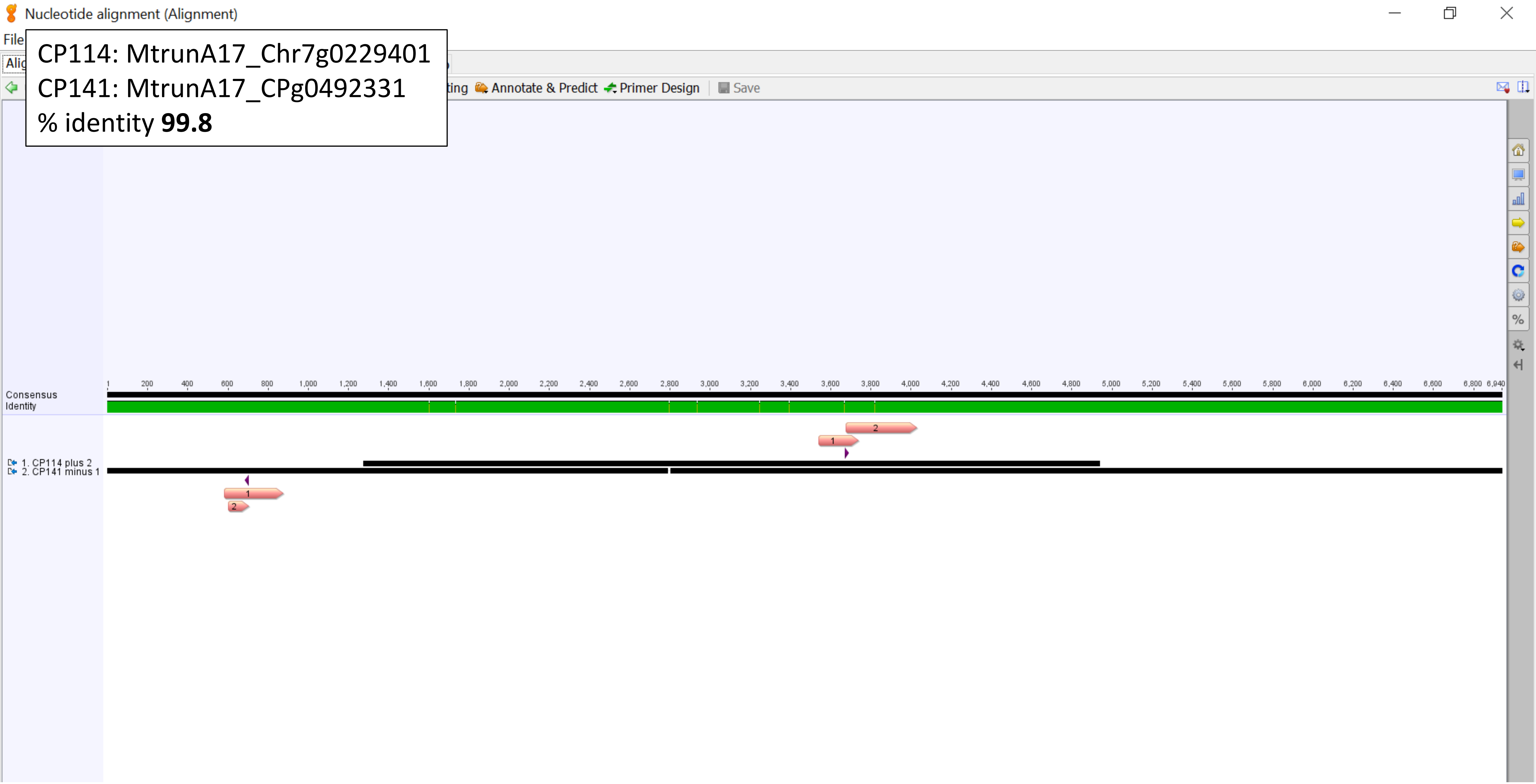

CP114: MtrunA17_Chr7g0229401
CP141: MtrunA17_CPg0492331
Zoom-in view, middle

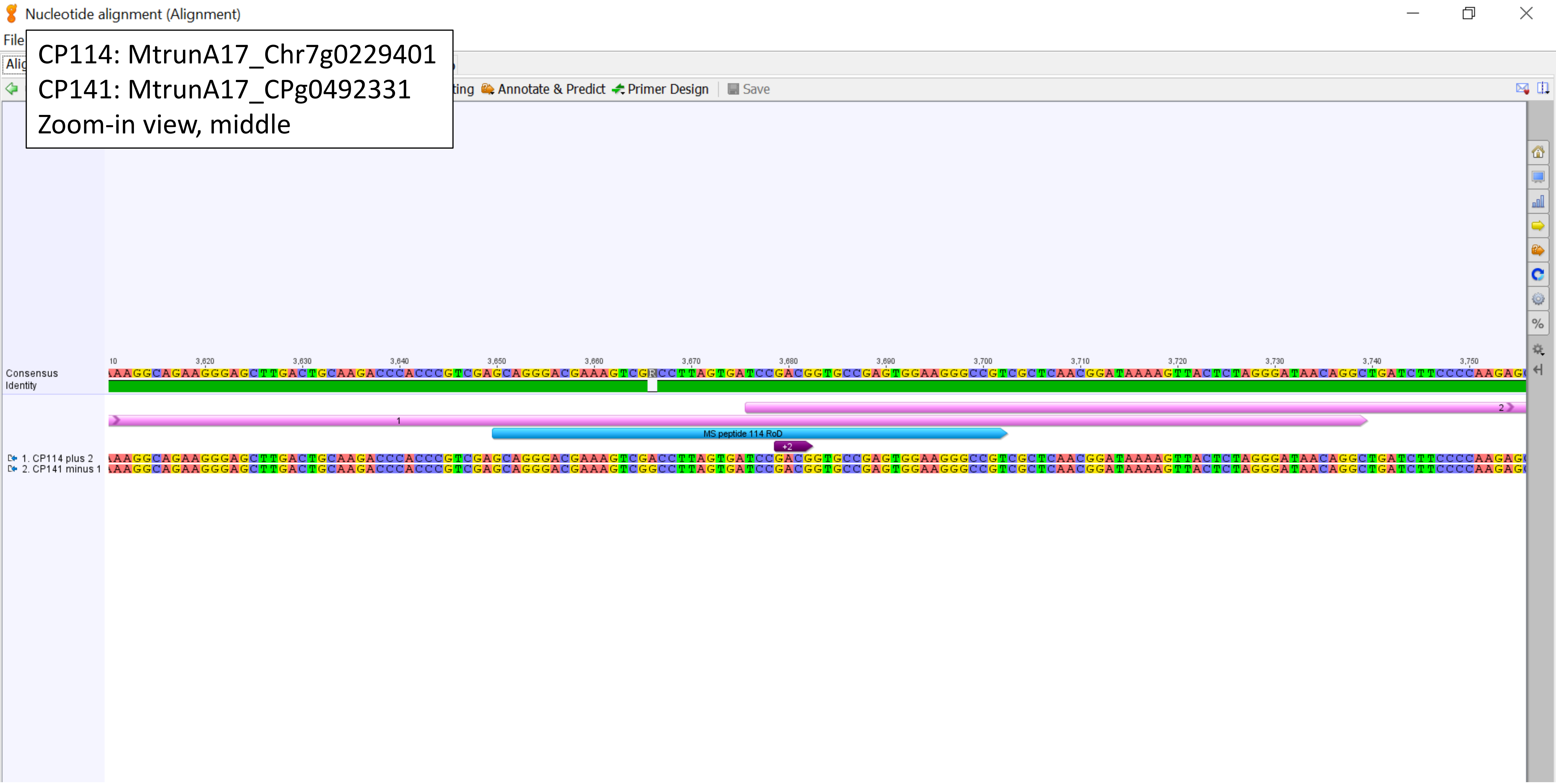

CP114: MtrunA17_Chr7g0229401
CP154: MtrunA17_MTg0491501
Geneious alignment, % identity **NA**

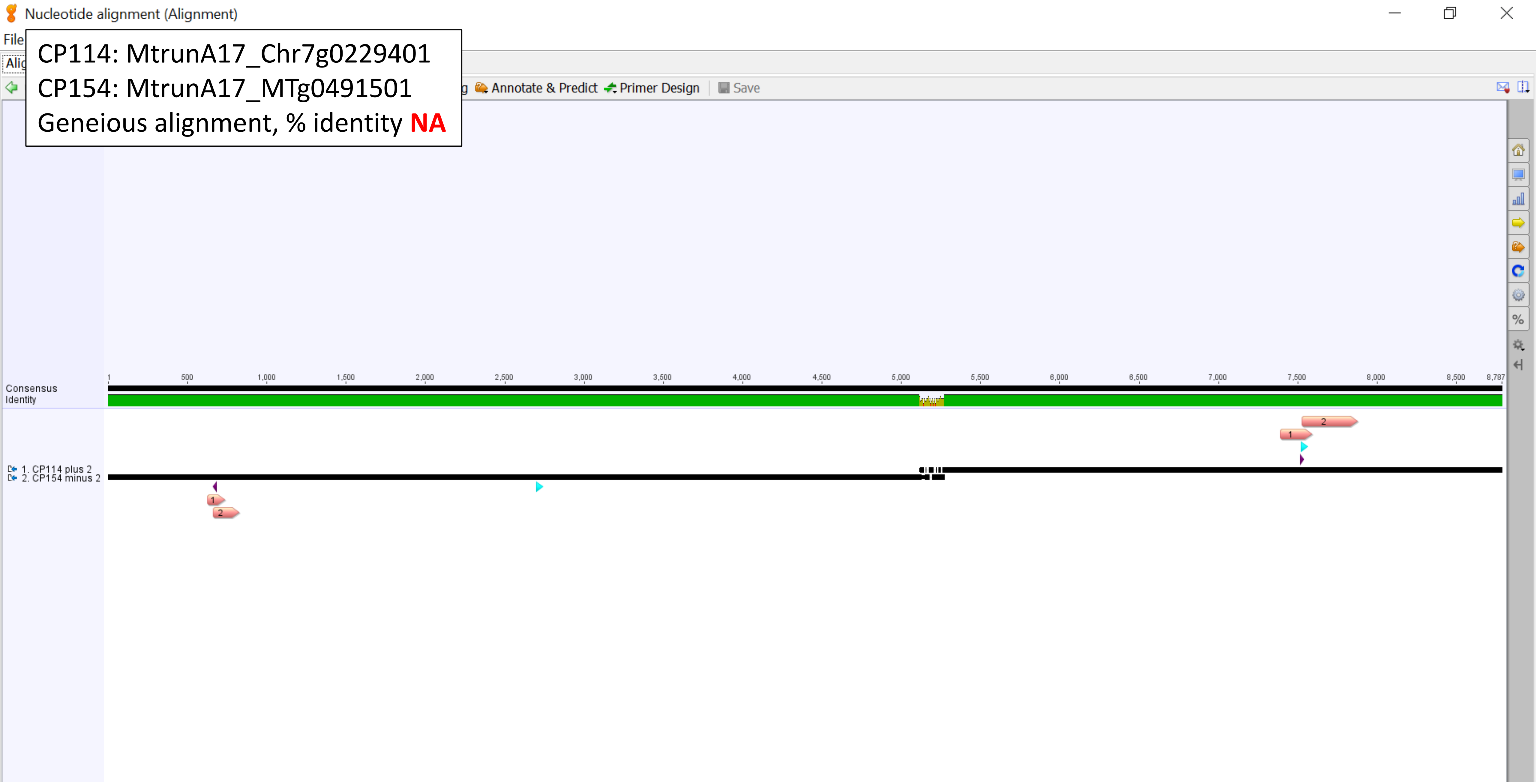

CP114: MtrunA17_Chr7g0229401
CP154: MtrunA17_MTg0491501
MUSCLE alignment, % identity **37.1**

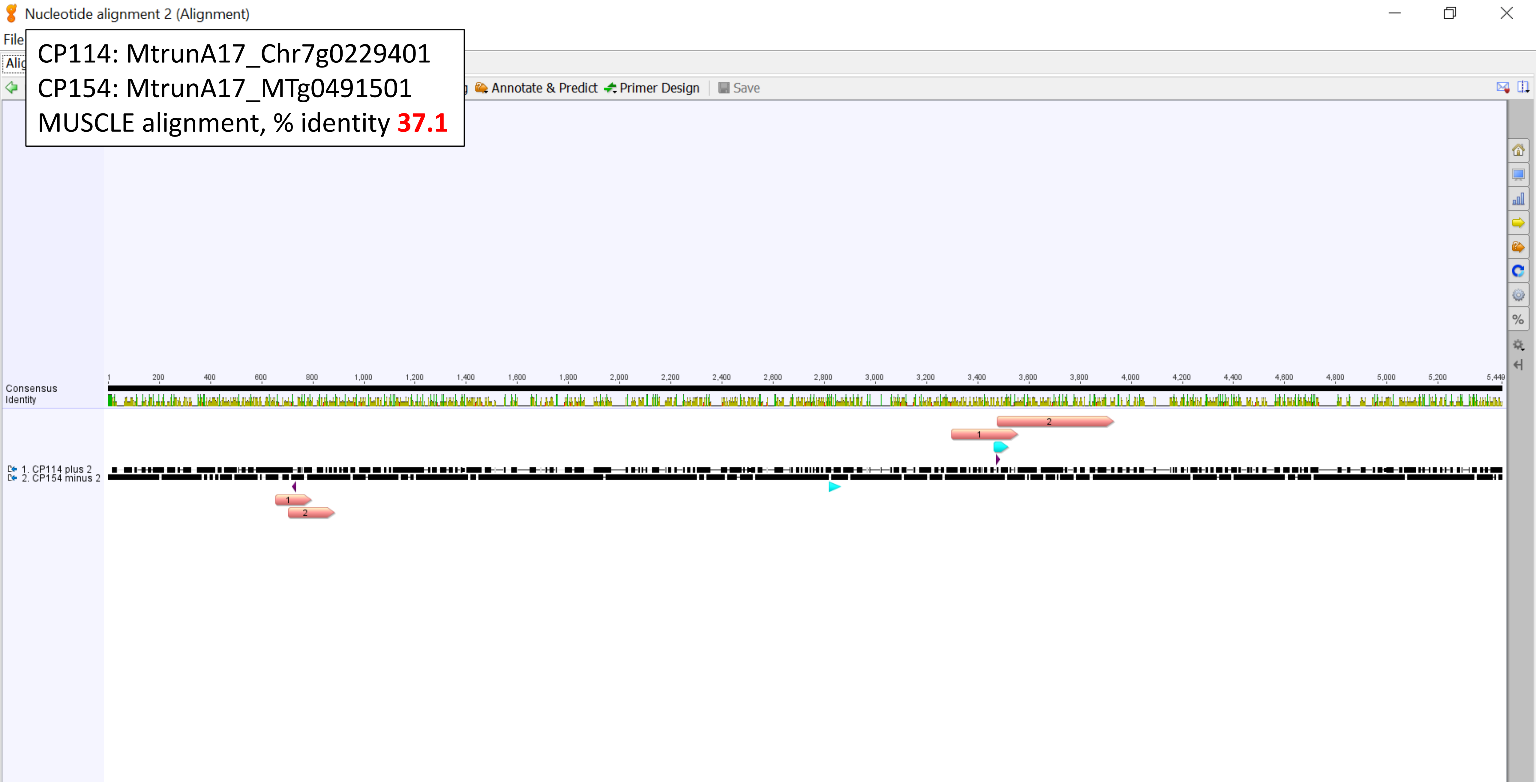

CP114: MtrunA17_Chr7g0229401
CP154: MtrunA17_MTg0491501
Homology region, % identity **90.9**

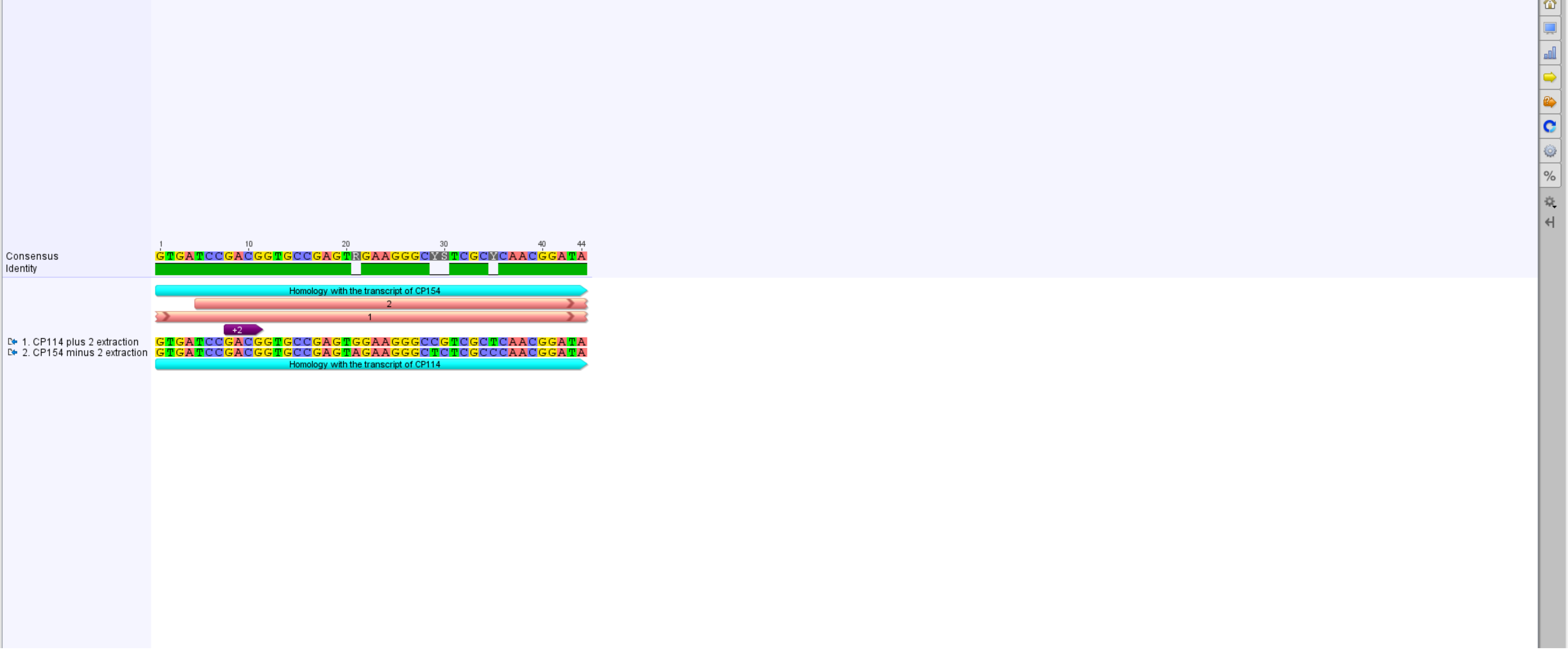

CP141: MtrunA17_CPg0492331
CP154: MtrunA17_MTg0491501
Overview

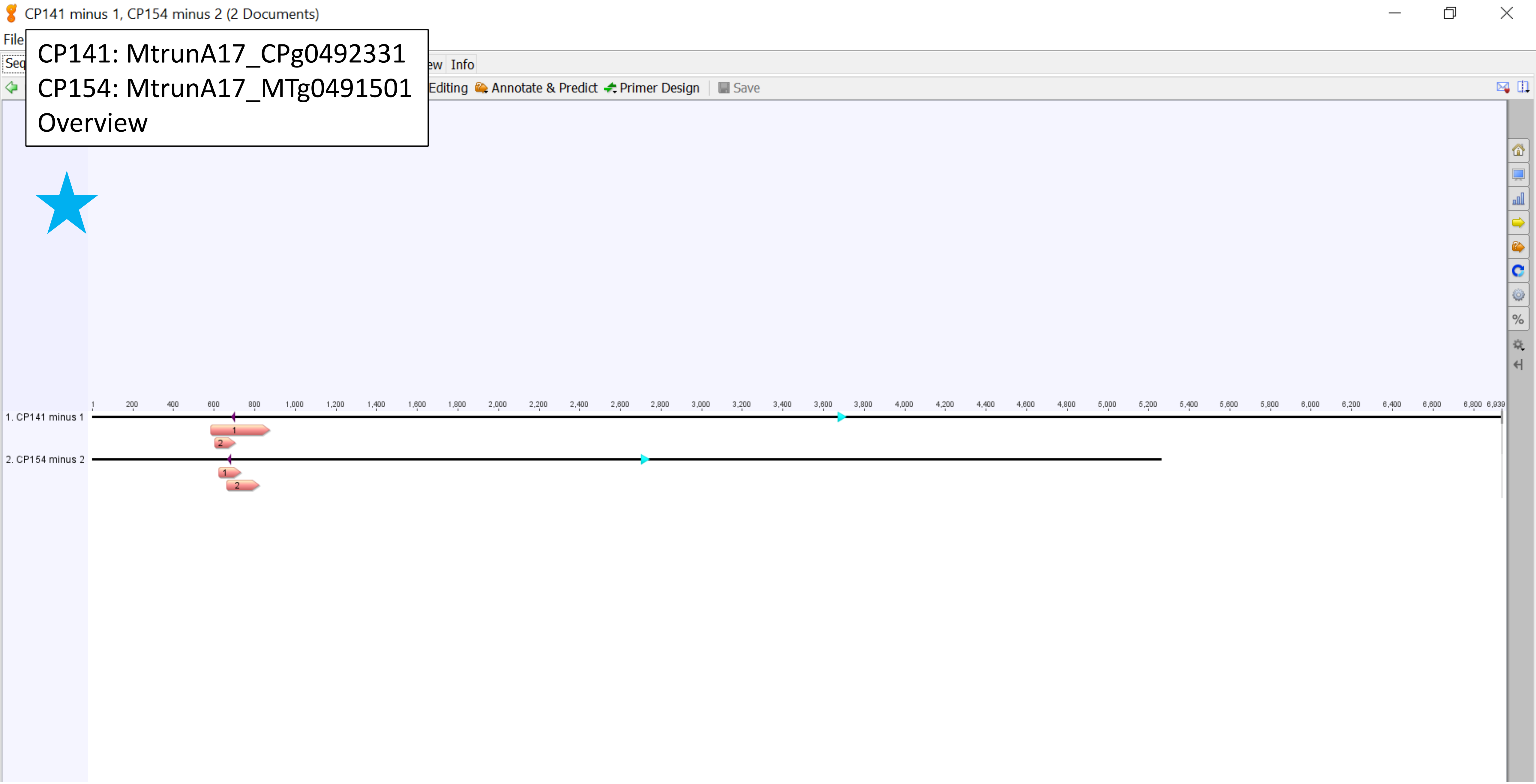

CP141: MtrunA17_CPg0492331
CP154: MtrunA17_MTg0491501
Geneious alignment, % identity **50.0**

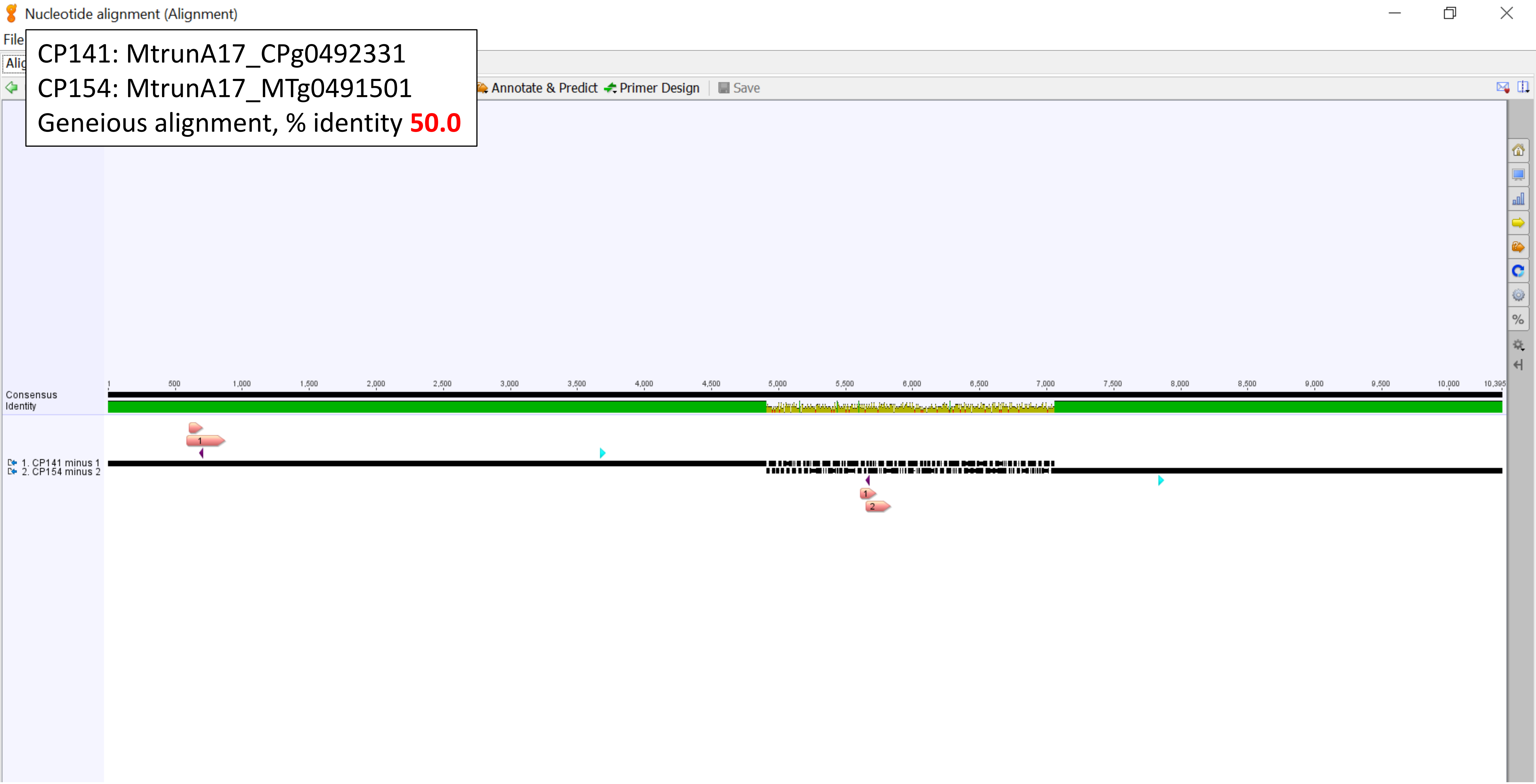

CP141: MtrunA17_CPg0492331
CP154: MtrunA17_MTg0491501
MUSCLE alignment, % identity **39.0**

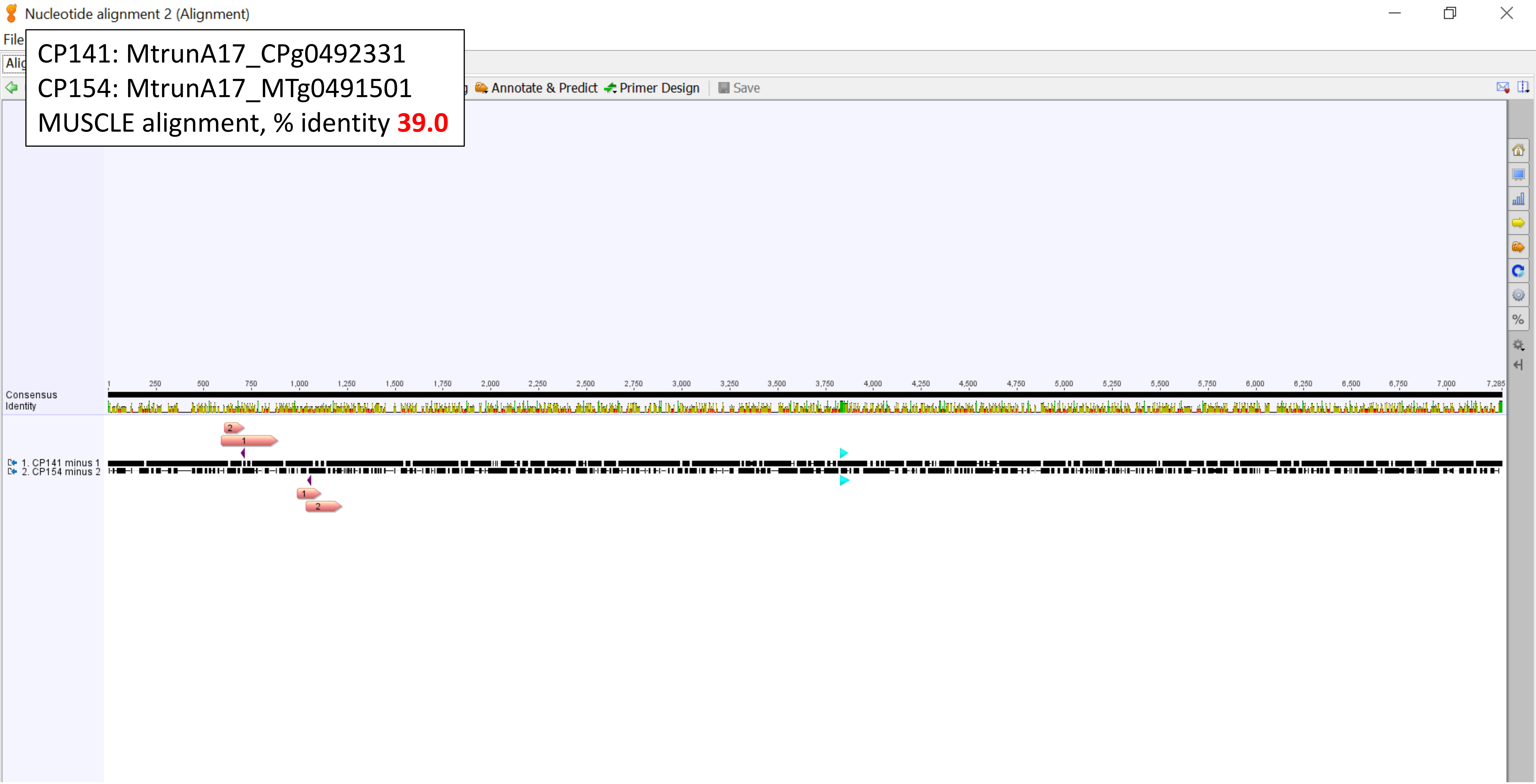

CP141: MtrunA17_CPg0492331
CP154: MtrunA17_MTg0491501
Homology region, % identity **90.9**

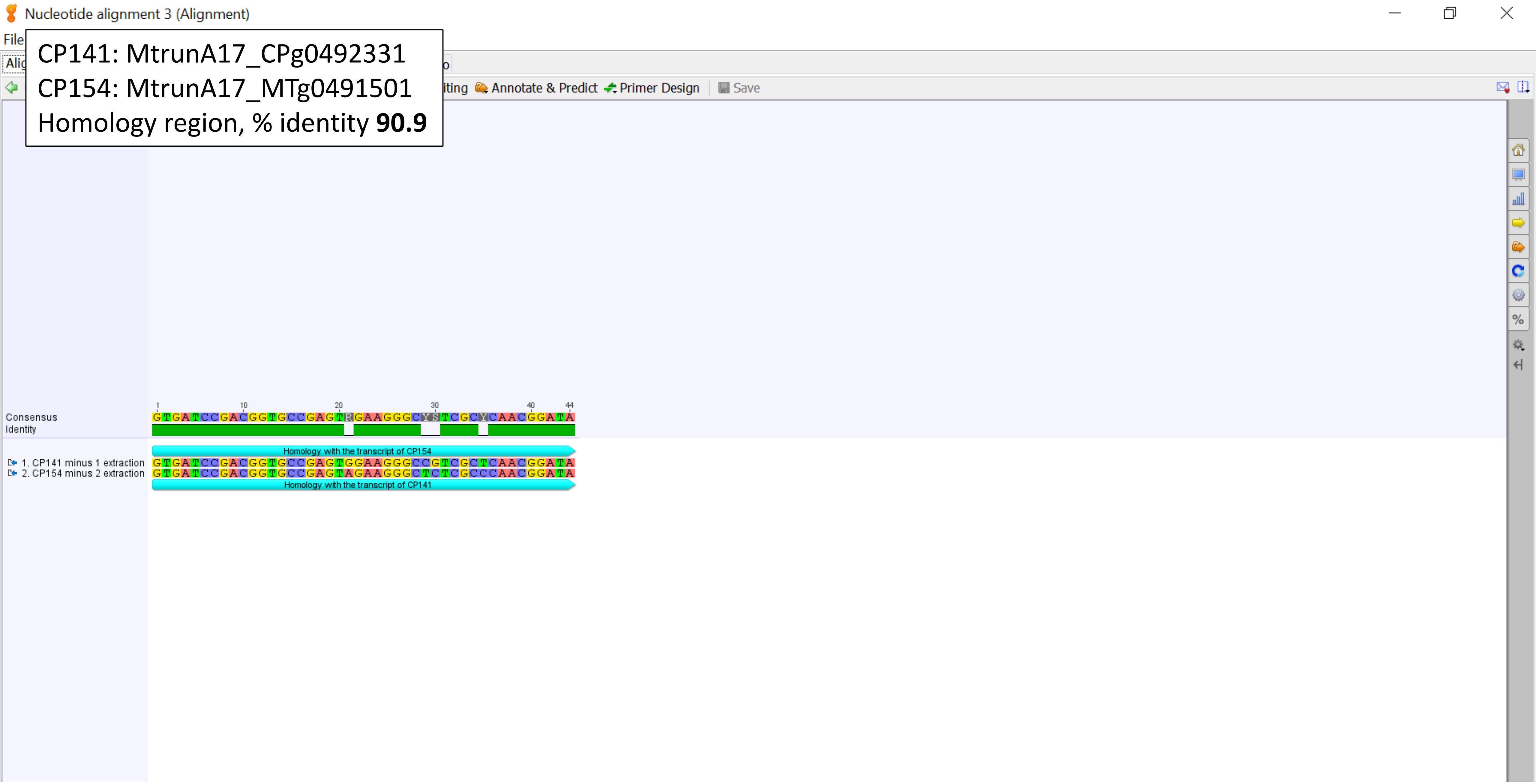

CP145: MtrunA17_CPg0492941
CP148_149: MtrunA17_MTg0490471
% identity **60.4**

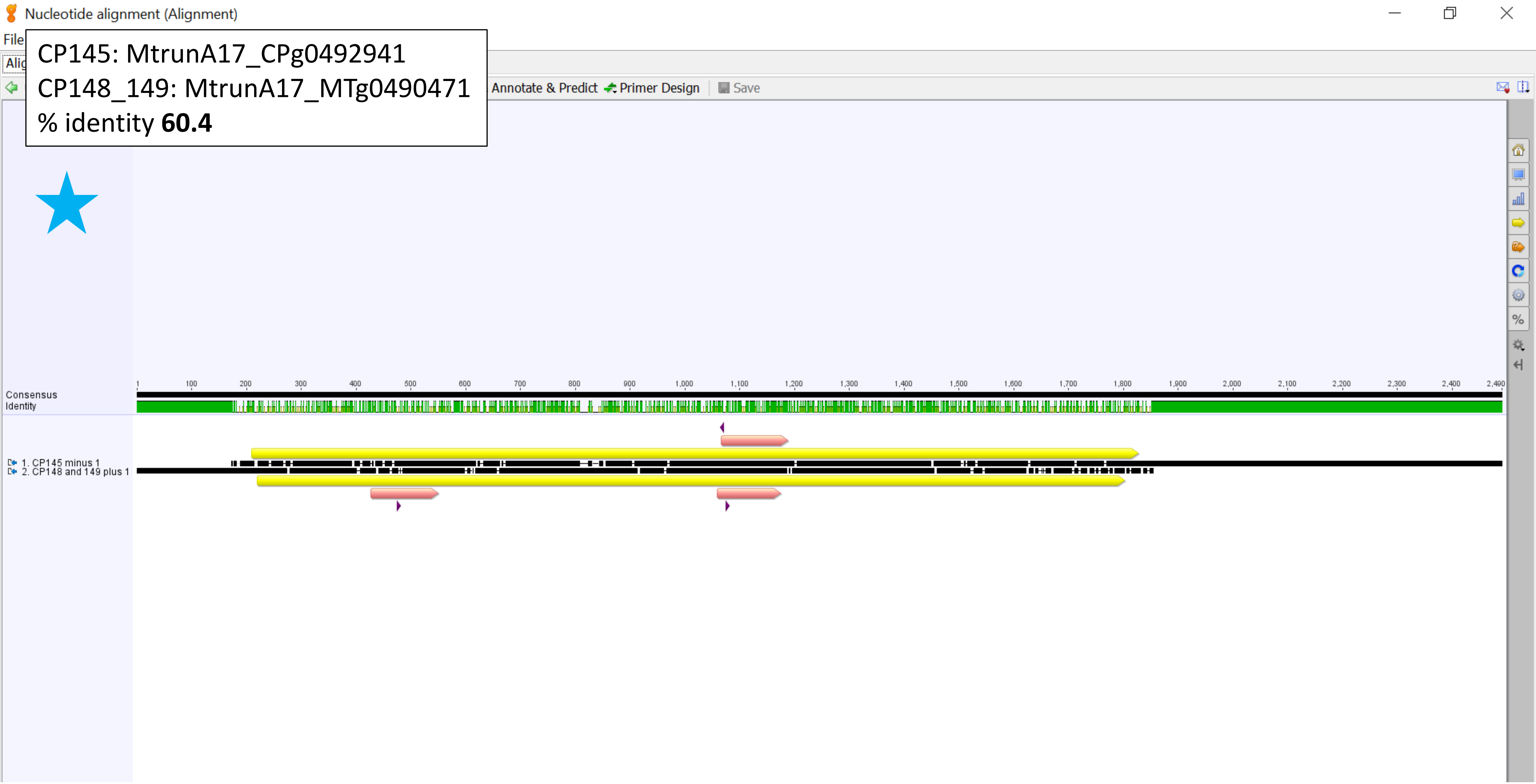

CP150_151: MtrunA17_MTg0490971
CP152: MtrunA17_MTg0491151
Geneious alignment, % identity **NA**

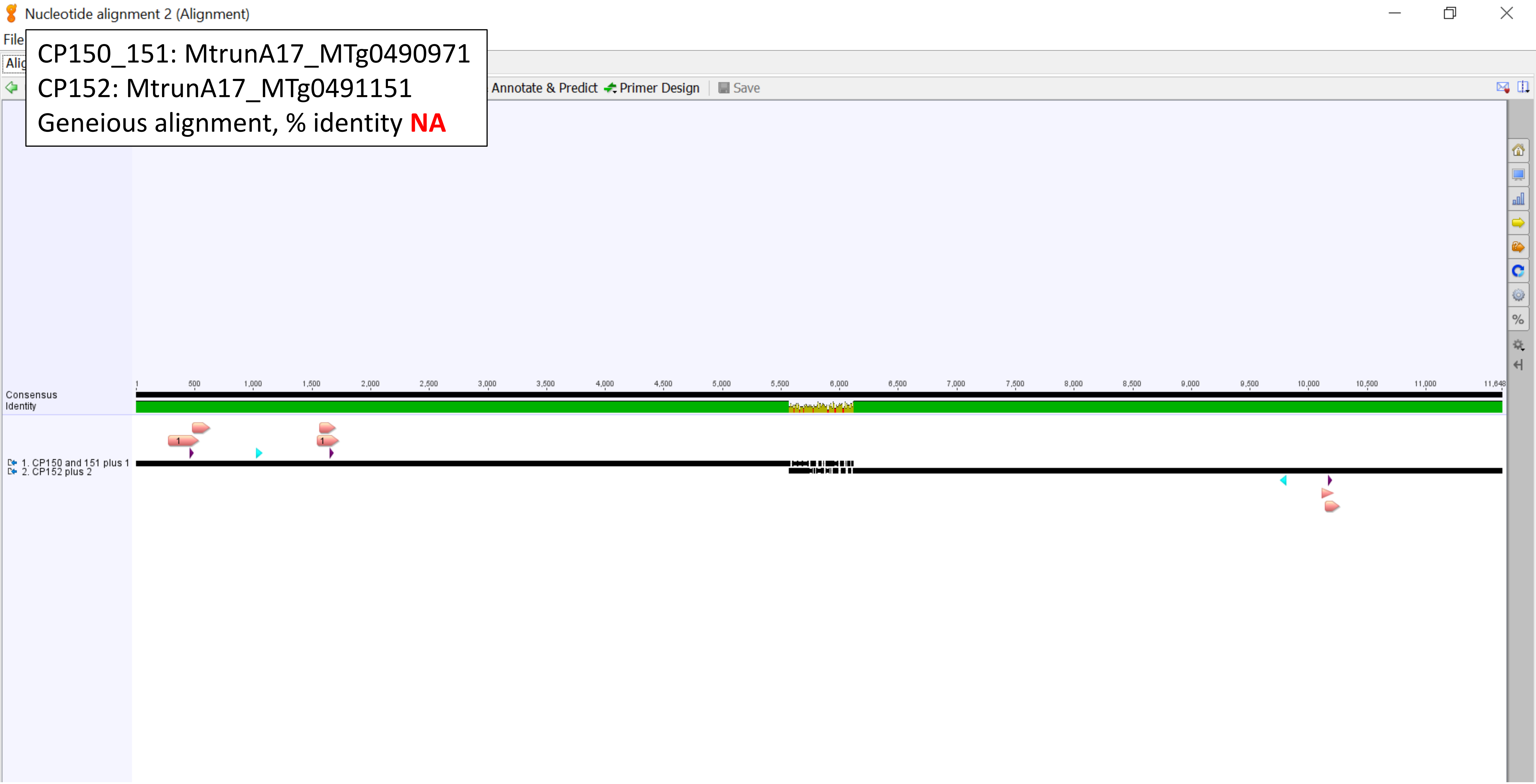

CP150_151: MtrunA17_MTg0490971
CP152: MtrunA17_MTg0491151
MUSCLE alignment, % identity **40.5**

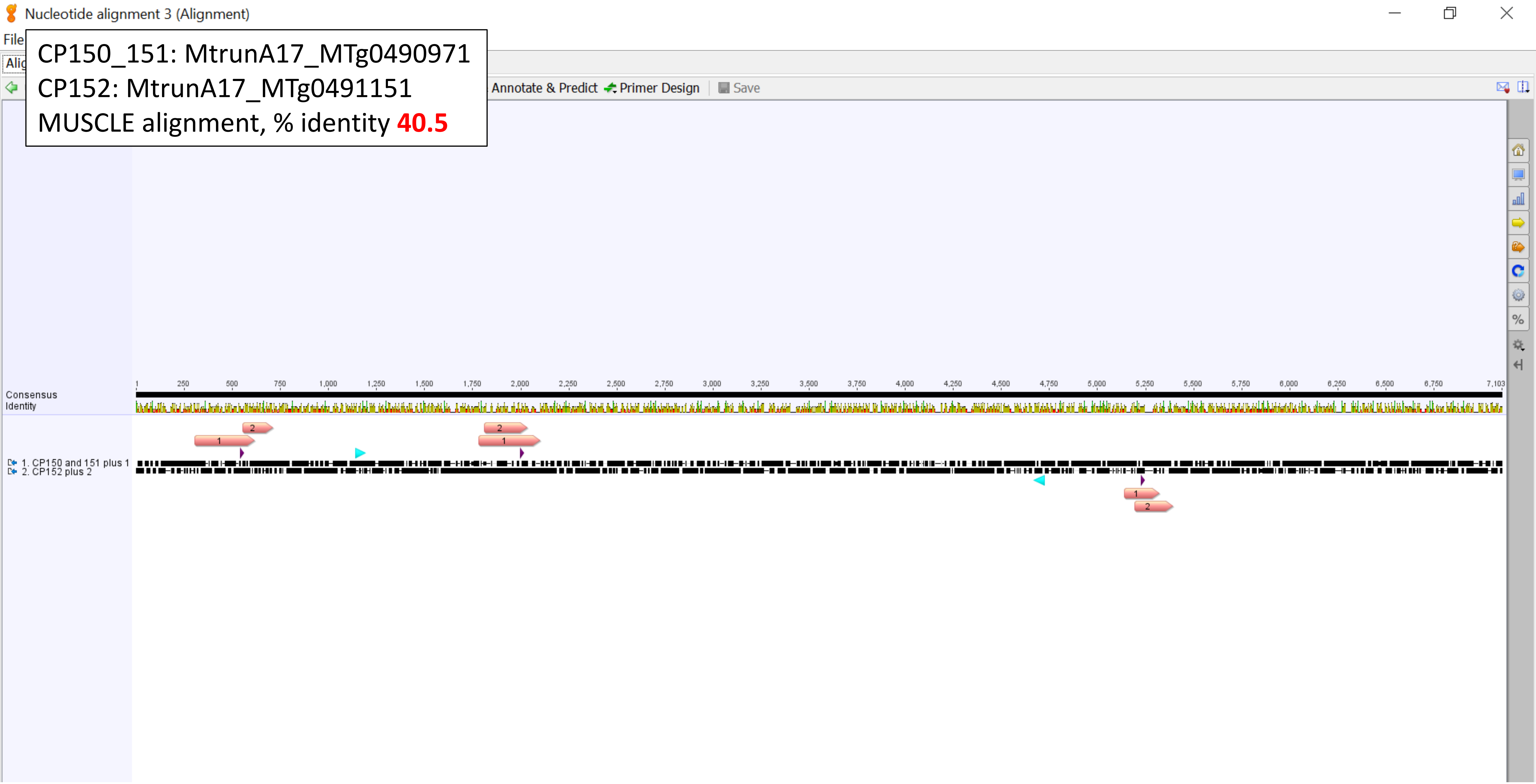

CP150_151: MtrunA17_MTg0490971
CP152: MtrunA17_MTg0491151
Homology region, % identity **100.0**

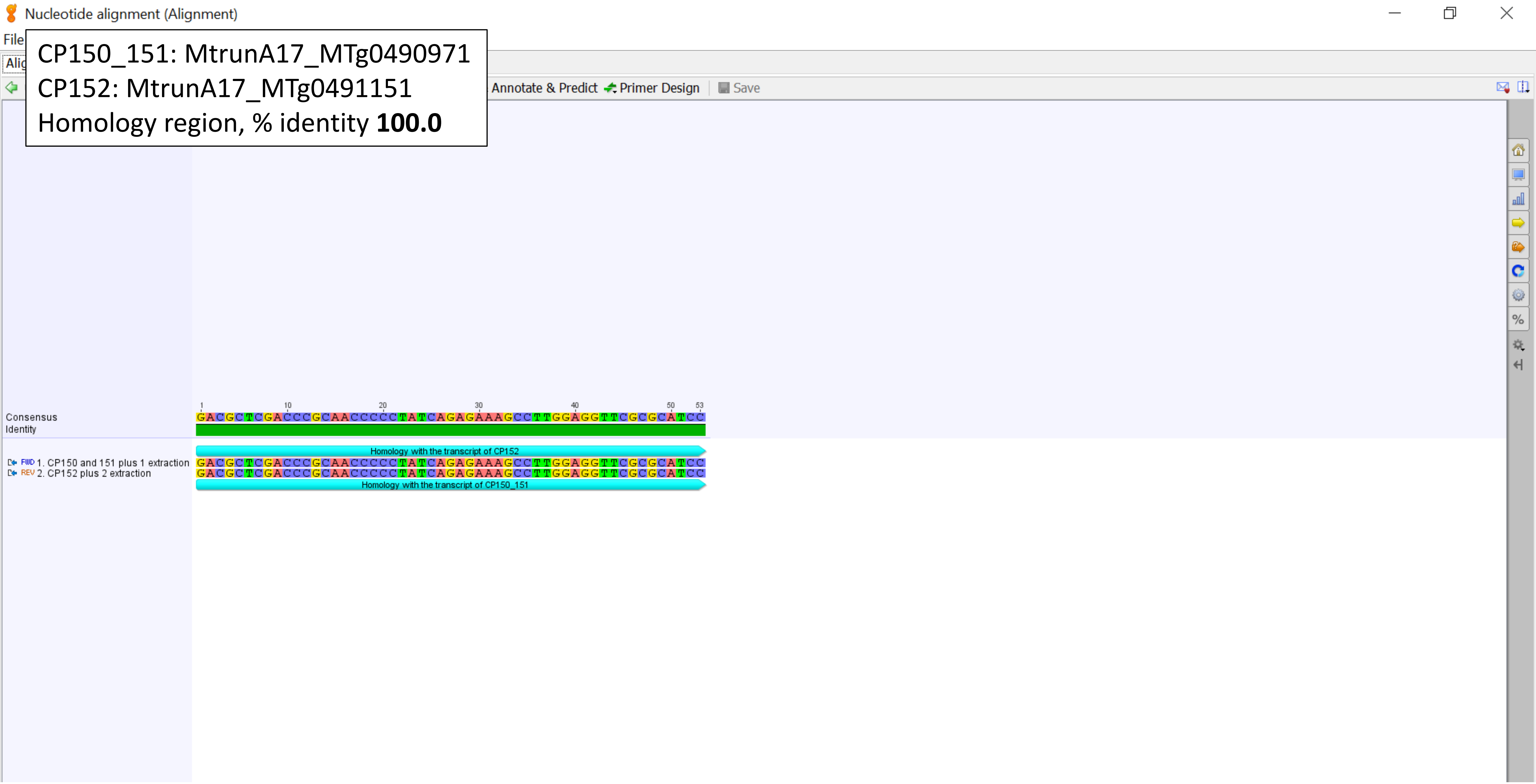

**Supplementary Dataset S14**. A graphical summary on nucleotide alignments among all 34 primary-source transcripts that have the global or the local similarity to each other. We discriminate between the alignments that concern the entire length of at least one transcript and those that concern only a part of the sequence. We refer to the latter ones as homology regions. They are highlighted with cyan in this dataset and with dark green in cells of Supplementary Dataset 15B. For alignments of the entire sequences, percent identity values are based on Geneious® alignments (Geneious® v. 7.1, Dotmatics Ltd., MA, USA, https://www.geneious.com). For alignments of the homology regions, percent identity values are based on BLASTN results (the *Medicago truncatula* genome browser v. 5.1.9). Reference ORFs (refORFs) are shown in yellow. Alternative ORFs (altORFs) are shown in pink. Numbers 1 and 2 on altORFs indicate the following. An altORF1 in this study is defined as the one that starts earlier than an altORF2. Switches from altORF2 to altORF1 are also found in this dataset but are five times less frequent (e.g. CP108 and four other CPs). Positions and the direction of programmed ribosomal frameshifting (PRF) events are labeled with deep purple arrowheads. Blue stars mark alignments with features of special interest. Screenshots are from Geneious® v. 7.1.
