## Supplementary material for "Discovery of diverse chimeric peptides in a eukaryotic proteome sets the stage for the experimental proof of the mosaic translation hypothesis": Supplementary Dataset S19 RNA-Seq read alignments.pdf

**CP31**, flowers:

MtrunA17_Chr1g0212961, primary source; 10-dpi nodules

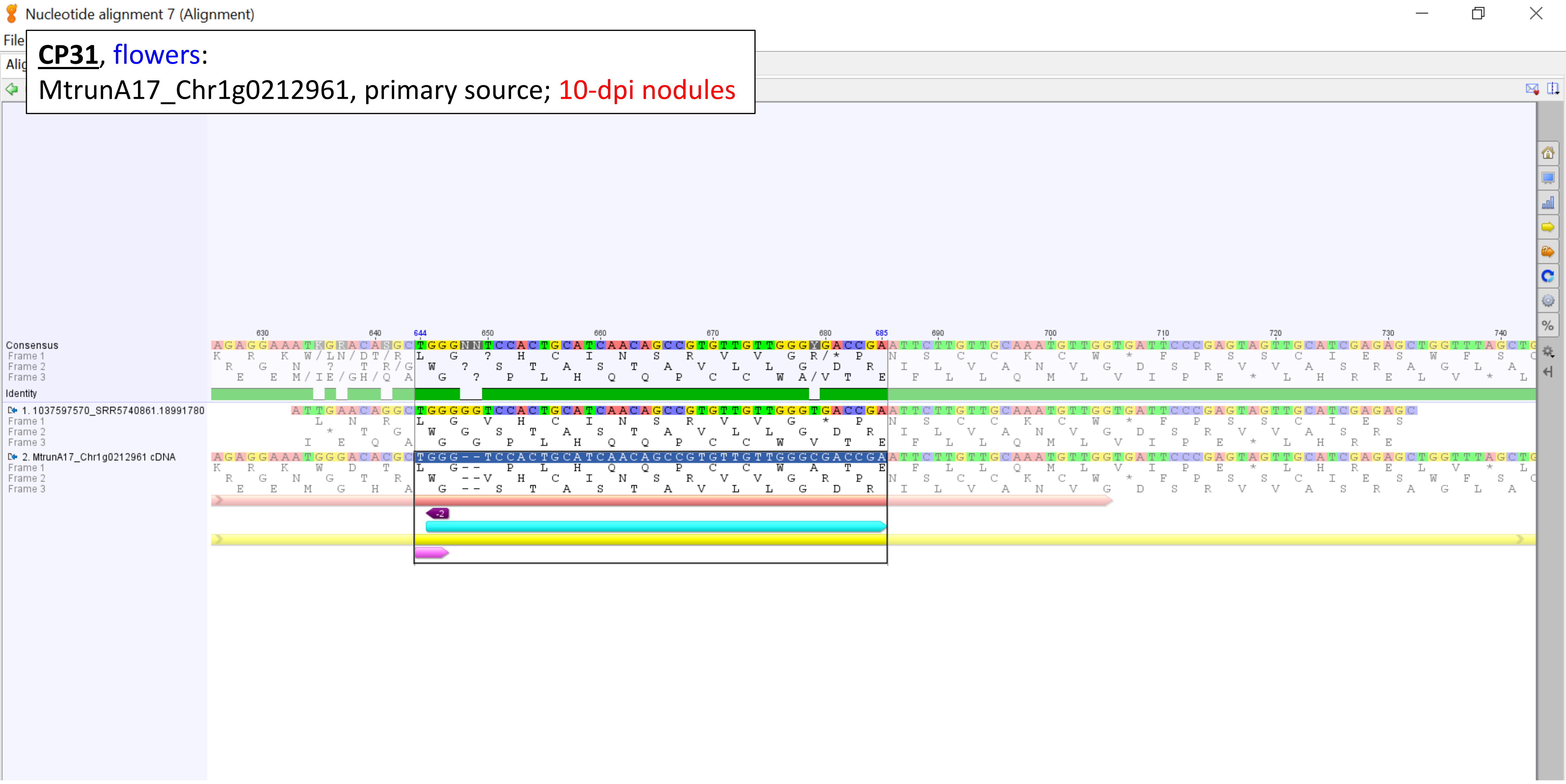

**CP35**, seeds:

Group 1, MtrunA17_Chr1g0203181, alternative source

roots (5), roots ribo-minus (19)

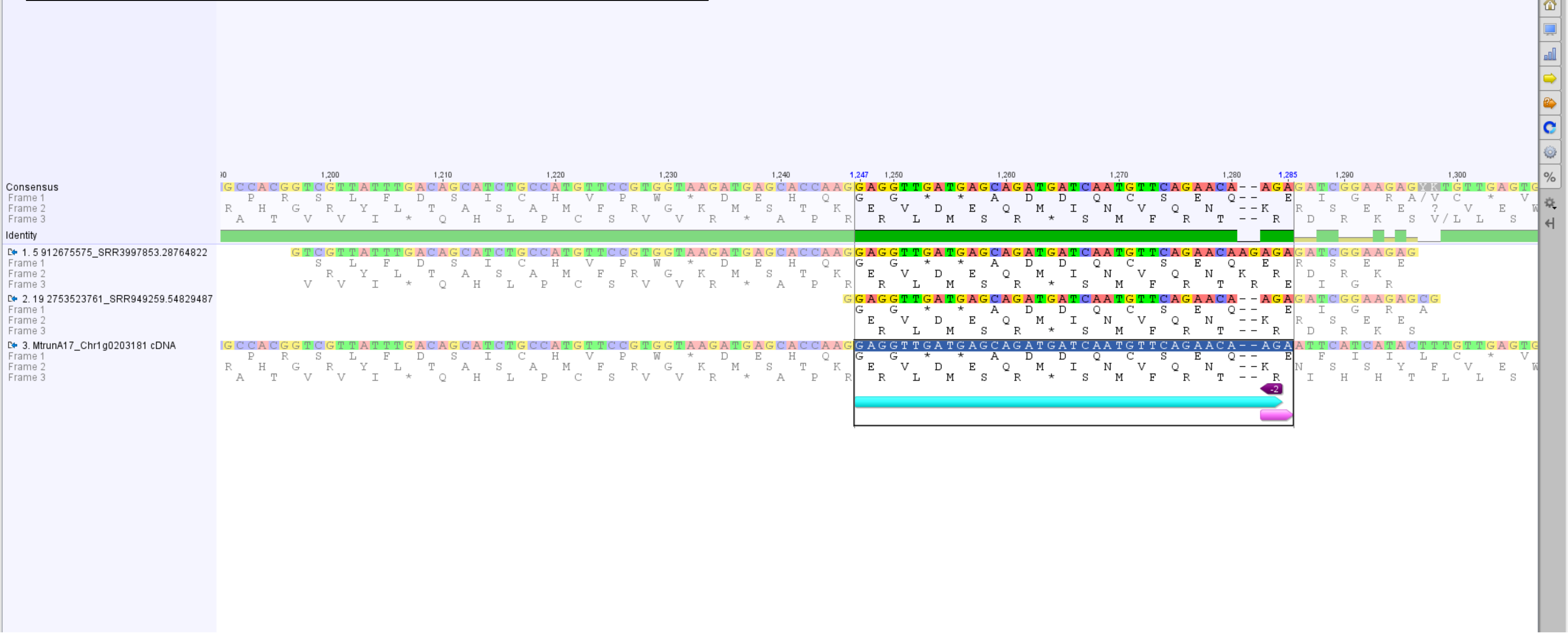

**CP35**, seeds:

Group 2, MtrunA17_Chr4g0008361, alternative source

seedlings (7, 15, 16), shoot apical buds (11)

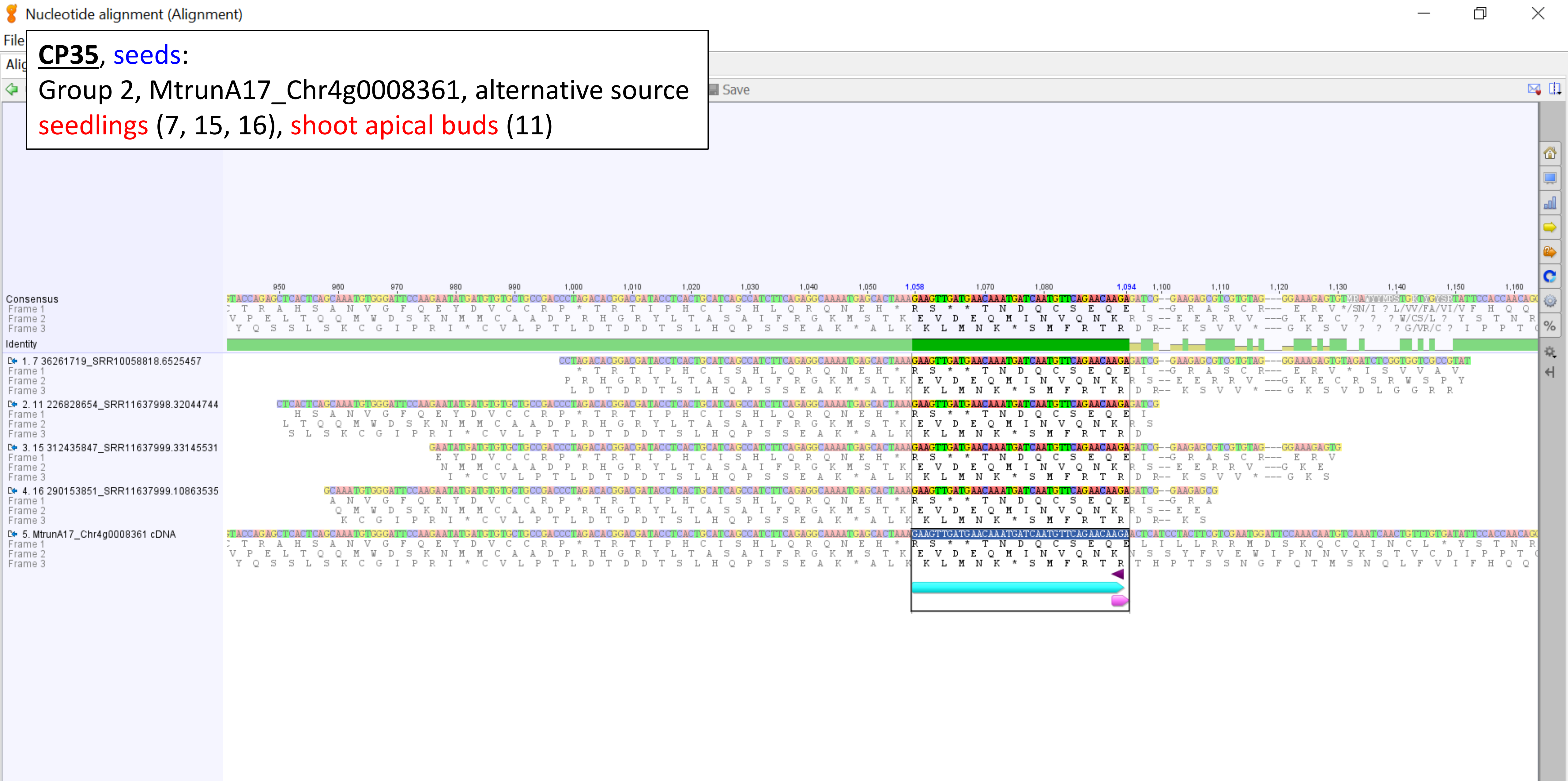

**CP35**, seeds:

Group 3, MtrunA17_Chr4g0008401, alternative source

roots (1, 2, 3, 4), petioles (9, 12, 18), seedlings (13, 14), shoot apical buds (6), roots ribo-minus (17)

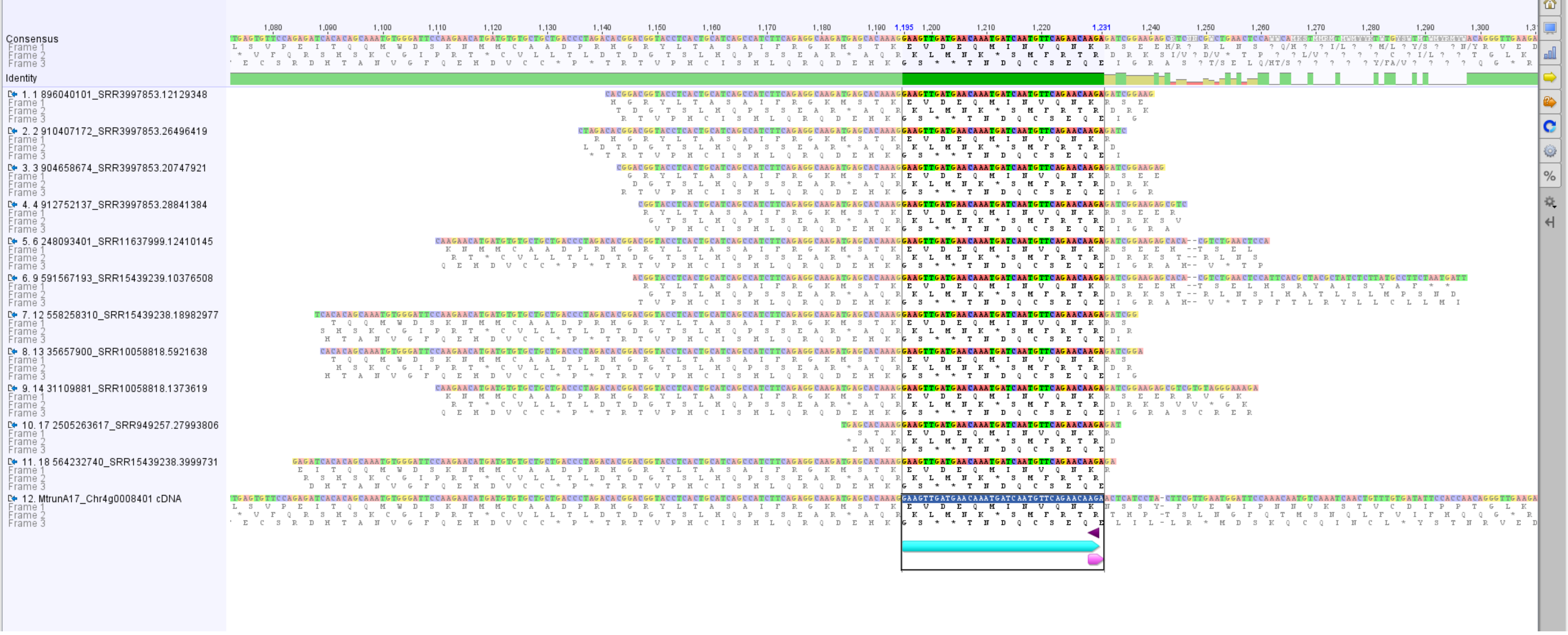

**<u>CP35</u>**, seeds:

Group 4, MtrunA17_Chr7g0255791, alternative source

seedlings (8), shoot apical buds (10)

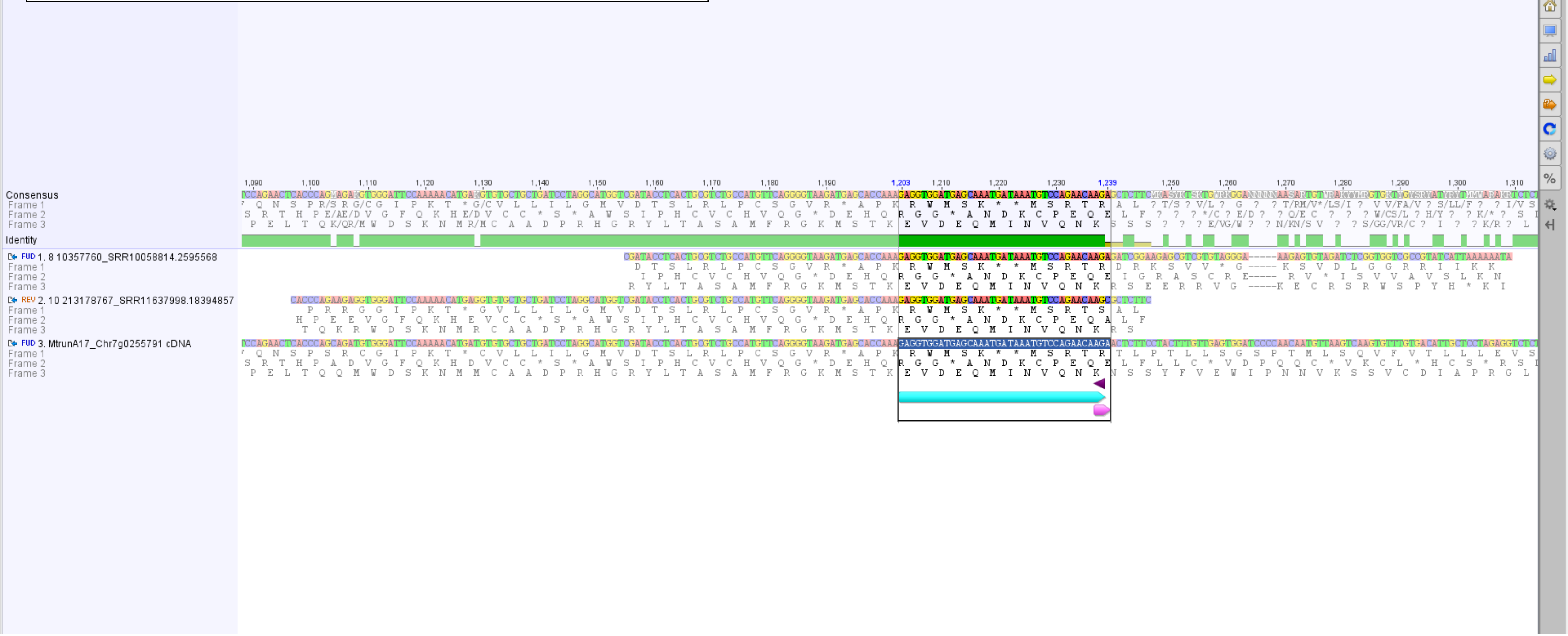

**CP54**, 14-dpi nodules, buds, seeds:

Overview, MtrunA17_Chr3g0105981, primary source

shoot apical buds (1, 2, 4, 5, 6, 11, 14, 15, 16, 20, 21, 23, 24, 25, 26, 27, 28, 29, 30, 32, 34, 35, 36, [37], 38, 39, 40, 42, 43, 44, 45, 46, 47, 48, 49, 51, 52, 53, 54, 55, 58, 68), 10-dpi nodules ([9], 10, [56], 57, 59, 60, 62, 63, 66, [67]), 14-dpi nodules (9, [10], 17, 19, [54], 61, 64, 65), petioles ([2], 8, 13, [24], [25], [26], 31, 33, 41, [43], 50), seedlings (7, 12, 18), shoots (3, 37, [40]), whole seedings (56, [66], 67), leaves (22)

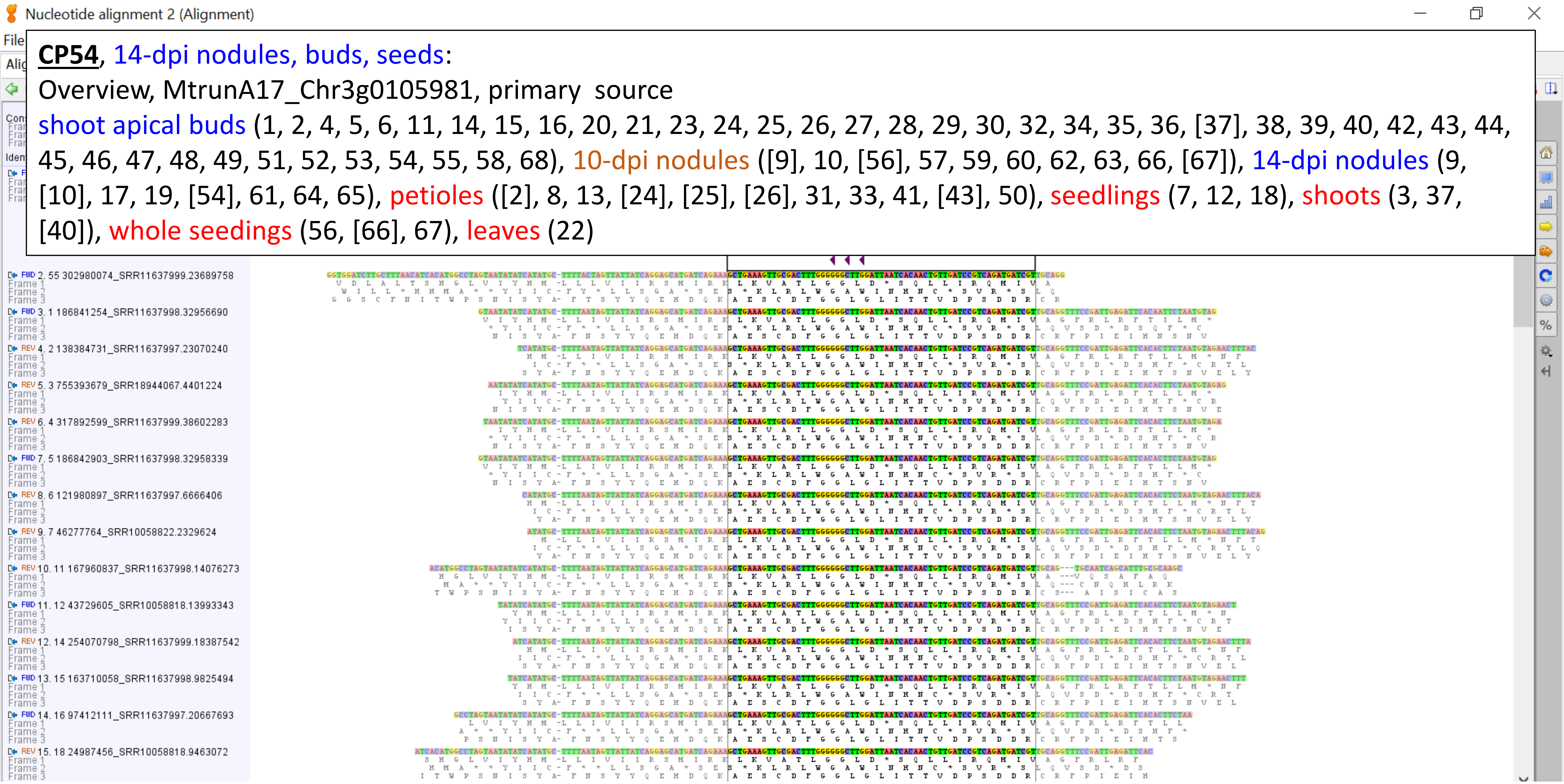

**CP54**, 14-dpi nodules, buds, seeds: Part 1, MtrunA17_Chr3g0105981, primary source

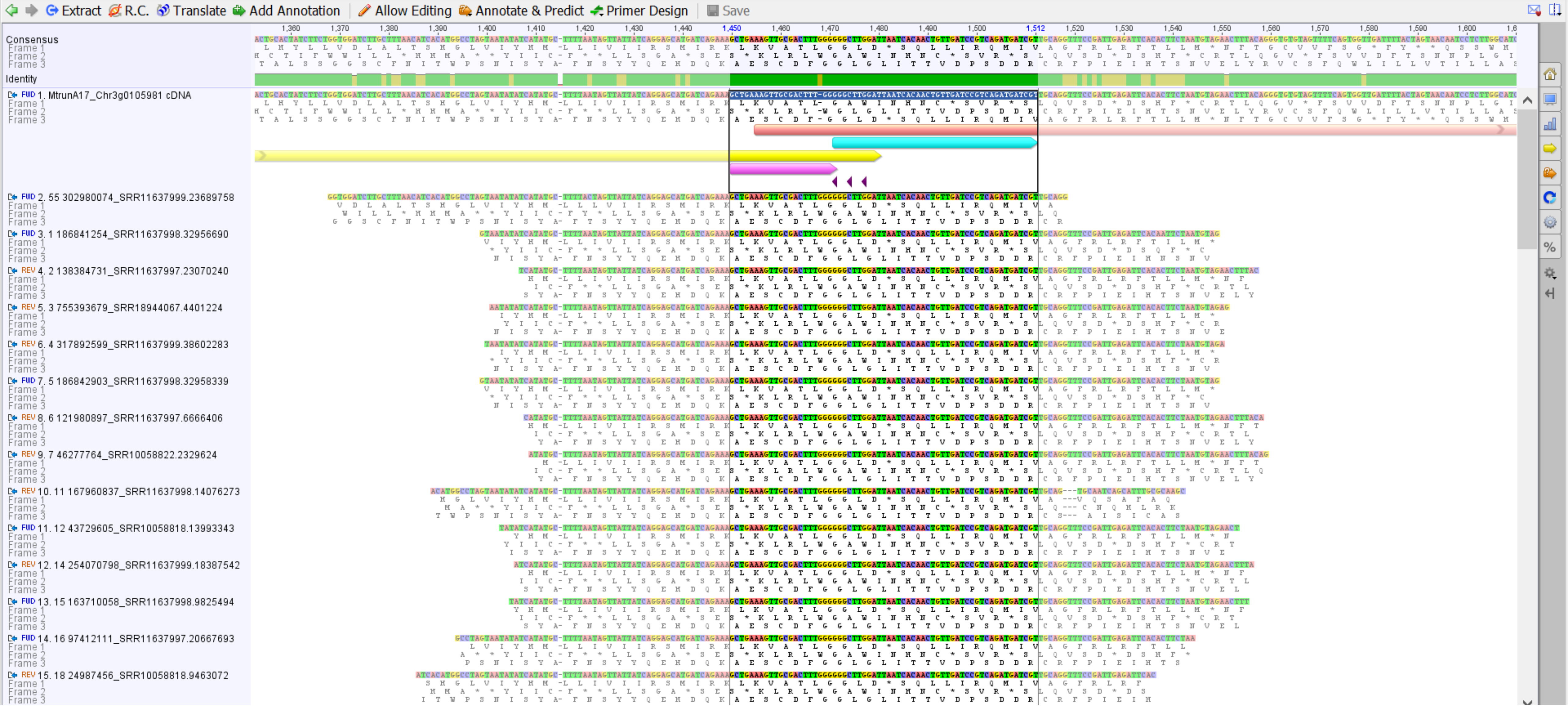

**CP54**, 14-dpi nodules, buds, seeds: Part 2, MtrunA17_Chr3g0105981, primary source

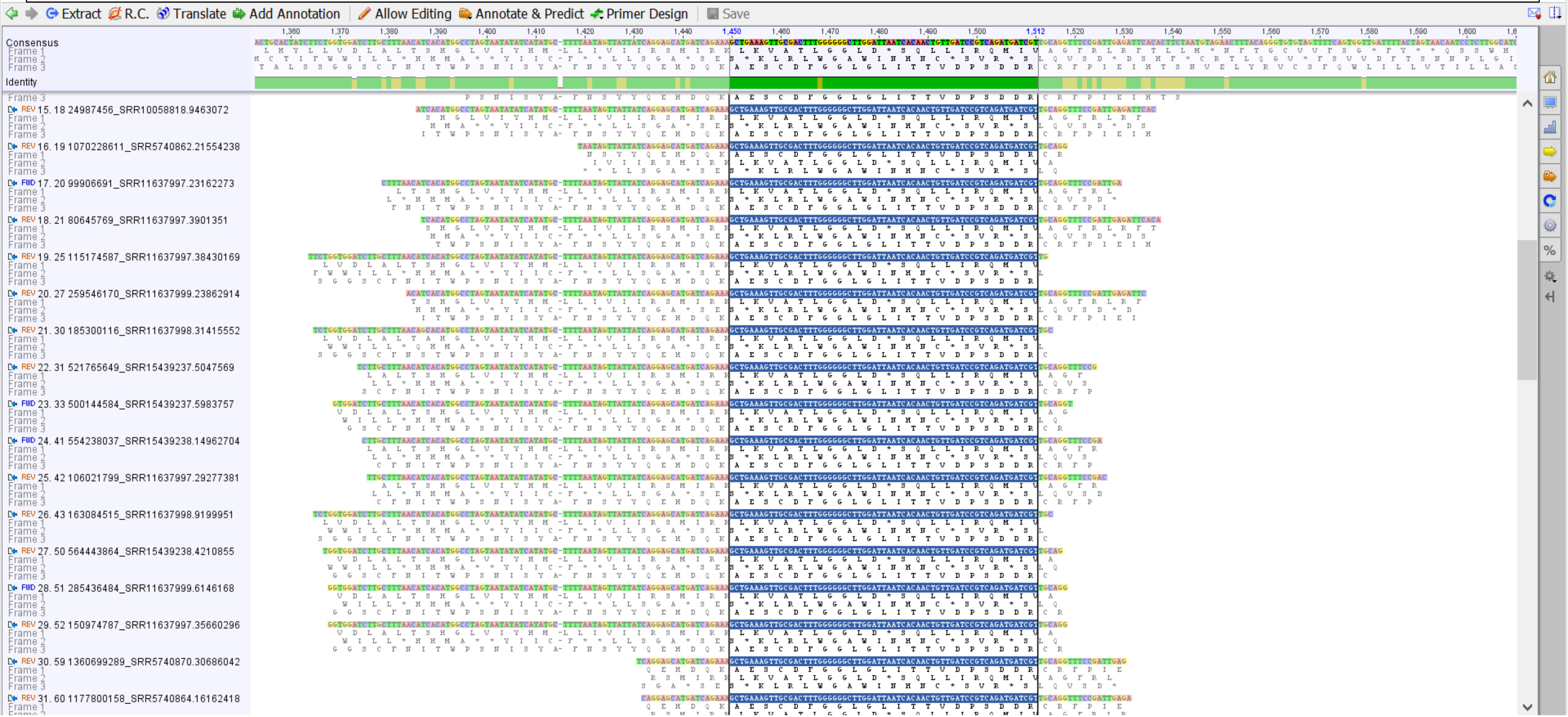

### **CP54**, 14-dpi nodules, buds, seeds: Part 3, MtrunA17_Chr3g0105981, primary source

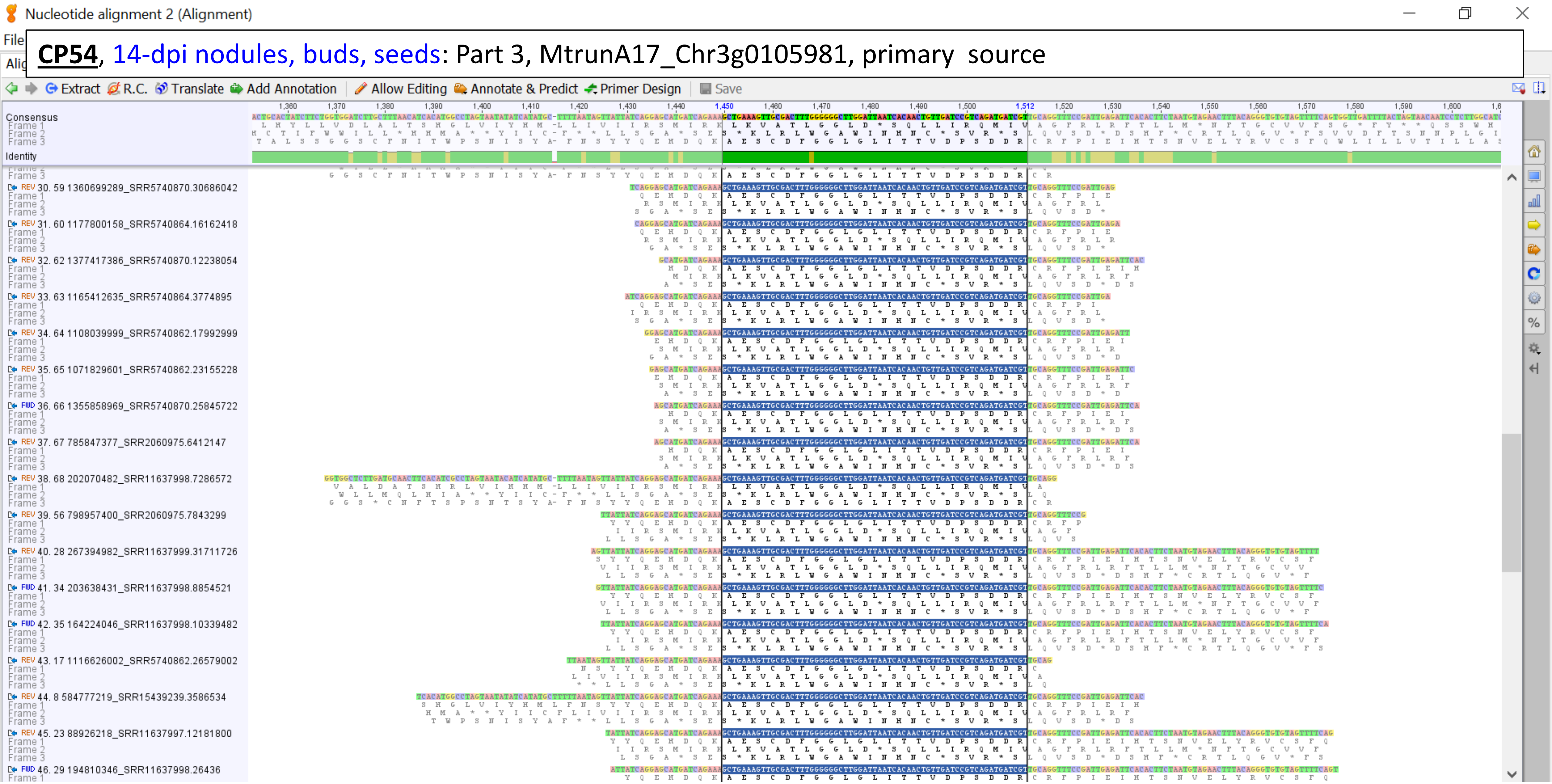

**CP54**, 14-dpi nodules, buds, seeds: Part 4, MtrunA17_Chr3g0105981, primary  source

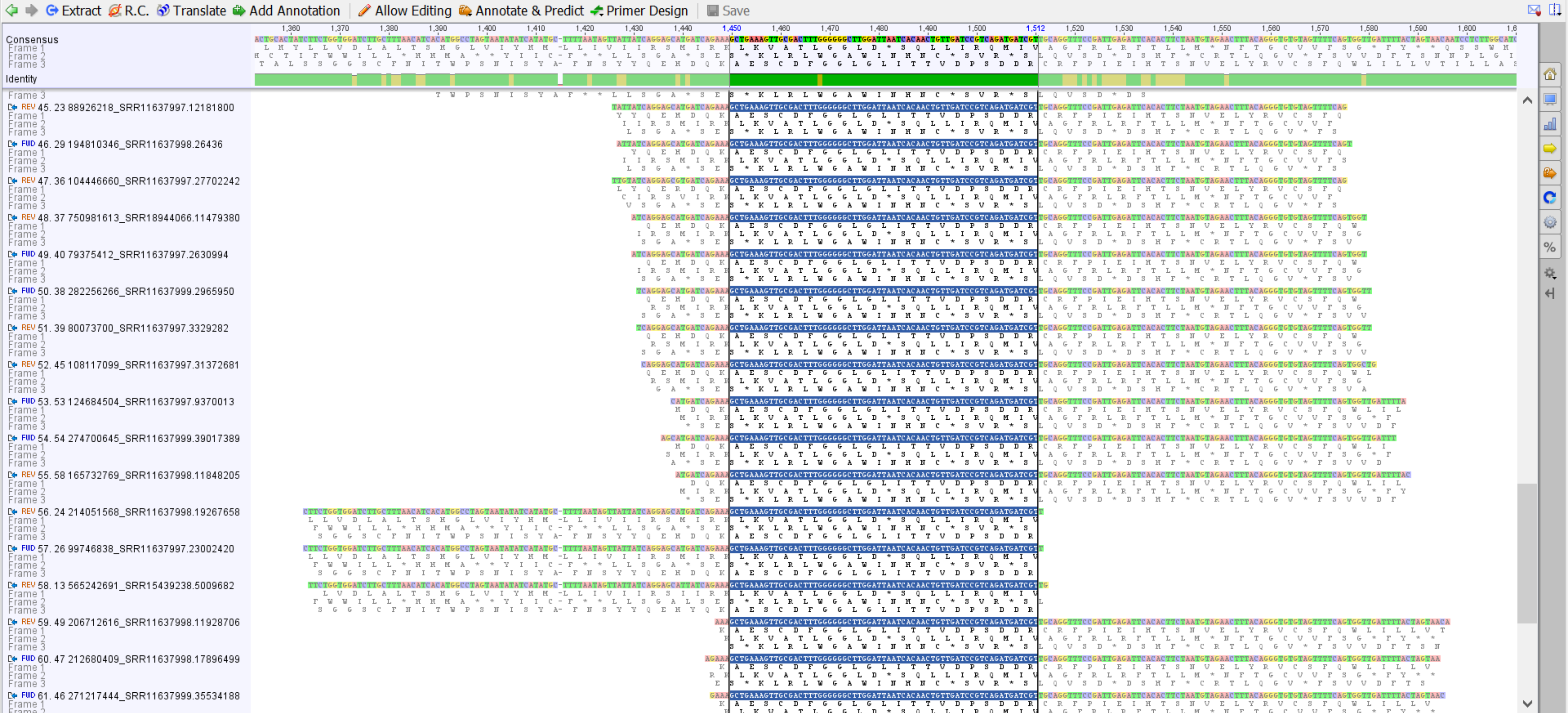

**CP54**, 14-dpi nodules, buds, seeds: Part 5, MtrunA17_Chr3g0105981, primary source

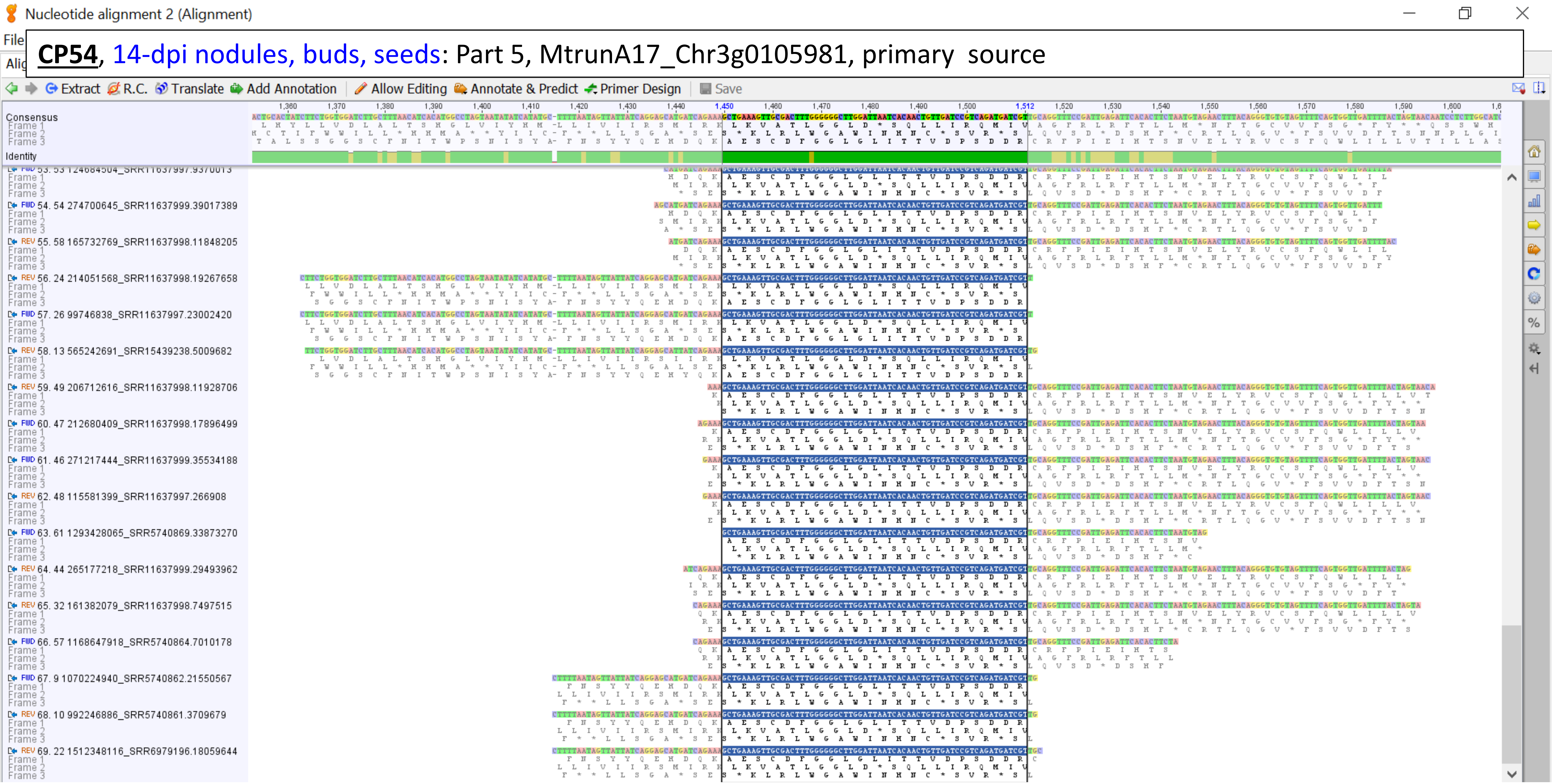

**CP73**, petioles under drought:
MtrunA17_Chr4g0034271, primary source
roots polyA

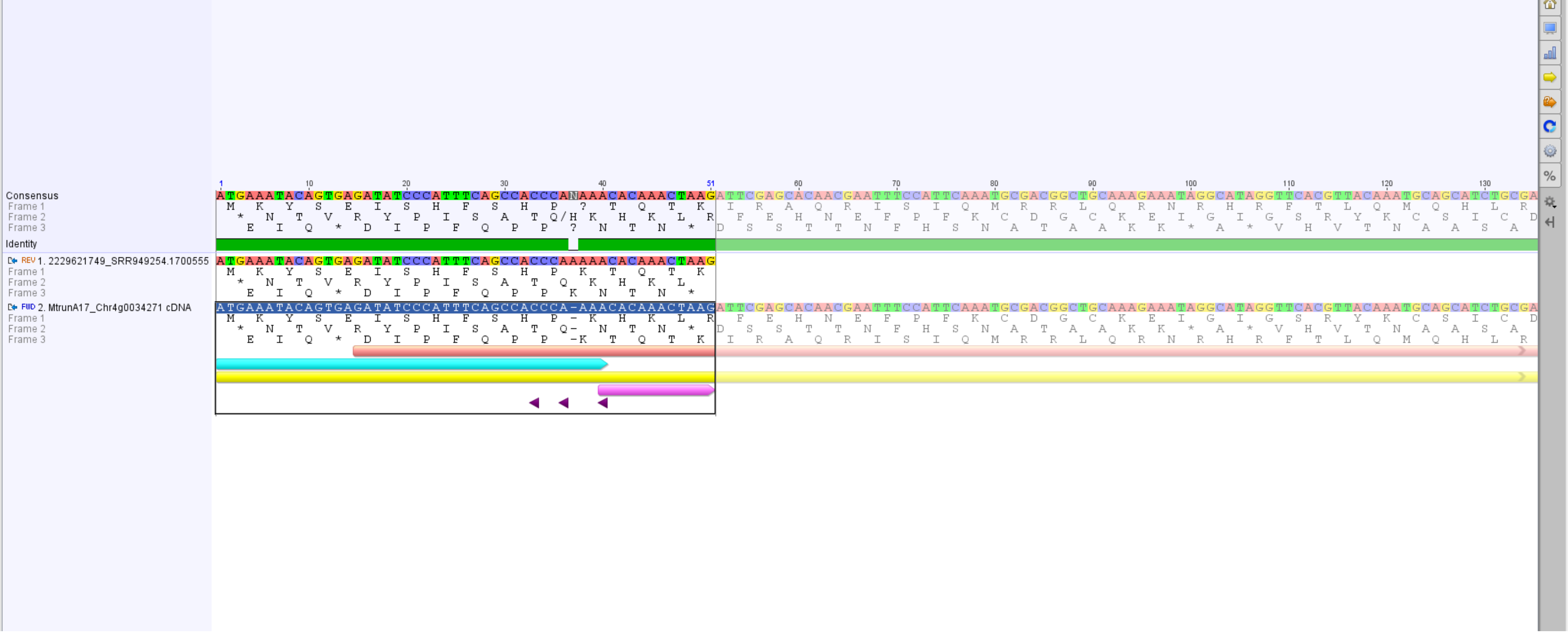

**CP74**, flowers:
MtrunA17_Chr4g0034491, primary source
shoot apical buds (1, 2)

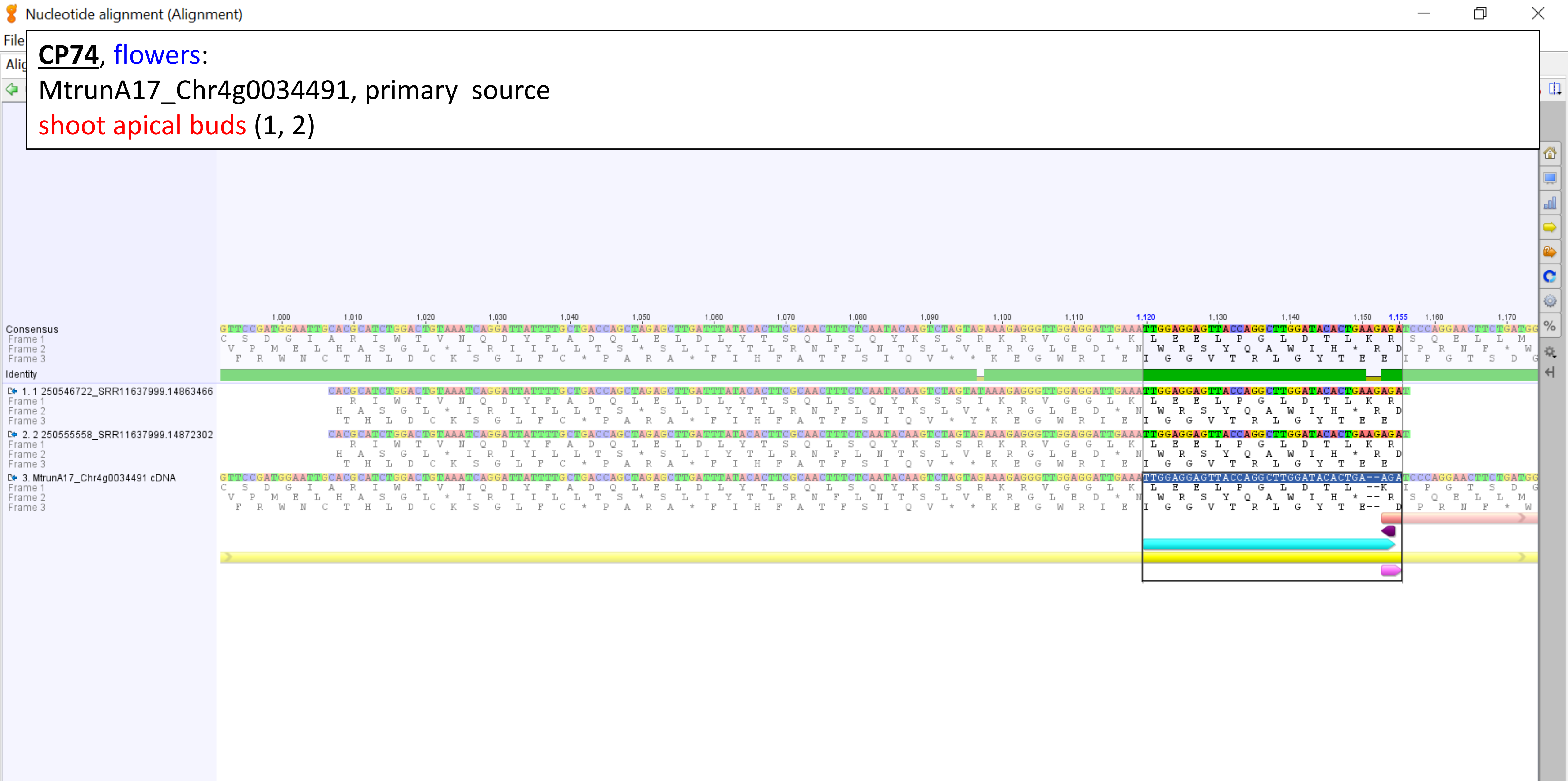

**CP92**, roots, stems:

MtrunA17_Chr5g0431401, primary source

14-dpi nodules (1, 2)

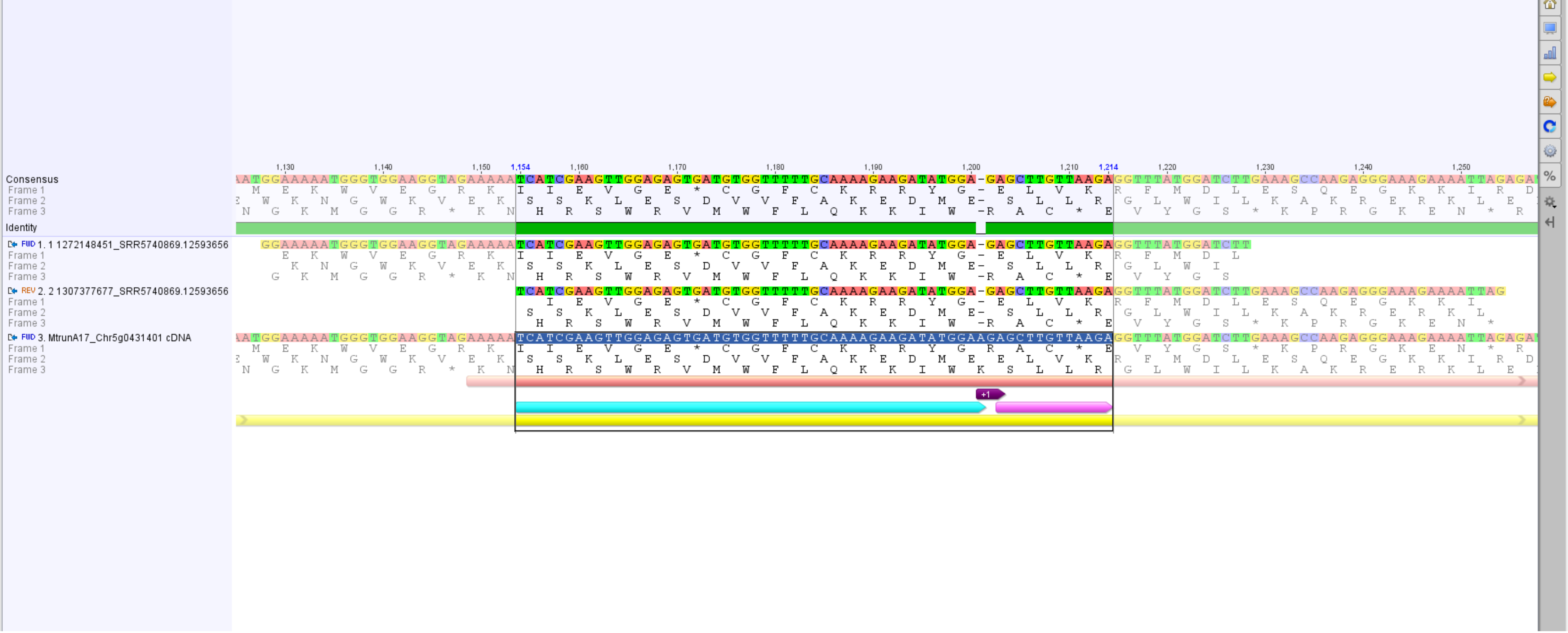

**CP93**, seeds:

MtrunA17_Chr5g0435191, primary source

10-dpi nodules (2, 3, 5), 14-dpi nodules (1, 6, 7), nodules polyA (8), roots (4), roots ribo-minus ([8], 9)

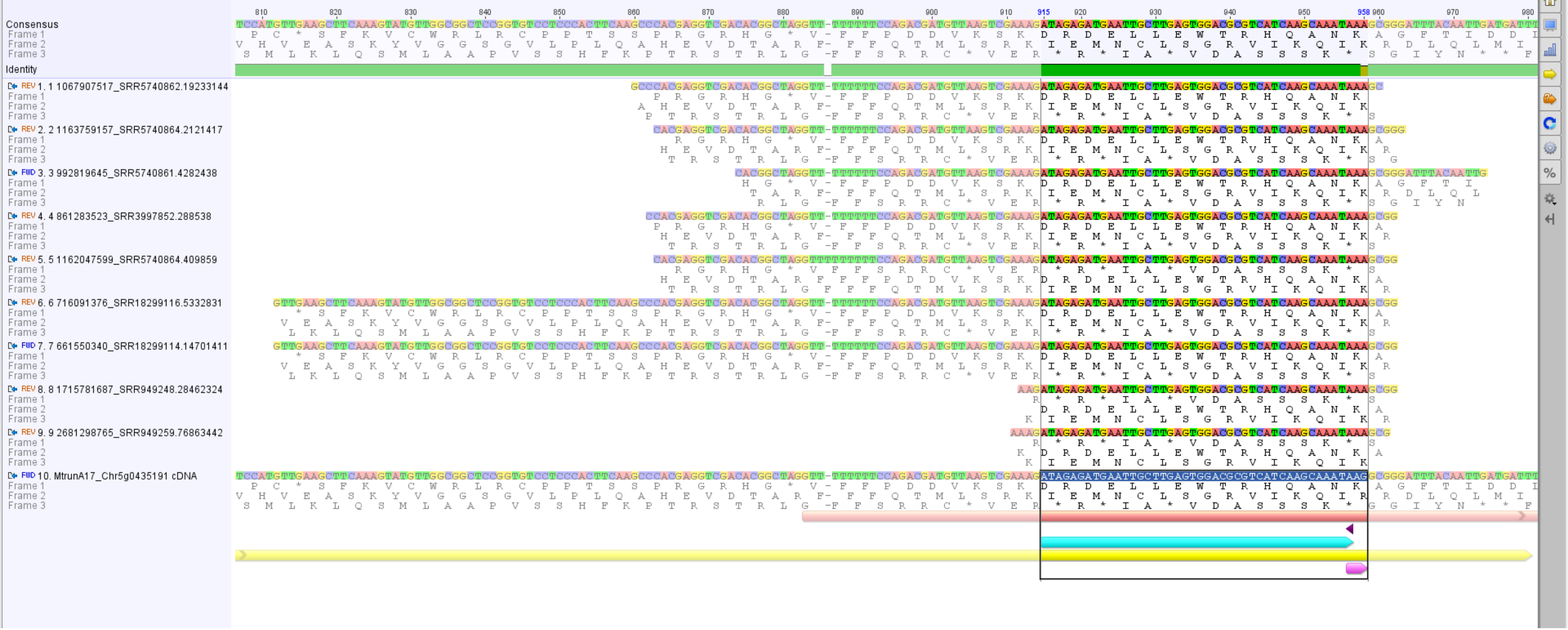

**<u>CP94</u>**, flowers:

MtrunA17_Chr5g0444231, primary source

petioles (1), 10-dpi nodules (3), 14-dpi nodules (2)

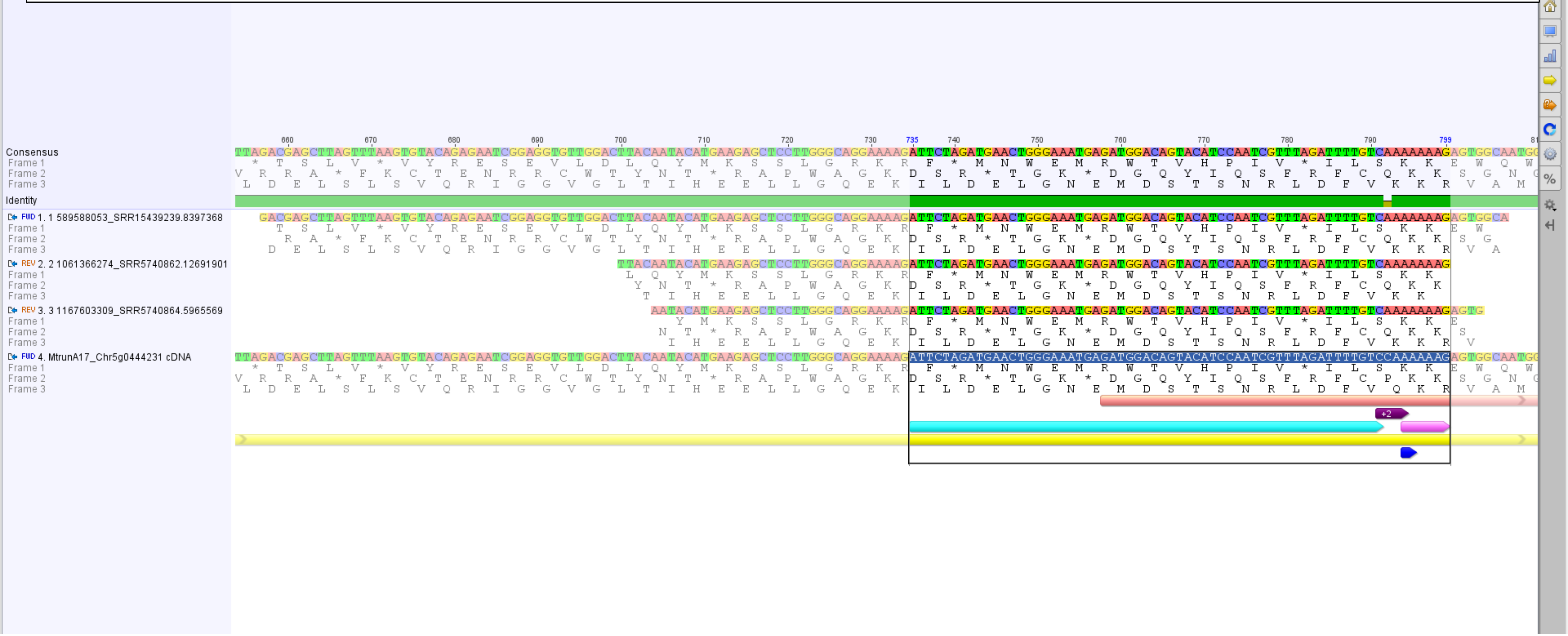

Nucleotide alignment 7 (Alignment)

**CP98**, buds:

MtrunA17_Chr6g0457471, alternative source

shoots

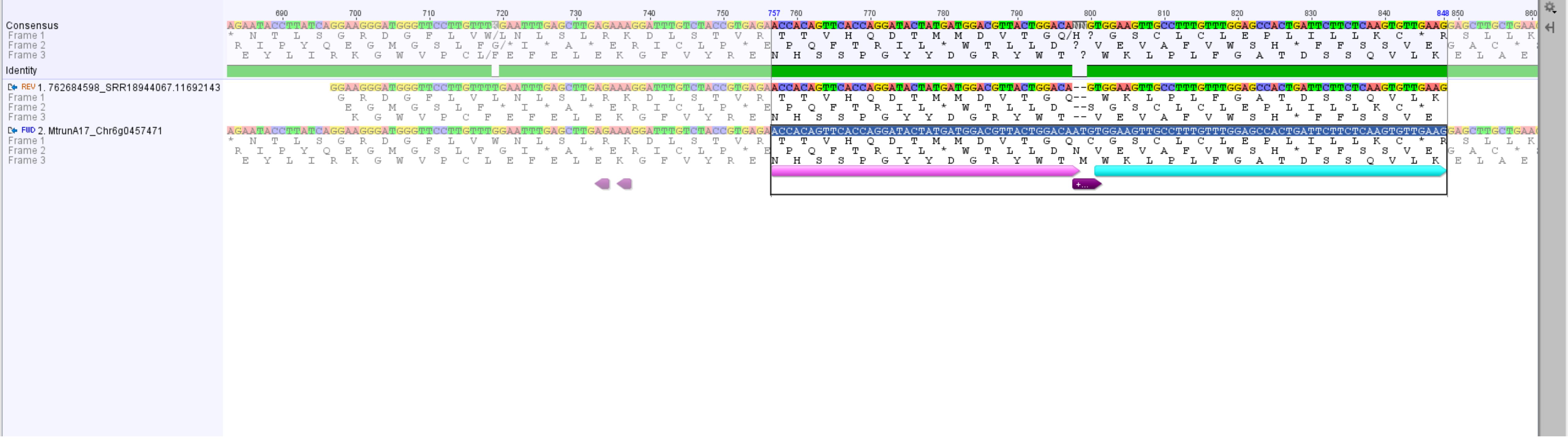

**CP100**, buds, whole plants, petioles, petioles under drought:
MtrunA17_Chr6g0457461, primary source
seedlings (1, 2)

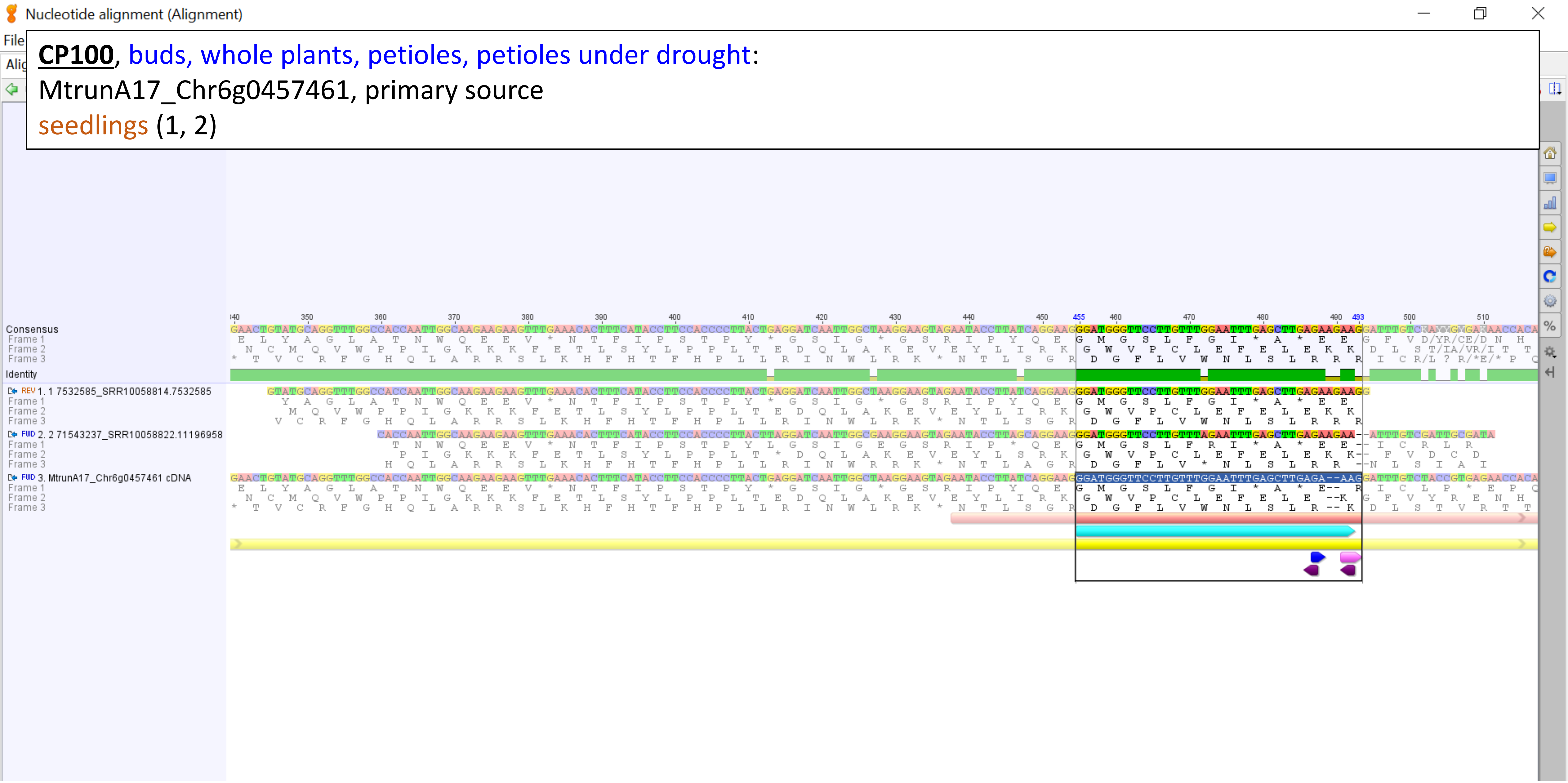

**CP110**, seeds:

MtrunA17_Chr7g0214741, primary source

shoots (1), 10-dpi nodules (2)

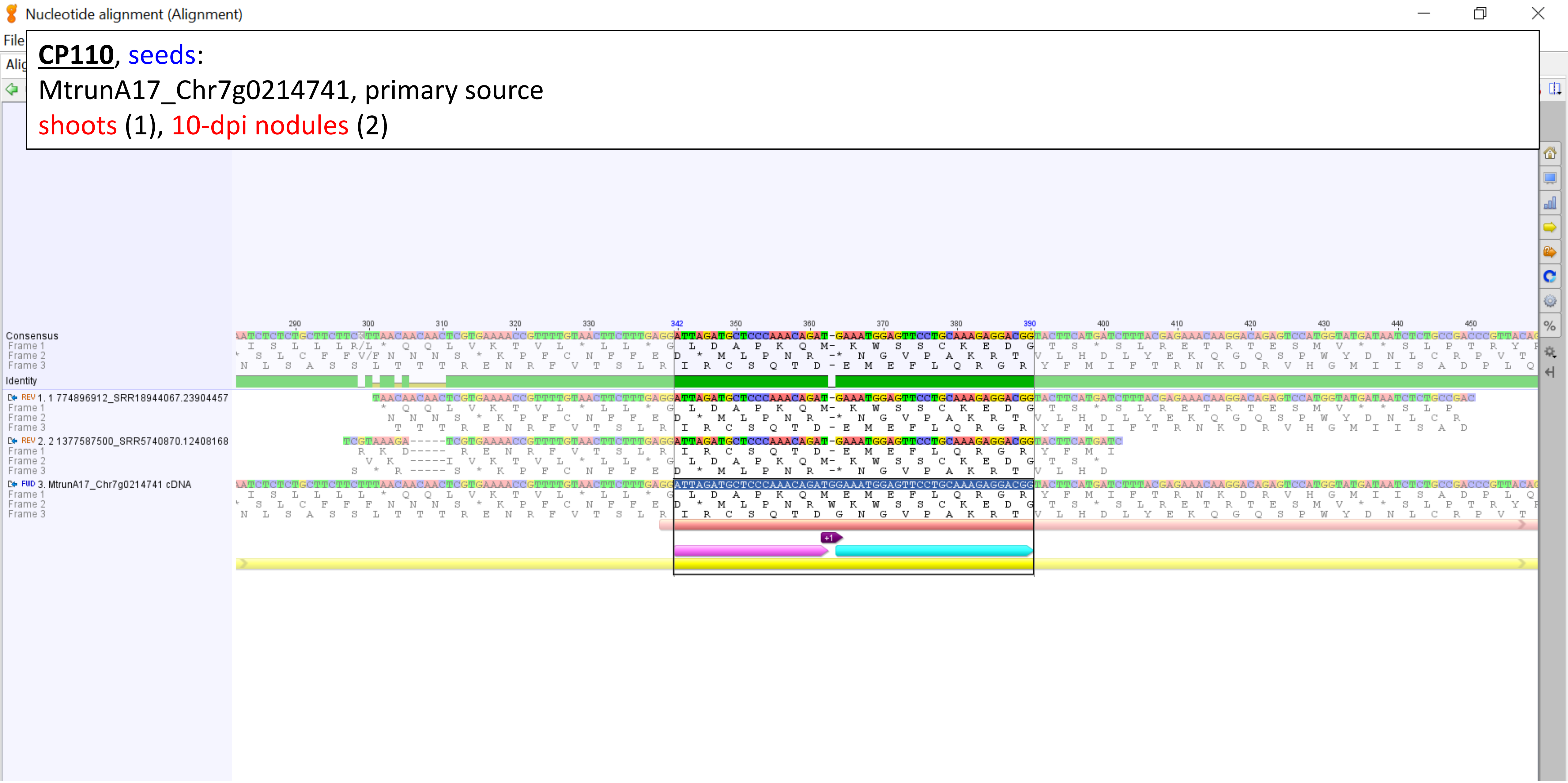

**<u>CP123</u>**, seeds:

Overview, MtrunA17_Chr8g0380331, alternative source

seedlings (1), shoot apical buds (2)

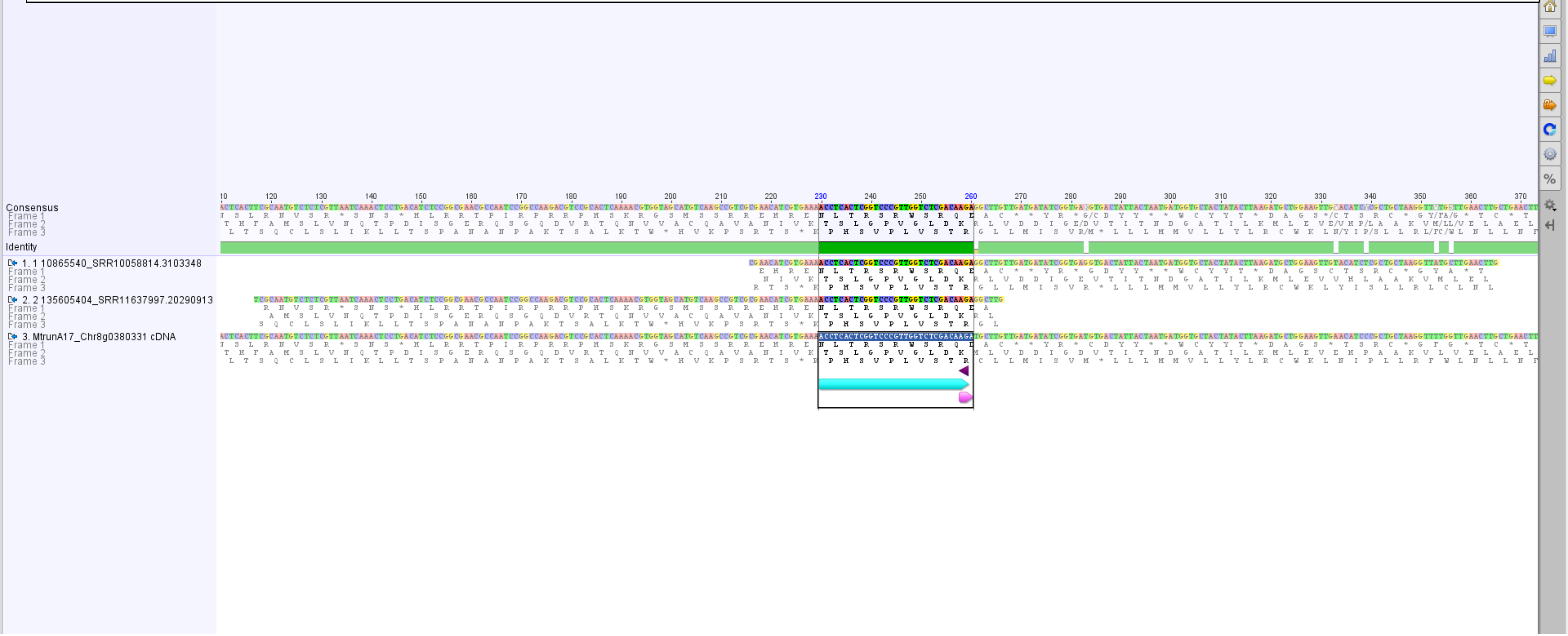

**CP123**, seeds:

Zoom-in view, MtrunA17_Chr8g0380331, alternative source

seedlings (1), shoot apical buds (2)

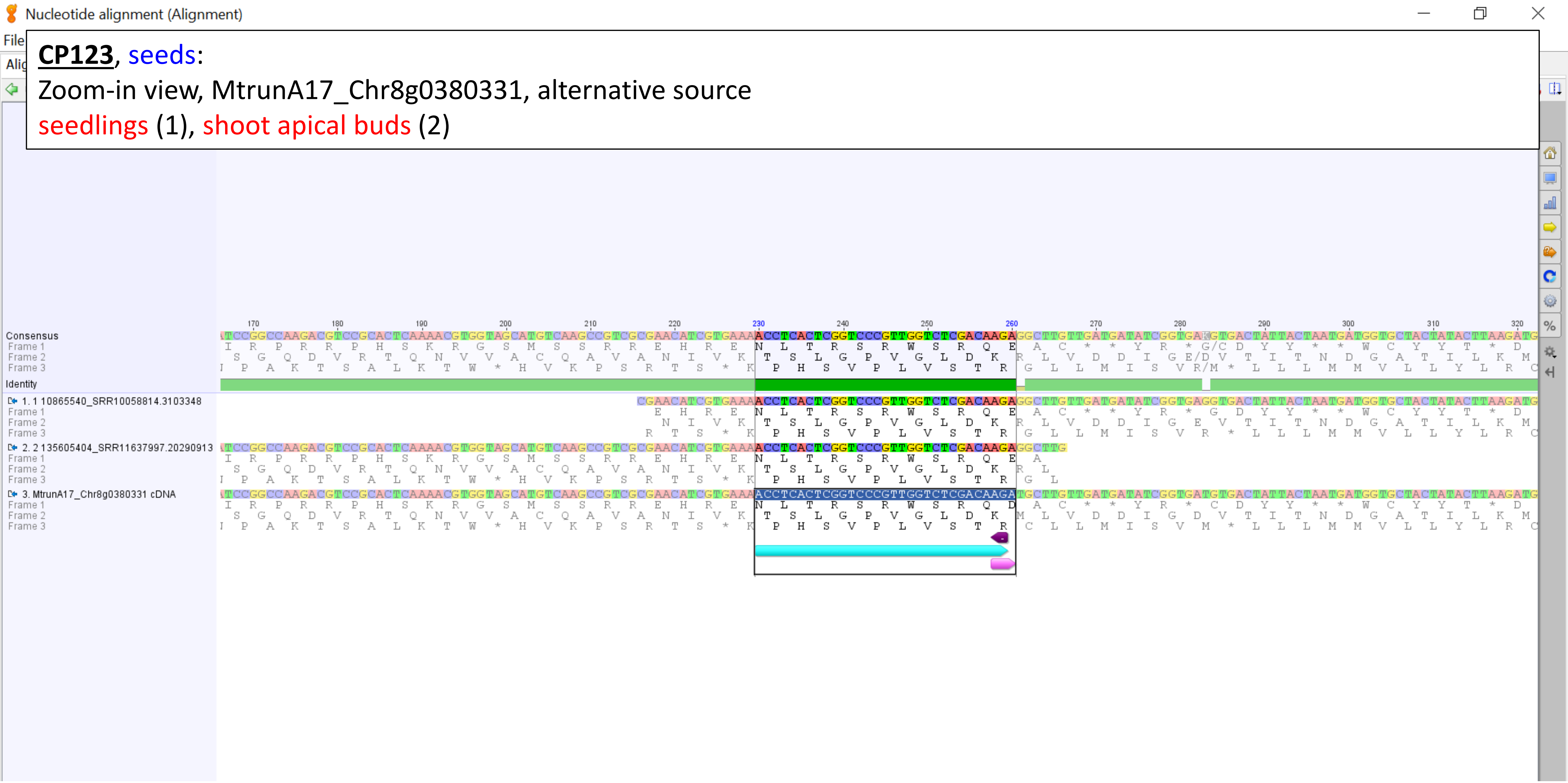

**CP140**, petioles under drought:

Group 1, overview, MtrunA17_Chr8g0392351, primary source

14-dpi nodules (7, 8, 9, 12, 13, 27), mature leaves (1, 2, 3, 4), nodules ribo-minus (14, 16, 19, 25), roots ribo-minus ([16], 18, [19], 20, 21, [25]), roots (5)

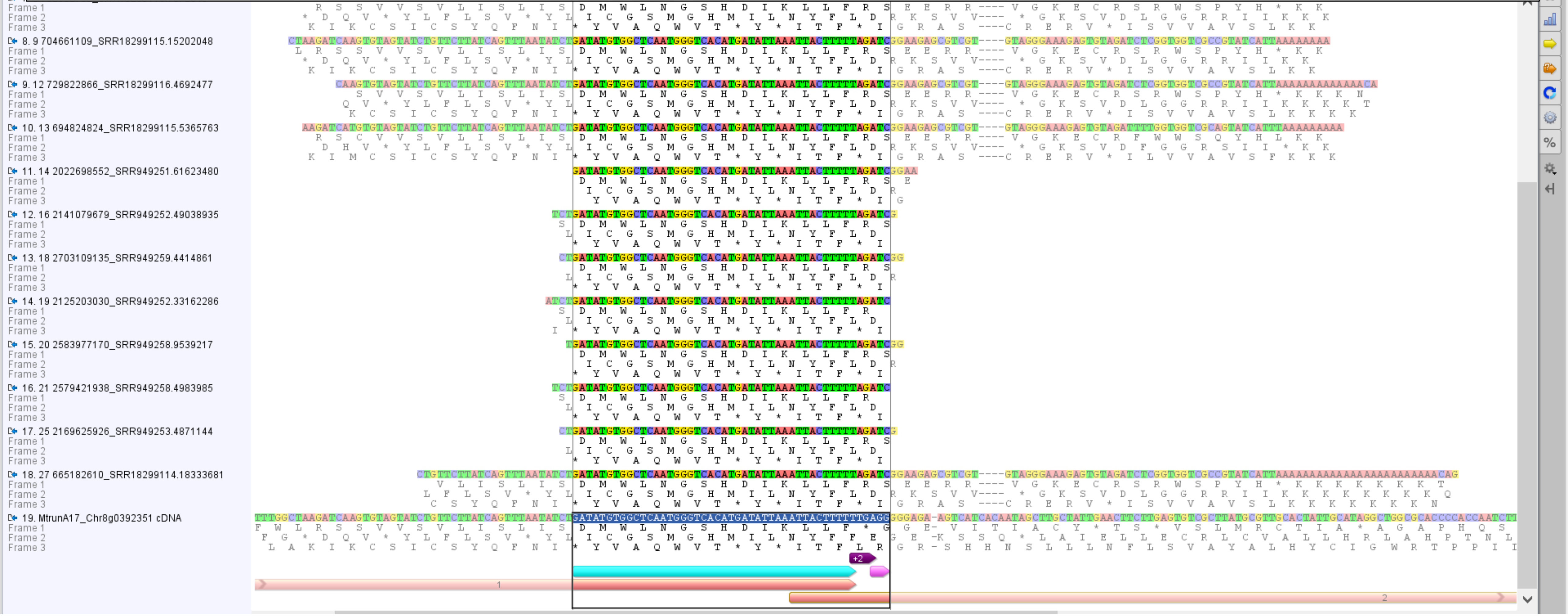

**CP140**, petioles under drought: Group 1, MtrunA17_Chr8g0392351, primary source

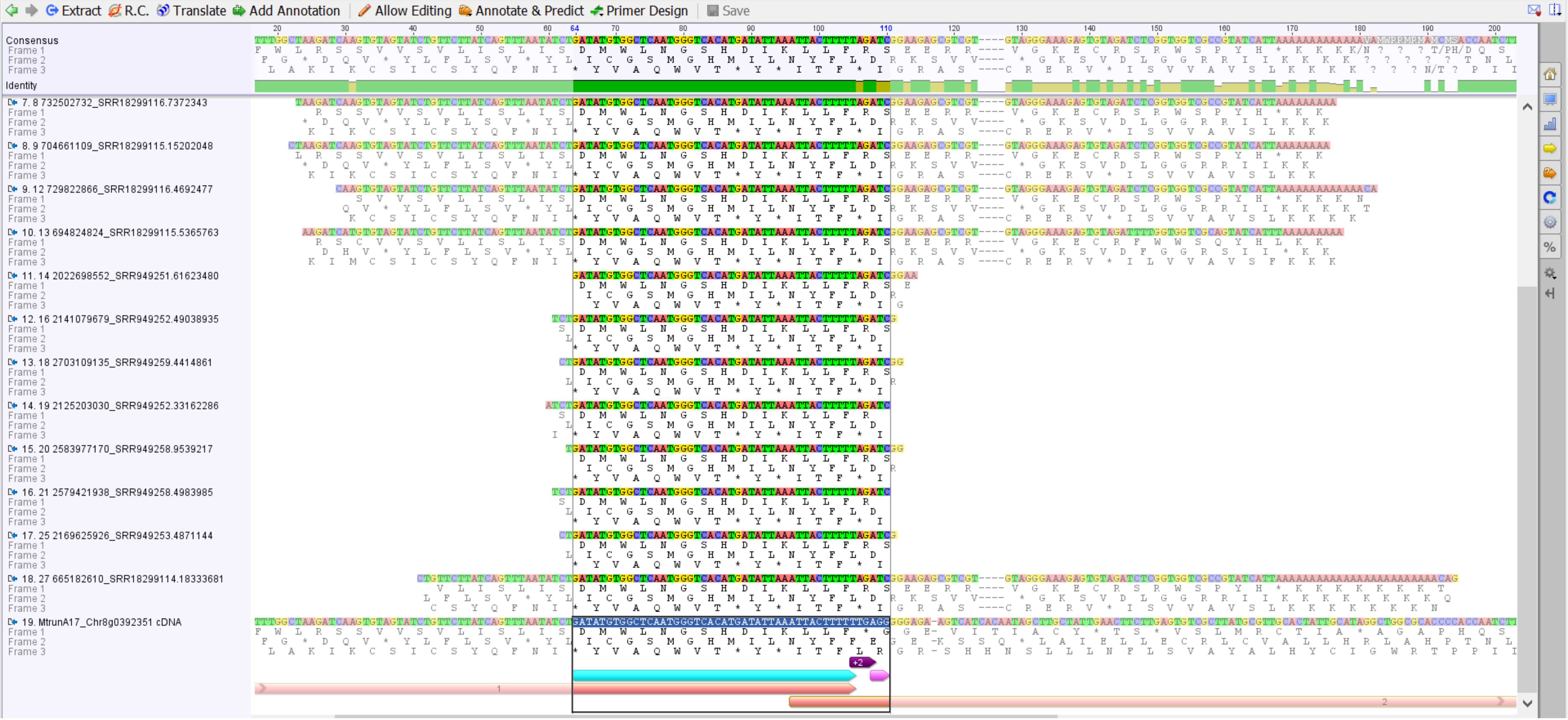

**<u>CP140</u>**, petioles under drought:

Group 2, MtrunA17_Chr8g0392351, primary source

roots ribo-minus (22, 23, 24, 28), 14-dpi nodules (11, 15), roots (26)

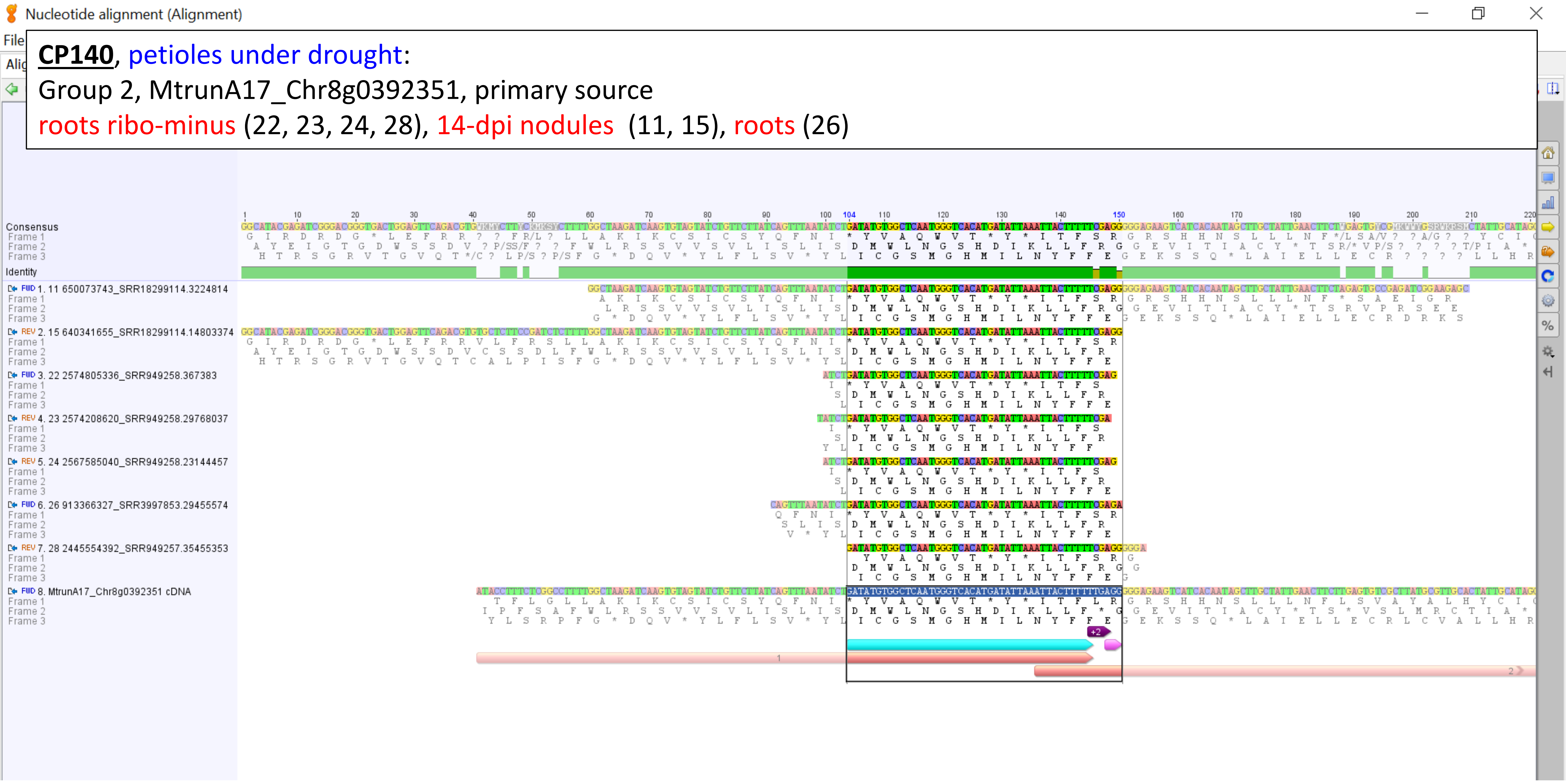

**CP140**, petioles under drought:

Group 3, MtrunA17_Chr8g0392351, primary source

nodules ribo-minus (29, 31), roots ribo-minus (30)

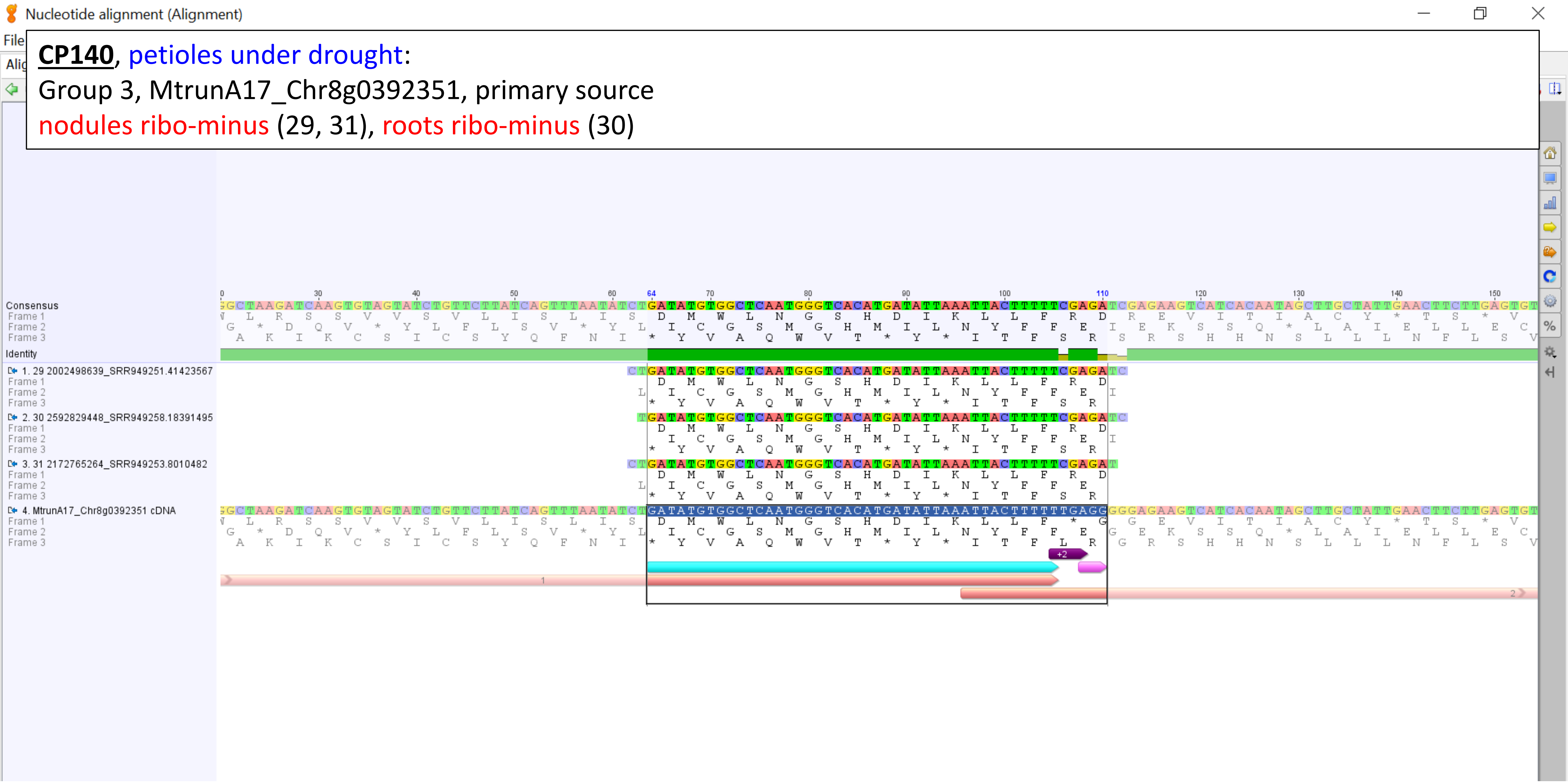

**CP140**, petioles under drought:

Group 4, MtrunA17_Chr8g0392351, primary source

14-dpi nodules (6, 10)

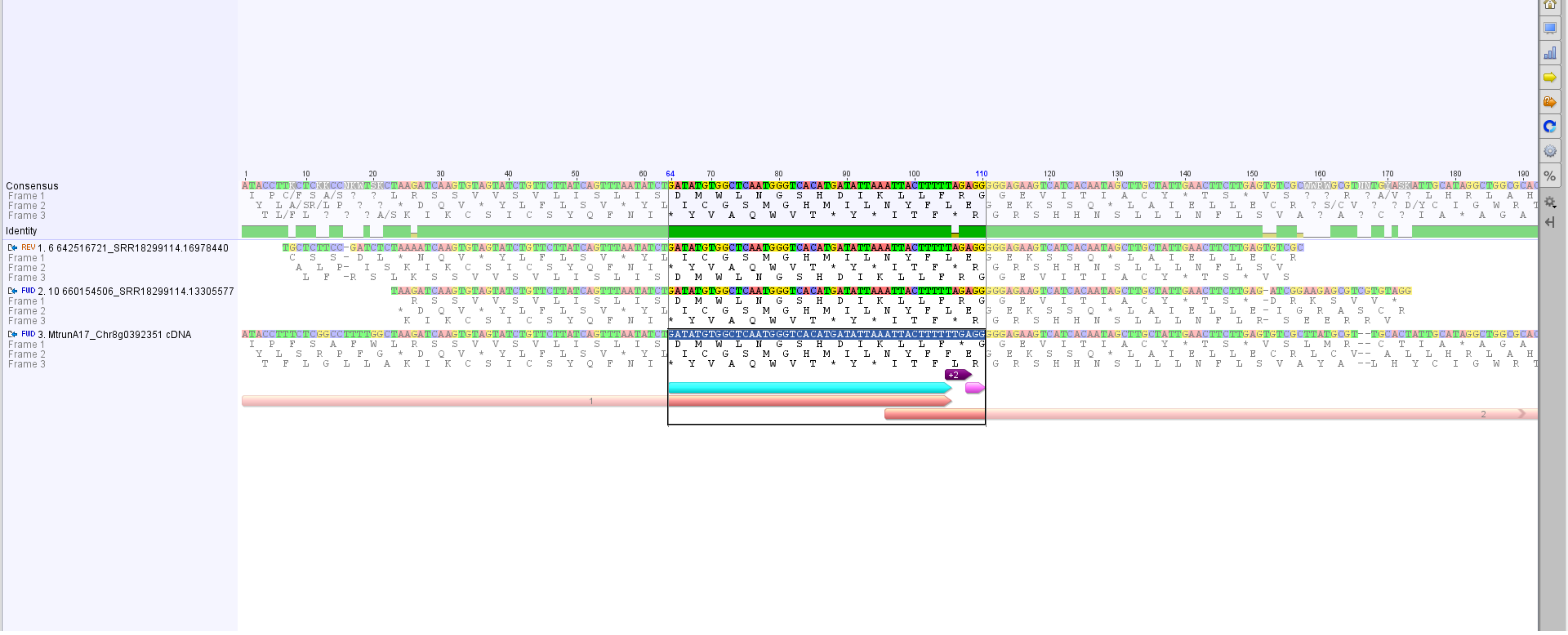

**CP140**, petioles under drought:

Group 5, MtrunA17_Chr8g0392351, primary source

14-dpi nodules (17)

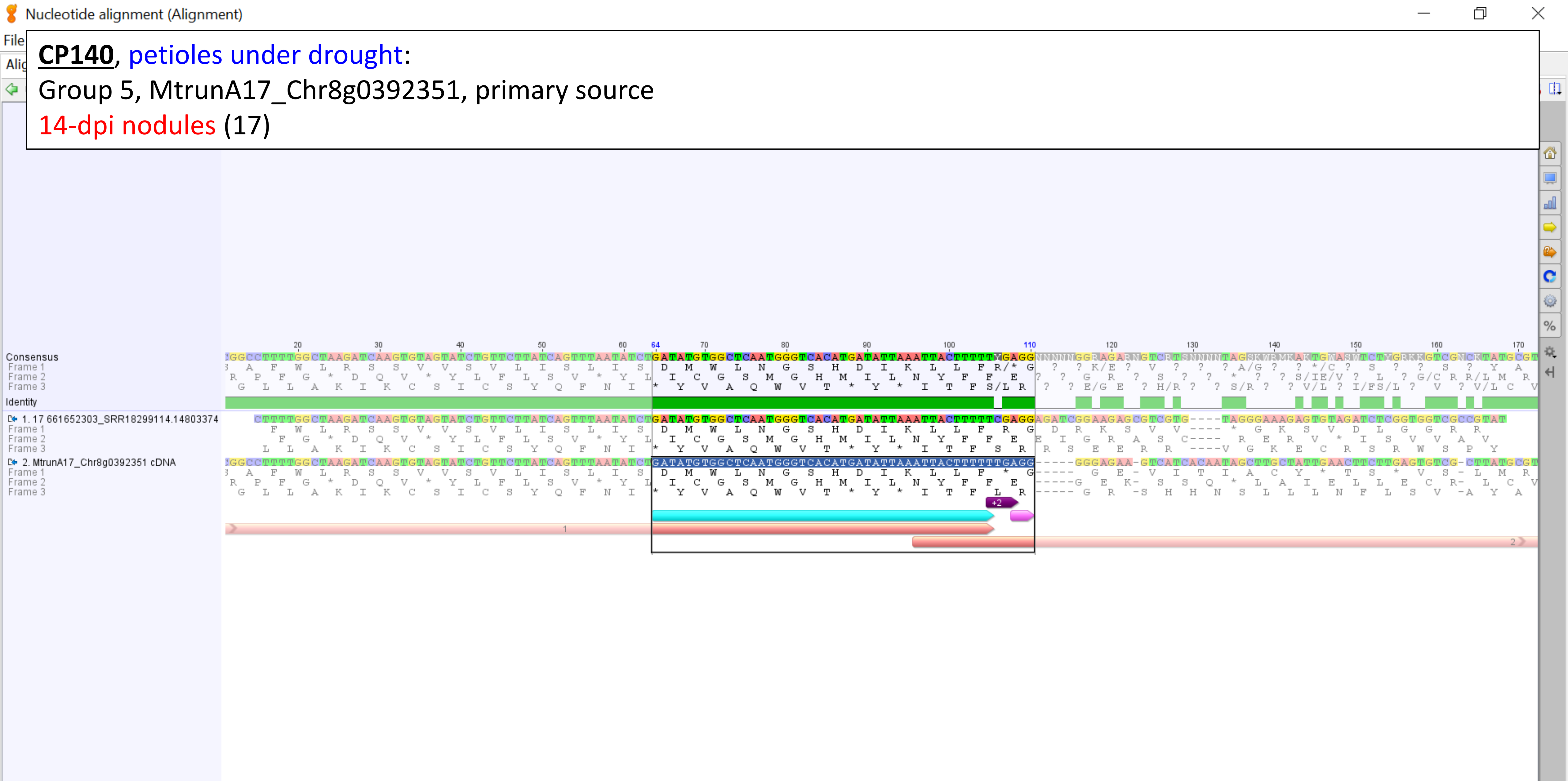

**CP148**, seeds:

Group 1, overview, MtrunA17_MTg0490471, primary source

14-dpi nodules (2, 3, 4, 5, 6, 7, 8, 9, 11, 12, 14, 15, 17, 18, 20, 21, 26, 27, 29, 30, 45, 46, 47), mature leaves (13, 16, 22, 23, 24, 25), nodules ribo-minus (31, 32, 34, 36, 38, 39, 44), roots ribo-minus ([31], [32], [34], 35, [36], 42), seedlings (10)

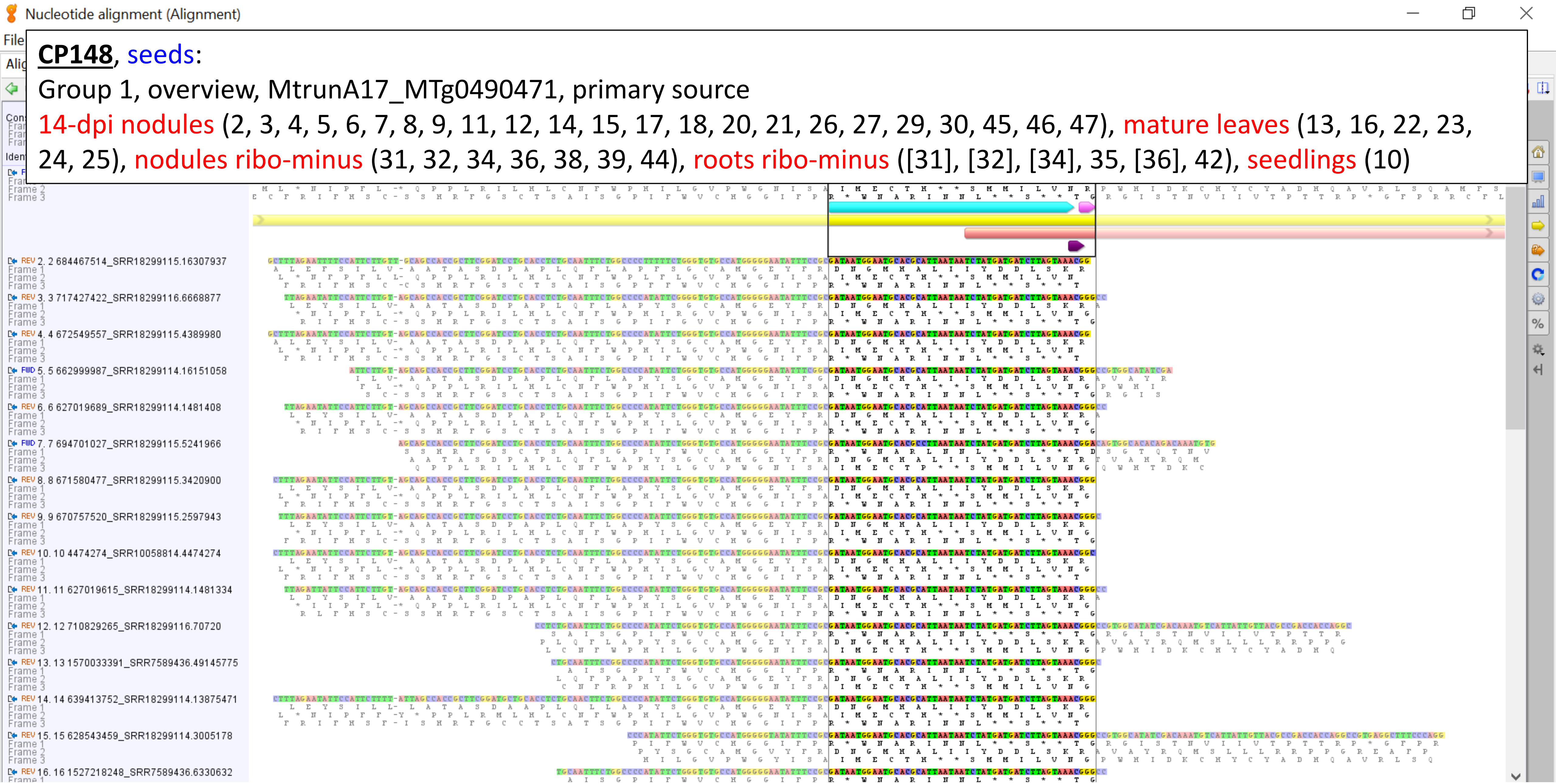

**CP148**, seeds:

Group 1, Part 1, MtrunA17_MTg0490471, primary source

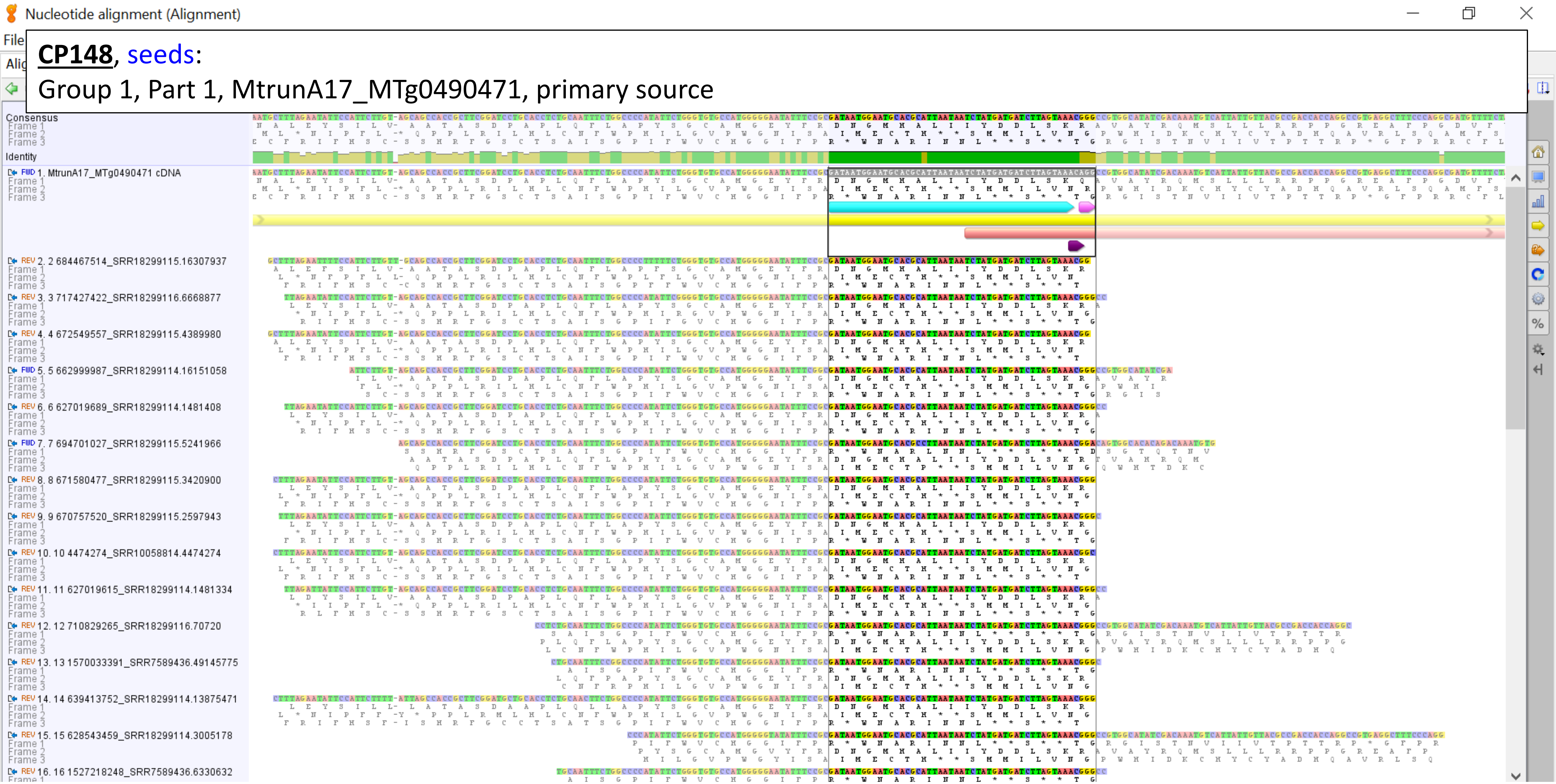

**CP148**, seeds:

Group 1, Part 2, MtrunA17_MTg0490471, primary source

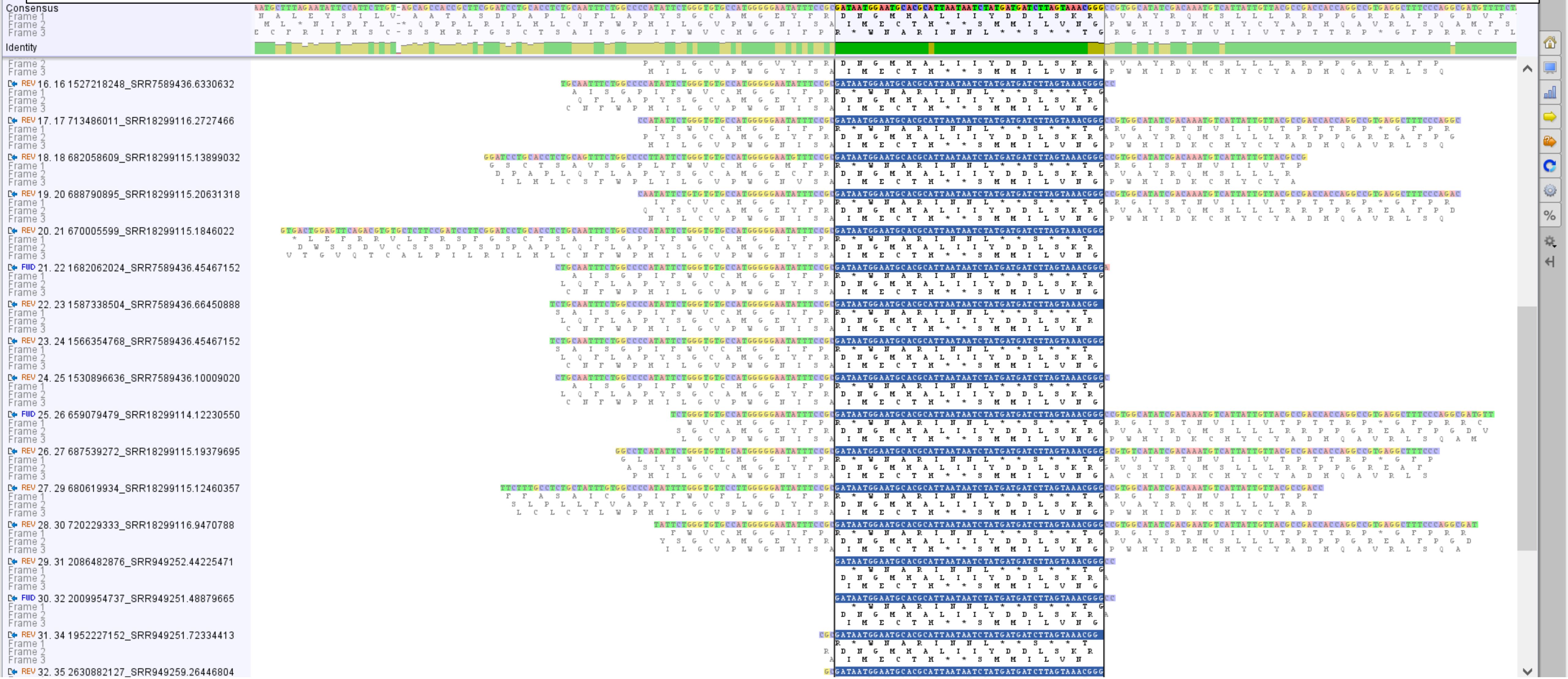

**CP148**, seeds:

Group 1, Part 3, MtrunA17_MTg0490471, primary source

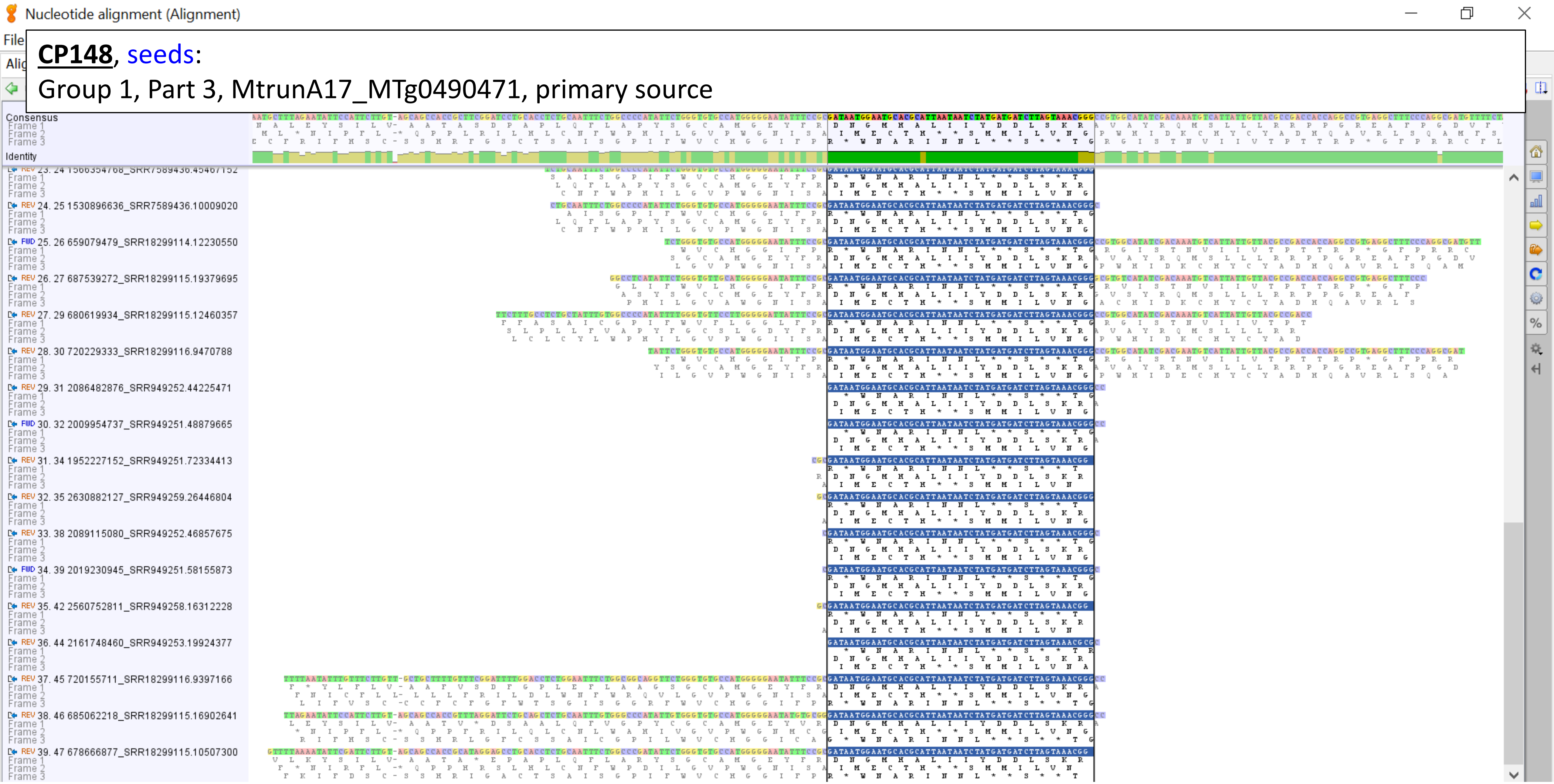

**CP148**, seeds:

Group 2, MtrunA17_MTg0490471, primary source

nodules ribo-minus (33, 36, 40, 43), 14-dpi nodules (1), roots ribo-minus (41), mature leaves (19)

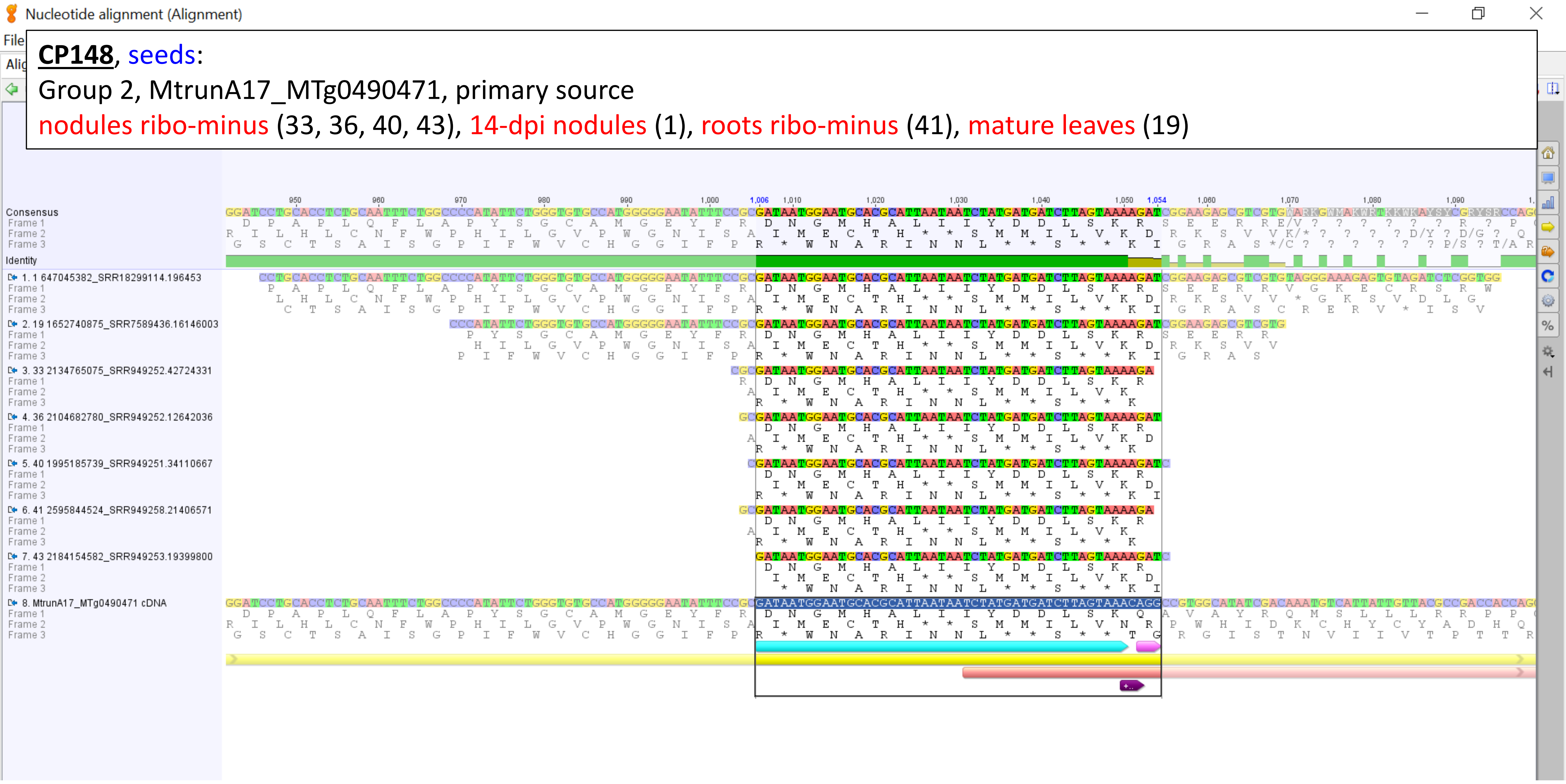

**CP148**, seeds:

Group 3, MtrunA17_MTg0490471, primary source

14-dpi nodules (28)

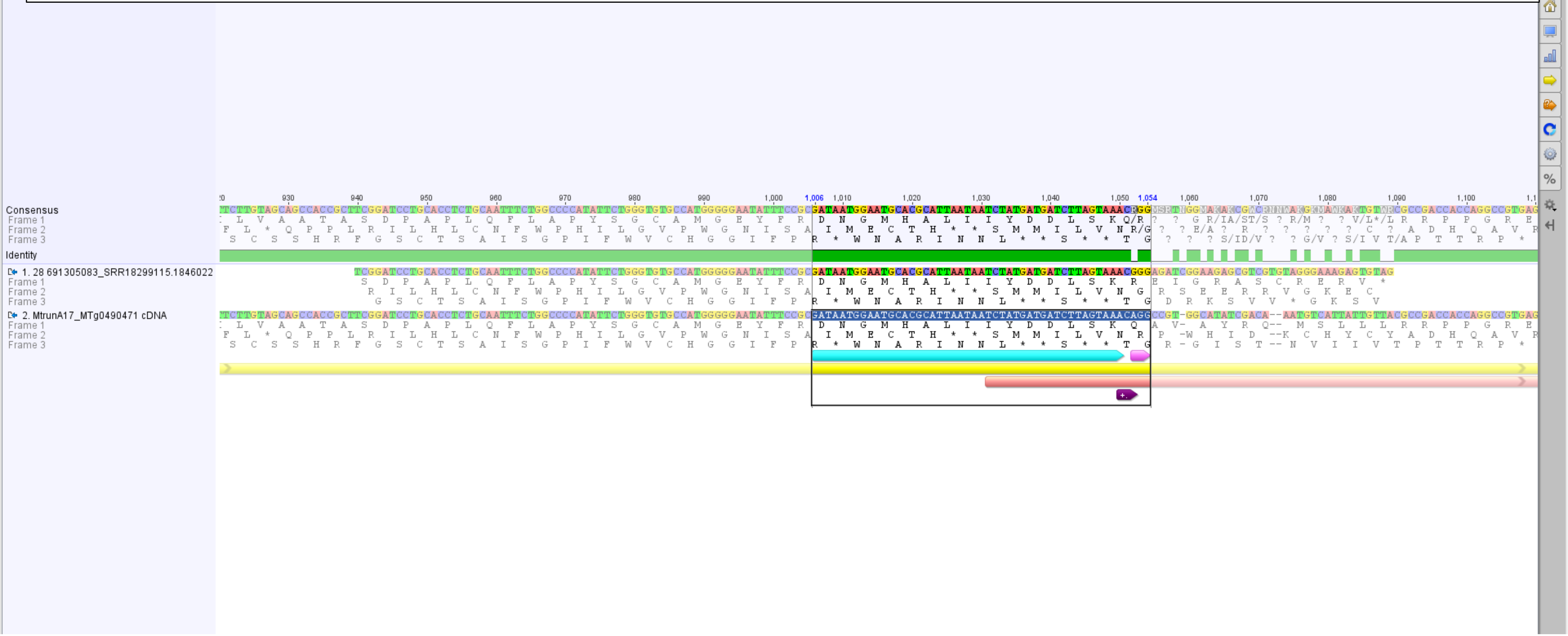

**CP148**, seeds:

Group 4, MtrunA17_MTg0490471, primary source

nodules ribo-minus (37)

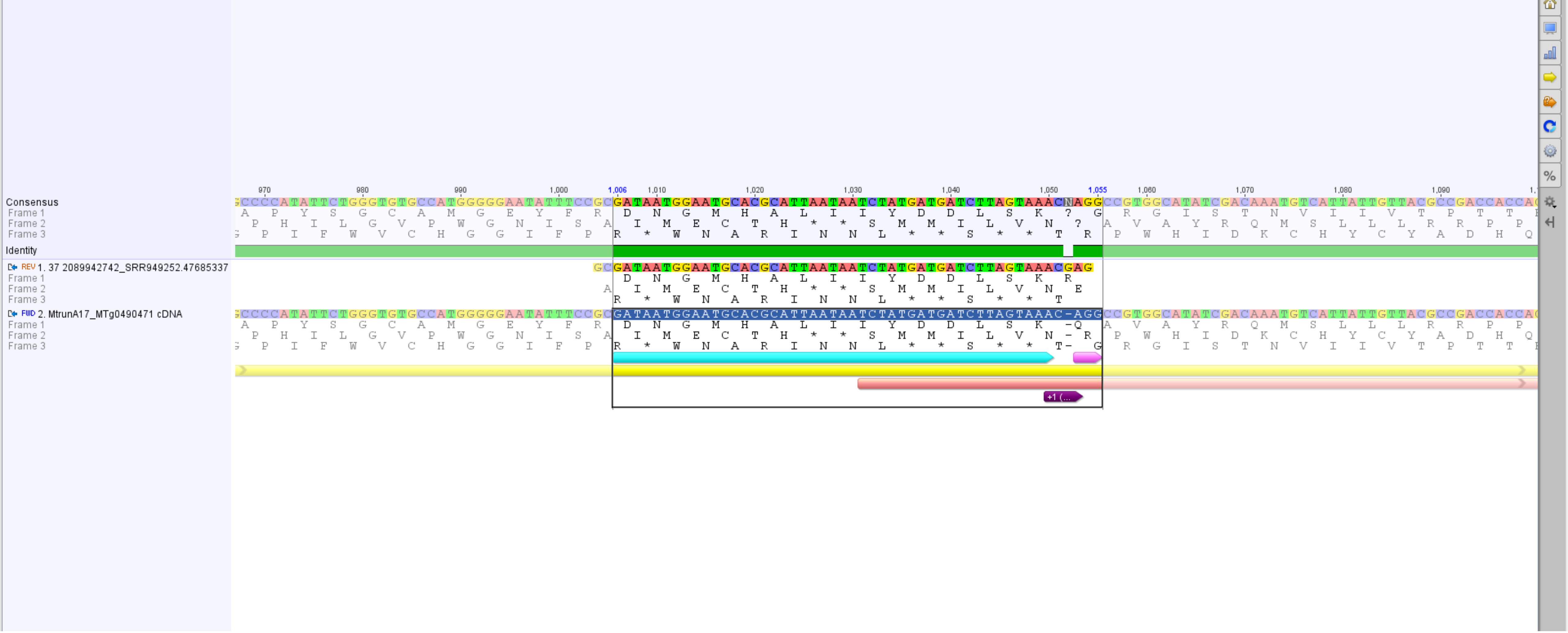

**<u>CP149</u>**, 10-dpi nodules, 14-dpi nodules, flowers, leaves, seeds:
MtrunA17_MTg0490471, primary source
14-dpi nodules (37)

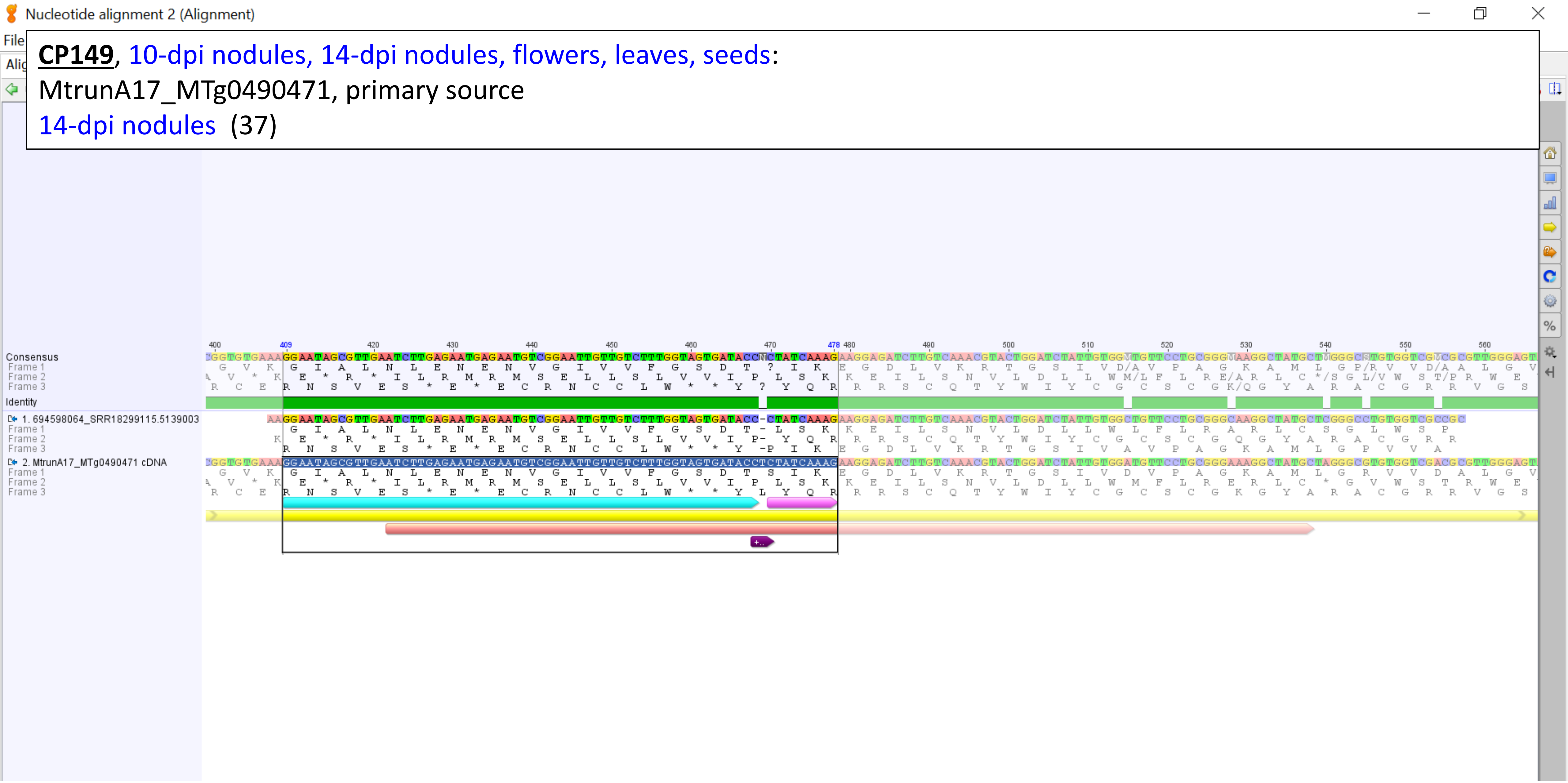

**Supplementary Dataset S19**. A graphical summary on alignments between transcripts of 15 MS-supported chimeric peptides and RNA-Seq reads from 50 selected RNA-Seq runs. RNA-Seq runs with names shown in blue match MS proteomic samples shown in the first line. Run names shown in red do not match corresponding MS samples. Run names shown in brown have partial match, for example, “10-dpi nodules” and “14-dpi nodules ”. Numbers between round brackets indicate read sequences that match a corresponding transcript. Numbers shown in square brackets indicate read sequences that are found in more than one run. For example, read 9 of CP54 is primarily found in a run from 14-dpi nodules. However, an identical sequence with a different ID is found also in a run from 10-dpi nodules. In each alignment, bases corresponding to the longer side of a chimeric peptide are annotated with cyan whereas bases of the shorter side are marked with purple. The PRF sites are annotated with deep purple. RefORFs and altORFs are labeled with yellow and pink, respectively. Each read number and ID along with the transcript ID are shown in alignment images. Screenshots are from Geneious® v. 7.1 (Dotmatics Ltd., MA, USA, https://www.geneious.com).
