## Supplementary material for "Discovery of diverse chimeric peptides in a eukaryotic proteome sets the stage for the experimental proof of the mosaic translation hypothesis": Supplementary Dataset S26 Expression profiles of non-overlapping repeat elements.pdf

INRAE/CNRS Medicago Express | Expression Atlas Server

lipm-browsers.toulouse.inra.fr/pub/expressionAtlas/app/v3/aa_reference_dataset/MtrunA17_Chr2R0106940

MtExpress V3 | Core (20220901)

! Switch to another dataset using the left menu

TMM | METADATA | SYNONYMOUS | ANNOTATION | GENOME PORTAL | LEGOO

repeat_region: MtrunA17_Chr2R0106940;

Log2 TMM Normalisation using EdgeR (Core [20220901])

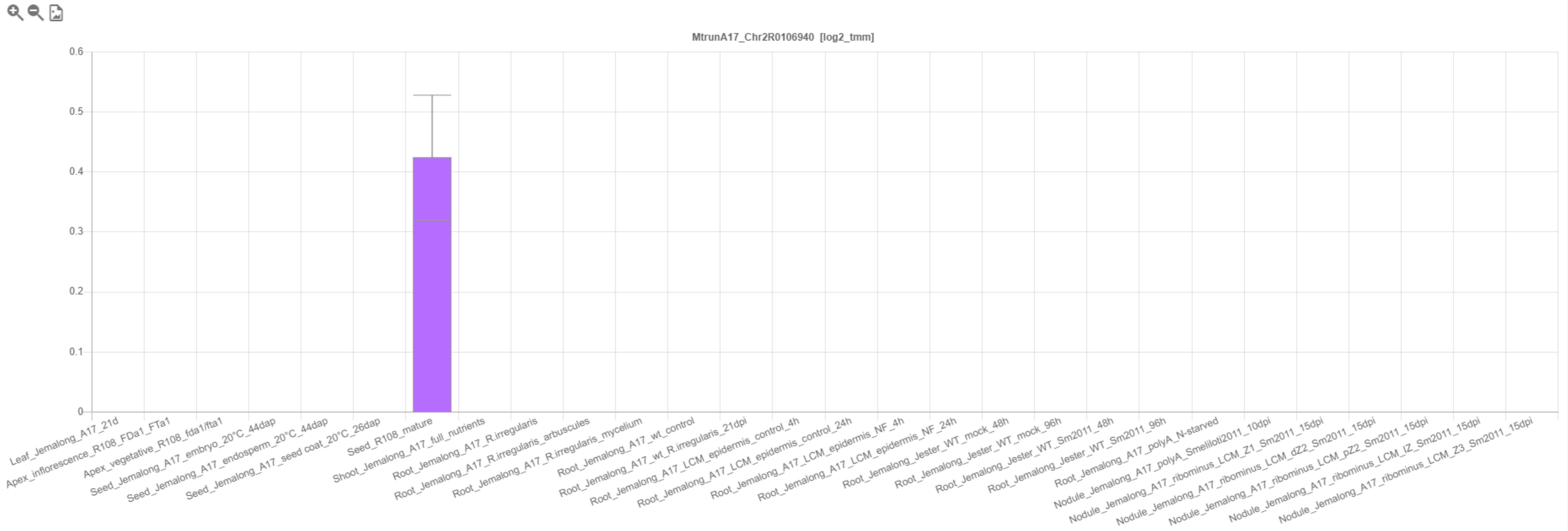

INRAE/CNRS Medicago Express | Expression Atlas Server

lipm-browsers.toulouse.inra.fr/pub/expressionAtlas/app/v3/aa_reference_dataset/MtrunA17_Chr3R0310480

MtExpress V3 | Core (20220901)

! Switch to another dataset using the left menu

TMM | METADATA | SYNONYMOUS | ANNOTATION | GENOME PORTAL | LEGOO

repeat_region: MtrunA17_Chr3R0310480;

Log2 TMM Normalisation using EdgeR (Core [20220901])

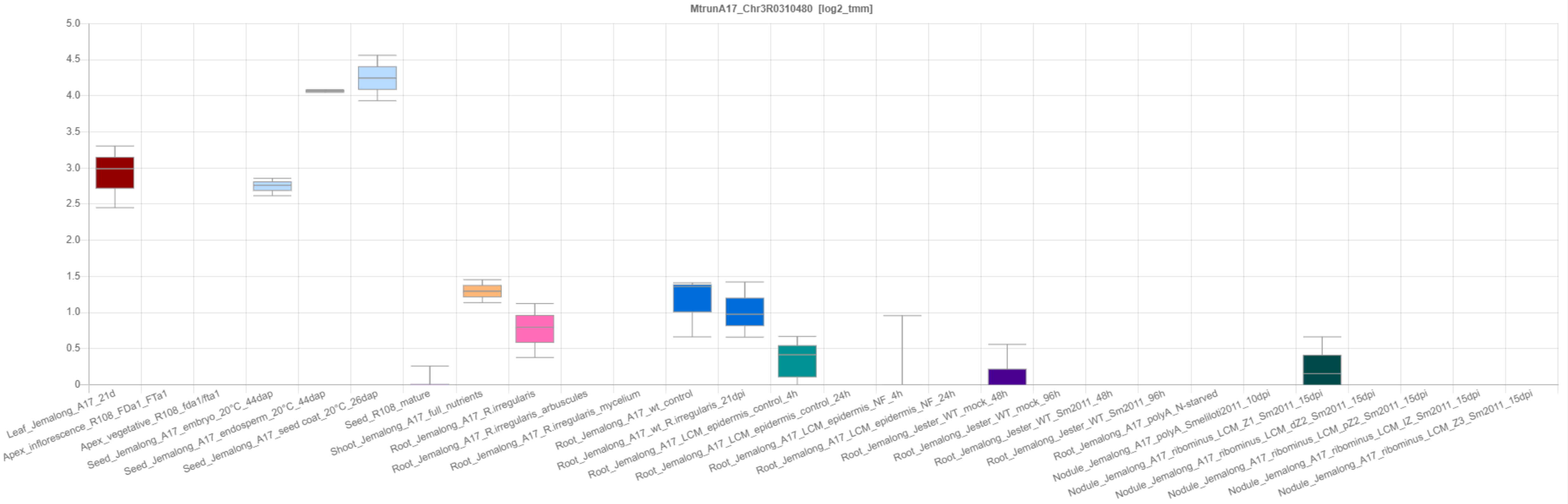

INRAE/CNRS Medicago Express | Expression Atlas Server

lipm-browsers.toulouse.inra.fr/pub/expressionAtlas/app/v3/aa_reference_dataset/MtrunA17_Chr4R0052500

MtExpress V3 | Core (20220901)

! Switch to another dataset using the left menu

TMM | METADATA | SYNONYMOUS | ANNOTATION | GENOME PORTAL | LEGOO

repeat_region: MtrunA17_Chr4R0052500;

Log2 TMM Normalisation using EdgeR (Core [20220901])

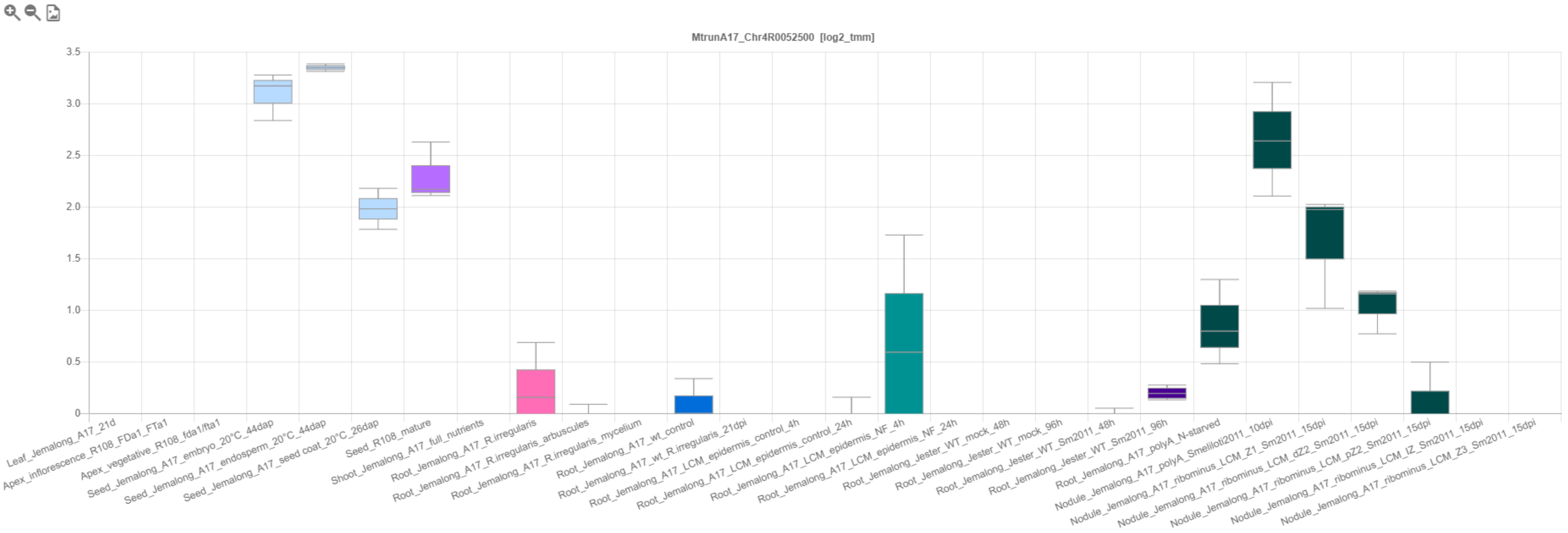

INRAE/CNRS Medicago Express | Expression Atlas Server

lipm-browsers.toulouse.inra.fr/pub/expressionAtlas/app/v3/aa_reference_dataset/MtrunA17_Chr1R0148110

MtExpress V3 | Core (20220901)

! Switch to another dataset using the left menu

TMM | METADATA | SYNONYMOUS | ANNOTATION | GENOME PORTAL | LEGOO

repeat_region: MtrunA17_Chr1R0148110;

Log2 TMM Normalisation using EdgeR (Core [20220901])

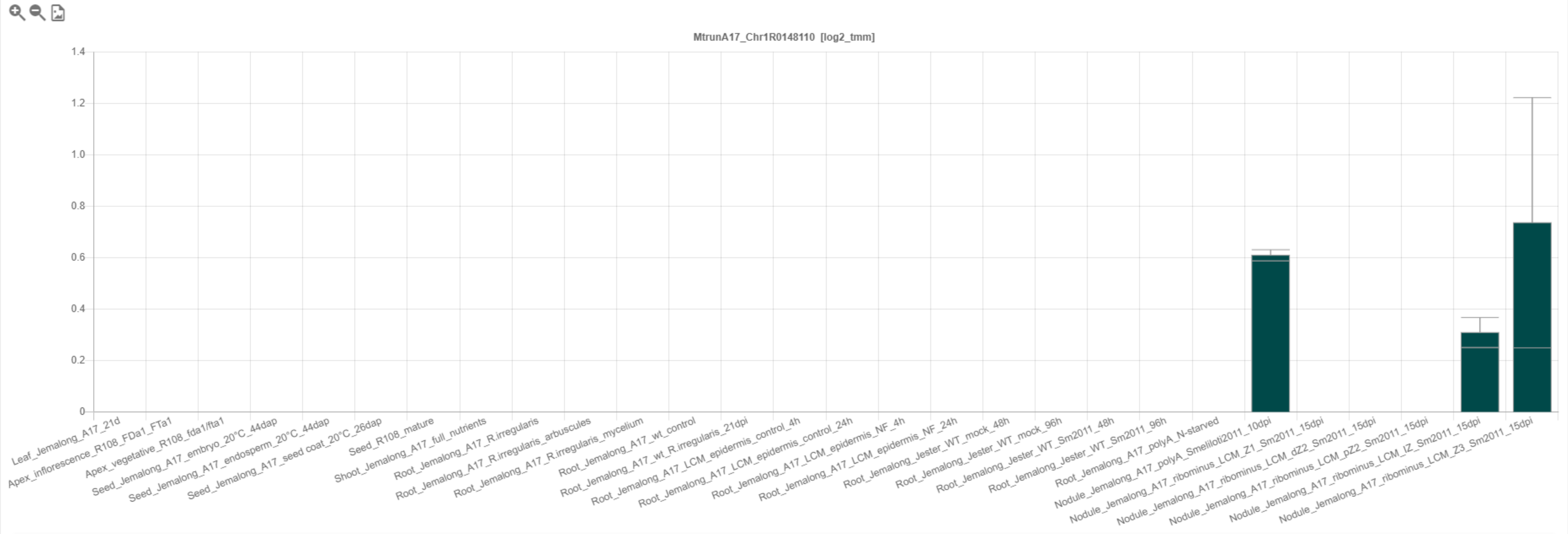

INRAE/CNRS Medicago Express | Expression Atlas Server

lipm-browsers.toulouse.inra.fr/pub/expressionAtlas/app/v3/aa_reference_dataset/MtrunA17_Chr1R0014150

MtExpress V3 | Core (20220901)

! Switch to another dataset using the left menu

lipme INRAE cnrs

TMM | METADATA | SYNONYMOUS | ANNOTATION | GENOME PORTAL | LEGOO

repeat_region: MtrunA17_Chr1R0014150;

Log2 TMM Normalisation using EdgeR (Core [20220901])

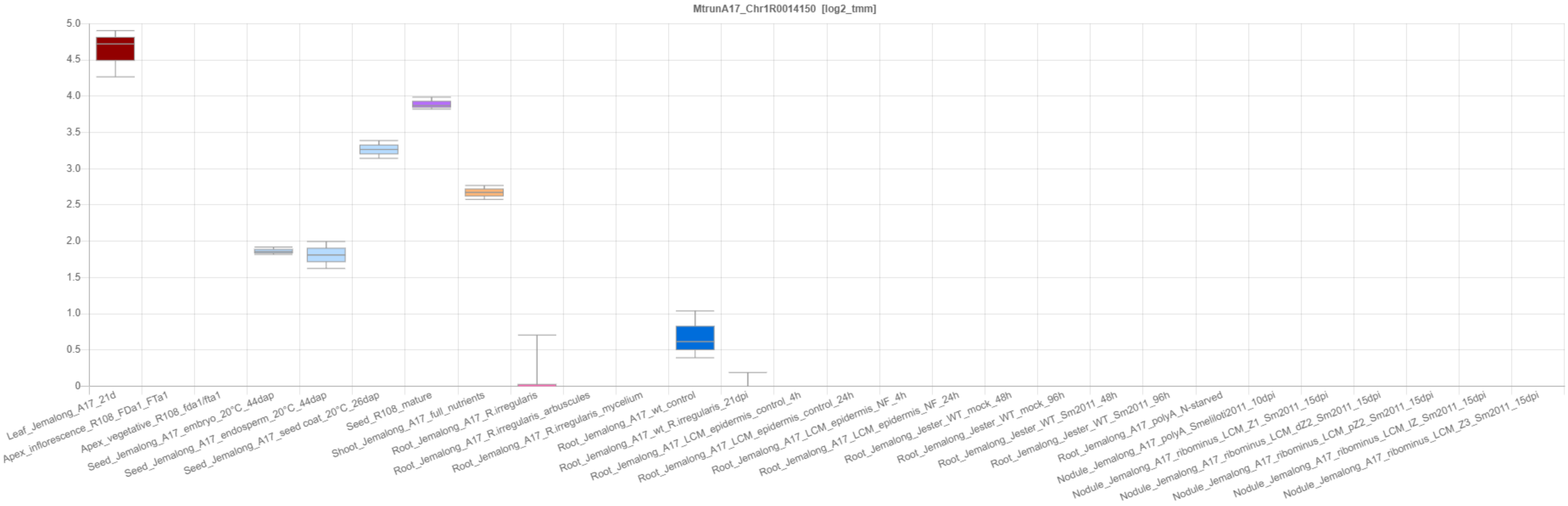

INRAE/CNRS Medicago Express | Expression Atlas Server

lipm-browsers.toulouse.inra.fr/pub/expressionAtlas/app/v3/aa_reference_dataset/MtrunA17_Chr7R0161930

MtExpress V3 | Core (20220901)

! Switch to another dataset using the left menu

TMM | METADATA | SYNONYMOUS | ANNOTATION | GENOME PORTAL | LEGOO

repeat_region: MtrunA17_Chr7R0161930;

Log2 TMM Normalisation using EdgeR (Core [20220901])

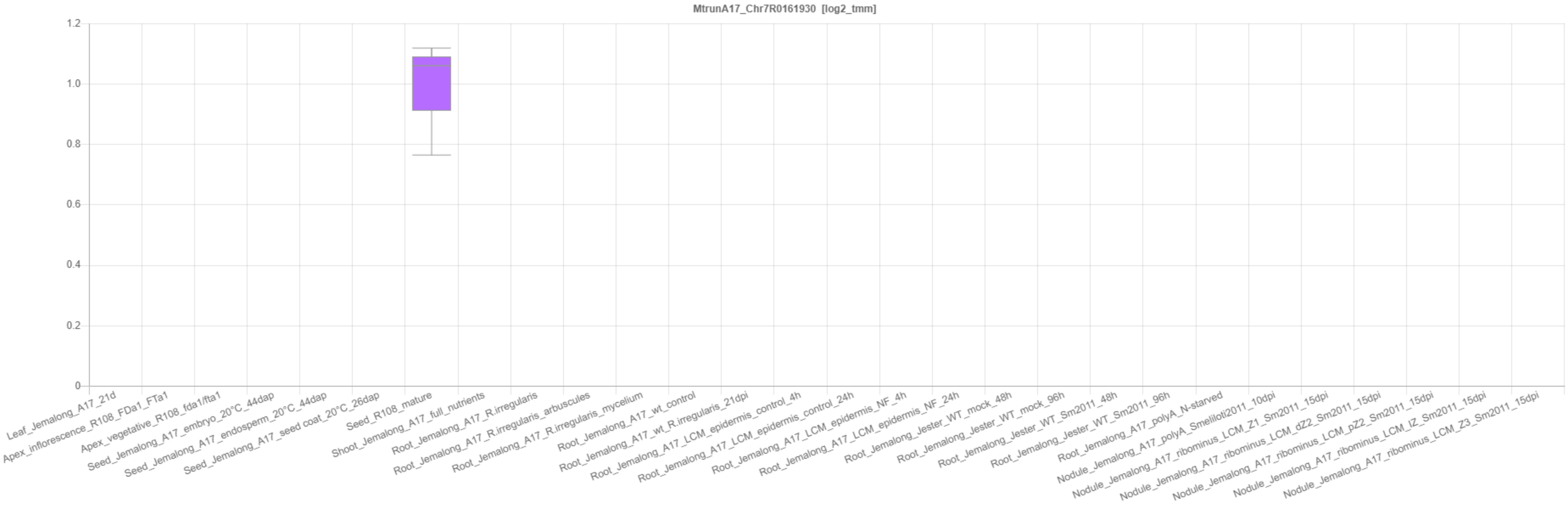

INRAE/CNRS Medicago Express | Expression Atlas Server

lipm-browsers.toulouse.inra.fr/pub/expressionAtlas/app/v3/aa_reference_dataset/MtrunA17_Chr3R0062580

MtExpress V3 | Core (20220901)

! Switch to another dataset using the left menu

lipme INRAE cnrs

TMM | METADATA | SYNONYMOUS | ANNOTATION | GENOME PORTAL | LEGOO

repeat_region: MtrunA17_Chr3R0062580;

Log2 TMM Normalisation using EdgeR (Core [20220901])

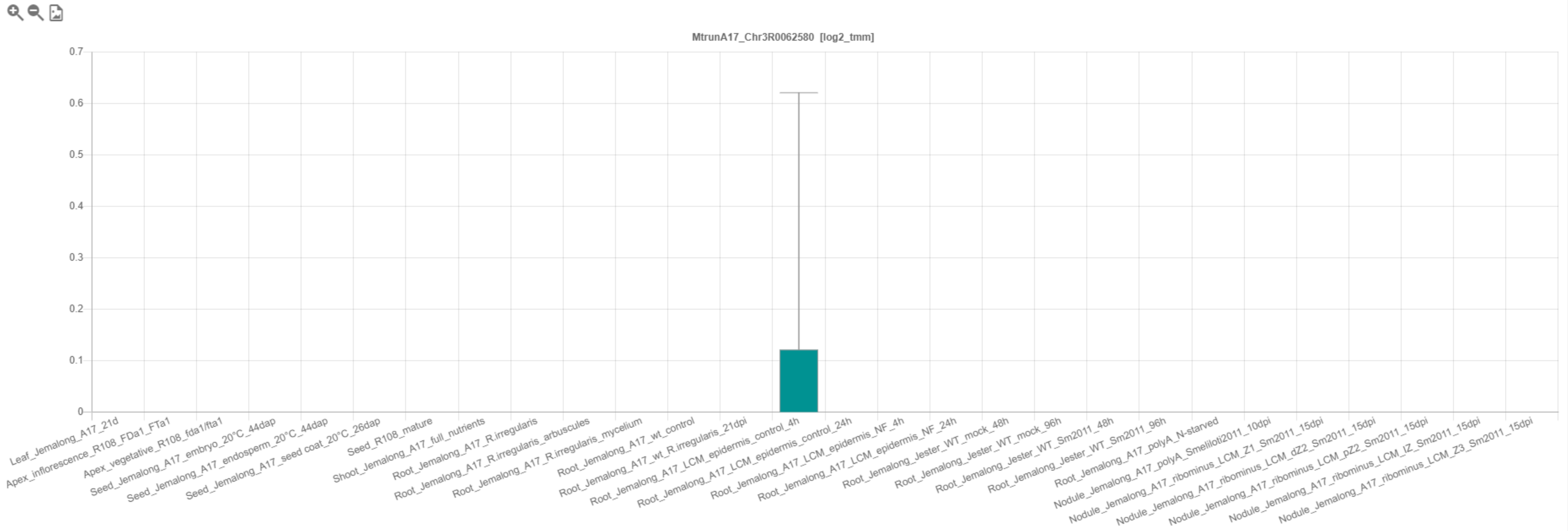

INRAE/CNRS Medicago Express | Expression Atlas Server

lipm-browsers.toulouse.inra.fr/pub/expressionAtlas/app/v3/aa_reference_dataset/MtrunA17_Chr0c04R0002520

MtExpress V3 | Core (20220901)

! Switch to another dataset using the left menu

TMM | METADATA | SYNONYMOUS | ANNOTATION | GENOME PORTAL | LEGOO

repeat_region: MtrunA17_Chr0c04R0002520;

Log2 TMM Normalisation using EdgeR (Core [20220901])

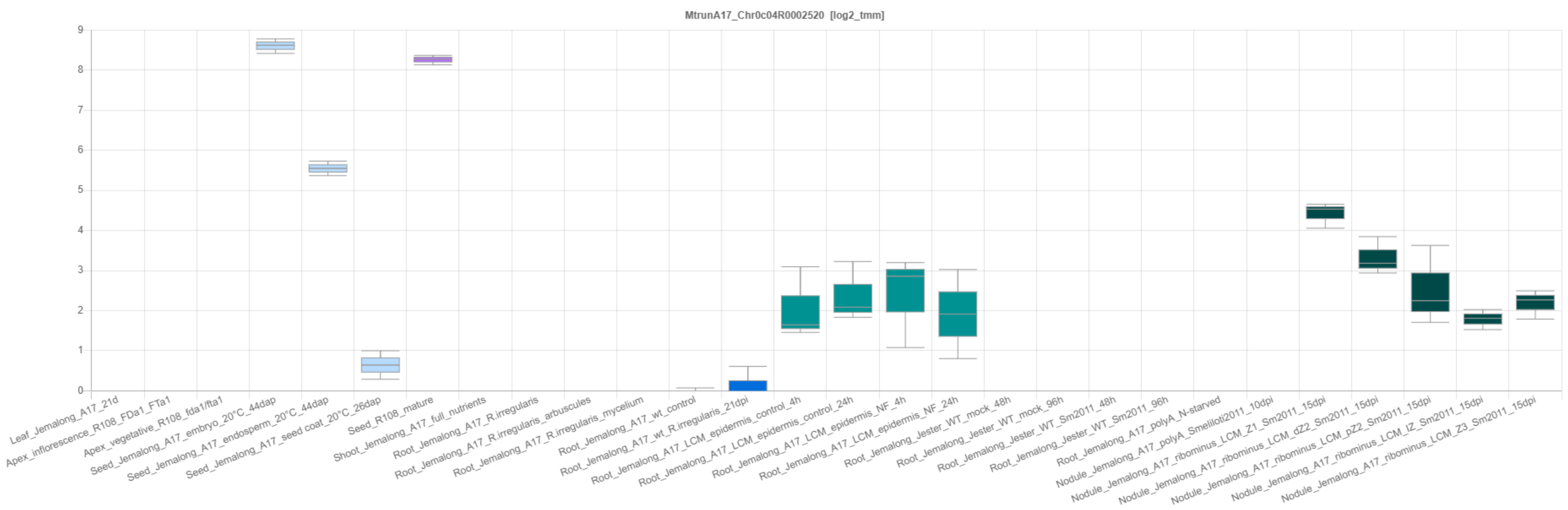

INRAE/CNRS Medicago Express | Expression Atlas Server

lipm-browsers.toulouse.inra.fr/pub/expressionAtlas/app/v3/aa_reference_dataset/MtrunA17_Chr3R0074220

MtExpress V3 | Core (20220901)

! Switch to another dataset using the left menu

TMM | METADATA | SYNONYMOUS | ANNOTATION | GENOME PORTAL | LEGOO

repeat_region: MtrunA17_Chr3R0074220;

Log2 TMM Normalisation using EdgeR (Core [20220901])

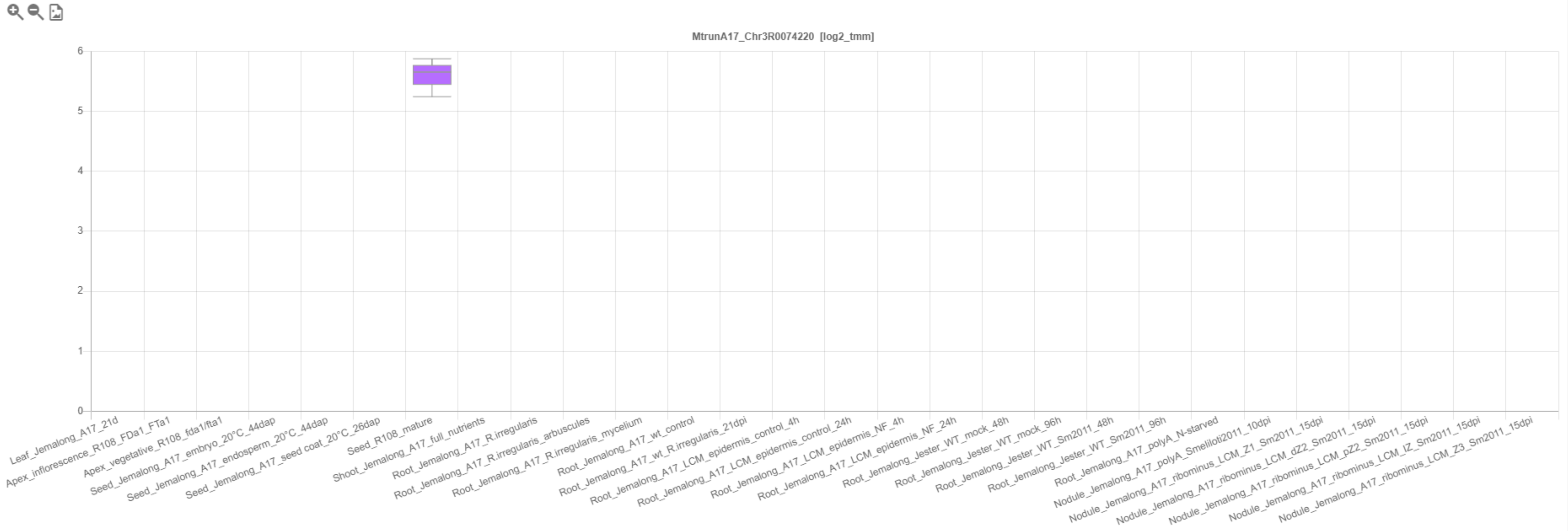

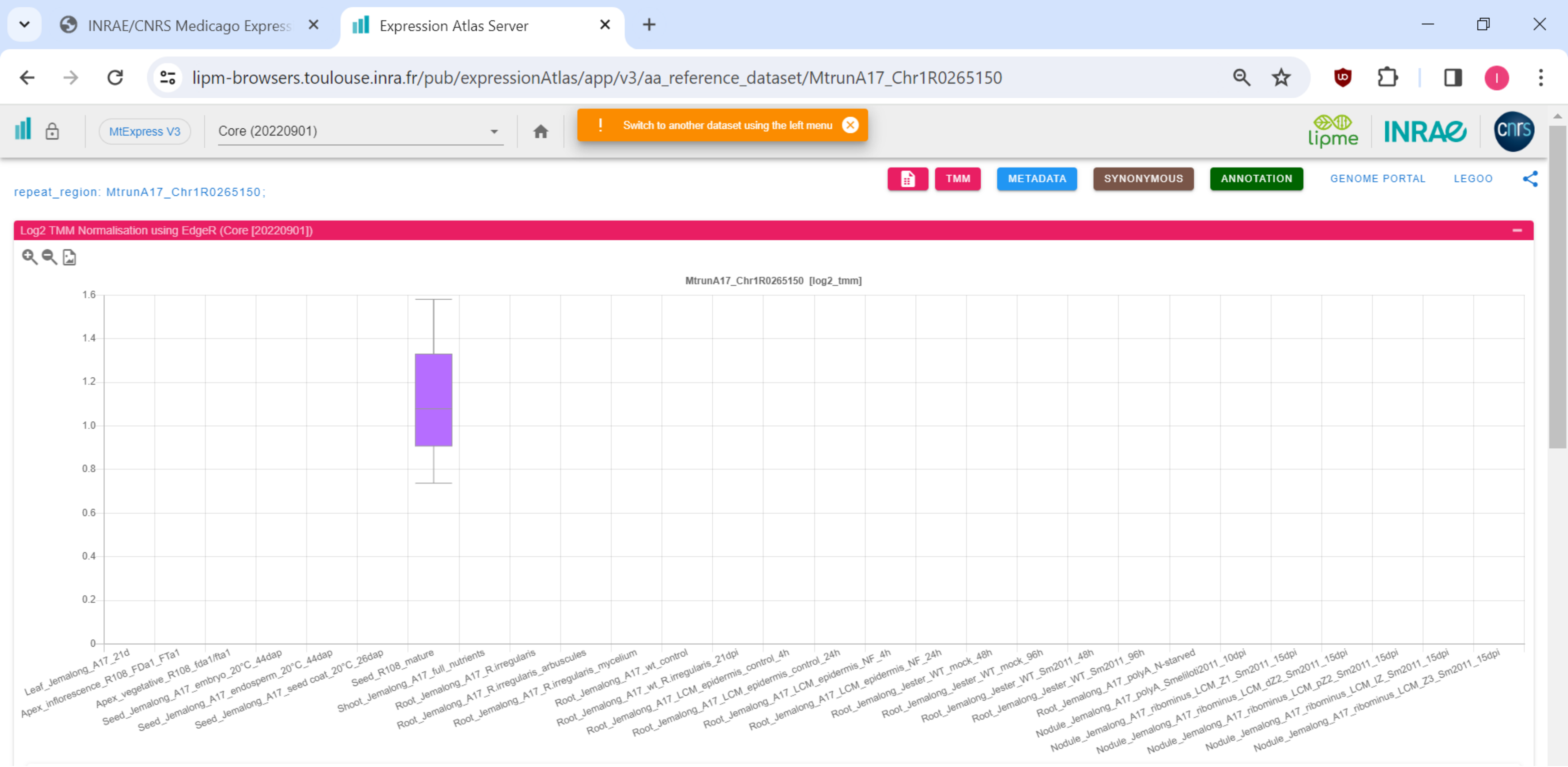
INRAE/CNRS Medicago Express
Expression Atlas Server
lipm-browsers.toulouse.inra.fr/pub/expressionAtlas/app/v3/aa_reference_dataset/MtrunA17_Chr1R0265150
MtExpress V3
Core (20220901)
Switch to another dataset using the left menu
TMM
METADATA
SYNONYMOUS
ANNOTATION
GENOME PORTAL
LEGOO
repeat_region: MtrunA17_Chr1R0265150;
Log2 TMM Normalisation using EdgeR (Core [20220901])
MtrunA17_Chr1R0265150 [log2_tmm]
1.6
1.4
1.2
1.0
0.8
0.6
0.4
0.2
0
Leaf_Jemalong_A17_21d
Apex_inflorescence_R108_FDa1_FTa1
Apex_vegetative_R108_fda1/fta1
Seed_Jemalong_A17_embryo_20°C_44dap
Seed_Jemalong_A17_endosperm_20°C_44dap
Seed_Jemalong_A17_seed coat_20°C_26dap
Seed_R108_mature
Shoot_Jemalong_A17_full_nutrients
Root_Jemalong_A17_R.irregularis
Root_Jemalong_A17_R.irregularis_arbuscules
Root_Jemalong_A17_R.irregularis_mycelium
Root_Jemalong_A17_wt_control
Root_Jemalong_A17_wt_R.irregularis_21dpi
Root_Jemalong_A17_LCM_epidermis_control_4h
Root_Jemalong_A17_LCM_epidermis_control_24h
Root_Jemalong_A17_LCM_epidermis_NF_4h
Root_Jemalong_A17_LCM_epidermis_NF_24h
Root_Jemalong_Jester_WT_mock_48h
Root_Jemalong_Jester_WT_mock_96h
Root_Jemalong_Jester_WT_Sm2011_48h
Root_Jemalong_Jester_WT_Sm2011_96h
Root_Jemalong_A17_polyA_N-starved
Nodule_Jemalong_A17_polyA_Smeliloti2011_10dpi
Nodule_Jemalong_A17_ribominus_LCM_Z1_Sm2011_15dpi
Nodule_Jemalong_A17_ribominus_LCM_dZ2_Sm2011_15dpi
Nodule_Jemalong_A17_ribominus_LCM_pZ2_Sm2011_15dpi
Nodule_Jemalong_A17_ribominus_LCM_IZ_Sm2011_15dpi
Nodule_Jemalong_A17_ribominus_LCM_Z3_Sm2011_15dpi

**Supplementary Dataset S26**. Expression profiles of ten non-overlapping repeat elements that have log2 TMM values above zero. Screenshots are from the RNA-Seq-based gene expression atlas of *Medicago truncatula* (MtExpress v. 3, https://medicago.toulouse.inrae.fr/GEA). The list corresponds to repeat elements marked with “Yes” in column B of Supplementary Dataset S25. For clarity and better visibility, only the core sample set is shown. Note that some repeat elements in this list have tissue- or sample-specific expression.
