## Supplementary material for "Discovery of diverse chimeric peptides in a eukaryotic proteome sets the stage for the experimental proof of the mosaic translation hypothesis": Supplementary Dataset S27 Expression profiles of 156 primary-source transcripts.pdf

CP1: MtrunA17_Chr0c01g0489091

INRAE/CNRS Medicago Expressi | Expression Atlas Server

...expressionAtlas/app/v3/aa_reference_dataset/MtrunA17_Chr0c01g0489091

MtExpress V3 | Core (20220901)

! Switch to another dataset using the left menu

lipme INRAE cnrs

TMM METADATA SYNONYMOUS ANNOTATION GENOME PORTAL LEGOO

ncRNA: MtrunA17_Chr0c01g0489091;

Log2 TMM Normalisation using EdgeR (Core [20220901])

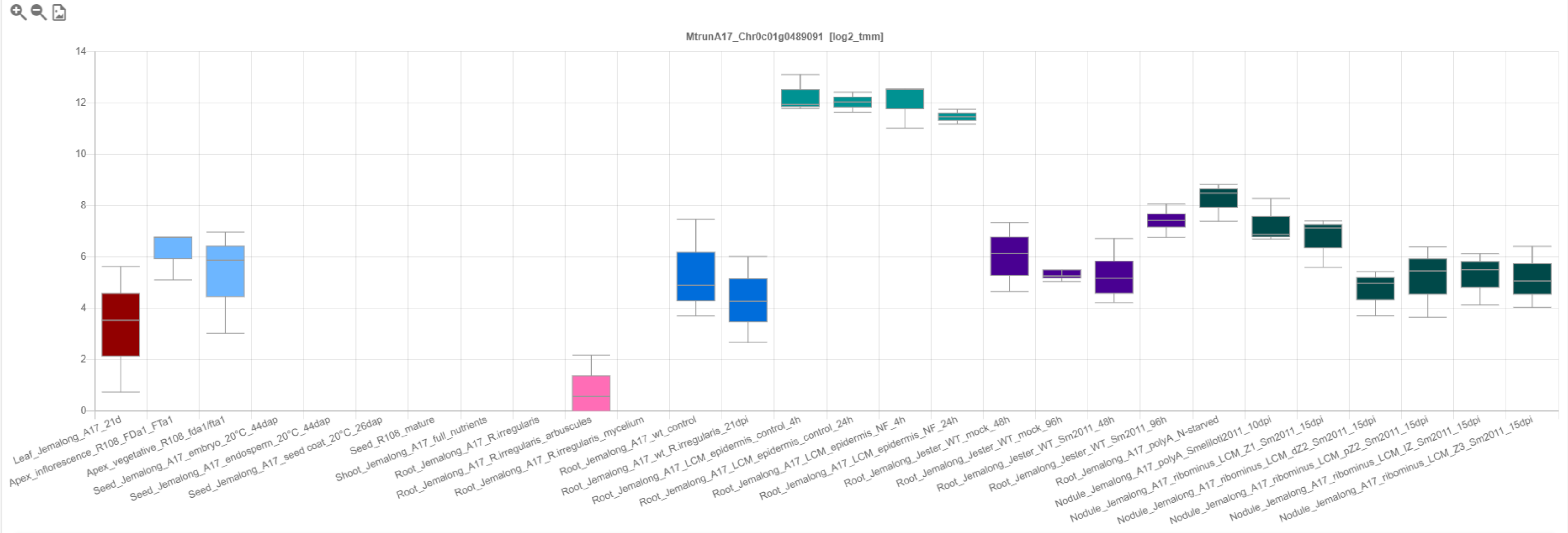

INRAE/CNRS Medicago Expressi | Expression Atlas Server

### CP2: MtrunA17_Chr0c28g0493951

[illegible]xpressionAtlas/app/v3/aa_reference_dataset/MtrunA17_Chr0c28g0493951

MtExpress V3 | Core (20220901)

! Switch to another dataset using the left menu

lipme INRAE cnrs

TMM METADATA SYNONYMOUS ANNOTATION GENOME PORTAL LEGOO

mRNA: MtrunA17_Chr0c28g0493951;

Log2 TMM Normalisation using EdgeR (Core [20220901])

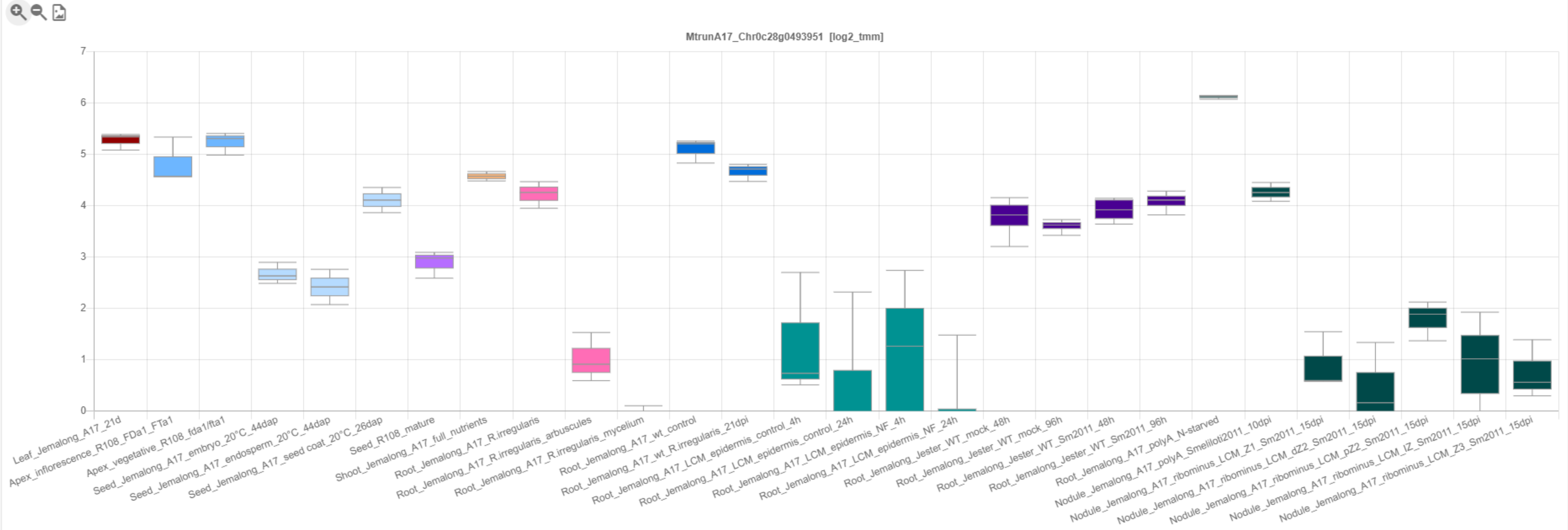

CP3: MtrunA17_Chr1g0148371

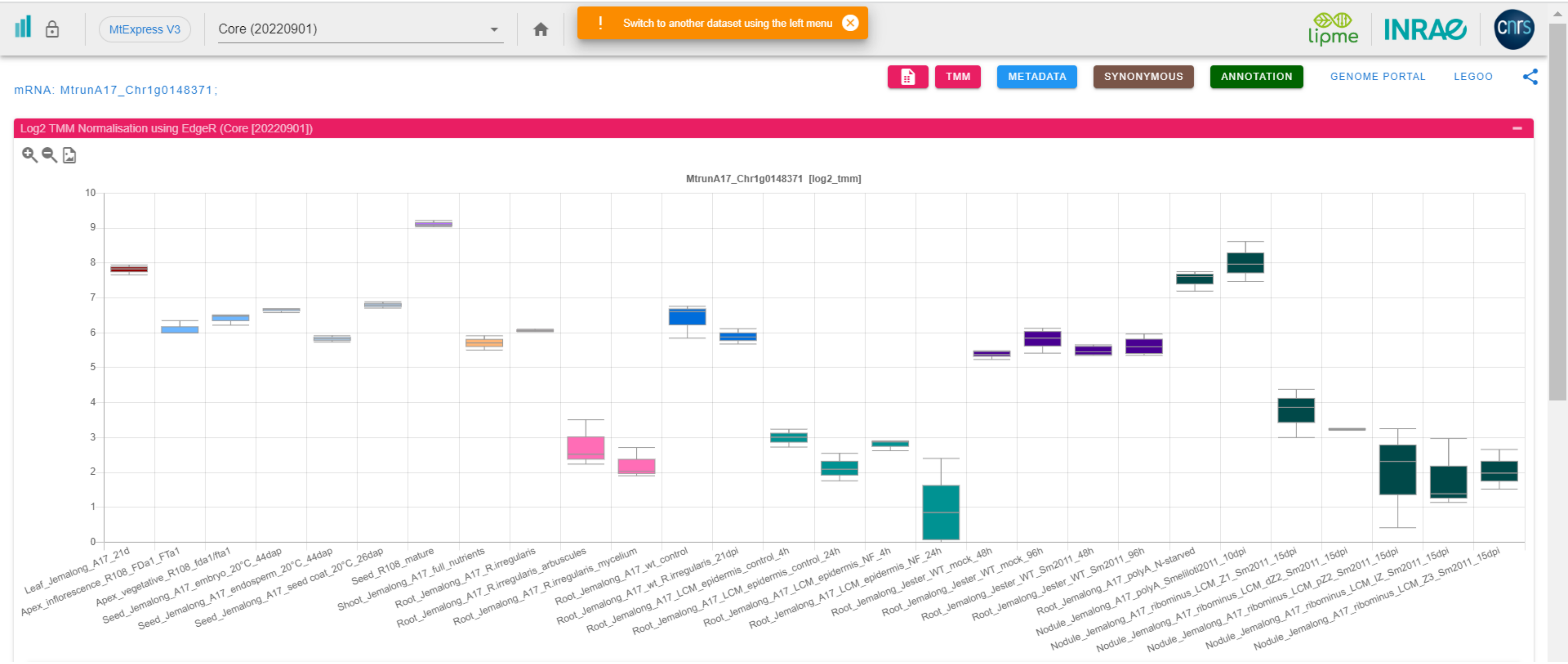

### CP4: MtrunA17_Chr1g0149991

INRAE/CNRS Medicago Expressi… | Expression Atlas Server

…pub/expressionAtlas/app/v3/aa_reference_dataset/MtrunA17_Chr1g0149991

MtExpress V3 | Core (20220901)

! Switch to another dataset using the left menu

TMM | METADATA | SYNONYMOUS | ANNOTATION | GENOME PORTAL | LEGOO

mRNA: MtrunA17_Chr1g0149991;

Log2 TMM Normalisation using EdgeR (Core [20220901])

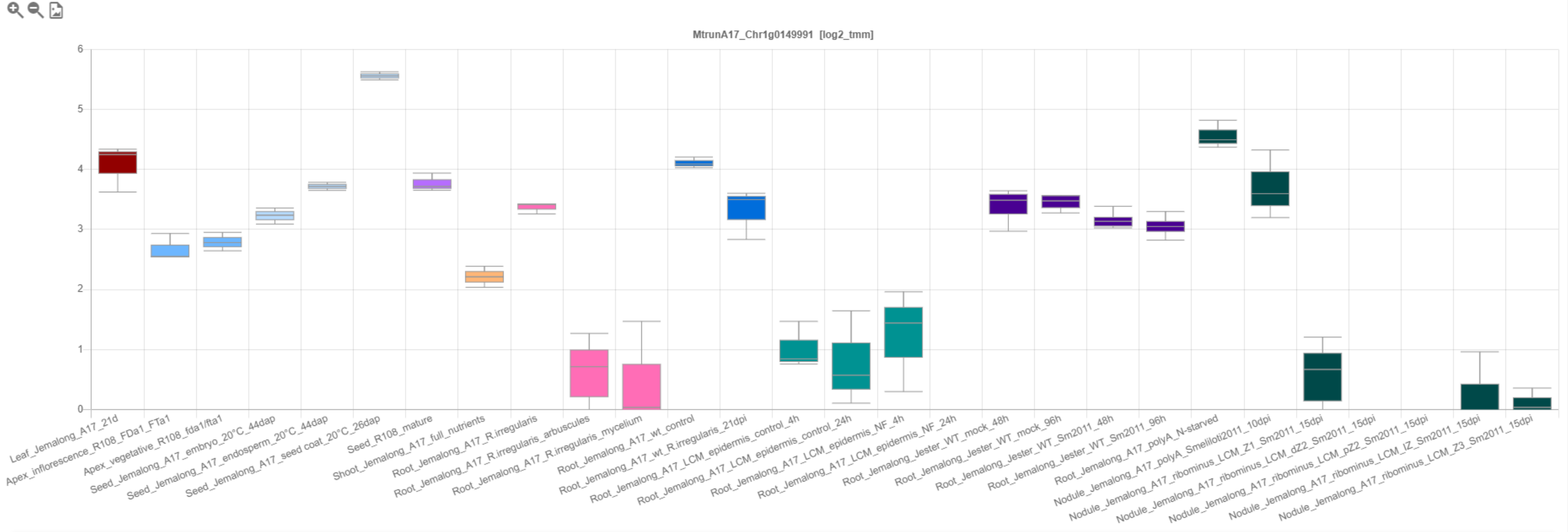

### CP5: MtrunA17_Chr1g0150571

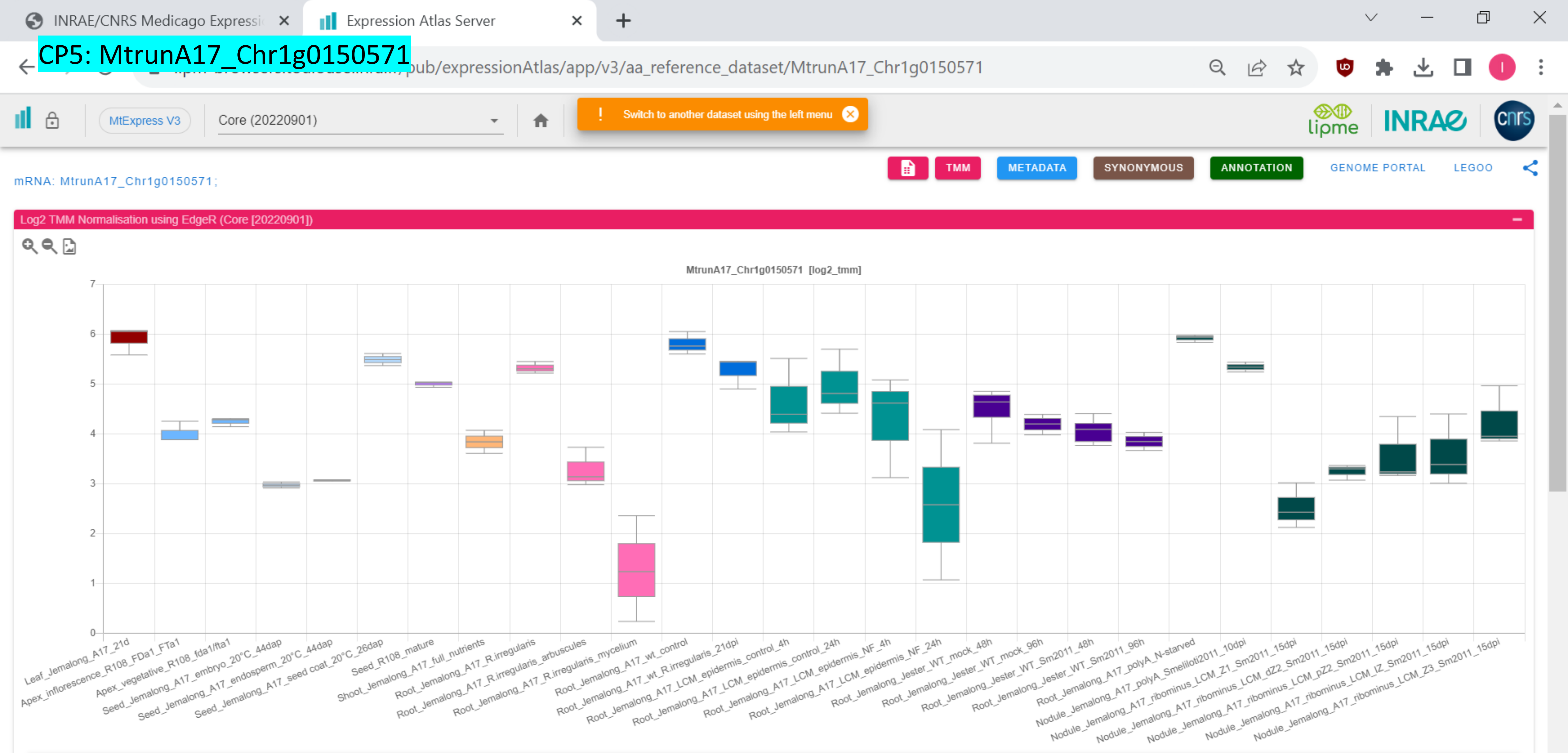

*CP6: MtrunA17_Chr1g0152521

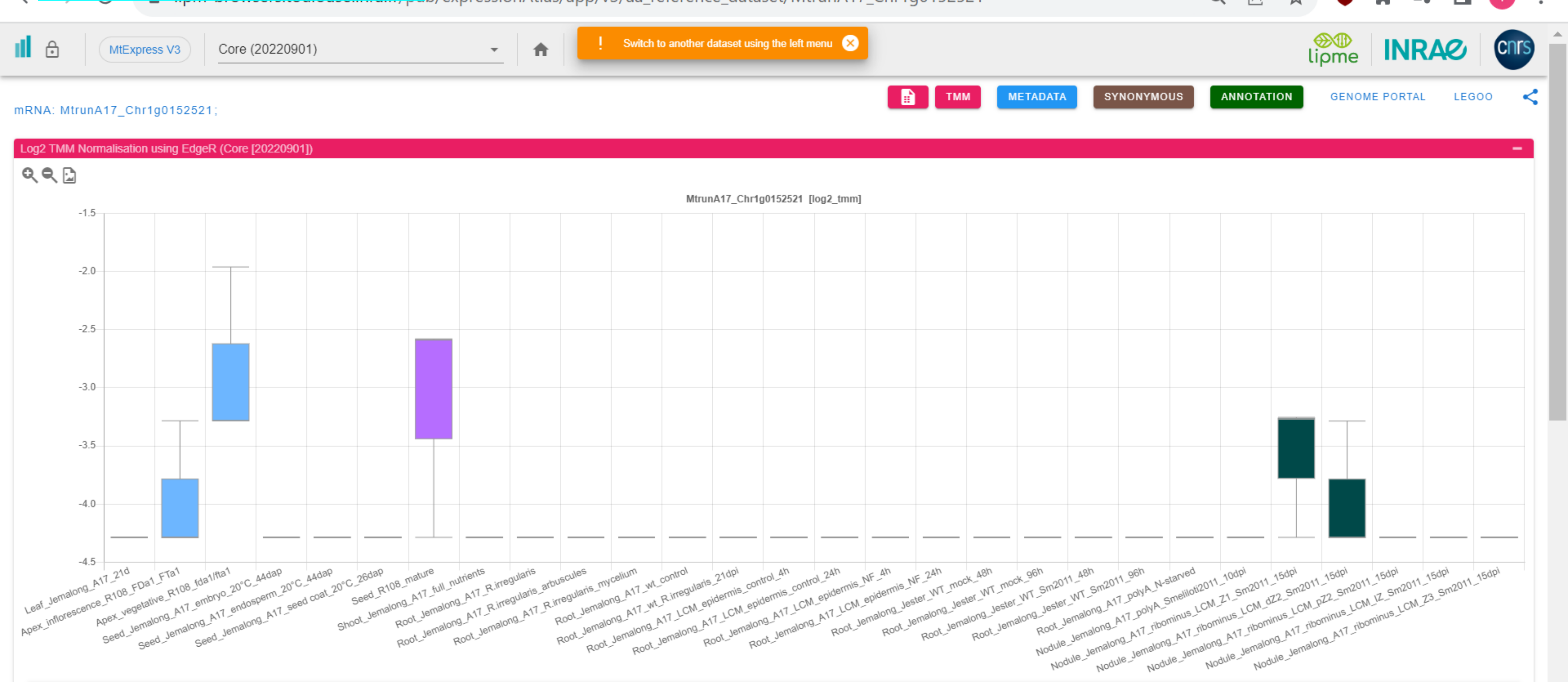

CP7: MtrunA17_Chr1g0153001

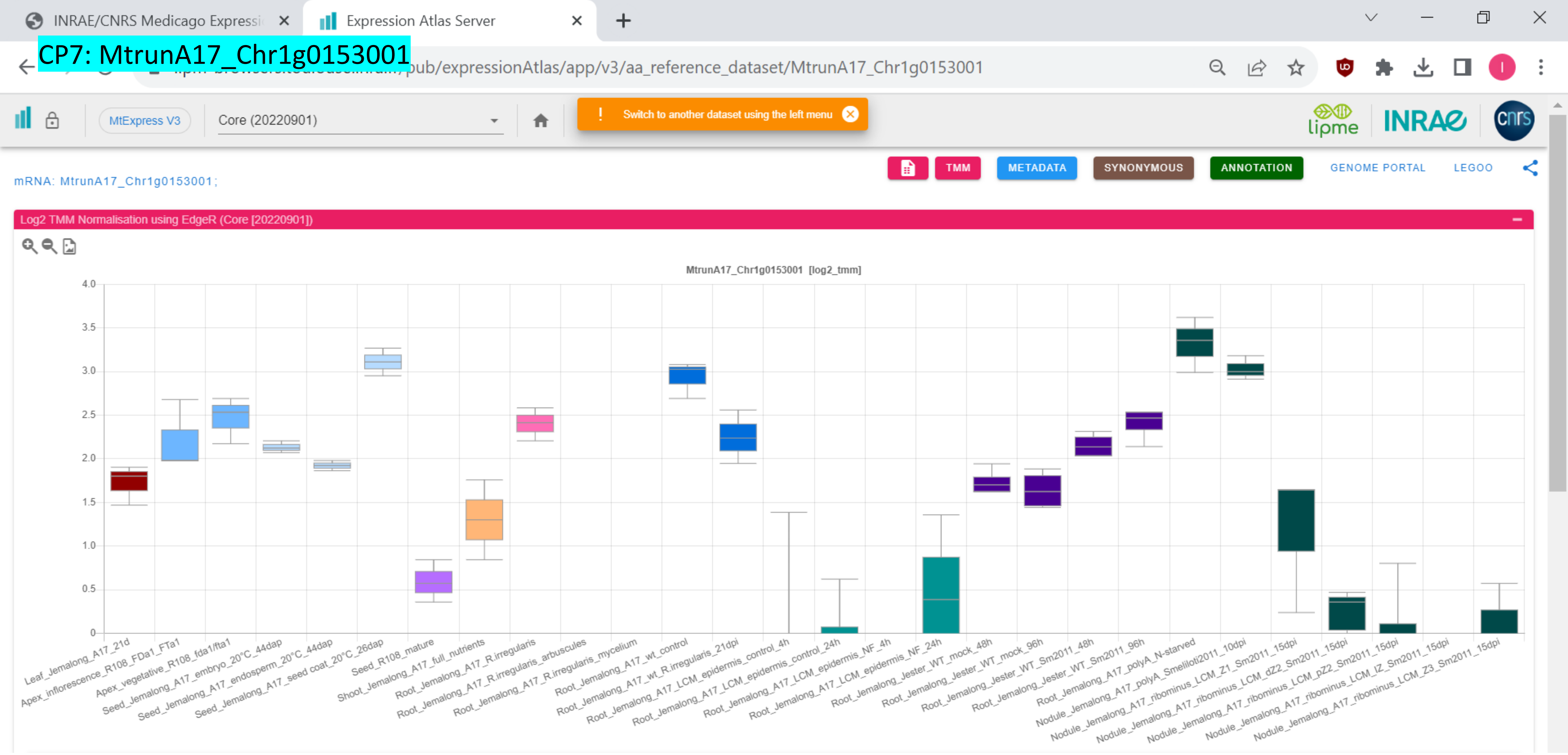

### CP8: MtrunA17_Chr1g0155251

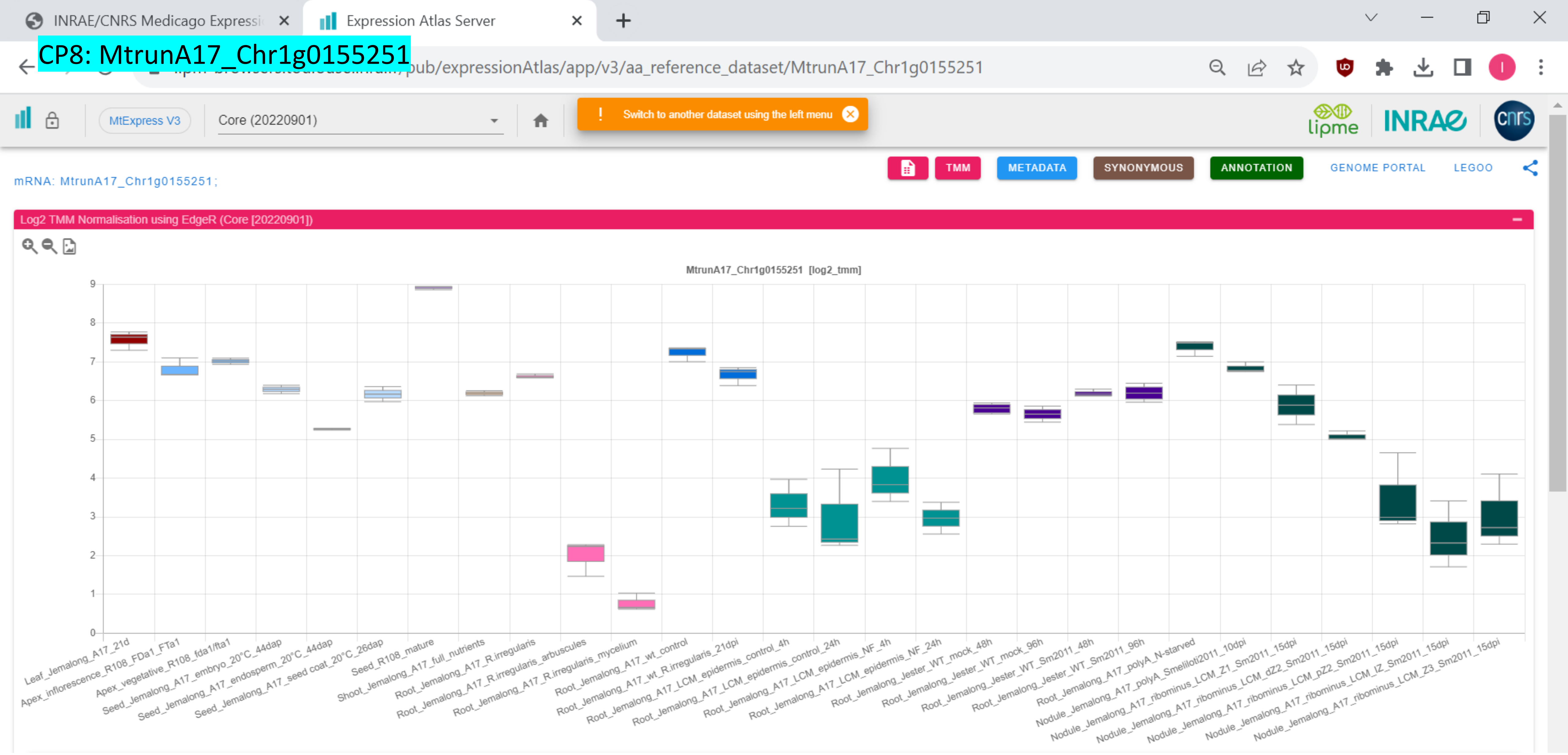

CP9: MtrunA17_Chr1g0158341

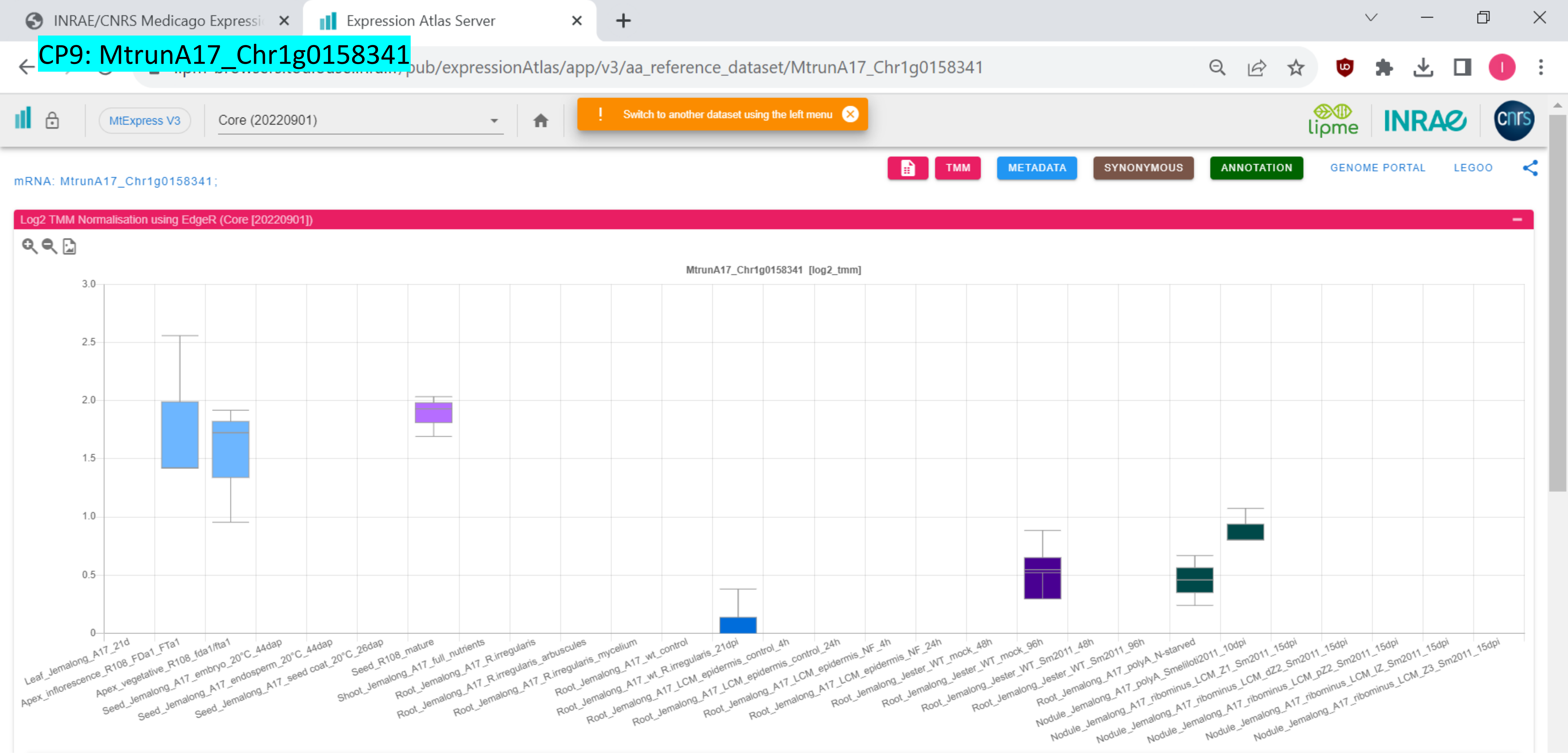

### CP10: MtrunA17_Chr1g0162101

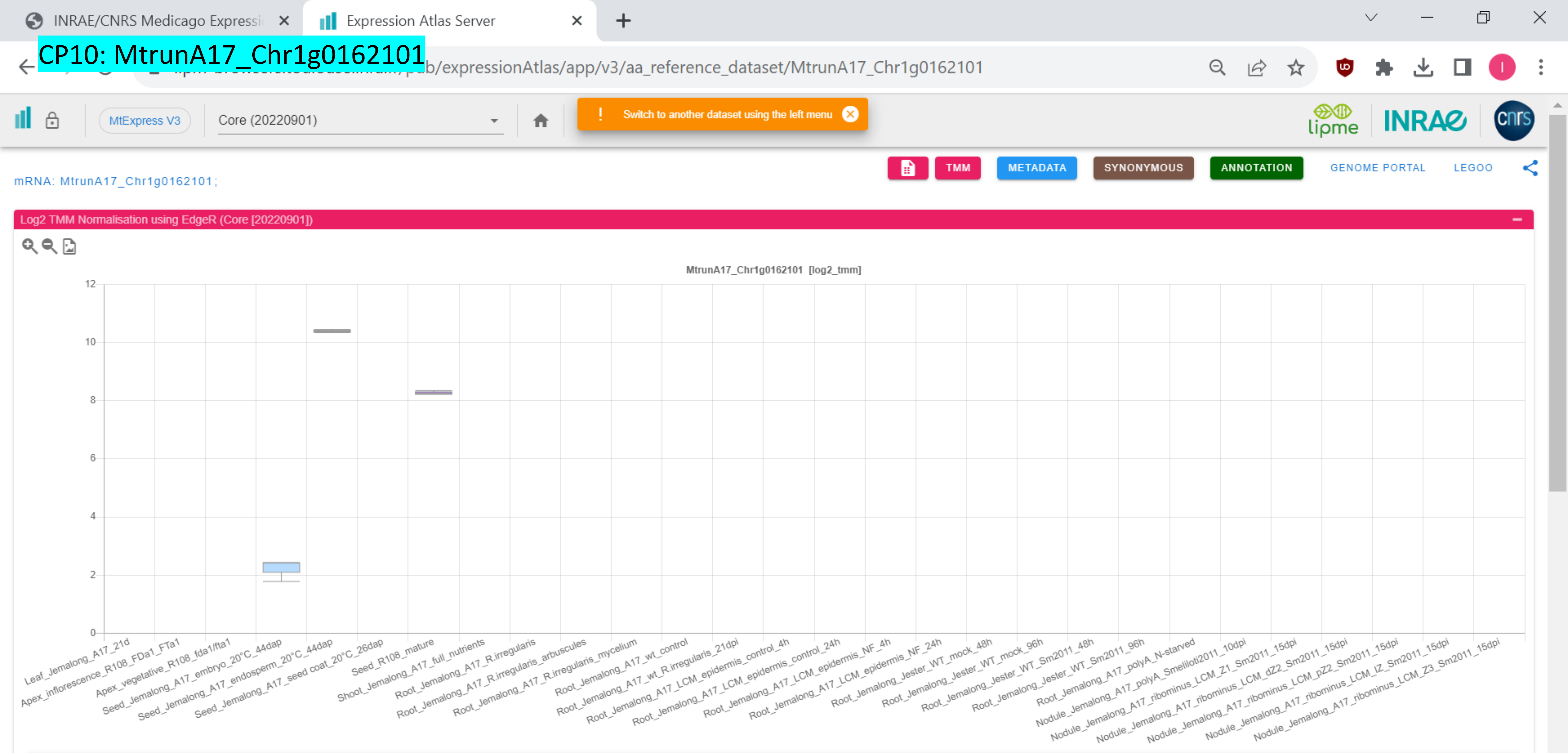

### CP11: MtrunA17_Chr1g0164591

INRAE/CNRS Medicago Expressi… | Expression Atlas Server

…pub/expressionAtlas/app/v3/aa_reference_dataset/MtrunA17_Chr1g0164591

MtExpress V3 | Core (20220901)

! Switch to another dataset using the left menu

TMM | METADATA | SYNONYMOUS | ANNOTATION | GENOME PORTAL | LEGOO

mRNA: MtrunA17_Chr1g0164591;

Log2 TMM Normalisation using EdgeR (Core [20220901])

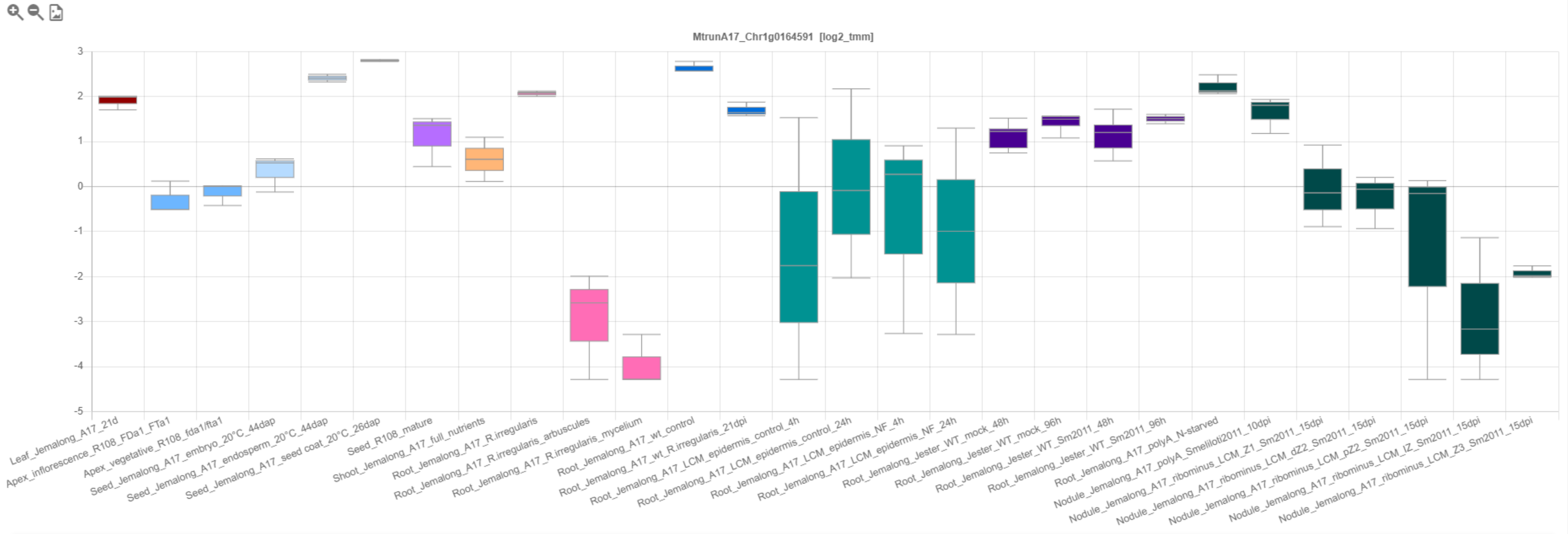

### CP12: MtrunA17_Chr1g0178361

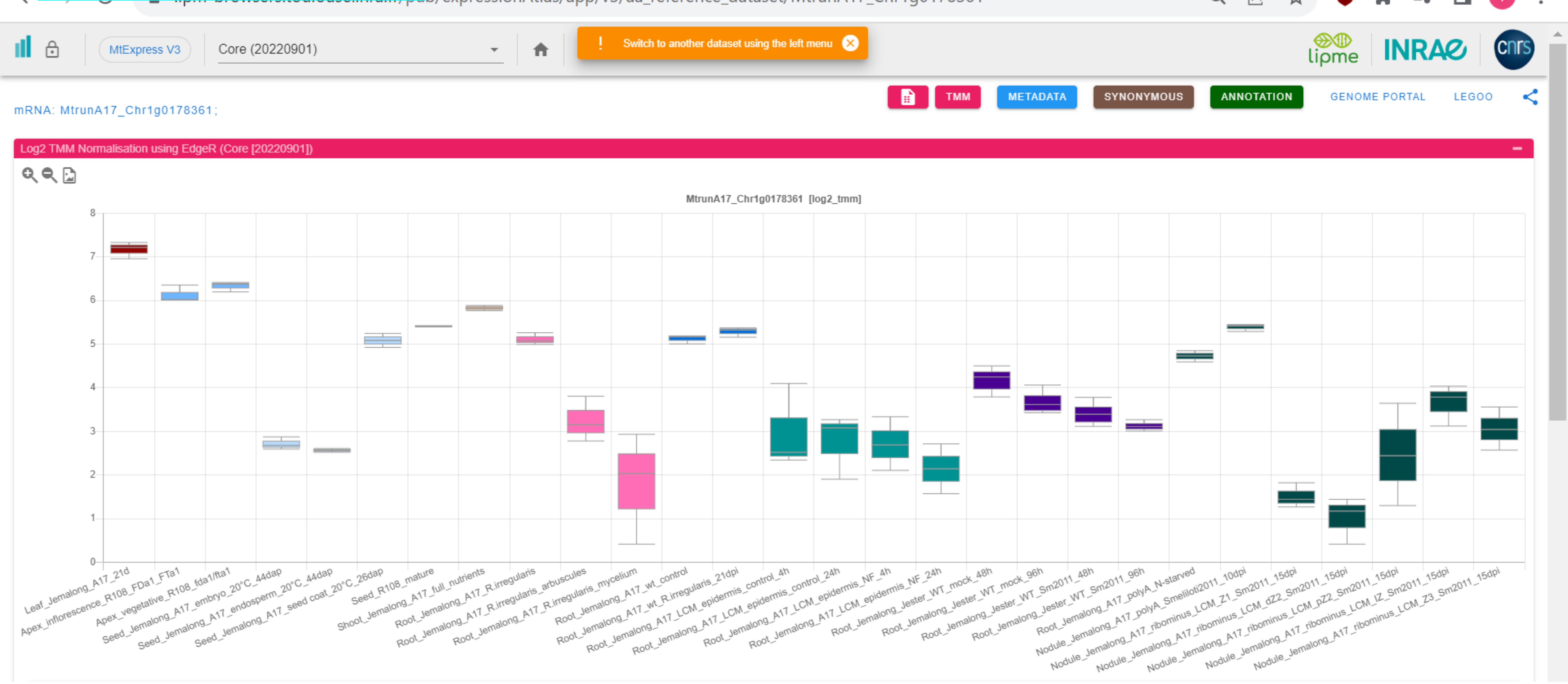

### CP13: MtrunA17_Chr1g0181761

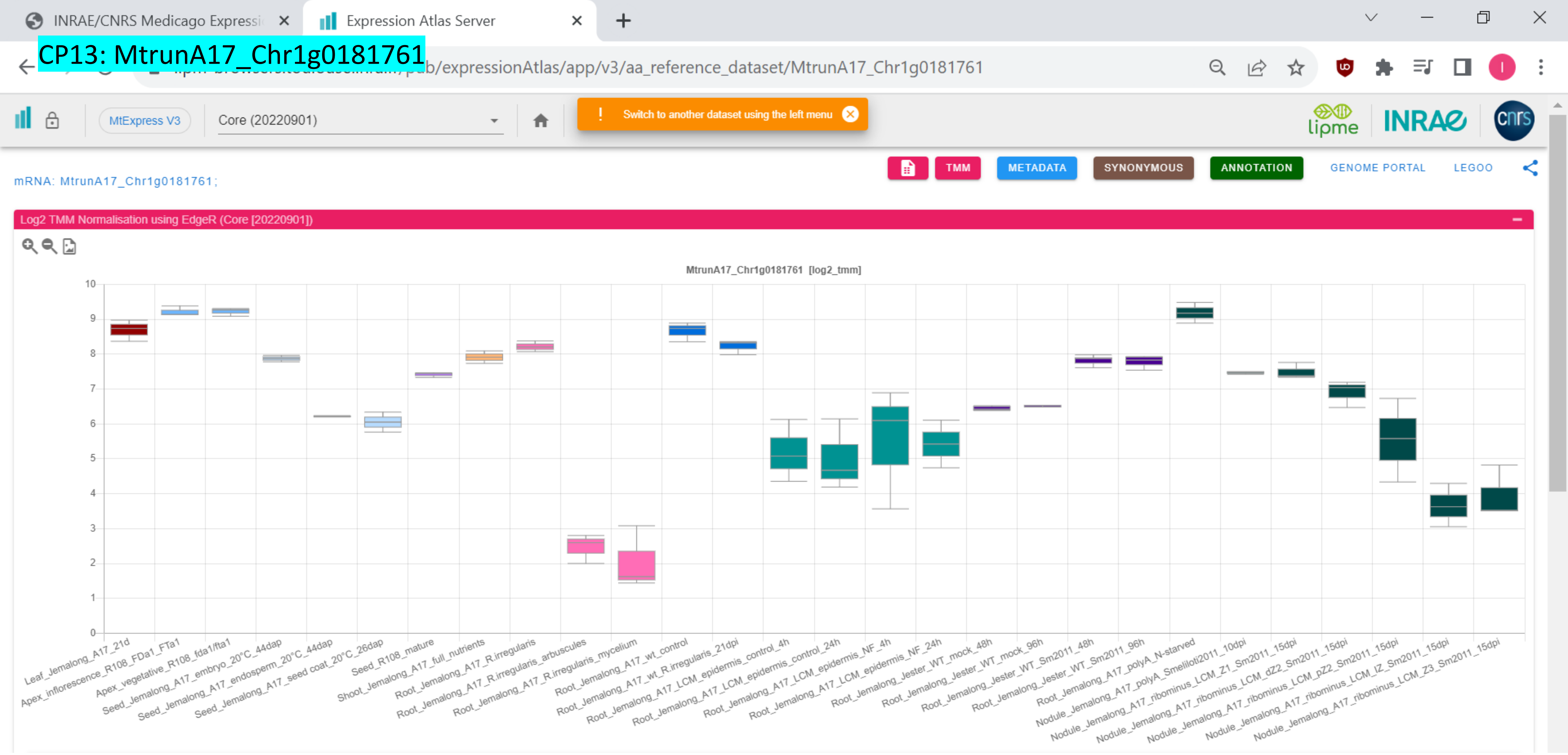

CP14: MtrunA17_Chr1g0182591

INRAE/CNRS Medicago Expressi… | Expression Atlas Server

…pub/expressionAtlas/app/v3/aa_reference_dataset/MtrunA17_Chr1g0182591

MtExpress V3 | Core (20220901)

! Switch to another dataset using the left menu

lipme | INRAE | cnrs

TMM | METADATA | SYNONYMOUS | ANNOTATION | GENOME PORTAL | LEGOO

mRNA: MtrunA17_Chr1g0182591;

Log2 TMM Normalisation using EdgeR (Core [20220901])

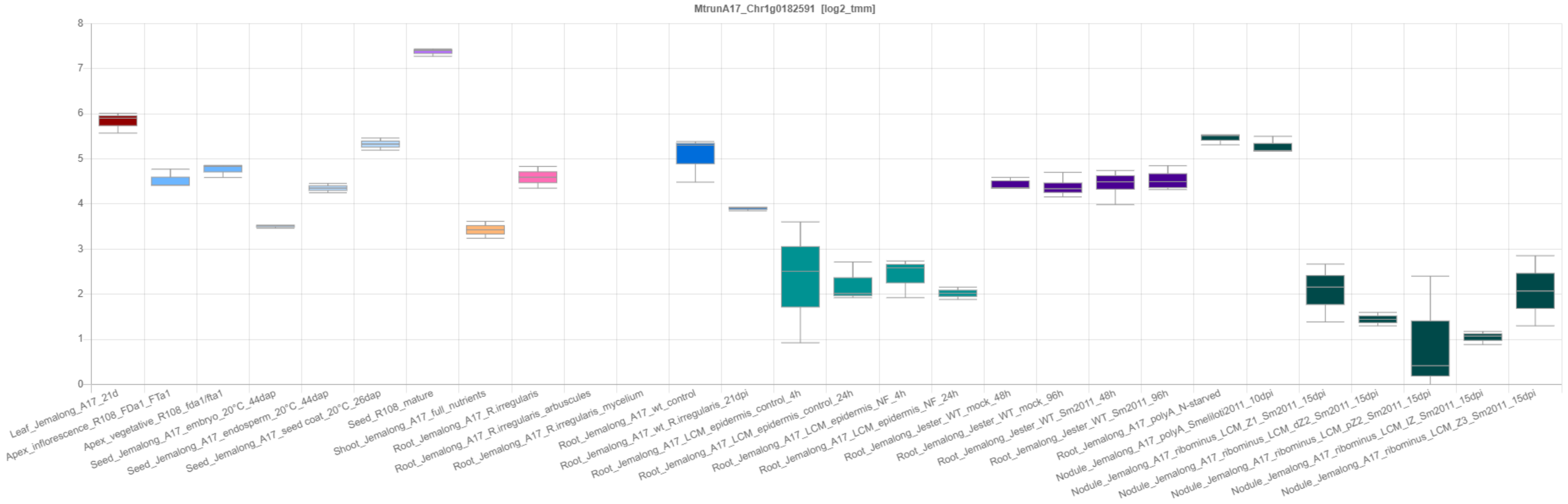

### CP15: MtrunA17_Chr1g0183001

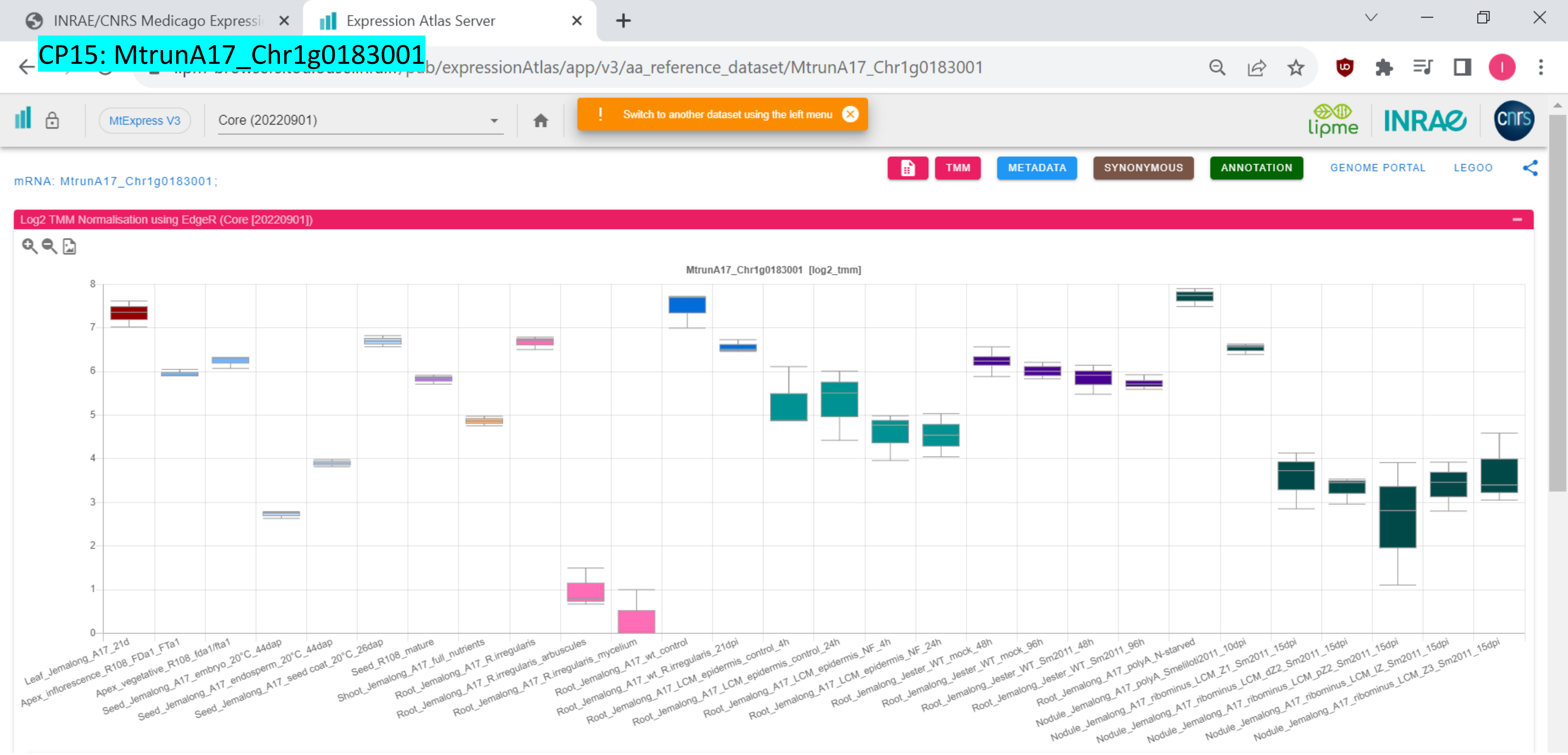

CP16: MtrunA17_Chr1g0185811

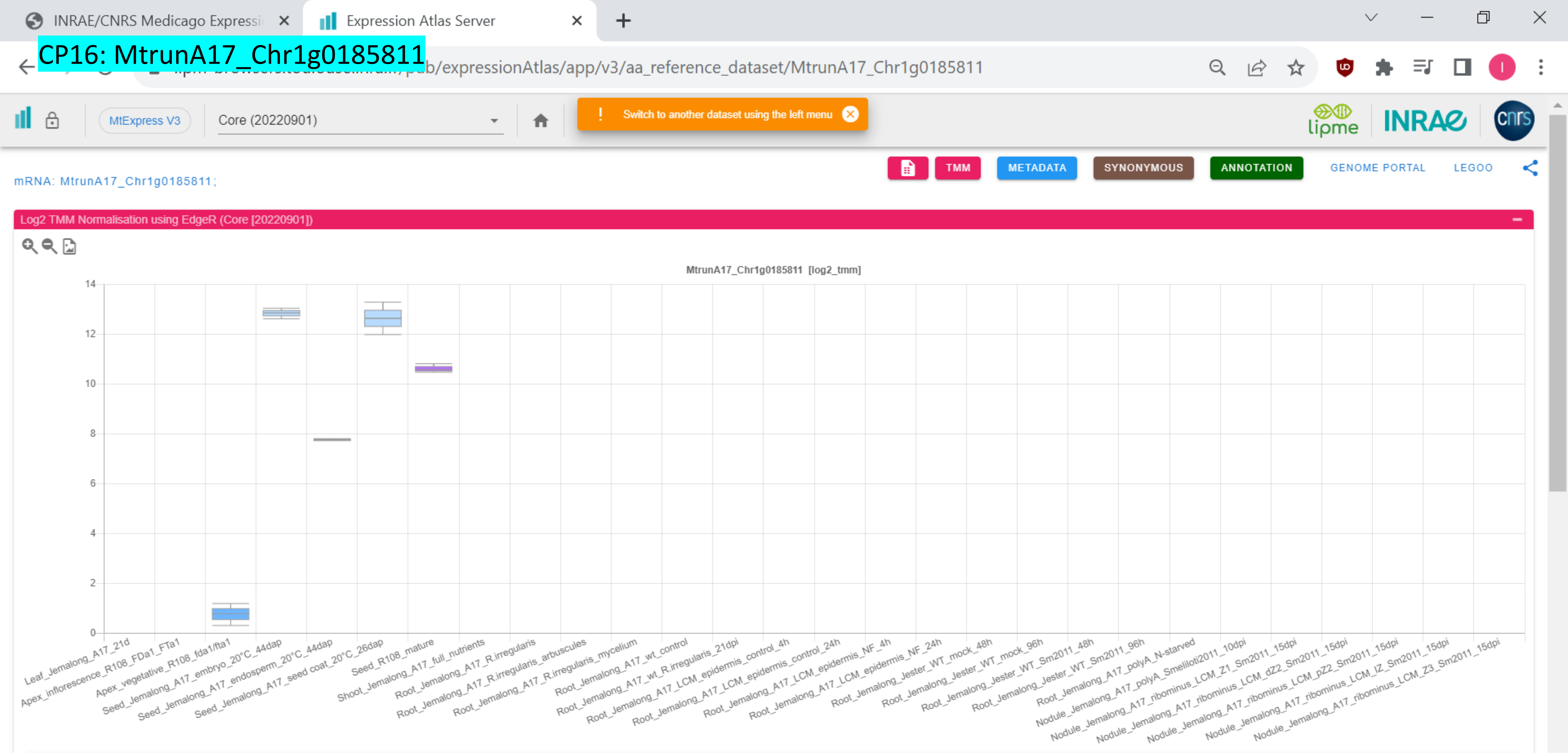

CP17: MtrunA17_Chr1g0185811

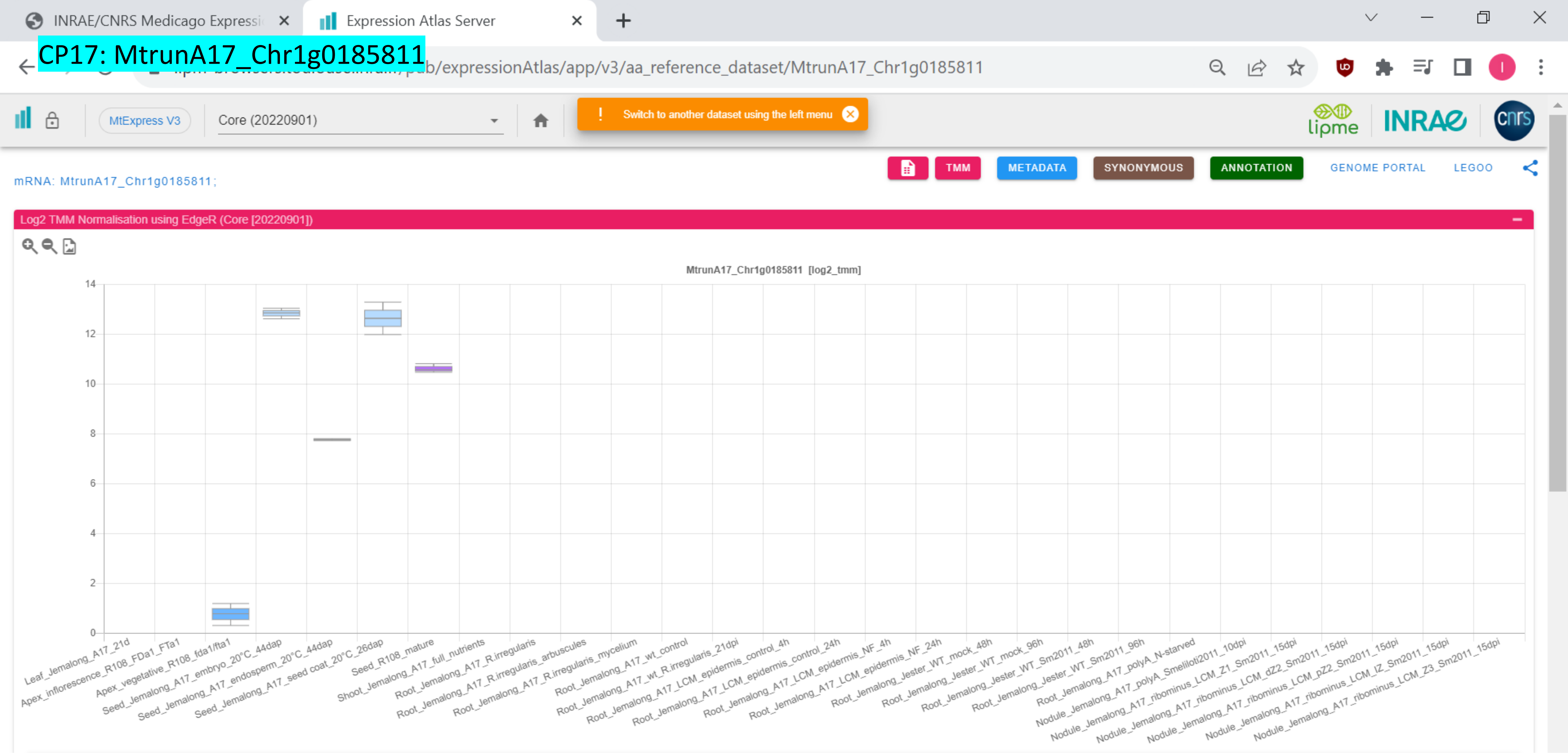

### CP18: MtrunA17_Chr1g0185811

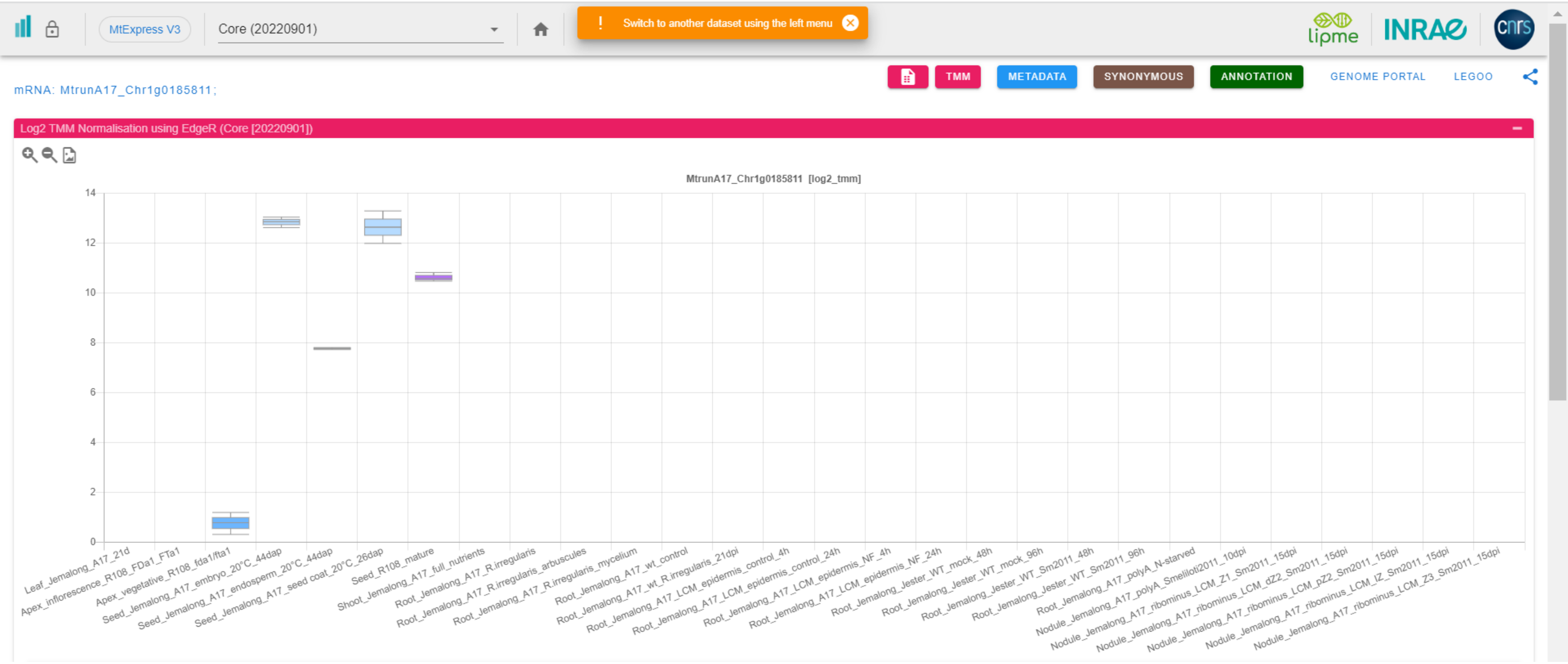

CP19: MtrunA17_Chr1g0185871

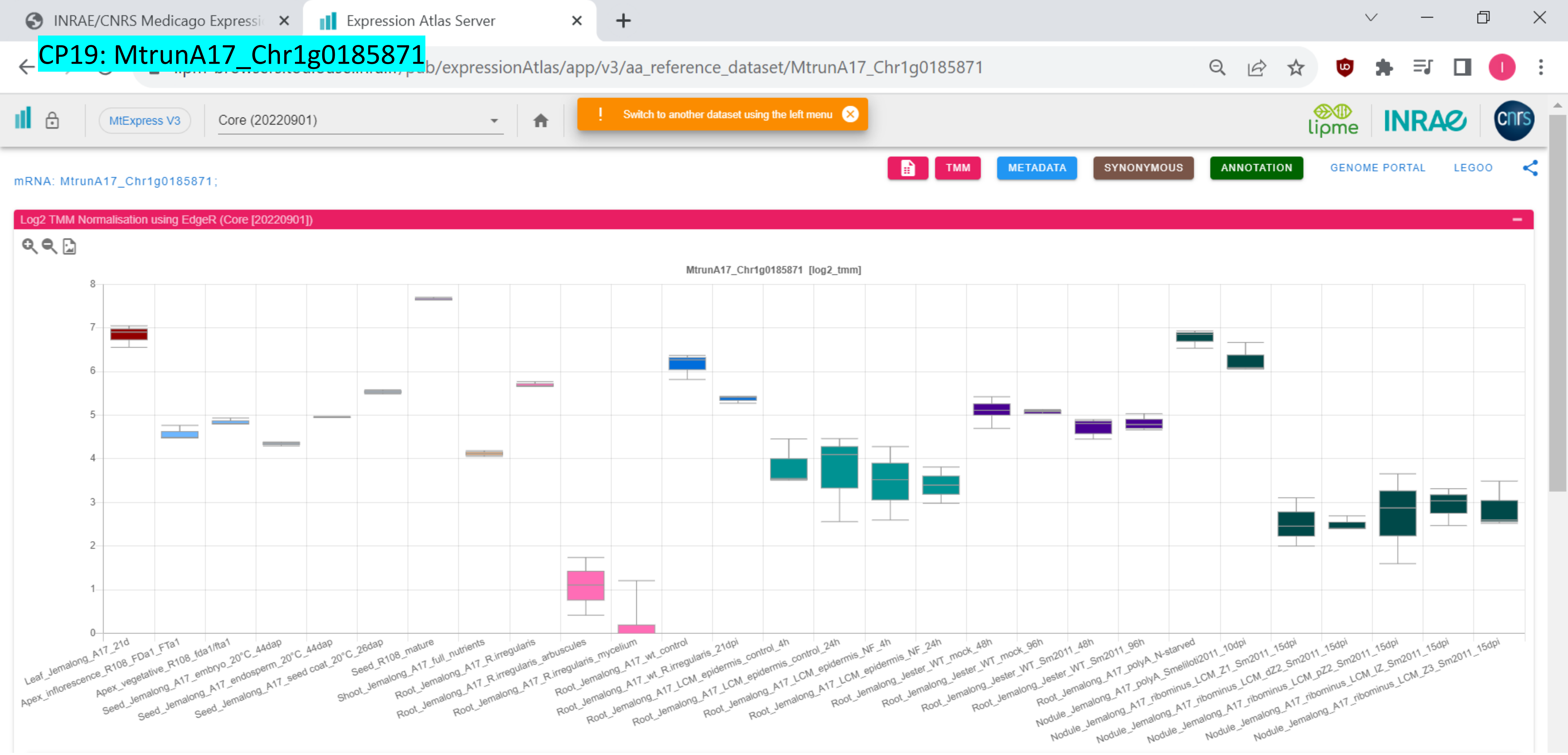

*CP20: MtrunA17_Chr1g0190571

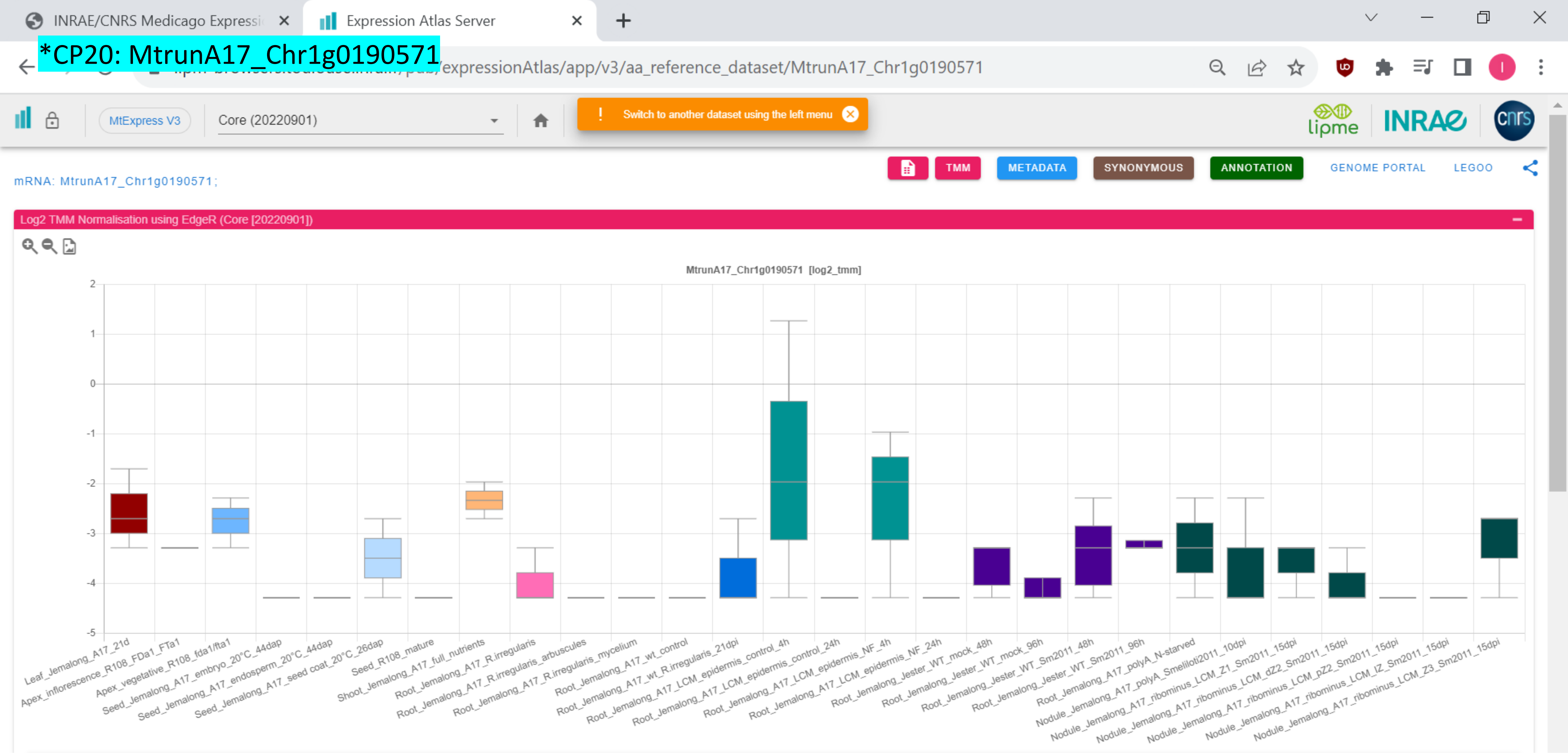

### CP21: MtrunA17_Chr1g0191411

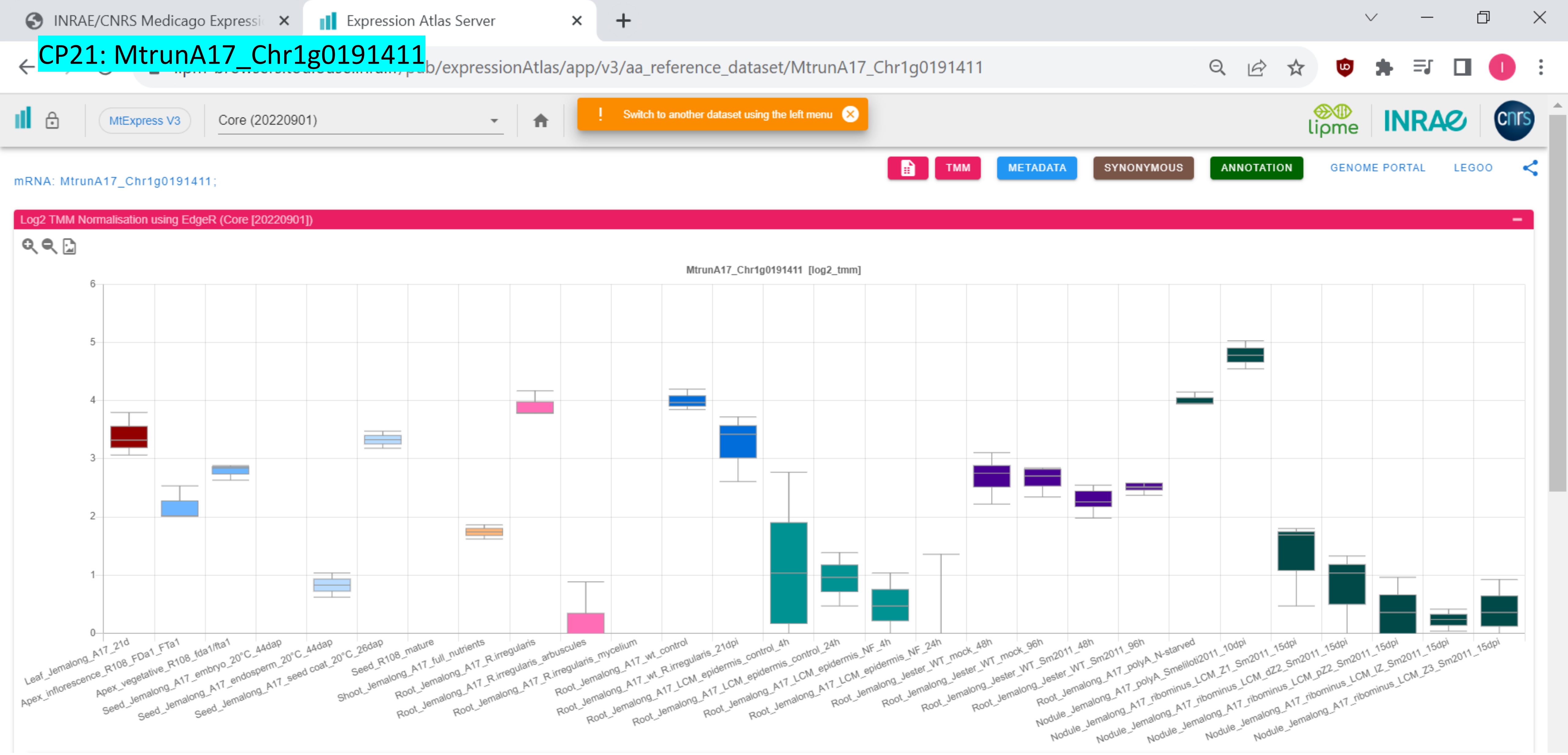

CP22: MtrunA17_Chr1g0198091

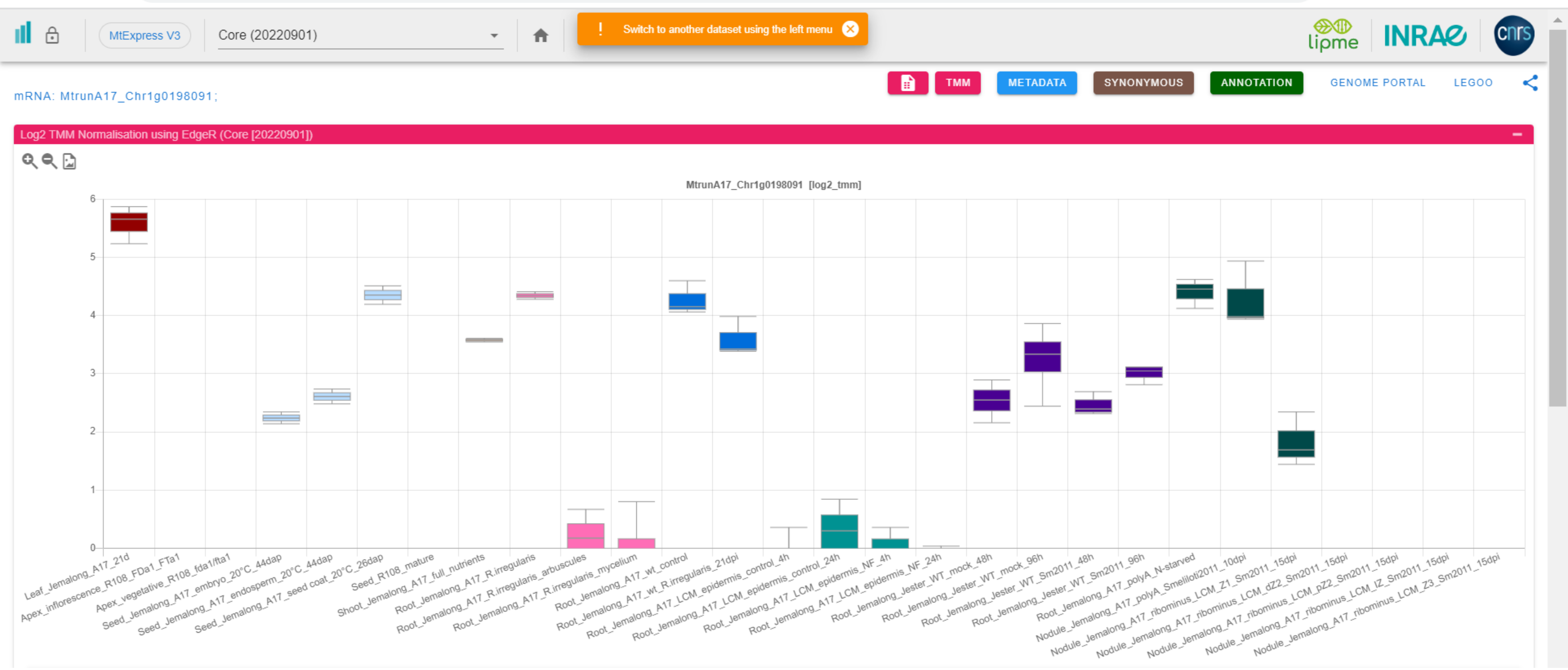

### CP23: MtrunA17_Chr1g0200071

mRNA: MtrunA17_Chr1g0200071;

Log2 TMM Normalisation using EdgeR (Core [20220901])

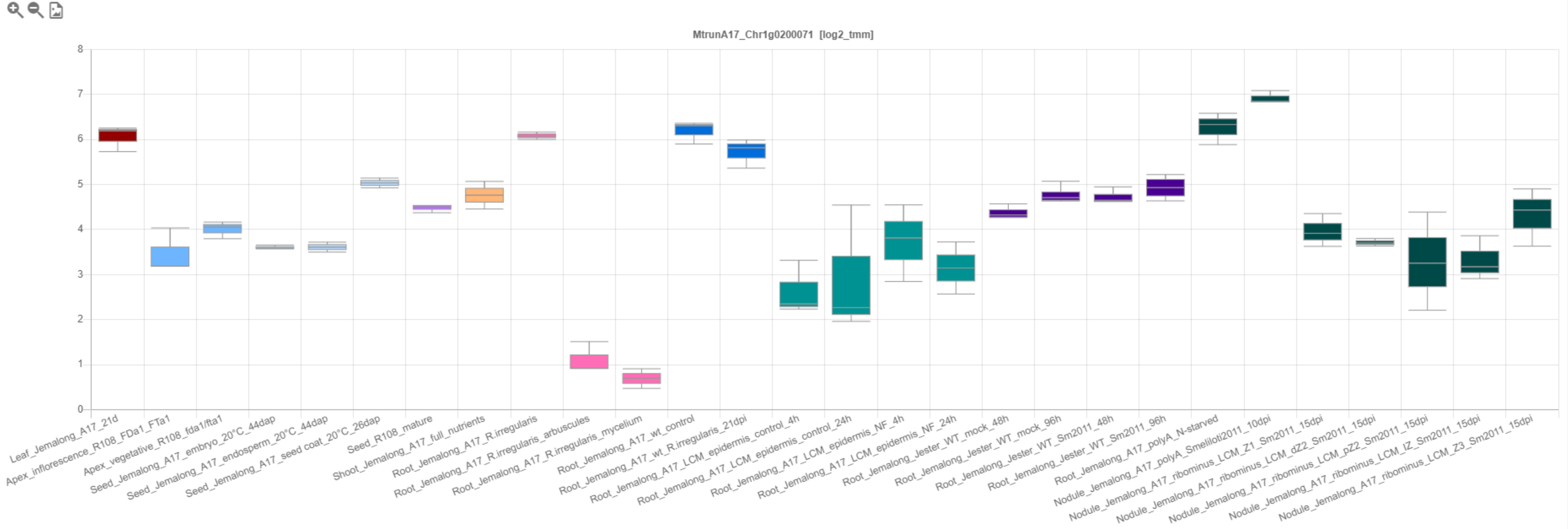

### CP24: MtrunA17_Chr1g0200071

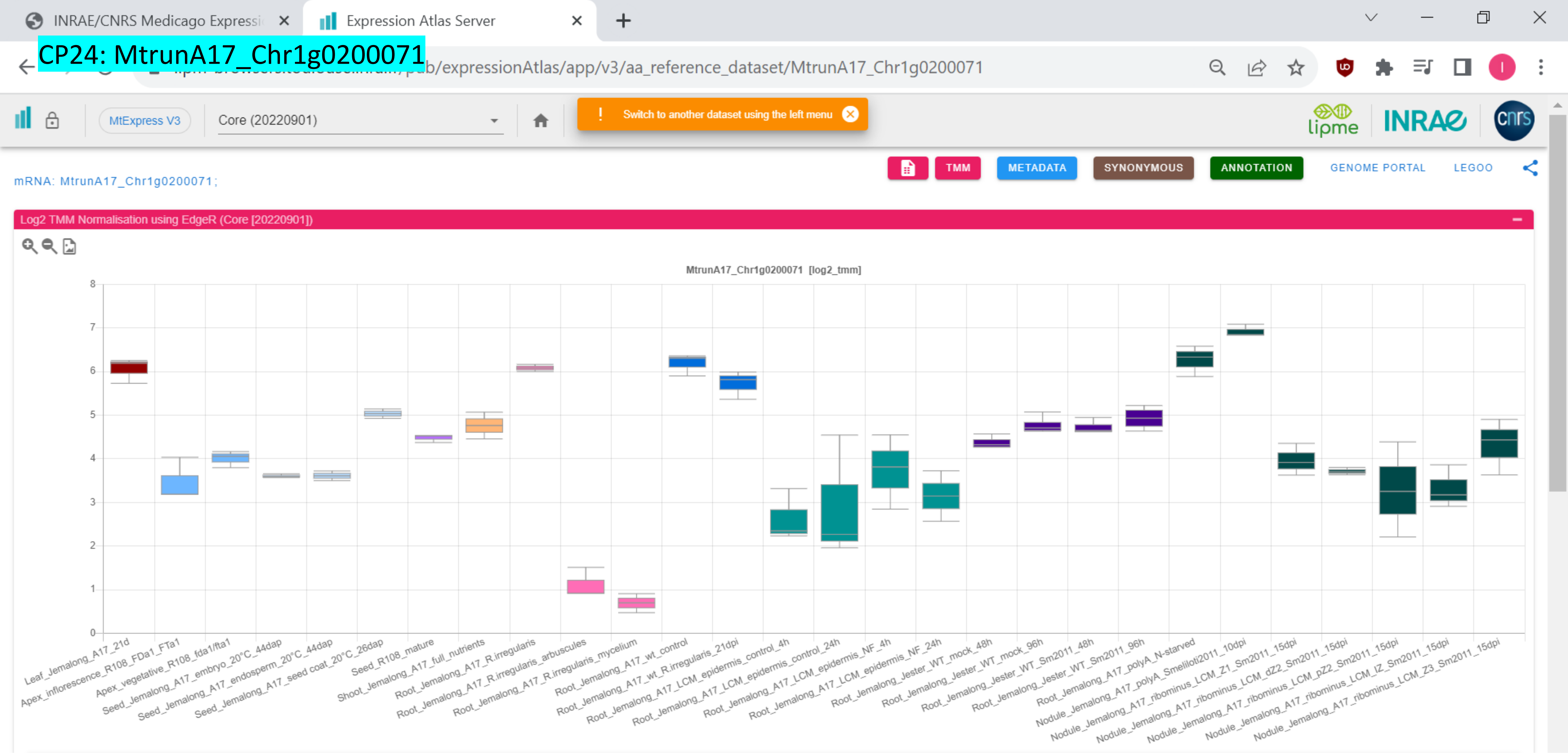

*CP25: MtrunA17_Chr1g0202001

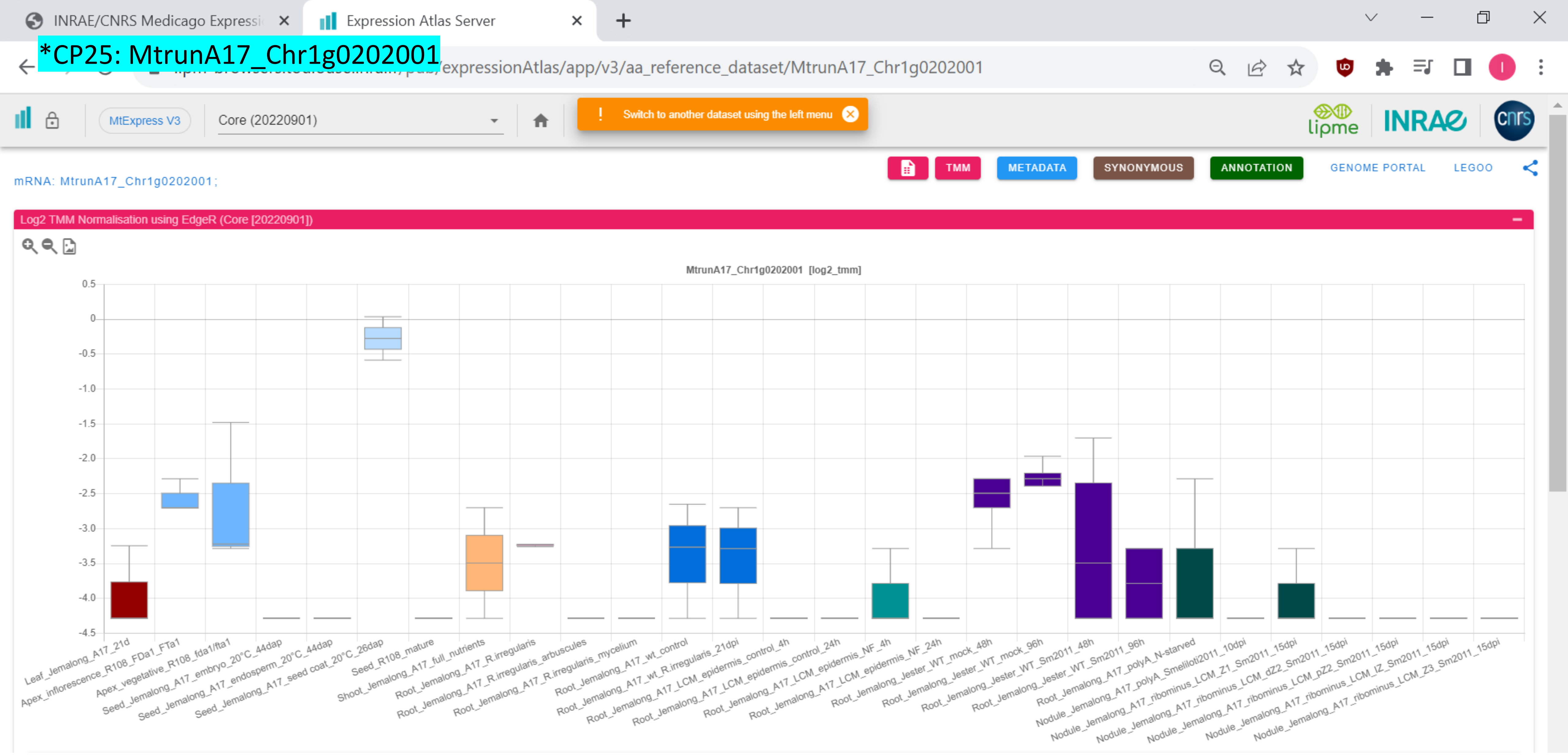

### CP26: MtrunA17_Chr1g0205601

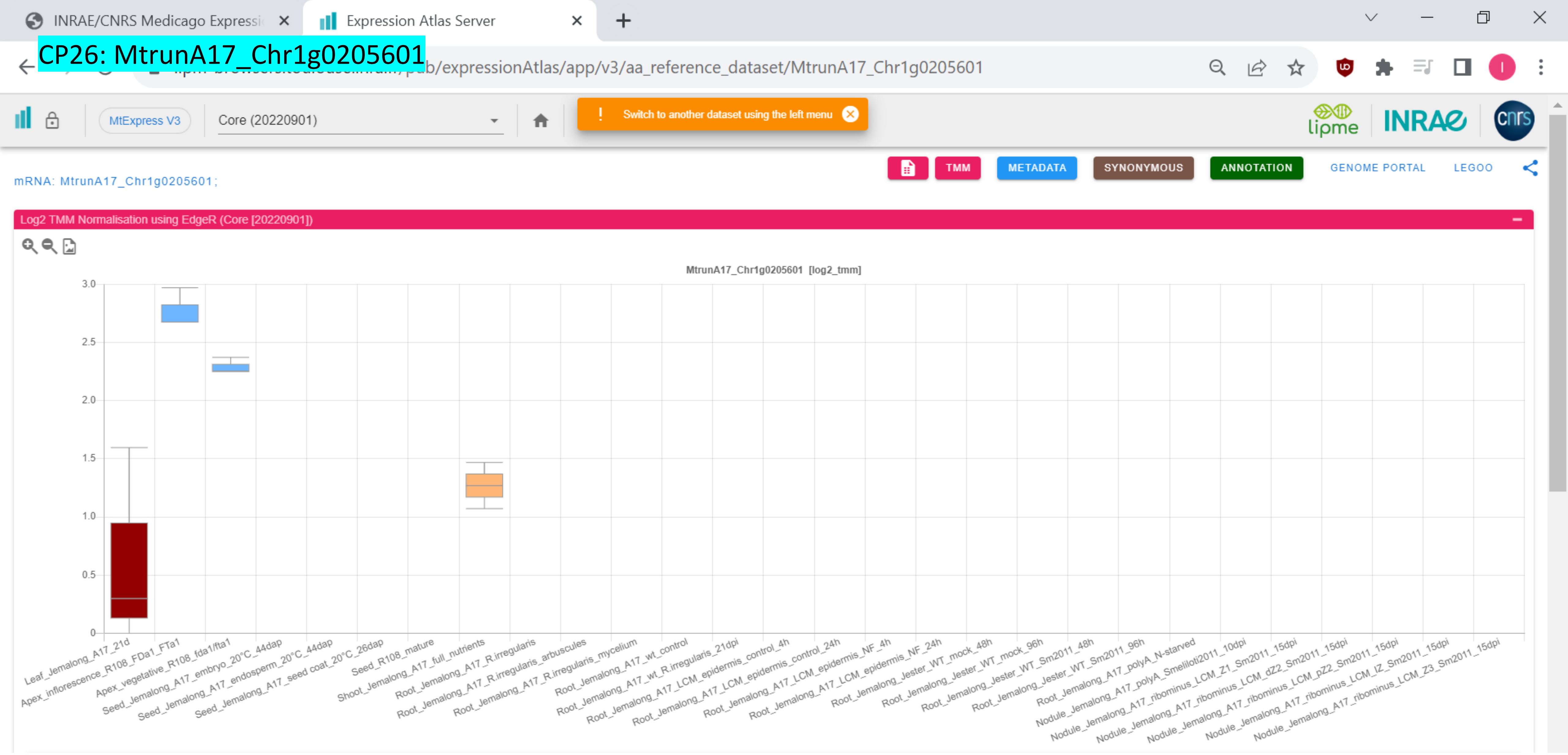

CP27: MtrunA17_Chr1g0207811

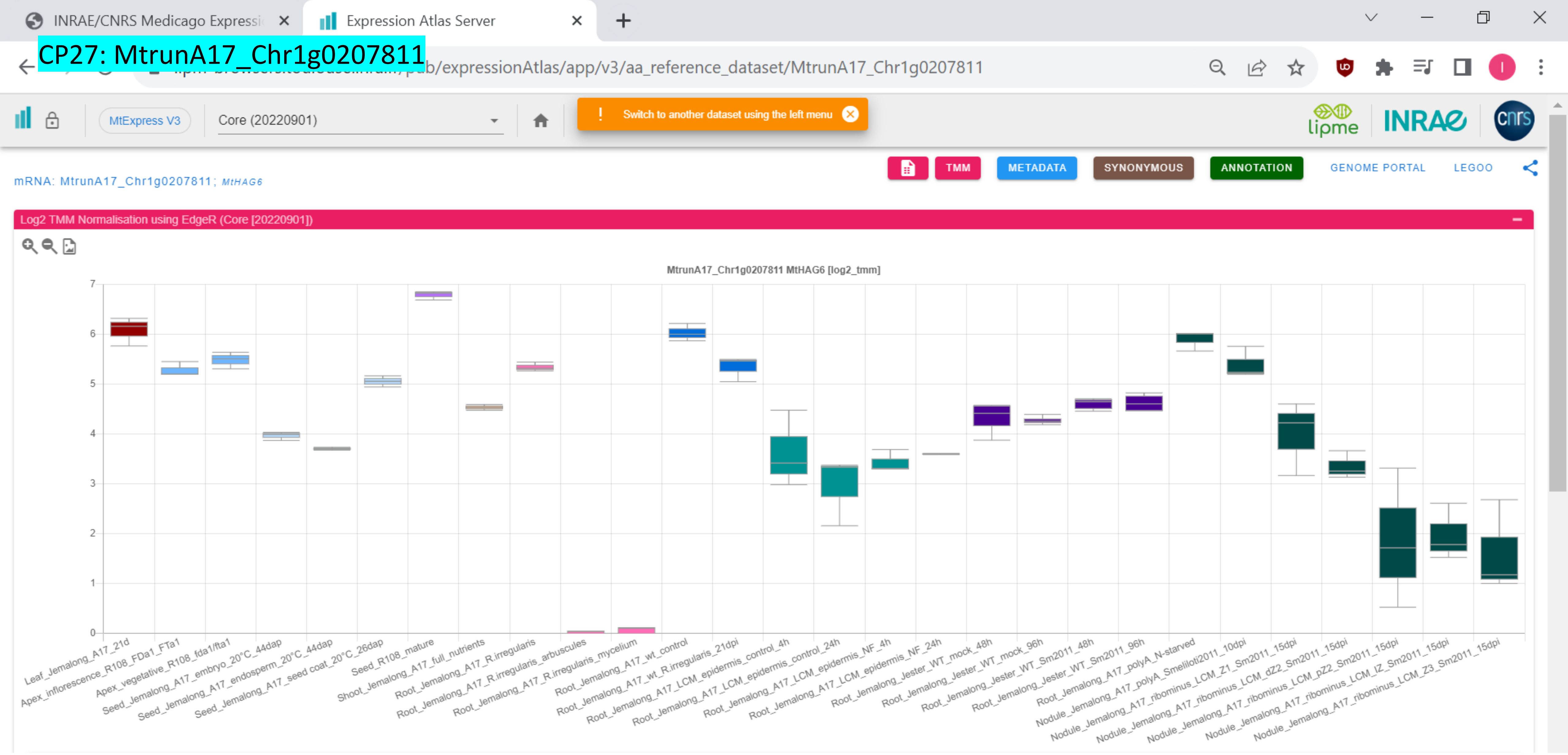

*CP28: MtrunA17_Chr1g0207921

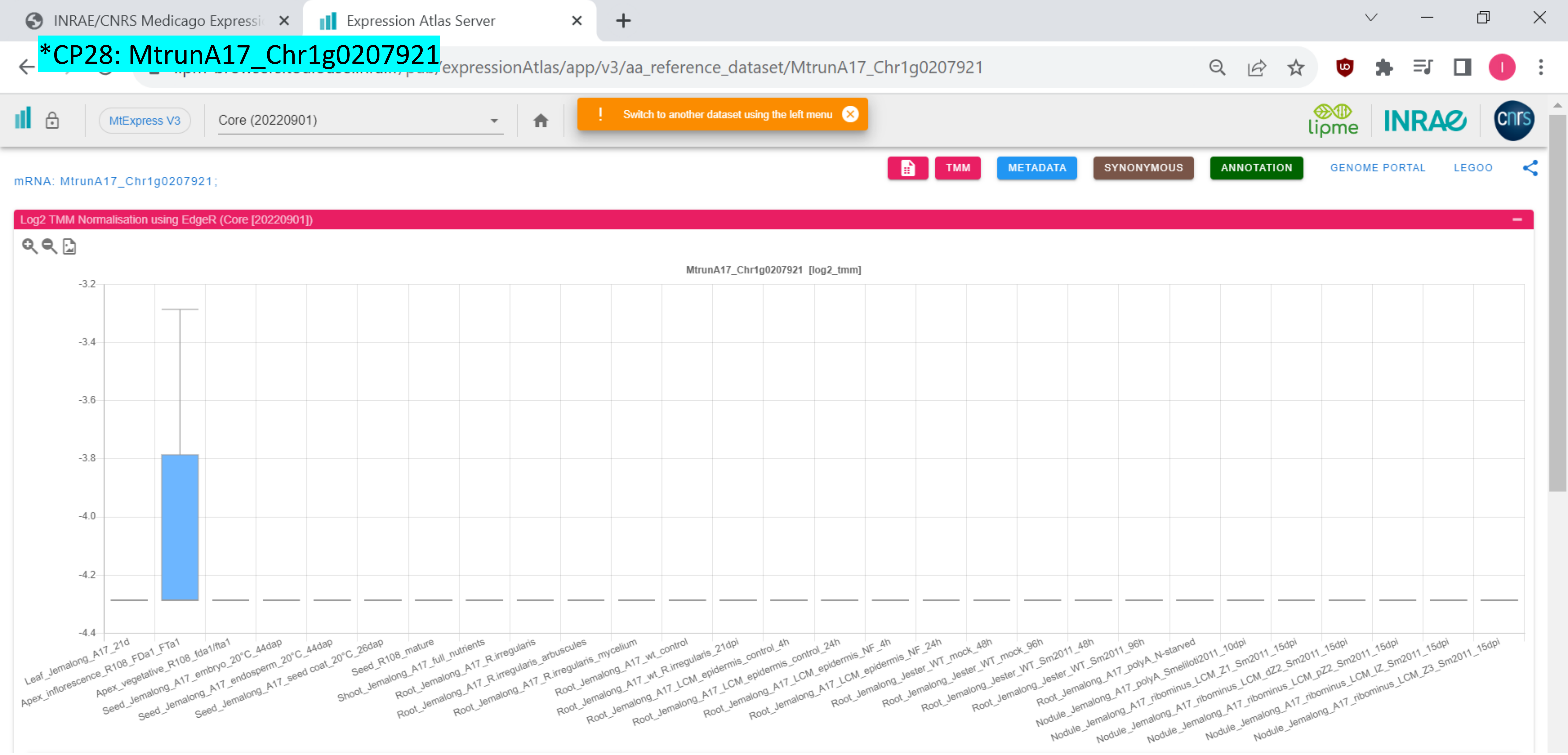

*CP29: MtrunA17_Chr1g0209791

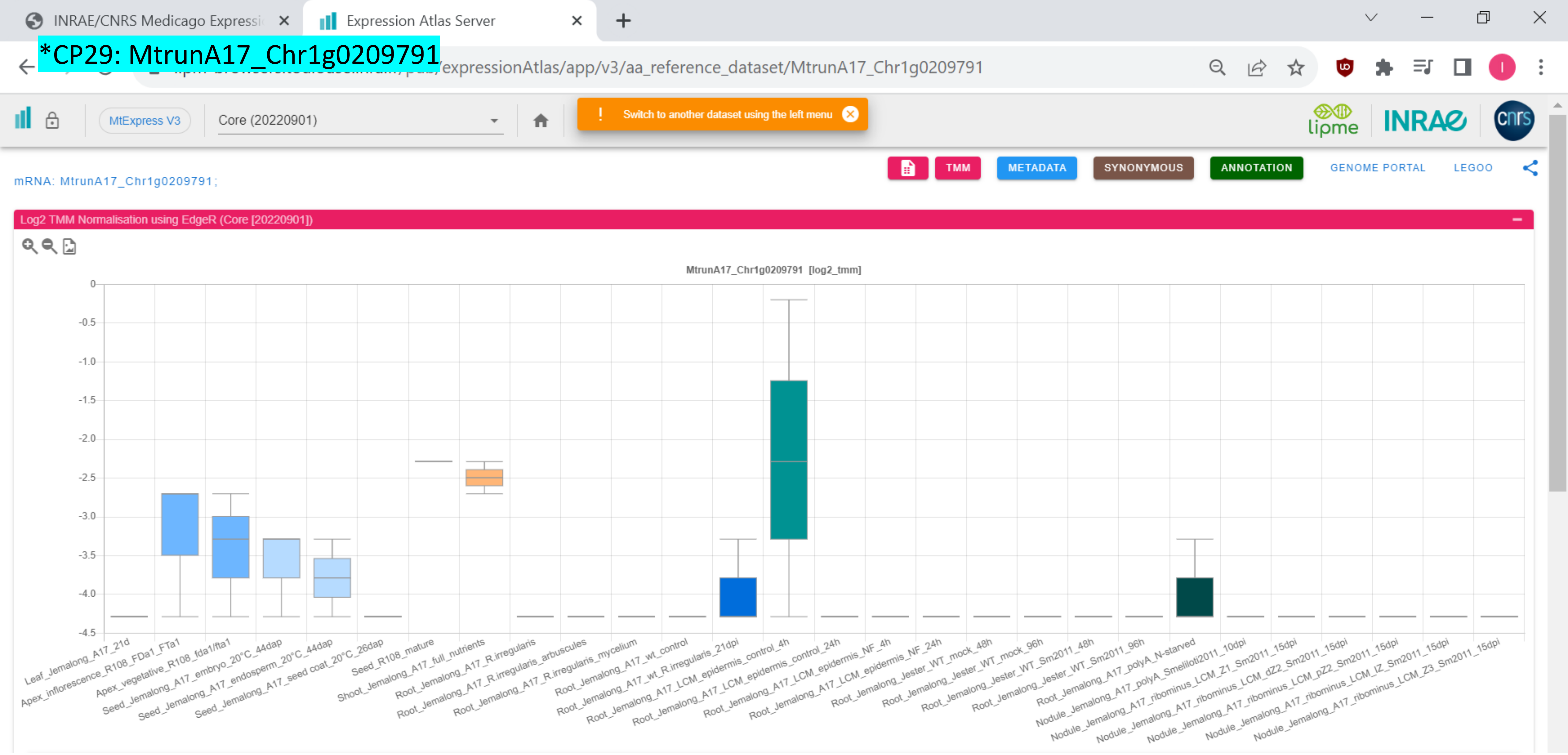

CP30: MtrunA17_Chr1g0210521

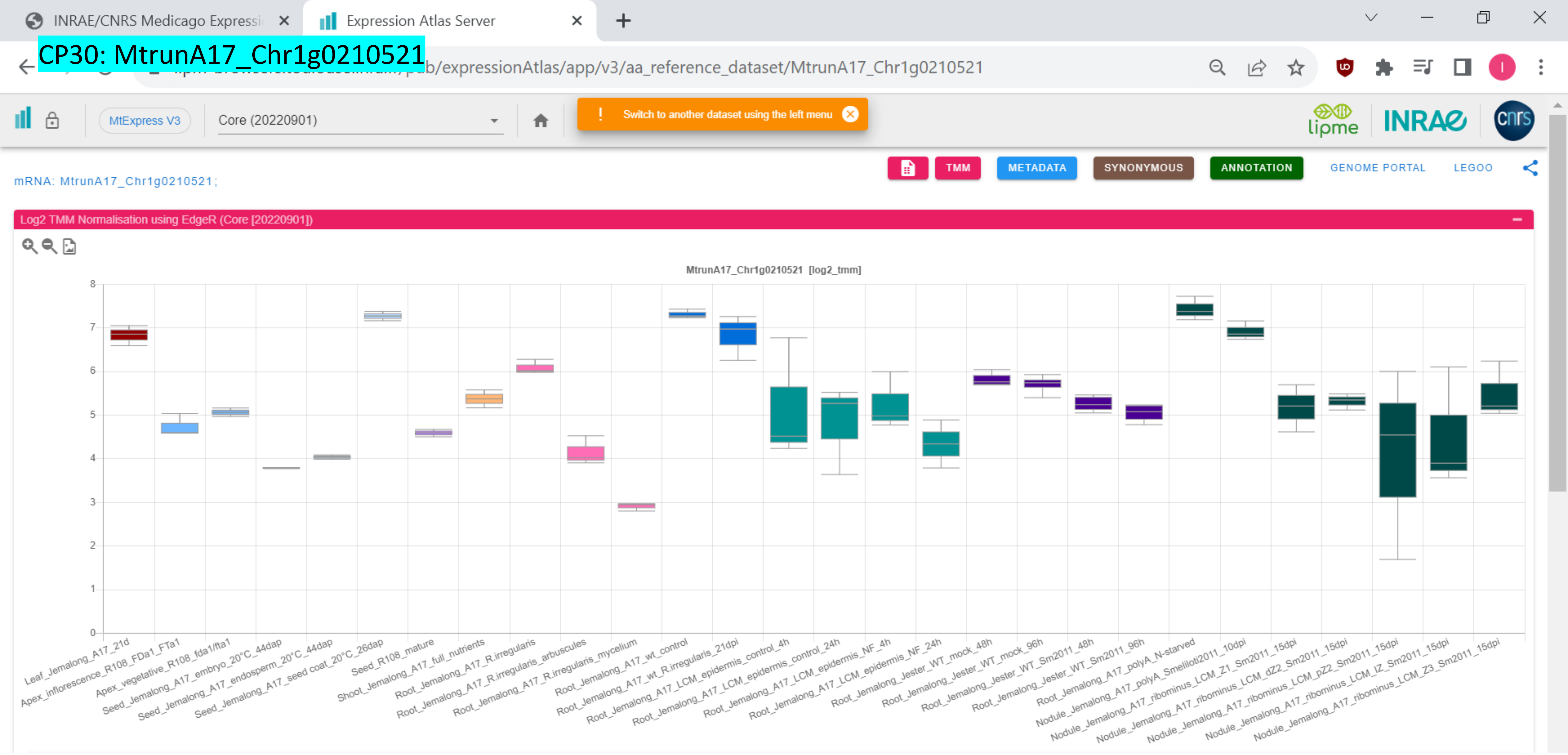

CP31: MtrunA17_Chr1g0212961

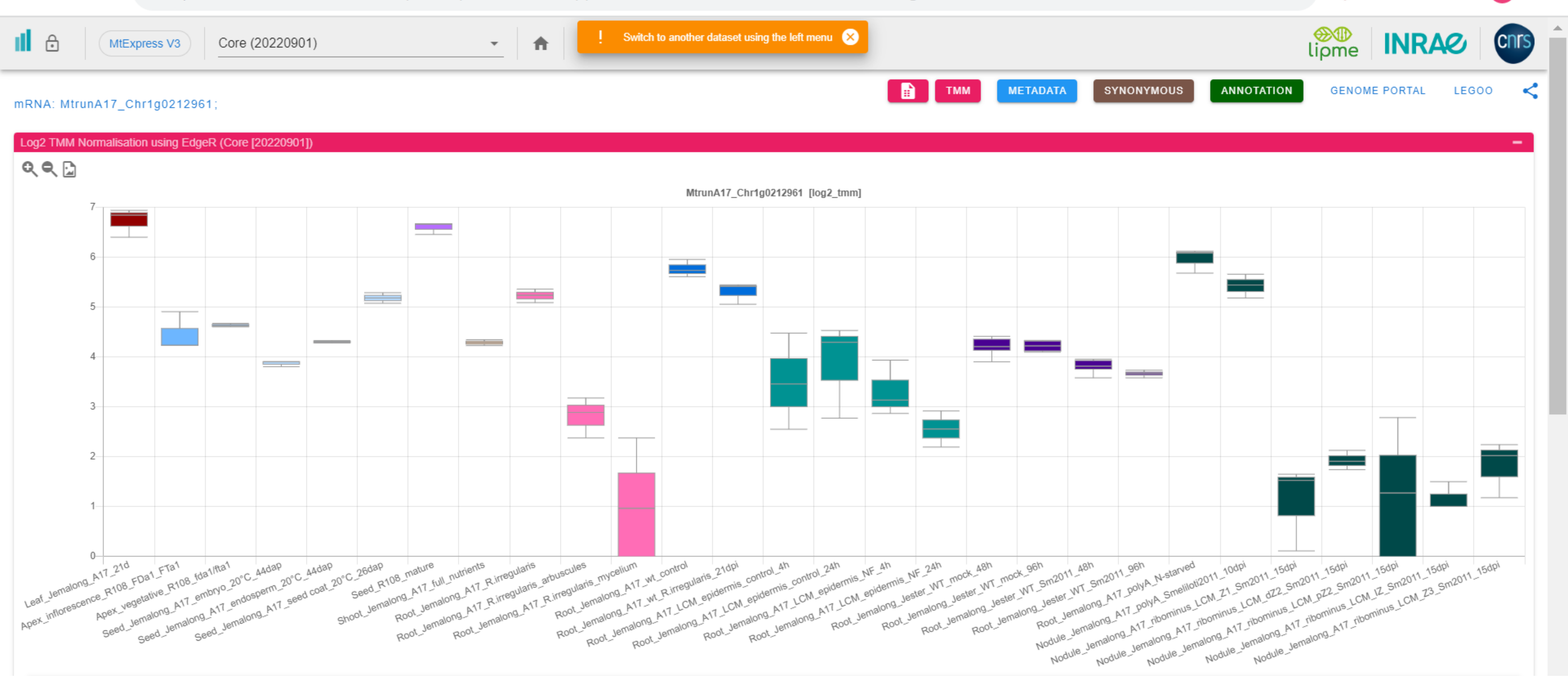

### CP32: MtrunA17_Chr1g1004575

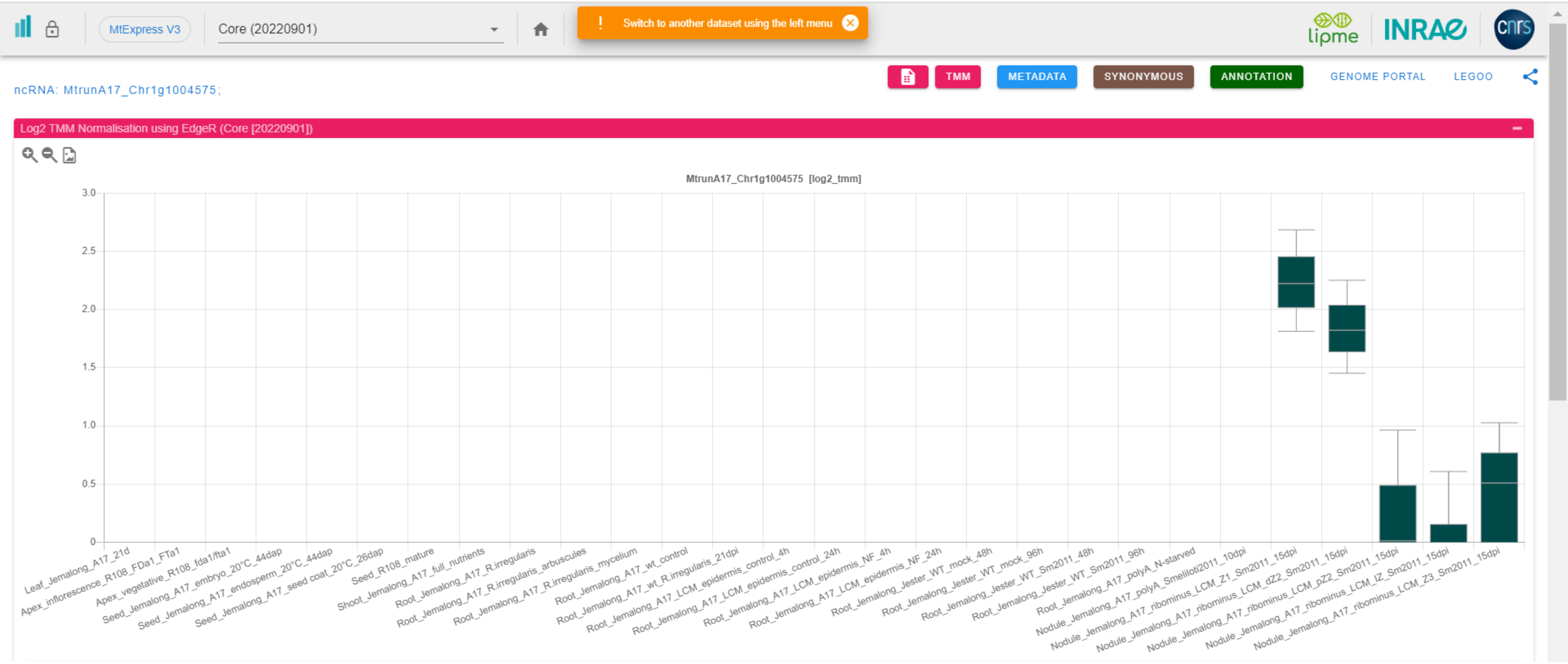

CP33: MtrunA17_Chr2g0283311

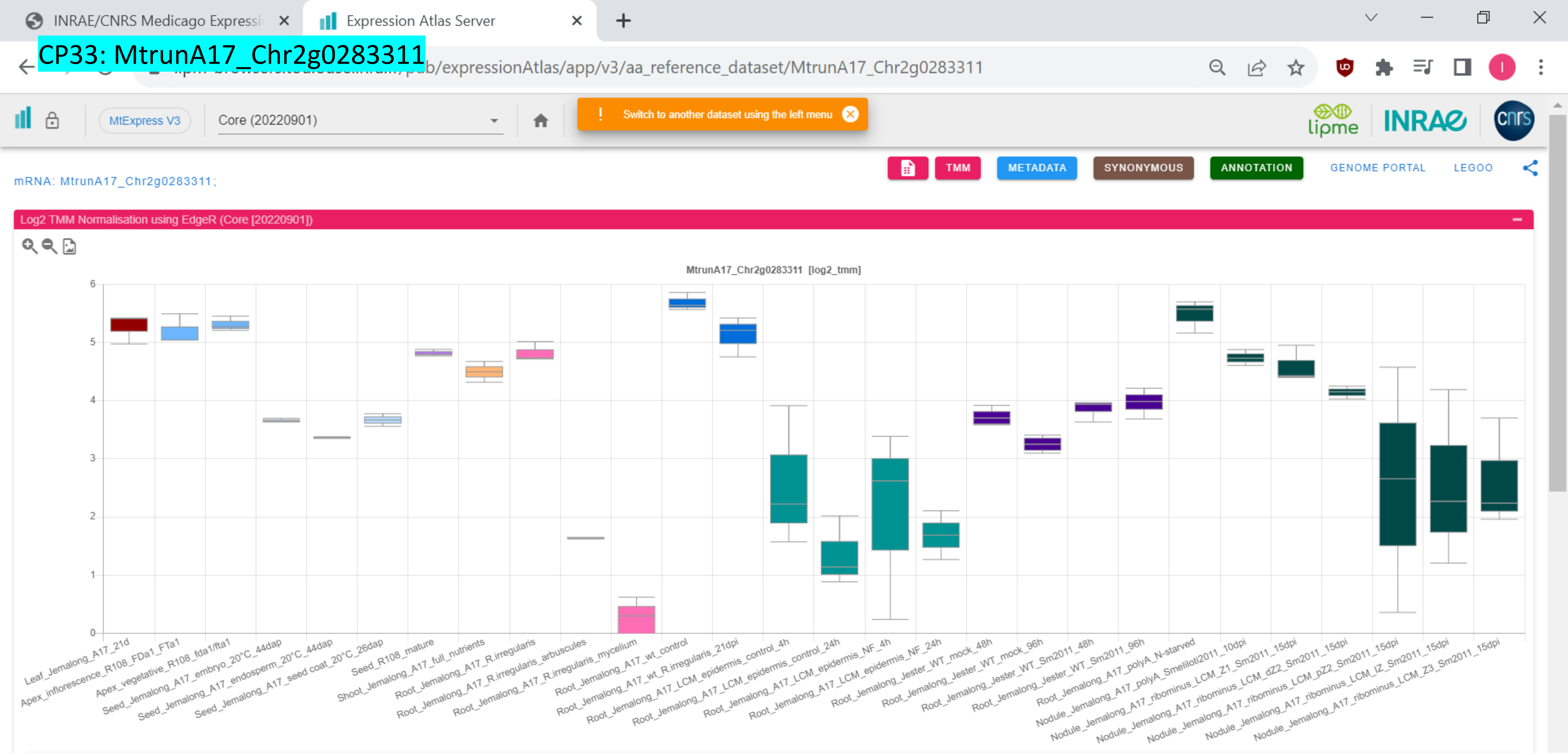

INRAE/CNRS Medicago Expressi
Expression Atlas Server

### CP34: MtrunA17_Chr2g0285461

…/expressionAtlas/app/v3/aa_reference_dataset/MtrunA17_Chr2g0285461

MtExpress V3 | Core (20220901)

Switch to another dataset using the left menu

lipme INRAE cnrs

TMM | METADATA | SYNONYMOUS | ANNOTATION | GENOME PORTAL | LEGOO

mRNA: MtrunA17_Chr2g0285461; *MtARF6|MtARF19a*

Log2 TMM Normalisation using EdgeR (Core [20220901])

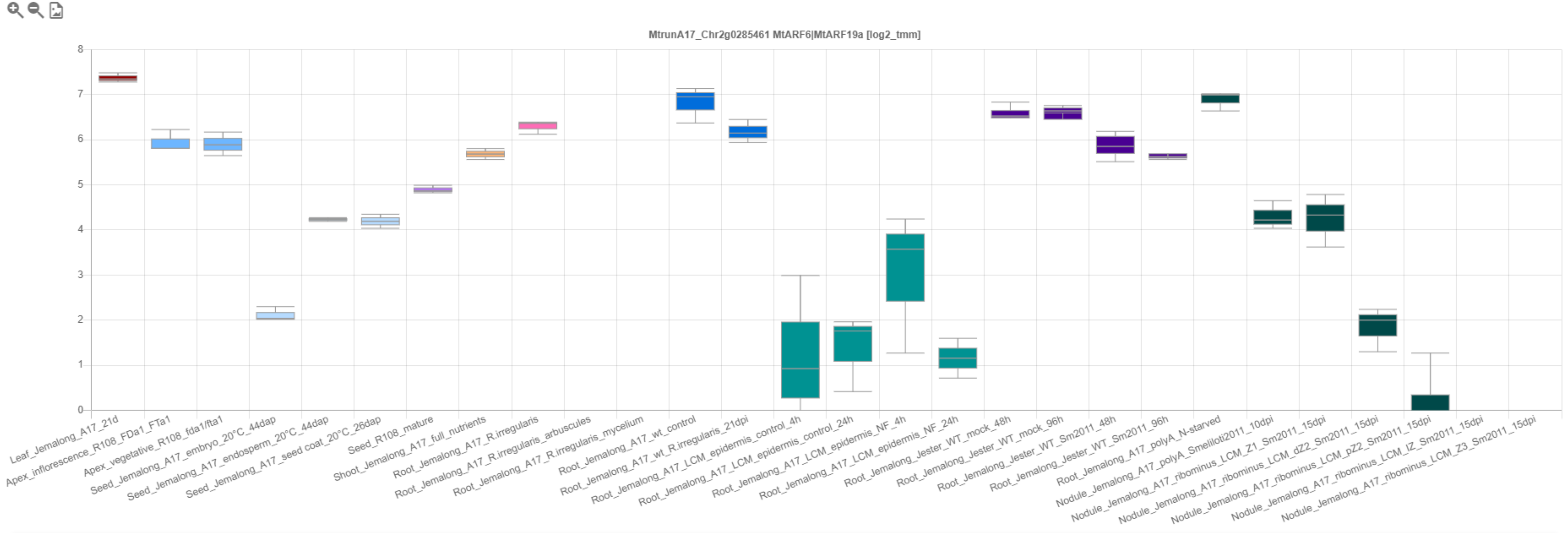

CP35: MtrunA17_Chr2g0292921

INRAE/CNRS Medicago Expressi | Expression Atlas Server

.../pub/expressionAtlas/app/v3/aa_reference_dataset/MtrunA17_Chr2g0292921

MtExpress V3 | Core (20220901) | Switch to another dataset using the left menu

TMM | METADATA | SYNONYMOUS | ANNOTATION | GENOME PORTAL | LEGOO

mRNA: MtrunA17_Chr2g0292921;

Log2 TMM Normalisation using EdgeR (Core [20220901])

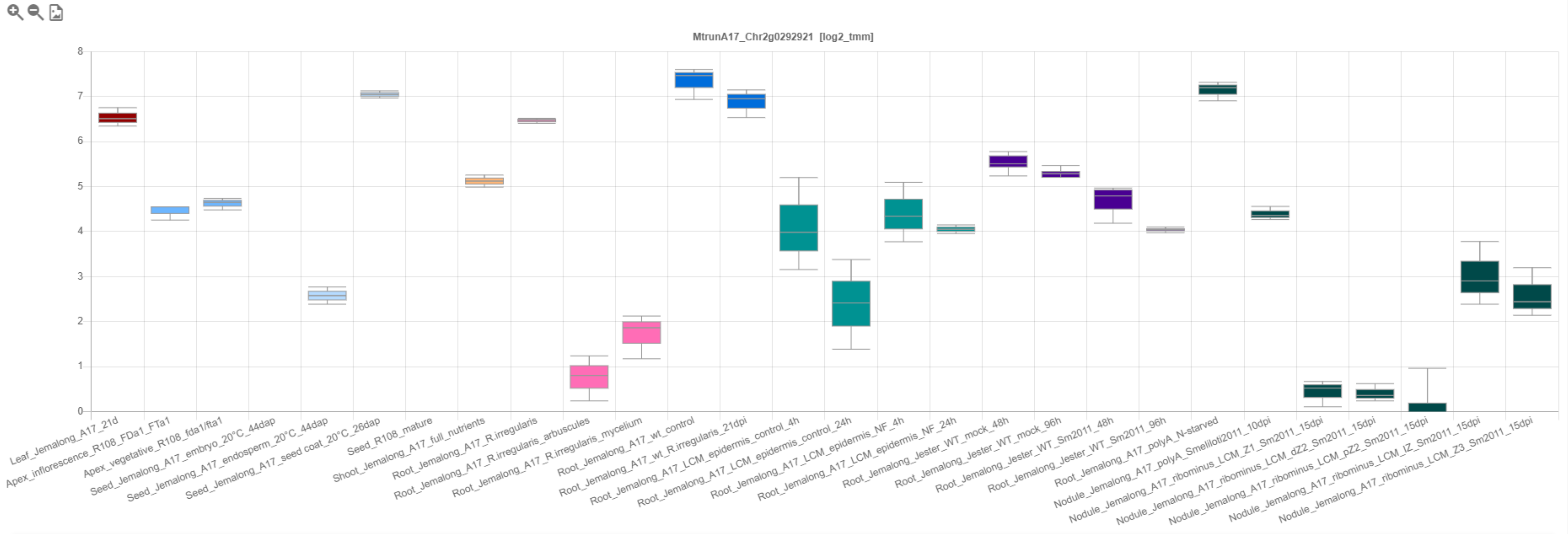

### CP36: MtrunA17_Chr2g0298731

INRAE/CNRS Medicago Expressi | Expression Atlas Server

.../pub/expressionAtlas/app/v3/aa_reference_dataset/MtrunA17_Chr2g0298731

MtExpress V3 | Core (20220901)

! Switch to another dataset using the left menu

TMM | METADATA | SYNONYMOUS | ANNOTATION | GENOME PORTAL | LEGOO

mRNA: MtrunA17_Chr2g0298731;

Log2 TMM Normalisation using EdgeR (Core [20220901])

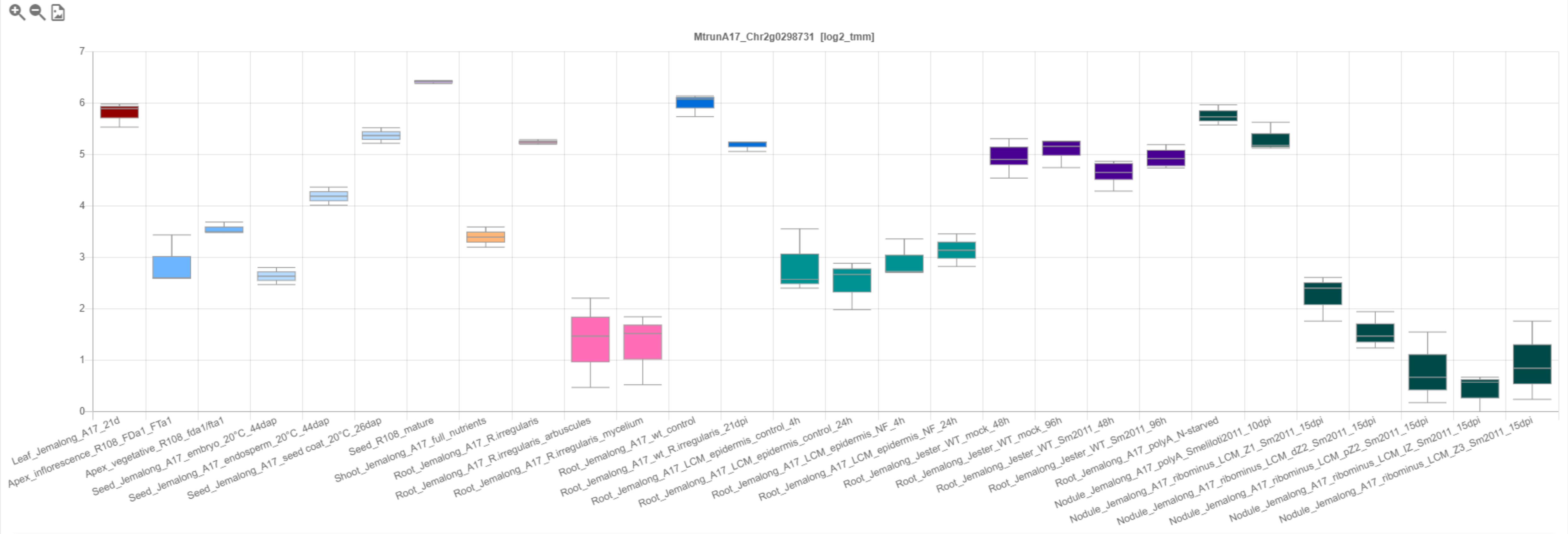

### CP37: MtrunA17_Chr2g0299561

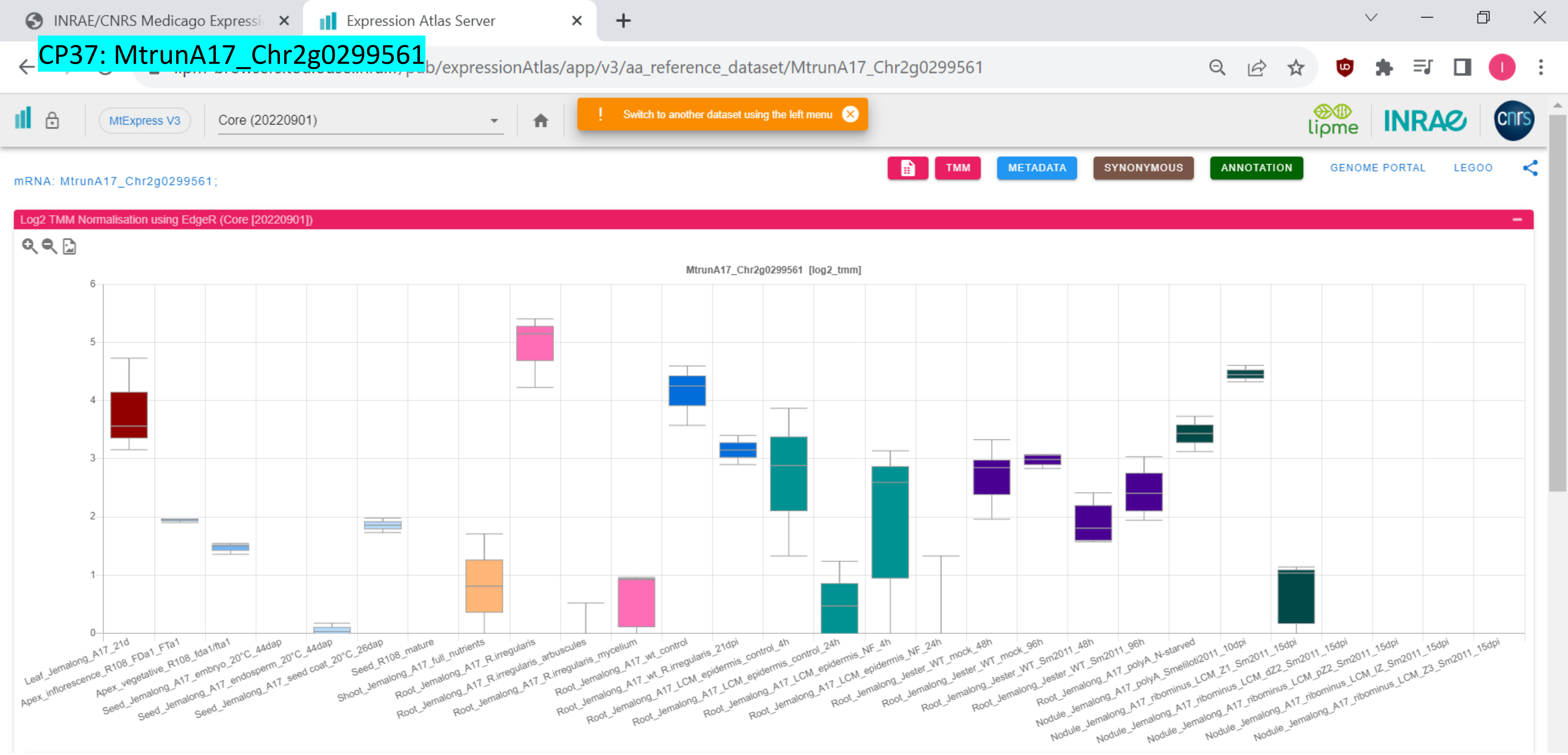

### CP38: MtrunA17_Chr2g0304891

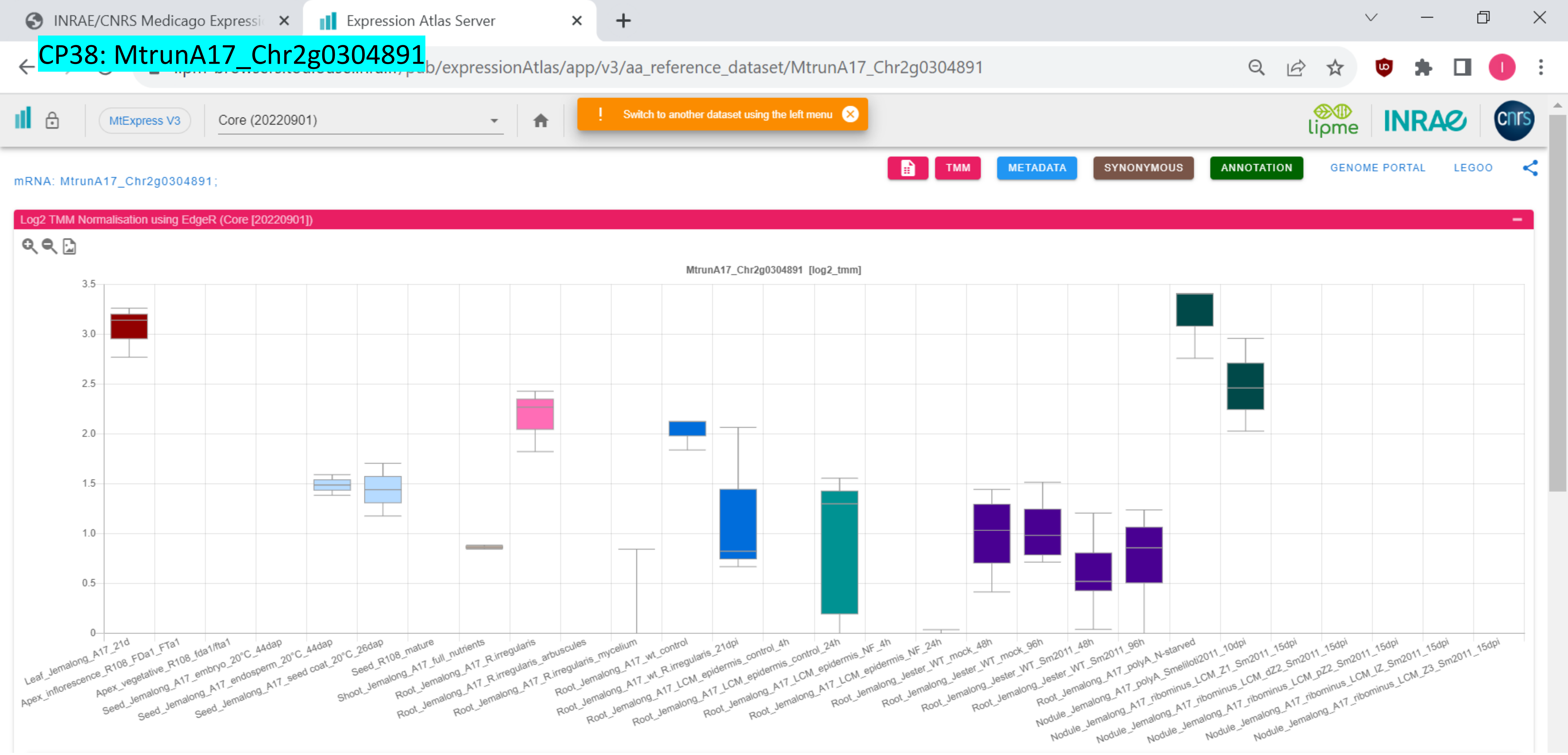

*CP39: MtrunA17_Chr2g0305951

INRAE/CNRS Medicago Expressi | Expression Atlas Server

.../expressionAtlas/app/v3/aa_reference_dataset/MtrunA17_Chr2g0305951

MtExpress V3 | Core (20220901) | ! Switch to another dataset using the left menu

lipme INRAE cnrs

TMM | METADATA | SYNONYMOUS | ANNOTATION | GENOME PORTAL | LEGOO

mRNA: MtrunA17_Chr2g0305951 ;

Log2 TMM Normalisation using EdgeR (Core [20220901])

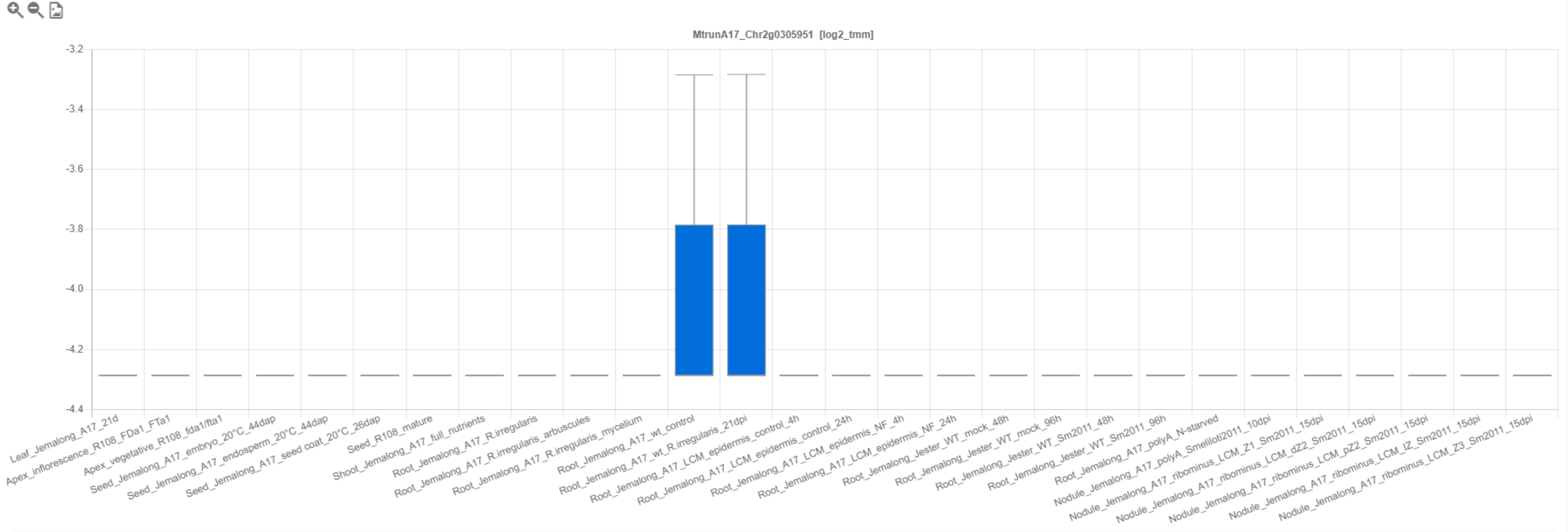

CP40: MtrunA17_Chr2g0309251

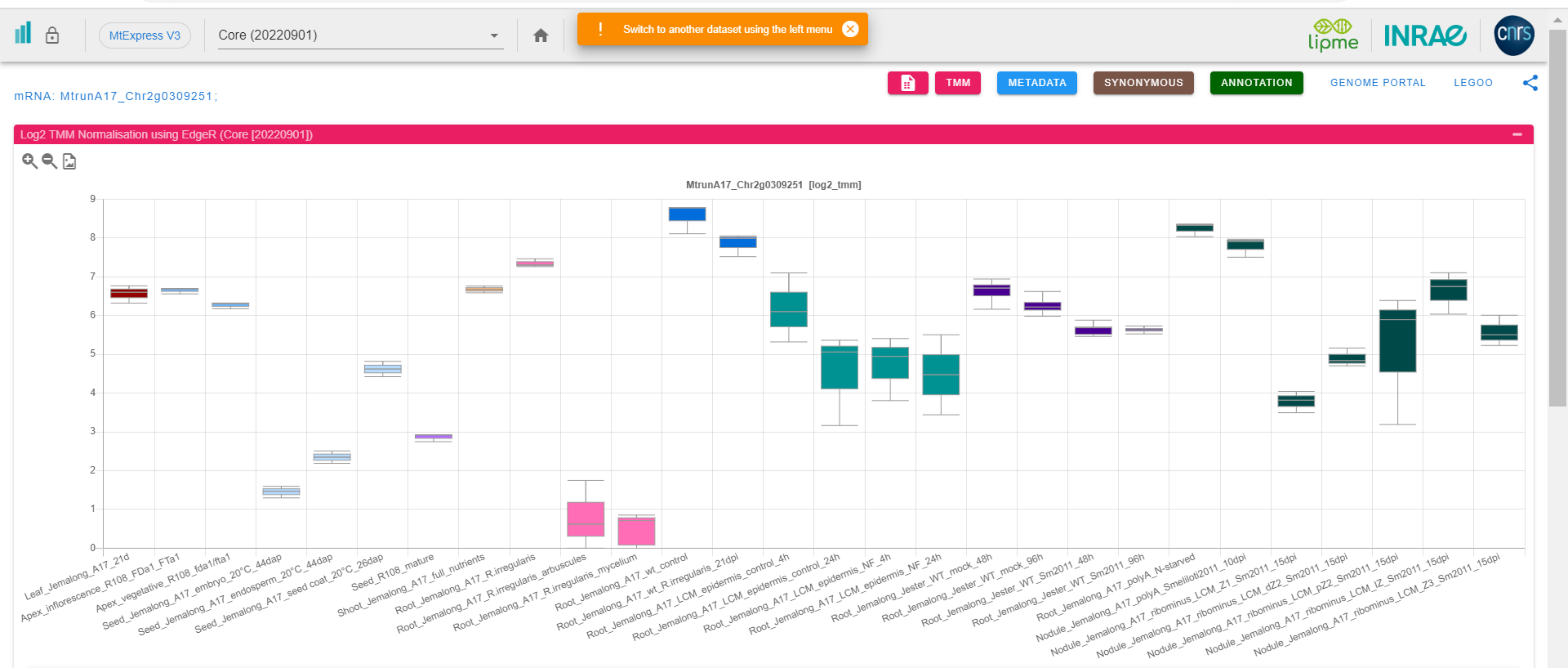

CP41: MtrunA17_Chr2g0310041

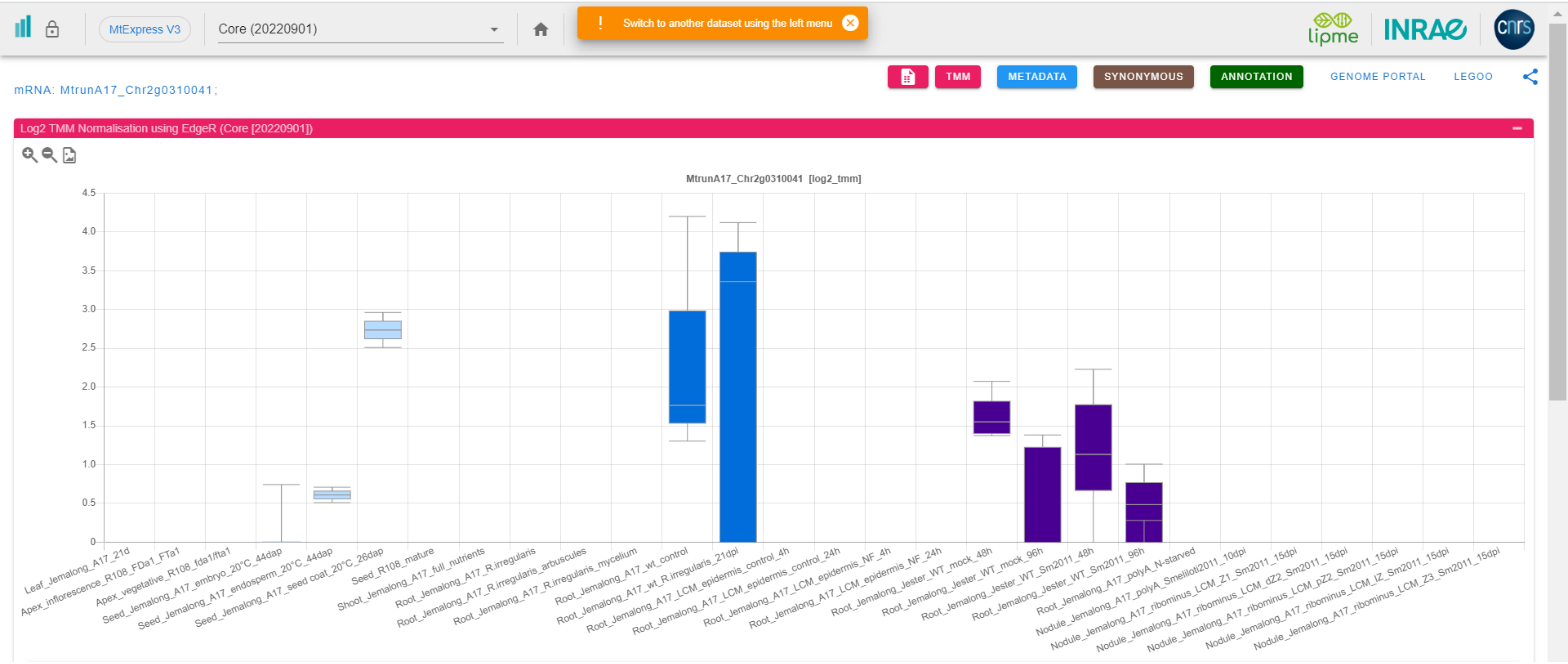

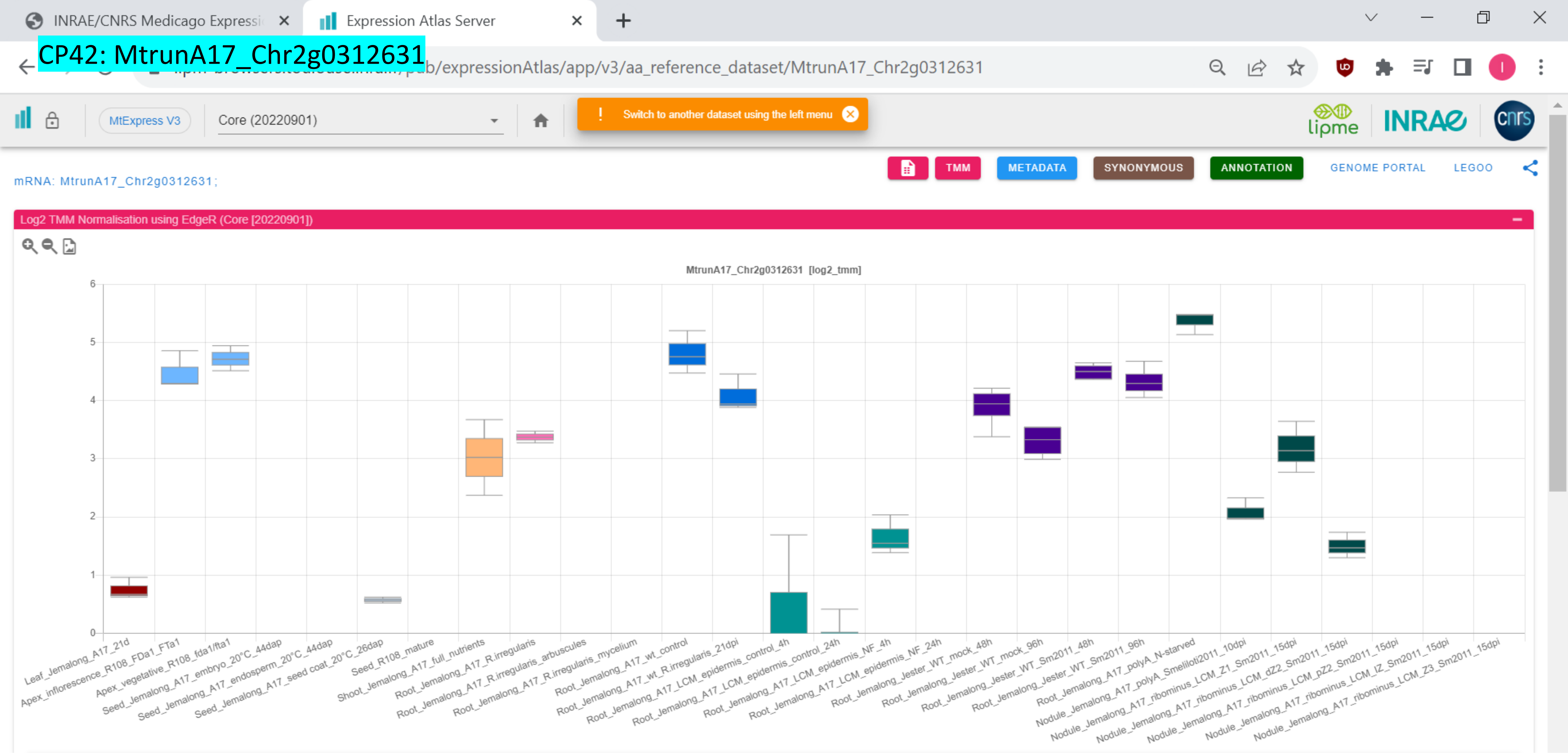
CP42: MtrunA17_Chr2g0312631
INRAE/CNRS Medicago Expressi
Expression Atlas Server
pub/expressionAtlas/app/v3/aa_reference_dataset/MtrunA17_Chr2g0312631
MtExpress V3
Core (20220901)
Switch to another dataset using the left menu
TMM
METADATA
SYNONYMOUS
ANNOTATION
GENOME PORTAL
LEGOO
mRNA: MtrunA17_Chr2g0312631;
Log2 TMM Normalisation using EdgeR (Core [20220901])
MtrunA17_Chr2g0312631 [log2_tmm]
6
5
4
3
2
1
0
Leaf_Jemalong_A17_21d
Apex_inflorescence_R108_FDa1_FTa1
Apex_vegetative_R108_fda1/fta1
Seed_Jemalong_A17_embryo_20°C_44dap
Seed_Jemalong_A17_endosperm_20°C_44dap
Seed_Jemalong_A17_seed coat_20°C_26dap
Seed_R108_mature
Shoot_Jemalong_A17_full_nutrients
Root_Jemalong_A17_R.irregularis
Root_Jemalong_A17_R.irregularis_arbuscules
Root_Jemalong_A17_R.irregularis_mycelium
Root_Jemalong_A17_wt_control
Root_Jemalong_A17_wt_R.irregularis_21dpi
Root_Jemalong_A17_LCM_epidermis_control_4h
Root_Jemalong_A17_LCM_epidermis_control_24h
Root_Jemalong_A17_LCM_epidermis_NF_4h
Root_Jemalong_A17_LCM_epidermis_NF_24h
Root_Jemalong_Jester_WT_mock_48h
Root_Jemalong_Jester_WT_mock_96h
Root_Jemalong_Jester_WT_Sm2011_48h
Root_Jemalong_Jester_WT_Sm2011_96h
Root_Jemalong_A17_polyA_N-starved
Nodule_Jemalong_A17_polyA_Smeliloti2011_10dpi
Nodule_Jemalong_A17_ribominus_LCM_Z1_Sm2011_15dpi
Nodule_Jemalong_A17_ribominus_LCM_dZ2_Sm2011_15dpi
Nodule_Jemalong_A17_ribominus_LCM_pZ2_Sm2011_15dpi
Nodule_Jemalong_A17_ribominus_LCM_IZ_Sm2011_15dpi
Nodule_Jemalong_A17_ribominus_LCM_Z3_Sm2011_15dpi

CP43: MtrunA17_Chr2g0316291

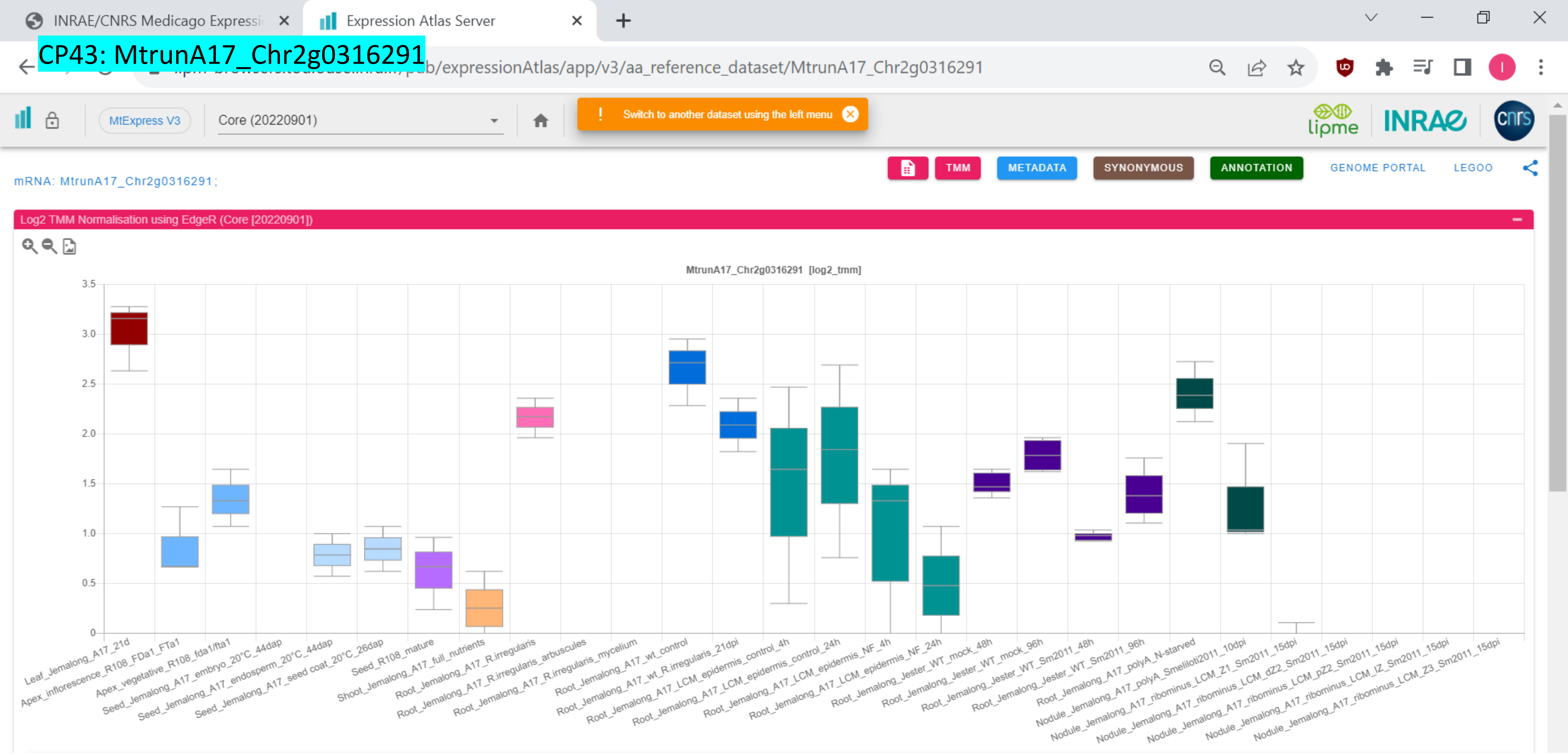

### CP44: MtrunA17_Chr2g0326801

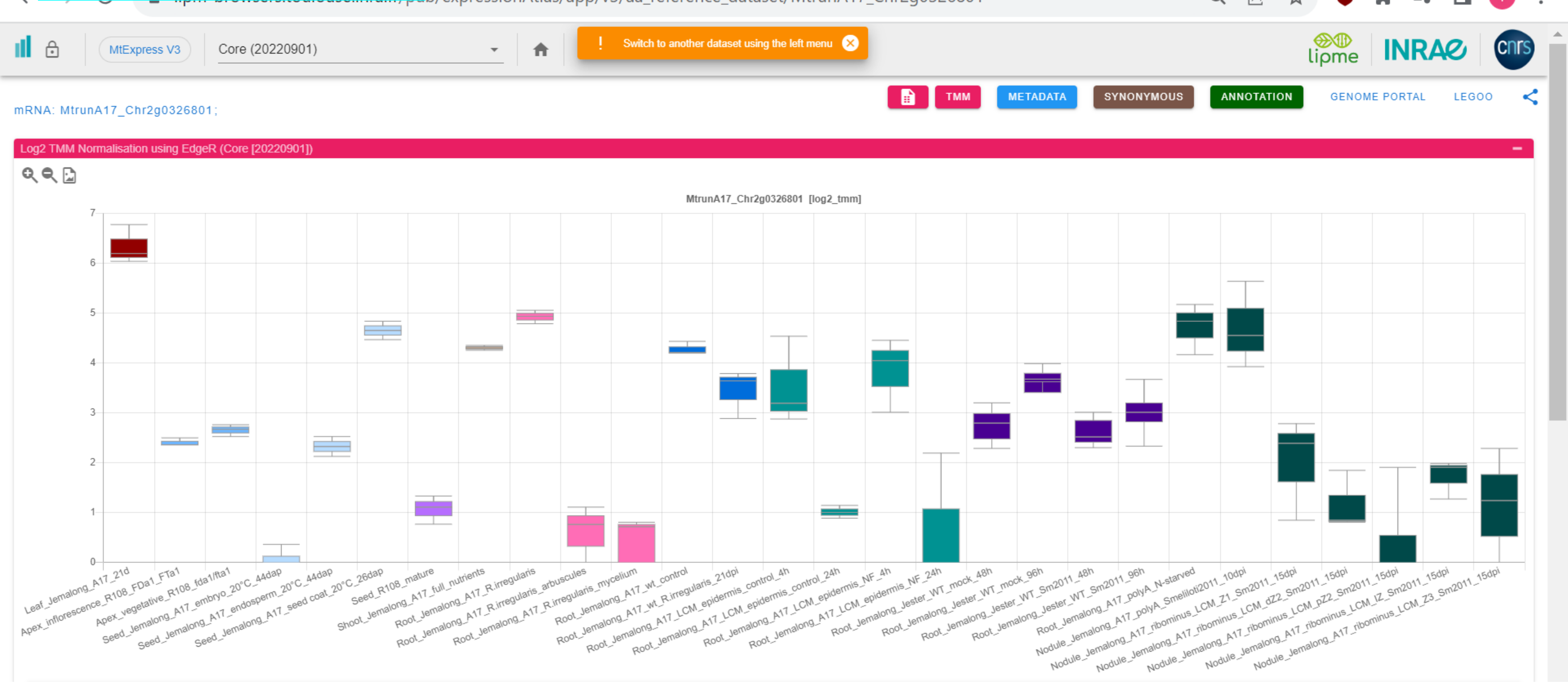

CP45: MtrunA17_Chr2g0328091

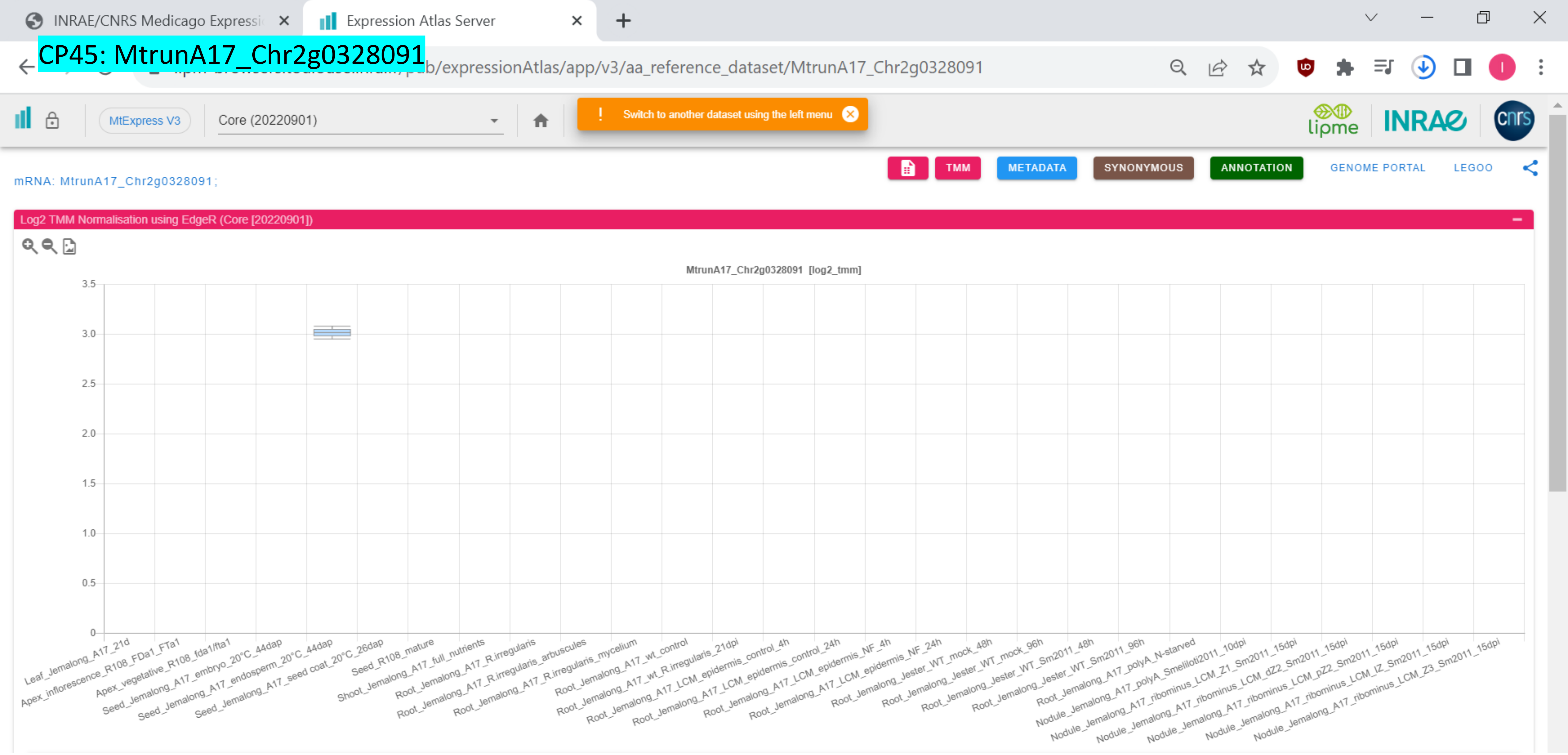

### CP46: MtrunA17_Chr2g0329031

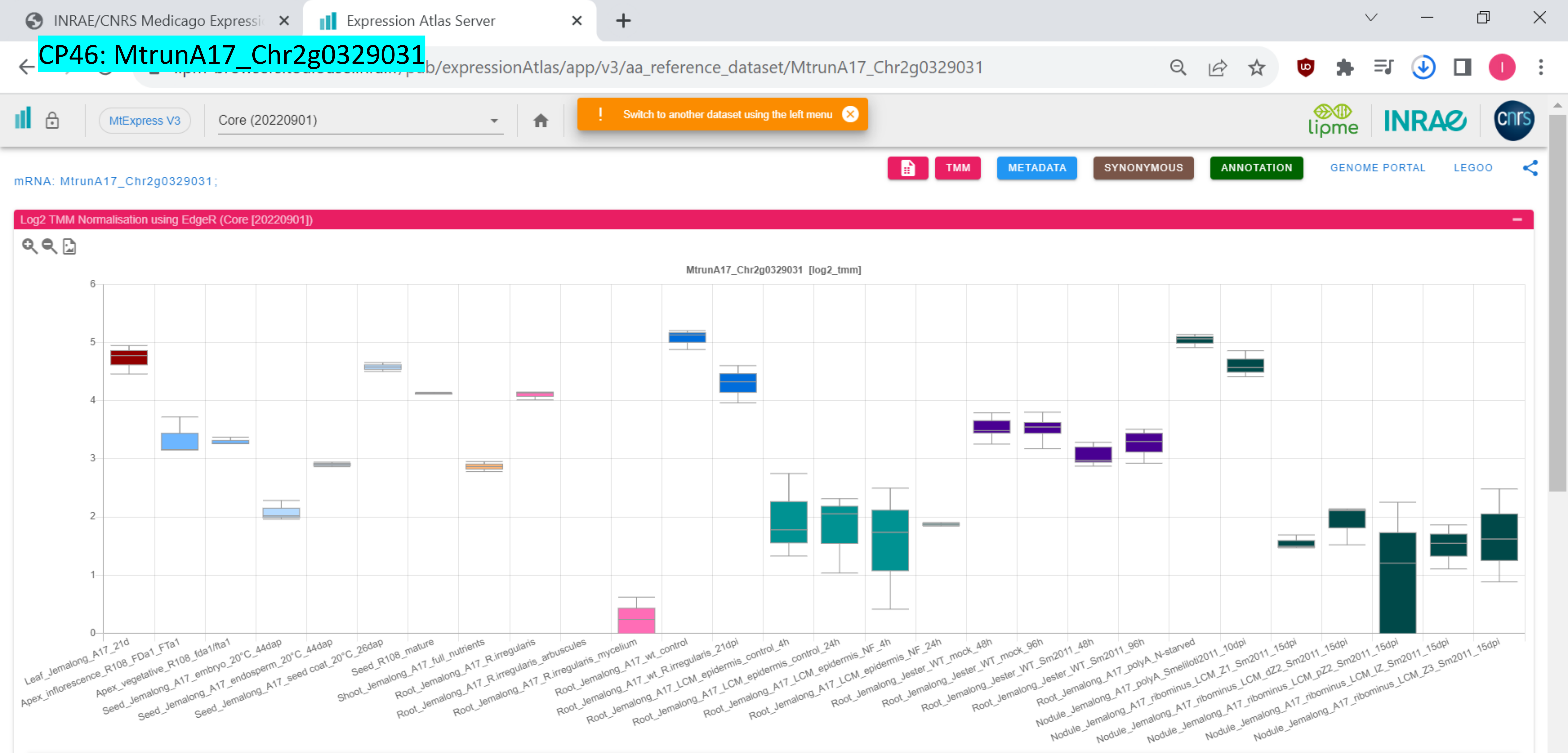

CP47: MtrunA17_Chr3g0079521

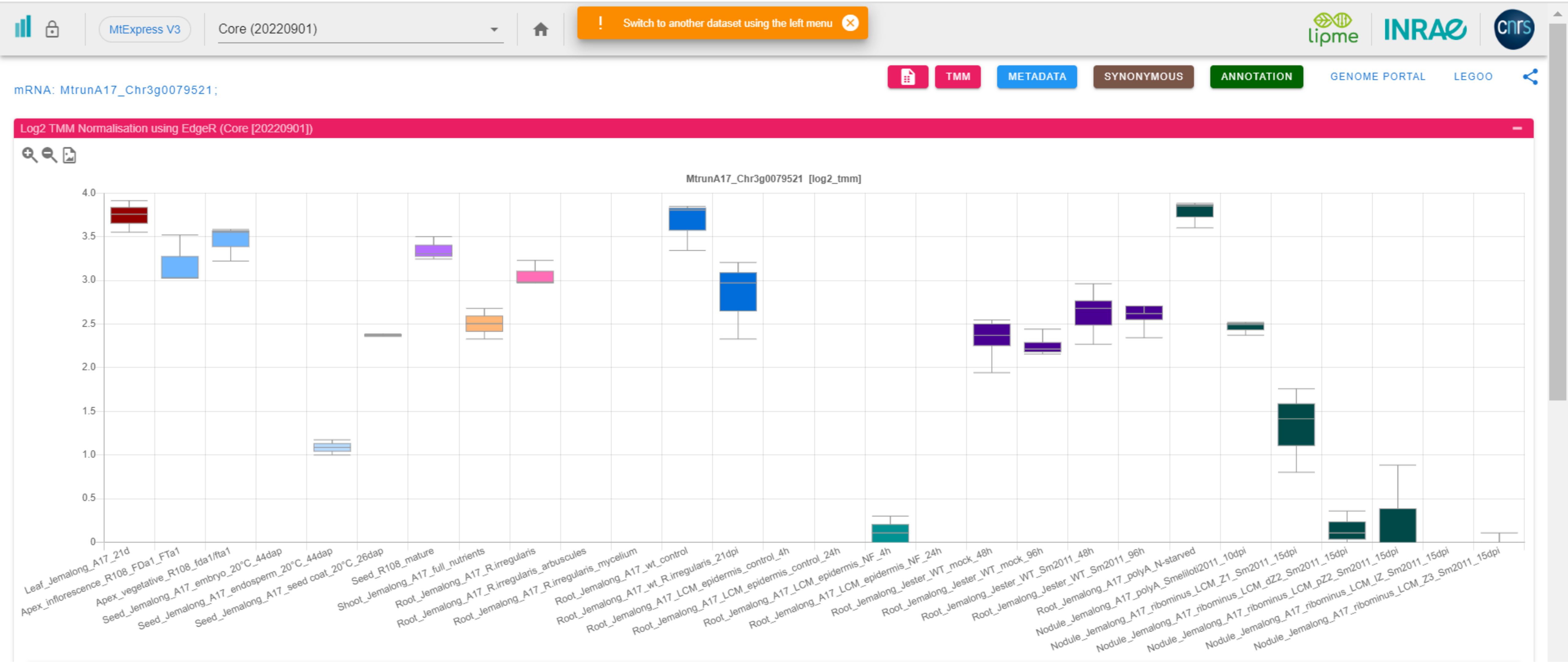

CP48: MtrunA17_Chr3g0083801

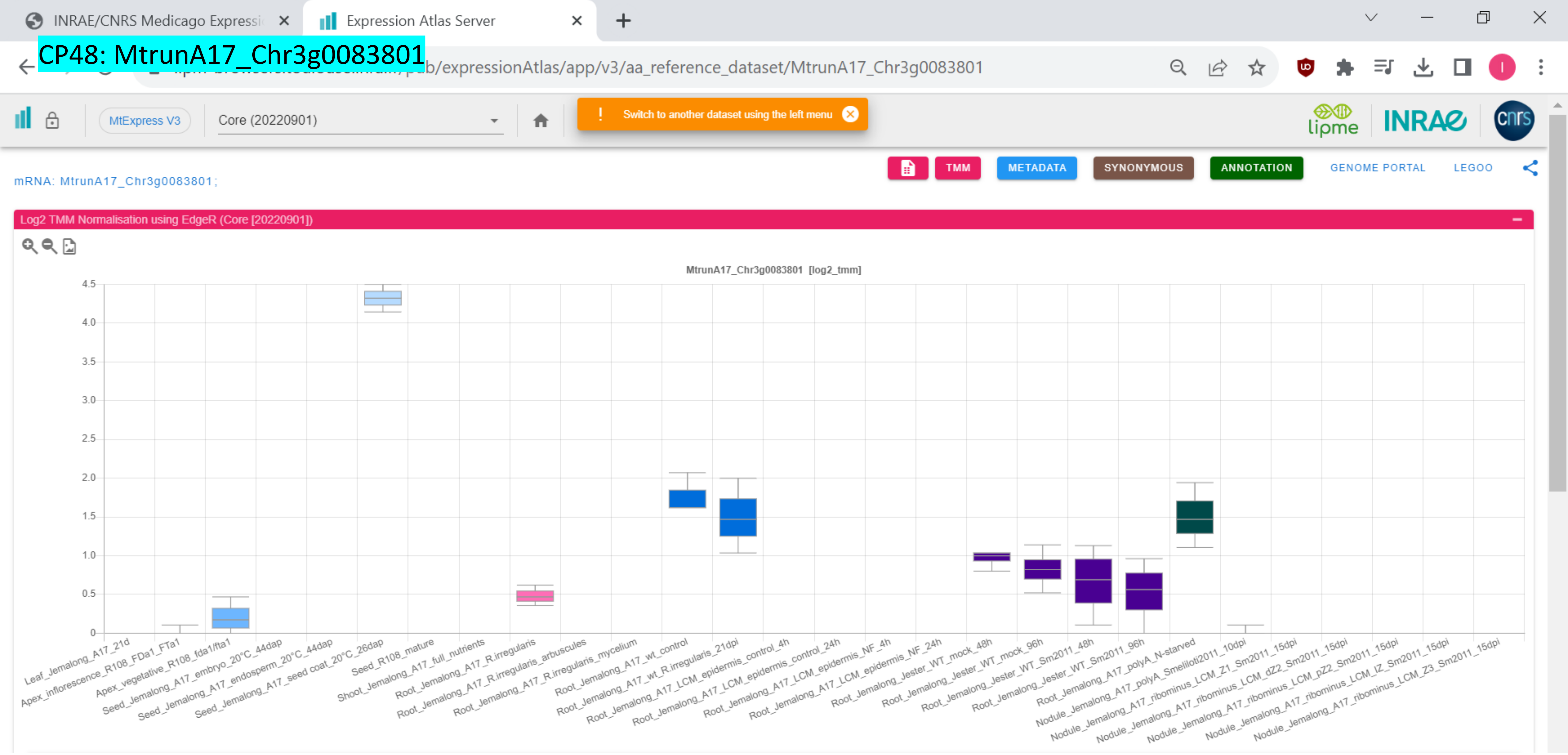

CP49: MtrunA17_Chr3g0091141

INRAE/CNRS Medicago Expressi | Expression Atlas Server

pub/expressionAtlas/app/v3/aa_reference_dataset/MtrunA17_Chr3g0091141

MtExpress V3 | Core (20220901)

! Switch to another dataset using the left menu

lipme INRAE cnrs

TMM | METADATA | SYNONYMOUS | ANNOTATION | GENOME PORTAL | LEGOO

mRNA: MtrunA17_Chr3g0091141; *MtCKX6*

Log2 TMM Normalisation using EdgeR (Core [20220901])

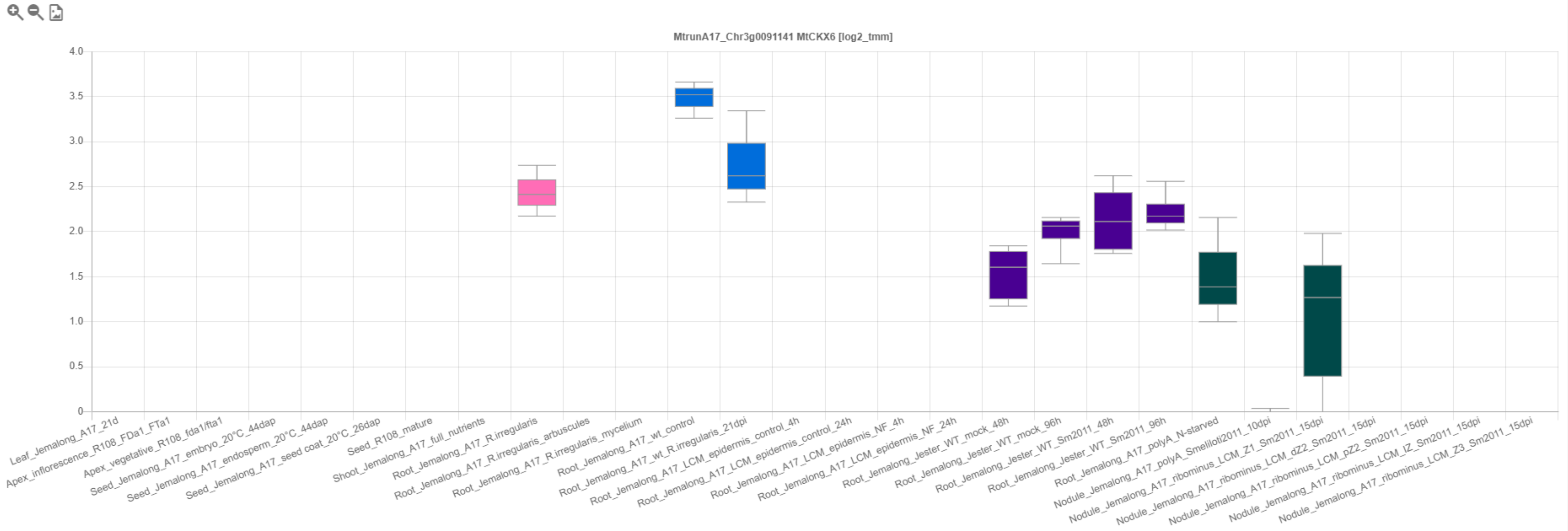

*CP50: MtrunA17_Chr3g0091671

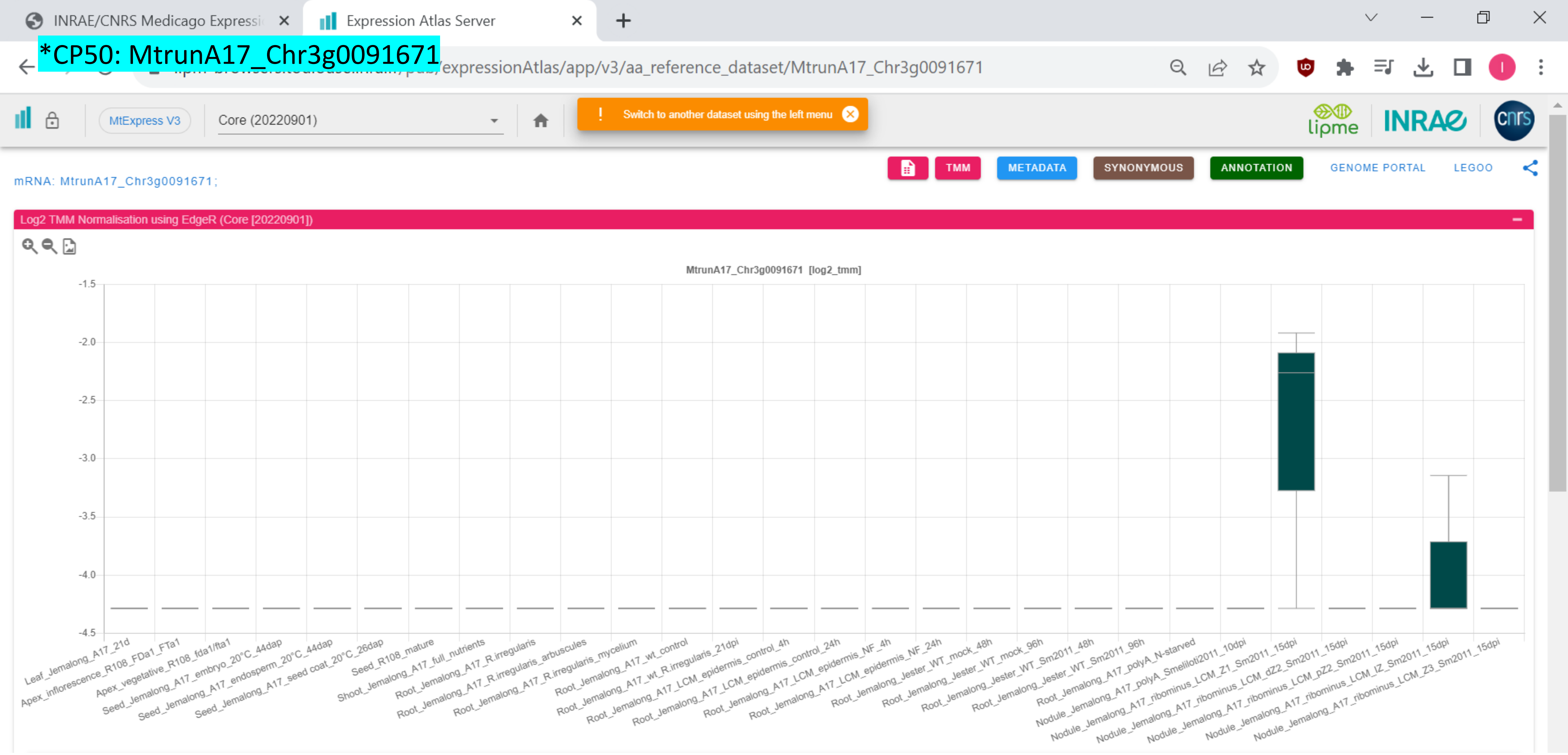

### CP51: MtrunA17_Chr3g0096421

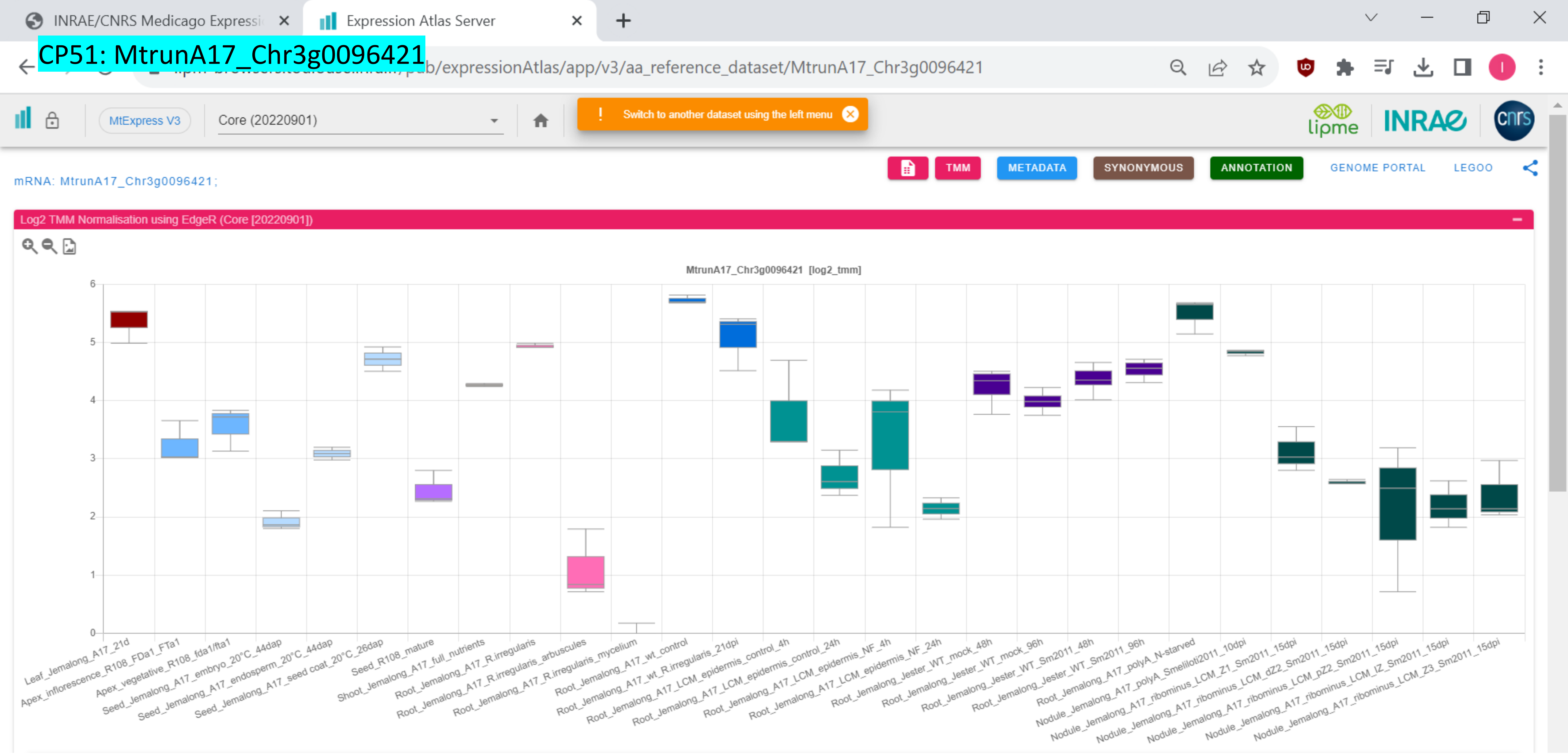

*CP52: MtrunA17_Chr3g0100221

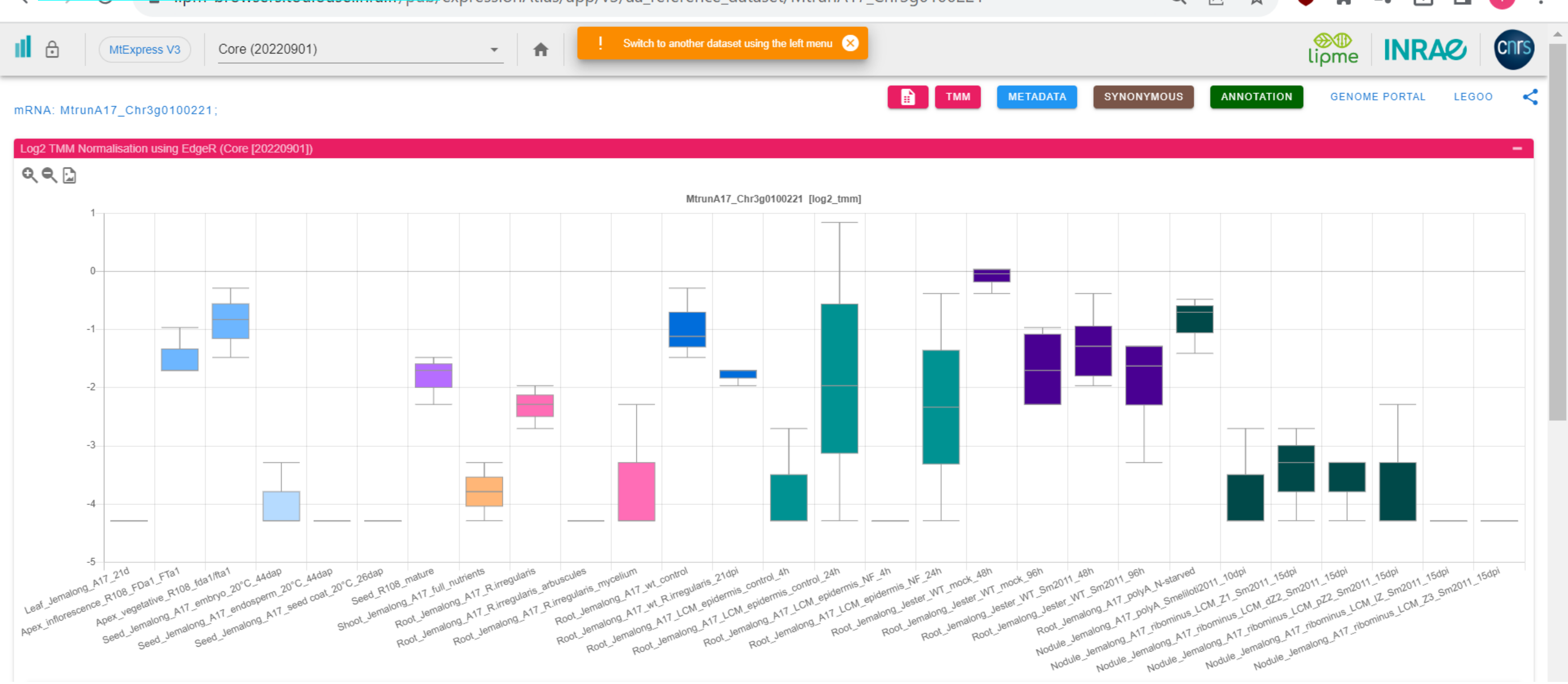

### CP53: MtrunA17_Chr3g0102171

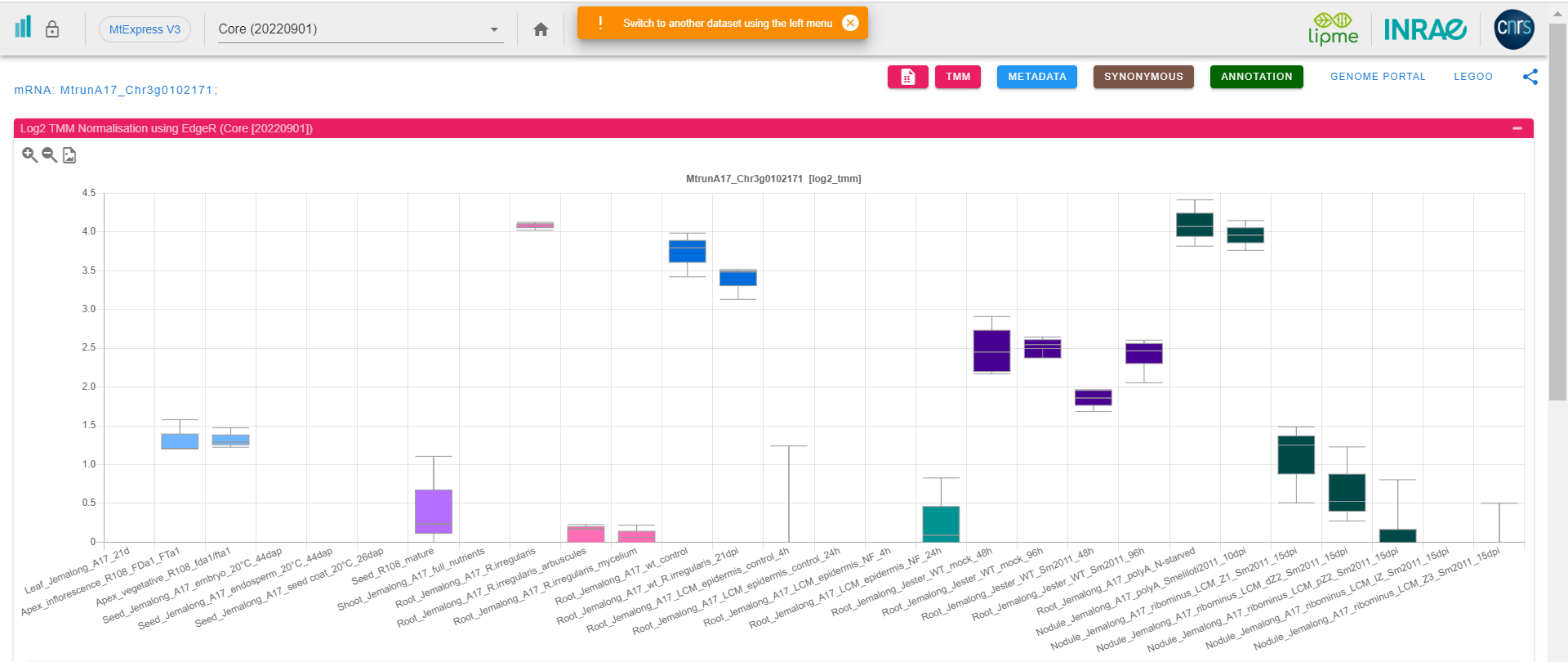

### CP54: MtrunA17_Chr3g0105981

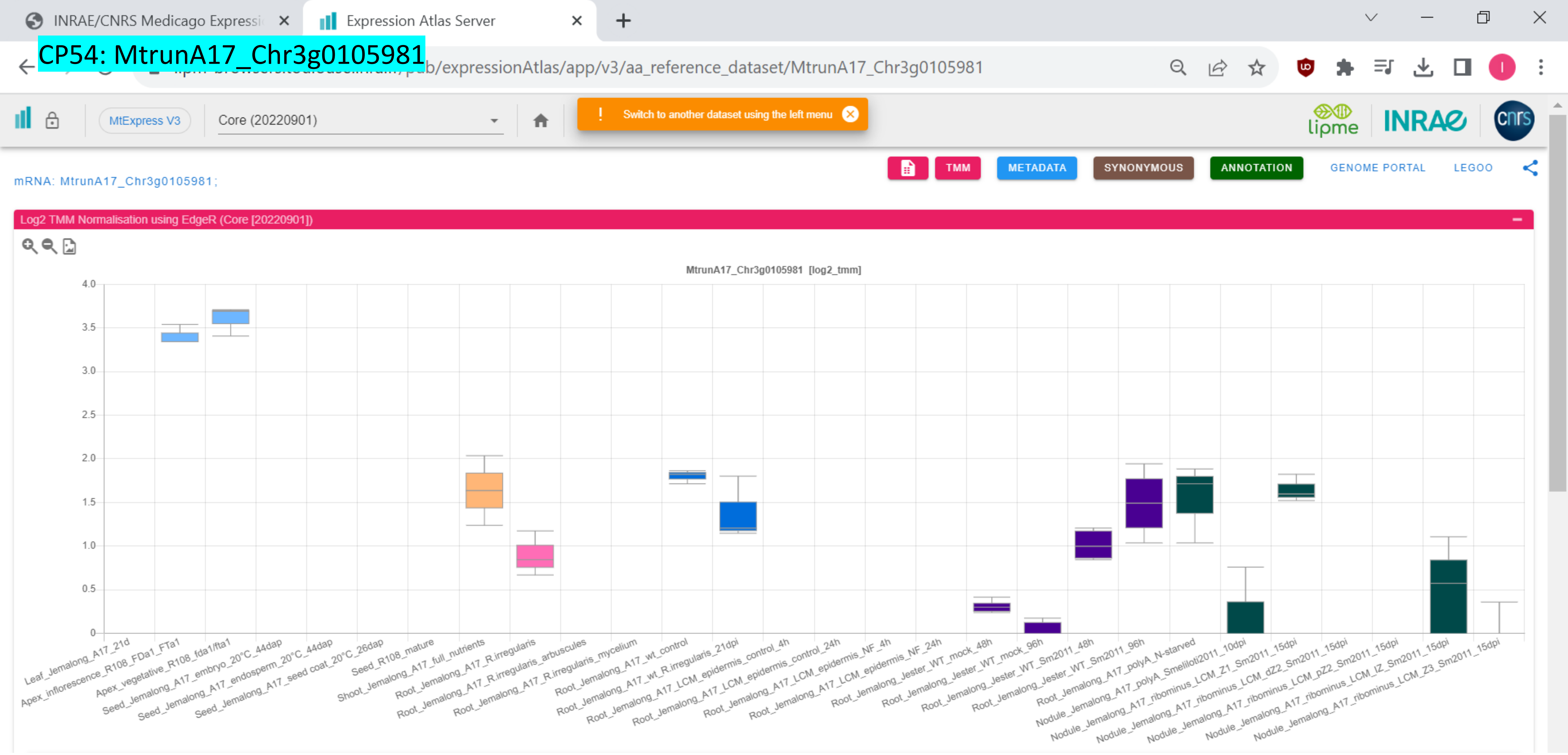

### CP55: MtrunA17_Chr3g0110451

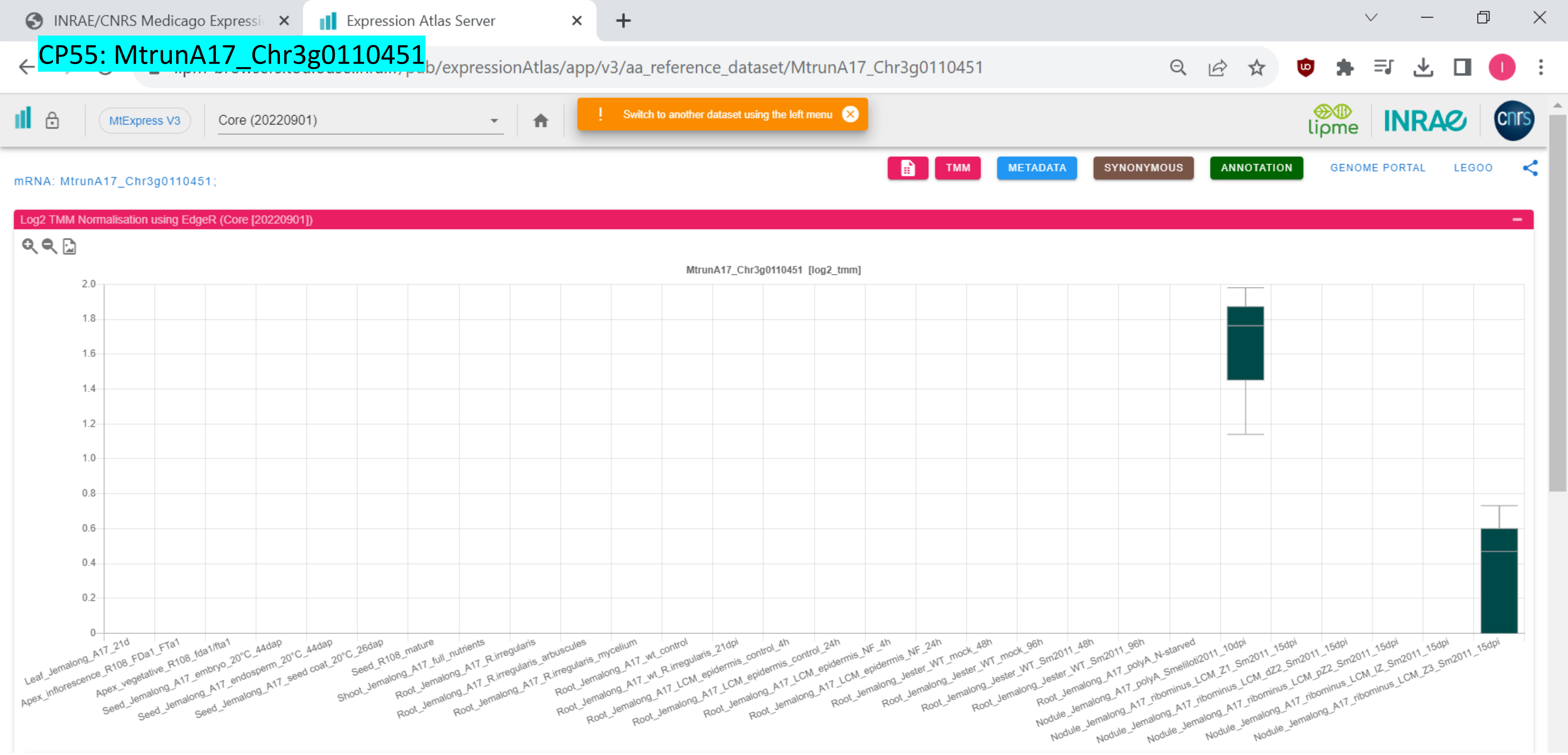

CP56: MtrunA17_Chr3g0113591

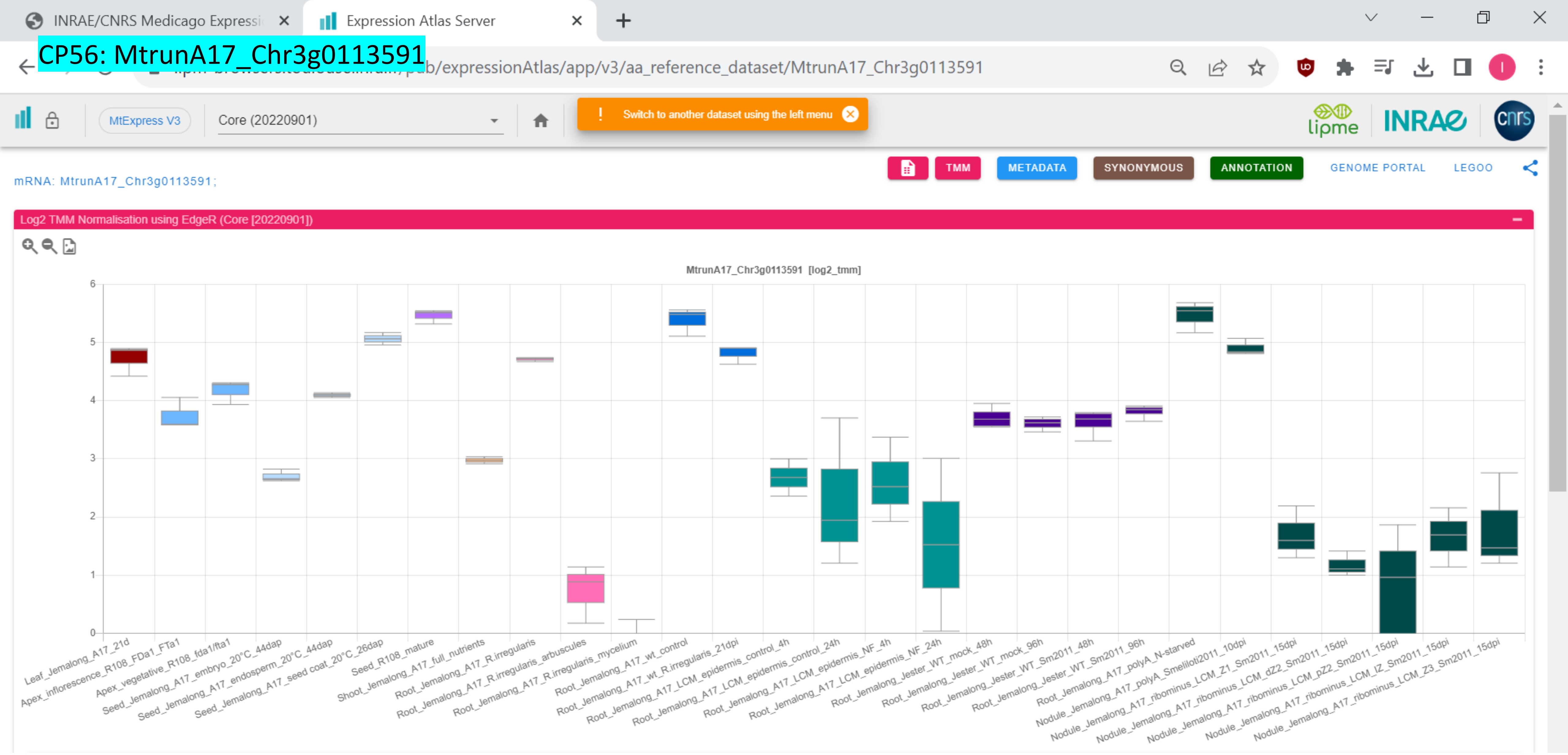

### CP57: MtrunA17_Chr3g0124631

INRAE/CNRS Medicago Expressi | Expression Atlas Server

[illegible]b/expressionAtlas/app/v3/aa_reference_dataset/MtrunA17_Chr3g0124631

MtExpress V3 | Core (20220901)

! Switch to another dataset using the left menu

lipme INRAE cnrs

TMM | METADATA | SYNONYMOUS | ANNOTATION | GENOME PORTAL | LEGOO

mRNA: MtrunA17_Chr3g0124631;

Log2 TMM Normalisation using EdgeR (Core [20220901])

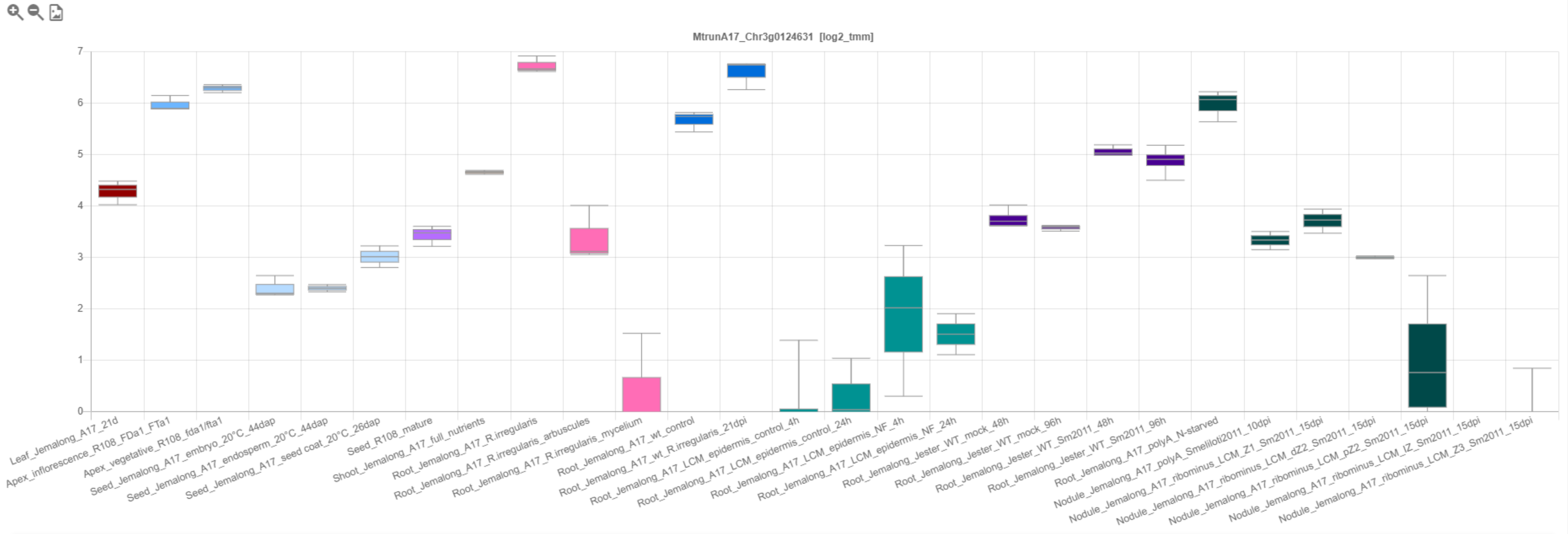

### *CP58: MtrunA17_Chr3g0127321

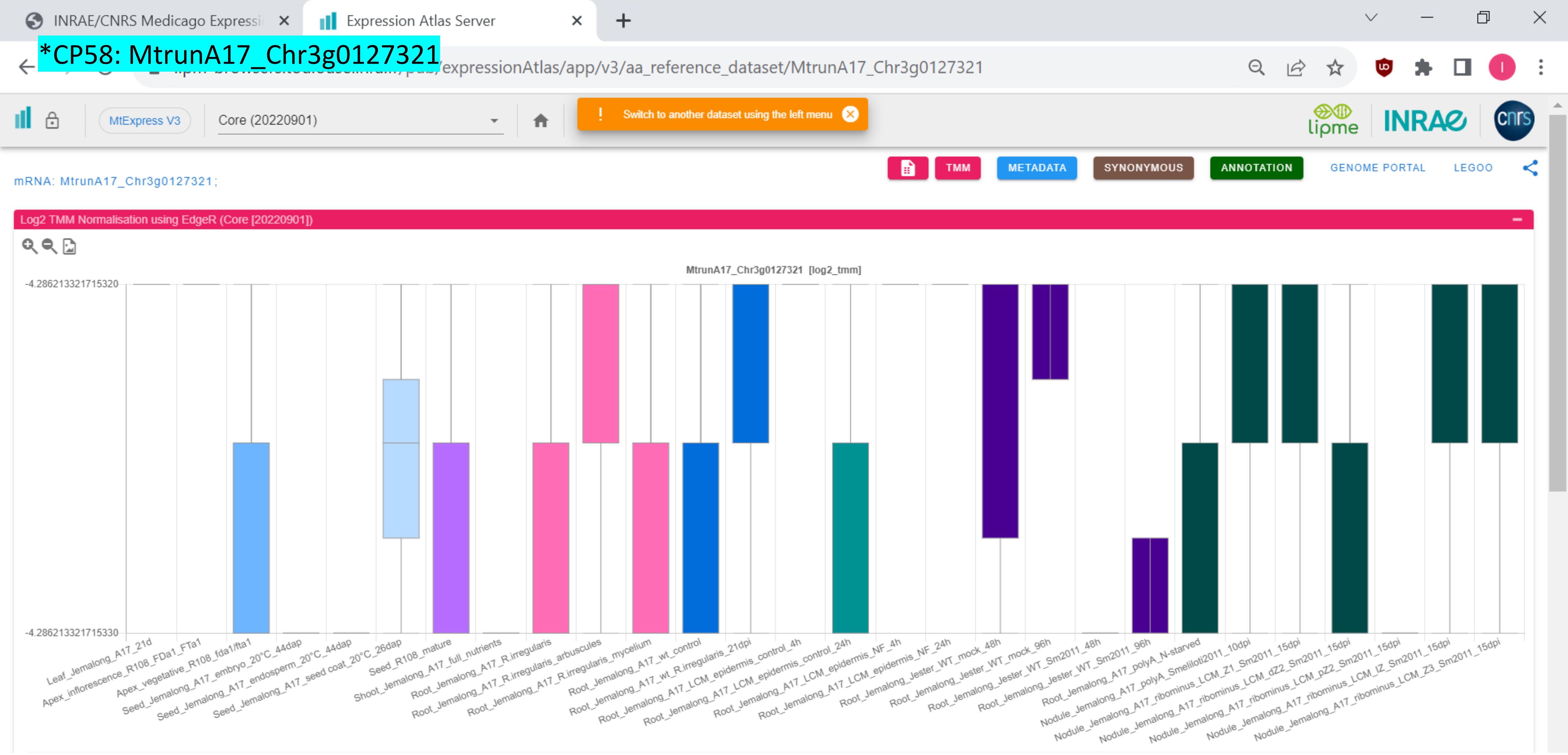

### CP59: MtrunA17_Chr3g0130671

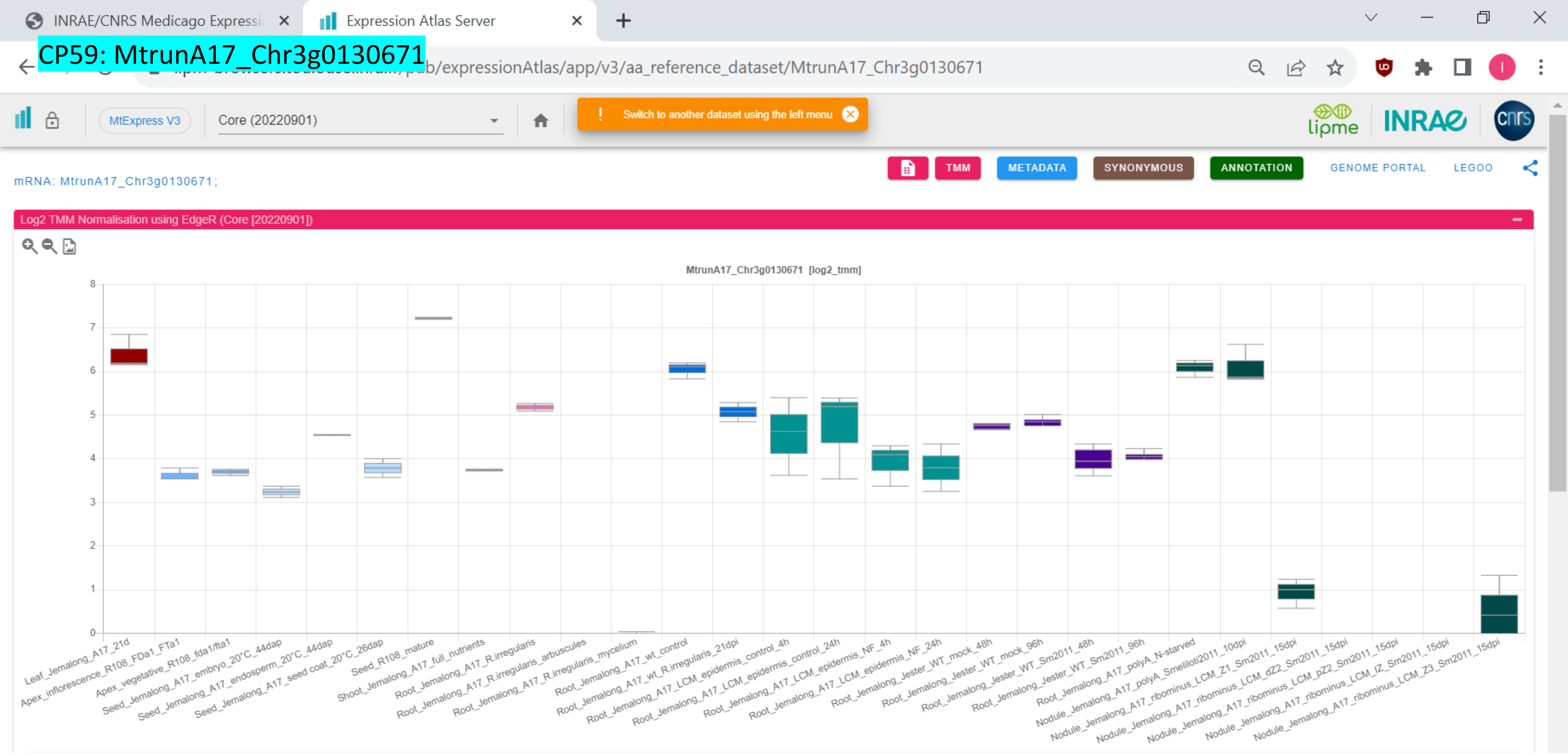

*CP60: MtrunA17_Chr3g0135761

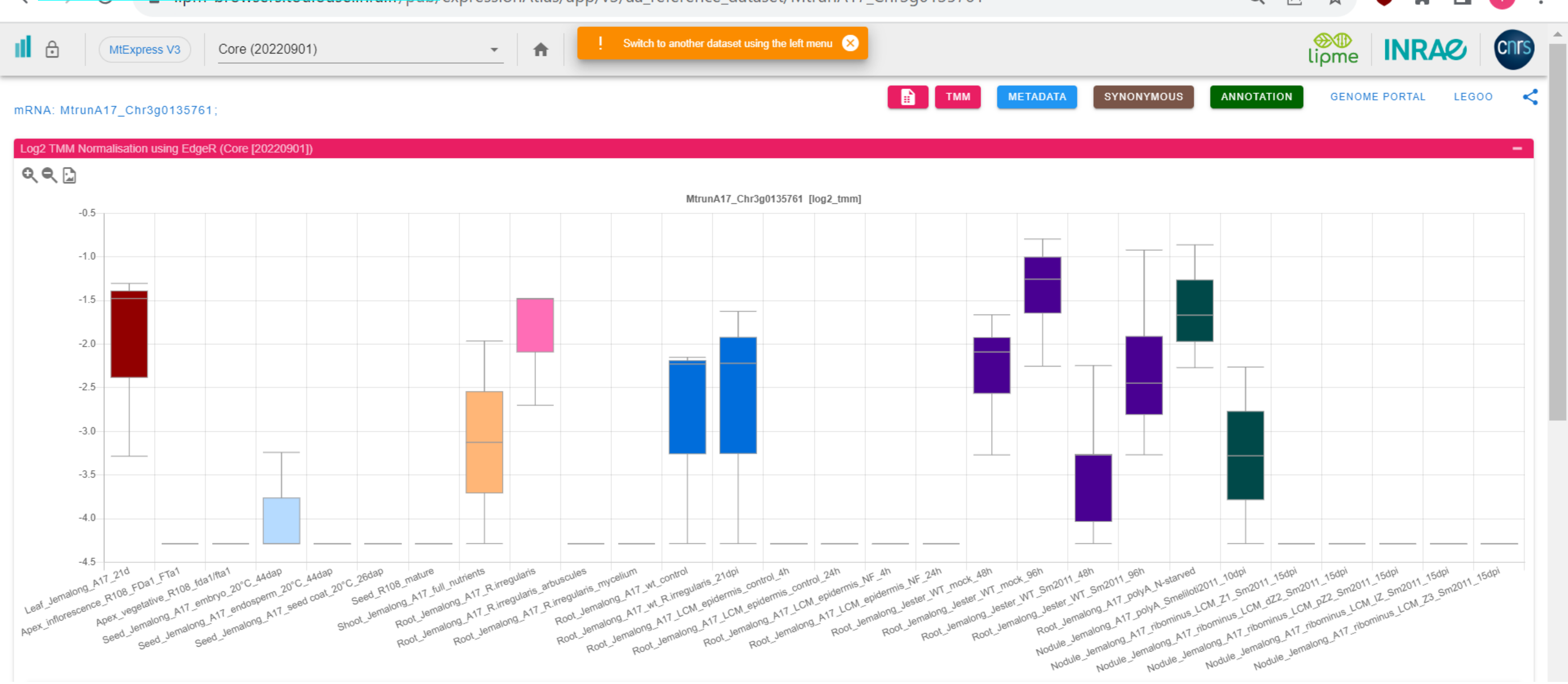

CP61: MtrunA17_Chr3g0137391

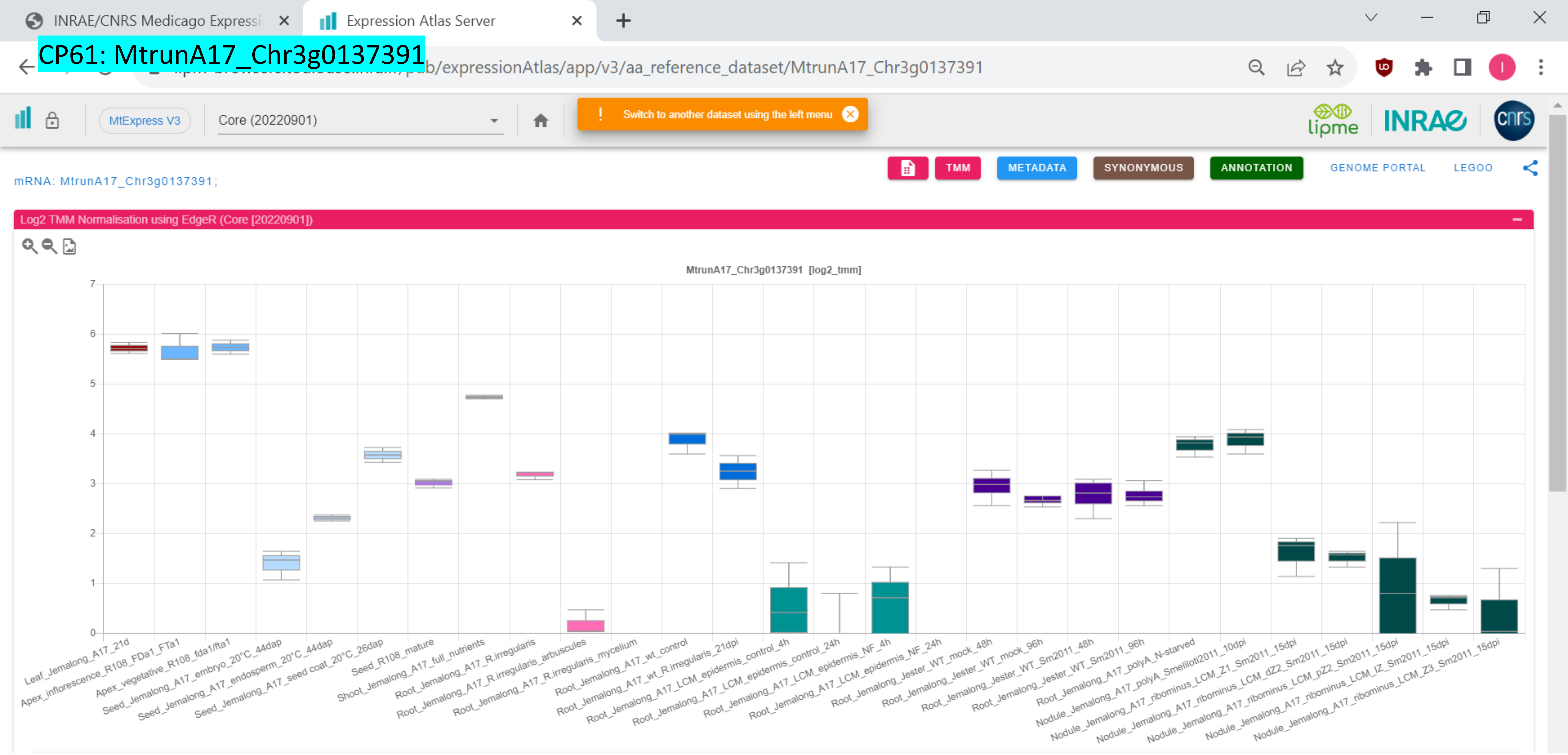

CP62: MtrunA17_Chr3g0141901

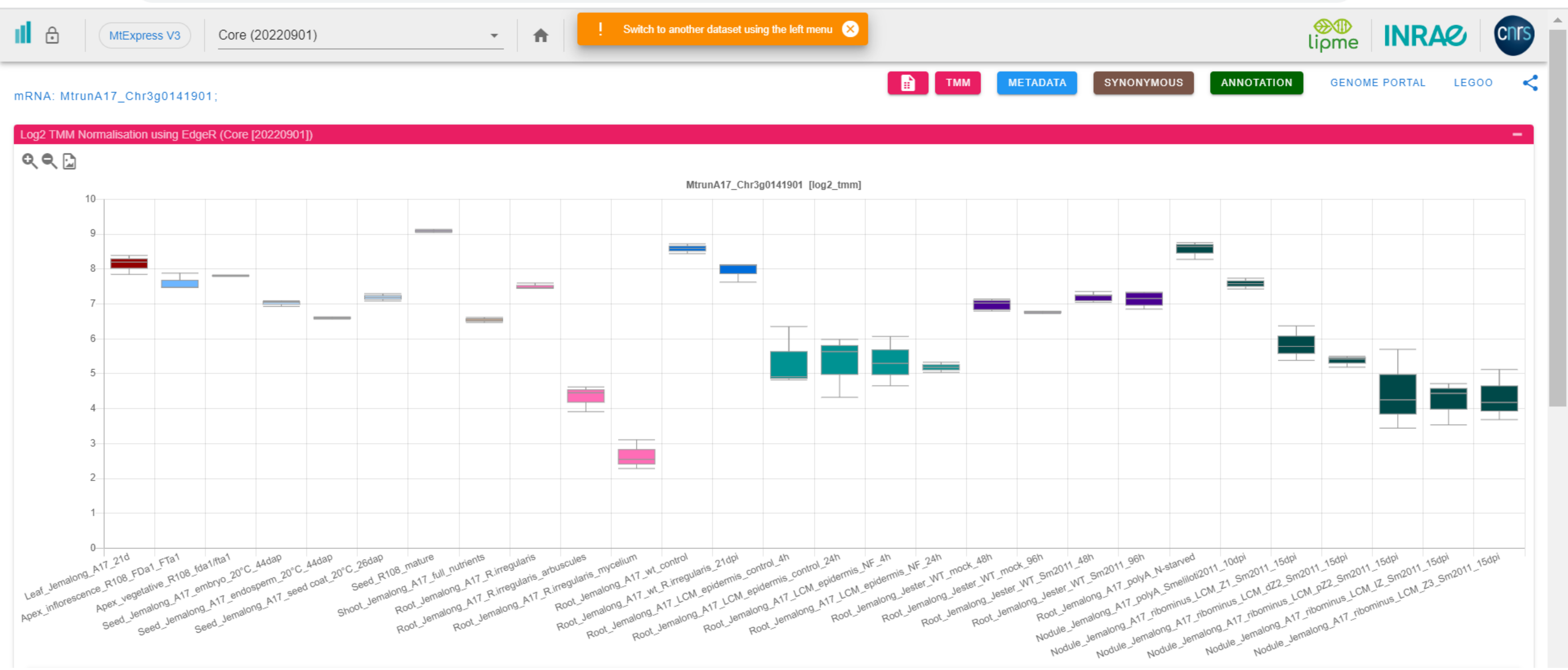

### CP63: MtrunA17_Chr3g0144151

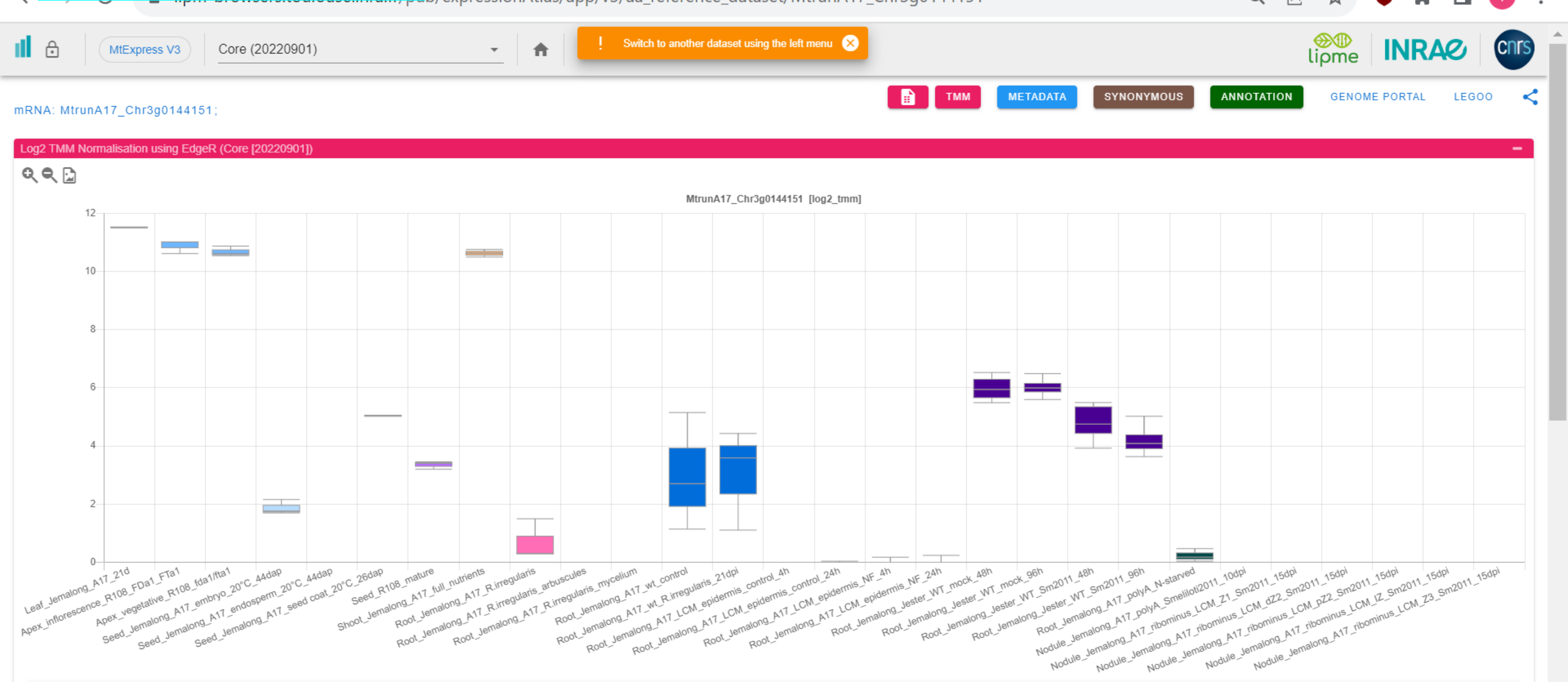

CP64: MtrunA17_Chr3g0144151

Expression Atlas Server

.../pub/expressionAtlas/app/v3/aa_reference_dataset/MtrunA17_Chr3g0144151

MtExpress V3 | Core (20220901)

Switch to another dataset using the left menu

TMM | METADATA | SYNONYMOUS | ANNOTATION | GENOME PORTAL | LEGOO

mRNA: MtrunA17_Chr3g0144151;

Log2 TMM Normalisation using EdgeR (Core [20220901])

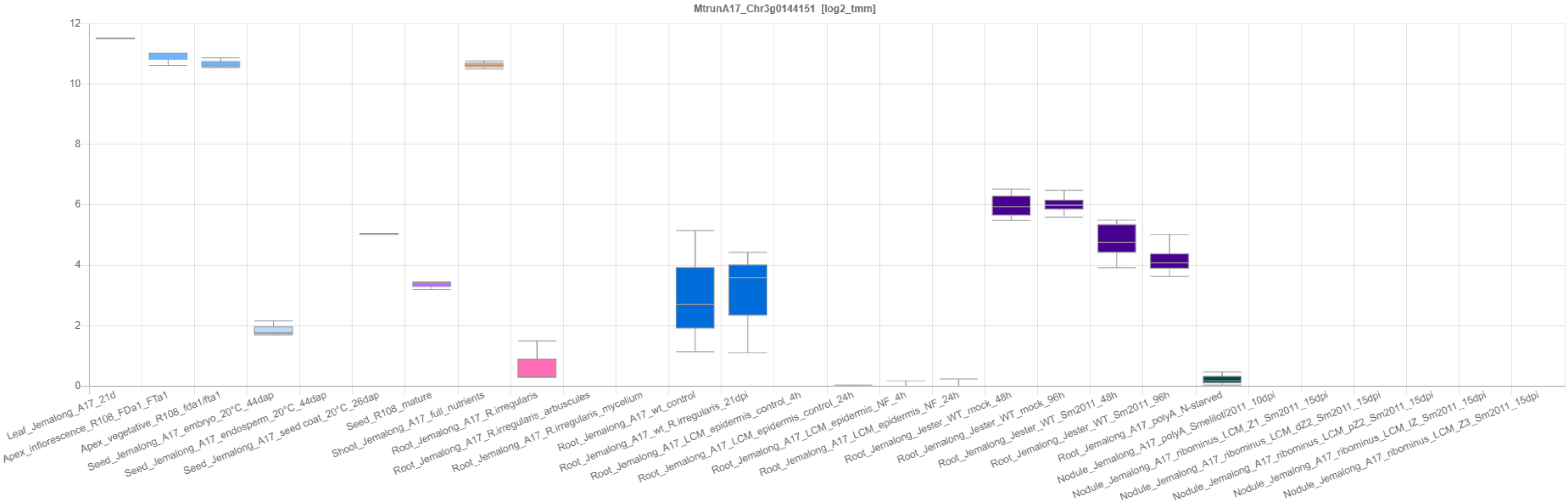

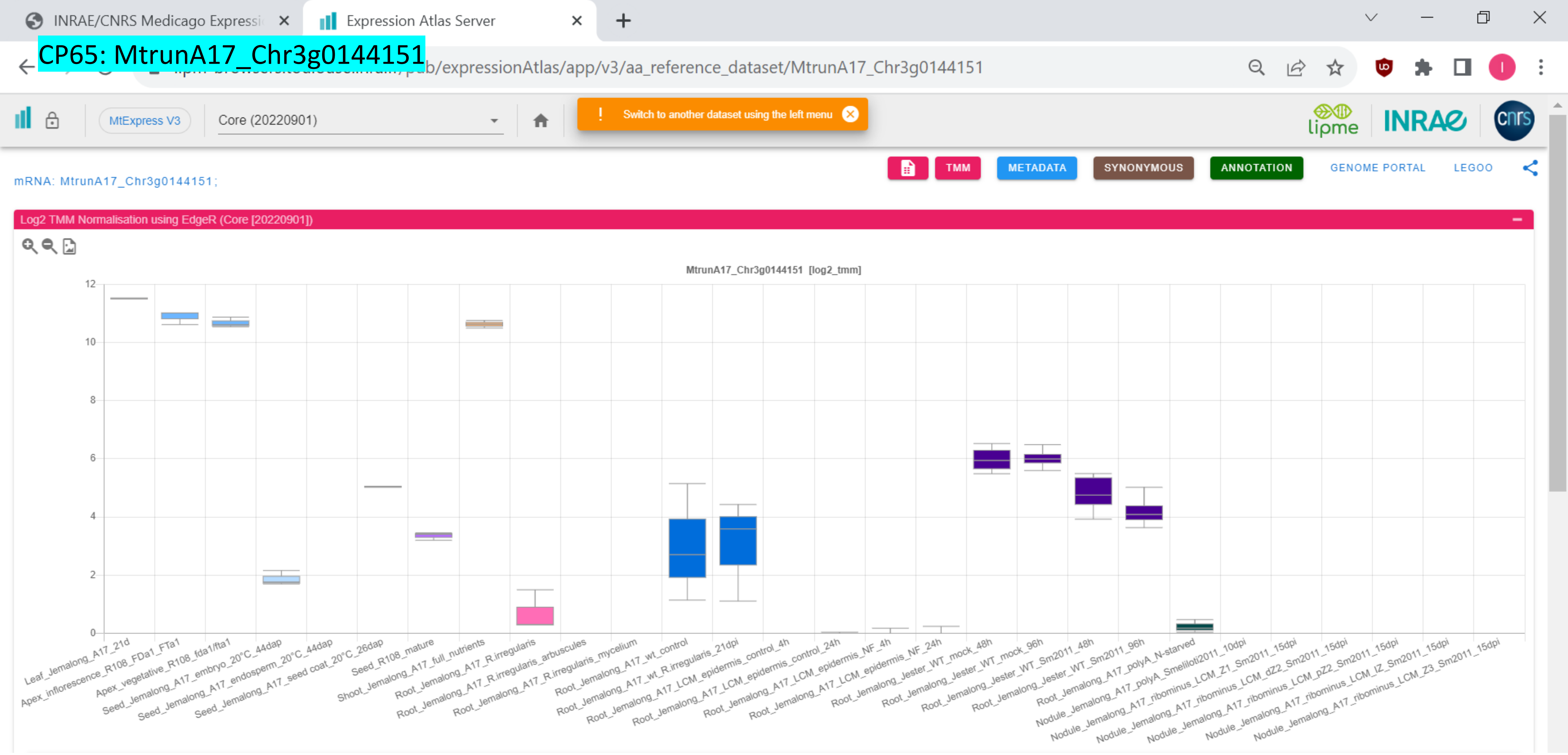

CP65: MtrunA17_Chr3g0144151
INRAE/CNRS Medicago Expressi
Expression Atlas Server
pub/expressionAtlas/app/v3/aa_reference_dataset/MtrunA17_Chr3g0144151
MtExpress V3
Core (20220901)
Switch to another dataset using the left menu
lipme
INRAE
cnrs
TMM
METADATA
SYNONYMOUS
ANNOTATION
GENOME PORTAL
LEGOO
mRNA: MtrunA17_Chr3g0144151;
Log2 TMM Normalisation using EdgeR (Core [20220901])
MtrunA17_Chr3g0144151 [log2_tmm]
12
10
8
6
4
2
0
Leaf_Jemalong_A17_21d
Apex_inflorescence_R108_FDa1_FTa1
Apex_vegetative_R108_fda1/fta1
Seed_Jemalong_A17_embryo_20°C_44dap
Seed_Jemalong_A17_endosperm_20°C_44dap
Seed_Jemalong_A17_seed coat_20°C_26dap
Seed_R108_mature
Shoot_Jemalong_A17_full_nutrients
Root_Jemalong_A17_full_nutrients
Root_Jemalong_A17_R.irregularis
Root_Jemalong_A17_R.irregularis_arbuscules
Root_Jemalong_A17_R.irregularis_mycelium
Root_Jemalong_A17_wt_control
Root_Jemalong_A17_wt_R.irregularis_21dpi
Root_Jemalong_A17_LCM_epidermis_control_4h
Root_Jemalong_A17_LCM_epidermis_control_24h
Root_Jemalong_A17_LCM_epidermis_NF_4h
Root_Jemalong_A17_LCM_epidermis_NF_24h
Root_Jemalong_Jester_WT_mock_48h
Root_Jemalong_Jester_WT_mock_96h
Root_Jemalong_Jester_WT_Sm2011_48h
Root_Jemalong_Jester_WT_Sm2011_96h
Root_Jemalong_A17_polyA_N-starved
Nodule_Jemalong_A17_polyA_Smeliloti2011_10dpi
Nodule_Jemalong_A17_ribominus_LCM_Z1_Sm2011_15dpi
Nodule_Jemalong_A17_ribominus_LCM_dZ2_Sm2011_15dpi
Nodule_Jemalong_A17_ribominus_LCM_pZ2_Sm2011_15dpi
Nodule_Jemalong_A17_ribominus_LCM_IZ_Sm2011_15dpi
Nodule_Jemalong_A17_ribominus_LCM_Z3_Sm2011_15dpi

*CP66: MtrunA17_Chr3g1011650

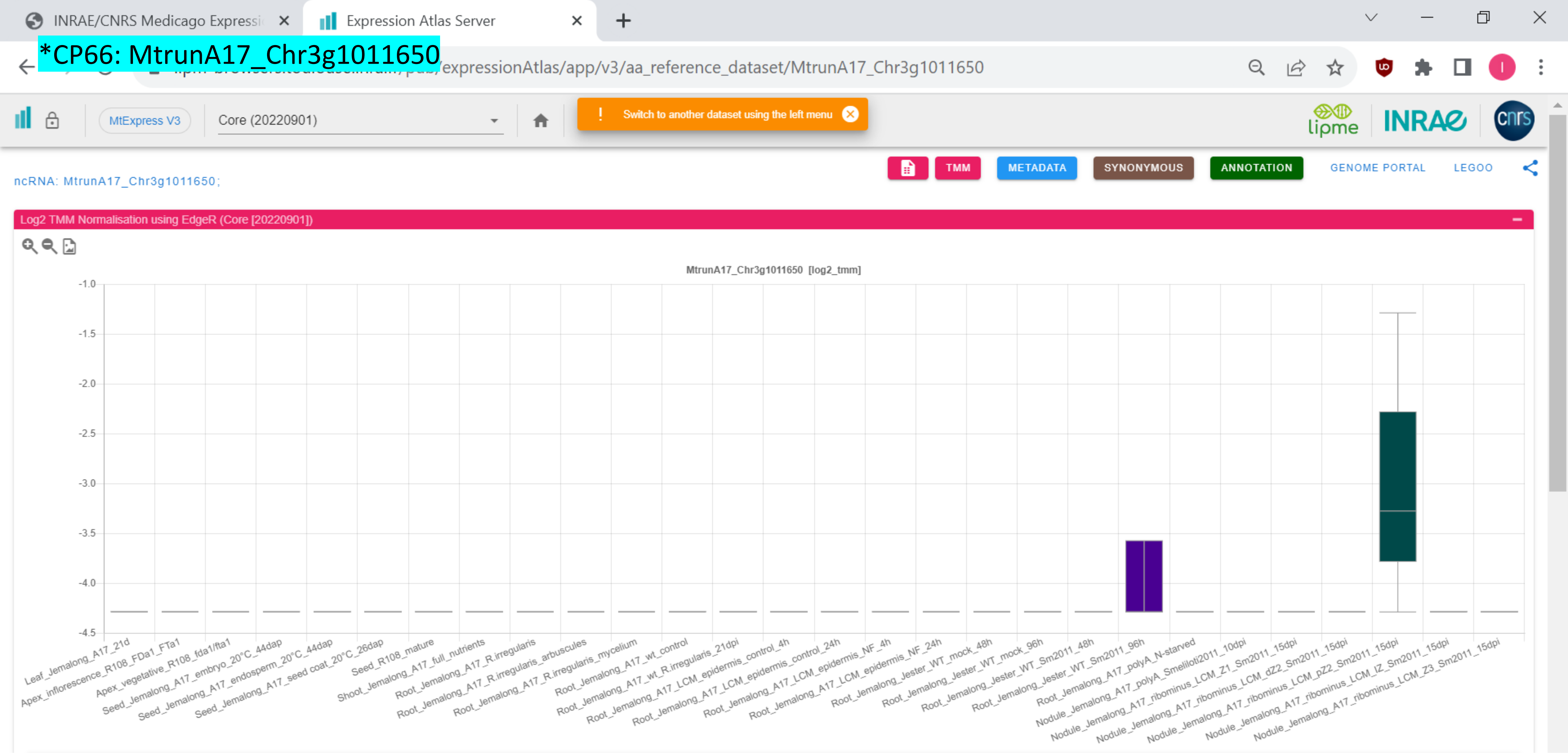

### CP67: MtrunA17_Chr4g0000131

INRAE/CNRS Medicago Expressi | Expression Atlas Server

.../pub/expressionAtlas/app/v3/aa_reference_dataset/MtrunA17_Chr4g0000131

MtExpress V3 | Core (20220901)

Switch to another dataset using the left menu

TMM | METADATA | SYNONYMOUS | ANNOTATION | GENOME PORTAL | LEGOO

mRNA: MtrunA17_Chr4g0000131;

Log2 TMM Normalisation using EdgeR (Core [20220901])

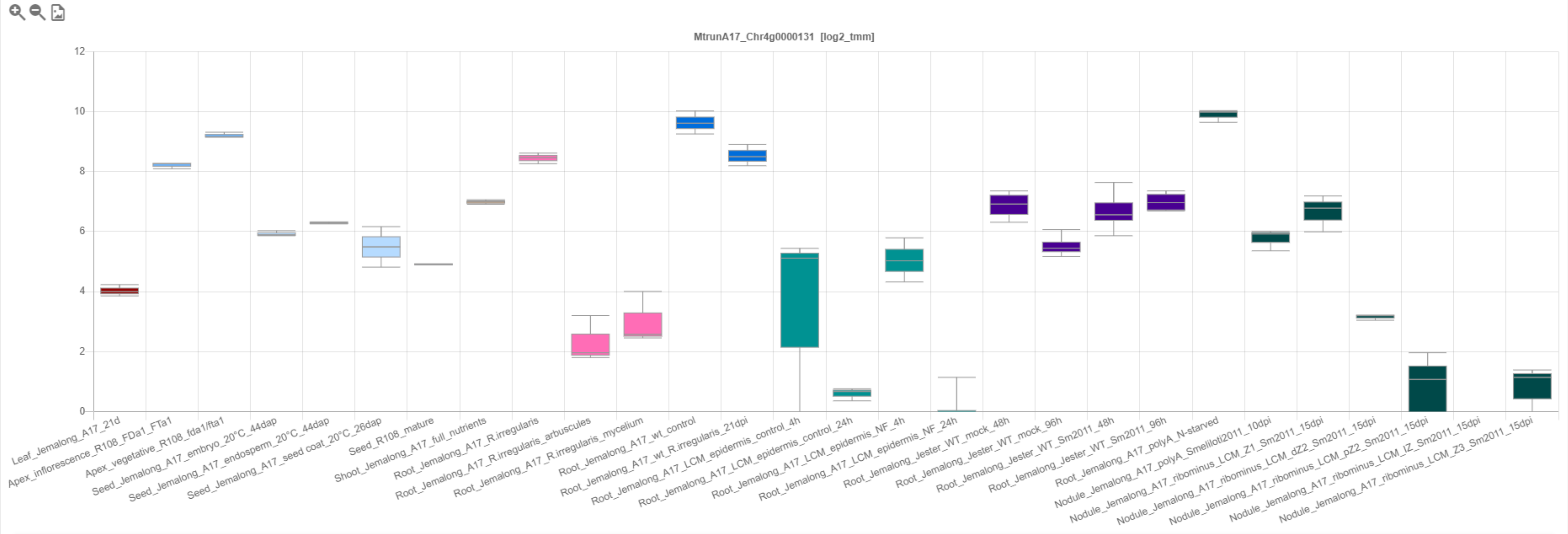

*CP68: MtrunA17_Chr4g0000891

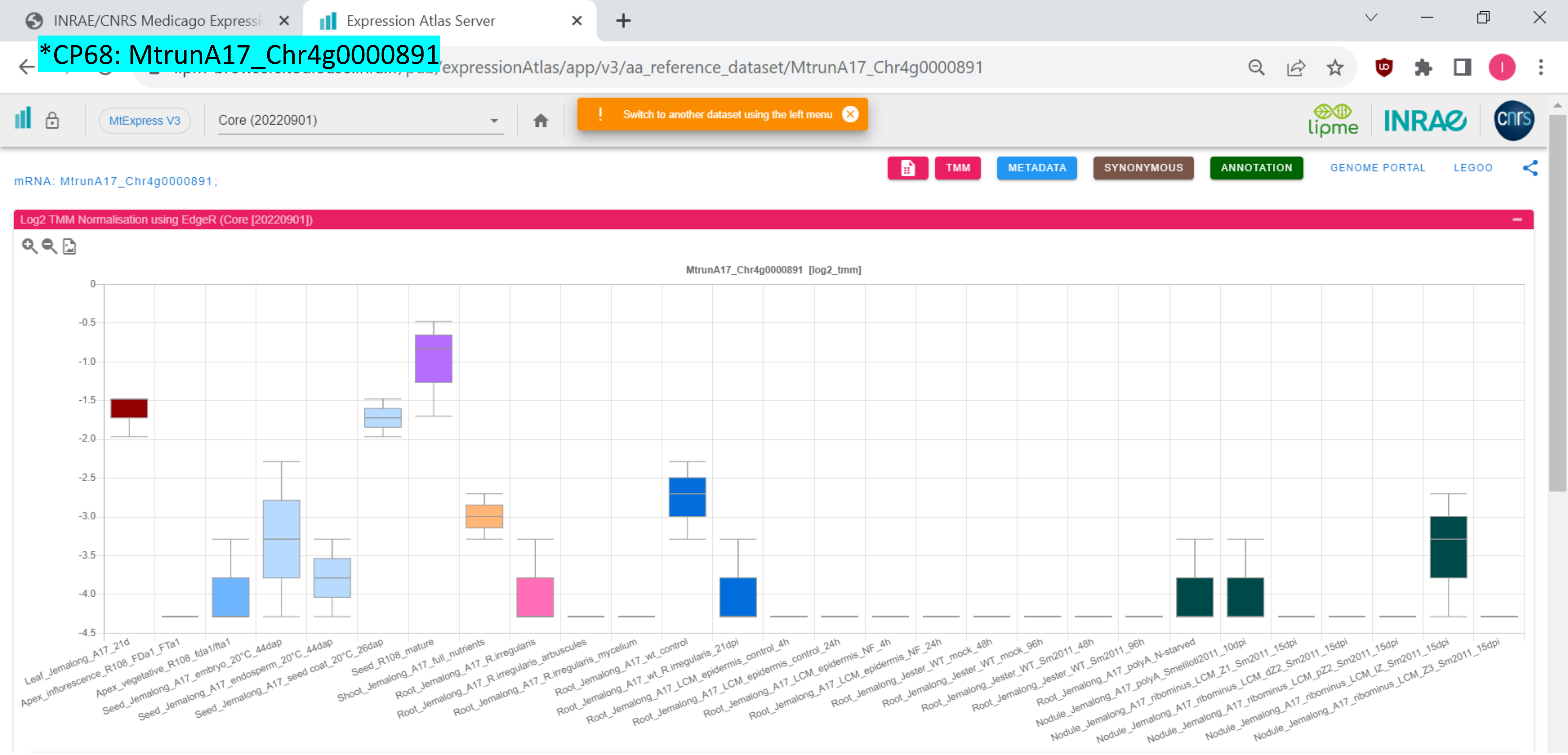

CP69: MtrunA17_Chr4g0004721

INRAE/CNRS Medicago Expressi | Expression Atlas Server

…pub/expressionAtlas/app/v3/aa_reference_dataset/MtrunA17_Chr4g0004721

MtExpress V3 | Core (20220901)

! Switch to another dataset using the left menu

TMM | METADATA | SYNONYMOUS | ANNOTATION | GENOME PORTAL | LEGOO

mRNA: MtrunA17_Chr4g0004721;

Log2 TMM Normalisation using EdgeR (Core [20220901])

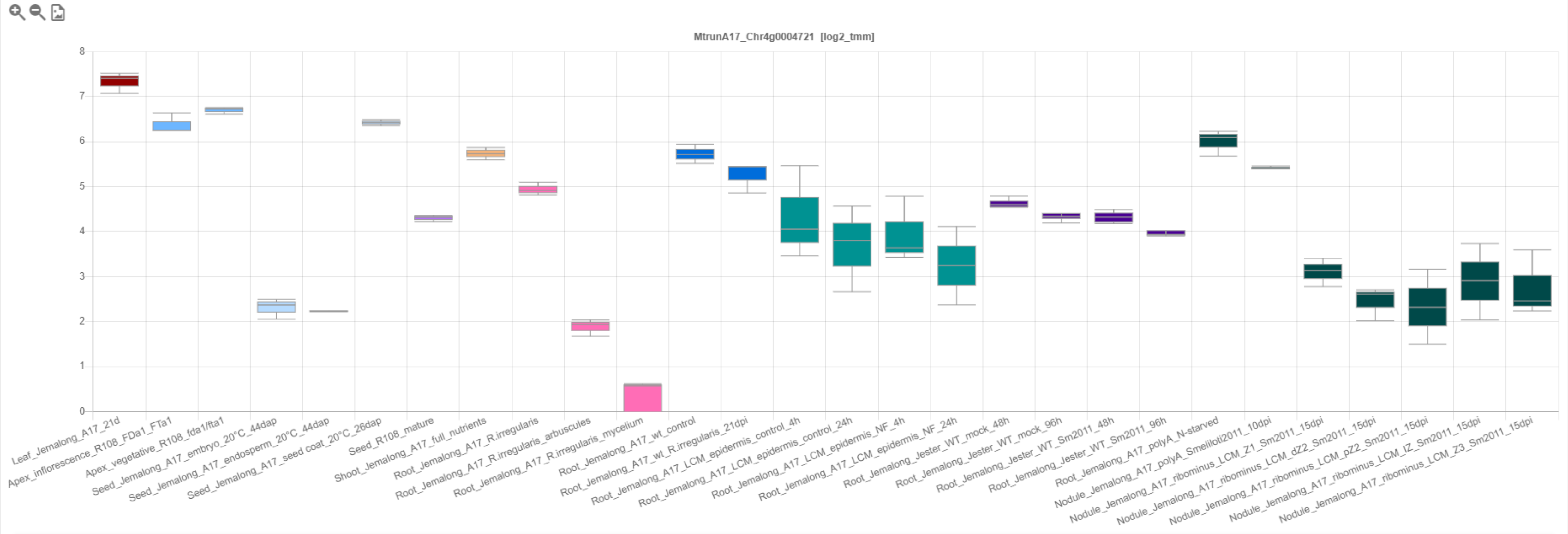

*CP70: MtrunA17_Chr4g0014251

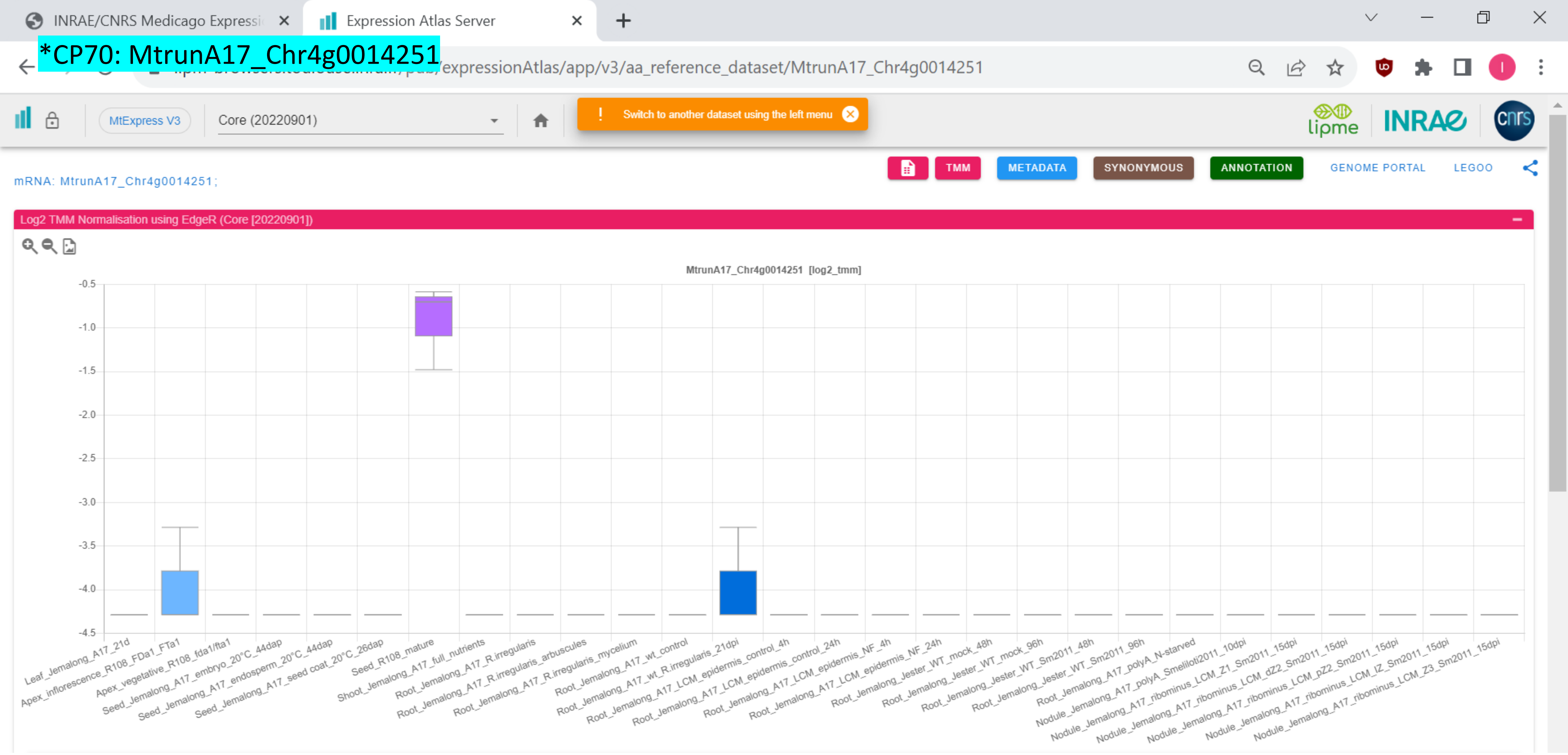

### CP71: MtrunA17_Chr4g0022491

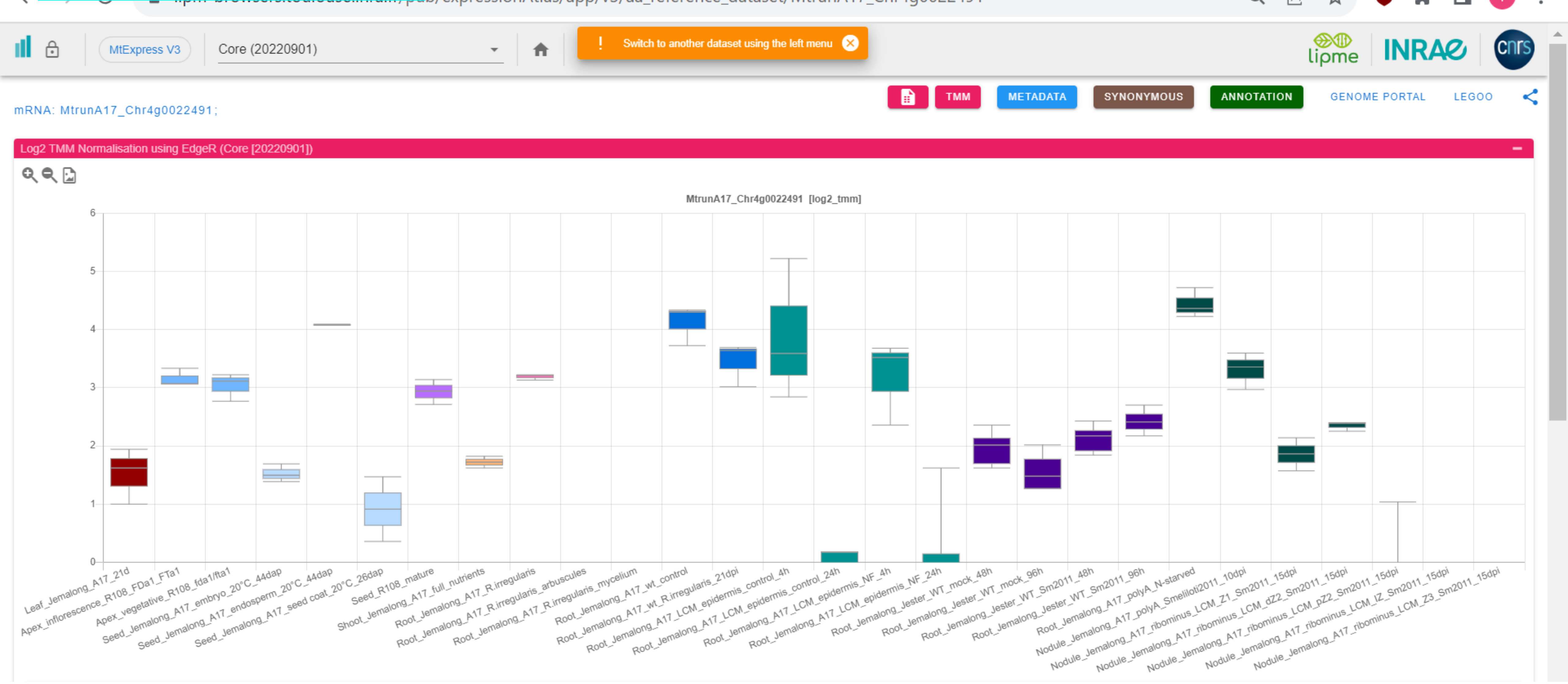

*CP72: MtrunA17_Chr4g0023791

INRAE/CNRS Medicago Expressi | Expression Atlas Server

.../expressionAtlas/app/v3/aa_reference_dataset/MtrunA17_Chr4g0023791

MtExpress V3 | Core (20220901)

! Switch to another dataset using the left menu

lipme INRAE cnrs

TMM | METADATA | SYNONYMOUS | ANNOTATION | GENOME PORTAL | LEGOO

tRNA: MtrunA17_Chr4g0023791;

Log2 TMM Normalisation using EdgeR (Core [20220901])

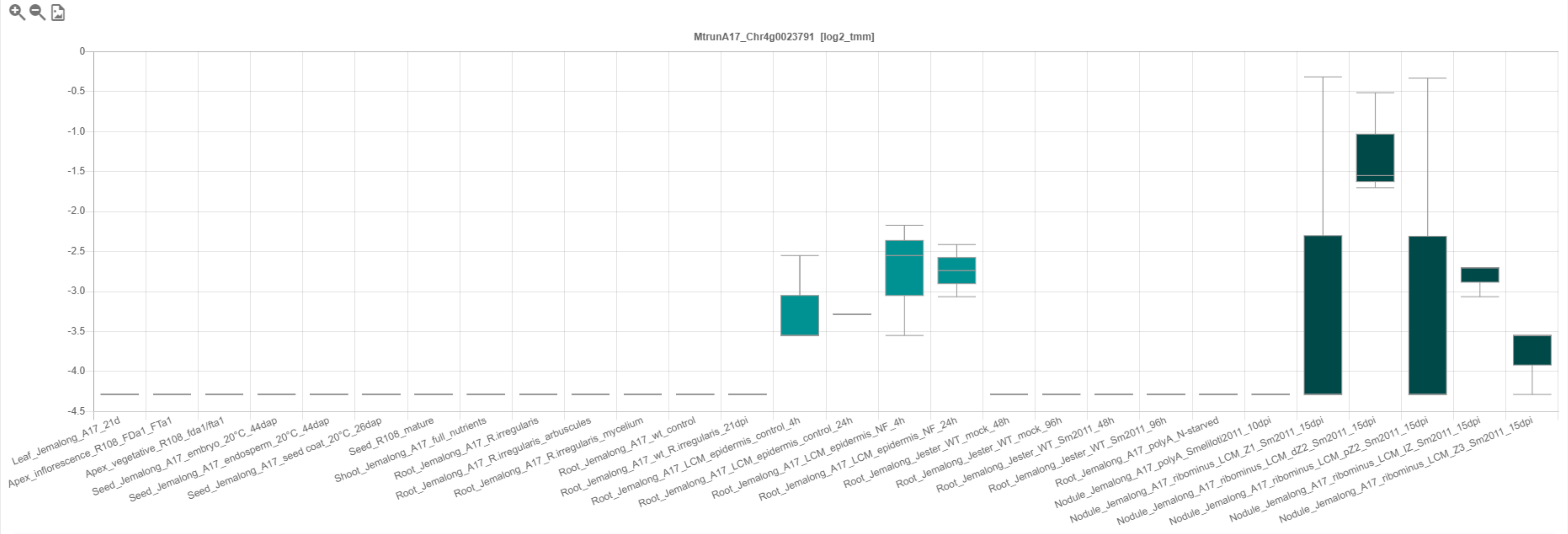

### CP73: MtrunA17_Chr4g0034271

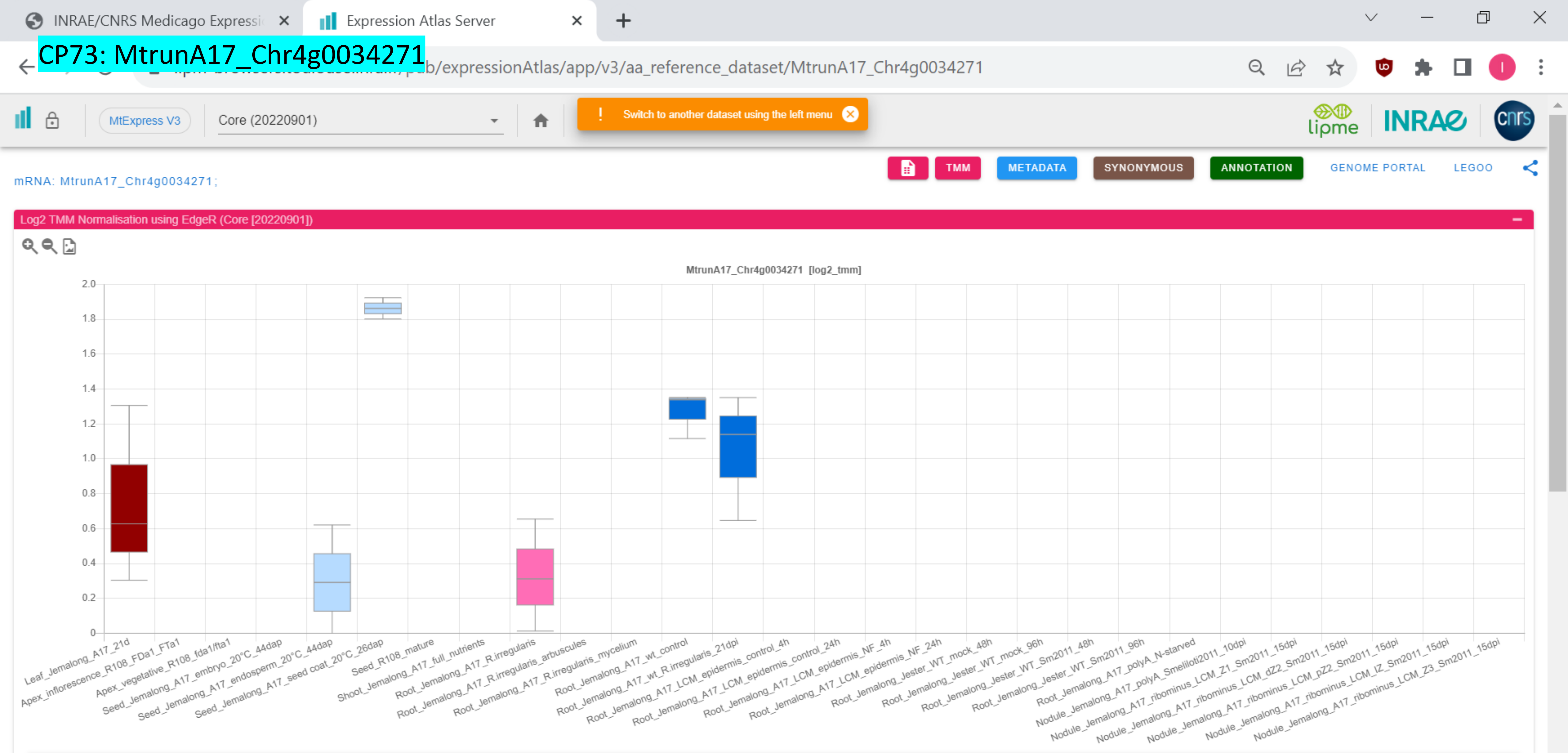

CP74: MtrunA17_Chr4g0034491

MtExpress V3

Core (20220901)

! Switch to another dataset using the left menu

TMM | METADATA | SYNONYMOUS | ANNOTATION | GENOME PORTAL | LEGOO

mRNA: MtrunA17_Chr4g0034491;

Log2 TMM Normalisation using EdgeR (Core [20220901])

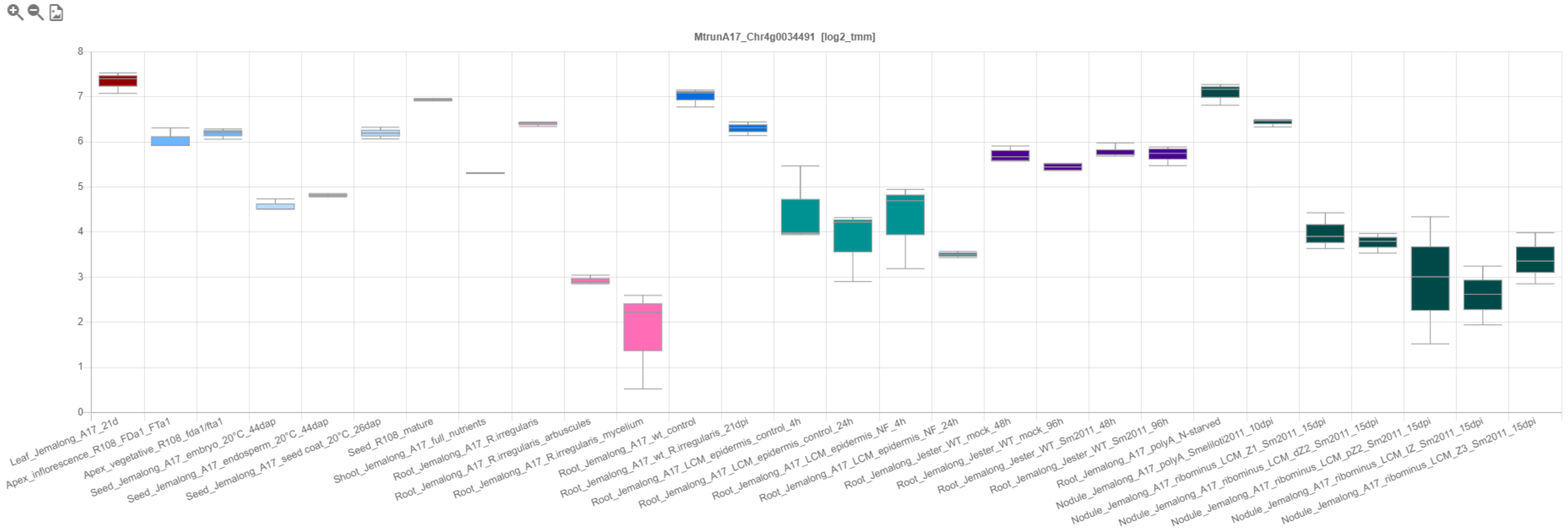

CP75: MtrunA17_Chr4g0037381

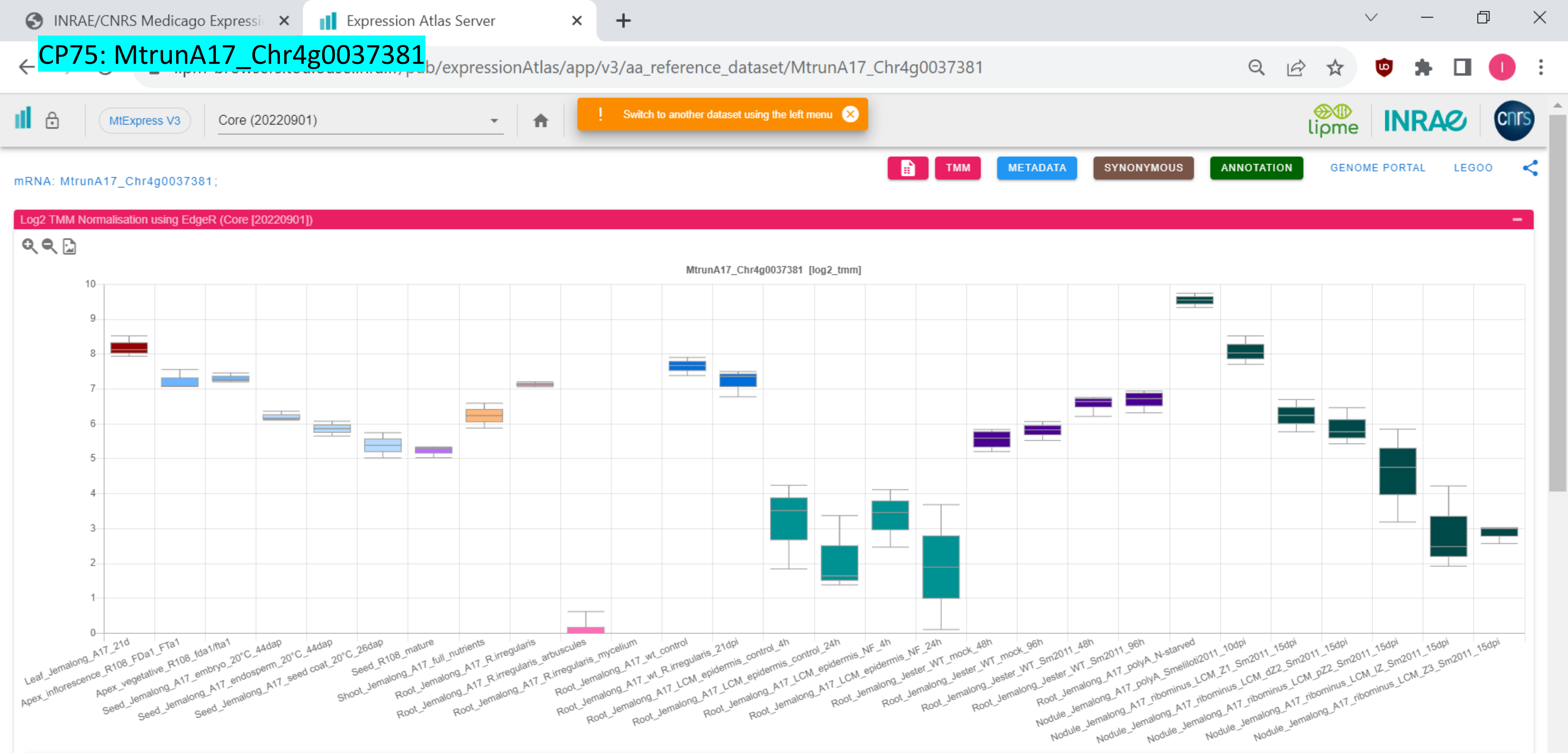

### CP76: MtrunA17_Chr4g0040471

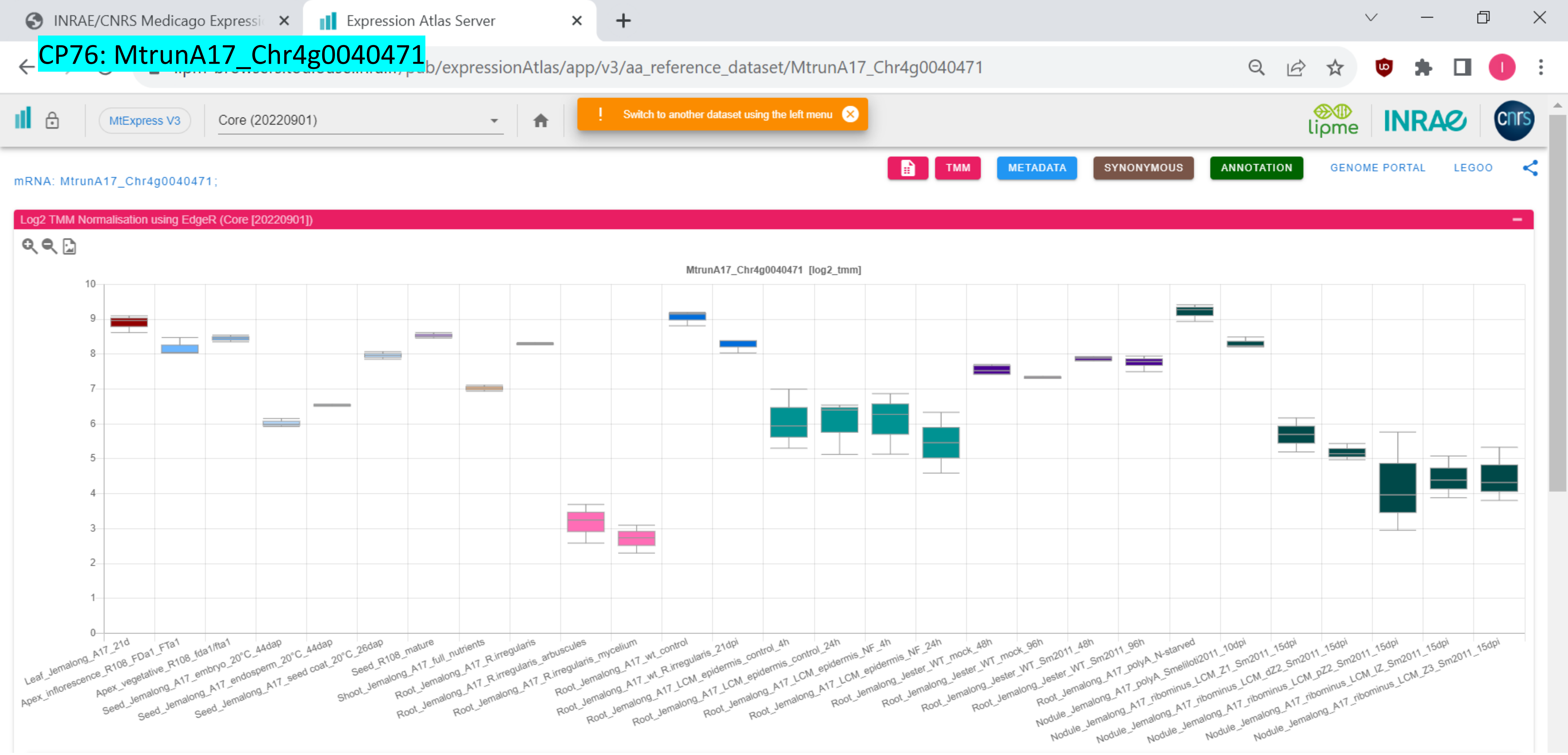

### CP77: MtrunA17_Chr4g0055331

INRAE/CNRS Medicago Expressi | Expression Atlas Server

…pub/expressionAtlas/app/v3/aa_reference_dataset/MtrunA17_Chr4g0055331

MtExpress V3 | Core (20220901) | ! Switch to another dataset using the left menu

TMM | METADATA | SYNONYMOUS | ANNOTATION | GENOME PORTAL | LEGOO

mRNA: MtrunA17_Chr4g0055331;

Log2 TMM Normalisation using EdgeR (Core [20220901])

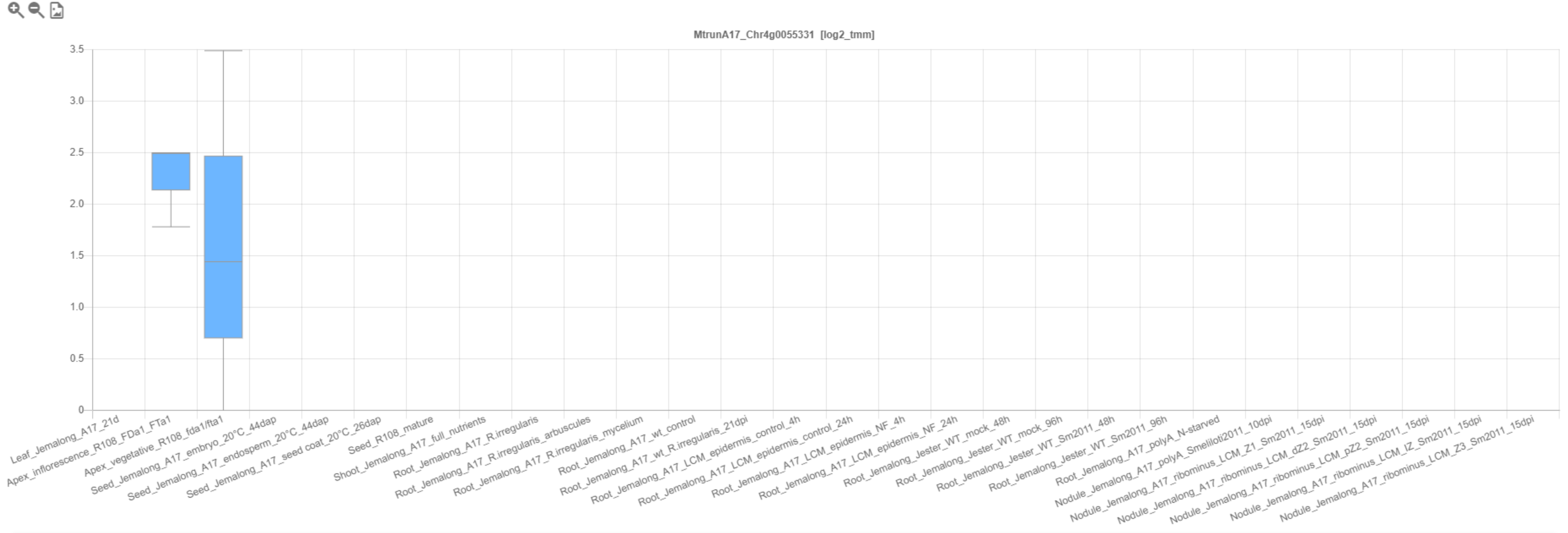

CP78: MtrunA17_Chr4g0059001

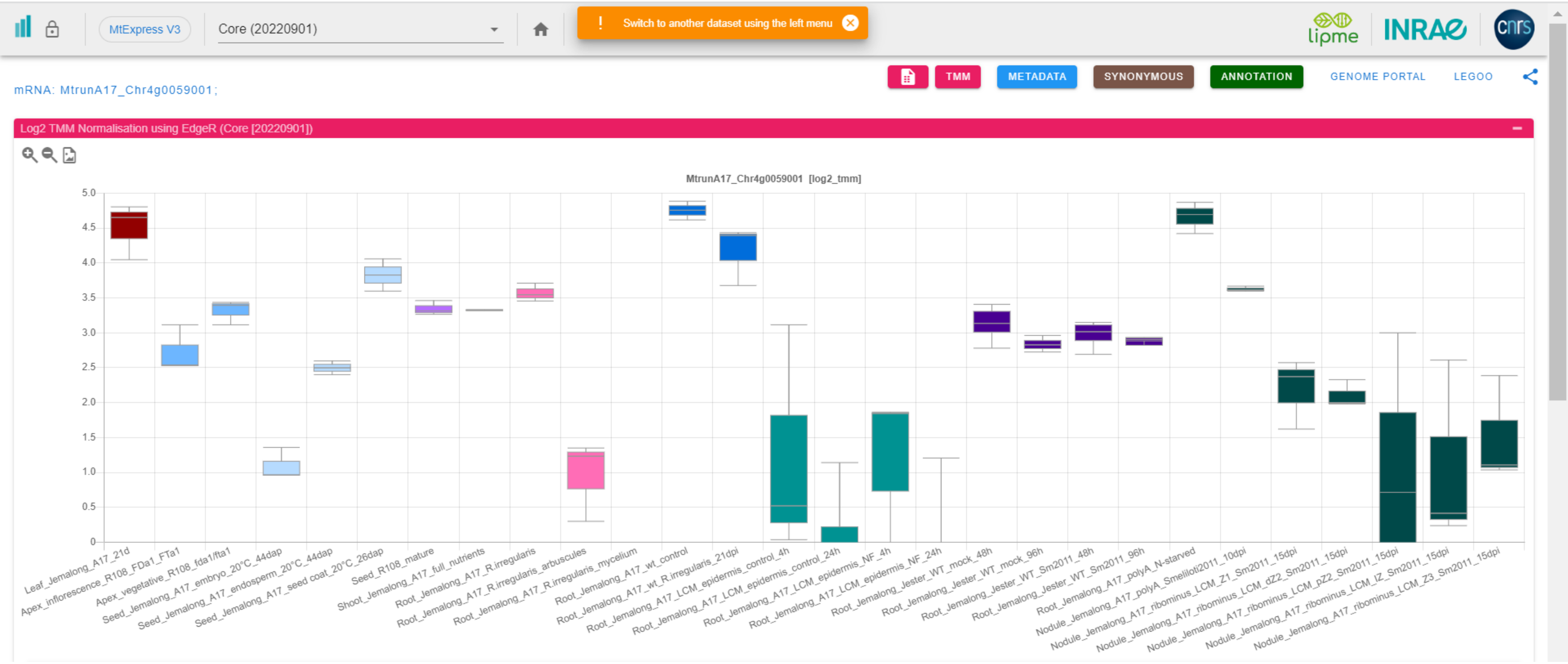

*CP79: MtrunA17_Chr4g0063201

INRAE/CNRS Medicago Expressi | Expression Atlas Server

expressionAtlas/app/v3/aa_reference_dataset/MtrunA17_Chr4g0063201

MtExpress V3 | Core (20220901)

! Switch to another dataset using the left menu

TMM | METADATA | SYNONYMOUS | ANNOTATION | GENOME PORTAL | LEGOO

mRNA: MtrunA17_Chr4g0063201 ;

Log2 TMM Normalisation using EdgeR (Core [20220901])

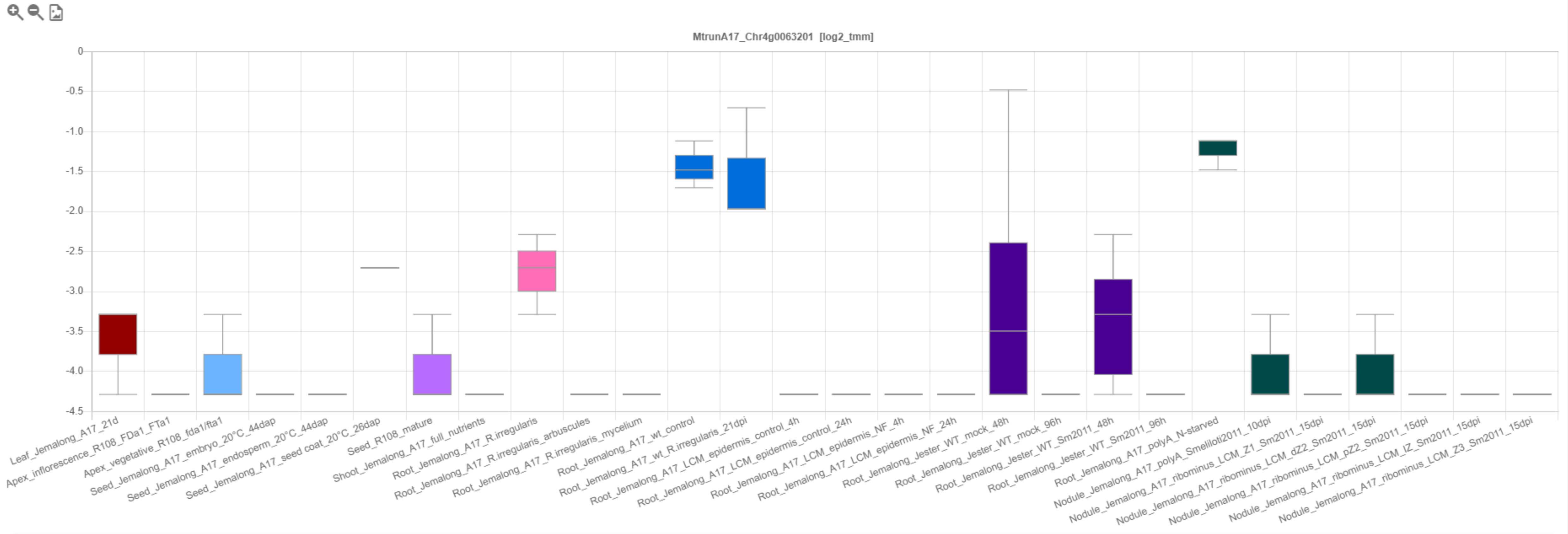

CP80: MtrunA17_Chr4g0070011

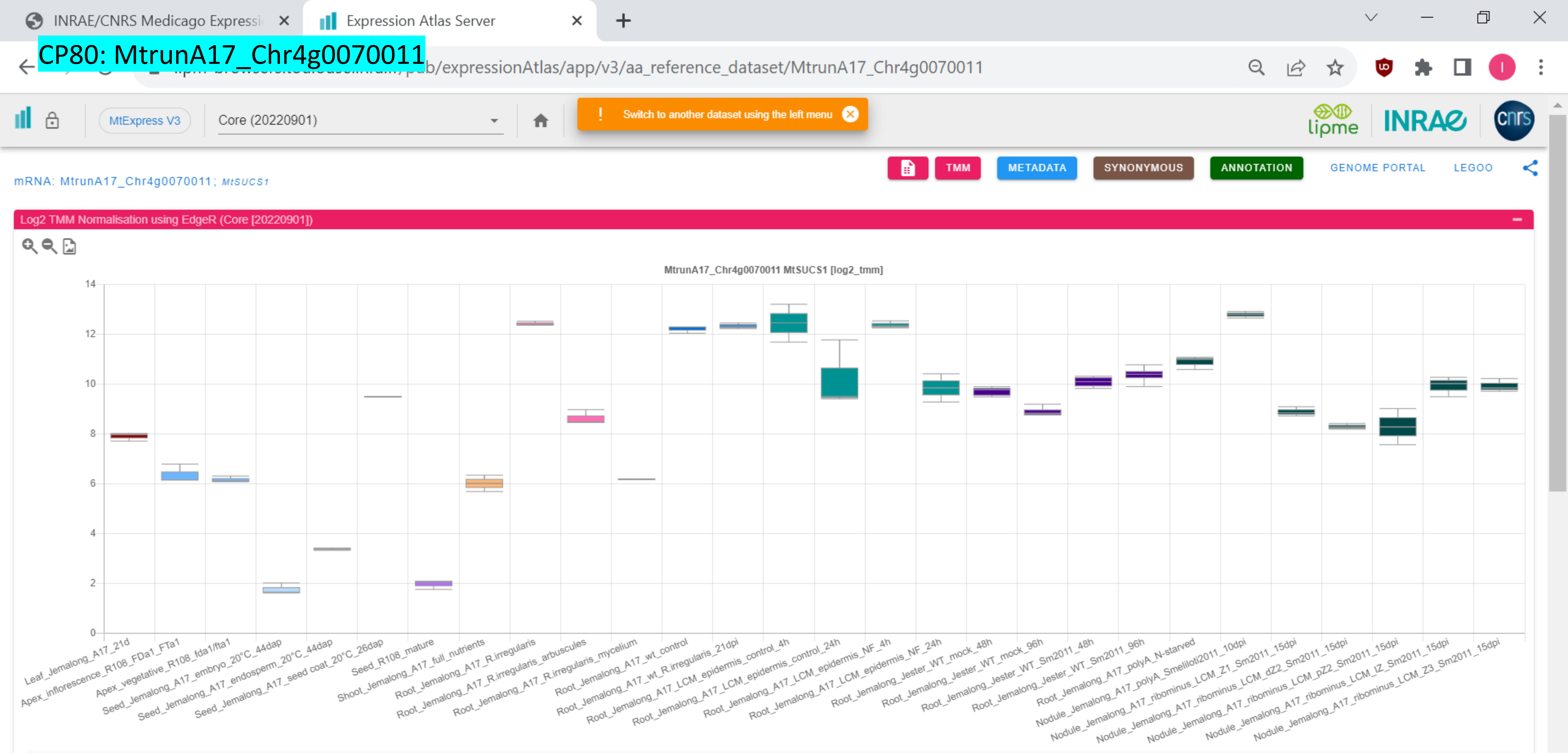

### CP81: MtrunA17_Chr5g0393401

INRAE/CNRS Medicago Expressi… | Expression Atlas Server

…pub/expressionAtlas/app/v3/aa_reference_dataset/MtrunA17_Chr5g0393401

MtExpress V3 | Core (20220901)

! Switch to another dataset using the left menu

TMM | METADATA | SYNONYMOUS | ANNOTATION | GENOME PORTAL | LEGOO

mRNA: MtrunA17_Chr5g0393401 ; *MtHDA9*

Log2 TMM Normalisation using EdgeR (Core [20220901])

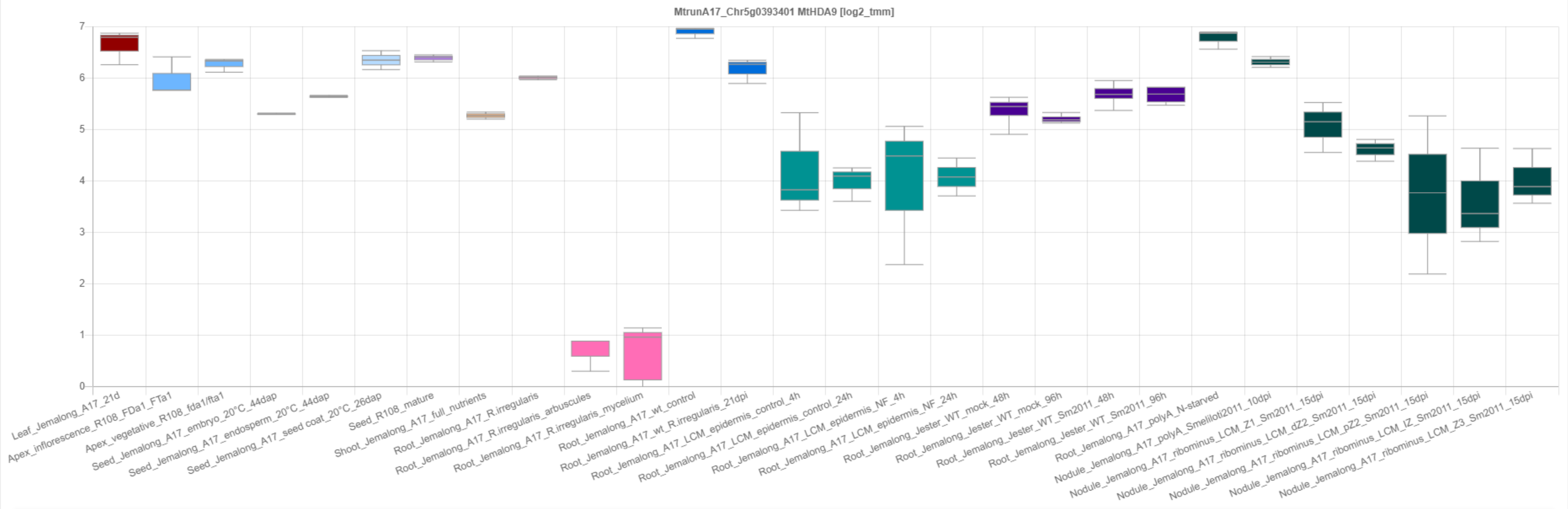

### CP82: MtrunA17_Chr5g0400181

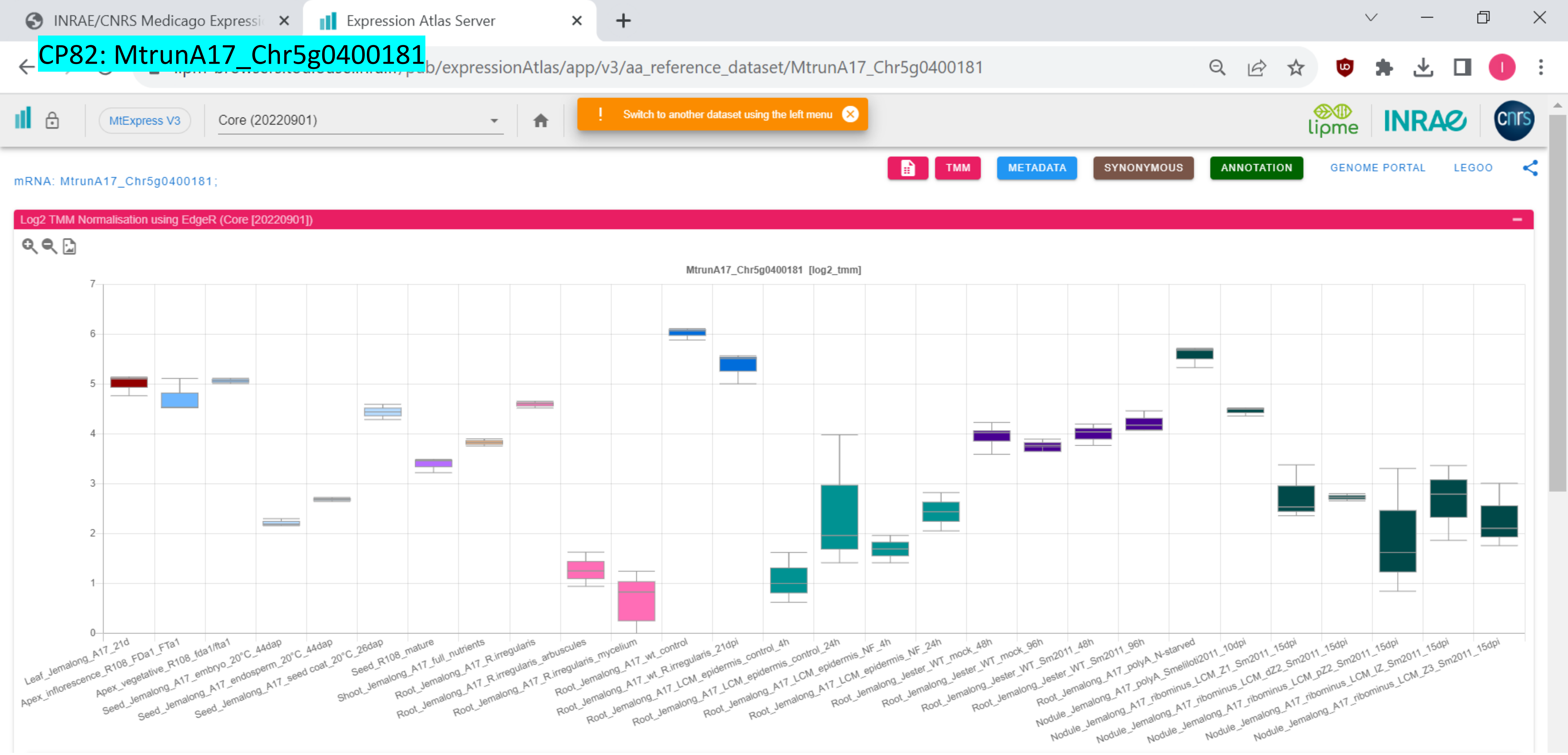

### CP83: MtrunA17_Chr5g0405061

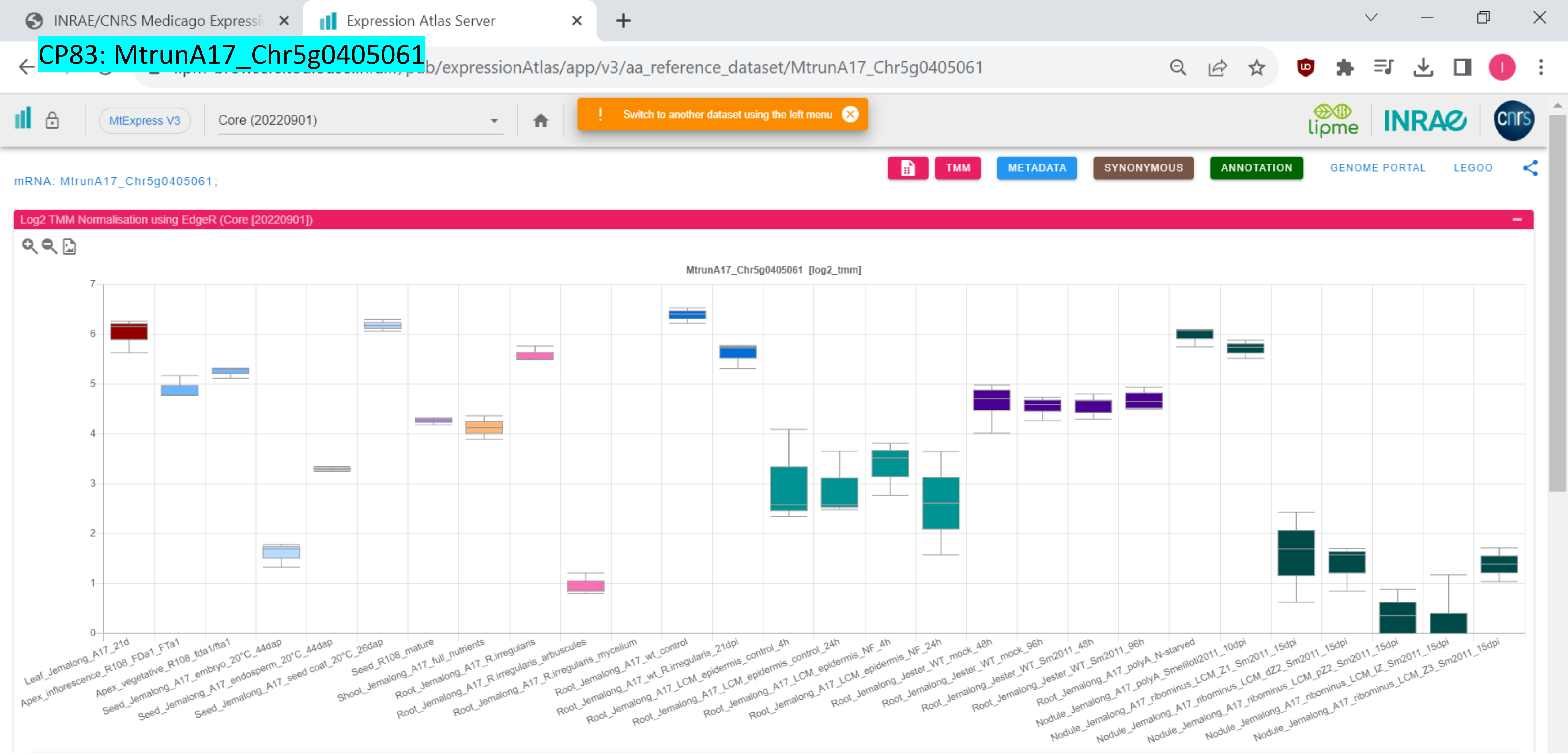

CP84: MtrunA17_Chr5g0408581

INRAE/CNRS Medicago Expressi… | Expression Atlas Server

…pub/expressionAtlas/app/v3/aa_reference_dataset/MtrunA17_Chr5g0408581

MtExpress V3 | Core (20220901)

! Switch to another dataset using the left menu

TMM | METADATA | SYNONYMOUS | ANNOTATION | GENOME PORTAL | LEGOO

mRNA: MtrunA17_Chr5g0408581;

Log2 TMM Normalisation using EdgeR (Core [20220901])

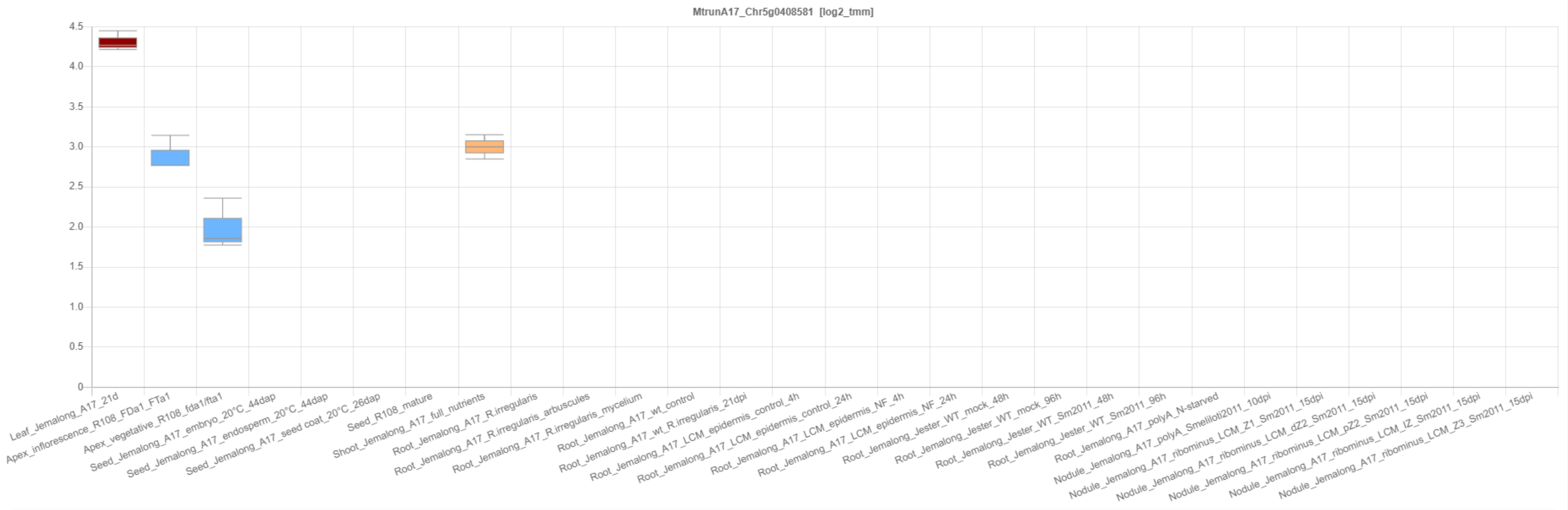

CP85: MtrunA17_Chr5g0415031

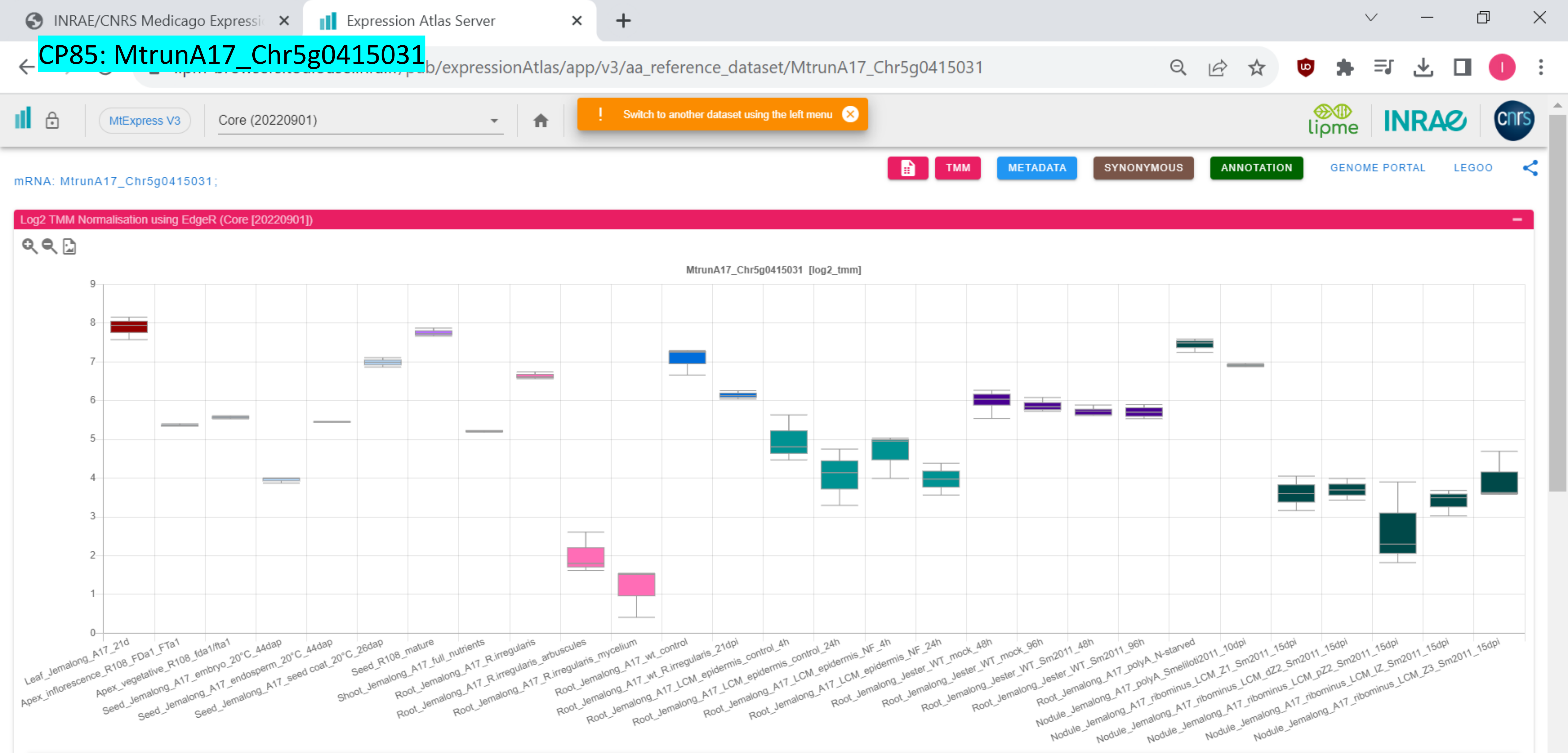

CP86: MtrunA17_Chr5g0415911

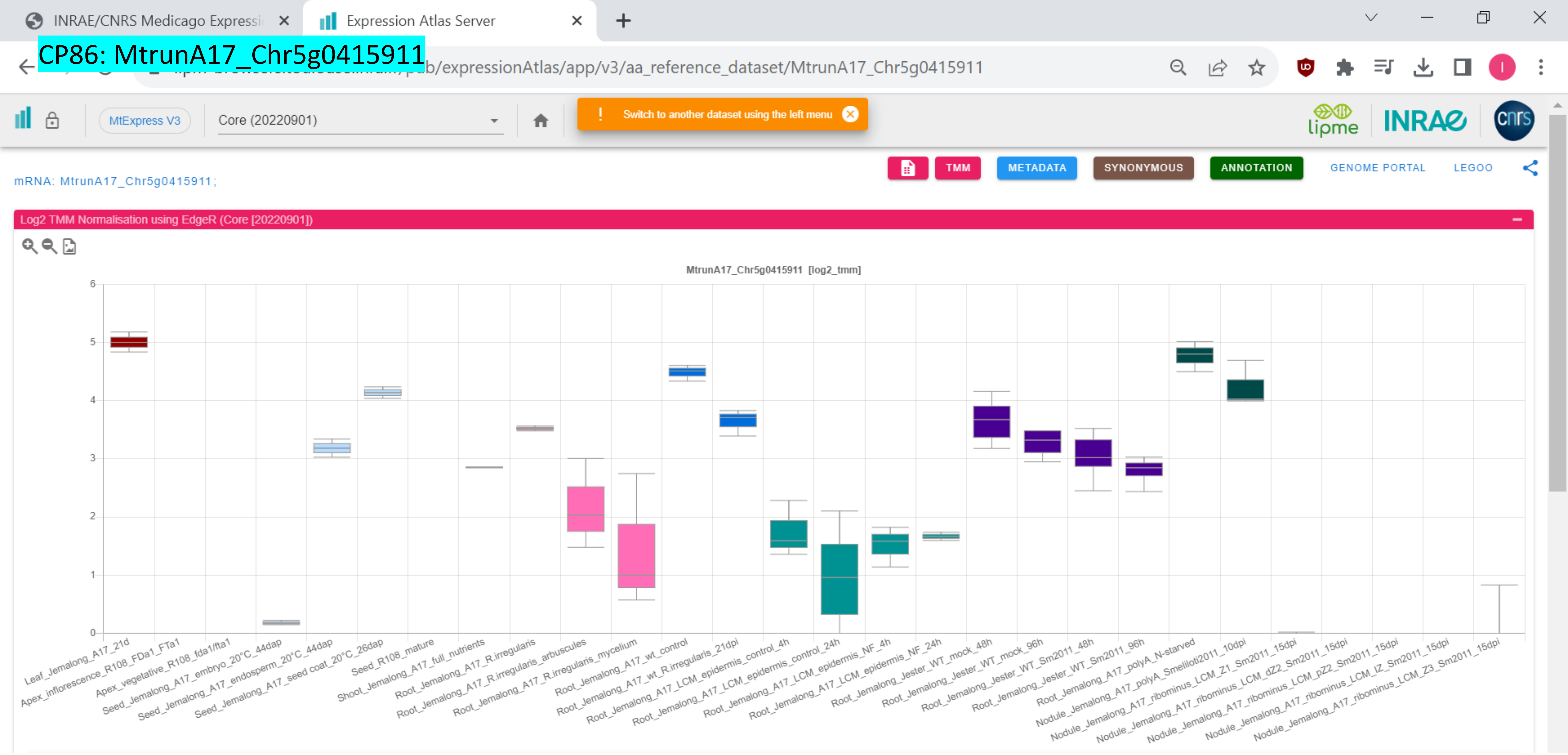

CP87: MtrunA17_Chr5g0421761

ncRNA: MtrunA17_Chr5g0421761;

Log2 TMM Normalisation using EdgeR (Core [20220901])

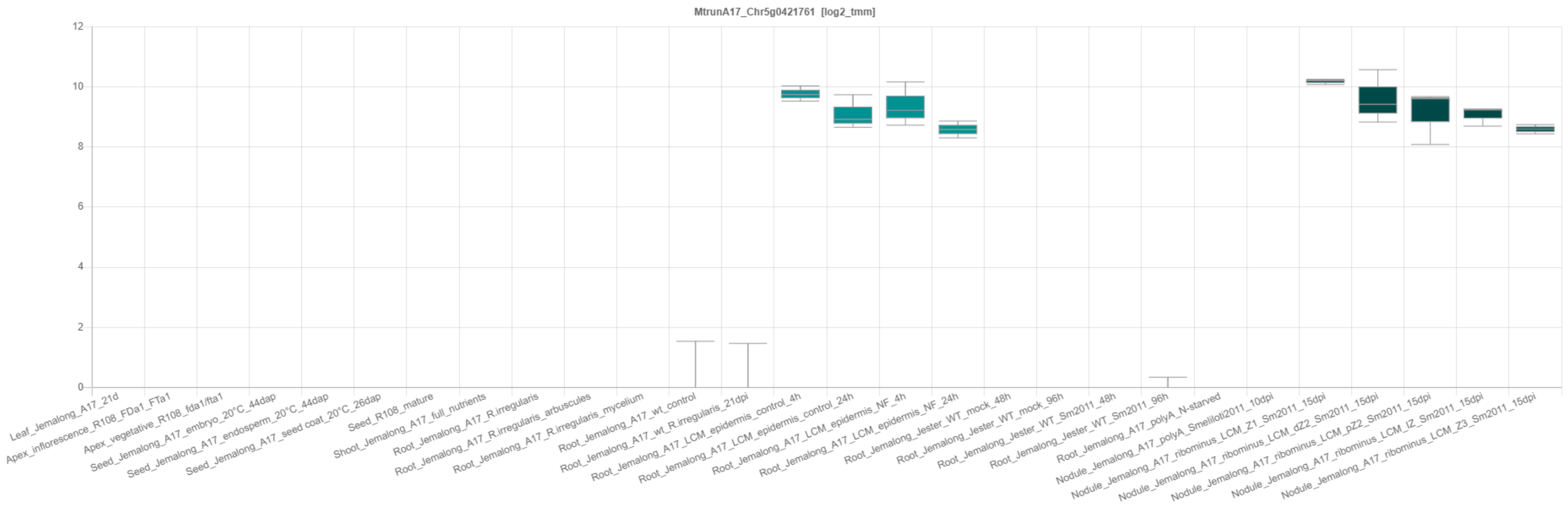

CP88: MtrunA17_Chr5g0422291

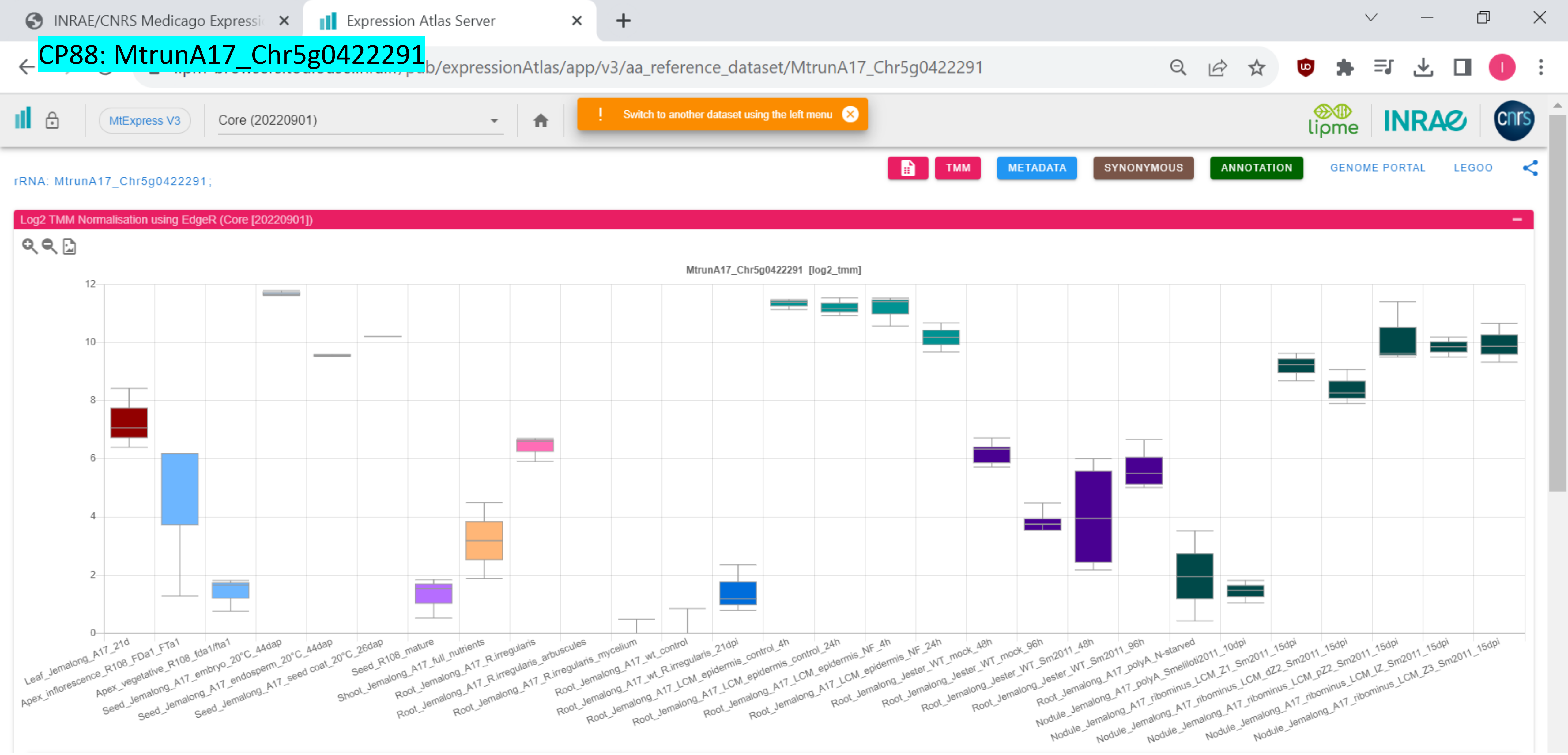

CP89: MtrunA17_Chr5g0422291

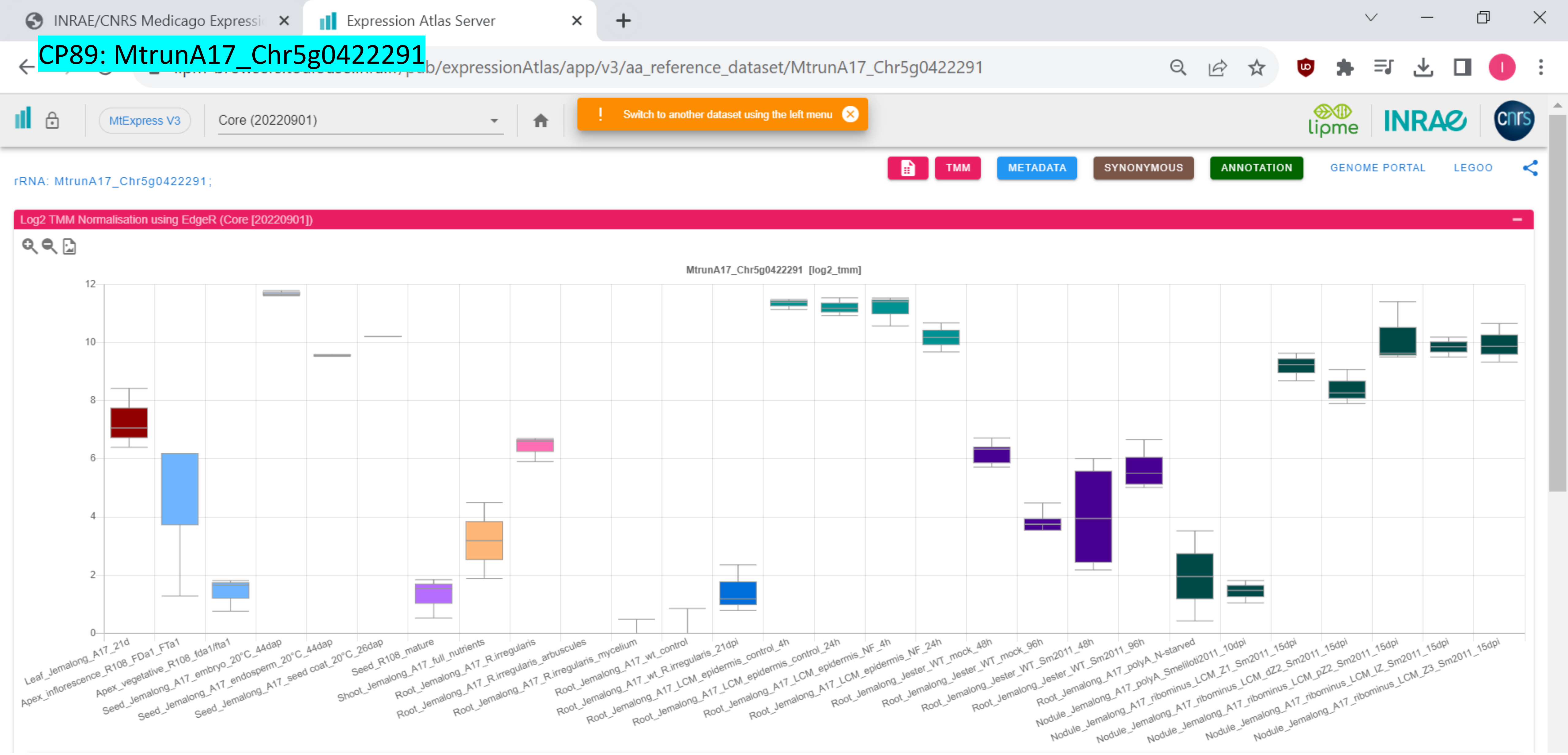

### CP90: MtrunA17_Chr5g0430341

INRAE/CNRS Medicago Expressi | Expression Athas Server

.../pub/expressionAtlas/app/v3/aa_reference_dataset/MtrunA17_Chr5g0430341

MtExpress V3 | Core (20220901) | ! Switch to another dataset using the left menu

lipme INRAE cnrs

TMM | METADATA | SYNONYMOUS | ANNOTATION | GENOME PORTAL | LEGOO

mRNA: MtrunA17_Chr5g0430341;

Log2 TMM Normalisation using EdgeR (Core [20220901])

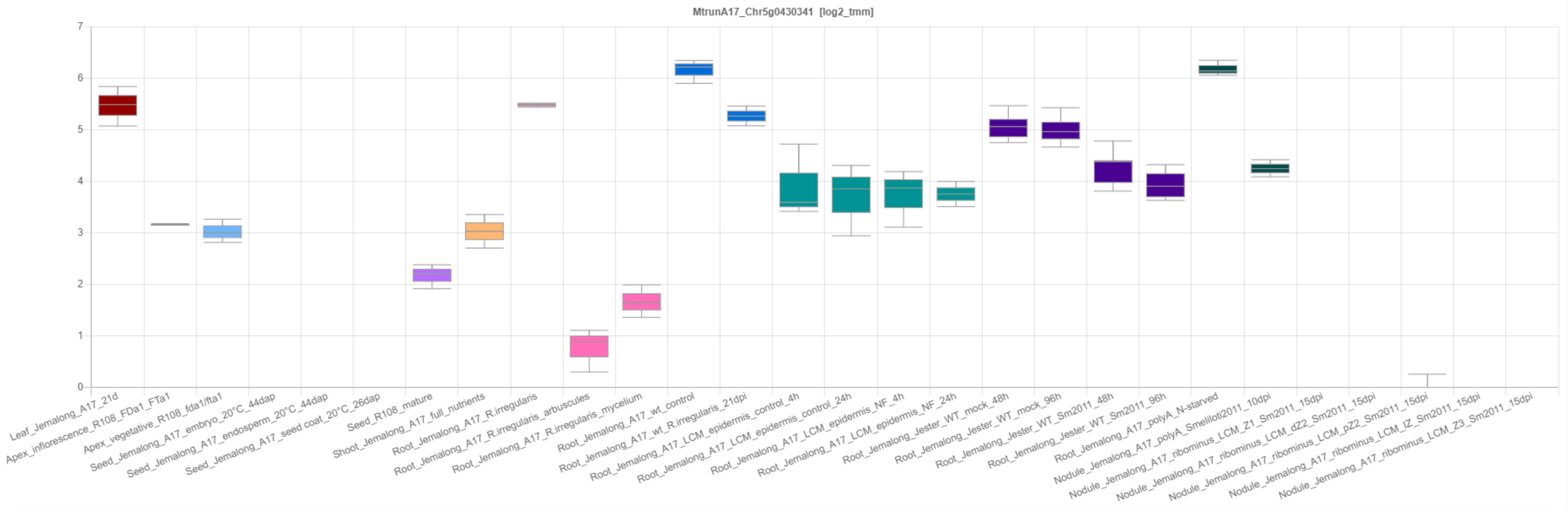

CP91: MtrunA17_Chr5g0430341

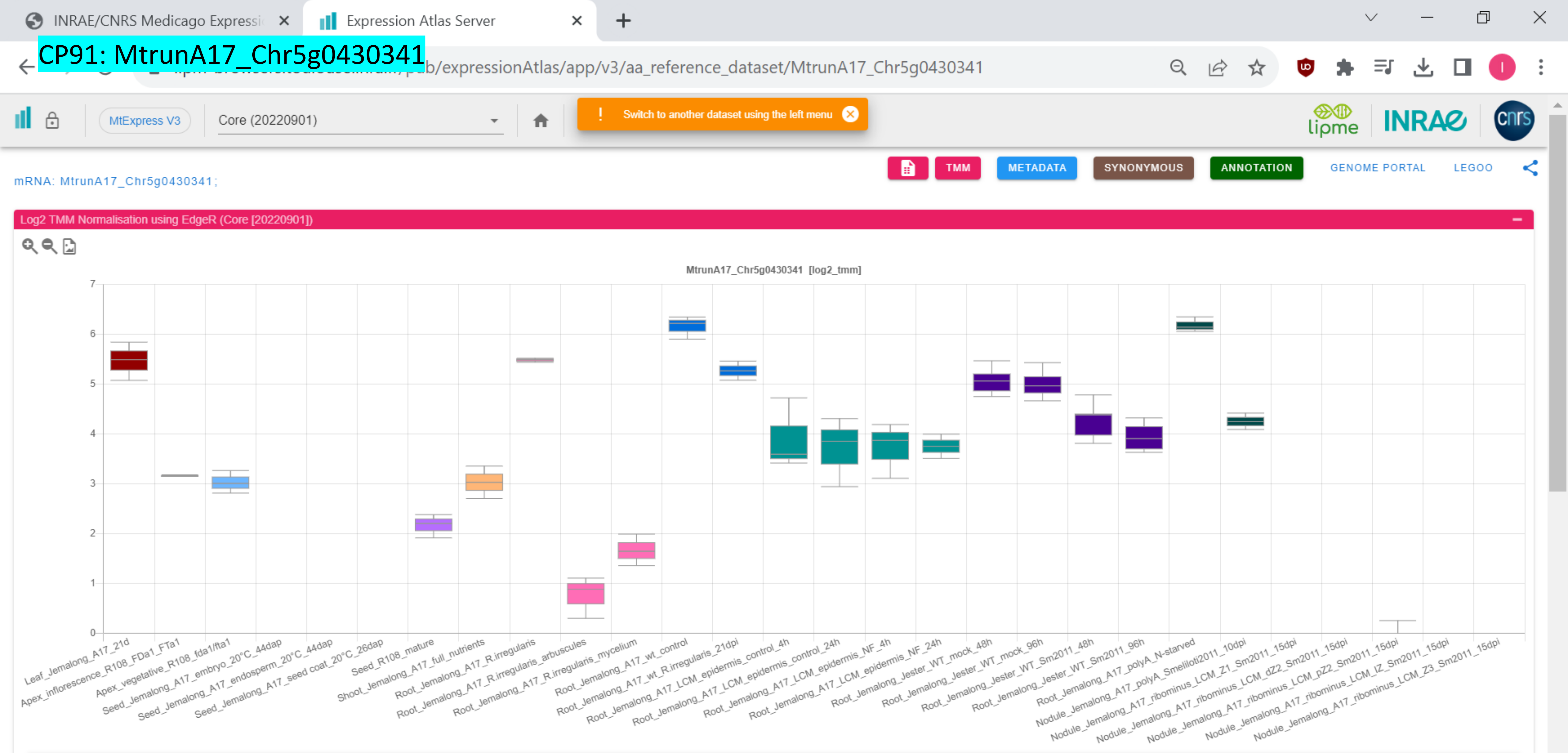

### CP92: MtrunA17_Chr5g0431401

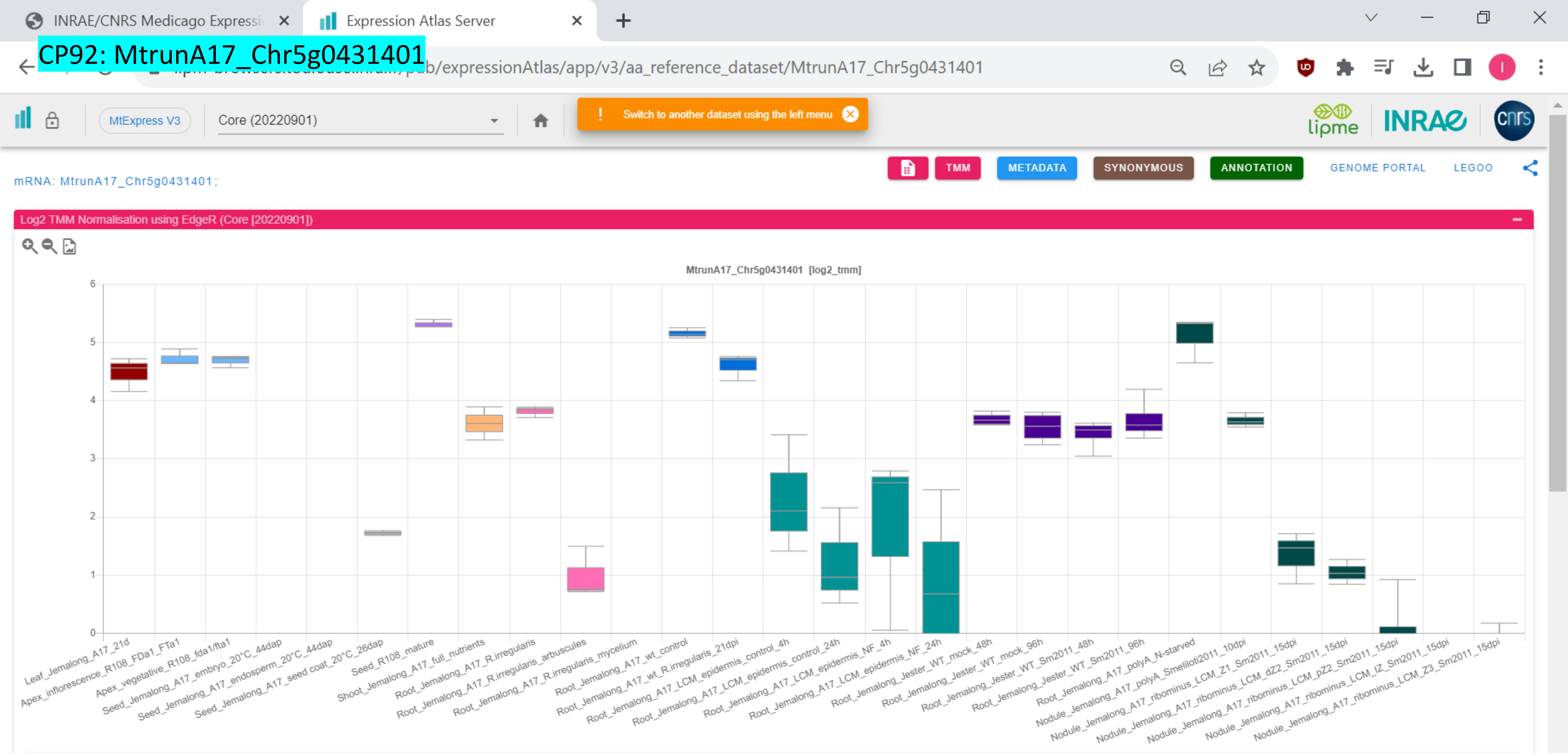

CP93: MtrunA17_Chr5g0435191

INRAE/CNRS Medicago Expressi | Expression Atlas Server

…pub/expressionAtlas/app/v3/aa_reference_dataset/MtrunA17_Chr5g0435191

MtExpress V3 | Core (20220901)

! Switch to another dataset using the left menu

TMM | METADATA | SYNONYMOUS | ANNOTATION | GENOME PORTAL | LEGOO

mRNA: MtrunA17_Chr5g0435191;

Log2 TMM Normalisation using EdgeR (Core [20220901])

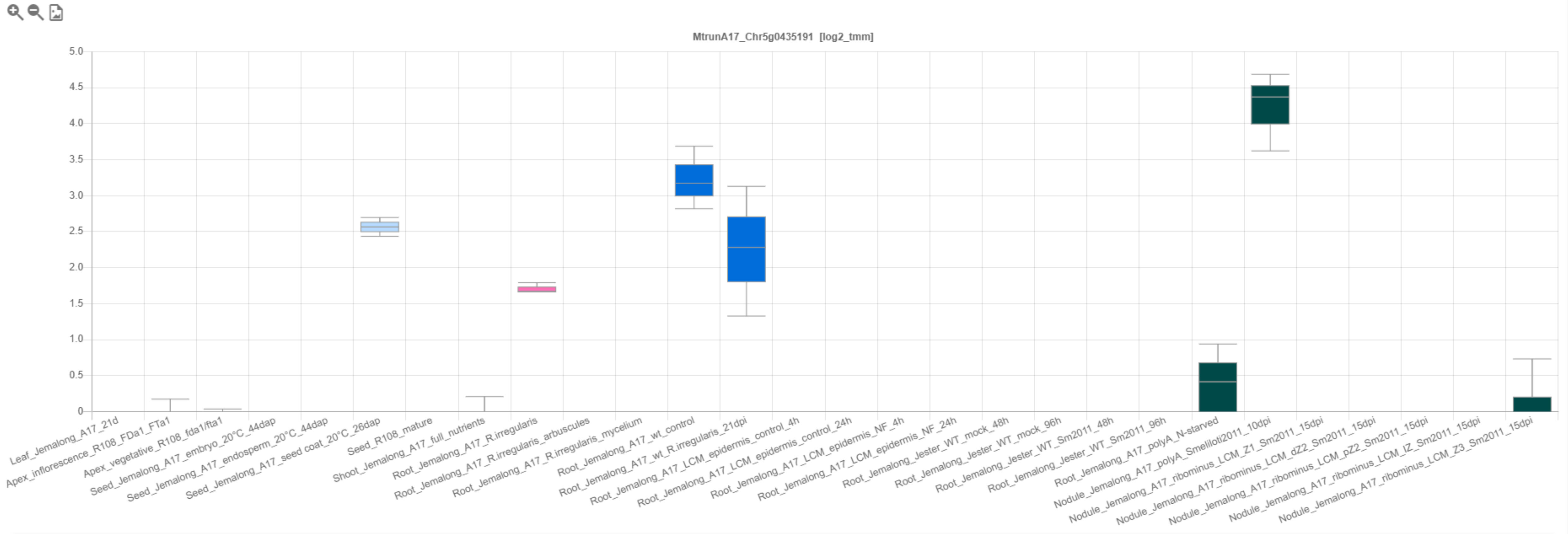

CP94: MtrunA17_Chr5g0444231

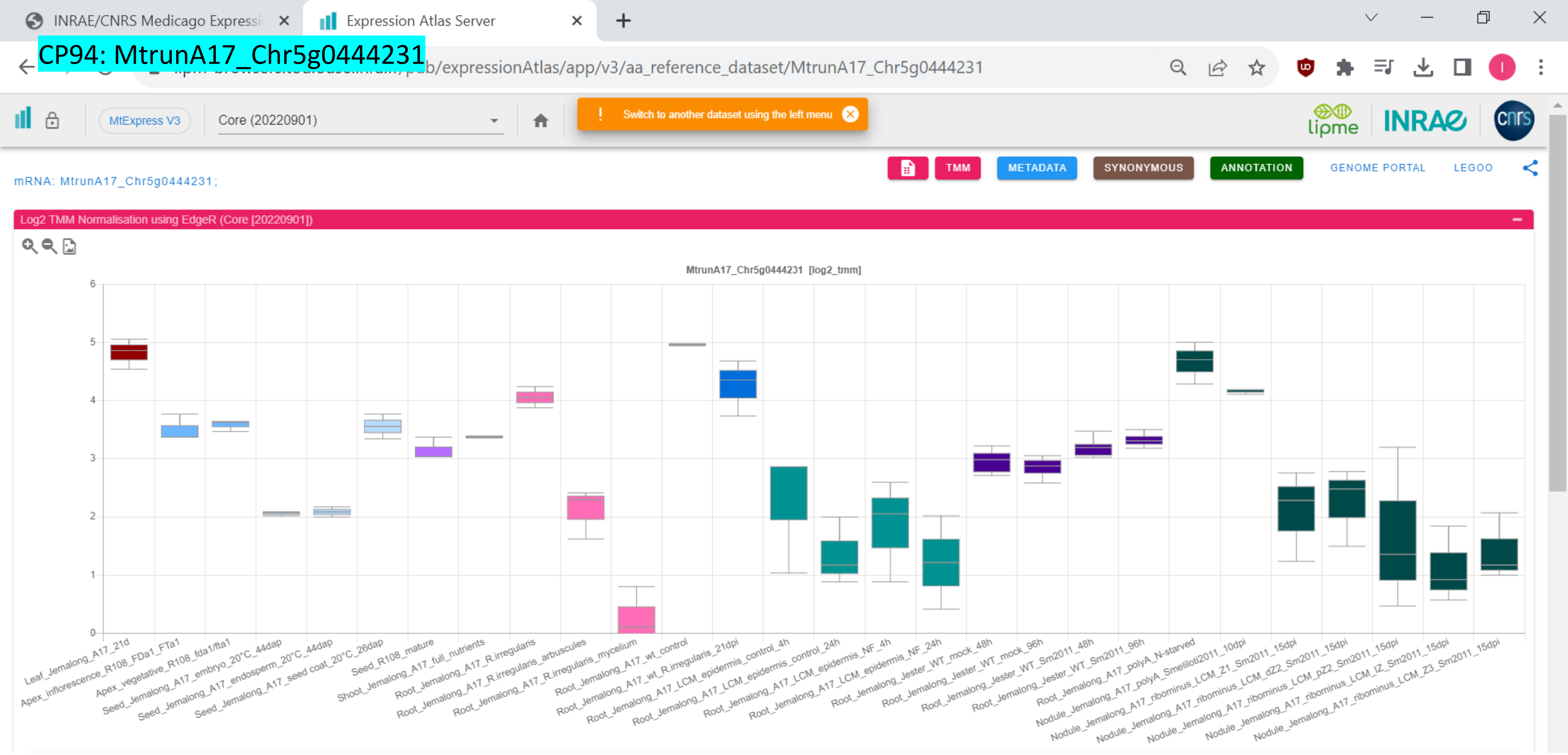

CP95: MtrunA17_Chr6g0451601

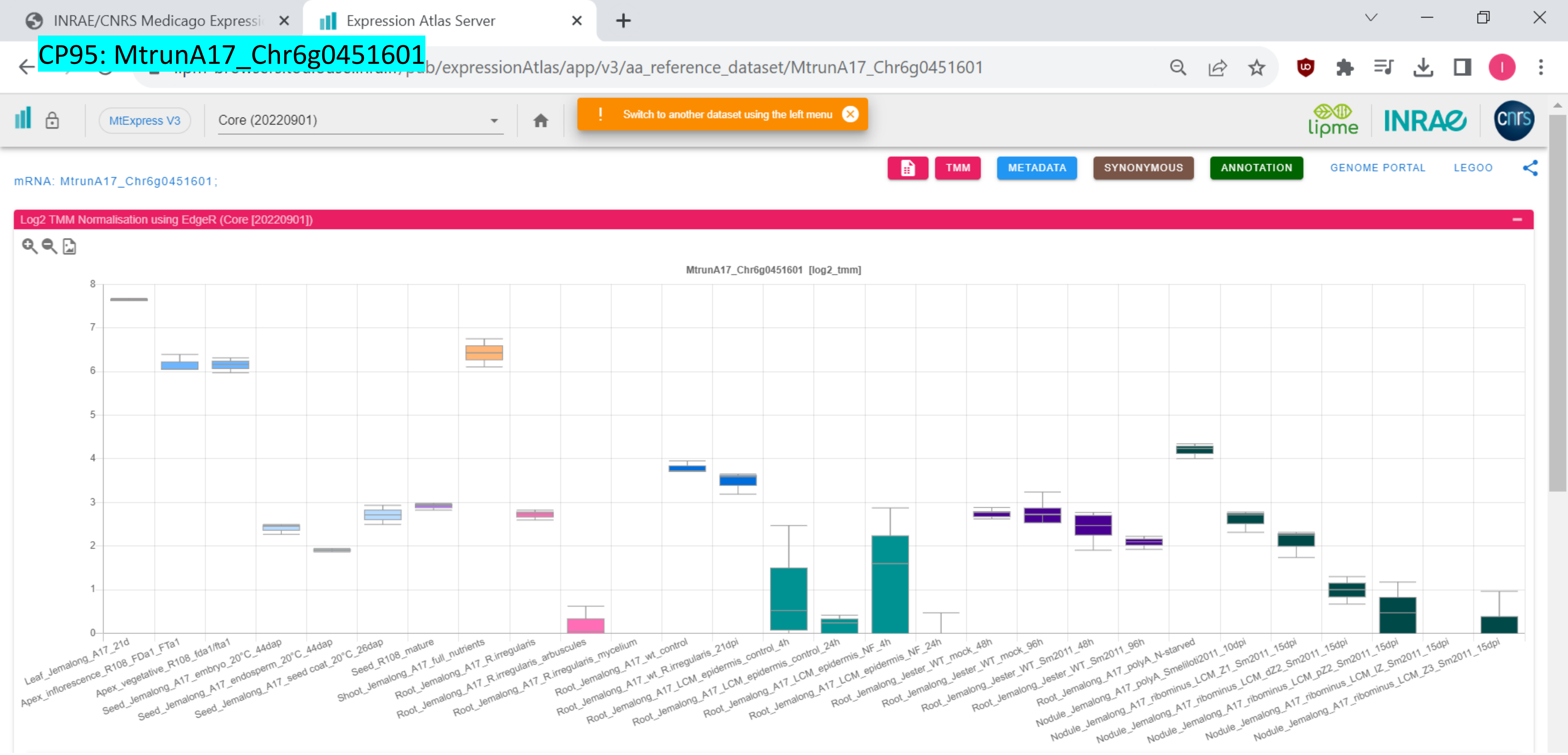

CP96: MtrunA17_Chr6g0452781

INRAE/CNRS Medicago Expressi | Expression Atlas Server

.../pub/expressionAtlas/app/v3/aa_reference_dataset/MtrunA17_Chr6g0452781

MtExpress V3 | Core (20220901)

! Switch to another dataset using the left menu

TMM | METADATA | SYNONYMOUS | ANNOTATION | GENOME PORTAL | LEGOO

mRNA: MtrunA17_Chr6g0452781;

Log2 TMM Normalisation using EdgeR (Core [20220901])

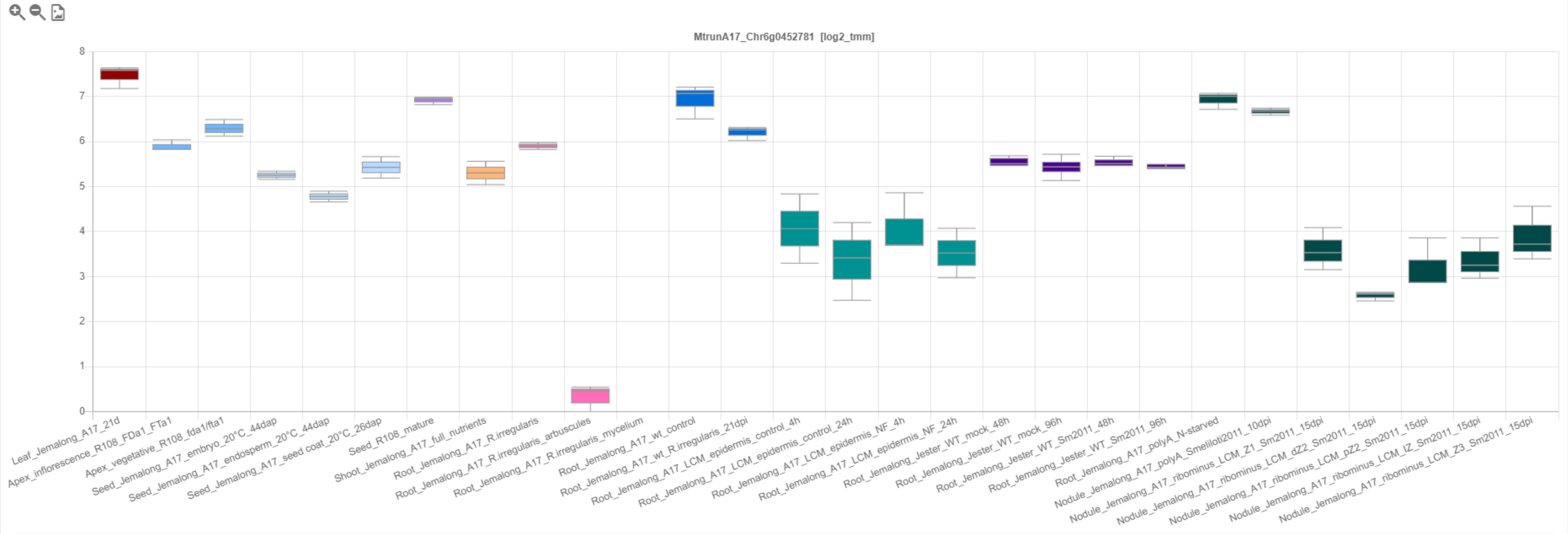

CP97: MtrunA17_Chr6g0457351

INRAE/CNRS Medicago Expressi | Expression Atlas Server

…pub/expressionAtlas/app/v3/aa_reference_dataset/MtrunA17_Chr6g0457351

MtExpress V3 | Core (20220901) | ! Switch to another dataset using the left menu

TMM | METADATA | SYNONYMOUS | ANNOTATION | GENOME PORTAL | LEGOO

mRNA: MtrunA17_Chr6g0457351;

Log2 TMM Normalisation using EdgeR (Core [20220901])

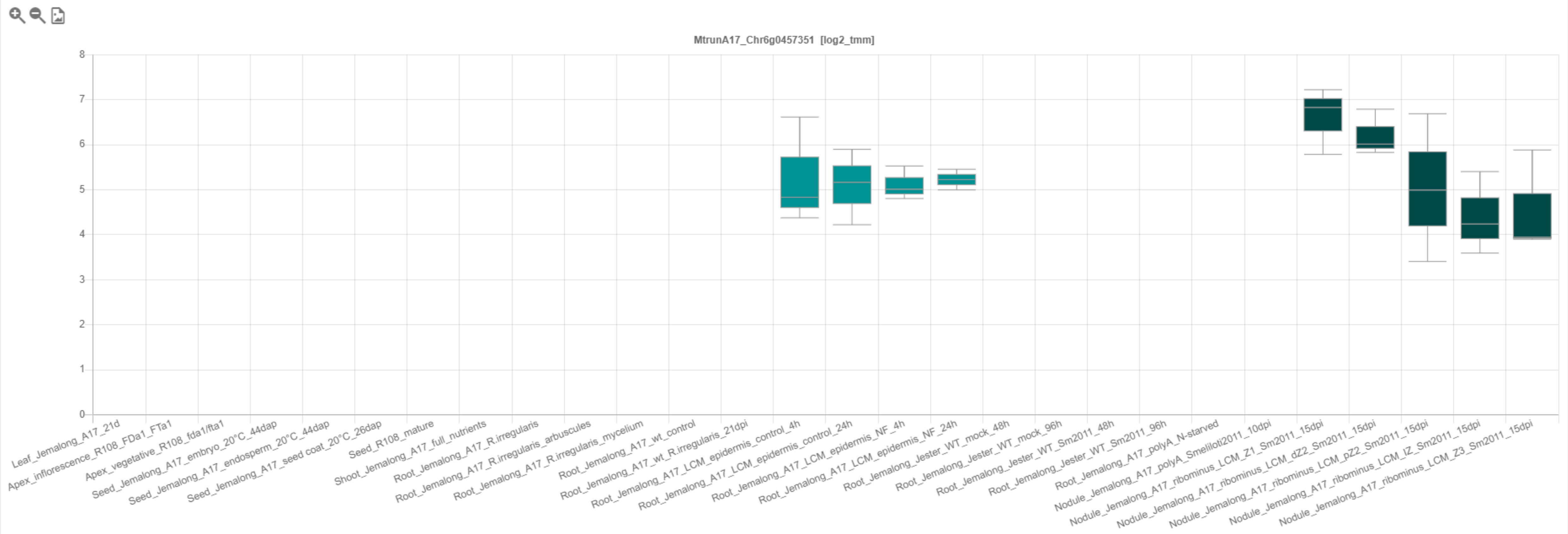

INRAE/CNRS Medicago Expressi

Expression Atlas Server

CP98: MtrunA17_Chr6g0457461

[illegible]b/expressionAtlas/app/v3/aa_reference_dataset/MtrunA17_Chr6g0457461

MtExpress V3

Core (20220901)

! Switch to another dataset using the left menu

TMM METADATA SYNONYMOUS ANNOTATION GENOME PORTAL LEGOO

mRNA: MtrunA17_Chr6g0457461 ;

Log2 TMM Normalisation using EdgeR (Core [20220901])

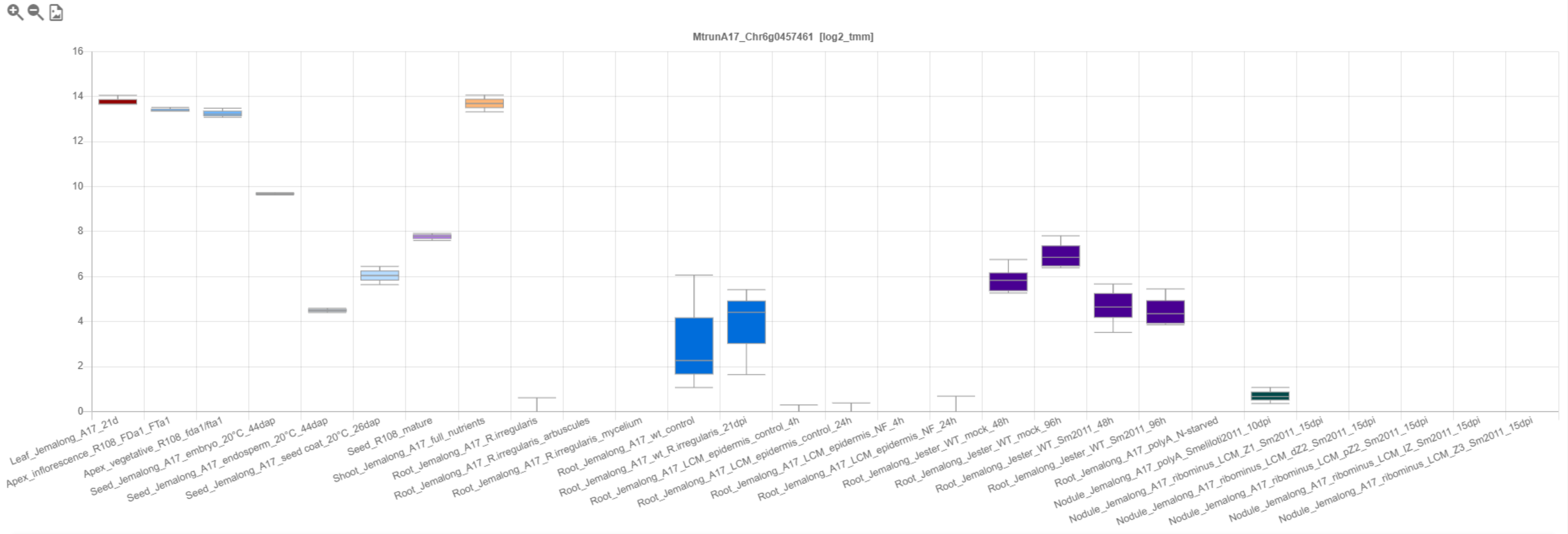

INRAE/CNRS Medicago Expressi | Expression Atlas Server

CP99: MtrunA17_Chr6g0457461

.../pub/expressionAtlas/app/v3/aa_reference_dataset/MtrunA17_Chr6g0457461

MtExpress V3 | Core (20220901)

! Switch to another dataset using the left menu

lipme INRAE cnrs

TMM | METADATA | SYNONYMOUS | ANNOTATION | GENOME PORTAL | LEGOO

mRNA: MtrunA17_Chr6g0457461;

Log2 TMM Normalisation using EdgeR (Core [20220901])

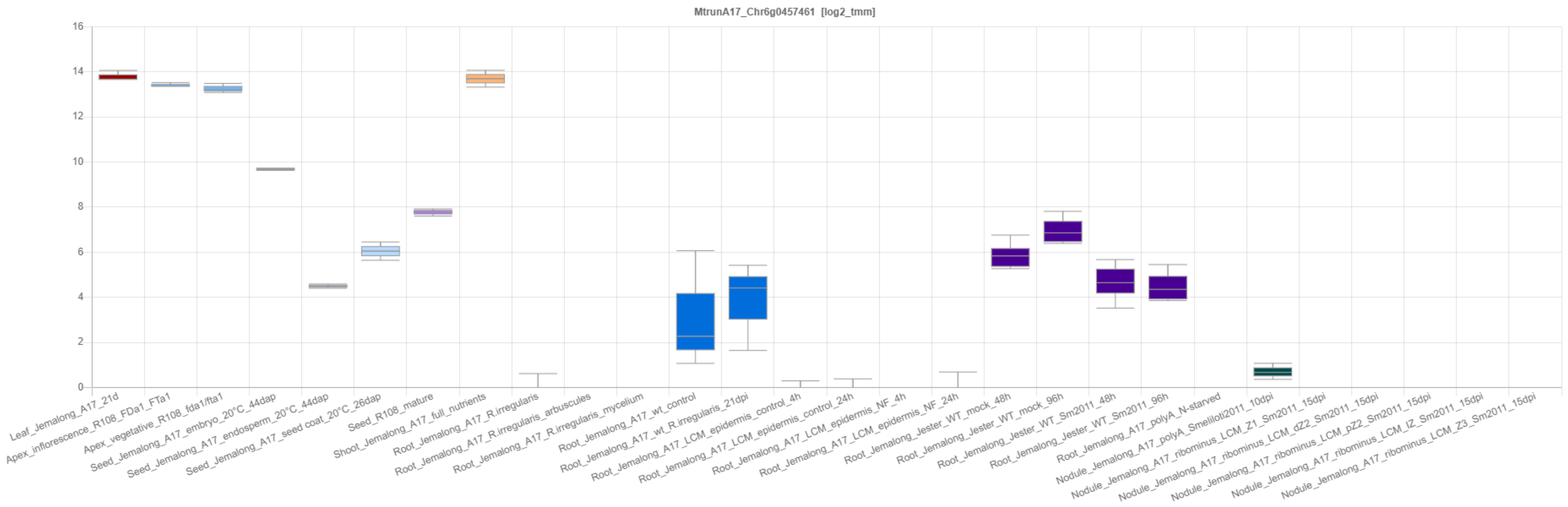

CP100: MtrunA17_Chr6g0457461

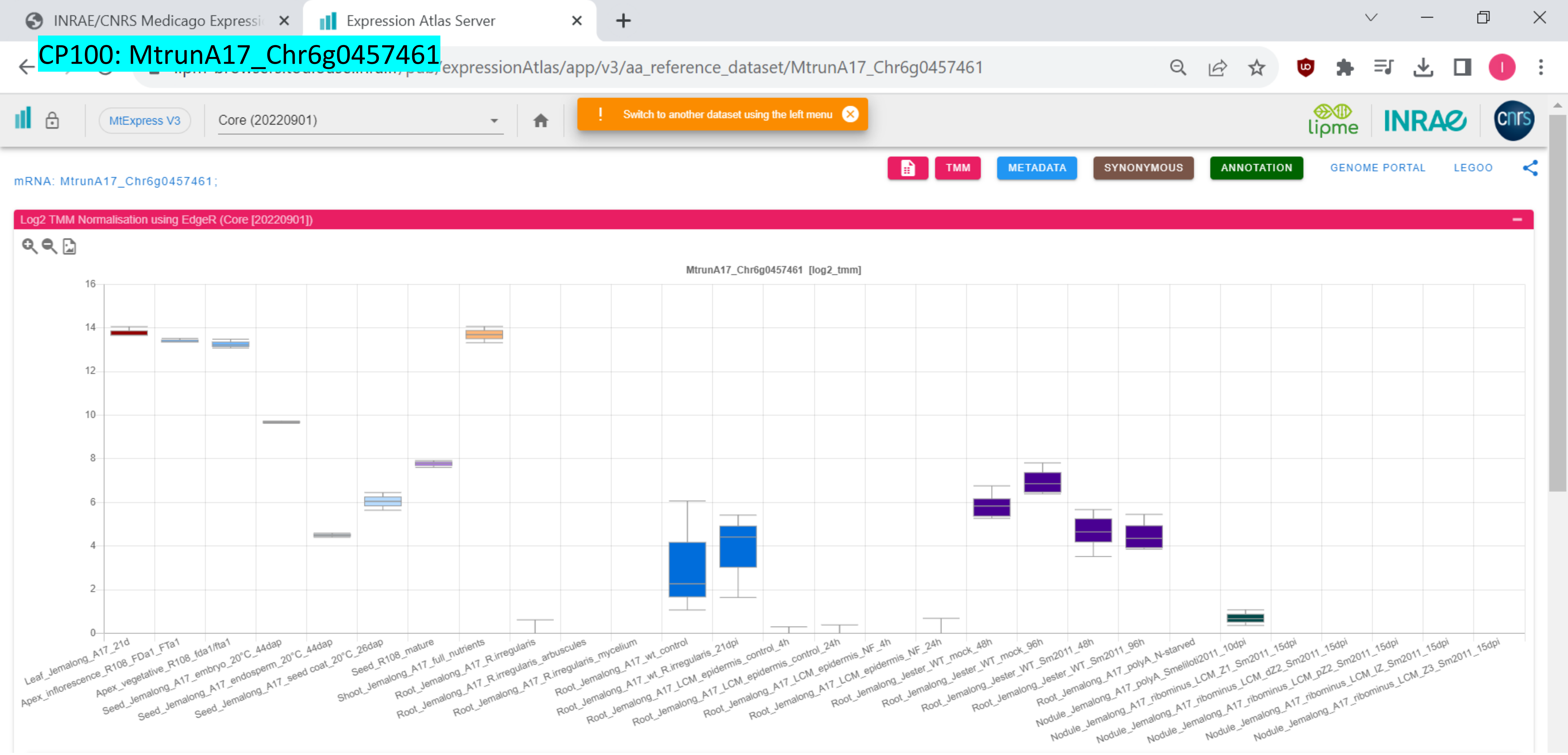

### CP101: MtrunA17_Chr6g0458091

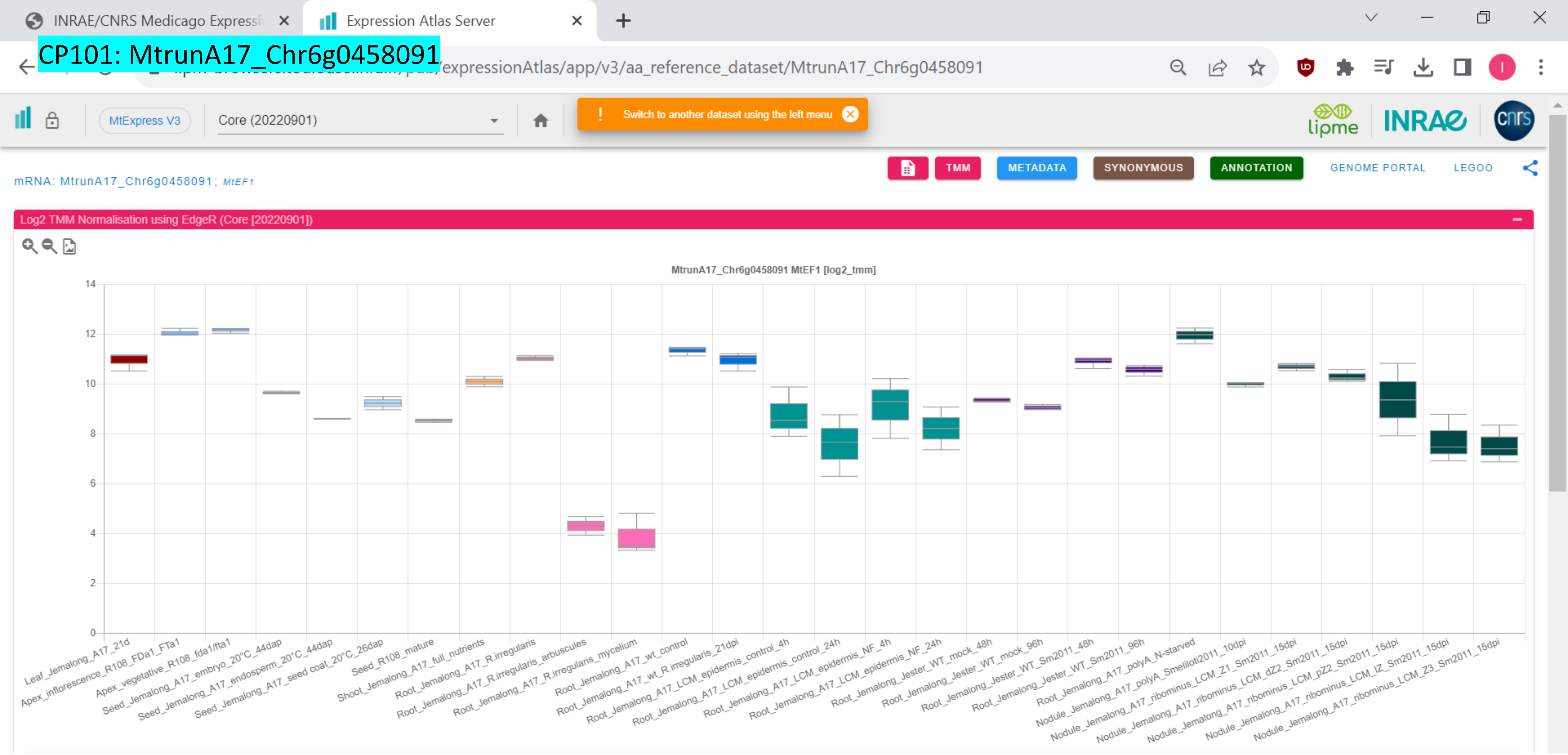

*CP102: MtrunA17_Chr6g0459481

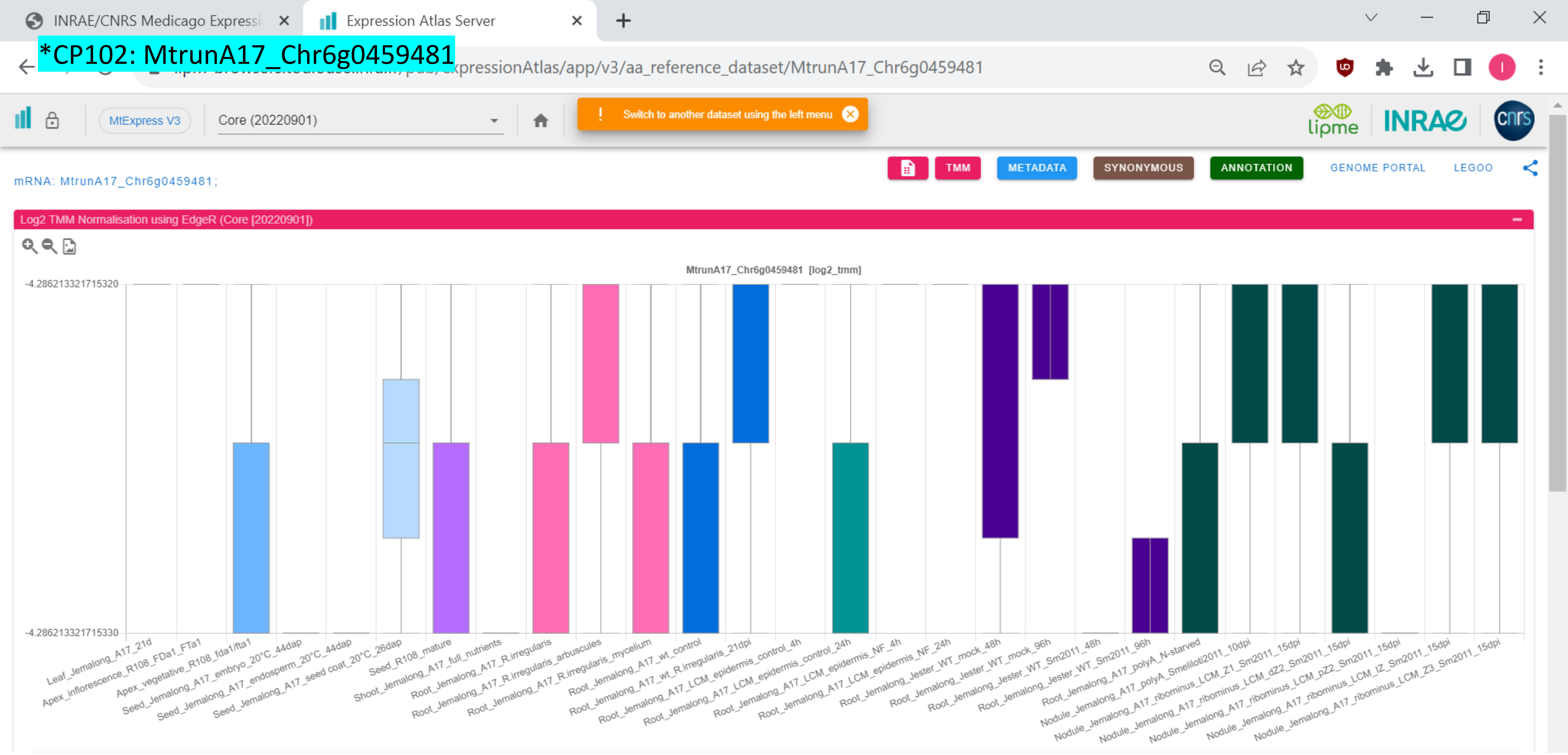

### CP103: MtrunA17_Chr6g0461931

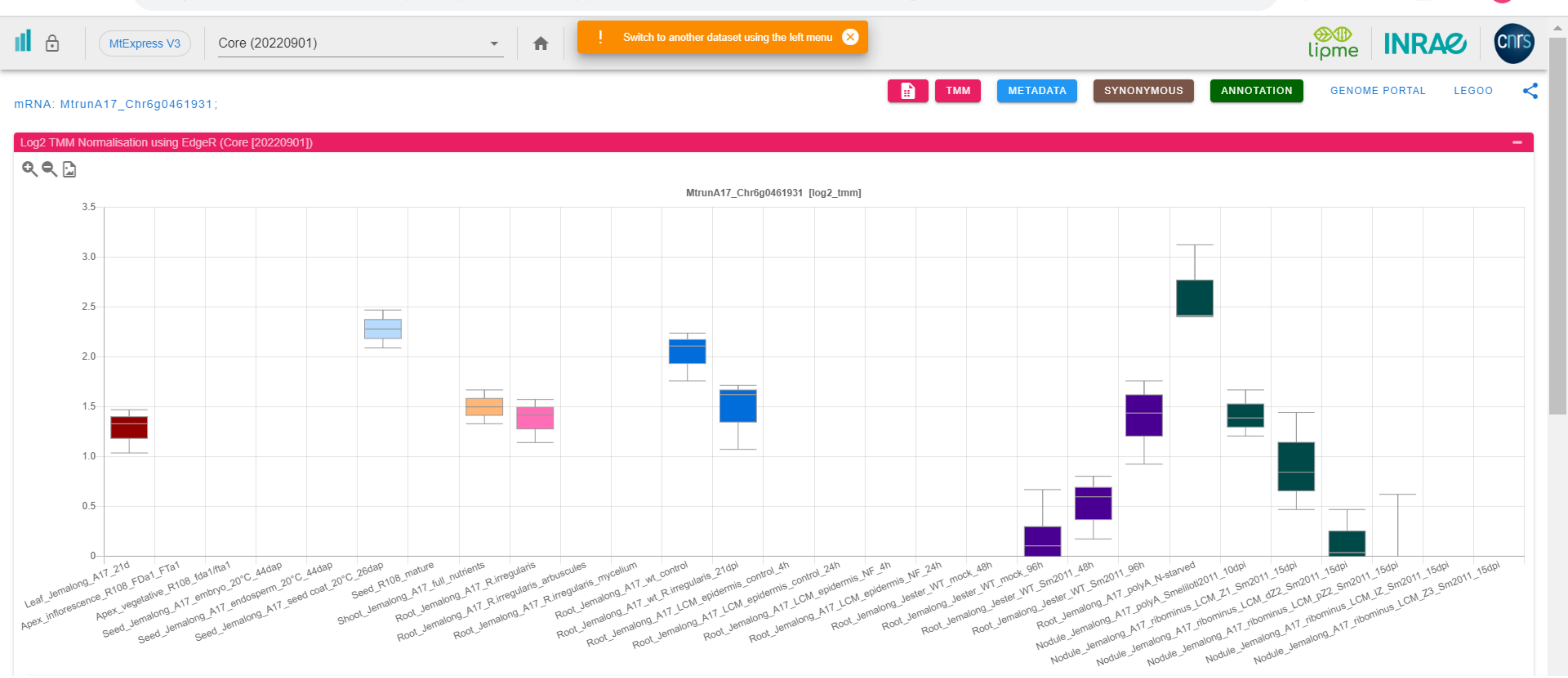

### CP104: MtrunA17_Chr6g0462271

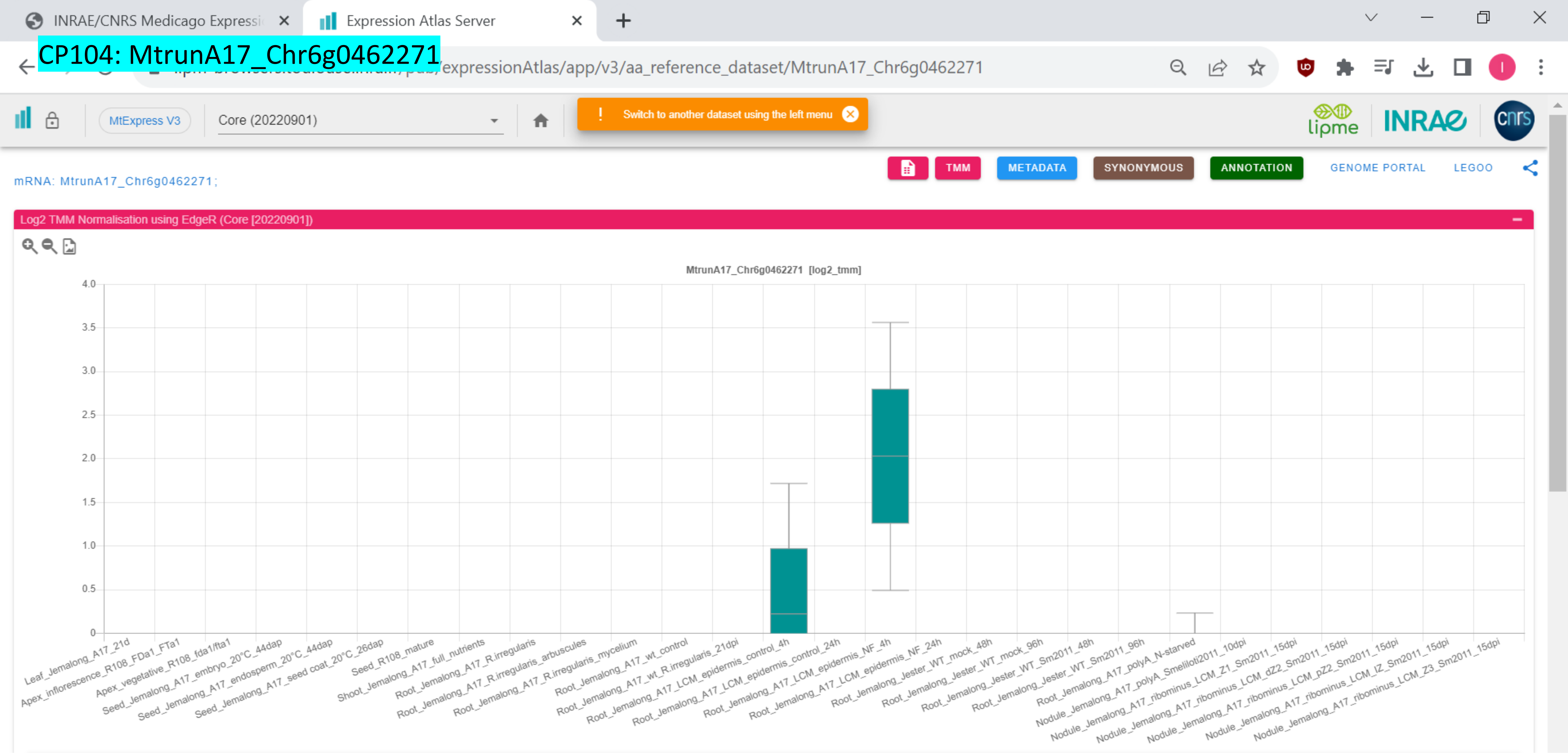

*CP105: MtrunA17_Chr6g0468411

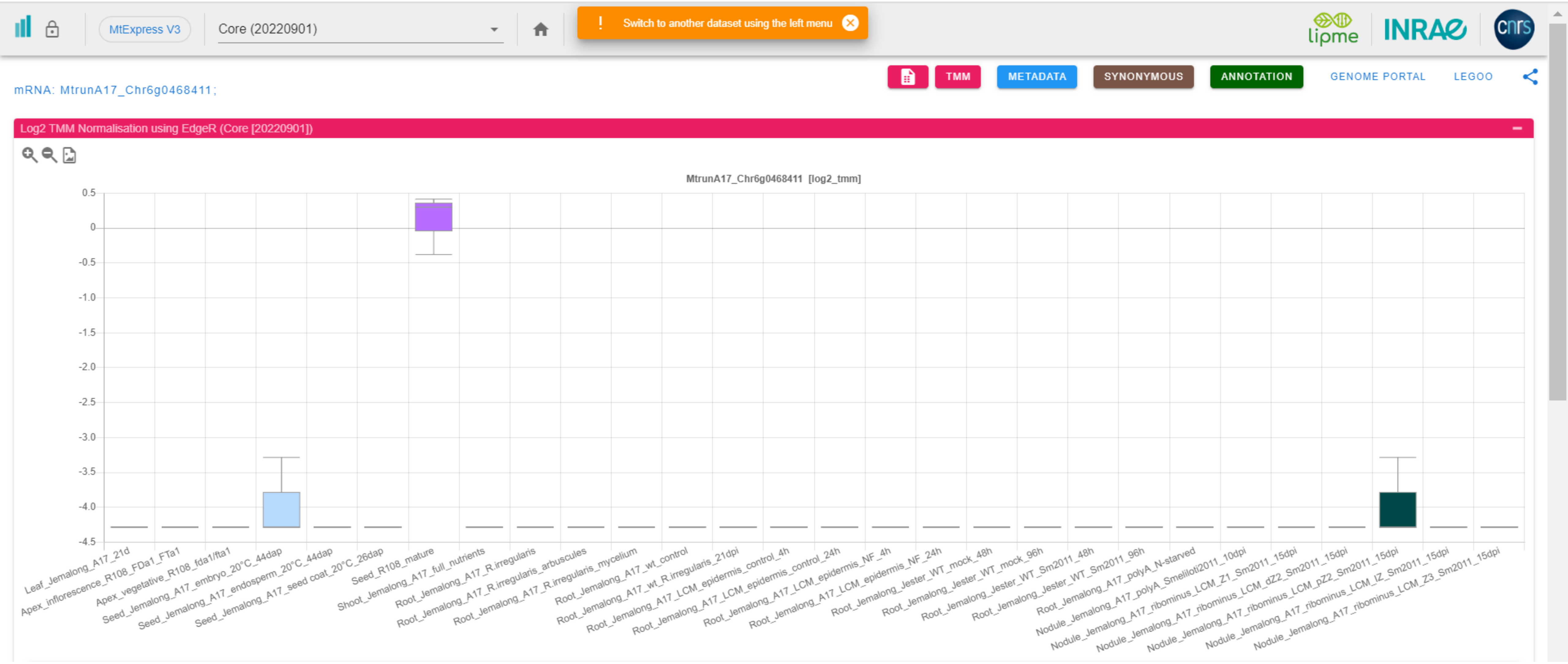

CP106: MtrunA17_Chr6g0476751

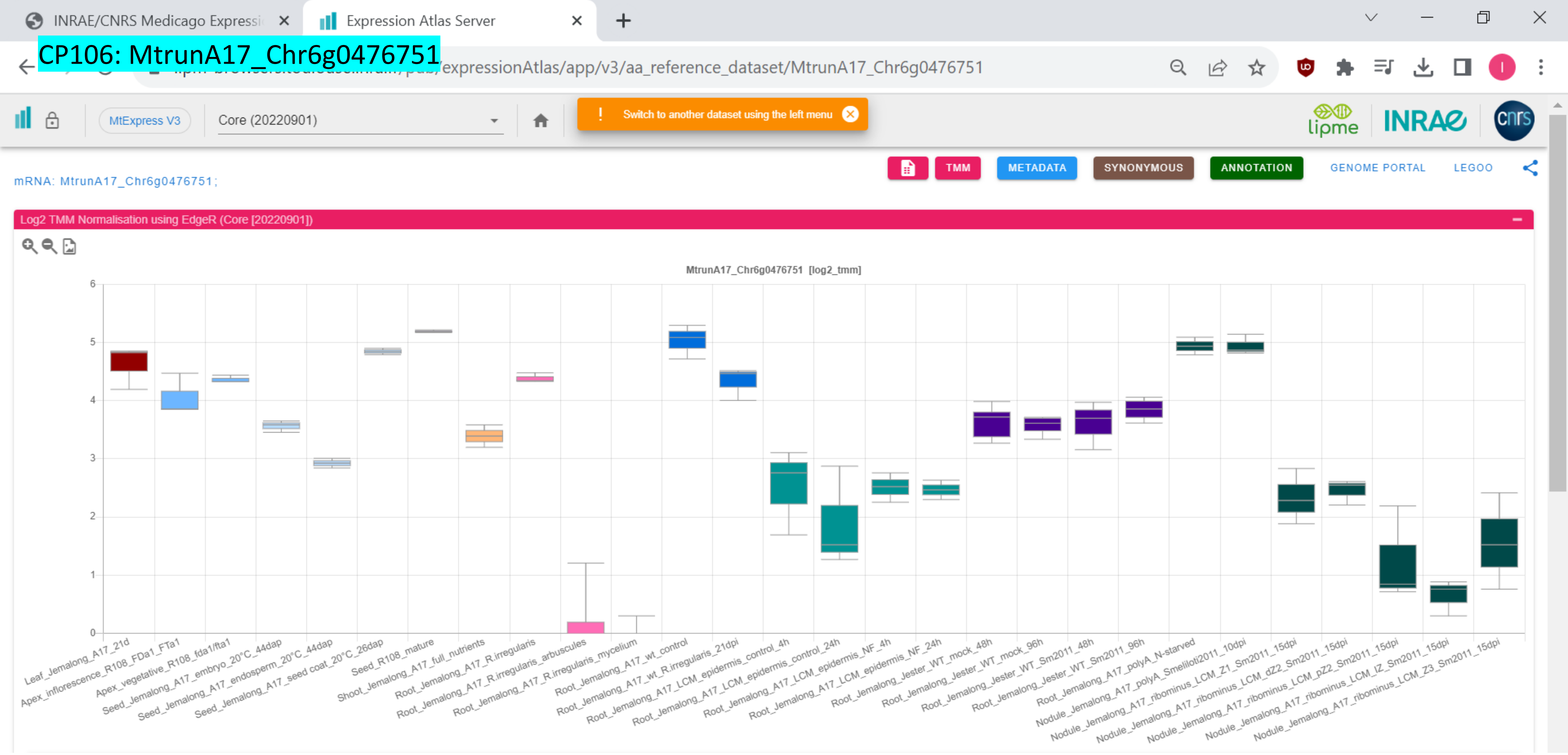

### CP107: MtrunA17_Chr6g0479001

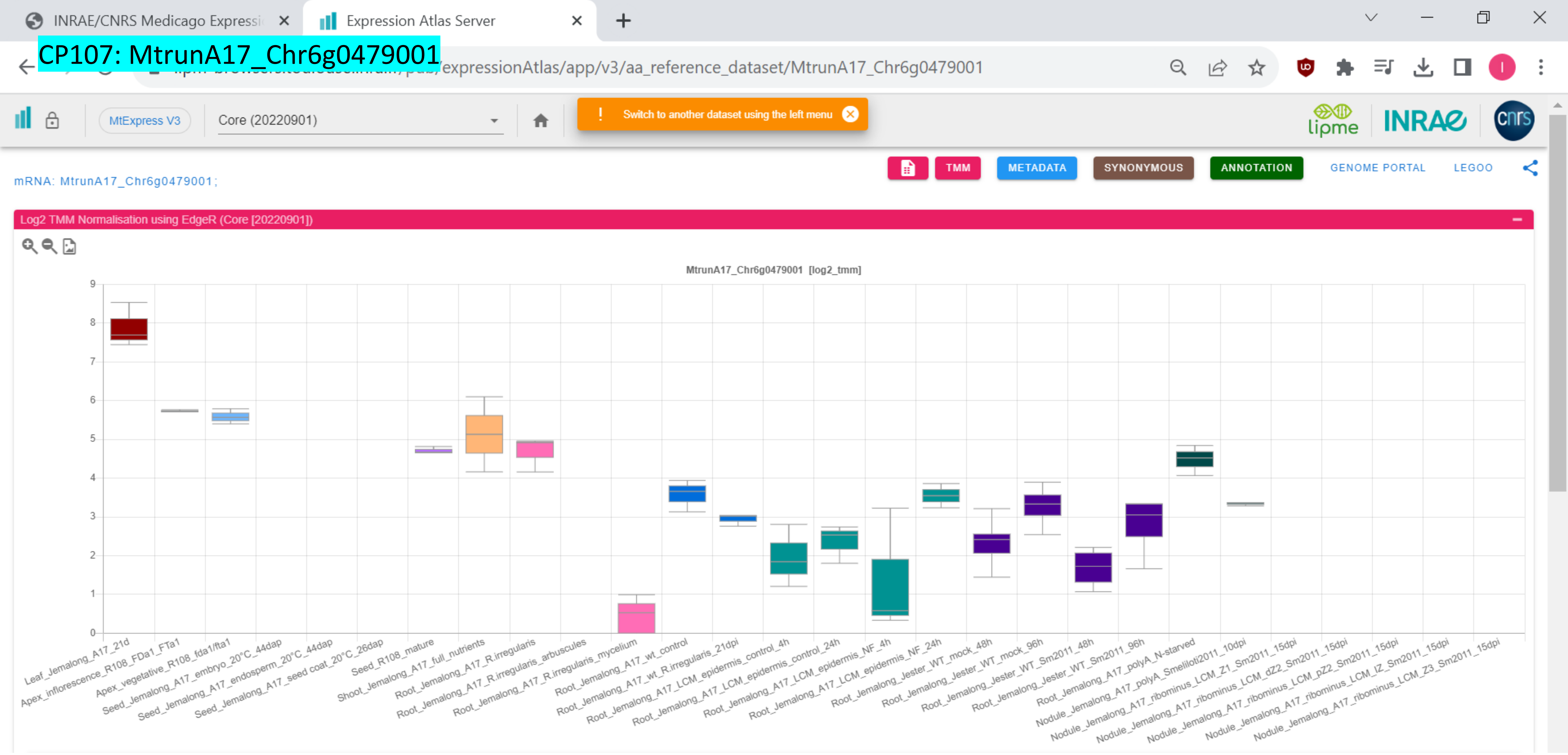

CP108: MtrunA17_Chr6g0485321

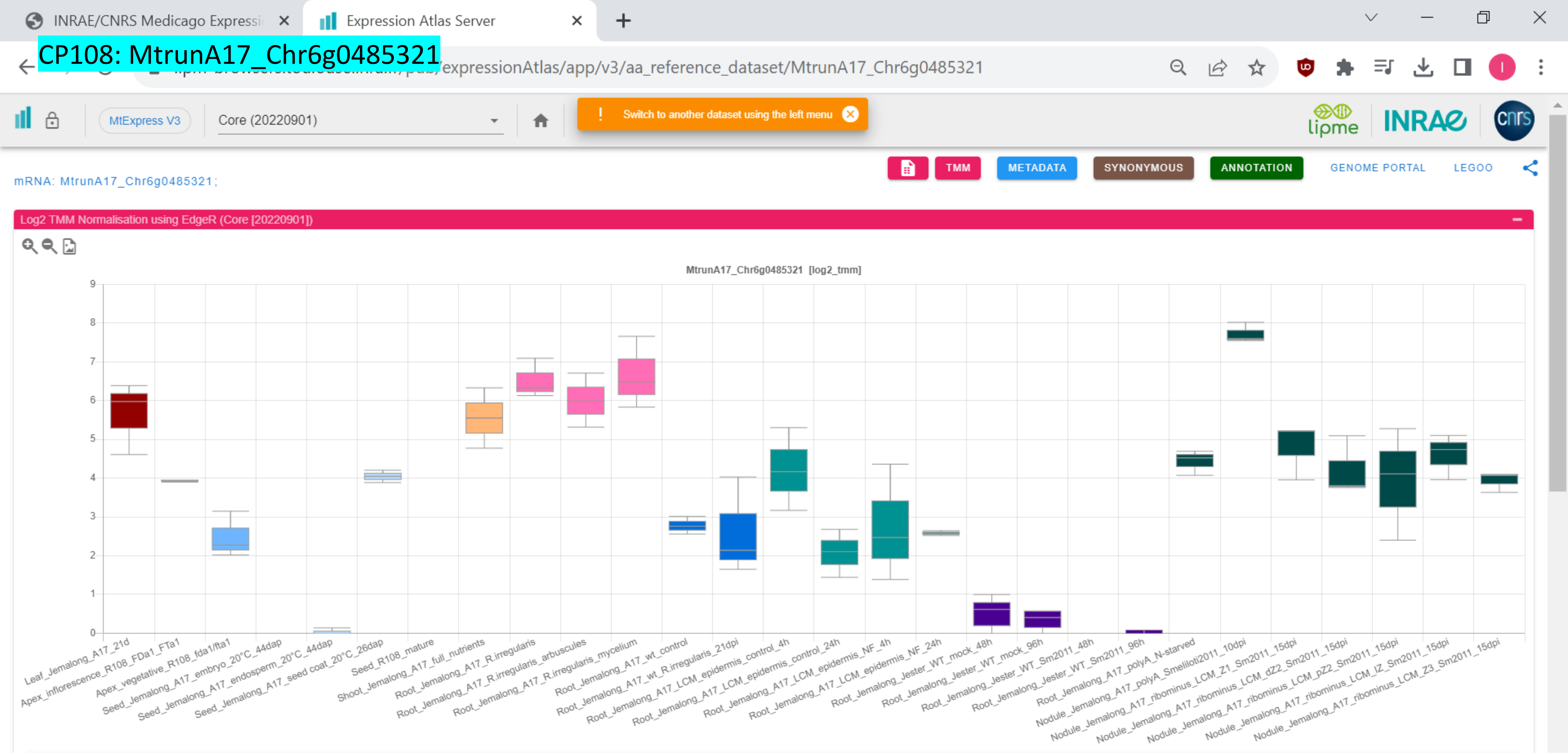

### CP109: MtrunA17_Chr6g0486961

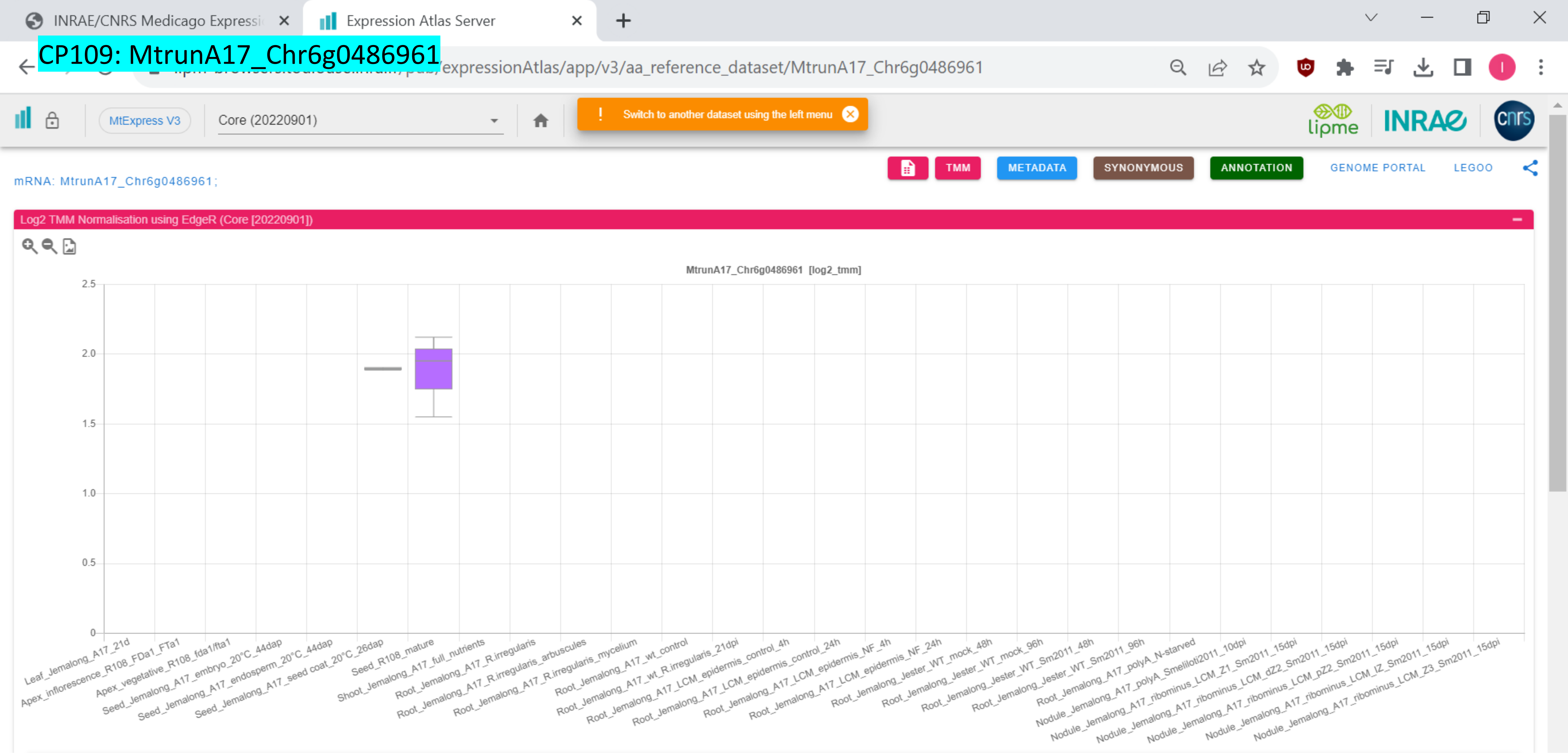

### CP110: MtrunA17_Chr7g0214741

INRAE/CNRS Medicago Expressi… | Expression Atlas Server

…/expressionAtlas/app/v3/aa_reference_dataset/MtrunA17_Chr7g0214741

MtExpress V3 | Core (20220901)

! Switch to another dataset using the left menu

TMM | METADATA | SYNONYMOUS | ANNOTATION | GENOME PORTAL | LEGOO

mRNA: MtrunA17_Chr7g0214741;

Log2 TMM Normalisation using EdgeR (Core [20220901])

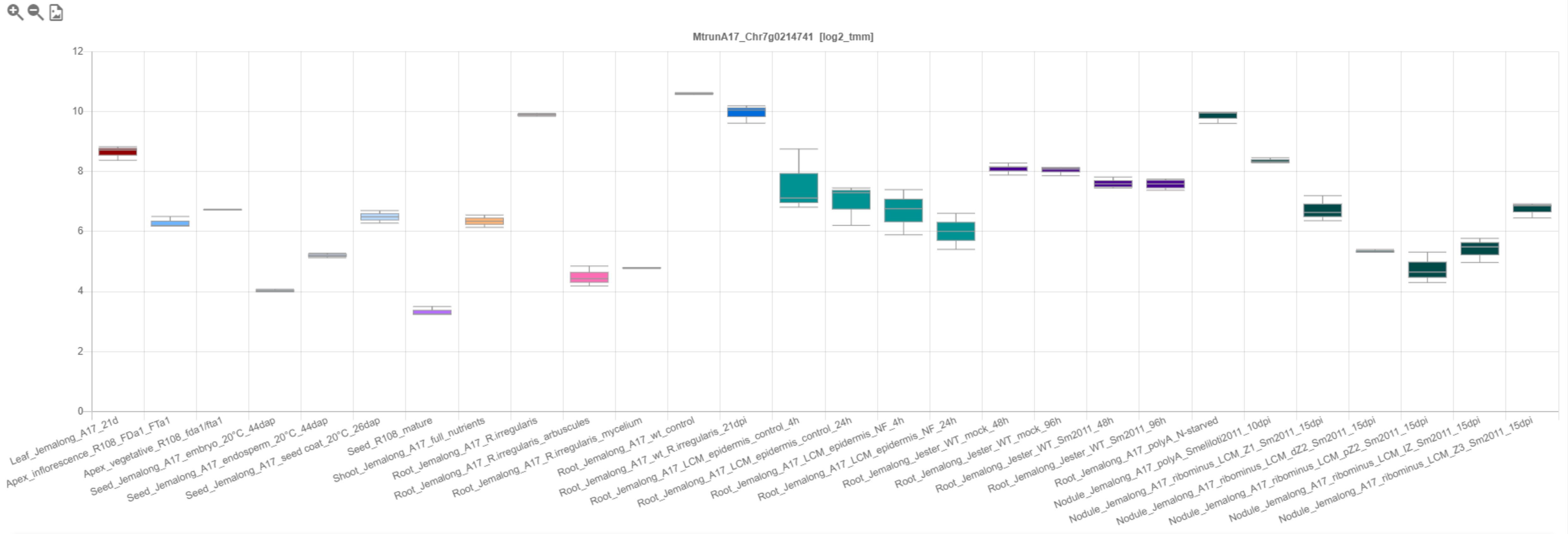

### CP111: MtrunA17_Chr7g0214911

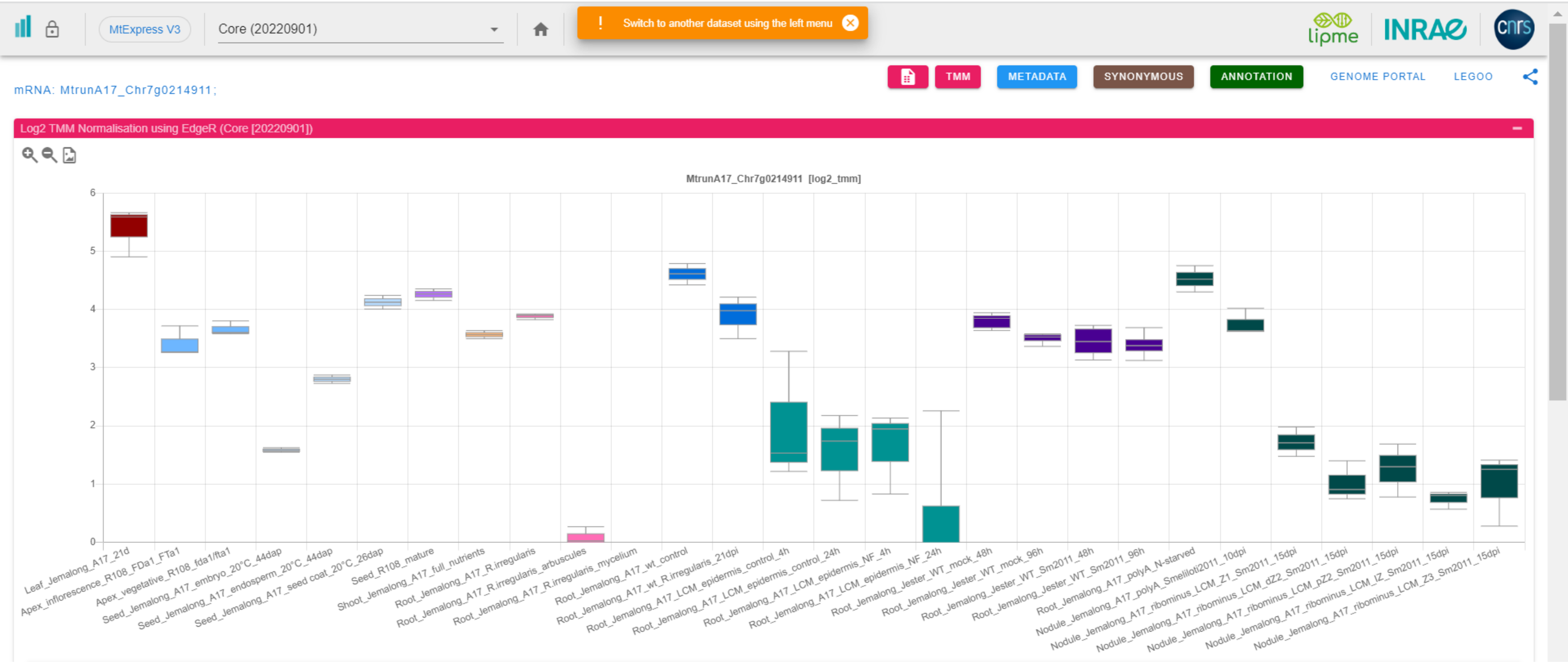

CP112: MtrunA17_Chr7g0219691

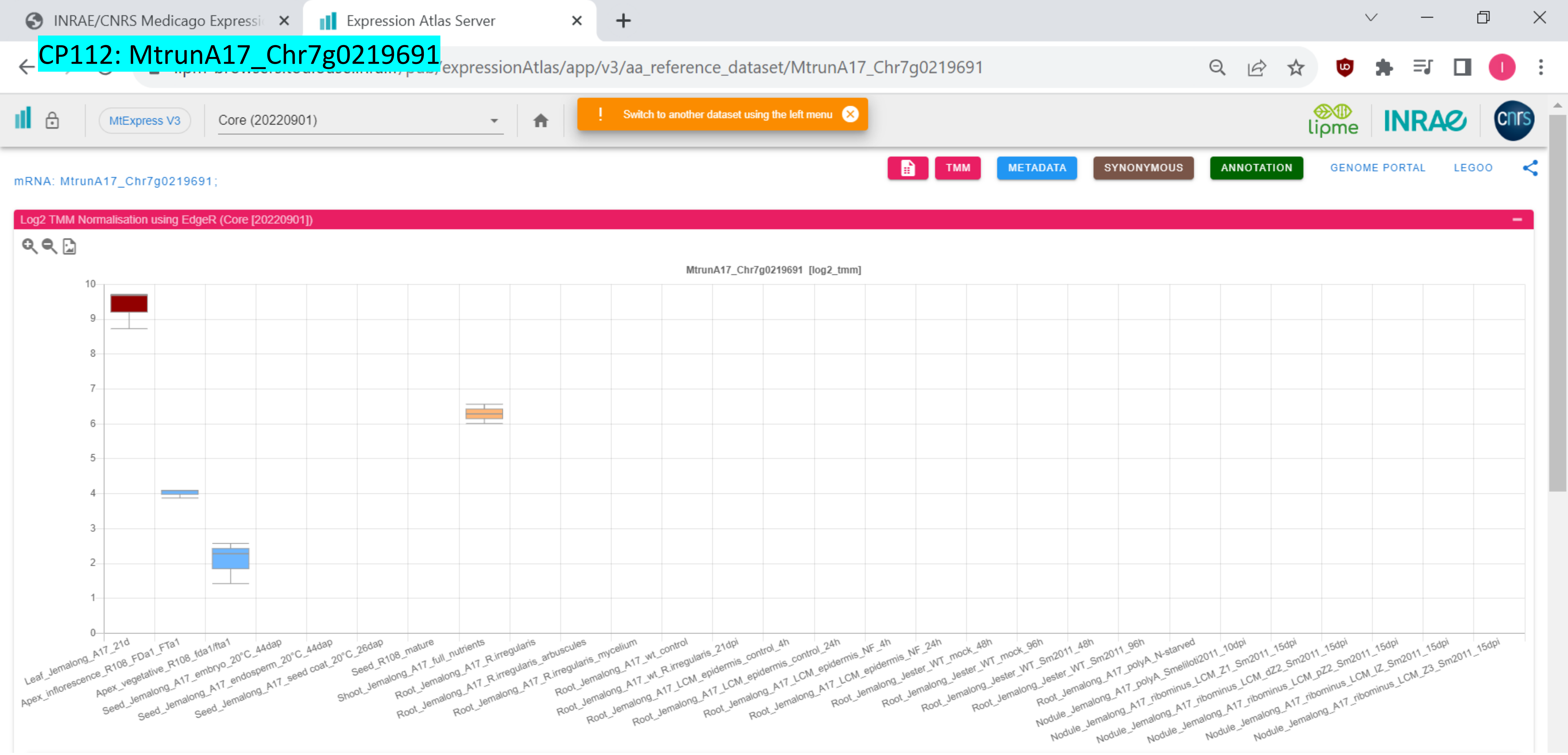

*CP113: MtrunA17_Chr7g0221631

INRAE/CNRS Medicago Expressi | Expression Atlas Server

...expressionAtlas/app/v3/aa_reference_dataset/MtrunA17_Chr7g0221631

MtExpress V3 | Core (20220901) | Switch to another dataset using the left menu

TMM | METADATA | SYNONYMOUS | ANNOTATION | GENOME PORTAL | LEGOO

mRNA: MtrunA17_Chr7g0221631;

Log2 TMM Normalisation using EdgeR (Core [20220901])

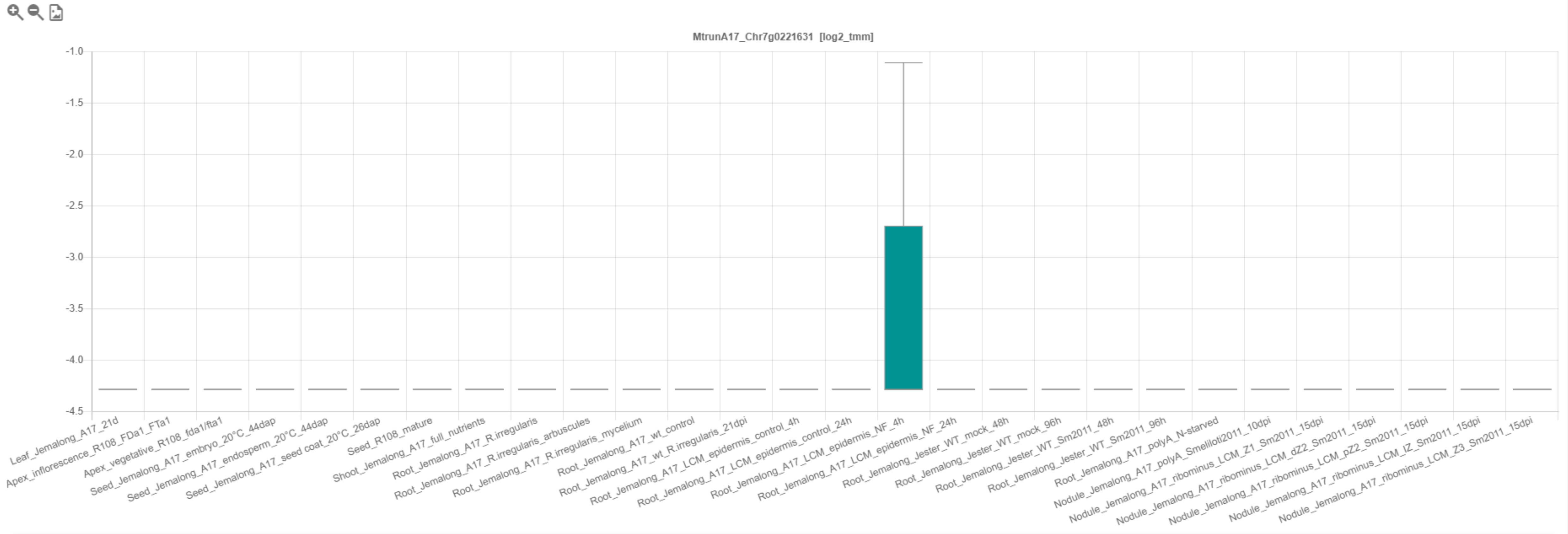

### CP114: MtrunA17_Chr7g0229401

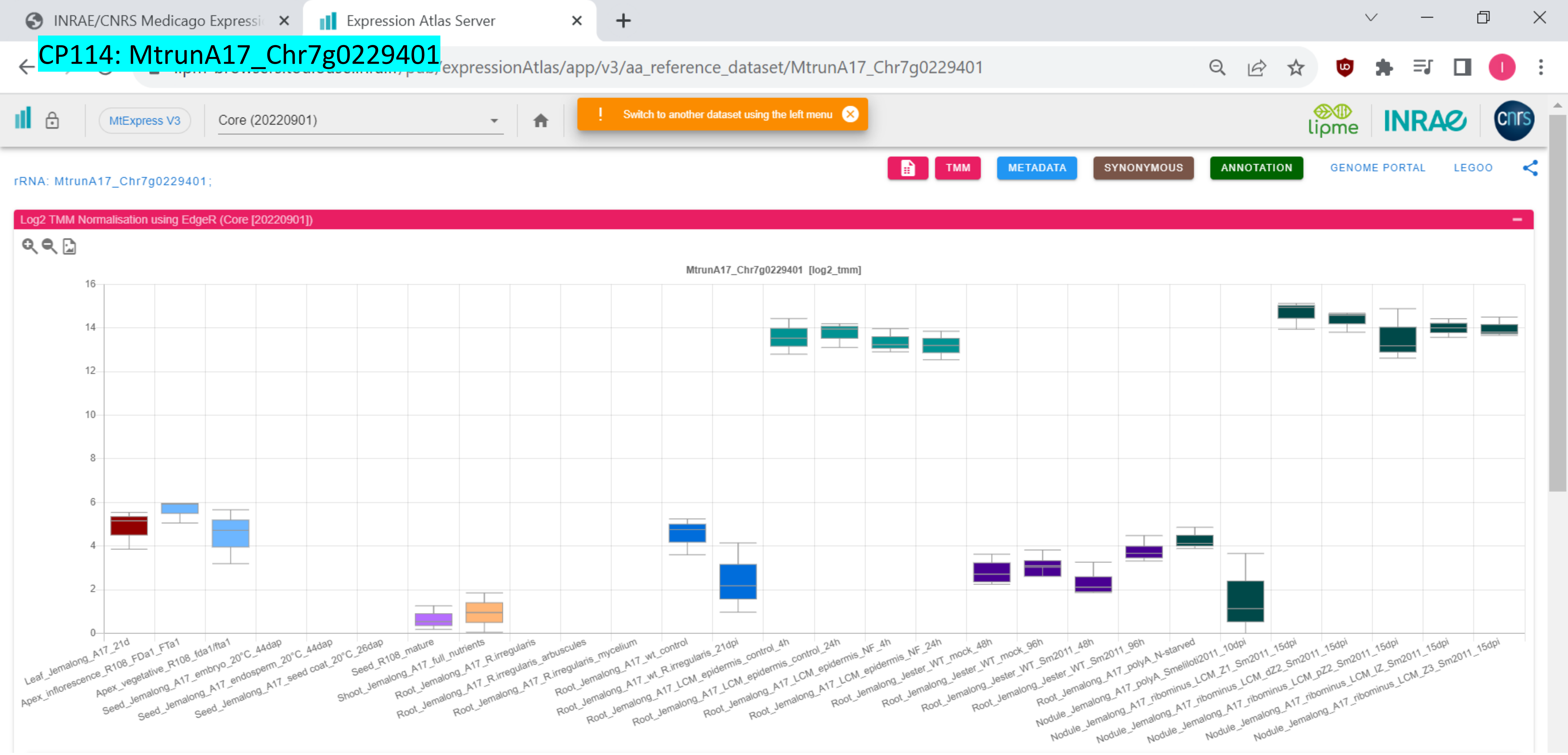

### CP115: MtrunA17_Chr7g0230341

INRAE/CNRS Medicago Expressi | Expression Atlas Server

.../expressionAtlas/app/v3/aa_reference_dataset/MtrunA17_Chr7g0230341

MtExpress V3 | Core (20220901)

! Switch to another dataset using the left menu

TMM | METADATA | SYNONYMOUS | ANNOTATION | GENOME PORTAL | LEGOO

lipme INRAE cnrs

mRNA: MtrunA17_Chr7g0230341;

Log2 TMM Normalisation using EdgeR (Core [20220901])

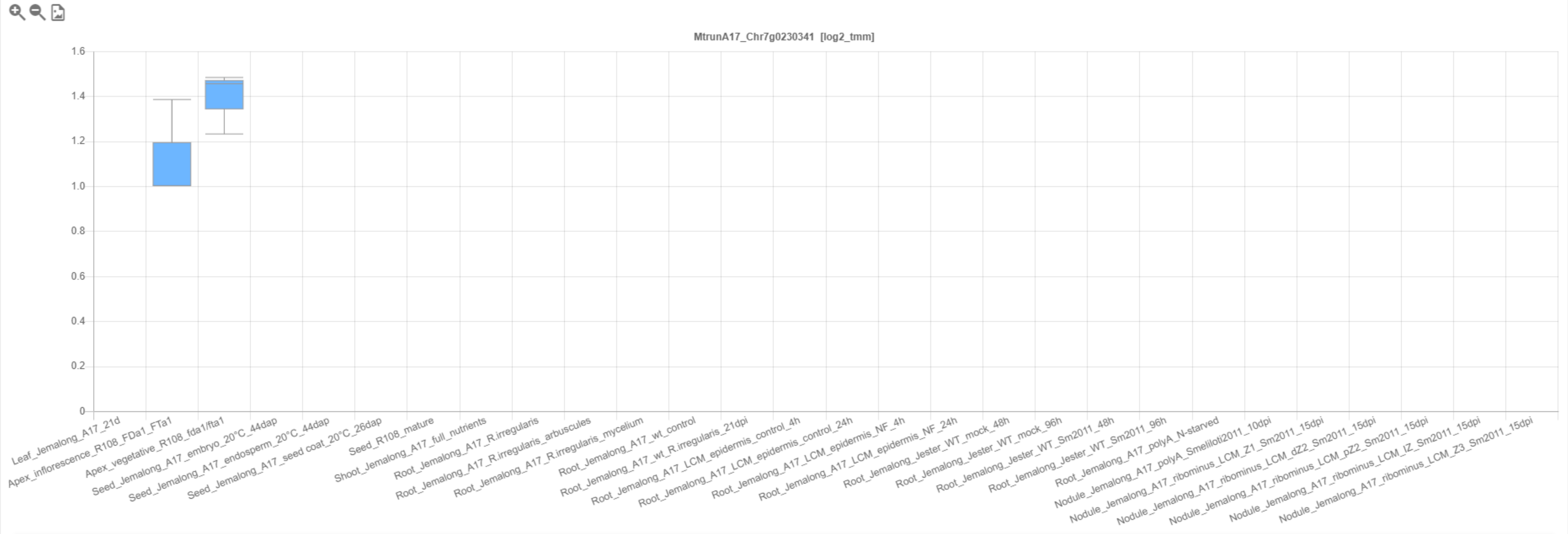

### CP116: MtrunA17_Chr7g0232851

INRAE/CNRS Medicago Expressi… | Expression Atlas Server

…/expressionAtlas/app/v3/aa_reference_dataset/MtrunA17_Chr7g0232851

MtExpress V3 | Core (20220901) | Switch to another dataset using the left menu

lipme INRAE cnrs

TMM | METADATA | SYNONYMOUS | ANNOTATION | GENOME PORTAL | LEGOO

mRNA: MtrunA17_Chr7g0232851 ; *MtSDG53*

Log2 TMM Normalisation using EdgeR (Core [20220901])

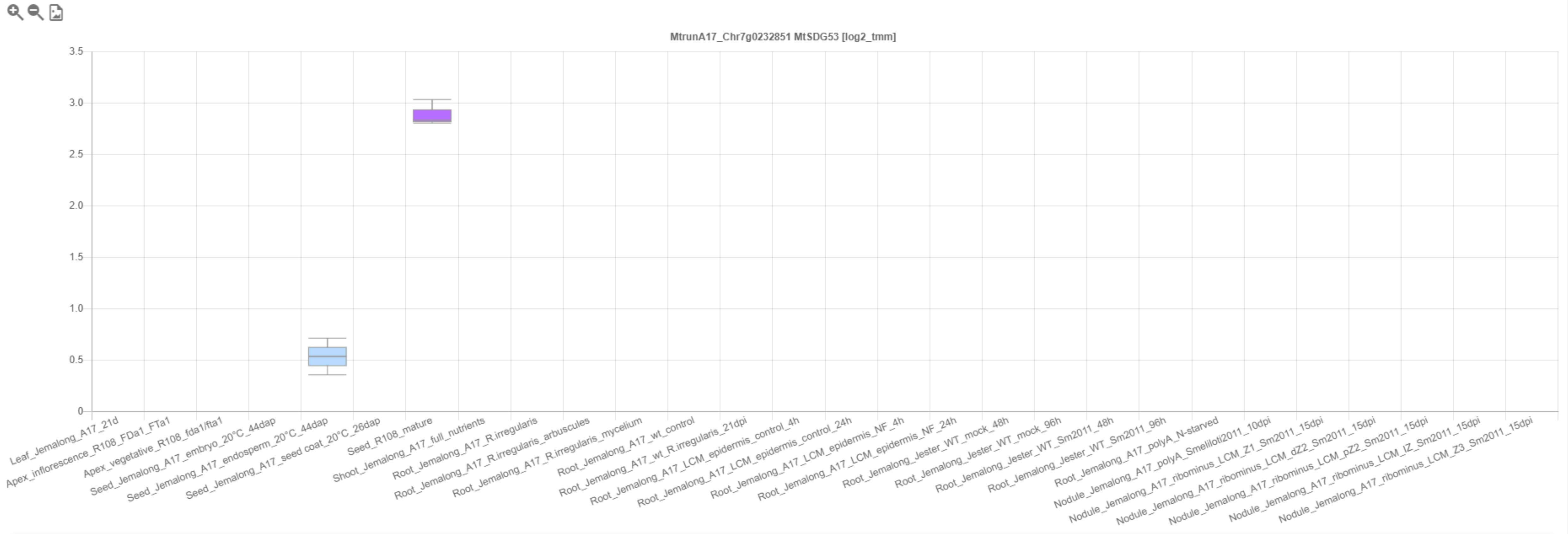

### CP117: MtrunA17_Chr7g0237331

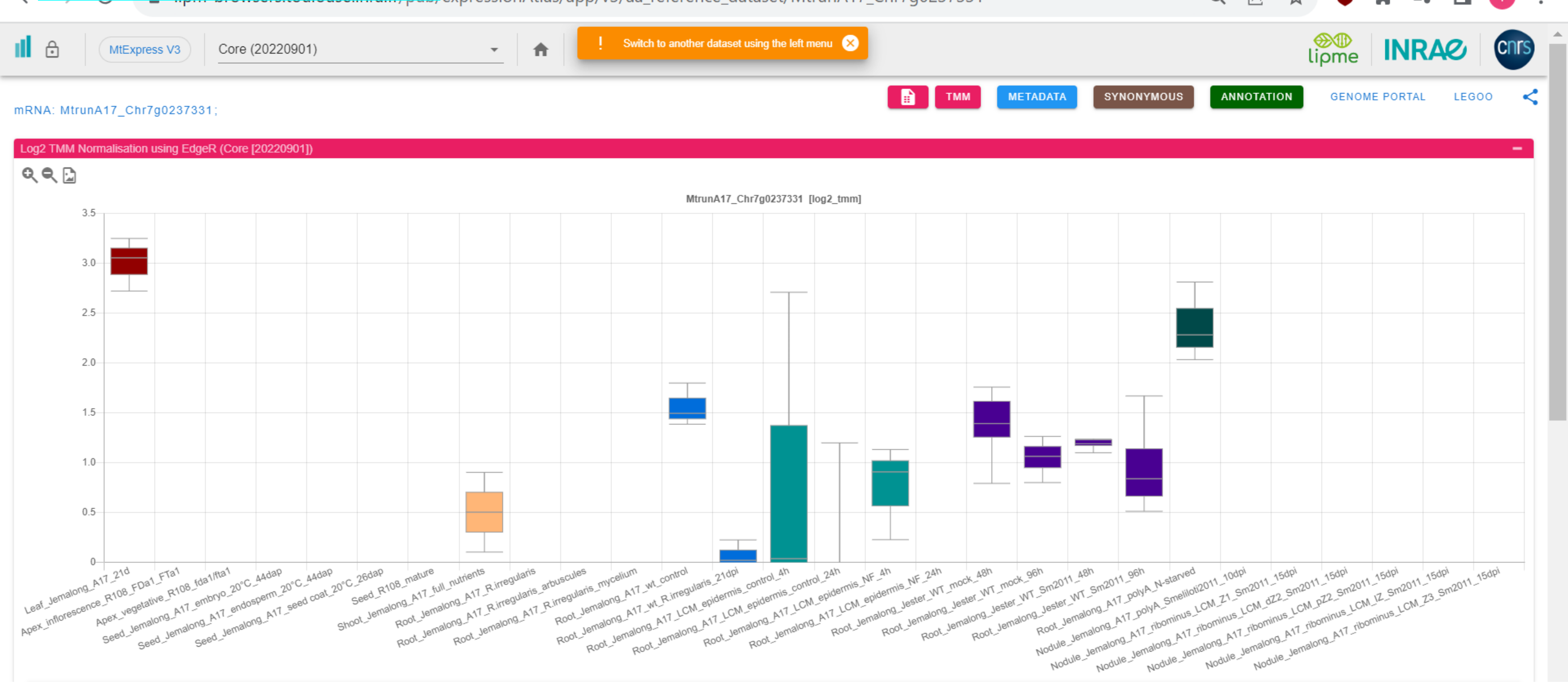

*CP118: MtrunA17_Chr7g0251971

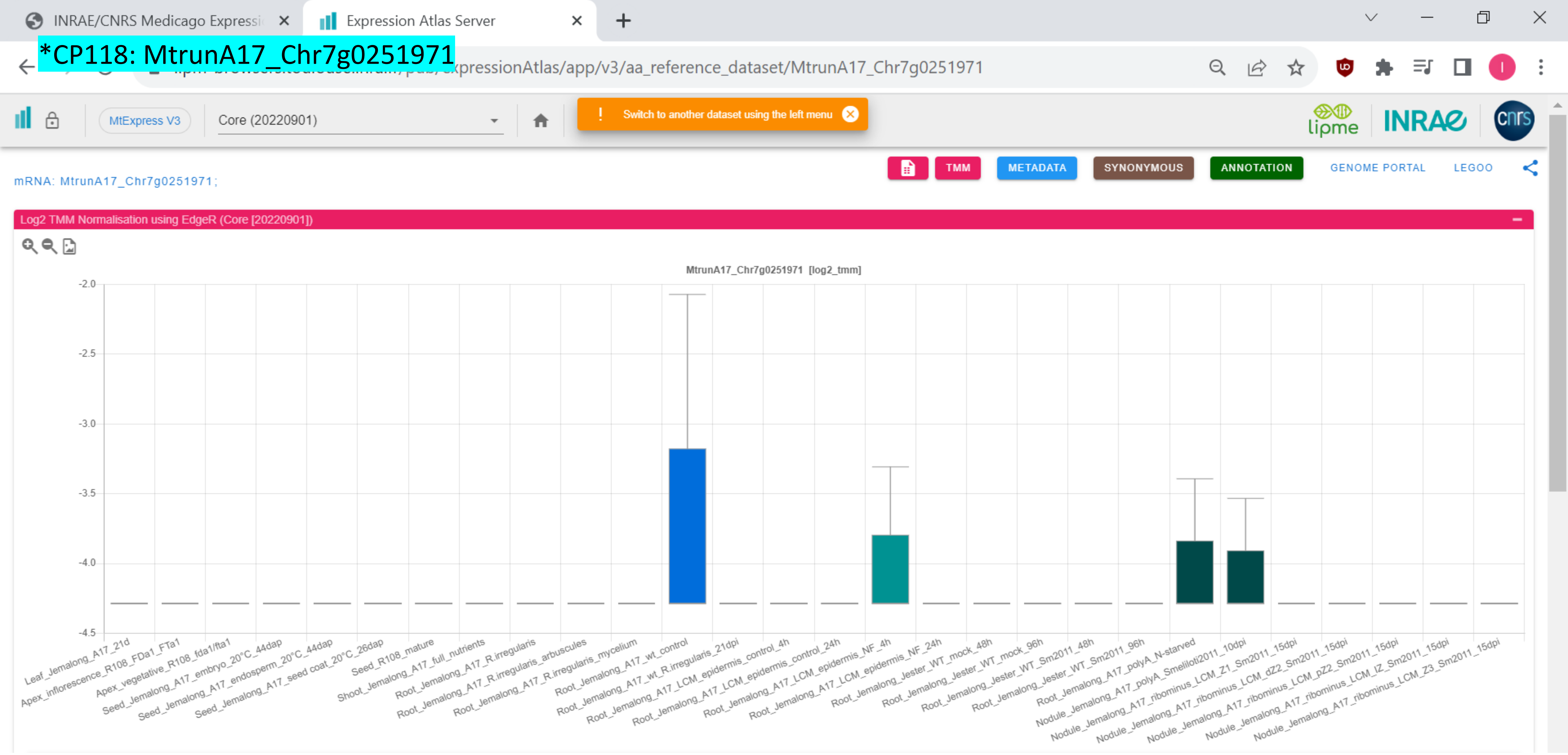

### CP119: MtrunA17_Chr7g0259611

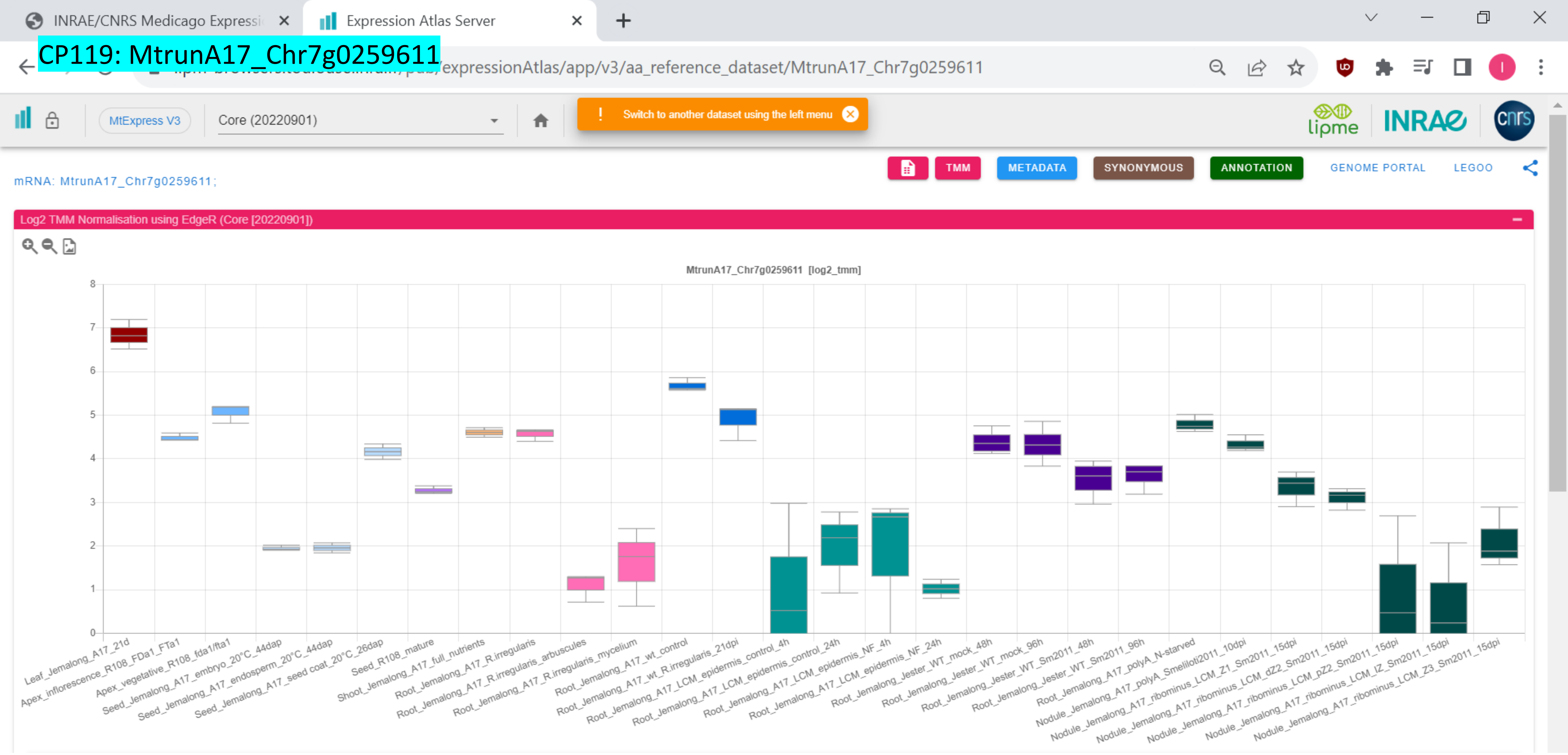

CP120: MtrunA17_Chr7g0262821

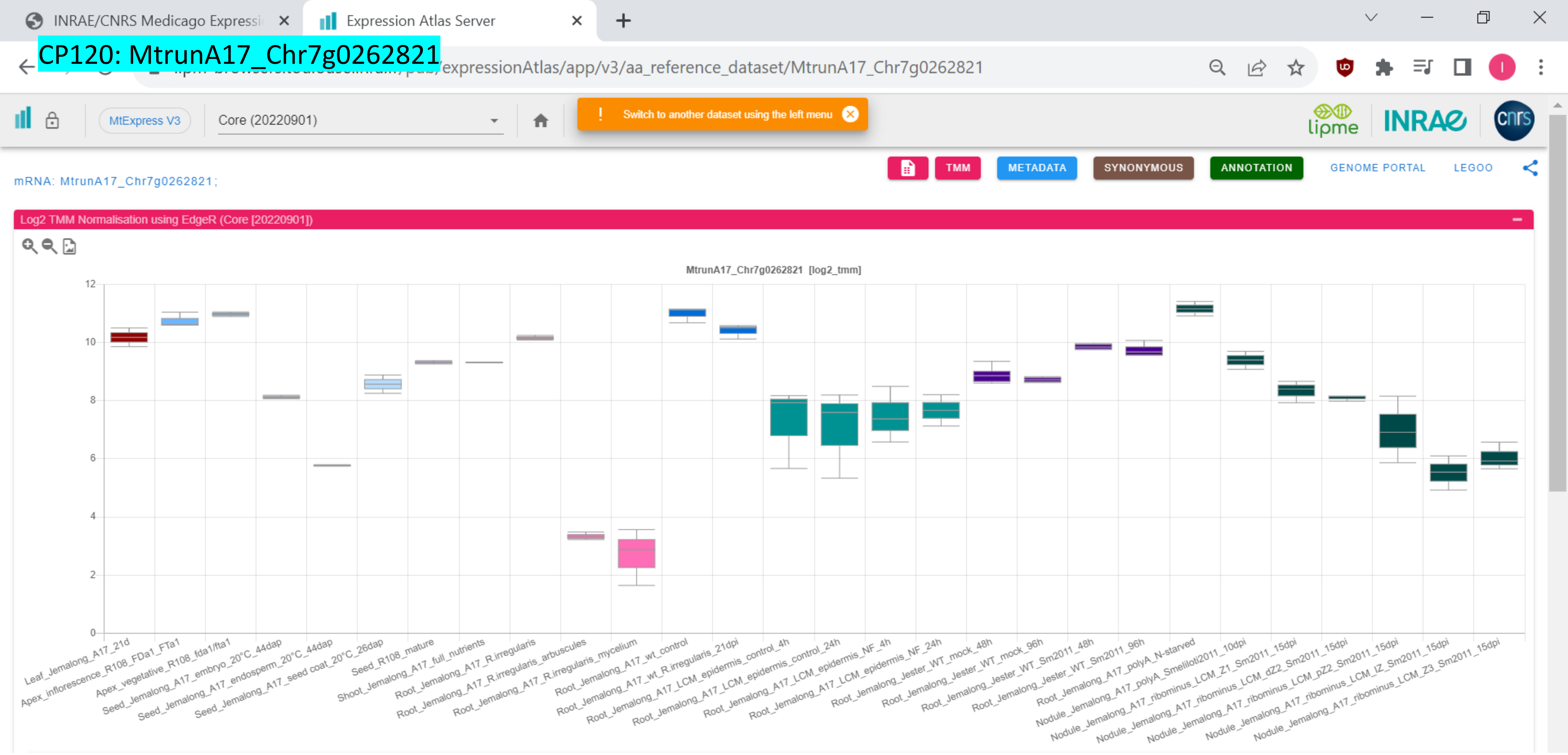

### CP121: MtrunA17_Chr7g0270811

mRNA: MtrunA17_Chr7g0270811;

Log2 TMM Normalisation using EdgeR (Core [20220901])

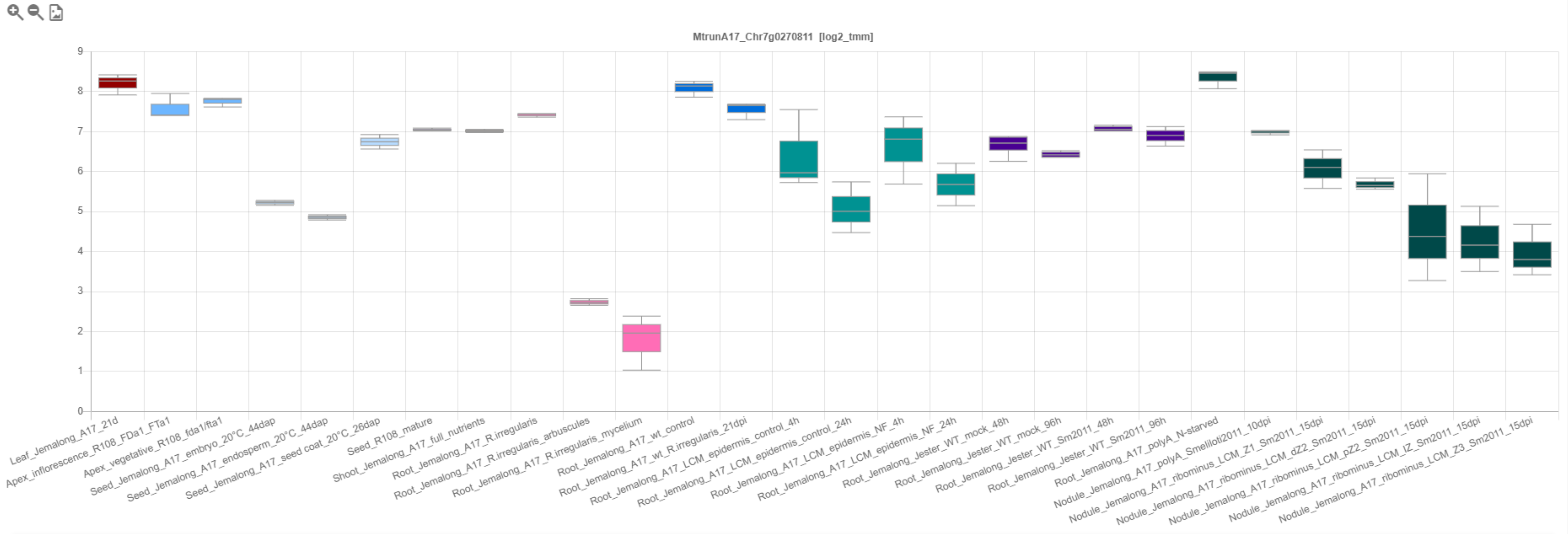

### CP122: MtrunA17_Chr7g1034306

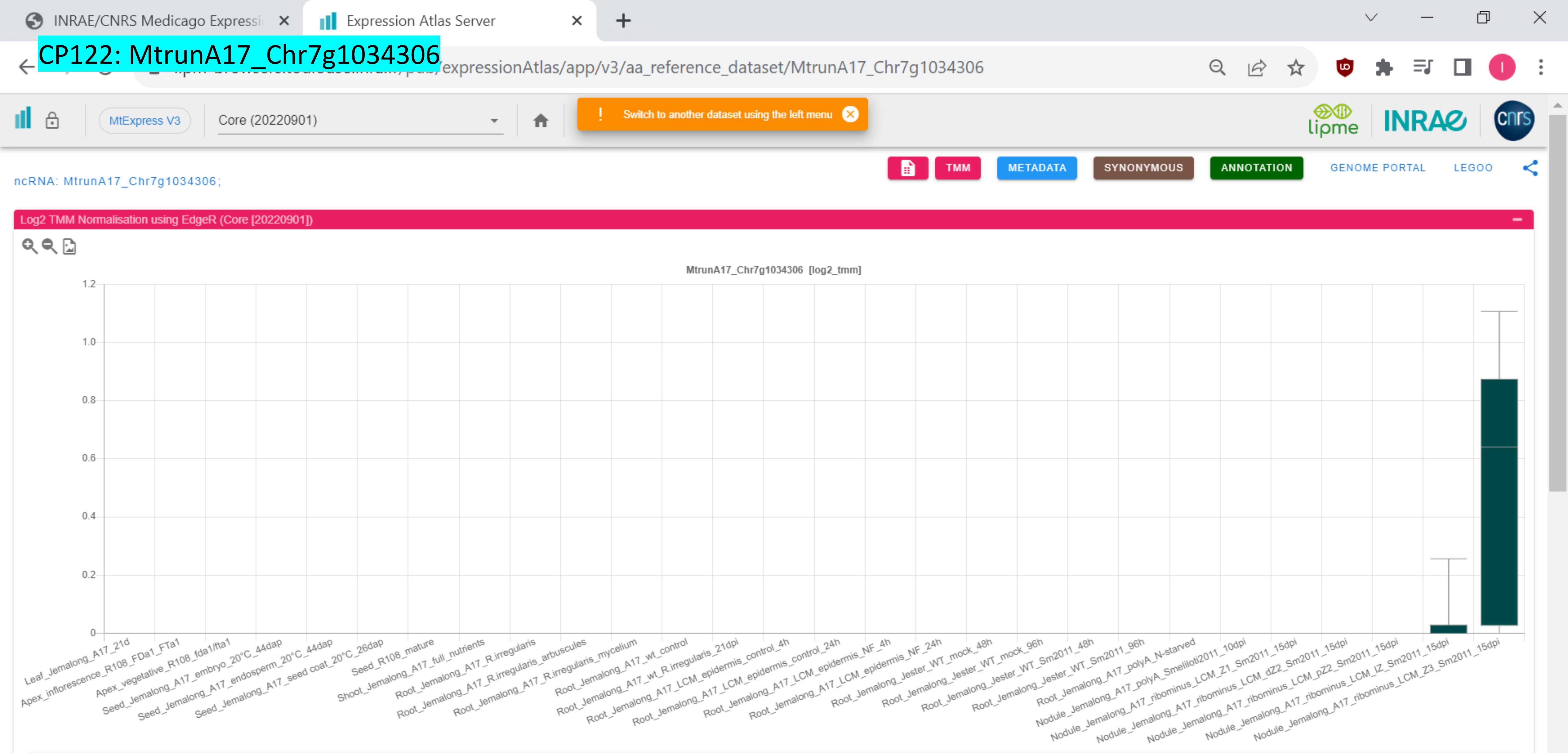

### CP123: MtrunA17_Chr8g0338111

INRAE/CNRS Medicago Expressi | Expression Atlas Server

…/expressionAtlas/app/v3/aa_reference_dataset/MtrunA17_Chr8g0338111

MtExpress V3 | Core (20220901)

! Switch to another dataset using the left menu

TMM | METADATA | SYNONYMOUS | ANNOTATION | GENOME PORTAL | LEGOO

mRNA: MtrunA17_Chr8g0338111;

Log2 TMM Normalisation using EdgeR (Core [20220901])

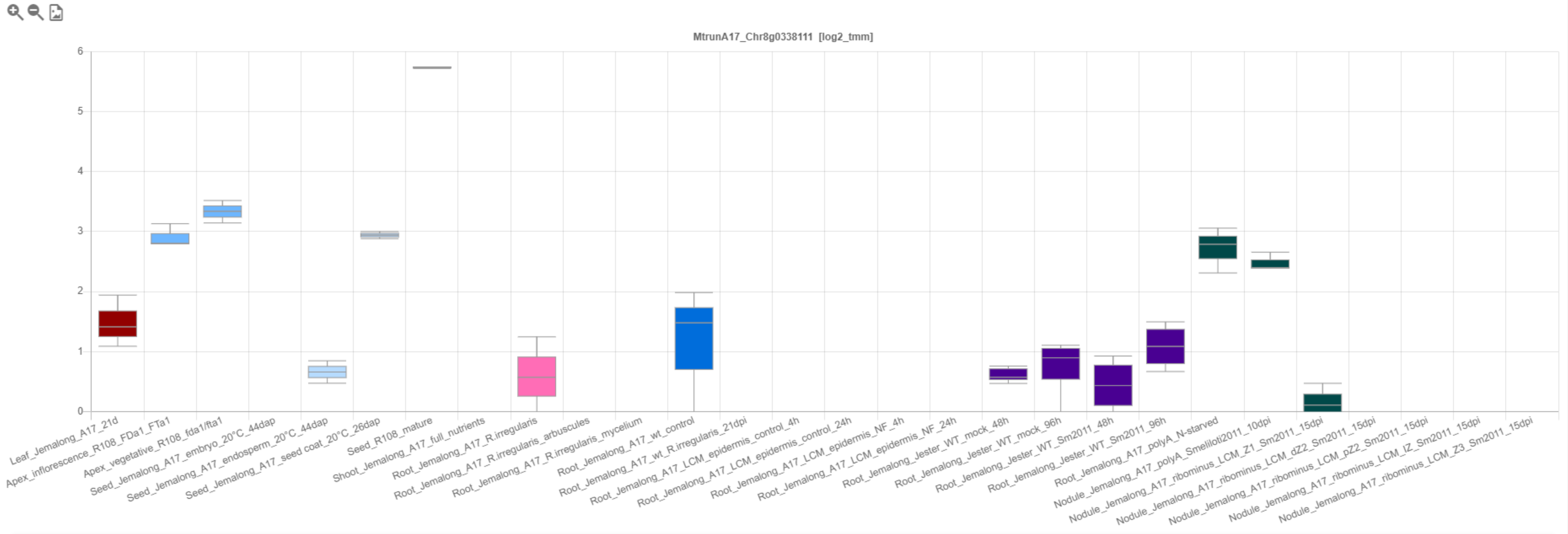

*CP124: MtrunA17_Chr8g0338301

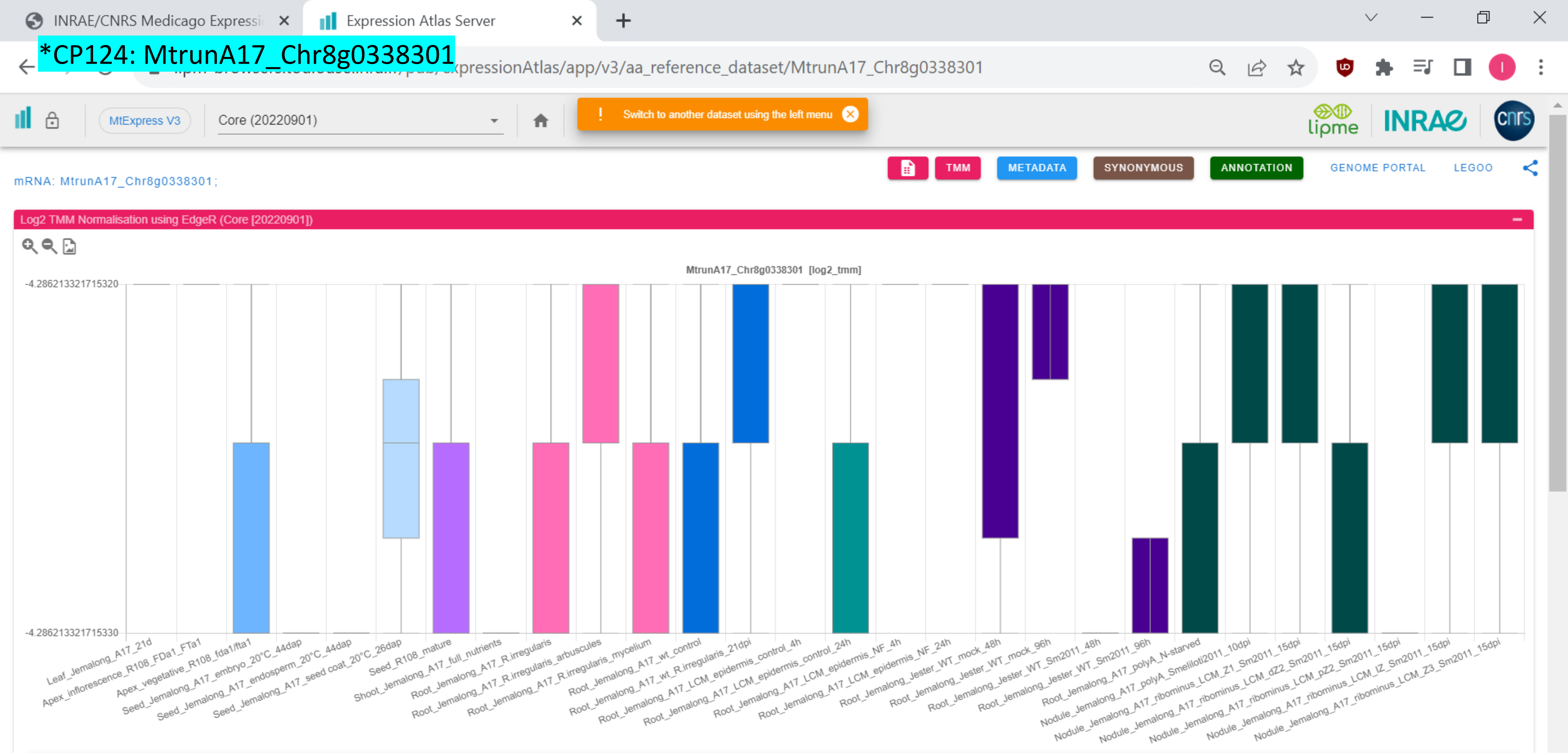

### CP125: MtrunA17_Chr8g0339891

INRAE/CNRS Medicago Expressi | Expression Atlas Server

.../expressionAtlas/app/v3/aa_reference_dataset/MtrunA17_Chr8g0339891

MtExpress V3 | Core (20220901)

! Switch to another dataset using the left menu

TMM | METADATA | SYNONYMOUS | ANNOTATION | GENOME PORTAL | LEGOO

mRNA: MtrunA17_Chr8g0339891;

Log2 TMM Normalisation using EdgeR (Core [20220901])

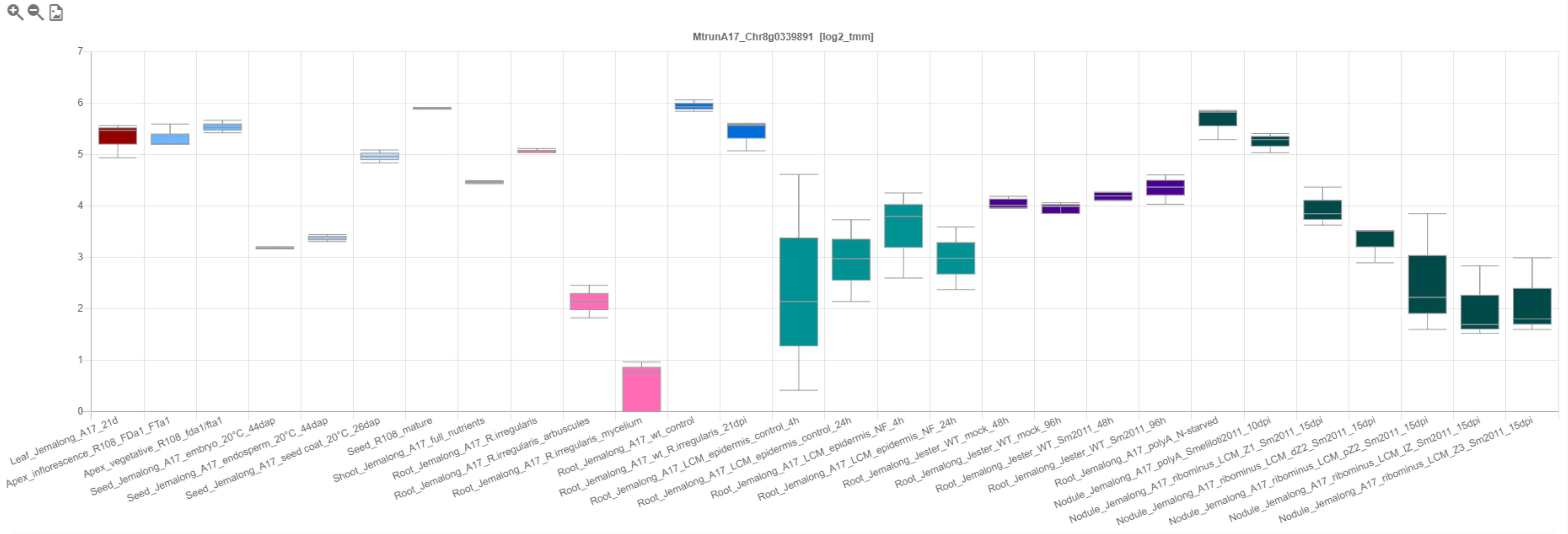

CP126: MtrunA17_Chr8g0342881

INRAE/CNRS Medicago Expressi | Expression Atlas Server

.../expressionAtlas/app/v3/aa_reference_dataset/MtrunA17_Chr8g0342881

MtExpress V3 | Core (20220901)

! Switch to another dataset using the left menu

TMM | METADATA | SYNONYMOUS | ANNOTATION | GENOME PORTAL | LEGOO

mRNA: MtrunA17_Chr8g0342881 ;

Log2 TMM Normalisation using EdgeR (Core [20220901])

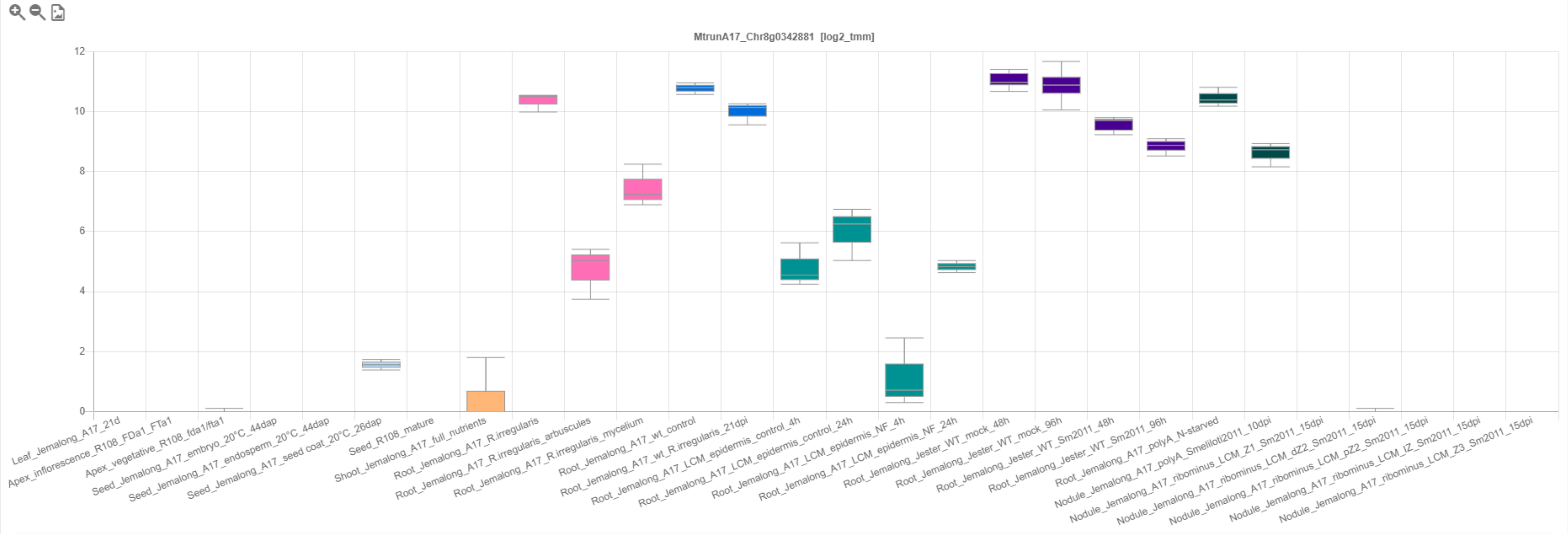

### CP127: MtrunA17_Chr8g0345421

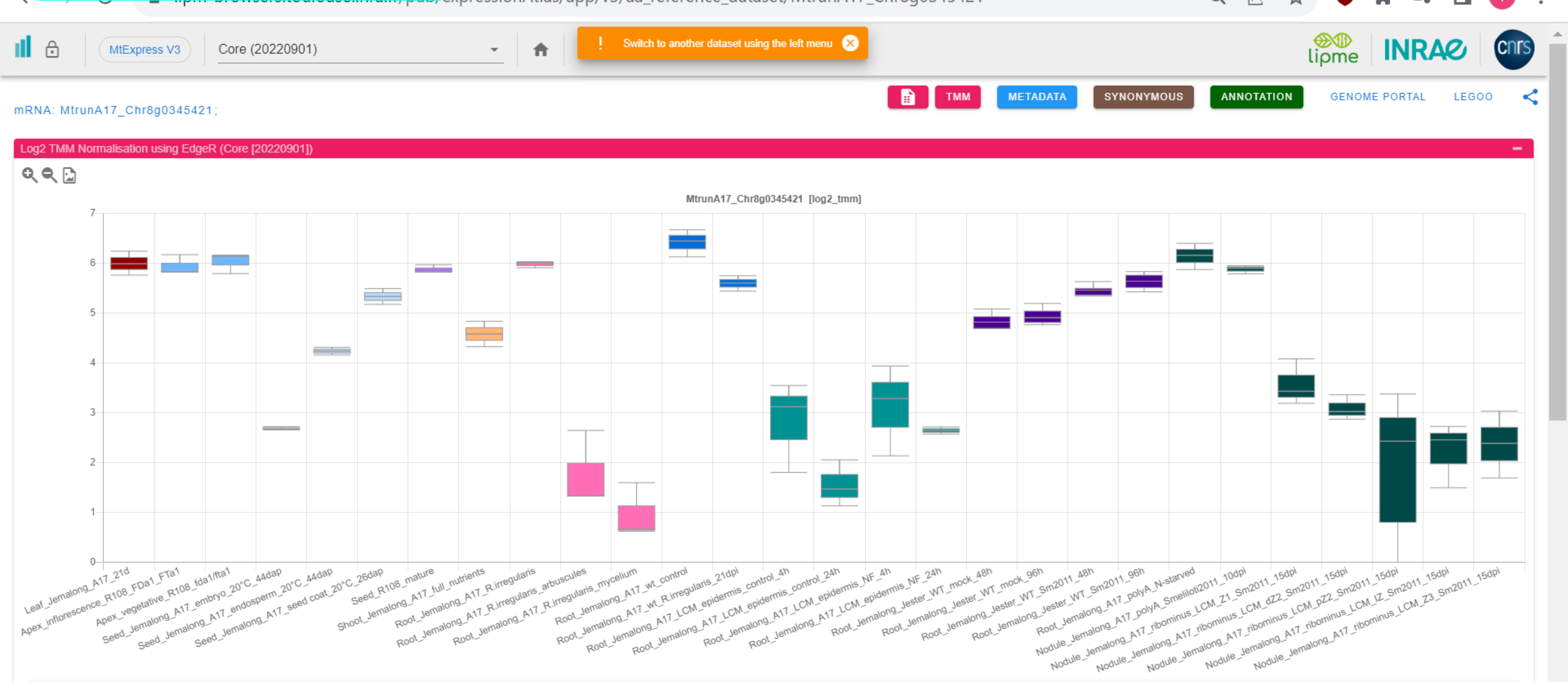

*CP128: MtrunA17_Chr8g0353711

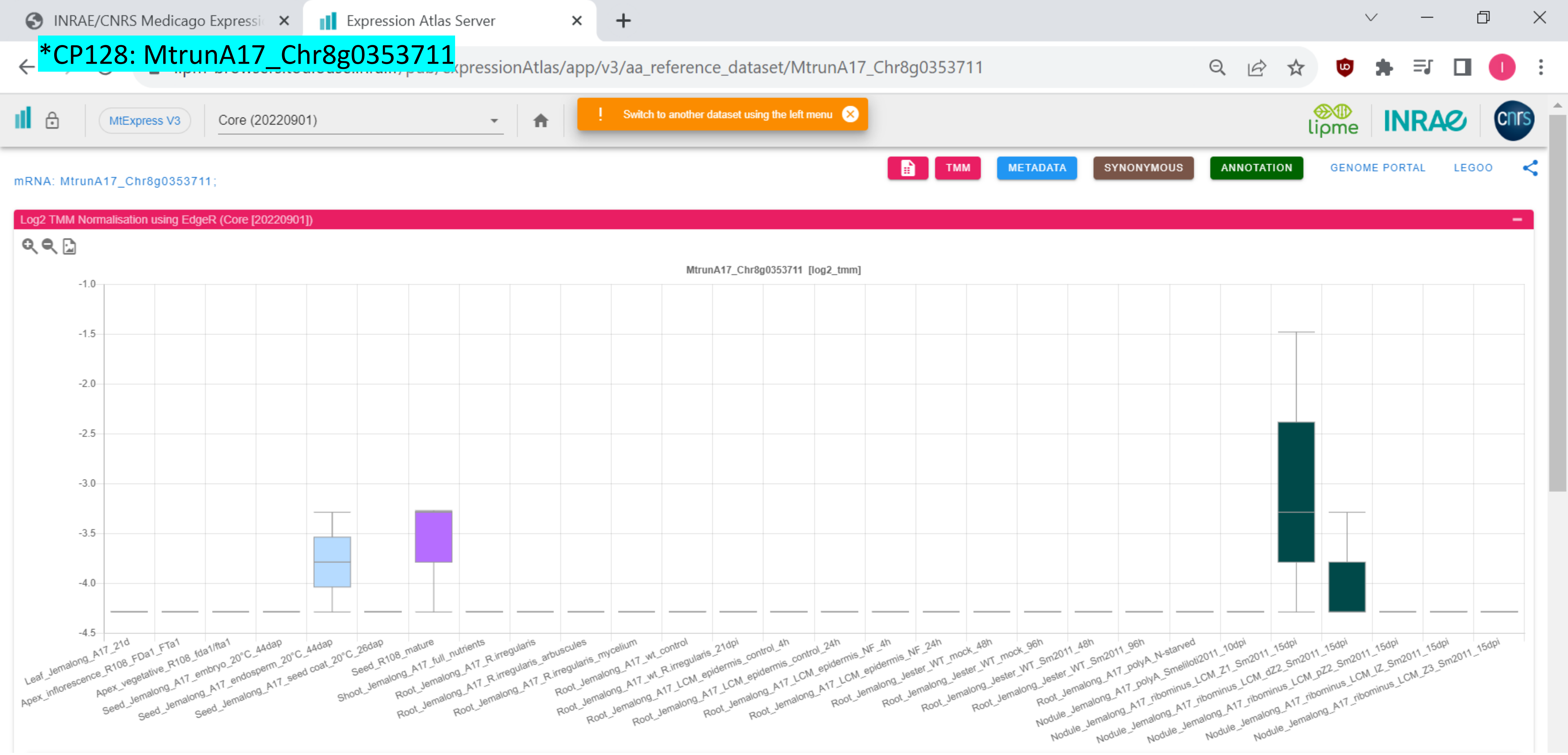

CP129: MtrunA17_Chr8g0355501

INRAE/CNRS Medicago Expressi | Expression Atlas Server

/expressionAtlas/app/v3/aa_reference_dataset/MtrunA17_Chr8g0355501

MtExpress V3 | Core (20220901) | Switch to another dataset using the left menu

TMM | METADATA | SYNONYMOUS | ANNOTATION | GENOME PORTAL | LEGOO

mRNA: MtrunA17_Chr8g0355501;

Log2 TMM Normalisation using EdgeR (Core [20220901])

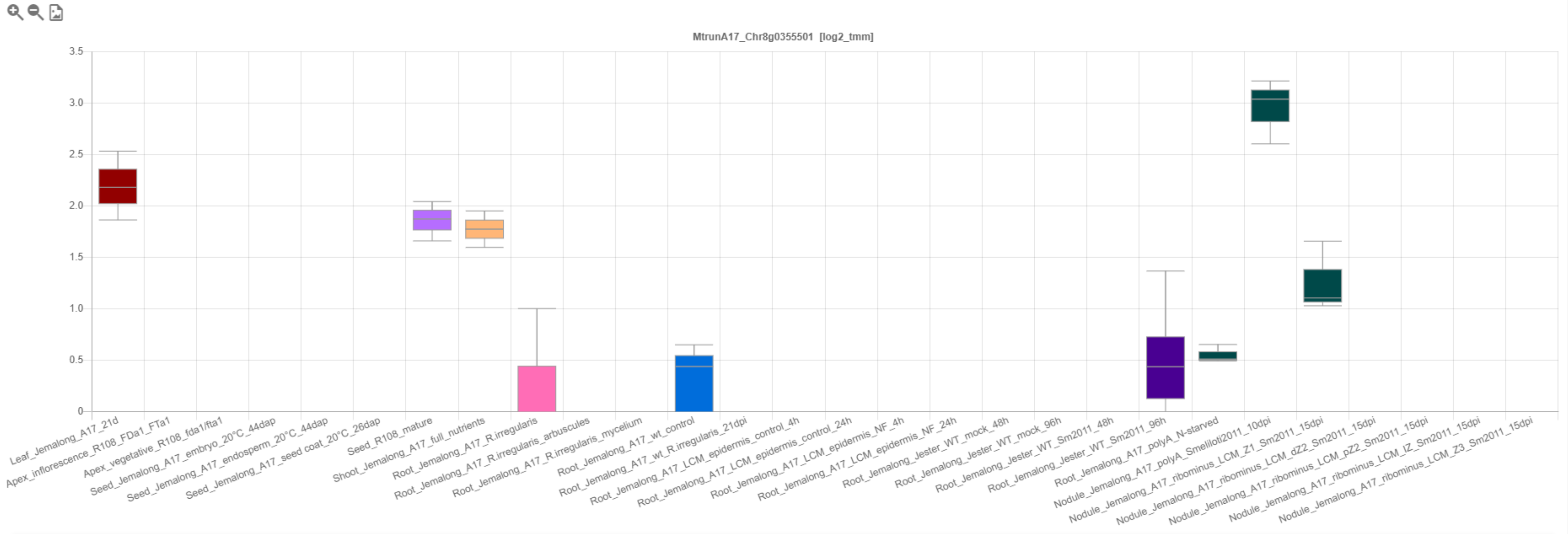

*CP130: MtrunA17_Chr8g0356581

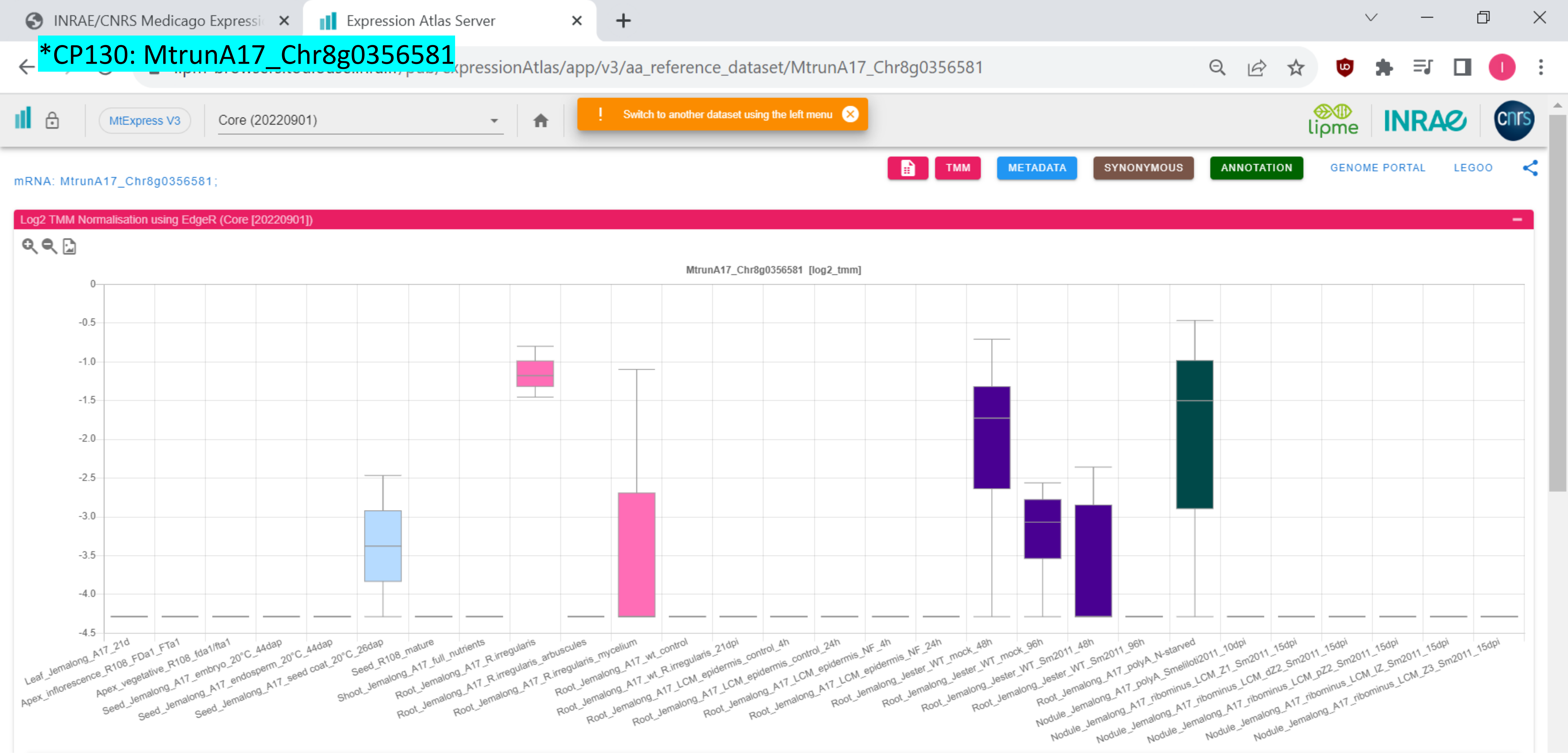

CP131: MtrunA17_Chr8g0365341

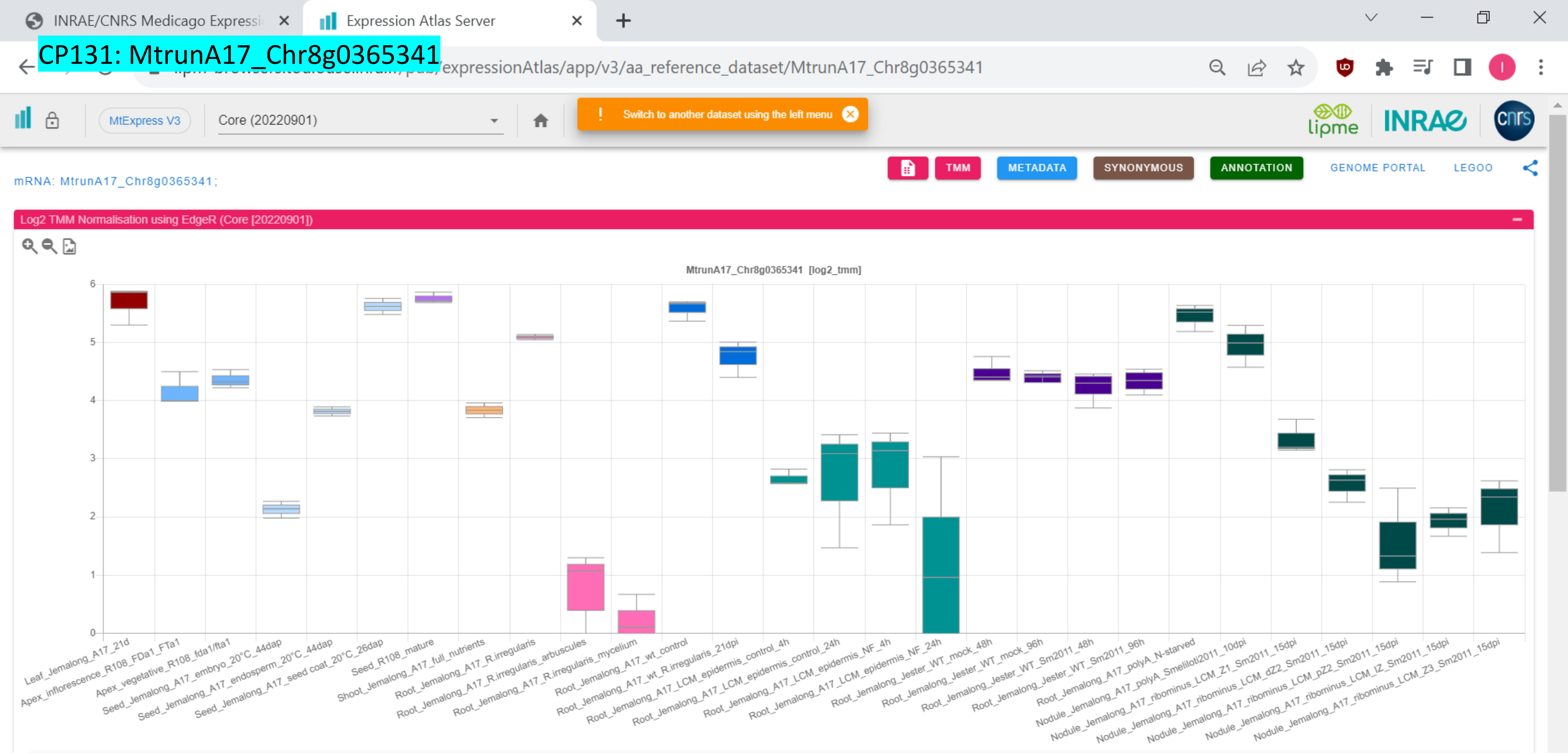

CP132: MtrunA17_Chr8g0368731

INRAE/CNRS Medicago Expressi | Expression Atlas Server

/expressionAtlas/app/v3/aa_reference_dataset/MtrunA17_Chr8g0368731

MtExpress V3 | Core (20220901) | Switch to another dataset using the left menu

TMM | METADATA | SYNONYMOUS | ANNOTATION | GENOME PORTAL | LEGOO

mRNA: MtrunA17_Chr8g0368731;

Log2 TMM Normalisation using EdgeR (Core [20220901])

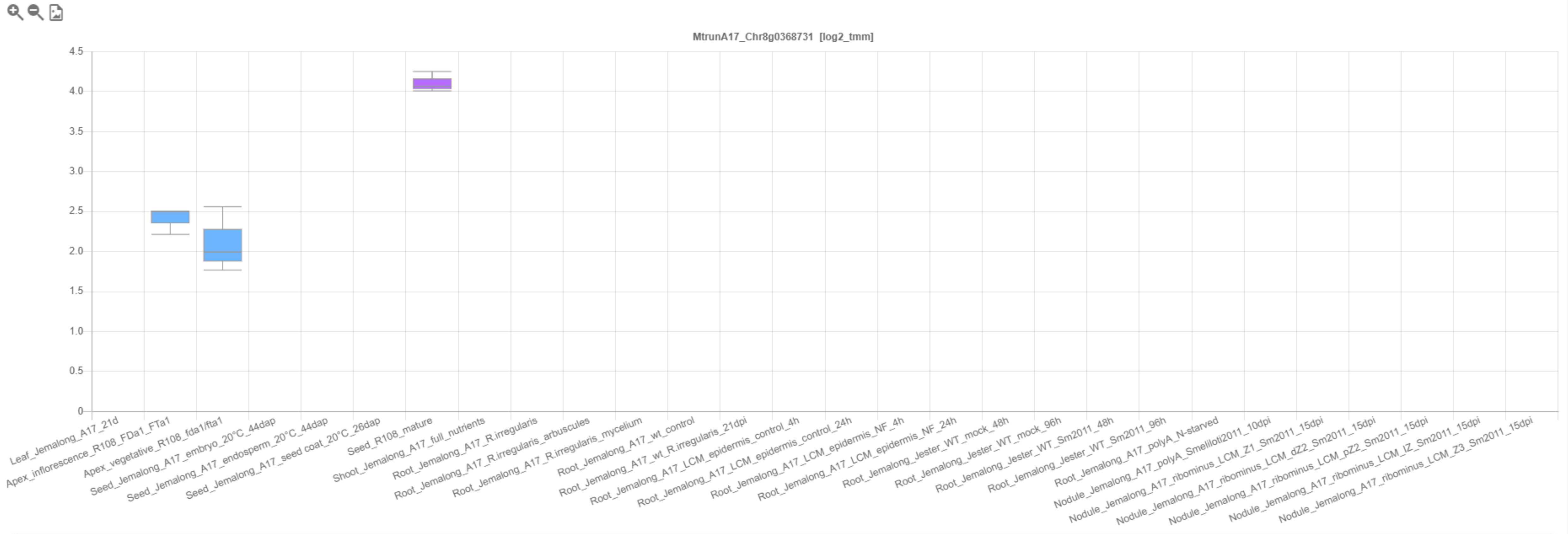

CP133: MtrunA17_Chr8g0368931

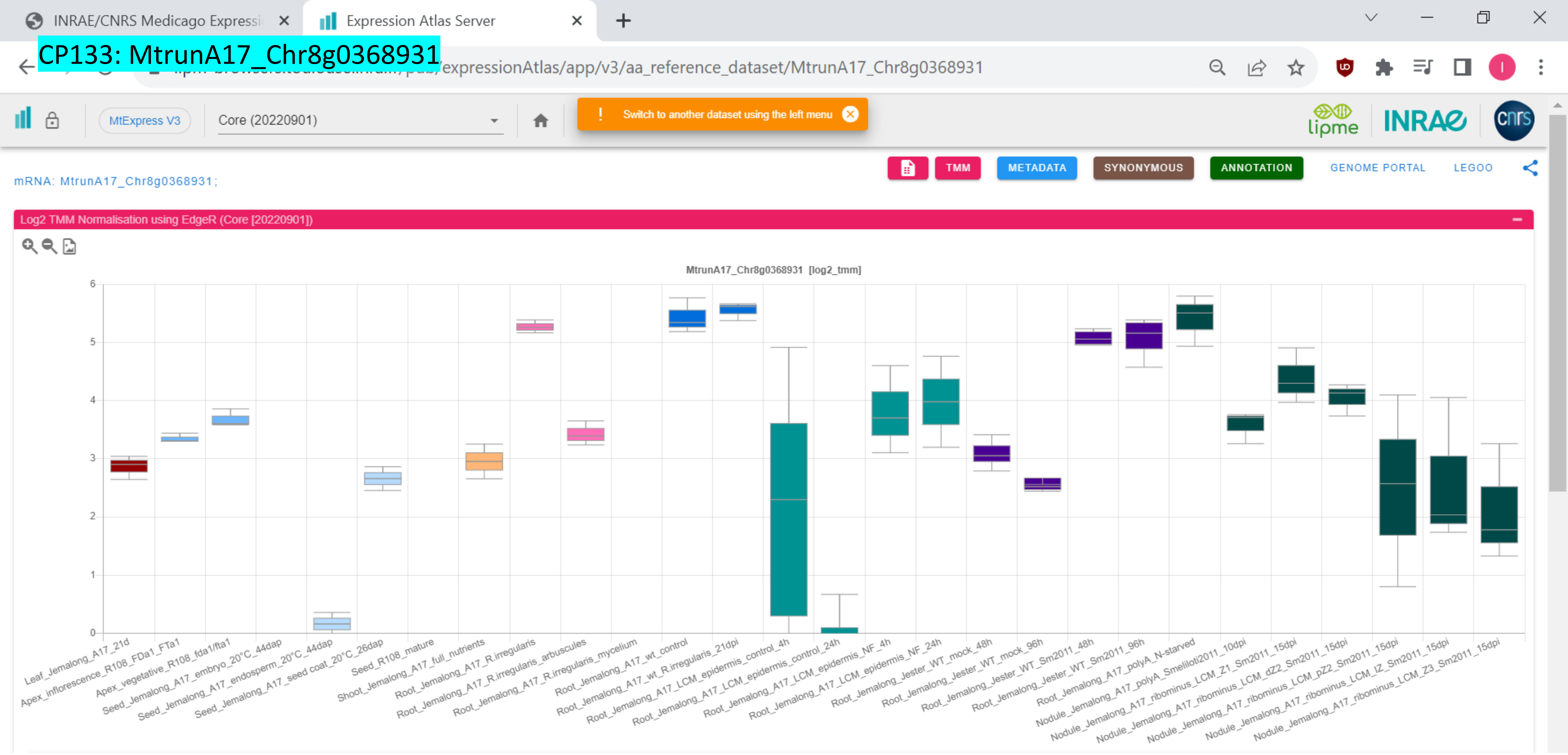

CP134: MtrunA17_Chr8g0371281

INRAE/CNRS Medicago Expressi | Expression Atlas Server

.../expressionAtlas/app/v3/aa_reference_dataset/MtrunA17_Chr8g0371281

MtExpress V3 | Core (20220901)

! Switch to another dataset using the left menu

TMM | METADATA | SYNONYMOUS | ANNOTATION | GENOME PORTAL | LEGOO

mRNA: MtrunA17_Chr8g0371281;

Log2 TMM Normalisation using EdgeR (Core [20220901])

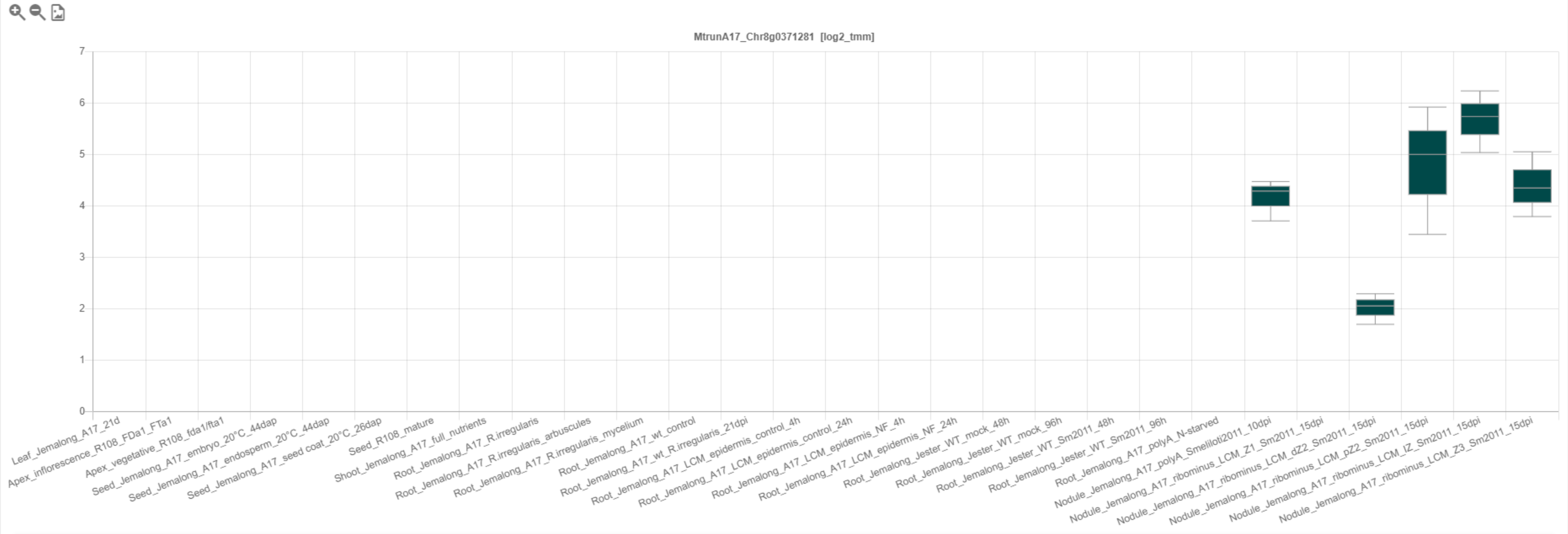

### CP135: MtrunA17_Chr8g0371741

INRAE/CNRS Medicago Expressi | Expression Atlas Server

…/expressionAtlas/app/v3/aa_reference_dataset/MtrunA17_Chr8g0371741

MtExpress V3 | Core (20220901)

! Switch to another dataset using the left menu

TMM | METADATA | SYNONYMOUS | ANNOTATION | GENOME PORTAL | LEGOO

lipme INRAE cnrs

mRNA: MtrunA17_Chr8g0371741;

Log2 TMM Normalisation using EdgeR (Core [20220901])

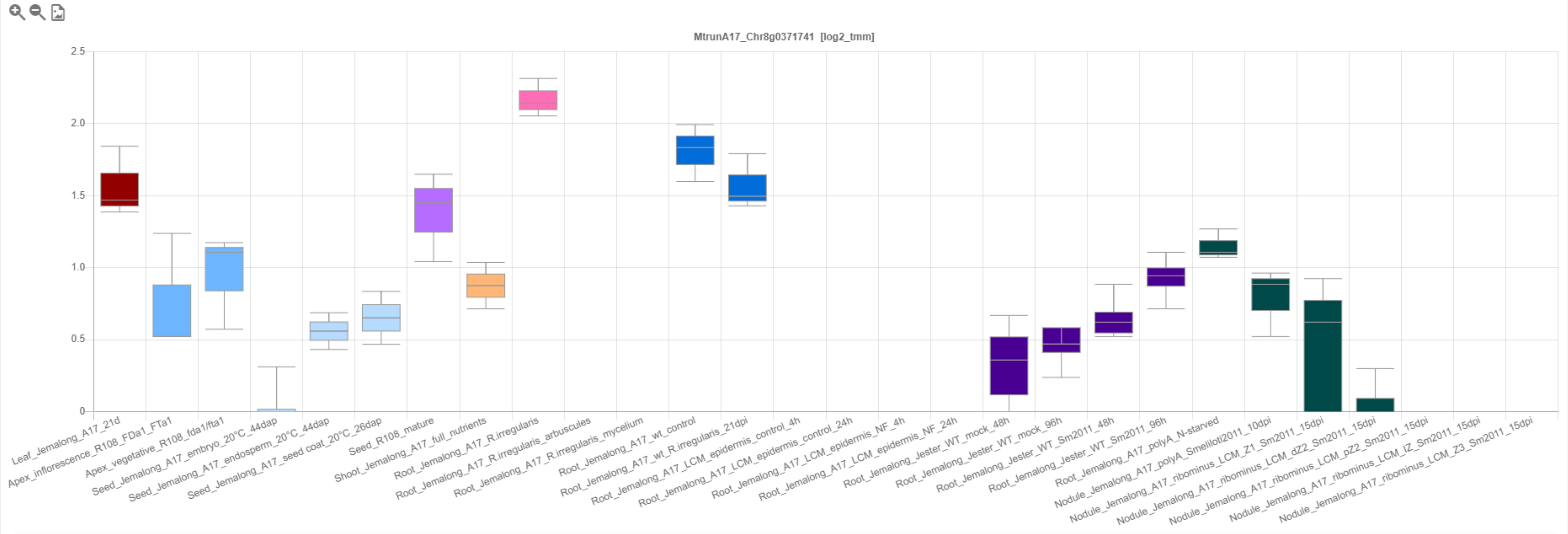

CP136: MtrunA17_Chr8g0373091

mRNA: MtrunA17_Chr8g0373091;

Log2 TMM Normalisation using EdgeR (Core [20220901])

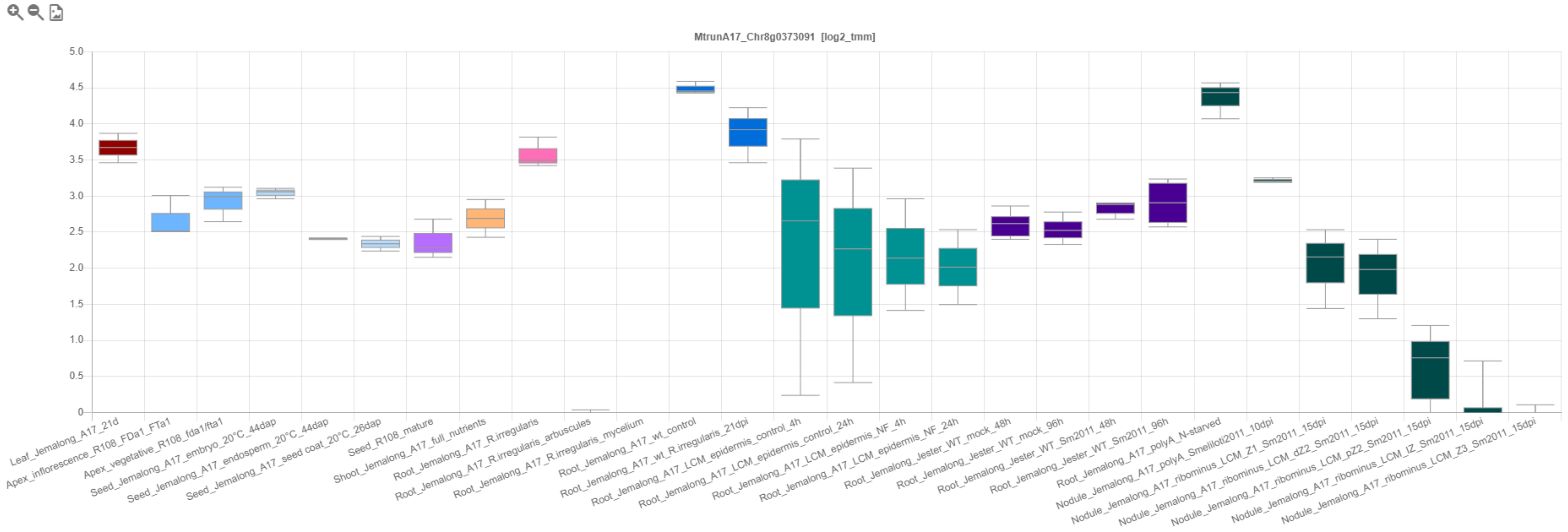

CP137: MtrunA17_Chr8g0376411

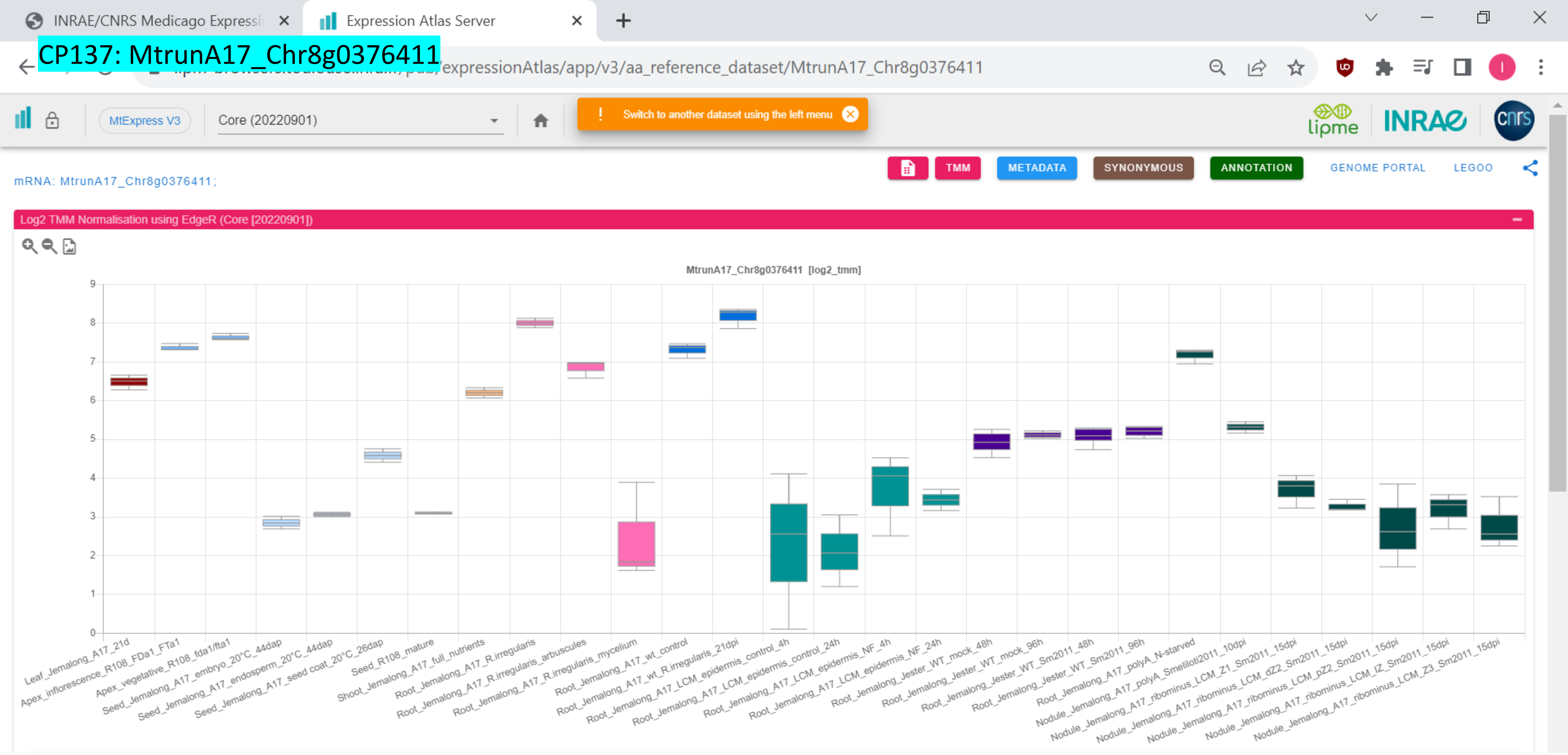

CP138: MtrunA17_Chr8g0377071

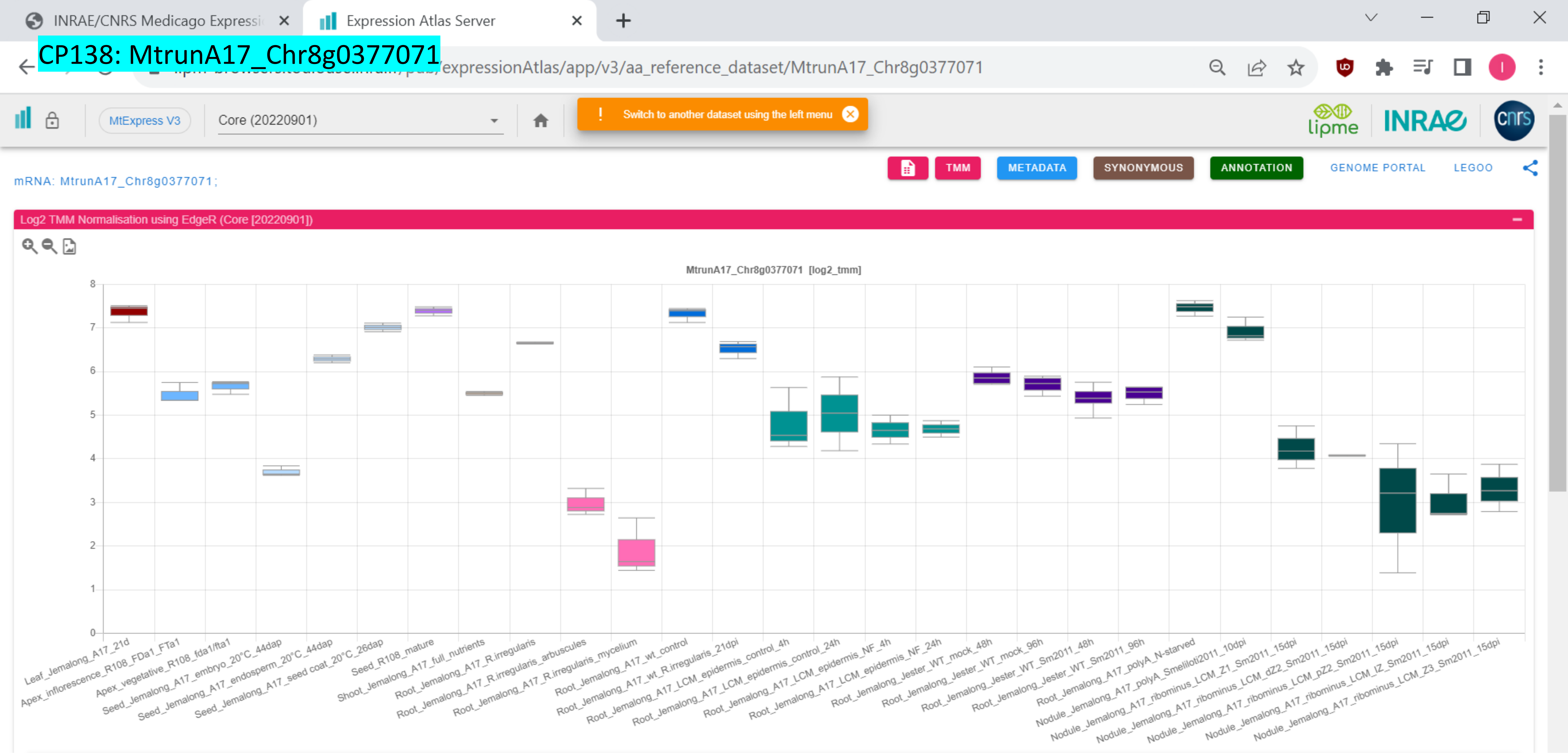

CP139: MtrunA17_Chr8g0385331

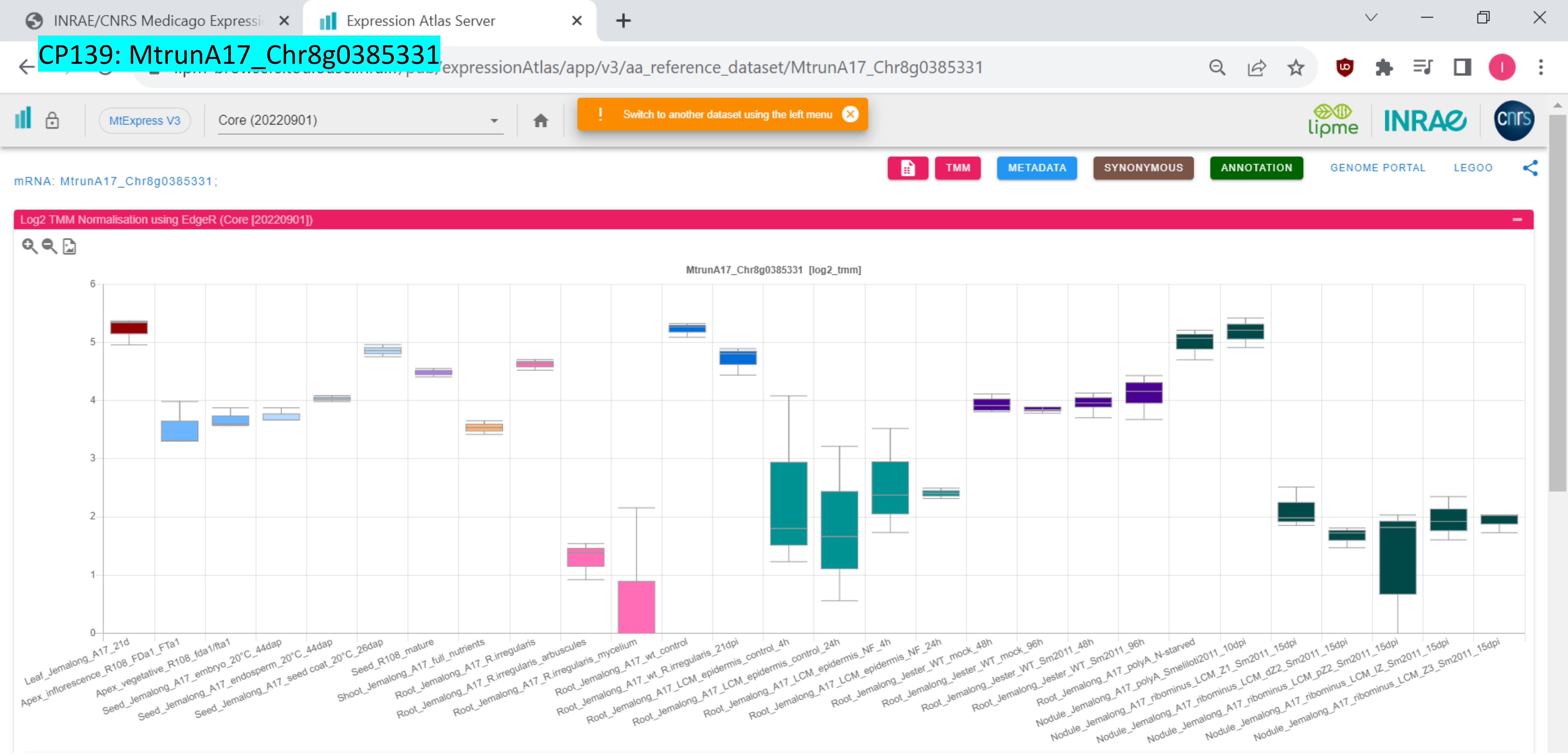

### CP140: MtrunA17_Chr8g0392351

INRAE/CNRS Medicago Expressi | Expression Atlas Server

/expressionAtlas/app/v3/aa_reference_dataset/MtrunA17_Chr8g0392351

MtExpress V3 | Core (20220901) | Switch to another dataset using the left menu

TMM | METADATA | SYNONYMOUS | ANNOTATION | GENOME PORTAL | LEGOO

ncRNA: MtrunA17_Chr8g0392351;

Log2 TMM Normalisation using EdgeR (Core [20220901])

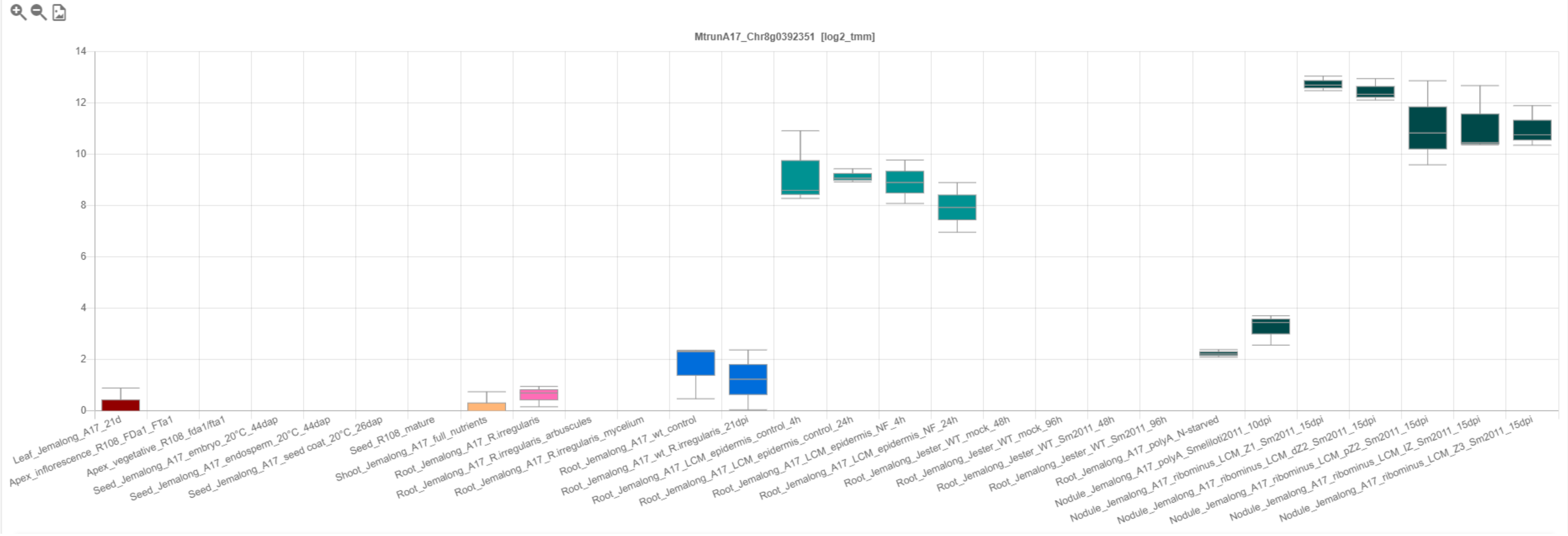

### CP141: MtrunA17_CPg0492331

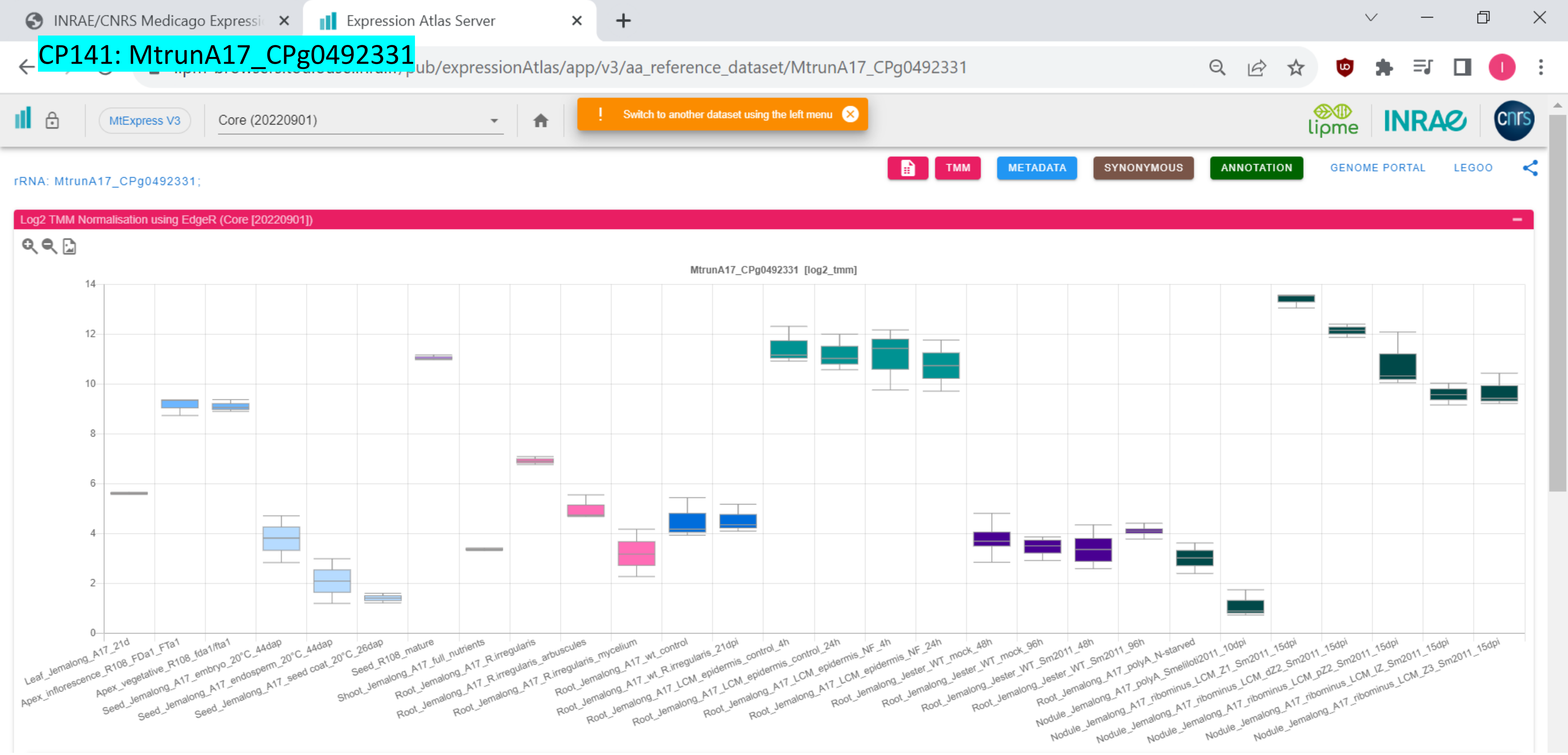

### CP142: MtrunA17_CPg0492381

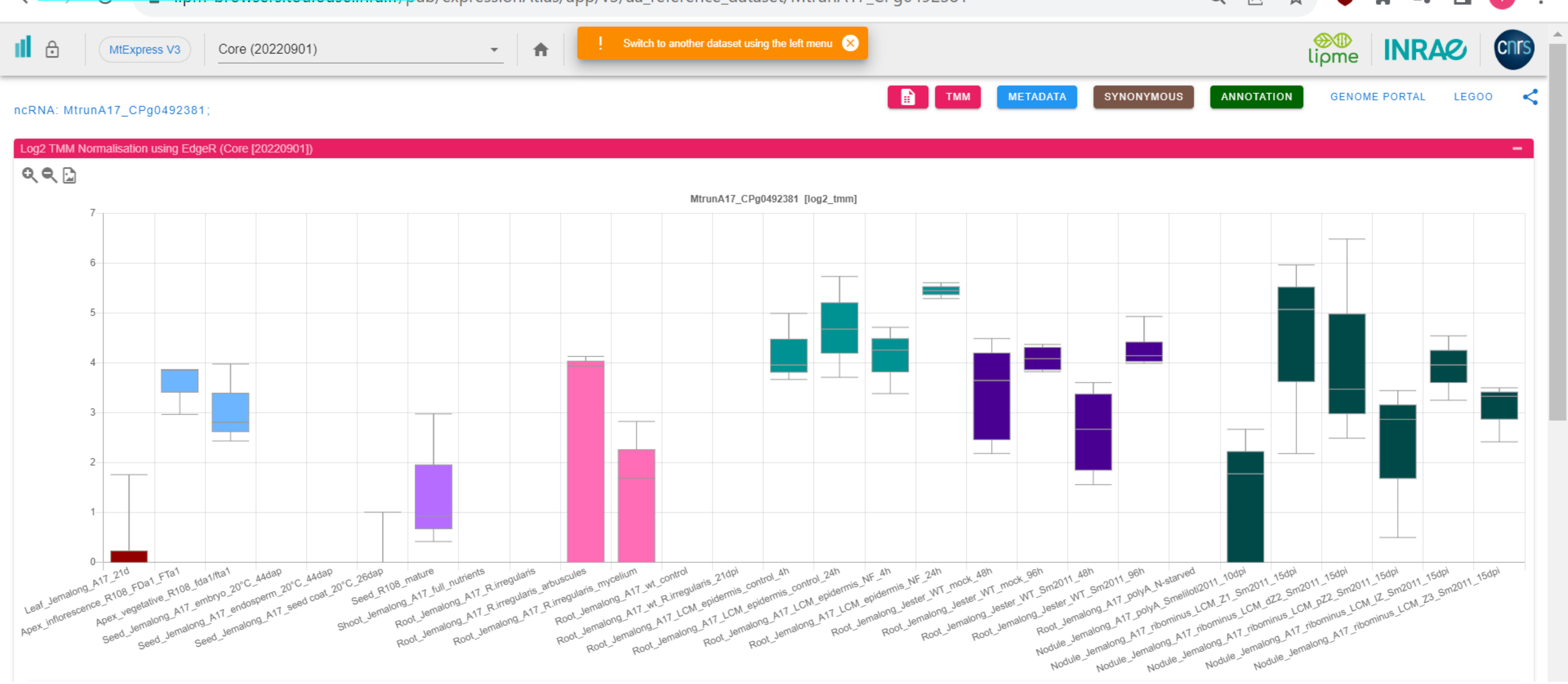

### CP143: MtrunA17_CPg0492461

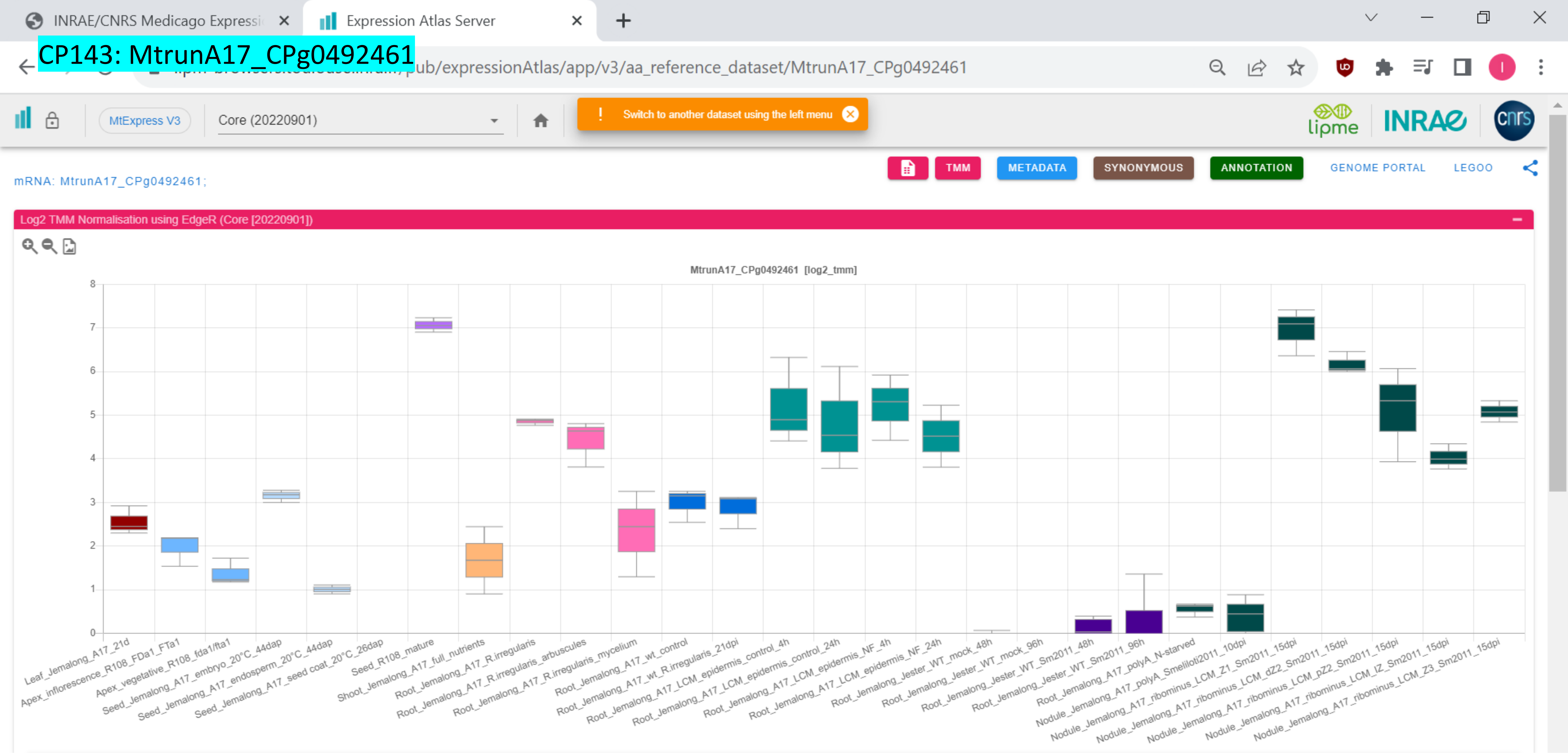

CP144: MtrunA17_CPg0492851

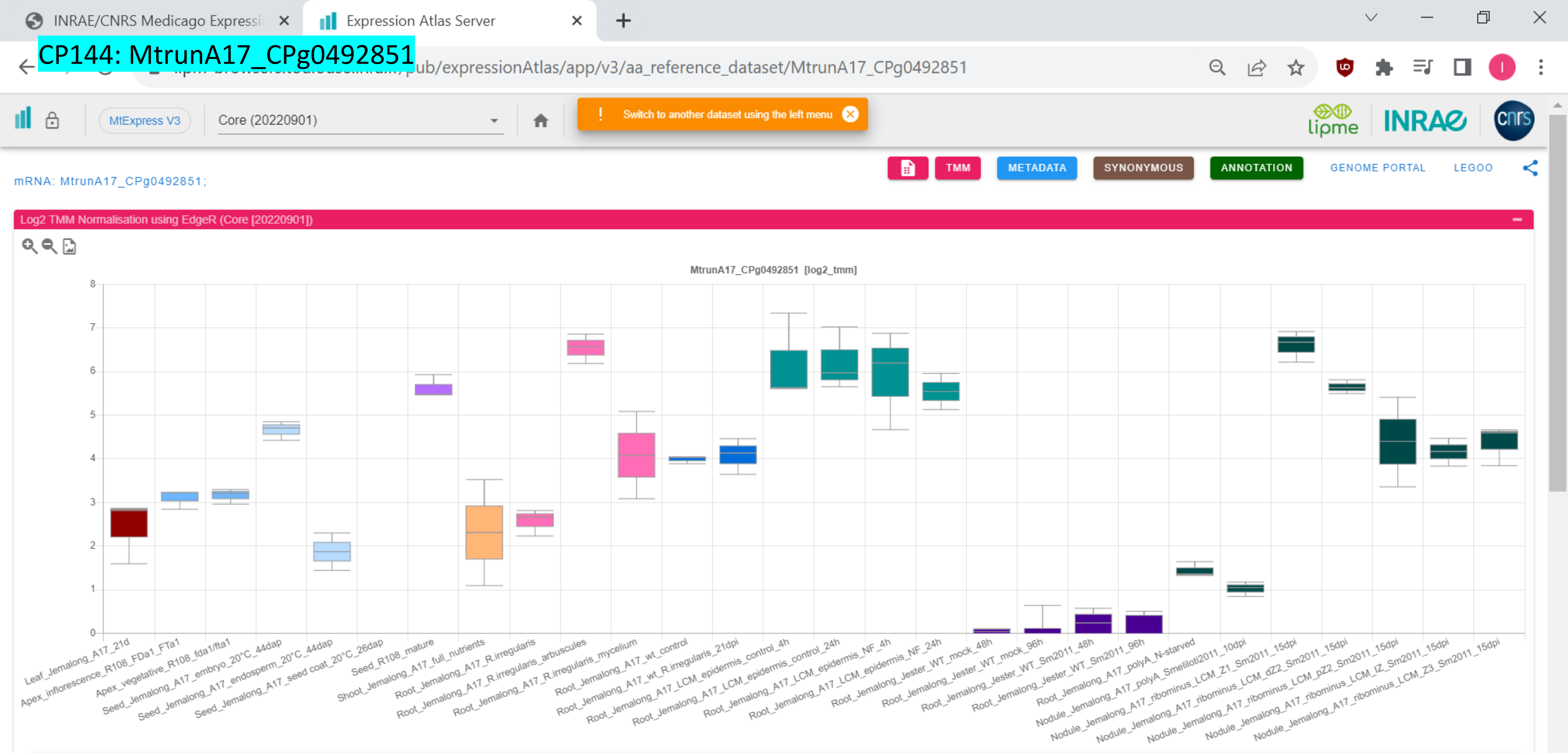

### CP145: MtrunA17_CPg0492941

Expression Atlas Server

pub/expressionAtlas/app/v3/aa_reference_dataset/MtrunA17_CPg0492941

MtExpress V3 | Core (20220901)

Switch to another dataset using the left menu

TMM | METADATA | SYNONYMOUS | ANNOTATION | GENOME PORTAL | LEGOO

mRNA: MtrunA17_CPg0492941;

Log2 TMM Normalisation using EdgeR (Core [20220901])

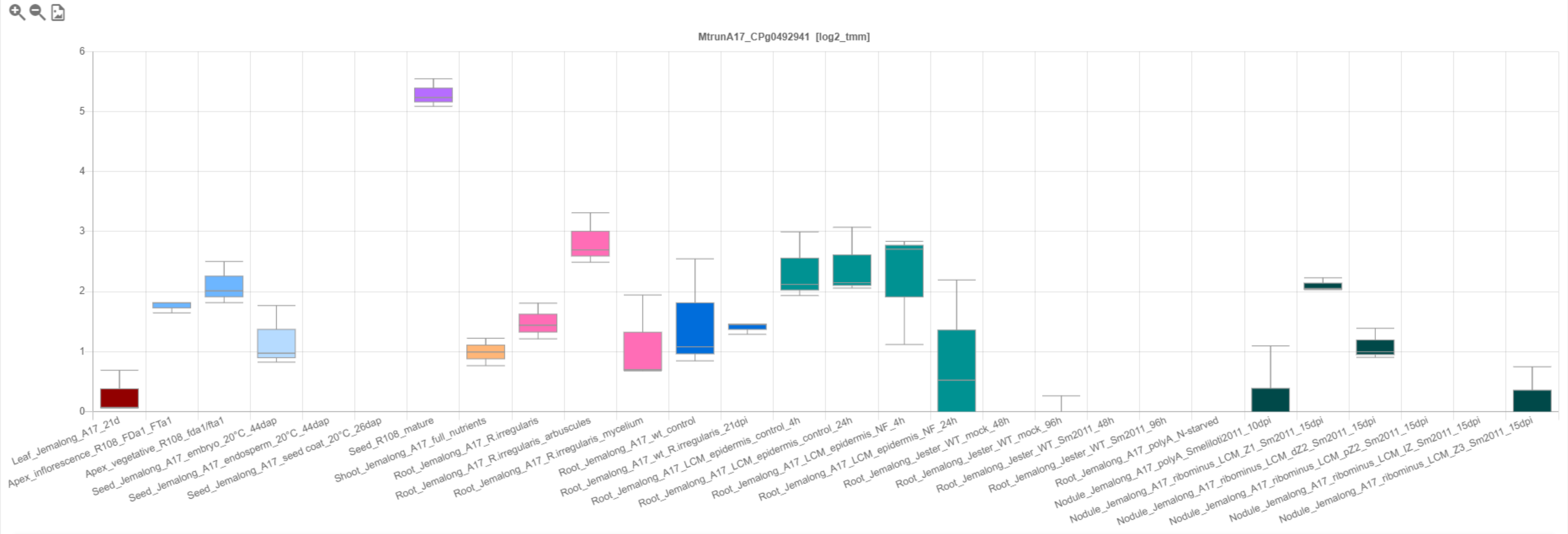

### CP146: MtrunA17_CPg0493291

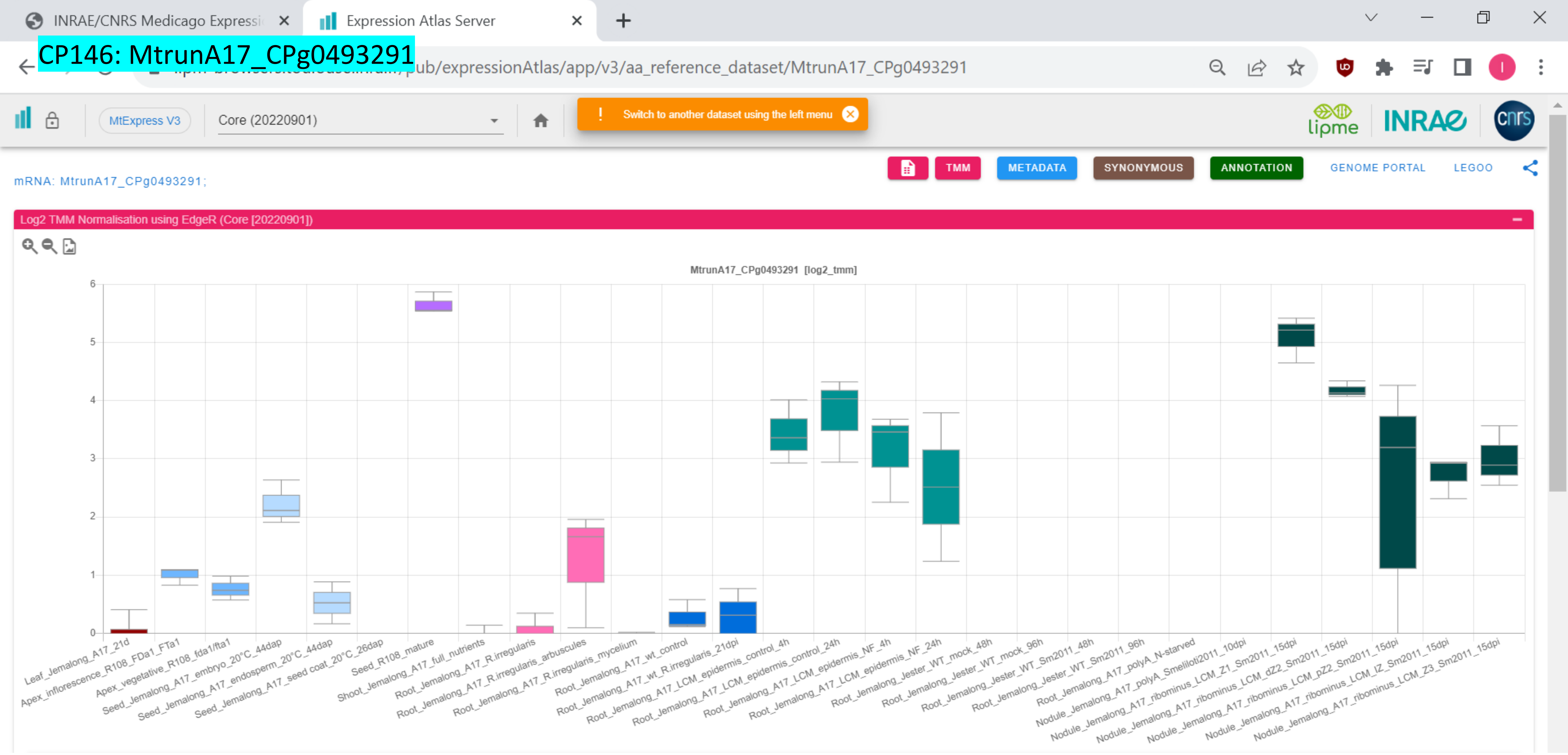

### CP147: MtrunA17_CPg0493401

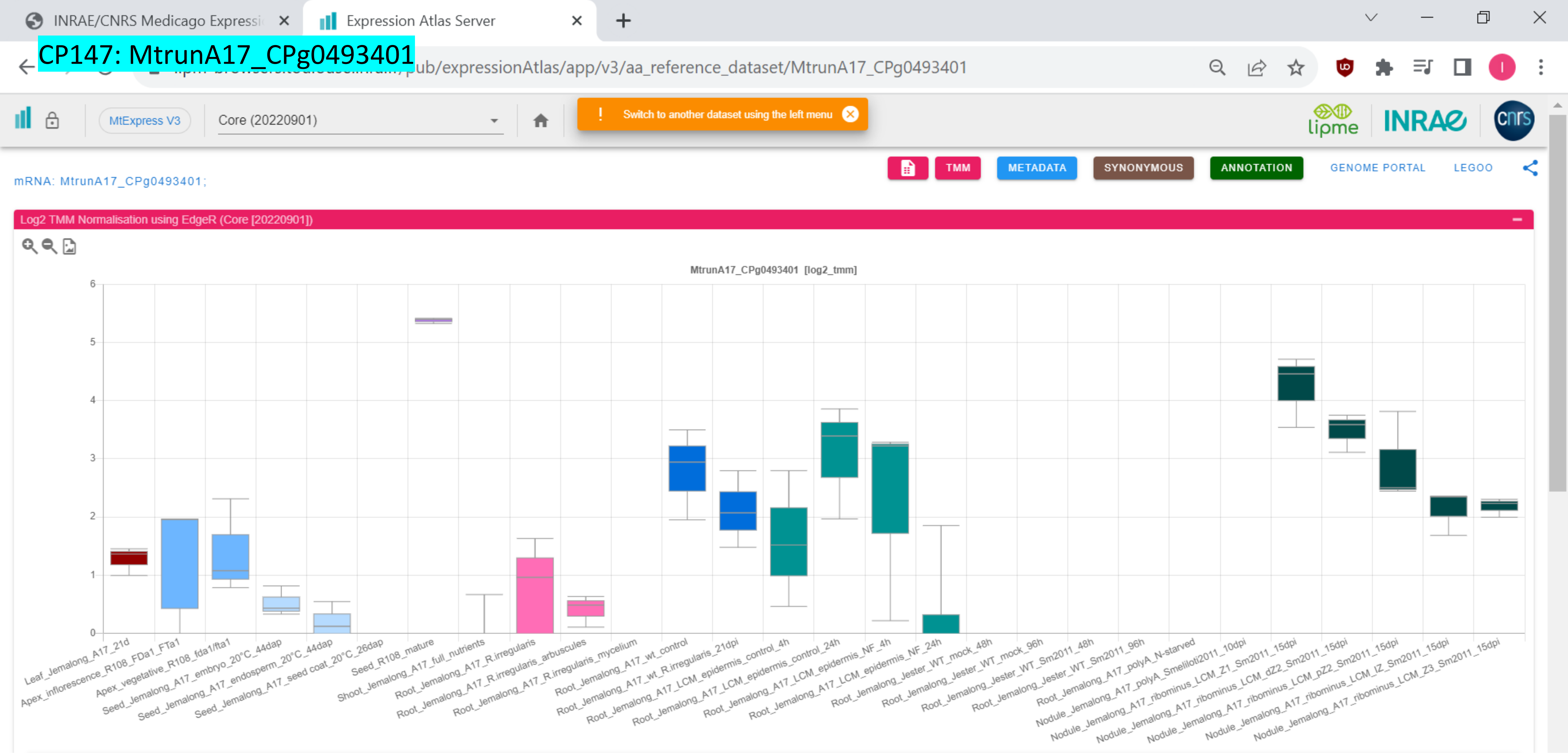

### CP148: MtrunA17_MTg0490471

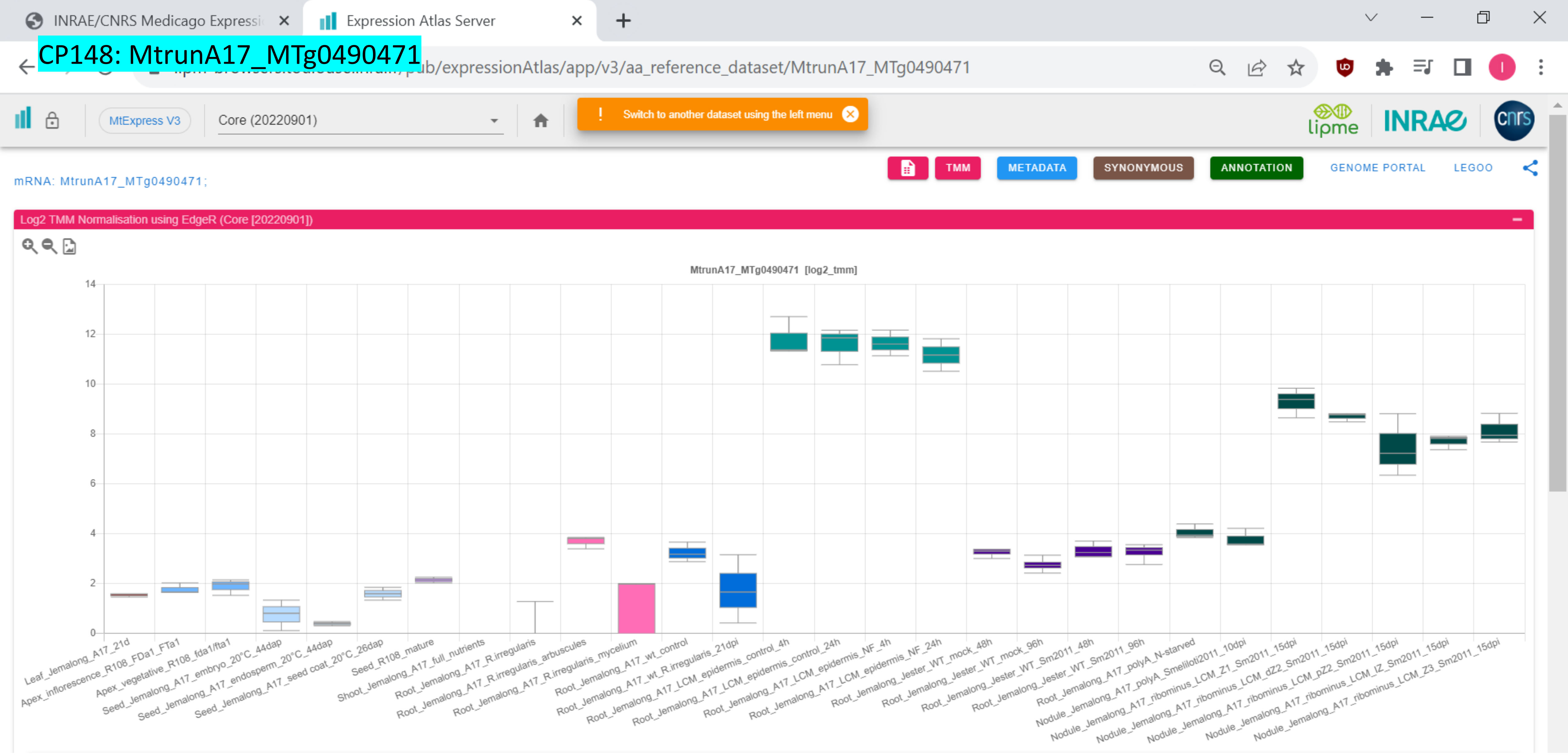

### CP149: MtrunA17_MTg0490471

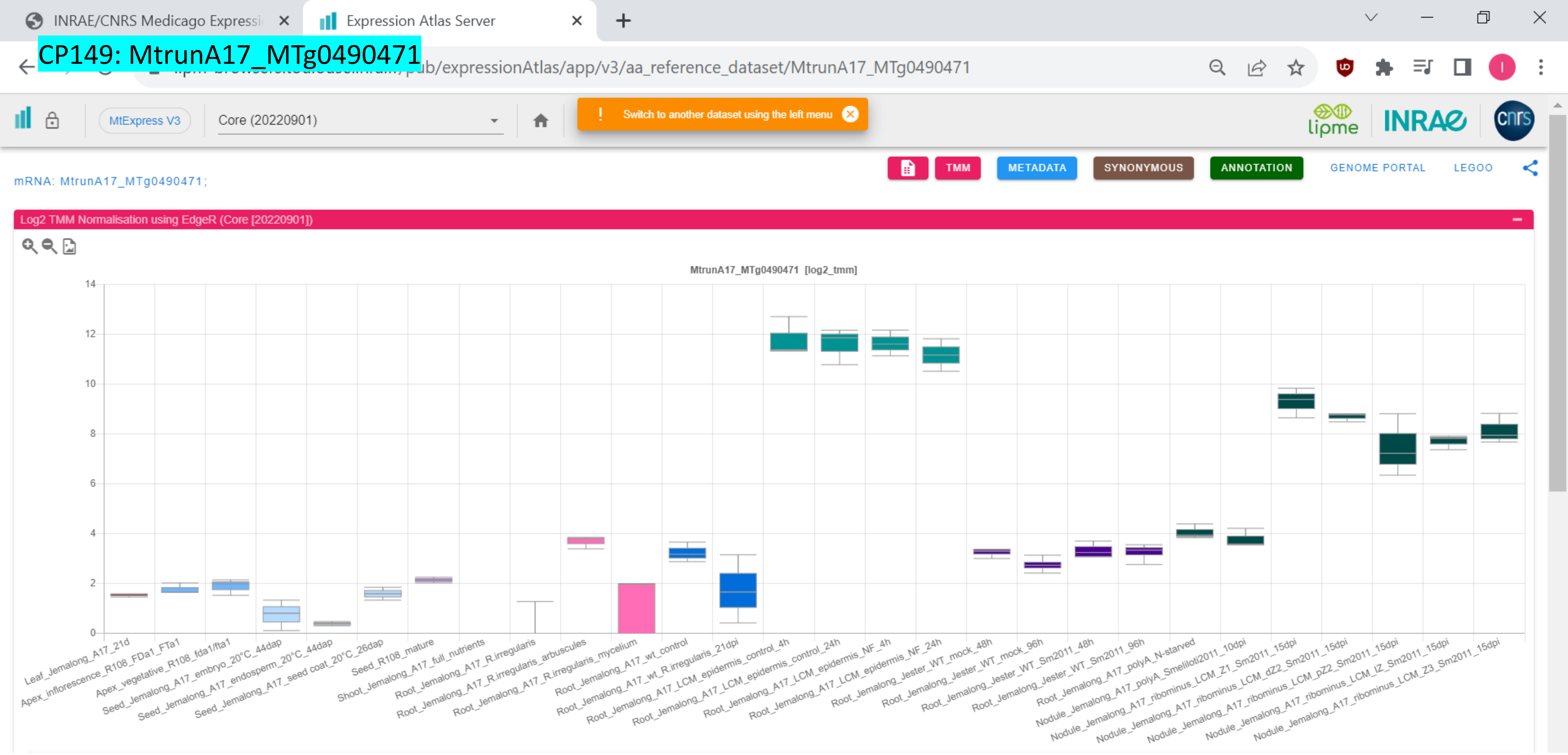

### CP150: MtrunA17_MTg0490971

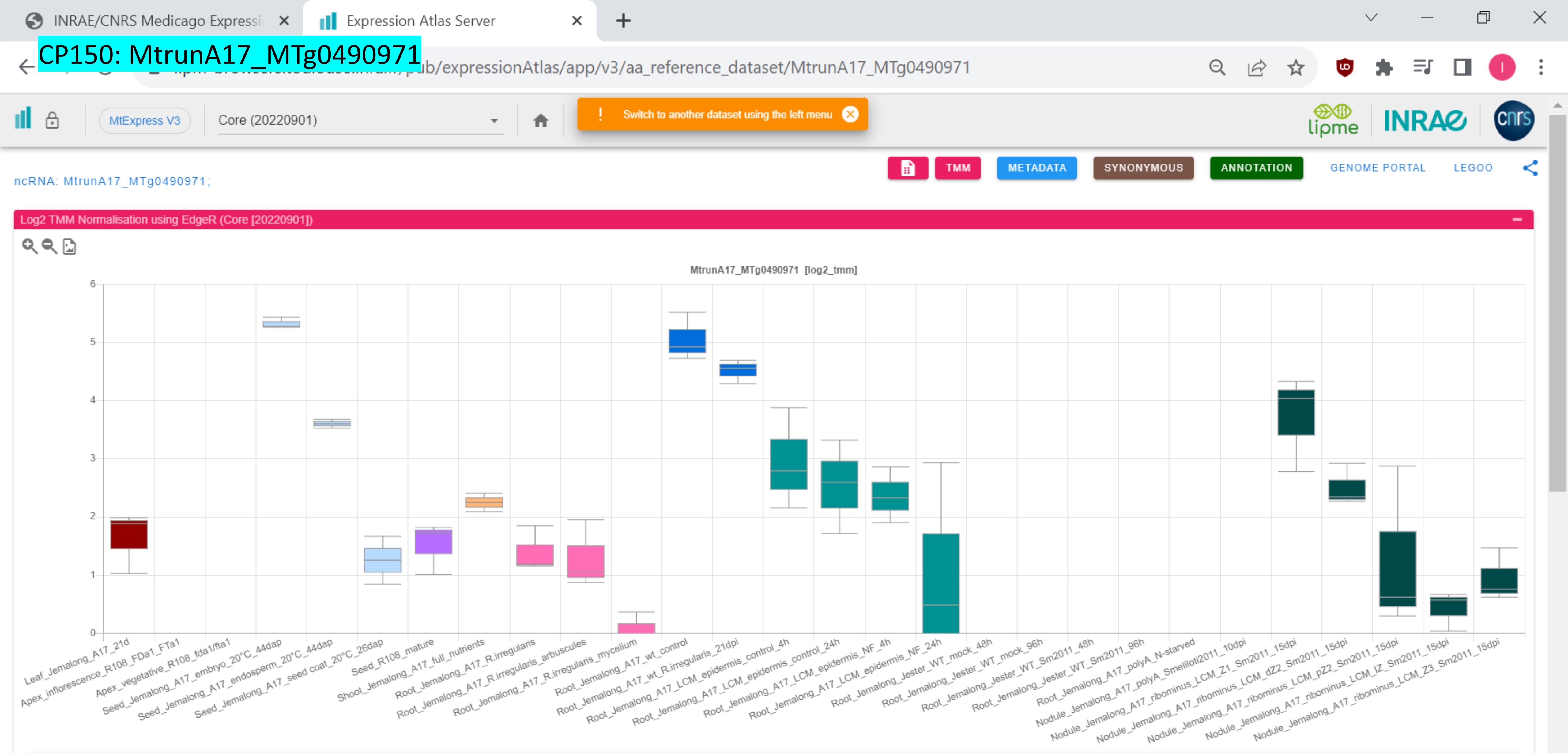

CP151: MtrunA17_MTg0490971

INRAE/CNRS Medicago Expressi | Expression Atlas Server

pub/expressionAtlas/app/v3/aa_reference_dataset/MtrunA17_MTg0490971

MtExpress V3 | Core (20220901)

Switch to another dataset using the left menu

TMM | METADATA | SYNONYMOUS | ANNOTATION | GENOME PORTAL | LEGOO

ncRNA: MtrunA17_MTg0490971;

Log2 TMM Normalisation using EdgeR (Core [20220901])

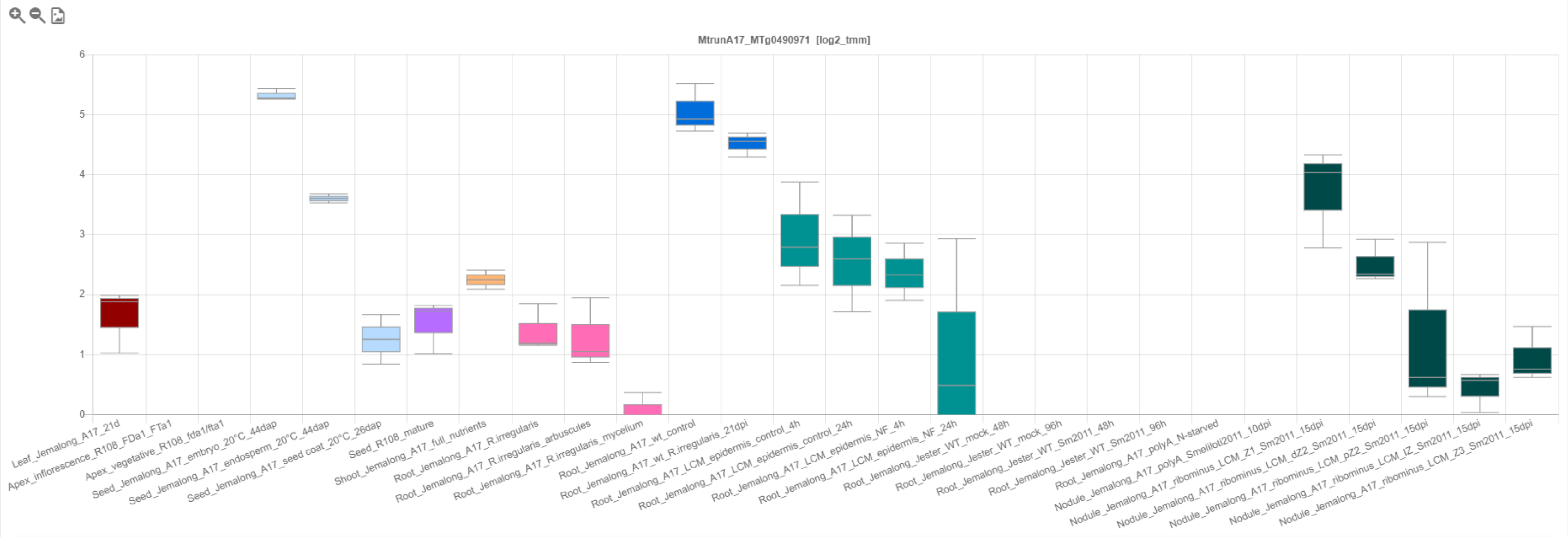

### CP152: MtrunA17_MTg0491151

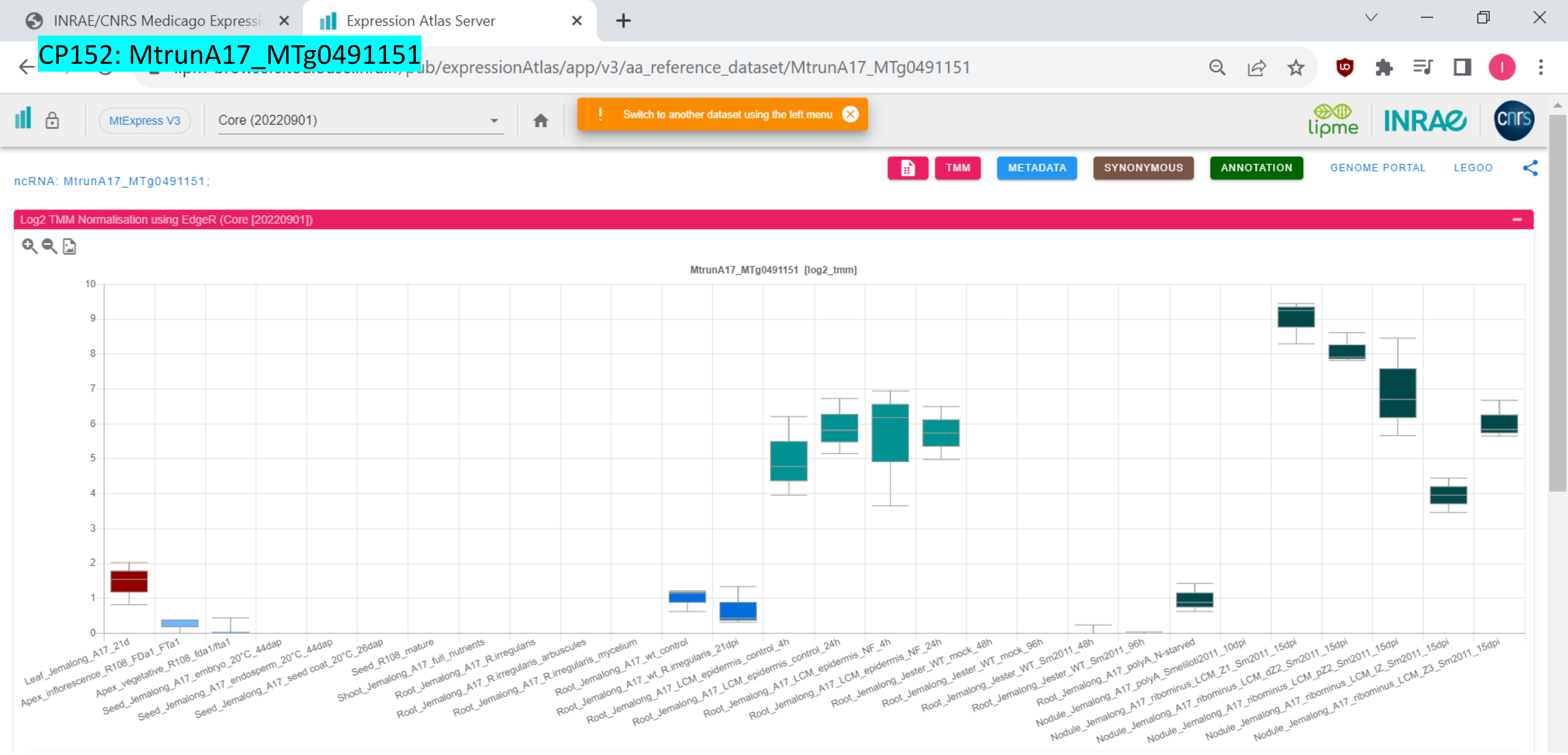

CP153: MtrunA17_MTg0491291

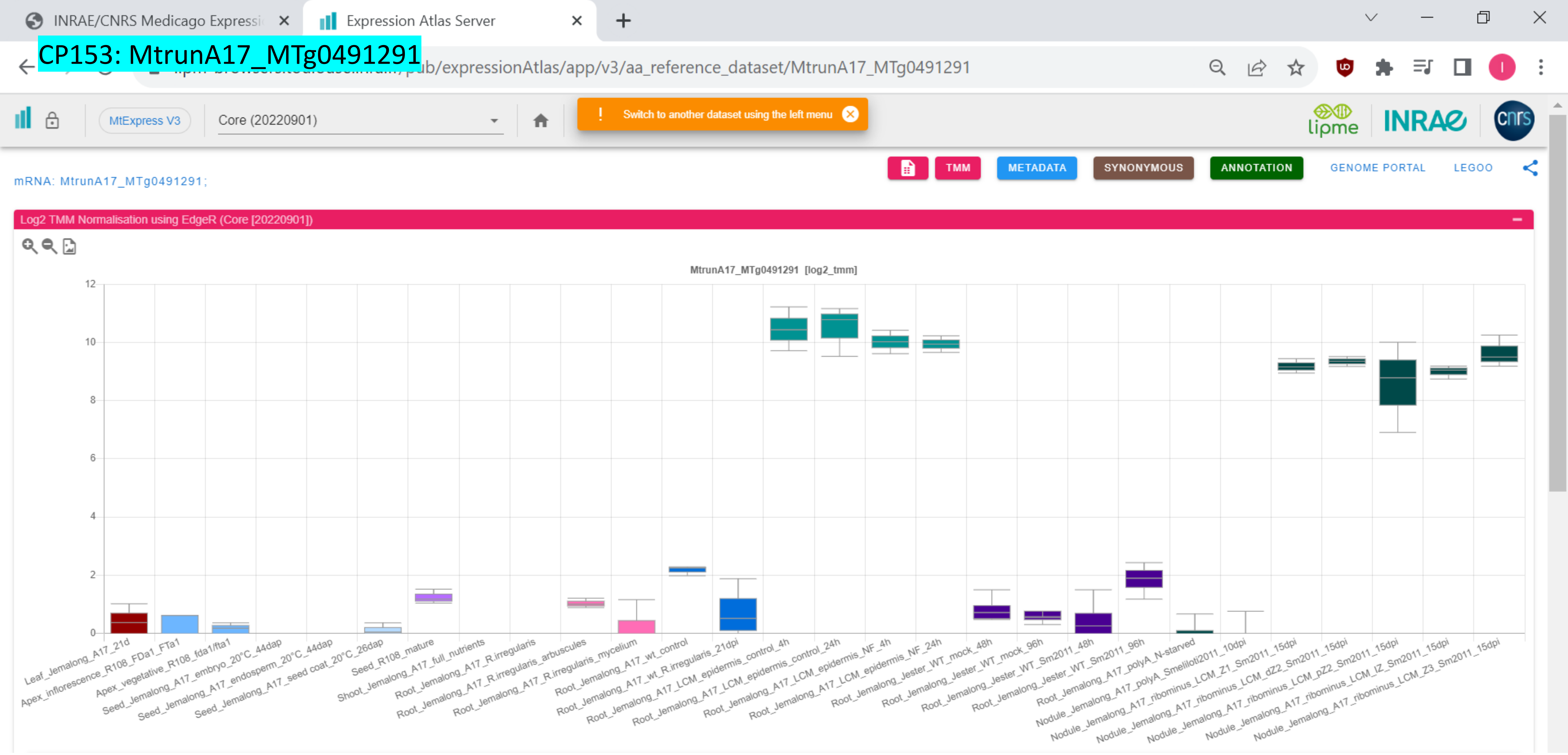

CP154: MtrunA17_MTg0491501

INRAE/CNRS Medicago Expressi | Expression Atlas Server

.../pub/expressionAtlas/app/v3/aa_reference_dataset/MtrunA17_MTg0491501

MtExpress V3 | Core (20220901) | ! Switch to another dataset using the left menu

TMM | METADATA | SYNONYMOUS | ANNOTATION | GENOME PORTAL | LEGOO

ncRNA: MtrunA17_MTg0491501;

Log2 TMM Normalisation using EdgeR (Core [20220901])

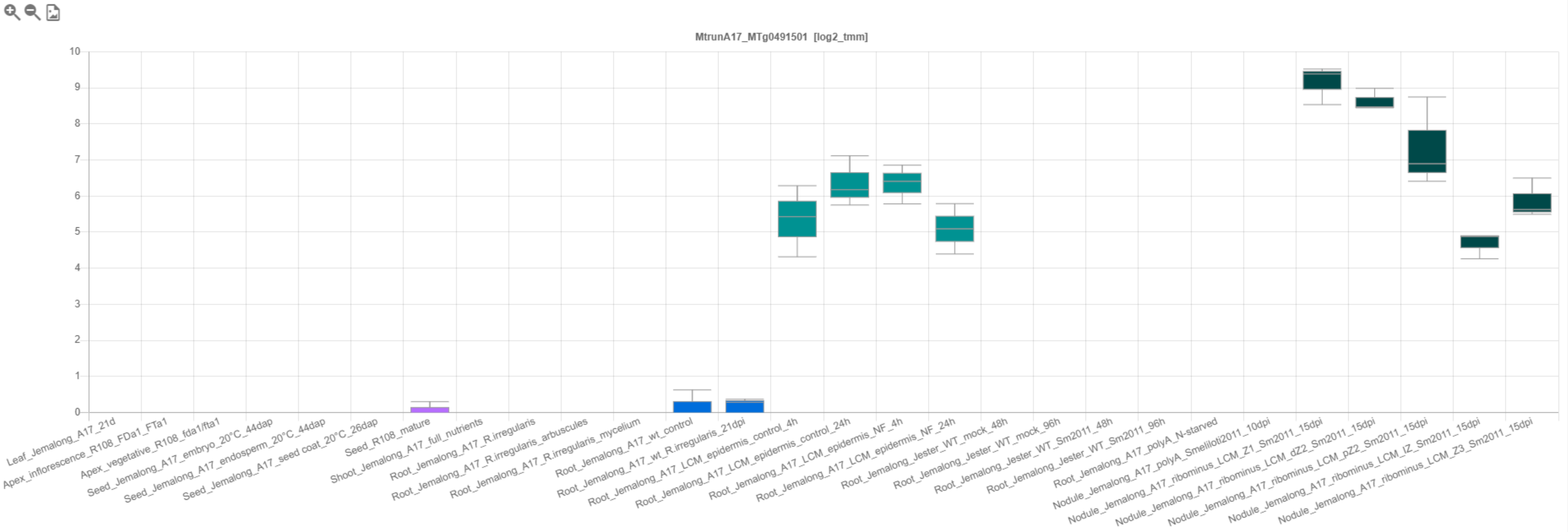

### CP155: MtrunA17_MTg0491621

INRAE/CNRS Medicago Expressi… | Expression Atlas Server

…pub/expressionAtlas/app/v3/aa_reference_dataset/MtrunA17_MTg0491621

MtExpress V3 | Core (20220901)

! Switch to another dataset using the left menu

TMM | METADATA | SYNONYMOUS | ANNOTATION | GENOME PORTAL | LEGOO

ncRNA: MtrunA17_MTg0491621;

Log2 TMM Normalisation using EdgeR (Core [20220901])

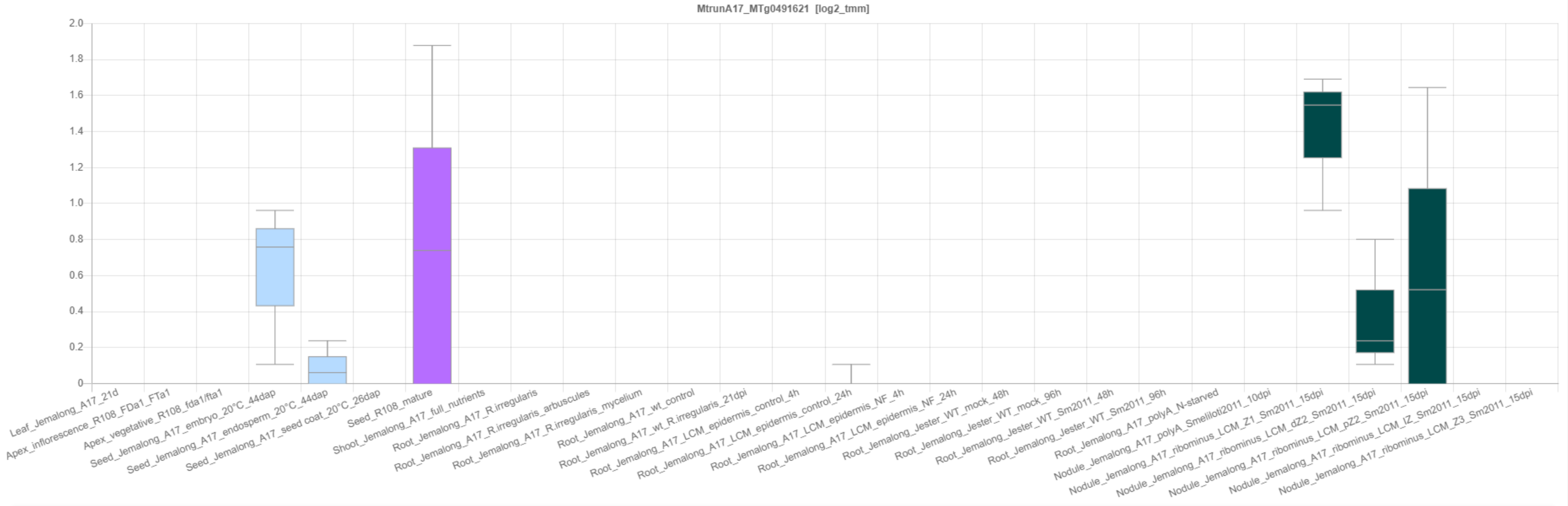

### CP156: MtrunA17_MTg0491711

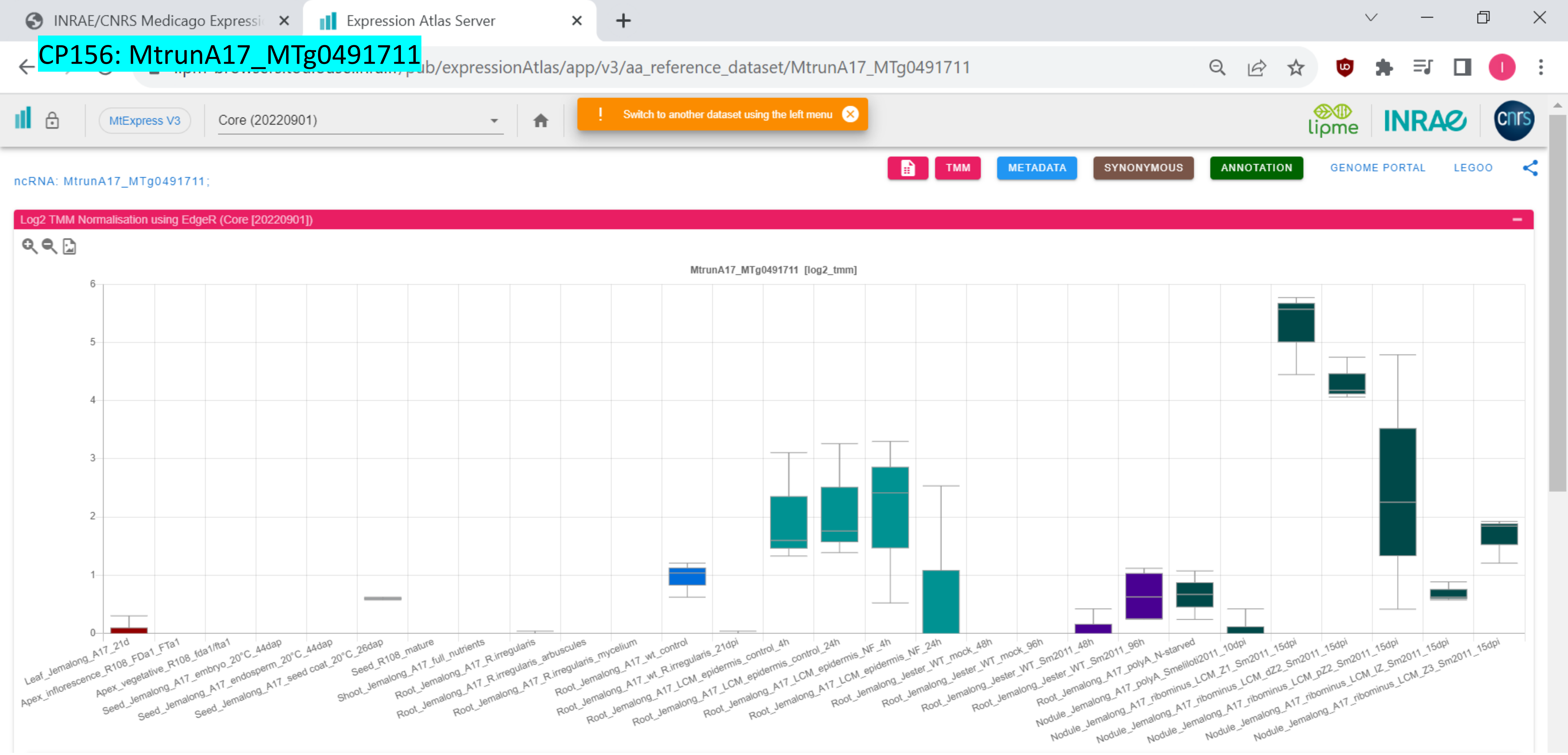

**Supplementary Dataset S27**. Expression profiles of 156 primary-source transcripts. Screenshots are from the RNA-Seq-based gene expression atlas of *Medicago truncatula* (MtExpress v. 3, https://medicago.toulouse.inrae.fr/GEA). For clarity and better visibility, only the core sample set is shown. Chimeric peptide identifiers that begin with an asterisk correspond to transcripts with log2 TMM values below zero.
