## Supplementary material for "Discovery of diverse chimeric peptides in a eukaryotic proteome sets the stage for the experimental proof of the mosaic translation hypothesis": Supplementary Dataset S28 Alternative-source transcripts associated with multiple PRF events.pdf

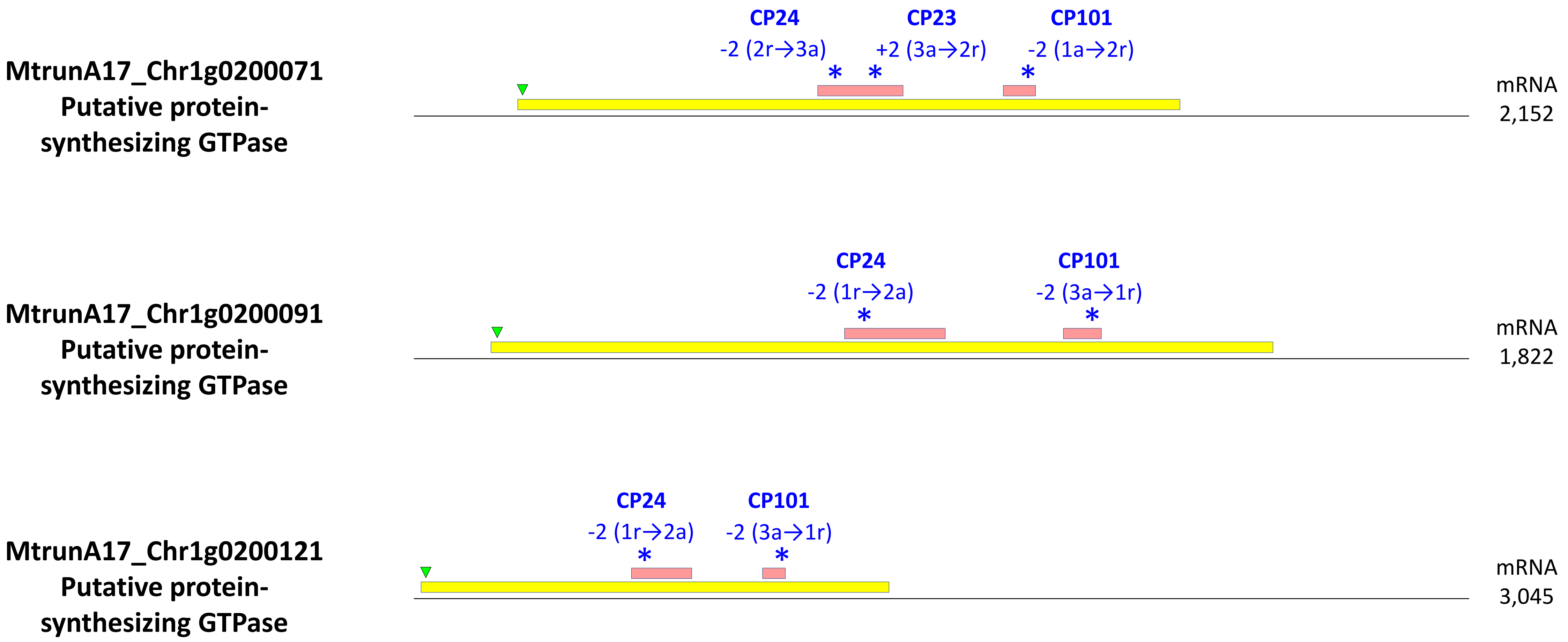

MtrunA17_Chr1g0200071
Putative protein-synthesizing GTPase
CP24
-2 (2r→3a)
CP23
+2 (3a→2r)
CP101
-2 (1a→2r)
mRNA
2,152
MtrunA17_Chr1g0200091
Putative protein-synthesizing GTPase
CP24
-2 (1r→2a)
CP101
-2 (3a→1r)
mRNA
1,822
MtrunA17_Chr1g0200121
Putative protein-synthesizing GTPase
CP24
-2 (1r→2a)
CP101
-2 (3a→1r)
mRNA
3,045

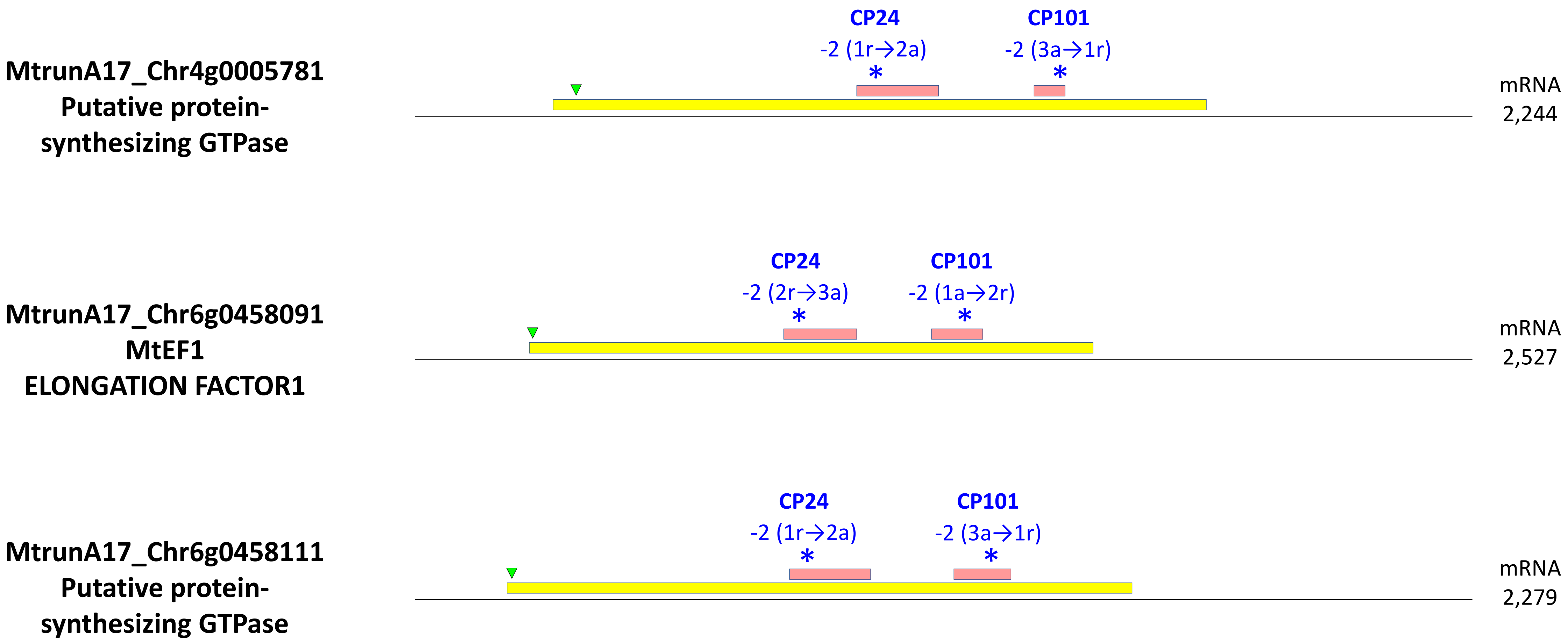
MtrunA17_Chr4g0005781
Putative protein-
synthesizing GTPase
CP24
-2 (1r→2a)
*
CP101
-2 (3a→1r)
*
mRNA
2,244
MtrunA17_Chr6g0458091
MtEF1
ELONGATION FACTOR1
CP24
-2 (2r→3a)
*
CP101
-2 (1a→2r)
*
mRNA
2,527
MtrunA17_Chr6g0458111
Putative protein-
synthesizing GTPase
CP24
-2 (1r→2a)
*
CP101
-2 (3a→1r)
*
mRNA
2,279

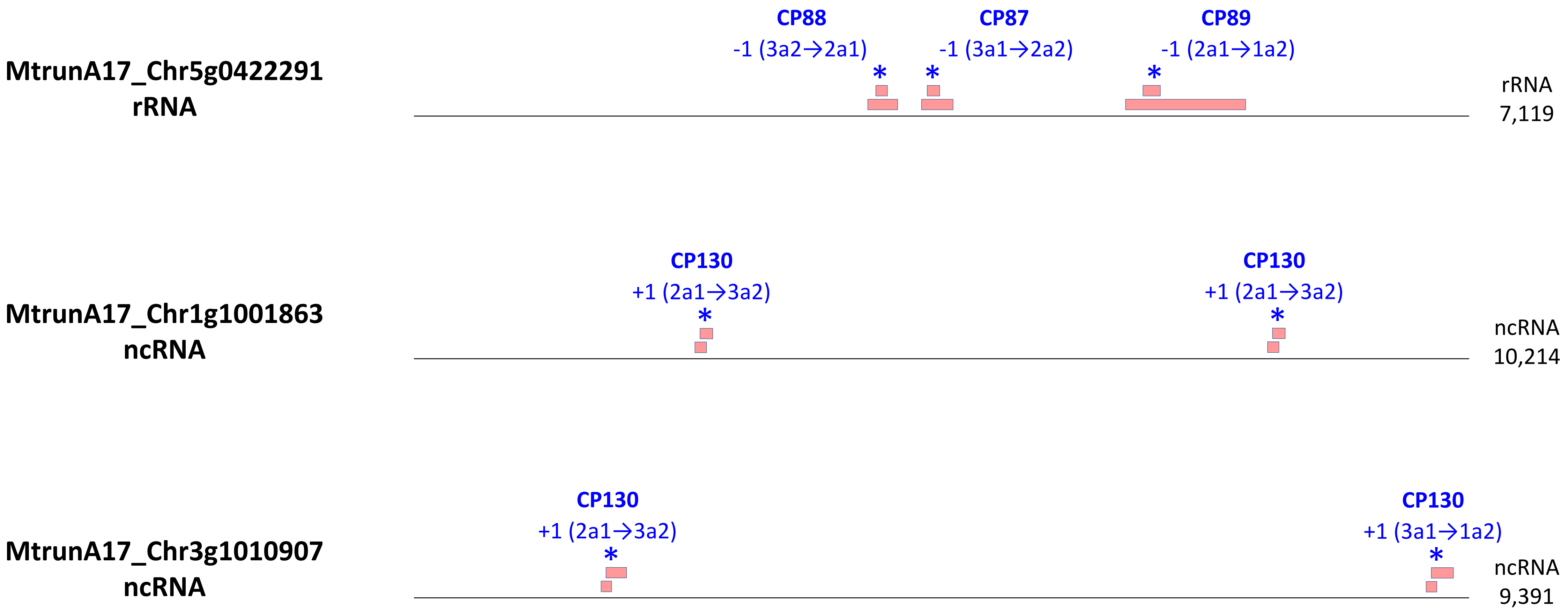

MtrunA17_Chr5g0422291
rRNA
CP88
-1 (3a2→2a1)
CP87
-1 (3a1→2a2)
CP89
-1 (2a1→1a2)
rRNA
7,119
MtrunA17_Chr1g1001863
ncRNA
CP130
+1 (2a1→3a2)
CP130
+1 (2a1→3a2)
ncRNA
10,214
MtrunA17_Chr3g1010907
ncRNA
CP130
+1 (2a1→3a2)
CP130
+1 (3a1→1a2)
ncRNA
9,391

**Supplementary Dataset S28**. Nine alternative-source transcripts associated with multiple PRF events. The images represent transcript models that feature the transcript type, length in nucleotides, and relative positions of ORFs (to scale) involved in the production of CPs. Reference ORFs (refORFs) are shown in yellow. Alternative ORFs (altORFs) are shown in pink. The first in-frame start codon (AUG) in each refORF is marked with a green triangle. Transcript models also show the positions and characteristics of PRF events, which are mapped with an asterisk. The description of a PRF event should be interpreted as follows. For example, CP88, -1 (3a2→2a1): a minus 1 frameshift changes the translation from frame 3 to frame 2, which corresponds to the change from altORF2 to altORF1. An altORF1 in this study is defined as the one that starts earlier than an altORF2. Note that three loci in this list, namely MtrunA17_Chr1g0200071, MtrunA17_Chr6g0458091, and MtrunA17_Chr5g0422291, can be regarded as primary sources for some CPs but as alternative sources for other CPs.
