## Supplementary material for "Discovery of diverse chimeric peptides in a eukaryotic proteome sets the stage for the experimental proof of the mosaic translation hypothesis": Supplementary Dataset S31 Alignments of rRNA transcripts with known rRNA-like mRNA transcripts.pdf

CP1: MtrunA17_Chr0c01g0489091

*Mus musculus* mRNA for testin2, X78990.1

58.6% in a region of 256 nt

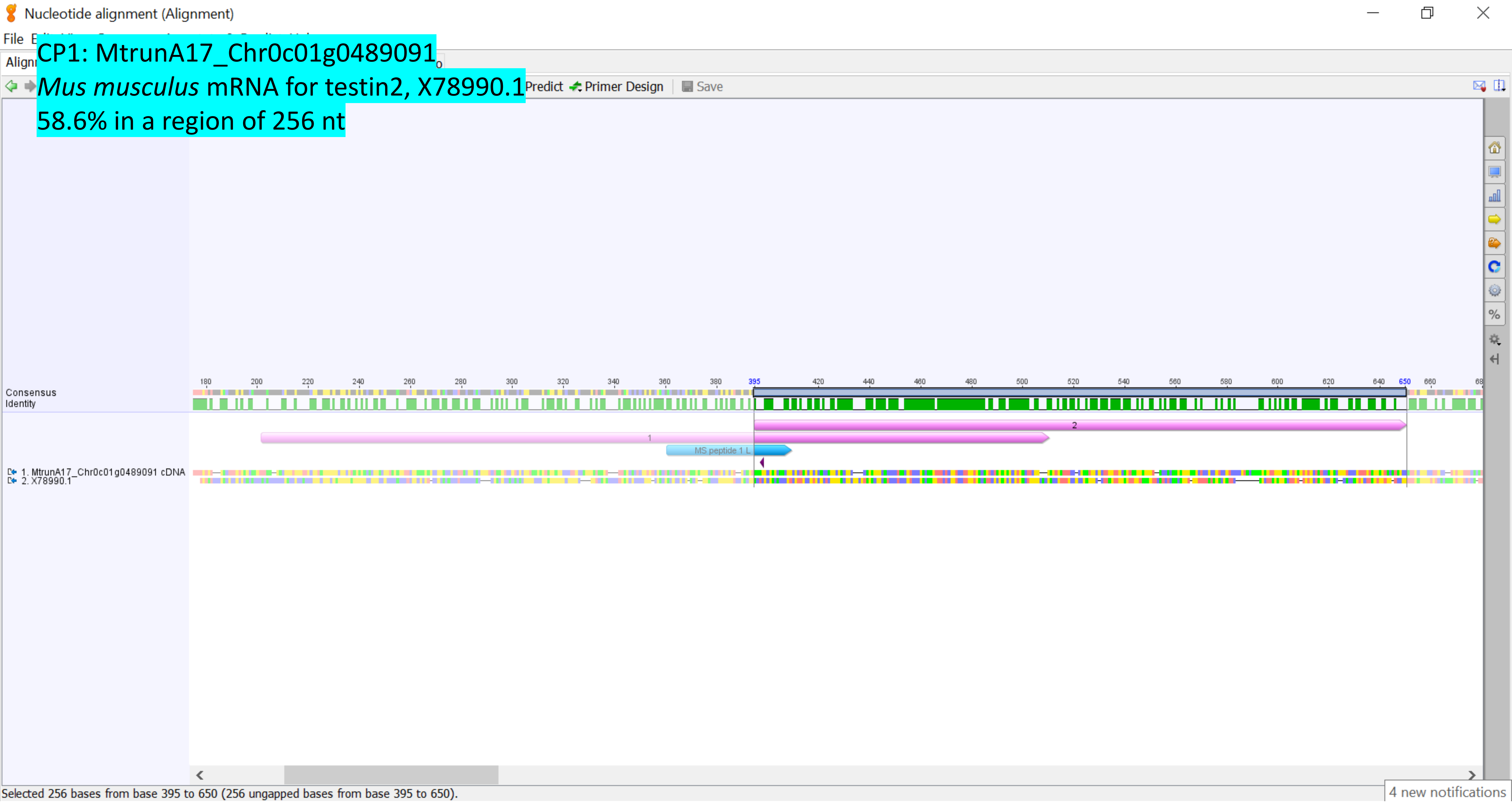

CP1: MtrunA17_Chr0c01g0489091, altORF2
*Mus musculus* testin2, X78990.1
32.9% in a region of 70 aa

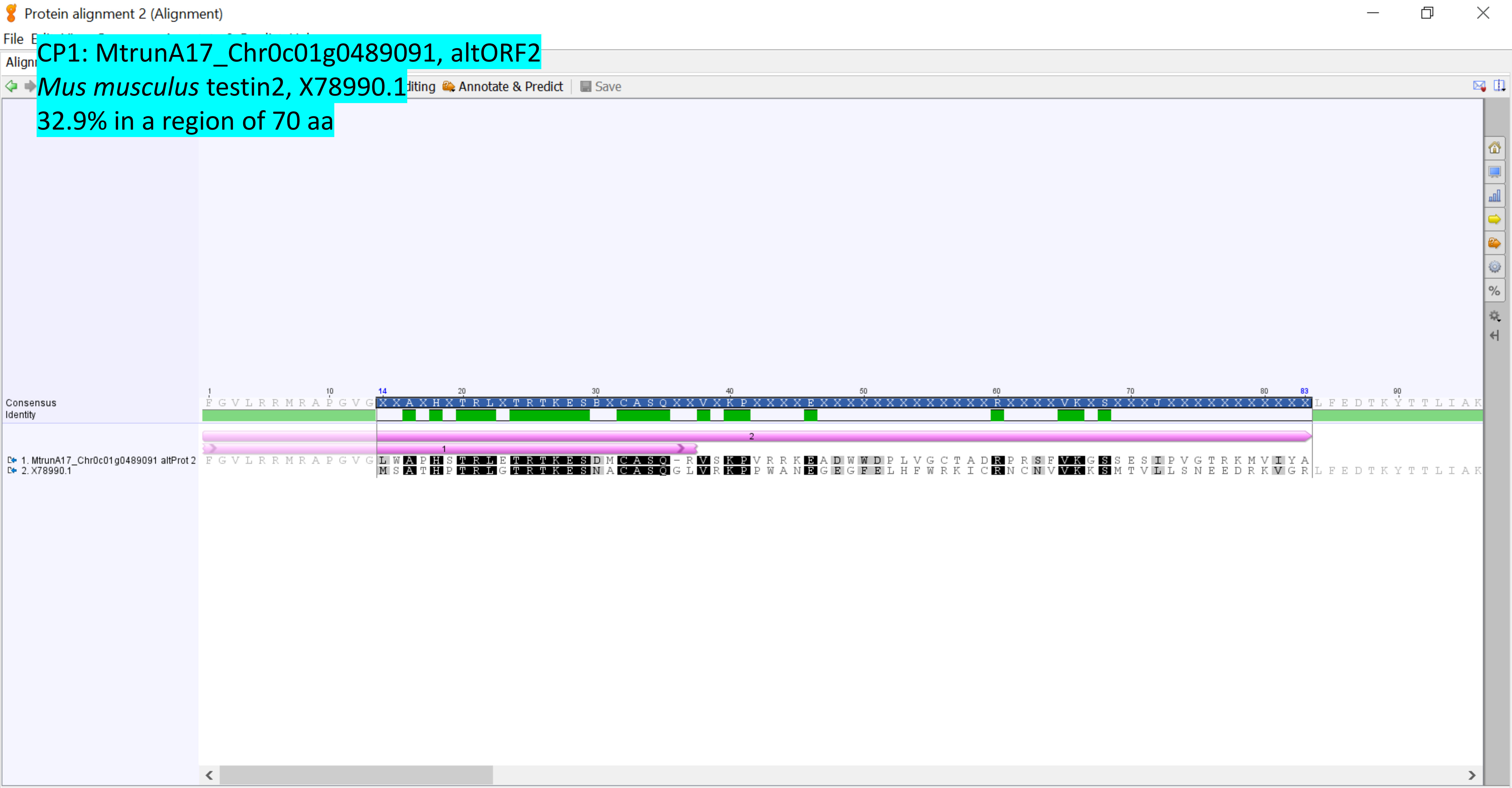

Selected 70 residues from residue 14 to 83 (70 ungapped residues from residue 14 to 83).

CP114: MtrunA17_Chr7g0229401

*Homo sapiens* Humanin (HN1) mRNA, complete cds, AY029066.1

67.4% in a region of 196 nt

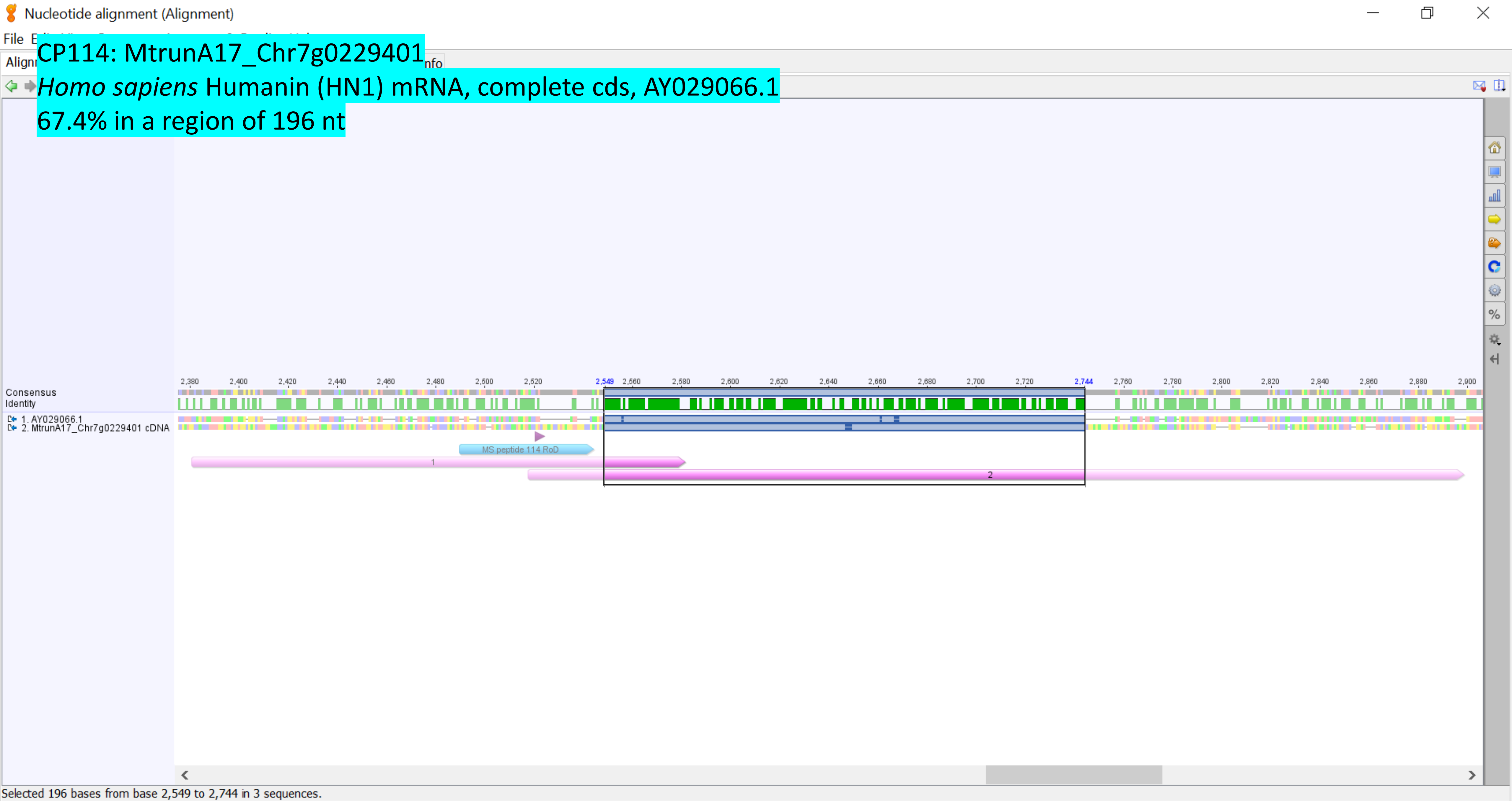

CP89: MtrunA17_Chr5g0422291
*Candida albicans* SC5314 Tar1p (TAR1), partial mRNA, XM_019475565.1
73.3% in a region of 285 nt

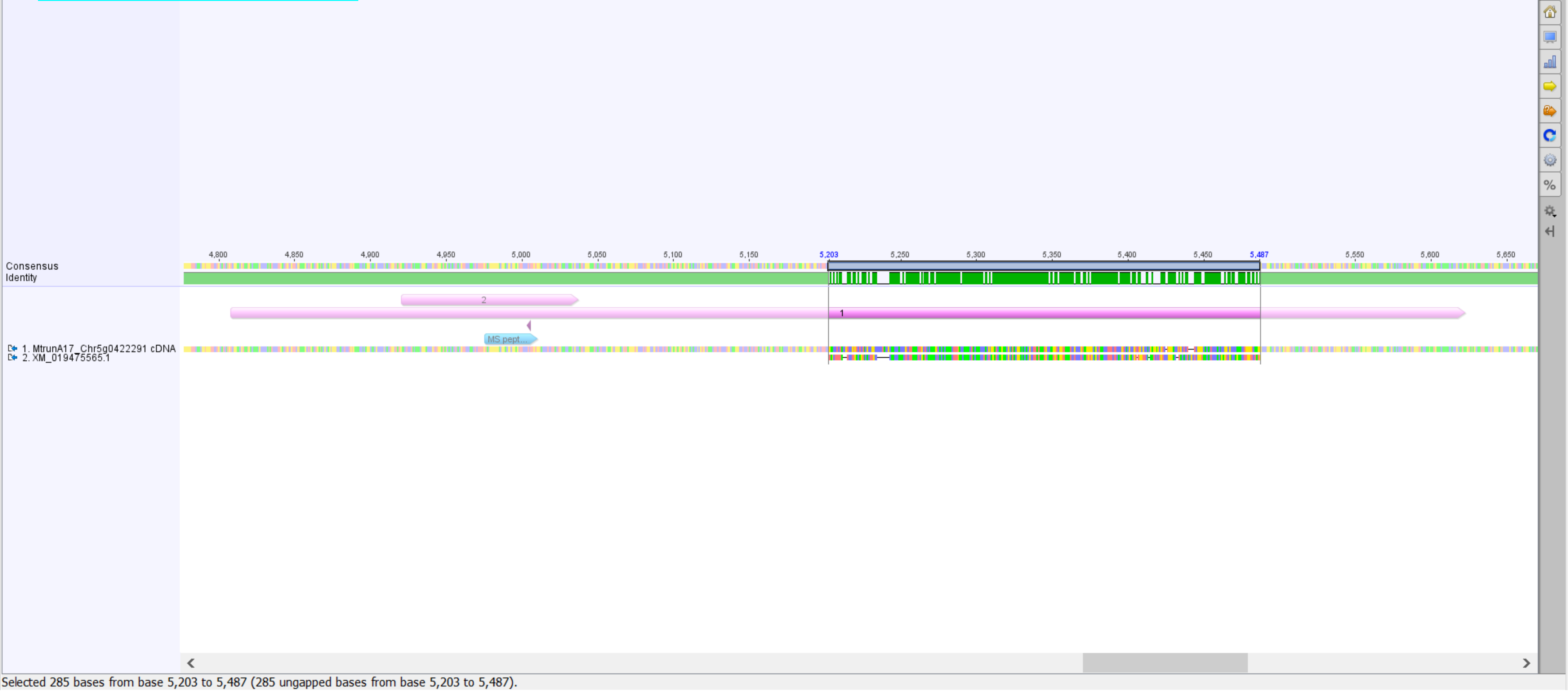

CP87: MtrunA17_Chr5g0421761

*Reticulitermes flavipes* rRNA-like mRNA, partial sequence, AY572860.1

80.5% in a region of 41 nt

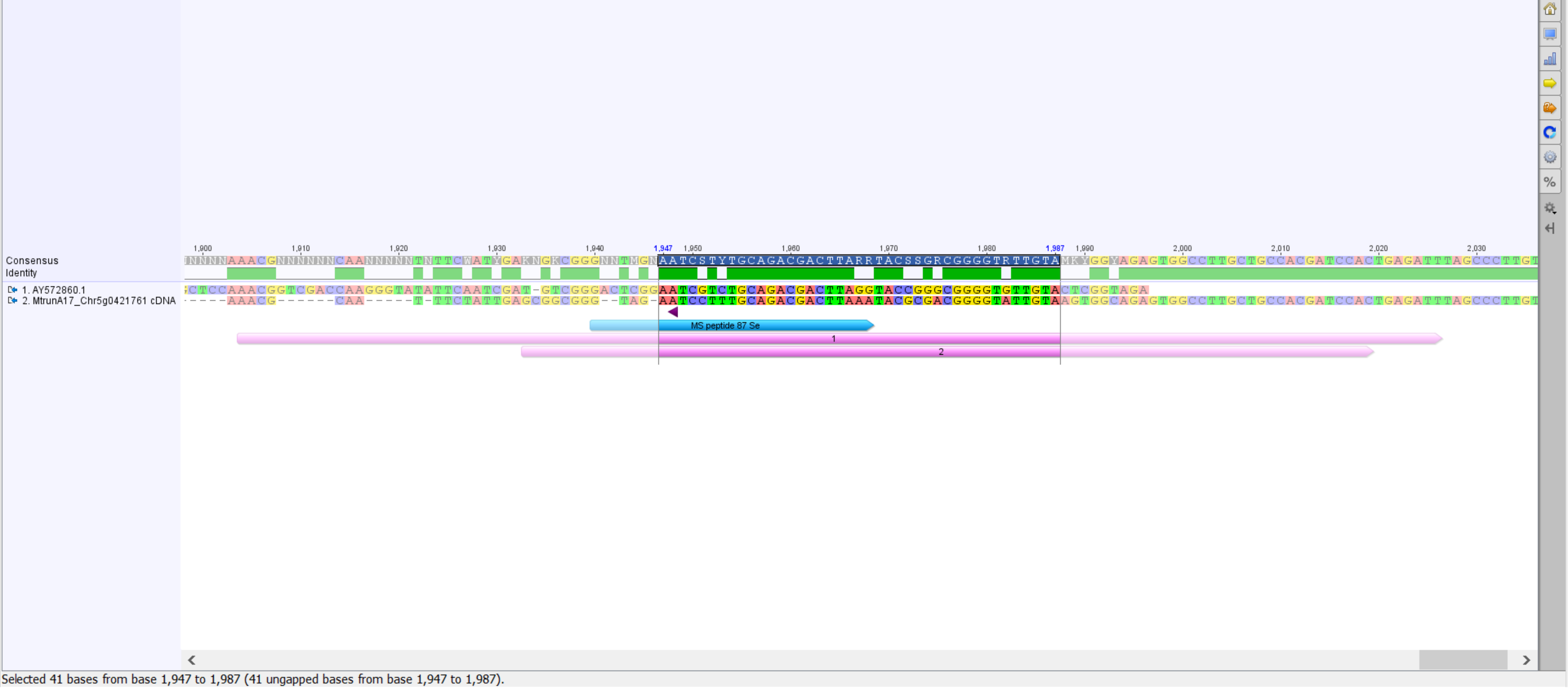

CP88: MtrunA17_Chr5g0422291
*Reticulitermes flavipes* rRNA-like mRNA, partial sequence, AY572860.1
88.3% in a region of 417 nt

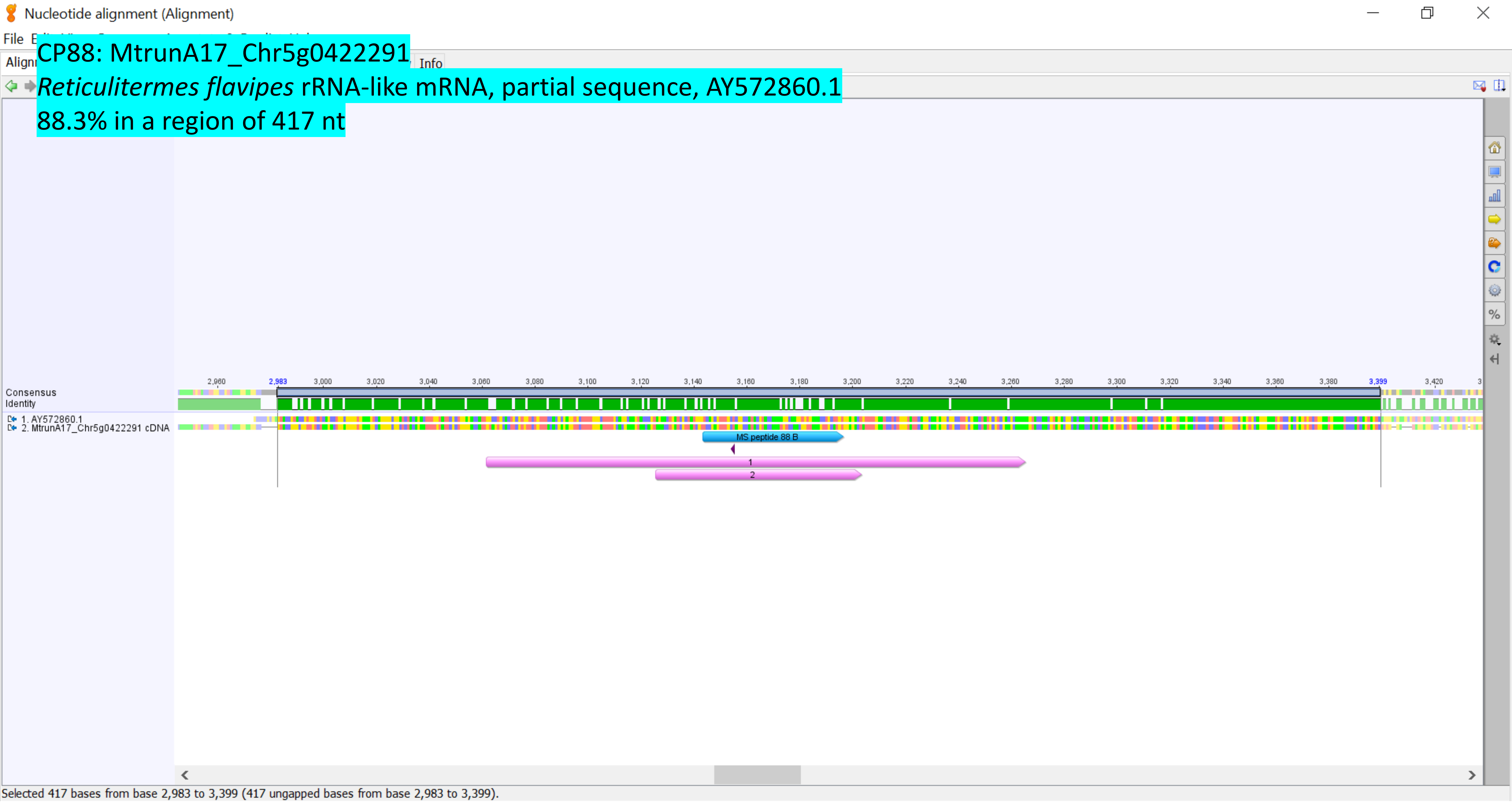

**Supplementary Dataset S31**. Alignments of four PRF-related rRNA transcripts with known rRNA-like mRNA transcripts. Percent identity values are based on Geneious® alignments (Geneious® v. 7.1, Dotmatics Ltd., MA, USA, https://www.geneious.com). Codes at the end of the annotated gene names are GenBank identifiers. Alternative ORFs (altORFs) are shown in pink. Numbers 1 and 2 on altORFs indicate the following. An altORF1 in this study is defined as the one that starts earlier than an altORF2. Switches from altORF2 to altORF1 are also found in this dataset but are five times less frequent (e.g. CP108 and four other CPs). Positions and the direction of programmed ribosomal frameshifting (PRF) events are labeled with deep purple arrowheads.
