## Supplementary material for "Discovery of diverse chimeric peptides in a eukaryotic proteome sets the stage for the experimental proof of the mosaic translation hypothesis": Supplementary figures.pdf

### Supplementary Figure S1

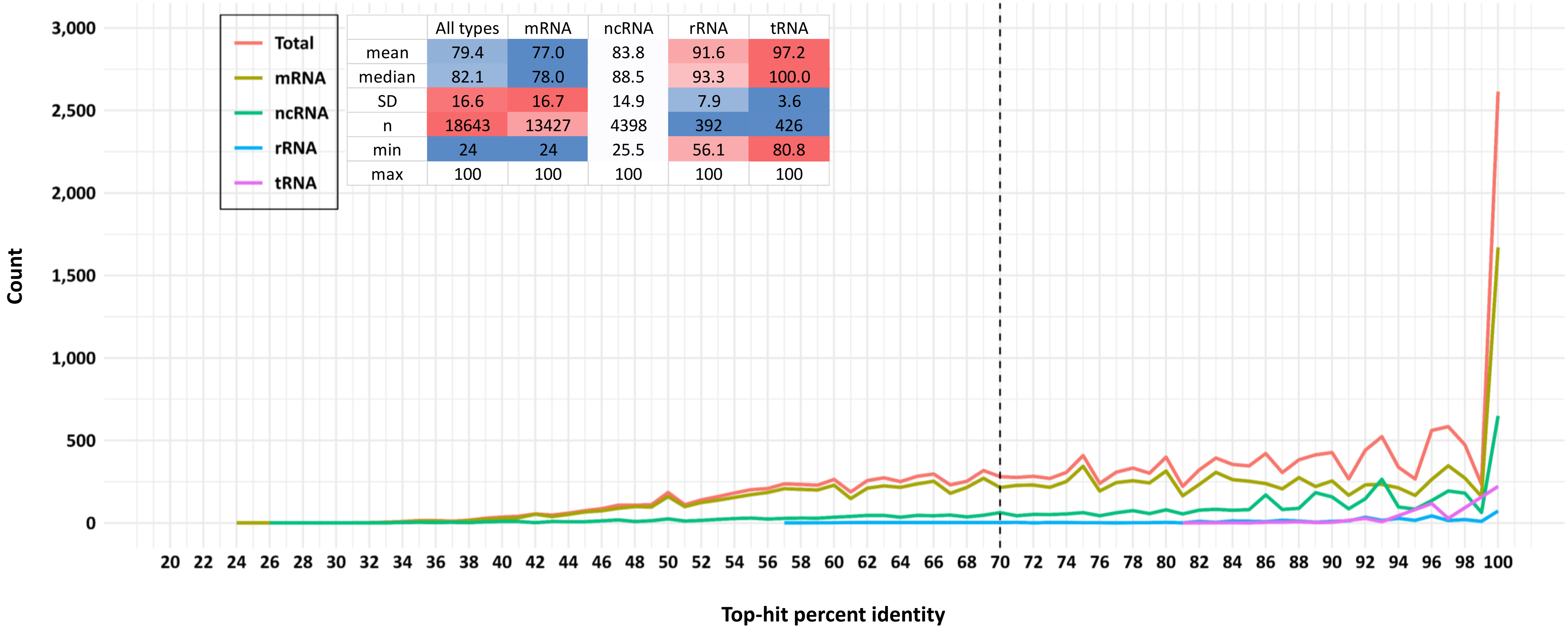


| | All types | mRNA | ncRNA | rRNA | tRNA |
|---|---|---|---|---|---|
| mean | 79.4 | 77.0 | 83.8 | 91.6 | 97.2 |
| median | 82.1 | 78.0 | 88.5 | 93.3 | 100.0 |
| SD | 16.6 | 16.7 | 14.9 | 7.9 | 3.6 |
| n | 18643 | 13427 | 4398 | 392 | 426 |
| min | 24 | 24 | 25.5 | 56.1 | 80.8 |
| max | 100 | 100 | 100 | 100 | 100 |

**Supplementary Figure S1**. Top-hit percent identity distributions of altProts from all RNA types (red) and separate RNA types (other colors). The threshold for the BLASTP filter: e-value ≤ 0.001. The bin size was set to one. The inset shows descriptive statistics of the data (SD, standard deviation; n, sample size). The X-axis reference line at 70.0 shows the threshold for robust conservation evidence, which is a proxy for translation evidence. The distributions are significantly different between different RNA types (Kolmogorov–Smirnov test, Bonferroni-adjusted p-values of all pairwise comparisons are close to zero). The image was generated in R.

### Supplementary Figure S2

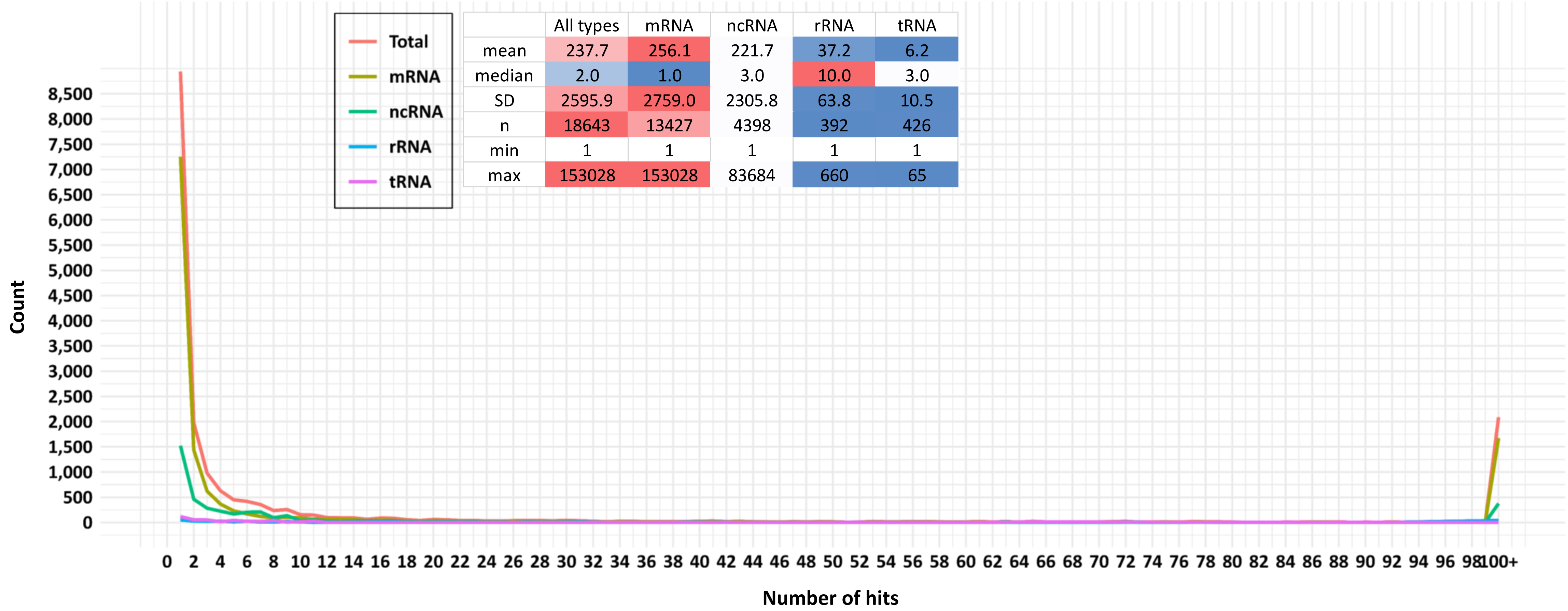


| | All types | mRNA | ncRNA | rRNA | tRNA |
|---|---|---|---|---|---|
| mean | 237.7 | 256.1 | 221.7 | 37.2 | 6.2 |
| median | 2.0 | 1.0 | 3.0 | 10.0 | 3.0 |
| SD | 2595.9 | 2759.0 | 2305.8 | 63.8 | 10.5 |
| n | 18643 | 13427 | 4398 | 392 | 426 |
| min | 1 | 1 | 1 | 1 | 1 |
| max | 153028 | 153028 | 83684 | 660 | 65 |

**Supplementary Figure S2**. Distributions of the number of hits per query of altProts from all RNA types (red) and separate RNA types (other colors) **before** the elimination of altProts with top-hit % identity below 70. The threshold for the BLASTP filter: e-value ≤ 0.001. The bin size was set to one. The inset shows descriptive statistics of the data (SD, standard deviation; n, sample size). The distributions are significantly different between different RNA types (Kolmogorov–Smirnov test, Bonferroni-adjusted p-values of all pairwise comparisons are close to zero, except for the comparison between ncRNA and tRNA, where p = 4.000E-11). The image was generated in R.

### Supplementary Figure S3

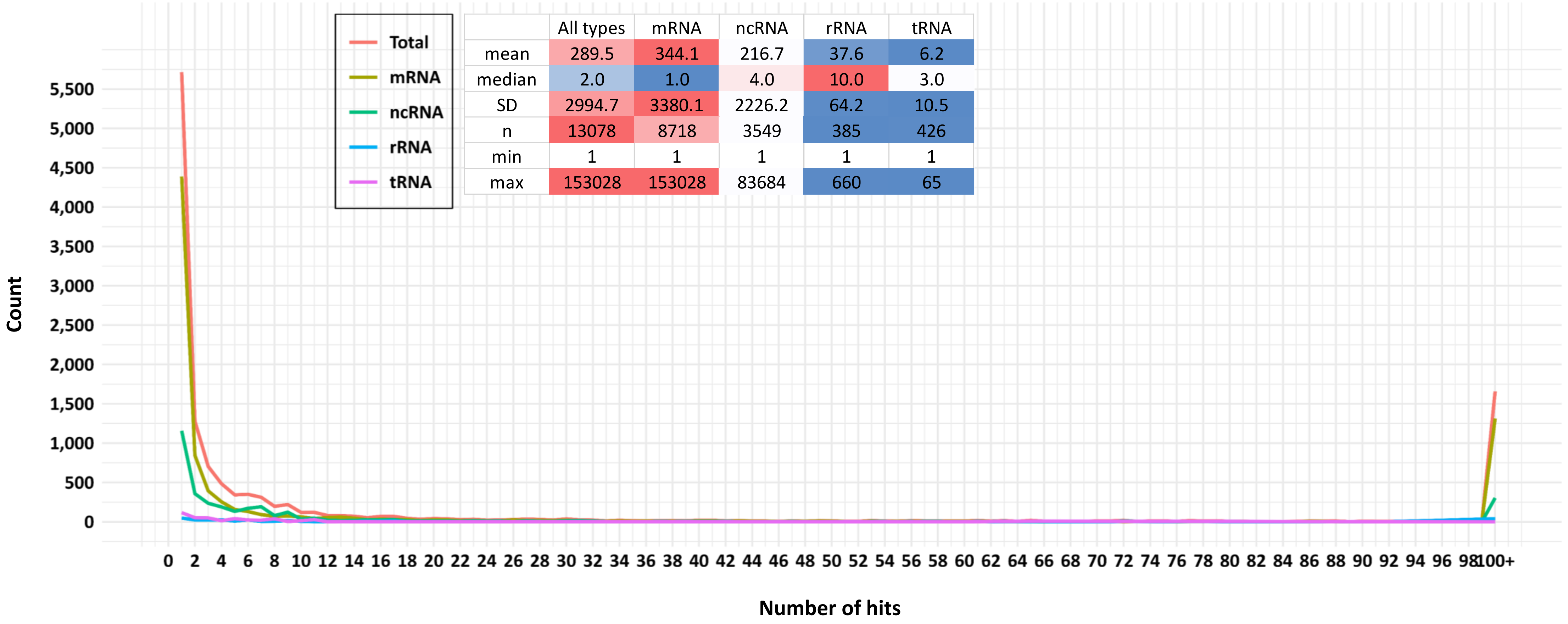


| | All types | mRNA | ncRNA | rRNA | tRNA |
|---|---|---|---|---|---|
| mean | 289.5 | 344.1 | 216.7 | 37.6 | 6.2 |
| median | 2.0 | 1.0 | 4.0 | 10.0 | 3.0 |
| SD | 2994.7 | 3380.1 | 2226.2 | 64.2 | 10.5 |
| n | 13078 | 8718 | 3549 | 385 | 426 |
| min | 1 | 1 | 1 | 1 | 1 |
| max | 153028 | 153028 | 83684 | 660 | 65 |

**Supplementary Figure S3**. Distributions of the number of hits per query of altProts from all RNA types (red) and separate RNA types (other colors) **after** the elimination of altProts with top-hit % identity below 70. The threshold for the BLASTP filter: e-value ≤ 0.001. The bin size was set to one. The inset shows descriptive statistics of the data (SD, standard deviation; n, sample size). The distributions are significantly different between different RNA types (Kolmogorov–Smirnov test, Bonferroni-adjusted p-values of all pairwise comparisons are close to zero, except for the comparison between ncRNA and tRNA, where $p = 3.160E-11$). The image was generated in R.

### Supplementary Figure S4

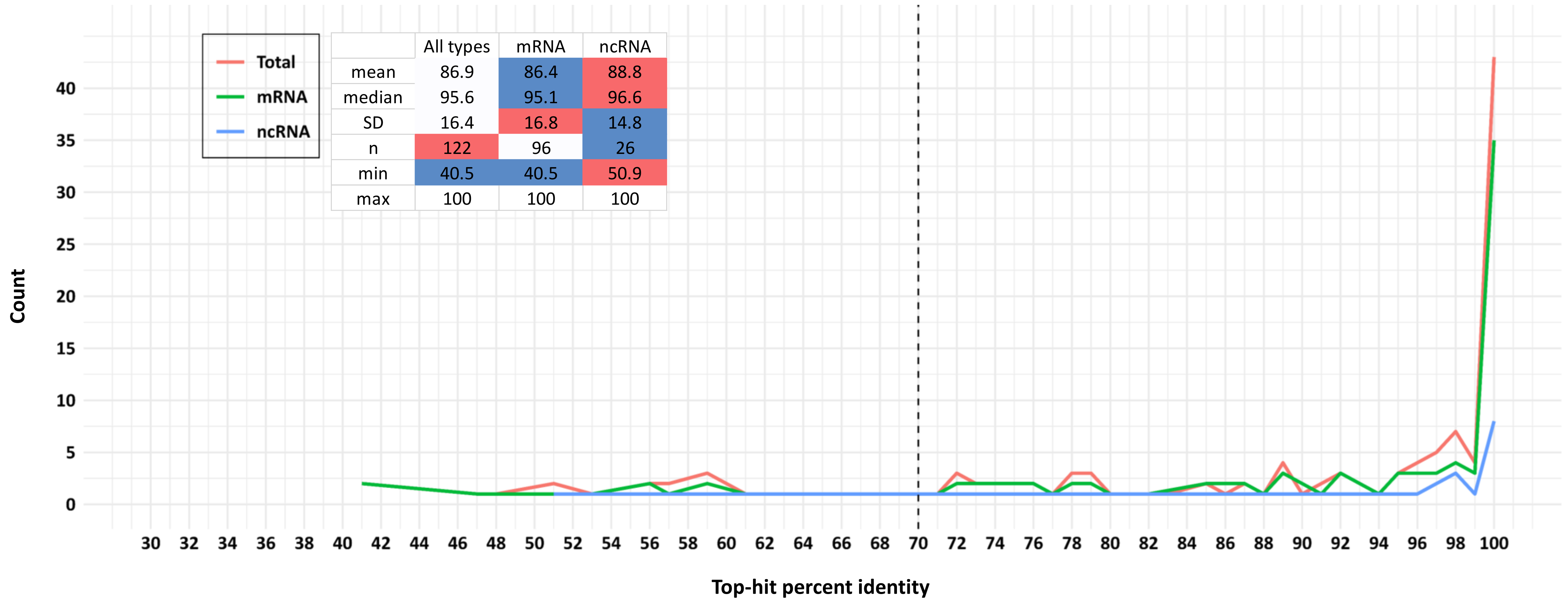


| | All types | mRNA | ncRNA |
|---|---|---|---|
| mean | 86.9 | 86.4 | 88.8 |
| median | 95.6 | 95.1 | 96.6 |
| SD | 16.4 | 16.8 | 14.8 |
| n | 122 | 96 | 26 |
| min | 40.5 | 40.5 | 50.9 |
| max | 100 | 100 | 100 |

**Supplementary Figure S4**. Top-hit percent identity distributions of 122 MS-supported conserved altProts from mRNA and ncRNA (green and blue, respectively) and the two types of RNA combined (red). The threshold for the BLASTP filter: e-value ≤ 0.001. The bin size was set to one. The inset shows descriptive statistics of the data (SD, standard deviation; n, sample size). The X-axis reference line at 70.0 shows the threshold for robust conservation evidence, which is a proxy for translation evidence. There are 103 MS-supported conserved altProts with percent identity values at least 70. The distributions are not significantly different from each other. The image was generated in R.

### Supplementary Figure S5

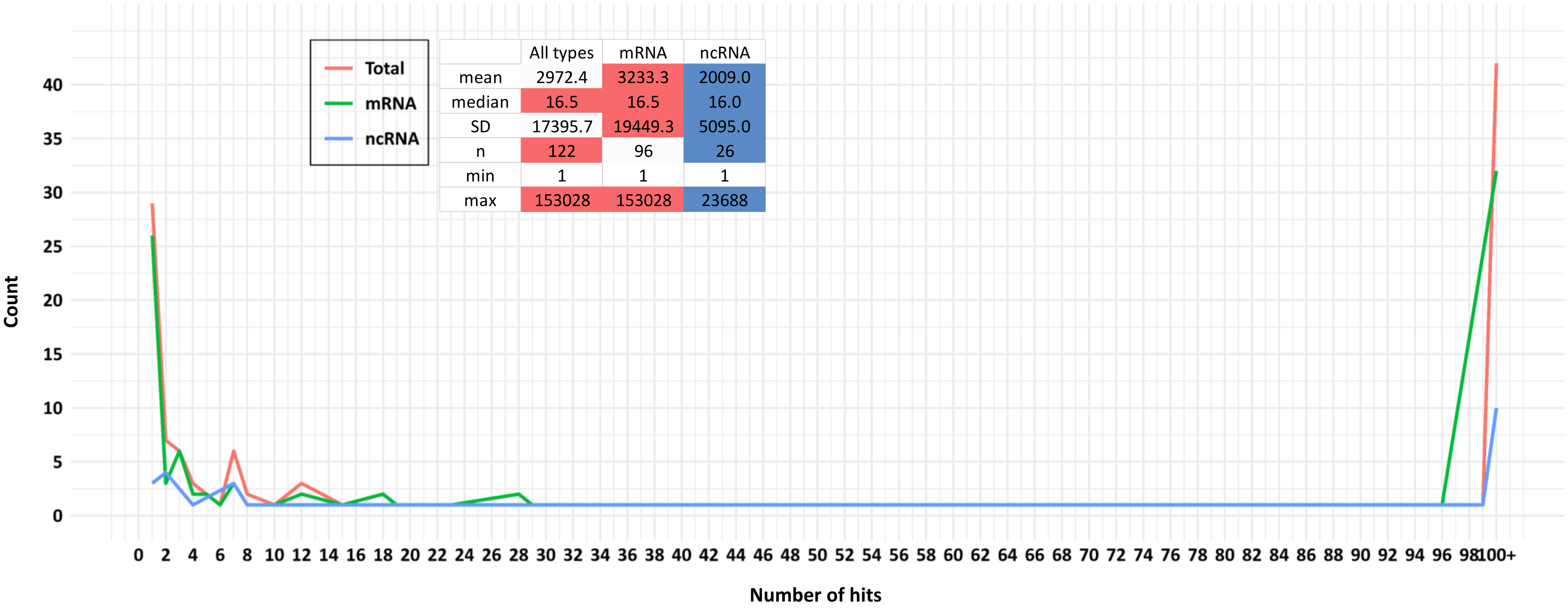


| | All types | mRNA | ncRNA |
|---|---|---|---|
| mean | 2972.4 | 3233.3 | 2009.0 |
| median | 16.5 | 16.5 | 16.0 |
| SD | 17395.7 | 19449.3 | 5095.0 |
| n | 122 | 96 | 26 |
| min | 1 | 1 | 1 |
| max | 153028 | 153028 | 23688 |

**Supplementary Figure S5**. Distributions of the number of hits per query of 122 MS-supported conserved altProts from mRNA and ncRNA (green and blue, respectively) and the two types of RNA combined (red). The threshold for the BLASTP filter: e-value ≤ 0.001. The bin size was set to one. The inset shows descriptive statistics of the data (SD, standard deviation; n, sample size). The distributions are not significantly different from each other. The image was generated in R.

### Supplementary Figure S6

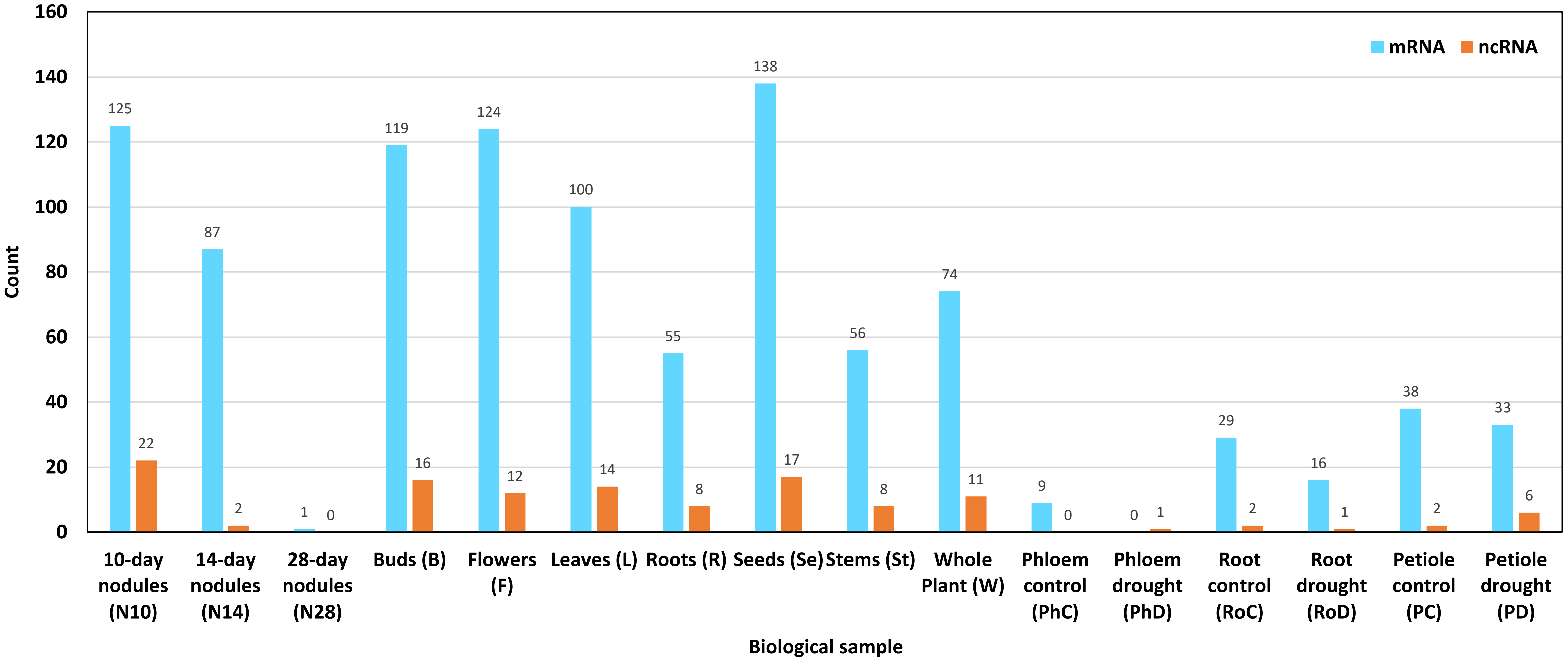


**Supplementary Figure S6**. Counts of 1,126 MS-validated altProts (805 unique sequences). The numbers of validated mRNA-derived altProts (1,004 in total; 720 unique) are shown from the second search of the two-step procedure, while the MS search for ncRNA-derived altProts (122 in total; 85 unique) consisted of a single step. No rRNA- and tRNA-derived non-chimeric altProts were validated by MS.

### Supplementary Figure S7

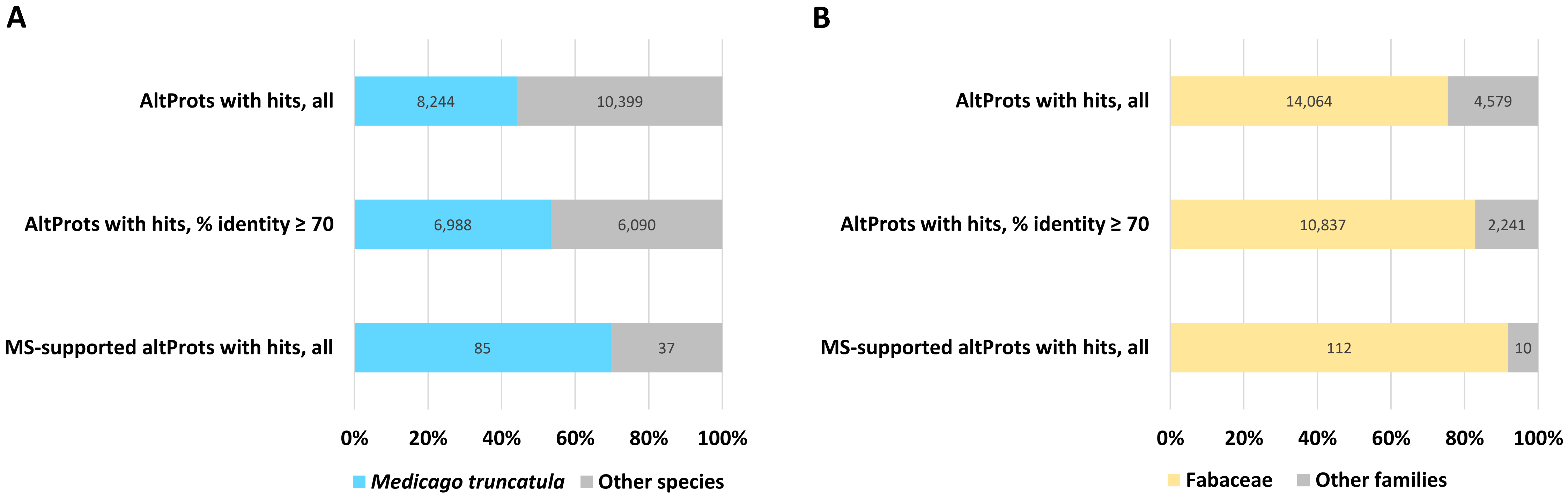


**Supplementary Figure S7**. The relative abundance of altProts with the top-BLASTP hits in *Medicago truncatula* and other species from the Fabaceae family. The proportions are shown for three altProt groups: altProts with at least one hit before the 70% identity filter (top), after the filter (middle), and MS-supported altProts with at least one hit regardless of the filter (bottom). A, altProts with *M. truncatula* hits vs altProts with hits in other species. B, altProts with Fabaceae hits vs altProts with hits in other families.

**Supplementary Figure S8**

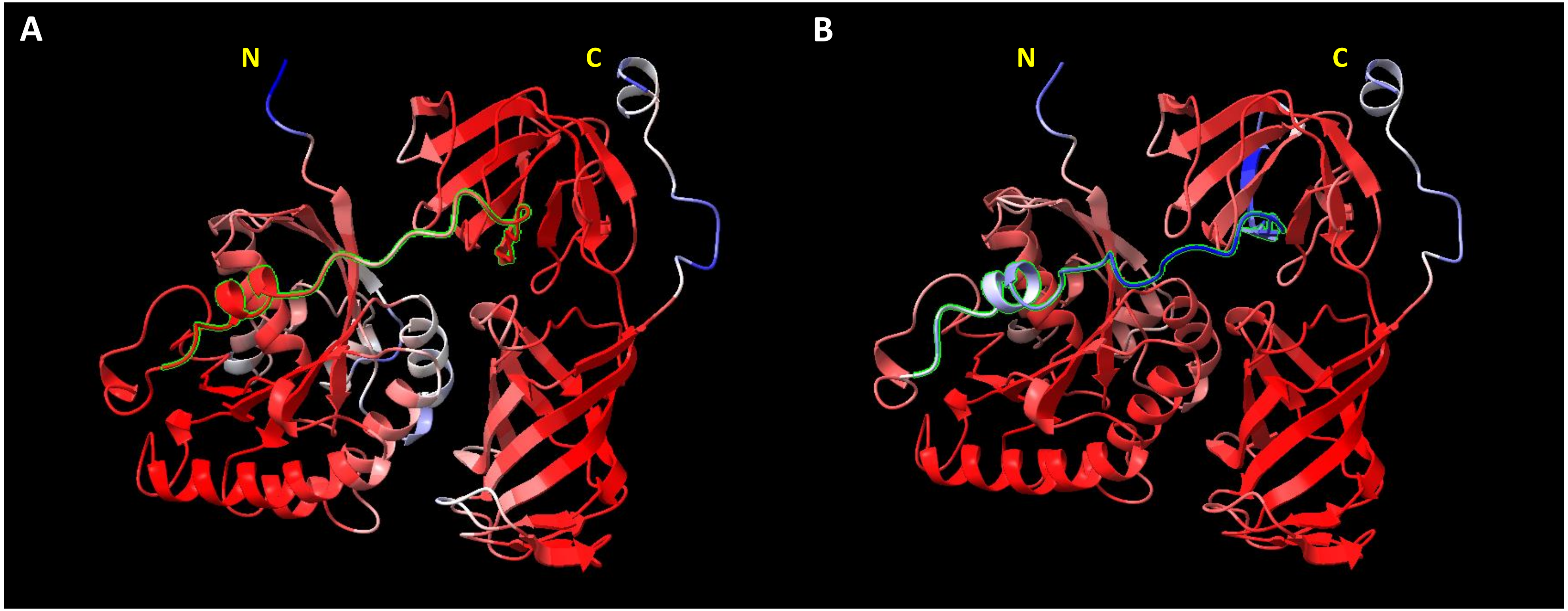


**Supplementary Figure S8**. Folding predictions for the putative protein-synthesizing GTPase encoded by transcript MtrunA17_Chr1g0200071. A, the annotated protein. B, the hypothetical mosaic protein 1 (Supplementary Dataset S5). Folding was predicted by ColabFold v. 1.5.2 and visualized with ChimeraX v. 1.6.1. The structure shown corresponds to the prediction with rank 1. Each structure was manually oriented in space to maximize the two-dimensional projection and to position the N-terminus (labeled with N in the figure) on the left-hand side. The blue-white-red gradient indicates the prediction confidence based on the range of b-factor values. Blue stands for the lowest and red stands for the highest values (the highest confidence). A sequence segment highlighted with lime delimits the portion that is different between the two proteins. Note that the introduction of a 28 amino acid-long segment from the altORF located in frame 3 does not change the overall structure of the protein.

**Supplementary Figure S9**

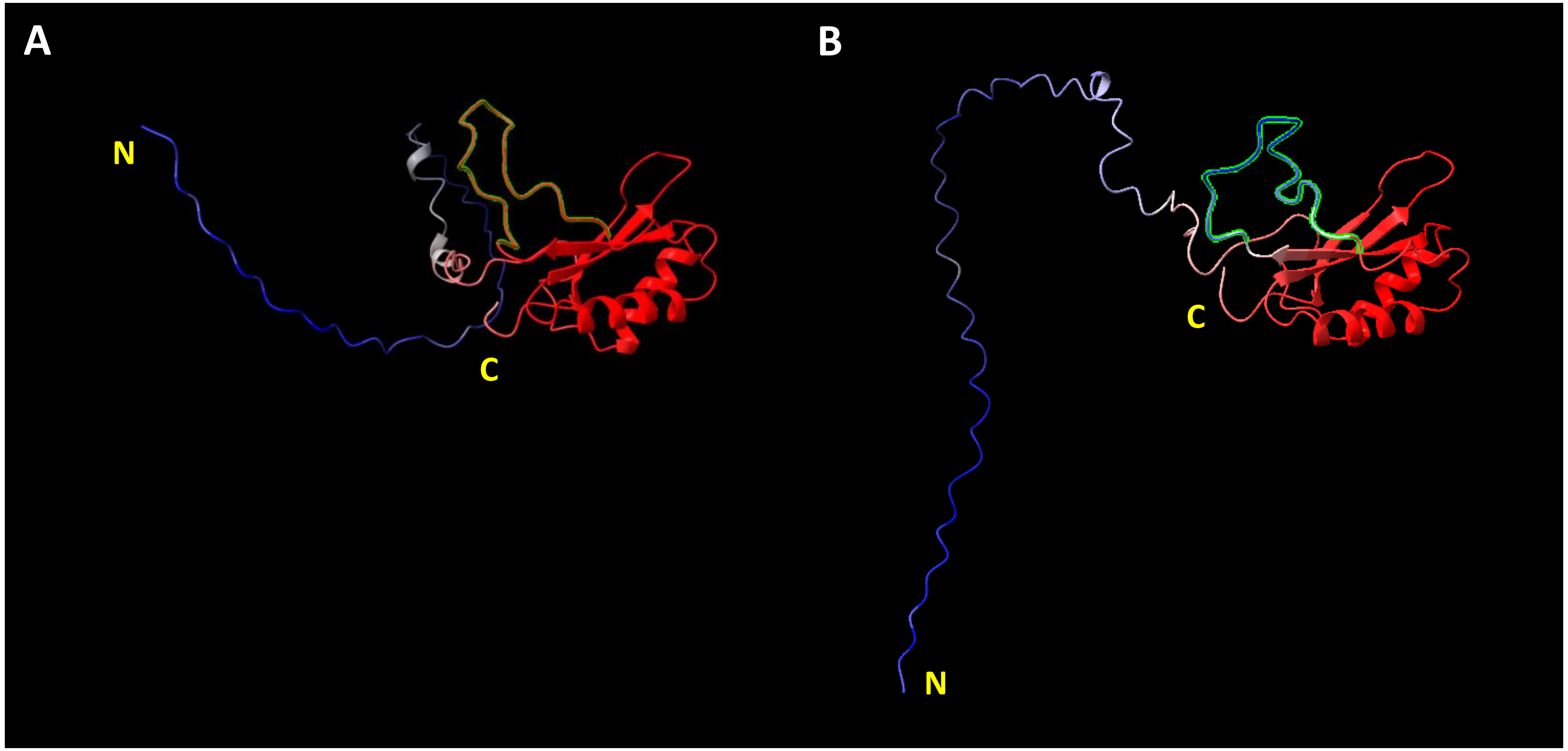


**Supplementary Figure S9**. Folding predictions for the putative ribulose-bisphosphate carboxylase (RuBisCo) encoded by transcript MtrunA17_Chr6g0457461. A, the annotated protein. B, the hypothetical mosaic protein 2 (Supplementary Dataset S5). Folding was predicted by ColabFold v. 1.5.2 and visualized with ChimeraX v. 1.6.1. The structure shown corresponds to the prediction with rank 1. Each structure was manually oriented in space to maximize the two-dimensional projection and to position the N-terminus (labeled with N in the figure) on the left-hand side. The blue-white-red gradient indicates the prediction confidence based on the range of b-factor values. Blue stands for the lowest and red stands for the highest values (the highest confidence). A sequence segment highlighted with lime delimits the portion that is different between the two proteins. Note that the introduction of a 22 amino acid-long segment from the altORF located in frame 3 does not change the overall structure of the protein.

**Supplementary Figure S10**

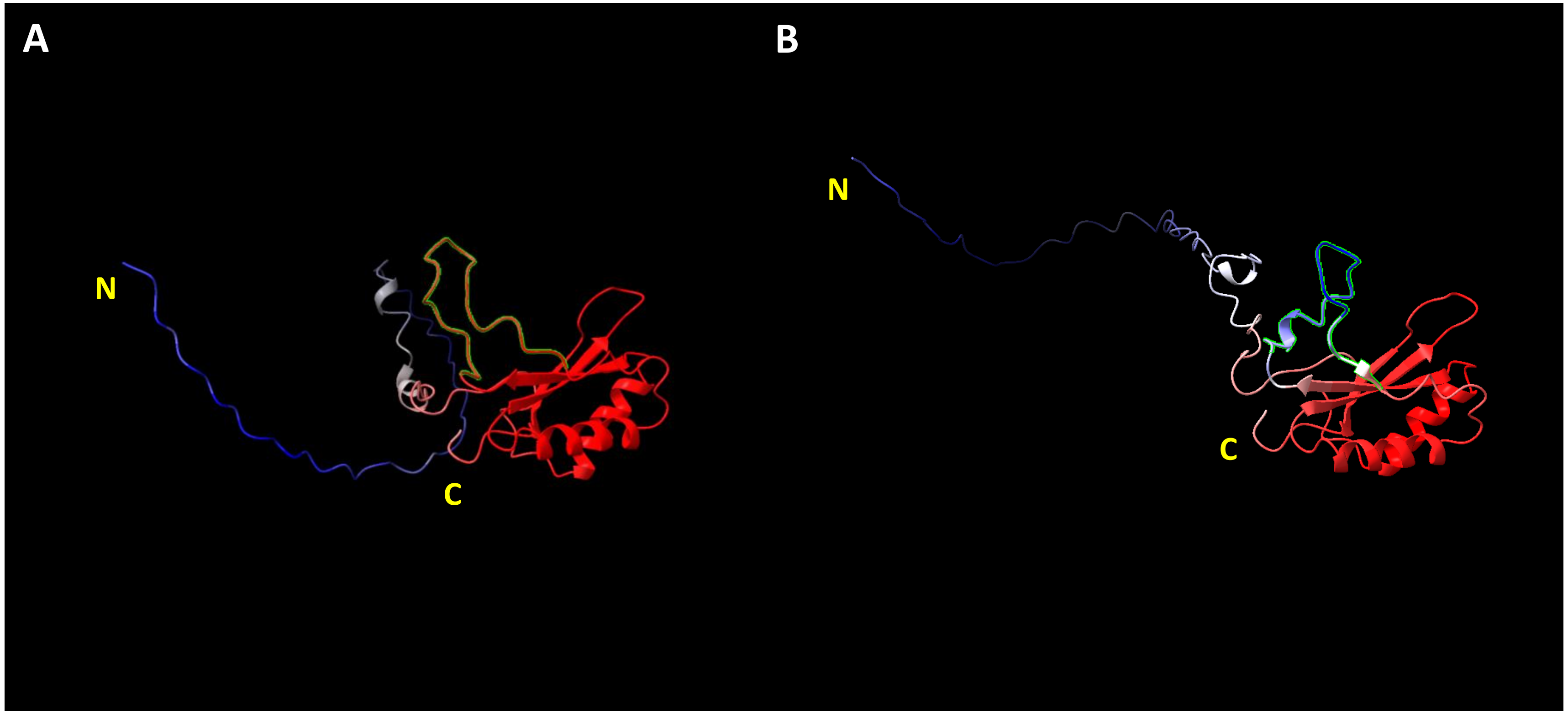


**Supplementary Figure S10**. Folding predictions for the putative ribulose-bisphosphate carboxylase (RuBisCo) encoded by transcript MtrunA17_Chr6g0457461. A, the annotated protein. B, the hypothetical mosaic protein 3 (Supplementary Dataset S5). Folding was predicted by ColabFold v. 1.5.2 and visualized with ChimeraX v. 1.6.1. The structure shown corresponds to the prediction with rank 1. Each structure was manually oriented in space to maximize the two-dimensional projection and to position the N-terminus (labeled with N in the figure) on the left-hand side. The blue-white-red gradient indicates the prediction confidence based on the range of b-factor values. Blue stands for the lowest and red stands for the highest values (the highest confidence). A sequence segment highlighted with lime delimits the portion that is different between the two proteins. Note that the introduction of a 21 amino acid-long segment from the altORF located in frame 3 does not change the overall structure of the protein.

**Supplementary Figure S11**

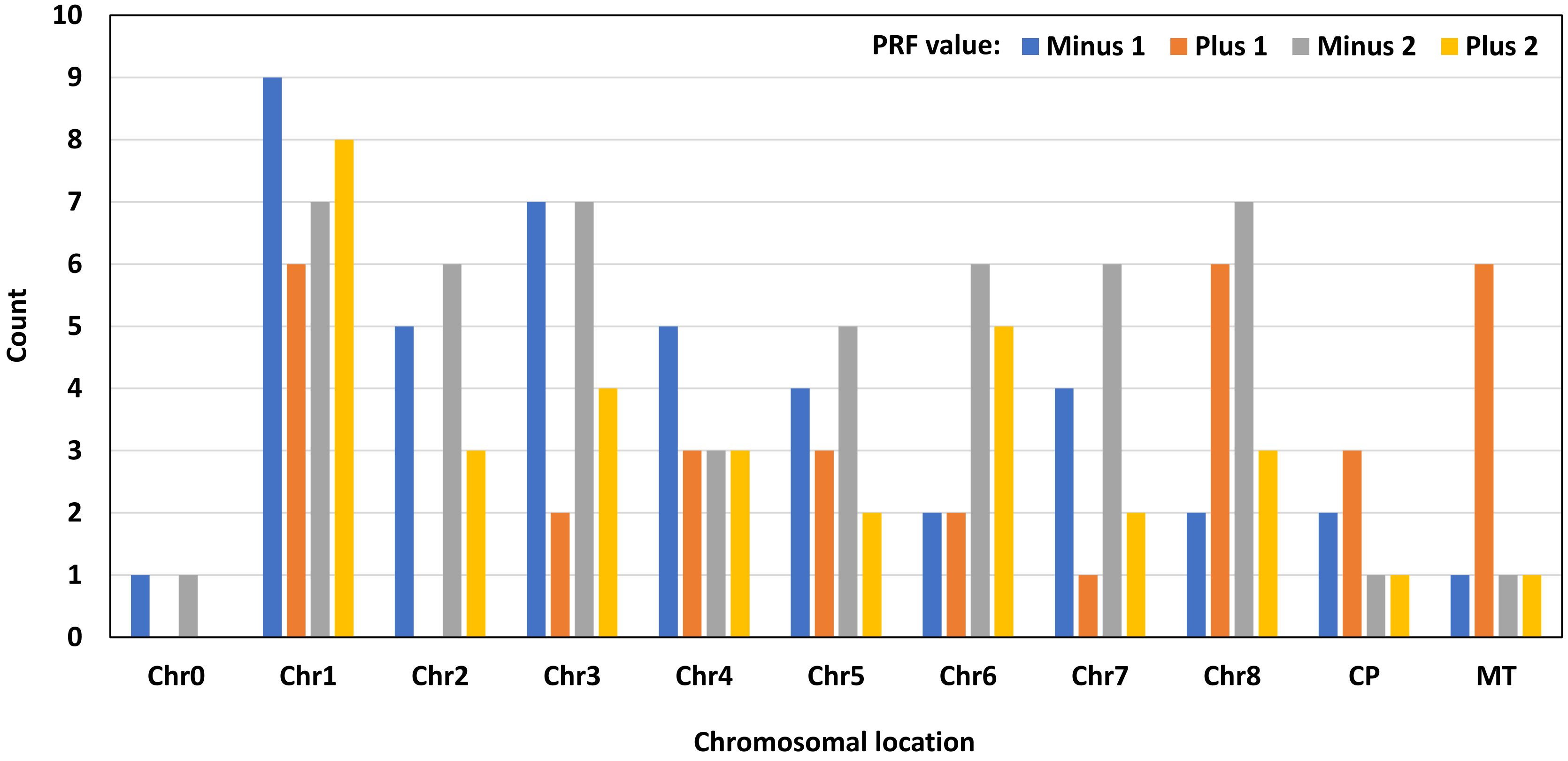


**Supplementary Figure S11**. Distribution of 156 chimeric MS peptides based on the chromosomal location of their primary-source genes and the PRF value. Chr0, unmapped loci. Chr1 to Chr8, Chromosome 1 to Chromosome 8. CP, chloroplasts. Mt, mitochondria. The Fisher's exact test with a simulated p-value shows significant association between the chromosomal location and the PRF value ($p = 0.003$). Within individual chromosomal locations, the distribution is not significantly different from the uniform distribution. Note the absence of chimeric peptides with PRF value plus 1 from Chromosome 2. Chloroplasts and mitochondria are the only two locations in which PRF value plus 1 is overrepresented.

**Supplementary Figure S12**

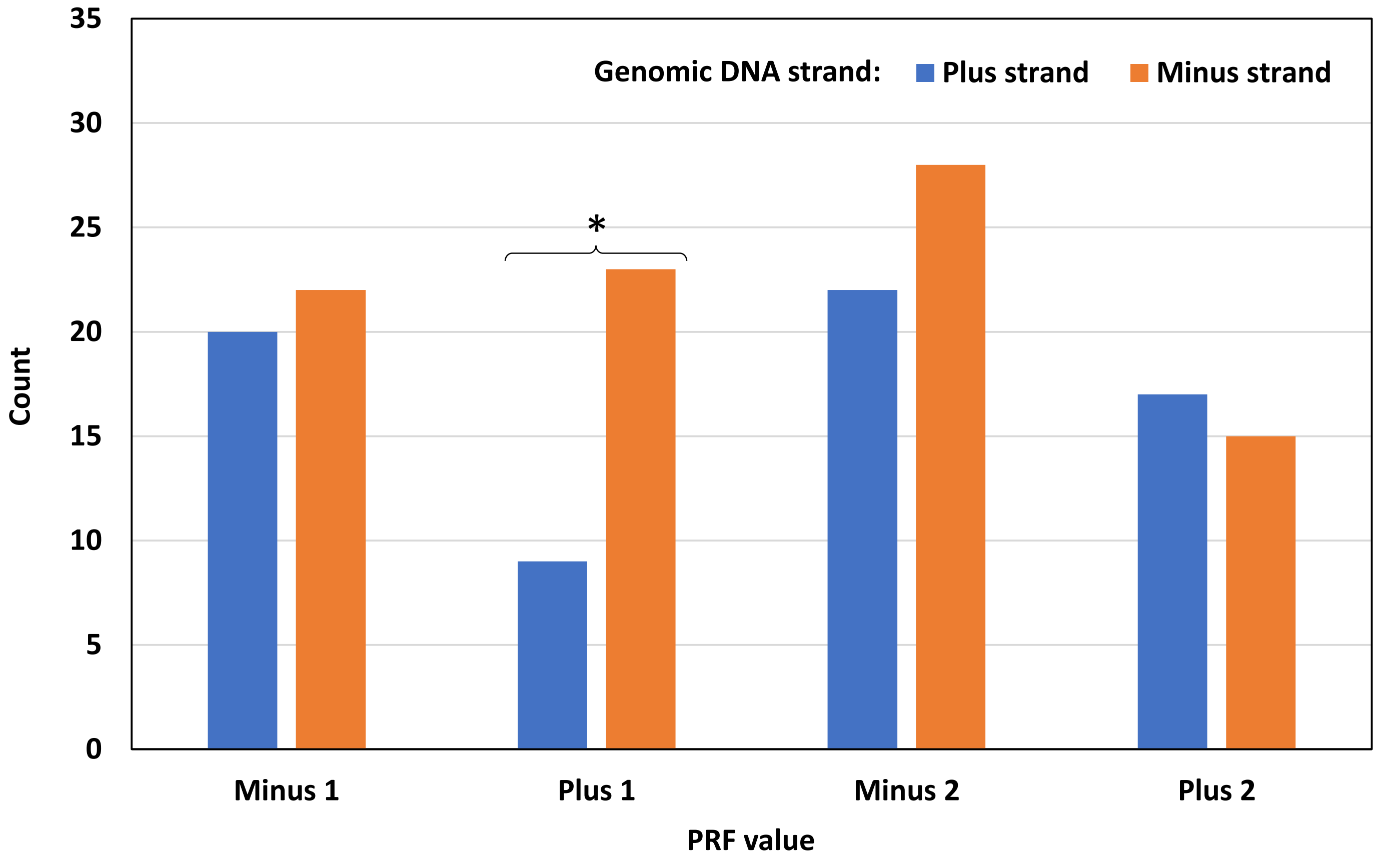


**Supplementary Figure S12**. Distribution of 156 chimeric MS peptides in different PRF categories depending on the genomic DNA strand of their primary-source genes. There is no significant association between the PRF value and the DNA strand. Within individual categories of PRF, the distribution is significantly different from the uniform distribution in one case indicated with an asterisk (the chi-square test for homogeneity, $p = 0.013$). The abundance of chimeric peptides with PRF value +1 on the plus strand of genomic DNA is less than half of their abundance on the minus strand.

**Supplementary Figure S13**

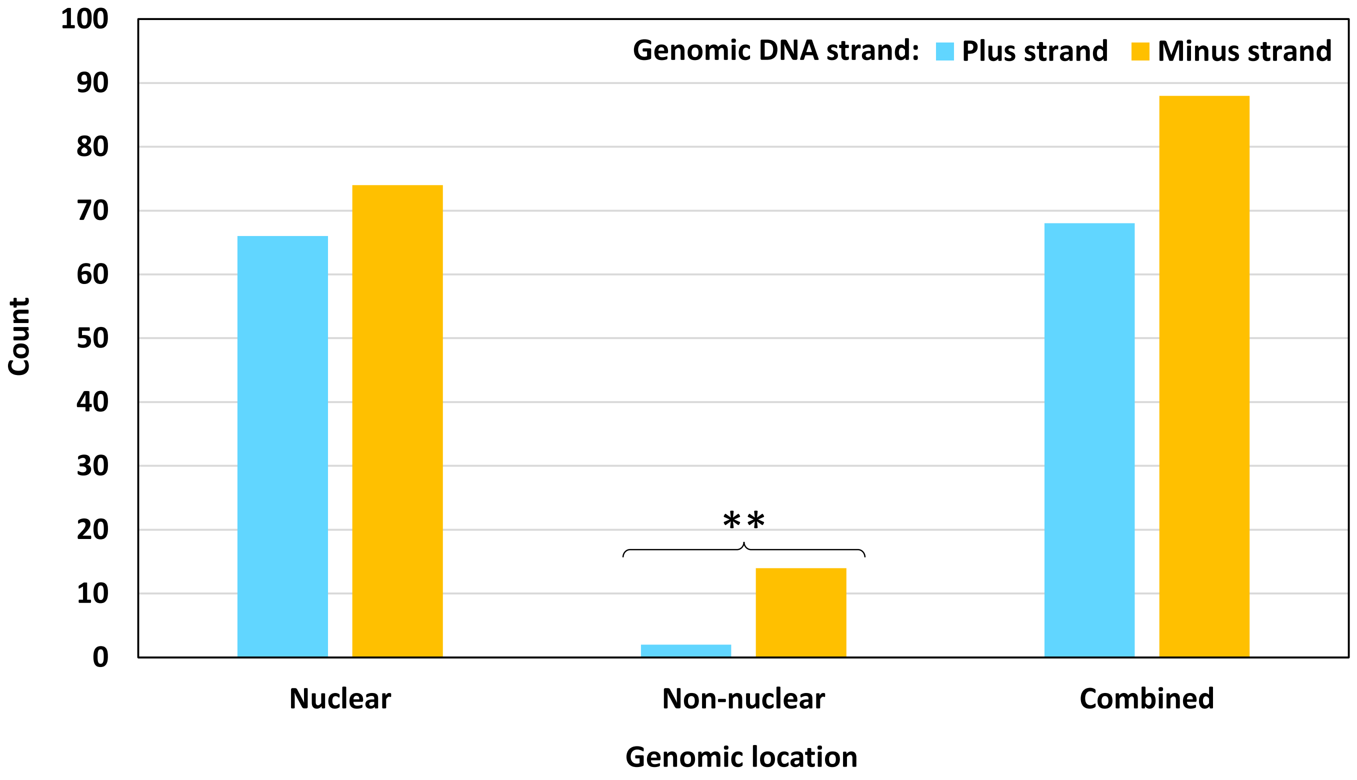


**Supplementary Figure S13**. Distribution of 156 chimeric MS peptides based on the genomic location and the genomic DNA strand of their primary-source genes. There is significant association between the genomic location and the DNA strand (the chi-square test for association, p = 0.008). Within individual categories of the genomic location, the distribution is significantly different from the uniform distribution in one case indicated with asterisks (the chi-square test for homogeneity, p = 0.003). The abundance of non-nuclear chimeric peptides is disproportionally lower on the plus strand of genomic DNA. Genes of all chloroplastic chimeric peptides and the majority of mitochondrial ones are located on the minus strand. This is not a general feature of the *M. truncatula* genome.

**Supplementary Figure S14**

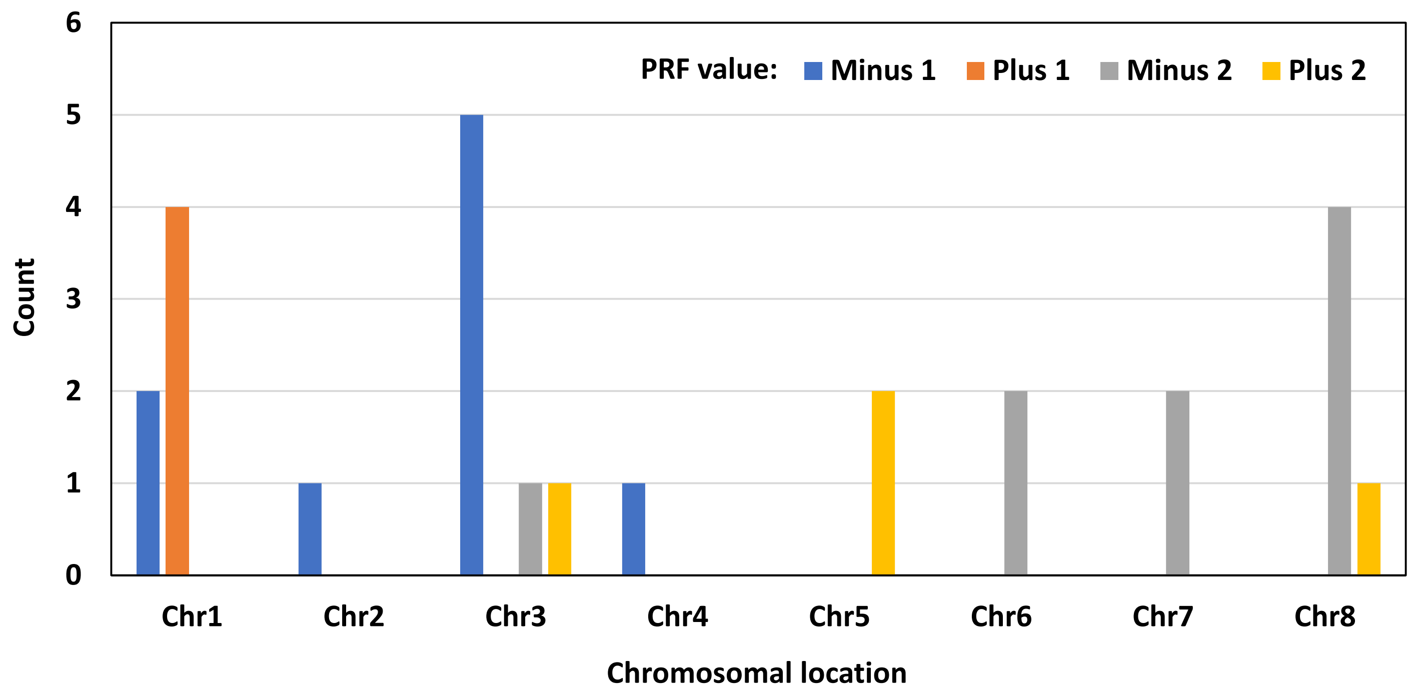


**Supplementary Figure S14**. Distribution of 26 chimeric MS peptides with PRF positions located close to a splicing site. The graph illustrates the counts of such peptides by different chromosomes from which they originate depending on the frameshift value. “Chr” stands for chromosome. There is significant association between the chromosomal location and the PRF value for this subset (the Fisher’s exact test, p = 7.718E-05). Within individual chromosomes, the distribution is not significantly different from the uniform distribution. In this subset of 26 chimeric peptides, the PRF sites were located within a few nucleotides to the left or to the right of the exon-exon borders or coincided with them so that the corresponding MS peptides covered the splicing sites.

**Supplementary Figure S15**

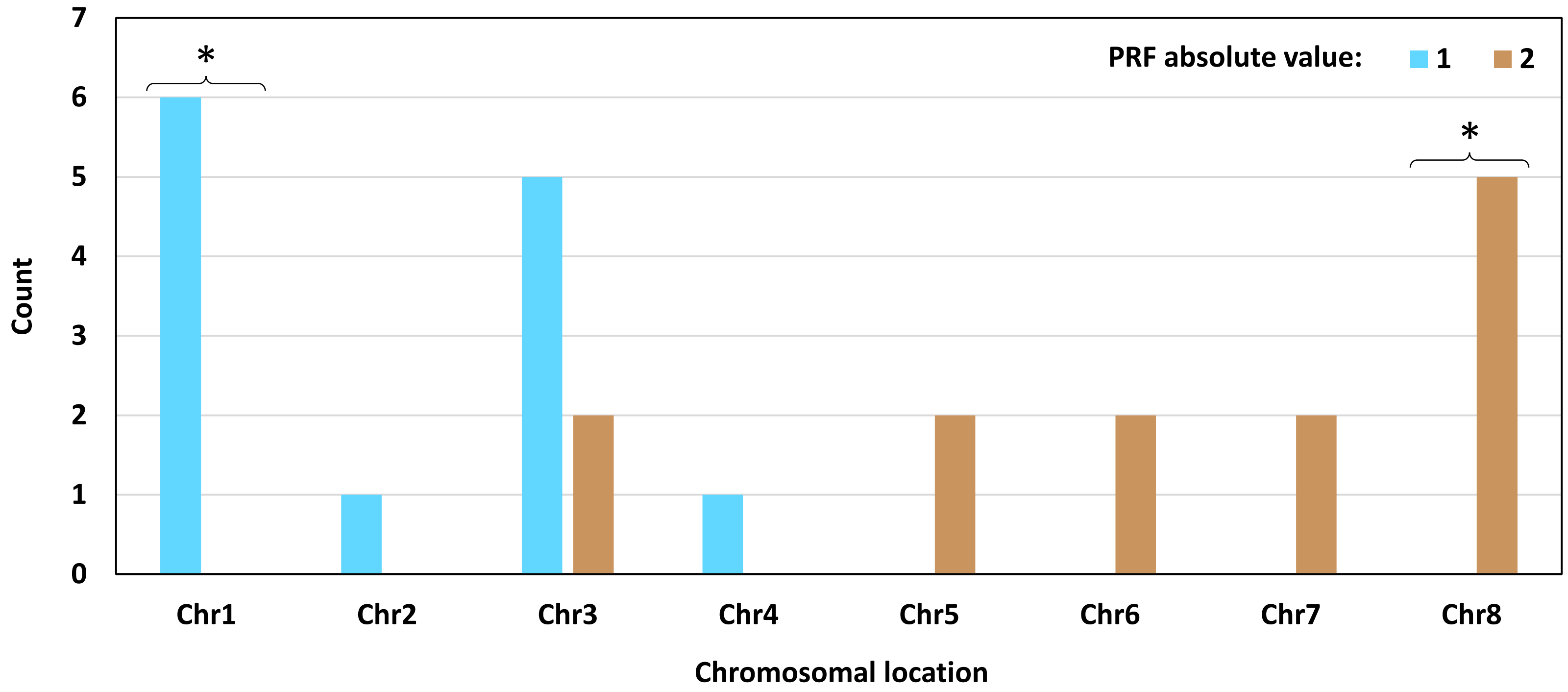


**Supplementary Figure S15**. Distribution of 26 chimeric MS peptides with PRF positions located close to a splicing site. The graph illustrates the counts of such peptides by different chromosomes from which they originate depending on the absolute value of PRF. “Chr” stands for chromosome. There is significant association between the chromosomal location and the PRF absolute value for this subset (the Fisher’s exact test, p = 6.161E-05). Within individual chromosomes, the distribution is significantly different from the uniform distribution in two cases indicated with asterisks (the Exact Binomial Test, Chr1, p-value = 0.016; Chr8, p = 0.031). Chromosomes 1 to 4 are enriched in the “short” PRF events, regardless of the sign. Chromosomes 5 to 8 are enriched in the “long” PRF events, regardless of the sign. In this subset of 26 chimeric peptides, the PRF sites were located within a few nucleotides to the left or to the right of the exon-exon borders or coincided with them so that the corresponding MS peptides covered the splicing sites.

**Supplementary Figure S16**

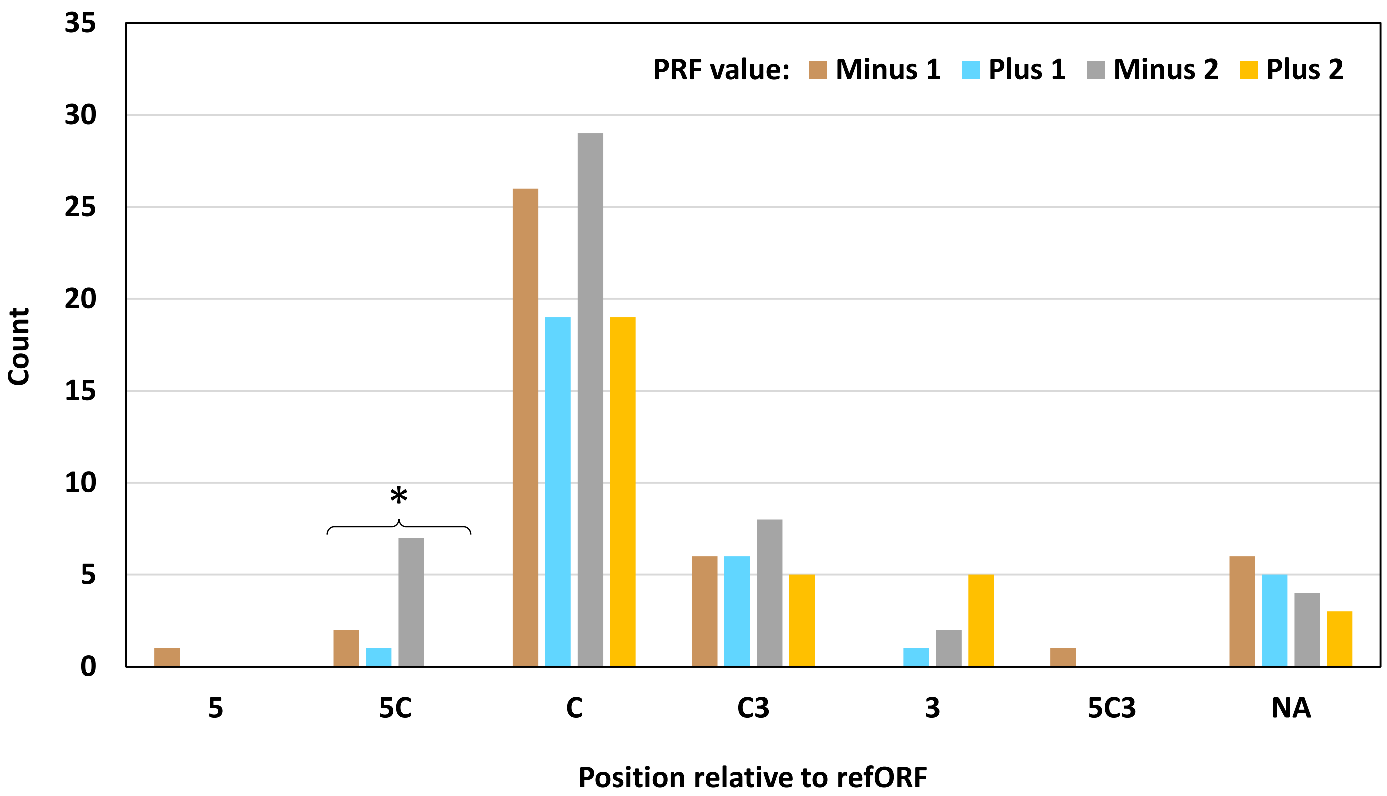


**Supplementary Figure S16**. Distribution of 156 chimeric MS peptides according to the position of an altORF relative to the refORF and the PRF value. 5, the 5'-untranslated region (UTR); 5C, the border between the 5'-UTR and the coding sequence (CDS); C, the CDS; C3, the border between the CDS and the 3'-UTR; 5C3, an altORF starts in the 5'-UTR and ends in the 3'-UTR; NA, all types of non-coding RNA, which have no refORF. There is no significant association between the altORF position and the PRF value. Within individual categories, the distribution of PRF values is significantly different from the uniform distribution in one case indicated with an asterisk (the Exact Multinomial Test, $p = 0.015$). PRF events with value +2 are absent from the following categories: 5, 5C, and 5C3.

**Supplementary Figure S17**

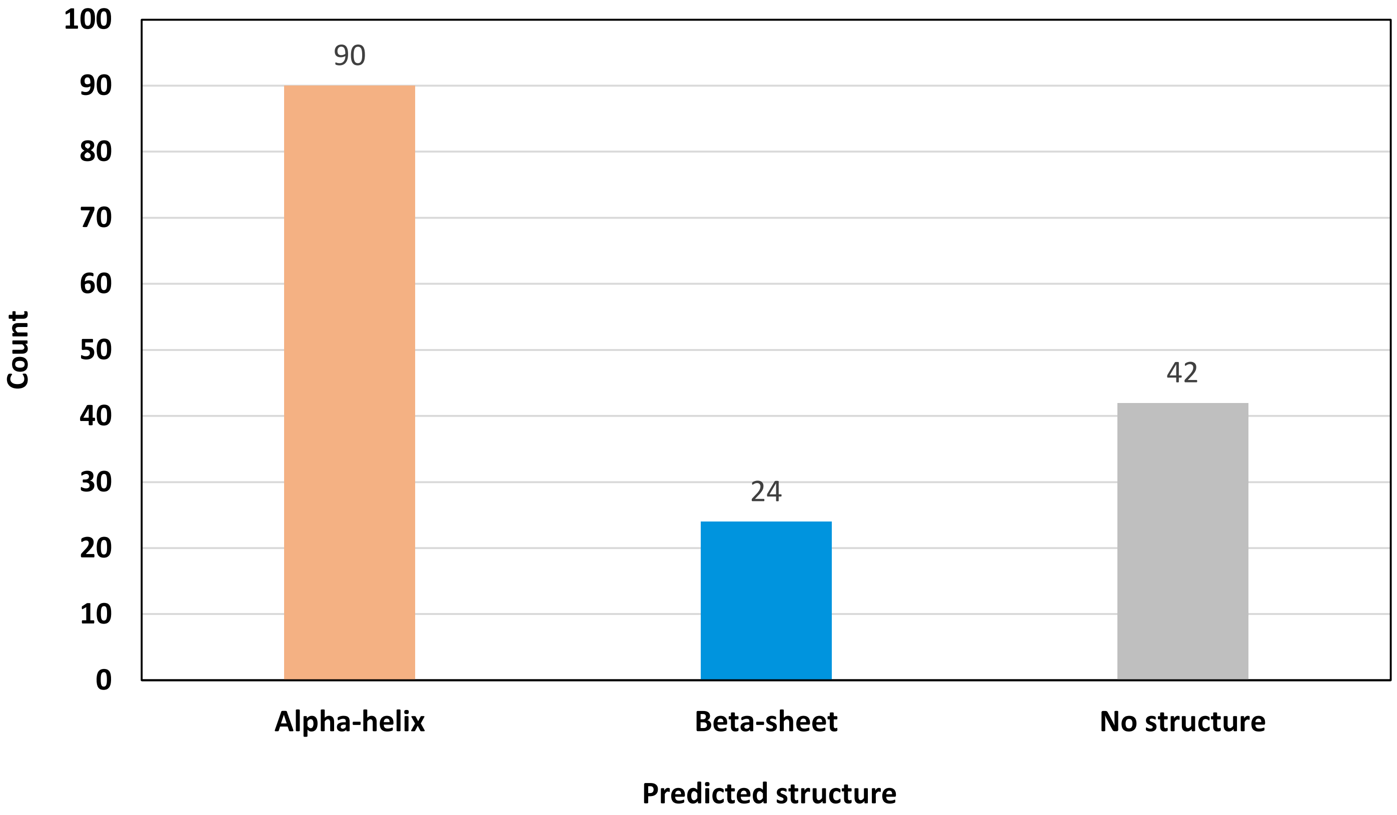


**Supplementary Figure S17**. Distribution of 156 MS-supported chimeric peptide models in three folding categories predicted by ColabFold v1.5.1. Alpha-helix, at least three full turns of the helix found in rank 1 predictions in the absence of a beta-sheet structure in any of the five predictions made for each chimeric model. Beta-sheet, beta-sheet structures are present in any of the five predictions regardless of the presence of alpha-helices. No structure, no beta-sheets and no alpha-helices with at least three full turns in predictions with rank 1. The categories do not take into account the PRF position relative to alpha-helices or beta-sheets. The distribution is significantly different from the uniform distribution (the chi-square test for homogeneity, $p = 1.899E-10$). Models with no structure constitute about one-quarter of the dataset (27%).

**Supplementary Figure S18**

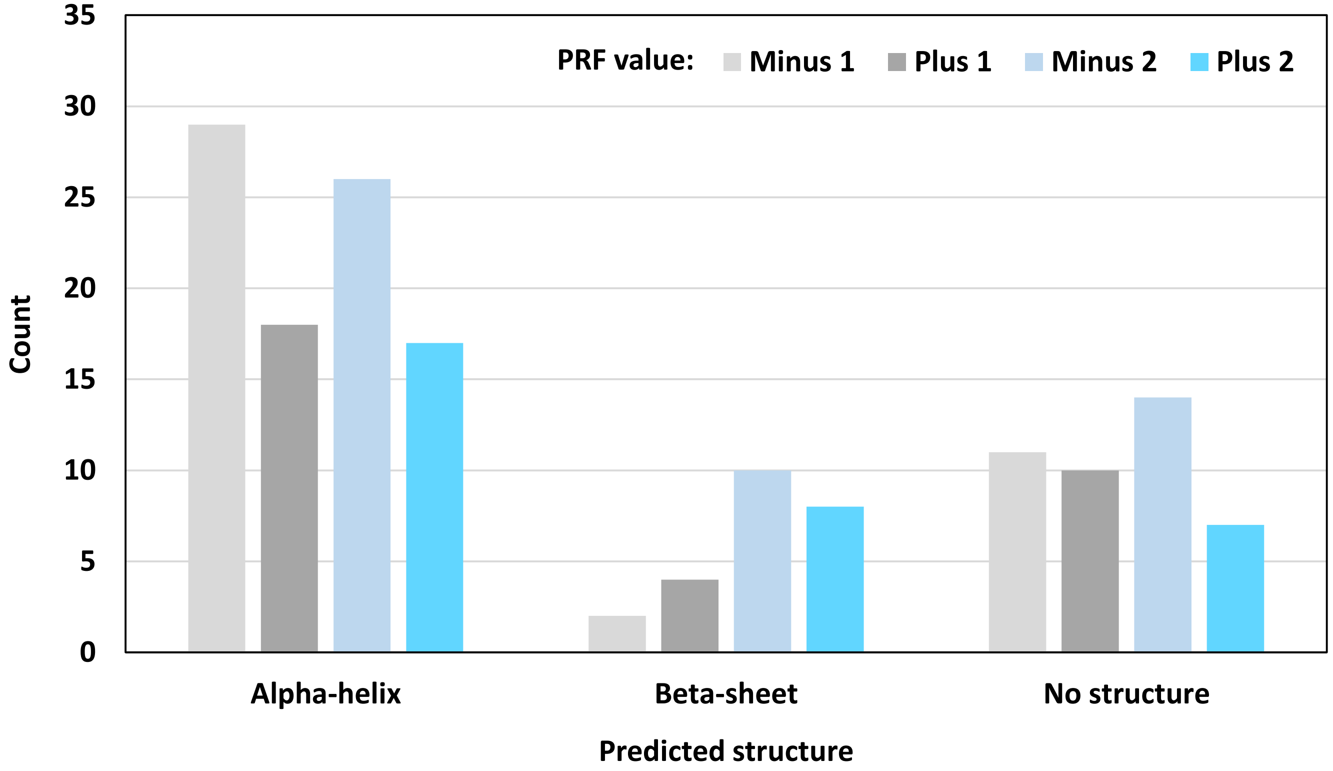


**Supplementary Figure S18**. PRF value-based distribution of 156 MS-supported chimeric peptide models in three folding categories described in the previous figure. There is no significant association between the folding category and the PRF value. Within individual folding categories, the distribution is not significantly different from the uniform distribution.

### Supplementary Figure S19

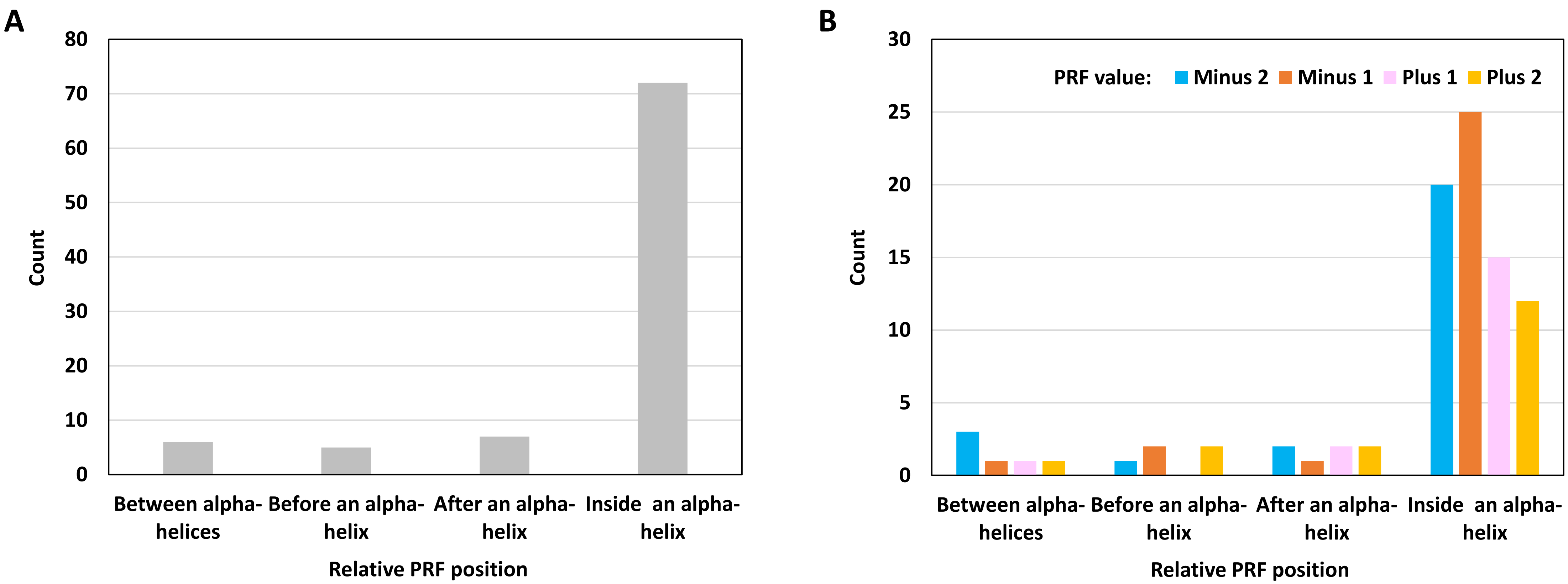


**Supplementary Figure S19**. Distribution of 90 alpha-helix-forming models of chimeric MS peptides in four categories based on the PRF position relative to the helices. A, groups with all PRF values combined. B, grouping by PRF values within each category. The distribution is significantly different from the uniform distribution in A (the chi-square test for homogeneity, p = 2.734E-31). There is no significant association between the PRF position relative to the alpha-helices and PRF value (B). The distribution of PRF values within each location group is not significantly different from the uniform distribution (B). The majority of the models with alpha-helices contain PRF sites within alpha-helices.

### Supplementary Figure S20

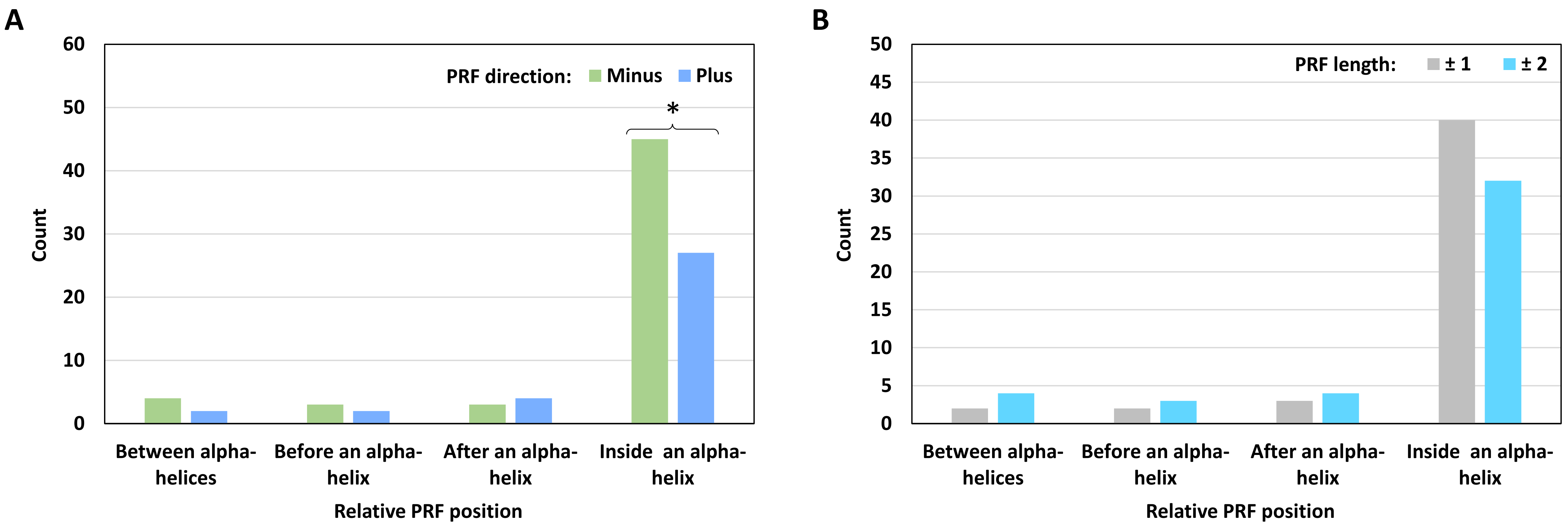


**Supplementary Figure S20**. Distribution of 90 alpha-helix-forming models of chimeric MS peptides in four categories based on the PRF position, direction, and length. A, grouping by the direction of PRF (minus, backward frameshifts; plus, forward frameshifts). B, grouping by the length of PRF (±1, "short" frameshifts; ±2, "long" frameshifts). There is significant association in A and B (the Fisher's exact test, p-values are 0.002 and 0.003, respectively). Within individual folding categories, the distribution is significantly different from the uniform distribution in one case indicated with an asterisk (the chi-square test for homogeneity, $p = 0.034$). Backward frameshifts are significantly more abundant in the models in which PRF sites are located inside alpha-helices.

### Supplementary Figure S21

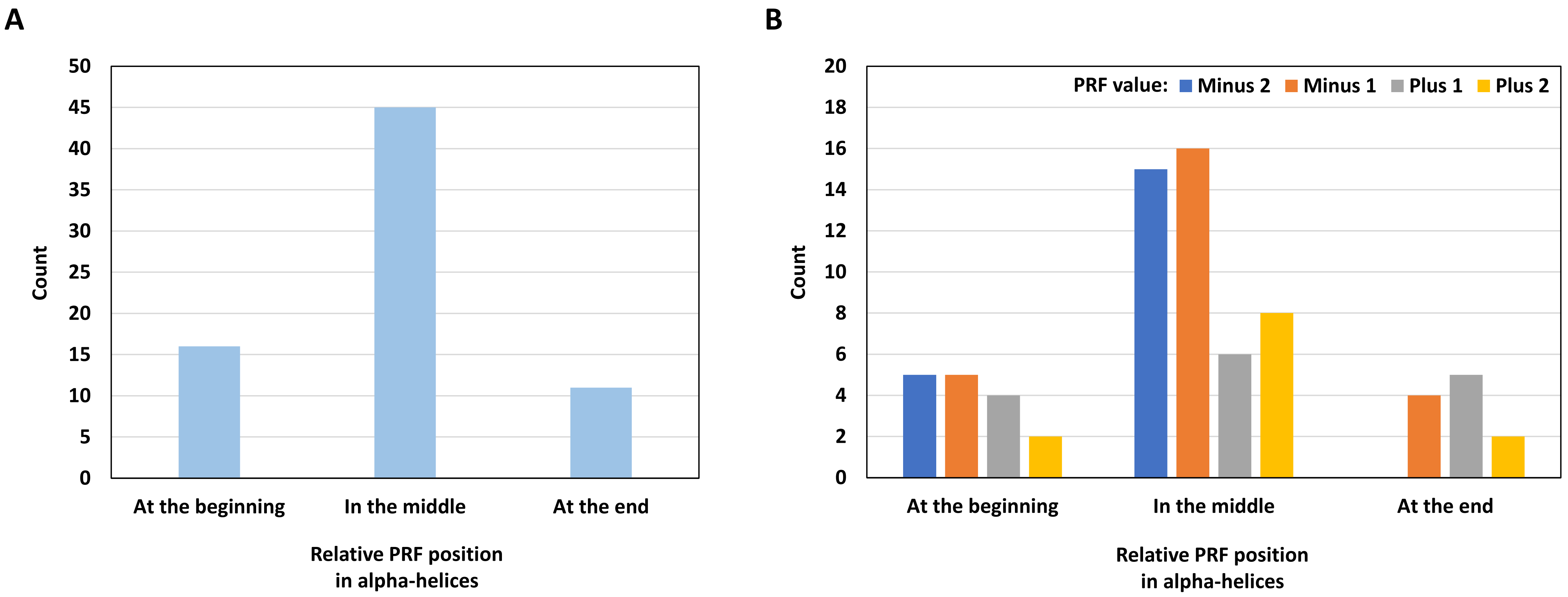


**Supplementary Figure S21**. Distribution of 72 MS-supported models based on the position of PRF sites inside alpha-helices. A, groups with all PRF values combined. B, grouping by PRF values within each category. The distribution is significantly different from the uniform distribution in A (the chi-square test for homogeneity, p = 7.976E-07). There is significant association between the PRF position inside the alpha-helix and the PRF value (the Fisher's exact test, p = 3.324E-05). The distribution within individual groups is not significantly different from the uniform distribution (B). The majority of the models with alpha-helices have PRF sites in the middle of alpha-helices.

### Supplementary Figure S22

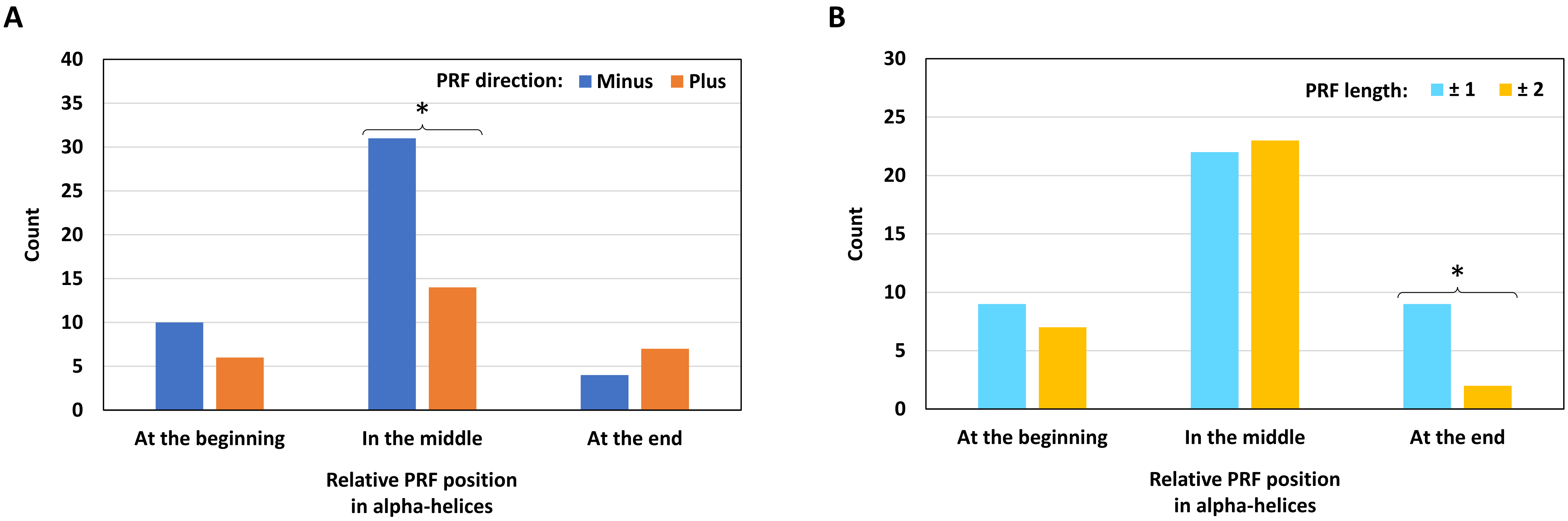


**Supplementary Figure S22**. Distribution of 72 MS-supported models based on the PRF position inside alpha-helices, PRF direction, and PRF length. A, grouping by the direction of PRF (minus, backward frameshifts; plus, forward frameshifts). B, grouping by the length of PRF (± 1, “short” frameshifts; ± 2, “long” frameshifts). There is no significant association in A and B. Within individual position categories, the distribution is significantly different from the uniform distribution in two cases indicated with asterisks (the chi-square test for homogeneity, A, $p = 0.011$; B, $p = 0.035$). Backward frameshifts are significantly more abundant in the models in which PRF sites are located in the middle of alpha-helices. “Long” frameshifts are significantly less abundant in the models in which PRF sites are located at the end of alpha-helices.

**Supplementary Figure S23**

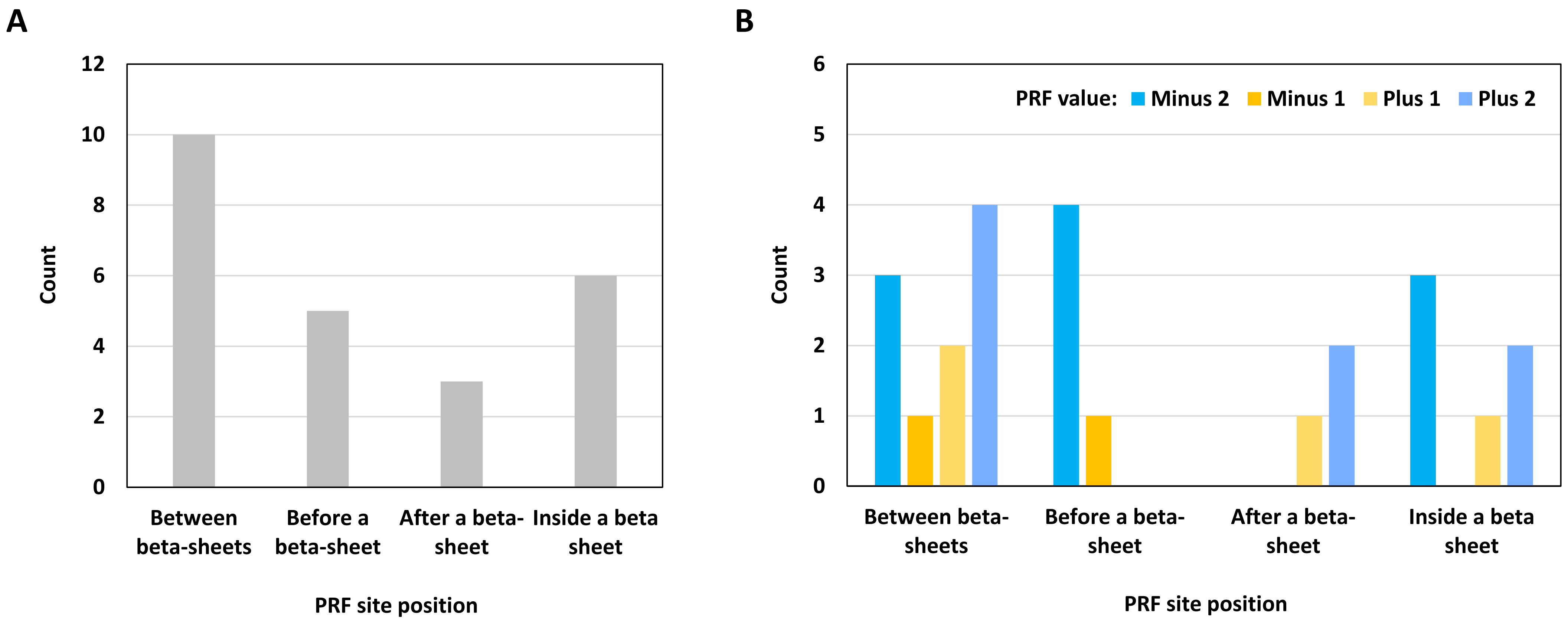


**Supplementary Figure S23**. Distribution of 24 MS-supported chimeric peptide models with predicted beta-sheets in four categories based on the position of PRF sites relative to the beta-sheets. A, combined counts regardless of the PRF value. B, counts with different PRF values shown separately. The distribution is not significantly different from the uniform distribution in A and in individual groups in B. There is no significant association between the PRF value and the category in B. However, the association is significant when the categories "before" and "after" are considered alone (the Fisher's exact test, p = 0.036). "Backward" and "forward" PRF values (minus and plus, respectively) are represented differentially before and after a beta-sheet.

**Supplementary Figure S24**

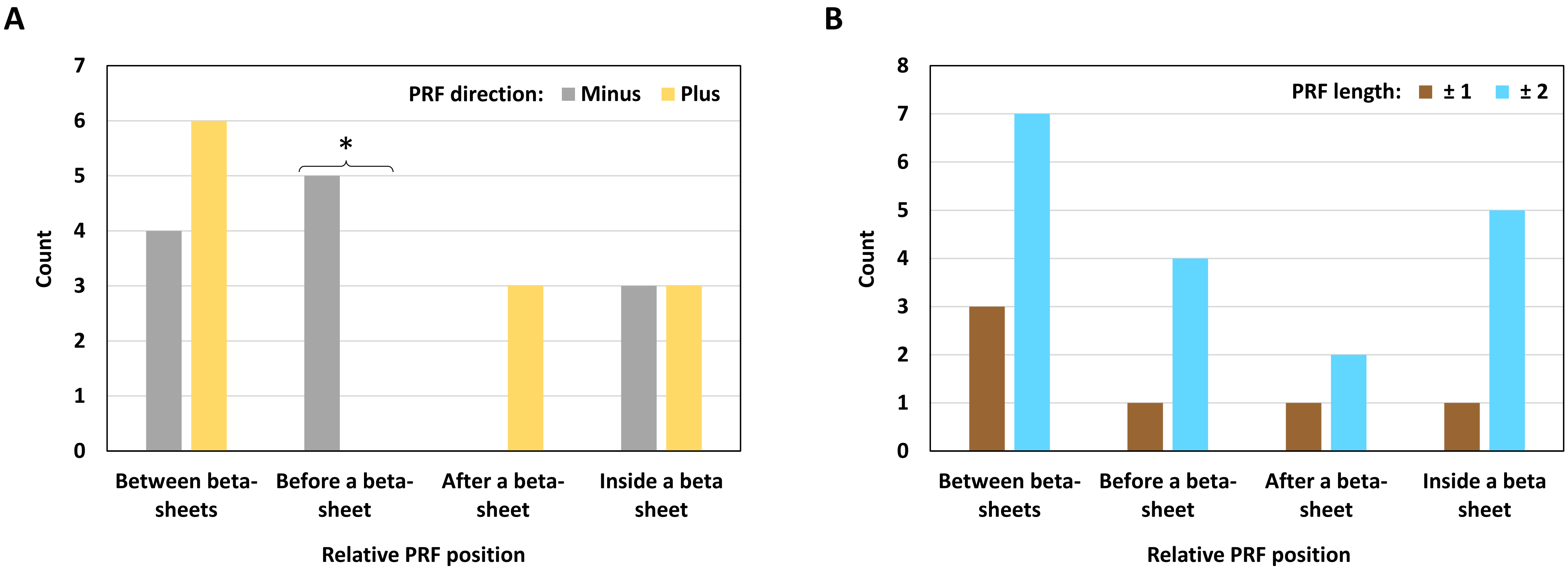


**Supplementary Figure S24**. Distribution of 24 MS-supported chimeric peptide models with predicted beta-sheets based on the PRF position, direction, and length. A, grouping by the direction of PRF (minus, backward frameshifts; plus, forward frameshifts). B, grouping by the length of PRF (± 1, "short" frameshifts; ± 2, "long" frameshifts). There is significant association in A (the Fisher's exact test, $p = 0.025$). Within individual position categories, the distribution is significantly different from the uniform distribution in one case indicated with an asterisk (the Exact Binomial Test, $p = 0.031$). All PRF sites located before beta-sheets have the negative value. All PRF sites located after beta-sheets have the positive value.

### Supplementary Figure S25

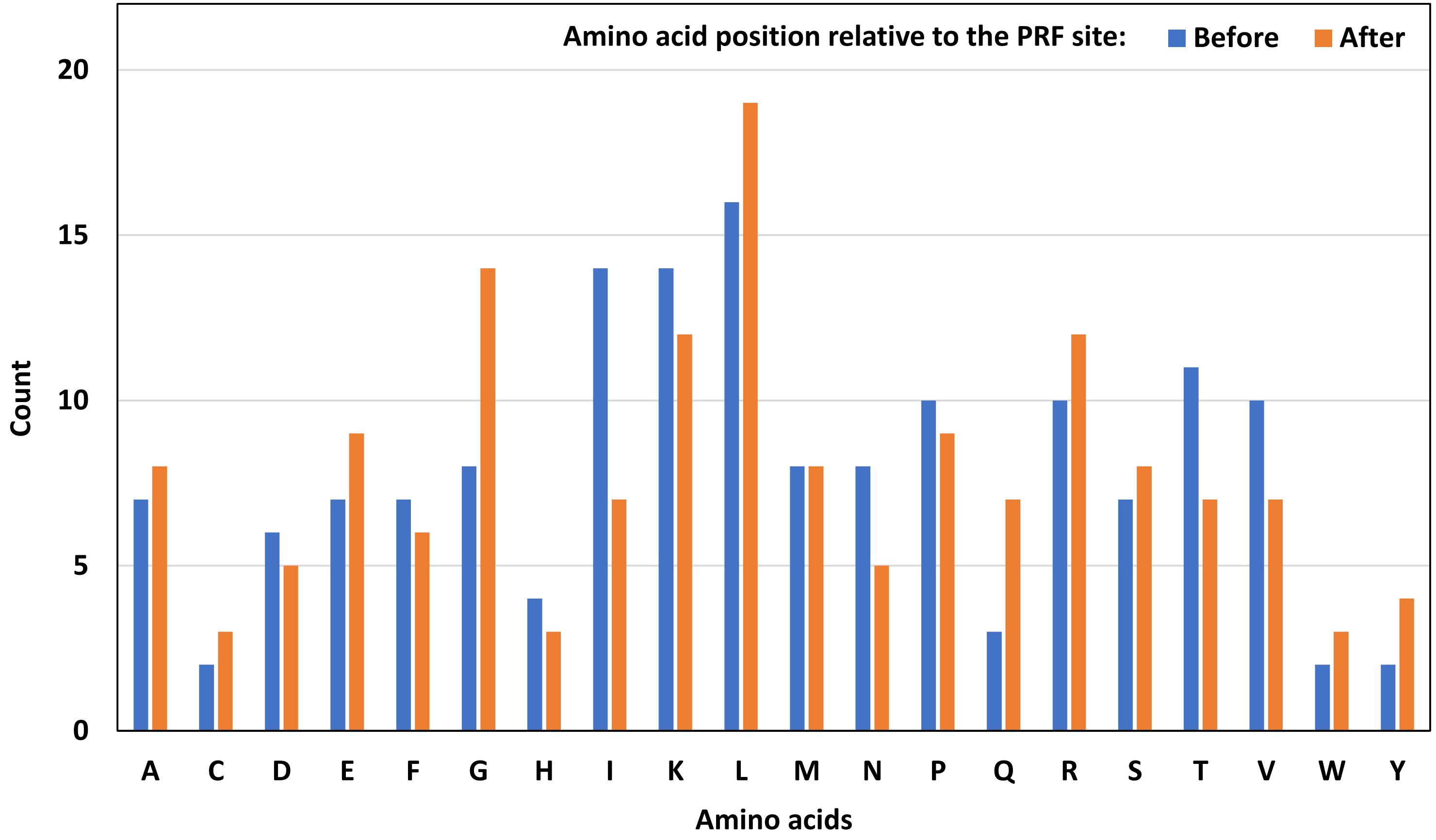


**Supplementary Figure S25**. The abundance of 20 amino acids found in 156 chimeric MS peptides immediately **before** and **after** the PRF site. There is no significant association between the amino acid and its position relative to the PRF site. For each amino acid, the distribution before and after is not significantly different from the uniform distribution.

### Supplementary Figure S26

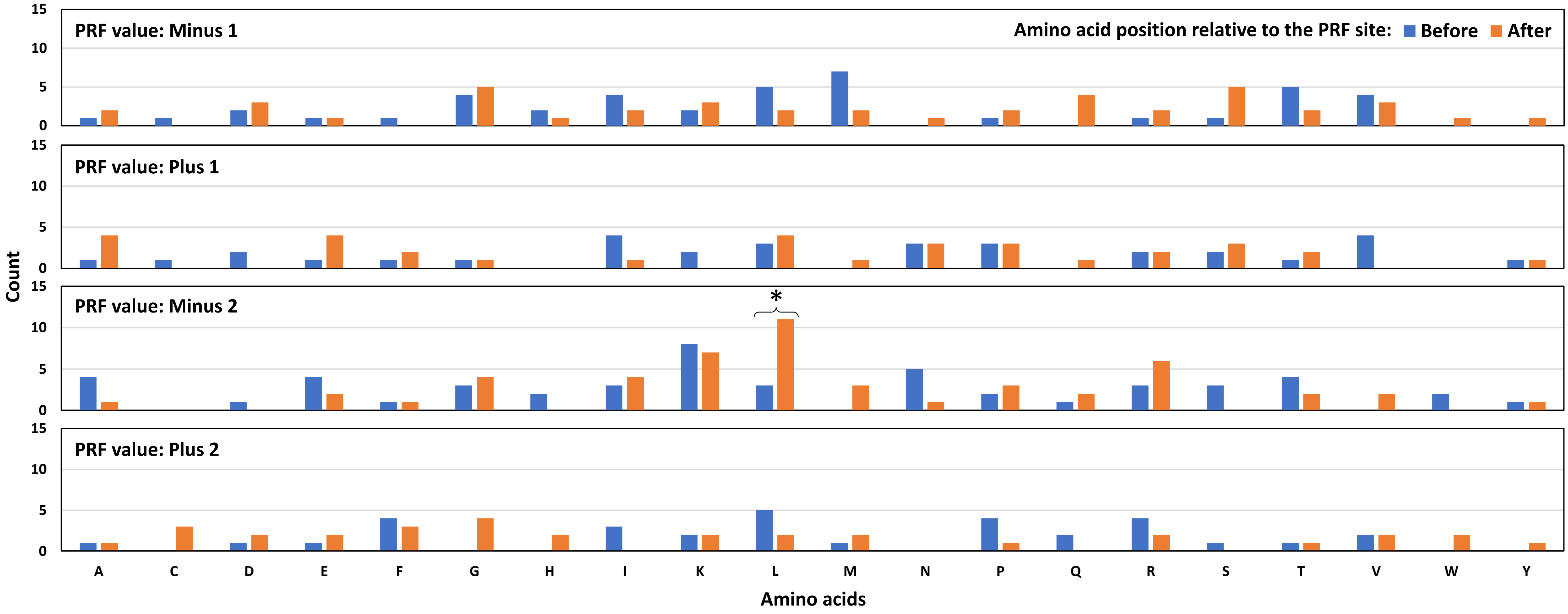


**Supplementary Figure S26**. The abundance of 20 amino acids found in 156 chimeric MS peptides immediately **before** and **after** the PRF site plotted by the PRF value. Within individual categories, there is significant association between the amino acid and the position relative to the PRF site in PRF categories Plus 1 and Minus 2 (the Fisher's exact test, p-values are 0.015 and 0.002, respectively). For individual amino acids, the distribution before and after is significantly different from the uniform distribution in one case indicated with an asterisk (the chi-square test for homogeneity, $p = 0.033$).

**Supplementary Figure S27**

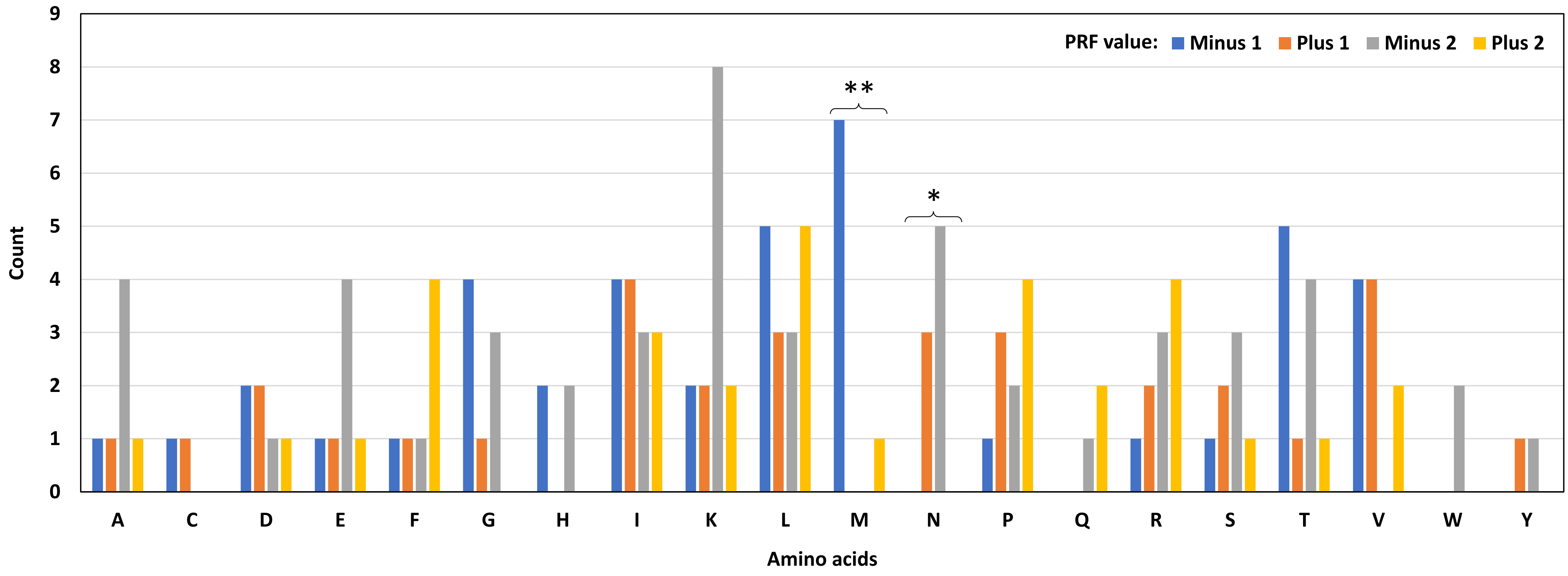


**Supplementary Figure S27**. Distribution of 20 amino acids found in 156 chimeric MS peptides immediately **before** the PRF site plotted by the PRF value. There is significant association between the amino acid and the PRF value (the Fisher's exact test with a simulated p-value, p = 0.003). For individual amino acids, the distribution in categories is significantly different from the uniform distribution in two cases indicated with asterisks (the Exact Multinomial Test, *p = 0.027, **p = 0.002). Nine amino acids are not found in certain PRF categories (C, G, H, M, N, Q, V, W, and Y), six of which are missing from the Plus 2 category.

**Supplementary Figure S28**

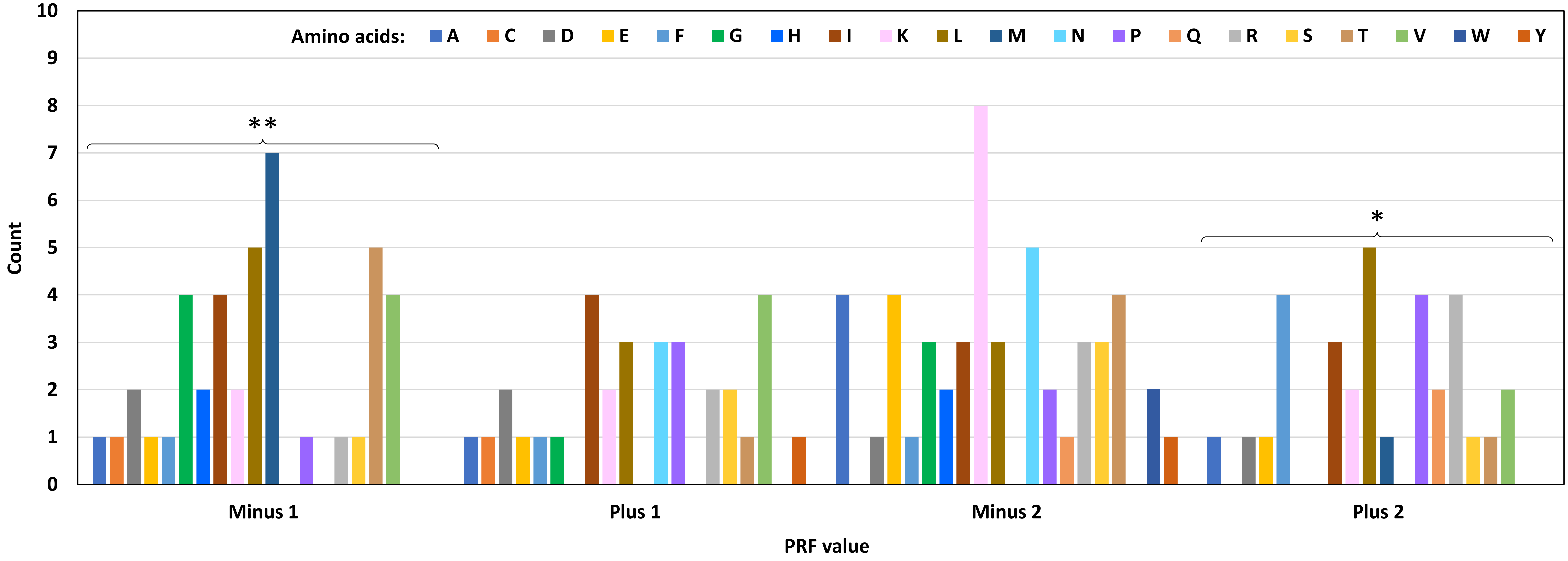


**Supplementary Figure S28**. Distribution of 20 amino acids found in 156 chimeric MS peptides immediately **before** the PRF site plotted by the PRF value. There is significant association between the amino acid and the PRF value (the Fisher's exact test with a simulated p-value, p = 0.003). For individual PRF categories, the distribution may be significantly different from the uniform distribution in two cases indicated with asterisks (the chi-square test for homogeneity, *p = 0.046, **p = 0.008). The Exact Multinomial Test cannot be used with these groups due to the large number of categories. Thus, the actual significance of the differences is unknown. Nine amino acids are not found in certain PRF categories (C, G, H, M, N, Q, V, W, and Y), six of which are missing from the Plus 2 category.

**Supplementary Figure S29**

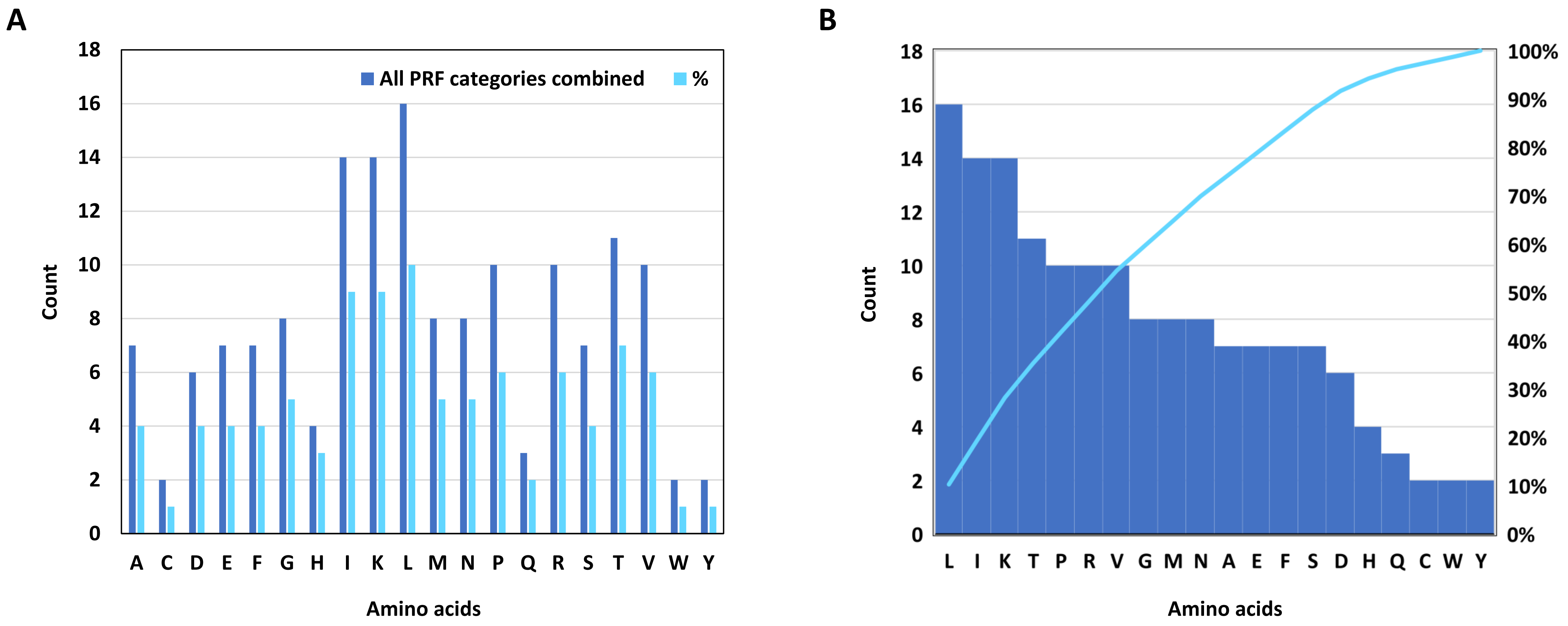


**Supplementary Figure S29**. The absolute and the relative abundance of 20 amino acids found in 156 chimeric MS peptides immediately **before** the PRF site. A, amino acids in the alphabetical order. B, a Pareto chart showing frequencies as dark blue bars and cumulative percentages out of 156 as a light blue curve. Frequencies of amino acids are distributed non-uniformly (the chi-square test for homogeneity, $p = 0.003$). Only three amino acids (L, I, and K) constitute 28% of the whole dataset. Seven amino acids (L, I, K, T, P, R, and V) constitute half of the dataset.

**Supplementary Figure S30**

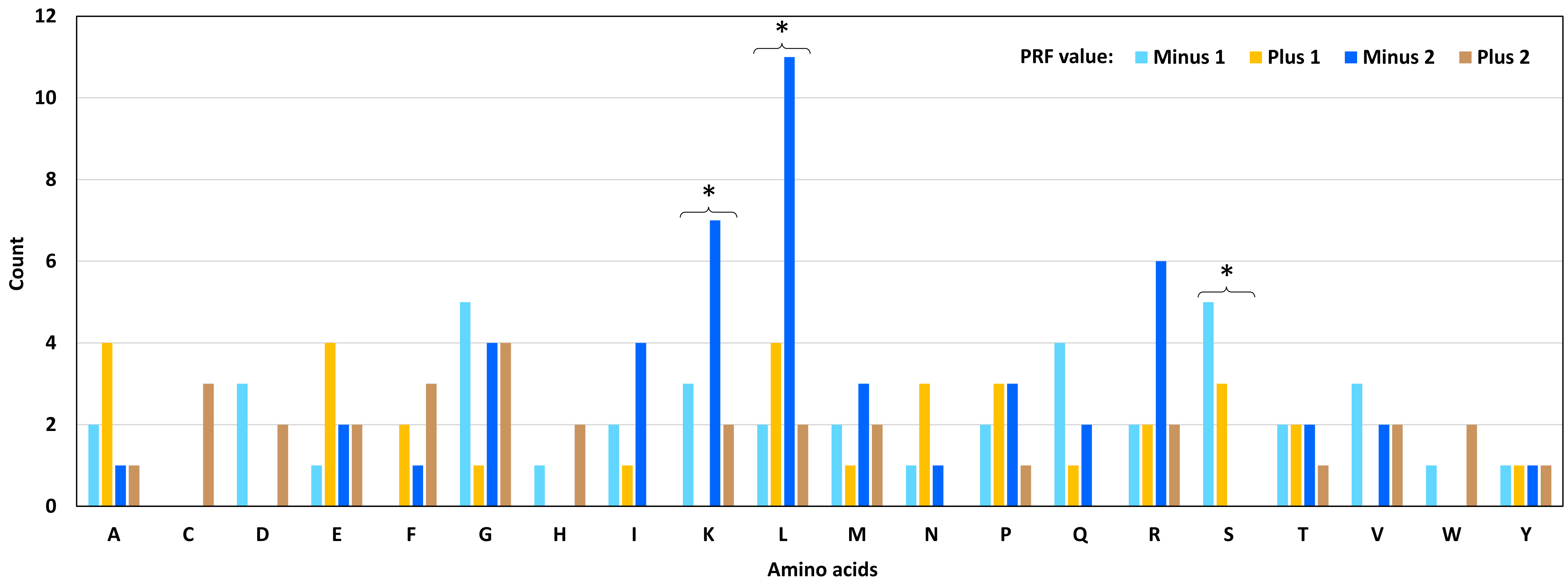


**Supplementary Figure S30**. Distribution of 20 amino acids found in 156 chimeric MS peptides immediately **after** the PRF site plotted by the PRF value. There is significant association between the amino acid and the PRF value (the Fisher's exact test with a simulated p-value, p = 0.041). For individual amino acids, the distribution in categories is significantly different from the uniform distribution in four cases indicated with asterisks. P-values of the Exact Multinomial Test are as follows: K, 0.037; L, 0.019; S, 0.027. Eleven amino acids are not found in certain PRF categories (C, D, F, H, I, K, N, Q, S, V, and W), six of which are missing from the Plus 1 category.

**Supplementary Figure S31**

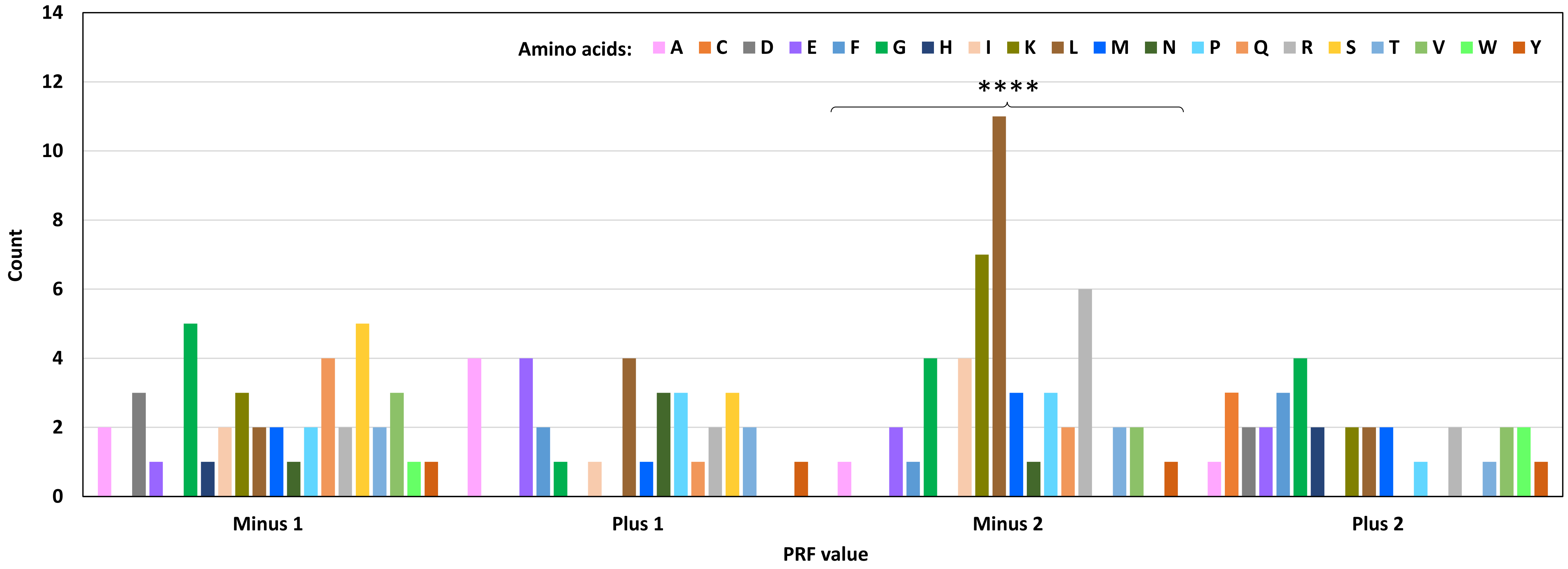


**Supplementary Figure S31**. Distribution of 20 amino acids found in 156 chimeric MS peptides immediately **after** the PRF site plotted by the PRF value. There is significant association between the amino acid and the PRF value (the Fisher's exact test with a simulated p-value, p = 0.041). For individual PRF categories, the distribution may be significantly different from the uniform distribution in one case indicated with asterisks (the chi-square test for homogeneity, p = 3.345E-06). The Exact Multinomial Test cannot be used with these groups due to the large number of categories. Thus, the actual significance of the differences is unknow. Eleven amino acids are not found in certain PRF categories (C, D, F, H, I, K, N, Q, S, V, and W), six of which are missing from the Plus 1 category.

**Supplementary Figure S32**

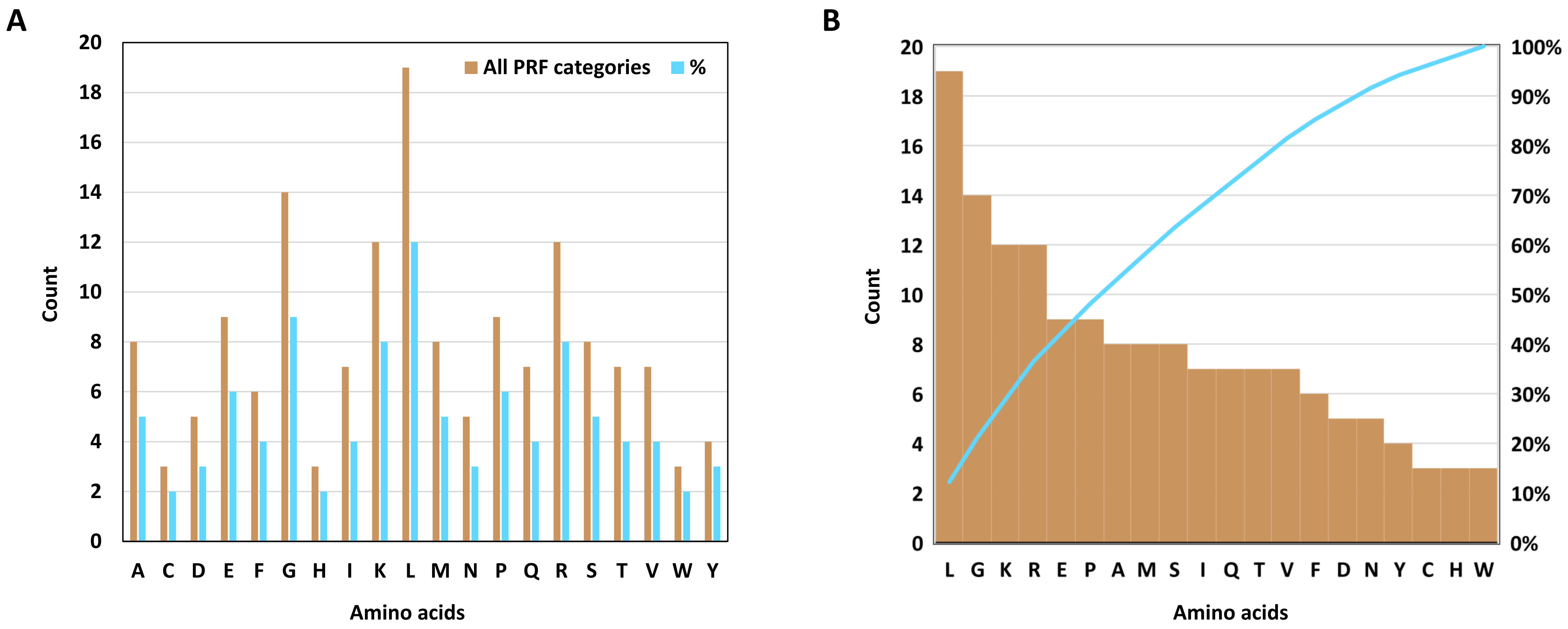


**Supplementary Figure S32**. The absolute and the relative abundance of 20 amino acids found in 156 chimeric MS peptides immediately **after** the PRF site. A, amino acids in the alphabetical order. B, a Pareto chart showing frequencies as brown bars and cumulative percentages out of 156 as a light blue curve. Frequencies of amino acids are distributed non-uniformly (the chi-square test for homogeneity, $p = 0.004$). Only three amino acids (L, G, and K) constitute 29% of the whole dataset. Six amino acids (L, G, K, R, E, and P) constitute half of the dataset.

**Supplementary Figure S33**

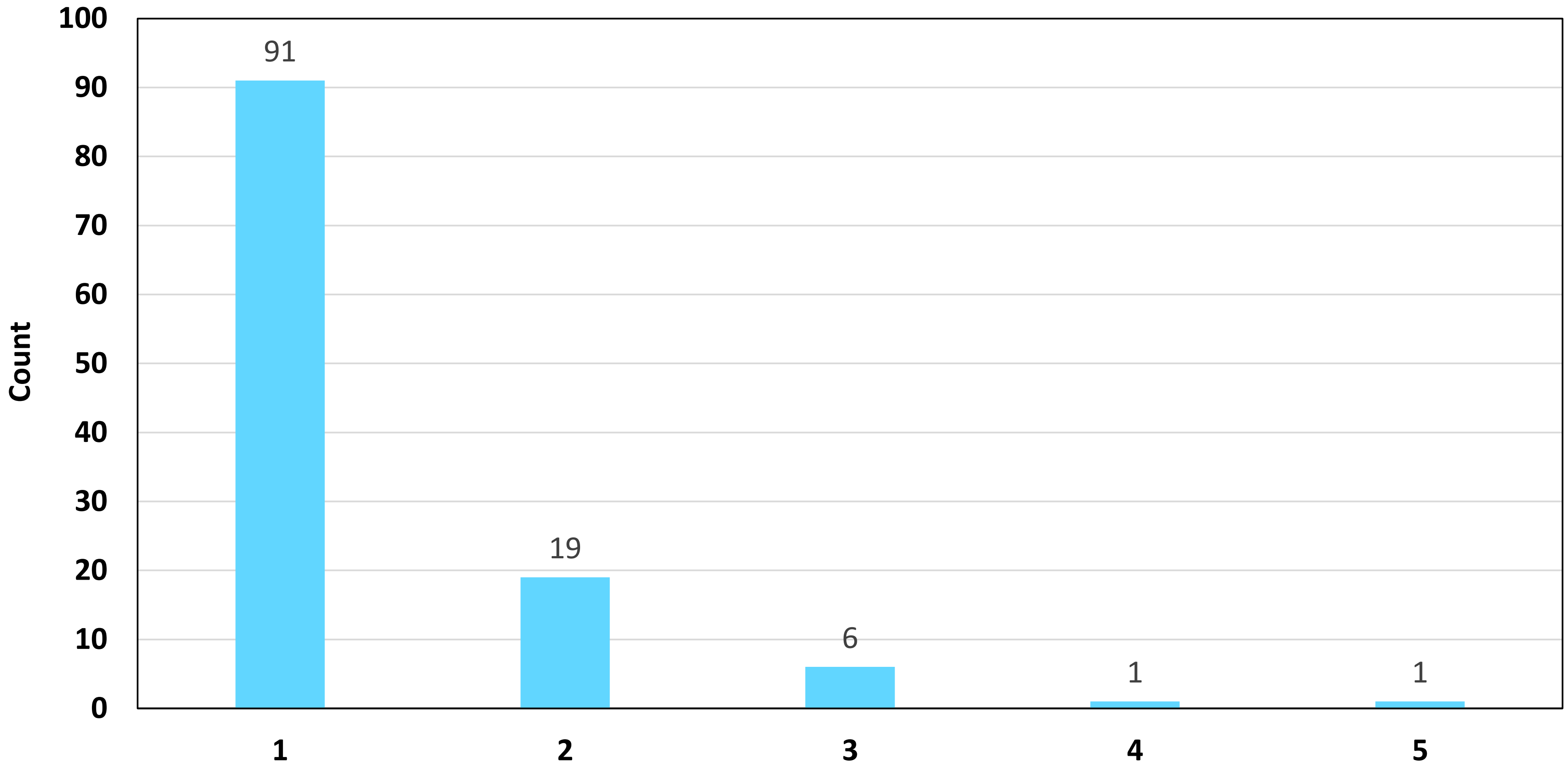


**Supplementary Figure S33**. Distribution of 118 pairs of amino acids found in 156 chimeric MS peptides immediately before and after the PRF site. Group one contains amino acid pairs found only once in the whole dataset; amino acid pairs found twice constitute group two, etc. Groups four and five contain only one amino acid pair each (AL and KR, respectively).

**Supplementary Figure S34**

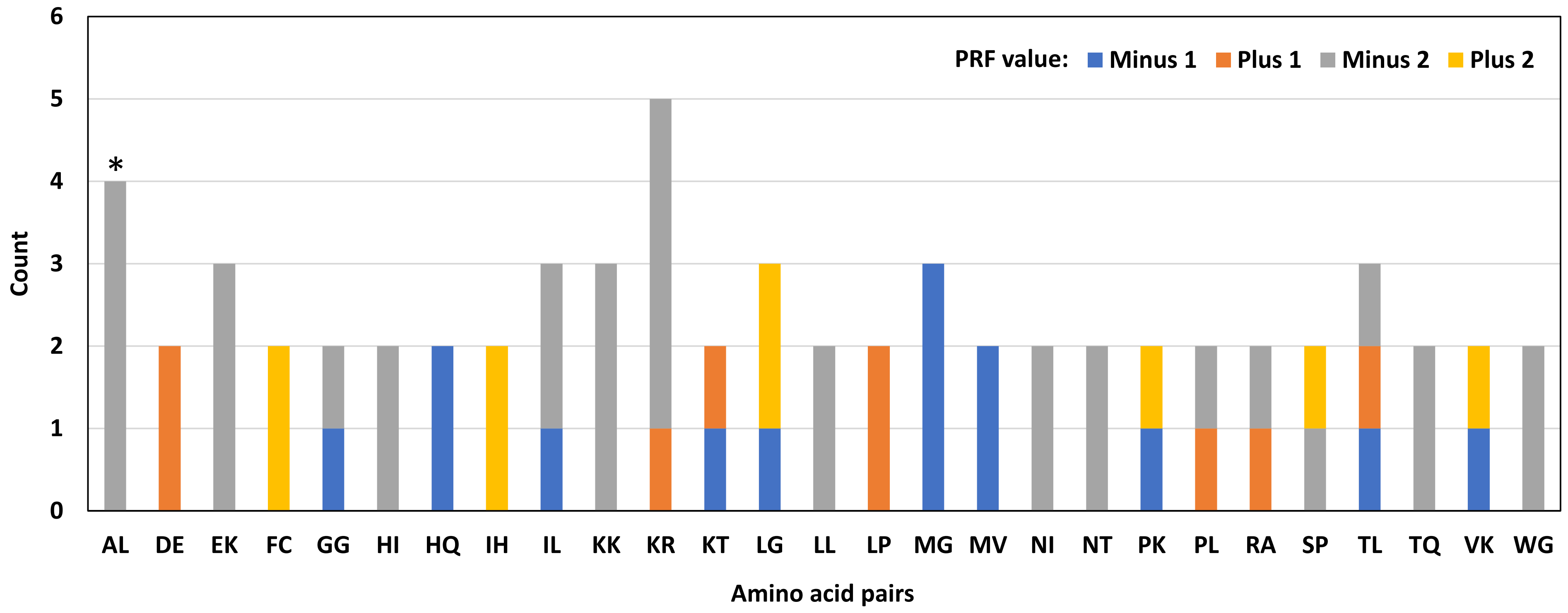


**Supplementary Figure S34**. Distribution of 27 pairs of amino acids found more than once in chimeric MS peptides immediately at the PRF site plotted by the PRF value. There is significant association between the amino acid pair and the PRF value (the Fisher's exact test with a simulated p-value, p = 0.002). Distribution of individual amino acid pairs in the PRF categories is significantly different from the uniform distribution in one case indicated with an asterisk (the Exact Multinomial Test, p = 0.016). Some amino acid pairs are unique to a specific PRF category (mostly, Minus 2).

### Supplementary Figure S35

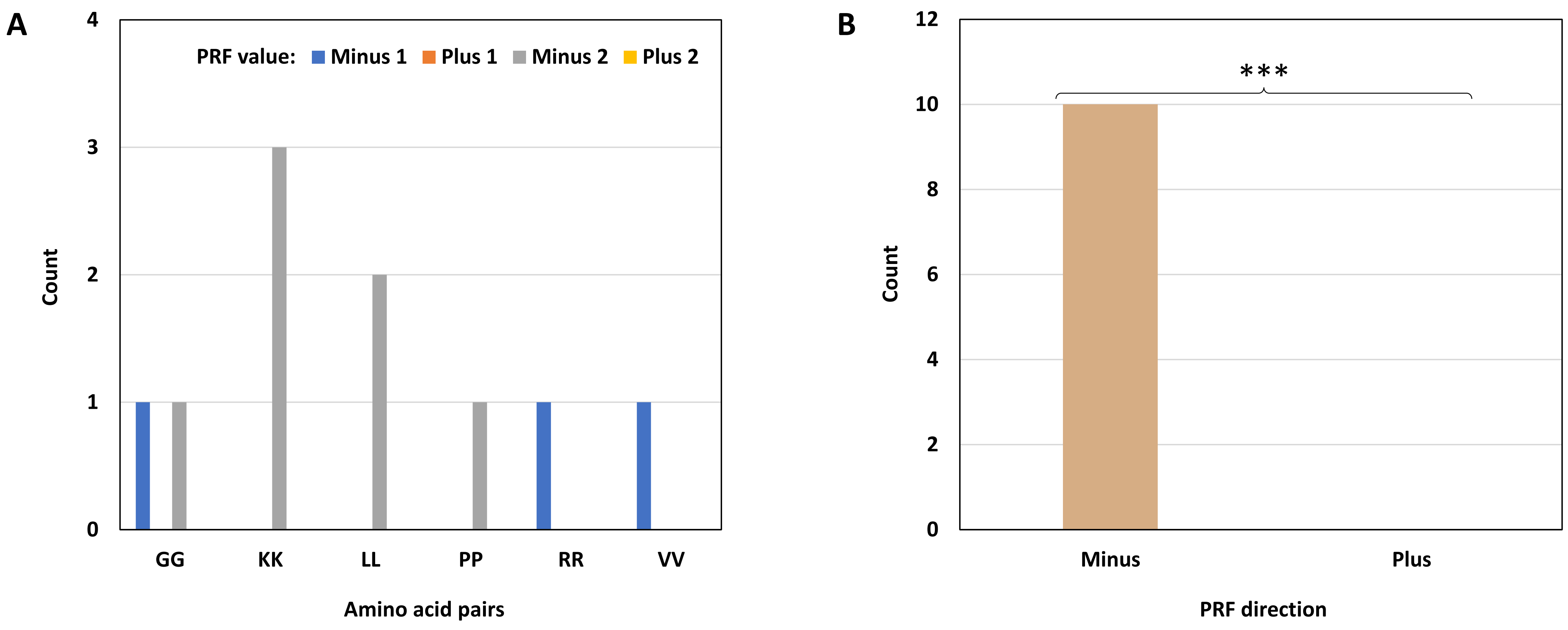


**Supplementary Figure S35**. Distribution of six pairs of repeated amino acids found in chimeric MS peptides immediately at the PRF site plotted by the PRF value. A, distribution of individual amino acid pairs by separate PRF categories. B, distribution of all pairs by combined PRF categories. The amino acid pair is not significantly associated with the PRF category in A. However, repeated amino acid pairs are present exclusively in chimeric MS peptides with backward PRF sites, which makes their distribution significantly different from the uniform distribution (the Exact Binomial Test, p = 9.766E-4).

**Supplementary Figure S36**

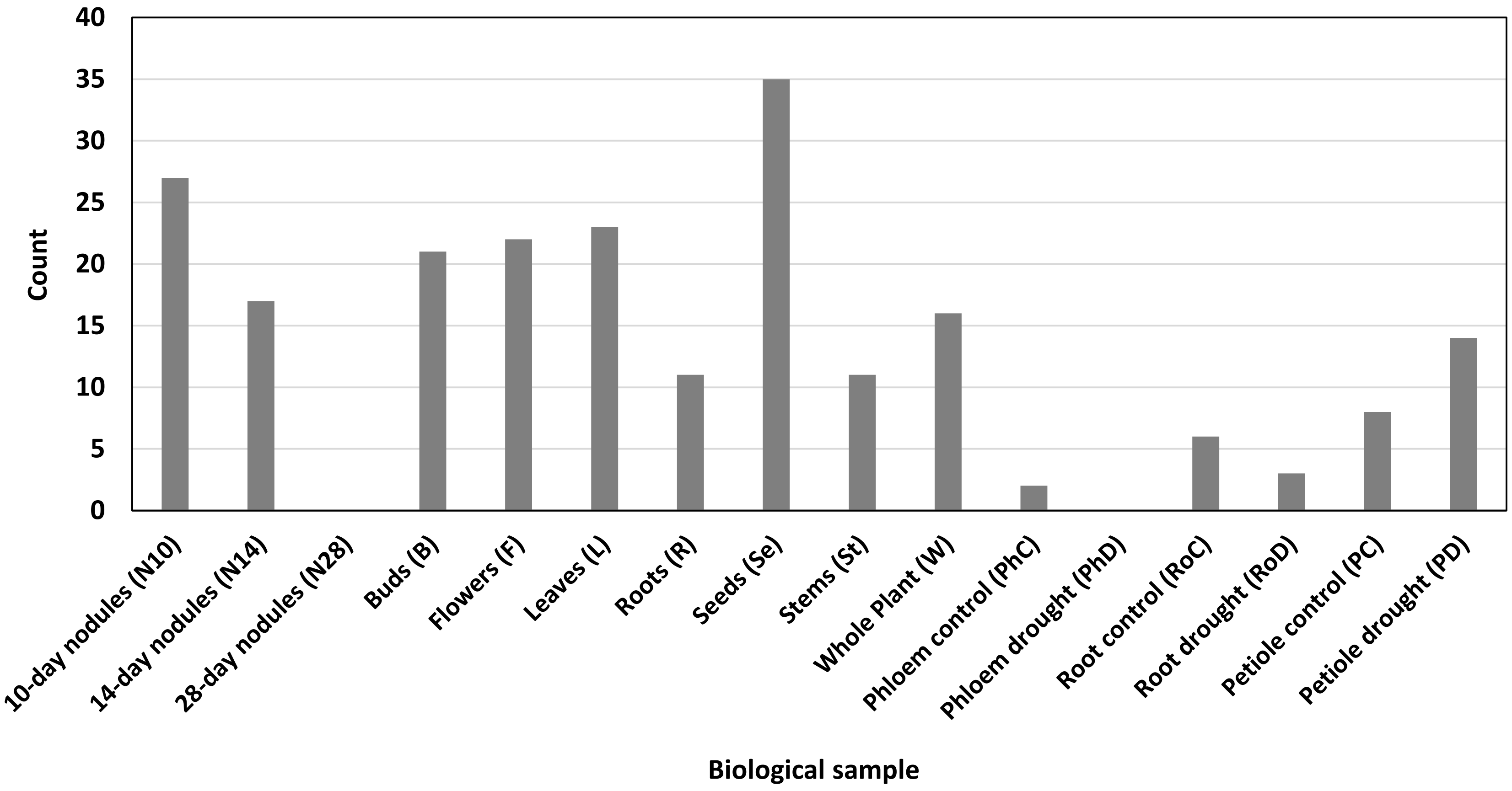


**Supplementary Figure S36**. Distribution of 156 chimeric MS peptides in 16 biological samples from three selected studies (Marx et al., 2016; Shin et al., 2021; Castañeda et al., 2021). The counts are not additive because some chimeric peptides were identified in multiple samples. No chimeric peptides were detected in 28-day nodules (N28) and the phloem under drought stress (PhD). The distribution of chimeric peptides is significantly different from the uniform distribution (the chi-square test for homogeneity, p = 5.45073E-18).

**Supplementary Figure S37**

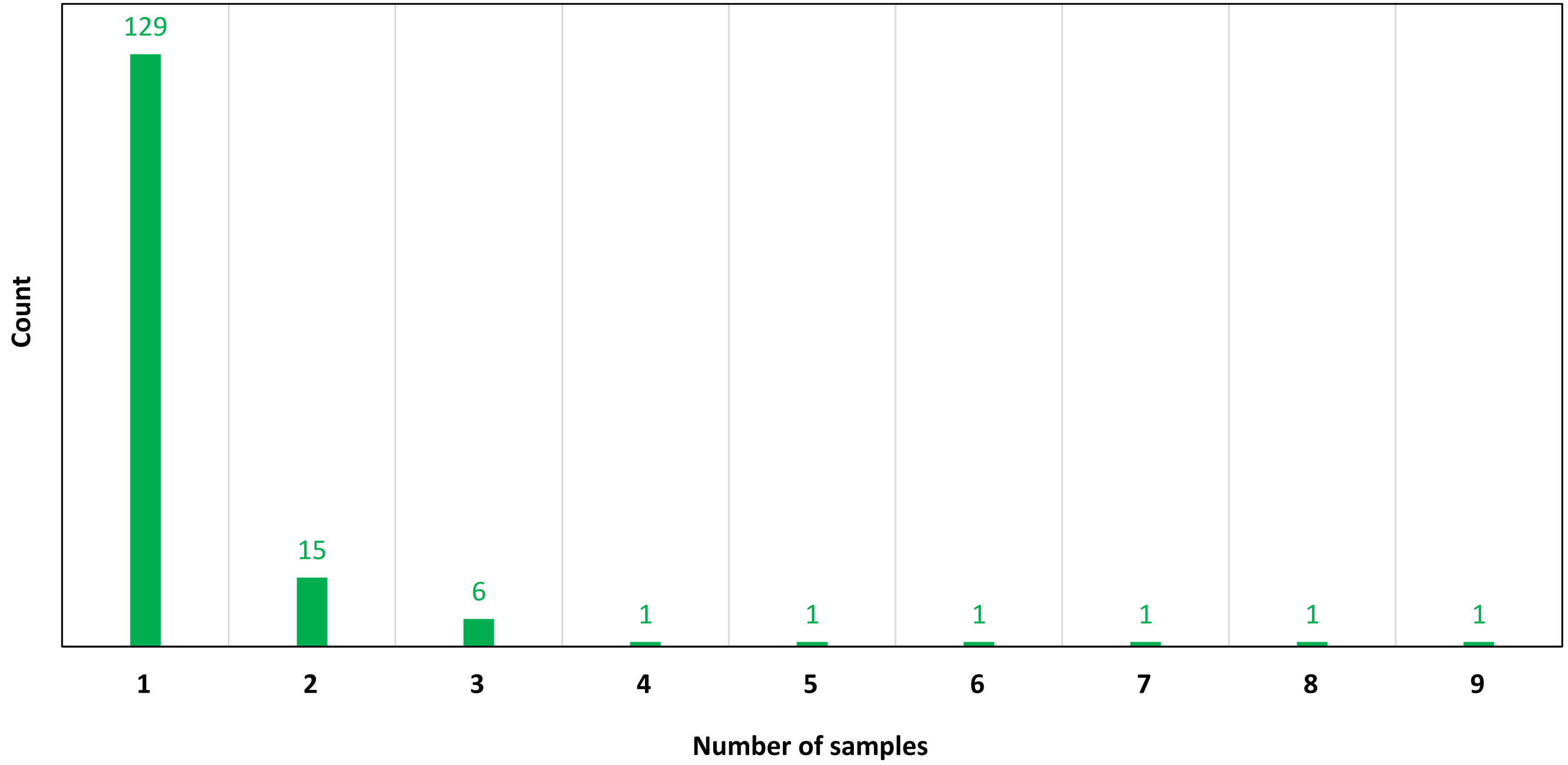


**Supplementary Figure S37**. Sample counts of 156 chimeric MS peptides identified in three selected studies (Marx et al., 2016; Shin et al., 2021; Castañeda et al., 2021). Group one contains chimeric peptides identified in a single sample (129 peptides); chimeric peptides identified in two samples constitute group two (15 peptides) etc. Groups four to nine contain only one peptide each.

### Supplementary Figure S38

| | 10-day nodules (N10) | 14-day nodules (N14) | 28-day nodules (N28) | Buds (B) | Flowers (F) | Leaves (L) | Roots (R) | Seeds (Se) | Stems (St) | Whole Plant (W) | Phloem control (PhC) | Phloem drought (PhD) | Root control (RoC) | Root drought (RoD) | Petiole control (PC) | Petiole drought (PD) |
|---|---|---|---|---|---|---|---|---|---|---|---|---|---|---|---|---|
| 10-day nodules (N10) | | 7 | nd | 1 | 4 | 2 | 1 | 4 | 1 | 2 | 0 | nd | 1 | 0 | 1 | 1 |
| 14-day nodules (N14) | 7 | | nd | 4 | 4 | 3 | 2 | 8 | 1 | 2 | 0 | nd | 1 | 0 | 2 | 2 |
| 28-day nodules (N28) | nd | nd | | nd | nd | nd | nd | nd | nd | nd | nd | nd | nd | nd | nd | nd |
| Buds (B) | 1 | 4 | nd | | 4 | 1 | 0 | 5 | 0 | 4 | 0 | nd | 1 | 0 | 3 | 3 |
| Flowers (F) | 4 | 4 | nd | 4 | | 5 | 2 | 4 | 2 | 3 | 0 | nd | 1 | 0 | 3 | 4 |
| Leaves (L) | 2 | 3 | nd | 1 | 5 | | 1 | 3 | 5 | 3 | 0 | nd | 0 | 0 | 2 | 2 |
| Roots (R) | 1 | 2 | nd | 0 | 2 | 1 | | 1 | 2 | 0 | 0 | nd | 0 | 0 | 0 | 0 |
| Seeds (Se) | 4 | 8 | nd | 5 | 4 | 3 | 1 | | 1 | 3 | 0 | nd | 1 | 0 | 2 | 2 |
| Stems (St) | 1 | 1 | nd | 0 | 2 | 5 | 2 | 1 | | 2 | 0 | nd | 0 | 0 | 1 | 1 |
| Whole Plant (W) | 2 | 2 | nd | 4 | 3 | 3 | 0 | 3 | 2 | | 0 | nd | 1 | 0 | 4 | 4 |
| Phloem control (PhC) | 0 | 0 | nd | 0 | 0 | 0 | 0 | 0 | 0 | 0 | | nd | 0 | 0 | 0 | 0 |
| Phloem drought (PhD) | nd | nd | nd | nd | nd | nd | nd | nd | nd | nd | nd | | nd | nd | nd | nd |
| Root control (RoC) | 1 | 1 | nd | 1 | 1 | 0 | 0 | 1 | 0 | 1 | 0 | nd | | 1 | 1 | 2 |
| Root drought (RoD) | 0 | 0 | nd | 0 | 0 | 0 | 0 | 0 | 0 | 0 | 0 | nd | 1 | | 0 | 1 |
| Petiole control (PC) | 1 | 2 | nd | 3 | 3 | 2 | 0 | 2 | 1 | 4 | 0 | nd | 1 | 0 | | 4 |
| Petiole drought (PD) | 1 | 2 | nd | 3 | 4 | 2 | 0 | 2 | 1 | 4 | 0 | nd | 2 | 1 | 4 | |
| Sum | 25 | 36 | nd | 26 | 36 | 27 | 9 | 34 | 16 | 28 | 0 | nd | 10 | 2 | 23 | 26 |

**Supplementary Figure S38**. Heatmap summary on the multi-sample occurrence of 27 putative chimeric peptides. 27 peptides out of 156 were found in more than one sample. Numbers indicate frequencies at which a given sample identifier is found together with identifiers of other samples in chimeric peptide names. Interpretation examples: (1) N14 x Se is 8, there are eight chimeric peptides found in 14-day nodules and seeds at the same time; (2) R x B is 0, chimeric peptides found in roots are never found in buds; (3) the sum of N14 is 36 and the sum of F is 36, chimeric peptides found in 14-day nodules and in flowers are most frequently found also in other samples, followed by seeds. Note that the sum row contains only cumulative frequencies, not the sums of chimeric peptides. Chimeric peptides found in root samples (R, RoC, and RoD) were the least frequently found in other samples. Sample PhC contained only chimeric peptides unique to that sample. No chimeric peptides were detected in samples N28 and PhD (marked with nd – not detected).

### Supplementary Figure S39

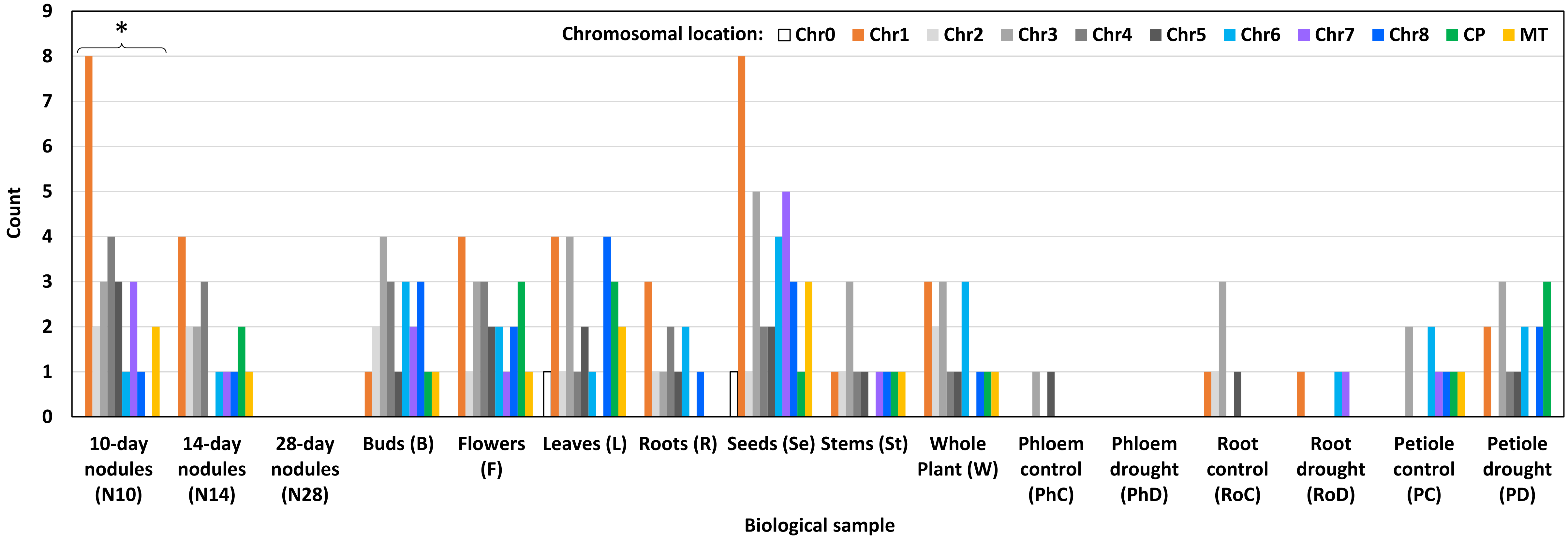


**Supplementary Figure S39**. Distribution of 156 chimeric MS peptides in 16 biological samples based on the chromosomal location of their primary-source loci. Chr0, unmapped loci. Chr1 to Chr8, Chromosome 1 to Chromosome 8. CP, chloroplasts. Mt, mitochondria. The counts are not additive because some chimeric peptides were identified in multiple samples. No chimeric peptides were detected in samples N28 and PhD. The chromosomal location is not associated with the sample. Within individual samples, the distribution of chromosomal locations may be significantly different from the uniform distribution in one sample indicated with an asterisk (the chi-square test for homogeneity, $p = 0.024$; the Exact Multinomial Test cannot be used for this group with many categories). Chloroplast-derived chimeric peptides are absent from samples N10, R, PhC, RoC, and RoD. Mitochondrial chimeric peptides are not found in samples R, PhC, RoC, RoD, and PD.

**Supplementary Figure S40**

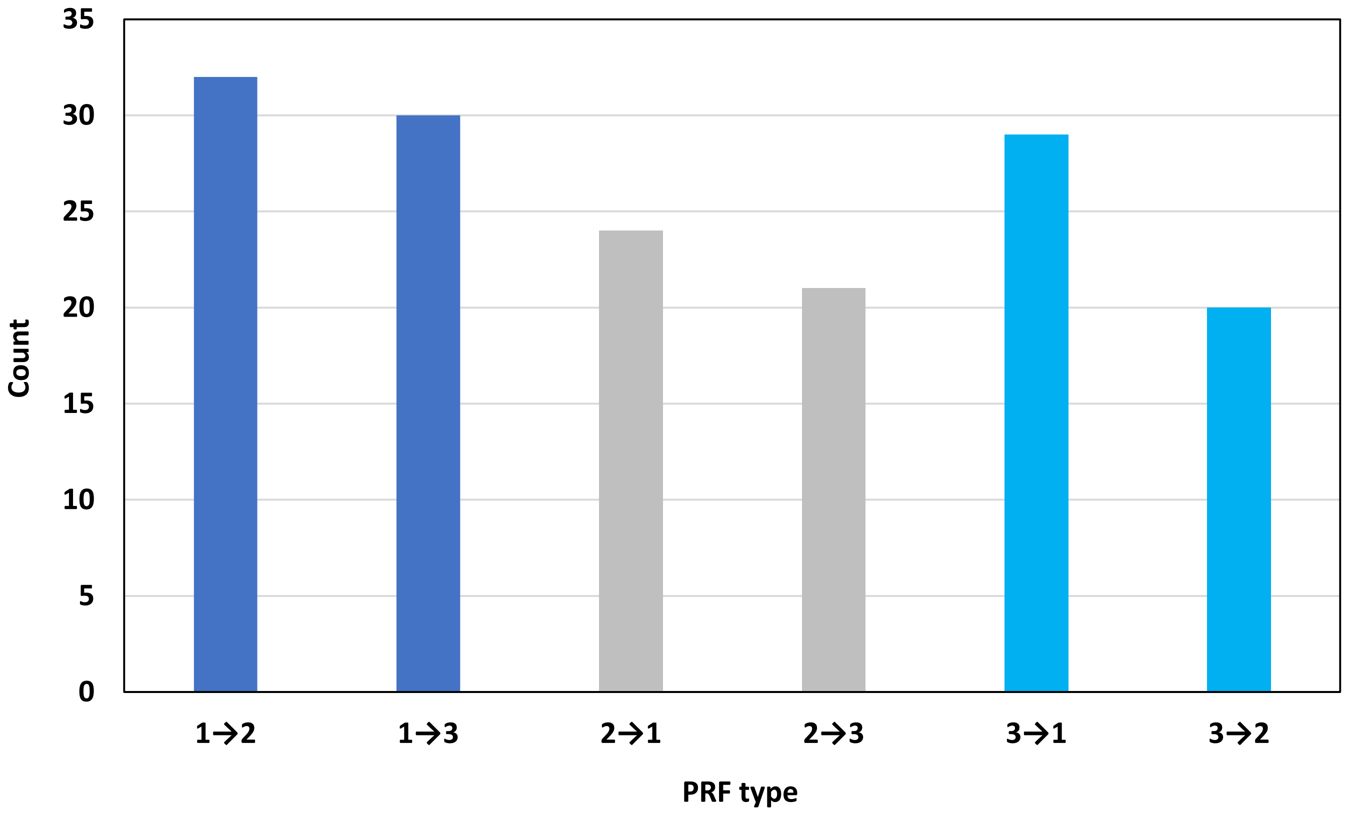


**Supplementary Figure S40**. Distribution of 156 chimeric MS peptides in six PRF types. The distribution is not significantly different from the uniform distribution.

### Supplementary Figure S41

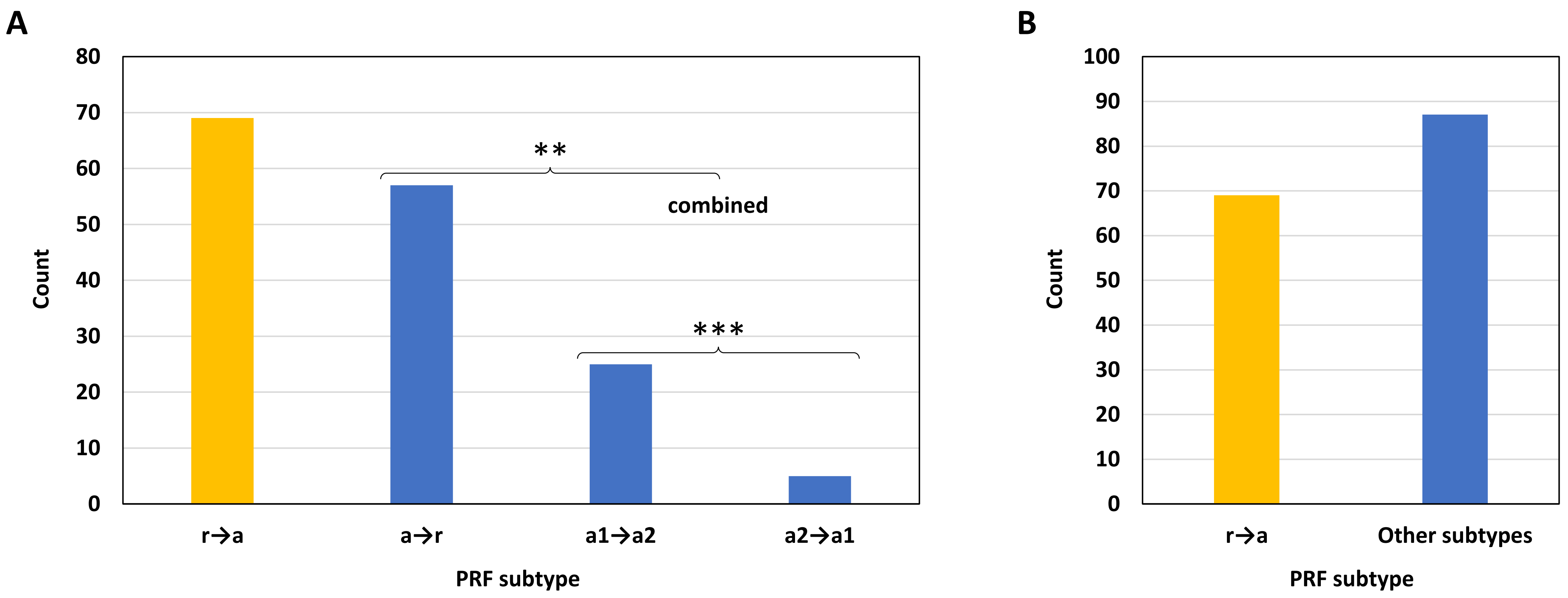


**Supplementary Figure S41**. Distribution of 156 chimeric MS peptides in different PRF subtypes. A, separate categories. B, PRF subtypes that start from altORFs are combined in one group. r→a, PRF from refORF to altORF; a→r, PRF from altORF to refORF; a1→a2, PRF from altORF1 to altORF2, where altORF1 starts upstream of altORF2; a2→a1, PRF from altORF2 to altORF1, where altORF1 starts upstream of altORF2. Proportion pairs in which the distribution is significantly different from the uniform distribution are marked with asterisks. P-values of the chi-square test for homogeneity of two proportions: **0.004; ***0.0003. PRF events that start from altORFs (blue bars) are statistically as abundant as PRF events that start from refORFs (B, p = 0.150).

**Supplementary Figure S42**

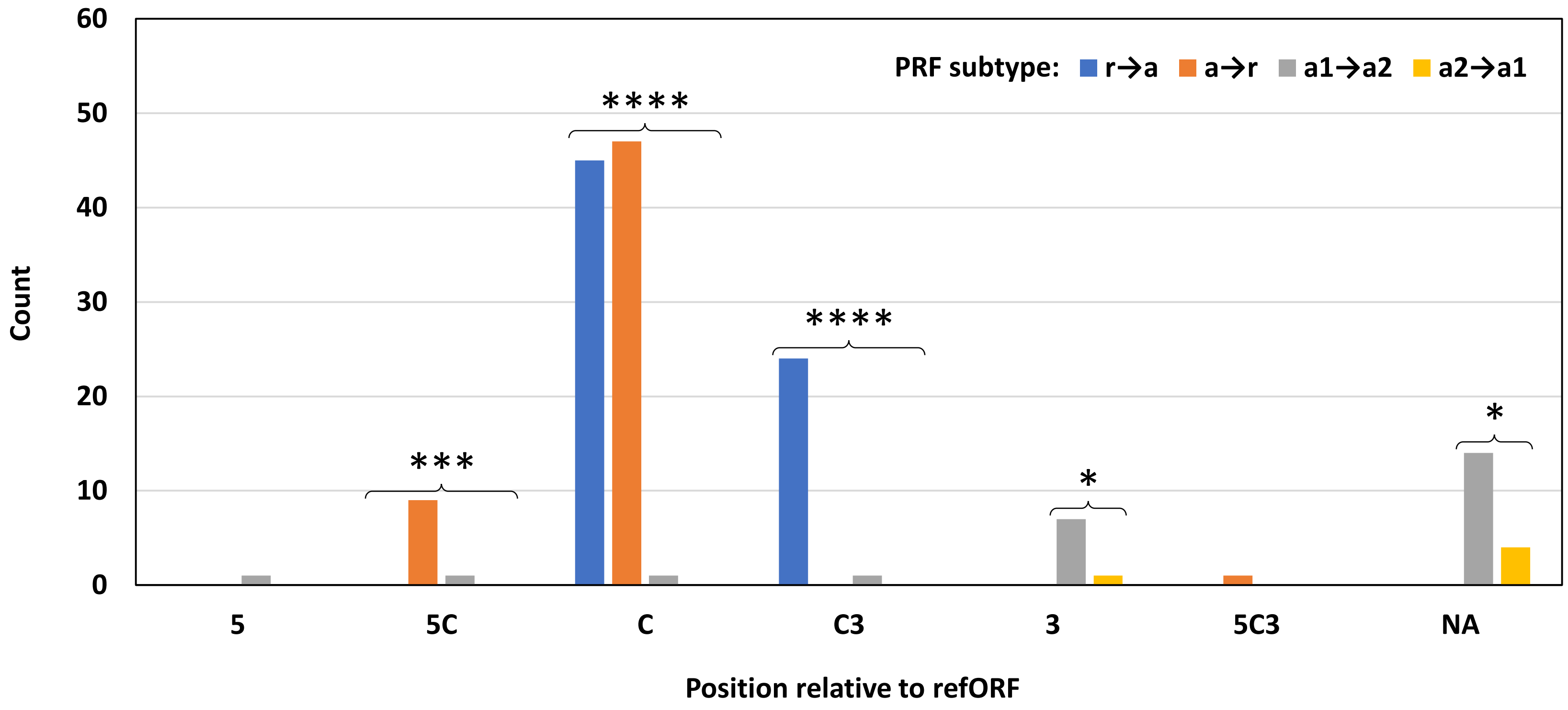


**Supplementary Figure S42**. Distribution of 156 chimeric MS peptides according to the position of an altORF relative to the refORF and the PRF subtype. 5, the 5'-untranslated region (UTR); 5C, the border between the 5'-UTR and the coding sequence (CDS); C, the CDS; C3, the border between the CDS and the 3'-UTR; 5C3, an altORF starts in the 5'-UTR and ends in the 3'-UTR; NA, all types of non-coding RNA, which have no refORF. Within individual categories, the distribution of PRF subtypes is significantly different from the uniform distribution in all cases except for 5 and 5C3 (the chi-square test for homogeneity for C and C3, the Exact Multinomial Test for 5C, and the Exact Binomial Test for 3 and NA, p-values < 0.05). Most observations in this graph are expected because of the nature of each category. The only striking observation here is approximately equal abundance of PRF subtypes r→a and a→r in PRF events located in the CDS, which is in line with the information presented in Supplementary Figure S41.

### Supplementary Figure S43

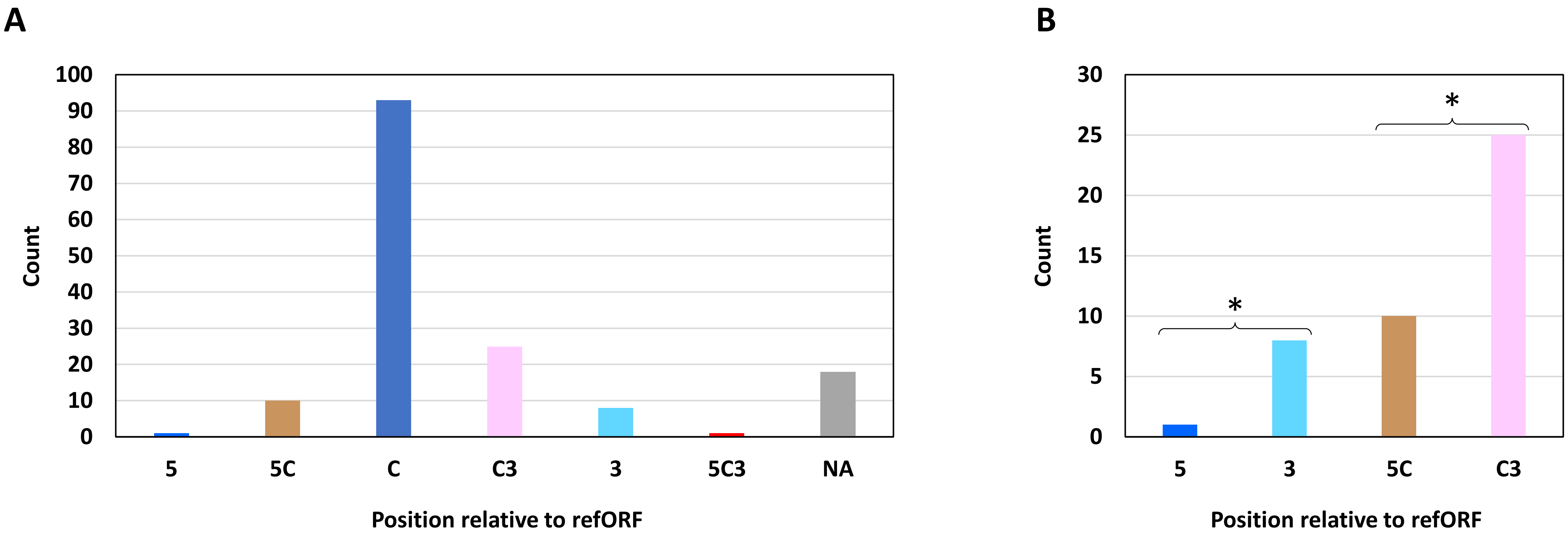


**Supplementary Figure S43**. Distribution of 156 chimeric MS peptides according to the position of an altORF involved in PRF relative to the refORF. A, all positions. B, selected positions for pairwise comparisons. Category names are the same as in Supplementary Figure S42. The distribution in A is significantly different from the uniform distribution (the chi-square test for homogeneity, p-value = 5.503E-58). The majority of altORFs are embedded in the CDS (dark blue bar in A). AltORFs located in the 3'-UTRs are significantly more frequent than altORFs located in the 5'-UTRs (B, the Exact Binomial Test, p = 0.020). The border between the CDS and the 3'-UTR has significantly more altORFs compared to the border between the 5'-UTR and the CDS (B, the chi-square test for homogeneity of two proportions, p = 0.011).

**Supplementary Figure S44**

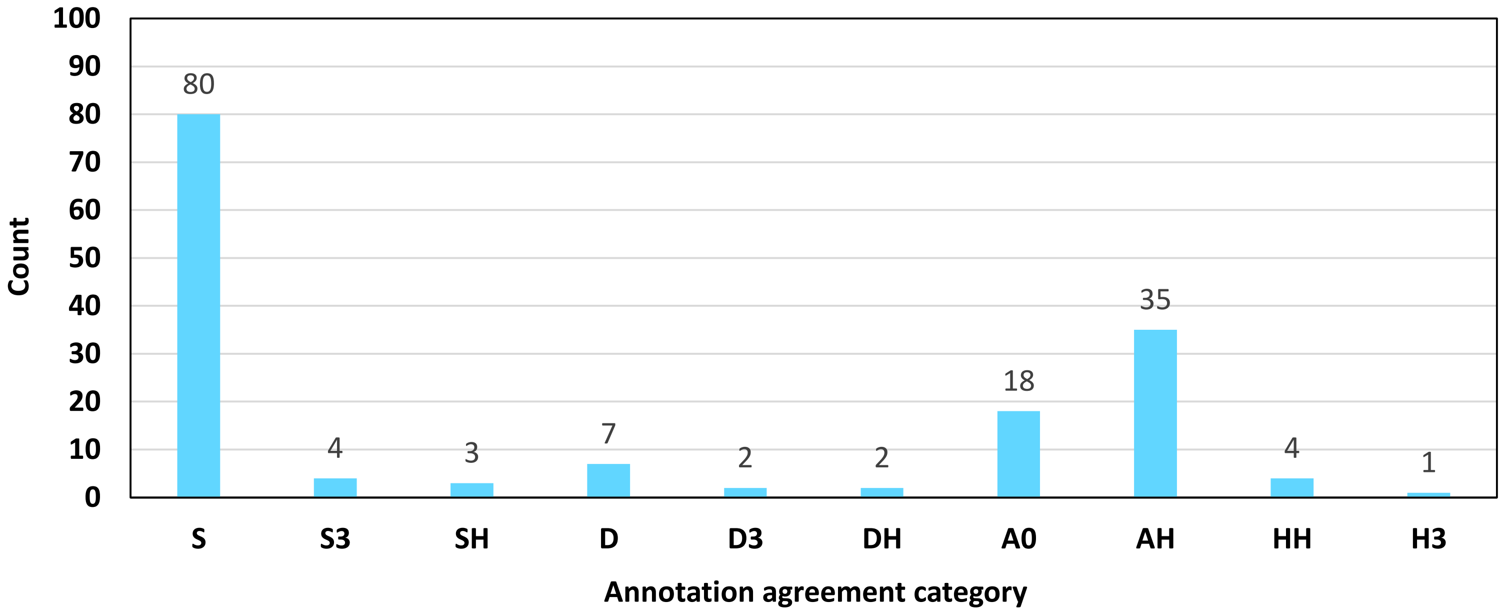


**Supplementary Figure S44**. Distribution of 156 chimeric MS peptides in ten categories of the agreement between the annotation of ORFs involved in PRF. The annotation is based on the global BLASTP search of each sequence using NCBI. S, the annotation is the same for proteins of both ORFs involved in PRF. S3, the same for proteins of two altORFs involved in PRF and the refORF of the transcript. SH, the same for two proteins, whereas the third one is annotated as a hypothetical or unknown protein. D, different for both proteins. D3, different for proteins of two altORFs involved in PRF and the refORF of the transcript. DH, different for two proteins, whereas the third one is annotated as a hypothetical or unknown protein. A0, one protein is clearly annotated but the other one has no significant similarity with any protein in the NCBI database. AH, one protein is clearly annotated but the other one is annotated as a hypothetical or unknown protein. HH, both proteins are annotated as hypothetical or unknown proteins. H3, all three proteins are annotated as hypothetical or unknown proteins. The distribution of chimeric peptides is significantly different from the uniform distribution (the chi-square test for homogeneity, p = 4.843E-72). More than half of altProts belong to the same family as their refProts and/or other altProts involved in PRF.

**Supplementary Figure S45**

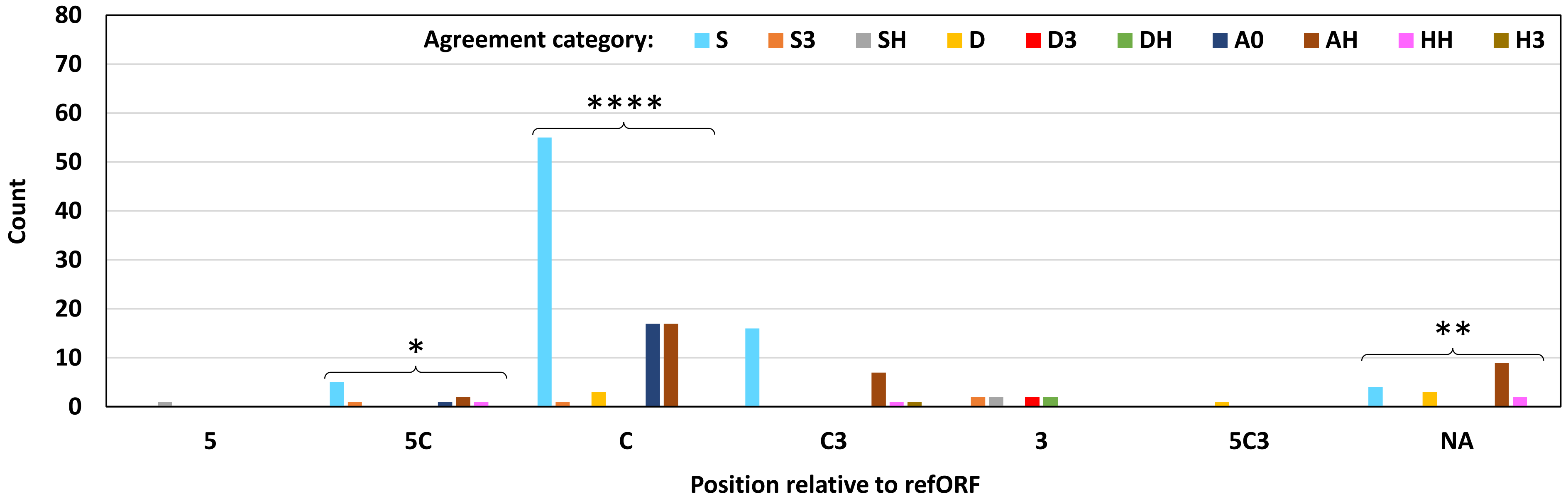


**Supplementary Figure S45**. Distribution of 156 chimeric MS peptides in ten annotation agreement categories grouped by altORF location relative to the refORF. Designation of categories is the same as in Supplementary Figure S44. There may be significant association between the agreement category and the location (the chi-square test for association, p = 1.262E-25, the Fisher's exact test is not applicable to these groups). Within individual location categories, the distribution is significantly different from the uniform distribution in 5C, C, and NA (the chi-square test for homogeneity for C, the Exact Multinomial Test for 5C and NA, p-values < 0.05). Agreement category S is predominant in C, C3, and 5C. This is in line with a hypothesis according to which most altORFs involved in PRF emerged by frameshifting mutations and only very few (agreement groups D, D3, and DH) are likely to result from DNA translocation events. The remaining categories possibly correspond to ORFs that have *de novo* origin (agreement groups A0, AH, HH, and H3).

**Supplementary Figure S46**

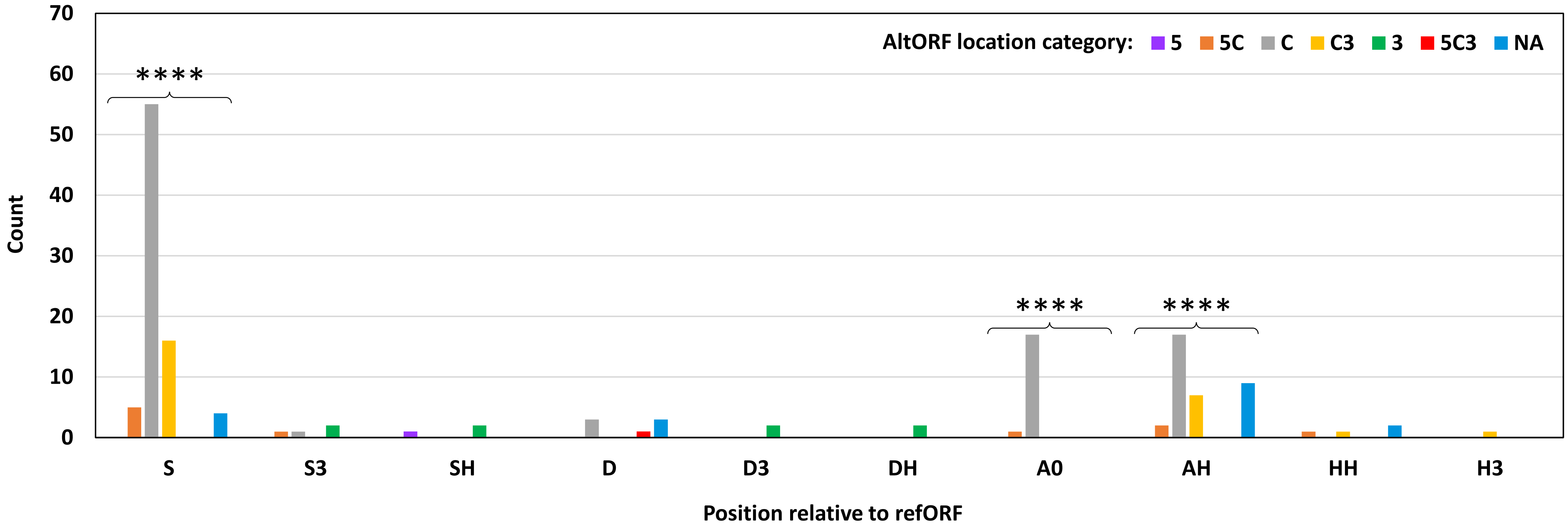


**Supplementary Figure S46**. Distribution of 156 chimeric MS peptides in seven altORF locations relative to the refORF grouped by annotation agreement categories. Designation of categories is the same as in Supplementary Figure S44. There may be significant association between the agreement category and the location (the chi-square test for association, p = 1.262E-25, the Fisher's exact test is not applicable to these groups). Within individual agreement categories, the distribution is significantly different from the uniform distribution in three cases indicated with asterisks (the chi-square test for homogeneity for S and AH, the Exact Multinomial Test for A0, p-values < 0.05). Agreement category S is predominant in C, C3, and 5C. Significantly more chimeric peptides of category S come from the CDS-3'-UTR border compared to the upstream border (the chi-square test for homogeneity, p = 0.016). Categories A0 and AH are abundant in chimeric peptides derived from altORFs located within the CDS (location C), which means altORFs with *de novo* origin are common in the CDSs.

**Supplementary Figure S47**

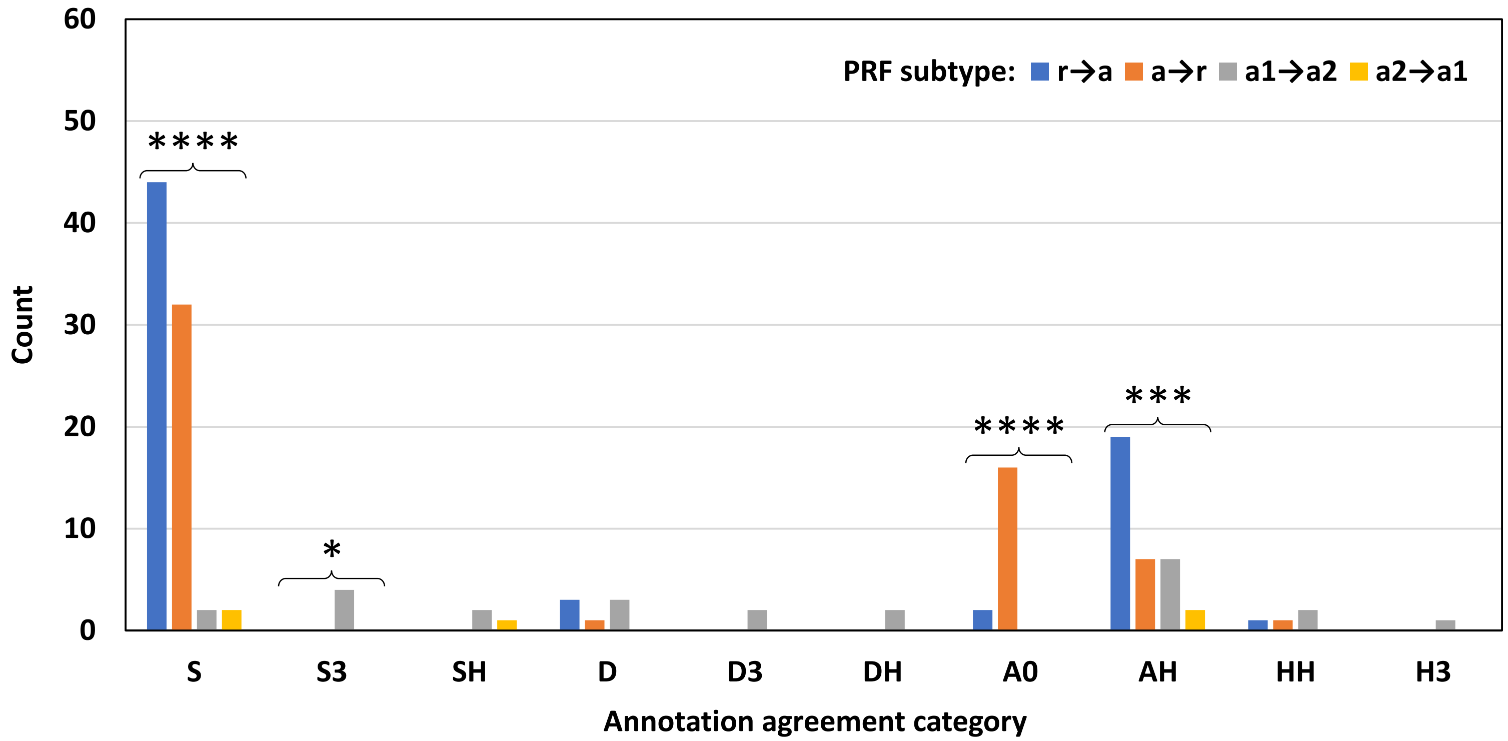


**Supplementary Figure S47**. Distribution of 156 chimeric MS peptides in ten categories of the agreement between the annotation of ORFs involved in PRF. The second analyzed factor in this graph is the PRF subtype. Designation of categories is the same as in Supplementary Figure S44. There <u>may be</u> significant association between the annotation agreement category and the PRF subtype (the chi-square test for association, p = 1.122E-11, the Fisher's exact test is not applicable to these groups). Within individual agreement categories, the distribution is significantly different from the uniform distribution in four cases indicated with asterisks (the chi-square test for homogeneity for S and AH, the Exact Multinomial Test for S3 and A0, p-values: S, 9.393E-15; S3, 0.016; A0, 2.259E-7; AH, 4.581E-4). The PRF subtypes r→a and a→r are differentially represented in the agreement categories A0 and AH.

**Supplementary Figure S48**

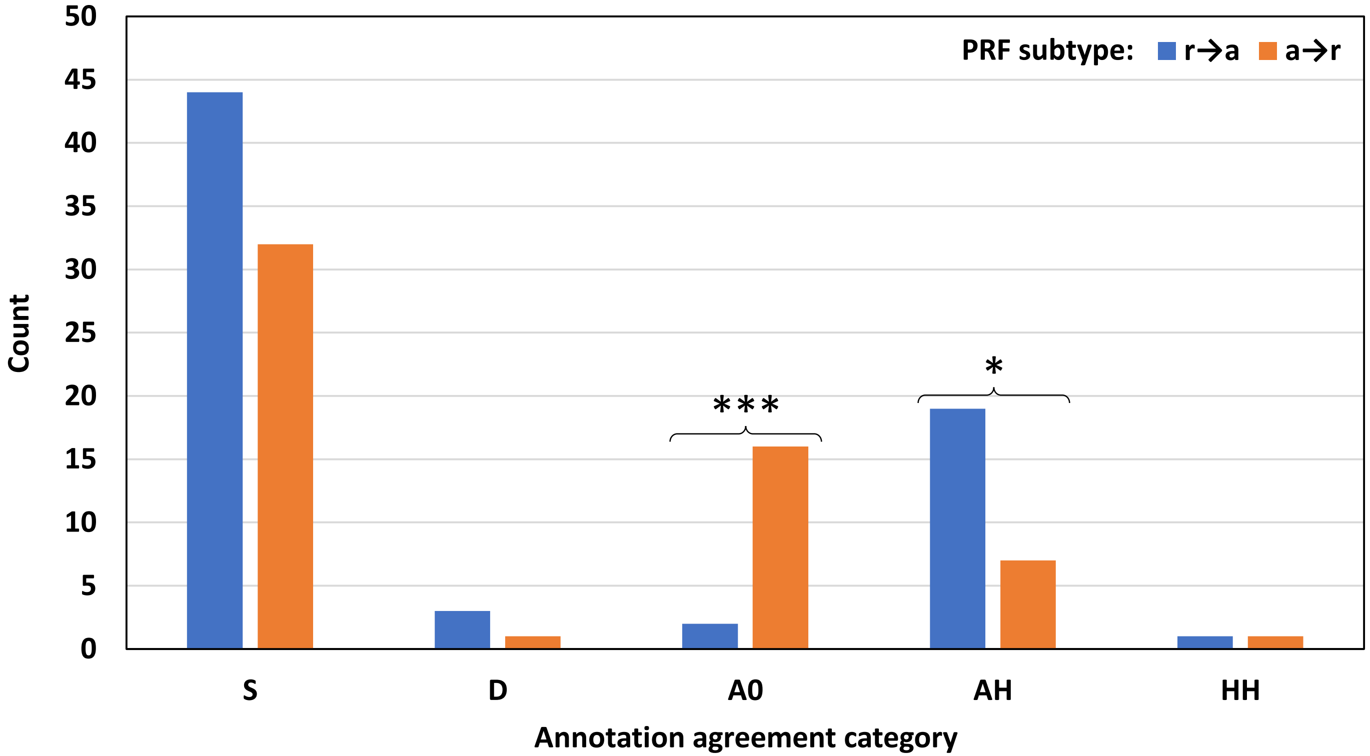


**Supplementary Figure S48**. Distribution of 126 chimeric MS peptides in five selected categories of the agreement between the annotation of ORFs involved in PRF. The second analyzed factor in this graph is the PRF subtype. Designation of categories is the same as in Supplementary Figure S44. There is significant association between the annotation agreement category and the PRF subtype (the chi-square test for association, p = 0.001). Within individual agreement categories, the distribution is significantly different from the uniform distribution in two cases indicated with asterisks. P-values of the chi-square test for homogeneity are as follows: A0, 9.670E-04; AH, 0.019. The PRF subtypes r→a and a→r are differentially represented in the agreement categories A0 and AH but not in the category S. All proteins with no significant similarity (A0) are altProts in mRNA transcripts.

**Supplementary Figure S49**

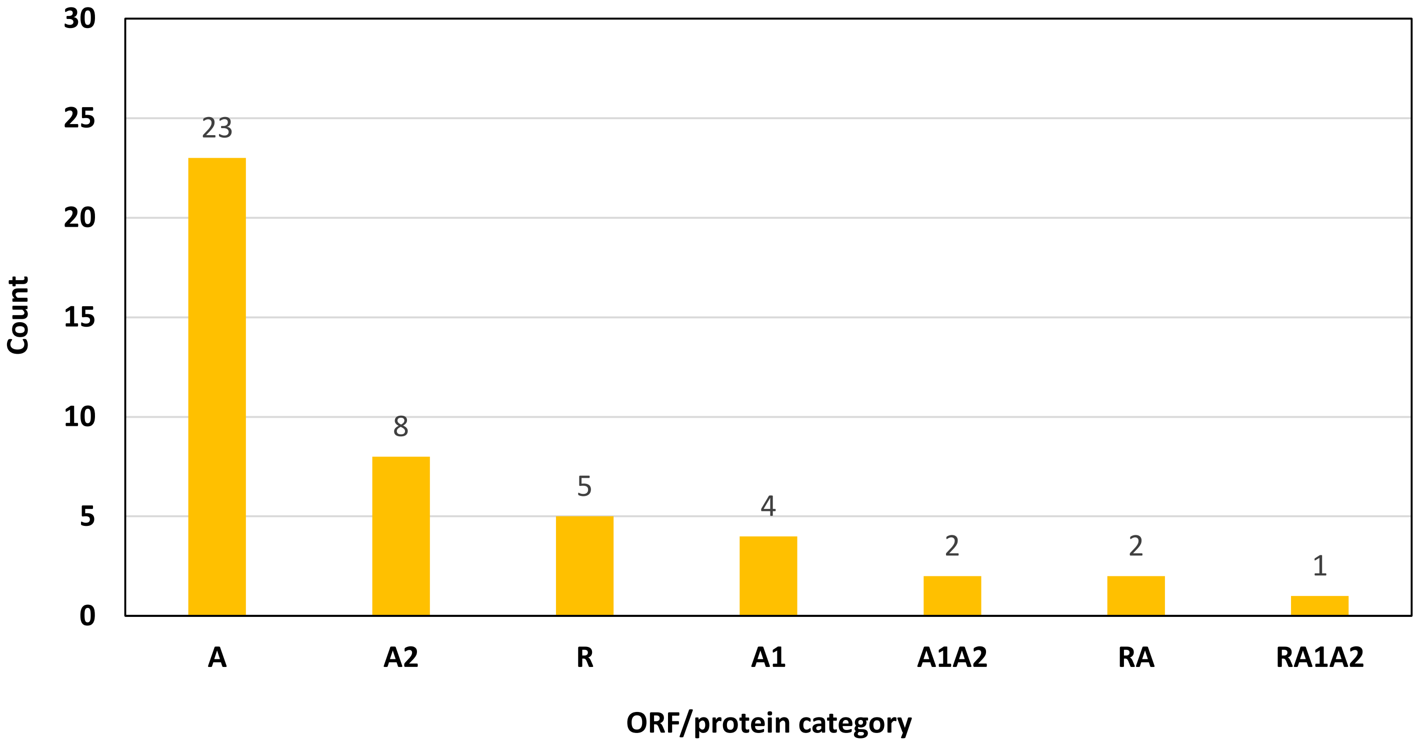


**Supplementary Figure S49**. Distribution of 45 chimeric MS peptides depending on the type of ORF products annotated as hypothetical or unknown. The categories indicate which ORF product is hypothetical or unknown. A, the altProt in the absence of the second altProt involved in PRF. A2, the second altProt regardless of the presence of the refProt (mRNA and non-mRNA transcripts). R, the refProt; A1, the first altProt in the absence of the refProt involved in PRF (non-mRNA transcripts only). A1A2, both altProts in the absence of the refProt involved in PRF (non-mRNA transcripts only). RA, both the refProt and the altProt. RA1A2, the refProt and two altProts. The distribution of chimeric peptides is significantly different from the uniform distribution (the chi-square test for homogeneity, $p = 4.588E-10$). The majority of proteins annotated as hypothetical or unknown are altProts. Half of them (23) are altProts of mRNA transcripts that have no second altProt involved in PRF.

### Supplementary Figure S50

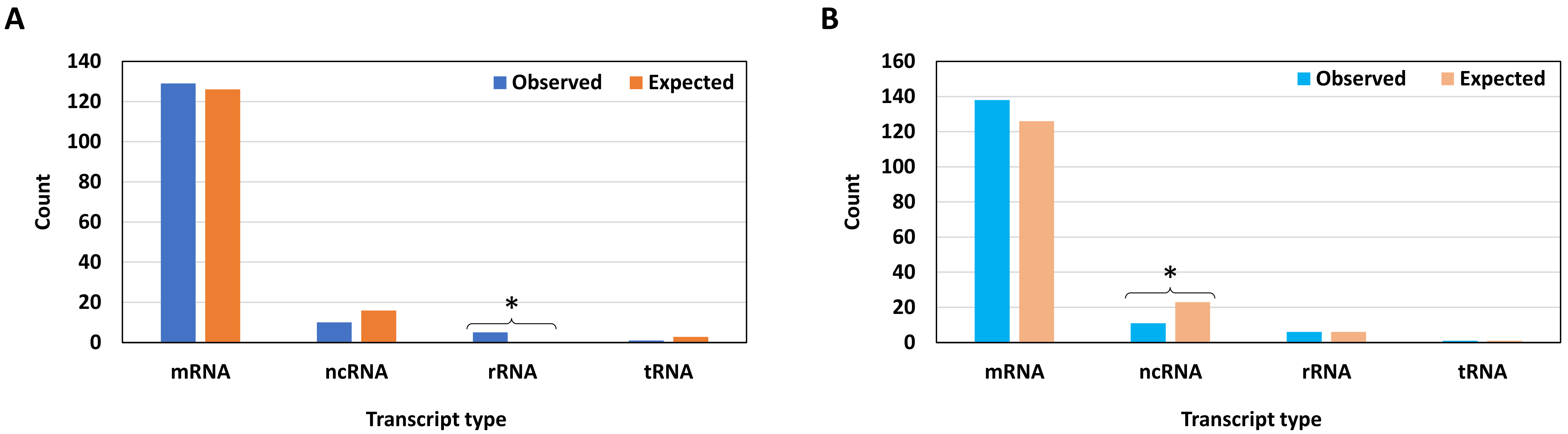


**Supplementary Figure S50**. Distribution of 145 transcripts (A) and their respective 156 chimeric MS peptides (B) in different groups of RNA. The basis for expected counts in A is the occurrence of corresponding transcript groups in the whole transcriptome (with the correction for two rRNA transcripts formerly annotated as ncRNA, MtrunA17_Chr0c01g0489091 and MtrunA17_Chr5g0421761). In B, the basis is the number of non-redundant chimeric protein models (Supplementary Table S9, 469,600 models in total). The Multinomial Exact Test of goodness-of-fit shows significant differences between the observed and expected proportions (A, p = 5.426E-06; B, p = 0.028). Within individual transcript categories, the distribution of observed and expected counts is significantly different from the uniform distribution in two cases indicated with asterisks (A, the Exact Binomial Test with expected value 0.2 rounded to zero, p = 0.032; B, the chi-square test for two proportions, p = 0.037). Ribosomal RNA transcripts are significantly overrepresented in the dataset (25-fold), evidently due to the generation of more models (note no significant difference for rRNA in B). This means rRNA transcripts stand out with regard to the higher number of conserved and/or translated overlapping ORFs compared to other RNA types. In contrast, ncRNA transcripts are distinguished by the detection of nearly two-fold fewer MS-supported CPs than could be expected based on the number of generated chimeric models (B).

**Supplementary Figure S51**

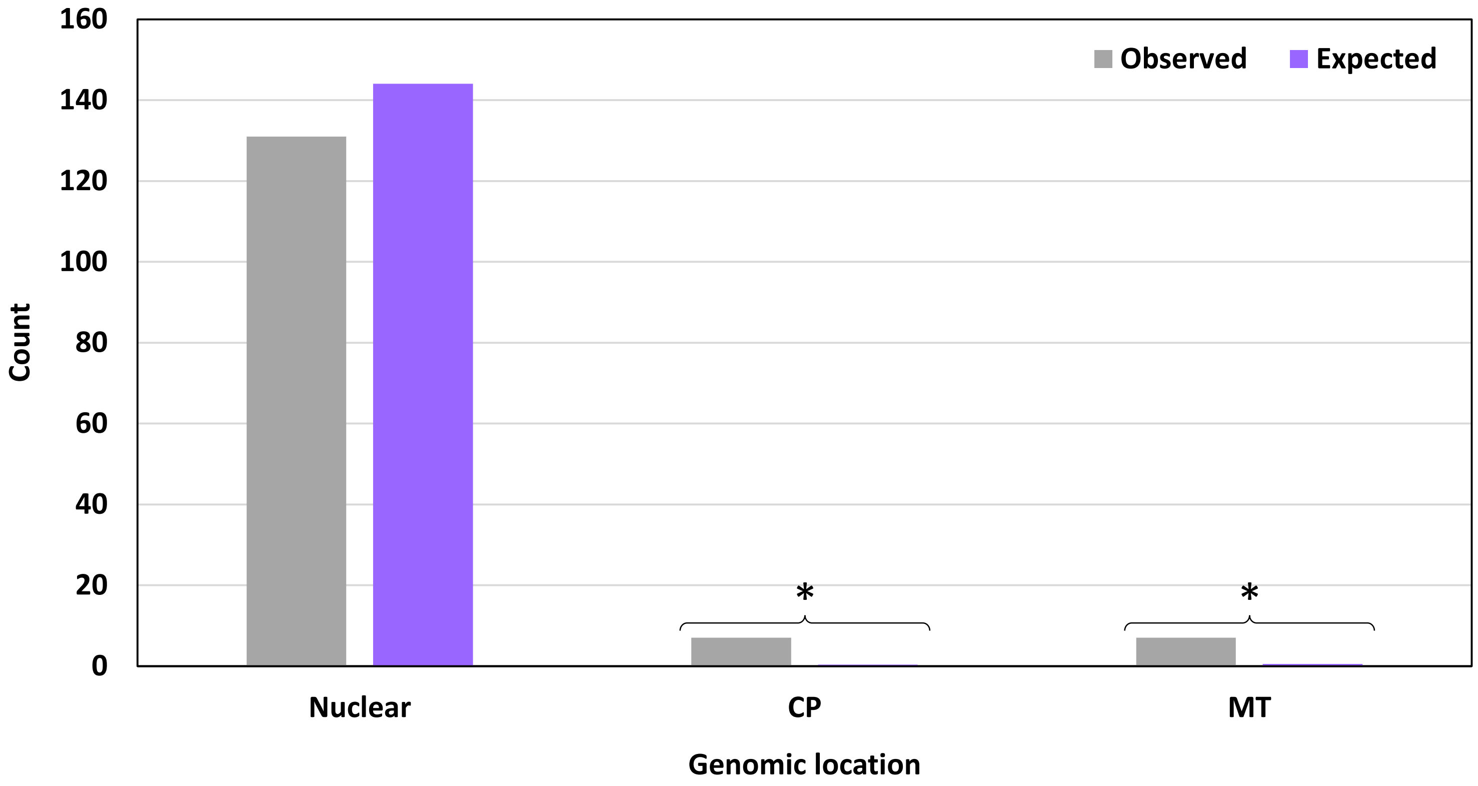


**Supplementary Figure S51**. Distribution of 145 chimeric MS peptide-producing transcripts in different genomic locations. Nuclear, Chromosome 0 (unmapped loci) to Chromosome 8. CP, chloroplasts. Mt, mitochondria. The basis for expected counts is the occurrence of corresponding transcript groups in the whole transcriptome. The Multinomial Exact Test of goodness-of-fit shows a significant difference between the observed and expected proportions (p = 5.422E-08). Within individual transcript categories, the distribution of counts is significantly different from the uniform distribution in two cases indicated with asterisks (the Exact Binomial Test with expected values 0.4 and 0.6 rounded to one, p = 0.035). Chloroplast and mitochondrial transcripts are significantly overrepresented in the dataset (19-fold and 12-fold, respectively). Five out of seven mitochondrial transcripts are ncRNAs.

### Supplementary Figure S52

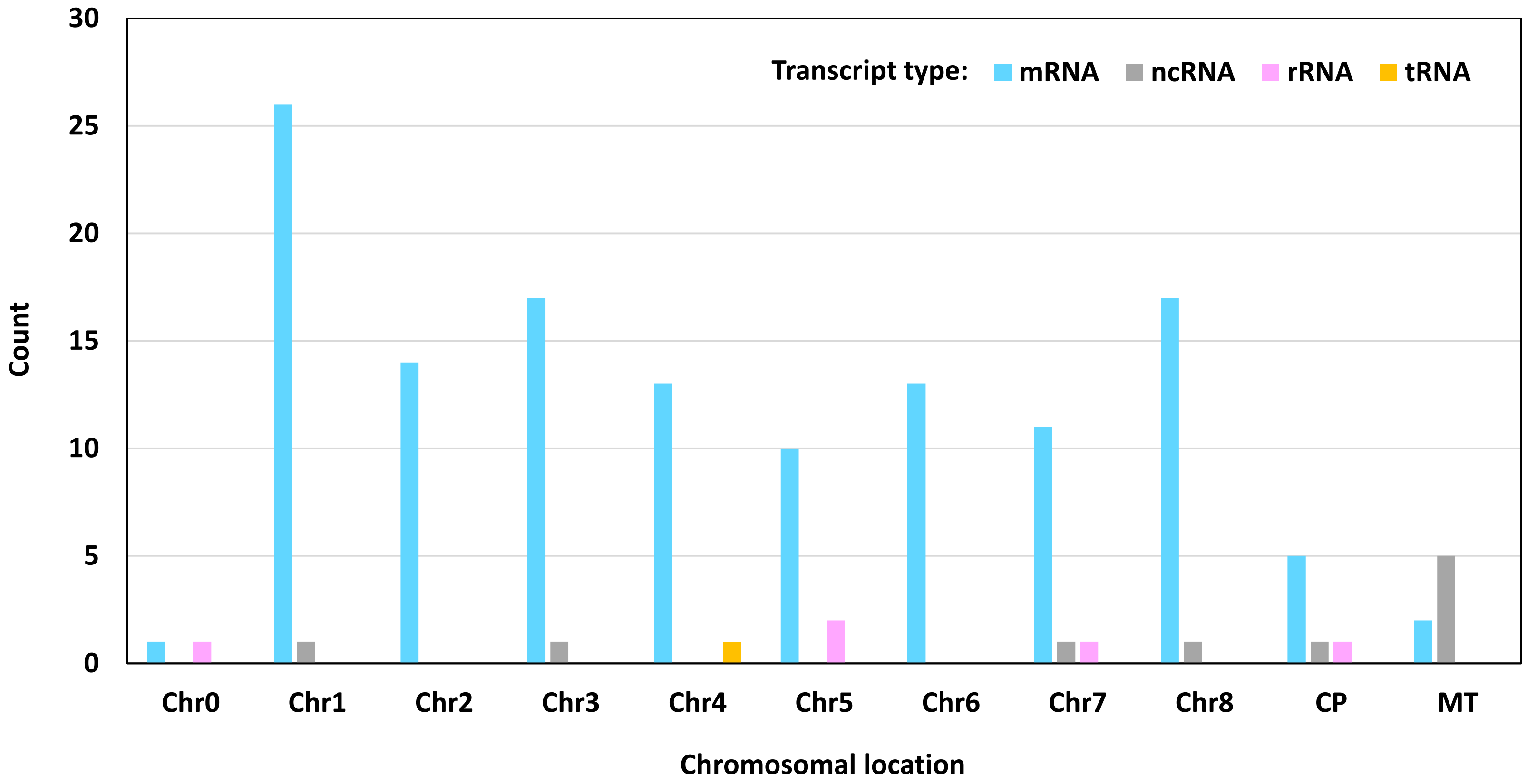


**Supplementary Figure S52**. Distribution of 145 chimeric MS peptide-producing transcripts in different chromosomal locations based on the transcript type. Chr0, unmapped loci. Chr1 to Chr8, Chromosome 1 to Chromosome 8. CP, chloroplasts. Mt, mitochondria. The Fisher's exact test with a simulated p-value shows significant association between the chromosomal location and the transcript type ($p = 4.998E-4$ with or without tRNA). Mitochondria is the only location in which ncRNA transcripts that produce chimeric peptides are more abundant than other RNA types. The high proportion of ncRNA in mitochondria is a general feature of the *M. truncatula* genome.

**Supplementary Figure S53**

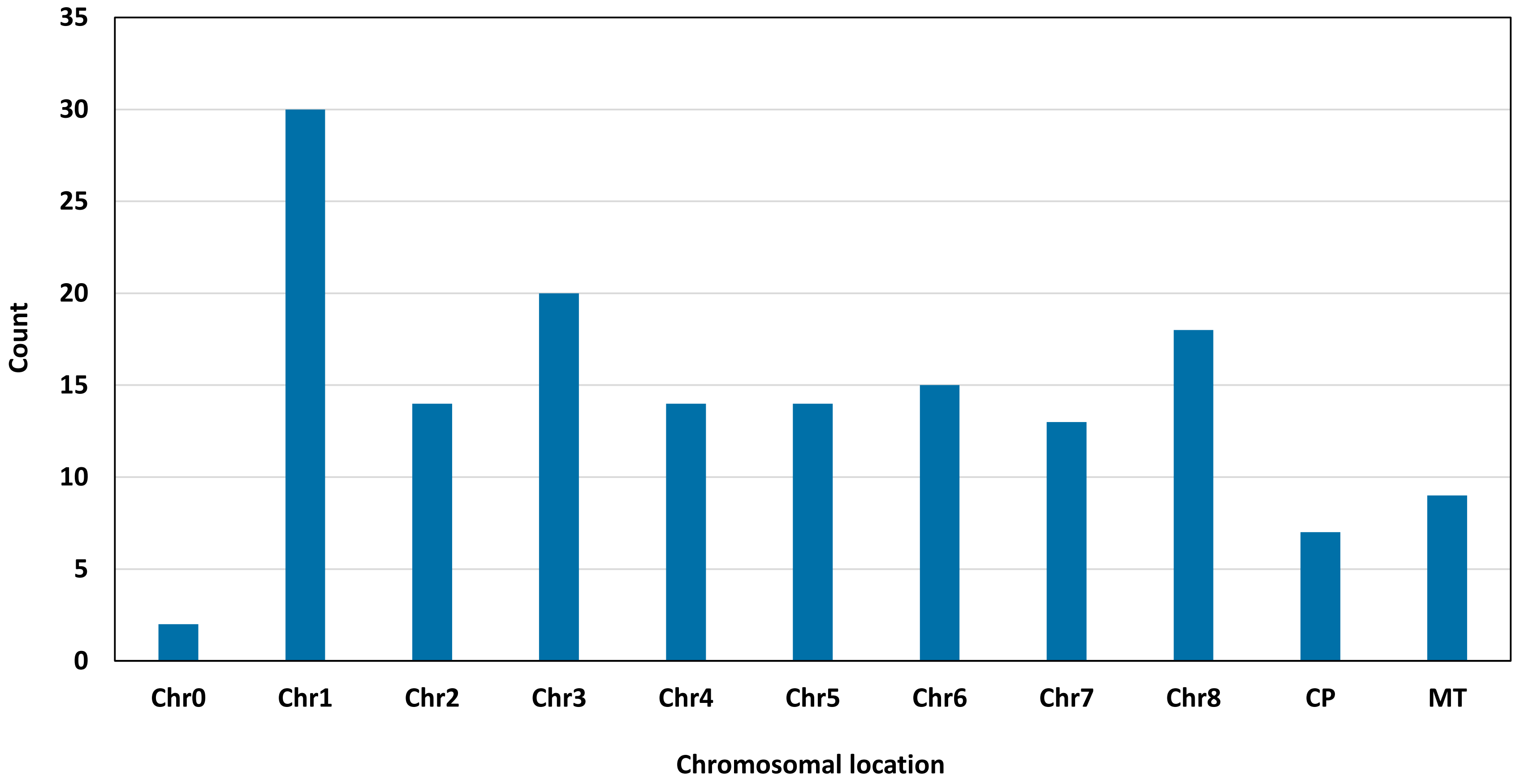


**Supplementary Figure S53**. Distribution of 156 chimeric MS peptides in different chromosomal locations. Chr0, unmapped loci. Chr1 to Chr8, Chromosome 1 to Chromosome 8. CP, chloroplasts. Mt, mitochondria. The distribution of chimeric peptides is significantly different from the uniform distribution (the chi-square test for homogeneity, p = 5.215E-05).

### Supplementary Figure S54

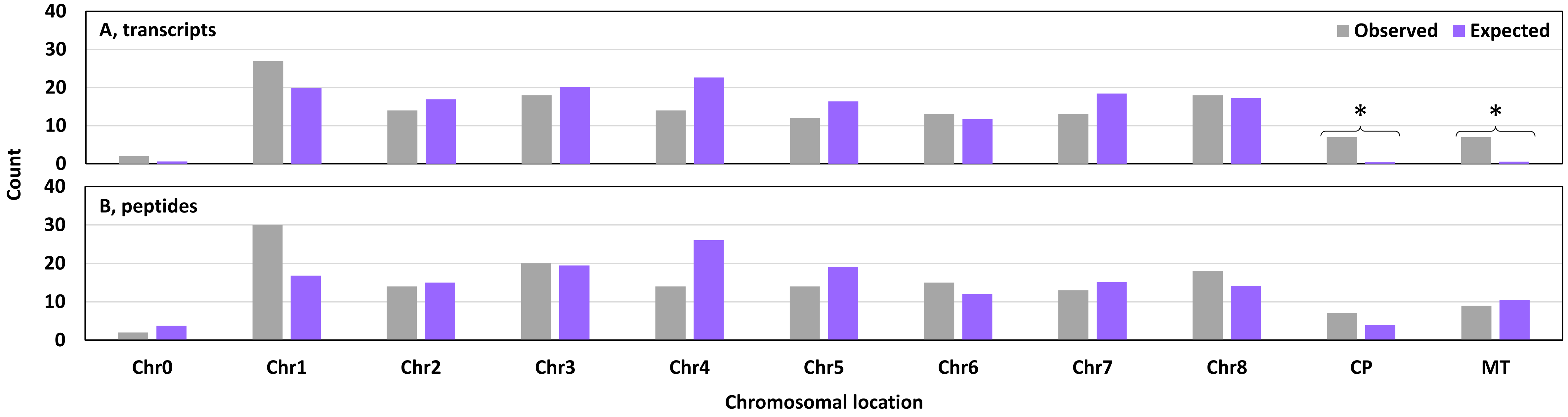


**Supplementary Figure S54**. Distribution of 145 transcripts (A) and their respective 156 chimeric MS peptides (B) in different chromosomal locations. Chr0, unmapped loci. Chr1 to Chr8, Chromosome 1 to Chromosome 8. CP, chloroplasts. Mt, mitochondria. The basis for expected counts in A is the occurrence of transcripts from different chromosomal locations in the whole transcriptome. In B, the basis is the non-redundant number of chimeric protein models associated with different chromosomal locations (Supplementary Figures S55 and S56, 469,600 models in total). The differences between the observed and expected proportions in A and B may be significant (the chi-square test for goodness-of-fit, A, p = 2.433E-39; B, p = 0.012). The Exact Multinomial Test cannot be used with these groups due to the large number of categories. Thus, the actual significance of the differences is unknow. Within individual chromosomal locations, the distribution of counts is significantly different from the uniform distribution in two cases indicated with asterisks (the Exact Binomial Test with expected values 0.4 and 0.6 rounded to one, p = 0.035). The many-fold overrepresentation of chloroplast and mitochondrial transcripts in our dataset may be due to the fact that transcripts from these locations had more conserved and/or MS-supported altORFs that overlapped with refORFs and/or each other. Thus, more chimeric protein models were produced from these transcripts (Supplementary Figures S55 and S56).

**Supplementary Figure S55**

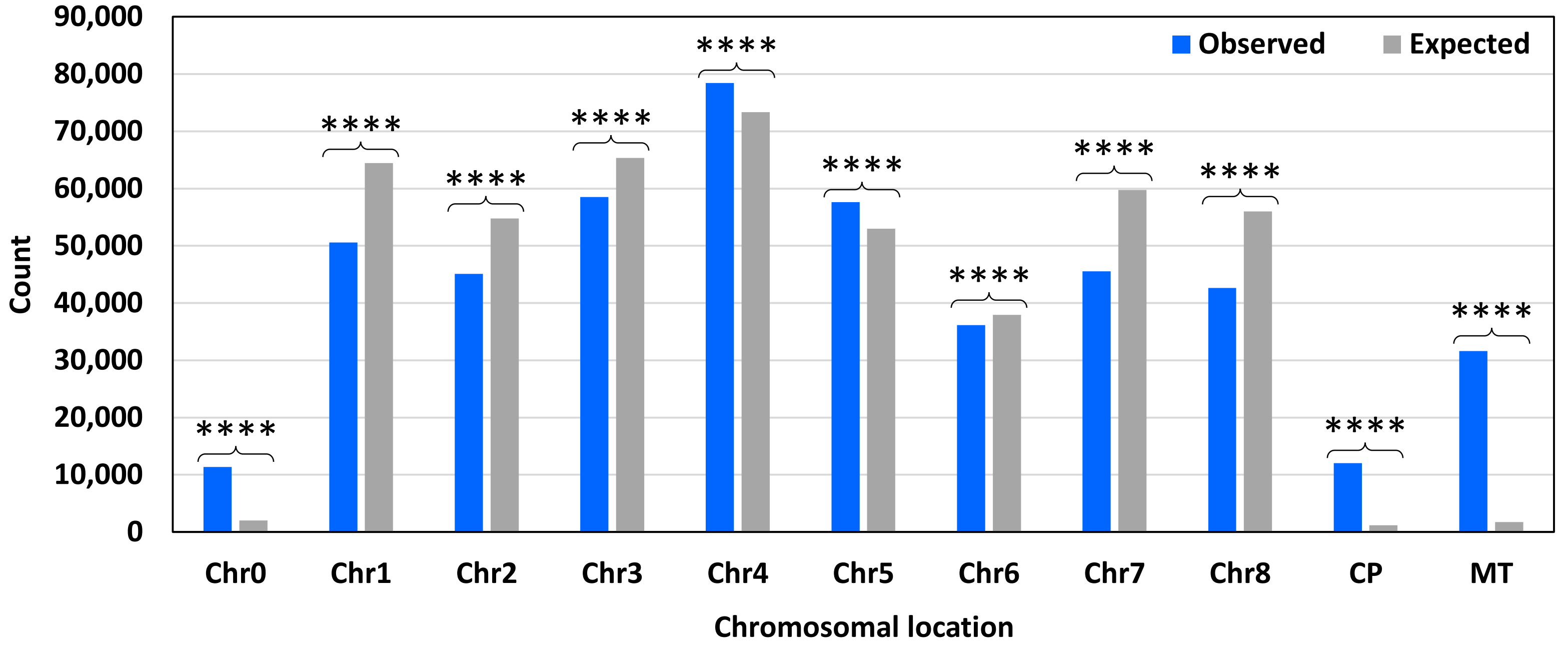


**Supplementary Figure S55**. Distribution of 469,600 non-redundant chimeric protein models in different chromosomal locations. Chr0, unmapped loci. Chr1 to Chr8, Chromosome 1 to Chromosome 8. CP, chloroplasts. Mt, mitochondria. The basis for expected counts is the occurrence of transcripts from different chromosomal locations in the whole transcriptome (the assumption of chimeric model counts being directly proportional to the number of transcripts). The chi-square test for goodness-of-fit shows a significant difference between the observed and expected proportions (p-value = 0). Within individual chromosomal locations, the distribution of counts is significantly different from the uniform distribution in all cases. P-values of the chi-square test for homogeneity are as follows: Chr0, Chr1, Chr7, Chr8, CP, and MT, p = 0; Chr2, p = 2.172E-206; Chr3, p = 3.075E-83; Chr4, p = 1.130E-38; Chr5, p = 1.041E-43; Chr6, p = 4.970E-11. Chimeric model counts depend not on the number of transcripts alone. Their counts are the function of the number of overlapping ORFs that have either conservation evidence or are MS-supported and also depend on the length of the overlapping region. The ratio of such overlapping ORFs relative to the expected counts is location-specific and is maximal in non-nuclear transcripts (CP and MT).

### Supplementary Figure S56

**A**

| A | B | C | D | E | F | G |
|---|---|---|---|---|---|---|
| Chromosomal location | Chimeric peptides | Chimeric protein models | Transcripts | Cumulative length of transcripts, nt | % (B/C*100) | Ratio (C/D) |
| Chr0 | 2 | 11,348 | 220 | 246,059 | 0.018 | 51.582 |
| Chr1 | 30 | 50,588 | 7,046 | 10,202,666 | 0.059 | 7.180 |
| Chr2 | 14 | 45,098 | 5,987 | 8,434,653 | 0.031 | 7.533 |
| Chr3 | 20 | 58,535 | 7,140 | 10,325,001 | 0.034 | 8.198 |
| Chr4 | 14 | 78,440 | 8,018 | 11,297,616 | 0.018 | 9.783 |
| Chr5 | 14 | 57,604 | 5,791 | 8,014,186 | 0.024 | 9.947 |
| Chr6 | 15 | 36,160 | 4,147 | 5,526,671 | 0.041 | 8.720 |
| Chr7 | 13 | 45,534 | 6,527 | 8,917,834 | 0.029 | 6.976 |
| Chr8 | 18 | 42,637 | 6,118 | 8,749,898 | 0.042 | 6.969 |
| CP | 7 | 12,034 | 130 | 144,093 | 0.058 | 92.569 |
| MT | 9 | 31,622 | 193 | 291,395 | 0.028 | 163.845 |
| Sum | 156 | 469,600 | 51,317 | 72,150,072 | 0.033 | 9.151 |

**B**

| Pair number | Column pair | Correlation test | Correlation coefficient | Sample size | T-test p-value |
|---|---|---|---|---|---|
| 1 | **BC** | Pearson | 0.597 | 11 | 0.053 |
| 2 | **BD** | Spearman | 0.670 | 11 | 0.024 |
| 3 | **BE** | Spearman | 0.688 | 11 | 0.019 |
| 4 | **BG** | Spearman | -0.661 | 11 | 0.027 |
| 5 | CD | Spearman | 0.873 | 11 | 0.001 |
| 6 | CE | Spearman | 0.891 | 11 | 0.000 |
| 7 | DE | Spearman | 0.991 | 11 | 0.000 |
| 8 | DF | Spearman | 0.018 | 11 | 0.958 |
| 9 | EF | Spearman | 0.045 | 11 | 0.894 |
| 10 | EG | Spearman | -0.636 | 11 | 0.035 |

**Supplementary Figure S56**. Distribution of 156 chimeric peptides, 469,600 chimeric protein models, and 51,317 non-microRNA transcripts in different chromosomal locations. A, heatmaps of the relationships between the distributions. Chr0, unmapped loci. Chr1 to Chr8, Chromosome 1 to Chromosome 8. CP, chloroplasts. Mt, mitochondria. Cells with blue fill show not sums but ratios of sums according to the formulas indicated in the header. B, a heatmap of correlation coefficients associated with the distributions shown in A. Data in columns B, C, and F are normally distributed. Columns D, E, and G contain skewed data. The relationships BC, BD, BE, and BG are non-monotonous, which makes their analysis with conventional tools unreliable. The t-test for a correlation coefficient was used to assess the significance of each coefficient. P-values above 0.05 are highlighted with light red. Surprisingly, there is no significant correlation between B and C. Likewise, the proportions of chimeric protein models that were supported with MS-peptides per chromosome (column F) do not correlate with any other independent column in this table. The data suggest the existence of location-specific determinants of chimeric protein abundance.

**Supplementary Figure S57**

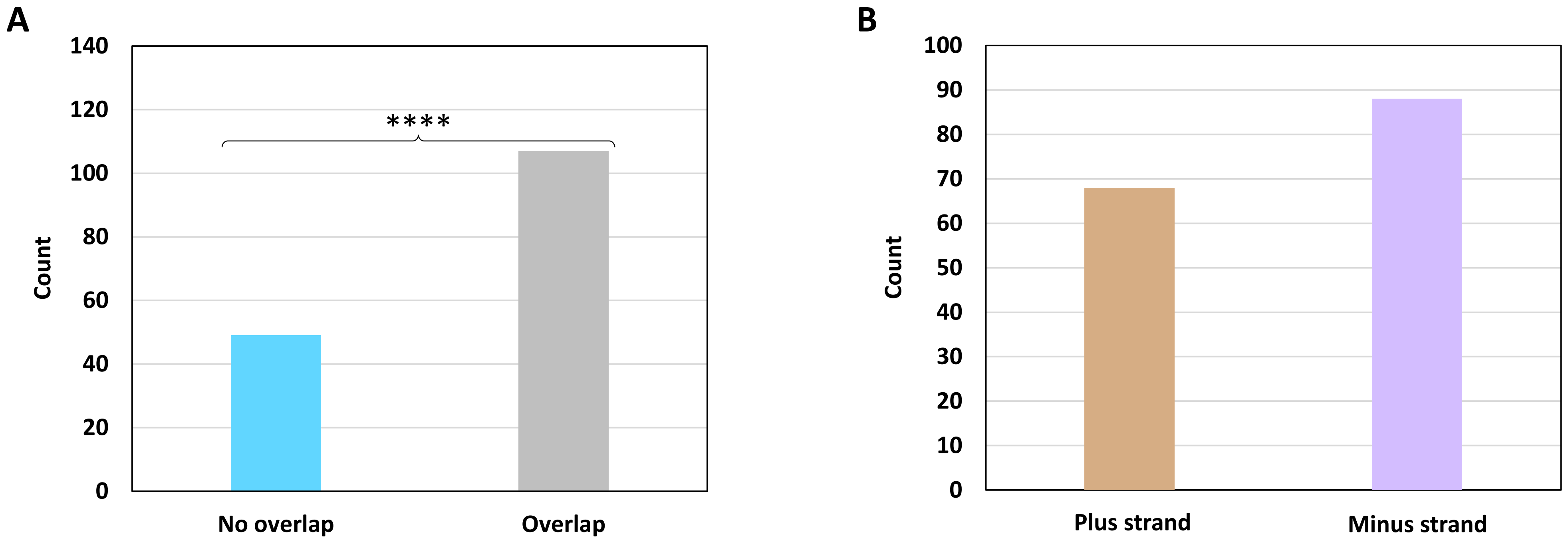


**Supplementary Figure S57**. Distribution of 156 chimeric MS peptides depending on whether their respective genes overlap with other genomic loci (A) or depending on the genomic DNA strand of the genes (B). The distribution is significantly different from the uniform distribution in A (the chi-square test for homogeneity, p = 3.422E-06). Only ca. one third of chimeric peptides correspond to genes that have no overlap with other loci. There is a trend toward more chimeric MS peptide-producing genes located on the minus strand.

### Supplementary Figure S58

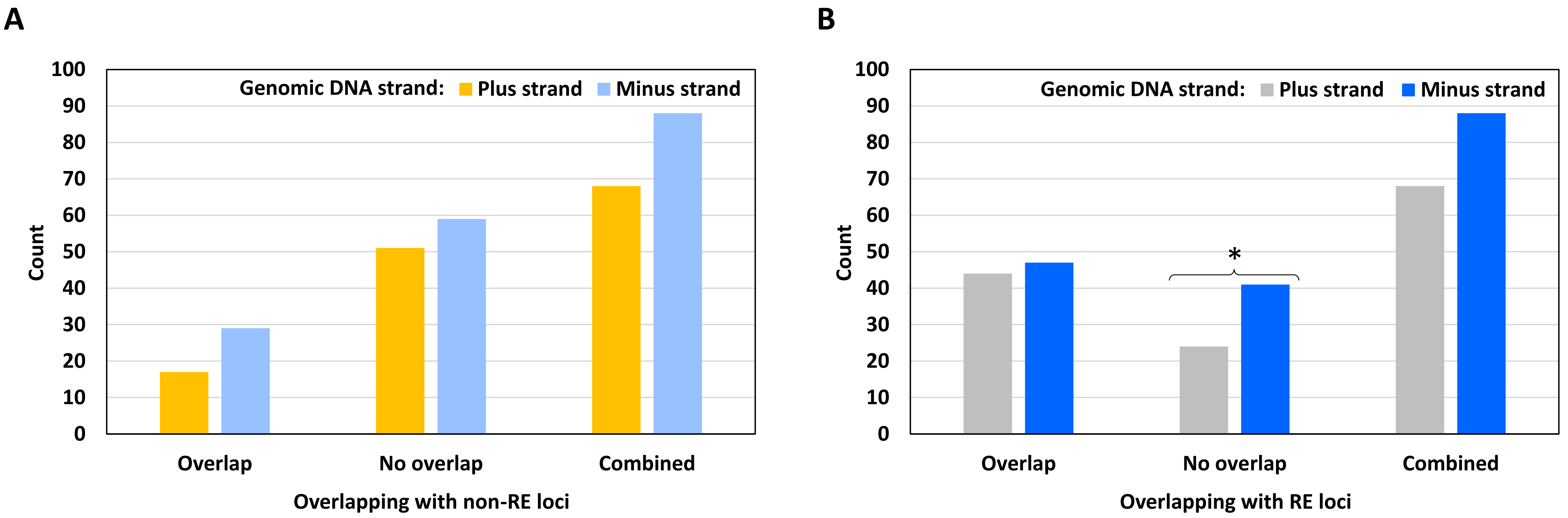


**Supplementary Figure S58**. Distribution of 156 chimeric MS peptides in different genomic DNA strands depending on whether their respective genes overlap with other loci. A, categories based on overlapping with loci not related to repeat elements (RE). B, categories based on overlapping with RE loci. There is no significant association between the DNA strand and the overlapping status. Within individual categories of overlapping, the distribution is significantly different from the uniform distribution in one case indicated with an asterisk (the chi-square test for homogeneity, $p = 0.035$). Significantly fewer chimeric peptides derive from genes that do not overlap with RE loci in the plus strand of genomic DNA (B). Note the opposite trend in the proportions of overlapping and non-overlapping genes in A and B.

### Supplementary Figure S59

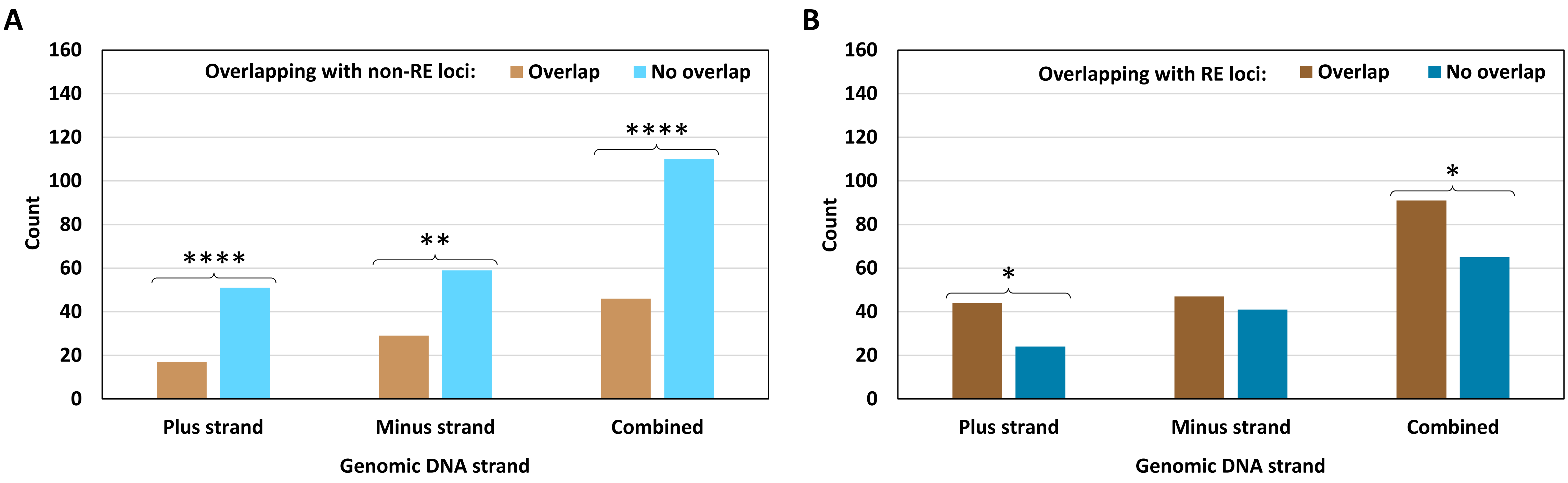


**Supplementary Figure S59**. Distribution of 156 chimeric MS peptides in different genomic DNA strands depending on whether their respective genes overlap with other loci. A, categories based on overlapping with loci not related to repeat elements (RE). B, categories based on overlapping with RE loci. There is no significant association between the DNA strand and the overlapping status. Within individual strands and for the strands combined, the distribution is significantly different from the uniform distribution in five cases indicated with asterisks. P-values of the chi-square test for homogeneity are as follows: A, plus strand, 3.738E-05; A, minus strand, 0.001; A, combined, 2.990E-07; B, plus strand, 0.015; B, combined, 0.037. A clear inverse relationship exists for chimeric peptides that come from genes overlapping with other loci: significantly fewer peptides correspond to genes that overlap with non-RE loci and more peptides derive from genes overlapping with RE loci, regardless of the DNA strand.

**Supplementary Figure S60**

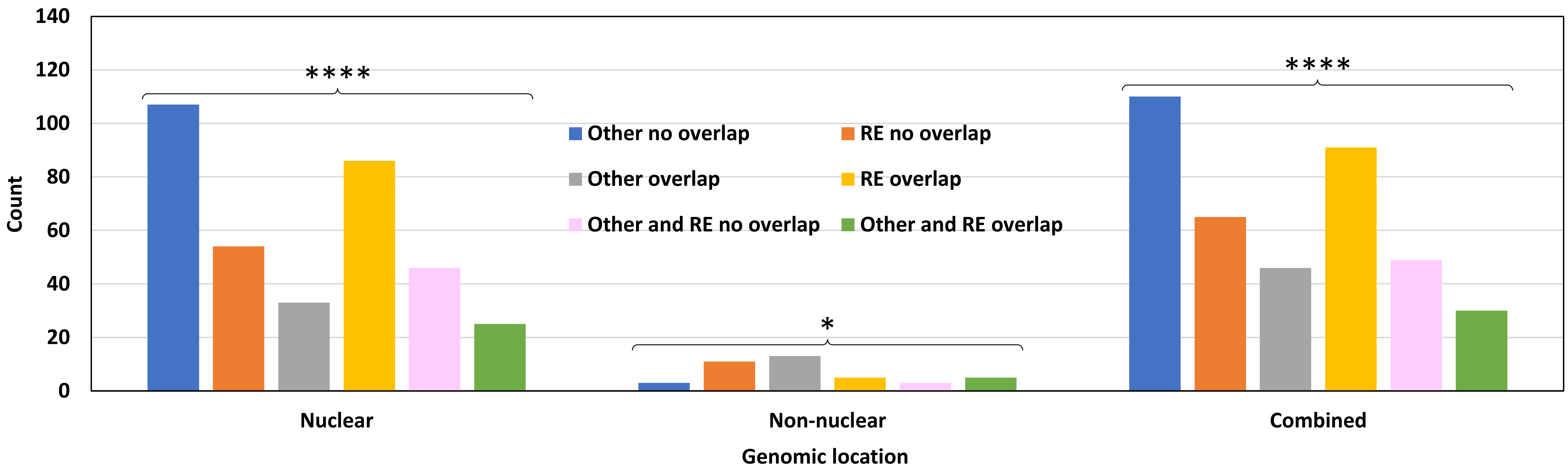


**Supplementary Figure S60**. Distribution of 156 chimeric MS peptides in two categories based on the genomic location of their genes and the overlap status. Per-category counts are not additive. “Other” refers to loci not related to repeat elements (RE). Interpretation example: “RE no overlap” combines chimeric peptides derived from genes that overlap with no RE loci regardless of the overlap with non-RE loci. “Nuclear”: Chromosome 0 (unmapped loci) to Chromosome 8. “Non-nuclear”: chloroplasts and mitochondria. “Combined”: nuclear and non-nuclear together. The chi-square test shows significant association between the genomic location and the overlapping status (p-value = 1.087E-05). Within individual genomic locations and the combined group, the distribution is significantly different from the uniform distribution in all three categories. P-values of the chi-square test for homogeneity are as follows: nuclear, 3.732E-17; non-nuclear, 0.018; combined, 1.177E-13. Nuclear genes mostly overlap with RE but not non-RE loci. Chloroplast and mitochondrial genes have the opposite trend: they mostly overlap with non-RE loci but not RE loci. This may be a general feature of the *M. truncatula* genome.

**Supplementary Figure S61**

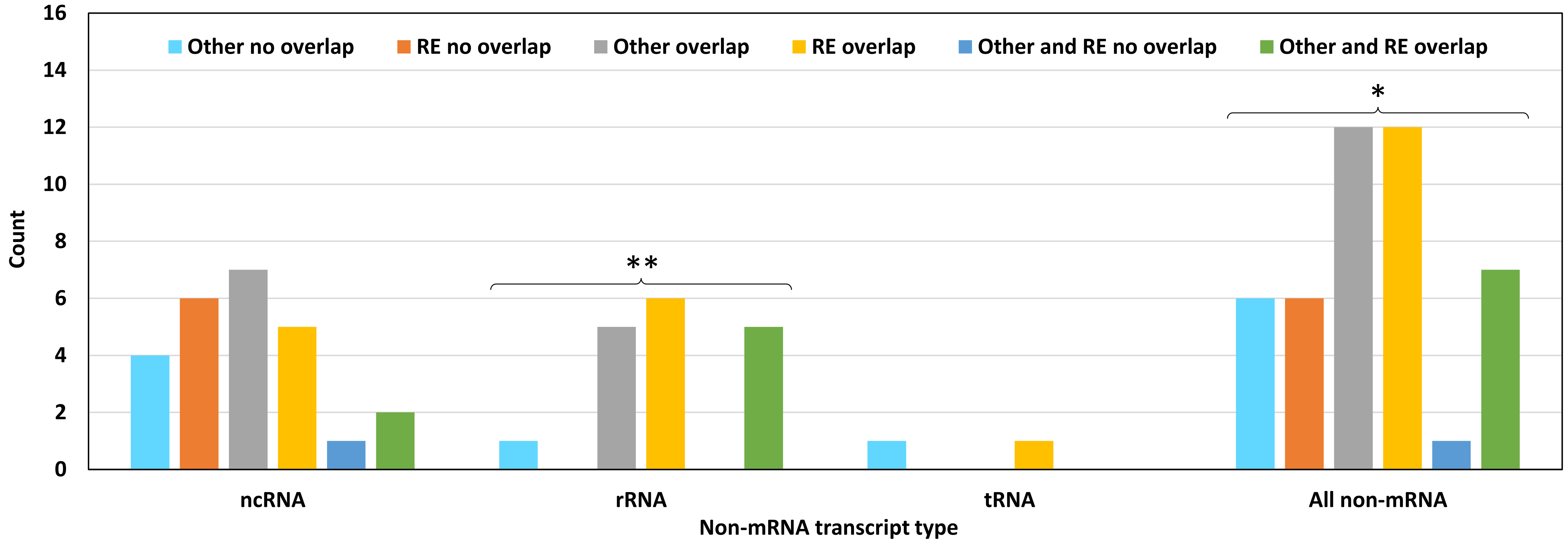


**Supplementary Figure S61**. Distribution of 18 chimeric MS peptides produced by non-mRNA transcripts in six categories depending on whether their genes overlap with different loci on genomic DNA. Per-category counts are not additive. “Other” refers to loci not related to repeat elements (RE). There is no significant association between the non-mRNA transcript type and the overlapping status. Within individual transcript types and for all non-mRNA types combined, the distribution is significantly different from the uniform distribution in two cases indicated with asterisks (rRNA, the Exact Multinomial Test, p = 0.006; combined, the chi-square test for homogeneity, p = 0.036). The dataset of 156 sequences contains six chimeric peptides derived from five rRNA genes. These five genes overlap with RE-like loci, and most of them also overlap with non-RE loci. This feature may be common in the *M. truncatula* genome, although rRNA loci with no overlap are present too.

### Supplementary Figure S62

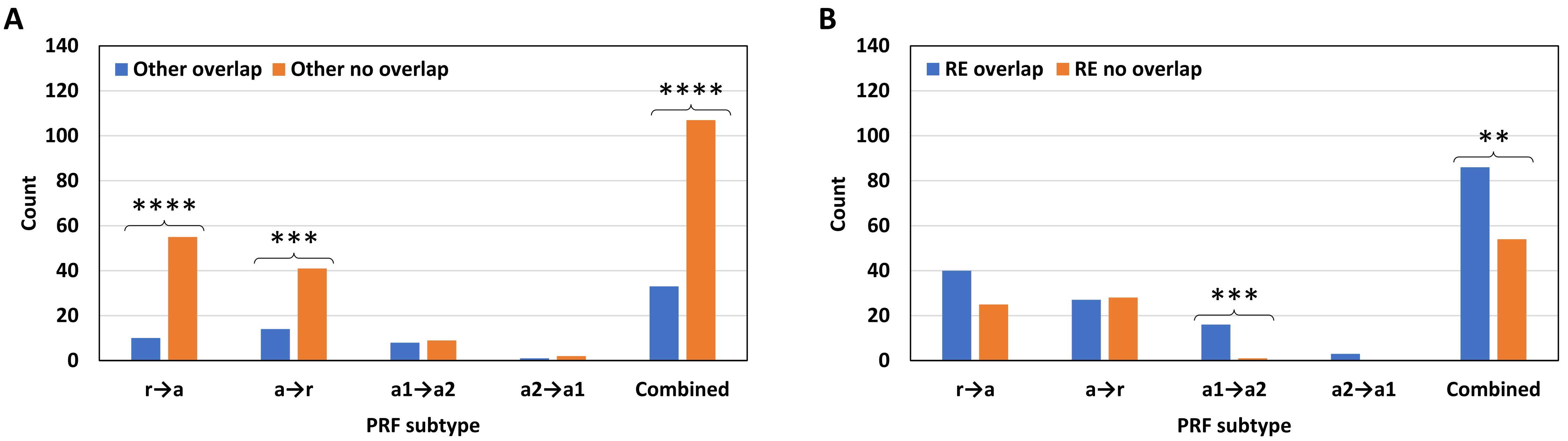


**Supplementary Figure S62**. Distribution of 140 chimeric MS peptides produced by nuclear transcripts in four categories of PRF depending on whether their genes overlap with different loci. "Other" refers to loci not related to repeat elements (RE). There is significant association between the PRF subtype and the overlapping status (the Fisher's exact test with "Combined" excluded, A, p = 0.033; B, p = 0.019). Within individual PRF subtypes and for all nuclear genes combined, the distribution is significantly different from the uniform distribution in five cases indicated with asterisks. P-values of the chi-square test for homogeneity are as follows: A, r→a, 2.384E-08; A, a→r, 0.0003; A, combined, 3.997E-10; B, a1→a2, 0.0003; B, combined, 0.007. The ratios are different for specific PRF subtypes and depend on the type of overlapping loci. Non-nuclear transcripts that produce putative chimeric proteins were excluded from this analysis because half of them are non-mRNA transcripts (no PRF subtypes r→a and a→r). Note that the general trends observed in A and B are absent from the PRF subtypes a1→a2 and a→r, respectively.

**Supplementary Figure S63**

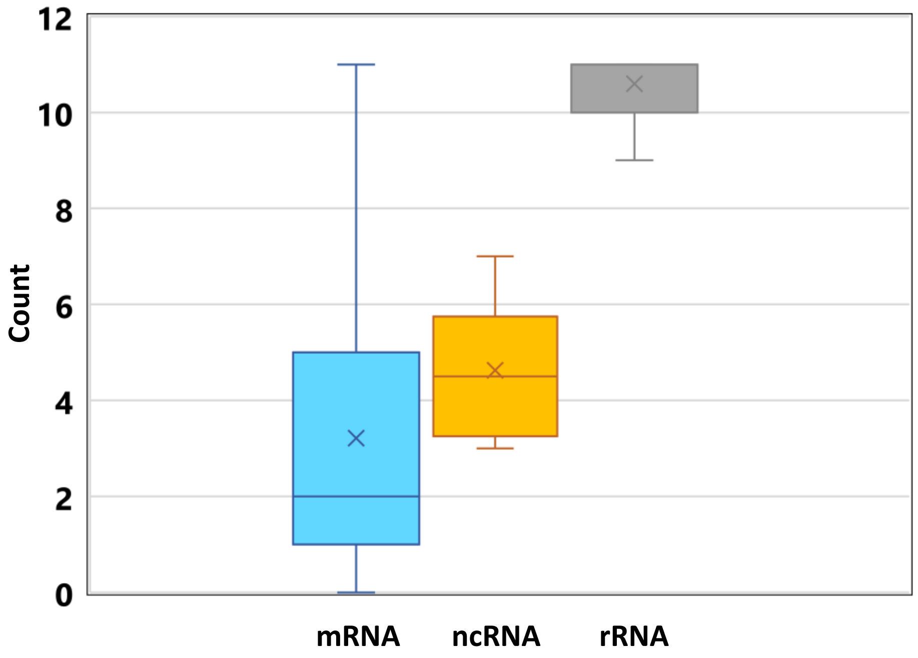


| | All types | mRNA | ncRNA | rRNA |
|---|---|---|---|---|
| mean | 4.0 | 3.2 | 4.6 | 10.6 |
| median | 3.0 | 2.0 | 4.5 | 11.0 |
| SD | 3.1 | 2.6 | 1.4 | 0.9 |
| n | 65.0 | 52.0 | 8.0 | 5.0 |
| min | 0.0 | 0.0 | 3.0 | 9.0 |
| max | 11.0 | 11.0 | 7.0 | 11.0 |

**Supplementary Figure S63**. Boxplots that summarize counts of sequences with shared subjects in the BLASTN analysis of 145 transcripts based on the *M. truncatula* transcriptome. tRNA is not included because there is only one tRNA sequence in the dataset. Interpretation example: Among 52 mRNA transcripts that have shared BLASTN subjects with other transcripts in the dataset, each mRNA transcript has shared subjects with 3.2 other sequences, on average. However, the maximal count of sequences with shared subjects is 11 for this RNA type and also for rRNA. Two mRNA transcripts in the dataset had no shared subjects with other sequences despite having meaningful alignments with other sequences (thus, zero is shown as a minimal value for mRNA). Visibly more sequences with shared subjects are found among rRNA transcripts, followed by ncRNA. The inset shows descriptive statistics of the data (SD, standard deviation; n, sample size). Cumulative distribution functions of the three samples are significantly different from each other. Bonferroni-adjusted p-values of Kolmogorov–Smirnov test are as follows: mRNA vs ncRNA, 0.016; mRNA vs rRNA, 1.505E-05; ncRNA vs rRNA, 0.002.

**Supplementary Figure S64**

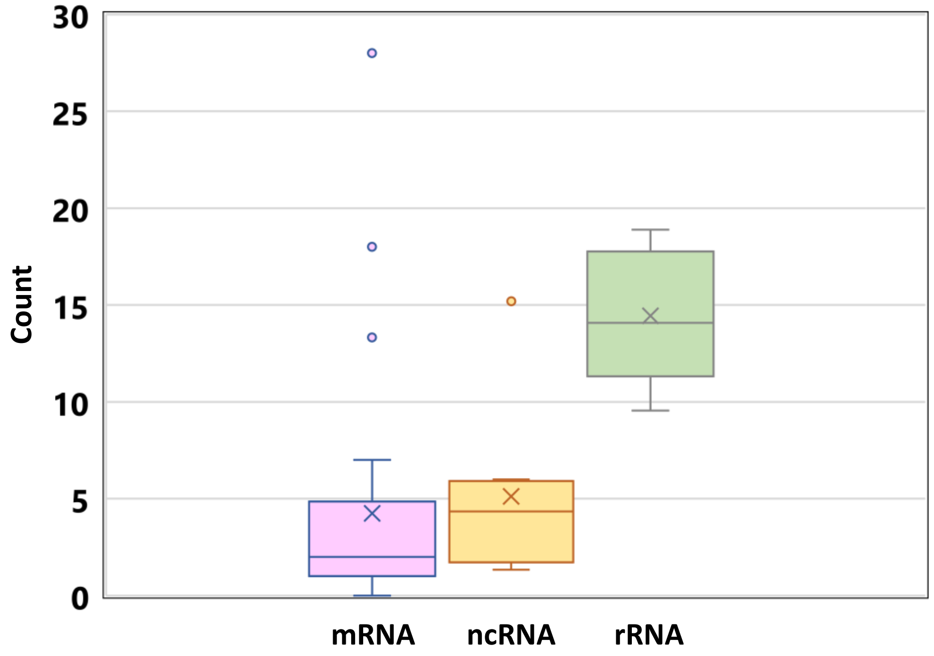


| | All types | mRNA | ncRNA | rRNA |
|---|---|---|---|---|
| mean | 5.1 | 4.3 | 5.1 | 14.4 |
| median | 3.0 | 2.0 | 4.3 | 14.1 |
| SD | 6.3 | 6.1 | 4.5 | 3.6 |
| n | 65.0 | 52.0 | 8.0 | 5.0 |
| min | 0.0 | 0.0 | 1.3 | 9.5 |
| max | 28.0 | 28.0 | 15.2 | 18.9 |

**Supplementary Figure S64**. Boxplots that summarize mean numbers of shared subjects in the BLASTN analysis of 145 transcripts based on the *M. truncatula* transcriptome. tRNA is not included because there is only one tRNA sequence in the dataset. Interpretation example: Among 52 mRNA transcripts that have shared BLASTN subjects with other transcripts in the dataset, each mRNA transcript has 4.3 shared subjects with other sequences, on average. However, the maximal number of shared subjects is 28 for this RNA type, which is higher than for rRNA and ncRNA. Two mRNA transcripts in the dataset had no shared subjects with other sequences despite having meaningful alignments with other sequences (thus, zero is shown as a minimal value for mRNA). rRNA transcripts have visibly more shared subjects, followed by ncRNA. The inset shows descriptive statistics of the data (SD, standard deviation; n, sample size). Cumulative distribution functions of the three samples are significantly different from each other in two cases. Bonferroni-adjusted p-values of Kolmogorov–Smirnov test are as follows: mRNA vs rRNA, 1.805E-04; ncRNA vs rRNA, 0.014.

**Supplementary Figure S65**

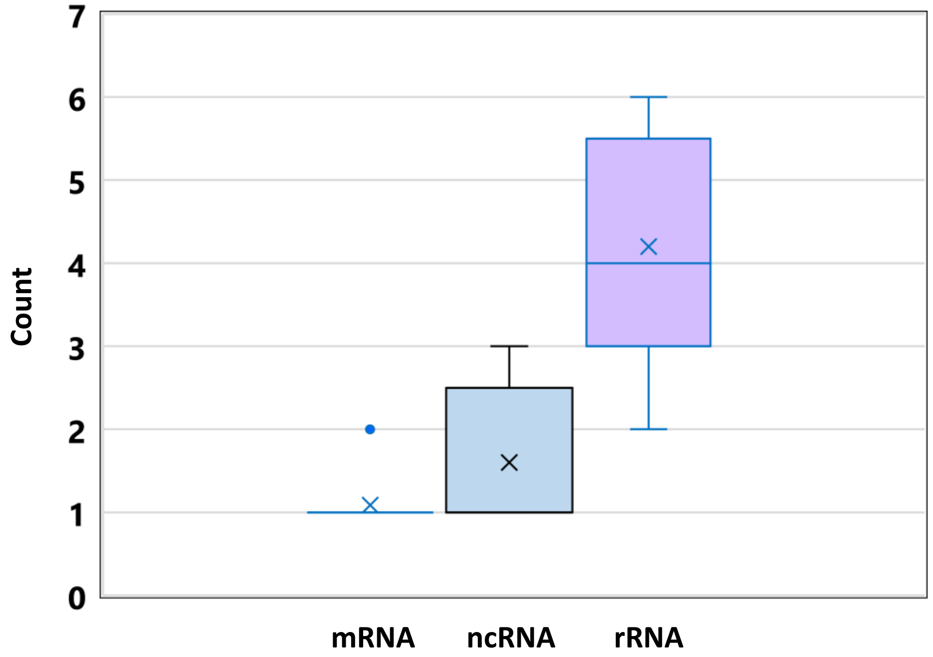


| | All types | mRNA | ncRNA | rRNA |
|---|---|---|---|---|
| mean | 1.6 | 1.1 | 1.6 | 4.2 |
| median | 1.0 | 1.0 | 1.0 | 4.0 |
| SD | 1.3 | 0.3 | 0.9 | 1.5 |
| n | 33.0 | 23.0 | 5.0 | 5.0 |
| min | 1.0 | 1.0 | 1.0 | 2.0 |
| max | 6.0 | 2.0 | 3.0 | 6.0 |

**Supplementary Figure S65**. Boxplots that summarize per-transcript counts of similar sequences in the dataset, based on meaningful pairwise nucleotide alignments. tRNA is not included because there is only one tRNA sequence in the dataset. Interpretation example: Among 23 mRNA transcripts that meaningfully align with other transcripts in the dataset, each mRNA transcript aligns with 1.1 other sequences, on average. The maximal per-transcript count of similar sequences is 2 for this RNA type, which is lower than for ncRNA and rRNA. rRNA transcripts have visibly more similar sequences, on average. The difference between mRNA and ncRNA is very small in this respect. The inset shows descriptive statistics of the data (SD, standard deviation; n, sample size). Cumulative distribution functions of the mRNA and the rRNA samples are significantly different from each other. Bonferroni-adjusted p-value of Kolmogorov–Smirnov test is 6.411E-4.

**Supplementary Figure S66**

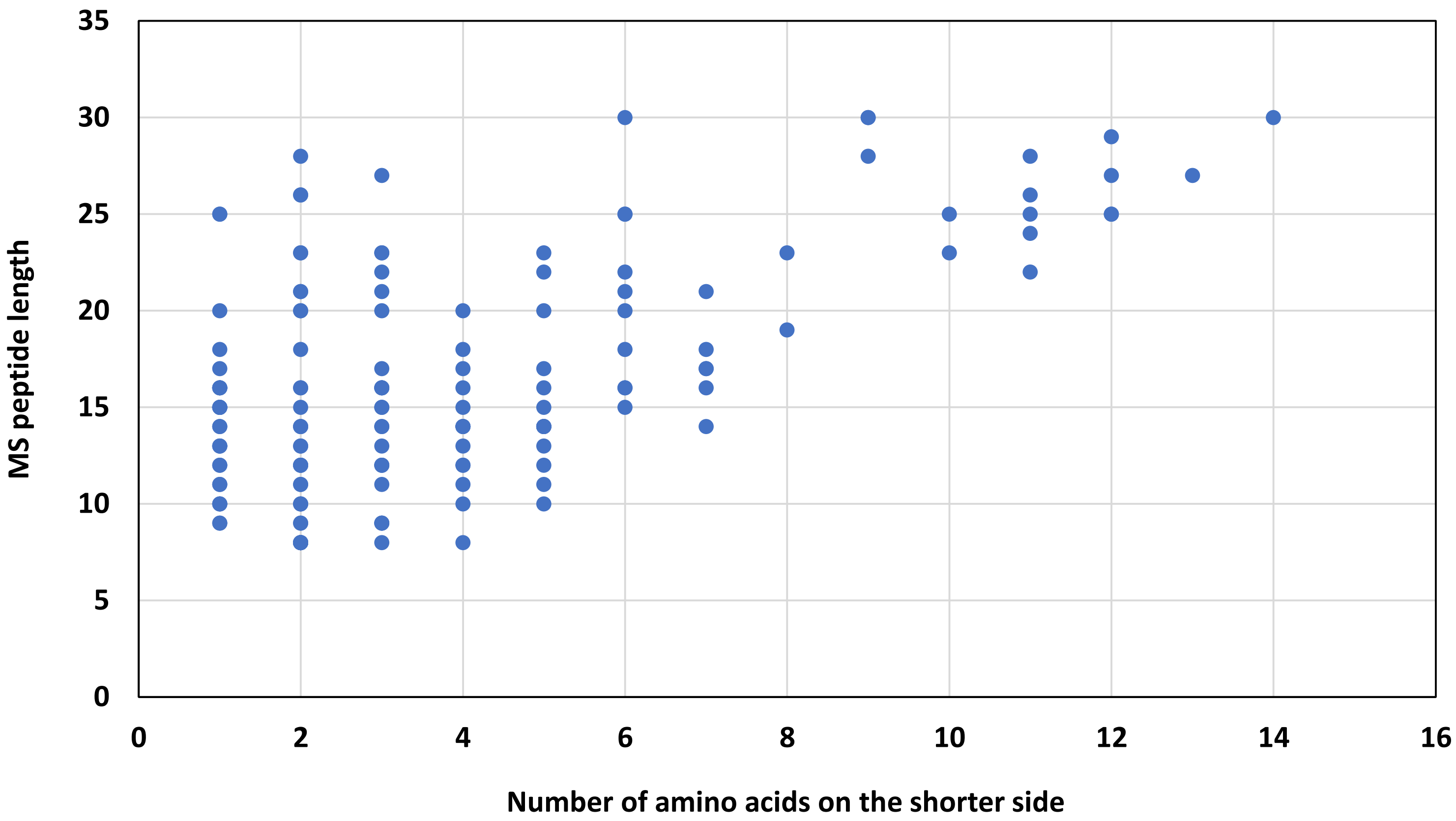


**Supplementary Figure S66**. A scatter chart of the relationship between the number of amino acids on the shorter side of 156 chimeric MS peptides and the length MS peptides. The graph illustrates that shorter peptides tend to have fewer amino acids on their shorter sides. Here, we define the shorter side of an MS peptide as a region that is aligned with fewer amino acids from a corresponding chimeric protein immediately upstream or downstream of the frameshifting site. The more amino acids are found on the shorter side of an MS peptide, the higher the chance that this MS peptide is specific to its chimeric protein.

**Supplementary Figure S67**

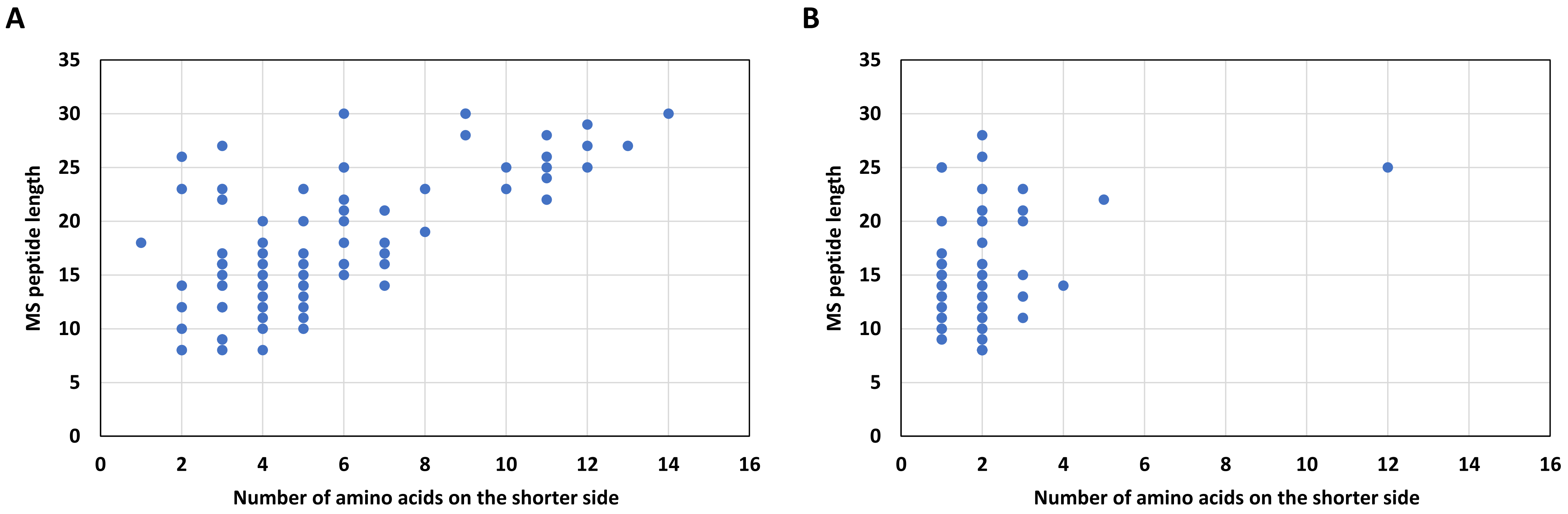


**Supplementary Figure S67**. Scatter charts of the relationship between the number of amino acids on the shorter side and the length of 156 chimeric MS peptides depending on whether the peptides could potentially originate from unknown splicing events. A, 87 MS peptides that could not result from uncharacterized splicing of their transcripts (peptides truly unique to PRF events). B, 69 MS peptides that could potentially originate from unknown splicing events that operate on their transcripts (peptides conditionally unique to PRF). Graph A illustrates that truly unique peptides can be of very short lengths (eight amino acids) and can have as few as one amino acid on their shorter sides. Graph B illustrates that even long peptides with more than four amino acids on their shorter sides can be conditionally unique.
