## Supplementary material for "Discovery of diverse chimeric peptides in a eukaryotic proteome sets the stage for the experimental proof of the mosaic translation hypothesis": Supplementary results.pdf

**4.5.1. The occurrence of PRF values is non-random across categories based on RNA type, chromosomal location, genomic location, DNA strand, splicing, PRF type, and PRF position relative to the CDS**

The proportions of observed PRF values -2, -1, +1, and +2 in the dataset are not significantly different from the expected proportions. However, a combined group of PRF values with a negative sign (-1 and -2) is significantly overrepresented in the detected chimeric peptides (Figure 2). PRF value -2 is the most abundant in the dataset, although its occurrence is not significantly different from the expected count.

With regard to RNA types, PRF value -1 is missing from the group of ncRNA transcripts but is significantly overrepresented in rRNA transcripts (five out of six). Likewise, a chimeric peptide derived from the only tRNA transcript in our data has PRF value -1, which is in line with the close evolutionary relationship between rRNA and tRNA, tRNA being a building block of rRNA (Root-Bernstein and Root-Bernstein, 2015, 2016, 2019; Caetano-Anollés and Caetano-Anollés, 2016; de Farias et al., 2016). The transcript type and the PRF value are significantly associated (Figure 3).

When chimeric peptides are grouped by the chromosomal location of their transcripts, we notice a significant association between the chromosomal location and the PRF value. PRF value +1 exhibits distinct occurrence. It is missing from transcripts on chromosome 2 and is visibly underrepresented in transcripts from chromosomes 1, 3, and 7. Chloroplasts and mitochondria are the only sources of transcripts in which PRF value +1 is overrepresented (Supplementary Figure S11). This is further supported by the grouping of transcripts into nuclear and non-nuclear ones. PRF events with value + 1 are significantly underrepresented in nuclear transcripts and visibly overrepresented in non-nuclear transcripts. This opposite trend means that transcripts with prokaryotic origin (chloroplasts and mitochondria) are different from truly eukaryotic transcripts in the distribution of PRF values and in the occurrence of PRF value +1 in particular (Figure 4).

The overrepresentation of PRF value +1 in non-nuclear transcripts is also evident from the following significant observation. The counts of chimeric peptides with PRF value +1 that originate from the plus strand of genomic DNA (nuclear and non-nuclear combined) is less than half of the counts from the minus strand (Supplementary Figure S12). This is probably because the abundance of non-nuclear chimeric peptides is significantly lower (seven-fold) for loci located on the plus strand of genomic DNA (Supplementary Figure S13). Genes of all chloroplast chimeric peptides and the majority of mitochondrial ones are located on the minus strand (Supplementary Dataset S1). This is not a common feature of the *M. truncatula* genome as can be judged from the relatively equal distribution of plus-strand and minus-strand genes in the non-nuclear genomes (the *M. truncatula* genome browser, Pecrix et al., 2018).

While studying the relationship between various parameters summarized in Supplementary Dataset S1, we have noticed a curious link between the PRF value, chromosomal location, and splicing. We found 26 chimeric models for which corresponding MS peptides cover annotated splicing sites. In other words, PRF sites in corresponding transcripts are located within a few nucleotides from splicing sites or coincide with them. Similar to the whole set of 156 chimeric peptides (Supplementary Figure S11), in this group, the association between the PRF value and the chromosome is highly significant, with a noticeable overrepresentation of PRF values -1 and -2 on chromosomes 3 and 8, respectively (Supplementary Figure S14). Furthermore, it turned out that “short” PRF events with absolute value 1 (-1 and +1) are clearly more frequent on chromosomes 1 to 4, while the remaining chromosomes 5 to 8 are associated exclusively with “long” PRF events, that is PRF values -2 and +2 (Supplementary Figure S15).

The next observation supporting the non-randomness of our dataset concerns highly significant association between PRF value and PRF type (Figure 5). The definition of forward frames (1 to 3) depends solely on the number of nucleotides between the beginning of a transcript and a feature of interest in its sequence. Because the 5’-UTRs are relatively tolerant to insertions and deletions compared to the CDSs, the frame is not a parameter expected to have any biological relevance. Surprisingly, our data clearly indicate that frame 3 stands out with regard to the PRF value. Namely, backward PRF events (-1 and -2) are significantly more frequent for the shift from any frame to frame 3. The fold-difference between forward and backward PRF events is numerically large (two and a half- to nearly four-fold), which makes it highly significant for individual PRF types 1→3 and 2→3 (Figure 5). To clarify if PRF value and type depend on the position of PRF sites, we correlated these parameters with distances of PRF sites from either end of a transcript, and also with transcript sizes. This analysis revealed only non-significant trends: (1) PRF sites of types 1→3 and 2→3 tend to be closer to the beginning of a transcript; (2) transcripts in which such PRF sites are found tend to be shorter (results not shown). Thus, it is difficult to explain the unusual features of PRF types 1→3 and 2→3.

Next, we looked into the distribution of PRF values within chimeric peptide categories based on the position of an involved altORF relative to a refORF of a transcript (if any). This analysis revealed significant predominance of PRF value -2 in one category: altORFs that start in the 5’-UTR and end in the refORF (Supplementary Figure S16). It may be remembered that PRF value -2 has the highest frequency in the whole dataset (Figure 2).

### 4.5.2. PRF values exhibit association with predicted folding structures and the PRF location relative to alpha-helices and beta-sheets

In the course of our analysis, we also predicted three-dimensional structures of the modeled portions of MS-supported chimeric peptides using ColabFold v1.5.2 (Mirdita et al., 2022). More than half of the models exhibit the presence of alpha-helices without beta-sheets (ca. 57%). Models with no structure constitute about one-quarter of the dataset (ca. 27%). A small group of models (ca. 15%) contain beta-sheets (Supplementary Figure S17). Different PRF values are approximately uniformly represented in these three folding groups (Supplementary Figure S18). However, when the counts of backward (-1 and -2) and forward (+1 and +2) PRF events are compared, significantly more PRF values with the negative sign are found in chimeric peptides that fold into alpha-helices (Figure 6A). Likewise, when PRF values are grouped into “short” (-1 and +1) and “long” (-2 and +2) categories, it turns out that beta-sheet structures are three times more abundant in the combined PRF category with the absolute value 2 (Figure 6B). This is in line with what is known about the determinants of PRF events (Riegger and Caliskan, 2022). Namely, apart from the well-characterized effect of RNA folding on the PRF definition, the protein folding or sequence features associated with specific folding structures, may also play role in the determination of PRF location and value.

Next, we took a close look into the location of PRF sites relative to alpha-helices and beta-sheets (Supplementary Figures S19-S24, Supplementary Datasets S7-S10). The majority of chimeric peptides that form alpha-helices (80%) harbor PRF sites within alpha-helices, regardless of the PRF value (Supplementary Figure S19). In line with the significantly higher proportion of backward PRF values in chimeric peptides with alpha-helices, PRF values with the negative sign are also significantly overrepresented in this majority-subgroup (Supplementary Figure S20). Further, we observed that ca. 63% of PRF sites located inside alpha-helices are in the middle of the structure, with ca. one-third of PRF sites occurring at either end of the helix, regardless of the PRF value (Supplementary Figure S21). Grouping of chimeric peptides in this category by PRF direction and length reveals that backward PRF events are significantly more abundant (ca. two-fold) in the middle of alpha-helices (Supplementary Figure S22A). In contrast, the proportion of “short” PRF events (-1 and +1) is significantly larger in chimeric peptides with PRF sites toward the end of an alpha-helix (Supplementary Figure S22B). Differential occurrence of PRF sites with contrasting characteristics in different locations points to a possibility that the relative position of PRF sites inside alpha-helices may have biological relevance.

In contrast to chimeric peptides with alpha-helices, nearly half of PRF sites in chimeric peptides with beta-sheets are located between beta-sheets (Supplementary Figure S23). There appears to be a significant association between the location of PRF sites and the direction of PRF when PRF values of the same sign are combined (Supplementary Figure S24A). All PRF sites located before beta-sheets have the negative sign whereas all PRF sites located after beta-sheets have the positive sign (Supplementary Figure S24A). These observations indicate that the ability of chimeric peptides to fold into alpha-helices or beta-sheets influences PRF characteristics differently.

In our dataset, 42 chimeric peptide models have no predicted structure (Supplementary Figure S17). This, however, does not imply the lack of functional relevance of this group. Intrinsically disordered proteins are ubiquitous in all organisms and may have important biological functions (Tesei et al., 2024). Although such proteins are known to have low conservation, in our study, chimeric models with no structure were generated from both MS-supported altORFs (eight) and conserved altORFs (34).

#### 4.5.3. PRF values depend on the diversity of amino acids found immediately before and after the PRF sites

Our next analytic domain focused on the search for relationships between amino acids immediately before and after the PRF site on one hand and PRF values on the other hand. The dataset contains chimeric peptides with all 20 usual proteinogenic amino acids flanking the PRF site. For each amino acid, the distribution before and after is not significantly different from the uniform distribution (Supplementary Figure S25). However, when chimeric peptides are grouped by the PRF value, the association is significant in two PRF categories: +1 and -2. The only amino acid significantly more abundant (nearly four-fold) immediately after the PRF site is leucine (L) in chimeric peptides with -2 frameshifts (Supplementary Figure S26).

In the position immediately before the PRF site, the distribution of two amino acids, polar methionine (M) and non-polar asparagine (N), is significantly different from the uniform distribution in four groups based on PRF values (Supplementary Figures S27 and S28). Namely, seven out of eight methionine amino acids in this position are found in -1 frameshifts. No methionine (M) is present before the PRF site in chimeric peptides with PRF values +1 and -2. The non-polar amino acid asparagine shows the opposite pattern: immediately before the PRF site, asparagine (N) is found exclusively in chimeric peptides with +1 and -2 frameshifts. Overall, there is significant association between the amino acid before the PRF site and the PRF value. Nine amino acids are not found in certain PRF categories (C, G, H, M, N, Q, V, W, and Y), six of which are missing from the chimeric peptides with PRF value +2 (Supplementary Figures S27 and S28). The distribution of 20 amino acids is visually unique for each PRF value and may be different from the uniform distribution for PRF categories -1 and +2. However, statistical significance of these differences is hard to estimate due to low counts and many categories (Supplementary Figure S28). The most abundant amino acids vary by PRF group: methionine (M) for -1, isoleucine (I) and valine (V) for +1, lysine (K) for -2, and leucine (L) for +2. Intriguingly, four out of these five amino acids are non-polar. It is interesting to compare these amino acids with the combined list of the most abundant amino acids in the position before the PRF site (Supplementary Figure S29). Only three amino acids (L, I, and K) constitute 28% of the whole dataset. Together with other four amino acids (T, P, R, and V), they comprise half of the dataset. Remarkably, methionine (M) and asparagine (N) prominent in certain PRF groups are not the most abundant amino acids upstream of the PRF site.

Similar significant observations were made on amino acids immediately after the PRF site. In this position, three amino acids, lysine (K), leucine (L), and serine (S), exhibit distribution significantly different from the uniform distribution in four PRF groups (Supplementary Figures S30 and S31). Lysine (K) is overrepresented in -2 frameshifts and absent in +1. Leucine (L) is mostly in -2 frameshifts, while serine (S) appears only in “short” frameshifts, namely, -1 and +1. Like in the case with the upstream position, there is significant association between the amino acid after the PRF site and the PRF value. Eleven amino acids are not found in certain PRF categories (C, D, F, H, I, K, N, Q, S, V, and W), six of which are missing from the PRF category with value +1 (Supplementary Figures S30 and S31). Each PRF value has a unique amino acid distribution, particularly -2, though exact p-values cannot be calculated due to low counts and many categories (Supplementary Figure S31). The most abundant amino acids were not the same for each PRF group when we considered the upstream position. This is somewhat different for the downstream position: polar amino acids glycine (G) and serine (S) are found in the largest number at this position in chimeric peptides with PRF value -1. Glycine (G) is also most frequent in the PRF category with value +2. Chimeric peptides with PRF value +1 have three equally abundant amino acids at this position: non-polar amino acids alanine (A) and leucine (L) together with a negatively charged glutamic acid (E). Leucine (L) is predominant also at the downstream position of chimeric peptides with PRF value -2. Leucine (L) has already been mentioned as outstanding in the comparison between the upstream and downstream positions, and also in the context of the non-uniform distribution of amino acids in the upstream position (Supplementary Figures S26-S28). Similar to amino acids in the position before the PRF site, at the downstream position, only three amino acids (L, G, and K) constitute 29% of the whole dataset. Together with three more amino acids (R, E, and P), they make up half of the dataset (Supplementary Figure S32). Thus, two non-polar amino acids leucine (L) and proline (P) together with two positively charged amino acids lysine (K) arginine (R), are the most commonly found amino acids at both positions flanking the PRF sites (compare Supplementary Figures S29 and S32). Leucine (L) is also the amino acid most commonly found in the reference proteome of *M. truncatula*, in its altProts, and in chimeric models (results not shown).

Naturally, in the next step, it was important to understand whether the amino acids commonly found immediately upstream or downstream of the PRF sites were also most abundant in *M. truncatula* proteins in general. To obtain a comprehensive view on frequencies of amino acids in this organism, we counted the occurrence of 20 usual proteinogenic amino acids in five different categories relevant to the goal of our study: (1) chimeric peptides, which are only the MS-supported parts of the modeled chimeric proteins; (2) modeled chimeric proteins, which include the MS-supported part but also a few more amino acids not covered by the MS peptide; (3) altProts involved in the formation of chimeric peptides; (4) refProts involved in the formation of chimeric peptides; and finally (5) the whole list of annotated proteins (refProts) found in *M. truncatula* regardless of their involvement in the production of chimeric peptides. This analysis showed that amino acids most abundant upstream or downstream of the PRF sites are not always found in the largest number in the annotated proteome. Amino acids C, H, W, and Y are consistently rare in all the five categories and also before and after the PRF sites. Leucine (L) is the most frequently found amino acid in all cases. In contrast, isoleucine (I) and lysine (K) have very high abundance in the upstream position but are moderately present in the remaining categories. However, according to the chi-square test for goodness-of-fit, the observed proportions of amino acids upstream or downstream of the PRF sites are not significantly different from the proportions expected based on the abundance in any of the five categories (results not shown). For individual amino acids, the differences are non-significant either (results not shown). This means that significantly non-uniform occurrence of some amino acids evident in Supplementary Figures S26-S28, S30, and S31 is associated only with different PRF categories, and there are no amino acids significantly overrepresented in all PRF categories combined.

As an extension of this experimental domain, we counted the occurrence of pairs formed by amino acids found immediately before and after the PRF sites. There are 118 such pairs. Most of them (ca. 77%) are found only once in the whole dataset. However, there are 19 amino acid pairs found twice and six pairs found three times each. Amino acid pairs AL and KR occur four and five times at the PRF sites, respectively (Supplementary Figure S33). We tested the distribution of 27 amino acid pairs found more than once in different categories based on the PRF values. This analysis revealed significant association between the amino acid pair and the PRF value (Supplementary Figure S34). The distribution of amino acid pair AL is significantly different from the uniform distribution. Four pairs, AL, EK, KK, and MG, are unique to a specific PRF category (found in at least three chimeric peptides each), and a different set of four, AL, EK, KK, and KR are unique to or most abundant in the PRF category -2 (Supplementary Figure S34).

Further, we noticed that six of 118 pairs are composed of repeated amino acids: GG, KK, LL, PP, RR, and VV. Intriguingly, all of them are found exclusively in backward PRF sites (-1 and -2). Three pairs, KK, LL, and PP are unique to PRF sites with value -2. Two pairs, RR and VV, are found only in -1 frameshifts. Amino acid pair GG is found once in a -1 PRF site and once in a -2 PRF site (Supplementary Figure S35A). The very low counts of these six repeated amino acid pairs speak against their importance in PRF events. However, their exclusive occurrence in backward PRF events may point to their biological relevance (Supplementary Figure S35B).

As it was the case with individual amino acids immediately before and after the PRF sites, we checked whether the frequencies of amino acid pairs were significantly different from the frequencies in the same five categories as described above: chimeric peptides, chimeric models, altProts and refProts involved in PRF, and all refProts regardless of their involvement in PRF. Out of 400 theoretically possible amino acid pairs, 282 are not found in the PRF sites but only 17 have zero-frequency in chimeric peptides. Only one amino acid pair, YW, is absent from all chimeric peptides and chimeric protein models. The remaining categories have non-zero occurrence of all 400 amino acids (results not shown). Surprisingly, 83 amino acid pairs are at least two-fold more abundant at PRF sites compared with their expected abundance based on frequencies in all refProts of *M. truncatula*. Moreover, 21 of them are at least five-fold overrepresented, and six pairs, FC, HQ, MG, MY, QW, and WG are overrepresented 10-19-fold compared to the expected counts based on frequencies in the annotated proteome. Despite these large fold-differences between observed and expected counts, none of them are statistically significant, probably because of the very small numerical values of observed and expected counts (results not shown).

Collectively, these observations point to a very large diversity of the immediate frameshifting sites with regard to their codon composition. As will be shown later, PRF sites in our dataset display almost no conservation signature within the *M. truncatula* transcriptome. At the same time, statistically significant deviations from the uniform occurrence of some amino acids in different PRF categories suggest the possibility of specific roles for certain codons and amino acids in the frameshifting process.

### 4.5.4. Chimeric peptides with specific PRF values have differential occurrence in 16 biological samples

We analyzed the distribution of 156 chimeric peptides in 16 biological samples from three MS-based studies (Marx et al., 2016; Shin et al., 2021; Castañeda et al., 2021) to determine correlations with different PRF categories. No chimeric peptides were detected in 28-day nodules and phloem under drought stress (Supplementary Figure S36). Seeds and 10-day nodules had the highest number of MS-supported chimeric peptides (35 and 27, respectively). Together with 17 chimeric peptides detected in 14-day nodules, they constitute 41% of the dataset. Nodule samples, combined, contributed 24% of chimeric peptides. Organs related to the reproduction and symbiotic nitrogen fixation are most represented in the dataset, which may point to the biological importance of chimeric peptides for these processes. Approximately 83% of chimeric peptides were found in a single sample. However, there are 15 chimeric peptides found in two samples, and 12 identified in three to nine samples (Supplementary Figure S37). Peptides in 10-day nodules often appeared in 14-day nodules, and those in 14-day nodules frequently occurred in seeds. Flowers also showed a notable overlap with nodules (Supplementary Figure S38). When examining PRF categories (Figure 7), the MS sample was found to be significantly associated with the PRF value. In seeds, buds, and root control, distribution of chimeric peptides based on PRF values is significantly different from the uniform distribution. In seed-derived chimeric peptides, PRF events with value -2 are more than two-fold overrepresented compared to -1 and +1 frameshifts and ca. three-fold more frequent compared to +2 PRF events. No chimeric peptides with PRF value +1 are found in buds. PRF events with value -1 are predominant in buds, flowers, leaves, stems, and 14-day nodules. Drought-stressed root samples contained only -2 and +2 PRF chimeric peptides, unlike other root samples. This observation probably reflects the specificity of certain chimeric peptides to particular environments. Namely, the same organ can presumably produce a different array of chimeric peptides under different conditions even when no stress is applied (compare sample R with sample RoC in Figure 7). Lastly, as an extension of this analysis, we followed the distribution of chimeric peptides in different samples depending on the chromosomal location of their respective genes. Chimeric peptides from chromosome 1 are predominant in nodules, seeds, flowers, and roots, suggesting "chimeric hotspots" on chromosome 1 involved in reproduction, nitrogen fixation, and root functions (Supplementary Figure S39).

### 4.5.5. Significant deviations from the randomness assumption with regard to PRF type, subtype, and location relative to the CDS

In our study, we refer to the type of shifting from one reading frame to another as PRF type. There are six possible PRF types, 1→2, 1→3, 2→1, 2→3, 3→1, and 3→2, all of which are found in approximately equal proportions in 156 chimeric peptides although numerically, shifts from frame 1 to other frames prevailed in the dataset (Supplementary Figure S40). Another PRF attribute, which we call PRF subtype here, indicates which ORFs are involved in the frameshifting. There are four possible PRF subtypes. PRF subtype r→a describes a frameshift that occurs from a refORF to an altORF. The opposite situation is designated as PRF subtype a→r. Two remaining PRF subtypes correspond to frameshifts between altORFs regardless of the presence of a refORF in a transcript. An altORF that starts earlier is called altORF1and the one that follows is referred to as altORF2. Accordingly, PRF subtype a1→a2 stands for a sequential frameshift from altORF1 to altORF2, whereas the opposite situation is designated as PRF subtype a2→a1. PRF subtypes r→a and a→r are approximately equally abundant in the dataset. PRF subtypes that involve no refORF are significantly less common, with PRF subtype a2→a1 being significantly less frequent compared to the sequential frameshift a1→a2 (Supplementary Figure S41A). Interestingly, PRF events that start from altORFs are statistically as abundant as those that start from refORFs (Supplementary Figure S41B). Since we do not know if chimeric peptides are fragments of longer mosaic proteins, it is not clear if this equality in abundance indicates the equality in translation initiation frequency at altORFs and refORFs.

We have noticed a very intriguing relationship between PRF type and PRF subtype in certain PRF categories (Figure 8). Although the overall association is not statistically significant (p-value = 0.055), in PRF type 1→2, PRF subtype r→a is significantly more frequent than PRF subtype a→r. Conversely, in PRF type 2→1, it is the other way around, which means refORFs and altORFs involved in PRF are associated with frame 1 and frame 2 in those two categories, respectively. There is a non-significant trend of the same kind in PRF types 2→3 and 3→2, where refORFs are associated with frame 2. These observations are very unexpected because, as we mentioned in Section 4.5.1, the frame definition depends solely on the number of nucleotides in the 5'-UTR of a transcript, which can tolerate insertions and deletions.

As an extension of this analysis, we also looked into the relationship between the PRF subtype and the altORF location relative to the CDS (Supplementary Figure S42). In line with the definition of PRF subtypes, the observed associations are quite expected, with the exception of the rare PRF subtype a2→a1, which is found only in the non-mRNA transcripts and in the 3'-UTR of one transcript. The approximately equal abundance of PRF subtypes r→a and a→r in PRF events located in the CDS is also remarkable and agrees with the information presented in Supplementary Figure S41.

Further, we noticed significant differences in the abundance of chimeric peptides derived from the 3'-UTRs and the 5'-UTRs, and also from the border regions (Supplementary Figure S43). Chimeric peptide-producing altORFs located in the 3'-UTRs are significantly more frequent than altORFs located in the 5'-UTRs. This observation agrees with a clear trend toward the longer 3-UTRs in our study (Supplementary Dataset S2). However, the difference in the UTR lengths does not account for the same direction of significant difference in chimeric peptides derived from the border regions. Namely, the border between the CDS and the 3'-UTR has significantly more altORFs involved in PRF compared to the border between the 5'-UTR and the CDS. Let us consider this observation in the context of altORF conservation. Traditionally, the 3'-UTRs are recommended for the design of qRT-PCR primers because they are much less conserved than the CDSs and the 5'-UTRs (Udvardi et al., 2008). Since the majority of our MS-supported chimeric peptides were modeled with conserved altORFs (Supplementary Tables S5 and S9), the low conservation of the 3'-UTRs makes the border with the CDS a very unexpected hotspot for chimeric peptides.

To understand the nature of the apparent paradox associated with the CDS-3'-UTR border, it may be useful to consider the evolutionary time passed since the emergence of a stop codon at the end of each CDS. If a stop codon is relatively "young", the downstream sequence is expected to retain functional features of the original protein-coding gene. Importantly, the 3'-UTR regions of such genes truncated by nonsense mutations in *M. truncatula* would resemble the CDSs of non-mutated members of the same protein class/family as in close relatives. Some of such nonsense mutations in *M. truncatula* probably coincided with or were followed by frameshifting mutations. In such transcripts, chimeric translation initiated within the CDS might simply "repair" the detrimental effect of a recent truncation/frameshift mutation. It may help restore the potential to synthesize a long protein. In the border between the CDS and the 5'-UTR, altORFs that resemble their CDSs are likely to emerge from mutations that disable the use of the normal translation initiation codon followed by a frameshift mutation. In this scenario, the beginning of the initial CDS becomes separated from the rest of the sequence. Like in the case of chimeric translation that starts at the end of the CDS, producing a frameshifted protein that initiates in the 5'-UTR may restore the ability to synthesize a longer amino acid sequence, which would have many amino acids at the same positions as in the ancestral longer form of the protein. This view on the potential origin of the UTR-altORFs is supported by our observations reported below. To assess the proportion of genes with the "young" UTR-altORFs derived from the formerly coding parts of the gene, it is important to reveal how many altORFs are related to refORFs and other altORFs in transcripts that produce chimeric peptides. To address this point, we conducted a comparative BLASTP analysis of refProts and altProts that are fused by PRF events to form chimeric peptides.

#### 4.5.6. Most ORFs involved in chimeric translation inside the CDSs and the CDS-UTR borders fall into the same annotation agreement category

To understand the extent of annotation agreement between refProts and altProts or among altProts involved in PRF within each transcript, we searched for sequence similarity using the global protein database (NCBI BLASTP tool). Based on this comparative analysis, 156 putative chimeric peptides were grouped in ten categories (Supplementary Dataset S1, Supplementary Figure S44). More than half of altProts that constitute chimeric peptides fall into the same functional group, class, or even family of proteins as corresponding refProts or other altProts associated with PRF on the same transcript (categories S, S3, and SH combined account for ca. 55.8% of the dataset). Naturally, we were curious to learn whether these high-agreement groups occur in different proportions in specific locations on a transcript (relative to the CDS). This way, we found association between the altORF location and the annotation agreement category (Supplementary Figures S45 and S46). However, statistical significance of this association cannot be determined accurately due to small counts and large number of groups. Category S prevails in chimeric peptides derived from the following locations: the CDS, the CDS-3'-UTR border, and the 5'-UTR-CDS border. This is in line with a hypothesis according to which most altORFs involved in PRF emerged by frameshifting mutations and only very few (agreement groups D, D3, and DH) are likely to result from DNA translocation events. The remaining categories possibly correspond to ORFs that have *de novo* origin (agreement groups A0, AH, HH, and H3). Significantly more chimeric peptides of category S come from the CDS-3'-UTR border compared to the upstream border (Supplementary Figures S45 and S46). Because upstream regions are less tolerant to mutations than downstream regions, the cause of this difference remains unclear.

#### 4.5.7. PRF subtype is associated with annotation agreement category of ORFs involved in chimeric translation

There may be a highly significant association between the annotation agreement category and the PRF subtype (Supplementary Figure S47). However, the statistical significance of this association cannot be determined accurately due to small counts and a large number of groups. Within individual agreement categories, the distribution is significantly different from the uniform distribution in four groups: S, S3, A0, and AH. In the agreement categories A0 and AH, PRF subtypes r→a and a→r occur differentially: eight times more chimeric peptides with PRF subtype a→r is found in category A0, and the opposite situation is observed in category AH. To assess the significance of these differences more accurately, we considered the distribution of two PRF subtypes r→a and a→r separately from other subtypes (Supplementary Figure S48). This analysis revealed significant association between the annotation agreement category and the PRF subtype and confirmed the differential representation of PRF subtypes r→a and a→r in the agreement categories A0 and AH. Interestingly, all 18 proteins with no significant similarity in the BLASTP searches (A0) are altProts in mRNA transcripts. Groups AH and HH contain chimeric peptides derived from both coding and non-coding transcript types (Supplementary Dataset S1). The difference in the definition of these groups is very subtle. Category A0 corresponds to refProt/altProt pairs in which a refProt is similar to an annotated protein but an altProt is not similar to any annotated protein. In this group, PRF mostly switches translation from an orphan (*de novo* evolved) altORF to a refORF. In group AH, the situation is different. The annotated part of the pair can be either an altProt or a refProt. The other part is not orphan but has similarity only with a hypothetical protein. In other words, it has limited conservation signature (evolutionarily younger than most refProts but older than a *de novo* emerged sequence). Interestingly, in five chimeric peptides out of 35 that belong to group AH (CP10, CP22, CP39, CP70, and CP118 in Supplementary Dataset S1), it is a refProt that has similarity with a hypothetical protein while an altProt is similar to annotated proteins. If the annotation of the corresponding genes was accurate and the refORF was defined correctly in the current genome release (v. 5.1.9, Pecrix et al., 2018), in this group of genes, the formerly functional refORF was "pushed" to a non-reference frame by a frameshift mutation while the formerly non-functional altORF with a very limited conservation signature acquired the main function. The *M. truncatula* genome annotation community might find it useful to dedicate special attention to these genes.

For more complete understanding of ORFs with limited conservation at the protein level, we plotted the distribution of 45 chimeric peptides from categories SH, DH, AH, HH, and H3 combined, depending on the type of ORF products annotated as hypothetical or unknown. In nearly half of the cases, such hypothetical/unknown ORFs correspond to altProts in the absence of the second altProt involved in frameshifting (group A, Supplementary Figure S49). The second largest group (A2) was composed of chimeric peptides in which the second altProt (derived from altORF2, see the definition of PRF subtypes) was similar to a hypothetical protein regardless of the presence of the refProt (mRNA and non-mRNA transcripts). Interestingly, there are two times fewer chimeric peptides in which the part similar to a hypothetical protein was not the second but the first altProt in the absence of the refProt involved in PRF (A1, non-mRNA transcripts only). Although the difference is not significant, numerically, the sum of groups A2, A1A2, and RA1A2 (11) is larger than the sum of groups A1, A1A2, and RA1A2 (7). This means that whenever two altORFs are involved, the upstream one tends to exhibit stronger conservation at the protein level. It would be useful to study this trend in a larger dataset of chimeric peptides, as it can help better understand the evolution of translated altORFs.

### 4.5.8. rRNA and non-nuclear transcripts are significantly overrepresented in the dataset

In Section 4.5.1, we have already mentioned significant unexpected observations associated with rRNA and non-nuclear transcripts. Namely, five out of six rRNA-derived chimeric peptides have PRF value -1 (Figure 3). Non-nuclear transcripts mostly produce chimeric peptides with PRF value +1, which is the least common among peptides derived from nuclear transcripts (Figure 4). In addition, it turns out that rRNA and non-nuclear transcripts stand out with regard to their strong overrepresentation in the dataset. To estimate the significance of these differences, we calculated expected proportions of four RNA types based on the occurrence of corresponding gene groups in the whole genome. BLASTN searches using the NCBI non-redundant database (Supplementary Dataset S1) and nucleotide alignments with annotated rRNA transcripts (homology group 1, Section 4.6.1) revealed that at least two transcripts annotated as ncRNA in the current *M. truncatula* genome (v. 5.1.9, Pecrix et al., 2018) are actually rRNA transcripts. These two transcripts, MtrunA17_Chr0c01g0489091 and MtrunA17_Chr5g0421761, are considered as rRNA throughout the manuscript except for supplementary tables, which are meant to show the "history" of their annotation status.

Statistical analysis of the difference between observed and expected proportions of four RNA types visualized in Supplementary Figure S50 incorporates the correction for the misannotation of those two rRNA transcripts. The difference between the observed and expected proportions is highly significant for the whole dataset. The observed number of rRNA transcripts is 25-fold higher than expected based on the number of rRNA loci in the whole genome (Supplementary Figure S50A). This is likely to be due to the generation of more models than expected (note no significant difference for rRNA in Supplementary Figure S50B). This means rRNA transcripts stand out with regard to the higher number of conserved and/or translated overlapping ORFs compared to other RNA types. In contrast, ncRNA transcripts are distinguished by the detection of nearly two-fold fewer MS-supported chimeric peptides than could be expected based on the number of generated chimeric models (Supplementary Figure S50B). This agrees with the classical definition of ncRNA as a non-translatable transcript type.

Next, we calculated expected proportions of nuclear, chloroplast, and mitochondrial transcripts based on the occurrence of corresponding gene groups in the whole genome. The difference between observed and expected proportions is highly significant (Supplementary Figure S51). Chloroplast and mitochondrial transcripts are significantly overrepresented in the dataset (19-fold and 12-fold, respectively). Interestingly, five out of seven mitochondrial transcripts are ncRNAs. The transcript type and the chromosomal location are significantly associated (Supplementary Figure S52). Mitochondria is the only location in which ncRNA transcripts that produce chimeric peptides are more abundant than other types. As can be seen in the current version of the *M. truncatula* genome browser (v. 5.1.9, Pecrix et al., 2018), the high proportion of ncRNA loci in mitochondria is a general feature. This may be true also for mitogenomes of other plant species although available reports on this topic do not specify if the highly abundant non-coding regions correspond to lncRNA-producing loci or other types of non-coding DNA (e.g., Morley et al., 2019; Anand and Pandi, 2021).

#### 4.5.9. Correlation analysis suggests the existence of chromosome-specific determinants of the chimeric peptide abundance

Visibly unequal counts of chimeric peptides in different chromosomal locations (Supplementary Figure S52) raise the following questions. Are the differences significant? If yes, are the counts proportional to the effective size of chromosomes and other relevant parameters? To address these questions, first, we looked into the distribution of chimeric peptide counts in different chromosomal locations regardless of the RNA type (Supplementary Figure S53). The distribution is non-uniform, with chromosomes 1, 3, and 8 producing the largest number of chimeric peptides. Expectedly, chloroplasts and mitochondria produce the smallest number of chimeric peptides (except for chromosome 0, which designates the minority of non-mapped genes) because non-nuclear genomes are smaller than any chromosome. However, chloroplasts and mitochondria have more chimeric peptide-producing transcripts than expected based on the total number of transcripts per location (Supplementary Figures S51 and S64A). Next, we wanted to learn if the number of chimeric peptides is proportional to the number of chimeric peptide models generated from each chromosomal location. This analysis revealed a non-significant but clear deviation (nearly two-fold) from the expected counts in chromosomes 1 and 4 but not in non-nuclear locations (Supplementary Figure S54B). The comparison between the two panels of Supplementary Figure S54 suggests the explanation for the many-fold overrepresentation of chloroplast and mitochondrial transcripts in our dataset. Transcripts from these locations have more conserved and/or MS-supported altORFs that overlap with refProts and/or each other. In addition, the overlapping regions between ORFs are probably longer. Thus, more chimeric protein models were generated from these transcripts.

In the next step of our analysis, we studied the relationship between the number of chimeric models and the total number of transcripts per location (Supplementary Figure S55). This way, we learned that observed counts are very significantly different from the expected counts for all locations. Chromosomes 0, 4, 5, chloroplasts, and mitochondria produce significantly more chimeric models than expected. For chromosomes 1, 2, 3, 7, and 8, the relationship is opposite. From this analysis, we conclude that chimeric model counts do not depend on the number of transcripts alone. Their counts are the function of the number of overlapping ORFs that have either conservation evidence for translation or are MS-supported and also depend on the length of the overlapping region. The proportion of such overlapping ORFs is location-specific and is maximal in non-nuclear transcripts.

To obtain more complete understanding of these relationships, we added one more factor to the analysis, cumulative length of transcripts per chromosome (Supplementary Figure S56A, column E), and calculated two ratios that we found illustrative for the purpose. The first ratio (Supplementary Figure S56A, column F) expresses the number of chimeric peptides per chimeric model, which is the proportion of chimeric models that received MS support. The second ratio (Supplementary Figure S56A, column G) reflects the number of chimeric models per transcript. It indicates the enrichment of transcripts with overlapping ORFs that are either conserved or MS-supported. This parameter also depends on the overlap length: the longer the overlapping region the more models are generated. Next, we conducted correlation analysis for all meaningful combinations of column factors. This analysis revealed a significant positive correlation in five cases and a significant negative correlation in two cases (Supplementary Figure S56B). Chimeric peptide numbers correlate positively with the numbers of transcripts and the cumulative lengths of transcripts. However, the relationships are non-monotonous, which makes the quantification of their significance difficult. At the same time, the non-monotonous nature of these relationships brings once again chromosomes 1 and 4 into the focus (compare with Supplementary Figure S54B). The number of chimeric models correlates positively and very strongly with the number of transcripts and the cumulative length of transcripts. This correlation contrasts with a very significant discrepancy between observed and expected counts of chimeric models per chromosome reported in Supplementary Figure S55. The maximal strength of a positive correlation is observed in the relationship between the number of transcripts and the cumulative length. A strong negative correlation (non-monotonous) exists between the number of chimeric peptides and the second ratio and between the cumulative length and the second ratio (Supplementary Figure S56B). The more chimeric models are generated per transcript per chromosome the fewer of them are supported with MS. The greater the cumulative length of transcripts per chromosome the fewer chimeric models are generated per transcript per chromosome. These two negative relationships can be partially explained by positive correlations among columns C, D, and E on one side and among B, D, and E on the other side (Supplementary Figure S56B). The absence of a significant correlation between the number of chimeric peptides (B) and the number of chimeric models (C) is surprising and probably contributes to the negative relationship with the second ratio. At the same time, it is possible that the first negative relationship (BG) can be attributed to the nature of MS proteomics as a method: the larger the search database the fewer peptides are validated.

Here are a few striking examples of this negative relationship. Only ca. seven chimeric models per transcript were generated from chromosome 1, which is the third lowest value in column G. However, the number of chimeric peptides validated with MS is the highest for this location (30, compare with Supplementary Figure S54B). In mitochondria, each transcript, on average, gave rise to the maximal number of models (ca. 164) but only nine chimeric peptides were validated with MS, which is the third lowest value in column B (Supplementary Figure S56A, compare with Supplementary Figure S54). A very similar situation is observed in chloroplasts, which gave on average ca. 93 models per transcript. More models generated per location means there are more overlapping ORFs that are conserved and/or MS-supported, and the overlap regions between them are longer. This notion is difficult to align with the overrepresentation of chimeric peptide-producing transcripts in non-nuclear locations (Supplementary Figures S51 and S54A). A caution should be taken with the interpretation of this correlation analysis because 11 is a very low sample size for accurate estimation of the true correlation coefficients. However, even with small sample sizes, strong and very strong correlations like those observed in our study can be estimated more accurately (Rousselet, 2018). Even with our small sample size conditioned by the genome structure of *M. truncatula* (eight chromosomes), it is evident that major differences with regard to the ability to produce chimeric peptides exist among different chromosomes and different genomic locations.

#### 4.5.10. The abundance of chimeric peptides depends on the overlapping of corresponding genes with repeat elements and other genetic loci

Many chimeric peptide-producing genes in our study overlap with other genetic loci. Some of them overlap with transposable elements or other repeated sequences (designated collectively as RE here). Others overlap with non-RE loci. Supplementary Dataset S11 contains screenshots from the *M. truncatula* genome browser that illustrate the overlapping landscape of each chimeric peptide-producing locus. We decided to investigate whether the fact of their overlapping with RE and non-RE loci is associated with other characteristics. To provide a more direct link with Supplementary Dataset S1 and to simplify the reproducibility of our statistical analysis, we based Supplementary Figures S57-S62 on chimeric peptide counts, not on gene counts (156 peptides originate from 145 genes in our dataset). This was also necessary for assessing the significance of differences in groups with low numbers of chimeric peptide-producing genes (non-mRNA transcripts). Thus, chimeric peptide counts serve as a proxy for gene counts in this analysis. Only ca. one third of chimeric peptides come from genes that have no overlap with other loci (Supplementary Figure S57A).

First, we looked into the relationship with the genomic DNA strand. More chimeric peptides correspond to genes that are located on the minus strand. However, this trend is not significant (Supplementary Figure S57B). Likewise, there is no significant difference in that respect between chimeric peptides from genes that overlap with non-RE loci and those that do not overlap with such loci (Supplementary Figure S58A). However, when we consider the overlap with RE-loci, we find that significantly fewer chimeric peptides come from non-overlapping genes located on the plus strand (Supplementary Figure S58B). A clear inverse relationship exists for chimeric peptides from genes that overlap with other loci: significantly fewer peptides derive from non-RE-overlapping loci (Supplementary Figure S59A) and more peptides correspond to genes that overlap with RE loci, regardless of the DNA strand (Supplementary Figure S59B). The predominance of genes that overlap with transposons and other RE loci may be a general feature of the *M. truncatula* genome or may be specific to our dataset. A dedicated study is required to ascertain this point.

Our next step was to understand whether overlapping is associated with the genomic location: nuclear genes vs. non-nuclear ones. Distribution of chimeric peptides in six overlap categories is non-uniform and depends on the location. Nuclear genes that produce chimeric peptides mostly overlap with RE but not non-RE loci, as can be judged from the chimeric peptide abundance. Chimeric peptide-producing chloroplast and mitochondrial genes have the opposite trend: they mostly overlap with non-RE loci but not RE loci (Supplementary Figure S60). This may also be a common characteristic of the *M. truncatula* genome, which deserves a separate study.

Further, we noticed significant deviations from the uniform distribution in chimeric peptides from non-mRNA transcripts when the same six overlap categories were considered. As indicated above, two-thirds of chimeric peptides come from genes that overlap with either a RE locus or a non-RE locus, or both (Supplementary Figure S57A). The predominance of peptides from overlapping loci is also significant in non-mRNA transcripts (Supplementary Figure S61). However, there is a striking difference in the proportion of chimeric peptides from non-overlapping loci between ncRNA and rRNA. Namely, five out of six rRNA-derived chimeric peptides come from genes that overlap with both RE loci and non-RE loci, and all six correspond to RE-overlapping loci. In contrast, only five out of 11 ncRNA-derived chimeric peptides correspond to RE-overlapping genes (Supplementary Figure S61). The trend of rRNA loci to overlap with other genes may be a common feature of the *M. truncatula* genome, although rRNA loci with no overlap are present too (the *M. truncatula* genome browser v. 5.1.9, Pecrix et al., 2018). The analysis presented here should motivate the *M. truncatula* genome annotation community for a large-scale study that addresses these fundamental but so far unclarified aspects. If the trends reported here are quantitatively different from the characteristics the whole genome, they may be revisited later and considered as PRF-specific features.

In the last analytical domain of this section, we looked into the relationship between the overlap status and the PRF subtype specifically in nuclear transcripts (Supplementary Figure S62). Non-nuclear transcripts that produce chimeric peptides were excluded from this analysis because half of them are non-mRNA transcripts (no PRF subtypes r→a and a→r). The overall trend of preferential overlapping with RE loci but not non-RE loci is evident in this group. The surprising message of Supplementary Figure S62 is the absence of this trend in PRF subtypes a1→a2 and a→r when they are considered in gene groups that overlap with non-RE and RE loci, respectively. These subtle differences can be indicators of processes specific to PRF, which we cannot characterize so far. Much more information is required for the accurate biological interpretation of these observations, as well as many other significant observations reported in Section 4.5.

### 4.6. Searches for sequence similarities associated with the ability to produce chimeric peptides

We extensively searched for similarities among nucleotide and protein sequences from 156 chimeric peptides to identify sequence determinants for different PRF categories. We extracted two sequences from each transcript: one with 50 nucleotides upstream and downstream of the PRF site and a shorter one with ten nucleotides from each side. Alignments using various algorithms, both as a whole group and within PRF value-based groups, revealed no significant sequence similarities. Using MEME software (Bailey and Elkan, 1994; Bailey et al., 2015), we also found no common motifs, indicating minimal conservation of PRF sites and flanking sequences at the transcript level. Similar results were obtained when analyzing amino acid sequences upstream and downstream of PRF sites.

Expanding our search to the whole transcript level, we explored secondary RNA structures like pseudoknots using several prediction algorithms. However, the predicted pseudoknots were too numerous and did not consistently align with PRF sites. These analyses suggest very little sequence conservation at and around PRF sites, both at the nucleotide and protein levels, making it challenging to identify common sequence determinants for PRF categories.

#### 4.6.1. One-sixth of chimeric peptide-producing transcripts have global homology with at least one other sequence in the collection

We searched for sequence similarities among all 145 transcripts that produce 156 chimeric peptides. Since pairwise alignments of these sequences would be difficult to inspect visually (many thousands of alignments), we employed the Clustal Omega alignment tool (Sievers et al., 2011) to align sequences grouped by PRF value. This way, we generated six multi-sequence alignments, which served as the primary source for further analysis. First, we focused on alignments that have reasonably high nucleotide percent identity over the whole length of transcripts. To select such alignments, we searched the literature for an established threshold above which two nucleotide sequences can be considered evolutionarily related. Finding such information turned out to be a challenge. Thus, we decided to estimate such a threshold empirically, on a small sample size. Using the *M. truncatula* genome browser, we randomly selected ten transcript sequences per RNA type. The random choice of loci had one element of a useful bias: we made sure that no sequences belong to the same closely related group, as can be seen from their annotations (Supplementary Dataset S12). Then we used the Geneious® alignment algorithm, a built-in option in the Geneious® software package v. 7.1, to learn percent nucleotide identity values for all 45 combinations of ten sequences per RNA type (100 minus ten divided by two). Geneious® alignment is very convenient for illustration purposes and is very similar to Clustal Omega alignment with regard to its outputs. To our surprise, none of the 45 values in the mRNA sample fell below 49.0%, whereas the maximal value was as high as 54.3%. The median 51.1% was very close to the mean value (51.3%). For the ncRNA and rRNA samples, the ranges were much larger (7.0-55.0% and 49.3-100%, respectively). However, the median values were very close to each other and to the median of the mRNA sample (ranged from 50.9% to 51.1%). With regard to this parameter, the tRNA sample was significantly different from other samples, with the median value of 55.8% and the range of 26.7-82.9%. Based on these estimates, we used an empirical threshold of 56% to decide which sequences have biologically relevant similarity. This threshold was backed up by the threshold of 50% identity at the protein level for the majority of alignments and 30% for three special cases where the similarity was obvious despite lower percent identity values (53%). This analysis revealed ten homology groups thoroughly described in Supplementary Dataset S13. Corresponding alignments are visualized in Supplementary Dataset S14.

Based on the annotation and the overall sequence similarity, two-thirds of 24 transcripts that constitute ten homology groups are involved in just two processes: transcription and translation. This is not very surprising because our dataset emerged from chimeric protein models generated from mostly conserved altORFs (Supplementary Tables S5 and S9). In at least 50% of cases, *in-silico* translation products of those altORFs had similarity with *M. truncatula* proteins (Supplementary Figure S7). Thus, our dataset derives from a "pre-selected" collection of sequences many of which can be expected to have some homology.

### 4.6.2. Conserved location of PRF sites in alignments is rarely associated with the conservation of PRF values

Homology groups 1 and 2 are composed of four sequences each. The remaining groups are pairs. Homology group 1 consists of three rRNA transcripts and the only tRNA transcript in our dataset. All four sequences are associated exclusively with PRF value -1. Maximal values of percent nucleotide identity (column G) vary from 87.5 to 58.8 in this group. In pairwise alignments between sequences of homology group 1, the distance between two closest PRF sites was generally large: at least 933 nt for high-percent-identity alignments and 220 for low-percent-identity alignments, which indicates no conservation of the PRF position despite high overall similarity and the same PRF value.

Homology group 2 consists of three putative disease resistance protein-coding transcripts and one ncRNA. This group is associated with PRF values -1, -2, and +2, but not +1. Here, maximal percent-identity values range from 59.9% to 93.0%. Remarkably, in the alignment between transcripts that produce CP53 (mRNA MtrunA17_Chr3g0102171) and CP66 (ncRNA MtrunA17_Chr3g1011650), PRF sites with value -2 were only 16 nt apart, which points to the conservation of both the PRF position and the PRF value in this pair.

Evidence for such conservation can also be found in homology group 6, where the PRF sites with value -2 are separated by 53 nt (Supplementary Datasets S13 and S14). This is highly unexpected given that MtrunA17_Chr3g0091671 (CP50) is ten times shorter than MtrunA17_Chr2g0283311 (CP33). Among the remaining seven groups, a few were associated with other outstanding features. For example, in homology groups 4 and 10, the PRF sites were only five and eight nucleotides apart, respectively. However, they had different PRF values: -1 and +2 in homology group 4, and -1 and +1 in homology group 10. This suggests a possibility that factors that define the PRF position and the PRF value may be different. The PRF values were conserved in homology group 8 (+1) and partially conserved in homology group 5 ("long" values -2 and +2), which contained one of the very likely candidates for mosaic translation described in Section 4.4 (MtrunA17_Chr1g0200071).

Another surprising observation was made in homology group 9, which is composed of two rRNA transcripts. The transcript that corresponds to CP114 (3,662 nt) is two times shorter than its matching transcript (transcript of CP141). However, it is nearly 100% identical to the transcript of CP141. Despite this extraordinary similarity, in the transcript of CP141, the PRF site with value -1 is outside the alignment region and is 2,979 nt upstream of the +2 PRF site in the other transcript (Supplementary Dataset S14). Interestingly, the region around the PRF site is conserved between the two transcripts, with only one mismatch, 12 nucleotides upstream of the site.

#### 4.6.3. Nearly half of the 145 transcripts form a network of small homology groups interconnected by shared homology to other sequences and/or direct similarity over short regions

Following this search for the global sequence similarity among 145 transcripts, we looked for more subtle similarities that could not be captured by the Clustal Omega alignment. We used two approaches to this task. One was based on the application of the MEME software to the transcript sequences rather than to extracted PRF-site containing regions (we conducted motif analysis). The other approach relied on the whole-transcriptome wide (*M. truncatula* only) BLASTN search for sequences that have shared subjects (query sequences similar to a shared subject with or without obvious similarity between query sequences). Both strategies revealed additional sequences in which high percent nucleotide identity concerned not the entire sequence but shorter regions (Supplementary Dataset S15). Supplementary Dataset S16 contains primary data from this BLASTN analysis. In total, 64 transcripts out of 145 (ca. 44%) have at least one shared subject with at least one other sequence in the dataset, according to the BLASTN analysis. This list of 64 transcripts contains all items from homology groups 1-10 except for transcripts that produce CP10 and CP33. These two transcripts have no shared subjects with any other sequence in the dataset despite having direct similarity with transcripts in homology groups 3 and 6 (Supplementary Dataset S13). Out of the combined 66 sequences with detectable similarity, direct alignments of various lengths and percent identity values were possible for 34 sequences, which is ca. one-quarter of the dataset (Supplementary Dataset S15). Fine details of all meaningful alignments are illustrated in Supplementary Dataset S14. In this analysis, we used MEME and BLASTN results as a guide and manually aligned corresponding local similarity regions for visualization.

Supplementary Figures S63-S65 summarize the data presented in Supplementary Dataset S15. From these summaries, we learn that rRNA transcripts, on average, have shared subjects with more other sequences in the dataset (10.6, row counts, column BQ in Supplementary Dataset S15A), have a larger mean number of shared subjects per transcript (14.5, row averages, column BR in Supplementary Dataset S15A), and a larger per-transcript mean count of similar sequences in the dataset, based on meaningful pairwise nucleotide alignments (4.2, row counts, column AK in Supplementary Dataset S15B). These values are consistently lower for ncRNA transcripts and are the lowest for mRNA transcripts, which further supports the idea about the ancestral role of rRNA-like molecules in genome evolution (Root-Bernstein and Root-Bernstein, 2015, 2016, 2019; Caetano-Anollés and Caetano-Anollés, 2016; de Farias et al., 2016). The highest values associated with shared subjects are found in both rRNA and mRNA transcripts. For example, rRNA MtrunA17_Chr0c01g0489091 (CP1) and mRNA MtrunA17_Chr2g0326801 (CP44) each have shared subjects with 11 other transcripts in the dataset. Per-transcript mean number of shared subjects is the highest in mRNA transcripts: sequences from homology group 4 (MtrunA17_Chr1g0190571, CP20, and MtrunA17_Chr1g0202001, CP25) have 28 shared subjects with each other but not with any other sequence in the dataset. However, the highest absolute number of shared subjects is found in rRNA transcripts: sequences from homology group 9, MtrunA17_Chr7g0229401 (CP114) and MtrunA17_CPg0492331 (CP141) have 68 shared subjects with each other. They also have shared subjects with nine and 11 other sequences, respectively, which makes their mean values lower compared to homology group 4 (Supplementary Dataset S15A).

In the analysis of direct sequence similarities (similarities evident from the pairwise alignment, not from the existence of shared subjects), we discriminate between the alignments that concern the entire length of at least one transcript and those that concern only a part of the sequence (identified with the MEME software). We refer to the latter ones as homology regions. They are highlighted with cyan in alignments of Supplementary Dataset S14 and with moss green in cells of Supplementary Dataset 15B. There are a few remarkable observations associated with homology regions. Among all alignments, homology regions included the PRF sites only in one pair of transcripts: rRNA MtrunA17_Chr7g0229401 (CP114) and ncRNA MtrunA17_MTg0491501 (CP154), with percent nucleotide identity ca. 90 over 44 nt (Supplementary Dataset S14). These transcripts cannot be meaningfully aligned over their entire lengths. When homology regions of ncRNA MtrunA17_MTg0491501 (CP154) and of a different rRNA transcript, MtrunA17_CPg0492331 (CP141), are aligned in an otherwise meaningless alignment (percent identity 39.0), the PRF sites with values -2 and -1, respectively, occur somewhat close to each other (338 ungapped bases), given the entire length of the transcripts. Surprisingly, these sites are almost equally far away from the 5'-end (675 and 697 nt, respectively), which may point to a link between the homology regions and the PRF sites. Ribosomal MtrunA17_Chr7g0229401 (CP114) makes a noteworthy alignment with another rRNA transcript, MtrunA17_Chr5g0422291 (CP88_89), which has two PRF sites. These sequences share four homology regions. One of them, region 4, is an inverted version of region 2 in MtrunA17_Chr7g0229401. The distance between regions 1 and 2 is conserved. Region 3 is located at nearly the same short distance downstream of the PRF sites. The overall percent identity of the alignment is only 51.4, which is below the threshold we mentioned earlier. This example illustrates how informative some of the low-percent-identity alignments can be, with local similarities non-detectable by Clustal Omega. In many alignments shown in Supplementary Dataset S14, a refORF and/or an altORF of one transcript align with such features in the matching transcript even when one of the transcripts is ten times shorter than the other one (e.g., MtrunA17_Chr2g0283311, CP33, and MtrunA17_Chr3g0091671, CP50; MtrunA17_Chr1g0162101, CP10, and MtrunA17_Chr4g0037381, CP75). This is true not only for homology groups 1-10, but also for some alignments with low percent nucleotide identity (e.g., MtrunA17_Chr1g0155251, CP8, and MtrunA17_Chr2g0326801, CP44).

The analysis of sequence similarities among 145 transcripts that constitute the dataset can be summarized as follows. Nearly 56% of sequences have no similarity with any other item in the collection, which means the ability to undergo productive frameshifting probably evolved independently in those sequences. The remaining 44% of transcripts do not constitute one homology group with one conserved PRF-related motif but form a network of interconnected small homology groups. This highlights very high diversity of sequences capable of producing chimeric peptides. The message we drew from this analysis has large practical importance. Namely, the strategy followed in our study, which is based on the deduction of the novel PRF sites directly from MS proteomic data, can conceptually be much more informative compared to any homology-based prediction strategy that relies upon similarities with the already known PRF sites. Furthermore, the absence of one or a few common frameshifting motifs is in line with what is known about the origin of plant genomes and the role of transposable elements in their evolution (Mhiri et al., 2022; Pulido and Casacuberta, 2023). Depending on a plant species, transposons may constitute up to 89% of the genome. In *M. truncatula*, their proportion is ca. 46% (Table S2 in Wang et al., 2021). They are closely related to viruses, which are entities known to use PRF for their activity (Mustafin, 2018; Hayward and Gilbert, 2022). In our dataset, many genes overlap with repeat elements including transposons. Because PRF sites in viruses are not conserved across different species (McNair et al., 2024), their relatives, transposons, may be expected to contain no "universal" frameshifting motifs. However, not all PRF sites in our dataset coincide with transposons. Thus, linking PRF in a plant genome to transposons alone would probably be incorrect. In Section 5.8, we explain our view on the contribution of non-viral sequences to the diversity of chimeric peptides. According to this view, the ability to undergo frameshifting is an ancient feature present in all genomes since the earliest times of their evolution.

### 4.7. Searches for alternative sources of amino acid sequences identical to MS-supported chimeric peptides

Given the short length of peptides deduced by MS proteomics (7-35 aa, Swaney et al., 2010), researchers should be aware of uncertainty about the true origin of each MS-validated sequence. Without dedicated additional analysis, one cannot even be certain about the chimeric origin of such peptides. Likewise, if the sequences are truly chimeric, the second concern is about the range of loci that can produce a given chimeric peptide. Due to the chimeric nature of such sequences, they can potentially be produced by more than one locus. We conducted comprehensive searches that address these concerns. Our analysis discriminated between sequences that can potentially have a non-PRF origin and truly chimeric sequences. It also showed which chimeric peptides have a single origin and which may have multiple origins. Furthermore, using next-generation transcriptomic data (RNA-Seq), we show which transcripts are the most likely sources of chimeric peptides detected by our approach. This information is crucial for the downstream functional analysis of these sequences. The details of analytical domains aimed at ascertaining the true origin of each chimeric peptide are presented below.

#### 4.7.1. Only one non-chimeric sequence identical to an MS-validated chimeric peptide can be encoded by the genome of *M. truncatula*

We searched for genomic locations that can potentially produce peptides that look like MS-validated chimeric sequences. This search concerned all DNA regions regardless of their annotation status. Only one chimeric peptide, CP80 (mRNA MtrunA17_Chr4g0070011), could potentially come from genomic DNA without any frameshift. Two RE loci contain nucleotide sequences that have potential to encode CP80: MtrunA17_Chr2R0265000 (frame 3F) and MtrunA17_Chr7R0255370 (frame 2R). This finding indicates that most peptides in our dataset are likely to be truly chimeric. We have also considered the genome of *Sinorhizobium meliloti* strain 2011 and its two plasmids as potential non-chimeric sources of MS peptides identified in nodules (Marx et al., 2016). Both “Standard” and “The Bacterial, Archaeal and Plant Plastid Code” tables were used (https://www.ncbi.nlm.nih.gov/Taxonomy/Utils/wprintgc.cgi). This analysis showed that none of nodule-detected MS peptides could be generated from the *M. truncatula* symbiont without PRF.

### 4.7.2. In theory, up to 44% of MS-validated chimeric peptides could have alternative origin from unknown exon skipping or intron retention events in their transcripts

Theoretically, translation of alternatively spliced forms of some transcripts detected in our study could mimic MS-validated chimeric peptides if exon skipping or intron retention events perfectly join two nucleotide sequences that correspond to the short arm and the long arm of a chimeric peptide (Supplementary Dataset S1). To address this ambiguity, we conducted a comprehensive search for duplicated regions that correspond to those two arms in each transcript. For the purpose of this analysis, the definition of arms is based on the PRF position in a chimeric peptide: the short arm is the portion of a chimeric peptide that contains fewer amino acids on either side of the frameshift. Accordingly, the long arm is the remaining portion. When two arms are of the same length, the left arm is depicted as “short” and the right arm as “long” in Supplementary Dataset S1. Supplementary Figure S66 illustrates that shorter peptides tend to have fewer amino acids in their short arms. Intuitively, the more amino acids are found in the short arm of an MS peptide, the higher the chance that this MS peptide is specific to its hypothetical chimeric protein. Out of 156 MS peptides, 107 (ca. 69%) contained between one and four amino acids in their short arms (Supplementary Figure S66), which makes it statistically likely to find exactly the same amino acid(s) more than once in three-frame translations of corresponding transcripts. The remaining 49 MS peptides (ca. 31%) contained between five and 14 amino acids in their short arms (Supplementary Figure S66). In the search for duplicated arm sequences, we counted the number of multiple occurrences on respective sides relative to the frameshift and recorded them in Supplementary Dataset S1. Surprisingly, 87 MS peptides (ca. 56%) could not result from exon skipping or intron retention events in their transcripts because their arm sequences are found only once (Supplementary Figure S67A). These peptides are likely to be truly chimeric. The remaining 69 MS peptides (ca. 44%) could potentially originate from exon skipping or intron retention events in their transcripts. These peptides are conditionally unique to the PRF events (Supplementary Figure S67B). Supplementary Figure S67A shows that chimeric peptides with no duplicated arms can be of very short lengths (eight amino acids) and can have as few as one amino acid in their short arms. Conversely, even long peptides with more than four amino acids in their short arms can be conditionally unique (Supplementary Figure 67B).

These theoretical considerations can be of value for the discrimination between truly chimeric peptides and chimeric-like sequences only in conjunction with high-quality transcriptomic data obtained from sequencing of RNA actually present in the same biological samples from which MS peptides were deduced. The following section describes the use of RNA-Seq data for this purpose.

### 4.7.3. Based on RNA-Seq reads from 50 runs, only four MS-validated peptides could potentially have non-chimeric origin; none of these peptides correspond to alternative splicing forms

Translation of alternatively spliced forms derived from exon skipping or intron retention can potentially mimic chimeric peptides detected in our study. Natural single nucleotide polymorphisms and RNA editing events could also produce sequences identical to chimeric peptides. Thus, it is very important to study actual transcript sequences of *M. truncatula*, which is best achieved using the next-generation RNA sequencing (RNA-Seq). RNA-Seq combines the quantitative power of qRT-PCR with the throughput of microarrays. In contrast to qRT-PCR and microarray analyses that measure the transcription levels of known transcripts only, RNA-Seq offers advantages previously unthinkable for a high-throughput method: it does not only quantify the transcripts; it reveals their nucleotide sequences in such a way that entirely novel transcripts can be detected (Wang et al., 2009). The NCBI SRA database (Leinonen et al., 2011) contains raw sequencing data for at least 4,000 of *M. truncatula* RNA-Seq runs (BioSample) that correspond to nearly all organs, tissues, and cell types under many normal and stress conditions. Because the size of raw data corresponding to each RNA-Seq run is very large, it was important to consider the computational feasibility of the analysis. We pre-selected 50 RNA-Seq runs (Supplementary Dataset S17) that have the best possible correspondence with the 16 samples of the three MS-based studies from which the chimeric peptides were deduced (Marx et al., 2016; Shin et al., 2021; Castañeda et al., 2021). These 50 runs contain data exclusively from the same ecotype of *M. truncatula* as was used for the sequencing of the *M. truncatula* genome (Jemalong A17, Pecrix et al., 2018). Also, the runs do not include any mutant or transgenic versions of ecotype A17. They come from wild-type samples. This is important because the three MS samples mentioned above also originate from the wild-type A17 plants.

To search for RNA-Seq reads that correspond exactly to our chimeric peptides, we considered each individual read in the RNA-Seq runs as a transcript sequence, with a unique identifier. The identifier contains a tag showing the RNA-Seq run from which the read came. We *in-silico* translated RNA-Seq reads in three forward reading frames and used the resulting sequences as a subject database to search for exact matches. Sequences of 156 chimeric peptides served as a query database. Upon the detection of exact matches, we manually aligned individual reads with corresponding spliced transcripts, which had the chimeric peptide encoding sites mapped on each sequence. In this procedure, we considered not only transcripts constituting the primary sources of chimeric peptides (transcripts from which the chimeric models were deduced) but also other transcripts that can potentially produce chimeric peptides via PRF. We refer to the latter transcripts as alternative sources (Section 4.7.4). Careful analysis of each sequence and each alignment permitted discrimination between the true chimeric sources of MS-validated peptides, potential non-chimeric native sources, and artifacts of RNA-Seq. The primary outputs of this analysis are shown in Supplementary Dataset S18. All relevant alignments are visualized in Supplementary Dataset S19. For additional information, Supplementary Dataset S20 lists RNA-Seq reads found across multiple runs.

In the course of this work, we found that only four MS-validated chimeric peptides could have non-chimeric origin supported by multiple RNA-Seq reads: CP54 (primary source: mRNA MtrunA17_Chr3g0105981), CP93 (mRNA MtrunA17_Chr5g0435191), CP140 (ncRNA MtrunA17_Chr8g0392351), and CP148 (mRNA MtrunA17_MTg0490471). Interestingly, although three peptides from this group were predicted to come possibly from exon skipping or intron retention events (Supplementary Dataset S1), RNA-Seq data clearly show that none of them can be produced via unconventional splicing of their transcripts. In all four cases, a few nucleotides different from the sequence of annotated transcripts were responsible for mimicking the chimeric peptides. This suggests that alternative splicing is unlikely to generate chimeric peptide-like sequences (Supplementary Dataset S19).

Several other chimeric peptides had RNA-Seq reads that mimic the frameshift products when translated in three forward frames. However, in each case, only a single read (CP31, CP73, CP98, and CP149), two reads (CP92, CP100, CP110, and CP123) or four reads (CP94) were present. One read out of three shown in the alignment for CP94 (589588053_

SRR15439239.8397368, Supplementary Datasets S18 and S19) can be found twice in the same run (SRR15439239, petioles, Supplementary Dataset S20), which makes the effective read number four. The remaining chimeric peptides mentioned above had reads present only once in any run. The very few reads found for these chimeric peptides are not sufficient for the solid support of their non-chimeric sources. Given the good average depth of sequencing (Supplementary Dataset S18), the scarcity of these reads makes them likely to contain sequencing errors that mimic chimeric peptides (Supplementary Dataset S19).

A large group of reads that mimic chimeric peptides (CP35, CP74, CP140, and CP148) originated from artifacts of the method. Adapters and primers commonly used in RNA-Seq were responsible for these artifacts. These sequences can be easily recognized as misaligned portions of the reads in Supplementary Dataset S19 and viewed in Supplementary Dataset S18. A comprehensive list of RNA-Seq adapters and primers downloaded from https://github.com/csf-ngs/fastqc/blob/master/Contaminants/contaminant_list.txt (Andrews, 2010) can be found in Supplementary Dataset S21. It should be noted that chimeric peptides are highly unique sequences that cannot be mimicked by pieces of artificial DNA such as adapters and primers alone. This is evident from the fact that in each of the four chimeric peptides mentioned above (CP35, CP74, CP140, and CP148), the short arm contains only one amino acid. This amino acid is R (arginine) in all four peptides (Supplementary Dataset S1). It is the only part contributed by the artificial DNA sequence in each case (AGA, AGG, and CGG). Given the total very large number of reads in each run and purely random start/end positions of each read, the coincidence of the artificial sequence with a PRF site is an event with non-zero probability.

Another important consideration associated with the use of RNA-Seq data for our purpose is the agreement between the source of the RNA-Seq run and the MS proteomic sample source. If an MS-validated chimeric peptide can indeed be mimicked by translation from unusual versions of native transcripts, one can reasonably expect that the RNA-Seq reads that support its non-chimeric translation should be present in a sufficiently large number (more than one count per million). Furthermore, these reads should come from the same source as the proteomic data from which the chimeric peptide was deduced. This agreement is found rarely in our analysis of RNA-Seq data (Supplementary Datasets S18 and S19). In fact, there are only two cases of an imperfect agreement (CP100 and CP140) and two cases of a perfect agreement (CP54 and CP149). Given that CP149 has only a single read, which corresponds to 0.047 counts per million, only CP54 is truly likely to have an alternative non-chimeric origin.

Based on the analysis of the genomic DNA described earlier, we predicted the alternative non-chimeric origin for CP80 (MtrunA17_Chr2R0265000 and MtrunA17_Chr7R0255370). Contrary to that prediction, our RNA-Seq analysis indicates that CP80 is likely to be truly chimeric because not a single read corresponding to those two loci was found in 50 representative RNA-Seq runs (Supplementary Dataset S18). This is further supported by the fact that these loci have a zero-transcription signal in the comprehensive RNA-Seq based gene expression atlas of *M. truncatula*, which includes data from 77 projects (MtExpress v. 3, Carrere et al., 2021). Repeat elements are commonly not expected to be transcribed and polyadenylated. This may seem to explain why MtrunA17_Chr2R0265000 and MtrunA17_Chr7R0255370 do not show up in the RNA-Seq analysis. However, not all RNA-Seq libraries among the selected 50 samples are constructed using the poly-A tail (Supplementary Dataset S17). CP80 was identified in 14 dpi nodules and roots. RNA-Seq libraries that correspond to 14 dpi nodules and roots were generated using random primers, which enables capturing of non-polyadenylated transcripts. Thus, truly transcribed repeat elements would be detected by RNA-Seq. Since MtrunA17_Chr2R0265000 and

MtrunA17_Chr7R0255370 are not among them, and they are also not expressed according to MtExpress v. 3, CP80 is likely to have truly chimeric nature.

Taken together, the evidence presented in this section rules out a possibility of non-chimeric origin for at least 152 MS-validated peptides out of 156 (ca. 97%), and only four peptides (CP54, CP100, CP140, and CP149) may have ambiguous origin.

#### 4.7.4. At least 58% of MS-supported chimeric peptides have unique sources, which simplifies their functional characterization

Our next goal was to identify all possible alternative chimeric sources of 156 chimeric peptides, which is crucial for their functional analysis. Classical sequence alignment tools such as TBLASTN are designed for non-chimeric query sequences, which makes it impossible to modify them for chimeric queries. For this reason, we designed a script that can address this challenging task. Because the potential field of application of this script reaches beyond the scope of our study, its details will be described in a separate publication (Çakır et al., preprint in preparation). The script can be used for the detection of alternative non-chimeric sources and chimeric sources that involve a wide range of PRF values. For consistency with the data on primary sources of chimeric peptides, in this study, we have limited the description of alternative sources to PRF values -2, -1, +1, and +2. Two different search databases were used for this analysis: (1) the entire transcriptome, which included mRNA, ncRNA, rRNA, and tRNA sequences, and (2) the entire set of repeat elements (RE), most of which are transposons (the *M. truncatula* genome portal MtrunA17r5.0-ANR, Pecrix et al., 2018; Wang et al., 2021b). The rationale behind the inclusion of the latter search database was the idea that transposons can be transcribed and possibly translated, and are likely to bear virus features associated with PRF (Mustafin, 2018; Hayward and Gilbert, 2022). Whereas non-RE transcripts can be reasonably assumed to undergo no reverse transcription and translation, reverse complements of transposons can potentially serve as a template for translation because a similar process is well-characterized for their virus counterparts (Varmus, 1987; Whitcomb and Hughes, 1992). For these reasons, we considered only forward reading frames for non-RE transcripts but both forward and reverse reading frames for RE sequences. The primary outputs of these searches can be found in Supplementary Datasets S22 and S23, for non-RE and RE sequences, respectively. Although our analysis was focused on PRF values -2, -1, +1, and +2, potential alternative sources that involve PRF values up to -29 and +29 are also listed in those datasets. All non-RE and RE alternative sources considered in this study are summarized in Supplementary Datasets S1 and S24. In Supplementary Dataset S1, we subdivided RE alternative sources into two categories: those that do not overlap with non-RE loci at the putative PRF sites and those that do. These data reveal very unexpected findings. First of all, despite the short nature of MS peptides, 91 out of 156 chimeric peptides (ca. 58%) have no alternative sources. Out of these chimeric peptides with unique sources, 13 (ca. 14%) have only one amino acid in their short arms, and 61 (ca. 67%) have between one and four amino acids in their short arms. This indicates that MS peptides are by far more unique than could be expected based on their limited length. Secondly, 115 (ca. 74%) of 156 chimeric peptides have no non-RE alternative sources. Furthermore, only five of these chimeric peptides have RE alternative sources that do not overlap with non-RE loci at the putative PRF sites. For 19 chimeric peptides, alternative sources are limited to RE sequences that overlap with non-RE loci in either orientation at the PRF sites. Given the symbiotic nature of nodule samples, we have also searched for loci that can potentially encode nodule-derived chimeric MS peptides in *S. meliloti* strain 2011 (Marx et al., 2016). This search revealed that only CP142 may be translated from rhizobial transcripts: three nearly identical loci with high similarity to 16S rRNA (GenBank accession NR_113670.1) can be found on the chromosome but not on the plasmids of that symbiotic organism, using the standard and prokaryotic codon tables

(Supplementary Dataset S1). In summary, chimeric peptides detected in nodules cannot be products of translation from non-chimeric loci of rhizobia.

### 4.7.5. Ten repeat-element loci are likely alternative sources for eight MS-supported chimeric peptides

When a putative non-RE chimeric peptide-producing locus overlaps with an RE locus, there is ambiguity about which gene produces a given chimeric peptide. To solve this ambiguity, it is essential to consider the transcription level of RE loci. In most cases, such loci have log2 TMM values below zero (MtExpress v. 3, Carrere et al., 2021), which means they are unlikely alternative sources in the presence of more likely primary or alternative non-RE sources. However, we found a few remarkable exceptions. Out of 70 alternative RE sources that overlap with no loci at the putative PRF sites, ten have log2 TMM values above zero (Supplementary Dataset S25). Details of their expression profiles in MtExpress v. 3 can be viewed in Supplementary Dataset S26. Four of these loci (MtrunA17_Chr1R0265150, MtrunA17_Chr3R0062580, MtrunA17_Chr3R0074220, and MtrunA17_Chr7R0161930) overlap with no other genes along the whole length of their sequences. Thus, ten RE loci can be regarded as likely alternative sources for the following eight chimeric peptides: CP24 and CP101 (MtrunA17_Chr1R0014150), CP28 (MtrunA17_Chr2R0106940, MtrunA17_Chr3R0310480, and MtrunA17_Chr4R0052500), CP37 (MtrunA17_Chr1R0148110), CP116 (MtrunA17_Chr7R0161930), CP122 (MtrunA17_Chr3R0062580), CP141 (MtrunA17_Chr0c04R0002520 and MtrunA17_Chr3R0074220), and CP143 (MtrunA17_Chr1R0265150). Among these RE loci, three can potentially produce chimeric peptides from their reverse complements: CP24 and CP101 (MtrunA17_Chr1R0014150), CP37 (MtrunA17_Chr1R0148110), and CP141 (MtrunA17_Chr3R0074220).

### 4.7.6. Two chimeric peptides are highly conserved among non-repeat alternative sources

Among 156 chimeric peptides, 37 can potentially come from non-RE alternative sources (Supplementary Dataset S1). Thirty-five of them have between one and five alternative sources. However, there are two highly-conserved chimeric peptides. CP24 has 11 alternative sources of non-repeat nature, all of which are annotated as putative protein-synthesizing GTPases, which is the same as the annotation of the primary source of CP24, CP23, and CP101. CP130 has nine alternative non-RE sources, among which there are four ncRNAs, two hypothetical transcripts, one transcript for transcription factor interactor and regulator, and two symbiotic nitrogen fixation-associated transcripts (MtNCR238, MtrunA17_Chr2g0301771, and leginsulin related MtN11/16/17 family member MtrunA17Chr3g0092704). The primary source of CP130 is annotated as putative reverse transcriptase, RNA-dependent DNA polymerase (Supplementary Dataset S1).

### 4.7.7. Four chimeric peptides are highly conserved among alternative sources of repeat nature and one is ultra-conserved

With regard to alternative RE sources that overlap with the regular (non-RE) loci at putative PRF sites, CP24 stands out because it has 11 alternative sources, which is the same as the number of non-RE alternative sources. When we consider the occurrence of alternative RE sources that do not overlap with the regular genes at the PRF sites, we observe a much stronger conservation. Whereas most chimeric peptides in this category have between one and seven alternative RE sources (Supplementary Dataset S1), there are two that have 15 sources (CP28 and CP122) and one with 20 such sources (CP37). Interestingly, primary sources of these three chimeric peptides are related to DNA or RNA binding, one of them being a helicase-encoding gene. The absolute record in the nomination for the strongest conservation belongs to CP130, which has 363 alternative RE sources regardless of their overlap status. This is intriguing because the primary source of CP130 is annotated as a locus related to the retrotransposon activity (reverse transcriptase).

### 4.7.8. PRF values and positions in MS peptides are highly conserved between the primary sources and the alternative sources

Theoretically, it is possible to produce exactly the same peptide sequence via frameshifting events of different values, with different positions of the frameshift along the peptide length. In view of the short length of MS peptides (8-30 aa in our study, Supplementary Dataset S1), one could expect the existence of multiple unrelated alternative sources of chimeric peptides, with different PRF values and positions involved in the frameshifting. The exhaustive detection of all alternative sources of 156 chimeric peptides with the aid of our in-house script demonstrates that this is partially true. Many chimeric peptides produced via conventional (well-studied) frameshifts with PRF values -2, -1, +1, and +2 according to their primary sources can also be products of much longer frameshifts (up to ±29), if alternative sources are considered (Supplementary Datasets S22 and S23). In contrast, if we follow the conservation of PRF values and positions within the conventional range of PRF values (-2, -1, +1, and +2), we find very few deviations from the PRF value and/or position of the primary source. Only four chimeric peptides, CP24, CP32, CP37, and CP80, feature such deviations. Among these peptides, only the last two have alternative sources in which the PRF position is different from that in the primary source. In this respect, CP37 stands out. It can be produced via frameshifting with three PRF values (-2, -1, and +2), which concerns three different positions in the peptide: 4, 6, and 7. LTR retrotransposon MtrunA17_Chr4R0290860 can produce CP37 from its reverse complement at two different positions: one involves a -2 frameshift and the other involves a -1 frameshift, both at position 7 in the peptide. Multiple frameshifting is not limited to non-RE loci, according to our data (Section 4.7.10).

### 4.7.9. Comparative analysis of expression patterns helps focus future functional characterization studies on the most likely sources of chimeric peptides

Whether a potential alternative source of a chimeric peptide is likely to be its true source depends on two factors: the magnitude of expression (relative transcript abundance) and the timing/location of expression. Using these two criteria, we compared the expression profiles of all primary (Supplementary Dataset S27) and alternative sources with the aid of MtExpress v. 3 (Carrere et al., 2021). This analysis revealed that 31 chimeric peptides can have alternative sources that are at least as likely or even more likely to be responsible for the production of chimeric peptides compared to the primary sources. This information is summarized in the last column of Supplementary Dataset S1.

### 4.7.10. The exhaustive search for alternative sources of chimeric peptides revealed 15 non-repeat and eight repeat loci potentially associated with multiple frameshifting

Following our exhaustive search for alternative sources of chimeric peptides, we realized that a primary source of peptide A can be an alternative source of peptide B, and the other way around. This consideration helped us identify additional sequences potentially associated with multiple frameshifting as well as additional frameshifting sites on the primary sources. The results of this analysis are depicted in Supplementary Dataset S28. Intriguingly, in all nine transcripts described in this dataset, the first frameshifting site has PRF value identical to the last frameshifting site. Moreover, all mRNA transcripts are associated with “long” frameshifts (mostly -2) while non-mRNA transcripts have “short” frameshifts (-1 and +1). It may be remembered that four out of eight primary sources associated with multiple frameshifting, which are described in Supplementary Dataset S4, also have repetitive PRF values (MtrunA17_Chr1g0185811, MtrunA17_Chr5g0422291, MtrunA17_MTg0490471, and MtrunA17_MTg0490971). It is useful to consider this observation together with another unusual recurring pattern: three mRNA transcripts associated with multiple frameshifting contain a sequence of PRF sites “-2 +2” (MtrunA17_Chr1g0200071, MtrunA17_Chr5g0430341, MtrunA17_Chr6g0457461, among which there are two candidates for mosaic translation). When we combine the information from Supplementary Datasets S4 and S28, we find that 15 non-RE transcripts can potentially be associated with multiple frameshifting in our study. This list can be extended with eight RE loci associated with multiple frameshifting: MtrunA17_Chr1R0250150 (CP23, CP24, and CP101), MtrunA17_Chr6R0039140, MtrunA17_Chr6R0039080, MtrunA17_Chr1R0250300, MtrunA17_Chr1R0014150 (CP24 and CP101), MtrunA17_Chr4R0035060 (CP35), MtrunA17_Chr4R0290860 (CP37), MtrunA17_Chr5R0130760 (CP87, CP88, and CP89). Unfortunately, none of these additional loci are as good candidates for mosaic translation as MtrunA17_Chr1g0200071 (protein-synthesizing GTPase) and MtrunA17_Chr6g0457461 (RuBisCo). Moreover, additional loci found for chimeric peptides primarily associated with MtrunA17_Chr1g0200071 and MtrunA17_Chr6g0457461 create the following ambiguity. It is possible that chimeric peptides originally deduced from these transcripts are expressed individually from different sources instead of being expressed from the same primary sources in the course of mosaic translation. Differential mutagenesis of corresponding loci, digestion of original MS samples with different proteases, and long-read proteomics are required to address this ambiguity.
