## Supplementary material for "Discovery of diverse chimeric peptides in a eukaryotic proteome sets the stage for the experimental proof of the mosaic translation hypothesis": Supplementary tables.pdf

**Supplementary Table S1**. Statistics on all ORFs 60 nt or longer grouped by RNA type

| Statistic | mRNA[1] | mRNA[2] | ncRNA | rRNA | tRNA |
|---|---|---|---|---|---|
| Number of transcripts | 44,624 | 44,624 | 5,657* | 62* | 974 |
| Median length of transcripts, nt | 1,280 | 1,280 | 413 | 120 | 75 |
| Number of ORFs[#] | 846,711 | 802,087 | 71,127 | 831 | 1,311 |
| ORFs per transcript | 19 | 18 | 13 | 13 | 1 |
| Combined length of ORFs, nt | 132,065,895 | 85,361,370 | 7,740,336 | 97,470 | 93,618 |
| Median length of ORFs, nt | 93 | 90 | 93 | 96 | 72 |
| Median length of refORFs, nt | 810 | - | - | - | - |

[1]Before the elimination of refORFs
[2]After the elimination of refORFs
*Blastn searches against the NCBI non-redundant database and nucleotide alignments with annotated rRNA transcripts (Homology Group 1 in this manuscript) revealed that at least two transcripts annotated as ncRNA in the current *Medicago truncatula* genome (1.9) are actually rRNA transcripts. These two transcripts, MtrunA17_Chr0c01g0489091 and MtrunA17_Chr5g0421761, are considered as rRNA throughout the manuscript except for supplementary tables, which are meant to show the “history” of their annotation status.
[#]There are 875,356 non-refORFs (altORFs) of the given length range in total.

**Supplementary Table S2**. Statistics on altProts with at least one hit in the global BLASTP analysis before the application of the filter

| Row | Statistic | mRNA | ncRNA | rRNA | tRNA |
|---|---|---|---|---|---|
| R1 | Number of transcripts | 44,624 | 5,657[#] | 62[#] | 974 |
| R2 | Number of altProts* | 802,087 | 71,127 | 831 | 1,311 |
| R3 | Number of altProts that have at least one hit | 13,427 | 4,398 | 392 | 426 |
| R4 | Percentage of altProts with at least one hit** | 1.67 | 6.18 | 47.17 | 32.49 |
| R5 | AltProts per transcript (R3/R1) | 0.30 | 0.78 | 6.32 | 0.44 |
| R6 | Transcripts per altProts (R1/R3) | 3.32 | 1.29 | 0.16 | 2.29 |
| R7 | Median length of altProts with hit(s), aa | 59.0 | 50.0 | 45.0 | 25.0 |
| R8 | Median length of alignment, aa | 44.0 | 39.0 | 40.0 | 24.0 |
| R9 | Median % identity | 78.0 | 88.5 | 93.3 | 100.0 |
| R10 | Median % query coverage | 82.4 | 84.6 | 94.4 | 100.0 |

[#]Blastn searches against the NCBI non-redundant database and nucleotide alignments with annotated rRNA transcripts (Homology Group 1 in this manuscript) revealed that at least two transcripts annotated as ncRNA in the current Medicago truncatula genome (1.9) are actually rRNA transcripts. These two transcripts, MtrunA17_Chr0c01g0489091 and MtrunA17_Chr5g0421761, are considered as rRNA throughout the manuscript except for supplementary tables, which are meant to show the “history” of their annotation status.
*All theoretically possible altProts regardless of BLASTP results
**Percentage of altProts that have at least one hit per all altProts in the group (R3/R2*100)

**Supplementary Table S3**. Numbers of altProts before and after the elimination of queries with less than 70% identity to annotated proteins

| Category | mRNA | ncRNA | rRNA | tRNA | Total |
|---|---|---|---|---|---|
| Total altProts with hits | 13,427 | 4,398 | 392 | 426 | 18,643 |
| Eliminated | 4,709 (35%) | 849 (19%) | 7 (2%) | 0 (0%) | 5,565 |
| Retained | 8,718 (65%) | 3,549 (81%) | 385 (98%) | 426 (100%) | 13,078 |

**Supplementary Table S4**. Distribution of transcripts in different categories based on RNA type, cellular localization, mass spectrometry (MS) support of altORFs, and conservation of altORF translation products

| Category | Total | mRNA | ncRNA | rRNA | tRNA | Nuclear transcripts | Mitochondrial transcripts | Chloroplast transcripts |
|---|---|---|---|---|---|---|---|---|
| Transcriptome | 51,317 | 44,624 | 5,657 | 62 | 974 | 50,994 | 193* | 130** |
| Transcripts with MS-supported altORFs | 797 | 712 | 85 | 0 | 0 | 785 | 6 | 6 |
| Transcripts with conserved altORFs (% identity ≥ 70) | 7,784 | 6,052 | 1,313 | 45 | 374 | 7,561 | 150 | 73 |
| Transcripts with conserved or MS-supported altORFs | 8,372 | 6,596 | 1,357 | 45 | 374 | 7,561 | 150 | 73 |
| Transcripts with conserved and MS-supported altORFs*** | 211 | 168 | 41 | 0 | 0 | 199 | 6 | 6 |
| Transcripts with modeled chimeric proteins | 5,502 | 5,036 | 392 | 22 | 52 | 5,325 | 119 | 58 |
| Transcripts with MS-supported chimeric proteins | 145 | 129 | 12 | 3 | 1 | 131 | 7 | 7 |

*73 mRNA and 120 ncRNA
**86 mRNA, 34 ncRNA, 1 rRNA, and 9 tRNA
***These transcripts are both conserved and have at least one MS-supported altORF.

**Supplementary Table S5**. Redundant counts of chimeric protein models, shown per altProt type, transcript type, and programmed ribosomal frameshifting (PRF) value

| PRF value | Modelled with MS-supported altProts | Modelled with conserved altProts | Sum | mRNA | ncRNA | rRNA | tRNA |
|---|---|---|---|---|---|---|---|
| +1 | 9,740 | 138,600 | 148,340 | 109,477 | 30,667 | 6,815 | 1,381 |
| +2 | 8,752 | 130,865 | 139,617 | 102,661 | 29,005 | 6,009 | 1,942 |
| -1 | 8,673 | 128,209 | 136,882 | 100,796 | 28,260 | 5,858 | 1,968 |
| -2 | 9,355 | 134,127 | 143,482 | 105,972 | 29,607 | 6,525 | 1,378 |
| Other (+1 to +10) # | 16 | 1,768 | 1,784 | 1,110 | 540 | 134 | 0 |
| Total | 36,536 | 533,569 | 570,105 | 420,016 | 118,079 | 25,341 | 6,669 |
| Total without other | 36,520 | 531,801 | 568,321 | 418,906 | 117,539 | 25,207 | 6,669 |

#Adjacent altORFs separated by 1-10 nucleotides that can potentially be joined by PRF events with values +1 to +10. Note that this category includes PRF values of +1 and +2 that are excluded from the four remaining categories.

**Supplementary Table S6**. Non-redundant counts of chimeric proteins modeled with altORFs that are MS-supported and conserved at the same time, shown per transcript type and PRF value

| PRF value | Modelled with MS-supported and conserved at the same time | mRNA [#] | ncRNA | rRNA | tRNA |
|---|---|---|---|---|---|
| +1 | 930 | 930 | 0 | 0 | 0 |
| +2 | 497 | 497 | 0 | 0 | 0 |
| -1 | 487 | 487 | 0 | 0 | 0 |
| -2 | 901 | 901 | 0 | 0 | 0 |
| Other (+1 to +10) [##] | 4 | 4 | 0 | 0 | 0 |
| Total | 2,819 | 2,819 | 0 | 0 | 0 |
| Total without other | 2,815 | 2,815 | 0 | 0 | 0 |

[#]All models in this table are restricted to mRNA because the definition of this category requires two overlapping or adjacent ORFs supported by MS. Since all canonical refORFs are assumed to be translated in mRNA by the genome annotation pipeline, there is a higher probability to find such models in the mRNA group.

[##]Adjacent altORFs separated by 1-10 nucleotides that can potentially be joined by PRF events with values +1 to +10. Note that this category includes PRF values of +1 and +2 that are excluded from the four remaining categories.

**Supplementary Table S7**. Non-redundant counts of chimeric proteins modeled with MS-supported altORFs, shown per proteomic study and PRF value

| PRF value | Modelled with MS-supported altProts (non-redundant) [#] | PXD002692 Marx et al., 2016 | PXD013606 Shin et al., 2021 | PXD022278 Castañeda et al., 2021 |
|---|---|---|---|---|
| +1 | 9,740 | 8,067 | 974 | 1,321 |
| +2 | 8,752 | 7,184 | 803 | 1,271 |
| -1 | 8,673 | 7,120 | 800 | 1,256 |
| -2 | 9,355 | 7,744 | 934 | 1,278 |
| Other (+1 to +10) [##] | 16 | 16 | 2 | 0 |
| Total | 36,536 | 30,131 | 3,513 | 5,126 |
| Total without other | 36,520 | 30,115 | 3,511 | 5,126 |

[#]All models in this table are restricted to mRNA because the definition of this category requires two overlapping or adjacent ORFs supported by MS. Since all canonical refORFs are assumed to be translated in mRNA by the genome annotation pipeline, there is a higher probability to find such models in the mRNA group.

[##]Adjacent altORFs separated by 1-10 nucleotides that can potentially be joined by PRF events with values +1 to +10. Note that this category includes PRF values of +1 and +2 that are excluded from the four remaining categories.

**Supplementary Table S8**. Non-redundant counts of chimeric proteins modeled with conserved altORFs, shown per transcript type and PRF value

| PRF value | mRNA | ncRNA | rRNA | tRNA |
|---|---|---|---|---|
| +1 | 90,089 | 17,889 | 4,574 | 815 |
| +2 | 84,660 | 17,491 | 4,140 | 1,064 |
| -1 | 83,100 | 17,087 | 4,046 | 1,090 |
| -2 | 87,360 | 17,288 | 4,384 | 818 |
| Other (+1 to +10) [#] | 765 | 297 | 76 | 0 |
| Total | 345,974 | 70,052 | 17,220 | 3,787 |
| Total without other | 345,209 | 69,755 | 17,144 | 3,787 |

[#]Adjacent altORFs separated by 1-10 nucleotides that can potentially be joined by PRF events with values +1 to +10. Note that this category includes PRF values of +1 and +2 that are excluded from the four remaining categories.

**Supplementary Table S9**. Non-redundant counts of chimeric protein models, shown per altProt type, transcript type, and PRF value

| PRF value | Modelled with MS-supported altProts | Modelled with conserved altProts [#] | Sum | mRNA | ncRNA | rRNA | tRNA |
|---|---|---|---|---|---|---|---|
| +1 | 9,740 | 113,367 | 122,177 | 98,899 | 17,889 | 4,574 | 815 |
| +2 | 8,752 | 107,355 | 115,610 | 92,915 | 17,491 | 4,140 | 1,064 |
| -1 | 8,673 | 105,323 | 113,509 | 91,286 | 17,087 | 4,046 | 1,090 |
| -2 | 9,355 | 109,850 | 118,304 | 95,814 | 17,288 | 4,384 | 818 |
| Other (+1 to +10) [##] | 16 | 1,138 | 1,150 | 777 | 297 | 76 | 0 |
| Total | 36,536 | 437,033 | 470,750 | 379,691 | 70,052 | 17,220 | 3,787 |
| Total without other | 36,520 | 435,895 | 469,600 | 378,914 | 69,755 | 17,144 | 3,787 |

[#] Counts of chimeric proteins modeled with altORFs that are MS-supported and conserved at the same time were subtracted from the “conserved” category. The redundancy due to altORF-altORF ambiguity (see the text in Section 4.3) concerns exclusively chimeric proteins modeled with conserved altORFs. This can be explained by the fact that the majority of PRF events that involve two altORFs instead of one altORF and one refORF are associated with non-mRNA transcript types, most of which have no MS-supported altORFs.

[##] Adjacent altORFs separated by 1-10 nucleotides that can potentially be joined by PRF events with values +1 to +10. Note that this category includes PRF values of +1 and +2 that are excluded from the four remaining categories.
